## Supplementary Tables for "Comparative developmental genomics of sex-biased gene expression in early embryogenesis across mammals"


| | |
|---|---|
| Supplementary Table S1. Transcriptomic datasets used across developmental time and sex | 2 |
| Supplementary Table S2. Mouse sex-differential expression results | 3 |
| Supplementary Table S3. Human sex-differential expression results | 52 |
| Supplementary Table S4. Mouse male GSEA of NMF clusters | 148 |
| Supplementary Table S5. Mouse female GSEA of NMF clusters | 283 |
| Supplementary Table S6. Human male GSEA of NMF clusters | 343 |
| Supplementary Table S7. Human female GSEA of NMF clusters | 426 |
| Supplementary Table S8. Mouse male NMF coefficient matrix | 557 |
| Supplementary Table S9. Mouse female NMF coefficient matrix | 787 |
| Supplementary Table S10. Human male NMF coefficient matrix | 1046 |
| Supplementary Table S11. Human female NMF coefficient matrix | 1500 |

**Supplementary Table S1.**
**Transcriptomic datasets used across developmental time and sex.**

| Mouse | E1.5 | E2 | E2.5 | E3 | E3.5 | E4 |
|---|---|---|---|---|---|---|
| Male | 3 | 2 | 6 | 4 | 12 | 8 |
| Female | 3 | 8 | 23 | 11 | 14 | 12 |

| Human | E3 | E4 | E5 | E6 | E7 |
|---|---|---|---|---|---|
| Male | 48 | 98 | 206 | 180 | 176 |
| Female | 33 | 92 | 171 | 235 | 290 |

### Supplementary Table S2

#### Mouse: differential expression results

| Gene | padj_E1.5 | log2fc_E1.5 | sex_E1.5 | padj_E2 | log2fc_E2 | sex_E2 | padj_E2.5 | log2fc_E2.5 | sex_E2.5 | padj_E3 | log2fc_E3 | sex_E3 | padj_E3.5 | log2fc_E3.5 | sex_E3.5 | padj_E4 | log2fc_E4 | sex_E4 |
|---|---|---|---|---|---|---|---|---|---|---|---|---|---|---|---|---|---|---|
| Fdxacb1 | | | | | | | | | | | | | | | | 0.00514 | -3.77291 | Male |
| Plekhh3 | | | | 0.00853 | -1.83615 | Male | | | | | | | | | | | | |
| Gm4737 | | | | | | | | | | 0.00493 | 2.27809 | Female | | | | | | |
| Ccdc15 | | | | | | | | | | 0.00000 | 1.43222 | Female | | | | | | |
| Sec61g | | | | | | | | | | 0.00000 | -5.41077 | Male | 0.00007 | 1.94508 | Female | | | |
| Usp5 | | | | | | | | | | 0.02467 | 1.69283 | Female | | | | | | |
| Mppe1 | | | | | | | | | | 0.00000 | 5.60231 | Female | | | | | | |
| Rwdd4a | | | | | | | | | | 0.00841 | -2.44859 | Male | | | | | | |
| Evi5l | | | | | | | | | | 0.02878 | -1.29386 | Male | | | | 0.02960 | -3.95567 | Male |
| Rpia | | | | | | | | | | 0.00000 | -5.30180 | Male | 0.00344 | 2.17864 | Female | | | |
| Sh2d5 | | | | | | | | | | 0.00000 | -5.19746 | Male | | | | | | |
| Xlr5a | | | | | | | | | | | | | 0.00222 | -3.50321 | Male | | | |
| Snhg6 | | | | | | | | | | 0.00008 | -3.54764 | Male | | | | | | |
| Lhfp | | | | | | | | | | 0.00224 | -1.91512 | Male | | | | | | |
| Mmrn2 | | | | 0.00004 | 0.93124 | Female | | | | | | | | | | | | |
| Slc10a6 | | | | | | | | | | | | | | | | 0.00001 | -6.05510 | Male |
| Rps19bp1 | | | | | | | | | | 0.00541 | -3.19644 | Male | | | | | | |
| Timm8b | | | | | | | | | | 0.00001 | -3.82171 | Male | 0.00948 | 1.86593 | Female | | | |
| Cdk14 | | | | | | | 0.00511 | 3.67960 | Female | | | | | | | | | |
| Gfpt2 | | | | | | | | | | | | | 0.00311 | 5.33344 | Female | | | |
| Efhb | | | | | | | | | | | | | 0.01147 | -3.44539 | Male | 0.00108 | 4.36913 | Female |
| Cndp1 | | | | | | | | | | 0.00285 | -1.29659 | Male | | | | 0.00000 | -1.25494 | Male |
| Eef2k | | | | | | | | | | | | | | | | 0.02610 | 3.59078 | Female |
| Alox8 | | | | | | | | | | 0.00386 | -2.52890 | Male | | | | | | |
| D2Wsu81e | | | | | | | | | | 0.04462 | 2.22777 | Female | | | | | | |
| Pramel5 | | | | | | | | | | | | | | | | 0.00332 | -1.53824 | Male |
| Rpl3 | | | | | | | | | | 0.02882 | 0.93406 | Female | | | | | | |
| St3gal2 | | | | | | | | | | | | | 0.00000 | 6.17789 | Female | | | |
| Sall1 | | | | 0.00004 | 4.16055 | Female | | | | | | | | | | | | |
| Ccng1 | | | | 0.02618 | 1.59409 | Female | | | | | | | | | | | | |
| Ube2e1 | | | | | | | | | | 0.00000 | 5.56333 | Female | | | | | | |
| Slc38a9 | | | | | | | | | | 0.00000 | 3.67057 | Female | | | | | | |
| Steap1 | | | | | | | | | | | | | 0.04457 | 1.88863 | Female | | | |
| Tmem108 | | | | | | | 0.00000 | 1.01149 | Female | | | | | | | | | |
| Atp6v0e2 | | | | 0.01890 | 0.98399 | Female | 0.00000 | -2.67621 | Male | | | | | | | | | |
| Pfn1 | | | | | | | | | | 0.00000 | 4.53822 | Female | | | | | | |
| Mbd6 | | | | | | | | | | | | | 0.04102 | 2.62668 | Female | | | |
| Slc35b3 | | | | | | | | | | 0.04958 | 2.59948 | Female | | | | | | |
| Rac1 | | | | | | | | | | 0.02251 | -1.30285 | Male | | | | | | |
| Galns | | | | | | | | | | | | | 0.01488 | 4.34897 | Female | | | |
| Khdrbs3 | 0.00000 | -2.56800 | Male | 0.00000 | 1.94152 | Female | | | | 0.00451 | -1.42879 | Male | 0.00000 | 1.57259 | Female | 0.00000 | -1.19161 | Male |
| Dnajc6 | 0.00007 | -2.23308 | Male | | | | | | | | | | 0.00225 | -5.39671 | Male | | | |
| Vipas39 | | | | | | | | | | 0.02105 | 2.69127 | Female | | | | | | |
| Nat8f1 | | | | | | | 0.00000 | 1.00208 | Female | 0.00013 | -1.16987 | Male | | | | | | |
| Tagln | | | | | | | | | | | | | | | | 0.00293 | -4.82329 | Male |
| Trmt12 | | | | | | | | | | | | | | | | 0.01925 | -3.26180 | Male |
| Gm12338 | | | | | | | | | | 0.00000 | -4.90393 | Male | 0.00106 | 1.71279 | Female | | | |
| Kat2b | | | | | | | | | | | | | | | | 0.02986 | 3.01617 | Female |
| Ifitm3 | | | | 0.00000 | 0.88794 | Female | | | | | | | | | | | | |
| Btg3 | | | | | | | 0.00969 | -3.97680 | Male | | | | | | | | | |
| Lrrc2 | | | | | | | | | | 0.04422 | -2.40942 | Male | | | | | | |
| Plag1 | | | | | | | | | | | | | | | | 0.03377 | -4.49298 | Male |
| Wdr45 | | | | | | | | | | 0.04923 | 1.48385 | Female | | | | | | |
| Poldip3 | | | | | | | | | | | | | 0.01233 | 1.01073 | Female | | | |
| Anxa5 | | | | | | | | | | 0.00000 | 1.34952 | Female | | | | | | |
| Zfp644 | | | | | | | | | | 0.01843 | -3.94087 | Male | 0.00000 | 5.23736 | Female | | | |
| Cep41 | | | | | | | | | | 0.00450 | 3.55418 | Female | | | | | | |
| Efcab5 | | | | | | | | | | | | | | | | 0.01058 | 4.00984 | Female |
| Usp26 | | | | | | | | | | 0.03259 | -1.00850 | Male | | | | | | |
| Naa25 | | | | | | | | | | 0.00152 | 2.45727 | Female | | | | | | |
| Ppp1r16b | | | | | | | | | | 0.00000 | -4.26653 | Male | 0.00025 | 2.00724 | Female | | | |
| Fbxw19 | | | | | | | | | | | | | 0.00007 | 4.89803 | Female | | | |
| Mfsd6 | | | | | | | | | | 0.00011 | -2.12729 | Male | | | | | | |
| Ythdf2 | | | | | | | | | | 0.00055 | -2.36489 | Male | 0.00364 | 1.51426 | Female | | | |
| Eif2s3y | 0.00028 | -6.14121 | Male | 0.00000 | -8.34606 | Male | 0.00000 | -8.11657 | Male | 0.00000 | -9.35471 | Male | 0.00000 | -9.08826 | Male | 0.00000 | -8.22968 | Male |
| Obox2 | | | | | | | | | | 0.02092 | -4.57991 | Male | | | | | | |
| Tnfsf13b | | | | | | | | | | | | | | | | 0.02309 | -2.15461 | Male |
| Icam4 | | | | 0.00000 | -2.61720 | Male | | | | | | | | | | | | |

| | | | | | | | | | | | | | | | | | | |
|---|---|---|---|---|---|---|---|---|---|---|---|---|---|---|---|---|---|---|
| Plk1 | | | | | | | | | | 0.02164 | 2.98153 | Female | | | | | | |
| Dnajb4 | | | | | | | | | | 0.00000 | -3.05312 | Male | | | | | | |
| Gm16157 | | | | | | | | | | | | | | | | 0.02396 | -4.83569 | Male |
| Srgn | | | | | | | | | | | | | | | | 0.01420 | 3.96057 | Female |
| Alg14 | | | | 0.00168 | 3.94545 | Female | | | | | | | | | | | | |
| Pitx1 | 0.00003 | 0.74747 | Female | | | | | | | | | | | | | | | |
| Wnt3a | | | | | | | 0.00000 | -5.08947 | Male | 0.00000 | 3.62858 | Female | | | | | | |
| Rpl38 | | | | | | | | | | 0.00000 | -5.52442 | Male | 0.00395 | 1.46699 | Female | | | |
| Acta1 | | | | | | | | | | 0.00000 | 6.16149 | Female | | | | | | |
| Gm10865 | | | | | | | | | | | | | | | | 0.01632 | -1.70022 | Male |
| Firre | | | | | | | 0.03472 | 3.49807 | Female | | | | | | | | | |
| Cox7b | | | | | | | 0.04109 | 1.46185 | Female | 0.00000 | -4.06131 | Male | 0.00007 | 1.89555 | Female | | | |
| Fgd4 | | | | | | | 0.00015 | 4.79156 | Female | | | | | | | 0.00115 | -4.14248 | Male |
| Krt7 | 0.00015 | 0.61925 | Female | 0.00072 | -1.64017 | Male | | | | | | | | | | | | |
| Gcm1 | | | | | | | | | | 0.00001 | -0.91730 | Male | | | | 0.00000 | -1.68549 | Male |
| Lrriq4 | | | | | | | | | | | | | 0.00003 | 1.01842 | Female | 0.00001 | 4.54247 | Female |
| Prr19 | 0.00193 | 1.25707 | Female | | | | | | | | | | | | | | | |
| Aarsd1 | | | | | | | | | | 0.04098 | 2.34308 | Female | | | | | | |
| Khdc3 | | | | | | | | | | 0.00002 | -4.29517 | Male | | | | | | |
| Bbs10 | 0.00025 | 1.05675 | Female | | | | 0.00000 | 1.78572 | Female | | | | 0.03302 | 3.07102 | Female | | | |
| Prss45 | | | | | | | | | | | | | 0.00000 | 3.25400 | Female | | | |
| Gm4262 | | | | | | | | | | 0.00866 | -1.88331 | Male | 0.02393 | 1.28021 | Female | | | |
| Rnf32 | 0.00000 | -1.11682 | Male | | | | | | | 0.00002 | -2.24375 | Male | | | | | | |
| Tenm3 | | | | | | | | | | | | | 0.00000 | -2.72038 | Male | | | |
| Ccdc177 | | | | | | | | | | 0.02243 | -1.24219 | Male | | | | | | |
| Rpa3 | | | | | | | | | | 0.01408 | -2.31230 | Male | | | | | | |
| Slc5a5 | | | | | | | | | | | | | | | | 0.00919 | 4.14089 | Female |
| C87436 | | | | | | | | | | | | | | | | 0.00380 | 3.07794 | Female |
| Gjb3 | | | | 0.00211 | 2.96843 | Female | | | | | | | | | | | | |
| Mtl5 | | | | | | | | | | 0.01679 | 2.87412 | Female | | | | 0.00265 | 0.64390 | Female |
| Pnp2 | | | | | | | | | | | | | | | | 0.01368 | -1.72884 | Male |
| Hdac2 | | | | | | | 0.04953 | 1.50779 | Female | | | | | | | | | |
| Phlda3 | | | | 0.00000 | -0.60011 | Male | | | | | | | | | | | | |
| Adamts10 | | | | 0.00212 | -1.47847 | Male | | | | | | | | | | | | |
| Atp8b1 | | | | | | | 0.00756 | 4.21216 | Female | | | | | | | | | |
| Slc25a5 | | | | | | | 0.01222 | 1.15322 | Female | | | | | | | | | |
| Ltf | | | | | | | | | | 0.00013 | 1.65296 | Female | | | | | | |
| Gm7325 | 0.00002 | 1.99378 | Female | | | | 0.00000 | -0.98662 | Male | | | | | | | | | |
| Prtg | | | | 0.00000 | -1.11132 | Male | 0.00000 | 2.44361 | Female | 0.00129 | -1.58329 | Male | | | | | | |
| Mogat2 | | | | 0.00886 | -2.09344 | Male | | | | | | | | | | 0.02170 | 4.78222 | Female |
| Msc | | | | | | | 0.01139 | 3.69329 | Female | | | | | | | | | |
| Peli2 | | | | | | | 0.00000 | -1.09913 | Male | 0.00303 | -1.52452 | Male | | | | | | |
| Apol6 | | | | | | | | | | 0.00133 | -3.61898 | Male | | | | | | |
| Zfp458 | | | | | | | | | | | | | | | | 0.00000 | 3.72621 | Female |
| Rab17 | | | | | | | 0.04109 | 2.06799 | Female | | | | | | | | | |
| Pih1d2 | | | | 0.00000 | 2.04613 | Female | | | | | | | | | | | | |
| Nab1 | | | | | | | | | | 0.00000 | -4.46070 | Male | | | | | | |
| LOC102634812 | | | | | | | 0.00004 | 2.28771 | Female | 0.00000 | -5.60601 | Male | 0.00000 | 2.52966 | Female | | | |
| Baz2b | | | | 0.02884 | 3.54772 | Female | | | | | | | | | | | | |
| Glrx2 | | | | | | | | | | 0.00001 | -3.52869 | Male | | | | | | |
| Hal | | | | 0.00000 | -2.60856 | Male | | | | 0.00000 | 0.58535 | Female | | | | | | |
| Cdk16 | | | | | | | | | | 0.00157 | 1.98727 | Female | | | | | | |
| Atp6v1e1 | | | | | | | | | | 0.00816 | 1.49817 | Female | | | | | | |
| Dbi | | | | | | | | | | 0.00006 | -3.83307 | Male | | | | | | |
| Snora69 | | | | | | | | | | | | | 0.02348 | 4.09287 | Female | | | |
| Vmn2r.ps129 | | | | | | | | | | 0.00000 | -5.56052 | Male | 0.00001 | 2.46528 | Female | | | |
| BC065397 | | | | | | | | | | | | | | | | 0.03087 | 3.83885 | Female |
| Hand1 | | | | | | | | | | | | | | | | 0.00657 | 3.09911 | Female |
| Shroom4 | | | | | | | | | | | | | 0.00085 | 4.98337 | Female | | | |
| Herpud1 | | | | 0.00316 | -2.17867 | Male | | | | 0.00000 | 3.21441 | Female | | | | | | |
| Smim24 | | | | | | | | | | | | | | | | 0.00000 | 1.59549 | Female |
| Csl | | | | | | | | | | 0.00341 | 1.73232 | Female | | | | | | |
| Src | | | | | | | | | | 0.00000 | 1.40145 | Female | | | | | | |
| Kbtbd13 | | | | 0.00001 | -0.67215 | Male | | | | | | | | | | | | |
| Ergic2 | | | | | | | | | | 0.02456 | 1.71453 | Female | | | | | | |
| Flt3l | | | | | | | | | | | | | | | | 0.00053 | -4.87323 | Male |
| Foxa1 | | | | | | | | | | | | | | | | 0.00000 | 4.63820 | Female |
| Tmem173 | | | | 0.00002 | 3.19806 | Female | | | | 0.00685 | -1.20603 | Male | | | | | | |
| Nmrk1 | | | | | | | | | | 0.00011 | -3.20294 | Male | | | | | | |
| Klf17 | | | | | | | | | | | | | | | | 0.00003 | -4.68710 | Male |

| | | | | | | | | | | | | | | | | | | |
|---|---|---|---|---|---|---|---|---|---|---|---|---|---|---|---|---|---|---|
| Pef1 | | | | | | | | | | 0.01118 | 1.98500 | Female | | | | | | |
| Emx2os | | | | | | | 0.01924 | -3.60325 | Male | | | | | | | | | |
| Dut | | | | | | | | | | 0.00000 | -3.82686 | Male | | | | | | |
| Med9 | | | | | | | | | | 0.01672 | -2.75621 | Male | | | | | | |
| Retsat | | | | 0.02785 | 1.91303 | Female | | | | | | | | | | | | |
| Tfe3 | | | | 0.00157 | 3.87909 | Female | | | | | | | | | | | | |
| Ano10 | | | | | | | | | | 0.00005 | -2.28747 | Male | | | | | | |
| Foxc1 | | | | 0.01121 | -2.06925 | Male | | | | | | | | | | | | |
| Mocs2 | | | | | | | 0.01325 | 4.05283 | Female | | | | | | | | | |
| Prrt3 | | | | | | | | | | | | | 0.00000 | -1.51487 | Male | | | |
| Ero1lb | | | | | | | | | | | | | | | | 0.00401 | -4.96907 | Male |
| Cdh23 | | | | | | | | | | | | | | | | 0.00000 | -1.84145 | Male |
| Kng2 | | | | | | | | | | | | | | | | 0.00207 | -3.50856 | Male |
| Llgl2 | | | | | | | | | | 0.00853 | 3.82171 | Female | | | | | | |
| Gm5148 | | | | | | | | | | 0.00000 | -3.67239 | Male | 0.00156 | 1.51817 | Female | | | |
| Thnsl1 | | | | | | | | | | 0.00000 | -0.73443 | Male | 0.00000 | 5.31457 | Female | | | |
| Eef1a1 | | | | | | | | | | 0.00248 | 1.05993 | Female | | | | | | |
| Gm19434 | | | | | | | | | | 0.00956 | -2.23278 | Male | | | | | | |
| Midn | | | | | | | | | | 0.04425 | -2.97189 | Male | | | | | | |
| Chrna10 | | | | | | | | | | | | | | | | 0.02986 | -4.01453 | Male |
| Psmg1 | | | | | | | | | | 0.02776 | 1.86284 | Female | | | | | | |
| Smtnl2 | | | | | | | | | | 0.00000 | -3.76921 | Male | 0.04102 | 1.34324 | Female | | | |
| Ankrd24 | | | | | | | 0.00000 | 1.91613 | Female | | | | | | | | | |
| Slc19a2 | | | | 0.00000 | 2.12996 | Female | | | | | | | | | | | | |
| Cast | | | | | | | | | | | | | | | | 0.00076 | -2.81264 | Male |
| Cd300lg | | | | 0.00000 | 2.50401 | Female | | | | | | | | | | 0.00000 | 0.94834 | Female |
| C1rl | | | | | | | 0.00834 | -4.01326 | Male | | | | | | | | | |
| Ninl | | | | | | | | | | | | | | | | 0.00000 | 5.53655 | Female |
| Hacd1 | | | | 0.03245 | -1.18456 | Male | | | | | | | | | | | | |
| Atp12a | | | | | | | 0.00000 | -1.98485 | Male | | | | | | | | | |
| AA619741 | | | | | | | | | | 0.02291 | -4.94562 | Male | | | | | | |
| Arid5b | | | | | | | | | | 0.00000 | 1.80734 | Female | | | | | | |
| Pdgfra | | | | | | | | | | 0.02682 | 3.28070 | Female | | | | | | |
| Miox | | | | | | | | | | | | | 0.04033 | -3.49798 | Male | | | |
| Hk1 | | | | | | | | | | | | | | | | 0.00958 | 3.02725 | Female |
| Chil5 | | | | 0.00851 | 0.79913 | Female | | | | | | | | | | | | |
| Plek | | | | | | | | | | 0.00090 | -4.78016 | Male | | | | | | |
| Ctnna1 | | | | | | | | | | 0.04871 | 1.09821 | Female | | | | | | |
| Nfatc1 | | | | | | | | | | | | | 0.00000 | 4.68039 | Female | | | |
| Cbfa2t2 | | | | | | | | | | | | | | | | 0.04412 | 1.64376 | Female |
| Klhdc3 | | | | | | | | | | 0.00000 | 4.81172 | Female | | | | | | |
| Nfatc2ip | | | | | | | | | | 0.00010 | -2.60777 | Male | | | | | | |
| Capn6 | | | | | | | | | | 0.00000 | 1.43578 | Female | | | | | | |
| Gsdmc4 | | | | | | | 0.02820 | 3.88482 | Female | | | | | | | 0.00054 | 5.09059 | Female |
| Tmem206 | | | | | | | | | | 0.00000 | -4.37510 | Male | | | | | | |
| Zfp316 | | | | | | | | | | 0.00000 | -2.88733 | Male | | | | | | |
| Rogdi | | | | | | | | | | 0.02933 | 2.04793 | Female | | | | | | |
| Rad23b | | | | | | | | | | 0.00553 | -1.39290 | Male | | | | | | |
| Ccdc58 | | | | | | | | | | 0.01973 | -1.43701 | Male | | | | | | |
| Jund | | | | | | | | | | 0.00059 | -4.45970 | Male | | | | | | |
| Il34 | | | | | | | | | | | | | 0.00083 | 5.57799 | Female | | | |
| Purg | | | | | | | 0.00000 | 0.70268 | Female | | | | | | | | | |
| Cntln | | | | | | | | | | | | | 0.01651 | 4.26157 | Female | | | |
| Mpdu1 | | | | | | | | | | 0.01369 | 1.95603 | Female | | | | | | |
| Magee1 | | | | | | | | | | | | | 0.00000 | 1.87103 | Female | | | |
| Gm10046 | | | | | | | | | | 0.00932 | -0.96461 | Male | | | | | | |
| Pip4k2c | | | | | | | | | | 0.02224 | 1.98694 | Female | | | | | | |
| Lta | | | | | | | | | | 0.01431 | -1.16353 | Male | | | | 0.00000 | -0.74065 | Male |
| Ppp1r14b | | | | | | | | | | 0.04535 | -2.21667 | Male | | | | | | |
| Fsd1l | | | | | | | 0.01448 | 3.38186 | Female | | | | | | | | | |
| Uhmk1 | | | | | | | | | | | | | | | | 0.00243 | 3.82479 | Female |
| Frk | | | | | | | | | | | | | 0.00000 | 6.34356 | Female | | | |
| Cdc20 | | | | | | | | | | 0.00570 | 1.85891 | Female | | | | | | |
| Maoa | | | | | | | | | | | | | 0.00237 | 3.31403 | Female | | | |
| Neu3 | | | | | | | | | | 0.00000 | -4.24098 | Male | | | | | | |
| Cpsf4 | | | | | | | | | | 0.00059 | 3.52305 | Female | | | | | | |
| Slc10a1 | | | | | | | | | | 0.02491 | -3.97601 | Male | | | | | | |
| Zbtb37 | | | | | | | | | | | | | | | | 0.00000 | 1.14491 | Female |
| Lhfpl5 | 0.00000 | -1.39584 | Male | | | | | | | | | | | | | | | |
| Slc44a3 | | | | | | | | | | | | | 0.02446 | 4.00732 | Female | | | |

| | | | | | | | | | | | | | | | | | | |
|---|---|---|---|---|---|---|---|---|---|---|---|---|---|---|---|---|---|---|
| Epb41l3 | | | | 0.00000 | 2.42946 | Female | | | | | | | | | | 0.00000 | 3.59365 | Female |
| Smim8 | | | | 0.01178 | 1.86348 | Female | | | | | | | | | | | | |
| Omd | | | | | | | 0.04281 | 1.80604 | Female | 0.00000 | -6.39173 | Male | 0.00536 | 1.83930 | Female | | | |
| Gm2837 | | | | | | | | | | | | | | | | 0.02200 | 3.81967 | Female |
| Dpm3 | | | | | | | | | | 0.02644 | -2.69261 | Male | | | | | | |
| Ecel1 | | | | | | | 0.00022 | -0.63435 | Male | | | | | | | | | |
| Gm20337 | | | | | | | 0.02585 | 2.12741 | Female | 0.00000 | -4.27736 | Male | | | | 0.01430 | -2.13152 | Male |
| Rpl32l | | | | | | | | | | 0.00000 | -4.29586 | Male | 0.00025 | 1.51050 | Female | | | |
| Nt5e | | | | | | | 0.00001 | 1.63449 | Female | | | | 0.00000 | 1.87843 | Female | | | |
| Kpna6 | | | | | | | | | | 0.00121 | 2.05206 | Female | | | | | | |
| Gm10548 | | | | | | | | | | 0.00464 | -2.71119 | Male | | | | | | |
| Gapt | | | | | | | | | | | | | | | | 0.00000 | -0.98692 | Male |
| Tcam1 | | | | | | | | | | | | | | | | 0.00014 | 4.63207 | Female |
| Tcstv3 | | | | | | | | | | | | | | | | 0.00000 | -6.51847 | Male |
| Nek9 | | | | 0.03790 | 2.94394 | Female | | | | | | | | | | | | |
| Cracr2a | | | | | | | | | | | | | | | | 0.00000 | -2.23258 | Male |
| Mysm1 | | | | | | | | | | | | | | | | 0.01683 | -1.52448 | Male |
| Ebpl | | | | | | | | | | 0.02925 | -3.98087 | Male | | | | | | |
| Brd4 | | | | | | | | | | 0.02764 | 1.67126 | Female | | | | | | |
| Rad1 | | | | | | | | | | | | | 0.00206 | 1.23502 | Female | | | |
| Ppl | | | | 0.00890 | -1.87736 | Male | | | | | | | | | | | | |
| Inadl | | | | | | | | | | 0.04777 | 3.12091 | Female | | | | | | |
| Urb1 | | | | | | | | | | | | | | | | 0.02592 | 2.07015 | Female |
| Dnajc11 | | | | | | | | | | 0.02045 | 1.69288 | Female | | | | | | |
| Tomm7 | | | | | | | | | | 0.00000 | -5.14391 | Male | 0.00195 | 1.45370 | Female | | | |
| Shcbp1l | | | | 0.00000 | 1.03556 | Female | | | | | | | | | | 0.00000 | -4.75315 | Male |
| Tmem30b | | | | | | | | | | | | | | | | 0.00182 | -5.15733 | Male |
| Gm17801 | | | | | | | | | | 0.00000 | -5.76689 | Male | | | | | | |
| Gm13078 | | | | | | | | | | 0.00000 | 3.74294 | Female | | | | 0.00927 | -1.82513 | Male |
| Manea | | | | | | | | | | | | | | | | 0.01310 | 3.53146 | Female |
| Syt12 | | | | | | | | | | 0.00003 | -3.39901 | Male | | | | | | |
| Rabgap1l | | | | 0.00000 | 1.02126 | Female | | | | 0.00002 | -2.66663 | Male | | | | 0.00000 | 2.17764 | Female |
| Sepp1 | | | | 0.00000 | 2.12798 | Female | | | | | | | | | | 0.00000 | 3.74677 | Female |
| Rasip1 | | | | 0.01003 | -1.53147 | Male | | | | 0.00000 | 2.63768 | Female | | | | | | |
| Nr3c1 | | | | | | | | | | | | | | | | 0.01925 | 4.56282 | Female |
| Rpgrip1l | | | | | | | | | | | | | | | | 0.00898 | 3.83739 | Female |
| Gzmm | | | | | | | | | | | | | 0.00001 | 5.57221 | Female | | | |
| Nsmce4a | | | | | | | | | | 0.00460 | -2.14869 | Male | | | | | | |
| Apod | | | | | | | | | | 0.04425 | -3.59747 | Male | | | | | | |
| Srek1 | | | | | | | | | | | | | | | | 0.00814 | 2.12831 | Female |
| Svbp | | | | | | | | | | | | | 0.04821 | 3.48128 | Female | | | |
| H2.T3 | | | | | | | | | | 0.00460 | -3.55126 | Male | | | | 0.00010 | -3.61662 | Male |
| Gm13498 | | | | | | | | | | 0.00000 | 3.96194 | Female | | | | | | |
| Asap2 | | | | | | | | | | | | | | | | 0.01490 | 2.82365 | Female |
| Grwd1 | | | | | | | | | | 0.01567 | 1.83976 | Female | | | | | | |
| Rrm1 | | | | | | | | | | 0.00556 | 1.27071 | Female | | | | | | |
| Cyp2b10 | | | | 0.00166 | 0.92526 | Female | | | | | | | | | | | | |
| Fkbp2 | | | | | | | | | | 0.00598 | -2.54158 | Male | | | | | | |
| Uqcrh | | | | | | | | | | 0.00000 | -4.46858 | Male | 0.00017 | 1.76873 | Female | | | |
| Krt19 | | | | | | | | | | 0.00001 | -3.15208 | Male | | | | | | |
| Gm11961 | | | | | | | | | | 0.00000 | -3.43855 | Male | | | | | | |
| Zfp937 | | | | | | | | | | | | | | | | 0.02476 | 2.00005 | Female |
| Dppa3 | | | | | | | | | | 0.01938 | -0.91937 | Male | | | | | | |
| Vasn | | | | | | | | | | | | | | | | 0.00000 | 3.26973 | Female |
| Cldn12 | | | | | | | | | | | | | | | | 0.03018 | -3.02657 | Male |
| Ptpn9 | | | | | | | | | | | | | | | | 0.00834 | 3.40619 | Female |
| Gm5113 | | | | | | | 0.01406 | 2.44254 | Female | | | | | | | | | |
| Mina | | | | | | | | | | 0.00027 | 3.90902 | Female | | | | | | |
| Serpinb6b | | | | | | | | | | | | | 0.04410 | 2.53972 | Female | | | |
| Acp7 | | | | | | | | | | | | | | | | 0.00000 | 2.98426 | Female |
| Myl6b | 0.02308 | 4.64816 | Female | | | | | | | | | | | | | | | |
| Ddx18 | | | | | | | | | | 0.00402 | 1.61139 | Female | | | | | | |
| Aph1c | | | | | | | | | | | | | 0.02267 | 3.53359 | Female | | | |
| Ier5 | | | | | | | | | | | | | 0.00000 | -0.58678 | Male | | | |
| Dennd5b | | | | | | | 0.00866 | 2.20824 | Female | 0.04371 | -2.00823 | Male | | | | | | |
| Fmo5 | | | | | | | | | | 0.00807 | -4.98537 | Male | | | | | | |
| Pdap1 | | | | | | | | | | 0.00000 | -3.73668 | Male | | | | | | |
| Pik3cb | | | | | | | | | | 0.04422 | -2.71645 | Male | | | | | | |
| Rps12 | | | | | | | | | | 0.00000 | -3.65519 | Male | 0.03290 | 1.26914 | Female | | | |
| Ndufa13 | | | | | | | | | | 0.00015 | -3.17128 | Male | 0.02665 | 1.59217 | Female | | | |

| | | | | | | | | | | | | | | | | | | |
|---|---|---|---|---|---|---|---|---|---|---|---|---|---|---|---|---|---|---|
| Casd1 | | | | 0.00545 | -0.69772 | Male | 0.00000 | 2.00729 | Female | | | | | | | 0.04721 | 3.99700 | Female |
| Mog | | | | | | | | | | | | | 0.00000 | 2.67401 | Female | | | |
| Ccdc7b | | | | | | | | | | 0.01661 | -5.05499 | Male | | | | | | |
| Mrps5 | 0.00005 | 5.45015 | Female | | | | | | | | | | | | | | | |
| Hapln4 | 0.00000 | -0.81156 | Male | | | | | | | | | | | | | | | |
| Tmem198 | | | | 0.02027 | -1.46130 | Male | | | | | | | | | | | | |
| Rel | | | | | | | 0.00000 | 2.75289 | Female | 0.00003 | 1.01418 | Female | | | | | | |
| Pld2 | | | | | | | | | | | | | 0.04642 | 2.65417 | Female | | | |
| Rbl2 | | | | | | | | | | 0.00000 | 2.65393 | Female | | | | | | |
| Coq3 | | | | | | | | | | | | | | | | 0.01639 | -1.66709 | Male |
| Zcchc17 | | | | | | | | | | 0.00333 | 2.97078 | Female | | | | | | |
| Ryr3 | | | | | | | | | | | | | 0.03811 | 4.69088 | Female | | | |
| Chac2 | | | | 0.00284 | 1.48131 | Female | | | | | | | | | | | | |
| Lysmd3 | | | | 0.00000 | 1.53543 | Female | | | | | | | | | | 0.00426 | -5.13692 | Male |
| Cox4i2 | | | | | | | | | | | | | 0.01621 | -3.35622 | Male | | | |
| Fam129a | | | | 0.00903 | 1.14211 | Female | | | | | | | | | | | | |
| Gm7534 | | | | | | | | | | | | | | | | 0.00000 | -0.98676 | Male |
| Zfp184 | | | | | | | | | | 0.01186 | 3.20760 | Female | | | | | | |
| Trim7 | | | | | | | | | | 0.00000 | 9.61748 | Female | | | | | | |
| Tmem45b | | | | | | | | | | 0.00051 | 4.57552 | Female | | | | | | |
| Drosha | | | | | | | | | | | | | | | | 0.03851 | -1.71703 | Male |
| Nudt16 | | | | | | | 0.02274 | -2.42447 | Male | | | | | | | | | |
| Bloc1s1 | 0.01889 | 3.80662 | Female | | | | | | | 0.00002 | -4.20511 | Male | | | | | | |
| Gpr157 | | | | | | | | | | 0.00341 | -1.85650 | Male | | | | | | |
| Gm5 | | | | | | | | | | 0.01588 | -3.77921 | Male | | | | | | |
| Lrrc31 | | | | | | | | | | 0.00000 | -4.45198 | Male | | | | | | |
| Fam69b | | | | 0.00021 | 1.34843 | Female | | | | | | | | | | | | |
| Dhps | | | | | | | 0.04338 | 1.85432 | Female | 0.02540 | -2.14596 | Male | | | | | | |
| Gm5039 | | | | | | | | | | 0.00525 | 2.51154 | Female | | | | | | |
| Romo1 | | | | | | | | | | 0.00107 | -3.25567 | Male | | | | | | |
| Tapbpl | | | | | | | | | | 0.00000 | -5.69813 | Male | 0.00051 | 2.17742 | Female | | | |
| Cnksr1 | | | | 0.00000 | 0.85412 | Female | | | | | | | 0.01272 | 4.12640 | Female | | | |
| Cat | | | | | | | 0.01182 | -3.36692 | Male | | | | | | | | | |
| Insig2 | | | | | | | | | | | | | | | | 0.01368 | 4.14387 | Female |
| Npr3 | 0.02417 | 0.91612 | Female | | | | | | | | | | | | | 0.00000 | 2.39946 | Female |
| AA792892 | | | | | | | | | | | | | | | | 0.04222 | -1.31952 | Male |
| Slc18b1 | | | | | | | | | | | | | | | | 0.00869 | 3.36203 | Female |
| Slc17a8 | | | | | | | | | | 0.00059 | -3.24359 | Male | | | | | | |
| H2.T24 | | | | | | | | | | 0.00000 | -4.82018 | Male | 0.03762 | 1.56804 | Female | | | |
| P3h3 | | | | | | | | | | 0.00000 | -6.14815 | Male | 0.01837 | 1.65264 | Female | | | |
| Ptbp1 | | | | | | | | | | 0.00668 | 1.36793 | Female | | | | | | |
| Anxa11 | | | | | | | | | | 0.04498 | -2.03979 | Male | | | | | | |
| Pag1 | | | | | | | | | | | | | | | | 0.03241 | 3.50257 | Female |
| Lbh | | | | 0.00000 | 1.34646 | Female | | | | | | | | | | 0.00000 | 5.75167 | Female |
| Usp29 | | | | | | | | | | 0.02265 | -3.18995 | Male | | | | | | |
| Prelid2 | | | | | | | | | | | | | | | | 0.00078 | -1.99170 | Male |
| Trappc1 | | | | 0.00075 | 3.98929 | Female | | | | | | | | | | | | |
| Zfp606 | | | | 0.00022 | -0.72603 | Male | | | | 0.00000 | 2.16074 | Female | | | | | | |
| Acyp1 | | | | | | | | | | 0.01183 | -3.23495 | Male | | | | 0.00372 | 3.05656 | Female |
| Fcho2 | | | | | | | | | | | | | 0.00082 | -2.81176 | Male | | | |
| Ptpn4 | | | | | | | | | | | | | | | | 0.01161 | 4.12950 | Female |
| Mmp17 | 0.00267 | 1.32744 | Female | | | | | | | | | | | | | 0.00000 | 2.84252 | Female |
| Pcbp2 | | | | | | | | | | 0.00724 | 1.76590 | Female | | | | | | |
| Pla2g12b | | | | | | | | | | | | | 0.00000 | 3.11887 | Female | | | |
| Rsrc2 | | | | | | | | | | | | | | | | 0.02694 | -1.11011 | Male |
| Psmc5 | | | | | | | | | | 0.01976 | 1.14517 | Female | | | | | | |
| Slc2a8 | | | | | | | | | | | | | | | | 0.04721 | -1.95855 | Male |
| Rn4.5s | | | | | | | | | | 0.00032 | -4.36233 | Male | | | | | | |
| Zfp503 | | | | | | | | | | 0.00022 | -2.36617 | Male | | | | | | |
| Sdr39u1 | | | | | | | | | | 0.00016 | 5.68025 | Female | | | | | | |
| Zfp341 | | | | | | | | | | | | | | | | 0.02986 | 3.39606 | Female |
| Mrgprb1 | | | | | | | | | | 0.01135 | -3.36892 | Male | | | | | | |
| Cobl | | | | 0.00000 | 3.43332 | Female | | | | | | | | | | | | |
| Wbscr25 | | | | | | | 0.00045 | -2.15392 | Male | | | | | | | | | |
| Stambpl1 | | | | 0.00173 | -0.62385 | Male | | | | 0.04411 | -0.99098 | Male | | | | | | |
| Cycs | | | | | | | | | | 0.04762 | -1.00670 | Male | | | | | | |
| Ica1l | | | | | | | | | | | | | 0.03392 | -3.76336 | Male | 0.00000 | 4.70515 | Female |
| Ubr5 | | | | | | | | | | | | | 0.01123 | -1.25284 | Male | | | |
| Vmp1 | | | | | | | | | | | | | | | | 0.02484 | -1.02125 | Male |
| Cd9 | | | | | | | | | | 0.00807 | 1.54067 | Female | | | | | | |

| | | | | | | | | | | | | | | | | | | |
|---|---|---|---|---|---|---|---|---|---|---|---|---|---|---|---|---|---|---|
| Cox19 | 0.01977 | -3.71392 | Male | | | | | | | | | | | | | | | |
| Nudt12 | | | | 0.00000 | 0.76686 | Female | 0.00000 | 4.86001 | Female | 0.00000 | 0.83194 | Female | | | | | | |
| Taco1 | | | | | | | | | | 0.01584 | 3.73669 | Female | | | | | | |
| Ccl9 | | | | | | | | | | | | | | | | 0.01180 | -3.03112 | Male |
| Lgals12 | | | | 0.03842 | -1.09920 | Male | | | | | | | | | | | | |
| Ppp1r26 | | | | 0.00822 | 3.21524 | Female | | | | | | | | | | | | |
| Prkcg | | | | | | | | | | | | | | | | 0.01925 | -2.42796 | Male |
| Stk25 | | | | | | | | | | 0.01785 | 2.31376 | Female | | | | | | |
| Ndrg2 | | | | 0.03818 | -1.50999 | Male | | | | | | | | | | | | |
| Gm6548 | | | | | | | | | | 0.00000 | 1.61925 | Female | | | | | | |
| Mir682 | | | | | | | | | | 0.00000 | -5.33800 | Male | | | | | | |
| Pex26 | | | | 0.02807 | 2.19265 | Female | 0.02064 | 3.28156 | Female | | | | | | | | | |
| Fbxl21 | | | | | | | 0.00202 | -0.95095 | Male | 0.03317 | -0.58887 | Male | | | | | | |
| Emg1 | | | | | | | | | | 0.00000 | 2.91767 | Female | | | | | | |
| Oscar | 0.00000 | -2.18917 | Male | | | | | | | | | | | | | | | |
| Csrnp2 | | | | | | | | | | | | | | | | 0.00542 | -3.12642 | Male |
| Epb41l1 | | | | 0.00000 | 0.73440 | Female | | | | | | | | | | | | |
| Galnt7 | | | | 0.00015 | 2.95637 | Female | | | | 0.02482 | -0.72283 | Male | | | | | | |
| Acta2 | | | | | | | | | | 0.00002 | 5.35068 | Female | | | | | | |
| Tsacc | | | | | | | | | | 0.04690 | -3.49793 | Male | | | | | | |
| Zfp940 | | | | 0.00368 | 0.60130 | Female | | | | 0.00002 | 0.79507 | Female | | | | | | |
| Agrp | 0.00000 | 2.73842 | Female | | | | | | | | | | | | | | | |
| Bbs12 | | | | | | | | | | 0.01411 | -1.19897 | Male | | | | | | |
| Zfp467 | | | | | | | | | | | | | 0.00000 | -1.45844 | Male | 0.00000 | -1.57971 | Male |
| Crtam | | | | | | | 0.00000 | 3.15926 | Female | | | | | | | | | |
| Fstl3 | | | | | | | | | | 0.00016 | 5.94318 | Female | | | | | | |
| Crxos | | | | | | | | | | 0.00000 | -2.39659 | Male | | | | | | |
| Mageb16.ps1 | | | | | | | | | | 0.00115 | 3.25692 | Female | | | | | | |
| Ddb2 | | | | | | | | | | 0.00002 | 5.20697 | Female | | | | | | |
| C3 | | | | | | | | | | | | | 0.00000 | -3.16136 | Male | | | |
| Tada3 | | | | | | | | | | | | | | | | 0.01977 | -2.13687 | Male |
| Gm561 | | | | | | | | | | 0.00000 | -5.21377 | Male | 0.00001 | 2.98591 | Female | | | |
| Ctsk | | | | 0.00004 | 2.03092 | Female | | | | | | | | | | | | |
| Prkx | | | | | | | 0.01016 | 3.31095 | Female | | | | | | | | | |
| Luc7l | | | | | | | | | | 0.01788 | 1.56621 | Female | | | | 0.00019 | -1.79954 | Male |
| Tex40 | 0.00001 | 1.37656 | Female | 0.00000 | -0.76392 | Male | | | | | | | | | | | | |
| Pou4f1 | | | | | | | | | | | | | 0.00000 | -2.75635 | Male | 0.00000 | -4.76786 | Male |
| Rn45s | | | | | | | | | | 0.00896 | 2.53498 | Female | | | | | | |
| Eif1 | | | | | | | | | | 0.00000 | -2.61084 | Male | | | | | | |
| Lsm5 | | | | | | | | | | 0.00000 | -3.74887 | Male | 0.01960 | 1.29019 | Female | | | |
| Chchd3 | | | | | | | | | | 0.00242 | 2.91667 | Female | | | | | | |
| Ifngr1 | | | | | | | | | | 0.02033 | -4.17162 | Male | 0.01646 | 3.86840 | Female | | | |
| Crym | | | | | | | | | | 0.00000 | 0.87848 | Female | | | | | | |
| Lrrc23 | | | | 0.00000 | -3.18166 | Male | | | | | | | 0.04703 | 1.12625 | Female | 0.01130 | 4.31276 | Female |
| Ifit1bl2 | | | | | | | | | | | | | 0.01508 | 2.53099 | Female | | | |
| Filip1l | | | | | | | | | | | | | | | | 0.00753 | -4.87449 | Male |
| Myo1d | | | | | | | | | | 0.01053 | 3.60706 | Female | | | | | | |
| Irgm1 | | | | | | | 0.04497 | 3.10797 | Female | 0.00000 | -5.72966 | Male | | | | | | |
| Acoxl | 0.00007 | 1.61879 | Female | | | | | | | | | | | | | | | |
| Gmds | | | | 0.00000 | 0.92976 | Female | | | | | | | | | | | | |
| Pcdh19 | | | | | | | | | | 0.00025 | -0.61378 | Male | 0.00000 | 3.60998 | Female | | | |
| Myo3b | | | | | | | | | | 0.00000 | -3.50508 | Male | | | | | | |
| Prom1 | | | | | | | | | | 0.00000 | 1.12579 | Female | | | | | | |
| Pfdn1 | | | | | | | | | | 0.04060 | -1.58303 | Male | | | | | | |
| Bik | | | | | | | | | | 0.00056 | -1.10189 | Male | | | | | | |
| Ddx60 | | | | | | | | | | | | | | | | 0.00000 | -0.60318 | Male |
| Rpl41 | | | | | | | | | | 0.00000 | -4.55243 | Male | 0.04893 | 1.11076 | Female | | | |
| Nsun3 | | | | | | | 0.00000 | 1.75009 | Female | 0.00204 | -0.61155 | Male | | | | | | |
| Daxx | | | | | | | | | | 0.00840 | 2.89737 | Female | | | | | | |
| Crls1 | | | | | | | | | | 0.03987 | -2.00547 | Male | | | | | | |
| Plekhg4 | | | | | | | | | | | | | 0.00667 | 0.92392 | Female | 0.00559 | 2.35886 | Female |
| Ostc | | | | | | | | | | 0.04776 | 1.20685 | Female | | | | | | |
| Serinc5 | | | | | | | | | | | | | | | | 0.00939 | 3.92812 | Female |
| Tom1l1 | | | | | | | | | | | | | | | | 0.02374 | -2.97980 | Male |
| Camk4 | | | | | | | | | | 0.02645 | -2.13104 | Male | | | | | | |
| Cpxm1 | 0.00000 | -0.67725 | Male | | | | | | | | | | | | | | | |
| Eif2a | | | | | | | | | | 0.02953 | 1.31551 | Female | | | | 0.01009 | 1.22215 | Female |
| Alas2 | | | | 0.00280 | -1.30672 | Male | | | | | | | | | | | | |
| Mib2 | | | | | | | | | | 0.00441 | 4.01289 | Female | | | | | | |
| Nedd8 | | | | | | | | | | 0.00570 | -1.80013 | Male | | | | | | |

| | | | | | | | | | | | | | | | | | | |
|---|---|---|---|---|---|---|---|---|---|---|---|---|---|---|---|---|---|---|
| Gng5 | | | | | | | | | | 0.00000 | -3.97865 | Male | 0.00919 | 1.82156 | Female | | | |
| Asb1 | 0.00000 | 0.85456 | Female | | | | | | | 0.00005 | -1.92925 | Male | | | | | | |
| Phf8 | | | | | | | 0.03684 | 2.50551 | Female | | | | | | | | | |
| Arpp19 | | | | | | | | | | 0.00000 | -3.53401 | Male | 0.00283 | 1.57634 | Female | | | |
| Smim13 | | | | | | | | | | | | | | | | 0.01937 | -1.67468 | Male |
| Slfn2 | | | | | | | | | | | | | 0.00079 | -1.63657 | Male | 0.00000 | 3.48426 | Female |
| Cox6a1 | | | | | | | | | | 0.00005 | -3.85348 | Male | | | | | | |
| Sumo1 | | | | | | | | | | | | | | | | 0.01529 | -0.99395 | Male |
| Nek4 | | | | | | | | | | 0.00005 | -4.37706 | Male | | | | | | |
| BC024386 | | | | | | | | | | 0.00000 | -6.16394 | Male | | | | | | |
| Zfp943 | | | | | | | | | | | | | | | | 0.00753 | 3.96407 | Female |
| Dnah9 | | | | 0.00000 | 3.11246 | Female | | | | | | | 0.00000 | -1.87255 | Male | | | |
| Smagp | | | | | | | | | | 0.00999 | -3.96964 | Male | | | | | | |
| Dand5 | | | | | | | | | | 0.00000 | -3.95391 | Male | | | | | | |
| Nup54 | | | | | | | | | | | | | 0.00964 | -1.52845 | Male | | | |
| Adck2 | | | | 0.00000 | 1.79559 | Female | | | | | | | | | | | | |
| Adam8 | | | | | | | | | | | | | 0.00000 | 1.61559 | Female | 0.00000 | 3.23140 | Female |
| Ndufs6 | | | | | | | | | | 0.00000 | -3.90509 | Male | 0.01737 | 1.61582 | Female | | | |
| Tbca | | | | | | | | | | 0.00000 | -4.05557 | Male | | | | | | |
| AY761184 | | | | | | | | | | 0.04952 | -4.23206 | Male | | | | | | |
| Ormdl2 | | | | | | | | | | 0.00002 | 3.05181 | Female | | | | | | |
| AI662270 | | | | | | | 0.00333 | 4.62375 | Female | | | | | | | | | |
| Morf4l1.ps1 | | | | | | | | | | 0.01785 | 1.47026 | Female | | | | | | |
| Clcnkb | | | | 0.00006 | -1.64596 | Male | 0.00000 | 0.91394 | Female | | | | | | | | | |
| Snx17 | | | | | | | | | | 0.02328 | 2.51115 | Female | | | | | | |
| U2af2 | | | | | | | | | | 0.00000 | -5.24699 | Male | 0.00000 | 2.21427 | Female | | | |
| Fbf1 | | | | 0.04835 | 1.84845 | Female | | | | | | | | | | | | |
| Dalrd3 | | | | | | | | | | 0.03457 | 2.35275 | Female | | | | | | |
| Itga11 | 0.00000 | -2.61824 | Male | | | | | | | | | | | | | | | |
| Nckap5 | | | | | | | | | | 0.04429 | -0.97433 | Male | | | | | | |
| Pop4 | | | | | | | | | | 0.03145 | 1.38495 | Female | | | | | | |
| Gbp9 | | | | | | | | | | | | | | | | 0.01438 | -4.17097 | Male |
| Gad2 | | | | | | | | | | 0.00350 | -2.22018 | Male | | | | | | |
| N4bp2 | | | | | | | | | | | | | | | | 0.01339 | 3.47311 | Female |
| Rpl27 | | | | | | | | | | 0.00007 | -2.64158 | Male | | | | | | |
| Fam151a | | | | | | | | | | 0.00000 | 7.56759 | Female | | | | | | |
| Gm14139 | | | | | | | 0.02606 | 2.92039 | Female | | | | | | | | | |
| Bgn | | | | | | | | | | 0.00017 | -1.93846 | Male | | | | 0.00000 | -5.28439 | Male |
| Ccne1 | | | | | | | | | | | | | | | | 0.02590 | -1.54120 | Male |
| Ctxn1 | | | | | | | | | | | | | 0.00018 | 5.18606 | Female | | | |
| Ube2f | 0.03154 | 3.32821 | Female | | | | | | | | | | | | | | | |
| Kri1 | | | | | | | | | | 0.00000 | 5.66460 | Female | | | | | | |
| Mir3091 | | | | | | | | | | 0.02009 | -4.28641 | Male | | | | | | |
| Nlrp5 | | | | | | | | | | | | | | | | 0.00008 | -5.05946 | Male |
| B3galt6 | | | | | | | 0.00796 | 4.29375 | Female | | | | | | | 0.00007 | -5.92067 | Male |
| Spcs1 | | | | | | | | | | 0.03334 | -1.14858 | Male | | | | | | |
| Trpm6 | | | | | | | 0.00000 | -3.17316 | Male | 0.00003 | -2.17291 | Male | | | | | | |
| Fgfbp1 | | | | | | | 0.00000 | 1.57270 | Female | | | | | | | | | |
| Il1rl1 | | | | | | | | | | 0.00007 | -4.84121 | Male | | | | | | |
| Kdelc1 | | | | | | | | | | 0.00000 | 1.18602 | Female | | | | | | |
| Rhag | | | | | | | | | | 0.04735 | -3.60648 | Male | | | | | | |
| Plp2 | | | | | | | | | | 0.00226 | 2.11038 | Female | | | | | | |
| Mbd3l2 | | | | | | | | | | | | | 0.00005 | -4.26991 | Male | | | |
| Plxna2 | | | | | | | | | | | | | 0.00000 | 2.62955 | Female | | | |
| Plcg1 | | | | | | | | | | 0.00016 | 5.70096 | Female | | | | | | |
| Egr1 | | | | | | | | | | | | | | | | 0.00000 | 7.56750 | Female |
| Sp9 | | | | | | | | | | | | | | | | 0.00069 | 3.25091 | Female |
| Uggt2 | | | | 0.00046 | 1.53046 | Female | | | | | | | | | | | | |
| Nudt16l1 | 0.02652 | 3.77261 | Female | | | | | | | | | | | | | | | |
| Nfib | | | | | | | | | | | | | 0.00000 | 2.10635 | Female | | | |
| Nfil3 | | | | | | | | | | | | | 0.00222 | 4.45876 | Female | | | |
| Cables1 | | | | | | | | | | | | | 0.04831 | 3.64524 | Female | | | |
| Rpl37 | | | | | | | | | | 0.00000 | -5.44875 | Male | 0.01542 | 1.25609 | Female | | | |
| Rom1 | | | | | | | | | | 0.00000 | 1.63023 | Female | 0.00023 | 4.81442 | Female | 0.00181 | 4.45603 | Female |
| Dennd4a | | | | | | | | | | | | | | | | 0.00000 | 4.11680 | Female |
| Gm4832 | | | | | | | | | | 0.00355 | -3.05493 | Male | 0.04089 | 1.93737 | Female | | | |
| Setdb2 | | | | | | | | | | 0.04010 | -4.54777 | Male | | | | | | |
| Obox5 | | | | | | | | | | 0.03149 | -3.83070 | Male | | | | 0.00395 | 4.01718 | Female |
| Dpysl2 | | | | | | | | | | 0.00000 | 0.69507 | Female | | | | | | |
| Agpat4 | | | | | | | | | | 0.00473 | -4.84132 | Male | | | | | | |

| | | | | | | | | | | | | | | | | | | |
|---|---|---|---|---|---|---|---|---|---|---|---|---|---|---|---|---|---|---|
| Dio1 | | | | | | | | | | | | | | | | 0.00300 | -4.77379 | Male |
| Kif5a | | | | 0.02695 | -0.91936 | Male | | | | | | | | | | | | |
| Sp100 | | | | | | | | | | 0.01770 | -2.46404 | Male | | | | | | |
| Smim4 | | | | | | | | | | 0.00590 | -3.62614 | Male | | | | | | |
| Poc1b | | | | | | | | | | | | | | | | 0.03714 | 3.35807 | Female |
| S100a16 | 0.00000 | -2.05502 | Male | | | | | | | | | | | | | | | |
| Gtsf1 | | | | | | | | | | | | | | | | 0.00050 | 4.50008 | Female |
| Gm12504 | | | | | | | | | | 0.00556 | 1.33103 | Female | | | | | | |
| Prkd3 | | | | | | | | | | | | | 0.00000 | -3.80720 | Male | 0.01543 | 4.25520 | Female |
| Ly6f | | | | 0.00000 | -2.66461 | Male | 0.00000 | -1.14648 | Male | | | | | | | | | |
| Lcn2 | | | | | | | | | | | | | | | | 0.00000 | 2.25304 | Female |
| Setx | | | | | | | | | | | | | | | | 0.01103 | 1.35533 | Female |
| Tmem14a | | | | | | | 0.00000 | -0.58195 | Male | | | | | | | | | |
| Clta | | | | | | | | | | 0.00001 | -3.24426 | Male | | | | | | |
| Lrrc46 | | | | | | | | | | | | | | | | 0.00791 | 4.15817 | Female |
| Pdlim3 | | | | | | | | | | 0.03265 | 4.20419 | Female | 0.00078 | 5.12270 | Female | | | |
| Rpl37rt | | | | | | | | | | 0.00000 | -5.25391 | Male | 0.01256 | 1.30046 | Female | | | |
| Pigx | | | | | | | | | | 0.00005 | -3.94966 | Male | | | | | | |
| Tc2n | | | | | | | | | | 0.00000 | -5.47817 | Male | | | | | | |
| Nutf2 | | | | | | | | | | 0.00231 | 4.46561 | Female | | | | | | |
| Snx25 | | | | 0.00000 | 1.30270 | Female | | | | | | | | | | | | |
| Cnr2 | | | | | | | | | | 0.02566 | -1.99460 | Male | | | | | | |
| Vps13c | | | | | | | | | | 0.00571 | 3.93656 | Female | | | | | | |
| Hmgb1 | | | | | | | | | | 0.04837 | 1.13206 | Female | | | | | | |
| Appbp2 | | | | | | | | | | 0.03063 | 2.13374 | Female | | | | | | |
| Snrpe | | | | | | | | | | 0.00000 | -4.83987 | Male | 0.00581 | 1.25952 | Female | | | |
| Fth1 | | | | | | | | | | 0.00370 | -2.75218 | Male | | | | | | |
| Arhgap18 | | | | | | | | | | | | | | | | 0.00000 | -4.16451 | Male |
| Ncam1 | | | | | | | | | | | | | | | | 0.04573 | 3.89964 | Female |
| Laptm5 | 0.01083 | 1.42788 | Female | | | | | | | | | | | | | | | |
| Gm6402 | | | | | | | | | | 0.00000 | -4.05335 | Male | 0.01524 | 1.12476 | Female | | | |
| Adcy3 | | | | | | | | | | | | | 0.03116 | -4.39448 | Male | | | |
| H2.T23 | | | | | | | | | | 0.00000 | -4.95971 | Male | 0.00246 | 1.65370 | Female | | | |
| Syt7 | | | | | | | | | | | | | | | | 0.00000 | 3.11809 | Female |
| B4galt7 | | | | | | | | | | 0.00017 | -2.92967 | Male | | | | | | |
| Fam98b | | | | | | | | | | 0.00000 | -4.43895 | Male | | | | | | |
| Prpf3 | | | | | | | | | | 0.03767 | 1.92675 | Female | | | | | | |
| Hif1a | | | | 0.01556 | 1.59140 | Female | 0.03112 | -3.94087 | Male | | | | | | | | | |
| Rtn1 | 0.00000 | 0.94029 | Female | | | | | | | | | | 0.00000 | 2.21129 | Female | | | |
| Mrps35 | | | | 0.00004 | 4.53528 | Female | | | | | | | | | | | | |
| Tat | 0.04473 | -3.06229 | Male | | | | | | | | | | | | | | | |
| Rnf165 | | | | | | | 0.04765 | 4.06354 | Female | | | | | | | | | |
| Adrb3 | | | | | | | 0.00000 | 2.83908 | Female | | | | | | | | | |
| Ptma | | | | | | | | | | 0.00942 | 1.24551 | Female | | | | | | |
| Asap3 | | | | | | | | | | | | | | | | 0.00000 | -4.22093 | Male |
| Cd209c | | | | | | | | | | | | | 0.01684 | 2.06353 | Female | | | |
| Xk | | | | | | | | | | | | | 0.01892 | 3.79770 | Female | | | |
| Slc9b2 | | | | | | | 0.00513 | -0.68794 | Male | | | | 0.00061 | 1.07324 | Female | | | |
| Setmar | | | | 0.00000 | 2.45953 | Female | | | | | | | 0.00000 | 5.71632 | Female | | | |
| Aco2 | | | | | | | | | | 0.01189 | 1.63238 | Female | | | | | | |
| Zfp692 | | | | | | | 0.02064 | 2.90142 | Female | | | | | | | | | |
| Asf1b | | | | | | | | | | 0.03219 | 1.77723 | Female | | | | | | |
| Necab2 | | | | | | | | | | | | | 0.00014 | 4.99460 | Female | | | |
| Hspa1b | | | | | | | | | | 0.04890 | -3.39838 | Male | | | | | | |
| Zp3 | | | | | | | | | | | | | | | | 0.00461 | 3.07838 | Female |
| LOC100505025 | | | | | | | | | | 0.00008 | -4.97330 | Male | | | | | | |
| Arhgap5 | | | | | | | 0.02631 | 3.18232 | Female | | | | | | | | | |
| Tspan9 | | | | | | | | | | | | | | | | 0.00121 | 4.49346 | Female |
| Vdac2 | | | | | | | | | | 0.00740 | 1.35046 | Female | | | | | | |
| Zfp865 | | | | | | | | | | | | | | | | 0.03020 | 3.43489 | Female |
| Mgst2 | | | | | | | 0.00000 | -2.99823 | Male | | | | | | | | | |
| Tmem232 | | | | | | | 0.00000 | 1.91832 | Female | | | | 0.00000 | -0.98662 | Male | | | |
| Serpinb1c | | | | 0.00000 | 2.53756 | Female | | | | | | | | | | | | |
| Pkd2l1 | | | | | | | 0.00004 | -0.68053 | Male | | | | | | | | | |
| Spcs3 | 0.00324 | 4.26602 | Female | 0.00739 | 3.35770 | Female | | | | | | | | | | | | |
| Zfp207 | | | | | | | | | | 0.00245 | 1.68633 | Female | | | | | | |
| Kpnb1 | | | | | | | | | | 0.00644 | 1.25004 | Female | | | | | | |
| Phactr4 | | | | | | | | | | | | | 0.00806 | -3.42686 | Male | | | |
| Chtf8 | | | | | | | | | | 0.00410 | 2.41795 | Female | | | | | | |
| Cbx7 | | | | | | | | | | 0.01294 | 2.18293 | Female | | | | | | |

| | | | | | | | | | | | | | | | | | | |
|---|---|---|---|---|---|---|---|---|---|---|---|---|---|---|---|---|---|---|
| Nrip2 | | | | | | | | | | | | | 0.00000 | -1.60863 | Male | | | |
| Tpd52l2 | | | | | | | | | | | | | | | | 0.01172 | -1.18099 | Male |
| Rab8a | | | | | | | | | | 0.04902 | 1.69224 | Female | | | | | | |
| Stard5 | | | | | | | | | | 0.00000 | -3.29843 | Male | | | | | | |
| Taf10 | | | | | | | | | | 0.00000 | -5.54993 | Male | | | | | | |
| Dpp10 | | | | 0.01235 | 2.79976 | Female | | | | | | | | | | | | |
| Yod1 | | | | | | | 0.00000 | 2.87290 | Female | 0.01002 | -1.33424 | Male | | | | | | |
| Ep300 | | | | | | | | | | 0.00002 | -5.90884 | Male | | | | | | |
| Pi4k2b | | | | | | | | | | | | | | | | 0.00225 | -2.18208 | Male |
| C1s2 | | | | | | | | | | | | | 0.03515 | 4.26202 | Female | | | |
| Phpt1 | | | | | | | | | | 0.03640 | -2.37789 | Male | | | | | | |
| Armc9 | | | | | | | | | | 0.03326 | 2.86824 | Female | | | | | | |
| Mfap2 | | | | 0.00000 | 1.23841 | Female | | | | 0.00001 | -2.98367 | Male | | | | | | |
| Atg9a | | | | | | | | | | | | | | | | 0.00652 | -1.73504 | Male |
| Zfp646 | | | | | | | | | | 0.00002 | 5.83143 | Female | | | | | | |
| Zfp612 | | | | | | | | | | | | | | | | 0.02738 | 3.52187 | Female |
| Fam214b | | | | | | | | | | | | | | | | 0.00001 | -3.06670 | Male |
| Ndufa11 | | | | | | | | | | 0.01727 | -2.10968 | Male | | | | | | |
| Ss18 | | | | | | | | | | 0.04902 | 1.07618 | Female | | | | | | |
| Thra | | | | | | | | | | | | | 0.00000 | 6.36616 | Female | | | |
| Csf3r | | | | | | | | | | 0.03682 | 2.18995 | Female | | | | | | |
| Foxl1 | | | | | | | | | | 0.00003 | -4.05525 | Male | | | | | | |
| Efr3b | | | | | | | | | | 0.04389 | -3.03019 | Male | | | | | | |
| Sema4a | | | | | | | | | | | | | 0.00354 | -4.75603 | Male | | | |
| Slc16a13 | | | | | | | 0.04296 | 2.93240 | Female | | | | | | | | | |
| Tmed6 | | | | | | | | | | | | | | | | 0.00292 | -4.49714 | Male |
| Rpl27a | | | | | | | | | | 0.00643 | -2.39215 | Male | | | | | | |
| Tfap2e | | | | | | | | | | | | | 0.04343 | -4.29755 | Male | | | |
| Plxdc2 | | | | | | | | | | 0.00036 | -0.92276 | Male | | | | | | |
| Zfp799 | | | | | | | | | | 0.00002 | -2.66262 | Male | 0.03040 | 1.20272 | Female | | | |
| Calr | | | | | | | | | | | | | 0.01580 | 1.07054 | Female | | | |
| Srek1ip1 | | | | 0.01796 | 3.52203 | Female | | | | 0.00015 | 4.82218 | Female | | | | | | |
| LOC106029237 | | | | | | | | | | | | | | | | 0.00044 | -5.86203 | Male |
| Trim62 | | | | | | | | | | 0.01561 | -0.90936 | Male | | | | | | |
| P2ry4 | | | | | | | | | | 0.00134 | -3.92231 | Male | 0.00464 | 2.59848 | Female | | | |
| Fxn | | | | | | | | | | 0.00780 | -3.80592 | Male | | | | | | |
| Ppp2r2b | | | | | | | | | | | | | 0.00206 | 4.31115 | Female | 0.00005 | -6.15749 | Male |
| Fut10 | | | | | | | | | | | | | | | | 0.00000 | 1.22345 | Female |
| Pctp | | | | | | | | | | | | | 0.03851 | 3.00000 | Female | | | |
| Ifrd2 | | | | | | | | | | 0.03707 | 2.16512 | Female | | | | | | |
| Fthl17f | | | | 0.01451 | 3.83223 | Female | 0.00000 | 5.63787 | Female | 0.00026 | 5.12400 | Female | 0.01249 | 3.06270 | Female | | | |
| Kit | | | | | | | | | | | | | 0.01125 | -4.04951 | Male | | | |
| Utp6 | | | | | | | | | | | | | | | | 0.00013 | -2.61978 | Male |
| Myoz3 | | | | | | | | | | 0.00299 | -3.67081 | Male | | | | | | |
| Rsl1d1 | | | | | | | | | | 0.01061 | 1.02298 | Female | | | | | | |
| Dnah2 | | | | | | | | | | 0.04733 | 3.43649 | Female | | | | | | |
| Nop56 | | | | | | | | | | 0.00987 | 0.88698 | Female | | | | | | |
| Ogfod1 | | | | | | | | | | | | | | | | 0.01859 | 2.20129 | Female |
| Grhl1 | | | | 0.00000 | 2.07708 | Female | | | | | | | | | | | | |
| Zfp202 | | | | | | | | | | 0.00011 | -4.62294 | Male | 0.00004 | 3.46806 | Female | | | |
| Rnf113a1 | | | | | | | | | | 0.02570 | 3.27924 | Female | | | | | | |
| Zfp106 | | | | | | | | | | | | | 0.04965 | -0.84795 | Male | | | |
| Trp73 | | | | 0.00900 | -1.60223 | Male | | | | | | | | | | | | |
| Nsf | | | | | | | | | | 0.02468 | 1.89884 | Female | | | | | | |
| Rnf185 | | | | | | | | | | | | | | | | 0.04406 | -1.60240 | Male |
| Ndufa8 | 0.00707 | 4.11402 | Female | | | | | | | | | | | | | | | |
| Ndufb4 | | | | | | | | | | 0.00000 | -3.77180 | Male | 0.00209 | 1.92846 | Female | | | |
| Nxpe4 | | | | | | | | | | 0.03708 | -0.86158 | Male | | | | | | |
| Guf1 | | | | | | | | | | | | | | | | 0.03813 | -3.54832 | Male |
| Uba1 | | | | | | | | | | 0.00000 | 2.06399 | Female | | | | | | |
| Pou6f1 | | | | | | | | | | | | | 0.02856 | 2.46661 | Female | | | |
| Magea5 | | | | | | | 0.02476 | 4.30822 | Female | 0.00000 | -1.53256 | Male | | | | | | |
| D5Ertd579e | | | | 0.03905 | 2.75936 | Female | | | | | | | | | | | | |
| Tmem81 | | | | | | | | | | 0.04120 | 2.99181 | Female | | | | | | |
| Ugt2b36 | | | | | | | | | | 0.00406 | -6.28266 | Male | | | | | | |
| Il1r1 | | | | | | | | | | 0.01649 | -1.15239 | Male | | | | | | |
| Alg3 | | | | 0.00732 | 2.94335 | Female | | | | | | | | | | | | |
| Nr2f2 | | | | | | | | | | | | | 0.03627 | 3.32955 | Female | | | |
| Exosc6 | | | | | | | | | | 0.00013 | -4.63509 | Male | | | | | | |
| Etaa1 | | | | | | | | | | | | | | | | 0.00028 | 2.91708 | Female |

| | | | | | | | | | | | | | | | | | | |
|---|---|---|---|---|---|---|---|---|---|---|---|---|---|---|---|---|---|---|
| Eif4g1 | | | | | | | | | | 0.00432 | 3.12923 | Female | | | | | | |
| Gngt1 | | | | | | | | | | 0.00001 | -4.47529 | Male | | | | | | |
| Fam65b | | | | | | | | | | 0.00022 | -3.82409 | Male | 0.01027 | 2.20226 | Female | | | |
| Cbx3 | | | | | | | | | | 0.00002 | -2.69131 | Male | | | | | | |
| Cox6b1 | | | | | | | | | | 0.00000 | -4.60905 | Male | 0.00106 | 1.52278 | Female | | | |
| Lrrc1 | | | | | | | | | | | | | 0.00001 | 0.88862 | Female | | | |
| Zfp82 | | | | | | | | | | 0.04923 | 3.02539 | Female | | | | | | |
| Actr3 | | | | | | | | | | 0.00100 | 1.62849 | Female | | | | | | |
| Ubb | | | | | | | | | | 0.00010 | 1.85566 | Female | | | | | | |
| Rab42 | | | | | | | | | | 0.00111 | 0.73413 | Female | | | | | | |
| Acads | | | | | | | | | | 0.01187 | 2.81366 | Female | | | | | | |
| Cnot9 | | | | | | | | | | 0.00033 | 1.93438 | Female | | | | | | |
| Ick | | | | | | | | | | 0.03109 | 3.02813 | Female | 0.03053 | 3.23934 | Female | | | |
| Cd8b1 | | | | | | | | | | 0.01658 | -1.76117 | Male | | | | | | |
| Lix1 | | | | | | | | | | 0.00010 | -2.73939 | Male | 0.04831 | 1.27204 | Female | | | |
| Ndufb6 | | | | | | | | | | 0.00012 | -3.10313 | Male | | | | | | |
| Hebp1 | | | | | | | | | | | | | 0.04102 | 4.11028 | Female | | | |
| Actb | | | | | | | | | | 0.00381 | -1.86233 | Male | 0.00000 | 1.97040 | Female | | | |
| Abca7 | | | | 0.00509 | -1.42962 | Male | | | | 0.00000 | 2.75148 | Female | | | | | | |
| Ggt5 | | | | | | | 0.00201 | -1.32654 | Male | | | | | | | 0.00000 | -4.40083 | Male |
| Cdc25a | | | | | | | | | | | | | | | | 0.02971 | -1.62168 | Male |
| Zfp39 | | | | | | | | | | 0.00000 | 1.06973 | Female | | | | | | |
| Itpka | 0.00000 | 2.02428 | Female | | | | | | | | | | | | | | | |
| Tm9sf1 | | | | | | | | | | | | | | | | 0.00069 | -2.90560 | Male |
| Cfap53 | | | | | | | | | | 0.00410 | -1.46497 | Male | | | | 0.00000 | 3.86357 | Female |
| Csnk1e | | | | | | | | | | 0.02482 | 1.47020 | Female | | | | | | |
| Syce3 | | | | | | | | | | 0.00909 | -5.38123 | Male | | | | | | |
| Tufm | | | | | | | | | | 0.00086 | 1.93177 | Female | | | | | | |
| Camsap1 | | | | | | | | | | | | | | | | 0.01375 | -1.47438 | Male |
| Pcnx3 | | | | | | | | | | 0.00001 | 5.90875 | Female | | | | | | |
| Phf7 | | | | | | | 0.04604 | -4.38249 | Male | | | | | | | 0.01375 | 4.10327 | Female |
| Bysl | | | | | | | | | | 0.00126 | 2.15551 | Female | | | | | | |
| Capn15 | | | | | | | | | | 0.00025 | 5.48475 | Female | | | | | | |
| Tor3a | | | | | | | | | | | | | | | | 0.00000 | 1.88766 | Female |
| Thbs1 | | | | | | | | | | | | | 0.00000 | 2.14951 | Female | 0.00001 | 1.31556 | Female |
| Xlr5b | | | | | | | | | | | | | 0.00089 | -3.58208 | Male | | | |
| Maneal | | | | | | | | | | | | | 0.00213 | 4.54662 | Female | | | |
| Mmachc | | | | | | | 0.02647 | 2.23532 | Female | | | | | | | | | |
| Mt2 | | | | | | | | | | 0.00000 | -6.58727 | Male | | | | | | |
| Slc31a2 | | | | | | | | | | 0.04814 | -4.34949 | Male | | | | | | |
| Cox6c | | | | | | | | | | 0.00000 | -5.25416 | Male | 0.00000 | 2.29033 | Female | | | |
| T2 | | | | | | | | | | 0.00001 | -5.29979 | Male | | | | | | |
| Snx21 | | | | | | | | | | | | | | | | 0.03255 | 4.69744 | Female |
| Uqcr10 | | | | | | | | | | 0.00000 | -4.46817 | Male | 0.00000 | 2.38102 | Female | | | |
| Fbxo46 | | | | | | | | | | | | | 0.00284 | 4.09081 | Female | | | |
| Cdh11 | | | | | | | | | | 0.04837 | 2.11740 | Female | | | | 0.03018 | -1.80093 | Male |
| Spef1 | | | | | | | | | | 0.00029 | -2.01615 | Male | | | | | | |
| Plekhf1 | | | | 0.00060 | 2.27351 | Female | | | | | | | | | | | | |
| Stim1 | | | | | | | | | | 0.04688 | 2.90391 | Female | | | | | | |
| Rbm11 | | | | | | | | | | 0.00000 | -4.41398 | Male | | | | | | |
| Yif1b | | | | | | | | | | 0.00194 | 3.88304 | Female | | | | | | |
| Akap10 | | | | | | | | | | | | | 0.00550 | -4.24871 | Male | 0.00384 | -4.23796 | Male |
| Ctf2 | | | | | | | | | | 0.01638 | 2.04209 | Female | | | | 0.00123 | 1.03566 | Female |
| Rpl10a | | | | | | | | | | 0.04498 | -1.43898 | Male | | | | | | |
| Imp3 | | | | | | | | | | 0.00075 | -3.50782 | Male | | | | | | |
| Nrg4 | | | | 0.00001 | 1.65025 | Female | | | | | | | | | | | | |
| Kansl1l | | | | | | | 0.02287 | 3.63318 | Female | | | | | | | | | |
| Pgp | | | | | | | | | | 0.00709 | -3.21443 | Male | | | | | | |
| Clu | | | | | | | | | | | | | | | | 0.00027 | 6.46424 | Female |
| Ssr4 | 0.00315 | 2.43496 | Female | | | | | | | | | | | | | | | |
| Htra2 | | | | | | | | | | 0.00008 | 3.55405 | Female | | | | 0.02200 | -1.89051 | Male |
| Cass4 | | | | | | | | | | 0.00007 | 0.67570 | Female | | | | | | |
| Lif | | | | | | | | | | 0.00297 | 4.46161 | Female | | | | | | |
| Supt20 | | | | | | | | | | 0.00088 | 3.25494 | Female | | | | | | |
| Rps13 | | | | | | | | | | 0.00000 | -3.51229 | Male | | | | | | |
| Tnnc2 | | | | | | | | | | | | | | | | 0.04792 | -3.98745 | Male |
| Homez | | | | | | | | | | 0.03585 | -1.77157 | Male | | | | | | |
| B3glct | 0.04177 | 3.24623 | Female | | | | | | | | | | | | | | | |
| Ccdc60 | 0.00166 | -1.94581 | Male | 0.03318 | -0.70926 | Male | | | | | | | | | | | | |
| Uck1 | | | | 0.00493 | 3.38780 | Female | | | | | | | | | | | | |

| | | | | | | | | | | | | | | | | | | |
|---|---|---|---|---|---|---|---|---|---|---|---|---|---|---|---|---|---|---|
| Tmem136 | | | | | | | | | | 0.00000 | -0.93329 | Male | | | | | | |
| Rps4x | | | | | | | 0.00139 | 0.92507 | Female | 0.03633 | 0.78822 | Female | 0.00020 | 0.85620 | Female | | | |
| Zscan22 | | | | | | | | | | 0.00582 | 3.95260 | Female | 0.01464 | 4.06473 | Female | | | |
| Tm4sf5 | | | | 0.00000 | 1.17357 | Female | | | | | | | 0.02069 | 4.28880 | Female | | | |
| Fthl17d | | | | 0.00181 | 4.93143 | Female | 0.00000 | 5.63447 | Female | 0.00211 | 4.63801 | Female | 0.00001 | 4.44212 | Female | | | |
| Nptx1 | | | | | | | | | | | | | | | | 0.00810 | 5.19007 | Female |
| Prmt5 | | | | | | | | | | 0.00454 | 1.55307 | Female | | | | | | |
| Ralgps1 | | | | | | | | | | | | | | | | 0.01430 | 3.26828 | Female |
| Pif1 | | | | 0.01346 | 2.16705 | Female | | | | | | | | | | | | |
| Csrnp3 | | | | | | | 0.00013 | 4.54791 | Female | 0.00340 | -4.12152 | Male | | | | | | |
| D17H6S53E | | | | | | | | | | 0.00000 | -4.86311 | Male | | | | | | |
| Creb3l3 | | | | | | | | | | | | | 0.00000 | -2.86202 | Male | 0.01483 | 3.31340 | Female |
| Azin2 | | | | | | | | | | 0.00038 | 1.63980 | Female | 0.00000 | 3.61945 | Female | | | |
| Pcbd2 | | | | | | | | | | 0.04660 | -4.02392 | Male | | | | | | |
| Zfp422 | | | | | | | | | | | | | 0.00070 | 2.63610 | Female | | | |
| Tmed7 | | | | 0.00000 | 5.16549 | Female | | | | | | | | | | | | |
| Pla2g4e | | | | | | | | | | | | | | | | 0.00001 | -5.68444 | Male |
| Ccdc124 | | | | | | | | | | 0.01219 | 3.11202 | Female | | | | | | |
| Nes | | | | | | | | | | 0.02257 | -1.24834 | Male | | | | | | |
| Rbpms | | | | | | | | | | 0.00346 | -1.92202 | Male | | | | | | |
| Hyal2 | | | | | | | 0.00634 | -4.89772 | Male | | | | | | | | | |
| Disp1 | | | | | | | | | | | | | | | | 0.00000 | 1.10951 | Female |
| Mpdz | | | | | | | | | | | | | | | | 0.00042 | 4.71556 | Female |
| Tnfrsf11a | | | | | | | | | | | | | | | | 0.02614 | -2.64309 | Male |
| Fbxl6 | | | | | | | | | | 0.00000 | 5.49555 | Female | | | | | | |
| Pld1 | | | | | | | | | | | | | 0.00000 | -3.30411 | Male | | | |
| Pla2g4c | | | | | | | | | | 0.03336 | -4.43069 | Male | 0.00073 | 4.77047 | Female | | | |
| Il17rb | | | | 0.00115 | -0.59662 | Male | | | | | | | | | | | | |
| Adgra2 | | | | | | | | | | | | | | | | 0.04062 | -2.70386 | Male |
| Dusp3 | | | | | | | | | | | | | | | | 0.00034 | -3.85993 | Male |
| Itpripl1 | | | | | | | | | | 0.00000 | 2.10512 | Female | | | | | | |
| D6Ertd527e | | | | | | | | | | | | | 0.00000 | -5.40200 | Male | | | |
| Hyal3 | | | | | | | | | | 0.01068 | -2.40999 | Male | | | | | | |
| Fn1 | | | | | | | | | | 0.03272 | 3.19702 | Female | | | | | | |
| Slco5a1 | | | | | | | | | | | | | 0.00000 | 0.98784 | Female | | | |
| Skida1 | | | | | | | | | | | | | 0.02149 | 4.40289 | Female | | | |
| Erh | | | | | | | | | | 0.00180 | -1.29269 | Male | | | | | | |
| Wdfy2 | | | | | | | | | | 0.04087 | 2.46063 | Female | | | | | | |
| Fus | | | | | | | | | | 0.00000 | -6.16046 | Male | | | | | | |
| Fam210a | | | | | | | | | | | | | | | | 0.01673 | 1.84945 | Female |
| Birc2 | | | | | | | | | | | | | | | | 0.03615 | 2.52207 | Female |
| Tbc1d13 | | | | | | | | | | | | | | | | 0.00453 | 2.06258 | Female |
| Daglb | | | | | | | | | | | | | 0.03894 | -3.70589 | Male | 0.00000 | 3.77486 | Female |
| Gm8624 | | | | | | | | | | 0.00000 | -4.43874 | Male | 0.00043 | 1.53111 | Female | | | |
| Pdha1 | | | | | | | 0.00201 | 1.16881 | Female | 0.00002 | 1.75351 | Female | | | | | | |
| Fbxw24 | | | | | | | | | | | | | | | | 0.00012 | -5.27778 | Male |
| S1pr5 | | | | | | | 0.00000 | 4.21860 | Female | | | | | | | | | |
| Rps21 | | | | | | | | | | 0.00000 | -5.44461 | Male | 0.00048 | 1.72274 | Female | | | |
| Ttf1 | | | | | | | | | | | | | 0.00103 | 1.84875 | Female | | | |
| Pik3r6 | | | | 0.00000 | -1.00483 | Male | | | | 0.01337 | 4.40380 | Female | | | | | | |
| Otud5 | | | | | | | | | | | | | | | | 0.02703 | 1.48341 | Female |
| Slc6a9 | | | | | | | | | | | | | 0.00000 | -3.27419 | Male | | | |
| Selplg | | | | | | | | | | 0.00632 | -1.97373 | Male | | | | | | |
| Gm9926 | | | | | | | | | | 0.01062 | -3.16195 | Male | | | | | | |
| Prex2 | | | | | | | | | | 0.00435 | -1.53342 | Male | | | | | | |
| Cnga3 | | | | | | | | | | | | | 0.00985 | 3.36609 | Female | | | |
| Ptk6 | | | | | | | | | | | | | 0.02209 | 2.74664 | Female | 0.00683 | 3.41139 | Female |
| Mrpl54 | | | | | | | | | | 0.00002 | -4.36985 | Male | | | | | | |
| Ndufa12 | | | | | | | | | | 0.00014 | -3.12484 | Male | | | | | | |
| Actg2 | | | | 0.00035 | 1.15379 | Female | | | | | | | | | | | | |
| Trim43c | | | | | | | | | | | | | | | | 0.04915 | -0.99092 | Male |
| Shc1 | | | | | | | | | | 0.01311 | 2.41574 | Female | | | | | | |
| Sft2d3 | | | | | | | | | | | | | 0.01127 | 5.24460 | Female | | | |
| Tchh | | | | 0.00000 | 4.53483 | Female | | | | | | | | | | | | |
| Mospd4 | | | | 0.00000 | -0.74637 | Male | 0.00000 | 2.58552 | Female | | | | | | | | | |
| Chga | | | | | | | | | | 0.00099 | 5.19588 | Female | | | | | | |
| Nanp | | | | | | | | | | 0.00000 | 1.75562 | Female | | | | | | |
| Tcl1b2 | | | | | | | | | | | | | | | | 0.00130 | -6.23177 | Male |
| Saxo1 | | | | 0.01080 | 2.50166 | Female | | | | 0.04981 | 1.69201 | Female | | | | | | |
| Kank4 | | | | | | | | | | 0.02452 | -3.48482 | Male | | | | | | |

| | | | | | | | | | | | | | | | | | | |
|---|---|---|---|---|---|---|---|---|---|---|---|---|---|---|---|---|---|---|
| Slc38a6 | | | | | | | | | | | | | | | | 0.02122 | -2.33147 | Male |
| Trim43b | | | | | | | | | | | | | | | | 0.03865 | -1.13370 | Male |
| BC055324 | | | | | | | | | | | | | | | | 0.00272 | 2.18347 | Female |
| Kctd2 | | | | | | | | | | 0.00658 | 3.44954 | Female | | | | | | |
| Klhl7 | | | | | | | | | | 0.04552 | -1.82585 | Male | | | | | | |
| Ttc39a | | | | 0.00005 | 0.81345 | Female | | | | | | | | | | | | |
| Teddm1b | | | | | | | | | | 0.00000 | -5.44439 | Male | 0.00078 | 1.75281 | Female | | | |
| Sgcb | | | | | | | | | | | | | 0.00000 | -4.31118 | Male | 0.00006 | -0.63811 | Male |
| Upk3b | | | | 0.01785 | -1.90488 | Male | | | | | | | | | | | | |
| Dennd1c | | | | | | | | | | | | | | | | 0.04334 | -3.75051 | Male |
| Oasl2 | | | | | | | | | | | | | | | | 0.00275 | 4.58033 | Female |
| Sox2 | | | | 0.00000 | -1.18891 | Male | | | | | | | | | | | | |
| Gm11127 | | | | | | | | | | 0.00703 | -1.88877 | Male | 0.03851 | 1.19436 | Female | | | |
| Exosc5 | | | | | | | | | | 0.01611 | 3.09397 | Female | | | | | | |
| Slc4a1ap | | | | | | | | | | | | | | | | 0.00002 | -2.93174 | Male |
| Mycn | | | | | | | 0.03588 | -5.41718 | Male | | | | | | | | | |
| Cyba | 0.00000 | 1.79552 | Female | | | | | | | | | | | | | 0.00135 | -4.71031 | Male |
| Thbs3 | | | | | | | | | | | | | 0.00000 | -3.43719 | Male | | | |
| Myo5b | | | | | | | | | | | | | 0.04617 | 2.95141 | Female | | | |
| Tfpi | | | | 0.00000 | 2.38298 | Female | | | | | | | | | | | | |
| Klhl8 | | | | | | | | | | | | | 0.00311 | 3.79520 | Female | 0.00096 | 4.11199 | Female |
| Cryba1 | 0.00458 | -1.73712 | Male | | | | | | | | | | | | | | | |
| Cd74 | | | | | | | | | | | | | | | | 0.00000 | -4.09045 | Male |
| Lcn9 | 0.00981 | 1.05594 | Female | | | | | | | | | | | | | | | |
| D1Pas1 | | | | | | | | | | | | | | | | 0.00000 | -1.07619 | Male |
| Umodl1 | | | | | | | | | | | | | | | | 0.04513 | -4.25919 | Male |
| Nars | | | | | | | | | | 0.00209 | 1.56485 | Female | | | | | | |
| Zfp974 | | | | | | | | | | | | | 0.00222 | 4.43554 | Female | | | |
| Bhmt2 | | | | | | | | | | | | | | | | 0.00542 | -2.17363 | Male |
| Gm805 | | | | 0.00021 | -0.86375 | Male | 0.00000 | -0.64439 | Male | | | | | | | | | |
| Mirt1 | | | | | | | | | | 0.00018 | -2.56939 | Male | | | | | | |
| Tbxa2r | | | | | | | | | | | | | 0.00797 | -4.27647 | Male | | | |
| Bcorl1 | | | | | | | | | | 0.04036 | 3.36114 | Female | | | | | | |
| Mia2 | | | | | | | | | | 0.00134 | -0.96927 | Male | | | | | | |
| Gpd1l | | | | | | | | | | | | | | | | 0.00031 | -1.69959 | Male |
| B3gnt6 | | | | | | | 0.00008 | -1.82751 | Male | 0.00001 | 2.36807 | Female | | | | | | |
| Osbpl2 | | | | | | | | | | | | | | | | 0.00422 | -2.28890 | Male |
| Rnd2 | | | | | | | | | | 0.00510 | 3.62446 | Female | | | | | | |
| Hdac10 | | | | 0.00001 | 0.58146 | Female | | | | | | | | | | | | |
| Hist1h1a | | | | 0.00000 | 1.11356 | Female | | | | | | | 0.00000 | 2.45704 | Female | | | |
| Cpe | | | | 0.00009 | -0.84862 | Male | | | | | | | 0.01816 | 3.90038 | Female | | | |
| Hmgcll1 | | | | | | | | | | 0.00001 | -2.56787 | Male | | | | | | |
| Tmem2 | | | | | | | 0.00031 | 5.06938 | Female | | | | | | | 0.00002 | -6.06453 | Male |
| Spag6l | | | | | | | | | | | | | | | | 0.00000 | 1.82934 | Female |
| Mrps34 | | | | | | | | | | 0.01230 | -3.23207 | Male | | | | | | |
| Inafm1 | | | | | | | | | | | | | | | | 0.00747 | -4.97469 | Male |
| Insm1 | | | | 0.03069 | 4.43684 | Female | | | | | | | | | | | | |
| Kdm4a | | | | | | | | | | 0.00001 | -5.73509 | Male | | | | | | |
| Prr11 | | | | | | | | | | 0.02209 | -2.64123 | Male | | | | | | |
| Rps6ka6 | | | | | | | | | | | | | | | | 0.00000 | 3.00531 | Female |
| Zfp873 | | | | | | | | | | | | | | | | 0.00017 | 4.80272 | Female |
| Ovol2 | | | | | | | | | | | | | | | | 0.00247 | -3.38714 | Male |
| Gpha2 | | | | | | | | | | | | | | | | 0.01009 | 4.37518 | Female |
| Zrsr2 | | | | | | | | | | | | | 0.00691 | 2.10437 | Female | | | |
| Mrpl47 | | | | | | | | | | | | | | | | 0.01478 | -1.06292 | Male |
| Trpm4 | | | | | | | 0.00749 | -5.04123 | Male | | | | | | | | | |
| BC021614 | | | | | | | | | | | | | | | | 0.02440 | 3.58812 | Female |
| Ccdc136 | | | | | | | | | | 0.00760 | -1.29877 | Male | | | | 0.00000 | -2.46636 | Male |
| Lrrc36 | | | | | | | 0.02720 | -2.53699 | Male | | | | | | | | | |
| Dgcr8 | | | | | | | | | | 0.03575 | 2.30427 | Female | | | | | | |
| Sil1 | | | | | | | | | | | | | | | | 0.00465 | -3.37091 | Male |
| Gng7 | 0.00440 | 0.98677 | Female | | | | | | | | | | | | | | | |
| Mapk8ip3 | | | | 0.00903 | 1.55477 | Female | | | | | | | | | | | | |
| Lig3 | | | | | | | | | | 0.00418 | 2.05335 | Female | | | | | | |
| Sepn1 | | | | 0.00343 | -1.47285 | Male | | | | | | | | | | | | |
| Abo | | | | | | | | | | 0.00032 | 0.96798 | Female | | | | 0.00000 | -2.90140 | Male |
| Eif3i | | | | | | | | | | 0.00000 | 1.76084 | Female | | | | | | |
| Rpl18a | | | | | | | | | | 0.00023 | -2.72421 | Male | | | | | | |
| Megf8 | | | | | | | | | | | | | | | | 0.02747 | 3.53112 | Female |
| Zfp991 | | | | | | | | | | | | | | | | 0.02818 | -1.91711 | Male |

| | | | | | | | | | | | | | | | | | | |
|---|---|---|---|---|---|---|---|---|---|---|---|---|---|---|---|---|---|---|
| Dpep1 | | | | | | | | | | | | | 0.00911 | 3.61932 | Female | 0.00000 | 3.74154 | Female |
| Gsdmc2 | | | | | | | | | | | | | | | | 0.00013 | 5.10245 | Female |
| Mep1b | | | | | | | | | | | | | | | | 0.00021 | 4.12151 | Female |
| Rdh9 | | | | | | | | | | | | | 0.03798 | 3.18664 | Female | | | |
| Gch1 | | | | | | | | | | 0.00808 | -2.39898 | Male | | | | | | |
| Rps28 | | | | | | | | | | 0.00000 | -5.67313 | Male | 0.00000 | 2.28773 | Female | | | |
| Klf6 | | | | | | | | | | 0.00175 | 4.72018 | Female | | | | | | |
| Sp3os | | | | | | | | | | 0.00000 | -4.88784 | Male | | | | | | |
| Morn1 | | | | | | | | | | | | | | | | 0.02284 | -4.23910 | Male |
| Nrap | | | | 0.01026 | 2.64181 | Female | | | | | | | | | | 0.03725 | 5.60874 | Female |
| Mtus1 | | | | | | | | | | | | | | | | 0.00311 | -2.19127 | Male |
| Dgkz | | | | | | | | | | 0.00056 | 3.13342 | Female | | | | | | |
| Oosp1 | | | | | | | | | | | | | 0.00515 | -3.78362 | Male | 0.01135 | -3.71347 | Male |
| Hsd3b1 | 0.02611 | -0.95814 | Male | | | | | | | | | | 0.00000 | 1.66209 | Female | | | |
| Trim66 | | | | | | | | | | | | | 0.00000 | -5.30157 | Male | 0.00000 | 2.04365 | Female |
| Krt18 | | | | 0.01084 | -6.09707 | Male | | | | | | | | | | | | |
| Asxl2 | | | | | | | | | | | | | | | | 0.00014 | -2.73610 | Male |
| Fam174b | | | | | | | | | | | | | 0.00003 | 1.08558 | Female | | | |
| Snord100 | | | | 0.00869 | 3.78730 | Female | | | | | | | | | | | | |
| AU015228 | | | | | | | | | | | | | 0.03040 | -3.14294 | Male | | | |
| Drd3 | | | | | | | | | | 0.00000 | 1.48695 | Female | | | | | | |
| Htatsf1 | | | | | | | 0.02866 | 1.86099 | Female | | | | | | | | | |
| Ube2n | | | | | | | | | | | | | | | | 0.04792 | -0.92877 | Male |
| Ooep | | | | | | | | | | 0.02224 | -2.64419 | Male | | | | | | |
| Adprm | | | | 0.01817 | 1.87456 | Female | 0.01571 | 3.45862 | Female | | | | | | | | | |
| Car8 | | | | | | | | | | 0.00016 | -4.42070 | Male | | | | | | |
| Arl6 | | | | | | | | | | 0.02188 | 3.07370 | Female | | | | | | |
| Mtx2 | | | | | | | | | | 0.02762 | -3.05102 | Male | | | | | | |
| Ptcra | | | | | | | | | | 0.00000 | -4.95646 | Male | 0.01731 | 1.65779 | Female | | | |
| Dtx3l | | | | | | | | | | 0.00173 | -3.78448 | Male | | | | | | |
| Arl13b | | | | | | | | | | | | | | | | 0.03731 | -3.24980 | Male |
| Nphp1 | | | | 0.00143 | 1.32581 | Female | | | | | | | | | | | | |
| Wdr73 | | | | | | | | | | | | | | | | 0.04930 | 1.46476 | Female |
| Nxpe3 | | | | | | | | | | 0.01893 | -5.01212 | Male | | | | | | |
| Golga4 | | | | | | | | | | 0.00194 | 3.15692 | Female | | | | | | |
| Slc39a11 | | | | | | | | | | | | | | | | 0.00191 | 4.59391 | Female |
| Rmdn2 | | | | | | | 0.00000 | 3.47719 | Female | | | | | | | | | |
| Erich1 | | | | | | | | | | | | | | | | 0.00606 | 2.90555 | Female |
| Capn1 | | | | | | | | | | 0.00383 | -3.37766 | Male | | | | | | |
| Ccdc78 | | | | | | | 0.02297 | -1.97729 | Male | 0.00396 | -2.73528 | Male | 0.00318 | 2.10860 | Female | | | |
| Aurka | | | | | | | | | | 0.01348 | 1.73466 | Female | | | | | | |
| Nectin2 | | | | 0.00362 | 1.98298 | Female | | | | | | | | | | | | |
| Chpf2 | | | | | | | | | | 0.02363 | 4.01443 | Female | | | | | | |
| Fgl1 | | | | | | | | | | | | | | | | 0.00014 | 1.81939 | Female |
| Mcrs1 | | | | | | | | | | 0.01702 | 1.93660 | Female | | | | | | |
| Ccr8 | | | | | | | 0.00000 | 4.17801 | Female | | | | | | | | | |
| Tfr2 | | | | | | | | | | | | | 0.00000 | -2.90021 | Male | | | |
| S100a14 | | | | | | | | | | 0.00000 | -4.05231 | Male | | | | | | |
| Arfgap3 | | | | | | | | | | 0.00004 | -2.25493 | Male | | | | | | |
| Rxrg | | | | | | | | | | | | | | | | 0.00821 | 3.27519 | Female |
| Hnf1b | | | | 0.00001 | -2.76609 | Male | | | | 0.00000 | 3.11720 | Female | | | | | | |
| Tcea1 | | | | | | | | | | | | | 0.00141 | -1.17921 | Male | | | |
| Ube2l6 | | | | | | | 0.03914 | -1.57943 | Male | | | | | | | 0.00040 | 2.10602 | Female |
| Foxc2 | | | | | | | | | | | | | | | | 0.00000 | -5.58371 | Male |
| Hmgcs1 | | | | | | | | | | | | | 0.00618 | 1.67314 | Female | | | |
| Pigp | | | | | | | | | | 0.00000 | -3.73791 | Male | 0.00112 | 1.35497 | Female | | | |
| Fmn1 | | | | | | | | | | 0.00000 | 0.89269 | Female | | | | | | |
| Ube2j2 | | | | | | | | | | 0.00000 | -2.63959 | Male | | | | | | |
| Krtcap2 | | | | | | | | | | 0.00945 | -2.46617 | Male | | | | | | |
| Psmd11 | | | | | | | | | | 0.00227 | 1.39397 | Female | | | | | | |
| Pigh | | | | | | | | | | | | | | | | 0.00109 | 5.03119 | Female |
| Terf2ip | | | | | | | | | | | | | | | | 0.00035 | 3.16757 | Female |
| Adprhl1 | 0.02693 | -0.64315 | Male | | | | 0.00000 | 1.26209 | Female | | | | | | | | | |
| Gstm1 | 0.00000 | -0.94518 | Male | | | | | | | 0.00000 | -1.55676 | Male | | | | | | |
| Amot | | | | | | | 0.00029 | 3.21206 | Female | | | | | | | | | |
| Tmem47 | | | | | | | | | | | | | | | | 0.03020 | 3.43826 | Female |
| Pigz | | | | 0.01419 | -1.53306 | Male | | | | | | | | | | | | |
| Neb | | | | | | | 0.01615 | -5.04453 | Male | | | | | | | | | |
| Scaper | | | | | | | | | | 0.03400 | 3.49928 | Female | | | | | | |
| Ppm1g | | | | | | | | | | 0.04196 | 1.30266 | Female | | | | | | |

| | | | | | | | | | | | | | | | | | | |
|---|---|---|---|---|---|---|---|---|---|---|---|---|---|---|---|---|---|---|
| Parp8 | | | | | | | | | | | | | 0.01878 | -1.50349 | Male | | | |
| Strn4 | | | | 0.04107 | 2.33245 | Female | | | | | | | | | | | | |
| C5ar2 | | | | | | | | | | 0.00014 | -4.19597 | Male | | | | | | |
| Wnt7b | | | | | | | | | | 0.02709 | 4.18252 | Female | | | | | | |
| Tmppe | | | | | | | | | | 0.03766 | -1.72024 | Male | | | | | | |
| Vprbp | | | | | | | | | | | | | 0.00052 | -2.55745 | Male | | | |
| Hdhd3 | 0.00045 | 1.28159 | Female | 0.00000 | 2.25557 | Female | 0.00000 | -3.56407 | Male | 0.00445 | -1.34922 | Male | | | | | | |
| Hibch | | | | | | | | | | 0.01795 | 2.49194 | Female | | | | | | |
| Rtn4r | | | | | | | | | | | | | | | | 0.00058 | -5.22263 | Male |
| Ahnak | | | | | | | | | | 0.00411 | -4.43929 | Male | 0.00062 | 3.69439 | Female | | | |
| Kpna4 | | | | | | | | | | 0.01441 | -1.92514 | Male | | | | | | |
| Ndst1 | | | | | | | | | | | | | 0.01487 | 2.87144 | Female | | | |
| Rnasel | | | | | | | | | | | | | | | | 0.04546 | 3.11045 | Female |
| Fam35a | | | | | | | | | | 0.03111 | 2.82144 | Female | | | | | | |
| Fam110c | | | | | | | | | | | | | | | | 0.00076 | 1.12260 | Female |
| Htr5b | | | | | | | | | | | | | | | | 0.00000 | 3.04466 | Female |
| Akna | | | | | | | | | | | | | | | | 0.00000 | 3.49230 | Female |
| Dpys | | | | 0.00144 | 1.13995 | Female | | | | | | | | | | | | |
| Kcp | 0.03593 | -1.05664 | Male | | | | 0.00002 | -1.76994 | Male | | | | | | | | | |
| Edn2 | | | | | | | | | | 0.02465 | 2.44955 | Female | | | | | | |
| Tcf24 | | | | 0.00222 | -2.13248 | Male | | | | | | | | | | | | |
| Elmod2 | | | | | | | | | | | | | | | | 0.01350 | -3.08476 | Male |
| Diexf | | | | | | | | | | 0.02682 | 2.26249 | Female | | | | | | |
| Arpin | | | | 0.00001 | 0.99013 | Female | | | | | | | 0.02822 | 3.93599 | Female | | | |
| Med30 | | | | | | | | | | 0.00027 | -6.09736 | Male | | | | | | |
| Syt9 | | | | | | | | | | 0.02385 | 3.72357 | Female | | | | | | |
| Tomm20 | | | | | | | | | | 0.00000 | -3.26891 | Male | | | | | | |
| Zfp523 | | | | | | | | | | 0.00000 | 3.32838 | Female | 0.00000 | -1.39169 | Male | | | |
| Mpp1 | | | | | | | | | | | | | 0.00052 | 1.51894 | Female | | | |
| Lnpep | | | | | | | 0.04416 | 4.37514 | Female | | | | | | | | | |
| Adtrp | | | | | | | | | | | | | | | | 0.00274 | -2.49132 | Male |
| Pggt1b | 0.00000 | 1.49106 | Female | | | | | | | | | | | | | | | |
| Catsper2 | | | | | | | | | | | | | | | | 0.01368 | -4.65355 | Male |
| Impdh2 | | | | | | | | | | | | | 0.02406 | -0.88626 | Male | | | |
| Lzts3 | | | | | | | | | | 0.03753 | -3.62920 | Male | | | | | | |
| Ccnt1 | | | | | | | | | | | | | | | | 0.04932 | -2.20948 | Male |
| Fhl3 | | | | | | | | | | 0.00165 | -5.28173 | Male | | | | | | |
| Slc25a26 | | | | | | | | | | | | | | | | 0.04387 | -2.39084 | Male |
| Arpc4 | | | | | | | | | | 0.00120 | 3.34100 | Female | | | | | | |
| Idi2 | | | | | | | | | | 0.00401 | 4.55564 | Female | | | | | | |
| Pnisr | | | | | | | | | | 0.00247 | 2.86422 | Female | | | | | | |
| Lrrc3 | | | | | | | | | | 0.00003 | -3.13651 | Male | | | | | | |
| Cpne9 | | | | | | | | | | | | | | | | 0.00946 | 3.58691 | Female |
| Ccl25 | 0.00000 | 1.08097 | Female | | | | | | | | | | | | | | | |
| Cfc1 | | | | | | | 0.00000 | -2.64789 | Male | | | | | | | | | |
| Shfm1 | | | | | | | | | | 0.00000 | -4.44805 | Male | 0.00000 | 2.37109 | Female | | | |
| Aif1l | | | | | | | | | | 0.02814 | -4.74745 | Male | | | | | | |
| Erdr1 | | | | | | | | | | 0.01912 | -2.67785 | Male | | | | | | |
| Ipo5 | | | | | | | | | | 0.04960 | 0.81033 | Female | | | | | | |
| Elovl3 | 0.04770 | 1.56531 | Female | | | | | | | | | | | | | | | |
| Timm10 | | | | | | | | | | 0.00840 | -1.89805 | Male | | | | | | |
| Zfp28 | 0.00000 | -1.22371 | Male | | | | | | | | | | | | | | | |
| Rpl4 | | | | | | | | | | 0.03218 | 0.80285 | Female | | | | | | |
| Abhd6 | | | | | | | | | | | | | | | | 0.00051 | -3.28655 | Male |
| Trps1 | | | | | | | | | | 0.00022 | -2.40182 | Male | 0.00000 | 0.88037 | Female | | | |
| D17Ertd648e | | | | | | | | | | | | | | | | 0.00958 | -1.47275 | Male |
| Mfsd4a | | | | | | | 0.04090 | 2.36846 | Female | 0.00274 | -3.19045 | Male | | | | | | |
| Dpysl4 | | | | 0.00071 | -2.67254 | Male | | | | | | | | | | | | |
| Net1 | | | | | | | | | | | | | | | | 0.04194 | -2.10813 | Male |
| Msrb1 | | | | | | | | | | 0.00003 | -3.85576 | Male | | | | | | |
| Cfap43 | | | | | | | 0.00000 | 5.59705 | Female | | | | | | | | | |
| Prrc2b | | | | 0.01041 | 3.09745 | Female | | | | | | | | | | | | |
| Adora1 | | | | | | | | | | 0.00001 | -1.10645 | Male | | | | | | |
| Xist | | | | 0.00003 | 5.13387 | Female | 0.00000 | 7.59985 | Female | 0.00000 | 5.55145 | Female | 0.00000 | 8.04782 | Female | 0.00000 | 7.38646 | Female |
| Actn2 | | | | | | | | | | | | | | | | 0.03198 | -2.07619 | Male |
| Sptssa | | | | | | | | | | 0.01890 | 1.42715 | Female | | | | | | |
| Psg16 | | | | | | | | | | 0.03207 | 1.53936 | Female | | | | | | |
| Lyrm9 | | | | | | | | | | 0.01895 | -3.20321 | Male | | | | | | |
| Dpy19l4 | | | | | | | | | | | | | | | | 0.00358 | -4.90775 | Male |
| Xpot | | | | | | | | | | | | | 0.00316 | -1.21172 | Male | | | |

| | | | | | | | | | | | | | | | | | | |
|---|---|---|---|---|---|---|---|---|---|---|---|---|---|---|---|---|---|---|
| Brip1os | | | | | | | | | | 0.00084 | -2.61769 | Male | | | | | | |
| Bex1 | | | | | | | | | | 0.02272 | 4.20701 | Female | | | | | | |
| Txn1 | | | | | | | | | | 0.00001 | -3.11379 | Male | 0.01140 | 1.46842 | Female | | | |
| Myoz1 | | | | | | | | | | 0.01068 | 2.27977 | Female | | | | | | |
| Prss41 | | | | | | | | | | | | | 0.00000 | 4.02721 | Female | | | |
| Igf1r | | | | | | | | | | | | | | | | 0.00000 | 2.82480 | Female |
| Snai2 | | | | | | | 0.00977 | -3.49010 | Male | | | | | | | | | |
| Ak4 | | | | | | | | | | 0.00033 | 2.06284 | Female | | | | | | |
| Kcnn4 | | | | 0.00002 | -2.34127 | Male | 0.01804 | -4.80920 | Male | 0.00000 | 3.53432 | Female | | | | | | |
| Ccdc181 | | | | | | | | | | | | | | | | 0.01178 | -5.27784 | Male |
| Apoe | | | | | | | | | | 0.01004 | -3.76743 | Male | | | | 0.00152 | 3.77981 | Female |
| Scml4 | | | | | | | | | | 0.04869 | -3.04073 | Male | | | | | | |
| Prl3b1 | | | | | | | | | | | | | | | | 0.00723 | -4.25803 | Male |
| Pomgnt2 | | | | 0.00340 | -0.80649 | Male | 0.00000 | 1.30704 | Female | | | | | | | | | |
| Ap3m2 | | | | | | | 0.00000 | 3.57698 | Female | | | | 0.00000 | 3.77661 | Female | | | |
| S100pbp | | | | | | | | | | | | | 0.00550 | 3.18554 | Female | | | |
| Hspe1 | | | | | | | | | | 0.00000 | -3.01664 | Male | 0.00000 | 1.95775 | Female | | | |
| Hpgds | | | | | | | | | | 0.00002 | -4.13297 | Male | | | | | | |
| Thg1l | | | | 0.02915 | 3.02308 | Female | | | | | | | | | | | | |
| Rab4a | | | | 0.00000 | -0.76277 | Male | | | | | | | | | | | | |
| Fam168a | | | | | | | | | | | | | | | | 0.00034 | 2.88112 | Female |
| Fam199x | | | | | | | | | | 0.00262 | -6.12352 | Male | | | | 0.02453 | -4.12826 | Male |
| Fcgbp | | | | | | | | | | | | | | | | 0.00753 | -5.13083 | Male |
| Cblc | | | | | | | | | | | | | | | | 0.01410 | 4.84727 | Female |
| Dcun1d5 | | | | | | | | | | 0.02440 | -1.82108 | Male | | | | | | |
| Abcc3 | | | | | | | | | | | | | | | | 0.00015 | 5.09256 | Female |
| Gm6525 | | | | | | | 0.00598 | 1.64061 | Female | 0.00000 | -4.77553 | Male | 0.00020 | 1.66824 | Female | | | |
| Rpl13 | | | | | | | | | | 0.00000 | -4.90932 | Male | | | | | | |
| Slirp | | | | | | | | | | 0.00000 | -5.42636 | Male | 0.00034 | 1.84790 | Female | | | |
| Dio3 | | | | | | | 0.00000 | 0.80920 | Female | 0.00020 | -2.15744 | Male | | | | 0.00000 | 2.75710 | Female |
| Psmd1 | | | | | | | | | | 0.00945 | 1.41184 | Female | | | | | | |
| Rbm20 | | | | 0.00000 | 0.99918 | Female | | | | | | | | | | | | |
| Clptm1 | | | | | | | | | | 0.01947 | 2.76364 | Female | | | | | | |
| Aox4 | 0.00438 | 0.80260 | Female | | | | | | | | | | | | | 0.00123 | -0.58628 | Male |
| Usf1 | | | | | | | | | | 0.00006 | 4.78162 | Female | | | | | | |
| Cpb2 | | | | | | | 0.00000 | -1.32918 | Male | | | | | | | | | |
| Ubl5 | | | | | | | | | | 0.00000 | -5.16160 | Male | | | | | | |
| Rps14 | | | | | | | | | | 0.00001 | -2.78685 | Male | 0.00989 | 1.31536 | Female | | | |
| Zbtb25 | | | | | | | | | | | | | 0.01565 | 4.51637 | Female | | | |
| Rcan3 | | | | | | | | | | | | | | | | 0.00251 | 3.77074 | Female |
| Apoa1 | | | | | | | | | | 0.00056 | 6.22215 | Female | | | | | | |
| Btbd3 | | | | | | | | | | 0.00000 | -3.04588 | Male | | | | 0.00791 | 4.61751 | Female |
| Elac2 | | | | | | | | | | 0.03125 | 2.28392 | Female | | | | | | |
| Sys1 | | | | | | | | | | 0.03143 | -2.29664 | Male | | | | | | |
| Mob3b | | | | | | | | | | 0.00000 | -5.21187 | Male | | | | | | |
| Prdx3 | | | | | | | | | | 0.00000 | 3.50572 | Female | | | | | | |
| Snrpb2 | | | | | | | 0.00435 | 1.23801 | Female | | | | | | | | | |
| Fmc1 | | | | | | | | | | | | | 0.01721 | 2.49675 | Female | | | |
| Tnfrsf25 | | | | 0.04154 | -0.68147 | Male | | | | 0.00005 | 2.58205 | Female | | | | 0.00000 | -4.09122 | Male |
| Sox2ot | | | | | | | | | | 0.03539 | -2.79234 | Male | | | | | | |
| Inpp4a | | | | | | | | | | | | | | | | 0.03731 | -4.08173 | Male |
| Cyld | | | | | | | | | | 0.00000 | -3.71244 | Male | 0.01542 | 1.42300 | Female | | | |
| Smyd1 | | | | 0.00000 | 0.95189 | Female | | | | | | | | | | | | |
| Zdhhc16 | | | | | | | | | | 0.02417 | 2.97399 | Female | | | | | | |
| AF067061 | | | | | | | | | | | | | | | | 0.00036 | -4.14294 | Male |
| Tnfaip6 | | | | | | | | | | 0.01624 | -3.18043 | Male | | | | | | |
| Wtap | | | | | | | | | | | | | | | | 0.01293 | -1.41479 | Male |
| Cald1 | | | | | | | | | | | | | 0.00000 | 3.56662 | Female | | | |
| Fam171a1 | | | | | | | | | | 0.00000 | 1.20480 | Female | | | | | | |
| Impg2 | | | | | | | | | | | | | 0.01374 | -4.73626 | Male | | | |
| Rpl35a | | | | | | | | | | 0.00000 | -4.55379 | Male | 0.00012 | 1.58643 | Female | | | |
| Lrmp | | | | | | | | | | | | | | | | 0.00002 | 0.91979 | Female |
| Rin3 | 0.00000 | -0.97951 | Male | | | | | | | | | | | | | | | |
| Ube2l3 | | | | | | | | | | 0.00631 | -1.17988 | Male | | | | | | |
| Otud3 | | | | | | | | | | | | | 0.03311 | 3.19569 | Female | | | |
| Ggt6 | | | | | | | | | | | | | | | | 0.00612 | 3.98016 | Female |
| Mei4 | | | | | | | | | | 0.00102 | -3.89608 | Male | | | | | | |
| Ranbp3l | | | | | | | | | | 0.00220 | -2.62125 | Male | 0.03851 | 1.44260 | Female | | | |
| Fhl1 | | | | 0.01587 | -0.91472 | Male | | | | | | | 0.00000 | -1.65848 | Male | 0.00000 | 1.66972 | Female |
| Rgs11 | | | | | | | | | | | | | | | | 0.04035 | -2.60654 | Male |

| | | | | | | | | | | | | | | | | | | |
|---|---|---|---|---|---|---|---|---|---|---|---|---|---|---|---|---|---|---|
| Fcgr1 | | | | 0.03492 | 1.87472 | Female | | | | | | | | | | 0.00011 | -0.75088 | Male |
| Bccip | | | | | | | | | | 0.00000 | 5.36605 | Female | | | | | | |
| Fdx1 | | | | | | | | | | 0.01940 | -2.66533 | Male | | | | | | |
| Abcc9 | | | | | | | | | | 0.00001 | -3.16429 | Male | | | | | | |
| Pdia3 | | | | | | | | | | 0.02060 | -2.21762 | Male | | | | | | |
| Myh15 | | | | | | | 0.00723 | 1.06328 | Female | 0.03365 | 1.42980 | Female | | | | | | |
| Egf | | | | | | | | | | | | | | | | 0.02824 | 3.52891 | Female |
| Tert | | | | | | | | | | 0.00000 | 3.41413 | Female | | | | | | |
| Aim2 | | | | | | | 0.03477 | 3.85219 | Female | | | | | | | | | |
| Oat | | | | | | | | | | 0.00081 | -5.50312 | Male | | | | | | |
| Saal1 | 0.03830 | 3.26349 | Female | | | | | | | | | | | | | | | |
| Slc50a1 | | | | | | | | | | 0.00144 | -0.94723 | Male | | | | | | |
| Erich5 | | | | | | | | | | 0.00000 | -5.30243 | Male | | | | | | |
| Hcfc2 | | | | | | | | | | 0.00030 | -1.95403 | Male | | | | | | |
| Elk4 | | | | | | | | | | | | | | | | 0.00217 | 4.05393 | Female |
| Tnfaip8l2 | | | | | | | | | | | | | | | | 0.02887 | 3.00388 | Female |
| Akap5 | | | | | | | | | | 0.00110 | -4.99053 | Male | | | | 0.02200 | 3.31185 | Female |
| Ndufa3 | | | | | | | | | | 0.00000 | -4.47148 | Male | | | | | | |
| Pax6 | | | | | | | | | | | | | 0.01038 | 3.57854 | Female | | | |
| Atp2a3 | | | | 0.00801 | -1.71477 | Male | 0.00000 | -1.09525 | Male | | | | | | | | | |
| Plscr1 | 0.00000 | -1.82470 | Male | | | | | | | | | | | | | | | |
| Slc30a3 | | | | 0.00009 | -0.68383 | Male | | | | | | | | | | | | |
| Dok1 | 0.00000 | 1.50949 | Female | | | | 0.00064 | 4.03773 | Female | | | | | | | | | |
| Eif3m | | | | | | | | | | 0.00176 | 1.13223 | Female | | | | | | |
| Inf2 | 0.00000 | 1.18635 | Female | | | | | | | | | | | | | | | |
| Ripply3 | | | | | | | | | | 0.00000 | -5.00587 | Male | 0.02612 | 1.55219 | Female | | | |
| Slc43a2 | | | | | | | | | | 0.01589 | -3.86036 | Male | | | | | | |
| Slc7a2 | | | | | | | | | | 0.00006 | -3.49606 | Male | | | | | | |
| Ciita | | | | | | | | | | | | | 0.00000 | -4.40902 | Male | | | |
| Idi1 | | | | | | | 0.00513 | 3.21856 | Female | | | | | | | | | |
| Ids | | | | 0.03951 | 3.15070 | Female | | | | | | | | | | | | |
| Chchd5 | | | | | | | | | | | | | | | | 0.00016 | -2.22975 | Male |
| Trp53i11 | | | | | | | 0.01012 | 3.90832 | Female | | | | | | | | | |
| Astl | | | | | | | | | | | | | 0.00674 | 4.01295 | Female | | | |
| Nupl2 | | | | | | | | | | | | | | | | 0.00000 | 4.53890 | Female |
| Sdf2l1 | | | | | | | | | | | | | | | | 0.02818 | 3.71992 | Female |
| Ccdc85b | | | | | | | | | | 0.00001 | -5.33963 | Male | | | | | | |
| Tmem64 | | | | | | | | | | | | | | | | 0.01716 | -2.20738 | Male |
| X3.Sep | | | | | | | | | | | | | | | | 0.00075 | 3.81940 | Female |
| Mrpl52 | | | | | | | | | | 0.03397 | -2.60380 | Male | | | | | | |
| Ttll4 | | | | | | | | | | 0.02732 | 2.90358 | Female | | | | | | |
| Mprip | | | | | | | | | | 0.04052 | 1.39745 | Female | | | | | | |
| Snhg15 | | | | | | | | | | 0.02511 | -2.52151 | Male | | | | | | |
| Gm6654 | | | | | | | | | | 0.00000 | -3.98004 | Male | 0.01711 | 1.38931 | Female | | | |
| Calm2 | | | | | | | | | | 0.00324 | 1.43769 | Female | | | | | | |
| Tbrg3 | | | | | | | | | | 0.02012 | -2.64703 | Male | | | | | | |
| Fanca | | | | | | | | | | 0.04981 | 3.04557 | Female | | | | | | |
| Actn1 | | | | | | | | | | 0.00051 | -3.83608 | Male | | | | | | |
| Gtf3a | | | | | | | | | | 0.02619 | 3.78463 | Female | | | | | | |
| Fam193a | | | | | | | | | | 0.00091 | 4.21681 | Female | | | | | | |
| Eftud2 | | | | | | | | | | 0.01592 | 1.65832 | Female | | | | | | |
| Kcnj2 | | | | | | | | | | 0.00185 | -5.21077 | Male | | | | | | |
| Rwdd1 | | | | | | | | | | 0.00152 | -2.32830 | Male | | | | | | |
| Gm2560 | | | | 0.03876 | 0.90465 | Female | | | | | | | 0.00000 | 0.74513 | Female | 0.00000 | -3.48719 | Male |
| Ppp1ca | | | | | | | | | | 0.01162 | 2.00843 | Female | | | | | | |
| Psg29 | | | | | | | | | | 0.00173 | -1.93422 | Male | 0.00985 | 1.29689 | Female | | | |
| Hnrnpf | | | | | | | | | | 0.00281 | 1.28900 | Female | | | | | | |
| Piwil1 | | | | | | | | | | | | | | | | 0.00491 | -4.99219 | Male |
| Apobr | | | | | | | | | | 0.00000 | 2.39186 | Female | | | | | | |
| Gm10012 | | | | | | | 0.00760 | 1.70295 | Female | 0.00000 | -6.07376 | Male | 0.00000 | 2.17363 | Female | | | |
| Etv1 | | | | | | | | | | 0.00241 | -1.58024 | Male | | | | | | |
| Naip5 | | | | | | | | | | 0.00001 | -3.33290 | Male | | | | | | |
| Wdr77 | | | | | | | | | | 0.00473 | 1.21446 | Female | | | | | | |
| Wbscr17 | | | | 0.00434 | 1.99249 | Female | | | | | | | | | | | | |
| Tm2d2 | | | | | | | | | | 0.01554 | 3.06236 | Female | | | | | | |
| Ankrd11 | | | | | | | | | | 0.00000 | 5.07561 | Female | | | | | | |
| Znrd1 | | | | | | | | | | 0.00704 | -1.80642 | Male | | | | | | |
| Dclre1c | | | | | | | | | | 0.00036 | -2.84711 | Male | | | | | | |
| Zcrb1 | | | | | | | | | | 0.00019 | -3.25478 | Male | | | | | | |
| Myeov2 | | | | | | | | | | 0.00000 | -6.91333 | Male | | | | | | |

| | | | | | | | | | | | | | | | | | | |
|---|---|---|---|---|---|---|---|---|---|---|---|---|---|---|---|---|---|---|
| Lrrc58 | | | | | | | | | | 0.00690 | -1.71312 | Male | | | | | | |
| Aard | | | | | | | 0.00000 | 1.81349 | Female | 0.00001 | -1.47158 | Male | | | | | | |
| Cdc16 | | | | | | | | | | | | | | | | 0.01179 | -1.58518 | Male |
| Rpl24 | | | | | | | | | | 0.00000 | -2.50288 | Male | 0.01636 | 1.04549 | Female | | | |
| Tmem88 | | | | 0.02039 | -1.07682 | Male | | | | 0.00000 | 1.71506 | Female | | | | | | |
| Mrpl23 | | | | | | | | | | 0.00000 | -4.08856 | Male | | | | | | |
| Cox5a | | | | | | | | | | 0.00000 | -5.74158 | Male | | | | | | |
| Nfkb1 | | | | | | | | | | 0.01320 | -1.13040 | Male | 0.00000 | 2.20132 | Female | | | |
| Lap3 | | | | | | | | | | 0.03633 | 1.30383 | Female | | | | | | |
| Rpl22l1 | | | | | | | | | | 0.00000 | -4.89446 | Male | 0.00550 | 1.47828 | Female | | | |
| Usp48 | | | | | | | | | | 0.03238 | 0.89636 | Female | | | | | | |
| Psd2 | | | | | | | | | | | | | 0.00000 | 1.02155 | Female | | | |
| Calml4 | | | | | | | | | | 0.00454 | -5.06879 | Male | | | | | | |
| Gpld1 | 0.00000 | 0.76780 | Female | | | | | | | 0.01728 | 3.35685 | Female | | | | | | |
| Mylk | | | | 0.00000 | 1.27292 | Female | | | | 0.00556 | -1.17482 | Male | | | | | | |
| Slc25a21 | | | | | | | | | | 0.00537 | 4.68219 | Female | | | | 0.00004 | -5.66013 | Male |
| Ciao1 | | | | | | | | | | | | | | | | 0.01159 | -1.33908 | Male |
| Umad1 | | | | 0.00000 | 1.31565 | Female | | | | | | | | | | | | |
| Slc26a8 | | | | | | | | | | 0.00006 | -0.80059 | Male | | | | | | |
| Sdk1 | | | | 0.00049 | -1.40301 | Male | | | | | | | | | | | | |
| Hypk | | | | | | | | | | 0.00000 | -4.19164 | Male | 0.00057 | 1.64101 | Female | | | |
| Irf5 | 0.00116 | 0.81327 | Female | | | | | | | | | | | | | 0.00600 | -6.32617 | Male |
| Atpif1 | | | | | | | | | | 0.00000 | -4.50595 | Male | 0.00014 | 1.82449 | Female | | | |
| Sap30l | | | | | | | | | | 0.00004 | -3.97696 | Male | | | | | | |
| Gm9895 | | | | | | | | | | | | | 0.01361 | 4.69221 | Female | | | |
| Rab27b | | | | | | | | | | 0.02110 | -3.17663 | Male | | | | | | |
| Myom1 | | | | 0.04200 | 2.39793 | Female | | | | | | | | | | | | |
| Mdh1 | | | | | | | | | | 0.00087 | 3.11411 | Female | | | | | | |
| Il5ra | | | | | | | | | | 0.00268 | -3.60730 | Male | | | | | | |
| Rpp38 | | | | | | | 0.01424 | 2.17352 | Female | | | | | | | | | |
| Pcnx2 | | | | | | | | | | 0.00016 | 2.80433 | Female | | | | 0.00000 | 1.79226 | Female |
| Upk1a | | | | | | | | | | | | | 0.00107 | 4.79215 | Female | | | |
| Bre | | | | 0.00434 | 2.55340 | Female | | | | | | | | | | | | |
| Rnaseh2b | 0.03878 | 3.01960 | Female | | | | | | | | | | | | | | | |
| Slc22a18 | | | | | | | 0.00114 | -2.23616 | Male | | | | | | | | | |
| Stub1 | | | | | | | | | | 0.00000 | 6.94877 | Female | | | | | | |
| Creb3l2 | | | | 0.00006 | -1.41669 | Male | | | | 0.00015 | -1.26425 | Male | | | | | | |
| Prune2 | | | | | | | | | | 0.00034 | -2.21509 | Male | 0.00000 | 1.32451 | Female | | | |
| Utp14a | | | | | | | | | | 0.00004 | 2.86149 | Female | | | | | | |
| Fam45a | | | | | | | | | | | | | | | | 0.02638 | -1.70212 | Male |
| Srsf4 | | | | | | | | | | 0.01828 | 4.73947 | Female | | | | | | |
| Fam65c | | | | | | | | | | 0.04530 | 5.05708 | Female | | | | | | |
| Flii | | | | | | | | | | 0.00207 | 2.96892 | Female | | | | 0.02410 | -1.91390 | Male |
| Aldoart1 | | | | | | | | | | 0.00000 | 3.27953 | Female | | | | | | |
| Rhbdl2 | | | | | | | | | | 0.02402 | 3.10504 | Female | | | | | | |
| Rbbp7 | | | | | | | 0.02207 | 1.15091 | Female | 0.00000 | 2.26104 | Female | | | | | | |
| Akap8l | | | | | | | | | | 0.01732 | 3.65937 | Female | | | | | | |
| Ccny | | | | | | | | | | 0.02569 | 3.30630 | Female | | | | | | |
| Stra13 | | | | | | | | | | 0.00003 | -3.21547 | Male | 0.00249 | 1.83844 | Female | | | |
| Fos | | | | | | | | | | 0.00000 | -2.77262 | Male | | | | | | |
| Ldhc | | | | | | | | | | | | | 0.03455 | -4.32241 | Male | 0.04387 | -4.14120 | Male |
| Tmc5 | | | | | | | | | | 0.00030 | -2.47775 | Male | | | | | | |
| Anapc13 | | | | | | | | | | 0.00000 | -4.76707 | Male | 0.00049 | 2.09099 | Female | | | |
| Copg2 | | | | | | | | | | 0.00252 | -5.60735 | Male | | | | | | |
| Sf3b3 | | | | | | | | | | 0.04380 | 1.23515 | Female | | | | | | |
| Mfsd4b1 | 0.00000 | -1.04633 | Male | | | | | | | | | | | | | | | |
| Ticam1 | | | | 0.02985 | 1.53237 | Female | | | | | | | | | | | | |
| Loxl2 | | | | | | | | | | 0.00000 | -6.07562 | Male | 0.00106 | 1.95095 | Female | | | |
| Itgb5 | | | | | | | | | | 0.02642 | 2.03541 | Female | | | | | | |
| Nubpl | | | | 0.00000 | 1.29839 | Female | | | | | | | | | | | | |
| Hus1b | | | | | | | 0.00000 | -1.28869 | Male | | | | | | | | | |
| Gtf2b | | | | | | | | | | 0.01407 | 1.98801 | Female | | | | | | |
| Patl2 | | | | | | | | | | 0.00000 | 1.32961 | Female | | | | | | |
| Alox12e | | | | | | | | | | | | | 0.00000 | 2.61267 | Female | | | |
| Lrp4 | | | | | | | | | | | | | | | | 0.00894 | 4.16413 | Female |
| Accsl | | | | | | | | | | | | | | | | 0.00568 | 3.73426 | Female |
| Upb1 | | | | | | | | | | | | | 0.02387 | 3.48192 | Female | | | |
| Ncbp3 | | | | | | | | | | 0.02882 | -1.54367 | Male | | | | | | |
| Xlr5c | | | | | | | | | | | | | 0.00243 | -3.46474 | Male | | | |
| Nol11 | | | | | | | | | | 0.00152 | 2.23476 | Female | | | | | | |

| | | | | | | | | | | | | | | | | | | |
|---|---|---|---|---|---|---|---|---|---|---|---|---|---|---|---|---|---|---|
| Rltpr | | | | | | | 0.00000 | 3.52778 | Female | | | | | | | | | |
| Lgr5 | | | | | | | 0.02677 | 2.70120 | Female | 0.00516 | -3.40032 | Male | | | | | | |
| Spink2 | | | | | | | | | | 0.00599 | -3.51946 | Male | | | | | | |
| Slc6a4 | | | | | | | | | | | | | 0.00703 | 3.80916 | Female | | | |
| Ptprn | | | | 0.00000 | -1.23620 | Male | 0.00000 | 3.79962 | Female | | | | | | | | | |
| Smu1 | | | | | | | | | | 0.01169 | 1.24372 | Female | | | | | | |
| Enpp3 | | | | | | | | | | | | | | | | 0.00224 | 3.95312 | Female |
| Mroh2b | | | | | | | | | | | | | | | | 0.02148 | 1.70137 | Female |
| Zfp345 | | | | | | | 0.00033 | 2.76658 | Female | | | | | | | | | |
| Gm5607 | | | | | | | | | | 0.00069 | -2.43125 | Male | | | | | | |
| Dhrs3 | | | | 0.00000 | -1.27641 | Male | | | | | | | | | | 0.00000 | 4.66698 | Female |
| C87414 | | | | | | | | | | | | | 0.01737 | -1.82203 | Male | | | |
| Pfas | | | | | | | | | | | | | | | | 0.02168 | 1.84193 | Female |
| Arrdc2 | | | | | | | | | | | | | | | | 0.00778 | 3.38820 | Female |
| Klhl41 | | | | | | | | | | 0.00000 | -5.10428 | Male | | | | | | |
| Trp53 | | | | | | | | | | 0.01156 | 3.08746 | Female | | | | | | |
| Nlrp12 | | | | | | | | | | | | | | | | 0.00754 | 3.75341 | Female |
| Ddx6 | | | | | | | | | | | | | 0.00195 | 2.21374 | Female | | | |
| Rbbp4 | | | | | | | | | | 0.00239 | -2.48755 | Male | | | | | | |
| Lats2 | | | | | | | | | | | | | | | | 0.00225 | -3.58674 | Male |
| Emb | | | | 0.00000 | 0.77275 | Female | | | | | | | | | | | | |
| Noc2l | | | | | | | | | | 0.00000 | 3.39166 | Female | | | | | | |
| Dph3 | | | | | | | | | | 0.00556 | -2.20821 | Male | | | | | | |
| Stra8 | | | | 0.00002 | -0.83668 | Male | | | | | | | | | | | | |
| Fam160b1 | | | | 0.01026 | 2.65635 | Female | | | | | | | | | | | | |
| Eed | | | | | | | | | | 0.03315 | 1.42811 | Female | | | | | | |
| Myom3 | | | | 0.00028 | 1.65114 | Female | | | | | | | | | | | | |
| Tacstd2 | | | | | | | | | | 0.04867 | -4.12544 | Male | | | | | | |
| Chm | | | | | | | 0.01289 | 3.99816 | Female | | | | | | | | | |
| Ebf3 | | | | | | | | | | 0.00044 | -1.04980 | Male | | | | | | |
| Il33 | | | | | | | | | | 0.00000 | 1.85050 | Female | | | | | | |
| Ttll9 | | | | | | | | | | 0.00000 | 0.80599 | Female | | | | 0.00000 | 1.10056 | Female |
| Atp5l | | | | | | | | | | 0.00000 | -5.16106 | Male | 0.00023 | 1.66496 | Female | | | |
| Fgd3 | | | | 0.00050 | 0.94843 | Female | | | | | | | | | | 0.00000 | 3.03920 | Female |
| Sash1 | | | | 0.00000 | 1.97787 | Female | | | | | | | | | | | | |
| Spns3 | | | | | | | 0.00001 | -1.98794 | Male | | | | | | | | | |
| Lmx1a | | | | 0.00000 | 2.76526 | Female | | | | | | | | | | | | |
| Bnc1 | | | | | | | | | | 0.00000 | -3.34499 | Male | | | | | | |
| Serf2 | | | | | | | | | | 0.00002 | -2.80371 | Male | | | | | | |
| Mgme1 | | | | | | | | | | 0.03807 | 2.07240 | Female | | | | | | |
| Exosc9 | | | | | | | | | | 0.04058 | 1.61650 | Female | | | | | | |
| Camk2b | | | | 0.00000 | -0.75275 | Male | | | | | | | | | | | | |
| Prcc | | | | | | | | | | 0.01624 | 3.59951 | Female | | | | | | |
| Map3k8 | | | | | | | 0.00005 | -3.41451 | Male | | | | | | | | | |
| Gstm7 | | | | 0.01168 | -1.44850 | Male | | | | 0.00000 | 1.69988 | Female | | | | | | |
| Cdc23 | | | | | | | | | | 0.01625 | 2.25856 | Female | | | | | | |
| Rps24 | | | | | | | | | | 0.00000 | -4.22623 | Male | 0.00207 | 1.24643 | Female | | | |
| Noa1 | | | | | | | | | | | | | | | | 0.00002 | 3.64080 | Female |
| Dnajc9 | | | | | | | | | | 0.00073 | -3.68373 | Male | | | | | | |
| App | 0.00000 | 1.13995 | Female | 0.00001 | 0.70822 | Female | | | | | | | | | | | | |
| Fgf13 | | | | | | | 0.00154 | 4.47044 | Female | | | | | | | | | |
| Adam28 | | | | | | | | | | 0.00000 | -3.44536 | Male | | | | | | |
| Ccdc152 | | | | 0.00258 | 0.90265 | Female | | | | | | | | | | | | |
| Rarb | | | | | | | | | | 0.00120 | 4.85845 | Female | | | | | | |
| Ech1 | | | | 0.00676 | 2.54155 | Female | | | | | | | | | | | | |
| Rnf41 | | | | | | | | | | | | | | | | 0.00581 | -2.19675 | Male |
| Ccdc88b | | | | | | | | | | 0.04886 | -0.97587 | Male | | | | | | |
| Snrpb | | | | | | | | | | 0.04425 | 2.24764 | Female | | | | | | |
| Cdo1 | | | | | | | | | | | | | 0.00000 | 2.04340 | Female | | | |
| Elovl5 | | | | | | | 0.00033 | 3.47474 | Female | | | | | | | | | |
| Prkra | | | | 0.00600 | -1.74882 | Male | | | | | | | | | | 0.00287 | -5.58293 | Male |
| Gtf2h4 | | | | | | | | | | 0.00000 | 6.93752 | Female | | | | | | |
| Elf3 | | | | | | | | | | 0.02926 | 2.31737 | Female | | | | | | |
| Ier3ip1 | | | | | | | | | | 0.02776 | -2.24269 | Male | | | | | | |
| Scpep1os | | | | | | | | | | 0.00000 | -3.46408 | Male | | | | | | |
| Carf | | | | | | | 0.03425 | 2.35465 | Female | | | | | | | | | |
| Arl10 | | | | | | | | | | | | | 0.00000 | -3.06252 | Male | | | |
| Rassf8 | 0.00000 | -1.47378 | Male | | | | | | | | | | | | | | | |
| Bnip3 | | | | | | | | | | 0.02475 | -2.28776 | Male | | | | | | |
| Flywch2 | | | | | | | | | | | | | 0.00691 | 4.30251 | Female | 0.00312 | -5.05810 | Male |

| | | | | | | | | | | | | | | | | | | |
|---|---|---|---|---|---|---|---|---|---|---|---|---|---|---|---|---|---|---|
| Cse1l | | | | | | | | | | | | | 0.04572 | -0.98025 | Male | | | |
| Fah | | | | 0.00001 | 2.77266 | Female | | | | | | | 0.00000 | 2.50506 | Female | | | |
| Pi4kb | | | | | | | 0.00575 | 4.44337 | Female | | | | | | | | | |
| Ddx3x | | | | | | | 0.01758 | 1.40820 | Female | | | | | | | | | |
| Olfr1372.ps1 | | | | | | | | | | | | | 0.02069 | 2.27467 | Female | | | |
| Klf7 | | | | | | | | | | | | | 0.00000 | -2.62698 | Male | | | |
| Zfp516 | 0.00000 | -1.05797 | Male | | | | | | | 0.00046 | -1.80781 | Male | 0.00513 | 5.15890 | Female | | | |
| Sdhd | | | | | | | 0.00999 | 2.45849 | Female | | | | | | | | | |
| Hcfc1r1 | | | | | | | | | | 0.04425 | 2.37409 | Female | | | | | | |
| Hdac9 | | | | | | | | | | 0.02817 | -0.58746 | Male | 0.00000 | -2.12554 | Male | | | |
| Syt15 | | | | | | | | | | 0.00000 | -4.71006 | Male | | | | | | |
| Sh3kbp1 | | | | | | | | | | | | | 0.04487 | 3.58396 | Female | | | |
| Eml6 | 0.00243 | -0.86405 | Male | 0.00001 | 2.13757 | Female | 0.00000 | 3.82153 | Female | | | | | | | 0.00000 | -1.86099 | Male |
| Nlrc4 | | | | 0.00010 | 0.83686 | Female | | | | | | | | | | 0.00000 | 2.46016 | Female |
| Slc22a17 | | | | 0.00005 | -2.48306 | Male | | | | | | | | | | | | |
| Smad7 | | | | | | | 0.00426 | 3.65207 | Female | | | | | | | | | |
| Abhd15 | | | | | | | 0.00092 | -3.37153 | Male | 0.00079 | -4.08974 | Male | | | | | | |
| Fbxl8 | | | | 0.00434 | -0.92634 | Male | | | | 0.01811 | -1.06081 | Male | | | | | | |
| Cbr4 | | | | 0.02199 | 2.41873 | Female | 0.01772 | 3.81304 | Female | 0.00350 | 4.19971 | Female | | | | | | |
| Blnk | | | | | | | 0.00000 | -4.23994 | Male | | | | | | | | | |
| Rdh11 | | | | | | | 0.04589 | 2.91823 | Female | | | | | | | | | |
| Dkk1 | | | | | | | | | | 0.00288 | -4.62722 | Male | | | | | | |
| Ncapd2 | | | | | | | | | | 0.00004 | 2.32593 | Female | | | | | | |
| Mak | | | | 0.00028 | 2.80838 | Female | | | | 0.02142 | -1.09383 | Male | | | | | | |
| Cenpv | | | | | | | | | | 0.01557 | -4.88312 | Male | | | | | | |
| Ms4a6c | | | | | | | | | | | | | | | | 0.00175 | 4.47798 | Female |
| Zfp647 | | | | | | | | | | | | | | | | 0.00000 | 2.43758 | Female |
| Osbpl5 | | | | 0.00027 | -0.58681 | Male | | | | | | | | | | | | |
| Mir3473f | | | | | | | | | | 0.00014 | -4.59252 | Male | | | | | | |
| Krcc1 | | | | | | | | | | | | | | | | 0.03415 | -2.72971 | Male |
| Psmg4 | | | | | | | | | | 0.02171 | -3.38279 | Male | | | | | | |
| Nyap1 | | | | | | | | | | | | | | | | 0.03477 | -4.32129 | Male |
| Rmi1 | | | | | | | | | | | | | | | | 0.00007 | -3.82189 | Male |
| Dnajc14 | | | | | | | | | | 0.00063 | 2.93366 | Female | | | | | | |
| Bhlhb9 | | | | | | | 0.00022 | 5.42556 | Female | | | | | | | 0.00003 | 5.79291 | Female |
| Ptpn2 | | | | | | | | | | | | | 0.04050 | -1.17905 | Male | | | |
| Btbd18 | | | | | | | | | | | | | | | | 0.00000 | -4.96604 | Male |
| Cox7a2l | | | | | | | | | | 0.03093 | 1.60746 | Female | | | | | | |
| Nagk | 0.00663 | 4.02191 | Female | | | | | | | | | | | | | | | |
| Rpl11 | | | | | | | | | | 0.00000 | -2.57758 | Male | 0.01508 | 1.15840 | Female | | | |
| Prob1 | | | | | | | | | | 0.00190 | -2.08352 | Male | | | | | | |
| Zap70 | 0.00000 | 2.21040 | Female | | | | | | | | | | | | | | | |
| Adap1 | | | | | | | | | | | | | | | | 0.04222 | 3.69659 | Female |
| Lrrn4cl | | | | | | | | | | 0.00000 | -4.06858 | Male | | | | | | |
| Foxo4 | | | | | | | | | | | | | 0.04100 | -3.67280 | Male | | | |
| Polr1c | | | | | | | | | | 0.00087 | 1.56057 | Female | | | | | | |
| Plk5 | | | | | | | | | | | | | | | | 0.00976 | -4.38070 | Male |
| Adamts4 | | | | | | | | | | 0.00000 | -5.50132 | Male | 0.00261 | 2.10389 | Female | | | |
| Clk3 | | | | 0.01412 | 3.14417 | Female | | | | | | | | | | | | |
| Polr1b | | | | | | | | | | 0.04867 | 1.41180 | Female | | | | | | |
| Asb4 | | | | | | | | | | | | | 0.03988 | 3.90226 | Female | | | |
| Gm14403 | | | | | | | | | | | | | | | | 0.00606 | -1.99832 | Male |
| Rbm22 | | | | | | | | | | 0.03336 | -1.43140 | Male | | | | | | |
| Cldn9 | | | | 0.00238 | -2.05751 | Male | | | | | | | | | | | | |
| Prss36 | | | | 0.02581 | 2.15915 | Female | | | | | | | | | | | | |
| Fam43a | | | | | | | 0.00000 | 2.98778 | Female | | | | | | | | | |
| Cox8a | | | | | | | | | | 0.00003 | -4.27761 | Male | | | | | | |
| Gtf2h5 | | | | | | | | | | 0.01289 | -1.64569 | Male | | | | | | |
| Sox17 | | | | | | | | | | 0.00001 | -0.77633 | Male | | | | 0.00123 | 6.30443 | Female |
| Fbxo3 | | | | 0.00000 | 1.04098 | Female | | | | 0.00070 | 4.84341 | Female | | | | | | |
| Atp1a1 | | | | | | | | | | 0.01874 | 1.80350 | Female | | | | | | |
| Abr | | | | 0.04808 | 2.65099 | Female | | | | | | | | | | | | |
| Snx11 | | | | | | | | | | 0.03800 | 3.71845 | Female | | | | 0.02519 | 3.27780 | Female |
| Zfand5 | | | | | | | | | | 0.01116 | -2.06992 | Male | | | | | | |
| Galntl6 | | | | | | | | | | 0.02112 | -2.90402 | Male | | | | | | |
| Fam73b | | | | | | | | | | | | | | | | 0.02876 | -3.56883 | Male |
| Npm3 | | | | | | | | | | 0.03895 | -2.98422 | Male | | | | | | |
| Rbfa | | | | | | | | | | 0.00075 | 5.45660 | Female | | | | | | |
| Mrpl42 | | | | | | | | | | 0.00000 | -2.48619 | Male | | | | | | |
| Lrch1 | | | | | | | | | | | | | 0.00608 | 4.09059 | Female | | | |

| | | | | | | | | | | | | | | | | | | |
|---|---|---|---|---|---|---|---|---|---|---|---|---|---|---|---|---|---|---|
| Sfswap | | | | | | | | | | 0.00218 | 3.53294 | Female | | | | | | |
| Gm13648 | | | | | | | | | | | | | | | | 0.04579 | 2.75627 | Female |
| Oplah | | | | | | | | | | 0.02858 | 4.03370 | Female | | | | | | |
| Coro1c | | | | | | | | | | 0.04522 | 1.73376 | Female | | | | | | |
| Gm10636 | | | | | | | | | | 0.00268 | -3.46767 | Male | | | | | | |
| B9d2 | | | | 0.00087 | -0.65646 | Male | | | | | | | | | | | | |
| Exoc3l4 | | | | 0.00686 | -1.14549 | Male | | | | | | | | | | | | |
| Cebpzos | | | | | | | | | | 0.00003 | -3.52758 | Male | | | | | | |
| Hspd1 | | | | | | | | | | 0.01973 | 1.15400 | Female | | | | | | |
| Acaa1a | | | | 0.03360 | 0.90742 | Female | | | | | | | | | | | | |
| Doc2g | | | | | | | | | | 0.03588 | 3.36597 | Female | | | | | | |
| Serpinf1 | | | | | | | | | | | | | 0.00000 | -2.12354 | Male | | | |
| Gm16702 | | | | | | | | | | 0.00000 | -5.46626 | Male | 0.00000 | 1.87275 | Female | | | |
| Rest | | | | | | | | | | 0.01406 | 3.46633 | Female | | | | | | |
| Kdsr | | | | | | | | | | | | | | | | 0.01134 | -2.41351 | Male |
| Brdt | | | | | | | | | | | | | 0.00392 | 3.73711 | Female | | | |
| Bud13 | | | | | | | 0.03732 | 2.02874 | Female | 0.00001 | 3.89750 | Female | | | | | | |
| Polr2i | | | | | | | | | | 0.03776 | -2.22292 | Male | | | | | | |
| Apoc1 | | | | | | | | | | 0.00000 | -4.68959 | Male | 0.01919 | 1.73956 | Female | | | |
| Cog8 | | | | | | | | | | 0.00396 | 4.57200 | Female | | | | | | |
| Ift22 | | | | 0.00001 | -1.43509 | Male | | | | | | | | | | | | |
| Epb42 | | | | | | | | | | 0.01209 | -3.45434 | Male | | | | | | |
| Wfdc2 | 0.00000 | 2.49662 | Female | 0.00000 | -1.13265 | Male | | | | | | | | | | | | |
| Gpr155 | | | | | | | | | | 0.00099 | -1.22510 | Male | 0.00000 | 3.11186 | Female | 0.00000 | 1.44541 | Female |
| Tmem145 | | | | | | | | | | | | | | | | 0.01178 | -4.37832 | Male |
| Zfp706 | | | | | | | | | | 0.00180 | -1.27362 | Male | | | | | | |
| Exoc6 | | | | | | | | | | | | | 0.00271 | -2.66966 | Male | 0.00078 | -3.26380 | Male |
| P2rx5 | | | | | | | | | | | | | 0.00000 | 1.76241 | Female | 0.00000 | -2.06668 | Male |
| Chmp4b | | | | | | | | | | 0.00000 | -5.29950 | Male | | | | | | |
| Cab39l | | | | | | | | | | | | | | | | 0.02284 | 1.60871 | Female |
| Vim | 0.00000 | 1.42790 | Female | | | | | | | | | | | | | | | |
| Dnajb11 | | | | | | | | | | 0.04916 | 1.55948 | Female | | | | | | |
| Ppp6r2 | | | | | | | | | | 0.00002 | -3.02614 | Male | | | | | | |
| Rpl39l | | | | | | | | | | 0.00000 | -3.23751 | Male | 0.02446 | 1.36066 | Female | | | |
| Ccdc22 | | | | | | | | | | 0.00000 | 6.80855 | Female | | | | | | |
| Pds5a | | | | | | | | | | 0.00000 | 4.26416 | Female | | | | | | |
| Ap2a1 | | | | | | | | | | 0.04064 | 3.14049 | Female | | | | | | |
| Mrps31 | 0.00147 | 4.01875 | Female | | | | | | | 0.04404 | 1.58073 | Female | | | | | | |
| Fpgt | | | | | | | | | | 0.01496 | -4.71710 | Male | | | | | | |
| Gal3st3 | | | | | | | | | | | | | | | | 0.00036 | -3.27949 | Male |
| Ffar2 | | | | | | | | | | | | | | | | 0.01632 | 0.65065 | Female |
| Rab4b | | | | | | | 0.01887 | -4.38853 | Male | | | | | | | | | |
| Trim71 | | | | | | | 0.00000 | 3.06179 | Female | 0.00001 | -1.61900 | Male | | | | | | |
| Iqgap2 | | | | | | | | | | | | | 0.00000 | -5.14337 | Male | 0.00000 | 4.09358 | Female |
| Zeb1 | | | | | | | | | | | | | 0.01947 | 3.51418 | Female | | | |
| Znrf2 | | | | | | | | | | | | | 0.04274 | -1.05973 | Male | | | |
| Fthl17b | | | | 0.00110 | 4.77047 | Female | 0.00000 | 5.51200 | Female | 0.00011 | 5.39636 | Female | 0.00066 | 3.72784 | Female | | | |
| Kdm7a | | | | | | | | | | | | | | | | 0.00003 | -3.04969 | Male |
| Gsk3a | | | | | | | | | | 0.00000 | -5.43927 | Male | | | | | | |
| AW209491 | | | | | | | | | | | | | | | | 0.02828 | 2.68050 | Female |
| Ranbp1 | | | | | | | | | | 0.02527 | -1.71054 | Male | | | | | | |
| Trappc4 | | | | | | | | | | | | | | | | 0.00171 | -1.47207 | Male |
| Siva1 | | | | | | | | | | 0.03651 | -2.21558 | Male | | | | | | |
| Naa15 | | | | | | | 0.04604 | 1.44646 | Female | | | | | | | | | |
| Taf11 | | | | | | | | | | 0.01068 | 1.77928 | Female | | | | | | |
| Gmfb | | | | | | | | | | | | | | | | 0.02238 | -2.40214 | Male |
| Vbp1 | | | | | | | 0.01314 | 1.69362 | Female | | | | | | | | | |
| Mfn1 | | | | | | | | | | 0.02245 | 1.33276 | Female | | | | | | |
| Rpl32 | | | | | | | | | | 0.00000 | -3.75954 | Male | 0.00039 | 1.49414 | Female | | | |
| Xlr3b | | | | | | | 0.01893 | 2.75915 | Female | | | | | | | | | |
| Dgat2 | | | | | | | | | | | | | | | | 0.00013 | 3.45975 | Female |
| Uqcrb | | | | | | | | | | 0.00000 | -4.99217 | Male | | | | | | |
| Itm2a | | | | | | | | | | | | | | | | 0.01005 | -4.02127 | Male |
| Itga5 | | | | 0.00352 | 1.83469 | Female | | | | | | | | | | | | |
| Ccne2 | | | | | | | | | | 0.01060 | 3.92612 | Female | | | | 0.04697 | -3.28215 | Male |
| Spesp1 | | | | 0.00005 | 1.38788 | Female | | | | | | | | | | | | |
| Tpm2 | 0.00000 | -2.05728 | Male | 0.00121 | -2.71596 | Male | | | | | | | | | | | | |
| Ly6g5b | | | | | | | | | | | | | | | | 0.00292 | 4.07829 | Female |
| Dusp6 | | | | 0.00000 | 2.83411 | Female | | | | | | | | | | | | |
| Nespas | | | | | | | 0.00000 | 1.50585 | Female | 0.00017 | 1.19907 | Female | | | | | | |

| | | | | | | | | | | | | | | | | | | |
|---|---|---|---|---|---|---|---|---|---|---|---|---|---|---|---|---|---|---|
| Mmel1 | | | | | | | 0.00000 | -1.31354 | Male | | | | | | | | | |
| Trmt13 | | | | | | | | | | 0.00006 | 5.70418 | Female | | | | | | |
| Clmp | | | | | | | | | | | | | | | | 0.00000 | -1.31708 | Male |
| Pak7 | | | | | | | | | | 0.00021 | -1.83260 | Male | | | | | | |
| Nmrk2 | | | | | | | | | | 0.00001 | -3.58251 | Male | | | | | | |
| Psmc2 | | | | | | | | | | 0.00000 | 1.79580 | Female | | | | | | |
| Rnf40 | | | | | | | | | | | | | | | | 0.00737 | 2.26931 | Female |
| Lrrc25 | | | | | | | | | | 0.00025 | 2.47344 | Female | | | | | | |
| Cyp2e1 | | | | | | | | | | 0.00000 | -5.27752 | Male | 0.00013 | 2.16697 | Female | | | |
| Platr14 | | | | | | | | | | 0.02698 | 2.72804 | Female | | | | | | |
| Polr2d | | | | | | | | | | 0.00000 | 3.54290 | Female | | | | | | |
| Hmgn1 | | | | | | | | | | 0.00003 | -2.58129 | Male | | | | | | |
| Capn2 | | | | 0.01061 | -2.12202 | Male | | | | 0.00000 | 2.03376 | Female | | | | | | |
| Cybrd1 | | | | | | | | | | 0.00605 | -2.95127 | Male | | | | | | |
| Gstm6 | 0.00031 | -0.95387 | Male | 0.00000 | 1.01458 | Female | 0.00002 | -1.50879 | Male | 0.00048 | -1.29876 | Male | | | | | | |
| Gk5 | | | | | | | | | | 0.00180 | 4.67155 | Female | | | | | | |
| Dsg1b | | | | | | | | | | 0.00003 | -4.02825 | Male | | | | | | |
| Pard3b | | | | 0.02695 | 5.94732 | Female | | | | | | | | | | | | |
| Zar1 | | | | | | | | | | 0.02965 | -5.35694 | Male | | | | | | |
| Arhgap30 | | | | | | | | | | | | | | | | 0.04580 | -3.62807 | Male |
| Ap1s2 | | | | | | | | | | | | | | | | 0.00000 | 4.44631 | Female |
| Adarb1 | | | | | | | | | | 0.02569 | -2.29676 | Male | | | | | | |
| Chpf | | | | | | | 0.01865 | 3.41618 | Female | | | | | | | | | |
| Cask | | | | | | | | | | | | | | | | 0.01375 | 3.72795 | Female |
| Cdc37 | | | | | | | | | | 0.03125 | 2.04513 | Female | | | | | | |
| Rfx2 | | | | | | | | | | 0.02805 | 2.97883 | Female | | | | | | |
| Zfp36l2 | | | | | | | | | | 0.00345 | -3.39956 | Male | | | | | | |
| Casc4 | | | | | | | | | | 0.00080 | -2.47048 | Male | | | | | | |
| Timp3 | | | | | | | | | | | | | 0.01560 | -4.70928 | Male | | | |
| Plekhb2 | | | | | | | | | | | | | | | | 0.00063 | -3.37869 | Male |
| BC051665 | | | | 0.00018 | 0.88250 | Female | | | | | | | | | | 0.03678 | 5.12562 | Female |
| Dvl3 | | | | | | | | | | 0.00584 | 4.09657 | Female | | | | | | |
| Nsun6 | | | | 0.01815 | 2.77310 | Female | | | | | | | | | | | | |
| Plod2 | | | | 0.00000 | 2.27615 | Female | | | | | | | | | | | | |
| Adcy7 | | | | | | | | | | | | | 0.00000 | 1.67924 | Female | | | |
| Fam46b | | | | | | | | | | | | | | | | 0.00181 | -4.43207 | Male |
| Tigd3 | | | | | | | | | | 0.00001 | -3.84814 | Male | | | | | | |
| Flcn | | | | | | | 0.02375 | 2.11969 | Female | | | | | | | 0.00013 | -2.94851 | Male |
| Btrc | | | | | | | | | | 0.00841 | 1.89534 | Female | | | | | | |
| Sec16b | | | | | | | | | | | | | 0.00000 | -2.18332 | Male | | | |
| Slc7a6os | | | | | | | | | | | | | | | | 0.00005 | 2.92150 | Female |
| Bola2 | | | | | | | | | | 0.00000 | -3.87800 | Male | | | | | | |
| Ccdc109b | | | | | | | 0.00000 | 1.54829 | Female | | | | | | | 0.00001 | 0.58695 | Female |
| Gm20939 | | | | | | | 0.03788 | 1.64436 | Female | | | | | | | | | |
| Neat1 | | | | | | | 0.00618 | 3.21864 | Female | | | | | | | | | |
| Atat1 | | | | | | | | | | | | | | | | 0.04462 | 3.58271 | Female |
| Surf2 | | | | | | | | | | 0.00004 | 2.62395 | Female | | | | | | |
| Cyp17a1 | | | | | | | | | | | | | | | | 0.00000 | -4.02484 | Male |
| Pus10 | | | | 0.00124 | 3.85665 | Female | | | | | | | | | | | | |
| Trim47 | 0.00000 | -0.63017 | Male | | | | | | | | | | | | | | | |
| Jun | | | | 0.00000 | -0.77022 | Male | | | | | | | | | | | | |
| Mfsd9 | | | | | | | | | | | | | | | | 0.02064 | -5.02187 | Male |
| Snord72 | | | | | | | | | | | | | | | | 0.03014 | 3.28193 | Female |
| Zdhhc15 | | | | | | | | | | | | | | | | 0.00001 | 1.36383 | Female |
| Ankrd45 | | | | | | | 0.00977 | 3.90555 | Female | | | | | | | | | |
| Cacna2d1 | | | | | | | | | | | | | | | | 0.00000 | -3.04037 | Male |
| Wdr6 | | | | 0.04612 | 3.16851 | Female | | | | 0.01450 | 2.98382 | Female | | | | | | |
| Gm7102 | | | | | | | | | | 0.00016 | -2.07815 | Male | | | | | | |
| Rab34 | | | | | | | | | | | | | | | | 0.02064 | 2.83459 | Female |
| Mmp1b | | | | | | | 0.00653 | -1.93773 | Male | 0.02502 | -0.69979 | Male | | | | | | |
| Tuba1c | | | | | | | | | | 0.00012 | 1.54219 | Female | 0.02418 | 0.75537 | Female | | | |
| Psmd13 | | | | | | | | | | 0.00000 | 2.05378 | Female | | | | | | |
| Oc90 | | | | | | | 0.00000 | 3.58260 | Female | 0.00000 | 4.37796 | Female | | | | 0.00000 | 2.01926 | Female |
| Ramp2 | | | | | | | | | | 0.01521 | -3.77552 | Male | | | | 0.00002 | 5.21776 | Female |
| Catsperd | | | | | | | | | | | | | | | | 0.00007 | -6.05338 | Male |
| Carm1 | | | | | | | | | | 0.00000 | 7.37471 | Female | | | | | | |
| Vamp2 | 0.00000 | 1.49169 | Female | | | | | | | | | | | | | 0.00730 | -4.74751 | Male |
| Fasn | | | | | | | | | | 0.04886 | 2.11586 | Female | | | | | | |
| Aldh9a1 | | | | | | | | | | | | | | | | 0.02105 | 2.06133 | Female |
| Rps19 | | | | | | | | | | 0.00000 | -3.23812 | Male | 0.01442 | 1.28894 | Female | | | |

| | | | | | | | | | | | | | | | | | | |
|---|---|---|---|---|---|---|---|---|---|---|---|---|---|---|---|---|---|---|
| Nrf1 | | | | | | | | | | 0.03746 | 1.95122 | Female | | | | | | |
| Lsm14b | | | | | | | | | | 0.03207 | -3.06068 | Male | | | | | | |
| Hspb2 | | | | | | | 0.00000 | -1.16493 | Male | | | | | | | | | |
| Cyp4f16 | | | | 0.00000 | -0.97820 | Male | 0.00000 | 1.56061 | Female | | | | | | | | | |
| Msantd3 | | | | | | | | | | 0.00000 | 1.70986 | Female | 0.00254 | 4.13538 | Female | | | |
| Zfp687 | | | | | | | | | | 0.03065 | 3.32689 | Female | | | | | | |
| Esr2 | | | | | | | | | | 0.00004 | -4.93441 | Male | | | | | | |
| Nlrp9c | | | | | | | | | | | | | 0.02387 | 4.62567 | Female | | | |
| Dph1 | | | | | | | | | | 0.00004 | 5.50976 | Female | | | | | | |
| Mvb12b | | | | | | | | | | | | | 0.01543 | -4.01521 | Male | 0.00542 | 4.24958 | Female |
| Cyp4f13 | | | | 0.00559 | -1.47041 | Male | 0.00000 | 0.64895 | Female | 0.00145 | -0.78801 | Male | | | | | | |
| Sesn2 | | | | | | | | | | 0.01010 | 3.18760 | Female | | | | | | |
| Susd2 | | | | | | | | | | 0.00000 | 2.45296 | Female | 0.02951 | 4.42462 | Female | | | |
| Gm4425 | 0.04632 | -3.93931 | Male | | | | | | | | | | | | | | | |
| Zfp976 | | | | | | | 0.02397 | 1.76380 | Female | | | | | | | | | |
| Gm11549 | | | | | | | | | | 0.00000 | -4.70250 | Male | | | | | | |
| Rps6kc1 | | | | | | | | | | | | | | | | 0.01805 | 3.70449 | Female |
| Dact3 | | | | 0.00000 | 0.60355 | Female | | | | 0.00000 | -4.20862 | Male | | | | | | |
| Cryl1 | | | | | | | | | | | | | | | | 0.00382 | 4.36913 | Female |
| Gas2 | 0.00007 | 2.14258 | Female | | | | | | | | | | | | | | | |
| BC002163 | | | | | | | | | | 0.00000 | -4.13769 | Male | 0.00049 | 2.01417 | Female | | | |
| Orai2 | | | | 0.00002 | 1.14004 | Female | | | | | | | | | | | | |
| Rpl36 | | | | | | | | | | 0.00019 | -3.43478 | Male | | | | | | |
| Chp1 | | | | | | | | | | | | | | | | 0.01368 | -1.39432 | Male |
| Cstb | | | | | | | | | | 0.00000 | -4.38263 | Male | 0.00210 | 1.72701 | Female | | | |
| Slc22a5 | | | | | | | 0.00000 | -7.22207 | Male | | | | | | | | | |
| Ttc8 | | | | | | | | | | | | | | | | 0.00003 | 1.22678 | Female |
| Cybb | | | | | | | | | | 0.00373 | 0.85682 | Female | | | | 0.00000 | 1.23652 | Female |
| Ppara | | | | 0.00110 | 1.35563 | Female | | | | | | | | | | 0.00000 | 1.87526 | Female |
| Zdhhc24 | | | | | | | 0.00723 | 3.84857 | Female | | | | | | | | | |
| Slc6a2 | | | | | | | | | | 0.00001 | -3.69141 | Male | 0.03361 | 1.73495 | Female | | | |
| Map6 | | | | | | | | | | | | | 0.00000 | -4.47637 | Male | 0.00000 | 2.11256 | Female |
| Gpr68 | | | | | | | | | | 0.03315 | -2.97432 | Male | | | | | | |
| Cep131 | | | | | | | | | | | | | | | | 0.00329 | 4.53398 | Female |
| Eps8l1 | | | | 0.04674 | -1.38727 | Male | | | | | | | | | | | | |
| BC028528 | | | | | | | 0.00668 | 3.24095 | Female | | | | | | | | | |
| Apobec3 | | | | | | | | | | | | | | | | 0.00001 | 5.03062 | Female |
| Pepd | 0.00391 | 4.32732 | Female | | | | | | | | | | | | | | | |
| Aifm3 | | | | | | | | | | | | | | | | 0.00458 | -4.76325 | Male |
| Orc3 | 0.00554 | -3.05594 | Male | | | | | | | | | | | | | | | |
| Mplkip | | | | | | | | | | 0.04588 | -2.31164 | Male | | | | | | |
| Trdmt1 | | | | | | | 0.02287 | 4.35685 | Female | | | | | | | | | |
| Rcn3 | | | | | | | | | | | | | | | | 0.02147 | 3.70540 | Female |
| Cox17 | | | | | | | | | | 0.00000 | -4.57302 | Male | 0.01589 | 1.75293 | Female | | | |
| Nav3 | | | | | | | | | | 0.00000 | 2.59997 | Female | | | | | | |
| Lsg1 | | | | | | | | | | 0.03421 | 2.17603 | Female | | | | | | |
| Myom2 | | | | | | | | | | | | | 0.00000 | 4.14248 | Female | 0.00000 | 0.81100 | Female |
| Pbld1 | | | | | | | | | | 0.00000 | -5.73353 | Male | 0.01532 | 1.46762 | Female | | | |
| Gcnt4 | | | | 0.00001 | 1.46614 | Female | | | | 0.00804 | -1.30277 | Male | | | | 0.00000 | 4.58966 | Female |
| Gm8234 | | | | | | | | | | 0.00254 | 1.57367 | Female | | | | | | |
| Ccdc150 | | | | | | | | | | 0.00000 | 2.95913 | Female | | | | | | |
| Gm16287 | | | | | | | | | | 0.00001 | -4.03076 | Male | | | | | | |
| Cilp2 | | | | | | | | | | | | | 0.00000 | -2.70926 | Male | | | |
| Nop10 | | | | | | | | | | 0.00000 | -4.17800 | Male | 0.00000 | 2.08994 | Female | | | |
| Abcd2 | | | | | | | | | | 0.00000 | -4.95965 | Male | | | | | | |
| Eci1 | | | | | | | | | | 0.03959 | -2.14054 | Male | | | | | | |
| Trpc5os | | | | 0.00081 | 1.42812 | Female | | | | | | | 0.00000 | 3.94133 | Female | 0.00000 | -2.84661 | Male |
| Rps19.ps3 | | | | | | | | | | 0.00000 | -3.02519 | Male | 0.01535 | 1.25689 | Female | | | |
| Phyhd1 | | | | | | | | | | | | | 0.00000 | 3.58847 | Female | | | |
| Kdm1b | | | | | | | | | | | | | | | | 0.02986 | 3.24689 | Female |
| Fmr1 | | | | 0.03048 | 2.91273 | Female | | | | | | | | | | | | |
| X6.Sep | 0.00000 | 2.65460 | Female | 0.00000 | 0.65376 | Female | | | | | | | | | | | | |
| Slc39a1 | | | | | | | | | | 0.00007 | -3.45362 | Male | 0.00094 | 2.18578 | Female | | | |
| Sycn | | | | | | | | | | 0.02955 | -3.19228 | Male | | | | | | |
| Med21 | | | | | | | | | | 0.00418 | -1.79943 | Male | | | | | | |
| Slc7a8 | | | | 0.00772 | 0.73369 | Female | 0.00000 | -0.68946 | Male | | | | | | | 0.00000 | 2.16773 | Female |
| Snx19 | | | | 0.00000 | 2.33822 | Female | | | | | | | 0.00193 | 4.43978 | Female | 0.00739 | -4.86206 | Male |
| Pdzd4 | | | | | | | | | | 0.03109 | -4.24060 | Male | | | | | | |
| Acbd4 | | | | 0.03638 | 2.29325 | Female | | | | | | | | | | | | |
| Oraov1 | | | | | | | | | | | | | | | | 0.01199 | -2.05442 | Male |

| | | | | | | | | | | | | | | | | | | |
|---|---|---|---|---|---|---|---|---|---|---|---|---|---|---|---|---|---|---|
| Gpr4 | | | | 0.00000 | -1.07747 | Male | | | | | | | | | | | | |
| Hmgb2 | | | | | | | | | | 0.00350 | 1.89889 | Female | | | | | | |
| Ogn | | | | | | | | | | 0.00001 | -4.05380 | Male | | | | | | |
| Pxmp2 | | | | | | | | | | 0.03205 | -2.43734 | Male | | | | | | |
| Fam96a | | | | | | | | | | 0.02468 | 1.29123 | Female | | | | | | |
| Cox7a1 | | | | | | | | | | 0.00307 | -5.04529 | Male | | | | | | |
| Mtss1l | | | | | | | | | | | | | | | | 0.00000 | 3.66415 | Female |
| Serpinh1 | | | | 0.00000 | -0.62282 | Male | | | | | | | | | | | | |
| BC051019 | | | | | | | | | | 0.00000 | 3.52681 | Female | | | | | | |
| Srsf1 | | | | | | | 0.00853 | 1.45260 | Female | | | | | | | | | |
| Dppa5a | | | | | | | | | | 0.00000 | -2.93566 | Male | 0.00380 | 1.30926 | Female | | | |
| Nfe2l1 | | | | | | | | | | 0.03082 | 1.78923 | Female | | | | | | |
| Ift27 | | | | | | | | | | | | | 0.03070 | -3.97234 | Male | | | |
| Zfp182 | | | | | | | | | | 0.00000 | 0.69827 | Female | | | | | | |
| Zc3h12a | | | | | | | | | | 0.02947 | -1.94215 | Male | | | | | | |
| Crx | | | | | | | | | | 0.00063 | -2.95014 | Male | | | | | | |
| Prpf19 | | | | | | | | | | 0.00000 | -3.59247 | Male | 0.03051 | 1.30734 | Female | | | |
| Btg4 | | | | | | | | | | | | | 0.00017 | -3.30770 | Male | | | |
| Btk | | | | | | | | | | | | | 0.00000 | -6.06134 | Male | | | |
| Cep57l1 | | | | | | | | | | | | | | | | 0.04873 | -1.82340 | Male |
| Gm15645 | | | | | | | | | | 0.02411 | -1.21430 | Male | | | | | | |
| Kng1 | | | | | | | | | | | | | | | | 0.00230 | -3.34041 | Male |
| Gm5577 | | | | | | | | | | | | | 0.02526 | -3.33084 | Male | | | |
| Eogt | | | | | | | | | | | | | | | | 0.01341 | 3.64542 | Female |
| Omt2a | | | | | | | | | | 0.00012 | -4.51060 | Male | 0.00008 | 3.75226 | Female | | | |
| Snhg1 | | | | | | | | | | 0.00000 | -3.38357 | Male | | | | 0.00976 | 1.55949 | Female |
| Chd1l | | | | 0.04154 | 2.41205 | Female | | | | | | | | | | | | |
| Pqlc1 | | | | | | | | | | | | | | | | 0.00247 | -2.66231 | Male |
| Cda | | | | | | | | | | | | | 0.00022 | 0.74217 | Female | | | |
| Rps15a | | | | | | | | | | 0.00000 | -4.57088 | Male | 0.01757 | 1.22559 | Female | | | |
| Psmd7 | | | | | | | | | | 0.00312 | 1.32658 | Female | | | | | | |
| Gm7616 | | | | | | | | | | 0.00000 | -5.38265 | Male | | | | | | |
| Jpx | | | | | | | | | | 0.00000 | -4.45096 | Male | | | | | | |
| Lrrc15 | | | | | | | | | | | | | 0.00000 | 5.13661 | Female | | | |
| B3gnt3 | 0.00019 | 2.25759 | Female | | | | 0.00000 | 1.03961 | Female | | | | | | | | | |
| Swi5 | | | | | | | | | | 0.00604 | -2.10187 | Male | | | | | | |
| Golt1a | 0.00054 | -0.81799 | Male | | | | | | | | | | 0.00000 | -4.79655 | Male | | | |
| Creb3l4 | | | | | | | | | | | | | | | | 0.00715 | 2.93280 | Female |
| Yipf2 | | | | | | | | | | 0.00669 | 3.91793 | Female | | | | | | |
| Thsd7b | | | | | | | 0.00854 | -0.63498 | Male | | | | | | | | | |
| Anks4b | | | | 0.04151 | 1.31495 | Female | | | | | | | | | | 0.00000 | 1.55855 | Female |
| Hdgfrp3 | | | | 0.02618 | -1.03208 | Male | | | | | | | | | | | | |
| AU016765 | | | | | | | 0.00000 | 2.40704 | Female | | | | | | | | | |
| Mdm1 | | | | | | | | | | 0.00167 | -5.30410 | Male | | | | | | |
| Serpinb1a | | | | | | | | | | 0.00009 | -1.85028 | Male | | | | | | |
| Mtcl1 | | | | 0.01893 | 1.74273 | Female | | | | 0.00740 | 3.86994 | Female | | | | | | |
| Duoxa1 | | | | | | | | | | | | | | | | 0.00396 | 2.53641 | Female |
| Ddx23 | | | | 0.00002 | -0.61652 | Male | | | | | | | | | | | | |
| Gm11544 | | | | | | | | | | | | | 0.04289 | -1.60683 | Male | | | |
| Reg1 | | | | | | | | | | | | | 0.01508 | -3.63956 | Male | 0.00002 | -0.75079 | Male |
| Clic1 | 0.00000 | 1.16519 | Female | 0.00000 | 1.01963 | Female | | | | | | | | | | | | |
| X6.Mar | | | | | | | | | | | | | | | | 0.00320 | -1.70671 | Male |
| Cflar | | | | | | | | | | 0.00000 | -5.68884 | Male | 0.00004 | 1.87638 | Female | | | |
| Khsrp | | | | | | | | | | 0.00112 | -2.48816 | Male | | | | | | |
| Tekt2 | | | | | | | | | | | | | | | | 0.00028 | 5.18841 | Female |
| Dhx9 | | | | | | | | | | 0.00000 | -4.80366 | Male | 0.00526 | 1.52524 | Female | | | |
| Rhox9 | | | | 0.00012 | 3.74271 | Female | | | | 0.00004 | 0.91563 | Female | 0.00000 | 4.78164 | Female | | | |
| Rps8 | | | | | | | | | | 0.02282 | -1.37993 | Male | | | | | | |
| Nmnat3 | | | | | | | 0.00000 | -2.41264 | Male | | | | | | | | | |
| Cth | | | | | | | | | | | | | | | | 0.00001 | -2.12577 | Male |
| Arrdc1 | | | | | | | | | | | | | | | | 0.00025 | 4.04988 | Female |
| Ttc26 | | | | | | | | | | 0.00015 | -3.69087 | Male | | | | 0.04362 | 1.92971 | Female |
| Gc | | | | | | | | | | 0.00001 | -7.62828 | Male | | | | | | |
| Huwe1 | | | | | | | 0.04289 | 1.22408 | Female | 0.02699 | 1.41454 | Female | | | | | | |
| Tcf15 | | | | | | | | | | 0.00226 | -4.87972 | Male | | | | | | |
| Furin | | | | | | | | | | | | | 0.03054 | 2.00237 | Female | | | |
| Tcp11 | | | | | | | | | | | | | 0.00000 | 2.84462 | Female | | | |
| Ganc | | | | | | | | | | | | | | | | 0.00030 | -6.13798 | Male |
| B3gnt5 | | | | | | | | | | 0.04408 | -4.66464 | Male | | | | | | |
| Rps11 | | | | | | | | | | 0.00000 | -3.35375 | Male | 0.00022 | 1.35928 | Female | | | |

| | | | | | | | | | | | | | | | | | | |
|---|---|---|---|---|---|---|---|---|---|---|---|---|---|---|---|---|---|---|
| Eif4e1b | | | | | | | | | | | | | | | | 0.01490 | 3.80391 | Female |
| Tcf7l1 | | | | 0.00018 | -1.76203 | Male | | | | | | | | | | | | |
| Chrnb4 | | | | | | | | | | | | | 0.00036 | 4.11256 | Female | | | |
| Ppib | | | | | | | | | | 0.00000 | 4.46323 | Female | | | | | | |
| Ppia | | | | | | | | | | 0.04970 | -1.02339 | Male | | | | | | |
| Tceal8 | | | | 0.00000 | 3.24440 | Female | | | | | | | 0.00004 | 5.69719 | Female | | | |
| Gsdmc3 | | | | | | | | | | | | | | | | 0.00834 | 3.91723 | Female |
| Muc13 | | | | | | | | | | | | | | | | 0.04753 | 4.47517 | Female |
| Gm13083 | | | | | | | | | | 0.00001 | 3.21397 | Female | | | | 0.00436 | -1.86074 | Male |
| Cd1d1 | | | | | | | | | | | | | 0.00000 | 2.23594 | Female | | | |
| Cyb5b | | | | | | | | | | | | | | | | 0.01243 | 2.61720 | Female |
| Mllt11 | | | | | | | | | | | | | | | | 0.01368 | 3.71380 | Female |
| Lsm6 | | | | | | | | | | | | | 0.03262 | 1.36209 | Female | | | |
| Nono | | | | | | | 0.00535 | 1.12623 | Female | | | | | | | | | |
| Zfp395 | | | | | | | | | | 0.03659 | 2.86014 | Female | | | | | | |
| H2afj | | | | | | | | | | | | | | | | 0.01139 | -2.95222 | Male |
| Spin2d | | | | | | | 0.00095 | 5.02628 | Female | | | | | | | | | |
| Rab20 | | | | | | | | | | 0.00075 | 3.03824 | Female | | | | | | |
| Set | | | | | | | | | | 0.00006 | 1.36376 | Female | | | | | | |
| Gm12250 | | | | | | | 0.02263 | 5.00251 | Female | | | | 0.00000 | 4.08463 | Female | | | |
| Efcab3 | | | | | | | | | | | | | 0.00000 | 2.24199 | Female | 0.00000 | -1.30991 | Male |
| Cdx1 | | | | | | | | | | | | | 0.00000 | 2.76202 | Female | | | |
| Amdhd2 | | | | 0.00268 | 2.25994 | Female | | | | | | | | | | | | |
| Arl1 | | | | | | | | | | 0.00082 | 2.61326 | Female | | | | | | |
| Rplp0 | | | | | | | | | | 0.03751 | 2.24538 | Female | | | | | | |
| Grm4 | | | | | | | 0.00000 | -1.85981 | Male | | | | | | | | | |
| Mnt | | | | | | | | | | | | | | | | 0.00069 | -3.34689 | Male |
| AY074887 | | | | | | | | | | 0.03763 | -1.58400 | Male | | | | | | |
| Pdha2 | | | | 0.00000 | 2.49929 | Female | 0.00511 | -1.41396 | Male | | | | | | | 0.00001 | 0.99797 | Female |
| Eif2s3x | | | | | | | 0.03818 | 1.13157 | Female | 0.04496 | 1.20253 | Female | | | | | | |
| Fbxo7 | | | | | | | | | | 0.00000 | 2.11393 | Female | | | | | | |
| Rpl39 | | | | | | | 0.00001 | 2.39503 | Female | 0.00000 | -5.78278 | Male | 0.00000 | 2.99210 | Female | | | |
| Ciart | | | | | | | | | | 0.01657 | 3.41814 | Female | | | | | | |
| Bcl3 | | | | 0.01292 | 1.77823 | Female | | | | | | | | | | | | |
| Afap1 | | | | | | | | | | 0.01467 | -3.23463 | Male | 0.00060 | 3.22107 | Female | | | |
| Gm6367 | | | | | | | | | | 0.02476 | -2.85390 | Male | | | | | | |
| Mroh8 | | | | | | | | | | | | | 0.02058 | -3.88238 | Male | | | |
| Cct5 | | | | | | | | | | 0.00268 | 1.16914 | Female | | | | | | |
| Id1 | | | | | | | | | | | | | 0.03197 | -3.09098 | Male | | | |
| Snord47 | | | | 0.03163 | 3.41663 | Female | | | | | | | | | | | | |
| Agbl5 | | | | | | | | | | | | | | | | 0.00000 | 4.00610 | Female |
| D5Ertd577e | | | | | | | | | | | | | | | | 0.00000 | -4.80607 | Male |
| Magoh | | | | | | | | | | 0.00000 | -2.62486 | Male | | | | | | |
| Gm6083 | | | | | | | | | | 0.00000 | -3.35856 | Male | 0.00229 | 1.36126 | Female | | | |
| Pawr | | | | | | | | | | 0.00350 | 4.00156 | Female | | | | | | |
| Pitpnc1 | | | | | | | 0.02728 | 2.77819 | Female | 0.00080 | 4.03039 | Female | | | | | | |
| Fam219a | | | | | | | 0.00000 | -1.26232 | Male | | | | | | | | | |
| Gjc3 | | | | | | | | | | 0.00007 | -3.04577 | Male | | | | | | |
| Pygo1 | | | | | | | | | | | | | | | | 0.00000 | -0.60929 | Male |
| S1pr4 | | | | | | | | | | 0.00015 | -4.31005 | Male | | | | | | |
| Neil1 | | | | | | | | | | 0.00722 | 4.92694 | Female | | | | | | |
| Mark2 | | | | | | | | | | 0.03588 | 2.06666 | Female | | | | | | |
| Kif2c | | | | | | | | | | | | | | | | 0.00002 | -3.04681 | Male |
| Lepr | | | | | | | | | | 0.00001 | -3.18273 | Male | | | | | | |
| Myo7a | 0.00002 | -1.16555 | Male | 0.00000 | 1.22320 | Female | | | | | | | | | | | | |
| Dapp1 | | | | | | | | | | 0.04183 | -3.79902 | Male | | | | | | |
| Ppp2r5a | | | | | | | | | | 0.00194 | 4.74331 | Female | | | | | | |
| Slamf8 | | | | | | | | | | 0.01183 | -3.35271 | Male | | | | | | |
| Grn | | | | | | | | | | | | | | | | 0.00959 | -1.59342 | Male |
| Nxn | | | | | | | | | | 0.00806 | 4.29011 | Female | | | | 0.00317 | 4.50242 | Female |
| Seh1l | | | | | | | | | | | | | 0.04617 | -0.99013 | Male | | | |
| Snhg9 | | | | | | | | | | | | | 0.01646 | 2.80881 | Female | | | |
| Pttg1 | | | | | | | | | | 0.00015 | -2.33813 | Male | | | | | | |
| Ilf2 | | | | | | | | | | 0.00229 | 2.31668 | Female | | | | | | |
| Mt1 | | | | | | | | | | 0.00057 | -4.01708 | Male | | | | | | |
| Cubn | | | | | | | | | | | | | | | | 0.00186 | 5.27981 | Female |
| Npepl1 | | | | 0.00000 | -1.85050 | Male | | | | | | | | | | | | |
| Liph | | | | | | | | | | | | | 0.00390 | 3.77526 | Female | | | |
| Prcp | 0.04461 | -3.77236 | Male | | | | | | | | | | | | | | | |
| Mrpl1 | | | | | | | | | | | | | | | | 0.00272 | -2.15782 | Male |

| | | | | | | | | | | | | | | | | | | |
|---|---|---|---|---|---|---|---|---|---|---|---|---|---|---|---|---|---|---|
| Rapgef4 | | | | | | | | | | 0.00000 | -4.60924 | Male | | | | | | |
| Rpl36a | | | | | | | 0.00301 | 1.61928 | Female | 0.00000 | -4.37088 | Male | 0.00008 | 1.65727 | Female | | | |
| Cacna1d | | | | 0.02354 | 1.15200 | Female | | | | 0.00519 | 0.80939 | Female | | | | | | |
| Ap2m1 | | | | | | | | | | | | | | | | 0.00086 | -1.31706 | Male |
| Txndc5 | | | | 0.00502 | 3.46929 | Female | | | | | | | | | | | | |
| Tmem63b | | | | | | | 0.02728 | 3.57825 | Female | | | | | | | | | |
| Gm6815 | | | | | | | | | | 0.00910 | 2.37626 | Female | | | | | | |
| Fsd1 | | | | | | | | | | 0.02690 | 4.74013 | Female | | | | | | |
| Arhgef2 | | | | | | | | | | 0.00000 | 1.73072 | Female | | | | | | |
| Gadd45gip1 | | | | | | | | | | 0.00001 | -4.65462 | Male | | | | | | |
| Ebp | | | | | | | | | | | | | 0.00109 | 1.82469 | Female | | | |
| Kank1 | | | | 0.02354 | 0.86710 | Female | | | | | | | | | | | | |
| Zfp972 | | | | | | | | | | | | | 0.00187 | 2.43023 | Female | 0.00000 | -4.85152 | Male |
| Parva | 0.00003 | -0.66695 | Male | 0.00000 | -0.68898 | Male | | | | 0.00000 | -0.71108 | Male | | | | | | |
| Cdk2 | | | | | | | | | | 0.00000 | 4.14766 | Female | | | | | | |
| Cdc42bpg | | | | | | | | | | 0.00002 | -5.69479 | Male | | | | | | |
| Cep164 | | | | | | | | | | 0.00185 | 3.89878 | Female | | | | | | |
| Arnt | | | | | | | | | | | | | | | | 0.02454 | -2.28703 | Male |
| Ddx26b | | | | | | | | | | | | | | | | 0.00000 | 4.96349 | Female |
| Gpc4 | | | | | | | | | | | | | | | | 0.00009 | 5.23292 | Female |
| Xdh | | | | 0.00110 | 0.59609 | Female | | | | | | | 0.00000 | -3.02347 | Male | | | |
| Rad9a | | | | | | | | | | 0.04074 | 2.35790 | Female | | | | | | |
| Morf4l2 | | | | | | | | | | 0.03848 | 1.39119 | Female | | | | | | |
| Kdm3b | | | | | | | | | | 0.00004 | 2.52148 | Female | | | | | | |
| Drc1 | | | | | | | | | | | | | | | | 0.00188 | -3.73702 | Male |
| Hnrnph2 | | | | | | | 0.03555 | 1.82076 | Female | | | | | | | | | |
| Jak3 | | | | 0.00022 | -1.35573 | Male | | | | | | | | | | | | |
| Fbxw13 | | | | | | | | | | 0.03076 | -5.15924 | Male | 0.00000 | 1.41077 | Female | | | |
| Zbed3 | | | | | | | | | | 0.00000 | -6.84021 | Male | | | | | | |
| Cul2 | | | | | | | | | | 0.01274 | 1.90022 | Female | | | | | | |
| Ndufab1 | | | | | | | | | | 0.00011 | -4.01008 | Male | | | | | | |
| Sbf1 | | | | | | | | | | 0.00179 | 4.49661 | Female | | | | | | |
| Sat2 | | | | | | | | | | | | | | | | 0.00315 | 3.05089 | Female |
| Themis3 | | | | | | | | | | 0.00001 | 2.97584 | Female | | | | | | |
| Wdr54 | | | | | | | 0.01056 | -5.96215 | Male | | | | | | | | | |
| Rps26 | | | | | | | | | | 0.00000 | -3.84468 | Male | 0.04266 | 1.25209 | Female | | | |
| Chac1 | | | | 0.00625 | 1.13567 | Female | | | | | | | | | | | | |
| Kcnv2 | | | | 0.02831 | 1.85758 | Female | 0.00000 | -2.12394 | Male | | | | | | | 0.00000 | 2.08342 | Female |
| Dap | | | | 0.01728 | 0.68462 | Female | | | | | | | | | | | | |
| Fbxo10 | | | | | | | | | | | | | | | | 0.03731 | 4.13439 | Female |
| Ireb2 | | | | | | | | | | | | | | | | 0.02986 | 1.86855 | Female |
| Grid2ip | | | | | | | | | | 0.00084 | 2.39138 | Female | 0.00000 | -2.95578 | Male | | | |
| Uba1y | | | | 0.00002 | -6.03973 | Male | 0.00000 | -5.10728 | Male | 0.00000 | -6.57654 | Male | 0.00035 | -3.02280 | Male | 0.00669 | -2.82586 | Male |
| BC017158 | | | | | | | | | | 0.00907 | 4.69578 | Female | | | | | | |
| Brd9 | | | | | | | | | | | | | | | | 0.03117 | -1.36244 | Male |
| Slc12a7 | | | | | | | | | | 0.04830 | 1.78644 | Female | | | | | | |
| Slc36a1os | | | | | | | | | | 0.00139 | -1.90856 | Male | 0.01123 | 1.20510 | Female | | | |
| Eno2 | | | | | | | | | | | | | | | | 0.02614 | 4.58283 | Female |
| Phf2 | | | | 0.00783 | -1.69543 | Male | | | | 0.00000 | 2.08076 | Female | | | | | | |
| Gm1110 | | | | | | | | | | | | | 0.01142 | -5.04329 | Male | | | |
| Kin | | | | | | | | | | 0.00740 | 2.88092 | Female | | | | | | |
| Tor4a | | | | | | | | | | 0.00000 | -4.39490 | Male | | | | | | |
| Slc6a8 | | | | | | | 0.01541 | 1.38320 | Female | 0.03055 | 1.44022 | Female | | | | | | |
| Irak1 | | | | | | | | | | 0.00000 | -5.47580 | Male | | | | | | |
| Mical2 | | | | | | | | | | 0.00016 | 2.39200 | Female | | | | 0.00000 | 1.38026 | Female |
| Paqr7 | | | | | | | | | | 0.04552 | -3.43062 | Male | | | | | | |
| Ulk3 | | | | | | | | | | 0.03767 | 3.49613 | Female | | | | | | |
| Klhl23 | | | | | | | | | | 0.00026 | -1.99765 | Male | | | | 0.00898 | -5.58448 | Male |
| Pth1r | | | | | | | | | | | | | | | | 0.00297 | 4.61127 | Female |
| Ddi2 | | | | | | | | | | 0.00033 | 5.68881 | Female | | | | | | |
| Irx2 | | | | | | | | | | 0.00102 | -1.80556 | Male | | | | | | |
| Mtmr9 | | | | 0.00000 | 1.19938 | Female | | | | | | | | | | | | |
| Fabp3 | | | | | | | | | | 0.01890 | -2.95793 | Male | | | | | | |
| Smtn | | | | | | | | | | 0.03488 | 3.59923 | Female | | | | | | |
| Idh3g | | | | | | | | | | 0.00000 | 1.94713 | Female | | | | 0.00305 | 1.10341 | Female |
| Ccno | | | | | | | | | | | | | | | | 0.04930 | 3.62696 | Female |
| Umps | | | | | | | | | | 0.00907 | 1.29137 | Female | | | | | | |
| Ogt | | | | | | | 0.02647 | 1.85252 | Female | | | | | | | | | |
| Pkd2l2 | | | | | | | | | | | | | 0.01336 | 4.10796 | Female | | | |
| Papss1 | | | | | | | | | | | | | | | | 0.00062 | -3.66739 | Male |

| | | | | | | | | | | | | | | | | | | |
|---|---|---|---|---|---|---|---|---|---|---|---|---|---|---|---|---|---|---|
| Sms | | | | | | | 0.00064 | 1.35330 | Female | | | | | | | 0.01902 | 0.98188 | Female |
| Scube1 | | | | | | | | | | 0.00000 | -5.99752 | Male | | | | | | |
| BC049715 | | | | | | | | | | | | | 0.01542 | -2.82765 | Male | | | |
| Lmo1 | | | | | | | | | | | | | 0.02748 | -4.62928 | Male | | | |
| Rtcb | | | | | | | | | | 0.02524 | 1.62459 | Female | | | | | | |
| Pdlim4 | 0.00001 | 2.20070 | Female | | | | | | | | | | | | | | | |
| Esrp2 | 0.00985 | 3.81520 | Female | | | | | | | 0.01041 | 2.84833 | Female | | | | | | |
| Mtrf1 | | | | 0.00000 | 1.18980 | Female | | | | | | | | | | | | |
| X7.Sep | | | | | | | | | | | | | 0.01233 | -2.00809 | Male | | | |
| Clpb | | | | | | | | | | | | | | | | 0.02218 | -2.22047 | Male |
| Trmt2b | | | | | | | 0.00643 | 4.65173 | Female | | | | | | | | | |
| Ehd2 | 0.00011 | 0.90509 | Female | | | | | | | | | | | | | 0.00000 | -1.32259 | Male |
| Frem2 | | | | | | | | | | 0.00000 | 1.78361 | Female | | | | | | |
| Uaca | 0.00000 | -0.64492 | Male | | | | | | | | | | | | | | | |
| Gm19583 | 0.01977 | 0.99125 | Female | | | | | | | | | | | | | | | |
| Ccdc114 | | | | | | | | | | 0.00014 | 1.94206 | Female | | | | 0.00000 | -3.08886 | Male |
| Rpl28 | | | | | | | | | | 0.00000 | -3.42045 | Male | 0.01698 | 1.40510 | Female | | | |
| Flnb | | | | | | | | | | 0.00082 | 2.33888 | Female | | | | | | |
| Arg1 | | | | | | | | | | | | | | | | 0.00226 | -4.37323 | Male |
| Svopl | | | | | | | | | | 0.01004 | -4.10461 | Male | | | | | | |
| Mlycd | 0.00004 | -1.26264 | Male | | | | | | | | | | | | | | | |
| Rab33b | | | | | | | | | | 0.00018 | -3.08689 | Male | | | | | | |
| Fbxw26 | | | | | | | | | | | | | 0.00228 | 4.94501 | Female | | | |
| Gbp4 | | | | | | | | | | | | | 0.01508 | -4.87146 | Male | | | |
| AU015836 | | | | | | | | | | | | | | | | 0.00020 | -5.62911 | Male |
| Snhg12 | | | | | | | | | | 0.02272 | -2.27603 | Male | | | | | | |
| Armc6 | | | | | | | | | | 0.02837 | 2.57192 | Female | | | | 0.04216 | -2.02227 | Male |
| Dnm3 | | | | | | | 0.00002 | 2.60450 | Female | | | | | | | | | |
| Rassf5 | | | | | | | | | | | | | | | | 0.00000 | 1.49834 | Female |
| Htatip2 | | | | 0.02878 | 1.89126 | Female | | | | | | | | | | | | |
| Rps2 | | | | | | | | | | 0.00000 | -3.43213 | Male | | | | | | |
| Mpc2 | | | | | | | | | | 0.00016 | -4.22145 | Male | | | | | | |
| Ndufa1 | | | | | | | | | | 0.00000 | -4.16300 | Male | 0.00001 | 2.43343 | Female | | | |
| Ppp2r1a | | | | | | | | | | 0.00002 | 2.13468 | Female | | | | | | |
| Chd3os | | | | | | | | | | | | | 0.00054 | 5.71448 | Female | | | |
| Rsrc1 | 0.00233 | -4.42714 | Male | | | | | | | | | | | | | | | |
| Pim2 | | | | | | | | | | | | | | | | 0.00000 | 5.53264 | Female |
| Glul | | | | | | | | | | | | | | | | 0.01746 | -1.15648 | Male |
| Scg3 | | | | | | | | | | | | | 0.00000 | 2.84901 | Female | | | |
| Ubox5 | | | | | | | | | | | | | | | | 0.01172 | 3.01933 | Female |
| Intu | | | | | | | | | | 0.00153 | -2.64659 | Male | | | | | | |
| BC039771 | | | | | | | | | | 0.00005 | -3.31207 | Male | | | | | | |
| Abcg1 | | | | 0.00002 | 1.15920 | Female | | | | | | | | | | 0.00999 | 2.23520 | Female |
| Kctd10 | | | | | | | | | | 0.00000 | 4.58765 | Female | | | | | | |
| Pus1 | | | | | | | | | | 0.00609 | 2.15677 | Female | | | | | | |
| Ano9 | | | | | | | | | | | | | | | | 0.00000 | -5.35433 | Male |
| Hoxb7 | | | | | | | | | | | | | | | | 0.00000 | 2.07587 | Female |
| Atg9b | | | | | | | 0.00000 | 0.80915 | Female | 0.00000 | 2.07521 | Female | | | | | | |
| Zfp407 | | | | | | | | | | | | | | | | 0.00045 | -3.96158 | Male |
| Fbxo43 | | | | | | | | | | 0.02847 | -3.35016 | Male | | | | | | |
| Ttll3 | | | | | | | | | | | | | | | | 0.02273 | -3.33458 | Male |
| Tspo | 0.00036 | 0.75263 | Female | | | | | | | 0.00001 | -0.65132 | Male | | | | | | |
| Hspa4l | | | | | | | | | | 0.00005 | -2.26269 | Male | | | | | | |
| Shkbp1 | | | | | | | | | | 0.01481 | 3.78956 | Female | | | | | | |
| Rps15 | | | | | | | | | | 0.00000 | -4.55735 | Male | 0.03379 | 1.32596 | Female | | | |
| Ccdc88c | | | | 0.00853 | 2.96714 | Female | | | | 0.00000 | 6.91284 | Female | | | | | | |
| Frmpd1 | | | | 0.00021 | 2.74456 | Female | | | | 0.04243 | 2.97934 | Female | | | | | | |
| Omt2b | | | | | | | | | | 0.00109 | -4.02775 | Male | 0.00024 | 3.46765 | Female | | | |
| Irs3 | | | | | | | | | | | | | 0.00066 | 0.93316 | Female | 0.00354 | 1.50108 | Female |
| Sapcd2 | | | | | | | | | | | | | | | | 0.01279 | -2.51738 | Male |
| Tmod2 | | | | | | | | | | | | | | | | 0.00013 | 4.79100 | Female |
| Med28 | | | | | | | | | | 0.03070 | -2.19798 | Male | | | | | | |
| Smco1 | | | | | | | | | | 0.00000 | -4.12456 | Male | | | | | | |
| Tmem256 | | | | | | | | | | 0.00335 | -4.42212 | Male | | | | | | |
| Pced1b | | | | | | | | | | 0.00081 | -3.92168 | Male | | | | 0.01126 | 2.70805 | Female |
| Prr14 | | | | | | | | | | 0.03079 | 2.41042 | Female | | | | | | |
| Mcts1 | | | | | | | 0.00087 | 2.02786 | Female | | | | | | | | | |
| Dmrt1 | 0.00061 | 0.91860 | Female | | | | | | | | | | | | | | | |
| Kcnj14 | | | | 0.00151 | 1.04538 | Female | | | | | | | 0.00000 | 4.42813 | Female | | | |
| Fam213b | | | | 0.00005 | -1.38680 | Male | 0.00000 | 0.85119 | Female | 0.00000 | -4.11689 | Male | | | | | | |

| | | | | | | | | | | | | | | | | | | |
|---|---|---|---|---|---|---|---|---|---|---|---|---|---|---|---|---|---|---|
| Ncln | | | | | | | | | | 0.00061 | 4.06760 | Female | | | | | | |
| Psmc3 | | | | | | | | | | 0.03240 | 1.18936 | Female | | | | | | |
| Hax1 | | | | | | | | | | | | | | | | 0.01210 | -1.10356 | Male |
| P3h1 | | | | | | | | | | | | | 0.02856 | 3.29271 | Female | | | |
| Gstm2 | 0.00000 | -0.84695 | Male | | | | | | | 0.00001 | -1.96343 | Male | | | | | | |
| Gm9376 | | | | | | | | | | 0.00000 | -3.55465 | Male | | | | | | |
| Tst | | | | | | | | | | 0.00000 | 1.69062 | Female | | | | | | |
| Hspa13 | | | | | | | | | | 0.00000 | 0.93371 | Female | | | | | | |
| Mex3a | | | | 0.00000 | 1.33509 | Female | 0.00712 | 3.67696 | Female | | | | | | | | | |
| Fam8a1 | | | | | | | | | | | | | | | | 0.00022 | -6.55832 | Male |
| Kdm6a | | | | | | | 0.01124 | 2.55206 | Female | 0.00183 | 3.22370 | Female | | | | | | |
| Zfp729b | | | | | | | | | | | | | | | | 0.04728 | -2.29112 | Male |
| Thoc2 | | | | | | | 0.00885 | 2.65998 | Female | | | | | | | | | |
| Parp14 | | | | | | | | | | 0.00030 | -2.27132 | Male | | | | | | |
| BB557941 | | | | | | | | | | | | | 0.00000 | 5.23475 | Female | | | |
| Sgsm1 | | | | | | | | | | | | | | | | 0.02280 | 2.82265 | Female |
| Rnf150 | | | | | | | | | | | | | | | | 0.00258 | 4.14651 | Female |
| Zfp941 | | | | | | | | | | 0.00006 | -4.31486 | Male | | | | | | |
| Prrg4 | | | | 0.02182 | 0.74182 | Female | | | | | | | | | | | | |
| Jmy | | | | | | | | | | | | | | | | 0.01746 | -2.36826 | Male |
| Ephx3 | 0.00066 | 3.24681 | Female | | | | 0.04379 | -4.98809 | Male | | | | | | | | | |
| Tppp3 | | | | | | | | | | | | | 0.02859 | 3.62316 | Female | | | |
| H2afv | | | | | | | | | | 0.00017 | -3.29820 | Male | | | | | | |
| Pld4 | | | | | | | 0.00024 | -3.34654 | Male | | | | | | | | | |
| Sh3bp5 | | | | 0.02307 | -1.19466 | Male | | | | | | | | | | | | |
| Stk11 | | | | | | | | | | 0.03037 | 2.57353 | Female | | | | | | |
| Zfp709 | | | | 0.00314 | 1.16986 | Female | 0.00000 | 1.63343 | Female | | | | | | | | | |
| Adgrf3 | 0.00000 | 1.88551 | Female | | | | | | | | | | 0.00010 | -5.28888 | Male | | | |
| Swap70 | | | | | | | | | | 0.04733 | 2.17834 | Female | | | | | | |
| Rps29 | | | | | | | | | | 0.00000 | -5.81626 | Male | 0.00001 | 2.19724 | Female | | | |
| Cav2 | | | | | | | | | | | | | 0.00000 | -0.67298 | Male | | | |
| Syngr3 | | | | | | | | | | | | | | | | 0.00349 | -4.96864 | Male |
| Snrpd1 | | | | | | | | | | 0.00000 | -2.93878 | Male | 0.02652 | 0.99348 | Female | | | |
| Dis3 | | | | | | | | | | | | | 0.00143 | -2.04936 | Male | | | |
| Tmem63a | | | | | | | | | | | | | | | | 0.00919 | 3.83025 | Female |
| Dmbx1 | | | | | | | | | | | | | 0.00004 | -5.35156 | Male | | | |
| Ptcd3 | | | | | | | | | | 0.04255 | 1.22953 | Female | | | | | | |
| Fcer2a | | | | | | | | | | 0.00000 | -3.92778 | Male | | | | | | |
| Map7 | | | | | | | | | | | | | | | | 0.00051 | -1.62249 | Male |
| Trip4 | | | | | | | | | | | | | | | | 0.01376 | 1.63667 | Female |
| Gm15698 | | | | | | | | | | | | | | | | 0.03936 | -1.24187 | Male |
| Psmd14 | | | | | | | | | | 0.03055 | 1.08893 | Female | | | | | | |
| Rps15a.ps4 | | | | | | | | | | 0.00000 | -4.76692 | Male | | | | | | |
| Ccpg1 | | | | | | | | | | | | | | | | 0.02150 | -1.55115 | Male |
| Adh1 | | | | | | | | | | 0.00005 | -2.01665 | Male | | | | | | |
| Phldb2 | 0.00000 | -1.18628 | Male | | | | | | | | | | | | | | | |
| Homer3 | | | | | | | | | | | | | | | | 0.00000 | -5.15409 | Male |
| Tma16 | | | | | | | | | | | | | | | | 0.04334 | -1.55151 | Male |
| Crip1 | | | | | | | | | | 0.00098 | -4.62323 | Male | | | | | | |
| Slc7a1 | | | | 0.00000 | 1.26327 | Female | | | | 0.00000 | 2.61798 | Female | | | | 0.01043 | 4.92636 | Female |
| Taf9b | | | | | | | | | | 0.00000 | -2.59728 | Male | 0.00000 | -2.58666 | Male | | | |
| Vcl | | | | 0.01016 | 3.40260 | Female | | | | | | | | | | | | |
| Herc3 | | | | | | | 0.00019 | -2.28488 | Male | | | | | | | | | |
| Cog6 | | | | | | | | | | | | | | | | 0.02448 | 3.45069 | Female |
| Lpl | | | | | | | 0.04770 | -3.06465 | Male | | | | | | | | | |
| Spag4 | | | | | | | | | | | | | | | | 0.03082 | 3.39320 | Female |
| Tubb3 | | | | | | | | | | | | | | | | 0.00158 | 4.14308 | Female |
| Med6 | | | | | | | | | | | | | | | | 0.00132 | -2.30939 | Male |
| Sowahb | | | | | | | 0.00037 | -1.76509 | Male | 0.00033 | 1.93396 | Female | | | | | | |
| Srp9 | | | | | | | | | | 0.00858 | -2.04017 | Male | | | | | | |
| Zfand2a | | | | | | | | | | | | | | | | 0.00000 | -2.37051 | Male |
| Prr18 | | | | | | | | | | 0.00000 | -1.97094 | Male | | | | | | |
| Pianp | | | | 0.00011 | -1.40417 | Male | | | | 0.00000 | 4.49359 | Female | | | | | | |
| Zfp113 | | | | | | | 0.01406 | 2.12311 | Female | 0.00280 | -2.73733 | Male | | | | | | |
| Pin4 | | | | | | | | | | 0.00000 | -4.23892 | Male | 0.00000 | 2.56559 | Female | | | |
| Bach2 | | | | | | | | | | 0.00311 | -3.45274 | Male | | | | | | |
| Gm1600 | | | | | | | | | | | | | | | | 0.01825 | -4.09340 | Male |
| Pip5k1a | | | | | | | | | | | | | 0.01508 | 2.29800 | Female | | | |
| Hsd17b2 | 0.00030 | 2.39164 | Female | | | | | | | | | | 0.00000 | 2.38120 | Female | | | |
| Npc1l1 | | | | | | | | | | | | | | | | 0.03777 | 2.66140 | Female |

| | | | | | | | | | | | | | | | | | | |
|---|---|---|---|---|---|---|---|---|---|---|---|---|---|---|---|---|---|---|
| Rasal2 | | | | | | | | | | | | | | | | 0.02042 | 3.18484 | Female |
| Acrbp | | | | | | | | | | | | | | | | 0.02614 | -3.59926 | Male |
| Gm29443 | | | | | | | | | | 0.00014 | -4.12214 | Male | | | | | | |
| Mpnd | | | | | | | 0.02728 | 3.44574 | Female | | | | | | | | | |
| Rpl12 | | | | | | | | | | 0.01241 | -2.05412 | Male | | | | | | |
| Tbc1d14 | | | | 0.00005 | 4.50728 | Female | | | | | | | | | | | | |
| Abhd4 | | | | 0.00190 | 2.00919 | Female | | | | | | | | | | | | |
| Eef1e1 | | | | | | | | | | | | | | | | 0.04546 | -0.94482 | Male |
| Mcm3 | | | | | | | | | | 0.01374 | 1.73411 | Female | | | | | | |
| Mef2d | | | | 0.00034 | 3.99606 | Female | | | | | | | | | | | | |
| Popdc3 | | | | | | | | | | | | | | | | 0.01818 | 4.38620 | Female |
| Ddx39 | | | | | | | | | | 0.02500 | 1.05268 | Female | | | | | | |
| Mdh2 | | | | | | | | | | 0.00468 | 1.71566 | Female | | | | | | |
| Taf7 | | | | | | | | | | 0.00000 | 5.40855 | Female | | | | | | |
| Gcat | 0.02674 | 3.52782 | Female | | | | | | | | | | | | | | | |
| Amn | | | | | | | | | | 0.03141 | -4.62970 | Male | | | | 0.01879 | 3.96676 | Female |
| Smad2 | | | | | | | | | | | | | | | | 0.01199 | 1.91192 | Female |
| Sod2 | | | | | | | | | | 0.00131 | -1.65041 | Male | | | | | | |
| Bri3 | | | | | | | | | | 0.00000 | -5.17423 | Male | | | | | | |
| Mir684.1 | | | | | | | | | | 0.00003 | -1.84676 | Male | | | | | | |
| Slc27a1 | | | | | | | | | | | | | | | | 0.00413 | -2.81339 | Male |
| Aldh1l1 | | | | | | | | | | | | | | | | 0.02410 | 4.12008 | Female |
| Tbx3 | | | | | | | | | | | | | | | | 0.03520 | 1.51906 | Female |
| Ndufa6 | | | | | | | | | | 0.04923 | -2.30260 | Male | | | | | | |
| Fthl17e | | | | 0.00131 | 4.90826 | Female | 0.00000 | 6.45251 | Female | 0.00204 | 4.38553 | Female | 0.00011 | 3.89300 | Female | | | |
| Guca1b | | | | 0.00030 | 0.72247 | Female | | | | | | | | | | | | |
| Myod1 | | | | 0.00062 | 0.95399 | Female | 0.00000 | -0.65521 | Male | | | | 0.00000 | 1.04838 | Female | | | |
| Hspb8 | | | | 0.04399 | -1.85033 | Male | | | | | | | | | | | | |
| Mbd5 | | | | | | | | | | 0.03530 | 2.87874 | Female | | | | | | |
| Arfgap2 | | | | 0.04211 | 2.32202 | Female | | | | | | | | | | | | |
| Tuba8 | | | | | | | | | | 0.00000 | 3.74089 | Female | 0.00364 | 5.07508 | Female | 0.00021 | -6.61241 | Male |
| Acmsd | | | | | | | | | | 0.01084 | -2.59730 | Male | | | | | | |
| Tsnaxip1 | | | | | | | | | | | | | 0.00081 | 4.08242 | Female | | | |
| As3mt | 0.00786 | -4.22613 | Male | 0.02226 | 2.09438 | Female | | | | | | | | | | | | |
| Pfdn5 | | | | | | | | | | 0.00000 | -3.31243 | Male | | | | | | |
| Abcb8 | | | | | | | 0.00013 | 4.17560 | Female | | | | | | | | | |
| Nfkbia | | | | | | | | | | | | | | | | 0.00174 | 4.40127 | Female |
| Ccdc94 | | | | | | | | | | 0.04464 | 2.89430 | Female | | | | 0.00813 | -2.92762 | Male |
| Zfp607a | | | | | | | | | | 0.00001 | -3.30454 | Male | 0.00669 | 1.66669 | Female | | | |
| Trnau1ap | | | | | | | | | | 0.02512 | -2.70297 | Male | | | | | | |
| Oas1b | | | | | | | 0.03414 | -2.42593 | Male | | | | | | | | | |
| Cd59b | | | | | | | | | | 0.01001 | -4.25347 | Male | | | | | | |
| Tbx15 | | | | 0.00006 | -1.37748 | Male | | | | 0.00013 | -1.53157 | Male | | | | | | |
| Zfp599 | | | | | | | 0.03797 | 2.95801 | Female | | | | | | | | | |
| Ndufv3 | | | | | | | | | | 0.00001 | -3.45375 | Male | 0.00026 | 2.16609 | Female | | | |
| Nfe2 | | | | | | | | | | | | | | | | 0.00542 | 4.33079 | Female |
| Ap2s1 | | | | | | | | | | 0.00168 | 4.30680 | Female | | | | | | |
| Hspb1 | | | | | | | | | | 0.00000 | -7.18998 | Male | | | | | | |
| Ftx | | | | | | | | | | | | | | | | 0.00159 | 4.17310 | Female |
| Ddx49 | | | | | | | | | | 0.00283 | 3.28897 | Female | | | | | | |
| Dad1 | | | | | | | | | | 0.01004 | -1.80261 | Male | | | | | | |
| Etv6 | | | | | | | | | | 0.03076 | 3.34789 | Female | | | | | | |
| Il6ra | | | | | | | | | | 0.00161 | 3.87411 | Female | | | | | | |
| Rbm3 | | | | | | | 0.04338 | 0.93144 | Female | | | | | | | 0.03858 | 0.85502 | Female |
| Tnk1 | | | | | | | | | | | | | 0.00000 | 1.48167 | Female | | | |
| Arl6ip5 | 0.00000 | -0.98798 | Male | | | | | | | 0.00055 | -2.02744 | Male | | | | | | |
| Palld | | | | | | | | | | | | | | | | 0.00002 | 4.88139 | Female |
| Hck | | | | | | | | | | 0.00000 | 1.82726 | Female | | | | | | |
| Fundc2 | | | | | | | | | | | | | 0.00002 | -5.65334 | Male | | | |
| Hint3 | | | | | | | | | | 0.00033 | -3.88702 | Male | | | | | | |
| Kdm5d | 0.00075 | -1.45859 | Male | 0.00000 | -1.18539 | Male | | | | 0.00004 | -3.82511 | Male | 0.00000 | -3.86405 | Male | 0.00000 | -5.11971 | Male |
| Tex19.1 | | | | | | | 0.03914 | 1.27864 | Female | 0.00158 | 1.91826 | Female | | | | | | |
| Irf1 | | | | | | | | | | | | | 0.00000 | -0.81227 | Male | 0.02247 | 3.79377 | Female |
| Ksr1 | 0.00174 | -0.87426 | Male | | | | | | | | | | | | | 0.00000 | -1.40205 | Male |
| Hspb6 | | | | | | | | | | 0.00000 | 0.87873 | Female | | | | | | |
| Fabp4 | | | | | | | | | | 0.02125 | -3.56619 | Male | | | | | | |
| Mlh3 | | | | | | | 0.00200 | 3.84158 | Female | | | | 0.01752 | 2.67365 | Female | | | |
| Hoxa10 | | | | | | | | | | 0.00798 | 1.42068 | Female | 0.00001 | 1.44273 | Female | | | |
| Pgap3 | | | | | | | | | | 0.00121 | 1.05095 | Female | | | | | | |
| Cfap74 | | | | | | | | | | | | | 0.04458 | 0.96142 | Female | | | |

| | | | | | | | | | | | | | | | | | | |
|---|---|---|---|---|---|---|---|---|---|---|---|---|---|---|---|---|---|---|
| Gorasp1 | | | | | | | | | | | | | | | | 0.04789 | 2.01253 | Female |
| Dcdc2c | | | | | | | | | | 0.00003 | -2.28513 | Male | | | | | | |
| Cops3 | | | | | | | | | | 0.00000 | 2.63223 | Female | | | | | | |
| Stx1a | | | | | | | | | | | | | | | | 0.00018 | -4.26593 | Male |
| Tnfrsf10b | | | | | | | | | | 0.00553 | -2.56848 | Male | | | | | | |
| Sgpp2 | | | | | | | | | | 0.00464 | -2.66286 | Male | | | | | | |
| Mtg2 | | | | | | | | | | 0.00039 | 5.24057 | Female | | | | | | |
| Gm14005 | | | | | | | | | | 0.00000 | -4.53308 | Male | 0.00165 | 1.95621 | Female | | | |
| Lgals1 | | | | | | | | | | 0.00194 | -4.99306 | Male | | | | | | |
| Fundc1 | | | | | | | | | | 0.01281 | 2.68229 | Female | | | | | | |
| Obsl1 | 0.00000 | -1.81988 | Male | 0.00000 | -1.08215 | Male | | | | | | | | | | | | |
| Nt5c | 0.00974 | 4.19316 | Female | | | | | | | | | | | | | | | |
| Dnajb3 | | | | | | | | | | | | | | | | 0.02808 | -4.73899 | Male |
| Fbxo36 | | | | | | | | | | | | | | | | 0.00000 | -5.46971 | Male |
| Cdh4 | | | | | | | | | | | | | 0.00000 | -3.78905 | Male | | | |
| Rps15a.ps6 | | | | | | | | | | 0.00000 | -4.68437 | Male | 0.04089 | 1.14174 | Female | | | |
| Spin2c | | | | | | | | | | | | | | | | 0.00113 | -6.03546 | Male |
| Stk35 | | | | | | | | | | | | | | | | 0.00500 | 2.85029 | Female |
| Nckap1l | 0.00499 | -1.42903 | Male | | | | | | | 0.01827 | -0.91486 | Male | | | | | | |
| Cpa1 | | | | | | | | | | | | | 0.00229 | -4.13230 | Male | 0.00335 | -4.28484 | Male |
| Atp5e | | | | | | | | | | 0.00000 | -4.78423 | Male | 0.03851 | 1.55447 | Female | | | |
| Pi4ka | | | | | | | | | | | | | | | | 0.00223 | 2.36373 | Female |
| Txn2 | | | | | | | | | | 0.00627 | 2.17368 | Female | | | | | | |
| Rimbp3 | | | | 0.00168 | 1.97392 | Female | 0.00000 | 0.81763 | Female | | | | | | | 0.00000 | -2.80560 | Male |
| Kcnd1 | | | | | | | 0.00000 | 2.27503 | Female | | | | | | | | | |
| Rnf13 | 0.00004 | 5.96659 | Female | | | | | | | | | | | | | | | |
| Atp5k | | | | | | | | | | 0.00000 | -5.23206 | Male | 0.00033 | 1.73469 | Female | | | |
| Atp11b | | | | | | | | | | | | | | | | 0.00063 | -3.54252 | Male |
| Toe1 | | | | | | | | | | 0.01241 | 1.85387 | Female | | | | | | |
| Tmem40 | | | | | | | | | | | | | | | | 0.04485 | -4.52642 | Male |
| Spry2 | | | | | | | | | | 0.00910 | -3.32535 | Male | | | | | | |
| Sfi1 | | | | | | | | | | | | | | | | 0.00217 | -2.17832 | Male |
| Mrpl44 | | | | | | | | | | 0.00741 | 1.60410 | Female | | | | | | |
| Ell3 | | | | | | | | | | | | | | | | 0.04222 | -2.66922 | Male |
| Aqp11 | | | | | | | | | | | | | 0.04811 | 3.02229 | Female | 0.00784 | 3.71252 | Female |
| Pcolce2 | 0.00867 | -1.19772 | Male | | | | | | | | | | 0.00000 | 1.98116 | Female | | | |
| Tmem218 | | | | | | | | | | | | | 0.00000 | 4.21276 | Female | | | |
| Ccnd1 | | | | | | | | | | | | | | | | 0.00004 | -5.95355 | Male |
| Faxc | | | | | | | | | | | | | | | | 0.00000 | -1.21057 | Male |
| Fbxo2 | | | | 0.00000 | -0.80163 | Male | | | | | | | | | | | | |
| Stk4 | | | | | | | | | | | | | | | | 0.00783 | -2.04865 | Male |
| Klhl17 | 0.00004 | 1.47797 | Female | | | | | | | | | | | | | 0.01545 | 3.98165 | Female |
| Lst1 | | | | | | | | | | 0.00000 | -7.25638 | Male | | | | 0.00366 | -3.72670 | Male |
| Thyn1 | | | | | | | | | | 0.00902 | 2.22540 | Female | | | | | | |
| Tmem258 | | | | | | | | | | 0.00000 | -5.38273 | Male | | | | | | |
| Pde4c | | | | | | | | | | 0.00050 | -2.90285 | Male | | | | | | |
| Tgfb2 | | | | | | | | | | | | | 0.02209 | -5.03503 | Male | | | |
| Pced1a | 0.00000 | 2.95797 | Female | | | | | | | | | | | | | | | |
| Nedd4l | | | | | | | | | | 0.00051 | 3.83249 | Female | | | | | | |
| Ost4 | | | | | | | | | | 0.00015 | -2.97751 | Male | | | | | | |
| Asprv1 | | | | | | | 0.00561 | 1.92543 | Female | | | | | | | | | |
| Smdt1 | | | | | | | | | | 0.00704 | -3.02588 | Male | | | | | | |
| Man1c1 | | | | | | | | | | | | | 0.00998 | 4.21165 | Female | | | |
| Psmb11 | | | | | | | | | | | | | | | | 0.00506 | -2.49302 | Male |
| Ndufa5 | 0.01390 | 3.28566 | Female | | | | | | | 0.00000 | -5.73748 | Male | 0.00000 | 2.87191 | Female | | | |
| Bin1 | | | | | | | | | | 0.00000 | 1.73298 | Female | | | | | | |
| Gm3139 | | | | | | | | | | | | | | | | 0.00622 | -1.27933 | Male |
| Marveld1 | 0.00004 | 1.84516 | Female | | | | | | | | | | 0.00000 | 1.24417 | Female | | | |
| Zfp423 | | | | | | | | | | 0.02188 | -1.08644 | Male | | | | | | |
| Ptch1 | | | | | | | 0.03472 | -4.65974 | Male | | | | | | | | | |
| Ptprh | | | | | | | | | | 0.04717 | -4.59780 | Male | | | | | | |
| Plxna3 | | | | 0.00854 | 1.64759 | Female | | | | | | | 0.00000 | 2.28939 | Female | | | |
| Prtn3 | 0.00001 | 1.20994 | Female | | | | | | | | | | | | | 0.02149 | -4.56276 | Male |
| Nub1 | | | | | | | | | | | | | | | | 0.00977 | -1.16290 | Male |
| Hif3a | | | | | | | | | | | | | 0.02522 | 3.90372 | Female | | | |
| Nbl1 | 0.00237 | 1.25013 | Female | 0.01135 | -1.39659 | Male | | | | | | | | | | | | |
| Idh3b | | | | | | | | | | 0.01135 | 1.43807 | Female | | | | | | |
| Eln | | | | | | | | | | | | | | | | 0.00000 | -0.59285 | Male |
| Cs | | | | | | | | | | 0.00004 | 1.81581 | Female | | | | | | |
| Fam135a | | | | | | | | | | | | | | | | 0.00108 | -4.91274 | Male |

| | | | | | | | | | | | | | | | | | | |
|---|---|---|---|---|---|---|---|---|---|---|---|---|---|---|---|---|---|---|
| Ilk | | | | | | | | | | | | | | | | 0.00009 | -3.02986 | Male |
| Fndc4 | 0.00014 | 1.28215 | Female | | | | | | | | | | | | | | | |
| Tuba1a | 0.03040 | -1.66131 | Male | | | | | | | 0.00019 | 1.51307 | Female | 0.01508 | 0.80046 | Female | | | |
| Iars | | | | | | | | | | | | | | | | 0.00167 | 1.63425 | Female |
| Milr1 | | | | | | | 0.01685 | 3.77690 | Female | | | | | | | 0.01728 | 3.46265 | Female |
| Mettl20 | 0.00000 | 1.47951 | Female | | | | | | | | | | | | | | | |
| Dleu2 | | | | 0.02896 | -1.53664 | Male | | | | | | | | | | | | |
| Ddx3y | | | | | | | 0.00000 | -7.41047 | Male | 0.00000 | -7.48924 | Male | 0.00000 | -8.94889 | Male | 0.00000 | -7.30054 | Male |
| Ube2h | | | | | | | | | | 0.02775 | 1.68128 | Female | | | | | | |
| Il6st | | | | | | | | | | 0.00000 | 0.91302 | Female | 0.01919 | 3.80364 | Female | | | |
| Cep126 | 0.00000 | -0.79742 | Male | 0.00000 | 1.25789 | Female | | | | | | | | | | | | |
| Gm7977 | | | | | | | | | | 0.00000 | 4.43525 | Female | | | | | | |
| Zfp938 | | | | | | | | | | | | | | | | 0.00000 | -6.07022 | Male |
| Minos1 | | | | | | | | | | 0.00000 | -4.61146 | Male | 0.01374 | 1.42755 | Female | | | |
| Uqcrfs1 | | | | | | | | | | 0.00798 | -2.07589 | Male | | | | | | |
| Abt1 | | | | | | | | | | 0.00013 | 2.96801 | Female | | | | | | |
| Tspan31 | | | | | | | | | | 0.02251 | 1.77835 | Female | | | | | | |
| Dync2h1 | | | | 0.01709 | 2.89765 | Female | | | | | | | | | | | | |
| Sirt4 | | | | | | | | | | 0.03205 | 3.90969 | Female | | | | 0.02775 | -4.19699 | Male |
| Gm6498 | | | | | | | | | | 0.00583 | 2.06804 | Female | | | | | | |
| Nme2 | | | | | | | | | | 0.00317 | -1.92133 | Male | | | | | | |
| H60b | | | | | | | | | | | | | 0.03851 | 4.33461 | Female | | | |
| Actn4 | | | | | | | | | | 0.00000 | 3.93987 | Female | | | | | | |
| Npy1r | | | | | | | | | | 0.00051 | -2.71495 | Male | | | | | | |
| Pnrc1 | | | | | | | | | | | | | 0.01140 | -3.95297 | Male | | | |
| Desi1 | | | | | | | | | | | | | | | | 0.01945 | 2.29734 | Female |
| Slc33a1 | | | | 0.01479 | 3.26583 | Female | | | | | | | | | | | | |
| Acadl | 0.00002 | -3.39687 | Male | | | | | | | | | | | | | | | |
| Gpr50 | | | | | | | | | | | | | | | | 0.00000 | -0.88266 | Male |
| Elf1 | | | | | | | | | | 0.04733 | -2.83783 | Male | | | | | | |
| Gpr146 | | | | 0.00043 | 2.32073 | Female | 0.00000 | 0.59287 | Female | | | | | | | 0.01859 | 4.26171 | Female |
| Chrna1 | | | | | | | | | | 0.00000 | -6.51781 | Male | 0.00000 | 2.50949 | Female | | | |
| Pipox | | | | 0.00011 | 1.67577 | Female | | | | | | | | | | | | |
| Ttyh3 | | | | | | | | | | | | | | | | 0.00226 | 4.42663 | Female |
| Mslnl | | | | | | | | | | | | | 0.02766 | 4.10452 | Female | | | |
| Cox7a2 | | | | | | | | | | 0.00000 | -5.36937 | Male | 0.00000 | 2.32044 | Female | | | |
| Dynlt1b | | | | | | | | | | 0.00568 | -1.77402 | Male | | | | | | |
| Taf4b | | | | | | | | | | | | | 0.00793 | -2.01429 | Male | | | |
| Mtif2 | | | | | | | 0.00073 | 2.35657 | Female | 0.02661 | -2.00166 | Male | | | | | | |
| Sfmbt2 | 0.00000 | 1.67045 | Female | 0.00000 | 1.72036 | Female | | | | 0.00377 | 3.89139 | Female | | | | | | |
| Cgnl1 | | | | | | | | | | 0.00055 | 3.59584 | Female | | | | | | |
| Uqcc2 | | | | | | | | | | 0.00008 | -4.05983 | Male | 0.02251 | 2.04768 | Female | | | |
| Rps16 | | | | | | | | | | 0.01912 | -2.01628 | Male | | | | | | |
| Zc3h7b | | | | | | | | | | | | | 0.00174 | 4.88141 | Female | | | |
| Hibadh | | | | 0.00073 | 3.05853 | Female | | | | | | | | | | | | |
| Myl6 | | | | | | | | | | 0.03133 | -1.49696 | Male | | | | | | |
| Arl14epl | 0.04499 | 3.72141 | Female | | | | | | | | | | | | | | | |
| Obox7 | | | | | | | | | | 0.01091 | -4.31587 | Male | 0.00464 | -3.82784 | Male | | | |
| Pom121 | | | | | | | | | | 0.03524 | 2.02379 | Female | | | | | | |
| Pdzk1ip1 | | | | | | | | | | 0.00000 | -0.87429 | Male | | | | | | |
| Slc35b1 | | | | | | | | | | 0.03334 | 1.48727 | Female | | | | | | |
| Greb1 | | | | | | | | | | 0.00000 | -0.58277 | Male | | | | | | |
| Iqcb1 | | | | | | | | | | | | | | | | 0.04717 | 5.42506 | Female |
| Stk17b | | | | | | | | | | | | | | | | 0.01508 | -2.92535 | Male |
| Cdc5l | | | | | | | | | | 0.01436 | 1.30286 | Female | | | | | | |
| Magi3 | | | | | | | | | | 0.03763 | -1.45958 | Male | | | | | | |
| Gm6710 | | | | | | | | | | | | | | | | 0.00141 | -2.67145 | Male |
| Dock2 | | | | | | | 0.00002 | -3.06822 | Male | | | | | | | | | |
| Vat1 | | | | | | | | | | 0.01308 | -2.83161 | Male | | | | 0.03168 | -2.25804 | Male |
| Zfp287 | | | | | | | 0.02631 | 3.35135 | Female | 0.04414 | -2.88947 | Male | | | | | | |
| Obox6 | | | | | | | | | | | | | | | | 0.00100 | -3.10345 | Male |
| Sepw1 | | | | | | | | | | 0.00856 | -2.26848 | Male | | | | | | |
| Dync1h1 | | | | | | | | | | 0.04890 | 1.96668 | Female | | | | | | |
| Fbxo32 | | | | | | | | | | 0.00000 | -4.95008 | Male | 0.02938 | 2.07233 | Female | | | |
| Map4k3 | | | | | | | 0.00614 | 2.77177 | Female | | | | 0.01721 | -2.10785 | Male | | | |
| Emilin2 | | | | | | | | | | | | | 0.02665 | -5.23748 | Male | | | |
| Lrrc59 | | | | | | | | | | 0.00106 | 1.56969 | Female | | | | | | |
| Gfra3 | | | | | | | | | | | | | | | | 0.02881 | 2.66334 | Female |
| Zfp398 | | | | | | | 0.00000 | 0.83740 | Female | | | | | | | | | |
| Med15 | | | | 0.00099 | 3.64919 | Female | | | | | | | | | | | | |

| | | | | | | | | | | | | | | | | | | |
|---|---|---|---|---|---|---|---|---|---|---|---|---|---|---|---|---|---|---|
| Scai | | | | | | | | | | | | | | | | 0.03770 | 2.99090 | Female |
| Slc30a7 | 0.01335 | -3.92928 | Male | | | | | | | | | | | | | | | |
| Smo | | | | 0.00945 | 0.79654 | Female | | | | 0.00054 | 1.38560 | Female | 0.00000 | -4.39828 | Male | | | |
| Snhg8 | | | | | | | | | | 0.02352 | -3.21683 | Male | | | | | | |
| Maml1 | | | | | | | | | | 0.03272 | 3.01441 | Female | | | | | | |
| Cnot3 | | | | | | | | | | 0.00771 | 3.11524 | Female | | | | | | |
| Clec2g | | | | | | | | | | | | | 0.00034 | 4.00657 | Female | | | |
| Copz1 | | | | | | | | | | 0.04944 | 1.02880 | Female | | | | | | |
| Pdlim2 | | | | | | | | | | | | | | | | 0.01824 | -4.90558 | Male |
| Nol7 | | | | | | | | | | 0.00000 | -3.90312 | Male | | | | | | |
| Gm1976 | | | | | | | | | | 0.04391 | -2.31555 | Male | | | | | | |
| Gm15421 | | | | | | | | | | 0.00000 | -5.12728 | Male | 0.00962 | 1.40849 | Female | | | |
| Nphp3 | | | | | | | | | | | | | | | | 0.03042 | 3.85826 | Female |
| Gpr137b | | | | | | | | | | | | | | | | 0.00976 | -1.39669 | Male |
| Evx1 | | | | | | | 0.00001 | 2.32493 | Female | | | | 0.00000 | 1.72037 | Female | | | |
| Evpl | | | | 0.00734 | -0.81822 | Male | | | | | | | | | | | | |
| Epb41 | | | | | | | | | | 0.00226 | 3.02613 | Female | | | | | | |
| Mrps24 | | | | | | | | | | 0.00000 | -5.81371 | Male | | | | | | |
| Zcchc6 | | | | | | | | | | 0.00001 | 5.03579 | Female | | | | 0.01724 | -2.59517 | Male |
| Cmip | | | | | | | | | | 0.00226 | 3.77339 | Female | | | | | | |
| Tceb1 | | | | | | | | | | 0.00002 | -2.11553 | Male | | | | 0.00731 | -1.23414 | Male |
| Cdnf | | | | | | | | | | | | | | | | 0.00013 | -3.43680 | Male |
| Itgb7 | | | | | | | | | | | | | | | | 0.01406 | 3.13165 | Female |
| Zp2 | | | | | | | | | | | | | | | | 0.00000 | 4.85828 | Female |
| Eif3e | | | | | | | | | | 0.01690 | 1.03764 | Female | | | | | | |
| Fhad1 | | | | | | | | | | | | | | | | 0.00009 | 1.47545 | Female |
| Snrnp27 | | | | | | | | | | 0.00395 | -1.93958 | Male | | | | | | |
| Stamos | | | | | | | 0.00000 | -1.86992 | Male | | | | | | | | | |
| Siae | 0.00031 | -0.88599 | Male | 0.00008 | 0.71216 | Female | | | | | | | | | | | | |
| Rrp15 | | | | | | | | | | 0.00869 | 3.59679 | Female | | | | | | |
| Panx1 | 0.00000 | -1.59029 | Male | | | | | | | | | | | | | | | |
| Fam19a5 | | | | 0.02896 | -1.56403 | Male | | | | 0.00000 | 2.22829 | Female | | | | | | |
| Cisd1 | | | | | | | | | | 0.02063 | -2.36713 | Male | | | | | | |
| Gm16513 | | | | | | | | | | 0.00525 | -2.08362 | Male | | | | | | |
| Map1a | | | | | | | | | | 0.00171 | -1.91367 | Male | | | | 0.00000 | 1.36423 | Female |
| Adamtsl1 | | | | | | | | | | 0.00000 | -3.38454 | Male | 0.00000 | -2.13787 | Male | | | |
| Ifngr2 | | | | | | | | | | | | | 0.00189 | 4.00724 | Female | | | |
| Lrrn4 | | | | | | | | | | | | | 0.03370 | 3.70158 | Female | 0.00558 | 4.60653 | Female |
| Trim14 | | | | | | | | | | | | | | | | 0.04172 | 2.47966 | Female |
| Lncenc1 | 0.00007 | -0.93671 | Male | 0.00001 | 0.97310 | Female | | | | | | | | | | | | |
| Fkbp1b | | | | | | | | | | 0.04762 | -3.95641 | Male | | | | | | |
| Fbxw21 | | | | | | | | | | | | | 0.00216 | 4.74281 | Female | | | |
| Ift74 | | | | | | | | | | | | | | | | 0.00495 | 4.01886 | Female |
| Wdr93 | | | | | | | | | | | | | | | | 0.00144 | -4.50594 | Male |
| Cox7c | | | | | | | 0.02952 | 1.52394 | Female | 0.00000 | -4.55599 | Male | 0.00019 | 1.80958 | Female | | | |
| Gpx6 | | | | | | | 0.00038 | 0.98074 | Female | | | | 0.00000 | 1.12607 | Female | | | |
| Cntn2 | | | | | | | | | | 0.00117 | -3.63164 | Male | | | | | | |
| Ing5 | | | | | | | | | | | | | | | | 0.00898 | -1.89853 | Male |
| Cd200 | | | | | | | | | | | | | | | | 0.00000 | 0.95999 | Female |
| Chil1 | | | | | | | | | | 0.00011 | 1.72141 | Female | | | | | | |
| Rpl9 | | | | | | | | | | 0.02754 | -1.11213 | Male | | | | | | |
| Kdm4b | | | | | | | 0.00122 | 4.01631 | Female | | | | | | | | | |
| Crlf1 | | | | | | | | | | | | | | | | 0.00022 | -0.91327 | Male |
| Fkbp10 | | | | | | | | | | | | | 0.02446 | -3.19349 | Male | | | |
| Cpne5 | | | | | | | | | | | | | | | | 0.00000 | 2.99269 | Female |
| Bfsp1 | | | | 0.00006 | 2.02862 | Female | 0.00001 | 1.03037 | Female | | | | 0.04674 | 4.62054 | Female | | | |
| Abtb2 | | | | | | | 0.00627 | 2.41574 | Female | | | | | | | | | |
| Mrps14 | | | | | | | | | | 0.00667 | -1.84090 | Male | | | | | | |
| Galnt16 | | | | | | | | | | | | | 0.04980 | 2.91373 | Female | | | |
| Tceanc | | | | | | | 0.00885 | 4.28255 | Female | | | | | | | | | |
| Tmem242 | | | | | | | | | | 0.02251 | 2.31091 | Female | | | | | | |
| Tpm3 | | | | | | | | | | 0.00439 | 1.29503 | Female | | | | | | |
| Naip2 | | | | | | | | | | 0.02212 | -2.95176 | Male | | | | | | |
| Pptc7 | | | | | | | | | | | | | 0.02058 | 2.63319 | Female | | | |
| Gm13125 | | | | 0.00001 | 2.05101 | Female | 0.00000 | 4.53706 | Female | 0.02081 | -0.84542 | Male | 0.00000 | 1.15704 | Female | | | |
| Prmt6 | | | | | | | | | | | | | | | | 0.00841 | 2.21999 | Female |
| Smarce1 | | | | | | | | | | 0.01765 | 1.57566 | Female | | | | 0.00611 | -1.44202 | Male |
| Malsu1 | | | | | | | | | | 0.00000 | -6.08102 | Male | | | | | | |
| Mgl2 | | | | | | | | | | | | | 0.03135 | 2.04827 | Female | 0.00052 | -3.21397 | Male |
| Oacyl | | | | | | | | | | | | | | | | 0.00034 | -0.63747 | Male |

| | | | | | | | | | | | | | | | | | | |
|---|---|---|---|---|---|---|---|---|---|---|---|---|---|---|---|---|---|---|
| Ntpcr | 0.00037 | -1.03056 | Male | 0.00210 | -1.22548 | Male | | | | 0.00000 | 1.18456 | Female | | | | | | |
| Colec12 | | | | | | | | | | 0.04365 | -1.59258 | Male | | | | | | |
| Parp12 | | | | | | | | | | | | | 0.00000 | -6.27793 | Male | | | |
| Ptpdc1 | | | | | | | | | | | | | 0.00012 | -5.59945 | Male | 0.00000 | -1.18430 | Male |
| Eno1 | | | | | | | | | | 0.02750 | 3.26389 | Female | | | | | | |
| Dcaf12l1 | | | | | | | 0.00215 | 3.85573 | Female | | | | | | | | | |
| Cdkl1 | | | | | | | | | | | | | 0.00007 | 0.86137 | Female | 0.00132 | -0.59352 | Male |
| Uqcr11 | | | | | | | | | | 0.00057 | -3.19844 | Male | | | | | | |
| Rab19 | | | | | | | | | | | | | 0.00000 | 0.73503 | Female | 0.00000 | 0.58654 | Female |
| Eomes | | | | 0.00493 | 2.67357 | Female | 0.00045 | 3.81038 | Female | | | | | | | | | |
| Gpsm1 | | | | | | | | | | | | | | | | 0.00000 | 5.25684 | Female |
| Snrpg | | | | | | | | | | 0.00000 | -5.62905 | Male | 0.00000 | 2.05797 | Female | | | |
| Vcp | | | | | | | | | | 0.01022 | 1.76301 | Female | | | | | | |
| Chd5 | | | | 0.00824 | -1.53897 | Male | | | | | | | | | | 0.00181 | -0.58610 | Male |
| Taf5l | | | | | | | | | | | | | 0.00937 | -1.78698 | Male | | | |
| Dynlrb1 | | | | | | | | | | 0.00000 | -2.77953 | Male | | | | | | |
| Phkg2 | | | | | | | | | | 0.00001 | 6.16665 | Female | | | | | | |
| Ramp1 | | | | | | | | | | | | | | | | 0.00000 | -1.07148 | Male |
| Glod5 | | | | | | | | | | | | | | | | 0.00960 | 4.61662 | Female |
| Ric8b | 0.00000 | 2.02973 | Female | | | | | | | | | | | | | | | |
| Dpcr1 | | | | | | | 0.04338 | -3.74022 | Male | | | | | | | | | |
| Chsy1 | | | | | | | | | | 0.01766 | -4.69000 | Male | 0.00011 | 5.05200 | Female | | | |
| Tmem29 | | | | | | | | | | 0.02692 | -2.82187 | Male | | | | 0.00888 | 2.67076 | Female |
| Megf6 | | | | | | | 0.01758 | 4.01972 | Female | | | | | | | | | |
| Chml | | | | | | | 0.00000 | 3.42248 | Female | 0.00014 | -2.18994 | Male | | | | | | |
| Nphs1 | | | | | | | | | | 0.01187 | -2.67213 | Male | | | | | | |
| Slc29a4 | 0.00001 | -2.44885 | Male | | | | | | | | | | | | | | | |
| Bcar3 | | | | 0.00000 | 2.14319 | Female | | | | | | | | | | | | |
| Apela | | | | 0.00014 | -1.28366 | Male | | | | | | | | | | | | |
| Ppp4c | | | | | | | | | | 0.00000 | 6.17599 | Female | | | | | | |
| Wwc1 | | | | | | | | | | 0.00016 | 5.69222 | Female | | | | | | |
| Atrnl1 | | | | | | | | | | | | | 0.03379 | 4.63977 | Female | | | |
| Ndufb7 | | | | | | | | | | 0.00000 | -4.12652 | Male | | | | | | |
| Ube2s | | | | | | | | | | 0.00464 | -2.33386 | Male | | | | | | |
| Rps17 | | | | | | | | | | 0.00000 | -4.37961 | Male | 0.00452 | 1.33334 | Female | | | |
| Stxbp4 | | | | | | | | | | 0.00027 | -2.93139 | Male | | | | | | |
| Gm5860 | | | | | | | | | | 0.00563 | -1.70818 | Male | | | | | | |
| Gm5127 | | | | | | | | | | | | | 0.00000 | 2.66080 | Female | | | |
| Csta1 | | | | | | | | | | 0.03143 | -3.64928 | Male | | | | | | |
| Taf2 | | | | | | | | | | | | | 0.00835 | -1.69691 | Male | | | |
| Mttp | | | | | | | | | | | | | | | | 0.00000 | -5.57816 | Male |
| Lemd2 | | | | 0.00000 | -0.97693 | Male | | | | | | | | | | | | |
| Fcgr2b | | | | | | | | | | 0.00000 | 7.21667 | Female | | | | 0.01728 | 2.49932 | Female |
| Amigo3 | | | | | | | | | | | | | | | | 0.01432 | -4.45209 | Male |
| Tdpoz1 | | | | | | | | | | | | | | | | 0.04385 | 2.49431 | Female |
| Ulk1 | | | | | | | | | | 0.03111 | 1.39247 | Female | | | | | | |
| Uqcc3 | | | | | | | | | | 0.02272 | -2.15902 | Male | | | | | | |
| Erg | | | | | | | | | | 0.00067 | -1.69416 | Male | | | | | | |
| Vmn1r65 | | | | | | | | | | | | | | | | 0.00573 | 2.98542 | Female |
| Lpin2 | | | | | | | | | | | | | 0.02139 | -3.72459 | Male | | | |
| Acvr1b | | | | 0.00000 | -1.50102 | Male | | | | 0.00000 | -2.66974 | Male | | | | | | |
| Crtc2 | | | | | | | 0.03552 | 3.18567 | Female | | | | | | | | | |
| Trappc2 | | | | 0.03142 | 2.62958 | Female | | | | | | | | | | | | |
| Pex6 | | | | 0.03048 | 0.69806 | Female | | | | | | | | | | | | |
| Gm17644 | | | | | | | | | | 0.00158 | 3.65493 | Female | | | | | | |
| Scand1 | | | | | | | | | | 0.00041 | -5.36301 | Male | | | | | | |
| Med31 | | | | | | | | | | 0.00510 | -2.45237 | Male | | | | | | |
| Snhg3 | | | | | | | | | | 0.00057 | -3.25418 | Male | | | | | | |
| Inhbb | | | | | | | | | | | | | 0.00000 | 1.83752 | Female | | | |
| Abhd8 | | | | 0.01597 | -1.55263 | Male | | | | | | | | | | | | |
| Atg4a | | | | 0.01815 | 2.28002 | Female | | | | 0.02090 | 2.85154 | Female | | | | | | |
| Tdrd3 | | | | 0.00560 | 3.56749 | Female | | | | | | | | | | | | |
| Nradd | | | | | | | | | | | | | | | | 0.00699 | 4.82835 | Female |
| Magea8 | 0.00000 | 1.80301 | Female | | | | | | | 0.00000 | -1.14576 | Male | | | | | | |
| Itga3 | | | | 0.03229 | 0.81900 | Female | | | | | | | | | | | | |
| Mbnl2 | | | | | | | | | | 0.00000 | 3.01679 | Female | | | | | | |
| Spon2 | | | | | | | | | | 0.00192 | 1.42034 | Female | | | | | | |
| AI854703 | | | | | | | | | | | | | 0.00000 | -5.37839 | Male | 0.00000 | 1.99130 | Female |
| Gstt2 | | | | | | | | | | 0.01066 | -1.00901 | Male | | | | | | |
| Lsm2 | | | | | | | | | | 0.01082 | -2.23363 | Male | | | | | | |

| | | | | | | | | | | | | | | | | | | |
|---|---|---|---|---|---|---|---|---|---|---|---|---|---|---|---|---|---|---|
| Nenf | | | | | | | | | | 0.00824 | -6.18776 | Male | | | | 0.03197 | 4.20276 | Female |
| Slc6a15 | | | | | | | | | | | | | 0.00179 | 3.25254 | Female | | | |
| Vwa5b2 | | | | | | | | | | | | | 0.00000 | -3.24677 | Male | | | |
| Aldh3b2 | | | | | | | | | | | | | 0.04667 | 4.76169 | Female | | | |
| Sp3 | | | | | | | 0.04676 | 2.66246 | Female | | | | | | | | | |
| Tulp1 | | | | | | | | | | | | | 0.00420 | -3.96768 | Male | | | |
| Sae1 | | | | | | | | | | 0.00952 | 1.47040 | Female | | | | | | |
| Nostrin | | | | | | | | | | | | | | | | 0.01028 | 4.56571 | Female |
| Cln6 | | | | 0.00001 | -1.44129 | Male | | | | | | | | | | | | |
| Map1lc3a | | | | | | | | | | 0.03395 | -2.24575 | Male | | | | | | |
| Coq9 | | | | | | | | | | | | | | | | 0.03669 | 1.70207 | Female |
| Ppie | | | | | | | | | | 0.00004 | 3.51426 | Female | | | | | | |
| Usmg5 | | | | | | | | | | 0.00000 | -6.27587 | Male | 0.00000 | 2.43228 | Female | | | |
| Triap1 | | | | | | | | | | 0.00000 | 2.50252 | Female | | | | 0.00146 | -1.45516 | Male |
| Pga5 | | | | | | | | | | | | | 0.00000 | 0.59535 | Female | | | |
| Luzp2 | 0.00378 | 2.06272 | Female | | | | | | | | | | | | | | | |
| Sbspon | | | | | | | | | | | | | 0.00000 | 0.87875 | Female | 0.00001 | 1.68069 | Female |
| Gm26705 | | | | | | | | | | | | | | | | 0.04728 | 2.86158 | Female |
| Mllt10 | | | | | | | | | | | | | | | | 0.03813 | -1.77792 | Male |
| Rpl37a | | | | | | | | | | 0.00000 | -4.94562 | Male | 0.00007 | 1.66772 | Female | | | |
| Kars | | | | | | | | | | 0.02699 | 1.38695 | Female | | | | | | |
| Chek2 | | | | | | | | | | | | | 0.00002 | -1.94784 | Male | | | |
| Nlrp5.ps | | | | | | | | | | | | | | | | 0.00002 | -5.57155 | Male |
| Gas2l3 | | | | | | | | | | 0.00004 | -3.22218 | Male | | | | | | |
| Obox3 | | | | | | | | | | 0.00000 | -6.00544 | Male | | | | | | |
| Slc27a3 | 0.00000 | -0.86850 | Male | | | | | | | | | | | | | | | |
| Lman1 | 0.03235 | 3.92329 | Female | | | | | | | | | | | | | | | |
| Cnga4 | | | | | | | 0.00000 | 2.47854 | Female | 0.02455 | -1.02196 | Male | | | | 0.00000 | 3.07516 | Female |
| Pomp | | | | | | | | | | 0.00214 | -1.43809 | Male | | | | 0.03900 | -0.89547 | Male |
| Slc28a3 | | | | | | | | | | 0.02482 | -2.03608 | Male | | | | | | |
| Panct2 | | | | | | | | | | | | | 0.00029 | 3.83548 | Female | | | |
| Zbed6 | | | | | | | | | | 0.00007 | -1.58719 | Male | 0.01404 | 3.61985 | Female | | | |
| Sfr1 | | | | | | | | | | 0.01455 | 1.43411 | Female | | | | | | |
| Ppt2 | | | | | | | | | | 0.00000 | -4.25813 | Male | | | | | | |
| Hsf5 | 0.02826 | -1.60465 | Male | 0.00045 | 0.77093 | Female | 0.01661 | -1.24229 | Male | | | | | | | 0.00051 | 0.93400 | Female |
| Ovgp1 | | | | 0.00006 | -1.29050 | Male | | | | | | | 0.00000 | 2.22655 | Female | 0.00000 | -0.61114 | Male |
| Aqp7 | | | | | | | | | | 0.04511 | -2.40388 | Male | | | | | | |
| Ankrd33b | | | | | | | | | | 0.00000 | 2.48665 | Female | | | | 0.00989 | 5.15915 | Female |
| Lmntd2 | 0.03022 | 0.84705 | Female | 0.00369 | -0.92357 | Male | 0.00000 | 1.64169 | Female | | | | | | | | | |
| Acin1 | | | | | | | | | | 0.00003 | 3.96252 | Female | | | | | | |
| Gm13830 | | | | | | | | | | | | | 0.03553 | 2.53230 | Female | | | |
| Slc7a11 | | | | | | | | | | 0.00022 | 1.90444 | Female | | | | | | |
| Snx24 | | | | | | | | | | | | | 0.04667 | 4.38218 | Female | | | |
| Art5 | | | | | | | | | | | | | | | | 0.00000 | -1.42034 | Male |
| Smim6 | 0.00107 | 1.53167 | Female | 0.00050 | 1.12777 | Female | | | | | | | | | | | | |
| Spice1 | 0.00000 | -2.25756 | Male | | | | | | | | | | | | | | | |
| Epha10 | | | | | | | | | | | | | | | | 0.00243 | 4.46207 | Female |
| Ubtfl1 | | | | | | | | | | 0.03831 | 2.78878 | Female | | | | | | |
| Lrrc56 | | | | | | | | | | | | | | | | 0.02362 | -4.44316 | Male |
| Gm15910 | | | | | | | | | | 0.00026 | -3.75086 | Male | | | | | | |
| Nol4l | | | | | | | | | | | | | | | | 0.00000 | -3.72824 | Male |
| Tefm | | | | | | | | | | | | | | | | 0.00413 | -2.05636 | Male |
| Dkc1 | | | | | | | | | | 0.00116 | 1.82677 | Female | | | | | | |
| Pou2f3 | | | | | | | | | | | | | 0.00000 | 5.51908 | Female | | | |
| Brip1 | | | | | | | | | | 0.00194 | -3.12945 | Male | | | | | | |
| Dcxr | 0.00000 | -1.30118 | Male | 0.00000 | -2.73201 | Male | | | | | | | | | | | | |
| B4galt1 | | | | 0.00890 | 2.87878 | Female | | | | | | | | | | | | |
| Ftsj1 | | | | | | | | | | 0.01103 | 1.81039 | Female | | | | | | |
| Ddx58 | | | | | | | | | | | | | | | | 0.00682 | 4.21205 | Female |
| Lemd3 | | | | | | | | | | | | | | | | 0.02505 | -2.95232 | Male |
| Scpep1 | | | | | | | | | | | | | | | | 0.00128 | -1.59267 | Male |
| BC005537 | | | | | | | | | | 0.00796 | 2.02902 | Female | | | | | | |
| Atxn7l1 | | | | 0.00000 | -1.88714 | Male | | | | 0.00000 | 3.05708 | Female | | | | | | |
| Ndufv1 | 0.03885 | 3.10800 | Female | | | | | | | | | | | | | | | |
| Anapc16 | | | | | | | | | | 0.00521 | -2.66136 | Male | | | | | | |
| Tuba1b | | | | | | | | | | 0.00718 | 1.15306 | Female | 0.03851 | 0.70588 | Female | | | |
| Cdc42ep3 | | | | 0.00000 | 1.15353 | Female | | | | | | | | | | | | |
| Cyp11a1 | | | | | | | | | | | | | 0.00000 | 2.32993 | Female | | | |
| Slc4a2 | | | | 0.00000 | -3.03052 | Male | | | | | | | | | | | | |
| Tspyl2 | | | | | | | | | | | | | | | | 0.03871 | -3.77955 | Male |

| | | | | | | | | | | | | | | | | | | |
|---|---|---|---|---|---|---|---|---|---|---|---|---|---|---|---|---|---|---|
| Gm20219 | | | | 0.00860 | 0.90709 | Female | | | | | | | | | | | | |
| Cnot1 | | | | | | | | | | 0.00007 | 1.75477 | Female | | | | | | |
| Rac2 | | | | | | | | | | 0.00160 | -3.12261 | Male | | | | | | |
| Armcx4 | | | | | | | 0.00458 | 2.74309 | Female | 0.00011 | -3.83083 | Male | | | | | | |
| X2.Mar | | | | | | | | | | | | | 0.03509 | 2.20107 | Female | | | |
| Spint5 | | | | | | | 0.00000 | 1.48257 | Female | | | | | | | | | |
| Tomm5 | | | | | | | | | | 0.00395 | -1.75876 | Male | | | | | | |
| Hey1 | | | | 0.00008 | 0.92956 | Female | | | | | | | | | | 0.00000 | -0.79213 | Male |
| Dexi | | | | 0.00002 | 0.87904 | Female | | | | | | | | | | | | |
| Fam195a | | | | | | | | | | 0.01737 | -3.19217 | Male | | | | | | |
| Wdr70 | | | | | | | | | | 0.03079 | 2.54026 | Female | | | | | | |
| Fthl17.ps1 | | | | 0.00102 | 4.82321 | Female | 0.00000 | 5.62317 | Female | 0.00160 | 4.62315 | Female | 0.00001 | 4.51635 | Female | | | |
| Hes1 | | | | | | | | | | 0.00022 | -5.23411 | Male | | | | | | |
| Cfap126 | | | | | | | | | | 0.00000 | 0.80390 | Female | | | | 0.00000 | 2.24337 | Female |
| Rpl22 | | | | | | | | | | 0.00000 | -4.64774 | Male | 0.00462 | 1.13036 | Female | | | |
| Dynll1 | | | | | | | | | | 0.02502 | -1.79867 | Male | | | | | | |
| Tnpo1 | | | | | | | | | | 0.00181 | -2.64507 | Male | | | | | | |
| Camk1 | | | | | | | 0.00327 | -4.22619 | Male | | | | | | | | | |
| Igfbp2 | | | | | | | | | | | | | | | | 0.02210 | 3.37869 | Female |
| Vstm5 | 0.00189 | 0.90527 | Female | 0.00024 | 0.76882 | Female | 0.00000 | -2.10228 | Male | 0.00000 | 2.10076 | Female | | | | | | |
| Nbas | | | | 0.02618 | 1.65774 | Female | | | | | | | | | | | | |
| Nudt7 | | | | | | | | | | | | | | | | 0.03239 | 4.04304 | Female |
| Pxdn | | | | | | | | | | 0.00130 | -0.89101 | Male | | | | 0.02737 | 5.19587 | Female |
| Ccdc153 | 0.00000 | 2.81209 | Female | | | | | | | | | | | | | | | |
| Pinlyp | | | | | | | | | | | | | | | | 0.00000 | -6.44535 | Male |
| Strip2 | | | | 0.00000 | 2.33596 | Female | | | | 0.00000 | -3.49614 | Male | | | | | | |
| Dimt1 | | | | | | | 0.00501 | 2.15820 | Female | | | | 0.02665 | -1.54768 | Male | 0.02448 | 1.72733 | Female |
| Cldn3 | | | | 0.00038 | -2.30537 | Male | | | | | | | | | | | | |
| Stx11 | | | | | | | 0.02003 | -0.65348 | Male | | | | | | | | | |
| Cav1 | | | | | | | | | | 0.00350 | 0.80670 | Female | | | | | | |
| Ctrl | | | | | | | | | | | | | | | | 0.00000 | -5.97194 | Male |
| St7l | | | | | | | | | | | | | | | | 0.00052 | -3.34762 | Male |
| Atp6v1c2 | | | | | | | | | | | | | | | | 0.00000 | 2.24698 | Female |
| Tpbgl | | | | | | | | | | 0.00647 | -1.60929 | Male | | | | | | |
| Mixl1 | | | | | | | | | | | | | 0.00000 | 0.71854 | Female | | | |
| Deb1 | | | | | | | | | | 0.00000 | -3.66688 | Male | 0.00174 | 1.82477 | Female | | | |
| Pla2g10os | | | | | | | | | | 0.04656 | -4.09930 | Male | | | | | | |
| Gm21284 | | | | | | | | | | 0.00000 | -4.40855 | Male | 0.01964 | 1.80147 | Female | | | |
| Zfp939 | | | | | | | | | | | | | | | | 0.00156 | -3.95123 | Male |
| Asb6 | | | | | | | | | | 0.01012 | 3.02904 | Female | | | | | | |
| Hmox2 | | | | | | | | | | 0.01769 | 2.04395 | Female | | | | | | |
| Epn1 | | | | | | | | | | 0.00000 | -4.18862 | Male | | | | 0.02365 | -2.04506 | Male |
| Emc2 | | | | 0.00000 | 4.74220 | Female | | | | | | | | | | | | |
| Nup93 | | | | | | | | | | 0.02500 | 1.80426 | Female | | | | | | |
| Cxcl13 | | | | | | | 0.00123 | -1.92795 | Male | | | | | | | | | |
| Lrrc75a | 0.00000 | -1.23038 | Male | | | | | | | 0.00004 | -2.27981 | Male | 0.01752 | 5.17048 | Female | 0.00731 | -6.23560 | Male |
| Rac3 | | | | | | | | | | | | | | | | 0.03691 | -3.34011 | Male |
| Wscd2 | | | | | | | 0.01058 | -4.75219 | Male | | | | | | | | | |
| Il11 | | | | | | | | | | 0.00000 | -4.71839 | Male | | | | | | |
| Kank3 | | | | 0.01113 | 2.31563 | Female | | | | | | | | | | | | |
| Bmp5 | | | | | | | | | | | | | 0.04780 | 3.08347 | Female | | | |
| Sult5a1 | | | | | | | | | | | | | | | | 0.04407 | 2.90208 | Female |
| Sppl3 | | | | | | | | | | 0.01241 | 1.86557 | Female | | | | | | |
| Lama4 | | | | | | | | | | 0.00000 | 0.62362 | Female | | | | | | |
| Setd1b | | | | | | | | | | 0.03395 | -2.40724 | Male | | | | | | |
| Gm7104 | | | | | | | | | | | | | 0.00004 | 4.62011 | Female | | | |
| Bmp8a | | | | | | | | | | 0.03336 | -0.61872 | Male | 0.00000 | 2.18008 | Female | | | |
| Allc | | | | | | | | | | | | | | | | 0.00000 | 5.00333 | Female |
| Ly6k | | | | | | | 0.00001 | -1.32892 | Male | | | | | | | | | |
| Snrpf | | | | | | | | | | 0.00036 | -3.83804 | Male | | | | | | |
| Got1 | | | | | | | | | | 0.01130 | 1.48160 | Female | | | | | | |
| LOC105242579 | | | | | | | | | | 0.00000 | -3.20214 | Male | | | | | | |
| Cilp | | | | | | | | | | | | | | | | 0.00000 | -5.89358 | Male |
| Dgka | | | | | | | | | | | | | | | | 0.00158 | -2.47985 | Male |
| Tmem69 | | | | | | | | | | 0.00000 | -4.63047 | Male | 0.00011 | 1.95354 | Female | | | |
| Park2 | | | | | | | 0.00282 | 2.65597 | Female | 0.00000 | -5.26255 | Male | 0.02954 | 1.81805 | Female | | | |
| Gm6578 | | | | 0.03915 | -2.45061 | Male | | | | | | | | | | | | |
| Cdx2 | | | | | | | 0.01758 | 3.23658 | Female | | | | | | | | | |
| Klf10 | | | | 0.00002 | 3.89391 | Female | | | | | | | | | | | | |
| Mrps21 | | | | | | | | | | 0.00000 | -4.20969 | Male | 0.00083 | 1.62450 | Female | | | |

| | | | | | | | | | | | | | | | | | | |
|---|---|---|---|---|---|---|---|---|---|---|---|---|---|---|---|---|---|---|
| Cdc37l1 | | | | | | | | | | | | | | | | 0.03787 | 2.51442 | Female |
| Abcg2 | 0.00017 | -0.81850 | Male | 0.00359 | -1.68703 | Male | | | | | | | | | | | | |
| Crebrf | | | | | | | | | | | | | | | | 0.00833 | -2.69753 | Male |
| Srd5a1 | | | | | | | | | | | | | | | | 0.03851 | -4.44724 | Male |
| Gapvd1 | | | | | | | | | | 0.02284 | 1.96451 | Female | | | | | | |
| Fau | | | | | | | | | | 0.00685 | -2.52960 | Male | | | | | | |
| Ppme1 | | | | | | | | | | 0.02797 | 2.91149 | Female | | | | | | |
| Gm5532 | | | | | | | | | | | | | 0.01370 | 3.91090 | Female | | | |
| Ist1 | | | | | | | | | | 0.03101 | 1.40844 | Female | | | | | | |
| Gm6639 | | | | | | | | | | | | | 0.00107 | 4.00160 | Female | | | |
| Dgkq | | | | | | | 0.03409 | 3.49218 | Female | | | | | | | | | |
| Oxa1l | | | | | | | | | | 0.00445 | 2.23844 | Female | | | | | | |
| Syt2 | | | | | | | | | | | | | | | | 0.00000 | 0.78795 | Female |
| C2cd2 | | | | | | | | | | 0.03959 | 2.90565 | Female | | | | | | |
| Scn10a | | | | | | | | | | 0.00435 | 1.75406 | Female | | | | | | |
| Nr1d1 | | | | | | | | | | | | | | | | 0.00108 | -4.87950 | Male |
| Mt4 | | | | | | | 0.00000 | 1.70650 | Female | | | | | | | | | |
| Igbp1 | | | | | | | | | | 0.00032 | 2.17804 | Female | | | | | | |
| Oosp3 | | | | | | | | | | 0.00000 | -6.07396 | Male | 0.00000 | -1.71503 | Male | | | |
| Aqp8 | | | | | | | | | | 0.00000 | 1.15814 | Female | | | | | | |
| Chchd2 | | | | | | | | | | 0.00245 | -3.46851 | Male | | | | | | |
| Usp49 | | | | | | | | | | | | | 0.00004 | 4.52866 | Female | | | |
| Foxo3 | 0.00000 | 1.72744 | Female | | | | | | | | | | | | | | | |
| Lta4h | 0.00000 | 1.43437 | Female | | | | | | | | | | | | | | | |
| Fut4 | | | | 0.00160 | 2.05294 | Female | | | | | | | | | | | | |
| Mlf1 | | | | | | | | | | 0.00030 | -1.63185 | Male | | | | | | |
| Piezo1 | | | | | | | | | | 0.00013 | 5.69843 | Female | | | | | | |
| Becn1 | | | | | | | | | | 0.00087 | 2.48755 | Female | | | | | | |
| Dstyk | | | | | | | | | | 0.03007 | -4.36990 | Male | | | | | | |
| Rbp7 | | | | | | | | | | | | | 0.00000 | 2.18818 | Female | | | |
| Ppargc1b | 0.00000 | -1.06446 | Male | 0.00000 | -0.85478 | Male | 0.00410 | -5.06668 | Male | 0.00336 | 4.23538 | Female | | | | | | |
| Ecd | | | | | | | | | | | | | | | | 0.02194 | -1.49308 | Male |
| Polr2j | | | | | | | | | | 0.00000 | -2.79546 | Male | 0.00115 | 1.51065 | Female | | | |
| Gsto1 | | | | 0.04153 | -3.69096 | Male | | | | | | | | | | | | |
| Rhox5 | | | | 0.00000 | 3.88092 | Female | | | | 0.00008 | 6.67684 | Female | 0.00000 | 9.49075 | Female | 0.00069 | 5.32766 | Female |
| Ptpru | 0.02483 | 0.61282 | Female | 0.04107 | -0.59342 | Male | 0.00000 | -3.89379 | Male | | | | | | | | | |
| Gfra1 | | | | | | | | | | 0.00000 | 1.19665 | Female | | | | | | |
| Ppef1 | | | | 0.03192 | 1.29326 | Female | | | | | | | 0.00000 | 1.49385 | Female | | | |
| Cdh16 | | | | | | | | | | 0.02208 | -3.10598 | Male | | | | | | |
| Anxa7 | | | | | | | | | | 0.00579 | 1.64726 | Female | | | | | | |
| Atg10 | | | | | | | | | | | | | | | | 0.03517 | 3.51722 | Female |
| Tmed9 | | | | | | | | | | | | | 0.00687 | 2.01872 | Female | | | |
| Zxda | | | | | | | | | | 0.01123 | -2.80236 | Male | 0.04588 | 1.81303 | Female | | | |
| Bend6 | | | | | | | | | | 0.00012 | -2.00250 | Male | | | | | | |
| Rps20 | | | | | | | | | | 0.00001 | -3.32314 | Male | | | | | | |
| Eif2ak1 | | | | | | | | | | | | | | | | 0.00722 | 2.57756 | Female |
| Irf2bp2 | | | | | | | | | | 0.00001 | -2.32918 | Male | 0.00513 | 4.33741 | Female | | | |
| Mrps16 | | | | | | | | | | 0.00000 | -3.68362 | Male | | | | | | |
| Mrpl33 | | | | | | | | | | 0.00000 | -4.99440 | Male | 0.00150 | 1.66089 | Female | | | |
| Npas2 | | | | | | | | | | 0.02443 | 3.83714 | Female | | | | | | |
| Lama5 | | | | 0.02045 | 1.08620 | Female | | | | | | | | | | | | |
| Lypla1 | | | | | | | | | | | | | 0.00063 | -1.54425 | Male | | | |
| Pik3c2b | | | | | | | | | | | | | 0.01565 | 2.41987 | Female | | | |
| Csf2rb | | | | | | | 0.00923 | -3.33735 | Male | 0.00714 | -4.12416 | Male | | | | | | |
| Oas1c | | | | | | | | | | | | | | | | 0.00061 | -4.69713 | Male |
| Rpl35 | | | | | | | | | | 0.00000 | -4.42468 | Male | 0.00009 | 1.68941 | Female | | | |
| Tppp | 0.00864 | 1.75536 | Female | | | | | | | | | | | | | | | |
| H2.DMb1 | 0.00704 | 0.61217 | Female | | | | | | | | | | | | | | | |
| Thap7 | | | | | | | | | | 0.04931 | 3.02119 | Female | | | | | | |
| Serpinb9b | | | | | | | | | | 0.00000 | 7.48603 | Female | | | | | | |
| Cebpz | | | | | | | | | | 0.04886 | 1.49580 | Female | | | | | | |
| Arxes2 | | | | | | | 0.00000 | 5.42911 | Female | | | | | | | | | |
| Dnajc19 | | | | | | | | | | 0.04616 | -2.45927 | Male | | | | | | |
| Dppa2 | | | | | | | | | | | | | 0.01593 | -1.42796 | Male | | | |
| Aebp1 | | | | | | | | | | | | | 0.00282 | 5.26652 | Female | | | |
| Cdc42ep2 | | | | | | | | | | | | | | | | 0.00000 | -6.16491 | Male |
| Pigb | | | | 0.00000 | 1.32919 | Female | | | | 0.00365 | 4.60522 | Female | | | | | | |
| Ramp3 | | | | | | | | | | | | | | | | 0.00000 | -1.98852 | Male |
| Crebbp | | | | | | | | | | 0.00003 | 5.39565 | Female | | | | 0.00009 | 4.42639 | Female |
| Cldn5 | | | | | | | | | | 0.00016 | -3.14020 | Male | | | | | | |

| | | | | | | | | | | | | | | | | | | |
|---|---|---|---|---|---|---|---|---|---|---|---|---|---|---|---|---|---|---|
| Epha2 | | | | 0.00409 | -1.74487 | Male | | | | | | | | | | | | |
| Slc25a35 | 0.00039 | -1.00358 | Male | | | | | | | | | | | | | | | |
| Gtpbp6 | | | | | | | | | | 0.00148 | 5.27272 | Female | | | | | | |
| Apof | | | | | | | | | | 0.02286 | -2.43199 | Male | | | | | | |
| Syvn1 | | | | | | | | | | 0.00999 | 3.24490 | Female | | | | | | |
| Lpar6 | | | | 0.02383 | 2.18160 | Female | | | | | | | | | | 0.00000 | 2.21832 | Female |
| Ndufc1 | | | | | | | | | | 0.00000 | -4.77690 | Male | | | | | | |
| Spon1 | | | | | | | | | | | | | | | | 0.00025 | -0.66432 | Male |
| Man2a1 | | | | 0.02312 | 1.28992 | Female | | | | | | | | | | | | |
| Mfsd7a | 0.00000 | 1.79902 | Female | | | | | | | | | | | | | 0.00000 | 0.75701 | Female |
| Rpl3l | | | | | | | | | | | | | | | | 0.00000 | -5.11366 | Male |
| Add3 | | | | | | | | | | 0.00418 | 4.43034 | Female | | | | | | |
| Fam13b | | | | | | | | | | 0.01556 | 3.56285 | Female | 0.01058 | -3.19476 | Male | | | |
| Tenm4 | | | | | | | | | | | | | 0.00000 | -1.10583 | Male | 0.00001 | 2.41244 | Female |
| Kcnf1 | | | | | | | | | | | | | | | | 0.00288 | -4.64910 | Male |
| Polr2c | | | | | | | | | | 0.04132 | 1.49523 | Female | | | | | | |
| Rcsd1 | | | | 0.00051 | 2.24596 | Female | | | | | | | 0.00000 | -4.14851 | Male | | | |
| Klhl26 | | | | 0.00005 | -1.67352 | Male | | | | | | | | | | | | |
| Smc1a | | | | | | | 0.00009 | 1.88399 | Female | 0.03182 | 1.40640 | Female | | | | | | |
| Cgref1 | | | | | | | | | | | | | | | | 0.02599 | 3.19102 | Female |
| BC037704 | | | | 0.00036 | -0.95804 | Male | | | | | | | | | | 0.00511 | -5.08356 | Male |
| Csnk1g2 | | | | | | | | | | 0.02822 | 3.09926 | Female | | | | | | |
| BC147527 | | | | | | | | | | | | | 0.00817 | 4.22890 | Female | 0.03928 | -4.01978 | Male |
| Psg23 | | | | | | | | | | 0.00268 | -1.83911 | Male | 0.00391 | 1.31526 | Female | | | |
| Slx1b | | | | | | | | | | 0.00007 | -2.42070 | Male | | | | | | |
| Gm4926 | | | | | | | | | | | | | 0.01532 | 2.31778 | Female | | | |
| Pclo | | | | | | | | | | 0.00082 | -3.48015 | Male | | | | | | |
| Dsg2 | | | | | | | 0.00138 | -3.69443 | Male | 0.00082 | 4.58233 | Female | | | | | | |
| Ccnk | | | | | | | | | | 0.00274 | -2.50534 | Male | | | | | | |
| Gm4951 | | | | | | | | | | 0.00001 | -4.21917 | Male | | | | | | |
| Olfr376 | | | | | | | 0.00076 | 1.75127 | Female | | | | | | | | | |
| Gm10653 | | | | | | | | | | 0.00000 | -3.69544 | Male | | | | | | |
| Cir1 | | | | | | | | | | | | | 0.01345 | -1.48186 | Male | | | |
| Klrb1a | | | | | | | | | | | | | | | | 0.00000 | -1.96999 | Male |
| Trpc5 | | | | | | | 0.02189 | 2.98681 | Female | 0.00256 | -4.05305 | Male | | | | 0.00869 | -2.97799 | Male |
| Gm12185 | | | | | | | | | | 0.01838 | -2.13212 | Male | | | | | | |
| Drg1 | | | | | | | | | | | | | | | | 0.03082 | 0.92650 | Female |
| Ccdc125 | | | | | | | | | | 0.00226 | -3.86045 | Male | | | | | | |
| Snrpd3 | | | | | | | | | | 0.00001 | -3.25298 | Male | | | | | | |
| Ildr1 | | | | 0.00000 | 0.88255 | Female | | | | 0.00000 | -1.10746 | Male | | | | | | |
| Sac3d1 | | | | | | | | | | 0.01279 | -3.08078 | Male | | | | | | |
| BC026585 | | | | | | | | | | 0.00577 | 3.36528 | Female | | | | | | |
| Slc22a15 | | | | | | | | | | 0.00028 | -3.43668 | Male | | | | | | |
| Gm38426 | | | | | | | | | | | | | 0.04089 | 4.98432 | Female | | | |
| Ssrp1 | | | | | | | | | | 0.02443 | 1.24185 | Female | | | | | | |
| Plce1 | | | | | | | | | | 0.00038 | -1.97574 | Male | | | | 0.00000 | 0.97439 | Female |
| Sema6d | | | | | | | | | | | | | | | | 0.01683 | 3.42067 | Female |
| Pfn2 | | | | 0.00000 | 1.17581 | Female | | | | | | | | | | | | |
| Zfp780b | | | | | | | | | | 0.02471 | -2.57980 | Male | | | | | | |
| Rangap1 | | | | | | | | | | 0.00000 | 1.98203 | Female | | | | | | |
| Alg9 | | | | 0.00158 | 2.23354 | Female | | | | | | | | | | | | |
| Txnrd3 | | | | | | | 0.00000 | 2.38410 | Female | | | | | | | 0.00000 | -2.76793 | Male |
| Ubtf | | | | | | | | | | 0.00907 | 1.84734 | Female | | | | | | |
| Pemt | | | | | | | | | | | | | | | | 0.00001 | -2.52029 | Male |
| Cpn1 | | | | | | | | | | 0.03079 | 2.57477 | Female | | | | | | |
| Paip1 | | | | | | | | | | 0.04082 | -3.18019 | Male | 0.02339 | 2.66658 | Female | | | |
| Rhbdl3 | | | | | | | | | | | | | | | | 0.00953 | 4.27003 | Female |
| Gm20767 | | | | | | | | | | | | | | | | 0.00201 | -1.80713 | Male |
| Gtf2h3 | | | | | | | | | | 0.04547 | 2.18400 | Female | | | | | | |
| Ttc38 | | | | | | | | | | 0.02171 | -2.66422 | Male | | | | | | |
| Lsm7 | | | | | | | | | | 0.00000 | -4.35280 | Male | 0.02460 | 1.59792 | Female | | | |
| Grb14 | | | | 0.01450 | 1.03099 | Female | | | | | | | | | | | | |
| Crlf2 | | | | | | | | | | 0.00239 | -5.75280 | Male | | | | | | |
| Becn2 | | | | | | | | | | | | | | | | 0.00042 | -2.38116 | Male |
| Akr1c21 | | | | 0.00000 | 1.89946 | Female | | | | | | | | | | | | |
| Stat6 | | | | 0.02801 | 1.00477 | Female | | | | | | | | | | | | |
| Zfp369 | | | | | | | | | | 0.03697 | 3.18255 | Female | | | | | | |
| Hdac3 | | | | | | | | | | 0.00006 | 5.49180 | Female | | | | | | |
| Atf7 | | | | | | | | | | | | | | | | 0.01338 | 3.18792 | Female |
| Ddr2 | | | | | | | | | | | | | | | | 0.01859 | 3.12543 | Female |

| | | | | | | | | | | | | | | | | | | |
|---|---|---|---|---|---|---|---|---|---|---|---|---|---|---|---|---|---|---|
| Rps7 | | | | | | | | | | 0.01270 | -1.48589 | Male | | | | | | |
| Abhd11os | | | | | | | | | | 0.01450 | -3.70864 | Male | | | | | | |
| Map7d2 | | | | | | | | | | | | | | | | 0.04915 | -2.66123 | Male |
| Snw1 | | | | | | | 0.00690 | 2.48745 | Female | | | | | | | | | |
| Nfatc4 | | | | 0.00067 | -2.12179 | Male | | | | | | | | | | | | |
| Ndst2 | 0.00648 | 0.60452 | Female | | | | | | | | | | | | | | | |
| Trpa1 | 0.03782 | -1.20857 | Male | | | | | | | | | | | | | | | |
| Frrs1l | | | | 0.00000 | -1.54698 | Male | 0.00000 | 4.15232 | Female | | | | | | | | | |
| Aamp | | | | | | | | | | 0.00066 | 2.12928 | Female | | | | | | |
| Setdb1 | | | | | | | | | | | | | | | | 0.00657 | -1.82374 | Male |
| Dip2a | | | | 0.00000 | 0.70064 | Female | | | | 0.00000 | -3.46933 | Male | | | | | | |
| Qars | | | | | | | | | | 0.01170 | 1.22773 | Female | | | | | | |
| Krt27 | | | | | | | | | | | | | | | | 0.00000 | -5.19515 | Male |
| Pcbp4 | | | | | | | | | | | | | 0.01696 | 4.29701 | Female | | | |
| Pelp1 | | | | | | | | | | 0.00435 | -2.77741 | Male | | | | | | |
| Trim54 | | | | 0.00020 | -0.87459 | Male | | | | | | | | | | | | |
| Fam83a | | | | | | | | | | 0.02417 | -2.98575 | Male | | | | | | |
| Pgam5 | | | | | | | | | | 0.03024 | 1.82241 | Female | | | | | | |
| Zfp558 | | | | | | | | | | | | | | | | 0.03584 | -3.59100 | Male |
| Tpgs2 | | | | | | | | | | 0.00541 | -3.95932 | Male | | | | | | |
| Mien1 | 0.00183 | 4.23268 | Female | | | | | | | | | | | | | | | |
| Atp5j | | | | | | | | | | 0.00000 | -4.45589 | Male | 0.00003 | 1.87595 | Female | | | |
| Rrn3 | | | | | | | | | | | | | | | | 0.00624 | 1.21864 | Female |
| Hprt | | | | | | | 0.00758 | 1.04152 | Female | | | | | | | 0.00867 | -0.95857 | Male |
| Pdcd5 | | | | | | | | | | 0.00000 | -4.83402 | Male | 0.04163 | 1.68206 | Female | | | |
| Tcerg1l | | | | | | | 0.00000 | -4.40469 | Male | 0.04469 | -0.71447 | Male | | | | | | |
| Arsk | | | | | | | | | | 0.00000 | -5.16665 | Male | | | | | | |
| Hoxb13 | 0.00000 | 0.80072 | Female | | | | 0.00000 | -2.75247 | Male | 0.00006 | -2.20853 | Male | | | | | | |
| Nr1h4 | | | | | | | | | | | | | | | | 0.00726 | 1.72170 | Female |
| Exd2 | | | | | | | | | | | | | | | | 0.00939 | 2.68469 | Female |
| Vamp8 | | | | | | | | | | 0.02724 | -1.72446 | Male | | | | 0.04915 | -1.34550 | Male |
| Maged1 | | | | | | | | | | 0.00025 | 3.92877 | Female | | | | | | |
| Plekhh2 | | | | | | | | | | | | | | | | 0.00319 | 4.02414 | Female |
| Sugct | | | | | | | 0.00310 | -0.81189 | Male | | | | | | | 0.00000 | 1.29983 | Female |
| Cfap45 | | | | | | | 0.00000 | -1.55985 | Male | 0.00002 | -2.45608 | Male | | | | | | |
| Erf | | | | | | | | | | 0.00149 | -4.08838 | Male | | | | | | |
| Snora41 | | | | | | | | | | | | | | | | 0.02218 | -3.50912 | Male |
| Cxcr2 | | | | | | | | | | 0.00000 | -5.89698 | Male | 0.00047 | 1.80264 | Female | | | |
| Aptx | | | | | | | | | | | | | | | | 0.00495 | -2.11171 | Male |
| Luc7l2 | | | | | | | | | | 0.01320 | -1.85144 | Male | | | | | | |
| Vsig10 | | | | | | | | | | | | | 0.00210 | 3.47654 | Female | | | |
| Mmgt2 | | | | | | | | | | | | | | | | 0.01375 | -3.81120 | Male |
| Cited4 | | | | | | | | | | 0.00000 | -2.87001 | Male | | | | | | |
| Lars | | | | | | | | | | | | | | | | 0.00879 | 1.65669 | Female |
| Meg3 | | | | | | | | | | | | | | | | 0.03944 | 3.89913 | Female |
| Shb | | | | | | | | | | | | | | | | 0.00158 | -5.23087 | Male |
| Bloc1s3 | | | | | | | | | | 0.00098 | -2.57108 | Male | | | | | | |
| Ccnb1ip1 | | | | | | | | | | | | | | | | 0.00128 | -2.65474 | Male |
| Gm715 | | | | | | | | | | 0.03219 | -1.67585 | Male | | | | | | |
| Anxa8 | | | | 0.00001 | 1.02642 | Female | | | | | | | | | | 0.00500 | -5.47841 | Male |
| Nudt4 | | | | | | | | | | 0.00469 | -1.36078 | Male | | | | | | |
| Rcn1 | | | | | | | | | | 0.00000 | -0.68456 | Male | | | | | | |
| Lgals8 | | | | | | | | | | | | | | | | 0.04753 | -1.35893 | Male |
| Cdkl3 | | | | | | | | | | | | | 0.01462 | -4.03064 | Male | | | |
| Sap130 | | | | | | | | | | | | | | | | 0.00061 | -2.51142 | Male |
| Cggbp1 | | | | | | | 0.00660 | 1.57221 | Female | | | | | | | | | |
| Sbds | | | | | | | | | | 0.01956 | 1.73401 | Female | | | | | | |
| Ephx1 | | | | | | | | | | 0.00180 | 4.80061 | Female | | | | | | |
| Ndufa4 | | | | | | | | | | 0.00002 | -3.04015 | Male | 0.00016 | 1.96678 | Female | | | |
| Ndufs4 | | | | | | | | | | 0.00406 | -2.84793 | Male | | | | | | |
| Prkaca | | | | | | | | | | 0.02669 | 2.99614 | Female | | | | | | |
| Dnajc5g | | | | | | | | | | | | | | | | 0.00000 | 4.41147 | Female |
| Fscn1 | | | | | | | | | | 0.00031 | -4.40478 | Male | | | | | | |
| Snord99 | | | | | | | | | | | | | | | | 0.00008 | 5.16779 | Female |
| Pank1 | | | | | | | 0.00536 | 4.21528 | Female | 0.00006 | -5.93855 | Male | | | | | | |
| Tmem8b | | | | | | | | | | | | | 0.00049 | 4.35212 | Female | 0.00493 | 4.27606 | Female |
| Ccdc166 | | | | | | | | | | | | | | | | 0.01602 | 3.57115 | Female |
| Trappc9 | | | | | | | | | | | | | | | | 0.00467 | -1.91552 | Male |
| Cd300ld | | | | | | | | | | | | | 0.00000 | 3.92333 | Female | 0.00000 | 2.20261 | Female |
| Suclg1 | | | | | | | | | | 0.03181 | 1.63126 | Female | | | | | | |

| | | | | | | | | | | | | | | | | | | |
|---|---|---|---|---|---|---|---|---|---|---|---|---|---|---|---|---|---|---|
| Pramef6 | | | | | | | | | | 0.00000 | 5.40959 | Female | | | | 0.00392 | -2.12604 | Male |
| Ccar1 | | | | | | | | | | | | | | | | 0.01328 | -1.45481 | Male |
| Taf1c | | | | | | | | | | 0.00004 | 4.63672 | Female | | | | | | |
| Pygb | | | | 0.00708 | 2.26033 | Female | | | | | | | | | | | | |
| Spta1 | | | | | | | 0.00205 | -3.42268 | Male | | | | | | | | | |
| Cand2 | | | | | | | | | | 0.01891 | -3.09655 | Male | | | | | | |
| Zfp36l1 | | | | 0.00000 | -1.44469 | Male | | | | 0.00000 | -0.98902 | Male | | | | | | |
| Tnrc6b | | | | | | | | | | | | | 0.00246 | 3.31116 | Female | | | |
| Mageb16 | | | | | | | 0.02682 | 2.14330 | Female | | | | | | | | | |
| Sugp2 | | | | | | | | | | | | | 0.01524 | 3.54623 | Female | 0.00450 | -4.36846 | Male |
| Gm6787 | | | | | | | | | | | | | | | | 0.02167 | -4.40426 | Male |
| Osr2 | | | | 0.00075 | -1.75695 | Male | | | | | | | | | | | | |
| Kank2 | | | | | | | | | | 0.03913 | -2.79648 | Male | | | | | | |
| Nptn | | | | | | | | | | | | | | | | 0.01255 | 3.03769 | Female |
| Gm1995 | | | | | | | | | | | | | 0.00146 | 2.97419 | Female | | | |
| Drg2 | | | | | | | | | | 0.00779 | 1.59836 | Female | | | | | | |
| Egflam | 0.01459 | -1.58323 | Male | | | | | | | | | | | | | | | |
| Dlgap4 | | | | | | | | | | 0.03336 | 2.76088 | Female | | | | | | |
| Yars | | | | | | | | | | 0.04371 | 1.33569 | Female | | | | | | |
| Tmcc3 | | | | | | | | | | | | | 0.03361 | -3.39911 | Male | | | |
| Col1a1 | | | | 0.00204 | 1.23360 | Female | | | | | | | | | | | | |
| Dynlt1a | | | | | | | | | | 0.01578 | -1.44092 | Male | | | | | | |
| Acsl4 | | | | | | | | | | | | | | | | 0.00000 | 4.18666 | Female |
| Ddx47 | | | | | | | | | | 0.01187 | 1.81961 | Female | | | | | | |
| Ikbkb | | | | 0.00000 | -0.81737 | Male | | | | | | | | | | | | |
| Pde4d | | | | | | | | | | 0.00000 | -3.38018 | Male | 0.00000 | -2.16944 | Male | | | |
| Plekhg1 | | | | | | | 0.03241 | -4.98447 | Male | | | | | | | | | |
| Gm14461 | | | | | | | | | | 0.00959 | -1.66893 | Male | | | | | | |
| Cept1 | | | | 0.00038 | 3.65240 | Female | 0.00055 | 3.56324 | Female | | | | | | | | | |
| Ppil3 | 0.01009 | 4.02737 | Female | | | | | | | | | | | | | | | |
| Suox | | | | | | | | | | 0.03049 | 3.47198 | Female | | | | | | |
| Lca5 | | | | 0.00629 | 0.68271 | Female | | | | | | | | | | | | |
| Nup98 | | | | | | | 0.00977 | 1.82830 | Female | 0.00097 | -2.48206 | Male | | | | | | |
| Atp2b1 | | | | | | | | | | 0.00057 | 3.81175 | Female | | | | | | |
| Fam168b | | | | | | | | | | 0.00932 | -2.01187 | Male | | | | | | |
| Sema3e | | | | | | | | | | 0.00000 | -4.32210 | Male | | | | | | |
| Tnpo2 | | | | | | | 0.00859 | 2.65683 | Female | 0.00223 | -3.25722 | Male | | | | | | |
| Gstm4 | 0.00002 | 1.28398 | Female | | | | 0.00000 | 2.46600 | Female | | | | | | | | | |
| Nkiras2 | | | | 0.00157 | 3.22173 | Female | | | | 0.00193 | 4.23435 | Female | | | | | | |
| Tmem241 | | | | | | | | | | 0.00001 | 1.45185 | Female | | | | | | |
| Pet2 | | | | | | | | | | | | | | | | 0.00000 | 1.01367 | Female |
| Lrrc8e | | | | | | | | | | | | | 0.03894 | 2.29135 | Female | | | |
| Gje1 | | | | | | | | | | 0.00000 | -4.15108 | Male | 0.00233 | 1.97293 | Female | | | |
| Glis3 | | | | | | | | | | 0.00000 | -3.22576 | Male | | | | 0.02740 | 4.05061 | Female |
| Ctrc | | | | 0.00393 | 1.18719 | Female | | | | | | | | | | | | |
| Pdcd11 | | | | | | | | | | 0.00003 | 4.01528 | Female | | | | | | |
| Zfp626 | | | | | | | | | | 0.00374 | -2.73787 | Male | | | | | | |
| Gpcpd1 | | | | | | | | | | | | | 0.02016 | -1.47116 | Male | | | |
| Tpt1 | | | | | | | | | | 0.00000 | -3.82438 | Male | | | | | | |
| Slc28a1 | | | | | | | | | | 0.00858 | -0.66626 | Male | | | | 0.00000 | 2.98817 | Female |
| Rps18 | | | | | | | | | | 0.00000 | -3.04460 | Male | 0.01374 | 1.37220 | Female | | | |
| Hnrnpl | | | | | | | | | | 0.01272 | -3.11222 | Male | | | | | | |
| Gm16381 | | | | | | | 0.03060 | 3.97280 | Female | | | | | | | | | |
| Alms1 | | | | | | | | | | 0.03395 | 3.27240 | Female | | | | | | |
| Adamts2 | | | | | | | | | | 0.00516 | -1.70951 | Male | | | | | | |
| Rhobtb1 | | | | | | | | | | | | | | | | 0.00000 | -1.38595 | Male |
| Sdcbp2 | | | | | | | | | | | | | | | | 0.00013 | 5.84344 | Female |
| Cnih4 | | | | | | | | | | 0.00000 | -2.97270 | Male | | | | | | |
| Slc35e2 | | | | | | | | | | 0.00002 | -3.09854 | Male | | | | | | |
| Gm15401 | | | | | | | | | | 0.00000 | -4.34968 | Male | | | | | | |
| Pde6g | | | | | | | | | | | | | | | | 0.04266 | 4.18954 | Female |
| Kcnip2 | | | | 0.00488 | 1.68210 | Female | | | | | | | | | | | | |
| Ccdc69 | | | | | | | | | | 0.01349 | -4.67735 | Male | | | | | | |
| Fgd1 | | | | | | | | | | 0.00000 | 2.79776 | Female | | | | | | |
| Sub1 | | | | | | | | | | 0.00000 | -3.07392 | Male | | | | | | |
| Hdlbp | | | | | | | | | | 0.00987 | 1.45183 | Female | | | | | | |
| Efcc1 | | | | | | | 0.02250 | 2.92162 | Female | 0.00000 | -6.14459 | Male | 0.04445 | 2.28480 | Female | | | |
| Trim63 | | | | | | | 0.00001 | 1.02027 | Female | | | | | | | | | |
| Ndufa2 | | | | | | | | | | 0.00000 | -4.33894 | Male | | | | | | |
| Spats2 | | | | | | | | | | 0.00009 | 5.17020 | Female | | | | | | |

| | | | | | | | | | | | | | | | | | | |
|---|---|---|---|---|---|---|---|---|---|---|---|---|---|---|---|---|---|---|
| AF067063 | | | | | | | | | | | | | | | | 0.00000 | -6.62289 | Male |
| Lilrb4a | | | | | | | 0.00030 | -3.76468 | Male | | | | | | | | | |
| Car2 | | | | | | | | | | | | | 0.00017 | -0.62322 | Male | 0.00000 | 2.73369 | Female |
| Spib | | | | | | | | | | 0.03679 | -3.63404 | Male | | | | | | |
| Rnf168 | | | | | | | | | | 0.00001 | -3.13258 | Male | | | | | | |
| Rpl21 | | | | | | | | | | 0.00000 | -3.25276 | Male | 0.01508 | 1.15197 | Female | | | |
| Grin3b | 0.03154 | -0.80980 | Male | 0.00004 | 0.73906 | Female | | | | | | | | | | | | |
| Gjb4 | | | | 0.00001 | -0.71737 | Male | | | | | | | | | | 0.00976 | 5.43551 | Female |
| Fastkd1 | | | | | | | 0.02336 | 3.25792 | Female | | | | | | | | | |
| Xrcc3 | | | | 0.01744 | 1.39852 | Female | | | | | | | | | | | | |
| Tex13 | 0.03830 | 2.88427 | Female | | | | | | | | | | | | | | | |
| Gsdma3 | | | | | | | | | | | | | | | | 0.00627 | -4.73217 | Male |
| Glrx | | | | | | | | | | 0.00221 | 3.86654 | Female | | | | | | |
| Mrps18a | 0.00010 | 5.24422 | Female | | | | | | | | | | | | | | | |
| Cct6a | | | | | | | | | | 0.00196 | 1.03032 | Female | | | | | | |
| Morf4l1 | | | | | | | | | | 0.02858 | 1.36930 | Female | | | | | | |
| Xlr3c | | | | | | | 0.00776 | 3.06706 | Female | | | | | | | | | |
| Gm13985 | | | | | | | | | | 0.00000 | -3.45842 | Male | | | | | | |
| Snrpd2 | | | | | | | | | | 0.00000 | -3.80067 | Male | 0.04168 | 1.34645 | Female | | | |
| Slc25a18 | 0.00003 | -2.05964 | Male | | | | | | | | | | 0.00000 | -1.36917 | Male | 0.00000 | -5.49529 | Male |
| Slfn5 | | | | | | | | | | 0.00000 | -5.96726 | Male | 0.00243 | 1.77773 | Female | | | |
| Smox | | | | 0.00003 | -1.00315 | Male | | | | | | | | | | | | |
| Entpd8 | | | | | | | 0.00058 | -3.76872 | Male | | | | | | | | | |
| Cyct | | | | | | | | | | 0.02640 | -4.67464 | Male | | | | | | |
| Aatf | | | | | | | | | | 0.00009 | 3.03963 | Female | | | | | | |
| Gm5069 | | | | | | | | | | 0.03057 | 1.77885 | Female | | | | | | |
| Meis1 | | | | | | | | | | | | | | | | 0.04462 | 4.30125 | Female |
| Rps23 | | | | | | | | | | 0.00000 | -3.76844 | Male | 0.00111 | 1.51454 | Female | | | |
| Gpr137b.ps | | | | | | | | | | | | | | | | 0.00001 | -1.93522 | Male |
| Slc22a13 | | | | | | | | | | 0.00004 | 5.02624 | Female | | | | | | |
| Rgl1 | | | | 0.00007 | 0.85339 | Female | | | | | | | | | | | | |
| Pramel4 | | | | | | | | | | | | | | | | 0.02730 | -1.25203 | Male |
| Dusp23 | | | | | | | 0.00000 | 3.30782 | Female | | | | 0.00000 | 2.75765 | Female | | | |
| Cap1 | | | | 0.00168 | 3.98405 | Female | | | | | | | | | | | | |
| Cct7 | | | | | | | | | | 0.00045 | 1.78712 | Female | | | | | | |
| Gm5803 | | | | | | | | | | 0.00168 | 2.83809 | Female | | | | | | |
| Lcn10 | | | | | | | | | | 0.00000 | 2.27022 | Female | | | | | | |
| Ppm1m | | | | | | | | | | 0.00000 | 2.52568 | Female | | | | | | |
| Pdia4 | | | | | | | | | | 0.00100 | 4.15872 | Female | | | | | | |
| Gm595 | | | | | | | | | | | | | 0.00000 | -2.77472 | Male | | | |
| Slain1 | | | | | | | | | | | | | 0.00000 | -6.20980 | Male | | | |
| Pld6 | | | | | | | | | | 0.03171 | -4.31244 | Male | | | | 0.00000 | -2.07344 | Male |
| Psmb10 | | | | | | | | | | 0.01716 | -2.92613 | Male | | | | | | |
| Calm3 | | | | | | | | | | 0.04126 | -2.17355 | Male | | | | | | |
| Alms1.ps2 | | | | | | | | | | 0.03124 | 3.15169 | Female | | | | | | |
| Wfdc3 | | | | 0.00019 | 2.06799 | Female | | | | | | | 0.01012 | -5.17504 | Male | | | |
| Akr1b10 | | | | 0.03438 | 3.51429 | Female | 0.00055 | -4.47978 | Male | 0.01819 | 3.87859 | Female | | | | | | |
| Rnf223 | | | | | | | | | | | | | | | | 0.01632 | -4.85776 | Male |
| Dync1i1 | 0.03296 | -0.89756 | Male | | | | | | | 0.01499 | -0.78828 | Male | | | | | | |
| Dmgdh | | | | | | | | | | | | | | | | 0.00991 | -1.68089 | Male |
| Krba1 | | | | | | | 0.04109 | 3.46580 | Female | | | | | | | | | |
| Mael | | | | | | | | | | | | | | | | 0.00000 | 1.14887 | Female |
| Sh3pxd2a | | | | | | | | | | | | | | | | 0.00320 | 4.18602 | Female |
| AI427809 | | | | | | | 0.00000 | -0.76242 | Male | | | | | | | | | |
| Adamtsl4 | | | | | | | | | | | | | | | | 0.02904 | 3.43995 | Female |
| Nolc1 | | | | | | | | | | 0.00027 | 1.69937 | Female | | | | | | |
| Gm20187 | | | | | | | | | | | | | | | | 0.01627 | 3.96498 | Female |
| Gm4285 | | | | | | | 0.00931 | 3.01540 | Female | | | | | | | | | |
| H2.D1 | | | | | | | | | | | | | | | | 0.02241 | 2.21061 | Female |
| Gm3286 | | | | | | | | | | | | | | | | 0.00000 | -2.45340 | Male |
| Bnip2 | | | | | | | | | | | | | | | | 0.00576 | -3.60210 | Male |
| Oxld1 | | | | | | | | | | | | | | | | 0.02122 | 4.06204 | Female |
| Gm2381 | | | | | | | 0.04247 | 1.91750 | Female | | | | | | | | | |
| Nobox | | | | | | | | | | | | | 0.00000 | -3.82514 | Male | | | |
| Slc38a2 | | | | | | | | | | | | | | | | 0.00044 | 5.62594 | Female |
| Ssb | | | | | | | | | | 0.00100 | 2.03541 | Female | | | | | | |
| Eif5a2 | | | | | | | | | | 0.00000 | -4.36485 | Male | | | | | | |
| Lysmd2 | | | | 0.00065 | -1.65282 | Male | | | | | | | | | | | | |
| Agl | | | | | | | | | | 0.00667 | -2.61247 | Male | | | | | | |
| Gm12253 | | | | | | | | | | 0.00205 | -3.80249 | Male | | | | | | |

| | | | | | | | | | | | | | | | | | | |
|---|---|---|---|---|---|---|---|---|---|---|---|---|---|---|---|---|---|---|
| Prox2 | | | | | | | | | | 0.00011 | -5.39193 | Male | | | | | | |
| Zfp979 | | | | 0.02602 | 3.01449 | Female | | | | | | | | | | | | |
| Rps27l | | | | | | | | | | 0.00000 | -4.53986 | Male | 0.00135 | 1.64140 | Female | | | |
| Tma7 | | | | | | | | | | 0.00000 | -3.48247 | Male | 0.00074 | 1.34941 | Female | | | |
| Emc3 | | | | | | | | | | | | | | | | 0.02614 | -1.06430 | Male |
| Abca3 | | | | | | | | | | 0.00180 | 4.13020 | Female | | | | | | |
| Slc11a1 | | | | | | | | | | | | | 0.00232 | 2.80825 | Female | | | |
| Nifk | | | | | | | | | | 0.02199 | 1.13750 | Female | | | | | | |
| B3gnt8 | | | | 0.00000 | -1.93018 | Male | | | | | | | | | | | | |
| Tpd52l1 | | | | 0.00004 | 1.53937 | Female | | | | | | | | | | | | |
| Pilra | | | | | | | | | | 0.00000 | -4.82687 | Male | 0.00641 | 2.45493 | Female | | | |
| Rpl23a | | | | | | | | | | 0.00000 | -3.15994 | Male | 0.03135 | 1.11879 | Female | | | |
| Lyrm1 | | | | | | | | | | | | | | | | 0.01484 | -3.07913 | Male |
| Lca5l | | | | | | | | | | | | | | | | 0.00008 | 0.79494 | Female |
| Rpa2 | | | | | | | | | | 0.00318 | 1.77353 | Female | | | | | | |
| Erap1 | | | | | | | | | | | | | 0.00000 | 3.29262 | Female | | | |
| Ngfrap1 | | | | | | | | | | 0.00000 | 3.67500 | Female | | | | 0.00236 | 5.14965 | Female |
| Trim23 | | | | | | | | | | 0.00295 | 4.50757 | Female | | | | | | |
| Samd10 | | | | | | | 0.00000 | -2.01505 | Male | | | | 0.00000 | -4.16175 | Male | | | |
| Atp1a2 | | | | | | | | | | 0.01704 | -3.65428 | Male | | | | | | |
| Traf5 | | | | | | | | | | 0.00147 | 3.19967 | Female | | | | | | |
| Majin | 0.00000 | 0.81113 | Female | | | | | | | | | | | | | | | |
| Gm14124 | | | | | | | | | | | | | | | | 0.00108 | -3.03951 | Male |
| Letm2 | | | | | | | | | | | | | | | | 0.00000 | -1.59762 | Male |
| Tpp1 | | | | | | | | | | 0.03400 | 2.82323 | Female | | | | | | |
| Acsm3 | | | | | | | | | | 0.04256 | 2.89184 | Female | | | | | | |
| Mrpl18 | | | | | | | | | | 0.01835 | 1.32276 | Female | | | | | | |
| Zhx3 | | | | | | | | | | | | | 0.00004 | 5.30071 | Female | | | |
| Ufsp1 | | | | 0.00021 | -1.35551 | Male | | | | | | | | | | | | |
| Ipo13 | | | | | | | | | | 0.03336 | 2.73876 | Female | | | | | | |
| Prkar2a | | | | | | | | | | 0.02908 | 2.28472 | Female | | | | | | |
| Mcm5 | | | | | | | | | | 0.01515 | 1.58608 | Female | | | | | | |
| Tbc1d16 | 0.00000 | -1.12391 | Male | | | | | | | 0.00000 | -3.51596 | Male | | | | | | |
| Fgf1 | | | | | | | | | | | | | | | | 0.00000 | -0.90159 | Male |
| Cecr2 | | | | | | | | | | | | | 0.04089 | -1.06656 | Male | | | |
| Fancb | | | | 0.04674 | 2.20331 | Female | | | | | | | | | | 0.00010 | 5.18589 | Female |
| Maged2 | | | | | | | | | | 0.00000 | 3.24126 | Female | 0.01721 | -4.17004 | Male | | | |
| Cep162 | | | | | | | | | | | | | | | | 0.00283 | 4.65000 | Female |
| Usp20 | | | | | | | | | | | | | | | | 0.03669 | -3.51135 | Male |
| Csrp1 | | | | 0.00379 | 2.76581 | Female | | | | | | | | | | | | |
| Fam49a | | | | | | | | | | 0.00004 | -2.27914 | Male | 0.00000 | 4.65364 | Female | 0.00000 | -1.50144 | Male |
| Tbl1x | | | | | | | 0.04071 | 2.27074 | Female | | | | | | | | | |
| Kif20a | | | | | | | | | | 0.00000 | 6.22460 | Female | | | | | | |
| Glg1 | | | | | | | | | | 0.00000 | 6.59347 | Female | | | | | | |
| Abcd3 | 0.01299 | 4.37850 | Female | 0.00071 | 3.74975 | Female | | | | | | | | | | | | |
| Clec2f | 0.00027 | -0.82119 | Male | 0.02226 | -1.58701 | Male | | | | | | | | | | | | |
| Ppm1k | | | | | | | | | | 0.00000 | -4.65320 | Male | | | | | | |
| Rab11a | | | | | | | | | | 0.00052 | -2.41626 | Male | | | | | | |
| Tcf19 | | | | | | | | | | 0.01562 | 3.25453 | Female | | | | | | |
| Dguok | | | | 0.00686 | 1.69184 | Female | | | | | | | | | | | | |
| Gm6086 | | | | | | | | | | | | | 0.03361 | 2.63050 | Female | | | |
| Spp1 | 0.00000 | -2.25428 | Male | | | | | | | 0.00000 | 1.52746 | Female | | | | | | |
| Zfp575 | | | | | | | | | | 0.01538 | -1.82720 | Male | | | | | | |
| Asz1 | | | | 0.02045 | 3.35625 | Female | | | | | | | | | | | | |
| Hnrnpul1 | | | | | | | | | | 0.00001 | 5.73036 | Female | | | | | | |
| Slc15a1 | | | | | | | | | | 0.04082 | -2.17573 | Male | | | | | | |
| Trim61 | | | | | | | | | | 0.00917 | -4.45364 | Male | 0.00005 | 5.30588 | Female | | | |
| Timd2 | | | | | | | | | | | | | | | | 0.00942 | -1.62738 | Male |
| Mmp2 | | | | | | | | | | 0.01300 | -0.76343 | Male | 0.00000 | 1.70071 | Female | | | |
| Nuak1 | | | | 0.00005 | -1.31242 | Male | | | | 0.00000 | 2.74564 | Female | | | | 0.03170 | 4.33680 | Female |
| Gm10324 | | | | | | | 0.00923 | 1.83467 | Female | | | | | | | | | |
| Gnpda1 | | | | | | | | | | | | | 0.01508 | -1.38985 | Male | | | |
| Ifitm6 | | | | 0.00316 | 1.70528 | Female | | | | 0.01673 | -1.11959 | Male | | | | | | |
| Card6 | | | | | | | | | | | | | | | | 0.03851 | 2.94473 | Female |
| Zfp454 | 0.00003 | 0.74002 | Female | | | | | | | | | | | | | 0.00000 | -0.59083 | Male |
| Gm4925 | | | | | | | | | | 0.02224 | -2.43293 | Male | | | | | | |
| Klhdc1 | | | | | | | | | | 0.00000 | 2.42782 | Female | | | | 0.00898 | -4.95814 | Male |
| Nxf7 | | | | | | | | | | | | | 0.00000 | 2.69711 | Female | | | |
| Psmb2 | | | | | | | | | | 0.00000 | 2.51841 | Female | | | | | | |
| Mrps18c | | | | | | | | | | 0.00000 | -4.94894 | Male | | | | | | |

| | | | | | | | | | | | | | | | | | | |
|---|---|---|---|---|---|---|---|---|---|---|---|---|---|---|---|---|---|---|
| Xcr1 | | | | | | | | | | | | | 0.00624 | 2.90197 | Female | | | |
| Rps27a | | | | | | | | | | 0.00000 | -3.06589 | Male | | | | | | |
| Eif3b | | | | | | | | | | 0.01412 | 1.88446 | Female | | | | | | |
| Fez1 | | | | | | | | | | | | | 0.00000 | 6.08447 | Female | | | |
| Ccdc173 | | | | | | | | | | | | | | | | 0.00002 | -5.77860 | Male |
| Dhx32 | | | | | | | | | | 0.00747 | 2.06286 | Female | | | | | | |
| Brms1l | | | | | | | | | | | | | | | | 0.01362 | -2.20357 | Male |
| Traf4 | | | | | | | | | | 0.03852 | 2.22521 | Female | | | | | | |
| Exd1 | | | | | | | | | | | | | | | | 0.00052 | -3.74828 | Male |
| Tktl1 | | | | | | | | | | | | | | | | 0.00010 | 3.59931 | Female |
| Dyx1c1 | 0.00373 | -0.97927 | Male | | | | 0.00000 | 4.38638 | Female | | | | | | | 0.00000 | -1.91906 | Male |
| Atp5g1 | | | | | | | | | | 0.00115 | -2.27051 | Male | 0.02626 | 1.28185 | Female | | | |
| Cela2a | | | | 0.00837 | -1.85800 | Male | | | | 0.00000 | 2.72644 | Female | | | | | | |
| Cul7 | | | | | | | | | | 0.00311 | 3.75284 | Female | | | | | | |
| Srrm1 | | | | | | | | | | 0.01141 | 1.39936 | Female | | | | | | |
| Sat1 | | | | | | | 0.00075 | 2.23345 | Female | 0.01679 | 2.00360 | Female | | | | | | |
| Tcstv1 | | | | | | | | | | | | | | | | 0.00052 | -5.32606 | Male |
| Ankrd44 | | | | | | | | | | | | | | | | 0.00000 | -5.90767 | Male |
| Gjb5 | | | | | | | | | | 0.00001 | 6.03737 | Female | | | | | | |
| Snord16a | | | | | | | | | | 0.00481 | 3.36172 | Female | | | | | | |
| Arpc3 | | | | | | | | | | | | | 0.02291 | -1.67526 | Male | | | |
| Reln | | | | | | | | | | | | | 0.00621 | -4.84052 | Male | | | |
| Bex4 | | | | | | | | | | 0.04637 | 4.47237 | Female | 0.02010 | 4.42274 | Female | | | |
| Gmeb1 | | | | | | | | | | 0.00445 | -2.14209 | Male | | | | | | |
| Dennd6a | | | | | | | | | | 0.04087 | -2.07263 | Male | | | | | | |
| Atox1 | | | | | | | | | | 0.00000 | -4.69788 | Male | 0.00006 | 2.15364 | Female | | | |
| Cnbd2 | | | | | | | | | | | | | | | | 0.02239 | -1.36025 | Male |
| Arfgef3 | | | | | | | 0.00005 | -2.16362 | Male | | | | | | | | | |
| Gm9776 | | | | | | | 0.01253 | 2.02120 | Female | 0.00000 | -5.51205 | Male | 0.00128 | 1.99915 | Female | | | |
| Rhou | | | | | | | | | | | | | 0.02251 | 3.55660 | Female | | | |
| Usp44 | | | | | | | | | | | | | | | | 0.00002 | 1.88065 | Female |
| Tmsb10 | | | | | | | | | | 0.00005 | -4.06328 | Male | | | | | | |
| Ctdp1 | | | | | | | | | | 0.00897 | 3.28898 | Female | | | | | | |
| Adk | | | | | | | | | | 0.02978 | 2.77251 | Female | | | | | | |
| Ddx24 | | | | | | | | | | 0.02985 | 1.38881 | Female | | | | | | |
| Icam2 | | | | | | | | | | 0.00164 | -1.42854 | Male | | | | | | |
| Snx5 | | | | | | | | | | | | | 0.01628 | -0.94129 | Male | | | |
| Dcbld1 | | | | | | | | | | | | | | | | 0.03311 | -1.80527 | Male |
| Ccdc43 | | | | 0.00006 | 4.22222 | Female | 0.02595 | 1.96501 | Female | | | | | | | | | |
| Nt5dc3 | | | | | | | | | | 0.00000 | -5.27847 | Male | | | | | | |
| Camkmt | | | | 0.00107 | -2.00837 | Male | | | | | | | | | | | | |
| Sft2d2 | 0.00003 | 0.74962 | Female | 0.00000 | -1.63808 | Male | 0.02243 | 3.76835 | Female | 0.00001 | -1.32607 | Male | | | | | | |
| Apom | | | | | | | | | | | | | | | | 0.00076 | 4.40154 | Female |
| Fam210b | | | | 0.00000 | 1.46729 | Female | | | | 0.00000 | 1.50598 | Female | | | | | | |
| Sod1 | | | | | | | | | | | | | 0.01443 | 1.42666 | Female | | | |
| Israa | | | | | | | | | | 0.00041 | -1.94928 | Male | | | | | | |
| Hcn3 | | | | | | | | | | | | | 0.00001 | -5.60213 | Male | 0.00009 | -5.64545 | Male |
| Rhoh | | | | | | | | | | 0.01354 | -3.49833 | Male | | | | | | |
| Gcn1l1 | | | | | | | | | | 0.00460 | 3.62354 | Female | | | | | | |
| Ints3 | | | | | | | | | | 0.01608 | 1.84966 | Female | | | | | | |
| Nlrp1a | | | | | | | | | | 0.02251 | -1.44681 | Male | | | | | | |
| Pisd.ps2 | | | | | | | | | | 0.00341 | 3.47092 | Female | | | | | | |
| Rmi2 | | | | | | | | | | 0.00000 | -5.05188 | Male | 0.00170 | 1.46204 | Female | | | |
| Pet100 | | | | | | | | | | 0.00000 | -5.90432 | Male | | | | | | |
| Abhd11 | | | | | | | | | | 0.01470 | 2.47652 | Female | | | | | | |
| Gm15800 | | | | | | | | | | 0.04276 | 2.46456 | Female | | | | | | |
| Ccdc39 | | | | | | | | | | 0.01062 | 1.03579 | Female | 0.00000 | -2.10066 | Male | 0.00000 | -1.37862 | Male |
| Setd7 | 0.00001 | 2.25463 | Female | 0.00050 | -0.76672 | Male | | | | | | | | | | | | |
| Pyy | | | | 0.00002 | -1.05680 | Male | | | | | | | | | | | | |
| Dnajc2 | | | | | | | | | | | | | | | | 0.00500 | -1.26703 | Male |
| BC025920 | | | | | | | | | | | | | | | | 0.00002 | -0.82343 | Male |
| Fam3b | 0.00000 | 0.84432 | Female | | | | | | | 0.00000 | -4.08393 | Male | | | | 0.00000 | 3.22624 | Female |
| Cdk5rap2 | | | | 0.00594 | 3.18650 | Female | | | | | | | | | | | | |
| Maea | | | | | | | | | | 0.04414 | 1.95566 | Female | | | | | | |
| Wdr81 | | | | | | | | | | | | | | | | 0.00106 | -2.97805 | Male |
| Wnt10b | | | | | | | | | | | | | 0.02604 | 4.27035 | Female | | | |
| Sh3gl3 | | | | | | | | | | 0.01134 | -1.20849 | Male | 0.00000 | 3.13272 | Female | | | |
| Reps2 | | | | | | | | | | | | | | | | 0.00009 | -4.53765 | Male |
| Ptpn7 | | | | | | | | | | 0.00001 | -3.55859 | Male | | | | | | |
| Etohd2 | | | | 0.00006 | 0.90426 | Female | | | | 0.00000 | 1.42641 | Female | | | | 0.00000 | -5.07336 | Male |

| | | | | | | | | | | | | | | | | | | |
|---|---|---|---|---|---|---|---|---|---|---|---|---|---|---|---|---|---|---|
| Vnn1 | | | | | | | | | | | | | | | | 0.04842 | -1.81991 | Male |
| Dbhos | | | | | | | | | | | | | 0.00557 | 1.98203 | Female | | | |
| Slc6a14 | | | | 0.00000 | 0.89453 | Female | | | | | | | | | | | | |
| Pla2g10 | | | | 0.04789 | 2.13078 | Female | | | | | | | | | | | | |
| Kif2a | | | | | | | | | | 0.03165 | 1.63729 | Female | | | | | | |
| Bzw2 | | | | | | | | | | 0.01138 | 1.48701 | Female | | | | | | |
| Dqx1 | | | | | | | | | | | | | | | | 0.00284 | -2.52732 | Male |
| Dync2li1 | | | | | | | | | | | | | 0.00000 | -1.68844 | Male | | | |
| Ppp3ca | | | | | | | | | | 0.03336 | 3.45019 | Female | | | | | | |
| Ube2e2 | | | | 0.00161 | 1.13146 | Female | | | | | | | | | | | | |
| H2.Q4 | | | | | | | | | | 0.00581 | 3.11406 | Female | | | | | | |
| Srsf2 | | | | | | | | | | 0.04947 | -1.05991 | Male | | | | | | |
| Wnt4 | | | | | | | | | | 0.00011 | -3.18384 | Male | | | | | | |
| Arhgap20os | | | | | | | 0.03559 | 2.96821 | Female | 0.04422 | -3.03894 | Male | | | | | | |
| Rpl30 | | | | | | | | | | 0.01027 | -2.27568 | Male | | | | | | |
| Ptov1 | | | | | | | | | | 0.01272 | 3.45343 | Female | | | | | | |
| Syna | | | | | | | | | | | | | 0.00000 | 2.92116 | Female | 0.00000 | -2.90164 | Male |
| Gm8267 | | | | | | | | | | 0.00000 | 2.26521 | Female | | | | | | |
| Prom2 | | | | | | | | | | | | | | | | 0.00556 | -4.79518 | Male |
| Mmp23 | | | | 0.03735 | -1.12237 | Male | | | | | | | | | | | | |
| Nr0b1 | | | | | | | 0.00054 | -0.74640 | Male | | | | | | | | | |
| Derl3 | | | | | | | | | | | | | | | | 0.00656 | 4.21465 | Female |
| Suz12 | | | | | | | | | | | | | 0.02031 | -1.06911 | Male | | | |
| Unc45b | | | | | | | | | | | | | 0.00000 | -5.86486 | Male | | | |
| Zkscan6 | | | | | | | 0.03513 | 3.40623 | Female | | | | | | | | | |
| Nob1 | | | | | | | | | | 0.01272 | 1.61532 | Female | | | | | | |
| Kat2a | | | | | | | | | | 0.03679 | 1.68708 | Female | | | | | | |
| Cercam | | | | | | | | | | 0.04203 | -2.55875 | Male | | | | | | |
| Polr2k | | | | | | | | | | 0.00000 | -6.52237 | Male | 0.02954 | 1.44749 | Female | | | |
| Ntf5 | | | | | | | | | | | | | | | | 0.00212 | 4.26235 | Female |
| Ptpn22 | | | | | | | | | | | | | | | | 0.04004 | -3.23470 | Male |
| Snord58b | | | | | | | | | | | | | | | | 0.00622 | -4.32357 | Male |
| Stard10 | | | | | | | | | | 0.00030 | 4.08282 | Female | | | | | | |
| Eapp | | | | | | | | | | | | | | | | 0.00288 | -1.42930 | Male |
| Stab1 | | | | 0.02363 | 1.54505 | Female | | | | | | | 0.00000 | -0.67503 | Male | | | |
| Chad | | | | | | | 0.00003 | -2.49841 | Male | | | | | | | | | |
| Mir17hg | 0.00000 | -1.82324 | Male | | | | | | | | | | | | | | | |
| Kcnk6 | | | | | | | | | | 0.01515 | -2.31145 | Male | | | | | | |
| Qrich1 | | | | | | | | | | 0.00121 | 3.20264 | Female | | | | | | |
| Slco3a1 | | | | | | | | | | | | | 0.02999 | 4.54186 | Female | | | |
| Smad9 | | | | | | | | | | 0.02415 | -3.53911 | Male | 0.00094 | 3.69481 | Female | | | |
| Adgrg1 | | | | | | | | | | | | | | | | 0.02908 | -3.23430 | Male |
| Wdr35 | | | | | | | | | | 0.00001 | -2.62012 | Male | | | | | | |
| Shbg | | | | | | | | | | | | | 0.01819 | -3.36196 | Male | | | |
| Lrig3 | | | | | | | | | | | | | 0.00000 | -2.85424 | Male | 0.00000 | 1.21828 | Female |
| Khdc1c | | | | | | | | | | 0.00086 | -3.54969 | Male | | | | | | |
| Ogdh | | | | | | | | | | 0.01744 | -1.24162 | Male | | | | | | |
| Snord96a | | | | | | | | | | 0.04388 | 3.01886 | Female | | | | | | |
| Mex3c | | | | | | | 0.02709 | 2.14658 | Female | | | | | | | | | |
| Hnrnpk | | | | | | | | | | 0.03815 | 1.04135 | Female | | | | | | |
| Rpl31 | | | | | | | | | | 0.00000 | -4.60588 | Male | 0.00004 | 1.66310 | Female | | | |
| Arl5b | | | | | | | | | | | | | | | | 0.00829 | 4.26555 | Female |
| Ankrd23 | | | | | | | | | | | | | | | | 0.03018 | 2.55817 | Female |
| Nrp2 | | | | | | | | | | | | | | | | 0.01992 | 3.92677 | Female |
| Myh11 | | | | 0.00002 | 2.69886 | Female | | | | | | | 0.00000 | -4.78778 | Male | | | |
| Oog3 | | | | | | | | | | | | | 0.03377 | -3.25092 | Male | 0.00226 | -4.54538 | Male |
| Iqcf1 | | | | | | | | | | | | | | | | 0.00000 | -3.09290 | Male |
| Arid3b | | | | | | | | | | 0.03209 | 2.64150 | Female | | | | | | |
| Cops5 | 0.02389 | 1.91064 | Female | | | | | | | | | | | | | | | |
| Gdi1 | | | | 0.00695 | 3.59819 | Female | | | | | | | | | | | | |
| Slc14a1 | | | | | | | | | | | | | 0.00004 | -1.13475 | Male | | | |
| Plxnb3 | | | | | | | | | | 0.04058 | -0.89757 | Male | | | | | | |
| Uba3 | | | | | | | | | | | | | 0.01299 | -1.99141 | Male | | | |
| Tnfsf10 | | | | | | | | | | 0.00000 | -4.40707 | Male | 0.03509 | 1.32469 | Female | | | |
| Eif3d | | | | | | | | | | 0.00509 | 1.41048 | Female | | | | | | |
| Gadd45g | | | | 0.00026 | -2.42934 | Male | | | | | | | | | | | | |
| Nek7 | | | | | | | | | | 0.02049 | -2.08214 | Male | | | | | | |
| Trim40 | | | | | | | | | | | | | 0.00000 | 3.18120 | Female | | | |
| St8sia1 | | | | | | | | | | | | | | | | 0.02096 | -3.32782 | Male |
| Adamtsl5 | | | | 0.00006 | -1.39929 | Male | | | | | | | | | | | | |

| | | | | | | | | | | | | | | | | | | |
|---|---|---|---|---|---|---|---|---|---|---|---|---|---|---|---|---|---|---|
| Mcm6 | | | | | | | | | | | | | | | | 0.01138 | 1.12298 | Female |
| Il2rg | | | | | | | | | | | | | | | | 0.00284 | 4.10068 | Female |
| Phlpp1 | | | | 0.00158 | -1.41952 | Male | | | | | | | | | | | | |
| Mmp12 | | | | | | | 0.00000 | -0.74192 | Male | 0.00000 | 1.93204 | Female | | | | | | |
| Bag1 | | | | | | | | | | | | | | | | 0.02505 | -1.55804 | Male |
| Chchd1 | | | | | | | | | | 0.00052 | -2.54436 | Male | | | | | | |
| Polg | | | | | | | | | | 0.01380 | 2.56893 | Female | | | | | | |
| Psma5 | | | | | | | | | | 0.01409 | 1.33851 | Female | | | | | | |
| Ctcfl | | | | | | | | | | 0.00000 | -4.00540 | Male | 0.04793 | 1.36232 | Female | | | |
| Psen2 | | | | | | | | | | | | | | | | 0.00391 | 5.24762 | Female |
| Ak3 | | | | | | | | | | | | | 0.00000 | 1.47143 | Female | | | |
| Dmkn | | | | 0.00783 | -1.72971 | Male | | | | | | | | | | | | |
| Brcc3 | | | | | | | 0.00923 | 3.28827 | Female | | | | | | | | | |
| Stip1 | | | | | | | | | | 0.00102 | 1.69009 | Female | | | | 0.01641 | -1.09663 | Male |
| Hist3h2a | | | | 0.00019 | -1.98842 | Male | | | | 0.03379 | 4.13176 | Female | | | | | | |
| Irf7 | | | | | | | | | | | | | 0.03886 | 4.47386 | Female | | | |
| Zkscan7 | | | | 0.02033 | 1.87371 | Female | 0.00000 | -1.18880 | Male | 0.00045 | 0.60829 | Female | | | | | | |
| Glmp | | | | | | | | | | | | | | | | 0.00538 | -1.49244 | Male |
| Igip | 0.00001 | 1.46287 | Female | | | | | | | 0.00000 | 1.82153 | Female | | | | | | |
| Zar1l | | | | | | | | | | | | | | | | 0.02374 | -4.63327 | Male |
| Gm13102 | | | | | | | | | | | | | | | | 0.03227 | -1.21545 | Male |
| C1qbp | | | | | | | | | | 0.00003 | -2.25761 | Male | | | | | | |
| Fbxo44 | | | | | | | | | | | | | 0.00237 | 4.82520 | Female | | | |
| Gigyf2 | | | | | | | | | | | | | | | | 0.03731 | 1.85452 | Female |
| Map2 | | | | | | | | | | | | | | | | 0.00000 | 1.14212 | Female |
| Cks2 | | | | | | | | | | 0.00000 | -3.18967 | Male | | | | | | |
| Mrps33 | | | | | | | | | | 0.02272 | -1.83883 | Male | | | | | | |
| Cdc42bpb | | | | | | | | | | 0.00027 | 4.48326 | Female | | | | | | |
| Tceb2 | | | | | | | | | | 0.00000 | -4.20991 | Male | 0.00336 | 1.93327 | Female | | | |
| Xrcc6bp1 | | | | | | | | | | 0.03987 | -3.81136 | Male | | | | | | |
| Mfap3l | | | | | | | 0.00000 | -1.33514 | Male | | | | | | | | | |
| Gin1 | | | | 0.00991 | 3.93150 | Female | | | | | | | | | | | | |
| Havcr2 | | | | | | | 0.01411 | -4.23472 | Male | | | | | | | | | |
| Vdr | | | | | | | | | | 0.00003 | 2.65214 | Female | | | | 0.00000 | -3.68265 | Male |
| Iscu | | | | | | | | | | 0.00454 | -2.79321 | Male | | | | | | |
| Def6 | | | | 0.00001 | -0.74564 | Male | | | | | | | | | | | | |
| Sh3bp2 | | | | | | | | | | | | | | | | 0.00183 | -4.99613 | Male |
| Rab3b | | | | | | | | | | | | | | | | 0.00633 | 4.35163 | Female |
| Mb21d1 | | | | | | | | | | 0.01557 | -2.80457 | Male | 0.02209 | 2.03104 | Female | | | |
| Adarb2 | | | | | | | | | | 0.03149 | -1.43907 | Male | | | | | | |
| Ctrcos | | | | | | | | | | 0.00013 | -4.20084 | Male | | | | | | |
| Arhgdib | | | | | | | | | | | | | 0.01722 | 4.21439 | Female | | | |
| Cd47 | | | | 0.00000 | 2.87802 | Female | | | | 0.00001 | -2.40641 | Male | | | | 0.02592 | 4.61323 | Female |
| Ifi35 | | | | | | | | | | | | | | | | 0.04462 | 2.45250 | Female |
| Dr1 | | | | | | | | | | | | | | | | 0.00688 | -2.14765 | Male |
| Kif3c | | | | | | | | | | | | | | | | 0.00958 | -3.47806 | Male |
| Mov10l1 | | | | | | | | | | | | | | | | 0.02170 | 3.22429 | Female |
| Ccdc112 | | | | 0.00005 | 1.58545 | Female | 0.00000 | 1.45238 | Female | | | | | | | | | |
| Abca1 | 0.00057 | 0.96321 | Female | | | | | | | | | | | | | | | |
| Oas1e | | | | | | | | | | | | | | | | 0.00012 | -5.70665 | Male |
| Rbx1 | | | | | | | | | | 0.00000 | -3.33912 | Male | 0.01921 | 0.94558 | Female | | | |
| Dnajc5 | | | | | | | | | | | | | 0.01436 | 1.42851 | Female | | | |
| Eepd1 | | | | | | | 0.00000 | 1.86210 | Female | | | | | | | | | |
| Hist1h2bc | | | | 0.00001 | -1.23866 | Male | | | | | | | | | | | | |
| S100a10 | | | | 0.00974 | -5.79702 | Male | | | | | | | | | | | | |
| Thop1 | | | | | | | | | | 0.01000 | 3.93179 | Female | | | | | | |
| Ddx31 | | | | | | | | | | | | | | | | 0.02986 | 2.26850 | Female |
| P4htm | 0.00000 | 3.60077 | Female | | | | | | | | | | | | | | | |
| Gpr3 | | | | | | | | | | 0.04612 | -3.44534 | Male | 0.00452 | -3.77916 | Male | | | |
| Invs | | | | | | | | | | 0.00235 | 1.41843 | Female | | | | | | |
| Kat6a | | | | | | | | | | 0.00027 | -3.97711 | Male | | | | | | |
| Tsr2 | | | | | | | | | | 0.00000 | 3.30587 | Female | | | | | | |
| Ccdc13 | 0.00039 | -0.86631 | Male | 0.00040 | -1.06658 | Male | | | | | | | | | | | | |
| Glrx3 | | | | | | | | | | 0.02822 | -1.31556 | Male | | | | | | |
| Mylip | | | | | | | 0.03368 | -4.64797 | Male | | | | | | | | | |
| Zfp385c | | | | | | | | | | 0.00000 | -4.03071 | Male | | | | | | |
| Akr1c14 | | | | | | | | | | | | | | | | 0.00000 | -2.10979 | Male |
| Gja4 | | | | | | | | | | | | | | | | 0.00000 | 3.58707 | Female |
| Nudcd3 | | | | 0.00095 | 3.50818 | Female | | | | | | | | | | | | |
| Hk2 | | | | | | | | | | 0.00571 | 3.72535 | Female | | | | | | |

| | | | | | | | | | | | | | | | | | | |
|---|---|---|---|---|---|---|---|---|---|---|---|---|---|---|---|---|---|---|
| Agpat5 | | | | | | | | | | | | | 0.01841 | -1.33957 | Male | | | |
| Ifnar1 | | | | | | | | | | 0.00000 | -3.06188 | Male | | | | 0.04682 | -0.94531 | Male |
| Higd1a | | | | | | | | | | 0.00000 | -3.98451 | Male | 0.02066 | 1.62645 | Female | | | |
| Lcor | | | | 0.00000 | 0.75528 | Female | | | | | | | | | | | | |
| Tspan3 | | | | | | | | | | 0.00015 | 5.22592 | Female | | | | | | |
| Arglu1 | | | | | | | | | | | | | 0.00082 | 1.51890 | Female | | | |
| Iars2 | | | | | | | | | | | | | 0.02961 | -2.09164 | Male | | | |
| Mfsd11 | | | | | | | | | | 0.00056 | -3.08063 | Male | | | | | | |
| Gata6 | 0.00275 | -3.66339 | Male | | | | | | | | | | | | | | | |
| Efhc2 | | | | | | | 0.00000 | 2.12247 | Female | 0.00140 | 0.88358 | Female | | | | | | |
| Txnip | | | | | | | | | | | | | | | | 0.00985 | 4.42884 | Female |
| Zfp811 | | | | | | | | | | 0.00001 | -0.78609 | Male | | | | | | |
| Olfr1055 | | | | | | | | | | 0.01625 | -1.81632 | Male | | | | | | |
| Rps25 | | | | | | | | | | 0.00000 | -4.87860 | Male | 0.00024 | 1.49045 | Female | | | |
| Trpc1 | | | | | | | | | | | | | 0.04050 | 2.01060 | Female | | | |
| LOC101056073 | | | | | | | | | | 0.01624 | 2.26273 | Female | | | | | | |
| Ncan | | | | | | | | | | | | | 0.00036 | 3.73096 | Female | | | |
| Pcgf5 | | | | | | | | | | | | | | | | 0.00753 | -2.45065 | Male |
| H6pd | 0.00997 | -1.22959 | Male | 0.00078 | -1.03715 | Male | | | | 0.03208 | -1.13141 | Male | | | | | | |
| Hint1 | | | | | | | | | | 0.00548 | -2.75429 | Male | | | | | | |
| Dpy30 | | | | | | | | | | 0.03509 | -1.56887 | Male | | | | | | |
| Id3 | | | | | | | | | | | | | | | | 0.03547 | 2.80959 | Female |
| Lat2 | | | | | | | | | | | | | | | | 0.00186 | -4.46558 | Male |
| Zfp708 | | | | | | | | | | | | | | | | 0.03197 | -3.54174 | Male |
| Mcc | | | | | | | | | | 0.00002 | -3.57782 | Male | 0.01954 | 1.65307 | Female | | | |
| Xkrx | | | | | | | 0.01527 | 3.75375 | Female | | | | | | | | | |
| Chd4 | | | | | | | | | | 0.02251 | 2.40810 | Female | | | | | | |
| Spry4 | | | | | | | | | | | | | | | | 0.00071 | 4.60284 | Female |
| Ptgfrn | | | | 0.00000 | 1.27481 | Female | | | | 0.00000 | 4.93986 | Female | | | | | | |
| Pard6a | | | | 0.01192 | 1.68638 | Female | | | | | | | | | | | | |
| Blcap | | | | | | | | | | | | | | | | 0.01775 | -2.39619 | Male |
| Cct3 | | | | | | | | | | 0.00001 | 1.32523 | Female | | | | | | |
| Traf2 | | | | | | | | | | 0.00171 | 3.30811 | Female | | | | | | |
| Gucy2e | | | | | | | | | | | | | | | | 0.00000 | -7.17421 | Male |
| P2rx4 | | | | | | | | | | | | | | | | 0.00054 | -3.43979 | Male |
| Dancr | | | | | | | | | | 0.00113 | -4.32362 | Male | | | | | | |
| Fam83b | 0.04038 | -1.54187 | Male | | | | | | | 0.00002 | 2.73556 | Female | 0.00000 | -4.93788 | Male | | | |
| Pars2 | | | | 0.00000 | 1.96708 | Female | | | | | | | | | | | | |
| Tomm6 | | | | | | | | | | 0.00000 | -3.68340 | Male | 0.01139 | 1.44257 | Female | | | |
| Gm14325 | | | | | | | | | | | | | | | | 0.00422 | -1.66483 | Male |
| Ankrd13d | | | | 0.01037 | -0.70707 | Male | | | | | | | | | | | | |
| Prps1 | | | | 0.01459 | 1.69731 | Female | 0.01436 | 1.03678 | Female | | | | | | | | | |
| Pfkfb4 | | | | | | | | | | | | | | | | 0.04434 | -3.44480 | Male |
| Sergef | | | | | | | | | | | | | | | | 0.03813 | 4.08947 | Female |
| Atp5j2 | | | | | | | | | | 0.00000 | -4.78137 | Male | 0.00004 | 1.81533 | Female | | | |
| Mylpf | | | | | | | | | | 0.00017 | -2.87654 | Male | | | | | | |
| Apcdd1 | | | | 0.00000 | 1.16853 | Female | | | | | | | | | | | | |
| Sspn | | | | | | | | | | 0.00008 | -2.04368 | Male | | | | | | |
| Msh5 | | | | | | | | | | 0.00000 | 1.21040 | Female | | | | | | |
| Rpl31.ps12 | | | | | | | | | | 0.00000 | -5.44792 | Male | 0.00002 | 1.73436 | Female | | | |
| Slc7a14 | | | | | | | | | | 0.00000 | -4.90548 | Male | 0.00155 | 1.57255 | Female | | | |
| Bex6 | | | | | | | | | | | | | | | | 0.03731 | -2.88197 | Male |
| Tbc1d24 | | | | | | | | | | 0.00930 | -2.81502 | Male | | | | | | |
| Vangl2 | | | | | | | | | | | | | 0.03610 | 3.86527 | Female | | | |
| Pgap1 | | | | 0.00001 | 2.21872 | Female | | | | | | | | | | | | |
| AV320801 | 0.02369 | 1.26054 | Female | 0.00002 | 3.35756 | Female | 0.00001 | 0.81469 | Female | 0.03820 | -0.77111 | Male | | | | 0.00000 | -4.66968 | Male |
| Tmem88b | | | | | | | | | | 0.00033 | -4.40472 | Male | | | | | | |
| Sf3b2 | | | | | | | | | | 0.04779 | 1.21119 | Female | | | | | | |
| Uxt | | | | | | | | | | 0.00102 | -2.52630 | Male | | | | | | |
| Zfyve19 | | | | | | | | | | 0.01441 | 3.12743 | Female | | | | | | |
| Dync1li1 | | | | | | | | | | 0.03867 | 1.94430 | Female | | | | | | |
| Mib1 | | | | | | | | | | | | | 0.00893 | -3.53429 | Male | | | |
| Gm6607 | | | | | | | | | | 0.00717 | -4.38655 | Male | | | | | | |
| Rpl17 | | | | | | | | | | 0.00001 | -2.19022 | Male | | | | | | |
| Nubp2 | | | | | | | | | | 0.01533 | 1.60505 | Female | | | | | | |
| Smap1 | | | | | | | | | | 0.00000 | 6.05170 | Female | | | | | | |
| Zbtb20 | | | | | | | | | | | | | 0.00000 | 3.11167 | Female | 0.00012 | 1.02435 | Female |
| Ldlrad2 | | | | | | | | | | 0.00000 | -3.81941 | Male | | | | | | |
| Fgf3 | | | | | | | | | | | | | 0.00008 | -4.93099 | Male | | | |
| AI661453 | | | | | | | | | | 0.00299 | -3.49174 | Male | 0.00983 | 2.41490 | Female | | | |

| | | | | | | | | | | | | | | | | | | |
|---|---|---|---|---|---|---|---|---|---|---|---|---|---|---|---|---|---|---|
| Mrps15 | | | | | | | | | | 0.00080 | -3.57996 | Male | | | | | | |
| Ccdc64b | | | | | | | | | | 0.00000 | 3.23201 | Female | | | | | | |
| Eif2ak4 | | | | | | | 0.00114 | -4.36553 | Male | | | | | | | | | |
| Il1f9 | | | | | | | | | | | | | 0.00173 | 2.83813 | Female | 0.00032 | -3.63694 | Male |
| Klf5 | | | | | | | | | | 0.00000 | 5.62447 | Female | | | | | | |
| Mettl1 | | | | | | | | | | 0.04779 | 2.20582 | Female | | | | | | |
| Mfap1b | | | | | | | | | | 0.00001 | 3.58696 | Female | | | | | | |
| Rspo2 | | | | | | | | | | 0.00000 | -3.84181 | Male | | | | 0.00000 | -1.42865 | Male |
| Srpr | | | | | | | | | | | | | | | | 0.04485 | -1.21002 | Male |
| Pdgfa | | | | | | | 0.00000 | 5.35796 | Female | | | | | | | | | |
| Ddx19b | | | | | | | | | | | | | | | | 0.00002 | -2.64823 | Male |
| Jakmip3 | | | | | | | 0.00013 | -3.20477 | Male | | | | | | | | | |
| Per2 | | | | | | | | | | | | | | | | 0.00002 | -0.68163 | Male |
| Gm13752 | | | | | | | | | | | | | | | | 0.00000 | -3.58289 | Male |
| Slc29a3 | | | | | | | | | | | | | | | | 0.00500 | -3.26699 | Male |
| Ppm1j | | | | | | | | | | 0.00000 | 2.67258 | Female | | | | | | |
| Col7a1 | | | | 0.00006 | -0.89479 | Male | | | | | | | | | | | | |
| Arf5 | | | | | | | | | | 0.00631 | 4.33166 | Female | | | | | | |
| Rph3a | | | | | | | 0.01483 | -3.32532 | Male | | | | | | | | | |
| Cd68 | | | | | | | 0.00332 | 4.64384 | Female | | | | | | | 0.00102 | 4.85894 | Female |
| Cyb5rl | | | | | | | | | | 0.00000 | 1.57475 | Female | | | | | | |
| Sec1 | | | | | | | | | | | | | | | | 0.00450 | 3.79439 | Female |
| Duoxa2 | | | | | | | 0.01182 | -1.61896 | Male | 0.00219 | 0.73871 | Female | | | | | | |
| Dusp26 | | | | | | | | | | | | | | | | 0.00898 | -4.43959 | Male |
| Churc1 | | | | | | | | | | 0.00000 | -4.38725 | Male | | | | | | |
| Fam214a | | | | | | | | | | | | | | | | 0.00000 | -5.32480 | Male |
| Flrt3 | | | | | | | | | | | | | | | | 0.01239 | 5.99879 | Female |
| Mbtps2 | | | | | | | | | | 0.01283 | 3.08970 | Female | | | | | | |
| Mospd2 | | | | | | | 0.00000 | 6.39885 | Female | | | | | | | | | |
| Thap2 | | | | | | | | | | | | | | | | 0.02241 | -3.20003 | Male |
| Ccser1 | | | | | | | 0.00000 | -1.46649 | Male | 0.00044 | -0.94798 | Male | | | | | | |
| Igsf23 | | | | | | | | | | | | | | | | 0.00001 | -2.47133 | Male |
| Ccdc42 | | | | | | | | | | | | | 0.02406 | 5.21441 | Female | | | |
| Gpr143 | | | | | | | 0.00000 | 3.24329 | Female | | | | | | | | | |
| Slc12a9 | | | | | | | 0.04294 | 3.25771 | Female | | | | | | | | | |
| A4galt | | | | | | | | | | 0.02165 | -2.36061 | Male | | | | | | |
| Skp1a | | | | | | | | | | 0.04300 | -1.23642 | Male | | | | | | |
| Sox15 | | | | | | | | | | | | | | | | 0.00443 | -2.77751 | Male |
| Fsbp | | | | | | | | | | | | | | | | 0.01450 | -4.46533 | Male |
| Rasa2 | | | | | | | | | | 0.00741 | -4.44420 | Male | | | | | | |
| Kdelc2 | | | | | | | | | | 0.00000 | -4.84028 | Male | | | | | | |
| Rpl23 | | | | | | | | | | 0.00000 | -3.38527 | Male | 0.00095 | 1.20203 | Female | | | |
| Dock1 | | | | 0.00000 | -0.93773 | Male | | | | | | | | | | | | |
| Ctps2 | | | | | | | | | | | | | | | | 0.00117 | 3.75980 | Female |
| Nop16 | | | | | | | | | | 0.01535 | 1.63619 | Female | | | | | | |
| Slc1a7 | | | | | | | | | | 0.00076 | -2.92262 | Male | 0.03191 | 1.63568 | Female | | | |
| Sirt5 | | | | | | | | | | | | | | | | 0.00167 | -4.74592 | Male |
| Fsd2 | | | | 0.00043 | 0.90387 | Female | | | | 0.00000 | 1.58447 | Female | | | | | | |
| Wdr19 | | | | | | | | | | | | | | | | 0.02988 | 4.54690 | Female |
| Akr1d1 | | | | | | | | | | 0.01468 | -3.93154 | Male | | | | | | |
| Crygc | | | | | | | 0.00000 | -2.61036 | Male | | | | | | | | | |
| Hint2 | | | | | | | | | | 0.00935 | -4.66299 | Male | | | | | | |
| Padi3 | | | | | | | 0.00076 | -1.74728 | Male | | | | | | | 0.00000 | 3.45493 | Female |
| Mtx1 | | | | | | | | | | 0.02048 | 2.84638 | Female | | | | | | |
| Gmfg | | | | | | | | | | 0.00025 | -6.80773 | Male | | | | | | |
| Pi16 | | | | | | | | | | | | | 0.00000 | 3.82566 | Female | 0.00000 | 1.36971 | Female |
| Dopey2 | | | | | | | | | | | | | | | | 0.04890 | 3.36979 | Female |
| Mdp1 | | | | | | | | | | | | | | | | 0.02238 | -1.96556 | Male |
| Mpv17l | | | | | | | | | | 0.00000 | -5.06489 | Male | 0.00377 | 1.70889 | Female | | | |
| Fbxo27 | 0.00003 | 1.14463 | Female | | | | | | | | | | | | | | | |
| Spryd4 | | | | 0.04422 | 2.97750 | Female | | | | | | | | | | | | |
| Ahsa1 | | | | | | | | | | 0.00940 | 1.49131 | Female | | | | | | |
| Cxxc1 | | | | | | | | | | | | | | | | 0.02986 | -1.88796 | Male |
| Rgs10 | | | | | | | | | | | | | | | | 0.00000 | -4.86897 | Male |
| Cgrrf1 | 0.00000 | -0.96662 | Male | | | | | | | | | | | | | | | |
| Zfp605 | | | | | | | 0.00756 | 2.75043 | Female | 0.01766 | -2.81267 | Male | | | | 0.02122 | -2.36126 | Male |
| Eny2 | | | | | | | | | | 0.00000 | -4.26139 | Male | 0.00098 | 1.71096 | Female | | | |
| Mrap | | | | | | | | | | 0.00000 | 1.71085 | Female | | | | 0.00000 | 1.56800 | Female |
| Smim11 | | | | | | | | | | 0.00006 | -5.09898 | Male | 0.02196 | 2.49037 | Female | | | |
| Dnajc1 | | | | | | | | | | | | | | | | 0.00015 | 3.83595 | Female |

| | | | | | | | | | | | | | | | | | | |
|---|---|---|---|---|---|---|---|---|---|---|---|---|---|---|---|---|---|---|
| Xaf1 | | | | | | | | | | 0.00000 | -5.07574 | Male | 0.00007 | 2.39185 | Female | | | |
| Arhgap26 | | | | 0.00000 | 2.02340 | Female | | | | | | | 0.00158 | 4.25039 | Female | | | |
| H2.Eb1 | | | | 0.00006 | -1.93870 | Male | | | | | | | | | | | | |
| Cox5b | | | | | | | | | | 0.00000 | -4.70022 | Male | | | | | | |
| Sema4b | | | | | | | | | | 0.00000 | -4.84313 | Male | | | | | | |
| Prokr2 | | | | | | | | | | | | | | | | 0.00834 | -4.75217 | Male |
| Fsip1 | | | | | | | | | | 0.00003 | 2.15036 | Female | 0.00000 | 1.95085 | Female | | | |
| Clca3a1 | | | | | | | | | | 0.01865 | -1.97450 | Male | | | | | | |
| Rps6ka5 | | | | | | | | | | | | | | | | 0.00525 | 3.72044 | Female |
| Gabarapl2 | | | | | | | | | | | | | 0.01695 | 0.66074 | Female | | | |
| Rfx4 | 0.00054 | -1.29235 | Male | | | | | | | 0.00000 | 1.12481 | Female | | | | | | |
| Epb41l4aos | | | | | | | | | | 0.00001 | -5.17735 | Male | | | | | | |
| Ccr4 | | | | | | | 0.02109 | 4.31974 | Female | | | | 0.00627 | 3.95618 | Female | | | |
| Clint1 | | | | | | | | | | 0.01953 | 2.38588 | Female | | | | | | |
| Lypd8 | | | | | | | 0.00029 | -1.67612 | Male | | | | | | | | | |
| Naf1 | | | | | | | | | | 0.01118 | -3.25658 | Male | | | | | | |
| Rpp21 | | | | | | | | | | 0.00003 | -3.17321 | Male | | | | | | |
| AU023762 | | | | | | | | | | 0.00044 | -3.29211 | Male | | | | | | |
| Egfl8 | | | | 0.00034 | -1.76545 | Male | | | | | | | 0.01123 | -4.37317 | Male | 0.00028 | -5.94410 | Male |
| Rsph9 | 0.00116 | 1.45757 | Female | | | | | | | | | | | | | | | |
| Mvd | | | | | | | | | | | | | | | | 0.01450 | 2.89136 | Female |
| H1foo | | | | | | | | | | | | | | | | 0.00006 | -5.75362 | Male |
| Gm5779 | | | | | | | | | | 0.00800 | 2.58437 | Female | | | | | | |
| Smim14 | | | | | | | | | | | | | | | | 0.03311 | -3.38504 | Male |
| Nrros | | | | | | | | | | | | | | | | 0.00000 | 2.50222 | Female |
| Prickle1 | | | | | | | | | | | | | | | | 0.04932 | 2.24698 | Female |
| Tmem260 | 0.00106 | -1.13333 | Male | | | | | | | 0.01004 | -1.43437 | Male | | | | | | |
| Tmem243 | | | | | | | | | | 0.00001 | 0.96656 | Female | | | | | | |
| Ucp2 | | | | | | | | | | | | | | | | 0.00001 | 5.97585 | Female |
| Rfesd | | | | | | | | | | 0.01395 | 3.33125 | Female | | | | | | |
| Gas5 | | | | | | | | | | 0.00002 | -2.35116 | Male | | | | | | |
| Gon4l | | | | | | | | | | 0.01281 | 1.84491 | Female | | | | | | |
| Prkag2os1 | | | | | | | | | | | | | 0.00000 | 1.33094 | Female | | | |
| Cel | | | | | | | 0.00222 | -3.17481 | Male | | | | | | | | | |
| B3galt4 | | | | | | | | | | | | | | | | 0.00000 | 1.99758 | Female |
| Calm5 | | | | | | | | | | | | | | | | 0.00000 | -1.71437 | Male |
| Carkd | 0.00329 | 4.71602 | Female | | | | | | | | | | | | | | | |
| BC029214 | | | | 0.00156 | -2.07721 | Male | | | | | | | | | | | | |
| Proc | | | | | | | | | | 0.00488 | -1.22402 | Male | 0.00000 | 1.63057 | Female | | | |
| Il10rb | 0.00020 | -1.98505 | Male | 0.00006 | -0.98613 | Male | | | | 0.00280 | -1.21508 | Male | | | | 0.00000 | 4.74496 | Female |
| Cdc42ep1 | | | | | | | 0.00041 | 4.15617 | Female | | | | | | | | | |
| Crabp2 | | | | | | | | | | | | | 0.00000 | -1.25469 | Male | | | |
| Dpep3 | | | | | | | | | | | | | | | | 0.03105 | -2.63596 | Male |
| Brk1 | | | | | | | | | | 0.04182 | 1.08452 | Female | | | | | | |
| Ly6e | | | | 0.00000 | 0.86688 | Female | | | | | | | | | | | | |
| Gspt2 | | | | | | | | | | | | | | | | 0.01240 | 3.77694 | Female |
| Sec24b | | | | | | | | | | | | | | | | 0.00587 | -2.25680 | Male |
| Enpep | | | | | | | | | | 0.00086 | 4.92655 | Female | | | | | | |
| Gab2 | | | | | | | | | | 0.00648 | 3.51106 | Female | | | | | | |
| Ptprt | | | | | | | 0.03477 | -4.63761 | Male | | | | | | | | | |
| Fcf1 | | | | | | | | | | 0.00315 | -1.83219 | Male | | | | | | |
| Qtrt1 | | | | | | | | | | 0.04890 | 2.16870 | Female | | | | | | |
| Snx7 | | | | | | | | | | | | | | | | 0.04153 | -5.00382 | Male |
| Clk2 | | | | | | | | | | | | | | | | 0.03377 | -2.04750 | Male |
| Cfi | | | | 0.01416 | 1.07407 | Female | | | | | | | 0.00000 | -1.45765 | Male | | | |
| Lpin1 | 0.00000 | -1.07371 | Male | | | | | | | | | | | | | | | |
| B4galt2 | | | | | | | | | | 0.00000 | 2.99034 | Female | | | | 0.00202 | 4.49081 | Female |
| Jph1 | | | | | | | 0.00712 | 5.36017 | Female | | | | | | | | | |
| Slc23a1 | | | | | | | | | | 0.00000 | -5.52324 | Male | 0.01750 | 1.48150 | Female | | | |
| Polr2e | | | | | | | | | | 0.04508 | 1.45973 | Female | | | | | | |
| Cpa2 | | | | | | | | | | 0.00000 | -3.19641 | Male | | | | | | |
| Slco2a1 | | | | | | | | | | 0.01493 | 3.68050 | Female | | | | | | |
| Mbl2 | | | | | | | | | | | | | | | | 0.00000 | 2.63123 | Female |
| Ak8 | | | | | | | | | | | | | 0.00156 | 5.31058 | Female | | | |
| Ldhb | | | | | | | | | | | | | | | | 0.00000 | -5.45576 | Male |
| Pomk | | | | | | | | | | 0.00000 | 2.79596 | Female | | | | | | |
| Pla2g7 | | | | 0.00016 | -0.95318 | Male | | | | | | | | | | | | |
| Map9 | 0.04829 | -0.85903 | Male | | | | | | | 0.04902 | -3.87005 | Male | | | | | | |
| Myl9 | | | | 0.00024 | -1.58884 | Male | | | | | | | | | | | | |
| Cyp2d9 | | | | | | | | | | 0.00275 | 6.38591 | Female | | | | | | |

| | | | | | | | | | | | | | | | | | | |
|---|---|---|---|---|---|---|---|---|---|---|---|---|---|---|---|---|---|---|
| Polr2l | | | | | | | | | | 0.00000 | -4.67031 | Male | 0.01232 | 1.43822 | Female | | | |
| Snhg20 | | | | | | | | | | 0.01455 | -4.01441 | Male | | | | | | |
| Trpc6 | | | | | | | | | | 0.00049 | -2.33927 | Male | | | | | | |
| Scn2b | | | | | | | | | | 0.00004 | -4.07402 | Male | | | | | | |
| Uba52 | | | | | | | | | | 0.00000 | -3.93836 | Male | 0.00069 | 1.56338 | Female | | | |
| Tigar | | | | | | | | | | 0.04733 | 1.26356 | Female | | | | 0.00660 | -1.30389 | Male |
| Naa38 | | | | | | | | | | 0.00790 | -2.66985 | Male | | | | | | |
| Muc1 | 0.00000 | -1.52713 | Male | 0.00321 | -2.39382 | Male | | | | 0.00000 | 3.74846 | Female | | | | | | |
| Tomm40l | | | | 0.00000 | -1.29306 | Male | | | | 0.00000 | -1.96199 | Male | | | | | | |
| Nucks1 | | | | | | | 0.03552 | 1.08312 | Female | | | | | | | | | |
| Lypd1 | | | | | | | | | | 0.00000 | -5.20912 | Male | 0.00446 | 1.64122 | Female | | | |
| Mirc35hg | | | | | | | | | | 0.00000 | -4.30334 | Male | | | | | | |
| Tcp1 | | | | | | | | | | 0.00000 | 1.88232 | Female | | | | | | |
| Galnt9 | | | | | | | | | | | | | 0.00550 | 5.36034 | Female | | | |
| Zbtb33 | | | | | | | | | | | | | 0.01638 | 4.79754 | Female | | | |
| Zfp667 | | | | 0.00000 | 1.48893 | Female | 0.00000 | -1.87104 | Male | | | | | | | | | |
| D3Ertd254e | | | | | | | 0.00211 | 2.77456 | Female | | | | | | | 0.00452 | -2.53787 | Male |
| Fam213a | | | | | | | 0.00018 | -4.66688 | Male | 0.01912 | 3.24334 | Female | | | | | | |
| Ing1 | | | | | | | | | | | | | 0.01263 | 1.27100 | Female | | | |
| Fam212b | | | | | | | | | | | | | | | | 0.01254 | -4.63077 | Male |
| Dhx37 | | | | | | | | | | 0.01034 | 3.19237 | Female | | | | | | |
| Itgal | | | | | | | | | | 0.00001 | -0.83940 | Male | 0.00000 | -5.20786 | Male | 0.00000 | -3.63552 | Male |
| Upp1 | | | | | | | | | | 0.04885 | 1.64697 | Female | | | | | | |
| Minpp1 | | | | | | | | | | 0.03334 | 2.66849 | Female | | | | | | |
| Pfkm | 0.00057 | 0.74551 | Female | | | | | | | | | | | | | | | |
| Duox1 | 0.00015 | -0.97103 | Male | | | | | | | | | | | | | | | |
| Ppp4r1l.ps | | | | | | | | | | | | | | | | 0.00781 | -4.97268 | Male |
| Ahsg | | | | | | | 0.00000 | -1.77586 | Male | | | | | | | | | |
| Cd93 | | | | | | | | | | 0.00000 | -4.29083 | Male | | | | | | |
| Prr15l | | | | | | | | | | | | | | | | 0.00449 | -5.05855 | Male |
| Arf1 | | | | | | | | | | 0.00308 | 2.13313 | Female | | | | | | |
| Rin1 | | | | | | | | | | 0.00182 | -3.37415 | Male | | | | | | |
| Gimap9 | | | | | | | | | | 0.00891 | -1.00100 | Male | | | | | | |
| Dusp22 | | | | | | | | | | | | | 0.04329 | 2.06187 | Female | | | |
| Eif2b1 | | | | | | | | | | 0.00024 | 2.13559 | Female | | | | | | |
| Rab2b | | | | | | | | | | | | | | | | 0.02280 | -3.05753 | Male |
| Gja1 | | | | | | | 0.00489 | 2.50286 | Female | | | | | | | | | |
| Bcl2l10 | | | | | | | | | | 0.00269 | -5.56647 | Male | | | | | | |
| Scaf4 | | | | | | | | | | 0.04789 | 2.42451 | Female | | | | | | |
| Scamp3 | | | | | | | | | | 0.00000 | 6.86772 | Female | | | | | | |
| Coq10b | | | | 0.00703 | 3.02400 | Female | | | | | | | | | | | | |
| Ndufb2 | | | | | | | | | | 0.00002 | -5.22281 | Male | | | | | | |
| Kdm5b | | | | | | | | | | | | | | | | 0.04574 | 1.66741 | Female |
| Ube3a | | | | | | | 0.01128 | 2.90563 | Female | | | | | | | | | |
| Gm1965 | | | | | | | | | | | | | 0.02612 | -4.53895 | Male | | | |
| Agpat9 | | | | | | | 0.03513 | 1.84010 | Female | | | | | | | | | |
| Tcf7l2 | | | | 0.00001 | -0.61156 | Male | | | | | | | | | | | | |
| Wdsub1 | | | | | | | | | | 0.03694 | 3.46568 | Female | | | | | | |
| Tpcn1 | | | | | | | | | | | | | | | | 0.01694 | 4.33558 | Female |
| Gpbp1l1 | | | | | | | | | | | | | | | | 0.03033 | 2.14106 | Female |
| Nol10 | | | | | | | | | | | | | | | | 0.00137 | 2.06384 | Female |
| Slc2a1 | | | | | | | 0.00380 | -3.74658 | Male | | | | | | | | | |
| Rab26os | | | | | | | | | | 0.01698 | -3.99777 | Male | | | | | | |
| Il17rc | 0.00241 | 1.35501 | Female | | | | | | | 0.00541 | -0.78649 | Male | | | | | | |
| Sapcd1 | 0.00396 | 1.69022 | Female | | | | | | | | | | | | | | | |
| Sptb | 0.00000 | 1.76626 | Female | | | | | | | | | | | | | | | |
| Rhbdd3 | | | | | | | 0.02016 | 3.11628 | Female | | | | | | | | | |
| Ndufa7 | | | | | | | | | | 0.00000 | -4.00912 | Male | 0.00002 | 1.94062 | Female | | | |
| Bmp15 | | | | | | | | | | | | | 0.02856 | 3.93043 | Female | | | |
| Itpr2 | | | | | | | | | | | | | 0.00071 | 4.31398 | Female | 0.00000 | -0.63322 | Male |
| Nxf2 | | | | | | | | | | 0.00000 | 3.01091 | Female | | | | | | |
| Mgst3 | 0.03830 | -4.54815 | Male | | | | | | | | | | | | | | | |
| Rnf128 | | | | | | | | | | 0.04690 | 1.82432 | Female | | | | | | |
| Pbdc1 | | | | | | | | | | 0.00027 | 2.07810 | Female | | | | | | |
| Lcmt1 | | | | 0.00199 | 3.38003 | Female | | | | | | | | | | | | |
| Ndufb3 | | | | | | | | | | 0.00000 | -3.34490 | Male | 0.00228 | 1.46736 | Female | | | |
| Gm11517 | | | | | | | | | | 0.00942 | -1.92964 | Male | | | | | | |
| Ociad2 | | | | | | | | | | 0.00004 | -3.63371 | Male | | | | | | |
| D6Wsu163e | | | | | | | | | | | | | | | | 0.01368 | -2.73153 | Male |
| Cdca3 | | | | | | | | | | 0.04141 | 1.37036 | Female | | | | | | |

| | | | | | | | | | | | | | | | | | | |
|---|---|---|---|---|---|---|---|---|---|---|---|---|---|---|---|---|---|---|
| Fbln7 | | | | | | | | | | | | | 0.00000 | 4.36891 | Female | | | |
| Zfp961 | | | | | | | | | | | | | | | | 0.02986 | 1.83459 | Female |
| Gramd1a | 0.00000 | -0.85796 | Male | | | | | | | | | | | | | | | |
| Aloxe3 | | | | | | | | | | | | | | | | 0.00265 | 4.07118 | Female |
| Dll3 | | | | | | | | | | 0.00074 | -5.76939 | Male | | | | | | |
| Oxct2a | | | | | | | | | | | | | | | | 0.00305 | 2.66676 | Female |
| Psma7 | | | | | | | | | | 0.00368 | 1.53973 | Female | | | | | | |
| Hmgn3 | | | | 0.00359 | -0.72766 | Male | | | | | | | | | | | | |
| Tyr | | | | | | | | | | 0.00000 | -4.88792 | Male | 0.01542 | 1.39370 | Female | | | |
| Phb | | | | | | | | | | 0.01091 | 2.82138 | Female | | | | | | |
| Mrpl45 | | | | | | | | | | 0.02816 | 1.67526 | Female | | | | | | |
| Ndc80 | | | | | | | | | | | | | | | | 0.00007 | -2.31335 | Male |
| Tfcp2l1 | | | | 0.02436 | 2.25613 | Female | 0.00000 | 3.12162 | Female | | | | | | | | | |
| Syncrip | | | | | | | 0.02243 | 1.07765 | Female | | | | | | | | | |
| Fam46a | 0.00027 | 1.27984 | Female | | | | | | | 0.00009 | -0.71023 | Male | | | | | | |
| Chchd10 | | | | | | | | | | 0.00129 | -3.96770 | Male | | | | | | |
| Sap18 | | | | | | | | | | 0.00054 | -2.97728 | Male | | | | | | |
| Plp1 | | | | | | | | | | | | | 0.01743 | -1.33325 | Male | | | |
| Ndufb11 | | | | | | | | | | | | | 0.00738 | 1.53053 | Female | | | |
| Zfp951 | | | | | | | | | | | | | | | | 0.00001 | -3.44937 | Male |
| Vps72 | | | | | | | | | | 0.03176 | 2.77541 | Female | | | | | | |
| Dnah17 | | | | | | | | | | | | | 0.00042 | -4.86646 | Male | | | |
| Cd84 | | | | | | | | | | 0.00000 | 2.92339 | Female | | | | | | |
| Eef1g | | | | | | | | | | 0.00002 | 1.46590 | Female | | | | | | |
| Agmat | | | | | | | | | | | | | | | | 0.01257 | -2.95200 | Male |
| Gm10190 | | | | | | | | | | 0.00814 | -3.25472 | Male | | | | | | |
| Pdk1 | | | | | | | | | | | | | 0.00093 | 4.63461 | Female | | | |
| Fndc10 | | | | | | | 0.00000 | -4.88522 | Male | | | | | | | | | |
| Rpl10 | | | | | | | 0.00222 | 1.02045 | Female | | | | 0.00010 | 1.00695 | Female | 0.00929 | 0.86087 | Female |
| Irf8 | | | | | | | | | | 0.03149 | -0.92345 | Male | | | | | | |
| Pja1 | | | | 0.00000 | 1.42735 | Female | | | | | | | | | | | | |
| Ttc9c | | | | | | | | | | | | | | | | 0.02464 | 2.44609 | Female |
| Acly | | | | | | | | | | 0.03586 | 1.37743 | Female | | | | | | |
| Mir703 | | | | | | | | | | 0.00000 | -1.64591 | Male | | | | | | |
| Tmc2 | | | | | | | | | | | | | | | | 0.00005 | -3.59487 | Male |
| Poli | | | | | | | | | | 0.00062 | -5.37255 | Male | | | | | | |
| Perp | | | | 0.00264 | 2.17134 | Female | | | | | | | | | | | | |
| Jmjd4 | | | | | | | 0.00095 | 3.70937 | Female | | | | | | | | | |
| Kctd14 | | | | | | | 0.04812 | -2.38603 | Male | 0.00870 | -3.39202 | Male | | | | | | |
| Ppm1e | | | | | | | | | | 0.00000 | 1.25153 | Female | | | | | | |
| Immp1l | | | | 0.00797 | 2.70436 | Female | | | | | | | | | | | | |
| Ptafr | | | | | | | | | | 0.00001 | -3.86809 | Male | | | | | | |
| Sox12 | | | | | | | | | | | | | | | | 0.00001 | 5.02657 | Female |
| Ltv1 | | | | | | | | | | | | | | | | 0.00226 | -1.00410 | Male |
| Acp6 | | | | 0.02436 | 1.11650 | Female | | | | | | | | | | | | |
| Hist1h2be | | | | 0.00001 | -1.14551 | Male | | | | 0.00001 | -0.93746 | Male | | | | | | |
| Lrrc14b | | | | | | | | | | 0.00333 | -2.96496 | Male | | | | | | |
| Ptrh2 | | | | | | | | | | | | | | | | 0.03188 | -1.41900 | Male |
| Arsa | 0.00031 | 0.84510 | Female | | | | | | | | | | | | | | | |
| Cbln3 | | | | | | | | | | 0.00000 | -3.54297 | Male | 0.02954 | 1.43190 | Female | | | |
| Ints6 | | | | | | | | | | | | | | | | 0.00306 | -3.23619 | Male |
| Arrdc3 | | | | | | | | | | 0.00000 | 2.16005 | Female | | | | 0.00970 | 5.04206 | Female |
| Zfp783 | 0.00663 | 2.41923 | Female | 0.00000 | 2.32510 | Female | | | | | | | | | | | | |
| Wdr36 | | | | | | | | | | | | | 0.03383 | -1.09785 | Male | | | |
| Orc6 | | | | | | | | | | 0.00628 | 1.91084 | Female | | | | 0.00944 | -1.53261 | Male |
| Fgr | | | | | | | | | | | | | 0.00000 | -5.52209 | Male | | | |
| Tnfsf12 | | | | | | | | | | | | | | | | 0.01199 | -3.36224 | Male |
| Topaz1 | | | | 0.03318 | 1.00788 | Female | | | | | | | | | | | | |
| Atp5h | | | | | | | | | | 0.00000 | -3.24627 | Male | 0.00917 | 1.09110 | Female | | | |
| Gmppb | 0.01788 | 3.65186 | Female | | | | | | | | | | | | | | | |
| Mrps30 | | | | | | | | | | 0.01188 | -2.34221 | Male | | | | | | |
| Phkg1 | | | | | | | | | | | | | | | | 0.01273 | 3.29507 | Female |
| Nicn1 | | | | | | | | | | 0.00724 | 3.15098 | Female | | | | | | |
| Phlda2 | | | | | | | | | | | | | 0.00032 | 5.09566 | Female | | | |
| Anxa3 | | | | 0.00036 | -3.36235 | Male | | | | | | | | | | | | |
| Vgll4 | | | | 0.00000 | -2.25446 | Male | | | | 0.00000 | 3.15926 | Female | | | | | | |
| Cpsf4l | | | | | | | | | | | | | | | | 0.00000 | -0.83870 | Male |
| Pop5 | | | | | | | | | | 0.00016 | -3.44114 | Male | | | | | | |
| Copg1 | | | | | | | | | | 0.00307 | 1.63946 | Female | | | | | | |
| Prps2 | | | | | | | 0.03693 | -4.76215 | Male | | | | | | | | | |

| | | | | | | | | | | | | | | | | | | |
|---|---|---|---|---|---|---|---|---|---|---|---|---|---|---|---|---|---|---|
| Rims3 | | | | | | | | | | | | | | | | 0.02284 | 3.24997 | Female |
| Ssr2 | | | | | | | | | | 0.02008 | 1.29703 | Female | | | | | | |
| Sh2d4b | 0.00796 | -1.57053 | Male | | | | 0.00000 | -2.74912 | Male | 0.00000 | 2.79001 | Female | | | | | | |
| Rpl36al | | | | | | | | | | 0.00000 | -3.81740 | Male | 0.00016 | 1.55227 | Female | | | |

### Supplementary Table S3

#### Human: differential expression results

| Gene | padj_E3 | log2fc_E3 | sex_E3 | padj_E4 | log2fc_E4 | sex_E4 | padj_E5 | log2fc_E5 | sex_E5 | padj_E6 | log2fc_E6 | sex_E6 | padj_E7 | log2fc_E7 | sex_E7 |
|---|---|---|---|---|---|---|---|---|---|---|---|---|---|---|---|
| ZNF791 | | | | | | | | | | 0.00203 | 0.58666 | Female | | | |
| INSR | | | | | | | | | | 0.00000 | 1.03876 | Female | | | |
| NCAM1 | | | | | | | 0.03580 | 0.74937 | Female | | | | 0.02441 | -0.75766 | Male |
| ZNF280B | | | | | | | | | | 0.00000 | 1.89189 | Female | | | |
| AGK | | | | | | | | | | 0.01101 | 0.66244 | Female | | | |
| PIH1D1 | 0.01276 | 0.90882 | Female | | | | | | | | | | | | |
| ZNF275 | 0.00081 | 4.74426 | Female | | | | 0.00266 | 2.38854 | Female | | | | | | |
| MEF2D | | | | | | | | | | | | | 0.03502 | 0.64670 | Female |
| MKRN2 | | | | | | | | | | | | | 0.00014 | 0.65549 | Female |
| MVB12B | | | | | | | | | | | | | 0.00539 | 1.92401 | Female |
| TXNRD2 | 0.00000 | 3.81244 | Female | 0.01406 | 1.34763 | Female | | | | | | | | | |
| BAX | | | | 0.00290 | 0.75400 | Female | | | | 0.00003 | -0.65178 | Male | | | |
| CCL15 | | | | | | | 0.00101 | -1.99953 | Male | | | | | | |
| COL9A3 | | | | | | | 0.01072 | -0.85559 | Male | | | | | | |
| TMEM256 | | | | | | | 0.00000 | -0.76163 | Male | | | | | | |
| ZNF672 | | | | | | | 0.00050 | -0.95802 | Male | 0.01492 | -0.67451 | Male | | | |
| METAP1 | 0.00211 | -0.91666 | Male | | | | | | | | | | 0.00000 | -0.59825 | Male |
| MIR4435-1HG | | | | 0.00236 | 1.28959 | Female | 0.00000 | 1.04438 | Female | | | | | | |
| PSMB1 | 0.00008 | -0.64094 | Male | | | | | | | | | | | | |
| SLC7A11 | | | | 0.00001 | 2.04664 | Female | | | | | | | 0.02382 | -0.71622 | Male |
| RARRES1 | | | | | | | 0.00001 | -2.23215 | Male | 0.01863 | -1.19473 | Male | | | |
| SNHG10 | | | | | | | 0.01293 | 0.58438 | Female | | | | | | |
| PRDM1 | 0.03623 | 2.90588 | Female | | | | | | | | | | | | |
| SNORA75 | | | | | | | | | | | | | 0.00879 | 1.15678 | Female |
| MINOS1 | 0.00000 | 1.02018 | Female | 0.00000 | 0.58473 | Female | | | | | | | | | |
| ALDH18A1 | | | | 0.01216 | 0.98415 | Female | | | | | | | | | |
| CNNM4 | | | | 0.03773 | 0.79253 | Female | | | | | | | | | |
| LSR | 0.01497 | 1.10240 | Female | | | | | | | | | | | | |
| PDLIM1 | 0.04221 | -0.79606 | Male | | | | | | | | | | | | |
| MAGI1 | | | | 0.00253 | 0.88093 | Female | | | | | | | | | |
| NSUN3 | | | | | | | 0.04408 | -0.97718 | Male | | | | | | |
| ATP6V0E1 | 0.00038 | 0.68717 | Female | | | | | | | | | | | | |
| KCNK6 | 0.00005 | 3.46718 | Female | | | | | | | | | | | | |
| HSDL1 | | | | | | | 0.02291 | -1.04942 | Male | | | | | | |
| RNASE1 | | | | | | | 0.04010 | 0.88301 | Female | | | | 0.04853 | -0.79742 | Male |
| TEX14 | | | | | | | | | | 0.04957 | -1.29640 | Male | | | |
| RIPK2 | | | | | | | | | | | | | 0.00452 | -0.60710 | Male |
| LINC00624 | | | | 0.04676 | -1.05204 | Male | | | | | | | | | |
| ANO1 | 0.00002 | 4.29809 | Female | 0.00152 | 2.21785 | Female | | | | | | | | | |
| ZNF37BP | 0.04403 | 1.70746 | Female | | | | 0.00784 | 0.88819 | Female | | | | | | |
| POLI | | | | | | | 0.01095 | -0.86898 | Male | | | | 0.01932 | -0.76098 | Male |
| SLC3A1 | | | | | | | | | | 0.01325 | 1.77448 | Female | | | |
| P2RX5 | | | | 0.00104 | 3.46192 | Female | 0.00000 | 3.65361 | Female | | | | | | |
| MBOAT1 | | | | | | | 0.00001 | 1.16507 | Female | 0.00949 | 0.73049 | Female | 0.00001 | 1.13808 | Female |
| C14orf159 | | | | | | | | | | 0.02744 | 0.89376 | Female | | | |
| SLC39A7 | | | | | | | | | | 0.00000 | -0.85315 | Male | | | |
| NPIPB5 | | | | | | | | | | | | | 0.00009 | 0.62310 | Female |
| TFCP2L1 | 0.00000 | -1.52471 | Male | 0.00236 | 0.61010 | Female | | | | | | | | | |
| ZNF286A | | | | 0.03246 | -0.59062 | Male | | | | | | | | | |

| | | | | | | | | | | | | | | | |
|---|---|---|---|---|---|---|---|---|---|---|---|---|---|---|---|
| CFTR | | | | | | | | | | 0.00073 | -3.49012 | Male | | | |
| BCL3 | | | | 0.00255 | 2.42034 | Female | | | | 0.00048 | 1.85951 | Female | | | |
| HAS2 | | | | | | | | | | 0.00000 | 2.79413 | Female | 0.01368 | -1.49878 | Male |
| PRKCH | | | | | | | | | | 0.03123 | -1.12677 | Male | | | |
| GAST | | | | | | | | | | 0.00927 | -1.48057 | Male | | | |
| GLTPD2 | | | | | | | | | | 0.01326 | 1.31632 | Female | | | |
| MPO | | | | 0.01420 | -2.92706 | Male | 0.02445 | 2.13603 | Female | | | | | | |
| CCDC66 | | | | 0.03537 | -0.88393 | Male | | | | | | | | | |
| HNF4A | | | | | | | | | | 0.00000 | 1.20521 | Female | 0.02052 | -0.63075 | Male |
| PBDC1 | | | | 0.00000 | 1.06665 | Female | | | | | | | | | |
| ABCC6P1 | | | | | | | 0.03999 | 1.42136 | Female | | | | 0.00269 | -1.89930 | Male |
| PNMA5 | 0.00076 | 3.90801 | Female | | | | | | | 0.00000 | 3.62065 | Female | | | |
| COPRS | | | | 0.01672 | 0.79571 | Female | | | | | | | | | |
| MAT2A | | | | | | | 0.00000 | -0.59575 | Male | | | | | | |
| THUMPD2 | | | | | | | | | | 0.00001 | -1.02470 | Male | | | |
| CD274 | 0.01042 | -2.28716 | Male | 0.00388 | 1.77977 | Female | | | | 0.00580 | -1.04423 | Male | | | |
| RTKN | | | | 0.00000 | 1.57683 | Female | | | | | | | | | |
| SLC16A4 | | | | | | | 0.02344 | -1.24733 | Male | | | | | | |
| RPL10L | | | | | | | | | | 0.00000 | -2.63195 | Male | 0.03236 | 0.80432 | Female |
| TXLNGY | | | | 0.00000 | -4.78953 | Male | 0.00000 | -5.99741 | Male | 0.00000 | -6.37012 | Male | 0.00000 | -5.59213 | Male |
| C9orf37 | | | | | | | 0.02258 | 0.66160 | Female | | | | 0.01610 | 0.64772 | Female |
| COX7B | 0.00022 | 0.61136 | Female | 0.00000 | 0.77757 | Female | 0.00000 | 0.66975 | Female | 0.00000 | 0.58748 | Female | | | |
| CRYBB2P1 | | | | | | | 0.00001 | 0.99795 | Female | | | | | | |
| ANKRD18B | | | | | | | | | | 0.03917 | 2.06242 | Female | | | |
| TMEM229B | | | | | | | | | | 0.00000 | 2.61210 | Female | | | |
| DENND2D | | | | | | | | | | | | | 0.00200 | 1.26810 | Female |
| PJA2 | | | | | | | 0.00000 | -1.06734 | Male | | | | | | |
| C5orf34 | | | | | | | 0.03814 | -0.66435 | Male | | | | | | |
| NELL2 | | | | | | | | | | 0.02678 | 1.50653 | Female | | | |
| ENOSF1 | | | | 0.01799 | 0.58392 | Female | | | | | | | | | |
| RAB8B | | | | | | | 0.00018 | -1.00284 | Male | | | | | | |
| DST | | | | 0.01816 | 0.85750 | Female | | | | 0.00007 | 0.85817 | Female | | | |
| UNC119B | | | | | | | | | | | | | 0.00241 | -0.96224 | Male |
| NDUFS2 | 0.01969 | -0.64880 | Male | | | | | | | | | | | | |
| ERVFRD-1 | | | | | | | 0.00013 | 2.17375 | Female | | | | 0.00054 | 1.90182 | Female |
| PPP1R16B | | | | | | | | | | 0.02062 | 2.51241 | Female | | | |
| PTGES | | | | 0.00000 | 1.80697 | Female | | | | | | | 0.00028 | 0.72578 | Female |
| TMEM67 | | | | 0.00656 | -1.81806 | Male | | | | | | | | | |
| OTX2 | | | | | | | | | | | | | 0.02745 | -1.16516 | Male |
| TMEM185A | | | | 0.03451 | 1.21180 | Female | | | | | | | 0.02711 | -0.81823 | Male |
| NANOGNB | | | | | | | | | | 0.00000 | 1.90535 | Female | 0.00000 | 2.17604 | Female |
| MAL | | | | | | | | | | 0.02716 | 2.43234 | Female | | | |
| GNAI1 | | | | | | | 0.02442 | -1.07653 | Male | | | | | | |
| DECR1 | | | | 0.00000 | -3.01803 | Male | 0.00547 | -1.16240 | Male | | | | | | |
| BNC1 | | | | | | | | | | 0.00395 | -2.53877 | Male | | | |
| FANCL | | | | | | | 0.00050 | -0.68722 | Male | | | | | | |
| NPIPA1 | | | | | | | | | | 0.01364 | 0.77016 | Female | 0.00104 | 0.94288 | Female |
| TPT1-AS1 | 0.04232 | 1.21869 | Female | | | | 0.00012 | -0.72479 | Male | | | | | | |
| PRKACB | | | | 0.00179 | 2.72591 | Female | | | | | | | | | |
| NRG1 | | | | | | | | | | 0.01098 | 1.72999 | Female | | | |
| TTTY15 | 0.00000 | -6.74729 | Male | 0.00000 | -6.63303 | Male | 0.00000 | -7.37300 | Male | 0.00000 | -7.22747 | Male | 0.00000 | -7.39977 | Male |

| | | | | | | | | | | | | | | | |
|---|---|---|---|---|---|---|---|---|---|---|---|---|---|---|---|
| FAM115A | | | | 0.00006 | -1.83872 | Male | 0.04080 | -0.72659 | Male | 0.02549 | -0.70431 | Male | 0.02002 | -0.69947 | Male |
| SETBP1 | | | | 0.02120 | -1.54782 | Male | | | | | | | | | |
| SLC8A3 | | | | | | | | | | | | | 0.04509 | -2.10976 | Male |
| BAG6 | | | | 0.00235 | 0.69048 | Female | | | | | | | | | |
| SBNO1 | 0.00149 | -0.65986 | Male | | | | | | | | | | | | |
| CD52 | | | | | | | 0.00019 | 2.16791 | Female | 0.01132 | 1.50627 | Female | 0.00000 | 3.36998 | Female |
| LRRC47 | | | | | | | 0.00000 | 0.71443 | Female | | | | | | |
| KCNQ1OT1 | | | | 0.00962 | -1.16366 | Male | | | | | | | | | |
| NIM1K | | | | 0.00050 | -2.37445 | Male | 0.01232 | -1.20211 | Male | 0.04375 | -0.93821 | Male | | | |
| LOC101927380 | | | | 0.04491 | -1.62601 | Male | | | | | | | | | |
| SMNDC1 | 0.04252 | -0.61759 | Male | | | | | | | | | | | | |
| LOC101928254 | | | | | | | | | | | | | 0.04712 | -1.34187 | Male |
| EYS | | | | | | | | | | | | | 0.00606 | -1.47950 | Male |
| FAM220A | | | | | | | 0.03627 | -0.74631 | Male | | | | | | |
| ZSWIM7 | | | | 0.00253 | 0.58636 | Female | | | | | | | | | |
| MYEF2 | | | | | | | 0.01537 | -1.76398 | Male | | | | | | |
| GABRQ | | | | 0.04306 | 2.90530 | Female | | | | | | | | | |
| FCGBP | | | | | | | 0.03678 | 1.30915 | Female | | | | | | |
| SCRN3 | | | | | | | 0.03438 | -0.82361 | Male | | | | | | |
| TRMT61A | | | | | | | 0.00054 | 0.60453 | Female | 0.00024 | -0.60755 | Male | | | |
| DLGAP1-AS2 | | | | | | | | | | 0.00894 | 1.04178 | Female | | | |
| ID4 | | | | | | | | | | | | | 0.00472 | -1.27504 | Male |
| FGF13 | | | | 0.02440 | 1.29487 | Female | | | | | | | | | |
| LOC284379 | | | | | | | 0.00555 | -1.73501 | Male | 0.02104 | -1.40637 | Male | | | |
| SFT2D3 | | | | | | | | | | 0.02956 | -0.94984 | Male | | | |
| FBXO44 | | | | | | | | | | 0.00016 | 2.37390 | Female | 0.00248 | 1.93983 | Female |
| HDDC3 | 0.01196 | -1.63230 | Male | | | | | | | | | | | | |
| DLX4 | | | | 0.00000 | 2.52118 | Female | | | | | | | | | |
| HPS3 | | | | 0.02848 | 1.14888 | Female | | | | | | | | | |
| DSTN | | | | | | | 0.00000 | -0.64117 | Male | | | | | | |
| HMGCL | | | | | | | | | | 0.03660 | 0.85133 | Female | | | |
| RAB40B | | | | | | | | | | 0.00011 | 1.76139 | Female | | | |
| SETD1A | | | | | | | | | | 0.00610 | 0.63681 | Female | 0.00108 | 0.72700 | Female |
| FCER1G | | | | | | | | | | 0.03191 | 1.53530 | Female | | | |
| NIT2 | | | | 0.00000 | 0.88947 | Female | | | | | | | | | |
| BACH2 | | | | | | | | | | | | | 0.00204 | -1.23456 | Male |
| COPS8 | | | | 0.00293 | 0.64570 | Female | | | | | | | | | |
| PKMYT1 | | | | | | | 0.00002 | -0.87981 | Male | | | | | | |
| PRDX2 | | | | | | | 0.00000 | 0.87532 | Female | | | | | | |
| NOVA1 | | | | 0.03859 | -1.11784 | Male | | | | 0.03562 | -0.73112 | Male | | | |
| FAM122C | 0.00280 | 2.43047 | Female | | | | | | | 0.00000 | 1.77959 | Female | | | |
| CCDC22 | | | | 0.00008 | 2.28898 | Female | 0.00960 | 1.11700 | Female | 0.01950 | 0.99026 | Female | | | |
| PI4K2B | | | | | | | | | | | | | 0.03917 | -0.87841 | Male |
| MAN2B2 | 0.00010 | 3.11095 | Female | | | | | | | | | | | | |
| SNAR-G1 | 0.00000 | -6.24438 | Male | | | | | | | | | | 0.00331 | 0.93880 | Female |
| PINK1-AS | | | | | | | | | | | | | 0.01605 | 0.59918 | Female |
| ARHGEF2 | 0.00031 | 3.08875 | Female | 0.00013 | -1.96965 | Male | 0.01204 | -0.93650 | Male | | | | | | |
| LOC100506675 | | | | | | | 0.00447 | 0.64873 | Female | 0.00002 | -0.89955 | Male | 0.00000 | -1.21733 | Male |
| SLC35F6 | | | | | | | 0.00000 | 0.77898 | Female | 0.00002 | 0.67010 | Female | | | |
| NR2F2 | | | | | | | | | | 0.03610 | -0.61933 | Male | | | |
| CUTC | 0.00074 | -1.31676 | Male | | | | | | | | | | | | |

| | | | | | | | | | | | | | | | |
|---|---|---|---|---|---|---|---|---|---|---|---|---|---|---|---|
| ATAD3C | | | | 0.00765 | 1.51258 | Female | | | | | | | | | |
| LINC00842 | | | | | | | | | | 0.01102 | -0.66787 | Male | | | |
| CASC2 | | | | | | | | | | | | | 0.01008 | 1.93492 | Female |
| KCND1 | 0.01337 | 3.40651 | Female | | | | | | | | | | | | |
| PUS1 | 0.00001 | -1.53493 | Male | | | | | | | | | | | | |
| PTBP3 | | | | 0.00002 | 0.91103 | Female | | | | | | | | | |
| FYTTD1 | | | | | | | | | | 0.01390 | 0.58467 | Female | | | |
| TMEM191B | | | | | | | 0.00004 | -1.14993 | Male | | | | | | |
| SORBS1 | | | | | | | | | | 0.00023 | 0.76318 | Female | | | |
| SPATS2L | | | | | | | | | | 0.01235 | 0.93334 | Female | | | |
| KIF14 | | | | | | | 0.01369 | -0.58344 | Male | | | | | | |
| TRIM77 | | | | | | | | | | | | | 0.00022 | -3.03583 | Male |
| LINC00116 | | | | 0.00160 | 0.76814 | Female | | | | | | | | | |
| LOC399715 | | | | | | | | | | 0.00048 | 1.05940 | Female | 0.04509 | 0.65858 | Female |
| EXOSC10 | 0.00266 | -0.77153 | Male | | | | | | | | | | | | |
| ART5 | | | | | | | | | | 0.01349 | 1.97489 | Female | | | |
| PDLIM3 | | | | | | | | | | | | | 0.00753 | -2.57999 | Male |
| EIF3M | 0.00011 | -0.64643 | Male | | | | | | | | | | | | |
| LRRC8A | | | | | | | 0.00001 | 0.67193 | Female | | | | | | |
| CASKIN1 | | | | | | | | | | 0.00351 | 1.88443 | Female | | | |
| MCM9 | | | | | | | 0.00545 | -0.91076 | Male | | | | | | |
| RAB18 | 0.00000 | -1.67025 | Male | | | | | | | | | | | | |
| C1D | | | | 0.00216 | -0.61788 | Male | | | | | | | 0.00000 | -0.82067 | Male |
| TRMU | | | | | | | 0.00000 | 0.78359 | Female | | | | | | |
| S100P | 0.00009 | 2.89802 | Female | | | | | | | 0.00092 | 0.88570 | Female | 0.00347 | 0.79428 | Female |
| CCT7 | 0.00029 | -0.76001 | Male | | | | | | | | | | | | |
| SLC25A16 | | | | | | | 0.00020 | -0.60378 | Male | 0.00000 | 0.73142 | Female | | | |
| CYB561A3 | | | | | | | | | | 0.01817 | -0.70757 | Male | | | |
| PLOD2 | | | | | | | | | | | | | 0.00088 | -0.94578 | Male |
| ZFP37 | | | | | | | 0.00154 | -1.57323 | Male | | | | | | |
| PER2 | | | | | | | 0.00066 | 0.73294 | Female | | | | | | |
| TMEM91 | | | | | | | | | | 0.03537 | -1.37904 | Male | | | |
| MINA | | | | | | | 0.03301 | -0.68709 | Male | | | | | | |
| MT1E | | | | | | | | | | | | | 0.00193 | 1.70150 | Female |
| TRPV1 | | | | | | | | | | | | | 0.03592 | 0.76319 | Female |
| LOC101928559 | | | | | | | | | | | | | 0.00001 | -5.61557 | Male |
| ALG6 | | | | | | | 0.00132 | -0.65956 | Male | | | | | | |
| HMGN2 | | | | | | | 0.00000 | 0.91059 | Female | | | | | | |
| NRGN | | | | | | | | | | 0.00000 | 3.87322 | Female | | | |
| TRPS1 | 0.00710 | 3.52169 | Female | | | | | | | | | | | | |
| VTI1B | | | | | | | 0.00000 | 0.60048 | Female | | | | | | |
| NTN4 | | | | | | | | | | 0.01635 | 1.70785 | Female | | | |
| ZNF480 | | | | | | | | | | 0.00001 | 0.60356 | Female | | | |
| DNAJB6 | 0.01091 | -0.66507 | Male | | | | | | | | | | | | |
| ANKRD19P | 0.00014 | 0.87745 | Female | | | | | | | | | | | | |
| GLRX | | | | 0.00054 | 1.93161 | Female | | | | | | | | | |
| PDCL3P4 | | | | | | | | | | 0.00019 | 0.75494 | Female | | | |
| LEFTY2 | | | | | | | 0.00018 | 0.74562 | Female | 0.04129 | 0.73288 | Female | | | |
| TBC1D25 | | | | | | | | | | 0.04756 | 0.65870 | Female | 0.04860 | 0.64806 | Female |
| FNBP1 | | | | | | | 0.00052 | 0.63287 | Female | 0.00091 | 0.58541 | Female | | | |
| DYRK1A | 0.00147 | -1.55470 | Male | | | | | | | | | | | | |

| | | | | | | | | | | | | | | | |
|---|---|---|---|---|---|---|---|---|---|---|---|---|---|---|---|
| TPTE2P6 | | | | | | | | | | 0.00225 | 1.61576 | Female | | | |
| HSP90AB4P | | | | | | | | | | 0.00000 | -1.50791 | Male | | | |
| EGR1 | | | | | | | | | | | | | 0.00018 | 1.67233 | Female |
| CLDN3 | | | | 0.00000 | 2.81681 | Female | 0.00003 | -1.34708 | Male | 0.00117 | -1.01062 | Male | | | |
| DDX43 | | | | | | | | | | 0.00355 | 0.69981 | Female | | | |
| LOC100996324 | | | | | | | | | | 0.04046 | 1.47660 | Female | | | |
| ADAMTS20 | | | | | | | | | | 0.00832 | -1.73507 | Male | | | |
| ACTA1 | | | | | | | | | | 0.00000 | -2.20342 | Male | | | |
| FBXO48 | | | | | | | | | | 0.01316 | 1.01817 | Female | | | |
| KLF17 | | | | | | | | | | 0.00000 | 2.75074 | Female | | | |
| NSUN5P1 | | | | | | | | | | | | | 0.00313 | 0.60767 | Female |
| PTCD1 | | | | | | | 0.00062 | -0.89241 | Male | | | | 0.00674 | 0.69617 | Female |
| FBN2 | | | | | | | | | | 0.01363 | -1.83664 | Male | | | |
| SNORA78 | | | | | | | | | | | | | 0.00964 | 0.88780 | Female |
| CYB5R1 | | | | | | | | | | 0.01446 | -0.60097 | Male | | | |
| LDHAL6A | | | | | | | | | | 0.01321 | -1.41164 | Male | | | |
| AURKA | 0.00939 | -0.77924 | Male | | | | | | | | | | | | |
| ZNF75D | | | | 0.00001 | 2.08534 | Female | 0.01094 | 0.93814 | Female | 0.01989 | 0.83570 | Female | | | |
| PGC | | | | | | | 0.01342 | 1.46898 | Female | | | | 0.03157 | 1.27032 | Female |
| LYAR | 0.00338 | -0.67760 | Male | | | | | | | | | | | | |
| OGFR | 0.04955 | -1.76441 | Male | | | | | | | | | | | | |
| RAB15 | | | | | | | | | | 0.00000 | 1.56130 | Female | | | |
| TGFBR2 | | | | | | | | | | 0.03489 | -1.14414 | Male | | | |
| USP25 | | | | | | | | | | | | | 0.00007 | -0.86445 | Male |
| FOXE1 | | | | 0.03388 | 1.88374 | Female | | | | | | | | | |
| TPST2 | | | | | | | | | | 0.00046 | 0.71152 | Female | | | |
| PCDH1 | | | | | | | | | | 0.00735 | 0.72445 | Female | | | |
| CCDC102A | | | | | | | | | | | | | 0.03242 | 1.21785 | Female |
| SERHL2 | | | | | | | | | | | | | 0.00000 | 0.96046 | Female |
| C15orf41 | | | | | | | 0.01991 | -0.59804 | Male | | | | | | |
| AUH | | | | | | | 0.00315 | -0.74865 | Male | | | | 0.00737 | -0.64470 | Male |
| MCAT | | | | | | | 0.00126 | -0.67413 | Male | 0.00177 | -0.62491 | Male | | | |
| TMEM106C | | | | | | | 0.00000 | -0.85231 | Male | 0.00000 | -1.03371 | Male | | | |
| ROGDI | | | | 0.00000 | 1.70101 | Female | | | | | | | | | |
| KCNC3 | | | | | | | | | | | | | 0.00053 | 0.94575 | Female |
| FKBP9 | 0.04437 | 1.94102 | Female | | | | | | | | | | | | |
| NFAT5 | | | | 0.04367 | 0.80122 | Female | | | | | | | | | |
| GEM | | | | | | | | | | | | | 0.00398 | 1.88854 | Female |
| SKAP2 | | | | 0.03670 | -0.67002 | Male | | | | | | | | | |
| PDLIM7 | | | | | | | 0.00761 | 0.58084 | Female | 0.00003 | 0.81915 | Female | | | |
| OSBPL1A | | | | | | | 0.00084 | -1.24549 | Male | | | | 0.01835 | -0.87287 | Male |
| RPL34-AS1 | | | | | | | 0.01304 | -1.28586 | Male | | | | | | |
| TENC1 | | | | | | | 0.03527 | -2.50078 | Male | | | | | | |
| CBFA2T2 | | | | | | | | | | 0.00000 | 1.08430 | Female | | | |
| ERF | | | | | | | | | | 0.00000 | 0.92054 | Female | | | |
| PDE4A | | | | | | | | | | 0.00175 | 1.50997 | Female | | | |
| ITGB3 | | | | | | | | | | | | | 0.02542 | -2.41590 | Male |
| DCAF10 | | | | 0.00183 | 1.79663 | Female | | | | | | | | | |
| FZD5 | | | | | | | 0.00000 | 0.95966 | Female | | | | | | |
| SNTB1 | | | | | | | | | | | | | 0.00000 | -3.36521 | Male |
| DIS3L | 0.01582 | -1.21070 | Male | | | | | | | | | | | | |

| | | | | | | | | | | | | | | | |
|---|---|---|---|---|---|---|---|---|---|---|---|---|---|---|---|
| LINC00987 | | | | | | | | | | 0.00003 | -0.61459 | Male | | | |
| MKL1 | | | | | | | 0.00559 | -0.62675 | Male | | | | | | |
| RNF139 | | | | | | | 0.00198 | -0.67701 | Male | | | | | | |
| AP2A1 | | | | | | | 0.00011 | -0.71310 | Male | 0.00067 | 0.61258 | Female | | | |
| KDM4B | | | | | | | | | | | | | 0.01777 | 0.79591 | Female |
| BPGM | | | | | | | | | | 0.00119 | -0.82973 | Male | | | |
| DDX60L | | | | | | | | | | | | | 0.00798 | 2.37705 | Female |
| POMT2 | | | | | | | 0.01010 | 0.65529 | Female | | | | | | |
| ARHGEF1 | | | | | | | 0.02425 | -0.66499 | Male | | | | | | |
| CFL1P1 | | | | | | | | | | 0.01132 | -1.36996 | Male | | | |
| GRK5 | | | | | | | | | | 0.00005 | 1.92005 | Female | | | |
| ACADM | | | | 0.01218 | -0.76063 | Male | | | | | | | | | |
| MAFG | 0.03750 | -1.06886 | Male | | | | | | | | | | | | |
| LAMA4 | | | | | | | | | | 0.00038 | 2.12061 | Female | | | |
| LINC00668 | | | | | | | | | | 0.00000 | -0.90123 | Male | | | |
| NMRK2 | | | | | | | | | | 0.00029 | 1.11174 | Female | 0.00047 | 1.06381 | Female |
| IFNGR2 | | | | | | | | | | | | | 0.00003 | -1.56165 | Male |
| IL6 | | | | | | | 0.03266 | 0.92102 | Female | 0.00112 | -1.24519 | Male | | | |
| TMEM65 | | | | | | | | | | | | | 0.00472 | -0.86960 | Male |
| PTPRZ1 | | | | | | | | | | 0.00080 | 2.19468 | Female | | | |
| DPF1 | | | | 0.04433 | 1.58873 | Female | | | | 0.00000 | 2.44583 | Female | | | |
| SCD5 | | | | | | | | | | 0.02560 | -1.11310 | Male | | | |
| MIR6847 | | | | | | | 0.00000 | 1.17564 | Female | | | | | | |
| CIPC | | | | 0.00234 | -1.09489 | Male | | | | | | | | | |
| ARL4C | | | | 0.00161 | 1.73339 | Female | | | | | | | | | |
| PGRMC1 | 0.00904 | 1.13676 | Female | | | | | | | 0.00009 | 0.63203 | Female | | | |
| APP | | | | 0.01431 | 0.90292 | Female | | | | 0.00000 | -1.00791 | Male | | | |
| TNFSF13B | | | | 0.00045 | -1.43621 | Male | | | | 0.00001 | -1.19088 | Male | | | |
| ARL8A | | | | | | | | | | 0.01037 | 0.83003 | Female | | | |
| CLCN6 | | | | 0.00083 | -1.41136 | Male | | | | | | | | | |
| HIST1H2AC | | | | | | | 0.00060 | 1.27224 | Female | | | | | | |
| OAF | | | | 0.00664 | -2.12335 | Male | | | | | | | | | |
| SLC23A2 | | | | | | | | | | 0.00022 | 0.75815 | Female | 0.00012 | 0.77144 | Female |
| TPM4 | | | | 0.00005 | 1.02255 | Female | | | | | | | | | |
| VASH1 | | | | | | | | | | 0.00024 | 1.89216 | Female | | | |
| ACTR6 | | | | | | | 0.00016 | -0.58397 | Male | | | | | | |
| FKBPL | | | | | | | | | | 0.00407 | -0.58714 | Male | | | |
| PIEZO1 | | | | 0.00049 | 1.27878 | Female | | | | | | | | | |
| PTN | | | | | | | 0.00009 | -0.89238 | Male | 0.00027 | -0.79523 | Male | | | |
| AGTRAP | | | | 0.02798 | 0.99722 | Female | | | | | | | | | |
| EPSTI1 | | | | 0.00677 | 1.43504 | Female | | | | | | | | | |
| CCDC181 | | | | | | | | | | 0.01944 | -1.41295 | Male | | | |
| CNNM3 | 0.00002 | 1.83651 | Female | | | | | | | | | | | | |
| LINC00479 | | | | 0.01120 | 2.03454 | Female | | | | | | | | | |
| NPTX1 | | | | 0.00013 | 2.67907 | Female | | | | 0.00497 | 1.44971 | Female | 0.00000 | 2.23873 | Female |
| CTDP1 | | | | | | | | | | 0.01113 | 0.64104 | Female | | | |
| PTS | | | | | | | | | | 0.00000 | -0.74990 | Male | | | |
| CITED4 | 0.00000 | 3.22419 | Female | 0.00006 | 1.51599 | Female | | | | | | | 0.00002 | 1.07075 | Female |
| GRK6 | 0.00000 | 3.31965 | Female | | | | | | | | | | | | |
| JAK1 | | | | 0.00420 | 1.03271 | Female | | | | | | | | | |
| CDKN1B | | | | 0.04806 | -1.35059 | Male | | | | 0.02888 | -0.91955 | Male | | | |

| | | | | | | | | | | | | | | | |
|---|---|---|---|---|---|---|---|---|---|---|---|---|---|---|---|
| SRSF12 | | | | 0.04563 | 1.35614 | Female | | | | | | | | | |
| MFN1 | | | | | | | 0.00000 | -0.71993 | Male | | | | | | |
| SKIL | | | | | | | | | | 0.00000 | 0.89026 | Female | | | |
| ATP5L2 | | | | | | | | | | | | | 0.00320 | 0.87538 | Female |
| DNAH12 | | | | | | | 0.02752 | -1.37478 | Male | | | | | | |
| S100A14 | 0.00064 | 2.51005 | Female | 0.00000 | 2.12827 | Female | | | | | | | | | |
| SLC35D1 | | | | 0.03101 | 1.05204 | Female | | | | | | | | | |
| CBX2 | | | | | | | | | | 0.00306 | -0.86584 | Male | | | |
| DLG3 | | | | 0.04476 | 0.73180 | Female | 0.00000 | 1.07878 | Female | 0.00000 | 1.05966 | Female | | | |
| RNF14 | 0.02490 | -1.07682 | Male | | | | | | | | | | | | |
| KIAA0020 | 0.00008 | -0.75943 | Male | | | | | | | | | | | | |
| SLC22A5 | | | | | | | | | | | | | 0.02351 | 0.80850 | Female |
| AP1G2 | | | | 0.01641 | 0.97694 | Female | | | | | | | | | |
| SNORD25 | | | | | | | | | | | | | 0.01816 | 0.69624 | Female |
| CHI3L1 | | | | | | | 0.02473 | 2.09446 | Female | | | | | | |
| LMOD2 | | | | | | | 0.01950 | 1.12736 | Female | | | | 0.02849 | 1.01459 | Female |
| RCN2 | | | | | | | 0.00009 | -0.72677 | Male | | | | 0.00000 | -0.87782 | Male |
| GNB4 | | | | | | | 0.00766 | -0.81101 | Male | 0.00005 | -1.09880 | Male | 0.00253 | -0.84017 | Male |
| RPS6KA5 | | | | | | | 0.01322 | 0.63050 | Female | | | | | | |
| PROCR | 0.00074 | 1.46372 | Female | | | | | | | | | | | | |
| ABCB6 | 0.00013 | 1.83011 | Female | | | | 0.00000 | -0.90676 | Male | 0.00003 | -0.79368 | Male | | | |
| CRYGB | | | | | | | | | | 0.00464 | 2.87308 | Female | | | |
| BICD1 | | | | 0.00831 | 0.95793 | Female | | | | 0.00000 | 1.01272 | Female | | | |
| PLXNA2 | | | | 0.01428 | 1.27939 | Female | | | | | | | | | |
| ZP4 | | | | | | | | | | 0.00000 | -2.73189 | Male | | | |
| KAZN | 0.02760 | 2.32670 | Female | | | | | | | 0.00036 | 1.18602 | Female | | | |
| STOM | | | | | | | | | | 0.00019 | 0.76693 | Female | | | |
| INPPL1 | 0.03750 | -1.34360 | Male | 0.02449 | 0.84908 | Female | | | | | | | | | |
| NTMT1 | | | | | | | 0.00000 | 0.77884 | Female | | | | | | |
| LGALS16 | | | | | | | | | | 0.00047 | -1.80054 | Male | | | |
| IGFBP3 | | | | | | | | | | | | | 0.00000 | -3.93043 | Male |
| TCTA | | | | | | | | | | | | | 0.01111 | 0.74937 | Female |
| CD81 | | | | 0.00000 | 1.57861 | Female | | | | 0.00056 | 0.58435 | Female | | | |
| MYO10 | | | | | | | | | | 0.00676 | -0.63813 | Male | | | |
| HPCAL1 | 0.00012 | 2.67574 | Female | 0.00001 | 1.78896 | Female | | | | | | | | | |
| OR51B5 | | | | | | | 0.02169 | -1.13465 | Male | | | | | | |
| DYSF | | | | 0.01279 | -0.94230 | Male | | | | | | | | | |
| ARPC4 | 0.01091 | -0.59554 | Male | | | | 0.00000 | 0.68538 | Female | | | | | | |
| KIAA0355 | | | | | | | | | | 0.00019 | -0.95460 | Male | | | |
| HEXB | | | | | | | 0.00000 | -0.77814 | Male | | | | | | |
| LLPH | 0.01497 | -0.58302 | Male | | | | | | | | | | | | |
| UBE2E2 | | | | | | | 0.00005 | -0.80997 | Male | | | | | | |
| PRSS3 | | | | | | | | | | 0.00000 | 4.08549 | Female | 0.01670 | -1.50691 | Male |
| NPTX2 | | | | | | | 0.00000 | 2.91338 | Female | | | | | | |
| SLC38A3 | 0.03308 | -1.83046 | Male | | | | | | | | | | 0.01552 | 1.73132 | Female |
| PPCS | 0.02265 | -0.89691 | Male | | | | | | | | | | | | |
| C21orf33 | | | | 0.00000 | 0.92607 | Female | | | | | | | | | |
| IFITM3 | | | | | | | | | | 0.00000 | 1.09687 | Female | 0.00000 | 1.14456 | Female |
| PREB | | | | | | | 0.00001 | -0.65440 | Male | | | | | | |
| FAM127B | 0.02218 | 1.35944 | Female | 0.00021 | 1.22658 | Female | 0.00005 | 0.92729 | Female | | | | | | |
| FGF23 | | | | 0.02600 | -0.95004 | Male | | | | 0.00746 | -0.76667 | Male | 0.00248 | -0.83098 | Male |

| | | | | | | | | | | | | | | | |
|---|---|---|---|---|---|---|---|---|---|---|---|---|---|---|---|
| OTUD5 | | | | 0.00010 | 1.05212 | Female | 0.00029 | 0.69068 | Female | | | | | | |
| HIST1H2BE | | | | | | | | | | 0.00215 | 1.63868 | Female | | | |
| HINT3 | 0.04081 | 1.25206 | Female | | | | | | | | | | | | |
| ZYG11A | | | | | | | | | | 0.00557 | 0.64298 | Female | | | |
| PHF2 | 0.01101 | -2.49582 | Male | | | | | | | | | | | | |
| FADS1 | | | | 0.03258 | -0.99029 | Male | | | | 0.00000 | -1.29066 | Male | | | |
| TSPYL4 | | | | | | | 0.00269 | -1.15191 | Male | | | | | | |
| RGS5 | | | | | | | | | | | | | 0.00000 | -1.86928 | Male |
| GPR176 | | | | | | | | | | 0.00026 | 1.73133 | Female | | | |
| CBFB | | | | | | | | | | | | | 0.00010 | -0.58733 | Male |
| KLRG2 | | | | 0.00035 | 1.15040 | Female | | | | 0.00060 | 0.74038 | Female | | | |
| SLC30A1 | | | | 0.00263 | 1.65528 | Female | | | | | | | | | |
| MAPRE2 | | | | | | | 0.00581 | 1.03931 | Female | | | | | | |
| FAM9B | | | | 0.00509 | 3.22259 | Female | | | | | | | | | |
| RHBDD1 | | | | | | | 0.00159 | -1.15155 | Male | | | | | | |
| OXA1L | 0.01471 | -0.58269 | Male | | | | | | | | | | | | |
| FAM120C | | | | 0.00062 | 1.40673 | Female | 0.00038 | 1.01799 | Female | | | | 0.01265 | 0.70448 | Female |
| SURF1 | | | | | | | | | | 0.01017 | -0.60909 | Male | | | |
| LGALS3BP | | | | | | | | | | | | | 0.01641 | 1.57708 | Female |
| EGFR | | | | | | | 0.00023 | -2.56109 | Male | | | | | | |
| CENPM | | | | | | | | | | 0.00529 | 0.63570 | Female | | | |
| DXO | | | | | | | | | | | | | 0.01340 | 0.71586 | Female |
| PPP1R35 | | | | | | | 0.00107 | -0.97617 | Male | | | | | | |
| CXorf38 | | | | | | | | | | 0.02200 | 0.60170 | Female | | | |
| SLC44A3 | | | | | | | 0.00145 | -2.21958 | Male | | | | | | |
| CD70 | | | | | | | | | | 0.00225 | 2.80962 | Female | | | |
| SPAG5-AS1 | | | | | | | 0.00381 | 0.80078 | Female | 0.01735 | -0.64935 | Male | 0.03310 | 0.58713 | Female |
| GPR39 | | | | | | | | | | | | | 0.00015 | -3.48925 | Male |
| CD151 | 0.02522 | -1.09646 | Male | 0.00000 | 1.34585 | Female | | | | | | | | | |
| CELF6 | | | | | | | | | | 0.02542 | -1.54951 | Male | | | |
| KLK7 | 0.00273 | 3.84075 | Female | | | | | | | | | | | | |
| EFNA4 | | | | | | | | | | 0.00953 | 0.58283 | Female | | | |
| SUSD5 | | | | | | | | | | 0.00514 | 2.67760 | Female | | | |
| C8orf31 | | | | | | | 0.01238 | 1.84152 | Female | | | | 0.00001 | 2.85295 | Female |
| IDO1 | | | | | | | 0.02534 | 2.24577 | Female | | | | | | |
| SNHG9 | | | | | | | 0.00000 | 0.93412 | Female | | | | 0.00000 | 0.75218 | Female |
| HAUS1 | 0.00178 | 1.13018 | Female | | | | | | | | | | | | |
| CGB | | | | | | | | | | 0.00000 | -2.91123 | Male | | | |
| S100A16 | 0.02761 | -0.89333 | Male | 0.00000 | 1.00986 | Female | | | | | | | | | |
| ASRGL1 | | | | | | | | | | 0.00000 | 0.77797 | Female | | | |
| CTBP1-AS | | | | | | | | | | | | | 0.01340 | -1.12171 | Male |
| THSD7A | | | | | | | | | | | | | 0.00004 | -3.71429 | Male |
| CCDC142 | | | | 0.00696 | -1.07916 | Male | | | | 0.02497 | -0.60061 | Male | | | |
| MTRNR2L8 | | | | | | | | | | 0.00000 | 1.32022 | Female | | | |
| IFT43 | | | | 0.01047 | 1.29270 | Female | | | | | | | | | |
| CDA | | | | | | | | | | 0.00000 | 3.40254 | Female | | | |
| ZNF160 | | | | | | | | | | 0.03814 | 0.60402 | Female | | | |
| DNMT3L | 0.00477 | -0.87300 | Male | 0.00000 | 1.26444 | Female | | | | 0.00000 | 0.73879 | Female | | | |
| TUBB2B | | | | 0.00252 | 1.63057 | Female | | | | 0.00000 | 1.47386 | Female | | | |
| SH3TC1 | | | | 0.00000 | 2.65476 | Female | | | | | | | | | |
| CAPZA2 | 0.00019 | 1.28785 | Female | | | | | | | | | | | | |

| | | | | | | | | | | | | | | | |
|---|---|---|---|---|---|---|---|---|---|---|---|---|---|---|---|
| C9orf135 | | | | | | | | | | 0.00338 | 1.75596 | Female | 0.03003 | -1.32456 | Male |
| ZNF350-AS1 | | | | | | | 0.00003 | 2.33626 | Female | | | | | | |
| NOA1 | | | | | | | | | | | | | 0.00001 | -0.60873 | Male |
| SPNS3 | | | | | | | 0.04443 | 1.84254 | Female | 0.00780 | 2.25262 | Female | | | |
| DENND6A | | | | 0.02089 | 1.35321 | Female | | | | | | | | | |
| LOC100294362 | | | | | | | | | | | | | 0.02250 | 1.30804 | Female |
| C11orf83 | | | | | | | 0.00000 | 0.72759 | Female | 0.00000 | 0.91117 | Female | | | |
| MDP1 | | | | | | | | | | 0.00072 | 1.11759 | Female | | | |
| RND3 | | | | | | | | | | 0.03357 | -1.99547 | Male | | | |
| ZBED6 | | | | | | | 0.00000 | -2.07381 | Male | | | | | | |
| SH3D19 | | | | 0.00702 | 1.38278 | Female | | | | | | | | | |
| ABCB10 | 0.00011 | -2.26911 | Male | | | | | | | | | | | | |
| DNAJC19 | 0.00015 | -0.90402 | Male | | | | | | | | | | | | |
| ERBB3 | 0.02274 | 1.18308 | Female | | | | | | | | | | | | |
| STAG2 | | | | 0.00008 | 0.70975 | Female | 0.00000 | 0.78938 | Female | | | | | | |
| DALRD3 | | | | | | | 0.01655 | -0.65457 | Male | | | | | | |
| HMGN3 | | | | 0.00107 | 1.17779 | Female | | | | | | | 0.00084 | -0.72985 | Male |
| FBXL3 | | | | | | | | | | | | | 0.00505 | -0.77708 | Male |
| MID1IP1 | | | | 0.00000 | 2.40293 | Female | | | | 0.02153 | 0.58964 | Female | | | |
| CDS2 | 0.01112 | -1.20864 | Male | | | | | | | 0.00053 | -0.63194 | Male | | | |
| EIF1AD | | | | | | | 0.00000 | 0.58930 | Female | | | | | | |
| EGLN3 | | | | 0.00000 | 1.27556 | Female | | | | 0.00012 | 0.73298 | Female | | | |
| NDUFAF6 | 0.03913 | 1.18124 | Female | 0.02402 | 0.68907 | Female | | | | | | | | | |
| MIR6764 | | | | | | | 0.00244 | 0.76216 | Female | | | | | | |
| ST6GALNAC3 | | | | 0.00033 | 3.19448 | Female | | | | 0.00074 | 2.33309 | Female | | | |
| GOLPH3 | | | | | | | | | | | | | 0.00000 | -0.59498 | Male |
| LOC100130093 | | | | | | | 0.00159 | 0.78964 | Female | | | | | | |
| MIR663A | | | | | | | | | | 0.00000 | 3.93251 | Female | | | |
| C10orf32 | | | | | | | | | | 0.00008 | -0.77097 | Male | | | |
| TOP1MT | 0.02366 | -0.97836 | Male | | | | | | | | | | | | |
| RSU1 | 0.00180 | -1.10921 | Male | | | | | | | | | | | | |
| FLNA | | | | 0.00000 | 1.91172 | Female | | | | | | | | | |
| DSC2 | 0.00001 | 1.91749 | Female | 0.00058 | 0.91522 | Female | | | | | | | | | |
| MIR17HG | | | | 0.00027 | -1.05873 | Male | | | | | | | | | |
| H2AFX | | | | 0.00392 | 0.69629 | Female | | | | | | | | | |
| EMC2 | | | | | | | 0.00001 | -0.69562 | Male | | | | | | |
| EMX1 | | | | | | | 0.03734 | 1.25958 | Female | | | | | | |
| RBM38 | | | | | | | 0.00000 | 0.66499 | Female | | | | | | |
| STARD4 | | | | | | | 0.03911 | -0.75220 | Male | | | | | | |
| PITPNM2 | | | | | | | | | | 0.04707 | 1.22785 | Female | | | |
| IQCH-AS1 | | | | 0.00528 | 1.28401 | Female | | | | | | | | | |
| IGSF9 | | | | | | | | | | | | | 0.04656 | 0.87687 | Female |
| LYPD3 | | | | 0.03947 | -1.27989 | Male | | | | 0.00000 | 1.86345 | Female | | | |
| CLIC4 | 0.00105 | -0.86126 | Male | | | | | | | | | | | | |
| S100A9 | | | | | | | | | | | | | 0.01663 | 2.29450 | Female |
| SPARC | | | | | | | | | | 0.00000 | 1.42447 | Female | | | |
| SREBF1 | | | | | | | 0.00993 | -0.71912 | Male | | | | | | |
| SUZ12P1 | | | | | | | | | | 0.01448 | 0.65387 | Female | | | |
| KLF8 | 0.00162 | 2.41552 | Female | 0.01200 | 1.18328 | Female | 0.02594 | 0.72744 | Female | 0.00005 | 1.16369 | Female | | | |
| PDZD11 | | | | 0.00000 | 1.01931 | Female | 0.00000 | 0.83731 | Female | 0.00000 | 0.74159 | Female | | | |
| LMO7 | | | | | | | | | | | | | 0.00393 | -1.24493 | Male |

| | | | | | | | | | | | | | | | |
|---|---|---|---|---|---|---|---|---|---|---|---|---|---|---|---|
| PCBP1 | 0.00006 | -0.85928 | Male | | | | | | | | | | | | |
| FZD7 | | | | | | | | | | 0.00000 | 4.08769 | Female | | | |
| PJA1 | | | | | | | | | | | | | 0.00003 | -1.63796 | Male |
| ATRX | | | | 0.00001 | 0.84558 | Female | | | | 0.00000 | 0.68471 | Female | | | |
| SNHG21 | | | | | | | | | | 0.01326 | 0.61066 | Female | | | |
| NDUFS8 | 0.03061 | 0.58521 | Female | | | | | | | | | | | | |
| MREG | 0.01700 | 2.17592 | Female | | | | 0.04823 | -0.74603 | Male | | | | | | |
| C12orf57 | | | | 0.00010 | -0.77847 | Male | | | | | | | | | |
| NHSL1 | | | | 0.00020 | 0.88830 | Female | | | | | | | | | |
| APEX2 | | | | 0.02610 | 0.62059 | Female | 0.00000 | 0.77040 | Female | | | | | | |
| ZNF138 | | | | | | | 0.00007 | -0.74190 | Male | | | | | | |
| CCDC57 | 0.01174 | -1.93576 | Male | 0.03859 | 1.24533 | Female | | | | | | | 0.01902 | 0.88357 | Female |
| E2F5 | | | | | | | | | | 0.04930 | -1.13272 | Male | | | |
| POLR2G | 0.00021 | 0.80558 | Female | | | | | | | | | | | | |
| CRYBA2 | | | | 0.02519 | 1.73437 | Female | 0.01639 | 1.25790 | Female | | | | 0.00043 | 1.65049 | Female |
| TNIP2 | | | | 0.00171 | 0.88725 | Female | | | | | | | | | |
| NOS2 | | | | 0.00584 | -2.23894 | Male | | | | | | | | | |
| ELOVL6 | | | | | | | 0.00000 | -0.83808 | Male | | | | | | |
| PDE1A | | | | | | | 0.00549 | -2.68535 | Male | | | | | | |
| TM4SF5 | | | | | | | | | | 0.00020 | 1.58299 | Female | 0.00096 | 1.43368 | Female |
| GATA3 | | | | 0.00018 | 1.05485 | Female | | | | | | | | | |
| RASSF8 | | | | | | | | | | | | | 0.01984 | -0.58158 | Male |
| AIF1L | 0.00057 | -1.06164 | Male | 0.00120 | 0.62365 | Female | | | | | | | | | |
| MAP1B | | | | | | | | | | 0.00058 | 1.45750 | Female | | | |
| CD19 | | | | | | | | | | 0.04871 | 1.38189 | Female | | | |
| KPNA5 | 0.01280 | -2.04727 | Male | | | | | | | | | | | | |
| PRAMEF19.uid-865 | | | | | | | | | | | | | 0.00002 | 0.83415 | Female |
| MOB1B | | | | | | | | | | 0.00006 | -0.73165 | Male | 0.00001 | -0.77202 | Male |
| FBXW5 | 0.00009 | -1.51511 | Male | | | | | | | | | | | | |
| FBXW4 | | | | 0.04634 | 1.21367 | Female | | | | | | | | | |
| ESD | | | | 0.00277 | 0.95283 | Female | | | | | | | | | |
| MIR4668 | | | | | | | 0.00222 | -0.69209 | Male | 0.01019 | 0.62088 | Female | | | |
| HIST1H2BF | | | | 0.00139 | -2.35547 | Male | | | | | | | | | |
| C19orf48 | 0.00497 | 0.67630 | Female | | | | | | | | | | | | |
| HMHA1 | | | | 0.01686 | 0.86458 | Female | | | | | | | 0.00265 | -0.65656 | Male |
| FAM174B | | | | | | | | | | 0.00498 | 1.88095 | Female | | | |
| SUN1 | | | | | | | | | | 0.00006 | 0.64877 | Female | | | |
| PPFIBP2 | | | | | | | | | | | | | 0.00006 | 0.71810 | Female |
| NCRNA00185 | | | | | | | 0.00073 | -3.00643 | Male | 0.00000 | -3.98578 | Male | 0.00000 | -4.28839 | Male |
| ARHGAP17 | | | | 0.01663 | 0.74860 | Female | | | | | | | | | |
| SGSH | | | | | | | 0.00313 | -1.16947 | Male | | | | | | |
| EIF4EBP2 | 0.00002 | -1.34657 | Male | | | | | | | | | | | | |
| SMIM1 | | | | 0.03651 | -1.57785 | Male | | | | | | | | | |
| CDCA7 | 0.00002 | -1.97614 | Male | | | | | | | | | | | | |
| C1orf64 | | | | | | | 0.02550 | 2.41118 | Female | 0.01399 | 2.33370 | Female | 0.00000 | -4.40564 | Male |
| ZDHHC2 | | | | | | | 0.04902 | -1.01410 | Male | | | | | | |
| GTF2IRD1 | | | | | | | | | | 0.00018 | 1.90494 | Female | | | |
| PSME2 | | | | 0.01663 | -0.58700 | Male | | | | | | | | | |
| GTPBP4 | 0.00032 | -0.76468 | Male | | | | | | | | | | | | |
| WNT7A | | | | 0.00000 | 1.03309 | Female | | | | | | | | | |
| LOC100506071 | | | | | | | | | | 0.00000 | -1.29662 | Male | 0.00922 | -0.73123 | Male |

| | | | | | | | | | | | | | | | |
|---|---|---|---|---|---|---|---|---|---|---|---|---|---|---|---|
| NMT2 | | | | | | | | | | 0.00000 | 1.39918 | Female | | | |
| LINC00643 | | | | 0.00000 | 0.94508 | Female | | | | | | | 0.01820 | -1.21481 | Male |
| FOXN4 | | | | | | | | | | | | | 0.01904 | 1.83369 | Female |
| HOXB7 | | | | | | | | | | | | | 0.01253 | -0.89585 | Male |
| LOC101927069 | | | | | | | | | | | | | 0.02836 | 1.42445 | Female |
| FOSB | | | | | | | 0.00000 | -3.09954 | Male | 0.00001 | -2.37949 | Male | | | |
| EIF5A2 | | | | 0.01500 | 0.85518 | Female | 0.00527 | 0.65561 | Female | | | | | | |
| ID3 | | | | | | | 0.00263 | -0.99972 | Male | 0.00664 | 0.87816 | Female | | | |
| LOC728024 | 0.01676 | 0.99187 | Female | 0.01372 | 0.64453 | Female | 0.00000 | 1.10005 | Female | 0.00000 | 0.98910 | Female | 0.00000 | 0.94540 | Female |
| APOOP5 | | | | 0.00004 | 1.11819 | Female | | | | | | | | | |
| NUDT10 | | | | | | | 0.00004 | 2.15311 | Female | 0.01401 | 0.99384 | Female | 0.00119 | 2.05808 | Female |
| C8orf37-AS1 | | | | | | | | | | 0.03032 | -1.08458 | Male | | | |
| FAM214B | | | | | | | | | | 0.00610 | 1.22775 | Female | | | |
| HRAS | | | | | | | | | | 0.00020 | -0.90440 | Male | | | |
| HOXD1 | | | | | | | | | | 0.00149 | 2.61843 | Female | | | |
| HIST1H1E | | | | 0.00032 | -1.83682 | Male | | | | 0.00397 | 1.02653 | Female | | | |
| LOC653513 | | | | | | | | | | 0.00190 | 1.38804 | Female | | | |
| COBLL1 | | | | | | | 0.01012 | -0.69602 | Male | | | | | | |
| KRT8 | | | | 0.00120 | 0.68320 | Female | | | | | | | | | |
| MUC4 | | | | | | | 0.03733 | 1.64197 | Female | | | | | | |
| SLC43A2 | | | | | | | 0.00011 | 0.98883 | Female | 0.00151 | 0.80150 | Female | | | |
| B3GNT7 | | | | 0.00000 | 1.88400 | Female | 0.00000 | 1.23933 | Female | 0.02898 | 0.58901 | Female | 0.00013 | 0.92577 | Female |
| AJAP1 | | | | | | | | | | | | | 0.00323 | 2.93403 | Female |
| ZYG11B | | | | | | | 0.00720 | -0.78916 | Male | | | | | | |
| BTN2A1 | | | | | | | 0.03503 | -0.96997 | Male | | | | | | |
| CAPN5 | | | | | | | | | | 0.04505 | 0.95608 | Female | | | |
| RPS28 | | | | | | | | | | | | | 0.00000 | -0.71416 | Male |
| LOC101928371 | | | | | | | 0.00012 | 2.19669 | Female | | | | | | |
| ZP2 | | | | | | | | | | | | | 0.00003 | -2.16639 | Male |
| DNAH3 | | | | | | | 0.00006 | 1.61788 | Female | | | | 0.00768 | 1.08738 | Female |
| PSMD7 | 0.00000 | -0.85967 | Male | | | | | | | | | | | | |
| PYCARD | | | | | | | | | | 0.00191 | 2.16374 | Female | | | |
| GLA | 0.00016 | 1.32140 | Female | 0.00010 | 0.85378 | Female | | | | | | | | | |
| MAP3K9 | | | | | | | | | | 0.04101 | 0.66730 | Female | | | |
| THOC2 | | | | 0.00000 | 0.94456 | Female | 0.00000 | 1.02230 | Female | 0.00000 | 0.80635 | Female | 0.00000 | 0.78369 | Female |
| RMRP | | | | | | | | | | 0.00002 | 0.88433 | Female | | | |
| TRIM46 | 0.00052 | 4.09968 | Female | | | | | | | | | | | | |
| LOC641367 | | | | | | | | | | 0.00533 | 1.30163 | Female | | | |
| IPP | | | | | | | 0.03602 | -0.58957 | Male | | | | | | |
| TMEM107 | | | | | | | 0.01176 | -0.95471 | Male | | | | | | |
| KMT2B | | | | | | | | | | | | | 0.00076 | 0.64935 | Female |
| FGFBP2 | | | | | | | | | | | | | 0.00858 | 1.82739 | Female |
| GAB2 | | | | | | | 0.00469 | -0.71643 | Male | | | | 0.00582 | 0.66417 | Female |
| OVOL1 | | | | 0.00000 | 1.77437 | Female | | | | | | | 0.02288 | 0.62603 | Female |
| ANKRD36 | | | | | | | | | | 0.00053 | 0.95998 | Female | | | |
| SLC15A4 | | | | 0.00562 | 1.07548 | Female | | | | | | | | | |
| DUSP4 | | | | | | | 0.03698 | 0.99053 | Female | | | | | | |
| PTGDS | | | | | | | 0.03742 | 1.47291 | Female | | | | | | |
| DEFB122 | | | | | | | | | | 0.00000 | 2.42756 | Female | | | |
| PRUNE2 | | | | | | | | | | 0.00345 | 1.49264 | Female | | | |
| BCL2L12 | 0.03391 | 0.65170 | Female | | | | | | | | | | | | |

| | | | | | | | | | | | | | | | |
|---|---|---|---|---|---|---|---|---|---|---|---|---|---|---|---|
| MSMB | | | | | | | | | | 0.00000 | 2.18016 | Female | 0.00113 | 1.55975 | Female |
| SH2D4A | | | | | | | | | | 0.00304 | -0.68004 | Male | | | |
| MIPOL1 | | | | | | | 0.03999 | -1.29220 | Male | | | | | | |
| MOCOS | | | | | | | 0.00694 | -1.77417 | Male | 0.00135 | -1.98832 | Male | | | |
| EPHB2 | | | | 0.00001 | 3.23612 | Female | | | | 0.03914 | 1.20177 | Female | | | |
| COMMD10 | 0.00979 | -1.28945 | Male | | | | | | | | | | | | |
| CITED2 | | | | | | | 0.00087 | 0.84607 | Female | 0.01997 | -0.59293 | Male | | | |
| LRAT | | | | | | | | | | | | | 0.00200 | -2.43670 | Male |
| PRKAR1B | | | | | | | 0.00116 | 0.85356 | Female | | | | | | |
| ZC3HAV1 | | | | | | | | | | 0.02297 | 0.73527 | Female | 0.04456 | -0.65327 | Male |
| FA2H | | | | | | | 0.00734 | -0.97414 | Male | | | | 0.00000 | -1.74918 | Male |
| GLRX3 | 0.00000 | -0.91739 | Male | | | | | | | | | | | | |
| LOC100507291 | | | | | | | | | | 0.04103 | 0.61949 | Female | | | |
| RSAD1 | | | | | | | | | | 0.04762 | 0.60464 | Female | | | |
| CHPT1 | | | | | | | 0.00001 | -0.75516 | Male | | | | | | |
| MRPL36 | 0.00100 | 0.79734 | Female | | | | 0.00000 | 0.65832 | Female | | | | | | |
| ATAT1 | 0.04446 | 1.42855 | Female | | | | | | | | | | | | |
| HAP1 | | | | | | | | | | 0.00152 | -1.42202 | Male | | | |
| SEC61A2 | | | | | | | | | | 0.01754 | 0.60244 | Female | 0.01187 | 0.61893 | Female |
| UCHL1 | | | | | | | | | | 0.00023 | -1.23993 | Male | | | |
| LOC100131564 | 0.04780 | -2.34327 | Male | | | | | | | | | | | | |
| MOSPD2 | | | | | | | | | | 0.00185 | 0.80704 | Female | | | |
| SLITRK1 | | | | | | | | | | 0.03659 | -1.56220 | Male | | | |
| PIGU | | | | | | | 0.00927 | -0.69377 | Male | | | | | | |
| RPS4X | | | | 0.00000 | 0.59722 | Female | | | | 0.00000 | 0.73626 | Female | 0.00000 | 0.90349 | Female |
| HMGN4 | | | | | | | | | | 0.00000 | -0.60727 | Male | | | |
| CMTM7 | 0.00477 | -1.69356 | Male | 0.00000 | 1.70383 | Female | | | | | | | | | |
| FAM13A | | | | | | | | | | 0.00116 | -1.71129 | Male | | | |
| BCKDHA | | | | | | | | | | | | | 0.00019 | 0.76174 | Female |
| PPP2R2A | 0.03273 | -0.72610 | Male | | | | | | | | | | | | |
| MKRN3 | | | | | | | | | | | | | 0.02428 | 0.68793 | Female |
| ACADVL | | | | 0.02039 | 0.75555 | Female | | | | | | | | | |
| RAPGEF1 | | | | 0.00004 | 1.37793 | Female | | | | | | | | | |
| PRCP | 0.00052 | 2.43290 | Female | | | | | | | | | | | | |
| DDAH2 | | | | | | | 0.00005 | -1.16973 | Male | | | | | | |
| AHNAK | | | | | | | 0.00021 | -0.67059 | Male | | | | | | |
| UTF1 | | | | | | | | | | 0.00017 | 2.47101 | Female | | | |
| CEACAM6 | | | | | | | | | | 0.00014 | 2.55502 | Female | | | |
| NTNG1 | | | | | | | | | | 0.00293 | 2.03108 | Female | | | |
| TOM1L2 | | | | | | | | | | | | | 0.02018 | 1.21745 | Female |
| TXNIP | 0.04780 | 1.62035 | Female | | | | | | | 0.03548 | 0.60242 | Female | | | |
| RAI14 | | | | 0.00021 | 1.02235 | Female | | | | | | | | | |
| BCLAF1 | 0.00155 | -0.66089 | Male | | | | | | | | | | | | |
| G3BP1 | 0.00105 | -0.73635 | Male | | | | | | | | | | | | |
| CEBPD | | | | | | | 0.00053 | -1.36605 | Male | | | | | | |
| SNORD58C | | | | | | | | | | 0.03948 | 0.67817 | Female | | | |
| BEX2 | 0.00012 | 2.78865 | Female | | | | | | | 0.00000 | -1.52764 | Male | | | |
| WNK3 | | | | 0.02326 | 1.29857 | Female | 0.00003 | 1.48407 | Female | 0.00000 | 1.67813 | Female | 0.00064 | 1.17916 | Female |
| GDAP1 | | | | | | | | | | 0.00583 | -1.28932 | Male | | | |
| ANKRD55 | | | | | | | | | | 0.00034 | 1.65701 | Female | | | |
| KLHL3 | | | | | | | | | | 0.01654 | 2.03392 | Female | | | |

| | | | | | | | | | | | | | | | |
|---|---|---|---|---|---|---|---|---|---|---|---|---|---|---|---|
| STIM1 | | | | | | | 0.01749 | -0.77950 | Male | | | | | | |
| STEAP3 | | | | 0.00188 | -1.64672 | Male | | | | | | | | | |
| DAZL | | | | | | | | | | 0.00129 | -2.21201 | Male | | | |
| LINC00707 | | | | | | | | | | | | | 0.04084 | 1.23488 | Female |
| UBASH3B | | | | | | | | | | 0.00003 | 1.04963 | Female | | | |
| POU6F1 | | | | | | | | | | 0.02518 | 1.55632 | Female | | | |
| LOC339166 | | | | | | | 0.00027 | 3.35654 | Female | | | | | | |
| PDPN | | | | | | | 0.01058 | -0.92146 | Male | 0.00498 | 0.95217 | Female | | | |
| ARFGAP3 | | | | | | | | | | 0.00030 | 0.90183 | Female | | | |
| MED12L | | | | | | | | | | 0.00000 | -0.88297 | Male | | | |
| CPT2 | | | | | | | 0.00081 | -1.07588 | Male | | | | | | |
| REST | | | | | | | | | | 0.00000 | 0.76459 | Female | | | |
| LILRB3 | 0.03671 | 2.57484 | Female | | | | | | | | | | | | |
| SCAMP3 | 0.00000 | 1.65787 | Female | 0.00085 | 0.73189 | Female | | | | | | | | | |
| POLM | | | | | | | 0.02188 | -0.74528 | Male | | | | | | |
| TGFB1 | | | | | | | | | | 0.02801 | 1.23623 | Female | | | |
| TMBIM1 | | | | | | | | | | 0.00000 | 1.30368 | Female | | | |
| HBA1 | | | | | | | 0.00256 | 1.02232 | Female | | | | | | |
| MAEL | | | | | | | | | | 0.00000 | 1.83231 | Female | 0.00051 | 0.99264 | Female |
| UBE3B | | | | | | | | | | | | | 0.00060 | -0.62859 | Male |
| TNNT1 | | | | | | | | | | 0.00049 | 1.60463 | Female | | | |
| SMAGP | | | | 0.00000 | 1.96964 | Female | | | | | | | | | |
| SHROOM2 | | | | 0.00000 | 2.35228 | Female | | | | | | | | | |
| FAM133A | | | | | | | 0.00000 | 3.06431 | Female | | | | | | |
| MICB | 0.00211 | -2.92662 | Male | | | | | | | | | | | | |
| KNG1 | | | | | | | 0.00000 | -1.47920 | Male | | | | | | |
| 1.00000 | 0.00001 | -1.04603 | Male | | | | | | | | | | | | |
| TARDBP | 0.00124 | 0.62340 | Female | | | | | | | | | | | | |
| PLOD1 | | | | | | | 0.00050 | -0.84867 | Male | | | | | | |
| ITPRIPL2 | | | | | | | | | | 0.02268 | -0.97147 | Male | | | |
| HIST1H2BG | | | | 0.00694 | -1.84447 | Male | 0.00000 | 2.23570 | Female | 0.02373 | 1.07397 | Female | | | |
| CYP19A1 | 0.04291 | -1.70028 | Male | | | | 0.02240 | 1.14577 | Female | | | | | | |
| DOK1 | | | | | | | 0.02234 | -1.53867 | Male | | | | | | |
| LPCAT1 | | | | 0.01441 | 0.91931 | Female | | | | | | | | | |
| NLRP8 | | | | | | | 0.02568 | 2.61490 | Female | | | | 0.01227 | 4.31542 | Female |
| PSKH1 | | | | | | | 0.01671 | -1.18803 | Male | | | | | | |
| UCHL5 | | | | 0.00059 | 0.71791 | Female | | | | | | | | | |
| PLD6 | | | | | | | 0.02187 | -1.28815 | Male | | | | | | |
| UPK2 | | | | | | | 0.00183 | -1.94915 | Male | | | | | | |
| ZNF385A | | | | | | | 0.00283 | 0.97624 | Female | 0.00890 | 0.83470 | Female | 0.02204 | 0.74350 | Female |
| SH3GLB2 | | | | 0.00000 | 2.16776 | Female | | | | 0.00003 | 1.01099 | Female | | | |
| NACC2 | | | | 0.02343 | 1.61010 | Female | | | | | | | | | |
| EPHA7 | | | | | | | | | | 0.03492 | 1.74487 | Female | | | |
| CD47 | | | | | | | 0.00000 | -1.34441 | Male | | | | | | |
| TTC38 | | | | | | | | | | 0.03480 | -0.95409 | Male | | | |
| GNL1 | 0.01314 | 0.86397 | Female | | | | | | | | | | | | |
| LIMK1 | | | | | | | 0.01204 | -0.93838 | Male | | | | | | |
| EEFSEC | | | | | | | | | | 0.02116 | 0.72499 | Female | | | |
| LOC101928378 | | | | 0.03835 | 1.18366 | Female | | | | | | | 0.04059 | 0.73119 | Female |
| SNORD32A | | | | | | | 0.00121 | 0.74896 | Female | 0.01112 | 0.59517 | Female | | | |
| MYOF | | | | 0.00000 | 1.21739 | Female | | | | | | | | | |

| | | | | | | | | | | | | | | | |
|---|---|---|---|---|---|---|---|---|---|---|---|---|---|---|---|
| CREG1 | | | | | | | 0.00000 | -1.35186 | Male | | | | | | |
| KIFC2 | | | | | | | | | | 0.04064 | 1.07769 | Female | | | |
| WDR47 | | | | | | | | | | 0.00305 | 0.63059 | Female | | | |
| KRT17 | | | | | | | | | | | | | 0.00000 | 4.60549 | Female |
| RAB13 | | | | | | | 0.00624 | -0.81174 | Male | | | | | | |
| BRD8 | | | | | | | | | | | | | 0.00000 | 0.70641 | Female |
| ANXA2P2 | 0.00217 | 0.99573 | Female | 0.00014 | -0.69763 | Male | | | | | | | | | |
| JAG1 | | | | | | | 0.01772 | 1.04914 | Female | 0.00013 | 1.49374 | Female | | | |
| EPHX1 | | | | | | | | | | 0.00001 | 1.27137 | Female | | | |
| TTC9 | | | | | | | | | | | | | 0.04989 | 0.88041 | Female |
| CCDC71 | | | | | | | 0.03429 | -1.46684 | Male | 0.00000 | -3.07586 | Male | | | |
| ANKS4B | | | | | | | | | | | | | 0.00652 | -2.33969 | Male |
| ABCD4 | | | | | | | | | | 0.00077 | 0.90502 | Female | 0.01383 | 0.68573 | Female |
| MYO1E | | | | | | | | | | 0.02740 | -0.58990 | Male | | | |
| GALNT8 | | | | | | | | | | 0.03918 | 1.20421 | Female | | | |
| TCF7L1 | | | | 0.00000 | 1.92499 | Female | | | | | | | 0.01843 | 0.60538 | Female |
| RASGRP2 | 0.00279 | -1.44274 | Male | | | | | | | | | | | | |
| KRT18 | | | | 0.00017 | 0.79724 | Female | | | | | | | | | |
| MAFA | | | | | | | 0.03094 | 1.02490 | Female | | | | | | |
| FAM149B1 | | | | | | | | | | | | | 0.00245 | -0.65323 | Male |
| LZTR1 | 0.00003 | -3.17266 | Male | | | | | | | | | | | | |
| TRABD2A | | | | | | | 0.00001 | 2.85949 | Female | | | | | | |
| EIF5AL1 | | | | | | | | | | | | | 0.00000 | -1.02198 | Male |
| LOC101927258 | | | | | | | | | | 0.00326 | 1.69742 | Female | | | |
| MAP1S | | | | | | | 0.00436 | -0.59018 | Male | | | | | | |
| DAZAP1 | 0.00222 | -1.13786 | Male | 0.00032 | 0.75981 | Female | | | | | | | | | |
| CES2 | | | | 0.00049 | 1.19401 | Female | | | | 0.00302 | -0.69013 | Male | 0.00440 | 0.66215 | Female |
| CRYM | | | | | | | | | | 0.00215 | 2.34948 | Female | | | |
| MUC21 | | | | | | | | | | 0.03808 | 1.78631 | Female | | | |
| SRSF8 | | | | 0.01231 | 1.22038 | Female | | | | | | | | | |
| PMEL | | | | 0.01284 | -1.48393 | Male | 0.03586 | -0.88336 | Male | | | | | | |
| C15orf65 | | | | | | | | | | 0.00731 | 1.36068 | Female | | | |
| ZNF782 | 0.02853 | -2.67295 | Male | | | | | | | | | | | | |
| GSE1 | | | | 0.00000 | 1.02404 | Female | | | | | | | | | |
| ODAM | | | | | | | | | | 0.00000 | -1.33628 | Male | 0.03051 | -0.64975 | Male |
| SCAND2P | | | | | | | 0.01396 | -0.86805 | Male | | | | | | |
| HLTF | | | | | | | | | | 0.00568 | 0.74915 | Female | | | |
| NPM2 | | | | | | | | | | | | | 0.01919 | -1.30003 | Male |
| KCMF1 | | | | 0.00000 | 1.35766 | Female | | | | | | | | | |
| MAP7 | | | | 0.01467 | 0.71677 | Female | | | | | | | | | |
| THY1 | | | | | | | 0.00049 | 2.90034 | Female | 0.00197 | 2.51496 | Female | | | |
| HOMEZ | | | | 0.00387 | -2.09670 | Male | | | | | | | | | |
| SOCS5 | | | | | | | 0.02081 | 0.86509 | Female | | | | | | |
| S100A6 | | | | 0.03442 | 0.81458 | Female | | | | | | | | | |
| POLR3G | | | | 0.03507 | -0.69038 | Male | 0.00533 | -0.58430 | Male | | | | 0.00027 | -0.68941 | Male |
| YIPF6 | | | | 0.00000 | 1.05689 | Female | 0.00000 | 0.74186 | Female | 0.00000 | 0.61585 | Female | 0.00000 | 0.61467 | Female |
| TOR3A | 0.00477 | -1.32719 | Male | | | | | | | | | | 0.00075 | -0.61710 | Male |
| TNRC6C | | | | | | | 0.02038 | 0.83536 | Female | | | | | | |
| CRIM1 | | | | | | | | | | 0.00008 | -0.81173 | Male | | | |
| COBL | | | | 0.00714 | 0.82799 | Female | | | | | | | | | |
| LUZP6+MTPN | | | | | | | | | | 0.00000 | -0.62038 | Male | | | |

| | | | | | | | | | | | | | | | |
|---|---|---|---|---|---|---|---|---|---|---|---|---|---|---|---|
| DPH3P1 | 0.03830 | -0.76511 | Male | | | | | | | | | | | | |
| TLE4 | | | | | | | | | | 0.00012 | -0.88449 | Male | | | |
| FAM91A1 | | | | | | | | | | | | | 0.00004 | -0.62001 | Male |
| HBA2 | | | | | | | 0.00002 | 1.93275 | Female | | | | | | |
| PIBF1 | | | | | | | | | | | | | 0.00000 | -1.02207 | Male |
| ALDH3A2 | | | | | | | | | | 0.00179 | 0.60340 | Female | | | |
| RDH10 | | | | 0.01307 | -0.76108 | Male | | | | | | | | | |
| SPTBN2 | | | | | | | | | | 0.00267 | -1.33625 | Male | | | |
| SEPT14 | | | | | | | | | | | | | 0.00522 | -2.27579 | Male |
| SAMD4A | | | | | | | | | | | | | 0.00899 | -2.05990 | Male |
| TPCN2 | 0.03519 | -1.65438 | Male | | | | 0.00805 | -0.82000 | Male | | | | | | |
| MIR492 | | | | | | | 0.00402 | 0.60705 | Female | | | | | | |
| H19 | | | | | | | 0.04069 | -0.77141 | Male | | | | | | |
| DYNC1LI1 | 0.00241 | -0.87494 | Male | | | | | | | | | | | | |
| ANTXR2 | | | | 0.03023 | 2.20420 | Female | | | | | | | | | |
| NEB | | | | | | | | | | 0.00392 | 0.89399 | Female | | | |
| SEPT12 | | | | | | | | | | 0.01138 | 1.24299 | Female | 0.00020 | 1.67913 | Female |
| CCDC87 | | | | | | | | | | | | | 0.04541 | 1.57738 | Female |
| ZSCAN22 | | | | | | | | | | | | | 0.01248 | 0.98880 | Female |
| SOX2-OT | | | | | | | | | | 0.01913 | 1.48075 | Female | | | |
| ATXN1 | | | | | | | | | | 0.02940 | 1.47581 | Female | | | |
| DHX33 | | | | | | | 0.00000 | 0.73186 | Female | | | | | | |
| ALDH1B1 | | | | 0.00000 | 3.02346 | Female | | | | | | | | | |
| SMS | | | | 0.01378 | 0.58053 | Female | | | | | | | | | |
| GPATCH3 | | | | 0.00028 | -0.86919 | Male | | | | | | | | | |
| C11orf70 | | | | | | | | | | 0.00142 | -1.59629 | Male | | | |
| ZDHHC11 | 0.01714 | 1.39007 | Female | | | | 0.00362 | 0.91635 | Female | | | | | | |
| FAM96B | | | | 0.00000 | 0.67486 | Female | | | | | | | | | |
| FAM120B | | | | | | | 0.03444 | -0.93126 | Male | | | | | | |
| GRIN2A | | | | | | | | | | 0.04305 | 1.56284 | Female | | | |
| SLCO4C1 | | | | | | | | | | | | | 0.00012 | -1.00636 | Male |
| MRS2 | 0.00000 | -1.81590 | Male | | | | | | | | | | | | |
| WBP5 | 0.00108 | 1.24363 | Female | 0.00000 | 1.09907 | Female | | | | 0.00000 | -0.93489 | Male | 0.00000 | -1.05267 | Male |
| NLRP7 | | | | 0.00000 | 0.75240 | Female | | | | | | | | | |
| PHF7 | | | | | | | | | | 0.04424 | -0.80685 | Male | | | |
| RGS3 | | | | 0.03817 | 1.42913 | Female | | | | | | | | | |
| KLF16 | | | | | | | 0.00447 | 0.60404 | Female | | | | | | |
| FRZB | | | | | | | 0.00106 | -4.11027 | Male | | | | | | |
| GADD45A | | | | | | | 0.00208 | 0.77376 | Female | | | | 0.00839 | 0.64020 | Female |
| SLC35A5 | | | | | | | 0.00003 | -0.82213 | Male | | | | | | |
| NFYC | 0.00014 | -1.00735 | Male | | | | | | | | | | | | |
| SLC8B1 | 0.00002 | 2.47399 | Female | 0.03859 | 0.77012 | Female | | | | 0.00001 | 0.94310 | Female | 0.00868 | 0.59383 | Female |
| PARD3B | | | | | | | 0.00000 | -2.89010 | Male | | | | | | |
| LRRC32 | | | | | | | | | | 0.00033 | 2.46855 | Female | | | |
| CELSR3 | | | | | | | 0.04075 | -1.00304 | Male | | | | | | |
| BTG1 | | | | | | | | | | 0.00003 | -0.64252 | Male | | | |
| LOC101929397 | | | | | | | | | | | | | 0.02119 | 1.39068 | Female |
| MTURN | | | | | | | 0.04261 | -1.61434 | Male | | | | | | |
| CCDC120 | | | | 0.00135 | 1.56293 | Female | 0.02953 | 0.78641 | Female | | | | | | |
| DERL3 | | | | | | | | | | 0.01584 | 1.12145 | Female | 0.01321 | 1.12445 | Female |
| LEUTX | | | | | | | | | | 0.00168 | 6.17071 | Female | | | |

| | | | | | | | | | | | | | | | |
|---|---|---|---|---|---|---|---|---|---|---|---|---|---|---|---|
| TOB1 | | | | | | | 0.00008 | 1.27151 | Female | | | | 0.01143 | 0.82282 | Female |
| SLC47A1 | | | | | | | | | | 0.00000 | 2.99780 | Female | | | |
| SLC25A30 | | | | | | | | | | | | | 0.02799 | 1.00233 | Female |
| NCMAP | | | | | | | | | | 0.00173 | -1.03656 | Male | 0.00000 | 1.47679 | Female |
| SLC12A3 | | | | 0.00001 | 1.40430 | Female | | | | 0.00200 | 0.69437 | Female | | | |
| BRD3 | | | | | | | 0.00000 | 0.83357 | Female | | | | | | |
| LOC101929718 | | | | | | | 0.00746 | 0.68172 | Female | | | | | | |
| TEAD3 | | | | | | | | | | 0.00204 | -0.81462 | Male | | | |
| TP53RK | | | | | | | | | | | | | 0.00000 | -0.64006 | Male |
| EEF1A2 | | | | | | | | | | | | | 0.00000 | 2.97390 | Female |
| TRMT6 | 0.00418 | -0.70372 | Male | | | | | | | | | | | | |
| BAAT | | | | | | | 0.02818 | -2.27453 | Male | | | | | | |
| CXADRP3 | | | | 0.00754 | 0.71369 | Female | | | | | | | | | |
| MGAT2 | | | | | | | 0.02320 | -0.61754 | Male | | | | | | |
| ZDHHC20 | | | | | | | 0.00013 | -0.64250 | Male | | | | | | |
| ADORA3 | | | | | | | | | | 0.03352 | 1.90203 | Female | | | |
| PLCB4 | | | | | | | | | | 0.00007 | 2.26740 | Female | | | |
| ZNF280A | | | | | | | | | | 0.00000 | 2.60259 | Female | 0.00000 | -1.68276 | Male |
| ECM1 | 0.00077 | 4.10138 | Female | 0.02674 | -1.98531 | Male | | | | 0.00015 | 2.06517 | Female | 0.00094 | 1.82616 | Female |
| ATP6V1B1 | | | | 0.00000 | 2.88888 | Female | | | | | | | | | |
| SLC7A4 | | | | 0.00000 | 3.45268 | Female | | | | | | | 0.00259 | 0.76905 | Female |
| SEPHS1 | 0.00651 | -0.72344 | Male | | | | | | | | | | | | |
| H1F0 | | | | 0.04536 | -0.58970 | Male | 0.00000 | 1.44727 | Female | | | | 0.00000 | 0.97052 | Female |
| PDE4C | | | | | | | | | | 0.00093 | 3.21629 | Female | | | |
| PRKAB2 | | | | | | | | | | 0.00036 | 0.96703 | Female | | | |
| TESK2 | | | | | | | | | | 0.01223 | 0.86991 | Female | | | |
| CCND2 | | | | | | | | | | | | | 0.00191 | -2.03534 | Male |
| KLHL5 | | | | | | | | | | | | | 0.01494 | -1.08713 | Male |
| RNF11 | | | | | | | | | | | | | 0.00027 | -0.63964 | Male |
| SUMO3 | 0.00013 | -1.02789 | Male | | | | | | | | | | | | |
| SLC27A4 | | | | 0.00067 | 0.82522 | Female | | | | | | | | | |
| TMEM42 | | | | | | | | | | 0.01537 | -1.37102 | Male | | | |
| SH3BP5-AS1 | | | | | | | 0.00002 | -1.80367 | Male | | | | | | |
| WDTC1 | | | | | | | | | | 0.00859 | 0.66624 | Female | | | |
| NAPRT1 | | | | | | | 0.00110 | -0.94296 | Male | | | | | | |
| CKMT1A | | | | | | | 0.00188 | 1.27187 | Female | | | | 0.00023 | 1.33167 | Female |
| RAD9A | | | | | | | | | | | | | 0.00452 | 0.66316 | Female |
| MIR3661 | | | | | | | 0.00017 | -0.79535 | Male | | | | | | |
| MOB3B | | | | 0.00000 | 1.27566 | Female | | | | | | | | | |
| RAB17 | | | | 0.00001 | 3.50244 | Female | | | | 0.00004 | -2.13889 | Male | | | |
| ZNF773 | | | | | | | | | | | | | 0.01794 | 0.66953 | Female |
| LINC00341 | | | | | | | | | | 0.01325 | -1.73696 | Male | | | |
| HENMT1 | | | | | | | 0.00062 | -0.67398 | Male | | | | | | |
| SLC44A1 | | | | 0.00000 | 1.57729 | Female | | | | | | | | | |
| GLS | | | | | | | | | | 0.00149 | -0.63609 | Male | | | |
| SCOC | | | | | | | 0.00039 | -0.63293 | Male | | | | 0.00002 | -0.69526 | Male |
| AP3B2 | | | | | | | | | | 0.03245 | -1.80978 | Male | 0.03127 | 1.77017 | Female |
| ARPC1A | 0.02391 | -0.61170 | Male | | | | | | | | | | | | |
| LOC729218 | | | | | | | | | | 0.04581 | 0.70736 | Female | | | |
| EPS8 | | | | | | | | | | | | | 0.04337 | -0.84743 | Male |
| N4BP2 | | | | | | | | | | 0.00000 | 0.70042 | Female | | | |

| | | | | | | | | | | | | | | | |
|---|---|---|---|---|---|---|---|---|---|---|---|---|---|---|---|
| NBPF10 | | | | | | | 0.00071 | 1.07950 | Female | | | | | | |
| FAM151A | | | | | | | 0.02363 | 0.74353 | Female | 0.00000 | 2.77893 | Female | | | |
| STRIP2 | | | | | | | | | | | | | 0.02190 | 0.65721 | Female |
| POLD4 | | | | | | | | | | 0.00518 | 1.06016 | Female | | | |
| HDAC1 | | | | 0.00002 | -0.64686 | Male | | | | | | | | | |
| TYSND1 | | | | | | | 0.01998 | -0.80176 | Male | | | | | | |
| ASPH | | | | | | | | | | | | | 0.00262 | -0.66156 | Male |
| RIC3 | | | | | | | | | | | | | 0.04523 | 1.01936 | Female |
| NXT2 | | | | | | | | | | 0.00746 | 0.71659 | Female | | | |
| HCAR2 | | | | | | | | | | 0.00614 | 1.70726 | Female | | | |
| SLC7A1 | 0.01011 | -3.14678 | Male | 0.00147 | -2.71952 | Male | 0.00415 | -1.75833 | Male | | | | | | |
| SPG20 | | | | 0.01181 | -0.79524 | Male | 0.00001 | -0.85488 | Male | 0.00036 | -0.68423 | Male | | | |
| RBM10 | | | | 0.00004 | 0.77545 | Female | | | | | | | | | |
| IRAK4 | 0.00064 | 3.01683 | Female | | | | 0.01447 | -0.98360 | Male | | | | | | |
| PAQR5 | | | | | | | 0.01049 | -1.52679 | Male | | | | | | |
| B2M | | | | | | | | | | 0.04926 | 0.90340 | Female | | | |
| CD55 | | | | | | | 0.00000 | -1.17545 | Male | | | | | | |
| FGFR4 | | | | 0.04036 | -0.69421 | Male | 0.00001 | -0.87134 | Male | | | | | | |
| SLC30A9 | | | | | | | | | | | | | 0.00060 | -0.60532 | Male |
| TRIM4 | | | | | | | 0.00748 | -1.14748 | Male | | | | | | |
| DPPA3 | | | | | | | 0.00000 | -0.58725 | Male | | | | | | |
| DYNC1H1 | 0.00027 | 1.15653 | Female | | | | | | | | | | | | |
| GNAI2 | | | | | | | | | | 0.00000 | 1.03890 | Female | | | |
| ZP3 | | | | | | | | | | 0.00000 | -1.77672 | Male | 0.00098 | 1.22059 | Female |
| DOCK3 | | | | | | | | | | 0.00138 | 1.12752 | Female | | | |
| ALOX12 | | | | | | | | | | 0.01608 | -1.03586 | Male | | | |
| LPHN1 | | | | | | | | | | 0.00003 | 1.10496 | Female | 0.00053 | 0.92981 | Female |
| CCRN4L | | | | | | | 0.00000 | 1.35609 | Female | | | | | | |
| MINOS1P1 | | | | | | | | | | 0.01140 | -1.09686 | Male | | | |
| OTUD1 | | | | | | | | | | 0.00036 | 1.24886 | Female | | | |
| ALPK3 | | | | | | | | | | | | | 0.02009 | 1.24507 | Female |
| C6orf132 | | | | 0.00067 | 0.83040 | Female | | | | | | | | | |
| TMEM133 | | | | 0.00089 | -2.49959 | Male | | | | | | | | | |
| THAP1 | | | | | | | | | | 0.00018 | -0.60746 | Male | | | |
| SNORA57 | | | | 0.00007 | -1.18117 | Male | | | | | | | | | |
| AIFM1 | | | | 0.00000 | 0.93080 | Female | 0.00000 | 0.83614 | Female | | | | | | |
| ARHGEF4 | | | | | | | 0.00687 | -2.09456 | Male | | | | | | |
| KATNB1 | | | | | | | | | | 0.01182 | 0.69079 | Female | | | |
| LOC646762 | | | | | | | | | | 0.03426 | -0.76185 | Male | | | |
| SUV420H2 | | | | 0.00016 | 2.03697 | Female | | | | | | | | | |
| DPP4 | | | | | | | | | | 0.00220 | 1.48231 | Female | | | |
| MRPL55 | | | | | | | 0.00000 | 0.59650 | Female | | | | | | |
| HOMER2 | | | | | | | | | | 0.00000 | 2.06388 | Female | | | |
| ARID1B | | | | 0.00024 | 1.00701 | Female | | | | | | | | | |
| BVES | | | | | | | 0.04699 | 2.05757 | Female | | | | | | |
| P2RX4 | | | | 0.00803 | 1.49891 | Female | | | | | | | | | |
| SENP3 | | | | | | | | | | | | | 0.00047 | -1.48627 | Male |
| NDST1 | | | | | | | 0.00168 | -0.67650 | Male | | | | | | |
| RNF207 | | | | 0.00536 | 1.78201 | Female | | | | | | | | | |
| KLF11 | | | | | | | | | | 0.00016 | 1.19464 | Female | 0.00014 | 1.17832 | Female |
| MMP9 | | | | | | | | | | 0.02373 | 1.79783 | Female | | | |

| | | | | | | | | | | | | | | | |
|---|---|---|---|---|---|---|---|---|---|---|---|---|---|---|---|
| TREML2 | | | | 0.00112 | -1.83727 | Male | | | | | | | | | |
| LRRC61 | | | | 0.02144 | 0.92920 | Female | | | | 0.00342 | -0.72435 | Male | | | |
| GBAS | | | | | | | 0.00559 | -0.71522 | Male | | | | | | |
| IRX4 | | | | | | | 0.02937 | -1.11054 | Male | | | | | | |
| TPTE2P1 | | | | | | | | | | 0.00007 | 1.96732 | Female | | | |
| NEFL | | | | | | | | | | 0.00964 | 1.36910 | Female | | | |
| LAS1L | 0.00023 | 1.03320 | Female | 0.00000 | 0.92379 | Female | 0.00000 | 1.70413 | Female | 0.00000 | 0.82325 | Female | 0.00000 | 1.06904 | Female |
| SNHG17 | | | | | | | | | | 0.00000 | 0.63402 | Female | | | |
| PKD1L2 | | | | 0.03743 | 1.42338 | Female | | | | | | | | | |
| VPS18 | | | | | | | 0.00143 | -0.70288 | Male | | | | | | |
| MED10 | | | | | | | | | | 0.00000 | -0.62422 | Male | | | |
| CYP27B1 | 0.00930 | -2.60524 | Male | | | | | | | | | | | | |
| TMEM8A | | | | | | | | | | 0.00042 | 1.01525 | Female | | | |
| LYPLA1 | | | | | | | | | | | | | 0.00000 | -0.59215 | Male |
| STAT3 | | | | | | | 0.00000 | 0.91717 | Female | | | | | | |
| LOC100128164 | | | | | | | | | | 0.04744 | -1.17714 | Male | | | |
| PAPOLG | | | | | | | 0.00166 | -0.73886 | Male | | | | | | |
| TMED1 | | | | | | | 0.00010 | 0.65401 | Female | | | | | | |
| CGA | | | | 0.00000 | -4.63811 | Male | 0.02006 | -1.19357 | Male | 0.00000 | -3.16567 | Male | 0.00883 | 1.39356 | Female |
| ALOXE3 | | | | | | | 0.00178 | 4.58746 | Female | | | | | | |
| PUS10 | | | | | | | | | | | | | 0.02952 | -0.75801 | Male |
| C5orf24 | | | | | | | | | | 0.00017 | -0.72536 | Male | | | |
| LAMB2 | | | | | | | 0.04835 | 0.96745 | Female | | | | | | |
| CBWD5.uid-11734 | | | | 0.00083 | 0.71166 | Female | 0.00000 | 0.79295 | Female | | | | | | |
| LARP4B | | | | | | | 0.00000 | 0.65316 | Female | | | | | | |
| RAB34 | | | | 0.03092 | -1.27361 | Male | 0.00000 | -2.16844 | Male | 0.00004 | -1.32565 | Male | | | |
| MICU2 | | | | | | | 0.00000 | -0.93676 | Male | | | | 0.00000 | -0.66309 | Male |
| PFKFB4 | | | | 0.04508 | 0.79550 | Female | | | | | | | | | |
| TRMT10B | | | | 0.03870 | 1.08167 | Female | | | | | | | | | |
| NOS3 | | | | | | | 0.00005 | 1.32173 | Female | 0.00475 | 0.98486 | Female | 0.00001 | 1.37778 | Female |
| PAX8-AS1.uid-3717 | | | | | | | 0.00478 | 0.73375 | Female | | | | | | |
| SWSAP1 | | | | | | | 0.04516 | 0.60312 | Female | | | | | | |
| MAPKAP1 | | | | 0.00234 | 0.78425 | Female | | | | | | | | | |
| IFT172 | | | | | | | | | | 0.00354 | 0.98736 | Female | | | |
| PEMT | 0.00170 | 1.64370 | Female | 0.00000 | 1.64122 | Female | | | | 0.00000 | 1.09077 | Female | | | |
| LOC101929694 | | | | | | | 0.02612 | 2.26663 | Female | | | | | | |
| RYR3 | | | | | | | 0.01538 | -1.19553 | Male | | | | | | |
| EDIL3 | | | | | | | | | | 0.04142 | -1.66071 | Male | | | |
| QTRT1 | | | | | | | 0.00102 | 0.58431 | Female | | | | | | |
| UBXN11 | | | | | | | | | | | | | 0.02135 | 0.65061 | Female |
| SNORD49A | | | | 0.01973 | -0.77689 | Male | | | | | | | | | |
| IL1R1 | | | | | | | 0.04912 | -1.24741 | Male | | | | | | |
| NDUFB2 | | | | 0.00000 | 0.79115 | Female | | | | | | | | | |
| TDH | | | | | | | | | | 0.00106 | -2.10192 | Male | | | |
| GRSF1 | | | | | | | 0.00000 | -0.58481 | Male | | | | | | |
| LOC100289230 | | | | | | | | | | | | | 0.00707 | 1.58282 | Female |
| PPP1R13L | 0.00290 | 1.99979 | Female | | | | | | | | | | | | |
| MROH6 | | | | | | | 0.00019 | 1.34657 | Female | | | | | | |
| HUS1 | | | | | | | | | | 0.00000 | 0.74581 | Female | | | |
| SLC35E2 | | | | | | | | | | 0.00544 | -0.83538 | Male | | | |
| AKT1S1 | 0.00733 | 1.02409 | Female | | | | | | | | | | | | |

| | | | | | | | | | | | | | | | |
|---|---|---|---|---|---|---|---|---|---|---|---|---|---|---|---|
| WWOX | | | | | | | 0.01914 | 0.70248 | Female | | | | | | |
| ARL8B | 0.03076 | -0.59245 | Male | | | | | | | | | | | | |
| RBMY2FP | 0.04045 | -3.61808 | Male | 0.00000 | -5.70147 | Male | 0.00049 | -2.90832 | Male | 0.00977 | -2.23021 | Male | 0.00001 | -3.55604 | Male |
| LRIG1 | | | | | | | | | | 0.00068 | 1.41836 | Female | | | |
| ZNF48 | | | | | | | | | | 0.00886 | 1.04855 | Female | 0.00855 | 1.03539 | Female |
| SPHK1 | | | | | | | 0.01271 | 1.00449 | Female | 0.00001 | 1.57861 | Female | | | |
| EMP2 | | | | | | | 0.00001 | -0.91992 | Male | 0.00000 | -1.00402 | Male | | | |
| FAM9C | | | | 0.02120 | 1.62968 | Female | | | | | | | | | |
| LEMD3 | | | | | | | 0.03608 | -0.64090 | Male | | | | | | |
| AQP6 | | | | | | | | | | | | | 0.02988 | 0.61426 | Female |
| TRERF1 | | | | | | | 0.01369 | -0.63439 | Male | | | | | | |
| MAGEA4 | | | | 0.02641 | 1.37498 | Female | 0.00012 | 1.47855 | Female | | | | 0.00000 | 2.22244 | Female |
| PRAMEF4 | | | | | | | | | | 0.00001 | 6.62271 | Female | | | |
| C19orf67 | | | | | | | | | | | | | 0.00488 | -2.75999 | Male |
| GLYATL1 | | | | | | | | | | | | | 0.00121 | -6.16333 | Male |
| ACACB | | | | | | | | | | 0.01895 | 1.64463 | Female | | | |
| OTUB2 | | | | | | | 0.02074 | 1.79359 | Female | | | | | | |
| SNORD16 | | | | 0.01189 | -1.17813 | Male | | | | | | | | | |
| VMA21 | | | | 0.00001 | 0.71230 | Female | 0.00000 | 0.58693 | Female | | | | | | |
| LOXL2 | | | | | | | | | | | | | 0.01369 | 1.25439 | Female |
| CTDSPL | | | | | | | | | | 0.00037 | -0.93438 | Male | | | |
| RPS6KL1 | | | | | | | | | | | | | 0.00005 | 1.56353 | Female |
| SCN2A | 0.01175 | 2.81435 | Female | | | | | | | | | | | | |
| C19orf44 | | | | | | | | | | 0.00566 | -1.10035 | Male | | | |
| DDX21 | 0.00002 | -0.74541 | Male | | | | | | | | | | | | |
| TRAM1L1 | | | | | | | | | | | | | 0.00189 | -0.77051 | Male |
| CYHR1 | | | | | | | | | | | | | 0.00147 | 0.78052 | Female |
| TMSB4X | | | | | | | | | | | | | 0.00000 | 0.58113 | Female |
| BMPR1A | | | | 0.02101 | 0.65670 | Female | 0.00002 | -0.73871 | Male | | | | 0.00000 | -0.79898 | Male |
| TUBA4B | | | | | | | | | | 0.04762 | -0.86885 | Male | 0.00997 | -1.04943 | Male |
| C14orf37 | | | | | | | | | | | | | 0.02975 | 1.32696 | Female |
| KIAA0825 | | | | 0.02725 | -1.03848 | Male | | | | 0.00138 | -0.93512 | Male | 0.00139 | -0.88537 | Male |
| FAM69B | | | | 0.01188 | 1.12664 | Female | 0.00680 | 0.79685 | Female | 0.00037 | 0.96572 | Female | | | |
| SLC38A8 | | | | | | | 0.00585 | 2.14436 | Female | | | | | | |
| ITPR2 | | | | | | | | | | 0.04356 | 1.02210 | Female | | | |
| PCNA | 0.00023 | -0.75403 | Male | | | | | | | | | | | | |
| MSX1 | | | | | | | 0.00501 | 1.68513 | Female | | | | 0.00004 | 2.20671 | Female |
| HSPA6 | | | | | | | | | | 0.00002 | -3.98997 | Male | | | |
| FLNB | | | | 0.00006 | 0.83643 | Female | | | | | | | | | |
| BAIAP2L1 | | | | | | | 0.00001 | -0.60110 | Male | | | | | | |
| WSB2 | | | | | | | 0.00025 | -0.73393 | Male | | | | | | |
| APOC1P1 | | | | | | | | | | 0.00001 | 1.01959 | Female | | | |
| BAG3 | | | | | | | | | | 0.00359 | -0.66955 | Male | | | |
| COL27A1 | | | | | | | | | | 0.00006 | 1.93680 | Female | | | |
| FAM83E | | | | | | | | | | 0.01177 | -1.55708 | Male | | | |
| LIN54 | | | | | | | | | | 0.01523 | -0.58158 | Male | | | |
| CSPG5 | | | | | | | | | | | | | 0.02803 | 1.51002 | Female |
| ANXA2 | 0.03460 | 0.66770 | Female | | | | | | | | | | | | |
| HBZ | | | | | | | | | | | | | 0.01713 | 1.61601 | Female |
| LGALS13 | | | | | | | | | | 0.00267 | -2.13361 | Male | | | |
| OGT | | | | 0.00015 | 0.70275 | Female | | | | 0.00000 | 0.95275 | Female | 0.00000 | 0.66294 | Female |

| | | | | | | | | | | | | | | | |
|---|---|---|---|---|---|---|---|---|---|---|---|---|---|---|---|
| CXCL14 | | | | | | | | | | | | | 0.01806 | -1.70949 | Male |
| TCF7L2 | | | | | | | | | | 0.00034 | -0.62362 | Male | | | |
| MCM3AP-AS1 | | | | | | | 0.02680 | -0.78027 | Male | | | | | | |
| EPB41L2 | | | | | | | | | | 0.00007 | 0.91523 | Female | | | |
| UST | | | | | | | | | | | | | 0.01254 | -1.05932 | Male |
| ZIC2 | | | | | | | | | | | | | 0.01900 | -1.68934 | Male |
| LOC101928099 | | | | | | | | | | 0.01034 | -1.06154 | Male | | | |
| CD44 | 0.00000 | -1.42113 | Male | | | | | | | | | | | | |
| HP1BP3 | 0.02265 | -0.75804 | Male | | | | | | | | | | | | |
| EPB41L5 | | | | | | | | | | 0.00000 | 0.79799 | Female | | | |
| PMPCA | | | | | | | 0.00000 | 0.67595 | Female | | | | | | |
| LAD1 | | | | 0.00199 | 1.15537 | Female | | | | | | | | | |
| C11orf95 | | | | | | | 0.00006 | 1.24913 | Female | 0.00656 | 0.86516 | Female | | | |
| RIMS2 | | | | | | | 0.00862 | 2.02336 | Female | | | | | | |
| PNCK | | | | | | | 0.00995 | 2.18043 | Female | | | | | | |
| EIF2S3 | 0.00054 | 0.70312 | Female | 0.00000 | 0.64675 | Female | 0.00000 | 1.07476 | Female | | | | | | |
| GABBR2 | | | | | | | | | | 0.01230 | 0.99137 | Female | | | |
| PTGR2 | | | | | | | | | | | | | 0.04643 | 0.75970 | Female |
| NOL4 | | | | | | | 0.03051 | 2.97679 | Female | | | | | | |
| CA2 | 0.00009 | -1.45726 | Male | 0.01433 | 0.62153 | Female | | | | | | | | | |
| SCPEP1 | | | | | | | | | | 0.00000 | 1.13300 | Female | | | |
| ATAD3A | | | | 0.00000 | 1.05149 | Female | | | | | | | | | |
| TCERG1 | 0.01539 | -0.59171 | Male | | | | | | | | | | | | |
| RHOF | | | | | | | 0.03784 | -0.86958 | Male | | | | | | |
| STRADB | | | | | | | 0.00165 | -0.76723 | Male | | | | | | |
| YY1 | | | | | | | 0.00000 | 0.63404 | Female | | | | | | |
| SMPDL3A | 0.01868 | 1.02817 | Female | | | | | | | | | | | | |
| LOC100506730 | | | | | | | 0.01432 | -1.58616 | Male | | | | | | |
| EML3 | | | | | | | 0.00040 | -0.90793 | Male | | | | | | |
| KLHL11 | | | | | | | | | | 0.00000 | 1.74394 | Female | | | |
| ZNF823 | | | | | | | 0.00006 | -0.92833 | Male | | | | | | |
| RSPH10B+RSPH10B2.uid-9484 | | | | | | | 0.04394 | -0.91502 | Male | | | | | | |
| SCO2.uid-2610 | 0.04168 | 1.15205 | Female | | | | | | | | | | | | |
| SCARB2 | | | | | | | | | | 0.00447 | 0.63742 | Female | | | |
| ENTPD5 | | | | | | | | | | | | | 0.00969 | 1.46164 | Female |
| GSTM3 | | | | | | | | | | | | | 0.00262 | 1.13186 | Female |
| CMTM3 | | | | 0.02562 | 1.84065 | Female | | | | | | | | | |
| UPP1 | | | | | | | 0.00000 | 0.87873 | Female | 0.00000 | 1.04249 | Female | 0.00000 | 0.76541 | Female |
| ZHX1 | | | | | | | | | | 0.01070 | 0.76399 | Female | | | |
| LBH | | | | 0.02865 | 1.64104 | Female | | | | 0.00273 | 1.39011 | Female | | | |
| NANOG | | | | | | | 0.03429 | 0.86979 | Female | 0.00000 | 2.22803 | Female | 0.00000 | 1.84032 | Female |
| NABP2 | | | | | | | | | | 0.00000 | -1.15588 | Male | | | |
| MX1 | | | | | | | | | | 0.00040 | 0.97803 | Female | | | |
| PEX5 | | | | | | | | | | | | | 0.00056 | 0.90316 | Female |
| TPD52L1 | | | | 0.04806 | 1.09286 | Female | 0.00310 | -0.91407 | Male | 0.00027 | -1.01940 | Male | 0.02005 | -0.68294 | Male |
| KLK13 | | | | | | | | | | 0.00140 | 2.40690 | Female | | | |
| LOC100507420.uid-10773 | | | | | | | | | | 0.01597 | 1.86434 | Female | 0.04963 | -1.56743 | Male |
| METTL12 | | | | 0.00008 | -0.86587 | Male | | | | | | | | | |
| CPPED1 | | | | | | | | | | 0.00486 | 0.61858 | Female | | | |
| SCAMP1 | | | | | | | | | | 0.00000 | 0.83331 | Female | | | |
| PARP12 | | | | 0.02171 | 2.38023 | Female | | | | 0.00025 | 2.36411 | Female | 0.00329 | 1.91987 | Female |

| | | | | | | | | | | | | | | | |
|---|---|---|---|---|---|---|---|---|---|---|---|---|---|---|---|
| CTAG2 | 0.03393 | 2.66150 | Female | | | | 0.02571 | 1.19646 | Female | | | | 0.00015 | 1.76054 | Female |
| ZNF841 | | | | | | | 0.04451 | 0.73095 | Female | | | | | | |
| HMGCR | | | | 0.00283 | -0.66383 | Male | | | | | | | | | |
| JPH3 | | | | | | | | | | 0.00020 | 1.68900 | Female | | | |
| SPOCK1 | | | | | | | | | | | | | 0.00145 | -2.58235 | Male |
| IQGAP3 | | | | 0.00000 | 1.31083 | Female | | | | 0.00001 | 0.73731 | Female | | | |
| ATG4C | | | | | | | 0.00675 | -0.61435 | Male | | | | | | |
| AQPEP | | | | | | | | | | 0.00056 | 1.80278 | Female | | | |
| REEP1 | 0.00200 | -1.47254 | Male | 0.00000 | 1.52920 | Female | 0.00001 | -0.76294 | Male | | | | | | |
| BRSK1 | | | | | | | 0.00004 | 1.13254 | Female | 0.00000 | -1.57841 | Male | | | |
| ZNF142 | | | | | | | | | | 0.00002 | 0.96404 | Female | | | |
| MCTP1 | | | | | | | | | | | | | 0.00125 | -1.54587 | Male |
| TSPY2 | | | | | | | 0.00000 | -3.57098 | Male | 0.00000 | -2.71574 | Male | 0.00000 | -4.11341 | Male |
| UBE2E4P | 0.00012 | -1.62506 | Male | | | | | | | | | | 0.00003 | -0.58999 | Male |
| MIR205HG | | | | | | | 0.00003 | 1.42232 | Female | | | | | | |
| PATL2 | | | | | | | | | | 0.02471 | -1.69106 | Male | | | |
| MOCS1 | | | | 0.02543 | 1.48569 | Female | | | | | | | | | |
| ALPL | | | | | | | | | | 0.00000 | 1.25925 | Female | 0.00262 | 0.62729 | Female |
| RFX1 | | | | | | | | | | | | | 0.00665 | 0.60621 | Female |
| LAMTOR4 | 0.00116 | 0.80126 | Female | | | | | | | | | | | | |
| NUDT21 | 0.00497 | -0.60365 | Male | | | | | | | | | | | | |
| GDF15 | | | | | | | 0.02291 | 1.03605 | Female | 0.04166 | -0.90783 | Male | | | |
| GTF2B | | | | | | | 0.00000 | -0.72573 | Male | | | | | | |
| MRPS31P5 | | | | | | | | | | 0.00004 | 1.04104 | Female | | | |
| SLC25A42 | | | | | | | | | | | | | 0.04219 | 1.20786 | Female |
| SNORD102 | | | | 0.00316 | -1.09589 | Male | | | | 0.00255 | 0.71566 | Female | | | |
| MBD3L2 | | | | | | | | | | 0.00000 | 7.05651 | Female | | | |
| CDKN1A | | | | | | | 0.02618 | 1.07485 | Female | | | | | | |
| ZFP42 | 0.00008 | -0.87707 | Male | | | | | | | | | | | | |
| SRD5A3 | | | | | | | 0.01463 | -0.68866 | Male | 0.01284 | -0.65976 | Male | 0.00138 | -0.81306 | Male |
| CASP8 | | | | | | | | | | 0.00005 | 1.56327 | Female | | | |
| RARRES2 | | | | | | | | | | 0.00032 | 1.78895 | Female | | | |
| EZH2 | 0.00082 | -0.97667 | Male | | | | | | | | | | | | |
| NRCAM | | | | | | | | | | 0.01341 | 1.68955 | Female | | | |
| DUSP9 | | | | | | | 0.02811 | 1.12540 | Female | | | | 0.00203 | -1.46881 | Male |
| AMOTL1 | | | | | | | 0.00099 | -1.50792 | Male | | | | | | |
| TDRP | | | | 0.00000 | 2.22765 | Female | | | | 0.00092 | -0.82461 | Male | 0.00113 | -0.79598 | Male |
| FYB | | | | | | | 0.00807 | -1.02958 | Male | 0.04179 | -0.74179 | Male | | | |
| FAM83F | | | | 0.03552 | 1.51184 | Female | 0.01512 | 1.10531 | Female | | | | | | |
| SNHG20 | | | | | | | 0.04892 | 0.92142 | Female | | | | | | |
| MAGEA10 | | | | | | | | | | | | | 0.00645 | 1.67071 | Female |
| TSIX | | | | | | | 0.00000 | 3.04874 | Female | 0.04595 | 0.89237 | Female | 0.00000 | 2.82697 | Female |
| WEE1 | 0.00000 | -2.62183 | Male | | | | | | | | | | | | |
| HGS | | | | 0.03372 | -0.58800 | Male | 0.00007 | -0.66559 | Male | | | | | | |
| FZD4 | | | | | | | | | | 0.00684 | 0.98773 | Female | | | |
| LOC441601 | | | | | | | | | | 0.00357 | -0.64181 | Male | | | |
| KDM5C | | | | 0.00001 | 1.29500 | Female | 0.00082 | 0.71006 | Female | 0.00000 | 1.05931 | Female | 0.00000 | 1.06142 | Female |
| FAM102A | | | | | | | 0.01761 | -0.64913 | Male | 0.02581 | 0.58660 | Female | | | |
| CSNK2A2 | | | | | | | 0.00117 | 0.74432 | Female | | | | | | |
| ACRC | 0.00000 | 4.99105 | Female | 0.00010 | 2.94109 | Female | 0.00000 | 2.81186 | Female | 0.00005 | 2.09331 | Female | | | |
| ZNF613 | | | | | | | 0.01810 | 0.99806 | Female | | | | | | |

| | | | | | | | | | | | | | | | |
|---|---|---|---|---|---|---|---|---|---|---|---|---|---|---|---|
| ABCC13 | | | | | | | | | | 0.00000 | 2.91687 | Female | | | |
| ARPC5 | 0.00268 | -0.77083 | Male | | | | | | | | | | | | |
| POU5F1B | 0.00999 | -1.59507 | Male | | | | | | | 0.00000 | 1.77920 | Female | | | |
| TAX1BP3 | | | | 0.00000 | 1.32084 | Female | | | | | | | | | |
| VASH2 | | | | | | | | | | 0.03544 | 1.35819 | Female | | | |
| MYLK-AS1 | | | | | | | 0.00302 | -1.55044 | Male | | | | | | |
| CNNM2 | | | | | | | 0.01722 | 0.83208 | Female | 0.01449 | -0.81218 | Male | | | |
| PWARSN | | | | | | | | | | 0.00003 | 1.04306 | Female | | | |
| ANKRD1 | | | | | | | | | | 0.02090 | 1.10763 | Female | | | |
| NDUFS6 | 0.00003 | 0.79610 | Female | | | | | | | | | | | | |
| LEPROTL1 | | | | | | | | | | | | | 0.00000 | -0.58268 | Male |
| FLJ37453 | | | | | | | 0.00354 | 1.21567 | Female | | | | | | |
| LINC00618 | | | | | | | | | | | | | 0.03770 | 1.78911 | Female |
| COMMD9 | 0.04397 | 0.75660 | Female | | | | | | | | | | | | |
| SPPL2B | | | | | | | 0.00001 | -1.26855 | Male | | | | | | |
| HIST2H2BE | | | | | | | | | | 0.00056 | 1.66811 | Female | | | |
| CEP192 | | | | | | | 0.00601 | 0.61869 | Female | | | | | | |
| ITGB4 | | | | | | | | | | 0.00142 | -1.45278 | Male | 0.01768 | 1.16041 | Female |
| PRC1 | | | | | | | 0.00000 | -0.65172 | Male | | | | | | |
| LOC391003.uid-852 | | | | | | | 0.00001 | 1.26755 | Female | 0.00000 | 1.53196 | Female | | | |
| CXXC5 | 0.03671 | 2.67896 | Female | | | | | | | | | | | | |
| CDC25C | | | | 0.04007 | -0.88468 | Male | 0.02972 | -0.62593 | Male | | | | | | |
| GAD1 | | | | | | | | | | | | | 0.03920 | -1.57975 | Male |
| GCSHP3 | 0.03076 | 1.58363 | Female | | | | | | | | | | | | |
| APOO | | | | 0.00124 | 1.01531 | Female | 0.00721 | 0.60379 | Female | | | | | | |
| CXCR6 | | | | | | | 0.03207 | -1.71642 | Male | | | | | | |
| AOC3 | | | | | | | | | | 0.03537 | -0.86547 | Male | | | |
| SLC44A2 | | | | 0.00332 | 0.60507 | Female | | | | | | | | | |
| ARRDC2 | | | | | | | | | | 0.01062 | 1.02977 | Female | | | |
| SEPT10 | | | | | | | 0.00001 | -1.07143 | Male | | | | | | |
| PSMG3 | 0.00337 | 0.78415 | Female | | | | 0.00000 | 0.79969 | Female | | | | | | |
| USP28 | | | | 0.02470 | 0.60989 | Female | | | | | | | | | |
| E2F6 | | | | | | | 0.00313 | -0.67759 | Male | | | | | | |
| EPAS1 | | | | | | | | | | | | | 0.00772 | 0.90454 | Female |
| NEK7 | | | | | | | 0.03184 | -0.61035 | Male | | | | 0.01581 | -0.62718 | Male |
| RPL13AP20 | | | | 0.00195 | -0.66507 | Male | | | | | | | | | |
| PARP16 | | | | | | | | | | 0.00191 | 1.41113 | Female | | | |
| AAMDC | 0.02218 | 1.32848 | Female | | | | 0.01419 | 0.59309 | Female | 0.00000 | 0.98690 | Female | | | |
| SHPK | | | | | | | | | | 0.00000 | 1.09311 | Female | | | |
| ARHGAP32 | | | | 0.00927 | 1.18754 | Female | | | | | | | | | |
| ZNF708 | | | | 0.01056 | -1.21059 | Male | 0.03703 | -0.70461 | Male | | | | | | |
| TM7SF3 | | | | | | | 0.00163 | -0.87369 | Male | | | | | | |
| PNPO | | | | | | | | | | 0.03920 | 0.74517 | Female | 0.01221 | 0.86200 | Female |
| BRCA2 | | | | | | | | | | | | | 0.00049 | -0.60380 | Male |
| ZSCAN5B | | | | | | | | | | | | | 0.00579 | 1.59029 | Female |
| C1orf56 | | | | 0.00627 | -1.32081 | Male | | | | | | | | | |
| KCNV1 | 0.01821 | 3.33147 | Female | | | | | | | | | | | | |
| FBXL21 | | | | 0.00492 | -2.13178 | Male | 0.00706 | -1.43040 | Male | | | | | | |
| ETNPPL | | | | | | | | | | 0.00000 | 3.20840 | Female | | | |
| IRX3 | | | | | | | | | | | | | 0.00001 | 1.99136 | Female |
| SLC52A2 | | | | 0.01479 | 0.61511 | Female | | | | | | | | | |

| | | | | | | | | | | | | | | | |
|---|---|---|---|---|---|---|---|---|---|---|---|---|---|---|---|
| TIMP3 | | | | | | | | | | | | | 0.02712 | -1.61370 | Male |
| LOC643387 | | | | | | | | | | 0.00000 | -0.59558 | Male | | | |
| ANKRD20A19P | 0.01592 | 3.23334 | Female | 0.04999 | 1.86891 | Female | | | | | | | | | |
| ZFP36L1 | 0.00000 | 2.41357 | Female | 0.04495 | -0.59667 | Male | | | | | | | | | |
| RMND5A | | | | | | | 0.00000 | 0.68222 | Female | | | | | | |
| CTNND1 | | | | | | | 0.00000 | -1.17668 | Male | 0.00004 | -0.69175 | Male | | | |
| HID1 | | | | 0.03815 | -2.28634 | Male | | | | | | | | | |
| GDPD5 | | | | | | | | | | 0.00918 | 0.96722 | Female | | | |
| SEMA4G | | | | | | | | | | 0.00573 | 1.25847 | Female | | | |
| CHIC1 | | | | | | | | | | | | | 0.00014 | -2.74972 | Male |
| RNF122 | | | | | | | | | | | | | 0.03370 | -0.71898 | Male |
| PKP3 | 0.02262 | -1.71508 | Male | | | | | | | | | | | | |
| FSTL1 | | | | 0.00004 | 2.17462 | Female | | | | | | | | | |
| UBAC2 | 0.00751 | -1.05892 | Male | | | | | | | | | | | | |
| LINC00673 | 0.03638 | -2.32022 | Male | | | | | | | 0.00001 | 2.88577 | Female | | | |
| RNF212 | | | | | | | 0.00348 | -0.83144 | Male | | | | 0.00001 | -1.11993 | Male |
| SUV39H1 | | | | | | | 0.00532 | 0.79437 | Female | | | | | | |
| RABAC1 | | | | | | | | | | 0.00000 | 0.62514 | Female | | | |
| SLC16A10 | | | | | | | | | | 0.04024 | 0.85716 | Female | | | |
| HPRT1 | | | | 0.00005 | 0.93186 | Female | | | | | | | 0.00000 | -0.93433 | Male |
| LASP1 | 0.01145 | 0.77782 | Female | | | | | | | | | | | | |
| LZIC | 0.02292 | -0.78594 | Male | | | | | | | | | | | | |
| CELA1 | | | | | | | 0.00594 | 1.29248 | Female | | | | 0.02737 | 1.01340 | Female |
| GLT1D1 | | | | 0.00005 | 2.70478 | Female | | | | 0.03801 | 1.00602 | Female | | | |
| ITM2B | | | | | | | 0.00000 | -0.78905 | Male | | | | 0.00000 | -0.81358 | Male |
| A4GALT | | | | 0.00005 | 0.82022 | Female | | | | | | | | | |
| ST6GAL1 | 0.00000 | -1.62190 | Male | | | | | | | | | | | | |
| EPB41L4A-AS1 | | | | 0.00654 | -0.60370 | Male | | | | | | | | | |
| GPM6A | | | | 0.00000 | -2.32608 | Male | 0.00000 | 1.52798 | Female | 0.01928 | -0.88724 | Male | | | |
| BATF3 | 0.03913 | -1.57718 | Male | | | | | | | 0.03074 | 0.65769 | Female | | | |
| RSRP1 | | | | | | | 0.00017 | 0.66077 | Female | | | | | | |
| ANXA8L1 | | | | | | | | | | 0.00012 | -1.03259 | Male | 0.00929 | 0.72373 | Female |
| BECN1P1 | | | | | | | | | | 0.00000 | 4.12122 | Female | 0.00000 | 1.08415 | Female |
| MMP14 | | | | | | | | | | 0.01634 | -0.98663 | Male | | | |
| EIF2AK2 | | | | | | | 0.00447 | -0.93460 | Male | | | | | | |
| ABLIM3 | | | | | | | | | | 0.01840 | -1.63833 | Male | | | |
| ACAA2 | 0.02987 | -0.77139 | Male | | | | | | | | | | | | |
| TFG | 0.00151 | -0.75998 | Male | | | | | | | | | | | | |
| PNMAL1 | | | | | | | | | | 0.00317 | 1.56331 | Female | | | |
| DYRK4 | | | | | | | | | | 0.00373 | 1.16805 | Female | | | |
| ROCK1P1 | | | | | | | | | | 0.00579 | -1.42279 | Male | 0.02787 | -1.17179 | Male |
| EBP | 0.00017 | 0.87855 | Female | 0.00000 | 0.87029 | Female | 0.00000 | 1.21493 | Female | 0.00000 | 1.03733 | Female | 0.00000 | 0.60136 | Female |
| ADAD1 | | | | | | | | | | 0.00235 | 1.98117 | Female | | | |
| C5orf66-AS1 | | | | | | | | | | 0.04712 | 1.55994 | Female | | | |
| CCDC122 | | | | | | | 0.02081 | -1.23165 | Male | | | | | | |
| FAM65B | | | | | | | | | | | | | 0.04135 | 1.04473 | Female |
| PCAT4 | | | | | | | | | | 0.00031 | 2.93679 | Female | | | |
| ZFP62 | | | | | | | | | | 0.00001 | -1.15971 | Male | | | |
| PAQR8 | | | | | | | 0.00241 | -1.32859 | Male | | | | | | |
| DIP2B | | | | | | | 0.00061 | -0.65521 | Male | 0.00060 | -0.62264 | Male | | | |
| SEPT6 | | | | | | | 0.03742 | 1.62315 | Female | 0.00016 | 2.58010 | Female | | | |

| | | | | | | | | | | | | | | | |
|---|---|---|---|---|---|---|---|---|---|---|---|---|---|---|---|
| ZNF398 | | | | | | | | | | 0.00028 | 0.96603 | Female | | | |
| EIF1AX | 0.00008 | 0.71965 | Female | 0.00000 | 0.86250 | Female | 0.00000 | 0.86788 | Female | 0.00000 | 0.58826 | Female | | | |
| STYK1 | | | | | | | | | | 0.00170 | 2.23771 | Female | | | |
| STK10 | 0.04221 | -1.11612 | Male | | | | | | | | | | 0.00268 | 0.59711 | Female |
| P2RY6 | 0.00057 | 2.65289 | Female | 0.00002 | -1.77736 | Male | | | | 0.03594 | -0.60404 | Male | | | |
| COL4A1 | | | | 0.00000 | 0.83107 | Female | 0.00000 | -3.00033 | Male | 0.00092 | 2.26047 | Female | | | |
| PTDSS2 | | | | 0.00004 | 1.14670 | Female | | | | | | | | | |
| ACE2 | | | | | | | | | | 0.03734 | 0.85007 | Female | | | |
| PSMB10 | 0.01224 | 1.32953 | Female | | | | | | | | | | | | |
| ARHGAP44 | | | | 0.04921 | 1.38226 | Female | | | | | | | | | |
| CAPN6 | | | | | | | | | | 0.00993 | -1.83462 | Male | | | |
| HIST1H2AH | | | | | | | | | | 0.00000 | 2.36239 | Female | | | |
| NUPR1 | | | | | | | | | | | | | 0.00000 | 2.89257 | Female |
| IGBP1 | | | | 0.00041 | 0.61578 | Female | | | | | | | | | |
| SNORA55 | | | | | | | | | | | | | 0.01104 | 1.45211 | Female |
| ZNF519 | | | | | | | | | | | | | 0.01741 | -0.77841 | Male |
| C16orf74 | | | | | | | 0.00155 | -1.12691 | Male | | | | | | |
| G6PC3 | | | | 0.00038 | 1.21099 | Female | | | | | | | | | |
| MYO1G | 0.00000 | 1.65785 | Female | | | | | | | 0.00077 | 2.57661 | Female | | | |
| MDK | | | | | | | | | | 0.00000 | 1.14141 | Female | | | |
| ITM2C | | | | | | | 0.00168 | -0.62621 | Male | 0.00000 | 1.49304 | Female | | | |
| SEC23A | | | | 0.01792 | -0.88200 | Male | | | | | | | | | |
| JUNB | | | | | | | 0.00355 | 0.80506 | Female | | | | | | |
| CSF3R | | | | 0.00000 | 3.35037 | Female | | | | | | | | | |
| CSRNP1 | | | | | | | | | | 0.01725 | 0.75578 | Female | 0.00681 | 0.82634 | Female |
| PARM1 | | | | | | | | | | 0.00115 | 2.14903 | Female | | | |
| LOC286437 | | | | | | | 0.01425 | 1.80694 | Female | | | | 0.03348 | 1.55613 | Female |
| HOXC10 | | | | | | | | | | | | | 0.00000 | 3.21491 | Female |
| CHODL | | | | | | | | | | 0.02039 | 2.04239 | Female | | | |
| PDSS1 | | | | | | | 0.00062 | -0.80659 | Male | | | | | | |
| TMEM31 | | | | | | | | | | 0.01155 | 1.26523 | Female | | | |
| PQLC1 | | | | | | | | | | 0.00000 | 1.88726 | Female | | | |
| ITPR3 | | | | | | | | | | | | | 0.04075 | -0.58653 | Male |
| AZIN2 | | | | | | | | | | 0.00610 | 1.59055 | Female | | | |
| ZRSR2 | | | | | | | 0.00313 | 0.70832 | Female | | | | | | |
| CYP1B1 | 0.00065 | -2.03450 | Male | | | | | | | 0.00242 | 1.18292 | Female | 0.00899 | 1.03569 | Female |
| MMP2 | | | | | | | | | | 0.00000 | 5.44101 | Female | | | |
| CASK | | | | 0.00027 | 1.62933 | Female | | | | | | | | | |
| FOXRED2 | | | | | | | | | | | | | 0.00218 | 0.79712 | Female |
| SNN | | | | | | | | | | | | | 0.00392 | 1.41695 | Female |
| STON2 | | | | 0.00628 | 0.81502 | Female | | | | | | | | | |
| REM1 | | | | | | | | | | 0.00019 | 2.47928 | Female | | | |
| GATA4 | | | | | | | 0.00646 | -2.08075 | Male | | | | | | |
| GSTM4 | 0.02554 | 2.73012 | Female | 0.00003 | 2.53909 | Female | | | | | | | | | |
| SF3A1 | 0.04441 | -0.60309 | Male | | | | | | | | | | | | |
| WDR73 | | | | 0.01654 | -0.62619 | Male | | | | | | | | | |
| OSTM1 | | | | | | | 0.00757 | -0.60080 | Male | | | | | | |
| TSPAN18 | | | | 0.03453 | 1.41975 | Female | | | | 0.00000 | 1.96966 | Female | | | |
| SIGMAR1 | 0.03152 | 1.40276 | Female | | | | | | | | | | | | |
| NNT-AS1 | 0.00000 | -3.20431 | Male | | | | | | | | | | | | |
| PSTPIP2 | | | | | | | | | | | | | 0.00309 | 1.67071 | Female |

| | | | | | | | | | | | | | | | |
|---|---|---|---|---|---|---|---|---|---|---|---|---|---|---|---|
| MAPT | | | | | | | | | | 0.02001 | -1.38991 | Male | | | |
| POU5F1P4 | 0.02391 | -1.31544 | Male | | | | | | | 0.00000 | 1.31947 | Female | | | |
| FMR1NB | | | | 0.00002 | 1.47030 | Female | 0.00003 | 1.02981 | Female | 0.00000 | 1.77483 | Female | 0.00680 | 0.67025 | Female |
| ABHD17C | 0.00603 | -1.47703 | Male | | | | | | | | | | 0.00066 | -0.66829 | Male |
| ALKBH7 | 0.01011 | 0.78668 | Female | | | | | | | | | | | | |
| COPS2 | 0.02338 | -0.68624 | Male | | | | | | | | | | | | |
| CCBL2 | | | | | | | 0.00110 | -0.78431 | Male | | | | | | |
| NME2.uid-2204 | 0.00796 | -0.61755 | Male | | | | | | | | | | | | |
| TMEM106B | | | | | | | 0.00125 | -0.88023 | Male | | | | | | |
| TXNDC12 | | | | 0.00019 | 0.60601 | Female | | | | | | | | | |
| ZDHHC14 | | | | 0.00000 | 3.01134 | Female | | | | | | | | | |
| LOC100506790 | | | | | | | 0.00002 | 2.03742 | Female | 0.00000 | 4.13009 | Female | | | |
| GPM6B | | | | | | | 0.00006 | 1.51374 | Female | | | | | | |
| FAM162A | 0.00010 | 1.06359 | Female | | | | | | | | | | | | |
| ZNF677 | | | | | | | 0.01683 | -0.60810 | Male | | | | | | |
| RNASEK.uid-21 | 0.00279 | 2.31777 | Female | | | | | | | | | | | | |
| NFKBIA | | | | | | | | | | 0.00000 | 0.67770 | Female | | | |
| TMEM92 | | | | | | | | | | 0.01899 | 1.02069 | Female | | | |
| MLK7-AS1 | | | | | | | 0.04599 | 1.05784 | Female | | | | | | |
| ZNF165 | | | | | | | 0.01712 | 0.60613 | Female | | | | | | |
| FAM124A | | | | | | | | | | 0.00000 | 2.67323 | Female | | | |
| CNN3 | 0.03671 | -0.69383 | Male | | | | | | | | | | | | |
| PRR13 | | | | | | | | | | 0.00000 | 0.92895 | Female | | | |
| RAB3D | | | | | | | | | | 0.00016 | 0.91866 | Female | 0.01523 | 0.62096 | Female |
| MUC5B | | | | | | | | | | 0.00890 | 2.02696 | Female | 0.01265 | 1.91079 | Female |
| FRMD4B | | | | | | | | | | | | | 0.00014 | 0.64815 | Female |
| LOC100129138 | | | | | | | 0.04306 | 0.69672 | Female | | | | 0.00079 | -0.95313 | Male |
| CXCL12 | | | | | | | | | | 0.01425 | 1.95673 | Female | | | |
| GNB3 | | | | | | | | | | | | | 0.02135 | 1.02538 | Female |
| PPP1CC | | | | | | | | | | | | | 0.00000 | -0.62360 | Male |
| DCTD | | | | | | | 0.00291 | -0.68718 | Male | 0.00154 | -0.68764 | Male | | | |
| C2CD5 | | | | | | | | | | 0.00627 | 0.86963 | Female | | | |
| SLC20A2 | | | | | | | | | | 0.00376 | -0.79778 | Male | | | |
| EVC2 | | | | | | | | | | | | | 0.03892 | 1.87096 | Female |
| EGFLAM | | | | | | | | | | 0.00000 | 1.80899 | Female | | | |
| DDX56 | 0.00794 | -0.64434 | Male | | | | | | | | | | | | |
| HSD17B3 | | | | | | | | | | 0.00411 | 2.53651 | Female | | | |
| RXRA | | | | | | | | | | 0.03448 | -0.66825 | Male | | | |
| EPM2AIP1 | | | | | | | | | | | | | 0.02415 | 0.63666 | Female |
| UFSP1 | | | | 0.01441 | 2.11938 | Female | | | | 0.03819 | -1.23539 | Male | | | |
| ADCY3 | | | | 0.00739 | -0.88754 | Male | | | | | | | | | |
| C22orf29 | | | | | | | | | | | | | 0.00000 | 1.06492 | Female |
| ID1 | | | | | | | | | | 0.00000 | 1.89915 | Female | 0.00339 | 0.78277 | Female |
| CTNNA2 | | | | | | | | | | | | | 0.00056 | -6.58023 | Male |
| RFC3 | 0.04026 | -0.67263 | Male | | | | | | | | | | | | |
| SDCBP | 0.00419 | 1.10173 | Female | 0.01400 | 0.60231 | Female | | | | | | | | | |
| DKC1 | | | | 0.00000 | 0.67080 | Female | 0.00000 | 0.76250 | Female | | | | 0.00000 | 0.68908 | Female |
| FUT1 | | | | | | | 0.04170 | -1.03975 | Male | | | | | | |
| CDX2 | | | | | | | | | | 0.02387 | 0.66398 | Female | | | |
| ALG13 | | | | 0.00001 | 0.81977 | Female | 0.00000 | 0.66479 | Female | | | | 0.00000 | 0.65955 | Female |
| SLK | 0.03947 | -0.90803 | Male | 0.01271 | 0.62621 | Female | | | | | | | | | |

| | | | | | | | | | | | | | | | |
|---|---|---|---|---|---|---|---|---|---|---|---|---|---|---|---|
| NQO2 | | | | 0.02534 | 0.73934 | Female | | | | 0.00000 | 1.26018 | Female | | | |
| CD38 | | | | | | | | | | 0.00000 | 3.19014 | Female | | | |
| PBX1 | | | | | | | | | | 0.00009 | 1.88351 | Female | | | |
| HTT | | | | | | | | | | 0.00187 | 0.65995 | Female | | | |
| ACKR2 | | | | 0.00475 | -1.48309 | Male | 0.00000 | -1.95086 | Male | | | | | | |
| RELL1 | 0.01948 | -0.79587 | Male | | | | | | | | | | | | |
| TAF12 | 0.02218 | -0.75649 | Male | | | | | | | | | | | | |
| SNAR-F | 0.00000 | -4.58776 | Male | 0.02370 | 1.31771 | Female | 0.00000 | 2.58889 | Female | 0.01098 | 0.94388 | Female | | | |
| CGB7 | | | | 0.04848 | -1.85775 | Male | | | | 0.00001 | -2.40895 | Male | | | |
| ERCC6L | 0.01036 | 1.25611 | Female | | | | 0.00000 | 0.87573 | Female | 0.00000 | 0.98921 | Female | 0.00200 | 0.58886 | Female |
| ARAP1 | | | | | | | 0.01913 | -0.60473 | Male | | | | | | |
| FGF4 | | | | | | | | | | 0.00851 | 2.44255 | Female | | | |
| HORMAD1 | | | | | | | | | | | | | 0.02456 | -0.66879 | Male |
| HECA | | | | | | | 0.00161 | -0.98986 | Male | | | | | | |
| TPM1 | | | | 0.00040 | 0.76786 | Female | 0.00000 | -0.95130 | Male | 0.00000 | -0.86314 | Male | | | |
| MARK1 | 0.03268 | 1.85006 | Female | | | | | | | | | | | | |
| CBS | 0.04717 | -1.46406 | Male | 0.00000 | -2.22041 | Male | 0.00001 | -1.10737 | Male | 0.00783 | -0.68342 | Male | | | |
| MAGI2-AS3 | | | | | | | 0.04019 | 1.96284 | Female | | | | | | |
| FRMD8 | | | | | | | 0.00000 | -1.05877 | Male | | | | | | |
| IL23R | | | | | | | | | | | | | 0.03770 | -1.67375 | Male |
| CPSF2 | 0.00603 | -0.94414 | Male | | | | | | | | | | | | |
| PITPNA-AS1 | | | | | | | | | | | | | 0.00203 | 0.64913 | Female |
| FADS2 | | | | 0.00000 | -2.45613 | Male | 0.00000 | -1.73408 | Male | 0.00059 | -1.03194 | Male | 0.01554 | -0.74745 | Male |
| FRRS1 | | | | | | | 0.04918 | -0.96632 | Male | | | | | | |
| MAZ | 0.00578 | -0.77476 | Male | | | | | | | | | | | | |
| LINC00909 | | | | | | | | | | 0.00664 | -0.97610 | Male | | | |
| SVOPL | | | | | | | | | | 0.03753 | 1.16636 | Female | | | |
| GAGE2A.uid-12821 | | | | 0.00033 | 2.79296 | Female | 0.00000 | 7.36993 | Female | | | | | | |
| KANTR | | | | | | | | | | 0.00000 | -1.62648 | Male | | | |
| LETMD1 | | | | | | | 0.00761 | -0.63571 | Male | | | | | | |
| FOXRED1 | 0.00000 | 1.79149 | Female | | | | | | | | | | | | |
| TMEM14A | | | | | | | | | | | | | 0.00000 | -0.87730 | Male |
| SMEK3P | | | | | | | 0.00922 | 1.82912 | Female | | | | | | |
| TLE6 | | | | | | | | | | | | | 0.00012 | 2.23114 | Female |
| NEK8 | | | | 0.00045 | 1.94006 | Female | | | | | | | | | |
| PCDH9 | | | | 0.03624 | -2.06493 | Male | | | | | | | | | |
| BOLA2+BOLA2B+LOC388242+LOC613038+SLX1A+SLX1A-SULT1A3+SLX1B+ | | | | | | | 0.00000 | 0.89701 | Female | | | | | | |
| VOPP1 | | | | | | | 0.00264 | -0.84827 | Male | | | | | | |
| ACTA2 | | | | | | | | | | 0.00000 | -2.11332 | Male | | | |
| IGFBP2 | | | | | | | | | | 0.00378 | 1.03595 | Female | | | |
| MS4A15 | | | | 0.00000 | 2.97443 | Female | | | | | | | | | |
| AARS2 | | | | | | | | | | 0.00011 | 0.82203 | Female | | | |
| TRMT2B | | | | 0.00027 | 1.58037 | Female | 0.02610 | 0.72442 | Female | | | | | | |
| LOC728323 | | | | | | | 0.00384 | -0.76605 | Male | | | | | | |
| ZNF589 | 0.00796 | 1.12691 | Female | | | | | | | | | | | | |
| GSTZ1 | 0.02218 | 1.25541 | Female | | | | | | | | | | | | |
| CD83 | | | | | | | | | | 0.00213 | 1.89575 | Female | | | |
| MARC2 | | | | | | | 0.00006 | -2.54394 | Male | 0.00319 | -1.86629 | Male | | | |
| TULP2 | | | | | | | | | | 0.00082 | 1.31106 | Female | | | |
| DOCK9 | | | | 0.01139 | 1.06340 | Female | | | | 0.00893 | -0.63623 | Male | | | |
| SEC24B | | | | | | | 0.00128 | -0.62146 | Male | | | | | | |

| | | | | | | | | | | | | | | | |
|---|---|---|---|---|---|---|---|---|---|---|---|---|---|---|---|
| IPO7 | | | | | | | | | | 0.00000 | -0.67413 | Male | | | |
| CACUL1 | | | | | | | 0.00000 | 0.80436 | Female | | | | | | |
| PCBP3 | | | | | | | | | | 0.03012 | 1.96651 | Female | 0.00306 | 2.44783 | Female |
| RRP15 | 0.00392 | -0.70666 | Male | | | | | | | | | | | | |
| HMBS | 0.00009 | 1.94241 | Female | | | | | | | | | | | | |
| ME1 | | | | | | | | | | 0.00885 | -1.75934 | Male | | | |
| CRKL | | | | | | | 0.00000 | 0.66403 | Female | | | | | | |
| ATAD3B | | | | | | | 0.00000 | 0.58970 | Female | | | | | | |
| NKX3-1 | | | | 0.02960 | 1.02341 | Female | 0.00000 | 1.91552 | Female | 0.01091 | -0.77504 | Male | | | |
| SPATA17 | | | | | | | | | | 0.01049 | 1.16353 | Female | | | |
| IFITM2 | | | | 0.00000 | 3.63427 | Female | 0.00024 | 0.92238 | Female | 0.00000 | -1.07632 | Male | 0.00969 | 0.65856 | Female |
| DAB2IP | | | | 0.00864 | 1.49501 | Female | | | | | | | | | |
| CHST2 | 0.01813 | 1.78555 | Female | | | | | | | 0.00000 | 1.32955 | Female | | | |
| SP7 | 0.03013 | 3.07047 | Female | | | | | | | | | | | | |
| DGKE | | | | | | | | | | 0.04229 | 1.46493 | Female | | | |
| ZSWIM4 | | | | | | | | | | 0.01818 | -0.94631 | Male | | | |
| PRAP1 | | | | 0.00000 | 1.86957 | Female | | | | 0.00009 | 0.75350 | Female | | | |
| RAB5A | 0.00089 | -0.93769 | Male | | | | | | | | | | | | |
| SYNJ2BP | | | | 0.03202 | 0.59358 | Female | 0.00000 | 0.95827 | Female | | | | | | |
| ANKRD37 | | | | | | | 0.00163 | 1.27826 | Female | | | | | | |
| PRAMEF11 | | | | | | | 0.04579 | 2.94680 | Female | 0.00082 | 2.74669 | Female | | | |
| PVRL3 | | | | | | | | | | 0.03885 | -0.84299 | Male | | | |
| HOXB4 | | | | | | | | | | 0.03366 | -0.75394 | Male | | | |
| PDPR | | | | | | | 0.02187 | -0.71488 | Male | | | | | | |
| FAM184A | | | | | | | 0.00000 | -1.60649 | Male | | | | 0.02603 | -0.72555 | Male |
| C1orf198 | | | | | | | | | | 0.00381 | -1.11398 | Male | | | |
| KDELC2 | | | | | | | 0.00606 | -1.08544 | Male | | | | | | |
| CDH3 | | | | | | | | | | | | | 0.01718 | 0.65681 | Female |
| MSL3 | | | | 0.04404 | 0.59955 | Female | 0.00102 | 0.60220 | Female | | | | | | |
| LOC729739 | | | | 0.03507 | -1.36309 | Male | | | | | | | 0.02440 | 0.92346 | Female |
| TIFA | | | | | | | | | | | | | 0.00143 | -4.25897 | Male |
| L1CAM | | | | 0.00059 | 3.14298 | Female | | | | 0.01972 | 1.62021 | Female | | | |
| RET | | | | | | | | | | | | | 0.00000 | 2.91567 | Female |
| LOC100130744 | | | | | | | | | | 0.04292 | -1.24223 | Male | | | |
| TRNP1 | | | | | | | | | | 0.04305 | 1.19552 | Female | | | |
| HCCS | | | | 0.00000 | 0.92967 | Female | 0.00000 | 1.16878 | Female | | | | | | |
| TCEAL8 | | | | 0.00002 | 1.34481 | Female | | | | | | | 0.00001 | -0.92308 | Male |
| BTBD2 | | | | | | | 0.02386 | -0.72698 | Male | | | | | | |
| ZNF680 | | | | | | | 0.00002 | -0.83075 | Male | | | | | | |
| YWHAG | | | | | | | | | | 0.00000 | -0.65631 | Male | | | |
| MPP7 | | | | | | | | | | | | | 0.00144 | -0.87510 | Male |
| SLC16A2 | 0.00000 | 2.94265 | Female | | | | | | | 0.00001 | 2.45905 | Female | | | |
| ZNF596 | | | | | | | 0.02126 | -1.34000 | Male | 0.02334 | 1.25958 | Female | | | |
| BTG2 | | | | 0.02983 | 1.19384 | Female | | | | 0.03709 | -0.70519 | Male | 0.03617 | 0.71513 | Female |
| CHCHD3 | | | | 0.00000 | 0.74234 | Female | | | | | | | | | |
| PCYT2 | | | | 0.04112 | 0.74839 | Female | | | | | | | | | |
| NCK2 | | | | | | | 0.00599 | 0.58937 | Female | | | | | | |
| KLC4 | | | | | | | | | | | | | 0.00481 | 0.85463 | Female |
| BRCC3 | 0.00000 | 2.85989 | Female | 0.00000 | 1.03153 | Female | | | | | | | | | |
| DUSP5 | | | | | | | 0.00010 | 1.02977 | Female | | | | | | |
| EDA | | | | | | | | | | 0.01231 | 1.42037 | Female | | | |

| | | | | | | | | | | | | | | | |
|---|---|---|---|---|---|---|---|---|---|---|---|---|---|---|---|
| HOXC13 | | | | | | | | | | | | | 0.02793 | 0.99135 | Female |
| ECD | 0.00732 | -1.19985 | Male | | | | | | | | | | | | |
| SLC36A2 | | | | | | | 0.02321 | 1.21974 | Female | | | | 0.03543 | 1.07490 | Female |
| ENSA | 0.00001 | 0.76718 | Female | | | | | | | | | | | | |
| MZT1 | | | | | | | 0.00000 | -0.73652 | Male | | | | | | |
| ABCB7 | | | | | | | 0.01537 | 0.58125 | Female | | | | | | |
| SLC26A11 | | | | 0.03454 | -1.08464 | Male | | | | | | | | | |
| BMPR2 | | | | | | | 0.01450 | -0.67577 | Male | | | | | | |
| FOXK2 | | | | | | | 0.00000 | 0.84406 | Female | | | | | | |
| DNAJC3 | | | | 0.00002 | 0.75140 | Female | | | | | | | | | |
| PDCL2 | | | | | | | 0.00009 | -1.25230 | Male | 0.01267 | 0.82731 | Female | | | |
| APOD | | | | | | | | | | 0.00026 | 1.92723 | Female | 0.00007 | 2.03616 | Female |
| NCR3LG1 | | | | | | | | | | 0.04279 | -2.09377 | Male | | | |
| ZNF516 | | | | | | | | | | 0.00872 | 1.18536 | Female | | | |
| C20orf194 | | | | | | | | | | | | | 0.00950 | 0.89786 | Female |
| KIF1A | | | | | | | | | | 0.02612 | 1.40568 | Female | | | |
| HIST1H1B | | | | | | | | | | 0.00031 | 1.21980 | Female | | | |
| MTRNR2L10 | | | | | | | | | | 0.00028 | 0.72664 | Female | | | |
| ZNF331 | | | | | | | | | | | | | 0.00029 | 0.67747 | Female |
| NES | | | | | | | | | | 0.00951 | 2.06513 | Female | | | |
| WDR11-AS1 | | | | | | | | | | 0.00000 | 2.82338 | Female | | | |
| SH2B2 | | | | | | | 0.00062 | -1.96021 | Male | | | | | | |
| CLN5 | | | | | | | | | | | | | 0.00001 | -0.81241 | Male |
| TTC28 | | | | 0.00903 | 1.44403 | Female | | | | | | | | | |
| SLC17A5 | | | | | | | 0.00241 | -0.64798 | Male | | | | | | |
| DYNLT1 | 0.04719 | -0.63198 | Male | | | | | | | | | | | | |
| ARHGAP9 | | | | | | | 0.02446 | 1.88816 | Female | | | | | | |
| COA3 | 0.00266 | 0.94370 | Female | | | | 0.00000 | 0.72682 | Female | | | | | | |
| FAM3A | | | | 0.03207 | 1.11174 | Female | | | | | | | 0.03807 | 0.69059 | Female |
| ZBTB33 | | | | 0.03386 | 0.88374 | Female | | | | | | | | | |
| C9orf142 | | | | | | | 0.00012 | -0.60224 | Male | | | | | | |
| BHMT | | | | | | | | | | 0.00846 | 1.64018 | Female | 0.00008 | -2.31244 | Male |
| DACH1 | | | | | | | | | | 0.03814 | 1.73923 | Female | | | |
| MATN2 | | | | | | | | | | 0.00294 | 2.03783 | Female | 0.02064 | 1.40330 | Female |
| ZNF664 | | | | | | | | | | 0.00000 | -0.88273 | Male | | | |
| IQSEC2 | | | | | | | | | | 0.00497 | 2.19265 | Female | | | |
| RPL41 | 0.00020 | 0.65821 | Female | | | | | | | | | | | | |
| PLK2 | | | | | | | 0.03454 | 1.03717 | Female | | | | | | |
| GPSM3 | | | | | | | | | | 0.00354 | 1.09650 | Female | | | |
| STARD9 | | | | | | | | | | | | | 0.02626 | 1.18785 | Female |
| LPCAT4 | | | | 0.00577 | 0.81713 | Female | | | | 0.00015 | 0.69645 | Female | | | |
| FAM120AOS | | | | | | | 0.00235 | -0.62791 | Male | | | | | | |
| BCL7C | | | | 0.03815 | 1.09534 | Female | | | | | | | | | |
| WNT3 | | | | | | | 0.00065 | 1.85423 | Female | 0.00000 | 2.59368 | Female | | | |
| STK32B | | | | | | | | | | | | | 0.03206 | 1.44447 | Female |
| TSPAN5 | | | | | | | | | | 0.00057 | -1.74436 | Male | | | |
| SNAR-H | | | | | | | 0.00808 | 0.79572 | Female | | | | | | |
| RCC2 | 0.00259 | -0.72944 | Male | | | | | | | | | | | | |
| GFER | | | | 0.03019 | 0.65268 | Female | | | | | | | | | |
| DUSP7 | | | | | | | 0.00002 | 1.12414 | Female | 0.02716 | 0.62015 | Female | 0.03211 | 0.59468 | Female |
| FREM2 | | | | | | | | | | 0.01866 | -1.30977 | Male | | | |

| | | | | | | | | | | | | | | | |
|---|---|---|---|---|---|---|---|---|---|---|---|---|---|---|---|
| PAGE1 | 0.00683 | 2.59844 | Female | 0.00002 | 2.41648 | Female | 0.00000 | 2.06920 | Female | 0.00006 | 1.53339 | Female | | | |
| LINC00936 | | | | 0.03998 | 0.62734 | Female | | | | | | | | | |
| PMEPA1 | | | | | | | | | | 0.00416 | 2.57561 | Female | | | |
| ARL5B | | | | | | | | | | | | | 0.00010 | -0.74850 | Male |
| FAM193B | | | | 0.00024 | -1.24330 | Male | | | | | | | | | |
| SNORA19 | | | | | | | 0.00816 | -0.93775 | Male | | | | | | |
| FBXL17 | | | | 0.02039 | 2.12230 | Female | | | | | | | | | |
| SLC6A6 | 0.03076 | -1.39331 | Male | | | | | | | | | | | | |
| RAB11FIP4 | | | | 0.00019 | 1.01101 | Female | | | | | | | | | |
| GABRB3 | | | | | | | 0.00036 | -1.08059 | Male | | | | | | |
| IDH1 | | | | 0.00010 | 0.71458 | Female | | | | 0.00000 | 0.58207 | Female | | | |
| SFT2D2 | | | | 0.04354 | 0.67789 | Female | | | | | | | | | |
| LINC00630 | | | | | | | 0.02447 | 0.98964 | Female | | | | | | |
| SMTN | | | | 0.00255 | 1.29910 | Female | | | | | | | | | |
| IFT57 | | | | | | | 0.01733 | 0.79868 | Female | | | | | | |
| GNPTAB | | | | | | | | | | 0.00140 | 0.59583 | Female | | | |
| MIR4461 | | | | | | | 0.01810 | -0.67527 | Male | 0.00939 | 0.73839 | Female | | | |
| RASSF1 | | | | | | | | | | 0.00316 | -0.64555 | Male | | | |
| CPOX | 0.00972 | -1.70678 | Male | | | | | | | | | | | | |
| LOC100129520 | | | | 0.02564 | 2.38613 | Female | | | | | | | 0.02945 | 1.57321 | Female |
| RTBDN | | | | | | | | | | 0.00232 | 1.49896 | Female | | | |
| DGKK | | | | | | | | | | 0.04038 | 3.02899 | Female | | | |
| C2CD4C | | | | | | | | | | | | | 0.02332 | 1.18292 | Female |
| GSN | | | | | | | | | | 0.00000 | 1.88546 | Female | | | |
| TMEM243 | | | | | | | 0.00201 | -1.21957 | Male | 0.00147 | -1.17104 | Male | 0.00027 | -1.27428 | Male |
| CROT | | | | | | | 0.04686 | 1.36960 | Female | 0.01983 | -1.48918 | Male | 0.02091 | -1.43676 | Male |
| HDGFL1 | | | | | | | 0.00042 | -1.61969 | Male | | | | | | |
| ATP1B2 | | | | | | | | | | 0.01173 | 1.36877 | Female | | | |
| ESRG | | | | | | | 0.00178 | -0.73892 | Male | 0.00000 | 1.10770 | Female | | | |
| SH3GL3 | | | | 0.00001 | 1.73790 | Female | 0.00001 | -1.09666 | Male | | | | | | |
| TUBB4A | 0.01330 | 1.23133 | Female | | | | | | | 0.00000 | 0.80307 | Female | | | |
| LTBR | | | | | | | 0.03455 | -0.95007 | Male | | | | | | |
| VEGFA | | | | 0.01488 | -1.42279 | Male | 0.04080 | -0.75084 | Male | | | | | | |
| EPHA2 | | | | | | | | | | 0.00451 | -1.75362 | Male | 0.01648 | 1.50409 | Female |
| JADE3 | | | | 0.04628 | 0.83375 | Female | | | | | | | | | |
| STAT6 | | | | | | | 0.03539 | 1.59580 | Female | | | | | | |
| CD2AP | | | | | | | | | | | | | 0.00001 | -0.59037 | Male |
| LY6E | | | | 0.01933 | 0.77715 | Female | | | | 0.00004 | 0.80318 | Female | 0.00001 | 0.86095 | Female |
| LGALS2 | | | | | | | | | | 0.00003 | 2.56310 | Female | | | |
| CCZ1B | | | | | | | | | | | | | 0.00162 | -0.61880 | Male |
| SAT1 | | | | 0.01188 | 1.21616 | Female | | | | 0.00000 | 2.18227 | Female | | | |
| VCX3A | | | | 0.00000 | 4.19069 | Female | 0.00000 | 2.61251 | Female | 0.00002 | 1.65742 | Female | 0.00000 | 2.57034 | Female |
| CLEC6A | | | | | | | | | | 0.00000 | 2.45809 | Female | 0.00000 | 2.71919 | Female |
| ZNF204P | 0.01032 | 3.66417 | Female | | | | | | | | | | | | |
| ING1 | | | | | | | 0.00340 | 0.65711 | Female | | | | | | |
| UNC45A | | | | | | | | | | 0.00285 | -0.70582 | Male | | | |
| PTCH1 | | | | | | | 0.01145 | -1.37938 | Male | | | | | | |
| YAE1D1 | | | | | | | | | | | | | 0.00156 | -0.64449 | Male |
| ACVR2B | | | | | | | | | | 0.00019 | -0.67990 | Male | | | |
| TLE1 | | | | 0.00000 | 1.64331 | Female | | | | | | | | | |
| RCN1 | | | | 0.00000 | -1.32744 | Male | | | | | | | | | |

| | | | | | | | | | | | | | | | |
|---|---|---|---|---|---|---|---|---|---|---|---|---|---|---|---|
| PRR9 | | | | 0.00000 | -1.53259 | Male | 0.00000 | -1.09060 | Male | | | | | | |
| EBNA1BP2 | | | | | | | 0.00000 | 0.58573 | Female | | | | | | |
| IMP3 | 0.00009 | 0.74245 | Female | | | | | | | | | | | | |
| ECHS1 | | | | 0.00000 | 0.77171 | Female | | | | | | | | | |
| DOCK5 | | | | | | | | | | 0.00858 | 1.47470 | Female | | | |
| FAM104B | | | | 0.00959 | 0.72163 | Female | | | | | | | | | |
| DSCR8 | | | | 0.02674 | 0.74536 | Female | | | | | | | | | |
| PITX2 | | | | | | | | | | 0.02824 | 1.22217 | Female | | | |
| RBM41 | | | | 0.00000 | 2.20301 | Female | 0.00041 | 0.91811 | Female | 0.00000 | 1.09221 | Female | | | |
| TRAPPC2 | | | | 0.04957 | 0.70373 | Female | 0.00005 | 0.86097 | Female | 0.00157 | 0.65833 | Female | 0.00023 | 0.73465 | Female |
| LOC202181 | | | | | | | | | | 0.03113 | 1.30621 | Female | | | |
| VCX2 | 0.00000 | 5.19443 | Female | 0.00000 | 3.99797 | Female | 0.00000 | 6.06016 | Female | 0.00000 | 4.38935 | Female | 0.00000 | 4.83440 | Female |
| GADD45G | | | | 0.00000 | 3.04060 | Female | | | | | | | | | |
| H1FOO | | | | | | | 0.04918 | 1.54691 | Female | 0.04059 | -1.59679 | Male | 0.00045 | -2.44088 | Male |
| SKP2 | 0.00139 | -1.06278 | Male | | | | | | | | | | | | |
| EPCAM | | | | 0.00000 | 1.14356 | Female | | | | | | | | | |
| CMC1 | 0.04271 | -0.91817 | Male | | | | | | | | | | | | |
| PDS5B | 0.00201 | -1.03332 | Male | | | | | | | | | | | | |
| MAN1C1 | | | | | | | 0.04894 | 1.11969 | Female | 0.00000 | 3.00890 | Female | | | |
| TOR2A | | | | | | | | | | 0.00091 | 0.96880 | Female | 0.00064 | 0.97659 | Female |
| PTPRB | | | | | | | | | | 0.00611 | 0.86071 | Female | 0.02424 | 0.71817 | Female |
| SLC7A3 | | | | | | | 0.01960 | 0.89997 | Female | 0.00000 | 1.75110 | Female | 0.00621 | 0.97857 | Female |
| SPIN2B | | | | | | | 0.00447 | 1.78290 | Female | | | | | | |
| HNF1B | | | | 0.00000 | 2.01882 | Female | | | | | | | | | |
| ATP6V0C | | | | 0.00007 | 0.72798 | Female | | | | | | | | | |
| LINC01030 | | | | | | | 0.01176 | 1.37888 | Female | 0.03339 | 1.15125 | Female | 0.01457 | 1.26573 | Female |
| TAGAP | | | | | | | | | | 0.01919 | -1.09511 | Male | | | |
| HS6ST2 | | | | | | | | | | 0.00339 | -1.38117 | Male | 0.00001 | -1.91228 | Male |
| NDUFA5 | | | | | | | 0.00000 | -0.61785 | Male | | | | | | |
| PRMT7 | | | | 0.04090 | 0.58045 | Female | | | | | | | | | |
| GLTSCR1L | | | | 0.00419 | 1.53107 | Female | | | | | | | | | |
| CYB561D2 | | | | | | | 0.00022 | 0.59222 | Female | | | | | | |
| PARD6A | | | | 0.01604 | 2.48667 | Female | | | | | | | | | |
| SLC4A8 | | | | | | | | | | 0.00028 | 1.35678 | Female | | | |
| DPF3 | | | | | | | 0.00089 | -1.02007 | Male | | | | 0.01739 | 0.72699 | Female |
| ITGAV | | | | | | | 0.01267 | -1.22693 | Male | | | | | | |
| MAP4K4 | | | | 0.00918 | 0.73229 | Female | | | | | | | | | |
| LINC01176 | | | | | | | | | | | | | 0.01417 | 1.74983 | Female |
| SMPD1 | | | | | | | | | | 0.04101 | 0.59884 | Female | | | |
| PXN-AS1 | | | | 0.01772 | -0.77579 | Male | | | | | | | | | |
| MORF4L2 | 0.00000 | 1.26249 | Female | 0.00027 | 0.59818 | Female | 0.00000 | 0.59570 | Female | | | | | | |
| TRPV2 | | | | | | | 0.00004 | 2.13130 | Female | | | | | | |
| HDGF | | | | | | | | | | 0.00001 | -0.61353 | Male | | | |
| NUDT7 | | | | | | | | | | | | | 0.01476 | -1.51390 | Male |
| HIST1H4C | 0.00000 | 1.77180 | Female | | | | 0.00000 | 0.70689 | Female | | | | | | |
| FAM46C | | | | | | | | | | 0.04219 | 1.06613 | Female | | | |
| UXT | | | | 0.00000 | 0.88883 | Female | 0.00000 | 0.75976 | Female | 0.00000 | 0.83842 | Female | 0.00000 | 0.84482 | Female |
| CALCOCO1 | | | | | | | | | | 0.00762 | -0.83326 | Male | 0.00016 | 1.09677 | Female |
| MIR8072 | | | | 0.02991 | -0.61047 | Male | | | | | | | | | |
| FAM117B | | | | | | | | | | | | | 0.00145 | -0.71187 | Male |
| MEIS2 | | | | 0.00416 | -2.54047 | Male | | | | | | | | | |

| | | | | | | | | | | | | | | | |
|---|---|---|---|---|---|---|---|---|---|---|---|---|---|---|---|
| TBL1XR1 | 0.00292 | -0.81580 | Male | | | | | | | | | | | | |
| SGPL1 | | | | | | | 0.00000 | 0.80693 | Female | | | | 0.00001 | 0.58322 | Female |
| GAL3ST4 | | | | | | | | | | | | | 0.02204 | 1.39313 | Female |
| LOC650368 | | | | | | | 0.01103 | 1.58126 | Female | 0.03704 | 1.28576 | Female | 0.01426 | 1.46470 | Female |
| KCNE3 | | | | | | | 0.00606 | 1.60194 | Female | | | | | | |
| ITGB7 | | | | | | | | | | | | | 0.00991 | 1.54340 | Female |
| IGSF1 | | | | | | | | | | 0.00852 | 2.10393 | Female | 0.00091 | 2.46820 | Female |
| EIF4E1B | | | | | | | 0.00364 | 2.25691 | Female | | | | | | |
| LOC102724814 | | | | 0.00001 | 1.93581 | Female | | | | | | | | | |
| SYNGR4 | | | | | | | 0.00117 | 2.26386 | Female | | | | 0.01823 | 1.59910 | Female |
| BRPF1 | | | | | | | 0.00045 | 0.75393 | Female | | | | | | |
| ARHGEF15 | | | | | | | 0.03269 | -2.32872 | Male | | | | | | |
| PIN4 | | | | 0.00000 | 1.09381 | Female | 0.00000 | 1.22479 | Female | 0.00000 | 0.88378 | Female | 0.00000 | 0.84722 | Female |
| PKD1 | | | | | | | | | | 0.00005 | 1.41296 | Female | | | |
| BCAP29 | | | | | | | 0.00000 | -0.98934 | Male | | | | | | |
| SP6 | | | | 0.00000 | -2.82669 | Male | 0.02474 | -0.72401 | Male | 0.02952 | -0.65802 | Male | | | |
| BLOC1S3 | | | | | | | 0.00000 | 1.04542 | Female | | | | 0.00008 | 0.75879 | Female |
| MZF1 | | | | | | | 0.00425 | 1.08428 | Female | | | | | | |
| STT3B | | | | | | | 0.00000 | -0.64517 | Male | | | | 0.00001 | -0.59038 | Male |
| GRB14 | | | | | | | | | | | | | 0.02070 | -1.39604 | Male |
| HNRNPCL2 | | | | | | | | | | 0.00000 | 2.83108 | Female | | | |
| BATF | | | | 0.00248 | 2.40378 | Female | | | | | | | | | |
| POT1 | | | | | | | 0.04594 | -0.64912 | Male | | | | | | |
| ZNF74 | 0.00457 | -2.43743 | Male | | | | | | | | | | | | |
| ATG10 | 0.00817 | -1.31759 | Male | | | | | | | | | | | | |
| C7orf13 | | | | | | | | | | 0.00001 | -2.64762 | Male | | | |
| AGBL5 | 0.00452 | 2.06775 | Female | | | | | | | 0.04305 | -0.64348 | Male | | | |
| NUDT16L1 | | | | | | | 0.00024 | 0.58100 | Female | | | | | | |
| DGKD | | | | | | | | | | 0.02136 | -0.62514 | Male | | | |
| PRSS41 | | | | | | | | | | 0.00006 | 1.77793 | Female | | | |
| SLC36A4 | | | | | | | | | | 0.00001 | -1.69514 | Male | 0.01793 | -0.93643 | Male |
| ZNF704 | | | | 0.00003 | 2.10511 | Female | | | | | | | | | |
| OSMR | | | | | | | | | | 0.04985 | 1.43479 | Female | | | |
| POLE4 | | | | | | | | | | 0.00003 | -0.78342 | Male | | | |
| CD97 | | | | 0.00138 | 1.36964 | Female | | | | | | | | | |
| UBTD1 | | | | | | | | | | 0.00001 | 1.56887 | Female | | | |
| SNORD88B | | | | | | | 0.00249 | 0.69240 | Female | | | | | | |
| ONECUT2 | | | | | | | | | | | | | 0.03039 | 2.65753 | Female |
| CEP170B | | | | 0.00000 | 2.53342 | Female | | | | | | | 0.03660 | 0.77901 | Female |
| CDH23 | | | | | | | 0.00003 | 2.06591 | Female | | | | | | |
| GDF3 | | | | | | | | | | 0.00014 | 1.82201 | Female | | | |
| FOSL1 | 0.00216 | 3.54401 | Female | | | | | | | 0.00016 | -1.87358 | Male | | | |
| NDUFB11 | 0.00000 | 1.14935 | Female | 0.00000 | 0.76340 | Female | 0.00000 | 0.69058 | Female | 0.00000 | 0.67624 | Female | 0.00000 | 0.63380 | Female |
| IRS2 | | | | | | | 0.00154 | 2.06749 | Female | | | | | | |
| EAF2 | | | | | | | | | | 0.01592 | -0.79077 | Male | | | |
| P2RY14 | | | | | | | | | | 0.00004 | -1.63919 | Male | | | |
| PHLDA1 | | | | | | | | | | 0.00000 | 2.06279 | Female | | | |
| SLC9A3 | | | | | | | | | | 0.00052 | 1.83405 | Female | | | |
| ELK3 | | | | | | | | | | 0.00259 | 1.19286 | Female | | | |
| DMC1 | | | | | | | | | | 0.01022 | 1.31143 | Female | 0.00979 | 1.29775 | Female |
| HEG1 | | | | | | | 0.00052 | -1.50156 | Male | | | | 0.02091 | 1.01836 | Female |

| | | | | | | | | | | | | | | | |
|---|---|---|---|---|---|---|---|---|---|---|---|---|---|---|---|
| CHAC2 | | | | 0.02900 | 1.20472 | Female | | | | | | | | | |
| DPY19L1 | | | | | | | | | | 0.00090 | 1.17402 | Female | | | |
| LARP6 | | | | | | | | | | 0.00000 | 2.16409 | Female | | | |
| COMMD5 | | | | | | | 0.00278 | 0.63788 | Female | | | | | | |
| PITX1 | | | | | | | | | | 0.02433 | 0.66723 | Female | | | |
| LINC00839 | | | | | | | | | | | | | 0.00665 | -2.31897 | Male |
| MACF1 | 0.01767 | -1.09962 | Male | | | | | | | | | | | | |
| CRNDE | | | | | | | | | | | | | 0.00867 | -1.20037 | Male |
| TMEM50B | | | | | | | | | | 0.00001 | 0.77716 | Female | | | |
| UHRF1BP1L | | | | | | | | | | 0.00070 | 0.85379 | Female | | | |
| SLC6A19 | | | | | | | | | | 0.00000 | 1.16966 | Female | | | |
| RBMX | 0.00000 | 1.16621 | Female | | | | | | | | | | | | |
| PRKAR2B | | | | | | | | | | 0.01456 | 1.17065 | Female | | | |
| LOC100420587 | | | | | | | | | | | | | 0.03210 | 1.66428 | Female |
| HDDC2 | 0.00000 | -1.08263 | Male | | | | | | | | | | | | |
| HYKK | | | | | | | | | | 0.04142 | 1.16602 | Female | | | |
| CNGB3 | | | | | | | | | | | | | 0.01357 | -5.11164 | Male |
| LMOD3 | | | | | | | 0.00000 | 1.89111 | Female | 0.02713 | -0.84195 | Male | | | |
| TARS2 | | | | | | | | | | 0.00110 | 0.78262 | Female | 0.00974 | 0.63423 | Female |
| HTATSF1 | | | | | | | 0.00002 | 0.59960 | Female | | | | | | |
| C16orf91 | | | | | | | 0.00001 | 0.62732 | Female | | | | | | |
| C1orf27 | 0.00516 | 1.41439 | Female | | | | | | | | | | | | |
| RTFDC1 | 0.00064 | -0.79218 | Male | | | | | | | | | | | | |
| PPIL1 | | | | | | | 0.00000 | -0.70310 | Male | | | | | | |
| C10orf35 | | | | | | | | | | 0.00000 | -1.67283 | Male | | | |
| TPM3P9 | | | | | | | | | | | | | 0.00005 | 0.76934 | Female |
| PAPSS2 | | | | | | | | | | 0.01326 | 1.69543 | Female | | | |
| DONSON | | | | | | | 0.00150 | -0.68128 | Male | | | | | | |
| HNRNPU | 0.00047 | -0.65885 | Male | | | | | | | | | | | | |
| SSTR5-AS1 | | | | | | | 0.01671 | -1.31253 | Male | 0.00027 | 1.79317 | Female | 0.00800 | -1.39319 | Male |
| C6orf201 | | | | | | | | | | 0.00054 | 0.99483 | Female | | | |
| UBE2D1 | | | | | | | 0.00763 | -0.58314 | Male | | | | | | |
| ZDBF2 | | | | | | | 0.00538 | -0.74194 | Male | | | | | | |
| SCARNA2 | | | | | | | | | | | | | 0.00407 | 1.89460 | Female |
| ZC4H2 | | | | 0.03101 | 1.41853 | Female | 0.04364 | 0.91327 | Female | | | | | | |
| FECH | | | | | | | 0.00002 | -1.26723 | Male | | | | | | |
| TTYH1 | | | | 0.01569 | 2.37921 | Female | | | | | | | | | |
| SLC4A7 | | | | | | | 0.01543 | -0.81532 | Male | | | | 0.04963 | -0.64597 | Male |
| ENO2 | | | | | | | | | | 0.00000 | 2.06329 | Female | | | |
| MGST2 | | | | | | | | | | 0.00012 | 0.83852 | Female | 0.00077 | -0.74021 | Male |
| MT1F | | | | | | | | | | 0.04605 | 1.22458 | Female | | | |
| WWC3 | | | | | | | | | | | | | 0.00530 | 0.89646 | Female |
| RTKN2 | | | | | | | 0.00226 | -0.68513 | Male | | | | | | |
| GRAMD1A | | | | | | | | | | 0.03745 | 0.94697 | Female | | | |
| PRIM2 | | | | | | | | | | | | | 0.00010 | -0.66346 | Male |
| TMEM45B | | | | | | | 0.02648 | -0.75715 | Male | | | | | | |
| SH3D21 | | | | 0.00797 | 1.58511 | Female | | | | | | | | | |
| ATP10D | | | | | | | | | | 0.02757 | -1.56050 | Male | | | |
| HAGH | | | | 0.00973 | 1.18510 | Female | | | | | | | | | |
| RASGEF1A | | | | | | | | | | 0.00000 | 1.73709 | Female | 0.01187 | 0.81779 | Female |
| KLC1 | | | | | | | 0.00008 | -0.83888 | Male | | | | | | |

| | | | | | | | | | | | | | | | |
|---|---|---|---|---|---|---|---|---|---|---|---|---|---|---|---|
| PLBD1 | | | | | | | 0.00074 | -0.61456 | Male | | | | | | |
| FAM127C | | | | | | | | | | | | | 0.00002 | -1.99924 | Male |
| CLPSL1 | | | | | | | | | | | | | 0.03153 | 1.22738 | Female |
| GAS7 | | | | | | | | | | | | | 0.00625 | 0.63383 | Female |
| MAD2L2 | | | | | | | | | | 0.00000 | 0.91485 | Female | | | |
| MRPL28 | | | | | | | 0.00000 | 0.76346 | Female | | | | | | |
| BIVM | | | | 0.00285 | 2.49506 | Female | | | | | | | | | |
| LOC728392 | | | | | | | 0.04738 | 2.11402 | Female | | | | | | |
| ACSF2 | | | | | | | | | | 0.04762 | 1.40308 | Female | | | |
| GMDS | | | | 0.01123 | 1.11074 | Female | | | | | | | | | |
| ATP2B4 | | | | 0.01874 | 0.96534 | Female | | | | | | | 0.00021 | 0.88435 | Female |
| HK1 | | | | | | | 0.00004 | 0.65018 | Female | | | | | | |
| APOA4 | | | | | | | | | | 0.00291 | 2.17975 | Female | | | |
| ABHD15 | 0.00034 | -3.05495 | Male | | | | | | | | | | | | |
| NFE2 | | | | 0.01139 | 2.14753 | Female | 0.00000 | 3.11750 | Female | | | | | | |
| COMMD1 | | | | 0.00021 | 0.99729 | Female | | | | | | | | | |
| ZNF593 | 0.00200 | -0.66610 | Male | | | | | | | | | | | | |
| EPHB3 | | | | | | | 0.00278 | 2.08926 | Female | | | | | | |
| N6AMT2 | | | | | | | | | | 0.00746 | -0.89235 | Male | | | |
| HEATR3 | | | | | | | 0.00663 | -0.79688 | Male | | | | | | |
| SERINC5 | | | | | | | 0.00113 | 0.62939 | Female | 0.00013 | 0.68979 | Female | | | |
| UROS | | | | | | | 0.04373 | -0.71098 | Male | | | | | | |
| HIST1H4J | | | | | | | | | | | | | 0.02371 | -0.73154 | Male |
| USE1 | | | | 0.00497 | -0.99926 | Male | | | | | | | | | |
| MIR372 | | | | | | | 0.00000 | 1.25276 | Female | | | | | | |
| CC2D1A | | | | | | | 0.00044 | 0.94584 | Female | | | | | | |
| TDRD3 | 0.02915 | -1.42458 | Male | | | | | | | | | | | | |
| HIST1H3G | | | | | | | 0.00343 | 1.16672 | Female | | | | | | |
| MIEF1 | | | | 0.00009 | -0.61353 | Male | | | | | | | | | |
| PNPLA4 | | | | 0.00000 | 1.49364 | Female | 0.00000 | 0.98398 | Female | 0.00013 | 0.76311 | Female | 0.00272 | 0.61186 | Female |
| ZNF98 | | | | | | | 0.03139 | 0.75116 | Female | | | | | | |
| NGEF | | | | | | | | | | 0.00000 | 2.40980 | Female | | | |
| RPL23AP7 | | | | | | | | | | 0.00475 | -0.65951 | Male | | | |
| GRHL2 | | | | 0.00000 | 2.04254 | Female | | | | | | | | | |
| MAPKAPK3 | | | | | | | 0.00000 | -1.78538 | Male | | | | | | |
| SFXN2 | 0.00039 | 1.43956 | Female | | | | | | | | | | | | |
| TRIM53AP.uid-15951 | | | | | | | | | | 0.00092 | 1.52771 | Female | | | |
| NCF2 | | | | | | | | | | | | | 0.02812 | 1.48404 | Female |
| OLFML1 | | | | | | | | | | | | | 0.01099 | 0.85045 | Female |
| ESAM | | | | | | | | | | 0.02281 | 1.80326 | Female | | | |
| SMO | | | | | | | 0.03660 | -2.44894 | Male | | | | | | |
| MAGEB16 | | | | | | | | | | | | | 0.02925 | 2.67994 | Female |
| CALML4 | 0.04719 | -1.30998 | Male | | | | | | | | | | | | |
| SLC1A5 | | | | 0.04271 | -0.67530 | Male | 0.00015 | -0.75813 | Male | | | | | | |
| UGP2 | 0.00074 | -0.84530 | Male | | | | | | | | | | | | |
| PLXNB2 | | | | 0.01410 | 1.06542 | Female | | | | 0.00344 | 0.80434 | Female | 0.02284 | 0.64412 | Female |
| C4orf48 | | | | 0.00880 | 0.78542 | Female | | | | | | | | | |
| SRSF6 | 0.00619 | -0.65613 | Male | | | | | | | | | | | | |
| ZC3H7B | | | | 0.00888 | 1.36867 | Female | | | | | | | | | |
| CEP55 | | | | | | | 0.02409 | -0.62095 | Male | 0.00238 | -0.75474 | Male | | | |
| MAGEB1 | | | | | | | 0.03641 | 1.05182 | Female | | | | | | |

| | | | | | | | | | | | | | | | |
|---|---|---|---|---|---|---|---|---|---|---|---|---|---|---|---|
| LYPD6B | 0.02263 | -2.23979 | Male | | | | | | | 0.00305 | 1.43964 | Female | | | |
| SCGB3A2 | | | | | | | | | | 0.00000 | 2.04432 | Female | | | |
| NFKBIB | | | | 0.02277 | -0.62497 | Male | | | | | | | | | |
| UPRT | | | | | | | | | | | | | 0.00664 | -1.16550 | Male |
| LINC01021 | | | | | | | 0.00601 | 1.13316 | Female | | | | | | |
| FAM89A | | | | | | | 0.00427 | -1.25115 | Male | 0.00052 | -1.38270 | Male | | | |
| IL36RN | | | | | | | | | | 0.00068 | -2.43418 | Male | | | |
| ACP5 | | | | | | | | | | 0.03636 | 0.59825 | Female | | | |
| EPHA1-AS1 | | | | | | | 0.04275 | 1.29922 | Female | | | | | | |
| DHCR7 | | | | | | | 0.00194 | -0.67015 | Male | | | | | | |
| HSD17B10 | 0.00000 | 1.21690 | Female | 0.00000 | 0.92630 | Female | 0.00000 | 0.87437 | Female | 0.00000 | 0.74702 | Female | 0.00000 | 1.24506 | Female |
| RPIA | | | | | | | | | | 0.00000 | -0.93906 | Male | | | |
| SLC9A2 | | | | | | | | | | | | | 0.00424 | -2.81156 | Male |
| WDR44 | | | | | | | 0.00385 | 0.79244 | Female | 0.00003 | 1.02963 | Female | | | |
| MPP1 | | | | 0.00612 | 1.03350 | Female | | | | | | | | | |
| MAGEB2 | | | | | | | 0.00000 | 1.34729 | Female | | | | 0.00063 | 0.77539 | Female |
| FIRRE | | | | | | | | | | 0.00540 | 0.72280 | Female | 0.00000 | -1.41878 | Male |
| DLG5 | | | | | | | | | | 0.00000 | -0.76748 | Male | | | |
| SPRYD3 | | | | 0.00387 | -1.45206 | Male | | | | | | | | | |
| TMEM99 | | | | | | | 0.00309 | -0.73858 | Male | 0.00397 | -0.69461 | Male | 0.00914 | -0.62965 | Male |
| ROBO3 | | | | | | | 0.00258 | -2.28182 | Male | | | | | | |
| PALM | | | | | | | | | | 0.02898 | -1.55706 | Male | | | |
| TWIST1 | | | | | | | | | | 0.03803 | 1.12975 | Female | | | |
| LOC440416 | | | | | | | | | | | | | 0.01251 | 0.73022 | Female |
| GNAL | | | | | | | 0.03241 | 0.77998 | Female | | | | | | |
| PROSER2-AS1 | | | | | | | | | | 0.00014 | 0.85251 | Female | | | |
| RILPL2 | | | | | | | 0.00469 | -1.33004 | Male | 0.00030 | -1.52569 | Male | | | |
| AGAP3 | | | | 0.00143 | 1.18286 | Female | | | | | | | | | |
| ALKBH6 | 0.00450 | 2.11891 | Female | | | | | | | | | | | | |
| BMS1 | | | | | | | 0.00000 | 0.59143 | Female | | | | | | |
| PRNCR1 | 0.00748 | -0.66495 | Male | | | | | | | | | | | | |
| TOMM20 | | | | 0.00000 | -0.59053 | Male | | | | | | | | | |
| SCN1A | | | | | | | | | | | | | 0.00331 | -3.31540 | Male |
| TMEM180 | | | | | | | 0.00733 | 0.60281 | Female | | | | | | |
| COLGALT1 | | | | | | | 0.00000 | 0.72272 | Female | | | | | | |
| OSGEPL1 | | | | | | | | | | | | | 0.02769 | -0.59301 | Male |
| PANK4 | | | | 0.00991 | 1.26591 | Female | 0.00882 | 0.86955 | Female | | | | | | |
| EGFL7 | 0.00086 | 2.27823 | Female | | | | | | | 0.01463 | 0.64648 | Female | 0.01237 | 0.65221 | Female |
| GPR75 | | | | | | | 0.00010 | -1.98341 | Male | | | | | | |
| MAGOHB | 0.00003 | -1.07942 | Male | | | | | | | | | | | | |
| TRIM61 | 0.00749 | -0.99341 | Male | | | | | | | 0.00000 | 0.83900 | Female | | | |
| ANO6 | | | | 0.00045 | 0.95496 | Female | | | | | | | | | |
| NEK6 | | | | 0.00001 | 1.48989 | Female | | | | | | | | | |
| DCAF11 | 0.02070 | 1.52310 | Female | | | | | | | | | | | | |
| SHMT2 | | | | 0.00001 | -0.78898 | Male | | | | | | | | | |
| MGC12916 | | | | | | | 0.04611 | 1.71791 | Female | | | | | | |
| MST4 | | | | | | | | | | 0.01623 | -0.82391 | Male | 0.00000 | -1.73566 | Male |
| CPTP | | | | | | | 0.03924 | 0.73602 | Female | | | | | | |
| CCNI2 | | | | | | | | | | | | | 0.00365 | 1.25227 | Female |
| FBRSL1 | | | | | | | | | | | | | 0.01108 | 0.75930 | Female |
| ZBTB20 | | | | | | | | | | | | | 0.00004 | -0.73359 | Male |

| | | | | | | | | | | | | | | | |
|---|---|---|---|---|---|---|---|---|---|---|---|---|---|---|---|
| DOCK8 | | | | | | | | | | 0.01090 | -1.48529 | Male | | | |
| RPL13AP3 | | | | | | | | | | 0.00000 | -1.30295 | Male | | | |
| HIST1H2BM | | | | | | | | | | | | | 0.01075 | -1.38727 | Male |
| MRPS26 | | | | 0.00000 | 1.12526 | Female | | | | | | | | | |
| DDIT4L | | | | | | | | | | | | | 0.00220 | -0.87663 | Male |
| IL13RA1 | | | | | | | | | | 0.00429 | 1.73639 | Female | | | |
| SOD2 | 0.00008 | -1.22171 | Male | 0.00058 | 0.67681 | Female | 0.00000 | 0.76818 | Female | | | | 0.00000 | 0.62322 | Female |
| MFSD2A | | | | | | | 0.02360 | -1.38889 | Male | 0.00000 | 3.55932 | Female | | | |
| FASN | | | | | | | 0.00291 | -0.62572 | Male | | | | | | |
| ACTN4 | 0.00925 | 0.84451 | Female | | | | | | | | | | | | |
| HIST1H4H | | | | | | | 0.02674 | 1.26197 | Female | 0.01107 | 1.34292 | Female | | | |
| HSDL2 | | | | | | | 0.00113 | -0.73213 | Male | | | | | | |
| HIST1H2BD | | | | | | | | | | 0.00020 | 1.59095 | Female | | | |
| NEFH | | | | | | | | | | 0.00011 | 0.83378 | Female | | | |
| ZMYM6NB | | | | | | | | | | 0.00001 | 1.50492 | Female | | | |
| ZNF729 | | | | | | | | | | 0.00132 | 0.79224 | Female | | | |
| JPX | | | | 0.00000 | 1.05019 | Female | | | | 0.00000 | 1.20135 | Female | | | |
| ZNF280D | | | | | | | 0.03249 | -1.30679 | Male | | | | | | |
| FAM49A | | | | | | | | | | 0.01632 | -1.19062 | Male | | | |
| SPOP | | | | | | | 0.00136 | 0.66940 | Female | | | | | | |
| WFDC2 | | | | | | | | | | | | | 0.00000 | 2.15822 | Female |
| ANKRD6 | | | | 0.00050 | -1.57327 | Male | | | | | | | | | |
| BDH1 | | | | 0.01930 | 2.41239 | Female | | | | | | | | | |
| KDM4E | | | | | | | | | | 0.00000 | 2.87887 | Female | | | |
| ZNF57 | | | | | | | | | | 0.00147 | 0.61040 | Female | | | |
| EXOSC6 | | | | 0.00000 | 1.13737 | Female | 0.00000 | 0.94659 | Female | | | | | | |
| POC1B | | | | 0.00182 | 1.35509 | Female | | | | | | | | | |
| MFAP2 | | | | | | | 0.00637 | 1.38927 | Female | 0.00001 | 1.99722 | Female | | | |
| LYN | 0.03485 | 1.47477 | Female | 0.00000 | 1.51511 | Female | | | | | | | | | |
| PDP1 | | | | | | | 0.00641 | 0.84387 | Female | | | | | | |
| SULT2B1 | | | | | | | 0.00035 | 2.29841 | Female | 0.00708 | 1.73342 | Female | | | |
| MT1M | | | | | | | | | | | | | 0.01634 | 1.68353 | Female |
| CTSD | | | | 0.01201 | 0.64776 | Female | 0.00001 | -0.71666 | Male | | | | | | |
| CASZ1 | | | | | | | | | | | | | 0.02561 | 1.78025 | Female |
| CTSE | 0.00000 | 1.19979 | Female | 0.00000 | 0.94255 | Female | | | | | | | | | |
| SEPT7 | 0.00224 | -0.67111 | Male | | | | | | | | | | | | |
| HSH2D | | | | 0.00045 | -1.08816 | Male | | | | | | | | | |
| SLC25A53 | | | | | | | 0.00123 | 1.04580 | Female | 0.00346 | 0.93181 | Female | 0.00004 | 1.16862 | Female |
| CCDC109B | | | | 0.00005 | 1.25995 | Female | 0.00900 | -0.60774 | Male | | | | | | |
| PDE6G | | | | | | | | | | 0.00001 | 2.44979 | Female | | | |
| FAM45A | | | | | | | 0.02617 | 0.64338 | Female | 0.02894 | 0.69156 | Female | | | |
| FPGS | | | | | | | 0.00455 | -0.60953 | Male | | | | | | |
| MMAB | | | | | | | | | | | | | 0.00857 | 0.61834 | Female |
| FOXK1 | | | | | | | 0.00107 | 0.58624 | Female | | | | 0.00000 | 0.78637 | Female |
| TMEM14C | | | | | | | 0.00000 | -0.69900 | Male | | | | | | |
| IQCG | | | | | | | | | | | | | 0.01051 | 0.86390 | Female |
| ADAMTSL4 | | | | 0.00036 | -1.87877 | Male | | | | | | | | | |
| RHOB | | | | | | | | | | | | | 0.00064 | 0.66087 | Female |
| TFAP2A | | | | | | | | | | 0.00009 | -0.79189 | Male | | | |
| FAM227B | | | | 0.01748 | -1.82267 | Male | | | | | | | | | |
| DCLRE1B | | | | | | | 0.00691 | -0.59703 | Male | | | | | | |

| | | | | | | | | | | | | | | | |
|---|---|---|---|---|---|---|---|---|---|---|---|---|---|---|---|
| HLA-C | | | | | | | | | | 0.04800 | -0.83116 | Male | | | |
| CENPI | | | | 0.02385 | 1.17886 | Female | | | | 0.02109 | 0.77846 | Female | | | |
| ZFP14 | | | | | | | 0.02111 | -1.36155 | Male | | | | | | |
| FMO2 | | | | | | | 0.00001 | -2.64114 | Male | | | | | | |
| LIAS | 0.03081 | -0.99733 | Male | | | | | | | | | | | | |
| NCOR2 | | | | | | | | | | 0.00000 | 1.22070 | Female | | | |
| LIG4 | | | | | | | | | | | | | 0.02750 | -0.76717 | Male |
| EPB41L3 | | | | 0.00085 | -1.49282 | Male | | | | | | | | | |
| C1orf106 | | | | 0.00000 | 1.41179 | Female | | | | | | | | | |
| PAGE5 | | | | 0.00298 | 1.52542 | Female | 0.00000 | 2.13505 | Female | 0.00000 | 2.17903 | Female | | | |
| GLRX5 | | | | | | | | | | 0.00000 | -0.66853 | Male | | | |
| TUSC1 | | | | | | | | | | 0.03369 | -0.74356 | Male | | | |
| FAM53C | | | | 0.01827 | -0.63845 | Male | | | | | | | | | |
| GATA2 | | | | 0.00000 | 2.53942 | Female | | | | | | | | | |
| RGS14 | | | | | | | | | | 0.01332 | 1.20080 | Female | | | |
| HLA-G | | | | 0.03197 | -2.27195 | Male | | | | | | | | | |
| FLJ32255 | | | | | | | | | | 0.04561 | -2.30523 | Male | | | |
| ZNF586 | | | | | | | | | | | | | 0.00212 | 0.61522 | Female |
| FGFR3 | | | | | | | | | | 0.00037 | 0.73830 | Female | 0.00432 | 0.60615 | Female |
| MRPS6 | | | | | | | 0.00000 | -0.62705 | Male | | | | | | |
| USP9Y | 0.00000 | -4.07012 | Male | 0.00000 | -7.32548 | Male | 0.00000 | -5.83612 | Male | 0.00000 | -6.08672 | Male | 0.00000 | -6.49154 | Male |
| TIMP2 | | | | | | | 0.00025 | -1.87394 | Male | | | | | | |
| C4orf19 | | | | | | | 0.00384 | -0.84382 | Male | 0.03750 | -0.60626 | Male | | | |
| DNAJB4 | | | | | | | 0.00140 | -0.80849 | Male | | | | | | |
| SEMA4B | | | | | | | | | | 0.00544 | 0.74816 | Female | | | |
| HSD17B1 | 0.00460 | 1.85061 | Female | | | | | | | | | | | | |
| ABCC5 | 0.01314 | 0.99128 | Female | | | | | | | | | | | | |
| VCAN | | | | | | | | | | 0.01946 | 1.23510 | Female | | | |
| SLFN13 | | | | | | | | | | | | | 0.03842 | 1.79405 | Female |
| FGFR1 | | | | | | | | | | 0.00000 | 1.59192 | Female | | | |
| COTL1 | | | | | | | 0.00087 | 1.12471 | Female | 0.00520 | 0.91938 | Female | 0.01818 | 0.79539 | Female |
| NFYB | | | | | | | 0.00000 | -0.66067 | Male | | | | | | |
| FAM162B | | | | | | | | | | 0.02256 | 1.10109 | Female | | | |
| TRIM56 | | | | | | | | | | 0.00422 | 1.35101 | Female | | | |
| MATK | | | | | | | | | | | | | 0.03636 | -1.95610 | Male |
| APOBEC3F | | | | | | | | | | 0.00314 | 1.73195 | Female | | | |
| MEIS3 | | | | | | | | | | | | | 0.00181 | 2.11559 | Female |
| C16orf62 | | | | | | | | | | 0.01165 | 0.62096 | Female | 0.00046 | 0.80746 | Female |
| CDK2 | 0.01101 | 1.12088 | Female | 0.01139 | -0.69597 | Male | | | | | | | | | |
| ZNF544 | | | | | | | | | | 0.00138 | -0.74887 | Male | | | |
| DNMT3B | | | | 0.00000 | 2.08602 | Female | | | | | | | | | |
| STX4 | | | | | | | 0.01862 | 0.67554 | Female | | | | | | |
| PHKA1 | 0.01630 | 1.51064 | Female | 0.00000 | 1.82137 | Female | 0.00000 | 1.03884 | Female | 0.00610 | 0.60012 | Female | | | |
| PITRM1-AS1 | | | | | | | | | | 0.02894 | -0.67993 | Male | | | |
| CORO2A | | | | 0.00014 | 1.19127 | Female | | | | | | | | | |
| NONO | 0.00033 | 0.65083 | Female | 0.00000 | 0.76484 | Female | 0.00000 | 0.72548 | Female | | | | | | |
| MTRNR2L1 | | | | 0.01834 | -1.29948 | Male | | | | 0.00001 | -1.47781 | Male | | | |
| NUDT4 | | | | | | | 0.01975 | 0.68105 | Female | | | | | | |
| BMP2K | | | | | | | | | | 0.03461 | 0.61513 | Female | | | |
| HRH1 | | | | 0.02089 | -2.06782 | Male | | | | | | | | | |
| SH2D1B | | | | | | | | | | 0.00438 | 2.43757 | Female | | | |

| | | | | | | | | | | | | | | | |
|---|---|---|---|---|---|---|---|---|---|---|---|---|---|---|---|
| TUSC3 | | | | | | | 0.00349 | -0.83806 | Male | | | | | | |
| ACCSL | | | | | | | 0.03858 | 1.82255 | Female | | | | 0.00000 | -4.19578 | Male |
| LAGE3 | | | | 0.00000 | 1.25951 | Female | | | | 0.00034 | 0.61159 | Female | | | |
| BRIX1 | 0.00052 | -0.81178 | Male | | | | | | | | | | | | |
| FBLIM1 | | | | 0.01898 | 1.82028 | Female | | | | | | | | | |
| NUP93 | | | | 0.00000 | 0.77769 | Female | | | | | | | | | |
| GSTA3 | | | | | | | 0.03767 | 1.05039 | Female | | | | | | |
| VAT1 | | | | | | | 0.00000 | -0.61606 | Male | | | | | | |
| ZAR1L | | | | | | | 0.03305 | 1.79115 | Female | | | | 0.00346 | -2.40498 | Male |
| GPRC5C | 0.00037 | -2.92719 | Male | | | | | | | 0.00000 | 1.87684 | Female | | | |
| LOC100287792 | | | | | | | | | | 0.00000 | -1.03239 | Male | 0.00330 | -0.63680 | Male |
| MLXIP | | | | | | | | | | 0.00001 | 0.86401 | Female | | | |
| FAM98B | | | | | | | | | | | | | 0.00661 | -0.60666 | Male |
| PRPF39 | | | | 0.00010 | 0.92106 | Female | | | | | | | | | |
| SOCS6 | | | | 0.01403 | 0.92649 | Female | | | | | | | | | |
| KIAA1671 | | | | | | | | | | 0.00068 | 1.07849 | Female | | | |
| FCHO1 | | | | | | | 0.00000 | 2.02267 | Female | | | | | | |
| CDR1 | | | | | | | 0.00144 | -0.86623 | Male | | | | 0.00583 | 0.72528 | Female |
| PIM1 | | | | | | | | | | | | | 0.00059 | 0.59967 | Female |
| FUZ | 0.01750 | -1.84067 | Male | | | | | | | 0.01472 | -0.76354 | Male | | | |
| UGCG | | | | | | | 0.00131 | -0.65448 | Male | | | | | | |
| ST8SIA1 | | | | | | | | | | 0.00637 | -2.47972 | Male | | | |
| PSMD5 | | | | | | | | | | | | | 0.01652 | -0.60173 | Male |
| CREBL2 | | | | | | | | | | 0.00013 | 1.13349 | Female | | | |
| ZSCAN16 | | | | | | | 0.04835 | -0.99785 | Male | | | | | | |
| ADA | | | | 0.00000 | 2.26907 | Female | | | | | | | | | |
| CSNK2A3 | 0.01160 | -0.76956 | Male | | | | | | | | | | | | |
| GAPDHS | | | | | | | | | | | | | 0.00131 | 1.85690 | Female |
| HIST1H1C | | | | 0.00113 | -1.31244 | Male | | | | | | | | | |
| NBN | | | | | | | | | | | | | 0.00083 | -0.58869 | Male |
| B4GALT1 | | | | 0.00000 | 1.27957 | Female | 0.00000 | 0.92796 | Female | | | | | | |
| CMTM8 | | | | | | | 0.02185 | -0.98692 | Male | | | | | | |
| ENTPD6 | 0.00000 | -2.50537 | Male | 0.00000 | 1.53813 | Female | | | | | | | | | |
| SERTAD4 | 0.02380 | -2.24661 | Male | | | | | | | 0.00049 | -1.52745 | Male | | | |
| HES4 | | | | | | | 0.00138 | -2.33218 | Male | | | | | | |
| TMEM170B | | | | | | | 0.01610 | -2.47956 | Male | | | | | | |
| WWC2 | 0.00834 | -1.15135 | Male | | | | | | | | | | | | |
| NREP | | | | 0.01284 | -1.94449 | Male | | | | | | | | | |
| CLCC1 | | | | | | | 0.00297 | -0.78093 | Male | | | | | | |
| MASP2 | | | | | | | 0.00000 | 1.56058 | Female | | | | 0.02863 | 0.69227 | Female |
| APOBEC3D | | | | | | | | | | 0.00064 | 2.21779 | Female | | | |
| PDHA1 | | | | 0.00000 | 0.94346 | Female | 0.00000 | 0.96411 | Female | 0.00000 | 0.67905 | Female | 0.00000 | 0.64429 | Female |
| FMR1 | | | | 0.00006 | 1.17155 | Female | | | | | | | 0.00313 | 0.59649 | Female |
| LINC00273 | | | | | | | 0.00006 | -1.07862 | Male | | | | | | |
| COQ5 | | | | | | | | | | 0.00008 | 0.90781 | Female | | | |
| SH3PXD2A | | | | | | | | | | 0.00000 | 2.00006 | Female | | | |
| STS | | | | 0.00000 | 1.87705 | Female | | | | | | | 0.00009 | 0.90126 | Female |
| NPNT | | | | 0.00008 | -2.87258 | Male | | | | | | | | | |
| DNAJC30 | 0.02203 | 1.37066 | Female | | | | | | | | | | | | |
| SOCS3 | 0.03152 | -1.98441 | Male | | | | 0.00000 | 1.76829 | Female | | | | | | |
| SMAD5-AS1 | | | | | | | | | | | | | 0.02330 | 1.10712 | Female |

| | | | | | | | | | | | | | | | |
|---|---|---|---|---|---|---|---|---|---|---|---|---|---|---|---|
| CDH11 | 0.00000 | 1.11363 | Female | 0.03769 | -0.68201 | Male | | | | | | | | | |
| SBSPON | | | | | | | 0.00818 | -1.25113 | Male | | | | | | |
| SAP30 | | | | | | | | | | | | | 0.00001 | -1.42698 | Male |
| SMLR1 | | | | | | | | | | 0.03462 | 1.79096 | Female | 0.01243 | -2.23889 | Male |
| CLDN19 | 0.00000 | 2.36421 | Female | | | | | | | 0.00000 | 2.21277 | Female | | | |
| TLE3 | | | | 0.00000 | 1.42824 | Female | | | | | | | | | |
| RMI2 | | | | | | | | | | | | | 0.04647 | -0.74675 | Male |
| SLC5A3 | | | | | | | 0.02269 | -0.58421 | Male | | | | | | |
| RNF187 | | | | 0.00000 | 1.12238 | Female | | | | | | | | | |
| SMPD4 | | | | | | | 0.00000 | -0.81345 | Male | | | | | | |
| TMTC4 | | | | | | | 0.00051 | -1.18289 | Male | | | | 0.04164 | -0.70789 | Male |
| OCA2 | | | | | | | | | | 0.00218 | 1.82452 | Female | | | |
| PTPLB | | | | | | | | | | 0.00649 | -0.73565 | Male | 0.01253 | -0.66819 | Male |
| ACOT8 | | | | | | | | | | 0.00039 | 0.79852 | Female | | | |
| RNF113A | 0.00007 | 1.43071 | Female | 0.00001 | 0.99373 | Female | 0.00000 | 1.23424 | Female | 0.00000 | 0.86450 | Female | 0.00000 | 0.89060 | Female |
| CASP10 | | | | | | | | | | 0.00156 | 1.71841 | Female | | | |
| SGCE | | | | | | | 0.00016 | 1.80652 | Female | | | | | | |
| SOGA1 | | | | | | | | | | | | | 0.02803 | 1.09571 | Female |
| DYNC2LI1 | | | | | | | | | | 0.04239 | 0.72702 | Female | | | |
| PINK1 | | | | | | | | | | 0.00000 | 1.61398 | Female | | | |
| KDM5D | 0.00000 | -3.65515 | Male | 0.00000 | -9.29016 | Male | 0.00000 | -8.54804 | Male | 0.00000 | -6.81853 | Male | 0.00000 | -6.13763 | Male |
| LMBR1 | | | | 0.03260 | 0.81860 | Female | 0.00266 | -0.72491 | Male | | | | | | |
| SIX5 | 0.04258 | -1.84949 | Male | 0.01799 | 1.32070 | Female | | | | | | | | | |
| GPR137C | | | | 0.01514 | 2.93608 | Female | | | | | | | | | |
| GRIPAP1 | | | | | | | 0.00002 | 0.87595 | Female | | | | | | |
| PITPNC1 | | | | 0.00000 | 1.49191 | Female | | | | | | | | | |
| GPR157 | | | | | | | 0.00174 | -2.07970 | Male | | | | | | |
| EEA1 | | | | | | | | | | 0.00000 | 0.63294 | Female | | | |
| BOK-AS1 | | | | 0.01069 | -2.18306 | Male | 0.03484 | 1.29955 | Female | | | | | | |
| DCBLD1 | | | | | | | 0.03958 | -0.88021 | Male | | | | | | |
| PCNT | | | | | | | | | | 0.00000 | 0.64294 | Female | | | |
| RGL4 | | | | | | | | | | | | | 0.01839 | 1.44048 | Female |
| ETHE1 | | | | 0.02758 | 0.77736 | Female | | | | 0.00000 | 1.21760 | Female | | | |
| PITHD1 | | | | | | | 0.00000 | -0.61941 | Male | | | | | | |
| DNAH11 | | | | | | | | | | | | | 0.00652 | -1.91477 | Male |
| LOC374443 | | | | | | | | | | | | | 0.01316 | -0.65043 | Male |
| MRAS | | | | | | | | | | 0.01323 | 1.49501 | Female | 0.02951 | 1.32359 | Female |
| MYCT1 | | | | | | | 0.04526 | -0.64241 | Male | | | | | | |
| HARS | | | | | | | 0.00000 | 0.62918 | Female | | | | | | |
| HAND1 | | | | 0.00000 | -2.13463 | Male | 0.00992 | -0.95309 | Male | | | | 0.00007 | 1.30134 | Female |
| XAGE3 | | | | 0.00000 | 2.46433 | Female | 0.00000 | 2.03920 | Female | | | | | | |
| AMPD2 | | | | | | | 0.01585 | -0.75570 | Male | | | | | | |
| NRXN2 | | | | | | | | | | 0.00723 | 0.92486 | Female | | | |
| TMSB4Y | | | | 0.00000 | -2.98467 | Male | 0.00000 | -2.68284 | Male | 0.00000 | -4.29379 | Male | 0.00000 | -5.03917 | Male |
| TRIM38 | | | | | | | | | | 0.00000 | 2.44214 | Female | | | |
| LOC101927873 | | | | | | | | | | 0.00813 | 1.82856 | Female | | | |
| SOX2 | | | | | | | | | | 0.00008 | 1.58045 | Female | | | |
| OPLAH | | | | | | | 0.00027 | -2.63837 | Male | | | | | | |
| SATB1 | | | | | | | 0.00000 | -3.45919 | Male | 0.00000 | -1.80558 | Male | | | |
| SNORD83B | | | | | | | 0.00996 | -0.93330 | Male | | | | | | |
| C3orf56 | | | | | | | | | | 0.00032 | -2.70965 | Male | | | |

| | | | | | | | | | | | | | | | |
|---|---|---|---|---|---|---|---|---|---|---|---|---|---|---|---|
| NKAPP1 | | | | | | | | | | 0.01295 | 1.32081 | Female | | | |
| GAGE12J | 0.01217 | 2.44617 | Female | 0.00001 | 2.23773 | Female | 0.00000 | 4.26257 | Female | | | | 0.00016 | -1.32364 | Male |
| HSD17B8 | | | | | | | 0.00008 | -2.15343 | Male | | | | | | |
| DPPA5 | | | | | | | | | | 0.00000 | 1.91754 | Female | | | |
| FGFR1OP2 | | | | 0.00098 | -0.76053 | Male | | | | | | | | | |
| TMEM209 | | | | | | | 0.00000 | -0.83899 | Male | | | | | | |
| ETV5 | | | | | | | | | | 0.03845 | 0.74955 | Female | | | |
| PTGR1 | | | | 0.00001 | 1.33524 | Female | | | | | | | | | |
| MYLK3 | | | | | | | | | | 0.01647 | -1.61182 | Male | | | |
| RPS6KA2 | | | | | | | | | | 0.00020 | 2.41150 | Female | | | |
| MOSPD3 | | | | | | | 0.00003 | -1.34726 | Male | | | | | | |
| MAGEC2 | 0.00003 | 5.25505 | Female | 0.00000 | 7.78050 | Female | 0.00000 | 7.25890 | Female | 0.00000 | 2.90823 | Female | 0.00000 | 7.28963 | Female |
| ARHGEF11 | | | | | | | | | | 0.00265 | 0.87295 | Female | | | |
| MKRN1 | | | | | | | | | | 0.00000 | 0.61839 | Female | | | |
| GDPD2 | 0.00000 | 0.76925 | Female | 0.00014 | 2.11075 | Female | 0.01023 | 1.05797 | Female | 0.00000 | 3.62059 | Female | 0.01102 | 1.00792 | Female |
| SNORD65 | | | | 0.03678 | -0.87671 | Male | 0.00000 | -1.22508 | Male | | | | | | |
| LOC646471 | 0.03979 | 1.81657 | Female | | | | | | | | | | | | |
| TMEM125 | | | | 0.03466 | -2.13931 | Male | | | | | | | | | |
| NT5DC2 | | | | 0.00000 | -2.86361 | Male | | | | 0.00096 | -1.12955 | Male | | | |
| UBA1 | 0.00001 | 0.94694 | Female | 0.00000 | 0.94345 | Female | 0.00000 | 0.84940 | Female | | | | 0.00000 | 0.66089 | Female |
| PTK2 | 0.00751 | -1.07256 | Male | | | | | | | | | | | | |
| CLEC11A | | | | | | | | | | | | | 0.00005 | 1.47012 | Female |
| BTBD18 | | | | | | | 0.04948 | 1.70783 | Female | | | | | | |
| POU2F3 | | | | 0.04722 | 2.10540 | Female | | | | | | | | | |
| ARSD | | | | 0.03353 | 2.31774 | Female | | | | | | | | | |
| CHMP4B | | | | 0.03004 | -2.63156 | Male | | | | 0.00000 | 4.70545 | Female | | | |
| AP4E1 | | | | | | | | | | 0.00300 | -0.63231 | Male | | | |
| RND2 | | | | | | | | | | 0.00000 | 1.42275 | Female | | | |
| NEURL1 | | | | | | | | | | 0.01473 | -1.11568 | Male | | | |
| LAMA1 | 0.00105 | 1.13328 | Female | 0.00130 | 0.66534 | Female | | | | | | | | | |
| MAP7D1 | | | | | | | | | | 0.04027 | -0.75519 | Male | | | |
| ST3GAL6 | 0.02380 | -1.79263 | Male | | | | | | | | | | | | |
| AVIL | | | | | | | 0.01463 | 1.61529 | Female | | | | | | |
| TST | | | | 0.00725 | 1.58431 | Female | | | | | | | | | |
| CCAR1 | 0.00331 | -0.61859 | Male | | | | | | | | | | | | |
| DEFB124 | | | | 0.03199 | 2.03027 | Female | | | | 0.00479 | 1.70395 | Female | | | |
| CCNG2 | | | | | | | | | | | | | 0.01270 | -0.95536 | Male |
| ADORA2B | 0.02826 | -2.05814 | Male | | | | | | | | | | | | |
| SNORA66 | | | | 0.03763 | -1.19911 | Male | | | | | | | 0.00981 | 0.89798 | Female |
| SNORD17 | | | | | | | 0.00000 | 0.76072 | Female | | | | | | |
| PLEKHM2 | | | | 0.03114 | 0.71261 | Female | | | | | | | | | |
| LRP5L | | | | 0.00367 | -2.74284 | Male | | | | | | | | | |
| ALDH7A1 | | | | 0.00103 | -0.84527 | Male | | | | | | | | | |
| MIR6817 | | | | | | | 0.00022 | 1.01993 | Female | 0.01204 | -0.75359 | Male | 0.00887 | 0.71880 | Female |
| FAM20B | 0.00011 | 1.95847 | Female | | | | | | | | | | | | |
| CAPG | | | | | | | | | | 0.00000 | 1.63520 | Female | | | |
| NAV2 | | | | | | | | | | | | | 0.01409 | 0.74767 | Female |
| HNRNPA1P33 | | | | 0.01676 | 1.17623 | Female | | | | 0.01230 | 0.74490 | Female | | | |
| MEX3A | | | | | | | | | | 0.00000 | -1.58806 | Male | | | |
| LINC00260 | | | | | | | | | | | | | 0.03592 | 1.54019 | Female |
| OSGIN1 | | | | | | | | | | 0.01980 | -2.11610 | Male | | | |

| | | | | | | | | | | | | | | | |
|---|---|---|---|---|---|---|---|---|---|---|---|---|---|---|---|
| MAP3K1 | | | | | | | | | | | | | 0.04737 | -0.63082 | Male |
| SMC1A | | | | 0.00000 | 0.97815 | Female | 0.00000 | 0.83991 | Female | 0.00000 | 0.90950 | Female | 0.00000 | 0.96211 | Female |
| APOBR | | | | | | | 0.00099 | 2.23150 | Female | 0.00000 | 3.07456 | Female | 0.01917 | 1.59173 | Female |
| MRPL23 | | | | 0.00943 | 0.83965 | Female | | | | | | | | | |
| PRAF2 | | | | | | | 0.00198 | 1.23674 | Female | | | | 0.03661 | 0.83581 | Female |
| NDUFAF3 | | | | | | | | | | 0.00076 | 0.68384 | Female | | | |
| PRR19 | | | | | | | | | | 0.02203 | -0.69963 | Male | 0.02699 | 0.67067 | Female |
| SNHG15 | | | | | | | | | | 0.00000 | 0.71458 | Female | | | |
| GPR1-AS | 0.00822 | 2.26289 | Female | | | | | | | | | | | | |
| RAB2A | | | | | | | | | | | | | 0.00000 | -0.69501 | Male |
| CCPG1 | | | | | | | | | | 0.00006 | 0.86832 | Female | | | |
| MTRNR2L2 | | | | | | | | | | 0.00000 | 0.90820 | Female | | | |
| PTGES2 | | | | | | | | | | 0.00000 | 0.96987 | Female | | | |
| OAZ2 | | | | | | | | | | 0.00500 | -0.64951 | Male | | | |
| PHACTR3 | | | | | | | 0.01919 | 1.69092 | Female | | | | | | |
| SHROOM3 | | | | 0.00146 | 0.99262 | Female | | | | | | | | | |
| MMACHC | | | | | | | | | | | | | 0.00204 | 0.73444 | Female |
| PHKA2 | | | | | | | 0.00000 | 1.49703 | Female | 0.01454 | 0.82449 | Female | | | |
| SYNE3 | | | | | | | 0.03018 | -1.97862 | Male | | | | | | |
| MPST | | | | | | | | | | 0.00001 | -0.85513 | Male | | | |
| SNORA76C | | | | | | | | | | | | | 0.01094 | 0.83518 | Female |
| KIF18B | | | | | | | 0.00393 | -0.66858 | Male | | | | | | |
| ELAVL1 | 0.00258 | -1.07741 | Male | | | | | | | | | | | | |
| MIIP | 0.03789 | -0.60417 | Male | | | | | | | | | | | | |
| SNX10 | | | | | | | 0.00016 | -1.32165 | Male | | | | | | |
| VPS51 | | | | | | | 0.00669 | -0.68534 | Male | | | | | | |
| SSX3 | | | | | | | | | | 0.00000 | 2.89104 | Female | | | |
| SCNN1A | | | | | | | | | | 0.00156 | 1.67242 | Female | | | |
| C7orf50 | 0.03493 | -0.73998 | Male | | | | | | | | | | | | |
| GSC | | | | | | | | | | 0.00000 | 3.20498 | Female | | | |
| MDH1B | | | | | | | | | | | | | 0.00139 | 1.94338 | Female |
| SLC31A1 | 0.00802 | 0.92088 | Female | | | | | | | | | | | | |
| SNORA70 | | | | | | | 0.00000 | 1.07480 | Female | 0.00000 | 1.26609 | Female | 0.00000 | 1.02506 | Female |
| PRAMEF2 | | | | | | | 0.00000 | 3.77385 | Female | 0.00000 | 5.95855 | Female | 0.00028 | 1.54098 | Female |
| TNFRSF25 | | | | | | | 0.02413 | 1.68495 | Female | 0.00000 | 2.97508 | Female | | | |
| WDR87 | | | | | | | 0.02374 | 0.65566 | Female | | | | | | |
| TRIM71 | | | | | | | 0.00072 | 0.58300 | Female | | | | | | |
| RAB3B | | | | 0.00001 | -1.70407 | Male | 0.00000 | -1.50159 | Male | | | | | | |
| LOC101929549 | | | | | | | | | | 0.00013 | 1.87640 | Female | | | |
| SLC16A9 | | | | | | | | | | 0.00000 | 2.36036 | Female | | | |
| TMED7 | | | | | | | 0.00000 | -0.74001 | Male | | | | | | |
| STXBP1 | | | | | | | | | | | | | 0.03236 | 1.03152 | Female |
| XIAP | | | | 0.00045 | 0.83645 | Female | 0.00000 | 0.82104 | Female | | | | | | |
| C1orf85 | | | | | | | 0.00184 | -0.80217 | Male | | | | | | |
| ANKMY1 | 0.04350 | 2.77950 | Female | | | | | | | | | | | | |
| KIT | | | | | | | | | | 0.00236 | 1.41549 | Female | | | |
| APBA3 | 0.00216 | 2.30209 | Female | | | | 0.01470 | -0.74450 | Male | | | | | | |
| TUBA3C | | | | | | | | | | 0.00000 | 2.97349 | Female | 0.01700 | 1.24072 | Female |
| SRRM5 | | | | | | | 0.03426 | 1.30300 | Female | | | | | | |
| MLYCD | | | | 0.04187 | 1.76546 | Female | | | | | | | | | |
| CREBRF | | | | | | | | | | | | | 0.00525 | 1.02410 | Female |

| | | | | | | | | | | | | | | | |
|---|---|---|---|---|---|---|---|---|---|---|---|---|---|---|---|
| UBLCP1 | | | | 0.01056 | -0.78369 | Male | 0.00001 | -0.86833 | Male | | | | 0.00024 | -0.67363 | Male |
| TARS | 0.00749 | -0.70396 | Male | | | | | | | | | | | | |
| ATF5 | | | | | | | 0.00000 | 0.85442 | Female | | | | | | |
| EBF2 | | | | | | | 0.04577 | 2.19163 | Female | | | | | | |
| ATG3 | | | | | | | | | | 0.00000 | 0.66812 | Female | | | |
| NSMAF | | | | 0.02987 | 0.65555 | Female | | | | | | | | | |
| PHLDA3 | | | | 0.00024 | 1.54433 | Female | 0.00579 | 0.84144 | Female | | | | | | |
| KMT2E-AS1 | | | | | | | 0.01470 | 0.60892 | Female | | | | 0.01289 | 0.58025 | Female |
| CAMSAP3 | | | | 0.00005 | 2.00998 | Female | | | | | | | | | |
| ZNF703 | | | | 0.00149 | 1.19434 | Female | | | | 0.00000 | 1.07526 | Female | 0.00000 | 1.24225 | Female |
| ABHD14A | 0.00132 | -3.21363 | Male | | | | | | | | | | | | |
| MAML1 | | | | 0.01400 | -0.79387 | Male | 0.00048 | -0.72056 | Male | | | | | | |
| POTEF | | | | | | | 0.01554 | 1.03359 | Female | 0.00000 | -2.58486 | Male | 0.00166 | 1.18985 | Female |
| SMAD9 | | | | | | | | | | 0.00363 | 0.80992 | Female | | | |
| UNC119 | | | | | | | | | | 0.02263 | 0.83950 | Female | | | |
| VTCN1 | | | | | | | | | | | | | 0.04923 | 1.10157 | Female |
| SNX33 | | | | | | | 0.01245 | -1.27375 | Male | | | | | | |
| ARSE | | | | | | | | | | 0.00355 | 1.58062 | Female | | | |
| MIR6790 | | | | 0.02091 | -0.63549 | Male | | | | | | | | | |
| TMEM74B | | | | | | | | | | 0.00222 | 1.92485 | Female | | | |
| GUCA1A | | | | | | | | | | 0.00028 | 2.14468 | Female | | | |
| MZT2B | | | | | | | | | | 0.00036 | -0.62842 | Male | | | |
| RAD54B | | | | | | | | | | | | | 0.00762 | -0.59330 | Male |
| IRF2BPL | | | | 0.02960 | 0.59534 | Female | | | | | | | | | |
| DENND2A | | | | | | | | | | 0.00000 | 2.71107 | Female | | | |
| RNPEPL1 | | | | 0.00023 | 1.93296 | Female | | | | | | | | | |
| ZZEF1 | | | | | | | 0.00242 | 0.73668 | Female | | | | 0.00080 | 0.75610 | Female |
| CDC26 | | | | 0.00179 | -0.59935 | Male | | | | | | | | | |
| ABCC6 | | | | | | | 0.00905 | 1.80109 | Female | | | | | | |
| PXDN | | | | | | | | | | 0.00094 | 2.02317 | Female | | | |
| BMP2 | | | | | | | | | | 0.00002 | 2.53132 | Female | | | |
| SP8 | | | | | | | | | | 0.00228 | 1.41972 | Female | | | |
| RAP2B | 0.03391 | -0.63500 | Male | | | | 0.00000 | 0.59302 | Female | | | | | | |
| LYPD2 | | | | | | | | | | 0.03710 | -2.20761 | Male | | | |
| AP4S1 | | | | | | | | | | 0.01740 | -0.84652 | Male | | | |
| SNAP29 | | | | 0.02029 | 0.70658 | Female | | | | | | | | | |
| RITA1 | 0.01749 | 1.18994 | Female | | | | | | | | | | | | |
| IQUB | | | | | | | | | | | | | 0.04141 | -1.58716 | Male |
| HLA-E | | | | | | | 0.00147 | 0.62617 | Female | | | | | | |
| TVP23C.uid-20973 | | | | | | | 0.01094 | -0.73558 | Male | 0.03050 | 0.64116 | Female | | | |
| SEMA3F | | | | | | | | | | | | | 0.00234 | 1.30900 | Female |
| IFNLR1 | | | | | | | | | | 0.03288 | 1.50388 | Female | | | |
| LY6G6C | | | | | | | 0.00606 | -1.53302 | Male | | | | | | |
| BOLA1 | | | | 0.01834 | -0.89785 | Male | 0.00614 | -0.68933 | Male | 0.00000 | -1.35994 | Male | 0.00154 | -0.73112 | Male |
| FBXO30 | | | | | | | | | | 0.00864 | -0.69167 | Male | | | |
| PHGDH | | | | 0.00000 | -2.03432 | Male | 0.00044 | -1.08084 | Male | 0.04863 | -0.63043 | Male | | | |
| RPS7P5 | | | | | | | | | | 0.00000 | -3.85426 | Male | | | |
| NHLRC2 | | | | | | | | | | 0.00348 | 0.66901 | Female | | | |
| NF2 | 0.00045 | -1.02999 | Male | | | | | | | | | | | | |
| WDR66 | | | | | | | | | | | | | 0.00076 | -0.60371 | Male |
| FILIP1 | | | | | | | 0.01789 | -0.60432 | Male | | | | | | |

| | | | | | | | | | | | | | | | |
|---|---|---|---|---|---|---|---|---|---|---|---|---|---|---|---|
| CCHCR1 | 0.01125 | -1.76082 | Male | | | | | | | | | | | | |
| SDHAP2 | | | | | | | | | | 0.00293 | 0.66791 | Female | 0.00198 | 0.66759 | Female |
| DENND3 | | | | | | | | | | | | | 0.02451 | 1.67172 | Female |
| TRAPPC5 | 0.00012 | 1.09720 | Female | | | | 0.00000 | 0.66296 | Female | | | | | | |
| NUDT2 | 0.00914 | 1.47097 | Female | | | | | | | | | | | | |
| NCBP2-AS2 | | | | | | | 0.00000 | 0.74189 | Female | | | | | | |
| PRTG | | | | | | | 0.00297 | -0.89691 | Male | | | | | | |
| PDCD7 | | | | | | | 0.00000 | 0.87454 | Female | | | | | | |
| DUSP19 | | | | | | | | | | | | | 0.00085 | 1.74105 | Female |
| ZSCAN16-AS1 | 0.00014 | 3.15673 | Female | | | | | | | | | | 0.02001 | 0.88180 | Female |
| TTC39B | | | | | | | | | | 0.00003 | 1.73056 | Female | | | |
| FGD4 | | | | | | | | | | 0.00093 | 0.88765 | Female | | | |
| LOC101927284 | | | | | | | | | | 0.00001 | -1.19800 | Male | 0.00004 | -1.05764 | Male |
| SLC35A1 | | | | 0.00038 | 1.18253 | Female | | | | 0.00484 | 0.64440 | Female | | | |
| LOC158960 | | | | | | | 0.00049 | 1.32419 | Female | 0.00195 | 1.15897 | Female | | | |
| SEZ6L2 | | | | | | | | | | 0.00037 | 0.84448 | Female | | | |
| S1PR3 | 0.00718 | 3.21334 | Female | | | | | | | | | | | | |
| IRAK2 | | | | | | | 0.01599 | -1.51632 | Male | | | | | | |
| FAM195B | | | | | | | 0.00003 | -1.05490 | Male | | | | | | |
| GLIPR1 | | | | | | | | | | 0.00038 | -0.97917 | Male | 0.00000 | 1.31533 | Female |
| NDRG2 | | | | | | | | | | 0.01718 | 1.18636 | Female | | | |
| LINC00657 | | | | 0.00641 | 0.66051 | Female | 0.00002 | 0.65486 | Female | | | | | | |
| HIST1H2BB | | | | | | | 0.01371 | 1.79784 | Female | | | | | | |
| LRG1 | 0.01072 | 3.64235 | Female | | | | | | | 0.00128 | 1.98613 | Female | | | |
| SGSM1 | | | | | | | | | | | | | 0.00060 | 2.48824 | Female |
| NEDD1 | | | | 0.01398 | -0.71905 | Male | | | | | | | | | |
| ZNF655 | | | | | | | 0.00083 | -0.76546 | Male | 0.00000 | -1.03551 | Male | | | |
| WBP1L | | | | | | | 0.00016 | 0.68007 | Female | | | | 0.00018 | 0.63485 | Female |
| ARFGEF1 | | | | | | | | | | 0.00004 | 0.59433 | Female | | | |
| CLCN4 | | | | | | | | | | 0.00436 | -2.30550 | Male | 0.00005 | -3.07360 | Male |
| KIAA1958 | | | | 0.03998 | 1.04049 | Female | | | | | | | | | |
| FAM216A | | | | | | | | | | 0.00042 | -1.06551 | Male | | | |
| KHDC1L | | | | | | | | | | 0.00000 | 1.19393 | Female | | | |
| DHRS3 | 0.00000 | 4.22044 | Female | | | | | | | 0.00195 | 1.04621 | Female | | | |
| ATL3 | | | | | | | | | | 0.00010 | 0.78304 | Female | | | |
| CACNA2D1 | | | | | | | | | | 0.00031 | 1.92498 | Female | | | |
| KSR1 | | | | | | | | | | | | | 0.02803 | 0.84290 | Female |
| ZFPM2 | | | | | | | | | | | | | 0.00096 | -2.24967 | Male |
| MARCH9 | 0.01568 | 2.76225 | Female | | | | | | | | | | | | |
| ACOT9 | | | | | | | 0.00558 | 0.97591 | Female | | | | | | |
| RNFT2 | 0.02115 | 2.15343 | Female | | | | | | | | | | | | |
| MGST1 | | | | | | | 0.00044 | -0.92054 | Male | | | | 0.00151 | -0.78758 | Male |
| MTRNR2L9 | | | | | | | | | | 0.00000 | 1.08303 | Female | | | |
| ZMAT2 | | | | | | | | | | 0.00000 | -0.61181 | Male | | | |
| FAR2 | | | | | | | | | | | | | 0.00769 | 2.02414 | Female |
| ATP13A2 | | | | 0.00127 | 0.94910 | Female | | | | 0.00002 | 0.78856 | Female | | | |
| LOC100506844 | | | | | | | 0.00497 | 2.05233 | Female | | | | | | |
| ADARB1 | | | | | | | | | | | | | 0.00033 | 1.20607 | Female |
| SPPL2A | | | | | | | | | | | | | 0.00000 | -0.62738 | Male |
| ECEL1P2 | | | | 0.00330 | 1.89021 | Female | 0.00000 | -2.74113 | Male | 0.00000 | -2.25895 | Male | | | |
| ZNF524 | | | | | | | 0.03191 | -1.06412 | Male | | | | | | |

| | | | | | | | | | | | | | | | |
|---|---|---|---|---|---|---|---|---|---|---|---|---|---|---|---|
| THOC7 | | | | | | | 0.00000 | -0.65559 | Male | | | | | | |
| HSPB1 | | | | | | | | | | 0.00416 | 0.73874 | Female | | | |
| AIFM2 | 0.02819 | -2.05874 | Male | 0.00068 | 1.91455 | Female | | | | | | | | | |
| SLC39A10 | | | | | | | | | | 0.00004 | 0.76024 | Female | | | |
| LINC00605 | | | | | | | | | | 0.00000 | 3.15241 | Female | | | |
| COA6 | 0.00241 | -0.68110 | Male | | | | | | | | | | | | |
| SNX12 | | | | 0.01646 | 0.90002 | Female | 0.02000 | 0.59454 | Female | | | | 0.00001 | 0.94103 | Female |
| GNAQ | 0.04481 | -1.14746 | Male | | | | | | | | | | | | |
| TBC1D4 | | | | | | | 0.02125 | 1.39800 | Female | | | | | | |
| MICU3 | | | | 0.03288 | -1.81220 | Male | | | | 0.02952 | -1.16692 | Male | 0.03418 | -1.11367 | Male |
| EPPK1 | | | | 0.00000 | 1.67623 | Female | 0.00000 | 1.12066 | Female | 0.00001 | 1.03864 | Female | | | |
| CCNY | | | | 0.00027 | 0.99966 | Female | | | | | | | | | |
| TAAR5 | | | | 0.00679 | -1.20999 | Male | 0.00622 | 0.85300 | Female | 0.00000 | -1.47562 | Male | 0.00000 | -3.42031 | Male |
| TMEM144 | | | | | | | | | | | | | 0.01552 | -1.12697 | Male |
| CRELD1 | | | | | | | | | | 0.00444 | 0.77187 | Female | | | |
| DNHD1 | | | | | | | | | | | | | 0.03892 | 0.71197 | Female |
| AIG1 | | | | | | | 0.03300 | -1.52724 | Male | | | | | | |
| SLC25A13 | | | | 0.00025 | 0.97831 | Female | | | | | | | | | |
| BAMBI | | | | | | | | | | 0.00000 | 2.62920 | Female | | | |
| LRRC37BP1 | | | | | | | | | | 0.02325 | 0.64867 | Female | 0.01020 | 0.70454 | Female |
| CYP26A1 | | | | | | | | | | | | | 0.00144 | 1.57427 | Female |
| LINC00200 | | | | | | | | | | | | | 0.02509 | 1.56643 | Female |
| IRAK1 | | | | | | | 0.00014 | 1.15070 | Female | 0.00456 | -0.85368 | Male | | | |
| EDNRB | 0.00010 | 2.61617 | Female | | | | | | | | | | | | |
| PLEC | | | | 0.00000 | 1.64802 | Female | | | | 0.00006 | 0.78265 | Female | | | |
| RHOV | 0.00000 | 0.84452 | Female | 0.02277 | 1.16289 | Female | | | | | | | | | |
| BAIAP2L2 | | | | | | | | | | 0.00000 | 1.91818 | Female | 0.00047 | 1.44980 | Female |
| FABP5 | | | | 0.00007 | 0.82310 | Female | | | | | | | | | |
| MFI2 | | | | | | | | | | 0.00500 | 2.37263 | Female | | | |
| TMTC1 | | | | | | | 0.01933 | -1.89142 | Male | | | | | | |
| DGKB | | | | | | | 0.04174 | -1.96088 | Male | | | | | | |
| EDEM1 | | | | | | | | | | 0.00595 | 0.63610 | Female | | | |
| SYCE1 | | | | | | | | | | 0.00486 | 1.28021 | Female | 0.00007 | 1.67737 | Female |
| MTHFS.uid-196 | 0.04306 | 1.14213 | Female | | | | | | | 0.00036 | 0.61083 | Female | | | |
| PPAPDC1B | 0.00349 | 2.62083 | Female | | | | | | | | | | | | |
| IPW | | | | | | | | | | 0.01610 | 0.75943 | Female | | | |
| TRIO | | | | | | | 0.00278 | 0.89041 | Female | | | | | | |
| DAGLA | | | | | | | | | | 0.00005 | 2.53503 | Female | | | |
| BCL6 | | | | | | | 0.00093 | -1.43167 | Male | 0.00011 | -1.53772 | Male | | | |
| SGMS1 | | | | | | | 0.00000 | -1.27759 | Male | | | | 0.00248 | -0.64957 | Male |
| SYAP1 | 0.00233 | 0.77579 | Female | 0.00002 | 0.62832 | Female | 0.00000 | 1.06123 | Female | 0.00000 | 0.65838 | Female | | | |
| MMP15 | | | | | | | | | | 0.00002 | 1.82553 | Female | | | |
| HOMER3 | | | | | | | | | | 0.00364 | 1.88699 | Female | | | |
| PP14571 | | | | | | | 0.03742 | -1.19849 | Male | | | | | | |
| CYP51A1-AS1 | | | | | | | | | | | | | 0.04287 | 0.81373 | Female |
| JADE2 | | | | 0.00000 | 2.99838 | Female | | | | 0.00000 | 1.91281 | Female | 0.01525 | 0.98341 | Female |
| MBTD1 | | | | | | | | | | 0.01467 | -0.64700 | Male | | | |
| LONRF2 | | | | | | | | | | | | | 0.04807 | 2.35242 | Female |
| PRR12 | | | | 0.04834 | -1.07565 | Male | | | | | | | | | |
| SCAMP5 | | | | | | | | | | | | | 0.01232 | 1.37447 | Female |
| ZNF608 | | | | | | | | | | 0.03462 | 1.24255 | Female | | | |

| | | | | | | | | | | | | | | | |
|---|---|---|---|---|---|---|---|---|---|---|---|---|---|---|---|
| HRASLS5 | | | | | | | | | | 0.00404 | 0.96172 | Female | 0.00068 | 1.09139 | Female |
| ZMIZ1 | 0.00516 | 1.40192 | Female | 0.00001 | 1.09544 | Female | | | | | | | | | |
| GABARAP | 0.00000 | 1.49995 | Female | 0.00006 | -0.65006 | Male | | | | | | | | | |
| TMEM184B | | | | | | | 0.00003 | -0.82795 | Male | | | | | | |
| TUBB2A | | | | | | | | | | 0.00000 | 1.38263 | Female | | | |
| TOP3A | | | | | | | 0.01862 | 0.59565 | Female | | | | | | |
| BCAR4 | | | | | | | | | | 0.00885 | -1.04300 | Male | | | |
| LOC401242 | | | | | | | | | | 0.00000 | -3.48675 | Male | | | |
| MXRA7 | 0.04508 | 1.57892 | Female | 0.01507 | 1.03043 | Female | 0.01275 | -0.70779 | Male | | | | | | |
| FBXO11 | | | | 0.00316 | 0.80448 | Female | | | | | | | | | |
| IGDCC3 | | | | 0.04563 | 1.41853 | Female | | | | | | | | | |
| RBPMS2 | | | | 0.00133 | 0.68203 | Female | | | | 0.00000 | 0.83535 | Female | | | |
| SFR1 | | | | | | | 0.01754 | -0.64320 | Male | | | | 0.00505 | -0.70113 | Male |
| XBP1 | 0.04126 | -1.01377 | Male | | | | | | | | | | | | |
| COL26A1 | | | | | | | | | | 0.04821 | 1.47293 | Female | | | |
| MED18 | | | | | | | | | | 0.00216 | -0.60447 | Male | | | |
| TPTEP1 | | | | | | | | | | 0.00000 | 0.63641 | Female | | | |
| ZFYVE1 | | | | | | | | | | | | | 0.00012 | 0.87075 | Female |
| HOXC13-AS | | | | 0.03291 | -2.41919 | Male | | | | | | | | | |
| TTN | | | | | | | 0.00246 | -0.66194 | Male | | | | | | |
| C1orf159 | 0.00200 | -2.15517 | Male | | | | | | | | | | | | |
| NMI | | | | 0.01664 | -1.36695 | Male | 0.02045 | 0.75482 | Female | | | | | | |
| ZCCHC11 | | | | 0.02046 | -0.91275 | Male | | | | | | | | | |
| DNALI1 | | | | | | | 0.00002 | -1.80268 | Male | 0.00528 | -1.16765 | Male | | | |
| TIGD5 | 0.04955 | 2.54259 | Female | 0.00204 | 2.11875 | Female | | | | | | | | | |
| NSUN7 | | | | | | | 0.03013 | -0.81019 | Male | 0.04690 | -0.72325 | Male | | | |
| PHF21B | | | | | | | 0.01824 | 1.26854 | Female | | | | 0.00130 | 1.56456 | Female |
| PIGM | | | | | | | 0.04399 | -0.78633 | Male | | | | | | |
| HIST1H3I | | | | | | | | | | 0.00317 | -1.14891 | Male | | | |
| KRT80 | | | | | | | | | | 0.00000 | -2.86760 | Male | | | |
| COMMD8 | | | | | | | | | | | | | 0.00004 | -0.66172 | Male |
| ARAP2 | | | | | | | | | | | | | 0.02724 | -1.72930 | Male |
| HCLS1 | | | | 0.02925 | -1.31325 | Male | | | | | | | | | |
| SORBS3 | | | | | | | | | | 0.03150 | 0.94626 | Female | | | |
| RAB11FIP3 | | | | | | | | | | 0.00388 | -0.81567 | Male | | | |
| TBX19 | | | | 0.00032 | 2.84414 | Female | | | | | | | | | |
| RHNO1 | | | | | | | 0.00137 | -0.72058 | Male | | | | | | |
| SH2B1 | | | | | | | | | | 0.00913 | 0.70030 | Female | 0.03304 | 0.58234 | Female |
| CARS2 | | | | 0.03930 | 1.00988 | Female | | | | | | | | | |
| FUOM | | | | 0.00000 | 1.62133 | Female | | | | | | | | | |
| HEY2 | | | | | | | | | | 0.00071 | 1.70687 | Female | 0.02990 | 1.14674 | Female |
| TUBA3D | | | | | | | 0.00028 | 1.13056 | Female | | | | 0.00000 | 1.52102 | Female |
| RB1 | | | | | | | | | | 0.03543 | -0.63652 | Male | | | |
| ANKRD9 | | | | 0.02425 | 1.30949 | Female | | | | | | | | | |
| ZNF311 | | | | | | | | | | 0.02116 | 1.50068 | Female | | | |
| KITLG | | | | | | | | | | | | | 0.01442 | -1.09152 | Male |
| DUXA | | | | | | | | | | 0.00000 | 2.65777 | Female | | | |
| TMEM63B | | | | | | | | | | 0.00395 | -0.76873 | Male | | | |
| WDR18 | 0.00388 | -0.88695 | Male | | | | 0.00000 | -0.58449 | Male | | | | | | |
| 15.00000 | 0.03996 | -1.12846 | Male | | | | | | | | | | 0.00000 | -0.85573 | Male |
| CLEC2D | | | | | | | 0.00034 | -0.75818 | Male | | | | 0.00004 | -0.75631 | Male |

| | | | | | | | | | | | | | | | |
|---|---|---|---|---|---|---|---|---|---|---|---|---|---|---|---|
| DBNDD2 | | | | | | | | | | 0.00486 | -0.73337 | Male | | | |
| CACNA2D2 | | | | | | | | | | 0.00001 | 1.74183 | Female | | | |
| ADAMTSL3 | | | | | | | 0.04130 | -1.34027 | Male | | | | | | |
| SYVN1 | | | | | | | 0.00015 | -0.82644 | Male | | | | | | |
| BTBD3 | | | | 0.02109 | -0.89067 | Male | | | | | | | | | |
| LCP1 | | | | | | | 0.00187 | -0.92010 | Male | | | | | | |
| OGDH | 0.02522 | 0.91448 | Female | | | | | | | | | | | | |
| ALG1L | | | | | | | | | | 0.04214 | 1.09695 | Female | | | |
| JUND | | | | 0.00595 | 0.83831 | Female | | | | | | | 0.00001 | 0.78663 | Female |
| CCDC171 | | | | | | | | | | 0.02045 | 1.41830 | Female | | | |
| PRAME | | | | | | | | | | 0.00004 | 1.97099 | Female | 0.00000 | 3.26499 | Female |
| MXRA8 | | | | | | | | | | | | | 0.02990 | -0.78772 | Male |
| TRG-AS1 | | | | | | | | | | | | | 0.04217 | 1.76225 | Female |
| PURA | | | | | | | | | | 0.00007 | -0.94826 | Male | | | |
| MTRF1 | | | | | | | | | | | | | 0.00140 | -0.62852 | Male |
| NRN1L | | | | 0.01967 | 1.58731 | Female | 0.00649 | 1.19885 | Female | | | | | | |
| HAUS6 | 0.00594 | -0.90979 | Male | | | | | | | | | | | | |
| TSPAN6 | 0.00053 | 1.29458 | Female | 0.00113 | 0.73138 | Female | | | | | | | | | |
| ZFY | 0.00006 | -3.42423 | Male | 0.00000 | -5.94546 | Male | 0.00000 | -6.65757 | Male | 0.00000 | -7.08029 | Male | 0.00000 | -7.27079 | Male |
| MEX3D | | | | | | | | | | 0.01725 | -0.60493 | Male | | | |
| CREB3L1 | | | | 0.00252 | 2.57730 | Female | | | | 0.00001 | 2.44555 | Female | | | |
| TPI1 | 0.00043 | 0.67091 | Female | 0.00000 | 0.89481 | Female | | | | | | | | | |
| CDK4 | 0.02160 | 0.60816 | Female | | | | | | | | | | | | |
| FN1 | | | | | | | | | | 0.04766 | 0.84140 | Female | 0.00310 | -1.15887 | Male |
| PGPEP1 | | | | | | | | | | | | | 0.00489 | 0.95387 | Female |
| SPINK1 | | | | | | | | | | 0.02485 | 1.34236 | Female | 0.00495 | -1.61269 | Male |
| KLF7 | | | | | | | | | | 0.00651 | 1.86659 | Female | | | |
| BTLA | | | | | | | | | | 0.01359 | 1.46545 | Female | | | |
| IFFO2 | | | | | | | | | | 0.03250 | -1.67147 | Male | | | |
| OCIAD2 | | | | 0.01307 | -2.53576 | Male | | | | | | | | | |
| PKIA | | | | 0.00184 | -2.25623 | Male | | | | 0.00018 | -1.70595 | Male | 0.01629 | -1.11241 | Male |
| KIF21A | | | | 0.00001 | 1.61332 | Female | | | | | | | | | |
| DENND1A | | | | 0.04772 | 0.94170 | Female | | | | | | | | | |
| ATF4 | | | | 0.00000 | -0.90521 | Male | | | | | | | | | |
| UTP20 | 0.02822 | -0.74291 | Male | | | | | | | | | | | | |
| MRPS35 | 0.02000 | -0.60555 | Male | | | | | | | | | | | | |
| ABHD13 | | | | | | | 0.01645 | -0.62727 | Male | | | | 0.01777 | -0.58117 | Male |
| MRPS10 | 0.00007 | -1.04179 | Male | | | | | | | | | | | | |
| DROSHA | | | | 0.00001 | 1.05271 | Female | | | | | | | | | |
| FOXD1 | | | | | | | 0.00965 | 1.97265 | Female | 0.02835 | -1.65832 | Male | | | |
| HSPB8 | 0.00000 | 1.24730 | Female | 0.00000 | -1.35724 | Male | | | | | | | | | |
| DUSP6 | 0.00147 | -2.31497 | Male | | | | | | | | | | | | |
| PSAT1 | | | | 0.01640 | -0.85876 | Male | 0.00000 | -1.10664 | Male | | | | | | |
| APTR | | | | | | | 0.00074 | -0.81728 | Male | | | | | | |
| LOC100505817 | | | | | | | 0.01271 | -0.85119 | Male | | | | 0.02715 | -0.72677 | Male |
| SSBP3 | | | | | | | 0.00000 | 0.78636 | Female | | | | | | |
| NPAS1 | | | | | | | | | | 0.01244 | 1.20613 | Female | | | |
| CAB39 | | | | 0.00000 | 1.38890 | Female | | | | | | | | | |
| ANPEP | 0.00000 | 2.07824 | Female | 0.00000 | 1.54957 | Female | | | | 0.00000 | 1.04063 | Female | | | |
| GPR37L1 | | | | | | | | | | 0.00213 | 1.53183 | Female | | | |
| CNN2 | | | | | | | | | | 0.00528 | -0.62680 | Male | | | |

| | | | | | | | | | | | | | | | |
|---|---|---|---|---|---|---|---|---|---|---|---|---|---|---|---|
| CNOT6L | | | | 0.00135 | 1.12989 | Female | | | | 0.00070 | -0.78396 | Male | | | |
| SLC30A2 | | | | 0.00017 | 2.72795 | Female | | | | 0.02391 | 1.20114 | Female | | | |
| PPL | | | | | | | | | | 0.03935 | -0.99192 | Male | | | |
| PLEKHG1 | | | | | | | | | | | | | 0.00966 | 1.89213 | Female |
| EN1 | | | | | | | | | | | | | 0.00000 | -1.64045 | Male |
| XYLT2 | | | | | | | | | | 0.02164 | 0.68064 | Female | | | |
| LOC729603 | | | | 0.04331 | 1.45633 | Female | | | | | | | | | |
| RARS | 0.00412 | -0.72964 | Male | | | | | | | | | | | | |
| SRC | | | | | | | 0.00001 | -1.65045 | Male | 0.01758 | -0.88880 | Male | | | |
| TEX11 | | | | | | | | | | 0.00000 | 1.96961 | Female | 0.00000 | 1.68319 | Female |
| TFF1 | | | | | | | | | | 0.03733 | 1.51526 | Female | | | |
| FAM127A | | | | 0.00022 | 1.07230 | Female | 0.00000 | 1.10141 | Female | | | | | | |
| ECHDC2 | | | | | | | | | | 0.00251 | -0.63312 | Male | | | |
| MEI1 | | | | | | | | | | | | | 0.01054 | 1.03364 | Female |
| OXR1 | | | | | | | | | | | | | 0.01289 | -0.71839 | Male |
| FAM60A | 0.01022 | -0.61663 | Male | | | | | | | | | | | | |
| CASP3 | 0.02822 | -0.84213 | Male | | | | | | | | | | | | |
| SLC30A6 | | | | | | | 0.00001 | -0.75557 | Male | | | | | | |
| CAPN13 | | | | | | | | | | 0.01456 | 1.65129 | Female | | | |
| CD200 | | | | | | | | | | 0.00984 | 1.85631 | Female | | | |
| FARP1 | | | | 0.00002 | 1.64183 | Female | | | | | | | | | |
| NUDT19 | | | | | | | 0.02183 | 1.11850 | Female | | | | | | |
| BAGE | | | | | | | | | | 0.00735 | -1.16773 | Male | | | |
| C10orf95 | | | | | | | 0.00099 | 0.75763 | Female | 0.00198 | 0.73022 | Female | 0.00162 | 0.69436 | Female |
| GPR160 | | | | | | | | | | 0.00000 | 1.16512 | Female | | | |
| FUCA1 | | | | | | | | | | 0.00001 | 0.84970 | Female | | | |
| FGF12 | | | | | | | | | | | | | 0.02325 | -1.17115 | Male |
| RAB20 | | | | 0.00000 | 2.03580 | Female | | | | | | | | | |
| LETM1 | 0.00828 | -0.78405 | Male | | | | 0.00000 | 0.64895 | Female | | | | | | |
| CSTA | | | | | | | 0.00316 | -1.96537 | Male | 0.01955 | 1.56029 | Female | | | |
| PHLDA2 | 0.00089 | -1.49988 | Male | | | | 0.00000 | 1.05439 | Female | 0.00000 | 0.84501 | Female | | | |
| ESYT3 | | | | | | | 0.03824 | -1.84773 | Male | | | | | | |
| GGCX | | | | 0.00000 | 1.54653 | Female | | | | | | | | | |
| ANKRD35 | | | | | | | 0.00000 | -4.07778 | Male | | | | | | |
| COLEC12 | | | | | | | 0.00054 | -0.88020 | Male | | | | | | |
| C6orf99 | | | | 0.00339 | 1.26098 | Female | | | | | | | 0.00232 | 0.81717 | Female |
| ZMYM4 | | | | 0.02983 | 0.71394 | Female | | | | | | | | | |
| ALX1 | | | | | | | | | | | | | 0.04137 | -1.18607 | Male |
| GOLGA8A | 0.04258 | 1.32417 | Female | | | | 0.02186 | -0.59416 | Male | | | | | | |
| PIN4P1 | | | | 0.00000 | 1.14841 | Female | 0.00000 | 1.48207 | Female | 0.00000 | 0.89177 | Female | 0.00000 | 0.78510 | Female |
| GADD45B | | | | | | | | | | | | | 0.02338 | 0.89669 | Female |
| NID2 | | | | | | | 0.00017 | -1.98402 | Male | | | | | | |
| SLC25A37 | | | | | | | | | | 0.00306 | -0.62818 | Male | | | |
| TAF1 | | | | | | | 0.00094 | 0.86570 | Female | 0.01230 | 0.65630 | Female | 0.00500 | 0.71224 | Female |
| B4GALT2 | | | | | | | 0.02381 | -0.92262 | Male | 0.03854 | -0.81222 | Male | | | |
| KLK8 | | | | 0.01816 | -1.01310 | Male | 0.00932 | 0.74270 | Female | 0.00000 | 1.48466 | Female | | | |
| SLC38A1 | | | | | | | 0.00278 | -0.87063 | Male | 0.00076 | -0.90442 | Male | | | |
| RUFY2 | | | | | | | 0.01551 | -0.63820 | Male | 0.00174 | 0.75783 | Female | | | |
| MTMR11 | | | | | | | | | | 0.03923 | 1.79559 | Female | | | |
| TEX40 | | | | | | | | | | 0.01590 | 1.02157 | Female | | | |
| ZSCAN26 | | | | | | | | | | 0.01892 | 1.45708 | Female | | | |

| | | | | | | | | | | | | | | | |
|---|---|---|---|---|---|---|---|---|---|---|---|---|---|---|---|
| SEC14L1P1 | | | | | | | | | | | | | 0.00654 | -0.90835 | Male |
| NEDD4L | | | | | | | | | | 0.03891 | 0.65667 | Female | | | |
| CDK5RAP3 | | | | | | | | | | 0.00494 | 0.62474 | Female | | | |
| MANF | | | | | | | | | | 0.00000 | -0.78100 | Male | | | |
| MAGEA12 | | | | | | | 0.04610 | 2.23485 | Female | | | | 0.00489 | 1.99260 | Female |
| MBTPS2 | | | | 0.03716 | 1.31942 | Female | 0.00278 | 1.16190 | Female | | | | | | |
| ATP1A1-AS1 | 0.02475 | -1.03704 | Male | | | | | | | | | | | | |
| HSD17B7 | | | | 0.00000 | -1.97216 | Male | | | | | | | | | |
| COCH | | | | 0.00016 | 1.81060 | Female | 0.00000 | -2.06396 | Male | 0.00003 | -1.26227 | Male | 0.01598 | -0.77068 | Male |
| WEE2 | | | | | | | 0.04534 | 1.51814 | Female | 0.00494 | -1.95250 | Male | 0.00018 | -2.46507 | Male |
| DENND2C | | | | | | | 0.03120 | -1.50885 | Male | | | | | | |
| GAGE2B | | | | | | | 0.00000 | 3.60793 | Female | | | | | | |
| MFSD3 | | | | | | | 0.00290 | -1.33290 | Male | | | | | | |
| C6orf57 | | | | | | | | | | | | | 0.00789 | -1.03429 | Male |
| TNFAIP2 | | | | | | | 0.00405 | 1.79639 | Female | | | | | | |
| MANBA | 0.02282 | 1.88786 | Female | | | | | | | | | | | | |
| RALGDS | | | | 0.00437 | 1.84977 | Female | | | | | | | | | |
| RPL13A | | | | | | | 0.00000 | -0.61038 | Male | | | | | | |
| CCDC80 | | | | | | | 0.01964 | 1.30450 | Female | 0.00000 | 2.34262 | Female | | | |
| MAN2B1 | | | | | | | 0.01518 | -0.76946 | Male | | | | | | |
| CHD7 | 0.04384 | -0.72110 | Male | | | | | | | | | | | | |
| ZNF34 | | | | | | | | | | 0.04414 | 1.01950 | Female | | | |
| PODXL | | | | | | | | | | 0.00654 | 0.84077 | Female | | | |
| DIAPH1 | 0.04441 | -0.82131 | Male | | | | | | | | | | | | |
| POLB | | | | | | | 0.00003 | -0.69559 | Male | | | | | | |
| APOC3 | | | | | | | | | | 0.00563 | 1.24704 | Female | 0.00002 | 1.77724 | Female |
| SLC1A3 | 0.00712 | -1.68389 | Male | | | | | | | | | | | | |
| NAALAD2 | | | | | | | 0.00032 | 0.94521 | Female | | | | | | |
| HBQ1 | | | | | | | | | | 0.03020 | -1.81345 | Male | | | |
| SYTL2 | | | | | | | | | | | | | 0.00511 | -1.89138 | Male |
| EPHA4 | | | | | | | | | | 0.00156 | 2.12743 | Female | | | |
| HAUS7 | | | | 0.00001 | 1.10838 | Female | 0.00000 | 1.45036 | Female | | | | 0.00000 | 0.79962 | Female |
| NME1 | 0.00088 | -0.59767 | Male | | | | | | | | | | | | |
| HPSE | | | | | | | | | | 0.00160 | 1.82086 | Female | | | |
| GMPR2 | 0.00264 | 1.34736 | Female | | | | | | | | | | | | |
| SLC22A11 | | | | 0.00061 | 2.58237 | Female | | | | | | | | | |
| ERICH1-AS1 | | | | | | | 0.00046 | -2.34046 | Male | | | | | | |
| KCNK12 | | | | | | | | | | 0.00994 | 2.19892 | Female | | | |
| TOMM40L | | | | 0.03537 | 1.23004 | Female | | | | | | | | | |
| ZNF440 | | | | 0.00004 | 2.05034 | Female | 0.02111 | -0.83544 | Male | | | | | | |
| DBN1 | | | | | | | 0.00732 | -0.61594 | Male | 0.00764 | 0.58705 | Female | | | |
| DEPTOR | | | | | | | | | | 0.00000 | 1.80300 | Female | | | |
| SLC7A11-AS1 | | | | | | | | | | 0.01249 | 1.56295 | Female | 0.00370 | -1.75109 | Male |
| TYK2 | 0.04423 | -1.38371 | Male | | | | | | | | | | | | |
| ASPHD1 | | | | | | | | | | | | | 0.00978 | 1.41654 | Female |
| BCAP31 | | | | 0.00000 | 0.95324 | Female | 0.00000 | 0.96767 | Female | | | | | | |
| GNG7 | | | | | | | | | | 0.01741 | 1.63857 | Female | | | |
| IMPAD1 | | | | | | | | | | | | | 0.00001 | -0.65135 | Male |
| SPRY4 | | | | | | | | | | 0.00000 | 1.88233 | Female | 0.04068 | 0.86478 | Female |
| MORN2 | | | | | | | 0.00107 | -0.67465 | Male | | | | | | |
| MYL12A | 0.00000 | -0.94118 | Male | | | | | | | | | | | | |

| | | | | | | | | | | | | | | | |
|---|---|---|---|---|---|---|---|---|---|---|---|---|---|---|---|
| ATP5G2 | | | | | | | 0.00000 | -0.61314 | Male | | | | | | |
| DGUOK-AS1 | 0.03121 | -1.29776 | Male | | | | | | | | | | | | |
| BCOR | | | | | | | 0.00340 | 0.80108 | Female | 0.00000 | 1.61311 | Female | | | |
| MND1 | | | | | | | 0.00001 | -0.76556 | Male | | | | | | |
| ASGR2 | | | | | | | | | | 0.03158 | 2.01909 | Female | | | |
| AASS | | | | | | | 0.01160 | -0.76130 | Male | 0.00251 | -0.83467 | Male | | | |
| TRIM54 | | | | | | | | | | | | | 0.00196 | 2.03536 | Female |
| IRF6 | | | | | | | 0.01338 | -0.93305 | Male | | | | | | |
| ACD | | | | | | | | | | | | | 0.00006 | 0.62669 | Female |
| MIAT | | | | | | | 0.00777 | 1.69421 | Female | 0.00004 | 2.24994 | Female | | | |
| DDIT3 | | | | | | | | | | | | | 0.00001 | 0.95776 | Female |
| HLA-F | | | | 0.01549 | 1.21879 | Female | | | | 0.00000 | 1.96216 | Female | 0.00140 | 0.98772 | Female |
| PRKY | | | | 0.00000 | -5.45828 | Male | 0.00000 | -6.00501 | Male | 0.00000 | -7.01340 | Male | 0.00000 | -7.13151 | Male |
| HN1L | | | | | | | | | | 0.00000 | -0.65321 | Male | | | |
| RASIP1 | | | | | | | 0.01776 | -1.54653 | Male | | | | | | |
| SCRN2 | | | | 0.04090 | -1.45130 | Male | | | | | | | | | |
| RHOC | | | | 0.00000 | 0.94790 | Female | | | | | | | | | |
| ZNF503 | | | | | | | 0.00032 | 2.53333 | Female | | | | | | |
| MED31 | | | | 0.00001 | -1.05998 | Male | | | | | | | | | |
| ZBTB6 | | | | | | | | | | 0.04968 | -0.71868 | Male | 0.04710 | -0.71392 | Male |
| DOT1L | | | | | | | | | | 0.00000 | 0.77329 | Female | | | |
| SMYD3 | | | | | | | | | | | | | 0.00574 | 0.72791 | Female |
| TPPP3 | | | | | | | | | | | | | 0.03059 | -2.13382 | Male |
| TMEM129 | 0.02488 | -2.12059 | Male | | | | | | | | | | | | |
| MTRNR2L3 | | | | 0.00056 | -1.27798 | Male | | | | 0.00894 | 0.72449 | Female | | | |
| SEH1L | 0.00828 | 0.59762 | Female | | | | | | | | | | | | |
| PRR14 | | | | 0.01275 | -0.79347 | Male | | | | | | | | | |
| SLC35E4 | | | | | | | | | | 0.00000 | 1.53152 | Female | | | |
| LRRC37A11P | | | | | | | | | | | | | 0.04953 | 0.98312 | Female |
| HIST1H2AI | | | | | | | | | | 0.00000 | -2.73474 | Male | | | |
| TUBD1 | 0.01855 | -1.23821 | Male | | | | | | | | | | | | |
| CABIN1 | 0.02109 | 1.24810 | Female | | | | 0.00484 | 0.60988 | Female | | | | | | |
| PRKX | | | | 0.00000 | 1.47020 | Female | 0.00000 | 1.42473 | Female | 0.00004 | 0.82006 | Female | 0.00000 | 1.05850 | Female |
| GRIN2D | | | | | | | 0.03670 | 1.01936 | Female | | | | 0.03266 | 0.97713 | Female |
| IFNAR2 | | | | | | | 0.00283 | -0.99219 | Male | | | | | | |
| BPNT1 | | | | | | | 0.00021 | -0.91211 | Male | | | | | | |
| REEP4 | | | | | | | 0.00066 | -0.58164 | Male | | | | | | |
| OCRL | | | | | | | | | | 0.00026 | 1.28437 | Female | | | |
| PROS1 | | | | | | | | | | 0.02894 | 1.21613 | Female | | | |
| FBXO36 | | | | | | | | | | 0.00191 | 0.83560 | Female | | | |
| WDR72 | | | | | | | 0.00704 | -1.09146 | Male | | | | 0.00001 | -1.56390 | Male |
| APOA2 | | | | | | | | | | 0.04410 | 1.04247 | Female | 0.01519 | 1.20440 | Female |
| ST14 | | | | 0.00000 | 1.69884 | Female | | | | | | | | | |
| NUCB1 | | | | 0.00000 | 0.87447 | Female | | | | 0.00000 | 0.59470 | Female | | | |
| MSH6 | | | | | | | | | | 0.00000 | 0.63795 | Female | | | |
| PHF5A | | | | 0.00000 | -0.97125 | Male | | | | | | | | | |
| PTPRD | | | | | | | | | | 0.00088 | 2.40184 | Female | | | |
| PHACTR1 | | | | | | | | | | 0.03845 | 1.10940 | Female | | | |
| CD276 | 0.00000 | -2.14791 | Male | | | | | | | | | | | | |
| CHMP4C | | | | 0.00007 | -2.19807 | Male | | | | | | | | | |
| ICOSLG | | | | | | | | | | 0.00239 | 0.88687 | Female | 0.00047 | 0.98256 | Female |

| | | | | | | | | | | | | | | | |
|---|---|---|---|---|---|---|---|---|---|---|---|---|---|---|---|
| PVR | | | | 0.00033 | 1.16908 | Female | | | | | | | | | |
| LOC101927151 | | | | | | | 0.00386 | 0.81477 | Female | 0.02231 | -0.64252 | Male | | | |
| FOS | | | | | | | | | | | | | 0.00000 | 2.00518 | Female |
| RPS6KA4 | 0.01192 | 2.10199 | Female | 0.00000 | 1.99562 | Female | | | | | | | 0.02536 | 0.60524 | Female |
| BAG1 | | | | 0.00000 | 1.06976 | Female | | | | | | | | | |
| HRH2 | | | | | | | 0.00039 | 2.71369 | Female | | | | | | |
| SLC37A4 | | | | | | | 0.00010 | -0.71265 | Male | | | | | | |
| SSR4 | | | | 0.00000 | 1.14034 | Female | 0.00000 | 0.91369 | Female | 0.00000 | 0.64060 | Female | | | |
| FASTKD2 | 0.00915 | -0.87378 | Male | | | | | | | | | | | | |
| LEP | | | | | | | 0.02817 | 1.18989 | Female | 0.00491 | 1.38775 | Female | 0.03279 | 1.10773 | Female |
| NDUFS5 | 0.00000 | 1.11903 | Female | | | | | | | | | | | | |
| TREX1 | | | | 0.04657 | 1.61892 | Female | | | | | | | | | |
| GGPS1 | | | | | | | 0.00002 | -0.77017 | Male | 0.00015 | -0.66842 | Male | | | |
| LOC388553 | | | | | | | 0.00777 | 1.61114 | Female | | | | 0.04229 | 1.22402 | Female |
| SLC12A9 | | | | 0.00564 | 1.22181 | Female | 0.00918 | -0.78578 | Male | | | | | | |
| DBP | | | | | | | | | | 0.00005 | 1.64686 | Female | | | |
| WDR34 | | | | | | | 0.00000 | -0.81255 | Male | | | | | | |
| PAGE2 | | | | 0.00000 | 2.44745 | Female | 0.00000 | 2.80843 | Female | 0.00002 | 1.08795 | Female | 0.01844 | 0.64185 | Female |
| SAYSD1 | | | | | | | 0.00113 | 0.87093 | Female | | | | | | |
| BCL11A | | | | 0.01569 | 1.30781 | Female | 0.01114 | 0.93055 | Female | 0.00060 | 1.16365 | Female | | | |
| HMGN5 | | | | | | | | | | 0.00377 | 0.79634 | Female | | | |
| SNX25 | | | | | | | | | | 0.01955 | 1.14743 | Female | | | |
| SLC6A9 | | | | | | | | | | | | | 0.02989 | 0.90683 | Female |
| SEPSECS | | | | | | | | | | | | | 0.02240 | -0.78431 | Male |
| TEX35 | | | | | | | | | | | | | 0.02553 | 1.91467 | Female |
| PLA2G10 | | | | | | | | | | 0.00000 | 1.79873 | Female | | | |
| FAM173A | 0.01700 | 1.64983 | Female | 0.00233 | 1.02742 | Female | | | | | | | | | |
| KIF19 | | | | | | | 0.01268 | -5.41309 | Male | | | | | | |
| CCDC84 | 0.00269 | 1.31404 | Female | | | | | | | | | | | | |
| PNRC1 | | | | | | | 0.00486 | 0.71112 | Female | | | | | | |
| DRAP1 | | | | 0.00202 | 0.63830 | Female | | | | 0.00000 | 0.67928 | Female | | | |
| CRTAP | | | | 0.00486 | 0.90956 | Female | | | | | | | | | |
| FUCA2 | | | | 0.00008 | 1.35023 | Female | | | | | | | | | |
| ACTG1P4 | | | | | | | 0.02704 | -0.67122 | Male | 0.00310 | -0.84027 | Male | | | |
| SRPK2 | 0.01507 | -0.65517 | Male | | | | | | | | | | | | |
| CGB5 | | | | 0.00005 | -3.03851 | Male | | | | 0.00000 | -3.01220 | Male | | | |
| OLR1 | | | | | | | 0.01953 | -0.66374 | Male | | | | | | |
| YPEL5 | | | | | | | | | | 0.00000 | -0.67431 | Male | | | |
| PAX5 | | | | | | | | | | 0.00432 | 2.03111 | Female | | | |
| VMO1 | | | | | | | | | | | | | 0.00421 | 1.39782 | Female |
| PLCB3 | | | | | | | | | | 0.00003 | 0.77382 | Female | | | |
| GRINA | | | | | | | | | | | | | 0.00005 | 0.84784 | Female |
| XAB2 | | | | | | | 0.00001 | -0.63371 | Male | | | | 0.00000 | 0.88002 | Female |
| FST | | | | 0.01943 | 1.86995 | Female | 0.00639 | -1.41947 | Male | | | | | | |
| LRP1 | | | | | | | | | | 0.00131 | 0.77476 | Female | 0.00516 | 0.68103 | Female |
| DYRK1B | | | | | | | | | | | | | 0.02715 | 0.96468 | Female |
| U2AF1L4 | | | | | | | 0.00000 | 1.05493 | Female | 0.00517 | 0.59889 | Female | | | |
| PRELP | | | | | | | | | | 0.01277 | 1.98931 | Female | 0.02750 | 0.86337 | Female |
| FBXL19 | | | | | | | 0.03519 | -0.95420 | Male | | | | | | |
| SEMA6D | | | | 0.01999 | 1.25495 | Female | | | | | | | 0.00393 | -0.92590 | Male |
| TENM1 | | | | | | | | | | | | | 0.00007 | 3.37879 | Female |

| | | | | | | | | | | | | | | | |
|---|---|---|---|---|---|---|---|---|---|---|---|---|---|---|---|
| DPEP1 | | | | | | | | | | | | | 0.00530 | 1.82939 | Female |
| CRIP1 | 0.00003 | 2.07415 | Female | 0.00000 | 1.74658 | Female | | | | | | | 0.00000 | 0.72321 | Female |
| IQSEC1 | | | | 0.00001 | 1.38626 | Female | | | | | | | | | |
| RNF149 | | | | | | | 0.00830 | -0.61680 | Male | | | | | | |
| ACOXL | | | | 0.00075 | 1.37234 | Female | | | | | | | | | |
| ACBD7 | | | | | | | | | | | | | 0.02460 | 1.54881 | Female |
| NLGN3 | | | | | | | 0.01022 | 2.16612 | Female | | | | | | |
| JOSD2 | | | | | | | 0.00100 | -0.94404 | Male | 0.01167 | -0.71452 | Male | | | |
| ARGFXP2 | | | | | | | | | | 0.00040 | 1.20483 | Female | 0.00030 | 1.20901 | Female |
| MBNL3 | 0.01211 | 1.71008 | Female | | | | | | | 0.00002 | -1.02729 | Male | | | |
| ABCA1 | | | | 0.00001 | 2.01531 | Female | | | | 0.00000 | 1.56885 | Female | | | |
| LOC100130899 | | | | 0.01367 | 1.85372 | Female | 0.00380 | 1.44143 | Female | | | | | | |
| HDAC10 | | | | | | | | | | | | | 0.03423 | 1.06118 | Female |
| PSD3 | | | | | | | | | | | | | 0.03252 | -0.90518 | Male |
| HESX1 | | | | | | | | | | 0.00455 | 1.74914 | Female | | | |
| SUN2 | | | | | | | 0.01501 | -0.73469 | Male | | | | | | |
| CDC42EP4 | | | | | | | | | | 0.00000 | 1.18698 | Female | | | |
| RPS26P11 | | | | | | | 0.00000 | 0.93173 | Female | | | | | | |
| SSBP3-AS1 | | | | | | | | | | 0.01507 | 1.18778 | Female | | | |
| LOC646214 | 0.00275 | 0.95435 | Female | | | | | | | | | | | | |
| ADPRH | | | | | | | | | | 0.01272 | -2.84054 | Male | | | |
| ETS1 | | | | | | | | | | 0.00057 | 2.26028 | Female | | | |
| ZNF518A | 0.02265 | -0.82319 | Male | | | | | | | | | | | | |
| FTSJ1 | | | | 0.00005 | 0.96030 | Female | 0.00001 | 0.70834 | Female | 0.00000 | 0.78038 | Female | 0.00010 | 0.60740 | Female |
| GALNT14 | | | | | | | 0.00791 | 1.84071 | Female | | | | | | |
| GUSBP5 | | | | | | | | | | 0.00578 | 1.44682 | Female | | | |
| ZCCHC3 | | | | | | | 0.00521 | 0.75051 | Female | | | | | | |
| ST5 | | | | 0.01569 | 1.26225 | Female | | | | | | | | | |
| FRMD8P1 | | | | | | | 0.00000 | -1.98648 | Male | | | | | | |
| RNF7 | 0.00508 | 0.71623 | Female | | | | | | | | | | | | |
| BBS5 | | | | | | | | | | | | | 0.02328 | 0.97637 | Female |
| ZNF689 | | | | | | | | | | | | | 0.00994 | 0.63084 | Female |
| PADI6 | | | | | | | 0.00305 | 2.16765 | Female | 0.00536 | -1.90773 | Male | 0.00002 | 2.63057 | Female |
| LOC101926892 | | | | | | | | | | | | | 0.02455 | 1.86423 | Female |
| ABHD12 | | | | 0.00027 | 0.93121 | Female | | | | | | | | | |
| CSGALNACT1 | | | | 0.01380 | 1.56915 | Female | | | | | | | | | |
| FAM101B | | | | 0.00000 | 1.98187 | Female | | | | | | | | | |
| LOC100506548 | | | | 0.00596 | -0.84631 | Male | 0.00315 | -0.61857 | Male | | | | | | |
| FLRT3 | | | | | | | 0.02895 | -2.56620 | Male | | | | | | |
| LOC100289019 | | | | | | | 0.03089 | 0.85380 | Female | | | | | | |
| LINC00501 | | | | | | | 0.03334 | 1.31869 | Female | | | | 0.00612 | -1.57134 | Male |
| COQ3 | | | | | | | | | | 0.02580 | 0.69917 | Female | | | |
| KIAA0368 | | | | 0.00278 | 0.70725 | Female | | | | | | | | | |
| SLC2A1 | 0.00000 | 1.71285 | Female | 0.01009 | 0.64856 | Female | | | | | | | | | |
| SMYD2 | | | | 0.04701 | 0.60115 | Female | 0.00001 | -0.78973 | Male | | | | | | |
| VPS13C | | | | | | | | | | 0.02397 | 0.88204 | Female | | | |
| VAMP2 | | | | | | | | | | 0.00044 | -0.76841 | Male | | | |
| DYM | | | | 0.03202 | 0.73027 | Female | | | | | | | | | |
| CRYBG3 | | | | | | | 0.00675 | -2.09331 | Male | | | | | | |
| PPFIA3 | | | | 0.01201 | 1.46486 | Female | | | | | | | | | |
| GATA6-AS1 | 0.00006 | -2.59968 | Male | | | | | | | | | | | | |

| | | | | | | | | | | | | | | | |
|---|---|---|---|---|---|---|---|---|---|---|---|---|---|---|---|
| LXN | 0.04089 | -2.63969 | Male | | | | | | | | | | | | |
| TCN2 | | | | | | | | | | 0.00614 | 1.62384 | Female | | | |
| ADAMTS5 | | | | | | | 0.00180 | 2.02940 | Female | | | | | | |
| NBAS | | | | | | | | | | 0.00029 | 0.74579 | Female | | | |
| ANGPT1 | | | | | | | | | | | | | 0.01943 | -0.64101 | Male |
| CDCA7L | | | | | | | | | | 0.03587 | 1.36680 | Female | | | |
| OFD1 | | | | 0.00001 | 0.68196 | Female | 0.00000 | 0.85297 | Female | 0.00000 | 0.68821 | Female | | | |
| TIPRL | 0.02237 | -0.67618 | Male | | | | | | | | | | | | |
| TYMSOS | | | | | | | | | | 0.00336 | 0.97931 | Female | | | |
| PI4KAP2 | | | | | | | 0.00000 | 0.82476 | Female | | | | 0.00000 | 0.76687 | Female |
| RRP36 | 0.04178 | -0.86954 | Male | | | | | | | | | | | | |
| EXTL2 | | | | | | | | | | | | | 0.00009 | -0.88353 | Male |
| HIST1H1D | 0.03963 | -1.70650 | Male | 0.00006 | -1.79849 | Male | | | | 0.00007 | 1.20378 | Female | | | |
| FAM208B | | | | 0.00153 | 0.84123 | Female | | | | | | | | | |
| C3orf33 | | | | | | | | | | | | | 0.01563 | -1.29455 | Male |
| DPP6 | | | | | | | 0.00000 | -2.39717 | Male | | | | | | |
| HIST2H2BA | | | | | | | 0.03091 | 1.43777 | Female | | | | | | |
| NLRC4 | | | | | | | 0.00099 | 1.73664 | Female | | | | | | |
| NSL1 | 0.04136 | -0.81348 | Male | | | | | | | | | | | | |
| DDB1 | | | | | | | | | | 0.00000 | -0.58304 | Male | | | |
| UBXN8 | 0.03706 | -1.06400 | Male | | | | 0.00050 | -0.67041 | Male | | | | 0.00001 | -0.78934 | Male |
| CDK16 | | | | 0.00000 | 1.09000 | Female | | | | | | | | | |
| PCDH17 | | | | 0.01407 | 2.17297 | Female | | | | | | | | | |
| ZNF787 | | | | 0.00397 | 1.26084 | Female | | | | | | | | | |
| PARP14 | | | | | | | | | | 0.00724 | 2.00399 | Female | | | |
| ALDH6A1 | 0.02942 | 1.21271 | Female | | | | | | | | | | | | |
| CDKN2AIPNL | | | | | | | | | | 0.00043 | 0.58694 | Female | | | |
| ATF3 | | | | 0.02695 | -0.87683 | Male | | | | | | | 0.00018 | 0.86404 | Female |
| MIR4458HG | 0.02523 | -2.01206 | Male | | | | | | | | | | | | |
| NSDHL | 0.02218 | 0.77575 | Female | 0.00000 | 0.96774 | Female | 0.00000 | 0.98937 | Female | 0.00000 | 0.94191 | Female | 0.00000 | 0.59874 | Female |
| ITGA6 | | | | 0.00244 | 1.02468 | Female | | | | 0.00000 | 1.07578 | Female | | | |
| LEF1 | | | | | | | 0.02663 | 2.96568 | Female | | | | 0.02252 | -4.86570 | Male |
| RAC2 | | | | | | | | | | 0.00315 | 1.74156 | Female | | | |
| PHF1 | | | | | | | 0.01446 | -0.96732 | Male | | | | | | |
| MTM1 | | | | 0.04513 | 1.13440 | Female | 0.02610 | 0.82996 | Female | 0.01869 | 0.82908 | Female | | | |
| SETD8 | | | | | | | 0.00000 | 0.73927 | Female | | | | | | |
| SYNPO2L | | | | | | | | | | 0.00668 | -1.98250 | Male | | | |
| OBSCN | | | | | | | 0.00221 | 2.89051 | Female | | | | | | |
| SLC35D2 | | | | | | | 0.01520 | -0.94405 | Male | | | | | | |
| GFRA3 | | | | | | | | | | 0.01634 | 1.97826 | Female | | | |
| THEM6 | | | | | | | | | | 0.00003 | -2.50896 | Male | | | |
| MPHOSPH6 | 0.02621 | -0.65303 | Male | | | | | | | | | | | | |
| EEPD1 | | | | | | | | | | 0.00315 | -2.03323 | Male | | | |
| LGALS1 | | | | | | | | | | 0.00000 | 1.33339 | Female | | | |
| VGLL1 | | | | 0.00000 | -0.67923 | Male | | | | 0.00118 | -0.86564 | Male | | | |
| FLJ42627 | | | | | | | 0.01667 | 1.15859 | Female | | | | | | |
| PLLP | | | | | | | | | | 0.00004 | 1.54523 | Female | | | |
| KIAA0930 | | | | 0.00002 | 1.48818 | Female | | | | | | | | | |
| PWWP2B | | | | 0.00125 | 1.51671 | Female | | | | 0.02260 | -0.64674 | Male | | | |
| ANKRD40 | | | | | | | 0.00000 | 0.75238 | Female | | | | | | |
| DPRX | | | | | | | | | | 0.01506 | 1.15694 | Female | 0.00166 | 1.40050 | Female |

| | | | | | | | | | | | | | | | |
|---|---|---|---|---|---|---|---|---|---|---|---|---|---|---|---|
| CERS5 | 0.00376 | -0.85169 | Male | | | | | | | | | | | | |
| ACTG2 | | | | 0.00106 | 1.80997 | Female | | | | | | | | | |
| L3HYPDH | | | | | | | | | | 0.00404 | -0.75066 | Male | | | |
| MARC1 | | | | | | | 0.04921 | -0.98791 | Male | | | | 0.04928 | -0.89645 | Male |
| MTOR | | | | 0.00048 | 0.74020 | Female | | | | | | | | | |
| SPP1 | 0.00000 | 2.24261 | Female | | | | | | | 0.00033 | 2.51087 | Female | | | |
| FMN1 | | | | | | | | | | | | | 0.03984 | -0.94830 | Male |
| PIM2 | 0.04148 | 1.37524 | Female | | | | 0.00000 | 1.10452 | Female | 0.00000 | 2.00017 | Female | | | |
| ANXA6 | | | | 0.00097 | -1.06673 | Male | | | | | | | | | |
| DOCK7 | | | | 0.01976 | 0.98790 | Female | | | | | | | | | |
| UBE2E3 | 0.00396 | -0.99529 | Male | | | | | | | | | | 0.00000 | -0.63396 | Male |
| PAQR4 | | | | 0.01188 | 0.99606 | Female | | | | | | | | | |
| SMEK2 | | | | 0.00148 | -0.59254 | Male | | | | | | | | | |
| LINC00261 | | | | | | | 0.00000 | -4.64942 | Male | | | | | | |
| EFNB1 | | | | | | | 0.04109 | 1.51195 | Female | | | | | | |
| ACER3 | | | | | | | | | | 0.01306 | -0.58668 | Male | | | |
| PQBP1 | 0.00619 | 0.69379 | Female | 0.00000 | 0.88048 | Female | 0.00000 | 0.87685 | Female | 0.00000 | 0.60670 | Female | | | |
| FOXN3 | | | | 0.00000 | 1.25706 | Female | | | | | | | | | |
| NDRG1 | | | | 0.01654 | 1.46069 | Female | | | | | | | | | |
| ELK1 | | | | | | | | | | 0.01905 | 0.63518 | Female | 0.00406 | 0.74061 | Female |
| PRSS16 | | | | | | | 0.00004 | -1.72161 | Male | | | | | | |
| KLF3 | 0.00010 | -1.79419 | Male | | | | | | | | | | 0.00068 | -0.66277 | Male |
| EBPL | | | | | | | 0.00000 | -1.04589 | Male | | | | 0.00004 | -0.71230 | Male |
| MSI2 | | | | | | | | | | 0.04707 | -0.80779 | Male | | | |
| PRKCQ-AS1 | | | | | | | 0.03218 | -0.94804 | Male | | | | | | |
| RORA | | | | | | | 0.03142 | 0.63956 | Female | 0.00000 | -1.22444 | Male | 0.00623 | -0.74197 | Male |
| ZNF24 | 0.00828 | -0.93887 | Male | | | | | | | | | | | | |
| SOAT2 | | | | | | | | | | 0.00036 | 1.17478 | Female | | | |
| ATP1A3 | | | | | | | | | | | | | 0.00009 | 2.36482 | Female |
| CLIC1 | 0.00000 | -0.72617 | Male | | | | | | | | | | | | |
| TRIM2 | 0.00036 | 2.76841 | Female | | | | | | | | | | 0.00010 | -1.03784 | Male |
| AOX1 | | | | | | | | | | 0.00162 | 1.83134 | Female | | | |
| RENBP | | | | | | | | | | 0.00741 | -2.25259 | Male | | | |
| SDF2 | | | | 0.00010 | -0.86247 | Male | | | | | | | | | |
| GPC4 | | | | 0.00213 | 2.99074 | Female | | | | | | | | | |
| MAGOH2 | 0.00963 | -0.58603 | Male | | | | | | | | | | | | |
| TDGF1P3 | | | | | | | | | | 0.00002 | 1.69590 | Female | 0.00099 | 1.34359 | Female |
| UROD | 0.00231 | 0.96013 | Female | | | | | | | | | | | | |
| SOX9 | | | | | | | | | | 0.00476 | 2.09199 | Female | | | |
| TNFSF9 | | | | | | | 0.00002 | 1.66737 | Female | 0.00001 | 1.70075 | Female | 0.00000 | 2.47222 | Female |
| CYP2F1 | | | | | | | | | | 0.00107 | 1.82407 | Female | 0.01212 | 1.43783 | Female |
| NAB1 | | | | | | | | | | | | | 0.00464 | -0.90450 | Male |
| SC5D | | | | | | | 0.00963 | -0.67150 | Male | | | | 0.00969 | -0.62488 | Male |
| IL17RC | | | | | | | | | | 0.00497 | 1.89461 | Female | | | |
| SELM | | | | | | | | | | 0.04808 | 0.68657 | Female | | | |
| ACN9 | 0.03519 | 0.69564 | Female | | | | | | | | | | | | |
| CENPH | | | | | | | 0.00002 | -0.77049 | Male | | | | | | |
| RPL36A.uid-13034 | | | | | | | 0.01063 | 0.80716 | Female | | | | 0.00781 | 0.78517 | Female |
| MTMR1 | | | | 0.01829 | 1.13932 | Female | | | | | | | | | |
| CHST12 | | | | | | | | | | 0.00000 | 1.09040 | Female | | | |
| C17orf75 | | | | | | | | | | 0.00711 | 0.74004 | Female | | | |

| | | | | | | | | | | | | | | | |
|---|---|---|---|---|---|---|---|---|---|---|---|---|---|---|---|
| DHCR24 | 0.00822 | 0.98926 | Female | | | | | | | | | | | | |
| KRTCAP3 | | | | | | | | | | 0.01880 | 0.60478 | Female | | | |
| MEAF6 | 0.03758 | -0.66720 | Male | | | | | | | | | | | | |
| FIBCD1 | | | | 0.00000 | 3.91663 | Female | | | | | | | | | |
| SUSD3 | | | | 0.00724 | 2.85349 | Female | | | | 0.00708 | 1.96956 | Female | | | |
| PLAC8 | 0.00166 | 2.07203 | Female | 0.01078 | 0.94996 | Female | | | | | | | | | |
| CEBPA | | | | 0.00000 | 1.67493 | Female | 0.00997 | 0.63796 | Female | | | | | | |
| C6orf147 | | | | | | | | | | 0.00407 | -0.74896 | Male | | | |
| REG1A | | | | | | | | | | 0.03859 | 2.36803 | Female | | | |
| CHCHD10 | | | | 0.00083 | 0.90469 | Female | | | | | | | | | |
| NCOA1 | | | | | | | | | | 0.00027 | 0.85855 | Female | | | |
| FAM222A | | | | | | | 0.00537 | 1.21488 | Female | | | | | | |
| SLC25A44 | | | | | | | | | | 0.00000 | 0.60245 | Female | | | |
| RFX2 | | | | | | | 0.02350 | -1.11224 | Male | | | | | | |
| PSD4 | | | | | | | | | | 0.04666 | -1.23322 | Male | | | |
| FXYD5 | 0.00113 | 2.70634 | Female | | | | 0.00045 | 1.10926 | Female | | | | | | |
| KIAA0226L | 0.01604 | -3.37964 | Male | | | | | | | | | | | | |
| PRPSAP2 | 0.00861 | -0.99575 | Male | | | | | | | | | | | | |
| TMEM159 | | | | | | | | | | 0.00000 | 2.07951 | Female | | | |
| FDXR | | | | | | | | | | | | | 0.00001 | 0.86940 | Female |
| DNASE1L3 | | | | | | | | | | 0.01989 | 2.34632 | Female | | | |
| TRIP6 | | | | | | | 0.00144 | -0.99580 | Male | 0.00097 | -0.95299 | Male | | | |
| ACTL8 | | | | | | | | | | 0.00555 | -1.81141 | Male | 0.00046 | -2.21113 | Male |
| ATXN7L2 | | | | | | | | | | | | | 0.02853 | 0.76409 | Female |
| MIR1292 | | | | | | | | | | | | | 0.00001 | 0.59070 | Female |
| RNF24 | | | | | | | | | | 0.00027 | 1.06963 | Female | | | |
| PPAP2A | | | | | | | | | | 0.00000 | 2.26898 | Female | 0.00596 | 0.82186 | Female |
| LTBP4 | | | | | | | | | | 0.00237 | 0.82829 | Female | | | |
| EPS8L2 | | | | 0.01320 | 1.15339 | Female | 0.02984 | -0.66942 | Male | | | | | | |
| C1orf61 | | | | 0.00019 | -1.43237 | Male | | | | | | | | | |
| GLI3 | | | | 0.00401 | 3.21107 | Female | | | | | | | | | |
| SIAE | | | | 0.00923 | 1.27517 | Female | | | | 0.04994 | 0.65834 | Female | | | |
| GREM2 | 0.00065 | 3.20853 | Female | | | | 0.00021 | -1.41277 | Male | | | | | | |
| PPP1R14D | | | | | | | | | | | | | 0.00495 | 1.48494 | Female |
| PFN2 | | | | | | | 0.00000 | -0.84827 | Male | 0.00000 | -1.07662 | Male | | | |
| SULT2A1 | | | | | | | 0.01295 | 1.76207 | Female | | | | | | |
| TP53 | | | | | | | | | | 0.00002 | -0.83030 | Male | | | |
| NOC4L | | | | 0.00001 | 0.98619 | Female | 0.00000 | 1.26750 | Female | | | | | | |
| PDCD11 | | | | | | | 0.00000 | 0.63892 | Female | | | | | | |
| DCUN1D1 | | | | | | | | | | 0.00008 | -0.58761 | Male | | | |
| PRPSAP1 | | | | | | | | | | 0.00000 | 0.86595 | Female | | | |
| DUSP1 | | | | | | | | | | | | | 0.04984 | 0.74449 | Female |
| MIR6834 | | | | | | | | | | 0.00003 | -1.06738 | Male | 0.00025 | 0.93879 | Female |
| FOXR1 | | | | | | | 0.00189 | 0.69435 | Female | 0.00000 | 1.04389 | Female | | | |
| HERC2P2 | | | | | | | | | | 0.00668 | 0.92750 | Female | | | |
| IDH2 | | | | | | | | | | 0.00094 | -1.06583 | Male | | | |
| DHRS11 | | | | | | | | | | | | | 0.01844 | 0.91946 | Female |
| UHMK1 | | | | | | | 0.00000 | 0.86077 | Female | | | | | | |
| EI24 | | | | 0.00110 | 0.59585 | Female | | | | | | | | | |
| ATP6AP1 | | | | 0.00000 | 1.18104 | Female | 0.00000 | 0.97668 | Female | 0.00000 | 0.64424 | Female | 0.00000 | 0.61299 | Female |
| PDP2 | | | | | | | | | | 0.00000 | -1.22373 | Male | | | |

| | | | | | | | | | | | | | | | |
|---|---|---|---|---|---|---|---|---|---|---|---|---|---|---|---|
| TRIM53AP.uid-15946 | | | | | | | | | | 0.00784 | 1.46138 | Female | | | |
| MYLK | | | | | | | 0.03826 | -1.05245 | Male | | | | | | |
| LOC643355 | | | | | | | | | | 0.00001 | -1.41518 | Male | 0.00304 | -0.96111 | Male |
| LGR5 | | | | | | | | | | | | | 0.04929 | -2.47594 | Male |
| CDH2 | | | | | | | | | | | | | 0.00193 | -1.63105 | Male |
| NENF | | | | | | | 0.00000 | -1.17842 | Male | 0.00914 | -0.61451 | Male | | | |
| PIDD1 | | | | | | | 0.00389 | -0.96968 | Male | | | | | | |
| EDA2R | 0.00003 | 4.03111 | Female | | | | 0.04159 | 0.95864 | Female | | | | | | |
| PI4KA | 0.00085 | 1.36158 | Female | | | | | | | | | | | | |
| FAM199X | | | | 0.00001 | 1.35239 | Female | | | | 0.00001 | 0.89025 | Female | | | |
| FOXD3-AS1 | | | | | | | | | | 0.02236 | 1.12313 | Female | | | |
| PARK2 | | | | | | | | | | 0.00000 | 2.22313 | Female | | | |
| PRR14L | | | | 0.00304 | -0.63811 | Male | | | | | | | | | |
| IVNS1ABP | | | | | | | | | | 0.00181 | -0.66266 | Male | | | |
| NGFRAP1 | 0.00000 | 2.26138 | Female | | | | | | | 0.00000 | -1.40130 | Male | 0.00099 | -0.63128 | Male |
| LINC00907 | | | | | | | | | | | | | 0.00825 | -1.71084 | Male |
| PGF | | | | | | | | | | 0.00003 | -1.14361 | Male | | | |
| ARHGEF16 | | | | | | | | | | 0.04240 | 0.71054 | Female | | | |
| FBXO6 | | | | | | | | | | 0.00000 | 2.51053 | Female | | | |
| XIST | | | | 0.02960 | 0.95411 | Female | 0.00000 | 2.12944 | Female | 0.00000 | 1.97173 | Female | 0.00000 | 3.34972 | Female |
| SLC34A2 | | | | 0.02051 | -0.96851 | Male | | | | 0.00256 | -0.78026 | Male | | | |
| NAT1 | | | | | | | | | | | | | 0.04548 | -0.83230 | Male |
| TLN1 | | | | | | | | | | | | | 0.01552 | 0.58412 | Female |
| YTHDF1 | 0.00330 | -0.81458 | Male | | | | | | | | | | | | |
| TUBA8 | | | | | | | | | | 0.01176 | -0.98552 | Male | 0.00203 | 1.16451 | Female |
| RASAL2 | | | | | | | 0.01845 | 0.91359 | Female | | | | | | |
| NLGN2 | | | | | | | 0.00398 | -1.38154 | Male | | | | | | |
| GBX2 | | | | | | | | | | 0.00224 | 1.37607 | Female | | | |
| GIN1 | | | | | | | 0.00120 | -1.17959 | Male | | | | | | |
| B3GNT2 | | | | | | | 0.00000 | -1.02302 | Male | 0.00190 | -0.58315 | Male | | | |
| CCDC106 | | | | | | | 0.00926 | -1.11569 | Male | | | | | | |
| LAPTM4B | | | | | | | | | | 0.00000 | 0.81473 | Female | | | |
| SLC52A1 | | | | | | | | | | | | | 0.01089 | 0.83235 | Female |
| PPA1 | 0.00044 | -0.71158 | Male | | | | | | | | | | | | |
| IL10RB | | | | | | | 0.00546 | -0.84261 | Male | | | | | | |
| TMEM183A | | | | | | | | | | | | | 0.00024 | -1.62682 | Male |
| SYNPO | | | | | | | 0.00579 | -1.22772 | Male | 0.02442 | -0.94054 | Male | 0.00511 | 1.15430 | Female |
| CLDN6 | | | | 0.00176 | 0.68238 | Female | | | | | | | | | |
| NEURL2 | | | | | | | | | | 0.04214 | 0.92947 | Female | | | |
| AP1S1 | | | | 0.02208 | 0.64029 | Female | | | | | | | | | |
| INMT | | | | | | | | | | | | | 0.03433 | 2.12665 | Female |
| LDLRAP1 | | | | | | | | | | 0.00002 | 1.27936 | Female | | | |
| NUDT22 | 0.01328 | -1.58816 | Male | | | | | | | | | | | | |
| CA5B | 0.01100 | 2.53602 | Female | 0.00000 | 4.01878 | Female | 0.00948 | 1.16324 | Female | 0.00000 | 2.17857 | Female | 0.00001 | 1.69674 | Female |
| NUAK2 | 0.01314 | 1.88134 | Female | | | | | | | | | | 0.00056 | 0.97842 | Female |
| CLTB | | | | | | | | | | 0.00160 | 0.61811 | Female | | | |
| HNRNPKP3 | | | | | | | | | | 0.00000 | -1.03734 | Male | | | |
| SALL3 | | | | | | | | | | 0.00111 | 1.65008 | Female | | | |
| CARD8 | | | | | | | | | | | | | 0.01119 | 0.94315 | Female |
| AGRN | | | | | | | | | | 0.04508 | 0.66340 | Female | 0.04920 | 0.64791 | Female |
| CKMT2-AS1 | | | | | | | | | | | | | 0.00454 | 0.88997 | Female |

| | | | | | | | | | | | | | | | |
|---|---|---|---|---|---|---|---|---|---|---|---|---|---|---|---|
| AKAP9 | 0.00122 | 1.16710 | Female | | | | | | | | | | | | |
| SNORA67 | | | | | | | 0.00004 | 0.75834 | Female | | | | 0.00032 | 0.63317 | Female |
| MGC72080 | | | | | | | | | | 0.03300 | -0.61972 | Male | | | |
| ARMCX6 | | | | | | | | | | | | | 0.02873 | -2.08223 | Male |
| DUS4L | | | | | | | 0.04477 | -0.59051 | Male | | | | | | |
| WTAP | 0.00015 | -0.82709 | Male | | | | | | | | | | | | |
| ING2 | 0.00306 | -1.37209 | Male | | | | | | | | | | | | |
| PRDM14 | | | | | | | | | | 0.00009 | 1.78776 | Female | | | |
| DTX2 | | | | | | | | | | 0.00143 | -0.72440 | Male | | | |
| HPS4 | | | | 0.04080 | -0.81434 | Male | | | | | | | | | |
| ZDHHC9 | | | | 0.00017 | 1.07192 | Female | 0.00261 | 0.61913 | Female | 0.00006 | 0.75618 | Female | | | |
| PDE5A | | | | | | | | | | 0.00030 | 1.17205 | Female | | | |
| PPARD | | | | | | | | | | 0.00011 | -0.69982 | Male | | | |
| KIAA1551 | | | | | | | | | | 0.00000 | 0.69608 | Female | | | |
| HUNK | | | | 0.00127 | 1.45472 | Female | | | | | | | | | |
| PACSIN1 | | | | | | | 0.01700 | -1.83722 | Male | | | | | | |
| OR5AU1 | | | | | | | 0.00000 | 2.48193 | Female | 0.01531 | -1.34116 | Male | | | |
| CYP1A1 | | | | | | | 0.00155 | -1.56938 | Male | | | | 0.00969 | 1.27731 | Female |
| SLC29A2 | | | | | | | 0.00064 | -0.79707 | Male | | | | | | |
| PTPRU | | | | 0.00000 | 1.79396 | Female | | | | | | | | | |
| RSG1 | 0.04508 | -2.81566 | Male | | | | | | | | | | | | |
| TNNC2 | | | | 0.01796 | 1.60961 | Female | | | | 0.00082 | 1.40598 | Female | 0.00452 | 1.21127 | Female |
| PQLC3 | | | | | | | | | | 0.00000 | 1.03464 | Female | | | |
| RTCB | 0.00124 | -0.81449 | Male | | | | | | | | | | | | |
| ZNF750 | | | | 0.00181 | -1.68986 | Male | | | | | | | | | |
| MSI1 | | | | | | | | | | | | | 0.01554 | -0.72284 | Male |
| WNT3A | | | | | | | | | | 0.04367 | 2.01604 | Female | | | |
| CEBPG | 0.02338 | -0.72702 | Male | | | | | | | | | | | | |
| YBX2 | | | | | | | | | | 0.00008 | 1.54468 | Female | | | |
| ZNF112 | 0.01358 | 3.32058 | Female | | | | | | | | | | | | |
| NEAT1 | | | | | | | 0.00239 | 0.86518 | Female | | | | | | |
| RNF103 | | | | | | | 0.04328 | 0.82019 | Female | | | | | | |
| CNTN6 | | | | | | | | | | 0.03989 | 0.92865 | Female | 0.00004 | -1.67750 | Male |
| ELOVL2 | | | | | | | | | | 0.03146 | -1.72099 | Male | | | |
| NCAM2 | | | | | | | 0.00424 | -2.81145 | Male | | | | | | |
| LINC01420 | | | | 0.01534 | 1.78122 | Female | | | | | | | | | |
| IGSF3 | | | | 0.00446 | 1.14805 | Female | | | | | | | | | |
| MIR3064 | | | | | | | 0.01160 | 0.65023 | Female | | | | | | |
| VAV2 | 0.04441 | -1.19940 | Male | 0.00010 | 1.17049 | Female | | | | | | | | | |
| ADNP2 | | | | | | | 0.00303 | 0.62251 | Female | | | | | | |
| NOS1 | | | | | | | 0.00001 | 1.77415 | Female | 0.00000 | -1.52667 | Male | | | |
| LRRC58 | 0.00053 | -2.12798 | Male | | | | | | | 0.00020 | -0.77273 | Male | | | |
| REXO4 | | | | | | | 0.00000 | 0.91607 | Female | | | | | | |
| LIME1 | | | | | | | 0.00459 | 1.18024 | Female | | | | | | |
| ZER1 | | | | | | | 0.02059 | 0.69703 | Female | | | | | | |
| ZNF316 | | | | | | | | | | 0.03461 | -0.91098 | Male | | | |
| EMG1 | 0.00019 | -0.78659 | Male | | | | | | | | | | | | |
| IL17RB | 0.00198 | 3.17429 | Female | | | | | | | | | | 0.02479 | -0.93582 | Male |
| RSPO3 | | | | 0.00000 | 1.02220 | Female | 0.00034 | -2.45246 | Male | 0.03300 | 1.61985 | Female | | | |
| ENG | | | | | | | 0.02543 | 1.59746 | Female | | | | | | |
| ABCC4 | | | | | | | | | | | | | 0.00000 | -1.22886 | Male |

| | | | | | | | | | | | | | | | |
|---|---|---|---|---|---|---|---|---|---|---|---|---|---|---|---|
| TTC3P1 | | | | 0.00405 | 1.29047 | Female | | | | | | | | | |
| HEXIM1 | | | | 0.00455 | -0.70201 | Male | | | | | | | | | |
| TYMS | 0.01125 | -1.09379 | Male | | | | 0.00000 | -0.81472 | Male | | | | | | |
| ZNF780B | | | | 0.04563 | -1.56237 | Male | | | | | | | | | |
| DDX1 | 0.00000 | -0.94020 | Male | | | | | | | | | | | | |
| CTBP1 | | | | 0.00651 | 0.60509 | Female | 0.00001 | 0.61851 | Female | | | | | | |
| TCEB2 | 0.00040 | -0.72157 | Male | | | | 0.00000 | 0.63279 | Female | | | | | | |
| WLS | | | | | | | 0.00000 | -1.80110 | Male | 0.00008 | -1.14411 | Male | 0.00684 | -0.81581 | Male |
| ZNF730 | | | | | | | 0.00895 | -0.74964 | Male | | | | | | |
| L1TD1 | | | | | | | | | | 0.00001 | 0.59140 | Female | | | |
| FUNDC2P2 | | | | | | | 0.00000 | 0.73815 | Female | | | | | | |
| NAV1 | | | | | | | | | | 0.03843 | 0.80376 | Female | | | |
| AJUBA | | | | | | | 0.00017 | -1.05957 | Male | 0.00281 | -0.82629 | Male | | | |
| OXLD1 | | | | | | | | | | 0.01419 | -0.70841 | Male | | | |
| SLC39A11 | | | | | | | | | | | | | 0.00080 | -0.99990 | Male |
| CSTF2 | | | | | | | 0.00000 | 1.36127 | Female | 0.00000 | 0.73659 | Female | 0.00000 | 0.88028 | Female |
| RRAD | | | | 0.00000 | 3.93504 | Female | 0.00027 | 1.62257 | Female | 0.00068 | 1.48182 | Female | | | |
| CHRNB1 | | | | | | | | | | 0.04150 | -0.68368 | Male | 0.02382 | 0.72725 | Female |
| SLC4A11 | | | | | | | | | | | | | 0.00000 | 0.91533 | Female |
| MIR614 | | | | | | | 0.00157 | 1.01242 | Female | | | | 0.00572 | 0.89340 | Female |
| CASP6 | 0.00008 | -1.60469 | Male | | | | | | | | | | | | |
| ARIH1 | | | | 0.00454 | 0.69839 | Female | | | | | | | | | |
| SLC6A8 | | | | 0.00000 | 1.34749 | Female | 0.00000 | 2.03541 | Female | 0.00000 | 1.29215 | Female | | | |
| ALPPL2 | | | | | | | | | | 0.00000 | 1.02489 | Female | 0.00000 | 0.87681 | Female |
| ANXA2P1 | | | | 0.00003 | -0.74485 | Male | | | | | | | | | |
| TTLL3 | | | | | | | 0.00075 | -1.26583 | Male | | | | | | |
| PELO | | | | | | | | | | 0.00003 | -0.75433 | Male | | | |
| DENND4B | | | | | | | | | | | | | 0.02643 | 0.79638 | Female |
| VWA5A | | | | | | | | | | 0.00000 | 3.19480 | Female | | | |
| BRI3 | | | | 0.03101 | 0.73192 | Female | | | | | | | | | |
| WNT6 | | | | | | | 0.00008 | -1.44781 | Male | | | | | | |
| PKIB | 0.02822 | -1.37703 | Male | | | | 0.00002 | -1.00569 | Male | | | | | | |
| FSCN1 | | | | 0.00069 | 0.79306 | Female | | | | | | | | | |
| FAM64A | | | | | | | | | | 0.00000 | 2.70617 | Female | 0.02208 | 1.43861 | Female |
| TPP1 | | | | | | | 0.00085 | -1.03800 | Male | | | | | | |
| METTL2B | 0.03283 | 0.76495 | Female | | | | | | | | | | | | |
| WWC1 | | | | | | | | | | | | | 0.00169 | 0.64387 | Female |
| PIGQ | | | | 0.01578 | 1.37478 | Female | | | | | | | | | |
| TNFRSF1A | | | | | | | | | | 0.00095 | 1.23730 | Female | | | |
| DYNLT3 | | | | | | | | | | | | | 0.00000 | -1.87321 | Male |
| SCML1 | | | | | | | | | | | | | 0.00133 | -1.35504 | Male |
| NFE2L3 | | | | 0.03113 | 0.64715 | Female | 0.00000 | -0.82416 | Male | | | | | | |
| MGAT4A | | | | | | | 0.00022 | -1.34210 | Male | | | | | | |
| LYG2 | | | | | | | 0.04542 | 1.16073 | Female | | | | | | |
| ZNF92 | | | | | | | 0.00003 | -0.72991 | Male | | | | | | |
| SAPCD2 | 0.00000 | -3.59182 | Male | 0.00000 | 4.48750 | Female | | | | | | | | | |
| APOL2 | | | | | | | 0.00000 | 2.90471 | Female | 0.03442 | 1.27682 | Female | | | |
| KBTBD4 | | | | | | | | | | 0.00006 | -1.11299 | Male | | | |
| PLXNA3 | | | | | | | | | | | | | 0.01489 | 0.98353 | Female |
| SNORD48 | | | | | | | | | | | | | 0.00305 | 0.81342 | Female |
| TP53I13 | 0.01969 | -1.88716 | Male | | | | 0.00030 | -1.15715 | Male | 0.01260 | -0.80412 | Male | 0.00601 | -0.86186 | Male |

| | | | | | | | | | | | | | | | |
|---|---|---|---|---|---|---|---|---|---|---|---|---|---|---|---|
| RHOT1 | | | | 0.02458 | 1.08099 | Female | 0.01112 | -0.80323 | Male | | | | | | |
| ASNS | | | | 0.00000 | -2.55705 | Male | 0.00001 | -1.15898 | Male | | | | | | |
| C1orf86 | | | | | | | 0.00278 | -0.79613 | Male | 0.02216 | -0.59681 | Male | | | |
| FRAT2 | | | | | | | 0.00013 | 0.62749 | Female | | | | | | |
| PIGO | | | | | | | 0.01050 | -0.73353 | Male | | | | | | |
| DDN | | | | | | | | | | 0.00458 | -2.26651 | Male | | | |
| LMBR1L | | | | 0.01937 | -0.68266 | Male | | | | | | | | | |
| MLLT11 | | | | | | | | | | 0.00251 | -1.20733 | Male | | | |
| PBX3 | | | | | | | | | | 0.00187 | 1.08579 | Female | | | |
| GSTP1 | | | | 0.00000 | 1.51136 | Female | | | | 0.00000 | 0.91471 | Female | | | |
| POLR2I | 0.00495 | 0.64197 | Female | | | | | | | | | | | | |
| SHISA2 | | | | | | | | | | 0.02546 | 1.65916 | Female | | | |
| TTTY14 | | | | | | | 0.00085 | -2.47344 | Male | 0.00000 | -3.13467 | Male | 0.00000 | -4.15901 | Male |
| ARGFX | | | | | | | | | | 0.00000 | 1.69603 | Female | | | |
| MED12 | | | | 0.00231 | 0.95817 | Female | 0.00268 | 0.65160 | Female | 0.00149 | 0.65563 | Female | 0.00007 | 0.78053 | Female |
| SLC38A9 | 0.00006 | -1.94675 | Male | | | | | | | | | | | | |
| LOC728613 | | | | 0.00007 | 1.45546 | Female | | | | | | | | | |
| AGPAT9 | | | | | | | 0.03709 | -0.78490 | Male | 0.02999 | -0.77570 | Male | 0.04610 | -0.73342 | Male |
| PRNP | | | | | | | | | | 0.03025 | 0.63139 | Female | | | |
| 20.00000 | 0.00340 | 3.17853 | Female | | | | | | | 0.00351 | 1.28526 | Female | | | |
| ICE1 | 0.04852 | -0.91080 | Male | | | | | | | | | | | | |
| FAM168B | 0.02349 | -0.80872 | Male | | | | | | | | | | | | |
| ATG16L1 | | | | | | | 0.00011 | 0.59565 | Female | | | | | | |
| BCORL1 | | | | 0.00102 | 1.18575 | Female | | | | | | | | | |
| PDHA2 | 0.00043 | -1.16231 | Male | 0.00000 | 0.88294 | Female | 0.00000 | 0.83101 | Female | | | | 0.00000 | 0.65613 | Female |
| RPS4Y2 | | | | | | | 0.00000 | -3.64839 | Male | 0.00012 | -3.35831 | Male | | | |
| MIR3648 | | | | | | | | | | 0.00000 | 2.84103 | Female | | | |
| CTB-113P19.1 | | | | | | | | | | 0.00054 | 0.95426 | Female | | | |
| LOC400043 | | | | | | | | | | 0.03659 | 1.22694 | Female | | | |
| C20orf96 | | | | | | | 0.00138 | -1.29939 | Male | | | | | | |
| DEDD2 | | | | | | | 0.00471 | -0.63751 | Male | 0.00007 | -0.80847 | Male | | | |
| LOC641746 | | | | | | | | | | | | | 0.02418 | -0.83088 | Male |
| SYCP1 | | | | | | | | | | | | | 0.00149 | -2.33862 | Male |
| CGREF1 | | | | | | | 0.00391 | 1.33796 | Female | 0.04309 | 0.95512 | Female | | | |
| TMX3 | | | | | | | 0.00053 | -0.71577 | Male | | | | 0.00033 | -0.68777 | Male |
| MOGS | | | | | | | 0.01052 | -0.58776 | Male | | | | | | |
| ZNF124 | 0.00492 | -1.61073 | Male | | | | | | | | | | | | |
| GPRC5B | | | | | | | | | | 0.00024 | 1.80598 | Female | | | |
| ZBTB34 | | | | | | | 0.02887 | 0.76754 | Female | | | | | | |
| CNBP | 0.00013 | -0.64531 | Male | | | | | | | | | | | | |
| TMEM205 | 0.00164 | -1.38999 | Male | | | | | | | | | | | | |
| PDGFA | | | | 0.00056 | 0.90829 | Female | 0.00009 | 0.69318 | Female | | | | 0.00036 | 0.60897 | Female |
| COX7B2 | | | | | | | 0.00000 | -0.77283 | Male | | | | | | |
| PIH1D2 | | | | | | | | | | 0.01495 | -1.44844 | Male | | | |
| SOX17 | | | | | | | | | | 0.00007 | 2.97450 | Female | | | |
| LMO1 | | | | | | | | | | | | | 0.00405 | 2.05647 | Female |
| SLC7A6OS | | | | | | | 0.00325 | 0.61895 | Female | | | | | | |
| RPL39 | 0.00102 | 0.83588 | Female | 0.00000 | 0.69687 | Female | 0.00000 | 0.62035 | Female | | | | | | |
| MYBBP1A | | | | | | | | | | 0.00009 | -0.62934 | Male | | | |
| SLC25A12 | | | | | | | | | | 0.02116 | 1.00870 | Female | | | |
| PPP1R14A | 0.01011 | -1.08578 | Male | 0.00000 | 1.45749 | Female | 0.00000 | 0.84348 | Female | | | | | | |

| | | | | | | | | | | | | | | | |
|---|---|---|---|---|---|---|---|---|---|---|---|---|---|---|---|
| TM7SF2 | | | | | | | 0.00000 | -2.45304 | Male | | | | | | |
| CKMT2 | | | | 0.03142 | 1.66862 | Female | | | | 0.03870 | 1.06556 | Female | | | |
| MOSPD1 | | | | 0.00120 | 0.87327 | Female | | | | 0.00069 | 0.60429 | Female | | | |
| SUPT20HL2 | | | | | | | 0.01274 | 2.58057 | Female | | | | | | |
| ETV1 | | | | | | | | | | 0.00854 | 1.82801 | Female | | | |
| GPC3 | | | | | | | | | | 0.02263 | 1.30187 | Female | | | |
| PURB | | | | | | | | | | 0.04717 | -0.63547 | Male | | | |
| CASP7 | | | | | | | | | | | | | 0.00021 | 1.94673 | Female |
| SNORD87 | | | | | | | | | | | | | 0.00612 | -0.62794 | Male |
| NTRK3 | | | | | | | | | | 0.03529 | 1.67656 | Female | | | |
| VENTX | 0.00015 | -2.13145 | Male | | | | | | | 0.00002 | 2.35907 | Female | | | |
| ZNF217 | | | | 0.03343 | -0.62649 | Male | | | | | | | | | |
| CLDN4 | | | | 0.00000 | 3.83957 | Female | | | | | | | | | |
| KLHL23 | | | | | | | | | | 0.00076 | -1.23230 | Male | | | |
| UACA | | | | 0.00008 | 1.27827 | Female | | | | 0.00000 | 1.58171 | Female | | | |
| CPSF6 | 0.00005 | -0.87863 | Male | | | | | | | | | | | | |
| TPM2 | | | | | | | 0.00198 | 0.63532 | Female | | | | | | |
| FABP7 | | | | 0.00816 | 1.60472 | Female | | | | 0.00856 | 0.92945 | Female | | | |
| VWA1 | | | | 0.00906 | 1.07943 | Female | | | | | | | | | |
| CLDN12 | | | | | | | 0.00104 | -0.75898 | Male | | | | | | |
| GAL | | | | | | | | | | 0.02852 | 0.71678 | Female | | | |
| LIPG | | | | | | | 0.04110 | 1.16700 | Female | | | | | | |
| NLRP12 | | | | | | | | | | 0.01082 | 0.79051 | Female | | | |
| BTBD1 | | | | | | | | | | | | | 0.00129 | -0.75962 | Male |
| CUEDC2 | 0.00027 | 2.69153 | Female | | | | | | | | | | | | |
| IKBKB | | | | | | | 0.00152 | 0.80116 | Female | | | | | | |
| IL17RE | | | | | | | 0.02125 | -1.46436 | Male | 0.02846 | -1.30186 | Male | | | |
| SRGAP2D.uid-421 | | | | | | | | | | 0.00033 | 0.68632 | Female | | | |
| SNORA84 | | | | | | | 0.03239 | 0.83715 | Female | | | | | | |
| UNC93A | | | | | | | | | | 0.00191 | 2.23673 | Female | | | |
| LOC100506178 | | | | 0.00169 | 1.76223 | Female | 0.00214 | 1.17513 | Female | | | | | | |
| TOB2 | 0.01330 | 0.83514 | Female | 0.00206 | -0.61476 | Male | | | | | | | | | |
| PIGY+PYURF | 0.02581 | 0.78755 | Female | | | | | | | | | | | | |
| JPH2 | | | | | | | | | | | | | 0.02277 | 2.80061 | Female |
| RTP1 | | | | | | | | | | | | | 0.00874 | -1.21333 | Male |
| SPATA13 | | | | | | | | | | 0.02008 | -0.87079 | Male | | | |
| EXOSC4 | | | | | | | 0.00000 | 0.64557 | Female | | | | | | |
| CCDC50 | | | | 0.00454 | 1.07026 | Female | | | | | | | | | |
| KIAA1804 | | | | | | | | | | 0.00380 | 1.53060 | Female | | | |
| PILRB | | | | | | | 0.00000 | 1.96068 | Female | | | | | | |
| RAD50 | | | | | | | | | | 0.00014 | -0.66743 | Male | | | |
| PRDX1 | 0.00000 | 0.77490 | Female | | | | | | | | | | | | |
| CDKN1C | | | | 0.00010 | -2.10275 | Male | 0.04005 | 0.85862 | Female | | | | 0.01828 | 0.90478 | Female |
| CCDC183-AS1 | | | | | | | 0.00125 | 2.05597 | Female | | | | | | |
| RGCC | | | | | | | | | | 0.00024 | 1.02066 | Female | | | |
| TPRX1 | | | | | | | | | | 0.00000 | 6.03232 | Female | | | |
| SHISA4 | | | | 0.00634 | -2.04887 | Male | | | | | | | | | |
| APOBEC3G | | | | | | | 0.02164 | -2.35228 | Male | | | | | | |
| NMB | | | | | | | | | | 0.00495 | -1.29439 | Male | | | |
| DDX3X | 0.00000 | 0.78957 | Female | | | | 0.00000 | 0.95360 | Female | 0.00000 | 0.61970 | Female | | | |
| PEG10 | | | | | | | 0.00049 | 1.21567 | Female | | | | 0.02328 | -0.77397 | Male |

| | | | | | | | | | | | | | | | |
|---|---|---|---|---|---|---|---|---|---|---|---|---|---|---|---|
| GGA2 | | | | 0.00611 | 0.63786 | Female | | | | | | | | | |
| SUN3 | | | | 0.00050 | 0.85190 | Female | 0.00007 | -0.64318 | Male | | | | | | |
| NT5C3B | | | | 0.00002 | 1.25052 | Female | | | | | | | | | |
| ZNF484 | | | | | | | 0.03230 | -1.09564 | Male | | | | | | |
| ZNF557 | | | | | | | 0.00003 | 0.78491 | Female | | | | | | |
| AKAP2.uid-11393 | | | | | | | 0.00838 | -0.95068 | Male | | | | | | |
| C5orf30 | | | | | | | | | | 0.00047 | -1.52743 | Male | | | |
| FSD2 | | | | | | | | | | 0.01827 | 0.77385 | Female | | | |
| SH3BGRL | | | | | | | 0.04199 | -2.03567 | Male | | | | | | |
| TPRXL | | | | | | | | | | 0.00140 | -1.65979 | Male | | | |
| NMNAT1 | | | | | | | | | | | | | 0.00803 | 0.73577 | Female |
| PRDX4 | 0.00025 | 1.47728 | Female | 0.00000 | 1.10733 | Female | | | | | | | | | |
| CCNK | | | | | | | 0.00000 | -0.64602 | Male | | | | | | |
| HIST1H2AM | | | | 0.00416 | -1.33898 | Male | | | | | | | | | |
| TGFBI | | | | | | | 0.02617 | -1.69938 | Male | | | | 0.00424 | 1.93665 | Female |
| CCDC88B | | | | | | | 0.01396 | 1.84598 | Female | | | | | | |
| OLAH | | | | | | | | | | 0.00694 | 1.59907 | Female | | | |
| FBLN7 | | | | | | | | | | | | | 0.00190 | -4.34915 | Male |
| FNTA | | | | 0.00122 | 0.75502 | Female | | | | | | | | | |
| TMEM109 | | | | 0.01568 | 0.62114 | Female | | | | | | | | | |
| FAM83H | | | | 0.00265 | 1.06538 | Female | | | | | | | | | |
| CTSH | | | | | | | | | | 0.03339 | 0.68597 | Female | 0.04223 | 0.65433 | Female |
| PLP1 | | | | | | | | | | 0.01693 | 2.37549 | Female | | | |
| TBC1D24 | | | | | | | | | | 0.00383 | -0.94668 | Male | | | |
| TESK1 | | | | | | | 0.00494 | 0.95317 | Female | | | | | | |
| TUBA3E | | | | | | | 0.02006 | 1.15977 | Female | 0.00008 | 1.75160 | Female | 0.02405 | 1.06151 | Female |
| CAPN2 | | | | 0.03870 | 0.77355 | Female | | | | | | | | | |
| FAM207A | | | | 0.00576 | 0.77142 | Female | | | | | | | | | |
| GAS6 | | | | 0.00120 | 1.04738 | Female | | | | | | | 0.00198 | -0.65300 | Male |
| ERGIC1 | | | | | | | | | | 0.00472 | -0.99867 | Male | | | |
| PLCD1 | | | | | | | | | | 0.01284 | 0.97932 | Female | | | |
| CHGA | | | | | | | 0.00091 | 1.66255 | Female | 0.00000 | 4.48667 | Female | | | |
| PRAMEF1 | | | | | | | | | | 0.00000 | 7.19203 | Female | | | |
| IRS1 | | | | | | | | | | 0.00166 | -1.26912 | Male | | | |
| GGT1 | | | | | | | | | | 0.00064 | 1.73178 | Female | | | |
| EFCAB2 | | | | | | | 0.00694 | -0.74366 | Male | | | | | | |
| AK8 | | | | | | | | | | 0.01332 | 1.36028 | Female | 0.01396 | -1.35374 | Male |
| CCDC58 | | | | | | | | | | | | | 0.00000 | -0.71785 | Male |
| ZNF462 | | | | | | | | | | 0.00320 | 1.84069 | Female | | | |
| ZDHHC7 | 0.01875 | -1.14622 | Male | | | | | | | | | | | | |
| EIF4E | | | | | | | | | | | | | 0.00000 | -0.67128 | Male |
| SNORD52 | | | | | | | | | | 0.00000 | 0.98005 | Female | | | |
| LOC100505666 | | | | | | | | | | | | | 0.03581 | 1.68759 | Female |
| TTC4 | | | | | | | | | | 0.00019 | 0.70742 | Female | | | |
| PALLD | | | | 0.04229 | 0.64207 | Female | | | | | | | | | |
| RPS10 | | | | | | | 0.00000 | 0.63793 | Female | | | | | | |
| TMEM86A | | | | | | | | | | 0.00001 | 2.35583 | Female | | | |
| ATG9B | | | | | | | 0.00004 | 1.08130 | Female | 0.00000 | 2.20551 | Female | 0.00000 | 1.60549 | Female |
| NDFIP2 | | | | | | | | | | | | | 0.00000 | -0.62210 | Male |
| GNB1L | 0.00092 | 1.74337 | Female | | | | | | | | | | | | |
| PPP2R4 | | | | 0.00906 | 0.65392 | Female | | | | | | | | | |

| | | | | | | | | | | | | | | | |
|---|---|---|---|---|---|---|---|---|---|---|---|---|---|---|---|
| ISG15 | | | | | | | | | | 0.00494 | 1.79328 | Female | | | |
| SESN1 | | | | | | | | | | 0.00555 | 0.69095 | Female | | | |
| ZNF529 | 0.01644 | 1.45954 | Female | | | | | | | | | | | | |
| SLC9A6 | | | | | | | 0.00184 | 0.90637 | Female | | | | | | |
| PPP1R8 | | | | | | | | | | 0.00000 | -1.30246 | Male | | | |
| MALT1 | | | | | | | 0.00436 | -1.02219 | Male | | | | | | |
| RELT | | | | 0.00157 | -1.93182 | Male | | | | | | | | | |
| ARPC3 | 0.00000 | -0.88441 | Male | | | | | | | | | | | | |
| HBP1 | | | | | | | | | | | | | 0.00215 | 0.63827 | Female |
| 1.00000 | | | | | | | 0.00419 | -0.59394 | Male | | | | | | |
| PLEKHF2 | 0.00788 | -1.57337 | Male | 0.04973 | 0.78917 | Female | | | | | | | | | |
| YES1 | 0.02948 | -0.79072 | Male | | | | 0.00000 | -0.62226 | Male | | | | | | |
| MEG3 | | | | | | | | | | 0.00168 | 0.94744 | Female | | | |
| PLEKHB1 | | | | | | | | | | 0.00961 | 1.58987 | Female | 0.01509 | 1.47637 | Female |
| LGALS3 | | | | | | | 0.00001 | -1.44756 | Male | 0.00000 | -1.88678 | Male | | | |
| SMPDL3B | | | | | | | | | | 0.00000 | -3.48437 | Male | | | |
| ELMOD2 | | | | | | | | | | | | | 0.00144 | -0.71130 | Male |
| IGSF10 | | | | | | | 0.04522 | -1.59154 | Male | | | | | | |
| OTUD3 | | | | | | | 0.00030 | 0.99571 | Female | | | | | | |
| RGS19 | | | | 0.00073 | -1.91106 | Male | | | | | | | | | |
| MAPK3 | 0.00166 | -1.50862 | Male | 0.00039 | 1.00189 | Female | | | | | | | 0.00012 | 0.70298 | Female |
| TNNC1 | | | | 0.03322 | 1.26247 | Female | | | | | | | | | |
| FAM219A | | | | | | | 0.04757 | 0.75755 | Female | | | | | | |
| NMT1 | | | | | | | 0.00000 | 0.78536 | Female | | | | | | |
| GPR114 | | | | | | | | | | 0.00161 | -1.63209 | Male | | | |
| SERTAD4-AS1 | | | | | | | | | | 0.02486 | -1.88060 | Male | | | |
| CTSF | | | | | | | | | | 0.00000 | 2.16345 | Female | | | |
| MAGT1 | | | | 0.02120 | 0.68275 | Female | | | | | | | | | |
| CPA1 | | | | | | | | | | 0.01936 | -2.21941 | Male | 0.00939 | 2.33186 | Female |
| GNPDA2 | | | | | | | | | | 0.00886 | -0.84873 | Male | | | |
| ALDH4A1 | | | | 0.00002 | 1.62950 | Female | 0.04614 | -0.59962 | Male | | | | | | |
| ANXA1 | 0.00015 | 1.92643 | Female | 0.00000 | -2.19766 | Male | | | | | | | | | |
| HPGD | | | | 0.00010 | 1.15638 | Female | | | | | | | | | |
| KAT7 | | | | 0.01795 | -0.64421 | Male | | | | | | | | | |
| TMPRSS2 | | | | | | | 0.00095 | -1.17736 | Male | | | | | | |
| COL6A1 | | | | | | | | | | 0.00003 | 1.92893 | Female | 0.00008 | 1.79146 | Female |
| TP53INP1 | | | | | | | | | | 0.00869 | -0.72058 | Male | | | |
| ZNF69 | | | | | | | 0.00006 | -1.72477 | Male | 0.02127 | -1.00276 | Male | | | |
| ZNF300 | | | | | | | | | | | | | 0.02585 | 0.80680 | Female |
| MIR6878 | | | | | | | | | | 0.04751 | 0.64863 | Female | 0.00680 | 0.83685 | Female |
| NPFFR2 | | | | | | | 0.02413 | -1.14999 | Male | | | | | | |
| VAMP8 | | | | 0.00002 | 0.73245 | Female | | | | | | | | | |
| PIP5K1P1 | | | | | | | | | | 0.00196 | -0.66621 | Male | | | |
| C5orf58 | | | | | | | | | | | | | 0.00140 | -1.28553 | Male |
| TMEM128 | | | | | | | | | | | | | 0.00001 | -0.82188 | Male |
| MRPS36 | 0.00471 | 0.79723 | Female | | | | | | | | | | | | |
| SWT1 | | | | | | | | | | 0.03484 | -0.58704 | Male | | | |
| CNST | | | | | | | | | | 0.00016 | -0.70633 | Male | | | |
| HNRNPA1L2 | | | | | | | | | | 0.00003 | 0.88221 | Female | 0.01045 | -0.58107 | Male |
| AVL9 | | | | | | | 0.00844 | -0.69708 | Male | | | | | | |
| SMDT1 | | | | | | | | | | 0.00018 | -0.93331 | Male | | | |

| | | | | | | | | | | | | | | | |
|---|---|---|---|---|---|---|---|---|---|---|---|---|---|---|---|
| LDOC1L | | | | | | | 0.00109 | -2.35323 | Male | | | | | | |
| WWP1 | | | | | | | | | | | | | 0.00196 | -0.58422 | Male |
| SDC4 | | | | 0.00003 | -0.66480 | Male | | | | | | | | | |
| RECK | | | | | | | | | | | | | 0.00593 | 1.42615 | Female |
| CCDC144A | | | | | | | 0.00419 | 1.59932 | Female | | | | | | |
| NYNRIN | | | | 0.03920 | -1.04391 | Male | | | | | | | | | |
| CFLAR | | | | | | | | | | 0.00000 | 2.09167 | Female | | | |
| SIGLEC10 | | | | | | | 0.03568 | -1.79420 | Male | | | | | | |
| GCNT2 | | | | 0.02868 | 1.07176 | Female | | | | | | | | | |
| MIR6857 | | | | | | | 0.00000 | 1.20518 | Female | 0.00000 | 1.08565 | Female | 0.00000 | 0.90911 | Female |
| ELF4 | | | | 0.03251 | 1.09910 | Female | | | | | | | | | |
| MPRIP | | | | 0.00016 | 0.90116 | Female | | | | | | | | | |
| NFKB2 | | | | 0.00715 | 0.91908 | Female | | | | 0.00000 | 1.21241 | Female | | | |
| AKIRIN1 | | | | | | | 0.00000 | -0.66906 | Male | | | | | | |
| HDHD1 | | | | | | | | | | 0.00003 | 0.79193 | Female | 0.00004 | 0.76748 | Female |
| CBWD1 | 0.04221 | -0.69254 | Male | | | | | | | | | | | | |
| SNHG4.uid-7434 | | | | 0.00828 | -1.08148 | Male | | | | | | | | | |
| CEACAM1 | | | | | | | | | | 0.00000 | 2.34641 | Female | 0.01599 | 1.25084 | Female |
| TALDO1 | | | | 0.00000 | 0.87859 | Female | | | | | | | | | |
| CDC42EP2 | | | | | | | | | | 0.00225 | -0.95522 | Male | 0.01389 | 0.79079 | Female |
| RASSF2 | 0.00417 | 2.55770 | Female | | | | | | | | | | | | |
| SAMHD1 | | | | | | | | | | 0.00162 | 0.74896 | Female | 0.01270 | 0.60633 | Female |
| TFE3 | | | | 0.03705 | 0.85979 | Female | 0.00928 | 0.67693 | Female | | | | | | |
| ASTN2 | | | | | | | | | | 0.00809 | 0.92337 | Female | | | |
| IFI6 | 0.02590 | 2.38199 | Female | 0.00005 | 2.39063 | Female | 0.02984 | 0.99567 | Female | | | | 0.00264 | 1.18660 | Female |
| LOC643623 | | | | | | | | | | 0.00000 | 3.16991 | Female | | | |
| VSIG10 | 0.01211 | -1.06972 | Male | | | | | | | 0.00000 | 0.97058 | Female | | | |
| C5orf63 | | | | | | | 0.00075 | -1.06372 | Male | | | | | | |
| ZRANB2 | 0.00094 | -0.77171 | Male | | | | | | | | | | | | |
| AFG3L1P | | | | | | | | | | 0.00070 | 0.91662 | Female | | | |
| DLST | | | | | | | | | | 0.00000 | -1.40176 | Male | | | |
| PLEKHA8 | | | | 0.00110 | 1.18457 | Female | | | | | | | | | |
| MGST3 | 0.00421 | 0.82510 | Female | | | | | | | | | | | | |
| GPR89B | 0.00007 | 2.14326 | Female | | | | | | | | | | | | |
| RP2 | | | | 0.01377 | 0.83757 | Female | 0.00001 | 0.93862 | Female | | | | | | |
| BCAM | | | | | | | 0.00010 | -1.60446 | Male | | | | | | |
| CHST8 | | | | | | | | | | 0.00009 | 1.84387 | Female | 0.02354 | 1.14378 | Female |
| AP5B1 | | | | 0.03381 | -1.17874 | Male | | | | | | | | | |
| NME1-NME2 | | | | | | | | | | 0.00917 | -0.62727 | Male | | | |
| OMA1 | 0.00550 | -1.56572 | Male | | | | 0.01757 | -0.58111 | Male | | | | | | |
| PDLIM2 | | | | | | | | | | 0.02997 | -1.04545 | Male | | | |
| ZNF668 | | | | | | | | | | 0.01859 | -0.60469 | Male | | | |
| HIP1 | | | | 0.01664 | 0.86780 | Female | | | | | | | | | |
| PTPN18 | | | | 0.03500 | 1.54757 | Female | | | | | | | | | |
| CCDC138 | | | | | | | 0.00768 | -0.65693 | Male | | | | | | |
| CETN2 | 0.00080 | 0.99575 | Female | 0.00002 | 0.75838 | Female | 0.00000 | 0.75882 | Female | 0.00000 | 0.64227 | Female | | | |
| IRX2 | 0.02684 | -1.76139 | Male | | | | 0.00921 | 0.86293 | Female | 0.00197 | -0.92951 | Male | | | |
| UBE2D3 | 0.00000 | -0.69400 | Male | | | | | | | | | | | | |
| HNRNPH2 | | | | 0.01062 | 0.60485 | Female | 0.00000 | 0.77935 | Female | 0.00000 | 0.67437 | Female | 0.00001 | 0.60655 | Female |
| CCIN | 0.00000 | -1.27162 | Male | | | | | | | | | | | | |
| ACKR3 | 0.00000 | 1.17307 | Female | 0.00189 | 1.38531 | Female | | | | | | | | | |

| | | | | | | | | | | | | | | | |
|---|---|---|---|---|---|---|---|---|---|---|---|---|---|---|---|
| TCF15 | | | | | | | 0.02287 | 1.40170 | Female | | | | | | |
| BCRP3 | | | | | | | | | | 0.01989 | 1.89772 | Female | | | |
| KCNJ14 | | | | | | | | | | 0.00170 | -2.00623 | Male | | | |
| HSPA1A | | | | | | | | | | | | | 0.00001 | 1.18948 | Female |
| CLCN5 | | | | 0.00419 | 1.17585 | Female | | | | 0.01387 | 0.67818 | Female | | | |
| VPS53 | | | | | | | 0.00000 | 0.72496 | Female | | | | | | |
| CA11 | | | | | | | | | | | | | 0.00795 | 1.96285 | Female |
| NOL12 | | | | | | | 0.00000 | 0.72534 | Female | | | | | | |
| SNHG6 | | | | 0.00000 | -0.71717 | Male | | | | | | | | | |
| PPP3R1 | | | | | | | | | | 0.00458 | -0.60878 | Male | | | |
| KIAA1462 | | | | | | | 0.03153 | -0.75497 | Male | | | | | | |
| MTAP | | | | | | | 0.01337 | -0.73020 | Male | 0.00832 | -0.72563 | Male | | | |
| ATP11A | 0.00004 | -1.45043 | Male | | | | | | | | | | | | |
| ELF3 | 0.00052 | -1.47918 | Male | | | | | | | 0.00001 | 0.66036 | Female | 0.00000 | 0.76542 | Female |
| HOXB3 | | | | | | | 0.00070 | -1.92293 | Male | | | | | | |
| NRIP3 | | | | | | | 0.00005 | 0.99394 | Female | 0.00000 | 1.57531 | Female | | | |
| OSBPL7 | | | | | | | | | | | | | 0.00488 | 2.60276 | Female |
| ZEB1 | | | | | | | | | | | | | 0.01519 | 1.94956 | Female |
| LOC100132352 | | | | | | | 0.00000 | -0.90126 | Male | | | | | | |
| CRMP1 | | | | | | | 0.01824 | 2.21332 | Female | | | | | | |
| CASC8 | | | | | | | | | | 0.01483 | 1.78587 | Female | | | |
| ACSS2 | | | | | | | 0.00000 | -1.36329 | Male | 0.00356 | 0.71096 | Female | | | |
| ACTC1 | | | | | | | | | | 0.00000 | -2.50845 | Male | 0.00023 | 1.71789 | Female |
| PFDN6 | | | | | | | | | | 0.00000 | -1.24077 | Male | 0.00002 | 0.69213 | Female |
| HTATIP2 | | | | | | | | | | | | | 0.00103 | -0.68142 | Male |
| GLIS2 | | | | | | | 0.03942 | 0.97385 | Female | 0.00035 | 1.48450 | Female | | | |
| HLA-F-AS1 | | | | | | | | | | 0.00082 | 1.79608 | Female | | | |
| DDHD2 | | | | | | | 0.01830 | -0.88791 | Male | | | | | | |
| ABHD2 | 0.01855 | -0.67716 | Male | | | | | | | | | | | | |
| FGD1 | | | | 0.03537 | 1.21894 | Female | 0.00001 | 1.57718 | Female | 0.04394 | 0.77830 | Female | | | |
| KIF26A | | | | 0.01189 | 1.29865 | Female | | | | | | | 0.00012 | 1.20481 | Female |
| BIN1 | 0.00598 | 1.90844 | Female | 0.00000 | 2.08089 | Female | | | | | | | | | |
| CHAMP1 | | | | | | | 0.01622 | -0.63434 | Male | | | | 0.00602 | -0.66192 | Male |
| NT5C | | | | 0.00923 | 0.60009 | Female | | | | | | | | | |
| DOCK6 | | | | | | | | | | 0.00004 | 1.08958 | Female | | | |
| HLA-DRB5 | | | | | | | | | | 0.00154 | -2.23502 | Male | | | |
| CHML | | | | | | | | | | | | | 0.00488 | -0.71075 | Male |
| GATA6 | 0.04947 | -0.85421 | Male | | | | | | | | | | | | |
| PRR5 | | | | 0.02090 | 0.70388 | Female | | | | | | | | | |
| MEF2BNB.uid-23522 | | | | | | | 0.00320 | 1.19100 | Female | | | | | | |
| ZNF622 | | | | | | | 0.00000 | 0.67083 | Female | | | | | | |
| ORMDL3 | | | | | | | 0.03228 | 0.61871 | Female | | | | | | |
| HAVCR2 | 0.00452 | 3.26731 | Female | 0.00935 | -1.86119 | Male | | | | | | | | | |
| TCEA1 | 0.03428 | -0.61949 | Male | | | | | | | | | | | | |
| CCDC112 | 0.00021 | -3.20488 | Male | | | | | | | | | | | | |
| GALNT12 | | | | | | | | | | 0.00717 | 2.05614 | Female | | | |
| NRIP1 | | | | | | | | | | | | | 0.04728 | -1.10427 | Male |
| HEY1 | | | | | | | 0.02932 | -1.65951 | Male | | | | | | |
| GRTP1 | | | | | | | | | | 0.00034 | 0.75123 | Female | 0.00217 | -0.64813 | Male |
| GSR | | | | | | | | | | 0.00129 | -0.74495 | Male | | | |
| WDR45 | 0.04664 | 0.78503 | Female | 0.00111 | 0.69576 | Female | 0.00000 | 1.58305 | Female | | | | 0.00000 | 0.87534 | Female |

| | | | | | | | | | | | | | | | |
|---|---|---|---|---|---|---|---|---|---|---|---|---|---|---|---|
| RASSF8-AS1 | | | | | | | | | | 0.00815 | 1.17351 | Female | | | |
| BNIP3L | | | | | | | | | | 0.00070 | -0.73817 | Male | | | |
| FOXI3 | | | | | | | 0.04019 | 0.86577 | Female | | | | 0.04928 | 0.79538 | Female |
| MARCKS | | | | 0.00043 | 1.01123 | Female | | | | | | | | | |
| CCDC64B | | | | | | | 0.00487 | -1.64226 | Male | | | | | | |
| LRRC8D | | | | | | | 0.00052 | -0.87985 | Male | | | | | | |
| NPEPPS | 0.00451 | -0.62543 | Male | | | | | | | | | | | | |
| PELI2 | | | | | | | | | | 0.00117 | 1.93225 | Female | | | |
| MT1HL1 | | | | | | | | | | | | | 0.00004 | 2.04353 | Female |
| ZBTB11-AS1 | | | | | | | | | | | | | 0.03254 | 0.60065 | Female |
| ARHGEF38 | | | | | | | | | | 0.00101 | 0.96323 | Female | | | |
| FNDC4 | | | | 0.03260 | -2.06019 | Male | 0.01795 | -2.02906 | Male | 0.00001 | 3.15846 | Female | | | |
| LINC00884 | | | | | | | 0.04689 | 1.64039 | Female | | | | 0.00044 | 2.43830 | Female |
| TMEM247 | | | | | | | 0.00215 | 1.64165 | Female | | | | | | |
| OCIAD1 | 0.01811 | -0.59274 | Male | | | | | | | | | | | | |
| UTP14A | | | | | | | 0.00000 | 0.73585 | Female | | | | | | |
| CCNJ | | | | | | | | | | 0.00080 | -0.70341 | Male | | | |
| CSAG1 | | | | 0.03693 | 2.01625 | Female | | | | | | | | | |
| SMG1P3 | | | | | | | | | | 0.00181 | 1.06034 | Female | | | |
| RPSAP58 | 0.00005 | 0.89086 | Female | | | | | | | | | | | | |
| PYROXD1 | 0.00028 | -1.76205 | Male | | | | | | | | | | | | |
| UAP1L1 | | | | 0.00005 | 1.70867 | Female | | | | 0.01218 | 0.76088 | Female | | | |
| KIF4B | | | | | | | 0.00002 | 0.68296 | Female | | | | | | |
| TAS1R1 | | | | | | | | | | | | | 0.00342 | 1.71984 | Female |
| CNIH4 | 0.04579 | 0.61244 | Female | | | | | | | | | | | | |
| C6orf211 | 0.02743 | -0.70987 | Male | | | | | | | | | | 0.00000 | -0.58678 | Male |
| MED14 | | | | 0.00080 | 1.01489 | Female | 0.00015 | 0.78474 | Female | | | | | | |
| TMEM164 | | | | 0.00001 | 1.57603 | Female | 0.00001 | 1.07327 | Female | 0.01074 | 0.65400 | Female | | | |
| PTPRS | | | | | | | 0.02348 | 0.71822 | Female | 0.00001 | 1.23234 | Female | | | |
| CENPT | 0.01091 | 0.92107 | Female | | | | | | | | | | | | |
| AFAP1-AS1 | | | | | | | | | | 0.01105 | 1.44542 | Female | 0.00404 | 1.55666 | Female |
| DNAJC22 | | | | | | | | | | 0.03097 | 1.99095 | Female | | | |
| DLEU2L | | | | | | | | | | 0.00002 | 0.89732 | Female | | | |
| RERE | | | | 0.00073 | 0.78806 | Female | | | | | | | | | |
| PHOSPHO1 | | | | | | | 0.00000 | 1.19052 | Female | 0.00000 | 2.68424 | Female | | | |
| PPIL4 | 0.03963 | -0.78445 | Male | | | | | | | | | | | | |
| SLC35G3 | 0.03835 | 3.45057 | Female | | | | | | | | | | | | |
| PI16 | | | | | | | | | | 0.00000 | 3.58505 | Female | | | |
| TPK1 | | | | | | | | | | 0.00000 | -1.45569 | Male | | | |
| TMEM160 | 0.00388 | 1.46949 | Female | | | | | | | 0.00000 | 0.89354 | Female | | | |
| LHX4 | | | | | | | | | | | | | 0.01358 | 1.59004 | Female |
| ANKRD45 | | | | | | | | | | 0.02428 | 1.77538 | Female | | | |
| DUS1L | | | | 0.03787 | 0.61563 | Female | | | | | | | | | |
| MTA2 | | | | 0.00002 | 0.76983 | Female | | | | | | | | | |
| TMEM52 | | | | | | | 0.00018 | -1.41554 | Male | | | | | | |
| LAMC3 | | | | 0.03653 | 1.78522 | Female | | | | | | | | | |
| ADI1 | 0.00002 | -1.23768 | Male | | | | | | | | | | | | |
| TIMM17B | | | | 0.00000 | 0.81518 | Female | 0.00000 | 0.64759 | Female | 0.00000 | 0.62960 | Female | 0.00000 | 0.63661 | Female |
| TRMT10A | | | | | | | | | | | | | 0.00000 | -0.73842 | Male |
| PRPS1L1 | | | | 0.00277 | 0.78159 | Female | 0.00000 | 1.24322 | Female | | | | | | |
| LINC01356 | | | | | | | 0.00000 | -1.33429 | Male | 0.01069 | 0.68368 | Female | | | |

| | | | | | | | | | | | | | | | |
|---|---|---|---|---|---|---|---|---|---|---|---|---|---|---|---|
| PCBD1 | | | | | | | 0.00113 | -0.88385 | Male | | | | | | |
| SNAPC2 | | | | | | | | | | 0.01234 | 0.70200 | Female | | | |
| LOC101927746 | | | | 0.01645 | 0.78246 | Female | | | | | | | | | |
| AMBN | | | | | | | 0.00036 | -1.26847 | Male | | | | | | |
| HIST1H2BN | | | | | | | 0.00996 | 1.66774 | Female | | | | | | |
| CLU | 0.03215 | -2.63384 | Male | | | | 0.00036 | 2.08930 | Female | | | | | | |
| STKLD1 | | | | 0.01747 | 2.19666 | Female | | | | | | | | | |
| TUBB8 | | | | | | | 0.03006 | 1.48713 | Female | 0.00003 | -2.53188 | Male | 0.01463 | -1.59244 | Male |
| CCDC39 | | | | | | | | | | | | | 0.00264 | -1.56765 | Male |
| NXN | | | | 0.04891 | 1.29075 | Female | | | | | | | | | |
| ORC3 | | | | | | | 0.00008 | -0.61772 | Male | | | | | | |
| ZNF439 | | | | | | | | | | 0.04934 | -1.28601 | Male | | | |
| CCDC8 | | | | 0.00943 | -2.29508 | Male | | | | | | | | | |
| EPHB1 | | | | | | | | | | 0.00060 | 1.95040 | Female | | | |
| MSL3P1 | | | | | | | 0.00184 | -0.88158 | Male | 0.02204 | -0.65965 | Male | 0.00257 | -0.81202 | Male |
| ERVMER34-1 | | | | | | | | | | 0.00759 | -0.90005 | Male | | | |
| ZNF205 | | | | | | | | | | 0.01853 | 1.09408 | Female | | | |
| ACOT1 | | | | | | | 0.00550 | -1.15246 | Male | 0.00029 | -1.37475 | Male | | | |
| HDGFRP3 | | | | | | | | | | 0.01002 | 1.02967 | Female | | | |
| CCND1 | | | | 0.00001 | 2.28150 | Female | 0.00000 | 1.65086 | Female | | | | 0.00000 | 1.89435 | Female |
| MSX2 | | | | | | | | | | 0.00071 | -0.63105 | Male | | | |
| M1AP | | | | | | | 0.02023 | -1.12176 | Male | | | | | | |
| SLC5A10 | | | | 0.00000 | 2.22944 | Female | | | | | | | | | |
| CCR8 | | | | | | | | | | 0.00000 | 3.45671 | Female | | | |
| ALDH1A1 | 0.00000 | -2.18180 | Male | | | | | | | 0.00000 | 3.17188 | Female | | | |
| SLC22A18 | | | | | | | | | | 0.00211 | 1.86365 | Female | | | |
| SH3BGRL2 | | | | | | | 0.01938 | -0.58599 | Male | 0.00004 | -0.88812 | Male | | | |
| HSPG2 | | | | | | | 0.00096 | -1.95392 | Male | | | | | | |
| RELL2 | | | | | | | 0.02829 | -1.48851 | Male | | | | | | |
| RPL23AP32 | | | | | | | | | | 0.00062 | -0.90785 | Male | 0.02283 | -0.62980 | Male |
| PLAU | | | | | | | | | | 0.00003 | 1.62566 | Female | | | |
| CACNB3 | | | | | | | | | | 0.00020 | 1.34739 | Female | | | |
| TREM1 | | | | | | | | | | 0.00233 | -1.87300 | Male | | | |
| KANK4 | | | | 0.00131 | 1.32277 | Female | | | | | | | | | |
| SPTAN1 | | | | 0.00834 | 0.58995 | Female | | | | | | | | | |
| GOLGA8B | | | | | | | | | | 0.00239 | -0.86290 | Male | | | |
| LHFP | | | | | | | 0.00183 | -1.64733 | Male | | | | | | |
| PAIP1 | | | | | | | 0.00000 | -0.61111 | Male | | | | | | |
| SULT1A1 | | | | | | | | | | 0.00324 | 1.47370 | Female | | | |
| LOC100507002 | 0.03493 | 2.82094 | Female | | | | | | | 0.00237 | 1.65365 | Female | | | |
| JAZF1 | | | | | | | | | | 0.01990 | 0.94090 | Female | | | |
| HIST3H2A | | | | 0.00235 | -1.82588 | Male | | | | 0.00000 | -2.13758 | Male | | | |
| SNX7 | 0.00053 | 2.92745 | Female | | | | | | | | | | | | |
| SLC15A3 | | | | | | | | | | 0.02525 | 1.85710 | Female | | | |
| CCNA1 | | | | | | | | | | 0.00000 | 1.99648 | Female | 0.01454 | 1.06499 | Female |
| WNT9A | | | | | | | | | | | | | 0.00000 | 2.88801 | Female |
| VWF | | | | | | | 0.03907 | 2.33401 | Female | | | | | | |
| PHACTR2 | 0.00000 | 3.02121 | Female | 0.02698 | 0.77868 | Female | 0.00000 | -1.03557 | Male | | | | | | |
| OLFML3 | 0.00216 | 3.22726 | Female | 0.00000 | 1.31822 | Female | | | | | | | 0.00021 | 2.44913 | Female |
| HNRNPD | 0.00012 | -0.92347 | Male | | | | | | | | | | | | |
| ZHX2 | | | | | | | 0.00400 | -2.02162 | Male | 0.01288 | -1.75727 | Male | | | |

| | | | | | | | | | | | | | | | |
|---|---|---|---|---|---|---|---|---|---|---|---|---|---|---|---|
| SPIN4 | 0.00000 | 5.55047 | Female | 0.01783 | 0.87204 | Female | 0.00217 | 0.72447 | Female | | | | | | |
| DCAF4L2 | | | | | | | | | | | | | 0.04744 | -0.65737 | Male |
| TCF12 | | | | 0.01834 | 0.88923 | Female | 0.00288 | -0.72157 | Male | | | | | | |
| SNORA27 | | | | 0.01012 | -0.97690 | Male | | | | | | | | | |
| H1FX | | | | | | | 0.00000 | 1.34104 | Female | | | | | | |
| NHEJ1 | 0.02329 | 1.49216 | Female | | | | | | | | | | | | |
| MIR5047 | | | | | | | 0.02363 | 0.60415 | Female | 0.00000 | 1.46237 | Female | 0.00838 | 0.64433 | Female |
| KAT2B | | | | | | | | | | 0.03843 | -0.74678 | Male | | | |
| DCXR | | | | 0.00053 | 0.83321 | Female | | | | | | | | | |
| SPIC | | | | | | | | | | 0.00000 | 3.09262 | Female | | | |
| HIST1H4F | | | | | | | | | | 0.00859 | 2.24655 | Female | | | |
| IFT140 | | | | | | | 0.03930 | 0.94540 | Female | | | | | | |
| MAPKAPK2 | | | | | | | 0.00181 | 0.83692 | Female | | | | | | |
| TMLHE | | | | | | | 0.02809 | 1.04895 | Female | | | | | | |
| LPGAT1 | | | | | | | | | | 0.01074 | 0.66808 | Female | | | |
| PLCG1 | | | | | | | | | | 0.00122 | -0.89587 | Male | | | |
| RAD23B | | | | | | | | | | 0.00000 | -1.16871 | Male | | | |
| GPR133 | | | | | | | | | | | | | 0.00766 | 1.56534 | Female |
| PP7080 | | | | | | | | | | 0.02011 | 0.62491 | Female | | | |
| SAC3D1 | | | | | | | | | | 0.01983 | -0.69163 | Male | | | |
| ZFX | | | | 0.00071 | 0.92923 | Female | 0.00000 | 0.84142 | Female | 0.00142 | 0.58534 | Female | | | |
| CATSPER2 | | | | | | | | | | | | | 0.02954 | 1.60311 | Female |
| PIP5KL1 | | | | | | | | | | | | | 0.00580 | 2.70568 | Female |
| SUMO4 | | | | | | | | | | 0.00004 | -1.67711 | Male | | | |
| POMC | | | | | | | | | | 0.00586 | 0.71425 | Female | 0.00004 | 0.98365 | Female |
| LIX1 | | | | | | | | | | | | | 0.00519 | 2.58371 | Female |
| CYP2S1 | | | | | | | | | | 0.00000 | 0.83800 | Female | | | |
| MACC1 | | | | 0.00060 | -2.60351 | Male | | | | 0.00182 | -1.72540 | Male | | | |
| HLA-DRB6 | | | | | | | | | | 0.00000 | -1.83240 | Male | | | |
| HIST1H3C | | | | 0.00044 | -1.82519 | Male | | | | 0.04891 | 0.76982 | Female | | | |
| NPC1 | | | | 0.00308 | -0.89021 | Male | | | | | | | | | |
| RMDN2 | | | | | | | 0.00963 | -1.41931 | Male | | | | 0.03585 | -1.14470 | Male |
| DPH5 | 0.00382 | 1.37789 | Female | | | | | | | | | | | | |
| FABP3 | 0.00000 | -2.71120 | Male | 0.00018 | 1.43954 | Female | 0.00337 | 0.78460 | Female | | | | | | |
| LOC728739 | | | | | | | 0.01738 | -0.60020 | Male | | | | | | |
| ARRDC3 | | | | | | | | | | 0.00528 | 0.63822 | Female | | | |
| C9orf84 | | | | | | | | | | 0.02263 | 1.30832 | Female | | | |
| FAM71E1 | | | | | | | | | | | | | 0.00107 | 0.78571 | Female |
| CACNA1H | | | | | | | | | | 0.00593 | 2.27757 | Female | | | |
| 8.00000 | | | | | | | 0.00000 | -0.63866 | Male | | | | | | |
| FAM76B | | | | | | | 0.00127 | -0.66891 | Male | | | | | | |
| S100Z | | | | | | | | | | 0.01094 | 1.81823 | Female | | | |
| SCARNA22 | | | | 0.01921 | 0.94627 | Female | | | | 0.00124 | 0.84472 | Female | 0.00685 | 0.69751 | Female |
| MRVI1 | | | | | | | | | | | | | 0.01888 | 1.62604 | Female |
| SEMA6A | | | | 0.00000 | 1.55524 | Female | | | | | | | | | |
| NELFCD | 0.00074 | -0.92316 | Male | | | | | | | | | | | | |
| ZNF667 | | | | | | | | | | | | | 0.03770 | 0.90487 | Female |
| MIER1 | 0.00855 | -0.86237 | Male | | | | | | | | | | | | |
| GRAMD1B | | | | | | | | | | 0.00003 | -2.54099 | Male | | | |
| SOX11 | | | | | | | | | | 0.00012 | -1.88650 | Male | | | |
| MYO5C | | | | | | | | | | | | | 0.02595 | -0.83645 | Male |

| | | | | | | | | | | | | | | | |
|---|---|---|---|---|---|---|---|---|---|---|---|---|---|---|---|
| HMGB3 | | | | 0.00000 | 1.00794 | Female | 0.00000 | 0.97645 | Female | 0.00000 | 1.33932 | Female | 0.00000 | 0.66574 | Female |
| RNF39 | | | | | | | | | | | | | 0.04844 | 1.02627 | Female |
| NFKBIL1 | | | | | | | 0.00028 | -1.14906 | Male | | | | | | |
| HERPUD1 | | | | 0.01372 | -0.80249 | Male | | | | | | | | | |
| DCAF8L1 | | | | | | | | | | 0.03250 | -1.15252 | Male | 0.01196 | 1.30976 | Female |
| CADM1 | | | | 0.00104 | 1.44478 | Female | | | | | | | 0.02282 | 0.69984 | Female |
| CYGB | | | | | | | 0.00184 | -1.58040 | Male | | | | 0.01811 | 1.20529 | Female |
| LOC100288778 | | | | | | | | | | 0.01735 | 0.90950 | Female | 0.00113 | -1.14383 | Male |
| NCEH1 | | | | | | | | | | 0.00009 | 0.90984 | Female | | | |
| MRPL32 | 0.00000 | -0.93057 | Male | | | | | | | | | | | | |
| PAMR1 | | | | | | | | | | 0.00045 | 2.32062 | Female | 0.00002 | 2.80848 | Female |
| PLAUR | | | | | | | | | | 0.00070 | 1.43045 | Female | | | |
| SIPA1L2 | | | | | | | | | | 0.00192 | 1.35244 | Female | | | |
| CLCN7 | | | | | | | | | | | | | 0.00413 | -0.92437 | Male |
| NPPB | | | | | | | | | | | | | 0.00000 | 2.34801 | Female |
| LPHN3 | | | | | | | | | | | | | 0.01750 | -1.08589 | Male |
| YWHAZ | 0.00004 | -0.69597 | Male | | | | | | | | | | | | |
| ABHD6 | | | | | | | 0.03051 | -0.69744 | Male | | | | | | |
| B3GNT5 | | | | | | | | | | 0.00725 | -0.97553 | Male | | | |
| IGF1 | | | | | | | 0.01170 | -1.67926 | Male | | | | | | |
| NDN | | | | | | | | | | 0.00004 | -1.46472 | Male | | | |
| STYXL1 | | | | | | | | | | 0.00351 | 0.83876 | Female | | | |
| RMDN1 | | | | 0.00924 | 0.66595 | Female | | | | | | | | | |
| LOC100131691 | | | | | | | | | | 0.00570 | -1.23502 | Male | | | |
| NT5E | | | | 0.01769 | 2.81559 | Female | | | | | | | | | |
| CCT4 | 0.00151 | -0.60730 | Male | | | | | | | | | | | | |
| CBLC | | | | 0.00048 | -2.63452 | Male | | | | | | | | | |
| ZBTB39 | | | | | | | | | | | | | 0.03300 | 0.89715 | Female |
| HSPA1B | | | | | | | | | | 0.00001 | -0.92203 | Male | | | |
| HOTAIRM1 | | | | | | | 0.01470 | 1.22885 | Female | | | | | | |
| C19orf60 | | | | 0.00002 | 1.05782 | Female | | | | 0.00007 | 0.66292 | Female | | | |
| ZNF496 | | | | 0.02394 | 1.22819 | Female | | | | 0.00542 | 0.94653 | Female | 0.02860 | 0.76372 | Female |
| NPIPA5 | | | | | | | 0.04195 | 0.68994 | Female | | | | 0.00632 | 0.84039 | Female |
| PLEK2 | | | | | | | 0.00526 | 1.18785 | Female | | | | 0.00772 | 1.06342 | Female |
| CRHR2 | | | | | | | | | | | | | 0.01289 | 2.09631 | Female |
| FANCA | | | | | | | | | | 0.00952 | 0.65552 | Female | | | |
| LIX1L | | | | | | | | | | 0.00194 | 2.26766 | Female | | | |
| LOC100506469 | | | | 0.01188 | -1.61711 | Male | 0.02723 | 0.99272 | Female | | | | 0.03370 | 0.90735 | Female |
| R3HCC1 | | | | | | | | | | 0.00029 | 1.07718 | Female | | | |
| PMS2P4 | | | | | | | | | | 0.00000 | 2.00900 | Female | | | |
| LARP1B | 0.01628 | -0.77837 | Male | | | | | | | | | | | | |
| IGSF8 | 0.00000 | -3.61432 | Male | | | | | | | | | | | | |
| LOC100129518 | | | | | | | | | | | | | 0.00495 | 1.33712 | Female |
| PAWR | | | | | | | 0.00984 | -0.59472 | Male | | | | | | |
| PAM | | | | | | | | | | | | | 0.02825 | -1.30297 | Male |
| SRGAP1 | | | | | | | | | | 0.00027 | 1.42006 | Female | | | |
| ANKRD36BP1 | | | | | | | | | | | | | 0.02215 | 0.85772 | Female |
| IL17RD | | | | | | | | | | 0.03772 | 1.43799 | Female | | | |
| BHLHE40 | | | | | | | 0.01855 | 0.93221 | Female | | | | | | |
| EXPH5 | 0.00067 | 2.15430 | Female | 0.01251 | 0.99851 | Female | 0.00124 | -0.83439 | Male | | | | | | |
| CUBN | | | | | | | 0.00663 | -2.76033 | Male | | | | | | |

| | | | | | | | | | | | | | | | |
|---|---|---|---|---|---|---|---|---|---|---|---|---|---|---|---|
| MAP3K8 | | | | | | | 0.00005 | -1.02432 | Male | 0.01754 | -0.61692 | Male | | | |
| XPR1 | | | | | | | 0.00004 | -1.03848 | Male | | | | | | |
| YPEL2 | | | | | | | 0.00000 | 0.87755 | Female | 0.00000 | 0.87258 | Female | 0.00012 | 0.62000 | Female |
| APOE | 0.03283 | -1.50467 | Male | 0.00000 | 1.93423 | Female | | | | 0.00000 | 1.71841 | Female | | | |
| COMMD4 | 0.02133 | 0.85841 | Female | | | | | | | | | | | | |
| PAK1 | | | | | | | | | | 0.00002 | 1.03582 | Female | | | |
| PLEKHA6 | | | | 0.00000 | -3.24069 | Male | | | | | | | | | |
| UBA2 | 0.04335 | -0.63731 | Male | | | | | | | | | | 0.00000 | -0.63587 | Male |
| FAM117A | | | | | | | | | | 0.03153 | 0.95171 | Female | | | |
| C17orf96 | | | | | | | | | | | | | 0.02194 | 1.00455 | Female |
| ZBTB17 | | | | 0.02448 | -0.94782 | Male | | | | | | | | | |
| NFATC1 | | | | | | | | | | 0.03610 | 1.35974 | Female | | | |
| PAK2 | | | | 0.00006 | 0.64546 | Female | | | | | | | | | |
| STK19.uid-8457 | | | | | | | | | | | | | 0.00521 | 0.71934 | Female |
| TMEM165 | | | | | | | 0.00000 | -1.05593 | Male | | | | 0.00000 | -0.88014 | Male |
| MTCL1 | | | | | | | | | | 0.00889 | 1.33144 | Female | | | |
| AIMP2 | 0.00061 | -1.05900 | Male | | | | | | | | | | | | |
| SLC25A5-AS1 | | | | | | | | | | 0.00498 | 1.80486 | Female | | | |
| SLC35A2 | | | | | | | | | | 0.00058 | 0.61568 | Female | 0.00005 | 0.69637 | Female |
| LECT2 | | | | | | | | | | 0.02955 | -1.65492 | Male | | | |
| INSIG2 | | | | | | | 0.01964 | -0.64781 | Male | | | | | | |
| BLVRB | | | | | | | | | | | | | 0.00354 | 0.62758 | Female |
| PPARG | 0.00707 | -1.91640 | Male | | | | 0.00008 | -1.12537 | Male | | | | | | |
| GNG10 | | | | 0.00000 | 1.01720 | Female | | | | | | | | | |
| AFAP1L1 | 0.03862 | 2.70844 | Female | | | | | | | | | | | | |
| SLC16A3 | | | | 0.00001 | 2.61462 | Female | | | | | | | | | |
| ANP32C | | | | | | | 0.00000 | -0.66555 | Male | | | | | | |
| LOC100507547 | | | | | | | | | | 0.03829 | 1.32500 | Female | | | |
| SCAPER | | | | | | | | | | | | | 0.04042 | -0.58872 | Male |
| RNASEH2B | | | | | | | 0.00000 | -0.91136 | Male | | | | 0.00123 | -0.62585 | Male |
| NCCRP1 | 0.00001 | 4.97866 | Female | | | | | | | | | | | | |
| KBTBD8 | | | | | | | | | | | | | 0.03236 | -0.72005 | Male |
| CCDC107 | | | | 0.01780 | -1.07935 | Male | | | | | | | | | |
| PTPRJ | | | | | | | | | | 0.04752 | 0.87218 | Female | | | |
| CTSA | 0.02113 | 0.91799 | Female | | | | | | | 0.00001 | 0.61570 | Female | | | |
| PACSIN2 | 0.01283 | -0.79409 | Male | | | | | | | | | | | | |
| SNX8 | | | | | | | | | | 0.00001 | 0.58720 | Female | | | |
| FAM83G | | | | 0.00000 | 2.80144 | Female | | | | | | | | | |
| FGF11 | | | | | | | | | | 0.03421 | -1.40975 | Male | | | |
| TET1 | | | | 0.00050 | 0.71044 | Female | | | | 0.00000 | 0.60007 | Female | | | |
| SUSD2 | | | | | | | | | | 0.00000 | 1.18878 | Female | 0.00706 | 0.69031 | Female |
| FHDC1 | | | | 0.00530 | -1.82401 | Male | | | | 0.02298 | -0.98345 | Male | | | |
| NKRF | | | | 0.00000 | 1.39988 | Female | 0.00002 | 0.84009 | Female | 0.00017 | 0.71371 | Female | | | |
| CEP72 | 0.00733 | -2.09812 | Male | | | | | | | | | | | | |
| SIX1 | | | | | | | | | | 0.01311 | 2.48425 | Female | 0.03851 | 2.09868 | Female |
| TAB3 | | | | | | | | | | 0.00950 | 0.71579 | Female | | | |
| TGFB3 | | | | | | | | | | 0.00006 | 2.04273 | Female | | | |
| ATP11C | | | | 0.00000 | 1.56747 | Female | 0.00000 | 0.81027 | Female | 0.00000 | 1.30391 | Female | | | |
| CIDEC | | | | | | | | | | | | | 0.00103 | 1.14007 | Female |
| KRT23 | | | | 0.00048 | -1.00041 | Male | | | | | | | | | |
| NFIL3 | | | | | | | | | | 0.00001 | -1.22328 | Male | | | |

| | | | | | | | | | | | | | | | |
|---|---|---|---|---|---|---|---|---|---|---|---|---|---|---|---|
| NRF1 | | | | 0.02386 | -0.74095 | Male | | | | | | | | | |
| ARL13B | | | | | | | 0.00007 | 1.14087 | Female | | | | | | |
| TIMM8A | 0.00008 | 0.91094 | Female | 0.00000 | 1.11093 | Female | 0.00000 | 1.11624 | Female | 0.00000 | 0.74008 | Female | | | |
| GCNT1 | | | | 0.01288 | -1.44136 | Male | 0.00225 | 1.16601 | Female | 0.00547 | 1.03020 | Female | | | |
| NRDE2 | | | | 0.02561 | -0.65788 | Male | | | | | | | | | |
| DLG2 | | | | | | | 0.00549 | 1.01363 | Female | | | | | | |
| CASP9 | | | | | | | | | | | | | 0.03087 | 0.78936 | Female |
| HDAC8 | | | | 0.00000 | 1.52203 | Female | 0.00000 | 1.23410 | Female | 0.00000 | 1.28746 | Female | 0.00002 | 0.92106 | Female |
| SP140L | | | | 0.01394 | 1.94448 | Female | | | | | | | 0.00926 | 1.33983 | Female |
| ISOC2 | | | | 0.01400 | 1.01055 | Female | 0.00707 | 0.73254 | Female | 0.00000 | 1.32746 | Female | 0.02314 | 0.60466 | Female |
| VEGFB | | | | | | | 0.00266 | -1.41034 | Male | 0.03328 | -0.99451 | Male | | | |
| ACSL4 | | | | | | | | | | 0.00558 | 0.61447 | Female | | | |
| REC114 | | | | | | | | | | | | | 0.00036 | -2.67818 | Male |
| SH3PXD2B | | | | | | | | | | 0.01437 | 1.21577 | Female | | | |
| GPANK1 | | | | 0.00000 | -0.77691 | Male | | | | | | | | | |
| TIMM8B | 0.00275 | 0.62447 | Female | | | | | | | | | | | | |
| SAP30L-AS1 | 0.00816 | 3.05252 | Female | | | | | | | | | | | | |
| TGIF1 | 0.01700 | -0.83631 | Male | | | | | | | | | | | | |
| LGALS4 | | | | | | | | | | 0.00001 | 1.88088 | Female | | | |
| CLPTM1L | | | | | | | 0.00000 | 0.74493 | Female | | | | | | |
| LPAR5 | | | | 0.02209 | 2.87641 | Female | | | | | | | | | |
| CGNL1 | | | | | | | | | | 0.00377 | 1.51625 | Female | | | |
| PLSCR1 | | | | | | | | | | | | | 0.02860 | -0.59135 | Male |
| CRHBP | | | | | | | | | | 0.04584 | 1.60837 | Female | 0.00124 | -2.43508 | Male |
| SLC35B4 | | | | 0.03165 | 0.61773 | Female | | | | | | | | | |
| LOC391322 | | | | | | | 0.02703 | -1.03221 | Male | | | | | | |
| ANKRD20A5P | | | | | | | | | | 0.00314 | 1.69239 | Female | 0.00857 | 1.51832 | Female |
| SNHG5 | | | | 0.00000 | -1.00010 | Male | | | | | | | 0.00000 | -0.90935 | Male |
| AUTS2 | | | | | | | | | | 0.00233 | 1.07915 | Female | | | |
| CBR3 | | | | | | | | | | 0.00000 | 2.31937 | Female | | | |
| CA5BP1 | | | | 0.00130 | 0.87572 | Female | 0.00000 | 1.65283 | Female | 0.00002 | 0.73714 | Female | 0.00005 | 0.69007 | Female |
| GCDH | | | | | | | 0.00000 | 0.83088 | Female | | | | | | |
| INA | | | | | | | 0.02348 | 0.75231 | Female | | | | | | |
| DHRS4-AS1 | | | | | | | | | | 0.03646 | -1.51519 | Male | | | |
| HMGN2P46 | | | | | | | | | | 0.00935 | -1.48382 | Male | | | |
| ARL4D | | | | | | | 0.00001 | 0.67867 | Female | | | | | | |
| CYFIP2 | | | | | | | | | | | | | 0.00870 | 1.44164 | Female |
| MT1G | | | | | | | | | | | | | 0.03748 | 1.03729 | Female |
| ZNF415 | | | | | | | | | | | | | 0.00001 | -1.36120 | Male |
| RAB9A | | | | 0.00034 | 0.76388 | Female | | | | | | | | | |
| PMAIP1 | | | | | | | | | | 0.00006 | 1.40022 | Female | | | |
| MIR22HG | 0.00613 | 1.51235 | Female | 0.01188 | -0.89488 | Male | | | | | | | | | |
| SNORD12C | | | | 0.00728 | -1.08386 | Male | | | | | | | | | |
| H3F3C | | | | | | | 0.00000 | 1.16492 | Female | | | | | | |
| LSS | | | | | | | 0.00000 | -1.12253 | Male | | | | | | |
| CRAT | | | | | | | 0.00444 | -1.82470 | Male | | | | | | |
| PFKFB2 | | | | | | | 0.04074 | -0.68384 | Male | | | | | | |
| BMF | | | | 0.00023 | -2.42615 | Male | | | | | | | | | |
| FASTK | | | | | | | 0.00996 | -0.58929 | Male | | | | | | |
| DARS | 0.01806 | 0.58448 | Female | | | | | | | | | | | | |
| LPIN3 | | | | | | | 0.00000 | 1.37343 | Female | | | | 0.00648 | 0.68188 | Female |

| | | | | | | | | | | | | | | | |
|---|---|---|---|---|---|---|---|---|---|---|---|---|---|---|---|
| PTRF | 0.01307 | 2.20629 | Female | 0.01306 | 1.40495 | Female | | | | | | | 0.01174 | 0.91649 | Female |
| SNAR-I | | | | 0.00002 | 1.51114 | Female | 0.00001 | 1.08758 | Female | 0.00003 | 1.01715 | Female | | | |
| DLGAP3 | | | | | | | | | | 0.00000 | 2.38659 | Female | | | |
| CYP51A1 | | | | | | | 0.00004 | -0.61969 | Male | | | | | | |
| TRIM43 | | | | | | | | | | 0.01955 | 6.25041 | Female | | | |
| C14orf80 | 0.01101 | -2.04335 | Male | | | | | | | | | | 0.00500 | 0.93459 | Female |
| TMCO6 | 0.01750 | 2.25656 | Female | | | | 0.00512 | -1.09636 | Male | | | | | | |
| MAK16 | 0.01581 | -0.72119 | Male | | | | | | | | | | | | |
| WDR13 | | | | | | | 0.00153 | 0.90218 | Female | | | | 0.03414 | 0.61220 | Female |
| NDUFA1 | 0.00000 | 1.13300 | Female | 0.00000 | 0.99716 | Female | 0.00000 | 0.80168 | Female | 0.00000 | 0.69878 | Female | 0.00000 | 0.87029 | Female |
| KCTD13 | | | | 0.04036 | 0.96152 | Female | | | | | | | | | |
| TRAF3IP1 | | | | | | | 0.00215 | 1.03222 | Female | | | | 0.00289 | 0.94503 | Female |
| IKBKAP | 0.00074 | 1.44003 | Female | | | | | | | | | | | | |
| CCDC170 | | | | | | | | | | 0.00153 | -2.58332 | Male | 0.00495 | -2.36789 | Male |
| APOOL | | | | 0.02725 | 0.58083 | Female | 0.00000 | 1.06849 | Female | | | | | | |
| MAP7D3 | | | | 0.03053 | 0.77457 | Female | 0.00136 | 0.72170 | Female | | | | | | |
| SESN3 | | | | 0.01069 | 1.44396 | Female | | | | 0.00000 | 1.57717 | Female | | | |
| GYG2 | | | | | | | | | | 0.03341 | 1.13747 | Female | | | |
| PLS3 | | | | 0.00000 | 1.19344 | Female | | | | | | | | | |
| CDX1 | | | | | | | 0.00000 | 1.66336 | Female | | | | | | |
| LINC01447 | | | | | | | 0.00426 | 2.38671 | Female | | | | | | |
| BUB1 | | | | | | | | | | 0.00000 | 1.58157 | Female | | | |
| OTUD6A | | | | | | | 0.04441 | 2.23510 | Female | | | | | | |
| EFR3B | | | | | | | 0.00249 | 0.81398 | Female | | | | 0.00014 | 0.93746 | Female |
| FAM107B | | | | | | | | | | 0.01067 | 0.59092 | Female | | | |
| FCHSD2 | 0.00685 | -2.44344 | Male | | | | | | | | | | | | |
| RPS4Y1 | 0.00000 | -9.31666 | Male | 0.00000 | -9.45687 | Male | 0.00000 | -11.05569 | Male | 0.00000 | -10.47092 | Male | 0.00000 | -8.98075 | Male |
| PPAP2B | | | | 0.00319 | 0.99723 | Female | | | | | | | | | |
| POLR2J4 | | | | | | | 0.02699 | 0.60996 | Female | | | | | | |
| DOHH | | | | | | | 0.00728 | -0.83806 | Male | | | | | | |
| ISYNA1 | 0.00013 | -2.09996 | Male | 0.00000 | 2.66713 | Female | | | | | | | | | |
| RPAP1 | | | | | | | 0.00000 | -0.91070 | Male | | | | | | |
| ALDH2 | | | | | | | | | | 0.00183 | 0.74232 | Female | | | |
| AGGF1 | | | | | | | 0.00001 | -0.74215 | Male | | | | | | |
| PGAP2 | | | | | | | 0.00000 | -0.95936 | Male | | | | | | |
| PGM1 | | | | | | | 0.00189 | -0.91422 | Male | 0.02030 | 0.68707 | Female | 0.00933 | -0.74068 | Male |
| HTRA2 | 0.00829 | 0.91723 | Female | | | | | | | | | | | | |
| MDFI | 0.00000 | 2.38708 | Female | 0.00000 | 2.75262 | Female | 0.03269 | 1.20167 | Female | | | | | | |
| GAGE1 | 0.00043 | 3.18219 | Female | | | | 0.00000 | 2.90962 | Female | 0.00000 | 2.48396 | Female | 0.00004 | 1.50252 | Female |
| AKT1 | | | | 0.01558 | 0.98531 | Female | | | | | | | | | |
| CERS6 | | | | | | | 0.03824 | -1.46930 | Male | | | | | | |
| SLC25A25 | | | | | | | 0.04745 | 0.61674 | Female | | | | | | |
| USP2 | | | | | | | 0.04118 | 0.77374 | Female | | | | | | |
| PPDPF | | | | | | | | | | 0.00000 | 1.18919 | Female | 0.00153 | 0.68931 | Female |
| GPKOW | | | | 0.00542 | 0.93171 | Female | 0.00003 | 0.90230 | Female | 0.00138 | 0.68825 | Female | | | |
| SNORA53 | | | | | | | | | | | | | 0.01504 | 1.50848 | Female |
| SYT2 | | | | | | | | | | 0.00002 | 1.38235 | Female | | | |
| STARD10 | | | | | | | 0.00242 | -0.72088 | Male | | | | | | |
| FDXACB1 | | | | | | | 0.04669 | -0.83473 | Male | | | | | | |
| RWDD2B | | | | | | | 0.03082 | -0.99060 | Male | | | | | | |
| ALYREF | | | | | | | | | | 0.00000 | -0.63373 | Male | | | |

| | | | | | | | | | | | | | | | |
|---|---|---|---|---|---|---|---|---|---|---|---|---|---|---|---|
| ZNF679 | | | | | | | | | | 0.00000 | -3.37744 | Male | | | |
| TMEM200A | | | | 0.01324 | 2.86442 | Female | | | | 0.00076 | -2.62803 | Male | | | |
| HS3ST3A1 | | | | | | | 0.02761 | 0.95207 | Female | | | | | | |
| RPAP2 | | | | | | | 0.01317 | -0.58402 | Male | | | | | | |
| TOP2B | | | | 0.00011 | 1.00669 | Female | | | | | | | | | |
| FAM83B | 0.01636 | 1.99258 | Female | | | | 0.00004 | -1.29238 | Male | | | | 0.04514 | -0.65867 | Male |
| UBL4A | 0.00000 | 3.25343 | Female | 0.00430 | 0.77639 | Female | | | | | | | | | |
| FTX | 0.01592 | 2.14135 | Female | | | | | | | 0.01368 | 0.87195 | Female | | | |
| LINC01278 | | | | 0.00183 | 1.34640 | Female | | | | | | | | | |
| TIGD2 | | | | | | | 0.00512 | -1.52765 | Male | | | | | | |
| LAMA2 | | | | | | | | | | 0.02791 | 1.26846 | Female | | | |
| FOXH1 | | | | 0.01035 | 1.14120 | Female | | | | 0.00138 | 0.89462 | Female | | | |
| NAA10 | | | | 0.00000 | 1.15899 | Female | 0.00000 | 0.75203 | Female | | | | | | |
| ZNF618 | | | | | | | | | | 0.00007 | 2.38323 | Female | 0.00102 | -2.00727 | Male |
| FAM8A1 | | | | | | | | | | | | | 0.04071 | -0.60677 | Male |
| NCBP2 | 0.00006 | -0.78032 | Male | | | | | | | | | | | | |
| SLBP | | | | | | | 0.00000 | -0.67248 | Male | | | | 0.00000 | -0.65595 | Male |
| LRP4 | | | | | | | | | | 0.00098 | 1.37332 | Female | | | |
| TRPM1 | | | | | | | | | | 0.02400 | -1.94775 | Male | | | |
| ZG16 | | | | | | | 0.00402 | 0.71091 | Female | | | | | | |
| AXL | | | | 0.00110 | -1.23194 | Male | | | | 0.00000 | -1.09037 | Male | 0.00096 | -0.81318 | Male |
| IMPA1 | | | | | | | 0.00103 | -0.71036 | Male | | | | | | |
| TLCD1 | | | | | | | 0.00074 | -0.78976 | Male | | | | | | |
| ETV4 | | | | | | | | | | 0.00000 | 2.02998 | Female | | | |
| RAP1GAP2 | | | | | | | | | | 0.00328 | 1.34358 | Female | 0.02542 | 1.05102 | Female |
| EFCAB10 | | | | | | | | | | | | | 0.01319 | 0.88748 | Female |
| RRAGB | | | | | | | 0.01217 | 1.07962 | Female | 0.00250 | 1.20666 | Female | | | |
| MYT1L | | | | | | | | | | 0.00560 | -1.93502 | Male | | | |
| ABAT | | | | | | | | | | 0.00003 | 2.41277 | Female | | | |
| KLHL2 | | | | | | | | | | | | | 0.01830 | -0.76044 | Male |
| MGC45800 | | | | | | | | | | 0.00458 | -1.92672 | Male | | | |
| FAM217B | 0.00860 | 1.92369 | Female | | | | | | | | | | | | |
| SEPHS2 | | | | 0.00148 | 0.83759 | Female | | | | | | | | | |
| FARSA | | | | | | | 0.00000 | 0.67909 | Female | | | | | | |
| POP1 | | | | | | | 0.00003 | 0.68774 | Female | | | | | | |
| HIVEP3 | | | | | | | | | | | | | 0.01885 | 0.85414 | Female |
| COLQ | | | | | | | 0.02542 | 1.79676 | Female | | | | | | |
| SFN | | | | | | | | | | 0.00262 | -0.70832 | Male | 0.00001 | 0.96607 | Female |
| WRB | | | | | | | 0.00080 | -0.74350 | Male | | | | | | |
| PLD1 | | | | | | | 0.00343 | -1.64093 | Male | | | | | | |
| TMEM18 | | | | 0.00098 | 0.79522 | Female | | | | | | | | | |
| SLC2A14 | 0.03414 | 1.39060 | Female | | | | | | | | | | 0.00869 | 0.65800 | Female |
| ACE | | | | | | | 0.02360 | 0.97070 | Female | 0.00000 | 2.96502 | Female | | | |
| KIAA1549L | | | | | | | | | | 0.01588 | 1.89326 | Female | | | |
| MAP3K15 | | | | | | | 0.02025 | 1.48614 | Female | | | | | | |
| PCSK7 | | | | 0.00691 | 0.89243 | Female | | | | | | | | | |
| MYLPF | | | | | | | | | | 0.00000 | -1.15531 | Male | | | |
| TP63 | | | | 0.01890 | 1.60036 | Female | 0.00535 | 1.29853 | Female | | | | | | |
| CCDC7 | | | | 0.00214 | -0.65693 | Male | | | | | | | | | |
| POMK | | | | | | | | | | 0.00394 | -0.88844 | Male | | | |
| ZNRF2 | | | | | | | | | | | | | 0.02138 | -0.91056 | Male |

| | | | | | | | | | | | | | | | |
|---|---|---|---|---|---|---|---|---|---|---|---|---|---|---|---|
| MAGEC1 | | | | 0.00827 | 2.74167 | Female | 0.01777 | 1.80980 | Female | | | | 0.00045 | 0.74968 | Female |
| F2RL1 | | | | 0.00147 | -0.74835 | Male | | | | | | | | | |
| MRPL44 | 0.00124 | -0.74270 | Male | | | | | | | | | | | | |
| FAM212B | | | | | | | | | | 0.01038 | 1.60052 | Female | | | |
| WDR6 | | | | | | | | | | 0.00818 | -0.58166 | Male | | | |
| LACTB2 | | | | | | | 0.01713 | -0.74104 | Male | | | | | | |
| HAPLN1 | | | | | | | | | | 0.00110 | -0.96541 | Male | 0.00111 | -0.94234 | Male |
| TINAGL1 | | | | | | | | | | 0.02579 | -0.77574 | Male | | | |
| PCLO | | | | | | | | | | 0.00228 | 1.66058 | Female | | | |
| WNK2 | | | | | | | | | | | | | 0.04594 | 0.90481 | Female |
| MED29 | | | | | | | | | | | | | 0.00000 | 0.62605 | Female |
| KIF4A | | | | | | | 0.00000 | 0.70709 | Female | | | | | | |
| QRICH2 | | | | | | | 0.00683 | 1.29493 | Female | | | | | | |
| GINS3 | | | | | | | 0.00000 | -0.82914 | Male | | | | 0.00008 | -0.61642 | Male |
| DEPDC7 | | | | | | | | | | 0.00076 | 1.53323 | Female | | | |
| GTF2I | | | | | | | | | | 0.00064 | 0.87093 | Female | | | |
| PLEKHH1 | | | | 0.00005 | -2.49317 | Male | | | | | | | | | |
| TRIM24 | | | | 0.03778 | 0.79157 | Female | | | | | | | | | |
| KATNBL1 | | | | 0.02452 | -0.77969 | Male | | | | | | | | | |
| AICDA | | | | | | | 0.01418 | 2.31250 | Female | 0.00000 | 3.19631 | Female | | | |
| SYNGR3 | | | | | | | 0.04796 | 0.63803 | Female | | | | | | |
| HES1 | | | | 0.00040 | -1.57283 | Male | | | | | | | | | |
| PLA2G4F | | | | | | | | | | 0.02504 | 1.08386 | Female | 0.01692 | 1.11602 | Female |
| FAM214A | | | | | | | | | | | | | 0.00000 | -1.50901 | Male |
| DPPA2 | | | | | | | | | | 0.00002 | 1.04334 | Female | | | |
| VAMP3 | 0.01042 | -0.83426 | Male | | | | | | | | | | | | |
| SPHK2 | | | | | | | 0.01824 | -0.61836 | Male | 0.00000 | -1.33341 | Male | | | |
| PRPS1 | | | | 0.00000 | 1.14281 | Female | 0.00000 | 1.32780 | Female | | | | | | |
| ANKRD11 | | | | 0.01005 | 0.60409 | Female | 0.00002 | 0.61891 | Female | | | | | | |
| UGDH | | | | | | | 0.01493 | -0.58505 | Male | | | | | | |
| FHL1 | | | | | | | | | | 0.00088 | -2.82594 | Male | | | |
| UBE2Q2P1 | | | | | | | | | | 0.04349 | 1.68983 | Female | | | |
| METTL2A | 0.00207 | 0.97265 | Female | | | | | | | | | | | | |
| HORMAD2-AS1 | | | | | | | | | | 0.02265 | 0.73020 | Female | | | |
| VCX3B | 0.00001 | 3.81261 | Female | 0.00000 | 3.41708 | Female | 0.00000 | 2.98189 | Female | 0.00076 | 1.36547 | Female | 0.00000 | 4.53617 | Female |
| PSMD10 | | | | 0.00010 | 0.73816 | Female | 0.00000 | 0.64899 | Female | 0.00000 | 0.59307 | Female | | | |
| SCMH1 | | | | | | | 0.02774 | -1.32200 | Male | | | | | | |
| NME4 | | | | | | | | | | 0.01142 | -0.72612 | Male | 0.01986 | -0.66501 | Male |
| CALB2 | | | | 0.02510 | -2.34201 | Male | | | | | | | 0.00000 | 3.59048 | Female |
| MFSD9 | 0.01779 | -1.71856 | Male | | | | | | | | | | | | |
| TPRG1L | | | | | | | 0.00679 | 0.69986 | Female | | | | | | |
| MANEA | | | | | | | 0.00001 | -1.13491 | Male | | | | 0.00717 | -0.68438 | Male |
| LOC645513 | | | | | | | | | | 0.00010 | 1.02572 | Female | | | |
| MT2A | | | | | | | | | | | | | 0.00008 | 1.41807 | Female |
| ARHGAP24 | | | | 0.04102 | 1.26103 | Female | 0.00003 | -1.56766 | Male | 0.02234 | -0.88000 | Male | | | |
| JAG2 | | | | 0.01402 | 1.31488 | Female | | | | 0.00076 | 1.12392 | Female | | | |
| LRRC8B | | | | | | | 0.00000 | -1.80421 | Male | 0.00008 | -1.27816 | Male | 0.00601 | -0.92285 | Male |
| CDC42 | | | | | | | | | | 0.00000 | -0.59064 | Male | | | |
| LINC01234 | | | | | | | | | | 0.04860 | 1.07896 | Female | 0.00008 | 1.88477 | Female |
| PMP22 | | | | | | | | | | 0.00000 | 1.93195 | Female | 0.00379 | 1.02363 | Female |
| BEX1 | | | | | | | | | | 0.00555 | -1.59459 | Male | 0.00505 | -1.51852 | Male |

| | | | | | | | | | | | | | | | |
|---|---|---|---|---|---|---|---|---|---|---|---|---|---|---|---|
| SLC30A5 | | | | | | | 0.00040 | -0.60438 | Male | | | | | | |
| PPME1 | | | | | | | | | | 0.00000 | -0.87609 | Male | | | |
| MYO9A | | | | 0.03064 | 0.75606 | Female | | | | 0.00164 | 0.66309 | Female | | | |
| WASH1 | | | | | | | | | | 0.01096 | -0.67879 | Male | 0.00047 | 0.88007 | Female |
| ACP6 | | | | | | | | | | 0.01954 | -0.98018 | Male | | | |
| RNF6 | | | | | | | | | | | | | 0.00070 | -0.59902 | Male |
| FAM50A | 0.00174 | -0.97019 | Male | 0.00000 | 1.11586 | Female | 0.00000 | 0.72210 | Female | | | | 0.00000 | 0.72195 | Female |
| MAGED2 | 0.00006 | 3.74677 | Female | | | | | | | | | | 0.00041 | -1.39739 | Male |
| GLE1 | 0.04947 | -1.06273 | Male | | | | | | | | | | | | |
| NAA11 | | | | | | | | | | | | | 0.00181 | -0.66750 | Male |
| C1orf233 | | | | | | | | | | | | | 0.02284 | 1.11320 | Female |
| HUWE1 | | | | 0.00000 | 1.18110 | Female | | | | 0.00000 | 0.91328 | Female | 0.00000 | 0.79153 | Female |
| SKA1 | | | | 0.00531 | -0.62796 | Male | | | | | | | | | |
| ZNF490 | | | | | | | | | | 0.01487 | 0.68043 | Female | | | |
| PPP2R5B | | | | | | | | | | 0.02256 | 1.05624 | Female | | | |
| MBD3L3 | | | | 0.02309 | -2.68636 | Male | | | | 0.01712 | 4.53425 | Female | | | |
| LRP2 | 0.00110 | -1.88583 | Male | | | | | | | | | | | | |
| SLC38A2 | | | | 0.00002 | -1.19606 | Male | 0.00001 | -0.85985 | Male | | | | | | |
| PTMS | | | | | | | | | | | | | 0.00013 | 0.80034 | Female |
| SUMO1P1 | | | | | | | | | | | | | 0.00833 | -2.27223 | Male |
| CCDC163P | | | | | | | | | | 0.00152 | 0.84558 | Female | | | |
| STMN2 | | | | | | | | | | 0.00049 | -4.64513 | Male | | | |
| RASL11A | | | | | | | | | | | | | 0.01001 | -1.12933 | Male |
| SFI1 | | | | 0.03476 | 1.28688 | Female | | | | | | | | | |
| DNPEP | | | | | | | 0.00527 | 0.80585 | Female | | | | 0.03103 | 0.61303 | Female |
| SNORD63 | | | | | | | | | | | | | 0.00494 | 0.66069 | Female |
| ZNF354A | | | | | | | | | | | | | 0.00639 | 0.87925 | Female |
| MOGAT3 | | | | | | | | | | 0.00000 | 3.23219 | Female | | | |
| ZNF277 | | | | | | | 0.00053 | -0.73457 | Male | 0.00100 | -0.66864 | Male | | | |
| TCL1B | | | | | | | | | | 0.00317 | 1.17337 | Female | | | |
| SNHG7 | | | | | | | | | | | | | 0.00001 | 0.70242 | Female |
| LY6G5C | | | | | | | 0.02875 | -2.20925 | Male | | | | | | |
| PGLS | 0.00105 | 1.63866 | Female | | | | | | | | | | | | |
| MYO15B | | | | | | | | | | | | | 0.00007 | 2.30291 | Female |
| AXIN1 | | | | 0.01245 | 0.89406 | Female | | | | | | | 0.00270 | 0.66221 | Female |
| SGPP2 | | | | | | | 0.01486 | -1.42594 | Male | | | | | | |
| BTN1A1 | | | | | | | | | | 0.00089 | -1.98550 | Male | | | |
| PPP1R32 | | | | | | | 0.03438 | -1.80736 | Male | | | | | | |
| ENOX2 | | | | | | | 0.00001 | 1.11169 | Female | | | | | | |
| ALPP | | | | | | | 0.00000 | -1.06652 | Male | | | | | | |
| MORN1 | | | | | | | | | | | | | 0.04006 | 1.02450 | Female |
| CIC | 0.01007 | -1.73504 | Male | | | | | | | | | | | | |
| DND1 | | | | | | | 0.01503 | 0.72209 | Female | 0.00021 | 0.98166 | Female | | | |
| AK9 | | | | 0.00503 | -2.42559 | Male | | | | | | | | | |
| KCNJ3 | | | | | | | | | | 0.00000 | -1.41488 | Male | 0.00000 | -1.73897 | Male |
| SEPW1 | | | | | | | | | | 0.00000 | 0.62127 | Female | | | |
| FAM129B | | | | | | | 0.00054 | 1.36512 | Female | | | | | | |
| TSPAN3 | | | | | | | 0.00085 | -0.62940 | Male | | | | | | |
| SSX6 | | | | | | | 0.00001 | 2.37482 | Female | 0.01104 | 1.42995 | Female | | | |
| TGDS | | | | | | | | | | | | | 0.00121 | -0.62000 | Male |
| TGM2 | 0.00000 | 2.49664 | Female | | | | | | | | | | 0.00021 | 1.29681 | Female |

| | | | | | | | | | | | | | | | |
|---|---|---|---|---|---|---|---|---|---|---|---|---|---|---|---|
| WNT7B | | | | 0.00241 | 1.87760 | Female | 0.01610 | -0.94412 | Male | | | | | | |
| CLIC3 | | | | | | | | | | 0.00028 | -1.22574 | Male | | | |
| LHFPL3-AS2 | | | | | | | | | | 0.00045 | 0.83444 | Female | | | |
| GLIPR2 | | | | 0.01937 | 0.58729 | Female | | | | | | | | | |
| ARL4A | | | | | | | | | | 0.02479 | 0.59452 | Female | | | |
| NOL9 | 0.01648 | -0.84873 | Male | | | | | | | | | | | | |
| CCDC124 | 0.01011 | -0.59982 | Male | | | | | | | | | | | | |
| TNS3 | | | | 0.00000 | 1.61782 | Female | | | | | | | | | |
| GYPC | | | | | | | | | | 0.00000 | 3.75179 | Female | | | |
| SNX29 | | | | 0.01288 | 1.05878 | Female | | | | | | | | | |
| NAGA | | | | | | | 0.04985 | 0.63290 | Female | 0.00005 | 1.11230 | Female | | | |
| BLOC1S2 | 0.03710 | 0.67660 | Female | | | | | | | | | | | | |
| SNORD104 | | | | | | | | | | | | | 0.00001 | 0.88360 | Female |
| SNORD10 | 0.00594 | 1.83060 | Female | | | | | | | 0.00726 | -0.70118 | Male | 0.00001 | 1.06633 | Female |
| NSUN6 | | | | | | | 0.04263 | -0.59710 | Male | | | | | | |
| TFPT | | | | 0.00003 | -0.94511 | Male | | | | | | | | | |
| CGB8 | | | | | | | | | | 0.00000 | -2.50946 | Male | | | |
| ACVR1B | | | | | | | | | | | | | 0.00969 | 0.79549 | Female |
| LOC102724153 | | | | | | | 0.00267 | 1.73695 | Female | 0.03952 | 1.22001 | Female | | | |
| GPR126 | | | | 0.03039 | 1.45050 | Female | | | | | | | 0.01405 | -0.97758 | Male |
| RANBP17 | | | | | | | | | | 0.02163 | 0.78672 | Female | | | |
| SHB | | | | | | | | | | 0.00184 | 0.91369 | Female | 0.00681 | 0.79923 | Female |
| TMCO4 | | | | | | | | | | 0.03336 | 1.19370 | Female | | | |
| RCBTB1 | | | | | | | 0.01394 | -0.63878 | Male | 0.00000 | 1.19022 | Female | | | |
| RPP25L | | | | | | | 0.01125 | 0.59029 | Female | | | | | | |
| NT5DC1 | | | | | | | 0.00200 | -0.75611 | Male | | | | | | |
| ACO1 | | | | | | | | | | 0.00027 | 0.76213 | Female | | | |
| CPZ | | | | | | | | | | | | | 0.00056 | 1.75379 | Female |
| AIRE | | | | 0.00009 | 2.86872 | Female | | | | 0.00000 | 3.23400 | Female | | | |
| LAMA5 | | | | 0.00009 | 1.18945 | Female | | | | | | | | | |
| MIDN | | | | 0.00000 | 1.01489 | Female | 0.00003 | 0.64542 | Female | | | | | | |
| TBCD | | | | | | | | | | 0.00000 | 1.36586 | Female | | | |
| WASF1 | | | | | | | 0.00000 | -1.16694 | Male | 0.00873 | -0.64018 | Male | | | |
| TMPPE | | | | | | | | | | | | | 0.04921 | 0.85065 | Female |
| NKAP | 0.01074 | 1.00068 | Female | 0.00017 | 0.82254 | Female | 0.00000 | 1.35248 | Female | 0.00000 | 1.14389 | Female | | | |
| CLEC4GP1 | | | | | | | 0.02694 | -1.34003 | Male | | | | | | |
| MORC4 | | | | 0.00312 | 1.20845 | Female | | | | 0.00326 | 0.78612 | Female | | | |
| SIRT2 | | | | | | | 0.04395 | 0.62804 | Female | | | | | | |
| SHD | | | | | | | | | | 0.00076 | 1.80654 | Female | | | |
| RHOU | 0.01191 | 1.91174 | Female | | | | | | | | | | | | |
| HEBP1 | | | | | | | 0.00232 | -1.05379 | Male | | | | | | |
| KIAA0895 | | | | | | | | | | 0.02342 | -1.48262 | Male | | | |
| TIGIT | | | | | | | | | | 0.00582 | -5.33296 | Male | | | |
| MED17 | | | | | | | 0.00101 | -0.61433 | Male | | | | | | |
| LINC01108 | | | | | | | | | | 0.00848 | 1.37517 | Female | | | |
| PPP1R26 | | | | | | | | | | 0.00022 | 2.13703 | Female | | | |
| LBR | | | | | | | 0.00000 | -0.58143 | Male | | | | | | |
| G6PD | | | | 0.01449 | 0.67563 | Female | | | | | | | | | |
| CNOT3 | | | | 0.02983 | 0.75037 | Female | | | | | | | | | |
| IZUMO4 | | | | | | | 0.04647 | -1.99548 | Male | | | | | | |
| FAM200A | | | | | | | 0.00953 | -0.98359 | Male | | | | | | |

| | | | | | | | | | | | | | | | |
|---|---|---|---|---|---|---|---|---|---|---|---|---|---|---|---|
| ZNF619 | | | | | | | | | | | | | 0.00203 | 1.94628 | Female |
| EIF1AY | 0.00000 | -6.27476 | Male | 0.00000 | -8.53148 | Male | 0.00000 | -9.75581 | Male | 0.00000 | -9.81738 | Male | 0.00000 | -9.01656 | Male |
| APEH | 0.04026 | 0.86475 | Female | | | | | | | | | | | | |
| HIST1H3H | | | | 0.00792 | -2.05326 | Male | | | | | | | | | |
| MIR1973 | | | | | | | | | | 0.00030 | 0.96313 | Female | | | |
| CT55 | | | | | | | 0.00024 | 2.01156 | Female | | | | | | |
| TMEM37 | | | | | | | 0.01165 | 0.81475 | Female | 0.00080 | 0.99833 | Female | | | |
| ZNF132 | | | | | | | | | | 0.02116 | -1.55161 | Male | | | |
| C16orf59 | | | | 0.00889 | -1.25555 | Male | 0.00278 | -0.95970 | Male | | | | | | |
| HIST1H4I | | | | | | | 0.00002 | -1.89327 | Male | 0.03110 | -0.98829 | Male | 0.03553 | -0.95233 | Male |
| NARS | 0.00003 | -0.76750 | Male | | | | | | | | | | | | |
| DMXL2 | | | | | | | 0.00246 | -1.19499 | Male | | | | | | |
| HIBADH | | | | | | | 0.00490 | -0.74529 | Male | | | | | | |
| PRAMEF19.uid-875 | | | | | | | 0.01808 | 0.85461 | Female | | | | | | |
| SNAR-E | | | | | | | 0.00000 | 1.06178 | Female | 0.00000 | 0.98568 | Female | | | |
| ACADS | | | | | | | | | | 0.00000 | 1.54492 | Female | 0.01649 | -0.83694 | Male |
| ARHGAP42 | | | | | | | | | | 0.01311 | 0.77220 | Female | | | |
| RFPL4B | | | | | | | | | | 0.02777 | -0.70651 | Male | | | |
| ADCK4 | | | | | | | 0.00975 | -0.95546 | Male | | | | | | |
| LINC00152 | | | | 0.00004 | 1.69556 | Female | 0.00001 | 1.00544 | Female | | | | 0.00948 | 0.59224 | Female |
| SELO | 0.02275 | -1.91694 | Male | 0.00637 | 1.39370 | Female | | | | | | | | | |
| SAA1 | | | | | | | | | | 0.03508 | -1.33796 | Male | | | |
| CNKSR1 | | | | | | | | | | 0.01693 | -0.86887 | Male | | | |
| PDE6C | | | | 0.00009 | 3.94779 | Female | | | | | | | | | |
| HPD | | | | | | | | | | | | | 0.04235 | 1.40027 | Female |
| SNORD50A | | | | | | | | | | 0.00000 | -0.93233 | Male | | | |
| HSPA12B | | | | | | | | | | | | | 0.02415 | 1.94244 | Female |
| UGGT2 | | | | | | | | | | | | | 0.00068 | -0.81210 | Male |
| PRMT5-AS1 | | | | | | | 0.00046 | 0.81138 | Female | | | | 0.00000 | 1.00838 | Female |
| TOPORS-AS1 | | | | | | | 0.00416 | 0.71600 | Female | | | | | | |
| LOC101928012 | | | | | | | | | | 0.01096 | 1.16896 | Female | | | |
| PAX8-AS1.uid-3716 | | | | | | | 0.01296 | 0.84730 | Female | | | | | | |
| SDHAP1 | 0.00322 | 1.04590 | Female | | | | 0.00000 | 0.77674 | Female | | | | | | |
| PYGB | | | | 0.00651 | 0.78147 | Female | | | | 0.00000 | 0.94626 | Female | | | |
| TSR2 | | | | 0.00002 | 0.85910 | Female | 0.00000 | 1.27730 | Female | 0.00000 | 0.80409 | Female | | | |
| EMB | | | | | | | | | | | | | 0.00422 | -2.60952 | Male |
| EFHD2 | | | | 0.00000 | 1.41174 | Female | | | | | | | | | |
| RPL10 | 0.00065 | 0.62077 | Female | 0.00000 | 0.65224 | Female | 0.00000 | 0.64325 | Female | 0.00000 | 0.77890 | Female | 0.00000 | 0.76620 | Female |
| NIPA2 | | | | | | | 0.00000 | -0.68519 | Male | | | | 0.00000 | -0.74635 | Male |
| PDCL3 | | | | | | | 0.00000 | 0.59220 | Female | | | | | | |
| WASH3P | | | | | | | 0.00073 | 1.06168 | Female | | | | | | |
| APOBEC3C | | | | | | | | | | 0.00084 | 2.48201 | Female | | | |
| PRSS22 | | | | | | | | | | | | | 0.00709 | -1.30769 | Male |
| XRRA1 | | | | | | | | | | | | | 0.00898 | 0.79066 | Female |
| SRXN1 | | | | 0.04772 | 0.72345 | Female | | | | | | | | | |
| SLC40A1 | 0.00311 | -2.43806 | Male | | | | | | | | | | | | |
| C4orf27 | 0.03287 | -0.80620 | Male | | | | | | | | | | 0.00000 | -0.65566 | Male |
| GPRC5A | | | | 0.00000 | 2.07147 | Female | | | | 0.01654 | 0.73625 | Female | | | |
| SLC15A2 | | | | | | | 0.01315 | -1.12652 | Male | | | | | | |
| PSORS1C2 | | | | | | | | | | 0.00000 | 2.67471 | Female | | | |
| ARID3B | | | | 0.00215 | -0.71241 | Male | | | | | | | | | |

| | | | | | | | | | | | | | | | |
|---|---|---|---|---|---|---|---|---|---|---|---|---|---|---|---|
| LOC100506655 | 0.00099 | 3.00384 | Female | | | | | | | | | | | | |
| TMEM62 | | | | | | | 0.00504 | -0.69504 | Male | | | | | | |
| SLC33A1 | | | | | | | | | | 0.00433 | 0.59949 | Female | | | |
| SEPT9 | | | | 0.00411 | 1.10444 | Female | 0.00125 | 0.82795 | Female | | | | | | |
| THOP1 | 0.00529 | -1.39245 | Male | | | | | | | | | | | | |
| LEFTY1 | 0.00000 | 0.89122 | Female | | | | | | | 0.00000 | 3.13977 | Female | | | |
| YJEFN3 | | | | | | | 0.00000 | 1.50948 | Female | 0.02675 | -0.64289 | Male | 0.00678 | 0.74295 | Female |
| CCDC178 | | | | | | | | | | | | | 0.00133 | 1.87294 | Female |
| BCL2L10 | | | | | | | | | | | | | 0.01104 | -2.59770 | Male |
| ELANE | | | | | | | 0.00046 | -2.67627 | Male | | | | | | |
| ABHD12B | | | | | | | | | | 0.00141 | 0.94391 | Female | | | |
| PLXNB3 | | | | | | | 0.00362 | 2.19508 | Female | 0.00005 | 2.77206 | Female | | | |
| IL11RA | 0.00574 | 2.89394 | Female | 0.02825 | -1.49666 | Male | | | | | | | | | |
| TENM3 | | | | 0.00888 | -1.26460 | Male | | | | 0.00785 | -0.79746 | Male | | | |
| KCNH2 | | | | | | | | | | 0.00723 | 2.16217 | Female | | | |
| APELA | | | | | | | | | | 0.02052 | 0.92683 | Female | | | |
| BRF1 | | | | | | | | | | | | | 0.03254 | 0.67424 | Female |
| EIF4EBP1 | | | | 0.00000 | 1.11508 | Female | | | | | | | | | |
| NR5A2 | | | | | | | | | | 0.00103 | 1.05344 | Female | | | |
| FAM45B.uid-12498 | | | | | | | | | | 0.00000 | 0.79355 | Female | | | |
| PKN3 | | | | 0.00069 | -1.79003 | Male | | | | | | | | | |
| IKBKG | | | | | | | 0.00449 | 0.74122 | Female | 0.02803 | 0.58245 | Female | 0.00042 | 0.83330 | Female |
| VBP1 | | | | 0.00021 | 0.59692 | Female | | | | 0.00000 | 0.59506 | Female | | | |
| FBP1 | | | | | | | | | | 0.00405 | 1.83261 | Female | | | |
| ARL17A.uid-21953 | | | | | | | 0.03584 | 1.67450 | Female | | | | | | |
| SEMA7A | | | | | | | 0.00012 | -1.72160 | Male | | | | | | |
| ATP9B | | | | | | | | | | 0.01542 | 0.61365 | Female | | | |
| HIPK2 | | | | | | | | | | 0.00020 | 1.70920 | Female | 0.01606 | -1.19933 | Male |
| HIST1H2BJ | | | | | | | 0.00573 | 1.29555 | Female | | | | | | |
| DHFR | | | | | | | | | | 0.00000 | -0.72905 | Male | | | |
| CAPN12 | | | | | | | | | | 0.00000 | 2.04383 | Female | 0.01696 | 1.10403 | Female |
| SH3BP5L | | | | | | | | | | 0.01429 | -0.79761 | Male | | | |
| ISL2 | | | | | | | | | | | | | 0.01108 | 2.01724 | Female |
| SNRPD1 | 0.00000 | -0.69480 | Male | | | | | | | | | | | | |
| ODC1 | 0.03307 | -0.69619 | Male | | | | | | | | | | | | |
| TXLNG | 0.04258 | 0.71360 | Female | 0.00000 | 1.15153 | Female | 0.00000 | 0.59777 | Female | 0.00000 | 0.65231 | Female | | | |
| FAM178B | | | | | | | 0.02356 | 1.02193 | Female | | | | | | |
| SLC23A1 | | | | | | | | | | | | | 0.01572 | 1.49103 | Female |
| SPSB1 | | | | | | | | | | | | | 0.01794 | 0.81656 | Female |
| SF3A2 | 0.02218 | -1.07467 | Male | | | | | | | | | | | | |
| IBTK | | | | | | | | | | | | | 0.00175 | -0.61303 | Male |
| TMPRSS4 | | | | | | | | | | | | | 0.01809 | -1.37136 | Male |
| CHRAC1 | | | | | | | 0.00000 | -0.76337 | Male | 0.00004 | -0.64850 | Male | | | |
| KRT18P55 | 0.00000 | -3.75021 | Male | | | | 0.00481 | 0.62835 | Female | | | | | | |
| PSMG3-AS1 | | | | 0.00584 | -1.68575 | Male | | | | | | | | | |
| CCKBR | 0.00000 | 2.89552 | Female | | | | 0.00000 | -1.22161 | Male | 0.00000 | -1.48136 | Male | | | |
| ACADSB | | | | 0.00045 | 1.25897 | Female | | | | | | | | | |
| KIAA0101 | | | | | | | 0.01108 | -0.63900 | Male | | | | 0.00543 | -0.63554 | Male |
| GTF2H3 | | | | | | | 0.00000 | -0.94624 | Male | | | | | | |
| ECHDC3 | | | | 0.00000 | 3.53977 | Female | 0.03866 | -1.26460 | Male | | | | | | |
| DCK | | | | | | | 0.00623 | -0.69189 | Male | | | | | | |

| | | | | | | | | | | | | | | | |
|---|---|---|---|---|---|---|---|---|---|---|---|---|---|---|---|
| COL18A1 | | | | | | | | | | 0.02488 | 1.67951 | Female | | | |
| CXorf57 | | | | | | | | | | | | | 0.03065 | -1.55725 | Male |
| GATAD2B | | | | | | | | | | | | | 0.00542 | 0.63330 | Female |
| KCNC4 | | | | | | | | | | 0.01097 | 1.21982 | Female | | | |
| DEPDC1B | | | | | | | 0.00066 | -0.74670 | Male | | | | | | |
| SHANK2 | | | | | | | 0.00579 | 1.38552 | Female | 0.00003 | 1.90425 | Female | | | |
| CDH8 | | | | 0.02716 | 2.01304 | Female | | | | 0.02092 | 1.39149 | Female | | | |
| BCL9 | | | | | | | 0.01834 | -0.68119 | Male | | | | | | |
| LIM2 | | | | | | | 0.03880 | 1.70603 | Female | 0.00019 | 2.65064 | Female | | | |
| RABEP2 | | | | | | | | | | 0.02893 | -0.76928 | Male | | | |
| RPSAP9 | | | | | | | | | | 0.02525 | -0.85641 | Male | | | |
| FCF1 | 0.00355 | -0.75506 | Male | | | | | | | | | | | | |
| DDHD1 | | | | 0.01229 | 0.67123 | Female | | | | | | | | | |
| GRAMD1C | | | | | | | | | | 0.04715 | 1.35921 | Female | | | |
| SYCP3 | | | | | | | | | | | | | 0.00000 | 0.62090 | Female |
| NIPSNAP1 | | | | | | | | | | | | | 0.04235 | 0.60545 | Female |
| HIST1H3B | | | | 0.00067 | -2.34961 | Male | | | | 0.01512 | 1.20872 | Female | | | |
| ABCA17P | | | | | | | 0.04682 | 0.99826 | Female | | | | | | |
| CDC42BPG | 0.02404 | 1.95255 | Female | | | | | | | | | | | | |
| DLX5 | | | | | | | 0.03060 | -0.90479 | Male | | | | | | |
| CAB39L | | | | | | | 0.02155 | -0.64707 | Male | 0.00077 | -0.84258 | Male | 0.00001 | -1.05109 | Male |
| IL6R | | | | | | | | | | 0.00000 | 1.92466 | Female | | | |
| LIMCH1 | | | | | | | | | | 0.00000 | 3.24333 | Female | | | |
| ZNF610 | | | | | | | | | | | | | 0.00316 | 1.46097 | Female |
| ENDOG | | | | 0.00006 | 1.60275 | Female | | | | | | | | | |
| SMARCD1 | | | | | | | 0.00163 | -0.58541 | Male | | | | | | |
| SKIDA1 | | | | 0.02543 | 0.97269 | Female | 0.02718 | 0.63761 | Female | | | | | | |
| KLF6 | | | | 0.03652 | 0.81942 | Female | 0.00016 | 0.87454 | Female | | | | 0.00000 | 1.03628 | Female |
| NRBP2 | | | | | | | | | | 0.00000 | 1.91582 | Female | 0.00982 | 1.02678 | Female |
| C15orf40 | 0.02835 | 1.13961 | Female | | | | | | | | | | | | |
| LAT | | | | | | | 0.04376 | 1.10715 | Female | | | | | | |
| SERPINB6 | | | | | | | | | | 0.00000 | 1.50878 | Female | | | |
| EMC1 | 0.00733 | 1.21204 | Female | | | | | | | | | | | | |
| URAD | | | | | | | 0.00072 | -1.81978 | Male | | | | | | |
| SNORA4 | | | | | | | | | | 0.00348 | 0.80874 | Female | | | |
| PTPN5 | | | | | | | 0.00015 | -2.24273 | Male | 0.00191 | 1.81736 | Female | | | |
| CELA3A | | | | | | | | | | 0.02739 | 1.10636 | Female | | | |
| JAKMIP1 | | | | | | | | | | 0.00477 | -0.86528 | Male | | | |
| CECR1 | | | | | | | | | | 0.00000 | 3.08528 | Female | | | |
| NT5C3A | | | | 0.00759 | 0.66112 | Female | | | | | | | | | |
| GCLM | | | | | | | | | | 0.00026 | -0.69593 | Male | | | |
| DOK2 | | | | | | | 0.03861 | 1.68698 | Female | | | | | | |
| DUSP10 | | | | | | | | | | 0.00817 | 1.91575 | Female | | | |
| GMFB | | | | | | | | | | 0.00000 | 1.23403 | Female | | | |
| SYNE4 | | | | | | | | | | 0.01461 | -1.57618 | Male | | | |
| ITLN2 | | | | | | | | | | 0.00498 | 1.23723 | Female | 0.00026 | 1.51276 | Female |
| CTHRC1 | | | | | | | 0.00891 | -1.74575 | Male | | | | | | |
| ASS1 | 0.00171 | 2.17773 | Female | | | | | | | | | | | | |
| ATP10A | | | | | | | | | | 0.00779 | 2.25438 | Female | | | |
| MMP24 | | | | | | | 0.04062 | 1.05195 | Female | 0.00500 | 1.33370 | Female | | | |
| ABRACL | 0.04324 | -1.01423 | Male | 0.00114 | 0.82661 | Female | 0.00061 | -0.59585 | Male | | | | 0.00007 | -0.63586 | Male |

| | | | | | | | | | | | | | | | |
|---|---|---|---|---|---|---|---|---|---|---|---|---|---|---|---|
| YEATS4 | | | | | | | 0.00000 | -0.60338 | Male | | | | | | |
| HSPA2 | | | | | | | | | | 0.00031 | -0.99790 | Male | | | |
| ANKRD46 | | | | | | | 0.01840 | -0.93599 | Male | | | | | | |
| TMEM219 | | | | | | | 0.00036 | -0.69708 | Male | | | | | | |
| TMEM261 | 0.00261 | 0.85839 | Female | 0.00000 | 0.86175 | Female | | | | | | | | | |
| CCR7 | | | | 0.00000 | -1.86286 | Male | | | | | | | | | |
| LRCH4 | | | | | | | | | | 0.01326 | 0.73492 | Female | | | |
| HDAC7 | | | | | | | | | | 0.00124 | 1.58170 | Female | | | |
| PEX12 | | | | | | | 0.01697 | -0.93692 | Male | | | | | | |
| PRELID2 | | | | | | | 0.02102 | -1.40178 | Male | | | | | | |
| ZFR2 | | | | | | | | | | 0.00373 | 1.74350 | Female | | | |
| UBXN6 | | | | | | | 0.00579 | 0.67757 | Female | | | | | | |
| SLC25A5 | 0.00000 | 1.39078 | Female | 0.00000 | 1.10371 | Female | 0.00000 | 1.15505 | Female | 0.00000 | 1.01317 | Female | 0.00000 | 0.89787 | Female |
| MID1 | | | | 0.04950 | 0.85729 | Female | | | | 0.00243 | -0.77985 | Male | | | |
| HIST1H3D | | | | 0.03369 | -1.55680 | Male | 0.01283 | 1.21386 | Female | | | | | | |
| 1.00000 | | | | 0.00000 | 1.15878 | Female | 0.00000 | 0.92934 | Female | | | | | | |
| ARL5A | | | | | | | | | | 0.00000 | -0.62838 | Male | | | |
| SLC27A2 | | | | | | | 0.00000 | -1.94373 | Male | 0.00000 | -1.48211 | Male | 0.00000 | -1.54956 | Male |
| SPDYC | | | | | | | | | | 0.04516 | 0.91502 | Female | | | |
| BLACE | | | | | | | | | | | | | 0.02135 | -1.20913 | Male |
| CCNT2-AS1 | | | | | | | | | | 0.00196 | -1.55424 | Male | | | |
| PER3 | | | | | | | | | | 0.00145 | -1.07439 | Male | | | |
| LOC100129534 | | | | 0.00447 | 0.98662 | Female | 0.00100 | 0.78634 | Female | | | | | | |
| KIAA0141 | | | | 0.03385 | -0.97026 | Male | 0.00120 | -0.91937 | Male | | | | | | |
| C5orf28 | | | | | | | 0.03056 | -0.70784 | Male | | | | | | |
| TSPYL6 | | | | | | | | | | 0.00936 | -1.25071 | Male | | | |
| TSPAN15 | | | | 0.00211 | -1.31685 | Male | | | | | | | | | |
| MAP2K4P1 | | | | | | | 0.01339 | -0.87453 | Male | | | | | | |
| MVP | | | | | | | 0.01755 | 0.82241 | Female | | | | | | |
| TMC6 | | | | | | | 0.02473 | -1.38626 | Male | | | | | | |
| TUBGCP5 | | | | | | | | | | | | | 0.00400 | -0.81831 | Male |
| FANCB | | | | 0.02277 | 0.76049 | Female | 0.00000 | 0.95097 | Female | | | | | | |
| S100A10 | 0.00000 | -1.90955 | Male | | | | | | | | | | | | |
| HIST1H2BH | 0.03092 | 1.99379 | Female | | | | 0.01180 | 1.03245 | Female | 0.03150 | 0.85952 | Female | | | |
| ZNF107 | | | | | | | 0.00281 | -0.68489 | Male | | | | | | |
| ANKRD20A9P | | | | | | | | | | 0.00030 | -1.31292 | Male | | | |
| ZNF667-AS1 | | | | | | | | | | | | | 0.00042 | 0.90488 | Female |
| METTL9 | | | | 0.02100 | 0.58749 | Female | | | | | | | 0.00007 | -0.58333 | Male |
| RHOBTB1 | | | | | | | | | | 0.00056 | -1.19137 | Male | | | |
| ABHD4 | 0.00002 | 3.21021 | Female | 0.00000 | 1.53935 | Female | 0.00077 | -1.14713 | Male | 0.01570 | -0.81768 | Male | | | |
| NLRP5 | | | | | | | | | | | | | 0.00000 | -4.02136 | Male |
| LDLR | | | | 0.00000 | 1.75844 | Female | 0.00692 | -0.71453 | Male | | | | 0.02460 | 0.58261 | Female |
| CBR4 | | | | | | | | | | | | | 0.00000 | -0.78844 | Male |
| RBBP7 | | | | 0.00009 | 0.69749 | Female | | | | | | | 0.00000 | -0.75616 | Male |
| NODAL | | | | | | | | | | 0.01230 | 1.60894 | Female | | | |
| MKRN9P | | | | | | | | | | | | | 0.01080 | -0.58921 | Male |
| GTF2H2 | | | | | | | 0.00185 | -0.61947 | Male | | | | | | |
| MMP24-AS1 | | | | | | | | | | 0.00091 | 1.26312 | Female | | | |
| CAMK2D | 0.00108 | 2.88268 | Female | | | | 0.00724 | -0.96645 | Male | | | | | | |
| EXT1 | | | | | | | | | | 0.03414 | 0.65586 | Female | | | |
| C1orf168 | | | | 0.03528 | 1.98662 | Female | | | | | | | | | |

| | | | | | | | | | | | | | | | |
|---|---|---|---|---|---|---|---|---|---|---|---|---|---|---|---|
| KLF4 | | | | | | | 0.01729 | 0.63049 | Female | 0.00000 | 1.79376 | Female | | | |
| SPATA22 | | | | | | | 0.02164 | 0.79135 | Female | | | | | | |
| C1QTNF9 | | | | | | | | | | 0.01366 | 2.19304 | Female | 0.00022 | -3.03984 | Male |
| SFXN1 | 0.00074 | -1.08873 | Male | | | | | | | | | | | | |
| HBM | | | | 0.02674 | -2.55662 | Male | | | | | | | | | |
| MYLIP | | | | | | | | | | 0.00700 | 0.74423 | Female | 0.01814 | 0.65838 | Female |
| DGAT2 | | | | | | | | | | 0.00993 | 1.91466 | Female | 0.03823 | 1.55834 | Female |
| RMI1 | | | | | | | 0.00006 | -0.79883 | Male | | | | | | |
| AK6 | 0.01593 | -0.72122 | Male | | | | | | | | | | | | |
| ZNF880 | | | | | | | | | | 0.00404 | -0.73526 | Male | | | |
| EPS15L1 | | | | 0.00209 | 0.78056 | Female | | | | | | | | | |
| DVL1 | | | | 0.03693 | 0.89435 | Female | | | | | | | | | |
| AMHR2 | | | | | | | | | | | | | 0.04522 | 1.39264 | Female |
| ZNF771 | | | | | | | | | | 0.01182 | -0.93288 | Male | | | |
| KIAA1161 | | | | 0.00000 | 2.57557 | Female | | | | | | | | | |
| TPD52 | | | | | | | 0.00001 | -0.93293 | Male | | | | | | |
| FZD6 | | | | | | | 0.02641 | 0.98601 | Female | | | | | | |
| GNG12 | | | | | | | | | | 0.01058 | -0.84861 | Male | | | |
| RAP2A | | | | | | | | | | | | | 0.00270 | -0.65296 | Male |
| ATP6AP2 | | | | 0.00000 | 0.91135 | Female | 0.00000 | 0.64940 | Female | 0.00000 | 0.62400 | Female | | | |
| TUBA4A | | | | | | | 0.00130 | 1.57636 | Female | 0.00000 | -3.27196 | Male | 0.00082 | -1.51885 | Male |
| EMP3 | | | | | | | | | | 0.00225 | 1.79368 | Female | | | |
| CACNA1A | | | | | | | | | | 0.00001 | -1.27479 | Male | 0.00166 | -0.92651 | Male |
| RBM46 | | | | | | | | | | 0.00000 | -1.96041 | Male | 0.00574 | -1.19977 | Male |
| APOC1 | 0.02275 | -0.63959 | Male | | | | | | | 0.00000 | 0.96251 | Female | | | |
| HSD3B1 | | | | | | | 0.03213 | -1.13326 | Male | | | | | | |
| TACC3 | 0.04397 | -0.64103 | Male | | | | | | | | | | | | |
| TXN | 0.00001 | -0.81623 | Male | | | | | | | | | | | | |
| MAFG-AS1 | | | | | | | 0.02025 | 1.62601 | Female | | | | | | |
| B3GAT3 | | | | | | | | | | 0.00975 | -0.71753 | Male | | | |
| C11orf96 | | | | | | | 0.00232 | 2.57012 | Female | 0.01610 | 2.08300 | Female | | | |
| TACSTD2 | | | | 0.03563 | -0.70162 | Male | 0.01450 | -0.72861 | Male | 0.00373 | -0.79076 | Male | | | |
| NOG | | | | | | | 0.00155 | 1.01953 | Female | | | | 0.00000 | -2.25544 | Male |
| RALGPS1 | | | | | | | 0.00714 | -1.15670 | Male | | | | | | |
| C6orf223 | | | | | | | 0.00000 | -1.87979 | Male | | | | | | |
| GUSBP1 | | | | | | | 0.00264 | 0.61291 | Female | | | | | | |
| DLEU1 | | | | | | | 0.00011 | -0.64875 | Male | | | | | | |
| FKBP8 | | | | | | | | | | 0.00532 | 0.62519 | Female | | | |
| RPS6KA3 | | | | | | | 0.03550 | 0.63478 | Female | | | | | | |
| MRPL38 | 0.00074 | -0.77699 | Male | | | | | | | | | | | | |
| TMX2 | 0.00068 | -0.67350 | Male | | | | | | | | | | | | |
| KBTBD3 | | | | | | | 0.02704 | -1.35480 | Male | | | | | | |
| FAR2P1 | | | | | | | | | | | | | 0.03321 | 1.35181 | Female |
| GSTM1 | | | | | | | | | | | | | 0.00347 | 1.68110 | Female |
| TRIM8 | | | | 0.01250 | 1.22178 | Female | 0.01501 | 0.80442 | Female | 0.00000 | 1.33519 | Female | | | |
| FER | | | | | | | 0.02433 | -0.60361 | Male | | | | | | |
| WWC2-AS2 | 0.03033 | 3.21808 | Female | | | | | | | | | | | | |
| ARHGAP30 | | | | | | | 0.00595 | 1.98426 | Female | | | | | | |
| PRAMEF12 | | | | | | | | | | 0.00885 | 2.14979 | Female | | | |
| TDRD6 | | | | | | | | | | | | | 0.01362 | -1.29652 | Male |
| LOC100133077 | | | | | | | 0.03032 | 2.04027 | Female | | | | | | |

| | | | | | | | | | | | | | | | |
|---|---|---|---|---|---|---|---|---|---|---|---|---|---|---|---|
| TOX2 | | | | | | | | | | | | | 0.00067 | 1.84616 | Female |
| CEPT1 | 0.00996 | -1.00874 | Male | | | | | | | | | | | | |
| IL13RA2 | | | | | | | | | | 0.01664 | 2.17348 | Female | | | |
| ATF6B | | | | | | | | | | 0.02453 | -0.68476 | Male | 0.03262 | 0.65017 | Female |
| HEATR6 | 0.00052 | 2.43862 | Female | | | | | | | | | | | | |
| COQ2 | | | | | | | 0.00000 | -1.24617 | Male | | | | 0.00124 | -0.78823 | Male |
| FAT2 | | | | | | | 0.00043 | 2.16047 | Female | | | | | | |
| KCNN1 | | | | | | | | | | 0.00000 | 2.86682 | Female | 0.00822 | 1.58898 | Female |
| PIF1 | | | | | | | | | | 0.02116 | 0.65527 | Female | | | |
| TJP2 | | | | 0.00000 | 1.99200 | Female | | | | | | | | | |
| LINC01118 | | | | | | | | | | | | | 0.00007 | -5.25767 | Male |
| GPR153 | | | | | | | | | | 0.00001 | 2.61639 | Female | 0.01426 | 1.54923 | Female |
| RAB31 | | | | 0.00001 | 1.78339 | Female | 0.00000 | -1.30753 | Male | 0.00007 | -1.02160 | Male | | | |
| SERPINB1 | | | | 0.00000 | 2.84665 | Female | | | | | | | | | |
| STK36 | | | | | | | | | | 0.00456 | 1.13506 | Female | | | |
| TMEM27 | | | | 0.02395 | 2.30323 | Female | | | | | | | | | |
| ZNF140 | | | | | | | | | | 0.04812 | 0.62605 | Female | | | |
| GOLGA3 | | | | | | | 0.00000 | 0.76638 | Female | | | | | | |
| ARAF | | | | | | | 0.00000 | 0.89193 | Female | 0.00095 | 0.59956 | Female | 0.00004 | 0.70758 | Female |
| ZNF276 | | | | | | | 0.04267 | 0.66021 | Female | | | | | | |
| FAM122B | 0.03152 | 1.44222 | Female | 0.00059 | 1.29161 | Female | 0.00001 | 1.08797 | Female | 0.00317 | 0.74795 | Female | | | |
| ZFAND2A | | | | | | | 0.00000 | 0.81123 | Female | 0.00000 | -1.04958 | Male | | | |
| UBALD2 | | | | | | | | | | | | | 0.00674 | 0.67988 | Female |
| NUP62 | | | | | | | 0.00000 | 0.61150 | Female | | | | | | |
| DCAF8L2 | | | | | | | | | | | | | 0.04739 | 0.97369 | Female |
| MAPKBP1 | | | | | | | 0.00074 | -1.12777 | Male | | | | | | |
| IDH3G | | | | 0.00005 | 0.87320 | Female | 0.00000 | 1.21235 | Female | 0.00000 | 0.83905 | Female | 0.00000 | 0.70751 | Female |
| EFNB3 | | | | | | | | | | 0.00052 | 1.65512 | Female | | | |
| ANXA2P3 | | | | 0.00000 | -0.82025 | Male | | | | | | | | | |
| MSMO1 | | | | 0.00043 | -0.85212 | Male | 0.00044 | -0.58883 | Male | | | | | | |
| BBC3 | | | | | | | 0.00098 | 1.32567 | Female | | | | | | |
| TTLL1 | | | | | | | 0.03125 | -1.69939 | Male | | | | | | |
| ZMYND11 | | | | | | | | | | | | | 0.00264 | -0.73160 | Male |
| RIBC1 | | | | | | | | | | | | | 0.00912 | 1.98299 | Female |
| ZNF789 | | | | | | | 0.00002 | -0.94133 | Male | | | | | | |
| VPS9D1-AS1 | | | | | | | | | | 0.00292 | 0.85897 | Female | | | |
| TMEM52B | 0.00000 | 3.53591 | Female | | | | | | | | | | | | |
| DPM3 | | | | | | | 0.00000 | 0.90943 | Female | | | | 0.00025 | 0.68573 | Female |
| TMPRSS13 | | | | | | | 0.03249 | -0.74655 | Male | 0.01304 | -0.78347 | Male | | | |
| HAS3 | | | | | | | | | | 0.00009 | -1.94036 | Male | | | |
| RIMKLB | | | | | | | | | | 0.00000 | 1.22855 | Female | 0.00004 | 0.75437 | Female |
| CS | 0.00158 | -0.62182 | Male | | | | | | | | | | | | |
| LOC100862671 | | | | | | | 0.03384 | 0.70709 | Female | | | | | | |
| ITPK1 | | | | 0.04090 | 0.65219 | Female | | | | | | | | | |
| CHMP7 | | | | | | | | | | 0.00001 | -0.80533 | Male | | | |
| ELOVL5 | | | | | | | 0.00000 | -0.88007 | Male | | | | | | |
| ZC3H12B | 0.00289 | 2.53801 | Female | | | | 0.00101 | 1.19296 | Female | | | | 0.00983 | 0.92311 | Female |
| PRSS12 | | | | 0.04165 | 1.36271 | Female | | | | | | | | | |
| STC2 | | | | 0.00000 | -3.47917 | Male | | | | 0.00355 | 0.76525 | Female | | | |
| ITGA2 | | | | | | | | | | 0.00867 | 1.64557 | Female | | | |
| HOXB9 | | | | | | | | | | 0.00288 | 2.56012 | Female | | | |

| | | | | | | | | | | | | | | | |
|---|---|---|---|---|---|---|---|---|---|---|---|---|---|---|---|
| SETD9 | 0.00411 | 2.42976 | Female | | | | 0.00384 | -0.97633 | Male | | | | | | |
| SHISA5 | | | | 0.00021 | 0.71502 | Female | | | | | | | | | |
| ANXA4 | | | | | | | | | | 0.00058 | 1.73998 | Female | | | |
| HIST2H2AC | | | | 0.04067 | -1.16483 | Male | | | | 0.00110 | 1.11638 | Female | | | |
| MAGED1 | 0.00032 | 2.14771 | Female | | | | | | | 0.00056 | -0.86042 | Male | 0.00005 | -0.96153 | Male |
| MPZL1 | | | | 0.00000 | 0.81241 | Female | | | | | | | | | |
| POLA1 | | | | 0.00066 | 0.99326 | Female | 0.00002 | 0.82817 | Female | | | | | | |
| UBE2A | | | | 0.00000 | 1.40334 | Female | 0.00000 | 0.89346 | Female | 0.00000 | 0.74638 | Female | 0.00000 | 0.85734 | Female |
| P4HA1 | | | | | | | 0.00000 | -1.44474 | Male | | | | 0.04573 | -0.58930 | Male |
| PLBD2 | | | | | | | 0.01470 | -0.82658 | Male | | | | | | |
| ZNF675 | | | | | | | 0.00246 | -0.69228 | Male | | | | | | |
| SLPI | | | | | | | | | | | | | 0.02518 | 1.15608 | Female |
| THNSL1 | | | | | | | 0.01678 | -1.60349 | Male | | | | | | |
| MSMP | | | | | | | 0.00279 | -1.62213 | Male | | | | | | |
| MYADML | | | | | | | | | | 0.00035 | -1.51132 | Male | | | |
| SLITRK2 | | | | 0.00201 | 1.88331 | Female | | | | | | | | | |
| COL7A1 | | | | | | | 0.00048 | 1.37071 | Female | | | | | | |
| USP9X | | | | 0.00000 | 1.21680 | Female | 0.00000 | 0.90991 | Female | 0.00000 | 0.81426 | Female | | | |
| SMOX | | | | 0.00263 | 0.68067 | Female | | | | | | | | | |
| RPS15AP10 | | | | | | | | | | 0.01704 | -1.16359 | Male | | | |
| MUC3A | | | | | | | | | | | | | 0.01928 | 1.00769 | Female |
| LINC00997 | | | | | | | | | | 0.00536 | -1.25886 | Male | | | |
| SPTBN1 | | | | 0.00228 | 0.63294 | Female | | | | | | | | | |
| SMIM14 | | | | | | | 0.01130 | -1.25079 | Male | | | | | | |
| GPR125 | | | | 0.00006 | 2.00039 | Female | | | | | | | | | |
| CAMKV | 0.00000 | -0.60420 | Male | | | | | | | | | | | | |
| CXorf40B | 0.02057 | 0.92177 | Female | | | | 0.00000 | 1.03277 | Female | 0.00000 | 0.75536 | Female | | | |
| SSX5 | | | | | | | 0.03649 | 1.91254 | Female | | | | | | |
| MRGPRX1 | | | | | | | 0.00652 | -1.12480 | Male | | | | | | |
| SNORA48 | | | | 0.02847 | -1.14104 | Male | 0.03142 | 0.77303 | Female | | | | | | |
| MYC | | | | 0.00244 | -0.78038 | Male | | | | | | | | | |
| RHBDL2 | | | | | | | 0.00966 | 2.19219 | Female | | | | | | |
| TRMT2A | | | | | | | 0.00050 | -0.65699 | Male | | | | | | |
| LOC100287225 | | | | | | | | | | | | | 0.00018 | -1.78606 | Male |
| NDUFB4 | 0.00000 | 0.82165 | Female | | | | | | | | | | | | |
| PM20D2 | | | | | | | | | | 0.00017 | -1.27349 | Male | | | |
| GYLTL1B | 0.00000 | 1.17156 | Female | 0.00000 | 1.90924 | Female | | | | | | | | | |
| DTWD1 | | | | | | | 0.00356 | -0.62105 | Male | | | | 0.00012 | -0.72766 | Male |
| FAM171A1 | | | | | | | | | | 0.03158 | 1.13577 | Female | | | |
| TET2 | | | | | | | 0.00045 | 0.60429 | Female | | | | | | |
| MYL9 | | | | | | | | | | 0.00000 | -1.99597 | Male | | | |
| MCTS1 | 0.00231 | 0.71671 | Female | 0.00000 | 1.04909 | Female | | | | 0.00000 | 0.82320 | Female | | | |
| MUTYH | | | | | | | | | | | | | 0.00389 | 0.81603 | Female |
| SLCO2A1 | | | | | | | 0.00120 | -1.56223 | Male | | | | | | |
| BIK | | | | | | | | | | 0.03440 | -0.61107 | Male | | | |
| PIK3R3 | | | | | | | | | | 0.00223 | 0.79075 | Female | | | |
| SNORA16A | 0.00322 | 2.43059 | Female | | | | 0.04359 | -0.72853 | Male | | | | | | |
| ATP6V0D2 | | | | | | | 0.02413 | -0.89768 | Male | 0.01107 | 0.94242 | Female | | | |
| TSNARE1 | | | | | | | | | | | | | 0.01345 | 1.37899 | Female |
| GNL3L | 0.00667 | 0.89624 | Female | 0.00000 | 0.87342 | Female | 0.00000 | 1.01263 | Female | | | | 0.00000 | 0.69863 | Female |
| CNIH3 | | | | 0.00552 | 2.23959 | Female | | | | 0.00348 | 2.45466 | Female | | | |

| | | | | | | | | | | | | | | | |
|---|---|---|---|---|---|---|---|---|---|---|---|---|---|---|---|
| RUNX1T1 | | | | | | | | | | 0.00000 | 1.91501 | Female | | | |
| RGS16 | | | | | | | 0.00355 | 0.81159 | Female | | | | | | |
| DDX3Y | 0.00000 | -9.41916 | Male | 0.00000 | -7.70717 | Male | 0.00000 | -8.05749 | Male | 0.00000 | -9.67669 | Male | 0.00000 | -7.93532 | Male |
| WBSCR16 | | | | 0.00009 | 1.29805 | Female | | | | | | | | | |
| GRIP1 | | | | | | | 0.00001 | -2.15350 | Male | | | | | | |
| FAM151B | | | | | | | 0.00784 | -1.29876 | Male | | | | | | |
| POU5F1 | | | | | | | | | | 0.00000 | 1.73161 | Female | | | |
| RPL13P5 | | | | | | | | | | 0.00191 | 1.05233 | Female | | | |
| AQP3 | | | | | | | | | | | | | 0.00026 | 0.70527 | Female |
| LINC00458 | | | | | | | | | | | | | 0.00000 | -2.39711 | Male |
| IGBP1P1 | 0.04402 | 0.75835 | Female | 0.00093 | 0.66369 | Female | 0.00000 | 0.67523 | Female | | | | | | |
| STX2 | | | | | | | | | | 0.01291 | 1.02103 | Female | | | |
| GABRE | 0.04804 | 3.45485 | Female | | | | | | | | | | | | |
| EEF1E1.uid-8163 | | | | | | | | | | | | | 0.00000 | -0.59710 | Male |
| NRARP | | | | 0.00833 | 2.72243 | Female | | | | | | | | | |
| AGPAT2 | | | | 0.00000 | 1.71098 | Female | | | | 0.00000 | 1.30572 | Female | | | |
| CKMT1B | | | | | | | | | | 0.04816 | 0.99578 | Female | | | |
| PQLC2 | | | | | | | | | | 0.00422 | -0.69924 | Male | | | |
| SLC8A2 | | | | | | | 0.00852 | 1.99921 | Female | | | | | | |
| MTMR10 | | | | | | | | | | 0.00209 | -0.78072 | Male | | | |
| ANXA8.uid-14184 | | | | | | | | | | 0.00909 | -0.90628 | Male | 0.00000 | 1.58382 | Female |
| KIAA1143 | | | | | | | | | | 0.00000 | -0.66393 | Male | | | |
| GIGYF1 | | | | | | | | | | | | | 0.00026 | 0.60827 | Female |
| FAM58A | | | | | | | 0.00000 | 1.37234 | Female | 0.00213 | 0.71980 | Female | 0.00304 | 0.68663 | Female |
| RAB11FIP2 | | | | | | | 0.00602 | 0.58316 | Female | | | | | | |
| ST6GALNAC1 | 0.00110 | 2.59941 | Female | | | | | | | | | | | | |
| TTPAL | | | | | | | | | | 0.01334 | -0.63070 | Male | | | |
| MYO1D | 0.01207 | 2.75591 | Female | | | | | | | | | | | | |
| TRIB3 | | | | | | | 0.03811 | -0.73604 | Male | | | | | | |
| CBLB | | | | | | | | | | 0.00195 | -0.80440 | Male | | | |
| GCNT3 | | | | | | | | | | 0.00000 | 2.65862 | Female | | | |
| ZBTB16 | | | | | | | | | | 0.02601 | -0.92989 | Male | 0.01104 | 3.78587 | Female |
| POGLUT1 | | | | 0.00004 | 2.51098 | Female | | | | | | | | | |
| ENTPD1 | | | | | | | | | | 0.00133 | -0.58811 | Male | | | |
| GJC1 | | | | | | | 0.00042 | -0.80830 | Male | 0.00275 | -0.66442 | Male | | | |
| RFFL | | | | | | | 0.00025 | -1.70582 | Male | | | | | | |
| ASAP2 | | | | 0.02120 | 0.91998 | Female | | | | | | | | | |
| CLEC10A | | | | | | | 0.02105 | 2.96350 | Female | | | | | | |
| FCER2 | | | | | | | 0.02703 | 1.13505 | Female | | | | | | |
| LINC00669 | | | | | | | 0.02543 | -1.01936 | Male | | | | | | |
| ABCE1 | | | | | | | | | | | | | 0.00000 | -0.58356 | Male |
| NMU | | | | | | | | | | | | | 0.00003 | -1.38216 | Male |
| PDXP | | | | | | | | | | | | | 0.01283 | 0.68849 | Female |
| ALS2CR11 | | | | | | | | | | 0.04521 | 1.21952 | Female | | | |
| POLQ | | | | | | | | | | 0.00919 | 0.74110 | Female | | | |
| CASC5 | | | | | | | | | | 0.00000 | 5.89009 | Female | | | |
| APOA1 | 0.00000 | -1.07822 | Male | | | | 0.00000 | -4.35021 | Male | | | | | | |
| RAB25 | | | | 0.00045 | 1.13477 | Female | 0.00000 | -1.09731 | Male | | | | | | |
| PRSS23 | | | | | | | 0.00557 | -0.61204 | Male | 0.00000 | 1.02456 | Female | | | |
| TMEM132B | | | | | | | 0.03429 | 1.65476 | Female | | | | | | |
| CD59 | | | | | | | | | | | | | 0.00000 | 0.71580 | Female |

| | | | | | | | | | | | | | | | |
|---|---|---|---|---|---|---|---|---|---|---|---|---|---|---|---|
| EIF2S2 | 0.00004 | -0.69450 | Male | | | | | | | | | | | | |
| RND1 | | | | | | | | | | | | | 0.00109 | -2.01253 | Male |
| NBR2 | 0.02158 | 1.67283 | Female | | | | | | | | | | | | |
| KRT10 | | | | | | | 0.00000 | -0.80141 | Male | 0.00000 | -0.80406 | Male | | | |
| CD37 | | | | | | | | | | 0.04505 | 0.98294 | Female | | | |
| CCM2 | | | | | | | 0.03698 | 0.60113 | Female | | | | | | |
| OPN3 | | | | | | | | | | 0.04145 | -1.04140 | Male | 0.03320 | -1.01566 | Male |
| SPC25 | 0.00012 | 1.51398 | Female | | | | | | | | | | | | |
| CTNS | 0.01439 | 1.30318 | Female | | | | | | | | | | | | |
| CRIP2 | | | | | | | 0.02330 | -1.56949 | Male | | | | | | |
| DPH6 | | | | | | | 0.00532 | -0.72038 | Male | | | | 0.01392 | -0.61541 | Male |
| CCDC117 | 0.00930 | -0.87110 | Male | | | | 0.00000 | -0.59198 | Male | | | | | | |
| MAN1A1 | | | | | | | 0.01337 | -1.13159 | Male | | | | 0.00970 | -1.09737 | Male |
| C2orf74 | | | | | | | 0.03093 | -0.67466 | Male | | | | | | |
| YWHAE | 0.00000 | -0.73397 | Male | | | | | | | | | | | | |
| SNORD69 | | | | | | | | | | 0.00003 | 1.19686 | Female | | | |
| EDAR | | | | | | | | | | | | | 0.04172 | -3.28575 | Male |
| LINC01012 | | | | 0.00013 | -2.51466 | Male | | | | | | | | | |
| CRIPAK | 0.02831 | 3.24271 | Female | | | | | | | | | | | | |
| PDXK | | | | 0.01747 | 0.65561 | Female | | | | | | | | | |
| SP100 | | | | 0.00542 | -1.31214 | Male | 0.00614 | 1.66309 | Female | | | | | | |
| ZNF428 | | | | | | | 0.00514 | 0.76530 | Female | 0.01110 | 0.67840 | Female | | | |
| STEAP1 | 0.00019 | 3.32622 | Female | | | | 0.00137 | -1.26819 | Male | | | | | | |
| ZNF800 | | | | | | | 0.00399 | -0.67326 | Male | | | | | | |
| CA4 | | | | | | | | | | 0.01719 | 2.44553 | Female | | | |
| CLEC4D | | | | | | | | | | 0.00000 | 2.84470 | Female | | | |
| NLRP13 | | | | | | | | | | 0.00001 | -2.94440 | Male | 0.00000 | -3.79474 | Male |
| ESYT1 | | | | | | | | | | | | | 0.00169 | 0.69585 | Female |
| MCMDC2 | | | | | | | 0.00754 | -1.72778 | Male | | | | | | |
| SVIL | | | | | | | | | | | | | 0.01732 | 0.65938 | Female |
| C4BPB | | | | | | | 0.00198 | -1.14304 | Male | | | | 0.01345 | -0.87563 | Male |
| COMT | | | | 0.04834 | -1.01311 | Male | | | | | | | | | |
| TRIM52 | | | | 0.02391 | -1.43338 | Male | | | | | | | | | |
| ZSCAN18 | | | | | | | | | | 0.03120 | -1.06700 | Male | 0.03002 | 1.02509 | Female |
| ABCA3 | | | | | | | 0.03305 | -1.56635 | Male | | | | | | |
| RIC8B | | | | | | | | | | 0.02805 | -0.77918 | Male | | | |
| SLC27A1 | | | | | | | | | | | | | 0.01080 | 1.49740 | Female |
| MFGE8 | | | | | | | | | | | | | 0.00143 | 1.36949 | Female |
| GRB7 | | | | | | | | | | 0.01745 | 0.68536 | Female | | | |
| LOC101928372 | | | | | | | | | | | | | 0.00650 | 1.06612 | Female |
| FBLN1 | | | | 0.00034 | -1.36260 | Male | | | | | | | 0.02793 | 0.58527 | Female |
| HECTD3 | | | | 0.02638 | 0.62225 | Female | | | | | | | | | |
| ACPP | 0.00000 | 2.51003 | Female | | | | | | | | | | | | |
| GSPT2 | 0.00001 | 1.99087 | Female | | | | 0.00278 | 0.64050 | Female | | | | 0.00003 | -0.78157 | Male |
| FAM154A | | | | | | | 0.02885 | 1.89795 | Female | | | | 0.00997 | -2.17117 | Male |
| MT1H | | | | | | | | | | 0.00622 | 1.52061 | Female | 0.00003 | 2.22852 | Female |
| ORAI1 | | | | | | | | | | 0.02580 | 0.60556 | Female | | | |
| DDX51 | | | | | | | 0.00033 | 0.74267 | Female | | | | | | |
| ITFG1 | | | | | | | | | | | | | 0.00003 | -0.76491 | Male |
| RNF219 | | | | | | | | | | | | | 0.00003 | -0.61416 | Male |
| USP22 | | | | | | | 0.00000 | 0.77216 | Female | | | | | | |

| | | | | | | | | | | | | | | | |
|---|---|---|---|---|---|---|---|---|---|---|---|---|---|---|---|
| MSX2P1 | | | | | | | 0.00000 | -1.33319 | Male | 0.00001 | -1.18162 | Male | | | |
| TBC1D9 | | | | | | | | | | | | | 0.01630 | -1.28567 | Male |
| SCARNA9 | | | | | | | | | | 0.00583 | 1.33594 | Female | | | |
| RNF115 | | | | | | | 0.00015 | -0.61877 | Male | | | | | | |
| CBR1 | 0.04508 | 1.35415 | Female | 0.00909 | 1.01781 | Female | 0.01712 | 0.64898 | Female | | | | | | |
| DNPH1 | | | | 0.00878 | 0.85519 | Female | | | | | | | | | |
| THG1L | | | | | | | | | | 0.02940 | 0.59414 | Female | | | |
| DSTNP2 | | | | | | | | | | 0.00001 | 0.72603 | Female | | | |
| KIAA1211 | | | | | | | | | | 0.01132 | -1.22493 | Male | 0.04124 | -1.00808 | Male |
| FOXP4 | | | | 0.01933 | 0.96805 | Female | | | | | | | | | |
| SMIM20 | | | | | | | | | | | | | 0.00014 | -0.66028 | Male |
| HIVEP2 | | | | | | | | | | | | | 0.01121 | 1.53392 | Female |
| PSORS1C1 | | | | | | | | | | 0.00359 | 1.83447 | Female | | | |
| AGPS | | | | 0.02448 | 0.60500 | Female | | | | 0.00000 | -0.70401 | Male | | | |
| UBE2E1 | | | | | | | 0.00194 | -0.58532 | Male | 0.00000 | -1.31741 | Male | | | |
| WNT10B | 0.04860 | -0.87678 | Male | | | | | | | | | | | | |
| TSHZ3 | | | | | | | 0.04040 | 2.13385 | Female | 0.04542 | 2.04725 | Female | | | |
| TMPRSS11E | | | | | | | | | | 0.00733 | 1.05040 | Female | | | |
| SNORD38A | | | | 0.01058 | -1.47581 | Male | | | | | | | | | |
| ATP8B2 | | | | | | | | | | 0.01964 | 1.24276 | Female | | | |
| VTRNA1-3 | | | | | | | | | | | | | 0.00000 | -1.85142 | Male |
| ERVV-2 | | | | | | | | | | 0.00587 | -1.18354 | Male | | | |
| METTL21A | | | | | | | 0.00009 | -0.90561 | Male | | | | | | |
| MRPL11 | 0.00064 | 0.65380 | Female | | | | | | | | | | | | |
| TPPP | | | | | | | 0.02359 | 1.31942 | Female | 0.01607 | 1.33931 | Female | 0.03272 | 1.17795 | Female |
| LGALS12 | | | | | | | | | | 0.00014 | 2.41142 | Female | | | |
| PERP | 0.00105 | -1.18405 | Male | | | | | | | | | | | | |
| ENPEP | | | | 0.00002 | -1.80293 | Male | 0.00628 | -0.73454 | Male | | | | 0.01504 | -0.61647 | Male |
| STAP2 | | | | | | | | | | 0.00259 | 1.77896 | Female | 0.00087 | -1.95852 | Male |
| BRWD3 | | | | 0.00001 | 1.22776 | Female | 0.00000 | 1.39046 | Female | 0.00000 | 1.30171 | Female | | | |
| SLC25A35 | | | | | | | | | | | | | 0.00203 | 0.59488 | Female |
| SDHAP3 | | | | | | | 0.01172 | 0.71695 | Female | 0.00001 | 1.13004 | Female | | | |
| GHITM | 0.00001 | -0.78390 | Male | | | | | | | | | | | | |
| CXorf65 | 0.03186 | 2.11383 | Female | 0.00122 | 1.46443 | Female | 0.04577 | 0.68363 | Female | | | | | | |
| ADAMTS2 | | | | 0.01255 | 2.75564 | Female | 0.04453 | 1.61190 | Female | 0.00956 | 1.91270 | Female | | | |
| ECI1 | | | | | | | 0.00014 | -0.67794 | Male | | | | | | |
| SBF2-AS1 | | | | | | | | | | | | | 0.00102 | 0.95735 | Female |
| FAM49B | | | | | | | 0.00000 | -0.59935 | Male | | | | | | |
| ZNF749 | | | | | | | | | | 0.01708 | 0.62775 | Female | | | |
| SCN8A | | | | | | | | | | | | | 0.02708 | 0.98844 | Female |
| FBN1 | | | | | | | 0.03709 | 1.87020 | Female | | | | | | |
| NLRP9 | | | | | | | | | | 0.00000 | 2.58257 | Female | | | |
| ADAMTS1 | | | | | | | 0.00074 | -1.48653 | Male | 0.00741 | -1.15899 | Male | 0.02860 | -0.95244 | Male |
| CTXN1 | | | | | | | 0.04040 | 1.10483 | Female | | | | 0.00650 | 1.34222 | Female |
| GDF9 | | | | | | | 0.00001 | 0.87135 | Female | | | | | | |
| RTN3 | | | | | | | 0.00000 | -0.62287 | Male | | | | | | |
| BOLL | | | | | | | | | | | | | 0.00540 | -2.02353 | Male |
| AIM1L | | | | | | | 0.02720 | -0.65170 | Male | | | | | | |
| MT1X | | | | | | | | | | 0.00000 | 1.80154 | Female | 0.01243 | 0.80538 | Female |
| COMMD7 | | | | 0.04635 | 0.59093 | Female | | | | | | | | | |
| ZNF449 | | | | | | | 0.02246 | 1.06422 | Female | | | | | | |

| | | | | | | | | | | | | | | | |
|---|---|---|---|---|---|---|---|---|---|---|---|---|---|---|---|
| AHRR | | | | | | | 0.00436 | 1.90502 | Female | | | | | | |
| NPB | | | | | | | | | | | | | 0.00544 | -1.80527 | Male |
| NEK4 | | | | | | | 0.04835 | -0.59266 | Male | | | | | | |
| ZBTB44 | | | | 0.04441 | 0.59226 | Female | | | | | | | | | |
| HDAC9 | | | | | | | | | | 0.01507 | 1.79734 | Female | | | |
| MARCH1 | | | | | | | | | | | | | 0.00461 | -1.33910 | Male |
| CLUHP3 | | | | 0.00427 | -2.23911 | Male | | | | | | | | | |
| ETFB | | | | | | | 0.00034 | 0.61286 | Female | | | | | | |
| TMEM150A | | | | 0.00002 | 1.60099 | Female | | | | | | | | | |
| SIGIRR | | | | 0.01188 | 0.81353 | Female | | | | | | | | | |
| SNORA24 | | | | 0.00946 | -1.14344 | Male | | | | | | | | | |
| ATP6V0A4 | | | | | | | 0.04130 | -0.59087 | Male | | | | | | |
| CACNA1B | | | | | | | 0.00001 | 2.44579 | Female | | | | 0.00000 | 2.43227 | Female |
| KLF9 | | | | | | | | | | 0.00116 | -1.17052 | Male | | | |
| TMEM123 | | | | | | | 0.00000 | -1.71510 | Male | | | | 0.00000 | -1.02885 | Male |
| TERC | | | | | | | | | | 0.00137 | 1.35256 | Female | 0.01272 | 1.06247 | Female |
| AIDA | | | | | | | | | | | | | 0.00011 | -0.68692 | Male |
| ALOX15 | | | | | | | | | | | | | 0.00665 | 2.06184 | Female |
| HAUS5 | | | | | | | | | | | | | 0.00297 | 0.66585 | Female |
| LGR4 | | | | | | | | | | | | | 0.00307 | -0.61224 | Male |
| IFI16 | | | | 0.03436 | 1.74731 | Female | | | | 0.00010 | 1.93137 | Female | | | |
| SLC22A20 | | | | | | | | | | 0.01183 | 1.28972 | Female | | | |
| CCDC144CP | | | | | | | 0.00103 | 0.96403 | Female | | | | | | |
| HADH | | | | 0.04198 | 0.70887 | Female | | | | | | | | | |
| KCNN4 | | | | | | | | | | 0.01284 | -0.89998 | Male | | | |
| OOSP2 | | | | | | | | | | | | | 0.00000 | -3.36799 | Male |
| PDCD4 | | | | 0.00492 | -3.08086 | Male | | | | 0.00000 | 4.63040 | Female | | | |
| EMD | | | | 0.00000 | 0.92597 | Female | 0.00000 | 0.79043 | Female | | | | | | |
| MAP7D2 | | | | | | | | | | 0.00091 | 1.15355 | Female | 0.00016 | 1.25760 | Female |
| TSPAN4 | | | | | | | | | | 0.03154 | -0.66109 | Male | | | |
| PLCD3 | | | | 0.02043 | 0.84698 | Female | | | | | | | | | |
| TPST1 | | | | | | | 0.01335 | -0.84040 | Male | | | | | | |
| IRF3 | 0.00005 | -1.39851 | Male | | | | | | | | | | | | |
| PRKCD | | | | 0.00153 | 1.33420 | Female | | | | | | | | | |
| CCDC152 | | | | | | | | | | | | | 0.00320 | -1.42826 | Male |
| HSPB3 | | | | 0.02752 | -2.42787 | Male | | | | | | | | | |
| HSD17B11 | 0.00955 | -1.68211 | Male | | | | | | | | | | | | |
| LURAP1L | | | | | | | | | | 0.02885 | -1.57400 | Male | | | |
| PROX2 | | | | | | | | | | | | | 0.02418 | -1.39978 | Male |
| UBR3 | | | | | | | | | | | | | 0.00829 | -0.58161 | Male |
| RGMA | | | | 0.00159 | 1.63415 | Female | | | | | | | | | |
| PLXDC1 | | | | | | | | | | 0.01275 | -0.88785 | Male | | | |
| ZIC5 | | | | | | | | | | | | | 0.02495 | -1.20136 | Male |
| NPPA | | | | | | | 0.00003 | -1.90112 | Male | | | | | | |
| METTL4 | | | | 0.02672 | -0.65303 | Male | | | | | | | | | |
| TMEM141 | | | | 0.00058 | 1.16588 | Female | | | | | | | | | |
| ADCY5 | | | | | | | 0.00427 | 1.81063 | Female | | | | | | |
| ZNF286B | | | | | | | 0.00706 | 0.62968 | Female | 0.00160 | -0.64736 | Male | | | |
| RASSF7 | 0.01725 | 1.72501 | Female | | | | 0.00004 | -0.90708 | Male | | | | | | |
| MVD | | | | | | | 0.00161 | -0.62940 | Male | | | | | | |
| CHRNA5 | | | | | | | | | | 0.00013 | -1.27573 | Male | | | |

| | | | | | | | | | | | | | | | |
|---|---|---|---|---|---|---|---|---|---|---|---|---|---|---|---|
| MRPL57 | 0.00129 | 0.62372 | Female | | | | | | | | | | | | |
| ACTBL2 | | | | | | | | | | 0.00000 | -2.75950 | Male | | | |
| INSL4 | | | | | | | 0.04179 | 1.54466 | Female | | | | | | |
| LZTS1 | | | | | | | | | | 0.02969 | 1.82529 | Female | | | |
| ZMAT5 | | | | | | | | | | 0.00011 | -0.63558 | Male | | | |
| RNF5P1.uid-10791 | | | | | | | | | | 0.00021 | -0.66744 | Male | | | |
| MOV10 | | | | | | | | | | | | | 0.00022 | 0.77407 | Female |
| MIR6869 | | | | | | | 0.00000 | 0.64559 | Female | | | | | | |
| DLX3 | | | | 0.00010 | -1.70653 | Male | | | | | | | | | |
| NBEAL2 | | | | 0.00549 | 1.49462 | Female | 0.00211 | -1.10983 | Male | | | | | | |
| FOXO4 | | | | | | | | | | | | | 0.04485 | 0.72191 | Female |
| MCPH1 | | | | 0.02385 | -0.59210 | Male | | | | | | | | | |
| PDE4D | | | | | | | 0.03187 | 0.88163 | Female | | | | | | |
| SNHG1 | 0.02562 | 0.60237 | Female | | | | | | | | | | | | |
| LINC00888 | | | | 0.03628 | 1.13145 | Female | | | | | | | | | |
| HIST1H2BC | | | | | | | 0.00187 | 1.21502 | Female | 0.00000 | 2.23780 | Female | | | |
| SERPINH1 | 0.00000 | -2.18232 | Male | | | | | | | 0.00000 | 2.05446 | Female | 0.02319 | -1.06067 | Male |
| VCX | 0.03059 | 2.09321 | Female | 0.00000 | 6.54785 | Female | 0.00000 | 4.13820 | Female | 0.00000 | 4.13908 | Female | 0.00000 | 3.35299 | Female |
| NUMA1 | | | | | | | 0.00415 | -0.59042 | Male | | | | | | |
| SLC25A24 | 0.00000 | -3.67632 | Male | | | | 0.00000 | -1.68033 | Male | 0.02624 | -0.82219 | Male | | | |
| RAB39A | | | | | | | | | | 0.00029 | -1.39612 | Male | | | |
| FUT3 | | | | | | | | | | | | | 0.00014 | 2.19456 | Female |
| ATPAF1 | | | | 0.03652 | 0.89806 | Female | | | | 0.01042 | 0.68047 | Female | | | |
| FOLR1 | | | | 0.00000 | 2.19677 | Female | | | | | | | | | |
| HLA-A | | | | | | | 0.02060 | -1.57269 | Male | | | | | | |
| B3GALT6 | | | | | | | | | | 0.01138 | -0.90883 | Male | | | |
| RBP7 | | | | | | | | | | 0.00001 | 1.73755 | Female | | | |
| PABPN1L | | | | | | | 0.00090 | 2.43310 | Female | | | | | | |
| COX20 | | | | 0.00000 | 1.28040 | Female | | | | | | | | | |
| SNORD56 | | | | | | | | | | | | | 0.03485 | 0.60770 | Female |
| TLE2 | | | | | | | | | | 0.02191 | 1.68555 | Female | | | |
| POLG2 | 0.00416 | 1.60097 | Female | | | | | | | | | | | | |
| ZIM3 | | | | | | | 0.00925 | -1.64532 | Male | | | | | | |
| MAL2 | | | | | | | 0.00068 | -1.63535 | Male | | | | | | |
| ULK2 | | | | | | | | | | 0.04932 | 0.79362 | Female | | | |
| SNORD34 | | | | 0.01898 | -1.11930 | Male | | | | | | | | | |
| TAF5 | | | | | | | | | | | | | 0.01787 | -0.58655 | Male |
| PRDM16 | | | | 0.01618 | 3.64570 | Female | 0.00000 | 4.63525 | Female | | | | | | |
| SERPINE1 | | | | 0.00213 | 1.98240 | Female | 0.00000 | 2.03960 | Female | 0.03402 | 0.97904 | Female | | | |
| MAP2K2 | | | | | | | 0.00008 | 0.63178 | Female | | | | | | |
| CLPS | | | | | | | | | | | | | 0.01055 | 1.52868 | Female |
| KIAA1430 | | | | | | | | | | 0.01017 | 0.95584 | Female | 0.00377 | -1.05652 | Male |
| LOC401127 | | | | | | | 0.00000 | -0.59034 | Male | | | | | | |
| CD3EAP | | | | | | | | | | 0.00000 | 0.62110 | Female | | | |
| PUS7L | | | | | | | | | | 0.00527 | -0.64252 | Male | 0.00193 | -0.68897 | Male |
| SEPT3 | | | | | | | | | | | | | 0.00765 | 2.00382 | Female |
| BSCL2.uid-15607 | | | | | | | 0.00060 | 0.69916 | Female | | | | | | |
| PKP2 | 0.00022 | -1.83658 | Male | 0.03947 | 0.70432 | Female | | | | | | | | | |
| AQP11 | | | | | | | | | | 0.00468 | 1.14543 | Female | | | |
| RAB43 | | | | | | | | | | 0.00015 | 1.31784 | Female | | | |
| SLC25A40 | | | | | | | 0.00038 | -1.26322 | Male | | | | | | |

| | | | | | | | | | | | | | | | |
|---|---|---|---|---|---|---|---|---|---|---|---|---|---|---|---|
| SVIP | | | | | | | 0.00000 | -0.98810 | Male | | | | | | |
| ICA1 | | | | 0.00771 | 2.32574 | Female | 0.00001 | -2.54569 | Male | | | | 0.00664 | -1.56906 | Male |
| ERMN | | | | | | | 0.02654 | -2.00140 | Male | 0.03036 | -1.88293 | Male | 0.00343 | 2.50158 | Female |
| TRIM14 | | | | | | | | | | 0.01480 | 1.12808 | Female | | | |
| SH2D3A | 0.04355 | 1.67424 | Female | | | | | | | | | | | | |
| MCTS2P | 0.00902 | 0.94603 | Female | | | | | | | | | | | | |
| SREK1IP1 | | | | 0.00093 | -0.62974 | Male | | | | | | | | | |
| TMEM220 | | | | | | | 0.01241 | -1.32870 | Male | | | | | | |
| FAM46B | | | | 0.01069 | 1.06716 | Female | 0.03875 | 0.60764 | Female | 0.00000 | 1.52383 | Female | 0.00530 | 0.73444 | Female |
| LDOC1 | | | | 0.00113 | 1.23700 | Female | 0.00185 | 0.82362 | Female | | | | 0.00004 | 0.97442 | Female |
| ERCC1 | | | | | | | | | | | | | 0.00096 | 0.66435 | Female |
| LOC100335030 | | | | | | | | | | | | | 0.01562 | -0.72741 | Male |
| GDI1 | | | | 0.00002 | 1.03869 | Female | | | | | | | 0.00002 | 0.68573 | Female |
| ANXA3 | | | | | | | 0.00000 | -0.87708 | Male | | | | | | |
| DLX2 | | | | | | | | | | 0.02050 | 1.93028 | Female | | | |
| GCM1 | | | | | | | | | | 0.00870 | -0.86482 | Male | | | |
| KIF16B | | | | | | | | | | 0.01234 | -0.64430 | Male | | | |
| FYN | | | | | | | | | | 0.00544 | 1.31016 | Female | | | |
| PTGFRN | | | | | | | | | | 0.00032 | 1.15776 | Female | 0.00000 | 1.52909 | Female |
| POU5F1P3 | | | | | | | | | | 0.00000 | 1.69331 | Female | | | |
| LOC643441 | | | | | | | | | | 0.00627 | -1.27942 | Male | | | |
| PAGE4 | | | | | | | | | | 0.00568 | -1.70745 | Male | 0.00070 | -1.96380 | Male |
| MFAP5 | | | | | | | | | | 0.01547 | 1.06999 | Female | | | |
| TMEM88 | 0.00074 | -3.25554 | Male | | | | 0.00235 | 1.39915 | Female | | | | | | |
| ATP7B | 0.00251 | -2.08883 | Male | | | | | | | | | | | | |
| DIAPH2 | | | | 0.00000 | 2.47273 | Female | 0.00063 | 1.33664 | Female | 0.00003 | 1.51499 | Female | | | |
| POU4F1 | | | | 0.04436 | 1.56666 | Female | 0.01071 | 1.30161 | Female | | | | | | |
| WNT11 | 0.00006 | -2.74840 | Male | | | | 0.00626 | 0.88410 | Female | | | | 0.00003 | 1.20507 | Female |
| UPF3B | 0.00003 | 1.18744 | Female | 0.00002 | 0.76420 | Female | 0.00000 | 0.74975 | Female | | | | | | |
| DSP | 0.01392 | -0.91382 | Male | | | | | | | | | | | | |
| ALG5 | | | | | | | | | | | | | 0.00000 | -0.83661 | Male |
| TAF1A | | | | | | | 0.00282 | -0.77413 | Male | | | | | | |
| ZNF780A | 0.01374 | 2.12443 | Female | | | | | | | | | | | | |
| SLC4A5 | | | | | | | | | | 0.00300 | -0.68825 | Male | | | |
| TDGF1 | | | | 0.00001 | 1.56705 | Female | | | | 0.00000 | 1.93119 | Female | | | |
| PPM1N | | | | | | | 0.00004 | 0.84017 | Female | | | | | | |
| ZNF22 | | | | | | | | | | 0.00776 | 0.68200 | Female | | | |
| SLC46A1 | 0.04026 | 2.22457 | Female | | | | | | | | | | | | |
| FBXL20 | | | | | | | | | | 0.00002 | 0.76628 | Female | | | |
| INE1 | | | | | | | 0.00070 | 2.23918 | Female | | | | | | |
| AFF2 | | | | | | | | | | 0.03273 | 2.16776 | Female | | | |
| MZT2A | | | | | | | | | | 0.00003 | -0.75375 | Male | | | |
| LOC100129148 | | | | | | | | | | | | | 0.00722 | 0.62526 | Female |
| PAGE2B | 0.00832 | 1.99583 | Female | | | | 0.00000 | 2.16843 | Female | 0.00000 | 1.90778 | Female | | | |
| BEX4 | 0.02948 | 1.33672 | Female | | | | | | | | | | | | |
| HSPH1 | | | | 0.00703 | 0.64885 | Female | | | | | | | | | |
| ACSBG1 | | | | | | | | | | | | | 0.02592 | 2.01490 | Female |
| NRSN2 | | | | 0.00577 | 1.41262 | Female | | | | | | | | | |
| RUNX1 | | | | 0.01377 | 1.13233 | Female | | | | | | | | | |
| MRFAP1 | 0.00001 | 0.69778 | Female | | | | | | | | | | | | |
| ZNF367 | | | | 0.00409 | 1.98663 | Female | | | | | | | | | |

| | | | | | | | | | | | | | | | |
|---|---|---|---|---|---|---|---|---|---|---|---|---|---|---|---|
| RHOBTB2 | | | | | | | | | | 0.01208 | -0.84940 | Male | | | |
| APH1B | | | | | | | 0.00155 | -2.12709 | Male | | | | | | |
| PNLDC1 | | | | | | | | | | 0.01074 | 1.33418 | Female | | | |
| CSMD3 | | | | 0.04765 | -0.95080 | Male | 0.04609 | -0.70075 | Male | 0.00023 | -1.10031 | Male | | | |
| MAP1LC3C | | | | 0.01457 | 2.98828 | Female | | | | 0.00031 | 2.83106 | Female | | | |
| GNRH2 | | | | | | | 0.00002 | 3.75390 | Female | | | | | | |
| NBEAP1 | | | | | | | 0.02479 | -1.59415 | Male | | | | | | |
| JUN | | | | | | | | | | | | | 0.00002 | 2.69327 | Female |
| MRM1 | | | | | | | | | | | | | 0.01733 | 0.67166 | Female |
| ANKHD1.uid-6876 | | | | | | | 0.00131 | 1.02350 | Female | | | | | | |
| NID1 | 0.02531 | -1.26529 | Male | | | | | | | | | | | | |
| CUL4B | | | | 0.00874 | 0.71192 | Female | | | | | | | | | |
| MBNL2 | | | | | | | 0.00004 | -2.17745 | Male | | | | | | |
| LOC729083 | | | | | | | 0.02868 | -1.77076 | Male | | | | | | |
| NR1D1 | | | | | | | 0.01245 | 1.32376 | Female | | | | | | |
| SLC28A3 | | | | | | | 0.01290 | -0.58025 | Male | | | | 0.00079 | 0.70319 | Female |
| TMEM135 | | | | | | | 0.00080 | -0.80424 | Male | | | | | | |
| PPAPDC3 | | | | 0.02046 | 2.88491 | Female | | | | | | | | | |
| HN1 | 0.00005 | -0.69888 | Male | | | | | | | | | | | | |
| MAOA | | | | 0.00204 | 2.63472 | Female | 0.00697 | 1.67659 | Female | | | | | | |
| CLDN11 | | | | 0.00000 | -1.09934 | Male | | | | | | | | | |
| COPS7A | | | | 0.00465 | -0.75657 | Male | | | | | | | | | |
| TFEB | | | | 0.01201 | 1.25532 | Female | | | | 0.04275 | -0.68309 | Male | | | |
| TMEM87B | | | | | | | | | | 0.02829 | 0.62217 | Female | | | |
| PAIP2B | | | | | | | | | | 0.00208 | -0.91243 | Male | 0.00179 | 0.91245 | Female |
| LINC00086 | | | | | | | | | | 0.00449 | 2.45825 | Female | 0.04996 | 1.78792 | Female |
| PRSS8 | 0.00002 | 2.71387 | Female | 0.00013 | 1.28702 | Female | | | | | | | | | |
| F3 | 0.04324 | -1.43313 | Male | | | | | | | | | | 0.00001 | 1.05239 | Female |
| BACE1 | | | | | | | 0.02618 | 0.81527 | Female | | | | | | |
| SNORD19 | | | | | | | | | | | | | 0.01254 | 1.21405 | Female |
| TFAP4 | | | | 0.00013 | 1.88162 | Female | 0.00000 | 1.68181 | Female | | | | | | |
| PHKA2-AS1 | | | | | | | 0.00036 | 1.70973 | Female | 0.00000 | 2.73932 | Female | | | |
| CELA3B | | | | | | | | | | 0.02162 | 1.14105 | Female | | | |
| HNRNPM | | | | | | | 0.00000 | 0.59862 | Female | | | | | | |
| FCHO2 | | | | | | | | | | 0.02344 | -0.66698 | Male | | | |
| PPP2R2D | | | | 0.01332 | 0.79414 | Female | | | | | | | | | |
| LPPR3 | | | | | | | | | | 0.00701 | -1.59155 | Male | | | |
| ZNF713 | | | | | | | | | | 0.03841 | -1.57694 | Male | | | |
| ZNF587B | | | | | | | 0.00004 | 0.82161 | Female | | | | | | |
| ERO1L | | | | | | | 0.00154 | -0.83506 | Male | | | | 0.00106 | -0.79622 | Male |
| SYT6 | | | | | | | | | | | | | 0.02055 | 2.75920 | Female |
| BMP15 | | | | | | | | | | | | | 0.00197 | 5.02944 | Female |
| BHMT2 | | | | | | | | | | 0.00242 | -2.13845 | Male | | | |
| CD9 | | | | | | | | | | 0.00000 | 1.13415 | Female | | | |
| ESRP2 | 0.00397 | -1.47791 | Male | | | | | | | | | | | | |
| GPR78 | | | | | | | | | | 0.03955 | -1.29066 | Male | | | |
| ARSI | | | | | | | | | | 0.03056 | 1.86127 | Female | | | |
| HOPX | | | | | | | | | | 0.04515 | -1.25026 | Male | | | |
| ZNF318 | | | | | | | | | | 0.00608 | 0.63707 | Female | | | |
| MYH13 | | | | 0.00161 | -2.89807 | Male | 0.01779 | -1.66764 | Male | | | | | | |
| SNORD89 | | | | | | | | | | | | | 0.04949 | 0.90213 | Female |

| | | | | | | | | | | | | | | | |
|---|---|---|---|---|---|---|---|---|---|---|---|---|---|---|---|
| GPHA2 | | | | | | | | | | 0.01635 | 1.76648 | Female | 0.00017 | -2.64868 | Male |
| RYR1 | | | | | | | | | | | | | 0.00158 | 1.01093 | Female |
| NCOA7 | | | | 0.01413 | 0.95838 | Female | | | | | | | | | |
| TERT | | | | | | | | | | | | | 0.00614 | -1.08634 | Male |
| CALD1 | 0.00071 | -1.89680 | Male | | | | 0.00033 | -0.81593 | Male | 0.00540 | -0.62300 | Male | | | |
| GSTO1 | 0.00120 | -1.79330 | Male | | | | | | | | | | 0.00000 | -1.09551 | Male |
| SLC35A3 | 0.00984 | -1.59104 | Male | | | | | | | | | | | | |
| KIAA1217 | 0.04136 | -1.55067 | Male | | | | | | | | | | | | |
| CGN | | | | 0.01188 | 1.04569 | Female | | | | | | | | | |
| AKT3 | | | | | | | 0.04076 | -0.58222 | Male | | | | | | |
| DDX55 | | | | | | | 0.00000 | 0.62045 | Female | | | | | | |
| 1.00000 | | | | | | | 0.00000 | 0.74002 | Female | | | | | | |
| ACOT11 | | | | | | | | | | 0.00100 | 1.47001 | Female | | | |
| BAZ2B.uid-2643 | | | | 0.03859 | -2.16731 | Male | | | | 0.02642 | -1.52430 | Male | | | |
| LOC441666 | | | | | | | | | | | | | 0.00000 | -2.26516 | Male |
| FOXP1 | | | | | | | 0.02622 | -0.75906 | Male | 0.00509 | 0.88093 | Female | | | |
| MGC32805 | | | | | | | | | | 0.00108 | 1.35976 | Female | | | |
| LAMP2 | | | | 0.00008 | 0.79571 | Female | 0.00000 | 0.97197 | Female | 0.00000 | 0.78427 | Female | | | |
| DCAF4 | | | | | | | | | | | | | 0.00000 | 0.70831 | Female |
| COL21A1 | | | | 0.00291 | -1.80413 | Male | | | | | | | | | |
| WNT1 | | | | | | | | | | | | | 0.00034 | 2.95530 | Female |
| KLF15 | | | | | | | | | | 0.00010 | -0.86744 | Male | | | |
| GALNT18 | | | | | | | | | | 0.01712 | 1.83713 | Female | | | |
| MEI4 | | | | | | | | | | | | | 0.04185 | -2.07625 | Male |
| LARP7 | | | | | | | 0.00000 | 0.82024 | Female | 0.00000 | 0.72614 | Female | | | |
| RIN2 | | | | 0.00664 | 1.21261 | Female | 0.03187 | 0.68138 | Female | | | | | | |
| RGL2 | | | | | | | 0.00000 | -1.16530 | Male | 0.00900 | -0.64884 | Male | | | |
| NEDD4 | | | | | | | 0.00767 | -1.22287 | Male | 0.02898 | -0.99025 | Male | | | |
| SPA17 | | | | | | | 0.00074 | -0.80128 | Male | | | | 0.01202 | -0.58275 | Male |
| SYBU | | | | 0.00007 | -2.29729 | Male | | | | | | | | | |
| PGBD1 | | | | | | | | | | 0.00237 | 1.26197 | Female | | | |
| QSOX1 | | | | | | | | | | | | | 0.00041 | 0.92402 | Female |
| SPATA6L | | | | | | | | | | | | | 0.02390 | -1.60969 | Male |
| KIF18A | | | | | | | 0.00487 | -0.67417 | Male | | | | | | |
| SERPINB9 | | | | 0.00001 | 1.66928 | Female | | | | | | | | | |
| ASXL3 | | | | | | | | | | 0.04595 | -1.88611 | Male | | | |
| FOXO3 | | | | | | | | | | 0.02612 | 0.63906 | Female | | | |
| OGFRL1 | | | | | | | | | | 0.00957 | 1.20249 | Female | | | |
| HMG20B | | | | | | | 0.00143 | -0.58365 | Male | | | | | | |
| TATDN1 | | | | | | | 0.00008 | -0.88969 | Male | | | | | | |
| CLMP | | | | | | | | | | 0.01745 | -1.33927 | Male | | | |
| GSTA2 | | | | | | | | | | 0.03012 | 2.38477 | Female | | | |
| MSN | | | | 0.00298 | 1.60620 | Female | 0.00013 | 1.36740 | Female | 0.03486 | 0.80633 | Female | | | |
| GK | | | | 0.01663 | 1.60287 | Female | 0.00003 | 1.73644 | Female | 0.00000 | 2.12479 | Female | | | |
| DPRXP4 | | | | | | | | | | 0.00746 | -1.29412 | Male | | | |
| GSTT1 | | | | | | | | | | 0.01972 | -0.79217 | Male | 0.00126 | 1.05788 | Female |
| MAN2C1 | 0.02275 | 1.97726 | Female | | | | | | | | | | | | |
| CDH4 | | | | | | | | | | | | | 0.00079 | 2.08044 | Female |
| SOHLH2 | | | | | | | | | | | | | 0.00008 | -1.30601 | Male |
| EFNB2 | | | | 0.01757 | 1.40254 | Female | 0.03524 | 0.85824 | Female | 0.00195 | 1.12863 | Female | | | |
| DDIT4 | | | | 0.00000 | -2.18939 | Male | | | | | | | 0.01911 | 0.75352 | Female |

| | | | | | | | | | | | | | | | |
|---|---|---|---|---|---|---|---|---|---|---|---|---|---|---|---|
| TUBB3 | 0.01821 | -1.46033 | Male | | | | 0.00001 | 1.03678 | Female | 0.00000 | 1.27454 | Female | | | |
| ANKRD30BL | | | | | | | | | | 0.02978 | 0.63616 | Female | | | |
| HSPA1L | | | | | | | 0.03162 | -1.89568 | Male | 0.00000 | -3.95501 | Male | | | |
| CDO1 | | | | | | | | | | 0.03529 | -0.94230 | Male | | | |
| SLC5A5 | | | | | | | | | | 0.00000 | 1.27139 | Female | | | |
| FAM132A | | | | | | | | | | 0.00228 | 1.91297 | Female | | | |
| LONRF1 | | | | | | | 0.03570 | -0.88256 | Male | | | | | | |
| JTB | | | | | | | 0.00000 | -0.63258 | Male | | | | | | |
| SUPT4H1 | | | | 0.00000 | -0.58052 | Male | | | | | | | | | |
| NME7 | | | | | | | | | | 0.00493 | -0.91376 | Male | | | |
| SLC25A43 | | | | | | | | | | 0.00286 | 0.92146 | Female | | | |
| CTCFL | | | | | | | | | | | | | 0.03067 | -1.60132 | Male |
| NDUFB2-AS1 | | | | | | | | | | | | | 0.01580 | 0.67297 | Female |
| SNORD76 | 0.02524 | 1.54619 | Female | | | | | | | | | | | | |
| RFPL4A | | | | | | | | | | 0.01603 | 4.55433 | Female | | | |
| LUCAT1 | | | | | | | | | | | | | 0.00043 | 1.64696 | Female |
| DDX53 | | | | | | | | | | | | | 0.00622 | 2.98975 | Female |
| TNFSF12 | | | | 0.03507 | 1.59490 | Female | | | | | | | | | |
| PI4K2A | | | | | | | 0.01283 | 0.67544 | Female | | | | | | |
| MGC27345 | 0.00184 | -2.03623 | Male | | | | | | | | | | | | |
| WDSUB1 | 0.00329 | -1.68392 | Male | | | | | | | | | | | | |
| TBX3 | | | | 0.00160 | 0.86623 | Female | | | | | | | | | |
| RAB3IP | | | | | | | 0.00001 | -0.74617 | Male | | | | | | |
| WDR24 | | | | | | | 0.03226 | -0.68726 | Male | | | | | | |
| EMC3-AS1 | | | | | | | | | | 0.00161 | -0.61413 | Male | | | |
| ATP2A1 | 0.04868 | 2.57303 | Female | | | | | | | | | | | | |
| COL4A2 | | | | 0.01361 | 1.91611 | Female | 0.01213 | -1.46947 | Male | 0.01365 | 1.37010 | Female | | | |
| C6orf52 | | | | | | | | | | | | | 0.03069 | 0.85504 | Female |
| RAB29 | | | | | | | 0.04030 | 0.74737 | Female | | | | | | |
| GRAMD2 | 0.00297 | -2.07813 | Male | 0.00001 | 1.87122 | Female | 0.04275 | -0.65962 | Male | 0.00037 | -0.99055 | Male | | | |
| NFATC2 | | | | | | | | | | 0.02577 | -1.35093 | Male | | | |
| PCDH15 | | | | | | | 0.00127 | -1.89377 | Male | 0.01623 | 1.42001 | Female | | | |
| LOC389906 | | | | 0.01676 | 1.10692 | Female | 0.00000 | 1.49478 | Female | 0.00555 | 0.81448 | Female | 0.00000 | 1.42997 | Female |
| IFITM1 | | | | 0.01387 | 1.24478 | Female | | | | 0.00000 | 2.27292 | Female | | | |
| NET1 | | | | | | | | | | 0.00009 | -0.64108 | Male | | | |
| ADM | 0.04208 | -1.20100 | Male | | | | | | | | | | | | |
| S1PR2 | | | | 0.02066 | -1.27923 | Male | | | | 0.00199 | -1.03710 | Male | | | |
| LTBP3 | | | | | | | 0.00098 | -1.09783 | Male | | | | | | |
| SLC9A1 | | | | | | | | | | 0.00023 | 0.60259 | Female | | | |
| TRPT1 | | | | | | | 0.00009 | -0.95060 | Male | | | | | | |
| FN3K | | | | | | | | | | 0.00000 | 2.35470 | Female | | | |
| PCSK5 | | | | | | | | | | | | | 0.01033 | 0.95862 | Female |
| ZNF578 | | | | | | | 0.01757 | 0.69854 | Female | | | | | | |
| NUTM1 | 0.01814 | 3.00937 | Female | | | | | | | | | | | | |
| MEST | | | | | | | 0.00000 | -0.75163 | Male | | | | 0.00003 | -0.58628 | Male |
| PNP | | | | | | | 0.00037 | 0.76366 | Female | | | | | | |
| RFNG | | | | | | | | | | | | | 0.01843 | 0.62823 | Female |
| TBC1D14 | | | | | | | 0.00532 | -0.83734 | Male | | | | | | |
| ZNF625 | | | | | | | 0.02583 | -0.98194 | Male | | | | | | |
| C15orf61 | | | | 0.00000 | 1.08799 | Female | | | | | | | | | |
| CORO1B | | | | | | | 0.00144 | -0.59980 | Male | | | | | | |

| | | | | | | | | | | | | | | | |
|---|---|---|---|---|---|---|---|---|---|---|---|---|---|---|---|
| KIFC1 | | | | | | | | | | 0.00001 | -0.86985 | Male | | | |
| GPX2 | | | | | | | | | | 0.00000 | 2.37714 | Female | | | |
| PCOLCE2 | | | | | | | 0.04436 | -1.47104 | Male | | | | | | |
| LINC01160 | 0.00120 | -2.44050 | Male | 0.01500 | 1.23750 | Female | | | | 0.00110 | 1.04040 | Female | 0.00096 | -1.02838 | Male |
| GNA12 | | | | | | | | | | 0.01570 | -0.65098 | Male | | | |
| ISCA2 | 0.00037 | 1.14243 | Female | | | | | | | | | | | | |
| PSME1 | | | | | | | 0.03598 | -0.62932 | Male | | | | | | |
| GAS2L1 | | | | | | | | | | 0.04773 | 0.70487 | Female | | | |
| FAM110B | 0.00000 | 4.35376 | Female | | | | | | | | | | | | |
| SRSF2 | | | | | | | 0.00000 | -0.67800 | Male | | | | | | |
| NSA2 | 0.00506 | -0.63286 | Male | | | | | | | | | | | | |
| PLOD3 | | | | | | | 0.00360 | -0.70111 | Male | | | | | | |
| B3GNT1 | | | | 0.03810 | -0.90061 | Male | | | | | | | | | |
| SCCPDH | | | | | | | 0.02078 | -1.02418 | Male | 0.00085 | -1.31674 | Male | | | |
| APRT | | | | 0.00006 | 0.59275 | Female | | | | | | | | | |
| ACTN1 | | | | 0.00037 | 0.79336 | Female | | | | | | | | | |
| RNPEP | | | | 0.00421 | 1.02643 | Female | | | | | | | | | |
| TMEM191C | | | | | | | 0.00000 | -1.20079 | Male | 0.00510 | -0.66145 | Male | | | |
| VPS9D1 | | | | | | | 0.00380 | 1.04301 | Female | 0.03370 | 0.79114 | Female | | | |
| ETV3 | | | | | | | | | | 0.00000 | 1.60720 | Female | | | |
| TULP1 | | | | | | | | | | 0.03237 | -1.42660 | Male | 0.03592 | 1.38616 | Female |
| UCP2 | | | | | | | | | | | | | 0.00383 | 0.79347 | Female |
| DZANK1 | | | | | | | | | | 0.01806 | 1.04033 | Female | | | |
| IL4R | | | | | | | | | | 0.01677 | 1.42762 | Female | | | |
| PROSER1 | | | | 0.01748 | 0.58450 | Female | 0.00004 | -0.61969 | Male | | | | | | |
| HS6ST1 | | | | 0.01933 | 0.83096 | Female | 0.00287 | 0.68136 | Female | 0.00000 | 1.06546 | Female | 0.00390 | 0.63880 | Female |
| PANK1 | | | | | | | 0.00127 | -1.28610 | Male | | | | | | |
| ARHGAP11A | | | | | | | 0.00001 | -0.64279 | Male | | | | 0.00002 | -0.58067 | Male |
| PTPLA | | | | | | | 0.00203 | -1.43392 | Male | | | | | | |
| TCL1A | | | | | | | | | | | | | 0.00000 | -2.90740 | Male |
| CCDC90B | | | | | | | 0.00010 | -0.63889 | Male | | | | | | |
| PTPMT1 | 0.00007 | 1.54948 | Female | | | | | | | | | | | | |
| BAIAP2 | | | | | | | 0.00017 | -0.62892 | Male | | | | | | |
| RBM3 | 0.00000 | 0.89402 | Female | 0.00000 | 0.82676 | Female | 0.00000 | 0.59626 | Female | 0.00000 | 0.69020 | Female | | | |
| PHPT1 | | | | 0.00000 | 0.74698 | Female | 0.00000 | 0.75461 | Female | | | | 0.00000 | 0.60628 | Female |
| CSNK1E | | | | | | | | | | | | | 0.00014 | 0.58424 | Female |
| ZFP57 | 0.00037 | -2.18484 | Male | | | | | | | | | | | | |
| C19orf70 | 0.01750 | 0.69301 | Female | | | | | | | | | | | | |
| OXT | | | | | | | | | | 0.00000 | 1.76863 | Female | 0.00011 | 0.93218 | Female |
| MIR373 | | | | | | | 0.00008 | 0.96442 | Female | | | | | | |
| PYY2 | | | | | | | | | | 0.00239 | 1.06742 | Female | | | |
| PEBP1 | 0.04026 | -0.64328 | Male | | | | | | | | | | | | |
| ZNF717 | | | | | | | | | | 0.02001 | -0.83231 | Male | | | |
| PVRL1 | | | | 0.01058 | 0.92916 | Female | 0.00254 | 0.71421 | Female | | | | 0.00000 | 1.08780 | Female |
| JUP | | | | 0.00000 | 1.24520 | Female | 0.00017 | -0.65362 | Male | | | | | | |
| LANCL2 | | | | 0.00194 | 1.09467 | Female | | | | | | | | | |
| IL2RB | | | | | | | 0.01783 | 1.56693 | Female | | | | | | |
| SENP5 | 0.02323 | 0.67664 | Female | | | | | | | | | | | | |
| YWHAEP1 | | | | | | | 0.00520 | -0.88994 | Male | | | | | | |
| WIPI1 | | | | | | | 0.02363 | -1.13657 | Male | | | | | | |
| ABCA13 | | | | | | | | | | 0.02937 | 1.02246 | Female | 0.02987 | -1.01557 | Male |

| | | | | | | | | | | | | | | | |
|---|---|---|---|---|---|---|---|---|---|---|---|---|---|---|---|
| PIGP | | | | | | | 0.00120 | -0.92345 | Male | | | | 0.02292 | -0.64774 | Male |
| TCEAL4 | | | | | | | | | | 0.02721 | 0.94404 | Female | 0.00403 | -1.15906 | Male |
| NETO2 | | | | | | | | | | | | | 0.01809 | -0.74221 | Male |
| SUMF2 | | | | | | | 0.00074 | -0.67940 | Male | | | | | | |
| TFDP3 | 0.00052 | 4.62559 | Female | 0.00000 | 4.71916 | Female | | | | | | | | | |
| CD320 | | | | 0.00000 | 1.51032 | Female | 0.00000 | 0.79204 | Female | | | | | | |
| PFKL | | | | 0.00000 | 1.01774 | Female | 0.00000 | -0.66429 | Male | | | | | | |
| FAM175A | | | | | | | | | | 0.00612 | -0.59634 | Male | 0.00000 | -0.94223 | Male |
| ZSCAN4 | | | | | | | | | | 0.00000 | 5.62373 | Female | | | |
| C11orf1 | 0.01969 | 2.23518 | Female | | | | | | | | | | | | |
| TBL1X | | | | 0.00531 | 1.17205 | Female | 0.00027 | 1.00750 | Female | | | | 0.00408 | 0.77612 | Female |
| PRKXP1 | | | | | | | 0.04084 | -0.89059 | Male | | | | | | |
| RBM15 | 0.00713 | -1.19243 | Male | 0.00671 | -0.74368 | Male | | | | | | | | | |
| DCAF4L1 | | | | 0.03451 | -0.87209 | Male | | | | | | | | | |
| PECAM1 | | | | | | | 0.02934 | 1.65411 | Female | | | | | | |
| RRP7B | | | | | | | 0.00000 | 0.78508 | Female | | | | | | |
| FAM86EP | | | | | | | | | | 0.00066 | 1.37485 | Female | | | |
| NSMCE4A | | | | | | | 0.00000 | -0.77024 | Male | | | | | | |
| MRI1 | | | | | | | | | | 0.00469 | -0.70670 | Male | | | |
| SLC6A3 | | | | | | | | | | 0.00003 | 2.36658 | Female | 0.00017 | 2.16628 | Female |
| KLF12 | | | | | | | | | | | | | 0.02527 | -1.57523 | Male |
| TKTL1 | | | | 0.01490 | 1.50066 | Female | 0.00000 | 1.86782 | Female | 0.00000 | 3.48605 | Female | 0.00268 | 1.14649 | Female |
| MTERFD1 | | | | | | | | | | | | | 0.00009 | -0.64427 | Male |
| SNORA9 | | | | | | | | | | | | | 0.03788 | 0.59396 | Female |
| ZNRD1-AS1 | | | | | | | | | | | | | 0.00895 | -1.66630 | Male |
| CHKA | | | | | | | | | | 0.00721 | -0.88649 | Male | | | |
| SLC25A10 | | | | | | | 0.02470 | -0.70859 | Male | | | | | | |
| SGK494 | | | | | | | 0.03429 | 0.64192 | Female | | | | | | |
| TRIML1 | | | | 0.04525 | -1.11616 | Male | | | | | | | | | |
| KDM6A | | | | | | | 0.00275 | 0.64423 | Female | 0.00000 | 1.08773 | Female | 0.00000 | 1.16022 | Female |
| SSX1 | | | | 0.00392 | 2.86658 | Female | | | | | | | | | |
| PML | | | | | | | 0.01610 | -0.86170 | Male | | | | | | |
| CAPNS1 | | | | 0.04397 | 0.58529 | Female | | | | 0.00019 | 0.62212 | Female | | | |
| TEAD2 | | | | | | | 0.00007 | 1.06041 | Female | | | | | | |
| LOC101927501 | 0.04208 | 2.88881 | Female | 0.00000 | 3.59877 | Female | 0.00000 | 3.14790 | Female | | | | | | |
| UNC5B | 0.00004 | 2.91642 | Female | | | | | | | 0.00002 | 2.23962 | Female | | | |
| NACAP1 | 0.00183 | -0.70836 | Male | | | | | | | | | | | | |
| PTK7 | | | | | | | 0.02096 | -0.80156 | Male | | | | | | |
| GNE | | | | | | | | | | 0.00000 | -2.40283 | Male | | | |
| YAP1 | | | | | | | | | | 0.00000 | -0.76251 | Male | | | |
| CDIP1 | | | | | | | 0.04688 | 0.82182 | Female | | | | | | |
| HDC | | | | | | | | | | | | | 0.00541 | -2.59301 | Male |
| GSK3B | | | | | | | | | | 0.00029 | -0.64743 | Male | | | |
| LMO2 | | | | | | | | | | 0.00001 | -2.70688 | Male | | | |
| TBL2 | | | | | | | 0.00365 | -0.73614 | Male | | | | | | |
| MUC15 | | | | | | | | | | 0.00727 | -1.12218 | Male | | | |
| CTTN | | | | | | | 0.00000 | 0.82427 | Female | | | | | | |
| HSPE1 | | | | 0.00000 | 0.95875 | Female | | | | | | | | | |
| TSPO | | | | | | | | | | 0.00005 | -1.73411 | Male | | | |
| IMPACT | | | | | | | 0.00051 | -0.76825 | Male | | | | 0.00001 | -0.88830 | Male |
| C1GALT1 | | | | | | | 0.00007 | -0.74266 | Male | | | | | | |

| | | | | | | | | | | | | | | | |
|---|---|---|---|---|---|---|---|---|---|---|---|---|---|---|---|
| TMX4 | | | | 0.00217 | 1.82621 | Female | | | | | | | | | |
| ZFYVE20 | 0.01885 | -1.23303 | Male | | | | | | | | | | | | |
| PPP1R9B | | | | | | | 0.00797 | 1.08518 | Female | | | | | | |
| RASD1 | | | | | | | | | | 0.00000 | 2.28732 | Female | | | |
| ABHD14B | | | | | | | 0.00051 | -1.03875 | Male | | | | | | |
| FUNDC2 | 0.02338 | 0.64937 | Female | 0.00001 | 0.66787 | Female | 0.00000 | 0.80060 | Female | | | | | | |
| DLX6 | | | | | | | | | | 0.00909 | -0.84754 | Male | 0.00625 | -0.86354 | Male |
| FOXA2 | | | | | | | | | | 0.03158 | 1.18161 | Female | | | |
| SLC43A3 | | | | | | | | | | | | | 0.00106 | -1.83242 | Male |
| ERCC2 | | | | | | | 0.00040 | -0.99755 | Male | | | | | | |
| SPTB | | | | | | | | | | | | | 0.00000 | 1.59070 | Female |
| SLCO3A1 | | | | | | | 0.02038 | 1.89885 | Female | | | | | | |
| ALG14 | 0.00001 | 2.45127 | Female | | | | | | | | | | | | |
| SLC9A7 | | | | | | | 0.03573 | 1.21818 | Female | 0.00860 | -1.40622 | Male | | | |
| LOC100130075 | | | | | | | | | | | | | 0.01648 | 0.73843 | Female |
| PLRG1 | 0.02742 | -0.62760 | Male | | | | | | | | | | | | |
| MAPK12 | | | | | | | | | | 0.00909 | 1.96370 | Female | | | |
| 1.00000 | | | | | | | | | | 0.02995 | 1.25112 | Female | | | |
| TANK | | | | | | | 0.00201 | -0.64923 | Male | | | | | | |
| CPNE8 | 0.02023 | -2.39407 | Male | | | | | | | | | | | | |
| UTY | 0.00000 | -6.03357 | Male | 0.00000 | -7.39858 | Male | 0.00000 | -6.61215 | Male | 0.00000 | -6.81694 | Male | 0.00000 | -7.02608 | Male |
| KATNBL1P6 | | | | 0.01401 | -0.87444 | Male | 0.00207 | -0.71205 | Male | | | | | | |
| ZNF649 | | | | 0.04487 | 1.04919 | Female | | | | | | | | | |
| GREB1 | | | | | | | 0.00156 | -2.08740 | Male | | | | | | |
| LRGUK | | | | | | | 0.03496 | -2.16723 | Male | | | | 0.03461 | 2.06515 | Female |
| LINC00665 | | | | | | | 0.00281 | 0.76122 | Female | 0.00066 | -0.79634 | Male | | | |
| COMTD1 | 0.00181 | 2.07670 | Female | | | | | | | | | | | | |
| MECP2 | | | | 0.00001 | 1.22860 | Female | 0.00000 | 1.27923 | Female | | | | | | |
| HSPBAP1 | | | | | | | 0.00001 | -1.16714 | Male | | | | | | |
| HOOK1 | | | | | | | 0.00852 | -0.88609 | Male | | | | | | |
| CHST4 | | | | | | | | | | | | | 0.00867 | -2.20504 | Male |
| FBXO41 | | | | | | | | | | | | | 0.00019 | 1.55737 | Female |
| AGPAT5 | | | | | | | | | | 0.00055 | -0.64643 | Male | 0.00016 | -0.68619 | Male |
| PAQR3 | 0.00093 | 2.03938 | Female | | | | | | | | | | 0.00325 | -0.67381 | Male |
| ERICH1 | 0.01793 | 0.80432 | Female | | | | | | | | | | | | |
| HDAC6 | 0.01118 | 1.81244 | Female | | | | | | | | | | | | |
| CNBD2 | | | | | | | 0.02429 | -1.75708 | Male | | | | | | |
| AGER | | | | | | | | | | | | | 0.00085 | 1.09801 | Female |
| EIF2B5 | | | | 0.03585 | 0.76065 | Female | | | | | | | | | |
| H2AFY2 | | | | | | | | | | 0.00083 | 1.54408 | Female | | | |
| DHX32 | | | | | | | | | | | | | 0.01959 | -0.66214 | Male |
| BCKDK | | | | 0.00006 | 1.38532 | Female | | | | | | | | | |
| ANKRD33B | | | | 0.00000 | 1.62935 | Female | | | | | | | 0.00616 | 0.68889 | Female |
| ENGASE | | | | 0.02431 | 1.97659 | Female | | | | | | | | | |
| FAM3C | | | | | | | | | | | | | 0.00001 | -0.67638 | Male |
| LOC101927027 | 0.01426 | -2.37809 | Male | | | | | | | | | | | | |
| CDR2L | | | | 0.01863 | 0.98070 | Female | 0.00527 | 0.75547 | Female | | | | 0.00432 | 0.73039 | Female |
| PLXNB1 | | | | | | | 0.00000 | -1.99598 | Male | 0.02525 | -0.75063 | Male | | | |
| NEK3 | | | | | | | | | | 0.01062 | 0.91625 | Female | | | |
| CACNG8 | | | | | | | 0.00004 | 3.28541 | Female | | | | | | |
| PDGFRA | | | | | | | | | | 0.00000 | 2.26425 | Female | | | |

| | | | | | | | | | | | | | | | |
|---|---|---|---|---|---|---|---|---|---|---|---|---|---|---|---|
| C16orf13 | | | | 0.00110 | 0.68738 | Female | | | | | | | | | |
| CXorf56 | 0.00231 | 0.92038 | Female | 0.00002 | 0.75220 | Female | 0.00000 | 1.03998 | Female | 0.00000 | 0.79820 | Female | 0.00000 | 0.81735 | Female |
| TEC | | | | | | | | | | 0.00233 | 1.24546 | Female | | | |
| SNAR-G2 | | | | | | | 0.00081 | 0.88937 | Female | 0.00174 | 0.82654 | Female | | | |
| GRM4 | | | | | | | | | | 0.00285 | 2.01632 | Female | 0.02443 | 1.55866 | Female |
| POTEE | | | | | | | 0.02703 | 1.05219 | Female | | | | 0.00776 | 1.12956 | Female |
| ARID3A | | | | | | | | | | 0.00000 | -0.87383 | Male | | | |
| KHDC3L | | | | | | | | | | 0.00008 | 1.84105 | Female | | | |
| C1orf131 | 0.00264 | -0.85617 | Male | | | | | | | | | | | | |
| SFXN5 | | | | | | | | | | 0.01628 | 0.68758 | Female | 0.00732 | 0.74301 | Female |
| REPS1 | 0.03312 | -0.96759 | Male | | | | | | | | | | | | |
| PTENP1 | | | | | | | 0.00000 | -1.07651 | Male | | | | | | |
| TSPAN1 | | | | 0.00004 | 3.01803 | Female | | | | | | | 0.00544 | 1.44046 | Female |
| ITGAM | | | | | | | | | | 0.00176 | 1.76084 | Female | | | |
| DSCC1 | | | | | | | 0.00012 | -0.62571 | Male | | | | | | |
| MIXL1 | | | | | | | | | | 0.00075 | 1.54964 | Female | 0.00000 | -2.00224 | Male |
| LAMC2 | | | | 0.00099 | 2.22496 | Female | | | | | | | | | |
| RPPH1 | | | | | | | 0.00000 | 1.51070 | Female | | | | | | |
| CDC14A | 0.03849 | 2.33813 | Female | | | | | | | | | | | | |
| ZC2HC1C | | | | | | | | | | | | | 0.01320 | 1.34108 | Female |
| DDAH1 | | | | 0.02311 | 2.34875 | Female | | | | | | | | | |
| CHI3L2 | 0.00169 | -1.80209 | Male | | | | | | | 0.00008 | 0.91998 | Female | 0.01246 | 0.62002 | Female |
| GHDC | | | | | | | 0.01203 | -1.56293 | Male | | | | | | |
| ERRFI1 | | | | 0.03431 | -0.67849 | Male | | | | | | | | | |
| HLA-DRB1 | | | | | | | | | | 0.00000 | -2.92991 | Male | | | |
| RNF10 | 0.02831 | -0.65012 | Male | | | | | | | | | | | | |
| SNTB2 | | | | | | | | | | 0.00027 | -0.69750 | Male | | | |
| FOXO3B | | | | | | | | | | 0.00065 | -1.17103 | Male | | | |
| PHTF1 | | | | | | | 0.00235 | -0.86002 | Male | 0.00906 | -0.71359 | Male | | | |
| UBALD1 | | | | 0.00792 | 0.96238 | Female | | | | | | | | | |
| LYPLAL1 | | | | | | | | | | | | | 0.00219 | -0.58322 | Male |
| FGFR2 | | | | 0.00006 | 1.52132 | Female | | | | | | | | | |
| SORT1 | | | | 0.00000 | 1.80142 | Female | | | | | | | | | |
| BAG2 | | | | | | | | | | 0.00000 | -1.74154 | Male | 0.00000 | -1.71801 | Male |
| BZRAP1-AS1 | | | | | | | | | | 0.03012 | -1.04593 | Male | 0.00016 | 1.60836 | Female |
| PPP1R14B | 0.00094 | -1.22374 | Male | | | | | | | | | | | | |
| PHF8 | | | | | | | | | | 0.00055 | 0.68521 | Female | | | |
| C5orf51 | | | | | | | 0.00000 | -0.82055 | Male | 0.00000 | -0.76684 | Male | | | |
| UBQLN2 | | | | | | | 0.00166 | 0.76689 | Female | | | | | | |
| PLEKHA5 | | | | | | | | | | 0.00046 | 0.71463 | Female | | | |
| SYDE2 | | | | 0.00000 | 3.41512 | Female | | | | | | | | | |
| SLC25A22 | | | | 0.00000 | 2.24070 | Female | | | | | | | | | |
| EPN1 | | | | 0.00846 | 0.88208 | Female | | | | | | | | | |
| LLGL2 | 0.02453 | -0.93804 | Male | | | | | | | | | | 0.00000 | 0.72078 | Female |
| SLC2A11 | | | | 0.04673 | -1.05316 | Male | | | | | | | 0.02371 | 0.73888 | Female |
| LOC101928259 | | | | | | | 0.03586 | 1.31679 | Female | | | | | | |
| LRBA | | | | 0.04505 | -0.79895 | Male | | | | | | | | | |
| CASC4 | | | | | | | 0.00039 | -1.08461 | Male | | | | | | |
| CD24 | | | | 0.00045 | 1.35062 | Female | 0.00009 | -0.97824 | Male | 0.00001 | -1.01100 | Male | | | |
| PPIC | | | | | | | 0.00050 | -0.67756 | Male | | | | | | |
| PPP1R15A | | | | | | | | | | | | | 0.00002 | 0.73525 | Female |

| | | | | | | | | | | | | | | | |
|---|---|---|---|---|---|---|---|---|---|---|---|---|---|---|---|
| AFMID | | | | 0.01514 | -0.67770 | Male | | | | | | | | | |
| ANKLE1 | | | | | | | | | | 0.00912 | -1.10977 | Male | | | |
| CRYZ | | | | | | | | | | | | | 0.00014 | -0.83613 | Male |
| ADCK1 | 0.00056 | -2.42060 | Male | | | | | | | | | | | | |
| FAM131A | | | | | | | 0.03965 | -1.23628 | Male | | | | | | |
| STAG3 | | | | | | | | | | | | | 0.00000 | 1.14501 | Female |
| SERAC1 | | | | 0.00831 | 0.99880 | Female | | | | | | | | | |
| CAV1 | 0.00330 | 1.94240 | Female | 0.00394 | -2.40086 | Male | 0.00764 | -1.70848 | Male | | | | 0.00089 | -1.89598 | Male |
| CHAC1 | | | | 0.02448 | -1.41494 | Male | 0.00443 | 1.18156 | Female | 0.00282 | 1.17288 | Female | | | |
| MRPL15 | 0.00652 | 0.60234 | Female | | | | | | | | | | | | |
| LINC00926 | | | | | | | | | | 0.03710 | -2.09112 | Male | | | |
| MIB1 | | | | | | | | | | | | | 0.02638 | -0.60951 | Male |
| ST3GAL2 | | | | | | | 0.00032 | 0.80012 | Female | | | | | | |
| LEPR | | | | | | | | | | 0.01721 | -1.22863 | Male | | | |
| IGF2BP1 | 0.00094 | -0.62781 | Male | | | | | | | | | | | | |
| GLDC | | | | 0.00840 | 0.84666 | Female | | | | | | | | | |
| NR2F6 | | | | 0.01279 | 0.73836 | Female | | | | | | | | | |
| METTL23 | | | | | | | 0.00239 | -0.87009 | Male | | | | | | |
| NRP1 | | | | | | | | | | 0.00021 | -1.71493 | Male | | | |
| MECOM | | | | | | | | | | 0.00000 | 2.16827 | Female | | | |
| DHODH | | | | | | | | | | 0.00984 | 0.62084 | Female | | | |
| GLS2 | | | | | | | | | | | | | 0.02127 | 1.71438 | Female |
| ABHD5 | | | | | | | 0.00045 | -0.59153 | Male | | | | | | |
| PI4KAP1 | | | | | | | 0.00015 | 0.75262 | Female | | | | | | |
| THRSP | | | | | | | | | | | | | 0.01449 | 1.66047 | Female |
| TSR3 | 0.00678 | -1.62186 | Male | | | | | | | | | | | | |
| EXOC6 | | | | | | | 0.00126 | -0.91727 | Male | | | | | | |
| RAB30-AS1 | | | | | | | | | | | | | 0.00228 | 0.59813 | Female |
| CSK | | | | | | | 0.00196 | -0.59370 | Male | | | | | | |
| MTSS1L | | | | | | | | | | 0.00124 | -1.68521 | Male | | | |
| TEX37 | | | | 0.04848 | -1.30848 | Male | 0.00170 | 1.31131 | Female | | | | | | |
| ANGPT4 | | | | | | | | | | | | | 0.03770 | 2.84185 | Female |
| PLEKHA2 | | | | | | | | | | | | | 0.00004 | 2.59894 | Female |
| TCERG1L | | | | 0.01840 | 1.36093 | Female | | | | 0.00382 | 1.01939 | Female | | | |
| KLHL22 | | | | | | | | | | 0.01110 | -1.21360 | Male | | | |
| CYB5R2 | | | | | | | 0.02229 | -0.89780 | Male | | | | 0.02410 | -0.82284 | Male |
| KRT19 | 0.00051 | -1.93054 | Male | | | | | | | | | | | | |
| LINC01091 | | | | | | | 0.02250 | -1.16557 | Male | | | | | | |
| LOC113230 | | | | | | | | | | | | | 0.00489 | 2.34202 | Female |
| SNORD79 | 0.04434 | 1.08099 | Female | | | | | | | | | | | | |
| MORC2-AS1 | | | | | | | 0.00074 | 0.79461 | Female | | | | | | |
| GLCE | | | | | | | 0.00403 | -0.89733 | Male | | | | | | |
| ZBTB25 | 0.02841 | 2.47100 | Female | | | | | | | | | | | | |
| TBC1D16 | | | | | | | | | | 0.04890 | 1.06358 | Female | | | |
| ZNF93 | | | | | | | 0.00499 | -0.58945 | Male | | | | | | |
| BTF3P11 | | | | | | | | | | 0.00018 | -1.44780 | Male | | | |
| ANKS6 | | | | | | | | | | | | | 0.02715 | 1.07614 | Female |
| ENTPD1-AS1 | | | | | | | | | | | | | 0.03756 | -1.07820 | Male |
| NKD2 | | | | 0.00649 | 1.01437 | Female | | | | | | | | | |
| LCMT2 | | | | | | | | | | 0.02417 | -0.77984 | Male | | | |
| MYCL | | | | | | | | | | 0.00000 | 1.48776 | Female | | | |

| | | | | | | | | | | | | | | | |
|---|---|---|---|---|---|---|---|---|---|---|---|---|---|---|---|
| AMMECR1 | 0.00395 | 1.83305 | Female | 0.01654 | 0.91538 | Female | | | | | | | | | |
| LMNB1 | | | | 0.00405 | -0.68403 | Male | | | | 0.00000 | -0.81594 | Male | | | |
| PRRT2 | | | | | | | | | | 0.00175 | -0.64519 | Male | | | |
| AKNA | | | | | | | | | | 0.04349 | 1.15980 | Female | | | |
| LRRC45 | | | | | | | 0.00480 | -1.06690 | Male | | | | | | |
| IPMK | 0.02921 | -1.40996 | Male | | | | | | | | | | | | |
| CLDN15 | | | | | | | | | | 0.02098 | -0.94688 | Male | | | |
| LOC100129935 | | | | | | | | | | 0.00003 | -2.10395 | Male | 0.04418 | 1.18359 | Female |
| KPNA7 | | | | | | | 0.00037 | 2.93562 | Female | | | | | | |
| LZTS2 | | | | | | | 0.00083 | -1.28074 | Male | | | | | | |
| LINC00974 | | | | 0.01150 | -2.46589 | Male | 0.04823 | 1.57947 | Female | | | | | | |
| UBE2J1 | | | | | | | 0.02035 | -0.69689 | Male | 0.00059 | -0.88874 | Male | | | |
| ZNF214 | | | | | | | 0.04380 | -1.23650 | Male | | | | | | |
| BUB1B | | | | 0.00224 | -1.01749 | Male | 0.00030 | -0.80636 | Male | 0.00000 | 2.21089 | Female | | | |
| ZNF814 | | | | | | | 0.00000 | 0.65144 | Female | | | | | | |
| TLN2 | | | | | | | | | | 0.01245 | 0.65350 | Female | | | |
| DGAT1 | | | | | | | | | | | | | 0.04555 | 0.67426 | Female |
| CPB2-AS1 | | | | | | | | | | | | | 0.00581 | 2.09276 | Female |
| MRPL3 | 0.00003 | -0.79278 | Male | | | | | | | | | | | | |
| SNORD116-26 | | | | | | | | | | 0.01645 | 1.03805 | Female | | | |
| ABCG2 | | | | 0.00001 | 1.75523 | Female | | | | | | | | | |
| PRR7 | | | | 0.03870 | 2.17174 | Female | | | | | | | | | |
| FUNDC1 | 0.00042 | 1.14693 | Female | 0.00010 | 0.77021 | Female | 0.00001 | 0.58960 | Female | 0.00000 | 1.13317 | Female | | | |
| PORCN | | | | 0.00866 | 1.77000 | Female | | | | 0.00000 | 2.04665 | Female | | | |
| RABL6 | | | | 0.00033 | 1.19023 | Female | | | | | | | | | |
| MRPS11 | 0.00000 | -1.14426 | Male | | | | | | | | | | | | |
| GNA11 | | | | | | | 0.00000 | 0.95753 | Female | | | | | | |
| LRP8 | | | | | | | 0.00034 | -1.56818 | Male | | | | | | |
| TBC1D23 | | | | | | | | | | 0.00002 | 0.59000 | Female | | | |
| HPDL | 0.00000 | 2.74396 | Female | | | | | | | | | | | | |
| GUCA1B | | | | | | | | | | | | | 0.02905 | 1.43431 | Female |
| DPH1 | 0.01713 | 1.94778 | Female | | | | 0.01919 | -0.77147 | Male | | | | | | |
| CREBBP | | | | | | | 0.00020 | 0.60773 | Female | 0.00011 | 0.60099 | Female | | | |
| TRIM43B | | | | | | | | | | 0.01857 | 4.27432 | Female | | | |
| RAB33B | | | | 0.00146 | -1.09897 | Male | | | | | | | | | |
| LINC00847 | | | | | | | | | | 0.04821 | -1.98458 | Male | | | |
| FAHD1 | | | | 0.00609 | 1.01515 | Female | | | | | | | 0.00336 | 0.68741 | Female |
| PPP2CA | | | | | | | 0.00000 | -0.68237 | Male | | | | | | |
| ZNF826P | | | | | | | | | | | | | 0.00142 | 0.65851 | Female |
| SMG1P7 | | | | | | | | | | 0.00010 | 1.20441 | Female | | | |
| ARSB | | | | | | | 0.02375 | -0.97462 | Male | | | | | | |
| EBLN1 | | | | | | | 0.03268 | 3.05477 | Female | | | | | | |
| SLC19A2 | | | | | | | 0.01438 | -0.67386 | Male | 0.00064 | -0.84098 | Male | 0.00262 | -0.74634 | Male |
| CORO1A | | | | | | | | | | 0.00000 | 1.77142 | Female | | | |
| LINC00339 | | | | | | | | | | 0.01838 | 0.91620 | Female | | | |
| LYZ | | | | | | | | | | 0.00000 | 2.99910 | Female | | | |
| ATP1B1 | | | | | | | | | | 0.00000 | 1.49558 | Female | | | |
| NOTCH1 | | | | | | | | | | | | | 0.00650 | 1.59701 | Female |
| ECHDC1 | | | | | | | 0.00083 | -0.73346 | Male | 0.00000 | -1.46478 | Male | 0.00001 | -0.85410 | Male |
| ZNF669 | | | | | | | 0.04573 | -0.81170 | Male | | | | | | |
| SLC35A4 | | | | | | | | | | 0.00000 | -0.72113 | Male | | | |

| | | | | | | | | | | | | | | | |
|---|---|---|---|---|---|---|---|---|---|---|---|---|---|---|---|
| KIF12 | | | | | | | | | | 0.00008 | 2.31752 | Female | | | |
| LRSAM1 | | | | | | | | | | 0.04106 | -0.65659 | Male | | | |
| H3F3B | | | | | | | 0.00000 | 0.63400 | Female | | | | | | |
| ANKRD36B | | | | | | | 0.00006 | 1.27830 | Female | 0.00527 | 0.89069 | Female | | | |
| EEF1A1 | | | | | | | 0.00000 | -0.68029 | Male | | | | | | |
| ZNF687 | | | | | | | 0.00000 | -1.74665 | Male | | | | | | |
| WDR81 | | | | | | | | | | 0.03568 | -1.06330 | Male | | | |
| TPI1P2 | | | | | | | 0.00000 | 0.59399 | Female | | | | | | |
| FBXO28 | 0.01002 | -0.92994 | Male | | | | | | | | | | | | |
| SMARCA1 | | | | | | | 0.02112 | -1.84922 | Male | | | | | | |
| VWA8 | | | | | | | | | | 0.02163 | 0.68157 | Female | | | |
| TCIRG1 | 0.04508 | -1.68826 | Male | | | | | | | | | | | | |
| SMCO4 | | | | | | | | | | 0.02094 | 0.96039 | Female | | | |
| RBMX2 | | | | 0.00002 | 0.75087 | Female | 0.00000 | 0.73784 | Female | | | | | | |
| SNORD45A | | | | | | | | | | | | | 0.02252 | -0.81773 | Male |
| ATG4A | | | | | | | | | | 0.01438 | 0.64788 | Female | | | |
| CIDECP | | | | | | | | | | | | | 0.00270 | 0.67195 | Female |

### Supplementary Table S4

#### Mouse male GSEA of NMF clusters

| **Cluster 1** | | | | | | |
|---|---|---|---|---|---|---|
| pathway | pval | padj | log2err | ES | NES | size |
| GO_ACTIVATION_OF_IMMUNE_RESPONSE | 0.00000 | 0.00000 | 0.78819 | 0.62851 | 1.39255 | 212 |
| GO_ACTIVATION_OF_INNATE_IMMUNE_RESPONSE | 0.00000 | 0.00000 | | 0.75054 | 1.62130 | 88 |
| GO_AEROBIC_RESPIRATION | 0.00001 | 0.00025 | 0.59333 | 0.69302 | 1.48168 | 65 |
| GO_ALTERNATIVE_MRNA_SPLICING_VIA_SPLICEOSOME | 0.00102 | 0.01782 | 0.45506 | 0.65168 | 1.38687 | 60 |
| GO_AMINE_METABOLIC_PROCESS | 0.00000 | 0.00000 | | 0.79189 | 1.70783 | 80 |
| GO_ANAPHASE_PROMOTING_COMPLEX_DEPENDENT_CATABOLIC_P | 0.00000 | 0.00000 | | 0.76611 | 1.65070 | 79 |
| GO_ANCHORING_JUNCTION | 0.00000 | 0.00000 | | 0.59615 | 1.33633 | 479 |
| GO_ANIMAL_ORGAN_REGENERATION | 0.00257 | 0.03716 | 0.43171 | 0.68240 | 1.41020 | 33 |
| GO_ANTIGEN_PROCESSING_AND_PRESENTATION | 0.00000 | 0.00000 | 0.70498 | 0.64776 | 1.42241 | 143 |
| GO_ANTIGEN_PROCESSING_AND_PRESENTATION_OF_EXOGENOUS_ | 0.00000 | 0.00000 | | 0.81051 | 1.72358 | 57 |
| GO_ANTIGEN_PROCESSING_AND_PRESENTATION_OF_PEPTIDE_ANTI | 0.00000 | 0.00000 | 0.70498 | 0.65912 | 1.44248 | 126 |
| GO_ANTIGEN_PROCESSING_AND_PRESENTATION_OF_PEPTIDE_ANTI | 0.00000 | 0.00000 | | 0.76925 | 1.65015 | 70 |
| GO_ANTIGEN_RECEPTOR_MEDIATED_SIGNALING_PATHWAY | 0.00000 | 0.00000 | 0.80122 | 0.67877 | 1.48641 | 127 |
| GO_APOPTOTIC_SIGNALING_PATHWAY | 0.00154 | 0.02464 | 0.45506 | 0.51561 | 1.15085 | 358 |
| GO_ATP_DEPENDENT_CHROMATIN_REMODELING | 0.00253 | 0.03681 | 0.43171 | 0.62124 | 1.32400 | 61 |
| GO_ATP_METABOLIC_PROCESS | 0.00000 | 0.00000 | 0.71951 | 0.62252 | 1.37693 | 196 |
| GO_ATP_SYNTHESIS_COUPLED_ELECTRON_TRANSPORT | 0.00005 | 0.00131 | 0.55733 | 0.65519 | 1.40723 | 74 |
| GO_BINDING_OF_SPERM_TO_ZONA_PELLUCIDA | 0.00000 | 0.00001 | 0.69013 | 0.89879 | 1.80846 | 18 |
| GO_CADHERIN_BINDING | 0.00000 | 0.00000 | | 0.63068 | 1.39961 | 236 |
| GO_CANONICAL_WNT_SIGNALING_PATHWAY | 0.00000 | 0.00003 | 0.65944 | 0.60816 | 1.34553 | 197 |
| GO_CATALYTIC_STEP_2_SPLICEOSOME | 0.00000 | 0.00002 | 0.65944 | 0.68068 | 1.46976 | 83 |
| GO_CELLULAR_AMINO_ACID_METABOLIC_PROCESS | 0.00000 | 0.00000 | 0.74774 | 0.61752 | 1.36824 | 216 |
| GO_CELLULAR_COMPONENT_DISASSEMBLY | 0.00000 | 0.00005 | 0.64355 | 0.55505 | 1.24197 | 401 |
| GO_CELLULAR_KETONE_METABOLIC_PROCESS | 0.00000 | 0.00000 | 0.77494 | 0.65955 | 1.44893 | 149 |
| GO_CELLULAR_NITROGEN_COMPOUND_CATABOLIC_PROCESS | 0.00000 | 0.00000 | | 0.74079 | 1.65848 | 436 |
| GO_CELLULAR_PROTEIN_COMPLEX_DISASSEMBLY | 0.00006 | 0.00139 | 0.55733 | 0.58222 | 1.28459 | 176 |
| GO_CELLULAR_RESPIRATION | 0.00000 | 0.00000 | 0.71951 | 0.64997 | 1.42492 | 136 |
| GO_CELLULAR_RESPONSE_TO_HEAT | 0.00018 | 0.00403 | 0.51885 | 0.61962 | 1.34016 | 91 |
| GO_CELLULAR_RESPONSE_TO_ORGANIC_CYCLIC_COMPOUND | 0.00302 | 0.04108 | 0.43171 | 0.52402 | 1.16585 | 283 |
| GO_CELLULAR_RESPONSE_TO_OXYGEN_LEVELS | 0.00000 | 0.00000 | 0.78819 | 0.65911 | 1.44882 | 157 |
| GO_CELL_ACTIVATION_INVOLVED_IN_IMMUNE_RESPONSE | 0.00000 | 0.00001 | 0.67496 | 0.57339 | 1.27913 | 341 |
| GO_CELL_ADHESION_MOLECULE_BINDING | 0.00000 | 0.00000 | 0.81404 | 0.60529 | 1.34788 | 308 |
| GO_CELL_CELL_RECOGNITION | 0.00000 | 0.00002 | 0.65944 | 0.83826 | 1.71003 | 26 |
| GO_CELL_CELL_SIGNALING_BY_WNT | 0.00000 | 0.00002 | 0.67496 | 0.57736 | 1.28559 | 299 |
| GO_CELL_CYCLE_G2_M_PHASE_TRANSITION | 0.00001 | 0.00024 | 0.59333 | 0.58034 | 1.28602 | 214 |
| GO_CELL_CYCLE_PHASE_TRANSITION | 0.00000 | 0.00004 | 0.64355 | 0.54319 | 1.21681 | 464 |
| GO_CELL_DIVISION | 0.00154 | 0.02464 | 0.45506 | 0.51551 | 1.15425 | 427 |
| GO_CELL_RECOGNITION | 0.00012 | 0.00270 | 0.53843 | 0.69554 | 1.46186 | 48 |
| GO_CELL_SUBSTRATE_JUNCTION | 0.00000 | 0.00000 | | 0.67751 | 1.50775 | 291 |
| GO_CELL_SURFACE_RECEPTOR_SIGNALING_PATHWAY_INVOLVED_IN | 0.00000 | 0.00003 | 0.65944 | 0.57035 | 1.27067 | 319 |
| GO_CHAPERONE_BINDING | 0.00007 | 0.00163 | 0.53843 | 0.64754 | 1.39253 | 76 |
| GO_CHAPERONE_COMPLEX | 0.00000 | 0.00004 | 0.64355 | 0.85718 | 1.73925 | 22 |
| GO_CHAPERONE_MEDIATED_PROTEIN_COMPLEX_ASSEMBLY | 0.00170 | 0.02662 | 0.45506 | 0.77423 | 1.54792 | 17 |
| GO_CHROMATIN_BINDING | 0.00071 | 0.01323 | 0.47727 | 0.52699 | 1.17680 | 369 |
| GO_CHROMOSOMAL_REGION | 0.00246 | 0.03601 | 0.43171 | 0.52919 | 1.17661 | 276 |
| GO_COP9_SIGNALOSOME | 0.00128 | 0.02141 | 0.45506 | 0.73616 | 1.49715 | 24 |
| GO_COTRANSLATIONAL_PROTEIN_TARGETING_TO_MEMBRANE | 0.00000 | 0.00000 | | 0.89471 | 1.93513 | 91 |
| GO_CYTOCHROME_COMPLEX | 0.00168 | 0.02641 | 0.45506 | 0.75311 | 1.52417 | 20 |
| GO_CYTOKINE_MEDIATED_SIGNALING_PATHWAY | 0.00000 | 0.00000 | 0.81404 | 0.59698 | 1.33170 | 339 |
| GO_CYTOPLASMIC_TRANSLATION | 0.00000 | 0.00000 | | 0.84753 | 1.83080 | 88 |
| GO_CYTOPLASMIC_TRANSLATIONAL_INITIATION | 0.00000 | 0.00000 | 0.74774 | 0.85688 | 1.76176 | 30 |
| GO_CYTOSOLIC_LARGE_RIBOSOMAL_SUBUNIT | 0.00000 | 0.00000 | | 0.91186 | 1.92988 | 53 |
| GO_CYTOSOLIC_RIBOSOME | 0.00000 | 0.00000 | | 0.89907 | 1.94710 | 95 |
| GO_CYTOSOLIC_SMALL_RIBOSOMAL_SUBUNIT | 0.00000 | 0.00000 | | 0.89114 | 1.85533 | 39 |
| GO_DEFENSE_RESPONSE_TO_OTHER_ORGANISM | 0.00004 | 0.00099 | 0.55733 | 0.53673 | 1.20127 | 415 |
| GO_DE_NOVO_PROTEIN_FOLDING | 0.00050 | 0.00978 | 0.47727 | 0.74232 | 1.51431 | 26 |
| GO_DISORDERED_DOMAIN_SPECIFIC_BINDING | 0.00161 | 0.02546 | 0.45506 | 0.71970 | 1.47197 | 27 |
| GO_DNA_BIOSYNTHETIC_PROCESS | 0.00000 | 0.00010 | 0.62726 | 0.62505 | 1.37017 | 134 |
| GO_DNA_CONFORMATION_CHANGE | 0.00027 | 0.00572 | 0.49849 | 0.55300 | 1.22527 | 216 |
| GO_DNA_DAMAGE_RESPONSE_DETECTION_OF_DNA_DAMAGE | 0.00198 | 0.03028 | 0.43171 | 0.67949 | 1.40669 | 35 |
| GO_DNA_GEOMETRIC_CHANGE | 0.00046 | 0.00925 | 0.49849 | 0.60807 | 1.31855 | 98 |
| GO_DNA_POLYMERASE_BINDING | 0.00268 | 0.03799 | 0.43171 | 0.76255 | 1.52457 | 17 |
| GO_DNA_REPAIR | 0.00265 | 0.03775 | 0.43171 | 0.51159 | 1.14547 | 427 |
| GO_DRUG_BINDING | 0.00176 | 0.02733 | 0.45506 | 0.64119 | 1.35096 | 49 |
| GO_ELECTRON_TRANSFER_ACTIVITY | 0.00001 | 0.00020 | 0.61053 | 0.67481 | 1.44938 | 74 |
| GO_ELECTRON_TRANSPORT_CHAIN | 0.00002 | 0.00053 | 0.57561 | 0.62257 | 1.36145 | 124 |
| GO_ENDOPEPTIDASE_ACTIVITY | 0.00124 | 0.02085 | 0.45506 | 0.54955 | 1.21076 | 171 |
| GO_ENDOPEPTIDASE_COMPLEX | 0.00000 | 0.00000 | | 0.80930 | 1.73234 | 66 |
| GO_ENERGY_DERIVATION_BY_OXIDATION_OF_ORGANIC_COMPOUN | 0.00000 | 0.00000 | 0.71951 | 0.63390 | 1.39707 | 172 |
| GO_ESTABLISHMENT_OF_PROTEIN_LOCALIZATION_TO_CHROMOSON | 0.00020 | 0.00432 | 0.51885 | 0.77339 | 1.57217 | 23 |
| GO_ESTABLISHMENT_OF_PROTEIN_LOCALIZATION_TO_ENDOPLASM | 0.00000 | 0.00000 | | 0.88026 | 1.91149 | 102 |
| GO_ESTABLISHMENT_OF_PROTEIN_LOCALIZATION_TO_MEMBRANE | 0.00000 | 0.00000 | | 0.74874 | 1.66164 | 240 |
| GO_ESTABLISHMENT_OF_PROTEIN_LOCALIZATION_TO_ORGANELLE | 0.00000 | 0.00000 | | 0.69519 | 1.55605 | 423 |

| | | | | | | |
|---|---|---|---|---|---|---|
| GO_ESTABLISHMENT_OF_PROTEIN_LOCALIZATION_TO_TELOMERE | 0.00006 | 0.00139 | 0.55733 | 0.83470 | 1.66881 | 17 |
| GO_ESTABLISHMENT_OF_RNA_LOCALIZATION | 0.00106 | 0.01819 | 0.45506 | 0.55577 | 1.22400 | 167 |
| GO_ESTABLISHMENT_OF_TISSUE_POLARITY | 0.00000 | 0.00000 | | 0.75051 | 1.62122 | 88 |
| GO_EUKARYOTIC_TRANSLATION_INITIATION_FACTOR_3_COMPLEX | 0.00003 | 0.00074 | 0.57561 | 0.86987 | 1.72555 | 15 |
| GO_EXOCYTOSIS | 0.00000 | 0.00015 | 0.61053 | 0.53905 | 1.20753 | 465 |
| GO_FC_EPSILON_RECEPTOR_SIGNALING_PATHWAY | 0.00000 | 0.00000 | | 0.75117 | 1.62103 | 82 |
| GO_FC_RECEPTOR_SIGNALING_PATHWAY | 0.00000 | 0.00000 | | 0.72054 | 1.57787 | 127 |
| GO_FERTILIZATION | 0.00089 | 0.01592 | 0.47727 | 0.63620 | 1.35888 | 64 |
| GO_FICOLIN_1_RICH_GRANULE | 0.00000 | 0.00000 | 0.78819 | 0.71716 | 1.55340 | 93 |
| GO_GENERATION_OF_PRECURSOR_METABOLITES_AND_ENERGY | 0.00000 | 0.00000 | 0.77494 | 0.59261 | 1.32026 | 320 |
| GO_GENE_SILENCING | 0.00261 | 0.03750 | 0.43171 | 0.55578 | 1.22252 | 161 |
| GO_GLAND_DEVELOPMENT | 0.00154 | 0.02464 | 0.45506 | 0.54117 | 1.19797 | 206 |
| GO_GLYCOSYL_COMPOUND_METABOLIC_PROCESS | 0.00071 | 0.01323 | 0.47727 | 0.62524 | 1.34204 | 71 |
| GO_GUANYL_NUCLEOTIDE_BINDING | 0.00095 | 0.01679 | 0.47727 | 0.54096 | 1.19861 | 216 |
| GO_HEAT_SHOCK_PROTEIN_BINDING | 0.00003 | 0.00092 | 0.55733 | 0.64369 | 1.39056 | 85 |
| GO_HEMATOPOIETIC_PROGENITOR_CELL_DIFFERENTIATION | 0.00000 | 0.00000 | 0.82666 | 0.70307 | 1.53239 | 113 |
| GO_HEMATOPOIETIC_STEM_CELL_DIFFERENTIATION | 0.00000 | 0.00000 | | 0.79776 | 1.71217 | 72 |
| GO_HSP70_PROTEIN_BINDING | 0.00228 | 0.03391 | 0.43171 | 0.71143 | 1.45504 | 27 |
| GO_IMMUNE_RESPONSE_REGULATING_SIGNALING_PATHWAY | 0.00000 | 0.00000 | 0.76146 | 0.63494 | 1.40480 | 197 |
| GO_INNATE_IMMUNE_RESPONSE | 0.00003 | 0.00071 | 0.57561 | 0.55173 | 1.23107 | 347 |
| GO_INNATE_IMMUNE_RESPONSE_ACTIVATING_SIGNAL_TRANSDUCT | 0.00000 | 0.00000 | | 0.79340 | 1.70281 | 72 |
| GO_INNER_MITOCHONDRIAL_MEMBRANE_PROTEIN_COMPLEX | 0.00002 | 0.00060 | 0.57561 | 0.65124 | 1.40678 | 88 |
| GO_INTERACTION_WITH_SYMBIONT | 0.00001 | 0.00026 | 0.59333 | 0.75084 | 1.56620 | 41 |
| GO_INTERLEUKIN_1_MEDIATED_SIGNALING_PATHWAY | 0.00000 | 0.00000 | | 0.81689 | 1.75671 | 76 |
| GO_INTRINSIC_APOPTOTIC_SIGNALING_PATHWAY | 0.00121 | 0.02029 | 0.45506 | 0.55118 | 1.21845 | 190 |
| GO_ION_CHANNEL_BINDING | 0.00087 | 0.01584 | 0.47727 | 0.63182 | 1.34683 | 62 |
| GO_ISOPEPTIDASE_ACTIVITY | 0.00139 | 0.02285 | 0.45506 | 0.78940 | 1.57097 | 16 |
| GO_LARGE_RIBOSOMAL_SUBUNIT | 0.00000 | 0.00000 | | 0.78135 | 1.70183 | 109 |
| GO_LEUKOCYTE_MEDIATED_IMMUNITY | 0.00000 | 0.00000 | 0.71951 | 0.57282 | 1.27895 | 364 |
| GO_MACROMOLECULE_TRANSMEMBRANE_TRANSPORTER_ACTIVITY | 0.00239 | 0.03510 | 0.43171 | 0.77734 | 1.54696 | 16 |
| GO_MAINTENANCE_OF_CELL_NUMBER | 0.00087 | 0.01584 | 0.47727 | 0.58944 | 1.28512 | 114 |
| GO_MATURATION_OF_LSU_RRNA | 0.00006 | 0.00151 | 0.53843 | 0.77606 | 1.58723 | 27 |
| GO_METHYLTRANSFERASE_COMPLEX | 0.00290 | 0.04014 | 0.43171 | 0.59476 | 1.28479 | 88 |
| GO_METHYL_CPG_BINDING | 0.00302 | 0.04108 | 0.43171 | 0.78053 | 1.54833 | 15 |
| GO_MIDBRAIN_DEVELOPMENT | 0.00021 | 0.00468 | 0.51885 | 0.68832 | 1.44457 | 47 |
| GO_MITOCHONDRIAL_ELECTRON_TRANSPORT_NADH_TO_UBIQUINO | 0.00331 | 0.04437 | 0.43171 | 0.65428 | 1.36724 | 42 |
| GO_MITOCHONDRIAL_GENE_EXPRESSION | 0.00048 | 0.00949 | 0.49849 | 0.57608 | 1.26525 | 152 |
| GO_MITOCHONDRIAL_LARGE_RIBOSOMAL_SUBUNIT | 0.00002 | 0.00050 | 0.57561 | 0.69789 | 1.48086 | 55 |
| GO_MITOCHONDRIAL_MATRIX | 0.00000 | 0.00002 | 0.65944 | 0.56045 | 1.25158 | 371 |
| GO_MITOCHONDRIAL_MEMBRANE_ORGANIZATION | 0.00361 | 0.04734 | 0.43171 | 0.59309 | 1.28094 | 86 |
| GO_MITOCHONDRIAL_PROTEIN_COMPLEX | 0.00000 | 0.00000 | 0.81404 | 0.63650 | 1.40931 | 208 |
| GO_MITOCHONDRIAL_TRANSLATION | 0.00004 | 0.00093 | 0.55733 | 0.61219 | 1.34135 | 129 |
| GO_MITOCHONDRIAL_TRANSLATIONAL_TERMINATION | 0.00000 | 0.00010 | 0.62726 | 0.66694 | 1.44128 | 87 |
| GO_MITOCHONDRION_ORGANIZATION | 0.00051 | 0.00993 | 0.47727 | 0.52284 | 1.16863 | 390 |
| GO_MODIFICATION_DEPENDENT_MACROMOLECULE_CATABOLIC_PR | 0.00002 | 0.00053 | 0.57561 | 0.53142 | 1.19156 | 486 |
| GO_MODULATION_BY_HOST_OF_SYMBIONT_PROCESS | 0.00015 | 0.00336 | 0.51885 | 0.72848 | 1.50542 | 33 |
| GO_MODULATION_OF_PROCESS_OF_OTHER_ORGANISM | 0.00019 | 0.00423 | 0.51885 | 0.66904 | 1.42286 | 59 |
| GO_MONOSACCHARIDE_BIOSYNTHETIC_PROCESS | 0.00335 | 0.04466 | 0.43171 | 0.61992 | 1.31781 | 58 |
| GO_MORPHOGENESIS_OF_AN_EPITHELIUM | 0.00006 | 0.00151 | 0.53843 | 0.55161 | 1.22755 | 290 |
| GO_MORPHOGENESIS_OF_A_POLARIZED_EPITHELIUM | 0.00000 | 0.00000 | | 0.72298 | 1.57057 | 103 |
| GO_MRNA_3_UTR_BINDING | 0.00003 | 0.00089 | 0.55733 | 0.67596 | 1.44827 | 68 |
| GO_MRNA_5_UTR_BINDING | 0.00006 | 0.00156 | 0.53843 | 0.80133 | 1.62592 | 22 |
| GO_MRNA_BINDING | 0.00000 | 0.00000 | | 0.66002 | 1.46353 | 222 |
| GO_MRNA_PROCESSING | 0.00000 | 0.00000 | 0.73376 | 0.56859 | 1.27295 | 418 |
| GO_MULTI_ORGANISM_LOCALIZATION | 0.00268 | 0.03799 | 0.43171 | 0.63245 | 1.33612 | 51 |
| GO_MULTI_ORGANISM_PROCESS | 0.00108 | 0.01839 | 0.45506 | 0.51114 | 1.14599 | 484 |
| GO_MYD88_DEPENDENT_TOLL_LIKE_RECEPTOR_SIGNALING_PATHW | 0.00002 | 0.00066 | 0.57561 | 0.85581 | 1.70312 | 16 |
| GO_MYELOID_LEUKOCYTE_ACTIVATION | 0.00000 | 0.00000 | 0.74774 | 0.58508 | 1.30479 | 327 |
| GO_MYELOID_LEUKOCYTE_MEDIATED_IMMUNITY | 0.00000 | 0.00000 | 0.76146 | 0.59841 | 1.33291 | 301 |
| GO_NADH_DEHYDROGENASE_ACTIVITY | 0.00383 | 0.04935 | 0.43171 | 0.66244 | 1.37139 | 35 |
| GO_NADH_DEHYDROGENASE_COMPLEX | 0.00376 | 0.04884 | 0.43171 | 0.65649 | 1.36421 | 38 |
| GO_NADH_METABOLIC_PROCESS | 0.00294 | 0.04036 | 0.43171 | 0.70642 | 1.44481 | 27 |
| GO_NCRNA_EXPORT_FROM_NUCLEUS | 0.00302 | 0.04108 | 0.43171 | 0.66820 | 1.38331 | 35 |
| GO_NCRNA_METABOLIC_PROCESS | 0.00000 | 0.00003 | 0.65944 | 0.55907 | 1.24907 | 381 |
| GO_NCRNA_PROCESSING | 0.00000 | 0.00003 | 0.65944 | 0.57175 | 1.27504 | 328 |
| GO_NEGATIVE_REGULATION_OF_CANONICAL_WNT_SIGNALING_PAT | 0.00000 | 0.00000 | | 0.71611 | 1.56038 | 111 |
| GO_NEGATIVE_REGULATION_OF_CELLULAR_AMIDE_METABOLIC_PRO | 0.00001 | 0.00020 | 0.61053 | 0.63031 | 1.37423 | 114 |
| GO_NEGATIVE_REGULATION_OF_CELL_CYCLE | 0.00003 | 0.00076 | 0.57561 | 0.53833 | 1.20513 | 425 |
| GO_NEGATIVE_REGULATION_OF_CELL_CYCLE_G2_M_PHASE_TRANS | 0.00000 | 0.00000 | 0.76146 | 0.71490 | 1.54439 | 85 |
| GO_NEGATIVE_REGULATION_OF_CELL_CYCLE_PHASE_TRANSITION | 0.00002 | 0.00060 | 0.57561 | 0.59041 | 1.30442 | 184 |
| GO_NEGATIVE_REGULATION_OF_CELL_CYCLE_PROCESS | 0.00000 | 0.00002 | 0.65944 | 0.58690 | 1.30374 | 249 |
| GO_NEGATIVE_REGULATION_OF_MITOTIC_CELL_CYCLE | 0.00004 | 0.00095 | 0.55733 | 0.56735 | 1.25902 | 232 |
| GO_NEGATIVE_REGULATION_OF_MRNA_METABOLIC_PROCESS | 0.00023 | 0.00491 | 0.51885 | 0.66670 | 1.41788 | 59 |
| GO_NEGATIVE_REGULATION_OF_MRNA_SPLICING_VIA_SPLICEOSOM | 0.00239 | 0.03510 | 0.43171 | 0.74857 | 1.51204 | 19 |
| GO_NEGATIVE_REGULATION_OF_RNA_SPLICING | 0.00029 | 0.00607 | 0.49849 | 0.76729 | 1.55976 | 23 |
| GO_NEGATIVE_REGULATION_OF_TRANSLATIONAL_INITIATION | 0.00018 | 0.00403 | 0.51885 | 0.83901 | 1.66434 | 15 |

| | | | | | | |
|---|---|---|---|---|---|---|
| GO_NEGATIVE_REGULATION_OF_WNT_SIGNALING_PATHWAY | 0.00000 | 0.00000 | | 0.69525 | 1.52249 | 127 |
| GO_NEURAL_NUCLEUS_DEVELOPMENT | 0.00001 | 0.00026 | 0.59333 | 0.79148 | 1.62187 | 28 |
| GO_NEURON_TO_NEURON_SYNAPSE | 0.00005 | 0.00134 | 0.55733 | 0.59636 | 1.31006 | 154 |
| GO_NIK_NF_KAPPAB_SIGNALING | 0.00000 | 0.00000 | | 0.72896 | 1.58354 | 103 |
| GO_NON_CANONICAL_WNT_SIGNALING_PATHWAY | 0.00000 | 0.00000 | | 0.72784 | 1.58023 | 101 |
| GO_NUCLEAR_EXPORT | 0.00031 | 0.00645 | 0.49849 | 0.56963 | 1.25542 | 172 |
| GO_NUCLEAR_TRANSCRIBED_MRNA_CATABOLIC_PROCESS | 0.00000 | 0.00000 | | 0.79179 | 1.74917 | 183 |
| GO_NUCLEAR_TRANSCRIBED_MRNA_CATABOLIC_PROCESS_NONSEN | 0.00000 | 0.00000 | | 0.89008 | 1.93909 | 110 |
| GO_NUCLEAR_TRANSPORT | 0.00008 | 0.00193 | 0.53843 | 0.55511 | 1.23408 | 273 |
| GO_NUCLEOBASE_CONTAINING_COMPOUND_TRANSPORT | 0.00146 | 0.02386 | 0.45506 | 0.55053 | 1.21842 | 202 |
| GO_NUCLEOBASE_CONTAINING_SMALL_MOLECULE_BIOSYNTHETIC_ | 0.00137 | 0.02261 | 0.45506 | 0.62749 | 1.34353 | 67 |
| GO_NUCLEOBASE_CONTAINING_SMALL_MOLECULE_INTERCONVERS | 0.00383 | 0.04935 | 0.43171 | 0.71259 | 1.44857 | 23 |
| GO_NUCLEOBASE_CONTAINING_SMALL_MOLECULE_METABOLIC_PR | 0.00320 | 0.04339 | 0.43171 | 0.51448 | 1.14825 | 352 |
| GO_NUCLEOSIDE_DIPHOSPHATE_METABOLIC_PROCESS | 0.00110 | 0.01865 | 0.45506 | 0.59637 | 1.29295 | 97 |
| GO_NUCLEOSIDE_METABOLIC_PROCESS | 0.00061 | 0.01169 | 0.47727 | 0.65315 | 1.38907 | 59 |
| GO_NUCLEOSIDE_MONOPHOSPHATE_BIOSYNTHETIC_PROCESS | 0.00250 | 0.03636 | 0.43171 | 0.68841 | 1.42033 | 32 |
| GO_NUCLEOSIDE_MONOPHOSPHATE_METABOLIC_PROCESS | 0.00152 | 0.02464 | 0.45506 | 0.64377 | 1.36002 | 51 |
| GO_NUCLEOSIDE_PHOSPHATE_BIOSYNTHETIC_PROCESS | 0.00043 | 0.00865 | 0.49849 | 0.58185 | 1.27831 | 148 |
| GO_NUCLEOSIDE_TRIPHOSPHATE_BIOSYNTHETIC_PROCESS | 0.00043 | 0.00865 | 0.49849 | 0.69332 | 1.44539 | 40 |
| GO_NUCLEOSIDE_TRIPHOSPHATE_METABOLIC_PROCESS | 0.00016 | 0.00360 | 0.51885 | 0.66788 | 1.41976 | 58 |
| GO_NUCLEOSOMAL_DNA_BINDING | 0.00023 | 0.00507 | 0.51885 | 0.79852 | 1.61801 | 21 |
| GO_NUCLEOSOME_BINDING | 0.00143 | 0.02332 | 0.45506 | 0.66132 | 1.38492 | 45 |
| GO_NUCLEOTIDE_EXCISION_REPAIR | 0.00004 | 0.00096 | 0.55733 | 0.63343 | 1.37355 | 98 |
| GO_NUCLEOTIDE_EXCISION_REPAIR_DNA_DAMAGE_RECOGNITION | 0.00015 | 0.00349 | 0.51885 | 0.78678 | 1.59640 | 22 |
| GO_NUCLEOTIDE_EXCISION_REPAIR_DNA_INCISION | 0.00313 | 0.04249 | 0.43171 | 0.66439 | 1.37898 | 36 |
| GO_NUCLEOTIDE_PHOSPHORYLATION | 0.00383 | 0.04935 | 0.43171 | 0.59002 | 1.27454 | 88 |
| GO_ORGANELLAR_RIBOSOME | 0.00000 | 0.00014 | 0.61053 | 0.66412 | 1.43471 | 85 |
| GO_ORGANELLE_INNER_MEMBRANE | 0.00000 | 0.00000 | 0.76146 | 0.57601 | 1.28697 | 378 |
| GO_ORGANIC_CYCLIC_COMPOUND_CATABOLIC_PROCESS | 0.00000 | 0.00000 | | 0.73350 | 1.64281 | 449 |
| GO_OXIDATIVE_PHOSPHORYLATION | 0.00001 | 0.00030 | 0.59333 | 0.64619 | 1.40027 | 96 |
| GO_OXIDOREDUCTASE_ACTIVITY | 0.00006 | 0.00139 | 0.55733 | 0.53544 | 1.19703 | 392 |
| GO_OXIDOREDUCTASE_COMPLEX | 0.00001 | 0.00027 | 0.59333 | 0.66015 | 1.42461 | 82 |
| GO_PEPTIDASE_COMPLEX | 0.00000 | 0.00000 | | 0.75988 | 1.63982 | 82 |
| GO_PIGMENT_GRANULE | 0.00003 | 0.00089 | 0.55733 | 0.66098 | 1.42344 | 78 |
| GO_POLYSOMAL_RIBOSOME | 0.00000 | 0.00000 | | 0.90624 | 1.86629 | 31 |
| GO_POLYSOME | 0.00000 | 0.00000 | | 0.82337 | 1.75481 | 61 |
| GO_POLY_PYRIMIDINE_TRACT_BINDING | 0.00327 | 0.04398 | 0.43171 | 0.75471 | 1.51855 | 18 |
| GO_POSITIVE_REGULATION_OF_APOPTOTIC_SIGNALING_PATHWAY | 0.00279 | 0.03889 | 0.43171 | 0.57666 | 1.25518 | 107 |
| GO_POSITIVE_REGULATION_OF_ATPASE_ACTIVITY | 0.00294 | 0.04036 | 0.43171 | 0.65446 | 1.36795 | 43 |
| GO_POSITIVE_REGULATION_OF_BINDING | 0.00088 | 0.01590 | 0.47727 | 0.60284 | 1.30556 | 95 |
| GO_POSITIVE_REGULATION_OF_CANONICAL_WNT_SIGNALING_PATH | 0.00000 | 0.00000 | 0.77494 | 0.70504 | 1.53160 | 103 |
| GO_POSITIVE_REGULATION_OF_CELLULAR_AMIDE_METABOLIC_PRO | 0.00003 | 0.00074 | 0.57561 | 0.62296 | 1.36083 | 118 |
| GO_POSITIVE_REGULATION_OF_CELLULAR_PROTEIN_LOCALIZATION | 0.00041 | 0.00840 | 0.49849 | 0.54915 | 1.21815 | 225 |
| GO_POSITIVE_REGULATION_OF_CELL_DEATH | 0.00069 | 0.01297 | 0.47727 | 0.52595 | 1.17423 | 363 |
| GO_POSITIVE_REGULATION_OF_CELL_POPULATION_PROLIFERATION | 0.00078 | 0.01455 | 0.47727 | 0.52449 | 1.17108 | 368 |
| GO_POSITIVE_REGULATION_OF_CHROMOSOME_ORGANIZATION | 0.00231 | 0.03437 | 0.43171 | 0.56990 | 1.24871 | 133 |
| GO_POSITIVE_REGULATION_OF_DEFENSE_RESPONSE | 0.00000 | 0.00003 | 0.65944 | 0.62175 | 1.36730 | 158 |
| GO_POSITIVE_REGULATION_OF_DNA_BIOSYNTHETIC_PROCESS | 0.00001 | 0.00041 | 0.59333 | 0.71929 | 1.52055 | 52 |
| GO_POSITIVE_REGULATION_OF_DNA_METABOLIC_PROCESS | 0.00364 | 0.04761 | 0.43171 | 0.56647 | 1.24189 | 135 |
| GO_POSITIVE_REGULATION_OF_DNA_TEMPLATED_TRANSCRIPTION_ | 0.00276 | 0.03885 | 0.43171 | 0.68932 | 1.41586 | 29 |
| GO_POSITIVE_REGULATION_OF_ESTABLISHMENT_OF_PROTEIN_LOCA | 0.00007 | 0.00168 | 0.53843 | 0.56461 | 1.25165 | 219 |
| GO_POSITIVE_REGULATION_OF_IMMUNE_RESPONSE | 0.00000 | 0.00001 | 0.69013 | 0.59037 | 1.31265 | 277 |
| GO_POSITIVE_REGULATION_OF_IMMUNE_SYSTEM_PROCESS | 0.00004 | 0.00110 | 0.55733 | 0.54018 | 1.20716 | 386 |
| GO_POSITIVE_REGULATION_OF_INNATE_IMMUNE_RESPONSE | 0.00000 | 0.00000 | 0.81404 | 0.69693 | 1.51947 | 114 |
| GO_POSITIVE_REGULATION_OF_INTRINSIC_APOPTOTIC_SIGNALING_ | 0.00104 | 0.01798 | 0.45506 | 0.69097 | 1.43578 | 37 |
| GO_POSITIVE_REGULATION_OF_ORGANELLE_ORGANIZATION | 0.00043 | 0.00865 | 0.49849 | 0.52093 | 1.16605 | 411 |
| GO_POSITIVE_REGULATION_OF_PROTEIN_LOCALIZATION_TO_NUCLE | 0.00001 | 0.00020 | 0.61053 | 0.69907 | 1.48990 | 61 |
| GO_POSITIVE_REGULATION_OF_RESPONSE_TO_BIOTIC_STIMULUS | 0.00000 | 0.00000 | 0.73376 | 0.66340 | 1.45310 | 128 |
| GO_POSITIVE_REGULATION_OF_RESPONSE_TO_EXTERNAL_STIMULU | 0.00002 | 0.00050 | 0.57561 | 0.58133 | 1.28681 | 201 |
| GO_POSITIVE_REGULATION_OF_TELOMERASE_ACTIVITY | 0.00061 | 0.01169 | 0.47727 | 0.73476 | 1.50565 | 28 |
| GO_POSITIVE_REGULATION_OF_TELOMERASE_RNA_LOCALIZATION_ | 0.00000 | 0.00001 | 0.69013 | 0.92451 | 1.83395 | 15 |
| GO_POSITIVE_REGULATION_OF_TELOMERE_MAINTENANCE_VIA_TE | 0.00130 | 0.02159 | 0.45506 | 0.71259 | 1.46508 | 30 |
| GO_POSITIVE_REGULATION_OF_TRANSCRIPTION_INITIATION_FROM_ | 0.00372 | 0.04847 | 0.43171 | 0.70671 | 1.43993 | 25 |
| GO_POSITIVE_REGULATION_OF_TRANSFERASE_ACTIVITY | 0.00174 | 0.02712 | 0.45506 | 0.51642 | 1.15220 | 345 |
| GO_POSITIVE_REGULATION_OF_TRANSLATION | 0.00001 | 0.00019 | 0.61053 | 0.64767 | 1.40419 | 97 |
| GO_POSITIVE_REGULATION_OF_VIRAL_GENOME_REPLICATION | 0.00287 | 0.03973 | 0.43171 | 0.68362 | 1.40783 | 31 |
| GO_POSITIVE_REGULATION_OF_VIRAL_PROCESS | 0.00028 | 0.00590 | 0.49849 | 0.62440 | 1.34741 | 81 |
| GO_POSITIVE_REGULATION_OF_WNT_SIGNALING_PATHWAY | 0.00000 | 0.00000 | 0.73376 | 0.66230 | 1.44763 | 122 |
| GO_POSTREPLICATION_REPAIR | 0.00024 | 0.00515 | 0.51885 | 0.69140 | 1.44518 | 43 |
| GO_POSTSYNAPSE | 0.00004 | 0.00109 | 0.55733 | 0.55969 | 1.24369 | 259 |
| GO_POSTSYNAPTIC_CYTOSOL | 0.00279 | 0.03889 | 0.43171 | 0.78295 | 1.55313 | 15 |
| GO_POSTTRANSCRIPTIONAL_REGULATION_OF_GENE_EXPRESSION | 0.00000 | 0.00000 | | 0.66656 | 1.49327 | 462 |
| GO_POST_TRANSLATIONAL_PROTEIN_MODIFICATION | 0.00002 | 0.00052 | 0.57561 | 0.57851 | 1.28412 | 229 |
| GO_PRECATALYTIC_SPLICEOSOME | 0.00085 | 0.01559 | 0.47727 | 0.65235 | 1.37448 | 49 |
| GO_PRERIBOSOME | 0.00087 | 0.01577 | 0.47727 | 0.62899 | 1.34772 | 69 |
| GO_PRERIBOSOME_LARGE_SUBUNIT_PRECURSOR | 0.00235 | 0.03474 | 0.43171 | 0.74079 | 1.50308 | 22 |

| | | | | | | |
|---|---|---|---|---|---|---|
| GO_PRE_MRNA_BINDING | 0.00133 | 0.02213 | 0.45506 | 0.72393 | 1.48062 | 27 |
| GO_PROTEASOMAL_PROTEIN_CATABOLIC_PROCESS | 0.00002 | 0.00044 | 0.59333 | 0.54166 | 1.21005 | 377 |
| GO_PROTEASOMAL_UBIQUITIN_INDEPENDENT_PROTEIN_CATABOLI( | 0.00000 | 0.00000 | 0.80122 | 0.92254 | 1.86931 | 21 |
| GO_PROTEASOME_ACCESSORY_COMPLEX | 0.00000 | 0.00001 | 0.67496 | 0.85464 | 1.73811 | 24 |
| GO_PROTEASOME_CORE_COMPLEX | 0.00000 | 0.00000 | | 0.94375 | 1.90631 | 19 |
| GO_PROTEIN_CONTAINING_COMPLEX_DISASSEMBLY | 0.00000 | 0.00007 | 0.62726 | 0.57810 | 1.28438 | 256 |
| GO_PROTEIN_CONTAINING_COMPLEX_LOCALIZATION | 0.00063 | 0.01197 | 0.47727 | 0.55335 | 1.22520 | 208 |
| GO_PROTEIN_DNA_COMPLEX_SUBUNIT_ORGANIZATION | 0.00001 | 0.00018 | 0.61053 | 0.60174 | 1.32491 | 165 |
| GO_PROTEIN_FOLDING | 0.00000 | 0.00002 | 0.65944 | 0.63113 | 1.38644 | 154 |
| GO_PROTEIN_FOLDING_CHAPERONE | 0.00135 | 0.02238 | 0.45506 | 0.77903 | 1.55752 | 17 |
| GO_PROTEIN_IMPORT | 0.00268 | 0.03799 | 0.43171 | 0.56317 | 1.23581 | 140 |
| GO_PROTEIN_LOCALIZATION_TO_CHROMOSOME | 0.00056 | 0.01096 | 0.47727 | 0.63523 | 1.36108 | 69 |
| GO_PROTEIN_LOCALIZATION_TO_CHROMOSOME_TELOMERIC_REGI( | 0.00335 | 0.04466 | 0.43171 | 0.70399 | 1.43984 | 27 |
| GO_PROTEIN_LOCALIZATION_TO_ENDOPLASMIC_RETICULUM | 0.00000 | 0.00000 | | 0.85521 | 1.87021 | 124 |
| GO_PROTEIN_LOCALIZATION_TO_MEMBRANE | 0.00000 | 0.00000 | | 0.65123 | 1.45714 | 406 |
| GO_PROTEIN_LOCALIZATION_TO_MITOCHONDRION | 0.00019 | 0.00420 | 0.51885 | 0.60773 | 1.32398 | 110 |
| GO_PROTEIN_LOCALIZATION_TO_NUCLEUS | 0.00002 | 0.00055 | 0.57561 | 0.58926 | 1.30220 | 187 |
| GO_PROTEIN_MODIFICATION_BY_SMALL_PROTEIN_REMOVAL | 0.00000 | 0.00000 | 0.71951 | 0.60745 | 1.34689 | 221 |
| GO_PROTEIN_POLYUBIQUITINATION | 0.00004 | 0.00106 | 0.55733 | 0.55874 | 1.24108 | 253 |
| GO_PROTEIN_REFOLDING | 0.00294 | 0.04036 | 0.43171 | 0.77039 | 1.53314 | 16 |
| GO_PROTEIN_STABILIZATION | 0.00001 | 0.00024 | 0.61053 | 0.62333 | 1.36586 | 130 |
| GO_PROTEIN_TARGETING | 0.00000 | 0.00000 | | 0.71181 | 1.58743 | 327 |
| GO_PROTEIN_TARGETING_TO_MEMBRANE | 0.00000 | 0.00000 | | 0.82370 | 1.80953 | 149 |
| GO_PROTEIN_TARGETING_TO_MITOCHONDRION | 0.00346 | 0.04593 | 0.43171 | 0.60408 | 1.30091 | 78 |
| GO_PROTEIN_TRANSMEMBRANE_TRANSPORT | 0.00265 | 0.03775 | 0.43171 | 0.64528 | 1.36171 | 50 |
| GO_PROTON_TRANSMEMBRANE_TRANSPORT | 0.00213 | 0.03229 | 0.43171 | 0.60815 | 1.30536 | 71 |
| GO_PURINE_CONTAINING_COMPOUND_BIOSYNTHETIC_PROCESS | 0.00246 | 0.03601 | 0.43171 | 0.57796 | 1.25936 | 111 |
| GO_PURINE_CONTAINING_COMPOUND_METABOLIC_PROCESS | 0.00100 | 0.01761 | 0.45506 | 0.53679 | 1.19242 | 255 |
| GO_PURINE_NUCLEOSIDE_TRIPHOSPHATE_METABOLIC_PROCESS | 0.00089 | 0.01592 | 0.47727 | 0.67157 | 1.40374 | 43 |
| GO_PYRIMIDINE_CONTAINING_COMPOUND_METABOLIC_PROCESS | 0.00353 | 0.04658 | 0.43171 | 0.64744 | 1.35878 | 47 |
| GO_REGENERATION | 0.00216 | 0.03242 | 0.43171 | 0.59987 | 1.29453 | 82 |
| GO_REGULATION_OF_ANIMAL_ORGAN_MORPHOGENESIS | 0.00000 | 0.00000 | | 0.69163 | 1.51541 | 133 |
| GO_REGULATION_OF_APOPTOTIC_SIGNALING_PATHWAY | 0.00031 | 0.00645 | 0.49849 | 0.55171 | 1.22506 | 247 |
| GO_REGULATION_OF_ATPASE_ACTIVITY | 0.00324 | 0.04369 | 0.43171 | 0.63031 | 1.34038 | 57 |
| GO_REGULATION_OF_ATP_METABOLIC_PROCESS | 0.00353 | 0.04658 | 0.43171 | 0.60176 | 1.29780 | 80 |
| GO_REGULATION_OF_BINDING | 0.00250 | 0.03636 | 0.43171 | 0.54476 | 1.20598 | 203 |
| GO_REGULATION_OF_CELLULAR_AMIDE_METABOLIC_PROCESS | 0.00000 | 0.00000 | | 0.62967 | 1.40226 | 311 |
| GO_REGULATION_OF_CELLULAR_AMINE_METABOLIC_PROCESS | 0.00000 | 0.00000 | | 0.82173 | 1.74682 | 58 |
| GO_REGULATION_OF_CELLULAR_AMINO_ACID_METABOLIC_PROCES | 0.00000 | 0.00000 | | 0.83776 | 1.77766 | 55 |
| GO_REGULATION_OF_CELLULAR_KETONE_METABOLIC_PROCESS | 0.00000 | 0.00000 | | 0.72641 | 1.58680 | 118 |
| GO_REGULATION_OF_CELLULAR_PROTEIN_LOCALIZATION | 0.00082 | 0.01513 | 0.47727 | 0.52542 | 1.17290 | 362 |
| GO_REGULATION_OF_CELLULAR_RESPONSE_TO_HEAT | 0.00276 | 0.03885 | 0.43171 | 0.61652 | 1.32004 | 67 |
| GO_REGULATION_OF_CELLULAR_RESPONSE_TO_STRESS | 0.00224 | 0.03344 | 0.43171 | 0.50545 | 1.13232 | 468 |
| GO_REGULATION_OF_CELL_CYCLE_G2_M_PHASE_TRANSITION | 0.00000 | 0.00000 | 0.71951 | 0.63230 | 1.39282 | 169 |
| GO_REGULATION_OF_CELL_CYCLE_PHASE_TRANSITION | 0.00001 | 0.00026 | 0.59333 | 0.55488 | 1.23783 | 337 |
| GO_REGULATION_OF_CHROMOSOME_ORGANIZATION | 0.00081 | 0.01501 | 0.47727 | 0.54255 | 1.20527 | 261 |
| GO_REGULATION_OF_CYSTEINE_TYPE_ENDOPEPTIDASE_ACTIVITY | 0.00202 | 0.03077 | 0.43171 | 0.57120 | 1.24837 | 121 |
| GO_REGULATION_OF_CYTOPLASMIC_TRANSLATION | 0.00130 | 0.02159 | 0.45506 | 0.73678 | 1.49773 | 23 |
| GO_REGULATION_OF_DEFENSE_RESPONSE | 0.00065 | 0.01237 | 0.47727 | 0.53633 | 1.19355 | 290 |
| GO_REGULATION_OF_DNA_BIOSYNTHETIC_PROCESS | 0.00016 | 0.00360 | 0.51885 | 0.64215 | 1.37923 | 74 |
| GO_REGULATION_OF_DNA_METABOLIC_PROCESS | 0.00069 | 0.01297 | 0.47727 | 0.54425 | 1.20775 | 237 |
| GO_REGULATION_OF_DNA_TEMPLATED_TRANSCRIPTION_INITIATIOI | 0.00163 | 0.02568 | 0.45506 | 0.68289 | 1.41372 | 35 |
| GO_REGULATION_OF_DNA_TEMPLATED_TRANSCRIPTION_IN_RESPO | 0.00000 | 0.00000 | | 0.73944 | 1.59800 | 89 |
| GO_REGULATION_OF_GENERATION_OF_PRECURSOR_METABOLITES_ | 0.00060 | 0.01164 | 0.47727 | 0.60476 | 1.31116 | 97 |
| GO_REGULATION_OF_HEMATOPOIETIC_PROGENITOR_CELL_DIFFERE | 0.00000 | 0.00000 | | 0.78717 | 1.69251 | 75 |
| GO_REGULATION_OF_HEMATOPOIETIC_STEM_CELL_DIFFERENTIATIC | 0.00000 | 0.00000 | | 0.81505 | 1.74256 | 65 |
| GO_REGULATION_OF_HEMOPOIESIS | 0.00000 | 0.00008 | 0.62726 | 0.58859 | 1.30634 | 228 |
| GO_REGULATION_OF_IMMUNE_RESPONSE | 0.00000 | 0.00016 | 0.61053 | 0.55362 | 1.23583 | 362 |
| GO_REGULATION_OF_INNATE_IMMUNE_RESPONSE | 0.00000 | 0.00000 | 0.71951 | 0.63883 | 1.40521 | 161 |
| GO_REGULATION_OF_INTRINSIC_APOPTOTIC_SIGNALING_PATHWAY | 0.00001 | 0.00039 | 0.59333 | 0.63478 | 1.38259 | 109 |
| GO_REGULATION_OF_MITOTIC_CELL_CYCLE | 0.00001 | 0.00018 | 0.61053 | 0.53899 | 1.20736 | 458 |
| GO_REGULATION_OF_MORPHOGENESIS_OF_AN_EPITHELIUM | 0.00000 | 0.00000 | | 0.72213 | 1.57284 | 109 |
| GO_REGULATION_OF_MRNA_CATABOLIC_PROCESS | 0.00000 | 0.00000 | | 0.69977 | 1.54127 | 168 |
| GO_REGULATION_OF_MRNA_METABOLIC_PROCESS | 0.00000 | 0.00000 | | 0.66528 | 1.47872 | 266 |
| GO_REGULATION_OF_MRNA_PROCESSING | 0.00004 | 0.00105 | 0.55733 | 0.61521 | 1.34090 | 113 |
| GO_REGULATION_OF_MRNA_SPLICING_VIA_SPLICEOSOME | 0.00001 | 0.00026 | 0.59333 | 0.67424 | 1.45411 | 80 |
| GO_REGULATION_OF_NUCLEASE_ACTIVITY | 0.00106 | 0.01819 | 0.45506 | 0.77918 | 1.56780 | 18 |
| GO_REGULATION_OF_OXIDATIVE_STRESS_INDUCED_CELL_DEATH | 0.00198 | 0.03028 | 0.43171 | 0.67287 | 1.40355 | 41 |
| GO_REGULATION_OF_PEPTIDASE_ACTIVITY | 0.00050 | 0.00978 | 0.47727 | 0.56122 | 1.24054 | 191 |
| GO_REGULATION_OF_POSTTRANSCRIPTIONAL_GENE_SILENCING | 0.00364 | 0.04761 | 0.43171 | 0.59824 | 1.28492 | 74 |
| GO_REGULATION_OF_PROTEIN_CATABOLIC_PROCESS | 0.00179 | 0.02783 | 0.45506 | 0.53057 | 1.17929 | 270 |
| GO_REGULATION_OF_PROTEIN_LOCALIZATION_TO_NUCLEUS | 0.00001 | 0.00042 | 0.59333 | 0.65505 | 1.41558 | 87 |
| GO_REGULATION_OF_PROTEIN_MODIFICATION_BY_SMALL_PROTEIN | 0.00025 | 0.00532 | 0.49849 | 0.57068 | 1.25684 | 167 |
| GO_REGULATION_OF_PROTEIN_STABILITY | 0.00093 | 0.01652 | 0.47727 | 0.55382 | 1.22571 | 202 |
| GO_REGULATION_OF_PROTEOLYSIS | 0.00013 | 0.00301 | 0.51885 | 0.52997 | 1.18392 | 377 |
| GO_REGULATION_OF_RESPONSE_TO_BIOTIC_STIMULUS | 0.00001 | 0.00020 | 0.61053 | 0.59049 | 1.30719 | 204 |

| | | | | | | |
|---|---|---|---|---|---|---|
| GO_REGULATION_OF_RNA_SPLICING | 0.00000 | 0.00001 | 0.69013 | 0.65896 | 1.43669 | 114 |
| GO_REGULATION_OF_SMALL_MOLECULE_METABOLIC_PROCESS | 0.00000 | 0.00000 | 0.71951 | 0.59134 | 1.31394 | 265 |
| GO_REGULATION_OF_STEM_CELL_DIFFERENTIATION | 0.00000 | 0.00000 | | 0.74049 | 1.60113 | 90 |
| GO_REGULATION_OF_SYMBIOTIC_PROCESS | 0.00006 | 0.00144 | 0.55733 | 0.60071 | 1.31682 | 134 |
| GO_REGULATION_OF_TELOMERASE_ACTIVITY | 0.00088 | 0.01590 | 0.47727 | 0.69161 | 1.43179 | 35 |
| GO_REGULATION_OF_TRANSCRIPTION_FROM_RNA_POLYMERASE_II | 0.00000 | 0.00000 | | 0.81194 | 1.73800 | 66 |
| GO_REGULATION_OF_TRANSCRIPTION_INITIATION_FROM_RNA_POL | 0.00143 | 0.02332 | 0.45506 | 0.70390 | 1.44582 | 29 |
| GO_REGULATION_OF_TRANSLATIONAL_INITIATION | 0.00002 | 0.00052 | 0.57561 | 0.68129 | 1.45518 | 64 |
| GO_REGULATION_OF_UBIQUITIN_PROTEIN_TRANSFERASE_ACTIVITY | 0.00061 | 0.01169 | 0.47727 | 0.69335 | 1.44627 | 41 |
| GO_REGULATION_OF_VIRAL_GENOME_REPLICATION | 0.00086 | 0.01566 | 0.47727 | 0.65316 | 1.38389 | 54 |
| GO_REGULATION_OF_VIRAL_TRANSCRIPTION | 0.00198 | 0.03028 | 0.43171 | 0.63932 | 1.34703 | 49 |
| GO_REGULATION_OF_WNT_SIGNALING_PATHWAY | 0.00000 | 0.00001 | 0.69013 | 0.60353 | 1.33816 | 220 |
| GO_RESPIRASOME | 0.00008 | 0.00181 | 0.53843 | 0.65420 | 1.40421 | 71 |
| GO_RESPIRATORY_CHAIN_COMPLEX | 0.00016 | 0.00353 | 0.51885 | 0.67626 | 1.43918 | 60 |
| GO_RESPIRATORY_ELECTRON_TRANSPORT_CHAIN | 0.00001 | 0.00025 | 0.59333 | 0.65819 | 1.42190 | 85 |
| GO_RESPONSE_TO_ALCOHOL | 0.00261 | 0.03750 | 0.43171 | 0.58804 | 1.27350 | 95 |
| GO_RESPONSE_TO_CADMIUM_ION | 0.00265 | 0.03775 | 0.43171 | 0.67227 | 1.39174 | 35 |
| GO_RESPONSE_TO_DRUG | 0.00104 | 0.01798 | 0.45506 | 0.55364 | 1.22258 | 180 |
| GO_RESPONSE_TO_ETHANOL | 0.00086 | 0.01566 | 0.47727 | 0.67309 | 1.40691 | 43 |
| GO_RESPONSE_TO_HEAT | 0.00012 | 0.00270 | 0.53843 | 0.61665 | 1.34221 | 107 |
| GO_RESPONSE_TO_HORMONE | 0.00048 | 0.00949 | 0.49849 | 0.51277 | 1.14814 | 453 |
| GO_RESPONSE_TO_HYDROGEN_PEROXIDE | 0.00257 | 0.03716 | 0.43171 | 0.60505 | 1.30368 | 79 |
| GO_RESPONSE_TO_INORGANIC_SUBSTANCE | 0.00004 | 0.00107 | 0.55733 | 0.55714 | 1.23870 | 275 |
| GO_RESPONSE_TO_INTERLEUKIN_1 | 0.00000 | 0.00000 | | 0.74844 | 1.62908 | 107 |
| GO_RESPONSE_TO_INTERLEUKIN_12 | 0.00012 | 0.00270 | 0.53843 | 0.72605 | 1.50694 | 36 |
| GO_RESPONSE_TO_INTERLEUKIN_4 | 0.00324 | 0.04369 | 0.43171 | 0.75812 | 1.51571 | 17 |
| GO_RESPONSE_TO_METAL_ION | 0.00030 | 0.00623 | 0.49849 | 0.56387 | 1.24233 | 171 |
| GO_RESPONSE_TO_ORGANIC_CYCLIC_COMPOUND | 0.00000 | 0.00016 | 0.61053 | 0.54603 | 1.22259 | 439 |
| GO_RESPONSE_TO_OXYGEN_LEVELS | 0.00000 | 0.00000 | 0.71951 | 0.60132 | 1.33438 | 233 |
| GO_RESPONSE_TO_STEROID_HORMONE | 0.00154 | 0.02464 | 0.45506 | 0.55514 | 1.22446 | 174 |
| GO_RESPONSE_TO_TEMPERATURE_STIMULUS | 0.00027 | 0.00572 | 0.49849 | 0.59216 | 1.29675 | 127 |
| GO_RESPONSE_TO_TUMOR_NECROSIS_FACTOR | 0.00000 | 0.00000 | 0.76146 | 0.65649 | 1.44236 | 151 |
| GO_RETINA_HOMEOSTASIS | 0.00104 | 0.01798 | 0.45506 | 0.73063 | 1.49432 | 27 |
| GO_RIBONUCLEOPROTEIN_COMPLEX_BINDING | 0.00000 | 0.00000 | 0.71951 | 0.66906 | 1.46050 | 117 |
| GO_RIBONUCLEOPROTEIN_COMPLEX_BIOGENESIS | 0.00000 | 0.00000 | | 0.66339 | 1.48264 | 389 |
| GO_RIBONUCLEOPROTEIN_COMPLEX_SUBUNIT_ORGANIZATION | 0.00000 | 0.00000 | | 0.69241 | 1.52681 | 173 |
| GO_RIBONUCLEOPROTEIN_GRANULE | 0.00000 | 0.00001 | 0.67496 | 0.61979 | 1.36366 | 163 |
| GO_RIBONUCLEOSIDE_TRIPHOSPHATE_BIOSYNTHETIC_PROCESS | 0.00041 | 0.00840 | 0.49849 | 0.71253 | 1.47248 | 33 |
| GO_RIBONUCLEOSIDE_TRIPHOSPHATE_METABOLIC_PROCESS | 0.00026 | 0.00550 | 0.49849 | 0.69681 | 1.45611 | 42 |
| GO_RIBOSE_PHOSPHATE_BIOSYNTHETIC_PROCESS | 0.00191 | 0.02939 | 0.45506 | 0.57932 | 1.26207 | 110 |
| GO_RIBOSE_PHOSPHATE_METABOLIC_PROCESS | 0.00108 | 0.01839 | 0.45506 | 0.53891 | 1.19687 | 245 |
| GO_RIBOSOMAL_LARGE_SUBUNIT_ASSEMBLY | 0.00000 | 0.00016 | 0.61053 | 0.80999 | 1.65663 | 27 |
| GO_RIBOSOMAL_LARGE_SUBUNIT_BIOGENESIS | 0.00000 | 0.00000 | | 0.77542 | 1.65982 | 66 |
| GO_RIBOSOMAL_SMALL_SUBUNIT_ASSEMBLY | 0.00023 | 0.00494 | 0.51885 | 0.81139 | 1.62221 | 17 |
| GO_RIBOSOMAL_SMALL_SUBUNIT_BINDING | 0.00383 | 0.04935 | 0.43171 | 0.75288 | 1.50523 | 17 |
| GO_RIBOSOMAL_SMALL_SUBUNIT_BIOGENESIS | 0.00001 | 0.00022 | 0.61053 | 0.69076 | 1.47997 | 68 |
| GO_RIBOSOMAL_SUBUNIT | 0.00000 | 0.00000 | | 0.77603 | 1.71031 | 172 |
| GO_RIBOSOME | 0.00000 | 0.00000 | | 0.76643 | 1.69659 | 206 |
| GO_RIBOSOME_ASSEMBLY | 0.00000 | 0.00000 | 0.73376 | 0.76173 | 1.61998 | 59 |
| GO_RIBOSOME_BINDING | 0.00019 | 0.00416 | 0.51885 | 0.68955 | 1.44715 | 47 |
| GO_RIBOSOME_BIOGENESIS | 0.00000 | 0.00000 | | 0.65877 | 1.46466 | 275 |
| GO_RNA_CATABOLIC_PROCESS | 0.00000 | 0.00000 | | 0.76774 | 1.71264 | 339 |
| GO_RNA_DEPENDENT_DNA_BIOSYNTHETIC_PROCESS | 0.00003 | 0.00076 | 0.57561 | 0.68926 | 1.46521 | 58 |
| GO_RNA_EXPORT_FROM_NUCLEUS | 0.00062 | 0.01183 | 0.47727 | 0.58378 | 1.27758 | 126 |
| GO_RNA_LOCALIZATION | 0.00000 | 0.00006 | 0.64355 | 0.59670 | 1.32020 | 198 |
| GO_RNA_POLYMERASE_II_PREINITIATION_COMPLEX_ASSEMBLY | 0.00089 | 0.01592 | 0.47727 | 0.77046 | 1.55626 | 19 |
| GO_RNA_SPLICING | 0.00000 | 0.00000 | 0.78819 | 0.59079 | 1.31927 | 369 |
| GO_RNA_SPLICING_VIA_TRANSESTERIFICATION_REACTIONS | 0.00000 | 0.00000 | | 0.60940 | 1.35714 | 300 |
| GO_RNA_STABILIZATION | 0.00065 | 0.01223 | 0.47727 | 0.69743 | 1.44384 | 35 |
| GO_ROUGH_ENDOPLASMIC_RETICULUM | 0.00001 | 0.00039 | 0.59333 | 0.71853 | 1.51629 | 50 |
| GO_RRNA_BINDING | 0.00000 | 0.00001 | 0.69013 | 0.74782 | 1.58865 | 56 |
| GO_RRNA_METABOLIC_PROCESS | 0.00000 | 0.00000 | 0.77494 | 0.63047 | 1.39562 | 206 |
| GO_SCF_DEPENDENT_PROTEASOMAL_UBIQUITIN_DEPENDENT_PRO | 0.00000 | 0.00000 | | 0.77527 | 1.67044 | 79 |
| GO_SECRETORY_GRANULE | 0.00000 | 0.00000 | 0.71951 | 0.56518 | 1.26461 | 399 |
| GO_SECRETORY_VESICLE | 0.00000 | 0.00002 | 0.67496 | 0.54866 | 1.22905 | 465 |
| GO_SIGNAL_TRANSDUCTION_BY_PROTEIN_PHOSPHORYLATION | 0.00004 | 0.00109 | 0.55733 | 0.53817 | 1.20474 | 424 |
| GO_SINGLE_FERTILIZATION | 0.00048 | 0.00949 | 0.49849 | 0.67414 | 1.42260 | 50 |
| GO_SINGLE_STRANDED_RNA_BINDING | 0.00039 | 0.00793 | 0.49849 | 0.66553 | 1.41527 | 57 |
| GO_SMALL_MOLECULE_BIOSYNTHETIC_PROCESS | 0.00216 | 0.03242 | 0.43171 | 0.51195 | 1.14522 | 397 |
| GO_SMALL_RIBOSOMAL_SUBUNIT | 0.00000 | 0.00000 | | 0.77648 | 1.66209 | 66 |
| GO_SM_LIKE_PROTEIN_FAMILY_COMPLEX | 0.00327 | 0.04398 | 0.43171 | 0.60468 | 1.29713 | 70 |
| GO_SPERM_EGG_RECOGNITION | 0.00000 | 0.00007 | 0.64355 | 0.85241 | 1.72955 | 22 |
| GO_SPLICEOSOMAL_COMPLEX | 0.00000 | 0.00003 | 0.65944 | 0.60939 | 1.34261 | 171 |
| GO_STEM_CELL_DIFFERENTIATION | 0.00000 | 0.00000 | 0.78819 | 0.66886 | 1.46930 | 147 |
| GO_STRUCTURAL_CONSTITUENT_OF_CYTOSKELETON | 0.00102 | 0.01782 | 0.45506 | 0.65913 | 1.39337 | 52 |
| GO_STRUCTURAL_CONSTITUENT_OF_RIBOSOME | 0.00000 | 0.00000 | | 0.79909 | 1.75541 | 150 |

| | | | | | | |
|---|---|---|---|---|---|---|
| GO_STRUCTURAL_MOLECULE_ACTIVITY | 0.00000 | 0.00000 | | 0.72651 | 1.61895 | 324 |
| GO_SUBSTANTIA_NIGRA_DEVELOPMENT | 0.00000 | 0.00013 | 0.62726 | 0.81622 | 1.66507 | 26 |
| GO_TELOMERASE_RNA_LOCALIZATION | 0.00001 | 0.00017 | 0.61053 | 0.85164 | 1.72025 | 19 |
| GO_TELOMERE_MAINTENANCE_VIA_TELOMERE_LENGTHENING | 0.00011 | 0.00250 | 0.53843 | 0.65659 | 1.40546 | 66 |
| GO_TELOMERE_ORGANIZATION | 0.00342 | 0.04554 | 0.43171 | 0.57000 | 1.24649 | 124 |
| GO_THREONINE_TYPE_ENDOPEPTIDASE_ACTIVITY | 0.00000 | 0.00000 | | 0.95179 | 1.92255 | 19 |
| GO_THREONINE_TYPE_PEPTIDASE_ACTIVITY | 0.00000 | 0.00000 | | 0.92199 | 1.87423 | 23 |
| GO_TISSUE_MORPHOGENESIS | 0.00097 | 0.01707 | 0.47727 | 0.52661 | 1.17402 | 326 |
| GO_TOXIN_TRANSPORT | 0.00113 | 0.01922 | 0.45506 | 0.72897 | 1.48708 | 26 |
| GO_TRANSCRIPTION_COUPLED_NUCLEOTIDE_EXCISION_REPAIR | 0.00001 | 0.00019 | 0.61053 | 0.68106 | 1.46186 | 71 |
| GO_TRANSCRIPTION_FACTOR_BINDING | 0.00043 | 0.00865 | 0.49849 | 0.52447 | 1.17376 | 409 |
| GO_TRANSCRIPTION_PREINITIATION_COMPLEX_ASSEMBLY | 0.00119 | 0.02004 | 0.45506 | 0.71399 | 1.46797 | 30 |
| GO_TRANSLATIONAL_ELONGATION | 0.00000 | 0.00000 | 0.71951 | 0.66087 | 1.44451 | 122 |
| GO_TRANSLATIONAL_INITIATION | 0.00000 | 0.00000 | | 0.84691 | 1.86515 | 166 |
| GO_TRANSLATIONAL_TERMINATION | 0.00000 | 0.00005 | 0.64355 | 0.65946 | 1.43060 | 99 |
| GO_TRANSLATION_ELONGATION_FACTOR_ACTIVITY | 0.00161 | 0.02546 | 0.45506 | 0.79860 | 1.58418 | 15 |
| GO_TRANSLATION_FACTOR_ACTIVITY_RNA_BINDING | 0.00000 | 0.00000 | | 0.79008 | 1.69288 | 69 |
| GO_TRANSLATION_INITIATION_FACTOR_ACTIVITY | 0.00000 | 0.00000 | 0.76146 | 0.80422 | 1.68417 | 45 |
| GO_TRANSLATION_INITIATION_FACTOR_BINDING | 0.00021 | 0.00468 | 0.51885 | 0.76186 | 1.55231 | 25 |
| GO_TRANSLATION_PREINITIATION_COMPLEX | 0.00000 | 0.00001 | 0.67496 | 0.90245 | 1.79595 | 16 |
| GO_TRANSLATION_REGULATOR_ACTIVITY | 0.00000 | 0.00000 | | 0.77663 | 1.68904 | 105 |
| GO_TRANSLATION_REGULATOR_ACTIVITY_NUCLEIC_ACID_BINDING | 0.00000 | 0.00000 | | 0.77150 | 1.66586 | 83 |
| GO_TRICARBOXYLIC_ACID_CYCLE | 0.00154 | 0.02464 | 0.45506 | 0.71854 | 1.47240 | 28 |
| GO_TUMOR_NECROSIS_FACTOR_MEDIATED_SIGNALING_PATHWAY | 0.00000 | 0.00000 | | 0.76277 | 1.65372 | 97 |
| GO_T_CELL_RECEPTOR_SIGNALING_PATHWAY | 0.00000 | 0.00000 | 0.81404 | 0.69994 | 1.52527 | 112 |
| GO_U12_TYPE_SPLICEOSOMAL_COMPLEX | 0.00279 | 0.03889 | 0.43171 | 0.70746 | 1.44693 | 27 |
| GO_U2_SNRNP | 0.00005 | 0.00134 | 0.55733 | 0.83081 | 1.67168 | 18 |
| GO_U2_TYPE_SPLICEOSOMAL_COMPLEX | 0.00010 | 0.00243 | 0.53843 | 0.63403 | 1.36970 | 85 |
| GO_UBIQUITIN_LIKE_PROTEIN_LIGASE_BINDING | 0.00000 | 0.00000 | 0.76146 | 0.62273 | 1.38051 | 219 |
| GO_UBIQUITIN_PROTEIN_TRANSFERASE_REGULATOR_ACTIVITY | 0.00165 | 0.02590 | 0.45506 | 0.79605 | 1.57911 | 15 |
| GO_UNFOLDED_PROTEIN_BINDING | 0.00000 | 0.00000 | 0.77494 | 0.72157 | 1.55881 | 85 |
| GO_VESICLE_LUMEN | 0.00000 | 0.00000 | 0.74774 | 0.64475 | 1.42008 | 168 |
| GO_VIRAL_GENE_EXPRESSION | 0.00000 | 0.00000 | | 0.80028 | 1.76317 | 171 |
| GO_VIRAL_GENOME_REPLICATION | 0.00100 | 0.01761 | 0.45506 | 0.61997 | 1.33078 | 73 |
| GO_VIRAL_LIFE_CYCLE | 0.00031 | 0.00640 | 0.49849 | 0.55767 | 1.23509 | 209 |
| HP_ABNORMALITY_OF_ACID_BASE_HOMEOSTASIS | 0.00216 | 0.03242 | 0.43171 | 0.53556 | 1.18846 | 232 |
| HP_ABNORMALITY_OF_CARDIOVASCULAR_SYSTEM_ELECTROPHYSIO | 0.00027 | 0.00572 | 0.49849 | 0.55832 | 1.23929 | 229 |
| HP_ABNORMALITY_OF_LOWER_LIP | 0.00181 | 0.02804 | 0.45506 | 0.56360 | 1.23779 | 145 |
| HP_ABNORMALITY_OF_THE_MITOCHONDRION | 0.00213 | 0.03229 | 0.43171 | 0.57831 | 1.26240 | 117 |
| HP_ABNORMALITY_OF_THE_NASAL_DORSUM | 0.00005 | 0.00116 | 0.55733 | 0.60288 | 1.32157 | 134 |
| HP_ABNORMAL_ELECTROPHYSIOLOGY_OF_SINOATRIAL_NODE_ORIG | 0.00279 | 0.03889 | 0.43171 | 0.68436 | 1.40934 | 31 |
| HP_ABNORMAL_ERYTHROCYTE_MORPHOLOGY | 0.00000 | 0.00015 | 0.61053 | 0.56049 | 1.25036 | 341 |
| HP_ABNORMAL_MEAN_CORPUSCULAR_VOLUME | 0.00357 | 0.04696 | 0.43171 | 0.75226 | 1.51363 | 18 |
| HP_ABNORMAL_RETICULOCYTE_MORPHOLOGY | 0.00064 | 0.01210 | 0.47727 | 0.71420 | 1.47081 | 31 |
| HP_ACUTE_LEUKEMIA | 0.00000 | 0.00000 | 0.73376 | 0.72786 | 1.56137 | 70 |
| HP_ANEMIA_OF_INADEQUATE_PRODUCTION | 0.00000 | 0.00008 | 0.62726 | 0.65593 | 1.42209 | 97 |
| HP_CONSTITUTIONAL_SYMPTOM | 0.00119 | 0.02004 | 0.45506 | 0.51427 | 1.15108 | 423 |
| HP_DELAYED_PUBERTY | 0.00005 | 0.00115 | 0.55733 | 0.62579 | 1.36211 | 107 |
| HP_DEPRESSED_NASAL_RIDGE | 0.00000 | 0.00011 | 0.62726 | 0.68731 | 1.47511 | 72 |
| HP_FATIGUE | 0.00039 | 0.00808 | 0.49849 | 0.56529 | 1.24923 | 187 |
| HP_HEADACHE | 0.00015 | 0.00340 | 0.51885 | 0.59065 | 1.29716 | 146 |
| HP_HEMATOLOGICAL_NEOPLASM | 0.00003 | 0.00078 | 0.57561 | 0.60108 | 1.32044 | 154 |
| HP_INCREASED_CSF_LACTATE | 0.00157 | 0.02509 | 0.45506 | 0.61622 | 1.31904 | 66 |
| HP_INCREASED_SERUM_LACTATE | 0.00216 | 0.03242 | 0.43171 | 0.56920 | 1.24724 | 130 |
| HP_LEUKEMIA | 0.00001 | 0.00026 | 0.59333 | 0.64877 | 1.40680 | 98 |
| HP_MACROCYTIC_ANEMIA | 0.00000 | 0.00000 | 0.74774 | 0.82468 | 1.71698 | 39 |
| HP_MIGRAINE | 0.00000 | 0.00000 | 0.71951 | 0.72105 | 1.54497 | 69 |
| HP_MYOTONIA | 0.00161 | 0.02546 | 0.45506 | 0.79820 | 1.58338 | 15 |
| HP_NEOPLASM | 0.00216 | 0.03242 | 0.43171 | 0.50820 | 1.13805 | 446 |
| HP_PALLOR | 0.00001 | 0.00040 | 0.59333 | 0.65169 | 1.41044 | 92 |
| HP_PUBERTY_AND_GONADAL_DISORDERS | 0.00287 | 0.03973 | 0.43171 | 0.51995 | 1.15843 | 322 |
| HP_RETICULOCYTOPENIA | 0.00004 | 0.00105 | 0.55733 | 0.86417 | 1.71425 | 15 |
| HP_RIMMED_VACUOLES | 0.00279 | 0.03889 | 0.43171 | 0.74033 | 1.49831 | 20 |
| HP_SHORT_NOSE | 0.00235 | 0.03474 | 0.43171 | 0.54919 | 1.21134 | 174 |
| HP_THICK_LOWER_LIP_VERMILION | 0.00048 | 0.00949 | 0.49849 | 0.60773 | 1.31617 | 94 |
| HP_THICK_VERMILION_BORDER | 0.00156 | 0.02486 | 0.45506 | 0.55983 | 1.23000 | 153 |
| HP_VENTRICULAR_ARRHYTHMIA | 0.00353 | 0.04658 | 0.43171 | 0.66198 | 1.37398 | 36 |
| **Cluster 1** | | | | | | |
| pathway | pval | padj | log2err | ES | NES | size |
| GO_ACTIVATION_OF_IMMUNE_RESPONSE | 0.00000 | 0.00000 | 0.78819 | 0.62851 | 1.39255 | 212 |
| GO_ACTIVATION_OF_INNATE_IMMUNE_RESPONSE | 0.00000 | 0.00000 | | 0.75054 | 1.62130 | 88 |
| GO_AEROBIC_RESPIRATION | 0.00001 | 0.00025 | 0.59333 | 0.69302 | 1.48168 | 65 |
| GO_ALTERNATIVE_MRNA_SPLICING_VIA_SPLICEOSOME | 0.00102 | 0.01782 | 0.45506 | 0.65168 | 1.38687 | 60 |
| GO_AMINE_METABOLIC_PROCESS | 0.00000 | 0.00000 | | 0.79189 | 1.70783 | 80 |
| GO_ANAPHASE_PROMOTING_COMPLEX_DEPENDENT_CATABOLIC_P | 0.00000 | 0.00000 | | 0.76611 | 1.65070 | 79 |
| GO_ANCHORING_JUNCTION | 0.00000 | 0.00000 | | 0.59615 | 1.33633 | 479 |

| | | | | | | |
|---|---|---|---|---|---|---|
| GO_ANIMAL_ORGAN_REGENERATION | 0.00257 | 0.03716 | 0.43171 | 0.68240 | 1.41020 | 33 |
| GO_ANTIGEN_PROCESSING_AND_PRESENTATION | 0.00000 | 0.00000 | 0.70498 | 0.64776 | 1.42241 | 143 |
| GO_ANTIGEN_PROCESSING_AND_PRESENTATION_OF_EXOGENOUS_ | 0.00000 | 0.00000 | | 0.81051 | 1.72358 | 57 |
| GO_ANTIGEN_PROCESSING_AND_PRESENTATION_OF_PEPTIDE_ANTI | 0.00000 | 0.00000 | 0.70498 | 0.65912 | 1.44248 | 126 |
| GO_ANTIGEN_PROCESSING_AND_PRESENTATION_OF_PEPTIDE_ANTI | 0.00000 | 0.00000 | | 0.76925 | 1.65015 | 70 |
| GO_ANTIGEN_RECEPTOR_MEDIATED_SIGNALING_PATHWAY | 0.00000 | 0.00000 | 0.80122 | 0.67877 | 1.48641 | 127 |
| GO_APOPTOTIC_SIGNALING_PATHWAY | 0.00154 | 0.02464 | 0.45506 | 0.51561 | 1.15085 | 358 |
| GO_ATP_DEPENDENT_CHROMATIN_REMODELING | 0.00253 | 0.03681 | 0.43171 | 0.62124 | 1.32400 | 61 |
| GO_ATP_METABOLIC_PROCESS | 0.00000 | 0.00000 | 0.71951 | 0.62252 | 1.37693 | 196 |
| GO_ATP_SYNTHESIS_COUPLED_ELECTRON_TRANSPORT | 0.00005 | 0.00131 | 0.55733 | 0.65519 | 1.40723 | 74 |
| GO_BINDING_OF_SPERM_TO_ZONA_PELLUCIDA | 0.00000 | 0.00001 | 0.69013 | 0.89879 | 1.80846 | 18 |
| GO_CADHERIN_BINDING | 0.00000 | 0.00000 | | 0.63068 | 1.39961 | 236 |
| GO_CANONICAL_WNT_SIGNALING_PATHWAY | 0.00000 | 0.00003 | 0.65944 | 0.60816 | 1.34553 | 197 |
| GO_CATALYTIC_STEP_2_SPLICEOSOME | 0.00000 | 0.00002 | 0.65944 | 0.68068 | 1.46976 | 83 |
| GO_CELLULAR_AMINO_ACID_METABOLIC_PROCESS | 0.00000 | 0.00000 | 0.74774 | 0.61752 | 1.36824 | 216 |
| GO_CELLULAR_COMPONENT_DISASSEMBLY | 0.00000 | 0.00005 | 0.64355 | 0.55505 | 1.24197 | 401 |
| GO_CELLULAR_KETONE_METABOLIC_PROCESS | 0.00000 | 0.00000 | 0.77494 | 0.65955 | 1.44893 | 149 |
| GO_CELLULAR_NITROGEN_COMPOUND_CATABOLIC_PROCESS | 0.00000 | 0.00000 | | 0.74079 | 1.65848 | 436 |
| GO_CELLULAR_PROTEIN_COMPLEX_DISASSEMBLY | 0.00006 | 0.00139 | 0.55733 | 0.58222 | 1.28459 | 176 |
| GO_CELLULAR_RESPIRATION | 0.00000 | 0.00000 | 0.71951 | 0.64997 | 1.42492 | 136 |
| GO_CELLULAR_RESPONSE_TO_HEAT | 0.00018 | 0.00403 | 0.51885 | 0.61962 | 1.34016 | 91 |
| GO_CELLULAR_RESPONSE_TO_ORGANIC_CYCLIC_COMPOUND | 0.00302 | 0.04108 | 0.43171 | 0.52402 | 1.16585 | 283 |
| GO_CELLULAR_RESPONSE_TO_OXYGEN_LEVELS | 0.00000 | 0.00000 | 0.78819 | 0.65911 | 1.44882 | 157 |
| GO_CELL_ACTIVATION_INVOLVED_IN_IMMUNE_RESPONSE | 0.00000 | 0.00001 | 0.67496 | 0.57339 | 1.27913 | 341 |
| GO_CELL_ADHESION_MOLECULE_BINDING | 0.00000 | 0.00000 | 0.81404 | 0.60529 | 1.34788 | 308 |
| GO_CELL_CELL_RECOGNITION | 0.00000 | 0.00002 | 0.65944 | 0.83826 | 1.71003 | 26 |
| GO_CELL_CELL_SIGNALING_BY_WNT | 0.00000 | 0.00002 | 0.67496 | 0.57736 | 1.28559 | 299 |
| GO_CELL_CYCLE_G2_M_PHASE_TRANSITION | 0.00001 | 0.00024 | 0.59333 | 0.58034 | 1.28602 | 214 |
| GO_CELL_CYCLE_PHASE_TRANSITION | 0.00000 | 0.00004 | 0.64355 | 0.54319 | 1.21681 | 464 |
| GO_CELL_DIVISION | 0.00154 | 0.02464 | 0.45506 | 0.51551 | 1.15425 | 427 |
| GO_CELL_RECOGNITION | 0.00012 | 0.00270 | 0.53843 | 0.69554 | 1.46186 | 48 |
| GO_CELL_SUBSTRATE_JUNCTION | 0.00000 | 0.00000 | | 0.67751 | 1.50775 | 291 |
| GO_CELL_SURFACE_RECEPTOR_SIGNALING_PATHWAY_INVOLVED_IN | 0.00000 | 0.00003 | 0.65944 | 0.57035 | 1.27067 | 319 |
| GO_CHAPERONE_BINDING | 0.00007 | 0.00163 | 0.53843 | 0.64754 | 1.39253 | 76 |
| GO_CHAPERONE_COMPLEX | 0.00000 | 0.00004 | 0.64355 | 0.85718 | 1.73925 | 22 |
| GO_CHAPERONE_MEDIATED_PROTEIN_COMPLEX_ASSEMBLY | 0.00170 | 0.02662 | 0.45506 | 0.77423 | 1.54792 | 17 |
| GO_CHROMATIN_BINDING | 0.00071 | 0.01323 | 0.47727 | 0.52699 | 1.17680 | 369 |
| GO_CHROMOSOMAL_REGION | 0.00246 | 0.03601 | 0.43171 | 0.52919 | 1.17661 | 276 |
| GO_COP9_SIGNALOSOME | 0.00128 | 0.02141 | 0.45506 | 0.73616 | 1.49715 | 24 |
| GO_COTRANSLATIONAL_PROTEIN_TARGETING_TO_MEMBRANE | 0.00000 | 0.00000 | | 0.89471 | 1.93513 | 91 |
| GO_CYTOCHROME_COMPLEX | 0.00168 | 0.02641 | 0.45506 | 0.75311 | 1.52417 | 20 |
| GO_CYTOKINE_MEDIATED_SIGNALING_PATHWAY | 0.00000 | 0.00000 | 0.81404 | 0.59698 | 1.33170 | 339 |
| GO_CYTOPLASMIC_TRANSLATION | 0.00000 | 0.00000 | | 0.84753 | 1.83080 | 88 |
| GO_CYTOPLASMIC_TRANSLATIONAL_INITIATION | 0.00000 | 0.00000 | 0.74774 | 0.85688 | 1.76176 | 30 |
| GO_CYTOSOLIC_LARGE_RIBOSOMAL_SUBUNIT | 0.00000 | 0.00000 | | 0.91186 | 1.92988 | 53 |
| GO_CYTOSOLIC_RIBOSOME | 0.00000 | 0.00000 | | 0.89907 | 1.94710 | 95 |
| GO_CYTOSOLIC_SMALL_RIBOSOMAL_SUBUNIT | 0.00000 | 0.00000 | | 0.89114 | 1.85533 | 39 |
| GO_DEFENSE_RESPONSE_TO_OTHER_ORGANISM | 0.00004 | 0.00099 | 0.55733 | 0.53673 | 1.20127 | 415 |
| GO_DE_NOVO_PROTEIN_FOLDING | 0.00050 | 0.00978 | 0.47727 | 0.74232 | 1.51431 | 26 |
| GO_DISORDERED_DOMAIN_SPECIFIC_BINDING | 0.00161 | 0.02546 | 0.45506 | 0.71970 | 1.47197 | 27 |
| GO_DNA_BIOSYNTHETIC_PROCESS | 0.00000 | 0.00010 | 0.62726 | 0.62505 | 1.37017 | 134 |
| GO_DNA_CONFORMATION_CHANGE | 0.00027 | 0.00572 | 0.49849 | 0.55300 | 1.22527 | 216 |
| GO_DNA_DAMAGE_RESPONSE_DETECTION_OF_DNA_DAMAGE | 0.00198 | 0.03028 | 0.43171 | 0.67949 | 1.40669 | 35 |
| GO_DNA_GEOMETRIC_CHANGE | 0.00046 | 0.00925 | 0.49849 | 0.60807 | 1.31855 | 98 |
| GO_DNA_POLYMERASE_BINDING | 0.00268 | 0.03799 | 0.43171 | 0.76255 | 1.52457 | 17 |
| GO_DNA_REPAIR | 0.00265 | 0.03775 | 0.43171 | 0.51159 | 1.14547 | 427 |
| GO_DRUG_BINDING | 0.00176 | 0.02733 | 0.45506 | 0.64119 | 1.35096 | 49 |
| GO_ELECTRON_TRANSFER_ACTIVITY | 0.00001 | 0.00020 | 0.61053 | 0.67481 | 1.44938 | 74 |
| GO_ELECTRON_TRANSPORT_CHAIN | 0.00002 | 0.00053 | 0.57561 | 0.62257 | 1.36145 | 124 |
| GO_ENDOPEPTIDASE_ACTIVITY | 0.00124 | 0.02085 | 0.45506 | 0.54955 | 1.21076 | 171 |
| GO_ENDOPEPTIDASE_COMPLEX | 0.00000 | 0.00000 | | 0.80930 | 1.73234 | 66 |
| GO_ENERGY_DERIVATION_BY_OXIDATION_OF_ORGANIC_COMPOUN | 0.00000 | 0.00000 | 0.71951 | 0.63390 | 1.39707 | 172 |
| GO_ESTABLISHMENT_OF_PROTEIN_LOCALIZATION_TO_CHROMOSON | 0.00020 | 0.00432 | 0.51885 | 0.77339 | 1.57217 | 23 |
| GO_ESTABLISHMENT_OF_PROTEIN_LOCALIZATION_TO_ENDOPLASM | 0.00000 | 0.00000 | | 0.88026 | 1.91149 | 102 |
| GO_ESTABLISHMENT_OF_PROTEIN_LOCALIZATION_TO_MEMBRANE | 0.00000 | 0.00000 | | 0.74874 | 1.66164 | 240 |
| GO_ESTABLISHMENT_OF_PROTEIN_LOCALIZATION_TO_ORGANELLE | 0.00000 | 0.00000 | | 0.69519 | 1.55605 | 423 |
| GO_ESTABLISHMENT_OF_PROTEIN_LOCALIZATION_TO_TELOMERE | 0.00006 | 0.00139 | 0.55733 | 0.83470 | 1.66881 | 17 |
| GO_ESTABLISHMENT_OF_RNA_LOCALIZATION | 0.00106 | 0.01819 | 0.45506 | 0.55577 | 1.22400 | 167 |
| GO_ESTABLISHMENT_OF_TISSUE_POLARITY | 0.00000 | 0.00000 | | 0.75051 | 1.62122 | 88 |
| GO_EUKARYOTIC_TRANSLATION_INITIATION_FACTOR_3_COMPLEX | 0.00003 | 0.00074 | 0.57561 | 0.86987 | 1.72555 | 15 |
| GO_EXOCYTOSIS | 0.00000 | 0.00015 | 0.61053 | 0.53905 | 1.20753 | 465 |
| GO_FC_EPSILON_RECEPTOR_SIGNALING_PATHWAY | 0.00000 | 0.00000 | | 0.75117 | 1.62103 | 82 |
| GO_FC_RECEPTOR_SIGNALING_PATHWAY | 0.00000 | 0.00000 | | 0.72054 | 1.57787 | 127 |
| GO_FERTILIZATION | 0.00089 | 0.01592 | 0.47727 | 0.63620 | 1.35888 | 64 |
| GO_FICOLIN_1_RICH_GRANULE | 0.00000 | 0.00000 | 0.78819 | 0.71716 | 1.55340 | 93 |
| GO_GENERATION_OF_PRECURSOR_METABOLITES_AND_ENERGY | 0.00000 | 0.00000 | 0.77494 | 0.59261 | 1.32026 | 320 |

| | | | | | | |
|---|---|---|---|---|---|---|
| GO_GENE_SILENCING | 0.00261 | 0.03750 | 0.43171 | 0.55578 | 1.22252 | 161 |
| GO_GLAND_DEVELOPMENT | 0.00154 | 0.02464 | 0.45506 | 0.54117 | 1.19797 | 206 |
| GO_GLYCOSYL_COMPOUND_METABOLIC_PROCESS | 0.00071 | 0.01323 | 0.47727 | 0.62524 | 1.34204 | 71 |
| GO_GUANYL_NUCLEOTIDE_BINDING | 0.00095 | 0.01679 | 0.47727 | 0.54096 | 1.19861 | 216 |
| GO_HEAT_SHOCK_PROTEIN_BINDING | 0.00003 | 0.00092 | 0.55733 | 0.64369 | 1.39056 | 85 |
| GO_HEMATOPOIETIC_PROGENITOR_CELL_DIFFERENTIATION | 0.00000 | 0.00000 | 0.82666 | 0.70307 | 1.53239 | 113 |
| GO_HEMATOPOIETIC_STEM_CELL_DIFFERENTIATION | 0.00000 | 0.00000 | | 0.79776 | 1.71217 | 72 |
| GO_HSP70_PROTEIN_BINDING | 0.00228 | 0.03391 | 0.43171 | 0.71143 | 1.45504 | 27 |
| GO_IMMUNE_RESPONSE_REGULATING_SIGNALING_PATHWAY | 0.00000 | 0.00000 | 0.76146 | 0.63494 | 1.40480 | 197 |
| GO_INNATE_IMMUNE_RESPONSE | 0.00003 | 0.00071 | 0.57561 | 0.55173 | 1.23107 | 347 |
| GO_INNATE_IMMUNE_RESPONSE_ACTIVATING_SIGNAL_TRANSDUCT | 0.00000 | 0.00000 | | 0.79340 | 1.70281 | 72 |
| GO_INNER_MITOCHONDRIAL_MEMBRANE_PROTEIN_COMPLEX | 0.00002 | 0.00060 | 0.57561 | 0.65124 | 1.40678 | 88 |
| GO_INTERACTION_WITH_SYMBIONT | 0.00001 | 0.00026 | 0.59333 | 0.75084 | 1.56620 | 41 |
| GO_INTERLEUKIN_1_MEDIATED_SIGNALING_PATHWAY | 0.00000 | 0.00000 | | 0.81689 | 1.75671 | 76 |
| GO_INTRINSIC_APOPTOTIC_SIGNALING_PATHWAY | 0.00121 | 0.02029 | 0.45506 | 0.55118 | 1.21845 | 190 |
| GO_ION_CHANNEL_BINDING | 0.00087 | 0.01584 | 0.47727 | 0.63182 | 1.34683 | 62 |
| GO_ISOPEPTIDASE_ACTIVITY | 0.00139 | 0.02285 | 0.45506 | 0.78940 | 1.57097 | 16 |
| GO_LARGE_RIBOSOMAL_SUBUNIT | 0.00000 | 0.00000 | | 0.78135 | 1.70183 | 109 |
| GO_LEUKOCYTE_MEDIATED_IMMUNITY | 0.00000 | 0.00000 | 0.71951 | 0.57282 | 1.27895 | 364 |
| GO_MACROMOLECULE_TRANSMEMBRANE_TRANSPORTER_ACTIVITY | 0.00239 | 0.03510 | 0.43171 | 0.77734 | 1.54696 | 16 |
| GO_MAINTENANCE_OF_CELL_NUMBER | 0.00087 | 0.01584 | 0.47727 | 0.58944 | 1.28512 | 114 |
| GO_MATURATION_OF_LSU_RRNA | 0.00006 | 0.00151 | 0.53843 | 0.77606 | 1.58723 | 27 |
| GO_METHYLTRANSFERASE_COMPLEX | 0.00290 | 0.04014 | 0.43171 | 0.59476 | 1.28479 | 88 |
| GO_METHYL_CPG_BINDING | 0.00302 | 0.04108 | 0.43171 | 0.78053 | 1.54833 | 15 |
| GO_MIDBRAIN_DEVELOPMENT | 0.00021 | 0.00468 | 0.51885 | 0.68832 | 1.44457 | 47 |
| GO_MITOCHONDRIAL_ELECTRON_TRANSPORT_NADH_TO_UBIQUIN | 0.00331 | 0.04437 | 0.43171 | 0.65428 | 1.36724 | 42 |
| GO_MITOCHONDRIAL_GENE_EXPRESSION | 0.00048 | 0.00949 | 0.49849 | 0.57608 | 1.26525 | 152 |
| GO_MITOCHONDRIAL_LARGE_RIBOSOMAL_SUBUNIT | 0.00002 | 0.00050 | 0.57561 | 0.69789 | 1.48086 | 55 |
| GO_MITOCHONDRIAL_MATRIX | 0.00000 | 0.00002 | 0.65944 | 0.56045 | 1.25158 | 371 |
| GO_MITOCHONDRIAL_MEMBRANE_ORGANIZATION | 0.00361 | 0.04734 | 0.43171 | 0.59309 | 1.28094 | 86 |
| GO_MITOCHONDRIAL_PROTEIN_COMPLEX | 0.00000 | 0.00000 | 0.81404 | 0.63650 | 1.40931 | 208 |
| GO_MITOCHONDRIAL_TRANSLATION | 0.00004 | 0.00093 | 0.55733 | 0.61219 | 1.34135 | 129 |
| GO_MITOCHONDRIAL_TRANSLATIONAL_TERMINATION | 0.00000 | 0.00010 | 0.62726 | 0.66694 | 1.44128 | 87 |
| GO_MITOCHONDRION_ORGANIZATION | 0.00051 | 0.00993 | 0.47727 | 0.52284 | 1.16863 | 390 |
| GO_MODIFICATION_DEPENDENT_MACROMOLECULE_CATABOLIC_PR | 0.00002 | 0.00053 | 0.57561 | 0.53142 | 1.19156 | 486 |
| GO_MODULATION_BY_HOST_OF_SYMBIONT_PROCESS | 0.00015 | 0.00336 | 0.51885 | 0.72848 | 1.50542 | 33 |
| GO_MODULATION_OF_PROCESS_OF_OTHER_ORGANISM | 0.00019 | 0.00423 | 0.51885 | 0.66904 | 1.42286 | 59 |
| GO_MONOSACCHARIDE_BIOSYNTHETIC_PROCESS | 0.00335 | 0.04466 | 0.43171 | 0.61992 | 1.31781 | 58 |
| GO_MORPHOGENESIS_OF_AN_EPITHELIUM | 0.00006 | 0.00151 | 0.53843 | 0.55161 | 1.22755 | 290 |
| GO_MORPHOGENESIS_OF_A_POLARIZED_EPITHELIUM | 0.00000 | 0.00000 | | 0.72298 | 1.57057 | 103 |
| GO_MRNA_3_UTR_BINDING | 0.00003 | 0.00089 | 0.55733 | 0.67596 | 1.44827 | 68 |
| GO_MRNA_5_UTR_BINDING | 0.00006 | 0.00156 | 0.53843 | 0.80133 | 1.62592 | 22 |
| GO_MRNA_BINDING | 0.00000 | 0.00000 | | 0.66002 | 1.46353 | 222 |
| GO_MRNA_PROCESSING | 0.00000 | 0.00000 | 0.73376 | 0.56859 | 1.27295 | 418 |
| GO_MULTI_ORGANISM_LOCALIZATION | 0.00268 | 0.03799 | 0.43171 | 0.63245 | 1.33612 | 51 |
| GO_MULTI_ORGANISM_PROCESS | 0.00108 | 0.01839 | 0.45506 | 0.51114 | 1.14599 | 484 |
| GO_MYD88_DEPENDENT_TOLL_LIKE_RECEPTOR_SIGNALING_PATHW | 0.00002 | 0.00066 | 0.57561 | 0.85581 | 1.70312 | 16 |
| GO_MYELOID_LEUKOCYTE_ACTIVATION | 0.00000 | 0.00000 | 0.74774 | 0.58508 | 1.30479 | 327 |
| GO_MYELOID_LEUKOCYTE_MEDIATED_IMMUNITY | 0.00000 | 0.00000 | 0.76146 | 0.59841 | 1.33291 | 301 |
| GO_NADH_DEHYDROGENASE_ACTIVITY | 0.00383 | 0.04935 | 0.43171 | 0.66244 | 1.37139 | 35 |
| GO_NADH_DEHYDROGENASE_COMPLEX | 0.00376 | 0.04884 | 0.43171 | 0.65649 | 1.36421 | 38 |
| GO_NADH_METABOLIC_PROCESS | 0.00294 | 0.04036 | 0.43171 | 0.70642 | 1.44481 | 27 |
| GO_NCRNA_EXPORT_FROM_NUCLEUS | 0.00302 | 0.04108 | 0.43171 | 0.66820 | 1.38331 | 35 |
| GO_NCRNA_METABOLIC_PROCESS | 0.00000 | 0.00003 | 0.65944 | 0.55907 | 1.24907 | 381 |
| GO_NCRNA_PROCESSING | 0.00000 | 0.00003 | 0.65944 | 0.57175 | 1.27504 | 328 |
| GO_NEGATIVE_REGULATION_OF_CANONICAL_WNT_SIGNALING_PAT | 0.00000 | 0.00000 | | 0.71611 | 1.56038 | 111 |
| GO_NEGATIVE_REGULATION_OF_CELLULAR_AMIDE_METABOLIC_PR | 0.00001 | 0.00020 | 0.61053 | 0.63031 | 1.37423 | 114 |
| GO_NEGATIVE_REGULATION_OF_CELL_CYCLE | 0.00003 | 0.00076 | 0.57561 | 0.53833 | 1.20513 | 425 |
| GO_NEGATIVE_REGULATION_OF_CELL_CYCLE_G2_M_PHASE_TRANS | 0.00000 | 0.00000 | 0.76146 | 0.71490 | 1.54439 | 85 |
| GO_NEGATIVE_REGULATION_OF_CELL_CYCLE_PHASE_TRANSITION | 0.00002 | 0.00060 | 0.57561 | 0.59041 | 1.30442 | 184 |
| GO_NEGATIVE_REGULATION_OF_CELL_CYCLE_PROCESS | 0.00000 | 0.00002 | 0.65944 | 0.58690 | 1.30374 | 249 |
| GO_NEGATIVE_REGULATION_OF_MITOTIC_CELL_CYCLE | 0.00004 | 0.00095 | 0.55733 | 0.56735 | 1.25902 | 232 |
| GO_NEGATIVE_REGULATION_OF_MRNA_METABOLIC_PROCESS | 0.00023 | 0.00491 | 0.51885 | 0.66670 | 1.41788 | 59 |
| GO_NEGATIVE_REGULATION_OF_MRNA_SPLICING_VIA_SPLICEOSOM | 0.00239 | 0.03510 | 0.43171 | 0.74857 | 1.51204 | 19 |
| GO_NEGATIVE_REGULATION_OF_RNA_SPLICING | 0.00029 | 0.00607 | 0.49849 | 0.76729 | 1.55976 | 23 |
| GO_NEGATIVE_REGULATION_OF_TRANSLATIONAL_INITIATION | 0.00018 | 0.00403 | 0.51885 | 0.83901 | 1.66434 | 15 |
| GO_NEGATIVE_REGULATION_OF_WNT_SIGNALING_PATHWAY | 0.00000 | 0.00000 | | 0.69525 | 1.52249 | 127 |
| GO_NEURAL_NUCLEUS_DEVELOPMENT | 0.00001 | 0.00026 | 0.59333 | 0.79148 | 1.62187 | 28 |
| GO_NEURON_TO_NEURON_SYNAPSE | 0.00005 | 0.00134 | 0.55733 | 0.59636 | 1.31006 | 154 |
| GO_NIK_NF_KAPPAB_SIGNALING | 0.00000 | 0.00000 | | 0.72896 | 1.58354 | 103 |
| GO_NON_CANONICAL_WNT_SIGNALING_PATHWAY | 0.00000 | 0.00000 | | 0.72784 | 1.58023 | 101 |
| GO_NUCLEAR_EXPORT | 0.00031 | 0.00645 | 0.49849 | 0.56963 | 1.25542 | 172 |
| GO_NUCLEAR_TRANSCRIBED_MRNA_CATABOLIC_PROCESS | 0.00000 | 0.00000 | | 0.79179 | 1.74917 | 183 |
| GO_NUCLEAR_TRANSCRIBED_MRNA_CATABOLIC_PROCESS_NONSEN | 0.00000 | 0.00000 | | 0.89008 | 1.93909 | 110 |
| GO_NUCLEAR_TRANSPORT | 0.00008 | 0.00193 | 0.53843 | 0.55511 | 1.23408 | 273 |
| GO_NUCLEOBASE_CONTAINING_COMPOUND_TRANSPORT | 0.00146 | 0.02386 | 0.45506 | 0.55053 | 1.21842 | 202 |

| | | | | | | |
|---|---|---|---|---|---|---|
| GO_NUCLEOBASE_CONTAINING_SMALL_MOLECULE_BIOSYNTHETIC_ | 0.00137 | 0.02261 | 0.45506 | 0.62749 | 1.34353 | 67 |
| GO_NUCLEOBASE_CONTAINING_SMALL_MOLECULE_INTERCONVERS | 0.00383 | 0.04935 | 0.43171 | 0.71259 | 1.44857 | 23 |
| GO_NUCLEOBASE_CONTAINING_SMALL_MOLECULE_METABOLIC_PR | 0.00320 | 0.04339 | 0.43171 | 0.51448 | 1.14825 | 352 |
| GO_NUCLEOSIDE_DIPHOSPHATE_METABOLIC_PROCESS | 0.00110 | 0.01865 | 0.45506 | 0.59637 | 1.29295 | 97 |
| GO_NUCLEOSIDE_METABOLIC_PROCESS | 0.00061 | 0.01169 | 0.47727 | 0.65315 | 1.38907 | 59 |
| GO_NUCLEOSIDE_MONOPHOSPHATE_BIOSYNTHETIC_PROCESS | 0.00250 | 0.03636 | 0.43171 | 0.68841 | 1.42033 | 32 |
| GO_NUCLEOSIDE_MONOPHOSPHATE_METABOLIC_PROCESS | 0.00152 | 0.02464 | 0.45506 | 0.64377 | 1.36002 | 51 |
| GO_NUCLEOSIDE_PHOSPHATE_BIOSYNTHETIC_PROCESS | 0.00043 | 0.00865 | 0.49849 | 0.58185 | 1.27831 | 148 |
| GO_NUCLEOSIDE_TRIPHOSPHATE_BIOSYNTHETIC_PROCESS | 0.00043 | 0.00865 | 0.49849 | 0.69332 | 1.44539 | 40 |
| GO_NUCLEOSIDE_TRIPHOSPHATE_METABOLIC_PROCESS | 0.00016 | 0.00360 | 0.51885 | 0.66788 | 1.41976 | 58 |
| GO_NUCLEOSOMAL_DNA_BINDING | 0.00023 | 0.00507 | 0.51885 | 0.79852 | 1.61801 | 21 |
| GO_NUCLEOSOME_BINDING | 0.00143 | 0.02332 | 0.45506 | 0.66132 | 1.38492 | 45 |
| GO_NUCLEOTIDE_EXCISION_REPAIR | 0.00004 | 0.00096 | 0.55733 | 0.63343 | 1.37355 | 98 |
| GO_NUCLEOTIDE_EXCISION_REPAIR_DNA_DAMAGE_RECOGNITION | 0.00015 | 0.00349 | 0.51885 | 0.78678 | 1.59640 | 22 |
| GO_NUCLEOTIDE_EXCISION_REPAIR_DNA_INCISION | 0.00313 | 0.04249 | 0.43171 | 0.66439 | 1.37898 | 36 |
| GO_NUCLEOTIDE_PHOSPHORYLATION | 0.00383 | 0.04935 | 0.43171 | 0.59002 | 1.27454 | 88 |
| GO_ORGANELLAR_RIBOSOME | 0.00000 | 0.00014 | 0.61053 | 0.66412 | 1.43471 | 85 |
| GO_ORGANELLE_INNER_MEMBRANE | 0.00000 | 0.00000 | 0.76146 | 0.57601 | 1.28697 | 378 |
| GO_ORGANIC_CYCLIC_COMPOUND_CATABOLIC_PROCESS | 0.00000 | 0.00000 | | 0.73350 | 1.64281 | 449 |
| GO_OXIDATIVE_PHOSPHORYLATION | 0.00001 | 0.00030 | 0.59333 | 0.64619 | 1.40027 | 96 |
| GO_OXIDOREDUCTASE_ACTIVITY | 0.00006 | 0.00139 | 0.55733 | 0.53544 | 1.19703 | 392 |
| GO_OXIDOREDUCTASE_COMPLEX | 0.00001 | 0.00027 | 0.59333 | 0.66015 | 1.42461 | 82 |
| GO_PEPTIDASE_COMPLEX | 0.00000 | 0.00000 | | 0.75988 | 1.63982 | 82 |
| GO_PIGMENT_GRANULE | 0.00003 | 0.00089 | 0.55733 | 0.66098 | 1.42344 | 78 |
| GO_POLYSOMAL_RIBOSOME | 0.00000 | 0.00000 | | 0.90624 | 1.86629 | 31 |
| GO_POLYSOME | 0.00000 | 0.00000 | | 0.82337 | 1.75481 | 61 |
| GO_POLY_PYRIMIDINE_TRACT_BINDING | 0.00327 | 0.04398 | 0.43171 | 0.75471 | 1.51855 | 18 |
| GO_POSITIVE_REGULATION_OF_APOPTOTIC_SIGNALING_PATHWAY | 0.00279 | 0.03889 | 0.43171 | 0.57666 | 1.25518 | 107 |
| GO_POSITIVE_REGULATION_OF_ATPASE_ACTIVITY | 0.00294 | 0.04036 | 0.43171 | 0.65446 | 1.36795 | 43 |
| GO_POSITIVE_REGULATION_OF_BINDING | 0.00088 | 0.01590 | 0.47727 | 0.60284 | 1.30556 | 95 |
| GO_POSITIVE_REGULATION_OF_CANONICAL_WNT_SIGNALING_PATH | 0.00000 | 0.00000 | 0.77494 | 0.70504 | 1.53160 | 103 |
| GO_POSITIVE_REGULATION_OF_CELLULAR_AMIDE_METABOLIC_PRO | 0.00003 | 0.00074 | 0.57561 | 0.62296 | 1.36083 | 118 |
| GO_POSITIVE_REGULATION_OF_CELLULAR_PROTEIN_LOCALIZATION | 0.00041 | 0.00840 | 0.49849 | 0.54915 | 1.21815 | 225 |
| GO_POSITIVE_REGULATION_OF_CELL_DEATH | 0.00069 | 0.01297 | 0.47727 | 0.52595 | 1.17423 | 363 |
| GO_POSITIVE_REGULATION_OF_CELL_POPULATION_PROLIFERATION | 0.00078 | 0.01455 | 0.47727 | 0.52449 | 1.17108 | 368 |
| GO_POSITIVE_REGULATION_OF_CHROMOSOME_ORGANIZATION | 0.00231 | 0.03437 | 0.43171 | 0.56990 | 1.24871 | 133 |
| GO_POSITIVE_REGULATION_OF_DEFENSE_RESPONSE | 0.00000 | 0.00003 | 0.65944 | 0.62175 | 1.36730 | 158 |
| GO_POSITIVE_REGULATION_OF_DNA_BIOSYNTHETIC_PROCESS | 0.00001 | 0.00041 | 0.59333 | 0.71929 | 1.52055 | 52 |
| GO_POSITIVE_REGULATION_OF_DNA_METABOLIC_PROCESS | 0.00364 | 0.04761 | 0.43171 | 0.56647 | 1.24189 | 135 |
| GO_POSITIVE_REGULATION_OF_DNA_TEMPLATED_TRANSCRIPTION_ | 0.00276 | 0.03885 | 0.43171 | 0.68932 | 1.41586 | 29 |
| GO_POSITIVE_REGULATION_OF_ESTABLISHMENT_OF_PROTEIN_LOCA | 0.00007 | 0.00168 | 0.53843 | 0.56461 | 1.25165 | 219 |
| GO_POSITIVE_REGULATION_OF_IMMUNE_RESPONSE | 0.00000 | 0.00001 | 0.69013 | 0.59037 | 1.31265 | 277 |
| GO_POSITIVE_REGULATION_OF_IMMUNE_SYSTEM_PROCESS | 0.00004 | 0.00110 | 0.55733 | 0.54018 | 1.20716 | 386 |
| GO_POSITIVE_REGULATION_OF_INNATE_IMMUNE_RESPONSE | 0.00000 | 0.00000 | 0.81404 | 0.69693 | 1.51947 | 114 |
| GO_POSITIVE_REGULATION_OF_INTRINSIC_APOPTOTIC_SIGNALING_ | 0.00104 | 0.01798 | 0.45506 | 0.69097 | 1.43578 | 37 |
| GO_POSITIVE_REGULATION_OF_ORGANELLE_ORGANIZATION | 0.00043 | 0.00865 | 0.49849 | 0.52093 | 1.16605 | 411 |
| GO_POSITIVE_REGULATION_OF_PROTEIN_LOCALIZATION_TO_NUCLE | 0.00001 | 0.00020 | 0.61053 | 0.69907 | 1.48990 | 61 |
| GO_POSITIVE_REGULATION_OF_RESPONSE_TO_BIOTIC_STIMULUS | 0.00000 | 0.00000 | 0.73376 | 0.66340 | 1.45310 | 128 |
| GO_POSITIVE_REGULATION_OF_RESPONSE_TO_EXTERNAL_STIMULU | 0.00002 | 0.00050 | 0.57561 | 0.58133 | 1.28681 | 201 |
| GO_POSITIVE_REGULATION_OF_TELOMERASE_ACTIVITY | 0.00061 | 0.01169 | 0.47727 | 0.73476 | 1.50565 | 28 |
| GO_POSITIVE_REGULATION_OF_TELOMERASE_RNA_LOCALIZATION_ | 0.00000 | 0.00001 | 0.69013 | 0.92451 | 1.83395 | 15 |
| GO_POSITIVE_REGULATION_OF_TELOMERE_MAINTENANCE_VIA_TE | 0.00130 | 0.02159 | 0.45506 | 0.71259 | 1.46508 | 30 |
| GO_POSITIVE_REGULATION_OF_TRANSCRIPTION_INITIATION_FROM | 0.00372 | 0.04847 | 0.43171 | 0.70671 | 1.43993 | 25 |
| GO_POSITIVE_REGULATION_OF_TRANSFERASE_ACTIVITY | 0.00174 | 0.02712 | 0.45506 | 0.51642 | 1.15220 | 345 |
| GO_POSITIVE_REGULATION_OF_TRANSLATION | 0.00001 | 0.00019 | 0.61053 | 0.64767 | 1.40419 | 97 |
| GO_POSITIVE_REGULATION_OF_VIRAL_GENOME_REPLICATION | 0.00287 | 0.03973 | 0.43171 | 0.68362 | 1.40783 | 31 |
| GO_POSITIVE_REGULATION_OF_VIRAL_PROCESS | 0.00028 | 0.00590 | 0.49849 | 0.62440 | 1.34741 | 81 |
| GO_POSITIVE_REGULATION_OF_WNT_SIGNALING_PATHWAY | 0.00000 | 0.00000 | 0.73376 | 0.66230 | 1.44763 | 122 |
| GO_POSTREPLICATION_REPAIR | 0.00024 | 0.00515 | 0.51885 | 0.69140 | 1.44518 | 43 |
| GO_POSTSYNAPSE | 0.00004 | 0.00109 | 0.55733 | 0.55969 | 1.24369 | 259 |
| GO_POSTSYNAPTIC_CYTOSOL | 0.00279 | 0.03889 | 0.43171 | 0.78295 | 1.55313 | 15 |
| GO_POSTTRANSCRIPTIONAL_REGULATION_OF_GENE_EXPRESSION | 0.00000 | 0.00000 | | 0.66656 | 1.49327 | 462 |
| GO_POST_TRANSLATIONAL_PROTEIN_MODIFICATION | 0.00002 | 0.00052 | 0.57561 | 0.57851 | 1.28412 | 229 |
| GO_PRECATALYTIC_SPLICEOSOME | 0.00085 | 0.01559 | 0.47727 | 0.65235 | 1.37448 | 49 |
| GO_PRERIBOSOME | 0.00087 | 0.01577 | 0.47727 | 0.62899 | 1.34772 | 69 |
| GO_PRERIBOSOME_LARGE_SUBUNIT_PRECURSOR | 0.00235 | 0.03474 | 0.43171 | 0.74079 | 1.50308 | 22 |
| GO_PRE_MRNA_BINDING | 0.00133 | 0.02213 | 0.45506 | 0.72393 | 1.48062 | 27 |
| GO_PROTEASOMAL_PROTEIN_CATABOLIC_PROCESS | 0.00002 | 0.00044 | 0.59333 | 0.54166 | 1.21005 | 377 |
| GO_PROTEASOMAL_UBIQUITIN_INDEPENDENT_PROTEIN_CATABOLI( | 0.00000 | 0.00000 | 0.80122 | 0.92254 | 1.86931 | 21 |
| GO_PROTEASOME_ACCESSORY_COMPLEX | 0.00000 | 0.00001 | 0.67496 | 0.85464 | 1.73811 | 24 |
| GO_PROTEASOME_CORE_COMPLEX | 0.00000 | 0.00000 | | 0.94375 | 1.90631 | 19 |
| GO_PROTEIN_CONTAINING_COMPLEX_DISASSEMBLY | 0.00000 | 0.00007 | 0.62726 | 0.57810 | 1.28438 | 256 |
| GO_PROTEIN_CONTAINING_COMPLEX_LOCALIZATION | 0.00063 | 0.01197 | 0.47727 | 0.55335 | 1.22520 | 208 |
| GO_PROTEIN_DNA_COMPLEX_SUBUNIT_ORGANIZATION | 0.00001 | 0.00018 | 0.61053 | 0.60174 | 1.32491 | 165 |
| GO_PROTEIN_FOLDING | 0.00000 | 0.00002 | 0.65944 | 0.63113 | 1.38644 | 154 |
| GO_PROTEIN_FOLDING_CHAPERONE | 0.00135 | 0.02238 | 0.45506 | 0.77903 | 1.55752 | 17 |

| | | | | | | |
|---|---|---|---|---|---|---|
| GO_PROTEIN_IMPORT | 0.00268 | 0.03799 | 0.43171 | 0.56317 | 1.23581 | 140 |
| GO_PROTEIN_LOCALIZATION_TO_CHROMOSOME | 0.00056 | 0.01096 | 0.47727 | 0.63523 | 1.36108 | 69 |
| GO_PROTEIN_LOCALIZATION_TO_CHROMOSOME_TELOMERIC_REGIO | 0.00335 | 0.04466 | 0.43171 | 0.70399 | 1.43984 | 27 |
| GO_PROTEIN_LOCALIZATION_TO_ENDOPLASMIC_RETICULUM | 0.00000 | 0.00000 | | 0.85521 | 1.87021 | 124 |
| GO_PROTEIN_LOCALIZATION_TO_MEMBRANE | 0.00000 | 0.00000 | | 0.65123 | 1.45714 | 406 |
| GO_PROTEIN_LOCALIZATION_TO_MITOCHONDRION | 0.00019 | 0.00420 | 0.51885 | 0.60773 | 1.32398 | 110 |
| GO_PROTEIN_LOCALIZATION_TO_NUCLEUS | 0.00002 | 0.00055 | 0.57561 | 0.58926 | 1.30220 | 187 |
| GO_PROTEIN_MODIFICATION_BY_SMALL_PROTEIN_REMOVAL | 0.00000 | 0.00000 | 0.71951 | 0.60745 | 1.34689 | 221 |
| GO_PROTEIN_POLYUBIQUITINATION | 0.00004 | 0.00106 | 0.55733 | 0.55874 | 1.24108 | 253 |
| GO_PROTEIN_REFOLDING | 0.00294 | 0.04036 | 0.43171 | 0.77039 | 1.53314 | 16 |
| GO_PROTEIN_STABILIZATION | 0.00001 | 0.00024 | 0.61053 | 0.62333 | 1.36586 | 130 |
| GO_PROTEIN_TARGETING | 0.00000 | 0.00000 | | 0.71181 | 1.58743 | 327 |
| GO_PROTEIN_TARGETING_TO_MEMBRANE | 0.00000 | 0.00000 | | 0.82370 | 1.80953 | 149 |
| GO_PROTEIN_TARGETING_TO_MITOCHONDRION | 0.00346 | 0.04593 | 0.43171 | 0.60408 | 1.30091 | 78 |
| GO_PROTEIN_TRANSMEMBRANE_TRANSPORT | 0.00265 | 0.03775 | 0.43171 | 0.64528 | 1.36171 | 50 |
| GO_PROTON_TRANSMEMBRANE_TRANSPORT | 0.00213 | 0.03229 | 0.43171 | 0.60815 | 1.30536 | 71 |
| GO_PURINE_CONTAINING_COMPOUND_BIOSYNTHETIC_PROCESS | 0.00246 | 0.03601 | 0.43171 | 0.57796 | 1.25936 | 111 |
| GO_PURINE_CONTAINING_COMPOUND_METABOLIC_PROCESS | 0.00100 | 0.01761 | 0.45506 | 0.53679 | 1.19242 | 255 |
| GO_PURINE_NUCLEOSIDE_TRIPHOSPHATE_METABOLIC_PROCESS | 0.00089 | 0.01592 | 0.47727 | 0.67157 | 1.40374 | 43 |
| GO_PYRIMIDINE_CONTAINING_COMPOUND_METABOLIC_PROCESS | 0.00353 | 0.04658 | 0.43171 | 0.64744 | 1.35878 | 47 |
| GO_REGENERATION | 0.00216 | 0.03242 | 0.43171 | 0.59987 | 1.29453 | 82 |
| GO_REGULATION_OF_ANIMAL_ORGAN_MORPHOGENESIS | 0.00000 | 0.00000 | | 0.69163 | 1.51541 | 133 |
| GO_REGULATION_OF_APOPTOTIC_SIGNALING_PATHWAY | 0.00031 | 0.00645 | 0.49849 | 0.55171 | 1.22506 | 247 |
| GO_REGULATION_OF_ATPASE_ACTIVITY | 0.00324 | 0.04369 | 0.43171 | 0.63031 | 1.34038 | 57 |
| GO_REGULATION_OF_ATP_METABOLIC_PROCESS | 0.00353 | 0.04658 | 0.43171 | 0.60176 | 1.29780 | 80 |
| GO_REGULATION_OF_BINDING | 0.00250 | 0.03636 | 0.43171 | 0.54476 | 1.20598 | 203 |
| GO_REGULATION_OF_CELLULAR_AMIDE_METABOLIC_PROCESS | 0.00000 | 0.00000 | | 0.62967 | 1.40226 | 311 |
| GO_REGULATION_OF_CELLULAR_AMINE_METABOLIC_PROCESS | 0.00000 | 0.00000 | | 0.82173 | 1.74682 | 58 |
| GO_REGULATION_OF_CELLULAR_AMINO_ACID_METABOLIC_PROCES | 0.00000 | 0.00000 | | 0.83776 | 1.77766 | 55 |
| GO_REGULATION_OF_CELLULAR_KETONE_METABOLIC_PROCESS | 0.00000 | 0.00000 | | 0.72641 | 1.58680 | 118 |
| GO_REGULATION_OF_CELLULAR_PROTEIN_LOCALIZATION | 0.00082 | 0.01513 | 0.47727 | 0.52542 | 1.17290 | 362 |
| GO_REGULATION_OF_CELLULAR_RESPONSE_TO_HEAT | 0.00276 | 0.03885 | 0.43171 | 0.61652 | 1.32004 | 67 |
| GO_REGULATION_OF_CELLULAR_RESPONSE_TO_STRESS | 0.00224 | 0.03344 | 0.43171 | 0.50545 | 1.13232 | 468 |
| GO_REGULATION_OF_CELL_CYCLE_G2_M_PHASE_TRANSITION | 0.00000 | 0.00000 | 0.71951 | 0.63230 | 1.39282 | 169 |
| GO_REGULATION_OF_CELL_CYCLE_PHASE_TRANSITION | 0.00001 | 0.00026 | 0.59333 | 0.55488 | 1.23783 | 337 |
| GO_REGULATION_OF_CHROMOSOME_ORGANIZATION | 0.00081 | 0.01501 | 0.47727 | 0.54255 | 1.20527 | 261 |
| GO_REGULATION_OF_CYSTEINE_TYPE_ENDOPEPTIDASE_ACTIVITY | 0.00202 | 0.03077 | 0.43171 | 0.57120 | 1.24837 | 121 |
| GO_REGULATION_OF_CYTOPLASMIC_TRANSLATION | 0.00130 | 0.02159 | 0.45506 | 0.73678 | 1.49773 | 23 |
| GO_REGULATION_OF_DEFENSE_RESPONSE | 0.00065 | 0.01237 | 0.47727 | 0.53633 | 1.19355 | 290 |
| GO_REGULATION_OF_DNA_BIOSYNTHETIC_PROCESS | 0.00016 | 0.00360 | 0.51885 | 0.64215 | 1.37923 | 74 |
| GO_REGULATION_OF_DNA_METABOLIC_PROCESS | 0.00069 | 0.01297 | 0.47727 | 0.54425 | 1.20775 | 237 |
| GO_REGULATION_OF_DNA_TEMPLATED_TRANSCRIPTION_INITIATIOI | 0.00163 | 0.02568 | 0.45506 | 0.68289 | 1.41372 | 35 |
| GO_REGULATION_OF_DNA_TEMPLATED_TRANSCRIPTION_IN_RESPO | 0.00000 | 0.00000 | | 0.73944 | 1.59800 | 89 |
| GO_REGULATION_OF_GENERATION_OF_PRECURSOR_METABOLITES_ | 0.00060 | 0.01164 | 0.47727 | 0.60476 | 1.31116 | 97 |
| GO_REGULATION_OF_HEMATOPOIETIC_PROGENITOR_CELL_DIFFERE | 0.00000 | 0.00000 | | 0.78717 | 1.69251 | 75 |
| GO_REGULATION_OF_HEMATOPOIETIC_STEM_CELL_DIFFERENTIATIO | 0.00000 | 0.00000 | | 0.81505 | 1.74256 | 65 |
| GO_REGULATION_OF_HEMOPOIESIS | 0.00000 | 0.00008 | 0.62726 | 0.58859 | 1.30634 | 228 |
| GO_REGULATION_OF_IMMUNE_RESPONSE | 0.00000 | 0.00016 | 0.61053 | 0.55362 | 1.23583 | 362 |
| GO_REGULATION_OF_INNATE_IMMUNE_RESPONSE | 0.00000 | 0.00000 | 0.71951 | 0.63883 | 1.40521 | 161 |
| GO_REGULATION_OF_INTRINSIC_APOPTOTIC_SIGNALING_PATHWAY | 0.00001 | 0.00039 | 0.59333 | 0.63478 | 1.38259 | 109 |
| GO_REGULATION_OF_MITOTIC_CELL_CYCLE | 0.00001 | 0.00018 | 0.61053 | 0.53899 | 1.20736 | 458 |
| GO_REGULATION_OF_MORPHOGENESIS_OF_AN_EPITHELIUM | 0.00000 | 0.00000 | | 0.72213 | 1.57284 | 109 |
| GO_REGULATION_OF_MRNA_CATABOLIC_PROCESS | 0.00000 | 0.00000 | | 0.69977 | 1.54127 | 168 |
| GO_REGULATION_OF_MRNA_METABOLIC_PROCESS | 0.00000 | 0.00000 | | 0.66528 | 1.47872 | 266 |
| GO_REGULATION_OF_MRNA_PROCESSING | 0.00004 | 0.00105 | 0.55733 | 0.61521 | 1.34090 | 113 |
| GO_REGULATION_OF_MRNA_SPLICING_VIA_SPLICEOSOME | 0.00001 | 0.00026 | 0.59333 | 0.67424 | 1.45411 | 80 |
| GO_REGULATION_OF_NUCLEASE_ACTIVITY | 0.00106 | 0.01819 | 0.45506 | 0.77918 | 1.56780 | 18 |
| GO_REGULATION_OF_OXIDATIVE_STRESS_INDUCED_CELL_DEATH | 0.00198 | 0.03028 | 0.43171 | 0.67287 | 1.40355 | 41 |
| GO_REGULATION_OF_PEPTIDASE_ACTIVITY | 0.00050 | 0.00978 | 0.47727 | 0.56122 | 1.24054 | 191 |
| GO_REGULATION_OF_POSTTRANSCRIPTIONAL_GENE_SILENCING | 0.00364 | 0.04761 | 0.43171 | 0.59824 | 1.28492 | 74 |
| GO_REGULATION_OF_PROTEIN_CATABOLIC_PROCESS | 0.00179 | 0.02783 | 0.45506 | 0.53057 | 1.17929 | 270 |
| GO_REGULATION_OF_PROTEIN_LOCALIZATION_TO_NUCLEUS | 0.00001 | 0.00042 | 0.59333 | 0.65505 | 1.41558 | 87 |
| GO_REGULATION_OF_PROTEIN_MODIFICATION_BY_SMALL_PROTEIN | 0.00025 | 0.00532 | 0.49849 | 0.57068 | 1.25684 | 167 |
| GO_REGULATION_OF_PROTEIN_STABILITY | 0.00093 | 0.01652 | 0.47727 | 0.55382 | 1.22571 | 202 |
| GO_REGULATION_OF_PROTEOLYSIS | 0.00013 | 0.00301 | 0.51885 | 0.52997 | 1.18392 | 377 |
| GO_REGULATION_OF_RESPONSE_TO_BIOTIC_STIMULUS | 0.00001 | 0.00020 | 0.61053 | 0.59049 | 1.30719 | 204 |
| GO_REGULATION_OF_RNA_SPLICING | 0.00000 | 0.00001 | 0.69013 | 0.65896 | 1.43669 | 114 |
| GO_REGULATION_OF_SMALL_MOLECULE_METABOLIC_PROCESS | 0.00000 | 0.00000 | 0.71951 | 0.59134 | 1.31394 | 265 |
| GO_REGULATION_OF_STEM_CELL_DIFFERENTIATION | 0.00000 | 0.00000 | | 0.74049 | 1.60113 | 90 |
| GO_REGULATION_OF_SYMBIOTIC_PROCESS | 0.00006 | 0.00144 | 0.55733 | 0.60071 | 1.31682 | 134 |
| GO_REGULATION_OF_TELOMERASE_ACTIVITY | 0.00088 | 0.01590 | 0.47727 | 0.69161 | 1.43179 | 35 |
| GO_REGULATION_OF_TRANSCRIPTION_FROM_RNA_POLYMERASE_II | 0.00000 | 0.00000 | | 0.81194 | 1.73800 | 66 |
| GO_REGULATION_OF_TRANSCRIPTION_INITIATION_FROM_RNA_POL | 0.00143 | 0.02332 | 0.45506 | 0.70390 | 1.44582 | 29 |
| GO_REGULATION_OF_TRANSLATIONAL_INITIATION | 0.00002 | 0.00052 | 0.57561 | 0.68129 | 1.45518 | 64 |
| GO_REGULATION_OF_UBIQUITIN_PROTEIN_TRANSFERASE_ACTIVITY | 0.00061 | 0.01169 | 0.47727 | 0.69335 | 1.44627 | 41 |
| GO_REGULATION_OF_VIRAL_GENOME_REPLICATION | 0.00086 | 0.01566 | 0.47727 | 0.65316 | 1.38389 | 54 |

| | | | | | | |
|---|---|---|---|---|---|---|
| GO_REGULATION_OF_VIRAL_TRANSCRIPTION | 0.00198 | 0.03028 | 0.43171 | 0.63932 | 1.34703 | 49 |
| GO_REGULATION_OF_WNT_SIGNALING_PATHWAY | 0.00000 | 0.00001 | 0.69013 | 0.60353 | 1.33816 | 220 |
| GO_RESPIRASOME | 0.00008 | 0.00181 | 0.53843 | 0.65420 | 1.40421 | 71 |
| GO_RESPIRATORY_CHAIN_COMPLEX | 0.00016 | 0.00353 | 0.51885 | 0.67626 | 1.43918 | 60 |
| GO_RESPIRATORY_ELECTRON_TRANSPORT_CHAIN | 0.00001 | 0.00025 | 0.59333 | 0.65819 | 1.42190 | 85 |
| GO_RESPONSE_TO_ALCOHOL | 0.00261 | 0.03750 | 0.43171 | 0.58804 | 1.27350 | 95 |
| GO_RESPONSE_TO_CADMIUM_ION | 0.00265 | 0.03775 | 0.43171 | 0.67227 | 1.39174 | 35 |
| GO_RESPONSE_TO_DRUG | 0.00104 | 0.01798 | 0.45506 | 0.55364 | 1.22258 | 180 |
| GO_RESPONSE_TO_ETHANOL | 0.00086 | 0.01566 | 0.47727 | 0.67309 | 1.40691 | 43 |
| GO_RESPONSE_TO_HEAT | 0.00012 | 0.00270 | 0.53843 | 0.61665 | 1.34221 | 107 |
| GO_RESPONSE_TO_HORMONE | 0.00048 | 0.00949 | 0.49849 | 0.51277 | 1.14814 | 453 |
| GO_RESPONSE_TO_HYDROGEN_PEROXIDE | 0.00257 | 0.03716 | 0.43171 | 0.60505 | 1.30368 | 79 |
| GO_RESPONSE_TO_INORGANIC_SUBSTANCE | 0.00004 | 0.00107 | 0.55733 | 0.55714 | 1.23870 | 275 |
| GO_RESPONSE_TO_INTERLEUKIN_1 | 0.00000 | 0.00000 | | 0.74844 | 1.62908 | 107 |
| GO_RESPONSE_TO_INTERLEUKIN_12 | 0.00012 | 0.00270 | 0.53843 | 0.72605 | 1.50694 | 36 |
| GO_RESPONSE_TO_INTERLEUKIN_4 | 0.00324 | 0.04369 | 0.43171 | 0.75812 | 1.51571 | 17 |
| GO_RESPONSE_TO_METAL_ION | 0.00030 | 0.00623 | 0.49849 | 0.56387 | 1.24233 | 171 |
| GO_RESPONSE_TO_ORGANIC_CYCLIC_COMPOUND | 0.00000 | 0.00016 | 0.61053 | 0.54603 | 1.22259 | 439 |
| GO_RESPONSE_TO_OXYGEN_LEVELS | 0.00000 | 0.00000 | 0.71951 | 0.60132 | 1.33438 | 233 |
| GO_RESPONSE_TO_STEROID_HORMONE | 0.00154 | 0.02464 | 0.45506 | 0.55514 | 1.22446 | 174 |
| GO_RESPONSE_TO_TEMPERATURE_STIMULUS | 0.00027 | 0.00572 | 0.49849 | 0.59216 | 1.29675 | 127 |
| GO_RESPONSE_TO_TUMOR_NECROSIS_FACTOR | 0.00000 | 0.00000 | 0.76146 | 0.65649 | 1.44236 | 151 |
| GO_RETINA_HOMEOSTASIS | 0.00104 | 0.01798 | 0.45506 | 0.73063 | 1.49432 | 27 |
| GO_RIBONUCLEOPROTEIN_COMPLEX_BINDING | 0.00000 | 0.00000 | 0.71951 | 0.66906 | 1.46050 | 117 |
| GO_RIBONUCLEOPROTEIN_COMPLEX_BIOGENESIS | 0.00000 | 0.00000 | | 0.66339 | 1.48264 | 389 |
| GO_RIBONUCLEOPROTEIN_COMPLEX_SUBUNIT_ORGANIZATION | 0.00000 | 0.00000 | | 0.69241 | 1.52681 | 173 |
| GO_RIBONUCLEOPROTEIN_GRANULE | 0.00000 | 0.00001 | 0.67496 | 0.61979 | 1.36366 | 163 |
| GO_RIBONUCLEOSIDE_TRIPHOSPHATE_BIOSYNTHETIC_PROCESS | 0.00041 | 0.00840 | 0.49849 | 0.71253 | 1.47248 | 33 |
| GO_RIBONUCLEOSIDE_TRIPHOSPHATE_METABOLIC_PROCESS | 0.00026 | 0.00550 | 0.49849 | 0.69681 | 1.45611 | 42 |
| GO_RIBOSE_PHOSPHATE_BIOSYNTHETIC_PROCESS | 0.00191 | 0.02939 | 0.45506 | 0.57932 | 1.26207 | 110 |
| GO_RIBOSE_PHOSPHATE_METABOLIC_PROCESS | 0.00108 | 0.01839 | 0.45506 | 0.53891 | 1.19687 | 245 |
| GO_RIBOSOMAL_LARGE_SUBUNIT_ASSEMBLY | 0.00000 | 0.00016 | 0.61053 | 0.80999 | 1.65663 | 27 |
| GO_RIBOSOMAL_LARGE_SUBUNIT_BIOGENESIS | 0.00000 | 0.00000 | | 0.77542 | 1.65982 | 66 |
| GO_RIBOSOMAL_SMALL_SUBUNIT_ASSEMBLY | 0.00023 | 0.00494 | 0.51885 | 0.81139 | 1.62221 | 17 |
| GO_RIBOSOMAL_SMALL_SUBUNIT_BINDING | 0.00383 | 0.04935 | 0.43171 | 0.75288 | 1.50523 | 17 |
| GO_RIBOSOMAL_SMALL_SUBUNIT_BIOGENESIS | 0.00001 | 0.00022 | 0.61053 | 0.69076 | 1.47997 | 68 |
| GO_RIBOSOMAL_SUBUNIT | 0.00000 | 0.00000 | | 0.77603 | 1.71031 | 172 |
| GO_RIBOSOME | 0.00000 | 0.00000 | | 0.76643 | 1.69659 | 206 |
| GO_RIBOSOME_ASSEMBLY | 0.00000 | 0.00000 | 0.73376 | 0.76173 | 1.61998 | 59 |
| GO_RIBOSOME_BINDING | 0.00019 | 0.00416 | 0.51885 | 0.68955 | 1.44715 | 47 |
| GO_RIBOSOME_BIOGENESIS | 0.00000 | 0.00000 | | 0.65877 | 1.46466 | 275 |
| GO_RNA_CATABOLIC_PROCESS | 0.00000 | 0.00000 | | 0.76774 | 1.71264 | 339 |
| GO_RNA_DEPENDENT_DNA_BIOSYNTHETIC_PROCESS | 0.00003 | 0.00076 | 0.57561 | 0.68926 | 1.46521 | 58 |
| GO_RNA_EXPORT_FROM_NUCLEUS | 0.00062 | 0.01183 | 0.47727 | 0.58378 | 1.27758 | 126 |
| GO_RNA_LOCALIZATION | 0.00000 | 0.00006 | 0.64355 | 0.59670 | 1.32020 | 198 |
| GO_RNA_POLYMERASE_II_PREINITIATION_COMPLEX_ASSEMBLY | 0.00089 | 0.01592 | 0.47727 | 0.77046 | 1.55626 | 19 |
| GO_RNA_SPLICING | 0.00000 | 0.00000 | 0.78819 | 0.59079 | 1.31927 | 369 |
| GO_RNA_SPLICING_VIA_TRANSESTERIFICATION_REACTIONS | 0.00000 | 0.00000 | | 0.60940 | 1.35714 | 300 |
| GO_RNA_STABILIZATION | 0.00065 | 0.01223 | 0.47727 | 0.69743 | 1.44384 | 35 |
| GO_ROUGH_ENDOPLASMIC_RETICULUM | 0.00001 | 0.00039 | 0.59333 | 0.71853 | 1.51629 | 50 |
| GO_RRNA_BINDING | 0.00000 | 0.00001 | 0.69013 | 0.74782 | 1.58865 | 56 |
| GO_RRNA_METABOLIC_PROCESS | 0.00000 | 0.00000 | 0.77494 | 0.63047 | 1.39562 | 206 |
| GO_SCF_DEPENDENT_PROTEASOMAL_UBIQUITIN_DEPENDENT_PRO | 0.00000 | 0.00000 | | 0.77527 | 1.67044 | 79 |
| GO_SECRETORY_GRANULE | 0.00000 | 0.00000 | 0.71951 | 0.56518 | 1.26461 | 399 |
| GO_SECRETORY_VESICLE | 0.00000 | 0.00002 | 0.67496 | 0.54866 | 1.22905 | 465 |
| GO_SIGNAL_TRANSDUCTION_BY_PROTEIN_PHOSPHORYLATION | 0.00004 | 0.00109 | 0.55733 | 0.53817 | 1.20474 | 424 |
| GO_SINGLE_FERTILIZATION | 0.00048 | 0.00949 | 0.49849 | 0.67414 | 1.42260 | 50 |
| GO_SINGLE_STRANDED_RNA_BINDING | 0.00039 | 0.00793 | 0.49849 | 0.66553 | 1.41527 | 57 |
| GO_SMALL_MOLECULE_BIOSYNTHETIC_PROCESS | 0.00216 | 0.03242 | 0.43171 | 0.51195 | 1.14522 | 397 |
| GO_SMALL_RIBOSOMAL_SUBUNIT | 0.00000 | 0.00000 | | 0.77648 | 1.66209 | 66 |
| GO_SM_LIKE_PROTEIN_FAMILY_COMPLEX | 0.00327 | 0.04398 | 0.43171 | 0.60468 | 1.29713 | 70 |
| GO_SPERM_EGG_RECOGNITION | 0.00000 | 0.00007 | 0.64355 | 0.85241 | 1.72955 | 22 |
| GO_SPLICEOSOMAL_COMPLEX | 0.00000 | 0.00003 | 0.65944 | 0.60939 | 1.34261 | 171 |
| GO_STEM_CELL_DIFFERENTIATION | 0.00000 | 0.00000 | 0.78819 | 0.66886 | 1.46930 | 147 |
| GO_STRUCTURAL_CONSTITUENT_OF_CYTOSKELETON | 0.00102 | 0.01782 | 0.45506 | 0.65913 | 1.39337 | 52 |
| GO_STRUCTURAL_CONSTITUENT_OF_RIBOSOME | 0.00000 | 0.00000 | | 0.79909 | 1.75541 | 150 |
| GO_STRUCTURAL_MOLECULE_ACTIVITY | 0.00000 | 0.00000 | | 0.72651 | 1.61895 | 324 |
| GO_SUBSTANTIA_NIGRA_DEVELOPMENT | 0.00000 | 0.00013 | 0.62726 | 0.81622 | 1.66507 | 26 |
| GO_TELOMERASE_RNA_LOCALIZATION | 0.00001 | 0.00017 | 0.61053 | 0.85164 | 1.72025 | 19 |
| GO_TELOMERE_MAINTENANCE_VIA_TELOMERE_LENGTHENING | 0.00011 | 0.00250 | 0.53843 | 0.65659 | 1.40546 | 66 |
| GO_TELOMERE_ORGANIZATION | 0.00342 | 0.04554 | 0.43171 | 0.57000 | 1.24649 | 124 |
| GO_THREONINE_TYPE_ENDOPEPTIDASE_ACTIVITY | 0.00000 | 0.00000 | | 0.95179 | 1.92255 | 19 |
| GO_THREONINE_TYPE_PEPTIDASE_ACTIVITY | 0.00000 | 0.00000 | | 0.92199 | 1.87423 | 23 |
| GO_TISSUE_MORPHOGENESIS | 0.00097 | 0.01707 | 0.47727 | 0.52661 | 1.17402 | 326 |
| GO_TOXIN_TRANSPORT | 0.00113 | 0.01922 | 0.45506 | 0.72897 | 1.48708 | 26 |
| GO_TRANSCRIPTION_COUPLED_NUCLEOTIDE_EXCISION_REPAIR | 0.00001 | 0.00019 | 0.61053 | 0.68106 | 1.46186 | 71 |

| | | | | | | |
|---|---|---|---|---|---|---|
| GO_TRANSCRIPTION_FACTOR_BINDING | 0.00043 | 0.00865 | 0.49849 | 0.52447 | 1.17376 | 409 |
| GO_TRANSCRIPTION_PREINITIATION_COMPLEX_ASSEMBLY | 0.00119 | 0.02004 | 0.45506 | 0.71399 | 1.46797 | 30 |
| GO_TRANSLATIONAL_ELONGATION | 0.00000 | 0.00000 | 0.71951 | 0.66087 | 1.44451 | 122 |
| GO_TRANSLATIONAL_INITIATION | 0.00000 | 0.00000 | | 0.84691 | 1.86515 | 166 |
| GO_TRANSLATIONAL_TERMINATION | 0.00000 | 0.00005 | 0.64355 | 0.65946 | 1.43060 | 99 |
| GO_TRANSLATION_ELONGATION_FACTOR_ACTIVITY | 0.00161 | 0.02546 | 0.45506 | 0.79860 | 1.58418 | 15 |
| GO_TRANSLATION_FACTOR_ACTIVITY_RNA_BINDING | 0.00000 | 0.00000 | | 0.79008 | 1.69288 | 69 |
| GO_TRANSLATION_INITIATION_FACTOR_ACTIVITY | 0.00000 | 0.00000 | 0.76146 | 0.80422 | 1.68417 | 45 |
| GO_TRANSLATION_INITIATION_FACTOR_BINDING | 0.00021 | 0.00468 | 0.51885 | 0.76186 | 1.55231 | 25 |
| GO_TRANSLATION_PREINITIATION_COMPLEX | 0.00000 | 0.00001 | 0.67496 | 0.90245 | 1.79595 | 16 |
| GO_TRANSLATION_REGULATOR_ACTIVITY | 0.00000 | 0.00000 | | 0.77663 | 1.68904 | 105 |
| GO_TRANSLATION_REGULATOR_ACTIVITY_NUCLEIC_ACID_BINDING | 0.00000 | 0.00000 | | 0.77150 | 1.66586 | 83 |
| GO_TRICARBOXYLIC_ACID_CYCLE | 0.00154 | 0.02464 | 0.45506 | 0.71854 | 1.47240 | 28 |
| GO_TUMOR_NECROSIS_FACTOR_MEDIATED_SIGNALING_PATHWAY | 0.00000 | 0.00000 | | 0.76277 | 1.65372 | 97 |
| GO_T_CELL_RECEPTOR_SIGNALING_PATHWAY | 0.00000 | 0.00000 | 0.81404 | 0.69994 | 1.52527 | 112 |
| GO_U12_TYPE_SPLICEOSOMAL_COMPLEX | 0.00279 | 0.03889 | 0.43171 | 0.70746 | 1.44693 | 27 |
| GO_U2_SNRNP | 0.00005 | 0.00134 | 0.55733 | 0.83081 | 1.67168 | 18 |
| GO_U2_TYPE_SPLICEOSOMAL_COMPLEX | 0.00010 | 0.00243 | 0.53843 | 0.63403 | 1.36970 | 85 |
| GO_UBIQUITIN_LIKE_PROTEIN_LIGASE_BINDING | 0.00000 | 0.00000 | 0.76146 | 0.62273 | 1.38051 | 219 |
| GO_UBIQUITIN_PROTEIN_TRANSFERASE_REGULATOR_ACTIVITY | 0.00165 | 0.02590 | 0.45506 | 0.79605 | 1.57911 | 15 |
| GO_UNFOLDED_PROTEIN_BINDING | 0.00000 | 0.00000 | 0.77494 | 0.72157 | 1.55881 | 85 |
| GO_VESICLE_LUMEN | 0.00000 | 0.00000 | 0.74774 | 0.64475 | 1.42008 | 168 |
| GO_VIRAL_GENE_EXPRESSION | 0.00000 | 0.00000 | | 0.80028 | 1.76317 | 171 |
| GO_VIRAL_GENOME_REPLICATION | 0.00100 | 0.01761 | 0.45506 | 0.61997 | 1.33078 | 73 |
| GO_VIRAL_LIFE_CYCLE | 0.00031 | 0.00640 | 0.49849 | 0.55767 | 1.23509 | 209 |
| HP_ABNORMALITY_OF_ACID_BASE_HOMEOSTASIS | 0.00216 | 0.03242 | 0.43171 | 0.53556 | 1.18846 | 232 |
| HP_ABNORMALITY_OF_CARDIOVASCULAR_SYSTEM_ELECTROPHYSIO | 0.00027 | 0.00572 | 0.49849 | 0.55832 | 1.23929 | 229 |
| HP_ABNORMALITY_OF_LOWER_LIP | 0.00181 | 0.02804 | 0.45506 | 0.56360 | 1.23779 | 145 |
| HP_ABNORMALITY_OF_THE_MITOCHONDRION | 0.00213 | 0.03229 | 0.43171 | 0.57831 | 1.26240 | 117 |
| HP_ABNORMALITY_OF_THE_NASAL_DORSUM | 0.00005 | 0.00116 | 0.55733 | 0.60288 | 1.32157 | 134 |
| HP_ABNORMAL_ELECTROPHYSIOLOGY_OF_SINOATRIAL_NODE_ORIG | 0.00279 | 0.03889 | 0.43171 | 0.68436 | 1.40934 | 31 |
| HP_ABNORMAL_ERYTHROCYTE_MORPHOLOGY | 0.00000 | 0.00015 | 0.61053 | 0.56049 | 1.25036 | 341 |
| HP_ABNORMAL_MEAN_CORPUSCULAR_VOLUME | 0.00357 | 0.04696 | 0.43171 | 0.75226 | 1.51363 | 18 |
| HP_ABNORMAL_RETICULOCYTE_MORPHOLOGY | 0.00064 | 0.01210 | 0.47727 | 0.71420 | 1.47081 | 31 |
| HP_ACUTE_LEUKEMIA | 0.00000 | 0.00000 | 0.73376 | 0.72786 | 1.56137 | 70 |
| HP_ANEMIA_OF_INADEQUATE_PRODUCTION | 0.00000 | 0.00008 | 0.62726 | 0.65593 | 1.42209 | 97 |
| HP_CONSTITUTIONAL_SYMPTOM | 0.00119 | 0.02004 | 0.45506 | 0.51427 | 1.15108 | 423 |
| HP_DELAYED_PUBERTY | 0.00005 | 0.00115 | 0.55733 | 0.62579 | 1.36211 | 107 |
| HP_DEPRESSED_NASAL_RIDGE | 0.00000 | 0.00011 | 0.62726 | 0.68731 | 1.47511 | 72 |
| HP_FATIGUE | 0.00039 | 0.00808 | 0.49849 | 0.56529 | 1.24923 | 187 |
| HP_HEADACHE | 0.00015 | 0.00340 | 0.51885 | 0.59065 | 1.29716 | 146 |
| HP_HEMATOLOGICAL_NEOPLASM | 0.00003 | 0.00078 | 0.57561 | 0.60108 | 1.32044 | 154 |
| HP_INCREASED_CSF_LACTATE | 0.00157 | 0.02509 | 0.45506 | 0.61622 | 1.31904 | 66 |
| HP_INCREASED_SERUM_LACTATE | 0.00216 | 0.03242 | 0.43171 | 0.56920 | 1.24724 | 130 |
| HP_LEUKEMIA | 0.00001 | 0.00026 | 0.59333 | 0.64877 | 1.40680 | 98 |
| HP_MACROCYTIC_ANEMIA | 0.00000 | 0.00000 | 0.74774 | 0.82468 | 1.71698 | 39 |
| HP_MIGRAINE | 0.00000 | 0.00000 | 0.71951 | 0.72105 | 1.54497 | 69 |
| HP_MYOTONIA | 0.00161 | 0.02546 | 0.45506 | 0.79820 | 1.58338 | 15 |
| HP_NEOPLASM | 0.00216 | 0.03242 | 0.43171 | 0.50820 | 1.13805 | 446 |
| HP_PALLOR | 0.00001 | 0.00040 | 0.59333 | 0.65169 | 1.41044 | 92 |
| HP_PUBERTY_AND_GONADAL_DISORDERS | 0.00287 | 0.03973 | 0.43171 | 0.51995 | 1.15843 | 322 |
| HP_RETICULOCYTOPENIA | 0.00004 | 0.00105 | 0.55733 | 0.86417 | 1.71425 | 15 |
| HP_RIMMED_VACUOLES | 0.00279 | 0.03889 | 0.43171 | 0.74033 | 1.49831 | 20 |
| HP_SHORT_NOSE | 0.00235 | 0.03474 | 0.43171 | 0.54919 | 1.21134 | 174 |
| HP_THICK_LOWER_LIP_VERMILION | 0.00048 | 0.00949 | 0.49849 | 0.60773 | 1.31617 | 94 |
| HP_THICK_VERMILION_BORDER | 0.00156 | 0.02486 | 0.45506 | 0.55983 | 1.23000 | 153 |
| HP_VENTRICULAR_ARRHYTHMIA | 0.00353 | 0.04658 | 0.43171 | 0.66198 | 1.37398 | 36 |
| **Cluster 1** | | | | | | |
| pathway | pval | padj | log2err | ES | NES | size |
| GO_ACTIVATION_OF_IMMUNE_RESPONSE | 0.00000 | 0.00000 | 0.78819 | 0.62851 | 1.39255 | 212 |
| GO_ACTIVATION_OF_INNATE_IMMUNE_RESPONSE | 0.00000 | 0.00000 | | 0.75054 | 1.62130 | 88 |
| GO_AEROBIC_RESPIRATION | 0.00001 | 0.00025 | 0.59333 | 0.69302 | 1.48168 | 65 |
| GO_ALTERNATIVE_MRNA_SPLICING_VIA_SPLICEOSOME | 0.00102 | 0.01782 | 0.45506 | 0.65168 | 1.38687 | 60 |
| GO_AMINE_METABOLIC_PROCESS | 0.00000 | 0.00000 | | 0.79189 | 1.70783 | 80 |
| GO_ANAPHASE_PROMOTING_COMPLEX_DEPENDENT_CATABOLIC_P | 0.00000 | 0.00000 | | 0.76611 | 1.65070 | 79 |
| GO_ANCHORING_JUNCTION | 0.00000 | 0.00000 | | 0.59615 | 1.33633 | 479 |
| GO_ANIMAL_ORGAN_REGENERATION | 0.00257 | 0.03716 | 0.43171 | 0.68240 | 1.41020 | 33 |
| GO_ANTIGEN_PROCESSING_AND_PRESENTATION | 0.00000 | 0.00000 | 0.70498 | 0.64776 | 1.42241 | 143 |
| GO_ANTIGEN_PROCESSING_AND_PRESENTATION_OF_EXOGENOUS_ | 0.00000 | 0.00000 | | 0.81051 | 1.72358 | 57 |
| GO_ANTIGEN_PROCESSING_AND_PRESENTATION_OF_PEPTIDE_ANTI | 0.00000 | 0.00000 | 0.70498 | 0.65912 | 1.44248 | 126 |
| GO_ANTIGEN_PROCESSING_AND_PRESENTATION_OF_PEPTIDE_ANTI | 0.00000 | 0.00000 | | 0.76925 | 1.65015 | 70 |
| GO_ANTIGEN_RECEPTOR_MEDIATED_SIGNALING_PATHWAY | 0.00000 | 0.00000 | 0.80122 | 0.67877 | 1.48641 | 127 |
| GO_APOPTOTIC_SIGNALING_PATHWAY | 0.00154 | 0.02464 | 0.45506 | 0.51561 | 1.15085 | 358 |
| GO_ATP_DEPENDENT_CHROMATIN_REMODELING | 0.00253 | 0.03681 | 0.43171 | 0.62124 | 1.32400 | 61 |
| GO_ATP_METABOLIC_PROCESS | 0.00000 | 0.00000 | 0.71951 | 0.62252 | 1.37693 | 196 |
| GO_ATP_SYNTHESIS_COUPLED_ELECTRON_TRANSPORT | 0.00005 | 0.00131 | 0.55733 | 0.65519 | 1.40723 | 74 |

| | | | | | | |
|---|---|---|---|---|---|---|
| GO_BINDING_OF_SPERM_TO_ZONA_PELLUCIDA | 0.00000 | 0.00001 | 0.69013 | 0.89879 | 1.80846 | 18 |
| GO_CADHERIN_BINDING | 0.00000 | 0.00000 | | 0.63068 | 1.39961 | 236 |
| GO_CANONICAL_WNT_SIGNALING_PATHWAY | 0.00000 | 0.00003 | 0.65944 | 0.60816 | 1.34553 | 197 |
| GO_CATALYTIC_STEP_2_SPLICEOSOME | 0.00000 | 0.00002 | 0.65944 | 0.68068 | 1.46976 | 83 |
| GO_CELLULAR_AMINO_ACID_METABOLIC_PROCESS | 0.00000 | 0.00000 | 0.74774 | 0.61752 | 1.36824 | 216 |
| GO_CELLULAR_COMPONENT_DISASSEMBLY | 0.00000 | 0.00005 | 0.64355 | 0.55505 | 1.24197 | 401 |
| GO_CELLULAR_KETONE_METABOLIC_PROCESS | 0.00000 | 0.00000 | 0.77494 | 0.65955 | 1.44893 | 149 |
| GO_CELLULAR_NITROGEN_COMPOUND_CATABOLIC_PROCESS | 0.00000 | 0.00000 | | 0.74079 | 1.65848 | 436 |
| GO_CELLULAR_PROTEIN_COMPLEX_DISASSEMBLY | 0.00006 | 0.00139 | 0.55733 | 0.58222 | 1.28459 | 176 |
| GO_CELLULAR_RESPIRATION | 0.00000 | 0.00000 | 0.71951 | 0.64997 | 1.42492 | 136 |
| GO_CELLULAR_RESPONSE_TO_HEAT | 0.00018 | 0.00403 | 0.51885 | 0.61962 | 1.34016 | 91 |
| GO_CELLULAR_RESPONSE_TO_ORGANIC_CYCLIC_COMPOUND | 0.00302 | 0.04108 | 0.43171 | 0.52402 | 1.16585 | 283 |
| GO_CELLULAR_RESPONSE_TO_OXYGEN_LEVELS | 0.00000 | 0.00000 | 0.78819 | 0.65911 | 1.44882 | 157 |
| GO_CELL_ACTIVATION_INVOLVED_IN_IMMUNE_RESPONSE | 0.00000 | 0.00001 | 0.67496 | 0.57339 | 1.27913 | 341 |
| GO_CELL_ADHESION_MOLECULE_BINDING | 0.00000 | 0.00000 | 0.81404 | 0.60529 | 1.34788 | 308 |
| GO_CELL_CELL_RECOGNITION | 0.00000 | 0.00002 | 0.65944 | 0.83826 | 1.71003 | 26 |
| GO_CELL_CELL_SIGNALING_BY_WNT | 0.00000 | 0.00002 | 0.67496 | 0.57736 | 1.28559 | 299 |
| GO_CELL_CYCLE_G2_M_PHASE_TRANSITION | 0.00001 | 0.00024 | 0.59333 | 0.58034 | 1.28602 | 214 |
| GO_CELL_CYCLE_PHASE_TRANSITION | 0.00000 | 0.00004 | 0.64355 | 0.54319 | 1.21681 | 464 |
| GO_CELL_DIVISION | 0.00154 | 0.02464 | 0.45506 | 0.51551 | 1.15425 | 427 |
| GO_CELL_RECOGNITION | 0.00012 | 0.00270 | 0.53843 | 0.69554 | 1.46186 | 48 |
| GO_CELL_SUBSTRATE_JUNCTION | 0.00000 | 0.00000 | | 0.67751 | 1.50775 | 291 |
| GO_CELL_SURFACE_RECEPTOR_SIGNALING_PATHWAY_INVOLVED_IN | 0.00000 | 0.00003 | 0.65944 | 0.57035 | 1.27067 | 319 |
| GO_CHAPERONE_BINDING | 0.00007 | 0.00163 | 0.53843 | 0.64754 | 1.39253 | 76 |
| GO_CHAPERONE_COMPLEX | 0.00000 | 0.00004 | 0.64355 | 0.85718 | 1.73925 | 22 |
| GO_CHAPERONE_MEDIATED_PROTEIN_COMPLEX_ASSEMBLY | 0.00170 | 0.02662 | 0.45506 | 0.77423 | 1.54792 | 17 |
| GO_CHROMATIN_BINDING | 0.00071 | 0.01323 | 0.47727 | 0.52699 | 1.17680 | 369 |
| GO_CHROMOSOMAL_REGION | 0.00246 | 0.03601 | 0.43171 | 0.52919 | 1.17661 | 276 |
| GO_COP9_SIGNALOSOME | 0.00128 | 0.02141 | 0.45506 | 0.73616 | 1.49715 | 24 |
| GO_COTRANSLATIONAL_PROTEIN_TARGETING_TO_MEMBRANE | 0.00000 | 0.00000 | | 0.89471 | 1.93513 | 91 |
| GO_CYTOCHROME_COMPLEX | 0.00168 | 0.02641 | 0.45506 | 0.75311 | 1.52417 | 20 |
| GO_CYTOKINE_MEDIATED_SIGNALING_PATHWAY | 0.00000 | 0.00000 | 0.81404 | 0.59698 | 1.33170 | 339 |
| GO_CYTOPLASMIC_TRANSLATION | 0.00000 | 0.00000 | | 0.84753 | 1.83080 | 88 |
| GO_CYTOPLASMIC_TRANSLATIONAL_INITIATION | 0.00000 | 0.00000 | 0.74774 | 0.85688 | 1.76176 | 30 |
| GO_CYTOSOLIC_LARGE_RIBOSOMAL_SUBUNIT | 0.00000 | 0.00000 | | 0.91186 | 1.92988 | 53 |
| GO_CYTOSOLIC_RIBOSOME | 0.00000 | 0.00000 | | 0.89907 | 1.94710 | 95 |
| GO_CYTOSOLIC_SMALL_RIBOSOMAL_SUBUNIT | 0.00000 | 0.00000 | | 0.89114 | 1.85533 | 39 |
| GO_DEFENSE_RESPONSE_TO_OTHER_ORGANISM | 0.00004 | 0.00099 | 0.55733 | 0.53673 | 1.20127 | 415 |
| GO_DE_NOVO_PROTEIN_FOLDING | 0.00050 | 0.00978 | 0.47727 | 0.74232 | 1.51431 | 26 |
| GO_DISORDERED_DOMAIN_SPECIFIC_BINDING | 0.00161 | 0.02546 | 0.45506 | 0.71970 | 1.47197 | 27 |
| GO_DNA_BIOSYNTHETIC_PROCESS | 0.00000 | 0.00010 | 0.62726 | 0.62505 | 1.37017 | 134 |
| GO_DNA_CONFORMATION_CHANGE | 0.00027 | 0.00572 | 0.49849 | 0.55300 | 1.22527 | 216 |
| GO_DNA_DAMAGE_RESPONSE_DETECTION_OF_DNA_DAMAGE | 0.00198 | 0.03028 | 0.43171 | 0.67949 | 1.40669 | 35 |
| GO_DNA_GEOMETRIC_CHANGE | 0.00046 | 0.00925 | 0.49849 | 0.60807 | 1.31855 | 98 |
| GO_DNA_POLYMERASE_BINDING | 0.00268 | 0.03799 | 0.43171 | 0.76255 | 1.52457 | 17 |
| GO_DNA_REPAIR | 0.00265 | 0.03775 | 0.43171 | 0.51159 | 1.14547 | 427 |
| GO_DRUG_BINDING | 0.00176 | 0.02733 | 0.45506 | 0.64119 | 1.35096 | 49 |
| GO_ELECTRON_TRANSFER_ACTIVITY | 0.00001 | 0.00020 | 0.61053 | 0.67481 | 1.44938 | 74 |
| GO_ELECTRON_TRANSPORT_CHAIN | 0.00002 | 0.00053 | 0.57561 | 0.62257 | 1.36145 | 124 |
| GO_ENDOPEPTIDASE_ACTIVITY | 0.00124 | 0.02085 | 0.45506 | 0.54955 | 1.21076 | 171 |
| GO_ENDOPEPTIDASE_COMPLEX | 0.00000 | 0.00000 | | 0.80930 | 1.73234 | 66 |
| GO_ENERGY_DERIVATION_BY_OXIDATION_OF_ORGANIC_COMPOUN | 0.00000 | 0.00000 | 0.71951 | 0.63390 | 1.39707 | 172 |
| GO_ESTABLISHMENT_OF_PROTEIN_LOCALIZATION_TO_CHROMOSON | 0.00020 | 0.00432 | 0.51885 | 0.77339 | 1.57217 | 23 |
| GO_ESTABLISHMENT_OF_PROTEIN_LOCALIZATION_TO_ENDOPLASM | 0.00000 | 0.00000 | | 0.88026 | 1.91149 | 102 |
| GO_ESTABLISHMENT_OF_PROTEIN_LOCALIZATION_TO_MEMBRANE | 0.00000 | 0.00000 | | 0.74874 | 1.66164 | 240 |
| GO_ESTABLISHMENT_OF_PROTEIN_LOCALIZATION_TO_ORGANELLE | 0.00000 | 0.00000 | | 0.69519 | 1.55605 | 423 |
| GO_ESTABLISHMENT_OF_PROTEIN_LOCALIZATION_TO_TELOMERE | 0.00006 | 0.00139 | 0.55733 | 0.83470 | 1.66881 | 17 |
| GO_ESTABLISHMENT_OF_RNA_LOCALIZATION | 0.00106 | 0.01819 | 0.45506 | 0.55577 | 1.22400 | 167 |
| GO_ESTABLISHMENT_OF_TISSUE_POLARITY | 0.00000 | 0.00000 | | 0.75051 | 1.62122 | 88 |
| GO_EUKARYOTIC_TRANSLATION_INITIATION_FACTOR_3_COMPLEX | 0.00003 | 0.00074 | 0.57561 | 0.86987 | 1.72555 | 15 |
| GO_EXOCYTOSIS | 0.00000 | 0.00015 | 0.61053 | 0.53905 | 1.20753 | 465 |
| GO_FC_EPSILON_RECEPTOR_SIGNALING_PATHWAY | 0.00000 | 0.00000 | | 0.75117 | 1.62103 | 82 |
| GO_FC_RECEPTOR_SIGNALING_PATHWAY | 0.00000 | 0.00000 | | 0.72054 | 1.57787 | 127 |
| GO_FERTILIZATION | 0.00089 | 0.01592 | 0.47727 | 0.63620 | 1.35888 | 64 |
| GO_FICOLIN_1_RICH_GRANULE | 0.00000 | 0.00000 | 0.78819 | 0.71716 | 1.55340 | 93 |
| GO_GENERATION_OF_PRECURSOR_METABOLITES_AND_ENERGY | 0.00000 | 0.00000 | 0.77494 | 0.59261 | 1.32026 | 320 |
| GO_GENE_SILENCING | 0.00261 | 0.03750 | 0.43171 | 0.55578 | 1.22252 | 161 |
| GO_GLAND_DEVELOPMENT | 0.00154 | 0.02464 | 0.45506 | 0.54117 | 1.19797 | 206 |
| GO_GLYCOSYL_COMPOUND_METABOLIC_PROCESS | 0.00071 | 0.01323 | 0.47727 | 0.62524 | 1.34204 | 71 |
| GO_GUANYL_NUCLEOTIDE_BINDING | 0.00095 | 0.01679 | 0.47727 | 0.54096 | 1.19861 | 216 |
| GO_HEAT_SHOCK_PROTEIN_BINDING | 0.00003 | 0.00092 | 0.55733 | 0.64369 | 1.39056 | 85 |
| GO_HEMATOPOIETIC_PROGENITOR_CELL_DIFFERENTIATION | 0.00000 | 0.00000 | 0.82666 | 0.70307 | 1.53239 | 113 |
| GO_HEMATOPOIETIC_STEM_CELL_DIFFERENTIATION | 0.00000 | 0.00000 | | 0.79776 | 1.71217 | 72 |
| GO_HSP70_PROTEIN_BINDING | 0.00228 | 0.03391 | 0.43171 | 0.71143 | 1.45504 | 27 |
| GO_IMMUNE_RESPONSE_REGULATING_SIGNALING_PATHWAY | 0.00000 | 0.00000 | 0.76146 | 0.63494 | 1.40480 | 197 |
| GO_INNATE_IMMUNE_RESPONSE | 0.00003 | 0.00071 | 0.57561 | 0.55173 | 1.23107 | 347 |

| | | | | | | |
|---|---|---|---|---|---|---|
| GO_INNATE_IMMUNE_RESPONSE_ACTIVATING_SIGNAL_TRANSDUCT | 0.00000 | 0.00000 | | 0.79340 | 1.70281 | 72 |
| GO_INNER_MITOCHONDRIAL_MEMBRANE_PROTEIN_COMPLEX | 0.00002 | 0.00060 | 0.57561 | 0.65124 | 1.40678 | 88 |
| GO_INTERACTION_WITH_SYMBIONT | 0.00001 | 0.00026 | 0.59333 | 0.75084 | 1.56620 | 41 |
| GO_INTERLEUKIN_1_MEDIATED_SIGNALING_PATHWAY | 0.00000 | 0.00000 | | 0.81689 | 1.75671 | 76 |
| GO_INTRINSIC_APOPTOTIC_SIGNALING_PATHWAY | 0.00121 | 0.02029 | 0.45506 | 0.55118 | 1.21845 | 190 |
| GO_ION_CHANNEL_BINDING | 0.00087 | 0.01584 | 0.47727 | 0.63182 | 1.34683 | 62 |
| GO_ISOPEPTIDASE_ACTIVITY | 0.00139 | 0.02285 | 0.45506 | 0.78940 | 1.57097 | 16 |
| GO_LARGE_RIBOSOMAL_SUBUNIT | 0.00000 | 0.00000 | | 0.78135 | 1.70183 | 109 |
| GO_LEUKOCYTE_MEDIATED_IMMUNITY | 0.00000 | 0.00000 | 0.71951 | 0.57282 | 1.27895 | 364 |
| GO_MACROMOLECULE_TRANSMEMBRANE_TRANSPORTER_ACTIVITY | 0.00239 | 0.03510 | 0.43171 | 0.77734 | 1.54696 | 16 |
| GO_MAINTENANCE_OF_CELL_NUMBER | 0.00087 | 0.01584 | 0.47727 | 0.58944 | 1.28512 | 114 |
| GO_MATURATION_OF_LSU_RRNA | 0.00006 | 0.00151 | 0.53843 | 0.77606 | 1.58723 | 27 |
| GO_METHYLTRANSFERASE_COMPLEX | 0.00290 | 0.04014 | 0.43171 | 0.59476 | 1.28479 | 88 |
| GO_METHYL_CPG_BINDING | 0.00302 | 0.04108 | 0.43171 | 0.78053 | 1.54833 | 15 |
| GO_MIDBRAIN_DEVELOPMENT | 0.00021 | 0.00468 | 0.51885 | 0.68832 | 1.44457 | 47 |
| GO_MITOCHONDRIAL_ELECTRON_TRANSPORT_NADH_TO_UBIQUINO | 0.00331 | 0.04437 | 0.43171 | 0.65428 | 1.36724 | 42 |
| GO_MITOCHONDRIAL_GENE_EXPRESSION | 0.00048 | 0.00949 | 0.49849 | 0.57608 | 1.26525 | 152 |
| GO_MITOCHONDRIAL_LARGE_RIBOSOMAL_SUBUNIT | 0.00002 | 0.00050 | 0.57561 | 0.69789 | 1.48086 | 55 |
| GO_MITOCHONDRIAL_MATRIX | 0.00000 | 0.00002 | 0.65944 | 0.56045 | 1.25158 | 371 |
| GO_MITOCHONDRIAL_MEMBRANE_ORGANIZATION | 0.00361 | 0.04734 | 0.43171 | 0.59309 | 1.28094 | 86 |
| GO_MITOCHONDRIAL_PROTEIN_COMPLEX | 0.00000 | 0.00000 | 0.81404 | 0.63650 | 1.40931 | 208 |
| GO_MITOCHONDRIAL_TRANSLATION | 0.00004 | 0.00093 | 0.55733 | 0.61219 | 1.34135 | 129 |
| GO_MITOCHONDRIAL_TRANSLATIONAL_TERMINATION | 0.00000 | 0.00010 | 0.62726 | 0.66694 | 1.44128 | 87 |
| GO_MITOCHONDRION_ORGANIZATION | 0.00051 | 0.00993 | 0.47727 | 0.52284 | 1.16863 | 390 |
| GO_MODIFICATION_DEPENDENT_MACROMOLECULE_CATABOLIC_PR | 0.00002 | 0.00053 | 0.57561 | 0.53142 | 1.19156 | 486 |
| GO_MODULATION_BY_HOST_OF_SYMBIONT_PROCESS | 0.00015 | 0.00336 | 0.51885 | 0.72848 | 1.50542 | 33 |
| GO_MODULATION_OF_PROCESS_OF_OTHER_ORGANISM | 0.00019 | 0.00423 | 0.51885 | 0.66904 | 1.42286 | 59 |
| GO_MONOSACCHARIDE_BIOSYNTHETIC_PROCESS | 0.00335 | 0.04466 | 0.43171 | 0.61992 | 1.31781 | 58 |
| GO_MORPHOGENESIS_OF_AN_EPITHELIUM | 0.00006 | 0.00151 | 0.53843 | 0.55161 | 1.22755 | 290 |
| GO_MORPHOGENESIS_OF_A_POLARIZED_EPITHELIUM | 0.00000 | 0.00000 | | 0.72298 | 1.57057 | 103 |
| GO_MRNA_3_UTR_BINDING | 0.00003 | 0.00089 | 0.55733 | 0.67596 | 1.44827 | 68 |
| GO_MRNA_5_UTR_BINDING | 0.00006 | 0.00156 | 0.53843 | 0.80133 | 1.62592 | 22 |
| GO_MRNA_BINDING | 0.00000 | 0.00000 | | 0.66002 | 1.46353 | 222 |
| GO_MRNA_PROCESSING | 0.00000 | 0.00000 | 0.73376 | 0.56859 | 1.27295 | 418 |
| GO_MULTI_ORGANISM_LOCALIZATION | 0.00268 | 0.03799 | 0.43171 | 0.63245 | 1.33612 | 51 |
| GO_MULTI_ORGANISM_PROCESS | 0.00108 | 0.01839 | 0.45506 | 0.51114 | 1.14599 | 484 |
| GO_MYD88_DEPENDENT_TOLL_LIKE_RECEPTOR_SIGNALING_PATHW | 0.00002 | 0.00066 | 0.57561 | 0.85581 | 1.70312 | 16 |
| GO_MYELOID_LEUKOCYTE_ACTIVATION | 0.00000 | 0.00000 | 0.74774 | 0.58508 | 1.30479 | 327 |
| GO_MYELOID_LEUKOCYTE_MEDIATED_IMMUNITY | 0.00000 | 0.00000 | 0.76146 | 0.59841 | 1.33291 | 301 |
| GO_NADH_DEHYDROGENASE_ACTIVITY | 0.00383 | 0.04935 | 0.43171 | 0.66244 | 1.37139 | 35 |
| GO_NADH_DEHYDROGENASE_COMPLEX | 0.00376 | 0.04884 | 0.43171 | 0.65649 | 1.36421 | 38 |
| GO_NADH_METABOLIC_PROCESS | 0.00294 | 0.04036 | 0.43171 | 0.70642 | 1.44481 | 27 |
| GO_NCRNA_EXPORT_FROM_NUCLEUS | 0.00302 | 0.04108 | 0.43171 | 0.66820 | 1.38331 | 35 |
| GO_NCRNA_METABOLIC_PROCESS | 0.00000 | 0.00003 | 0.65944 | 0.55907 | 1.24907 | 381 |
| GO_NCRNA_PROCESSING | 0.00000 | 0.00003 | 0.65944 | 0.57175 | 1.27504 | 328 |
| GO_NEGATIVE_REGULATION_OF_CANONICAL_WNT_SIGNALING_PAT | 0.00000 | 0.00000 | | 0.71611 | 1.56038 | 111 |
| GO_NEGATIVE_REGULATION_OF_CELLULAR_AMIDE_METABOLIC_PRO | 0.00001 | 0.00020 | 0.61053 | 0.63031 | 1.37423 | 114 |
| GO_NEGATIVE_REGULATION_OF_CELL_CYCLE | 0.00003 | 0.00076 | 0.57561 | 0.53833 | 1.20513 | 425 |
| GO_NEGATIVE_REGULATION_OF_CELL_CYCLE_G2_M_PHASE_TRANS | 0.00000 | 0.00000 | 0.76146 | 0.71490 | 1.54439 | 85 |
| GO_NEGATIVE_REGULATION_OF_CELL_CYCLE_PHASE_TRANSITION | 0.00002 | 0.00060 | 0.57561 | 0.59041 | 1.30442 | 184 |
| GO_NEGATIVE_REGULATION_OF_CELL_CYCLE_PROCESS | 0.00000 | 0.00002 | 0.65944 | 0.58690 | 1.30374 | 249 |
| GO_NEGATIVE_REGULATION_OF_MITOTIC_CELL_CYCLE | 0.00004 | 0.00095 | 0.55733 | 0.56735 | 1.25902 | 232 |
| GO_NEGATIVE_REGULATION_OF_MRNA_METABOLIC_PROCESS | 0.00023 | 0.00491 | 0.51885 | 0.66670 | 1.41788 | 59 |
| GO_NEGATIVE_REGULATION_OF_MRNA_SPLICING_VIA_SPLICEOSOM | 0.00239 | 0.03510 | 0.43171 | 0.74857 | 1.51204 | 19 |
| GO_NEGATIVE_REGULATION_OF_RNA_SPLICING | 0.00029 | 0.00607 | 0.49849 | 0.76729 | 1.55976 | 23 |
| GO_NEGATIVE_REGULATION_OF_TRANSLATIONAL_INITIATION | 0.00018 | 0.00403 | 0.51885 | 0.83901 | 1.66434 | 15 |
| GO_NEGATIVE_REGULATION_OF_WNT_SIGNALING_PATHWAY | 0.00000 | 0.00000 | | 0.69525 | 1.52249 | 127 |
| GO_NEURAL_NUCLEUS_DEVELOPMENT | 0.00001 | 0.00026 | 0.59333 | 0.79148 | 1.62187 | 28 |
| GO_NEURON_TO_NEURON_SYNAPSE | 0.00005 | 0.00134 | 0.55733 | 0.59636 | 1.31006 | 154 |
| GO_NIK_NF_KAPPAB_SIGNALING | 0.00000 | 0.00000 | | 0.72896 | 1.58354 | 103 |
| GO_NON_CANONICAL_WNT_SIGNALING_PATHWAY | 0.00000 | 0.00000 | | 0.72784 | 1.58023 | 101 |
| GO_NUCLEAR_EXPORT | 0.00031 | 0.00645 | 0.49849 | 0.56963 | 1.25542 | 172 |
| GO_NUCLEAR_TRANSCRIBED_MRNA_CATABOLIC_PROCESS | 0.00000 | 0.00000 | | 0.79179 | 1.74917 | 183 |
| GO_NUCLEAR_TRANSCRIBED_MRNA_CATABOLIC_PROCESS_NONSEN | 0.00000 | 0.00000 | | 0.89008 | 1.93909 | 110 |
| GO_NUCLEAR_TRANSPORT | 0.00008 | 0.00193 | 0.53843 | 0.55511 | 1.23408 | 273 |
| GO_NUCLEOBASE_CONTAINING_COMPOUND_TRANSPORT | 0.00146 | 0.02386 | 0.45506 | 0.55053 | 1.21842 | 202 |
| GO_NUCLEOBASE_CONTAINING_SMALL_MOLECULE_BIOSYNTHETIC_ | 0.00137 | 0.02261 | 0.45506 | 0.62749 | 1.34353 | 67 |
| GO_NUCLEOBASE_CONTAINING_SMALL_MOLECULE_INTERCONVERS | 0.00383 | 0.04935 | 0.43171 | 0.71259 | 1.44857 | 23 |
| GO_NUCLEOBASE_CONTAINING_SMALL_MOLECULE_METABOLIC_PR | 0.00320 | 0.04339 | 0.43171 | 0.51448 | 1.14825 | 352 |
| GO_NUCLEOSIDE_DIPHOSPHATE_METABOLIC_PROCESS | 0.00110 | 0.01865 | 0.45506 | 0.59637 | 1.29295 | 97 |
| GO_NUCLEOSIDE_METABOLIC_PROCESS | 0.00061 | 0.01169 | 0.47727 | 0.65315 | 1.38907 | 59 |
| GO_NUCLEOSIDE_MONOPHOSPHATE_BIOSYNTHETIC_PROCESS | 0.00250 | 0.03636 | 0.43171 | 0.68841 | 1.42033 | 32 |
| GO_NUCLEOSIDE_MONOPHOSPHATE_METABOLIC_PROCESS | 0.00152 | 0.02464 | 0.45506 | 0.64377 | 1.36002 | 51 |
| GO_NUCLEOSIDE_PHOSPHATE_BIOSYNTHETIC_PROCESS | 0.00043 | 0.00865 | 0.49849 | 0.58185 | 1.27831 | 148 |
| GO_NUCLEOSIDE_TRIPHOSPHATE_BIOSYNTHETIC_PROCESS | 0.00043 | 0.00865 | 0.49849 | 0.69332 | 1.44539 | 40 |
| GO_NUCLEOSIDE_TRIPHOSPHATE_METABOLIC_PROCESS | 0.00016 | 0.00360 | 0.51885 | 0.66788 | 1.41976 | 58 |

| | | | | | | |
|---|---|---|---|---|---|---|
| GO_NUCLEOSOMAL_DNA_BINDING | 0.00023 | 0.00507 | 0.51885 | 0.79852 | 1.61801 | 21 |
| GO_NUCLEOSOME_BINDING | 0.00143 | 0.02332 | 0.45506 | 0.66132 | 1.38492 | 45 |
| GO_NUCLEOTIDE_EXCISION_REPAIR | 0.00004 | 0.00096 | 0.55733 | 0.63343 | 1.37355 | 98 |
| GO_NUCLEOTIDE_EXCISION_REPAIR_DNA_DAMAGE_RECOGNITION | 0.00015 | 0.00349 | 0.51885 | 0.78678 | 1.59640 | 22 |
| GO_NUCLEOTIDE_EXCISION_REPAIR_DNA_INCISION | 0.00313 | 0.04249 | 0.43171 | 0.66439 | 1.37898 | 36 |
| GO_NUCLEOTIDE_PHOSPHORYLATION | 0.00383 | 0.04935 | 0.43171 | 0.59002 | 1.27454 | 88 |
| GO_ORGANELLAR_RIBOSOME | 0.00000 | 0.00014 | 0.61053 | 0.66412 | 1.43471 | 85 |
| GO_ORGANELLE_INNER_MEMBRANE | 0.00000 | 0.00000 | 0.76146 | 0.57601 | 1.28697 | 378 |
| GO_ORGANIC_CYCLIC_COMPOUND_CATABOLIC_PROCESS | 0.00000 | 0.00000 | | 0.73350 | 1.64281 | 449 |
| GO_OXIDATIVE_PHOSPHORYLATION | 0.00001 | 0.00030 | 0.59333 | 0.64619 | 1.40027 | 96 |
| GO_OXIDOREDUCTASE_ACTIVITY | 0.00006 | 0.00139 | 0.55733 | 0.53544 | 1.19703 | 392 |
| GO_OXIDOREDUCTASE_COMPLEX | 0.00001 | 0.00027 | 0.59333 | 0.66015 | 1.42461 | 82 |
| GO_PEPTIDASE_COMPLEX | 0.00000 | 0.00000 | | 0.75988 | 1.63982 | 82 |
| GO_PIGMENT_GRANULE | 0.00003 | 0.00089 | 0.55733 | 0.66098 | 1.42344 | 78 |
| GO_POLYSOMAL_RIBOSOME | 0.00000 | 0.00000 | | 0.90624 | 1.86629 | 31 |
| GO_POLYSOME | 0.00000 | 0.00000 | | 0.82337 | 1.75481 | 61 |
| GO_POLY_PYRIMIDINE_TRACT_BINDING | 0.00327 | 0.04398 | 0.43171 | 0.75471 | 1.51855 | 18 |
| GO_POSITIVE_REGULATION_OF_APOPTOTIC_SIGNALING_PATHWAY | 0.00279 | 0.03889 | 0.43171 | 0.57666 | 1.25518 | 107 |
| GO_POSITIVE_REGULATION_OF_ATPASE_ACTIVITY | 0.00294 | 0.04036 | 0.43171 | 0.65446 | 1.36795 | 43 |
| GO_POSITIVE_REGULATION_OF_BINDING | 0.00088 | 0.01590 | 0.47727 | 0.60284 | 1.30556 | 95 |
| GO_POSITIVE_REGULATION_OF_CANONICAL_WNT_SIGNALING_PATH | 0.00000 | 0.00000 | 0.77494 | 0.70504 | 1.53160 | 103 |
| GO_POSITIVE_REGULATION_OF_CELLULAR_AMIDE_METABOLIC_PRC | 0.00003 | 0.00074 | 0.57561 | 0.62296 | 1.36083 | 118 |
| GO_POSITIVE_REGULATION_OF_CELLULAR_PROTEIN_LOCALIZATION | 0.00041 | 0.00840 | 0.49849 | 0.54915 | 1.21815 | 225 |
| GO_POSITIVE_REGULATION_OF_CELL_DEATH | 0.00069 | 0.01297 | 0.47727 | 0.52595 | 1.17423 | 363 |
| GO_POSITIVE_REGULATION_OF_CELL_POPULATION_PROLIFERATION | 0.00078 | 0.01455 | 0.47727 | 0.52449 | 1.17108 | 368 |
| GO_POSITIVE_REGULATION_OF_CHROMOSOME_ORGANIZATION | 0.00231 | 0.03437 | 0.43171 | 0.56990 | 1.24871 | 133 |
| GO_POSITIVE_REGULATION_OF_DEFENSE_RESPONSE | 0.00000 | 0.00003 | 0.65944 | 0.62175 | 1.36730 | 158 |
| GO_POSITIVE_REGULATION_OF_DNA_BIOSYNTHETIC_PROCESS | 0.00001 | 0.00041 | 0.59333 | 0.71929 | 1.52055 | 52 |
| GO_POSITIVE_REGULATION_OF_DNA_METABOLIC_PROCESS | 0.00364 | 0.04761 | 0.43171 | 0.56647 | 1.24189 | 135 |
| GO_POSITIVE_REGULATION_OF_DNA_TEMPLATED_TRANSCRIPTION_ | 0.00276 | 0.03885 | 0.43171 | 0.68932 | 1.41586 | 29 |
| GO_POSITIVE_REGULATION_OF_ESTABLISHMENT_OF_PROTEIN_LOCA | 0.00007 | 0.00168 | 0.53843 | 0.56461 | 1.25165 | 219 |
| GO_POSITIVE_REGULATION_OF_IMMUNE_RESPONSE | 0.00000 | 0.00001 | 0.69013 | 0.59037 | 1.31265 | 277 |
| GO_POSITIVE_REGULATION_OF_IMMUNE_SYSTEM_PROCESS | 0.00004 | 0.00110 | 0.55733 | 0.54018 | 1.20716 | 386 |
| GO_POSITIVE_REGULATION_OF_INNATE_IMMUNE_RESPONSE | 0.00000 | 0.00000 | 0.81404 | 0.69693 | 1.51947 | 114 |
| GO_POSITIVE_REGULATION_OF_INTRINSIC_APOPTOTIC_SIGNALING_ | 0.00104 | 0.01798 | 0.45506 | 0.69097 | 1.43578 | 37 |
| GO_POSITIVE_REGULATION_OF_ORGANELLE_ORGANIZATION | 0.00043 | 0.00865 | 0.49849 | 0.52093 | 1.16605 | 411 |
| GO_POSITIVE_REGULATION_OF_PROTEIN_LOCALIZATION_TO_NUCLE | 0.00001 | 0.00020 | 0.61053 | 0.69907 | 1.48990 | 61 |
| GO_POSITIVE_REGULATION_OF_RESPONSE_TO_BIOTIC_STIMULUS | 0.00000 | 0.00000 | 0.73376 | 0.66340 | 1.45310 | 128 |
| GO_POSITIVE_REGULATION_OF_RESPONSE_TO_EXTERNAL_STIMULU | 0.00002 | 0.00050 | 0.57561 | 0.58133 | 1.28681 | 201 |
| GO_POSITIVE_REGULATION_OF_TELOMERASE_ACTIVITY | 0.00061 | 0.01169 | 0.47727 | 0.73476 | 1.50565 | 28 |
| GO_POSITIVE_REGULATION_OF_TELOMERASE_RNA_LOCALIZATION_ | 0.00000 | 0.00001 | 0.69013 | 0.92451 | 1.83395 | 15 |
| GO_POSITIVE_REGULATION_OF_TELOMERE_MAINTENANCE_VIA_TE | 0.00130 | 0.02159 | 0.45506 | 0.71259 | 1.46508 | 30 |
| GO_POSITIVE_REGULATION_OF_TRANSCRIPTION_INITIATION_FROM | 0.00372 | 0.04847 | 0.43171 | 0.70671 | 1.43993 | 25 |
| GO_POSITIVE_REGULATION_OF_TRANSFERASE_ACTIVITY | 0.00174 | 0.02712 | 0.45506 | 0.51642 | 1.15220 | 345 |
| GO_POSITIVE_REGULATION_OF_TRANSLATION | 0.00001 | 0.00019 | 0.61053 | 0.64767 | 1.40419 | 97 |
| GO_POSITIVE_REGULATION_OF_VIRAL_GENOME_REPLICATION | 0.00287 | 0.03973 | 0.43171 | 0.68362 | 1.40783 | 31 |
| GO_POSITIVE_REGULATION_OF_VIRAL_PROCESS | 0.00028 | 0.00590 | 0.49849 | 0.62440 | 1.34741 | 81 |
| GO_POSITIVE_REGULATION_OF_WNT_SIGNALING_PATHWAY | 0.00000 | 0.00000 | 0.73376 | 0.66230 | 1.44763 | 122 |
| GO_POSTREPLICATION_REPAIR | 0.00024 | 0.00515 | 0.51885 | 0.69140 | 1.44518 | 43 |
| GO_POSTSYNAPSE | 0.00004 | 0.00109 | 0.55733 | 0.55969 | 1.24369 | 259 |
| GO_POSTSYNAPTIC_CYTOSOL | 0.00279 | 0.03889 | 0.43171 | 0.78295 | 1.55313 | 15 |
| GO_POSTTRANSCRIPTIONAL_REGULATION_OF_GENE_EXPRESSION | 0.00000 | 0.00000 | | 0.66656 | 1.49327 | 462 |
| GO_POST_TRANSLATIONAL_PROTEIN_MODIFICATION | 0.00002 | 0.00052 | 0.57561 | 0.57851 | 1.28412 | 229 |
| GO_PRECATALYTIC_SPLICEOSOME | 0.00085 | 0.01559 | 0.47727 | 0.65235 | 1.37448 | 49 |
| GO_PRERIBOSOME | 0.00087 | 0.01577 | 0.47727 | 0.62899 | 1.34772 | 69 |
| GO_PRERIBOSOME_LARGE_SUBUNIT_PRECURSOR | 0.00235 | 0.03474 | 0.43171 | 0.74079 | 1.50308 | 22 |
| GO_PRE_MRNA_BINDING | 0.00133 | 0.02213 | 0.45506 | 0.72393 | 1.48062 | 27 |
| GO_PROTEASOMAL_PROTEIN_CATABOLIC_PROCESS | 0.00002 | 0.00044 | 0.59333 | 0.54166 | 1.21005 | 377 |
| GO_PROTEASOMAL_UBIQUITIN_INDEPENDENT_PROTEIN_CATABOLI | 0.00000 | 0.00000 | 0.80122 | 0.92254 | 1.86931 | 21 |
| GO_PROTEASOME_ACCESSORY_COMPLEX | 0.00000 | 0.00001 | 0.67496 | 0.85464 | 1.73811 | 24 |
| GO_PROTEASOME_CORE_COMPLEX | 0.00000 | 0.00000 | | 0.94375 | 1.90631 | 19 |
| GO_PROTEIN_CONTAINING_COMPLEX_DISASSEMBLY | 0.00000 | 0.00007 | 0.62726 | 0.57810 | 1.28438 | 256 |
| GO_PROTEIN_CONTAINING_COMPLEX_LOCALIZATION | 0.00063 | 0.01197 | 0.47727 | 0.55335 | 1.22520 | 208 |
| GO_PROTEIN_DNA_COMPLEX_SUBUNIT_ORGANIZATION | 0.00001 | 0.00018 | 0.61053 | 0.60174 | 1.32491 | 165 |
| GO_PROTEIN_FOLDING | 0.00000 | 0.00002 | 0.65944 | 0.63113 | 1.38644 | 154 |
| GO_PROTEIN_FOLDING_CHAPERONE | 0.00135 | 0.02238 | 0.45506 | 0.77903 | 1.55752 | 17 |
| GO_PROTEIN_IMPORT | 0.00268 | 0.03799 | 0.43171 | 0.56317 | 1.23581 | 140 |
| GO_PROTEIN_LOCALIZATION_TO_CHROMOSOME | 0.00056 | 0.01096 | 0.47727 | 0.63523 | 1.36108 | 69 |
| GO_PROTEIN_LOCALIZATION_TO_CHROMOSOME_TELOMERIC_REGI | 0.00335 | 0.04466 | 0.43171 | 0.70399 | 1.43984 | 27 |
| GO_PROTEIN_LOCALIZATION_TO_ENDOPLASMIC_RETICULUM | 0.00000 | 0.00000 | | 0.85521 | 1.87021 | 124 |
| GO_PROTEIN_LOCALIZATION_TO_MEMBRANE | 0.00000 | 0.00000 | | 0.65123 | 1.45714 | 406 |
| GO_PROTEIN_LOCALIZATION_TO_MITOCHONDRION | 0.00019 | 0.00420 | 0.51885 | 0.60773 | 1.32398 | 110 |
| GO_PROTEIN_LOCALIZATION_TO_NUCLEUS | 0.00002 | 0.00055 | 0.57561 | 0.58926 | 1.30220 | 187 |
| GO_PROTEIN_MODIFICATION_BY_SMALL_PROTEIN_REMOVAL | 0.00000 | 0.00000 | 0.71951 | 0.60745 | 1.34689 | 221 |
| GO_PROTEIN_POLYUBIQUITINATION | 0.00004 | 0.00106 | 0.55733 | 0.55874 | 1.24108 | 253 |
| GO_PROTEIN_REFOLDING | 0.00294 | 0.04036 | 0.43171 | 0.77039 | 1.53314 | 16 |

| | | | | | | |
|---|---|---|---|---|---|---|
| GO_PROTEIN_STABILIZATION | 0.00001 | 0.00024 | 0.61053 | 0.62333 | 1.36586 | 130 |
| GO_PROTEIN_TARGETING | 0.00000 | 0.00000 | | 0.71181 | 1.58743 | 327 |
| GO_PROTEIN_TARGETING_TO_MEMBRANE | 0.00000 | 0.00000 | | 0.82370 | 1.80953 | 149 |
| GO_PROTEIN_TARGETING_TO_MITOCHONDRION | 0.00346 | 0.04593 | 0.43171 | 0.60408 | 1.30091 | 78 |
| GO_PROTEIN_TRANSMEMBRANE_TRANSPORT | 0.00265 | 0.03775 | 0.43171 | 0.64528 | 1.36171 | 50 |
| GO_PROTON_TRANSMEMBRANE_TRANSPORT | 0.00213 | 0.03229 | 0.43171 | 0.60815 | 1.30536 | 71 |
| GO_PURINE_CONTAINING_COMPOUND_BIOSYNTHETIC_PROCESS | 0.00246 | 0.03601 | 0.43171 | 0.57796 | 1.25936 | 111 |
| GO_PURINE_CONTAINING_COMPOUND_METABOLIC_PROCESS | 0.00100 | 0.01761 | 0.45506 | 0.53679 | 1.19242 | 255 |
| GO_PURINE_NUCLEOSIDE_TRIPHOSPHATE_METABOLIC_PROCESS | 0.00089 | 0.01592 | 0.47727 | 0.67157 | 1.40374 | 43 |
| GO_PYRIMIDINE_CONTAINING_COMPOUND_METABOLIC_PROCESS | 0.00353 | 0.04658 | 0.43171 | 0.64744 | 1.35878 | 47 |
| GO_REGENERATION | 0.00216 | 0.03242 | 0.43171 | 0.59987 | 1.29453 | 82 |
| GO_REGULATION_OF_ANIMAL_ORGAN_MORPHOGENESIS | 0.00000 | 0.00000 | | 0.69163 | 1.51541 | 133 |
| GO_REGULATION_OF_APOPTOTIC_SIGNALING_PATHWAY | 0.00031 | 0.00645 | 0.49849 | 0.55171 | 1.22506 | 247 |
| GO_REGULATION_OF_ATPASE_ACTIVITY | 0.00324 | 0.04369 | 0.43171 | 0.63031 | 1.34038 | 57 |
| GO_REGULATION_OF_ATP_METABOLIC_PROCESS | 0.00353 | 0.04658 | 0.43171 | 0.60176 | 1.29780 | 80 |
| GO_REGULATION_OF_BINDING | 0.00250 | 0.03636 | 0.43171 | 0.54476 | 1.20598 | 203 |
| GO_REGULATION_OF_CELLULAR_AMIDE_METABOLIC_PROCESS | 0.00000 | 0.00000 | | 0.62967 | 1.40226 | 311 |
| GO_REGULATION_OF_CELLULAR_AMINE_METABOLIC_PROCESS | 0.00000 | 0.00000 | | 0.82173 | 1.74682 | 58 |
| GO_REGULATION_OF_CELLULAR_AMINO_ACID_METABOLIC_PROCES | 0.00000 | 0.00000 | | 0.83776 | 1.77766 | 55 |
| GO_REGULATION_OF_CELLULAR_KETONE_METABOLIC_PROCESS | 0.00000 | 0.00000 | | 0.72641 | 1.58680 | 118 |
| GO_REGULATION_OF_CELLULAR_PROTEIN_LOCALIZATION | 0.00082 | 0.01513 | 0.47727 | 0.52542 | 1.17290 | 362 |
| GO_REGULATION_OF_CELLULAR_RESPONSE_TO_HEAT | 0.00276 | 0.03885 | 0.43171 | 0.61652 | 1.32004 | 67 |
| GO_REGULATION_OF_CELLULAR_RESPONSE_TO_STRESS | 0.00224 | 0.03344 | 0.43171 | 0.50545 | 1.13232 | 468 |
| GO_REGULATION_OF_CELL_CYCLE_G2_M_PHASE_TRANSITION | 0.00000 | 0.00000 | 0.71951 | 0.63230 | 1.39282 | 169 |
| GO_REGULATION_OF_CELL_CYCLE_PHASE_TRANSITION | 0.00001 | 0.00026 | 0.59333 | 0.55488 | 1.23783 | 337 |
| GO_REGULATION_OF_CHROMOSOME_ORGANIZATION | 0.00081 | 0.01501 | 0.47727 | 0.54255 | 1.20527 | 261 |
| GO_REGULATION_OF_CYSTEINE_TYPE_ENDOPEPTIDASE_ACTIVITY | 0.00202 | 0.03077 | 0.43171 | 0.57120 | 1.24837 | 121 |
| GO_REGULATION_OF_CYTOPLASMIC_TRANSLATION | 0.00130 | 0.02159 | 0.45506 | 0.73678 | 1.49773 | 23 |
| GO_REGULATION_OF_DEFENSE_RESPONSE | 0.00065 | 0.01237 | 0.47727 | 0.53633 | 1.19355 | 290 |
| GO_REGULATION_OF_DNA_BIOSYNTHETIC_PROCESS | 0.00016 | 0.00360 | 0.51885 | 0.64215 | 1.37923 | 74 |
| GO_REGULATION_OF_DNA_METABOLIC_PROCESS | 0.00069 | 0.01297 | 0.47727 | 0.54425 | 1.20775 | 237 |
| GO_REGULATION_OF_DNA_TEMPLATED_TRANSCRIPTION_INITIATIOI | 0.00163 | 0.02568 | 0.45506 | 0.68289 | 1.41372 | 35 |
| GO_REGULATION_OF_DNA_TEMPLATED_TRANSCRIPTION_IN_RESPO | 0.00000 | 0.00000 | | 0.73944 | 1.59800 | 89 |
| GO_REGULATION_OF_GENERATION_OF_PRECURSOR_METABOLITES_ | 0.00060 | 0.01164 | 0.47727 | 0.60476 | 1.31116 | 97 |
| GO_REGULATION_OF_HEMATOPOIETIC_PROGENITOR_CELL_DIFFERE | 0.00000 | 0.00000 | | 0.78717 | 1.69251 | 75 |
| GO_REGULATION_OF_HEMATOPOIETIC_STEM_CELL_DIFFERENTIATIC | 0.00000 | 0.00000 | | 0.81505 | 1.74256 | 65 |
| GO_REGULATION_OF_HEMOPOIESIS | 0.00000 | 0.00008 | 0.62726 | 0.58859 | 1.30634 | 228 |
| GO_REGULATION_OF_IMMUNE_RESPONSE | 0.00000 | 0.00016 | 0.61053 | 0.55362 | 1.23583 | 362 |
| GO_REGULATION_OF_INNATE_IMMUNE_RESPONSE | 0.00000 | 0.00000 | 0.71951 | 0.63883 | 1.40521 | 161 |
| GO_REGULATION_OF_INTRINSIC_APOPTOTIC_SIGNALING_PATHWAY | 0.00001 | 0.00039 | 0.59333 | 0.63478 | 1.38259 | 109 |
| GO_REGULATION_OF_MITOTIC_CELL_CYCLE | 0.00001 | 0.00018 | 0.61053 | 0.53899 | 1.20736 | 458 |
| GO_REGULATION_OF_MORPHOGENESIS_OF_AN_EPITHELIUM | 0.00000 | 0.00000 | | 0.72213 | 1.57284 | 109 |
| GO_REGULATION_OF_MRNA_CATABOLIC_PROCESS | 0.00000 | 0.00000 | | 0.69977 | 1.54127 | 168 |
| GO_REGULATION_OF_MRNA_METABOLIC_PROCESS | 0.00000 | 0.00000 | | 0.66528 | 1.47872 | 266 |
| GO_REGULATION_OF_MRNA_PROCESSING | 0.00004 | 0.00105 | 0.55733 | 0.61521 | 1.34090 | 113 |
| GO_REGULATION_OF_MRNA_SPLICING_VIA_SPLICEOSOME | 0.00001 | 0.00026 | 0.59333 | 0.67424 | 1.45411 | 80 |
| GO_REGULATION_OF_NUCLEASE_ACTIVITY | 0.00106 | 0.01819 | 0.45506 | 0.77918 | 1.56780 | 18 |
| GO_REGULATION_OF_OXIDATIVE_STRESS_INDUCED_CELL_DEATH | 0.00198 | 0.03028 | 0.43171 | 0.67287 | 1.40355 | 41 |
| GO_REGULATION_OF_PEPTIDASE_ACTIVITY | 0.00050 | 0.00978 | 0.47727 | 0.56122 | 1.24054 | 191 |
| GO_REGULATION_OF_POSTTRANSCRIPTIONAL_GENE_SILENCING | 0.00364 | 0.04761 | 0.43171 | 0.59824 | 1.28492 | 74 |
| GO_REGULATION_OF_PROTEIN_CATABOLIC_PROCESS | 0.00179 | 0.02783 | 0.45506 | 0.53057 | 1.17929 | 270 |
| GO_REGULATION_OF_PROTEIN_LOCALIZATION_TO_NUCLEUS | 0.00001 | 0.00042 | 0.59333 | 0.65505 | 1.41558 | 87 |
| GO_REGULATION_OF_PROTEIN_MODIFICATION_BY_SMALL_PROTEIN | 0.00025 | 0.00532 | 0.49849 | 0.57068 | 1.25684 | 167 |
| GO_REGULATION_OF_PROTEIN_STABILITY | 0.00093 | 0.01652 | 0.47727 | 0.55382 | 1.22571 | 202 |
| GO_REGULATION_OF_PROTEOLYSIS | 0.00013 | 0.00301 | 0.51885 | 0.52997 | 1.18392 | 377 |
| GO_REGULATION_OF_RESPONSE_TO_BIOTIC_STIMULUS | 0.00001 | 0.00020 | 0.61053 | 0.59049 | 1.30719 | 204 |
| GO_REGULATION_OF_RNA_SPLICING | 0.00000 | 0.00001 | 0.69013 | 0.65896 | 1.43669 | 114 |
| GO_REGULATION_OF_SMALL_MOLECULE_METABOLIC_PROCESS | 0.00000 | 0.00000 | 0.71951 | 0.59134 | 1.31394 | 265 |
| GO_REGULATION_OF_STEM_CELL_DIFFERENTIATION | 0.00000 | 0.00000 | | 0.74049 | 1.60113 | 90 |
| GO_REGULATION_OF_SYMBIOTIC_PROCESS | 0.00006 | 0.00144 | 0.55733 | 0.60071 | 1.31682 | 134 |
| GO_REGULATION_OF_TELOMERASE_ACTIVITY | 0.00088 | 0.01590 | 0.47727 | 0.69161 | 1.43179 | 35 |
| GO_REGULATION_OF_TRANSCRIPTION_FROM_RNA_POLYMERASE_II | 0.00000 | 0.00000 | | 0.81194 | 1.73800 | 66 |
| GO_REGULATION_OF_TRANSCRIPTION_INITIATION_FROM_RNA_POL | 0.00143 | 0.02332 | 0.45506 | 0.70390 | 1.44582 | 29 |
| GO_REGULATION_OF_TRANSLATIONAL_INITIATION | 0.00002 | 0.00052 | 0.57561 | 0.68129 | 1.45518 | 64 |
| GO_REGULATION_OF_UBIQUITIN_PROTEIN_TRANSFERASE_ACTIVITY | 0.00061 | 0.01169 | 0.47727 | 0.69335 | 1.44627 | 41 |
| GO_REGULATION_OF_VIRAL_GENOME_REPLICATION | 0.00086 | 0.01566 | 0.47727 | 0.65316 | 1.38389 | 54 |
| GO_REGULATION_OF_VIRAL_TRANSCRIPTION | 0.00198 | 0.03028 | 0.43171 | 0.63932 | 1.34703 | 49 |
| GO_REGULATION_OF_WNT_SIGNALING_PATHWAY | 0.00000 | 0.00001 | 0.69013 | 0.60353 | 1.33816 | 220 |
| GO_RESPIRASOME | 0.00008 | 0.00181 | 0.53843 | 0.65420 | 1.40421 | 71 |
| GO_RESPIRATORY_CHAIN_COMPLEX | 0.00016 | 0.00353 | 0.51885 | 0.67626 | 1.43918 | 60 |
| GO_RESPIRATORY_ELECTRON_TRANSPORT_CHAIN | 0.00001 | 0.00025 | 0.59333 | 0.65819 | 1.42190 | 85 |
| GO_RESPONSE_TO_ALCOHOL | 0.00261 | 0.03750 | 0.43171 | 0.58804 | 1.27350 | 95 |
| GO_RESPONSE_TO_CADMIUM_ION | 0.00265 | 0.03775 | 0.43171 | 0.67227 | 1.39174 | 35 |
| GO_RESPONSE_TO_DRUG | 0.00104 | 0.01798 | 0.45506 | 0.55364 | 1.22258 | 180 |
| GO_RESPONSE_TO_ETHANOL | 0.00086 | 0.01566 | 0.47727 | 0.67309 | 1.40691 | 43 |
| GO_RESPONSE_TO_HEAT | 0.00012 | 0.00270 | 0.53843 | 0.61665 | 1.34221 | 107 |

| | | | | | | |
|---|---|---|---|---|---|---|
| GO_RESPONSE_TO_HORMONE | 0.00048 | 0.00949 | 0.49849 | 0.51277 | 1.14814 | 453 |
| GO_RESPONSE_TO_HYDROGEN_PEROXIDE | 0.00257 | 0.03716 | 0.43171 | 0.60505 | 1.30368 | 79 |
| GO_RESPONSE_TO_INORGANIC_SUBSTANCE | 0.00004 | 0.00107 | 0.55733 | 0.55714 | 1.23870 | 275 |
| GO_RESPONSE_TO_INTERLEUKIN_1 | 0.00000 | 0.00000 | | 0.74844 | 1.62908 | 107 |
| GO_RESPONSE_TO_INTERLEUKIN_12 | 0.00012 | 0.00270 | 0.53843 | 0.72605 | 1.50694 | 36 |
| GO_RESPONSE_TO_INTERLEUKIN_4 | 0.00324 | 0.04369 | 0.43171 | 0.75812 | 1.51571 | 17 |
| GO_RESPONSE_TO_METAL_ION | 0.00030 | 0.00623 | 0.49849 | 0.56387 | 1.24233 | 171 |
| GO_RESPONSE_TO_ORGANIC_CYCLIC_COMPOUND | 0.00000 | 0.00016 | 0.61053 | 0.54603 | 1.22259 | 439 |
| GO_RESPONSE_TO_OXYGEN_LEVELS | 0.00000 | 0.00000 | 0.71951 | 0.60132 | 1.33438 | 233 |
| GO_RESPONSE_TO_STEROID_HORMONE | 0.00154 | 0.02464 | 0.45506 | 0.55514 | 1.22446 | 174 |
| GO_RESPONSE_TO_TEMPERATURE_STIMULUS | 0.00027 | 0.00572 | 0.49849 | 0.59216 | 1.29675 | 127 |
| GO_RESPONSE_TO_TUMOR_NECROSIS_FACTOR | 0.00000 | 0.00000 | 0.76146 | 0.65649 | 1.44236 | 151 |
| GO_RETINA_HOMEOSTASIS | 0.00104 | 0.01798 | 0.45506 | 0.73063 | 1.49432 | 27 |
| GO_RIBONUCLEOPROTEIN_COMPLEX_BINDING | 0.00000 | 0.00000 | 0.71951 | 0.66906 | 1.46050 | 117 |
| GO_RIBONUCLEOPROTEIN_COMPLEX_BIOGENESIS | 0.00000 | 0.00000 | | 0.66339 | 1.48264 | 389 |
| GO_RIBONUCLEOPROTEIN_COMPLEX_SUBUNIT_ORGANIZATION | 0.00000 | 0.00000 | | 0.69241 | 1.52681 | 173 |
| GO_RIBONUCLEOPROTEIN_GRANULE | 0.00000 | 0.00001 | 0.67496 | 0.61979 | 1.36366 | 163 |
| GO_RIBONUCLEOSIDE_TRIPHOSPHATE_BIOSYNTHETIC_PROCESS | 0.00041 | 0.00840 | 0.49849 | 0.71253 | 1.47248 | 33 |
| GO_RIBONUCLEOSIDE_TRIPHOSPHATE_METABOLIC_PROCESS | 0.00026 | 0.00550 | 0.49849 | 0.69681 | 1.45611 | 42 |
| GO_RIBOSE_PHOSPHATE_BIOSYNTHETIC_PROCESS | 0.00191 | 0.02939 | 0.45506 | 0.57932 | 1.26207 | 110 |
| GO_RIBOSE_PHOSPHATE_METABOLIC_PROCESS | 0.00108 | 0.01839 | 0.45506 | 0.53891 | 1.19687 | 245 |
| GO_RIBOSOMAL_LARGE_SUBUNIT_ASSEMBLY | 0.00000 | 0.00016 | 0.61053 | 0.80999 | 1.65663 | 27 |
| GO_RIBOSOMAL_LARGE_SUBUNIT_BIOGENESIS | 0.00000 | 0.00000 | | 0.77542 | 1.65982 | 66 |
| GO_RIBOSOMAL_SMALL_SUBUNIT_ASSEMBLY | 0.00023 | 0.00494 | 0.51885 | 0.81139 | 1.62221 | 17 |
| GO_RIBOSOMAL_SMALL_SUBUNIT_BINDING | 0.00383 | 0.04935 | 0.43171 | 0.75288 | 1.50523 | 17 |
| GO_RIBOSOMAL_SMALL_SUBUNIT_BIOGENESIS | 0.00001 | 0.00022 | 0.61053 | 0.69076 | 1.47997 | 68 |
| GO_RIBOSOMAL_SUBUNIT | 0.00000 | 0.00000 | | 0.77603 | 1.71031 | 172 |
| GO_RIBOSOME | 0.00000 | 0.00000 | | 0.76643 | 1.69659 | 206 |
| GO_RIBOSOME_ASSEMBLY | 0.00000 | 0.00000 | 0.73376 | 0.76173 | 1.61998 | 59 |
| GO_RIBOSOME_BINDING | 0.00019 | 0.00416 | 0.51885 | 0.68955 | 1.44715 | 47 |
| GO_RIBOSOME_BIOGENESIS | 0.00000 | 0.00000 | | 0.65877 | 1.46466 | 275 |
| GO_RNA_CATABOLIC_PROCESS | 0.00000 | 0.00000 | | 0.76774 | 1.71264 | 339 |
| GO_RNA_DEPENDENT_DNA_BIOSYNTHETIC_PROCESS | 0.00003 | 0.00076 | 0.57561 | 0.68926 | 1.46521 | 58 |
| GO_RNA_EXPORT_FROM_NUCLEUS | 0.00062 | 0.01183 | 0.47727 | 0.58378 | 1.27758 | 126 |
| GO_RNA_LOCALIZATION | 0.00000 | 0.00006 | 0.64355 | 0.59670 | 1.32020 | 198 |
| GO_RNA_POLYMERASE_II_PREINITIATION_COMPLEX_ASSEMBLY | 0.00089 | 0.01592 | 0.47727 | 0.77046 | 1.55626 | 19 |
| GO_RNA_SPLICING | 0.00000 | 0.00000 | 0.78819 | 0.59079 | 1.31927 | 369 |
| GO_RNA_SPLICING_VIA_TRANSESTERIFICATION_REACTIONS | 0.00000 | 0.00000 | | 0.60940 | 1.35714 | 300 |
| GO_RNA_STABILIZATION | 0.00065 | 0.01223 | 0.47727 | 0.69743 | 1.44384 | 35 |
| GO_ROUGH_ENDOPLASMIC_RETICULUM | 0.00001 | 0.00039 | 0.59333 | 0.71853 | 1.51629 | 50 |
| GO_RRNA_BINDING | 0.00000 | 0.00001 | 0.69013 | 0.74782 | 1.58865 | 56 |
| GO_RRNA_METABOLIC_PROCESS | 0.00000 | 0.00000 | 0.77494 | 0.63047 | 1.39562 | 206 |
| GO_SCF_DEPENDENT_PROTEASOMAL_UBIQUITIN_DEPENDENT_PRO | 0.00000 | 0.00000 | | 0.77527 | 1.67044 | 79 |
| GO_SECRETORY_GRANULE | 0.00000 | 0.00000 | 0.71951 | 0.56518 | 1.26461 | 399 |
| GO_SECRETORY_VESICLE | 0.00000 | 0.00002 | 0.67496 | 0.54866 | 1.22905 | 465 |
| GO_SIGNAL_TRANSDUCTION_BY_PROTEIN_PHOSPHORYLATION | 0.00004 | 0.00109 | 0.55733 | 0.53817 | 1.20474 | 424 |
| GO_SINGLE_FERTILIZATION | 0.00048 | 0.00949 | 0.49849 | 0.67414 | 1.42260 | 50 |
| GO_SINGLE_STRANDED_RNA_BINDING | 0.00039 | 0.00793 | 0.49849 | 0.66553 | 1.41527 | 57 |
| GO_SMALL_MOLECULE_BIOSYNTHETIC_PROCESS | 0.00216 | 0.03242 | 0.43171 | 0.51195 | 1.14522 | 397 |
| GO_SMALL_RIBOSOMAL_SUBUNIT | 0.00000 | 0.00000 | | 0.77648 | 1.66209 | 66 |
| GO_SM_LIKE_PROTEIN_FAMILY_COMPLEX | 0.00327 | 0.04398 | 0.43171 | 0.60468 | 1.29713 | 70 |
| GO_SPERM_EGG_RECOGNITION | 0.00000 | 0.00007 | 0.64355 | 0.85241 | 1.72955 | 22 |
| GO_SPLICEOSOMAL_COMPLEX | 0.00000 | 0.00003 | 0.65944 | 0.60939 | 1.34261 | 171 |
| GO_STEM_CELL_DIFFERENTIATION | 0.00000 | 0.00000 | 0.78819 | 0.66886 | 1.46930 | 147 |
| GO_STRUCTURAL_CONSTITUENT_OF_CYTOSKELETON | 0.00102 | 0.01782 | 0.45506 | 0.65913 | 1.39337 | 52 |
| GO_STRUCTURAL_CONSTITUENT_OF_RIBOSOME | 0.00000 | 0.00000 | | 0.79909 | 1.75541 | 150 |
| GO_STRUCTURAL_MOLECULE_ACTIVITY | 0.00000 | 0.00000 | | 0.72651 | 1.61895 | 324 |
| GO_SUBSTANTIA_NIGRA_DEVELOPMENT | 0.00000 | 0.00013 | 0.62726 | 0.81622 | 1.66507 | 26 |
| GO_TELOMERASE_RNA_LOCALIZATION | 0.00001 | 0.00017 | 0.61053 | 0.85164 | 1.72025 | 19 |
| GO_TELOMERE_MAINTENANCE_VIA_TELOMERE_LENGTHENING | 0.00011 | 0.00250 | 0.53843 | 0.65659 | 1.40546 | 66 |
| GO_TELOMERE_ORGANIZATION | 0.00342 | 0.04554 | 0.43171 | 0.57000 | 1.24649 | 124 |
| GO_THREONINE_TYPE_ENDOPEPTIDASE_ACTIVITY | 0.00000 | 0.00000 | | 0.95179 | 1.92255 | 19 |
| GO_THREONINE_TYPE_PEPTIDASE_ACTIVITY | 0.00000 | 0.00000 | | 0.92199 | 1.87423 | 23 |
| GO_TISSUE_MORPHOGENESIS | 0.00097 | 0.01707 | 0.47727 | 0.52661 | 1.17402 | 326 |
| GO_TOXIN_TRANSPORT | 0.00113 | 0.01922 | 0.45506 | 0.72897 | 1.48708 | 26 |
| GO_TRANSCRIPTION_COUPLED_NUCLEOTIDE_EXCISION_REPAIR | 0.00001 | 0.00019 | 0.61053 | 0.68106 | 1.46186 | 71 |
| GO_TRANSCRIPTION_FACTOR_BINDING | 0.00043 | 0.00865 | 0.49849 | 0.52447 | 1.17376 | 409 |
| GO_TRANSCRIPTION_PREINITIATION_COMPLEX_ASSEMBLY | 0.00119 | 0.02004 | 0.45506 | 0.71399 | 1.46797 | 30 |
| GO_TRANSLATIONAL_ELONGATION | 0.00000 | 0.00000 | 0.71951 | 0.66087 | 1.44451 | 122 |
| GO_TRANSLATIONAL_INITIATION | 0.00000 | 0.00000 | | 0.84691 | 1.86515 | 166 |
| GO_TRANSLATIONAL_TERMINATION | 0.00000 | 0.00005 | 0.64355 | 0.65946 | 1.43060 | 99 |
| GO_TRANSLATION_ELONGATION_FACTOR_ACTIVITY | 0.00161 | 0.02546 | 0.45506 | 0.79860 | 1.58418 | 15 |
| GO_TRANSLATION_FACTOR_ACTIVITY_RNA_BINDING | 0.00000 | 0.00000 | | 0.79008 | 1.69288 | 69 |
| GO_TRANSLATION_INITIATION_FACTOR_ACTIVITY | 0.00000 | 0.00000 | 0.76146 | 0.80422 | 1.68417 | 45 |
| GO_TRANSLATION_INITIATION_FACTOR_BINDING | 0.00021 | 0.00468 | 0.51885 | 0.76186 | 1.55231 | 25 |
| GO_TRANSLATION_PREINITIATION_COMPLEX | 0.00000 | 0.00001 | 0.67496 | 0.90245 | 1.79595 | 16 |

| | | | | | | |
|---|---|---|---|---|---|---|
| GO_TRANSLATION_REGULATOR_ACTIVITY | 0.00000 | 0.00000 | | 0.77663 | 1.68904 | 105 |
| GO_TRANSLATION_REGULATOR_ACTIVITY_NUCLEIC_ACID_BINDING | 0.00000 | 0.00000 | | 0.77150 | 1.66586 | 83 |
| GO_TRICARBOXYLIC_ACID_CYCLE | 0.00154 | 0.02464 | 0.45506 | 0.71854 | 1.47240 | 28 |
| GO_TUMOR_NECROSIS_FACTOR_MEDIATED_SIGNALING_PATHWAY | 0.00000 | 0.00000 | | 0.76277 | 1.65372 | 97 |
| GO_T_CELL_RECEPTOR_SIGNALING_PATHWAY | 0.00000 | 0.00000 | 0.81404 | 0.69994 | 1.52527 | 112 |
| GO_U12_TYPE_SPLICEOSOMAL_COMPLEX | 0.00279 | 0.03889 | 0.43171 | 0.70746 | 1.44693 | 27 |
| GO_U2_SNRNP | 0.00005 | 0.00134 | 0.55733 | 0.83081 | 1.67168 | 18 |
| GO_U2_TYPE_SPLICEOSOMAL_COMPLEX | 0.00010 | 0.00243 | 0.53843 | 0.63403 | 1.36970 | 85 |
| GO_UBIQUITIN_LIKE_PROTEIN_LIGASE_BINDING | 0.00000 | 0.00000 | 0.76146 | 0.62273 | 1.38051 | 219 |
| GO_UBIQUITIN_PROTEIN_TRANSFERASE_REGULATOR_ACTIVITY | 0.00165 | 0.02590 | 0.45506 | 0.79605 | 1.57911 | 15 |
| GO_UNFOLDED_PROTEIN_BINDING | 0.00000 | 0.00000 | 0.77494 | 0.72157 | 1.55881 | 85 |
| GO_VESICLE_LUMEN | 0.00000 | 0.00000 | 0.74774 | 0.64475 | 1.42008 | 168 |
| GO_VIRAL_GENE_EXPRESSION | 0.00000 | 0.00000 | | 0.80028 | 1.76317 | 171 |
| GO_VIRAL_GENOME_REPLICATION | 0.00100 | 0.01761 | 0.45506 | 0.61997 | 1.33078 | 73 |
| GO_VIRAL_LIFE_CYCLE | 0.00031 | 0.00640 | 0.49849 | 0.55767 | 1.23509 | 209 |
| HP_ABNORMALITY_OF_ACID_BASE_HOMEOSTASIS | 0.00216 | 0.03242 | 0.43171 | 0.53556 | 1.18846 | 232 |
| HP_ABNORMALITY_OF_CARDIOVASCULAR_SYSTEM_ELECTROPHYSIC | 0.00027 | 0.00572 | 0.49849 | 0.55832 | 1.23929 | 229 |
| HP_ABNORMALITY_OF_LOWER_LIP | 0.00181 | 0.02804 | 0.45506 | 0.56360 | 1.23779 | 145 |
| HP_ABNORMALITY_OF_THE_MITOCHONDRION | 0.00213 | 0.03229 | 0.43171 | 0.57831 | 1.26240 | 117 |
| HP_ABNORMALITY_OF_THE_NASAL_DORSUM | 0.00005 | 0.00116 | 0.55733 | 0.60288 | 1.32157 | 134 |
| HP_ABNORMAL_ELECTROPHYSIOLOGY_OF_SINOATRIAL_NODE_ORIG | 0.00279 | 0.03889 | 0.43171 | 0.68436 | 1.40934 | 31 |
| HP_ABNORMAL_ERYTHROCYTE_MORPHOLOGY | 0.00000 | 0.00015 | 0.61053 | 0.56049 | 1.25036 | 341 |
| HP_ABNORMAL_MEAN_CORPUSCULAR_VOLUME | 0.00357 | 0.04696 | 0.43171 | 0.75226 | 1.51363 | 18 |
| HP_ABNORMAL_RETICULOCYTE_MORPHOLOGY | 0.00064 | 0.01210 | 0.47727 | 0.71420 | 1.47081 | 31 |
| HP_ACUTE_LEUKEMIA | 0.00000 | 0.00000 | 0.73376 | 0.72786 | 1.56137 | 70 |
| HP_ANEMIA_OF_INADEQUATE_PRODUCTION | 0.00000 | 0.00008 | 0.62726 | 0.65593 | 1.42209 | 97 |
| HP_CONSTITUTIONAL_SYMPTOM | 0.00119 | 0.02004 | 0.45506 | 0.51427 | 1.15108 | 423 |
| HP_DELAYED_PUBERTY | 0.00005 | 0.00115 | 0.55733 | 0.62579 | 1.36211 | 107 |
| HP_DEPRESSED_NASAL_RIDGE | 0.00000 | 0.00011 | 0.62726 | 0.68731 | 1.47511 | 72 |
| HP_FATIGUE | 0.00039 | 0.00808 | 0.49849 | 0.56529 | 1.24923 | 187 |
| HP_HEADACHE | 0.00015 | 0.00340 | 0.51885 | 0.59065 | 1.29716 | 146 |
| HP_HEMATOLOGICAL_NEOPLASM | 0.00003 | 0.00078 | 0.57561 | 0.60108 | 1.32044 | 154 |
| HP_INCREASED_CSF_LACTATE | 0.00157 | 0.02509 | 0.45506 | 0.61622 | 1.31904 | 66 |
| HP_INCREASED_SERUM_LACTATE | 0.00216 | 0.03242 | 0.43171 | 0.56920 | 1.24724 | 130 |
| HP_LEUKEMIA | 0.00001 | 0.00026 | 0.59333 | 0.64877 | 1.40680 | 98 |
| HP_MACROCYTIC_ANEMIA | 0.00000 | 0.00000 | 0.74774 | 0.82468 | 1.71698 | 39 |
| HP_MIGRAINE | 0.00000 | 0.00000 | 0.71951 | 0.72105 | 1.54497 | 69 |
| HP_MYOTONIA | 0.00161 | 0.02546 | 0.45506 | 0.79820 | 1.58338 | 15 |
| HP_NEOPLASM | 0.00216 | 0.03242 | 0.43171 | 0.50820 | 1.13805 | 446 |
| HP_PALLOR | 0.00001 | 0.00040 | 0.59333 | 0.65169 | 1.41044 | 92 |
| HP_PUBERTY_AND_GONADAL_DISORDERS | 0.00287 | 0.03973 | 0.43171 | 0.51995 | 1.15843 | 322 |
| HP_RETICULOCYTOPENIA | 0.00004 | 0.00105 | 0.55733 | 0.86417 | 1.71425 | 15 |
| HP_RIMMED_VACUOLES | 0.00279 | 0.03889 | 0.43171 | 0.74033 | 1.49831 | 20 |
| HP_SHORT_NOSE | 0.00235 | 0.03474 | 0.43171 | 0.54919 | 1.21134 | 174 |
| HP_THICK_LOWER_LIP_VERMILION | 0.00048 | 0.00949 | 0.49849 | 0.60773 | 1.31617 | 94 |
| HP_THICK_VERMILION_BORDER | 0.00156 | 0.02486 | 0.45506 | 0.55983 | 1.23000 | 153 |
| HP_VENTRICULAR_ARRHYTHMIA | 0.00353 | 0.04658 | 0.43171 | 0.66198 | 1.37398 | 36 |
| **Cluster 1** | | | | | | |
| pathway | pval | padj | log2err | ES | NES | size |
| GO_ACTIVATION_OF_IMMUNE_RESPONSE | 0.00000 | 0.00000 | 0.78819 | 0.62851 | 1.39255 | 212 |
| GO_ACTIVATION_OF_INNATE_IMMUNE_RESPONSE | 0.00000 | 0.00000 | | 0.75054 | 1.62130 | 88 |
| GO_AEROBIC_RESPIRATION | 0.00001 | 0.00025 | 0.59333 | 0.69302 | 1.48168 | 65 |
| GO_ALTERNATIVE_MRNA_SPLICING_VIA_SPLICEOSOME | 0.00102 | 0.01782 | 0.45506 | 0.65168 | 1.38687 | 60 |
| GO_AMINE_METABOLIC_PROCESS | 0.00000 | 0.00000 | | 0.79189 | 1.70783 | 80 |
| GO_ANAPHASE_PROMOTING_COMPLEX_DEPENDENT_CATABOLIC_P | 0.00000 | 0.00000 | | 0.76611 | 1.65070 | 79 |
| GO_ANCHORING_JUNCTION | 0.00000 | 0.00000 | | 0.59615 | 1.33633 | 479 |
| GO_ANIMAL_ORGAN_REGENERATION | 0.00257 | 0.03716 | 0.43171 | 0.68240 | 1.41020 | 33 |
| GO_ANTIGEN_PROCESSING_AND_PRESENTATION | 0.00000 | 0.00000 | 0.70498 | 0.64776 | 1.42241 | 143 |
| GO_ANTIGEN_PROCESSING_AND_PRESENTATION_OF_EXOGENOUS_ | 0.00000 | 0.00000 | | 0.81051 | 1.72358 | 57 |
| GO_ANTIGEN_PROCESSING_AND_PRESENTATION_OF_PEPTIDE_ANTI | 0.00000 | 0.00000 | 0.70498 | 0.65912 | 1.44248 | 126 |
| GO_ANTIGEN_PROCESSING_AND_PRESENTATION_OF_PEPTIDE_ANTI | 0.00000 | 0.00000 | | 0.76925 | 1.65015 | 70 |
| GO_ANTIGEN_RECEPTOR_MEDIATED_SIGNALING_PATHWAY | 0.00000 | 0.00000 | 0.80122 | 0.67877 | 1.48641 | 127 |
| GO_APOPTOTIC_SIGNALING_PATHWAY | 0.00154 | 0.02464 | 0.45506 | 0.51561 | 1.15085 | 358 |
| GO_ATP_DEPENDENT_CHROMATIN_REMODELING | 0.00253 | 0.03681 | 0.43171 | 0.62124 | 1.32400 | 61 |
| GO_ATP_METABOLIC_PROCESS | 0.00000 | 0.00000 | 0.71951 | 0.62252 | 1.37693 | 196 |
| GO_ATP_SYNTHESIS_COUPLED_ELECTRON_TRANSPORT | 0.00005 | 0.00131 | 0.55733 | 0.65519 | 1.40723 | 74 |
| GO_BINDING_OF_SPERM_TO_ZONA_PELLUCIDA | 0.00000 | 0.00001 | 0.69013 | 0.89879 | 1.80846 | 18 |
| GO_CADHERIN_BINDING | 0.00000 | 0.00000 | | 0.63068 | 1.39961 | 236 |
| GO_CANONICAL_WNT_SIGNALING_PATHWAY | 0.00000 | 0.00003 | 0.65944 | 0.60816 | 1.34553 | 197 |
| GO_CATALYTIC_STEP_2_SPLICEOSOME | 0.00000 | 0.00002 | 0.65944 | 0.68068 | 1.46976 | 83 |
| GO_CELLULAR_AMINO_ACID_METABOLIC_PROCESS | 0.00000 | 0.00000 | 0.74774 | 0.61752 | 1.36824 | 216 |
| GO_CELLULAR_COMPONENT_DISASSEMBLY | 0.00000 | 0.00005 | 0.64355 | 0.55505 | 1.24197 | 401 |
| GO_CELLULAR_KETONE_METABOLIC_PROCESS | 0.00000 | 0.00000 | 0.77494 | 0.65955 | 1.44893 | 149 |
| GO_CELLULAR_NITROGEN_COMPOUND_CATABOLIC_PROCESS | 0.00000 | 0.00000 | | 0.74079 | 1.65848 | 436 |
| GO_CELLULAR_PROTEIN_COMPLEX_DISASSEMBLY | 0.00006 | 0.00139 | 0.55733 | 0.58222 | 1.28459 | 176 |
| GO_CELLULAR_RESPIRATION | 0.00000 | 0.00000 | 0.71951 | 0.64997 | 1.42492 | 136 |

| | | | | | | |
|---|---|---|---|---|---|---|
| GO_CELLULAR_RESPONSE_TO_HEAT | 0.00018 | 0.00403 | 0.51885 | 0.61962 | 1.34016 | 91 |
| GO_CELLULAR_RESPONSE_TO_ORGANIC_CYCLIC_COMPOUND | 0.00302 | 0.04108 | 0.43171 | 0.52402 | 1.16585 | 283 |
| GO_CELLULAR_RESPONSE_TO_OXYGEN_LEVELS | 0.00000 | 0.00000 | 0.78819 | 0.65911 | 1.44882 | 157 |
| GO_CELL_ACTIVATION_INVOLVED_IN_IMMUNE_RESPONSE | 0.00000 | 0.00001 | 0.67496 | 0.57339 | 1.27913 | 341 |
| GO_CELL_ADHESION_MOLECULE_BINDING | 0.00000 | 0.00000 | 0.81404 | 0.60529 | 1.34788 | 308 |
| GO_CELL_CELL_RECOGNITION | 0.00000 | 0.00002 | 0.65944 | 0.83826 | 1.71003 | 26 |
| GO_CELL_CELL_SIGNALING_BY_WNT | 0.00000 | 0.00002 | 0.67496 | 0.57736 | 1.28559 | 299 |
| GO_CELL_CYCLE_G2_M_PHASE_TRANSITION | 0.00001 | 0.00024 | 0.59333 | 0.58034 | 1.28602 | 214 |
| GO_CELL_CYCLE_PHASE_TRANSITION | 0.00000 | 0.00004 | 0.64355 | 0.54319 | 1.21681 | 464 |
| GO_CELL_DIVISION | 0.00154 | 0.02464 | 0.45506 | 0.51551 | 1.15425 | 427 |
| GO_CELL_RECOGNITION | 0.00012 | 0.00270 | 0.53843 | 0.69554 | 1.46186 | 48 |
| GO_CELL_SUBSTRATE_JUNCTION | 0.00000 | 0.00000 | | 0.67751 | 1.50775 | 291 |
| GO_CELL_SURFACE_RECEPTOR_SIGNALING_PATHWAY_INVOLVED_IN | 0.00000 | 0.00003 | 0.65944 | 0.57035 | 1.27067 | 319 |
| GO_CHAPERONE_BINDING | 0.00007 | 0.00163 | 0.53843 | 0.64754 | 1.39253 | 76 |
| GO_CHAPERONE_COMPLEX | 0.00000 | 0.00004 | 0.64355 | 0.85718 | 1.73925 | 22 |
| GO_CHAPERONE_MEDIATED_PROTEIN_COMPLEX_ASSEMBLY | 0.00170 | 0.02662 | 0.45506 | 0.77423 | 1.54792 | 17 |
| GO_CHROMATIN_BINDING | 0.00071 | 0.01323 | 0.47727 | 0.52699 | 1.17680 | 369 |
| GO_CHROMOSOMAL_REGION | 0.00246 | 0.03601 | 0.43171 | 0.52919 | 1.17661 | 276 |
| GO_COP9_SIGNALOSOME | 0.00128 | 0.02141 | 0.45506 | 0.73616 | 1.49715 | 24 |
| GO_COTRANSLATIONAL_PROTEIN_TARGETING_TO_MEMBRANE | 0.00000 | 0.00000 | | 0.89471 | 1.93513 | 91 |
| GO_CYTOCHROME_COMPLEX | 0.00168 | 0.02641 | 0.45506 | 0.75311 | 1.52417 | 20 |
| GO_CYTOKINE_MEDIATED_SIGNALING_PATHWAY | 0.00000 | 0.00000 | 0.81404 | 0.59698 | 1.33170 | 339 |
| GO_CYTOPLASMIC_TRANSLATION | 0.00000 | 0.00000 | | 0.84753 | 1.83080 | 88 |
| GO_CYTOPLASMIC_TRANSLATIONAL_INITIATION | 0.00000 | 0.00000 | 0.74774 | 0.85688 | 1.76176 | 30 |
| GO_CYTOSOLIC_LARGE_RIBOSOMAL_SUBUNIT | 0.00000 | 0.00000 | | 0.91186 | 1.92988 | 53 |
| GO_CYTOSOLIC_RIBOSOME | 0.00000 | 0.00000 | | 0.89907 | 1.94710 | 95 |
| GO_CYTOSOLIC_SMALL_RIBOSOMAL_SUBUNIT | 0.00000 | 0.00000 | | 0.89114 | 1.85533 | 39 |
| GO_DEFENSE_RESPONSE_TO_OTHER_ORGANISM | 0.00004 | 0.00099 | 0.55733 | 0.53673 | 1.20127 | 415 |
| GO_DE_NOVO_PROTEIN_FOLDING | 0.00050 | 0.00978 | 0.47727 | 0.74232 | 1.51431 | 26 |
| GO_DISORDERED_DOMAIN_SPECIFIC_BINDING | 0.00161 | 0.02546 | 0.45506 | 0.71970 | 1.47197 | 27 |
| GO_DNA_BIOSYNTHETIC_PROCESS | 0.00000 | 0.00010 | 0.62726 | 0.62505 | 1.37017 | 134 |
| GO_DNA_CONFORMATION_CHANGE | 0.00027 | 0.00572 | 0.49849 | 0.55300 | 1.22527 | 216 |
| GO_DNA_DAMAGE_RESPONSE_DETECTION_OF_DNA_DAMAGE | 0.00198 | 0.03028 | 0.43171 | 0.67949 | 1.40669 | 35 |
| GO_DNA_GEOMETRIC_CHANGE | 0.00046 | 0.00925 | 0.49849 | 0.60807 | 1.31855 | 98 |
| GO_DNA_POLYMERASE_BINDING | 0.00268 | 0.03799 | 0.43171 | 0.76255 | 1.52457 | 17 |
| GO_DNA_REPAIR | 0.00265 | 0.03775 | 0.43171 | 0.51159 | 1.14547 | 427 |
| GO_DRUG_BINDING | 0.00176 | 0.02733 | 0.45506 | 0.64119 | 1.35096 | 49 |
| GO_ELECTRON_TRANSFER_ACTIVITY | 0.00001 | 0.00020 | 0.61053 | 0.67481 | 1.44938 | 74 |
| GO_ELECTRON_TRANSPORT_CHAIN | 0.00002 | 0.00053 | 0.57561 | 0.62257 | 1.36145 | 124 |
| GO_ENDOPEPTIDASE_ACTIVITY | 0.00124 | 0.02085 | 0.45506 | 0.54955 | 1.21076 | 171 |
| GO_ENDOPEPTIDASE_COMPLEX | 0.00000 | 0.00000 | | 0.80930 | 1.73234 | 66 |
| GO_ENERGY_DERIVATION_BY_OXIDATION_OF_ORGANIC_COMPOUN | 0.00000 | 0.00000 | 0.71951 | 0.63390 | 1.39707 | 172 |
| GO_ESTABLISHMENT_OF_PROTEIN_LOCALIZATION_TO_CHROMOSON | 0.00020 | 0.00432 | 0.51885 | 0.77339 | 1.57217 | 23 |
| GO_ESTABLISHMENT_OF_PROTEIN_LOCALIZATION_TO_ENDOPLASM | 0.00000 | 0.00000 | | 0.88026 | 1.91149 | 102 |
| GO_ESTABLISHMENT_OF_PROTEIN_LOCALIZATION_TO_MEMBRANE | 0.00000 | 0.00000 | | 0.74874 | 1.66164 | 240 |
| GO_ESTABLISHMENT_OF_PROTEIN_LOCALIZATION_TO_ORGANELLE | 0.00000 | 0.00000 | | 0.69519 | 1.55605 | 423 |
| GO_ESTABLISHMENT_OF_PROTEIN_LOCALIZATION_TO_TELOMERE | 0.00006 | 0.00139 | 0.55733 | 0.83470 | 1.66881 | 17 |
| GO_ESTABLISHMENT_OF_RNA_LOCALIZATION | 0.00106 | 0.01819 | 0.45506 | 0.55577 | 1.22400 | 167 |
| GO_ESTABLISHMENT_OF_TISSUE_POLARITY | 0.00000 | 0.00000 | | 0.75051 | 1.62122 | 88 |
| GO_EUKARYOTIC_TRANSLATION_INITIATION_FACTOR_3_COMPLEX | 0.00003 | 0.00074 | 0.57561 | 0.86987 | 1.72555 | 15 |
| GO_EXOCYTOSIS | 0.00000 | 0.00015 | 0.61053 | 0.53905 | 1.20753 | 465 |
| GO_FC_EPSILON_RECEPTOR_SIGNALING_PATHWAY | 0.00000 | 0.00000 | | 0.75117 | 1.62103 | 82 |
| GO_FC_RECEPTOR_SIGNALING_PATHWAY | 0.00000 | 0.00000 | | 0.72054 | 1.57787 | 127 |
| GO_FERTILIZATION | 0.00089 | 0.01592 | 0.47727 | 0.63620 | 1.35888 | 64 |
| GO_FICOLIN_1_RICH_GRANULE | 0.00000 | 0.00000 | 0.78819 | 0.71716 | 1.55340 | 93 |
| GO_GENERATION_OF_PRECURSOR_METABOLITES_AND_ENERGY | 0.00000 | 0.00000 | 0.77494 | 0.59261 | 1.32026 | 320 |
| GO_GENE_SILENCING | 0.00261 | 0.03750 | 0.43171 | 0.55578 | 1.22252 | 161 |
| GO_GLAND_DEVELOPMENT | 0.00154 | 0.02464 | 0.45506 | 0.54117 | 1.19797 | 206 |
| GO_GLYCOSYL_COMPOUND_METABOLIC_PROCESS | 0.00071 | 0.01323 | 0.47727 | 0.62524 | 1.34204 | 71 |
| GO_GUANYL_NUCLEOTIDE_BINDING | 0.00095 | 0.01679 | 0.47727 | 0.54096 | 1.19861 | 216 |
| GO_HEAT_SHOCK_PROTEIN_BINDING | 0.00003 | 0.00092 | 0.55733 | 0.64369 | 1.39056 | 85 |
| GO_HEMATOPOIETIC_PROGENITOR_CELL_DIFFERENTIATION | 0.00000 | 0.00000 | 0.82666 | 0.70307 | 1.53239 | 113 |
| GO_HEMATOPOIETIC_STEM_CELL_DIFFERENTIATION | 0.00000 | 0.00000 | | 0.79776 | 1.71217 | 72 |
| GO_HSP70_PROTEIN_BINDING | 0.00228 | 0.03391 | 0.43171 | 0.71143 | 1.45504 | 27 |
| GO_IMMUNE_RESPONSE_REGULATING_SIGNALING_PATHWAY | 0.00000 | 0.00000 | 0.76146 | 0.63494 | 1.40480 | 197 |
| GO_INNATE_IMMUNE_RESPONSE | 0.00003 | 0.00071 | 0.57561 | 0.55173 | 1.23107 | 347 |
| GO_INNATE_IMMUNE_RESPONSE_ACTIVATING_SIGNAL_TRANSDUCT | 0.00000 | 0.00000 | | 0.79340 | 1.70281 | 72 |
| GO_INNER_MITOCHONDRIAL_MEMBRANE_PROTEIN_COMPLEX | 0.00002 | 0.00060 | 0.57561 | 0.65124 | 1.40678 | 88 |
| GO_INTERACTION_WITH_SYMBIONT | 0.00001 | 0.00026 | 0.59333 | 0.75084 | 1.56620 | 41 |
| GO_INTERLEUKIN_1_MEDIATED_SIGNALING_PATHWAY | 0.00000 | 0.00000 | | 0.81689 | 1.75671 | 76 |
| GO_INTRINSIC_APOPTOTIC_SIGNALING_PATHWAY | 0.00121 | 0.02029 | 0.45506 | 0.55118 | 1.21845 | 190 |
| GO_ION_CHANNEL_BINDING | 0.00087 | 0.01584 | 0.47727 | 0.63182 | 1.34683 | 62 |
| GO_ISOPEPTIDASE_ACTIVITY | 0.00139 | 0.02285 | 0.45506 | 0.78940 | 1.57097 | 16 |
| GO_LARGE_RIBOSOMAL_SUBUNIT | 0.00000 | 0.00000 | | 0.78135 | 1.70183 | 109 |
| GO_LEUKOCYTE_MEDIATED_IMMUNITY | 0.00000 | 0.00000 | 0.71951 | 0.57282 | 1.27895 | 364 |
| GO_MACROMOLECULE_TRANSMEMBRANE_TRANSPORTER_ACTIVITY | 0.00239 | 0.03510 | 0.43171 | 0.77734 | 1.54696 | 16 |

| | | | | | | |
|---|---|---|---|---|---|---|
| GO_MAINTENANCE_OF_CELL_NUMBER | 0.00087 | 0.01584 | 0.47727 | 0.58944 | 1.28512 | 114 |
| GO_MATURATION_OF_LSU_RRNA | 0.00006 | 0.00151 | 0.53843 | 0.77606 | 1.58723 | 27 |
| GO_METHYLTRANSFERASE_COMPLEX | 0.00290 | 0.04014 | 0.43171 | 0.59476 | 1.28479 | 88 |
| GO_METHYL_CPG_BINDING | 0.00302 | 0.04108 | 0.43171 | 0.78053 | 1.54833 | 15 |
| GO_MIDBRAIN_DEVELOPMENT | 0.00021 | 0.00468 | 0.51885 | 0.68832 | 1.44457 | 47 |
| GO_MITOCHONDRIAL_ELECTRON_TRANSPORT_NADH_TO_UBIQUIN( | 0.00331 | 0.04437 | 0.43171 | 0.65428 | 1.36724 | 42 |
| GO_MITOCHONDRIAL_GENE_EXPRESSION | 0.00048 | 0.00949 | 0.49849 | 0.57608 | 1.26525 | 152 |
| GO_MITOCHONDRIAL_LARGE_RIBOSOMAL_SUBUNIT | 0.00002 | 0.00050 | 0.57561 | 0.69789 | 1.48086 | 55 |
| GO_MITOCHONDRIAL_MATRIX | 0.00000 | 0.00002 | 0.65944 | 0.56045 | 1.25158 | 371 |
| GO_MITOCHONDRIAL_MEMBRANE_ORGANIZATION | 0.00361 | 0.04734 | 0.43171 | 0.59309 | 1.28094 | 86 |
| GO_MITOCHONDRIAL_PROTEIN_COMPLEX | 0.00000 | 0.00000 | 0.81404 | 0.63650 | 1.40931 | 208 |
| GO_MITOCHONDRIAL_TRANSLATION | 0.00004 | 0.00093 | 0.55733 | 0.61219 | 1.34135 | 129 |
| GO_MITOCHONDRIAL_TRANSLATIONAL_TERMINATION | 0.00000 | 0.00010 | 0.62726 | 0.66694 | 1.44128 | 87 |
| GO_MITOCHONDRION_ORGANIZATION | 0.00051 | 0.00993 | 0.47727 | 0.52284 | 1.16863 | 390 |
| GO_MODIFICATION_DEPENDENT_MACROMOLECULE_CATABOLIC_PR | 0.00002 | 0.00053 | 0.57561 | 0.53142 | 1.19156 | 486 |
| GO_MODULATION_BY_HOST_OF_SYMBIONT_PROCESS | 0.00015 | 0.00336 | 0.51885 | 0.72848 | 1.50542 | 33 |
| GO_MODULATION_OF_PROCESS_OF_OTHER_ORGANISM | 0.00019 | 0.00423 | 0.51885 | 0.66904 | 1.42286 | 59 |
| GO_MONOSACCHARIDE_BIOSYNTHETIC_PROCESS | 0.00335 | 0.04466 | 0.43171 | 0.61992 | 1.31781 | 58 |
| GO_MORPHOGENESIS_OF_AN_EPITHELIUM | 0.00006 | 0.00151 | 0.53843 | 0.55161 | 1.22755 | 290 |
| GO_MORPHOGENESIS_OF_A_POLARIZED_EPITHELIUM | 0.00000 | 0.00000 | | 0.72298 | 1.57057 | 103 |
| GO_MRNA_3_UTR_BINDING | 0.00003 | 0.00089 | 0.55733 | 0.67596 | 1.44827 | 68 |
| GO_MRNA_5_UTR_BINDING | 0.00006 | 0.00156 | 0.53843 | 0.80133 | 1.62592 | 22 |
| GO_MRNA_BINDING | 0.00000 | 0.00000 | | 0.66002 | 1.46353 | 222 |
| GO_MRNA_PROCESSING | 0.00000 | 0.00000 | 0.73376 | 0.56859 | 1.27295 | 418 |
| GO_MULTI_ORGANISM_LOCALIZATION | 0.00268 | 0.03799 | 0.43171 | 0.63245 | 1.33612 | 51 |
| GO_MULTI_ORGANISM_PROCESS | 0.00108 | 0.01839 | 0.45506 | 0.51114 | 1.14599 | 484 |
| GO_MYD88_DEPENDENT_TOLL_LIKE_RECEPTOR_SIGNALING_PATHW | 0.00002 | 0.00066 | 0.57561 | 0.85581 | 1.70312 | 16 |
| GO_MYELOID_LEUKOCYTE_ACTIVATION | 0.00000 | 0.00000 | 0.74774 | 0.58508 | 1.30479 | 327 |
| GO_MYELOID_LEUKOCYTE_MEDIATED_IMMUNITY | 0.00000 | 0.00000 | 0.76146 | 0.59841 | 1.33291 | 301 |
| GO_NADH_DEHYDROGENASE_ACTIVITY | 0.00383 | 0.04935 | 0.43171 | 0.66244 | 1.37139 | 35 |
| GO_NADH_DEHYDROGENASE_COMPLEX | 0.00376 | 0.04884 | 0.43171 | 0.65649 | 1.36421 | 38 |
| GO_NADH_METABOLIC_PROCESS | 0.00294 | 0.04036 | 0.43171 | 0.70642 | 1.44481 | 27 |
| GO_NCRNA_EXPORT_FROM_NUCLEUS | 0.00302 | 0.04108 | 0.43171 | 0.66820 | 1.38331 | 35 |
| GO_NCRNA_METABOLIC_PROCESS | 0.00000 | 0.00003 | 0.65944 | 0.55907 | 1.24907 | 381 |
| GO_NCRNA_PROCESSING | 0.00000 | 0.00003 | 0.65944 | 0.57175 | 1.27504 | 328 |
| GO_NEGATIVE_REGULATION_OF_CANONICAL_WNT_SIGNALING_PAT | 0.00000 | 0.00000 | | 0.71611 | 1.56038 | 111 |
| GO_NEGATIVE_REGULATION_OF_CELLULAR_AMIDE_METABOLIC_PR( | 0.00001 | 0.00020 | 0.61053 | 0.63031 | 1.37423 | 114 |
| GO_NEGATIVE_REGULATION_OF_CELL_CYCLE | 0.00003 | 0.00076 | 0.57561 | 0.53833 | 1.20513 | 425 |
| GO_NEGATIVE_REGULATION_OF_CELL_CYCLE_G2_M_PHASE_TRANS | 0.00000 | 0.00000 | 0.76146 | 0.71490 | 1.54439 | 85 |
| GO_NEGATIVE_REGULATION_OF_CELL_CYCLE_PHASE_TRANSITION | 0.00002 | 0.00060 | 0.57561 | 0.59041 | 1.30442 | 184 |
| GO_NEGATIVE_REGULATION_OF_CELL_CYCLE_PROCESS | 0.00000 | 0.00002 | 0.65944 | 0.58690 | 1.30374 | 249 |
| GO_NEGATIVE_REGULATION_OF_MITOTIC_CELL_CYCLE | 0.00004 | 0.00095 | 0.55733 | 0.56735 | 1.25902 | 232 |
| GO_NEGATIVE_REGULATION_OF_MRNA_METABOLIC_PROCESS | 0.00023 | 0.00491 | 0.51885 | 0.66670 | 1.41788 | 59 |
| GO_NEGATIVE_REGULATION_OF_MRNA_SPLICING_VIA_SPLICEOSOM | 0.00239 | 0.03510 | 0.43171 | 0.74857 | 1.51204 | 19 |
| GO_NEGATIVE_REGULATION_OF_RNA_SPLICING | 0.00029 | 0.00607 | 0.49849 | 0.76729 | 1.55976 | 23 |
| GO_NEGATIVE_REGULATION_OF_TRANSLATIONAL_INITIATION | 0.00018 | 0.00403 | 0.51885 | 0.83901 | 1.66434 | 15 |
| GO_NEGATIVE_REGULATION_OF_WNT_SIGNALING_PATHWAY | 0.00000 | 0.00000 | | 0.69525 | 1.52249 | 127 |
| GO_NEURAL_NUCLEUS_DEVELOPMENT | 0.00001 | 0.00026 | 0.59333 | 0.79148 | 1.62187 | 28 |
| GO_NEURON_TO_NEURON_SYNAPSE | 0.00005 | 0.00134 | 0.55733 | 0.59636 | 1.31006 | 154 |
| GO_NIK_NF_KAPPAB_SIGNALING | 0.00000 | 0.00000 | | 0.72896 | 1.58354 | 103 |
| GO_NON_CANONICAL_WNT_SIGNALING_PATHWAY | 0.00000 | 0.00000 | | 0.72784 | 1.58023 | 101 |
| GO_NUCLEAR_EXPORT | 0.00031 | 0.00645 | 0.49849 | 0.56963 | 1.25542 | 172 |
| GO_NUCLEAR_TRANSCRIBED_MRNA_CATABOLIC_PROCESS | 0.00000 | 0.00000 | | 0.79179 | 1.74917 | 183 |
| GO_NUCLEAR_TRANSCRIBED_MRNA_CATABOLIC_PROCESS_NONSEN | 0.00000 | 0.00000 | | 0.89008 | 1.93909 | 110 |
| GO_NUCLEAR_TRANSPORT | 0.00008 | 0.00193 | 0.53843 | 0.55511 | 1.23408 | 273 |
| GO_NUCLEOBASE_CONTAINING_COMPOUND_TRANSPORT | 0.00146 | 0.02386 | 0.45506 | 0.55053 | 1.21842 | 202 |
| GO_NUCLEOBASE_CONTAINING_SMALL_MOLECULE_BIOSYNTHETIC_ | 0.00137 | 0.02261 | 0.45506 | 0.62749 | 1.34353 | 67 |
| GO_NUCLEOBASE_CONTAINING_SMALL_MOLECULE_INTERCONVERS | 0.00383 | 0.04935 | 0.43171 | 0.71259 | 1.44857 | 23 |
| GO_NUCLEOBASE_CONTAINING_SMALL_MOLECULE_METABOLIC_PR | 0.00320 | 0.04339 | 0.43171 | 0.51448 | 1.14825 | 352 |
| GO_NUCLEOSIDE_DIPHOSPHATE_METABOLIC_PROCESS | 0.00110 | 0.01865 | 0.45506 | 0.59637 | 1.29295 | 97 |
| GO_NUCLEOSIDE_METABOLIC_PROCESS | 0.00061 | 0.01169 | 0.47727 | 0.65315 | 1.38907 | 59 |
| GO_NUCLEOSIDE_MONOPHOSPHATE_BIOSYNTHETIC_PROCESS | 0.00250 | 0.03636 | 0.43171 | 0.68841 | 1.42033 | 32 |
| GO_NUCLEOSIDE_MONOPHOSPHATE_METABOLIC_PROCESS | 0.00152 | 0.02464 | 0.45506 | 0.64377 | 1.36002 | 51 |
| GO_NUCLEOSIDE_PHOSPHATE_BIOSYNTHETIC_PROCESS | 0.00043 | 0.00865 | 0.49849 | 0.58185 | 1.27831 | 148 |
| GO_NUCLEOSIDE_TRIPHOSPHATE_BIOSYNTHETIC_PROCESS | 0.00043 | 0.00865 | 0.49849 | 0.69332 | 1.44539 | 40 |
| GO_NUCLEOSIDE_TRIPHOSPHATE_METABOLIC_PROCESS | 0.00016 | 0.00360 | 0.51885 | 0.66788 | 1.41976 | 58 |
| GO_NUCLEOSOMAL_DNA_BINDING | 0.00023 | 0.00507 | 0.51885 | 0.79852 | 1.61801 | 21 |
| GO_NUCLEOSOME_BINDING | 0.00143 | 0.02332 | 0.45506 | 0.66132 | 1.38492 | 45 |
| GO_NUCLEOTIDE_EXCISION_REPAIR | 0.00004 | 0.00096 | 0.55733 | 0.63343 | 1.37355 | 98 |
| GO_NUCLEOTIDE_EXCISION_REPAIR_DNA_DAMAGE_RECOGNITION | 0.00015 | 0.00349 | 0.51885 | 0.78678 | 1.59640 | 22 |
| GO_NUCLEOTIDE_EXCISION_REPAIR_DNA_INCISION | 0.00313 | 0.04249 | 0.43171 | 0.66439 | 1.37898 | 36 |
| GO_NUCLEOTIDE_PHOSPHORYLATION | 0.00383 | 0.04935 | 0.43171 | 0.59002 | 1.27454 | 88 |
| GO_ORGANELLAR_RIBOSOME | 0.00000 | 0.00014 | 0.61053 | 0.66412 | 1.43471 | 85 |
| GO_ORGANELLE_INNER_MEMBRANE | 0.00000 | 0.00000 | 0.76146 | 0.57601 | 1.28697 | 378 |
| GO_ORGANIC_CYCLIC_COMPOUND_CATABOLIC_PROCESS | 0.00000 | 0.00000 | | 0.73350 | 1.64281 | 449 |
| GO_OXIDATIVE_PHOSPHORYLATION | 0.00001 | 0.00030 | 0.59333 | 0.64619 | 1.40027 | 96 |

| | | | | | | |
|---|---|---|---|---|---|---|
| GO_OXIDOREDUCTASE_ACTIVITY | 0.00006 | 0.00139 | 0.55733 | 0.53544 | 1.19703 | 392 |
| GO_OXIDOREDUCTASE_COMPLEX | 0.00001 | 0.00027 | 0.59333 | 0.66015 | 1.42461 | 82 |
| GO_PEPTIDASE_COMPLEX | 0.00000 | 0.00000 | | 0.75988 | 1.63982 | 82 |
| GO_PIGMENT_GRANULE | 0.00003 | 0.00089 | 0.55733 | 0.66098 | 1.42344 | 78 |
| GO_POLYSOMAL_RIBOSOME | 0.00000 | 0.00000 | | 0.90624 | 1.86629 | 31 |
| GO_POLYSOME | 0.00000 | 0.00000 | | 0.82337 | 1.75481 | 61 |
| GO_POLY_PYRIMIDINE_TRACT_BINDING | 0.00327 | 0.04398 | 0.43171 | 0.75471 | 1.51855 | 18 |
| GO_POSITIVE_REGULATION_OF_APOPTOTIC_SIGNALING_PATHWAY | 0.00279 | 0.03889 | 0.43171 | 0.57666 | 1.25518 | 107 |
| GO_POSITIVE_REGULATION_OF_ATPASE_ACTIVITY | 0.00294 | 0.04036 | 0.43171 | 0.65446 | 1.36795 | 43 |
| GO_POSITIVE_REGULATION_OF_BINDING | 0.00088 | 0.01590 | 0.47727 | 0.60284 | 1.30556 | 95 |
| GO_POSITIVE_REGULATION_OF_CANONICAL_WNT_SIGNALING_PATH | 0.00000 | 0.00000 | 0.77494 | 0.70504 | 1.53160 | 103 |
| GO_POSITIVE_REGULATION_OF_CELLULAR_AMIDE_METABOLIC_PRC | 0.00003 | 0.00074 | 0.57561 | 0.62296 | 1.36083 | 118 |
| GO_POSITIVE_REGULATION_OF_CELLULAR_PROTEIN_LOCALIZATION | 0.00041 | 0.00840 | 0.49849 | 0.54915 | 1.21815 | 225 |
| GO_POSITIVE_REGULATION_OF_CELL_DEATH | 0.00069 | 0.01297 | 0.47727 | 0.52595 | 1.17423 | 363 |
| GO_POSITIVE_REGULATION_OF_CELL_POPULATION_PROLIFERATION | 0.00078 | 0.01455 | 0.47727 | 0.52449 | 1.17108 | 368 |
| GO_POSITIVE_REGULATION_OF_CHROMOSOME_ORGANIZATION | 0.00231 | 0.03437 | 0.43171 | 0.56990 | 1.24871 | 133 |
| GO_POSITIVE_REGULATION_OF_DEFENSE_RESPONSE | 0.00000 | 0.00003 | 0.65944 | 0.62175 | 1.36730 | 158 |
| GO_POSITIVE_REGULATION_OF_DNA_BIOSYNTHETIC_PROCESS | 0.00001 | 0.00041 | 0.59333 | 0.71929 | 1.52055 | 52 |
| GO_POSITIVE_REGULATION_OF_DNA_METABOLIC_PROCESS | 0.00364 | 0.04761 | 0.43171 | 0.56647 | 1.24189 | 135 |
| GO_POSITIVE_REGULATION_OF_DNA_TEMPLATED_TRANSCRIPTION_ | 0.00276 | 0.03885 | 0.43171 | 0.68932 | 1.41586 | 29 |
| GO_POSITIVE_REGULATION_OF_ESTABLISHMENT_OF_PROTEIN_LOC/ | 0.00007 | 0.00168 | 0.53843 | 0.56461 | 1.25165 | 219 |
| GO_POSITIVE_REGULATION_OF_IMMUNE_RESPONSE | 0.00000 | 0.00001 | 0.69013 | 0.59037 | 1.31265 | 277 |
| GO_POSITIVE_REGULATION_OF_IMMUNE_SYSTEM_PROCESS | 0.00004 | 0.00110 | 0.55733 | 0.54018 | 1.20716 | 386 |
| GO_POSITIVE_REGULATION_OF_INNATE_IMMUNE_RESPONSE | 0.00000 | 0.00000 | 0.81404 | 0.69693 | 1.51947 | 114 |
| GO_POSITIVE_REGULATION_OF_INTRINSIC_APOPTOTIC_SIGNALING_ | 0.00104 | 0.01798 | 0.45506 | 0.69097 | 1.43578 | 37 |
| GO_POSITIVE_REGULATION_OF_ORGANELLE_ORGANIZATION | 0.00043 | 0.00865 | 0.49849 | 0.52093 | 1.16605 | 411 |
| GO_POSITIVE_REGULATION_OF_PROTEIN_LOCALIZATION_TO_NUCLE | 0.00001 | 0.00020 | 0.61053 | 0.69907 | 1.48990 | 61 |
| GO_POSITIVE_REGULATION_OF_RESPONSE_TO_BIOTIC_STIMULUS | 0.00000 | 0.00000 | 0.73376 | 0.66340 | 1.45310 | 128 |
| GO_POSITIVE_REGULATION_OF_RESPONSE_TO_EXTERNAL_STIMULU | 0.00002 | 0.00050 | 0.57561 | 0.58133 | 1.28681 | 201 |
| GO_POSITIVE_REGULATION_OF_TELOMERASE_ACTIVITY | 0.00061 | 0.01169 | 0.47727 | 0.73476 | 1.50565 | 28 |
| GO_POSITIVE_REGULATION_OF_TELOMERASE_RNA_LOCALIZATION_ | 0.00000 | 0.00001 | 0.69013 | 0.92451 | 1.83395 | 15 |
| GO_POSITIVE_REGULATION_OF_TELOMERE_MAINTENANCE_VIA_TE | 0.00130 | 0.02159 | 0.45506 | 0.71259 | 1.46508 | 30 |
| GO_POSITIVE_REGULATION_OF_TRANSCRIPTION_INITIATION_FROM_ | 0.00372 | 0.04847 | 0.43171 | 0.70671 | 1.43993 | 25 |
| GO_POSITIVE_REGULATION_OF_TRANSFERASE_ACTIVITY | 0.00174 | 0.02712 | 0.45506 | 0.51642 | 1.15220 | 345 |
| GO_POSITIVE_REGULATION_OF_TRANSLATION | 0.00001 | 0.00019 | 0.61053 | 0.64767 | 1.40419 | 97 |
| GO_POSITIVE_REGULATION_OF_VIRAL_GENOME_REPLICATION | 0.00287 | 0.03973 | 0.43171 | 0.68362 | 1.40783 | 31 |
| GO_POSITIVE_REGULATION_OF_VIRAL_PROCESS | 0.00028 | 0.00590 | 0.49849 | 0.62440 | 1.34741 | 81 |
| GO_POSITIVE_REGULATION_OF_WNT_SIGNALING_PATHWAY | 0.00000 | 0.00000 | 0.73376 | 0.66230 | 1.44763 | 122 |
| GO_POSTREPLICATION_REPAIR | 0.00024 | 0.00515 | 0.51885 | 0.69140 | 1.44518 | 43 |
| GO_POSTSYNAPSE | 0.00004 | 0.00109 | 0.55733 | 0.55969 | 1.24369 | 259 |
| GO_POSTSYNAPTIC_CYTOSOL | 0.00279 | 0.03889 | 0.43171 | 0.78295 | 1.55313 | 15 |
| GO_POSTTRANSCRIPTIONAL_REGULATION_OF_GENE_EXPRESSION | 0.00000 | 0.00000 | | 0.66656 | 1.49327 | 462 |
| GO_POST_TRANSLATIONAL_PROTEIN_MODIFICATION | 0.00002 | 0.00052 | 0.57561 | 0.57851 | 1.28412 | 229 |
| GO_PRECATALYTIC_SPLICEOSOME | 0.00085 | 0.01559 | 0.47727 | 0.65235 | 1.37448 | 49 |
| GO_PRERIBOSOME | 0.00087 | 0.01577 | 0.47727 | 0.62899 | 1.34772 | 69 |
| GO_PRERIBOSOME_LARGE_SUBUNIT_PRECURSOR | 0.00235 | 0.03474 | 0.43171 | 0.74079 | 1.50308 | 22 |
| GO_PRE_MRNA_BINDING | 0.00133 | 0.02213 | 0.45506 | 0.72393 | 1.48062 | 27 |
| GO_PROTEASOMAL_PROTEIN_CATABOLIC_PROCESS | 0.00002 | 0.00044 | 0.59333 | 0.54166 | 1.21005 | 377 |
| GO_PROTEASOMAL_UBIQUITIN_INDEPENDENT_PROTEIN_CATABOLI( | 0.00000 | 0.00000 | 0.80122 | 0.92254 | 1.86931 | 21 |
| GO_PROTEASOME_ACCESSORY_COMPLEX | 0.00000 | 0.00001 | 0.67496 | 0.85464 | 1.73811 | 24 |
| GO_PROTEASOME_CORE_COMPLEX | 0.00000 | 0.00000 | | 0.94375 | 1.90631 | 19 |
| GO_PROTEIN_CONTAINING_COMPLEX_DISASSEMBLY | 0.00000 | 0.00007 | 0.62726 | 0.57810 | 1.28438 | 256 |
| GO_PROTEIN_CONTAINING_COMPLEX_LOCALIZATION | 0.00063 | 0.01197 | 0.47727 | 0.55335 | 1.22520 | 208 |
| GO_PROTEIN_DNA_COMPLEX_SUBUNIT_ORGANIZATION | 0.00001 | 0.00018 | 0.61053 | 0.60174 | 1.32491 | 165 |
| GO_PROTEIN_FOLDING | 0.00000 | 0.00002 | 0.65944 | 0.63113 | 1.38644 | 154 |
| GO_PROTEIN_FOLDING_CHAPERONE | 0.00135 | 0.02238 | 0.45506 | 0.77903 | 1.55752 | 17 |
| GO_PROTEIN_IMPORT | 0.00268 | 0.03799 | 0.43171 | 0.56317 | 1.23581 | 140 |
| GO_PROTEIN_LOCALIZATION_TO_CHROMOSOME | 0.00056 | 0.01096 | 0.47727 | 0.63523 | 1.36108 | 69 |
| GO_PROTEIN_LOCALIZATION_TO_CHROMOSOME_TELOMERIC_REGI( | 0.00335 | 0.04466 | 0.43171 | 0.70399 | 1.43984 | 27 |
| GO_PROTEIN_LOCALIZATION_TO_ENDOPLASMIC_RETICULUM | 0.00000 | 0.00000 | | 0.85521 | 1.87021 | 124 |
| GO_PROTEIN_LOCALIZATION_TO_MEMBRANE | 0.00000 | 0.00000 | | 0.65123 | 1.45714 | 406 |
| GO_PROTEIN_LOCALIZATION_TO_MITOCHONDRION | 0.00019 | 0.00420 | 0.51885 | 0.60773 | 1.32398 | 110 |
| GO_PROTEIN_LOCALIZATION_TO_NUCLEUS | 0.00002 | 0.00055 | 0.57561 | 0.58926 | 1.30220 | 187 |
| GO_PROTEIN_MODIFICATION_BY_SMALL_PROTEIN_REMOVAL | 0.00000 | 0.00000 | 0.71951 | 0.60745 | 1.34689 | 221 |
| GO_PROTEIN_POLYUBIQUITINATION | 0.00004 | 0.00106 | 0.55733 | 0.55874 | 1.24108 | 253 |
| GO_PROTEIN_REFOLDING | 0.00294 | 0.04036 | 0.43171 | 0.77039 | 1.53314 | 16 |
| GO_PROTEIN_STABILIZATION | 0.00001 | 0.00024 | 0.61053 | 0.62333 | 1.36586 | 130 |
| GO_PROTEIN_TARGETING | 0.00000 | 0.00000 | | 0.71181 | 1.58743 | 327 |
| GO_PROTEIN_TARGETING_TO_MEMBRANE | 0.00000 | 0.00000 | | 0.82370 | 1.80953 | 149 |
| GO_PROTEIN_TARGETING_TO_MITOCHONDRION | 0.00346 | 0.04593 | 0.43171 | 0.60408 | 1.30091 | 78 |
| GO_PROTEIN_TRANSMEMBRANE_TRANSPORT | 0.00265 | 0.03775 | 0.43171 | 0.64528 | 1.36171 | 50 |
| GO_PROTON_TRANSMEMBRANE_TRANSPORT | 0.00213 | 0.03229 | 0.43171 | 0.60815 | 1.30536 | 71 |
| GO_PURINE_CONTAINING_COMPOUND_BIOSYNTHETIC_PROCESS | 0.00246 | 0.03601 | 0.43171 | 0.57796 | 1.25936 | 111 |
| GO_PURINE_CONTAINING_COMPOUND_METABOLIC_PROCESS | 0.00100 | 0.01761 | 0.45506 | 0.53679 | 1.19242 | 255 |
| GO_PURINE_NUCLEOSIDE_TRIPHOSPHATE_METABOLIC_PROCESS | 0.00089 | 0.01592 | 0.47727 | 0.67157 | 1.40374 | 43 |
| GO_PYRIMIDINE_CONTAINING_COMPOUND_METABOLIC_PROCESS | 0.00353 | 0.04658 | 0.43171 | 0.64744 | 1.35878 | 47 |

| | | | | | | |
|---|---|---|---|---|---|---|
| GO_REGENERATION | 0.00216 | 0.03242 | 0.43171 | 0.59987 | 1.29453 | 82 |
| GO_REGULATION_OF_ANIMAL_ORGAN_MORPHOGENESIS | 0.00000 | 0.00000 | | 0.69163 | 1.51541 | 133 |
| GO_REGULATION_OF_APOPTOTIC_SIGNALING_PATHWAY | 0.00031 | 0.00645 | 0.49849 | 0.55171 | 1.22506 | 247 |
| GO_REGULATION_OF_ATPASE_ACTIVITY | 0.00324 | 0.04369 | 0.43171 | 0.63031 | 1.34038 | 57 |
| GO_REGULATION_OF_ATP_METABOLIC_PROCESS | 0.00353 | 0.04658 | 0.43171 | 0.60176 | 1.29780 | 80 |
| GO_REGULATION_OF_BINDING | 0.00250 | 0.03636 | 0.43171 | 0.54476 | 1.20598 | 203 |
| GO_REGULATION_OF_CELLULAR_AMIDE_METABOLIC_PROCESS | 0.00000 | 0.00000 | | 0.62967 | 1.40226 | 311 |
| GO_REGULATION_OF_CELLULAR_AMINE_METABOLIC_PROCESS | 0.00000 | 0.00000 | | 0.82173 | 1.74682 | 58 |
| GO_REGULATION_OF_CELLULAR_AMINO_ACID_METABOLIC_PROCES | 0.00000 | 0.00000 | | 0.83776 | 1.77766 | 55 |
| GO_REGULATION_OF_CELLULAR_KETONE_METABOLIC_PROCESS | 0.00000 | 0.00000 | | 0.72641 | 1.58680 | 118 |
| GO_REGULATION_OF_CELLULAR_PROTEIN_LOCALIZATION | 0.00082 | 0.01513 | 0.47727 | 0.52542 | 1.17290 | 362 |
| GO_REGULATION_OF_CELLULAR_RESPONSE_TO_HEAT | 0.00276 | 0.03885 | 0.43171 | 0.61652 | 1.32004 | 67 |
| GO_REGULATION_OF_CELLULAR_RESPONSE_TO_STRESS | 0.00224 | 0.03344 | 0.43171 | 0.50545 | 1.13232 | 468 |
| GO_REGULATION_OF_CELL_CYCLE_G2_M_PHASE_TRANSITION | 0.00000 | 0.00000 | 0.71951 | 0.63230 | 1.39282 | 169 |
| GO_REGULATION_OF_CELL_CYCLE_PHASE_TRANSITION | 0.00001 | 0.00026 | 0.59333 | 0.55488 | 1.23783 | 337 |
| GO_REGULATION_OF_CHROMOSOME_ORGANIZATION | 0.00081 | 0.01501 | 0.47727 | 0.54255 | 1.20527 | 261 |
| GO_REGULATION_OF_CYSTEINE_TYPE_ENDOPEPTIDASE_ACTIVITY | 0.00202 | 0.03077 | 0.43171 | 0.57120 | 1.24837 | 121 |
| GO_REGULATION_OF_CYTOPLASMIC_TRANSLATION | 0.00130 | 0.02159 | 0.45506 | 0.73678 | 1.49773 | 23 |
| GO_REGULATION_OF_DEFENSE_RESPONSE | 0.00065 | 0.01237 | 0.47727 | 0.53633 | 1.19355 | 290 |
| GO_REGULATION_OF_DNA_BIOSYNTHETIC_PROCESS | 0.00016 | 0.00360 | 0.51885 | 0.64215 | 1.37923 | 74 |
| GO_REGULATION_OF_DNA_METABOLIC_PROCESS | 0.00069 | 0.01297 | 0.47727 | 0.54425 | 1.20775 | 237 |
| GO_REGULATION_OF_DNA_TEMPLATED_TRANSCRIPTION_INITIATIOI | 0.00163 | 0.02568 | 0.45506 | 0.68289 | 1.41372 | 35 |
| GO_REGULATION_OF_DNA_TEMPLATED_TRANSCRIPTION_IN_RESPO | 0.00000 | 0.00000 | | 0.73944 | 1.59800 | 89 |
| GO_REGULATION_OF_GENERATION_OF_PRECURSOR_METABOLITES_ | 0.00060 | 0.01164 | 0.47727 | 0.60476 | 1.31116 | 97 |
| GO_REGULATION_OF_HEMATOPOIETIC_PROGENITOR_CELL_DIFFERE | 0.00000 | 0.00000 | | 0.78717 | 1.69251 | 75 |
| GO_REGULATION_OF_HEMATOPOIETIC_STEM_CELL_DIFFERENTIATIC | 0.00000 | 0.00000 | | 0.81505 | 1.74256 | 65 |
| GO_REGULATION_OF_HEMOPOIESIS | 0.00000 | 0.00008 | 0.62726 | 0.58859 | 1.30634 | 228 |
| GO_REGULATION_OF_IMMUNE_RESPONSE | 0.00000 | 0.00016 | 0.61053 | 0.55362 | 1.23583 | 362 |
| GO_REGULATION_OF_INNATE_IMMUNE_RESPONSE | 0.00000 | 0.00000 | 0.71951 | 0.63883 | 1.40521 | 161 |
| GO_REGULATION_OF_INTRINSIC_APOPTOTIC_SIGNALING_PATHWAY | 0.00001 | 0.00039 | 0.59333 | 0.63478 | 1.38259 | 109 |
| GO_REGULATION_OF_MITOTIC_CELL_CYCLE | 0.00001 | 0.00018 | 0.61053 | 0.53899 | 1.20736 | 458 |
| GO_REGULATION_OF_MORPHOGENESIS_OF_AN_EPITHELIUM | 0.00000 | 0.00000 | | 0.72213 | 1.57284 | 109 |
| GO_REGULATION_OF_MRNA_CATABOLIC_PROCESS | 0.00000 | 0.00000 | | 0.69977 | 1.54127 | 168 |
| GO_REGULATION_OF_MRNA_METABOLIC_PROCESS | 0.00000 | 0.00000 | | 0.66528 | 1.47872 | 266 |
| GO_REGULATION_OF_MRNA_PROCESSING | 0.00004 | 0.00105 | 0.55733 | 0.61521 | 1.34090 | 113 |
| GO_REGULATION_OF_MRNA_SPLICING_VIA_SPLICEOSOME | 0.00001 | 0.00026 | 0.59333 | 0.67424 | 1.45411 | 80 |
| GO_REGULATION_OF_NUCLEASE_ACTIVITY | 0.00106 | 0.01819 | 0.45506 | 0.77918 | 1.56780 | 18 |
| GO_REGULATION_OF_OXIDATIVE_STRESS_INDUCED_CELL_DEATH | 0.00198 | 0.03028 | 0.43171 | 0.67287 | 1.40355 | 41 |
| GO_REGULATION_OF_PEPTIDASE_ACTIVITY | 0.00050 | 0.00978 | 0.47727 | 0.56122 | 1.24054 | 191 |
| GO_REGULATION_OF_POSTTRANSCRIPTIONAL_GENE_SILENCING | 0.00364 | 0.04761 | 0.43171 | 0.59824 | 1.28492 | 74 |
| GO_REGULATION_OF_PROTEIN_CATABOLIC_PROCESS | 0.00179 | 0.02783 | 0.45506 | 0.53057 | 1.17929 | 270 |
| GO_REGULATION_OF_PROTEIN_LOCALIZATION_TO_NUCLEUS | 0.00001 | 0.00042 | 0.59333 | 0.65505 | 1.41558 | 87 |
| GO_REGULATION_OF_PROTEIN_MODIFICATION_BY_SMALL_PROTEIN | 0.00025 | 0.00532 | 0.49849 | 0.57068 | 1.25684 | 167 |
| GO_REGULATION_OF_PROTEIN_STABILITY | 0.00093 | 0.01652 | 0.47727 | 0.55382 | 1.22571 | 202 |
| GO_REGULATION_OF_PROTEOLYSIS | 0.00013 | 0.00301 | 0.51885 | 0.52997 | 1.18392 | 377 |
| GO_REGULATION_OF_RESPONSE_TO_BIOTIC_STIMULUS | 0.00001 | 0.00020 | 0.61053 | 0.59049 | 1.30719 | 204 |
| GO_REGULATION_OF_RNA_SPLICING | 0.00000 | 0.00001 | 0.69013 | 0.65896 | 1.43669 | 114 |
| GO_REGULATION_OF_SMALL_MOLECULE_METABOLIC_PROCESS | 0.00000 | 0.00000 | 0.71951 | 0.59134 | 1.31394 | 265 |
| GO_REGULATION_OF_STEM_CELL_DIFFERENTIATION | 0.00000 | 0.00000 | | 0.74049 | 1.60113 | 90 |
| GO_REGULATION_OF_SYMBIOTIC_PROCESS | 0.00006 | 0.00144 | 0.55733 | 0.60071 | 1.31682 | 134 |
| GO_REGULATION_OF_TELOMERASE_ACTIVITY | 0.00088 | 0.01590 | 0.47727 | 0.69161 | 1.43179 | 35 |
| GO_REGULATION_OF_TRANSCRIPTION_FROM_RNA_POLYMERASE_II | 0.00000 | 0.00000 | | 0.81194 | 1.73800 | 66 |
| GO_REGULATION_OF_TRANSCRIPTION_INITIATION_FROM_RNA_POL | 0.00143 | 0.02332 | 0.45506 | 0.70390 | 1.44582 | 29 |
| GO_REGULATION_OF_TRANSLATIONAL_INITIATION | 0.00002 | 0.00052 | 0.57561 | 0.68129 | 1.45518 | 64 |
| GO_REGULATION_OF_UBIQUITIN_PROTEIN_TRANSFERASE_ACTIVITY | 0.00061 | 0.01169 | 0.47727 | 0.69335 | 1.44627 | 41 |
| GO_REGULATION_OF_VIRAL_GENOME_REPLICATION | 0.00086 | 0.01566 | 0.47727 | 0.65316 | 1.38389 | 54 |
| GO_REGULATION_OF_VIRAL_TRANSCRIPTION | 0.00198 | 0.03028 | 0.43171 | 0.63932 | 1.34703 | 49 |
| GO_REGULATION_OF_WNT_SIGNALING_PATHWAY | 0.00000 | 0.00001 | 0.69013 | 0.60353 | 1.33816 | 220 |
| GO_RESPIRASOME | 0.00008 | 0.00181 | 0.53843 | 0.65420 | 1.40421 | 71 |
| GO_RESPIRATORY_CHAIN_COMPLEX | 0.00016 | 0.00353 | 0.51885 | 0.67626 | 1.43918 | 60 |
| GO_RESPIRATORY_ELECTRON_TRANSPORT_CHAIN | 0.00001 | 0.00025 | 0.59333 | 0.65819 | 1.42190 | 85 |
| GO_RESPONSE_TO_ALCOHOL | 0.00261 | 0.03750 | 0.43171 | 0.58804 | 1.27350 | 95 |
| GO_RESPONSE_TO_CADMIUM_ION | 0.00265 | 0.03775 | 0.43171 | 0.67227 | 1.39174 | 35 |
| GO_RESPONSE_TO_DRUG | 0.00104 | 0.01798 | 0.45506 | 0.55364 | 1.22258 | 180 |
| GO_RESPONSE_TO_ETHANOL | 0.00086 | 0.01566 | 0.47727 | 0.67309 | 1.40691 | 43 |
| GO_RESPONSE_TO_HEAT | 0.00012 | 0.00270 | 0.53843 | 0.61665 | 1.34221 | 107 |
| GO_RESPONSE_TO_HORMONE | 0.00048 | 0.00949 | 0.49849 | 0.51277 | 1.14814 | 453 |
| GO_RESPONSE_TO_HYDROGEN_PEROXIDE | 0.00257 | 0.03716 | 0.43171 | 0.60505 | 1.30368 | 79 |
| GO_RESPONSE_TO_INORGANIC_SUBSTANCE | 0.00004 | 0.00107 | 0.55733 | 0.55714 | 1.23870 | 275 |
| GO_RESPONSE_TO_INTERLEUKIN_1 | 0.00000 | 0.00000 | | 0.74844 | 1.62908 | 107 |
| GO_RESPONSE_TO_INTERLEUKIN_12 | 0.00012 | 0.00270 | 0.53843 | 0.72605 | 1.50694 | 36 |
| GO_RESPONSE_TO_INTERLEUKIN_4 | 0.00324 | 0.04369 | 0.43171 | 0.75812 | 1.51571 | 17 |
| GO_RESPONSE_TO_METAL_ION | 0.00030 | 0.00623 | 0.49849 | 0.56387 | 1.24233 | 171 |
| GO_RESPONSE_TO_ORGANIC_CYCLIC_COMPOUND | 0.00000 | 0.00016 | 0.61053 | 0.54603 | 1.22259 | 439 |
| GO_RESPONSE_TO_OXYGEN_LEVELS | 0.00000 | 0.00000 | 0.71951 | 0.60132 | 1.33438 | 233 |
| GO_RESPONSE_TO_STEROID_HORMONE | 0.00154 | 0.02464 | 0.45506 | 0.55514 | 1.22446 | 174 |

| | | | | | | |
|---|---|---|---|---|---|---|
| GO_RESPONSE_TO_TEMPERATURE_STIMULUS | 0.00027 | 0.00572 | 0.49849 | 0.59216 | 1.29675 | 127 |
| GO_RESPONSE_TO_TUMOR_NECROSIS_FACTOR | 0.00000 | 0.00000 | 0.76146 | 0.65649 | 1.44236 | 151 |
| GO_RETINA_HOMEOSTASIS | 0.00104 | 0.01798 | 0.45506 | 0.73063 | 1.49432 | 27 |
| GO_RIBONUCLEOPROTEIN_COMPLEX_BINDING | 0.00000 | 0.00000 | 0.71951 | 0.66906 | 1.46050 | 117 |
| GO_RIBONUCLEOPROTEIN_COMPLEX_BIOGENESIS | 0.00000 | 0.00000 | | 0.66339 | 1.48264 | 389 |
| GO_RIBONUCLEOPROTEIN_COMPLEX_SUBUNIT_ORGANIZATION | 0.00000 | 0.00000 | | 0.69241 | 1.52681 | 173 |
| GO_RIBONUCLEOPROTEIN_GRANULE | 0.00000 | 0.00001 | 0.67496 | 0.61979 | 1.36366 | 163 |
| GO_RIBONUCLEOSIDE_TRIPHOSPHATE_BIOSYNTHETIC_PROCESS | 0.00041 | 0.00840 | 0.49849 | 0.71253 | 1.47248 | 33 |
| GO_RIBONUCLEOSIDE_TRIPHOSPHATE_METABOLIC_PROCESS | 0.00026 | 0.00550 | 0.49849 | 0.69681 | 1.45611 | 42 |
| GO_RIBOSE_PHOSPHATE_BIOSYNTHETIC_PROCESS | 0.00191 | 0.02939 | 0.45506 | 0.57932 | 1.26207 | 110 |
| GO_RIBOSE_PHOSPHATE_METABOLIC_PROCESS | 0.00108 | 0.01839 | 0.45506 | 0.53891 | 1.19687 | 245 |
| GO_RIBOSOMAL_LARGE_SUBUNIT_ASSEMBLY | 0.00000 | 0.00016 | 0.61053 | 0.80999 | 1.65663 | 27 |
| GO_RIBOSOMAL_LARGE_SUBUNIT_BIOGENESIS | 0.00000 | 0.00000 | | 0.77542 | 1.65982 | 66 |
| GO_RIBOSOMAL_SMALL_SUBUNIT_ASSEMBLY | 0.00023 | 0.00494 | 0.51885 | 0.81139 | 1.62221 | 17 |
| GO_RIBOSOMAL_SMALL_SUBUNIT_BINDING | 0.00383 | 0.04935 | 0.43171 | 0.75288 | 1.50523 | 17 |
| GO_RIBOSOMAL_SMALL_SUBUNIT_BIOGENESIS | 0.00001 | 0.00022 | 0.61053 | 0.69076 | 1.47997 | 68 |
| GO_RIBOSOMAL_SUBUNIT | 0.00000 | 0.00000 | | 0.77603 | 1.71031 | 172 |
| GO_RIBOSOME | 0.00000 | 0.00000 | | 0.76643 | 1.69659 | 206 |
| GO_RIBOSOME_ASSEMBLY | 0.00000 | 0.00000 | 0.73376 | 0.76173 | 1.61998 | 59 |
| GO_RIBOSOME_BINDING | 0.00019 | 0.00416 | 0.51885 | 0.68955 | 1.44715 | 47 |
| GO_RIBOSOME_BIOGENESIS | 0.00000 | 0.00000 | | 0.65877 | 1.46466 | 275 |
| GO_RNA_CATABOLIC_PROCESS | 0.00000 | 0.00000 | | 0.76774 | 1.71264 | 339 |
| GO_RNA_DEPENDENT_DNA_BIOSYNTHETIC_PROCESS | 0.00003 | 0.00076 | 0.57561 | 0.68926 | 1.46521 | 58 |
| GO_RNA_EXPORT_FROM_NUCLEUS | 0.00062 | 0.01183 | 0.47727 | 0.58378 | 1.27758 | 126 |
| GO_RNA_LOCALIZATION | 0.00000 | 0.00006 | 0.64355 | 0.59670 | 1.32020 | 198 |
| GO_RNA_POLYMERASE_II_PREINITIATION_COMPLEX_ASSEMBLY | 0.00089 | 0.01592 | 0.47727 | 0.77046 | 1.55626 | 19 |
| GO_RNA_SPLICING | 0.00000 | 0.00000 | 0.78819 | 0.59079 | 1.31927 | 369 |
| GO_RNA_SPLICING_VIA_TRANSESTERIFICATION_REACTIONS | 0.00000 | 0.00000 | | 0.60940 | 1.35714 | 300 |
| GO_RNA_STABILIZATION | 0.00065 | 0.01223 | 0.47727 | 0.69743 | 1.44384 | 35 |
| GO_ROUGH_ENDOPLASMIC_RETICULUM | 0.00001 | 0.00039 | 0.59333 | 0.71853 | 1.51629 | 50 |
| GO_RRNA_BINDING | 0.00000 | 0.00001 | 0.69013 | 0.74782 | 1.58865 | 56 |
| GO_RRNA_METABOLIC_PROCESS | 0.00000 | 0.00000 | 0.77494 | 0.63047 | 1.39562 | 206 |
| GO_SCF_DEPENDENT_PROTEASOMAL_UBIQUITIN_DEPENDENT_PRO | 0.00000 | 0.00000 | | 0.77527 | 1.67044 | 79 |
| GO_SECRETORY_GRANULE | 0.00000 | 0.00000 | 0.71951 | 0.56518 | 1.26461 | 399 |
| GO_SECRETORY_VESICLE | 0.00000 | 0.00002 | 0.67496 | 0.54866 | 1.22905 | 465 |
| GO_SIGNAL_TRANSDUCTION_BY_PROTEIN_PHOSPHORYLATION | 0.00004 | 0.00109 | 0.55733 | 0.53817 | 1.20474 | 424 |
| GO_SINGLE_FERTILIZATION | 0.00048 | 0.00949 | 0.49849 | 0.67414 | 1.42260 | 50 |
| GO_SINGLE_STRANDED_RNA_BINDING | 0.00039 | 0.00793 | 0.49849 | 0.66553 | 1.41527 | 57 |
| GO_SMALL_MOLECULE_BIOSYNTHETIC_PROCESS | 0.00216 | 0.03242 | 0.43171 | 0.51195 | 1.14522 | 397 |
| GO_SMALL_RIBOSOMAL_SUBUNIT | 0.00000 | 0.00000 | | 0.77648 | 1.66209 | 66 |
| GO_SM_LIKE_PROTEIN_FAMILY_COMPLEX | 0.00327 | 0.04398 | 0.43171 | 0.60468 | 1.29713 | 70 |
| GO_SPERM_EGG_RECOGNITION | 0.00000 | 0.00007 | 0.64355 | 0.85241 | 1.72955 | 22 |
| GO_SPLICEOSOMAL_COMPLEX | 0.00000 | 0.00003 | 0.65944 | 0.60939 | 1.34261 | 171 |
| GO_STEM_CELL_DIFFERENTIATION | 0.00000 | 0.00000 | 0.78819 | 0.66886 | 1.46930 | 147 |
| GO_STRUCTURAL_CONSTITUENT_OF_CYTOSKELETON | 0.00102 | 0.01782 | 0.45506 | 0.65913 | 1.39337 | 52 |
| GO_STRUCTURAL_CONSTITUENT_OF_RIBOSOME | 0.00000 | 0.00000 | | 0.79909 | 1.75541 | 150 |
| GO_STRUCTURAL_MOLECULE_ACTIVITY | 0.00000 | 0.00000 | | 0.72651 | 1.61895 | 324 |
| GO_SUBSTANTIA_NIGRA_DEVELOPMENT | 0.00000 | 0.00013 | 0.62726 | 0.81622 | 1.66507 | 26 |
| GO_TELOMERASE_RNA_LOCALIZATION | 0.00001 | 0.00017 | 0.61053 | 0.85164 | 1.72025 | 19 |
| GO_TELOMERE_MAINTENANCE_VIA_TELOMERE_LENGTHENING | 0.00011 | 0.00250 | 0.53843 | 0.65659 | 1.40546 | 66 |
| GO_TELOMERE_ORGANIZATION | 0.00342 | 0.04554 | 0.43171 | 0.57000 | 1.24649 | 124 |
| GO_THREONINE_TYPE_ENDOPEPTIDASE_ACTIVITY | 0.00000 | 0.00000 | | 0.95179 | 1.92255 | 19 |
| GO_THREONINE_TYPE_PEPTIDASE_ACTIVITY | 0.00000 | 0.00000 | | 0.92199 | 1.87423 | 23 |
| GO_TISSUE_MORPHOGENESIS | 0.00097 | 0.01707 | 0.47727 | 0.52661 | 1.17402 | 326 |
| GO_TOXIN_TRANSPORT | 0.00113 | 0.01922 | 0.45506 | 0.72897 | 1.48708 | 26 |
| GO_TRANSCRIPTION_COUPLED_NUCLEOTIDE_EXCISION_REPAIR | 0.00001 | 0.00019 | 0.61053 | 0.68106 | 1.46186 | 71 |
| GO_TRANSCRIPTION_FACTOR_BINDING | 0.00043 | 0.00865 | 0.49849 | 0.52447 | 1.17376 | 409 |
| GO_TRANSCRIPTION_PREINITIATION_COMPLEX_ASSEMBLY | 0.00119 | 0.02004 | 0.45506 | 0.71399 | 1.46797 | 30 |
| GO_TRANSLATIONAL_ELONGATION | 0.00000 | 0.00000 | 0.71951 | 0.66087 | 1.44451 | 122 |
| GO_TRANSLATIONAL_INITIATION | 0.00000 | 0.00000 | | 0.84691 | 1.86515 | 166 |
| GO_TRANSLATIONAL_TERMINATION | 0.00000 | 0.00005 | 0.64355 | 0.65946 | 1.43060 | 99 |
| GO_TRANSLATION_ELONGATION_FACTOR_ACTIVITY | 0.00161 | 0.02546 | 0.45506 | 0.79860 | 1.58418 | 15 |
| GO_TRANSLATION_FACTOR_ACTIVITY_RNA_BINDING | 0.00000 | 0.00000 | | 0.79008 | 1.69288 | 69 |
| GO_TRANSLATION_INITIATION_FACTOR_ACTIVITY | 0.00000 | 0.00000 | 0.76146 | 0.80422 | 1.68417 | 45 |
| GO_TRANSLATION_INITIATION_FACTOR_BINDING | 0.00021 | 0.00468 | 0.51885 | 0.76186 | 1.55231 | 25 |
| GO_TRANSLATION_PREINITIATION_COMPLEX | 0.00000 | 0.00001 | 0.67496 | 0.90245 | 1.79595 | 16 |
| GO_TRANSLATION_REGULATOR_ACTIVITY | 0.00000 | 0.00000 | | 0.77663 | 1.68904 | 105 |
| GO_TRANSLATION_REGULATOR_ACTIVITY_NUCLEIC_ACID_BINDING | 0.00000 | 0.00000 | | 0.77150 | 1.66586 | 83 |
| GO_TRICARBOXYLIC_ACID_CYCLE | 0.00154 | 0.02464 | 0.45506 | 0.71854 | 1.47240 | 28 |
| GO_TUMOR_NECROSIS_FACTOR_MEDIATED_SIGNALING_PATHWAY | 0.00000 | 0.00000 | | 0.76277 | 1.65372 | 97 |
| GO_T_CELL_RECEPTOR_SIGNALING_PATHWAY | 0.00000 | 0.00000 | 0.81404 | 0.69994 | 1.52527 | 112 |
| GO_U12_TYPE_SPLICEOSOMAL_COMPLEX | 0.00279 | 0.03889 | 0.43171 | 0.70746 | 1.44693 | 27 |
| GO_U2_SNRNP | 0.00005 | 0.00134 | 0.55733 | 0.83081 | 1.67168 | 18 |
| GO_U2_TYPE_SPLICEOSOMAL_COMPLEX | 0.00010 | 0.00243 | 0.53843 | 0.63403 | 1.36970 | 85 |
| GO_UBIQUITIN_LIKE_PROTEIN_LIGASE_BINDING | 0.00000 | 0.00000 | 0.76146 | 0.62273 | 1.38051 | 219 |
| GO_UBIQUITIN_PROTEIN_TRANSFERASE_REGULATOR_ACTIVITY | 0.00165 | 0.02590 | 0.45506 | 0.79605 | 1.57911 | 15 |

| | | | | | | |
|---|---|---|---|---|---|---|
| GO_UNFOLDED_PROTEIN_BINDING | 0.00000 | 0.00000 | 0.77494 | 0.72157 | 1.55881 | 85 |
| GO_VESICLE_LUMEN | 0.00000 | 0.00000 | 0.74774 | 0.64475 | 1.42008 | 168 |
| GO_VIRAL_GENE_EXPRESSION | 0.00000 | 0.00000 | | 0.80028 | 1.76317 | 171 |
| GO_VIRAL_GENOME_REPLICATION | 0.00100 | 0.01761 | 0.45506 | 0.61997 | 1.33078 | 73 |
| GO_VIRAL_LIFE_CYCLE | 0.00031 | 0.00640 | 0.49849 | 0.55767 | 1.23509 | 209 |
| HP_ABNORMALITY_OF_ACID_BASE_HOMEOSTASIS | 0.00216 | 0.03242 | 0.43171 | 0.53556 | 1.18846 | 232 |
| HP_ABNORMALITY_OF_CARDIOVASCULAR_SYSTEM_ELECTROPHYSIC | 0.00027 | 0.00572 | 0.49849 | 0.55832 | 1.23929 | 229 |
| HP_ABNORMALITY_OF_LOWER_LIP | 0.00181 | 0.02804 | 0.45506 | 0.56360 | 1.23779 | 145 |
| HP_ABNORMALITY_OF_THE_MITOCHONDRION | 0.00213 | 0.03229 | 0.43171 | 0.57831 | 1.26240 | 117 |
| HP_ABNORMALITY_OF_THE_NASAL_DORSUM | 0.00005 | 0.00116 | 0.55733 | 0.60288 | 1.32157 | 134 |
| HP_ABNORMAL_ELECTROPHYSIOLOGY_OF_SINOATRIAL_NODE_ORIG | 0.00279 | 0.03889 | 0.43171 | 0.68436 | 1.40934 | 31 |
| HP_ABNORMAL_ERYTHROCYTE_MORPHOLOGY | 0.00000 | 0.00015 | 0.61053 | 0.56049 | 1.25036 | 341 |
| HP_ABNORMAL_MEAN_CORPUSCULAR_VOLUME | 0.00357 | 0.04696 | 0.43171 | 0.75226 | 1.51363 | 18 |
| HP_ABNORMAL_RETICULOCYTE_MORPHOLOGY | 0.00064 | 0.01210 | 0.47727 | 0.71420 | 1.47081 | 31 |
| HP_ACUTE_LEUKEMIA | 0.00000 | 0.00000 | 0.73376 | 0.72786 | 1.56137 | 70 |
| HP_ANEMIA_OF_INADEQUATE_PRODUCTION | 0.00000 | 0.00008 | 0.62726 | 0.65593 | 1.42209 | 97 |
| HP_CONSTITUTIONAL_SYMPTOM | 0.00119 | 0.02004 | 0.45506 | 0.51427 | 1.15108 | 423 |
| HP_DELAYED_PUBERTY | 0.00005 | 0.00115 | 0.55733 | 0.62579 | 1.36211 | 107 |
| HP_DEPRESSED_NASAL_RIDGE | 0.00000 | 0.00011 | 0.62726 | 0.68731 | 1.47511 | 72 |
| HP_FATIGUE | 0.00039 | 0.00808 | 0.49849 | 0.56529 | 1.24923 | 187 |
| HP_HEADACHE | 0.00015 | 0.00340 | 0.51885 | 0.59065 | 1.29716 | 146 |
| HP_HEMATOLOGICAL_NEOPLASM | 0.00003 | 0.00078 | 0.57561 | 0.60108 | 1.32044 | 154 |
| HP_INCREASED_CSF_LACTATE | 0.00157 | 0.02509 | 0.45506 | 0.61622 | 1.31904 | 66 |
| HP_INCREASED_SERUM_LACTATE | 0.00216 | 0.03242 | 0.43171 | 0.56920 | 1.24724 | 130 |
| HP_LEUKEMIA | 0.00001 | 0.00026 | 0.59333 | 0.64877 | 1.40680 | 98 |
| HP_MACROCYTIC_ANEMIA | 0.00000 | 0.00000 | 0.74774 | 0.82468 | 1.71698 | 39 |
| HP_MIGRAINE | 0.00000 | 0.00000 | 0.71951 | 0.72105 | 1.54497 | 69 |
| HP_MYOTONIA | 0.00161 | 0.02546 | 0.45506 | 0.79820 | 1.58338 | 15 |
| HP_NEOPLASM | 0.00216 | 0.03242 | 0.43171 | 0.50820 | 1.13805 | 446 |
| HP_PALLOR | 0.00001 | 0.00040 | 0.59333 | 0.65169 | 1.41044 | 92 |
| HP_PUBERTY_AND_GONADAL_DISORDERS | 0.00287 | 0.03973 | 0.43171 | 0.51995 | 1.15843 | 322 |
| HP_RETICULOCYTOPENIA | 0.00004 | 0.00105 | 0.55733 | 0.86417 | 1.71425 | 15 |
| HP_RIMMED_VACUOLES | 0.00279 | 0.03889 | 0.43171 | 0.74033 | 1.49831 | 20 |
| HP_SHORT_NOSE | 0.00235 | 0.03474 | 0.43171 | 0.54919 | 1.21134 | 174 |
| HP_THICK_LOWER_LIP_VERMILION | 0.00048 | 0.00949 | 0.49849 | 0.60773 | 1.31617 | 94 |
| HP_THICK_VERMILION_BORDER | 0.00156 | 0.02486 | 0.45506 | 0.55983 | 1.23000 | 153 |
| HP_VENTRICULAR_ARRHYTHMIA | 0.00353 | 0.04658 | 0.43171 | 0.66198 | 1.37398 | 36 |
| **Cluster 1** | | | | | | |
| pathway | pval | padj | log2err | ES | NES | size |
| GO_ACTIVATION_OF_IMMUNE_RESPONSE | 0.00000 | 0.00000 | 0.78819 | 0.62851 | 1.39255 | 212 |
| GO_ACTIVATION_OF_INNATE_IMMUNE_RESPONSE | 0.00000 | 0.00000 | | 0.75054 | 1.62130 | 88 |
| GO_AEROBIC_RESPIRATION | 0.00001 | 0.00025 | 0.59333 | 0.69302 | 1.48168 | 65 |
| GO_ALTERNATIVE_MRNA_SPLICING_VIA_SPLICEOSOME | 0.00102 | 0.01782 | 0.45506 | 0.65168 | 1.38687 | 60 |
| GO_AMINE_METABOLIC_PROCESS | 0.00000 | 0.00000 | | 0.79189 | 1.70783 | 80 |
| GO_ANAPHASE_PROMOTING_COMPLEX_DEPENDENT_CATABOLIC_P | 0.00000 | 0.00000 | | 0.76611 | 1.65070 | 79 |
| GO_ANCHORING_JUNCTION | 0.00000 | 0.00000 | | 0.59615 | 1.33633 | 479 |
| GO_ANIMAL_ORGAN_REGENERATION | 0.00257 | 0.03716 | 0.43171 | 0.68240 | 1.41020 | 33 |
| GO_ANTIGEN_PROCESSING_AND_PRESENTATION | 0.00000 | 0.00000 | 0.70498 | 0.64776 | 1.42241 | 143 |
| GO_ANTIGEN_PROCESSING_AND_PRESENTATION_OF_EXOGENOUS_ | 0.00000 | 0.00000 | | 0.81051 | 1.72358 | 57 |
| GO_ANTIGEN_PROCESSING_AND_PRESENTATION_OF_PEPTIDE_ANTI | 0.00000 | 0.00000 | 0.70498 | 0.65912 | 1.44248 | 126 |
| GO_ANTIGEN_PROCESSING_AND_PRESENTATION_OF_PEPTIDE_ANTI | 0.00000 | 0.00000 | | 0.76925 | 1.65015 | 70 |
| GO_ANTIGEN_RECEPTOR_MEDIATED_SIGNALING_PATHWAY | 0.00000 | 0.00000 | 0.80122 | 0.67877 | 1.48641 | 127 |
| GO_APOPTOTIC_SIGNALING_PATHWAY | 0.00154 | 0.02464 | 0.45506 | 0.51561 | 1.15085 | 358 |
| GO_ATP_DEPENDENT_CHROMATIN_REMODELING | 0.00253 | 0.03681 | 0.43171 | 0.62124 | 1.32400 | 61 |
| GO_ATP_METABOLIC_PROCESS | 0.00000 | 0.00000 | 0.71951 | 0.62252 | 1.37693 | 196 |
| GO_ATP_SYNTHESIS_COUPLED_ELECTRON_TRANSPORT | 0.00005 | 0.00131 | 0.55733 | 0.65519 | 1.40723 | 74 |
| GO_BINDING_OF_SPERM_TO_ZONA_PELLUCIDA | 0.00000 | 0.00001 | 0.69013 | 0.89879 | 1.80846 | 18 |
| GO_CADHERIN_BINDING | 0.00000 | 0.00000 | | 0.63068 | 1.39961 | 236 |
| GO_CANONICAL_WNT_SIGNALING_PATHWAY | 0.00000 | 0.00003 | 0.65944 | 0.60816 | 1.34553 | 197 |
| GO_CATALYTIC_STEP_2_SPLICEOSOME | 0.00000 | 0.00002 | 0.65944 | 0.68068 | 1.46976 | 83 |
| GO_CELLULAR_AMINO_ACID_METABOLIC_PROCESS | 0.00000 | 0.00000 | 0.74774 | 0.61752 | 1.36824 | 216 |
| GO_CELLULAR_COMPONENT_DISASSEMBLY | 0.00000 | 0.00005 | 0.64355 | 0.55505 | 1.24197 | 401 |
| GO_CELLULAR_KETONE_METABOLIC_PROCESS | 0.00000 | 0.00000 | 0.77494 | 0.65955 | 1.44893 | 149 |
| GO_CELLULAR_NITROGEN_COMPOUND_CATABOLIC_PROCESS | 0.00000 | 0.00000 | | 0.74079 | 1.65848 | 436 |
| GO_CELLULAR_PROTEIN_COMPLEX_DISASSEMBLY | 0.00006 | 0.00139 | 0.55733 | 0.58222 | 1.28459 | 176 |
| GO_CELLULAR_RESPIRATION | 0.00000 | 0.00000 | 0.71951 | 0.64997 | 1.42492 | 136 |
| GO_CELLULAR_RESPONSE_TO_HEAT | 0.00018 | 0.00403 | 0.51885 | 0.61962 | 1.34016 | 91 |
| GO_CELLULAR_RESPONSE_TO_ORGANIC_CYCLIC_COMPOUND | 0.00302 | 0.04108 | 0.43171 | 0.52402 | 1.16585 | 283 |
| GO_CELLULAR_RESPONSE_TO_OXYGEN_LEVELS | 0.00000 | 0.00000 | 0.78819 | 0.65911 | 1.44882 | 157 |
| GO_CELL_ACTIVATION_INVOLVED_IN_IMMUNE_RESPONSE | 0.00000 | 0.00001 | 0.67496 | 0.57339 | 1.27913 | 341 |
| GO_CELL_ADHESION_MOLECULE_BINDING | 0.00000 | 0.00000 | 0.81404 | 0.60529 | 1.34788 | 308 |
| GO_CELL_CELL_RECOGNITION | 0.00000 | 0.00002 | 0.65944 | 0.83826 | 1.71003 | 26 |
| GO_CELL_CELL_SIGNALING_BY_WNT | 0.00000 | 0.00002 | 0.67496 | 0.57736 | 1.28559 | 299 |
| GO_CELL_CYCLE_G2_M_PHASE_TRANSITION | 0.00001 | 0.00024 | 0.59333 | 0.58034 | 1.28602 | 214 |
| GO_CELL_CYCLE_PHASE_TRANSITION | 0.00000 | 0.00004 | 0.64355 | 0.54319 | 1.21681 | 464 |
| GO_CELL_DIVISION | 0.00154 | 0.02464 | 0.45506 | 0.51551 | 1.15425 | 427 |

| | | | | | | |
|---|---|---|---|---|---|---|
| GO_CELL_RECOGNITION | 0.00012 | 0.00270 | 0.53843 | 0.69554 | 1.46186 | 48 |
| GO_CELL_SUBSTRATE_JUNCTION | 0.00000 | 0.00000 | | 0.67751 | 1.50775 | 291 |
| GO_CELL_SURFACE_RECEPTOR_SIGNALING_PATHWAY_INVOLVED_IN | 0.00000 | 0.00003 | 0.65944 | 0.57035 | 1.27067 | 319 |
| GO_CHAPERONE_BINDING | 0.00007 | 0.00163 | 0.53843 | 0.64754 | 1.39253 | 76 |
| GO_CHAPERONE_COMPLEX | 0.00000 | 0.00004 | 0.64355 | 0.85718 | 1.73925 | 22 |
| GO_CHAPERONE_MEDIATED_PROTEIN_COMPLEX_ASSEMBLY | 0.00170 | 0.02662 | 0.45506 | 0.77423 | 1.54792 | 17 |
| GO_CHROMATIN_BINDING | 0.00071 | 0.01323 | 0.47727 | 0.52699 | 1.17680 | 369 |
| GO_CHROMOSOMAL_REGION | 0.00246 | 0.03601 | 0.43171 | 0.52919 | 1.17661 | 276 |
| GO_COP9_SIGNALOSOME | 0.00128 | 0.02141 | 0.45506 | 0.73616 | 1.49715 | 24 |
| GO_COTRANSLATIONAL_PROTEIN_TARGETING_TO_MEMBRANE | 0.00000 | 0.00000 | | 0.89471 | 1.93513 | 91 |
| GO_CYTOCHROME_COMPLEX | 0.00168 | 0.02641 | 0.45506 | 0.75311 | 1.52417 | 20 |
| GO_CYTOKINE_MEDIATED_SIGNALING_PATHWAY | 0.00000 | 0.00000 | 0.81404 | 0.59698 | 1.33170 | 339 |
| GO_CYTOPLASMIC_TRANSLATION | 0.00000 | 0.00000 | | 0.84753 | 1.83080 | 88 |
| GO_CYTOPLASMIC_TRANSLATIONAL_INITIATION | 0.00000 | 0.00000 | 0.74774 | 0.85688 | 1.76176 | 30 |
| GO_CYTOSOLIC_LARGE_RIBOSOMAL_SUBUNIT | 0.00000 | 0.00000 | | 0.91186 | 1.92988 | 53 |
| GO_CYTOSOLIC_RIBOSOME | 0.00000 | 0.00000 | | 0.89907 | 1.94710 | 95 |
| GO_CYTOSOLIC_SMALL_RIBOSOMAL_SUBUNIT | 0.00000 | 0.00000 | | 0.89114 | 1.85533 | 39 |
| GO_DEFENSE_RESPONSE_TO_OTHER_ORGANISM | 0.00004 | 0.00099 | 0.55733 | 0.53673 | 1.20127 | 415 |
| GO_DE_NOVO_PROTEIN_FOLDING | 0.00050 | 0.00978 | 0.47727 | 0.74232 | 1.51431 | 26 |
| GO_DISORDERED_DOMAIN_SPECIFIC_BINDING | 0.00161 | 0.02546 | 0.45506 | 0.71970 | 1.47197 | 27 |
| GO_DNA_BIOSYNTHETIC_PROCESS | 0.00000 | 0.00010 | 0.62726 | 0.62505 | 1.37017 | 134 |
| GO_DNA_CONFORMATION_CHANGE | 0.00027 | 0.00572 | 0.49849 | 0.55300 | 1.22527 | 216 |
| GO_DNA_DAMAGE_RESPONSE_DETECTION_OF_DNA_DAMAGE | 0.00198 | 0.03028 | 0.43171 | 0.67949 | 1.40669 | 35 |
| GO_DNA_GEOMETRIC_CHANGE | 0.00046 | 0.00925 | 0.49849 | 0.60807 | 1.31855 | 98 |
| GO_DNA_POLYMERASE_BINDING | 0.00268 | 0.03799 | 0.43171 | 0.76255 | 1.52457 | 17 |
| GO_DNA_REPAIR | 0.00265 | 0.03775 | 0.43171 | 0.51159 | 1.14547 | 427 |
| GO_DRUG_BINDING | 0.00176 | 0.02733 | 0.45506 | 0.64119 | 1.35096 | 49 |
| GO_ELECTRON_TRANSFER_ACTIVITY | 0.00001 | 0.00020 | 0.61053 | 0.67481 | 1.44938 | 74 |
| GO_ELECTRON_TRANSPORT_CHAIN | 0.00002 | 0.00053 | 0.57561 | 0.62257 | 1.36145 | 124 |
| GO_ENDOPEPTIDASE_ACTIVITY | 0.00124 | 0.02085 | 0.45506 | 0.54955 | 1.21076 | 171 |
| GO_ENDOPEPTIDASE_COMPLEX | 0.00000 | 0.00000 | | 0.80930 | 1.73234 | 66 |
| GO_ENERGY_DERIVATION_BY_OXIDATION_OF_ORGANIC_COMPOUN | 0.00000 | 0.00000 | 0.71951 | 0.63390 | 1.39707 | 172 |
| GO_ESTABLISHMENT_OF_PROTEIN_LOCALIZATION_TO_CHROMOSON | 0.00020 | 0.00432 | 0.51885 | 0.77339 | 1.57217 | 23 |
| GO_ESTABLISHMENT_OF_PROTEIN_LOCALIZATION_TO_ENDOPLASM | 0.00000 | 0.00000 | | 0.88026 | 1.91149 | 102 |
| GO_ESTABLISHMENT_OF_PROTEIN_LOCALIZATION_TO_MEMBRANE | 0.00000 | 0.00000 | | 0.74874 | 1.66164 | 240 |
| GO_ESTABLISHMENT_OF_PROTEIN_LOCALIZATION_TO_ORGANELLE | 0.00000 | 0.00000 | | 0.69519 | 1.55605 | 423 |
| GO_ESTABLISHMENT_OF_PROTEIN_LOCALIZATION_TO_TELOMERE | 0.00006 | 0.00139 | 0.55733 | 0.83470 | 1.66881 | 17 |
| GO_ESTABLISHMENT_OF_RNA_LOCALIZATION | 0.00106 | 0.01819 | 0.45506 | 0.55577 | 1.22400 | 167 |
| GO_ESTABLISHMENT_OF_TISSUE_POLARITY | 0.00000 | 0.00000 | | 0.75051 | 1.62122 | 88 |
| GO_EUKARYOTIC_TRANSLATION_INITIATION_FACTOR_3_COMPLEX | 0.00003 | 0.00074 | 0.57561 | 0.86987 | 1.72555 | 15 |
| GO_EXOCYTOSIS | 0.00000 | 0.00015 | 0.61053 | 0.53905 | 1.20753 | 465 |
| GO_FC_EPSILON_RECEPTOR_SIGNALING_PATHWAY | 0.00000 | 0.00000 | | 0.75117 | 1.62103 | 82 |
| GO_FC_RECEPTOR_SIGNALING_PATHWAY | 0.00000 | 0.00000 | | 0.72054 | 1.57787 | 127 |
| GO_FERTILIZATION | 0.00089 | 0.01592 | 0.47727 | 0.63620 | 1.35888 | 64 |
| GO_FICOLIN_1_RICH_GRANULE | 0.00000 | 0.00000 | 0.78819 | 0.71716 | 1.55340 | 93 |
| GO_GENERATION_OF_PRECURSOR_METABOLITES_AND_ENERGY | 0.00000 | 0.00000 | 0.77494 | 0.59261 | 1.32026 | 320 |
| GO_GENE_SILENCING | 0.00261 | 0.03750 | 0.43171 | 0.55578 | 1.22252 | 161 |
| GO_GLAND_DEVELOPMENT | 0.00154 | 0.02464 | 0.45506 | 0.54117 | 1.19797 | 206 |
| GO_GLYCOSYL_COMPOUND_METABOLIC_PROCESS | 0.00071 | 0.01323 | 0.47727 | 0.62524 | 1.34204 | 71 |
| GO_GUANYL_NUCLEOTIDE_BINDING | 0.00095 | 0.01679 | 0.47727 | 0.54096 | 1.19861 | 216 |
| GO_HEAT_SHOCK_PROTEIN_BINDING | 0.00003 | 0.00092 | 0.55733 | 0.64369 | 1.39056 | 85 |
| GO_HEMATOPOIETIC_PROGENITOR_CELL_DIFFERENTIATION | 0.00000 | 0.00000 | 0.82666 | 0.70307 | 1.53239 | 113 |
| GO_HEMATOPOIETIC_STEM_CELL_DIFFERENTIATION | 0.00000 | 0.00000 | | 0.79776 | 1.71217 | 72 |
| GO_HSP70_PROTEIN_BINDING | 0.00228 | 0.03391 | 0.43171 | 0.71143 | 1.45504 | 27 |
| GO_IMMUNE_RESPONSE_REGULATING_SIGNALING_PATHWAY | 0.00000 | 0.00000 | 0.76146 | 0.63494 | 1.40480 | 197 |
| GO_INNATE_IMMUNE_RESPONSE | 0.00003 | 0.00071 | 0.57561 | 0.55173 | 1.23107 | 347 |
| GO_INNATE_IMMUNE_RESPONSE_ACTIVATING_SIGNAL_TRANSDUCT | 0.00000 | 0.00000 | | 0.79340 | 1.70281 | 72 |
| GO_INNER_MITOCHONDRIAL_MEMBRANE_PROTEIN_COMPLEX | 0.00002 | 0.00060 | 0.57561 | 0.65124 | 1.40678 | 88 |
| GO_INTERACTION_WITH_SYMBIONT | 0.00001 | 0.00026 | 0.59333 | 0.75084 | 1.56620 | 41 |
| GO_INTERLEUKIN_1_MEDIATED_SIGNALING_PATHWAY | 0.00000 | 0.00000 | | 0.81689 | 1.75671 | 76 |
| GO_INTRINSIC_APOPTOTIC_SIGNALING_PATHWAY | 0.00121 | 0.02029 | 0.45506 | 0.55118 | 1.21845 | 190 |
| GO_ION_CHANNEL_BINDING | 0.00087 | 0.01584 | 0.47727 | 0.63182 | 1.34683 | 62 |
| GO_ISOPEPTIDASE_ACTIVITY | 0.00139 | 0.02285 | 0.45506 | 0.78940 | 1.57097 | 16 |
| GO_LARGE_RIBOSOMAL_SUBUNIT | 0.00000 | 0.00000 | | 0.78135 | 1.70183 | 109 |
| GO_LEUKOCYTE_MEDIATED_IMMUNITY | 0.00000 | 0.00000 | 0.71951 | 0.57282 | 1.27895 | 364 |
| GO_MACROMOLECULE_TRANSMEMBRANE_TRANSPORTER_ACTIVITY | 0.00239 | 0.03510 | 0.43171 | 0.77734 | 1.54696 | 16 |
| GO_MAINTENANCE_OF_CELL_NUMBER | 0.00087 | 0.01584 | 0.47727 | 0.58944 | 1.28512 | 114 |
| GO_MATURATION_OF_LSU_RRNA | 0.00006 | 0.00151 | 0.53843 | 0.77606 | 1.58723 | 27 |
| GO_METHYLTRANSFERASE_COMPLEX | 0.00290 | 0.04014 | 0.43171 | 0.59476 | 1.28479 | 88 |
| GO_METHYL_CPG_BINDING | 0.00302 | 0.04108 | 0.43171 | 0.78053 | 1.54833 | 15 |
| GO_MIDBRAIN_DEVELOPMENT | 0.00021 | 0.00468 | 0.51885 | 0.68832 | 1.44457 | 47 |
| GO_MITOCHONDRIAL_ELECTRON_TRANSPORT_NADH_TO_UBIQUINO | 0.00331 | 0.04437 | 0.43171 | 0.65428 | 1.36724 | 42 |
| GO_MITOCHONDRIAL_GENE_EXPRESSION | 0.00048 | 0.00949 | 0.49849 | 0.57608 | 1.26525 | 152 |
| GO_MITOCHONDRIAL_LARGE_RIBOSOMAL_SUBUNIT | 0.00002 | 0.00050 | 0.57561 | 0.69789 | 1.48086 | 55 |
| GO_MITOCHONDRIAL_MATRIX | 0.00000 | 0.00002 | 0.65944 | 0.56045 | 1.25158 | 371 |
| GO_MITOCHONDRIAL_MEMBRANE_ORGANIZATION | 0.00361 | 0.04734 | 0.43171 | 0.59309 | 1.28094 | 86 |

| | | | | | | |
|---|---|---|---|---|---|---|
| GO_MITOCHONDRIAL_PROTEIN_COMPLEX | 0.00000 | 0.00000 | 0.81404 | 0.63650 | 1.40931 | 208 |
| GO_MITOCHONDRIAL_TRANSLATION | 0.00004 | 0.00093 | 0.55733 | 0.61219 | 1.34135 | 129 |
| GO_MITOCHONDRIAL_TRANSLATIONAL_TERMINATION | 0.00000 | 0.00010 | 0.62726 | 0.66694 | 1.44128 | 87 |
| GO_MITOCHONDRION_ORGANIZATION | 0.00051 | 0.00993 | 0.47727 | 0.52284 | 1.16863 | 390 |
| GO_MODIFICATION_DEPENDENT_MACROMOLECULE_CATABOLIC_PR | 0.00002 | 0.00053 | 0.57561 | 0.53142 | 1.19156 | 486 |
| GO_MODULATION_BY_HOST_OF_SYMBIONT_PROCESS | 0.00015 | 0.00336 | 0.51885 | 0.72848 | 1.50542 | 33 |
| GO_MODULATION_OF_PROCESS_OF_OTHER_ORGANISM | 0.00019 | 0.00423 | 0.51885 | 0.66904 | 1.42286 | 59 |
| GO_MONOSACCHARIDE_BIOSYNTHETIC_PROCESS | 0.00335 | 0.04466 | 0.43171 | 0.61992 | 1.31781 | 58 |
| GO_MORPHOGENESIS_OF_AN_EPITHELIUM | 0.00006 | 0.00151 | 0.53843 | 0.55161 | 1.22755 | 290 |
| GO_MORPHOGENESIS_OF_A_POLARIZED_EPITHELIUM | 0.00000 | 0.00000 | | 0.72298 | 1.57057 | 103 |
| GO_MRNA_3_UTR_BINDING | 0.00003 | 0.00089 | 0.55733 | 0.67596 | 1.44827 | 68 |
| GO_MRNA_5_UTR_BINDING | 0.00006 | 0.00156 | 0.53843 | 0.80133 | 1.62592 | 22 |
| GO_MRNA_BINDING | 0.00000 | 0.00000 | | 0.66002 | 1.46353 | 222 |
| GO_MRNA_PROCESSING | 0.00000 | 0.00000 | 0.73376 | 0.56859 | 1.27295 | 418 |
| GO_MULTI_ORGANISM_LOCALIZATION | 0.00268 | 0.03799 | 0.43171 | 0.63245 | 1.33612 | 51 |
| GO_MULTI_ORGANISM_PROCESS | 0.00108 | 0.01839 | 0.45506 | 0.51114 | 1.14599 | 484 |
| GO_MYD88_DEPENDENT_TOLL_LIKE_RECEPTOR_SIGNALING_PATHW | 0.00002 | 0.00066 | 0.57561 | 0.85581 | 1.70312 | 16 |
| GO_MYELOID_LEUKOCYTE_ACTIVATION | 0.00000 | 0.00000 | 0.74774 | 0.58508 | 1.30479 | 327 |
| GO_MYELOID_LEUKOCYTE_MEDIATED_IMMUNITY | 0.00000 | 0.00000 | 0.76146 | 0.59841 | 1.33291 | 301 |
| GO_NADH_DEHYDROGENASE_ACTIVITY | 0.00383 | 0.04935 | 0.43171 | 0.66244 | 1.37139 | 35 |
| GO_NADH_DEHYDROGENASE_COMPLEX | 0.00376 | 0.04884 | 0.43171 | 0.65649 | 1.36421 | 38 |
| GO_NADH_METABOLIC_PROCESS | 0.00294 | 0.04036 | 0.43171 | 0.70642 | 1.44481 | 27 |
| GO_NCRNA_EXPORT_FROM_NUCLEUS | 0.00302 | 0.04108 | 0.43171 | 0.66820 | 1.38331 | 35 |
| GO_NCRNA_METABOLIC_PROCESS | 0.00000 | 0.00003 | 0.65944 | 0.55907 | 1.24907 | 381 |
| GO_NCRNA_PROCESSING | 0.00000 | 0.00003 | 0.65944 | 0.57175 | 1.27504 | 328 |
| GO_NEGATIVE_REGULATION_OF_CANONICAL_WNT_SIGNALING_PAT | 0.00000 | 0.00000 | | 0.71611 | 1.56038 | 111 |
| GO_NEGATIVE_REGULATION_OF_CELLULAR_AMIDE_METABOLIC_PR | 0.00001 | 0.00020 | 0.61053 | 0.63031 | 1.37423 | 114 |
| GO_NEGATIVE_REGULATION_OF_CELL_CYCLE | 0.00003 | 0.00076 | 0.57561 | 0.53833 | 1.20513 | 425 |
| GO_NEGATIVE_REGULATION_OF_CELL_CYCLE_G2_M_PHASE_TRANS | 0.00000 | 0.00000 | 0.76146 | 0.71490 | 1.54439 | 85 |
| GO_NEGATIVE_REGULATION_OF_CELL_CYCLE_PHASE_TRANSITION | 0.00002 | 0.00060 | 0.57561 | 0.59041 | 1.30442 | 184 |
| GO_NEGATIVE_REGULATION_OF_CELL_CYCLE_PROCESS | 0.00000 | 0.00002 | 0.65944 | 0.58690 | 1.30374 | 249 |
| GO_NEGATIVE_REGULATION_OF_MITOTIC_CELL_CYCLE | 0.00004 | 0.00095 | 0.55733 | 0.56735 | 1.25902 | 232 |
| GO_NEGATIVE_REGULATION_OF_MRNA_METABOLIC_PROCESS | 0.00023 | 0.00491 | 0.51885 | 0.66670 | 1.41788 | 59 |
| GO_NEGATIVE_REGULATION_OF_MRNA_SPLICING_VIA_SPLICEOSOM | 0.00239 | 0.03510 | 0.43171 | 0.74857 | 1.51204 | 19 |
| GO_NEGATIVE_REGULATION_OF_RNA_SPLICING | 0.00029 | 0.00607 | 0.49849 | 0.76729 | 1.55976 | 23 |
| GO_NEGATIVE_REGULATION_OF_TRANSLATIONAL_INITIATION | 0.00018 | 0.00403 | 0.51885 | 0.83901 | 1.66434 | 15 |
| GO_NEGATIVE_REGULATION_OF_WNT_SIGNALING_PATHWAY | 0.00000 | 0.00000 | | 0.69525 | 1.52249 | 127 |
| GO_NEURAL_NUCLEUS_DEVELOPMENT | 0.00001 | 0.00026 | 0.59333 | 0.79148 | 1.62187 | 28 |
| GO_NEURON_TO_NEURON_SYNAPSE | 0.00005 | 0.00134 | 0.55733 | 0.59636 | 1.31006 | 154 |
| GO_NIK_NF_KAPPAB_SIGNALING | 0.00000 | 0.00000 | | 0.72896 | 1.58354 | 103 |
| GO_NON_CANONICAL_WNT_SIGNALING_PATHWAY | 0.00000 | 0.00000 | | 0.72784 | 1.58023 | 101 |
| GO_NUCLEAR_EXPORT | 0.00031 | 0.00645 | 0.49849 | 0.56963 | 1.25542 | 172 |
| GO_NUCLEAR_TRANSCRIBED_MRNA_CATABOLIC_PROCESS | 0.00000 | 0.00000 | | 0.79179 | 1.74917 | 183 |
| GO_NUCLEAR_TRANSCRIBED_MRNA_CATABOLIC_PROCESS_NONSEN | 0.00000 | 0.00000 | | 0.89008 | 1.93909 | 110 |
| GO_NUCLEAR_TRANSPORT | 0.00008 | 0.00193 | 0.53843 | 0.55511 | 1.23408 | 273 |
| GO_NUCLEOBASE_CONTAINING_COMPOUND_TRANSPORT | 0.00146 | 0.02386 | 0.45506 | 0.55053 | 1.21842 | 202 |
| GO_NUCLEOBASE_CONTAINING_SMALL_MOLECULE_BIOSYNTHETIC_ | 0.00137 | 0.02261 | 0.45506 | 0.62749 | 1.34353 | 67 |
| GO_NUCLEOBASE_CONTAINING_SMALL_MOLECULE_INTERCONVERS | 0.00383 | 0.04935 | 0.43171 | 0.71259 | 1.44857 | 23 |
| GO_NUCLEOBASE_CONTAINING_SMALL_MOLECULE_METABOLIC_PR | 0.00320 | 0.04339 | 0.43171 | 0.51448 | 1.14825 | 352 |
| GO_NUCLEOSIDE_DIPHOSPHATE_METABOLIC_PROCESS | 0.00110 | 0.01865 | 0.45506 | 0.59637 | 1.29295 | 97 |
| GO_NUCLEOSIDE_METABOLIC_PROCESS | 0.00061 | 0.01169 | 0.47727 | 0.65315 | 1.38907 | 59 |
| GO_NUCLEOSIDE_MONOPHOSPHATE_BIOSYNTHETIC_PROCESS | 0.00250 | 0.03636 | 0.43171 | 0.68841 | 1.42033 | 32 |
| GO_NUCLEOSIDE_MONOPHOSPHATE_METABOLIC_PROCESS | 0.00152 | 0.02464 | 0.45506 | 0.64377 | 1.36002 | 51 |
| GO_NUCLEOSIDE_PHOSPHATE_BIOSYNTHETIC_PROCESS | 0.00043 | 0.00865 | 0.49849 | 0.58185 | 1.27831 | 148 |
| GO_NUCLEOSIDE_TRIPHOSPHATE_BIOSYNTHETIC_PROCESS | 0.00043 | 0.00865 | 0.49849 | 0.69332 | 1.44539 | 40 |
| GO_NUCLEOSIDE_TRIPHOSPHATE_METABOLIC_PROCESS | 0.00016 | 0.00360 | 0.51885 | 0.66788 | 1.41976 | 58 |
| GO_NUCLEOSOMAL_DNA_BINDING | 0.00023 | 0.00507 | 0.51885 | 0.79852 | 1.61801 | 21 |
| GO_NUCLEOSOME_BINDING | 0.00143 | 0.02332 | 0.45506 | 0.66132 | 1.38492 | 45 |
| GO_NUCLEOTIDE_EXCISION_REPAIR | 0.00004 | 0.00096 | 0.55733 | 0.63343 | 1.37355 | 98 |
| GO_NUCLEOTIDE_EXCISION_REPAIR_DNA_DAMAGE_RECOGNITION | 0.00015 | 0.00349 | 0.51885 | 0.78678 | 1.59640 | 22 |
| GO_NUCLEOTIDE_EXCISION_REPAIR_DNA_INCISION | 0.00313 | 0.04249 | 0.43171 | 0.66439 | 1.37898 | 36 |
| GO_NUCLEOTIDE_PHOSPHORYLATION | 0.00383 | 0.04935 | 0.43171 | 0.59002 | 1.27454 | 88 |
| GO_ORGANELLAR_RIBOSOME | 0.00000 | 0.00014 | 0.61053 | 0.66412 | 1.43471 | 85 |
| GO_ORGANELLE_INNER_MEMBRANE | 0.00000 | 0.00000 | 0.76146 | 0.57601 | 1.28697 | 378 |
| GO_ORGANIC_CYCLIC_COMPOUND_CATABOLIC_PROCESS | 0.00000 | 0.00000 | | 0.73350 | 1.64281 | 449 |
| GO_OXIDATIVE_PHOSPHORYLATION | 0.00001 | 0.00030 | 0.59333 | 0.64619 | 1.40027 | 96 |
| GO_OXIDOREDUCTASE_ACTIVITY | 0.00006 | 0.00139 | 0.55733 | 0.53544 | 1.19703 | 392 |
| GO_OXIDOREDUCTASE_COMPLEX | 0.00001 | 0.00027 | 0.59333 | 0.66015 | 1.42461 | 82 |
| GO_PEPTIDASE_COMPLEX | 0.00000 | 0.00000 | | 0.75988 | 1.63982 | 82 |
| GO_PIGMENT_GRANULE | 0.00003 | 0.00089 | 0.55733 | 0.66098 | 1.42344 | 78 |
| GO_POLYSOMAL_RIBOSOME | 0.00000 | 0.00000 | | 0.90624 | 1.86629 | 31 |
| GO_POLYSOME | 0.00000 | 0.00000 | | 0.82337 | 1.75481 | 61 |
| GO_POLY_PYRIMIDINE_TRACT_BINDING | 0.00327 | 0.04398 | 0.43171 | 0.75471 | 1.51855 | 18 |
| GO_POSITIVE_REGULATION_OF_APOPTOTIC_SIGNALING_PATHWAY | 0.00279 | 0.03889 | 0.43171 | 0.57666 | 1.25518 | 107 |
| GO_POSITIVE_REGULATION_OF_ATPASE_ACTIVITY | 0.00294 | 0.04036 | 0.43171 | 0.65446 | 1.36795 | 43 |
| GO_POSITIVE_REGULATION_OF_BINDING | 0.00088 | 0.01590 | 0.47727 | 0.60284 | 1.30556 | 95 |

| | | | | | | |
|---|---|---|---|---|---|---|
| GO_POSITIVE_REGULATION_OF_CANONICAL_WNT_SIGNALING_PATH | 0.00000 | 0.00000 | 0.77494 | 0.70504 | 1.53160 | 103 |
| GO_POSITIVE_REGULATION_OF_CELLULAR_AMIDE_METABOLIC_PRC | 0.00003 | 0.00074 | 0.57561 | 0.62296 | 1.36083 | 118 |
| GO_POSITIVE_REGULATION_OF_CELLULAR_PROTEIN_LOCALIZATION | 0.00041 | 0.00840 | 0.49849 | 0.54915 | 1.21815 | 225 |
| GO_POSITIVE_REGULATION_OF_CELL_DEATH | 0.00069 | 0.01297 | 0.47727 | 0.52595 | 1.17423 | 363 |
| GO_POSITIVE_REGULATION_OF_CELL_POPULATION_PROLIFERATION | 0.00078 | 0.01455 | 0.47727 | 0.52449 | 1.17108 | 368 |
| GO_POSITIVE_REGULATION_OF_CHROMOSOME_ORGANIZATION | 0.00231 | 0.03437 | 0.43171 | 0.56990 | 1.24871 | 133 |
| GO_POSITIVE_REGULATION_OF_DEFENSE_RESPONSE | 0.00000 | 0.00003 | 0.65944 | 0.62175 | 1.36730 | 158 |
| GO_POSITIVE_REGULATION_OF_DNA_BIOSYNTHETIC_PROCESS | 0.00001 | 0.00041 | 0.59333 | 0.71929 | 1.52055 | 52 |
| GO_POSITIVE_REGULATION_OF_DNA_METABOLIC_PROCESS | 0.00364 | 0.04761 | 0.43171 | 0.56647 | 1.24189 | 135 |
| GO_POSITIVE_REGULATION_OF_DNA_TEMPLATED_TRANSCRIPTION_ | 0.00276 | 0.03885 | 0.43171 | 0.68932 | 1.41586 | 29 |
| GO_POSITIVE_REGULATION_OF_ESTABLISHMENT_OF_PROTEIN_LOCA | 0.00007 | 0.00168 | 0.53843 | 0.56461 | 1.25165 | 219 |
| GO_POSITIVE_REGULATION_OF_IMMUNE_RESPONSE | 0.00000 | 0.00001 | 0.69013 | 0.59037 | 1.31265 | 277 |
| GO_POSITIVE_REGULATION_OF_IMMUNE_SYSTEM_PROCESS | 0.00004 | 0.00110 | 0.55733 | 0.54018 | 1.20716 | 386 |
| GO_POSITIVE_REGULATION_OF_INNATE_IMMUNE_RESPONSE | 0.00000 | 0.00000 | 0.81404 | 0.69693 | 1.51947 | 114 |
| GO_POSITIVE_REGULATION_OF_INTRINSIC_APOPTOTIC_SIGNALING_ | 0.00104 | 0.01798 | 0.45506 | 0.69097 | 1.43578 | 37 |
| GO_POSITIVE_REGULATION_OF_ORGANELLE_ORGANIZATION | 0.00043 | 0.00865 | 0.49849 | 0.52093 | 1.16605 | 411 |
| GO_POSITIVE_REGULATION_OF_PROTEIN_LOCALIZATION_TO_NUCLE | 0.00001 | 0.00020 | 0.61053 | 0.69907 | 1.48990 | 61 |
| GO_POSITIVE_REGULATION_OF_RESPONSE_TO_BIOTIC_STIMULUS | 0.00000 | 0.00000 | 0.73376 | 0.66340 | 1.45310 | 128 |
| GO_POSITIVE_REGULATION_OF_RESPONSE_TO_EXTERNAL_STIMULU | 0.00002 | 0.00050 | 0.57561 | 0.58133 | 1.28681 | 201 |
| GO_POSITIVE_REGULATION_OF_TELOMERASE_ACTIVITY | 0.00061 | 0.01169 | 0.47727 | 0.73476 | 1.50565 | 28 |
| GO_POSITIVE_REGULATION_OF_TELOMERASE_RNA_LOCALIZATION_ | 0.00000 | 0.00001 | 0.69013 | 0.92451 | 1.83395 | 15 |
| GO_POSITIVE_REGULATION_OF_TELOMERE_MAINTENANCE_VIA_TE | 0.00130 | 0.02159 | 0.45506 | 0.71259 | 1.46508 | 30 |
| GO_POSITIVE_REGULATION_OF_TRANSCRIPTION_INITIATION_FROM | 0.00372 | 0.04847 | 0.43171 | 0.70671 | 1.43993 | 25 |
| GO_POSITIVE_REGULATION_OF_TRANSFERASE_ACTIVITY | 0.00174 | 0.02712 | 0.45506 | 0.51642 | 1.15220 | 345 |
| GO_POSITIVE_REGULATION_OF_TRANSLATION | 0.00001 | 0.00019 | 0.61053 | 0.64767 | 1.40419 | 97 |
| GO_POSITIVE_REGULATION_OF_VIRAL_GENOME_REPLICATION | 0.00287 | 0.03973 | 0.43171 | 0.68362 | 1.40783 | 31 |
| GO_POSITIVE_REGULATION_OF_VIRAL_PROCESS | 0.00028 | 0.00590 | 0.49849 | 0.62440 | 1.34741 | 81 |
| GO_POSITIVE_REGULATION_OF_WNT_SIGNALING_PATHWAY | 0.00000 | 0.00000 | 0.73376 | 0.66230 | 1.44763 | 122 |
| GO_POSTREPLICATION_REPAIR | 0.00024 | 0.00515 | 0.51885 | 0.69140 | 1.44518 | 43 |
| GO_POSTSYNAPSE | 0.00004 | 0.00109 | 0.55733 | 0.55969 | 1.24369 | 259 |
| GO_POSTSYNAPTIC_CYTOSOL | 0.00279 | 0.03889 | 0.43171 | 0.78295 | 1.55313 | 15 |
| GO_POSTTRANSCRIPTIONAL_REGULATION_OF_GENE_EXPRESSION | 0.00000 | 0.00000 | | 0.66656 | 1.49327 | 462 |
| GO_POST_TRANSLATIONAL_PROTEIN_MODIFICATION | 0.00002 | 0.00052 | 0.57561 | 0.57851 | 1.28412 | 229 |
| GO_PRECATALYTIC_SPLICEOSOME | 0.00085 | 0.01559 | 0.47727 | 0.65235 | 1.37448 | 49 |
| GO_PRERIBOSOME | 0.00087 | 0.01577 | 0.47727 | 0.62899 | 1.34772 | 69 |
| GO_PRERIBOSOME_LARGE_SUBUNIT_PRECURSOR | 0.00235 | 0.03474 | 0.43171 | 0.74079 | 1.50308 | 22 |
| GO_PRE_MRNA_BINDING | 0.00133 | 0.02213 | 0.45506 | 0.72393 | 1.48062 | 27 |
| GO_PROTEASOMAL_PROTEIN_CATABOLIC_PROCESS | 0.00002 | 0.00044 | 0.59333 | 0.54166 | 1.21005 | 377 |
| GO_PROTEASOMAL_UBIQUITIN_INDEPENDENT_PROTEIN_CATABOLI | 0.00000 | 0.00000 | 0.80122 | 0.92254 | 1.86931 | 21 |
| GO_PROTEASOME_ACCESSORY_COMPLEX | 0.00000 | 0.00001 | 0.67496 | 0.85464 | 1.73811 | 24 |
| GO_PROTEASOME_CORE_COMPLEX | 0.00000 | 0.00000 | | 0.94375 | 1.90631 | 19 |
| GO_PROTEIN_CONTAINING_COMPLEX_DISASSEMBLY | 0.00000 | 0.00007 | 0.62726 | 0.57810 | 1.28438 | 256 |
| GO_PROTEIN_CONTAINING_COMPLEX_LOCALIZATION | 0.00063 | 0.01197 | 0.47727 | 0.55335 | 1.22520 | 208 |
| GO_PROTEIN_DNA_COMPLEX_SUBUNIT_ORGANIZATION | 0.00001 | 0.00018 | 0.61053 | 0.60174 | 1.32491 | 165 |
| GO_PROTEIN_FOLDING | 0.00000 | 0.00002 | 0.65944 | 0.63113 | 1.38644 | 154 |
| GO_PROTEIN_FOLDING_CHAPERONE | 0.00135 | 0.02238 | 0.45506 | 0.77903 | 1.55752 | 17 |
| GO_PROTEIN_IMPORT | 0.00268 | 0.03799 | 0.43171 | 0.56317 | 1.23581 | 140 |
| GO_PROTEIN_LOCALIZATION_TO_CHROMOSOME | 0.00056 | 0.01096 | 0.47727 | 0.63523 | 1.36108 | 69 |
| GO_PROTEIN_LOCALIZATION_TO_CHROMOSOME_TELOMERIC_REGI | 0.00335 | 0.04466 | 0.43171 | 0.70399 | 1.43984 | 27 |
| GO_PROTEIN_LOCALIZATION_TO_ENDOPLASMIC_RETICULUM | 0.00000 | 0.00000 | | 0.85521 | 1.87021 | 124 |
| GO_PROTEIN_LOCALIZATION_TO_MEMBRANE | 0.00000 | 0.00000 | | 0.65123 | 1.45714 | 406 |
| GO_PROTEIN_LOCALIZATION_TO_MITOCHONDRION | 0.00019 | 0.00420 | 0.51885 | 0.60773 | 1.32398 | 110 |
| GO_PROTEIN_LOCALIZATION_TO_NUCLEUS | 0.00002 | 0.00055 | 0.57561 | 0.58926 | 1.30220 | 187 |
| GO_PROTEIN_MODIFICATION_BY_SMALL_PROTEIN_REMOVAL | 0.00000 | 0.00000 | 0.71951 | 0.60745 | 1.34689 | 221 |
| GO_PROTEIN_POLYUBIQUITINATION | 0.00004 | 0.00106 | 0.55733 | 0.55874 | 1.24108 | 253 |
| GO_PROTEIN_REFOLDING | 0.00294 | 0.04036 | 0.43171 | 0.77039 | 1.53314 | 16 |
| GO_PROTEIN_STABILIZATION | 0.00001 | 0.00024 | 0.61053 | 0.62333 | 1.36586 | 130 |
| GO_PROTEIN_TARGETING | 0.00000 | 0.00000 | | 0.71181 | 1.58743 | 327 |
| GO_PROTEIN_TARGETING_TO_MEMBRANE | 0.00000 | 0.00000 | | 0.82370 | 1.80953 | 149 |
| GO_PROTEIN_TARGETING_TO_MITOCHONDRION | 0.00346 | 0.04593 | 0.43171 | 0.60408 | 1.30091 | 78 |
| GO_PROTEIN_TRANSMEMBRANE_TRANSPORT | 0.00265 | 0.03775 | 0.43171 | 0.64528 | 1.36171 | 50 |
| GO_PROTON_TRANSMEMBRANE_TRANSPORT | 0.00213 | 0.03229 | 0.43171 | 0.60815 | 1.30536 | 71 |
| GO_PURINE_CONTAINING_COMPOUND_BIOSYNTHETIC_PROCESS | 0.00246 | 0.03601 | 0.43171 | 0.57796 | 1.25936 | 111 |
| GO_PURINE_CONTAINING_COMPOUND_METABOLIC_PROCESS | 0.00100 | 0.01761 | 0.45506 | 0.53679 | 1.19242 | 255 |
| GO_PURINE_NUCLEOSIDE_TRIPHOSPHATE_METABOLIC_PROCESS | 0.00089 | 0.01592 | 0.47727 | 0.67157 | 1.40374 | 43 |
| GO_PYRIMIDINE_CONTAINING_COMPOUND_METABOLIC_PROCESS | 0.00353 | 0.04658 | 0.43171 | 0.64744 | 1.35878 | 47 |
| GO_REGENERATION | 0.00216 | 0.03242 | 0.43171 | 0.59987 | 1.29453 | 82 |
| GO_REGULATION_OF_ANIMAL_ORGAN_MORPHOGENESIS | 0.00000 | 0.00000 | | 0.69163 | 1.51541 | 133 |
| GO_REGULATION_OF_APOPTOTIC_SIGNALING_PATHWAY | 0.00031 | 0.00645 | 0.49849 | 0.55171 | 1.22506 | 247 |
| GO_REGULATION_OF_ATPASE_ACTIVITY | 0.00324 | 0.04369 | 0.43171 | 0.63031 | 1.34038 | 57 |
| GO_REGULATION_OF_ATP_METABOLIC_PROCESS | 0.00353 | 0.04658 | 0.43171 | 0.60176 | 1.29780 | 80 |
| GO_REGULATION_OF_BINDING | 0.00250 | 0.03636 | 0.43171 | 0.54476 | 1.20598 | 203 |
| GO_REGULATION_OF_CELLULAR_AMIDE_METABOLIC_PROCESS | 0.00000 | 0.00000 | | 0.62967 | 1.40226 | 311 |
| GO_REGULATION_OF_CELLULAR_AMINE_METABOLIC_PROCESS | 0.00000 | 0.00000 | | 0.82173 | 1.74682 | 58 |
| GO_REGULATION_OF_CELLULAR_AMINO_ACID_METABOLIC_PROCES | 0.00000 | 0.00000 | | 0.83776 | 1.77766 | 55 |
| GO_REGULATION_OF_CELLULAR_KETONE_METABOLIC_PROCESS | 0.00000 | 0.00000 | | 0.72641 | 1.58680 | 118 |

| | | | | | | |
|---|---|---|---|---|---|---|
| GO_REGULATION_OF_CELLULAR_PROTEIN_LOCALIZATION | 0.00082 | 0.01513 | 0.47727 | 0.52542 | 1.17290 | 362 |
| GO_REGULATION_OF_CELLULAR_RESPONSE_TO_HEAT | 0.00276 | 0.03885 | 0.43171 | 0.61652 | 1.32004 | 67 |
| GO_REGULATION_OF_CELLULAR_RESPONSE_TO_STRESS | 0.00224 | 0.03344 | 0.43171 | 0.50545 | 1.13232 | 468 |
| GO_REGULATION_OF_CELL_CYCLE_G2_M_PHASE_TRANSITION | 0.00000 | 0.00000 | 0.71951 | 0.63230 | 1.39282 | 169 |
| GO_REGULATION_OF_CELL_CYCLE_PHASE_TRANSITION | 0.00001 | 0.00026 | 0.59333 | 0.55488 | 1.23783 | 337 |
| GO_REGULATION_OF_CHROMOSOME_ORGANIZATION | 0.00081 | 0.01501 | 0.47727 | 0.54255 | 1.20527 | 261 |
| GO_REGULATION_OF_CYSTEINE_TYPE_ENDOPEPTIDASE_ACTIVITY | 0.00202 | 0.03077 | 0.43171 | 0.57120 | 1.24837 | 121 |
| GO_REGULATION_OF_CYTOPLASMIC_TRANSLATION | 0.00130 | 0.02159 | 0.45506 | 0.73678 | 1.49773 | 23 |
| GO_REGULATION_OF_DEFENSE_RESPONSE | 0.00065 | 0.01237 | 0.47727 | 0.53633 | 1.19355 | 290 |
| GO_REGULATION_OF_DNA_BIOSYNTHETIC_PROCESS | 0.00016 | 0.00360 | 0.51885 | 0.64215 | 1.37923 | 74 |
| GO_REGULATION_OF_DNA_METABOLIC_PROCESS | 0.00069 | 0.01297 | 0.47727 | 0.54425 | 1.20775 | 237 |
| GO_REGULATION_OF_DNA_TEMPLATED_TRANSCRIPTION_INITIATIOI | 0.00163 | 0.02568 | 0.45506 | 0.68289 | 1.41372 | 35 |
| GO_REGULATION_OF_DNA_TEMPLATED_TRANSCRIPTION_IN_RESPO | 0.00000 | 0.00000 | | 0.73944 | 1.59800 | 89 |
| GO_REGULATION_OF_GENERATION_OF_PRECURSOR_METABOLITES_ | 0.00060 | 0.01164 | 0.47727 | 0.60476 | 1.31116 | 97 |
| GO_REGULATION_OF_HEMATOPOIETIC_PROGENITOR_CELL_DIFFERE | 0.00000 | 0.00000 | | 0.78717 | 1.69251 | 75 |
| GO_REGULATION_OF_HEMATOPOIETIC_STEM_CELL_DIFFERENTIATIC | 0.00000 | 0.00000 | | 0.81505 | 1.74256 | 65 |
| GO_REGULATION_OF_HEMOPOIESIS | 0.00000 | 0.00008 | 0.62726 | 0.58859 | 1.30634 | 228 |
| GO_REGULATION_OF_IMMUNE_RESPONSE | 0.00000 | 0.00016 | 0.61053 | 0.55362 | 1.23583 | 362 |
| GO_REGULATION_OF_INNATE_IMMUNE_RESPONSE | 0.00000 | 0.00000 | 0.71951 | 0.63883 | 1.40521 | 161 |
| GO_REGULATION_OF_INTRINSIC_APOPTOTIC_SIGNALING_PATHWAY | 0.00001 | 0.00039 | 0.59333 | 0.63478 | 1.38259 | 109 |
| GO_REGULATION_OF_MITOTIC_CELL_CYCLE | 0.00001 | 0.00018 | 0.61053 | 0.53899 | 1.20736 | 458 |
| GO_REGULATION_OF_MORPHOGENESIS_OF_AN_EPITHELIUM | 0.00000 | 0.00000 | | 0.72213 | 1.57284 | 109 |
| GO_REGULATION_OF_MRNA_CATABOLIC_PROCESS | 0.00000 | 0.00000 | | 0.69977 | 1.54127 | 168 |
| GO_REGULATION_OF_MRNA_METABOLIC_PROCESS | 0.00000 | 0.00000 | | 0.66528 | 1.47872 | 266 |
| GO_REGULATION_OF_MRNA_PROCESSING | 0.00004 | 0.00105 | 0.55733 | 0.61521 | 1.34090 | 113 |
| GO_REGULATION_OF_MRNA_SPLICING_VIA_SPLICEOSOME | 0.00001 | 0.00026 | 0.59333 | 0.67424 | 1.45411 | 80 |
| GO_REGULATION_OF_NUCLEASE_ACTIVITY | 0.00106 | 0.01819 | 0.45506 | 0.77918 | 1.56780 | 18 |
| GO_REGULATION_OF_OXIDATIVE_STRESS_INDUCED_CELL_DEATH | 0.00198 | 0.03028 | 0.43171 | 0.67287 | 1.40355 | 41 |
| GO_REGULATION_OF_PEPTIDASE_ACTIVITY | 0.00050 | 0.00978 | 0.47727 | 0.56122 | 1.24054 | 191 |
| GO_REGULATION_OF_POSTTRANSCRIPTIONAL_GENE_SILENCING | 0.00364 | 0.04761 | 0.43171 | 0.59824 | 1.28492 | 74 |
| GO_REGULATION_OF_PROTEIN_CATABOLIC_PROCESS | 0.00179 | 0.02783 | 0.45506 | 0.53057 | 1.17929 | 270 |
| GO_REGULATION_OF_PROTEIN_LOCALIZATION_TO_NUCLEUS | 0.00001 | 0.00042 | 0.59333 | 0.65505 | 1.41558 | 87 |
| GO_REGULATION_OF_PROTEIN_MODIFICATION_BY_SMALL_PROTEIN | 0.00025 | 0.00532 | 0.49849 | 0.57068 | 1.25684 | 167 |
| GO_REGULATION_OF_PROTEIN_STABILITY | 0.00093 | 0.01652 | 0.47727 | 0.55382 | 1.22571 | 202 |
| GO_REGULATION_OF_PROTEOLYSIS | 0.00013 | 0.00301 | 0.51885 | 0.52997 | 1.18392 | 377 |
| GO_REGULATION_OF_RESPONSE_TO_BIOTIC_STIMULUS | 0.00001 | 0.00020 | 0.61053 | 0.59049 | 1.30719 | 204 |
| GO_REGULATION_OF_RNA_SPLICING | 0.00000 | 0.00001 | 0.69013 | 0.65896 | 1.43669 | 114 |
| GO_REGULATION_OF_SMALL_MOLECULE_METABOLIC_PROCESS | 0.00000 | 0.00000 | 0.71951 | 0.59134 | 1.31394 | 265 |
| GO_REGULATION_OF_STEM_CELL_DIFFERENTIATION | 0.00000 | 0.00000 | | 0.74049 | 1.60113 | 90 |
| GO_REGULATION_OF_SYMBIOTIC_PROCESS | 0.00006 | 0.00144 | 0.55733 | 0.60071 | 1.31682 | 134 |
| GO_REGULATION_OF_TELOMERASE_ACTIVITY | 0.00088 | 0.01590 | 0.47727 | 0.69161 | 1.43179 | 35 |
| GO_REGULATION_OF_TRANSCRIPTION_FROM_RNA_POLYMERASE_II | 0.00000 | 0.00000 | | 0.81194 | 1.73800 | 66 |
| GO_REGULATION_OF_TRANSCRIPTION_INITIATION_FROM_RNA_POL | 0.00143 | 0.02332 | 0.45506 | 0.70390 | 1.44582 | 29 |
| GO_REGULATION_OF_TRANSLATIONAL_INITIATION | 0.00002 | 0.00052 | 0.57561 | 0.68129 | 1.45518 | 64 |
| GO_REGULATION_OF_UBIQUITIN_PROTEIN_TRANSFERASE_ACTIVITY | 0.00061 | 0.01169 | 0.47727 | 0.69335 | 1.44627 | 41 |
| GO_REGULATION_OF_VIRAL_GENOME_REPLICATION | 0.00086 | 0.01566 | 0.47727 | 0.65316 | 1.38389 | 54 |
| GO_REGULATION_OF_VIRAL_TRANSCRIPTION | 0.00198 | 0.03028 | 0.43171 | 0.63932 | 1.34703 | 49 |
| GO_REGULATION_OF_WNT_SIGNALING_PATHWAY | 0.00000 | 0.00001 | 0.69013 | 0.60353 | 1.33816 | 220 |
| GO_RESPIRASOME | 0.00008 | 0.00181 | 0.53843 | 0.65420 | 1.40421 | 71 |
| GO_RESPIRATORY_CHAIN_COMPLEX | 0.00016 | 0.00353 | 0.51885 | 0.67626 | 1.43918 | 60 |
| GO_RESPIRATORY_ELECTRON_TRANSPORT_CHAIN | 0.00001 | 0.00025 | 0.59333 | 0.65819 | 1.42190 | 85 |
| GO_RESPONSE_TO_ALCOHOL | 0.00261 | 0.03750 | 0.43171 | 0.58804 | 1.27350 | 95 |
| GO_RESPONSE_TO_CADMIUM_ION | 0.00265 | 0.03775 | 0.43171 | 0.67227 | 1.39174 | 35 |
| GO_RESPONSE_TO_DRUG | 0.00104 | 0.01798 | 0.45506 | 0.55364 | 1.22258 | 180 |
| GO_RESPONSE_TO_ETHANOL | 0.00086 | 0.01566 | 0.47727 | 0.67309 | 1.40691 | 43 |
| GO_RESPONSE_TO_HEAT | 0.00012 | 0.00270 | 0.53843 | 0.61665 | 1.34221 | 107 |
| GO_RESPONSE_TO_HORMONE | 0.00048 | 0.00949 | 0.49849 | 0.51277 | 1.14814 | 453 |
| GO_RESPONSE_TO_HYDROGEN_PEROXIDE | 0.00257 | 0.03716 | 0.43171 | 0.60505 | 1.30368 | 79 |
| GO_RESPONSE_TO_INORGANIC_SUBSTANCE | 0.00004 | 0.00107 | 0.55733 | 0.55714 | 1.23870 | 275 |
| GO_RESPONSE_TO_INTERLEUKIN_1 | 0.00000 | 0.00000 | | 0.74844 | 1.62908 | 107 |
| GO_RESPONSE_TO_INTERLEUKIN_12 | 0.00012 | 0.00270 | 0.53843 | 0.72605 | 1.50694 | 36 |
| GO_RESPONSE_TO_INTERLEUKIN_4 | 0.00324 | 0.04369 | 0.43171 | 0.75812 | 1.51571 | 17 |
| GO_RESPONSE_TO_METAL_ION | 0.00030 | 0.00623 | 0.49849 | 0.56387 | 1.24233 | 171 |
| GO_RESPONSE_TO_ORGANIC_CYCLIC_COMPOUND | 0.00000 | 0.00016 | 0.61053 | 0.54603 | 1.22259 | 439 |
| GO_RESPONSE_TO_OXYGEN_LEVELS | 0.00000 | 0.00000 | 0.71951 | 0.60132 | 1.33438 | 233 |
| GO_RESPONSE_TO_STEROID_HORMONE | 0.00154 | 0.02464 | 0.45506 | 0.55514 | 1.22446 | 174 |
| GO_RESPONSE_TO_TEMPERATURE_STIMULUS | 0.00027 | 0.00572 | 0.49849 | 0.59216 | 1.29675 | 127 |
| GO_RESPONSE_TO_TUMOR_NECROSIS_FACTOR | 0.00000 | 0.00000 | 0.76146 | 0.65649 | 1.44236 | 151 |
| GO_RETINA_HOMEOSTASIS | 0.00104 | 0.01798 | 0.45506 | 0.73063 | 1.49432 | 27 |
| GO_RIBONUCLEOPROTEIN_COMPLEX_BINDING | 0.00000 | 0.00000 | 0.71951 | 0.66906 | 1.46050 | 117 |
| GO_RIBONUCLEOPROTEIN_COMPLEX_BIOGENESIS | 0.00000 | 0.00000 | | 0.66339 | 1.48264 | 389 |
| GO_RIBONUCLEOPROTEIN_COMPLEX_SUBUNIT_ORGANIZATION | 0.00000 | 0.00000 | | 0.69241 | 1.52681 | 173 |
| GO_RIBONUCLEOPROTEIN_GRANULE | 0.00000 | 0.00001 | 0.67496 | 0.61979 | 1.36366 | 163 |
| GO_RIBONUCLEOSIDE_TRIPHOSPHATE_BIOSYNTHETIC_PROCESS | 0.00041 | 0.00840 | 0.49849 | 0.71253 | 1.47248 | 33 |
| GO_RIBONUCLEOSIDE_TRIPHOSPHATE_METABOLIC_PROCESS | 0.00026 | 0.00550 | 0.49849 | 0.69681 | 1.45611 | 42 |
| GO_RIBOSE_PHOSPHATE_BIOSYNTHETIC_PROCESS | 0.00191 | 0.02939 | 0.45506 | 0.57932 | 1.26207 | 110 |

| | | | | | | |
|---|---|---|---|---|---|---|
| GO_RIBOSE_PHOSPHATE_METABOLIC_PROCESS | 0.00108 | 0.01839 | 0.45506 | 0.53891 | 1.19687 | 245 |
| GO_RIBOSOMAL_LARGE_SUBUNIT_ASSEMBLY | 0.00000 | 0.00016 | 0.61053 | 0.80999 | 1.65663 | 27 |
| GO_RIBOSOMAL_LARGE_SUBUNIT_BIOGENESIS | 0.00000 | 0.00000 | | 0.77542 | 1.65982 | 66 |
| GO_RIBOSOMAL_SMALL_SUBUNIT_ASSEMBLY | 0.00023 | 0.00494 | 0.51885 | 0.81139 | 1.62221 | 17 |
| GO_RIBOSOMAL_SMALL_SUBUNIT_BINDING | 0.00383 | 0.04935 | 0.43171 | 0.75288 | 1.50523 | 17 |
| GO_RIBOSOMAL_SMALL_SUBUNIT_BIOGENESIS | 0.00001 | 0.00022 | 0.61053 | 0.69076 | 1.47997 | 68 |
| GO_RIBOSOMAL_SUBUNIT | 0.00000 | 0.00000 | | 0.77603 | 1.71031 | 172 |
| GO_RIBOSOME | 0.00000 | 0.00000 | | 0.76643 | 1.69659 | 206 |
| GO_RIBOSOME_ASSEMBLY | 0.00000 | 0.00000 | 0.73376 | 0.76173 | 1.61998 | 59 |
| GO_RIBOSOME_BINDING | 0.00019 | 0.00416 | 0.51885 | 0.68955 | 1.44715 | 47 |
| GO_RIBOSOME_BIOGENESIS | 0.00000 | 0.00000 | | 0.65877 | 1.46466 | 275 |
| GO_RNA_CATABOLIC_PROCESS | 0.00000 | 0.00000 | | 0.76774 | 1.71264 | 339 |
| GO_RNA_DEPENDENT_DNA_BIOSYNTHETIC_PROCESS | 0.00003 | 0.00076 | 0.57561 | 0.68926 | 1.46521 | 58 |
| GO_RNA_EXPORT_FROM_NUCLEUS | 0.00062 | 0.01183 | 0.47727 | 0.58378 | 1.27758 | 126 |
| GO_RNA_LOCALIZATION | 0.00000 | 0.00006 | 0.64355 | 0.59670 | 1.32020 | 198 |
| GO_RNA_POLYMERASE_II_PREINITIATION_COMPLEX_ASSEMBLY | 0.00089 | 0.01592 | 0.47727 | 0.77046 | 1.55626 | 19 |
| GO_RNA_SPLICING | 0.00000 | 0.00000 | 0.78819 | 0.59079 | 1.31927 | 369 |
| GO_RNA_SPLICING_VIA_TRANSESTERIFICATION_REACTIONS | 0.00000 | 0.00000 | | 0.60940 | 1.35714 | 300 |
| GO_RNA_STABILIZATION | 0.00065 | 0.01223 | 0.47727 | 0.69743 | 1.44384 | 35 |
| GO_ROUGH_ENDOPLASMIC_RETICULUM | 0.00001 | 0.00039 | 0.59333 | 0.71853 | 1.51629 | 50 |
| GO_RRNA_BINDING | 0.00000 | 0.00001 | 0.69013 | 0.74782 | 1.58865 | 56 |
| GO_RRNA_METABOLIC_PROCESS | 0.00000 | 0.00000 | 0.77494 | 0.63047 | 1.39562 | 206 |
| GO_SCF_DEPENDENT_PROTEASOMAL_UBIQUITIN_DEPENDENT_PRO | 0.00000 | 0.00000 | | 0.77527 | 1.67044 | 79 |
| GO_SECRETORY_GRANULE | 0.00000 | 0.00000 | 0.71951 | 0.56518 | 1.26461 | 399 |
| GO_SECRETORY_VESICLE | 0.00000 | 0.00002 | 0.67496 | 0.54866 | 1.22905 | 465 |
| GO_SIGNAL_TRANSDUCTION_BY_PROTEIN_PHOSPHORYLATION | 0.00004 | 0.00109 | 0.55733 | 0.53817 | 1.20474 | 424 |
| GO_SINGLE_FERTILIZATION | 0.00048 | 0.00949 | 0.49849 | 0.67414 | 1.42260 | 50 |
| GO_SINGLE_STRANDED_RNA_BINDING | 0.00039 | 0.00793 | 0.49849 | 0.66553 | 1.41527 | 57 |
| GO_SMALL_MOLECULE_BIOSYNTHETIC_PROCESS | 0.00216 | 0.03242 | 0.43171 | 0.51195 | 1.14522 | 397 |
| GO_SMALL_RIBOSOMAL_SUBUNIT | 0.00000 | 0.00000 | | 0.77648 | 1.66209 | 66 |
| GO_SM_LIKE_PROTEIN_FAMILY_COMPLEX | 0.00327 | 0.04398 | 0.43171 | 0.60468 | 1.29713 | 70 |
| GO_SPERM_EGG_RECOGNITION | 0.00000 | 0.00007 | 0.64355 | 0.85241 | 1.72955 | 22 |
| GO_SPLICEOSOMAL_COMPLEX | 0.00000 | 0.00003 | 0.65944 | 0.60939 | 1.34261 | 171 |
| GO_STEM_CELL_DIFFERENTIATION | 0.00000 | 0.00000 | 0.78819 | 0.66886 | 1.46930 | 147 |
| GO_STRUCTURAL_CONSTITUENT_OF_CYTOSKELETON | 0.00102 | 0.01782 | 0.45506 | 0.65913 | 1.39337 | 52 |
| GO_STRUCTURAL_CONSTITUENT_OF_RIBOSOME | 0.00000 | 0.00000 | | 0.79909 | 1.75541 | 150 |
| GO_STRUCTURAL_MOLECULE_ACTIVITY | 0.00000 | 0.00000 | | 0.72651 | 1.61895 | 324 |
| GO_SUBSTANTIA_NIGRA_DEVELOPMENT | 0.00000 | 0.00013 | 0.62726 | 0.81622 | 1.66507 | 26 |
| GO_TELOMERASE_RNA_LOCALIZATION | 0.00001 | 0.00017 | 0.61053 | 0.85164 | 1.72025 | 19 |
| GO_TELOMERE_MAINTENANCE_VIA_TELOMERE_LENGTHENING | 0.00011 | 0.00250 | 0.53843 | 0.65659 | 1.40546 | 66 |
| GO_TELOMERE_ORGANIZATION | 0.00342 | 0.04554 | 0.43171 | 0.57000 | 1.24649 | 124 |
| GO_THREONINE_TYPE_ENDOPEPTIDASE_ACTIVITY | 0.00000 | 0.00000 | | 0.95179 | 1.92255 | 19 |
| GO_THREONINE_TYPE_PEPTIDASE_ACTIVITY | 0.00000 | 0.00000 | | 0.92199 | 1.87423 | 23 |
| GO_TISSUE_MORPHOGENESIS | 0.00097 | 0.01707 | 0.47727 | 0.52661 | 1.17402 | 326 |
| GO_TOXIN_TRANSPORT | 0.00113 | 0.01922 | 0.45506 | 0.72897 | 1.48708 | 26 |
| GO_TRANSCRIPTION_COUPLED_NUCLEOTIDE_EXCISION_REPAIR | 0.00001 | 0.00019 | 0.61053 | 0.68106 | 1.46186 | 71 |
| GO_TRANSCRIPTION_FACTOR_BINDING | 0.00043 | 0.00865 | 0.49849 | 0.52447 | 1.17376 | 409 |
| GO_TRANSCRIPTION_PREINITIATION_COMPLEX_ASSEMBLY | 0.00119 | 0.02004 | 0.45506 | 0.71399 | 1.46797 | 30 |
| GO_TRANSLATIONAL_ELONGATION | 0.00000 | 0.00000 | 0.71951 | 0.66087 | 1.44451 | 122 |
| GO_TRANSLATIONAL_INITIATION | 0.00000 | 0.00000 | | 0.84691 | 1.86515 | 166 |
| GO_TRANSLATIONAL_TERMINATION | 0.00000 | 0.00005 | 0.64355 | 0.65946 | 1.43060 | 99 |
| GO_TRANSLATION_ELONGATION_FACTOR_ACTIVITY | 0.00161 | 0.02546 | 0.45506 | 0.79860 | 1.58418 | 15 |
| GO_TRANSLATION_FACTOR_ACTIVITY_RNA_BINDING | 0.00000 | 0.00000 | | 0.79008 | 1.69288 | 69 |
| GO_TRANSLATION_INITIATION_FACTOR_ACTIVITY | 0.00000 | 0.00000 | 0.76146 | 0.80422 | 1.68417 | 45 |
| GO_TRANSLATION_INITIATION_FACTOR_BINDING | 0.00021 | 0.00468 | 0.51885 | 0.76186 | 1.55231 | 25 |
| GO_TRANSLATION_PREINITIATION_COMPLEX | 0.00000 | 0.00001 | 0.67496 | 0.90245 | 1.79595 | 16 |
| GO_TRANSLATION_REGULATOR_ACTIVITY | 0.00000 | 0.00000 | | 0.77663 | 1.68904 | 105 |
| GO_TRANSLATION_REGULATOR_ACTIVITY_NUCLEIC_ACID_BINDING | 0.00000 | 0.00000 | | 0.77150 | 1.66586 | 83 |
| GO_TRICARBOXYLIC_ACID_CYCLE | 0.00154 | 0.02464 | 0.45506 | 0.71854 | 1.47240 | 28 |
| GO_TUMOR_NECROSIS_FACTOR_MEDIATED_SIGNALING_PATHWAY | 0.00000 | 0.00000 | | 0.76277 | 1.65372 | 97 |
| GO_T_CELL_RECEPTOR_SIGNALING_PATHWAY | 0.00000 | 0.00000 | 0.81404 | 0.69994 | 1.52527 | 112 |
| GO_U12_TYPE_SPLICEOSOMAL_COMPLEX | 0.00279 | 0.03889 | 0.43171 | 0.70746 | 1.44693 | 27 |
| GO_U2_SNRNP | 0.00005 | 0.00134 | 0.55733 | 0.83081 | 1.67168 | 18 |
| GO_U2_TYPE_SPLICEOSOMAL_COMPLEX | 0.00010 | 0.00243 | 0.53843 | 0.63403 | 1.36970 | 85 |
| GO_UBIQUITIN_LIKE_PROTEIN_LIGASE_BINDING | 0.00000 | 0.00000 | 0.76146 | 0.62273 | 1.38051 | 219 |
| GO_UBIQUITIN_PROTEIN_TRANSFERASE_REGULATOR_ACTIVITY | 0.00165 | 0.02590 | 0.45506 | 0.79605 | 1.57911 | 15 |
| GO_UNFOLDED_PROTEIN_BINDING | 0.00000 | 0.00000 | 0.77494 | 0.72157 | 1.55881 | 85 |
| GO_VESICLE_LUMEN | 0.00000 | 0.00000 | 0.74774 | 0.64475 | 1.42008 | 168 |
| GO_VIRAL_GENE_EXPRESSION | 0.00000 | 0.00000 | | 0.80028 | 1.76317 | 171 |
| GO_VIRAL_GENOME_REPLICATION | 0.00100 | 0.01761 | 0.45506 | 0.61997 | 1.33078 | 73 |
| GO_VIRAL_LIFE_CYCLE | 0.00031 | 0.00640 | 0.49849 | 0.55767 | 1.23509 | 209 |
| HP_ABNORMALITY_OF_ACID_BASE_HOMEOSTASIS | 0.00216 | 0.03242 | 0.43171 | 0.53556 | 1.18846 | 232 |
| HP_ABNORMALITY_OF_CARDIOVASCULAR_SYSTEM_ELECTROPHYSIO | 0.00027 | 0.00572 | 0.49849 | 0.55832 | 1.23929 | 229 |
| HP_ABNORMALITY_OF_LOWER_LIP | 0.00181 | 0.02804 | 0.45506 | 0.56360 | 1.23779 | 145 |
| HP_ABNORMALITY_OF_THE_MITOCHONDRION | 0.00213 | 0.03229 | 0.43171 | 0.57831 | 1.26240 | 117 |
| HP_ABNORMALITY_OF_THE_NASAL_DORSUM | 0.00005 | 0.00116 | 0.55733 | 0.60288 | 1.32157 | 134 |

| | | | | | | |
|---|---|---|---|---|---|---|
| HP_ABNORMAL_ELECTROPHYSIOLOGY_OF_SINOATRIAL_NODE_ORIG | 0.00279 | 0.03889 | 0.43171 | 0.68436 | 1.40934 | 31 |
| HP_ABNORMAL_ERYTHROCYTE_MORPHOLOGY | 0.00000 | 0.00015 | 0.61053 | 0.56049 | 1.25036 | 341 |
| HP_ABNORMAL_MEAN_CORPUSCULAR_VOLUME | 0.00357 | 0.04696 | 0.43171 | 0.75226 | 1.51363 | 18 |
| HP_ABNORMAL_RETICULOCYTE_MORPHOLOGY | 0.00064 | 0.01210 | 0.47727 | 0.71420 | 1.47081 | 31 |
| HP_ACUTE_LEUKEMIA | 0.00000 | 0.00000 | 0.73376 | 0.72786 | 1.56137 | 70 |
| HP_ANEMIA_OF_INADEQUATE_PRODUCTION | 0.00000 | 0.00008 | 0.62726 | 0.65593 | 1.42209 | 97 |
| HP_CONSTITUTIONAL_SYMPTOM | 0.00119 | 0.02004 | 0.45506 | 0.51427 | 1.15108 | 423 |
| HP_DELAYED_PUBERTY | 0.00005 | 0.00115 | 0.55733 | 0.62579 | 1.36211 | 107 |
| HP_DEPRESSED_NASAL_RIDGE | 0.00000 | 0.00011 | 0.62726 | 0.68731 | 1.47511 | 72 |
| HP_FATIGUE | 0.00039 | 0.00808 | 0.49849 | 0.56529 | 1.24923 | 187 |
| HP_HEADACHE | 0.00015 | 0.00340 | 0.51885 | 0.59065 | 1.29716 | 146 |
| HP_HEMATOLOGICAL_NEOPLASM | 0.00003 | 0.00078 | 0.57561 | 0.60108 | 1.32044 | 154 |
| HP_INCREASED_CSF_LACTATE | 0.00157 | 0.02509 | 0.45506 | 0.61622 | 1.31904 | 66 |
| HP_INCREASED_SERUM_LACTATE | 0.00216 | 0.03242 | 0.43171 | 0.56920 | 1.24724 | 130 |
| HP_LEUKEMIA | 0.00001 | 0.00026 | 0.59333 | 0.64877 | 1.40680 | 98 |
| HP_MACROCYTIC_ANEMIA | 0.00000 | 0.00000 | 0.74774 | 0.82468 | 1.71698 | 39 |
| HP_MIGRAINE | 0.00000 | 0.00000 | 0.71951 | 0.72105 | 1.54497 | 69 |
| HP_MYOTONIA | 0.00161 | 0.02546 | 0.45506 | 0.79820 | 1.58338 | 15 |
| HP_NEOPLASM | 0.00216 | 0.03242 | 0.43171 | 0.50820 | 1.13805 | 446 |
| HP_PALLOR | 0.00001 | 0.00040 | 0.59333 | 0.65169 | 1.41044 | 92 |
| HP_PUBERTY_AND_GONADAL_DISORDERS | 0.00287 | 0.03973 | 0.43171 | 0.51995 | 1.15843 | 322 |
| HP_RETICULOCYTOPENIA | 0.00004 | 0.00105 | 0.55733 | 0.86417 | 1.71425 | 15 |
| HP_RIMMED_VACUOLES | 0.00279 | 0.03889 | 0.43171 | 0.74033 | 1.49831 | 20 |
| HP_SHORT_NOSE | 0.00235 | 0.03474 | 0.43171 | 0.54919 | 1.21134 | 174 |
| HP_THICK_LOWER_LIP_VERMILION | 0.00048 | 0.00949 | 0.49849 | 0.60773 | 1.31617 | 94 |
| HP_THICK_VERMILION_BORDER | 0.00156 | 0.02486 | 0.45506 | 0.55983 | 1.23000 | 153 |
| HP_VENTRICULAR_ARRHYTHMIA | 0.00353 | 0.04658 | 0.43171 | 0.66198 | 1.37398 | 36 |
| **Cluster 1** | | | | | | |
| pathway | pval | padj | log2err | ES | NES | size |
| GO_ACTIVATION_OF_IMMUNE_RESPONSE | 0.00000 | 0.00000 | 0.78819 | 0.62851 | 1.39255 | 212 |
| GO_ACTIVATION_OF_INNATE_IMMUNE_RESPONSE | 0.00000 | 0.00000 | | 0.75054 | 1.62130 | 88 |
| GO_AEROBIC_RESPIRATION | 0.00001 | 0.00025 | 0.59333 | 0.69302 | 1.48168 | 65 |
| GO_ALTERNATIVE_MRNA_SPLICING_VIA_SPLICEOSOME | 0.00102 | 0.01782 | 0.45506 | 0.65168 | 1.38687 | 60 |
| GO_AMINE_METABOLIC_PROCESS | 0.00000 | 0.00000 | | 0.79189 | 1.70783 | 80 |
| GO_ANAPHASE_PROMOTING_COMPLEX_DEPENDENT_CATABOLIC_P | 0.00000 | 0.00000 | | 0.76611 | 1.65070 | 79 |
| GO_ANCHORING_JUNCTION | 0.00000 | 0.00000 | | 0.59615 | 1.33633 | 479 |
| GO_ANIMAL_ORGAN_REGENERATION | 0.00257 | 0.03716 | 0.43171 | 0.68240 | 1.41020 | 33 |
| GO_ANTIGEN_PROCESSING_AND_PRESENTATION | 0.00000 | 0.00000 | 0.70498 | 0.64776 | 1.42241 | 143 |
| GO_ANTIGEN_PROCESSING_AND_PRESENTATION_OF_EXOGENOUS_ | 0.00000 | 0.00000 | | 0.81051 | 1.72358 | 57 |
| GO_ANTIGEN_PROCESSING_AND_PRESENTATION_OF_PEPTIDE_ANTI | 0.00000 | 0.00000 | 0.70498 | 0.65912 | 1.44248 | 126 |
| GO_ANTIGEN_PROCESSING_AND_PRESENTATION_OF_PEPTIDE_ANTI | 0.00000 | 0.00000 | | 0.76925 | 1.65015 | 70 |
| GO_ANTIGEN_RECEPTOR_MEDIATED_SIGNALING_PATHWAY | 0.00000 | 0.00000 | 0.80122 | 0.67877 | 1.48641 | 127 |
| GO_APOPTOTIC_SIGNALING_PATHWAY | 0.00154 | 0.02464 | 0.45506 | 0.51561 | 1.15085 | 358 |
| GO_ATP_DEPENDENT_CHROMATIN_REMODELING | 0.00253 | 0.03681 | 0.43171 | 0.62124 | 1.32400 | 61 |
| GO_ATP_METABOLIC_PROCESS | 0.00000 | 0.00000 | 0.71951 | 0.62252 | 1.37693 | 196 |
| GO_ATP_SYNTHESIS_COUPLED_ELECTRON_TRANSPORT | 0.00005 | 0.00131 | 0.55733 | 0.65519 | 1.40723 | 74 |
| GO_BINDING_OF_SPERM_TO_ZONA_PELLUCIDA | 0.00000 | 0.00001 | 0.69013 | 0.89879 | 1.80846 | 18 |
| GO_CADHERIN_BINDING | 0.00000 | 0.00000 | | 0.63068 | 1.39961 | 236 |
| GO_CANONICAL_WNT_SIGNALING_PATHWAY | 0.00000 | 0.00003 | 0.65944 | 0.60816 | 1.34553 | 197 |
| GO_CATALYTIC_STEP_2_SPLICEOSOME | 0.00000 | 0.00002 | 0.65944 | 0.68068 | 1.46976 | 83 |
| GO_CELLULAR_AMINO_ACID_METABOLIC_PROCESS | 0.00000 | 0.00000 | 0.74774 | 0.61752 | 1.36824 | 216 |
| GO_CELLULAR_COMPONENT_DISASSEMBLY | 0.00000 | 0.00005 | 0.64355 | 0.55505 | 1.24197 | 401 |
| GO_CELLULAR_KETONE_METABOLIC_PROCESS | 0.00000 | 0.00000 | 0.77494 | 0.65955 | 1.44893 | 149 |
| GO_CELLULAR_NITROGEN_COMPOUND_CATABOLIC_PROCESS | 0.00000 | 0.00000 | | 0.74079 | 1.65848 | 436 |
| GO_CELLULAR_PROTEIN_COMPLEX_DISASSEMBLY | 0.00006 | 0.00139 | 0.55733 | 0.58222 | 1.28459 | 176 |
| GO_CELLULAR_RESPIRATION | 0.00000 | 0.00000 | 0.71951 | 0.64997 | 1.42492 | 136 |
| GO_CELLULAR_RESPONSE_TO_HEAT | 0.00018 | 0.00403 | 0.51885 | 0.61962 | 1.34016 | 91 |
| GO_CELLULAR_RESPONSE_TO_ORGANIC_CYCLIC_COMPOUND | 0.00302 | 0.04108 | 0.43171 | 0.52402 | 1.16585 | 283 |
| GO_CELLULAR_RESPONSE_TO_OXYGEN_LEVELS | 0.00000 | 0.00000 | 0.78819 | 0.65911 | 1.44882 | 157 |
| GO_CELL_ACTIVATION_INVOLVED_IN_IMMUNE_RESPONSE | 0.00000 | 0.00001 | 0.67496 | 0.57339 | 1.27913 | 341 |
| GO_CELL_ADHESION_MOLECULE_BINDING | 0.00000 | 0.00000 | 0.81404 | 0.60529 | 1.34788 | 308 |
| GO_CELL_CELL_RECOGNITION | 0.00000 | 0.00002 | 0.65944 | 0.83826 | 1.71003 | 26 |
| GO_CELL_CELL_SIGNALING_BY_WNT | 0.00000 | 0.00002 | 0.67496 | 0.57736 | 1.28559 | 299 |
| GO_CELL_CYCLE_G2_M_PHASE_TRANSITION | 0.00001 | 0.00024 | 0.59333 | 0.58034 | 1.28602 | 214 |
| GO_CELL_CYCLE_PHASE_TRANSITION | 0.00000 | 0.00004 | 0.64355 | 0.54319 | 1.21681 | 464 |
| GO_CELL_DIVISION | 0.00154 | 0.02464 | 0.45506 | 0.51551 | 1.15425 | 427 |
| GO_CELL_RECOGNITION | 0.00012 | 0.00270 | 0.53843 | 0.69554 | 1.46186 | 48 |
| GO_CELL_SUBSTRATE_JUNCTION | 0.00000 | 0.00000 | | 0.67751 | 1.50775 | 291 |
| GO_CELL_SURFACE_RECEPTOR_SIGNALING_PATHWAY_INVOLVED_IN | 0.00000 | 0.00003 | 0.65944 | 0.57035 | 1.27067 | 319 |
| GO_CHAPERONE_BINDING | 0.00007 | 0.00163 | 0.53843 | 0.64754 | 1.39253 | 76 |
| GO_CHAPERONE_COMPLEX | 0.00000 | 0.00004 | 0.64355 | 0.85718 | 1.73925 | 22 |
| GO_CHAPERONE_MEDIATED_PROTEIN_COMPLEX_ASSEMBLY | 0.00170 | 0.02662 | 0.45506 | 0.77423 | 1.54792 | 17 |
| GO_CHROMATIN_BINDING | 0.00071 | 0.01323 | 0.47727 | 0.52699 | 1.17680 | 369 |
| GO_CHROMOSOMAL_REGION | 0.00246 | 0.03601 | 0.43171 | 0.52919 | 1.17661 | 276 |
| GO_COP9_SIGNALOSOME | 0.00128 | 0.02141 | 0.45506 | 0.73616 | 1.49715 | 24 |
| GO_COTRANSLATIONAL_PROTEIN_TARGETING_TO_MEMBRANE | 0.00000 | 0.00000 | | 0.89471 | 1.93513 | 91 |

| | | | | | | |
|---|---|---|---|---|---|---|
| GO_CYTOCHROME_COMPLEX | 0.00168 | 0.02641 | 0.45506 | 0.75311 | 1.52417 | 20 |
| GO_CYTOKINE_MEDIATED_SIGNALING_PATHWAY | 0.00000 | 0.00000 | 0.81404 | 0.59698 | 1.33170 | 339 |
| GO_CYTOPLASMIC_TRANSLATION | 0.00000 | 0.00000 | | 0.84753 | 1.83080 | 88 |
| GO_CYTOPLASMIC_TRANSLATIONAL_INITIATION | 0.00000 | 0.00000 | 0.74774 | 0.85688 | 1.76176 | 30 |
| GO_CYTOSOLIC_LARGE_RIBOSOMAL_SUBUNIT | 0.00000 | 0.00000 | | 0.91186 | 1.92988 | 53 |
| GO_CYTOSOLIC_RIBOSOME | 0.00000 | 0.00000 | | 0.89907 | 1.94710 | 95 |
| GO_CYTOSOLIC_SMALL_RIBOSOMAL_SUBUNIT | 0.00000 | 0.00000 | | 0.89114 | 1.85533 | 39 |
| GO_DEFENSE_RESPONSE_TO_OTHER_ORGANISM | 0.00004 | 0.00099 | 0.55733 | 0.53673 | 1.20127 | 415 |
| GO_DE_NOVO_PROTEIN_FOLDING | 0.00050 | 0.00978 | 0.47727 | 0.74232 | 1.51431 | 26 |
| GO_DISORDERED_DOMAIN_SPECIFIC_BINDING | 0.00161 | 0.02546 | 0.45506 | 0.71970 | 1.47197 | 27 |
| GO_DNA_BIOSYNTHETIC_PROCESS | 0.00000 | 0.00010 | 0.62726 | 0.62505 | 1.37017 | 134 |
| GO_DNA_CONFORMATION_CHANGE | 0.00027 | 0.00572 | 0.49849 | 0.55300 | 1.22527 | 216 |
| GO_DNA_DAMAGE_RESPONSE_DETECTION_OF_DNA_DAMAGE | 0.00198 | 0.03028 | 0.43171 | 0.67949 | 1.40669 | 35 |
| GO_DNA_GEOMETRIC_CHANGE | 0.00046 | 0.00925 | 0.49849 | 0.60807 | 1.31855 | 98 |
| GO_DNA_POLYMERASE_BINDING | 0.00268 | 0.03799 | 0.43171 | 0.76255 | 1.52457 | 17 |
| GO_DNA_REPAIR | 0.00265 | 0.03775 | 0.43171 | 0.51159 | 1.14547 | 427 |
| GO_DRUG_BINDING | 0.00176 | 0.02733 | 0.45506 | 0.64119 | 1.35096 | 49 |
| GO_ELECTRON_TRANSFER_ACTIVITY | 0.00001 | 0.00020 | 0.61053 | 0.67481 | 1.44938 | 74 |
| GO_ELECTRON_TRANSPORT_CHAIN | 0.00002 | 0.00053 | 0.57561 | 0.62257 | 1.36145 | 124 |
| GO_ENDOPEPTIDASE_ACTIVITY | 0.00124 | 0.02085 | 0.45506 | 0.54955 | 1.21076 | 171 |
| GO_ENDOPEPTIDASE_COMPLEX | 0.00000 | 0.00000 | | 0.80930 | 1.73234 | 66 |
| GO_ENERGY_DERIVATION_BY_OXIDATION_OF_ORGANIC_COMPOUN | 0.00000 | 0.00000 | 0.71951 | 0.63390 | 1.39707 | 172 |
| GO_ESTABLISHMENT_OF_PROTEIN_LOCALIZATION_TO_CHROMOSOM | 0.00020 | 0.00432 | 0.51885 | 0.77339 | 1.57217 | 23 |
| GO_ESTABLISHMENT_OF_PROTEIN_LOCALIZATION_TO_ENDOPLASM | 0.00000 | 0.00000 | | 0.88026 | 1.91149 | 102 |
| GO_ESTABLISHMENT_OF_PROTEIN_LOCALIZATION_TO_MEMBRANE | 0.00000 | 0.00000 | | 0.74874 | 1.66164 | 240 |
| GO_ESTABLISHMENT_OF_PROTEIN_LOCALIZATION_TO_ORGANELLE | 0.00000 | 0.00000 | | 0.69519 | 1.55605 | 423 |
| GO_ESTABLISHMENT_OF_PROTEIN_LOCALIZATION_TO_TELOMERE | 0.00006 | 0.00139 | 0.55733 | 0.83470 | 1.66881 | 17 |
| GO_ESTABLISHMENT_OF_RNA_LOCALIZATION | 0.00106 | 0.01819 | 0.45506 | 0.55577 | 1.22400 | 167 |
| GO_ESTABLISHMENT_OF_TISSUE_POLARITY | 0.00000 | 0.00000 | | 0.75051 | 1.62122 | 88 |
| GO_EUKARYOTIC_TRANSLATION_INITIATION_FACTOR_3_COMPLEX | 0.00003 | 0.00074 | 0.57561 | 0.86987 | 1.72555 | 15 |
| GO_EXOCYTOSIS | 0.00000 | 0.00015 | 0.61053 | 0.53905 | 1.20753 | 465 |
| GO_FC_EPSILON_RECEPTOR_SIGNALING_PATHWAY | 0.00000 | 0.00000 | | 0.75117 | 1.62103 | 82 |
| GO_FC_RECEPTOR_SIGNALING_PATHWAY | 0.00000 | 0.00000 | | 0.72054 | 1.57787 | 127 |
| GO_FERTILIZATION | 0.00089 | 0.01592 | 0.47727 | 0.63620 | 1.35888 | 64 |
| GO_FICOLIN_1_RICH_GRANULE | 0.00000 | 0.00000 | 0.78819 | 0.71716 | 1.55340 | 93 |
| GO_GENERATION_OF_PRECURSOR_METABOLITES_AND_ENERGY | 0.00000 | 0.00000 | 0.77494 | 0.59261 | 1.32026 | 320 |
| GO_GENE_SILENCING | 0.00261 | 0.03750 | 0.43171 | 0.55578 | 1.22252 | 161 |
| GO_GLAND_DEVELOPMENT | 0.00154 | 0.02464 | 0.45506 | 0.54117 | 1.19797 | 206 |
| GO_GLYCOSYL_COMPOUND_METABOLIC_PROCESS | 0.00071 | 0.01323 | 0.47727 | 0.62524 | 1.34204 | 71 |
| GO_GUANYL_NUCLEOTIDE_BINDING | 0.00095 | 0.01679 | 0.47727 | 0.54096 | 1.19861 | 216 |
| GO_HEAT_SHOCK_PROTEIN_BINDING | 0.00003 | 0.00092 | 0.55733 | 0.64369 | 1.39056 | 85 |
| GO_HEMATOPOIETIC_PROGENITOR_CELL_DIFFERENTIATION | 0.00000 | 0.00000 | 0.82666 | 0.70307 | 1.53239 | 113 |
| GO_HEMATOPOIETIC_STEM_CELL_DIFFERENTIATION | 0.00000 | 0.00000 | | 0.79776 | 1.71217 | 72 |
| GO_HSP70_PROTEIN_BINDING | 0.00228 | 0.03391 | 0.43171 | 0.71143 | 1.45504 | 27 |
| GO_IMMUNE_RESPONSE_REGULATING_SIGNALING_PATHWAY | 0.00000 | 0.00000 | 0.76146 | 0.63494 | 1.40480 | 197 |
| GO_INNATE_IMMUNE_RESPONSE | 0.00003 | 0.00071 | 0.57561 | 0.55173 | 1.23107 | 347 |
| GO_INNATE_IMMUNE_RESPONSE_ACTIVATING_SIGNAL_TRANSDUCT | 0.00000 | 0.00000 | | 0.79340 | 1.70281 | 72 |
| GO_INNER_MITOCHONDRIAL_MEMBRANE_PROTEIN_COMPLEX | 0.00002 | 0.00060 | 0.57561 | 0.65124 | 1.40678 | 88 |
| GO_INTERACTION_WITH_SYMBIONT | 0.00001 | 0.00026 | 0.59333 | 0.75084 | 1.56620 | 41 |
| GO_INTERLEUKIN_1_MEDIATED_SIGNALING_PATHWAY | 0.00000 | 0.00000 | | 0.81689 | 1.75671 | 76 |
| GO_INTRINSIC_APOPTOTIC_SIGNALING_PATHWAY | 0.00121 | 0.02029 | 0.45506 | 0.55118 | 1.21845 | 190 |
| GO_ION_CHANNEL_BINDING | 0.00087 | 0.01584 | 0.47727 | 0.63182 | 1.34683 | 62 |
| GO_ISOPEPTIDASE_ACTIVITY | 0.00139 | 0.02285 | 0.45506 | 0.78940 | 1.57097 | 16 |
| GO_LARGE_RIBOSOMAL_SUBUNIT | 0.00000 | 0.00000 | | 0.78135 | 1.70183 | 109 |
| GO_LEUKOCYTE_MEDIATED_IMMUNITY | 0.00000 | 0.00000 | 0.71951 | 0.57282 | 1.27895 | 364 |
| GO_MACROMOLECULE_TRANSMEMBRANE_TRANSPORTER_ACTIVITY | 0.00239 | 0.03510 | 0.43171 | 0.77734 | 1.54696 | 16 |
| GO_MAINTENANCE_OF_CELL_NUMBER | 0.00087 | 0.01584 | 0.47727 | 0.58944 | 1.28512 | 114 |
| GO_MATURATION_OF_LSU_RRNA | 0.00006 | 0.00151 | 0.53843 | 0.77606 | 1.58723 | 27 |
| GO_METHYLTRANSFERASE_COMPLEX | 0.00290 | 0.04014 | 0.43171 | 0.59476 | 1.28479 | 88 |
| GO_METHYL_CPG_BINDING | 0.00302 | 0.04108 | 0.43171 | 0.78053 | 1.54833 | 15 |
| GO_MIDBRAIN_DEVELOPMENT | 0.00021 | 0.00468 | 0.51885 | 0.68832 | 1.44457 | 47 |
| GO_MITOCHONDRIAL_ELECTRON_TRANSPORT_NADH_TO_UBIQUIN | 0.00331 | 0.04437 | 0.43171 | 0.65428 | 1.36724 | 42 |
| GO_MITOCHONDRIAL_GENE_EXPRESSION | 0.00048 | 0.00949 | 0.49849 | 0.57608 | 1.26525 | 152 |
| GO_MITOCHONDRIAL_LARGE_RIBOSOMAL_SUBUNIT | 0.00002 | 0.00050 | 0.57561 | 0.69789 | 1.48086 | 55 |
| GO_MITOCHONDRIAL_MATRIX | 0.00000 | 0.00002 | 0.65944 | 0.56045 | 1.25158 | 371 |
| GO_MITOCHONDRIAL_MEMBRANE_ORGANIZATION | 0.00361 | 0.04734 | 0.43171 | 0.59309 | 1.28094 | 86 |
| GO_MITOCHONDRIAL_PROTEIN_COMPLEX | 0.00000 | 0.00000 | 0.81404 | 0.63650 | 1.40931 | 208 |
| GO_MITOCHONDRIAL_TRANSLATION | 0.00004 | 0.00093 | 0.55733 | 0.61219 | 1.34135 | 129 |
| GO_MITOCHONDRIAL_TRANSLATIONAL_TERMINATION | 0.00000 | 0.00010 | 0.62726 | 0.66694 | 1.44128 | 87 |
| GO_MITOCHONDRION_ORGANIZATION | 0.00051 | 0.00993 | 0.47727 | 0.52284 | 1.16863 | 390 |
| GO_MODIFICATION_DEPENDENT_MACROMOLECULE_CATABOLIC_PR | 0.00002 | 0.00053 | 0.57561 | 0.53142 | 1.19156 | 486 |
| GO_MODULATION_BY_HOST_OF_SYMBIONT_PROCESS | 0.00015 | 0.00336 | 0.51885 | 0.72848 | 1.50542 | 33 |
| GO_MODULATION_OF_PROCESS_OF_OTHER_ORGANISM | 0.00019 | 0.00423 | 0.51885 | 0.66904 | 1.42286 | 59 |
| GO_MONOSACCHARIDE_BIOSYNTHETIC_PROCESS | 0.00335 | 0.04466 | 0.43171 | 0.61992 | 1.31781 | 58 |
| GO_MORPHOGENESIS_OF_AN_EPITHELIUM | 0.00006 | 0.00151 | 0.53843 | 0.55161 | 1.22755 | 290 |
| GO_MORPHOGENESIS_OF_A_POLARIZED_EPITHELIUM | 0.00000 | 0.00000 | | 0.72298 | 1.57057 | 103 |

| | | | | | | |
|---|---|---|---|---|---|---|
| GO_MRNA_3_UTR_BINDING | 0.00003 | 0.00089 | 0.55733 | 0.67596 | 1.44827 | 68 |
| GO_MRNA_5_UTR_BINDING | 0.00006 | 0.00156 | 0.53843 | 0.80133 | 1.62592 | 22 |
| GO_MRNA_BINDING | 0.00000 | 0.00000 | | 0.66002 | 1.46353 | 222 |
| GO_MRNA_PROCESSING | 0.00000 | 0.00000 | 0.73376 | 0.56859 | 1.27295 | 418 |
| GO_MULTI_ORGANISM_LOCALIZATION | 0.00268 | 0.03799 | 0.43171 | 0.63245 | 1.33612 | 51 |
| GO_MULTI_ORGANISM_PROCESS | 0.00108 | 0.01839 | 0.45506 | 0.51114 | 1.14599 | 484 |
| GO_MYD88_DEPENDENT_TOLL_LIKE_RECEPTOR_SIGNALING_PATHW | 0.00002 | 0.00066 | 0.57561 | 0.85581 | 1.70312 | 16 |
| GO_MYELOID_LEUKOCYTE_ACTIVATION | 0.00000 | 0.00000 | 0.74774 | 0.58508 | 1.30479 | 327 |
| GO_MYELOID_LEUKOCYTE_MEDIATED_IMMUNITY | 0.00000 | 0.00000 | 0.76146 | 0.59841 | 1.33291 | 301 |
| GO_NADH_DEHYDROGENASE_ACTIVITY | 0.00383 | 0.04935 | 0.43171 | 0.66244 | 1.37139 | 35 |
| GO_NADH_DEHYDROGENASE_COMPLEX | 0.00376 | 0.04884 | 0.43171 | 0.65649 | 1.36421 | 38 |
| GO_NADH_METABOLIC_PROCESS | 0.00294 | 0.04036 | 0.43171 | 0.70642 | 1.44481 | 27 |
| GO_NCRNA_EXPORT_FROM_NUCLEUS | 0.00302 | 0.04108 | 0.43171 | 0.66820 | 1.38331 | 35 |
| GO_NCRNA_METABOLIC_PROCESS | 0.00000 | 0.00003 | 0.65944 | 0.55907 | 1.24907 | 381 |
| GO_NCRNA_PROCESSING | 0.00000 | 0.00003 | 0.65944 | 0.57175 | 1.27504 | 328 |
| GO_NEGATIVE_REGULATION_OF_CANONICAL_WNT_SIGNALING_PAT | 0.00000 | 0.00000 | | 0.71611 | 1.56038 | 111 |
| GO_NEGATIVE_REGULATION_OF_CELLULAR_AMIDE_METABOLIC_PR | 0.00001 | 0.00020 | 0.61053 | 0.63031 | 1.37423 | 114 |
| GO_NEGATIVE_REGULATION_OF_CELL_CYCLE | 0.00003 | 0.00076 | 0.57561 | 0.53833 | 1.20513 | 425 |
| GO_NEGATIVE_REGULATION_OF_CELL_CYCLE_G2_M_PHASE_TRANS | 0.00000 | 0.00000 | 0.76146 | 0.71490 | 1.54439 | 85 |
| GO_NEGATIVE_REGULATION_OF_CELL_CYCLE_PHASE_TRANSITION | 0.00002 | 0.00060 | 0.57561 | 0.59041 | 1.30442 | 184 |
| GO_NEGATIVE_REGULATION_OF_CELL_CYCLE_PROCESS | 0.00000 | 0.00002 | 0.65944 | 0.58690 | 1.30374 | 249 |
| GO_NEGATIVE_REGULATION_OF_MITOTIC_CELL_CYCLE | 0.00004 | 0.00095 | 0.55733 | 0.56735 | 1.25902 | 232 |
| GO_NEGATIVE_REGULATION_OF_MRNA_METABOLIC_PROCESS | 0.00023 | 0.00491 | 0.51885 | 0.66670 | 1.41788 | 59 |
| GO_NEGATIVE_REGULATION_OF_MRNA_SPLICING_VIA_SPLICEOSOM | 0.00239 | 0.03510 | 0.43171 | 0.74857 | 1.51204 | 19 |
| GO_NEGATIVE_REGULATION_OF_RNA_SPLICING | 0.00029 | 0.00607 | 0.49849 | 0.76729 | 1.55976 | 23 |
| GO_NEGATIVE_REGULATION_OF_TRANSLATIONAL_INITIATION | 0.00018 | 0.00403 | 0.51885 | 0.83901 | 1.66434 | 15 |
| GO_NEGATIVE_REGULATION_OF_WNT_SIGNALING_PATHWAY | 0.00000 | 0.00000 | | 0.69525 | 1.52249 | 127 |
| GO_NEURAL_NUCLEUS_DEVELOPMENT | 0.00001 | 0.00026 | 0.59333 | 0.79148 | 1.62187 | 28 |
| GO_NEURON_TO_NEURON_SYNAPSE | 0.00005 | 0.00134 | 0.55733 | 0.59636 | 1.31006 | 154 |
| GO_NIK_NF_KAPPAB_SIGNALING | 0.00000 | 0.00000 | | 0.72896 | 1.58354 | 103 |
| GO_NON_CANONICAL_WNT_SIGNALING_PATHWAY | 0.00000 | 0.00000 | | 0.72784 | 1.58023 | 101 |
| GO_NUCLEAR_EXPORT | 0.00031 | 0.00645 | 0.49849 | 0.56963 | 1.25542 | 172 |
| GO_NUCLEAR_TRANSCRIBED_MRNA_CATABOLIC_PROCESS | 0.00000 | 0.00000 | | 0.79179 | 1.74917 | 183 |
| GO_NUCLEAR_TRANSCRIBED_MRNA_CATABOLIC_PROCESS_NONSEN | 0.00000 | 0.00000 | | 0.89008 | 1.93909 | 110 |
| GO_NUCLEAR_TRANSPORT | 0.00008 | 0.00193 | 0.53843 | 0.55511 | 1.23408 | 273 |
| GO_NUCLEOBASE_CONTAINING_COMPOUND_TRANSPORT | 0.00146 | 0.02386 | 0.45506 | 0.55053 | 1.21842 | 202 |
| GO_NUCLEOBASE_CONTAINING_SMALL_MOLECULE_BIOSYNTHETIC_ | 0.00137 | 0.02261 | 0.45506 | 0.62749 | 1.34353 | 67 |
| GO_NUCLEOBASE_CONTAINING_SMALL_MOLECULE_INTERCONVERS | 0.00383 | 0.04935 | 0.43171 | 0.71259 | 1.44857 | 23 |
| GO_NUCLEOBASE_CONTAINING_SMALL_MOLECULE_METABOLIC_PR | 0.00320 | 0.04339 | 0.43171 | 0.51448 | 1.14825 | 352 |
| GO_NUCLEOSIDE_DIPHOSPHATE_METABOLIC_PROCESS | 0.00110 | 0.01865 | 0.45506 | 0.59637 | 1.29295 | 97 |
| GO_NUCLEOSIDE_METABOLIC_PROCESS | 0.00061 | 0.01169 | 0.47727 | 0.65315 | 1.38907 | 59 |
| GO_NUCLEOSIDE_MONOPHOSPHATE_BIOSYNTHETIC_PROCESS | 0.00250 | 0.03636 | 0.43171 | 0.68841 | 1.42033 | 32 |
| GO_NUCLEOSIDE_MONOPHOSPHATE_METABOLIC_PROCESS | 0.00152 | 0.02464 | 0.45506 | 0.64377 | 1.36002 | 51 |
| GO_NUCLEOSIDE_PHOSPHATE_BIOSYNTHETIC_PROCESS | 0.00043 | 0.00865 | 0.49849 | 0.58185 | 1.27831 | 148 |
| GO_NUCLEOSIDE_TRIPHOSPHATE_BIOSYNTHETIC_PROCESS | 0.00043 | 0.00865 | 0.49849 | 0.69332 | 1.44539 | 40 |
| GO_NUCLEOSIDE_TRIPHOSPHATE_METABOLIC_PROCESS | 0.00016 | 0.00360 | 0.51885 | 0.66788 | 1.41976 | 58 |
| GO_NUCLEOSOMAL_DNA_BINDING | 0.00023 | 0.00507 | 0.51885 | 0.79852 | 1.61801 | 21 |
| GO_NUCLEOSOME_BINDING | 0.00143 | 0.02332 | 0.45506 | 0.66132 | 1.38492 | 45 |
| GO_NUCLEOTIDE_EXCISION_REPAIR | 0.00004 | 0.00096 | 0.55733 | 0.63343 | 1.37355 | 98 |
| GO_NUCLEOTIDE_EXCISION_REPAIR_DNA_DAMAGE_RECOGNITION | 0.00015 | 0.00349 | 0.51885 | 0.78678 | 1.59640 | 22 |
| GO_NUCLEOTIDE_EXCISION_REPAIR_DNA_INCISION | 0.00313 | 0.04249 | 0.43171 | 0.66439 | 1.37898 | 36 |
| GO_NUCLEOTIDE_PHOSPHORYLATION | 0.00383 | 0.04935 | 0.43171 | 0.59002 | 1.27454 | 88 |
| GO_ORGANELLAR_RIBOSOME | 0.00000 | 0.00014 | 0.61053 | 0.66412 | 1.43471 | 85 |
| GO_ORGANELLE_INNER_MEMBRANE | 0.00000 | 0.00000 | 0.76146 | 0.57601 | 1.28697 | 378 |
| GO_ORGANIC_CYCLIC_COMPOUND_CATABOLIC_PROCESS | 0.00000 | 0.00000 | | 0.73350 | 1.64281 | 449 |
| GO_OXIDATIVE_PHOSPHORYLATION | 0.00001 | 0.00030 | 0.59333 | 0.64619 | 1.40027 | 96 |
| GO_OXIDOREDUCTASE_ACTIVITY | 0.00006 | 0.00139 | 0.55733 | 0.53544 | 1.19703 | 392 |
| GO_OXIDOREDUCTASE_COMPLEX | 0.00001 | 0.00027 | 0.59333 | 0.66015 | 1.42461 | 82 |
| GO_PEPTIDASE_COMPLEX | 0.00000 | 0.00000 | | 0.75988 | 1.63982 | 82 |
| GO_PIGMENT_GRANULE | 0.00003 | 0.00089 | 0.55733 | 0.66098 | 1.42344 | 78 |
| GO_POLYSOMAL_RIBOSOME | 0.00000 | 0.00000 | | 0.90624 | 1.86629 | 31 |
| GO_POLYSOME | 0.00000 | 0.00000 | | 0.82337 | 1.75481 | 61 |
| GO_POLY_PYRIMIDINE_TRACT_BINDING | 0.00327 | 0.04398 | 0.43171 | 0.75471 | 1.51855 | 18 |
| GO_POSITIVE_REGULATION_OF_APOPTOTIC_SIGNALING_PATHWAY | 0.00279 | 0.03889 | 0.43171 | 0.57666 | 1.25518 | 107 |
| GO_POSITIVE_REGULATION_OF_ATPASE_ACTIVITY | 0.00294 | 0.04036 | 0.43171 | 0.65446 | 1.36795 | 43 |
| GO_POSITIVE_REGULATION_OF_BINDING | 0.00088 | 0.01590 | 0.47727 | 0.60284 | 1.30556 | 95 |
| GO_POSITIVE_REGULATION_OF_CANONICAL_WNT_SIGNALING_PATH | 0.00000 | 0.00000 | 0.77494 | 0.70504 | 1.53160 | 103 |
| GO_POSITIVE_REGULATION_OF_CELLULAR_AMIDE_METABOLIC_PRO | 0.00003 | 0.00074 | 0.57561 | 0.62296 | 1.36083 | 118 |
| GO_POSITIVE_REGULATION_OF_CELLULAR_PROTEIN_LOCALIZATION | 0.00041 | 0.00840 | 0.49849 | 0.54915 | 1.21815 | 225 |
| GO_POSITIVE_REGULATION_OF_CELL_DEATH | 0.00069 | 0.01297 | 0.47727 | 0.52595 | 1.17423 | 363 |
| GO_POSITIVE_REGULATION_OF_CELL_POPULATION_PROLIFERATION | 0.00078 | 0.01455 | 0.47727 | 0.52449 | 1.17108 | 368 |
| GO_POSITIVE_REGULATION_OF_CHROMOSOME_ORGANIZATION | 0.00231 | 0.03437 | 0.43171 | 0.56990 | 1.24871 | 133 |
| GO_POSITIVE_REGULATION_OF_DEFENSE_RESPONSE | 0.00000 | 0.00003 | 0.65944 | 0.62175 | 1.36730 | 158 |
| GO_POSITIVE_REGULATION_OF_DNA_BIOSYNTHETIC_PROCESS | 0.00001 | 0.00041 | 0.59333 | 0.71929 | 1.52055 | 52 |
| GO_POSITIVE_REGULATION_OF_DNA_METABOLIC_PROCESS | 0.00364 | 0.04761 | 0.43171 | 0.56647 | 1.24189 | 135 |
| GO_POSITIVE_REGULATION_OF_DNA_TEMPLATED_TRANSCRIPTION_ | 0.00276 | 0.03885 | 0.43171 | 0.68932 | 1.41586 | 29 |

| | | | | | | |
|---|---|---|---|---|---|---|
| GO_POSITIVE_REGULATION_OF_ESTABLISHMENT_OF_PROTEIN_LOC/ | 0.00007 | 0.00168 | 0.53843 | 0.56461 | 1.25165 | 219 |
| GO_POSITIVE_REGULATION_OF_IMMUNE_RESPONSE | 0.00000 | 0.00001 | 0.69013 | 0.59037 | 1.31265 | 277 |
| GO_POSITIVE_REGULATION_OF_IMMUNE_SYSTEM_PROCESS | 0.00004 | 0.00110 | 0.55733 | 0.54018 | 1.20716 | 386 |
| GO_POSITIVE_REGULATION_OF_INNATE_IMMUNE_RESPONSE | 0.00000 | 0.00000 | 0.81404 | 0.69693 | 1.51947 | 114 |
| GO_POSITIVE_REGULATION_OF_INTRINSIC_APOPTOTIC_SIGNALING_ | 0.00104 | 0.01798 | 0.45506 | 0.69097 | 1.43578 | 37 |
| GO_POSITIVE_REGULATION_OF_ORGANELLE_ORGANIZATION | 0.00043 | 0.00865 | 0.49849 | 0.52093 | 1.16605 | 411 |
| GO_POSITIVE_REGULATION_OF_PROTEIN_LOCALIZATION_TO_NUCLE | 0.00001 | 0.00020 | 0.61053 | 0.69907 | 1.48990 | 61 |
| GO_POSITIVE_REGULATION_OF_RESPONSE_TO_BIOTIC_STIMULUS | 0.00000 | 0.00000 | 0.73376 | 0.66340 | 1.45310 | 128 |
| GO_POSITIVE_REGULATION_OF_RESPONSE_TO_EXTERNAL_STIMULU | 0.00002 | 0.00050 | 0.57561 | 0.58133 | 1.28681 | 201 |
| GO_POSITIVE_REGULATION_OF_TELOMERASE_ACTIVITY | 0.00061 | 0.01169 | 0.47727 | 0.73476 | 1.50565 | 28 |
| GO_POSITIVE_REGULATION_OF_TELOMERASE_RNA_LOCALIZATION_ | 0.00000 | 0.00001 | 0.69013 | 0.92451 | 1.83395 | 15 |
| GO_POSITIVE_REGULATION_OF_TELOMERE_MAINTENANCE_VIA_TE | 0.00130 | 0.02159 | 0.45506 | 0.71259 | 1.46508 | 30 |
| GO_POSITIVE_REGULATION_OF_TRANSCRIPTION_INITIATION_FROM | 0.00372 | 0.04847 | 0.43171 | 0.70671 | 1.43993 | 25 |
| GO_POSITIVE_REGULATION_OF_TRANSFERASE_ACTIVITY | 0.00174 | 0.02712 | 0.45506 | 0.51642 | 1.15220 | 345 |
| GO_POSITIVE_REGULATION_OF_TRANSLATION | 0.00001 | 0.00019 | 0.61053 | 0.64767 | 1.40419 | 97 |
| GO_POSITIVE_REGULATION_OF_VIRAL_GENOME_REPLICATION | 0.00287 | 0.03973 | 0.43171 | 0.68362 | 1.40783 | 31 |
| GO_POSITIVE_REGULATION_OF_VIRAL_PROCESS | 0.00028 | 0.00590 | 0.49849 | 0.62440 | 1.34741 | 81 |
| GO_POSITIVE_REGULATION_OF_WNT_SIGNALING_PATHWAY | 0.00000 | 0.00000 | 0.73376 | 0.66230 | 1.44763 | 122 |
| GO_POSTREPLICATION_REPAIR | 0.00024 | 0.00515 | 0.51885 | 0.69140 | 1.44518 | 43 |
| GO_POSTSYNAPSE | 0.00004 | 0.00109 | 0.55733 | 0.55969 | 1.24369 | 259 |
| GO_POSTSYNAPTIC_CYTOSOL | 0.00279 | 0.03889 | 0.43171 | 0.78295 | 1.55313 | 15 |
| GO_POSTTRANSCRIPTIONAL_REGULATION_OF_GENE_EXPRESSION | 0.00000 | 0.00000 | | 0.66656 | 1.49327 | 462 |
| GO_POST_TRANSLATIONAL_PROTEIN_MODIFICATION | 0.00002 | 0.00052 | 0.57561 | 0.57851 | 1.28412 | 229 |
| GO_PRECATALYTIC_SPLICEOSOME | 0.00085 | 0.01559 | 0.47727 | 0.65235 | 1.37448 | 49 |
| GO_PRERIBOSOME | 0.00087 | 0.01577 | 0.47727 | 0.62899 | 1.34772 | 69 |
| GO_PRERIBOSOME_LARGE_SUBUNIT_PRECURSOR | 0.00235 | 0.03474 | 0.43171 | 0.74079 | 1.50308 | 22 |
| GO_PRE_MRNA_BINDING | 0.00133 | 0.02213 | 0.45506 | 0.72393 | 1.48062 | 27 |
| GO_PROTEASOMAL_PROTEIN_CATABOLIC_PROCESS | 0.00002 | 0.00044 | 0.59333 | 0.54166 | 1.21005 | 377 |
| GO_PROTEASOMAL_UBIQUITIN_INDEPENDENT_PROTEIN_CATABOLI( | 0.00000 | 0.00000 | 0.80122 | 0.92254 | 1.86931 | 21 |
| GO_PROTEASOME_ACCESSORY_COMPLEX | 0.00000 | 0.00001 | 0.67496 | 0.85464 | 1.73811 | 24 |
| GO_PROTEASOME_CORE_COMPLEX | 0.00000 | 0.00000 | | 0.94375 | 1.90631 | 19 |
| GO_PROTEIN_CONTAINING_COMPLEX_DISASSEMBLY | 0.00000 | 0.00007 | 0.62726 | 0.57810 | 1.28438 | 256 |
| GO_PROTEIN_CONTAINING_COMPLEX_LOCALIZATION | 0.00063 | 0.01197 | 0.47727 | 0.55335 | 1.22520 | 208 |
| GO_PROTEIN_DNA_COMPLEX_SUBUNIT_ORGANIZATION | 0.00001 | 0.00018 | 0.61053 | 0.60174 | 1.32491 | 165 |
| GO_PROTEIN_FOLDING | 0.00000 | 0.00002 | 0.65944 | 0.63113 | 1.38644 | 154 |
| GO_PROTEIN_FOLDING_CHAPERONE | 0.00135 | 0.02238 | 0.45506 | 0.77903 | 1.55752 | 17 |
| GO_PROTEIN_IMPORT | 0.00268 | 0.03799 | 0.43171 | 0.56317 | 1.23581 | 140 |
| GO_PROTEIN_LOCALIZATION_TO_CHROMOSOME | 0.00056 | 0.01096 | 0.47727 | 0.63523 | 1.36108 | 69 |
| GO_PROTEIN_LOCALIZATION_TO_CHROMOSOME_TELOMERIC_REGI( | 0.00335 | 0.04466 | 0.43171 | 0.70399 | 1.43984 | 27 |
| GO_PROTEIN_LOCALIZATION_TO_ENDOPLASMIC_RETICULUM | 0.00000 | 0.00000 | | 0.85521 | 1.87021 | 124 |
| GO_PROTEIN_LOCALIZATION_TO_MEMBRANE | 0.00000 | 0.00000 | | 0.65123 | 1.45714 | 406 |
| GO_PROTEIN_LOCALIZATION_TO_MITOCHONDRION | 0.00019 | 0.00420 | 0.51885 | 0.60773 | 1.32398 | 110 |
| GO_PROTEIN_LOCALIZATION_TO_NUCLEUS | 0.00002 | 0.00055 | 0.57561 | 0.58926 | 1.30220 | 187 |
| GO_PROTEIN_MODIFICATION_BY_SMALL_PROTEIN_REMOVAL | 0.00000 | 0.00000 | 0.71951 | 0.60745 | 1.34689 | 221 |
| GO_PROTEIN_POLYUBIQUITINATION | 0.00004 | 0.00106 | 0.55733 | 0.55874 | 1.24108 | 253 |
| GO_PROTEIN_REFOLDING | 0.00294 | 0.04036 | 0.43171 | 0.77039 | 1.53314 | 16 |
| GO_PROTEIN_STABILIZATION | 0.00001 | 0.00024 | 0.61053 | 0.62333 | 1.36586 | 130 |
| GO_PROTEIN_TARGETING | 0.00000 | 0.00000 | | 0.71181 | 1.58743 | 327 |
| GO_PROTEIN_TARGETING_TO_MEMBRANE | 0.00000 | 0.00000 | | 0.82370 | 1.80953 | 149 |
| GO_PROTEIN_TARGETING_TO_MITOCHONDRION | 0.00346 | 0.04593 | 0.43171 | 0.60408 | 1.30091 | 78 |
| GO_PROTEIN_TRANSMEMBRANE_TRANSPORT | 0.00265 | 0.03775 | 0.43171 | 0.64528 | 1.36171 | 50 |
| GO_PROTON_TRANSMEMBRANE_TRANSPORT | 0.00213 | 0.03229 | 0.43171 | 0.60815 | 1.30536 | 71 |
| GO_PURINE_CONTAINING_COMPOUND_BIOSYNTHETIC_PROCESS | 0.00246 | 0.03601 | 0.43171 | 0.57796 | 1.25936 | 111 |
| GO_PURINE_CONTAINING_COMPOUND_METABOLIC_PROCESS | 0.00100 | 0.01761 | 0.45506 | 0.53679 | 1.19242 | 255 |
| GO_PURINE_NUCLEOSIDE_TRIPHOSPHATE_METABOLIC_PROCESS | 0.00089 | 0.01592 | 0.47727 | 0.67157 | 1.40374 | 43 |
| GO_PYRIMIDINE_CONTAINING_COMPOUND_METABOLIC_PROCESS | 0.00353 | 0.04658 | 0.43171 | 0.64744 | 1.35878 | 47 |
| GO_REGENERATION | 0.00216 | 0.03242 | 0.43171 | 0.59987 | 1.29453 | 82 |
| GO_REGULATION_OF_ANIMAL_ORGAN_MORPHOGENESIS | 0.00000 | 0.00000 | | 0.69163 | 1.51541 | 133 |
| GO_REGULATION_OF_APOPTOTIC_SIGNALING_PATHWAY | 0.00031 | 0.00645 | 0.49849 | 0.55171 | 1.22506 | 247 |
| GO_REGULATION_OF_ATPASE_ACTIVITY | 0.00324 | 0.04369 | 0.43171 | 0.63031 | 1.34038 | 57 |
| GO_REGULATION_OF_ATP_METABOLIC_PROCESS | 0.00353 | 0.04658 | 0.43171 | 0.60176 | 1.29780 | 80 |
| GO_REGULATION_OF_BINDING | 0.00250 | 0.03636 | 0.43171 | 0.54476 | 1.20598 | 203 |
| GO_REGULATION_OF_CELLULAR_AMIDE_METABOLIC_PROCESS | 0.00000 | 0.00000 | | 0.62967 | 1.40226 | 311 |
| GO_REGULATION_OF_CELLULAR_AMINE_METABOLIC_PROCESS | 0.00000 | 0.00000 | | 0.82173 | 1.74682 | 58 |
| GO_REGULATION_OF_CELLULAR_AMINO_ACID_METABOLIC_PROCES | 0.00000 | 0.00000 | | 0.83776 | 1.77766 | 55 |
| GO_REGULATION_OF_CELLULAR_KETONE_METABOLIC_PROCESS | 0.00000 | 0.00000 | | 0.72641 | 1.58680 | 118 |
| GO_REGULATION_OF_CELLULAR_PROTEIN_LOCALIZATION | 0.00082 | 0.01513 | 0.47727 | 0.52542 | 1.17290 | 362 |
| GO_REGULATION_OF_CELLULAR_RESPONSE_TO_HEAT | 0.00276 | 0.03885 | 0.43171 | 0.61652 | 1.32004 | 67 |
| GO_REGULATION_OF_CELLULAR_RESPONSE_TO_STRESS | 0.00224 | 0.03344 | 0.43171 | 0.50545 | 1.13232 | 468 |
| GO_REGULATION_OF_CELL_CYCLE_G2_M_PHASE_TRANSITION | 0.00000 | 0.00000 | 0.71951 | 0.63230 | 1.39282 | 169 |
| GO_REGULATION_OF_CELL_CYCLE_PHASE_TRANSITION | 0.00001 | 0.00026 | 0.59333 | 0.55488 | 1.23783 | 337 |
| GO_REGULATION_OF_CHROMOSOME_ORGANIZATION | 0.00081 | 0.01501 | 0.47727 | 0.54255 | 1.20527 | 261 |
| GO_REGULATION_OF_CYSTEINE_TYPE_ENDOPEPTIDASE_ACTIVITY | 0.00202 | 0.03077 | 0.43171 | 0.57120 | 1.24837 | 121 |
| GO_REGULATION_OF_CYTOPLASMIC_TRANSLATION | 0.00130 | 0.02159 | 0.45506 | 0.73678 | 1.49773 | 23 |
| GO_REGULATION_OF_DEFENSE_RESPONSE | 0.00065 | 0.01237 | 0.47727 | 0.53633 | 1.19355 | 290 |
| GO_REGULATION_OF_DNA_BIOSYNTHETIC_PROCESS | 0.00016 | 0.00360 | 0.51885 | 0.64215 | 1.37923 | 74 |

| | | | | | | |
|---|---|---|---|---|---|---|
| GO_REGULATION_OF_DNA_METABOLIC_PROCESS | 0.00069 | 0.01297 | 0.47727 | 0.54425 | 1.20775 | 237 |
| GO_REGULATION_OF_DNA_TEMPLATED_TRANSCRIPTION_INITIATIO | 0.00163 | 0.02568 | 0.45506 | 0.68289 | 1.41372 | 35 |
| GO_REGULATION_OF_DNA_TEMPLATED_TRANSCRIPTION_IN_RESPO | 0.00000 | 0.00000 | | 0.73944 | 1.59800 | 89 |
| GO_REGULATION_OF_GENERATION_OF_PRECURSOR_METABOLITES_ | 0.00060 | 0.01164 | 0.47727 | 0.60476 | 1.31116 | 97 |
| GO_REGULATION_OF_HEMATOPOIETIC_PROGENITOR_CELL_DIFFERE | 0.00000 | 0.00000 | | 0.78717 | 1.69251 | 75 |
| GO_REGULATION_OF_HEMATOPOIETIC_STEM_CELL_DIFFERENTIATIO | 0.00000 | 0.00000 | | 0.81505 | 1.74256 | 65 |
| GO_REGULATION_OF_HEMOPOIESIS | 0.00000 | 0.00008 | 0.62726 | 0.58859 | 1.30634 | 228 |
| GO_REGULATION_OF_IMMUNE_RESPONSE | 0.00000 | 0.00016 | 0.61053 | 0.55362 | 1.23583 | 362 |
| GO_REGULATION_OF_INNATE_IMMUNE_RESPONSE | 0.00000 | 0.00000 | 0.71951 | 0.63883 | 1.40521 | 161 |
| GO_REGULATION_OF_INTRINSIC_APOPTOTIC_SIGNALING_PATHWAY | 0.00001 | 0.00039 | 0.59333 | 0.63478 | 1.38259 | 109 |
| GO_REGULATION_OF_MITOTIC_CELL_CYCLE | 0.00001 | 0.00018 | 0.61053 | 0.53899 | 1.20736 | 458 |
| GO_REGULATION_OF_MORPHOGENESIS_OF_AN_EPITHELIUM | 0.00000 | 0.00000 | | 0.72213 | 1.57284 | 109 |
| GO_REGULATION_OF_MRNA_CATABOLIC_PROCESS | 0.00000 | 0.00000 | | 0.69977 | 1.54127 | 168 |
| GO_REGULATION_OF_MRNA_METABOLIC_PROCESS | 0.00000 | 0.00000 | | 0.66528 | 1.47872 | 266 |
| GO_REGULATION_OF_MRNA_PROCESSING | 0.00004 | 0.00105 | 0.55733 | 0.61521 | 1.34090 | 113 |
| GO_REGULATION_OF_MRNA_SPLICING_VIA_SPLICEOSOME | 0.00001 | 0.00026 | 0.59333 | 0.67424 | 1.45411 | 80 |
| GO_REGULATION_OF_NUCLEASE_ACTIVITY | 0.00106 | 0.01819 | 0.45506 | 0.77918 | 1.56780 | 18 |
| GO_REGULATION_OF_OXIDATIVE_STRESS_INDUCED_CELL_DEATH | 0.00198 | 0.03028 | 0.43171 | 0.67287 | 1.40355 | 41 |
| GO_REGULATION_OF_PEPTIDASE_ACTIVITY | 0.00050 | 0.00978 | 0.47727 | 0.56122 | 1.24054 | 191 |
| GO_REGULATION_OF_POSTTRANSCRIPTIONAL_GENE_SILENCING | 0.00364 | 0.04761 | 0.43171 | 0.59824 | 1.28492 | 74 |
| GO_REGULATION_OF_PROTEIN_CATABOLIC_PROCESS | 0.00179 | 0.02783 | 0.45506 | 0.53057 | 1.17929 | 270 |
| GO_REGULATION_OF_PROTEIN_LOCALIZATION_TO_NUCLEUS | 0.00001 | 0.00042 | 0.59333 | 0.65505 | 1.41558 | 87 |
| GO_REGULATION_OF_PROTEIN_MODIFICATION_BY_SMALL_PROTEIN | 0.00025 | 0.00532 | 0.49849 | 0.57068 | 1.25684 | 167 |
| GO_REGULATION_OF_PROTEIN_STABILITY | 0.00093 | 0.01652 | 0.47727 | 0.55382 | 1.22571 | 202 |
| GO_REGULATION_OF_PROTEOLYSIS | 0.00013 | 0.00301 | 0.51885 | 0.52997 | 1.18392 | 377 |
| GO_REGULATION_OF_RESPONSE_TO_BIOTIC_STIMULUS | 0.00001 | 0.00020 | 0.61053 | 0.59049 | 1.30719 | 204 |
| GO_REGULATION_OF_RNA_SPLICING | 0.00000 | 0.00001 | 0.69013 | 0.65896 | 1.43669 | 114 |
| GO_REGULATION_OF_SMALL_MOLECULE_METABOLIC_PROCESS | 0.00000 | 0.00000 | 0.71951 | 0.59134 | 1.31394 | 265 |
| GO_REGULATION_OF_STEM_CELL_DIFFERENTIATION | 0.00000 | 0.00000 | | 0.74049 | 1.60113 | 90 |
| GO_REGULATION_OF_SYMBIOTIC_PROCESS | 0.00006 | 0.00144 | 0.55733 | 0.60071 | 1.31682 | 134 |
| GO_REGULATION_OF_TELOMERASE_ACTIVITY | 0.00088 | 0.01590 | 0.47727 | 0.69161 | 1.43179 | 35 |
| GO_REGULATION_OF_TRANSCRIPTION_FROM_RNA_POLYMERASE_II | 0.00000 | 0.00000 | | 0.81194 | 1.73800 | 66 |
| GO_REGULATION_OF_TRANSCRIPTION_INITIATION_FROM_RNA_POL | 0.00143 | 0.02332 | 0.45506 | 0.70390 | 1.44582 | 29 |
| GO_REGULATION_OF_TRANSLATIONAL_INITIATION | 0.00002 | 0.00052 | 0.57561 | 0.68129 | 1.45518 | 64 |
| GO_REGULATION_OF_UBIQUITIN_PROTEIN_TRANSFERASE_ACTIVITY | 0.00061 | 0.01169 | 0.47727 | 0.69335 | 1.44627 | 41 |
| GO_REGULATION_OF_VIRAL_GENOME_REPLICATION | 0.00086 | 0.01566 | 0.47727 | 0.65316 | 1.38389 | 54 |
| GO_REGULATION_OF_VIRAL_TRANSCRIPTION | 0.00198 | 0.03028 | 0.43171 | 0.63932 | 1.34703 | 49 |
| GO_REGULATION_OF_WNT_SIGNALING_PATHWAY | 0.00000 | 0.00001 | 0.69013 | 0.60353 | 1.33816 | 220 |
| GO_RESPIRASOME | 0.00008 | 0.00181 | 0.53843 | 0.65420 | 1.40421 | 71 |
| GO_RESPIRATORY_CHAIN_COMPLEX | 0.00016 | 0.00353 | 0.51885 | 0.67626 | 1.43918 | 60 |
| GO_RESPIRATORY_ELECTRON_TRANSPORT_CHAIN | 0.00001 | 0.00025 | 0.59333 | 0.65819 | 1.42190 | 85 |
| GO_RESPONSE_TO_ALCOHOL | 0.00261 | 0.03750 | 0.43171 | 0.58804 | 1.27350 | 95 |
| GO_RESPONSE_TO_CADMIUM_ION | 0.00265 | 0.03775 | 0.43171 | 0.67227 | 1.39174 | 35 |
| GO_RESPONSE_TO_DRUG | 0.00104 | 0.01798 | 0.45506 | 0.55364 | 1.22258 | 180 |
| GO_RESPONSE_TO_ETHANOL | 0.00086 | 0.01566 | 0.47727 | 0.67309 | 1.40691 | 43 |
| GO_RESPONSE_TO_HEAT | 0.00012 | 0.00270 | 0.53843 | 0.61665 | 1.34221 | 107 |
| GO_RESPONSE_TO_HORMONE | 0.00048 | 0.00949 | 0.49849 | 0.51277 | 1.14814 | 453 |
| GO_RESPONSE_TO_HYDROGEN_PEROXIDE | 0.00257 | 0.03716 | 0.43171 | 0.60505 | 1.30368 | 79 |
| GO_RESPONSE_TO_INORGANIC_SUBSTANCE | 0.00004 | 0.00107 | 0.55733 | 0.55714 | 1.23870 | 275 |
| GO_RESPONSE_TO_INTERLEUKIN_1 | 0.00000 | 0.00000 | | 0.74844 | 1.62908 | 107 |
| GO_RESPONSE_TO_INTERLEUKIN_12 | 0.00012 | 0.00270 | 0.53843 | 0.72605 | 1.50694 | 36 |
| GO_RESPONSE_TO_INTERLEUKIN_4 | 0.00324 | 0.04369 | 0.43171 | 0.75812 | 1.51571 | 17 |
| GO_RESPONSE_TO_METAL_ION | 0.00030 | 0.00623 | 0.49849 | 0.56387 | 1.24233 | 171 |
| GO_RESPONSE_TO_ORGANIC_CYCLIC_COMPOUND | 0.00000 | 0.00016 | 0.61053 | 0.54603 | 1.22259 | 439 |
| GO_RESPONSE_TO_OXYGEN_LEVELS | 0.00000 | 0.00000 | 0.71951 | 0.60132 | 1.33438 | 233 |
| GO_RESPONSE_TO_STEROID_HORMONE | 0.00154 | 0.02464 | 0.45506 | 0.55514 | 1.22446 | 174 |
| GO_RESPONSE_TO_TEMPERATURE_STIMULUS | 0.00027 | 0.00572 | 0.49849 | 0.59216 | 1.29675 | 127 |
| GO_RESPONSE_TO_TUMOR_NECROSIS_FACTOR | 0.00000 | 0.00000 | 0.76146 | 0.65649 | 1.44236 | 151 |
| GO_RETINA_HOMEOSTASIS | 0.00104 | 0.01798 | 0.45506 | 0.73063 | 1.49432 | 27 |
| GO_RIBONUCLEOPROTEIN_COMPLEX_BINDING | 0.00000 | 0.00000 | 0.71951 | 0.66906 | 1.46050 | 117 |
| GO_RIBONUCLEOPROTEIN_COMPLEX_BIOGENESIS | 0.00000 | 0.00000 | | 0.66339 | 1.48264 | 389 |
| GO_RIBONUCLEOPROTEIN_COMPLEX_SUBUNIT_ORGANIZATION | 0.00000 | 0.00000 | | 0.69241 | 1.52681 | 173 |
| GO_RIBONUCLEOPROTEIN_GRANULE | 0.00000 | 0.00001 | 0.67496 | 0.61979 | 1.36366 | 163 |
| GO_RIBONUCLEOSIDE_TRIPHOSPHATE_BIOSYNTHETIC_PROCESS | 0.00041 | 0.00840 | 0.49849 | 0.71253 | 1.47248 | 33 |
| GO_RIBONUCLEOSIDE_TRIPHOSPHATE_METABOLIC_PROCESS | 0.00026 | 0.00550 | 0.49849 | 0.69681 | 1.45611 | 42 |
| GO_RIBOSE_PHOSPHATE_BIOSYNTHETIC_PROCESS | 0.00191 | 0.02939 | 0.45506 | 0.57932 | 1.26207 | 110 |
| GO_RIBOSE_PHOSPHATE_METABOLIC_PROCESS | 0.00108 | 0.01839 | 0.45506 | 0.53891 | 1.19687 | 245 |
| GO_RIBOSOMAL_LARGE_SUBUNIT_ASSEMBLY | 0.00000 | 0.00016 | 0.61053 | 0.80999 | 1.65663 | 27 |
| GO_RIBOSOMAL_LARGE_SUBUNIT_BIOGENESIS | 0.00000 | 0.00000 | | 0.77542 | 1.65982 | 66 |
| GO_RIBOSOMAL_SMALL_SUBUNIT_ASSEMBLY | 0.00023 | 0.00494 | 0.51885 | 0.81139 | 1.62221 | 17 |
| GO_RIBOSOMAL_SMALL_SUBUNIT_BINDING | 0.00383 | 0.04935 | 0.43171 | 0.75288 | 1.50523 | 17 |
| GO_RIBOSOMAL_SMALL_SUBUNIT_BIOGENESIS | 0.00001 | 0.00022 | 0.61053 | 0.69076 | 1.47997 | 68 |
| GO_RIBOSOMAL_SUBUNIT | 0.00000 | 0.00000 | | 0.77603 | 1.71031 | 172 |
| GO_RIBOSOME | 0.00000 | 0.00000 | | 0.76643 | 1.69659 | 206 |
| GO_RIBOSOME_ASSEMBLY | 0.00000 | 0.00000 | 0.73376 | 0.76173 | 1.61998 | 59 |
| GO_RIBOSOME_BINDING | 0.00019 | 0.00416 | 0.51885 | 0.68955 | 1.44715 | 47 |

| | | | | | | |
|---|---|---|---|---|---|---|
| GO_RIBOSOME_BIOGENESIS | 0.00000 | 0.00000 | | 0.65877 | 1.46466 | 275 |
| GO_RNA_CATABOLIC_PROCESS | 0.00000 | 0.00000 | | 0.76774 | 1.71264 | 339 |
| GO_RNA_DEPENDENT_DNA_BIOSYNTHETIC_PROCESS | 0.00003 | 0.00076 | 0.57561 | 0.68926 | 1.46521 | 58 |
| GO_RNA_EXPORT_FROM_NUCLEUS | 0.00062 | 0.01183 | 0.47727 | 0.58378 | 1.27758 | 126 |
| GO_RNA_LOCALIZATION | 0.00000 | 0.00006 | 0.64355 | 0.59670 | 1.32020 | 198 |
| GO_RNA_POLYMERASE_II_PREINITIATION_COMPLEX_ASSEMBLY | 0.00089 | 0.01592 | 0.47727 | 0.77046 | 1.55626 | 19 |
| GO_RNA_SPLICING | 0.00000 | 0.00000 | 0.78819 | 0.59079 | 1.31927 | 369 |
| GO_RNA_SPLICING_VIA_TRANSESTERIFICATION_REACTIONS | 0.00000 | 0.00000 | | 0.60940 | 1.35714 | 300 |
| GO_RNA_STABILIZATION | 0.00065 | 0.01223 | 0.47727 | 0.69743 | 1.44384 | 35 |
| GO_ROUGH_ENDOPLASMIC_RETICULUM | 0.00001 | 0.00039 | 0.59333 | 0.71853 | 1.51629 | 50 |
| GO_RRNA_BINDING | 0.00000 | 0.00001 | 0.69013 | 0.74782 | 1.58865 | 56 |
| GO_RRNA_METABOLIC_PROCESS | 0.00000 | 0.00000 | 0.77494 | 0.63047 | 1.39562 | 206 |
| GO_SCF_DEPENDENT_PROTEASOMAL_UBIQUITIN_DEPENDENT_PRO | 0.00000 | 0.00000 | | 0.77527 | 1.67044 | 79 |
| GO_SECRETORY_GRANULE | 0.00000 | 0.00000 | 0.71951 | 0.56518 | 1.26461 | 399 |
| GO_SECRETORY_VESICLE | 0.00000 | 0.00002 | 0.67496 | 0.54866 | 1.22905 | 465 |
| GO_SIGNAL_TRANSDUCTION_BY_PROTEIN_PHOSPHORYLATION | 0.00004 | 0.00109 | 0.55733 | 0.53817 | 1.20474 | 424 |
| GO_SINGLE_FERTILIZATION | 0.00048 | 0.00949 | 0.49849 | 0.67414 | 1.42260 | 50 |
| GO_SINGLE_STRANDED_RNA_BINDING | 0.00039 | 0.00793 | 0.49849 | 0.66553 | 1.41527 | 57 |
| GO_SMALL_MOLECULE_BIOSYNTHETIC_PROCESS | 0.00216 | 0.03242 | 0.43171 | 0.51195 | 1.14522 | 397 |
| GO_SMALL_RIBOSOMAL_SUBUNIT | 0.00000 | 0.00000 | | 0.77648 | 1.66209 | 66 |
| GO_SM_LIKE_PROTEIN_FAMILY_COMPLEX | 0.00327 | 0.04398 | 0.43171 | 0.60468 | 1.29713 | 70 |
| GO_SPERM_EGG_RECOGNITION | 0.00000 | 0.00007 | 0.64355 | 0.85241 | 1.72955 | 22 |
| GO_SPLICEOSOMAL_COMPLEX | 0.00000 | 0.00003 | 0.65944 | 0.60939 | 1.34261 | 171 |
| GO_STEM_CELL_DIFFERENTIATION | 0.00000 | 0.00000 | 0.78819 | 0.66886 | 1.46930 | 147 |
| GO_STRUCTURAL_CONSTITUENT_OF_CYTOSKELETON | 0.00102 | 0.01782 | 0.45506 | 0.65913 | 1.39337 | 52 |
| GO_STRUCTURAL_CONSTITUENT_OF_RIBOSOME | 0.00000 | 0.00000 | | 0.79909 | 1.75541 | 150 |
| GO_STRUCTURAL_MOLECULE_ACTIVITY | 0.00000 | 0.00000 | | 0.72651 | 1.61895 | 324 |
| GO_SUBSTANTIA_NIGRA_DEVELOPMENT | 0.00000 | 0.00013 | 0.62726 | 0.81622 | 1.66507 | 26 |
| GO_TELOMERASE_RNA_LOCALIZATION | 0.00001 | 0.00017 | 0.61053 | 0.85164 | 1.72025 | 19 |
| GO_TELOMERE_MAINTENANCE_VIA_TELOMERE_LENGTHENING | 0.00011 | 0.00250 | 0.53843 | 0.65659 | 1.40546 | 66 |
| GO_TELOMERE_ORGANIZATION | 0.00342 | 0.04554 | 0.43171 | 0.57000 | 1.24649 | 124 |
| GO_THREONINE_TYPE_ENDOPEPTIDASE_ACTIVITY | 0.00000 | 0.00000 | | 0.95179 | 1.92255 | 19 |
| GO_THREONINE_TYPE_PEPTIDASE_ACTIVITY | 0.00000 | 0.00000 | | 0.92199 | 1.87423 | 23 |
| GO_TISSUE_MORPHOGENESIS | 0.00097 | 0.01707 | 0.47727 | 0.52661 | 1.17402 | 326 |
| GO_TOXIN_TRANSPORT | 0.00113 | 0.01922 | 0.45506 | 0.72897 | 1.48708 | 26 |
| GO_TRANSCRIPTION_COUPLED_NUCLEOTIDE_EXCISION_REPAIR | 0.00001 | 0.00019 | 0.61053 | 0.68106 | 1.46186 | 71 |
| GO_TRANSCRIPTION_FACTOR_BINDING | 0.00043 | 0.00865 | 0.49849 | 0.52447 | 1.17376 | 409 |
| GO_TRANSCRIPTION_PREINITIATION_COMPLEX_ASSEMBLY | 0.00119 | 0.02004 | 0.45506 | 0.71399 | 1.46797 | 30 |
| GO_TRANSLATIONAL_ELONGATION | 0.00000 | 0.00000 | 0.71951 | 0.66087 | 1.44451 | 122 |
| GO_TRANSLATIONAL_INITIATION | 0.00000 | 0.00000 | | 0.84691 | 1.86515 | 166 |
| GO_TRANSLATIONAL_TERMINATION | 0.00000 | 0.00005 | 0.64355 | 0.65946 | 1.43060 | 99 |
| GO_TRANSLATION_ELONGATION_FACTOR_ACTIVITY | 0.00161 | 0.02546 | 0.45506 | 0.79860 | 1.58418 | 15 |
| GO_TRANSLATION_FACTOR_ACTIVITY_RNA_BINDING | 0.00000 | 0.00000 | | 0.79008 | 1.69288 | 69 |
| GO_TRANSLATION_INITIATION_FACTOR_ACTIVITY | 0.00000 | 0.00000 | 0.76146 | 0.80422 | 1.68417 | 45 |
| GO_TRANSLATION_INITIATION_FACTOR_BINDING | 0.00021 | 0.00468 | 0.51885 | 0.76186 | 1.55231 | 25 |
| GO_TRANSLATION_PREINITIATION_COMPLEX | 0.00000 | 0.00001 | 0.67496 | 0.90245 | 1.79595 | 16 |
| GO_TRANSLATION_REGULATOR_ACTIVITY | 0.00000 | 0.00000 | | 0.77663 | 1.68904 | 105 |
| GO_TRANSLATION_REGULATOR_ACTIVITY_NUCLEIC_ACID_BINDING | 0.00000 | 0.00000 | | 0.77150 | 1.66586 | 83 |
| GO_TRICARBOXYLIC_ACID_CYCLE | 0.00154 | 0.02464 | 0.45506 | 0.71854 | 1.47240 | 28 |
| GO_TUMOR_NECROSIS_FACTOR_MEDIATED_SIGNALING_PATHWAY | 0.00000 | 0.00000 | | 0.76277 | 1.65372 | 97 |
| GO_T_CELL_RECEPTOR_SIGNALING_PATHWAY | 0.00000 | 0.00000 | 0.81404 | 0.69994 | 1.52527 | 112 |
| GO_U12_TYPE_SPLICEOSOMAL_COMPLEX | 0.00279 | 0.03889 | 0.43171 | 0.70746 | 1.44693 | 27 |
| GO_U2_SNRNP | 0.00005 | 0.00134 | 0.55733 | 0.83081 | 1.67168 | 18 |
| GO_U2_TYPE_SPLICEOSOMAL_COMPLEX | 0.00010 | 0.00243 | 0.53843 | 0.63403 | 1.36970 | 85 |
| GO_UBIQUITIN_LIKE_PROTEIN_LIGASE_BINDING | 0.00000 | 0.00000 | 0.76146 | 0.62273 | 1.38051 | 219 |
| GO_UBIQUITIN_PROTEIN_TRANSFERASE_REGULATOR_ACTIVITY | 0.00165 | 0.02590 | 0.45506 | 0.79605 | 1.57911 | 15 |
| GO_UNFOLDED_PROTEIN_BINDING | 0.00000 | 0.00000 | 0.77494 | 0.72157 | 1.55881 | 85 |
| GO_VESICLE_LUMEN | 0.00000 | 0.00000 | 0.74774 | 0.64475 | 1.42008 | 168 |
| GO_VIRAL_GENE_EXPRESSION | 0.00000 | 0.00000 | | 0.80028 | 1.76317 | 171 |
| GO_VIRAL_GENOME_REPLICATION | 0.00100 | 0.01761 | 0.45506 | 0.61997 | 1.33078 | 73 |
| GO_VIRAL_LIFE_CYCLE | 0.00031 | 0.00640 | 0.49849 | 0.55767 | 1.23509 | 209 |
| HP_ABNORMALITY_OF_ACID_BASE_HOMEOSTASIS | 0.00216 | 0.03242 | 0.43171 | 0.53556 | 1.18846 | 232 |
| HP_ABNORMALITY_OF_CARDIOVASCULAR_SYSTEM_ELECTROPHYSIO | 0.00027 | 0.00572 | 0.49849 | 0.55832 | 1.23929 | 229 |
| HP_ABNORMALITY_OF_LOWER_LIP | 0.00181 | 0.02804 | 0.45506 | 0.56360 | 1.23779 | 145 |
| HP_ABNORMALITY_OF_THE_MITOCHONDRION | 0.00213 | 0.03229 | 0.43171 | 0.57831 | 1.26240 | 117 |
| HP_ABNORMALITY_OF_THE_NASAL_DORSUM | 0.00005 | 0.00116 | 0.55733 | 0.60288 | 1.32157 | 134 |
| HP_ABNORMAL_ELECTROPHYSIOLOGY_OF_SINOATRIAL_NODE_ORIG | 0.00279 | 0.03889 | 0.43171 | 0.68436 | 1.40934 | 31 |
| HP_ABNORMAL_ERYTHROCYTE_MORPHOLOGY | 0.00000 | 0.00015 | 0.61053 | 0.56049 | 1.25036 | 341 |
| HP_ABNORMAL_MEAN_CORPUSCULAR_VOLUME | 0.00357 | 0.04696 | 0.43171 | 0.75226 | 1.51363 | 18 |
| HP_ABNORMAL_RETICULOCYTE_MORPHOLOGY | 0.00064 | 0.01210 | 0.47727 | 0.71420 | 1.47081 | 31 |
| HP_ACUTE_LEUKEMIA | 0.00000 | 0.00000 | 0.73376 | 0.72786 | 1.56137 | 70 |
| HP_ANEMIA_OF_INADEQUATE_PRODUCTION | 0.00000 | 0.00008 | 0.62726 | 0.65593 | 1.42209 | 97 |
| HP_CONSTITUTIONAL_SYMPTOM | 0.00119 | 0.02004 | 0.45506 | 0.51427 | 1.15108 | 423 |
| HP_DELAYED_PUBERTY | 0.00005 | 0.00115 | 0.55733 | 0.62579 | 1.36211 | 107 |
| HP_DEPRESSED_NASAL_RIDGE | 0.00000 | 0.00011 | 0.62726 | 0.68731 | 1.47511 | 72 |
| HP_FATIGUE | 0.00039 | 0.00808 | 0.49849 | 0.56529 | 1.24923 | 187 |

| | | | | | | |
|---|---|---|---|---|---|---|
| HP_HEADACHE | 0.00015 | 0.00340 | 0.51885 | 0.59065 | 1.29716 | 146 |
| HP_HEMATOLOGICAL_NEOPLASM | 0.00003 | 0.00078 | 0.57561 | 0.60108 | 1.32044 | 154 |
| HP_INCREASED_CSF_LACTATE | 0.00157 | 0.02509 | 0.45506 | 0.61622 | 1.31904 | 66 |
| HP_INCREASED_SERUM_LACTATE | 0.00216 | 0.03242 | 0.43171 | 0.56920 | 1.24724 | 130 |
| HP_LEUKEMIA | 0.00001 | 0.00026 | 0.59333 | 0.64877 | 1.40680 | 98 |
| HP_MACROCYTIC_ANEMIA | 0.00000 | 0.00000 | 0.74774 | 0.82468 | 1.71698 | 39 |
| HP_MIGRAINE | 0.00000 | 0.00000 | 0.71951 | 0.72105 | 1.54497 | 69 |
| HP_MYOTONIA | 0.00161 | 0.02546 | 0.45506 | 0.79820 | 1.58338 | 15 |
| HP_NEOPLASM | 0.00216 | 0.03242 | 0.43171 | 0.50820 | 1.13805 | 446 |
| HP_PALLOR | 0.00001 | 0.00040 | 0.59333 | 0.65169 | 1.41044 | 92 |
| HP_PUBERTY_AND_GONADAL_DISORDERS | 0.00287 | 0.03973 | 0.43171 | 0.51995 | 1.15843 | 322 |
| HP_RETICULOCYTOPENIA | 0.00004 | 0.00105 | 0.55733 | 0.86417 | 1.71425 | 15 |
| HP_RIMMED_VACUOLES | 0.00279 | 0.03889 | 0.43171 | 0.74033 | 1.49831 | 20 |
| HP_SHORT_NOSE | 0.00235 | 0.03474 | 0.43171 | 0.54919 | 1.21134 | 174 |
| HP_THICK_LOWER_LIP_VERMILION | 0.00048 | 0.00949 | 0.49849 | 0.60773 | 1.31617 | 94 |
| HP_THICK_VERMILION_BORDER | 0.00156 | 0.02486 | 0.45506 | 0.55983 | 1.23000 | 153 |
| HP_VENTRICULAR_ARRHYTHMIA | 0.00353 | 0.04658 | 0.43171 | 0.66198 | 1.37398 | 36 |
| **Cluster 1** | | | | | | |
| pathway | pval | padj | log2err | ES | NES | size |
| GO_ACTIVATION_OF_IMMUNE_RESPONSE | 0.00000 | 0.00000 | 0.78819 | 0.62851 | 1.39255 | 212 |
| GO_ACTIVATION_OF_INNATE_IMMUNE_RESPONSE | 0.00000 | 0.00000 | | 0.75054 | 1.62130 | 88 |
| GO_AEROBIC_RESPIRATION | 0.00001 | 0.00025 | 0.59333 | 0.69302 | 1.48168 | 65 |
| GO_ALTERNATIVE_MRNA_SPLICING_VIA_SPLICEOSOME | 0.00102 | 0.01782 | 0.45506 | 0.65168 | 1.38687 | 60 |
| GO_AMINE_METABOLIC_PROCESS | 0.00000 | 0.00000 | | 0.79189 | 1.70783 | 80 |
| GO_ANAPHASE_PROMOTING_COMPLEX_DEPENDENT_CATABOLIC_P | 0.00000 | 0.00000 | | 0.76611 | 1.65070 | 79 |
| GO_ANCHORING_JUNCTION | 0.00000 | 0.00000 | | 0.59615 | 1.33633 | 479 |
| GO_ANIMAL_ORGAN_REGENERATION | 0.00257 | 0.03716 | 0.43171 | 0.68240 | 1.41020 | 33 |
| GO_ANTIGEN_PROCESSING_AND_PRESENTATION | 0.00000 | 0.00000 | 0.70498 | 0.64776 | 1.42241 | 143 |
| GO_ANTIGEN_PROCESSING_AND_PRESENTATION_OF_EXOGENOUS_ | 0.00000 | 0.00000 | | 0.81051 | 1.72358 | 57 |
| GO_ANTIGEN_PROCESSING_AND_PRESENTATION_OF_PEPTIDE_ANTI | 0.00000 | 0.00000 | 0.70498 | 0.65912 | 1.44248 | 126 |
| GO_ANTIGEN_PROCESSING_AND_PRESENTATION_OF_PEPTIDE_ANTI | 0.00000 | 0.00000 | | 0.76925 | 1.65015 | 70 |
| GO_ANTIGEN_RECEPTOR_MEDIATED_SIGNALING_PATHWAY | 0.00000 | 0.00000 | 0.80122 | 0.67877 | 1.48641 | 127 |
| GO_APOPTOTIC_SIGNALING_PATHWAY | 0.00154 | 0.02464 | 0.45506 | 0.51561 | 1.15085 | 358 |
| GO_ATP_DEPENDENT_CHROMATIN_REMODELING | 0.00253 | 0.03681 | 0.43171 | 0.62124 | 1.32400 | 61 |
| GO_ATP_METABOLIC_PROCESS | 0.00000 | 0.00000 | 0.71951 | 0.62252 | 1.37693 | 196 |
| GO_ATP_SYNTHESIS_COUPLED_ELECTRON_TRANSPORT | 0.00005 | 0.00131 | 0.55733 | 0.65519 | 1.40723 | 74 |
| GO_BINDING_OF_SPERM_TO_ZONA_PELLUCIDA | 0.00000 | 0.00001 | 0.69013 | 0.89879 | 1.80846 | 18 |
| GO_CADHERIN_BINDING | 0.00000 | 0.00000 | | 0.63068 | 1.39961 | 236 |
| GO_CANONICAL_WNT_SIGNALING_PATHWAY | 0.00000 | 0.00003 | 0.65944 | 0.60816 | 1.34553 | 197 |
| GO_CATALYTIC_STEP_2_SPLICEOSOME | 0.00000 | 0.00002 | 0.65944 | 0.68068 | 1.46976 | 83 |
| GO_CELLULAR_AMINO_ACID_METABOLIC_PROCESS | 0.00000 | 0.00000 | 0.74774 | 0.61752 | 1.36824 | 216 |
| GO_CELLULAR_COMPONENT_DISASSEMBLY | 0.00000 | 0.00005 | 0.64355 | 0.55505 | 1.24197 | 401 |
| GO_CELLULAR_KETONE_METABOLIC_PROCESS | 0.00000 | 0.00000 | 0.77494 | 0.65955 | 1.44893 | 149 |
| GO_CELLULAR_NITROGEN_COMPOUND_CATABOLIC_PROCESS | 0.00000 | 0.00000 | | 0.74079 | 1.65848 | 436 |
| GO_CELLULAR_PROTEIN_COMPLEX_DISASSEMBLY | 0.00006 | 0.00139 | 0.55733 | 0.58222 | 1.28459 | 176 |
| GO_CELLULAR_RESPIRATION | 0.00000 | 0.00000 | 0.71951 | 0.64997 | 1.42492 | 136 |
| GO_CELLULAR_RESPONSE_TO_HEAT | 0.00018 | 0.00403 | 0.51885 | 0.61962 | 1.34016 | 91 |
| GO_CELLULAR_RESPONSE_TO_ORGANIC_CYCLIC_COMPOUND | 0.00302 | 0.04108 | 0.43171 | 0.52402 | 1.16585 | 283 |
| GO_CELLULAR_RESPONSE_TO_OXYGEN_LEVELS | 0.00000 | 0.00000 | 0.78819 | 0.65911 | 1.44882 | 157 |
| GO_CELL_ACTIVATION_INVOLVED_IN_IMMUNE_RESPONSE | 0.00000 | 0.00001 | 0.67496 | 0.57339 | 1.27913 | 341 |
| GO_CELL_ADHESION_MOLECULE_BINDING | 0.00000 | 0.00000 | 0.81404 | 0.60529 | 1.34788 | 308 |
| GO_CELL_CELL_RECOGNITION | 0.00000 | 0.00002 | 0.65944 | 0.83826 | 1.71003 | 26 |
| GO_CELL_CELL_SIGNALING_BY_WNT | 0.00000 | 0.00002 | 0.67496 | 0.57736 | 1.28559 | 299 |
| GO_CELL_CYCLE_G2_M_PHASE_TRANSITION | 0.00001 | 0.00024 | 0.59333 | 0.58034 | 1.28602 | 214 |
| GO_CELL_CYCLE_PHASE_TRANSITION | 0.00000 | 0.00004 | 0.64355 | 0.54319 | 1.21681 | 464 |
| GO_CELL_DIVISION | 0.00154 | 0.02464 | 0.45506 | 0.51551 | 1.15425 | 427 |
| GO_CELL_RECOGNITION | 0.00012 | 0.00270 | 0.53843 | 0.69554 | 1.46186 | 48 |
| GO_CELL_SUBSTRATE_JUNCTION | 0.00000 | 0.00000 | | 0.67751 | 1.50775 | 291 |
| GO_CELL_SURFACE_RECEPTOR_SIGNALING_PATHWAY_INVOLVED_IN | 0.00000 | 0.00003 | 0.65944 | 0.57035 | 1.27067 | 319 |
| GO_CHAPERONE_BINDING | 0.00007 | 0.00163 | 0.53843 | 0.64754 | 1.39253 | 76 |
| GO_CHAPERONE_COMPLEX | 0.00000 | 0.00004 | 0.64355 | 0.85718 | 1.73925 | 22 |
| GO_CHAPERONE_MEDIATED_PROTEIN_COMPLEX_ASSEMBLY | 0.00170 | 0.02662 | 0.45506 | 0.77423 | 1.54792 | 17 |
| GO_CHROMATIN_BINDING | 0.00071 | 0.01323 | 0.47727 | 0.52699 | 1.17680 | 369 |
| GO_CHROMOSOMAL_REGION | 0.00246 | 0.03601 | 0.43171 | 0.52919 | 1.17661 | 276 |
| GO_COP9_SIGNALOSOME | 0.00128 | 0.02141 | 0.45506 | 0.73616 | 1.49715 | 24 |
| GO_COTRANSLATIONAL_PROTEIN_TARGETING_TO_MEMBRANE | 0.00000 | 0.00000 | | 0.89471 | 1.93513 | 91 |
| GO_CYTOCHROME_COMPLEX | 0.00168 | 0.02641 | 0.45506 | 0.75311 | 1.52417 | 20 |
| GO_CYTOKINE_MEDIATED_SIGNALING_PATHWAY | 0.00000 | 0.00000 | 0.81404 | 0.59698 | 1.33170 | 339 |
| GO_CYTOPLASMIC_TRANSLATION | 0.00000 | 0.00000 | | 0.84753 | 1.83080 | 88 |
| GO_CYTOPLASMIC_TRANSLATIONAL_INITIATION | 0.00000 | 0.00000 | 0.74774 | 0.85688 | 1.76176 | 30 |
| GO_CYTOSOLIC_LARGE_RIBOSOMAL_SUBUNIT | 0.00000 | 0.00000 | | 0.91186 | 1.92988 | 53 |
| GO_CYTOSOLIC_RIBOSOME | 0.00000 | 0.00000 | | 0.89907 | 1.94710 | 95 |
| GO_CYTOSOLIC_SMALL_RIBOSOMAL_SUBUNIT | 0.00000 | 0.00000 | | 0.89114 | 1.85533 | 39 |
| GO_DEFENSE_RESPONSE_TO_OTHER_ORGANISM | 0.00004 | 0.00099 | 0.55733 | 0.53673 | 1.20127 | 415 |
| GO_DE_NOVO_PROTEIN_FOLDING | 0.00050 | 0.00978 | 0.47727 | 0.74232 | 1.51431 | 26 |
| GO_DISORDERED_DOMAIN_SPECIFIC_BINDING | 0.00161 | 0.02546 | 0.45506 | 0.71970 | 1.47197 | 27 |

| | | | | | | |
|---|---|---|---|---|---|---|
| GO_DNA_BIOSYNTHETIC_PROCESS | 0.00000 | 0.00010 | 0.62726 | 0.62505 | 1.37017 | 134 |
| GO_DNA_CONFORMATION_CHANGE | 0.00027 | 0.00572 | 0.49849 | 0.55300 | 1.22527 | 216 |
| GO_DNA_DAMAGE_RESPONSE_DETECTION_OF_DNA_DAMAGE | 0.00198 | 0.03028 | 0.43171 | 0.67949 | 1.40669 | 35 |
| GO_DNA_GEOMETRIC_CHANGE | 0.00046 | 0.00925 | 0.49849 | 0.60807 | 1.31855 | 98 |
| GO_DNA_POLYMERASE_BINDING | 0.00268 | 0.03799 | 0.43171 | 0.76255 | 1.52457 | 17 |
| GO_DNA_REPAIR | 0.00265 | 0.03775 | 0.43171 | 0.51159 | 1.14547 | 427 |
| GO_DRUG_BINDING | 0.00176 | 0.02733 | 0.45506 | 0.64119 | 1.35096 | 49 |
| GO_ELECTRON_TRANSFER_ACTIVITY | 0.00001 | 0.00020 | 0.61053 | 0.67481 | 1.44938 | 74 |
| GO_ELECTRON_TRANSPORT_CHAIN | 0.00002 | 0.00053 | 0.57561 | 0.62257 | 1.36145 | 124 |
| GO_ENDOPEPTIDASE_ACTIVITY | 0.00124 | 0.02085 | 0.45506 | 0.54955 | 1.21076 | 171 |
| GO_ENDOPEPTIDASE_COMPLEX | 0.00000 | 0.00000 | | 0.80930 | 1.73234 | 66 |
| GO_ENERGY_DERIVATION_BY_OXIDATION_OF_ORGANIC_COMPOUN | 0.00000 | 0.00000 | 0.71951 | 0.63390 | 1.39707 | 172 |
| GO_ESTABLISHMENT_OF_PROTEIN_LOCALIZATION_TO_CHROMOSOM | 0.00020 | 0.00432 | 0.51885 | 0.77339 | 1.57217 | 23 |
| GO_ESTABLISHMENT_OF_PROTEIN_LOCALIZATION_TO_ENDOPLASM | 0.00000 | 0.00000 | | 0.88026 | 1.91149 | 102 |
| GO_ESTABLISHMENT_OF_PROTEIN_LOCALIZATION_TO_MEMBRANE | 0.00000 | 0.00000 | | 0.74874 | 1.66164 | 240 |
| GO_ESTABLISHMENT_OF_PROTEIN_LOCALIZATION_TO_ORGANELLE | 0.00000 | 0.00000 | | 0.69519 | 1.55605 | 423 |
| GO_ESTABLISHMENT_OF_PROTEIN_LOCALIZATION_TO_TELOMERE | 0.00006 | 0.00139 | 0.55733 | 0.83470 | 1.66881 | 17 |
| GO_ESTABLISHMENT_OF_RNA_LOCALIZATION | 0.00106 | 0.01819 | 0.45506 | 0.55577 | 1.22400 | 167 |
| GO_ESTABLISHMENT_OF_TISSUE_POLARITY | 0.00000 | 0.00000 | | 0.75051 | 1.62122 | 88 |
| GO_EUKARYOTIC_TRANSLATION_INITIATION_FACTOR_3_COMPLEX | 0.00003 | 0.00074 | 0.57561 | 0.86987 | 1.72555 | 15 |
| GO_EXOCYTOSIS | 0.00000 | 0.00015 | 0.61053 | 0.53905 | 1.20753 | 465 |
| GO_FC_EPSILON_RECEPTOR_SIGNALING_PATHWAY | 0.00000 | 0.00000 | | 0.75117 | 1.62103 | 82 |
| GO_FC_RECEPTOR_SIGNALING_PATHWAY | 0.00000 | 0.00000 | | 0.72054 | 1.57787 | 127 |
| GO_FERTILIZATION | 0.00089 | 0.01592 | 0.47727 | 0.63620 | 1.35888 | 64 |
| GO_FICOLIN_1_RICH_GRANULE | 0.00000 | 0.00000 | 0.78819 | 0.71716 | 1.55340 | 93 |
| GO_GENERATION_OF_PRECURSOR_METABOLITES_AND_ENERGY | 0.00000 | 0.00000 | 0.77494 | 0.59261 | 1.32026 | 320 |
| GO_GENE_SILENCING | 0.00261 | 0.03750 | 0.43171 | 0.55578 | 1.22252 | 161 |
| GO_GLAND_DEVELOPMENT | 0.00154 | 0.02464 | 0.45506 | 0.54117 | 1.19797 | 206 |
| GO_GLYCOSYL_COMPOUND_METABOLIC_PROCESS | 0.00071 | 0.01323 | 0.47727 | 0.62524 | 1.34204 | 71 |
| GO_GUANYL_NUCLEOTIDE_BINDING | 0.00095 | 0.01679 | 0.47727 | 0.54096 | 1.19861 | 216 |
| GO_HEAT_SHOCK_PROTEIN_BINDING | 0.00003 | 0.00092 | 0.55733 | 0.64369 | 1.39056 | 85 |
| GO_HEMATOPOIETIC_PROGENITOR_CELL_DIFFERENTIATION | 0.00000 | 0.00000 | 0.82666 | 0.70307 | 1.53239 | 113 |
| GO_HEMATOPOIETIC_STEM_CELL_DIFFERENTIATION | 0.00000 | 0.00000 | | 0.79776 | 1.71217 | 72 |
| GO_HSP70_PROTEIN_BINDING | 0.00228 | 0.03391 | 0.43171 | 0.71143 | 1.45504 | 27 |
| GO_IMMUNE_RESPONSE_REGULATING_SIGNALING_PATHWAY | 0.00000 | 0.00000 | 0.76146 | 0.63494 | 1.40480 | 197 |
| GO_INNATE_IMMUNE_RESPONSE | 0.00003 | 0.00071 | 0.57561 | 0.55173 | 1.23107 | 347 |
| GO_INNATE_IMMUNE_RESPONSE_ACTIVATING_SIGNAL_TRANSDUCT | 0.00000 | 0.00000 | | 0.79340 | 1.70281 | 72 |
| GO_INNER_MITOCHONDRIAL_MEMBRANE_PROTEIN_COMPLEX | 0.00002 | 0.00060 | 0.57561 | 0.65124 | 1.40678 | 88 |
| GO_INTERACTION_WITH_SYMBIONT | 0.00001 | 0.00026 | 0.59333 | 0.75084 | 1.56620 | 41 |
| GO_INTERLEUKIN_1_MEDIATED_SIGNALING_PATHWAY | 0.00000 | 0.00000 | | 0.81689 | 1.75671 | 76 |
| GO_INTRINSIC_APOPTOTIC_SIGNALING_PATHWAY | 0.00121 | 0.02029 | 0.45506 | 0.55118 | 1.21845 | 190 |
| GO_ION_CHANNEL_BINDING | 0.00087 | 0.01584 | 0.47727 | 0.63182 | 1.34683 | 62 |
| GO_ISOPEPTIDASE_ACTIVITY | 0.00139 | 0.02285 | 0.45506 | 0.78940 | 1.57097 | 16 |
| GO_LARGE_RIBOSOMAL_SUBUNIT | 0.00000 | 0.00000 | | 0.78135 | 1.70183 | 109 |
| GO_LEUKOCYTE_MEDIATED_IMMUNITY | 0.00000 | 0.00000 | 0.71951 | 0.57282 | 1.27895 | 364 |
| GO_MACROMOLECULE_TRANSMEMBRANE_TRANSPORTER_ACTIVITY | 0.00239 | 0.03510 | 0.43171 | 0.77734 | 1.54696 | 16 |
| GO_MAINTENANCE_OF_CELL_NUMBER | 0.00087 | 0.01584 | 0.47727 | 0.58944 | 1.28512 | 114 |
| GO_MATURATION_OF_LSU_RRNA | 0.00006 | 0.00151 | 0.53843 | 0.77606 | 1.58723 | 27 |
| GO_METHYLTRANSFERASE_COMPLEX | 0.00290 | 0.04014 | 0.43171 | 0.59476 | 1.28479 | 88 |
| GO_METHYL_CPG_BINDING | 0.00302 | 0.04108 | 0.43171 | 0.78053 | 1.54833 | 15 |
| GO_MIDBRAIN_DEVELOPMENT | 0.00021 | 0.00468 | 0.51885 | 0.68832 | 1.44457 | 47 |
| GO_MITOCHONDRIAL_ELECTRON_TRANSPORT_NADH_TO_UBIQUIN | 0.00331 | 0.04437 | 0.43171 | 0.65428 | 1.36724 | 42 |
| GO_MITOCHONDRIAL_GENE_EXPRESSION | 0.00048 | 0.00949 | 0.49849 | 0.57608 | 1.26525 | 152 |
| GO_MITOCHONDRIAL_LARGE_RIBOSOMAL_SUBUNIT | 0.00002 | 0.00050 | 0.57561 | 0.69789 | 1.48086 | 55 |
| GO_MITOCHONDRIAL_MATRIX | 0.00000 | 0.00002 | 0.65944 | 0.56045 | 1.25158 | 371 |
| GO_MITOCHONDRIAL_MEMBRANE_ORGANIZATION | 0.00361 | 0.04734 | 0.43171 | 0.59309 | 1.28094 | 86 |
| GO_MITOCHONDRIAL_PROTEIN_COMPLEX | 0.00000 | 0.00000 | 0.81404 | 0.63650 | 1.40931 | 208 |
| GO_MITOCHONDRIAL_TRANSLATION | 0.00004 | 0.00093 | 0.55733 | 0.61219 | 1.34135 | 129 |
| GO_MITOCHONDRIAL_TRANSLATIONAL_TERMINATION | 0.00000 | 0.00010 | 0.62726 | 0.66694 | 1.44128 | 87 |
| GO_MITOCHONDRION_ORGANIZATION | 0.00051 | 0.00993 | 0.47727 | 0.52284 | 1.16863 | 390 |
| GO_MODIFICATION_DEPENDENT_MACROMOLECULE_CATABOLIC_PR | 0.00002 | 0.00053 | 0.57561 | 0.53142 | 1.19156 | 486 |
| GO_MODULATION_BY_HOST_OF_SYMBIONT_PROCESS | 0.00015 | 0.00336 | 0.51885 | 0.72848 | 1.50542 | 33 |
| GO_MODULATION_OF_PROCESS_OF_OTHER_ORGANISM | 0.00019 | 0.00423 | 0.51885 | 0.66904 | 1.42286 | 59 |
| GO_MONOSACCHARIDE_BIOSYNTHETIC_PROCESS | 0.00335 | 0.04466 | 0.43171 | 0.61992 | 1.31781 | 58 |
| GO_MORPHOGENESIS_OF_AN_EPITHELIUM | 0.00006 | 0.00151 | 0.53843 | 0.55161 | 1.22755 | 290 |
| GO_MORPHOGENESIS_OF_A_POLARIZED_EPITHELIUM | 0.00000 | 0.00000 | | 0.72298 | 1.57057 | 103 |
| GO_MRNA_3_UTR_BINDING | 0.00003 | 0.00089 | 0.55733 | 0.67596 | 1.44827 | 68 |
| GO_MRNA_5_UTR_BINDING | 0.00006 | 0.00156 | 0.53843 | 0.80133 | 1.62592 | 22 |
| GO_MRNA_BINDING | 0.00000 | 0.00000 | | 0.66002 | 1.46353 | 222 |
| GO_MRNA_PROCESSING | 0.00000 | 0.00000 | 0.73376 | 0.56859 | 1.27295 | 418 |
| GO_MULTI_ORGANISM_LOCALIZATION | 0.00268 | 0.03799 | 0.43171 | 0.63245 | 1.33612 | 51 |
| GO_MULTI_ORGANISM_PROCESS | 0.00108 | 0.01839 | 0.45506 | 0.51114 | 1.14599 | 484 |
| GO_MYD88_DEPENDENT_TOLL_LIKE_RECEPTOR_SIGNALING_PATHW | 0.00002 | 0.00066 | 0.57561 | 0.85581 | 1.70312 | 16 |
| GO_MYELOID_LEUKOCYTE_ACTIVATION | 0.00000 | 0.00000 | 0.74774 | 0.58508 | 1.30479 | 327 |
| GO_MYELOID_LEUKOCYTE_MEDIATED_IMMUNITY | 0.00000 | 0.00000 | 0.76146 | 0.59841 | 1.33291 | 301 |
| GO_NADH_DEHYDROGENASE_ACTIVITY | 0.00383 | 0.04935 | 0.43171 | 0.66244 | 1.37139 | 35 |

| | | | | | | |
|---|---|---|---|---|---|---|
| GO_NADH_DEHYDROGENASE_COMPLEX | 0.00376 | 0.04884 | 0.43171 | 0.65649 | 1.36421 | 38 |
| GO_NADH_METABOLIC_PROCESS | 0.00294 | 0.04036 | 0.43171 | 0.70642 | 1.44481 | 27 |
| GO_NCRNA_EXPORT_FROM_NUCLEUS | 0.00302 | 0.04108 | 0.43171 | 0.66820 | 1.38331 | 35 |
| GO_NCRNA_METABOLIC_PROCESS | 0.00000 | 0.00003 | 0.65944 | 0.55907 | 1.24907 | 381 |
| GO_NCRNA_PROCESSING | 0.00000 | 0.00003 | 0.65944 | 0.57175 | 1.27504 | 328 |
| GO_NEGATIVE_REGULATION_OF_CANONICAL_WNT_SIGNALING_PAT | 0.00000 | 0.00000 | | 0.71611 | 1.56038 | 111 |
| GO_NEGATIVE_REGULATION_OF_CELLULAR_AMIDE_METABOLIC_PR | 0.00001 | 0.00020 | 0.61053 | 0.63031 | 1.37423 | 114 |
| GO_NEGATIVE_REGULATION_OF_CELL_CYCLE | 0.00003 | 0.00076 | 0.57561 | 0.53833 | 1.20513 | 425 |
| GO_NEGATIVE_REGULATION_OF_CELL_CYCLE_G2_M_PHASE_TRANS | 0.00000 | 0.00000 | 0.76146 | 0.71490 | 1.54439 | 85 |
| GO_NEGATIVE_REGULATION_OF_CELL_CYCLE_PHASE_TRANSITION | 0.00002 | 0.00060 | 0.57561 | 0.59041 | 1.30442 | 184 |
| GO_NEGATIVE_REGULATION_OF_CELL_CYCLE_PROCESS | 0.00000 | 0.00002 | 0.65944 | 0.58690 | 1.30374 | 249 |
| GO_NEGATIVE_REGULATION_OF_MITOTIC_CELL_CYCLE | 0.00004 | 0.00095 | 0.55733 | 0.56735 | 1.25902 | 232 |
| GO_NEGATIVE_REGULATION_OF_MRNA_METABOLIC_PROCESS | 0.00023 | 0.00491 | 0.51885 | 0.66670 | 1.41788 | 59 |
| GO_NEGATIVE_REGULATION_OF_MRNA_SPLICING_VIA_SPLICEOSOM | 0.00239 | 0.03510 | 0.43171 | 0.74857 | 1.51204 | 19 |
| GO_NEGATIVE_REGULATION_OF_RNA_SPLICING | 0.00029 | 0.00607 | 0.49849 | 0.76729 | 1.55976 | 23 |
| GO_NEGATIVE_REGULATION_OF_TRANSLATIONAL_INITIATION | 0.00018 | 0.00403 | 0.51885 | 0.83901 | 1.66434 | 15 |
| GO_NEGATIVE_REGULATION_OF_WNT_SIGNALING_PATHWAY | 0.00000 | 0.00000 | | 0.69525 | 1.52249 | 127 |
| GO_NEURAL_NUCLEUS_DEVELOPMENT | 0.00001 | 0.00026 | 0.59333 | 0.79148 | 1.62187 | 28 |
| GO_NEURON_TO_NEURON_SYNAPSE | 0.00005 | 0.00134 | 0.55733 | 0.59636 | 1.31006 | 154 |
| GO_NIK_NF_KAPPAB_SIGNALING | 0.00000 | 0.00000 | | 0.72896 | 1.58354 | 103 |
| GO_NON_CANONICAL_WNT_SIGNALING_PATHWAY | 0.00000 | 0.00000 | | 0.72784 | 1.58023 | 101 |
| GO_NUCLEAR_EXPORT | 0.00031 | 0.00645 | 0.49849 | 0.56963 | 1.25542 | 172 |
| GO_NUCLEAR_TRANSCRIBED_MRNA_CATABOLIC_PROCESS | 0.00000 | 0.00000 | | 0.79179 | 1.74917 | 183 |
| GO_NUCLEAR_TRANSCRIBED_MRNA_CATABOLIC_PROCESS_NONSEN | 0.00000 | 0.00000 | | 0.89008 | 1.93909 | 110 |
| GO_NUCLEAR_TRANSPORT | 0.00008 | 0.00193 | 0.53843 | 0.55511 | 1.23408 | 273 |
| GO_NUCLEOBASE_CONTAINING_COMPOUND_TRANSPORT | 0.00146 | 0.02386 | 0.45506 | 0.55053 | 1.21842 | 202 |
| GO_NUCLEOBASE_CONTAINING_SMALL_MOLECULE_BIOSYNTHETIC_ | 0.00137 | 0.02261 | 0.45506 | 0.62749 | 1.34353 | 67 |
| GO_NUCLEOBASE_CONTAINING_SMALL_MOLECULE_INTERCONVERS | 0.00383 | 0.04935 | 0.43171 | 0.71259 | 1.44857 | 23 |
| GO_NUCLEOBASE_CONTAINING_SMALL_MOLECULE_METABOLIC_PR | 0.00320 | 0.04339 | 0.43171 | 0.51448 | 1.14825 | 352 |
| GO_NUCLEOSIDE_DIPHOSPHATE_METABOLIC_PROCESS | 0.00110 | 0.01865 | 0.45506 | 0.59637 | 1.29295 | 97 |
| GO_NUCLEOSIDE_METABOLIC_PROCESS | 0.00061 | 0.01169 | 0.47727 | 0.65315 | 1.38907 | 59 |
| GO_NUCLEOSIDE_MONOPHOSPHATE_BIOSYNTHETIC_PROCESS | 0.00250 | 0.03636 | 0.43171 | 0.68841 | 1.42033 | 32 |
| GO_NUCLEOSIDE_MONOPHOSPHATE_METABOLIC_PROCESS | 0.00152 | 0.02464 | 0.45506 | 0.64377 | 1.36002 | 51 |
| GO_NUCLEOSIDE_PHOSPHATE_BIOSYNTHETIC_PROCESS | 0.00043 | 0.00865 | 0.49849 | 0.58185 | 1.27831 | 148 |
| GO_NUCLEOSIDE_TRIPHOSPHATE_BIOSYNTHETIC_PROCESS | 0.00043 | 0.00865 | 0.49849 | 0.69332 | 1.44539 | 40 |
| GO_NUCLEOSIDE_TRIPHOSPHATE_METABOLIC_PROCESS | 0.00016 | 0.00360 | 0.51885 | 0.66788 | 1.41976 | 58 |
| GO_NUCLEOSOMAL_DNA_BINDING | 0.00023 | 0.00507 | 0.51885 | 0.79852 | 1.61801 | 21 |
| GO_NUCLEOSOME_BINDING | 0.00143 | 0.02332 | 0.45506 | 0.66132 | 1.38492 | 45 |
| GO_NUCLEOTIDE_EXCISION_REPAIR | 0.00004 | 0.00096 | 0.55733 | 0.63343 | 1.37355 | 98 |
| GO_NUCLEOTIDE_EXCISION_REPAIR_DNA_DAMAGE_RECOGNITION | 0.00015 | 0.00349 | 0.51885 | 0.78678 | 1.59640 | 22 |
| GO_NUCLEOTIDE_EXCISION_REPAIR_DNA_INCISION | 0.00313 | 0.04249 | 0.43171 | 0.66439 | 1.37898 | 36 |
| GO_NUCLEOTIDE_PHOSPHORYLATION | 0.00383 | 0.04935 | 0.43171 | 0.59002 | 1.27454 | 88 |
| GO_ORGANELLAR_RIBOSOME | 0.00000 | 0.00014 | 0.61053 | 0.66412 | 1.43471 | 85 |
| GO_ORGANELLE_INNER_MEMBRANE | 0.00000 | 0.00000 | 0.76146 | 0.57601 | 1.28697 | 378 |
| GO_ORGANIC_CYCLIC_COMPOUND_CATABOLIC_PROCESS | 0.00000 | 0.00000 | | 0.73350 | 1.64281 | 449 |
| GO_OXIDATIVE_PHOSPHORYLATION | 0.00001 | 0.00030 | 0.59333 | 0.64619 | 1.40027 | 96 |
| GO_OXIDOREDUCTASE_ACTIVITY | 0.00006 | 0.00139 | 0.55733 | 0.53544 | 1.19703 | 392 |
| GO_OXIDOREDUCTASE_COMPLEX | 0.00001 | 0.00027 | 0.59333 | 0.66015 | 1.42461 | 82 |
| GO_PEPTIDASE_COMPLEX | 0.00000 | 0.00000 | | 0.75988 | 1.63982 | 82 |
| GO_PIGMENT_GRANULE | 0.00003 | 0.00089 | 0.55733 | 0.66098 | 1.42344 | 78 |
| GO_POLYSOMAL_RIBOSOME | 0.00000 | 0.00000 | | 0.90624 | 1.86629 | 31 |
| GO_POLYSOME | 0.00000 | 0.00000 | | 0.82337 | 1.75481 | 61 |
| GO_POLY_PYRIMIDINE_TRACT_BINDING | 0.00327 | 0.04398 | 0.43171 | 0.75471 | 1.51855 | 18 |
| GO_POSITIVE_REGULATION_OF_APOPTOTIC_SIGNALING_PATHWAY | 0.00279 | 0.03889 | 0.43171 | 0.57666 | 1.25518 | 107 |
| GO_POSITIVE_REGULATION_OF_ATPASE_ACTIVITY | 0.00294 | 0.04036 | 0.43171 | 0.65446 | 1.36795 | 43 |
| GO_POSITIVE_REGULATION_OF_BINDING | 0.00088 | 0.01590 | 0.47727 | 0.60284 | 1.30556 | 95 |
| GO_POSITIVE_REGULATION_OF_CANONICAL_WNT_SIGNALING_PATH | 0.00000 | 0.00000 | 0.77494 | 0.70504 | 1.53160 | 103 |
| GO_POSITIVE_REGULATION_OF_CELLULAR_AMIDE_METABOLIC_PRO | 0.00003 | 0.00074 | 0.57561 | 0.62296 | 1.36083 | 118 |
| GO_POSITIVE_REGULATION_OF_CELLULAR_PROTEIN_LOCALIZATION | 0.00041 | 0.00840 | 0.49849 | 0.54915 | 1.21815 | 225 |
| GO_POSITIVE_REGULATION_OF_CELL_DEATH | 0.00069 | 0.01297 | 0.47727 | 0.52595 | 1.17423 | 363 |
| GO_POSITIVE_REGULATION_OF_CELL_POPULATION_PROLIFERATION | 0.00078 | 0.01455 | 0.47727 | 0.52449 | 1.17108 | 368 |
| GO_POSITIVE_REGULATION_OF_CHROMOSOME_ORGANIZATION | 0.00231 | 0.03437 | 0.43171 | 0.56990 | 1.24871 | 133 |
| GO_POSITIVE_REGULATION_OF_DEFENSE_RESPONSE | 0.00000 | 0.00003 | 0.65944 | 0.62175 | 1.36730 | 158 |
| GO_POSITIVE_REGULATION_OF_DNA_BIOSYNTHETIC_PROCESS | 0.00001 | 0.00041 | 0.59333 | 0.71929 | 1.52055 | 52 |
| GO_POSITIVE_REGULATION_OF_DNA_METABOLIC_PROCESS | 0.00364 | 0.04761 | 0.43171 | 0.56647 | 1.24189 | 135 |
| GO_POSITIVE_REGULATION_OF_DNA_TEMPLATED_TRANSCRIPTION_ | 0.00276 | 0.03885 | 0.43171 | 0.68932 | 1.41586 | 29 |
| GO_POSITIVE_REGULATION_OF_ESTABLISHMENT_OF_PROTEIN_LOC | 0.00007 | 0.00168 | 0.53843 | 0.56461 | 1.25165 | 219 |
| GO_POSITIVE_REGULATION_OF_IMMUNE_RESPONSE | 0.00000 | 0.00001 | 0.69013 | 0.59037 | 1.31265 | 277 |
| GO_POSITIVE_REGULATION_OF_IMMUNE_SYSTEM_PROCESS | 0.00004 | 0.00110 | 0.55733 | 0.54018 | 1.20716 | 386 |
| GO_POSITIVE_REGULATION_OF_INNATE_IMMUNE_RESPONSE | 0.00000 | 0.00000 | 0.81404 | 0.69693 | 1.51947 | 114 |
| GO_POSITIVE_REGULATION_OF_INTRINSIC_APOPTOTIC_SIGNALING_ | 0.00104 | 0.01798 | 0.45506 | 0.69097 | 1.43578 | 37 |
| GO_POSITIVE_REGULATION_OF_ORGANELLE_ORGANIZATION | 0.00043 | 0.00865 | 0.49849 | 0.52093 | 1.16605 | 411 |
| GO_POSITIVE_REGULATION_OF_PROTEIN_LOCALIZATION_TO_NUCLE | 0.00001 | 0.00020 | 0.61053 | 0.69907 | 1.48990 | 61 |
| GO_POSITIVE_REGULATION_OF_RESPONSE_TO_BIOTIC_STIMULUS | 0.00000 | 0.00000 | 0.73376 | 0.66340 | 1.45310 | 128 |
| GO_POSITIVE_REGULATION_OF_RESPONSE_TO_EXTERNAL_STIMULU | 0.00002 | 0.00050 | 0.57561 | 0.58133 | 1.28681 | 201 |
| GO_POSITIVE_REGULATION_OF_TELOMERASE_ACTIVITY | 0.00061 | 0.01169 | 0.47727 | 0.73476 | 1.50565 | 28 |

| | | | | | | |
|---|---|---|---|---|---|---|
| GO_POSITIVE_REGULATION_OF_TELOMERASE_RNA_LOCALIZATION_ | 0.00000 | 0.00001 | 0.69013 | 0.92451 | 1.83395 | 15 |
| GO_POSITIVE_REGULATION_OF_TELOMERE_MAINTENANCE_VIA_TE | 0.00130 | 0.02159 | 0.45506 | 0.71259 | 1.46508 | 30 |
| GO_POSITIVE_REGULATION_OF_TRANSCRIPTION_INITIATION_FROM | 0.00372 | 0.04847 | 0.43171 | 0.70671 | 1.43993 | 25 |
| GO_POSITIVE_REGULATION_OF_TRANSFERASE_ACTIVITY | 0.00174 | 0.02712 | 0.45506 | 0.51642 | 1.15220 | 345 |
| GO_POSITIVE_REGULATION_OF_TRANSLATION | 0.00001 | 0.00019 | 0.61053 | 0.64767 | 1.40419 | 97 |
| GO_POSITIVE_REGULATION_OF_VIRAL_GENOME_REPLICATION | 0.00287 | 0.03973 | 0.43171 | 0.68362 | 1.40783 | 31 |
| GO_POSITIVE_REGULATION_OF_VIRAL_PROCESS | 0.00028 | 0.00590 | 0.49849 | 0.62440 | 1.34741 | 81 |
| GO_POSITIVE_REGULATION_OF_WNT_SIGNALING_PATHWAY | 0.00000 | 0.00000 | 0.73376 | 0.66230 | 1.44763 | 122 |
| GO_POSTREPLICATION_REPAIR | 0.00024 | 0.00515 | 0.51885 | 0.69140 | 1.44518 | 43 |
| GO_POSTSYNAPSE | 0.00004 | 0.00109 | 0.55733 | 0.55969 | 1.24369 | 259 |
| GO_POSTSYNAPTIC_CYTOSOL | 0.00279 | 0.03889 | 0.43171 | 0.78295 | 1.55313 | 15 |
| GO_POSTTRANSCRIPTIONAL_REGULATION_OF_GENE_EXPRESSION | 0.00000 | 0.00000 | | 0.66656 | 1.49327 | 462 |
| GO_POST_TRANSLATIONAL_PROTEIN_MODIFICATION | 0.00002 | 0.00052 | 0.57561 | 0.57851 | 1.28412 | 229 |
| GO_PRECATALYTIC_SPLICEOSOME | 0.00085 | 0.01559 | 0.47727 | 0.65235 | 1.37448 | 49 |
| GO_PRERIBOSOME | 0.00087 | 0.01577 | 0.47727 | 0.62899 | 1.34772 | 69 |
| GO_PRERIBOSOME_LARGE_SUBUNIT_PRECURSOR | 0.00235 | 0.03474 | 0.43171 | 0.74079 | 1.50308 | 22 |
| GO_PRE_MRNA_BINDING | 0.00133 | 0.02213 | 0.45506 | 0.72393 | 1.48062 | 27 |
| GO_PROTEASOMAL_PROTEIN_CATABOLIC_PROCESS | 0.00002 | 0.00044 | 0.59333 | 0.54166 | 1.21005 | 377 |
| GO_PROTEASOMAL_UBIQUITIN_INDEPENDENT_PROTEIN_CATABOLIC | 0.00000 | 0.00000 | 0.80122 | 0.92254 | 1.86931 | 21 |
| GO_PROTEASOME_ACCESSORY_COMPLEX | 0.00000 | 0.00001 | 0.67496 | 0.85464 | 1.73811 | 24 |
| GO_PROTEASOME_CORE_COMPLEX | 0.00000 | 0.00000 | | 0.94375 | 1.90631 | 19 |
| GO_PROTEIN_CONTAINING_COMPLEX_DISASSEMBLY | 0.00000 | 0.00007 | 0.62726 | 0.57810 | 1.28438 | 256 |
| GO_PROTEIN_CONTAINING_COMPLEX_LOCALIZATION | 0.00063 | 0.01197 | 0.47727 | 0.55335 | 1.22520 | 208 |
| GO_PROTEIN_DNA_COMPLEX_SUBUNIT_ORGANIZATION | 0.00001 | 0.00018 | 0.61053 | 0.60174 | 1.32491 | 165 |
| GO_PROTEIN_FOLDING | 0.00000 | 0.00002 | 0.65944 | 0.63113 | 1.38644 | 154 |
| GO_PROTEIN_FOLDING_CHAPERONE | 0.00135 | 0.02238 | 0.45506 | 0.77903 | 1.55752 | 17 |
| GO_PROTEIN_IMPORT | 0.00268 | 0.03799 | 0.43171 | 0.56317 | 1.23581 | 140 |
| GO_PROTEIN_LOCALIZATION_TO_CHROMOSOME | 0.00056 | 0.01096 | 0.47727 | 0.63523 | 1.36108 | 69 |
| GO_PROTEIN_LOCALIZATION_TO_CHROMOSOME_TELOMERIC_REGIO | 0.00335 | 0.04466 | 0.43171 | 0.70399 | 1.43984 | 27 |
| GO_PROTEIN_LOCALIZATION_TO_ENDOPLASMIC_RETICULUM | 0.00000 | 0.00000 | | 0.85521 | 1.87021 | 124 |
| GO_PROTEIN_LOCALIZATION_TO_MEMBRANE | 0.00000 | 0.00000 | | 0.65123 | 1.45714 | 406 |
| GO_PROTEIN_LOCALIZATION_TO_MITOCHONDRION | 0.00019 | 0.00420 | 0.51885 | 0.60773 | 1.32398 | 110 |
| GO_PROTEIN_LOCALIZATION_TO_NUCLEUS | 0.00002 | 0.00055 | 0.57561 | 0.58926 | 1.30220 | 187 |
| GO_PROTEIN_MODIFICATION_BY_SMALL_PROTEIN_REMOVAL | 0.00000 | 0.00000 | 0.71951 | 0.60745 | 1.34689 | 221 |
| GO_PROTEIN_POLYUBIQUITINATION | 0.00004 | 0.00106 | 0.55733 | 0.55874 | 1.24108 | 253 |
| GO_PROTEIN_REFOLDING | 0.00294 | 0.04036 | 0.43171 | 0.77039 | 1.53314 | 16 |
| GO_PROTEIN_STABILIZATION | 0.00001 | 0.00024 | 0.61053 | 0.62333 | 1.36586 | 130 |
| GO_PROTEIN_TARGETING | 0.00000 | 0.00000 | | 0.71181 | 1.58743 | 327 |
| GO_PROTEIN_TARGETING_TO_MEMBRANE | 0.00000 | 0.00000 | | 0.82370 | 1.80953 | 149 |
| GO_PROTEIN_TARGETING_TO_MITOCHONDRION | 0.00346 | 0.04593 | 0.43171 | 0.60408 | 1.30091 | 78 |
| GO_PROTEIN_TRANSMEMBRANE_TRANSPORT | 0.00265 | 0.03775 | 0.43171 | 0.64528 | 1.36171 | 50 |
| GO_PROTON_TRANSMEMBRANE_TRANSPORT | 0.00213 | 0.03229 | 0.43171 | 0.60815 | 1.30536 | 71 |
| GO_PURINE_CONTAINING_COMPOUND_BIOSYNTHETIC_PROCESS | 0.00246 | 0.03601 | 0.43171 | 0.57796 | 1.25936 | 111 |
| GO_PURINE_CONTAINING_COMPOUND_METABOLIC_PROCESS | 0.00100 | 0.01761 | 0.45506 | 0.53679 | 1.19242 | 255 |
| GO_PURINE_NUCLEOSIDE_TRIPHOSPHATE_METABOLIC_PROCESS | 0.00089 | 0.01592 | 0.47727 | 0.67157 | 1.40374 | 43 |
| GO_PYRIMIDINE_CONTAINING_COMPOUND_METABOLIC_PROCESS | 0.00353 | 0.04658 | 0.43171 | 0.64744 | 1.35878 | 47 |
| GO_REGENERATION | 0.00216 | 0.03242 | 0.43171 | 0.59987 | 1.29453 | 82 |
| GO_REGULATION_OF_ANIMAL_ORGAN_MORPHOGENESIS | 0.00000 | 0.00000 | | 0.69163 | 1.51541 | 133 |
| GO_REGULATION_OF_APOPTOTIC_SIGNALING_PATHWAY | 0.00031 | 0.00645 | 0.49849 | 0.55171 | 1.22506 | 247 |
| GO_REGULATION_OF_ATPASE_ACTIVITY | 0.00324 | 0.04369 | 0.43171 | 0.63031 | 1.34038 | 57 |
| GO_REGULATION_OF_ATP_METABOLIC_PROCESS | 0.00353 | 0.04658 | 0.43171 | 0.60176 | 1.29780 | 80 |
| GO_REGULATION_OF_BINDING | 0.00250 | 0.03636 | 0.43171 | 0.54476 | 1.20598 | 203 |
| GO_REGULATION_OF_CELLULAR_AMIDE_METABOLIC_PROCESS | 0.00000 | 0.00000 | | 0.62967 | 1.40226 | 311 |
| GO_REGULATION_OF_CELLULAR_AMINE_METABOLIC_PROCESS | 0.00000 | 0.00000 | | 0.82173 | 1.74682 | 58 |
| GO_REGULATION_OF_CELLULAR_AMINO_ACID_METABOLIC_PROCES | 0.00000 | 0.00000 | | 0.83776 | 1.77766 | 55 |
| GO_REGULATION_OF_CELLULAR_KETONE_METABOLIC_PROCESS | 0.00000 | 0.00000 | | 0.72641 | 1.58680 | 118 |
| GO_REGULATION_OF_CELLULAR_PROTEIN_LOCALIZATION | 0.00082 | 0.01513 | 0.47727 | 0.52542 | 1.17290 | 362 |
| GO_REGULATION_OF_CELLULAR_RESPONSE_TO_HEAT | 0.00276 | 0.03885 | 0.43171 | 0.61652 | 1.32004 | 67 |
| GO_REGULATION_OF_CELLULAR_RESPONSE_TO_STRESS | 0.00224 | 0.03344 | 0.43171 | 0.50545 | 1.13232 | 468 |
| GO_REGULATION_OF_CELL_CYCLE_G2_M_PHASE_TRANSITION | 0.00000 | 0.00000 | 0.71951 | 0.63230 | 1.39282 | 169 |
| GO_REGULATION_OF_CELL_CYCLE_PHASE_TRANSITION | 0.00001 | 0.00026 | 0.59333 | 0.55488 | 1.23783 | 337 |
| GO_REGULATION_OF_CHROMOSOME_ORGANIZATION | 0.00081 | 0.01501 | 0.47727 | 0.54255 | 1.20527 | 261 |
| GO_REGULATION_OF_CYSTEINE_TYPE_ENDOPEPTIDASE_ACTIVITY | 0.00202 | 0.03077 | 0.43171 | 0.57120 | 1.24837 | 121 |
| GO_REGULATION_OF_CYTOPLASMIC_TRANSLATION | 0.00130 | 0.02159 | 0.45506 | 0.73678 | 1.49773 | 23 |
| GO_REGULATION_OF_DEFENSE_RESPONSE | 0.00065 | 0.01237 | 0.47727 | 0.53633 | 1.19355 | 290 |
| GO_REGULATION_OF_DNA_BIOSYNTHETIC_PROCESS | 0.00016 | 0.00360 | 0.51885 | 0.64215 | 1.37923 | 74 |
| GO_REGULATION_OF_DNA_METABOLIC_PROCESS | 0.00069 | 0.01297 | 0.47727 | 0.54425 | 1.20775 | 237 |
| GO_REGULATION_OF_DNA_TEMPLATED_TRANSCRIPTION_INITIATIO | 0.00163 | 0.02568 | 0.45506 | 0.68289 | 1.41372 | 35 |
| GO_REGULATION_OF_DNA_TEMPLATED_TRANSCRIPTION_IN_RESPO | 0.00000 | 0.00000 | | 0.73944 | 1.59800 | 89 |
| GO_REGULATION_OF_GENERATION_OF_PRECURSOR_METABOLITES_ | 0.00060 | 0.01164 | 0.47727 | 0.60476 | 1.31116 | 97 |
| GO_REGULATION_OF_HEMATOPOIETIC_PROGENITOR_CELL_DIFFERE | 0.00000 | 0.00000 | | 0.78717 | 1.69251 | 75 |
| GO_REGULATION_OF_HEMATOPOIETIC_STEM_CELL_DIFFERENTIATIO | 0.00000 | 0.00000 | | 0.81505 | 1.74256 | 65 |
| GO_REGULATION_OF_HEMOPOIESIS | 0.00000 | 0.00008 | 0.62726 | 0.58859 | 1.30634 | 228 |
| GO_REGULATION_OF_IMMUNE_RESPONSE | 0.00000 | 0.00016 | 0.61053 | 0.55362 | 1.23583 | 362 |
| GO_REGULATION_OF_INNATE_IMMUNE_RESPONSE | 0.00000 | 0.00000 | 0.71951 | 0.63883 | 1.40521 | 161 |
| GO_REGULATION_OF_INTRINSIC_APOPTOTIC_SIGNALING_PATHWAY | 0.00001 | 0.00039 | 0.59333 | 0.63478 | 1.38259 | 109 |

| | | | | | | |
|---|---|---|---|---|---|---|
| GO_REGULATION_OF_MITOTIC_CELL_CYCLE | 0.00001 | 0.00018 | 0.61053 | 0.53899 | 1.20736 | 458 |
| GO_REGULATION_OF_MORPHOGENESIS_OF_AN_EPITHELIUM | 0.00000 | 0.00000 | | 0.72213 | 1.57284 | 109 |
| GO_REGULATION_OF_MRNA_CATABOLIC_PROCESS | 0.00000 | 0.00000 | | 0.69977 | 1.54127 | 168 |
| GO_REGULATION_OF_MRNA_METABOLIC_PROCESS | 0.00000 | 0.00000 | | 0.66528 | 1.47872 | 266 |
| GO_REGULATION_OF_MRNA_PROCESSING | 0.00004 | 0.00105 | 0.55733 | 0.61521 | 1.34090 | 113 |
| GO_REGULATION_OF_MRNA_SPLICING_VIA_SPLICEOSOME | 0.00001 | 0.00026 | 0.59333 | 0.67424 | 1.45411 | 80 |
| GO_REGULATION_OF_NUCLEASE_ACTIVITY | 0.00106 | 0.01819 | 0.45506 | 0.77918 | 1.56780 | 18 |
| GO_REGULATION_OF_OXIDATIVE_STRESS_INDUCED_CELL_DEATH | 0.00198 | 0.03028 | 0.43171 | 0.67287 | 1.40355 | 41 |
| GO_REGULATION_OF_PEPTIDASE_ACTIVITY | 0.00050 | 0.00978 | 0.47727 | 0.56122 | 1.24054 | 191 |
| GO_REGULATION_OF_POSTTRANSCRIPTIONAL_GENE_SILENCING | 0.00364 | 0.04761 | 0.43171 | 0.59824 | 1.28492 | 74 |
| GO_REGULATION_OF_PROTEIN_CATABOLIC_PROCESS | 0.00179 | 0.02783 | 0.45506 | 0.53057 | 1.17929 | 270 |
| GO_REGULATION_OF_PROTEIN_LOCALIZATION_TO_NUCLEUS | 0.00001 | 0.00042 | 0.59333 | 0.65505 | 1.41558 | 87 |
| GO_REGULATION_OF_PROTEIN_MODIFICATION_BY_SMALL_PROTEIN | 0.00025 | 0.00532 | 0.49849 | 0.57068 | 1.25684 | 167 |
| GO_REGULATION_OF_PROTEIN_STABILITY | 0.00093 | 0.01652 | 0.47727 | 0.55382 | 1.22571 | 202 |
| GO_REGULATION_OF_PROTEOLYSIS | 0.00013 | 0.00301 | 0.51885 | 0.52997 | 1.18392 | 377 |
| GO_REGULATION_OF_RESPONSE_TO_BIOTIC_STIMULUS | 0.00001 | 0.00020 | 0.61053 | 0.59049 | 1.30719 | 204 |
| GO_REGULATION_OF_RNA_SPLICING | 0.00000 | 0.00001 | 0.69013 | 0.65896 | 1.43669 | 114 |
| GO_REGULATION_OF_SMALL_MOLECULE_METABOLIC_PROCESS | 0.00000 | 0.00000 | 0.71951 | 0.59134 | 1.31394 | 265 |
| GO_REGULATION_OF_STEM_CELL_DIFFERENTIATION | 0.00000 | 0.00000 | | 0.74049 | 1.60113 | 90 |
| GO_REGULATION_OF_SYMBIOTIC_PROCESS | 0.00006 | 0.00144 | 0.55733 | 0.60071 | 1.31682 | 134 |
| GO_REGULATION_OF_TELOMERASE_ACTIVITY | 0.00088 | 0.01590 | 0.47727 | 0.69161 | 1.43179 | 35 |
| GO_REGULATION_OF_TRANSCRIPTION_FROM_RNA_POLYMERASE_II | 0.00000 | 0.00000 | | 0.81194 | 1.73800 | 66 |
| GO_REGULATION_OF_TRANSCRIPTION_INITIATION_FROM_RNA_POL | 0.00143 | 0.02332 | 0.45506 | 0.70390 | 1.44582 | 29 |
| GO_REGULATION_OF_TRANSLATIONAL_INITIATION | 0.00002 | 0.00052 | 0.57561 | 0.68129 | 1.45518 | 64 |
| GO_REGULATION_OF_UBIQUITIN_PROTEIN_TRANSFERASE_ACTIVITY | 0.00061 | 0.01169 | 0.47727 | 0.69335 | 1.44627 | 41 |
| GO_REGULATION_OF_VIRAL_GENOME_REPLICATION | 0.00086 | 0.01566 | 0.47727 | 0.65316 | 1.38389 | 54 |
| GO_REGULATION_OF_VIRAL_TRANSCRIPTION | 0.00198 | 0.03028 | 0.43171 | 0.63932 | 1.34703 | 49 |
| GO_REGULATION_OF_WNT_SIGNALING_PATHWAY | 0.00000 | 0.00001 | 0.69013 | 0.60353 | 1.33816 | 220 |
| GO_RESPIRASOME | 0.00008 | 0.00181 | 0.53843 | 0.65420 | 1.40421 | 71 |
| GO_RESPIRATORY_CHAIN_COMPLEX | 0.00016 | 0.00353 | 0.51885 | 0.67626 | 1.43918 | 60 |
| GO_RESPIRATORY_ELECTRON_TRANSPORT_CHAIN | 0.00001 | 0.00025 | 0.59333 | 0.65819 | 1.42190 | 85 |
| GO_RESPONSE_TO_ALCOHOL | 0.00261 | 0.03750 | 0.43171 | 0.58804 | 1.27350 | 95 |
| GO_RESPONSE_TO_CADMIUM_ION | 0.00265 | 0.03775 | 0.43171 | 0.67227 | 1.39174 | 35 |
| GO_RESPONSE_TO_DRUG | 0.00104 | 0.01798 | 0.45506 | 0.55364 | 1.22258 | 180 |
| GO_RESPONSE_TO_ETHANOL | 0.00086 | 0.01566 | 0.47727 | 0.67309 | 1.40691 | 43 |
| GO_RESPONSE_TO_HEAT | 0.00012 | 0.00270 | 0.53843 | 0.61665 | 1.34221 | 107 |
| GO_RESPONSE_TO_HORMONE | 0.00048 | 0.00949 | 0.49849 | 0.51277 | 1.14814 | 453 |
| GO_RESPONSE_TO_HYDROGEN_PEROXIDE | 0.00257 | 0.03716 | 0.43171 | 0.60505 | 1.30368 | 79 |
| GO_RESPONSE_TO_INORGANIC_SUBSTANCE | 0.00004 | 0.00107 | 0.55733 | 0.55714 | 1.23870 | 275 |
| GO_RESPONSE_TO_INTERLEUKIN_1 | 0.00000 | 0.00000 | | 0.74844 | 1.62908 | 107 |
| GO_RESPONSE_TO_INTERLEUKIN_12 | 0.00012 | 0.00270 | 0.53843 | 0.72605 | 1.50694 | 36 |
| GO_RESPONSE_TO_INTERLEUKIN_4 | 0.00324 | 0.04369 | 0.43171 | 0.75812 | 1.51571 | 17 |
| GO_RESPONSE_TO_METAL_ION | 0.00030 | 0.00623 | 0.49849 | 0.56387 | 1.24233 | 171 |
| GO_RESPONSE_TO_ORGANIC_CYCLIC_COMPOUND | 0.00000 | 0.00016 | 0.61053 | 0.54603 | 1.22259 | 439 |
| GO_RESPONSE_TO_OXYGEN_LEVELS | 0.00000 | 0.00000 | 0.71951 | 0.60132 | 1.33438 | 233 |
| GO_RESPONSE_TO_STEROID_HORMONE | 0.00154 | 0.02464 | 0.45506 | 0.55514 | 1.22446 | 174 |
| GO_RESPONSE_TO_TEMPERATURE_STIMULUS | 0.00027 | 0.00572 | 0.49849 | 0.59216 | 1.29675 | 127 |
| GO_RESPONSE_TO_TUMOR_NECROSIS_FACTOR | 0.00000 | 0.00000 | 0.76146 | 0.65649 | 1.44236 | 151 |
| GO_RETINA_HOMEOSTASIS | 0.00104 | 0.01798 | 0.45506 | 0.73063 | 1.49432 | 27 |
| GO_RIBONUCLEOPROTEIN_COMPLEX_BINDING | 0.00000 | 0.00000 | 0.71951 | 0.66906 | 1.46050 | 117 |
| GO_RIBONUCLEOPROTEIN_COMPLEX_BIOGENESIS | 0.00000 | 0.00000 | | 0.66339 | 1.48264 | 389 |
| GO_RIBONUCLEOPROTEIN_COMPLEX_SUBUNIT_ORGANIZATION | 0.00000 | 0.00000 | | 0.69241 | 1.52681 | 173 |
| GO_RIBONUCLEOPROTEIN_GRANULE | 0.00000 | 0.00001 | 0.67496 | 0.61979 | 1.36366 | 163 |
| GO_RIBONUCLEOSIDE_TRIPHOSPHATE_BIOSYNTHETIC_PROCESS | 0.00041 | 0.00840 | 0.49849 | 0.71253 | 1.47248 | 33 |
| GO_RIBONUCLEOSIDE_TRIPHOSPHATE_METABOLIC_PROCESS | 0.00026 | 0.00550 | 0.49849 | 0.69681 | 1.45611 | 42 |
| GO_RIBOSE_PHOSPHATE_BIOSYNTHETIC_PROCESS | 0.00191 | 0.02939 | 0.45506 | 0.57932 | 1.26207 | 110 |
| GO_RIBOSE_PHOSPHATE_METABOLIC_PROCESS | 0.00108 | 0.01839 | 0.45506 | 0.53891 | 1.19687 | 245 |
| GO_RIBOSOMAL_LARGE_SUBUNIT_ASSEMBLY | 0.00000 | 0.00016 | 0.61053 | 0.80999 | 1.65663 | 27 |
| GO_RIBOSOMAL_LARGE_SUBUNIT_BIOGENESIS | 0.00000 | 0.00000 | | 0.77542 | 1.65982 | 66 |
| GO_RIBOSOMAL_SMALL_SUBUNIT_ASSEMBLY | 0.00023 | 0.00494 | 0.51885 | 0.81139 | 1.62221 | 17 |
| GO_RIBOSOMAL_SMALL_SUBUNIT_BINDING | 0.00383 | 0.04935 | 0.43171 | 0.75288 | 1.50523 | 17 |
| GO_RIBOSOMAL_SMALL_SUBUNIT_BIOGENESIS | 0.00001 | 0.00022 | 0.61053 | 0.69076 | 1.47997 | 68 |
| GO_RIBOSOMAL_SUBUNIT | 0.00000 | 0.00000 | | 0.77603 | 1.71031 | 172 |
| GO_RIBOSOME | 0.00000 | 0.00000 | | 0.76643 | 1.69659 | 206 |
| GO_RIBOSOME_ASSEMBLY | 0.00000 | 0.00000 | 0.73376 | 0.76173 | 1.61998 | 59 |
| GO_RIBOSOME_BINDING | 0.00019 | 0.00416 | 0.51885 | 0.68955 | 1.44715 | 47 |
| GO_RIBOSOME_BIOGENESIS | 0.00000 | 0.00000 | | 0.65877 | 1.46466 | 275 |
| GO_RNA_CATABOLIC_PROCESS | 0.00000 | 0.00000 | | 0.76774 | 1.71264 | 339 |
| GO_RNA_DEPENDENT_DNA_BIOSYNTHETIC_PROCESS | 0.00003 | 0.00076 | 0.57561 | 0.68926 | 1.46521 | 58 |
| GO_RNA_EXPORT_FROM_NUCLEUS | 0.00062 | 0.01183 | 0.47727 | 0.58378 | 1.27758 | 126 |
| GO_RNA_LOCALIZATION | 0.00000 | 0.00006 | 0.64355 | 0.59670 | 1.32020 | 198 |
| GO_RNA_POLYMERASE_II_PREINITIATION_COMPLEX_ASSEMBLY | 0.00089 | 0.01592 | 0.47727 | 0.77046 | 1.55626 | 19 |
| GO_RNA_SPLICING | 0.00000 | 0.00000 | 0.78819 | 0.59079 | 1.31927 | 369 |
| GO_RNA_SPLICING_VIA_TRANSESTERIFICATION_REACTIONS | 0.00000 | 0.00000 | | 0.60940 | 1.35714 | 300 |
| GO_RNA_STABILIZATION | 0.00065 | 0.01223 | 0.47727 | 0.69743 | 1.44384 | 35 |
| GO_ROUGH_ENDOPLASMIC_RETICULUM | 0.00001 | 0.00039 | 0.59333 | 0.71853 | 1.51629 | 50 |

| | | | | | | |
|---|---|---|---|---|---|---|
| GO_RRNA_BINDING | 0.00000 | 0.00001 | 0.69013 | 0.74782 | 1.58865 | 56 |
| GO_RRNA_METABOLIC_PROCESS | 0.00000 | 0.00000 | 0.77494 | 0.63047 | 1.39562 | 206 |
| GO_SCF_DEPENDENT_PROTEASOMAL_UBIQUITIN_DEPENDENT_PRO | 0.00000 | 0.00000 | | 0.77527 | 1.67044 | 79 |
| GO_SECRETORY_GRANULE | 0.00000 | 0.00000 | 0.71951 | 0.56518 | 1.26461 | 399 |
| GO_SECRETORY_VESICLE | 0.00000 | 0.00002 | 0.67496 | 0.54866 | 1.22905 | 465 |
| GO_SIGNAL_TRANSDUCTION_BY_PROTEIN_PHOSPHORYLATION | 0.00004 | 0.00109 | 0.55733 | 0.53817 | 1.20474 | 424 |
| GO_SINGLE_FERTILIZATION | 0.00048 | 0.00949 | 0.49849 | 0.67414 | 1.42260 | 50 |
| GO_SINGLE_STRANDED_RNA_BINDING | 0.00039 | 0.00793 | 0.49849 | 0.66553 | 1.41527 | 57 |
| GO_SMALL_MOLECULE_BIOSYNTHETIC_PROCESS | 0.00216 | 0.03242 | 0.43171 | 0.51195 | 1.14522 | 397 |
| GO_SMALL_RIBOSOMAL_SUBUNIT | 0.00000 | 0.00000 | | 0.77648 | 1.66209 | 66 |
| GO_SM_LIKE_PROTEIN_FAMILY_COMPLEX | 0.00327 | 0.04398 | 0.43171 | 0.60468 | 1.29713 | 70 |
| GO_SPERM_EGG_RECOGNITION | 0.00000 | 0.00007 | 0.64355 | 0.85241 | 1.72955 | 22 |
| GO_SPLICEOSOMAL_COMPLEX | 0.00000 | 0.00003 | 0.65944 | 0.60939 | 1.34261 | 171 |
| GO_STEM_CELL_DIFFERENTIATION | 0.00000 | 0.00000 | 0.78819 | 0.66886 | 1.46930 | 147 |
| GO_STRUCTURAL_CONSTITUENT_OF_CYTOSKELETON | 0.00102 | 0.01782 | 0.45506 | 0.65913 | 1.39337 | 52 |
| GO_STRUCTURAL_CONSTITUENT_OF_RIBOSOME | 0.00000 | 0.00000 | | 0.79909 | 1.75541 | 150 |
| GO_STRUCTURAL_MOLECULE_ACTIVITY | 0.00000 | 0.00000 | | 0.72651 | 1.61895 | 324 |
| GO_SUBSTANTIA_NIGRA_DEVELOPMENT | 0.00000 | 0.00013 | 0.62726 | 0.81622 | 1.66507 | 26 |
| GO_TELOMERASE_RNA_LOCALIZATION | 0.00001 | 0.00017 | 0.61053 | 0.85164 | 1.72025 | 19 |
| GO_TELOMERE_MAINTENANCE_VIA_TELOMERE_LENGTHENING | 0.00011 | 0.00250 | 0.53843 | 0.65659 | 1.40546 | 66 |
| GO_TELOMERE_ORGANIZATION | 0.00342 | 0.04554 | 0.43171 | 0.57000 | 1.24649 | 124 |
| GO_THREONINE_TYPE_ENDOPEPTIDASE_ACTIVITY | 0.00000 | 0.00000 | | 0.95179 | 1.92255 | 19 |
| GO_THREONINE_TYPE_PEPTIDASE_ACTIVITY | 0.00000 | 0.00000 | | 0.92199 | 1.87423 | 23 |
| GO_TISSUE_MORPHOGENESIS | 0.00097 | 0.01707 | 0.47727 | 0.52661 | 1.17402 | 326 |
| GO_TOXIN_TRANSPORT | 0.00113 | 0.01922 | 0.45506 | 0.72897 | 1.48708 | 26 |
| GO_TRANSCRIPTION_COUPLED_NUCLEOTIDE_EXCISION_REPAIR | 0.00001 | 0.00019 | 0.61053 | 0.68106 | 1.46186 | 71 |
| GO_TRANSCRIPTION_FACTOR_BINDING | 0.00043 | 0.00865 | 0.49849 | 0.52447 | 1.17376 | 409 |
| GO_TRANSCRIPTION_PREINITIATION_COMPLEX_ASSEMBLY | 0.00119 | 0.02004 | 0.45506 | 0.71399 | 1.46797 | 30 |
| GO_TRANSLATIONAL_ELONGATION | 0.00000 | 0.00000 | 0.71951 | 0.66087 | 1.44451 | 122 |
| GO_TRANSLATIONAL_INITIATION | 0.00000 | 0.00000 | | 0.84691 | 1.86515 | 166 |
| GO_TRANSLATIONAL_TERMINATION | 0.00000 | 0.00005 | 0.64355 | 0.65946 | 1.43060 | 99 |
| GO_TRANSLATION_ELONGATION_FACTOR_ACTIVITY | 0.00161 | 0.02546 | 0.45506 | 0.79860 | 1.58418 | 15 |
| GO_TRANSLATION_FACTOR_ACTIVITY_RNA_BINDING | 0.00000 | 0.00000 | | 0.79008 | 1.69288 | 69 |
| GO_TRANSLATION_INITIATION_FACTOR_ACTIVITY | 0.00000 | 0.00000 | 0.76146 | 0.80422 | 1.68417 | 45 |
| GO_TRANSLATION_INITIATION_FACTOR_BINDING | 0.00021 | 0.00468 | 0.51885 | 0.76186 | 1.55231 | 25 |
| GO_TRANSLATION_PREINITIATION_COMPLEX | 0.00000 | 0.00001 | 0.67496 | 0.90245 | 1.79595 | 16 |
| GO_TRANSLATION_REGULATOR_ACTIVITY | 0.00000 | 0.00000 | | 0.77663 | 1.68904 | 105 |
| GO_TRANSLATION_REGULATOR_ACTIVITY_NUCLEIC_ACID_BINDING | 0.00000 | 0.00000 | | 0.77150 | 1.66586 | 83 |
| GO_TRICARBOXYLIC_ACID_CYCLE | 0.00154 | 0.02464 | 0.45506 | 0.71854 | 1.47240 | 28 |
| GO_TUMOR_NECROSIS_FACTOR_MEDIATED_SIGNALING_PATHWAY | 0.00000 | 0.00000 | | 0.76277 | 1.65372 | 97 |
| GO_T_CELL_RECEPTOR_SIGNALING_PATHWAY | 0.00000 | 0.00000 | 0.81404 | 0.69994 | 1.52527 | 112 |
| GO_U12_TYPE_SPLICEOSOMAL_COMPLEX | 0.00279 | 0.03889 | 0.43171 | 0.70746 | 1.44693 | 27 |
| GO_U2_SNRNP | 0.00005 | 0.00134 | 0.55733 | 0.83081 | 1.67168 | 18 |
| GO_U2_TYPE_SPLICEOSOMAL_COMPLEX | 0.00010 | 0.00243 | 0.53843 | 0.63403 | 1.36970 | 85 |
| GO_UBIQUITIN_LIKE_PROTEIN_LIGASE_BINDING | 0.00000 | 0.00000 | 0.76146 | 0.62273 | 1.38051 | 219 |
| GO_UBIQUITIN_PROTEIN_TRANSFERASE_REGULATOR_ACTIVITY | 0.00165 | 0.02590 | 0.45506 | 0.79605 | 1.57911 | 15 |
| GO_UNFOLDED_PROTEIN_BINDING | 0.00000 | 0.00000 | 0.77494 | 0.72157 | 1.55881 | 85 |
| GO_VESICLE_LUMEN | 0.00000 | 0.00000 | 0.74774 | 0.64475 | 1.42008 | 168 |
| GO_VIRAL_GENE_EXPRESSION | 0.00000 | 0.00000 | | 0.80028 | 1.76317 | 171 |
| GO_VIRAL_GENOME_REPLICATION | 0.00100 | 0.01761 | 0.45506 | 0.61997 | 1.33078 | 73 |
| GO_VIRAL_LIFE_CYCLE | 0.00031 | 0.00640 | 0.49849 | 0.55767 | 1.23509 | 209 |
| HP_ABNORMALITY_OF_ACID_BASE_HOMEOSTASIS | 0.00216 | 0.03242 | 0.43171 | 0.53556 | 1.18846 | 232 |
| HP_ABNORMALITY_OF_CARDIOVASCULAR_SYSTEM_ELECTROPHYSIO | 0.00027 | 0.00572 | 0.49849 | 0.55832 | 1.23929 | 229 |
| HP_ABNORMALITY_OF_LOWER_LIP | 0.00181 | 0.02804 | 0.45506 | 0.56360 | 1.23779 | 145 |
| HP_ABNORMALITY_OF_THE_MITOCHONDRION | 0.00213 | 0.03229 | 0.43171 | 0.57831 | 1.26240 | 117 |
| HP_ABNORMALITY_OF_THE_NASAL_DORSUM | 0.00005 | 0.00116 | 0.55733 | 0.60288 | 1.32157 | 134 |
| HP_ABNORMAL_ELECTROPHYSIOLOGY_OF_SINOATRIAL_NODE_ORIG | 0.00279 | 0.03889 | 0.43171 | 0.68436 | 1.40934 | 31 |
| HP_ABNORMAL_ERYTHROCYTE_MORPHOLOGY | 0.00000 | 0.00015 | 0.61053 | 0.56049 | 1.25036 | 341 |
| HP_ABNORMAL_MEAN_CORPUSCULAR_VOLUME | 0.00357 | 0.04696 | 0.43171 | 0.75226 | 1.51363 | 18 |
| HP_ABNORMAL_RETICULOCYTE_MORPHOLOGY | 0.00064 | 0.01210 | 0.47727 | 0.71420 | 1.47081 | 31 |
| HP_ACUTE_LEUKEMIA | 0.00000 | 0.00000 | 0.73376 | 0.72786 | 1.56137 | 70 |
| HP_ANEMIA_OF_INADEQUATE_PRODUCTION | 0.00000 | 0.00008 | 0.62726 | 0.65593 | 1.42209 | 97 |
| HP_CONSTITUTIONAL_SYMPTOM | 0.00119 | 0.02004 | 0.45506 | 0.51427 | 1.15108 | 423 |
| HP_DELAYED_PUBERTY | 0.00005 | 0.00115 | 0.55733 | 0.62579 | 1.36211 | 107 |
| HP_DEPRESSED_NASAL_RIDGE | 0.00000 | 0.00011 | 0.62726 | 0.68731 | 1.47511 | 72 |
| HP_FATIGUE | 0.00039 | 0.00808 | 0.49849 | 0.56529 | 1.24923 | 187 |
| HP_HEADACHE | 0.00015 | 0.00340 | 0.51885 | 0.59065 | 1.29716 | 146 |
| HP_HEMATOLOGICAL_NEOPLASM | 0.00003 | 0.00078 | 0.57561 | 0.60108 | 1.32044 | 154 |
| HP_INCREASED_CSF_LACTATE | 0.00157 | 0.02509 | 0.45506 | 0.61622 | 1.31904 | 66 |
| HP_INCREASED_SERUM_LACTATE | 0.00216 | 0.03242 | 0.43171 | 0.56920 | 1.24724 | 130 |
| HP_LEUKEMIA | 0.00001 | 0.00026 | 0.59333 | 0.64877 | 1.40680 | 98 |
| HP_MACROCYTIC_ANEMIA | 0.00000 | 0.00000 | 0.74774 | 0.82468 | 1.71698 | 39 |
| HP_MIGRAINE | 0.00000 | 0.00000 | 0.71951 | 0.72105 | 1.54497 | 69 |
| HP_MYOTONIA | 0.00161 | 0.02546 | 0.45506 | 0.79820 | 1.58338 | 15 |
| HP_NEOPLASM | 0.00216 | 0.03242 | 0.43171 | 0.50820 | 1.13805 | 446 |
| HP_PALLOR | 0.00001 | 0.00040 | 0.59333 | 0.65169 | 1.41044 | 92 |

| | | | | | | |
|---|---|---|---|---|---|---|
| HP_PUBERTY_AND_GONADAL_DISORDERS | 0.00287 | 0.03973 | 0.43171 | 0.51995 | 1.15843 | 322 |
| HP_RETICULOCYTOPENIA | 0.00004 | 0.00105 | 0.55733 | 0.86417 | 1.71425 | 15 |
| HP_RIMMED_VACUOLES | 0.00279 | 0.03889 | 0.43171 | 0.74033 | 1.49831 | 20 |
| HP_SHORT_NOSE | 0.00235 | 0.03474 | 0.43171 | 0.54919 | 1.21134 | 174 |
| HP_THICK_LOWER_LIP_VERMILION | 0.00048 | 0.00949 | 0.49849 | 0.60773 | 1.31617 | 94 |
| HP_THICK_VERMILION_BORDER | 0.00156 | 0.02486 | 0.45506 | 0.55983 | 1.23000 | 153 |
| HP_VENTRICULAR_ARRHYTHMIA | 0.00353 | 0.04658 | 0.43171 | 0.66198 | 1.37398 | 36 |
| **Cluster 1** | | | | | | |
| pathway | pval | padj | log2err | ES | NES | size |
| GO_ACTIVATION_OF_IMMUNE_RESPONSE | 0.00000 | 0.00000 | 0.78819 | 0.62851 | 1.39255 | 212 |
| GO_ACTIVATION_OF_INNATE_IMMUNE_RESPONSE | 0.00000 | 0.00000 | | 0.75054 | 1.62130 | 88 |
| GO_AEROBIC_RESPIRATION | 0.00001 | 0.00025 | 0.59333 | 0.69302 | 1.48168 | 65 |
| GO_ALTERNATIVE_MRNA_SPLICING_VIA_SPLICEOSOME | 0.00102 | 0.01782 | 0.45506 | 0.65168 | 1.38687 | 60 |
| GO_AMINE_METABOLIC_PROCESS | 0.00000 | 0.00000 | | 0.79189 | 1.70783 | 80 |
| GO_ANAPHASE_PROMOTING_COMPLEX_DEPENDENT_CATABOLIC_P | 0.00000 | 0.00000 | | 0.76611 | 1.65070 | 79 |
| GO_ANCHORING_JUNCTION | 0.00000 | 0.00000 | | 0.59615 | 1.33633 | 479 |
| GO_ANIMAL_ORGAN_REGENERATION | 0.00257 | 0.03716 | 0.43171 | 0.68240 | 1.41020 | 33 |
| GO_ANTIGEN_PROCESSING_AND_PRESENTATION | 0.00000 | 0.00000 | 0.70498 | 0.64776 | 1.42241 | 143 |
| GO_ANTIGEN_PROCESSING_AND_PRESENTATION_OF_EXOGENOUS_ | 0.00000 | 0.00000 | | 0.81051 | 1.72358 | 57 |
| GO_ANTIGEN_PROCESSING_AND_PRESENTATION_OF_PEPTIDE_ANTI | 0.00000 | 0.00000 | 0.70498 | 0.65912 | 1.44248 | 126 |
| GO_ANTIGEN_PROCESSING_AND_PRESENTATION_OF_PEPTIDE_ANTI | 0.00000 | 0.00000 | | 0.76925 | 1.65015 | 70 |
| GO_ANTIGEN_RECEPTOR_MEDIATED_SIGNALING_PATHWAY | 0.00000 | 0.00000 | 0.80122 | 0.67877 | 1.48641 | 127 |
| GO_APOPTOTIC_SIGNALING_PATHWAY | 0.00154 | 0.02464 | 0.45506 | 0.51561 | 1.15085 | 358 |
| GO_ATP_DEPENDENT_CHROMATIN_REMODELING | 0.00253 | 0.03681 | 0.43171 | 0.62124 | 1.32400 | 61 |
| GO_ATP_METABOLIC_PROCESS | 0.00000 | 0.00000 | 0.71951 | 0.62252 | 1.37693 | 196 |
| GO_ATP_SYNTHESIS_COUPLED_ELECTRON_TRANSPORT | 0.00005 | 0.00131 | 0.55733 | 0.65519 | 1.40723 | 74 |
| GO_BINDING_OF_SPERM_TO_ZONA_PELLUCIDA | 0.00000 | 0.00001 | 0.69013 | 0.89879 | 1.80846 | 18 |
| GO_CADHERIN_BINDING | 0.00000 | 0.00000 | | 0.63068 | 1.39961 | 236 |
| GO_CANONICAL_WNT_SIGNALING_PATHWAY | 0.00000 | 0.00003 | 0.65944 | 0.60816 | 1.34553 | 197 |
| GO_CATALYTIC_STEP_2_SPLICEOSOME | 0.00000 | 0.00002 | 0.65944 | 0.68068 | 1.46976 | 83 |
| GO_CELLULAR_AMINO_ACID_METABOLIC_PROCESS | 0.00000 | 0.00000 | 0.74774 | 0.61752 | 1.36824 | 216 |
| GO_CELLULAR_COMPONENT_DISASSEMBLY | 0.00000 | 0.00005 | 0.64355 | 0.55505 | 1.24197 | 401 |
| GO_CELLULAR_KETONE_METABOLIC_PROCESS | 0.00000 | 0.00000 | 0.77494 | 0.65955 | 1.44893 | 149 |
| GO_CELLULAR_NITROGEN_COMPOUND_CATABOLIC_PROCESS | 0.00000 | 0.00000 | | 0.74079 | 1.65848 | 436 |
| GO_CELLULAR_PROTEIN_COMPLEX_DISASSEMBLY | 0.00006 | 0.00139 | 0.55733 | 0.58222 | 1.28459 | 176 |
| GO_CELLULAR_RESPIRATION | 0.00000 | 0.00000 | 0.71951 | 0.64997 | 1.42492 | 136 |
| GO_CELLULAR_RESPONSE_TO_HEAT | 0.00018 | 0.00403 | 0.51885 | 0.61962 | 1.34016 | 91 |
| GO_CELLULAR_RESPONSE_TO_ORGANIC_CYCLIC_COMPOUND | 0.00302 | 0.04108 | 0.43171 | 0.52402 | 1.16585 | 283 |
| GO_CELLULAR_RESPONSE_TO_OXYGEN_LEVELS | 0.00000 | 0.00000 | 0.78819 | 0.65911 | 1.44882 | 157 |
| GO_CELL_ACTIVATION_INVOLVED_IN_IMMUNE_RESPONSE | 0.00000 | 0.00001 | 0.67496 | 0.57339 | 1.27913 | 341 |
| GO_CELL_ADHESION_MOLECULE_BINDING | 0.00000 | 0.00000 | 0.81404 | 0.60529 | 1.34788 | 308 |
| GO_CELL_CELL_RECOGNITION | 0.00000 | 0.00002 | 0.65944 | 0.83826 | 1.71003 | 26 |
| GO_CELL_CELL_SIGNALING_BY_WNT | 0.00000 | 0.00002 | 0.67496 | 0.57736 | 1.28559 | 299 |
| GO_CELL_CYCLE_G2_M_PHASE_TRANSITION | 0.00001 | 0.00024 | 0.59333 | 0.58034 | 1.28602 | 214 |
| GO_CELL_CYCLE_PHASE_TRANSITION | 0.00000 | 0.00004 | 0.64355 | 0.54319 | 1.21681 | 464 |
| GO_CELL_DIVISION | 0.00154 | 0.02464 | 0.45506 | 0.51551 | 1.15425 | 427 |
| GO_CELL_RECOGNITION | 0.00012 | 0.00270 | 0.53843 | 0.69554 | 1.46186 | 48 |
| GO_CELL_SUBSTRATE_JUNCTION | 0.00000 | 0.00000 | | 0.67751 | 1.50775 | 291 |
| GO_CELL_SURFACE_RECEPTOR_SIGNALING_PATHWAY_INVOLVED_IN | 0.00000 | 0.00003 | 0.65944 | 0.57035 | 1.27067 | 319 |
| GO_CHAPERONE_BINDING | 0.00007 | 0.00163 | 0.53843 | 0.64754 | 1.39253 | 76 |
| GO_CHAPERONE_COMPLEX | 0.00000 | 0.00004 | 0.64355 | 0.85718 | 1.73925 | 22 |
| GO_CHAPERONE_MEDIATED_PROTEIN_COMPLEX_ASSEMBLY | 0.00170 | 0.02662 | 0.45506 | 0.77423 | 1.54792 | 17 |
| GO_CHROMATIN_BINDING | 0.00071 | 0.01323 | 0.47727 | 0.52699 | 1.17680 | 369 |
| GO_CHROMOSOMAL_REGION | 0.00246 | 0.03601 | 0.43171 | 0.52919 | 1.17661 | 276 |
| GO_COP9_SIGNALOSOME | 0.00128 | 0.02141 | 0.45506 | 0.73616 | 1.49715 | 24 |
| GO_COTRANSLATIONAL_PROTEIN_TARGETING_TO_MEMBRANE | 0.00000 | 0.00000 | | 0.89471 | 1.93513 | 91 |
| GO_CYTOCHROME_COMPLEX | 0.00168 | 0.02641 | 0.45506 | 0.75311 | 1.52417 | 20 |
| GO_CYTOKINE_MEDIATED_SIGNALING_PATHWAY | 0.00000 | 0.00000 | 0.81404 | 0.59698 | 1.33170 | 339 |
| GO_CYTOPLASMIC_TRANSLATION | 0.00000 | 0.00000 | | 0.84753 | 1.83080 | 88 |
| GO_CYTOPLASMIC_TRANSLATIONAL_INITIATION | 0.00000 | 0.00000 | 0.74774 | 0.85688 | 1.76176 | 30 |
| GO_CYTOSOLIC_LARGE_RIBOSOMAL_SUBUNIT | 0.00000 | 0.00000 | | 0.91186 | 1.92988 | 53 |
| GO_CYTOSOLIC_RIBOSOME | 0.00000 | 0.00000 | | 0.89907 | 1.94710 | 95 |
| GO_CYTOSOLIC_SMALL_RIBOSOMAL_SUBUNIT | 0.00000 | 0.00000 | | 0.89114 | 1.85533 | 39 |
| GO_DEFENSE_RESPONSE_TO_OTHER_ORGANISM | 0.00004 | 0.00099 | 0.55733 | 0.53673 | 1.20127 | 415 |
| GO_DE_NOVO_PROTEIN_FOLDING | 0.00050 | 0.00978 | 0.47727 | 0.74232 | 1.51431 | 26 |
| GO_DISORDERED_DOMAIN_SPECIFIC_BINDING | 0.00161 | 0.02546 | 0.45506 | 0.71970 | 1.47197 | 27 |
| GO_DNA_BIOSYNTHETIC_PROCESS | 0.00000 | 0.00010 | 0.62726 | 0.62505 | 1.37017 | 134 |
| GO_DNA_CONFORMATION_CHANGE | 0.00027 | 0.00572 | 0.49849 | 0.55300 | 1.22527 | 216 |
| GO_DNA_DAMAGE_RESPONSE_DETECTION_OF_DNA_DAMAGE | 0.00198 | 0.03028 | 0.43171 | 0.67949 | 1.40669 | 35 |
| GO_DNA_GEOMETRIC_CHANGE | 0.00046 | 0.00925 | 0.49849 | 0.60807 | 1.31855 | 98 |
| GO_DNA_POLYMERASE_BINDING | 0.00268 | 0.03799 | 0.43171 | 0.76255 | 1.52457 | 17 |
| GO_DNA_REPAIR | 0.00265 | 0.03775 | 0.43171 | 0.51159 | 1.14547 | 427 |
| GO_DRUG_BINDING | 0.00176 | 0.02733 | 0.45506 | 0.64119 | 1.35096 | 49 |
| GO_ELECTRON_TRANSFER_ACTIVITY | 0.00001 | 0.00020 | 0.61053 | 0.67481 | 1.44938 | 74 |
| GO_ELECTRON_TRANSPORT_CHAIN | 0.00002 | 0.00053 | 0.57561 | 0.62257 | 1.36145 | 124 |
| GO_ENDOPEPTIDASE_ACTIVITY | 0.00124 | 0.02085 | 0.45506 | 0.54955 | 1.21076 | 171 |

| | | | | | | |
|---|---|---|---|---|---|---|
| GO_ENDOPEPTIDASE_COMPLEX | 0.00000 | 0.00000 | | 0.80930 | 1.73234 | 66 |
| GO_ENERGY_DERIVATION_BY_OXIDATION_OF_ORGANIC_COMPOUN | 0.00000 | 0.00000 | 0.71951 | 0.63390 | 1.39707 | 172 |
| GO_ESTABLISHMENT_OF_PROTEIN_LOCALIZATION_TO_CHROMOSOM | 0.00020 | 0.00432 | 0.51885 | 0.77339 | 1.57217 | 23 |
| GO_ESTABLISHMENT_OF_PROTEIN_LOCALIZATION_TO_ENDOPLASM | 0.00000 | 0.00000 | | 0.88026 | 1.91149 | 102 |
| GO_ESTABLISHMENT_OF_PROTEIN_LOCALIZATION_TO_MEMBRANE | 0.00000 | 0.00000 | | 0.74874 | 1.66164 | 240 |
| GO_ESTABLISHMENT_OF_PROTEIN_LOCALIZATION_TO_ORGANELLE | 0.00000 | 0.00000 | | 0.69519 | 1.55605 | 423 |
| GO_ESTABLISHMENT_OF_PROTEIN_LOCALIZATION_TO_TELOMERE | 0.00006 | 0.00139 | 0.55733 | 0.83470 | 1.66881 | 17 |
| GO_ESTABLISHMENT_OF_RNA_LOCALIZATION | 0.00106 | 0.01819 | 0.45506 | 0.55577 | 1.22400 | 167 |
| GO_ESTABLISHMENT_OF_TISSUE_POLARITY | 0.00000 | 0.00000 | | 0.75051 | 1.62122 | 88 |
| GO_EUKARYOTIC_TRANSLATION_INITIATION_FACTOR_3_COMPLEX | 0.00003 | 0.00074 | 0.57561 | 0.86987 | 1.72555 | 15 |
| GO_EXOCYTOSIS | 0.00000 | 0.00015 | 0.61053 | 0.53905 | 1.20753 | 465 |
| GO_FC_EPSILON_RECEPTOR_SIGNALING_PATHWAY | 0.00000 | 0.00000 | | 0.75117 | 1.62103 | 82 |
| GO_FC_RECEPTOR_SIGNALING_PATHWAY | 0.00000 | 0.00000 | | 0.72054 | 1.57787 | 127 |
| GO_FERTILIZATION | 0.00089 | 0.01592 | 0.47727 | 0.63620 | 1.35888 | 64 |
| GO_FICOLIN_1_RICH_GRANULE | 0.00000 | 0.00000 | 0.78819 | 0.71716 | 1.55340 | 93 |
| GO_GENERATION_OF_PRECURSOR_METABOLITES_AND_ENERGY | 0.00000 | 0.00000 | 0.77494 | 0.59261 | 1.32026 | 320 |
| GO_GENE_SILENCING | 0.00261 | 0.03750 | 0.43171 | 0.55578 | 1.22252 | 161 |
| GO_GLAND_DEVELOPMENT | 0.00154 | 0.02464 | 0.45506 | 0.54117 | 1.19797 | 206 |
| GO_GLYCOSYL_COMPOUND_METABOLIC_PROCESS | 0.00071 | 0.01323 | 0.47727 | 0.62524 | 1.34204 | 71 |
| GO_GUANYL_NUCLEOTIDE_BINDING | 0.00095 | 0.01679 | 0.47727 | 0.54096 | 1.19861 | 216 |
| GO_HEAT_SHOCK_PROTEIN_BINDING | 0.00003 | 0.00092 | 0.55733 | 0.64369 | 1.39056 | 85 |
| GO_HEMATOPOIETIC_PROGENITOR_CELL_DIFFERENTIATION | 0.00000 | 0.00000 | 0.82666 | 0.70307 | 1.53239 | 113 |
| GO_HEMATOPOIETIC_STEM_CELL_DIFFERENTIATION | 0.00000 | 0.00000 | | 0.79776 | 1.71217 | 72 |
| GO_HSP70_PROTEIN_BINDING | 0.00228 | 0.03391 | 0.43171 | 0.71143 | 1.45504 | 27 |
| GO_IMMUNE_RESPONSE_REGULATING_SIGNALING_PATHWAY | 0.00000 | 0.00000 | 0.76146 | 0.63494 | 1.40480 | 197 |
| GO_INNATE_IMMUNE_RESPONSE | 0.00003 | 0.00071 | 0.57561 | 0.55173 | 1.23107 | 347 |
| GO_INNATE_IMMUNE_RESPONSE_ACTIVATING_SIGNAL_TRANSDUCT | 0.00000 | 0.00000 | | 0.79340 | 1.70281 | 72 |
| GO_INNER_MITOCHONDRIAL_MEMBRANE_PROTEIN_COMPLEX | 0.00002 | 0.00060 | 0.57561 | 0.65124 | 1.40678 | 88 |
| GO_INTERACTION_WITH_SYMBIONT | 0.00001 | 0.00026 | 0.59333 | 0.75084 | 1.56620 | 41 |
| GO_INTERLEUKIN_1_MEDIATED_SIGNALING_PATHWAY | 0.00000 | 0.00000 | | 0.81689 | 1.75671 | 76 |
| GO_INTRINSIC_APOPTOTIC_SIGNALING_PATHWAY | 0.00121 | 0.02029 | 0.45506 | 0.55118 | 1.21845 | 190 |
| GO_ION_CHANNEL_BINDING | 0.00087 | 0.01584 | 0.47727 | 0.63182 | 1.34683 | 62 |
| GO_ISOPEPTIDASE_ACTIVITY | 0.00139 | 0.02285 | 0.45506 | 0.78940 | 1.57097 | 16 |
| GO_LARGE_RIBOSOMAL_SUBUNIT | 0.00000 | 0.00000 | | 0.78135 | 1.70183 | 109 |
| GO_LEUKOCYTE_MEDIATED_IMMUNITY | 0.00000 | 0.00000 | 0.71951 | 0.57282 | 1.27895 | 364 |
| GO_MACROMOLECULE_TRANSMEMBRANE_TRANSPORTER_ACTIVITY | 0.00239 | 0.03510 | 0.43171 | 0.77734 | 1.54696 | 16 |
| GO_MAINTENANCE_OF_CELL_NUMBER | 0.00087 | 0.01584 | 0.47727 | 0.58944 | 1.28512 | 114 |
| GO_MATURATION_OF_LSU_RRNA | 0.00006 | 0.00151 | 0.53843 | 0.77606 | 1.58723 | 27 |
| GO_METHYLTRANSFERASE_COMPLEX | 0.00290 | 0.04014 | 0.43171 | 0.59476 | 1.28479 | 88 |
| GO_METHYL_CPG_BINDING | 0.00302 | 0.04108 | 0.43171 | 0.78053 | 1.54833 | 15 |
| GO_MIDBRAIN_DEVELOPMENT | 0.00021 | 0.00468 | 0.51885 | 0.68832 | 1.44457 | 47 |
| GO_MITOCHONDRIAL_ELECTRON_TRANSPORT_NADH_TO_UBIQUIN | 0.00331 | 0.04437 | 0.43171 | 0.65428 | 1.36724 | 42 |
| GO_MITOCHONDRIAL_GENE_EXPRESSION | 0.00048 | 0.00949 | 0.49849 | 0.57608 | 1.26525 | 152 |
| GO_MITOCHONDRIAL_LARGE_RIBOSOMAL_SUBUNIT | 0.00002 | 0.00050 | 0.57561 | 0.69789 | 1.48086 | 55 |
| GO_MITOCHONDRIAL_MATRIX | 0.00000 | 0.00002 | 0.65944 | 0.56045 | 1.25158 | 371 |
| GO_MITOCHONDRIAL_MEMBRANE_ORGANIZATION | 0.00361 | 0.04734 | 0.43171 | 0.59309 | 1.28094 | 86 |
| GO_MITOCHONDRIAL_PROTEIN_COMPLEX | 0.00000 | 0.00000 | 0.81404 | 0.63650 | 1.40931 | 208 |
| GO_MITOCHONDRIAL_TRANSLATION | 0.00004 | 0.00093 | 0.55733 | 0.61219 | 1.34135 | 129 |
| GO_MITOCHONDRIAL_TRANSLATIONAL_TERMINATION | 0.00000 | 0.00010 | 0.62726 | 0.66694 | 1.44128 | 87 |
| GO_MITOCHONDRION_ORGANIZATION | 0.00051 | 0.00993 | 0.47727 | 0.52284 | 1.16863 | 390 |
| GO_MODIFICATION_DEPENDENT_MACROMOLECULE_CATABOLIC_PR | 0.00002 | 0.00053 | 0.57561 | 0.53142 | 1.19156 | 486 |
| GO_MODULATION_BY_HOST_OF_SYMBIONT_PROCESS | 0.00015 | 0.00336 | 0.51885 | 0.72848 | 1.50542 | 33 |
| GO_MODULATION_OF_PROCESS_OF_OTHER_ORGANISM | 0.00019 | 0.00423 | 0.51885 | 0.66904 | 1.42286 | 59 |
| GO_MONOSACCHARIDE_BIOSYNTHETIC_PROCESS | 0.00335 | 0.04466 | 0.43171 | 0.61992 | 1.31781 | 58 |
| GO_MORPHOGENESIS_OF_AN_EPITHELIUM | 0.00006 | 0.00151 | 0.53843 | 0.55161 | 1.22755 | 290 |
| GO_MORPHOGENESIS_OF_A_POLARIZED_EPITHELIUM | 0.00000 | 0.00000 | | 0.72298 | 1.57057 | 103 |
| GO_MRNA_3_UTR_BINDING | 0.00003 | 0.00089 | 0.55733 | 0.67596 | 1.44827 | 68 |
| GO_MRNA_5_UTR_BINDING | 0.00006 | 0.00156 | 0.53843 | 0.80133 | 1.62592 | 22 |
| GO_MRNA_BINDING | 0.00000 | 0.00000 | | 0.66002 | 1.46353 | 222 |
| GO_MRNA_PROCESSING | 0.00000 | 0.00000 | 0.73376 | 0.56859 | 1.27295 | 418 |
| GO_MULTI_ORGANISM_LOCALIZATION | 0.00268 | 0.03799 | 0.43171 | 0.63245 | 1.33612 | 51 |
| GO_MULTI_ORGANISM_PROCESS | 0.00108 | 0.01839 | 0.45506 | 0.51114 | 1.14599 | 484 |
| GO_MYD88_DEPENDENT_TOLL_LIKE_RECEPTOR_SIGNALING_PATHW | 0.00002 | 0.00066 | 0.57561 | 0.85581 | 1.70312 | 16 |
| GO_MYELOID_LEUKOCYTE_ACTIVATION | 0.00000 | 0.00000 | 0.74774 | 0.58508 | 1.30479 | 327 |
| GO_MYELOID_LEUKOCYTE_MEDIATED_IMMUNITY | 0.00000 | 0.00000 | 0.76146 | 0.59841 | 1.33291 | 301 |
| GO_NADH_DEHYDROGENASE_ACTIVITY | 0.00383 | 0.04935 | 0.43171 | 0.66244 | 1.37139 | 35 |
| GO_NADH_DEHYDROGENASE_COMPLEX | 0.00376 | 0.04884 | 0.43171 | 0.65649 | 1.36421 | 38 |
| GO_NADH_METABOLIC_PROCESS | 0.00294 | 0.04036 | 0.43171 | 0.70642 | 1.44481 | 27 |
| GO_NCRNA_EXPORT_FROM_NUCLEUS | 0.00302 | 0.04108 | 0.43171 | 0.66820 | 1.38331 | 35 |
| GO_NCRNA_METABOLIC_PROCESS | 0.00000 | 0.00003 | 0.65944 | 0.55907 | 1.24907 | 381 |
| GO_NCRNA_PROCESSING | 0.00000 | 0.00003 | 0.65944 | 0.57175 | 1.27504 | 328 |
| GO_NEGATIVE_REGULATION_OF_CANONICAL_WNT_SIGNALING_PAT | 0.00000 | 0.00000 | | 0.71611 | 1.56038 | 111 |
| GO_NEGATIVE_REGULATION_OF_CELLULAR_AMIDE_METABOLIC_PR | 0.00001 | 0.00020 | 0.61053 | 0.63031 | 1.37423 | 114 |
| GO_NEGATIVE_REGULATION_OF_CELL_CYCLE | 0.00003 | 0.00076 | 0.57561 | 0.53833 | 1.20513 | 425 |
| GO_NEGATIVE_REGULATION_OF_CELL_CYCLE_G2_M_PHASE_TRANS | 0.00000 | 0.00000 | 0.76146 | 0.71490 | 1.54439 | 85 |
| GO_NEGATIVE_REGULATION_OF_CELL_CYCLE_PHASE_TRANSITION | 0.00002 | 0.00060 | 0.57561 | 0.59041 | 1.30442 | 184 |

| | | | | | | |
|---|---|---|---|---|---|---|
| GO_NEGATIVE_REGULATION_OF_CELL_CYCLE_PROCESS | 0.00000 | 0.00002 | 0.65944 | 0.58690 | 1.30374 | 249 |
| GO_NEGATIVE_REGULATION_OF_MITOTIC_CELL_CYCLE | 0.00004 | 0.00095 | 0.55733 | 0.56735 | 1.25902 | 232 |
| GO_NEGATIVE_REGULATION_OF_MRNA_METABOLIC_PROCESS | 0.00023 | 0.00491 | 0.51885 | 0.66670 | 1.41788 | 59 |
| GO_NEGATIVE_REGULATION_OF_MRNA_SPLICING_VIA_SPLICEOSOM | 0.00239 | 0.03510 | 0.43171 | 0.74857 | 1.51204 | 19 |
| GO_NEGATIVE_REGULATION_OF_RNA_SPLICING | 0.00029 | 0.00607 | 0.49849 | 0.76729 | 1.55976 | 23 |
| GO_NEGATIVE_REGULATION_OF_TRANSLATIONAL_INITIATION | 0.00018 | 0.00403 | 0.51885 | 0.83901 | 1.66434 | 15 |
| GO_NEGATIVE_REGULATION_OF_WNT_SIGNALING_PATHWAY | 0.00000 | 0.00000 | | 0.69525 | 1.52249 | 127 |
| GO_NEURAL_NUCLEUS_DEVELOPMENT | 0.00001 | 0.00026 | 0.59333 | 0.79148 | 1.62187 | 28 |
| GO_NEURON_TO_NEURON_SYNAPSE | 0.00005 | 0.00134 | 0.55733 | 0.59636 | 1.31006 | 154 |
| GO_NIK_NF_KAPPAB_SIGNALING | 0.00000 | 0.00000 | | 0.72896 | 1.58354 | 103 |
| GO_NON_CANONICAL_WNT_SIGNALING_PATHWAY | 0.00000 | 0.00000 | | 0.72784 | 1.58023 | 101 |
| GO_NUCLEAR_EXPORT | 0.00031 | 0.00645 | 0.49849 | 0.56963 | 1.25542 | 172 |
| GO_NUCLEAR_TRANSCRIBED_MRNA_CATABOLIC_PROCESS | 0.00000 | 0.00000 | | 0.79179 | 1.74917 | 183 |
| GO_NUCLEAR_TRANSCRIBED_MRNA_CATABOLIC_PROCESS_NONSEN | 0.00000 | 0.00000 | | 0.89008 | 1.93909 | 110 |
| GO_NUCLEAR_TRANSPORT | 0.00008 | 0.00193 | 0.53843 | 0.55511 | 1.23408 | 273 |
| GO_NUCLEOBASE_CONTAINING_COMPOUND_TRANSPORT | 0.00146 | 0.02386 | 0.45506 | 0.55053 | 1.21842 | 202 |
| GO_NUCLEOBASE_CONTAINING_SMALL_MOLECULE_BIOSYNTHETIC_ | 0.00137 | 0.02261 | 0.45506 | 0.62749 | 1.34353 | 67 |
| GO_NUCLEOBASE_CONTAINING_SMALL_MOLECULE_INTERCONVERS | 0.00383 | 0.04935 | 0.43171 | 0.71259 | 1.44857 | 23 |
| GO_NUCLEOBASE_CONTAINING_SMALL_MOLECULE_METABOLIC_PR | 0.00320 | 0.04339 | 0.43171 | 0.51448 | 1.14825 | 352 |
| GO_NUCLEOSIDE_DIPHOSPHATE_METABOLIC_PROCESS | 0.00110 | 0.01865 | 0.45506 | 0.59637 | 1.29295 | 97 |
| GO_NUCLEOSIDE_METABOLIC_PROCESS | 0.00061 | 0.01169 | 0.47727 | 0.65315 | 1.38907 | 59 |
| GO_NUCLEOSIDE_MONOPHOSPHATE_BIOSYNTHETIC_PROCESS | 0.00250 | 0.03636 | 0.43171 | 0.68841 | 1.42033 | 32 |
| GO_NUCLEOSIDE_MONOPHOSPHATE_METABOLIC_PROCESS | 0.00152 | 0.02464 | 0.45506 | 0.64377 | 1.36002 | 51 |
| GO_NUCLEOSIDE_PHOSPHATE_BIOSYNTHETIC_PROCESS | 0.00043 | 0.00865 | 0.49849 | 0.58185 | 1.27831 | 148 |
| GO_NUCLEOSIDE_TRIPHOSPHATE_BIOSYNTHETIC_PROCESS | 0.00043 | 0.00865 | 0.49849 | 0.69332 | 1.44539 | 40 |
| GO_NUCLEOSIDE_TRIPHOSPHATE_METABOLIC_PROCESS | 0.00016 | 0.00360 | 0.51885 | 0.66788 | 1.41976 | 58 |
| GO_NUCLEOSOMAL_DNA_BINDING | 0.00023 | 0.00507 | 0.51885 | 0.79852 | 1.61801 | 21 |
| GO_NUCLEOSOME_BINDING | 0.00143 | 0.02332 | 0.45506 | 0.66132 | 1.38492 | 45 |
| GO_NUCLEOTIDE_EXCISION_REPAIR | 0.00004 | 0.00096 | 0.55733 | 0.63343 | 1.37355 | 98 |
| GO_NUCLEOTIDE_EXCISION_REPAIR_DNA_DAMAGE_RECOGNITION | 0.00015 | 0.00349 | 0.51885 | 0.78678 | 1.59640 | 22 |
| GO_NUCLEOTIDE_EXCISION_REPAIR_DNA_INCISION | 0.00313 | 0.04249 | 0.43171 | 0.66439 | 1.37898 | 36 |
| GO_NUCLEOTIDE_PHOSPHORYLATION | 0.00383 | 0.04935 | 0.43171 | 0.59002 | 1.27454 | 88 |
| GO_ORGANELLAR_RIBOSOME | 0.00000 | 0.00014 | 0.61053 | 0.66412 | 1.43471 | 85 |
| GO_ORGANELLE_INNER_MEMBRANE | 0.00000 | 0.00000 | 0.76146 | 0.57601 | 1.28697 | 378 |
| GO_ORGANIC_CYCLIC_COMPOUND_CATABOLIC_PROCESS | 0.00000 | 0.00000 | | 0.73350 | 1.64281 | 449 |
| GO_OXIDATIVE_PHOSPHORYLATION | 0.00001 | 0.00030 | 0.59333 | 0.64619 | 1.40027 | 96 |
| GO_OXIDOREDUCTASE_ACTIVITY | 0.00006 | 0.00139 | 0.55733 | 0.53544 | 1.19703 | 392 |
| GO_OXIDOREDUCTASE_COMPLEX | 0.00001 | 0.00027 | 0.59333 | 0.66015 | 1.42461 | 82 |
| GO_PEPTIDASE_COMPLEX | 0.00000 | 0.00000 | | 0.75988 | 1.63982 | 82 |
| GO_PIGMENT_GRANULE | 0.00003 | 0.00089 | 0.55733 | 0.66098 | 1.42344 | 78 |
| GO_POLYSOMAL_RIBOSOME | 0.00000 | 0.00000 | | 0.90624 | 1.86629 | 31 |
| GO_POLYSOME | 0.00000 | 0.00000 | | 0.82337 | 1.75481 | 61 |
| GO_POLY_PYRIMIDINE_TRACT_BINDING | 0.00327 | 0.04398 | 0.43171 | 0.75471 | 1.51855 | 18 |
| GO_POSITIVE_REGULATION_OF_APOPTOTIC_SIGNALING_PATHWAY | 0.00279 | 0.03889 | 0.43171 | 0.57666 | 1.25518 | 107 |
| GO_POSITIVE_REGULATION_OF_ATPASE_ACTIVITY | 0.00294 | 0.04036 | 0.43171 | 0.65446 | 1.36795 | 43 |
| GO_POSITIVE_REGULATION_OF_BINDING | 0.00088 | 0.01590 | 0.47727 | 0.60284 | 1.30556 | 95 |
| GO_POSITIVE_REGULATION_OF_CANONICAL_WNT_SIGNALING_PATH | 0.00000 | 0.00000 | 0.77494 | 0.70504 | 1.53160 | 103 |
| GO_POSITIVE_REGULATION_OF_CELLULAR_AMIDE_METABOLIC_PRO | 0.00003 | 0.00074 | 0.57561 | 0.62296 | 1.36083 | 118 |
| GO_POSITIVE_REGULATION_OF_CELLULAR_PROTEIN_LOCALIZATION | 0.00041 | 0.00840 | 0.49849 | 0.54915 | 1.21815 | 225 |
| GO_POSITIVE_REGULATION_OF_CELL_DEATH | 0.00069 | 0.01297 | 0.47727 | 0.52595 | 1.17423 | 363 |
| GO_POSITIVE_REGULATION_OF_CELL_POPULATION_PROLIFERATION | 0.00078 | 0.01455 | 0.47727 | 0.52449 | 1.17108 | 368 |
| GO_POSITIVE_REGULATION_OF_CHROMOSOME_ORGANIZATION | 0.00231 | 0.03437 | 0.43171 | 0.56990 | 1.24871 | 133 |
| GO_POSITIVE_REGULATION_OF_DEFENSE_RESPONSE | 0.00000 | 0.00003 | 0.65944 | 0.62175 | 1.36730 | 158 |
| GO_POSITIVE_REGULATION_OF_DNA_BIOSYNTHETIC_PROCESS | 0.00001 | 0.00041 | 0.59333 | 0.71929 | 1.52055 | 52 |
| GO_POSITIVE_REGULATION_OF_DNA_METABOLIC_PROCESS | 0.00364 | 0.04761 | 0.43171 | 0.56647 | 1.24189 | 135 |
| GO_POSITIVE_REGULATION_OF_DNA_TEMPLATED_TRANSCRIPTION_ | 0.00276 | 0.03885 | 0.43171 | 0.68932 | 1.41586 | 29 |
| GO_POSITIVE_REGULATION_OF_ESTABLISHMENT_OF_PROTEIN_LOC/ | 0.00007 | 0.00168 | 0.53843 | 0.56461 | 1.25165 | 219 |
| GO_POSITIVE_REGULATION_OF_IMMUNE_RESPONSE | 0.00000 | 0.00001 | 0.69013 | 0.59037 | 1.31265 | 277 |
| GO_POSITIVE_REGULATION_OF_IMMUNE_SYSTEM_PROCESS | 0.00004 | 0.00110 | 0.55733 | 0.54018 | 1.20716 | 386 |
| GO_POSITIVE_REGULATION_OF_INNATE_IMMUNE_RESPONSE | 0.00000 | 0.00000 | 0.81404 | 0.69693 | 1.51947 | 114 |
| GO_POSITIVE_REGULATION_OF_INTRINSIC_APOPTOTIC_SIGNALING_ | 0.00104 | 0.01798 | 0.45506 | 0.69097 | 1.43578 | 37 |
| GO_POSITIVE_REGULATION_OF_ORGANELLE_ORGANIZATION | 0.00043 | 0.00865 | 0.49849 | 0.52093 | 1.16605 | 411 |
| GO_POSITIVE_REGULATION_OF_PROTEIN_LOCALIZATION_TO_NUCLE | 0.00001 | 0.00020 | 0.61053 | 0.69907 | 1.48990 | 61 |
| GO_POSITIVE_REGULATION_OF_RESPONSE_TO_BIOTIC_STIMULUS | 0.00000 | 0.00000 | 0.73376 | 0.66340 | 1.45310 | 128 |
| GO_POSITIVE_REGULATION_OF_RESPONSE_TO_EXTERNAL_STIMULU | 0.00002 | 0.00050 | 0.57561 | 0.58133 | 1.28681 | 201 |
| GO_POSITIVE_REGULATION_OF_TELOMERASE_ACTIVITY | 0.00061 | 0.01169 | 0.47727 | 0.73476 | 1.50565 | 28 |
| GO_POSITIVE_REGULATION_OF_TELOMERASE_RNA_LOCALIZATION_ | 0.00000 | 0.00001 | 0.69013 | 0.92451 | 1.83395 | 15 |
| GO_POSITIVE_REGULATION_OF_TELOMERE_MAINTENANCE_VIA_TE | 0.00130 | 0.02159 | 0.45506 | 0.71259 | 1.46508 | 30 |
| GO_POSITIVE_REGULATION_OF_TRANSCRIPTION_INITIATION_FROM | 0.00372 | 0.04847 | 0.43171 | 0.70671 | 1.43993 | 25 |
| GO_POSITIVE_REGULATION_OF_TRANSFERASE_ACTIVITY | 0.00174 | 0.02712 | 0.45506 | 0.51642 | 1.15220 | 345 |
| GO_POSITIVE_REGULATION_OF_TRANSLATION | 0.00001 | 0.00019 | 0.61053 | 0.64767 | 1.40419 | 97 |
| GO_POSITIVE_REGULATION_OF_VIRAL_GENOME_REPLICATION | 0.00287 | 0.03973 | 0.43171 | 0.68362 | 1.40783 | 31 |
| GO_POSITIVE_REGULATION_OF_VIRAL_PROCESS | 0.00028 | 0.00590 | 0.49849 | 0.62440 | 1.34741 | 81 |
| GO_POSITIVE_REGULATION_OF_WNT_SIGNALING_PATHWAY | 0.00000 | 0.00000 | 0.73376 | 0.66230 | 1.44763 | 122 |
| GO_POSTREPLICATION_REPAIR | 0.00024 | 0.00515 | 0.51885 | 0.69140 | 1.44518 | 43 |
| GO_POSTSYNAPSE | 0.00004 | 0.00109 | 0.55733 | 0.55969 | 1.24369 | 259 |

| | | | | | | |
|---|---|---|---|---|---|---|
| GO_POSTSYNAPTIC_CYTOSOL | 0.00279 | 0.03889 | 0.43171 | 0.78295 | 1.55313 | 15 |
| GO_POSTTRANSCRIPTIONAL_REGULATION_OF_GENE_EXPRESSION | 0.00000 | 0.00000 | | 0.66656 | 1.49327 | 462 |
| GO_POST_TRANSLATIONAL_PROTEIN_MODIFICATION | 0.00002 | 0.00052 | 0.57561 | 0.57851 | 1.28412 | 229 |
| GO_PRECATALYTIC_SPLICEOSOME | 0.00085 | 0.01559 | 0.47727 | 0.65235 | 1.37448 | 49 |
| GO_PRERIBOSOME | 0.00087 | 0.01577 | 0.47727 | 0.62899 | 1.34772 | 69 |
| GO_PRERIBOSOME_LARGE_SUBUNIT_PRECURSOR | 0.00235 | 0.03474 | 0.43171 | 0.74079 | 1.50308 | 22 |
| GO_PRE_MRNA_BINDING | 0.00133 | 0.02213 | 0.45506 | 0.72393 | 1.48062 | 27 |
| GO_PROTEASOMAL_PROTEIN_CATABOLIC_PROCESS | 0.00002 | 0.00044 | 0.59333 | 0.54166 | 1.21005 | 377 |
| GO_PROTEASOMAL_UBIQUITIN_INDEPENDENT_PROTEIN_CATABOLI( | 0.00000 | 0.00000 | 0.80122 | 0.92254 | 1.86931 | 21 |
| GO_PROTEASOME_ACCESSORY_COMPLEX | 0.00000 | 0.00001 | 0.67496 | 0.85464 | 1.73811 | 24 |
| GO_PROTEASOME_CORE_COMPLEX | 0.00000 | 0.00000 | | 0.94375 | 1.90631 | 19 |
| GO_PROTEIN_CONTAINING_COMPLEX_DISASSEMBLY | 0.00000 | 0.00007 | 0.62726 | 0.57810 | 1.28438 | 256 |
| GO_PROTEIN_CONTAINING_COMPLEX_LOCALIZATION | 0.00063 | 0.01197 | 0.47727 | 0.55335 | 1.22520 | 208 |
| GO_PROTEIN_DNA_COMPLEX_SUBUNIT_ORGANIZATION | 0.00001 | 0.00018 | 0.61053 | 0.60174 | 1.32491 | 165 |
| GO_PROTEIN_FOLDING | 0.00000 | 0.00002 | 0.65944 | 0.63113 | 1.38644 | 154 |
| GO_PROTEIN_FOLDING_CHAPERONE | 0.00135 | 0.02238 | 0.45506 | 0.77903 | 1.55752 | 17 |
| GO_PROTEIN_IMPORT | 0.00268 | 0.03799 | 0.43171 | 0.56317 | 1.23581 | 140 |
| GO_PROTEIN_LOCALIZATION_TO_CHROMOSOME | 0.00056 | 0.01096 | 0.47727 | 0.63523 | 1.36108 | 69 |
| GO_PROTEIN_LOCALIZATION_TO_CHROMOSOME_TELOMERIC_REGI( | 0.00335 | 0.04466 | 0.43171 | 0.70399 | 1.43984 | 27 |
| GO_PROTEIN_LOCALIZATION_TO_ENDOPLASMIC_RETICULUM | 0.00000 | 0.00000 | | 0.85521 | 1.87021 | 124 |
| GO_PROTEIN_LOCALIZATION_TO_MEMBRANE | 0.00000 | 0.00000 | | 0.65123 | 1.45714 | 406 |
| GO_PROTEIN_LOCALIZATION_TO_MITOCHONDRION | 0.00019 | 0.00420 | 0.51885 | 0.60773 | 1.32398 | 110 |
| GO_PROTEIN_LOCALIZATION_TO_NUCLEUS | 0.00002 | 0.00055 | 0.57561 | 0.58926 | 1.30220 | 187 |
| GO_PROTEIN_MODIFICATION_BY_SMALL_PROTEIN_REMOVAL | 0.00000 | 0.00000 | 0.71951 | 0.60745 | 1.34689 | 221 |
| GO_PROTEIN_POLYUBIQUITINATION | 0.00004 | 0.00106 | 0.55733 | 0.55874 | 1.24108 | 253 |
| GO_PROTEIN_REFOLDING | 0.00294 | 0.04036 | 0.43171 | 0.77039 | 1.53314 | 16 |
| GO_PROTEIN_STABILIZATION | 0.00001 | 0.00024 | 0.61053 | 0.62333 | 1.36586 | 130 |
| GO_PROTEIN_TARGETING | 0.00000 | 0.00000 | | 0.71181 | 1.58743 | 327 |
| GO_PROTEIN_TARGETING_TO_MEMBRANE | 0.00000 | 0.00000 | | 0.82370 | 1.80953 | 149 |
| GO_PROTEIN_TARGETING_TO_MITOCHONDRION | 0.00346 | 0.04593 | 0.43171 | 0.60408 | 1.30091 | 78 |
| GO_PROTEIN_TRANSMEMBRANE_TRANSPORT | 0.00265 | 0.03775 | 0.43171 | 0.64528 | 1.36171 | 50 |
| GO_PROTON_TRANSMEMBRANE_TRANSPORT | 0.00213 | 0.03229 | 0.43171 | 0.60815 | 1.30536 | 71 |
| GO_PURINE_CONTAINING_COMPOUND_BIOSYNTHETIC_PROCESS | 0.00246 | 0.03601 | 0.43171 | 0.57796 | 1.25936 | 111 |
| GO_PURINE_CONTAINING_COMPOUND_METABOLIC_PROCESS | 0.00100 | 0.01761 | 0.45506 | 0.53679 | 1.19242 | 255 |
| GO_PURINE_NUCLEOSIDE_TRIPHOSPHATE_METABOLIC_PROCESS | 0.00089 | 0.01592 | 0.47727 | 0.67157 | 1.40374 | 43 |
| GO_PYRIMIDINE_CONTAINING_COMPOUND_METABOLIC_PROCESS | 0.00353 | 0.04658 | 0.43171 | 0.64744 | 1.35878 | 47 |
| GO_REGENERATION | 0.00216 | 0.03242 | 0.43171 | 0.59987 | 1.29453 | 82 |
| GO_REGULATION_OF_ANIMAL_ORGAN_MORPHOGENESIS | 0.00000 | 0.00000 | | 0.69163 | 1.51541 | 133 |
| GO_REGULATION_OF_APOPTOTIC_SIGNALING_PATHWAY | 0.00031 | 0.00645 | 0.49849 | 0.55171 | 1.22506 | 247 |
| GO_REGULATION_OF_ATPASE_ACTIVITY | 0.00324 | 0.04369 | 0.43171 | 0.63031 | 1.34038 | 57 |
| GO_REGULATION_OF_ATP_METABOLIC_PROCESS | 0.00353 | 0.04658 | 0.43171 | 0.60176 | 1.29780 | 80 |
| GO_REGULATION_OF_BINDING | 0.00250 | 0.03636 | 0.43171 | 0.54476 | 1.20598 | 203 |
| GO_REGULATION_OF_CELLULAR_AMIDE_METABOLIC_PROCESS | 0.00000 | 0.00000 | | 0.62967 | 1.40226 | 311 |
| GO_REGULATION_OF_CELLULAR_AMINE_METABOLIC_PROCESS | 0.00000 | 0.00000 | | 0.82173 | 1.74682 | 58 |
| GO_REGULATION_OF_CELLULAR_AMINO_ACID_METABOLIC_PROCES | 0.00000 | 0.00000 | | 0.83776 | 1.77766 | 55 |
| GO_REGULATION_OF_CELLULAR_KETONE_METABOLIC_PROCESS | 0.00000 | 0.00000 | | 0.72641 | 1.58680 | 118 |
| GO_REGULATION_OF_CELLULAR_PROTEIN_LOCALIZATION | 0.00082 | 0.01513 | 0.47727 | 0.52542 | 1.17290 | 362 |
| GO_REGULATION_OF_CELLULAR_RESPONSE_TO_HEAT | 0.00276 | 0.03885 | 0.43171 | 0.61652 | 1.32004 | 67 |
| GO_REGULATION_OF_CELLULAR_RESPONSE_TO_STRESS | 0.00224 | 0.03344 | 0.43171 | 0.50545 | 1.13232 | 468 |
| GO_REGULATION_OF_CELL_CYCLE_G2_M_PHASE_TRANSITION | 0.00000 | 0.00000 | 0.71951 | 0.63230 | 1.39282 | 169 |
| GO_REGULATION_OF_CELL_CYCLE_PHASE_TRANSITION | 0.00001 | 0.00026 | 0.59333 | 0.55488 | 1.23783 | 337 |
| GO_REGULATION_OF_CHROMOSOME_ORGANIZATION | 0.00081 | 0.01501 | 0.47727 | 0.54255 | 1.20527 | 261 |
| GO_REGULATION_OF_CYSTEINE_TYPE_ENDOPEPTIDASE_ACTIVITY | 0.00202 | 0.03077 | 0.43171 | 0.57120 | 1.24837 | 121 |
| GO_REGULATION_OF_CYTOPLASMIC_TRANSLATION | 0.00130 | 0.02159 | 0.45506 | 0.73678 | 1.49773 | 23 |
| GO_REGULATION_OF_DEFENSE_RESPONSE | 0.00065 | 0.01237 | 0.47727 | 0.53633 | 1.19355 | 290 |
| GO_REGULATION_OF_DNA_BIOSYNTHETIC_PROCESS | 0.00016 | 0.00360 | 0.51885 | 0.64215 | 1.37923 | 74 |
| GO_REGULATION_OF_DNA_METABOLIC_PROCESS | 0.00069 | 0.01297 | 0.47727 | 0.54425 | 1.20775 | 237 |
| GO_REGULATION_OF_DNA_TEMPLATED_TRANSCRIPTION_INITIATIO | 0.00163 | 0.02568 | 0.45506 | 0.68289 | 1.41372 | 35 |
| GO_REGULATION_OF_DNA_TEMPLATED_TRANSCRIPTION_IN_RESPO | 0.00000 | 0.00000 | | 0.73944 | 1.59800 | 89 |
| GO_REGULATION_OF_GENERATION_OF_PRECURSOR_METABOLITES_ | 0.00060 | 0.01164 | 0.47727 | 0.60476 | 1.31116 | 97 |
| GO_REGULATION_OF_HEMATOPOIETIC_PROGENITOR_CELL_DIFFERE | 0.00000 | 0.00000 | | 0.78717 | 1.69251 | 75 |
| GO_REGULATION_OF_HEMATOPOIETIC_STEM_CELL_DIFFERENTIATIO | 0.00000 | 0.00000 | | 0.81505 | 1.74256 | 65 |
| GO_REGULATION_OF_HEMOPOIESIS | 0.00000 | 0.00008 | 0.62726 | 0.58859 | 1.30634 | 228 |
| GO_REGULATION_OF_IMMUNE_RESPONSE | 0.00000 | 0.00016 | 0.61053 | 0.55362 | 1.23583 | 362 |
| GO_REGULATION_OF_INNATE_IMMUNE_RESPONSE | 0.00000 | 0.00000 | 0.71951 | 0.63883 | 1.40521 | 161 |
| GO_REGULATION_OF_INTRINSIC_APOPTOTIC_SIGNALING_PATHWAY | 0.00001 | 0.00039 | 0.59333 | 0.63478 | 1.38259 | 109 |
| GO_REGULATION_OF_MITOTIC_CELL_CYCLE | 0.00001 | 0.00018 | 0.61053 | 0.53899 | 1.20736 | 458 |
| GO_REGULATION_OF_MORPHOGENESIS_OF_AN_EPITHELIUM | 0.00000 | 0.00000 | | 0.72213 | 1.57284 | 109 |
| GO_REGULATION_OF_MRNA_CATABOLIC_PROCESS | 0.00000 | 0.00000 | | 0.69977 | 1.54127 | 168 |
| GO_REGULATION_OF_MRNA_METABOLIC_PROCESS | 0.00000 | 0.00000 | | 0.66528 | 1.47872 | 266 |
| GO_REGULATION_OF_MRNA_PROCESSING | 0.00004 | 0.00105 | 0.55733 | 0.61521 | 1.34090 | 113 |
| GO_REGULATION_OF_MRNA_SPLICING_VIA_SPLICEOSOME | 0.00001 | 0.00026 | 0.59333 | 0.67424 | 1.45411 | 80 |
| GO_REGULATION_OF_NUCLEASE_ACTIVITY | 0.00106 | 0.01819 | 0.45506 | 0.77918 | 1.56780 | 18 |
| GO_REGULATION_OF_OXIDATIVE_STRESS_INDUCED_CELL_DEATH | 0.00198 | 0.03028 | 0.43171 | 0.67287 | 1.40355 | 41 |
| GO_REGULATION_OF_PEPTIDASE_ACTIVITY | 0.00050 | 0.00978 | 0.47727 | 0.56122 | 1.24054 | 191 |
| GO_REGULATION_OF_POSTTRANSCRIPTIONAL_GENE_SILENCING | 0.00364 | 0.04761 | 0.43171 | 0.59824 | 1.28492 | 74 |

| | | | | | | |
|---|---|---|---|---|---|---|
| GO_REGULATION_OF_PROTEIN_CATABOLIC_PROCESS | 0.00179 | 0.02783 | 0.45506 | 0.53057 | 1.17929 | 270 |
| GO_REGULATION_OF_PROTEIN_LOCALIZATION_TO_NUCLEUS | 0.00001 | 0.00042 | 0.59333 | 0.65505 | 1.41558 | 87 |
| GO_REGULATION_OF_PROTEIN_MODIFICATION_BY_SMALL_PROTEIN | 0.00025 | 0.00532 | 0.49849 | 0.57068 | 1.25684 | 167 |
| GO_REGULATION_OF_PROTEIN_STABILITY | 0.00093 | 0.01652 | 0.47727 | 0.55382 | 1.22571 | 202 |
| GO_REGULATION_OF_PROTEOLYSIS | 0.00013 | 0.00301 | 0.51885 | 0.52997 | 1.18392 | 377 |
| GO_REGULATION_OF_RESPONSE_TO_BIOTIC_STIMULUS | 0.00001 | 0.00020 | 0.61053 | 0.59049 | 1.30719 | 204 |
| GO_REGULATION_OF_RNA_SPLICING | 0.00000 | 0.00001 | 0.69013 | 0.65896 | 1.43669 | 114 |
| GO_REGULATION_OF_SMALL_MOLECULE_METABOLIC_PROCESS | 0.00000 | 0.00000 | 0.71951 | 0.59134 | 1.31394 | 265 |
| GO_REGULATION_OF_STEM_CELL_DIFFERENTIATION | 0.00000 | 0.00000 | | 0.74049 | 1.60113 | 90 |
| GO_REGULATION_OF_SYMBIOTIC_PROCESS | 0.00006 | 0.00144 | 0.55733 | 0.60071 | 1.31682 | 134 |
| GO_REGULATION_OF_TELOMERASE_ACTIVITY | 0.00088 | 0.01590 | 0.47727 | 0.69161 | 1.43179 | 35 |
| GO_REGULATION_OF_TRANSCRIPTION_FROM_RNA_POLYMERASE_II | 0.00000 | 0.00000 | | 0.81194 | 1.73800 | 66 |
| GO_REGULATION_OF_TRANSCRIPTION_INITIATION_FROM_RNA_POL | 0.00143 | 0.02332 | 0.45506 | 0.70390 | 1.44582 | 29 |
| GO_REGULATION_OF_TRANSLATIONAL_INITIATION | 0.00002 | 0.00052 | 0.57561 | 0.68129 | 1.45518 | 64 |
| GO_REGULATION_OF_UBIQUITIN_PROTEIN_TRANSFERASE_ACTIVITY | 0.00061 | 0.01169 | 0.47727 | 0.69335 | 1.44627 | 41 |
| GO_REGULATION_OF_VIRAL_GENOME_REPLICATION | 0.00086 | 0.01566 | 0.47727 | 0.65316 | 1.38389 | 54 |
| GO_REGULATION_OF_VIRAL_TRANSCRIPTION | 0.00198 | 0.03028 | 0.43171 | 0.63932 | 1.34703 | 49 |
| GO_REGULATION_OF_WNT_SIGNALING_PATHWAY | 0.00000 | 0.00001 | 0.69013 | 0.60353 | 1.33816 | 220 |
| GO_RESPIRASOME | 0.00008 | 0.00181 | 0.53843 | 0.65420 | 1.40421 | 71 |
| GO_RESPIRATORY_CHAIN_COMPLEX | 0.00016 | 0.00353 | 0.51885 | 0.67626 | 1.43918 | 60 |
| GO_RESPIRATORY_ELECTRON_TRANSPORT_CHAIN | 0.00001 | 0.00025 | 0.59333 | 0.65819 | 1.42190 | 85 |
| GO_RESPONSE_TO_ALCOHOL | 0.00261 | 0.03750 | 0.43171 | 0.58804 | 1.27350 | 95 |
| GO_RESPONSE_TO_CADMIUM_ION | 0.00265 | 0.03775 | 0.43171 | 0.67227 | 1.39174 | 35 |
| GO_RESPONSE_TO_DRUG | 0.00104 | 0.01798 | 0.45506 | 0.55364 | 1.22258 | 180 |
| GO_RESPONSE_TO_ETHANOL | 0.00086 | 0.01566 | 0.47727 | 0.67309 | 1.40691 | 43 |
| GO_RESPONSE_TO_HEAT | 0.00012 | 0.00270 | 0.53843 | 0.61665 | 1.34221 | 107 |
| GO_RESPONSE_TO_HORMONE | 0.00048 | 0.00949 | 0.49849 | 0.51277 | 1.14814 | 453 |
| GO_RESPONSE_TO_HYDROGEN_PEROXIDE | 0.00257 | 0.03716 | 0.43171 | 0.60505 | 1.30368 | 79 |
| GO_RESPONSE_TO_INORGANIC_SUBSTANCE | 0.00004 | 0.00107 | 0.55733 | 0.55714 | 1.23870 | 275 |
| GO_RESPONSE_TO_INTERLEUKIN_1 | 0.00000 | 0.00000 | | 0.74844 | 1.62908 | 107 |
| GO_RESPONSE_TO_INTERLEUKIN_12 | 0.00012 | 0.00270 | 0.53843 | 0.72605 | 1.50694 | 36 |
| GO_RESPONSE_TO_INTERLEUKIN_4 | 0.00324 | 0.04369 | 0.43171 | 0.75812 | 1.51571 | 17 |
| GO_RESPONSE_TO_METAL_ION | 0.00030 | 0.00623 | 0.49849 | 0.56387 | 1.24233 | 171 |
| GO_RESPONSE_TO_ORGANIC_CYCLIC_COMPOUND | 0.00000 | 0.00016 | 0.61053 | 0.54603 | 1.22259 | 439 |
| GO_RESPONSE_TO_OXYGEN_LEVELS | 0.00000 | 0.00000 | 0.71951 | 0.60132 | 1.33438 | 233 |
| GO_RESPONSE_TO_STEROID_HORMONE | 0.00154 | 0.02464 | 0.45506 | 0.55514 | 1.22446 | 174 |
| GO_RESPONSE_TO_TEMPERATURE_STIMULUS | 0.00027 | 0.00572 | 0.49849 | 0.59216 | 1.29675 | 127 |
| GO_RESPONSE_TO_TUMOR_NECROSIS_FACTOR | 0.00000 | 0.00000 | 0.76146 | 0.65649 | 1.44236 | 151 |
| GO_RETINA_HOMEOSTASIS | 0.00104 | 0.01798 | 0.45506 | 0.73063 | 1.49432 | 27 |
| GO_RIBONUCLEOPROTEIN_COMPLEX_BINDING | 0.00000 | 0.00000 | 0.71951 | 0.66906 | 1.46050 | 117 |
| GO_RIBONUCLEOPROTEIN_COMPLEX_BIOGENESIS | 0.00000 | 0.00000 | | 0.66339 | 1.48264 | 389 |
| GO_RIBONUCLEOPROTEIN_COMPLEX_SUBUNIT_ORGANIZATION | 0.00000 | 0.00000 | | 0.69241 | 1.52681 | 173 |
| GO_RIBONUCLEOPROTEIN_GRANULE | 0.00000 | 0.00001 | 0.67496 | 0.61979 | 1.36366 | 163 |
| GO_RIBONUCLEOSIDE_TRIPHOSPHATE_BIOSYNTHETIC_PROCESS | 0.00041 | 0.00840 | 0.49849 | 0.71253 | 1.47248 | 33 |
| GO_RIBONUCLEOSIDE_TRIPHOSPHATE_METABOLIC_PROCESS | 0.00026 | 0.00550 | 0.49849 | 0.69681 | 1.45611 | 42 |
| GO_RIBOSE_PHOSPHATE_BIOSYNTHETIC_PROCESS | 0.00191 | 0.02939 | 0.45506 | 0.57932 | 1.26207 | 110 |
| GO_RIBOSE_PHOSPHATE_METABOLIC_PROCESS | 0.00108 | 0.01839 | 0.45506 | 0.53891 | 1.19687 | 245 |
| GO_RIBOSOMAL_LARGE_SUBUNIT_ASSEMBLY | 0.00000 | 0.00016 | 0.61053 | 0.80999 | 1.65663 | 27 |
| GO_RIBOSOMAL_LARGE_SUBUNIT_BIOGENESIS | 0.00000 | 0.00000 | | 0.77542 | 1.65982 | 66 |
| GO_RIBOSOMAL_SMALL_SUBUNIT_ASSEMBLY | 0.00023 | 0.00494 | 0.51885 | 0.81139 | 1.62221 | 17 |
| GO_RIBOSOMAL_SMALL_SUBUNIT_BINDING | 0.00383 | 0.04935 | 0.43171 | 0.75288 | 1.50523 | 17 |
| GO_RIBOSOMAL_SMALL_SUBUNIT_BIOGENESIS | 0.00001 | 0.00022 | 0.61053 | 0.69076 | 1.47997 | 68 |
| GO_RIBOSOMAL_SUBUNIT | 0.00000 | 0.00000 | | 0.77603 | 1.71031 | 172 |
| GO_RIBOSOME | 0.00000 | 0.00000 | | 0.76643 | 1.69659 | 206 |
| GO_RIBOSOME_ASSEMBLY | 0.00000 | 0.00000 | 0.73376 | 0.76173 | 1.61998 | 59 |
| GO_RIBOSOME_BINDING | 0.00019 | 0.00416 | 0.51885 | 0.68955 | 1.44715 | 47 |
| GO_RIBOSOME_BIOGENESIS | 0.00000 | 0.00000 | | 0.65877 | 1.46466 | 275 |
| GO_RNA_CATABOLIC_PROCESS | 0.00000 | 0.00000 | | 0.76774 | 1.71264 | 339 |
| GO_RNA_DEPENDENT_DNA_BIOSYNTHETIC_PROCESS | 0.00003 | 0.00076 | 0.57561 | 0.68926 | 1.46521 | 58 |
| GO_RNA_EXPORT_FROM_NUCLEUS | 0.00062 | 0.01183 | 0.47727 | 0.58378 | 1.27758 | 126 |
| GO_RNA_LOCALIZATION | 0.00000 | 0.00006 | 0.64355 | 0.59670 | 1.32020 | 198 |
| GO_RNA_POLYMERASE_II_PREINITIATION_COMPLEX_ASSEMBLY | 0.00089 | 0.01592 | 0.47727 | 0.77046 | 1.55626 | 19 |
| GO_RNA_SPLICING | 0.00000 | 0.00000 | 0.78819 | 0.59079 | 1.31927 | 369 |
| GO_RNA_SPLICING_VIA_TRANSESTERIFICATION_REACTIONS | 0.00000 | 0.00000 | | 0.60940 | 1.35714 | 300 |
| GO_RNA_STABILIZATION | 0.00065 | 0.01223 | 0.47727 | 0.69743 | 1.44384 | 35 |
| GO_ROUGH_ENDOPLASMIC_RETICULUM | 0.00001 | 0.00039 | 0.59333 | 0.71853 | 1.51629 | 50 |
| GO_RRNA_BINDING | 0.00000 | 0.00001 | 0.69013 | 0.74782 | 1.58865 | 56 |
| GO_RRNA_METABOLIC_PROCESS | 0.00000 | 0.00000 | 0.77494 | 0.63047 | 1.39562 | 206 |
| GO_SCF_DEPENDENT_PROTEASOMAL_UBIQUITIN_DEPENDENT_PRO | 0.00000 | 0.00000 | | 0.77527 | 1.67044 | 79 |
| GO_SECRETORY_GRANULE | 0.00000 | 0.00000 | 0.71951 | 0.56518 | 1.26461 | 399 |
| GO_SECRETORY_VESICLE | 0.00000 | 0.00002 | 0.67496 | 0.54866 | 1.22905 | 465 |
| GO_SIGNAL_TRANSDUCTION_BY_PROTEIN_PHOSPHORYLATION | 0.00004 | 0.00109 | 0.55733 | 0.53817 | 1.20474 | 424 |
| GO_SINGLE_FERTILIZATION | 0.00048 | 0.00949 | 0.49849 | 0.67414 | 1.42260 | 50 |
| GO_SINGLE_STRANDED_RNA_BINDING | 0.00039 | 0.00793 | 0.49849 | 0.66553 | 1.41527 | 57 |
| GO_SMALL_MOLECULE_BIOSYNTHETIC_PROCESS | 0.00216 | 0.03242 | 0.43171 | 0.51195 | 1.14522 | 397 |
| GO_SMALL_RIBOSOMAL_SUBUNIT | 0.00000 | 0.00000 | | 0.77648 | 1.66209 | 66 |

| | | | | | | |
|---|---|---|---|---|---|---|
| GO_SM_LIKE_PROTEIN_FAMILY_COMPLEX | 0.00327 | 0.04398 | 0.43171 | 0.60468 | 1.29713 | 70 |
| GO_SPERM_EGG_RECOGNITION | 0.00000 | 0.00007 | 0.64355 | 0.85241 | 1.72955 | 22 |
| GO_SPLICEOSOMAL_COMPLEX | 0.00000 | 0.00003 | 0.65944 | 0.60939 | 1.34261 | 171 |
| GO_STEM_CELL_DIFFERENTIATION | 0.00000 | 0.00000 | 0.78819 | 0.66886 | 1.46930 | 147 |
| GO_STRUCTURAL_CONSTITUENT_OF_CYTOSKELETON | 0.00102 | 0.01782 | 0.45506 | 0.65913 | 1.39337 | 52 |
| GO_STRUCTURAL_CONSTITUENT_OF_RIBOSOME | 0.00000 | 0.00000 | | 0.79909 | 1.75541 | 150 |
| GO_STRUCTURAL_MOLECULE_ACTIVITY | 0.00000 | 0.00000 | | 0.72651 | 1.61895 | 324 |
| GO_SUBSTANTIA_NIGRA_DEVELOPMENT | 0.00000 | 0.00013 | 0.62726 | 0.81622 | 1.66507 | 26 |
| GO_TELOMERASE_RNA_LOCALIZATION | 0.00001 | 0.00017 | 0.61053 | 0.85164 | 1.72025 | 19 |
| GO_TELOMERE_MAINTENANCE_VIA_TELOMERE_LENGTHENING | 0.00011 | 0.00250 | 0.53843 | 0.65659 | 1.40546 | 66 |
| GO_TELOMERE_ORGANIZATION | 0.00342 | 0.04554 | 0.43171 | 0.57000 | 1.24649 | 124 |
| GO_THREONINE_TYPE_ENDOPEPTIDASE_ACTIVITY | 0.00000 | 0.00000 | | 0.95179 | 1.92255 | 19 |
| GO_THREONINE_TYPE_PEPTIDASE_ACTIVITY | 0.00000 | 0.00000 | | 0.92199 | 1.87423 | 23 |
| GO_TISSUE_MORPHOGENESIS | 0.00097 | 0.01707 | 0.47727 | 0.52661 | 1.17402 | 326 |
| GO_TOXIN_TRANSPORT | 0.00113 | 0.01922 | 0.45506 | 0.72897 | 1.48708 | 26 |
| GO_TRANSCRIPTION_COUPLED_NUCLEOTIDE_EXCISION_REPAIR | 0.00001 | 0.00019 | 0.61053 | 0.68106 | 1.46186 | 71 |
| GO_TRANSCRIPTION_FACTOR_BINDING | 0.00043 | 0.00865 | 0.49849 | 0.52447 | 1.17376 | 409 |
| GO_TRANSCRIPTION_PREINITIATION_COMPLEX_ASSEMBLY | 0.00119 | 0.02004 | 0.45506 | 0.71399 | 1.46797 | 30 |
| GO_TRANSLATIONAL_ELONGATION | 0.00000 | 0.00000 | 0.71951 | 0.66087 | 1.44451 | 122 |
| GO_TRANSLATIONAL_INITIATION | 0.00000 | 0.00000 | | 0.84691 | 1.86515 | 166 |
| GO_TRANSLATIONAL_TERMINATION | 0.00000 | 0.00005 | 0.64355 | 0.65946 | 1.43060 | 99 |
| GO_TRANSLATION_ELONGATION_FACTOR_ACTIVITY | 0.00161 | 0.02546 | 0.45506 | 0.79860 | 1.58418 | 15 |
| GO_TRANSLATION_FACTOR_ACTIVITY_RNA_BINDING | 0.00000 | 0.00000 | | 0.79008 | 1.69288 | 69 |
| GO_TRANSLATION_INITIATION_FACTOR_ACTIVITY | 0.00000 | 0.00000 | 0.76146 | 0.80422 | 1.68417 | 45 |
| GO_TRANSLATION_INITIATION_FACTOR_BINDING | 0.00021 | 0.00468 | 0.51885 | 0.76186 | 1.55231 | 25 |
| GO_TRANSLATION_PREINITIATION_COMPLEX | 0.00000 | 0.00001 | 0.67496 | 0.90245 | 1.79595 | 16 |
| GO_TRANSLATION_REGULATOR_ACTIVITY | 0.00000 | 0.00000 | | 0.77663 | 1.68904 | 105 |
| GO_TRANSLATION_REGULATOR_ACTIVITY_NUCLEIC_ACID_BINDING | 0.00000 | 0.00000 | | 0.77150 | 1.66586 | 83 |
| GO_TRICARBOXYLIC_ACID_CYCLE | 0.00154 | 0.02464 | 0.45506 | 0.71854 | 1.47240 | 28 |
| GO_TUMOR_NECROSIS_FACTOR_MEDIATED_SIGNALING_PATHWAY | 0.00000 | 0.00000 | | 0.76277 | 1.65372 | 97 |
| GO_T_CELL_RECEPTOR_SIGNALING_PATHWAY | 0.00000 | 0.00000 | 0.81404 | 0.69994 | 1.52527 | 112 |
| GO_U12_TYPE_SPLICEOSOMAL_COMPLEX | 0.00279 | 0.03889 | 0.43171 | 0.70746 | 1.44693 | 27 |
| GO_U2_SNRNP | 0.00005 | 0.00134 | 0.55733 | 0.83081 | 1.67168 | 18 |
| GO_U2_TYPE_SPLICEOSOMAL_COMPLEX | 0.00010 | 0.00243 | 0.53843 | 0.63403 | 1.36970 | 85 |
| GO_UBIQUITIN_LIKE_PROTEIN_LIGASE_BINDING | 0.00000 | 0.00000 | 0.76146 | 0.62273 | 1.38051 | 219 |
| GO_UBIQUITIN_PROTEIN_TRANSFERASE_REGULATOR_ACTIVITY | 0.00165 | 0.02590 | 0.45506 | 0.79605 | 1.57911 | 15 |
| GO_UNFOLDED_PROTEIN_BINDING | 0.00000 | 0.00000 | 0.77494 | 0.72157 | 1.55881 | 85 |
| GO_VESICLE_LUMEN | 0.00000 | 0.00000 | 0.74774 | 0.64475 | 1.42008 | 168 |
| GO_VIRAL_GENE_EXPRESSION | 0.00000 | 0.00000 | | 0.80028 | 1.76317 | 171 |
| GO_VIRAL_GENOME_REPLICATION | 0.00100 | 0.01761 | 0.45506 | 0.61997 | 1.33078 | 73 |
| GO_VIRAL_LIFE_CYCLE | 0.00031 | 0.00640 | 0.49849 | 0.55767 | 1.23509 | 209 |
| HP_ABNORMALITY_OF_ACID_BASE_HOMEOSTASIS | 0.00216 | 0.03242 | 0.43171 | 0.53556 | 1.18846 | 232 |
| HP_ABNORMALITY_OF_CARDIOVASCULAR_SYSTEM_ELECTROPHYSIO | 0.00027 | 0.00572 | 0.49849 | 0.55832 | 1.23929 | 229 |
| HP_ABNORMALITY_OF_LOWER_LIP | 0.00181 | 0.02804 | 0.45506 | 0.56360 | 1.23779 | 145 |
| HP_ABNORMALITY_OF_THE_MITOCHONDRION | 0.00213 | 0.03229 | 0.43171 | 0.57831 | 1.26240 | 117 |
| HP_ABNORMALITY_OF_THE_NASAL_DORSUM | 0.00005 | 0.00116 | 0.55733 | 0.60288 | 1.32157 | 134 |
| HP_ABNORMAL_ELECTROPHYSIOLOGY_OF_SINOATRIAL_NODE_ORIG | 0.00279 | 0.03889 | 0.43171 | 0.68436 | 1.40934 | 31 |
| HP_ABNORMAL_ERYTHROCYTE_MORPHOLOGY | 0.00000 | 0.00015 | 0.61053 | 0.56049 | 1.25036 | 341 |
| HP_ABNORMAL_MEAN_CORPUSCULAR_VOLUME | 0.00357 | 0.04696 | 0.43171 | 0.75226 | 1.51363 | 18 |
| HP_ABNORMAL_RETICULOCYTE_MORPHOLOGY | 0.00064 | 0.01210 | 0.47727 | 0.71420 | 1.47081 | 31 |
| HP_ACUTE_LEUKEMIA | 0.00000 | 0.00000 | 0.73376 | 0.72786 | 1.56137 | 70 |
| HP_ANEMIA_OF_INADEQUATE_PRODUCTION | 0.00000 | 0.00008 | 0.62726 | 0.65593 | 1.42209 | 97 |
| HP_CONSTITUTIONAL_SYMPTOM | 0.00119 | 0.02004 | 0.45506 | 0.51427 | 1.15108 | 423 |
| HP_DELAYED_PUBERTY | 0.00005 | 0.00115 | 0.55733 | 0.62579 | 1.36211 | 107 |
| HP_DEPRESSED_NASAL_RIDGE | 0.00000 | 0.00011 | 0.62726 | 0.68731 | 1.47511 | 72 |
| HP_FATIGUE | 0.00039 | 0.00808 | 0.49849 | 0.56529 | 1.24923 | 187 |
| HP_HEADACHE | 0.00015 | 0.00340 | 0.51885 | 0.59065 | 1.29716 | 146 |
| HP_HEMATOLOGICAL_NEOPLASM | 0.00003 | 0.00078 | 0.57561 | 0.60108 | 1.32044 | 154 |
| HP_INCREASED_CSF_LACTATE | 0.00157 | 0.02509 | 0.45506 | 0.61622 | 1.31904 | 66 |
| HP_INCREASED_SERUM_LACTATE | 0.00216 | 0.03242 | 0.43171 | 0.56920 | 1.24724 | 130 |
| HP_LEUKEMIA | 0.00001 | 0.00026 | 0.59333 | 0.64877 | 1.40680 | 98 |
| HP_MACROCYTIC_ANEMIA | 0.00000 | 0.00000 | 0.74774 | 0.82468 | 1.71698 | 39 |
| HP_MIGRAINE | 0.00000 | 0.00000 | 0.71951 | 0.72105 | 1.54497 | 69 |
| HP_MYOTONIA | 0.00161 | 0.02546 | 0.45506 | 0.79820 | 1.58338 | 15 |
| HP_NEOPLASM | 0.00216 | 0.03242 | 0.43171 | 0.50820 | 1.13805 | 446 |
| HP_PALLOR | 0.00001 | 0.00040 | 0.59333 | 0.65169 | 1.41044 | 92 |
| HP_PUBERTY_AND_GONADAL_DISORDERS | 0.00287 | 0.03973 | 0.43171 | 0.51995 | 1.15843 | 322 |
| HP_RETICULOCYTOPENIA | 0.00004 | 0.00105 | 0.55733 | 0.86417 | 1.71425 | 15 |
| HP_RIMMED_VACUOLES | 0.00279 | 0.03889 | 0.43171 | 0.74033 | 1.49831 | 20 |
| HP_SHORT_NOSE | 0.00235 | 0.03474 | 0.43171 | 0.54919 | 1.21134 | 174 |
| HP_THICK_LOWER_LIP_VERMILION | 0.00048 | 0.00949 | 0.49849 | 0.60773 | 1.31617 | 94 |
| HP_THICK_VERMILION_BORDER | 0.00156 | 0.02486 | 0.45506 | 0.55983 | 1.23000 | 153 |
| HP_VENTRICULAR_ARRHYTHMIA | 0.00353 | 0.04658 | 0.43171 | 0.66198 | 1.37398 | 36 |
| **Cluster 1** | | | | | | |
| pathway | pval | padj | log2err | ES | NES | size |
| GO_ACTIVATION_OF_IMMUNE_RESPONSE | 0.00000 | 0.00000 | 0.78819 | 0.62851 | 1.39255 | 212 |

| | | | | | | |
|---|---|---|---|---|---|---|
| GO_ACTIVATION_OF_INNATE_IMMUNE_RESPONSE | 0.00000 | 0.00000 | | 0.75054 | 1.62130 | 88 |
| GO_AEROBIC_RESPIRATION | 0.00001 | 0.00025 | 0.59333 | 0.69302 | 1.48168 | 65 |
| GO_ALTERNATIVE_MRNA_SPLICING_VIA_SPLICEOSOME | 0.00102 | 0.01782 | 0.45506 | 0.65168 | 1.38687 | 60 |
| GO_AMINE_METABOLIC_PROCESS | 0.00000 | 0.00000 | | 0.79189 | 1.70783 | 80 |
| GO_ANAPHASE_PROMOTING_COMPLEX_DEPENDENT_CATABOLIC_P | 0.00000 | 0.00000 | | 0.76611 | 1.65070 | 79 |
| GO_ANCHORING_JUNCTION | 0.00000 | 0.00000 | | 0.59615 | 1.33633 | 479 |
| GO_ANIMAL_ORGAN_REGENERATION | 0.00257 | 0.03716 | 0.43171 | 0.68240 | 1.41020 | 33 |
| GO_ANTIGEN_PROCESSING_AND_PRESENTATION | 0.00000 | 0.00000 | 0.70498 | 0.64776 | 1.42241 | 143 |
| GO_ANTIGEN_PROCESSING_AND_PRESENTATION_OF_EXOGENOUS_ | 0.00000 | 0.00000 | | 0.81051 | 1.72358 | 57 |
| GO_ANTIGEN_PROCESSING_AND_PRESENTATION_OF_PEPTIDE_ANTI | 0.00000 | 0.00000 | 0.70498 | 0.65912 | 1.44248 | 126 |
| GO_ANTIGEN_PROCESSING_AND_PRESENTATION_OF_PEPTIDE_ANTI | 0.00000 | 0.00000 | | 0.76925 | 1.65015 | 70 |
| GO_ANTIGEN_RECEPTOR_MEDIATED_SIGNALING_PATHWAY | 0.00000 | 0.00000 | 0.80122 | 0.67877 | 1.48641 | 127 |
| GO_APOPTOTIC_SIGNALING_PATHWAY | 0.00154 | 0.02464 | 0.45506 | 0.51561 | 1.15085 | 358 |
| GO_ATP_DEPENDENT_CHROMATIN_REMODELING | 0.00253 | 0.03681 | 0.43171 | 0.62124 | 1.32400 | 61 |
| GO_ATP_METABOLIC_PROCESS | 0.00000 | 0.00000 | 0.71951 | 0.62252 | 1.37693 | 196 |
| GO_ATP_SYNTHESIS_COUPLED_ELECTRON_TRANSPORT | 0.00005 | 0.00131 | 0.55733 | 0.65519 | 1.40723 | 74 |
| GO_BINDING_OF_SPERM_TO_ZONA_PELLUCIDA | 0.00000 | 0.00001 | 0.69013 | 0.89879 | 1.80846 | 18 |
| GO_CADHERIN_BINDING | 0.00000 | 0.00000 | | 0.63068 | 1.39961 | 236 |
| GO_CANONICAL_WNT_SIGNALING_PATHWAY | 0.00000 | 0.00003 | 0.65944 | 0.60816 | 1.34553 | 197 |
| GO_CATALYTIC_STEP_2_SPLICEOSOME | 0.00000 | 0.00002 | 0.65944 | 0.68068 | 1.46976 | 83 |
| GO_CELLULAR_AMINO_ACID_METABOLIC_PROCESS | 0.00000 | 0.00000 | 0.74774 | 0.61752 | 1.36824 | 216 |
| GO_CELLULAR_COMPONENT_DISASSEMBLY | 0.00000 | 0.00005 | 0.64355 | 0.55505 | 1.24197 | 401 |
| GO_CELLULAR_KETONE_METABOLIC_PROCESS | 0.00000 | 0.00000 | 0.77494 | 0.65955 | 1.44893 | 149 |
| GO_CELLULAR_NITROGEN_COMPOUND_CATABOLIC_PROCESS | 0.00000 | 0.00000 | | 0.74079 | 1.65848 | 436 |
| GO_CELLULAR_PROTEIN_COMPLEX_DISASSEMBLY | 0.00006 | 0.00139 | 0.55733 | 0.58222 | 1.28459 | 176 |
| GO_CELLULAR_RESPIRATION | 0.00000 | 0.00000 | 0.71951 | 0.64997 | 1.42492 | 136 |
| GO_CELLULAR_RESPONSE_TO_HEAT | 0.00018 | 0.00403 | 0.51885 | 0.61962 | 1.34016 | 91 |
| GO_CELLULAR_RESPONSE_TO_ORGANIC_CYCLIC_COMPOUND | 0.00302 | 0.04108 | 0.43171 | 0.52402 | 1.16585 | 283 |
| GO_CELLULAR_RESPONSE_TO_OXYGEN_LEVELS | 0.00000 | 0.00000 | 0.78819 | 0.65911 | 1.44882 | 157 |
| GO_CELL_ACTIVATION_INVOLVED_IN_IMMUNE_RESPONSE | 0.00000 | 0.00001 | 0.67496 | 0.57339 | 1.27913 | 341 |
| GO_CELL_ADHESION_MOLECULE_BINDING | 0.00000 | 0.00000 | 0.81404 | 0.60529 | 1.34788 | 308 |
| GO_CELL_CELL_RECOGNITION | 0.00000 | 0.00002 | 0.65944 | 0.83826 | 1.71003 | 26 |
| GO_CELL_CELL_SIGNALING_BY_WNT | 0.00000 | 0.00002 | 0.67496 | 0.57736 | 1.28559 | 299 |
| GO_CELL_CYCLE_G2_M_PHASE_TRANSITION | 0.00001 | 0.00024 | 0.59333 | 0.58034 | 1.28602 | 214 |
| GO_CELL_CYCLE_PHASE_TRANSITION | 0.00000 | 0.00004 | 0.64355 | 0.54319 | 1.21681 | 464 |
| GO_CELL_DIVISION | 0.00154 | 0.02464 | 0.45506 | 0.51551 | 1.15425 | 427 |
| GO_CELL_RECOGNITION | 0.00012 | 0.00270 | 0.53843 | 0.69554 | 1.46186 | 48 |
| GO_CELL_SUBSTRATE_JUNCTION | 0.00000 | 0.00000 | | 0.67751 | 1.50775 | 291 |
| GO_CELL_SURFACE_RECEPTOR_SIGNALING_PATHWAY_INVOLVED_IN | 0.00000 | 0.00003 | 0.65944 | 0.57035 | 1.27067 | 319 |
| GO_CHAPERONE_BINDING | 0.00007 | 0.00163 | 0.53843 | 0.64754 | 1.39253 | 76 |
| GO_CHAPERONE_COMPLEX | 0.00000 | 0.00004 | 0.64355 | 0.85718 | 1.73925 | 22 |
| GO_CHAPERONE_MEDIATED_PROTEIN_COMPLEX_ASSEMBLY | 0.00170 | 0.02662 | 0.45506 | 0.77423 | 1.54792 | 17 |
| GO_CHROMATIN_BINDING | 0.00071 | 0.01323 | 0.47727 | 0.52699 | 1.17680 | 369 |
| GO_CHROMOSOMAL_REGION | 0.00246 | 0.03601 | 0.43171 | 0.52919 | 1.17661 | 276 |
| GO_COP9_SIGNALOSOME | 0.00128 | 0.02141 | 0.45506 | 0.73616 | 1.49715 | 24 |
| GO_COTRANSLATIONAL_PROTEIN_TARGETING_TO_MEMBRANE | 0.00000 | 0.00000 | | 0.89471 | 1.93513 | 91 |
| GO_CYTOCHROME_COMPLEX | 0.00168 | 0.02641 | 0.45506 | 0.75311 | 1.52417 | 20 |
| GO_CYTOKINE_MEDIATED_SIGNALING_PATHWAY | 0.00000 | 0.00000 | 0.81404 | 0.59698 | 1.33170 | 339 |
| GO_CYTOPLASMIC_TRANSLATION | 0.00000 | 0.00000 | | 0.84753 | 1.83080 | 88 |
| GO_CYTOPLASMIC_TRANSLATIONAL_INITIATION | 0.00000 | 0.00000 | 0.74774 | 0.85688 | 1.76176 | 30 |
| GO_CYTOSOLIC_LARGE_RIBOSOMAL_SUBUNIT | 0.00000 | 0.00000 | | 0.91186 | 1.92988 | 53 |
| GO_CYTOSOLIC_RIBOSOME | 0.00000 | 0.00000 | | 0.89907 | 1.94710 | 95 |
| GO_CYTOSOLIC_SMALL_RIBOSOMAL_SUBUNIT | 0.00000 | 0.00000 | | 0.89114 | 1.85533 | 39 |
| GO_DEFENSE_RESPONSE_TO_OTHER_ORGANISM | 0.00004 | 0.00099 | 0.55733 | 0.53673 | 1.20127 | 415 |
| GO_DE_NOVO_PROTEIN_FOLDING | 0.00050 | 0.00978 | 0.47727 | 0.74232 | 1.51431 | 26 |
| GO_DISORDERED_DOMAIN_SPECIFIC_BINDING | 0.00161 | 0.02546 | 0.45506 | 0.71970 | 1.47197 | 27 |
| GO_DNA_BIOSYNTHETIC_PROCESS | 0.00000 | 0.00010 | 0.62726 | 0.62505 | 1.37017 | 134 |
| GO_DNA_CONFORMATION_CHANGE | 0.00027 | 0.00572 | 0.49849 | 0.55300 | 1.22527 | 216 |
| GO_DNA_DAMAGE_RESPONSE_DETECTION_OF_DNA_DAMAGE | 0.00198 | 0.03028 | 0.43171 | 0.67949 | 1.40669 | 35 |
| GO_DNA_GEOMETRIC_CHANGE | 0.00046 | 0.00925 | 0.49849 | 0.60807 | 1.31855 | 98 |
| GO_DNA_POLYMERASE_BINDING | 0.00268 | 0.03799 | 0.43171 | 0.76255 | 1.52457 | 17 |
| GO_DNA_REPAIR | 0.00265 | 0.03775 | 0.43171 | 0.51159 | 1.14547 | 427 |
| GO_DRUG_BINDING | 0.00176 | 0.02733 | 0.45506 | 0.64119 | 1.35096 | 49 |
| GO_ELECTRON_TRANSFER_ACTIVITY | 0.00001 | 0.00020 | 0.61053 | 0.67481 | 1.44938 | 74 |
| GO_ELECTRON_TRANSPORT_CHAIN | 0.00002 | 0.00053 | 0.57561 | 0.62257 | 1.36145 | 124 |
| GO_ENDOPEPTIDASE_ACTIVITY | 0.00124 | 0.02085 | 0.45506 | 0.54955 | 1.21076 | 171 |
| GO_ENDOPEPTIDASE_COMPLEX | 0.00000 | 0.00000 | | 0.80930 | 1.73234 | 66 |
| GO_ENERGY_DERIVATION_BY_OXIDATION_OF_ORGANIC_COMPOUN | 0.00000 | 0.00000 | 0.71951 | 0.63390 | 1.39707 | 172 |
| GO_ESTABLISHMENT_OF_PROTEIN_LOCALIZATION_TO_CHROMOSON | 0.00020 | 0.00432 | 0.51885 | 0.77339 | 1.57217 | 23 |
| GO_ESTABLISHMENT_OF_PROTEIN_LOCALIZATION_TO_ENDOPLASM | 0.00000 | 0.00000 | | 0.88026 | 1.91149 | 102 |
| GO_ESTABLISHMENT_OF_PROTEIN_LOCALIZATION_TO_MEMBRANE | 0.00000 | 0.00000 | | 0.74874 | 1.66164 | 240 |
| GO_ESTABLISHMENT_OF_PROTEIN_LOCALIZATION_TO_ORGANELLE | 0.00000 | 0.00000 | | 0.69519 | 1.55605 | 423 |
| GO_ESTABLISHMENT_OF_PROTEIN_LOCALIZATION_TO_TELOMERE | 0.00006 | 0.00139 | 0.55733 | 0.83470 | 1.66881 | 17 |
| GO_ESTABLISHMENT_OF_RNA_LOCALIZATION | 0.00106 | 0.01819 | 0.45506 | 0.55577 | 1.22400 | 167 |
| GO_ESTABLISHMENT_OF_TISSUE_POLARITY | 0.00000 | 0.00000 | | 0.75051 | 1.62122 | 88 |
| GO_EUKARYOTIC_TRANSLATION_INITIATION_FACTOR_3_COMPLEX | 0.00003 | 0.00074 | 0.57561 | 0.86987 | 1.72555 | 15 |

| | | | | | | |
|---|---|---|---|---|---|---|
| GO_EXOCYTOSIS | 0.00000 | 0.00015 | 0.61053 | 0.53905 | 1.20753 | 465 |
| GO_FC_EPSILON_RECEPTOR_SIGNALING_PATHWAY | 0.00000 | 0.00000 | | 0.75117 | 1.62103 | 82 |
| GO_FC_RECEPTOR_SIGNALING_PATHWAY | 0.00000 | 0.00000 | | 0.72054 | 1.57787 | 127 |
| GO_FERTILIZATION | 0.00089 | 0.01592 | 0.47727 | 0.63620 | 1.35888 | 64 |
| GO_FICOLIN_1_RICH_GRANULE | 0.00000 | 0.00000 | 0.78819 | 0.71716 | 1.55340 | 93 |
| GO_GENERATION_OF_PRECURSOR_METABOLITES_AND_ENERGY | 0.00000 | 0.00000 | 0.77494 | 0.59261 | 1.32026 | 320 |
| GO_GENE_SILENCING | 0.00261 | 0.03750 | 0.43171 | 0.55578 | 1.22252 | 161 |
| GO_GLAND_DEVELOPMENT | 0.00154 | 0.02464 | 0.45506 | 0.54117 | 1.19797 | 206 |
| GO_GLYCOSYL_COMPOUND_METABOLIC_PROCESS | 0.00071 | 0.01323 | 0.47727 | 0.62524 | 1.34204 | 71 |
| GO_GUANYL_NUCLEOTIDE_BINDING | 0.00095 | 0.01679 | 0.47727 | 0.54096 | 1.19861 | 216 |
| GO_HEAT_SHOCK_PROTEIN_BINDING | 0.00003 | 0.00092 | 0.55733 | 0.64369 | 1.39056 | 85 |
| GO_HEMATOPOIETIC_PROGENITOR_CELL_DIFFERENTIATION | 0.00000 | 0.00000 | 0.82666 | 0.70307 | 1.53239 | 113 |
| GO_HEMATOPOIETIC_STEM_CELL_DIFFERENTIATION | 0.00000 | 0.00000 | | 0.79776 | 1.71217 | 72 |
| GO_HSP70_PROTEIN_BINDING | 0.00228 | 0.03391 | 0.43171 | 0.71143 | 1.45504 | 27 |
| GO_IMMUNE_RESPONSE_REGULATING_SIGNALING_PATHWAY | 0.00000 | 0.00000 | 0.76146 | 0.63494 | 1.40480 | 197 |
| GO_INNATE_IMMUNE_RESPONSE | 0.00003 | 0.00071 | 0.57561 | 0.55173 | 1.23107 | 347 |
| GO_INNATE_IMMUNE_RESPONSE_ACTIVATING_SIGNAL_TRANSDUCT | 0.00000 | 0.00000 | | 0.79340 | 1.70281 | 72 |
| GO_INNER_MITOCHONDRIAL_MEMBRANE_PROTEIN_COMPLEX | 0.00002 | 0.00060 | 0.57561 | 0.65124 | 1.40678 | 88 |
| GO_INTERACTION_WITH_SYMBIONT | 0.00001 | 0.00026 | 0.59333 | 0.75084 | 1.56620 | 41 |
| GO_INTERLEUKIN_1_MEDIATED_SIGNALING_PATHWAY | 0.00000 | 0.00000 | | 0.81689 | 1.75671 | 76 |
| GO_INTRINSIC_APOPTOTIC_SIGNALING_PATHWAY | 0.00121 | 0.02029 | 0.45506 | 0.55118 | 1.21845 | 190 |
| GO_ION_CHANNEL_BINDING | 0.00087 | 0.01584 | 0.47727 | 0.63182 | 1.34683 | 62 |
| GO_ISOPEPTIDASE_ACTIVITY | 0.00139 | 0.02285 | 0.45506 | 0.78940 | 1.57097 | 16 |
| GO_LARGE_RIBOSOMAL_SUBUNIT | 0.00000 | 0.00000 | | 0.78135 | 1.70183 | 109 |
| GO_LEUKOCYTE_MEDIATED_IMMUNITY | 0.00000 | 0.00000 | 0.71951 | 0.57282 | 1.27895 | 364 |
| GO_MACROMOLECULE_TRANSMEMBRANE_TRANSPORTER_ACTIVITY | 0.00239 | 0.03510 | 0.43171 | 0.77734 | 1.54696 | 16 |
| GO_MAINTENANCE_OF_CELL_NUMBER | 0.00087 | 0.01584 | 0.47727 | 0.58944 | 1.28512 | 114 |
| GO_MATURATION_OF_LSU_RRNA | 0.00006 | 0.00151 | 0.53843 | 0.77606 | 1.58723 | 27 |
| GO_METHYLTRANSFERASE_COMPLEX | 0.00290 | 0.04014 | 0.43171 | 0.59476 | 1.28479 | 88 |
| GO_METHYL_CPG_BINDING | 0.00302 | 0.04108 | 0.43171 | 0.78053 | 1.54833 | 15 |
| GO_MIDBRAIN_DEVELOPMENT | 0.00021 | 0.00468 | 0.51885 | 0.68832 | 1.44457 | 47 |
| GO_MITOCHONDRIAL_ELECTRON_TRANSPORT_NADH_TO_UBIQUINO | 0.00331 | 0.04437 | 0.43171 | 0.65428 | 1.36724 | 42 |
| GO_MITOCHONDRIAL_GENE_EXPRESSION | 0.00048 | 0.00949 | 0.49849 | 0.57608 | 1.26525 | 152 |
| GO_MITOCHONDRIAL_LARGE_RIBOSOMAL_SUBUNIT | 0.00002 | 0.00050 | 0.57561 | 0.69789 | 1.48086 | 55 |
| GO_MITOCHONDRIAL_MATRIX | 0.00000 | 0.00002 | 0.65944 | 0.56045 | 1.25158 | 371 |
| GO_MITOCHONDRIAL_MEMBRANE_ORGANIZATION | 0.00361 | 0.04734 | 0.43171 | 0.59309 | 1.28094 | 86 |
| GO_MITOCHONDRIAL_PROTEIN_COMPLEX | 0.00000 | 0.00000 | 0.81404 | 0.63650 | 1.40931 | 208 |
| GO_MITOCHONDRIAL_TRANSLATION | 0.00004 | 0.00093 | 0.55733 | 0.61219 | 1.34135 | 129 |
| GO_MITOCHONDRIAL_TRANSLATIONAL_TERMINATION | 0.00000 | 0.00010 | 0.62726 | 0.66694 | 1.44128 | 87 |
| GO_MITOCHONDRION_ORGANIZATION | 0.00051 | 0.00993 | 0.47727 | 0.52284 | 1.16863 | 390 |
| GO_MODIFICATION_DEPENDENT_MACROMOLECULE_CATABOLIC_PR | 0.00002 | 0.00053 | 0.57561 | 0.53142 | 1.19156 | 486 |
| GO_MODULATION_BY_HOST_OF_SYMBIONT_PROCESS | 0.00015 | 0.00336 | 0.51885 | 0.72848 | 1.50542 | 33 |
| GO_MODULATION_OF_PROCESS_OF_OTHER_ORGANISM | 0.00019 | 0.00423 | 0.51885 | 0.66904 | 1.42286 | 59 |
| GO_MONOSACCHARIDE_BIOSYNTHETIC_PROCESS | 0.00335 | 0.04466 | 0.43171 | 0.61992 | 1.31781 | 58 |
| GO_MORPHOGENESIS_OF_AN_EPITHELIUM | 0.00006 | 0.00151 | 0.53843 | 0.55161 | 1.22755 | 290 |
| GO_MORPHOGENESIS_OF_A_POLARIZED_EPITHELIUM | 0.00000 | 0.00000 | | 0.72298 | 1.57057 | 103 |
| GO_MRNA_3_UTR_BINDING | 0.00003 | 0.00089 | 0.55733 | 0.67596 | 1.44827 | 68 |
| GO_MRNA_5_UTR_BINDING | 0.00006 | 0.00156 | 0.53843 | 0.80133 | 1.62592 | 22 |
| GO_MRNA_BINDING | 0.00000 | 0.00000 | | 0.66002 | 1.46353 | 222 |
| GO_MRNA_PROCESSING | 0.00000 | 0.00000 | 0.73376 | 0.56859 | 1.27295 | 418 |
| GO_MULTI_ORGANISM_LOCALIZATION | 0.00268 | 0.03799 | 0.43171 | 0.63245 | 1.33612 | 51 |
| GO_MULTI_ORGANISM_PROCESS | 0.00108 | 0.01839 | 0.45506 | 0.51114 | 1.14599 | 484 |
| GO_MYD88_DEPENDENT_TOLL_LIKE_RECEPTOR_SIGNALING_PATHW | 0.00002 | 0.00066 | 0.57561 | 0.85581 | 1.70312 | 16 |
| GO_MYELOID_LEUKOCYTE_ACTIVATION | 0.00000 | 0.00000 | 0.74774 | 0.58508 | 1.30479 | 327 |
| GO_MYELOID_LEUKOCYTE_MEDIATED_IMMUNITY | 0.00000 | 0.00000 | 0.76146 | 0.59841 | 1.33291 | 301 |
| GO_NADH_DEHYDROGENASE_ACTIVITY | 0.00383 | 0.04935 | 0.43171 | 0.66244 | 1.37139 | 35 |
| GO_NADH_DEHYDROGENASE_COMPLEX | 0.00376 | 0.04884 | 0.43171 | 0.65649 | 1.36421 | 38 |
| GO_NADH_METABOLIC_PROCESS | 0.00294 | 0.04036 | 0.43171 | 0.70642 | 1.44481 | 27 |
| GO_NCRNA_EXPORT_FROM_NUCLEUS | 0.00302 | 0.04108 | 0.43171 | 0.66820 | 1.38331 | 35 |
| GO_NCRNA_METABOLIC_PROCESS | 0.00000 | 0.00003 | 0.65944 | 0.55907 | 1.24907 | 381 |
| GO_NCRNA_PROCESSING | 0.00000 | 0.00003 | 0.65944 | 0.57175 | 1.27504 | 328 |
| GO_NEGATIVE_REGULATION_OF_CANONICAL_WNT_SIGNALING_PAT | 0.00000 | 0.00000 | | 0.71611 | 1.56038 | 111 |
| GO_NEGATIVE_REGULATION_OF_CELLULAR_AMIDE_METABOLIC_PRO | 0.00001 | 0.00020 | 0.61053 | 0.63031 | 1.37423 | 114 |
| GO_NEGATIVE_REGULATION_OF_CELL_CYCLE | 0.00003 | 0.00076 | 0.57561 | 0.53833 | 1.20513 | 425 |
| GO_NEGATIVE_REGULATION_OF_CELL_CYCLE_G2_M_PHASE_TRANS | 0.00000 | 0.00000 | 0.76146 | 0.71490 | 1.54439 | 85 |
| GO_NEGATIVE_REGULATION_OF_CELL_CYCLE_PHASE_TRANSITION | 0.00002 | 0.00060 | 0.57561 | 0.59041 | 1.30442 | 184 |
| GO_NEGATIVE_REGULATION_OF_CELL_CYCLE_PROCESS | 0.00000 | 0.00002 | 0.65944 | 0.58690 | 1.30374 | 249 |
| GO_NEGATIVE_REGULATION_OF_MITOTIC_CELL_CYCLE | 0.00004 | 0.00095 | 0.55733 | 0.56735 | 1.25902 | 232 |
| GO_NEGATIVE_REGULATION_OF_MRNA_METABOLIC_PROCESS | 0.00023 | 0.00491 | 0.51885 | 0.66670 | 1.41788 | 59 |
| GO_NEGATIVE_REGULATION_OF_MRNA_SPLICING_VIA_SPLICEOSOM | 0.00239 | 0.03510 | 0.43171 | 0.74857 | 1.51204 | 19 |
| GO_NEGATIVE_REGULATION_OF_RNA_SPLICING | 0.00029 | 0.00607 | 0.49849 | 0.76729 | 1.55976 | 23 |
| GO_NEGATIVE_REGULATION_OF_TRANSLATIONAL_INITIATION | 0.00018 | 0.00403 | 0.51885 | 0.83901 | 1.66434 | 15 |
| GO_NEGATIVE_REGULATION_OF_WNT_SIGNALING_PATHWAY | 0.00000 | 0.00000 | | 0.69525 | 1.52249 | 127 |
| GO_NEURAL_NUCLEUS_DEVELOPMENT | 0.00001 | 0.00026 | 0.59333 | 0.79148 | 1.62187 | 28 |
| GO_NEURON_TO_NEURON_SYNAPSE | 0.00005 | 0.00134 | 0.55733 | 0.59636 | 1.31006 | 154 |
| GO_NIK_NF_KAPPAB_SIGNALING | 0.00000 | 0.00000 | | 0.72896 | 1.58354 | 103 |

| | | | | | | |
|---|---|---|---|---|---|---|
| GO_NON_CANONICAL_WNT_SIGNALING_PATHWAY | 0.00000 | 0.00000 | | 0.72784 | 1.58023 | 101 |
| GO_NUCLEAR_EXPORT | 0.00031 | 0.00645 | 0.49849 | 0.56963 | 1.25542 | 172 |
| GO_NUCLEAR_TRANSCRIBED_MRNA_CATABOLIC_PROCESS | 0.00000 | 0.00000 | | 0.79179 | 1.74917 | 183 |
| GO_NUCLEAR_TRANSCRIBED_MRNA_CATABOLIC_PROCESS_NONSEN | 0.00000 | 0.00000 | | 0.89008 | 1.93909 | 110 |
| GO_NUCLEAR_TRANSPORT | 0.00008 | 0.00193 | 0.53843 | 0.55511 | 1.23408 | 273 |
| GO_NUCLEOBASE_CONTAINING_COMPOUND_TRANSPORT | 0.00146 | 0.02386 | 0.45506 | 0.55053 | 1.21842 | 202 |
| GO_NUCLEOBASE_CONTAINING_SMALL_MOLECULE_BIOSYNTHETIC_ | 0.00137 | 0.02261 | 0.45506 | 0.62749 | 1.34353 | 67 |
| GO_NUCLEOBASE_CONTAINING_SMALL_MOLECULE_INTERCONVERS | 0.00383 | 0.04935 | 0.43171 | 0.71259 | 1.44857 | 23 |
| GO_NUCLEOBASE_CONTAINING_SMALL_MOLECULE_METABOLIC_PR | 0.00320 | 0.04339 | 0.43171 | 0.51448 | 1.14825 | 352 |
| GO_NUCLEOSIDE_DIPHOSPHATE_METABOLIC_PROCESS | 0.00110 | 0.01865 | 0.45506 | 0.59637 | 1.29295 | 97 |
| GO_NUCLEOSIDE_METABOLIC_PROCESS | 0.00061 | 0.01169 | 0.47727 | 0.65315 | 1.38907 | 59 |
| GO_NUCLEOSIDE_MONOPHOSPHATE_BIOSYNTHETIC_PROCESS | 0.00250 | 0.03636 | 0.43171 | 0.68841 | 1.42033 | 32 |
| GO_NUCLEOSIDE_MONOPHOSPHATE_METABOLIC_PROCESS | 0.00152 | 0.02464 | 0.45506 | 0.64377 | 1.36002 | 51 |
| GO_NUCLEOSIDE_PHOSPHATE_BIOSYNTHETIC_PROCESS | 0.00043 | 0.00865 | 0.49849 | 0.58185 | 1.27831 | 148 |
| GO_NUCLEOSIDE_TRIPHOSPHATE_BIOSYNTHETIC_PROCESS | 0.00043 | 0.00865 | 0.49849 | 0.69332 | 1.44539 | 40 |
| GO_NUCLEOSIDE_TRIPHOSPHATE_METABOLIC_PROCESS | 0.00016 | 0.00360 | 0.51885 | 0.66788 | 1.41976 | 58 |
| GO_NUCLEOSOMAL_DNA_BINDING | 0.00023 | 0.00507 | 0.51885 | 0.79852 | 1.61801 | 21 |
| GO_NUCLEOSOME_BINDING | 0.00143 | 0.02332 | 0.45506 | 0.66132 | 1.38492 | 45 |
| GO_NUCLEOTIDE_EXCISION_REPAIR | 0.00004 | 0.00096 | 0.55733 | 0.63343 | 1.37355 | 98 |
| GO_NUCLEOTIDE_EXCISION_REPAIR_DNA_DAMAGE_RECOGNITION | 0.00015 | 0.00349 | 0.51885 | 0.78678 | 1.59640 | 22 |
| GO_NUCLEOTIDE_EXCISION_REPAIR_DNA_INCISION | 0.00313 | 0.04249 | 0.43171 | 0.66439 | 1.37898 | 36 |
| GO_NUCLEOTIDE_PHOSPHORYLATION | 0.00383 | 0.04935 | 0.43171 | 0.59002 | 1.27454 | 88 |
| GO_ORGANELLAR_RIBOSOME | 0.00000 | 0.00014 | 0.61053 | 0.66412 | 1.43471 | 85 |
| GO_ORGANELLE_INNER_MEMBRANE | 0.00000 | 0.00000 | 0.76146 | 0.57601 | 1.28697 | 378 |
| GO_ORGANIC_CYCLIC_COMPOUND_CATABOLIC_PROCESS | 0.00000 | 0.00000 | | 0.73350 | 1.64281 | 449 |
| GO_OXIDATIVE_PHOSPHORYLATION | 0.00001 | 0.00030 | 0.59333 | 0.64619 | 1.40027 | 96 |
| GO_OXIDOREDUCTASE_ACTIVITY | 0.00006 | 0.00139 | 0.55733 | 0.53544 | 1.19703 | 392 |
| GO_OXIDOREDUCTASE_COMPLEX | 0.00001 | 0.00027 | 0.59333 | 0.66015 | 1.42461 | 82 |
| GO_PEPTIDASE_COMPLEX | 0.00000 | 0.00000 | | 0.75988 | 1.63982 | 82 |
| GO_PIGMENT_GRANULE | 0.00003 | 0.00089 | 0.55733 | 0.66098 | 1.42344 | 78 |
| GO_POLYSOMAL_RIBOSOME | 0.00000 | 0.00000 | | 0.90624 | 1.86629 | 31 |
| GO_POLYSOME | 0.00000 | 0.00000 | | 0.82337 | 1.75481 | 61 |
| GO_POLY_PYRIMIDINE_TRACT_BINDING | 0.00327 | 0.04398 | 0.43171 | 0.75471 | 1.51855 | 18 |
| GO_POSITIVE_REGULATION_OF_APOPTOTIC_SIGNALING_PATHWAY | 0.00279 | 0.03889 | 0.43171 | 0.57666 | 1.25518 | 107 |
| GO_POSITIVE_REGULATION_OF_ATPASE_ACTIVITY | 0.00294 | 0.04036 | 0.43171 | 0.65446 | 1.36795 | 43 |
| GO_POSITIVE_REGULATION_OF_BINDING | 0.00088 | 0.01590 | 0.47727 | 0.60284 | 1.30556 | 95 |
| GO_POSITIVE_REGULATION_OF_CANONICAL_WNT_SIGNALING_PATH | 0.00000 | 0.00000 | 0.77494 | 0.70504 | 1.53160 | 103 |
| GO_POSITIVE_REGULATION_OF_CELLULAR_AMIDE_METABOLIC_PRO | 0.00003 | 0.00074 | 0.57561 | 0.62296 | 1.36083 | 118 |
| GO_POSITIVE_REGULATION_OF_CELLULAR_PROTEIN_LOCALIZATION | 0.00041 | 0.00840 | 0.49849 | 0.54915 | 1.21815 | 225 |
| GO_POSITIVE_REGULATION_OF_CELL_DEATH | 0.00069 | 0.01297 | 0.47727 | 0.52595 | 1.17423 | 363 |
| GO_POSITIVE_REGULATION_OF_CELL_POPULATION_PROLIFERATION | 0.00078 | 0.01455 | 0.47727 | 0.52449 | 1.17108 | 368 |
| GO_POSITIVE_REGULATION_OF_CHROMOSOME_ORGANIZATION | 0.00231 | 0.03437 | 0.43171 | 0.56990 | 1.24871 | 133 |
| GO_POSITIVE_REGULATION_OF_DEFENSE_RESPONSE | 0.00000 | 0.00003 | 0.65944 | 0.62175 | 1.36730 | 158 |
| GO_POSITIVE_REGULATION_OF_DNA_BIOSYNTHETIC_PROCESS | 0.00001 | 0.00041 | 0.59333 | 0.71929 | 1.52055 | 52 |
| GO_POSITIVE_REGULATION_OF_DNA_METABOLIC_PROCESS | 0.00364 | 0.04761 | 0.43171 | 0.56647 | 1.24189 | 135 |
| GO_POSITIVE_REGULATION_OF_DNA_TEMPLATED_TRANSCRIPTION_ | 0.00276 | 0.03885 | 0.43171 | 0.68932 | 1.41586 | 29 |
| GO_POSITIVE_REGULATION_OF_ESTABLISHMENT_OF_PROTEIN_LOC/ | 0.00007 | 0.00168 | 0.53843 | 0.56461 | 1.25165 | 219 |
| GO_POSITIVE_REGULATION_OF_IMMUNE_RESPONSE | 0.00000 | 0.00001 | 0.69013 | 0.59037 | 1.31265 | 277 |
| GO_POSITIVE_REGULATION_OF_IMMUNE_SYSTEM_PROCESS | 0.00004 | 0.00110 | 0.55733 | 0.54018 | 1.20716 | 386 |
| GO_POSITIVE_REGULATION_OF_INNATE_IMMUNE_RESPONSE | 0.00000 | 0.00000 | 0.81404 | 0.69693 | 1.51947 | 114 |
| GO_POSITIVE_REGULATION_OF_INTRINSIC_APOPTOTIC_SIGNALING_ | 0.00104 | 0.01798 | 0.45506 | 0.69097 | 1.43578 | 37 |
| GO_POSITIVE_REGULATION_OF_ORGANELLE_ORGANIZATION | 0.00043 | 0.00865 | 0.49849 | 0.52093 | 1.16605 | 411 |
| GO_POSITIVE_REGULATION_OF_PROTEIN_LOCALIZATION_TO_NUCLE | 0.00001 | 0.00020 | 0.61053 | 0.69907 | 1.48990 | 61 |
| GO_POSITIVE_REGULATION_OF_RESPONSE_TO_BIOTIC_STIMULUS | 0.00000 | 0.00000 | 0.73376 | 0.66340 | 1.45310 | 128 |
| GO_POSITIVE_REGULATION_OF_RESPONSE_TO_EXTERNAL_STIMULU | 0.00002 | 0.00050 | 0.57561 | 0.58133 | 1.28681 | 201 |
| GO_POSITIVE_REGULATION_OF_TELOMERASE_ACTIVITY | 0.00061 | 0.01169 | 0.47727 | 0.73476 | 1.50565 | 28 |
| GO_POSITIVE_REGULATION_OF_TELOMERASE_RNA_LOCALIZATION_ | 0.00000 | 0.00001 | 0.69013 | 0.92451 | 1.83395 | 15 |
| GO_POSITIVE_REGULATION_OF_TELOMERE_MAINTENANCE_VIA_TE | 0.00130 | 0.02159 | 0.45506 | 0.71259 | 1.46508 | 30 |
| GO_POSITIVE_REGULATION_OF_TRANSCRIPTION_INITIATION_FROM | 0.00372 | 0.04847 | 0.43171 | 0.70671 | 1.43993 | 25 |
| GO_POSITIVE_REGULATION_OF_TRANSFERASE_ACTIVITY | 0.00174 | 0.02712 | 0.45506 | 0.51642 | 1.15220 | 345 |
| GO_POSITIVE_REGULATION_OF_TRANSLATION | 0.00001 | 0.00019 | 0.61053 | 0.64767 | 1.40419 | 97 |
| GO_POSITIVE_REGULATION_OF_VIRAL_GENOME_REPLICATION | 0.00287 | 0.03973 | 0.43171 | 0.68362 | 1.40783 | 31 |
| GO_POSITIVE_REGULATION_OF_VIRAL_PROCESS | 0.00028 | 0.00590 | 0.49849 | 0.62440 | 1.34741 | 81 |
| GO_POSITIVE_REGULATION_OF_WNT_SIGNALING_PATHWAY | 0.00000 | 0.00000 | 0.73376 | 0.66230 | 1.44763 | 122 |
| GO_POSTREPLICATION_REPAIR | 0.00024 | 0.00515 | 0.51885 | 0.69140 | 1.44518 | 43 |
| GO_POSTSYNAPSE | 0.00004 | 0.00109 | 0.55733 | 0.55969 | 1.24369 | 259 |
| GO_POSTSYNAPTIC_CYTOSOL | 0.00279 | 0.03889 | 0.43171 | 0.78295 | 1.55313 | 15 |
| GO_POSTTRANSCRIPTIONAL_REGULATION_OF_GENE_EXPRESSION | 0.00000 | 0.00000 | | 0.66656 | 1.49327 | 462 |
| GO_POST_TRANSLATIONAL_PROTEIN_MODIFICATION | 0.00002 | 0.00052 | 0.57561 | 0.57851 | 1.28412 | 229 |
| GO_PRECATALYTIC_SPLICEOSOME | 0.00085 | 0.01559 | 0.47727 | 0.65235 | 1.37448 | 49 |
| GO_PRERIBOSOME | 0.00087 | 0.01577 | 0.47727 | 0.62899 | 1.34772 | 69 |
| GO_PRERIBOSOME_LARGE_SUBUNIT_PRECURSOR | 0.00235 | 0.03474 | 0.43171 | 0.74079 | 1.50308 | 22 |
| GO_PRE_MRNA_BINDING | 0.00133 | 0.02213 | 0.45506 | 0.72393 | 1.48062 | 27 |
| GO_PROTEASOMAL_PROTEIN_CATABOLIC_PROCESS | 0.00002 | 0.00044 | 0.59333 | 0.54166 | 1.21005 | 377 |
| GO_PROTEASOMAL_UBIQUITIN_INDEPENDENT_PROTEIN_CATABOLI( | 0.00000 | 0.00000 | 0.80122 | 0.92254 | 1.86931 | 21 |
| GO_PROTEASOME_ACCESSORY_COMPLEX | 0.00000 | 0.00001 | 0.67496 | 0.85464 | 1.73811 | 24 |

| | | | | | | |
|---|---|---|---|---|---|---|
| GO_PROTEASOME_CORE_COMPLEX | 0.00000 | 0.00000 | | 0.94375 | 1.90631 | 19 |
| GO_PROTEIN_CONTAINING_COMPLEX_DISASSEMBLY | 0.00000 | 0.00007 | 0.62726 | 0.57810 | 1.28438 | 256 |
| GO_PROTEIN_CONTAINING_COMPLEX_LOCALIZATION | 0.00063 | 0.01197 | 0.47727 | 0.55335 | 1.22520 | 208 |
| GO_PROTEIN_DNA_COMPLEX_SUBUNIT_ORGANIZATION | 0.00001 | 0.00018 | 0.61053 | 0.60174 | 1.32491 | 165 |
| GO_PROTEIN_FOLDING | 0.00000 | 0.00002 | 0.65944 | 0.63113 | 1.38644 | 154 |
| GO_PROTEIN_FOLDING_CHAPERONE | 0.00135 | 0.02238 | 0.45506 | 0.77903 | 1.55752 | 17 |
| GO_PROTEIN_IMPORT | 0.00268 | 0.03799 | 0.43171 | 0.56317 | 1.23581 | 140 |
| GO_PROTEIN_LOCALIZATION_TO_CHROMOSOME | 0.00056 | 0.01096 | 0.47727 | 0.63523 | 1.36108 | 69 |
| GO_PROTEIN_LOCALIZATION_TO_CHROMOSOME_TELOMERIC_REGI( | 0.00335 | 0.04466 | 0.43171 | 0.70399 | 1.43984 | 27 |
| GO_PROTEIN_LOCALIZATION_TO_ENDOPLASMIC_RETICULUM | 0.00000 | 0.00000 | | 0.85521 | 1.87021 | 124 |
| GO_PROTEIN_LOCALIZATION_TO_MEMBRANE | 0.00000 | 0.00000 | | 0.65123 | 1.45714 | 406 |
| GO_PROTEIN_LOCALIZATION_TO_MITOCHONDRION | 0.00019 | 0.00420 | 0.51885 | 0.60773 | 1.32398 | 110 |
| GO_PROTEIN_LOCALIZATION_TO_NUCLEUS | 0.00002 | 0.00055 | 0.57561 | 0.58926 | 1.30220 | 187 |
| GO_PROTEIN_MODIFICATION_BY_SMALL_PROTEIN_REMOVAL | 0.00000 | 0.00000 | 0.71951 | 0.60745 | 1.34689 | 221 |
| GO_PROTEIN_POLYUBIQUITINATION | 0.00004 | 0.00106 | 0.55733 | 0.55874 | 1.24108 | 253 |
| GO_PROTEIN_REFOLDING | 0.00294 | 0.04036 | 0.43171 | 0.77039 | 1.53314 | 16 |
| GO_PROTEIN_STABILIZATION | 0.00001 | 0.00024 | 0.61053 | 0.62333 | 1.36586 | 130 |
| GO_PROTEIN_TARGETING | 0.00000 | 0.00000 | | 0.71181 | 1.58743 | 327 |
| GO_PROTEIN_TARGETING_TO_MEMBRANE | 0.00000 | 0.00000 | | 0.82370 | 1.80953 | 149 |
| GO_PROTEIN_TARGETING_TO_MITOCHONDRION | 0.00346 | 0.04593 | 0.43171 | 0.60408 | 1.30091 | 78 |
| GO_PROTEIN_TRANSMEMBRANE_TRANSPORT | 0.00265 | 0.03775 | 0.43171 | 0.64528 | 1.36171 | 50 |
| GO_PROTON_TRANSMEMBRANE_TRANSPORT | 0.00213 | 0.03229 | 0.43171 | 0.60815 | 1.30536 | 71 |
| GO_PURINE_CONTAINING_COMPOUND_BIOSYNTHETIC_PROCESS | 0.00246 | 0.03601 | 0.43171 | 0.57796 | 1.25936 | 111 |
| GO_PURINE_CONTAINING_COMPOUND_METABOLIC_PROCESS | 0.00100 | 0.01761 | 0.45506 | 0.53679 | 1.19242 | 255 |
| GO_PURINE_NUCLEOSIDE_TRIPHOSPHATE_METABOLIC_PROCESS | 0.00089 | 0.01592 | 0.47727 | 0.67157 | 1.40374 | 43 |
| GO_PYRIMIDINE_CONTAINING_COMPOUND_METABOLIC_PROCESS | 0.00353 | 0.04658 | 0.43171 | 0.64744 | 1.35878 | 47 |
| GO_REGENERATION | 0.00216 | 0.03242 | 0.43171 | 0.59987 | 1.29453 | 82 |
| GO_REGULATION_OF_ANIMAL_ORGAN_MORPHOGENESIS | 0.00000 | 0.00000 | | 0.69163 | 1.51541 | 133 |
| GO_REGULATION_OF_APOPTOTIC_SIGNALING_PATHWAY | 0.00031 | 0.00645 | 0.49849 | 0.55171 | 1.22506 | 247 |
| GO_REGULATION_OF_ATPASE_ACTIVITY | 0.00324 | 0.04369 | 0.43171 | 0.63031 | 1.34038 | 57 |
| GO_REGULATION_OF_ATP_METABOLIC_PROCESS | 0.00353 | 0.04658 | 0.43171 | 0.60176 | 1.29780 | 80 |
| GO_REGULATION_OF_BINDING | 0.00250 | 0.03636 | 0.43171 | 0.54476 | 1.20598 | 203 |
| GO_REGULATION_OF_CELLULAR_AMIDE_METABOLIC_PROCESS | 0.00000 | 0.00000 | | 0.62967 | 1.40226 | 311 |
| GO_REGULATION_OF_CELLULAR_AMINE_METABOLIC_PROCESS | 0.00000 | 0.00000 | | 0.82173 | 1.74682 | 58 |
| GO_REGULATION_OF_CELLULAR_AMINO_ACID_METABOLIC_PROCES | 0.00000 | 0.00000 | | 0.83776 | 1.77766 | 55 |
| GO_REGULATION_OF_CELLULAR_KETONE_METABOLIC_PROCESS | 0.00000 | 0.00000 | | 0.72641 | 1.58680 | 118 |
| GO_REGULATION_OF_CELLULAR_PROTEIN_LOCALIZATION | 0.00082 | 0.01513 | 0.47727 | 0.52542 | 1.17290 | 362 |
| GO_REGULATION_OF_CELLULAR_RESPONSE_TO_HEAT | 0.00276 | 0.03885 | 0.43171 | 0.61652 | 1.32004 | 67 |
| GO_REGULATION_OF_CELLULAR_RESPONSE_TO_STRESS | 0.00224 | 0.03344 | 0.43171 | 0.50545 | 1.13232 | 468 |
| GO_REGULATION_OF_CELL_CYCLE_G2_M_PHASE_TRANSITION | 0.00000 | 0.00000 | 0.71951 | 0.63230 | 1.39282 | 169 |
| GO_REGULATION_OF_CELL_CYCLE_PHASE_TRANSITION | 0.00001 | 0.00026 | 0.59333 | 0.55488 | 1.23783 | 337 |
| GO_REGULATION_OF_CHROMOSOME_ORGANIZATION | 0.00081 | 0.01501 | 0.47727 | 0.54255 | 1.20527 | 261 |
| GO_REGULATION_OF_CYSTEINE_TYPE_ENDOPEPTIDASE_ACTIVITY | 0.00202 | 0.03077 | 0.43171 | 0.57120 | 1.24837 | 121 |
| GO_REGULATION_OF_CYTOPLASMIC_TRANSLATION | 0.00130 | 0.02159 | 0.45506 | 0.73678 | 1.49773 | 23 |
| GO_REGULATION_OF_DEFENSE_RESPONSE | 0.00065 | 0.01237 | 0.47727 | 0.53633 | 1.19355 | 290 |
| GO_REGULATION_OF_DNA_BIOSYNTHETIC_PROCESS | 0.00016 | 0.00360 | 0.51885 | 0.64215 | 1.37923 | 74 |
| GO_REGULATION_OF_DNA_METABOLIC_PROCESS | 0.00069 | 0.01297 | 0.47727 | 0.54425 | 1.20775 | 237 |
| GO_REGULATION_OF_DNA_TEMPLATED_TRANSCRIPTION_INITIATIOI | 0.00163 | 0.02568 | 0.45506 | 0.68289 | 1.41372 | 35 |
| GO_REGULATION_OF_DNA_TEMPLATED_TRANSCRIPTION_IN_RESPO | 0.00000 | 0.00000 | | 0.73944 | 1.59800 | 89 |
| GO_REGULATION_OF_GENERATION_OF_PRECURSOR_METABOLITES_ | 0.00060 | 0.01164 | 0.47727 | 0.60476 | 1.31116 | 97 |
| GO_REGULATION_OF_HEMATOPOIETIC_PROGENITOR_CELL_DIFFERE | 0.00000 | 0.00000 | | 0.78717 | 1.69251 | 75 |
| GO_REGULATION_OF_HEMATOPOIETIC_STEM_CELL_DIFFERENTIATIC | 0.00000 | 0.00000 | | 0.81505 | 1.74256 | 65 |
| GO_REGULATION_OF_HEMOPOIESIS | 0.00000 | 0.00008 | 0.62726 | 0.58859 | 1.30634 | 228 |
| GO_REGULATION_OF_IMMUNE_RESPONSE | 0.00000 | 0.00016 | 0.61053 | 0.55362 | 1.23583 | 362 |
| GO_REGULATION_OF_INNATE_IMMUNE_RESPONSE | 0.00000 | 0.00000 | 0.71951 | 0.63883 | 1.40521 | 161 |
| GO_REGULATION_OF_INTRINSIC_APOPTOTIC_SIGNALING_PATHWAY | 0.00001 | 0.00039 | 0.59333 | 0.63478 | 1.38259 | 109 |
| GO_REGULATION_OF_MITOTIC_CELL_CYCLE | 0.00001 | 0.00018 | 0.61053 | 0.53899 | 1.20736 | 458 |
| GO_REGULATION_OF_MORPHOGENESIS_OF_AN_EPITHELIUM | 0.00000 | 0.00000 | | 0.72213 | 1.57284 | 109 |
| GO_REGULATION_OF_MRNA_CATABOLIC_PROCESS | 0.00000 | 0.00000 | | 0.69977 | 1.54127 | 168 |
| GO_REGULATION_OF_MRNA_METABOLIC_PROCESS | 0.00000 | 0.00000 | | 0.66528 | 1.47872 | 266 |
| GO_REGULATION_OF_MRNA_PROCESSING | 0.00004 | 0.00105 | 0.55733 | 0.61521 | 1.34090 | 113 |
| GO_REGULATION_OF_MRNA_SPLICING_VIA_SPLICEOSOME | 0.00001 | 0.00026 | 0.59333 | 0.67424 | 1.45411 | 80 |
| GO_REGULATION_OF_NUCLEASE_ACTIVITY | 0.00106 | 0.01819 | 0.45506 | 0.77918 | 1.56780 | 18 |
| GO_REGULATION_OF_OXIDATIVE_STRESS_INDUCED_CELL_DEATH | 0.00198 | 0.03028 | 0.43171 | 0.67287 | 1.40355 | 41 |
| GO_REGULATION_OF_PEPTIDASE_ACTIVITY | 0.00050 | 0.00978 | 0.47727 | 0.56122 | 1.24054 | 191 |
| GO_REGULATION_OF_POSTTRANSCRIPTIONAL_GENE_SILENCING | 0.00364 | 0.04761 | 0.43171 | 0.59824 | 1.28492 | 74 |
| GO_REGULATION_OF_PROTEIN_CATABOLIC_PROCESS | 0.00179 | 0.02783 | 0.45506 | 0.53057 | 1.17929 | 270 |
| GO_REGULATION_OF_PROTEIN_LOCALIZATION_TO_NUCLEUS | 0.00001 | 0.00042 | 0.59333 | 0.65505 | 1.41558 | 87 |
| GO_REGULATION_OF_PROTEIN_MODIFICATION_BY_SMALL_PROTEIN | 0.00025 | 0.00532 | 0.49849 | 0.57068 | 1.25684 | 167 |
| GO_REGULATION_OF_PROTEIN_STABILITY | 0.00093 | 0.01652 | 0.47727 | 0.55382 | 1.22571 | 202 |
| GO_REGULATION_OF_PROTEOLYSIS | 0.00013 | 0.00301 | 0.51885 | 0.52997 | 1.18392 | 377 |
| GO_REGULATION_OF_RESPONSE_TO_BIOTIC_STIMULUS | 0.00001 | 0.00020 | 0.61053 | 0.59049 | 1.30719 | 204 |
| GO_REGULATION_OF_RNA_SPLICING | 0.00000 | 0.00001 | 0.69013 | 0.65896 | 1.43669 | 114 |
| GO_REGULATION_OF_SMALL_MOLECULE_METABOLIC_PROCESS | 0.00000 | 0.00000 | 0.71951 | 0.59134 | 1.31394 | 265 |
| GO_REGULATION_OF_STEM_CELL_DIFFERENTIATION | 0.00000 | 0.00000 | | 0.74049 | 1.60113 | 90 |
| GO_REGULATION_OF_SYMBIOTIC_PROCESS | 0.00006 | 0.00144 | 0.55733 | 0.60071 | 1.31682 | 134 |

| | | | | | | |
|---|---|---|---|---|---|---|
| GO_REGULATION_OF_TELOMERASE_ACTIVITY | 0.00088 | 0.01590 | 0.47727 | 0.69161 | 1.43179 | 35 |
| GO_REGULATION_OF_TRANSCRIPTION_FROM_RNA_POLYMERASE_II | 0.00000 | 0.00000 | | 0.81194 | 1.73800 | 66 |
| GO_REGULATION_OF_TRANSCRIPTION_INITIATION_FROM_RNA_POL | 0.00143 | 0.02332 | 0.45506 | 0.70390 | 1.44582 | 29 |
| GO_REGULATION_OF_TRANSLATIONAL_INITIATION | 0.00002 | 0.00052 | 0.57561 | 0.68129 | 1.45518 | 64 |
| GO_REGULATION_OF_UBIQUITIN_PROTEIN_TRANSFERASE_ACTIVITY | 0.00061 | 0.01169 | 0.47727 | 0.69335 | 1.44627 | 41 |
| GO_REGULATION_OF_VIRAL_GENOME_REPLICATION | 0.00086 | 0.01566 | 0.47727 | 0.65316 | 1.38389 | 54 |
| GO_REGULATION_OF_VIRAL_TRANSCRIPTION | 0.00198 | 0.03028 | 0.43171 | 0.63932 | 1.34703 | 49 |
| GO_REGULATION_OF_WNT_SIGNALING_PATHWAY | 0.00000 | 0.00001 | 0.69013 | 0.60353 | 1.33816 | 220 |
| GO_RESPIRASOME | 0.00008 | 0.00181 | 0.53843 | 0.65420 | 1.40421 | 71 |
| GO_RESPIRATORY_CHAIN_COMPLEX | 0.00016 | 0.00353 | 0.51885 | 0.67626 | 1.43918 | 60 |
| GO_RESPIRATORY_ELECTRON_TRANSPORT_CHAIN | 0.00001 | 0.00025 | 0.59333 | 0.65819 | 1.42190 | 85 |
| GO_RESPONSE_TO_ALCOHOL | 0.00261 | 0.03750 | 0.43171 | 0.58804 | 1.27350 | 95 |
| GO_RESPONSE_TO_CADMIUM_ION | 0.00265 | 0.03775 | 0.43171 | 0.67227 | 1.39174 | 35 |
| GO_RESPONSE_TO_DRUG | 0.00104 | 0.01798 | 0.45506 | 0.55364 | 1.22258 | 180 |
| GO_RESPONSE_TO_ETHANOL | 0.00086 | 0.01566 | 0.47727 | 0.67309 | 1.40691 | 43 |
| GO_RESPONSE_TO_HEAT | 0.00012 | 0.00270 | 0.53843 | 0.61665 | 1.34221 | 107 |
| GO_RESPONSE_TO_HORMONE | 0.00048 | 0.00949 | 0.49849 | 0.51277 | 1.14814 | 453 |
| GO_RESPONSE_TO_HYDROGEN_PEROXIDE | 0.00257 | 0.03716 | 0.43171 | 0.60505 | 1.30368 | 79 |
| GO_RESPONSE_TO_INORGANIC_SUBSTANCE | 0.00004 | 0.00107 | 0.55733 | 0.55714 | 1.23870 | 275 |
| GO_RESPONSE_TO_INTERLEUKIN_1 | 0.00000 | 0.00000 | | 0.74844 | 1.62908 | 107 |
| GO_RESPONSE_TO_INTERLEUKIN_12 | 0.00012 | 0.00270 | 0.53843 | 0.72605 | 1.50694 | 36 |
| GO_RESPONSE_TO_INTERLEUKIN_4 | 0.00324 | 0.04369 | 0.43171 | 0.75812 | 1.51571 | 17 |
| GO_RESPONSE_TO_METAL_ION | 0.00030 | 0.00623 | 0.49849 | 0.56387 | 1.24233 | 171 |
| GO_RESPONSE_TO_ORGANIC_CYCLIC_COMPOUND | 0.00000 | 0.00016 | 0.61053 | 0.54603 | 1.22259 | 439 |
| GO_RESPONSE_TO_OXYGEN_LEVELS | 0.00000 | 0.00000 | 0.71951 | 0.60132 | 1.33438 | 233 |
| GO_RESPONSE_TO_STEROID_HORMONE | 0.00154 | 0.02464 | 0.45506 | 0.55514 | 1.22446 | 174 |
| GO_RESPONSE_TO_TEMPERATURE_STIMULUS | 0.00027 | 0.00572 | 0.49849 | 0.59216 | 1.29675 | 127 |
| GO_RESPONSE_TO_TUMOR_NECROSIS_FACTOR | 0.00000 | 0.00000 | 0.76146 | 0.65649 | 1.44236 | 151 |
| GO_RETINA_HOMEOSTASIS | 0.00104 | 0.01798 | 0.45506 | 0.73063 | 1.49432 | 27 |
| GO_RIBONUCLEOPROTEIN_COMPLEX_BINDING | 0.00000 | 0.00000 | 0.71951 | 0.66906 | 1.46050 | 117 |
| GO_RIBONUCLEOPROTEIN_COMPLEX_BIOGENESIS | 0.00000 | 0.00000 | | 0.66339 | 1.48264 | 389 |
| GO_RIBONUCLEOPROTEIN_COMPLEX_SUBUNIT_ORGANIZATION | 0.00000 | 0.00000 | | 0.69241 | 1.52681 | 173 |
| GO_RIBONUCLEOPROTEIN_GRANULE | 0.00000 | 0.00001 | 0.67496 | 0.61979 | 1.36366 | 163 |
| GO_RIBONUCLEOSIDE_TRIPHOSPHATE_BIOSYNTHETIC_PROCESS | 0.00041 | 0.00840 | 0.49849 | 0.71253 | 1.47248 | 33 |
| GO_RIBONUCLEOSIDE_TRIPHOSPHATE_METABOLIC_PROCESS | 0.00026 | 0.00550 | 0.49849 | 0.69681 | 1.45611 | 42 |
| GO_RIBOSE_PHOSPHATE_BIOSYNTHETIC_PROCESS | 0.00191 | 0.02939 | 0.45506 | 0.57932 | 1.26207 | 110 |
| GO_RIBOSE_PHOSPHATE_METABOLIC_PROCESS | 0.00108 | 0.01839 | 0.45506 | 0.53891 | 1.19687 | 245 |
| GO_RIBOSOMAL_LARGE_SUBUNIT_ASSEMBLY | 0.00000 | 0.00016 | 0.61053 | 0.80999 | 1.65663 | 27 |
| GO_RIBOSOMAL_LARGE_SUBUNIT_BIOGENESIS | 0.00000 | 0.00000 | | 0.77542 | 1.65982 | 66 |
| GO_RIBOSOMAL_SMALL_SUBUNIT_ASSEMBLY | 0.00023 | 0.00494 | 0.51885 | 0.81139 | 1.62221 | 17 |
| GO_RIBOSOMAL_SMALL_SUBUNIT_BINDING | 0.00383 | 0.04935 | 0.43171 | 0.75288 | 1.50523 | 17 |
| GO_RIBOSOMAL_SMALL_SUBUNIT_BIOGENESIS | 0.00001 | 0.00022 | 0.61053 | 0.69076 | 1.47997 | 68 |
| GO_RIBOSOMAL_SUBUNIT | 0.00000 | 0.00000 | | 0.77603 | 1.71031 | 172 |
| GO_RIBOSOME | 0.00000 | 0.00000 | | 0.76643 | 1.69659 | 206 |
| GO_RIBOSOME_ASSEMBLY | 0.00000 | 0.00000 | 0.73376 | 0.76173 | 1.61998 | 59 |
| GO_RIBOSOME_BINDING | 0.00019 | 0.00416 | 0.51885 | 0.68955 | 1.44715 | 47 |
| GO_RIBOSOME_BIOGENESIS | 0.00000 | 0.00000 | | 0.65877 | 1.46466 | 275 |
| GO_RNA_CATABOLIC_PROCESS | 0.00000 | 0.00000 | | 0.76774 | 1.71264 | 339 |
| GO_RNA_DEPENDENT_DNA_BIOSYNTHETIC_PROCESS | 0.00003 | 0.00076 | 0.57561 | 0.68926 | 1.46521 | 58 |
| GO_RNA_EXPORT_FROM_NUCLEUS | 0.00062 | 0.01183 | 0.47727 | 0.58378 | 1.27758 | 126 |
| GO_RNA_LOCALIZATION | 0.00000 | 0.00006 | 0.64355 | 0.59670 | 1.32020 | 198 |
| GO_RNA_POLYMERASE_II_PREINITIATION_COMPLEX_ASSEMBLY | 0.00089 | 0.01592 | 0.47727 | 0.77046 | 1.55626 | 19 |
| GO_RNA_SPLICING | 0.00000 | 0.00000 | 0.78819 | 0.59079 | 1.31927 | 369 |
| GO_RNA_SPLICING_VIA_TRANSESTERIFICATION_REACTIONS | 0.00000 | 0.00000 | | 0.60940 | 1.35714 | 300 |
| GO_RNA_STABILIZATION | 0.00065 | 0.01223 | 0.47727 | 0.69743 | 1.44384 | 35 |
| GO_ROUGH_ENDOPLASMIC_RETICULUM | 0.00001 | 0.00039 | 0.59333 | 0.71853 | 1.51629 | 50 |
| GO_RRNA_BINDING | 0.00000 | 0.00001 | 0.69013 | 0.74782 | 1.58865 | 56 |
| GO_RRNA_METABOLIC_PROCESS | 0.00000 | 0.00000 | 0.77494 | 0.63047 | 1.39562 | 206 |
| GO_SCF_DEPENDENT_PROTEASOMAL_UBIQUITIN_DEPENDENT_PRO | 0.00000 | 0.00000 | | 0.77527 | 1.67044 | 79 |
| GO_SECRETORY_GRANULE | 0.00000 | 0.00000 | 0.71951 | 0.56518 | 1.26461 | 399 |
| GO_SECRETORY_VESICLE | 0.00000 | 0.00002 | 0.67496 | 0.54866 | 1.22905 | 465 |
| GO_SIGNAL_TRANSDUCTION_BY_PROTEIN_PHOSPHORYLATION | 0.00004 | 0.00109 | 0.55733 | 0.53817 | 1.20474 | 424 |
| GO_SINGLE_FERTILIZATION | 0.00048 | 0.00949 | 0.49849 | 0.67414 | 1.42260 | 50 |
| GO_SINGLE_STRANDED_RNA_BINDING | 0.00039 | 0.00793 | 0.49849 | 0.66553 | 1.41527 | 57 |
| GO_SMALL_MOLECULE_BIOSYNTHETIC_PROCESS | 0.00216 | 0.03242 | 0.43171 | 0.51195 | 1.14522 | 397 |
| GO_SMALL_RIBOSOMAL_SUBUNIT | 0.00000 | 0.00000 | | 0.77648 | 1.66209 | 66 |
| GO_SM_LIKE_PROTEIN_FAMILY_COMPLEX | 0.00327 | 0.04398 | 0.43171 | 0.60468 | 1.29713 | 70 |
| GO_SPERM_EGG_RECOGNITION | 0.00000 | 0.00007 | 0.64355 | 0.85241 | 1.72955 | 22 |
| GO_SPLICEOSOMAL_COMPLEX | 0.00000 | 0.00003 | 0.65944 | 0.60939 | 1.34261 | 171 |
| GO_STEM_CELL_DIFFERENTIATION | 0.00000 | 0.00000 | 0.78819 | 0.66886 | 1.46930 | 147 |
| GO_STRUCTURAL_CONSTITUENT_OF_CYTOSKELETON | 0.00102 | 0.01782 | 0.45506 | 0.65913 | 1.39337 | 52 |
| GO_STRUCTURAL_CONSTITUENT_OF_RIBOSOME | 0.00000 | 0.00000 | | 0.79909 | 1.75541 | 150 |
| GO_STRUCTURAL_MOLECULE_ACTIVITY | 0.00000 | 0.00000 | | 0.72651 | 1.61895 | 324 |
| GO_SUBSTANTIA_NIGRA_DEVELOPMENT | 0.00000 | 0.00013 | 0.62726 | 0.81622 | 1.66507 | 26 |
| GO_TELOMERASE_RNA_LOCALIZATION | 0.00001 | 0.00017 | 0.61053 | 0.85164 | 1.72025 | 19 |
| GO_TELOMERE_MAINTENANCE_VIA_TELOMERE_LENGTHENING | 0.00011 | 0.00250 | 0.53843 | 0.65659 | 1.40546 | 66 |

| | | | | | | |
|---|---|---|---|---|---|---|
| GO_TELOMERE_ORGANIZATION | 0.00342 | 0.04554 | 0.43171 | 0.57000 | 1.24649 | 124 |
| GO_THREONINE_TYPE_ENDOPEPTIDASE_ACTIVITY | 0.00000 | 0.00000 | | 0.95179 | 1.92255 | 19 |
| GO_THREONINE_TYPE_PEPTIDASE_ACTIVITY | 0.00000 | 0.00000 | | 0.92199 | 1.87423 | 23 |
| GO_TISSUE_MORPHOGENESIS | 0.00097 | 0.01707 | 0.47727 | 0.52661 | 1.17402 | 326 |
| GO_TOXIN_TRANSPORT | 0.00113 | 0.01922 | 0.45506 | 0.72897 | 1.48708 | 26 |
| GO_TRANSCRIPTION_COUPLED_NUCLEOTIDE_EXCISION_REPAIR | 0.00001 | 0.00019 | 0.61053 | 0.68106 | 1.46186 | 71 |
| GO_TRANSCRIPTION_FACTOR_BINDING | 0.00043 | 0.00865 | 0.49849 | 0.52447 | 1.17376 | 409 |
| GO_TRANSCRIPTION_PREINITIATION_COMPLEX_ASSEMBLY | 0.00119 | 0.02004 | 0.45506 | 0.71399 | 1.46797 | 30 |
| GO_TRANSLATIONAL_ELONGATION | 0.00000 | 0.00000 | 0.71951 | 0.66087 | 1.44451 | 122 |
| GO_TRANSLATIONAL_INITIATION | 0.00000 | 0.00000 | | 0.84691 | 1.86515 | 166 |
| GO_TRANSLATIONAL_TERMINATION | 0.00000 | 0.00005 | 0.64355 | 0.65946 | 1.43060 | 99 |
| GO_TRANSLATION_ELONGATION_FACTOR_ACTIVITY | 0.00161 | 0.02546 | 0.45506 | 0.79860 | 1.58418 | 15 |
| GO_TRANSLATION_FACTOR_ACTIVITY_RNA_BINDING | 0.00000 | 0.00000 | | 0.79008 | 1.69288 | 69 |
| GO_TRANSLATION_INITIATION_FACTOR_ACTIVITY | 0.00000 | 0.00000 | 0.76146 | 0.80422 | 1.68417 | 45 |
| GO_TRANSLATION_INITIATION_FACTOR_BINDING | 0.00021 | 0.00468 | 0.51885 | 0.76186 | 1.55231 | 25 |
| GO_TRANSLATION_PREINITIATION_COMPLEX | 0.00000 | 0.00001 | 0.67496 | 0.90245 | 1.79595 | 16 |
| GO_TRANSLATION_REGULATOR_ACTIVITY | 0.00000 | 0.00000 | | 0.77663 | 1.68904 | 105 |
| GO_TRANSLATION_REGULATOR_ACTIVITY_NUCLEIC_ACID_BINDING | 0.00000 | 0.00000 | | 0.77150 | 1.66586 | 83 |
| GO_TRICARBOXYLIC_ACID_CYCLE | 0.00154 | 0.02464 | 0.45506 | 0.71854 | 1.47240 | 28 |
| GO_TUMOR_NECROSIS_FACTOR_MEDIATED_SIGNALING_PATHWAY | 0.00000 | 0.00000 | | 0.76277 | 1.65372 | 97 |
| GO_T_CELL_RECEPTOR_SIGNALING_PATHWAY | 0.00000 | 0.00000 | 0.81404 | 0.69994 | 1.52527 | 112 |
| GO_U12_TYPE_SPLICEOSOMAL_COMPLEX | 0.00279 | 0.03889 | 0.43171 | 0.70746 | 1.44693 | 27 |
| GO_U2_SNRNP | 0.00005 | 0.00134 | 0.55733 | 0.83081 | 1.67168 | 18 |
| GO_U2_TYPE_SPLICEOSOMAL_COMPLEX | 0.00010 | 0.00243 | 0.53843 | 0.63403 | 1.36970 | 85 |
| GO_UBIQUITIN_LIKE_PROTEIN_LIGASE_BINDING | 0.00000 | 0.00000 | 0.76146 | 0.62273 | 1.38051 | 219 |
| GO_UBIQUITIN_PROTEIN_TRANSFERASE_REGULATOR_ACTIVITY | 0.00165 | 0.02590 | 0.45506 | 0.79605 | 1.57911 | 15 |
| GO_UNFOLDED_PROTEIN_BINDING | 0.00000 | 0.00000 | 0.77494 | 0.72157 | 1.55881 | 85 |
| GO_VESICLE_LUMEN | 0.00000 | 0.00000 | 0.74774 | 0.64475 | 1.42008 | 168 |
| GO_VIRAL_GENE_EXPRESSION | 0.00000 | 0.00000 | | 0.80028 | 1.76317 | 171 |
| GO_VIRAL_GENOME_REPLICATION | 0.00100 | 0.01761 | 0.45506 | 0.61997 | 1.33078 | 73 |
| GO_VIRAL_LIFE_CYCLE | 0.00031 | 0.00640 | 0.49849 | 0.55767 | 1.23509 | 209 |
| HP_ABNORMALITY_OF_ACID_BASE_HOMEOSTASIS | 0.00216 | 0.03242 | 0.43171 | 0.53556 | 1.18846 | 232 |
| HP_ABNORMALITY_OF_CARDIOVASCULAR_SYSTEM_ELECTROPHYSIO | 0.00027 | 0.00572 | 0.49849 | 0.55832 | 1.23929 | 229 |
| HP_ABNORMALITY_OF_LOWER_LIP | 0.00181 | 0.02804 | 0.45506 | 0.56360 | 1.23779 | 145 |
| HP_ABNORMALITY_OF_THE_MITOCHONDRION | 0.00213 | 0.03229 | 0.43171 | 0.57831 | 1.26240 | 117 |
| HP_ABNORMALITY_OF_THE_NASAL_DORSUM | 0.00005 | 0.00116 | 0.55733 | 0.60288 | 1.32157 | 134 |
| HP_ABNORMAL_ELECTROPHYSIOLOGY_OF_SINOATRIAL_NODE_ORIG | 0.00279 | 0.03889 | 0.43171 | 0.68436 | 1.40934 | 31 |
| HP_ABNORMAL_ERYTHROCYTE_MORPHOLOGY | 0.00000 | 0.00015 | 0.61053 | 0.56049 | 1.25036 | 341 |
| HP_ABNORMAL_MEAN_CORPUSCULAR_VOLUME | 0.00357 | 0.04696 | 0.43171 | 0.75226 | 1.51363 | 18 |
| HP_ABNORMAL_RETICULOCYTE_MORPHOLOGY | 0.00064 | 0.01210 | 0.47727 | 0.71420 | 1.47081 | 31 |
| HP_ACUTE_LEUKEMIA | 0.00000 | 0.00000 | 0.73376 | 0.72786 | 1.56137 | 70 |
| HP_ANEMIA_OF_INADEQUATE_PRODUCTION | 0.00000 | 0.00008 | 0.62726 | 0.65593 | 1.42209 | 97 |
| HP_CONSTITUTIONAL_SYMPTOM | 0.00119 | 0.02004 | 0.45506 | 0.51427 | 1.15108 | 423 |
| HP_DELAYED_PUBERTY | 0.00005 | 0.00115 | 0.55733 | 0.62579 | 1.36211 | 107 |
| HP_DEPRESSED_NASAL_RIDGE | 0.00000 | 0.00011 | 0.62726 | 0.68731 | 1.47511 | 72 |
| HP_FATIGUE | 0.00039 | 0.00808 | 0.49849 | 0.56529 | 1.24923 | 187 |
| HP_HEADACHE | 0.00015 | 0.00340 | 0.51885 | 0.59065 | 1.29716 | 146 |
| HP_HEMATOLOGICAL_NEOPLASM | 0.00003 | 0.00078 | 0.57561 | 0.60108 | 1.32044 | 154 |
| HP_INCREASED_CSF_LACTATE | 0.00157 | 0.02509 | 0.45506 | 0.61622 | 1.31904 | 66 |
| HP_INCREASED_SERUM_LACTATE | 0.00216 | 0.03242 | 0.43171 | 0.56920 | 1.24724 | 130 |
| HP_LEUKEMIA | 0.00001 | 0.00026 | 0.59333 | 0.64877 | 1.40680 | 98 |
| HP_MACROCYTIC_ANEMIA | 0.00000 | 0.00000 | 0.74774 | 0.82468 | 1.71698 | 39 |
| HP_MIGRAINE | 0.00000 | 0.00000 | 0.71951 | 0.72105 | 1.54497 | 69 |
| HP_MYOTONIA | 0.00161 | 0.02546 | 0.45506 | 0.79820 | 1.58338 | 15 |
| HP_NEOPLASM | 0.00216 | 0.03242 | 0.43171 | 0.50820 | 1.13805 | 446 |
| HP_PALLOR | 0.00001 | 0.00040 | 0.59333 | 0.65169 | 1.41044 | 92 |
| HP_PUBERTY_AND_GONADAL_DISORDERS | 0.00287 | 0.03973 | 0.43171 | 0.51995 | 1.15843 | 322 |
| HP_RETICULOCYTOPENIA | 0.00004 | 0.00105 | 0.55733 | 0.86417 | 1.71425 | 15 |
| HP_RIMMED_VACUOLES | 0.00279 | 0.03889 | 0.43171 | 0.74033 | 1.49831 | 20 |
| HP_SHORT_NOSE | 0.00235 | 0.03474 | 0.43171 | 0.54919 | 1.21134 | 174 |
| HP_THICK_LOWER_LIP_VERMILION | 0.00048 | 0.00949 | 0.49849 | 0.60773 | 1.31617 | 94 |
| HP_THICK_VERMILION_BORDER | 0.00156 | 0.02486 | 0.45506 | 0.55983 | 1.23000 | 153 |
| HP_VENTRICULAR_ARRHYTHMIA | 0.00353 | 0.04658 | 0.43171 | 0.66198 | 1.37398 | 36 |
| **Cluster 1** | | | | | | |
| pathway | pval | padj | log2err | ES | NES | size |
| GO_ACTIVATION_OF_IMMUNE_RESPONSE | 0.00000 | 0.00000 | 0.78819 | 0.62851 | 1.39255 | 212 |
| GO_ACTIVATION_OF_INNATE_IMMUNE_RESPONSE | 0.00000 | 0.00000 | | 0.75054 | 1.62130 | 88 |
| GO_AEROBIC_RESPIRATION | 0.00001 | 0.00025 | 0.59333 | 0.69302 | 1.48168 | 65 |
| GO_ALTERNATIVE_MRNA_SPLICING_VIA_SPLICEOSOME | 0.00102 | 0.01782 | 0.45506 | 0.65168 | 1.38687 | 60 |
| GO_AMINE_METABOLIC_PROCESS | 0.00000 | 0.00000 | | 0.79189 | 1.70783 | 80 |
| GO_ANAPHASE_PROMOTING_COMPLEX_DEPENDENT_CATABOLIC_P | 0.00000 | 0.00000 | | 0.76611 | 1.65070 | 79 |
| GO_ANCHORING_JUNCTION | 0.00000 | 0.00000 | | 0.59615 | 1.33633 | 479 |
| GO_ANIMAL_ORGAN_REGENERATION | 0.00257 | 0.03716 | 0.43171 | 0.68240 | 1.41020 | 33 |
| GO_ANTIGEN_PROCESSING_AND_PRESENTATION | 0.00000 | 0.00000 | 0.70498 | 0.64776 | 1.42241 | 143 |
| GO_ANTIGEN_PROCESSING_AND_PRESENTATION_OF_EXOGENOUS_ | 0.00000 | 0.00000 | | 0.81051 | 1.72358 | 57 |
| GO_ANTIGEN_PROCESSING_AND_PRESENTATION_OF_PEPTIDE_ANTI | 0.00000 | 0.00000 | 0.70498 | 0.65912 | 1.44248 | 126 |

| | | | | | | |
|---|---|---|---|---|---|---|
| GO_ANTIGEN_PROCESSING_AND_PRESENTATION_OF_PEPTIDE_ANTI | 0.00000 | 0.00000 | | 0.76925 | 1.65015 | 70 |
| GO_ANTIGEN_RECEPTOR_MEDIATED_SIGNALING_PATHWAY | 0.00000 | 0.00000 | 0.80122 | 0.67877 | 1.48641 | 127 |
| GO_APOPTOTIC_SIGNALING_PATHWAY | 0.00154 | 0.02464 | 0.45506 | 0.51561 | 1.15085 | 358 |
| GO_ATP_DEPENDENT_CHROMATIN_REMODELING | 0.00253 | 0.03681 | 0.43171 | 0.62124 | 1.32400 | 61 |
| GO_ATP_METABOLIC_PROCESS | 0.00000 | 0.00000 | 0.71951 | 0.62252 | 1.37693 | 196 |
| GO_ATP_SYNTHESIS_COUPLED_ELECTRON_TRANSPORT | 0.00005 | 0.00131 | 0.55733 | 0.65519 | 1.40723 | 74 |
| GO_BINDING_OF_SPERM_TO_ZONA_PELLUCIDA | 0.00000 | 0.00001 | 0.69013 | 0.89879 | 1.80846 | 18 |
| GO_CADHERIN_BINDING | 0.00000 | 0.00000 | | 0.63068 | 1.39961 | 236 |
| GO_CANONICAL_WNT_SIGNALING_PATHWAY | 0.00000 | 0.00003 | 0.65944 | 0.60816 | 1.34553 | 197 |
| GO_CATALYTIC_STEP_2_SPLICEOSOME | 0.00000 | 0.00002 | 0.65944 | 0.68068 | 1.46976 | 83 |
| GO_CELLULAR_AMINO_ACID_METABOLIC_PROCESS | 0.00000 | 0.00000 | 0.74774 | 0.61752 | 1.36824 | 216 |
| GO_CELLULAR_COMPONENT_DISASSEMBLY | 0.00000 | 0.00005 | 0.64355 | 0.55505 | 1.24197 | 401 |
| GO_CELLULAR_KETONE_METABOLIC_PROCESS | 0.00000 | 0.00000 | 0.77494 | 0.65955 | 1.44893 | 149 |
| GO_CELLULAR_NITROGEN_COMPOUND_CATABOLIC_PROCESS | 0.00000 | 0.00000 | | 0.74079 | 1.65848 | 436 |
| GO_CELLULAR_PROTEIN_COMPLEX_DISASSEMBLY | 0.00006 | 0.00139 | 0.55733 | 0.58222 | 1.28459 | 176 |
| GO_CELLULAR_RESPIRATION | 0.00000 | 0.00000 | 0.71951 | 0.64997 | 1.42492 | 136 |
| GO_CELLULAR_RESPONSE_TO_HEAT | 0.00018 | 0.00403 | 0.51885 | 0.61962 | 1.34016 | 91 |
| GO_CELLULAR_RESPONSE_TO_ORGANIC_CYCLIC_COMPOUND | 0.00302 | 0.04108 | 0.43171 | 0.52402 | 1.16585 | 283 |
| GO_CELLULAR_RESPONSE_TO_OXYGEN_LEVELS | 0.00000 | 0.00000 | 0.78819 | 0.65911 | 1.44882 | 157 |
| GO_CELL_ACTIVATION_INVOLVED_IN_IMMUNE_RESPONSE | 0.00000 | 0.00001 | 0.67496 | 0.57339 | 1.27913 | 341 |
| GO_CELL_ADHESION_MOLECULE_BINDING | 0.00000 | 0.00000 | 0.81404 | 0.60529 | 1.34788 | 308 |
| GO_CELL_CELL_RECOGNITION | 0.00000 | 0.00002 | 0.65944 | 0.83826 | 1.71003 | 26 |
| GO_CELL_CELL_SIGNALING_BY_WNT | 0.00000 | 0.00002 | 0.67496 | 0.57736 | 1.28559 | 299 |
| GO_CELL_CYCLE_G2_M_PHASE_TRANSITION | 0.00001 | 0.00024 | 0.59333 | 0.58034 | 1.28602 | 214 |
| GO_CELL_CYCLE_PHASE_TRANSITION | 0.00000 | 0.00004 | 0.64355 | 0.54319 | 1.21681 | 464 |
| GO_CELL_DIVISION | 0.00154 | 0.02464 | 0.45506 | 0.51551 | 1.15425 | 427 |
| GO_CELL_RECOGNITION | 0.00012 | 0.00270 | 0.53843 | 0.69554 | 1.46186 | 48 |
| GO_CELL_SUBSTRATE_JUNCTION | 0.00000 | 0.00000 | | 0.67751 | 1.50775 | 291 |
| GO_CELL_SURFACE_RECEPTOR_SIGNALING_PATHWAY_INVOLVED_IN | 0.00000 | 0.00003 | 0.65944 | 0.57035 | 1.27067 | 319 |
| GO_CHAPERONE_BINDING | 0.00007 | 0.00163 | 0.53843 | 0.64754 | 1.39253 | 76 |
| GO_CHAPERONE_COMPLEX | 0.00000 | 0.00004 | 0.64355 | 0.85718 | 1.73925 | 22 |
| GO_CHAPERONE_MEDIATED_PROTEIN_COMPLEX_ASSEMBLY | 0.00170 | 0.02662 | 0.45506 | 0.77423 | 1.54792 | 17 |
| GO_CHROMATIN_BINDING | 0.00071 | 0.01323 | 0.47727 | 0.52699 | 1.17680 | 369 |
| GO_CHROMOSOMAL_REGION | 0.00246 | 0.03601 | 0.43171 | 0.52919 | 1.17661 | 276 |
| GO_COP9_SIGNALOSOME | 0.00128 | 0.02141 | 0.45506 | 0.73616 | 1.49715 | 24 |
| GO_COTRANSLATIONAL_PROTEIN_TARGETING_TO_MEMBRANE | 0.00000 | 0.00000 | | 0.89471 | 1.93513 | 91 |
| GO_CYTOCHROME_COMPLEX | 0.00168 | 0.02641 | 0.45506 | 0.75311 | 1.52417 | 20 |
| GO_CYTOKINE_MEDIATED_SIGNALING_PATHWAY | 0.00000 | 0.00000 | 0.81404 | 0.59698 | 1.33170 | 339 |
| GO_CYTOPLASMIC_TRANSLATION | 0.00000 | 0.00000 | | 0.84753 | 1.83080 | 88 |
| GO_CYTOPLASMIC_TRANSLATIONAL_INITIATION | 0.00000 | 0.00000 | 0.74774 | 0.85688 | 1.76176 | 30 |
| GO_CYTOSOLIC_LARGE_RIBOSOMAL_SUBUNIT | 0.00000 | 0.00000 | | 0.91186 | 1.92988 | 53 |
| GO_CYTOSOLIC_RIBOSOME | 0.00000 | 0.00000 | | 0.89907 | 1.94710 | 95 |
| GO_CYTOSOLIC_SMALL_RIBOSOMAL_SUBUNIT | 0.00000 | 0.00000 | | 0.89114 | 1.85533 | 39 |
| GO_DEFENSE_RESPONSE_TO_OTHER_ORGANISM | 0.00004 | 0.00099 | 0.55733 | 0.53673 | 1.20127 | 415 |
| GO_DE_NOVO_PROTEIN_FOLDING | 0.00050 | 0.00978 | 0.47727 | 0.74232 | 1.51431 | 26 |
| GO_DISORDERED_DOMAIN_SPECIFIC_BINDING | 0.00161 | 0.02546 | 0.45506 | 0.71970 | 1.47197 | 27 |
| GO_DNA_BIOSYNTHETIC_PROCESS | 0.00000 | 0.00010 | 0.62726 | 0.62505 | 1.37017 | 134 |
| GO_DNA_CONFORMATION_CHANGE | 0.00027 | 0.00572 | 0.49849 | 0.55300 | 1.22527 | 216 |
| GO_DNA_DAMAGE_RESPONSE_DETECTION_OF_DNA_DAMAGE | 0.00198 | 0.03028 | 0.43171 | 0.67949 | 1.40669 | 35 |
| GO_DNA_GEOMETRIC_CHANGE | 0.00046 | 0.00925 | 0.49849 | 0.60807 | 1.31855 | 98 |
| GO_DNA_POLYMERASE_BINDING | 0.00268 | 0.03799 | 0.43171 | 0.76255 | 1.52457 | 17 |
| GO_DNA_REPAIR | 0.00265 | 0.03775 | 0.43171 | 0.51159 | 1.14547 | 427 |
| GO_DRUG_BINDING | 0.00176 | 0.02733 | 0.45506 | 0.64119 | 1.35096 | 49 |
| GO_ELECTRON_TRANSFER_ACTIVITY | 0.00001 | 0.00020 | 0.61053 | 0.67481 | 1.44938 | 74 |
| GO_ELECTRON_TRANSPORT_CHAIN | 0.00002 | 0.00053 | 0.57561 | 0.62257 | 1.36145 | 124 |
| GO_ENDOPEPTIDASE_ACTIVITY | 0.00124 | 0.02085 | 0.45506 | 0.54955 | 1.21076 | 171 |
| GO_ENDOPEPTIDASE_COMPLEX | 0.00000 | 0.00000 | | 0.80930 | 1.73234 | 66 |
| GO_ENERGY_DERIVATION_BY_OXIDATION_OF_ORGANIC_COMPOUN | 0.00000 | 0.00000 | 0.71951 | 0.63390 | 1.39707 | 172 |
| GO_ESTABLISHMENT_OF_PROTEIN_LOCALIZATION_TO_CHROMOSOM | 0.00020 | 0.00432 | 0.51885 | 0.77339 | 1.57217 | 23 |
| GO_ESTABLISHMENT_OF_PROTEIN_LOCALIZATION_TO_ENDOPLASM | 0.00000 | 0.00000 | | 0.88026 | 1.91149 | 102 |
| GO_ESTABLISHMENT_OF_PROTEIN_LOCALIZATION_TO_MEMBRANE | 0.00000 | 0.00000 | | 0.74874 | 1.66164 | 240 |
| GO_ESTABLISHMENT_OF_PROTEIN_LOCALIZATION_TO_ORGANELLE | 0.00000 | 0.00000 | | 0.69519 | 1.55605 | 423 |
| GO_ESTABLISHMENT_OF_PROTEIN_LOCALIZATION_TO_TELOMERE | 0.00006 | 0.00139 | 0.55733 | 0.83470 | 1.66881 | 17 |
| GO_ESTABLISHMENT_OF_RNA_LOCALIZATION | 0.00106 | 0.01819 | 0.45506 | 0.55577 | 1.22400 | 167 |
| GO_ESTABLISHMENT_OF_TISSUE_POLARITY | 0.00000 | 0.00000 | | 0.75051 | 1.62122 | 88 |
| GO_EUKARYOTIC_TRANSLATION_INITIATION_FACTOR_3_COMPLEX | 0.00003 | 0.00074 | 0.57561 | 0.86987 | 1.72555 | 15 |
| GO_EXOCYTOSIS | 0.00000 | 0.00015 | 0.61053 | 0.53905 | 1.20753 | 465 |
| GO_FC_EPSILON_RECEPTOR_SIGNALING_PATHWAY | 0.00000 | 0.00000 | | 0.75117 | 1.62103 | 82 |
| GO_FC_RECEPTOR_SIGNALING_PATHWAY | 0.00000 | 0.00000 | | 0.72054 | 1.57787 | 127 |
| GO_FERTILIZATION | 0.00089 | 0.01592 | 0.47727 | 0.63620 | 1.35888 | 64 |
| GO_FICOLIN_1_RICH_GRANULE | 0.00000 | 0.00000 | 0.78819 | 0.71716 | 1.55340 | 93 |
| GO_GENERATION_OF_PRECURSOR_METABOLITES_AND_ENERGY | 0.00000 | 0.00000 | 0.77494 | 0.59261 | 1.32026 | 320 |
| GO_GENE_SILENCING | 0.00261 | 0.03750 | 0.43171 | 0.55578 | 1.22252 | 161 |
| GO_GLAND_DEVELOPMENT | 0.00154 | 0.02464 | 0.45506 | 0.54117 | 1.19797 | 206 |
| GO_GLYCOSYL_COMPOUND_METABOLIC_PROCESS | 0.00071 | 0.01323 | 0.47727 | 0.62524 | 1.34204 | 71 |
| GO_GUANYL_NUCLEOTIDE_BINDING | 0.00095 | 0.01679 | 0.47727 | 0.54096 | 1.19861 | 216 |

| | | | | | | |
|---|---|---|---|---|---|---|
| GO_HEAT_SHOCK_PROTEIN_BINDING | 0.00003 | 0.00092 | 0.55733 | 0.64369 | 1.39056 | 85 |
| GO_HEMATOPOIETIC_PROGENITOR_CELL_DIFFERENTIATION | 0.00000 | 0.00000 | 0.82666 | 0.70307 | 1.53239 | 113 |
| GO_HEMATOPOIETIC_STEM_CELL_DIFFERENTIATION | 0.00000 | 0.00000 | | 0.79776 | 1.71217 | 72 |
| GO_HSP70_PROTEIN_BINDING | 0.00228 | 0.03391 | 0.43171 | 0.71143 | 1.45504 | 27 |
| GO_IMMUNE_RESPONSE_REGULATING_SIGNALING_PATHWAY | 0.00000 | 0.00000 | 0.76146 | 0.63494 | 1.40480 | 197 |
| GO_INNATE_IMMUNE_RESPONSE | 0.00003 | 0.00071 | 0.57561 | 0.55173 | 1.23107 | 347 |
| GO_INNATE_IMMUNE_RESPONSE_ACTIVATING_SIGNAL_TRANSDUCT | 0.00000 | 0.00000 | | 0.79340 | 1.70281 | 72 |
| GO_INNER_MITOCHONDRIAL_MEMBRANE_PROTEIN_COMPLEX | 0.00002 | 0.00060 | 0.57561 | 0.65124 | 1.40678 | 88 |
| GO_INTERACTION_WITH_SYMBIONT | 0.00001 | 0.00026 | 0.59333 | 0.75084 | 1.56620 | 41 |
| GO_INTERLEUKIN_1_MEDIATED_SIGNALING_PATHWAY | 0.00000 | 0.00000 | | 0.81689 | 1.75671 | 76 |
| GO_INTRINSIC_APOPTOTIC_SIGNALING_PATHWAY | 0.00121 | 0.02029 | 0.45506 | 0.55118 | 1.21845 | 190 |
| GO_ION_CHANNEL_BINDING | 0.00087 | 0.01584 | 0.47727 | 0.63182 | 1.34683 | 62 |
| GO_ISOPEPTIDASE_ACTIVITY | 0.00139 | 0.02285 | 0.45506 | 0.78940 | 1.57097 | 16 |
| GO_LARGE_RIBOSOMAL_SUBUNIT | 0.00000 | 0.00000 | | 0.78135 | 1.70183 | 109 |
| GO_LEUKOCYTE_MEDIATED_IMMUNITY | 0.00000 | 0.00000 | 0.71951 | 0.57282 | 1.27895 | 364 |
| GO_MACROMOLECULE_TRANSMEMBRANE_TRANSPORTER_ACTIVITY | 0.00239 | 0.03510 | 0.43171 | 0.77734 | 1.54696 | 16 |
| GO_MAINTENANCE_OF_CELL_NUMBER | 0.00087 | 0.01584 | 0.47727 | 0.58944 | 1.28512 | 114 |
| GO_MATURATION_OF_LSU_RRNA | 0.00006 | 0.00151 | 0.53843 | 0.77606 | 1.58723 | 27 |
| GO_METHYLTRANSFERASE_COMPLEX | 0.00290 | 0.04014 | 0.43171 | 0.59476 | 1.28479 | 88 |
| GO_METHYL_CPG_BINDING | 0.00302 | 0.04108 | 0.43171 | 0.78053 | 1.54833 | 15 |
| GO_MIDBRAIN_DEVELOPMENT | 0.00021 | 0.00468 | 0.51885 | 0.68832 | 1.44457 | 47 |
| GO_MITOCHONDRIAL_ELECTRON_TRANSPORT_NADH_TO_UBIQUIN | 0.00331 | 0.04437 | 0.43171 | 0.65428 | 1.36724 | 42 |
| GO_MITOCHONDRIAL_GENE_EXPRESSION | 0.00048 | 0.00949 | 0.49849 | 0.57608 | 1.26525 | 152 |
| GO_MITOCHONDRIAL_LARGE_RIBOSOMAL_SUBUNIT | 0.00002 | 0.00050 | 0.57561 | 0.69789 | 1.48086 | 55 |
| GO_MITOCHONDRIAL_MATRIX | 0.00000 | 0.00002 | 0.65944 | 0.56045 | 1.25158 | 371 |
| GO_MITOCHONDRIAL_MEMBRANE_ORGANIZATION | 0.00361 | 0.04734 | 0.43171 | 0.59309 | 1.28094 | 86 |
| GO_MITOCHONDRIAL_PROTEIN_COMPLEX | 0.00000 | 0.00000 | 0.81404 | 0.63650 | 1.40931 | 208 |
| GO_MITOCHONDRIAL_TRANSLATION | 0.00004 | 0.00093 | 0.55733 | 0.61219 | 1.34135 | 129 |
| GO_MITOCHONDRIAL_TRANSLATIONAL_TERMINATION | 0.00000 | 0.00010 | 0.62726 | 0.66694 | 1.44128 | 87 |
| GO_MITOCHONDRION_ORGANIZATION | 0.00051 | 0.00993 | 0.47727 | 0.52284 | 1.16863 | 390 |
| GO_MODIFICATION_DEPENDENT_MACROMOLECULE_CATABOLIC_PR | 0.00002 | 0.00053 | 0.57561 | 0.53142 | 1.19156 | 486 |
| GO_MODULATION_BY_HOST_OF_SYMBIONT_PROCESS | 0.00015 | 0.00336 | 0.51885 | 0.72848 | 1.50542 | 33 |
| GO_MODULATION_OF_PROCESS_OF_OTHER_ORGANISM | 0.00019 | 0.00423 | 0.51885 | 0.66904 | 1.42286 | 59 |
| GO_MONOSACCHARIDE_BIOSYNTHETIC_PROCESS | 0.00335 | 0.04466 | 0.43171 | 0.61992 | 1.31781 | 58 |
| GO_MORPHOGENESIS_OF_AN_EPITHELIUM | 0.00006 | 0.00151 | 0.53843 | 0.55161 | 1.22755 | 290 |
| GO_MORPHOGENESIS_OF_A_POLARIZED_EPITHELIUM | 0.00000 | 0.00000 | | 0.72298 | 1.57057 | 103 |
| GO_MRNA_3_UTR_BINDING | 0.00003 | 0.00089 | 0.55733 | 0.67596 | 1.44827 | 68 |
| GO_MRNA_5_UTR_BINDING | 0.00006 | 0.00156 | 0.53843 | 0.80133 | 1.62592 | 22 |
| GO_MRNA_BINDING | 0.00000 | 0.00000 | | 0.66002 | 1.46353 | 222 |
| GO_MRNA_PROCESSING | 0.00000 | 0.00000 | 0.73376 | 0.56859 | 1.27295 | 418 |
| GO_MULTI_ORGANISM_LOCALIZATION | 0.00268 | 0.03799 | 0.43171 | 0.63245 | 1.33612 | 51 |
| GO_MULTI_ORGANISM_PROCESS | 0.00108 | 0.01839 | 0.45506 | 0.51114 | 1.14599 | 484 |
| GO_MYD88_DEPENDENT_TOLL_LIKE_RECEPTOR_SIGNALING_PATHW | 0.00002 | 0.00066 | 0.57561 | 0.85581 | 1.70312 | 16 |
| GO_MYELOID_LEUKOCYTE_ACTIVATION | 0.00000 | 0.00000 | 0.74774 | 0.58508 | 1.30479 | 327 |
| GO_MYELOID_LEUKOCYTE_MEDIATED_IMMUNITY | 0.00000 | 0.00000 | 0.76146 | 0.59841 | 1.33291 | 301 |
| GO_NADH_DEHYDROGENASE_ACTIVITY | 0.00383 | 0.04935 | 0.43171 | 0.66244 | 1.37139 | 35 |
| GO_NADH_DEHYDROGENASE_COMPLEX | 0.00376 | 0.04884 | 0.43171 | 0.65649 | 1.36421 | 38 |
| GO_NADH_METABOLIC_PROCESS | 0.00294 | 0.04036 | 0.43171 | 0.70642 | 1.44481 | 27 |
| GO_NCRNA_EXPORT_FROM_NUCLEUS | 0.00302 | 0.04108 | 0.43171 | 0.66820 | 1.38331 | 35 |
| GO_NCRNA_METABOLIC_PROCESS | 0.00000 | 0.00003 | 0.65944 | 0.55907 | 1.24907 | 381 |
| GO_NCRNA_PROCESSING | 0.00000 | 0.00003 | 0.65944 | 0.57175 | 1.27504 | 328 |
| GO_NEGATIVE_REGULATION_OF_CANONICAL_WNT_SIGNALING_PAT | 0.00000 | 0.00000 | | 0.71611 | 1.56038 | 111 |
| GO_NEGATIVE_REGULATION_OF_CELLULAR_AMIDE_METABOLIC_PR | 0.00001 | 0.00020 | 0.61053 | 0.63031 | 1.37423 | 114 |
| GO_NEGATIVE_REGULATION_OF_CELL_CYCLE | 0.00003 | 0.00076 | 0.57561 | 0.53833 | 1.20513 | 425 |
| GO_NEGATIVE_REGULATION_OF_CELL_CYCLE_G2_M_PHASE_TRANS | 0.00000 | 0.00000 | 0.76146 | 0.71490 | 1.54439 | 85 |
| GO_NEGATIVE_REGULATION_OF_CELL_CYCLE_PHASE_TRANSITION | 0.00002 | 0.00060 | 0.57561 | 0.59041 | 1.30442 | 184 |
| GO_NEGATIVE_REGULATION_OF_CELL_CYCLE_PROCESS | 0.00000 | 0.00002 | 0.65944 | 0.58690 | 1.30374 | 249 |
| GO_NEGATIVE_REGULATION_OF_MITOTIC_CELL_CYCLE | 0.00004 | 0.00095 | 0.55733 | 0.56735 | 1.25902 | 232 |
| GO_NEGATIVE_REGULATION_OF_MRNA_METABOLIC_PROCESS | 0.00023 | 0.00491 | 0.51885 | 0.66670 | 1.41788 | 59 |
| GO_NEGATIVE_REGULATION_OF_MRNA_SPLICING_VIA_SPLICEOSOM | 0.00239 | 0.03510 | 0.43171 | 0.74857 | 1.51204 | 19 |
| GO_NEGATIVE_REGULATION_OF_RNA_SPLICING | 0.00029 | 0.00607 | 0.49849 | 0.76729 | 1.55976 | 23 |
| GO_NEGATIVE_REGULATION_OF_TRANSLATIONAL_INITIATION | 0.00018 | 0.00403 | 0.51885 | 0.83901 | 1.66434 | 15 |
| GO_NEGATIVE_REGULATION_OF_WNT_SIGNALING_PATHWAY | 0.00000 | 0.00000 | | 0.69525 | 1.52249 | 127 |
| GO_NEURAL_NUCLEUS_DEVELOPMENT | 0.00001 | 0.00026 | 0.59333 | 0.79148 | 1.62187 | 28 |
| GO_NEURON_TO_NEURON_SYNAPSE | 0.00005 | 0.00134 | 0.55733 | 0.59636 | 1.31006 | 154 |
| GO_NIK_NF_KAPPAB_SIGNALING | 0.00000 | 0.00000 | | 0.72896 | 1.58354 | 103 |
| GO_NON_CANONICAL_WNT_SIGNALING_PATHWAY | 0.00000 | 0.00000 | | 0.72784 | 1.58023 | 101 |
| GO_NUCLEAR_EXPORT | 0.00031 | 0.00645 | 0.49849 | 0.56963 | 1.25542 | 172 |
| GO_NUCLEAR_TRANSCRIBED_MRNA_CATABOLIC_PROCESS | 0.00000 | 0.00000 | | 0.79179 | 1.74917 | 183 |
| GO_NUCLEAR_TRANSCRIBED_MRNA_CATABOLIC_PROCESS_NONSEN | 0.00000 | 0.00000 | | 0.89008 | 1.93909 | 110 |
| GO_NUCLEAR_TRANSPORT | 0.00008 | 0.00193 | 0.53843 | 0.55511 | 1.23408 | 273 |
| GO_NUCLEOBASE_CONTAINING_COMPOUND_TRANSPORT | 0.00146 | 0.02386 | 0.45506 | 0.55053 | 1.21842 | 202 |
| GO_NUCLEOBASE_CONTAINING_SMALL_MOLECULE_BIOSYNTHETIC_ | 0.00137 | 0.02261 | 0.45506 | 0.62749 | 1.34353 | 67 |
| GO_NUCLEOBASE_CONTAINING_SMALL_MOLECULE_INTERCONVERS | 0.00383 | 0.04935 | 0.43171 | 0.71259 | 1.44857 | 23 |
| GO_NUCLEOBASE_CONTAINING_SMALL_MOLECULE_METABOLIC_PR | 0.00320 | 0.04339 | 0.43171 | 0.51448 | 1.14825 | 352 |
| GO_NUCLEOSIDE_DIPHOSPHATE_METABOLIC_PROCESS | 0.00110 | 0.01865 | 0.45506 | 0.59637 | 1.29295 | 97 |

| | | | | | | |
|---|---|---|---|---|---|---|
| GO_NUCLEOSIDE_METABOLIC_PROCESS | 0.00061 | 0.01169 | 0.47727 | 0.65315 | 1.38907 | 59 |
| GO_NUCLEOSIDE_MONOPHOSPHATE_BIOSYNTHETIC_PROCESS | 0.00250 | 0.03636 | 0.43171 | 0.68841 | 1.42033 | 32 |
| GO_NUCLEOSIDE_MONOPHOSPHATE_METABOLIC_PROCESS | 0.00152 | 0.02464 | 0.45506 | 0.64377 | 1.36002 | 51 |
| GO_NUCLEOSIDE_PHOSPHATE_BIOSYNTHETIC_PROCESS | 0.00043 | 0.00865 | 0.49849 | 0.58185 | 1.27831 | 148 |
| GO_NUCLEOSIDE_TRIPHOSPHATE_BIOSYNTHETIC_PROCESS | 0.00043 | 0.00865 | 0.49849 | 0.69332 | 1.44539 | 40 |
| GO_NUCLEOSIDE_TRIPHOSPHATE_METABOLIC_PROCESS | 0.00016 | 0.00360 | 0.51885 | 0.66788 | 1.41976 | 58 |
| GO_NUCLEOSOMAL_DNA_BINDING | 0.00023 | 0.00507 | 0.51885 | 0.79852 | 1.61801 | 21 |
| GO_NUCLEOSOME_BINDING | 0.00143 | 0.02332 | 0.45506 | 0.66132 | 1.38492 | 45 |
| GO_NUCLEOTIDE_EXCISION_REPAIR | 0.00004 | 0.00096 | 0.55733 | 0.63343 | 1.37355 | 98 |
| GO_NUCLEOTIDE_EXCISION_REPAIR_DNA_DAMAGE_RECOGNITION | 0.00015 | 0.00349 | 0.51885 | 0.78678 | 1.59640 | 22 |
| GO_NUCLEOTIDE_EXCISION_REPAIR_DNA_INCISION | 0.00313 | 0.04249 | 0.43171 | 0.66439 | 1.37898 | 36 |
| GO_NUCLEOTIDE_PHOSPHORYLATION | 0.00383 | 0.04935 | 0.43171 | 0.59002 | 1.27454 | 88 |
| GO_ORGANELLAR_RIBOSOME | 0.00000 | 0.00014 | 0.61053 | 0.66412 | 1.43471 | 85 |
| GO_ORGANELLE_INNER_MEMBRANE | 0.00000 | 0.00000 | 0.76146 | 0.57601 | 1.28697 | 378 |
| GO_ORGANIC_CYCLIC_COMPOUND_CATABOLIC_PROCESS | 0.00000 | 0.00000 | | 0.73350 | 1.64281 | 449 |
| GO_OXIDATIVE_PHOSPHORYLATION | 0.00001 | 0.00030 | 0.59333 | 0.64619 | 1.40027 | 96 |
| GO_OXIDOREDUCTASE_ACTIVITY | 0.00006 | 0.00139 | 0.55733 | 0.53544 | 1.19703 | 392 |
| GO_OXIDOREDUCTASE_COMPLEX | 0.00001 | 0.00027 | 0.59333 | 0.66015 | 1.42461 | 82 |
| GO_PEPTIDASE_COMPLEX | 0.00000 | 0.00000 | | 0.75988 | 1.63982 | 82 |
| GO_PIGMENT_GRANULE | 0.00003 | 0.00089 | 0.55733 | 0.66098 | 1.42344 | 78 |
| GO_POLYSOMAL_RIBOSOME | 0.00000 | 0.00000 | | 0.90624 | 1.86629 | 31 |
| GO_POLYSOME | 0.00000 | 0.00000 | | 0.82337 | 1.75481 | 61 |
| GO_POLY_PYRIMIDINE_TRACT_BINDING | 0.00327 | 0.04398 | 0.43171 | 0.75471 | 1.51855 | 18 |
| GO_POSITIVE_REGULATION_OF_APOPTOTIC_SIGNALING_PATHWAY | 0.00279 | 0.03889 | 0.43171 | 0.57666 | 1.25518 | 107 |
| GO_POSITIVE_REGULATION_OF_ATPASE_ACTIVITY | 0.00294 | 0.04036 | 0.43171 | 0.65446 | 1.36795 | 43 |
| GO_POSITIVE_REGULATION_OF_BINDING | 0.00088 | 0.01590 | 0.47727 | 0.60284 | 1.30556 | 95 |
| GO_POSITIVE_REGULATION_OF_CANONICAL_WNT_SIGNALING_PATH | 0.00000 | 0.00000 | 0.77494 | 0.70504 | 1.53160 | 103 |
| GO_POSITIVE_REGULATION_OF_CELLULAR_AMIDE_METABOLIC_PRO | 0.00003 | 0.00074 | 0.57561 | 0.62296 | 1.36083 | 118 |
| GO_POSITIVE_REGULATION_OF_CELLULAR_PROTEIN_LOCALIZATION | 0.00041 | 0.00840 | 0.49849 | 0.54915 | 1.21815 | 225 |
| GO_POSITIVE_REGULATION_OF_CELL_DEATH | 0.00069 | 0.01297 | 0.47727 | 0.52595 | 1.17423 | 363 |
| GO_POSITIVE_REGULATION_OF_CELL_POPULATION_PROLIFERATION | 0.00078 | 0.01455 | 0.47727 | 0.52449 | 1.17108 | 368 |
| GO_POSITIVE_REGULATION_OF_CHROMOSOME_ORGANIZATION | 0.00231 | 0.03437 | 0.43171 | 0.56990 | 1.24871 | 133 |
| GO_POSITIVE_REGULATION_OF_DEFENSE_RESPONSE | 0.00000 | 0.00003 | 0.65944 | 0.62175 | 1.36730 | 158 |
| GO_POSITIVE_REGULATION_OF_DNA_BIOSYNTHETIC_PROCESS | 0.00001 | 0.00041 | 0.59333 | 0.71929 | 1.52055 | 52 |
| GO_POSITIVE_REGULATION_OF_DNA_METABOLIC_PROCESS | 0.00364 | 0.04761 | 0.43171 | 0.56647 | 1.24189 | 135 |
| GO_POSITIVE_REGULATION_OF_DNA_TEMPLATED_TRANSCRIPTION_ | 0.00276 | 0.03885 | 0.43171 | 0.68932 | 1.41586 | 29 |
| GO_POSITIVE_REGULATION_OF_ESTABLISHMENT_OF_PROTEIN_LOC/ | 0.00007 | 0.00168 | 0.53843 | 0.56461 | 1.25165 | 219 |
| GO_POSITIVE_REGULATION_OF_IMMUNE_RESPONSE | 0.00000 | 0.00001 | 0.69013 | 0.59037 | 1.31265 | 277 |
| GO_POSITIVE_REGULATION_OF_IMMUNE_SYSTEM_PROCESS | 0.00004 | 0.00110 | 0.55733 | 0.54018 | 1.20716 | 386 |
| GO_POSITIVE_REGULATION_OF_INNATE_IMMUNE_RESPONSE | 0.00000 | 0.00000 | 0.81404 | 0.69693 | 1.51947 | 114 |
| GO_POSITIVE_REGULATION_OF_INTRINSIC_APOPTOTIC_SIGNALING_ | 0.00104 | 0.01798 | 0.45506 | 0.69097 | 1.43578 | 37 |
| GO_POSITIVE_REGULATION_OF_ORGANELLE_ORGANIZATION | 0.00043 | 0.00865 | 0.49849 | 0.52093 | 1.16605 | 411 |
| GO_POSITIVE_REGULATION_OF_PROTEIN_LOCALIZATION_TO_NUCLE | 0.00001 | 0.00020 | 0.61053 | 0.69907 | 1.48990 | 61 |
| GO_POSITIVE_REGULATION_OF_RESPONSE_TO_BIOTIC_STIMULUS | 0.00000 | 0.00000 | 0.73376 | 0.66340 | 1.45310 | 128 |
| GO_POSITIVE_REGULATION_OF_RESPONSE_TO_EXTERNAL_STIMULU | 0.00002 | 0.00050 | 0.57561 | 0.58133 | 1.28681 | 201 |
| GO_POSITIVE_REGULATION_OF_TELOMERASE_ACTIVITY | 0.00061 | 0.01169 | 0.47727 | 0.73476 | 1.50565 | 28 |
| GO_POSITIVE_REGULATION_OF_TELOMERASE_RNA_LOCALIZATION_ | 0.00000 | 0.00001 | 0.69013 | 0.92451 | 1.83395 | 15 |
| GO_POSITIVE_REGULATION_OF_TELOMERE_MAINTENANCE_VIA_TE | 0.00130 | 0.02159 | 0.45506 | 0.71259 | 1.46508 | 30 |
| GO_POSITIVE_REGULATION_OF_TRANSCRIPTION_INITIATION_FROM | 0.00372 | 0.04847 | 0.43171 | 0.70671 | 1.43993 | 25 |
| GO_POSITIVE_REGULATION_OF_TRANSFERASE_ACTIVITY | 0.00174 | 0.02712 | 0.45506 | 0.51642 | 1.15220 | 345 |
| GO_POSITIVE_REGULATION_OF_TRANSLATION | 0.00001 | 0.00019 | 0.61053 | 0.64767 | 1.40419 | 97 |
| GO_POSITIVE_REGULATION_OF_VIRAL_GENOME_REPLICATION | 0.00287 | 0.03973 | 0.43171 | 0.68362 | 1.40783 | 31 |
| GO_POSITIVE_REGULATION_OF_VIRAL_PROCESS | 0.00028 | 0.00590 | 0.49849 | 0.62440 | 1.34741 | 81 |
| GO_POSITIVE_REGULATION_OF_WNT_SIGNALING_PATHWAY | 0.00000 | 0.00000 | 0.73376 | 0.66230 | 1.44763 | 122 |
| GO_POSTREPLICATION_REPAIR | 0.00024 | 0.00515 | 0.51885 | 0.69140 | 1.44518 | 43 |
| GO_POSTSYNAPSE | 0.00004 | 0.00109 | 0.55733 | 0.55969 | 1.24369 | 259 |
| GO_POSTSYNAPTIC_CYTOSOL | 0.00279 | 0.03889 | 0.43171 | 0.78295 | 1.55313 | 15 |
| GO_POSTTRANSCRIPTIONAL_REGULATION_OF_GENE_EXPRESSION | 0.00000 | 0.00000 | | 0.66656 | 1.49327 | 462 |
| GO_POST_TRANSLATIONAL_PROTEIN_MODIFICATION | 0.00002 | 0.00052 | 0.57561 | 0.57851 | 1.28412 | 229 |
| GO_PRECATALYTIC_SPLICEOSOME | 0.00085 | 0.01559 | 0.47727 | 0.65235 | 1.37448 | 49 |
| GO_PRERIBOSOME | 0.00087 | 0.01577 | 0.47727 | 0.62899 | 1.34772 | 69 |
| GO_PRERIBOSOME_LARGE_SUBUNIT_PRECURSOR | 0.00235 | 0.03474 | 0.43171 | 0.74079 | 1.50308 | 22 |
| GO_PRE_MRNA_BINDING | 0.00133 | 0.02213 | 0.45506 | 0.72393 | 1.48062 | 27 |
| GO_PROTEASOMAL_PROTEIN_CATABOLIC_PROCESS | 0.00002 | 0.00044 | 0.59333 | 0.54166 | 1.21005 | 377 |
| GO_PROTEASOMAL_UBIQUITIN_INDEPENDENT_PROTEIN_CATABOLI( | 0.00000 | 0.00000 | 0.80122 | 0.92254 | 1.86931 | 21 |
| GO_PROTEASOME_ACCESSORY_COMPLEX | 0.00000 | 0.00001 | 0.67496 | 0.85464 | 1.73811 | 24 |
| GO_PROTEASOME_CORE_COMPLEX | 0.00000 | 0.00000 | | 0.94375 | 1.90631 | 19 |
| GO_PROTEIN_CONTAINING_COMPLEX_DISASSEMBLY | 0.00000 | 0.00007 | 0.62726 | 0.57810 | 1.28438 | 256 |
| GO_PROTEIN_CONTAINING_COMPLEX_LOCALIZATION | 0.00063 | 0.01197 | 0.47727 | 0.55335 | 1.22520 | 208 |
| GO_PROTEIN_DNA_COMPLEX_SUBUNIT_ORGANIZATION | 0.00001 | 0.00018 | 0.61053 | 0.60174 | 1.32491 | 165 |
| GO_PROTEIN_FOLDING | 0.00000 | 0.00002 | 0.65944 | 0.63113 | 1.38644 | 154 |
| GO_PROTEIN_FOLDING_CHAPERONE | 0.00135 | 0.02238 | 0.45506 | 0.77903 | 1.55752 | 17 |
| GO_PROTEIN_IMPORT | 0.00268 | 0.03799 | 0.43171 | 0.56317 | 1.23581 | 140 |
| GO_PROTEIN_LOCALIZATION_TO_CHROMOSOME | 0.00056 | 0.01096 | 0.47727 | 0.63523 | 1.36108 | 69 |
| GO_PROTEIN_LOCALIZATION_TO_CHROMOSOME_TELOMERIC_REGI( | 0.00335 | 0.04466 | 0.43171 | 0.70399 | 1.43984 | 27 |
| GO_PROTEIN_LOCALIZATION_TO_ENDOPLASMIC_RETICULUM | 0.00000 | 0.00000 | | 0.85521 | 1.87021 | 124 |

| | | | | | | |
|---|---|---|---|---|---|---|
| GO_PROTEIN_LOCALIZATION_TO_MEMBRANE | 0.00000 | 0.00000 | | 0.65123 | 1.45714 | 406 |
| GO_PROTEIN_LOCALIZATION_TO_MITOCHONDRION | 0.00019 | 0.00420 | 0.51885 | 0.60773 | 1.32398 | 110 |
| GO_PROTEIN_LOCALIZATION_TO_NUCLEUS | 0.00002 | 0.00055 | 0.57561 | 0.58926 | 1.30220 | 187 |
| GO_PROTEIN_MODIFICATION_BY_SMALL_PROTEIN_REMOVAL | 0.00000 | 0.00000 | 0.71951 | 0.60745 | 1.34689 | 221 |
| GO_PROTEIN_POLYUBIQUITINATION | 0.00004 | 0.00106 | 0.55733 | 0.55874 | 1.24108 | 253 |
| GO_PROTEIN_REFOLDING | 0.00294 | 0.04036 | 0.43171 | 0.77039 | 1.53314 | 16 |
| GO_PROTEIN_STABILIZATION | 0.00001 | 0.00024 | 0.61053 | 0.62333 | 1.36586 | 130 |
| GO_PROTEIN_TARGETING | 0.00000 | 0.00000 | | 0.71181 | 1.58743 | 327 |
| GO_PROTEIN_TARGETING_TO_MEMBRANE | 0.00000 | 0.00000 | | 0.82370 | 1.80953 | 149 |
| GO_PROTEIN_TARGETING_TO_MITOCHONDRION | 0.00346 | 0.04593 | 0.43171 | 0.60408 | 1.30091 | 78 |
| GO_PROTEIN_TRANSMEMBRANE_TRANSPORT | 0.00265 | 0.03775 | 0.43171 | 0.64528 | 1.36171 | 50 |
| GO_PROTON_TRANSMEMBRANE_TRANSPORT | 0.00213 | 0.03229 | 0.43171 | 0.60815 | 1.30536 | 71 |
| GO_PURINE_CONTAINING_COMPOUND_BIOSYNTHETIC_PROCESS | 0.00246 | 0.03601 | 0.43171 | 0.57796 | 1.25936 | 111 |
| GO_PURINE_CONTAINING_COMPOUND_METABOLIC_PROCESS | 0.00100 | 0.01761 | 0.45506 | 0.53679 | 1.19242 | 255 |
| GO_PURINE_NUCLEOSIDE_TRIPHOSPHATE_METABOLIC_PROCESS | 0.00089 | 0.01592 | 0.47727 | 0.67157 | 1.40374 | 43 |
| GO_PYRIMIDINE_CONTAINING_COMPOUND_METABOLIC_PROCESS | 0.00353 | 0.04658 | 0.43171 | 0.64744 | 1.35878 | 47 |
| GO_REGENERATION | 0.00216 | 0.03242 | 0.43171 | 0.59987 | 1.29453 | 82 |
| GO_REGULATION_OF_ANIMAL_ORGAN_MORPHOGENESIS | 0.00000 | 0.00000 | | 0.69163 | 1.51541 | 133 |
| GO_REGULATION_OF_APOPTOTIC_SIGNALING_PATHWAY | 0.00031 | 0.00645 | 0.49849 | 0.55171 | 1.22506 | 247 |
| GO_REGULATION_OF_ATPASE_ACTIVITY | 0.00324 | 0.04369 | 0.43171 | 0.63031 | 1.34038 | 57 |
| GO_REGULATION_OF_ATP_METABOLIC_PROCESS | 0.00353 | 0.04658 | 0.43171 | 0.60176 | 1.29780 | 80 |
| GO_REGULATION_OF_BINDING | 0.00250 | 0.03636 | 0.43171 | 0.54476 | 1.20598 | 203 |
| GO_REGULATION_OF_CELLULAR_AMIDE_METABOLIC_PROCESS | 0.00000 | 0.00000 | | 0.62967 | 1.40226 | 311 |
| GO_REGULATION_OF_CELLULAR_AMINE_METABOLIC_PROCESS | 0.00000 | 0.00000 | | 0.82173 | 1.74682 | 58 |
| GO_REGULATION_OF_CELLULAR_AMINO_ACID_METABOLIC_PROCES | 0.00000 | 0.00000 | | 0.83776 | 1.77766 | 55 |
| GO_REGULATION_OF_CELLULAR_KETONE_METABOLIC_PROCESS | 0.00000 | 0.00000 | | 0.72641 | 1.58680 | 118 |
| GO_REGULATION_OF_CELLULAR_PROTEIN_LOCALIZATION | 0.00082 | 0.01513 | 0.47727 | 0.52542 | 1.17290 | 362 |
| GO_REGULATION_OF_CELLULAR_RESPONSE_TO_HEAT | 0.00276 | 0.03885 | 0.43171 | 0.61652 | 1.32004 | 67 |
| GO_REGULATION_OF_CELLULAR_RESPONSE_TO_STRESS | 0.00224 | 0.03344 | 0.43171 | 0.50545 | 1.13232 | 468 |
| GO_REGULATION_OF_CELL_CYCLE_G2_M_PHASE_TRANSITION | 0.00000 | 0.00000 | 0.71951 | 0.63230 | 1.39282 | 169 |
| GO_REGULATION_OF_CELL_CYCLE_PHASE_TRANSITION | 0.00001 | 0.00026 | 0.59333 | 0.55488 | 1.23783 | 337 |
| GO_REGULATION_OF_CHROMOSOME_ORGANIZATION | 0.00081 | 0.01501 | 0.47727 | 0.54255 | 1.20527 | 261 |
| GO_REGULATION_OF_CYSTEINE_TYPE_ENDOPEPTIDASE_ACTIVITY | 0.00202 | 0.03077 | 0.43171 | 0.57120 | 1.24837 | 121 |
| GO_REGULATION_OF_CYTOPLASMIC_TRANSLATION | 0.00130 | 0.02159 | 0.45506 | 0.73678 | 1.49773 | 23 |
| GO_REGULATION_OF_DEFENSE_RESPONSE | 0.00065 | 0.01237 | 0.47727 | 0.53633 | 1.19355 | 290 |
| GO_REGULATION_OF_DNA_BIOSYNTHETIC_PROCESS | 0.00016 | 0.00360 | 0.51885 | 0.64215 | 1.37923 | 74 |
| GO_REGULATION_OF_DNA_METABOLIC_PROCESS | 0.00069 | 0.01297 | 0.47727 | 0.54425 | 1.20775 | 237 |
| GO_REGULATION_OF_DNA_TEMPLATED_TRANSCRIPTION_INITIATIOI | 0.00163 | 0.02568 | 0.45506 | 0.68289 | 1.41372 | 35 |
| GO_REGULATION_OF_DNA_TEMPLATED_TRANSCRIPTION_IN_RESPO | 0.00000 | 0.00000 | | 0.73944 | 1.59800 | 89 |
| GO_REGULATION_OF_GENERATION_OF_PRECURSOR_METABOLITES_ | 0.00060 | 0.01164 | 0.47727 | 0.60476 | 1.31116 | 97 |
| GO_REGULATION_OF_HEMATOPOIETIC_PROGENITOR_CELL_DIFFERE | 0.00000 | 0.00000 | | 0.78717 | 1.69251 | 75 |
| GO_REGULATION_OF_HEMATOPOIETIC_STEM_CELL_DIFFERENTIATIC | 0.00000 | 0.00000 | | 0.81505 | 1.74256 | 65 |
| GO_REGULATION_OF_HEMOPOIESIS | 0.00000 | 0.00008 | 0.62726 | 0.58859 | 1.30634 | 228 |
| GO_REGULATION_OF_IMMUNE_RESPONSE | 0.00000 | 0.00016 | 0.61053 | 0.55362 | 1.23583 | 362 |
| GO_REGULATION_OF_INNATE_IMMUNE_RESPONSE | 0.00000 | 0.00000 | 0.71951 | 0.63883 | 1.40521 | 161 |
| GO_REGULATION_OF_INTRINSIC_APOPTOTIC_SIGNALING_PATHWAY | 0.00001 | 0.00039 | 0.59333 | 0.63478 | 1.38259 | 109 |
| GO_REGULATION_OF_MITOTIC_CELL_CYCLE | 0.00001 | 0.00018 | 0.61053 | 0.53899 | 1.20736 | 458 |
| GO_REGULATION_OF_MORPHOGENESIS_OF_AN_EPITHELIUM | 0.00000 | 0.00000 | | 0.72213 | 1.57284 | 109 |
| GO_REGULATION_OF_MRNA_CATABOLIC_PROCESS | 0.00000 | 0.00000 | | 0.69977 | 1.54127 | 168 |
| GO_REGULATION_OF_MRNA_METABOLIC_PROCESS | 0.00000 | 0.00000 | | 0.66528 | 1.47872 | 266 |
| GO_REGULATION_OF_MRNA_PROCESSING | 0.00004 | 0.00105 | 0.55733 | 0.61521 | 1.34090 | 113 |
| GO_REGULATION_OF_MRNA_SPLICING_VIA_SPLICEOSOME | 0.00001 | 0.00026 | 0.59333 | 0.67424 | 1.45411 | 80 |
| GO_REGULATION_OF_NUCLEASE_ACTIVITY | 0.00106 | 0.01819 | 0.45506 | 0.77918 | 1.56780 | 18 |
| GO_REGULATION_OF_OXIDATIVE_STRESS_INDUCED_CELL_DEATH | 0.00198 | 0.03028 | 0.43171 | 0.67287 | 1.40355 | 41 |
| GO_REGULATION_OF_PEPTIDASE_ACTIVITY | 0.00050 | 0.00978 | 0.47727 | 0.56122 | 1.24054 | 191 |
| GO_REGULATION_OF_POSTTRANSCRIPTIONAL_GENE_SILENCING | 0.00364 | 0.04761 | 0.43171 | 0.59824 | 1.28492 | 74 |
| GO_REGULATION_OF_PROTEIN_CATABOLIC_PROCESS | 0.00179 | 0.02783 | 0.45506 | 0.53057 | 1.17929 | 270 |
| GO_REGULATION_OF_PROTEIN_LOCALIZATION_TO_NUCLEUS | 0.00001 | 0.00042 | 0.59333 | 0.65505 | 1.41558 | 87 |
| GO_REGULATION_OF_PROTEIN_MODIFICATION_BY_SMALL_PROTEIN | 0.00025 | 0.00532 | 0.49849 | 0.57068 | 1.25684 | 167 |
| GO_REGULATION_OF_PROTEIN_STABILITY | 0.00093 | 0.01652 | 0.47727 | 0.55382 | 1.22571 | 202 |
| GO_REGULATION_OF_PROTEOLYSIS | 0.00013 | 0.00301 | 0.51885 | 0.52997 | 1.18392 | 377 |
| GO_REGULATION_OF_RESPONSE_TO_BIOTIC_STIMULUS | 0.00001 | 0.00020 | 0.61053 | 0.59049 | 1.30719 | 204 |
| GO_REGULATION_OF_RNA_SPLICING | 0.00000 | 0.00001 | 0.69013 | 0.65896 | 1.43669 | 114 |
| GO_REGULATION_OF_SMALL_MOLECULE_METABOLIC_PROCESS | 0.00000 | 0.00000 | 0.71951 | 0.59134 | 1.31394 | 265 |
| GO_REGULATION_OF_STEM_CELL_DIFFERENTIATION | 0.00000 | 0.00000 | | 0.74049 | 1.60113 | 90 |
| GO_REGULATION_OF_SYMBIOTIC_PROCESS | 0.00006 | 0.00144 | 0.55733 | 0.60071 | 1.31682 | 134 |
| GO_REGULATION_OF_TELOMERASE_ACTIVITY | 0.00088 | 0.01590 | 0.47727 | 0.69161 | 1.43179 | 35 |
| GO_REGULATION_OF_TRANSCRIPTION_FROM_RNA_POLYMERASE_II | 0.00000 | 0.00000 | | 0.81194 | 1.73800 | 66 |
| GO_REGULATION_OF_TRANSCRIPTION_INITIATION_FROM_RNA_POL | 0.00143 | 0.02332 | 0.45506 | 0.70390 | 1.44582 | 29 |
| GO_REGULATION_OF_TRANSLATIONAL_INITIATION | 0.00002 | 0.00052 | 0.57561 | 0.68129 | 1.45518 | 64 |
| GO_REGULATION_OF_UBIQUITIN_PROTEIN_TRANSFERASE_ACTIVITY | 0.00061 | 0.01169 | 0.47727 | 0.69335 | 1.44627 | 41 |
| GO_REGULATION_OF_VIRAL_GENOME_REPLICATION | 0.00086 | 0.01566 | 0.47727 | 0.65316 | 1.38389 | 54 |
| GO_REGULATION_OF_VIRAL_TRANSCRIPTION | 0.00198 | 0.03028 | 0.43171 | 0.63932 | 1.34703 | 49 |
| GO_REGULATION_OF_WNT_SIGNALING_PATHWAY | 0.00000 | 0.00001 | 0.69013 | 0.60353 | 1.33816 | 220 |
| GO_RESPIRASOME | 0.00008 | 0.00181 | 0.53843 | 0.65420 | 1.40421 | 71 |
| GO_RESPIRATORY_CHAIN_COMPLEX | 0.00016 | 0.00353 | 0.51885 | 0.67626 | 1.43918 | 60 |

| | | | | | | |
|---|---|---|---|---|---|---|
| GO_RESPIRATORY_ELECTRON_TRANSPORT_CHAIN | 0.00001 | 0.00025 | 0.59333 | 0.65819 | 1.42190 | 85 |
| GO_RESPONSE_TO_ALCOHOL | 0.00261 | 0.03750 | 0.43171 | 0.58804 | 1.27350 | 95 |
| GO_RESPONSE_TO_CADMIUM_ION | 0.00265 | 0.03775 | 0.43171 | 0.67227 | 1.39174 | 35 |
| GO_RESPONSE_TO_DRUG | 0.00104 | 0.01798 | 0.45506 | 0.55364 | 1.22258 | 180 |
| GO_RESPONSE_TO_ETHANOL | 0.00086 | 0.01566 | 0.47727 | 0.67309 | 1.40691 | 43 |
| GO_RESPONSE_TO_HEAT | 0.00012 | 0.00270 | 0.53843 | 0.61665 | 1.34221 | 107 |
| GO_RESPONSE_TO_HORMONE | 0.00048 | 0.00949 | 0.49849 | 0.51277 | 1.14814 | 453 |
| GO_RESPONSE_TO_HYDROGEN_PEROXIDE | 0.00257 | 0.03716 | 0.43171 | 0.60505 | 1.30368 | 79 |
| GO_RESPONSE_TO_INORGANIC_SUBSTANCE | 0.00004 | 0.00107 | 0.55733 | 0.55714 | 1.23870 | 275 |
| GO_RESPONSE_TO_INTERLEUKIN_1 | 0.00000 | 0.00000 | | 0.74844 | 1.62908 | 107 |
| GO_RESPONSE_TO_INTERLEUKIN_12 | 0.00012 | 0.00270 | 0.53843 | 0.72605 | 1.50694 | 36 |
| GO_RESPONSE_TO_INTERLEUKIN_4 | 0.00324 | 0.04369 | 0.43171 | 0.75812 | 1.51571 | 17 |
| GO_RESPONSE_TO_METAL_ION | 0.00030 | 0.00623 | 0.49849 | 0.56387 | 1.24233 | 171 |
| GO_RESPONSE_TO_ORGANIC_CYCLIC_COMPOUND | 0.00000 | 0.00016 | 0.61053 | 0.54603 | 1.22259 | 439 |
| GO_RESPONSE_TO_OXYGEN_LEVELS | 0.00000 | 0.00000 | 0.71951 | 0.60132 | 1.33438 | 233 |
| GO_RESPONSE_TO_STEROID_HORMONE | 0.00154 | 0.02464 | 0.45506 | 0.55514 | 1.22446 | 174 |
| GO_RESPONSE_TO_TEMPERATURE_STIMULUS | 0.00027 | 0.00572 | 0.49849 | 0.59216 | 1.29675 | 127 |
| GO_RESPONSE_TO_TUMOR_NECROSIS_FACTOR | 0.00000 | 0.00000 | 0.76146 | 0.65649 | 1.44236 | 151 |
| GO_RETINA_HOMEOSTASIS | 0.00104 | 0.01798 | 0.45506 | 0.73063 | 1.49432 | 27 |
| GO_RIBONUCLEOPROTEIN_COMPLEX_BINDING | 0.00000 | 0.00000 | 0.71951 | 0.66906 | 1.46050 | 117 |
| GO_RIBONUCLEOPROTEIN_COMPLEX_BIOGENESIS | 0.00000 | 0.00000 | | 0.66339 | 1.48264 | 389 |
| GO_RIBONUCLEOPROTEIN_COMPLEX_SUBUNIT_ORGANIZATION | 0.00000 | 0.00000 | | 0.69241 | 1.52681 | 173 |
| GO_RIBONUCLEOPROTEIN_GRANULE | 0.00000 | 0.00001 | 0.67496 | 0.61979 | 1.36366 | 163 |
| GO_RIBONUCLEOSIDE_TRIPHOSPHATE_BIOSYNTHETIC_PROCESS | 0.00041 | 0.00840 | 0.49849 | 0.71253 | 1.47248 | 33 |
| GO_RIBONUCLEOSIDE_TRIPHOSPHATE_METABOLIC_PROCESS | 0.00026 | 0.00550 | 0.49849 | 0.69681 | 1.45611 | 42 |
| GO_RIBOSE_PHOSPHATE_BIOSYNTHETIC_PROCESS | 0.00191 | 0.02939 | 0.45506 | 0.57932 | 1.26207 | 110 |
| GO_RIBOSE_PHOSPHATE_METABOLIC_PROCESS | 0.00108 | 0.01839 | 0.45506 | 0.53891 | 1.19687 | 245 |
| GO_RIBOSOMAL_LARGE_SUBUNIT_ASSEMBLY | 0.00000 | 0.00016 | 0.61053 | 0.80999 | 1.65663 | 27 |
| GO_RIBOSOMAL_LARGE_SUBUNIT_BIOGENESIS | 0.00000 | 0.00000 | | 0.77542 | 1.65982 | 66 |
| GO_RIBOSOMAL_SMALL_SUBUNIT_ASSEMBLY | 0.00023 | 0.00494 | 0.51885 | 0.81139 | 1.62221 | 17 |
| GO_RIBOSOMAL_SMALL_SUBUNIT_BINDING | 0.00383 | 0.04935 | 0.43171 | 0.75288 | 1.50523 | 17 |
| GO_RIBOSOMAL_SMALL_SUBUNIT_BIOGENESIS | 0.00001 | 0.00022 | 0.61053 | 0.69076 | 1.47997 | 68 |
| GO_RIBOSOMAL_SUBUNIT | 0.00000 | 0.00000 | | 0.77603 | 1.71031 | 172 |
| GO_RIBOSOME | 0.00000 | 0.00000 | | 0.76643 | 1.69659 | 206 |
| GO_RIBOSOME_ASSEMBLY | 0.00000 | 0.00000 | 0.73376 | 0.76173 | 1.61998 | 59 |
| GO_RIBOSOME_BINDING | 0.00019 | 0.00416 | 0.51885 | 0.68955 | 1.44715 | 47 |
| GO_RIBOSOME_BIOGENESIS | 0.00000 | 0.00000 | | 0.65877 | 1.46466 | 275 |
| GO_RNA_CATABOLIC_PROCESS | 0.00000 | 0.00000 | | 0.76774 | 1.71264 | 339 |
| GO_RNA_DEPENDENT_DNA_BIOSYNTHETIC_PROCESS | 0.00003 | 0.00076 | 0.57561 | 0.68926 | 1.46521 | 58 |
| GO_RNA_EXPORT_FROM_NUCLEUS | 0.00062 | 0.01183 | 0.47727 | 0.58378 | 1.27758 | 126 |
| GO_RNA_LOCALIZATION | 0.00000 | 0.00006 | 0.64355 | 0.59670 | 1.32020 | 198 |
| GO_RNA_POLYMERASE_II_PREINITIATION_COMPLEX_ASSEMBLY | 0.00089 | 0.01592 | 0.47727 | 0.77046 | 1.55626 | 19 |
| GO_RNA_SPLICING | 0.00000 | 0.00000 | 0.78819 | 0.59079 | 1.31927 | 369 |
| GO_RNA_SPLICING_VIA_TRANSESTERIFICATION_REACTIONS | 0.00000 | 0.00000 | | 0.60940 | 1.35714 | 300 |
| GO_RNA_STABILIZATION | 0.00065 | 0.01223 | 0.47727 | 0.69743 | 1.44384 | 35 |
| GO_ROUGH_ENDOPLASMIC_RETICULUM | 0.00001 | 0.00039 | 0.59333 | 0.71853 | 1.51629 | 50 |
| GO_RRNA_BINDING | 0.00000 | 0.00001 | 0.69013 | 0.74782 | 1.58865 | 56 |
| GO_RRNA_METABOLIC_PROCESS | 0.00000 | 0.00000 | 0.77494 | 0.63047 | 1.39562 | 206 |
| GO_SCF_DEPENDENT_PROTEASOMAL_UBIQUITIN_DEPENDENT_PRO | 0.00000 | 0.00000 | | 0.77527 | 1.67044 | 79 |
| GO_SECRETORY_GRANULE | 0.00000 | 0.00000 | 0.71951 | 0.56518 | 1.26461 | 399 |
| GO_SECRETORY_VESICLE | 0.00000 | 0.00002 | 0.67496 | 0.54866 | 1.22905 | 465 |
| GO_SIGNAL_TRANSDUCTION_BY_PROTEIN_PHOSPHORYLATION | 0.00004 | 0.00109 | 0.55733 | 0.53817 | 1.20474 | 424 |
| GO_SINGLE_FERTILIZATION | 0.00048 | 0.00949 | 0.49849 | 0.67414 | 1.42260 | 50 |
| GO_SINGLE_STRANDED_RNA_BINDING | 0.00039 | 0.00793 | 0.49849 | 0.66553 | 1.41527 | 57 |
| GO_SMALL_MOLECULE_BIOSYNTHETIC_PROCESS | 0.00216 | 0.03242 | 0.43171 | 0.51195 | 1.14522 | 397 |
| GO_SMALL_RIBOSOMAL_SUBUNIT | 0.00000 | 0.00000 | | 0.77648 | 1.66209 | 66 |
| GO_SM_LIKE_PROTEIN_FAMILY_COMPLEX | 0.00327 | 0.04398 | 0.43171 | 0.60468 | 1.29713 | 70 |
| GO_SPERM_EGG_RECOGNITION | 0.00000 | 0.00007 | 0.64355 | 0.85241 | 1.72955 | 22 |
| GO_SPLICEOSOMAL_COMPLEX | 0.00000 | 0.00003 | 0.65944 | 0.60939 | 1.34261 | 171 |
| GO_STEM_CELL_DIFFERENTIATION | 0.00000 | 0.00000 | 0.78819 | 0.66886 | 1.46930 | 147 |
| GO_STRUCTURAL_CONSTITUENT_OF_CYTOSKELETON | 0.00102 | 0.01782 | 0.45506 | 0.65913 | 1.39337 | 52 |
| GO_STRUCTURAL_CONSTITUENT_OF_RIBOSOME | 0.00000 | 0.00000 | | 0.79909 | 1.75541 | 150 |
| GO_STRUCTURAL_MOLECULE_ACTIVITY | 0.00000 | 0.00000 | | 0.72651 | 1.61895 | 324 |
| GO_SUBSTANTIA_NIGRA_DEVELOPMENT | 0.00000 | 0.00013 | 0.62726 | 0.81622 | 1.66507 | 26 |
| GO_TELOMERASE_RNA_LOCALIZATION | 0.00001 | 0.00017 | 0.61053 | 0.85164 | 1.72025 | 19 |
| GO_TELOMERE_MAINTENANCE_VIA_TELOMERE_LENGTHENING | 0.00011 | 0.00250 | 0.53843 | 0.65659 | 1.40546 | 66 |
| GO_TELOMERE_ORGANIZATION | 0.00342 | 0.04554 | 0.43171 | 0.57000 | 1.24649 | 124 |
| GO_THREONINE_TYPE_ENDOPEPTIDASE_ACTIVITY | 0.00000 | 0.00000 | | 0.95179 | 1.92255 | 19 |
| GO_THREONINE_TYPE_PEPTIDASE_ACTIVITY | 0.00000 | 0.00000 | | 0.92199 | 1.87423 | 23 |
| GO_TISSUE_MORPHOGENESIS | 0.00097 | 0.01707 | 0.47727 | 0.52661 | 1.17402 | 326 |
| GO_TOXIN_TRANSPORT | 0.00113 | 0.01922 | 0.45506 | 0.72897 | 1.48708 | 26 |
| GO_TRANSCRIPTION_COUPLED_NUCLEOTIDE_EXCISION_REPAIR | 0.00001 | 0.00019 | 0.61053 | 0.68106 | 1.46186 | 71 |
| GO_TRANSCRIPTION_FACTOR_BINDING | 0.00043 | 0.00865 | 0.49849 | 0.52447 | 1.17376 | 409 |
| GO_TRANSCRIPTION_PREINITIATION_COMPLEX_ASSEMBLY | 0.00119 | 0.02004 | 0.45506 | 0.71399 | 1.46797 | 30 |
| GO_TRANSLATIONAL_ELONGATION | 0.00000 | 0.00000 | 0.71951 | 0.66087 | 1.44451 | 122 |
| GO_TRANSLATIONAL_INITIATION | 0.00000 | 0.00000 | | 0.84691 | 1.86515 | 166 |

| | | | | | | |
|---|---|---|---|---|---|---|
| GO_TRANSLATIONAL_TERMINATION | 0.00000 | 0.00005 | 0.64355 | 0.65946 | 1.43060 | 99 |
| GO_TRANSLATION_ELONGATION_FACTOR_ACTIVITY | 0.00161 | 0.02546 | 0.45506 | 0.79860 | 1.58418 | 15 |
| GO_TRANSLATION_FACTOR_ACTIVITY_RNA_BINDING | 0.00000 | 0.00000 | | 0.79008 | 1.69288 | 69 |
| GO_TRANSLATION_INITIATION_FACTOR_ACTIVITY | 0.00000 | 0.00000 | 0.76146 | 0.80422 | 1.68417 | 45 |
| GO_TRANSLATION_INITIATION_FACTOR_BINDING | 0.00021 | 0.00468 | 0.51885 | 0.76186 | 1.55231 | 25 |
| GO_TRANSLATION_PREINITIATION_COMPLEX | 0.00000 | 0.00001 | 0.67496 | 0.90245 | 1.79595 | 16 |
| GO_TRANSLATION_REGULATOR_ACTIVITY | 0.00000 | 0.00000 | | 0.77663 | 1.68904 | 105 |
| GO_TRANSLATION_REGULATOR_ACTIVITY_NUCLEIC_ACID_BINDING | 0.00000 | 0.00000 | | 0.77150 | 1.66586 | 83 |
| GO_TRICARBOXYLIC_ACID_CYCLE | 0.00154 | 0.02464 | 0.45506 | 0.71854 | 1.47240 | 28 |
| GO_TUMOR_NECROSIS_FACTOR_MEDIATED_SIGNALING_PATHWAY | 0.00000 | 0.00000 | | 0.76277 | 1.65372 | 97 |
| GO_T_CELL_RECEPTOR_SIGNALING_PATHWAY | 0.00000 | 0.00000 | 0.81404 | 0.69994 | 1.52527 | 112 |
| GO_U12_TYPE_SPLICEOSOMAL_COMPLEX | 0.00279 | 0.03889 | 0.43171 | 0.70746 | 1.44693 | 27 |
| GO_U2_SNRNP | 0.00005 | 0.00134 | 0.55733 | 0.83081 | 1.67168 | 18 |
| GO_U2_TYPE_SPLICEOSOMAL_COMPLEX | 0.00010 | 0.00243 | 0.53843 | 0.63403 | 1.36970 | 85 |
| GO_UBIQUITIN_LIKE_PROTEIN_LIGASE_BINDING | 0.00000 | 0.00000 | 0.76146 | 0.62273 | 1.38051 | 219 |
| GO_UBIQUITIN_PROTEIN_TRANSFERASE_REGULATOR_ACTIVITY | 0.00165 | 0.02590 | 0.45506 | 0.79605 | 1.57911 | 15 |
| GO_UNFOLDED_PROTEIN_BINDING | 0.00000 | 0.00000 | 0.77494 | 0.72157 | 1.55881 | 85 |
| GO_VESICLE_LUMEN | 0.00000 | 0.00000 | 0.74774 | 0.64475 | 1.42008 | 168 |
| GO_VIRAL_GENE_EXPRESSION | 0.00000 | 0.00000 | | 0.80028 | 1.76317 | 171 |
| GO_VIRAL_GENOME_REPLICATION | 0.00100 | 0.01761 | 0.45506 | 0.61997 | 1.33078 | 73 |
| GO_VIRAL_LIFE_CYCLE | 0.00031 | 0.00640 | 0.49849 | 0.55767 | 1.23509 | 209 |
| HP_ABNORMALITY_OF_ACID_BASE_HOMEOSTASIS | 0.00216 | 0.03242 | 0.43171 | 0.53556 | 1.18846 | 232 |
| HP_ABNORMALITY_OF_CARDIOVASCULAR_SYSTEM_ELECTROPHYSIO | 0.00027 | 0.00572 | 0.49849 | 0.55832 | 1.23929 | 229 |
| HP_ABNORMALITY_OF_LOWER_LIP | 0.00181 | 0.02804 | 0.45506 | 0.56360 | 1.23779 | 145 |
| HP_ABNORMALITY_OF_THE_MITOCHONDRION | 0.00213 | 0.03229 | 0.43171 | 0.57831 | 1.26240 | 117 |
| HP_ABNORMALITY_OF_THE_NASAL_DORSUM | 0.00005 | 0.00116 | 0.55733 | 0.60288 | 1.32157 | 134 |
| HP_ABNORMAL_ELECTROPHYSIOLOGY_OF_SINOATRIAL_NODE_ORIG | 0.00279 | 0.03889 | 0.43171 | 0.68436 | 1.40934 | 31 |
| HP_ABNORMAL_ERYTHROCYTE_MORPHOLOGY | 0.00000 | 0.00015 | 0.61053 | 0.56049 | 1.25036 | 341 |
| HP_ABNORMAL_MEAN_CORPUSCULAR_VOLUME | 0.00357 | 0.04696 | 0.43171 | 0.75226 | 1.51363 | 18 |
| HP_ABNORMAL_RETICULOCYTE_MORPHOLOGY | 0.00064 | 0.01210 | 0.47727 | 0.71420 | 1.47081 | 31 |
| HP_ACUTE_LEUKEMIA | 0.00000 | 0.00000 | 0.73376 | 0.72786 | 1.56137 | 70 |
| HP_ANEMIA_OF_INADEQUATE_PRODUCTION | 0.00000 | 0.00008 | 0.62726 | 0.65593 | 1.42209 | 97 |
| HP_CONSTITUTIONAL_SYMPTOM | 0.00119 | 0.02004 | 0.45506 | 0.51427 | 1.15108 | 423 |
| HP_DELAYED_PUBERTY | 0.00005 | 0.00115 | 0.55733 | 0.62579 | 1.36211 | 107 |
| HP_DEPRESSED_NASAL_RIDGE | 0.00000 | 0.00011 | 0.62726 | 0.68731 | 1.47511 | 72 |
| HP_FATIGUE | 0.00039 | 0.00808 | 0.49849 | 0.56529 | 1.24923 | 187 |
| HP_HEADACHE | 0.00015 | 0.00340 | 0.51885 | 0.59065 | 1.29716 | 146 |
| HP_HEMATOLOGICAL_NEOPLASM | 0.00003 | 0.00078 | 0.57561 | 0.60108 | 1.32044 | 154 |
| HP_INCREASED_CSF_LACTATE | 0.00157 | 0.02509 | 0.45506 | 0.61622 | 1.31904 | 66 |
| HP_INCREASED_SERUM_LACTATE | 0.00216 | 0.03242 | 0.43171 | 0.56920 | 1.24724 | 130 |
| HP_LEUKEMIA | 0.00001 | 0.00026 | 0.59333 | 0.64877 | 1.40680 | 98 |
| HP_MACROCYTIC_ANEMIA | 0.00000 | 0.00000 | 0.74774 | 0.82468 | 1.71698 | 39 |
| HP_MIGRAINE | 0.00000 | 0.00000 | 0.71951 | 0.72105 | 1.54497 | 69 |
| HP_MYOTONIA | 0.00161 | 0.02546 | 0.45506 | 0.79820 | 1.58338 | 15 |
| HP_NEOPLASM | 0.00216 | 0.03242 | 0.43171 | 0.50820 | 1.13805 | 446 |
| HP_PALLOR | 0.00001 | 0.00040 | 0.59333 | 0.65169 | 1.41044 | 92 |
| HP_PUBERTY_AND_GONADAL_DISORDERS | 0.00287 | 0.03973 | 0.43171 | 0.51995 | 1.15843 | 322 |
| HP_RETICULOCYTOPENIA | 0.00004 | 0.00105 | 0.55733 | 0.86417 | 1.71425 | 15 |
| HP_RIMMED_VACUOLES | 0.00279 | 0.03889 | 0.43171 | 0.74033 | 1.49831 | 20 |
| HP_SHORT_NOSE | 0.00235 | 0.03474 | 0.43171 | 0.54919 | 1.21134 | 174 |
| HP_THICK_LOWER_LIP_VERMILION | 0.00048 | 0.00949 | 0.49849 | 0.60773 | 1.31617 | 94 |
| HP_THICK_VERMILION_BORDER | 0.00156 | 0.02486 | 0.45506 | 0.55983 | 1.23000 | 153 |
| HP_VENTRICULAR_ARRHYTHMIA | 0.00353 | 0.04658 | 0.43171 | 0.66198 | 1.37398 | 36 |
| **Cluster 1** | | | | | | |
| pathway | pval | padj | log2err | ES | NES | size |
| GO_ACTIVATION_OF_IMMUNE_RESPONSE | 0.00000 | 0.00000 | 0.78819 | 0.62851 | 1.39255 | 212 |
| GO_ACTIVATION_OF_INNATE_IMMUNE_RESPONSE | 0.00000 | 0.00000 | | 0.75054 | 1.62130 | 88 |
| GO_AEROBIC_RESPIRATION | 0.00001 | 0.00025 | 0.59333 | 0.69302 | 1.48168 | 65 |
| GO_ALTERNATIVE_MRNA_SPLICING_VIA_SPLICEOSOME | 0.00102 | 0.01782 | 0.45506 | 0.65168 | 1.38687 | 60 |
| GO_AMINE_METABOLIC_PROCESS | 0.00000 | 0.00000 | | 0.79189 | 1.70783 | 80 |
| GO_ANAPHASE_PROMOTING_COMPLEX_DEPENDENT_CATABOLIC_P | 0.00000 | 0.00000 | | 0.76611 | 1.65070 | 79 |
| GO_ANCHORING_JUNCTION | 0.00000 | 0.00000 | | 0.59615 | 1.33633 | 479 |
| GO_ANIMAL_ORGAN_REGENERATION | 0.00257 | 0.03716 | 0.43171 | 0.68240 | 1.41020 | 33 |
| GO_ANTIGEN_PROCESSING_AND_PRESENTATION | 0.00000 | 0.00000 | 0.70498 | 0.64776 | 1.42241 | 143 |
| GO_ANTIGEN_PROCESSING_AND_PRESENTATION_OF_EXOGENOUS_ | 0.00000 | 0.00000 | | 0.81051 | 1.72358 | 57 |
| GO_ANTIGEN_PROCESSING_AND_PRESENTATION_OF_PEPTIDE_ANTI | 0.00000 | 0.00000 | 0.70498 | 0.65912 | 1.44248 | 126 |
| GO_ANTIGEN_PROCESSING_AND_PRESENTATION_OF_PEPTIDE_ANTI | 0.00000 | 0.00000 | | 0.76925 | 1.65015 | 70 |
| GO_ANTIGEN_RECEPTOR_MEDIATED_SIGNALING_PATHWAY | 0.00000 | 0.00000 | 0.80122 | 0.67877 | 1.48641 | 127 |
| GO_APOPTOTIC_SIGNALING_PATHWAY | 0.00154 | 0.02464 | 0.45506 | 0.51561 | 1.15085 | 358 |
| GO_ATP_DEPENDENT_CHROMATIN_REMODELING | 0.00253 | 0.03681 | 0.43171 | 0.62124 | 1.32400 | 61 |
| GO_ATP_METABOLIC_PROCESS | 0.00000 | 0.00000 | 0.71951 | 0.62252 | 1.37693 | 196 |
| GO_ATP_SYNTHESIS_COUPLED_ELECTRON_TRANSPORT | 0.00005 | 0.00131 | 0.55733 | 0.65519 | 1.40723 | 74 |
| GO_BINDING_OF_SPERM_TO_ZONA_PELLUCIDA | 0.00000 | 0.00001 | 0.69013 | 0.89879 | 1.80846 | 18 |
| GO_CADHERIN_BINDING | 0.00000 | 0.00000 | | 0.63068 | 1.39961 | 236 |
| GO_CANONICAL_WNT_SIGNALING_PATHWAY | 0.00000 | 0.00003 | 0.65944 | 0.60816 | 1.34553 | 197 |
| GO_CATALYTIC_STEP_2_SPLICEOSOME | 0.00000 | 0.00002 | 0.65944 | 0.68068 | 1.46976 | 83 |

| | | | | | | |
|---|---|---|---|---|---|---|
| GO_CELLULAR_AMINO_ACID_METABOLIC_PROCESS | 0.00000 | 0.00000 | 0.74774 | 0.61752 | 1.36824 | 216 |
| GO_CELLULAR_COMPONENT_DISASSEMBLY | 0.00000 | 0.00005 | 0.64355 | 0.55505 | 1.24197 | 401 |
| GO_CELLULAR_KETONE_METABOLIC_PROCESS | 0.00000 | 0.00000 | 0.77494 | 0.65955 | 1.44893 | 149 |
| GO_CELLULAR_NITROGEN_COMPOUND_CATABOLIC_PROCESS | 0.00000 | 0.00000 | | 0.74079 | 1.65848 | 436 |
| GO_CELLULAR_PROTEIN_COMPLEX_DISASSEMBLY | 0.00006 | 0.00139 | 0.55733 | 0.58222 | 1.28459 | 176 |
| GO_CELLULAR_RESPIRATION | 0.00000 | 0.00000 | 0.71951 | 0.64997 | 1.42492 | 136 |
| GO_CELLULAR_RESPONSE_TO_HEAT | 0.00018 | 0.00403 | 0.51885 | 0.61962 | 1.34016 | 91 |
| GO_CELLULAR_RESPONSE_TO_ORGANIC_CYCLIC_COMPOUND | 0.00302 | 0.04108 | 0.43171 | 0.52402 | 1.16585 | 283 |
| GO_CELLULAR_RESPONSE_TO_OXYGEN_LEVELS | 0.00000 | 0.00000 | 0.78819 | 0.65911 | 1.44882 | 157 |
| GO_CELL_ACTIVATION_INVOLVED_IN_IMMUNE_RESPONSE | 0.00000 | 0.00001 | 0.67496 | 0.57339 | 1.27913 | 341 |
| GO_CELL_ADHESION_MOLECULE_BINDING | 0.00000 | 0.00000 | 0.81404 | 0.60529 | 1.34788 | 308 |
| GO_CELL_CELL_RECOGNITION | 0.00000 | 0.00002 | 0.65944 | 0.83826 | 1.71003 | 26 |
| GO_CELL_CELL_SIGNALING_BY_WNT | 0.00000 | 0.00002 | 0.67496 | 0.57736 | 1.28559 | 299 |
| GO_CELL_CYCLE_G2_M_PHASE_TRANSITION | 0.00001 | 0.00024 | 0.59333 | 0.58034 | 1.28602 | 214 |
| GO_CELL_CYCLE_PHASE_TRANSITION | 0.00000 | 0.00004 | 0.64355 | 0.54319 | 1.21681 | 464 |
| GO_CELL_DIVISION | 0.00154 | 0.02464 | 0.45506 | 0.51551 | 1.15425 | 427 |
| GO_CELL_RECOGNITION | 0.00012 | 0.00270 | 0.53843 | 0.69554 | 1.46186 | 48 |
| GO_CELL_SUBSTRATE_JUNCTION | 0.00000 | 0.00000 | | 0.67751 | 1.50775 | 291 |
| GO_CELL_SURFACE_RECEPTOR_SIGNALING_PATHWAY_INVOLVED_IN | 0.00000 | 0.00003 | 0.65944 | 0.57035 | 1.27067 | 319 |
| GO_CHAPERONE_BINDING | 0.00007 | 0.00163 | 0.53843 | 0.64754 | 1.39253 | 76 |
| GO_CHAPERONE_COMPLEX | 0.00000 | 0.00004 | 0.64355 | 0.85718 | 1.73925 | 22 |
| GO_CHAPERONE_MEDIATED_PROTEIN_COMPLEX_ASSEMBLY | 0.00170 | 0.02662 | 0.45506 | 0.77423 | 1.54792 | 17 |
| GO_CHROMATIN_BINDING | 0.00071 | 0.01323 | 0.47727 | 0.52699 | 1.17680 | 369 |
| GO_CHROMOSOMAL_REGION | 0.00246 | 0.03601 | 0.43171 | 0.52919 | 1.17661 | 276 |
| GO_COP9_SIGNALOSOME | 0.00128 | 0.02141 | 0.45506 | 0.73616 | 1.49715 | 24 |
| GO_COTRANSLATIONAL_PROTEIN_TARGETING_TO_MEMBRANE | 0.00000 | 0.00000 | | 0.89471 | 1.93513 | 91 |
| GO_CYTOCHROME_COMPLEX | 0.00168 | 0.02641 | 0.45506 | 0.75311 | 1.52417 | 20 |
| GO_CYTOKINE_MEDIATED_SIGNALING_PATHWAY | 0.00000 | 0.00000 | 0.81404 | 0.59698 | 1.33170 | 339 |
| GO_CYTOPLASMIC_TRANSLATION | 0.00000 | 0.00000 | | 0.84753 | 1.83080 | 88 |
| GO_CYTOPLASMIC_TRANSLATIONAL_INITIATION | 0.00000 | 0.00000 | 0.74774 | 0.85688 | 1.76176 | 30 |
| GO_CYTOSOLIC_LARGE_RIBOSOMAL_SUBUNIT | 0.00000 | 0.00000 | | 0.91186 | 1.92988 | 53 |
| GO_CYTOSOLIC_RIBOSOME | 0.00000 | 0.00000 | | 0.89907 | 1.94710 | 95 |
| GO_CYTOSOLIC_SMALL_RIBOSOMAL_SUBUNIT | 0.00000 | 0.00000 | | 0.89114 | 1.85533 | 39 |
| GO_DEFENSE_RESPONSE_TO_OTHER_ORGANISM | 0.00004 | 0.00099 | 0.55733 | 0.53673 | 1.20127 | 415 |
| GO_DE_NOVO_PROTEIN_FOLDING | 0.00050 | 0.00978 | 0.47727 | 0.74232 | 1.51431 | 26 |
| GO_DISORDERED_DOMAIN_SPECIFIC_BINDING | 0.00161 | 0.02546 | 0.45506 | 0.71970 | 1.47197 | 27 |
| GO_DNA_BIOSYNTHETIC_PROCESS | 0.00000 | 0.00010 | 0.62726 | 0.62505 | 1.37017 | 134 |
| GO_DNA_CONFORMATION_CHANGE | 0.00027 | 0.00572 | 0.49849 | 0.55300 | 1.22527 | 216 |
| GO_DNA_DAMAGE_RESPONSE_DETECTION_OF_DNA_DAMAGE | 0.00198 | 0.03028 | 0.43171 | 0.67949 | 1.40669 | 35 |
| GO_DNA_GEOMETRIC_CHANGE | 0.00046 | 0.00925 | 0.49849 | 0.60807 | 1.31855 | 98 |
| GO_DNA_POLYMERASE_BINDING | 0.00268 | 0.03799 | 0.43171 | 0.76255 | 1.52457 | 17 |
| GO_DNA_REPAIR | 0.00265 | 0.03775 | 0.43171 | 0.51159 | 1.14547 | 427 |
| GO_DRUG_BINDING | 0.00176 | 0.02733 | 0.45506 | 0.64119 | 1.35096 | 49 |
| GO_ELECTRON_TRANSFER_ACTIVITY | 0.00001 | 0.00020 | 0.61053 | 0.67481 | 1.44938 | 74 |
| GO_ELECTRON_TRANSPORT_CHAIN | 0.00002 | 0.00053 | 0.57561 | 0.62257 | 1.36145 | 124 |
| GO_ENDOPEPTIDASE_ACTIVITY | 0.00124 | 0.02085 | 0.45506 | 0.54955 | 1.21076 | 171 |
| GO_ENDOPEPTIDASE_COMPLEX | 0.00000 | 0.00000 | | 0.80930 | 1.73234 | 66 |
| GO_ENERGY_DERIVATION_BY_OXIDATION_OF_ORGANIC_COMPOUN | 0.00000 | 0.00000 | 0.71951 | 0.63390 | 1.39707 | 172 |
| GO_ESTABLISHMENT_OF_PROTEIN_LOCALIZATION_TO_CHROMOSON | 0.00020 | 0.00432 | 0.51885 | 0.77339 | 1.57217 | 23 |
| GO_ESTABLISHMENT_OF_PROTEIN_LOCALIZATION_TO_ENDOPLASM | 0.00000 | 0.00000 | | 0.88026 | 1.91149 | 102 |
| GO_ESTABLISHMENT_OF_PROTEIN_LOCALIZATION_TO_MEMBRANE | 0.00000 | 0.00000 | | 0.74874 | 1.66164 | 240 |
| GO_ESTABLISHMENT_OF_PROTEIN_LOCALIZATION_TO_ORGANELLE | 0.00000 | 0.00000 | | 0.69519 | 1.55605 | 423 |
| GO_ESTABLISHMENT_OF_PROTEIN_LOCALIZATION_TO_TELOMERE | 0.00006 | 0.00139 | 0.55733 | 0.83470 | 1.66881 | 17 |
| GO_ESTABLISHMENT_OF_RNA_LOCALIZATION | 0.00106 | 0.01819 | 0.45506 | 0.55577 | 1.22400 | 167 |
| GO_ESTABLISHMENT_OF_TISSUE_POLARITY | 0.00000 | 0.00000 | | 0.75051 | 1.62122 | 88 |
| GO_EUKARYOTIC_TRANSLATION_INITIATION_FACTOR_3_COMPLEX | 0.00003 | 0.00074 | 0.57561 | 0.86987 | 1.72555 | 15 |
| GO_EXOCYTOSIS | 0.00000 | 0.00015 | 0.61053 | 0.53905 | 1.20753 | 465 |
| GO_FC_EPSILON_RECEPTOR_SIGNALING_PATHWAY | 0.00000 | 0.00000 | | 0.75117 | 1.62103 | 82 |
| GO_FC_RECEPTOR_SIGNALING_PATHWAY | 0.00000 | 0.00000 | | 0.72054 | 1.57787 | 127 |
| GO_FERTILIZATION | 0.00089 | 0.01592 | 0.47727 | 0.63620 | 1.35888 | 64 |
| GO_FICOLIN_1_RICH_GRANULE | 0.00000 | 0.00000 | 0.78819 | 0.71716 | 1.55340 | 93 |
| GO_GENERATION_OF_PRECURSOR_METABOLITES_AND_ENERGY | 0.00000 | 0.00000 | 0.77494 | 0.59261 | 1.32026 | 320 |
| GO_GENE_SILENCING | 0.00261 | 0.03750 | 0.43171 | 0.55578 | 1.22252 | 161 |
| GO_GLAND_DEVELOPMENT | 0.00154 | 0.02464 | 0.45506 | 0.54117 | 1.19797 | 206 |
| GO_GLYCOSYL_COMPOUND_METABOLIC_PROCESS | 0.00071 | 0.01323 | 0.47727 | 0.62524 | 1.34204 | 71 |
| GO_GUANYL_NUCLEOTIDE_BINDING | 0.00095 | 0.01679 | 0.47727 | 0.54096 | 1.19861 | 216 |
| GO_HEAT_SHOCK_PROTEIN_BINDING | 0.00003 | 0.00092 | 0.55733 | 0.64369 | 1.39056 | 85 |
| GO_HEMATOPOIETIC_PROGENITOR_CELL_DIFFERENTIATION | 0.00000 | 0.00000 | 0.82666 | 0.70307 | 1.53239 | 113 |
| GO_HEMATOPOIETIC_STEM_CELL_DIFFERENTIATION | 0.00000 | 0.00000 | | 0.79776 | 1.71217 | 72 |
| GO_HSP70_PROTEIN_BINDING | 0.00228 | 0.03391 | 0.43171 | 0.71143 | 1.45504 | 27 |
| GO_IMMUNE_RESPONSE_REGULATING_SIGNALING_PATHWAY | 0.00000 | 0.00000 | 0.76146 | 0.63494 | 1.40480 | 197 |
| GO_INNATE_IMMUNE_RESPONSE | 0.00003 | 0.00071 | 0.57561 | 0.55173 | 1.23107 | 347 |
| GO_INNATE_IMMUNE_RESPONSE_ACTIVATING_SIGNAL_TRANSDUCT | 0.00000 | 0.00000 | | 0.79340 | 1.70281 | 72 |
| GO_INNER_MITOCHONDRIAL_MEMBRANE_PROTEIN_COMPLEX | 0.00002 | 0.00060 | 0.57561 | 0.65124 | 1.40678 | 88 |
| GO_INTERACTION_WITH_SYMBIONT | 0.00001 | 0.00026 | 0.59333 | 0.75084 | 1.56620 | 41 |
| GO_INTERLEUKIN_1_MEDIATED_SIGNALING_PATHWAY | 0.00000 | 0.00000 | | 0.81689 | 1.75671 | 76 |

| | | | | | | |
|---|---|---|---|---|---|---|
| GO_INTRINSIC_APOPTOTIC_SIGNALING_PATHWAY | 0.00121 | 0.02029 | 0.45506 | 0.55118 | 1.21845 | 190 |
| GO_ION_CHANNEL_BINDING | 0.00087 | 0.01584 | 0.47727 | 0.63182 | 1.34683 | 62 |
| GO_ISOPEPTIDASE_ACTIVITY | 0.00139 | 0.02285 | 0.45506 | 0.78940 | 1.57097 | 16 |
| GO_LARGE_RIBOSOMAL_SUBUNIT | 0.00000 | 0.00000 | | 0.78135 | 1.70183 | 109 |
| GO_LEUKOCYTE_MEDIATED_IMMUNITY | 0.00000 | 0.00000 | 0.71951 | 0.57282 | 1.27895 | 364 |
| GO_MACROMOLECULE_TRANSMEMBRANE_TRANSPORTER_ACTIVITY | 0.00239 | 0.03510 | 0.43171 | 0.77734 | 1.54696 | 16 |
| GO_MAINTENANCE_OF_CELL_NUMBER | 0.00087 | 0.01584 | 0.47727 | 0.58944 | 1.28512 | 114 |
| GO_MATURATION_OF_LSU_RRNA | 0.00006 | 0.00151 | 0.53843 | 0.77606 | 1.58723 | 27 |
| GO_METHYLTRANSFERASE_COMPLEX | 0.00290 | 0.04014 | 0.43171 | 0.59476 | 1.28479 | 88 |
| GO_METHYL_CPG_BINDING | 0.00302 | 0.04108 | 0.43171 | 0.78053 | 1.54833 | 15 |
| GO_MIDBRAIN_DEVELOPMENT | 0.00021 | 0.00468 | 0.51885 | 0.68832 | 1.44457 | 47 |
| GO_MITOCHONDRIAL_ELECTRON_TRANSPORT_NADH_TO_UBIQUIN | 0.00331 | 0.04437 | 0.43171 | 0.65428 | 1.36724 | 42 |
| GO_MITOCHONDRIAL_GENE_EXPRESSION | 0.00048 | 0.00949 | 0.49849 | 0.57608 | 1.26525 | 152 |
| GO_MITOCHONDRIAL_LARGE_RIBOSOMAL_SUBUNIT | 0.00002 | 0.00050 | 0.57561 | 0.69789 | 1.48086 | 55 |
| GO_MITOCHONDRIAL_MATRIX | 0.00000 | 0.00002 | 0.65944 | 0.56045 | 1.25158 | 371 |
| GO_MITOCHONDRIAL_MEMBRANE_ORGANIZATION | 0.00361 | 0.04734 | 0.43171 | 0.59309 | 1.28094 | 86 |
| GO_MITOCHONDRIAL_PROTEIN_COMPLEX | 0.00000 | 0.00000 | 0.81404 | 0.63650 | 1.40931 | 208 |
| GO_MITOCHONDRIAL_TRANSLATION | 0.00004 | 0.00093 | 0.55733 | 0.61219 | 1.34135 | 129 |
| GO_MITOCHONDRIAL_TRANSLATIONAL_TERMINATION | 0.00000 | 0.00010 | 0.62726 | 0.66694 | 1.44128 | 87 |
| GO_MITOCHONDRION_ORGANIZATION | 0.00051 | 0.00993 | 0.47727 | 0.52284 | 1.16863 | 390 |
| GO_MODIFICATION_DEPENDENT_MACROMOLECULE_CATABOLIC_PR | 0.00002 | 0.00053 | 0.57561 | 0.53142 | 1.19156 | 486 |
| GO_MODULATION_BY_HOST_OF_SYMBIONT_PROCESS | 0.00015 | 0.00336 | 0.51885 | 0.72848 | 1.50542 | 33 |
| GO_MODULATION_OF_PROCESS_OF_OTHER_ORGANISM | 0.00019 | 0.00423 | 0.51885 | 0.66904 | 1.42286 | 59 |
| GO_MONOSACCHARIDE_BIOSYNTHETIC_PROCESS | 0.00335 | 0.04466 | 0.43171 | 0.61992 | 1.31781 | 58 |
| GO_MORPHOGENESIS_OF_AN_EPITHELIUM | 0.00006 | 0.00151 | 0.53843 | 0.55161 | 1.22755 | 290 |
| GO_MORPHOGENESIS_OF_A_POLARIZED_EPITHELIUM | 0.00000 | 0.00000 | | 0.72298 | 1.57057 | 103 |
| GO_MRNA_3_UTR_BINDING | 0.00003 | 0.00089 | 0.55733 | 0.67596 | 1.44827 | 68 |
| GO_MRNA_5_UTR_BINDING | 0.00006 | 0.00156 | 0.53843 | 0.80133 | 1.62592 | 22 |
| GO_MRNA_BINDING | 0.00000 | 0.00000 | | 0.66002 | 1.46353 | 222 |
| GO_MRNA_PROCESSING | 0.00000 | 0.00000 | 0.73376 | 0.56859 | 1.27295 | 418 |
| GO_MULTI_ORGANISM_LOCALIZATION | 0.00268 | 0.03799 | 0.43171 | 0.63245 | 1.33612 | 51 |
| GO_MULTI_ORGANISM_PROCESS | 0.00108 | 0.01839 | 0.45506 | 0.51114 | 1.14599 | 484 |
| GO_MYD88_DEPENDENT_TOLL_LIKE_RECEPTOR_SIGNALING_PATHW | 0.00002 | 0.00066 | 0.57561 | 0.85581 | 1.70312 | 16 |
| GO_MYELOID_LEUKOCYTE_ACTIVATION | 0.00000 | 0.00000 | 0.74774 | 0.58508 | 1.30479 | 327 |
| GO_MYELOID_LEUKOCYTE_MEDIATED_IMMUNITY | 0.00000 | 0.00000 | 0.76146 | 0.59841 | 1.33291 | 301 |
| GO_NADH_DEHYDROGENASE_ACTIVITY | 0.00383 | 0.04935 | 0.43171 | 0.66244 | 1.37139 | 35 |
| GO_NADH_DEHYDROGENASE_COMPLEX | 0.00376 | 0.04884 | 0.43171 | 0.65649 | 1.36421 | 38 |
| GO_NADH_METABOLIC_PROCESS | 0.00294 | 0.04036 | 0.43171 | 0.70642 | 1.44481 | 27 |
| GO_NCRNA_EXPORT_FROM_NUCLEUS | 0.00302 | 0.04108 | 0.43171 | 0.66820 | 1.38331 | 35 |
| GO_NCRNA_METABOLIC_PROCESS | 0.00000 | 0.00003 | 0.65944 | 0.55907 | 1.24907 | 381 |
| GO_NCRNA_PROCESSING | 0.00000 | 0.00003 | 0.65944 | 0.57175 | 1.27504 | 328 |
| GO_NEGATIVE_REGULATION_OF_CANONICAL_WNT_SIGNALING_PAT | 0.00000 | 0.00000 | | 0.71611 | 1.56038 | 111 |
| GO_NEGATIVE_REGULATION_OF_CELLULAR_AMIDE_METABOLIC_PR | 0.00001 | 0.00020 | 0.61053 | 0.63031 | 1.37423 | 114 |
| GO_NEGATIVE_REGULATION_OF_CELL_CYCLE | 0.00003 | 0.00076 | 0.57561 | 0.53833 | 1.20513 | 425 |
| GO_NEGATIVE_REGULATION_OF_CELL_CYCLE_G2_M_PHASE_TRANS | 0.00000 | 0.00000 | 0.76146 | 0.71490 | 1.54439 | 85 |
| GO_NEGATIVE_REGULATION_OF_CELL_CYCLE_PHASE_TRANSITION | 0.00002 | 0.00060 | 0.57561 | 0.59041 | 1.30442 | 184 |
| GO_NEGATIVE_REGULATION_OF_CELL_CYCLE_PROCESS | 0.00000 | 0.00002 | 0.65944 | 0.58690 | 1.30374 | 249 |
| GO_NEGATIVE_REGULATION_OF_MITOTIC_CELL_CYCLE | 0.00004 | 0.00095 | 0.55733 | 0.56735 | 1.25902 | 232 |
| GO_NEGATIVE_REGULATION_OF_MRNA_METABOLIC_PROCESS | 0.00023 | 0.00491 | 0.51885 | 0.66670 | 1.41788 | 59 |
| GO_NEGATIVE_REGULATION_OF_MRNA_SPLICING_VIA_SPLICEOSOM | 0.00239 | 0.03510 | 0.43171 | 0.74857 | 1.51204 | 19 |
| GO_NEGATIVE_REGULATION_OF_RNA_SPLICING | 0.00029 | 0.00607 | 0.49849 | 0.76729 | 1.55976 | 23 |
| GO_NEGATIVE_REGULATION_OF_TRANSLATIONAL_INITIATION | 0.00018 | 0.00403 | 0.51885 | 0.83901 | 1.66434 | 15 |
| GO_NEGATIVE_REGULATION_OF_WNT_SIGNALING_PATHWAY | 0.00000 | 0.00000 | | 0.69525 | 1.52249 | 127 |
| GO_NEURAL_NUCLEUS_DEVELOPMENT | 0.00001 | 0.00026 | 0.59333 | 0.79148 | 1.62187 | 28 |
| GO_NEURON_TO_NEURON_SYNAPSE | 0.00005 | 0.00134 | 0.55733 | 0.59636 | 1.31006 | 154 |
| GO_NIK_NF_KAPPAB_SIGNALING | 0.00000 | 0.00000 | | 0.72896 | 1.58354 | 103 |
| GO_NON_CANONICAL_WNT_SIGNALING_PATHWAY | 0.00000 | 0.00000 | | 0.72784 | 1.58023 | 101 |
| GO_NUCLEAR_EXPORT | 0.00031 | 0.00645 | 0.49849 | 0.56963 | 1.25542 | 172 |
| GO_NUCLEAR_TRANSCRIBED_MRNA_CATABOLIC_PROCESS | 0.00000 | 0.00000 | | 0.79179 | 1.74917 | 183 |
| GO_NUCLEAR_TRANSCRIBED_MRNA_CATABOLIC_PROCESS_NONSEN | 0.00000 | 0.00000 | | 0.89008 | 1.93909 | 110 |
| GO_NUCLEAR_TRANSPORT | 0.00008 | 0.00193 | 0.53843 | 0.55511 | 1.23408 | 273 |
| GO_NUCLEOBASE_CONTAINING_COMPOUND_TRANSPORT | 0.00146 | 0.02386 | 0.45506 | 0.55053 | 1.21842 | 202 |
| GO_NUCLEOBASE_CONTAINING_SMALL_MOLECULE_BIOSYNTHETIC_ | 0.00137 | 0.02261 | 0.45506 | 0.62749 | 1.34353 | 67 |
| GO_NUCLEOBASE_CONTAINING_SMALL_MOLECULE_INTERCONVERS | 0.00383 | 0.04935 | 0.43171 | 0.71259 | 1.44857 | 23 |
| GO_NUCLEOBASE_CONTAINING_SMALL_MOLECULE_METABOLIC_PR | 0.00320 | 0.04339 | 0.43171 | 0.51448 | 1.14825 | 352 |
| GO_NUCLEOSIDE_DIPHOSPHATE_METABOLIC_PROCESS | 0.00110 | 0.01865 | 0.45506 | 0.59637 | 1.29295 | 97 |
| GO_NUCLEOSIDE_METABOLIC_PROCESS | 0.00061 | 0.01169 | 0.47727 | 0.65315 | 1.38907 | 59 |
| GO_NUCLEOSIDE_MONOPHOSPHATE_BIOSYNTHETIC_PROCESS | 0.00250 | 0.03636 | 0.43171 | 0.68841 | 1.42033 | 32 |
| GO_NUCLEOSIDE_MONOPHOSPHATE_METABOLIC_PROCESS | 0.00152 | 0.02464 | 0.45506 | 0.64377 | 1.36002 | 51 |
| GO_NUCLEOSIDE_PHOSPHATE_BIOSYNTHETIC_PROCESS | 0.00043 | 0.00865 | 0.49849 | 0.58185 | 1.27831 | 148 |
| GO_NUCLEOSIDE_TRIPHOSPHATE_BIOSYNTHETIC_PROCESS | 0.00043 | 0.00865 | 0.49849 | 0.69332 | 1.44539 | 40 |
| GO_NUCLEOSIDE_TRIPHOSPHATE_METABOLIC_PROCESS | 0.00016 | 0.00360 | 0.51885 | 0.66788 | 1.41976 | 58 |
| GO_NUCLEOSOMAL_DNA_BINDING | 0.00023 | 0.00507 | 0.51885 | 0.79852 | 1.61801 | 21 |
| GO_NUCLEOSOME_BINDING | 0.00143 | 0.02332 | 0.45506 | 0.66132 | 1.38492 | 45 |
| GO_NUCLEOTIDE_EXCISION_REPAIR | 0.00004 | 0.00096 | 0.55733 | 0.63343 | 1.37355 | 98 |
| GO_NUCLEOTIDE_EXCISION_REPAIR_DNA_DAMAGE_RECOGNITION | 0.00015 | 0.00349 | 0.51885 | 0.78678 | 1.59640 | 22 |

| | | | | | | |
|---|---|---|---|---|---|---|
| GO_NUCLEOTIDE_EXCISION_REPAIR_DNA_INCISION | 0.00313 | 0.04249 | 0.43171 | 0.66439 | 1.37898 | 36 |
| GO_NUCLEOTIDE_PHOSPHORYLATION | 0.00383 | 0.04935 | 0.43171 | 0.59002 | 1.27454 | 88 |
| GO_ORGANELLAR_RIBOSOME | 0.00000 | 0.00014 | 0.61053 | 0.66412 | 1.43471 | 85 |
| GO_ORGANELLE_INNER_MEMBRANE | 0.00000 | 0.00000 | 0.76146 | 0.57601 | 1.28697 | 378 |
| GO_ORGANIC_CYCLIC_COMPOUND_CATABOLIC_PROCESS | 0.00000 | 0.00000 | | 0.73350 | 1.64281 | 449 |
| GO_OXIDATIVE_PHOSPHORYLATION | 0.00001 | 0.00030 | 0.59333 | 0.64619 | 1.40027 | 96 |
| GO_OXIDOREDUCTASE_ACTIVITY | 0.00006 | 0.00139 | 0.55733 | 0.53544 | 1.19703 | 392 |
| GO_OXIDOREDUCTASE_COMPLEX | 0.00001 | 0.00027 | 0.59333 | 0.66015 | 1.42461 | 82 |
| GO_PEPTIDASE_COMPLEX | 0.00000 | 0.00000 | | 0.75988 | 1.63982 | 82 |
| GO_PIGMENT_GRANULE | 0.00003 | 0.00089 | 0.55733 | 0.66098 | 1.42344 | 78 |
| GO_POLYSOMAL_RIBOSOME | 0.00000 | 0.00000 | | 0.90624 | 1.86629 | 31 |
| GO_POLYSOME | 0.00000 | 0.00000 | | 0.82337 | 1.75481 | 61 |
| GO_POLY_PYRIMIDINE_TRACT_BINDING | 0.00327 | 0.04398 | 0.43171 | 0.75471 | 1.51855 | 18 |
| GO_POSITIVE_REGULATION_OF_APOPTOTIC_SIGNALING_PATHWAY | 0.00279 | 0.03889 | 0.43171 | 0.57666 | 1.25518 | 107 |
| GO_POSITIVE_REGULATION_OF_ATPASE_ACTIVITY | 0.00294 | 0.04036 | 0.43171 | 0.65446 | 1.36795 | 43 |
| GO_POSITIVE_REGULATION_OF_BINDING | 0.00088 | 0.01590 | 0.47727 | 0.60284 | 1.30556 | 95 |
| GO_POSITIVE_REGULATION_OF_CANONICAL_WNT_SIGNALING_PATH | 0.00000 | 0.00000 | 0.77494 | 0.70504 | 1.53160 | 103 |
| GO_POSITIVE_REGULATION_OF_CELLULAR_AMIDE_METABOLIC_PRO | 0.00003 | 0.00074 | 0.57561 | 0.62296 | 1.36083 | 118 |
| GO_POSITIVE_REGULATION_OF_CELLULAR_PROTEIN_LOCALIZATION | 0.00041 | 0.00840 | 0.49849 | 0.54915 | 1.21815 | 225 |
| GO_POSITIVE_REGULATION_OF_CELL_DEATH | 0.00069 | 0.01297 | 0.47727 | 0.52595 | 1.17423 | 363 |
| GO_POSITIVE_REGULATION_OF_CELL_POPULATION_PROLIFERATION | 0.00078 | 0.01455 | 0.47727 | 0.52449 | 1.17108 | 368 |
| GO_POSITIVE_REGULATION_OF_CHROMOSOME_ORGANIZATION | 0.00231 | 0.03437 | 0.43171 | 0.56990 | 1.24871 | 133 |
| GO_POSITIVE_REGULATION_OF_DEFENSE_RESPONSE | 0.00000 | 0.00003 | 0.65944 | 0.62175 | 1.36730 | 158 |
| GO_POSITIVE_REGULATION_OF_DNA_BIOSYNTHETIC_PROCESS | 0.00001 | 0.00041 | 0.59333 | 0.71929 | 1.52055 | 52 |
| GO_POSITIVE_REGULATION_OF_DNA_METABOLIC_PROCESS | 0.00364 | 0.04761 | 0.43171 | 0.56647 | 1.24189 | 135 |
| GO_POSITIVE_REGULATION_OF_DNA_TEMPLATED_TRANSCRIPTION_ | 0.00276 | 0.03885 | 0.43171 | 0.68932 | 1.41586 | 29 |
| GO_POSITIVE_REGULATION_OF_ESTABLISHMENT_OF_PROTEIN_LOC | 0.00007 | 0.00168 | 0.53843 | 0.56461 | 1.25165 | 219 |
| GO_POSITIVE_REGULATION_OF_IMMUNE_RESPONSE | 0.00000 | 0.00001 | 0.69013 | 0.59037 | 1.31265 | 277 |
| GO_POSITIVE_REGULATION_OF_IMMUNE_SYSTEM_PROCESS | 0.00004 | 0.00110 | 0.55733 | 0.54018 | 1.20716 | 386 |
| GO_POSITIVE_REGULATION_OF_INNATE_IMMUNE_RESPONSE | 0.00000 | 0.00000 | 0.81404 | 0.69693 | 1.51947 | 114 |
| GO_POSITIVE_REGULATION_OF_INTRINSIC_APOPTOTIC_SIGNALING_ | 0.00104 | 0.01798 | 0.45506 | 0.69097 | 1.43578 | 37 |
| GO_POSITIVE_REGULATION_OF_ORGANELLE_ORGANIZATION | 0.00043 | 0.00865 | 0.49849 | 0.52093 | 1.16605 | 411 |
| GO_POSITIVE_REGULATION_OF_PROTEIN_LOCALIZATION_TO_NUCLE | 0.00001 | 0.00020 | 0.61053 | 0.69907 | 1.48990 | 61 |
| GO_POSITIVE_REGULATION_OF_RESPONSE_TO_BIOTIC_STIMULUS | 0.00000 | 0.00000 | 0.73376 | 0.66340 | 1.45310 | 128 |
| GO_POSITIVE_REGULATION_OF_RESPONSE_TO_EXTERNAL_STIMULU | 0.00002 | 0.00050 | 0.57561 | 0.58133 | 1.28681 | 201 |
| GO_POSITIVE_REGULATION_OF_TELOMERASE_ACTIVITY | 0.00061 | 0.01169 | 0.47727 | 0.73476 | 1.50565 | 28 |
| GO_POSITIVE_REGULATION_OF_TELOMERASE_RNA_LOCALIZATION_ | 0.00000 | 0.00001 | 0.69013 | 0.92451 | 1.83395 | 15 |
| GO_POSITIVE_REGULATION_OF_TELOMERE_MAINTENANCE_VIA_TE | 0.00130 | 0.02159 | 0.45506 | 0.71259 | 1.46508 | 30 |
| GO_POSITIVE_REGULATION_OF_TRANSCRIPTION_INITIATION_FROM_ | 0.00372 | 0.04847 | 0.43171 | 0.70671 | 1.43993 | 25 |
| GO_POSITIVE_REGULATION_OF_TRANSFERASE_ACTIVITY | 0.00174 | 0.02712 | 0.45506 | 0.51642 | 1.15220 | 345 |
| GO_POSITIVE_REGULATION_OF_TRANSLATION | 0.00001 | 0.00019 | 0.61053 | 0.64767 | 1.40419 | 97 |
| GO_POSITIVE_REGULATION_OF_VIRAL_GENOME_REPLICATION | 0.00287 | 0.03973 | 0.43171 | 0.68362 | 1.40783 | 31 |
| GO_POSITIVE_REGULATION_OF_VIRAL_PROCESS | 0.00028 | 0.00590 | 0.49849 | 0.62440 | 1.34741 | 81 |
| GO_POSITIVE_REGULATION_OF_WNT_SIGNALING_PATHWAY | 0.00000 | 0.00000 | 0.73376 | 0.66230 | 1.44763 | 122 |
| GO_POSTREPLICATION_REPAIR | 0.00024 | 0.00515 | 0.51885 | 0.69140 | 1.44518 | 43 |
| GO_POSTSYNAPSE | 0.00004 | 0.00109 | 0.55733 | 0.55969 | 1.24369 | 259 |
| GO_POSTSYNAPTIC_CYTOSOL | 0.00279 | 0.03889 | 0.43171 | 0.78295 | 1.55313 | 15 |
| GO_POSTTRANSCRIPTIONAL_REGULATION_OF_GENE_EXPRESSION | 0.00000 | 0.00000 | | 0.66656 | 1.49327 | 462 |
| GO_POST_TRANSLATIONAL_PROTEIN_MODIFICATION | 0.00002 | 0.00052 | 0.57561 | 0.57851 | 1.28412 | 229 |
| GO_PRECATALYTIC_SPLICEOSOME | 0.00085 | 0.01559 | 0.47727 | 0.65235 | 1.37448 | 49 |
| GO_PRERIBOSOME | 0.00087 | 0.01577 | 0.47727 | 0.62899 | 1.34772 | 69 |
| GO_PRERIBOSOME_LARGE_SUBUNIT_PRECURSOR | 0.00235 | 0.03474 | 0.43171 | 0.74079 | 1.50308 | 22 |
| GO_PRE_MRNA_BINDING | 0.00133 | 0.02213 | 0.45506 | 0.72393 | 1.48062 | 27 |
| GO_PROTEASOMAL_PROTEIN_CATABOLIC_PROCESS | 0.00002 | 0.00044 | 0.59333 | 0.54166 | 1.21005 | 377 |
| GO_PROTEASOMAL_UBIQUITIN_INDEPENDENT_PROTEIN_CATABOLI | 0.00000 | 0.00000 | 0.80122 | 0.92254 | 1.86931 | 21 |
| GO_PROTEASOME_ACCESSORY_COMPLEX | 0.00000 | 0.00001 | 0.67496 | 0.85464 | 1.73811 | 24 |
| GO_PROTEASOME_CORE_COMPLEX | 0.00000 | 0.00000 | | 0.94375 | 1.90631 | 19 |
| GO_PROTEIN_CONTAINING_COMPLEX_DISASSEMBLY | 0.00000 | 0.00007 | 0.62726 | 0.57810 | 1.28438 | 256 |
| GO_PROTEIN_CONTAINING_COMPLEX_LOCALIZATION | 0.00063 | 0.01197 | 0.47727 | 0.55335 | 1.22520 | 208 |
| GO_PROTEIN_DNA_COMPLEX_SUBUNIT_ORGANIZATION | 0.00001 | 0.00018 | 0.61053 | 0.60174 | 1.32491 | 165 |
| GO_PROTEIN_FOLDING | 0.00000 | 0.00002 | 0.65944 | 0.63113 | 1.38644 | 154 |
| GO_PROTEIN_FOLDING_CHAPERONE | 0.00135 | 0.02238 | 0.45506 | 0.77903 | 1.55752 | 17 |
| GO_PROTEIN_IMPORT | 0.00268 | 0.03799 | 0.43171 | 0.56317 | 1.23581 | 140 |
| GO_PROTEIN_LOCALIZATION_TO_CHROMOSOME | 0.00056 | 0.01096 | 0.47727 | 0.63523 | 1.36108 | 69 |
| GO_PROTEIN_LOCALIZATION_TO_CHROMOSOME_TELOMERIC_REGI | 0.00335 | 0.04466 | 0.43171 | 0.70399 | 1.43984 | 27 |
| GO_PROTEIN_LOCALIZATION_TO_ENDOPLASMIC_RETICULUM | 0.00000 | 0.00000 | | 0.85521 | 1.87021 | 124 |
| GO_PROTEIN_LOCALIZATION_TO_MEMBRANE | 0.00000 | 0.00000 | | 0.65123 | 1.45714 | 406 |
| GO_PROTEIN_LOCALIZATION_TO_MITOCHONDRION | 0.00019 | 0.00420 | 0.51885 | 0.60773 | 1.32398 | 110 |
| GO_PROTEIN_LOCALIZATION_TO_NUCLEUS | 0.00002 | 0.00055 | 0.57561 | 0.58926 | 1.30220 | 187 |
| GO_PROTEIN_MODIFICATION_BY_SMALL_PROTEIN_REMOVAL | 0.00000 | 0.00000 | 0.71951 | 0.60745 | 1.34689 | 221 |
| GO_PROTEIN_POLYUBIQUITINATION | 0.00004 | 0.00106 | 0.55733 | 0.55874 | 1.24108 | 253 |
| GO_PROTEIN_REFOLDING | 0.00294 | 0.04036 | 0.43171 | 0.77039 | 1.53314 | 16 |
| GO_PROTEIN_STABILIZATION | 0.00001 | 0.00024 | 0.61053 | 0.62333 | 1.36586 | 130 |
| GO_PROTEIN_TARGETING | 0.00000 | 0.00000 | | 0.71181 | 1.58743 | 327 |
| GO_PROTEIN_TARGETING_TO_MEMBRANE | 0.00000 | 0.00000 | | 0.82370 | 1.80953 | 149 |
| GO_PROTEIN_TARGETING_TO_MITOCHONDRION | 0.00346 | 0.04593 | 0.43171 | 0.60408 | 1.30091 | 78 |

| | | | | | | |
|---|---|---|---|---|---|---|
| GO_PROTEIN_TRANSMEMBRANE_TRANSPORT | 0.00265 | 0.03775 | 0.43171 | 0.64528 | 1.36171 | 50 |
| GO_PROTON_TRANSMEMBRANE_TRANSPORT | 0.00213 | 0.03229 | 0.43171 | 0.60815 | 1.30536 | 71 |
| GO_PURINE_CONTAINING_COMPOUND_BIOSYNTHETIC_PROCESS | 0.00246 | 0.03601 | 0.43171 | 0.57796 | 1.25936 | 111 |
| GO_PURINE_CONTAINING_COMPOUND_METABOLIC_PROCESS | 0.00100 | 0.01761 | 0.45506 | 0.53679 | 1.19242 | 255 |
| GO_PURINE_NUCLEOSIDE_TRIPHOSPHATE_METABOLIC_PROCESS | 0.00089 | 0.01592 | 0.47727 | 0.67157 | 1.40374 | 43 |
| GO_PYRIMIDINE_CONTAINING_COMPOUND_METABOLIC_PROCESS | 0.00353 | 0.04658 | 0.43171 | 0.64744 | 1.35878 | 47 |
| GO_REGENERATION | 0.00216 | 0.03242 | 0.43171 | 0.59987 | 1.29453 | 82 |
| GO_REGULATION_OF_ANIMAL_ORGAN_MORPHOGENESIS | 0.00000 | 0.00000 | | 0.69163 | 1.51541 | 133 |
| GO_REGULATION_OF_APOPTOTIC_SIGNALING_PATHWAY | 0.00031 | 0.00645 | 0.49849 | 0.55171 | 1.22506 | 247 |
| GO_REGULATION_OF_ATPASE_ACTIVITY | 0.00324 | 0.04369 | 0.43171 | 0.63031 | 1.34038 | 57 |
| GO_REGULATION_OF_ATP_METABOLIC_PROCESS | 0.00353 | 0.04658 | 0.43171 | 0.60176 | 1.29780 | 80 |
| GO_REGULATION_OF_BINDING | 0.00250 | 0.03636 | 0.43171 | 0.54476 | 1.20598 | 203 |
| GO_REGULATION_OF_CELLULAR_AMIDE_METABOLIC_PROCESS | 0.00000 | 0.00000 | | 0.62967 | 1.40226 | 311 |
| GO_REGULATION_OF_CELLULAR_AMINE_METABOLIC_PROCESS | 0.00000 | 0.00000 | | 0.82173 | 1.74682 | 58 |
| GO_REGULATION_OF_CELLULAR_AMINO_ACID_METABOLIC_PROCES | 0.00000 | 0.00000 | | 0.83776 | 1.77766 | 55 |
| GO_REGULATION_OF_CELLULAR_KETONE_METABOLIC_PROCESS | 0.00000 | 0.00000 | | 0.72641 | 1.58680 | 118 |
| GO_REGULATION_OF_CELLULAR_PROTEIN_LOCALIZATION | 0.00082 | 0.01513 | 0.47727 | 0.52542 | 1.17290 | 362 |
| GO_REGULATION_OF_CELLULAR_RESPONSE_TO_HEAT | 0.00276 | 0.03885 | 0.43171 | 0.61652 | 1.32004 | 67 |
| GO_REGULATION_OF_CELLULAR_RESPONSE_TO_STRESS | 0.00224 | 0.03344 | 0.43171 | 0.50545 | 1.13232 | 468 |
| GO_REGULATION_OF_CELL_CYCLE_G2_M_PHASE_TRANSITION | 0.00000 | 0.00000 | 0.71951 | 0.63230 | 1.39282 | 169 |
| GO_REGULATION_OF_CELL_CYCLE_PHASE_TRANSITION | 0.00001 | 0.00026 | 0.59333 | 0.55488 | 1.23783 | 337 |
| GO_REGULATION_OF_CHROMOSOME_ORGANIZATION | 0.00081 | 0.01501 | 0.47727 | 0.54255 | 1.20527 | 261 |
| GO_REGULATION_OF_CYSTEINE_TYPE_ENDOPEPTIDASE_ACTIVITY | 0.00202 | 0.03077 | 0.43171 | 0.57120 | 1.24837 | 121 |
| GO_REGULATION_OF_CYTOPLASMIC_TRANSLATION | 0.00130 | 0.02159 | 0.45506 | 0.73678 | 1.49773 | 23 |
| GO_REGULATION_OF_DEFENSE_RESPONSE | 0.00065 | 0.01237 | 0.47727 | 0.53633 | 1.19355 | 290 |
| GO_REGULATION_OF_DNA_BIOSYNTHETIC_PROCESS | 0.00016 | 0.00360 | 0.51885 | 0.64215 | 1.37923 | 74 |
| GO_REGULATION_OF_DNA_METABOLIC_PROCESS | 0.00069 | 0.01297 | 0.47727 | 0.54425 | 1.20775 | 237 |
| GO_REGULATION_OF_DNA_TEMPLATED_TRANSCRIPTION_INITIATIOI | 0.00163 | 0.02568 | 0.45506 | 0.68289 | 1.41372 | 35 |
| GO_REGULATION_OF_DNA_TEMPLATED_TRANSCRIPTION_IN_RESPO | 0.00000 | 0.00000 | | 0.73944 | 1.59800 | 89 |
| GO_REGULATION_OF_GENERATION_OF_PRECURSOR_METABOLITES_ | 0.00060 | 0.01164 | 0.47727 | 0.60476 | 1.31116 | 97 |
| GO_REGULATION_OF_HEMATOPOIETIC_PROGENITOR_CELL_DIFFERE | 0.00000 | 0.00000 | | 0.78717 | 1.69251 | 75 |
| GO_REGULATION_OF_HEMATOPOIETIC_STEM_CELL_DIFFERENTIATIC | 0.00000 | 0.00000 | | 0.81505 | 1.74256 | 65 |
| GO_REGULATION_OF_HEMOPOIESIS | 0.00000 | 0.00008 | 0.62726 | 0.58859 | 1.30634 | 228 |
| GO_REGULATION_OF_IMMUNE_RESPONSE | 0.00000 | 0.00016 | 0.61053 | 0.55362 | 1.23583 | 362 |
| GO_REGULATION_OF_INNATE_IMMUNE_RESPONSE | 0.00000 | 0.00000 | 0.71951 | 0.63883 | 1.40521 | 161 |
| GO_REGULATION_OF_INTRINSIC_APOPTOTIC_SIGNALING_PATHWAY | 0.00001 | 0.00039 | 0.59333 | 0.63478 | 1.38259 | 109 |
| GO_REGULATION_OF_MITOTIC_CELL_CYCLE | 0.00001 | 0.00018 | 0.61053 | 0.53899 | 1.20736 | 458 |
| GO_REGULATION_OF_MORPHOGENESIS_OF_AN_EPITHELIUM | 0.00000 | 0.00000 | | 0.72213 | 1.57284 | 109 |
| GO_REGULATION_OF_MRNA_CATABOLIC_PROCESS | 0.00000 | 0.00000 | | 0.69977 | 1.54127 | 168 |
| GO_REGULATION_OF_MRNA_METABOLIC_PROCESS | 0.00000 | 0.00000 | | 0.66528 | 1.47872 | 266 |
| GO_REGULATION_OF_MRNA_PROCESSING | 0.00004 | 0.00105 | 0.55733 | 0.61521 | 1.34090 | 113 |
| GO_REGULATION_OF_MRNA_SPLICING_VIA_SPLICEOSOME | 0.00001 | 0.00026 | 0.59333 | 0.67424 | 1.45411 | 80 |
| GO_REGULATION_OF_NUCLEASE_ACTIVITY | 0.00106 | 0.01819 | 0.45506 | 0.77918 | 1.56780 | 18 |
| GO_REGULATION_OF_OXIDATIVE_STRESS_INDUCED_CELL_DEATH | 0.00198 | 0.03028 | 0.43171 | 0.67287 | 1.40355 | 41 |
| GO_REGULATION_OF_PEPTIDASE_ACTIVITY | 0.00050 | 0.00978 | 0.47727 | 0.56122 | 1.24054 | 191 |
| GO_REGULATION_OF_POSTTRANSCRIPTIONAL_GENE_SILENCING | 0.00364 | 0.04761 | 0.43171 | 0.59824 | 1.28492 | 74 |
| GO_REGULATION_OF_PROTEIN_CATABOLIC_PROCESS | 0.00179 | 0.02783 | 0.45506 | 0.53057 | 1.17929 | 270 |
| GO_REGULATION_OF_PROTEIN_LOCALIZATION_TO_NUCLEUS | 0.00001 | 0.00042 | 0.59333 | 0.65505 | 1.41558 | 87 |
| GO_REGULATION_OF_PROTEIN_MODIFICATION_BY_SMALL_PROTEIN | 0.00025 | 0.00532 | 0.49849 | 0.57068 | 1.25684 | 167 |
| GO_REGULATION_OF_PROTEIN_STABILITY | 0.00093 | 0.01652 | 0.47727 | 0.55382 | 1.22571 | 202 |
| GO_REGULATION_OF_PROTEOLYSIS | 0.00013 | 0.00301 | 0.51885 | 0.52997 | 1.18392 | 377 |
| GO_REGULATION_OF_RESPONSE_TO_BIOTIC_STIMULUS | 0.00001 | 0.00020 | 0.61053 | 0.59049 | 1.30719 | 204 |
| GO_REGULATION_OF_RNA_SPLICING | 0.00000 | 0.00001 | 0.69013 | 0.65896 | 1.43669 | 114 |
| GO_REGULATION_OF_SMALL_MOLECULE_METABOLIC_PROCESS | 0.00000 | 0.00000 | 0.71951 | 0.59134 | 1.31394 | 265 |
| GO_REGULATION_OF_STEM_CELL_DIFFERENTIATION | 0.00000 | 0.00000 | | 0.74049 | 1.60113 | 90 |
| GO_REGULATION_OF_SYMBIOTIC_PROCESS | 0.00006 | 0.00144 | 0.55733 | 0.60071 | 1.31682 | 134 |
| GO_REGULATION_OF_TELOMERASE_ACTIVITY | 0.00088 | 0.01590 | 0.47727 | 0.69161 | 1.43179 | 35 |
| GO_REGULATION_OF_TRANSCRIPTION_FROM_RNA_POLYMERASE_II | 0.00000 | 0.00000 | | 0.81194 | 1.73800 | 66 |
| GO_REGULATION_OF_TRANSCRIPTION_INITIATION_FROM_RNA_POL | 0.00143 | 0.02332 | 0.45506 | 0.70390 | 1.44582 | 29 |
| GO_REGULATION_OF_TRANSLATIONAL_INITIATION | 0.00002 | 0.00052 | 0.57561 | 0.68129 | 1.45518 | 64 |
| GO_REGULATION_OF_UBIQUITIN_PROTEIN_TRANSFERASE_ACTIVITY | 0.00061 | 0.01169 | 0.47727 | 0.69335 | 1.44627 | 41 |
| GO_REGULATION_OF_VIRAL_GENOME_REPLICATION | 0.00086 | 0.01566 | 0.47727 | 0.65316 | 1.38389 | 54 |
| GO_REGULATION_OF_VIRAL_TRANSCRIPTION | 0.00198 | 0.03028 | 0.43171 | 0.63932 | 1.34703 | 49 |
| GO_REGULATION_OF_WNT_SIGNALING_PATHWAY | 0.00000 | 0.00001 | 0.69013 | 0.60353 | 1.33816 | 220 |
| GO_RESPIRASOME | 0.00008 | 0.00181 | 0.53843 | 0.65420 | 1.40421 | 71 |
| GO_RESPIRATORY_CHAIN_COMPLEX | 0.00016 | 0.00353 | 0.51885 | 0.67626 | 1.43918 | 60 |
| GO_RESPIRATORY_ELECTRON_TRANSPORT_CHAIN | 0.00001 | 0.00025 | 0.59333 | 0.65819 | 1.42190 | 85 |
| GO_RESPONSE_TO_ALCOHOL | 0.00261 | 0.03750 | 0.43171 | 0.58804 | 1.27350 | 95 |
| GO_RESPONSE_TO_CADMIUM_ION | 0.00265 | 0.03775 | 0.43171 | 0.67227 | 1.39174 | 35 |
| GO_RESPONSE_TO_DRUG | 0.00104 | 0.01798 | 0.45506 | 0.55364 | 1.22258 | 180 |
| GO_RESPONSE_TO_ETHANOL | 0.00086 | 0.01566 | 0.47727 | 0.67309 | 1.40691 | 43 |
| GO_RESPONSE_TO_HEAT | 0.00012 | 0.00270 | 0.53843 | 0.61665 | 1.34221 | 107 |
| GO_RESPONSE_TO_HORMONE | 0.00048 | 0.00949 | 0.49849 | 0.51277 | 1.14814 | 453 |
| GO_RESPONSE_TO_HYDROGEN_PEROXIDE | 0.00257 | 0.03716 | 0.43171 | 0.60505 | 1.30368 | 79 |
| GO_RESPONSE_TO_INORGANIC_SUBSTANCE | 0.00004 | 0.00107 | 0.55733 | 0.55714 | 1.23870 | 275 |
| GO_RESPONSE_TO_INTERLEUKIN_1 | 0.00000 | 0.00000 | | 0.74844 | 1.62908 | 107 |

| | | | | | | |
|---|---|---|---|---|---|---|
| GO_RESPONSE_TO_INTERLEUKIN_12 | 0.00012 | 0.00270 | 0.53843 | 0.72605 | 1.50694 | 36 |
| GO_RESPONSE_TO_INTERLEUKIN_4 | 0.00324 | 0.04369 | 0.43171 | 0.75812 | 1.51571 | 17 |
| GO_RESPONSE_TO_METAL_ION | 0.00030 | 0.00623 | 0.49849 | 0.56387 | 1.24233 | 171 |
| GO_RESPONSE_TO_ORGANIC_CYCLIC_COMPOUND | 0.00000 | 0.00016 | 0.61053 | 0.54603 | 1.22259 | 439 |
| GO_RESPONSE_TO_OXYGEN_LEVELS | 0.00000 | 0.00000 | 0.71951 | 0.60132 | 1.33438 | 233 |
| GO_RESPONSE_TO_STEROID_HORMONE | 0.00154 | 0.02464 | 0.45506 | 0.55514 | 1.22446 | 174 |
| GO_RESPONSE_TO_TEMPERATURE_STIMULUS | 0.00027 | 0.00572 | 0.49849 | 0.59216 | 1.29675 | 127 |
| GO_RESPONSE_TO_TUMOR_NECROSIS_FACTOR | 0.00000 | 0.00000 | 0.76146 | 0.65649 | 1.44236 | 151 |
| GO_RETINA_HOMEOSTASIS | 0.00104 | 0.01798 | 0.45506 | 0.73063 | 1.49432 | 27 |
| GO_RIBONUCLEOPROTEIN_COMPLEX_BINDING | 0.00000 | 0.00000 | 0.71951 | 0.66906 | 1.46050 | 117 |
| GO_RIBONUCLEOPROTEIN_COMPLEX_BIOGENESIS | 0.00000 | 0.00000 | | 0.66339 | 1.48264 | 389 |
| GO_RIBONUCLEOPROTEIN_COMPLEX_SUBUNIT_ORGANIZATION | 0.00000 | 0.00000 | | 0.69241 | 1.52681 | 173 |
| GO_RIBONUCLEOPROTEIN_GRANULE | 0.00000 | 0.00001 | 0.67496 | 0.61979 | 1.36366 | 163 |
| GO_RIBONUCLEOSIDE_TRIPHOSPHATE_BIOSYNTHETIC_PROCESS | 0.00041 | 0.00840 | 0.49849 | 0.71253 | 1.47248 | 33 |
| GO_RIBONUCLEOSIDE_TRIPHOSPHATE_METABOLIC_PROCESS | 0.00026 | 0.00550 | 0.49849 | 0.69681 | 1.45611 | 42 |
| GO_RIBOSE_PHOSPHATE_BIOSYNTHETIC_PROCESS | 0.00191 | 0.02939 | 0.45506 | 0.57932 | 1.26207 | 110 |
| GO_RIBOSE_PHOSPHATE_METABOLIC_PROCESS | 0.00108 | 0.01839 | 0.45506 | 0.53891 | 1.19687 | 245 |
| GO_RIBOSOMAL_LARGE_SUBUNIT_ASSEMBLY | 0.00000 | 0.00016 | 0.61053 | 0.80999 | 1.65663 | 27 |
| GO_RIBOSOMAL_LARGE_SUBUNIT_BIOGENESIS | 0.00000 | 0.00000 | | 0.77542 | 1.65982 | 66 |
| GO_RIBOSOMAL_SMALL_SUBUNIT_ASSEMBLY | 0.00023 | 0.00494 | 0.51885 | 0.81139 | 1.62221 | 17 |
| GO_RIBOSOMAL_SMALL_SUBUNIT_BINDING | 0.00383 | 0.04935 | 0.43171 | 0.75288 | 1.50523 | 17 |
| GO_RIBOSOMAL_SMALL_SUBUNIT_BIOGENESIS | 0.00001 | 0.00022 | 0.61053 | 0.69076 | 1.47997 | 68 |
| GO_RIBOSOMAL_SUBUNIT | 0.00000 | 0.00000 | | 0.77603 | 1.71031 | 172 |
| GO_RIBOSOME | 0.00000 | 0.00000 | | 0.76643 | 1.69659 | 206 |
| GO_RIBOSOME_ASSEMBLY | 0.00000 | 0.00000 | 0.73376 | 0.76173 | 1.61998 | 59 |
| GO_RIBOSOME_BINDING | 0.00019 | 0.00416 | 0.51885 | 0.68955 | 1.44715 | 47 |
| GO_RIBOSOME_BIOGENESIS | 0.00000 | 0.00000 | | 0.65877 | 1.46466 | 275 |
| GO_RNA_CATABOLIC_PROCESS | 0.00000 | 0.00000 | | 0.76774 | 1.71264 | 339 |
| GO_RNA_DEPENDENT_DNA_BIOSYNTHETIC_PROCESS | 0.00003 | 0.00076 | 0.57561 | 0.68926 | 1.46521 | 58 |
| GO_RNA_EXPORT_FROM_NUCLEUS | 0.00062 | 0.01183 | 0.47727 | 0.58378 | 1.27758 | 126 |
| GO_RNA_LOCALIZATION | 0.00000 | 0.00006 | 0.64355 | 0.59670 | 1.32020 | 198 |
| GO_RNA_POLYMERASE_II_PREINITIATION_COMPLEX_ASSEMBLY | 0.00089 | 0.01592 | 0.47727 | 0.77046 | 1.55626 | 19 |
| GO_RNA_SPLICING | 0.00000 | 0.00000 | 0.78819 | 0.59079 | 1.31927 | 369 |
| GO_RNA_SPLICING_VIA_TRANSESTERIFICATION_REACTIONS | 0.00000 | 0.00000 | | 0.60940 | 1.35714 | 300 |
| GO_RNA_STABILIZATION | 0.00065 | 0.01223 | 0.47727 | 0.69743 | 1.44384 | 35 |
| GO_ROUGH_ENDOPLASMIC_RETICULUM | 0.00001 | 0.00039 | 0.59333 | 0.71853 | 1.51629 | 50 |
| GO_RRNA_BINDING | 0.00000 | 0.00001 | 0.69013 | 0.74782 | 1.58865 | 56 |
| GO_RRNA_METABOLIC_PROCESS | 0.00000 | 0.00000 | 0.77494 | 0.63047 | 1.39562 | 206 |
| GO_SCF_DEPENDENT_PROTEASOMAL_UBIQUITIN_DEPENDENT_PRO | 0.00000 | 0.00000 | | 0.77527 | 1.67044 | 79 |
| GO_SECRETORY_GRANULE | 0.00000 | 0.00000 | 0.71951 | 0.56518 | 1.26461 | 399 |
| GO_SECRETORY_VESICLE | 0.00000 | 0.00002 | 0.67496 | 0.54866 | 1.22905 | 465 |
| GO_SIGNAL_TRANSDUCTION_BY_PROTEIN_PHOSPHORYLATION | 0.00004 | 0.00109 | 0.55733 | 0.53817 | 1.20474 | 424 |
| GO_SINGLE_FERTILIZATION | 0.00048 | 0.00949 | 0.49849 | 0.67414 | 1.42260 | 50 |
| GO_SINGLE_STRANDED_RNA_BINDING | 0.00039 | 0.00793 | 0.49849 | 0.66553 | 1.41527 | 57 |
| GO_SMALL_MOLECULE_BIOSYNTHETIC_PROCESS | 0.00216 | 0.03242 | 0.43171 | 0.51195 | 1.14522 | 397 |
| GO_SMALL_RIBOSOMAL_SUBUNIT | 0.00000 | 0.00000 | | 0.77648 | 1.66209 | 66 |
| GO_SM_LIKE_PROTEIN_FAMILY_COMPLEX | 0.00327 | 0.04398 | 0.43171 | 0.60468 | 1.29713 | 70 |
| GO_SPERM_EGG_RECOGNITION | 0.00000 | 0.00007 | 0.64355 | 0.85241 | 1.72955 | 22 |
| GO_SPLICEOSOMAL_COMPLEX | 0.00000 | 0.00003 | 0.65944 | 0.60939 | 1.34261 | 171 |
| GO_STEM_CELL_DIFFERENTIATION | 0.00000 | 0.00000 | 0.78819 | 0.66886 | 1.46930 | 147 |
| GO_STRUCTURAL_CONSTITUENT_OF_CYTOSKELETON | 0.00102 | 0.01782 | 0.45506 | 0.65913 | 1.39337 | 52 |
| GO_STRUCTURAL_CONSTITUENT_OF_RIBOSOME | 0.00000 | 0.00000 | | 0.79909 | 1.75541 | 150 |
| GO_STRUCTURAL_MOLECULE_ACTIVITY | 0.00000 | 0.00000 | | 0.72651 | 1.61895 | 324 |
| GO_SUBSTANTIA_NIGRA_DEVELOPMENT | 0.00000 | 0.00013 | 0.62726 | 0.81622 | 1.66507 | 26 |
| GO_TELOMERASE_RNA_LOCALIZATION | 0.00001 | 0.00017 | 0.61053 | 0.85164 | 1.72025 | 19 |
| GO_TELOMERE_MAINTENANCE_VIA_TELOMERE_LENGTHENING | 0.00011 | 0.00250 | 0.53843 | 0.65659 | 1.40546 | 66 |
| GO_TELOMERE_ORGANIZATION | 0.00342 | 0.04554 | 0.43171 | 0.57000 | 1.24649 | 124 |
| GO_THREONINE_TYPE_ENDOPEPTIDASE_ACTIVITY | 0.00000 | 0.00000 | | 0.95179 | 1.92255 | 19 |
| GO_THREONINE_TYPE_PEPTIDASE_ACTIVITY | 0.00000 | 0.00000 | | 0.92199 | 1.87423 | 23 |
| GO_TISSUE_MORPHOGENESIS | 0.00097 | 0.01707 | 0.47727 | 0.52661 | 1.17402 | 326 |
| GO_TOXIN_TRANSPORT | 0.00113 | 0.01922 | 0.45506 | 0.72897 | 1.48708 | 26 |
| GO_TRANSCRIPTION_COUPLED_NUCLEOTIDE_EXCISION_REPAIR | 0.00001 | 0.00019 | 0.61053 | 0.68106 | 1.46186 | 71 |
| GO_TRANSCRIPTION_FACTOR_BINDING | 0.00043 | 0.00865 | 0.49849 | 0.52447 | 1.17376 | 409 |
| GO_TRANSCRIPTION_PREINITIATION_COMPLEX_ASSEMBLY | 0.00119 | 0.02004 | 0.45506 | 0.71399 | 1.46797 | 30 |
| GO_TRANSLATIONAL_ELONGATION | 0.00000 | 0.00000 | 0.71951 | 0.66087 | 1.44451 | 122 |
| GO_TRANSLATIONAL_INITIATION | 0.00000 | 0.00000 | | 0.84691 | 1.86515 | 166 |
| GO_TRANSLATIONAL_TERMINATION | 0.00000 | 0.00005 | 0.64355 | 0.65946 | 1.43060 | 99 |
| GO_TRANSLATION_ELONGATION_FACTOR_ACTIVITY | 0.00161 | 0.02546 | 0.45506 | 0.79860 | 1.58418 | 15 |
| GO_TRANSLATION_FACTOR_ACTIVITY_RNA_BINDING | 0.00000 | 0.00000 | | 0.79008 | 1.69288 | 69 |
| GO_TRANSLATION_INITIATION_FACTOR_ACTIVITY | 0.00000 | 0.00000 | 0.76146 | 0.80422 | 1.68417 | 45 |
| GO_TRANSLATION_INITIATION_FACTOR_BINDING | 0.00021 | 0.00468 | 0.51885 | 0.76186 | 1.55231 | 25 |
| GO_TRANSLATION_PREINITIATION_COMPLEX | 0.00000 | 0.00001 | 0.67496 | 0.90245 | 1.79595 | 16 |
| GO_TRANSLATION_REGULATOR_ACTIVITY | 0.00000 | 0.00000 | | 0.77663 | 1.68904 | 105 |
| GO_TRANSLATION_REGULATOR_ACTIVITY_NUCLEIC_ACID_BINDING | 0.00000 | 0.00000 | | 0.77150 | 1.66586 | 83 |
| GO_TRICARBOXYLIC_ACID_CYCLE | 0.00154 | 0.02464 | 0.45506 | 0.71854 | 1.47240 | 28 |
| GO_TUMOR_NECROSIS_FACTOR_MEDIATED_SIGNALING_PATHWAY | 0.00000 | 0.00000 | | 0.76277 | 1.65372 | 97 |

| | | | | | | |
|---|---|---|---|---|---|---|
| GO_T_CELL_RECEPTOR_SIGNALING_PATHWAY | 0.00000 | 0.00000 | 0.81404 | 0.69994 | 1.52527 | 112 |
| GO_U12_TYPE_SPLICEOSOMAL_COMPLEX | 0.00279 | 0.03889 | 0.43171 | 0.70746 | 1.44693 | 27 |
| GO_U2_SNRNP | 0.00005 | 0.00134 | 0.55733 | 0.83081 | 1.67168 | 18 |
| GO_U2_TYPE_SPLICEOSOMAL_COMPLEX | 0.00010 | 0.00243 | 0.53843 | 0.63403 | 1.36970 | 85 |
| GO_UBIQUITIN_LIKE_PROTEIN_LIGASE_BINDING | 0.00000 | 0.00000 | 0.76146 | 0.62273 | 1.38051 | 219 |
| GO_UBIQUITIN_PROTEIN_TRANSFERASE_REGULATOR_ACTIVITY | 0.00165 | 0.02590 | 0.45506 | 0.79605 | 1.57911 | 15 |
| GO_UNFOLDED_PROTEIN_BINDING | 0.00000 | 0.00000 | 0.77494 | 0.72157 | 1.55881 | 85 |
| GO_VESICLE_LUMEN | 0.00000 | 0.00000 | 0.74774 | 0.64475 | 1.42008 | 168 |
| GO_VIRAL_GENE_EXPRESSION | 0.00000 | 0.00000 | | 0.80028 | 1.76317 | 171 |
| GO_VIRAL_GENOME_REPLICATION | 0.00100 | 0.01761 | 0.45506 | 0.61997 | 1.33078 | 73 |
| GO_VIRAL_LIFE_CYCLE | 0.00031 | 0.00640 | 0.49849 | 0.55767 | 1.23509 | 209 |
| HP_ABNORMALITY_OF_ACID_BASE_HOMEOSTASIS | 0.00216 | 0.03242 | 0.43171 | 0.53556 | 1.18846 | 232 |
| HP_ABNORMALITY_OF_CARDIOVASCULAR_SYSTEM_ELECTROPHYSIO | 0.00027 | 0.00572 | 0.49849 | 0.55832 | 1.23929 | 229 |
| HP_ABNORMALITY_OF_LOWER_LIP | 0.00181 | 0.02804 | 0.45506 | 0.56360 | 1.23779 | 145 |
| HP_ABNORMALITY_OF_THE_MITOCHONDRION | 0.00213 | 0.03229 | 0.43171 | 0.57831 | 1.26240 | 117 |
| HP_ABNORMALITY_OF_THE_NASAL_DORSUM | 0.00005 | 0.00116 | 0.55733 | 0.60288 | 1.32157 | 134 |
| HP_ABNORMAL_ELECTROPHYSIOLOGY_OF_SINOATRIAL_NODE_ORIG | 0.00279 | 0.03889 | 0.43171 | 0.68436 | 1.40934 | 31 |
| HP_ABNORMAL_ERYTHROCYTE_MORPHOLOGY | 0.00000 | 0.00015 | 0.61053 | 0.56049 | 1.25036 | 341 |
| HP_ABNORMAL_MEAN_CORPUSCULAR_VOLUME | 0.00357 | 0.04696 | 0.43171 | 0.75226 | 1.51363 | 18 |
| HP_ABNORMAL_RETICULOCYTE_MORPHOLOGY | 0.00064 | 0.01210 | 0.47727 | 0.71420 | 1.47081 | 31 |
| HP_ACUTE_LEUKEMIA | 0.00000 | 0.00000 | 0.73376 | 0.72786 | 1.56137 | 70 |
| HP_ANEMIA_OF_INADEQUATE_PRODUCTION | 0.00000 | 0.00008 | 0.62726 | 0.65593 | 1.42209 | 97 |
| HP_CONSTITUTIONAL_SYMPTOM | 0.00119 | 0.02004 | 0.45506 | 0.51427 | 1.15108 | 423 |
| HP_DELAYED_PUBERTY | 0.00005 | 0.00115 | 0.55733 | 0.62579 | 1.36211 | 107 |
| HP_DEPRESSED_NASAL_RIDGE | 0.00000 | 0.00011 | 0.62726 | 0.68731 | 1.47511 | 72 |
| HP_FATIGUE | 0.00039 | 0.00808 | 0.49849 | 0.56529 | 1.24923 | 187 |
| HP_HEADACHE | 0.00015 | 0.00340 | 0.51885 | 0.59065 | 1.29716 | 146 |
| HP_HEMATOLOGICAL_NEOPLASM | 0.00003 | 0.00078 | 0.57561 | 0.60108 | 1.32044 | 154 |
| HP_INCREASED_CSF_LACTATE | 0.00157 | 0.02509 | 0.45506 | 0.61622 | 1.31904 | 66 |
| HP_INCREASED_SERUM_LACTATE | 0.00216 | 0.03242 | 0.43171 | 0.56920 | 1.24724 | 130 |
| HP_LEUKEMIA | 0.00001 | 0.00026 | 0.59333 | 0.64877 | 1.40680 | 98 |
| HP_MACROCYTIC_ANEMIA | 0.00000 | 0.00000 | 0.74774 | 0.82468 | 1.71698 | 39 |
| HP_MIGRAINE | 0.00000 | 0.00000 | 0.71951 | 0.72105 | 1.54497 | 69 |
| HP_MYOTONIA | 0.00161 | 0.02546 | 0.45506 | 0.79820 | 1.58338 | 15 |
| HP_NEOPLASM | 0.00216 | 0.03242 | 0.43171 | 0.50820 | 1.13805 | 446 |
| HP_PALLOR | 0.00001 | 0.00040 | 0.59333 | 0.65169 | 1.41044 | 92 |
| HP_PUBERTY_AND_GONADAL_DISORDERS | 0.00287 | 0.03973 | 0.43171 | 0.51995 | 1.15843 | 322 |
| HP_RETICULOCYTOPENIA | 0.00004 | 0.00105 | 0.55733 | 0.86417 | 1.71425 | 15 |
| HP_RIMMED_VACUOLES | 0.00279 | 0.03889 | 0.43171 | 0.74033 | 1.49831 | 20 |
| HP_SHORT_NOSE | 0.00235 | 0.03474 | 0.43171 | 0.54919 | 1.21134 | 174 |
| HP_THICK_LOWER_LIP_VERMILION | 0.00048 | 0.00949 | 0.49849 | 0.60773 | 1.31617 | 94 |
| HP_THICK_VERMILION_BORDER | 0.00156 | 0.02486 | 0.45506 | 0.55983 | 1.23000 | 153 |
| HP_VENTRICULAR_ARRHYTHMIA | 0.00353 | 0.04658 | 0.43171 | 0.66198 | 1.37398 | 36 |

**Cluster 1**

| pathway | pval | padj | log2err | ES | NES | size |
|---|---|---|---|---|---|---|
| GO_ACTIVATION_OF_IMMUNE_RESPONSE | 0.00000 | 0.00000 | 0.78819 | 0.62851 | 1.39255 | 212 |
| GO_ACTIVATION_OF_INNATE_IMMUNE_RESPONSE | 0.00000 | 0.00000 | | 0.75054 | 1.62130 | 88 |
| GO_AEROBIC_RESPIRATION | 0.00001 | 0.00025 | 0.59333 | 0.69302 | 1.48168 | 65 |
| GO_ALTERNATIVE_MRNA_SPLICING_VIA_SPLICEOSOME | 0.00102 | 0.01782 | 0.45506 | 0.65168 | 1.38687 | 60 |
| GO_AMINE_METABOLIC_PROCESS | 0.00000 | 0.00000 | | 0.79189 | 1.70783 | 80 |
| GO_ANAPHASE_PROMOTING_COMPLEX_DEPENDENT_CATABOLIC_P | 0.00000 | 0.00000 | | 0.76611 | 1.65070 | 79 |
| GO_ANCHORING_JUNCTION | 0.00000 | 0.00000 | | 0.59615 | 1.33633 | 479 |
| GO_ANIMAL_ORGAN_REGENERATION | 0.00257 | 0.03716 | 0.43171 | 0.68240 | 1.41020 | 33 |
| GO_ANTIGEN_PROCESSING_AND_PRESENTATION | 0.00000 | 0.00000 | 0.70498 | 0.64776 | 1.42241 | 143 |
| GO_ANTIGEN_PROCESSING_AND_PRESENTATION_OF_EXOGENOUS_ | 0.00000 | 0.00000 | | 0.81051 | 1.72358 | 57 |
| GO_ANTIGEN_PROCESSING_AND_PRESENTATION_OF_PEPTIDE_ANTI | 0.00000 | 0.00000 | 0.70498 | 0.65912 | 1.44248 | 126 |
| GO_ANTIGEN_PROCESSING_AND_PRESENTATION_OF_PEPTIDE_ANTI | 0.00000 | 0.00000 | | 0.76925 | 1.65015 | 70 |
| GO_ANTIGEN_RECEPTOR_MEDIATED_SIGNALING_PATHWAY | 0.00000 | 0.00000 | 0.80122 | 0.67877 | 1.48641 | 127 |
| GO_APOPTOTIC_SIGNALING_PATHWAY | 0.00154 | 0.02464 | 0.45506 | 0.51561 | 1.15085 | 358 |
| GO_ATP_DEPENDENT_CHROMATIN_REMODELING | 0.00253 | 0.03681 | 0.43171 | 0.62124 | 1.32400 | 61 |
| GO_ATP_METABOLIC_PROCESS | 0.00000 | 0.00000 | 0.71951 | 0.62252 | 1.37693 | 196 |
| GO_ATP_SYNTHESIS_COUPLED_ELECTRON_TRANSPORT | 0.00005 | 0.00131 | 0.55733 | 0.65519 | 1.40723 | 74 |
| GO_BINDING_OF_SPERM_TO_ZONA_PELLUCIDA | 0.00000 | 0.00001 | 0.69013 | 0.89879 | 1.80846 | 18 |
| GO_CADHERIN_BINDING | 0.00000 | 0.00000 | | 0.63068 | 1.39961 | 236 |
| GO_CANONICAL_WNT_SIGNALING_PATHWAY | 0.00000 | 0.00003 | 0.65944 | 0.60816 | 1.34553 | 197 |
| GO_CATALYTIC_STEP_2_SPLICEOSOME | 0.00000 | 0.00002 | 0.65944 | 0.68068 | 1.46976 | 83 |
| GO_CELLULAR_AMINO_ACID_METABOLIC_PROCESS | 0.00000 | 0.00000 | 0.74774 | 0.61752 | 1.36824 | 216 |
| GO_CELLULAR_COMPONENT_DISASSEMBLY | 0.00000 | 0.00005 | 0.64355 | 0.55505 | 1.24197 | 401 |
| GO_CELLULAR_KETONE_METABOLIC_PROCESS | 0.00000 | 0.00000 | 0.77494 | 0.65955 | 1.44893 | 149 |
| GO_CELLULAR_NITROGEN_COMPOUND_CATABOLIC_PROCESS | 0.00000 | 0.00000 | | 0.74079 | 1.65848 | 436 |
| GO_CELLULAR_PROTEIN_COMPLEX_DISASSEMBLY | 0.00006 | 0.00139 | 0.55733 | 0.58222 | 1.28459 | 176 |
| GO_CELLULAR_RESPIRATION | 0.00000 | 0.00000 | 0.71951 | 0.64997 | 1.42492 | 136 |
| GO_CELLULAR_RESPONSE_TO_HEAT | 0.00018 | 0.00403 | 0.51885 | 0.61962 | 1.34016 | 91 |
| GO_CELLULAR_RESPONSE_TO_ORGANIC_CYCLIC_COMPOUND | 0.00302 | 0.04108 | 0.43171 | 0.52402 | 1.16585 | 283 |
| GO_CELLULAR_RESPONSE_TO_OXYGEN_LEVELS | 0.00000 | 0.00000 | 0.78819 | 0.65911 | 1.44882 | 157 |
| GO_CELL_ACTIVATION_INVOLVED_IN_IMMUNE_RESPONSE | 0.00000 | 0.00001 | 0.67496 | 0.57339 | 1.27913 | 341 |

| | | | | | | |
|---|---|---|---|---|---|---|
| GO_CELL_ADHESION_MOLECULE_BINDING | 0.00000 | 0.00000 | 0.81404 | 0.60529 | 1.34788 | 308 |
| GO_CELL_CELL_RECOGNITION | 0.00000 | 0.00002 | 0.65944 | 0.83826 | 1.71003 | 26 |
| GO_CELL_CELL_SIGNALING_BY_WNT | 0.00000 | 0.00002 | 0.67496 | 0.57736 | 1.28559 | 299 |
| GO_CELL_CYCLE_G2_M_PHASE_TRANSITION | 0.00001 | 0.00024 | 0.59333 | 0.58034 | 1.28602 | 214 |
| GO_CELL_CYCLE_PHASE_TRANSITION | 0.00000 | 0.00004 | 0.64355 | 0.54319 | 1.21681 | 464 |
| GO_CELL_DIVISION | 0.00154 | 0.02464 | 0.45506 | 0.51551 | 1.15425 | 427 |
| GO_CELL_RECOGNITION | 0.00012 | 0.00270 | 0.53843 | 0.69554 | 1.46186 | 48 |
| GO_CELL_SUBSTRATE_JUNCTION | 0.00000 | 0.00000 | | 0.67751 | 1.50775 | 291 |
| GO_CELL_SURFACE_RECEPTOR_SIGNALING_PATHWAY_INVOLVED_IN | 0.00000 | 0.00003 | 0.65944 | 0.57035 | 1.27067 | 319 |
| GO_CHAPERONE_BINDING | 0.00007 | 0.00163 | 0.53843 | 0.64754 | 1.39253 | 76 |
| GO_CHAPERONE_COMPLEX | 0.00000 | 0.00004 | 0.64355 | 0.85718 | 1.73925 | 22 |
| GO_CHAPERONE_MEDIATED_PROTEIN_COMPLEX_ASSEMBLY | 0.00170 | 0.02662 | 0.45506 | 0.77423 | 1.54792 | 17 |
| GO_CHROMATIN_BINDING | 0.00071 | 0.01323 | 0.47727 | 0.52699 | 1.17680 | 369 |
| GO_CHROMOSOMAL_REGION | 0.00246 | 0.03601 | 0.43171 | 0.52919 | 1.17661 | 276 |
| GO_COP9_SIGNALOSOME | 0.00128 | 0.02141 | 0.45506 | 0.73616 | 1.49715 | 24 |
| GO_COTRANSLATIONAL_PROTEIN_TARGETING_TO_MEMBRANE | 0.00000 | 0.00000 | | 0.89471 | 1.93513 | 91 |
| GO_CYTOCHROME_COMPLEX | 0.00168 | 0.02641 | 0.45506 | 0.75311 | 1.52417 | 20 |
| GO_CYTOKINE_MEDIATED_SIGNALING_PATHWAY | 0.00000 | 0.00000 | 0.81404 | 0.59698 | 1.33170 | 339 |
| GO_CYTOPLASMIC_TRANSLATION | 0.00000 | 0.00000 | | 0.84753 | 1.83080 | 88 |
| GO_CYTOPLASMIC_TRANSLATIONAL_INITIATION | 0.00000 | 0.00000 | 0.74774 | 0.85688 | 1.76176 | 30 |
| GO_CYTOSOLIC_LARGE_RIBOSOMAL_SUBUNIT | 0.00000 | 0.00000 | | 0.91186 | 1.92988 | 53 |
| GO_CYTOSOLIC_RIBOSOME | 0.00000 | 0.00000 | | 0.89907 | 1.94710 | 95 |
| GO_CYTOSOLIC_SMALL_RIBOSOMAL_SUBUNIT | 0.00000 | 0.00000 | | 0.89114 | 1.85533 | 39 |
| GO_DEFENSE_RESPONSE_TO_OTHER_ORGANISM | 0.00004 | 0.00099 | 0.55733 | 0.53673 | 1.20127 | 415 |
| GO_DE_NOVO_PROTEIN_FOLDING | 0.00050 | 0.00978 | 0.47727 | 0.74232 | 1.51431 | 26 |
| GO_DISORDERED_DOMAIN_SPECIFIC_BINDING | 0.00161 | 0.02546 | 0.45506 | 0.71970 | 1.47197 | 27 |
| GO_DNA_BIOSYNTHETIC_PROCESS | 0.00000 | 0.00010 | 0.62726 | 0.62505 | 1.37017 | 134 |
| GO_DNA_CONFORMATION_CHANGE | 0.00027 | 0.00572 | 0.49849 | 0.55300 | 1.22527 | 216 |
| GO_DNA_DAMAGE_RESPONSE_DETECTION_OF_DNA_DAMAGE | 0.00198 | 0.03028 | 0.43171 | 0.67949 | 1.40669 | 35 |
| GO_DNA_GEOMETRIC_CHANGE | 0.00046 | 0.00925 | 0.49849 | 0.60807 | 1.31855 | 98 |
| GO_DNA_POLYMERASE_BINDING | 0.00268 | 0.03799 | 0.43171 | 0.76255 | 1.52457 | 17 |
| GO_DNA_REPAIR | 0.00265 | 0.03775 | 0.43171 | 0.51159 | 1.14547 | 427 |
| GO_DRUG_BINDING | 0.00176 | 0.02733 | 0.45506 | 0.64119 | 1.35096 | 49 |
| GO_ELECTRON_TRANSFER_ACTIVITY | 0.00001 | 0.00020 | 0.61053 | 0.67481 | 1.44938 | 74 |
| GO_ELECTRON_TRANSPORT_CHAIN | 0.00002 | 0.00053 | 0.57561 | 0.62257 | 1.36145 | 124 |
| GO_ENDOPEPTIDASE_ACTIVITY | 0.00124 | 0.02085 | 0.45506 | 0.54955 | 1.21076 | 171 |
| GO_ENDOPEPTIDASE_COMPLEX | 0.00000 | 0.00000 | | 0.80930 | 1.73234 | 66 |
| GO_ENERGY_DERIVATION_BY_OXIDATION_OF_ORGANIC_COMPOUN | 0.00000 | 0.00000 | 0.71951 | 0.63390 | 1.39707 | 172 |
| GO_ESTABLISHMENT_OF_PROTEIN_LOCALIZATION_TO_CHROMOSON | 0.00020 | 0.00432 | 0.51885 | 0.77339 | 1.57217 | 23 |
| GO_ESTABLISHMENT_OF_PROTEIN_LOCALIZATION_TO_ENDOPLASM | 0.00000 | 0.00000 | | 0.88026 | 1.91149 | 102 |
| GO_ESTABLISHMENT_OF_PROTEIN_LOCALIZATION_TO_MEMBRANE | 0.00000 | 0.00000 | | 0.74874 | 1.66164 | 240 |
| GO_ESTABLISHMENT_OF_PROTEIN_LOCALIZATION_TO_ORGANELLE | 0.00000 | 0.00000 | | 0.69519 | 1.55605 | 423 |
| GO_ESTABLISHMENT_OF_PROTEIN_LOCALIZATION_TO_TELOMERE | 0.00006 | 0.00139 | 0.55733 | 0.83470 | 1.66881 | 17 |
| GO_ESTABLISHMENT_OF_RNA_LOCALIZATION | 0.00106 | 0.01819 | 0.45506 | 0.55577 | 1.22400 | 167 |
| GO_ESTABLISHMENT_OF_TISSUE_POLARITY | 0.00000 | 0.00000 | | 0.75051 | 1.62122 | 88 |
| GO_EUKARYOTIC_TRANSLATION_INITIATION_FACTOR_3_COMPLEX | 0.00003 | 0.00074 | 0.57561 | 0.86987 | 1.72555 | 15 |
| GO_EXOCYTOSIS | 0.00000 | 0.00015 | 0.61053 | 0.53905 | 1.20753 | 465 |
| GO_FC_EPSILON_RECEPTOR_SIGNALING_PATHWAY | 0.00000 | 0.00000 | | 0.75117 | 1.62103 | 82 |
| GO_FC_RECEPTOR_SIGNALING_PATHWAY | 0.00000 | 0.00000 | | 0.72054 | 1.57787 | 127 |
| GO_FERTILIZATION | 0.00089 | 0.01592 | 0.47727 | 0.63620 | 1.35888 | 64 |
| GO_FICOLIN_1_RICH_GRANULE | 0.00000 | 0.00000 | 0.78819 | 0.71716 | 1.55340 | 93 |
| GO_GENERATION_OF_PRECURSOR_METABOLITES_AND_ENERGY | 0.00000 | 0.00000 | 0.77494 | 0.59261 | 1.32026 | 320 |
| GO_GENE_SILENCING | 0.00261 | 0.03750 | 0.43171 | 0.55578 | 1.22252 | 161 |
| GO_GLAND_DEVELOPMENT | 0.00154 | 0.02464 | 0.45506 | 0.54117 | 1.19797 | 206 |
| GO_GLYCOSYL_COMPOUND_METABOLIC_PROCESS | 0.00071 | 0.01323 | 0.47727 | 0.62524 | 1.34204 | 71 |
| GO_GUANYL_NUCLEOTIDE_BINDING | 0.00095 | 0.01679 | 0.47727 | 0.54096 | 1.19861 | 216 |
| GO_HEAT_SHOCK_PROTEIN_BINDING | 0.00003 | 0.00092 | 0.55733 | 0.64369 | 1.39056 | 85 |
| GO_HEMATOPOIETIC_PROGENITOR_CELL_DIFFERENTIATION | 0.00000 | 0.00000 | 0.82666 | 0.70307 | 1.53239 | 113 |
| GO_HEMATOPOIETIC_STEM_CELL_DIFFERENTIATION | 0.00000 | 0.00000 | | 0.79776 | 1.71217 | 72 |
| GO_HSP70_PROTEIN_BINDING | 0.00228 | 0.03391 | 0.43171 | 0.71143 | 1.45504 | 27 |
| GO_IMMUNE_RESPONSE_REGULATING_SIGNALING_PATHWAY | 0.00000 | 0.00000 | 0.76146 | 0.63494 | 1.40480 | 197 |
| GO_INNATE_IMMUNE_RESPONSE | 0.00003 | 0.00071 | 0.57561 | 0.55173 | 1.23107 | 347 |
| GO_INNATE_IMMUNE_RESPONSE_ACTIVATING_SIGNAL_TRANSDUCT | 0.00000 | 0.00000 | | 0.79340 | 1.70281 | 72 |
| GO_INNER_MITOCHONDRIAL_MEMBRANE_PROTEIN_COMPLEX | 0.00002 | 0.00060 | 0.57561 | 0.65124 | 1.40678 | 88 |
| GO_INTERACTION_WITH_SYMBIONT | 0.00001 | 0.00026 | 0.59333 | 0.75084 | 1.56620 | 41 |
| GO_INTERLEUKIN_1_MEDIATED_SIGNALING_PATHWAY | 0.00000 | 0.00000 | | 0.81689 | 1.75671 | 76 |
| GO_INTRINSIC_APOPTOTIC_SIGNALING_PATHWAY | 0.00121 | 0.02029 | 0.45506 | 0.55118 | 1.21845 | 190 |
| GO_ION_CHANNEL_BINDING | 0.00087 | 0.01584 | 0.47727 | 0.63182 | 1.34683 | 62 |
| GO_ISOPEPTIDASE_ACTIVITY | 0.00139 | 0.02285 | 0.45506 | 0.78940 | 1.57097 | 16 |
| GO_LARGE_RIBOSOMAL_SUBUNIT | 0.00000 | 0.00000 | | 0.78135 | 1.70183 | 109 |
| GO_LEUKOCYTE_MEDIATED_IMMUNITY | 0.00000 | 0.00000 | 0.71951 | 0.57282 | 1.27895 | 364 |
| GO_MACROMOLECULE_TRANSMEMBRANE_TRANSPORTER_ACTIVITY | 0.00239 | 0.03510 | 0.43171 | 0.77734 | 1.54696 | 16 |
| GO_MAINTENANCE_OF_CELL_NUMBER | 0.00087 | 0.01584 | 0.47727 | 0.58944 | 1.28512 | 114 |
| GO_MATURATION_OF_LSU_RRNA | 0.00006 | 0.00151 | 0.53843 | 0.77606 | 1.58723 | 27 |
| GO_METHYLTRANSFERASE_COMPLEX | 0.00290 | 0.04014 | 0.43171 | 0.59476 | 1.28479 | 88 |
| GO_METHYL_CPG_BINDING | 0.00302 | 0.04108 | 0.43171 | 0.78053 | 1.54833 | 15 |

| | | | | | | |
|---|---|---|---|---|---|---|
| GO_MIDBRAIN_DEVELOPMENT | 0.00021 | 0.00468 | 0.51885 | 0.68832 | 1.44457 | 47 |
| GO_MITOCHONDRIAL_ELECTRON_TRANSPORT_NADH_TO_UBIQUINO | 0.00331 | 0.04437 | 0.43171 | 0.65428 | 1.36724 | 42 |
| GO_MITOCHONDRIAL_GENE_EXPRESSION | 0.00048 | 0.00949 | 0.49849 | 0.57608 | 1.26525 | 152 |
| GO_MITOCHONDRIAL_LARGE_RIBOSOMAL_SUBUNIT | 0.00002 | 0.00050 | 0.57561 | 0.69789 | 1.48086 | 55 |
| GO_MITOCHONDRIAL_MATRIX | 0.00000 | 0.00002 | 0.65944 | 0.56045 | 1.25158 | 371 |
| GO_MITOCHONDRIAL_MEMBRANE_ORGANIZATION | 0.00361 | 0.04734 | 0.43171 | 0.59309 | 1.28094 | 86 |
| GO_MITOCHONDRIAL_PROTEIN_COMPLEX | 0.00000 | 0.00000 | 0.81404 | 0.63650 | 1.40931 | 208 |
| GO_MITOCHONDRIAL_TRANSLATION | 0.00004 | 0.00093 | 0.55733 | 0.61219 | 1.34135 | 129 |
| GO_MITOCHONDRIAL_TRANSLATIONAL_TERMINATION | 0.00000 | 0.00010 | 0.62726 | 0.66694 | 1.44128 | 87 |
| GO_MITOCHONDRION_ORGANIZATION | 0.00051 | 0.00993 | 0.47727 | 0.52284 | 1.16863 | 390 |
| GO_MODIFICATION_DEPENDENT_MACROMOLECULE_CATABOLIC_PR | 0.00002 | 0.00053 | 0.57561 | 0.53142 | 1.19156 | 486 |
| GO_MODULATION_BY_HOST_OF_SYMBIONT_PROCESS | 0.00015 | 0.00336 | 0.51885 | 0.72848 | 1.50542 | 33 |
| GO_MODULATION_OF_PROCESS_OF_OTHER_ORGANISM | 0.00019 | 0.00423 | 0.51885 | 0.66904 | 1.42286 | 59 |
| GO_MONOSACCHARIDE_BIOSYNTHETIC_PROCESS | 0.00335 | 0.04466 | 0.43171 | 0.61992 | 1.31781 | 58 |
| GO_MORPHOGENESIS_OF_AN_EPITHELIUM | 0.00006 | 0.00151 | 0.53843 | 0.55161 | 1.22755 | 290 |
| GO_MORPHOGENESIS_OF_A_POLARIZED_EPITHELIUM | 0.00000 | 0.00000 | | 0.72298 | 1.57057 | 103 |
| GO_MRNA_3_UTR_BINDING | 0.00003 | 0.00089 | 0.55733 | 0.67596 | 1.44827 | 68 |
| GO_MRNA_5_UTR_BINDING | 0.00006 | 0.00156 | 0.53843 | 0.80133 | 1.62592 | 22 |
| GO_MRNA_BINDING | 0.00000 | 0.00000 | | 0.66002 | 1.46353 | 222 |
| GO_MRNA_PROCESSING | 0.00000 | 0.00000 | 0.73376 | 0.56859 | 1.27295 | 418 |
| GO_MULTI_ORGANISM_LOCALIZATION | 0.00268 | 0.03799 | 0.43171 | 0.63245 | 1.33612 | 51 |
| GO_MULTI_ORGANISM_PROCESS | 0.00108 | 0.01839 | 0.45506 | 0.51114 | 1.14599 | 484 |
| GO_MYD88_DEPENDENT_TOLL_LIKE_RECEPTOR_SIGNALING_PATHW | 0.00002 | 0.00066 | 0.57561 | 0.85581 | 1.70312 | 16 |
| GO_MYELOID_LEUKOCYTE_ACTIVATION | 0.00000 | 0.00000 | 0.74774 | 0.58508 | 1.30479 | 327 |
| GO_MYELOID_LEUKOCYTE_MEDIATED_IMMUNITY | 0.00000 | 0.00000 | 0.76146 | 0.59841 | 1.33291 | 301 |
| GO_NADH_DEHYDROGENASE_ACTIVITY | 0.00383 | 0.04935 | 0.43171 | 0.66244 | 1.37139 | 35 |
| GO_NADH_DEHYDROGENASE_COMPLEX | 0.00376 | 0.04884 | 0.43171 | 0.65649 | 1.36421 | 38 |
| GO_NADH_METABOLIC_PROCESS | 0.00294 | 0.04036 | 0.43171 | 0.70642 | 1.44481 | 27 |
| GO_NCRNA_EXPORT_FROM_NUCLEUS | 0.00302 | 0.04108 | 0.43171 | 0.66820 | 1.38331 | 35 |
| GO_NCRNA_METABOLIC_PROCESS | 0.00000 | 0.00003 | 0.65944 | 0.55907 | 1.24907 | 381 |
| GO_NCRNA_PROCESSING | 0.00000 | 0.00003 | 0.65944 | 0.57175 | 1.27504 | 328 |
| GO_NEGATIVE_REGULATION_OF_CANONICAL_WNT_SIGNALING_PAT | 0.00000 | 0.00000 | | 0.71611 | 1.56038 | 111 |
| GO_NEGATIVE_REGULATION_OF_CELLULAR_AMIDE_METABOLIC_PRO | 0.00001 | 0.00020 | 0.61053 | 0.63031 | 1.37423 | 114 |
| GO_NEGATIVE_REGULATION_OF_CELL_CYCLE | 0.00003 | 0.00076 | 0.57561 | 0.53833 | 1.20513 | 425 |
| GO_NEGATIVE_REGULATION_OF_CELL_CYCLE_G2_M_PHASE_TRANS | 0.00000 | 0.00000 | 0.76146 | 0.71490 | 1.54439 | 85 |
| GO_NEGATIVE_REGULATION_OF_CELL_CYCLE_PHASE_TRANSITION | 0.00002 | 0.00060 | 0.57561 | 0.59041 | 1.30442 | 184 |
| GO_NEGATIVE_REGULATION_OF_CELL_CYCLE_PROCESS | 0.00000 | 0.00002 | 0.65944 | 0.58690 | 1.30374 | 249 |
| GO_NEGATIVE_REGULATION_OF_MITOTIC_CELL_CYCLE | 0.00004 | 0.00095 | 0.55733 | 0.56735 | 1.25902 | 232 |
| GO_NEGATIVE_REGULATION_OF_MRNA_METABOLIC_PROCESS | 0.00023 | 0.00491 | 0.51885 | 0.66670 | 1.41788 | 59 |
| GO_NEGATIVE_REGULATION_OF_MRNA_SPLICING_VIA_SPLICEOSOM | 0.00239 | 0.03510 | 0.43171 | 0.74857 | 1.51204 | 19 |
| GO_NEGATIVE_REGULATION_OF_RNA_SPLICING | 0.00029 | 0.00607 | 0.49849 | 0.76729 | 1.55976 | 23 |
| GO_NEGATIVE_REGULATION_OF_TRANSLATIONAL_INITIATION | 0.00018 | 0.00403 | 0.51885 | 0.83901 | 1.66434 | 15 |
| GO_NEGATIVE_REGULATION_OF_WNT_SIGNALING_PATHWAY | 0.00000 | 0.00000 | | 0.69525 | 1.52249 | 127 |
| GO_NEURAL_NUCLEUS_DEVELOPMENT | 0.00001 | 0.00026 | 0.59333 | 0.79148 | 1.62187 | 28 |
| GO_NEURON_TO_NEURON_SYNAPSE | 0.00005 | 0.00134 | 0.55733 | 0.59636 | 1.31006 | 154 |
| GO_NIK_NF_KAPPAB_SIGNALING | 0.00000 | 0.00000 | | 0.72896 | 1.58354 | 103 |
| GO_NON_CANONICAL_WNT_SIGNALING_PATHWAY | 0.00000 | 0.00000 | | 0.72784 | 1.58023 | 101 |
| GO_NUCLEAR_EXPORT | 0.00031 | 0.00645 | 0.49849 | 0.56963 | 1.25542 | 172 |
| GO_NUCLEAR_TRANSCRIBED_MRNA_CATABOLIC_PROCESS | 0.00000 | 0.00000 | | 0.79179 | 1.74917 | 183 |
| GO_NUCLEAR_TRANSCRIBED_MRNA_CATABOLIC_PROCESS_NONSEN | 0.00000 | 0.00000 | | 0.89008 | 1.93909 | 110 |
| GO_NUCLEAR_TRANSPORT | 0.00008 | 0.00193 | 0.53843 | 0.55511 | 1.23408 | 273 |
| GO_NUCLEOBASE_CONTAINING_COMPOUND_TRANSPORT | 0.00146 | 0.02386 | 0.45506 | 0.55053 | 1.21842 | 202 |
| GO_NUCLEOBASE_CONTAINING_SMALL_MOLECULE_BIOSYNTHETIC_ | 0.00137 | 0.02261 | 0.45506 | 0.62749 | 1.34353 | 67 |
| GO_NUCLEOBASE_CONTAINING_SMALL_MOLECULE_INTERCONVERS | 0.00383 | 0.04935 | 0.43171 | 0.71259 | 1.44857 | 23 |
| GO_NUCLEOBASE_CONTAINING_SMALL_MOLECULE_METABOLIC_PR | 0.00320 | 0.04339 | 0.43171 | 0.51448 | 1.14825 | 352 |
| GO_NUCLEOSIDE_DIPHOSPHATE_METABOLIC_PROCESS | 0.00110 | 0.01865 | 0.45506 | 0.59637 | 1.29295 | 97 |
| GO_NUCLEOSIDE_METABOLIC_PROCESS | 0.00061 | 0.01169 | 0.47727 | 0.65315 | 1.38907 | 59 |
| GO_NUCLEOSIDE_MONOPHOSPHATE_BIOSYNTHETIC_PROCESS | 0.00250 | 0.03636 | 0.43171 | 0.68841 | 1.42033 | 32 |
| GO_NUCLEOSIDE_MONOPHOSPHATE_METABOLIC_PROCESS | 0.00152 | 0.02464 | 0.45506 | 0.64377 | 1.36002 | 51 |
| GO_NUCLEOSIDE_PHOSPHATE_BIOSYNTHETIC_PROCESS | 0.00043 | 0.00865 | 0.49849 | 0.58185 | 1.27831 | 148 |
| GO_NUCLEOSIDE_TRIPHOSPHATE_BIOSYNTHETIC_PROCESS | 0.00043 | 0.00865 | 0.49849 | 0.69332 | 1.44539 | 40 |
| GO_NUCLEOSIDE_TRIPHOSPHATE_METABOLIC_PROCESS | 0.00016 | 0.00360 | 0.51885 | 0.66788 | 1.41976 | 58 |
| GO_NUCLEOSOMAL_DNA_BINDING | 0.00023 | 0.00507 | 0.51885 | 0.79852 | 1.61801 | 21 |
| GO_NUCLEOSOME_BINDING | 0.00143 | 0.02332 | 0.45506 | 0.66132 | 1.38492 | 45 |
| GO_NUCLEOTIDE_EXCISION_REPAIR | 0.00004 | 0.00096 | 0.55733 | 0.63343 | 1.37355 | 98 |
| GO_NUCLEOTIDE_EXCISION_REPAIR_DNA_DAMAGE_RECOGNITION | 0.00015 | 0.00349 | 0.51885 | 0.78678 | 1.59640 | 22 |
| GO_NUCLEOTIDE_EXCISION_REPAIR_DNA_INCISION | 0.00313 | 0.04249 | 0.43171 | 0.66439 | 1.37898 | 36 |
| GO_NUCLEOTIDE_PHOSPHORYLATION | 0.00383 | 0.04935 | 0.43171 | 0.59002 | 1.27454 | 88 |
| GO_ORGANELLAR_RIBOSOME | 0.00000 | 0.00014 | 0.61053 | 0.66412 | 1.43471 | 85 |
| GO_ORGANELLE_INNER_MEMBRANE | 0.00000 | 0.00000 | 0.76146 | 0.57601 | 1.28697 | 378 |
| GO_ORGANIC_CYCLIC_COMPOUND_CATABOLIC_PROCESS | 0.00000 | 0.00000 | | 0.73350 | 1.64281 | 449 |
| GO_OXIDATIVE_PHOSPHORYLATION | 0.00001 | 0.00030 | 0.59333 | 0.64619 | 1.40027 | 96 |
| GO_OXIDOREDUCTASE_ACTIVITY | 0.00006 | 0.00139 | 0.55733 | 0.53544 | 1.19703 | 392 |
| GO_OXIDOREDUCTASE_COMPLEX | 0.00001 | 0.00027 | 0.59333 | 0.66015 | 1.42461 | 82 |
| GO_PEPTIDASE_COMPLEX | 0.00000 | 0.00000 | | 0.75988 | 1.63982 | 82 |
| GO_PIGMENT_GRANULE | 0.00003 | 0.00089 | 0.55733 | 0.66098 | 1.42344 | 78 |

| | | | | | | |
|---|---|---|---|---|---|---|
| GO_POLYSOMAL_RIBOSOME | 0.00000 | 0.00000 | | 0.90624 | 1.86629 | 31 |
| GO_POLYSOME | 0.00000 | 0.00000 | | 0.82337 | 1.75481 | 61 |
| GO_POLY_PYRIMIDINE_TRACT_BINDING | 0.00327 | 0.04398 | 0.43171 | 0.75471 | 1.51855 | 18 |
| GO_POSITIVE_REGULATION_OF_APOPTOTIC_SIGNALING_PATHWAY | 0.00279 | 0.03889 | 0.43171 | 0.57666 | 1.25518 | 107 |
| GO_POSITIVE_REGULATION_OF_ATPASE_ACTIVITY | 0.00294 | 0.04036 | 0.43171 | 0.65446 | 1.36795 | 43 |
| GO_POSITIVE_REGULATION_OF_BINDING | 0.00088 | 0.01590 | 0.47727 | 0.60284 | 1.30556 | 95 |
| GO_POSITIVE_REGULATION_OF_CANONICAL_WNT_SIGNALING_PATH | 0.00000 | 0.00000 | 0.77494 | 0.70504 | 1.53160 | 103 |
| GO_POSITIVE_REGULATION_OF_CELLULAR_AMIDE_METABOLIC_PRC | 0.00003 | 0.00074 | 0.57561 | 0.62296 | 1.36083 | 118 |
| GO_POSITIVE_REGULATION_OF_CELLULAR_PROTEIN_LOCALIZATION | 0.00041 | 0.00840 | 0.49849 | 0.54915 | 1.21815 | 225 |
| GO_POSITIVE_REGULATION_OF_CELL_DEATH | 0.00069 | 0.01297 | 0.47727 | 0.52595 | 1.17423 | 363 |
| GO_POSITIVE_REGULATION_OF_CELL_POPULATION_PROLIFERATION | 0.00078 | 0.01455 | 0.47727 | 0.52449 | 1.17108 | 368 |
| GO_POSITIVE_REGULATION_OF_CHROMOSOME_ORGANIZATION | 0.00231 | 0.03437 | 0.43171 | 0.56990 | 1.24871 | 133 |
| GO_POSITIVE_REGULATION_OF_DEFENSE_RESPONSE | 0.00000 | 0.00003 | 0.65944 | 0.62175 | 1.36730 | 158 |
| GO_POSITIVE_REGULATION_OF_DNA_BIOSYNTHETIC_PROCESS | 0.00001 | 0.00041 | 0.59333 | 0.71929 | 1.52055 | 52 |
| GO_POSITIVE_REGULATION_OF_DNA_METABOLIC_PROCESS | 0.00364 | 0.04761 | 0.43171 | 0.56647 | 1.24189 | 135 |
| GO_POSITIVE_REGULATION_OF_DNA_TEMPLATED_TRANSCRIPTION_ | 0.00276 | 0.03885 | 0.43171 | 0.68932 | 1.41586 | 29 |
| GO_POSITIVE_REGULATION_OF_ESTABLISHMENT_OF_PROTEIN_LOC/ | 0.00007 | 0.00168 | 0.53843 | 0.56461 | 1.25165 | 219 |
| GO_POSITIVE_REGULATION_OF_IMMUNE_RESPONSE | 0.00000 | 0.00001 | 0.69013 | 0.59037 | 1.31265 | 277 |
| GO_POSITIVE_REGULATION_OF_IMMUNE_SYSTEM_PROCESS | 0.00004 | 0.00110 | 0.55733 | 0.54018 | 1.20716 | 386 |
| GO_POSITIVE_REGULATION_OF_INNATE_IMMUNE_RESPONSE | 0.00000 | 0.00000 | 0.81404 | 0.69693 | 1.51947 | 114 |
| GO_POSITIVE_REGULATION_OF_INTRINSIC_APOPTOTIC_SIGNALING_ | 0.00104 | 0.01798 | 0.45506 | 0.69097 | 1.43578 | 37 |
| GO_POSITIVE_REGULATION_OF_ORGANELLE_ORGANIZATION | 0.00043 | 0.00865 | 0.49849 | 0.52093 | 1.16605 | 411 |
| GO_POSITIVE_REGULATION_OF_PROTEIN_LOCALIZATION_TO_NUCLE | 0.00001 | 0.00020 | 0.61053 | 0.69907 | 1.48990 | 61 |
| GO_POSITIVE_REGULATION_OF_RESPONSE_TO_BIOTIC_STIMULUS | 0.00000 | 0.00000 | 0.73376 | 0.66340 | 1.45310 | 128 |
| GO_POSITIVE_REGULATION_OF_RESPONSE_TO_EXTERNAL_STIMULU | 0.00002 | 0.00050 | 0.57561 | 0.58133 | 1.28681 | 201 |
| GO_POSITIVE_REGULATION_OF_TELOMERASE_ACTIVITY | 0.00061 | 0.01169 | 0.47727 | 0.73476 | 1.50565 | 28 |
| GO_POSITIVE_REGULATION_OF_TELOMERASE_RNA_LOCALIZATION_ | 0.00000 | 0.00001 | 0.69013 | 0.92451 | 1.83395 | 15 |
| GO_POSITIVE_REGULATION_OF_TELOMERE_MAINTENANCE_VIA_TE | 0.00130 | 0.02159 | 0.45506 | 0.71259 | 1.46508 | 30 |
| GO_POSITIVE_REGULATION_OF_TRANSCRIPTION_INITIATION_FROM | 0.00372 | 0.04847 | 0.43171 | 0.70671 | 1.43993 | 25 |
| GO_POSITIVE_REGULATION_OF_TRANSFERASE_ACTIVITY | 0.00174 | 0.02712 | 0.45506 | 0.51642 | 1.15220 | 345 |
| GO_POSITIVE_REGULATION_OF_TRANSLATION | 0.00001 | 0.00019 | 0.61053 | 0.64767 | 1.40419 | 97 |
| GO_POSITIVE_REGULATION_OF_VIRAL_GENOME_REPLICATION | 0.00287 | 0.03973 | 0.43171 | 0.68362 | 1.40783 | 31 |
| GO_POSITIVE_REGULATION_OF_VIRAL_PROCESS | 0.00028 | 0.00590 | 0.49849 | 0.62440 | 1.34741 | 81 |
| GO_POSITIVE_REGULATION_OF_WNT_SIGNALING_PATHWAY | 0.00000 | 0.00000 | 0.73376 | 0.66230 | 1.44763 | 122 |
| GO_POSTREPLICATION_REPAIR | 0.00024 | 0.00515 | 0.51885 | 0.69140 | 1.44518 | 43 |
| GO_POSTSYNAPSE | 0.00004 | 0.00109 | 0.55733 | 0.55969 | 1.24369 | 259 |
| GO_POSTSYNAPTIC_CYTOSOL | 0.00279 | 0.03889 | 0.43171 | 0.78295 | 1.55313 | 15 |
| GO_POSTTRANSCRIPTIONAL_REGULATION_OF_GENE_EXPRESSION | 0.00000 | 0.00000 | | 0.66656 | 1.49327 | 462 |
| GO_POST_TRANSLATIONAL_PROTEIN_MODIFICATION | 0.00002 | 0.00052 | 0.57561 | 0.57851 | 1.28412 | 229 |
| GO_PRECATALYTIC_SPLICEOSOME | 0.00085 | 0.01559 | 0.47727 | 0.65235 | 1.37448 | 49 |
| GO_PRERIBOSOME | 0.00087 | 0.01577 | 0.47727 | 0.62899 | 1.34772 | 69 |
| GO_PRERIBOSOME_LARGE_SUBUNIT_PRECURSOR | 0.00235 | 0.03474 | 0.43171 | 0.74079 | 1.50308 | 22 |
| GO_PRE_MRNA_BINDING | 0.00133 | 0.02213 | 0.45506 | 0.72393 | 1.48062 | 27 |
| GO_PROTEASOMAL_PROTEIN_CATABOLIC_PROCESS | 0.00002 | 0.00044 | 0.59333 | 0.54166 | 1.21005 | 377 |
| GO_PROTEASOMAL_UBIQUITIN_INDEPENDENT_PROTEIN_CATABOLI( | 0.00000 | 0.00000 | 0.80122 | 0.92254 | 1.86931 | 21 |
| GO_PROTEASOME_ACCESSORY_COMPLEX | 0.00000 | 0.00001 | 0.67496 | 0.85464 | 1.73811 | 24 |
| GO_PROTEASOME_CORE_COMPLEX | 0.00000 | 0.00000 | | 0.94375 | 1.90631 | 19 |
| GO_PROTEIN_CONTAINING_COMPLEX_DISASSEMBLY | 0.00000 | 0.00007 | 0.62726 | 0.57810 | 1.28438 | 256 |
| GO_PROTEIN_CONTAINING_COMPLEX_LOCALIZATION | 0.00063 | 0.01197 | 0.47727 | 0.55335 | 1.22520 | 208 |
| GO_PROTEIN_DNA_COMPLEX_SUBUNIT_ORGANIZATION | 0.00001 | 0.00018 | 0.61053 | 0.60174 | 1.32491 | 165 |
| GO_PROTEIN_FOLDING | 0.00000 | 0.00002 | 0.65944 | 0.63113 | 1.38644 | 154 |
| GO_PROTEIN_FOLDING_CHAPERONE | 0.00135 | 0.02238 | 0.45506 | 0.77903 | 1.55752 | 17 |
| GO_PROTEIN_IMPORT | 0.00268 | 0.03799 | 0.43171 | 0.56317 | 1.23581 | 140 |
| GO_PROTEIN_LOCALIZATION_TO_CHROMOSOME | 0.00056 | 0.01096 | 0.47727 | 0.63523 | 1.36108 | 69 |
| GO_PROTEIN_LOCALIZATION_TO_CHROMOSOME_TELOMERIC_REGI( | 0.00335 | 0.04466 | 0.43171 | 0.70399 | 1.43984 | 27 |
| GO_PROTEIN_LOCALIZATION_TO_ENDOPLASMIC_RETICULUM | 0.00000 | 0.00000 | | 0.85521 | 1.87021 | 124 |
| GO_PROTEIN_LOCALIZATION_TO_MEMBRANE | 0.00000 | 0.00000 | | 0.65123 | 1.45714 | 406 |
| GO_PROTEIN_LOCALIZATION_TO_MITOCHONDRION | 0.00019 | 0.00420 | 0.51885 | 0.60773 | 1.32398 | 110 |
| GO_PROTEIN_LOCALIZATION_TO_NUCLEUS | 0.00002 | 0.00055 | 0.57561 | 0.58926 | 1.30220 | 187 |
| GO_PROTEIN_MODIFICATION_BY_SMALL_PROTEIN_REMOVAL | 0.00000 | 0.00000 | 0.71951 | 0.60745 | 1.34689 | 221 |
| GO_PROTEIN_POLYUBIQUITINATION | 0.00004 | 0.00106 | 0.55733 | 0.55874 | 1.24108 | 253 |
| GO_PROTEIN_REFOLDING | 0.00294 | 0.04036 | 0.43171 | 0.77039 | 1.53314 | 16 |
| GO_PROTEIN_STABILIZATION | 0.00001 | 0.00024 | 0.61053 | 0.62333 | 1.36586 | 130 |
| GO_PROTEIN_TARGETING | 0.00000 | 0.00000 | | 0.71181 | 1.58743 | 327 |
| GO_PROTEIN_TARGETING_TO_MEMBRANE | 0.00000 | 0.00000 | | 0.82370 | 1.80953 | 149 |
| GO_PROTEIN_TARGETING_TO_MITOCHONDRION | 0.00346 | 0.04593 | 0.43171 | 0.60408 | 1.30091 | 78 |
| GO_PROTEIN_TRANSMEMBRANE_TRANSPORT | 0.00265 | 0.03775 | 0.43171 | 0.64528 | 1.36171 | 50 |
| GO_PROTON_TRANSMEMBRANE_TRANSPORT | 0.00213 | 0.03229 | 0.43171 | 0.60815 | 1.30536 | 71 |
| GO_PURINE_CONTAINING_COMPOUND_BIOSYNTHETIC_PROCESS | 0.00246 | 0.03601 | 0.43171 | 0.57796 | 1.25936 | 111 |
| GO_PURINE_CONTAINING_COMPOUND_METABOLIC_PROCESS | 0.00100 | 0.01761 | 0.45506 | 0.53679 | 1.19242 | 255 |
| GO_PURINE_NUCLEOSIDE_TRIPHOSPHATE_METABOLIC_PROCESS | 0.00089 | 0.01592 | 0.47727 | 0.67157 | 1.40374 | 43 |
| GO_PYRIMIDINE_CONTAINING_COMPOUND_METABOLIC_PROCESS | 0.00353 | 0.04658 | 0.43171 | 0.64744 | 1.35878 | 47 |
| GO_REGENERATION | 0.00216 | 0.03242 | 0.43171 | 0.59987 | 1.29453 | 82 |
| GO_REGULATION_OF_ANIMAL_ORGAN_MORPHOGENESIS | 0.00000 | 0.00000 | | 0.69163 | 1.51541 | 133 |
| GO_REGULATION_OF_APOPTOTIC_SIGNALING_PATHWAY | 0.00031 | 0.00645 | 0.49849 | 0.55171 | 1.22506 | 247 |
| GO_REGULATION_OF_ATPASE_ACTIVITY | 0.00324 | 0.04369 | 0.43171 | 0.63031 | 1.34038 | 57 |

| | | | | | | |
|---|---|---|---|---|---|---|
| GO_REGULATION_OF_ATP_METABOLIC_PROCESS | 0.00353 | 0.04658 | 0.43171 | 0.60176 | 1.29780 | 80 |
| GO_REGULATION_OF_BINDING | 0.00250 | 0.03636 | 0.43171 | 0.54476 | 1.20598 | 203 |
| GO_REGULATION_OF_CELLULAR_AMIDE_METABOLIC_PROCESS | 0.00000 | 0.00000 | | 0.62967 | 1.40226 | 311 |
| GO_REGULATION_OF_CELLULAR_AMINE_METABOLIC_PROCESS | 0.00000 | 0.00000 | | 0.82173 | 1.74682 | 58 |
| GO_REGULATION_OF_CELLULAR_AMINO_ACID_METABOLIC_PROCES | 0.00000 | 0.00000 | | 0.83776 | 1.77766 | 55 |
| GO_REGULATION_OF_CELLULAR_KETONE_METABOLIC_PROCESS | 0.00000 | 0.00000 | | 0.72641 | 1.58680 | 118 |
| GO_REGULATION_OF_CELLULAR_PROTEIN_LOCALIZATION | 0.00082 | 0.01513 | 0.47727 | 0.52542 | 1.17290 | 362 |
| GO_REGULATION_OF_CELLULAR_RESPONSE_TO_HEAT | 0.00276 | 0.03885 | 0.43171 | 0.61652 | 1.32004 | 67 |
| GO_REGULATION_OF_CELLULAR_RESPONSE_TO_STRESS | 0.00224 | 0.03344 | 0.43171 | 0.50545 | 1.13232 | 468 |
| GO_REGULATION_OF_CELL_CYCLE_G2_M_PHASE_TRANSITION | 0.00000 | 0.00000 | 0.71951 | 0.63230 | 1.39282 | 169 |
| GO_REGULATION_OF_CELL_CYCLE_PHASE_TRANSITION | 0.00001 | 0.00026 | 0.59333 | 0.55488 | 1.23783 | 337 |
| GO_REGULATION_OF_CHROMOSOME_ORGANIZATION | 0.00081 | 0.01501 | 0.47727 | 0.54255 | 1.20527 | 261 |
| GO_REGULATION_OF_CYSTEINE_TYPE_ENDOPEPTIDASE_ACTIVITY | 0.00202 | 0.03077 | 0.43171 | 0.57120 | 1.24837 | 121 |
| GO_REGULATION_OF_CYTOPLASMIC_TRANSLATION | 0.00130 | 0.02159 | 0.45506 | 0.73678 | 1.49773 | 23 |
| GO_REGULATION_OF_DEFENSE_RESPONSE | 0.00065 | 0.01237 | 0.47727 | 0.53633 | 1.19355 | 290 |
| GO_REGULATION_OF_DNA_BIOSYNTHETIC_PROCESS | 0.00016 | 0.00360 | 0.51885 | 0.64215 | 1.37923 | 74 |
| GO_REGULATION_OF_DNA_METABOLIC_PROCESS | 0.00069 | 0.01297 | 0.47727 | 0.54425 | 1.20775 | 237 |
| GO_REGULATION_OF_DNA_TEMPLATED_TRANSCRIPTION_INITIATIOI | 0.00163 | 0.02568 | 0.45506 | 0.68289 | 1.41372 | 35 |
| GO_REGULATION_OF_DNA_TEMPLATED_TRANSCRIPTION_IN_RESPO | 0.00000 | 0.00000 | | 0.73944 | 1.59800 | 89 |
| GO_REGULATION_OF_GENERATION_OF_PRECURSOR_METABOLITES_ | 0.00060 | 0.01164 | 0.47727 | 0.60476 | 1.31116 | 97 |
| GO_REGULATION_OF_HEMATOPOIETIC_PROGENITOR_CELL_DIFFERE | 0.00000 | 0.00000 | | 0.78717 | 1.69251 | 75 |
| GO_REGULATION_OF_HEMATOPOIETIC_STEM_CELL_DIFFERENTIATIC | 0.00000 | 0.00000 | | 0.81505 | 1.74256 | 65 |
| GO_REGULATION_OF_HEMOPOIESIS | 0.00000 | 0.00008 | 0.62726 | 0.58859 | 1.30634 | 228 |
| GO_REGULATION_OF_IMMUNE_RESPONSE | 0.00000 | 0.00016 | 0.61053 | 0.55362 | 1.23583 | 362 |
| GO_REGULATION_OF_INNATE_IMMUNE_RESPONSE | 0.00000 | 0.00000 | 0.71951 | 0.63883 | 1.40521 | 161 |
| GO_REGULATION_OF_INTRINSIC_APOPTOTIC_SIGNALING_PATHWAY | 0.00001 | 0.00039 | 0.59333 | 0.63478 | 1.38259 | 109 |
| GO_REGULATION_OF_MITOTIC_CELL_CYCLE | 0.00001 | 0.00018 | 0.61053 | 0.53899 | 1.20736 | 458 |
| GO_REGULATION_OF_MORPHOGENESIS_OF_AN_EPITHELIUM | 0.00000 | 0.00000 | | 0.72213 | 1.57284 | 109 |
| GO_REGULATION_OF_MRNA_CATABOLIC_PROCESS | 0.00000 | 0.00000 | | 0.69977 | 1.54127 | 168 |
| GO_REGULATION_OF_MRNA_METABOLIC_PROCESS | 0.00000 | 0.00000 | | 0.66528 | 1.47872 | 266 |
| GO_REGULATION_OF_MRNA_PROCESSING | 0.00004 | 0.00105 | 0.55733 | 0.61521 | 1.34090 | 113 |
| GO_REGULATION_OF_MRNA_SPLICING_VIA_SPLICEOSOME | 0.00001 | 0.00026 | 0.59333 | 0.67424 | 1.45411 | 80 |
| GO_REGULATION_OF_NUCLEASE_ACTIVITY | 0.00106 | 0.01819 | 0.45506 | 0.77918 | 1.56780 | 18 |
| GO_REGULATION_OF_OXIDATIVE_STRESS_INDUCED_CELL_DEATH | 0.00198 | 0.03028 | 0.43171 | 0.67287 | 1.40355 | 41 |
| GO_REGULATION_OF_PEPTIDASE_ACTIVITY | 0.00050 | 0.00978 | 0.47727 | 0.56122 | 1.24054 | 191 |
| GO_REGULATION_OF_POSTTRANSCRIPTIONAL_GENE_SILENCING | 0.00364 | 0.04761 | 0.43171 | 0.59824 | 1.28492 | 74 |
| GO_REGULATION_OF_PROTEIN_CATABOLIC_PROCESS | 0.00179 | 0.02783 | 0.45506 | 0.53057 | 1.17929 | 270 |
| GO_REGULATION_OF_PROTEIN_LOCALIZATION_TO_NUCLEUS | 0.00001 | 0.00042 | 0.59333 | 0.65505 | 1.41558 | 87 |
| GO_REGULATION_OF_PROTEIN_MODIFICATION_BY_SMALL_PROTEIN | 0.00025 | 0.00532 | 0.49849 | 0.57068 | 1.25684 | 167 |
| GO_REGULATION_OF_PROTEIN_STABILITY | 0.00093 | 0.01652 | 0.47727 | 0.55382 | 1.22571 | 202 |
| GO_REGULATION_OF_PROTEOLYSIS | 0.00013 | 0.00301 | 0.51885 | 0.52997 | 1.18392 | 377 |
| GO_REGULATION_OF_RESPONSE_TO_BIOTIC_STIMULUS | 0.00001 | 0.00020 | 0.61053 | 0.59049 | 1.30719 | 204 |
| GO_REGULATION_OF_RNA_SPLICING | 0.00000 | 0.00001 | 0.69013 | 0.65896 | 1.43669 | 114 |
| GO_REGULATION_OF_SMALL_MOLECULE_METABOLIC_PROCESS | 0.00000 | 0.00000 | 0.71951 | 0.59134 | 1.31394 | 265 |
| GO_REGULATION_OF_STEM_CELL_DIFFERENTIATION | 0.00000 | 0.00000 | | 0.74049 | 1.60113 | 90 |
| GO_REGULATION_OF_SYMBIOTIC_PROCESS | 0.00006 | 0.00144 | 0.55733 | 0.60071 | 1.31682 | 134 |
| GO_REGULATION_OF_TELOMERASE_ACTIVITY | 0.00088 | 0.01590 | 0.47727 | 0.69161 | 1.43179 | 35 |
| GO_REGULATION_OF_TRANSCRIPTION_FROM_RNA_POLYMERASE_II | 0.00000 | 0.00000 | | 0.81194 | 1.73800 | 66 |
| GO_REGULATION_OF_TRANSCRIPTION_INITIATION_FROM_RNA_POL | 0.00143 | 0.02332 | 0.45506 | 0.70390 | 1.44582 | 29 |
| GO_REGULATION_OF_TRANSLATIONAL_INITIATION | 0.00002 | 0.00052 | 0.57561 | 0.68129 | 1.45518 | 64 |
| GO_REGULATION_OF_UBIQUITIN_PROTEIN_TRANSFERASE_ACTIVITY | 0.00061 | 0.01169 | 0.47727 | 0.69335 | 1.44627 | 41 |
| GO_REGULATION_OF_VIRAL_GENOME_REPLICATION | 0.00086 | 0.01566 | 0.47727 | 0.65316 | 1.38389 | 54 |
| GO_REGULATION_OF_VIRAL_TRANSCRIPTION | 0.00198 | 0.03028 | 0.43171 | 0.63932 | 1.34703 | 49 |
| GO_REGULATION_OF_WNT_SIGNALING_PATHWAY | 0.00000 | 0.00001 | 0.69013 | 0.60353 | 1.33816 | 220 |
| GO_RESPIRASOME | 0.00008 | 0.00181 | 0.53843 | 0.65420 | 1.40421 | 71 |
| GO_RESPIRATORY_CHAIN_COMPLEX | 0.00016 | 0.00353 | 0.51885 | 0.67626 | 1.43918 | 60 |
| GO_RESPIRATORY_ELECTRON_TRANSPORT_CHAIN | 0.00001 | 0.00025 | 0.59333 | 0.65819 | 1.42190 | 85 |
| GO_RESPONSE_TO_ALCOHOL | 0.00261 | 0.03750 | 0.43171 | 0.58804 | 1.27350 | 95 |
| GO_RESPONSE_TO_CADMIUM_ION | 0.00265 | 0.03775 | 0.43171 | 0.67227 | 1.39174 | 35 |
| GO_RESPONSE_TO_DRUG | 0.00104 | 0.01798 | 0.45506 | 0.55364 | 1.22258 | 180 |
| GO_RESPONSE_TO_ETHANOL | 0.00086 | 0.01566 | 0.47727 | 0.67309 | 1.40691 | 43 |
| GO_RESPONSE_TO_HEAT | 0.00012 | 0.00270 | 0.53843 | 0.61665 | 1.34221 | 107 |
| GO_RESPONSE_TO_HORMONE | 0.00048 | 0.00949 | 0.49849 | 0.51277 | 1.14814 | 453 |
| GO_RESPONSE_TO_HYDROGEN_PEROXIDE | 0.00257 | 0.03716 | 0.43171 | 0.60505 | 1.30368 | 79 |
| GO_RESPONSE_TO_INORGANIC_SUBSTANCE | 0.00004 | 0.00107 | 0.55733 | 0.55714 | 1.23870 | 275 |
| GO_RESPONSE_TO_INTERLEUKIN_1 | 0.00000 | 0.00000 | | 0.74844 | 1.62908 | 107 |
| GO_RESPONSE_TO_INTERLEUKIN_12 | 0.00012 | 0.00270 | 0.53843 | 0.72605 | 1.50694 | 36 |
| GO_RESPONSE_TO_INTERLEUKIN_4 | 0.00324 | 0.04369 | 0.43171 | 0.75812 | 1.51571 | 17 |
| GO_RESPONSE_TO_METAL_ION | 0.00030 | 0.00623 | 0.49849 | 0.56387 | 1.24233 | 171 |
| GO_RESPONSE_TO_ORGANIC_CYCLIC_COMPOUND | 0.00000 | 0.00016 | 0.61053 | 0.54603 | 1.22259 | 439 |
| GO_RESPONSE_TO_OXYGEN_LEVELS | 0.00000 | 0.00000 | 0.71951 | 0.60132 | 1.33438 | 233 |
| GO_RESPONSE_TO_STEROID_HORMONE | 0.00154 | 0.02464 | 0.45506 | 0.55514 | 1.22446 | 174 |
| GO_RESPONSE_TO_TEMPERATURE_STIMULUS | 0.00027 | 0.00572 | 0.49849 | 0.59216 | 1.29675 | 127 |
| GO_RESPONSE_TO_TUMOR_NECROSIS_FACTOR | 0.00000 | 0.00000 | 0.76146 | 0.65649 | 1.44236 | 151 |
| GO_RETINA_HOMEOSTASIS | 0.00104 | 0.01798 | 0.45506 | 0.73063 | 1.49432 | 27 |
| GO_RIBONUCLEOPROTEIN_COMPLEX_BINDING | 0.00000 | 0.00000 | 0.71951 | 0.66906 | 1.46050 | 117 |

| | | | | | | |
|---|---|---|---|---|---|---|
| GO_RIBONUCLEOPROTEIN_COMPLEX_BIOGENESIS | 0.00000 | 0.00000 | | 0.66339 | 1.48264 | 389 |
| GO_RIBONUCLEOPROTEIN_COMPLEX_SUBUNIT_ORGANIZATION | 0.00000 | 0.00000 | | 0.69241 | 1.52681 | 173 |
| GO_RIBONUCLEOPROTEIN_GRANULE | 0.00000 | 0.00001 | 0.67496 | 0.61979 | 1.36366 | 163 |
| GO_RIBONUCLEOSIDE_TRIPHOSPHATE_BIOSYNTHETIC_PROCESS | 0.00041 | 0.00840 | 0.49849 | 0.71253 | 1.47248 | 33 |
| GO_RIBONUCLEOSIDE_TRIPHOSPHATE_METABOLIC_PROCESS | 0.00026 | 0.00550 | 0.49849 | 0.69681 | 1.45611 | 42 |
| GO_RIBOSE_PHOSPHATE_BIOSYNTHETIC_PROCESS | 0.00191 | 0.02939 | 0.45506 | 0.57932 | 1.26207 | 110 |
| GO_RIBOSE_PHOSPHATE_METABOLIC_PROCESS | 0.00108 | 0.01839 | 0.45506 | 0.53891 | 1.19687 | 245 |
| GO_RIBOSOMAL_LARGE_SUBUNIT_ASSEMBLY | 0.00000 | 0.00016 | 0.61053 | 0.80999 | 1.65663 | 27 |
| GO_RIBOSOMAL_LARGE_SUBUNIT_BIOGENESIS | 0.00000 | 0.00000 | | 0.77542 | 1.65982 | 66 |
| GO_RIBOSOMAL_SMALL_SUBUNIT_ASSEMBLY | 0.00023 | 0.00494 | 0.51885 | 0.81139 | 1.62221 | 17 |
| GO_RIBOSOMAL_SMALL_SUBUNIT_BINDING | 0.00383 | 0.04935 | 0.43171 | 0.75288 | 1.50523 | 17 |
| GO_RIBOSOMAL_SMALL_SUBUNIT_BIOGENESIS | 0.00001 | 0.00022 | 0.61053 | 0.69076 | 1.47997 | 68 |
| GO_RIBOSOMAL_SUBUNIT | 0.00000 | 0.00000 | | 0.77603 | 1.71031 | 172 |
| GO_RIBOSOME | 0.00000 | 0.00000 | | 0.76643 | 1.69659 | 206 |
| GO_RIBOSOME_ASSEMBLY | 0.00000 | 0.00000 | 0.73376 | 0.76173 | 1.61998 | 59 |
| GO_RIBOSOME_BINDING | 0.00019 | 0.00416 | 0.51885 | 0.68955 | 1.44715 | 47 |
| GO_RIBOSOME_BIOGENESIS | 0.00000 | 0.00000 | | 0.65877 | 1.46466 | 275 |
| GO_RNA_CATABOLIC_PROCESS | 0.00000 | 0.00000 | | 0.76774 | 1.71264 | 339 |
| GO_RNA_DEPENDENT_DNA_BIOSYNTHETIC_PROCESS | 0.00003 | 0.00076 | 0.57561 | 0.68926 | 1.46521 | 58 |
| GO_RNA_EXPORT_FROM_NUCLEUS | 0.00062 | 0.01183 | 0.47727 | 0.58378 | 1.27758 | 126 |
| GO_RNA_LOCALIZATION | 0.00000 | 0.00006 | 0.64355 | 0.59670 | 1.32020 | 198 |
| GO_RNA_POLYMERASE_II_PREINITIATION_COMPLEX_ASSEMBLY | 0.00089 | 0.01592 | 0.47727 | 0.77046 | 1.55626 | 19 |
| GO_RNA_SPLICING | 0.00000 | 0.00000 | 0.78819 | 0.59079 | 1.31927 | 369 |
| GO_RNA_SPLICING_VIA_TRANSESTERIFICATION_REACTIONS | 0.00000 | 0.00000 | | 0.60940 | 1.35714 | 300 |
| GO_RNA_STABILIZATION | 0.00065 | 0.01223 | 0.47727 | 0.69743 | 1.44384 | 35 |
| GO_ROUGH_ENDOPLASMIC_RETICULUM | 0.00001 | 0.00039 | 0.59333 | 0.71853 | 1.51629 | 50 |
| GO_RRNA_BINDING | 0.00000 | 0.00001 | 0.69013 | 0.74782 | 1.58865 | 56 |
| GO_RRNA_METABOLIC_PROCESS | 0.00000 | 0.00000 | 0.77494 | 0.63047 | 1.39562 | 206 |
| GO_SCF_DEPENDENT_PROTEASOMAL_UBIQUITIN_DEPENDENT_PRO | 0.00000 | 0.00000 | | 0.77527 | 1.67044 | 79 |
| GO_SECRETORY_GRANULE | 0.00000 | 0.00000 | 0.71951 | 0.56518 | 1.26461 | 399 |
| GO_SECRETORY_VESICLE | 0.00000 | 0.00002 | 0.67496 | 0.54866 | 1.22905 | 465 |
| GO_SIGNAL_TRANSDUCTION_BY_PROTEIN_PHOSPHORYLATION | 0.00004 | 0.00109 | 0.55733 | 0.53817 | 1.20474 | 424 |
| GO_SINGLE_FERTILIZATION | 0.00048 | 0.00949 | 0.49849 | 0.67414 | 1.42260 | 50 |
| GO_SINGLE_STRANDED_RNA_BINDING | 0.00039 | 0.00793 | 0.49849 | 0.66553 | 1.41527 | 57 |
| GO_SMALL_MOLECULE_BIOSYNTHETIC_PROCESS | 0.00216 | 0.03242 | 0.43171 | 0.51195 | 1.14522 | 397 |
| GO_SMALL_RIBOSOMAL_SUBUNIT | 0.00000 | 0.00000 | | 0.77648 | 1.66209 | 66 |
| GO_SM_LIKE_PROTEIN_FAMILY_COMPLEX | 0.00327 | 0.04398 | 0.43171 | 0.60468 | 1.29713 | 70 |
| GO_SPERM_EGG_RECOGNITION | 0.00000 | 0.00007 | 0.64355 | 0.85241 | 1.72955 | 22 |
| GO_SPLICEOSOMAL_COMPLEX | 0.00000 | 0.00003 | 0.65944 | 0.60939 | 1.34261 | 171 |
| GO_STEM_CELL_DIFFERENTIATION | 0.00000 | 0.00000 | 0.78819 | 0.66886 | 1.46930 | 147 |
| GO_STRUCTURAL_CONSTITUENT_OF_CYTOSKELETON | 0.00102 | 0.01782 | 0.45506 | 0.65913 | 1.39337 | 52 |
| GO_STRUCTURAL_CONSTITUENT_OF_RIBOSOME | 0.00000 | 0.00000 | | 0.79909 | 1.75541 | 150 |
| GO_STRUCTURAL_MOLECULE_ACTIVITY | 0.00000 | 0.00000 | | 0.72651 | 1.61895 | 324 |
| GO_SUBSTANTIA_NIGRA_DEVELOPMENT | 0.00000 | 0.00013 | 0.62726 | 0.81622 | 1.66507 | 26 |
| GO_TELOMERASE_RNA_LOCALIZATION | 0.00001 | 0.00017 | 0.61053 | 0.85164 | 1.72025 | 19 |
| GO_TELOMERE_MAINTENANCE_VIA_TELOMERE_LENGTHENING | 0.00011 | 0.00250 | 0.53843 | 0.65659 | 1.40546 | 66 |
| GO_TELOMERE_ORGANIZATION | 0.00342 | 0.04554 | 0.43171 | 0.57000 | 1.24649 | 124 |
| GO_THREONINE_TYPE_ENDOPEPTIDASE_ACTIVITY | 0.00000 | 0.00000 | | 0.95179 | 1.92255 | 19 |
| GO_THREONINE_TYPE_PEPTIDASE_ACTIVITY | 0.00000 | 0.00000 | | 0.92199 | 1.87423 | 23 |
| GO_TISSUE_MORPHOGENESIS | 0.00097 | 0.01707 | 0.47727 | 0.52661 | 1.17402 | 326 |
| GO_TOXIN_TRANSPORT | 0.00113 | 0.01922 | 0.45506 | 0.72897 | 1.48708 | 26 |
| GO_TRANSCRIPTION_COUPLED_NUCLEOTIDE_EXCISION_REPAIR | 0.00001 | 0.00019 | 0.61053 | 0.68106 | 1.46186 | 71 |
| GO_TRANSCRIPTION_FACTOR_BINDING | 0.00043 | 0.00865 | 0.49849 | 0.52447 | 1.17376 | 409 |
| GO_TRANSCRIPTION_PREINITIATION_COMPLEX_ASSEMBLY | 0.00119 | 0.02004 | 0.45506 | 0.71399 | 1.46797 | 30 |
| GO_TRANSLATIONAL_ELONGATION | 0.00000 | 0.00000 | 0.71951 | 0.66087 | 1.44451 | 122 |
| GO_TRANSLATIONAL_INITIATION | 0.00000 | 0.00000 | | 0.84691 | 1.86515 | 166 |
| GO_TRANSLATIONAL_TERMINATION | 0.00000 | 0.00005 | 0.64355 | 0.65946 | 1.43060 | 99 |
| GO_TRANSLATION_ELONGATION_FACTOR_ACTIVITY | 0.00161 | 0.02546 | 0.45506 | 0.79860 | 1.58418 | 15 |
| GO_TRANSLATION_FACTOR_ACTIVITY_RNA_BINDING | 0.00000 | 0.00000 | | 0.79008 | 1.69288 | 69 |
| GO_TRANSLATION_INITIATION_FACTOR_ACTIVITY | 0.00000 | 0.00000 | 0.76146 | 0.80422 | 1.68417 | 45 |
| GO_TRANSLATION_INITIATION_FACTOR_BINDING | 0.00021 | 0.00468 | 0.51885 | 0.76186 | 1.55231 | 25 |
| GO_TRANSLATION_PREINITIATION_COMPLEX | 0.00000 | 0.00001 | 0.67496 | 0.90245 | 1.79595 | 16 |
| GO_TRANSLATION_REGULATOR_ACTIVITY | 0.00000 | 0.00000 | | 0.77663 | 1.68904 | 105 |
| GO_TRANSLATION_REGULATOR_ACTIVITY_NUCLEIC_ACID_BINDING | 0.00000 | 0.00000 | | 0.77150 | 1.66586 | 83 |
| GO_TRICARBOXYLIC_ACID_CYCLE | 0.00154 | 0.02464 | 0.45506 | 0.71854 | 1.47240 | 28 |
| GO_TUMOR_NECROSIS_FACTOR_MEDIATED_SIGNALING_PATHWAY | 0.00000 | 0.00000 | | 0.76277 | 1.65372 | 97 |
| GO_T_CELL_RECEPTOR_SIGNALING_PATHWAY | 0.00000 | 0.00000 | 0.81404 | 0.69994 | 1.52527 | 112 |
| GO_U12_TYPE_SPLICEOSOMAL_COMPLEX | 0.00279 | 0.03889 | 0.43171 | 0.70746 | 1.44693 | 27 |
| GO_U2_SNRNP | 0.00005 | 0.00134 | 0.55733 | 0.83081 | 1.67168 | 18 |
| GO_U2_TYPE_SPLICEOSOMAL_COMPLEX | 0.00010 | 0.00243 | 0.53843 | 0.63403 | 1.36970 | 85 |
| GO_UBIQUITIN_LIKE_PROTEIN_LIGASE_BINDING | 0.00000 | 0.00000 | 0.76146 | 0.62273 | 1.38051 | 219 |
| GO_UBIQUITIN_PROTEIN_TRANSFERASE_REGULATOR_ACTIVITY | 0.00165 | 0.02590 | 0.45506 | 0.79605 | 1.57911 | 15 |
| GO_UNFOLDED_PROTEIN_BINDING | 0.00000 | 0.00000 | 0.77494 | 0.72157 | 1.55881 | 85 |
| GO_VESICLE_LUMEN | 0.00000 | 0.00000 | 0.74774 | 0.64475 | 1.42008 | 168 |
| GO_VIRAL_GENE_EXPRESSION | 0.00000 | 0.00000 | | 0.80028 | 1.76317 | 171 |
| GO_VIRAL_GENOME_REPLICATION | 0.00100 | 0.01761 | 0.45506 | 0.61997 | 1.33078 | 73 |

| | | | | | | |
|---|---|---|---|---|---|---|
| GO_VIRAL_LIFE_CYCLE | 0.00031 | 0.00640 | 0.49849 | 0.55767 | 1.23509 | 209 |
| HP_ABNORMALITY_OF_ACID_BASE_HOMEOSTASIS | 0.00216 | 0.03242 | 0.43171 | 0.53556 | 1.18846 | 232 |
| HP_ABNORMALITY_OF_CARDIOVASCULAR_SYSTEM_ELECTROPHYSIO | 0.00027 | 0.00572 | 0.49849 | 0.55832 | 1.23929 | 229 |
| HP_ABNORMALITY_OF_LOWER_LIP | 0.00181 | 0.02804 | 0.45506 | 0.56360 | 1.23779 | 145 |
| HP_ABNORMALITY_OF_THE_MITOCHONDRION | 0.00213 | 0.03229 | 0.43171 | 0.57831 | 1.26240 | 117 |
| HP_ABNORMALITY_OF_THE_NASAL_DORSUM | 0.00005 | 0.00116 | 0.55733 | 0.60288 | 1.32157 | 134 |
| HP_ABNORMAL_ELECTROPHYSIOLOGY_OF_SINOATRIAL_NODE_ORIG | 0.00279 | 0.03889 | 0.43171 | 0.68436 | 1.40934 | 31 |
| HP_ABNORMAL_ERYTHROCYTE_MORPHOLOGY | 0.00000 | 0.00015 | 0.61053 | 0.56049 | 1.25036 | 341 |
| HP_ABNORMAL_MEAN_CORPUSCULAR_VOLUME | 0.00357 | 0.04696 | 0.43171 | 0.75226 | 1.51363 | 18 |
| HP_ABNORMAL_RETICULOCYTE_MORPHOLOGY | 0.00064 | 0.01210 | 0.47727 | 0.71420 | 1.47081 | 31 |
| HP_ACUTE_LEUKEMIA | 0.00000 | 0.00000 | 0.73376 | 0.72786 | 1.56137 | 70 |
| HP_ANEMIA_OF_INADEQUATE_PRODUCTION | 0.00000 | 0.00008 | 0.62726 | 0.65593 | 1.42209 | 97 |
| HP_CONSTITUTIONAL_SYMPTOM | 0.00119 | 0.02004 | 0.45506 | 0.51427 | 1.15108 | 423 |
| HP_DELAYED_PUBERTY | 0.00005 | 0.00115 | 0.55733 | 0.62579 | 1.36211 | 107 |
| HP_DEPRESSED_NASAL_RIDGE | 0.00000 | 0.00011 | 0.62726 | 0.68731 | 1.47511 | 72 |
| HP_FATIGUE | 0.00039 | 0.00808 | 0.49849 | 0.56529 | 1.24923 | 187 |
| HP_HEADACHE | 0.00015 | 0.00340 | 0.51885 | 0.59065 | 1.29716 | 146 |
| HP_HEMATOLOGICAL_NEOPLASM | 0.00003 | 0.00078 | 0.57561 | 0.60108 | 1.32044 | 154 |
| HP_INCREASED_CSF_LACTATE | 0.00157 | 0.02509 | 0.45506 | 0.61622 | 1.31904 | 66 |
| HP_INCREASED_SERUM_LACTATE | 0.00216 | 0.03242 | 0.43171 | 0.56920 | 1.24724 | 130 |
| HP_LEUKEMIA | 0.00001 | 0.00026 | 0.59333 | 0.64877 | 1.40680 | 98 |
| HP_MACROCYTIC_ANEMIA | 0.00000 | 0.00000 | 0.74774 | 0.82468 | 1.71698 | 39 |
| HP_MIGRAINE | 0.00000 | 0.00000 | 0.71951 | 0.72105 | 1.54497 | 69 |
| HP_MYOTONIA | 0.00161 | 0.02546 | 0.45506 | 0.79820 | 1.58338 | 15 |
| HP_NEOPLASM | 0.00216 | 0.03242 | 0.43171 | 0.50820 | 1.13805 | 446 |
| HP_PALLOR | 0.00001 | 0.00040 | 0.59333 | 0.65169 | 1.41044 | 92 |
| HP_PUBERTY_AND_GONADAL_DISORDERS | 0.00287 | 0.03973 | 0.43171 | 0.51995 | 1.15843 | 322 |
| HP_RETICULOCYTOPENIA | 0.00004 | 0.00105 | 0.55733 | 0.86417 | 1.71425 | 15 |
| HP_RIMMED_VACUOLES | 0.00279 | 0.03889 | 0.43171 | 0.74033 | 1.49831 | 20 |
| HP_SHORT_NOSE | 0.00235 | 0.03474 | 0.43171 | 0.54919 | 1.21134 | 174 |
| HP_THICK_LOWER_LIP_VERMILION | 0.00048 | 0.00949 | 0.49849 | 0.60773 | 1.31617 | 94 |
| HP_THICK_VERMILION_BORDER | 0.00156 | 0.02486 | 0.45506 | 0.55983 | 1.23000 | 153 |
| HP_VENTRICULAR_ARRHYTHMIA | 0.00353 | 0.04658 | 0.43171 | 0.66198 | 1.37398 | 36 |
| **Cluster 1** | | | | | | |
| pathway | pval | padj | log2err | ES | NES | size |
| GO_ACTIVATION_OF_IMMUNE_RESPONSE | 0.00000 | 0.00000 | 0.78819 | 0.62851 | 1.39255 | 212 |
| GO_ACTIVATION_OF_INNATE_IMMUNE_RESPONSE | 0.00000 | 0.00000 | | 0.75054 | 1.62130 | 88 |
| GO_AEROBIC_RESPIRATION | 0.00001 | 0.00025 | 0.59333 | 0.69302 | 1.48168 | 65 |
| GO_ALTERNATIVE_MRNA_SPLICING_VIA_SPLICEOSOME | 0.00102 | 0.01782 | 0.45506 | 0.65168 | 1.38687 | 60 |
| GO_AMINE_METABOLIC_PROCESS | 0.00000 | 0.00000 | | 0.79189 | 1.70783 | 80 |
| GO_ANAPHASE_PROMOTING_COMPLEX_DEPENDENT_CATABOLIC_P | 0.00000 | 0.00000 | | 0.76611 | 1.65070 | 79 |
| GO_ANCHORING_JUNCTION | 0.00000 | 0.00000 | | 0.59615 | 1.33633 | 479 |
| GO_ANIMAL_ORGAN_REGENERATION | 0.00257 | 0.03716 | 0.43171 | 0.68240 | 1.41020 | 33 |
| GO_ANTIGEN_PROCESSING_AND_PRESENTATION | 0.00000 | 0.00000 | 0.70498 | 0.64776 | 1.42241 | 143 |
| GO_ANTIGEN_PROCESSING_AND_PRESENTATION_OF_EXOGENOUS_ | 0.00000 | 0.00000 | | 0.81051 | 1.72358 | 57 |
| GO_ANTIGEN_PROCESSING_AND_PRESENTATION_OF_PEPTIDE_ANTI | 0.00000 | 0.00000 | 0.70498 | 0.65912 | 1.44248 | 126 |
| GO_ANTIGEN_PROCESSING_AND_PRESENTATION_OF_PEPTIDE_ANTI | 0.00000 | 0.00000 | | 0.76925 | 1.65015 | 70 |
| GO_ANTIGEN_RECEPTOR_MEDIATED_SIGNALING_PATHWAY | 0.00000 | 0.00000 | 0.80122 | 0.67877 | 1.48641 | 127 |
| GO_APOPTOTIC_SIGNALING_PATHWAY | 0.00154 | 0.02464 | 0.45506 | 0.51561 | 1.15085 | 358 |
| GO_ATP_DEPENDENT_CHROMATIN_REMODELING | 0.00253 | 0.03681 | 0.43171 | 0.62124 | 1.32400 | 61 |
| GO_ATP_METABOLIC_PROCESS | 0.00000 | 0.00000 | 0.71951 | 0.62252 | 1.37693 | 196 |
| GO_ATP_SYNTHESIS_COUPLED_ELECTRON_TRANSPORT | 0.00005 | 0.00131 | 0.55733 | 0.65519 | 1.40723 | 74 |
| GO_BINDING_OF_SPERM_TO_ZONA_PELLUCIDA | 0.00000 | 0.00001 | 0.69013 | 0.89879 | 1.80846 | 18 |
| GO_CADHERIN_BINDING | 0.00000 | 0.00000 | | 0.63068 | 1.39961 | 236 |
| GO_CANONICAL_WNT_SIGNALING_PATHWAY | 0.00000 | 0.00003 | 0.65944 | 0.60816 | 1.34553 | 197 |
| GO_CATALYTIC_STEP_2_SPLICEOSOME | 0.00000 | 0.00002 | 0.65944 | 0.68068 | 1.46976 | 83 |
| GO_CELLULAR_AMINO_ACID_METABOLIC_PROCESS | 0.00000 | 0.00000 | 0.74774 | 0.61752 | 1.36824 | 216 |
| GO_CELLULAR_COMPONENT_DISASSEMBLY | 0.00000 | 0.00005 | 0.64355 | 0.55505 | 1.24197 | 401 |
| GO_CELLULAR_KETONE_METABOLIC_PROCESS | 0.00000 | 0.00000 | 0.77494 | 0.65955 | 1.44893 | 149 |
| GO_CELLULAR_NITROGEN_COMPOUND_CATABOLIC_PROCESS | 0.00000 | 0.00000 | | 0.74079 | 1.65848 | 436 |
| GO_CELLULAR_PROTEIN_COMPLEX_DISASSEMBLY | 0.00006 | 0.00139 | 0.55733 | 0.58222 | 1.28459 | 176 |
| GO_CELLULAR_RESPIRATION | 0.00000 | 0.00000 | 0.71951 | 0.64997 | 1.42492 | 136 |
| GO_CELLULAR_RESPONSE_TO_HEAT | 0.00018 | 0.00403 | 0.51885 | 0.61962 | 1.34016 | 91 |
| GO_CELLULAR_RESPONSE_TO_ORGANIC_CYCLIC_COMPOUND | 0.00302 | 0.04108 | 0.43171 | 0.52402 | 1.16585 | 283 |
| GO_CELLULAR_RESPONSE_TO_OXYGEN_LEVELS | 0.00000 | 0.00000 | 0.78819 | 0.65911 | 1.44882 | 157 |
| GO_CELL_ACTIVATION_INVOLVED_IN_IMMUNE_RESPONSE | 0.00000 | 0.00001 | 0.67496 | 0.57339 | 1.27913 | 341 |
| GO_CELL_ADHESION_MOLECULE_BINDING | 0.00000 | 0.00000 | 0.81404 | 0.60529 | 1.34788 | 308 |
| GO_CELL_CELL_RECOGNITION | 0.00000 | 0.00002 | 0.65944 | 0.83826 | 1.71003 | 26 |
| GO_CELL_CELL_SIGNALING_BY_WNT | 0.00000 | 0.00002 | 0.67496 | 0.57736 | 1.28559 | 299 |
| GO_CELL_CYCLE_G2_M_PHASE_TRANSITION | 0.00001 | 0.00024 | 0.59333 | 0.58034 | 1.28602 | 214 |
| GO_CELL_CYCLE_PHASE_TRANSITION | 0.00000 | 0.00004 | 0.64355 | 0.54319 | 1.21681 | 464 |
| GO_CELL_DIVISION | 0.00154 | 0.02464 | 0.45506 | 0.51551 | 1.15425 | 427 |
| GO_CELL_RECOGNITION | 0.00012 | 0.00270 | 0.53843 | 0.69554 | 1.46186 | 48 |
| GO_CELL_SUBSTRATE_JUNCTION | 0.00000 | 0.00000 | | 0.67751 | 1.50775 | 291 |
| GO_CELL_SURFACE_RECEPTOR_SIGNALING_PATHWAY_INVOLVED_IN | 0.00000 | 0.00003 | 0.65944 | 0.57035 | 1.27067 | 319 |
| GO_CHAPERONE_BINDING | 0.00007 | 0.00163 | 0.53843 | 0.64754 | 1.39253 | 76 |

| | | | | | | |
|---|---|---|---|---|---|---|
| GO_CHAPERONE_COMPLEX | 0.00000 | 0.00004 | 0.64355 | 0.85718 | 1.73925 | 22 |
| GO_CHAPERONE_MEDIATED_PROTEIN_COMPLEX_ASSEMBLY | 0.00170 | 0.02662 | 0.45506 | 0.77423 | 1.54792 | 17 |
| GO_CHROMATIN_BINDING | 0.00071 | 0.01323 | 0.47727 | 0.52699 | 1.17680 | 369 |
| GO_CHROMOSOMAL_REGION | 0.00246 | 0.03601 | 0.43171 | 0.52919 | 1.17661 | 276 |
| GO_COP9_SIGNALOSOME | 0.00128 | 0.02141 | 0.45506 | 0.73616 | 1.49715 | 24 |
| GO_COTRANSLATIONAL_PROTEIN_TARGETING_TO_MEMBRANE | 0.00000 | 0.00000 | | 0.89471 | 1.93513 | 91 |
| GO_CYTOCHROME_COMPLEX | 0.00168 | 0.02641 | 0.45506 | 0.75311 | 1.52417 | 20 |
| GO_CYTOKINE_MEDIATED_SIGNALING_PATHWAY | 0.00000 | 0.00000 | 0.81404 | 0.59698 | 1.33170 | 339 |
| GO_CYTOPLASMIC_TRANSLATION | 0.00000 | 0.00000 | | 0.84753 | 1.83080 | 88 |
| GO_CYTOPLASMIC_TRANSLATIONAL_INITIATION | 0.00000 | 0.00000 | 0.74774 | 0.85688 | 1.76176 | 30 |
| GO_CYTOSOLIC_LARGE_RIBOSOMAL_SUBUNIT | 0.00000 | 0.00000 | | 0.91186 | 1.92988 | 53 |
| GO_CYTOSOLIC_RIBOSOME | 0.00000 | 0.00000 | | 0.89907 | 1.94710 | 95 |
| GO_CYTOSOLIC_SMALL_RIBOSOMAL_SUBUNIT | 0.00000 | 0.00000 | | 0.89114 | 1.85533 | 39 |
| GO_DEFENSE_RESPONSE_TO_OTHER_ORGANISM | 0.00004 | 0.00099 | 0.55733 | 0.53673 | 1.20127 | 415 |
| GO_DE_NOVO_PROTEIN_FOLDING | 0.00050 | 0.00978 | 0.47727 | 0.74232 | 1.51431 | 26 |
| GO_DISORDERED_DOMAIN_SPECIFIC_BINDING | 0.00161 | 0.02546 | 0.45506 | 0.71970 | 1.47197 | 27 |
| GO_DNA_BIOSYNTHETIC_PROCESS | 0.00000 | 0.00010 | 0.62726 | 0.62505 | 1.37017 | 134 |
| GO_DNA_CONFORMATION_CHANGE | 0.00027 | 0.00572 | 0.49849 | 0.55300 | 1.22527 | 216 |
| GO_DNA_DAMAGE_RESPONSE_DETECTION_OF_DNA_DAMAGE | 0.00198 | 0.03028 | 0.43171 | 0.67949 | 1.40669 | 35 |
| GO_DNA_GEOMETRIC_CHANGE | 0.00046 | 0.00925 | 0.49849 | 0.60807 | 1.31855 | 98 |
| GO_DNA_POLYMERASE_BINDING | 0.00268 | 0.03799 | 0.43171 | 0.76255 | 1.52457 | 17 |
| GO_DNA_REPAIR | 0.00265 | 0.03775 | 0.43171 | 0.51159 | 1.14547 | 427 |
| GO_DRUG_BINDING | 0.00176 | 0.02733 | 0.45506 | 0.64119 | 1.35096 | 49 |
| GO_ELECTRON_TRANSFER_ACTIVITY | 0.00001 | 0.00020 | 0.61053 | 0.67481 | 1.44938 | 74 |
| GO_ELECTRON_TRANSPORT_CHAIN | 0.00002 | 0.00053 | 0.57561 | 0.62257 | 1.36145 | 124 |
| GO_ENDOPEPTIDASE_ACTIVITY | 0.00124 | 0.02085 | 0.45506 | 0.54955 | 1.21076 | 171 |
| GO_ENDOPEPTIDASE_COMPLEX | 0.00000 | 0.00000 | | 0.80930 | 1.73234 | 66 |
| GO_ENERGY_DERIVATION_BY_OXIDATION_OF_ORGANIC_COMPOUN | 0.00000 | 0.00000 | 0.71951 | 0.63390 | 1.39707 | 172 |
| GO_ESTABLISHMENT_OF_PROTEIN_LOCALIZATION_TO_CHROMOSON | 0.00020 | 0.00432 | 0.51885 | 0.77339 | 1.57217 | 23 |
| GO_ESTABLISHMENT_OF_PROTEIN_LOCALIZATION_TO_ENDOPLASM | 0.00000 | 0.00000 | | 0.88026 | 1.91149 | 102 |
| GO_ESTABLISHMENT_OF_PROTEIN_LOCALIZATION_TO_MEMBRANE | 0.00000 | 0.00000 | | 0.74874 | 1.66164 | 240 |
| GO_ESTABLISHMENT_OF_PROTEIN_LOCALIZATION_TO_ORGANELLE | 0.00000 | 0.00000 | | 0.69519 | 1.55605 | 423 |
| GO_ESTABLISHMENT_OF_PROTEIN_LOCALIZATION_TO_TELOMERE | 0.00006 | 0.00139 | 0.55733 | 0.83470 | 1.66881 | 17 |
| GO_ESTABLISHMENT_OF_RNA_LOCALIZATION | 0.00106 | 0.01819 | 0.45506 | 0.55577 | 1.22400 | 167 |
| GO_ESTABLISHMENT_OF_TISSUE_POLARITY | 0.00000 | 0.00000 | | 0.75051 | 1.62122 | 88 |
| GO_EUKARYOTIC_TRANSLATION_INITIATION_FACTOR_3_COMPLEX | 0.00003 | 0.00074 | 0.57561 | 0.86987 | 1.72555 | 15 |
| GO_EXOCYTOSIS | 0.00000 | 0.00015 | 0.61053 | 0.53905 | 1.20753 | 465 |
| GO_FC_EPSILON_RECEPTOR_SIGNALING_PATHWAY | 0.00000 | 0.00000 | | 0.75117 | 1.62103 | 82 |
| GO_FC_RECEPTOR_SIGNALING_PATHWAY | 0.00000 | 0.00000 | | 0.72054 | 1.57787 | 127 |
| GO_FERTILIZATION | 0.00089 | 0.01592 | 0.47727 | 0.63620 | 1.35888 | 64 |
| GO_FICOLIN_1_RICH_GRANULE | 0.00000 | 0.00000 | 0.78819 | 0.71716 | 1.55340 | 93 |
| GO_GENERATION_OF_PRECURSOR_METABOLITES_AND_ENERGY | 0.00000 | 0.00000 | 0.77494 | 0.59261 | 1.32026 | 320 |
| GO_GENE_SILENCING | 0.00261 | 0.03750 | 0.43171 | 0.55578 | 1.22252 | 161 |
| GO_GLAND_DEVELOPMENT | 0.00154 | 0.02464 | 0.45506 | 0.54117 | 1.19797 | 206 |
| GO_GLYCOSYL_COMPOUND_METABOLIC_PROCESS | 0.00071 | 0.01323 | 0.47727 | 0.62524 | 1.34204 | 71 |
| GO_GUANYL_NUCLEOTIDE_BINDING | 0.00095 | 0.01679 | 0.47727 | 0.54096 | 1.19861 | 216 |
| GO_HEAT_SHOCK_PROTEIN_BINDING | 0.00003 | 0.00092 | 0.55733 | 0.64369 | 1.39056 | 85 |
| GO_HEMATOPOIETIC_PROGENITOR_CELL_DIFFERENTIATION | 0.00000 | 0.00000 | 0.82666 | 0.70307 | 1.53239 | 113 |
| GO_HEMATOPOIETIC_STEM_CELL_DIFFERENTIATION | 0.00000 | 0.00000 | | 0.79776 | 1.71217 | 72 |
| GO_HSP70_PROTEIN_BINDING | 0.00228 | 0.03391 | 0.43171 | 0.71143 | 1.45504 | 27 |
| GO_IMMUNE_RESPONSE_REGULATING_SIGNALING_PATHWAY | 0.00000 | 0.00000 | 0.76146 | 0.63494 | 1.40480 | 197 |
| GO_INNATE_IMMUNE_RESPONSE | 0.00003 | 0.00071 | 0.57561 | 0.55173 | 1.23107 | 347 |
| GO_INNATE_IMMUNE_RESPONSE_ACTIVATING_SIGNAL_TRANSDUCT | 0.00000 | 0.00000 | | 0.79340 | 1.70281 | 72 |
| GO_INNER_MITOCHONDRIAL_MEMBRANE_PROTEIN_COMPLEX | 0.00002 | 0.00060 | 0.57561 | 0.65124 | 1.40678 | 88 |
| GO_INTERACTION_WITH_SYMBIONT | 0.00001 | 0.00026 | 0.59333 | 0.75084 | 1.56620 | 41 |
| GO_INTERLEUKIN_1_MEDIATED_SIGNALING_PATHWAY | 0.00000 | 0.00000 | | 0.81689 | 1.75671 | 76 |
| GO_INTRINSIC_APOPTOTIC_SIGNALING_PATHWAY | 0.00121 | 0.02029 | 0.45506 | 0.55118 | 1.21845 | 190 |
| GO_ION_CHANNEL_BINDING | 0.00087 | 0.01584 | 0.47727 | 0.63182 | 1.34683 | 62 |
| GO_ISOPEPTIDASE_ACTIVITY | 0.00139 | 0.02285 | 0.45506 | 0.78940 | 1.57097 | 16 |
| GO_LARGE_RIBOSOMAL_SUBUNIT | 0.00000 | 0.00000 | | 0.78135 | 1.70183 | 109 |
| GO_LEUKOCYTE_MEDIATED_IMMUNITY | 0.00000 | 0.00000 | 0.71951 | 0.57282 | 1.27895 | 364 |
| GO_MACROMOLECULE_TRANSMEMBRANE_TRANSPORTER_ACTIVITY | 0.00239 | 0.03510 | 0.43171 | 0.77734 | 1.54696 | 16 |
| GO_MAINTENANCE_OF_CELL_NUMBER | 0.00087 | 0.01584 | 0.47727 | 0.58944 | 1.28512 | 114 |
| GO_MATURATION_OF_LSU_RRNA | 0.00006 | 0.00151 | 0.53843 | 0.77606 | 1.58723 | 27 |
| GO_METHYLTRANSFERASE_COMPLEX | 0.00290 | 0.04014 | 0.43171 | 0.59476 | 1.28479 | 88 |
| GO_METHYL_CPG_BINDING | 0.00302 | 0.04108 | 0.43171 | 0.78053 | 1.54833 | 15 |
| GO_MIDBRAIN_DEVELOPMENT | 0.00021 | 0.00468 | 0.51885 | 0.68832 | 1.44457 | 47 |
| GO_MITOCHONDRIAL_ELECTRON_TRANSPORT_NADH_TO_UBIQUIN | 0.00331 | 0.04437 | 0.43171 | 0.65428 | 1.36724 | 42 |
| GO_MITOCHONDRIAL_GENE_EXPRESSION | 0.00048 | 0.00949 | 0.49849 | 0.57608 | 1.26525 | 152 |
| GO_MITOCHONDRIAL_LARGE_RIBOSOMAL_SUBUNIT | 0.00002 | 0.00050 | 0.57561 | 0.69789 | 1.48086 | 55 |
| GO_MITOCHONDRIAL_MATRIX | 0.00000 | 0.00002 | 0.65944 | 0.56045 | 1.25158 | 371 |
| GO_MITOCHONDRIAL_MEMBRANE_ORGANIZATION | 0.00361 | 0.04734 | 0.43171 | 0.59309 | 1.28094 | 86 |
| GO_MITOCHONDRIAL_PROTEIN_COMPLEX | 0.00000 | 0.00000 | 0.81404 | 0.63650 | 1.40931 | 208 |
| GO_MITOCHONDRIAL_TRANSLATION | 0.00004 | 0.00093 | 0.55733 | 0.61219 | 1.34135 | 129 |
| GO_MITOCHONDRIAL_TRANSLATIONAL_TERMINATION | 0.00000 | 0.00010 | 0.62726 | 0.66694 | 1.44128 | 87 |
| GO_MITOCHONDRION_ORGANIZATION | 0.00051 | 0.00993 | 0.47727 | 0.52284 | 1.16863 | 390 |

| | | | | | | |
|---|---|---|---|---|---|---|
| GO_MODIFICATION_DEPENDENT_MACROMOLECULE_CATABOLIC_PR | 0.00002 | 0.00053 | 0.57561 | 0.53142 | 1.19156 | 486 |
| GO_MODULATION_BY_HOST_OF_SYMBIONT_PROCESS | 0.00015 | 0.00336 | 0.51885 | 0.72848 | 1.50542 | 33 |
| GO_MODULATION_OF_PROCESS_OF_OTHER_ORGANISM | 0.00019 | 0.00423 | 0.51885 | 0.66904 | 1.42286 | 59 |
| GO_MONOSACCHARIDE_BIOSYNTHETIC_PROCESS | 0.00335 | 0.04466 | 0.43171 | 0.61992 | 1.31781 | 58 |
| GO_MORPHOGENESIS_OF_AN_EPITHELIUM | 0.00006 | 0.00151 | 0.53843 | 0.55161 | 1.22755 | 290 |
| GO_MORPHOGENESIS_OF_A_POLARIZED_EPITHELIUM | 0.00000 | 0.00000 | | 0.72298 | 1.57057 | 103 |
| GO_MRNA_3_UTR_BINDING | 0.00003 | 0.00089 | 0.55733 | 0.67596 | 1.44827 | 68 |
| GO_MRNA_5_UTR_BINDING | 0.00006 | 0.00156 | 0.53843 | 0.80133 | 1.62592 | 22 |
| GO_MRNA_BINDING | 0.00000 | 0.00000 | | 0.66002 | 1.46353 | 222 |
| GO_MRNA_PROCESSING | 0.00000 | 0.00000 | 0.73376 | 0.56859 | 1.27295 | 418 |
| GO_MULTI_ORGANISM_LOCALIZATION | 0.00268 | 0.03799 | 0.43171 | 0.63245 | 1.33612 | 51 |
| GO_MULTI_ORGANISM_PROCESS | 0.00108 | 0.01839 | 0.45506 | 0.51114 | 1.14599 | 484 |
| GO_MYD88_DEPENDENT_TOLL_LIKE_RECEPTOR_SIGNALING_PATHW | 0.00002 | 0.00066 | 0.57561 | 0.85581 | 1.70312 | 16 |
| GO_MYELOID_LEUKOCYTE_ACTIVATION | 0.00000 | 0.00000 | 0.74774 | 0.58508 | 1.30479 | 327 |
| GO_MYELOID_LEUKOCYTE_MEDIATED_IMMUNITY | 0.00000 | 0.00000 | 0.76146 | 0.59841 | 1.33291 | 301 |
| GO_NADH_DEHYDROGENASE_ACTIVITY | 0.00383 | 0.04935 | 0.43171 | 0.66244 | 1.37139 | 35 |
| GO_NADH_DEHYDROGENASE_COMPLEX | 0.00376 | 0.04884 | 0.43171 | 0.65649 | 1.36421 | 38 |
| GO_NADH_METABOLIC_PROCESS | 0.00294 | 0.04036 | 0.43171 | 0.70642 | 1.44481 | 27 |
| GO_NCRNA_EXPORT_FROM_NUCLEUS | 0.00302 | 0.04108 | 0.43171 | 0.66820 | 1.38331 | 35 |
| GO_NCRNA_METABOLIC_PROCESS | 0.00000 | 0.00003 | 0.65944 | 0.55907 | 1.24907 | 381 |
| GO_NCRNA_PROCESSING | 0.00000 | 0.00003 | 0.65944 | 0.57175 | 1.27504 | 328 |
| GO_NEGATIVE_REGULATION_OF_CANONICAL_WNT_SIGNALING_PAT | 0.00000 | 0.00000 | | 0.71611 | 1.56038 | 111 |
| GO_NEGATIVE_REGULATION_OF_CELLULAR_AMIDE_METABOLIC_PR | 0.00001 | 0.00020 | 0.61053 | 0.63031 | 1.37423 | 114 |
| GO_NEGATIVE_REGULATION_OF_CELL_CYCLE | 0.00003 | 0.00076 | 0.57561 | 0.53833 | 1.20513 | 425 |
| GO_NEGATIVE_REGULATION_OF_CELL_CYCLE_G2_M_PHASE_TRANS | 0.00000 | 0.00000 | 0.76146 | 0.71490 | 1.54439 | 85 |
| GO_NEGATIVE_REGULATION_OF_CELL_CYCLE_PHASE_TRANSITION | 0.00002 | 0.00060 | 0.57561 | 0.59041 | 1.30442 | 184 |
| GO_NEGATIVE_REGULATION_OF_CELL_CYCLE_PROCESS | 0.00000 | 0.00002 | 0.65944 | 0.58690 | 1.30374 | 249 |
| GO_NEGATIVE_REGULATION_OF_MITOTIC_CELL_CYCLE | 0.00004 | 0.00095 | 0.55733 | 0.56735 | 1.25902 | 232 |
| GO_NEGATIVE_REGULATION_OF_MRNA_METABOLIC_PROCESS | 0.00023 | 0.00491 | 0.51885 | 0.66670 | 1.41788 | 59 |
| GO_NEGATIVE_REGULATION_OF_MRNA_SPLICING_VIA_SPLICEOSOM | 0.00239 | 0.03510 | 0.43171 | 0.74857 | 1.51204 | 19 |
| GO_NEGATIVE_REGULATION_OF_RNA_SPLICING | 0.00029 | 0.00607 | 0.49849 | 0.76729 | 1.55976 | 23 |
| GO_NEGATIVE_REGULATION_OF_TRANSLATIONAL_INITIATION | 0.00018 | 0.00403 | 0.51885 | 0.83901 | 1.66434 | 15 |
| GO_NEGATIVE_REGULATION_OF_WNT_SIGNALING_PATHWAY | 0.00000 | 0.00000 | | 0.69525 | 1.52249 | 127 |
| GO_NEURAL_NUCLEUS_DEVELOPMENT | 0.00001 | 0.00026 | 0.59333 | 0.79148 | 1.62187 | 28 |
| GO_NEURON_TO_NEURON_SYNAPSE | 0.00005 | 0.00134 | 0.55733 | 0.59636 | 1.31006 | 154 |
| GO_NIK_NF_KAPPAB_SIGNALING | 0.00000 | 0.00000 | | 0.72896 | 1.58354 | 103 |
| GO_NON_CANONICAL_WNT_SIGNALING_PATHWAY | 0.00000 | 0.00000 | | 0.72784 | 1.58023 | 101 |
| GO_NUCLEAR_EXPORT | 0.00031 | 0.00645 | 0.49849 | 0.56963 | 1.25542 | 172 |
| GO_NUCLEAR_TRANSCRIBED_MRNA_CATABOLIC_PROCESS | 0.00000 | 0.00000 | | 0.79179 | 1.74917 | 183 |
| GO_NUCLEAR_TRANSCRIBED_MRNA_CATABOLIC_PROCESS_NONSEN | 0.00000 | 0.00000 | | 0.89008 | 1.93909 | 110 |
| GO_NUCLEAR_TRANSPORT | 0.00008 | 0.00193 | 0.53843 | 0.55511 | 1.23408 | 273 |
| GO_NUCLEOBASE_CONTAINING_COMPOUND_TRANSPORT | 0.00146 | 0.02386 | 0.45506 | 0.55053 | 1.21842 | 202 |
| GO_NUCLEOBASE_CONTAINING_SMALL_MOLECULE_BIOSYNTHETIC_ | 0.00137 | 0.02261 | 0.45506 | 0.62749 | 1.34353 | 67 |
| GO_NUCLEOBASE_CONTAINING_SMALL_MOLECULE_INTERCONVERS | 0.00383 | 0.04935 | 0.43171 | 0.71259 | 1.44857 | 23 |
| GO_NUCLEOBASE_CONTAINING_SMALL_MOLECULE_METABOLIC_PR | 0.00320 | 0.04339 | 0.43171 | 0.51448 | 1.14825 | 352 |
| GO_NUCLEOSIDE_DIPHOSPHATE_METABOLIC_PROCESS | 0.00110 | 0.01865 | 0.45506 | 0.59637 | 1.29295 | 97 |
| GO_NUCLEOSIDE_METABOLIC_PROCESS | 0.00061 | 0.01169 | 0.47727 | 0.65315 | 1.38907 | 59 |
| GO_NUCLEOSIDE_MONOPHOSPHATE_BIOSYNTHETIC_PROCESS | 0.00250 | 0.03636 | 0.43171 | 0.68841 | 1.42033 | 32 |
| GO_NUCLEOSIDE_MONOPHOSPHATE_METABOLIC_PROCESS | 0.00152 | 0.02464 | 0.45506 | 0.64377 | 1.36002 | 51 |
| GO_NUCLEOSIDE_PHOSPHATE_BIOSYNTHETIC_PROCESS | 0.00043 | 0.00865 | 0.49849 | 0.58185 | 1.27831 | 148 |
| GO_NUCLEOSIDE_TRIPHOSPHATE_BIOSYNTHETIC_PROCESS | 0.00043 | 0.00865 | 0.49849 | 0.69332 | 1.44539 | 40 |
| GO_NUCLEOSIDE_TRIPHOSPHATE_METABOLIC_PROCESS | 0.00016 | 0.00360 | 0.51885 | 0.66788 | 1.41976 | 58 |
| GO_NUCLEOSOMAL_DNA_BINDING | 0.00023 | 0.00507 | 0.51885 | 0.79852 | 1.61801 | 21 |
| GO_NUCLEOSOME_BINDING | 0.00143 | 0.02332 | 0.45506 | 0.66132 | 1.38492 | 45 |
| GO_NUCLEOTIDE_EXCISION_REPAIR | 0.00004 | 0.00096 | 0.55733 | 0.63343 | 1.37355 | 98 |
| GO_NUCLEOTIDE_EXCISION_REPAIR_DNA_DAMAGE_RECOGNITION | 0.00015 | 0.00349 | 0.51885 | 0.78678 | 1.59640 | 22 |
| GO_NUCLEOTIDE_EXCISION_REPAIR_DNA_INCISION | 0.00313 | 0.04249 | 0.43171 | 0.66439 | 1.37898 | 36 |
| GO_NUCLEOTIDE_PHOSPHORYLATION | 0.00383 | 0.04935 | 0.43171 | 0.59002 | 1.27454 | 88 |
| GO_ORGANELLAR_RIBOSOME | 0.00000 | 0.00014 | 0.61053 | 0.66412 | 1.43471 | 85 |
| GO_ORGANELLE_INNER_MEMBRANE | 0.00000 | 0.00000 | 0.76146 | 0.57601 | 1.28697 | 378 |
| GO_ORGANIC_CYCLIC_COMPOUND_CATABOLIC_PROCESS | 0.00000 | 0.00000 | | 0.73350 | 1.64281 | 449 |
| GO_OXIDATIVE_PHOSPHORYLATION | 0.00001 | 0.00030 | 0.59333 | 0.64619 | 1.40027 | 96 |
| GO_OXIDOREDUCTASE_ACTIVITY | 0.00006 | 0.00139 | 0.55733 | 0.53544 | 1.19703 | 392 |
| GO_OXIDOREDUCTASE_COMPLEX | 0.00001 | 0.00027 | 0.59333 | 0.66015 | 1.42461 | 82 |
| GO_PEPTIDASE_COMPLEX | 0.00000 | 0.00000 | | 0.75988 | 1.63982 | 82 |
| GO_PIGMENT_GRANULE | 0.00003 | 0.00089 | 0.55733 | 0.66098 | 1.42344 | 78 |
| GO_POLYSOMAL_RIBOSOME | 0.00000 | 0.00000 | | 0.90624 | 1.86629 | 31 |
| GO_POLYSOME | 0.00000 | 0.00000 | | 0.82337 | 1.75481 | 61 |
| GO_POLY_PYRIMIDINE_TRACT_BINDING | 0.00327 | 0.04398 | 0.43171 | 0.75471 | 1.51855 | 18 |
| GO_POSITIVE_REGULATION_OF_APOPTOTIC_SIGNALING_PATHWAY | 0.00279 | 0.03889 | 0.43171 | 0.57666 | 1.25518 | 107 |
| GO_POSITIVE_REGULATION_OF_ATPASE_ACTIVITY | 0.00294 | 0.04036 | 0.43171 | 0.65446 | 1.36795 | 43 |
| GO_POSITIVE_REGULATION_OF_BINDING | 0.00088 | 0.01590 | 0.47727 | 0.60284 | 1.30556 | 95 |
| GO_POSITIVE_REGULATION_OF_CANONICAL_WNT_SIGNALING_PATH | 0.00000 | 0.00000 | 0.77494 | 0.70504 | 1.53160 | 103 |
| GO_POSITIVE_REGULATION_OF_CELLULAR_AMIDE_METABOLIC_PRO | 0.00003 | 0.00074 | 0.57561 | 0.62296 | 1.36083 | 118 |
| GO_POSITIVE_REGULATION_OF_CELLULAR_PROTEIN_LOCALIZATION | 0.00041 | 0.00840 | 0.49849 | 0.54915 | 1.21815 | 225 |
| GO_POSITIVE_REGULATION_OF_CELL_DEATH | 0.00069 | 0.01297 | 0.47727 | 0.52595 | 1.17423 | 363 |

| | | | | | | |
|---|---|---|---|---|---|---|
| GO_POSITIVE_REGULATION_OF_CELL_POPULATION_PROLIFERATION | 0.00078 | 0.01455 | 0.47727 | 0.52449 | 1.17108 | 368 |
| GO_POSITIVE_REGULATION_OF_CHROMOSOME_ORGANIZATION | 0.00231 | 0.03437 | 0.43171 | 0.56990 | 1.24871 | 133 |
| GO_POSITIVE_REGULATION_OF_DEFENSE_RESPONSE | 0.00000 | 0.00003 | 0.65944 | 0.62175 | 1.36730 | 158 |
| GO_POSITIVE_REGULATION_OF_DNA_BIOSYNTHETIC_PROCESS | 0.00001 | 0.00041 | 0.59333 | 0.71929 | 1.52055 | 52 |
| GO_POSITIVE_REGULATION_OF_DNA_METABOLIC_PROCESS | 0.00364 | 0.04761 | 0.43171 | 0.56647 | 1.24189 | 135 |
| GO_POSITIVE_REGULATION_OF_DNA_TEMPLATED_TRANSCRIPTION_ | 0.00276 | 0.03885 | 0.43171 | 0.68932 | 1.41586 | 29 |
| GO_POSITIVE_REGULATION_OF_ESTABLISHMENT_OF_PROTEIN_LOCA | 0.00007 | 0.00168 | 0.53843 | 0.56461 | 1.25165 | 219 |
| GO_POSITIVE_REGULATION_OF_IMMUNE_RESPONSE | 0.00000 | 0.00001 | 0.69013 | 0.59037 | 1.31265 | 277 |
| GO_POSITIVE_REGULATION_OF_IMMUNE_SYSTEM_PROCESS | 0.00004 | 0.00110 | 0.55733 | 0.54018 | 1.20716 | 386 |
| GO_POSITIVE_REGULATION_OF_INNATE_IMMUNE_RESPONSE | 0.00000 | 0.00000 | 0.81404 | 0.69693 | 1.51947 | 114 |
| GO_POSITIVE_REGULATION_OF_INTRINSIC_APOPTOTIC_SIGNALING_ | 0.00104 | 0.01798 | 0.45506 | 0.69097 | 1.43578 | 37 |
| GO_POSITIVE_REGULATION_OF_ORGANELLE_ORGANIZATION | 0.00043 | 0.00865 | 0.49849 | 0.52093 | 1.16605 | 411 |
| GO_POSITIVE_REGULATION_OF_PROTEIN_LOCALIZATION_TO_NUCLE | 0.00001 | 0.00020 | 0.61053 | 0.69907 | 1.48990 | 61 |
| GO_POSITIVE_REGULATION_OF_RESPONSE_TO_BIOTIC_STIMULUS | 0.00000 | 0.00000 | 0.73376 | 0.66340 | 1.45310 | 128 |
| GO_POSITIVE_REGULATION_OF_RESPONSE_TO_EXTERNAL_STIMULU | 0.00002 | 0.00050 | 0.57561 | 0.58133 | 1.28681 | 201 |
| GO_POSITIVE_REGULATION_OF_TELOMERASE_ACTIVITY | 0.00061 | 0.01169 | 0.47727 | 0.73476 | 1.50565 | 28 |
| GO_POSITIVE_REGULATION_OF_TELOMERASE_RNA_LOCALIZATION_ | 0.00000 | 0.00001 | 0.69013 | 0.92451 | 1.83395 | 15 |
| GO_POSITIVE_REGULATION_OF_TELOMERE_MAINTENANCE_VIA_TE | 0.00130 | 0.02159 | 0.45506 | 0.71259 | 1.46508 | 30 |
| GO_POSITIVE_REGULATION_OF_TRANSCRIPTION_INITIATION_FROM | 0.00372 | 0.04847 | 0.43171 | 0.70671 | 1.43993 | 25 |
| GO_POSITIVE_REGULATION_OF_TRANSFERASE_ACTIVITY | 0.00174 | 0.02712 | 0.45506 | 0.51642 | 1.15220 | 345 |
| GO_POSITIVE_REGULATION_OF_TRANSLATION | 0.00001 | 0.00019 | 0.61053 | 0.64767 | 1.40419 | 97 |
| GO_POSITIVE_REGULATION_OF_VIRAL_GENOME_REPLICATION | 0.00287 | 0.03973 | 0.43171 | 0.68362 | 1.40783 | 31 |
| GO_POSITIVE_REGULATION_OF_VIRAL_PROCESS | 0.00028 | 0.00590 | 0.49849 | 0.62440 | 1.34741 | 81 |
| GO_POSITIVE_REGULATION_OF_WNT_SIGNALING_PATHWAY | 0.00000 | 0.00000 | 0.73376 | 0.66230 | 1.44763 | 122 |
| GO_POSTREPLICATION_REPAIR | 0.00024 | 0.00515 | 0.51885 | 0.69140 | 1.44518 | 43 |
| GO_POSTSYNAPSE | 0.00004 | 0.00109 | 0.55733 | 0.55969 | 1.24369 | 259 |
| GO_POSTSYNAPTIC_CYTOSOL | 0.00279 | 0.03889 | 0.43171 | 0.78295 | 1.55313 | 15 |
| GO_POSTTRANSCRIPTIONAL_REGULATION_OF_GENE_EXPRESSION | 0.00000 | 0.00000 | | 0.66656 | 1.49327 | 462 |
| GO_POST_TRANSLATIONAL_PROTEIN_MODIFICATION | 0.00002 | 0.00052 | 0.57561 | 0.57851 | 1.28412 | 229 |
| GO_PRECATALYTIC_SPLICEOSOME | 0.00085 | 0.01559 | 0.47727 | 0.65235 | 1.37448 | 49 |
| GO_PRERIBOSOME | 0.00087 | 0.01577 | 0.47727 | 0.62899 | 1.34772 | 69 |
| GO_PRERIBOSOME_LARGE_SUBUNIT_PRECURSOR | 0.00235 | 0.03474 | 0.43171 | 0.74079 | 1.50308 | 22 |
| GO_PRE_MRNA_BINDING | 0.00133 | 0.02213 | 0.45506 | 0.72393 | 1.48062 | 27 |
| GO_PROTEASOMAL_PROTEIN_CATABOLIC_PROCESS | 0.00002 | 0.00044 | 0.59333 | 0.54166 | 1.21005 | 377 |
| GO_PROTEASOMAL_UBIQUITIN_INDEPENDENT_PROTEIN_CATABOLI | 0.00000 | 0.00000 | 0.80122 | 0.92254 | 1.86931 | 21 |
| GO_PROTEASOME_ACCESSORY_COMPLEX | 0.00000 | 0.00001 | 0.67496 | 0.85464 | 1.73811 | 24 |
| GO_PROTEASOME_CORE_COMPLEX | 0.00000 | 0.00000 | | 0.94375 | 1.90631 | 19 |
| GO_PROTEIN_CONTAINING_COMPLEX_DISASSEMBLY | 0.00000 | 0.00007 | 0.62726 | 0.57810 | 1.28438 | 256 |
| GO_PROTEIN_CONTAINING_COMPLEX_LOCALIZATION | 0.00063 | 0.01197 | 0.47727 | 0.55335 | 1.22520 | 208 |
| GO_PROTEIN_DNA_COMPLEX_SUBUNIT_ORGANIZATION | 0.00001 | 0.00018 | 0.61053 | 0.60174 | 1.32491 | 165 |
| GO_PROTEIN_FOLDING | 0.00000 | 0.00002 | 0.65944 | 0.63113 | 1.38644 | 154 |
| GO_PROTEIN_FOLDING_CHAPERONE | 0.00135 | 0.02238 | 0.45506 | 0.77903 | 1.55752 | 17 |
| GO_PROTEIN_IMPORT | 0.00268 | 0.03799 | 0.43171 | 0.56317 | 1.23581 | 140 |
| GO_PROTEIN_LOCALIZATION_TO_CHROMOSOME | 0.00056 | 0.01096 | 0.47727 | 0.63523 | 1.36108 | 69 |
| GO_PROTEIN_LOCALIZATION_TO_CHROMOSOME_TELOMERIC_REGI | 0.00335 | 0.04466 | 0.43171 | 0.70399 | 1.43984 | 27 |
| GO_PROTEIN_LOCALIZATION_TO_ENDOPLASMIC_RETICULUM | 0.00000 | 0.00000 | | 0.85521 | 1.87021 | 124 |
| GO_PROTEIN_LOCALIZATION_TO_MEMBRANE | 0.00000 | 0.00000 | | 0.65123 | 1.45714 | 406 |
| GO_PROTEIN_LOCALIZATION_TO_MITOCHONDRION | 0.00019 | 0.00420 | 0.51885 | 0.60773 | 1.32398 | 110 |
| GO_PROTEIN_LOCALIZATION_TO_NUCLEUS | 0.00002 | 0.00055 | 0.57561 | 0.58926 | 1.30220 | 187 |
| GO_PROTEIN_MODIFICATION_BY_SMALL_PROTEIN_REMOVAL | 0.00000 | 0.00000 | 0.71951 | 0.60745 | 1.34689 | 221 |
| GO_PROTEIN_POLYUBIQUITINATION | 0.00004 | 0.00106 | 0.55733 | 0.55874 | 1.24108 | 253 |
| GO_PROTEIN_REFOLDING | 0.00294 | 0.04036 | 0.43171 | 0.77039 | 1.53314 | 16 |
| GO_PROTEIN_STABILIZATION | 0.00001 | 0.00024 | 0.61053 | 0.62333 | 1.36586 | 130 |
| GO_PROTEIN_TARGETING | 0.00000 | 0.00000 | | 0.71181 | 1.58743 | 327 |
| GO_PROTEIN_TARGETING_TO_MEMBRANE | 0.00000 | 0.00000 | | 0.82370 | 1.80953 | 149 |
| GO_PROTEIN_TARGETING_TO_MITOCHONDRION | 0.00346 | 0.04593 | 0.43171 | 0.60408 | 1.30091 | 78 |
| GO_PROTEIN_TRANSMEMBRANE_TRANSPORT | 0.00265 | 0.03775 | 0.43171 | 0.64528 | 1.36171 | 50 |
| GO_PROTON_TRANSMEMBRANE_TRANSPORT | 0.00213 | 0.03229 | 0.43171 | 0.60815 | 1.30536 | 71 |
| GO_PURINE_CONTAINING_COMPOUND_BIOSYNTHETIC_PROCESS | 0.00246 | 0.03601 | 0.43171 | 0.57796 | 1.25936 | 111 |
| GO_PURINE_CONTAINING_COMPOUND_METABOLIC_PROCESS | 0.00100 | 0.01761 | 0.45506 | 0.53679 | 1.19242 | 255 |
| GO_PURINE_NUCLEOSIDE_TRIPHOSPHATE_METABOLIC_PROCESS | 0.00089 | 0.01592 | 0.47727 | 0.67157 | 1.40374 | 43 |
| GO_PYRIMIDINE_CONTAINING_COMPOUND_METABOLIC_PROCESS | 0.00353 | 0.04658 | 0.43171 | 0.64744 | 1.35878 | 47 |
| GO_REGENERATION | 0.00216 | 0.03242 | 0.43171 | 0.59987 | 1.29453 | 82 |
| GO_REGULATION_OF_ANIMAL_ORGAN_MORPHOGENESIS | 0.00000 | 0.00000 | | 0.69163 | 1.51541 | 133 |
| GO_REGULATION_OF_APOPTOTIC_SIGNALING_PATHWAY | 0.00031 | 0.00645 | 0.49849 | 0.55171 | 1.22506 | 247 |
| GO_REGULATION_OF_ATPASE_ACTIVITY | 0.00324 | 0.04369 | 0.43171 | 0.63031 | 1.34038 | 57 |
| GO_REGULATION_OF_ATP_METABOLIC_PROCESS | 0.00353 | 0.04658 | 0.43171 | 0.60176 | 1.29780 | 80 |
| GO_REGULATION_OF_BINDING | 0.00250 | 0.03636 | 0.43171 | 0.54476 | 1.20598 | 203 |
| GO_REGULATION_OF_CELLULAR_AMIDE_METABOLIC_PROCESS | 0.00000 | 0.00000 | | 0.62967 | 1.40226 | 311 |
| GO_REGULATION_OF_CELLULAR_AMINE_METABOLIC_PROCESS | 0.00000 | 0.00000 | | 0.82173 | 1.74682 | 58 |
| GO_REGULATION_OF_CELLULAR_AMINO_ACID_METABOLIC_PROCES | 0.00000 | 0.00000 | | 0.83776 | 1.77766 | 55 |
| GO_REGULATION_OF_CELLULAR_KETONE_METABOLIC_PROCESS | 0.00000 | 0.00000 | | 0.72641 | 1.58680 | 118 |
| GO_REGULATION_OF_CELLULAR_PROTEIN_LOCALIZATION | 0.00082 | 0.01513 | 0.47727 | 0.52542 | 1.17290 | 362 |
| GO_REGULATION_OF_CELLULAR_RESPONSE_TO_HEAT | 0.00276 | 0.03885 | 0.43171 | 0.61652 | 1.32004 | 67 |
| GO_REGULATION_OF_CELLULAR_RESPONSE_TO_STRESS | 0.00224 | 0.03344 | 0.43171 | 0.50545 | 1.13232 | 468 |
| GO_REGULATION_OF_CELL_CYCLE_G2_M_PHASE_TRANSITION | 0.00000 | 0.00000 | 0.71951 | 0.63230 | 1.39282 | 169 |

| | | | | | | |
|---|---|---|---|---|---|---|
| GO_REGULATION_OF_CELL_CYCLE_PHASE_TRANSITION | 0.00001 | 0.00026 | 0.59333 | 0.55488 | 1.23783 | 337 |
| GO_REGULATION_OF_CHROMOSOME_ORGANIZATION | 0.00081 | 0.01501 | 0.47727 | 0.54255 | 1.20527 | 261 |
| GO_REGULATION_OF_CYSTEINE_TYPE_ENDOPEPTIDASE_ACTIVITY | 0.00202 | 0.03077 | 0.43171 | 0.57120 | 1.24837 | 121 |
| GO_REGULATION_OF_CYTOPLASMIC_TRANSLATION | 0.00130 | 0.02159 | 0.45506 | 0.73678 | 1.49773 | 23 |
| GO_REGULATION_OF_DEFENSE_RESPONSE | 0.00065 | 0.01237 | 0.47727 | 0.53633 | 1.19355 | 290 |
| GO_REGULATION_OF_DNA_BIOSYNTHETIC_PROCESS | 0.00016 | 0.00360 | 0.51885 | 0.64215 | 1.37923 | 74 |
| GO_REGULATION_OF_DNA_METABOLIC_PROCESS | 0.00069 | 0.01297 | 0.47727 | 0.54425 | 1.20775 | 237 |
| GO_REGULATION_OF_DNA_TEMPLATED_TRANSCRIPTION_INITIATIOI | 0.00163 | 0.02568 | 0.45506 | 0.68289 | 1.41372 | 35 |
| GO_REGULATION_OF_DNA_TEMPLATED_TRANSCRIPTION_IN_RESPO | 0.00000 | 0.00000 | | 0.73944 | 1.59800 | 89 |
| GO_REGULATION_OF_GENERATION_OF_PRECURSOR_METABOLITES_ | 0.00060 | 0.01164 | 0.47727 | 0.60476 | 1.31116 | 97 |
| GO_REGULATION_OF_HEMATOPOIETIC_PROGENITOR_CELL_DIFFERE | 0.00000 | 0.00000 | | 0.78717 | 1.69251 | 75 |
| GO_REGULATION_OF_HEMATOPOIETIC_STEM_CELL_DIFFERENTIATIC | 0.00000 | 0.00000 | | 0.81505 | 1.74256 | 65 |
| GO_REGULATION_OF_HEMOPOIESIS | 0.00000 | 0.00008 | 0.62726 | 0.58859 | 1.30634 | 228 |
| GO_REGULATION_OF_IMMUNE_RESPONSE | 0.00000 | 0.00016 | 0.61053 | 0.55362 | 1.23583 | 362 |
| GO_REGULATION_OF_INNATE_IMMUNE_RESPONSE | 0.00000 | 0.00000 | 0.71951 | 0.63883 | 1.40521 | 161 |
| GO_REGULATION_OF_INTRINSIC_APOPTOTIC_SIGNALING_PATHWAY | 0.00001 | 0.00039 | 0.59333 | 0.63478 | 1.38259 | 109 |
| GO_REGULATION_OF_MITOTIC_CELL_CYCLE | 0.00001 | 0.00018 | 0.61053 | 0.53899 | 1.20736 | 458 |
| GO_REGULATION_OF_MORPHOGENESIS_OF_AN_EPITHELIUM | 0.00000 | 0.00000 | | 0.72213 | 1.57284 | 109 |
| GO_REGULATION_OF_MRNA_CATABOLIC_PROCESS | 0.00000 | 0.00000 | | 0.69977 | 1.54127 | 168 |
| GO_REGULATION_OF_MRNA_METABOLIC_PROCESS | 0.00000 | 0.00000 | | 0.66528 | 1.47872 | 266 |
| GO_REGULATION_OF_MRNA_PROCESSING | 0.00004 | 0.00105 | 0.55733 | 0.61521 | 1.34090 | 113 |
| GO_REGULATION_OF_MRNA_SPLICING_VIA_SPLICEOSOME | 0.00001 | 0.00026 | 0.59333 | 0.67424 | 1.45411 | 80 |
| GO_REGULATION_OF_NUCLEASE_ACTIVITY | 0.00106 | 0.01819 | 0.45506 | 0.77918 | 1.56780 | 18 |
| GO_REGULATION_OF_OXIDATIVE_STRESS_INDUCED_CELL_DEATH | 0.00198 | 0.03028 | 0.43171 | 0.67287 | 1.40355 | 41 |
| GO_REGULATION_OF_PEPTIDASE_ACTIVITY | 0.00050 | 0.00978 | 0.47727 | 0.56122 | 1.24054 | 191 |
| GO_REGULATION_OF_POSTTRANSCRIPTIONAL_GENE_SILENCING | 0.00364 | 0.04761 | 0.43171 | 0.59824 | 1.28492 | 74 |
| GO_REGULATION_OF_PROTEIN_CATABOLIC_PROCESS | 0.00179 | 0.02783 | 0.45506 | 0.53057 | 1.17929 | 270 |
| GO_REGULATION_OF_PROTEIN_LOCALIZATION_TO_NUCLEUS | 0.00001 | 0.00042 | 0.59333 | 0.65505 | 1.41558 | 87 |
| GO_REGULATION_OF_PROTEIN_MODIFICATION_BY_SMALL_PROTEIN | 0.00025 | 0.00532 | 0.49849 | 0.57068 | 1.25684 | 167 |
| GO_REGULATION_OF_PROTEIN_STABILITY | 0.00093 | 0.01652 | 0.47727 | 0.55382 | 1.22571 | 202 |
| GO_REGULATION_OF_PROTEOLYSIS | 0.00013 | 0.00301 | 0.51885 | 0.52997 | 1.18392 | 377 |
| GO_REGULATION_OF_RESPONSE_TO_BIOTIC_STIMULUS | 0.00001 | 0.00020 | 0.61053 | 0.59049 | 1.30719 | 204 |
| GO_REGULATION_OF_RNA_SPLICING | 0.00000 | 0.00001 | 0.69013 | 0.65896 | 1.43669 | 114 |
| GO_REGULATION_OF_SMALL_MOLECULE_METABOLIC_PROCESS | 0.00000 | 0.00000 | 0.71951 | 0.59134 | 1.31394 | 265 |
| GO_REGULATION_OF_STEM_CELL_DIFFERENTIATION | 0.00000 | 0.00000 | | 0.74049 | 1.60113 | 90 |
| GO_REGULATION_OF_SYMBIOTIC_PROCESS | 0.00006 | 0.00144 | 0.55733 | 0.60071 | 1.31682 | 134 |
| GO_REGULATION_OF_TELOMERASE_ACTIVITY | 0.00088 | 0.01590 | 0.47727 | 0.69161 | 1.43179 | 35 |
| GO_REGULATION_OF_TRANSCRIPTION_FROM_RNA_POLYMERASE_II | 0.00000 | 0.00000 | | 0.81194 | 1.73800 | 66 |
| GO_REGULATION_OF_TRANSCRIPTION_INITIATION_FROM_RNA_POL | 0.00143 | 0.02332 | 0.45506 | 0.70390 | 1.44582 | 29 |
| GO_REGULATION_OF_TRANSLATIONAL_INITIATION | 0.00002 | 0.00052 | 0.57561 | 0.68129 | 1.45518 | 64 |
| GO_REGULATION_OF_UBIQUITIN_PROTEIN_TRANSFERASE_ACTIVITY | 0.00061 | 0.01169 | 0.47727 | 0.69335 | 1.44627 | 41 |
| GO_REGULATION_OF_VIRAL_GENOME_REPLICATION | 0.00086 | 0.01566 | 0.47727 | 0.65316 | 1.38389 | 54 |
| GO_REGULATION_OF_VIRAL_TRANSCRIPTION | 0.00198 | 0.03028 | 0.43171 | 0.63932 | 1.34703 | 49 |
| GO_REGULATION_OF_WNT_SIGNALING_PATHWAY | 0.00000 | 0.00001 | 0.69013 | 0.60353 | 1.33816 | 220 |
| GO_RESPIRASOME | 0.00008 | 0.00181 | 0.53843 | 0.65420 | 1.40421 | 71 |
| GO_RESPIRATORY_CHAIN_COMPLEX | 0.00016 | 0.00353 | 0.51885 | 0.67626 | 1.43918 | 60 |
| GO_RESPIRATORY_ELECTRON_TRANSPORT_CHAIN | 0.00001 | 0.00025 | 0.59333 | 0.65819 | 1.42190 | 85 |
| GO_RESPONSE_TO_ALCOHOL | 0.00261 | 0.03750 | 0.43171 | 0.58804 | 1.27350 | 95 |
| GO_RESPONSE_TO_CADMIUM_ION | 0.00265 | 0.03775 | 0.43171 | 0.67227 | 1.39174 | 35 |
| GO_RESPONSE_TO_DRUG | 0.00104 | 0.01798 | 0.45506 | 0.55364 | 1.22258 | 180 |
| GO_RESPONSE_TO_ETHANOL | 0.00086 | 0.01566 | 0.47727 | 0.67309 | 1.40691 | 43 |
| GO_RESPONSE_TO_HEAT | 0.00012 | 0.00270 | 0.53843 | 0.61665 | 1.34221 | 107 |
| GO_RESPONSE_TO_HORMONE | 0.00048 | 0.00949 | 0.49849 | 0.51277 | 1.14814 | 453 |
| GO_RESPONSE_TO_HYDROGEN_PEROXIDE | 0.00257 | 0.03716 | 0.43171 | 0.60505 | 1.30368 | 79 |
| GO_RESPONSE_TO_INORGANIC_SUBSTANCE | 0.00004 | 0.00107 | 0.55733 | 0.55714 | 1.23870 | 275 |
| GO_RESPONSE_TO_INTERLEUKIN_1 | 0.00000 | 0.00000 | | 0.74844 | 1.62908 | 107 |
| GO_RESPONSE_TO_INTERLEUKIN_12 | 0.00012 | 0.00270 | 0.53843 | 0.72605 | 1.50694 | 36 |
| GO_RESPONSE_TO_INTERLEUKIN_4 | 0.00324 | 0.04369 | 0.43171 | 0.75812 | 1.51571 | 17 |
| GO_RESPONSE_TO_METAL_ION | 0.00030 | 0.00623 | 0.49849 | 0.56387 | 1.24233 | 171 |
| GO_RESPONSE_TO_ORGANIC_CYCLIC_COMPOUND | 0.00000 | 0.00016 | 0.61053 | 0.54603 | 1.22259 | 439 |
| GO_RESPONSE_TO_OXYGEN_LEVELS | 0.00000 | 0.00000 | 0.71951 | 0.60132 | 1.33438 | 233 |
| GO_RESPONSE_TO_STEROID_HORMONE | 0.00154 | 0.02464 | 0.45506 | 0.55514 | 1.22446 | 174 |
| GO_RESPONSE_TO_TEMPERATURE_STIMULUS | 0.00027 | 0.00572 | 0.49849 | 0.59216 | 1.29675 | 127 |
| GO_RESPONSE_TO_TUMOR_NECROSIS_FACTOR | 0.00000 | 0.00000 | 0.76146 | 0.65649 | 1.44236 | 151 |
| GO_RETINA_HOMEOSTASIS | 0.00104 | 0.01798 | 0.45506 | 0.73063 | 1.49432 | 27 |
| GO_RIBONUCLEOPROTEIN_COMPLEX_BINDING | 0.00000 | 0.00000 | 0.71951 | 0.66906 | 1.46050 | 117 |
| GO_RIBONUCLEOPROTEIN_COMPLEX_BIOGENESIS | 0.00000 | 0.00000 | | 0.66339 | 1.48264 | 389 |
| GO_RIBONUCLEOPROTEIN_COMPLEX_SUBUNIT_ORGANIZATION | 0.00000 | 0.00000 | | 0.69241 | 1.52681 | 173 |
| GO_RIBONUCLEOPROTEIN_GRANULE | 0.00000 | 0.00001 | 0.67496 | 0.61979 | 1.36366 | 163 |
| GO_RIBONUCLEOSIDE_TRIPHOSPHATE_BIOSYNTHETIC_PROCESS | 0.00041 | 0.00840 | 0.49849 | 0.71253 | 1.47248 | 33 |
| GO_RIBONUCLEOSIDE_TRIPHOSPHATE_METABOLIC_PROCESS | 0.00026 | 0.00550 | 0.49849 | 0.69681 | 1.45611 | 42 |
| GO_RIBOSE_PHOSPHATE_BIOSYNTHETIC_PROCESS | 0.00191 | 0.02939 | 0.45506 | 0.57932 | 1.26207 | 110 |
| GO_RIBOSE_PHOSPHATE_METABOLIC_PROCESS | 0.00108 | 0.01839 | 0.45506 | 0.53891 | 1.19687 | 245 |
| GO_RIBOSOMAL_LARGE_SUBUNIT_ASSEMBLY | 0.00000 | 0.00016 | 0.61053 | 0.80999 | 1.65663 | 27 |
| GO_RIBOSOMAL_LARGE_SUBUNIT_BIOGENESIS | 0.00000 | 0.00000 | | 0.77542 | 1.65982 | 66 |
| GO_RIBOSOMAL_SMALL_SUBUNIT_ASSEMBLY | 0.00023 | 0.00494 | 0.51885 | 0.81139 | 1.62221 | 17 |

| | | | | | | |
|---|---|---|---|---|---|---|
| GO_RIBOSOMAL_SMALL_SUBUNIT_BINDING | 0.00383 | 0.04935 | 0.43171 | 0.75288 | 1.50523 | 17 |
| GO_RIBOSOMAL_SMALL_SUBUNIT_BIOGENESIS | 0.00001 | 0.00022 | 0.61053 | 0.69076 | 1.47997 | 68 |
| GO_RIBOSOMAL_SUBUNIT | 0.00000 | 0.00000 | | 0.77603 | 1.71031 | 172 |
| GO_RIBOSOME | 0.00000 | 0.00000 | | 0.76643 | 1.69659 | 206 |
| GO_RIBOSOME_ASSEMBLY | 0.00000 | 0.00000 | 0.73376 | 0.76173 | 1.61998 | 59 |
| GO_RIBOSOME_BINDING | 0.00019 | 0.00416 | 0.51885 | 0.68955 | 1.44715 | 47 |
| GO_RIBOSOME_BIOGENESIS | 0.00000 | 0.00000 | | 0.65877 | 1.46466 | 275 |
| GO_RNA_CATABOLIC_PROCESS | 0.00000 | 0.00000 | | 0.76774 | 1.71264 | 339 |
| GO_RNA_DEPENDENT_DNA_BIOSYNTHETIC_PROCESS | 0.00003 | 0.00076 | 0.57561 | 0.68926 | 1.46521 | 58 |
| GO_RNA_EXPORT_FROM_NUCLEUS | 0.00062 | 0.01183 | 0.47727 | 0.58378 | 1.27758 | 126 |
| GO_RNA_LOCALIZATION | 0.00000 | 0.00006 | 0.64355 | 0.59670 | 1.32020 | 198 |
| GO_RNA_POLYMERASE_II_PREINITIATION_COMPLEX_ASSEMBLY | 0.00089 | 0.01592 | 0.47727 | 0.77046 | 1.55626 | 19 |
| GO_RNA_SPLICING | 0.00000 | 0.00000 | 0.78819 | 0.59079 | 1.31927 | 369 |
| GO_RNA_SPLICING_VIA_TRANSESTERIFICATION_REACTIONS | 0.00000 | 0.00000 | | 0.60940 | 1.35714 | 300 |
| GO_RNA_STABILIZATION | 0.00065 | 0.01223 | 0.47727 | 0.69743 | 1.44384 | 35 |
| GO_ROUGH_ENDOPLASMIC_RETICULUM | 0.00001 | 0.00039 | 0.59333 | 0.71853 | 1.51629 | 50 |
| GO_RRNA_BINDING | 0.00000 | 0.00001 | 0.69013 | 0.74782 | 1.58865 | 56 |
| GO_RRNA_METABOLIC_PROCESS | 0.00000 | 0.00000 | 0.77494 | 0.63047 | 1.39562 | 206 |
| GO_SCF_DEPENDENT_PROTEASOMAL_UBIQUITIN_DEPENDENT_PRO | 0.00000 | 0.00000 | | 0.77527 | 1.67044 | 79 |
| GO_SECRETORY_GRANULE | 0.00000 | 0.00000 | 0.71951 | 0.56518 | 1.26461 | 399 |
| GO_SECRETORY_VESICLE | 0.00000 | 0.00002 | 0.67496 | 0.54866 | 1.22905 | 465 |
| GO_SIGNAL_TRANSDUCTION_BY_PROTEIN_PHOSPHORYLATION | 0.00004 | 0.00109 | 0.55733 | 0.53817 | 1.20474 | 424 |
| GO_SINGLE_FERTILIZATION | 0.00048 | 0.00949 | 0.49849 | 0.67414 | 1.42260 | 50 |
| GO_SINGLE_STRANDED_RNA_BINDING | 0.00039 | 0.00793 | 0.49849 | 0.66553 | 1.41527 | 57 |
| **Cluster 2** | | | | | | |
| pathway | pval | padj | log2err | ES | NES | size |
| GO_ACTIVATION_OF_IMMUNE_RESPONSE | 0.00000 | 0.00000 | | 0.61836 | 1.52110 | 161 |
| GO_ACTIVATION_OF_INNATE_IMMUNE_RESPONSE | 0.00000 | 0.00000 | | 0.72021 | 1.71579 | 74 |
| GO_AEROBIC_RESPIRATION | 0.00198 | 0.02746 | 0.43171 | 0.57301 | 1.34854 | 58 |
| GO_ALTERNATIVE_MRNA_SPLICING_VIA_SPLICEOSOME | 0.00012 | 0.00256 | 0.53843 | 0.61615 | 1.45008 | 58 |
| GO_AMINE_METABOLIC_PROCESS | 0.00000 | 0.00000 | | 0.73026 | 1.73280 | 68 |
| GO_ANAPHASE_PROMOTING_COMPLEX_DEPENDENT_CATABOLIC_P | 0.00000 | 0.00000 | 0.78819 | 0.69443 | 1.65713 | 76 |
| GO_ANCHORING_JUNCTION | 0.00000 | 0.00000 | 0.73376 | 0.52693 | 1.32550 | 331 |
| GO_ANTIGEN_PROCESSING_AND_PRESENTATION | 0.00000 | 0.00000 | 0.73376 | 0.61865 | 1.50246 | 120 |
| GO_ANTIGEN_PROCESSING_AND_PRESENTATION_OF_EXOGENOUS_ | 0.00000 | 0.00000 | 0.74774 | 0.72805 | 1.70185 | 51 |
| GO_ANTIGEN_PROCESSING_AND_PRESENTATION_OF_PEPTIDE_ANTI | 0.00000 | 0.00001 | 0.69013 | 0.61884 | 1.49517 | 108 |
| GO_ANTIGEN_PROCESSING_AND_PRESENTATION_OF_PEPTIDE_ANTI | 0.00000 | 0.00000 | 0.71951 | 0.69233 | 1.63347 | 61 |
| GO_ANTIGEN_RECEPTOR_MEDIATED_SIGNALING_PATHWAY | 0.00000 | 0.00000 | 0.77494 | 0.65065 | 1.56590 | 99 |
| GO_APOPTOTIC_SIGNALING_PATHWAY | 0.00139 | 0.02067 | 0.45506 | 0.47751 | 1.19634 | 273 |
| GO_ATPASE_ACTIVITY | 0.00002 | 0.00043 | 0.57561 | 0.52146 | 1.30108 | 226 |
| GO_ATP_BIOSYNTHETIC_PROCESS | 0.00346 | 0.04447 | 0.43171 | 0.69466 | 1.52399 | 19 |
| GO_AZUROPHIL_GRANULE_LUMEN | 0.00181 | 0.02583 | 0.45506 | 0.61277 | 1.41812 | 42 |
| GO_CADHERIN_BINDING | 0.00000 | 0.00000 | | 0.60276 | 1.49846 | 206 |
| GO_CANONICAL_WNT_SIGNALING_PATHWAY | 0.00000 | 0.00005 | 0.64355 | 0.57217 | 1.39950 | 140 |
| GO_CARBOHYDRATE_BIOSYNTHETIC_PROCESS | 0.00119 | 0.01811 | 0.45506 | 0.57137 | 1.35924 | 72 |
| GO_CATALYTIC_STEP_2_SPLICEOSOME | 0.00002 | 0.00057 | 0.57561 | 0.61752 | 1.47361 | 76 |
| GO_CELLULAR_AMINO_ACID_METABOLIC_PROCESS | 0.00001 | 0.00034 | 0.59333 | 0.53669 | 1.32659 | 179 |
| GO_CELLULAR_CARBOHYDRATE_CATABOLIC_PROCESS | 0.00018 | 0.00367 | 0.51885 | 0.74026 | 1.63837 | 21 |
| GO_CELLULAR_KETONE_METABOLIC_PROCESS | 0.00000 | 0.00003 | 0.65944 | 0.59461 | 1.44409 | 120 |
| GO_CELLULAR_NITROGEN_COMPOUND_CATABOLIC_PROCESS | 0.00000 | 0.00000 | | 0.62805 | 1.58298 | 360 |
| GO_CELLULAR_PROTEIN_CATABOLIC_PROCESS | 0.00000 | 0.00001 | 0.67496 | 0.49851 | 1.26261 | 451 |
| GO_CELLULAR_RESPONSE_TO_ENDOGENOUS_STIMULUS | 0.00327 | 0.04244 | 0.43171 | 0.45146 | 1.14333 | 473 |
| GO_CELLULAR_RESPONSE_TO_HEAT | 0.00033 | 0.00619 | 0.49849 | 0.57966 | 1.38219 | 75 |
| GO_CELLULAR_RESPONSE_TO_OXYGEN_LEVELS | 0.00000 | 0.00000 | 0.73376 | 0.60768 | 1.48228 | 130 |
| GO_CELL_ACTIVATION | 0.00001 | 0.00032 | 0.59333 | 0.48387 | 1.22400 | 441 |
| GO_CELL_ACTIVATION_INVOLVED_IN_IMMUNE_RESPONSE | 0.00000 | 0.00000 | 0.70498 | 0.54068 | 1.35291 | 267 |
| GO_CELL_ADHESION_MOLECULE_BINDING | 0.00000 | 0.00000 | | 0.58388 | 1.45785 | 245 |
| GO_CELL_BODY | 0.00055 | 0.00974 | 0.47727 | 0.52017 | 1.28198 | 166 |
| GO_CELL_CELL_RECOGNITION | 0.00001 | 0.00021 | 0.59333 | 0.82905 | 1.78510 | 16 |
| GO_CELL_CELL_SIGNALING | 0.00003 | 0.00064 | 0.57561 | 0.48078 | 1.21744 | 455 |
| GO_CELL_CELL_SIGNALING_BY_WNT | 0.00000 | 0.00004 | 0.64355 | 0.54536 | 1.35753 | 214 |
| GO_CELL_CYCLE_G2_M_PHASE_TRANSITION | 0.00000 | 0.00004 | 0.64355 | 0.55209 | 1.36648 | 185 |
| GO_CELL_CYCLE_PHASE_TRANSITION | 0.00000 | 0.00004 | 0.64355 | 0.50359 | 1.27185 | 391 |
| GO_CELL_RECOGNITION | 0.00002 | 0.00061 | 0.57561 | 0.71147 | 1.62893 | 33 |
| GO_CELL_SUBSTRATE_JUNCTION | 0.00000 | 0.00000 | | 0.59447 | 1.47979 | 214 |
| GO_CELL_SURFACE_RECEPTOR_SIGNALING_PATHWAY_INVOLVED_IN | 0.00000 | 0.00003 | 0.65944 | 0.54307 | 1.35384 | 221 |
| GO_CHAPERONE_COMPLEX | 0.00000 | 0.00001 | 0.67496 | 0.84612 | 1.86334 | 20 |
| GO_CIRCADIAN_RHYTHM | 0.00031 | 0.00581 | 0.49849 | 0.57877 | 1.38121 | 79 |
| GO_COTRANSLATIONAL_PROTEIN_TARGETING_TO_MEMBRANE | 0.00000 | 0.00000 | 0.83909 | 0.68632 | 1.64421 | 87 |
| GO_CYTOKINE_MEDIATED_SIGNALING_PATHWAY | 0.00000 | 0.00000 | 0.80122 | 0.57707 | 1.44125 | 233 |
| GO_CYTOPLASMIC_TRANSLATION | 0.00000 | 0.00000 | | 0.70520 | 1.68683 | 84 |
| GO_CYTOPLASMIC_TRANSLATIONAL_INITIATION | 0.00000 | 0.00002 | 0.65944 | 0.76149 | 1.73775 | 31 |
| GO_CYTOSOLIC_LARGE_RIBOSOMAL_SUBUNIT | 0.00000 | 0.00000 | 0.71951 | 0.71458 | 1.67037 | 51 |
| GO_CYTOSOLIC_RIBOSOME | 0.00000 | 0.00000 | 0.82666 | 0.67980 | 1.63204 | 91 |
| GO_CYTOSOLIC_SMALL_RIBOSOMAL_SUBUNIT | 0.00048 | 0.00864 | 0.49849 | 0.65096 | 1.50040 | 39 |
| GO_DEFENSE_RESPONSE | 0.00000 | 0.00000 | 0.70498 | 0.51236 | 1.29450 | 403 |

| | | | | | | |
|---|---|---|---|---|---|---|
| GO_DEFENSE_RESPONSE_TO_OTHER_ORGANISM | 0.00000 | 0.00000 | 0.81404 | 0.55597 | 1.39440 | 293 |
| GO_DENDRITIC_TREE | 0.00383 | 0.04845 | 0.43171 | 0.48355 | 1.19649 | 184 |
| GO_DISORDERED_DOMAIN_SPECIFIC_BINDING | 0.00017 | 0.00352 | 0.51885 | 0.74171 | 1.64158 | 21 |
| GO_DNA_BIOSYNTHETIC_PROCESS | 0.00030 | 0.00575 | 0.49849 | 0.53930 | 1.30913 | 119 |
| GO_DNA_GEOMETRIC_CHANGE | 0.00268 | 0.03555 | 0.43171 | 0.54024 | 1.29425 | 87 |
| GO_DNA_POLYMERASE_BINDING | 0.00191 | 0.02659 | 0.45506 | 0.70733 | 1.54195 | 18 |
| GO_DNA_REPLICATION_ORIGIN_BINDING | 0.00228 | 0.03102 | 0.43171 | 0.68984 | 1.52680 | 21 |
| GO_DNA_TEMPLATED_TRANSCRIPTION_ELONGATION | 0.00139 | 0.02067 | 0.45506 | 0.54915 | 1.31559 | 87 |
| GO_DNA_TEMPLATED_TRANSCRIPTION_INITIATION | 0.00185 | 0.02604 | 0.45506 | 0.51435 | 1.25676 | 135 |
| GO_DNA_TEMPLATED_TRANSCRIPTION_TERMINATION | 0.00017 | 0.00352 | 0.51885 | 0.60744 | 1.43155 | 60 |
| GO_DOUBLE_STRANDED_RNA_BINDING | 0.00154 | 0.02251 | 0.45506 | 0.60959 | 1.41900 | 47 |
| GO_DRUG_BINDING | 0.00228 | 0.03102 | 0.43171 | 0.60191 | 1.39426 | 43 |
| GO_ENDOPEPTIDASE_COMPLEX | 0.00000 | 0.00000 | 0.78819 | 0.72384 | 1.70782 | 61 |
| GO_EPITHELIUM_DEVELOPMENT | 0.00018 | 0.00375 | 0.51885 | 0.47541 | 1.20006 | 386 |
| GO_ESTABLISHMENT_OF_PROTEIN_LOCALIZATION_TO_CHROMOSON | 0.00000 | 0.00014 | 0.61053 | 0.81240 | 1.78909 | 20 |
| GO_ESTABLISHMENT_OF_PROTEIN_LOCALIZATION_TO_ENDOPLASM | 0.00000 | 0.00000 | 0.80122 | 0.67307 | 1.61918 | 95 |
| GO_ESTABLISHMENT_OF_PROTEIN_LOCALIZATION_TO_MEMBRANE | 0.00000 | 0.00000 | 0.78819 | 0.58093 | 1.44079 | 198 |
| GO_ESTABLISHMENT_OF_PROTEIN_LOCALIZATION_TO_ORGANELLE | 0.00000 | 0.00000 | | 0.56436 | 1.42151 | 349 |
| GO_ESTABLISHMENT_OF_PROTEIN_LOCALIZATION_TO_TELOMERE | 0.00000 | 0.00000 | 0.70498 | 0.89287 | 1.91153 | 15 |
| GO_ESTABLISHMENT_OF_RNA_LOCALIZATION | 0.00000 | 0.00001 | 0.69013 | 0.58397 | 1.43049 | 147 |
| GO_ESTABLISHMENT_OF_TISSUE_POLARITY | 0.00000 | 0.00000 | 0.77494 | 0.69731 | 1.65581 | 69 |
| GO_EUKARYOTIC_TRANSLATION_INITIATION_FACTOR_3_COMPLEX | 0.00000 | 0.00013 | 0.61053 | 0.84890 | 1.81740 | 15 |
| GO_EXOCYTOSIS | 0.00006 | 0.00134 | 0.55733 | 0.48712 | 1.22779 | 359 |
| GO_FC_EPSILON_RECEPTOR_SIGNALING_PATHWAY | 0.00000 | 0.00000 | 0.77494 | 0.69219 | 1.64667 | 72 |
| GO_FC_RECEPTOR_SIGNALING_PATHWAY | 0.00000 | 0.00000 | 0.78819 | 0.65081 | 1.57004 | 105 |
| GO_FERTILIZATION | 0.00040 | 0.00733 | 0.49849 | 0.63948 | 1.47722 | 41 |
| GO_FIBRILLAR_CENTER | 0.00108 | 0.01675 | 0.45506 | 0.56319 | 1.34404 | 78 |
| GO_FICOLIN_1_RICH_GRANULE | 0.00000 | 0.00000 | | 0.72613 | 1.73790 | 85 |
| GO_GENERATION_OF_PRECURSOR_METABOLITES_AND_ENERGY | 0.00135 | 0.02038 | 0.45506 | 0.48154 | 1.20486 | 263 |
| GO_GLUCOSE_CATABOLIC_PROCESS | 0.00054 | 0.00965 | 0.47727 | 0.75975 | 1.64714 | 17 |
| GO_GLYCOSYL_COMPOUND_METABOLIC_PROCESS | 0.00055 | 0.00974 | 0.47727 | 0.59221 | 1.39901 | 62 |
| GO_GTPASE_ACTIVITY | 0.00298 | 0.03892 | 0.43171 | 0.51028 | 1.23965 | 121 |
| GO_GUANYL_NUCLEOTIDE_BINDING | 0.00110 | 0.01687 | 0.45506 | 0.50917 | 1.25153 | 158 |
| GO_HEAT_SHOCK_PROTEIN_BINDING | 0.00001 | 0.00015 | 0.61053 | 0.64645 | 1.52716 | 62 |
| GO_HEMATOPOIETIC_PROGENITOR_CELL_DIFFERENTIATION | 0.00000 | 0.00001 | 0.69013 | 0.63799 | 1.53167 | 91 |
| GO_HEMATOPOIETIC_STEM_CELL_DIFFERENTIATION | 0.00000 | 0.00000 | 0.77494 | 0.71081 | 1.67920 | 62 |
| GO_HEXOSE_CATABOLIC_PROCESS | 0.00023 | 0.00458 | 0.51885 | 0.69218 | 1.55844 | 26 |
| GO_HSP70_PROTEIN_BINDING | 0.00037 | 0.00675 | 0.49849 | 0.73742 | 1.60755 | 18 |
| GO_HSP90_PROTEIN_BINDING | 0.00253 | 0.03387 | 0.43171 | 0.68760 | 1.52184 | 21 |
| GO_HYDROLASE_ACTIVITY_ACTING_ON_ACID_ANHYDRIDES | 0.00001 | 0.00027 | 0.59333 | 0.49585 | 1.25132 | 384 |
| GO_IMMUNE_EFFECTOR_PROCESS | 0.00001 | 0.00017 | 0.61053 | 0.49513 | 1.25086 | 399 |
| GO_IMMUNE_RESPONSE_REGULATING_SIGNALING_PATHWAY | 0.00000 | 0.00000 | 0.74774 | 0.60625 | 1.48638 | 151 |
| GO_INNATE_IMMUNE_RESPONSE | 0.00000 | 0.00000 | | 0.58531 | 1.46184 | 247 |
| GO_INNATE_IMMUNE_RESPONSE_ACTIVATING_SIGNAL_TRANSDUCT | 0.00000 | 0.00000 | 0.80122 | 0.73289 | 1.72917 | 61 |
| GO_INTERACTION_WITH_SYMBIONT | 0.00257 | 0.03417 | 0.43171 | 0.62867 | 1.43937 | 33 |
| GO_INTERLEUKIN_1_MEDIATED_SIGNALING_PATHWAY | 0.00000 | 0.00000 | 0.81404 | 0.72257 | 1.70862 | 64 |
| GO_INTRINSIC_APOPTOTIC_SIGNALING_PATHWAY | 0.00106 | 0.01657 | 0.45506 | 0.50960 | 1.24629 | 139 |
| GO_ION_CHANNEL_BINDING | 0.00141 | 0.02081 | 0.45506 | 0.60909 | 1.41090 | 43 |
| GO_LARGE_RIBOSOMAL_SUBUNIT | 0.00003 | 0.00067 | 0.57561 | 0.58260 | 1.40290 | 102 |
| GO_LEUKOCYTE_MEDIATED_IMMUNITY | 0.00000 | 0.00001 | 0.69013 | 0.53684 | 1.34525 | 279 |
| GO_MAINTENANCE_OF_CELL_NUMBER | 0.00383 | 0.04845 | 0.43171 | 0.52528 | 1.26264 | 94 |
| GO_MATURATION_OF_LSU_RRNA | 0.00045 | 0.00810 | 0.49849 | 0.70646 | 1.58678 | 25 |
| GO_MEMBRANE_ORGANIZATION | 0.00250 | 0.03356 | 0.43171 | 0.46144 | 1.16395 | 376 |
| GO_MITOCHONDRIAL_MEMBRANE_ORGANIZATION | 0.00130 | 0.01967 | 0.45506 | 0.57169 | 1.35653 | 68 |
| GO_MITOCHONDRIAL_TRANSPORT | 0.00095 | 0.01519 | 0.47727 | 0.52631 | 1.28279 | 129 |
| GO_MODIFICATION_DEPENDENT_MACROMOLECULE_CATABOLIC_PR | 0.00000 | 0.00005 | 0.64355 | 0.50309 | 1.27019 | 388 |
| GO_MONOSACCHARIDE_BIOSYNTHETIC_PROCESS | 0.00161 | 0.02344 | 0.45506 | 0.61711 | 1.43322 | 44 |
| GO_MONOSACCHARIDE_CATABOLIC_PROCESS | 0.00016 | 0.00334 | 0.51885 | 0.68560 | 1.56966 | 32 |
| GO_MORPHOGENESIS_OF_AN_EPITHELIUM | 0.00017 | 0.00347 | 0.51885 | 0.50700 | 1.26076 | 208 |
| GO_MORPHOGENESIS_OF_A_POLARIZED_EPITHELIUM | 0.00000 | 0.00000 | 0.77494 | 0.67783 | 1.61926 | 80 |
| GO_MRNA_3_END_PROCESSING | 0.00028 | 0.00536 | 0.49849 | 0.58020 | 1.38463 | 79 |
| GO_MRNA_3_UTR_BINDING | 0.00003 | 0.00079 | 0.55733 | 0.63895 | 1.49993 | 54 |
| GO_MRNA_5_UTR_BINDING | 0.00054 | 0.00965 | 0.47727 | 0.70467 | 1.56707 | 22 |
| GO_MRNA_BINDING | 0.00000 | 0.00000 | | 0.61483 | 1.52423 | 196 |
| GO_MRNA_CIS_SPLICING_VIA_SPLICEOSOME | 0.00235 | 0.03175 | 0.43171 | 0.67982 | 1.52206 | 24 |
| GO_MRNA_EXPORT_FROM_NUCLEUS | 0.00007 | 0.00160 | 0.53843 | 0.58238 | 1.39670 | 89 |
| GO_MRNA_PROCESSING | 0.00000 | 0.00000 | | 0.55586 | 1.40117 | 369 |
| GO_MRNA_TRANSPORT | 0.00006 | 0.00141 | 0.53843 | 0.56697 | 1.37061 | 110 |
| GO_MYELOID_LEUKOCYTE_ACTIVATION | 0.00000 | 0.00000 | 0.76146 | 0.56021 | 1.39922 | 244 |
| GO_MYELOID_LEUKOCYTE_MEDIATED_IMMUNITY | 0.00000 | 0.00000 | 0.77494 | 0.56220 | 1.40395 | 238 |
| GO_NADH_METABOLIC_PROCESS | 0.00033 | 0.00613 | 0.49849 | 0.75388 | 1.65393 | 19 |
| GO_NAD_BINDING | 0.00011 | 0.00232 | 0.53843 | 0.68881 | 1.57705 | 33 |
| GO_NAD_METABOLIC_PROCESS | 0.00002 | 0.00039 | 0.59333 | 0.77006 | 1.71248 | 22 |
| GO_NCRNA_METABOLIC_PROCESS | 0.00021 | 0.00414 | 0.51885 | 0.48338 | 1.21635 | 334 |
| GO_NCRNA_PROCESSING | 0.00016 | 0.00331 | 0.51885 | 0.49354 | 1.23811 | 295 |
| GO_NEGATIVE_REGULATION_OF_CANONICAL_WNT_SIGNALING_PAT | 0.00000 | 0.00001 | 0.67496 | 0.63605 | 1.52191 | 86 |

| | | | | | | |
|---|---|---|---|---|---|---|
| GO_NEGATIVE_REGULATION_OF_CELLULAR_AMIDE_METABOLIC_PR( | 0.00021 | 0.00416 | 0.51885 | 0.57304 | 1.37283 | 87 |
| GO_NEGATIVE_REGULATION_OF_CELLULAR_CATABOLIC_PROCESS | 0.00170 | 0.02440 | 0.45506 | 0.51867 | 1.26683 | 133 |
| GO_NEGATIVE_REGULATION_OF_CELL_CYCLE | 0.00014 | 0.00305 | 0.51885 | 0.48699 | 1.22597 | 340 |
| GO_NEGATIVE_REGULATION_OF_CELL_CYCLE_G2_M_PHASE_TRANS | 0.00000 | 0.00000 | 0.70498 | 0.66009 | 1.57529 | 78 |
| GO_NEGATIVE_REGULATION_OF_CELL_CYCLE_PHASE_TRANSITION | 0.00000 | 0.00002 | 0.67496 | 0.57442 | 1.40939 | 153 |
| GO_NEGATIVE_REGULATION_OF_CELL_CYCLE_PROCESS | 0.00000 | 0.00004 | 0.64355 | 0.54413 | 1.35298 | 207 |
| GO_NEGATIVE_REGULATION_OF_CELL_DEATH | 0.00014 | 0.00297 | 0.51885 | 0.47932 | 1.21055 | 391 |
| GO_NEGATIVE_REGULATION_OF_MITOTIC_CELL_CYCLE | 0.00000 | 0.00011 | 0.62726 | 0.54201 | 1.34164 | 188 |
| GO_NEGATIVE_REGULATION_OF_MRNA_METABOLIC_PROCESS | 0.00022 | 0.00431 | 0.51885 | 0.63634 | 1.48435 | 49 |
| GO_NEGATIVE_REGULATION_OF_MRNA_SPLICING_VIA_SPLICEOSOM | 0.00390 | 0.04926 | 0.43171 | 0.70832 | 1.52515 | 16 |
| GO_NEGATIVE_REGULATION_OF_PROTEIN_MODIFICATION_BY_SMA | 0.00231 | 0.03134 | 0.43171 | 0.60106 | 1.39662 | 45 |
| GO_NEGATIVE_REGULATION_OF_RNA_CATABOLIC_PROCESS | 0.00072 | 0.01223 | 0.47727 | 0.65419 | 1.49795 | 34 |
| GO_NEGATIVE_REGULATION_OF_UBIQUITIN_DEPENDENT_PROTEIN_ | 0.00092 | 0.01485 | 0.47727 | 0.65361 | 1.49647 | 33 |
| GO_NEGATIVE_REGULATION_OF_WNT_SIGNALING_PATHWAY | 0.00000 | 0.00002 | 0.65944 | 0.62144 | 1.49289 | 93 |
| GO_NEURON_TO_NEURON_SYNAPSE | 0.00000 | 0.00006 | 0.64355 | 0.59882 | 1.44462 | 105 |
| GO_NIK_NF_KAPPAB_SIGNALING | 0.00000 | 0.00000 | | 0.69691 | 1.67090 | 88 |
| GO_NON_CANONICAL_WNT_SIGNALING_PATHWAY | 0.00000 | 0.00000 | 0.73376 | 0.66996 | 1.59884 | 78 |
| GO_NUCLEAR_BODY | 0.00085 | 0.01400 | 0.47727 | 0.46231 | 1.17038 | 459 |
| GO_NUCLEAR_EXPORT | 0.00001 | 0.00015 | 0.61053 | 0.56161 | 1.37772 | 152 |
| GO_NUCLEAR_MATRIX | 0.00183 | 0.02586 | 0.45506 | 0.56546 | 1.34519 | 72 |
| GO_NUCLEAR_PERIPHERY | 0.00331 | 0.04280 | 0.43171 | 0.53750 | 1.28907 | 89 |
| GO_NUCLEAR_SPECK | 0.00000 | 0.00005 | 0.64355 | 0.53105 | 1.32688 | 242 |
| GO_NUCLEAR_TRANSCRIBED_MRNA_CATABOLIC_PROCESS | 0.00000 | 0.00000 | | 0.63119 | 1.55463 | 164 |
| GO_NUCLEAR_TRANSCRIBED_MRNA_CATABOLIC_PROCESS_NONSEN | 0.00000 | 0.00000 | | 0.69649 | 1.68024 | 105 |
| GO_NUCLEAR_TRANSPORT | 0.00001 | 0.00029 | 0.59333 | 0.52583 | 1.31310 | 232 |
| GO_NUCLEOBASE_CONTAINING_COMPOUND_TRANSPORT | 0.00000 | 0.00001 | 0.69013 | 0.57837 | 1.42543 | 166 |
| GO_NUCLEOSIDE_METABOLIC_PROCESS | 0.00034 | 0.00643 | 0.49849 | 0.61108 | 1.42843 | 51 |
| GO_NUCLEOTIDE_EXCISION_REPAIR | 0.00035 | 0.00649 | 0.49849 | 0.57103 | 1.36614 | 82 |
| GO_NUCLEOTIDE_EXCISION_REPAIR_DNA_DAMAGE_RECOGNITION | 0.00097 | 0.01539 | 0.47727 | 0.71975 | 1.58504 | 20 |
| GO_NUCLEOTIDE_EXCISION_REPAIR_DNA_DUPLEX_UNWINDING | 0.00353 | 0.04518 | 0.43171 | 0.68869 | 1.50132 | 18 |
| GO_NUCLEOTIDE_EXCISION_REPAIR_DNA_INCISION | 0.00335 | 0.04316 | 0.43171 | 0.62857 | 1.43443 | 31 |
| GO_ORGANIC_ACID_METABOLIC_PROCESS | 0.00019 | 0.00378 | 0.51885 | 0.46738 | 1.18407 | 480 |
| GO_ORGANIC_CYCLIC_COMPOUND_CATABOLIC_PROCESS | 0.00000 | 0.00000 | | 0.62293 | 1.57002 | 366 |
| GO_PEPTIDASE_ACTIVITY | 0.00194 | 0.02702 | 0.45506 | 0.49345 | 1.22163 | 189 |
| GO_PEPTIDASE_COMPLEX | 0.00000 | 0.00000 | 0.73376 | 0.67313 | 1.60505 | 75 |
| GO_PEPTIDE_BINDING | 0.00098 | 0.01563 | 0.45506 | 0.56349 | 1.34364 | 75 |
| GO_PEPTIDE_BIOSYNTHETIC_PROCESS | 0.00000 | 0.00000 | | 0.56912 | 1.44147 | 451 |
| GO_PERINUCLEAR_REGION_OF_CYTOPLASM | 0.00078 | 0.01313 | 0.47727 | 0.47619 | 1.19564 | 303 |
| GO_PIGMENT_GRANULE | 0.00019 | 0.00390 | 0.51885 | 0.60935 | 1.44158 | 65 |
| GO_POLYSOMAL_RIBOSOME | 0.00078 | 0.01313 | 0.47727 | 0.66427 | 1.51283 | 30 |
| GO_POLYSOME | 0.00001 | 0.00030 | 0.59333 | 0.64974 | 1.52730 | 56 |
| GO_POLY_PYRIMIDINE_TRACT_BINDING | 0.00020 | 0.00402 | 0.51885 | 0.78820 | 1.68744 | 15 |
| GO_POSITIVE_REGULATION_OF_APOPTOTIC_SIGNALING_PATHWAY | 0.00272 | 0.03574 | 0.43171 | 0.55355 | 1.32096 | 76 |
| GO_POSITIVE_REGULATION_OF_CANONICAL_WNT_SIGNALING_PATH | 0.00000 | 0.00000 | 0.76146 | 0.67175 | 1.60474 | 80 |
| GO_POSITIVE_REGULATION_OF_CELLULAR_AMIDE_METABOLIC_PRC | 0.00007 | 0.00147 | 0.53843 | 0.58310 | 1.39843 | 89 |
| GO_POSITIVE_REGULATION_OF_CELLULAR_COMPONENT_ORGANIZA | 0.00002 | 0.00053 | 0.57561 | 0.47528 | 1.20411 | 485 |
| GO_POSITIVE_REGULATION_OF_CELLULAR_PROTEIN_LOCALIZATION | 0.00007 | 0.00149 | 0.53843 | 0.52892 | 1.30666 | 177 |
| GO_POSITIVE_REGULATION_OF_CELL_DEATH | 0.00167 | 0.02395 | 0.45506 | 0.48259 | 1.20685 | 261 |
| GO_POSITIVE_REGULATION_OF_CHROMOSOME_ORGANIZATION | 0.00028 | 0.00536 | 0.49849 | 0.54034 | 1.31167 | 119 |
| GO_POSITIVE_REGULATION_OF_DEFENSE_RESPONSE | 0.00000 | 0.00000 | 0.78819 | 0.63592 | 1.54368 | 119 |
| GO_POSITIVE_REGULATION_OF_DNA_BIOSYNTHETIC_PROCESS | 0.00000 | 0.00002 | 0.67496 | 0.72003 | 1.67308 | 45 |
| GO_POSITIVE_REGULATION_OF_DNA_METABOLIC_PROCESS | 0.00002 | 0.00052 | 0.57561 | 0.58703 | 1.41354 | 98 |
| GO_POSITIVE_REGULATION_OF_DNA_TEMPLATED_TRANSCRIPTION_ | 0.00006 | 0.00130 | 0.55733 | 0.76073 | 1.68368 | 21 |
| GO_POSITIVE_REGULATION_OF_ESTABLISHMENT_OF_PROTEIN_LOC/ | 0.00000 | 0.00009 | 0.62726 | 0.55563 | 1.37055 | 169 |
| GO_POSITIVE_REGULATION_OF_IMMUNE_RESPONSE | 0.00000 | 0.00000 | 0.82666 | 0.58635 | 1.45859 | 211 |
| GO_POSITIVE_REGULATION_OF_IMMUNE_SYSTEM_PROCESS | 0.00000 | 0.00000 | 0.71951 | 0.53816 | 1.34953 | 284 |
| GO_POSITIVE_REGULATION_OF_INNATE_IMMUNE_RESPONSE | 0.00000 | 0.00000 | | 0.67965 | 1.63371 | 94 |
| GO_POSITIVE_REGULATION_OF_INTRACELLULAR_TRANSPORT | 0.00133 | 0.02016 | 0.45506 | 0.52415 | 1.27236 | 119 |
| GO_POSITIVE_REGULATION_OF_INTRINSIC_APOPTOTIC_SIGNALING_ | 0.00183 | 0.02586 | 0.45506 | 0.65444 | 1.47349 | 26 |
| GO_POSITIVE_REGULATION_OF_MRNA_PROCESSING | 0.00052 | 0.00923 | 0.47727 | 0.70549 | 1.56889 | 22 |
| GO_POSITIVE_REGULATION_OF_MRNA_SPLICING_VIA_SPLICEOSOM | 0.00002 | 0.00045 | 0.57561 | 0.82833 | 1.77335 | 15 |
| GO_POSITIVE_REGULATION_OF_NF_KAPPAB_TRANSCRIPTION_FACT( | 0.00191 | 0.02659 | 0.45506 | 0.57891 | 1.36090 | 57 |
| GO_POSITIVE_REGULATION_OF_ORGANELLE_ORGANIZATION | 0.00106 | 0.01657 | 0.45506 | 0.47219 | 1.18632 | 315 |
| GO_POSITIVE_REGULATION_OF_PROTEIN_LOCALIZATION_TO_NUCLE | 0.00001 | 0.00016 | 0.61053 | 0.67876 | 1.57758 | 46 |
| GO_POSITIVE_REGULATION_OF_RESPONSE_TO_BIOTIC_STIMULUS | 0.00000 | 0.00000 | 0.82666 | 0.65649 | 1.58458 | 106 |
| GO_POSITIVE_REGULATION_OF_RESPONSE_TO_EXTERNAL_STIMULU | 0.00000 | 0.00000 | 0.74774 | 0.60641 | 1.48558 | 145 |
| GO_POSITIVE_REGULATION_OF_RNA_SPLICING | 0.00191 | 0.02659 | 0.45506 | 0.67848 | 1.50883 | 22 |
| GO_POSITIVE_REGULATION_OF_TELOMERASE_ACTIVITY | 0.00001 | 0.00030 | 0.59333 | 0.74807 | 1.68429 | 26 |
| GO_POSITIVE_REGULATION_OF_TELOMERASE_RNA_LOCALIZATION_ | 0.00000 | 0.00014 | 0.61053 | 0.84636 | 1.81195 | 15 |
| GO_POSITIVE_REGULATION_OF_TELOMERE_MAINTENANCE | 0.00000 | 0.00001 | 0.67496 | 0.75415 | 1.72681 | 35 |
| GO_POSITIVE_REGULATION_OF_TELOMERE_MAINTENANCE_VIA_TE | 0.00001 | 0.00018 | 0.61053 | 0.75825 | 1.71107 | 27 |
| GO_POSITIVE_REGULATION_OF_TRANSCRIPTION_INITIATION_FROM_ | 0.00036 | 0.00663 | 0.49849 | 0.73891 | 1.61079 | 18 |
| GO_POSITIVE_REGULATION_OF_TRANSLATION | 0.00004 | 0.00091 | 0.55733 | 0.60471 | 1.44313 | 78 |
| GO_POSITIVE_REGULATION_OF_TRANSPORT | 0.00015 | 0.00313 | 0.51885 | 0.48952 | 1.22884 | 301 |
| GO_POSITIVE_REGULATION_OF_VIRAL_GENOME_REPLICATION | 0.00383 | 0.04845 | 0.43171 | 0.63529 | 1.43988 | 29 |
| GO_POSITIVE_REGULATION_OF_VIRAL_PROCESS | 0.00097 | 0.01539 | 0.47727 | 0.57106 | 1.36274 | 76 |

| | | | | | | |
|---|---|---|---|---|---|---|
| GO_POSITIVE_REGULATION_OF_WNT_SIGNALING_PATHWAY | 0.00000 | 0.00001 | 0.67496 | 0.62997 | 1.51272 | 92 |
| GO_POSTSYNAPSE | 0.00085 | 0.01400 | 0.47727 | 0.50374 | 1.24334 | 174 |
| GO_POSTTRANSCRIPTIONAL_REGULATION_OF_GENE_EXPRESSION | 0.00000 | 0.00000 | | 0.60715 | 1.53240 | 385 |
| GO_POST_TRANSLATIONAL_PROTEIN_MODIFICATION | 0.00000 | 0.00002 | 0.65944 | 0.56161 | 1.38647 | 175 |
| GO_PRERIBOSOME | 0.00089 | 0.01451 | 0.47727 | 0.57795 | 1.36840 | 66 |
| GO_PRE_MRNA_BINDING | 0.00078 | 0.01313 | 0.47727 | 0.67301 | 1.51529 | 26 |
| GO_PROTEASOMAL_PROTEIN_CATABOLIC_PROCESS | 0.00000 | 0.00001 | 0.69013 | 0.52876 | 1.32646 | 296 |
| GO_PROTEASOMAL_UBIQUITIN_INDEPENDENT_PROTEIN_CATABOLI( | 0.00002 | 0.00043 | 0.57561 | 0.79049 | 1.74084 | 20 |
| GO_PROTEASOME_ACCESSORY_COMPLEX | 0.00001 | 0.00019 | 0.61053 | 0.78074 | 1.74801 | 24 |
| GO_PROTEASOME_CORE_COMPLEX | 0.00000 | 0.00005 | 0.64355 | 0.83051 | 1.81046 | 18 |
| GO_PROTEIN_CONTAINING_COMPLEX_LOCALIZATION | 0.00017 | 0.00355 | 0.51885 | 0.52970 | 1.30547 | 166 |
| GO_PROTEIN_DNA_COMPLEX_SUBUNIT_ORGANIZATION | 0.00102 | 0.01610 | 0.45506 | 0.51714 | 1.26682 | 146 |
| GO_PROTEIN_DOMAIN_SPECIFIC_BINDING | 0.00137 | 0.02059 | 0.45506 | 0.47457 | 1.19140 | 302 |
| GO_PROTEIN_FOLDING | 0.00005 | 0.00105 | 0.55733 | 0.55492 | 1.35536 | 133 |
| GO_PROTEIN_LOCALIZATION_TO_CHROMOSOME_TELOMERIC_REGI( | 0.00001 | 0.00033 | 0.59333 | 0.77159 | 1.72750 | 24 |
| GO_PROTEIN_LOCALIZATION_TO_ENDOPLASMIC_RETICULUM | 0.00000 | 0.00000 | 0.82666 | 0.65652 | 1.58730 | 111 |
| GO_PROTEIN_LOCALIZATION_TO_MEMBRANE | 0.00001 | 0.00021 | 0.59333 | 0.50579 | 1.27052 | 309 |
| GO_PROTEIN_LOCALIZATION_TO_MITOCHONDRION | 0.00009 | 0.00195 | 0.53843 | 0.58052 | 1.39225 | 89 |
| GO_PROTEIN_LOCALIZATION_TO_NUCLEUS | 0.00079 | 0.01323 | 0.47727 | 0.51503 | 1.26367 | 153 |
| GO_PROTEIN_MODIFICATION_BY_SMALL_PROTEIN_REMOVAL | 0.00001 | 0.00017 | 0.61053 | 0.54766 | 1.35380 | 181 |
| GO_PROTEIN_POLYUBIQUITINATION | 0.00000 | 0.00010 | 0.62726 | 0.54151 | 1.34439 | 202 |
| GO_PROTEIN_STABILIZATION | 0.00043 | 0.00771 | 0.49849 | 0.55882 | 1.34561 | 98 |
| GO_PROTEIN_TARGETING | 0.00000 | 0.00000 | | 0.56997 | 1.42619 | 267 |
| GO_PROTEIN_TARGETING_TO_MEMBRANE | 0.00000 | 0.00000 | 0.82666 | 0.63793 | 1.55389 | 127 |
| GO_PROTEIN_TARGETING_TO_MITOCHONDRION | 0.00294 | 0.03855 | 0.43171 | 0.57117 | 1.35124 | 65 |
| GO_PROTEIN_TRANSMEMBRANE_TRANSPORT | 0.00091 | 0.01476 | 0.47727 | 0.64932 | 1.48679 | 34 |
| GO_PURINE_NUCLEOSIDE_METABOLIC_PROCESS | 0.00159 | 0.02324 | 0.45506 | 0.64314 | 1.46769 | 31 |
| GO_PURINE_NUCLEOSIDE_TRIPHOSPHATE_METABOLIC_PROCESS | 0.00213 | 0.02925 | 0.43171 | 0.62519 | 1.44101 | 39 |
| GO_REGULATION_OF_ANATOMICAL_STRUCTURE_MORPHOGENESIS | 0.00059 | 0.01030 | 0.47727 | 0.47306 | 1.19129 | 342 |
| GO_REGULATION_OF_ANIMAL_ORGAN_MORPHOGENESIS | 0.00000 | 0.00000 | 0.73376 | 0.64866 | 1.56046 | 95 |
| GO_REGULATION_OF_APOPTOTIC_SIGNALING_PATHWAY | 0.00027 | 0.00518 | 0.49849 | 0.51323 | 1.26912 | 183 |
| GO_REGULATION_OF_BINDING | 0.00068 | 0.01173 | 0.47727 | 0.52094 | 1.27611 | 148 |
| GO_REGULATION_OF_CELLULAR_AMIDE_METABOLIC_PROCESS | 0.00000 | 0.00000 | | 0.58019 | 1.44861 | 246 |
| GO_REGULATION_OF_CELLULAR_AMINE_METABOLIC_PROCESS | 0.00000 | 0.00000 | | 0.77336 | 1.81788 | 56 |
| GO_REGULATION_OF_CELLULAR_AMINO_ACID_METABOLIC_PROCES | 0.00000 | 0.00000 | | 0.77476 | 1.81481 | 53 |
| GO_REGULATION_OF_CELLULAR_CATABOLIC_PROCESS | 0.00000 | 0.00000 | 0.76146 | 0.51426 | 1.30259 | 475 |
| GO_REGULATION_OF_CELLULAR_KETONE_METABOLIC_PROCESS | 0.00000 | 0.00000 | 0.73376 | 0.64294 | 1.54547 | 94 |
| GO_REGULATION_OF_CELLULAR_LOCALIZATION | 0.00113 | 0.01738 | 0.45506 | 0.46436 | 1.17325 | 400 |
| GO_REGULATION_OF_CELLULAR_PROTEIN_CATABOLIC_PROCESS | 0.00039 | 0.00722 | 0.49849 | 0.53145 | 1.30097 | 143 |
| GO_REGULATION_OF_CELLULAR_PROTEIN_LOCALIZATION | 0.00051 | 0.00910 | 0.47727 | 0.48678 | 1.21980 | 275 |
| GO_REGULATION_OF_CELL_CYCLE_G2_M_PHASE_TRANSITION | 0.00000 | 0.00005 | 0.64355 | 0.56492 | 1.38649 | 154 |
| GO_REGULATION_OF_CELL_CYCLE_PHASE_TRANSITION | 0.00000 | 0.00012 | 0.61053 | 0.51532 | 1.29233 | 287 |
| GO_REGULATION_OF_CELL_CYCLE_PROCESS | 0.00004 | 0.00090 | 0.55733 | 0.48436 | 1.22544 | 444 |
| GO_REGULATION_OF_DEFENSE_RESPONSE | 0.00000 | 0.00009 | 0.62726 | 0.53545 | 1.33172 | 209 |
| GO_REGULATION_OF_DNA_BIOSYNTHETIC_PROCESS | 0.00000 | 0.00007 | 0.62726 | 0.64700 | 1.53189 | 66 |
| GO_REGULATION_OF_DNA_METABOLIC_PROCESS | 0.00081 | 0.01349 | 0.47727 | 0.50261 | 1.24430 | 189 |
| GO_REGULATION_OF_DNA_TEMPLATED_TRANSCRIPTION_INITIATIOI | 0.00011 | 0.00228 | 0.53843 | 0.73687 | 1.65508 | 25 |
| GO_REGULATION_OF_DNA_TEMPLATED_TRANSCRIPTION_IN_RESPO | 0.00000 | 0.00000 | 0.78819 | 0.69293 | 1.65309 | 77 |
| GO_REGULATION_OF_ESTABLISHMENT_OF_PROTEIN_LOCALIZATION_ | 0.00145 | 0.02129 | 0.45506 | 0.60895 | 1.41057 | 43 |
| GO_REGULATION_OF_HEMATOPOIETIC_PROGENITOR_CELL_DIFFERE | 0.00000 | 0.00000 | 0.74774 | 0.69539 | 1.64647 | 66 |
| GO_REGULATION_OF_HEMATOPOIETIC_STEM_CELL_DIFFERENTIATIC | 0.00000 | 0.00000 | 0.78819 | 0.72523 | 1.70678 | 58 |
| GO_REGULATION_OF_HEMOPOIESIS | 0.00003 | 0.00070 | 0.57561 | 0.53713 | 1.32586 | 173 |
| GO_REGULATION_OF_IMMUNE_RESPONSE | 0.00000 | 0.00001 | 0.69013 | 0.53708 | 1.34382 | 263 |
| GO_REGULATION_OF_IMMUNE_SYSTEM_PROCESS | 0.00056 | 0.00986 | 0.47727 | 0.46699 | 1.18123 | 431 |
| GO_REGULATION_OF_INNATE_IMMUNE_RESPONSE | 0.00000 | 0.00000 | 0.80122 | 0.63526 | 1.54719 | 125 |
| GO_REGULATION_OF_INTRACELLULAR_TRANSPORT | 0.00324 | 0.04208 | 0.43171 | 0.49270 | 1.21741 | 178 |
| GO_REGULATION_OF_MITOTIC_CELL_CYCLE | 0.00002 | 0.00047 | 0.57561 | 0.49039 | 1.23633 | 372 |
| GO_REGULATION_OF_MORPHOGENESIS_OF_AN_EPITHELIUM | 0.00000 | 0.00000 | 0.74774 | 0.66796 | 1.59630 | 81 |
| GO_REGULATION_OF_MRNA_CATABOLIC_PROCESS | 0.00000 | 0.00000 | | 0.64645 | 1.58117 | 140 |
| GO_REGULATION_OF_MRNA_METABOLIC_PROCESS | 0.00000 | 0.00000 | | 0.63618 | 1.58853 | 231 |
| GO_REGULATION_OF_MRNA_PROCESSING | 0.00000 | 0.00000 | 0.71951 | 0.63159 | 1.52087 | 102 |
| GO_REGULATION_OF_MRNA_SPLICING_VIA_SPLICEOSOME | 0.00000 | 0.00001 | 0.69013 | 0.66357 | 1.57914 | 73 |
| GO_REGULATION_OF_NUCLEOCYTOPLASMIC_TRANSPORT | 0.00231 | 0.03134 | 0.43171 | 0.56922 | 1.34484 | 63 |
| GO_REGULATION_OF_PROTEASOMAL_PROTEIN_CATABOLIC_PROCES | 0.00110 | 0.01687 | 0.45506 | 0.53216 | 1.28821 | 113 |
| GO_REGULATION_OF_PROTEIN_CATABOLIC_PROCESS | 0.00003 | 0.00081 | 0.55733 | 0.52691 | 1.30990 | 206 |
| GO_REGULATION_OF_PROTEIN_CONTAINING_COMPLEX_ASSEMBLY | 0.00069 | 0.01184 | 0.47727 | 0.50771 | 1.25510 | 182 |
| GO_REGULATION_OF_PROTEIN_LOCALIZATION_TO_NUCLEUS | 0.00004 | 0.00096 | 0.55733 | 0.61908 | 1.46579 | 66 |
| GO_REGULATION_OF_PROTEIN_STABILITY | 0.00139 | 0.02067 | 0.45506 | 0.50752 | 1.24815 | 160 |
| GO_REGULATION_OF_PROTEOLYSIS | 0.00146 | 0.02149 | 0.45506 | 0.47765 | 1.19706 | 278 |
| GO_REGULATION_OF_RESPONSE_TO_BIOTIC_STIMULUS | 0.00000 | 0.00000 | 0.76146 | 0.59818 | 1.46991 | 156 |
| GO_REGULATION_OF_RESPONSE_TO_EXTERNAL_STIMULUS | 0.00115 | 0.01761 | 0.45506 | 0.47821 | 1.19919 | 288 |
| GO_REGULATION_OF_RNA_SPLICING | 0.00000 | 0.00001 | 0.67496 | 0.62939 | 1.51411 | 95 |
| GO_REGULATION_OF_SMALL_MOLECULE_METABOLIC_PROCESS | 0.00000 | 0.00002 | 0.67496 | 0.55107 | 1.36813 | 201 |
| GO_REGULATION_OF_STEM_CELL_DIFFERENTIATION | 0.00000 | 0.00000 | 0.76146 | 0.67943 | 1.62145 | 78 |
| GO_REGULATION_OF_SYMBIOTIC_PROCESS | 0.00006 | 0.00145 | 0.53843 | 0.56140 | 1.35731 | 111 |
| GO_REGULATION_OF_TELOMERASE_ACTIVITY | 0.00006 | 0.00136 | 0.55733 | 0.70522 | 1.61459 | 32 |

| | | | | | | |
|---|---|---|---|---|---|---|
| GO_REGULATION_OF_TELOMERE_MAINTENANCE | 0.00001 | 0.00031 | 0.59333 | 0.65202 | 1.53277 | 57 |
| GO_REGULATION_OF_TELOMERE_MAINTENANCE_VIA_TELOMERE_L | 0.00000 | 0.00014 | 0.61053 | 0.69530 | 1.61481 | 44 |
| GO_REGULATION_OF_TRANSCRIPTION_FROM_RNA_POLYMERASE_II | 0.00000 | 0.00000 | 0.81404 | 0.73605 | 1.73464 | 60 |
| GO_REGULATION_OF_TRANSCRIPTION_INITIATION_FROM_RNA_POL | 0.00027 | 0.00523 | 0.49849 | 0.74567 | 1.64214 | 20 |
| GO_REGULATION_OF_TRANSLATIONAL_INITIATION | 0.00005 | 0.00119 | 0.55733 | 0.63333 | 1.48673 | 54 |
| GO_REGULATION_OF_UBIQUITIN_DEPENDENT_PROTEIN_CATABOLIC | 0.00110 | 0.01687 | 0.45506 | 0.54074 | 1.29981 | 94 |
| GO_REGULATION_OF_VIRAL_GENOME_REPLICATION | 0.00253 | 0.03387 | 0.43171 | 0.60769 | 1.41133 | 44 |
| GO_REGULATION_OF_VIRAL_LIFE_CYCLE | 0.00122 | 0.01861 | 0.45506 | 0.56752 | 1.34761 | 69 |
| GO_REGULATION_OF_WNT_SIGNALING_PATHWAY | 0.00002 | 0.00039 | 0.59333 | 0.54784 | 1.34658 | 157 |
| GO_RESPONSE_TO_BIOTIC_STIMULUS | 0.00000 | 0.00000 | 0.80122 | 0.52941 | 1.33782 | 413 |
| GO_RESPONSE_TO_CYTOKINE | 0.00000 | 0.00000 | | 0.57213 | 1.44599 | 414 |
| GO_RESPONSE_TO_DOPAMINE | 0.00257 | 0.03417 | 0.43171 | 0.69687 | 1.51915 | 18 |
| GO_RESPONSE_TO_HEAT | 0.00008 | 0.00176 | 0.53843 | 0.58045 | 1.39236 | 90 |
| GO_RESPONSE_TO_INTERFERON_GAMMA | 0.00027 | 0.00523 | 0.49849 | 0.63146 | 1.46766 | 46 |
| GO_RESPONSE_TO_INTERLEUKIN_1 | 0.00000 | 0.00000 | 0.80122 | 0.67726 | 1.62424 | 89 |
| GO_RESPONSE_TO_INTERLEUKIN_12 | 0.00071 | 0.01211 | 0.47727 | 0.65969 | 1.51034 | 32 |
| GO_RESPONSE_TO_INTERLEUKIN_4 | 0.00272 | 0.03574 | 0.43171 | 0.72199 | 1.55459 | 16 |
| GO_RESPONSE_TO_LEUKEMIA_INHIBITORY_FACTOR | 0.00063 | 0.01090 | 0.47727 | 0.58082 | 1.37342 | 64 |
| GO_RESPONSE_TO_OXYGEN_LEVELS | 0.00000 | 0.00004 | 0.64355 | 0.55489 | 1.37107 | 178 |
| GO_RESPONSE_TO_TEMPERATURE_STIMULUS | 0.00022 | 0.00431 | 0.51885 | 0.55470 | 1.33521 | 101 |
| GO_RESPONSE_TO_TUMOR_NECROSIS_FACTOR | 0.00000 | 0.00000 | 0.73376 | 0.62911 | 1.52465 | 115 |
| GO_RHYTHMIC_PROCESS | 0.00030 | 0.00569 | 0.49849 | 0.55984 | 1.34808 | 102 |
| GO_RIBONUCLEOPROTEIN_COMPLEX_BINDING | 0.00000 | 0.00002 | 0.67496 | 0.61567 | 1.48253 | 102 |
| GO_RIBONUCLEOPROTEIN_COMPLEX_BIOGENESIS | 0.00000 | 0.00000 | | 0.59359 | 1.49554 | 354 |
| GO_RIBONUCLEOPROTEIN_COMPLEX_SUBUNIT_ORGANIZATION | 0.00000 | 0.00000 | 0.76146 | 0.60319 | 1.48263 | 157 |
| GO_RIBONUCLEOPROTEIN_GRANULE | 0.00001 | 0.00015 | 0.61053 | 0.57244 | 1.39816 | 133 |
| GO_RIBONUCLEOSIDE_METABOLIC_PROCESS | 0.00205 | 0.02840 | 0.43171 | 0.63998 | 1.46048 | 31 |
| GO_RIBONUCLEOSIDE_TRIPHOSPHATE_BIOSYNTHETIC_PROCESS | 0.00068 | 0.01173 | 0.47727 | 0.66692 | 1.51887 | 30 |
| GO_RIBONUCLEOSIDE_TRIPHOSPHATE_METABOLIC_PROCESS | 0.00087 | 0.01426 | 0.47727 | 0.64092 | 1.47726 | 39 |
| GO_RIBOSOMAL_LARGE_SUBUNIT_ASSEMBLY | 0.00002 | 0.00056 | 0.57561 | 0.76123 | 1.70431 | 24 |
| GO_RIBOSOMAL_LARGE_SUBUNIT_BIOGENESIS | 0.00000 | 0.00000 | 0.80122 | 0.72446 | 1.71143 | 62 |
| GO_RIBOSOMAL_SMALL_SUBUNIT_BIOGENESIS | 0.00309 | 0.04026 | 0.43171 | 0.56979 | 1.34797 | 65 |
| GO_RIBOSOMAL_SUBUNIT | 0.00000 | 0.00001 | 0.69013 | 0.57331 | 1.41106 | 163 |
| GO_RIBOSOME | 0.00000 | 0.00000 | 0.70498 | 0.56671 | 1.40393 | 192 |
| GO_RIBOSOME_ASSEMBLY | 0.00004 | 0.00096 | 0.55733 | 0.63652 | 1.49422 | 54 |
| GO_RIBOSOME_BINDING | 0.00017 | 0.00352 | 0.51885 | 0.65585 | 1.51324 | 40 |
| GO_RIBOSOME_BIOGENESIS | 0.00000 | 0.00000 | | 0.59237 | 1.47987 | 251 |
| GO_RNA_CAPPING | 0.00064 | 0.01102 | 0.47727 | 0.67529 | 1.53054 | 29 |
| GO_RNA_CATABOLIC_PROCESS | 0.00000 | 0.00000 | | 0.65338 | 1.63873 | 293 |
| GO_RNA_DEPENDENT_DNA_BIOSYNTHETIC_PROCESS | 0.00000 | 0.00003 | 0.65944 | 0.68625 | 1.60749 | 53 |
| GO_RNA_EXPORT_FROM_NUCLEUS | 0.00000 | 0.00008 | 0.62726 | 0.59326 | 1.43776 | 115 |
| GO_RNA_HELICASE_ACTIVITY | 0.00213 | 0.02925 | 0.43171 | 0.58082 | 1.36052 | 53 |
| GO_RNA_LOCALIZATION | 0.00000 | 0.00000 | 0.82666 | 0.60521 | 1.49539 | 178 |
| GO_RNA_POLYMERASE_II_PREINITIATION_COMPLEX_ASSEMBLY | 0.00272 | 0.03574 | 0.43171 | 0.71997 | 1.55022 | 16 |
| GO_RNA_SPLICING | 0.00000 | 0.00000 | | 0.55935 | 1.40640 | 322 |
| GO_RNA_SPLICING_VIA_TRANSESTERIFICATION_REACTIONS | 0.00000 | 0.00000 | | 0.57313 | 1.43491 | 269 |
| GO_RNA_STABILIZATION | 0.00074 | 0.01249 | 0.47727 | 0.67242 | 1.52405 | 29 |
| GO_RRNA_BINDING | 0.00000 | 0.00008 | 0.62726 | 0.67221 | 1.57133 | 51 |
| GO_RRNA_METABOLIC_PROCESS | 0.00000 | 0.00004 | 0.64355 | 0.55423 | 1.37210 | 189 |
| GO_SCF_DEPENDENT_PROTEASOMAL_UBIQUITIN_DEPENDENT_PRO | 0.00000 | 0.00000 | 0.74774 | 0.69972 | 1.65299 | 62 |
| GO_SECRETORY_GRANULE | 0.00000 | 0.00001 | 0.69013 | 0.52468 | 1.31740 | 303 |
| GO_SECRETORY_VESICLE | 0.00001 | 0.00023 | 0.59333 | 0.50063 | 1.26100 | 348 |
| GO_SIGNAL_TRANSDUCTION_BY_PROTEIN_PHOSPHORYLATION | 0.00013 | 0.00274 | 0.51885 | 0.49121 | 1.23227 | 296 |
| GO_SINGLE_FERTILIZATION | 0.00108 | 0.01675 | 0.45506 | 0.65197 | 1.49271 | 33 |
| GO_SINGLE_STRANDED_RNA_BINDING | 0.00007 | 0.00156 | 0.53843 | 0.65270 | 1.52253 | 49 |
| GO_SMALL_RIBOSOMAL_SUBUNIT | 0.00088 | 0.01441 | 0.47727 | 0.57722 | 1.36490 | 64 |
| GO_SNRNA_BINDING | 0.00165 | 0.02376 | 0.45506 | 0.63435 | 1.45251 | 34 |
| GO_SOMATODENDRITIC_COMPARTMENT | 0.00163 | 0.02363 | 0.45506 | 0.48653 | 1.21466 | 237 |
| GO_SPLICEOSOMAL_COMPLEX | 0.00000 | 0.00007 | 0.62726 | 0.56910 | 1.39530 | 151 |
| GO_STEM_CELL_DIFFERENTIATION | 0.00000 | 0.00000 | 0.73376 | 0.61581 | 1.49486 | 119 |
| GO_STRUCTURAL_CONSTITUENT_OF_RIBOSOME | 0.00000 | 0.00001 | 0.67496 | 0.58607 | 1.43468 | 143 |
| GO_STRUCTURAL_MOLECULE_ACTIVITY | 0.00000 | 0.00003 | 0.65944 | 0.52876 | 1.32300 | 263 |
| GO_SUPRAMOLECULAR_COMPLEX | 0.00001 | 0.00025 | 0.59333 | 0.48154 | 1.22083 | 496 |
| GO_SYNAPSE | 0.00000 | 0.00000 | 0.70498 | 0.51291 | 1.29344 | 374 |
| GO_TELOMERASE_RNA_LOCALIZATION | 0.00001 | 0.00036 | 0.59333 | 0.80465 | 1.76531 | 19 |
| GO_TELOMERE_MAINTENANCE_VIA_TELOMERE_LENGTHENING | 0.00000 | 0.00003 | 0.65944 | 0.67023 | 1.57733 | 58 |
| GO_TELOMERE_ORGANIZATION | 0.00040 | 0.00728 | 0.49849 | 0.54924 | 1.32571 | 106 |
| GO_TERMINATION_OF_RNA_POLYMERASE_I_TRANSCRIPTION | 0.00165 | 0.02376 | 0.45506 | 0.67373 | 1.52034 | 27 |
| GO_THREONINE_TYPE_ENDOPEPTIDASE_ACTIVITY | 0.00000 | 0.00014 | 0.61053 | 0.84076 | 1.82279 | 17 |
| GO_THREONINE_TYPE_PEPTIDASE_ACTIVITY | 0.00000 | 0.00008 | 0.62726 | 0.82241 | 1.79282 | 18 |
| GO_TISSUE_MORPHOGENESIS | 0.00087 | 0.01431 | 0.47727 | 0.49095 | 1.22563 | 234 |
| GO_TOXIN_TRANSPORT | 0.00000 | 0.00014 | 0.61053 | 0.82038 | 1.79982 | 19 |
| GO_TRANSCRIPTION_BY_RNA_POLYMERASE_I | 0.00095 | 0.01519 | 0.47727 | 0.61899 | 1.44088 | 47 |
| GO_TRANSCRIPTION_COUPLED_NUCLEOTIDE_EXCISION_REPAIR | 0.00004 | 0.00097 | 0.55733 | 0.61903 | 1.46378 | 64 |
| GO_TRANSCRIPTION_ELONGATION_FROM_RNA_POLYMERASE_II_PR | 0.00183 | 0.02586 | 0.45506 | 0.56543 | 1.33702 | 64 |
| GO_TRANSCRIPTION_ELONGATION_FROM_RNA_POLYMERASE_I_PR | 0.00102 | 0.01610 | 0.45506 | 0.66718 | 1.50217 | 26 |

| | | | | | | |
|---|---|---|---|---|---|---|
| GO_TRANSCRIPTION_FACTOR_BINDING | 0.00357 | 0.04553 | 0.43171 | 0.46976 | 1.17924 | 300 |
| GO_TRANSCRIPTION_INITIATION_FROM_RNA_POLYMERASE_II_PROI | 0.00174 | 0.02486 | 0.45506 | 0.52471 | 1.26584 | 105 |
| GO_TRANSLATIONAL_INITIATION | 0.00000 | 0.00000 | | 0.68616 | 1.68357 | 153 |
| GO_TRANSLATION_FACTOR_ACTIVITY_RNA_BINDING | 0.00000 | 0.00000 | | 0.75002 | 1.77437 | 65 |
| GO_TRANSLATION_INITIATION_FACTOR_ACTIVITY | 0.00000 | 0.00000 | 0.71951 | 0.73959 | 1.71767 | 44 |
| GO_TRANSLATION_PREINITIATION_COMPLEX | 0.00023 | 0.00458 | 0.51885 | 0.78478 | 1.68012 | 15 |
| GO_TRANSLATION_REGULATOR_ACTIVITY | 0.00000 | 0.00000 | | 0.69857 | 1.68054 | 95 |
| GO_TRANSLATION_REGULATOR_ACTIVITY_NUCLEIC_ACID_BINDING | 0.00000 | 0.00000 | | 0.71374 | 1.70333 | 78 |
| GO_TRANSMEMBRANE_TRANSPORT | 0.00001 | 0.00017 | 0.61053 | 0.48874 | 1.23549 | 421 |
| GO_TUMOR_NECROSIS_FACTOR_MEDIATED_SIGNALING_PATHWAY | 0.00000 | 0.00000 | | 0.71615 | 1.70612 | 74 |
| GO_T_CELL_RECEPTOR_SIGNALING_PATHWAY | 0.00000 | 0.00000 | 0.80122 | 0.67631 | 1.62198 | 89 |
| GO_U2_TYPE_SPLICEOSOMAL_COMPLEX | 0.00350 | 0.04483 | 0.43171 | 0.55165 | 1.31603 | 77 |
| GO_UBIQUITIN_LIKE_PROTEIN_LIGASE_BINDING | 0.00000 | 0.00000 | 0.74774 | 0.58402 | 1.44098 | 170 |
| GO_UNFOLDED_PROTEIN_BINDING | 0.00000 | 0.00000 | 0.74774 | 0.68130 | 1.61882 | 70 |
| GO_VACUOLAR_LUMEN | 0.00087 | 0.01431 | 0.47727 | 0.57747 | 1.36550 | 64 |
| GO_VESICLE_LUMEN | 0.00000 | 0.00000 | | 0.66852 | 1.63518 | 140 |
| GO_VIRAL_GENE_EXPRESSION | 0.00000 | 0.00000 | | 0.63328 | 1.55658 | 158 |
| GO_VIRAL_GENOME_REPLICATION | 0.00141 | 0.02081 | 0.45506 | 0.57593 | 1.36070 | 63 |
| GO_VIRAL_LIFE_CYCLE | 0.00004 | 0.00103 | 0.55733 | 0.53558 | 1.32062 | 168 |
| HP_ABNORMALITIES_OF_PLACENTA_OR_UMBILICAL_CORD | 0.00224 | 0.03069 | 0.43171 | 0.72858 | 1.57957 | 17 |
| HP_BULBAR_PALSY | 0.00037 | 0.00675 | 0.49849 | 0.77671 | 1.66283 | 15 |
| HP_RIMMED_VACUOLES | 0.00239 | 0.03216 | 0.43171 | 0.72637 | 1.56402 | 16 |
| **Cluster 3** | | | | | | |
| pathway | pval | padj | log2err | ES | NES | size |
| GO_ACTIVATION_OF_IMMUNE_RESPONSE | 0.00000 | 0.00001 | 0.69013 | 0.61965 | 1.32882 | 212 |
| GO_ACTIVATION_OF_INNATE_IMMUNE_RESPONSE | 0.00000 | 0.00000 | 0.76146 | 0.73203 | 1.52688 | 84 |
| GO_AEROBIC_RESPIRATION | 0.00001 | 0.00040 | 0.59333 | 0.68868 | 1.42420 | 67 |
| GO_AMIDE_BINDING | 0.00154 | 0.02627 | 0.45506 | 0.57331 | 1.21953 | 149 |
| GO_AMINE_METABOLIC_PROCESS | 0.00000 | 0.00000 | | 0.78474 | 1.63440 | 81 |
| GO_ANAPHASE_PROMOTING_COMPLEX_DEPENDENT_CATABOLIC_P | 0.00000 | 0.00000 | 0.78819 | 0.75491 | 1.56748 | 75 |
| GO_ANCHORING_JUNCTION | 0.00000 | 0.00000 | | 0.60858 | 1.32376 | 460 |
| GO_ANTIGEN_PROCESSING_AND_PRESENTATION | 0.00000 | 0.00000 | 0.73376 | 0.66495 | 1.41090 | 137 |
| GO_ANTIGEN_PROCESSING_AND_PRESENTATION_OF_EXOGENOUS_ | 0.00000 | 0.00000 | 0.78819 | 0.79988 | 1.63811 | 55 |
| GO_ANTIGEN_PROCESSING_AND_PRESENTATION_OF_PEPTIDE_ANTI | 0.00000 | 0.00000 | 0.76146 | 0.68701 | 1.45323 | 118 |
| GO_ANTIGEN_PROCESSING_AND_PRESENTATION_OF_PEPTIDE_ANTI | 0.00000 | 0.00000 | 0.81404 | 0.77851 | 1.60811 | 66 |
| GO_ANTIGEN_RECEPTOR_MEDIATED_SIGNALING_PATHWAY | 0.00000 | 0.00000 | 0.71951 | 0.67541 | 1.42999 | 129 |
| GO_ATP_METABOLIC_PROCESS | 0.00040 | 0.00879 | 0.49849 | 0.56924 | 1.21865 | 195 |
| GO_AZUROPHIL_GRANULE | 0.00287 | 0.04393 | 0.43171 | 0.59549 | 1.25036 | 93 |
| GO_BINDING_OF_SPERM_TO_ZONA_PELLUCIDA | 0.00000 | 0.00005 | 0.64355 | 0.90412 | 1.73107 | 16 |
| GO_BLOOD_MICROPARTICLE | 0.00045 | 0.00992 | 0.49849 | 0.75869 | 1.47975 | 24 |
| GO_CADHERIN_BINDING | 0.00000 | 0.00000 | | 0.65082 | 1.39782 | 235 |
| GO_CANONICAL_WNT_SIGNALING_PATHWAY | 0.00000 | 0.00004 | 0.65944 | 0.62399 | 1.33196 | 172 |
| GO_CATALYTIC_STEP_2_SPLICEOSOME | 0.00028 | 0.00670 | 0.49849 | 0.63766 | 1.32806 | 81 |
| GO_CELLULAR_AMINO_ACID_METABOLIC_PROCESS | 0.00000 | 0.00000 | 0.77494 | 0.63706 | 1.36641 | 214 |
| GO_CELLULAR_COMPONENT_DISASSEMBLY | 0.00072 | 0.01449 | 0.47727 | 0.53572 | 1.16183 | 371 |
| GO_CELLULAR_KETONE_METABOLIC_PROCESS | 0.00000 | 0.00011 | 0.62726 | 0.63031 | 1.34079 | 149 |
| GO_CELLULAR_NITROGEN_COMPOUND_CATABOLIC_PROCESS | 0.00000 | 0.00000 | | 0.71616 | 1.55568 | 425 |
| GO_CELLULAR_RESPIRATION | 0.00075 | 0.01478 | 0.47727 | 0.58820 | 1.24822 | 138 |
| GO_CELLULAR_RESPONSE_TO_DRUG | 0.00253 | 0.04003 | 0.43171 | 0.69231 | 1.37297 | 31 |
| GO_CELLULAR_RESPONSE_TO_HEAT | 0.00019 | 0.00464 | 0.51885 | 0.63952 | 1.33547 | 85 |
| GO_CELLULAR_RESPONSE_TO_OXYGEN_LEVELS | 0.00000 | 0.00000 | 0.78819 | 0.67246 | 1.43140 | 151 |
| GO_CELL_ACTIVATION_INVOLVED_IN_IMMUNE_RESPONSE | 0.00000 | 0.00000 | 0.74774 | 0.59190 | 1.28304 | 346 |
| GO_CELL_ADHESION_MOLECULE_BINDING | 0.00000 | 0.00000 | | 0.62847 | 1.35916 | 307 |
| GO_CELL_BODY | 0.00056 | 0.01173 | 0.47727 | 0.55497 | 1.19367 | 248 |
| GO_CELL_CELL_RECOGNITION | 0.00000 | 0.00000 | 0.74774 | 0.88625 | 1.73260 | 25 |
| GO_CELL_CELL_SIGNALING_BY_WNT | 0.00000 | 0.00001 | 0.69013 | 0.60066 | 1.29442 | 272 |
| GO_CELL_CYCLE_DNA_REPLICATION | 0.00156 | 0.02642 | 0.45506 | 0.65089 | 1.33271 | 54 |
| GO_CELL_CYCLE_G2_M_PHASE_TRANSITION | 0.00000 | 0.00014 | 0.62726 | 0.60123 | 1.28852 | 207 |
| GO_CELL_CYCLE_PHASE_TRANSITION | 0.00000 | 0.00001 | 0.67496 | 0.56656 | 1.23136 | 446 |
| GO_CELL_DEATH_IN_RESPONSE_TO_OXIDATIVE_STRESS | 0.00076 | 0.01491 | 0.47727 | 0.67965 | 1.38102 | 47 |
| GO_CELL_RECOGNITION | 0.00000 | 0.00011 | 0.62726 | 0.73506 | 1.49933 | 50 |
| GO_CELL_SUBSTRATE_JUNCTION | 0.00000 | 0.00000 | | 0.67163 | 1.44854 | 279 |
| GO_CELL_SURFACE_RECEPTOR_SIGNALING_PATHWAY_INVOLVED_IN | 0.00000 | 0.00001 | 0.67496 | 0.59085 | 1.27679 | 299 |
| GO_CEREBRAL_CORTEX_CELL_MIGRATION | 0.00213 | 0.03447 | 0.43171 | 0.75392 | 1.46314 | 19 |
| GO_CHAPERONE_BINDING | 0.00003 | 0.00084 | 0.57561 | 0.66959 | 1.38917 | 73 |
| GO_CHAPERONE_COFACTOR_DEPENDENT_PROTEIN_REFOLDING | 0.00202 | 0.03286 | 0.43171 | 0.75603 | 1.46723 | 19 |
| GO_CHAPERONE_COMPLEX | 0.00000 | 0.00009 | 0.62726 | 0.85489 | 1.66412 | 22 |
| GO_COP9_SIGNALOSOME | 0.00053 | 0.01122 | 0.47727 | 0.77057 | 1.49960 | 23 |
| GO_COTRANSLATIONAL_PROTEIN_TARGETING_TO_MEMBRANE | 0.00000 | 0.00000 | | 0.88708 | 1.86061 | 91 |
| GO_CYTOKINE_MEDIATED_SIGNALING_PATHWAY | 0.00000 | 0.00000 | | 0.62955 | 1.36196 | 314 |
| GO_CYTOPLASMIC_TRANSLATION | 0.00000 | 0.00000 | | 0.83616 | 1.75038 | 88 |
| GO_CYTOPLASMIC_TRANSLATIONAL_INITIATION | 0.00000 | 0.00009 | 0.62726 | 0.81481 | 1.61336 | 30 |
| GO_CYTOSOLIC_LARGE_RIBOSOMAL_SUBUNIT | 0.00000 | 0.00000 | | 0.89402 | 1.83052 | 54 |
| GO_CYTOSOLIC_RIBOSOME | 0.00000 | 0.00000 | | 0.88808 | 1.86634 | 96 |
| GO_CYTOSOLIC_SMALL_RIBOSOMAL_SUBUNIT | 0.00000 | 0.00000 | | 0.88673 | 1.78736 | 39 |
| GO_DEFENSE_RESPONSE_TO_OTHER_ORGANISM | 0.00000 | 0.00005 | 0.64355 | 0.56387 | 1.22473 | 412 |

| | | | | | | |
|---|---|---|---|---|---|---|
| GO_DE_NOVO_PROTEIN_FOLDING | 0.00143 | 0.02476 | 0.45506 | 0.73867 | 1.44784 | 26 |
| GO_DNA_BIOSYNTHETIC_PROCESS | 0.00016 | 0.00396 | 0.51885 | 0.60777 | 1.28897 | 135 |
| GO_DNA_DAMAGE_RESPONSE_DETECTION_OF_DNA_DAMAGE | 0.00053 | 0.01122 | 0.47727 | 0.70658 | 1.41020 | 34 |
| GO_DOUBLE_STRANDED_RNA_BINDING | 0.00276 | 0.04258 | 0.43171 | 0.64626 | 1.32217 | 53 |
| GO_DRUG_BINDING | 0.00067 | 0.01371 | 0.47727 | 0.67472 | 1.37290 | 48 |
| GO_ENDOPEPTIDASE_ACTIVITY | 0.00130 | 0.02302 | 0.45506 | 0.56733 | 1.20938 | 165 |
| GO_ENDOPEPTIDASE_COMPLEX | 0.00000 | 0.00001 | 0.69013 | 0.74915 | 1.54473 | 62 |
| GO_ENERGY_DERIVATION_BY_OXIDATION_OF_ORGANIC_COMPOUN | 0.00113 | 0.02108 | 0.45506 | 0.56693 | 1.21151 | 177 |
| GO_ERROR_PRONE_TRANSLESION_SYNTHESIS | 0.00265 | 0.04143 | 0.43171 | 0.76820 | 1.48751 | 18 |
| GO_ESTABLISHMENT_OF_PROTEIN_LOCALIZATION_TO_CHROMOSOM | 0.00012 | 0.00312 | 0.51885 | 0.80665 | 1.57209 | 21 |
| GO_ESTABLISHMENT_OF_PROTEIN_LOCALIZATION_TO_ENDOPLASM | 0.00000 | 0.00000 | | 0.86564 | 1.82201 | 101 |
| GO_ESTABLISHMENT_OF_PROTEIN_LOCALIZATION_TO_MEMBRANE | 0.00000 | 0.00000 | | 0.73931 | 1.58825 | 239 |
| GO_ESTABLISHMENT_OF_PROTEIN_LOCALIZATION_TO_ORGANELLE | 0.00000 | 0.00000 | | 0.68779 | 1.49321 | 404 |
| GO_ESTABLISHMENT_OF_PROTEIN_LOCALIZATION_TO_TELOMERE | 0.00010 | 0.00260 | 0.53843 | 0.84929 | 1.62608 | 16 |
| GO_ESTABLISHMENT_OF_RNA_LOCALIZATION | 0.00030 | 0.00696 | 0.49849 | 0.58379 | 1.24292 | 155 |
| GO_ESTABLISHMENT_OF_TISSUE_POLARITY | 0.00000 | 0.00000 | 0.76146 | 0.73944 | 1.53916 | 80 |
| GO_EUKARYOTIC_TRANSLATION_INITIATION_FACTOR_3_COMPLEX | 0.00002 | 0.00056 | 0.57561 | 0.87816 | 1.67568 | 15 |
| GO_EXOCYTOSIS | 0.00000 | 0.00000 | 0.71951 | 0.57464 | 1.25038 | 467 |
| GO_FATTY_ACID_BINDING | 0.00309 | 0.04657 | 0.43171 | 0.78231 | 1.49784 | 16 |
| GO_FC_EPSILON_RECEPTOR_SIGNALING_PATHWAY | 0.00000 | 0.00000 | 0.78819 | 0.74181 | 1.54729 | 84 |
| GO_FC_RECEPTOR_SIGNALING_PATHWAY | 0.00000 | 0.00000 | | 0.71132 | 1.50604 | 124 |
| GO_FERTILIZATION | 0.00014 | 0.00354 | 0.51885 | 0.66356 | 1.37066 | 66 |
| GO_FICOLIN_1_RICH_GRANULE | 0.00000 | 0.00000 | 0.80122 | 0.73005 | 1.53335 | 95 |
| GO_GENERATION_OF_PRECURSOR_METABOLITES_AND_ENERGY | 0.00012 | 0.00312 | 0.51885 | 0.55199 | 1.19421 | 321 |
| GO_GLUTAMATERGIC_SYNAPSE | 0.00110 | 0.02046 | 0.45506 | 0.58246 | 1.23655 | 139 |
| GO_GLYCOSAMINOGLYCAN_BINDING | 0.00242 | 0.03849 | 0.43171 | 0.66016 | 1.33957 | 46 |
| GO_GLYCOSYL_COMPOUND_METABOLIC_PROCESS | 0.00009 | 0.00248 | 0.53843 | 0.65587 | 1.36071 | 73 |
| GO_GUANYL_NUCLEOTIDE_BINDING | 0.00176 | 0.02925 | 0.45506 | 0.55366 | 1.18594 | 204 |
| GO_HEAT_SHOCK_PROTEIN_BINDING | 0.00010 | 0.00249 | 0.53843 | 0.64735 | 1.35182 | 85 |
| GO_HEMATOPOIETIC_PROGENITOR_CELL_DIFFERENTIATION | 0.00000 | 0.00000 | 0.71951 | 0.70033 | 1.47542 | 104 |
| GO_HEMATOPOIETIC_STEM_CELL_DIFFERENTIATION | 0.00000 | 0.00000 | | 0.78490 | 1.62385 | 69 |
| GO_HEPARIN_BINDING | 0.00261 | 0.04108 | 0.43171 | 0.68093 | 1.36690 | 37 |
| GO_HSP90_PROTEIN_BINDING | 0.00141 | 0.02467 | 0.45506 | 0.71957 | 1.41792 | 28 |
| GO_IMMUNE_RESPONSE_REGULATING_SIGNALING_PATHWAY | 0.00000 | 0.00000 | 0.70498 | 0.62924 | 1.34736 | 198 |
| GO_INFLAMMATORY_RESPONSE_TO_ANTIGENIC_STIMULUS | 0.00268 | 0.04167 | 0.43171 | 0.76996 | 1.48559 | 17 |
| GO_INNATE_IMMUNE_RESPONSE | 0.00000 | 0.00004 | 0.65944 | 0.58139 | 1.25992 | 338 |
| GO_INNATE_IMMUNE_RESPONSE_ACTIVATING_SIGNAL_TRANSDUCT | 0.00000 | 0.00000 | | 0.78447 | 1.62296 | 69 |
| GO_INTERACTION_WITH_SYMBIONT | 0.00117 | 0.02148 | 0.45506 | 0.67801 | 1.37096 | 42 |
| GO_INTERLEUKIN_1_MEDIATED_SIGNALING_PATHWAY | 0.00000 | 0.00000 | | 0.79806 | 1.65211 | 71 |
| GO_INTRINSIC_APOPTOTIC_SIGNALING_PATHWAY | 0.00161 | 0.02720 | 0.45506 | 0.56659 | 1.21110 | 181 |
| GO_ION_CHANNEL_BINDING | 0.00054 | 0.01124 | 0.47727 | 0.64389 | 1.32707 | 61 |
| GO_LARGE_RIBOSOMAL_SUBUNIT | 0.00000 | 0.00000 | | 0.76449 | 1.61367 | 108 |
| GO_LEUKOCYTE_MEDIATED_IMMUNITY | 0.00000 | 0.00000 | 0.70498 | 0.58229 | 1.26294 | 372 |
| GO_LYSOSOMAL_LUMEN | 0.00048 | 0.01048 | 0.49849 | 0.68248 | 1.38216 | 44 |
| GO_MATURATION_OF_LSU_RRNA | 0.00008 | 0.00206 | 0.53843 | 0.79914 | 1.56230 | 25 |
| GO_MATURATION_OF_LSU_RRNA_FROM_TRICISTRONIC_RRNA_TRA | 0.00126 | 0.02258 | 0.45506 | 0.80660 | 1.53914 | 15 |
| GO_MIDBRAIN_DEVELOPMENT | 0.00198 | 0.03245 | 0.43171 | 0.67047 | 1.35568 | 43 |
| GO_MITOCHONDRIAL_PROTEIN_COMPLEX | 0.00167 | 0.02788 | 0.45506 | 0.55664 | 1.19208 | 199 |
| GO_MODIFICATION_DEPENDENT_MACROMOLECULE_CATABOLIC_PR | 0.00000 | 0.00003 | 0.65944 | 0.56173 | 1.22204 | 461 |
| GO_MONOCARBOXYLIC_ACID_BINDING | 0.00102 | 0.01947 | 0.45506 | 0.74527 | 1.45357 | 24 |
| GO_MONOSACCHARIDE_BIOSYNTHETIC_PROCESS | 0.00117 | 0.02148 | 0.45506 | 0.65010 | 1.33303 | 57 |
| GO_MORPHOGENESIS_OF_AN_EPITHELIUM | 0.00004 | 0.00107 | 0.55733 | 0.57616 | 1.24075 | 265 |
| GO_MORPHOGENESIS_OF_A_POLARIZED_EPITHELIUM | 0.00000 | 0.00000 | | 0.74951 | 1.57160 | 90 |
| GO_MRNA_3_UTR_BINDING | 0.00028 | 0.00651 | 0.49849 | 0.66335 | 1.36781 | 62 |
| GO_MRNA_5_UTR_BINDING | 0.00143 | 0.02476 | 0.45506 | 0.75711 | 1.47379 | 22 |
| GO_MRNA_BINDING | 0.00000 | 0.00000 | | 0.65463 | 1.40527 | 222 |
| GO_MRNA_EXPORT_FROM_NUCLEUS | 0.00253 | 0.04003 | 0.43171 | 0.60313 | 1.26677 | 95 |
| GO_MRNA_PROCESSING | 0.00002 | 0.00068 | 0.57561 | 0.55302 | 1.20098 | 407 |
| GO_MRNA_TRANSPORT | 0.00205 | 0.03337 | 0.43171 | 0.58859 | 1.24544 | 119 |
| GO_MULTI_ORGANISM_PROCESS | 0.00228 | 0.03645 | 0.43171 | 0.51951 | 1.13072 | 476 |
| GO_MYD88_DEPENDENT_TOLL_LIKE_RECEPTOR_SIGNALING_PATHW | 0.00007 | 0.00197 | 0.53843 | 0.84983 | 1.63970 | 17 |
| GO_MYD88_INDEPENDENT_TOLL_LIKE_RECEPTOR_SIGNALING_PATH | 0.00130 | 0.02302 | 0.45506 | 0.80446 | 1.54025 | 16 |
| GO_MYELOID_LEUKOCYTE_ACTIVATION | 0.00000 | 0.00000 | 0.80122 | 0.60456 | 1.30920 | 330 |
| GO_MYELOID_LEUKOCYTE_MEDIATED_IMMUNITY | 0.00000 | 0.00000 | 0.77494 | 0.61172 | 1.32245 | 302 |
| GO_NADH_METABOLIC_PROCESS | 0.00084 | 0.01625 | 0.47727 | 0.75645 | 1.47885 | 25 |
| GO_NAD_BINDING | 0.00054 | 0.01124 | 0.47727 | 0.69962 | 1.41190 | 41 |
| GO_NAD_METABOLIC_PROCESS | 0.00287 | 0.04393 | 0.43171 | 0.70793 | 1.40174 | 30 |
| GO_NCRNA_METABOLIC_PROCESS | 0.00003 | 0.00080 | 0.57561 | 0.55651 | 1.20682 | 366 |
| GO_NCRNA_PROCESSING | 0.00001 | 0.00028 | 0.59333 | 0.57202 | 1.23751 | 314 |
| GO_NEGATIVE_REGULATION_OF_CALCIUM_ION_TRANSMEMBRANE_ | 0.00276 | 0.04258 | 0.43171 | 0.78792 | 1.50859 | 16 |
| GO_NEGATIVE_REGULATION_OF_CANONICAL_WNT_SIGNALING_PAT | 0.00000 | 0.00000 | 0.76146 | 0.71326 | 1.49906 | 98 |
| GO_NEGATIVE_REGULATION_OF_CATION_CHANNEL_ACTIVITY | 0.00202 | 0.03286 | 0.43171 | 0.79457 | 1.51618 | 15 |
| GO_NEGATIVE_REGULATION_OF_CELLULAR_AMIDE_METABOLIC_PR | 0.00005 | 0.00148 | 0.55733 | 0.62708 | 1.32472 | 112 |
| GO_NEGATIVE_REGULATION_OF_CELL_CYCLE | 0.00014 | 0.00358 | 0.51885 | 0.54493 | 1.18282 | 397 |
| GO_NEGATIVE_REGULATION_OF_CELL_CYCLE_G2_M_PHASE_TRANS | 0.00000 | 0.00002 | 0.65944 | 0.69672 | 1.45324 | 84 |
| GO_NEGATIVE_REGULATION_OF_CELL_CYCLE_PHASE_TRANSITION | 0.00006 | 0.00159 | 0.55733 | 0.59307 | 1.26738 | 177 |

| | | | | | | |
|---|---|---|---|---|---|---|
| GO_NEGATIVE_REGULATION_OF_CELL_CYCLE_PROCESS | 0.00001 | 0.00021 | 0.61053 | 0.58804 | 1.26334 | 237 |
| GO_NEGATIVE_REGULATION_OF_CELL_DEATH | 0.00000 | 0.00015 | 0.61053 | 0.55108 | 1.19971 | 485 |
| GO_NEGATIVE_REGULATION_OF_MITOTIC_CELL_CYCLE | 0.00003 | 0.00080 | 0.57561 | 0.58607 | 1.25706 | 214 |
| GO_NEGATIVE_REGULATION_OF_PROTEIN_MODIFICATION_BY_SMA | 0.00143 | 0.02476 | 0.45506 | 0.64216 | 1.31865 | 58 |
| GO_NEGATIVE_REGULATION_OF_TRANSLATIONAL_INITIATION | 0.00290 | 0.04438 | 0.43171 | 0.76785 | 1.48153 | 17 |
| GO_NEGATIVE_REGULATION_OF_WNT_SIGNALING_PATHWAY | 0.00000 | 0.00000 | 0.80122 | 0.70434 | 1.48794 | 112 |
| GO_NEURON_DEATH | 0.00316 | 0.04718 | 0.43171 | 0.55906 | 1.19300 | 169 |
| GO_NEURON_TO_NEURON_SYNAPSE | 0.00002 | 0.00063 | 0.57561 | 0.61701 | 1.31421 | 156 |
| GO_NIK_NF_KAPPAB_SIGNALING | 0.00000 | 0.00000 | | 0.74751 | 1.57103 | 98 |
| GO_NON_CANONICAL_WNT_SIGNALING_PATHWAY | 0.00000 | 0.00000 | 0.81404 | 0.74944 | 1.56884 | 88 |
| GO_NUCLEAR_EXPORT | 0.00040 | 0.00886 | 0.49849 | 0.58373 | 1.24430 | 163 |
| GO_NUCLEAR_TRANSCRIBED_MRNA_CATABOLIC_PROCESS | 0.00000 | 0.00000 | | 0.78371 | 1.67509 | 179 |
| GO_NUCLEAR_TRANSCRIBED_MRNA_CATABOLIC_PROCESS_NONSEN | 0.00000 | 0.00000 | | 0.85538 | 1.80619 | 110 |
| GO_NUCLEAR_TRANSPORT | 0.00004 | 0.00125 | 0.55733 | 0.57550 | 1.23827 | 254 |
| GO_NUCLEOBASE_CONTAINING_COMPOUND_TRANSPORT | 0.00032 | 0.00730 | 0.49849 | 0.57343 | 1.22760 | 192 |
| GO_NUCLEOBASE_CONTAINING_SMALL_MOLECULE_METABOLIC_PR | 0.00054 | 0.01124 | 0.47727 | 0.53917 | 1.16884 | 356 |
| GO_NUCLEOSIDE_DIPHOSPHATE_METABOLIC_PROCESS | 0.00302 | 0.04570 | 0.43171 | 0.59164 | 1.24564 | 103 |
| GO_NUCLEOSIDE_METABOLIC_PROCESS | 0.00009 | 0.00237 | 0.53843 | 0.66915 | 1.37913 | 61 |
| GO_NUCLEOSIDE_PHOSPHATE_BIOSYNTHETIC_PROCESS | 0.00110 | 0.02046 | 0.45506 | 0.58457 | 1.24220 | 145 |
| GO_NUCLEOSOMAL_DNA_BINDING | 0.00335 | 0.04955 | 0.43171 | 0.73315 | 1.42883 | 21 |
| GO_NUCLEOTIDE_EXCISION_REPAIR_DNA_DAMAGE_RECOGNITION | 0.00028 | 0.00651 | 0.49849 | 0.79196 | 1.54163 | 22 |
| GO_NUCLEOTIDE_EXCISION_REPAIR_DNA_INCISION | 0.00316 | 0.04718 | 0.43171 | 0.67470 | 1.35440 | 37 |
| GO_ORGANIC_CYCLIC_COMPOUND_CATABOLIC_PROCESS | 0.00000 | 0.00000 | | 0.70564 | 1.53343 | 435 |
| GO_PEPTIDASE_COMPLEX | 0.00000 | 0.00002 | 0.67496 | 0.70547 | 1.46703 | 77 |
| GO_PIGMENT_GRANULE | 0.00000 | 0.00008 | 0.62726 | 0.69784 | 1.44833 | 74 |
| GO_POLYSOMAL_RIBOSOME | 0.00000 | 0.00000 | 0.80122 | 0.88261 | 1.74762 | 30 |
| GO_POLYSOME | 0.00000 | 0.00000 | | 0.81168 | 1.66436 | 57 |
| GO_POSITIVE_REGULATION_OF_APOPTOTIC_SIGNALING_PATHWAY | 0.00016 | 0.00405 | 0.51885 | 0.62289 | 1.31341 | 106 |
| GO_POSITIVE_REGULATION_OF_ATP_METABOLIC_PROCESS | 0.00183 | 0.03029 | 0.45506 | 0.73405 | 1.43878 | 26 |
| GO_POSITIVE_REGULATION_OF_CANONICAL_WNT_SIGNALING_PATH | 0.00000 | 0.00000 | 0.74774 | 0.71815 | 1.50441 | 89 |
| GO_POSITIVE_REGULATION_OF_CELLULAR_AMIDE_METABOLIC_PRO | 0.00034 | 0.00767 | 0.49849 | 0.61023 | 1.28964 | 116 |
| GO_POSITIVE_REGULATION_OF_CELLULAR_PROTEIN_LOCALIZATION | 0.00001 | 0.00032 | 0.59333 | 0.59474 | 1.27661 | 223 |
| GO_POSITIVE_REGULATION_OF_CELL_DEATH | 0.00002 | 0.00056 | 0.57561 | 0.56037 | 1.21482 | 354 |
| GO_POSITIVE_REGULATION_OF_DEFENSE_RESPONSE | 0.00000 | 0.00001 | 0.69013 | 0.63657 | 1.35683 | 161 |
| GO_POSITIVE_REGULATION_OF_DNA_BIOSYNTHETIC_PROCESS | 0.00006 | 0.00161 | 0.55733 | 0.70731 | 1.44441 | 51 |
| GO_POSITIVE_REGULATION_OF_DNA_TEMPLATED_TRANSCRIPTION_ | 0.00194 | 0.03194 | 0.45506 | 0.71575 | 1.40550 | 27 |
| GO_POSITIVE_REGULATION_OF_ESTABLISHMENT_OF_PROTEIN_LOC/ | 0.00000 | 0.00002 | 0.67496 | 0.61458 | 1.31795 | 212 |
| GO_POSITIVE_REGULATION_OF_IMMUNE_RESPONSE | 0.00000 | 0.00008 | 0.62726 | 0.58407 | 1.25931 | 275 |
| GO_POSITIVE_REGULATION_OF_IMMUNE_SYSTEM_PROCESS | 0.00001 | 0.00024 | 0.61053 | 0.55797 | 1.21065 | 382 |
| GO_POSITIVE_REGULATION_OF_INNATE_IMMUNE_RESPONSE | 0.00000 | 0.00000 | 0.71951 | 0.68190 | 1.44022 | 111 |
| GO_POSITIVE_REGULATION_OF_INTRACELLULAR_PROTEIN_TRANSPO | 0.00011 | 0.00293 | 0.53843 | 0.61404 | 1.30020 | 123 |
| GO_POSITIVE_REGULATION_OF_INTRACELLULAR_TRANSPORT | 0.00031 | 0.00722 | 0.49849 | 0.58834 | 1.25233 | 151 |
| GO_POSITIVE_REGULATION_OF_INTRINSIC_APOPTOTIC_SIGNALING_ | 0.00078 | 0.01529 | 0.47727 | 0.70313 | 1.41147 | 37 |
| GO_POSITIVE_REGULATION_OF_MRNA_PROCESSING | 0.00121 | 0.02180 | 0.45506 | 0.74988 | 1.46600 | 25 |
| GO_POSITIVE_REGULATION_OF_MRNA_SPLICING_VIA_SPLICEOSOM | 0.00078 | 0.01529 | 0.47727 | 0.81599 | 1.55705 | 15 |
| GO_POSITIVE_REGULATION_OF_NITRIC_OXIDE_METABOLIC_PROCES | 0.00135 | 0.02385 | 0.45506 | 0.78786 | 1.52012 | 17 |
| GO_POSITIVE_REGULATION_OF_PROTEIN_IMPORT | 0.00176 | 0.02925 | 0.45506 | 0.71764 | 1.40921 | 27 |
| GO_POSITIVE_REGULATION_OF_PROTEIN_LOCALIZATION_TO_NUCLE | 0.00000 | 0.00001 | 0.69013 | 0.74630 | 1.53249 | 58 |
| GO_POSITIVE_REGULATION_OF_RESPONSE_TO_BIOTIC_STIMULUS | 0.00000 | 0.00001 | 0.69013 | 0.66569 | 1.40908 | 126 |
| GO_POSITIVE_REGULATION_OF_RESPONSE_TO_EXTERNAL_STIMULU | 0.00001 | 0.00024 | 0.61053 | 0.59897 | 1.28273 | 202 |
| GO_POSITIVE_REGULATION_OF_TELOMERASE_RNA_LOCALIZATION_ | 0.00002 | 0.00058 | 0.57561 | 0.87739 | 1.67422 | 15 |
| GO_POSITIVE_REGULATION_OF_TRANSLATION | 0.00007 | 0.00186 | 0.53843 | 0.63890 | 1.34277 | 98 |
| GO_POSITIVE_REGULATION_OF_TRANSPORT | 0.00294 | 0.04470 | 0.43171 | 0.52690 | 1.14440 | 409 |
| GO_POSITIVE_REGULATION_OF_VIRAL_GENOME_REPLICATION | 0.00033 | 0.00752 | 0.49849 | 0.73437 | 1.45637 | 31 |
| GO_POSITIVE_REGULATION_OF_VIRAL_LIFE_CYCLE | 0.00156 | 0.02642 | 0.45506 | 0.66828 | 1.35792 | 47 |
| GO_POSITIVE_REGULATION_OF_VIRAL_PROCESS | 0.00121 | 0.02180 | 0.45506 | 0.62119 | 1.29421 | 82 |
| GO_POSITIVE_REGULATION_OF_WNT_SIGNALING_PATHWAY | 0.00000 | 0.00000 | 0.73376 | 0.69153 | 1.45812 | 106 |
| GO_POSTREPLICATION_REPAIR | 0.00104 | 0.01969 | 0.45506 | 0.68163 | 1.37825 | 43 |
| GO_POSTSYNAPSE | 0.00000 | 0.00006 | 0.64355 | 0.59474 | 1.27934 | 249 |
| GO_POSTTRANSCRIPTIONAL_REGULATION_OF_GENE_EXPRESSION | 0.00000 | 0.00000 | | 0.65164 | 1.41662 | 448 |
| GO_POST_TRANSLATIONAL_PROTEIN_MODIFICATION | 0.00001 | 0.00020 | 0.61053 | 0.59432 | 1.27508 | 216 |
| GO_PROCESS_UTILIZING_AUTOPHAGIC_MECHANISM | 0.00316 | 0.04718 | 0.43171 | 0.52446 | 1.13739 | 370 |
| GO_PROTEASOMAL_PROTEIN_CATABOLIC_PROCESS | 0.00000 | 0.00003 | 0.65944 | 0.57518 | 1.24688 | 350 |
| GO_PROTEASOMAL_UBIQUITIN_INDEPENDENT_PROTEIN_CATABOLI( | 0.00000 | 0.00000 | 0.77494 | 0.92516 | 1.79546 | 19 |
| GO_PROTEASOME_ACCESSORY_COMPLEX | 0.00005 | 0.00134 | 0.55733 | 0.79801 | 1.55645 | 24 |
| GO_PROTEASOME_CORE_COMPLEX | 0.00000 | 0.00000 | 0.77494 | 0.93882 | 1.81140 | 17 |
| GO_PROTEIN_CONTAINING_COMPLEX_DISASSEMBLY | 0.00054 | 0.01124 | 0.47727 | 0.55917 | 1.20142 | 240 |
| GO_PROTEIN_CONTAINING_COMPLEX_LOCALIZATION | 0.00053 | 0.01122 | 0.47727 | 0.56704 | 1.21372 | 194 |
| GO_PROTEIN_DNA_COMPLEX_SUBUNIT_ORGANIZATION | 0.00097 | 0.01848 | 0.47727 | 0.57696 | 1.22932 | 159 |
| GO_PROTEIN_FOLDING | 0.00000 | 0.00000 | 0.73376 | 0.65390 | 1.39326 | 159 |
| GO_PROTEIN_IMPORT | 0.00075 | 0.01478 | 0.47727 | 0.59382 | 1.25704 | 125 |
| GO_PROTEIN_LOCALIZATION_TO_ENDOPLASMIC_RETICULUM | 0.00000 | 0.00000 | | 0.84159 | 1.78152 | 125 |
| GO_PROTEIN_LOCALIZATION_TO_MEMBRANE | 0.00000 | 0.00000 | | 0.64490 | 1.39982 | 397 |
| GO_PROTEIN_LOCALIZATION_TO_MITOCHONDRION | 0.00034 | 0.00767 | 0.49849 | 0.61453 | 1.29763 | 110 |
| GO_PROTEIN_LOCALIZATION_TO_NUCLEUS | 0.00000 | 0.00011 | 0.62726 | 0.61725 | 1.31850 | 174 |
| GO_PROTEIN_MODIFICATION_BY_SMALL_PROTEIN_REMOVAL | 0.00000 | 0.00002 | 0.65944 | 0.61241 | 1.31319 | 211 |

| | | | | | | |
|---|---|---|---|---|---|---|
| GO_PROTEIN_POLYUBIQUITINATION | 0.00018 | 0.00444 | 0.51885 | 0.56575 | 1.21602 | 244 |
| GO_PROTEIN_REFOLDING | 0.00167 | 0.02788 | 0.45506 | 0.79845 | 1.52875 | 16 |
| GO_PROTEIN_STABILIZATION | 0.00004 | 0.00123 | 0.55733 | 0.62551 | 1.32379 | 127 |
| GO_PROTEIN_TARGETING | 0.00000 | 0.00000 | | 0.70306 | 1.52157 | 326 |
| GO_PROTEIN_TARGETING_TO_MEMBRANE | 0.00000 | 0.00000 | | 0.82410 | 1.75363 | 150 |
| GO_PURINE_CONTAINING_COMPOUND_METABOLIC_PROCESS | 0.00065 | 0.01339 | 0.47727 | 0.55331 | 1.19089 | 260 |
| GO_PURINE_NUCLEOSIDE_METABOLIC_PROCESS | 0.00052 | 0.01119 | 0.47727 | 0.70586 | 1.42279 | 39 |
| GO_REGULATION_OF_ANIMAL_ORGAN_MORPHOGENESIS | 0.00000 | 0.00000 | 0.77494 | 0.70281 | 1.48505 | 114 |
| GO_REGULATION_OF_APOPTOTIC_SIGNALING_PATHWAY | 0.00002 | 0.00062 | 0.57561 | 0.58196 | 1.24980 | 233 |
| GO_REGULATION_OF_CELLULAR_AMIDE_METABOLIC_PROCESS | 0.00000 | 0.00000 | 0.82666 | 0.62228 | 1.34527 | 302 |
| GO_REGULATION_OF_CELLULAR_AMINE_METABOLIC_PROCESS | 0.00000 | 0.00000 | | 0.83142 | 1.70484 | 57 |
| GO_REGULATION_OF_CELLULAR_AMINO_ACID_METABOLIC_PROCES | 0.00000 | 0.00000 | | 0.84425 | 1.72863 | 54 |
| GO_REGULATION_OF_CELLULAR_KETONE_METABOLIC_PROCESS | 0.00000 | 0.00000 | 0.78819 | 0.70510 | 1.48999 | 113 |
| GO_REGULATION_OF_CELLULAR_PROTEIN_CATABOLIC_PROCESS | 0.00068 | 0.01385 | 0.47727 | 0.57435 | 1.22696 | 175 |
| GO_REGULATION_OF_CELLULAR_PROTEIN_LOCALIZATION | 0.00001 | 0.00029 | 0.59333 | 0.56551 | 1.22579 | 353 |
| GO_REGULATION_OF_CELLULAR_RESPONSE_TO_HEAT | 0.00157 | 0.02666 | 0.45506 | 0.64093 | 1.32395 | 64 |
| GO_REGULATION_OF_CELLULAR_RESPONSE_TO_TRANSFORMING_G | 0.00075 | 0.01478 | 0.47727 | 0.66516 | 1.36083 | 53 |
| GO_REGULATION_OF_CELL_CYCLE_G2_M_PHASE_TRANSITION | 0.00000 | 0.00007 | 0.64355 | 0.62790 | 1.33942 | 168 |
| GO_REGULATION_OF_CELL_CYCLE_PHASE_TRANSITION | 0.00000 | 0.00018 | 0.61053 | 0.57104 | 1.23586 | 326 |
| GO_REGULATION_OF_DEFENSE_RESPONSE | 0.00005 | 0.00147 | 0.55733 | 0.56577 | 1.22159 | 291 |
| GO_REGULATION_OF_DNA_BIOSYNTHETIC_PROCESS | 0.00148 | 0.02564 | 0.45506 | 0.62212 | 1.29177 | 75 |
| GO_REGULATION_OF_DNA_TEMPLATED_TRANSCRIPTION_INITIATIOI | 0.00079 | 0.01541 | 0.47727 | 0.71675 | 1.42458 | 32 |
| GO_REGULATION_OF_DNA_TEMPLATED_TRANSCRIPTION_IN_RESPO | 0.00000 | 0.00000 | 0.81404 | 0.75304 | 1.57492 | 87 |
| GO_REGULATION_OF_GENE_EXPRESSION_EPIGENETIC | 0.00216 | 0.03487 | 0.43171 | 0.58785 | 1.24220 | 115 |
| GO_REGULATION_OF_HEMATOPOIETIC_PROGENITOR_CELL_DIFFERE | 0.00000 | 0.00000 | 0.81404 | 0.77248 | 1.59886 | 70 |
| GO_REGULATION_OF_HEMATOPOIETIC_STEM_CELL_DIFFERENTIATIC | 0.00000 | 0.00000 | 0.82666 | 0.79534 | 1.63997 | 62 |
| GO_REGULATION_OF_HEMOPOIESIS | 0.00002 | 0.00058 | 0.57561 | 0.58647 | 1.25825 | 216 |
| GO_REGULATION_OF_IMMUNE_RESPONSE | 0.00009 | 0.00247 | 0.53843 | 0.55078 | 1.19421 | 363 |
| GO_REGULATION_OF_INNATE_IMMUNE_RESPONSE | 0.00000 | 0.00009 | 0.62726 | 0.62485 | 1.33136 | 159 |
| GO_REGULATION_OF_INTRACELLULAR_PROTEIN_TRANSPORT | 0.00002 | 0.00062 | 0.57561 | 0.59972 | 1.28057 | 173 |
| GO_REGULATION_OF_INTRACELLULAR_TRANSPORT | 0.00119 | 0.02161 | 0.45506 | 0.55547 | 1.19307 | 229 |
| GO_REGULATION_OF_INTRINSIC_APOPTOTIC_SIGNALING_PATHWAY | 0.00000 | 0.00010 | 0.62726 | 0.66083 | 1.39130 | 103 |
| GO_REGULATION_OF_MITOTIC_CELL_CYCLE | 0.00000 | 0.00008 | 0.62726 | 0.56084 | 1.21841 | 431 |
| GO_REGULATION_OF_MORPHOGENESIS_OF_AN_EPITHELIUM | 0.00000 | 0.00000 | 0.78819 | 0.72547 | 1.52373 | 95 |
| GO_REGULATION_OF_MRNA_CATABOLIC_PROCESS | 0.00000 | 0.00000 | | 0.68524 | 1.45966 | 157 |
| GO_REGULATION_OF_MRNA_METABOLIC_PROCESS | 0.00000 | 0.00000 | | 0.65890 | 1.41807 | 257 |
| GO_REGULATION_OF_MRNA_PROCESSING | 0.00087 | 0.01690 | 0.47727 | 0.60534 | 1.27911 | 114 |
| GO_REGULATION_OF_MRNA_SPLICING_VIA_SPLICEOSOME | 0.00006 | 0.00181 | 0.53843 | 0.65970 | 1.37318 | 80 |
| GO_REGULATION_OF_NUCLEOCYTOPLASMIC_TRANSPORT | 0.00023 | 0.00559 | 0.51885 | 0.65154 | 1.34878 | 71 |
| GO_REGULATION_OF_OXIDATIVE_STRESS_INDUCED_CELL_DEATH | 0.00013 | 0.00339 | 0.51885 | 0.73712 | 1.47226 | 35 |
| GO_REGULATION_OF_OXIDATIVE_STRESS_INDUCED_INTRINSIC_APO | 0.00150 | 0.02588 | 0.45506 | 0.74869 | 1.45703 | 23 |
| GO_REGULATION_OF_PEPTIDE_TRANSPORT | 0.00002 | 0.00066 | 0.57561 | 0.55788 | 1.20885 | 339 |
| GO_REGULATION_OF_PROTEASOMAL_PROTEIN_CATABOLIC_PROCES | 0.00035 | 0.00778 | 0.49849 | 0.59406 | 1.25862 | 133 |
| GO_REGULATION_OF_PROTEIN_CATABOLIC_PROCESS | 0.00006 | 0.00175 | 0.53843 | 0.57253 | 1.23187 | 254 |
| GO_REGULATION_OF_PROTEIN_CONTAINING_COMPLEX_ASSEMBLY | 0.00283 | 0.04360 | 0.43171 | 0.54592 | 1.17461 | 252 |
| GO_REGULATION_OF_PROTEIN_IMPORT | 0.00011 | 0.00284 | 0.53843 | 0.73264 | 1.46798 | 36 |
| GO_REGULATION_OF_PROTEIN_LOCALIZATION_TO_NUCLEUS | 0.00000 | 0.00001 | 0.69013 | 0.71294 | 1.48538 | 82 |
| GO_REGULATION_OF_PROTEIN_MODIFICATION_BY_SMALL_PROTEIN | 0.00009 | 0.00229 | 0.53843 | 0.59616 | 1.27081 | 163 |
| GO_REGULATION_OF_PROTEIN_STABILITY | 0.00003 | 0.00098 | 0.55733 | 0.58759 | 1.25835 | 199 |
| GO_REGULATION_OF_PROTEOLYSIS | 0.00061 | 0.01254 | 0.47727 | 0.54057 | 1.17225 | 362 |
| GO_REGULATION_OF_RESPONSE_TO_BIOTIC_STIMULUS | 0.00001 | 0.00022 | 0.61053 | 0.59892 | 1.28262 | 199 |
| GO_REGULATION_OF_RESPONSE_TO_OXIDATIVE_STRESS | 0.00228 | 0.03645 | 0.43171 | 0.65981 | 1.34070 | 47 |
| GO_REGULATION_OF_RNA_SPLICING | 0.00003 | 0.00083 | 0.57561 | 0.63786 | 1.34789 | 113 |
| GO_REGULATION_OF_SIGNAL_TRANSDUCTION_BY_P53_CLASS_MED | 0.00119 | 0.02161 | 0.45506 | 0.58909 | 1.24693 | 126 |
| GO_REGULATION_OF_SMALL_MOLECULE_METABOLIC_PROCESS | 0.00000 | 0.00001 | 0.70498 | 0.60596 | 1.30406 | 256 |
| GO_REGULATION_OF_STEM_CELL_DIFFERENTIATION | 0.00000 | 0.00000 | 0.81404 | 0.75434 | 1.56913 | 79 |
| GO_REGULATION_OF_SYMBIOTIC_PROCESS | 0.00032 | 0.00734 | 0.49849 | 0.59649 | 1.26582 | 138 |
| GO_REGULATION_OF_TRANSCRIPTION_FROM_RNA_POLYMERASE_II | 0.00000 | 0.00000 | | 0.79840 | 1.64627 | 62 |
| GO_REGULATION_OF_TRANSLATIONAL_INITIATION | 0.00010 | 0.00262 | 0.53843 | 0.66729 | 1.37997 | 67 |
| GO_REGULATION_OF_TRANSLATION_IN_RESPONSE_TO_STRESS | 0.00130 | 0.02302 | 0.45506 | 0.78414 | 1.51838 | 18 |
| GO_REGULATION_OF_VIRAL_GENOME_REPLICATION | 0.00009 | 0.00242 | 0.53843 | 0.68669 | 1.40699 | 56 |
| GO_REGULATION_OF_VIRAL_LIFE_CYCLE | 0.00095 | 0.01819 | 0.47727 | 0.62201 | 1.30089 | 87 |
| GO_REGULATION_OF_WNT_SIGNALING_PATHWAY | 0.00000 | 0.00000 | 0.71951 | 0.62970 | 1.34809 | 195 |
| GO_RESPONSE_TO_ALCOHOL | 0.00179 | 0.02977 | 0.45506 | 0.59719 | 1.25616 | 99 |
| GO_RESPONSE_TO_DRUG | 0.00000 | 0.00006 | 0.64355 | 0.61995 | 1.32447 | 176 |
| GO_RESPONSE_TO_ESTRADIOL | 0.00294 | 0.04470 | 0.43171 | 0.63628 | 1.30470 | 57 |
| GO_RESPONSE_TO_ETHANOL | 0.00095 | 0.01819 | 0.47727 | 0.67771 | 1.37519 | 46 |
| GO_RESPONSE_TO_HEAT | 0.00004 | 0.00123 | 0.55733 | 0.64699 | 1.36306 | 104 |
| GO_RESPONSE_TO_INTERFERON_GAMMA | 0.00122 | 0.02206 | 0.45506 | 0.62650 | 1.30084 | 75 |
| GO_RESPONSE_TO_INTERLEUKIN_1 | 0.00000 | 0.00000 | | 0.73620 | 1.55099 | 104 |
| GO_RESPONSE_TO_INTERLEUKIN_12 | 0.00012 | 0.00312 | 0.51885 | 0.75106 | 1.49627 | 33 |
| GO_RESPONSE_TO_INTERLEUKIN_4 | 0.00052 | 0.01119 | 0.47727 | 0.77692 | 1.51413 | 21 |
| GO_RESPONSE_TO_LEUKEMIA_INHIBITORY_FACTOR | 0.00017 | 0.00409 | 0.51885 | 0.65876 | 1.36348 | 70 |
| GO_RESPONSE_TO_ORGANIC_CYCLIC_COMPOUND | 0.00075 | 0.01478 | 0.47727 | 0.52796 | 1.14688 | 427 |
| GO_RESPONSE_TO_OXYGEN_LEVELS | 0.00000 | 0.00000 | 0.77494 | 0.63653 | 1.36711 | 228 |
| GO_RESPONSE_TO_TEMPERATURE_STIMULUS | 0.00017 | 0.00418 | 0.51885 | 0.61058 | 1.29273 | 129 |

| | | | | | | |
|---|---|---|---|---|---|---|
| GO_RESPONSE_TO_TOPOLOGICALLY_INCORRECT_PROTEIN | 0.00023 | 0.00559 | 0.51885 | 0.59541 | 1.26437 | 140 |
| GO_RESPONSE_TO_TUMOR_NECROSIS_FACTOR | 0.00000 | 0.00000 | 0.71951 | 0.65746 | 1.39672 | 143 |
| GO_RETINA_HOMEOSTASIS | 0.00154 | 0.02627 | 0.45506 | 0.75321 | 1.46793 | 21 |
| GO_RIBONUCLEOPROTEIN_COMPLEX_BINDING | 0.00000 | 0.00002 | 0.65944 | 0.66907 | 1.41384 | 113 |
| GO_RIBONUCLEOPROTEIN_COMPLEX_BIOGENESIS | 0.00000 | 0.00000 | | 0.65700 | 1.42554 | 377 |
| GO_RIBONUCLEOPROTEIN_COMPLEX_SUBUNIT_ORGANIZATION | 0.00000 | 0.00000 | | 0.67650 | 1.44391 | 170 |
| GO_RIBONUCLEOPROTEIN_GRANULE | 0.00001 | 0.00049 | 0.59333 | 0.60587 | 1.29154 | 165 |
| GO_RIBONUCLEOSIDE_METABOLIC_PROCESS | 0.00054 | 0.01139 | 0.47727 | 0.70341 | 1.41785 | 39 |
| GO_RIBONUCLEOSIDE_TRIPHOSPHATE_METABOLIC_PROCESS | 0.00313 | 0.04700 | 0.43171 | 0.66174 | 1.33803 | 43 |
| GO_RIBOSE_PHOSPHATE_METABOLIC_PROCESS | 0.00106 | 0.01991 | 0.45506 | 0.55008 | 1.18317 | 248 |
| GO_RIBOSOMAL_LARGE_SUBUNIT_ASSEMBLY | 0.00006 | 0.00173 | 0.53843 | 0.77198 | 1.52119 | 28 |
| GO_RIBOSOMAL_LARGE_SUBUNIT_BIOGENESIS | 0.00000 | 0.00000 | 0.77494 | 0.76877 | 1.58789 | 65 |
| GO_RIBOSOMAL_SMALL_SUBUNIT_ASSEMBLY | 0.00216 | 0.03487 | 0.43171 | 0.79448 | 1.52116 | 16 |
| GO_RIBOSOMAL_SMALL_SUBUNIT_BIOGENESIS | 0.00034 | 0.00774 | 0.49849 | 0.65362 | 1.35013 | 66 |
| GO_RIBOSOMAL_SUBUNIT | 0.00000 | 0.00000 | | 0.76254 | 1.62661 | 168 |
| GO_RIBOSOME | 0.00000 | 0.00000 | | 0.74023 | 1.58510 | 200 |
| GO_RIBOSOME_ASSEMBLY | 0.00000 | 0.00000 | 0.70498 | 0.75308 | 1.54644 | 58 |
| GO_RIBOSOME_BINDING | 0.00117 | 0.02148 | 0.45506 | 0.67449 | 1.36865 | 46 |
| GO_RIBOSOME_BIOGENESIS | 0.00000 | 0.00000 | | 0.65348 | 1.40727 | 265 |
| GO_RNA_CATABOLIC_PROCESS | 0.00000 | 0.00000 | | 0.74803 | 1.61889 | 326 |
| GO_RNA_DEPENDENT_DNA_BIOSYNTHETIC_PROCESS | 0.00187 | 0.03081 | 0.45506 | 0.64026 | 1.31669 | 59 |
| GO_RNA_EXPORT_FROM_NUCLEUS | 0.00064 | 0.01306 | 0.47727 | 0.60143 | 1.27334 | 121 |
| GO_RNA_LOCALIZATION | 0.00001 | 0.00038 | 0.59333 | 0.60784 | 1.30042 | 185 |
| GO_RNA_SPLICING | 0.00000 | 0.00006 | 0.64355 | 0.57634 | 1.24968 | 358 |
| GO_RNA_SPLICING_VIA_TRANSESTERIFICATION_REACTIONS | 0.00000 | 0.00001 | 0.67496 | 0.59096 | 1.27640 | 293 |
| GO_ROUGH_ENDOPLASMIC_RETICULUM | 0.00000 | 0.00003 | 0.65944 | 0.74560 | 1.52664 | 54 |
| GO_ROUGH_ENDOPLASMIC_RETICULUM_MEMBRANE | 0.00268 | 0.04167 | 0.43171 | 0.72712 | 1.41818 | 24 |
| GO_RRNA_BINDING | 0.00000 | 0.00002 | 0.67496 | 0.75574 | 1.54614 | 53 |
| GO_RRNA_METABOLIC_PROCESS | 0.00000 | 0.00000 | 0.71951 | 0.62741 | 1.34365 | 199 |
| GO_SCF_DEPENDENT_PROTEASOMAL_UBIQUITIN_DEPENDENT_PRO | 0.00000 | 0.00000 | 0.74774 | 0.74177 | 1.54019 | 75 |
| GO_SECRETORY_GRANULE | 0.00000 | 0.00000 | 0.80122 | 0.59417 | 1.28978 | 393 |
| GO_SECRETORY_VESICLE | 0.00000 | 0.00000 | 0.70498 | 0.57264 | 1.24609 | 464 |
| GO_SIGNAL_TRANSDUCTION_BY_PROTEIN_PHOSPHORYLATION | 0.00002 | 0.00061 | 0.57561 | 0.55370 | 1.20208 | 400 |
| GO_SINGLE_FERTILIZATION | 0.00002 | 0.00058 | 0.57561 | 0.71342 | 1.45519 | 50 |
| GO_SINGLE_STRANDED_DNA_BINDING | 0.00106 | 0.01991 | 0.45506 | 0.60062 | 1.26138 | 92 |
| GO_SINGLE_STRANDED_RNA_BINDING | 0.00124 | 0.02232 | 0.45506 | 0.65449 | 1.34007 | 54 |
| GO_SMALL_MOLECULE_BIOSYNTHETIC_PROCESS | 0.00072 | 0.01449 | 0.47727 | 0.53330 | 1.15709 | 381 |
| GO_SMALL_RIBOSOMAL_SUBUNIT | 0.00000 | 0.00000 | 0.82666 | 0.79153 | 1.63395 | 63 |
| GO_SPERM_EGG_RECOGNITION | 0.00000 | 0.00000 | 0.73376 | 0.89737 | 1.74681 | 22 |
| GO_SPLICEOSOMAL_COMPLEX | 0.00008 | 0.00207 | 0.53843 | 0.59716 | 1.27293 | 163 |
| GO_STEM_CELL_DIFFERENTIATION | 0.00000 | 0.00000 | | 0.70718 | 1.49814 | 131 |
| GO_STRUCTURAL_CONSTITUENT_OF_RIBOSOME | 0.00000 | 0.00000 | | 0.77387 | 1.64675 | 150 |
| GO_STRUCTURAL_MOLECULE_ACTIVITY | 0.00000 | 0.00000 | | 0.69839 | 1.51147 | 326 |
| GO_SUBSTANTIA_NIGRA_DEVELOPMENT | 0.00137 | 0.02410 | 0.45506 | 0.74006 | 1.44342 | 24 |
| GO_TELOMERASE_RNA_LOCALIZATION | 0.00029 | 0.00689 | 0.49849 | 0.80097 | 1.55444 | 19 |
| GO_THREONINE_TYPE_ENDOPEPTIDASE_ACTIVITY | 0.00000 | 0.00000 | 0.80122 | 0.94827 | 1.82964 | 17 |
| GO_THREONINE_TYPE_PEPTIDASE_ACTIVITY | 0.00000 | 0.00000 | 0.70498 | 0.89775 | 1.74228 | 20 |
| GO_TISSUE_MORPHOGENESIS | 0.00053 | 0.01122 | 0.47727 | 0.54807 | 1.18450 | 300 |
| GO_TOLL_LIKE_RECEPTOR_SIGNALING_PATHWAY | 0.00059 | 0.01221 | 0.47727 | 0.65427 | 1.35139 | 65 |
| GO_TOXIN_TRANSPORT | 0.00011 | 0.00291 | 0.53843 | 0.79554 | 1.54820 | 23 |
| GO_TRANSCRIPTION_COUPLED_NUCLEOTIDE_EXCISION_REPAIR | 0.00020 | 0.00490 | 0.51885 | 0.65744 | 1.36016 | 69 |
| GO_TRANSCRIPTION_FACTOR_BINDING | 0.00075 | 0.01478 | 0.47727 | 0.53508 | 1.16092 | 383 |
| GO_TRANSFORMING_GROWTH_FACTOR_BETA_RECEPTOR_SIGNALIN | 0.00231 | 0.03694 | 0.43171 | 0.58995 | 1.23897 | 92 |
| GO_TRANSLATIONAL_ELONGATION | 0.00039 | 0.00873 | 0.49849 | 0.60785 | 1.28446 | 115 |
| GO_TRANSLATIONAL_INITIATION | 0.00000 | 0.00000 | | 0.82013 | 1.74947 | 168 |
| GO_TRANSLATION_ELONGATION_FACTOR_ACTIVITY | 0.00001 | 0.00038 | 0.59333 | 0.88345 | 1.68578 | 15 |
| GO_TRANSLATION_FACTOR_ACTIVITY_RNA_BINDING | 0.00000 | 0.00000 | | 0.79975 | 1.65389 | 67 |
| GO_TRANSLATION_INITIATION_FACTOR_ACTIVITY | 0.00000 | 0.00000 | 0.71951 | 0.80067 | 1.61895 | 43 |
| GO_TRANSLATION_INITIATION_FACTOR_BINDING | 0.00008 | 0.00212 | 0.53843 | 0.76624 | 1.50989 | 28 |
| GO_TRANSLATION_PREINITIATION_COMPLEX | 0.00008 | 0.00207 | 0.53843 | 0.85357 | 1.63429 | 16 |
| GO_TRANSLATION_REGULATOR_ACTIVITY | 0.00000 | 0.00000 | | 0.74955 | 1.58011 | 105 |
| GO_TRANSLATION_REGULATOR_ACTIVITY_NUCLEIC_ACID_BINDING | 0.00000 | 0.00000 | | 0.77015 | 1.60401 | 81 |
| GO_TRICARBOXYLIC_ACID_CYCLE | 0.00167 | 0.02788 | 0.45506 | 0.72672 | 1.43623 | 29 |
| GO_TUMOR_NECROSIS_FACTOR_MEDIATED_SIGNALING_PATHWAY | 0.00000 | 0.00000 | 0.77494 | 0.72617 | 1.52120 | 89 |
| GO_T_CELL_RECEPTOR_SIGNALING_PATHWAY | 0.00000 | 0.00000 | 0.76146 | 0.69615 | 1.47122 | 116 |
| GO_U12_TYPE_SPLICEOSOMAL_COMPLEX | 0.00268 | 0.04167 | 0.43171 | 0.74522 | 1.45065 | 22 |
| GO_U2_SNRNP | 0.00324 | 0.04816 | 0.43171 | 0.78005 | 1.49352 | 16 |
| GO_UBIQUITIN_LIKE_PROTEIN_LIGASE_BINDING | 0.00000 | 0.00000 | 0.74774 | 0.63237 | 1.35547 | 208 |
| GO_UNFOLDED_PROTEIN_BINDING | 0.00000 | 0.00000 | 0.73376 | 0.72549 | 1.50983 | 78 |
| GO_VACUOLAR_LUMEN | 0.00009 | 0.00248 | 0.53843 | 0.65214 | 1.36025 | 84 |
| GO_VACUOLE | 0.00031 | 0.00722 | 0.49849 | 0.53125 | 1.15539 | 458 |
| GO_VESICLE_LUMEN | 0.00000 | 0.00000 | | 0.68820 | 1.46705 | 165 |
| GO_VIRAL_GENE_EXPRESSION | 0.00000 | 0.00000 | | 0.77984 | 1.66412 | 169 |
| GO_VIRAL_GENOME_REPLICATION | 0.00119 | 0.02161 | 0.45506 | 0.62701 | 1.30192 | 75 |
| GO_VIRAL_LIFE_CYCLE | 0.00002 | 0.00074 | 0.57561 | 0.58760 | 1.25998 | 211 |
| HP_ABNORMALITY_OF_CARDIOVASCULAR_SYSTEM_ELECTROPHYSIO | 0.00078 | 0.01529 | 0.47727 | 0.55708 | 1.19647 | 230 |

| | | | | | | |
|---|---|---|---|---|---|---|
| HP_ABNORMALITY_OF_LOWER_LIP | 0.00235 | 0.03742 | 0.43171 | 0.57156 | 1.21249 | 136 |
| HP_ABNORMALITY_OF_THE_NASAL_DORSUM | 0.00004 | 0.00123 | 0.55733 | 0.61752 | 1.30791 | 130 |
| HP_ABNORMAL_ERYTHROCYTE_MORPHOLOGY | 0.00004 | 0.00125 | 0.55733 | 0.55881 | 1.21031 | 333 |
| HP_ACUTE_LEUKEMIA | 0.00000 | 0.00000 | 0.74774 | 0.75462 | 1.56103 | 68 |
| HP_ANEMIA_OF_INADEQUATE_PRODUCTION | 0.00002 | 0.00069 | 0.57561 | 0.65138 | 1.37230 | 104 |
| HP_APHASIA | 0.00331 | 0.04913 | 0.43171 | 0.72276 | 1.41665 | 26 |
| HP_DELAYED_PUBERTY | 0.00001 | 0.00020 | 0.61053 | 0.65457 | 1.37786 | 102 |
| HP_DEPRESSED_NASAL_RIDGE | 0.00000 | 0.00008 | 0.62726 | 0.69490 | 1.44169 | 73 |
| HP_FATIGUE | 0.00037 | 0.00825 | 0.49849 | 0.58034 | 1.23965 | 174 |
| HP_HEMATOLOGICAL_NEOPLASM | 0.00032 | 0.00734 | 0.49849 | 0.58789 | 1.25138 | 151 |
| HP_JUVENILE_ASEPTIC_NECROSIS | 0.00265 | 0.04143 | 0.43171 | 0.78762 | 1.50292 | 15 |
| HP_LEUKEMIA | 0.00000 | 0.00011 | 0.62726 | 0.66699 | 1.40097 | 94 |
| HP_MACROCYTIC_ANEMIA | 0.00000 | 0.00000 | 0.73376 | 0.80941 | 1.63666 | 42 |
| HP_MACROGYRIA | 0.00104 | 0.01969 | 0.45506 | 0.80768 | 1.54642 | 16 |
| HP_MIGRAINE | 0.00000 | 0.00001 | 0.69013 | 0.74080 | 1.52924 | 63 |
| HP_MUTISM | 0.00305 | 0.04614 | 0.43171 | 0.71103 | 1.40524 | 29 |
| HP_PALLOR | 0.00022 | 0.00537 | 0.51885 | 0.62987 | 1.31947 | 89 |
| HP_RETICULOCYTOPENIA | 0.00005 | 0.00143 | 0.55733 | 0.86555 | 1.65163 | 15 |
| HP_SHORT_NOSE | 0.00154 | 0.02627 | 0.45506 | 0.57325 | 1.22146 | 160 |
| HP_SUPRANUCLEAR_GAZE_PALSY | 0.00133 | 0.02360 | 0.45506 | 0.75096 | 1.46145 | 23 |
| HP_THICK_LOWER_LIP_VERMILION | 0.00035 | 0.00778 | 0.49849 | 0.62275 | 1.30804 | 94 |
| HP_THICK_VERMILION_BORDER | 0.00004 | 0.00105 | 0.55733 | 0.61047 | 1.29690 | 143 |
| HP_THROMBOCYTOSIS | 0.00115 | 0.02135 | 0.45506 | 0.75943 | 1.48006 | 21 |
| HP_TRANSIENT_ISCHEMIC_ATTACK | 0.00058 | 0.01207 | 0.47727 | 0.76677 | 1.49222 | 23 |
| **Cluster 4** | | | | | | |
| pathway | pval | padj | log2err | ES | NES | size |
| GO_ALTERNATIVE_MRNA_SPLICING_VIA_SPLICEOSOME | 0.00009 | 0.00564 | 0.53843 | 0.65857 | 1.36993 | 66 |
| GO_ANAPHASE_PROMOTING_COMPLEX_DEPENDENT_CATABOLIC_P | 0.00055 | 0.02404 | 0.47727 | 0.62567 | 1.31041 | 75 |
| GO_APOPTOTIC_SIGNALING_PATHWAY | 0.00022 | 0.01142 | 0.51885 | 0.52793 | 1.15063 | 400 |
| GO_CATALYTIC_STEP_2_SPLICEOSOME | 0.00130 | 0.04542 | 0.45506 | 0.61141 | 1.28259 | 79 |
| GO_CELLULAR_NITROGEN_COMPOUND_CATABOLIC_PROCESS | 0.00000 | 0.00000 | | 0.61051 | 1.33409 | 451 |
| GO_CELLULAR_RESPONSE_TO_OXYGEN_LEVELS | 0.00030 | 0.01505 | 0.49849 | 0.57213 | 1.22462 | 160 |
| GO_CELL_CELL_SIGNALING_BY_WNT | 0.00133 | 0.04619 | 0.45506 | 0.52340 | 1.13818 | 340 |
| GO_CELL_CYCLE_PHASE_TRANSITION | 0.00004 | 0.00303 | 0.55733 | 0.52836 | 1.15628 | 498 |
| GO_CELL_DIVISION | 0.00000 | 0.00029 | 0.62726 | 0.54063 | 1.18182 | 459 |
| GO_CELL_SUBSTRATE_JUNCTION | 0.00002 | 0.00146 | 0.57561 | 0.55652 | 1.20830 | 307 |
| GO_CHROMATIN_ASSEMBLY_OR_DISASSEMBLY | 0.00122 | 0.04436 | 0.45506 | 0.57354 | 1.21631 | 117 |
| GO_CHROMATIN_BINDING | 0.00005 | 0.00361 | 0.55733 | 0.53762 | 1.17292 | 417 |
| GO_CHROMOSOMAL_REGION | 0.00001 | 0.00118 | 0.59333 | 0.55978 | 1.21367 | 292 |
| GO_CHROMOSOME_CENTROMERIC_REGION | 0.00005 | 0.00389 | 0.55733 | 0.58057 | 1.24462 | 168 |
| GO_CHROMOSOME_SEGREGATION | 0.00049 | 0.02219 | 0.47727 | 0.54272 | 1.17425 | 265 |
| GO_COLLAGEN_CATABOLIC_PROCESS | 0.00010 | 0.00622 | 0.53843 | 0.80574 | 1.56920 | 20 |
| GO_CONDENSED_CHROMOSOME | 0.00011 | 0.00657 | 0.53843 | 0.57934 | 1.24273 | 173 |
| GO_CONDENSED_CHROMOSOME_CENTROMERIC_REGION | 0.00068 | 0.02859 | 0.47727 | 0.59956 | 1.26809 | 98 |
| GO_COTRANSLATIONAL_PROTEIN_TARGETING_TO_MEMBRANE | 0.00000 | 0.00000 | | 0.78240 | 1.64917 | 90 |
| GO_COVALENT_CHROMATIN_MODIFICATION | 0.00000 | 0.00034 | 0.62726 | 0.55371 | 1.20497 | 359 |
| GO_CYTOPLASMIC_STRESS_GRANULE | 0.00019 | 0.01063 | 0.51885 | 0.65196 | 1.35122 | 61 |
| GO_CYTOPLASMIC_TRANSLATION | 0.00000 | 0.00000 | 0.78819 | 0.72025 | 1.51755 | 89 |
| GO_CYTOSOLIC_LARGE_RIBOSOMAL_SUBUNIT | 0.00000 | 0.00000 | | 0.80263 | 1.64865 | 51 |
| GO_CYTOSOLIC_RIBOSOME | 0.00000 | 0.00000 | | 0.81123 | 1.71392 | 94 |
| GO_CYTOSOLIC_SMALL_RIBOSOMAL_SUBUNIT | 0.00000 | 0.00000 | | 0.85220 | 1.72880 | 40 |
| GO_DNA_BINDING_TRANSCRIPTION_FACTOR_BINDING | 0.00008 | 0.00532 | 0.53843 | 0.55927 | 1.20774 | 242 |
| GO_DNA_CONFORMATION_CHANGE | 0.00012 | 0.00732 | 0.51885 | 0.55749 | 1.20254 | 228 |
| GO_DNA_DAMAGE_RESPONSE_SIGNAL_TRANSDUCTION_BY_P53_CL | 0.00082 | 0.03216 | 0.47727 | 0.60943 | 1.28207 | 85 |
| GO_DNA_INTEGRITY_CHECKPOINT | 0.00084 | 0.03247 | 0.47727 | 0.57642 | 1.22538 | 127 |
| GO_DNA_PACKAGING | 0.00141 | 0.04664 | 0.45506 | 0.56892 | 1.20969 | 128 |
| GO_DNA_REPAIR | 0.00013 | 0.00738 | 0.51885 | 0.52990 | 1.15748 | 439 |
| GO_DNA_REPLICATION | 0.00115 | 0.04244 | 0.45506 | 0.53728 | 1.15942 | 231 |
| GO_DNA_TEMPLATED_TRANSCRIPTION_ELONGATION | 0.00013 | 0.00750 | 0.51885 | 0.62097 | 1.31187 | 96 |
| GO_DNA_TEMPLATED_TRANSCRIPTION_INITIATION | 0.00007 | 0.00493 | 0.53843 | 0.57831 | 1.24097 | 177 |
| GO_DOUBLE_STRANDED_RNA_BINDING | 0.00081 | 0.03201 | 0.47727 | 0.63162 | 1.30750 | 60 |
| GO_ESTABLISHMENT_OF_PROTEIN_LOCALIZATION_TO_ENDOPLASM | 0.00000 | 0.00000 | | 0.77124 | 1.63106 | 99 |
| GO_ESTABLISHMENT_OF_PROTEIN_LOCALIZATION_TO_MEMBRANE | 0.00000 | 0.00000 | | 0.63813 | 1.37965 | 256 |
| GO_ESTABLISHMENT_OF_PROTEIN_LOCALIZATION_TO_ORGANELLE | 0.00000 | 0.00000 | | 0.58791 | 1.28419 | 439 |
| GO_ESTABLISHMENT_OF_RNA_LOCALIZATION | 0.00000 | 0.00016 | 0.64355 | 0.61208 | 1.31107 | 166 |
| GO_FICOLIN_1_RICH_GRANULE | 0.00137 | 0.04642 | 0.45506 | 0.59477 | 1.25652 | 96 |
| GO_GLOBAL_GENOME_NUCLEOTIDE_EXCISION_REPAIR | 0.00122 | 0.04436 | 0.45506 | 0.73910 | 1.45568 | 23 |
| GO_HETEROCHROMATIN | 0.00152 | 0.04976 | 0.45506 | 0.63285 | 1.30931 | 59 |
| GO_HISTONE_BINDING | 0.00051 | 0.02253 | 0.47727 | 0.56195 | 1.20697 | 187 |
| GO_HISTONE_H2A_UBIQUITINATION | 0.00042 | 0.01946 | 0.49849 | 0.76139 | 1.49315 | 22 |
| GO_HISTONE_METHYLTRANSFERASE_COMPLEX | 0.00064 | 0.02686 | 0.47727 | 0.62774 | 1.30696 | 68 |
| GO_HISTONE_UBIQUITINATION | 0.00028 | 0.01427 | 0.49849 | 0.69307 | 1.40909 | 42 |
| GO_IMPORT_INTO_NUCLEUS | 0.00009 | 0.00564 | 0.53843 | 0.59778 | 1.26926 | 122 |
| GO_INTRINSIC_APOPTOTIC_SIGNALING_PATHWAY | 0.00001 | 0.00111 | 0.59333 | 0.57958 | 1.24690 | 203 |
| GO_INTRINSIC_APOPTOTIC_SIGNALING_PATHWAY_BY_P53_CLASS_M | 0.00021 | 0.01133 | 0.51885 | 0.66127 | 1.36259 | 54 |
| GO_KINETOCHORE | 0.00030 | 0.01505 | 0.49849 | 0.58521 | 1.24111 | 118 |

| | | | | | | |
|---|---|---|---|---|---|---|
| GO_LARGE_RIBOSOMAL_SUBUNIT | 0.00000 | 0.00000 | 0.74774 | 0.68359 | 1.44689 | 104 |
| GO_MATURATION_OF_LSU_RRNA | 0.00029 | 0.01461 | 0.49849 | 0.75565 | 1.49635 | 26 |
| GO_METHYLTRANSFERASE_COMPLEX | 0.00003 | 0.00221 | 0.57561 | 0.63976 | 1.35165 | 94 |
| GO_MITOTIC_NUCLEAR_DIVISION | 0.00023 | 0.01205 | 0.51885 | 0.55705 | 1.19984 | 215 |
| GO_MITOTIC_SPINDLE | 0.00137 | 0.04642 | 0.45506 | 0.58151 | 1.23176 | 108 |
| GO_MRNA_3_UTR_BINDING | 0.00098 | 0.03745 | 0.45506 | 0.62514 | 1.30456 | 70 |
| GO_MRNA_5_UTR_BINDING | 0.00008 | 0.00542 | 0.53843 | 0.78789 | 1.54513 | 22 |
| GO_MRNA_BINDING | 0.00000 | 0.00000 | | 0.64899 | 1.40127 | 239 |
| GO_MRNA_EXPORT_FROM_NUCLEUS | 0.00009 | 0.00564 | 0.53843 | 0.61781 | 1.30657 | 99 |
| GO_MRNA_PROCESSING | 0.00000 | 0.00000 | 0.80122 | 0.57837 | 1.26180 | 419 |
| GO_MRNA_TRANSPORT | 0.00004 | 0.00348 | 0.55733 | 0.60580 | 1.28745 | 126 |
| GO_NCRNA_METABOLIC_PROCESS | 0.00137 | 0.04642 | 0.45506 | 0.52064 | 1.13402 | 384 |
| GO_NCRNA_PROCESSING | 0.00006 | 0.00426 | 0.55733 | 0.54502 | 1.18432 | 320 |
| GO_NCRNA_TRANSCRIPTION | 0.00036 | 0.01738 | 0.49849 | 0.61278 | 1.29491 | 95 |
| GO_NEGATIVE_REGULATION_OF_CELLULAR_AMIDE_METABOLIC_PR | 0.00021 | 0.01131 | 0.51885 | 0.59118 | 1.25471 | 120 |
| GO_NEGATIVE_REGULATION_OF_CELL_CYCLE | 0.00000 | 0.00042 | 0.62726 | 0.54118 | 1.18276 | 456 |
| GO_NEGATIVE_REGULATION_OF_CELL_CYCLE_PHASE_TRANSITION | 0.00128 | 0.04529 | 0.45506 | 0.54539 | 1.17255 | 197 |
| GO_NEGATIVE_REGULATION_OF_CELL_CYCLE_PROCESS | 0.00001 | 0.00103 | 0.59333 | 0.56639 | 1.22543 | 264 |
| GO_NUCLEAR_ENVELOPE | 0.00000 | 0.00002 | 0.70498 | 0.57236 | 1.24483 | 338 |
| GO_NUCLEAR_EXPORT | 0.00000 | 0.00052 | 0.61053 | 0.60264 | 1.29286 | 175 |
| GO_NUCLEAR_MEMBRANE | 0.00000 | 0.00036 | 0.62726 | 0.58367 | 1.25749 | 217 |
| GO_NUCLEAR_SPECK | 0.00000 | 0.00052 | 0.61053 | 0.56347 | 1.22270 | 302 |
| GO_NUCLEAR_TRANSCRIBED_MRNA_CATABOLIC_PROCESS | 0.00000 | 0.00000 | | 0.68474 | 1.47071 | 187 |
| GO_NUCLEAR_TRANSCRIBED_MRNA_CATABOLIC_PROCESS_NONSEN | 0.00000 | 0.00000 | | 0.77353 | 1.63940 | 113 |
| GO_NUCLEAR_TRANSPORT | 0.00000 | 0.00000 | 0.74774 | 0.59281 | 1.28469 | 281 |
| GO_NUCLEOBASE_CONTAINING_COMPOUND_TRANSPORT | 0.00001 | 0.00130 | 0.59333 | 0.57897 | 1.24622 | 207 |
| GO_NUCLEUS_ORGANIZATION | 0.00130 | 0.04542 | 0.45506 | 0.58640 | 1.24118 | 103 |
| GO_ORGANELLE_FISSION | 0.00072 | 0.02953 | 0.47727 | 0.52815 | 1.14857 | 348 |
| GO_ORGANIC_CYCLIC_COMPOUND_CATABOLIC_PROCESS | 0.00000 | 0.00000 | | 0.60373 | 1.31971 | 472 |
| GO_P53_BINDING | 0.00141 | 0.04664 | 0.45506 | 0.64893 | 1.33147 | 49 |
| GO_PCG_PROTEIN_COMPLEX | 0.00070 | 0.02916 | 0.47727 | 0.68066 | 1.38157 | 41 |
| GO_PEPTIDYL_LYSINE_MODIFICATION | 0.00009 | 0.00560 | 0.53843 | 0.54800 | 1.18892 | 300 |
| GO_POLYSOMAL_RIBOSOME | 0.00000 | 0.00005 | 0.67496 | 0.82912 | 1.65224 | 29 |
| GO_POLYSOME | 0.00000 | 0.00002 | 0.70498 | 0.72868 | 1.50844 | 60 |
| GO_POSITIVE_REGULATION_OF_CELLULAR_AMIDE_METABOLIC_PRC | 0.00059 | 0.02545 | 0.47727 | 0.58160 | 1.23585 | 124 |
| GO_POSITIVE_REGULATION_OF_CELL_CYCLE | 0.00040 | 0.01875 | 0.49849 | 0.54205 | 1.17387 | 271 |
| GO_POSITIVE_REGULATION_OF_INTRACELLULAR_PROTEIN_TRANSPC | 0.00080 | 0.03185 | 0.47727 | 0.57269 | 1.21874 | 133 |
| GO_POSITIVE_REGULATION_OF_NUCLEOCYTOPLASMIC_TRANSPORT | 0.00005 | 0.00389 | 0.55733 | 0.70109 | 1.43291 | 47 |
| GO_POSITIVE_REGULATION_OF_ORGANELLE_ORGANIZATION | 0.00074 | 0.03005 | 0.47727 | 0.51755 | 1.13079 | 447 |
| GO_POSITIVE_REGULATION_OF_PROTEIN_MODIFICATION_BY_SMAL | 0.00128 | 0.04529 | 0.45506 | 0.59057 | 1.25139 | 110 |
| GO_POSITIVE_REGULATION_OF_TRANSLATION | 0.00012 | 0.00725 | 0.53843 | 0.60939 | 1.28983 | 104 |
| GO_POSITIVE_REGULATION_OF_VIRAL_GENOME_REPLICATION | 0.00100 | 0.03745 | 0.45506 | 0.70122 | 1.40389 | 31 |
| GO_POSITIVE_REGULATION_OF_VIRAL_PROCESS | 0.00055 | 0.02404 | 0.47727 | 0.61069 | 1.28415 | 84 |
| GO_POSTTRANSCRIPTIONAL_REGULATION_OF_GENE_EXPRESSION | 0.00000 | 0.00000 | | 0.58728 | 1.28507 | 485 |
| GO_PRERIBOSOME | 0.00022 | 0.01144 | 0.51885 | 0.64153 | 1.33567 | 68 |
| GO_PROCESS_UTILIZING_AUTOPHAGIC_MECHANISM | 0.00124 | 0.04450 | 0.45506 | 0.52075 | 1.13508 | 403 |
| GO_PROTEASOMAL_PROTEIN_CATABOLIC_PROCESS | 0.00005 | 0.00361 | 0.55733 | 0.53567 | 1.16669 | 393 |
| GO_PROTEIN_CONTAINING_COMPLEX_LOCALIZATION | 0.00009 | 0.00565 | 0.53843 | 0.56446 | 1.21637 | 220 |
| GO_PROTEIN_DNA_COMPLEX_SUBUNIT_ORGANIZATION | 0.00001 | 0.00079 | 0.59333 | 0.59564 | 1.27749 | 172 |
| GO_PROTEIN_IMPORT | 0.00075 | 0.03005 | 0.47727 | 0.57410 | 1.22389 | 144 |
| GO_PROTEIN_LOCALIZATION_TO_ENDOPLASMIC_RETICULUM | 0.00000 | 0.00000 | | 0.75326 | 1.59940 | 122 |
| GO_PROTEIN_LOCALIZATION_TO_MEMBRANE | 0.00001 | 0.00052 | 0.61053 | 0.53869 | 1.17751 | 464 |
| GO_PROTEIN_LOCALIZATION_TO_NUCLEUS | 0.00000 | 0.00020 | 0.64355 | 0.59502 | 1.28012 | 205 |
| GO_PROTEIN_MODIFICATION_BY_SMALL_PROTEIN_REMOVAL | 0.00022 | 0.01142 | 0.51885 | 0.55190 | 1.19112 | 234 |
| GO_PROTEIN_POLYUBIQUITINATION | 0.00050 | 0.02244 | 0.47727 | 0.54255 | 1.17389 | 265 |
| GO_PROTEIN_TARGETING | 0.00000 | 0.00000 | | 0.60669 | 1.31928 | 346 |
| GO_PROTEIN_TARGETING_TO_MEMBRANE | 0.00000 | 0.00000 | | 0.72999 | 1.56030 | 154 |
| GO_REGULATION_OF_APOPTOTIC_SIGNALING_PATHWAY | 0.00141 | 0.04664 | 0.45506 | 0.53127 | 1.15080 | 274 |
| GO_REGULATION_OF_CELLULAR_AMIDE_METABOLIC_PROCESS | 0.00000 | 0.00000 | 0.73376 | 0.58439 | 1.26999 | 328 |
| GO_REGULATION_OF_CELL_CYCLE_PHASE_TRANSITION | 0.00046 | 0.02090 | 0.49849 | 0.52915 | 1.15142 | 360 |
| GO_REGULATION_OF_CHROMOSOME_ORGANIZATION | 0.00043 | 0.01994 | 0.49849 | 0.53943 | 1.16875 | 279 |
| GO_REGULATION_OF_CHROMOSOME_SEGREGATION | 0.00135 | 0.04642 | 0.45506 | 0.59731 | 1.26197 | 94 |
| GO_REGULATION_OF_DNA_METABOLIC_PROCESS | 0.00044 | 0.02042 | 0.49849 | 0.54388 | 1.17588 | 256 |
| GO_REGULATION_OF_GENE_EXPRESSION_EPIGENETIC | 0.00007 | 0.00493 | 0.53843 | 0.60144 | 1.27821 | 126 |
| GO_REGULATION_OF_INTRINSIC_APOPTOTIC_SIGNALING_PATHWAY | 0.00007 | 0.00493 | 0.53843 | 0.60459 | 1.28163 | 116 |
| GO_REGULATION_OF_MITOTIC_CELL_CYCLE | 0.00031 | 0.01538 | 0.49849 | 0.51940 | 1.13652 | 491 |
| GO_REGULATION_OF_MRNA_CATABOLIC_PROCESS | 0.00005 | 0.00382 | 0.55733 | 0.58299 | 1.25035 | 172 |
| GO_REGULATION_OF_MRNA_METABOLIC_PROCESS | 0.00000 | 0.00000 | 0.78819 | 0.60311 | 1.30671 | 279 |
| GO_REGULATION_OF_MRNA_PROCESSING | 0.00001 | 0.00117 | 0.59333 | 0.62012 | 1.31638 | 121 |
| GO_REGULATION_OF_MRNA_SPLICING_VIA_SPLICEOSOME | 0.00000 | 0.00042 | 0.62726 | 0.65761 | 1.38510 | 87 |
| GO_REGULATION_OF_NUCLEOCYTOPLASMIC_TRANSPORT | 0.00021 | 0.01136 | 0.51885 | 0.63236 | 1.32654 | 79 |
| GO_REGULATION_OF_PROTEASOMAL_UBIQUITIN_DEPENDENT_PRO | 0.00061 | 0.02605 | 0.47727 | 0.58952 | 1.24676 | 100 |
| GO_REGULATION_OF_PROTEIN_MODIFICATION_BY_SMALL_PROTEIN | 0.00002 | 0.00154 | 0.57561 | 0.59039 | 1.26832 | 188 |
| GO_REGULATION_OF_RESPONSE_TO_DNA_DAMAGE_STIMULUS | 0.00037 | 0.01738 | 0.49849 | 0.56982 | 1.22186 | 171 |
| GO_REGULATION_OF_RNA_SPLICING | 0.00000 | 0.00013 | 0.65944 | 0.63383 | 1.34581 | 122 |
| GO_REGULATION_OF_SIGNAL_TRANSDUCTION_BY_P53_CLASS_MED | 0.00001 | 0.00066 | 0.61053 | 0.61706 | 1.31456 | 141 |

| | | | | | | |
|---|---|---|---|---|---|---|
| GO_REGULATION_OF_SYMBIOTIC_PROCESS | 0.00071 | 0.02935 | 0.47727 | 0.56979 | 1.21429 | 143 |
| GO_REGULATION_OF_TRANSLATIONAL_INITIATION | 0.00020 | 0.01104 | 0.51885 | 0.64316 | 1.33906 | 68 |
| GO_REGULATION_OF_UBIQUITIN_DEPENDENT_PROTEIN_CATABOLIC | 0.00139 | 0.04664 | 0.45506 | 0.57415 | 1.21856 | 120 |
| GO_REGULATION_OF_UBIQUITIN_PROTEIN_TRANSFERASE_ACTIVITY | 0.00133 | 0.04619 | 0.45506 | 0.65155 | 1.33451 | 48 |
| GO_RESPONSE_TO_GROWTH_FACTOR | 0.00087 | 0.03347 | 0.47727 | 0.52124 | 1.13625 | 406 |
| GO_RESPONSE_TO_OXYGEN_LEVELS | 0.00013 | 0.00750 | 0.51885 | 0.55106 | 1.19076 | 251 |
| GO_RIBONUCLEOPROTEIN_COMPLEX_BIOGENESIS | 0.00000 | 0.00000 | | 0.62619 | 1.36391 | 384 |
| GO_RIBONUCLEOPROTEIN_COMPLEX_SUBUNIT_ORGANIZATION | 0.00000 | 0.00000 | 0.83909 | 0.65459 | 1.40453 | 178 |
| GO_RIBONUCLEOPROTEIN_GRANULE | 0.00000 | 0.00038 | 0.62726 | 0.59626 | 1.28067 | 187 |
| GO_RIBOSOMAL_LARGE_SUBUNIT_ASSEMBLY | 0.00006 | 0.00437 | 0.55733 | 0.76427 | 1.51820 | 28 |
| GO_RIBOSOMAL_LARGE_SUBUNIT_BIOGENESIS | 0.00000 | 0.00001 | 0.71951 | 0.72936 | 1.51561 | 65 |
| GO_RIBOSOMAL_SMALL_SUBUNIT_BIOGENESIS | 0.00077 | 0.03096 | 0.47727 | 0.63125 | 1.31172 | 65 |
| GO_RIBOSOMAL_SUBUNIT | 0.00000 | 0.00000 | | 0.70078 | 1.50108 | 166 |
| GO_RIBOSOME | 0.00000 | 0.00000 | | 0.67921 | 1.46057 | 198 |
| GO_RIBOSOME_ASSEMBLY | 0.00000 | 0.00020 | 0.64355 | 0.71839 | 1.48426 | 58 |
| GO_RIBOSOME_BIOGENESIS | 0.00000 | 0.00000 | | 0.61715 | 1.33564 | 267 |
| GO_RNA_CATABOLIC_PROCESS | 0.00000 | 0.00000 | | 0.64003 | 1.39194 | 345 |
| GO_RNA_EXPORT_FROM_NUCLEUS | 0.00001 | 0.00111 | 0.59333 | 0.61616 | 1.30948 | 126 |
| GO_RNA_LOCALIZATION | 0.00000 | 0.00006 | 0.67496 | 0.60483 | 1.30036 | 197 |
| GO_RNA_SPLICING | 0.00000 | 0.00000 | 0.77494 | 0.58403 | 1.27129 | 369 |
| GO_RNA_SPLICING_VIA_TRANSESTERIFICATION_REACTIONS | 0.00000 | 0.00001 | 0.73376 | 0.58584 | 1.27124 | 301 |
| GO_RRNA_BINDING | 0.00000 | 0.00039 | 0.62726 | 0.71355 | 1.46950 | 53 |
| GO_RRNA_METABOLIC_PROCESS | 0.00000 | 0.00002 | 0.70498 | 0.60693 | 1.30554 | 202 |
| GO_SIGNAL_TRANSDUCTION_BY_P53_CLASS_MEDIATOR | 0.00000 | 0.00028 | 0.62726 | 0.59241 | 1.27530 | 210 |
| GO_SINGLE_STRANDED_RNA_BINDING | 0.00003 | 0.00239 | 0.57561 | 0.66904 | 1.39025 | 65 |
| GO_SMALL_RIBOSOMAL_SUBUNIT | 0.00000 | 0.00000 | 0.81404 | 0.75868 | 1.57654 | 65 |
| GO_SNRNA_TRANSCRIPTION | 0.00124 | 0.04450 | 0.45506 | 0.61671 | 1.28535 | 69 |
| GO_SPINDLE | 0.00007 | 0.00493 | 0.53843 | 0.54922 | 1.19156 | 300 |
| GO_SPLICEOSOMAL_COMPLEX | 0.00003 | 0.00259 | 0.55733 | 0.58891 | 1.26052 | 164 |
| GO_SPLICEOSOMAL_COMPLEX_ASSEMBLY | 0.00084 | 0.03247 | 0.47727 | 0.65712 | 1.34828 | 49 |
| GO_STRUCTURAL_CONSTITUENT_OF_RIBOSOME | 0.00000 | 0.00000 | | 0.70322 | 1.50007 | 145 |
| GO_STRUCTURAL_MOLECULE_ACTIVITY | 0.00000 | 0.00004 | 0.67496 | 0.56438 | 1.22844 | 365 |
| GO_TISSUE_REGENERATION | 0.00100 | 0.03745 | 0.45506 | 0.68295 | 1.37956 | 36 |
| GO_TRANSCRIPTION_COREGULATOR_ACTIVITY | 0.00009 | 0.00560 | 0.53843 | 0.53521 | 1.16554 | 375 |
| GO_TRANSCRIPTION_COREPRESSOR_ACTIVITY | 0.00075 | 0.03005 | 0.47727 | 0.57121 | 1.21939 | 149 |
| GO_TRANSCRIPTION_ELONGATION_FROM_RNA_POLYMERASE_II_PR | 0.00006 | 0.00451 | 0.53843 | 0.65082 | 1.35932 | 72 |
| GO_TRANSCRIPTION_FACTOR_BINDING | 0.00001 | 0.00052 | 0.61053 | 0.54188 | 1.18409 | 452 |
| GO_TRANSCRIPTION_INITIATION_FROM_RNA_POLYMERASE_II_PROI | 0.00062 | 0.02626 | 0.47727 | 0.57004 | 1.21377 | 138 |
| GO_TRANSCRIPTION_REGULATOR_COMPLEX | 0.00015 | 0.00844 | 0.51885 | 0.54782 | 1.18703 | 280 |
| GO_TRANSLATIONAL_INITIATION | 0.00000 | 0.00000 | | 0.74227 | 1.59123 | 169 |
| GO_TRANSLATION_FACTOR_ACTIVITY_RNA_BINDING | 0.00002 | 0.00200 | 0.57561 | 0.66231 | 1.38275 | 71 |
| GO_TRANSLATION_INITIATION_FACTOR_ACTIVITY | 0.00013 | 0.00750 | 0.51885 | 0.69406 | 1.41602 | 46 |
| GO_TRANSLATION_REGULATOR_ACTIVITY | 0.00000 | 0.00029 | 0.62726 | 0.63526 | 1.34636 | 113 |
| GO_TRANSLATION_REGULATOR_ACTIVITY_NUCLEIC_ACID_BINDING | 0.00001 | 0.00064 | 0.61053 | 0.65318 | 1.37577 | 87 |
| GO_UBIQUITIN_LIGASE_COMPLEX | 0.00000 | 0.00021 | 0.64355 | 0.58219 | 1.25648 | 234 |
| GO_UBIQUITIN_LIKE_PROTEIN_BINDING | 0.00051 | 0.02253 | 0.47727 | 0.62070 | 1.30262 | 80 |
| GO_UBIQUITIN_LIKE_PROTEIN_LIGASE_ACTIVITY | 0.00037 | 0.01738 | 0.49849 | 0.56167 | 1.20773 | 196 |
| GO_UBIQUITIN_LIKE_PROTEIN_LIGASE_BINDING | 0.00013 | 0.00750 | 0.51885 | 0.55764 | 1.20460 | 244 |
| GO_UBIQUITIN_LIKE_PROTEIN_TRANSFERASE_ACTIVITY | 0.00003 | 0.00284 | 0.55733 | 0.55227 | 1.19928 | 311 |
| GO_UNFOLDED_PROTEIN_BINDING | 0.00012 | 0.00725 | 0.53843 | 0.63349 | 1.33269 | 85 |
| GO_VIRAL_GENE_EXPRESSION | 0.00000 | 0.00000 | | 0.71093 | 1.52444 | 171 |
| GO_VIRAL_LIFE_CYCLE | 0.00121 | 0.04421 | 0.45506 | 0.54839 | 1.18174 | 218 |
| GO_ZINC_ION_BINDING | 0.00148 | 0.04881 | 0.45506 | 0.51424 | 1.12409 | 467 |
| HP_ABNORMALITY_OF_LOWER_LIP | 0.00052 | 0.02277 | 0.47727 | 0.56616 | 1.21401 | 171 |
| HP_ABNORMALITY_OF_THE_NASAL_DORSUM | 0.00037 | 0.01738 | 0.49849 | 0.57156 | 1.22209 | 156 |
| HP_ACUTE_LEUKEMIA | 0.00022 | 0.01142 | 0.51885 | 0.63747 | 1.33089 | 71 |
| HP_DELAYED_PUBERTY | 0.00085 | 0.03262 | 0.47727 | 0.57400 | 1.21734 | 118 |
| HP_DEPRESSED_NASAL_RIDGE | 0.00005 | 0.00361 | 0.55733 | 0.64375 | 1.35428 | 85 |
| HP_LEUKEMIA | 0.00100 | 0.03745 | 0.45506 | 0.58774 | 1.24366 | 102 |
| HP_MACROCYTIC_ANEMIA | 0.00006 | 0.00448 | 0.53843 | 0.71337 | 1.44549 | 39 |
| HP_MIGRAINE | 0.00104 | 0.03860 | 0.45506 | 0.61737 | 1.28945 | 72 |
| HP_THICK_LOWER_LIP_VERMILION | 0.00037 | 0.01738 | 0.49849 | 0.59324 | 1.25731 | 115 |
| **Cluster 5** | | | | | | |
| pathway | pval | padj | log2err | ES | NES | size |
| GO_ACTIVATION_OF_IMMUNE_RESPONSE | 0.00000 | 0.00000 | 0.70498 | 0.58095 | 1.45882 | 160 |
| GO_ACTIVATION_OF_INNATE_IMMUNE_RESPONSE | 0.00000 | 0.00000 | 0.78819 | 0.69769 | 1.70168 | 73 |
| GO_ADP_BINDING | 0.00110 | 0.01910 | 0.45506 | 0.71568 | 1.58916 | 20 |
| GO_AEROBIC_RESPIRATION | 0.00000 | 0.00001 | 0.69013 | 0.67944 | 1.63526 | 59 |
| GO_ALPHA_AMINO_ACID_BIOSYNTHETIC_PROCESS | 0.00309 | 0.04513 | 0.43171 | 0.61109 | 1.42282 | 37 |
| GO_AMINE_METABOLIC_PROCESS | 0.00000 | 0.00000 | | 0.76193 | 1.84706 | 66 |
| GO_ANAPHASE_PROMOTING_COMPLEX_DEPENDENT_CATABOLIC_P | 0.00000 | 0.00000 | | 0.70579 | 1.72296 | 74 |
| GO_ANCHORING_JUNCTION | 0.00000 | 0.00000 | | 0.56907 | 1.45662 | 353 |
| GO_ANIMAL_ORGAN_REGENERATION | 0.00331 | 0.04779 | 0.43171 | 0.63462 | 1.44782 | 29 |
| GO_ANTIGEN_PROCESSING_AND_PRESENTATION | 0.00000 | 0.00008 | 0.62726 | 0.58379 | 1.45035 | 119 |
| GO_ANTIGEN_PROCESSING_AND_PRESENTATION_OF_EXOGENOUS_ | 0.00000 | 0.00000 | | 0.79724 | 1.89352 | 48 |
| GO_ANTIGEN_PROCESSING_AND_PRESENTATION_OF_PEPTIDE_ANTI | 0.00000 | 0.00001 | 0.69013 | 0.61157 | 1.51476 | 107 |

| | | | | | | |
|---|---|---|---|---|---|---|
| GO_ANTIGEN_PROCESSING_AND_PRESENTATION_OF_PEPTIDE_ANTI | 0.00000 | 0.00000 | | 0.74205 | 1.78593 | 59 |
| GO_ANTIGEN_RECEPTOR_MEDIATED_SIGNALING_PATHWAY | 0.00000 | 0.00008 | 0.62726 | 0.60625 | 1.49261 | 95 |
| GO_ANTIOXIDANT_ACTIVITY | 0.00205 | 0.03325 | 0.43171 | 0.67915 | 1.53107 | 23 |
| GO_APOPTOTIC_SIGNALING_PATHWAY | 0.00202 | 0.03288 | 0.43171 | 0.47555 | 1.21270 | 261 |
| GO_ATPASE_ACTIVITY | 0.00017 | 0.00370 | 0.51885 | 0.50427 | 1.27724 | 213 |
| GO_ATP_METABOLIC_PROCESS | 0.00040 | 0.00790 | 0.49849 | 0.51104 | 1.28514 | 169 |
| GO_ATP_SYNTHESIS_COUPLED_ELECTRON_TRANSPORT | 0.00064 | 0.01174 | 0.47727 | 0.57724 | 1.40541 | 71 |
| GO_BLOOD_MICROPARTICLE | 0.00268 | 0.04120 | 0.43171 | 0.72215 | 1.56989 | 16 |
| GO_CADHERIN_BINDING | 0.00000 | 0.00000 | | 0.59392 | 1.50312 | 208 |
| GO_CANONICAL_WNT_SIGNALING_PATHWAY | 0.00000 | 0.00000 | 0.73376 | 0.60755 | 1.51816 | 138 |
| GO_CATALYTIC_STEP_2_SPLICEOSOME | 0.00043 | 0.00824 | 0.49849 | 0.56688 | 1.38763 | 78 |
| GO_CELLULAR_AMINO_ACID_METABOLIC_PROCESS | 0.00000 | 0.00000 | 0.81404 | 0.60592 | 1.52355 | 167 |
| GO_CELLULAR_KETONE_METABOLIC_PROCESS | 0.00000 | 0.00000 | 0.76146 | 0.64483 | 1.59621 | 104 |
| GO_CELLULAR_NITROGEN_COMPOUND_CATABOLIC_PROCESS | 0.00000 | 0.00000 | | 0.70423 | 1.80206 | 355 |
| GO_CELLULAR_OXIDANT_DETOXIFICATION | 0.00257 | 0.03988 | 0.43171 | 0.63707 | 1.45853 | 30 |
| GO_CELLULAR_PROTEIN_CATABOLIC_PROCESS | 0.00003 | 0.00078 | 0.57561 | 0.47646 | 1.22618 | 453 |
| GO_CELLULAR_RESPIRATION | 0.00000 | 0.00000 | 0.71951 | 0.60617 | 1.50741 | 124 |
| GO_CELLULAR_RESPONSE_TO_CHEMICAL_STRESS | 0.00205 | 0.03325 | 0.43171 | 0.50799 | 1.27130 | 146 |
| GO_CELLULAR_RESPONSE_TO_OXYGEN_LEVELS | 0.00000 | 0.00000 | 0.73376 | 0.61276 | 1.52381 | 124 |
| GO_CELLULAR_RESPONSE_TO_TOXIC_SUBSTANCE | 0.00079 | 0.01428 | 0.47727 | 0.63789 | 1.48521 | 37 |
| GO_CELL_ACTIVATION | 0.00009 | 0.00221 | 0.53843 | 0.47063 | 1.21116 | 453 |
| GO_CELL_ACTIVATION_INVOLVED_IN_IMMUNE_RESPONSE | 0.00001 | 0.00036 | 0.59333 | 0.51264 | 1.30764 | 265 |
| GO_CELL_ADHESION_MOLECULE_BINDING | 0.00000 | 0.00000 | | 0.57740 | 1.47147 | 250 |
| GO_CELL_BODY | 0.00024 | 0.00513 | 0.51885 | 0.51240 | 1.29059 | 179 |
| GO_CELL_CELL_RECOGNITION | 0.00000 | 0.00002 | 0.67496 | 0.87353 | 1.89898 | 16 |
| GO_CELL_CELL_SIGNALING_BY_WNT | 0.00000 | 0.00000 | 0.70498 | 0.55700 | 1.41166 | 216 |
| GO_CELL_CYCLE_G2_M_PHASE_TRANSITION | 0.00003 | 0.00097 | 0.55733 | 0.53274 | 1.34084 | 175 |
| GO_CELL_CYCLE_PHASE_TRANSITION | 0.00005 | 0.00136 | 0.55733 | 0.48259 | 1.23639 | 377 |
| GO_CELL_RECOGNITION | 0.00007 | 0.00173 | 0.53843 | 0.71365 | 1.62814 | 29 |
| GO_CELL_SUBSTRATE_JUNCTION | 0.00000 | 0.00000 | | 0.65550 | 1.66470 | 230 |
| GO_CELL_SURFACE_RECEPTOR_SIGNALING_PATHWAY_INVOLVED_IN | 0.00000 | 0.00001 | 0.67496 | 0.53850 | 1.36793 | 233 |
| GO_CHAPERONE_COMPLEX | 0.00000 | 0.00012 | 0.62726 | 0.81445 | 1.80848 | 20 |
| GO_COTRANSLATIONAL_PROTEIN_TARGETING_TO_MEMBRANE | 0.00000 | 0.00000 | | 0.85574 | 2.10386 | 91 |
| GO_CYTOKINE_MEDIATED_SIGNALING_PATHWAY | 0.00000 | 0.00000 | | 0.57596 | 1.46518 | 241 |
| GO_CYTOPLASMIC_STRESS_GRANULE | 0.00009 | 0.00208 | 0.53843 | 0.64811 | 1.53479 | 46 |
| GO_CYTOPLASMIC_TRANSLATION | 0.00000 | 0.00000 | | 0.78766 | 1.92900 | 81 |
| GO_CYTOPLASMIC_TRANSLATIONAL_INITIATION | 0.00000 | 0.00000 | 0.70498 | 0.80535 | 1.83230 | 28 |
| GO_CYTOSOLIC_LARGE_RIBOSOMAL_SUBUNIT | 0.00000 | 0.00000 | | 0.86821 | 2.07468 | 53 |
| GO_CYTOSOLIC_RIBOSOME | 0.00000 | 0.00000 | | 0.86974 | 2.14116 | 94 |
| GO_CYTOSOLIC_SMALL_RIBOSOMAL_SUBUNIT | 0.00000 | 0.00000 | | 0.88443 | 2.06389 | 39 |
| GO_DEFENSE_RESPONSE | 0.00002 | 0.00065 | 0.57561 | 0.48098 | 1.23518 | 412 |
| GO_DEFENSE_RESPONSE_TO_OTHER_ORGANISM | 0.00000 | 0.00002 | 0.67496 | 0.52222 | 1.33266 | 294 |
| GO_DETOXIFICATION | 0.00298 | 0.04432 | 0.43171 | 0.61630 | 1.43699 | 38 |
| GO_DISORDERED_DOMAIN_SPECIFIC_BINDING | 0.00046 | 0.00882 | 0.49849 | 0.73400 | 1.62984 | 20 |
| GO_DNA_BIOSYNTHETIC_PROCESS | 0.00006 | 0.00161 | 0.53843 | 0.55812 | 1.38339 | 114 |
| GO_DNA_DAMAGE_RESPONSE_DETECTION_OF_DNA_DAMAGE | 0.00163 | 0.02750 | 0.45506 | 0.63495 | 1.46761 | 34 |
| GO_DNA_GEOMETRIC_CHANGE | 0.00350 | 0.04971 | 0.43171 | 0.54022 | 1.32115 | 77 |
| GO_DNA_POLYMERASE_BINDING | 0.00283 | 0.04265 | 0.43171 | 0.73755 | 1.59237 | 15 |
| GO_DOUBLE_STRANDED_RNA_BINDING | 0.00093 | 0.01639 | 0.47727 | 0.62915 | 1.47589 | 41 |
| GO_ELECTRON_TRANSFER_ACTIVITY | 0.00024 | 0.00513 | 0.51885 | 0.60211 | 1.45741 | 64 |
| GO_ELECTRON_TRANSPORT_CHAIN | 0.00006 | 0.00163 | 0.53843 | 0.56344 | 1.39479 | 110 |
| GO_ENDOPEPTIDASE_ACTIVITY | 0.00004 | 0.00110 | 0.55733 | 0.56318 | 1.39595 | 114 |
| GO_ENDOPEPTIDASE_COMPLEX | 0.00000 | 0.00000 | | 0.77516 | 1.86056 | 57 |
| GO_ENERGY_DERIVATION_BY_OXIDATION_OF_ORGANIC_COMPOUN | 0.00000 | 0.00005 | 0.64355 | 0.56209 | 1.41039 | 155 |
| GO_EPITHELIUM_DEVELOPMENT | 0.00268 | 0.04120 | 0.43171 | 0.45623 | 1.16991 | 391 |
| GO_ESTABLISHMENT_OF_PROTEIN_LOCALIZATION_TO_CHROMOSOM | 0.00003 | 0.00081 | 0.57561 | 0.78575 | 1.74474 | 20 |
| GO_ESTABLISHMENT_OF_PROTEIN_LOCALIZATION_TO_ENDOPLASM | 0.00000 | 0.00000 | | 0.84091 | 2.07814 | 101 |
| GO_ESTABLISHMENT_OF_PROTEIN_LOCALIZATION_TO_MEMBRANE | 0.00000 | 0.00000 | | 0.69865 | 1.76815 | 208 |
| GO_ESTABLISHMENT_OF_PROTEIN_LOCALIZATION_TO_ORGANELLE | 0.00000 | 0.00000 | | 0.62010 | 1.58840 | 370 |
| GO_ESTABLISHMENT_OF_PROTEIN_LOCALIZATION_TO_TELOMERE | 0.00005 | 0.00141 | 0.55733 | 0.80528 | 1.75060 | 16 |
| GO_ESTABLISHMENT_OF_TISSUE_POLARITY | 0.00000 | 0.00000 | | 0.74459 | 1.80990 | 68 |
| GO_EUKARYOTIC_TRANSLATION_INITIATION_FACTOR_3_COMPLEX | 0.00000 | 0.00015 | 0.61053 | 0.85479 | 1.84549 | 15 |
| GO_EXOCYTOSIS | 0.00055 | 0.01046 | 0.47727 | 0.47084 | 1.20498 | 356 |
| GO_FC_EPSILON_RECEPTOR_SIGNALING_PATHWAY | 0.00000 | 0.00000 | | 0.74014 | 1.79149 | 64 |
| GO_FC_RECEPTOR_MEDIATED_STIMULATORY_SIGNALING_PATHWAY | 0.00346 | 0.04963 | 0.43171 | 0.60915 | 1.42898 | 41 |
| GO_FC_RECEPTOR_SIGNALING_PATHWAY | 0.00000 | 0.00000 | | 0.70613 | 1.74094 | 96 |
| GO_FICOLIN_1_RICH_GRANULE | 0.00000 | 0.00000 | | 0.68593 | 1.68609 | 88 |
| GO_GENERATION_OF_PRECURSOR_METABOLITES_AND_ENERGY | 0.00000 | 0.00002 | 0.65944 | 0.52336 | 1.33489 | 274 |
| GO_GLYCOLYTIC_PROCESS_THROUGH_FRUCTOSE_6_PHOSPHATE | 0.00087 | 0.01565 | 0.47727 | 0.75277 | 1.63646 | 16 |
| GO_HEMATOPOIETIC_PROGENITOR_CELL_DIFFERENTIATION | 0.00000 | 0.00000 | 0.81404 | 0.67720 | 1.66461 | 88 |
| GO_HEMATOPOIETIC_STEM_CELL_DIFFERENTIATION | 0.00000 | 0.00000 | | 0.76499 | 1.84115 | 59 |
| GO_HYDROLASE_ACTIVITY_ACTING_ON_ACID_ANHYDRIDES | 0.00021 | 0.00446 | 0.51885 | 0.47370 | 1.21429 | 386 |
| GO_IMMUNE_EFFECTOR_PROCESS | 0.00004 | 0.00100 | 0.55733 | 0.48369 | 1.24158 | 405 |
| GO_IMMUNE_RESPONSE_REGULATING_SIGNALING_PATHWAY | 0.00000 | 0.00001 | 0.69013 | 0.58117 | 1.45592 | 150 |
| GO_INNATE_IMMUNE_RESPONSE | 0.00000 | 0.00000 | 0.77494 | 0.55831 | 1.42029 | 241 |
| GO_INNATE_IMMUNE_RESPONSE_ACTIVATING_SIGNAL_TRANSDUCT | 0.00000 | 0.00000 | | 0.74219 | 1.79355 | 62 |

| | | | | | | |
|---|---|---|---|---|---|---|
| GO_INNER_MITOCHONDRIAL_MEMBRANE_PROTEIN_COMPLEX | 0.00068 | 0.01254 | 0.47727 | 0.55495 | 1.35961 | 83 |
| GO_INTERACTION_WITH_SYMBIONT | 0.00002 | 0.00050 | 0.57561 | 0.70769 | 1.65008 | 38 |
| GO_INTERLEUKIN_1_MEDIATED_SIGNALING_PATHWAY | 0.00000 | 0.00000 | | 0.75262 | 1.82077 | 63 |
| GO_INTRINSIC_APOPTOTIC_SIGNALING_PATHWAY | 0.00302 | 0.04446 | 0.43171 | 0.50351 | 1.25753 | 136 |
| GO_LARGE_RIBOSOMAL_SUBUNIT | 0.00000 | 0.00000 | | 0.68688 | 1.70074 | 106 |
| GO_LEUKOCYTE_MEDIATED_IMMUNITY | 0.00000 | 0.00006 | 0.64355 | 0.51610 | 1.31637 | 274 |
| GO_MATURATION_OF_LSU_RRNA | 0.00130 | 0.02238 | 0.45506 | 0.67063 | 1.52055 | 27 |
| GO_MEMBRANE_PROTEIN_COMPLEX | 0.00213 | 0.03399 | 0.43171 | 0.45034 | 1.15662 | 413 |
| GO_MITOCHONDRIAL_ENVELOPE | 0.00028 | 0.00575 | 0.49849 | 0.46252 | 1.18914 | 435 |
| GO_MITOCHONDRIAL_PROTEIN_COMPLEX | 0.00008 | 0.00198 | 0.53843 | 0.51623 | 1.30382 | 192 |
| GO_MODIFICATION_DEPENDENT_MACROMOLECULE_CATABOLIC_PR | 0.00007 | 0.00175 | 0.53843 | 0.47881 | 1.22781 | 391 |
| GO_MODULATION_BY_HOST_OF_SYMBIONT_PROCESS | 0.00057 | 0.01072 | 0.47727 | 0.66712 | 1.53028 | 31 |
| GO_MODULATION_OF_PROCESS_OF_OTHER_ORGANISM | 0.00070 | 0.01282 | 0.47727 | 0.60853 | 1.44997 | 51 |
| GO_MORPHOGENESIS_OF_AN_EPITHELIUM | 0.00000 | 0.00015 | 0.61053 | 0.53405 | 1.34963 | 199 |
| GO_MORPHOGENESIS_OF_A_POLARIZED_EPITHELIUM | 0.00000 | 0.00000 | | 0.73525 | 1.79927 | 79 |
| GO_MRNA_3_UTR_BINDING | 0.00041 | 0.00804 | 0.49849 | 0.60308 | 1.44754 | 57 |
| GO_MRNA_5_UTR_BINDING | 0.00003 | 0.00083 | 0.57561 | 0.77667 | 1.73311 | 21 |
| GO_MRNA_BINDING | 0.00000 | 0.00000 | | 0.61177 | 1.54500 | 189 |
| GO_MRNA_PROCESSING | 0.00037 | 0.00736 | 0.49849 | 0.47409 | 1.21375 | 359 |
| GO_MYELOID_LEUKOCYTE_ACTIVATION | 0.00000 | 0.00013 | 0.62726 | 0.52371 | 1.33264 | 244 |
| GO_MYELOID_LEUKOCYTE_MEDIATED_IMMUNITY | 0.00000 | 0.00003 | 0.65944 | 0.53576 | 1.36070 | 231 |
| GO_NADH_METABOLIC_PROCESS | 0.00029 | 0.00601 | 0.49849 | 0.73578 | 1.64185 | 21 |
| GO_NAD_BINDING | 0.00024 | 0.00510 | 0.51885 | 0.66616 | 1.53974 | 34 |
| GO_NCRNA_METABOLIC_PROCESS | 0.00257 | 0.03988 | 0.43171 | 0.46330 | 1.18377 | 318 |
| GO_NCRNA_PROCESSING | 0.00042 | 0.00819 | 0.49849 | 0.48556 | 1.23871 | 279 |
| GO_NEGATIVE_REGULATION_OF_CANONICAL_WNT_SIGNALING_PAT | 0.00000 | 0.00000 | | 0.70146 | 1.71809 | 82 |
| GO_NEGATIVE_REGULATION_OF_CELLULAR_AMIDE_METABOLIC_PR( | 0.00010 | 0.00245 | 0.53843 | 0.57965 | 1.42376 | 87 |
| GO_NEGATIVE_REGULATION_OF_CELL_CYCLE | 0.00098 | 0.01723 | 0.45506 | 0.46775 | 1.19686 | 343 |
| GO_NEGATIVE_REGULATION_OF_CELL_CYCLE_G2_M_PHASE_TRANS | 0.00000 | 0.00000 | 0.74774 | 0.68261 | 1.66488 | 73 |
| GO_NEGATIVE_REGULATION_OF_CELL_CYCLE_PHASE_TRANSITION | 0.00004 | 0.00101 | 0.55733 | 0.54406 | 1.36360 | 152 |
| GO_NEGATIVE_REGULATION_OF_CELL_CYCLE_PROCESS | 0.00001 | 0.00020 | 0.61053 | 0.52928 | 1.33951 | 208 |
| GO_NEGATIVE_REGULATION_OF_CELL_DEATH | 0.00011 | 0.00267 | 0.53843 | 0.48050 | 1.23054 | 365 |
| GO_NEGATIVE_REGULATION_OF_MITOTIC_CELL_CYCLE | 0.00012 | 0.00279 | 0.53843 | 0.51800 | 1.30836 | 190 |
| GO_NEGATIVE_REGULATION_OF_MRNA_METABOLIC_PROCESS | 0.00320 | 0.04647 | 0.43171 | 0.58718 | 1.39700 | 50 |
| GO_NEGATIVE_REGULATION_OF_MRNA_SPLICING_VIA_SPLICEOSOM | 0.00191 | 0.03150 | 0.45506 | 0.72734 | 1.59749 | 18 |
| GO_NEGATIVE_REGULATION_OF_RNA_SPLICING | 0.00009 | 0.00208 | 0.53843 | 0.75696 | 1.68913 | 21 |
| GO_NEGATIVE_REGULATION_OF_VIRAL_PROCESS | 0.00231 | 0.03646 | 0.43171 | 0.63103 | 1.45604 | 33 |
| GO_NEGATIVE_REGULATION_OF_WNT_SIGNALING_PATHWAY | 0.00000 | 0.00000 | | 0.68552 | 1.68709 | 93 |
| GO_NEURAL_NUCLEUS_DEVELOPMENT | 0.00018 | 0.00397 | 0.51885 | 0.73484 | 1.65661 | 23 |
| GO_NEURON_TO_NEURON_SYNAPSE | 0.00001 | 0.00036 | 0.59333 | 0.57525 | 1.42584 | 114 |
| GO_NIK_NF_KAPPAB_SIGNALING | 0.00000 | 0.00000 | | 0.72152 | 1.76700 | 81 |
| GO_NON_CANONICAL_WNT_SIGNALING_PATHWAY | 0.00000 | 0.00000 | | 0.73271 | 1.79048 | 75 |
| GO_NUCLEAR_TRANSCRIBED_MRNA_CATABOLIC_PROCESS | 0.00000 | 0.00000 | | 0.76636 | 1.92701 | 168 |
| GO_NUCLEAR_TRANSCRIBED_MRNA_CATABOLIC_PROCESS_NONSEN | 0.00000 | 0.00000 | | 0.86116 | 2.13296 | 107 |
| GO_NUCLEAR_TRANSPORT | 0.00119 | 0.02063 | 0.45506 | 0.48296 | 1.22708 | 235 |
| GO_NUCLEOSIDE_PHOSPHATE_BIOSYNTHETIC_PROCESS | 0.00198 | 0.03249 | 0.43171 | 0.52737 | 1.30606 | 109 |
| GO_NUCLEOSIDE_TRIPHOSPHATE_METABOLIC_PROCESS | 0.00276 | 0.04193 | 0.43171 | 0.60155 | 1.42090 | 45 |
| GO_NUCLEOTIDE_EXCISION_REPAIR_DNA_DAMAGE_RECOGNITION | 0.00072 | 0.01311 | 0.47727 | 0.72391 | 1.60744 | 20 |
| GO_ORGANELLE_INNER_MEMBRANE | 0.00163 | 0.02750 | 0.45506 | 0.46511 | 1.18822 | 316 |
| GO_ORGANIC_ACID_METABOLIC_PROCESS | 0.00001 | 0.00024 | 0.61053 | 0.47924 | 1.23407 | 467 |
| GO_ORGANIC_CYCLIC_COMPOUND_CATABOLIC_PROCESS | 0.00000 | 0.00000 | | 0.69939 | 1.79091 | 361 |
| GO_OXIDATION_REDUCTION_PROCESS | 0.00033 | 0.00672 | 0.49849 | 0.46549 | 1.19689 | 430 |
| GO_OXIDATIVE_PHOSPHORYLATION | 0.00055 | 0.01046 | 0.47727 | 0.56785 | 1.39378 | 85 |
| GO_OXIDOREDUCTASE_ACTIVITY | 0.00092 | 0.01629 | 0.47727 | 0.47693 | 1.21745 | 297 |
| GO_OXIDOREDUCTASE_ACTIVITY_ACTING_ON_CH_OH_GROUP_OF_ | 0.00257 | 0.03988 | 0.43171 | 0.57889 | 1.38081 | 52 |
| GO_OXIDOREDUCTASE_ACTIVITY_ACTING_ON_THE_ALDEHYDE_OR_ | 0.00058 | 0.01085 | 0.47727 | 0.76117 | 1.65472 | 16 |
| GO_OXIDOREDUCTASE_COMPLEX | 0.00057 | 0.01072 | 0.47727 | 0.57548 | 1.40628 | 75 |
| GO_PEPTIDASE_ACTIVITY | 0.00082 | 0.01472 | 0.47727 | 0.50299 | 1.26860 | 183 |
| GO_PEPTIDASE_COMPLEX | 0.00000 | 0.00000 | 0.80122 | 0.70401 | 1.71421 | 72 |
| GO_PEPTIDE_BIOSYNTHETIC_PROCESS | 0.00000 | 0.00000 | | 0.63809 | 1.64206 | 452 |
| GO_PIGMENT_GRANULE | 0.00302 | 0.04446 | 0.43171 | 0.55820 | 1.35532 | 67 |
| GO_POLYSOMAL_RIBOSOME | 0.00000 | 0.00000 | 0.78819 | 0.82421 | 1.88697 | 30 |
| GO_POLYSOME | 0.00000 | 0.00000 | | 0.76864 | 1.84202 | 56 |
| GO_POLY_PYRIMIDINE_TRACT_BINDING | 0.00098 | 0.01723 | 0.45506 | 0.74631 | 1.62240 | 16 |
| GO_POSITIVE_REGULATION_OF_BINDING | 0.00228 | 0.03611 | 0.43171 | 0.54521 | 1.33540 | 82 |
| GO_POSITIVE_REGULATION_OF_CANONICAL_WNT_SIGNALING_PATH | 0.00000 | 0.00000 | 0.81404 | 0.70495 | 1.72264 | 75 |
| GO_POSITIVE_REGULATION_OF_CELLULAR_AMIDE_METABOLIC_PRC | 0.00033 | 0.00678 | 0.49849 | 0.56803 | 1.39490 | 86 |
| GO_POSITIVE_REGULATION_OF_CELLULAR_PROTEIN_LOCALIZATION | 0.00075 | 0.01351 | 0.47727 | 0.50146 | 1.26343 | 180 |
| GO_POSITIVE_REGULATION_OF_CELL_DEATH | 0.00004 | 0.00120 | 0.55733 | 0.49882 | 1.27168 | 260 |
| GO_POSITIVE_REGULATION_OF_DEFENSE_RESPONSE | 0.00000 | 0.00015 | 0.61053 | 0.58129 | 1.44631 | 125 |
| GO_POSITIVE_REGULATION_OF_DNA_BIOSYNTHETIC_PROCESS | 0.00005 | 0.00139 | 0.55733 | 0.65562 | 1.55257 | 46 |
| GO_POSITIVE_REGULATION_OF_DNA_METABOLIC_PROCESS | 0.00061 | 0.01128 | 0.47727 | 0.54229 | 1.34192 | 103 |
| GO_POSITIVE_REGULATION_OF_ESTABLISHMENT_OF_PROTEIN_LOC/ | 0.00031 | 0.00640 | 0.49849 | 0.51061 | 1.28464 | 170 |
| GO_POSITIVE_REGULATION_OF_IMMUNE_RESPONSE | 0.00000 | 0.00001 | 0.69013 | 0.54719 | 1.38534 | 210 |
| GO_POSITIVE_REGULATION_OF_IMMUNE_SYSTEM_PROCESS | 0.00021 | 0.00459 | 0.51885 | 0.48688 | 1.24249 | 289 |
| GO_POSITIVE_REGULATION_OF_INNATE_IMMUNE_RESPONSE | 0.00000 | 0.00000 | 0.74774 | 0.64407 | 1.58562 | 94 |

| | | | | | | |
|---|---|---|---|---|---|---|
| GO_POSITIVE_REGULATION_OF_PROTEIN_LOCALIZATION_TO_NUCLE | 0.00019 | 0.00416 | 0.51885 | 0.61913 | 1.47680 | 52 |
| GO_POSITIVE_REGULATION_OF_RESPONSE_TO_BIOTIC_STIMULUS | 0.00000 | 0.00000 | 0.74774 | 0.62905 | 1.55806 | 107 |
| GO_POSITIVE_REGULATION_OF_RESPONSE_TO_EXTERNAL_STIMULU | 0.00015 | 0.00346 | 0.51885 | 0.52777 | 1.32399 | 156 |
| GO_POSITIVE_REGULATION_OF_TELOMERASE_RNA_LOCALIZATION_ | 0.00000 | 0.00003 | 0.65944 | 0.87774 | 1.89503 | 15 |
| GO_POSITIVE_REGULATION_OF_TELOMERE_MAINTENANCE | 0.00055 | 0.01046 | 0.47727 | 0.64953 | 1.51447 | 38 |
| GO_POSITIVE_REGULATION_OF_TELOMERE_MAINTENANCE_VIA_TE | 0.00018 | 0.00405 | 0.51885 | 0.69492 | 1.58541 | 29 |
| GO_POSITIVE_REGULATION_OF_TRANSLATION | 0.00004 | 0.00107 | 0.55733 | 0.60919 | 1.48582 | 73 |
| GO_POSITIVE_REGULATION_OF_VIRAL_GENOME_REPLICATION | 0.00185 | 0.03079 | 0.45506 | 0.66330 | 1.50391 | 27 |
| GO_POSITIVE_REGULATION_OF_WNT_SIGNALING_PATHWAY | 0.00000 | 0.00000 | 0.76146 | 0.66585 | 1.63432 | 85 |
| GO_POSTSYNAPSE | 0.00007 | 0.00180 | 0.53843 | 0.52199 | 1.31844 | 190 |
| GO_POSTTRANSCRIPTIONAL_REGULATION_OF_GENE_EXPRESSION | 0.00000 | 0.00000 | | 0.58292 | 1.49426 | 386 |
| GO_POST_TRANSLATIONAL_PROTEIN_MODIFICATION | 0.00059 | 0.01098 | 0.47727 | 0.50058 | 1.26429 | 192 |
| GO_PRECATALYTIC_SPLICEOSOME | 0.00283 | 0.04265 | 0.43171 | 0.59303 | 1.40436 | 46 |
| GO_PROTEASOMAL_PROTEIN_CATABOLIC_PROCESS | 0.00009 | 0.00208 | 0.53843 | 0.48707 | 1.24362 | 301 |
| GO_PROTEASOMAL_UBIQUITIN_INDEPENDENT_PROTEIN_CATABOLIC | 0.00000 | 0.00000 | 0.73376 | 0.90205 | 1.96097 | 16 |
| GO_PROTEASOME_ACCESSORY_COMPLEX | 0.00001 | 0.00022 | 0.61053 | 0.79068 | 1.78471 | 24 |
| GO_PROTEASOME_CORE_COMPLEX | 0.00000 | 0.00000 | 0.77494 | 0.92477 | 1.99656 | 15 |
| GO_PROTEIN_DNA_COMPLEX_SUBUNIT_ORGANIZATION | 0.00305 | 0.04473 | 0.43171 | 0.50207 | 1.25573 | 143 |
| GO_PROTEIN_DOMAIN_SPECIFIC_BINDING | 0.00002 | 0.00070 | 0.57561 | 0.49590 | 1.26614 | 301 |
| GO_PROTEIN_FOLDING | 0.00000 | 0.00015 | 0.61053 | 0.57111 | 1.42505 | 132 |
| GO_PROTEIN_KINASE_C_BINDING | 0.00156 | 0.02644 | 0.45506 | 0.72590 | 1.58795 | 17 |
| GO_PROTEIN_LOCALIZATION_TO_CHROMOSOME | 0.00242 | 0.03796 | 0.43171 | 0.57768 | 1.38658 | 57 |
| GO_PROTEIN_LOCALIZATION_TO_CHROMOSOME_TELOMERIC_REGI | 0.00121 | 0.02087 | 0.45506 | 0.69520 | 1.56921 | 24 |
| GO_PROTEIN_LOCALIZATION_TO_ENDOPLASMIC_RETICULUM | 0.00000 | 0.00000 | | 0.81782 | 2.02898 | 116 |
| GO_PROTEIN_LOCALIZATION_TO_MEMBRANE | 0.00000 | 0.00000 | | 0.59983 | 1.53390 | 326 |
| GO_PROTEIN_LOCALIZATION_TO_NUCLEUS | 0.00335 | 0.04818 | 0.43171 | 0.49578 | 1.24200 | 150 |
| GO_PROTEIN_MODIFICATION_BY_SMALL_PROTEIN_REMOVAL | 0.00001 | 0.00025 | 0.59333 | 0.53764 | 1.35780 | 189 |
| GO_PROTEIN_POLYUBIQUITINATION | 0.00005 | 0.00136 | 0.55733 | 0.51434 | 1.30170 | 208 |
| GO_PROTEIN_STABILIZATION | 0.00283 | 0.04265 | 0.43171 | 0.52106 | 1.29059 | 107 |
| GO_PROTEIN_TARGETING | 0.00000 | 0.00000 | | 0.64974 | 1.65796 | 286 |
| GO_PROTEIN_TARGETING_TO_MEMBRANE | 0.00000 | 0.00000 | | 0.77354 | 1.93194 | 136 |
| GO_PURINE_CONTAINING_COMPOUND_BIOSYNTHETIC_PROCESS | 0.00198 | 0.03249 | 0.43171 | 0.55033 | 1.34673 | 79 |
| GO_PURINE_NUCLEOSIDE_TRIPHOSPHATE_METABOLIC_PROCESS | 0.00090 | 0.01608 | 0.47727 | 0.64956 | 1.50471 | 35 |
| GO_REGULATION_OF_ANIMAL_ORGAN_MORPHOGENESIS | 0.00000 | 0.00000 | | 0.68116 | 1.67707 | 95 |
| GO_REGULATION_OF_APOPTOTIC_SIGNALING_PATHWAY | 0.00014 | 0.00322 | 0.51885 | 0.51728 | 1.30171 | 172 |
| GO_REGULATION_OF_CELLULAR_AMIDE_METABOLIC_PROCESS | 0.00000 | 0.00000 | | 0.57124 | 1.45396 | 245 |
| GO_REGULATION_OF_CELLULAR_AMINE_METABOLIC_PROCESS | 0.00000 | 0.00000 | | 0.79694 | 1.89890 | 51 |
| GO_REGULATION_OF_CELLULAR_AMINO_ACID_METABOLIC_PROCES | 0.00000 | 0.00000 | | 0.80867 | 1.92066 | 48 |
| GO_REGULATION_OF_CELLULAR_CATABOLIC_PROCESS | 0.00000 | 0.00001 | 0.70498 | 0.49392 | 1.27281 | 486 |
| GO_REGULATION_OF_CELLULAR_KETONE_METABOLIC_PROCESS | 0.00000 | 0.00000 | | 0.69634 | 1.70534 | 81 |
| GO_REGULATION_OF_CELLULAR_PROTEIN_LOCALIZATION | 0.00272 | 0.04150 | 0.43171 | 0.47170 | 1.20336 | 279 |
| GO_REGULATION_OF_CELL_CYCLE_G2_M_PHASE_TRANSITION | 0.00000 | 0.00004 | 0.64355 | 0.57946 | 1.44890 | 141 |
| GO_REGULATION_OF_CELL_CYCLE_PHASE_TRANSITION | 0.00014 | 0.00322 | 0.51885 | 0.49080 | 1.25206 | 277 |
| GO_REGULATION_OF_CELL_CYCLE_PROCESS | 0.00030 | 0.00624 | 0.49849 | 0.46516 | 1.19588 | 429 |
| GO_REGULATION_OF_DEFENSE_RESPONSE | 0.00034 | 0.00684 | 0.49849 | 0.49885 | 1.26327 | 212 |
| GO_REGULATION_OF_DNA_BIOSYNTHETIC_PROCESS | 0.00002 | 0.00054 | 0.57561 | 0.62328 | 1.51093 | 66 |
| GO_REGULATION_OF_DNA_METABOLIC_PROCESS | 0.00006 | 0.00161 | 0.53843 | 0.51781 | 1.30783 | 192 |
| GO_REGULATION_OF_DNA_TEMPLATED_TRANSCRIPTION_IN_RESPO | 0.00000 | 0.00000 | 0.80122 | 0.68663 | 1.67921 | 77 |
| GO_REGULATION_OF_HEMATOPOIETIC_PROGENITOR_CELL_DIFFERE | 0.00000 | 0.00000 | | 0.74134 | 1.79347 | 63 |
| GO_REGULATION_OF_HEMATOPOIETIC_STEM_CELL_DIFFERENTIATIO | 0.00000 | 0.00000 | | 0.77847 | 1.86376 | 55 |
| GO_REGULATION_OF_HEMOPOIESIS | 0.00004 | 0.00110 | 0.55733 | 0.53399 | 1.34272 | 168 |
| GO_REGULATION_OF_IMMUNE_RESPONSE | 0.00000 | 0.00008 | 0.62726 | 0.51736 | 1.31978 | 267 |
| GO_REGULATION_OF_INNATE_IMMUNE_RESPONSE | 0.00000 | 0.00003 | 0.65944 | 0.58253 | 1.45187 | 128 |
| GO_REGULATION_OF_INTRINSIC_APOPTOTIC_SIGNALING_PATHWAY | 0.00005 | 0.00136 | 0.55733 | 0.59089 | 1.44639 | 78 |
| GO_REGULATION_OF_MITOTIC_CELL_CYCLE | 0.00014 | 0.00325 | 0.51885 | 0.47529 | 1.21730 | 366 |
| GO_REGULATION_OF_MORPHOGENESIS_OF_AN_EPITHELIUM | 0.00000 | 0.00000 | | 0.71616 | 1.75291 | 80 |
| GO_REGULATION_OF_MRNA_CATABOLIC_PROCESS | 0.00000 | 0.00000 | | 0.64380 | 1.60875 | 138 |
| GO_REGULATION_OF_MRNA_METABOLIC_PROCESS | 0.00000 | 0.00000 | | 0.59965 | 1.52158 | 228 |
| GO_REGULATION_OF_MRNA_PROCESSING | 0.00231 | 0.03646 | 0.43171 | 0.52268 | 1.29243 | 102 |
| GO_REGULATION_OF_MRNA_SPLICING_VIA_SPLICEOSOME | 0.00005 | 0.00135 | 0.55733 | 0.59684 | 1.45699 | 74 |
| GO_REGULATION_OF_PEPTIDASE_ACTIVITY | 0.00294 | 0.04391 | 0.43171 | 0.49740 | 1.24606 | 150 |
| GO_REGULATION_OF_PROTEIN_IMPORT | 0.00287 | 0.04294 | 0.43171 | 0.62643 | 1.44542 | 33 |
| GO_REGULATION_OF_PROTEIN_LOCALIZATION_TO_NUCLEUS | 0.00141 | 0.02403 | 0.45506 | 0.56306 | 1.37101 | 72 |
| GO_REGULATION_OF_PROTEIN_MODIFICATION_BY_SMALL_PROTEIN | 0.00191 | 0.03150 | 0.45506 | 0.51508 | 1.28192 | 126 |
| GO_REGULATION_OF_PROTEOLYSIS | 0.00139 | 0.02380 | 0.45506 | 0.47021 | 1.20010 | 291 |
| GO_REGULATION_OF_RESPONSE_TO_BIOTIC_STIMULUS | 0.00000 | 0.00014 | 0.61053 | 0.55528 | 1.39379 | 158 |
| GO_REGULATION_OF_RNA_SPLICING | 0.00008 | 0.00198 | 0.53843 | 0.57022 | 1.40587 | 96 |
| GO_REGULATION_OF_SMALL_MOLECULE_METABOLIC_PROCESS | 0.00001 | 0.00018 | 0.61053 | 0.53268 | 1.34564 | 194 |
| GO_REGULATION_OF_STEM_CELL_DIFFERENTIATION | 0.00000 | 0.00000 | 0.76146 | 0.68824 | 1.68183 | 75 |
| GO_REGULATION_OF_SYMBIOTIC_PROCESS | 0.00185 | 0.03079 | 0.45506 | 0.52773 | 1.30638 | 110 |
| GO_REGULATION_OF_TELOMERE_MAINTENANCE | 0.00053 | 0.01006 | 0.47727 | 0.58524 | 1.40854 | 59 |
| GO_REGULATION_OF_TELOMERE_MAINTENANCE_VIA_TELOMERE_L | 0.00004 | 0.00099 | 0.55733 | 0.66063 | 1.55731 | 44 |
| GO_REGULATION_OF_TRANSCRIPTION_FROM_RNA_POLYMERASE_II | 0.00000 | 0.00000 | | 0.75638 | 1.82043 | 59 |
| GO_REGULATION_OF_TRANSFERASE_ACTIVITY | 0.00183 | 0.03070 | 0.45506 | 0.45550 | 1.16861 | 401 |
| GO_REGULATION_OF_UBIQUITIN_PROTEIN_LIGASE_ACTIVITY | 0.00350 | 0.04971 | 0.43171 | 0.71134 | 1.56234 | 18 |
| GO_REGULATION_OF_VIRAL_GENOME_REPLICATION | 0.00027 | 0.00559 | 0.49849 | 0.64474 | 1.51561 | 42 |

| | | | | | | |
|---|---|---|---|---|---|---|
| GO_REGULATION_OF_VIRAL_LIFE_CYCLE | 0.00302 | 0.04446 | 0.43171 | 0.56827 | 1.37549 | 64 |
| GO_REGULATION_OF_WNT_SIGNALING_PATHWAY | 0.00000 | 0.00000 | 0.76146 | 0.60441 | 1.51400 | 151 |
| GO_RESPIRATORY_CHAIN_COMPLEX | 0.00132 | 0.02262 | 0.45506 | 0.57975 | 1.39618 | 60 |
| GO_RESPIRATORY_ELECTRON_TRANSPORT_CHAIN | 0.00016 | 0.00367 | 0.51885 | 0.58768 | 1.43843 | 80 |
| GO_RESPONSE_TO_ABIOTIC_STIMULUS | 0.00001 | 0.00022 | 0.61053 | 0.48250 | 1.24339 | 491 |
| GO_RESPONSE_TO_BIOTIC_STIMULUS | 0.00000 | 0.00000 | 0.71951 | 0.50930 | 1.30864 | 420 |
| GO_RESPONSE_TO_CYTOKINE | 0.00000 | 0.00000 | | 0.54183 | 1.39301 | 427 |
| GO_RESPONSE_TO_DRUG | 0.00305 | 0.04473 | 0.43171 | 0.50721 | 1.26760 | 139 |
| GO_RESPONSE_TO_INORGANIC_SUBSTANCE | 0.00091 | 0.01618 | 0.47727 | 0.49498 | 1.25054 | 196 |
| GO_RESPONSE_TO_INTERFERON_GAMMA | 0.00242 | 0.03796 | 0.43171 | 0.59103 | 1.40616 | 50 |
| GO_RESPONSE_TO_INTERLEUKIN_1 | 0.00000 | 0.00000 | | 0.70311 | 1.72262 | 83 |
| GO_RESPONSE_TO_INTERLEUKIN_12 | 0.00000 | 0.00000 | 0.70498 | 0.78040 | 1.79014 | 31 |
| GO_RESPONSE_TO_ORGANIC_CYCLIC_COMPOUND | 0.00014 | 0.00329 | 0.51885 | 0.48117 | 1.23135 | 331 |
| GO_RESPONSE_TO_OXIDATIVE_STRESS | 0.00004 | 0.00114 | 0.55733 | 0.52263 | 1.31753 | 182 |
| GO_RESPONSE_TO_OXYGEN_LEVELS | 0.00000 | 0.00002 | 0.65944 | 0.55259 | 1.39441 | 185 |
| GO_RESPONSE_TO_TOXIC_SUBSTANCE | 0.00040 | 0.00784 | 0.49849 | 0.57269 | 1.40146 | 79 |
| GO_RESPONSE_TO_TUMOR_NECROSIS_FACTOR | 0.00000 | 0.00000 | 0.78819 | 0.63147 | 1.56959 | 120 |
| GO_RIBONUCLEOPROTEIN_COMPLEX_BINDING | 0.00000 | 0.00005 | 0.64355 | 0.60570 | 1.49334 | 96 |
| GO_RIBONUCLEOPROTEIN_COMPLEX_BIOGENESIS | 0.00000 | 0.00000 | | 0.59145 | 1.51382 | 351 |
| GO_RIBONUCLEOPROTEIN_COMPLEX_SUBUNIT_ORGANIZATION | 0.00000 | 0.00000 | | 0.63308 | 1.58906 | 158 |
| GO_RIBONUCLEOPROTEIN_GRANULE | 0.00000 | 0.00004 | 0.65944 | 0.57610 | 1.43828 | 134 |
| GO_RIBONUCLEOSIDE_TRIPHOSPHATE_BIOSYNTHETIC_PROCESS | 0.00324 | 0.04686 | 0.43171 | 0.64609 | 1.46996 | 28 |
| GO_RIBONUCLEOSIDE_TRIPHOSPHATE_METABOLIC_PROCESS | 0.00050 | 0.00958 | 0.47727 | 0.65991 | 1.52867 | 35 |
| GO_RIBOSOMAL_LARGE_SUBUNIT_ASSEMBLY | 0.00001 | 0.00031 | 0.59333 | 0.76196 | 1.72401 | 26 |
| GO_RIBOSOMAL_LARGE_SUBUNIT_BIOGENESIS | 0.00000 | 0.00000 | 0.71951 | 0.69617 | 1.68233 | 62 |
| GO_RIBOSOMAL_SMALL_SUBUNIT_ASSEMBLY | 0.00181 | 0.03050 | 0.45506 | 0.75185 | 1.62324 | 15 |
| GO_RIBOSOMAL_SMALL_SUBUNIT_BIOGENESIS | 0.00032 | 0.00647 | 0.49849 | 0.59239 | 1.43154 | 62 |
| GO_RIBOSOMAL_SUBUNIT | 0.00000 | 0.00000 | | 0.70400 | 1.77041 | 169 |
| GO_RIBOSOME | 0.00000 | 0.00000 | | 0.67681 | 1.71109 | 200 |
| GO_RIBOSOME_ASSEMBLY | 0.00000 | 0.00000 | 0.74774 | 0.72461 | 1.73335 | 54 |
| GO_RIBOSOME_BINDING | 0.00202 | 0.03288 | 0.43171 | 0.62513 | 1.45759 | 38 |
| GO_RIBOSOME_BIOGENESIS | 0.00000 | 0.00000 | | 0.58688 | 1.49478 | 248 |
| GO_RNA_CATABOLIC_PROCESS | 0.00000 | 0.00000 | | 0.73302 | 1.87060 | 293 |
| GO_RNA_DEPENDENT_DNA_BIOSYNTHETIC_PROCESS | 0.00001 | 0.00023 | 0.61053 | 0.65771 | 1.56881 | 52 |
| GO_RNA_LOCALIZATION | 0.00019 | 0.00413 | 0.51885 | 0.51856 | 1.30515 | 175 |
| GO_RNA_SPLICING | 0.00000 | 0.00015 | 0.61053 | 0.50493 | 1.29052 | 321 |
| GO_RNA_SPLICING_VIA_TRANSESTERIFICATION_REACTIONS | 0.00001 | 0.00022 | 0.61053 | 0.51245 | 1.30733 | 270 |
| GO_RRNA_BINDING | 0.00000 | 0.00000 | 0.70498 | 0.71497 | 1.69988 | 49 |
| GO_RRNA_METABOLIC_PROCESS | 0.00000 | 0.00013 | 0.62726 | 0.54775 | 1.38179 | 184 |
| GO_SCF_DEPENDENT_PROTEASOMAL_UBIQUITIN_DEPENDENT_PRO | 0.00000 | 0.00000 | 0.82666 | 0.72747 | 1.76184 | 65 |
| GO_SECRETORY_GRANULE | 0.00000 | 0.00011 | 0.62726 | 0.51298 | 1.30931 | 295 |
| GO_SECRETORY_VESICLE | 0.00001 | 0.00022 | 0.61053 | 0.49581 | 1.26858 | 336 |
| GO_SIGNAL_TRANSDUCTION_BY_PROTEIN_PHOSPHORYLATION | 0.00005 | 0.00141 | 0.55733 | 0.49597 | 1.26600 | 296 |
| GO_SINGLE_FERTILIZATION | 0.00213 | 0.03399 | 0.43171 | 0.63283 | 1.46018 | 33 |
| GO_SINGLE_STRANDED_RNA_BINDING | 0.00024 | 0.00513 | 0.51885 | 0.61608 | 1.46952 | 52 |
| GO_SMALL_MOLECULE_BIOSYNTHETIC_PROCESS | 0.00209 | 0.03374 | 0.43171 | 0.46663 | 1.19080 | 289 |
| GO_SMALL_RIBOSOMAL_SUBUNIT | 0.00000 | 0.00000 | | 0.75303 | 1.82546 | 66 |
| GO_SPERM_EGG_RECOGNITION | 0.00000 | 0.00001 | 0.69013 | 0.88775 | 1.91664 | 15 |
| GO_SPLICEOSOMAL_COMPLEX | 0.00008 | 0.00191 | 0.53843 | 0.53752 | 1.34722 | 152 |
| GO_STEM_CELL_DIFFERENTIATION | 0.00000 | 0.00001 | 0.67496 | 0.60696 | 1.50583 | 116 |
| GO_STRUCTURAL_CONSTITUENT_OF_CYTOSKELETON | 0.00213 | 0.03399 | 0.43171 | 0.62386 | 1.45461 | 38 |
| GO_STRUCTURAL_CONSTITUENT_OF_RIBOSOME | 0.00000 | 0.00000 | | 0.72183 | 1.80798 | 148 |
| GO_STRUCTURAL_MOLECULE_ACTIVITY | 0.00000 | 0.00000 | | 0.63881 | 1.62974 | 271 |
| GO_SUBSTANTIA_NIGRA_DEVELOPMENT | 0.00013 | 0.00314 | 0.51885 | 0.74885 | 1.67103 | 21 |
| GO_SUPRAMOLECULAR_COMPLEX | 0.00038 | 0.00751 | 0.49849 | 0.45616 | 1.17551 | 495 |
| GO_SYNAPSE | 0.00000 | 0.00000 | 0.76146 | 0.52127 | 1.33684 | 392 |
| GO_TELOMERASE_RNA_LOCALIZATION | 0.00000 | 0.00009 | 0.62726 | 0.82548 | 1.82418 | 19 |
| GO_TELOMERE_MAINTENANCE_VIA_TELOMERE_LENGTHENING | 0.00007 | 0.00185 | 0.53843 | 0.63140 | 1.51780 | 58 |
| GO_THREONINE_TYPE_PEPTIDASE_ACTIVITY | 0.00000 | 0.00000 | 0.78819 | 0.92121 | 2.00262 | 16 |
| GO_TISSUE_MORPHOGENESIS | 0.00030 | 0.00624 | 0.49849 | 0.49552 | 1.25681 | 223 |
| GO_TOXIN_TRANSPORT | 0.00016 | 0.00367 | 0.51885 | 0.74617 | 1.66505 | 21 |
| GO_TRANSLATIONAL_ELONGATION | 0.00272 | 0.04150 | 0.43171 | 0.51935 | 1.28847 | 116 |
| GO_TRANSLATIONAL_INITIATION | 0.00000 | 0.00000 | | 0.81200 | 2.03419 | 150 |
| GO_TRANSLATION_FACTOR_ACTIVITY_RNA_BINDING | 0.00000 | 0.00000 | 0.80122 | 0.72696 | 1.75867 | 63 |
| GO_TRANSLATION_INITIATION_FACTOR_ACTIVITY | 0.00000 | 0.00003 | 0.65944 | 0.70666 | 1.66145 | 43 |
| GO_TRANSLATION_PREINITIATION_COMPLEX | 0.00001 | 0.00025 | 0.59333 | 0.83927 | 1.82449 | 16 |
| GO_TRANSLATION_REGULATOR_ACTIVITY | 0.00000 | 0.00000 | | 0.70073 | 1.72277 | 91 |
| GO_TRANSLATION_REGULATOR_ACTIVITY_NUCLEIC_ACID_BINDING | 0.00000 | 0.00000 | | 0.71146 | 1.73680 | 74 |
| GO_TRANSMEMBRANE_TRANSPORT | 0.00037 | 0.00730 | 0.49849 | 0.46340 | 1.19160 | 432 |
| GO_TRICARBOXYLIC_ACID_CYCLE | 0.00004 | 0.00120 | 0.55733 | 0.73357 | 1.66325 | 27 |
| GO_TUMOR_NECROSIS_FACTOR_MEDIATED_SIGNALING_PATHWAY | 0.00000 | 0.00000 | | 0.71379 | 1.74709 | 80 |
| GO_T_CELL_RECEPTOR_SIGNALING_PATHWAY | 0.00000 | 0.00003 | 0.65944 | 0.63113 | 1.54910 | 85 |
| GO_U2_SNRNP | 0.00313 | 0.04553 | 0.43171 | 0.71395 | 1.56809 | 18 |
| GO_UBIQUITIN_LIKE_PROTEIN_LIGASE_BINDING | 0.00000 | 0.00000 | 0.70498 | 0.57276 | 1.44133 | 172 |
| GO_UNFOLDED_PROTEIN_BINDING | 0.00000 | 0.00000 | 0.71951 | 0.68148 | 1.65760 | 69 |
| GO_VESICLE_LUMEN | 0.00000 | 0.00000 | 0.74774 | 0.60549 | 1.51321 | 139 |

| | | | | | | |
|---|---|---|---|---|---|---|
| GO_VIRAL_GENE_EXPRESSION | 0.00000 | 0.00000 | | 0.73976 | 1.85799 | 163 |
| GO_VIRAL_GENOME_REPLICATION | 0.00037 | 0.00730 | 0.49849 | 0.61131 | 1.46498 | 56 |
| GO_VIRAL_LIFE_CYCLE | 0.00019 | 0.00419 | 0.51885 | 0.52750 | 1.32460 | 160 |
| HP_ABNORMALITY_OF_LOWER_LIP | 0.00287 | 0.04294 | 0.43171 | 0.52003 | 1.28738 | 105 |
| HP_ABNORMAL_ERYTHROCYTE_MORPHOLOGY | 0.00018 | 0.00397 | 0.51885 | 0.49459 | 1.25885 | 245 |
| HP_ABNORMAL_RETICULOCYTE_MORPHOLOGY | 0.00074 | 0.01339 | 0.47727 | 0.73586 | 1.62615 | 19 |
| HP_ACUTE_LEUKEMIA | 0.00002 | 0.00070 | 0.57561 | 0.64355 | 1.54700 | 58 |
| HP_ANEMIA_OF_INADEQUATE_PRODUCTION | 0.00002 | 0.00049 | 0.57561 | 0.62675 | 1.52594 | 71 |
| HP_DELAYED_PUBERTY | 0.00001 | 0.00031 | 0.59333 | 0.60946 | 1.49663 | 86 |
| HP_DEPRESSED_NASAL_RIDGE | 0.00010 | 0.00228 | 0.53843 | 0.61043 | 1.47362 | 61 |
| HP_FATIGUE | 0.00008 | 0.00187 | 0.53843 | 0.55212 | 1.37768 | 133 |
| HP_HEADACHE | 0.00016 | 0.00350 | 0.51885 | 0.55705 | 1.37742 | 102 |
| HP_LARGE_HANDS | 0.00228 | 0.03611 | 0.43171 | 0.68670 | 1.53233 | 21 |
| HP_LEUKEMIA | 0.00015 | 0.00333 | 0.51885 | 0.58382 | 1.42778 | 77 |
| HP_MACROCYTIC_ANEMIA | 0.00000 | 0.00000 | 0.71951 | 0.79100 | 1.82515 | 33 |
| HP_MIGRAINE | 0.00000 | 0.00000 | 0.78819 | 0.74652 | 1.77609 | 50 |
| HP_PALLOR | 0.00011 | 0.00255 | 0.53843 | 0.58654 | 1.43186 | 74 |
| HP_SHORT_NOSE | 0.00034 | 0.00691 | 0.49849 | 0.53452 | 1.33375 | 132 |
| HP_THICK_LOWER_LIP_VERMILION | 0.00350 | 0.04971 | 0.43171 | 0.54390 | 1.32776 | 74 |
| HP_THICK_VERMILION_BORDER | 0.00265 | 0.04090 | 0.43171 | 0.51592 | 1.27880 | 114 |
| **Cluster 6** | | | | | | |
| pathway | pval | padj | log2err | ES | NES | size |
| GO_ACTIN_BINDING | 0.00029 | 0.00634 | 0.49849 | 0.50262 | 1.32114 | 149 |
| GO_ACTIN_CYTOSKELETON | 0.00004 | 0.00101 | 0.55733 | 0.49871 | 1.32321 | 189 |
| GO_ACTIN_FILAMENT_BASED_PROCESS | 0.00331 | 0.04212 | 0.43171 | 0.44190 | 1.18898 | 280 |
| GO_ACTIN_FILAMENT_BINDING | 0.00001 | 0.00017 | 0.61053 | 0.59100 | 1.51324 | 84 |
| GO_ACTIN_FILAMENT_ORGANIZATION | 0.00213 | 0.03041 | 0.43171 | 0.47525 | 1.25490 | 167 |
| GO_ACTIVATION_OF_IMMUNE_RESPONSE | 0.00000 | 0.00001 | 0.69013 | 0.54678 | 1.44186 | 164 |
| GO_ACTIVATION_OF_INNATE_IMMUNE_RESPONSE | 0.00000 | 0.00001 | 0.69013 | 0.63155 | 1.60742 | 74 |
| GO_AEROBIC_RESPIRATION | 0.00257 | 0.03437 | 0.43171 | 0.53837 | 1.35789 | 63 |
| GO_AMIDE_BINDING | 0.00302 | 0.03877 | 0.43171 | 0.48640 | 1.26925 | 127 |
| GO_AMINE_METABOLIC_PROCESS | 0.00000 | 0.00000 | 0.74774 | 0.66908 | 1.69628 | 69 |
| GO_ANAPHASE_PROMOTING_COMPLEX_DEPENDENT_CATABOLIC_P | 0.00000 | 0.00000 | 0.78819 | 0.67022 | 1.70585 | 74 |
| GO_ANCHORING_JUNCTION | 0.00000 | 0.00000 | | 0.57523 | 1.55510 | 353 |
| GO_ANIMAL_ORGAN_MORPHOGENESIS | 0.00003 | 0.00073 | 0.57561 | 0.47433 | 1.27759 | 301 |
| GO_ANTIGEN_PROCESSING_AND_PRESENTATION | 0.00000 | 0.00000 | | 0.62975 | 1.63781 | 119 |
| GO_ANTIGEN_PROCESSING_AND_PRESENTATION_OF_EXOGENOUS_ | 0.00000 | 0.00000 | 0.78819 | 0.70883 | 1.76639 | 53 |
| GO_ANTIGEN_PROCESSING_AND_PRESENTATION_OF_PEPTIDE_ANTI | 0.00000 | 0.00000 | 0.82666 | 0.64069 | 1.65799 | 105 |
| GO_ANTIGEN_PROCESSING_AND_PRESENTATION_OF_PEPTIDE_ANTI | 0.00000 | 0.00000 | 0.74774 | 0.67846 | 1.71029 | 62 |
| GO_ANTIGEN_PROCESSING_AND_PRESENTATION_OF_PEPTIDE_OR_ | 0.00054 | 0.01051 | 0.47727 | 0.58434 | 1.45232 | 51 |
| GO_ANTIGEN_RECEPTOR_MEDIATED_SIGNALING_PATHWAY | 0.00000 | 0.00009 | 0.62726 | 0.57741 | 1.49392 | 103 |
| GO_APOPTOTIC_MITOCHONDRIAL_CHANGES | 0.00146 | 0.02307 | 0.45506 | 0.56231 | 1.40374 | 55 |
| GO_APOPTOTIC_SIGNALING_PATHWAY | 0.00000 | 0.00001 | 0.69013 | 0.50912 | 1.36727 | 262 |
| GO_AXONAL_GROWTH_CONE | 0.00398 | 0.04755 | 0.40702 | 0.69424 | 1.57458 | 16 |
| GO_AZUROPHIL_GRANULE | 0.00122 | 0.02003 | 0.45506 | 0.53251 | 1.36347 | 84 |
| GO_AZUROPHIL_GRANULE_LUMEN | 0.00154 | 0.02383 | 0.45506 | 0.57225 | 1.41697 | 48 |
| GO_BLOOD_MICROPARTICLE | 0.00316 | 0.04046 | 0.43171 | 0.69492 | 1.58056 | 17 |
| GO_BRUSH_BORDER | 0.00002 | 0.00049 | 0.57561 | 0.65779 | 1.60597 | 40 |
| GO_CADHERIN_BINDING | 0.00000 | 0.00000 | | 0.59836 | 1.59541 | 216 |
| GO_CANONICAL_WNT_SIGNALING_PATHWAY | 0.00001 | 0.00040 | 0.59333 | 0.53217 | 1.39732 | 147 |
| GO_CARBOHYDRATE_BIOSYNTHETIC_PROCESS | 0.00009 | 0.00231 | 0.53843 | 0.57677 | 1.46930 | 75 |
| GO_CELLULAR_AMINO_ACID_METABOLIC_PROCESS | 0.00001 | 0.00036 | 0.59333 | 0.51692 | 1.36280 | 162 |
| GO_CELLULAR_CARBOHYDRATE_CATABOLIC_PROCESS | 0.00048 | 0.00967 | 0.49849 | 0.70334 | 1.62282 | 20 |
| GO_CELLULAR_COMPONENT_ASSEMBLY_INVOLVED_IN_MORPHOGE | 0.00272 | 0.03604 | 0.43171 | 0.59437 | 1.44079 | 37 |
| GO_CELLULAR_KETONE_METABOLIC_PROCESS | 0.00000 | 0.00000 | 0.71951 | 0.58683 | 1.52543 | 117 |
| GO_CELLULAR_NITROGEN_COMPOUND_CATABOLIC_PROCESS | 0.00000 | 0.00000 | | 0.59556 | 1.60961 | 350 |
| GO_CELLULAR_PROTEIN_CATABOLIC_PROCESS | 0.00001 | 0.00021 | 0.61053 | 0.46184 | 1.25549 | 441 |
| GO_CELLULAR_RESPONSE_TO_ENDOGENOUS_STIMULUS | 0.00205 | 0.02971 | 0.43171 | 0.42546 | 1.15783 | 480 |
| GO_CELLULAR_RESPONSE_TO_ORGANIC_CYCLIC_COMPOUND | 0.00006 | 0.00164 | 0.53843 | 0.49220 | 1.30909 | 202 |
| GO_CELLULAR_RESPONSE_TO_OXYGEN_LEVELS | 0.00000 | 0.00000 | | 0.63450 | 1.65574 | 127 |
| GO_CELLULAR_RESPONSE_TO_TOPOLOGICALLY_INCORRECT_PROTEI | 0.00294 | 0.03813 | 0.43171 | 0.51911 | 1.32918 | 84 |
| GO_CELL_ACTIVATION | 0.00000 | 0.00000 | | 0.51868 | 1.41102 | 456 |
| GO_CELL_ACTIVATION_INVOLVED_IN_IMMUNE_RESPONSE | 0.00000 | 0.00000 | | 0.56526 | 1.52005 | 276 |
| GO_CELL_ADHESION_MEDIATOR_ACTIVITY | 0.00231 | 0.03199 | 0.43171 | 0.65900 | 1.52053 | 20 |
| GO_CELL_ADHESION_MOLECULE_BINDING | 0.00000 | 0.00000 | | 0.59238 | 1.59106 | 263 |
| GO_CELL_BODY | 0.00357 | 0.04458 | 0.43171 | 0.45695 | 1.21395 | 193 |
| GO_CELL_CELL_ADHESION | 0.00023 | 0.00512 | 0.51885 | 0.47827 | 1.27174 | 197 |
| GO_CELL_CELL_JUNCTION | 0.00045 | 0.00919 | 0.49849 | 0.48905 | 1.29418 | 173 |
| GO_CELL_CELL_RECOGNITION | 0.00000 | 0.00003 | 0.65944 | 0.81922 | 1.88160 | 19 |
| GO_CELL_CELL_SIGNALING | 0.00001 | 0.00018 | 0.61053 | 0.45919 | 1.24927 | 469 |
| GO_CELL_CELL_SIGNALING_BY_WNT | 0.00000 | 0.00002 | 0.67496 | 0.51654 | 1.37800 | 220 |
| GO_CELL_CORTEX | 0.00357 | 0.04458 | 0.43171 | 0.48601 | 1.26335 | 117 |
| GO_CELL_CYCLE_G2_M_PHASE_TRANSITION | 0.00239 | 0.03282 | 0.43171 | 0.47123 | 1.24772 | 177 |
| GO_CELL_LEADING_EDGE | 0.00098 | 0.01706 | 0.45506 | 0.47205 | 1.25006 | 181 |
| GO_CELL_RECOGNITION | 0.00003 | 0.00075 | 0.57561 | 0.69697 | 1.67209 | 31 |
| GO_CELL_REDOX_HOMEOSTASIS | 0.00342 | 0.04319 | 0.43171 | 0.59931 | 1.44228 | 32 |

| | | | | | | |
|---|---|---|---|---|---|---|
| GO_CELL_SUBSTRATE_JUNCTION | 0.00000 | 0.00000 | | 0.61689 | 1.64848 | 230 |
| GO_CELL_SURFACE | 0.00361 | 0.04481 | 0.43171 | 0.47322 | 1.24176 | 144 |
| GO_CELL_SURFACE_RECEPTOR_SIGNALING_PATHWAY_INVOLVED_IN | 0.00000 | 0.00001 | 0.69013 | 0.51824 | 1.38534 | 232 |
| GO_CHAPERONE_BINDING | 0.00004 | 0.00115 | 0.55733 | 0.61871 | 1.53773 | 51 |
| GO_CHAPERONE_COMPLEX | 0.00000 | 0.00013 | 0.61053 | 0.79606 | 1.82840 | 19 |
| GO_CHAPERONE_MEDIATED_PROTEIN_FOLDING | 0.00111 | 0.01848 | 0.45506 | 0.60941 | 1.47505 | 36 |
| GO_CLATHRIN_DEPENDENT_ENDOCYTOSIS | 0.00364 | 0.04515 | 0.43171 | 0.61822 | 1.47436 | 28 |
| GO_CLUSTER_OF_ACTIN_BASED_CELL_PROJECTIONS | 0.00106 | 0.01787 | 0.45506 | 0.56738 | 1.41640 | 55 |
| GO_CONTRACTILE_FIBER | 0.00104 | 0.01775 | 0.45506 | 0.53617 | 1.36637 | 77 |
| GO_CORTICAL_CYTOSKELETON | 0.00250 | 0.03376 | 0.43171 | 0.56798 | 1.40251 | 46 |
| GO_COTRANSLATIONAL_PROTEIN_TARGETING_TO_MEMBRANE | 0.00000 | 0.00000 | | 0.73280 | 1.88229 | 89 |
| GO_CYTOKINE_MEDIATED_SIGNALING_PATHWAY | 0.00000 | 0.00000 | | 0.58253 | 1.55855 | 236 |
| GO_CYTOPLASMIC_STRESS_GRANULE | 0.00167 | 0.02534 | 0.45506 | 0.56645 | 1.40599 | 50 |
| GO_CYTOPLASMIC_TRANSLATION | 0.00000 | 0.00000 | | 0.72303 | 1.84476 | 79 |
| GO_CYTOPLASMIC_TRANSLATIONAL_INITIATION | 0.00000 | 0.00000 | 0.76146 | 0.80600 | 1.92370 | 29 |
| GO_CYTOSKELETAL_PROTEIN_BINDING | 0.00016 | 0.00380 | 0.51885 | 0.45597 | 1.23664 | 387 |
| GO_CYTOSOLIC_LARGE_RIBOSOMAL_SUBUNIT | 0.00000 | 0.00000 | 0.82666 | 0.74454 | 1.85233 | 52 |
| GO_CYTOSOLIC_RIBOSOME | 0.00000 | 0.00000 | | 0.73786 | 1.89767 | 92 |
| GO_CYTOSOLIC_SMALL_RIBOSOMAL_SUBUNIT | 0.00000 | 0.00000 | 0.74774 | 0.73892 | 1.80406 | 40 |
| GO_DEFENSE_RESPONSE | 0.00000 | 0.00000 | 0.70498 | 0.48701 | 1.32182 | 398 |
| GO_DEFENSE_RESPONSE_TO_OTHER_ORGANISM | 0.00000 | 0.00004 | 0.64355 | 0.49228 | 1.32502 | 288 |
| GO_DE_NOVO_PROTEIN_FOLDING | 0.00067 | 0.01258 | 0.47727 | 0.68208 | 1.58740 | 22 |
| GO_DISORDERED_DOMAIN_SPECIFIC_BINDING | 0.00006 | 0.00153 | 0.55733 | 0.74771 | 1.73149 | 21 |
| GO_ENDOPEPTIDASE_ACTIVITY | 0.00073 | 0.01339 | 0.47727 | 0.50610 | 1.31558 | 117 |
| GO_ENDOPEPTIDASE_COMPLEX | 0.00000 | 0.00000 | 0.82666 | 0.72560 | 1.82129 | 58 |
| GO_ENERGY_DERIVATION_BY_OXIDATION_OF_ORGANIC_COMPOUN | 0.00209 | 0.02997 | 0.43171 | 0.47961 | 1.26334 | 158 |
| GO_ENTRY_INTO_HOST | 0.00220 | 0.03081 | 0.43171 | 0.55535 | 1.39124 | 57 |
| GO_EPIDERMAL_CELL_DIFFERENTIATION | 0.00287 | 0.03748 | 0.43171 | 0.52958 | 1.34376 | 70 |
| GO_EPIDERMIS_DEVELOPMENT | 0.00124 | 0.02012 | 0.45506 | 0.51407 | 1.32500 | 96 |
| GO_EPITHELIAL_CELL_DIFFERENTIATION | 0.00065 | 0.01233 | 0.47727 | 0.47697 | 1.26867 | 198 |
| GO_EPITHELIUM_DEVELOPMENT | 0.00000 | 0.00000 | 0.80122 | 0.50793 | 1.37763 | 388 |
| GO_ESTABLISHMENT_OF_PROTEIN_LOCALIZATION_TO_CHROMOSON | 0.00194 | 0.02862 | 0.45506 | 0.67523 | 1.55088 | 19 |
| GO_ESTABLISHMENT_OF_PROTEIN_LOCALIZATION_TO_ENDOPLASM | 0.00000 | 0.00000 | | 0.72258 | 1.86387 | 98 |
| GO_ESTABLISHMENT_OF_PROTEIN_LOCALIZATION_TO_MEMBRANE | 0.00000 | 0.00000 | | 0.64144 | 1.70619 | 199 |
| GO_ESTABLISHMENT_OF_PROTEIN_LOCALIZATION_TO_ORGANELLE | 0.00000 | 0.00000 | | 0.58579 | 1.58158 | 338 |
| GO_ESTABLISHMENT_OF_PROTEIN_LOCALIZATION_TO_TELOMERE | 0.00069 | 0.01277 | 0.47727 | 0.75320 | 1.69192 | 15 |
| GO_ESTABLISHMENT_OF_TISSUE_POLARITY | 0.00000 | 0.00000 | | 0.71132 | 1.80337 | 69 |
| GO_EUKARYOTIC_TRANSLATION_INITIATION_FACTOR_3_COMPLEX | 0.00001 | 0.00044 | 0.59333 | 0.82820 | 1.86040 | 15 |
| GO_EXOCYTOSIS | 0.00000 | 0.00000 | | 0.54340 | 1.46802 | 344 |
| GO_FC_EPSILON_RECEPTOR_SIGNALING_PATHWAY | 0.00000 | 0.00001 | 0.69013 | 0.63132 | 1.60686 | 74 |
| GO_FC_RECEPTOR_MEDIATED_STIMULATORY_SIGNALING_PATHWAY | 0.00342 | 0.04319 | 0.43171 | 0.56764 | 1.39254 | 42 |
| GO_FC_RECEPTOR_SIGNALING_PATHWAY | 0.00000 | 0.00000 | 0.80122 | 0.63189 | 1.63522 | 105 |
| GO_FICOLIN_1_RICH_GRANULE | 0.00000 | 0.00000 | | 0.67530 | 1.72786 | 83 |
| GO_FICOLIN_1_RICH_GRANULE_MEMBRANE | 0.00148 | 0.02328 | 0.45506 | 0.67131 | 1.54891 | 20 |
| GO_GENERATION_OF_PRECURSOR_METABOLITES_AND_ENERGY | 0.00368 | 0.04537 | 0.43171 | 0.44642 | 1.20000 | 270 |
| GO_GLIOGENESIS | 0.00220 | 0.03081 | 0.43171 | 0.51235 | 1.32049 | 95 |
| GO_GLUCOSE_METABOLIC_PROCESS | 0.00008 | 0.00201 | 0.53843 | 0.57356 | 1.46242 | 78 |
| GO_GLUTAMATERGIC_SYNAPSE | 0.00032 | 0.00677 | 0.49849 | 0.52921 | 1.36630 | 101 |
| GO_HEAT_SHOCK_PROTEIN_BINDING | 0.00139 | 0.02206 | 0.45506 | 0.54894 | 1.38780 | 65 |
| GO_HEMATOPOIETIC_PROGENITOR_CELL_DIFFERENTIATION | 0.00000 | 0.00000 | 0.71951 | 0.62862 | 1.60842 | 83 |
| GO_HEMATOPOIETIC_STEM_CELL_DIFFERENTIATION | 0.00000 | 0.00000 | 0.80122 | 0.70723 | 1.77811 | 59 |
| GO_HORMONE_BINDING | 0.00074 | 0.01351 | 0.47727 | 0.75181 | 1.68880 | 15 |
| GO_IMMUNE_EFFECTOR_PROCESS | 0.00000 | 0.00000 | | 0.52987 | 1.43818 | 403 |
| GO_IMMUNE_RESPONSE_REGULATING_SIGNALING_PATHWAY | 0.00000 | 0.00001 | 0.70498 | 0.55890 | 1.47170 | 157 |
| GO_IMMUNE_SYSTEM_DEVELOPMENT | 0.00413 | 0.04908 | 0.40702 | 0.43339 | 1.17009 | 338 |
| GO_INFLAMMATORY_RESPONSE | 0.00156 | 0.02404 | 0.45506 | 0.48564 | 1.27083 | 137 |
| GO_INNATE_IMMUNE_RESPONSE | 0.00000 | 0.00003 | 0.65944 | 0.50827 | 1.36021 | 238 |
| GO_INNATE_IMMUNE_RESPONSE_ACTIVATING_SIGNAL_TRANSDUCT | 0.00000 | 0.00000 | 0.76146 | 0.68291 | 1.72149 | 62 |
| GO_INORGANIC_ANION_TRANSPORT | 0.00115 | 0.01889 | 0.45506 | 0.61660 | 1.48928 | 35 |
| GO_INTERACTION_WITH_HOST | 0.00011 | 0.00275 | 0.53843 | 0.54196 | 1.39687 | 96 |
| GO_INTERACTION_WITH_SYMBIONT | 0.00172 | 0.02609 | 0.45506 | 0.60973 | 1.47152 | 34 |
| GO_INTERLEUKIN_1_MEDIATED_SIGNALING_PATHWAY | 0.00000 | 0.00000 | | 0.71475 | 1.80276 | 63 |
| GO_INTRINSIC_APOPTOTIC_SIGNALING_PATHWAY | 0.00026 | 0.00576 | 0.49849 | 0.50725 | 1.32813 | 138 |
| GO_ION_CHANNEL_BINDING | 0.00135 | 0.02162 | 0.45506 | 0.59593 | 1.45897 | 41 |
| GO_KERATINOCYTE_DIFFERENTIATION | 0.00039 | 0.00814 | 0.49849 | 0.59690 | 1.47800 | 48 |
| GO_LARGE_RIBOSOMAL_SUBUNIT | 0.00000 | 0.00000 | 0.76146 | 0.61347 | 1.58456 | 102 |
| GO_LEUKOCYTE_MEDIATED_IMMUNITY | 0.00000 | 0.00000 | | 0.55602 | 1.49692 | 289 |
| GO_LYSOSOMAL_LUMEN | 0.00003 | 0.00093 | 0.55733 | 0.64938 | 1.58545 | 40 |
| GO_MEMBRANE_ORGANIZATION | 0.00020 | 0.00461 | 0.51885 | 0.44711 | 1.21346 | 397 |
| GO_MESENCHYMAL_CELL_DIFFERENTIATION | 0.00202 | 0.02926 | 0.43171 | 0.54899 | 1.38200 | 60 |
| GO_MONOSACCHARIDE_BIOSYNTHETIC_PROCESS | 0.00003 | 0.00089 | 0.55733 | 0.64298 | 1.58750 | 45 |
| GO_MONOSACCHARIDE_CATABOLIC_PROCESS | 0.00302 | 0.03877 | 0.43171 | 0.63919 | 1.51048 | 26 |
| GO_MONOSACCHARIDE_METABOLIC_PROCESS | 0.00001 | 0.00031 | 0.59333 | 0.55679 | 1.44274 | 109 |
| GO_MORPHOGENESIS_OF_AN_EPITHELIUM | 0.00000 | 0.00000 | 0.73376 | 0.54271 | 1.44538 | 208 |
| GO_MORPHOGENESIS_OF_A_POLARIZED_EPITHELIUM | 0.00000 | 0.00000 | | 0.69857 | 1.78513 | 81 |
| GO_MOVEMENT_IN_HOST_ENVIRONMENT | 0.00060 | 0.01152 | 0.47727 | 0.55442 | 1.40561 | 69 |

| | | | | | | |
|---|---|---|---|---|---|---|
| GO_MRNA_3_UTR_BINDING | 0.00077 | 0.01407 | 0.47727 | 0.56039 | 1.39648 | 53 |
| GO_MRNA_5_UTR_BINDING | 0.00084 | 0.01506 | 0.47727 | 0.69146 | 1.60123 | 21 |
| GO_MRNA_BINDING | 0.00000 | 0.00000 | 0.70498 | 0.53769 | 1.42665 | 189 |
| GO_MUSCLE_CELL_DIFFERENTIATION | 0.00113 | 0.01872 | 0.45506 | 0.50634 | 1.31342 | 111 |
| GO_MUSCLE_STRUCTURE_DEVELOPMENT | 0.00086 | 0.01533 | 0.47727 | 0.47446 | 1.26149 | 196 |
| GO_MYELOID_LEUKOCYTE_ACTIVATION | 0.00000 | 0.00000 | | 0.57599 | 1.54593 | 259 |
| GO_MYELOID_LEUKOCYTE_MEDIATED_IMMUNITY | 0.00000 | 0.00000 | | 0.58817 | 1.57527 | 242 |
| GO_NADH_METABOLIC_PROCESS | 0.00064 | 0.01208 | 0.47727 | 0.73617 | 1.66967 | 16 |
| GO_NAD_BINDING | 0.00065 | 0.01221 | 0.47727 | 0.62853 | 1.51688 | 34 |
| GO_NAD_METABOLIC_PROCESS | 0.00216 | 0.03066 | 0.43171 | 0.66719 | 1.54502 | 21 |
| GO_NEGATIVE_REGULATION_OF_CANONICAL_WNT_SIGNALING_PAT | 0.00000 | 0.00000 | 0.73376 | 0.63874 | 1.63547 | 84 |
| GO_NEGATIVE_REGULATION_OF_CELLULAR_AMIDE_METABOLIC_PR | 0.00002 | 0.00058 | 0.57561 | 0.57068 | 1.46377 | 87 |
| GO_NEGATIVE_REGULATION_OF_CELLULAR_PROTEIN_CATABOLIC_P | 0.00092 | 0.01618 | 0.47727 | 0.55760 | 1.38953 | 53 |
| GO_NEGATIVE_REGULATION_OF_CELL_CYCLE_G2_M_PHASE_TRANS | 0.00000 | 0.00006 | 0.64355 | 0.62210 | 1.57986 | 71 |
| GO_NEGATIVE_REGULATION_OF_CELL_CYCLE_PHASE_TRANSITION | 0.00009 | 0.00235 | 0.53843 | 0.51377 | 1.35046 | 149 |
| GO_NEGATIVE_REGULATION_OF_CELL_CYCLE_PROCESS | 0.00054 | 0.01064 | 0.47727 | 0.47729 | 1.26957 | 199 |
| GO_NEGATIVE_REGULATION_OF_CELL_DEATH | 0.00000 | 0.00003 | 0.65944 | 0.48350 | 1.30861 | 362 |
| GO_NEGATIVE_REGULATION_OF_MITOTIC_CELL_CYCLE | 0.00095 | 0.01648 | 0.47727 | 0.47486 | 1.25819 | 184 |
| GO_NEGATIVE_REGULATION_OF_PROTEIN_METABOLIC_PROCESS | 0.00027 | 0.00578 | 0.49849 | 0.44576 | 1.21173 | 437 |
| GO_NEGATIVE_REGULATION_OF_PROTEOLYSIS | 0.00093 | 0.01622 | 0.47727 | 0.50736 | 1.31435 | 108 |
| GO_NEGATIVE_REGULATION_OF_RNA_SPLICING | 0.00130 | 0.02095 | 0.45506 | 0.67496 | 1.55734 | 20 |
| GO_NEGATIVE_REGULATION_OF_SIGNALING | 0.00001 | 0.00038 | 0.59333 | 0.45520 | 1.23910 | 490 |
| GO_NEGATIVE_REGULATION_OF_TRANSMEMBRANE_RECEPTOR_PR | 0.00420 | 0.04947 | 0.40702 | 0.59385 | 1.43319 | 34 |
| GO_NEGATIVE_REGULATION_OF_UBIQUITIN_DEPENDENT_PROTEIN_ | 0.00290 | 0.03776 | 0.43171 | 0.60291 | 1.45093 | 32 |
| GO_NEGATIVE_REGULATION_OF_WNT_SIGNALING_PATHWAY | 0.00000 | 0.00000 | 0.77494 | 0.63802 | 1.63922 | 91 |
| GO_NEURAL_NUCLEUS_DEVELOPMENT | 0.00111 | 0.01848 | 0.45506 | 0.66445 | 1.57018 | 26 |
| GO_NEURON_TO_NEURON_SYNAPSE | 0.00088 | 0.01565 | 0.47727 | 0.50971 | 1.32075 | 109 |
| GO_NIK_NF_KAPPAB_SIGNALING | 0.00000 | 0.00000 | | 0.67859 | 1.73751 | 84 |
| GO_NON_CANONICAL_WNT_SIGNALING_PATHWAY | 0.00000 | 0.00000 | | 0.71041 | 1.81009 | 76 |
| GO_NUCLEAR_TRANSCRIBED_MRNA_CATABOLIC_PROCESS | 0.00000 | 0.00000 | | 0.64191 | 1.69123 | 159 |
| GO_NUCLEAR_TRANSCRIBED_MRNA_CATABOLIC_PROCESS_NONSEN | 0.00000 | 0.00000 | | 0.72147 | 1.86702 | 105 |
| GO_ORGANELLE_ASSEMBLY | 0.00253 | 0.03416 | 0.43171 | 0.43253 | 1.17418 | 410 |
| GO_ORGANIC_ACID_METABOLIC_PROCESS | 0.00001 | 0.00033 | 0.59333 | 0.45637 | 1.24153 | 470 |
| GO_ORGANIC_CYCLIC_COMPOUND_CATABOLIC_PROCESS | 0.00000 | 0.00000 | | 0.58902 | 1.59285 | 355 |
| GO_OXIDATION_REDUCTION_PROCESS | 0.00124 | 0.02012 | 0.45506 | 0.43204 | 1.17323 | 419 |
| GO_PEPTIDASE_ACTIVITY | 0.00167 | 0.02534 | 0.45506 | 0.46653 | 1.23784 | 189 |
| GO_PEPTIDASE_COMPLEX | 0.00000 | 0.00000 | 0.73376 | 0.65477 | 1.66279 | 72 |
| GO_PEPTIDE_BINDING | 0.00383 | 0.04659 | 0.43171 | 0.50634 | 1.30007 | 88 |
| GO_PEPTIDE_BIOSYNTHETIC_PROCESS | 0.00000 | 0.00000 | | 0.57197 | 1.55541 | 443 |
| GO_PEPTIDE_METABOLIC_PROCESS | 0.00000 | 0.00000 | | 0.55743 | 1.51834 | 500 |
| GO_PIGMENT_GRANULE | 0.00001 | 0.00034 | 0.59333 | 0.60361 | 1.52712 | 67 |
| GO_PLATELET_DEGRANULATION | 0.00420 | 0.04947 | 0.40702 | 0.59444 | 1.43101 | 33 |
| GO_POLYSOMAL_RIBOSOME | 0.00017 | 0.00412 | 0.51885 | 0.67541 | 1.61690 | 30 |
| GO_POLYSOME | 0.00001 | 0.00021 | 0.61053 | 0.63750 | 1.59449 | 56 |
| GO_POSITIVE_REGULATION_OF_CANONICAL_WNT_SIGNALING_PATH | 0.00000 | 0.00004 | 0.64355 | 0.60256 | 1.54507 | 85 |
| GO_POSITIVE_REGULATION_OF_CELLULAR_AMIDE_METABOLIC_PRC | 0.00115 | 0.01889 | 0.45506 | 0.51740 | 1.32947 | 90 |
| GO_POSITIVE_REGULATION_OF_CELLULAR_COMPONENT_ORGANIZA | 0.00009 | 0.00233 | 0.53843 | 0.44336 | 1.20755 | 499 |
| GO_POSITIVE_REGULATION_OF_CELLULAR_PROTEIN_CATABOLIC_PR | 0.00042 | 0.00866 | 0.49849 | 0.53672 | 1.37153 | 81 |
| GO_POSITIVE_REGULATION_OF_CELLULAR_PROTEIN_LOCALIZATION | 0.00064 | 0.01208 | 0.47727 | 0.48247 | 1.27684 | 174 |
| GO_POSITIVE_REGULATION_OF_CELL_DEATH | 0.00124 | 0.02012 | 0.45506 | 0.46045 | 1.23633 | 261 |
| GO_POSITIVE_REGULATION_OF_CELL_POPULATION_PROLIFERATION | 0.00220 | 0.03081 | 0.43171 | 0.45032 | 1.20936 | 262 |
| GO_POSITIVE_REGULATION_OF_DEFENSE_RESPONSE | 0.00001 | 0.00021 | 0.61053 | 0.55034 | 1.43058 | 117 |
| GO_POSITIVE_REGULATION_OF_DNA_BIOSYNTHETIC_PROCESS | 0.00043 | 0.00871 | 0.49849 | 0.61070 | 1.50119 | 43 |
| GO_POSITIVE_REGULATION_OF_DNA_METABOLIC_PROCESS | 0.00165 | 0.02522 | 0.45506 | 0.51818 | 1.33553 | 95 |
| GO_POSITIVE_REGULATION_OF_DNA_TEMPLATED_TRANSCRIPTION_ | 0.00216 | 0.03066 | 0.43171 | 0.66753 | 1.54581 | 21 |
| GO_POSITIVE_REGULATION_OF_ESTABLISHMENT_OF_PROTEIN_LOC | 0.00030 | 0.00641 | 0.49849 | 0.49403 | 1.30449 | 167 |
| GO_POSITIVE_REGULATION_OF_IMMUNE_RESPONSE | 0.00000 | 0.00008 | 0.62726 | 0.51608 | 1.37333 | 205 |
| GO_POSITIVE_REGULATION_OF_IMMUNE_SYSTEM_PROCESS | 0.00040 | 0.00826 | 0.49849 | 0.46254 | 1.24500 | 281 |
| GO_POSITIVE_REGULATION_OF_INNATE_IMMUNE_RESPONSE | 0.00000 | 0.00010 | 0.62726 | 0.58749 | 1.50940 | 91 |
| GO_POSITIVE_REGULATION_OF_ION_TRANSPORT | 0.00398 | 0.04755 | 0.40702 | 0.52697 | 1.33473 | 68 |
| GO_POSITIVE_REGULATION_OF_NEURON_APOPTOTIC_PROCESS | 0.00239 | 0.03282 | 0.43171 | 0.64710 | 1.51597 | 24 |
| GO_POSITIVE_REGULATION_OF_NEURON_DEATH | 0.00320 | 0.04082 | 0.43171 | 0.59591 | 1.43929 | 35 |
| GO_POSITIVE_REGULATION_OF_PROTEIN_CONTAINING_COMPLEX_ | 0.00106 | 0.01787 | 0.45506 | 0.49735 | 1.29407 | 121 |
| GO_POSITIVE_REGULATION_OF_PROTEIN_LOCALIZATION_TO_NUCLE | 0.00023 | 0.00512 | 0.51885 | 0.59954 | 1.48812 | 50 |
| GO_POSITIVE_REGULATION_OF_RESPONSE_TO_BIOTIC_STIMULUS | 0.00000 | 0.00011 | 0.62726 | 0.57878 | 1.49393 | 100 |
| GO_POSITIVE_REGULATION_OF_RESPONSE_TO_EXTERNAL_STIMULU | 0.00016 | 0.00388 | 0.51885 | 0.50551 | 1.32684 | 146 |
| GO_POSITIVE_REGULATION_OF_TELOMERE_MAINTENANCE_VIA_TE | 0.00093 | 0.01622 | 0.47727 | 0.66903 | 1.56734 | 24 |
| GO_POSITIVE_REGULATION_OF_TRANSCRIPTION_INITIATION_FROM | 0.00209 | 0.02997 | 0.43171 | 0.69160 | 1.58341 | 18 |
| GO_POSITIVE_REGULATION_OF_TRANSLATION | 0.00108 | 0.01806 | 0.45506 | 0.53232 | 1.35902 | 80 |
| GO_POSITIVE_REGULATION_OF_TRANSPORT | 0.00257 | 0.03437 | 0.43171 | 0.44125 | 1.18932 | 309 |
| GO_POSITIVE_REGULATION_OF_VIRAL_LIFE_CYCLE | 0.00246 | 0.03355 | 0.43171 | 0.58615 | 1.42788 | 39 |
| GO_POSITIVE_REGULATION_OF_WNT_SIGNALING_PATHWAY | 0.00000 | 0.00005 | 0.64355 | 0.58583 | 1.50856 | 94 |
| GO_POSTSYNAPSE | 0.00000 | 0.00013 | 0.62726 | 0.51931 | 1.37598 | 184 |
| GO_POSTSYNAPSE_ORGANIZATION | 0.00361 | 0.04481 | 0.43171 | 0.55386 | 1.37656 | 51 |
| GO_POSTTRANSCRIPTIONAL_REGULATION_OF_GENE_EXPRESSION | 0.00000 | 0.00000 | | 0.53698 | 1.45369 | 369 |
| GO_POST_TRANSLATIONAL_PROTEIN_MODIFICATION | 0.00006 | 0.00153 | 0.55733 | 0.50173 | 1.32865 | 178 |

| | | | | | | |
|---|---|---|---|---|---|---|
| GO_PROTEASOMAL_PROTEIN_CATABOLIC_PROCESS | 0.00013 | 0.00332 | 0.51885 | 0.46903 | 1.26284 | 294 |
| GO_PROTEASOMAL_UBIQUITIN_INDEPENDENT_PROTEIN_CATABOLI( | 0.00000 | 0.00002 | 0.65944 | 0.84370 | 1.93164 | 18 |
| GO_PROTEASOME_ACCESSORY_COMPLEX | 0.00006 | 0.00152 | 0.55733 | 0.72641 | 1.70177 | 24 |
| GO_PROTEASOME_CORE_COMPLEX | 0.00000 | 0.00002 | 0.67496 | 0.85863 | 1.94743 | 16 |
| GO_PROTEIN_DOMAIN_SPECIFIC_BINDING | 0.00052 | 0.01036 | 0.47727 | 0.45179 | 1.21885 | 319 |
| GO_PROTEIN_FOLDING | 0.00000 | 0.00001 | 0.70498 | 0.56869 | 1.49100 | 141 |
| GO_PROTEIN_IMPORT | 0.00027 | 0.00578 | 0.49849 | 0.53142 | 1.37557 | 106 |
| GO_PROTEIN_LOCALIZATION_TO_ENDOPLASMIC_RETICULUM | 0.00000 | 0.00000 | | 0.70095 | 1.81959 | 114 |
| GO_PROTEIN_LOCALIZATION_TO_MEMBRANE | 0.00000 | 0.00000 | | 0.55923 | 1.50865 | 325 |
| GO_PROTEIN_LOCALIZATION_TO_NUCLEUS | 0.00061 | 0.01165 | 0.47727 | 0.50004 | 1.31041 | 140 |
| GO_PROTEIN_MODIFICATION_BY_SMALL_PROTEIN_CONJUGATION | 0.00100 | 0.01725 | 0.45506 | 0.43033 | 1.17181 | 495 |
| GO_PROTEIN_MODIFICATION_BY_SMALL_PROTEIN_REMOVAL | 0.00014 | 0.00341 | 0.51885 | 0.49530 | 1.31160 | 178 |
| GO_PROTEIN_POLYUBIQUITINATION | 0.00004 | 0.00117 | 0.55733 | 0.49720 | 1.32142 | 195 |
| GO_PROTEIN_STABILIZATION | 0.00000 | 0.00002 | 0.65944 | 0.58525 | 1.51098 | 101 |
| GO_PROTEIN_TARGETING | 0.00000 | 0.00000 | | 0.60940 | 1.63825 | 271 |
| GO_PROTEIN_TARGETING_TO_MEMBRANE | 0.00000 | 0.00000 | | 0.69403 | 1.81435 | 131 |
| GO_PURINE_CONTAINING_COMPOUND_METABOLIC_PROCESS | 0.00313 | 0.04009 | 0.43171 | 0.45846 | 1.21796 | 193 |
| GO_PURINE_NUCLEOSIDE_METABOLIC_PROCESS | 0.00283 | 0.03720 | 0.43171 | 0.61495 | 1.46773 | 29 |
| GO_PURINE_NUCLEOSIDE_TRIPHOSPHATE_METABOLIC_PROCESS | 0.00298 | 0.03851 | 0.43171 | 0.59206 | 1.43709 | 38 |
| GO_REACTIVE_OXYGEN_SPECIES_METABOLIC_PROCESS | 0.00139 | 0.02206 | 0.45506 | 0.52438 | 1.34494 | 86 |
| GO_RECEPTOR_MEDIATED_ENDOCYTOSIS | 0.00000 | 0.00010 | 0.62726 | 0.58245 | 1.49661 | 90 |
| GO_REGULATION_OF_ANATOMICAL_STRUCTURE_MORPHOGENESIS | 0.00054 | 0.01051 | 0.47727 | 0.44715 | 1.21023 | 362 |
| GO_REGULATION_OF_ANIMAL_ORGAN_MORPHOGENESIS | 0.00000 | 0.00000 | 0.78819 | 0.63367 | 1.63388 | 97 |
| GO_REGULATION_OF_APOPTOTIC_SIGNALING_PATHWAY | 0.00001 | 0.00041 | 0.59333 | 0.51387 | 1.36051 | 176 |
| GO_REGULATION_OF_CELLULAR_AMIDE_METABOLIC_PROCESS | 0.00000 | 0.00000 | 0.74774 | 0.53377 | 1.42764 | 235 |
| GO_REGULATION_OF_CELLULAR_AMINE_METABOLIC_PROCESS | 0.00000 | 0.00000 | | 0.72712 | 1.81197 | 53 |
| GO_REGULATION_OF_CELLULAR_AMINO_ACID_METABOLIC_PROCES | 0.00000 | 0.00000 | 0.81404 | 0.73621 | 1.82737 | 50 |
| GO_REGULATION_OF_CELLULAR_CATABOLIC_PROCESS | 0.00000 | 0.00000 | 0.73376 | 0.48378 | 1.31656 | 481 |
| GO_REGULATION_OF_CELLULAR_KETONE_METABOLIC_PROCESS | 0.00000 | 0.00000 | 0.73376 | 0.62551 | 1.60708 | 91 |
| GO_REGULATION_OF_CELLULAR_LOCALIZATION | 0.00082 | 0.01479 | 0.47727 | 0.44122 | 1.19793 | 413 |
| GO_REGULATION_OF_CELLULAR_PROTEIN_CATABOLIC_PROCESS | 0.00001 | 0.00039 | 0.59333 | 0.52885 | 1.39009 | 149 |
| GO_REGULATION_OF_CELLULAR_PROTEIN_LOCALIZATION | 0.00012 | 0.00301 | 0.53843 | 0.46856 | 1.26002 | 274 |
| GO_REGULATION_OF_CELLULAR_RESPONSE_TO_GROWTH_FACTOR_ | 0.00165 | 0.02522 | 0.45506 | 0.53465 | 1.36321 | 78 |
| GO_REGULATION_OF_CELLULAR_RESPONSE_TO_TRANSFORMING_G | 0.00108 | 0.01806 | 0.45506 | 0.60011 | 1.47825 | 44 |
| GO_REGULATION_OF_CELL_CYCLE_G2_M_PHASE_TRANSITION | 0.00000 | 0.00011 | 0.62726 | 0.53894 | 1.41376 | 143 |
| GO_REGULATION_OF_CELL_CYCLE_PHASE_TRANSITION | 0.00179 | 0.02661 | 0.45506 | 0.45080 | 1.21221 | 273 |
| GO_REGULATION_OF_CELL_CYCLE_PROCESS | 0.00209 | 0.02997 | 0.43171 | 0.43425 | 1.17978 | 425 |
| GO_REGULATION_OF_CELL_POPULATION_PROLIFERATION | 0.00150 | 0.02342 | 0.45506 | 0.43226 | 1.17659 | 489 |
| GO_REGULATION_OF_CYTOPLASMIC_TRANSLATION | 0.00042 | 0.00866 | 0.49849 | 0.74417 | 1.69257 | 17 |
| GO_REGULATION_OF_DEFENSE_RESPONSE | 0.00004 | 0.00100 | 0.55733 | 0.49926 | 1.32796 | 198 |
| GO_REGULATION_OF_DNA_TEMPLATED_TRANSCRIPTION_INITIATIOI | 0.00390 | 0.04714 | 0.43171 | 0.63358 | 1.49723 | 26 |
| GO_REGULATION_OF_DNA_TEMPLATED_TRANSCRIPTION_IN_RESPO | 0.00000 | 0.00000 | 0.82666 | 0.69204 | 1.75746 | 71 |
| GO_REGULATION_OF_HEMATOPOIETIC_PROGENITOR_CELL_DIFFERE | 0.00000 | 0.00000 | 0.78819 | 0.69133 | 1.74273 | 62 |
| GO_REGULATION_OF_HEMATOPOIETIC_STEM_CELL_DIFFERENTIATI( | 0.00000 | 0.00000 | 0.78819 | 0.71220 | 1.78132 | 56 |
| GO_REGULATION_OF_HEMOPOIESIS | 0.00012 | 0.00301 | 0.53843 | 0.49851 | 1.31729 | 169 |
| GO_REGULATION_OF_IMMUNE_RESPONSE | 0.00018 | 0.00413 | 0.51885 | 0.47483 | 1.27493 | 261 |
| GO_REGULATION_OF_IMMUNE_SYSTEM_PROCESS | 0.00059 | 0.01144 | 0.47727 | 0.43900 | 1.19281 | 428 |
| GO_REGULATION_OF_INNATE_IMMUNE_RESPONSE | 0.00001 | 0.00021 | 0.61053 | 0.55564 | 1.44619 | 120 |
| GO_REGULATION_OF_INTRINSIC_APOPTOTIC_SIGNALING_PATHWAY | 0.00015 | 0.00371 | 0.51885 | 0.55715 | 1.42489 | 82 |
| GO_REGULATION_OF_MITOTIC_CELL_CYCLE | 0.00183 | 0.02707 | 0.45506 | 0.43512 | 1.17726 | 360 |
| GO_REGULATION_OF_MORPHOGENESIS_OF_AN_EPITHELIUM | 0.00000 | 0.00000 | | 0.68810 | 1.75446 | 78 |
| GO_REGULATION_OF_MRNA_CATABOLIC_PROCESS | 0.00000 | 0.00000 | 0.81404 | 0.60617 | 1.58623 | 137 |
| GO_REGULATION_OF_MRNA_METABOLIC_PROCESS | 0.00000 | 0.00000 | 0.81404 | 0.55402 | 1.47801 | 220 |
| GO_REGULATION_OF_MUSCLE_SYSTEM_PROCESS | 0.00287 | 0.03748 | 0.43171 | 0.54251 | 1.36569 | 60 |
| GO_REGULATION_OF_OXIDATIVE_STRESS_INDUCED_CELL_DEATH | 0.00279 | 0.03682 | 0.43171 | 0.61983 | 1.48385 | 30 |
| GO_REGULATION_OF_PLASMA_LIPOPROTEIN_PARTICLE_LEVELS | 0.00376 | 0.04593 | 0.43171 | 0.61322 | 1.46802 | 30 |
| GO_REGULATION_OF_PROTEASOMAL_PROTEIN_CATABOLIC_PROCES | 0.00202 | 0.02926 | 0.43171 | 0.49720 | 1.29088 | 112 |
| GO_REGULATION_OF_PROTEASOMAL_UBIQUITIN_DEPENDENT_PRO | 0.00413 | 0.04908 | 0.40702 | 0.51780 | 1.31955 | 77 |
| GO_REGULATION_OF_PROTEIN_CATABOLIC_PROCESS | 0.00011 | 0.00270 | 0.53843 | 0.48570 | 1.29504 | 214 |
| GO_REGULATION_OF_PROTEIN_CONTAINING_COMPLEX_ASSEMBLY | 0.00011 | 0.00279 | 0.53843 | 0.48880 | 1.30039 | 203 |
| GO_REGULATION_OF_PROTEIN_IMPORT | 0.00268 | 0.03565 | 0.43171 | 0.62058 | 1.48564 | 30 |
| GO_REGULATION_OF_PROTEIN_LOCALIZATION | 0.00017 | 0.00403 | 0.51885 | 0.45010 | 1.22114 | 393 |
| GO_REGULATION_OF_PROTEIN_LOCALIZATION_TO_NUCLEUS | 0.00021 | 0.00475 | 0.51885 | 0.57553 | 1.45480 | 66 |
| GO_REGULATION_OF_PROTEIN_MODIFICATION_BY_SMALL_PROTEIN | 0.00159 | 0.02453 | 0.45506 | 0.49548 | 1.29296 | 127 |
| GO_REGULATION_OF_PROTEIN_STABILITY | 0.00019 | 0.00437 | 0.51885 | 0.49477 | 1.30441 | 162 |
| GO_REGULATION_OF_PROTEOLYSIS | 0.00021 | 0.00481 | 0.51885 | 0.46618 | 1.25481 | 281 |
| GO_REGULATION_OF_REACTIVE_OXYGEN_SPECIES_METABOLIC_PRC | 0.00179 | 0.02661 | 0.45506 | 0.54234 | 1.36694 | 61 |
| GO_REGULATION_OF_RESPONSE_TO_BIOTIC_STIMULUS | 0.00004 | 0.00109 | 0.55733 | 0.51839 | 1.36064 | 146 |
| GO_REGULATION_OF_RESPONSE_TO_EXTERNAL_STIMULUS | 0.00002 | 0.00046 | 0.57561 | 0.47846 | 1.28780 | 282 |
| GO_REGULATION_OF_SMALL_MOLECULE_METABOLIC_PROCESS | 0.00001 | 0.00037 | 0.59333 | 0.50509 | 1.34409 | 205 |
| GO_REGULATION_OF_STEM_CELL_DIFFERENTIATION | 0.00000 | 0.00003 | 0.65944 | 0.62941 | 1.59840 | 71 |
| GO_REGULATION_OF_SYNAPSE_STRUCTURE_OR_ACTIVITY | 0.00198 | 0.02899 | 0.43171 | 0.55340 | 1.38101 | 54 |
| GO_REGULATION_OF_TRANSCRIPTION_FROM_RNA_POLYMERASE_II | 0.00000 | 0.00000 | 0.83909 | 0.72933 | 1.83064 | 58 |
| GO_REGULATION_OF_TRANSCRIPTION_INITIATION_FROM_RNA_POI | 0.00246 | 0.03355 | 0.43171 | 0.66231 | 1.53371 | 21 |
| GO_REGULATION_OF_TRANSLATIONAL_INITIATION | 0.00008 | 0.00209 | 0.53843 | 0.60767 | 1.51697 | 55 |
| GO_REGULATION_OF_UBIQUITIN_DEPENDENT_PROTEIN_CATABOLIC | 0.00027 | 0.00578 | 0.49849 | 0.53470 | 1.37390 | 90 |

| | | | | | | |
|---|---|---|---|---|---|---|
| GO_REGULATION_OF_VIRAL_LIFE_CYCLE | 0.00135 | 0.02162 | 0.45506 | 0.54272 | 1.37463 | 68 |
| GO_REGULATION_OF_WNT_SIGNALING_PATHWAY | 0.00000 | 0.00015 | 0.61053 | 0.53132 | 1.39883 | 156 |
| GO_RESPONSE_TO_ABIOTIC_STIMULUS | 0.00000 | 0.00000 | 0.73376 | 0.48017 | 1.30780 | 499 |
| GO_RESPONSE_TO_ALCOHOL | 0.00014 | 0.00350 | 0.51885 | 0.56340 | 1.43576 | 77 |
| GO_RESPONSE_TO_BIOTIC_STIMULUS | 0.00000 | 0.00000 | 0.71951 | 0.48507 | 1.31685 | 409 |
| GO_RESPONSE_TO_CYTOKINE | 0.00000 | 0.00000 | | 0.54767 | 1.48687 | 411 |
| GO_RESPONSE_TO_DRUG | 0.00398 | 0.04755 | 0.40702 | 0.47729 | 1.24894 | 136 |
| GO_RESPONSE_TO_ENDOPLASMIC_RETICULUM_STRESS | 0.00250 | 0.03376 | 0.43171 | 0.47719 | 1.25631 | 156 |
| GO_RESPONSE_TO_ESTRADIOL | 0.00342 | 0.04319 | 0.43171 | 0.59345 | 1.43336 | 35 |
| GO_RESPONSE_TO_ETHANOL | 0.00250 | 0.03376 | 0.43171 | 0.59573 | 1.44409 | 37 |
| GO_RESPONSE_TO_HEAT | 0.00420 | 0.04947 | 0.40702 | 0.49512 | 1.27339 | 92 |
| GO_RESPONSE_TO_INORGANIC_SUBSTANCE | 0.00000 | 0.00013 | 0.62726 | 0.50477 | 1.34221 | 197 |
| GO_RESPONSE_TO_INTERLEUKIN_1 | 0.00000 | 0.00000 | 0.76146 | 0.64865 | 1.66086 | 84 |
| GO_RESPONSE_TO_INTERLEUKIN_12 | 0.00000 | 0.00011 | 0.62726 | 0.71641 | 1.72898 | 34 |
| GO_RESPONSE_TO_INTERLEUKIN_4 | 0.00060 | 0.01152 | 0.47727 | 0.72088 | 1.65045 | 18 |
| GO_RESPONSE_TO_METAL_ION | 0.00132 | 0.02117 | 0.45506 | 0.50701 | 1.31861 | 119 |
| GO_RESPONSE_TO_NITROGEN_COMPOUND | 0.00174 | 0.02629 | 0.45506 | 0.43152 | 1.17149 | 412 |
| GO_RESPONSE_TO_ORGANIC_CYCLIC_COMPOUND | 0.00001 | 0.00029 | 0.59333 | 0.47722 | 1.28740 | 318 |
| GO_RESPONSE_TO_OXYGEN_LEVELS | 0.00000 | 0.00000 | | 0.58178 | 1.54150 | 184 |
| GO_RESPONSE_TO_TOPOLOGICALLY_INCORRECT_PROTEIN | 0.00025 | 0.00558 | 0.49849 | 0.52339 | 1.35470 | 107 |
| GO_RESPONSE_TO_TUMOR_NECROSIS_FACTOR | 0.00000 | 0.00000 | 0.80122 | 0.62032 | 1.61113 | 115 |
| GO_RESPONSE_TO_XENOBIOTIC_STIMULUS | 0.00021 | 0.00478 | 0.51885 | 0.66735 | 1.59277 | 29 |
| GO_RIBONUCLEOPROTEIN_COMPLEX_BINDING | 0.00390 | 0.04714 | 0.43171 | 0.50079 | 1.29264 | 100 |
| GO_RIBONUCLEOPROTEIN_COMPLEX_BIOGENESIS | 0.00000 | 0.00001 | 0.69013 | 0.49682 | 1.34028 | 318 |
| GO_RIBONUCLEOPROTEIN_COMPLEX_SUBUNIT_ORGANIZATION | 0.00000 | 0.00008 | 0.62726 | 0.54155 | 1.42419 | 150 |
| GO_RIBONUCLEOPROTEIN_GRANULE | 0.00025 | 0.00551 | 0.49849 | 0.51325 | 1.34174 | 132 |
| GO_RIBONUCLEOSIDE_METABOLIC_PROCESS | 0.00176 | 0.02631 | 0.45506 | 0.63233 | 1.50800 | 28 |
| GO_RIBONUCLEOSIDE_TRIPHOSPHATE_METABOLIC_PROCESS | 0.00154 | 0.02383 | 0.45506 | 0.60571 | 1.46829 | 37 |
| GO_RIBOSE_PHOSPHATE_METABOLIC_PROCESS | 0.00383 | 0.04659 | 0.43171 | 0.46106 | 1.22164 | 184 |
| GO_RIBOSOMAL_LARGE_SUBUNIT_ASSEMBLY | 0.00067 | 0.01258 | 0.47727 | 0.67416 | 1.59312 | 26 |
| GO_RIBOSOMAL_LARGE_SUBUNIT_BIOGENESIS | 0.00002 | 0.00058 | 0.57561 | 0.62435 | 1.56713 | 58 |
| GO_RIBOSOMAL_SUBUNIT | 0.00000 | 0.00000 | | 0.60649 | 1.60004 | 165 |
| GO_RIBOSOME | 0.00000 | 0.00000 | | 0.60057 | 1.59582 | 194 |
| GO_RIBOSOME_ASSEMBLY | 0.00080 | 0.01451 | 0.47727 | 0.57230 | 1.42867 | 55 |
| GO_RIBOSOME_BIOGENESIS | 0.00023 | 0.00512 | 0.51885 | 0.47660 | 1.27224 | 222 |
| GO_RNA_CATABOLIC_PROCESS | 0.00000 | 0.00000 | | 0.62639 | 1.68579 | 285 |
| GO_RNA_DEPENDENT_DNA_BIOSYNTHETIC_PROCESS | 0.00224 | 0.03124 | 0.43171 | 0.57461 | 1.42023 | 47 |
| GO_RNA_LOCALIZATION | 0.00141 | 0.02227 | 0.45506 | 0.48086 | 1.26725 | 161 |
| GO_RNA_POLYMERASE_II_PREINITIATION_COMPLEX_ASSEMBLY | 0.00179 | 0.02661 | 0.45506 | 0.71457 | 1.62069 | 16 |
| GO_RNA_SPLICING | 0.00040 | 0.00826 | 0.49849 | 0.45391 | 1.22307 | 303 |
| GO_RNA_SPLICING_VIA_TRANSESTERIFICATION_REACTIONS | 0.00176 | 0.02631 | 0.45506 | 0.45583 | 1.22189 | 248 |
| GO_ROUGH_ENDOPLASMIC_RETICULUM | 0.00420 | 0.04947 | 0.40702 | 0.57122 | 1.39461 | 40 |
| GO_RRNA_BINDING | 0.00000 | 0.00009 | 0.62726 | 0.66183 | 1.63877 | 48 |
| GO_SCAFFOLD_PROTEIN_BINDING | 0.00023 | 0.00512 | 0.51885 | 0.70744 | 1.64642 | 22 |
| GO_SCF_DEPENDENT_PROTEASOMAL_UBIQUITIN_DEPENDENT_PRO | 0.00000 | 0.00001 | 0.67496 | 0.65303 | 1.64789 | 64 |
| GO_SECRETORY_GRANULE | 0.00000 | 0.00000 | | 0.57804 | 1.55635 | 295 |
| GO_SECRETORY_GRANULE_MEMBRANE | 0.00033 | 0.00713 | 0.49849 | 0.53069 | 1.36981 | 100 |
| GO_SECRETORY_VESICLE | 0.00000 | 0.00000 | | 0.56279 | 1.52006 | 341 |
| GO_SIGNALING_RECEPTOR_BINDING | 0.00246 | 0.03355 | 0.43171 | 0.43376 | 1.17426 | 369 |
| GO_SIGNAL_TRANSDUCTION_BY_PROTEIN_PHOSPHORYLATION | 0.00000 | 0.00008 | 0.62726 | 0.48382 | 1.30369 | 303 |
| GO_SINGLE_STRANDED_RNA_BINDING | 0.00068 | 0.01270 | 0.47727 | 0.58822 | 1.45699 | 49 |
| GO_SKIN_DEVELOPMENT | 0.00202 | 0.02926 | 0.43171 | 0.52489 | 1.34398 | 84 |
| GO_SMALL_MOLECULE_BIOSYNTHETIC_PROCESS | 0.00350 | 0.04389 | 0.43171 | 0.44280 | 1.19179 | 283 |
| GO_SMALL_RIBOSOMAL_SUBUNIT | 0.00000 | 0.00016 | 0.61053 | 0.61907 | 1.56484 | 66 |
| GO_SPERM_EGG_RECOGNITION | 0.00000 | 0.00003 | 0.65944 | 0.84946 | 1.93204 | 17 |
| GO_SPINDLE_MIDZONE | 0.00346 | 0.04354 | 0.43171 | 0.64635 | 1.50426 | 22 |
| GO_SPLICEOSOMAL_COMPLEX | 0.00018 | 0.00413 | 0.51885 | 0.51198 | 1.33939 | 135 |
| GO_STEM_CELL_DIFFERENTIATION | 0.00000 | 0.00000 | 0.73376 | 0.60513 | 1.56799 | 109 |
| GO_STRUCTURAL_CONSTITUENT_OF_CYTOSKELETON | 0.00001 | 0.00033 | 0.59333 | 0.66427 | 1.62180 | 40 |
| GO_STRUCTURAL_CONSTITUENT_OF_RIBOSOME | 0.00000 | 0.00000 | | 0.64142 | 1.68420 | 147 |
| GO_STRUCTURAL_MOLECULE_ACTIVITY | 0.00000 | 0.00000 | | 0.58981 | 1.58694 | 280 |
| GO_SUBSTANTIA_NIGRA_DEVELOPMENT | 0.00053 | 0.01046 | 0.47727 | 0.68351 | 1.60125 | 24 |
| GO_SUPRAMOLECULAR_FIBER_ORGANIZATION | 0.00025 | 0.00551 | 0.49849 | 0.46695 | 1.25403 | 262 |
| GO_SYNAPSE | 0.00000 | 0.00000 | 0.78819 | 0.50131 | 1.36042 | 396 |
| GO_TBP_CLASS_PROTEIN_BINDING | 0.00110 | 0.01830 | 0.45506 | 0.70697 | 1.61861 | 18 |
| GO_TELOMERASE_RNA_LOCALIZATION | 0.00047 | 0.00952 | 0.49849 | 0.74444 | 1.68845 | 16 |
| GO_TERTIARY_GRANULE | 0.00053 | 0.01046 | 0.47727 | 0.59333 | 1.47614 | 52 |
| GO_THREONINE_TYPE_ENDOPEPTIDASE_ACTIVITY | 0.00000 | 0.00002 | 0.67496 | 0.85863 | 1.94743 | 16 |
| GO_THREONINE_TYPE_PEPTIDASE_ACTIVITY | 0.00000 | 0.00003 | 0.65944 | 0.83817 | 1.91900 | 18 |
| GO_TISSUE_MORPHOGENESIS | 0.00000 | 0.00000 | 0.74774 | 0.52996 | 1.41685 | 234 |
| GO_TOXIN_TRANSPORT | 0.00025 | 0.00551 | 0.49849 | 0.71626 | 1.65263 | 20 |
| GO_TRANSLATIONAL_INITIATION | 0.00000 | 0.00000 | | 0.71098 | 1.87182 | 156 |
| GO_TRANSLATION_FACTOR_ACTIVITY_RNA_BINDING | 0.00000 | 0.00000 | 0.80122 | 0.70693 | 1.77957 | 60 |
| GO_TRANSLATION_INITIATION_FACTOR_ACTIVITY | 0.00000 | 0.00002 | 0.67496 | 0.70577 | 1.73490 | 43 |
| GO_TRANSLATION_INITIATION_FACTOR_BINDING | 0.00398 | 0.04755 | 0.40702 | 0.63999 | 1.49459 | 23 |
| GO_TRANSLATION_PREINITIATION_COMPLEX | 0.00001 | 0.00034 | 0.59333 | 0.81602 | 1.85079 | 16 |

| | | | | | | |
|---|---|---|---|---|---|---|
| GO_TRANSLATION_REGULATOR_ACTIVITY | 0.00000 | 0.00000 | 0.80122 | 0.64883 | 1.66872 | 92 |
| GO_TRANSLATION_REGULATOR_ACTIVITY_NUCLEIC_ACID_BINDING | 0.00000 | 0.00000 | 0.80122 | 0.67718 | 1.72357 | 74 |
| GO_TRANSMEMBRANE_TRANSPORT | 0.00000 | 0.00016 | 0.61053 | 0.46171 | 1.25649 | 481 |
| GO_TRICARBOXYLIC_ACID_CYCLE | 0.00035 | 0.00738 | 0.49849 | 0.67297 | 1.59795 | 27 |
| GO_TUMOR_NECROSIS_FACTOR_MEDIATED_SIGNALING_PATHWAY | 0.00000 | 0.00000 | | 0.69105 | 1.76042 | 75 |
| GO_T_CELL_RECEPTOR_SIGNALING_PATHWAY | 0.00000 | 0.00003 | 0.65944 | 0.59306 | 1.52852 | 95 |
| GO_UBIQUITIN_LIKE_PROTEIN_LIGASE_BINDING | 0.00000 | 0.00002 | 0.65944 | 0.54124 | 1.42914 | 167 |
| GO_UNFOLDED_PROTEIN_BINDING | 0.00000 | 0.00007 | 0.64355 | 0.61557 | 1.56324 | 72 |
| GO_VACUOLAR_LUMEN | 0.00000 | 0.00008 | 0.62726 | 0.61247 | 1.56055 | 76 |
| GO_VACUOLAR_MEMBRANE | 0.00000 | 0.00003 | 0.65944 | 0.51296 | 1.36991 | 228 |
| GO_VACUOLE | 0.00000 | 0.00000 | 0.71951 | 0.49224 | 1.33278 | 371 |
| GO_VESICLE_LUMEN | 0.00000 | 0.00000 | 0.82666 | 0.60456 | 1.58433 | 140 |
| GO_VESICLE_MEMBRANE | 0.00087 | 0.01544 | 0.47727 | 0.45207 | 1.21676 | 282 |
| GO_VIRAL_GENE_EXPRESSION | 0.00000 | 0.00000 | | 0.63587 | 1.67574 | 160 |
| GO_VIRAL_LIFE_CYCLE | 0.00100 | 0.01725 | 0.45506 | 0.47877 | 1.26512 | 169 |
| HP_ABNORMALITY_OF_BLOOD_AND_BLOOD_FORMING_TISSUES | 0.00198 | 0.02899 | 0.43171 | 0.42984 | 1.16988 | 483 |
| HP_ABNORMALITY_OF_CARDIOVASCULAR_SYSTEM_ELECTROPHYSIO | 0.00004 | 0.00097 | 0.55733 | 0.51370 | 1.35525 | 165 |
| HP_ABNORMALITY_OF_SKIN_ADNEXA_PHYSIOLOGY | 0.00228 | 0.03157 | 0.43171 | 0.54015 | 1.36557 | 65 |
| HP_ABNORMALITY_OF_THE_NASAL_DORSUM | 0.00390 | 0.04714 | 0.43171 | 0.49046 | 1.26683 | 102 |
| HP_ABNORMAL_CARDIAC_TEST | 0.00088 | 0.01565 | 0.47727 | 0.65559 | 1.55669 | 27 |
| HP_ABNORMAL_ELECTROPHYSIOLOGY_OF_SINOATRIAL_NODE_ORIG | 0.00002 | 0.00060 | 0.57561 | 0.73430 | 1.72987 | 25 |
| HP_ABNORMAL_ENZYME_COENZYME_ACTIVITY | 0.00276 | 0.03643 | 0.43171 | 0.51477 | 1.31996 | 85 |
| HP_ABNORMAL_ERYTHROCYTE_MORPHOLOGY | 0.00039 | 0.00812 | 0.49849 | 0.46928 | 1.25762 | 247 |
| HP_ABNORMAL_QT_INTERVAL | 0.00054 | 0.01051 | 0.47727 | 0.72458 | 1.65892 | 18 |
| HP_ACUTE_LEUKEMIA | 0.00216 | 0.03066 | 0.43171 | 0.55430 | 1.38639 | 56 |
| HP_ANEMIA_OF_INADEQUATE_PRODUCTION | 0.00013 | 0.00328 | 0.51885 | 0.55982 | 1.42923 | 80 |
| HP_ARTERIAL_STENOSIS | 0.00150 | 0.02342 | 0.45506 | 0.58330 | 1.43096 | 42 |
| HP_ARTERIOSCLEROSIS | 0.00257 | 0.03437 | 0.43171 | 0.66003 | 1.52844 | 21 |
| HP_CEREBRAL_ISCHEMIA | 0.00290 | 0.03776 | 0.43171 | 0.62232 | 1.48414 | 28 |
| HP_CONSTITUTIONAL_SYMPTOM | 0.00265 | 0.03526 | 0.43171 | 0.44209 | 1.19171 | 306 |
| HP_DELAYED_PUBERTY | 0.00176 | 0.02631 | 0.45506 | 0.52163 | 1.33466 | 83 |
| HP_DEPRESSED_NASAL_RIDGE | 0.00106 | 0.01787 | 0.45506 | 0.56825 | 1.42358 | 57 |
| HP_DERMATOLOGICAL_MANIFESTATIONS_OF_SYSTEMIC_DISORDERS | 0.00220 | 0.03081 | 0.43171 | 0.51514 | 1.32133 | 87 |
| HP_FATIGUE | 0.00069 | 0.01277 | 0.47727 | 0.49481 | 1.29482 | 137 |
| HP_GLOBAL_BRAIN_ATROPHY | 0.00076 | 0.01396 | 0.47727 | 0.67098 | 1.58561 | 26 |
| HP_HEADACHE | 0.00090 | 0.01591 | 0.47727 | 0.51745 | 1.33877 | 103 |
| HP_HEMATOLOGICAL_NEOPLASM | 0.00372 | 0.04571 | 0.43171 | 0.49268 | 1.27662 | 109 |
| HP_LEUKEMIA | 0.00228 | 0.03157 | 0.43171 | 0.52895 | 1.34631 | 74 |
| HP_MACROCYTIC_ANEMIA | 0.00059 | 0.01144 | 0.47727 | 0.62198 | 1.50773 | 37 |
| HP_MIGRAINE | 0.00039 | 0.00814 | 0.49849 | 0.57342 | 1.42895 | 53 |
| HP_NEOPLASM | 0.00376 | 0.04593 | 0.43171 | 0.43492 | 1.17315 | 313 |
| HP_PALLOR | 0.00005 | 0.00136 | 0.55733 | 0.58180 | 1.47747 | 72 |
| HP_SINUS_TACHYCARDIA | 0.00044 | 0.00896 | 0.49849 | 0.74315 | 1.69025 | 17 |
| HP_THROMBOCYTOSIS | 0.00104 | 0.01775 | 0.45506 | 0.74256 | 1.66801 | 15 |
| HP_WEIGHT_LOSS | 0.00368 | 0.04537 | 0.43171 | 0.48917 | 1.26611 | 107 |
| **Cluster 7** | | | | | | |
| pathway | pval | padj | log2err | ES | NES | size |
| GO_ACTIN_BINDING | 0.00097 | 0.01519 | 0.47727 | 0.50277 | 1.25212 | 164 |
| GO_ACTIN_FILAMENT_BINDING | 0.00001 | 0.00031 | 0.59333 | 0.60863 | 1.46905 | 82 |
| GO_ACTIVATION_OF_IMMUNE_RESPONSE | 0.00000 | 0.00000 | 0.70498 | 0.56999 | 1.42611 | 183 |
| GO_ACTIVATION_OF_INNATE_IMMUNE_RESPONSE | 0.00000 | 0.00000 | 0.77494 | 0.69058 | 1.65994 | 77 |
| GO_AEROBIC_RESPIRATION | 0.00012 | 0.00283 | 0.53843 | 0.61541 | 1.46291 | 59 |
| GO_ALPHA_AMINO_ACID_METABOLIC_PROCESS | 0.00298 | 0.03823 | 0.43171 | 0.51721 | 1.26238 | 104 |
| GO_AMIDE_BINDING | 0.00023 | 0.00508 | 0.51885 | 0.53094 | 1.31048 | 131 |
| GO_AMINE_METABOLIC_PROCESS | 0.00000 | 0.00000 | | 0.72592 | 1.73686 | 70 |
| GO_ANAPHASE_PROMOTING_COMPLEX_DEPENDENT_CATABOLIC_P | 0.00000 | 0.00000 | 0.73376 | 0.67466 | 1.61992 | 76 |
| GO_ANATOMICAL_STRUCTURE_HOMEOSTASIS | 0.00368 | 0.04379 | 0.43171 | 0.48242 | 1.20991 | 203 |
| GO_ANCHORING_JUNCTION | 0.00000 | 0.00000 | | 0.56184 | 1.43173 | 391 |
| GO_ANIMAL_ORGAN_MORPHOGENESIS | 0.00239 | 0.03201 | 0.43171 | 0.46188 | 1.17302 | 338 |
| GO_ANTIGEN_PROCESSING_AND_PRESENTATION | 0.00000 | 0.00002 | 0.67496 | 0.58965 | 1.45473 | 130 |
| GO_ANTIGEN_PROCESSING_AND_PRESENTATION_OF_EXOGENOUS_ | 0.00000 | 0.00000 | 0.82666 | 0.76362 | 1.80152 | 51 |
| GO_ANTIGEN_PROCESSING_AND_PRESENTATION_OF_PEPTIDE_ANTI | 0.00000 | 0.00000 | 0.73376 | 0.62422 | 1.52982 | 113 |
| GO_ANTIGEN_PROCESSING_AND_PRESENTATION_OF_PEPTIDE_ANTI | 0.00000 | 0.00000 | 0.82666 | 0.73786 | 1.75698 | 63 |
| GO_ANTIGEN_RECEPTOR_MEDIATED_SIGNALING_PATHWAY | 0.00000 | 0.00000 | 0.73376 | 0.62508 | 1.53192 | 113 |
| GO_ATPASE_ACTIVITY | 0.00152 | 0.02227 | 0.45506 | 0.47853 | 1.20639 | 246 |
| GO_ATPASE_BINDING | 0.00287 | 0.03708 | 0.43171 | 0.58870 | 1.38100 | 47 |
| GO_ATP_METABOLIC_PROCESS | 0.00065 | 0.01115 | 0.47727 | 0.51157 | 1.27691 | 174 |
| GO_AUTOPHAGOSOME | 0.00413 | 0.04753 | 0.40702 | 0.57176 | 1.35670 | 56 |
| GO_AUTOPHAGOSOME_MEMBRANE | 0.00113 | 0.01725 | 0.45506 | 0.71155 | 1.56654 | 20 |
| GO_AXONAL_GROWTH_CONE | 0.00339 | 0.04125 | 0.43171 | 0.73366 | 1.56948 | 15 |
| GO_AZUROPHIL_GRANULE | 0.00121 | 0.01821 | 0.45506 | 0.55085 | 1.33072 | 83 |
| GO_AZUROPHIL_GRANULE_LUMEN | 0.00191 | 0.02655 | 0.45506 | 0.60991 | 1.42593 | 45 |
| GO_BASE_EXCISION_REPAIR | 0.00198 | 0.02728 | 0.43171 | 0.65479 | 1.48101 | 28 |
| GO_BLOOD_MICROPARTICLE | 0.00413 | 0.04753 | 0.40702 | 0.68588 | 1.50818 | 19 |
| GO_CADHERIN_BINDING | 0.00000 | 0.00000 | | 0.60438 | 1.51916 | 222 |
| GO_CANONICAL_WNT_SIGNALING_PATHWAY | 0.00000 | 0.00005 | 0.64355 | 0.56093 | 1.39258 | 151 |

| | | | | | | |
|---|---|---|---|---|---|---|
| GO_CARBOHYDRATE_BIOSYNTHETIC_PROCESS | 0.00165 | 0.02374 | 0.45506 | 0.54358 | 1.31892 | 89 |
| GO_CARBOHYDRATE_CATABOLIC_PROCESS | 0.00092 | 0.01460 | 0.47727 | 0.53259 | 1.29929 | 103 |
| GO_CARBOHYDRATE_DERIVATIVE_METABOLIC_PROCESS | 0.00179 | 0.02543 | 0.45506 | 0.45254 | 1.15619 | 484 |
| GO_CARBOHYDRATE_METABOLIC_PROCESS | 0.00024 | 0.00521 | 0.51885 | 0.48644 | 1.22868 | 272 |
| GO_CARBON_CARBON_LYASE_ACTIVITY | 0.00316 | 0.03949 | 0.43171 | 0.66142 | 1.48474 | 25 |
| GO_CARBON_OXYGEN_LYASE_ACTIVITY | 0.00305 | 0.03859 | 0.43171 | 0.64492 | 1.45868 | 28 |
| GO_CATALYTIC_STEP_2_SPLICEOSOME | 0.00242 | 0.03234 | 0.43171 | 0.55292 | 1.32907 | 77 |
| GO_CELLULAR_AMINO_ACID_METABOLIC_PROCESS | 0.00000 | 0.00001 | 0.67496 | 0.56476 | 1.41261 | 181 |
| GO_CELLULAR_CARBOHYDRATE_CATABOLIC_PROCESS | 0.00361 | 0.04332 | 0.43171 | 0.67681 | 1.49005 | 20 |
| GO_CELLULAR_HOMEOSTASIS | 0.00185 | 0.02599 | 0.45506 | 0.46659 | 1.18521 | 340 |
| GO_CELLULAR_KETONE_METABOLIC_PROCESS | 0.00000 | 0.00002 | 0.67496 | 0.59384 | 1.46395 | 128 |
| GO_CELLULAR_MODIFIED_AMINO_ACID_METABOLIC_PROCESS | 0.00302 | 0.03831 | 0.43171 | 0.53278 | 1.29541 | 93 |
| GO_CELLULAR_NITROGEN_COMPOUND_CATABOLIC_PROCESS | 0.00000 | 0.00000 | | 0.61816 | 1.57346 | 375 |
| GO_CELLULAR_PROTEIN_CATABOLIC_PROCESS | 0.00000 | 0.00014 | 0.61053 | 0.47996 | 1.22756 | 500 |
| GO_CELLULAR_RESPIRATION | 0.00049 | 0.00892 | 0.47727 | 0.52811 | 1.30215 | 129 |
| GO_CELLULAR_RESPONSE_TO_DRUG | 0.00043 | 0.00810 | 0.49849 | 0.67704 | 1.53901 | 30 |
| GO_CELLULAR_RESPONSE_TO_HEAT | 0.00088 | 0.01424 | 0.47727 | 0.56069 | 1.35333 | 82 |
| GO_CELLULAR_RESPONSE_TO_ORGANIC_CYCLIC_COMPOUND | 0.00117 | 0.01771 | 0.45506 | 0.48921 | 1.23018 | 226 |
| GO_CELLULAR_RESPONSE_TO_OXYGEN_LEVELS | 0.00000 | 0.00000 | 0.76146 | 0.60824 | 1.50419 | 138 |
| GO_CELL_ACTIVATION | 0.00000 | 0.00013 | 0.62726 | 0.48674 | 1.24382 | 487 |
| GO_CELL_ACTIVATION_INVOLVED_IN_IMMUNE_RESPONSE | 0.00000 | 0.00000 | 0.76146 | 0.54278 | 1.37292 | 287 |
| GO_CELL_ADHESION_MOLECULE_BINDING | 0.00000 | 0.00000 | | 0.57394 | 1.45068 | 280 |
| GO_CELL_BODY | 0.00014 | 0.00319 | 0.51885 | 0.51843 | 1.29615 | 180 |
| GO_CELL_CELL_RECOGNITION | 0.00000 | 0.00016 | 0.61053 | 0.85349 | 1.82582 | 15 |
| GO_CELL_CELL_SIGNALING_BY_WNT | 0.00000 | 0.00001 | 0.69013 | 0.54130 | 1.36251 | 234 |
| GO_CELL_CYCLE_G2_M_PHASE_TRANSITION | 0.00008 | 0.00201 | 0.53843 | 0.51746 | 1.29769 | 195 |
| GO_CELL_CYCLE_PHASE_TRANSITION | 0.00048 | 0.00882 | 0.49849 | 0.46830 | 1.19377 | 409 |
| GO_CELL_KILLING | 0.00194 | 0.02692 | 0.45506 | 0.62622 | 1.44040 | 35 |
| GO_CELL_RECOGNITION | 0.00008 | 0.00199 | 0.53843 | 0.70693 | 1.60694 | 30 |
| GO_CELL_SUBSTRATE_JUNCTION | 0.00000 | 0.00000 | | 0.62734 | 1.57985 | 240 |
| GO_CELL_SURFACE_RECEPTOR_SIGNALING_PATHWAY_INVOLVED_IN | 0.00000 | 0.00001 | 0.69013 | 0.53601 | 1.35200 | 248 |
| GO_CHAPERONE_BINDING | 0.00420 | 0.04805 | 0.40702 | 0.54895 | 1.30872 | 64 |
| GO_CHAPERONE_COMPLEX | 0.00000 | 0.00004 | 0.65944 | 0.82598 | 1.81846 | 20 |
| GO_CORTICAL_CYTOSKELETON | 0.00104 | 0.01604 | 0.45506 | 0.61837 | 1.43813 | 42 |
| GO_COTRANSLATIONAL_PROTEIN_TARGETING_TO_MEMBRANE | 0.00000 | 0.00000 | | 0.78229 | 1.90090 | 92 |
| GO_CYTOKINE_MEDIATED_SIGNALING_PATHWAY | 0.00000 | 0.00000 | 0.81404 | 0.56576 | 1.42719 | 255 |
| GO_CYTOPLASMIC_TRANSLATION | 0.00000 | 0.00000 | | 0.74956 | 1.80920 | 82 |
| GO_CYTOPLASMIC_TRANSLATIONAL_INITIATION | 0.00000 | 0.00000 | 0.73376 | 0.80409 | 1.82472 | 29 |
| GO_CYTOSKELETAL_PROTEIN_BINDING | 0.00183 | 0.02581 | 0.45506 | 0.45802 | 1.16800 | 417 |
| GO_CYTOSOLIC_LARGE_RIBOSOMAL_SUBUNIT | 0.00000 | 0.00000 | | 0.76181 | 1.80179 | 53 |
| GO_CYTOSOLIC_RIBOSOME | 0.00000 | 0.00000 | | 0.75168 | 1.82893 | 94 |
| GO_CYTOSOLIC_SMALL_RIBOSOMAL_SUBUNIT | 0.00000 | 0.00000 | 0.71951 | 0.75353 | 1.74620 | 40 |
| GO_DEFENSE_RESPONSE | 0.00025 | 0.00529 | 0.49849 | 0.46481 | 1.18660 | 461 |
| GO_DEFENSE_RESPONSE_TO_OTHER_ORGANISM | 0.00000 | 0.00015 | 0.61053 | 0.50465 | 1.27914 | 319 |
| GO_DETECTION_OF_STIMULUS | 0.00066 | 0.01139 | 0.47727 | 0.56553 | 1.35651 | 74 |
| GO_DISORDERED_DOMAIN_SPECIFIC_BINDING | 0.00073 | 0.01224 | 0.47727 | 0.70672 | 1.56749 | 22 |
| GO_DNA_BIOSYNTHETIC_PROCESS | 0.00137 | 0.02029 | 0.45506 | 0.51489 | 1.27031 | 130 |
| GO_DNA_CONFORMATION_CHANGE | 0.00331 | 0.04054 | 0.43171 | 0.48632 | 1.21830 | 192 |
| GO_DNA_DAMAGE_RESPONSE_DETECTION_OF_DNA_DAMAGE | 0.00007 | 0.00190 | 0.53843 | 0.70040 | 1.59776 | 31 |
| GO_DNA_POLYMERASE_BINDING | 0.00209 | 0.02834 | 0.43171 | 0.72453 | 1.57496 | 17 |
| GO_DNA_REPAIR | 0.00179 | 0.02543 | 0.45506 | 0.46218 | 1.17640 | 376 |
| GO_DOUBLE_STRANDED_RNA_BINDING | 0.00320 | 0.03966 | 0.43171 | 0.60174 | 1.40683 | 45 |
| GO_DRUG_BINDING | 0.00038 | 0.00738 | 0.49849 | 0.63439 | 1.47366 | 41 |
| GO_EMBRYO_DEVELOPMENT | 0.00033 | 0.00676 | 0.49849 | 0.47059 | 1.19960 | 409 |
| GO_EMBRYO_DEVELOPMENT_ENDING_IN_BIRTH_OR_EGG_HATCHIN | 0.00072 | 0.01213 | 0.47727 | 0.48256 | 1.22012 | 284 |
| GO_ENDOPEPTIDASE_ACTIVITY | 0.00004 | 0.00117 | 0.55733 | 0.54583 | 1.34724 | 131 |
| GO_ENDOPEPTIDASE_COMPLEX | 0.00000 | 0.00000 | 0.71951 | 0.69270 | 1.64767 | 61 |
| GO_ENERGY_DERIVATION_BY_OXIDATION_OF_ORGANIC_COMPOUN | 0.00102 | 0.01586 | 0.45506 | 0.50809 | 1.26488 | 163 |
| GO_EPITHELIAL_TO_MESENCHYMAL_TRANSITION | 0.00019 | 0.00427 | 0.51885 | 0.62436 | 1.47383 | 52 |
| GO_EPITHELIUM_DEVELOPMENT | 0.00002 | 0.00049 | 0.57561 | 0.48087 | 1.22656 | 433 |
| GO_ESTABLISHMENT_OF_PROTEIN_LOCALIZATION_TO_CHROMOSON | 0.00014 | 0.00319 | 0.51885 | 0.74002 | 1.64895 | 23 |
| GO_ESTABLISHMENT_OF_PROTEIN_LOCALIZATION_TO_ENDOPLASM | 0.00000 | 0.00000 | | 0.75963 | 1.85221 | 102 |
| GO_ESTABLISHMENT_OF_PROTEIN_LOCALIZATION_TO_MEMBRANE | 0.00000 | 0.00000 | | 0.63190 | 1.58847 | 221 |
| GO_ESTABLISHMENT_OF_PROTEIN_LOCALIZATION_TO_ORGANELLE | 0.00000 | 0.00000 | | 0.58407 | 1.48738 | 380 |
| GO_ESTABLISHMENT_OF_PROTEIN_LOCALIZATION_TO_TELOMERE | 0.00044 | 0.00822 | 0.49849 | 0.75757 | 1.64679 | 17 |
| GO_ESTABLISHMENT_OF_RNA_LOCALIZATION | 0.00018 | 0.00414 | 0.51885 | 0.52450 | 1.30214 | 154 |
| GO_ESTABLISHMENT_OF_TISSUE_POLARITY | 0.00000 | 0.00000 | 0.83909 | 0.69584 | 1.67660 | 81 |
| GO_EUKARYOTIC_TRANSLATION_INITIATION_FACTOR_3_COMPLEX | 0.00000 | 0.00002 | 0.67496 | 0.88152 | 1.88578 | 15 |
| GO_EXOCYTOSIS | 0.00000 | 0.00000 | 0.73376 | 0.51630 | 1.31533 | 385 |
| GO_FATTY_ACID_BINDING | 0.00122 | 0.01832 | 0.45506 | 0.75683 | 1.61904 | 15 |
| GO_FC_EPSILON_RECEPTOR_SIGNALING_PATHWAY | 0.00000 | 0.00000 | 0.80122 | 0.69464 | 1.66620 | 74 |
| GO_FC_RECEPTOR_SIGNALING_PATHWAY | 0.00000 | 0.00000 | 0.81404 | 0.65435 | 1.60239 | 111 |
| GO_FERTILIZATION | 0.00039 | 0.00753 | 0.49849 | 0.61347 | 1.44454 | 50 |
| GO_FICOLIN_1_RICH_GRANULE | 0.00000 | 0.00000 | | 0.69326 | 1.68012 | 87 |
| GO_GENERATION_OF_PRECURSOR_METABOLITES_AND_ENERGY | 0.00000 | 0.00006 | 0.64355 | 0.51945 | 1.31339 | 284 |
| GO_GENE_SILENCING | 0.00205 | 0.02792 | 0.43171 | 0.50651 | 1.25423 | 143 |

| | | | | | | |
|---|---|---|---|---|---|---|
| GO_GLIAL_CELL_MIGRATION | 0.00368 | 0.04379 | 0.43171 | 0.71126 | 1.55226 | 18 |
| GO_GLIOGENESIS | 0.00246 | 0.03274 | 0.43171 | 0.53230 | 1.29611 | 97 |
| GO_GLUCOSE_CATABOLIC_PROCESS | 0.00342 | 0.04160 | 0.43171 | 0.71313 | 1.55634 | 18 |
| GO_GLUCOSE_METABOLIC_PROCESS | 0.00088 | 0.01424 | 0.47727 | 0.53341 | 1.30191 | 104 |
| GO_GLUTAMATERGIC_SYNAPSE | 0.00165 | 0.02374 | 0.45506 | 0.52356 | 1.28265 | 112 |
| GO_GLYCOSAMINOGLYCAN_BINDING | 0.00167 | 0.02387 | 0.45506 | 0.64429 | 1.46208 | 29 |
| GO_GLYCOSYL_COMPOUND_METABOLIC_PROCESS | 0.00000 | 0.00008 | 0.62726 | 0.65507 | 1.55921 | 62 |
| GO_GTP_METABOLIC_PROCESS | 0.00019 | 0.00424 | 0.51885 | 0.77290 | 1.68010 | 17 |
| GO_GUANOSINE_CONTAINING_COMPOUND_METABOLIC_PROCESS | 0.00057 | 0.01006 | 0.47727 | 0.69556 | 1.56139 | 25 |
| GO_GUANYL_NUCLEOTIDE_BINDING | 0.00069 | 0.01170 | 0.47727 | 0.50155 | 1.25284 | 177 |
| GO_G_PROTEIN_COUPLED_RECEPTOR_BINDING | 0.00065 | 0.01115 | 0.47727 | 0.59354 | 1.41094 | 59 |
| GO_HEAT_SHOCK_PROTEIN_BINDING | 0.00069 | 0.01170 | 0.47727 | 0.57536 | 1.38052 | 75 |
| GO_HEMATOPOIETIC_PROGENITOR_CELL_DIFFERENTIATION | 0.00000 | 0.00000 | 0.76146 | 0.65655 | 1.59807 | 96 |
| GO_HEMATOPOIETIC_STEM_CELL_DIFFERENTIATION | 0.00000 | 0.00000 | | 0.74410 | 1.77114 | 62 |
| GO_HSP70_PROTEIN_BINDING | 0.00364 | 0.04366 | 0.43171 | 0.67420 | 1.48976 | 21 |
| GO_HYDROLASE_ACTIVITY_ACTING_ON_ACID_ANHYDRIDES | 0.00049 | 0.00892 | 0.47727 | 0.46020 | 1.17385 | 437 |
| GO_IMMUNE_EFFECTOR_PROCESS | 0.00000 | 0.00001 | 0.69013 | 0.49713 | 1.26810 | 441 |
| GO_IMMUNE_RESPONSE_REGULATING_SIGNALING_PATHWAY | 0.00000 | 0.00001 | 0.69013 | 0.56991 | 1.42254 | 174 |
| GO_IMMUNE_SYSTEM_DEVELOPMENT | 0.00036 | 0.00713 | 0.49849 | 0.46834 | 1.19266 | 380 |
| GO_INCLUSION_BODY | 0.00331 | 0.04054 | 0.43171 | 0.59102 | 1.37619 | 43 |
| GO_INNATE_IMMUNE_RESPONSE | 0.00000 | 0.00001 | 0.70498 | 0.53702 | 1.35560 | 261 |
| GO_INNATE_IMMUNE_RESPONSE_ACTIVATING_SIGNAL_TRANSDUCT | 0.00000 | 0.00000 | | 0.74486 | 1.77364 | 63 |
| GO_INTERACTION_WITH_HOST | 0.00372 | 0.04392 | 0.43171 | 0.51449 | 1.25573 | 104 |
| GO_INTERACTION_WITH_SYMBIONT | 0.00029 | 0.00603 | 0.49849 | 0.66048 | 1.52189 | 36 |
| GO_INTERLEUKIN_1_MEDIATED_SIGNALING_PATHWAY | 0.00000 | 0.00000 | | 0.74294 | 1.77390 | 67 |
| GO_ION_CHANNEL_BINDING | 0.00041 | 0.00779 | 0.49849 | 0.63347 | 1.47504 | 43 |
| GO_LARGE_RIBOSOMAL_SUBUNIT | 0.00000 | 0.00000 | 0.73376 | 0.63261 | 1.54529 | 106 |
| GO_LEUKOCYTE_MEDIATED_IMMUNITY | 0.00000 | 0.00000 | 0.77494 | 0.54053 | 1.36925 | 303 |
| GO_LYASE_ACTIVITY | 0.00010 | 0.00254 | 0.53843 | 0.59529 | 1.42575 | 71 |
| GO_LYSOSOMAL_LUMEN | 0.00039 | 0.00759 | 0.49849 | 0.63469 | 1.47788 | 43 |
| GO_MACROAUTOPHAGY | 0.00216 | 0.02919 | 0.43171 | 0.48383 | 1.21473 | 211 |
| GO_MEMBRANE_ORGANIZATION | 0.00163 | 0.02361 | 0.45506 | 0.45633 | 1.16358 | 418 |
| GO_MESENCHYMAL_CELL_DIFFERENTIATION | 0.00038 | 0.00732 | 0.49849 | 0.58433 | 1.39810 | 70 |
| GO_MESENCHYME_DEVELOPMENT | 0.00079 | 0.01314 | 0.47727 | 0.55708 | 1.34698 | 84 |
| GO_MIDBRAIN_DEVELOPMENT | 0.00283 | 0.03670 | 0.43171 | 0.60249 | 1.39627 | 38 |
| GO_MODIFICATION_DEPENDENT_MACROMOLECULE_CATABOLIC_PR | 0.00011 | 0.00256 | 0.53843 | 0.47368 | 1.20790 | 424 |
| GO_MODULATION_BY_HOST_OF_SYMBIONT_PROCESS | 0.00309 | 0.03876 | 0.43171 | 0.63352 | 1.43764 | 29 |
| GO_MONOCARBOXYLIC_ACID_BINDING | 0.00065 | 0.01127 | 0.47727 | 0.71716 | 1.58470 | 21 |
| GO_MONOCARBOXYLIC_ACID_METABOLIC_PROCESS | 0.00353 | 0.04274 | 0.43171 | 0.46814 | 1.18329 | 281 |
| GO_MONOSACCHARIDE_BIOSYNTHETIC_PROCESS | 0.00053 | 0.00945 | 0.47727 | 0.61041 | 1.44091 | 52 |
| GO_MONOSACCHARIDE_CATABOLIC_PROCESS | 0.00390 | 0.04569 | 0.43171 | 0.62108 | 1.42026 | 32 |
| GO_MONOSACCHARIDE_METABOLIC_PROCESS | 0.00055 | 0.00984 | 0.47727 | 0.52672 | 1.30294 | 139 |
| GO_MORPHOGENESIS_OF_AN_EPITHELIUM | 0.00001 | 0.00027 | 0.59333 | 0.52452 | 1.31966 | 230 |
| GO_MORPHOGENESIS_OF_A_POLARIZED_EPITHELIUM | 0.00000 | 0.00000 | | 0.69355 | 1.68441 | 91 |
| GO_MRNA_3_UTR_BINDING | 0.00010 | 0.00247 | 0.53843 | 0.62809 | 1.49120 | 57 |
| GO_MRNA_5_UTR_BINDING | 0.00011 | 0.00257 | 0.53843 | 0.74403 | 1.65025 | 22 |
| GO_MRNA_BINDING | 0.00000 | 0.00000 | | 0.59530 | 1.49349 | 204 |
| GO_MRNA_PROCESSING | 0.00002 | 0.00072 | 0.57561 | 0.48422 | 1.23255 | 375 |
| GO_MULTI_ORGANISM_LOCALIZATION | 0.00405 | 0.04710 | 0.40702 | 0.58155 | 1.36608 | 49 |
| GO_MULTI_ORGANISM_PROCESS | 0.00025 | 0.00532 | 0.49849 | 0.46896 | 1.19369 | 375 |
| GO_MYELOID_LEUKOCYTE_ACTIVATION | 0.00000 | 0.00000 | 0.76146 | 0.55220 | 1.39491 | 270 |
| GO_MYELOID_LEUKOCYTE_MEDIATED_IMMUNITY | 0.00000 | 0.00000 | 0.82666 | 0.56641 | 1.42828 | 253 |
| GO_NADH_METABOLIC_PROCESS | 0.00002 | 0.00074 | 0.57561 | 0.76787 | 1.71100 | 23 |
| GO_NADP_METABOLIC_PROCESS | 0.00276 | 0.03611 | 0.43171 | 0.68796 | 1.51459 | 20 |
| GO_NAD_BINDING | 0.00045 | 0.00838 | 0.49849 | 0.66374 | 1.51783 | 32 |
| GO_NAD_METABOLIC_PROCESS | 0.00047 | 0.00868 | 0.49849 | 0.70521 | 1.57428 | 24 |
| GO_NCRNA_METABOLIC_PROCESS | 0.00097 | 0.01519 | 0.47727 | 0.47357 | 1.20175 | 331 |
| GO_NCRNA_PROCESSING | 0.00031 | 0.00643 | 0.49849 | 0.48504 | 1.22706 | 289 |
| GO_NEGATIVE_REGULATION_OF_CANONICAL_WNT_SIGNALING_PAT | 0.00000 | 0.00001 | 0.69013 | 0.63922 | 1.55246 | 91 |
| GO_NEGATIVE_REGULATION_OF_CATABOLIC_PROCESS | 0.00242 | 0.03234 | 0.43171 | 0.49336 | 1.22867 | 164 |
| GO_NEGATIVE_REGULATION_OF_CELLULAR_AMIDE_METABOLIC_PR( | 0.00008 | 0.00209 | 0.53843 | 0.58181 | 1.41304 | 91 |
| GO_NEGATIVE_REGULATION_OF_CELLULAR_CATABOLIC_PROCESS | 0.00079 | 0.01314 | 0.47727 | 0.51670 | 1.27901 | 142 |
| GO_NEGATIVE_REGULATION_OF_CELLULAR_PROTEIN_CATABOLIC_PI | 0.00126 | 0.01876 | 0.45506 | 0.59569 | 1.41073 | 55 |
| GO_NEGATIVE_REGULATION_OF_CELL_CYCLE | 0.00017 | 0.00392 | 0.51885 | 0.48168 | 1.22564 | 358 |
| GO_NEGATIVE_REGULATION_OF_CELL_CYCLE_G2_M_PHASE_TRANS | 0.00000 | 0.00000 | 0.71951 | 0.65896 | 1.58519 | 78 |
| GO_NEGATIVE_REGULATION_OF_CELL_CYCLE_PHASE_TRANSITION | 0.00115 | 0.01748 | 0.45506 | 0.50581 | 1.25932 | 161 |
| GO_NEGATIVE_REGULATION_OF_CELL_CYCLE_PROCESS | 0.00021 | 0.00470 | 0.51885 | 0.49983 | 1.25646 | 224 |
| GO_NEGATIVE_REGULATION_OF_CELL_DEATH | 0.00001 | 0.00042 | 0.59333 | 0.48878 | 1.24596 | 409 |
| GO_NEGATIVE_REGULATION_OF_MITOTIC_CELL_CYCLE | 0.00042 | 0.00796 | 0.49849 | 0.50327 | 1.26253 | 197 |
| GO_NEGATIVE_REGULATION_OF_MRNA_METABOLIC_PROCESS | 0.00082 | 0.01347 | 0.47727 | 0.60602 | 1.43359 | 54 |
| GO_NEGATIVE_REGULATION_OF_PROTEASOMAL_PROTEIN_CATABOL | 0.00302 | 0.03831 | 0.43171 | 0.62053 | 1.42984 | 36 |
| GO_NEGATIVE_REGULATION_OF_PROTEASOMAL_UBIQUITIN_DEPEN | 0.00302 | 0.03831 | 0.43171 | 0.66002 | 1.47340 | 24 |
| GO_NEGATIVE_REGULATION_OF_PROTEIN_CONTAINING_COMPLEX_ | 0.00309 | 0.03876 | 0.43171 | 0.55758 | 1.33309 | 68 |
| GO_NEGATIVE_REGULATION_OF_PROTEIN_METABOLIC_PROCESS | 0.00057 | 0.01006 | 0.47727 | 0.46089 | 1.17693 | 470 |
| GO_NEGATIVE_REGULATION_OF_PROTEIN_MODIFICATION_BY_SMA | 0.00435 | 0.04964 | 0.40702 | 0.57677 | 1.35811 | 50 |
| GO_NEGATIVE_REGULATION_OF_RNA_CATABOLIC_PROCESS | 0.00026 | 0.00548 | 0.49849 | 0.65301 | 1.50871 | 37 |

| | | | | | | |
|---|---|---|---|---|---|---|
| GO_NEGATIVE_REGULATION_OF_UBIQUITIN_DEPENDENT_PROTEIN_ | 0.00098 | 0.01538 | 0.45506 | 0.64268 | 1.47828 | 35 |
| GO_NEGATIVE_REGULATION_OF_WNT_SIGNALING_PATHWAY | 0.00000 | 0.00000 | 0.73376 | 0.63503 | 1.54761 | 100 |
| GO_NEURAL_NUCLEUS_DEVELOPMENT | 0.00202 | 0.02771 | 0.43171 | 0.66640 | 1.48764 | 24 |
| GO_NEURON_DEATH | 0.00420 | 0.04805 | 0.40702 | 0.49876 | 1.23576 | 144 |
| GO_NEURON_DEATH_IN_RESPONSE_TO_OXIDATIVE_STRESS | 0.00045 | 0.00838 | 0.49849 | 0.75716 | 1.64589 | 17 |
| GO_NEURON_TO_NEURON_SYNAPSE | 0.00000 | 0.00013 | 0.62726 | 0.57838 | 1.42014 | 117 |
| GO_NIK_NF_KAPPAB_SIGNALING | 0.00000 | 0.00000 | | 0.70613 | 1.71584 | 92 |
| GO_NON_CANONICAL_WNT_SIGNALING_PATHWAY | 0.00000 | 0.00000 | | 0.71124 | 1.72340 | 86 |
| GO_NUCLEAR_ENVELOPE | 0.00083 | 0.01357 | 0.47727 | 0.47704 | 1.20420 | 266 |
| GO_NUCLEAR_EXPORT | 0.00006 | 0.00170 | 0.53843 | 0.53579 | 1.33135 | 156 |
| GO_NUCLEAR_PERIPHERY | 0.00194 | 0.02692 | 0.45506 | 0.53471 | 1.29761 | 90 |
| GO_NUCLEAR_TRANSCRIBED_MRNA_CATABOLIC_PROCESS | 0.00000 | 0.00000 | | 0.66286 | 1.65470 | 170 |
| GO_NUCLEAR_TRANSCRIBED_MRNA_CATABOLIC_PROCESS_NONSEN | 0.00000 | 0.00000 | | 0.73498 | 1.79892 | 110 |
| GO_NUCLEAR_TRANSPORT | 0.00004 | 0.00120 | 0.55733 | 0.50888 | 1.28126 | 237 |
| GO_NUCLEOBASE_CONTAINING_COMPOUND_TRANSPORT | 0.00032 | 0.00644 | 0.49849 | 0.51090 | 1.27713 | 179 |
| GO_NUCLEOBASE_CONTAINING_SMALL_MOLECULE_METABOLIC_PR | 0.00001 | 0.00024 | 0.61053 | 0.50503 | 1.27851 | 293 |
| GO_NUCLEOSIDE_DIPHOSPHATE_METABOLIC_PROCESS | 0.00091 | 0.01455 | 0.47727 | 0.55139 | 1.33607 | 86 |
| GO_NUCLEOSIDE_METABOLIC_PROCESS | 0.00000 | 0.00007 | 0.62726 | 0.67673 | 1.59654 | 51 |
| GO_NUCLEOSIDE_MONOPHOSPHATE_BIOSYNTHETIC_PROCESS | 0.00346 | 0.04194 | 0.43171 | 0.65357 | 1.47349 | 27 |
| GO_NUCLEOSIDE_PHOSPHATE_BIOSYNTHETIC_PROCESS | 0.00023 | 0.00490 | 0.51885 | 0.53878 | 1.32450 | 122 |
| GO_NUCLEOSIDE_TRIPHOSPHATE_BIOSYNTHETIC_PROCESS | 0.00108 | 0.01651 | 0.45506 | 0.64432 | 1.47931 | 34 |
| GO_NUCLEOSIDE_TRIPHOSPHATE_METABOLIC_PROCESS | 0.00036 | 0.00717 | 0.49849 | 0.61809 | 1.45192 | 49 |
| GO_NUCLEOTIDE_EXCISION_REPAIR | 0.00054 | 0.00958 | 0.47727 | 0.55345 | 1.34129 | 87 |
| GO_NUCLEOTIDE_EXCISION_REPAIR_DNA_DAMAGE_RECOGNITION | 0.00253 | 0.03355 | 0.43171 | 0.68632 | 1.51654 | 21 |
| GO_NUCLEOTIDE_EXCISION_REPAIR_DNA_GAP_FILLING | 0.00092 | 0.01460 | 0.47727 | 0.70871 | 1.56603 | 21 |
| GO_NUCLEOTIDE_EXCISION_REPAIR_DNA_INCISION | 0.00327 | 0.04028 | 0.43171 | 0.62578 | 1.43101 | 32 |
| GO_NUCLEOTIDE_EXCISION_REPAIR_PREINCISION_COMPLEX_ASSEM | 0.00279 | 0.03631 | 0.43171 | 0.65641 | 1.47705 | 26 |
| GO_NUCLEOTIDE_PHOSPHORYLATION | 0.00320 | 0.03966 | 0.43171 | 0.54807 | 1.31741 | 77 |
| GO_ORGANIC_ACID_BINDING | 0.00038 | 0.00732 | 0.49849 | 0.58538 | 1.39770 | 67 |
| GO_ORGANIC_CYCLIC_COMPOUND_CATABOLIC_PROCESS | 0.00000 | 0.00000 | | 0.60818 | 1.54933 | 384 |
| GO_ORGANOPHOSPHATE_METABOLIC_PROCESS | 0.00279 | 0.03631 | 0.43171 | 0.44682 | 1.14258 | 496 |
| GO_OXIDATION_REDUCTION_PROCESS | 0.00205 | 0.02792 | 0.43171 | 0.45109 | 1.15202 | 465 |
| GO_OXIDOREDUCTASE_ACTIVITY_ACTING_ON_CH_OH_GROUP_OF_ | 0.00113 | 0.01725 | 0.45506 | 0.59696 | 1.40917 | 52 |
| GO_PEPTIDASE_ACTIVITY | 0.00061 | 0.01059 | 0.47727 | 0.49566 | 1.24467 | 212 |
| GO_PEPTIDASE_COMPLEX | 0.00000 | 0.00003 | 0.65944 | 0.64838 | 1.55682 | 76 |
| GO_PEPTIDE_BINDING | 0.00074 | 0.01236 | 0.47727 | 0.55352 | 1.34122 | 86 |
| GO_PEPTIDE_BIOSYNTHETIC_PROCESS | 0.00000 | 0.00000 | | 0.60224 | 1.53770 | 468 |
| GO_PIGMENT_GRANULE | 0.00000 | 0.00001 | 0.69013 | 0.66873 | 1.60004 | 70 |
| GO_POLYSOMAL_RIBOSOME | 0.00000 | 0.00014 | 0.61053 | 0.73791 | 1.68333 | 31 |
| GO_POLYSOME | 0.00000 | 0.00004 | 0.64355 | 0.68252 | 1.61637 | 55 |
| GO_POLY_PURINE_TRACT_BINDING | 0.00357 | 0.04298 | 0.43171 | 0.67687 | 1.49017 | 20 |
| GO_POLY_PYRIMIDINE_TRACT_BINDING | 0.00191 | 0.02655 | 0.45506 | 0.72918 | 1.57316 | 16 |
| GO_POSITIVE_REGULATION_OF_CANONICAL_WNT_SIGNALING_PATH | 0.00000 | 0.00000 | 0.77494 | 0.66773 | 1.61309 | 83 |
| GO_POSITIVE_REGULATION_OF_CELLULAR_AMIDE_METABOLIC_PRO | 0.00009 | 0.00230 | 0.53843 | 0.56743 | 1.38288 | 99 |
| GO_POSITIVE_REGULATION_OF_CELLULAR_PROTEIN_LOCALIZATION | 0.00015 | 0.00346 | 0.51885 | 0.50934 | 1.27730 | 196 |
| GO_POSITIVE_REGULATION_OF_DEFENSE_RESPONSE | 0.00000 | 0.00006 | 0.64355 | 0.57870 | 1.42690 | 129 |
| GO_POSITIVE_REGULATION_OF_DNA_BIOSYNTHETIC_PROCESS | 0.00091 | 0.01455 | 0.47727 | 0.60752 | 1.42710 | 49 |
| GO_POSITIVE_REGULATION_OF_DNA_METABOLIC_PROCESS | 0.00040 | 0.00773 | 0.49849 | 0.54130 | 1.32611 | 112 |
| GO_POSITIVE_REGULATION_OF_ESTABLISHMENT_OF_PROTEIN_LOC/ | 0.00009 | 0.00220 | 0.53843 | 0.52659 | 1.31777 | 186 |
| GO_POSITIVE_REGULATION_OF_IMMUNE_RESPONSE | 0.00001 | 0.00044 | 0.59333 | 0.51775 | 1.30316 | 232 |
| GO_POSITIVE_REGULATION_OF_IMMUNE_SYSTEM_PROCESS | 0.00014 | 0.00326 | 0.51885 | 0.48605 | 1.23189 | 312 |
| GO_POSITIVE_REGULATION_OF_INNATE_IMMUNE_RESPONSE | 0.00000 | 0.00000 | 0.70498 | 0.63767 | 1.55275 | 98 |
| GO_POSITIVE_REGULATION_OF_INTRACELLULAR_TRANSPORT | 0.00335 | 0.04089 | 0.43171 | 0.50277 | 1.24308 | 137 |
| GO_POSITIVE_REGULATION_OF_INTRINSIC_APOPTOTIC_SIGNALING_ | 0.00405 | 0.04710 | 0.40702 | 0.62462 | 1.41745 | 29 |
| GO_POSITIVE_REGULATION_OF_MRNA_METABOLIC_PROCESS | 0.00102 | 0.01586 | 0.45506 | 0.59926 | 1.41459 | 52 |
| GO_POSITIVE_REGULATION_OF_MRNA_PROCESSING | 0.00170 | 0.02433 | 0.45506 | 0.68883 | 1.53489 | 23 |
| GO_POSITIVE_REGULATION_OF_MRNA_SPLICING_VIA_SPLICEOSOM | 0.00006 | 0.00161 | 0.55733 | 0.81613 | 1.74590 | 15 |
| GO_POSITIVE_REGULATION_OF_NUCLEOCYTOPLASMIC_TRANSPORT | 0.00294 | 0.03785 | 0.43171 | 0.59385 | 1.38277 | 43 |
| GO_POSITIVE_REGULATION_OF_ORGANELLE_ORGANIZATION | 0.00108 | 0.01651 | 0.45506 | 0.46875 | 1.19167 | 349 |
| GO_POSITIVE_REGULATION_OF_PROTEIN_CONTAINING_COMPLEX_/ | 0.00257 | 0.03395 | 0.43171 | 0.50538 | 1.24740 | 131 |
| GO_POSITIVE_REGULATION_OF_PROTEIN_LOCALIZATION_TO_NUCLE | 0.00001 | 0.00033 | 0.59333 | 0.65142 | 1.54070 | 53 |
| GO_POSITIVE_REGULATION_OF_RESPONSE_TO_BIOTIC_STIMULUS | 0.00000 | 0.00001 | 0.69013 | 0.61388 | 1.50327 | 111 |
| GO_POSITIVE_REGULATION_OF_RESPONSE_TO_EXTERNAL_STIMULU | 0.00009 | 0.00225 | 0.53843 | 0.52994 | 1.31939 | 161 |
| GO_POSITIVE_REGULATION_OF_RNA_SPLICING | 0.00294 | 0.03785 | 0.43171 | 0.67695 | 1.50841 | 23 |
| GO_POSITIVE_REGULATION_OF_TELOMERASE_RNA_LOCALIZATION_ | 0.00000 | 0.00002 | 0.65944 | 0.87989 | 1.88230 | 15 |
| GO_POSITIVE_REGULATION_OF_TELOMERE_MAINTENANCE | 0.00320 | 0.03966 | 0.43171 | 0.60048 | 1.39162 | 39 |
| GO_POSITIVE_REGULATION_OF_TELOMERE_MAINTENANCE_VIA_TE | 0.00420 | 0.04805 | 0.40702 | 0.63047 | 1.43314 | 30 |
| GO_POSITIVE_REGULATION_OF_TRANSLATION | 0.00003 | 0.00097 | 0.55733 | 0.59015 | 1.42949 | 85 |
| GO_POSITIVE_REGULATION_OF_VIRAL_GENOME_REPLICATION | 0.00017 | 0.00389 | 0.51885 | 0.70827 | 1.59681 | 27 |
| GO_POSITIVE_REGULATION_OF_VIRAL_LIFE_CYCLE | 0.00013 | 0.00306 | 0.51885 | 0.65181 | 1.51590 | 42 |
| GO_POSITIVE_REGULATION_OF_VIRAL_PROCESS | 0.00357 | 0.04298 | 0.43171 | 0.55317 | 1.32727 | 75 |
| GO_POSITIVE_REGULATION_OF_WNT_SIGNALING_PATHWAY | 0.00000 | 0.00001 | 0.69013 | 0.62644 | 1.52532 | 97 |
| GO_POSTREPLICATION_REPAIR | 0.00372 | 0.04392 | 0.43171 | 0.58855 | 1.37045 | 43 |
| GO_POSTSYNAPSE | 0.00000 | 0.00003 | 0.65944 | 0.54852 | 1.37605 | 197 |
| GO_POSTTRANSCRIPTIONAL_REGULATION_OF_GENE_EXPRESSION | 0.00000 | 0.00000 | | 0.60289 | 1.53646 | 401 |
| GO_POST_TRANSLATIONAL_PROTEIN_MODIFICATION | 0.00000 | 0.00001 | 0.69013 | 0.56069 | 1.40483 | 193 |

| | | | | | | |
|---|---|---|---|---|---|---|
| GO_PROCESS_UTILIZING_AUTOPHAGIC_MECHANISM | 0.00176 | 0.02505 | 0.45506 | 0.46914 | 1.18914 | 320 |
| GO_PROTEASOMAL_PROTEIN_CATABOLIC_PROCESS | 0.00004 | 0.00110 | 0.55733 | 0.49128 | 1.24578 | 326 |
| GO_PROTEASOMAL_UBIQUITIN_INDEPENDENT_PROTEIN_CATABOLIC | 0.00000 | 0.00000 | 0.71951 | 0.86468 | 1.90134 | 19 |
| GO_PROTEASOME_ACCESSORY_COMPLEX | 0.00000 | 0.00008 | 0.62726 | 0.77614 | 1.73261 | 24 |
| GO_PROTEASOME_CORE_COMPLEX | 0.00000 | 0.00000 | 0.77494 | 0.90397 | 1.96501 | 17 |
| GO_PROTEIN_CONTAINING_COMPLEX_LOCALIZATION | 0.00163 | 0.02361 | 0.45506 | 0.49847 | 1.24449 | 175 |
| GO_PROTEIN_DNA_COMPLEX_SUBUNIT_ORGANIZATION | 0.00022 | 0.00483 | 0.51885 | 0.51836 | 1.28600 | 150 |
| GO_PROTEIN_DOMAIN_SPECIFIC_BINDING | 0.00056 | 0.00997 | 0.47727 | 0.47396 | 1.20409 | 342 |
| GO_PROTEIN_FOLDING | 0.00000 | 0.00001 | 0.69013 | 0.58703 | 1.45214 | 139 |
| GO_PROTEIN_LOCALIZATION_TO_CHROMOSOME_TELOMERIC_REGIO | 0.00205 | 0.02792 | 0.43171 | 0.66406 | 1.49428 | 26 |
| GO_PROTEIN_LOCALIZATION_TO_ENDOPLASMIC_RETICULUM | 0.00000 | 0.00000 | | 0.74049 | 1.82035 | 121 |
| GO_PROTEIN_LOCALIZATION_TO_MEMBRANE | 0.00000 | 0.00000 | | 0.54526 | 1.38646 | 350 |
| GO_PROTEIN_LOCALIZATION_TO_NUCLEUS | 0.00004 | 0.00114 | 0.55733 | 0.53714 | 1.33627 | 158 |
| GO_PROTEIN_MODIFICATION_BY_SMALL_PROTEIN_REMOVAL | 0.00000 | 0.00005 | 0.64355 | 0.55118 | 1.38077 | 192 |
| GO_PROTEIN_POLYUBIQUITINATION | 0.00006 | 0.00163 | 0.55733 | 0.51252 | 1.28880 | 226 |
| GO_PROTEIN_STABILIZATION | 0.00002 | 0.00058 | 0.57561 | 0.57270 | 1.40127 | 109 |
| GO_PROTEIN_TARGETING | 0.00000 | 0.00000 | | 0.59889 | 1.51699 | 298 |
| GO_PROTEIN_TARGETING_TO_MEMBRANE | 0.00000 | 0.00000 | | 0.69737 | 1.72686 | 143 |
| GO_PURINE_CONTAINING_COMPOUND_BIOSYNTHETIC_PROCESS | 0.00055 | 0.00984 | 0.47727 | 0.55543 | 1.35047 | 93 |
| GO_PURINE_CONTAINING_COMPOUND_METABOLIC_PROCESS | 0.00004 | 0.00120 | 0.55733 | 0.51698 | 1.29954 | 220 |
| GO_PURINE_NUCLEOSIDE_METABOLIC_PROCESS | 0.00012 | 0.00293 | 0.51885 | 0.69106 | 1.58236 | 33 |
| GO_PURINE_NUCLEOSIDE_TRIPHOSPHATE_METABOLIC_PROCESS | 0.00033 | 0.00660 | 0.49849 | 0.65086 | 1.50835 | 38 |
| GO_PYRUVATE_METABOLIC_PROCESS | 0.00413 | 0.04753 | 0.40702 | 0.53268 | 1.28798 | 84 |
| GO_REGENERATION | 0.00163 | 0.02361 | 0.45506 | 0.56672 | 1.35494 | 68 |
| GO_REGULATION_OF_ANATOMICAL_STRUCTURE_MORPHOGENESIS | 0.00036 | 0.00717 | 0.49849 | 0.46916 | 1.19534 | 398 |
| GO_REGULATION_OF_ANIMAL_ORGAN_MORPHOGENESIS | 0.00000 | 0.00000 | 0.81404 | 0.65839 | 1.60826 | 106 |
| GO_REGULATION_OF_ATP_METABOLIC_PROCESS | 0.00095 | 0.01499 | 0.47727 | 0.57618 | 1.37778 | 69 |
| GO_REGULATION_OF_CELLULAR_AMIDE_METABOLIC_PROCESS | 0.00000 | 0.00000 | | 0.59027 | 1.49001 | 262 |
| GO_REGULATION_OF_CELLULAR_AMINE_METABOLIC_PROCESS | 0.00000 | 0.00000 | | 0.79894 | 1.88996 | 54 |
| GO_REGULATION_OF_CELLULAR_AMINO_ACID_METABOLIC_PROCES | 0.00000 | 0.00000 | | 0.82021 | 1.93502 | 51 |
| GO_REGULATION_OF_CELLULAR_KETONE_METABOLIC_PROCESS | 0.00000 | 0.00000 | 0.76146 | 0.65199 | 1.58895 | 99 |
| GO_REGULATION_OF_CELLULAR_LOCALIZATION | 0.00024 | 0.00521 | 0.51885 | 0.46425 | 1.18531 | 462 |
| GO_REGULATION_OF_CELLULAR_PROTEIN_CATABOLIC_PROCESS | 0.00012 | 0.00283 | 0.53843 | 0.52866 | 1.31277 | 155 |
| GO_REGULATION_OF_CELLULAR_PROTEIN_LOCALIZATION | 0.00002 | 0.00070 | 0.57561 | 0.49802 | 1.26218 | 309 |
| GO_REGULATION_OF_CELLULAR_RESPONSE_TO_HEAT | 0.00305 | 0.03859 | 0.43171 | 0.56535 | 1.34443 | 60 |
| GO_REGULATION_OF_CELLULAR_RESPONSE_TO_STRESS | 0.00024 | 0.00516 | 0.51885 | 0.47117 | 1.19988 | 380 |
| GO_REGULATION_OF_CELLULAR_RESPONSE_TO_TRANSFORMING_G | 0.00383 | 0.04503 | 0.43171 | 0.58686 | 1.37393 | 46 |
| GO_REGULATION_OF_CELL_CYCLE_G2_M_PHASE_TRANSITION | 0.00000 | 0.00015 | 0.61053 | 0.54988 | 1.36875 | 160 |
| GO_REGULATION_OF_CELL_CYCLE_PHASE_TRANSITION | 0.00126 | 0.01876 | 0.45506 | 0.46906 | 1.18818 | 303 |
| GO_REGULATION_OF_DEFENSE_RESPONSE | 0.00078 | 0.01308 | 0.47727 | 0.48689 | 1.22556 | 235 |
| GO_REGULATION_OF_DNA_BIOSYNTHETIC_PROCESS | 0.00413 | 0.04753 | 0.40702 | 0.55556 | 1.32847 | 69 |
| GO_REGULATION_OF_DNA_METABOLIC_PROCESS | 0.00253 | 0.03355 | 0.43171 | 0.48620 | 1.21939 | 203 |
| GO_REGULATION_OF_DNA_TEMPLATED_TRANSCRIPTION_IN_RESPO | 0.00000 | 0.00000 | | 0.70380 | 1.70175 | 84 |
| GO_REGULATION_OF_GENERATION_OF_PRECURSOR_METABOLITES_ | 0.00036 | 0.00713 | 0.49849 | 0.55823 | 1.35288 | 87 |
| GO_REGULATION_OF_HEMATOPOIETIC_PROGENITOR_CELL_DIFFERE | 0.00000 | 0.00000 | | 0.73385 | 1.74953 | 64 |
| GO_REGULATION_OF_HEMATOPOIETIC_STEM_CELL_DIFFERENTIATIO | 0.00000 | 0.00000 | | 0.75659 | 1.79731 | 58 |
| GO_REGULATION_OF_HEMOPOIESIS | 0.00018 | 0.00411 | 0.51885 | 0.51642 | 1.29113 | 180 |
| GO_REGULATION_OF_IMMUNE_RESPONSE | 0.00016 | 0.00365 | 0.51885 | 0.48970 | 1.23885 | 289 |
| GO_REGULATION_OF_IMMUNE_SYSTEM_PROCESS | 0.00268 | 0.03533 | 0.43171 | 0.44819 | 1.14449 | 475 |
| GO_REGULATION_OF_INNATE_IMMUNE_RESPONSE | 0.00000 | 0.00003 | 0.65944 | 0.58096 | 1.43638 | 136 |
| GO_REGULATION_OF_INTRACELLULAR_TRANSPORT | 0.00372 | 0.04392 | 0.43171 | 0.48211 | 1.20954 | 204 |
| GO_REGULATION_OF_INTRINSIC_APOPTOTIC_SIGNALING_PATHWAY | 0.00069 | 0.01170 | 0.47727 | 0.54881 | 1.33183 | 90 |
| GO_REGULATION_OF_MITOTIC_CELL_CYCLE | 0.00001 | 0.00039 | 0.59333 | 0.48457 | 1.23474 | 395 |
| GO_REGULATION_OF_MORPHOGENESIS_OF_AN_EPITHELIUM | 0.00000 | 0.00000 | | 0.68147 | 1.65378 | 90 |
| GO_REGULATION_OF_MRNA_CATABOLIC_PROCESS | 0.00000 | 0.00000 | | 0.64316 | 1.59163 | 140 |
| GO_REGULATION_OF_MRNA_METABOLIC_PROCESS | 0.00000 | 0.00000 | | 0.59710 | 1.50338 | 237 |
| GO_REGULATION_OF_MRNA_PROCESSING | 0.00010 | 0.00237 | 0.53843 | 0.55807 | 1.36547 | 109 |
| GO_REGULATION_OF_MRNA_SPLICING_VIA_SPLICEOSOME | 0.00006 | 0.00168 | 0.53843 | 0.60357 | 1.45082 | 77 |
| GO_REGULATION_OF_NUCLEOCYTOPLASMIC_TRANSPORT | 0.00081 | 0.01340 | 0.47727 | 0.57829 | 1.38282 | 69 |
| GO_REGULATION_OF_OXIDATIVE_PHOSPHORYLATION | 0.00152 | 0.02227 | 0.45506 | 0.73435 | 1.60264 | 18 |
| GO_REGULATION_OF_PROTEASOMAL_PROTEIN_CATABOLIC_PROCES | 0.00272 | 0.03572 | 0.43171 | 0.51665 | 1.26857 | 117 |
| GO_REGULATION_OF_PROTEIN_CATABOLIC_PROCESS | 0.00001 | 0.00029 | 0.59333 | 0.52606 | 1.32285 | 226 |
| GO_REGULATION_OF_PROTEIN_CONTAINING_COMPLEX_ASSEMBLY | 0.00013 | 0.00306 | 0.51885 | 0.50734 | 1.27525 | 223 |
| GO_REGULATION_OF_PROTEIN_IMPORT | 0.00224 | 0.03011 | 0.43171 | 0.62335 | 1.43380 | 35 |
| GO_REGULATION_OF_PROTEIN_LOCALIZATION | 0.00044 | 0.00822 | 0.49849 | 0.46428 | 1.18456 | 450 |
| GO_REGULATION_OF_PROTEIN_LOCALIZATION_TO_NUCLEUS | 0.00001 | 0.00024 | 0.61053 | 0.62295 | 1.49424 | 74 |
| GO_REGULATION_OF_PROTEIN_MODIFICATION_BY_SMALL_PROTEIN | 0.00010 | 0.00238 | 0.53843 | 0.53866 | 1.33462 | 144 |
| GO_REGULATION_OF_PROTEIN_STABILITY | 0.00302 | 0.03831 | 0.43171 | 0.49158 | 1.22903 | 180 |
| GO_REGULATION_OF_PROTEOLYSIS | 0.00051 | 0.00922 | 0.47727 | 0.48419 | 1.22687 | 308 |
| GO_REGULATION_OF_RESPONSE_TO_BIOTIC_STIMULUS | 0.00005 | 0.00146 | 0.55733 | 0.53578 | 1.33634 | 169 |
| GO_REGULATION_OF_RNA_SPLICING | 0.00004 | 0.00123 | 0.55733 | 0.57516 | 1.40243 | 102 |
| GO_REGULATION_OF_SMALL_MOLECULE_METABOLIC_PROCESS | 0.00000 | 0.00000 | 0.78819 | 0.56466 | 1.42014 | 229 |
| GO_REGULATION_OF_STEM_CELL_DIFFERENTIATION | 0.00000 | 0.00000 | 0.76146 | 0.68580 | 1.64666 | 76 |
| GO_REGULATION_OF_SYMBIOTIC_PROCESS | 0.00157 | 0.02301 | 0.45506 | 0.52922 | 1.29595 | 111 |
| GO_REGULATION_OF_TRANSCRIPTION_FROM_RNA_POLYMERASE_II | 0.00000 | 0.00000 | | 0.77207 | 1.83604 | 60 |
| GO_REGULATION_OF_TRANSLATIONAL_INITIATION | 0.00001 | 0.00046 | 0.59333 | 0.65025 | 1.54295 | 56 |

| | | | | | | |
|---|---|---|---|---|---|---|
| GO_REGULATION_OF_VIRAL_GENOME_REPLICATION | 0.00009 | 0.00223 | 0.53843 | 0.65654 | 1.52874 | 43 |
| GO_REGULATION_OF_VIRAL_LIFE_CYCLE | 0.00038 | 0.00732 | 0.49849 | 0.58945 | 1.40608 | 66 |
| GO_REGULATION_OF_WNT_SIGNALING_PATHWAY | 0.00000 | 0.00007 | 0.64355 | 0.55963 | 1.39538 | 168 |
| GO_RESPONSE_TO_ALCOHOL | 0.00205 | 0.02792 | 0.43171 | 0.54926 | 1.32574 | 82 |
| GO_RESPONSE_TO_BIOTIC_STIMULUS | 0.00000 | 0.00001 | 0.69013 | 0.49911 | 1.27396 | 455 |
| GO_RESPONSE_TO_CYTOKINE | 0.00000 | 0.00000 | | 0.55391 | 1.41310 | 448 |
| GO_RESPONSE_TO_DRUG | 0.00000 | 0.00011 | 0.62726 | 0.56034 | 1.38958 | 148 |
| GO_RESPONSE_TO_ETHANOL | 0.00213 | 0.02877 | 0.43171 | 0.60786 | 1.40873 | 39 |
| GO_RESPONSE_TO_GROWTH_FACTOR | 0.00191 | 0.02655 | 0.45506 | 0.47701 | 1.20449 | 269 |
| GO_RESPONSE_TO_HEAT | 0.00029 | 0.00610 | 0.49849 | 0.55989 | 1.36333 | 98 |
| GO_RESPONSE_TO_HORMONE | 0.00052 | 0.00932 | 0.47727 | 0.46830 | 1.19192 | 373 |
| GO_RESPONSE_TO_INTERFERON_GAMMA | 0.00041 | 0.00785 | 0.49849 | 0.61593 | 1.44684 | 49 |
| GO_RESPONSE_TO_INTERLEUKIN_1 | 0.00000 | 0.00000 | 0.80122 | 0.65874 | 1.60278 | 94 |
| GO_RESPONSE_TO_INTERLEUKIN_12 | 0.00122 | 0.01832 | 0.45506 | 0.65453 | 1.49313 | 31 |
| GO_RESPONSE_TO_INTERLEUKIN_4 | 0.00008 | 0.00216 | 0.53843 | 0.78661 | 1.70991 | 17 |
| GO_RESPONSE_TO_NITROGEN_COMPOUND | 0.00010 | 0.00246 | 0.53843 | 0.47139 | 1.20326 | 460 |
| GO_RESPONSE_TO_ORGANIC_CYCLIC_COMPOUND | 0.00006 | 0.00154 | 0.55733 | 0.48954 | 1.24452 | 349 |
| GO_RESPONSE_TO_OXYGEN_LEVELS | 0.00000 | 0.00000 | 0.70498 | 0.55549 | 1.39393 | 208 |
| GO_RESPONSE_TO_PEPTIDE_HORMONE | 0.00327 | 0.04028 | 0.43171 | 0.48635 | 1.21698 | 184 |
| GO_RESPONSE_TO_TEMPERATURE_STIMULUS | 0.00032 | 0.00644 | 0.49849 | 0.54106 | 1.32792 | 116 |
| GO_RESPONSE_TO_TOPOLOGICALLY_INCORRECT_PROTEIN | 0.00104 | 0.01604 | 0.45506 | 0.52339 | 1.28663 | 121 |
| GO_RESPONSE_TO_TUMOR_NECROSIS_FACTOR | 0.00000 | 0.00002 | 0.65944 | 0.58739 | 1.44917 | 130 |
| GO_RHYTHMIC_PROCESS | 0.00085 | 0.01383 | 0.47727 | 0.52994 | 1.30033 | 115 |
| GO_RIBONUCLEOPROTEIN_COMPLEX_BINDING | 0.00000 | 0.00001 | 0.69013 | 0.61833 | 1.50767 | 102 |
| GO_RIBONUCLEOPROTEIN_COMPLEX_BIOGENESIS | 0.00000 | 0.00000 | | 0.57486 | 1.46285 | 363 |
| GO_RIBONUCLEOPROTEIN_COMPLEX_SUBUNIT_ORGANIZATION | 0.00000 | 0.00000 | 0.76146 | 0.58739 | 1.46396 | 167 |
| GO_RIBONUCLEOPROTEIN_GRANULE | 0.00008 | 0.00197 | 0.53843 | 0.53509 | 1.32705 | 147 |
| GO_RIBONUCLEOSIDE_METABOLIC_PROCESS | 0.00014 | 0.00326 | 0.51885 | 0.68742 | 1.57405 | 33 |
| GO_RIBONUCLEOSIDE_MONOPHOSPHATE_BIOSYNTHETIC_PROCESS | 0.00122 | 0.01832 | 0.45506 | 0.70507 | 1.55798 | 21 |
| GO_RIBONUCLEOSIDE_MONOPHOSPHATE_METABOLIC_PROCESS | 0.00390 | 0.04569 | 0.43171 | 0.63547 | 1.44451 | 30 |
| GO_RIBONUCLEOSIDE_TRIPHOSPHATE_BIOSYNTHETIC_PROCESS | 0.00058 | 0.01019 | 0.47727 | 0.68350 | 1.54594 | 28 |
| GO_RIBONUCLEOSIDE_TRIPHOSPHATE_METABOLIC_PROCESS | 0.00006 | 0.00161 | 0.55733 | 0.67733 | 1.56488 | 37 |
| GO_RIBOSE_PHOSPHATE_BIOSYNTHETIC_PROCESS | 0.00042 | 0.00796 | 0.49849 | 0.56347 | 1.36848 | 91 |
| GO_RIBOSE_PHOSPHATE_METABOLIC_PROCESS | 0.00004 | 0.00105 | 0.55733 | 0.51874 | 1.30173 | 208 |
| GO_RIBOSOMAL_LARGE_SUBUNIT_ASSEMBLY | 0.00000 | 0.00003 | 0.65944 | 0.78578 | 1.77156 | 27 |
| GO_RIBOSOMAL_LARGE_SUBUNIT_BIOGENESIS | 0.00000 | 0.00000 | 0.73376 | 0.70148 | 1.67036 | 63 |
| GO_RIBOSOMAL_SMALL_SUBUNIT_BINDING | 0.00061 | 0.01059 | 0.47727 | 0.77154 | 1.65052 | 15 |
| GO_RIBOSOMAL_SMALL_SUBUNIT_BIOGENESIS | 0.00069 | 0.01170 | 0.47727 | 0.57520 | 1.37168 | 65 |
| GO_RIBOSOMAL_SUBUNIT | 0.00000 | 0.00000 | | 0.64641 | 1.61228 | 169 |
| GO_RIBOSOME | 0.00000 | 0.00000 | | 0.62658 | 1.57121 | 201 |
| GO_RIBOSOME_ASSEMBLY | 0.00000 | 0.00001 | 0.70498 | 0.68845 | 1.63545 | 58 |
| GO_RIBOSOME_BINDING | 0.00004 | 0.00103 | 0.55733 | 0.66941 | 1.55501 | 41 |
| GO_RIBOSOME_BIOGENESIS | 0.00000 | 0.00000 | 0.82666 | 0.56939 | 1.43630 | 256 |
| GO_RNA_CATABOLIC_PROCESS | 0.00000 | 0.00000 | | 0.65273 | 1.65313 | 300 |
| GO_RNA_DEPENDENT_DNA_BIOSYNTHETIC_PROCESS | 0.00025 | 0.00532 | 0.49849 | 0.61570 | 1.46097 | 56 |
| GO_RNA_EXPORT_FROM_NUCLEUS | 0.00009 | 0.00222 | 0.53843 | 0.54183 | 1.33194 | 120 |
| GO_RNA_LOCALIZATION | 0.00000 | 0.00009 | 0.62726 | 0.54514 | 1.36408 | 184 |
| GO_RNA_SPLICING | 0.00000 | 0.00004 | 0.64355 | 0.50850 | 1.28943 | 326 |
| GO_RNA_SPLICING_VIA_TRANSESTERIFICATION_REACTIONS | 0.00000 | 0.00006 | 0.64355 | 0.51940 | 1.31266 | 276 |
| GO_RNA_STABILIZATION | 0.00019 | 0.00424 | 0.51885 | 0.68349 | 1.55919 | 31 |
| GO_ROUGH_ENDOPLASMIC_RETICULUM | 0.00082 | 0.01347 | 0.47727 | 0.61157 | 1.43651 | 48 |
| GO_RRNA_BINDING | 0.00000 | 0.00000 | 0.80122 | 0.74557 | 1.75559 | 50 |
| GO_RRNA_METABOLIC_PROCESS | 0.00004 | 0.00107 | 0.55733 | 0.52714 | 1.32056 | 192 |
| GO_SCF_DEPENDENT_PROTEASOMAL_UBIQUITIN_DEPENDENT_PRO | 0.00000 | 0.00000 | 0.77494 | 0.70611 | 1.68437 | 66 |
| GO_SECRETORY_GRANULE | 0.00000 | 0.00000 | 0.82666 | 0.54875 | 1.39178 | 328 |
| GO_SECRETORY_VESICLE | 0.00000 | 0.00000 | 0.76146 | 0.52150 | 1.32743 | 375 |
| GO_SIDE_OF_MEMBRANE | 0.00398 | 0.04645 | 0.40702 | 0.50000 | 1.23651 | 138 |
| GO_SIGNAL_SEQUENCE_BINDING | 0.00031 | 0.00636 | 0.49849 | 0.69979 | 1.58278 | 28 |
| GO_SIGNAL_TRANSDUCTION_BY_PROTEIN_PHOSPHORYLATION | 0.00035 | 0.00701 | 0.49849 | 0.47753 | 1.21314 | 342 |
| GO_SINGLE_FERTILIZATION | 0.00009 | 0.00220 | 0.53843 | 0.67686 | 1.55964 | 36 |
| GO_SINGLE_STRANDED_RNA_BINDING | 0.00007 | 0.00192 | 0.53843 | 0.62660 | 1.48201 | 53 |
| GO_SMALL_MOLECULE_BIOSYNTHETIC_PROCESS | 0.00001 | 0.00040 | 0.59333 | 0.49679 | 1.26008 | 329 |
| GO_SMALL_RIBOSOMAL_SUBUNIT | 0.00000 | 0.00003 | 0.65944 | 0.66037 | 1.57525 | 66 |
| GO_SNRNA_BINDING | 0.00198 | 0.02728 | 0.43171 | 0.62769 | 1.44633 | 36 |
| GO_SPLICEOSOMAL_COMPLEX | 0.00037 | 0.00723 | 0.49849 | 0.51968 | 1.29046 | 155 |
| GO_STEM_CELL_DIFFERENTIATION | 0.00000 | 0.00000 | 0.80122 | 0.63220 | 1.55410 | 120 |
| GO_STRUCTURAL_CONSTITUENT_OF_RIBOSOME | 0.00000 | 0.00000 | | 0.64473 | 1.59952 | 150 |
| GO_STRUCTURAL_MOLECULE_ACTIVITY | 0.00000 | 0.00000 | | 0.57099 | 1.44354 | 285 |
| GO_SUBSTANTIA_NIGRA_DEVELOPMENT | 0.00183 | 0.02581 | 0.45506 | 0.68726 | 1.53137 | 23 |
| GO_SYNAPSE | 0.00000 | 0.00000 | 0.80122 | 0.52429 | 1.33704 | 430 |
| GO_TELOMERASE_RNA_LOCALIZATION | 0.00002 | 0.00060 | 0.57561 | 0.79154 | 1.74050 | 19 |
| GO_TELOMERE_MAINTENANCE_VIA_TELOMERE_LENGTHENING | 0.00167 | 0.02387 | 0.45506 | 0.57646 | 1.37211 | 62 |
| GO_TELOMERE_ORGANIZATION | 0.00309 | 0.03876 | 0.43171 | 0.51939 | 1.27369 | 114 |
| GO_THREONINE_TYPE_ENDOPEPTIDASE_ACTIVITY | 0.00000 | 0.00000 | 0.81404 | 0.91849 | 1.98160 | 16 |
| GO_THREONINE_TYPE_PEPTIDASE_ACTIVITY | 0.00000 | 0.00000 | 0.71951 | 0.88236 | 1.92566 | 18 |
| GO_TISSUE_MORPHOGENESIS | 0.00006 | 0.00154 | 0.55733 | 0.50379 | 1.27153 | 260 |

| | | | | | | |
|---|---|---|---|---|---|---|
| GO_TOLL_LIKE_RECEPTOR_SIGNALING_PATHWAY | 0.00313 | 0.03913 | 0.43171 | 0.57403 | 1.35768 | 53 |
| GO_TOXIN_TRANSPORT | 0.00025 | 0.00532 | 0.49849 | 0.72441 | 1.60674 | 22 |
| GO_TRANSCRIPTION_COUPLED_NUCLEOTIDE_EXCISION_REPAIR | 0.00012 | 0.00280 | 0.53843 | 0.61072 | 1.45425 | 63 |
| GO_TRANSLATIONAL_INITIATION | 0.00000 | 0.00000 | | 0.72823 | 1.81056 | 157 |
| GO_TRANSLATION_FACTOR_ACTIVITY_RNA_BINDING | 0.00000 | 0.00000 | 0.81404 | 0.72729 | 1.73111 | 62 |
| GO_TRANSLATION_INITIATION_FACTOR_ACTIVITY | 0.00000 | 0.00001 | 0.67496 | 0.72478 | 1.68559 | 42 |
| GO_TRANSLATION_PREINITIATION_COMPLEX | 0.00001 | 0.00024 | 0.61053 | 0.84715 | 1.81226 | 15 |
| GO_TRANSLATION_REGULATOR_ACTIVITY | 0.00000 | 0.00000 | | 0.68863 | 1.67435 | 93 |
| GO_TRANSLATION_REGULATOR_ACTIVITY_NUCLEIC_ACID_BINDING | 0.00000 | 0.00000 | 0.77494 | 0.69195 | 1.66144 | 76 |
| GO_TRANSMEMBRANE_TRANSPORT | 0.00001 | 0.00037 | 0.59333 | 0.47774 | 1.22154 | 495 |
| GO_TRICARBOXYLIC_ACID_CYCLE | 0.00098 | 0.01538 | 0.45506 | 0.68110 | 1.53262 | 26 |
| GO_TUMOR_NECROSIS_FACTOR_MEDIATED_SIGNALING_PATHWAY | 0.00000 | 0.00000 | 0.76146 | 0.67215 | 1.61692 | 78 |
| GO_T_CELL_RECEPTOR_SIGNALING_PATHWAY | 0.00000 | 0.00000 | 0.77494 | 0.64540 | 1.57451 | 103 |
| GO_UBIQUITIN_LIKE_PROTEIN_LIGASE_BINDING | 0.00004 | 0.00123 | 0.55733 | 0.52372 | 1.31085 | 187 |
| GO_UNFOLDED_PROTEIN_BINDING | 0.00001 | 0.00022 | 0.61053 | 0.62667 | 1.50363 | 75 |
| GO_VACUOLAR_LUMEN | 0.00001 | 0.00028 | 0.59333 | 0.61613 | 1.48287 | 79 |
| GO_VACUOLAR_MEMBRANE | 0.00052 | 0.00932 | 0.47727 | 0.49332 | 1.24236 | 241 |
| GO_VACUOLE | 0.00006 | 0.00163 | 0.55733 | 0.47940 | 1.22166 | 402 |
| GO_VESICLE_LUMEN | 0.00000 | 0.00000 | | 0.63803 | 1.58224 | 148 |
| GO_VIRAL_GENE_EXPRESSION | 0.00000 | 0.00000 | | 0.67780 | 1.68801 | 164 |
| GO_VIRAL_GENOME_REPLICATION | 0.00032 | 0.00644 | 0.49849 | 0.60095 | 1.42943 | 61 |
| GO_VIRAL_LIFE_CYCLE | 0.00001 | 0.00032 | 0.59333 | 0.54012 | 1.34919 | 177 |
| HP_ABNORMALITY_OF_THE_ANKLES | 0.00019 | 0.00417 | 0.51885 | 0.65849 | 1.52136 | 37 |
| HP_ABNORMAL_AUTONOMIC_NERVOUS_SYSTEM_PHYSIOLOGY | 0.00045 | 0.00835 | 0.49849 | 0.63223 | 1.47215 | 43 |
| HP_ACUTE_LEUKEMIA | 0.00327 | 0.04028 | 0.43171 | 0.58029 | 1.37770 | 57 |
| HP_BILATERAL_SENSORINEURAL_HEARING_IMPAIRMENT | 0.00279 | 0.03631 | 0.43171 | 0.67487 | 1.49685 | 22 |
| HP_DELAYED_CRANIAL_SUTURE_CLOSURE | 0.00087 | 0.01419 | 0.47727 | 0.65943 | 1.49643 | 29 |
| HP_DELAYED_PUBERTY | 0.00368 | 0.04379 | 0.43171 | 0.52978 | 1.28812 | 93 |
| HP_HEMATOLOGICAL_NEOPLASM | 0.00086 | 0.01393 | 0.47727 | 0.52498 | 1.29369 | 127 |
| HP_INTERSTITIAL_PULMONARY_ABNORMALITY | 0.00383 | 0.04503 | 0.43171 | 0.61308 | 1.41018 | 35 |
| HP_LEUKEMIA | 0.00143 | 0.02104 | 0.45506 | 0.55714 | 1.34089 | 79 |
| HP_MACROCYTIC_ANEMIA | 0.00132 | 0.01953 | 0.45506 | 0.61932 | 1.43529 | 39 |
| HP_SYNOSTOSIS_INVOLVING_BONES_OF_THE_HAND | 0.00090 | 0.01449 | 0.47727 | 0.76211 | 1.63034 | 15 |
| **Cluster 8** | | | | | | |
| pathway | pval | padj | log2err | ES | NES | size |
| GO_ACTIVATION_OF_IMMUNE_RESPONSE | 0.00000 | 0.00000 | 0.77494 | 0.63539 | 1.35136 | 213 |
| GO_ACTIVATION_OF_INNATE_IMMUNE_RESPONSE | 0.00000 | 0.00000 | 0.77494 | 0.73890 | 1.52868 | 86 |
| GO_AEROBIC_RESPIRATION | 0.00213 | 0.04005 | 0.43171 | 0.63506 | 1.30108 | 63 |
| GO_ALTERNATIVE_MRNA_SPLICING_VIA_SPLICEOSOME | 0.00242 | 0.04413 | 0.43171 | 0.63421 | 1.29816 | 62 |
| GO_AMIDE_TRANSMEMBRANE_TRANSPORTER_ACTIVITY | 0.00019 | 0.00540 | 0.51885 | 0.75374 | 1.48375 | 30 |
| GO_AMINE_METABOLIC_PROCESS | 0.00000 | 0.00000 | | 0.77048 | 1.58991 | 76 |
| GO_ANAPHASE_PROMOTING_COMPLEX_DEPENDENT_CATABOLIC_P | 0.00000 | 0.00000 | 0.74774 | 0.73995 | 1.52690 | 76 |
| GO_ANCHORED_COMPONENT_OF_PLASMA_MEMBRANE | 0.00202 | 0.03848 | 0.43171 | 0.79521 | 1.49431 | 15 |
| GO_ANCHORING_JUNCTION | 0.00000 | 0.00003 | 0.65944 | 0.56232 | 1.21139 | 452 |
| GO_ANIMAL_ORGAN_REGENERATION | 0.00097 | 0.02163 | 0.47727 | 0.71264 | 1.41768 | 34 |
| GO_ANTIGEN_PROCESSING_AND_PRESENTATION | 0.00000 | 0.00002 | 0.67496 | 0.64740 | 1.36447 | 140 |
| GO_ANTIGEN_PROCESSING_AND_PRESENTATION_OF_EXOGENOUS_ | 0.00000 | 0.00000 | 0.80122 | 0.78879 | 1.60618 | 56 |
| GO_ANTIGEN_PROCESSING_AND_PRESENTATION_OF_PEPTIDE_ANTI | 0.00000 | 0.00001 | 0.70498 | 0.66670 | 1.39770 | 123 |
| GO_ANTIGEN_PROCESSING_AND_PRESENTATION_OF_PEPTIDE_ANTI | 0.00000 | 0.00000 | 0.70498 | 0.74020 | 1.52040 | 67 |
| GO_ANTIGEN_RECEPTOR_MEDIATED_SIGNALING_PATHWAY | 0.00000 | 0.00001 | 0.69013 | 0.66251 | 1.39021 | 127 |
| GO_ATPASE_ACTIVITY | 0.00205 | 0.03905 | 0.43171 | 0.54216 | 1.15993 | 281 |
| GO_ATPASE_COMPLEX | 0.00242 | 0.04413 | 0.43171 | 0.63434 | 1.29842 | 62 |
| GO_AUTOPHAGOSOME_ORGANIZATION | 0.00183 | 0.03619 | 0.45506 | 0.62268 | 1.28399 | 75 |
| GO_AZUROPHIL_GRANULE_LUMEN | 0.00044 | 0.01119 | 0.49849 | 0.66849 | 1.36200 | 57 |
| GO_BINDING_OF_SPERM_TO_ZONA_PELLUCIDA | 0.00000 | 0.00017 | 0.61053 | 0.86278 | 1.64952 | 19 |
| GO_BRUSH_BORDER | 0.00102 | 0.02250 | 0.45506 | 0.65794 | 1.33973 | 56 |
| GO_CADHERIN_BINDING | 0.00000 | 0.00001 | 0.69013 | 0.61303 | 1.30689 | 236 |
| GO_CANONICAL_WNT_SIGNALING_PATHWAY | 0.00000 | 0.00012 | 0.62726 | 0.61780 | 1.30991 | 179 |
| GO_CATALYTIC_STEP_2_SPLICEOSOME | 0.00001 | 0.00031 | 0.59333 | 0.67764 | 1.39932 | 79 |
| GO_CELLULAR_AMINO_ACID_METABOLIC_PROCESS | 0.00000 | 0.00000 | 0.71951 | 0.62865 | 1.33572 | 204 |
| GO_CELLULAR_COMPONENT_DISASSEMBLY | 0.00013 | 0.00382 | 0.51885 | 0.54151 | 1.16465 | 381 |
| GO_CELLULAR_HOMEOSTASIS | 0.00202 | 0.03848 | 0.43171 | 0.52800 | 1.13640 | 406 |
| GO_CELLULAR_KETONE_METABOLIC_PROCESS | 0.00001 | 0.00037 | 0.59333 | 0.62383 | 1.31501 | 145 |
| GO_CELLULAR_METABOLIC_COMPOUND_SALVAGE | 0.00064 | 0.01522 | 0.47727 | 0.78267 | 1.50464 | 21 |
| GO_CELLULAR_NITROGEN_COMPOUND_CATABOLIC_PROCESS | 0.00000 | 0.00000 | | 0.66928 | 1.44067 | 420 |
| GO_CELLULAR_RESPONSE_TO_DRUG | 0.00087 | 0.01973 | 0.47727 | 0.71675 | 1.42586 | 34 |
| GO_CELLULAR_RESPONSE_TO_HEAT | 0.00246 | 0.04451 | 0.43171 | 0.61169 | 1.26866 | 92 |
| GO_CELLULAR_RESPONSE_TO_OXYGEN_LEVELS | 0.00000 | 0.00003 | 0.65944 | 0.64441 | 1.35967 | 149 |
| GO_CELL_ACTIVATION_INVOLVED_IN_IMMUNE_RESPONSE | 0.00003 | 0.00095 | 0.57561 | 0.56059 | 1.20379 | 351 |
| GO_CELL_ADHESION_MOLECULE_BINDING | 0.00000 | 0.00000 | 0.73376 | 0.60033 | 1.28629 | 304 |
| GO_CELL_BODY | 0.00039 | 0.01010 | 0.49849 | 0.56423 | 1.20336 | 244 |
| GO_CELL_CELL_RECOGNITION | 0.00000 | 0.00000 | 0.76146 | 0.86638 | 1.69754 | 28 |
| GO_CELL_CELL_SIGNALING_BY_WNT | 0.00001 | 0.00052 | 0.59333 | 0.57927 | 1.23980 | 286 |
| GO_CELL_CYCLE_G2_M_PHASE_TRANSITION | 0.00001 | 0.00033 | 0.59333 | 0.59213 | 1.26072 | 219 |
| GO_CELL_CYCLE_PHASE_TRANSITION | 0.00000 | 0.00002 | 0.67496 | 0.56104 | 1.20877 | 455 |
| GO_CELL_DEATH_IN_RESPONSE_TO_OXIDATIVE_STRESS | 0.00086 | 0.01948 | 0.47727 | 0.66160 | 1.34422 | 52 |

| | | | | | | |
|---|---|---|---|---|---|---|
| GO_CELL_DIVISION | 0.00070 | 0.01654 | 0.47727 | 0.52991 | 1.14062 | 408 |
| GO_CELL_RECOGNITION | 0.00005 | 0.00147 | 0.55733 | 0.69762 | 1.41863 | 53 |
| GO_CELL_SUBSTRATE_JUNCTION | 0.00000 | 0.00000 | 0.82666 | 0.62835 | 1.34433 | 279 |
| GO_CELL_SURFACE_RECEPTOR_SIGNALING_PATHWAY_INVOLVED_IN | 0.00002 | 0.00082 | 0.57561 | 0.56860 | 1.21850 | 305 |
| GO_CHAPERONE_COMPLEX | 0.00002 | 0.00062 | 0.57561 | 0.84078 | 1.61636 | 21 |
| GO_CHROMATIN_BINDING | 0.00030 | 0.00804 | 0.49849 | 0.54494 | 1.17109 | 358 |
| GO_COTRANSLATIONAL_PROTEIN_TARGETING_TO_MEMBRANE | 0.00000 | 0.00000 | | 0.77805 | 1.61369 | 92 |
| GO_CYTOKINE_MEDIATED_SIGNALING_PATHWAY | 0.00000 | 0.00003 | 0.65944 | 0.58117 | 1.24707 | 323 |
| GO_CYTOPLASMIC_TRANSLATION | 0.00000 | 0.00000 | | 0.77469 | 1.60642 | 91 |
| GO_CYTOPLASMIC_TRANSLATIONAL_INITIATION | 0.00000 | 0.00002 | 0.67496 | 0.82457 | 1.62778 | 31 |
| GO_CYTOSOLIC_LARGE_RIBOSOMAL_SUBUNIT | 0.00000 | 0.00000 | 0.77494 | 0.79127 | 1.60906 | 53 |
| GO_CYTOSOLIC_RIBOSOME | 0.00000 | 0.00000 | | 0.78441 | 1.62859 | 94 |
| GO_CYTOSOLIC_SMALL_RIBOSOMAL_SUBUNIT | 0.00000 | 0.00004 | 0.65944 | 0.79093 | 1.58692 | 40 |
| GO_DEFENSE_RESPONSE_TO_OTHER_ORGANISM | 0.00000 | 0.00000 | 0.73376 | 0.58521 | 1.25890 | 386 |
| GO_DNA_BINDING_TRANSCRIPTION_FACTOR_BINDING | 0.00042 | 0.01076 | 0.49849 | 0.57061 | 1.21318 | 212 |
| GO_DNA_BIOSYNTHETIC_PROCESS | 0.00141 | 0.02917 | 0.45506 | 0.58884 | 1.24015 | 137 |
| GO_DNA_CONFORMATION_CHANGE | 0.00276 | 0.04938 | 0.43171 | 0.55564 | 1.18075 | 207 |
| GO_DNA_POLYMERASE_BINDING | 0.00130 | 0.02750 | 0.45506 | 0.80162 | 1.51346 | 16 |
| GO_DOUBLE_STRANDED_RNA_BINDING | 0.00157 | 0.03201 | 0.45506 | 0.65267 | 1.32497 | 51 |
| GO_EMBRYO_IMPLANTATION | 0.00235 | 0.04335 | 0.43171 | 0.71871 | 1.40116 | 26 |
| GO_ENDOPEPTIDASE_ACTIVITY | 0.00054 | 0.01332 | 0.47727 | 0.58232 | 1.23139 | 159 |
| GO_ENDOPEPTIDASE_COMPLEX | 0.00000 | 0.00000 | 0.76146 | 0.76268 | 1.56253 | 63 |
| GO_ESTABLISHMENT_OF_PROTEIN_LOCALIZATION_TO_ENDOPLASM | 0.00000 | 0.00000 | | 0.75824 | 1.57983 | 102 |
| GO_ESTABLISHMENT_OF_PROTEIN_LOCALIZATION_TO_MEMBRANE | 0.00000 | 0.00000 | | 0.66876 | 1.42588 | 237 |
| GO_ESTABLISHMENT_OF_PROTEIN_LOCALIZATION_TO_ORGANELLE | 0.00000 | 0.00000 | | 0.62783 | 1.35139 | 405 |
| GO_ESTABLISHMENT_OF_PROTEIN_LOCALIZATION_TO_TELOMERE | 0.00191 | 0.03712 | 0.45506 | 0.79289 | 1.49697 | 16 |
| GO_ESTABLISHMENT_OF_RNA_LOCALIZATION | 0.00002 | 0.00059 | 0.57561 | 0.61027 | 1.29184 | 165 |
| GO_ESTABLISHMENT_OF_TISSUE_POLARITY | 0.00000 | 0.00000 | 0.74774 | 0.71868 | 1.48986 | 89 |
| GO_EUKARYOTIC_TRANSLATION_INITIATION_FACTOR_3_COMPLEX | 0.00002 | 0.00065 | 0.57561 | 0.88422 | 1.66157 | 15 |
| GO_EXOCYTOSIS | 0.00015 | 0.00435 | 0.51885 | 0.53844 | 1.16028 | 468 |
| GO_FC_EPSILON_RECEPTOR_SIGNALING_PATHWAY | 0.00000 | 0.00000 | 0.77494 | 0.73033 | 1.50958 | 84 |
| GO_FC_RECEPTOR_SIGNALING_PATHWAY | 0.00000 | 0.00000 | 0.80122 | 0.69149 | 1.45195 | 128 |
| GO_FERTILIZATION | 0.00209 | 0.03962 | 0.43171 | 0.63567 | 1.30603 | 68 |
| GO_FICOLIN_1_RICH_GRANULE | 0.00000 | 0.00000 | 0.78819 | 0.72951 | 1.51356 | 93 |
| GO_GLYCOSYL_COMPOUND_BIOSYNTHETIC_PROCESS | 0.00141 | 0.02917 | 0.45506 | 0.73027 | 1.42779 | 27 |
| GO_GLYCOSYL_COMPOUND_METABOLIC_PROCESS | 0.00004 | 0.00123 | 0.55733 | 0.68223 | 1.40167 | 68 |
| GO_GTP_METABOLIC_PROCESS | 0.00028 | 0.00746 | 0.49849 | 0.81160 | 1.54654 | 18 |
| GO_HEMATOPOIETIC_PROGENITOR_CELL_DIFFERENTIATION | 0.00000 | 0.00019 | 0.61053 | 0.66143 | 1.38015 | 104 |
| GO_HEMATOPOIETIC_STEM_CELL_DIFFERENTIATION | 0.00000 | 0.00000 | 0.82666 | 0.78177 | 1.60619 | 68 |
| GO_IMMUNE_RESPONSE_REGULATING_SIGNALING_PATHWAY | 0.00000 | 0.00001 | 0.70498 | 0.62859 | 1.33499 | 199 |
| GO_IMMUNE_SYSTEM_DEVELOPMENT | 0.00054 | 0.01332 | 0.47727 | 0.53120 | 1.14463 | 461 |
| GO_INNATE_IMMUNE_RESPONSE | 0.00000 | 0.00000 | 0.77494 | 0.60493 | 1.29764 | 318 |
| GO_INNATE_IMMUNE_RESPONSE_ACTIVATING_SIGNAL_TRANSDUCT | 0.00000 | 0.00000 | | 0.78085 | 1.60705 | 70 |
| GO_INTERACTION_WITH_SYMBIONT | 0.00124 | 0.02673 | 0.45506 | 0.68361 | 1.37295 | 41 |
| GO_INTERLEUKIN_1_MEDIATED_SIGNALING_PATHWAY | 0.00000 | 0.00000 | 0.80122 | 0.77371 | 1.58963 | 68 |
| GO_INTRINSIC_APOPTOTIC_SIGNALING_PATHWAY | 0.00062 | 0.01491 | 0.47727 | 0.57345 | 1.21551 | 180 |
| GO_ION_CHANNEL_BINDING | 0.00005 | 0.00143 | 0.55733 | 0.70009 | 1.42640 | 57 |
| GO_LARGE_RIBOSOMAL_SUBUNIT | 0.00000 | 0.00001 | 0.69013 | 0.68742 | 1.43439 | 104 |
| GO_LEUKOCYTE_MEDIATED_IMMUNITY | 0.00002 | 0.00059 | 0.57561 | 0.55932 | 1.20279 | 377 |
| GO_LIVER_REGENERATION | 0.00089 | 0.02006 | 0.47727 | 0.80683 | 1.52328 | 16 |
| GO_MATURATION_OF_LSU_RRNA | 0.00157 | 0.03201 | 0.45506 | 0.72854 | 1.42442 | 27 |
| GO_MITOCHONDRIAL_PROTEIN_COMPLEX | 0.00070 | 0.01654 | 0.47727 | 0.56892 | 1.20801 | 192 |
| GO_MITOCHONDRION_ORGANIZATION | 0.00062 | 0.01491 | 0.47727 | 0.53980 | 1.15980 | 356 |
| GO_MODIFICATION_DEPENDENT_MACROMOLECULE_CATABOLIC_PR | 0.00000 | 0.00017 | 0.61053 | 0.55356 | 1.19286 | 479 |
| GO_MODULATION_OF_PROCESS_OF_OTHER_ORGANISM | 0.00228 | 0.04212 | 0.43171 | 0.64010 | 1.30245 | 55 |
| GO_MORPHOGENESIS_OF_AN_EPITHELIUM | 0.00154 | 0.03160 | 0.45506 | 0.54951 | 1.17560 | 278 |
| GO_MORPHOGENESIS_OF_A_POLARIZED_EPITHELIUM | 0.00000 | 0.00000 | 0.73376 | 0.69470 | 1.44699 | 100 |
| GO_MRNA_3_UTR_BINDING | 0.00082 | 0.01887 | 0.47727 | 0.64373 | 1.32063 | 65 |
| GO_MRNA_5_UTR_BINDING | 0.00008 | 0.00235 | 0.53843 | 0.80287 | 1.54854 | 22 |
| GO_MRNA_BINDING | 0.00000 | 0.00000 | 0.78819 | 0.63511 | 1.35265 | 225 |
| GO_MRNA_EXPORT_FROM_NUCLEUS | 0.00018 | 0.00516 | 0.51885 | 0.62315 | 1.29797 | 100 |
| GO_MRNA_PROCESSING | 0.00000 | 0.00000 | 0.73376 | 0.58353 | 1.25593 | 406 |
| GO_MRNA_TRANSPORT | 0.00026 | 0.00712 | 0.49849 | 0.60619 | 1.27134 | 125 |
| GO_MYELOID_LEUKOCYTE_ACTIVATION | 0.00000 | 0.00003 | 0.65944 | 0.58531 | 1.25635 | 330 |
| GO_MYELOID_LEUKOCYTE_MEDIATED_IMMUNITY | 0.00000 | 0.00001 | 0.70498 | 0.59385 | 1.27261 | 305 |
| GO_NCRNA_METABOLIC_PROCESS | 0.00000 | 0.00004 | 0.65944 | 0.57310 | 1.23269 | 380 |
| GO_NCRNA_PROCESSING | 0.00000 | 0.00000 | 0.71951 | 0.59050 | 1.26716 | 327 |
| GO_NEGATIVE_REGULATION_OF_CANONICAL_WNT_SIGNALING_PAT | 0.00000 | 0.00000 | 0.78819 | 0.71325 | 1.48563 | 100 |
| GO_NEGATIVE_REGULATION_OF_CELLULAR_AMIDE_METABOLIC_PR | 0.00033 | 0.00852 | 0.49849 | 0.61667 | 1.29153 | 114 |
| GO_NEGATIVE_REGULATION_OF_CELL_CYCLE | 0.00001 | 0.00034 | 0.59333 | 0.55187 | 1.18796 | 419 |
| GO_NEGATIVE_REGULATION_OF_CELL_CYCLE_G2_M_PHASE_TRANS | 0.00000 | 0.00000 | 0.73376 | 0.71722 | 1.48351 | 85 |
| GO_NEGATIVE_REGULATION_OF_CELL_CYCLE_PHASE_TRANSITION | 0.00000 | 0.00018 | 0.61053 | 0.61242 | 1.29813 | 180 |
| GO_NEGATIVE_REGULATION_OF_CELL_CYCLE_PROCESS | 0.00000 | 0.00007 | 0.64355 | 0.59835 | 1.27643 | 245 |
| GO_NEGATIVE_REGULATION_OF_CELL_DEATH | 0.00003 | 0.00100 | 0.57561 | 0.54182 | 1.16802 | 485 |
| GO_NEGATIVE_REGULATION_OF_MITOTIC_CELL_CYCLE | 0.00001 | 0.00038 | 0.59333 | 0.59044 | 1.25721 | 227 |
| GO_NEGATIVE_REGULATION_OF_WNT_SIGNALING_PATHWAY | 0.00000 | 0.00000 | 0.78819 | 0.70575 | 1.47598 | 110 |

| | | | | | | |
|---|---|---|---|---|---|---|
| GO_NEURON_TO_NEURON_SYNAPSE | 0.00191 | 0.03712 | 0.45506 | 0.57105 | 1.20758 | 158 |
| GO_NIK_NF_KAPPAB_SIGNALING | 0.00000 | 0.00000 | | 0.72766 | 1.51565 | 100 |
| GO_NON_CANONICAL_WNT_SIGNALING_PATHWAY | 0.00000 | 0.00000 | 0.80122 | 0.71948 | 1.49892 | 101 |
| GO_NUCLEAR_EXPORT | 0.00003 | 0.00087 | 0.57561 | 0.60962 | 1.29187 | 173 |
| GO_NUCLEAR_PERIPHERY | 0.00224 | 0.04171 | 0.43171 | 0.59698 | 1.24344 | 100 |
| GO_NUCLEAR_SPECK | 0.00100 | 0.02218 | 0.45506 | 0.54371 | 1.16369 | 289 |
| GO_NUCLEAR_TRANSCRIBED_MRNA_CATABOLIC_PROCESS | 0.00000 | 0.00000 | | 0.69635 | 1.47683 | 184 |
| GO_NUCLEAR_TRANSCRIBED_MRNA_CATABOLIC_PROCESS_NONSEN | 0.00000 | 0.00000 | | 0.76570 | 1.60309 | 113 |
| GO_NUCLEAR_TRANSPORT | 0.00000 | 0.00002 | 0.67496 | 0.60113 | 1.28494 | 266 |
| GO_NUCLEOBASE_CONTAINING_COMPOUND_TRANSPORT | 0.00015 | 0.00435 | 0.51885 | 0.58392 | 1.24015 | 201 |
| GO_NUCLEOBASE_CONTAINING_SMALL_MOLECULE_BIOSYNTHETIC_ | 0.00170 | 0.03413 | 0.45506 | 0.64761 | 1.32429 | 61 |
| GO_NUCLEOSIDE_METABOLIC_PROCESS | 0.00000 | 0.00010 | 0.62726 | 0.72677 | 1.47882 | 55 |
| GO_NUCLEOSIDE_MONOPHOSPHATE_BIOSYNTHETIC_PROCESS | 0.00179 | 0.03559 | 0.45506 | 0.71973 | 1.41321 | 29 |
| GO_NUCLEOSIDE_TRIPHOSPHATE_METABOLIC_PROCESS | 0.00198 | 0.03816 | 0.43171 | 0.64554 | 1.31681 | 58 |
| GO_NUCLEOSOMAL_DNA_BINDING | 0.00179 | 0.03559 | 0.45506 | 0.76345 | 1.46771 | 21 |
| GO_ORGANIC_CYCLIC_COMPOUND_CATABOLIC_PROCESS | 0.00000 | 0.00000 | | 0.66119 | 1.42335 | 428 |
| GO_PEPTIDASE_COMPLEX | 0.00000 | 0.00002 | 0.67496 | 0.70647 | 1.45886 | 79 |
| GO_PEPTIDE_TRANSMEMBRANE_TRANSPORTER_ACTIVITY | 0.00048 | 0.01212 | 0.49849 | 0.80155 | 1.52740 | 18 |
| GO_PIGMENT_GRANULE | 0.00110 | 0.02393 | 0.45506 | 0.63058 | 1.30028 | 75 |
| GO_POLYSOMAL_RIBOSOME | 0.00004 | 0.00129 | 0.55733 | 0.77137 | 1.52277 | 31 |
| GO_POLYSOME | 0.00000 | 0.00000 | 0.71951 | 0.75732 | 1.54484 | 58 |
| GO_POLY_PYRIMIDINE_TRACT_BINDING | 0.00010 | 0.00299 | 0.53843 | 0.82847 | 1.57869 | 18 |
| GO_POSITIVE_REGULATION_OF_CANONICAL_WNT_SIGNALING_PATH | 0.00000 | 0.00000 | 0.77494 | 0.72056 | 1.49736 | 95 |
| GO_POSITIVE_REGULATION_OF_CELLULAR_AMIDE_METABOLIC_PRC | 0.00003 | 0.00113 | 0.55733 | 0.63446 | 1.32540 | 108 |
| GO_POSITIVE_REGULATION_OF_CELLULAR_PROTEIN_CATABOLIC_PR | 0.00132 | 0.02767 | 0.45506 | 0.60499 | 1.26039 | 101 |
| GO_POSITIVE_REGULATION_OF_CELLULAR_PROTEIN_LOCALIZATION | 0.00043 | 0.01089 | 0.49849 | 0.56341 | 1.19979 | 221 |
| GO_POSITIVE_REGULATION_OF_CELL_DEATH | 0.00121 | 0.02604 | 0.45506 | 0.53675 | 1.15326 | 356 |
| GO_POSITIVE_REGULATION_OF_DEFENSE_RESPONSE | 0.00000 | 0.00001 | 0.67496 | 0.63542 | 1.34386 | 160 |
| GO_POSITIVE_REGULATION_OF_DNA_BIOSYNTHETIC_PROCESS | 0.00042 | 0.01082 | 0.49849 | 0.67049 | 1.36229 | 52 |
| GO_POSITIVE_REGULATION_OF_ESTABLISHMENT_OF_PROTEIN_LOC/ | 0.00015 | 0.00435 | 0.51885 | 0.57633 | 1.22643 | 215 |
| GO_POSITIVE_REGULATION_OF_IMMUNE_RESPONSE | 0.00000 | 0.00018 | 0.61053 | 0.58426 | 1.25009 | 275 |
| GO_POSITIVE_REGULATION_OF_IMMUNE_SYSTEM_PROCESS | 0.00054 | 0.01321 | 0.47727 | 0.53963 | 1.16081 | 383 |
| GO_POSITIVE_REGULATION_OF_INNATE_IMMUNE_RESPONSE | 0.00000 | 0.00000 | 0.77494 | 0.69752 | 1.45922 | 111 |
| GO_POSITIVE_REGULATION_OF_MRNA_PROCESSING | 0.00250 | 0.04503 | 0.43171 | 0.72194 | 1.41150 | 27 |
| GO_POSITIVE_REGULATION_OF_MRNA_SPLICING_VIA_SPLICEOSOM | 0.00080 | 0.01853 | 0.47727 | 0.80311 | 1.52701 | 17 |
| GO_POSITIVE_REGULATION_OF_PROTEIN_LOCALIZATION_TO_NUCLE | 0.00008 | 0.00251 | 0.53843 | 0.68690 | 1.40602 | 62 |
| GO_POSITIVE_REGULATION_OF_RESPONSE_TO_BIOTIC_STIMULUS | 0.00000 | 0.00000 | 0.71951 | 0.67566 | 1.41741 | 126 |
| GO_POSITIVE_REGULATION_OF_RESPONSE_TO_EXTERNAL_STIMULU | 0.00002 | 0.00067 | 0.57561 | 0.59030 | 1.25375 | 197 |
| GO_POSITIVE_REGULATION_OF_TELOMERASE_RNA_LOCALIZATION_ | 0.00000 | 0.00010 | 0.62726 | 0.90826 | 1.70676 | 15 |
| GO_POSITIVE_REGULATION_OF_TRANSLATION | 0.00001 | 0.00047 | 0.59333 | 0.66039 | 1.36939 | 91 |
| GO_POSITIVE_REGULATION_OF_VIRAL_PROCESS | 0.00086 | 0.01948 | 0.47727 | 0.63553 | 1.31144 | 76 |
| GO_POSITIVE_REGULATION_OF_WNT_SIGNALING_PATHWAY | 0.00000 | 0.00000 | 0.76146 | 0.70149 | 1.46611 | 109 |
| GO_POSTSYNAPSE | 0.00060 | 0.01460 | 0.47727 | 0.55298 | 1.18055 | 257 |
| GO_POSTTRANSCRIPTIONAL_REGULATION_OF_GENE_EXPRESSION | 0.00000 | 0.00000 | | 0.64783 | 1.39537 | 448 |
| GO_POST_TRANSLATIONAL_PROTEIN_MODIFICATION | 0.00007 | 0.00226 | 0.53843 | 0.57454 | 1.22314 | 224 |
| GO_PRERIBOSOME | 0.00001 | 0.00031 | 0.59333 | 0.69153 | 1.42154 | 69 |
| GO_PRE_MRNA_BINDING | 0.00224 | 0.04171 | 0.43171 | 0.71024 | 1.39160 | 28 |
| GO_PROTEASOMAL_PROTEIN_CATABOLIC_PROCESS | 0.00001 | 0.00025 | 0.61053 | 0.56338 | 1.21151 | 375 |
| GO_PROTEASOMAL_UBIQUITIN_INDEPENDENT_PROTEIN_CATABOLI( | 0.00000 | 0.00007 | 0.64355 | 0.87516 | 1.67319 | 19 |
| GO_PROTEASOME_ACCESSORY_COMPLEX | 0.00000 | 0.00004 | 0.65944 | 0.84871 | 1.64698 | 24 |
| GO_PROTEASOME_CORE_COMPLEX | 0.00001 | 0.00023 | 0.61053 | 0.88018 | 1.67356 | 17 |
| GO_PROTEIN_CONTAINING_COMPLEX_DISASSEMBLY | 0.00028 | 0.00767 | 0.49849 | 0.55997 | 1.19455 | 247 |
| GO_PROTEIN_CONTAINING_COMPLEX_LOCALIZATION | 0.00054 | 0.01321 | 0.47727 | 0.57413 | 1.21934 | 201 |
| GO_PROTEIN_DNA_COMPLEX_SUBUNIT_ORGANIZATION | 0.00030 | 0.00800 | 0.49849 | 0.58315 | 1.23470 | 168 |
| GO_PROTEIN_FOLDING | 0.00000 | 0.00007 | 0.64355 | 0.63015 | 1.33287 | 161 |
| GO_PROTEIN_IMPORT | 0.00019 | 0.00540 | 0.51885 | 0.60827 | 1.27571 | 125 |
| GO_PROTEIN_LOCALIZATION_TO_CHROMOSOME | 0.00076 | 0.01776 | 0.47727 | 0.65067 | 1.33684 | 68 |
| GO_PROTEIN_LOCALIZATION_TO_ENDOPLASMIC_RETICULUM | 0.00000 | 0.00000 | | 0.73383 | 1.53844 | 123 |
| GO_PROTEIN_LOCALIZATION_TO_MEMBRANE | 0.00000 | 0.00000 | 0.71951 | 0.58090 | 1.25009 | 396 |
| GO_PROTEIN_LOCALIZATION_TO_MITOCHONDRION | 0.00052 | 0.01293 | 0.47727 | 0.61042 | 1.27370 | 105 |
| GO_PROTEIN_LOCALIZATION_TO_NUCLEUS | 0.00000 | 0.00019 | 0.61053 | 0.61011 | 1.29366 | 178 |
| GO_PROTEIN_MODIFICATION_BY_SMALL_PROTEIN_REMOVAL | 0.00001 | 0.00050 | 0.59333 | 0.59102 | 1.25812 | 218 |
| GO_PROTEIN_POLYUBIQUITINATION | 0.00048 | 0.01212 | 0.49849 | 0.56288 | 1.20031 | 240 |
| GO_PROTEIN_REFOLDING | 0.00194 | 0.03758 | 0.45506 | 0.76564 | 1.46380 | 19 |
| GO_PROTEIN_STABILIZATION | 0.00087 | 0.01960 | 0.47727 | 0.59140 | 1.24248 | 130 |
| GO_PROTEIN_TARGETING | 0.00000 | 0.00000 | | 0.62787 | 1.34680 | 319 |
| GO_PROTEIN_TARGETING_TO_MEMBRANE | 0.00000 | 0.00000 | | 0.70426 | 1.48760 | 152 |
| GO_PURINE_NUCLEOSIDE_METABOLIC_PROCESS | 0.00019 | 0.00533 | 0.51885 | 0.73456 | 1.46536 | 35 |
| GO_PURINE_NUCLEOSIDE_TRIPHOSPHATE_METABOLIC_PROCESS | 0.00228 | 0.04212 | 0.43171 | 0.67215 | 1.35401 | 43 |
| GO_REGENERATION | 0.00010 | 0.00301 | 0.53843 | 0.65986 | 1.36239 | 81 |
| GO_REGULATION_OF_ANIMAL_ORGAN_MORPHOGENESIS | 0.00000 | 0.00000 | 0.78819 | 0.68685 | 1.44053 | 124 |
| GO_REGULATION_OF_APOPTOTIC_SIGNALING_PATHWAY | 0.00272 | 0.04888 | 0.43171 | 0.54886 | 1.16848 | 224 |
| GO_REGULATION_OF_CELLULAR_AMIDE_METABOLIC_PROCESS | 0.00000 | 0.00000 | | 0.62318 | 1.33432 | 300 |
| GO_REGULATION_OF_CELLULAR_AMINE_METABOLIC_PROCESS | 0.00000 | 0.00000 | | 0.81263 | 1.65473 | 56 |
| GO_REGULATION_OF_CELLULAR_AMINO_ACID_METABOLIC_PROCES | 0.00000 | 0.00000 | | 0.82581 | 1.67787 | 52 |
| GO_REGULATION_OF_CELLULAR_KETONE_METABOLIC_PROCESS | 0.00000 | 0.00001 | 0.67496 | 0.66983 | 1.40330 | 115 |

| | | | | | | |
|---|---|---|---|---|---|---|
| GO_REGULATION_OF_CELLULAR_PROTEIN_CATABOLIC_PROCESS | 0.00017 | 0.00485 | 0.51885 | 0.58451 | 1.23945 | 185 |
| GO_REGULATION_OF_CELLULAR_PROTEIN_LOCALIZATION | 0.00003 | 0.00095 | 0.57561 | 0.56060 | 1.20382 | 351 |
| GO_REGULATION_OF_CELL_CYCLE_G2_M_PHASE_TRANSITION | 0.00000 | 0.00000 | 0.71951 | 0.63645 | 1.34927 | 175 |
| GO_REGULATION_OF_CELL_CYCLE_PHASE_TRANSITION | 0.00003 | 0.00112 | 0.55733 | 0.56381 | 1.21020 | 330 |
| GO_REGULATION_OF_CHROMOSOME_ORGANIZATION | 0.00157 | 0.03201 | 0.45506 | 0.54604 | 1.16573 | 257 |
| GO_REGULATION_OF_DEFENSE_RESPONSE | 0.00032 | 0.00832 | 0.49849 | 0.55348 | 1.18423 | 283 |
| GO_REGULATION_OF_DNA_TEMPLATED_TRANSCRIPTION_IN_RESPO | 0.00000 | 0.00000 | 0.78819 | 0.72815 | 1.50990 | 91 |
| GO_REGULATION_OF_HEMATOPOIETIC_PROGENITOR_CELL_DIFFERE | 0.00000 | 0.00000 | 0.76146 | 0.74664 | 1.53917 | 72 |
| GO_REGULATION_OF_HEMATOPOIETIC_STEM_CELL_DIFFERENTIATIC | 0.00000 | 0.00000 | 0.81404 | 0.79075 | 1.61858 | 62 |
| GO_REGULATION_OF_HEMOPOIESIS | 0.00005 | 0.00155 | 0.55733 | 0.57825 | 1.23105 | 224 |
| GO_REGULATION_OF_IMMUNE_RESPONSE | 0.00030 | 0.00804 | 0.49849 | 0.54624 | 1.17428 | 363 |
| GO_REGULATION_OF_INNATE_IMMUNE_RESPONSE | 0.00000 | 0.00000 | 0.76146 | 0.66017 | 1.39519 | 156 |
| GO_REGULATION_OF_INTRINSIC_APOPTOTIC_SIGNALING_PATHWAY | 0.00067 | 0.01603 | 0.47727 | 0.61401 | 1.28146 | 106 |
| GO_REGULATION_OF_MITOTIC_CELL_CYCLE | 0.00001 | 0.00048 | 0.59333 | 0.55125 | 1.18734 | 448 |
| GO_REGULATION_OF_MORPHOGENESIS_OF_AN_EPITHELIUM | 0.00000 | 0.00000 | 0.71951 | 0.69509 | 1.45110 | 107 |
| GO_REGULATION_OF_MRNA_CATABOLIC_PROCESS | 0.00000 | 0.00000 | | 0.68990 | 1.45887 | 159 |
| GO_REGULATION_OF_MRNA_METABOLIC_PROCESS | 0.00000 | 0.00000 | | 0.65454 | 1.39735 | 257 |
| GO_REGULATION_OF_MRNA_PROCESSING | 0.00005 | 0.00146 | 0.55733 | 0.62371 | 1.30482 | 111 |
| GO_REGULATION_OF_MRNA_SPLICING_VIA_SPLICEOSOME | 0.00011 | 0.00313 | 0.53843 | 0.65769 | 1.35772 | 78 |
| GO_REGULATION_OF_PEPTIDE_TRANSPORT | 0.00100 | 0.02218 | 0.45506 | 0.53892 | 1.15735 | 343 |
| GO_REGULATION_OF_PROTEASOMAL_PROTEIN_CATABOLIC_PROCES | 0.00191 | 0.03712 | 0.45506 | 0.57718 | 1.21626 | 144 |
| GO_REGULATION_OF_PROTEIN_CATABOLIC_PROCESS | 0.00001 | 0.00033 | 0.59333 | 0.58001 | 1.24018 | 270 |
| GO_REGULATION_OF_PROTEIN_LOCALIZATION_TO_NUCLEUS | 0.00003 | 0.00088 | 0.57561 | 0.65523 | 1.35434 | 84 |
| GO_REGULATION_OF_PROTEIN_STABILITY | 0.00224 | 0.04171 | 0.43171 | 0.55758 | 1.18487 | 207 |
| GO_REGULATION_OF_PROTEOLYSIS | 0.00011 | 0.00328 | 0.53843 | 0.54849 | 1.17936 | 369 |
| GO_REGULATION_OF_RESPONSE_TO_BIOTIC_STIMULUS | 0.00000 | 0.00003 | 0.65944 | 0.61854 | 1.31372 | 195 |
| GO_REGULATION_OF_RNA_SPLICING | 0.00004 | 0.00125 | 0.55733 | 0.63900 | 1.33401 | 107 |
| GO_REGULATION_OF_SMALL_MOLECULE_METABOLIC_PROCESS | 0.00001 | 0.00033 | 0.59333 | 0.58627 | 1.25291 | 264 |
| GO_REGULATION_OF_STEM_CELL_DIFFERENTIATION | 0.00000 | 0.00000 | 0.78819 | 0.73685 | 1.52305 | 84 |
| GO_REGULATION_OF_SYMBIOTIC_PROCESS | 0.00031 | 0.00824 | 0.49849 | 0.61161 | 1.28191 | 119 |
| GO_REGULATION_OF_TRANSCRIPTION_FROM_RNA_POLYMERASE_II | 0.00000 | 0.00000 | | 0.79026 | 1.61905 | 63 |
| GO_REGULATION_OF_TRANSLATIONAL_INITIATION | 0.00000 | 0.00012 | 0.62726 | 0.71481 | 1.46045 | 60 |
| GO_REGULATION_OF_TRANSLATION_IN_RESPONSE_TO_STRESS | 0.00194 | 0.03758 | 0.45506 | 0.78670 | 1.49580 | 17 |
| GO_REGULATION_OF_VIRAL_GENOME_REPLICATION | 0.00246 | 0.04451 | 0.43171 | 0.66846 | 1.35180 | 46 |
| GO_REGULATION_OF_VIRAL_LIFE_CYCLE | 0.00202 | 0.03848 | 0.43171 | 0.62782 | 1.29211 | 70 |
| GO_REGULATION_OF_WNT_SIGNALING_PATHWAY | 0.00000 | 0.00001 | 0.67496 | 0.62176 | 1.32029 | 193 |
| GO_RESPONSE_TO_DRUG | 0.00001 | 0.00053 | 0.59333 | 0.60112 | 1.27415 | 181 |
| GO_RESPONSE_TO_HEAT | 0.00035 | 0.00908 | 0.49849 | 0.60967 | 1.27562 | 112 |
| GO_RESPONSE_TO_INTERLEUKIN_1 | 0.00000 | 0.00000 | 0.80122 | 0.71880 | 1.49750 | 101 |
| GO_RESPONSE_TO_INTERLEUKIN_12 | 0.00085 | 0.01943 | 0.47727 | 0.71727 | 1.42689 | 34 |
| GO_RESPONSE_TO_INTERLEUKIN_4 | 0.00003 | 0.00086 | 0.57561 | 0.83786 | 1.60912 | 20 |
| GO_RESPONSE_TO_LEUKEMIA_INHIBITORY_FACTOR | 0.00080 | 0.01853 | 0.47727 | 0.63572 | 1.30986 | 73 |
| GO_RESPONSE_TO_ORGANIC_CYCLIC_COMPOUND | 0.00242 | 0.04413 | 0.43171 | 0.52454 | 1.12939 | 436 |
| GO_RESPONSE_TO_OXYGEN_LEVELS | 0.00001 | 0.00027 | 0.61053 | 0.58836 | 1.25321 | 229 |
| GO_RESPONSE_TO_TEMPERATURE_STIMULUS | 0.00167 | 0.03363 | 0.45506 | 0.59020 | 1.24094 | 133 |
| GO_RESPONSE_TO_TUMOR_NECROSIS_FACTOR | 0.00001 | 0.00025 | 0.61053 | 0.61272 | 1.29583 | 160 |
| GO_RIBONUCLEOPROTEIN_COMPLEX_BINDING | 0.00000 | 0.00006 | 0.64355 | 0.65634 | 1.37308 | 111 |
| GO_RIBONUCLEOPROTEIN_COMPLEX_BIOGENESIS | 0.00000 | 0.00000 | | 0.66654 | 1.43387 | 386 |
| GO_RIBONUCLEOPROTEIN_COMPLEX_SUBUNIT_ORGANIZATION | 0.00000 | 0.00000 | | 0.69056 | 1.46313 | 172 |
| GO_RIBONUCLEOPROTEIN_GRANULE | 0.00004 | 0.00142 | 0.55733 | 0.60664 | 1.28351 | 163 |
| GO_RIBONUCLEOSIDE_METABOLIC_PROCESS | 0.00009 | 0.00277 | 0.53843 | 0.74139 | 1.48045 | 36 |
| GO_RIBONUCLEOSIDE_MONOPHOSPHATE_BIOSYNTHETIC_PROCESS | 0.00213 | 0.04005 | 0.43171 | 0.75989 | 1.46085 | 21 |
| GO_RIBONUCLEOSIDE_TRIPHOSPHATE_METABOLIC_PROCESS | 0.00132 | 0.02767 | 0.45506 | 0.68134 | 1.36969 | 42 |
| GO_RIBOSOMAL_LARGE_SUBUNIT_ASSEMBLY | 0.00025 | 0.00678 | 0.49849 | 0.76151 | 1.48887 | 27 |
| GO_RIBOSOMAL_LARGE_SUBUNIT_BIOGENESIS | 0.00000 | 0.00001 | 0.69013 | 0.73948 | 1.51708 | 65 |
| GO_RIBOSOMAL_SMALL_SUBUNIT_BIOGENESIS | 0.00017 | 0.00496 | 0.51885 | 0.66140 | 1.35961 | 69 |
| GO_RIBOSOMAL_SUBUNIT | 0.00000 | 0.00000 | | 0.68625 | 1.45287 | 166 |
| GO_RIBOSOME | 0.00000 | 0.00000 | | 0.68345 | 1.45153 | 198 |
| GO_RIBOSOME_ASSEMBLY | 0.00000 | 0.00005 | 0.64355 | 0.73147 | 1.49319 | 59 |
| GO_RIBOSOME_BINDING | 0.00130 | 0.02750 | 0.45506 | 0.67935 | 1.37382 | 46 |
| GO_RIBOSOME_BIOGENESIS | 0.00000 | 0.00000 | | 0.65208 | 1.39492 | 274 |
| GO_RNA_CATABOLIC_PROCESS | 0.00000 | 0.00000 | | 0.69254 | 1.48642 | 329 |
| GO_RNA_DEPENDENT_DNA_BIOSYNTHETIC_PROCESS | 0.00179 | 0.03559 | 0.45506 | 0.64629 | 1.31932 | 59 |
| GO_RNA_EXPORT_FROM_NUCLEUS | 0.00002 | 0.00067 | 0.57561 | 0.63094 | 1.32360 | 126 |
| GO_RNA_LOCALIZATION | 0.00000 | 0.00000 | 0.71951 | 0.62878 | 1.33577 | 196 |
| GO_RNA_POLYMERASE_II_SPECIFIC_DNA_BINDING_TRANSCRIPTION | 0.00133 | 0.02796 | 0.45506 | 0.57341 | 1.21448 | 169 |
| GO_RNA_SPLICING | 0.00000 | 0.00000 | 0.81404 | 0.60493 | 1.29974 | 356 |
| GO_RNA_SPLICING_VIA_TRANSESTERIFICATION_REACTIONS | 0.00000 | 0.00000 | 0.83909 | 0.62464 | 1.33691 | 289 |
| GO_ROUGH_ENDOPLASMIC_RETICULUM | 0.00051 | 0.01276 | 0.47727 | 0.66367 | 1.34959 | 53 |
| GO_RRNA_BINDING | 0.00000 | 0.00016 | 0.62726 | 0.72680 | 1.47796 | 53 |
| GO_RRNA_METABOLIC_PROCESS | 0.00000 | 0.00000 | 0.77494 | 0.64000 | 1.36013 | 208 |
| GO_SCF_DEPENDENT_PROTEASOMAL_UBIQUITIN_DEPENDENT_PRO | 0.00000 | 0.00000 | 0.80122 | 0.75877 | 1.56420 | 72 |
| GO_SECRETORY_GRANULE | 0.00000 | 0.00001 | 0.69013 | 0.57393 | 1.23535 | 402 |
| GO_SECRETORY_VESICLE | 0.00000 | 0.00006 | 0.64355 | 0.55854 | 1.20347 | 463 |
| GO_SERINE_FAMILY_AMINO_ACID_METABOLIC_PROCESS | 0.00168 | 0.03388 | 0.45506 | 0.75855 | 1.46649 | 23 |
| GO_SINGLE_FERTILIZATION | 0.00024 | 0.00669 | 0.51885 | 0.67704 | 1.37862 | 56 |

| | | | | | | |
|---|---|---|---|---|---|---|
| GO_SINGLE_STRANDED_RNA_BINDING | 0.00018 | 0.00516 | 0.51885 | 0.68121 | 1.38712 | 56 |
| GO_SMALL_RIBOSOMAL_SUBUNIT | 0.00004 | 0.00127 | 0.55733 | 0.68455 | 1.40439 | 65 |
| GO_SMALL_SUBUNIT_PROCESSOME | 0.00242 | 0.04413 | 0.43171 | 0.70756 | 1.40088 | 32 |
| GO_SPERM_EGG_RECOGNITION | 0.00000 | 0.00000 | 0.73376 | 0.86796 | 1.69212 | 26 |
| GO_SPINDLE_MIDZONE | 0.00135 | 0.02824 | 0.45506 | 0.73086 | 1.42895 | 27 |
| GO_SPLICEOSOMAL_COMPLEX | 0.00000 | 0.00016 | 0.62726 | 0.62239 | 1.31683 | 163 |
| GO_STEM_CELL_DIFFERENTIATION | 0.00000 | 0.00000 | 0.83909 | 0.69112 | 1.45661 | 140 |
| GO_STRUCTURAL_CONSTITUENT_OF_RIBOSOME | 0.00000 | 0.00000 | | 0.69183 | 1.45937 | 148 |
| GO_STRUCTURAL_MOLECULE_ACTIVITY | 0.00000 | 0.00000 | | 0.63791 | 1.36788 | 315 |
| GO_TELOMERASE_RNA_LOCALIZATION | 0.00001 | 0.00042 | 0.59333 | 0.85107 | 1.62714 | 19 |
| GO_TELOMERE_MAINTENANCE_VIA_TELOMERE_LENGTHENING | 0.00145 | 0.02982 | 0.45506 | 0.63567 | 1.30569 | 67 |
| GO_THREONINE_TYPE_ENDOPEPTIDASE_ACTIVITY | 0.00000 | 0.00012 | 0.62726 | 0.88878 | 1.68990 | 17 |
| GO_THREONINE_TYPE_PEPTIDASE_ACTIVITY | 0.00001 | 0.00021 | 0.61053 | 0.85571 | 1.64507 | 21 |
| GO_TRANSCRIPTION_COREGULATOR_ACTIVITY | 0.00165 | 0.03338 | 0.45506 | 0.53903 | 1.15664 | 322 |
| GO_TRANSCRIPTION_FACTOR_BINDING | 0.00001 | 0.00038 | 0.59333 | 0.55943 | 1.20374 | 393 |
| GO_TRANSLATIONAL_ELONGATION | 0.00130 | 0.02750 | 0.45506 | 0.59838 | 1.25417 | 119 |
| GO_TRANSLATIONAL_INITIATION | 0.00000 | 0.00000 | | 0.77499 | 1.63935 | 162 |
| GO_TRANSLATION_ELONGATION_FACTOR_ACTIVITY | 0.00006 | 0.00180 | 0.55733 | 0.86819 | 1.63145 | 15 |
| GO_TRANSLATION_FACTOR_ACTIVITY_RNA_BINDING | 0.00000 | 0.00000 | | 0.80800 | 1.66007 | 68 |
| GO_TRANSLATION_INITIATION_FACTOR_ACTIVITY | 0.00000 | 0.00000 | 0.76146 | 0.81749 | 1.64681 | 43 |
| GO_TRANSLATION_INITIATION_FACTOR_BINDING | 0.00021 | 0.00581 | 0.51885 | 0.75853 | 1.48938 | 29 |
| GO_TRANSLATION_PREINITIATION_COMPLEX | 0.00001 | 0.00038 | 0.59333 | 0.88066 | 1.66268 | 16 |
| GO_TRANSLATION_REGULATOR_ACTIVITY | 0.00000 | 0.00000 | | 0.76829 | 1.60154 | 103 |
| GO_TRANSLATION_REGULATOR_ACTIVITY_NUCLEIC_ACID_BINDING | 0.00000 | 0.00000 | | 0.78830 | 1.62757 | 81 |
| GO_TUMOR_NECROSIS_FACTOR_MEDIATED_SIGNALING_PATHWAY | 0.00000 | 0.00000 | 0.71951 | 0.69228 | 1.44225 | 101 |
| GO_T_CELL_RECEPTOR_SIGNALING_PATHWAY | 0.00000 | 0.00000 | 0.73376 | 0.68231 | 1.42850 | 113 |
| GO_U2_TYPE_SPLICEOSOMAL_COMPLEX | 0.00126 | 0.02702 | 0.45506 | 0.61958 | 1.27954 | 82 |
| GO_UBIQUITIN_LIKE_PROTEIN_LIGASE_BINDING | 0.00002 | 0.00080 | 0.57561 | 0.58939 | 1.25269 | 210 |
| GO_UNFOLDED_PROTEIN_BINDING | 0.00000 | 0.00006 | 0.64355 | 0.68758 | 1.42102 | 83 |
| GO_VACUOLAR_LUMEN | 0.00007 | 0.00207 | 0.53843 | 0.64725 | 1.34503 | 95 |
| GO_VACUOLE_ORGANIZATION | 0.00023 | 0.00647 | 0.51885 | 0.60263 | 1.26338 | 123 |
| GO_VESICLE_LUMEN | 0.00000 | 0.00000 | | 0.69418 | 1.47187 | 177 |
| GO_VIRAL_GENE_EXPRESSION | 0.00000 | 0.00000 | | 0.71733 | 1.51880 | 168 |
| GO_VIRAL_GENOME_REPLICATION | 0.00119 | 0.02574 | 0.45506 | 0.63706 | 1.30696 | 65 |
| GO_VIRAL_LIFE_CYCLE | 0.00053 | 0.01310 | 0.47727 | 0.57367 | 1.21837 | 194 |
| HP_ABNORMALITY_OF_THE_AUDITORY_CANAL | 0.00185 | 0.03643 | 0.45506 | 0.69752 | 1.39146 | 35 |
| HP_ABNORMAL_CIRCULATING_SERINE_FAMILY_AMINO_ACID_CONC | 0.00098 | 0.02195 | 0.45506 | 0.80510 | 1.52003 | 16 |
| HP_ACUTE_LEUKEMIA | 0.00111 | 0.02424 | 0.45506 | 0.64224 | 1.31881 | 66 |
| HP_ATRESIA_OF_THE_EXTERNAL_AUDITORY_CANAL | 0.00080 | 0.01853 | 0.47727 | 0.75347 | 1.46569 | 25 |
| HP_DEPRESSED_NASAL_RIDGE | 0.00063 | 0.01507 | 0.47727 | 0.63704 | 1.31324 | 72 |
| HP_MACROCYTIC_ANEMIA | 0.00110 | 0.02393 | 0.45506 | 0.68374 | 1.37452 | 42 |
| HP_MACROGYRIA | 0.00080 | 0.01853 | 0.47727 | 0.80248 | 1.52582 | 17 |
| **Cluster 9** | | | | | | |
| pathway | pval | padj | log2err | ES | NES | size |
| GO_ALTERNATIVE_MRNA_SPLICING_VIA_SPLICEOSOME | 0.00002 | 0.00124 | 0.57561 | 0.69102 | 1.37373 | 62 |
| GO_ANAPHASE_PROMOTING_COMPLEX_DEPENDENT_CATABOLIC_P | 0.00010 | 0.00546 | 0.53843 | 0.67042 | 1.33839 | 73 |
| GO_ANCHORING_JUNCTION | 0.00018 | 0.00847 | 0.51885 | 0.54451 | 1.13569 | 498 |
| GO_CATALYTIC_STEP_2_SPLICEOSOME | 0.00005 | 0.00333 | 0.55733 | 0.67454 | 1.35199 | 78 |
| GO_CELLULAR_NITROGEN_COMPOUND_CATABOLIC_PROCESS | 0.00000 | 0.00000 | | 0.67366 | 1.40290 | 435 |
| GO_CELLULAR_RESPONSE_TO_HEAT | 0.00156 | 0.04394 | 0.45506 | 0.62128 | 1.25035 | 87 |
| GO_CELLULAR_RESPONSE_TO_OXYGEN_LEVELS | 0.00122 | 0.03764 | 0.45506 | 0.58776 | 1.20167 | 152 |
| GO_CELL_CYCLE_CHECKPOINT | 0.00006 | 0.00340 | 0.55733 | 0.60413 | 1.23963 | 174 |
| GO_CELL_CYCLE_G1_S_PHASE_TRANSITION | 0.00106 | 0.03434 | 0.45506 | 0.57088 | 1.17451 | 212 |
| GO_CELL_CYCLE_PHASE_TRANSITION | 0.00000 | 0.00010 | 0.64355 | 0.56669 | 1.18106 | 479 |
| GO_CELL_DIVISION | 0.00000 | 0.00005 | 0.67496 | 0.57391 | 1.19519 | 439 |
| GO_CELL_SUBSTRATE_JUNCTION | 0.00000 | 0.00000 | 0.80122 | 0.62343 | 1.29022 | 290 |
| GO_CHROMATIN_BINDING | 0.00014 | 0.00664 | 0.51885 | 0.55841 | 1.16066 | 377 |
| GO_CHROMOSOMAL_REGION | 0.00073 | 0.02559 | 0.47727 | 0.56160 | 1.16174 | 281 |
| GO_CHROMOSOME_CENTROMERIC_REGION | 0.00080 | 0.02719 | 0.47727 | 0.59280 | 1.21509 | 166 |
| GO_CHROMOSOME_SEGREGATION | 0.00000 | 0.00036 | 0.61053 | 0.59425 | 1.22693 | 264 |
| GO_COLLAGEN_CATABOLIC_PROCESS | 0.00013 | 0.00664 | 0.51885 | 0.82641 | 1.53931 | 18 |
| GO_CONDENSED_CHROMOSOME | 0.00033 | 0.01454 | 0.49849 | 0.58821 | 1.20651 | 170 |
| GO_COTRANSLATIONAL_PROTEIN_TARGETING_TO_MEMBRANE | 0.00000 | 0.00000 | | 0.87723 | 1.76838 | 91 |
| GO_COVALENT_CHROMATIN_MODIFICATION | 0.00030 | 0.01305 | 0.49849 | 0.55753 | 1.15669 | 349 |
| GO_CYTOKINE_MEDIATED_SIGNALING_PATHWAY | 0.00050 | 0.02061 | 0.47727 | 0.55140 | 1.14573 | 369 |
| GO_CYTOPLASMIC_STRESS_GRANULE | 0.00026 | 0.01172 | 0.49849 | 0.67135 | 1.32997 | 59 |
| GO_CYTOPLASMIC_TRANSLATION | 0.00000 | 0.00000 | | 0.75709 | 1.52367 | 87 |
| GO_CYTOSOLIC_LARGE_RIBOSOMAL_SUBUNIT | 0.00000 | 0.00000 | | 0.89387 | 1.76498 | 52 |
| GO_CYTOSOLIC_RIBOSOME | 0.00000 | 0.00000 | | 0.89827 | 1.81511 | 96 |
| GO_CYTOSOLIC_SMALL_RIBOSOMAL_SUBUNIT | 0.00000 | 0.00000 | | 0.91979 | 1.79600 | 41 |
| GO_DNA_BINDING_TRANSCRIPTION_FACTOR_BINDING | 0.00139 | 0.04062 | 0.45506 | 0.56441 | 1.16344 | 225 |
| GO_DNA_CONFORMATION_CHANGE | 0.00052 | 0.02107 | 0.47727 | 0.57678 | 1.18809 | 220 |
| GO_DNA_DAMAGE_RESPONSE_DETECTION_OF_DNA_DAMAGE | 0.00170 | 0.04764 | 0.45506 | 0.69847 | 1.35229 | 34 |
| GO_DNA_INTEGRITY_CHECKPOINT | 0.00176 | 0.04873 | 0.45506 | 0.59395 | 1.21100 | 126 |
| GO_DNA_PACKAGING | 0.00088 | 0.02948 | 0.47727 | 0.60214 | 1.22750 | 125 |
| GO_DNA_REPAIR | 0.00141 | 0.04076 | 0.45506 | 0.53837 | 1.12082 | 427 |

| | | | | | | |
|---|---|---|---|---|---|---|
| GO_DNA_REPLICATION | 0.00137 | 0.04028 | 0.45506 | 0.56454 | 1.16287 | 221 |
| GO_DNA_TEMPLATED_TRANSCRIPTION_ELONGATION | 0.00024 | 0.01078 | 0.51885 | 0.64240 | 1.29754 | 95 |
| GO_DNA_TEMPLATED_TRANSCRIPTION_INITIATION | 0.00003 | 0.00199 | 0.57561 | 0.60764 | 1.24683 | 174 |
| GO_DNA_TEMPLATED_TRANSCRIPTION_TERMINATION | 0.00041 | 0.01745 | 0.49849 | 0.65816 | 1.31114 | 70 |
| GO_ESTABLISHMENT_OF_PROTEIN_LOCALIZATION_TO_ENDOPLASM | 0.00000 | 0.00000 | | 0.86045 | 1.74199 | 102 |
| GO_ESTABLISHMENT_OF_PROTEIN_LOCALIZATION_TO_MEMBRANE | 0.00000 | 0.00000 | | 0.69871 | 1.44170 | 244 |
| GO_ESTABLISHMENT_OF_PROTEIN_LOCALIZATION_TO_ORGANELLE | 0.00000 | 0.00000 | | 0.63830 | 1.32891 | 417 |
| GO_ESTABLISHMENT_OF_RNA_LOCALIZATION | 0.00000 | 0.00002 | 0.69013 | 0.64230 | 1.31679 | 167 |
| GO_FIBRILLAR_CENTER | 0.00070 | 0.02492 | 0.47727 | 0.62052 | 1.25398 | 97 |
| GO_FICOLIN_1_RICH_GRANULE | 0.00000 | 0.00002 | 0.69013 | 0.70030 | 1.41379 | 94 |
| GO_HEAT_SHOCK_PROTEIN_BINDING | 0.00065 | 0.02325 | 0.47727 | 0.62844 | 1.26606 | 89 |
| GO_HISTONE_BINDING | 0.00135 | 0.03994 | 0.45506 | 0.57866 | 1.18770 | 179 |
| GO_IMPORT_INTO_NUCLEUS | 0.00100 | 0.03309 | 0.45506 | 0.60684 | 1.23462 | 118 |
| GO_INTRINSIC_APOPTOTIC_SIGNALING_PATHWAY_BY_P53_CLASS_M | 0.00124 | 0.03781 | 0.45506 | 0.67068 | 1.32454 | 50 |
| GO_LARGE_RIBOSOMAL_SUBUNIT | 0.00000 | 0.00000 | | 0.76188 | 1.54387 | 105 |
| GO_LEUKOCYTE_MEDIATED_IMMUNITY | 0.00152 | 0.04311 | 0.45506 | 0.54290 | 1.12920 | 393 |
| GO_MATURATION_OF_LSU_RRNA | 0.00013 | 0.00658 | 0.51885 | 0.76449 | 1.45464 | 27 |
| GO_MATURATION_OF_SSU_RRNA_FROM_TRICISTRONIC_RRNA_TRA | 0.00122 | 0.03764 | 0.45506 | 0.70404 | 1.36308 | 34 |
| GO_METAPHASE_ANAPHASE_TRANSITION_OF_CELL_CYCLE | 0.00023 | 0.01032 | 0.51885 | 0.69177 | 1.36591 | 52 |
| GO_MITOTIC_CELL_CYCLE_CHECKPOINT | 0.00053 | 0.02115 | 0.47727 | 0.60858 | 1.24282 | 133 |
| GO_MITOTIC_NUCLEAR_DIVISION | 0.00012 | 0.00634 | 0.51885 | 0.59169 | 1.21694 | 207 |
| GO_MITOTIC_SISTER_CHROMATID_SEGREGATION | 0.00007 | 0.00410 | 0.53843 | 0.62053 | 1.26726 | 134 |
| GO_MITOTIC_SPINDLE | 0.00113 | 0.03594 | 0.45506 | 0.61148 | 1.23782 | 101 |
| GO_MODIFICATION_DEPENDENT_MACROMOLECULE_CATABOLIC_PR | 0.00001 | 0.00042 | 0.61053 | 0.55923 | 1.16623 | 495 |
| GO_MRNA_3_END_PROCESSING | 0.00013 | 0.00664 | 0.51885 | 0.64488 | 1.29846 | 88 |
| GO_MRNA_5_UTR_BINDING | 0.00016 | 0.00745 | 0.51885 | 0.79669 | 1.49790 | 22 |
| GO_MRNA_BINDING | 0.00000 | 0.00000 | | 0.66367 | 1.36890 | 233 |
| GO_MRNA_CIS_SPLICING_VIA_SPLICEOSOME | 0.00170 | 0.04764 | 0.45506 | 0.74587 | 1.41523 | 25 |
| GO_MRNA_EXPORT_FROM_NUCLEUS | 0.00011 | 0.00591 | 0.53843 | 0.64068 | 1.29637 | 99 |
| GO_MRNA_PROCESSING | 0.00000 | 0.00000 | | 0.63110 | 1.31343 | 414 |
| GO_MRNA_TRANSPORT | 0.00002 | 0.00114 | 0.59333 | 0.63165 | 1.28787 | 126 |
| GO_MYELOID_LEUKOCYTE_ACTIVATION | 0.00018 | 0.00873 | 0.51885 | 0.55584 | 1.15393 | 352 |
| GO_MYELOID_LEUKOCYTE_MEDIATED_IMMUNITY | 0.00011 | 0.00571 | 0.53843 | 0.56927 | 1.17891 | 312 |
| GO_NCRNA_METABOLIC_PROCESS | 0.00000 | 0.00001 | 0.70498 | 0.58858 | 1.22345 | 376 |
| GO_NCRNA_PROCESSING | 0.00000 | 0.00000 | 0.76146 | 0.60869 | 1.26119 | 320 |
| GO_NCRNA_TRANSCRIPTION | 0.00004 | 0.00236 | 0.55733 | 0.66478 | 1.34183 | 93 |
| GO_NEGATIVE_REGULATION_OF_CELLULAR_AMIDE_METABOLIC_PR | 0.00077 | 0.02657 | 0.47727 | 0.60931 | 1.23927 | 117 |
| GO_NEGATIVE_REGULATION_OF_CELL_CYCLE | 0.00000 | 0.00001 | 0.70498 | 0.57942 | 1.20639 | 433 |
| GO_NEGATIVE_REGULATION_OF_CELL_CYCLE_G2_M_PHASE_TRANS | 0.00141 | 0.04076 | 0.45506 | 0.62639 | 1.25901 | 85 |
| GO_NEGATIVE_REGULATION_OF_CELL_CYCLE_PHASE_TRANSITION | 0.00001 | 0.00060 | 0.61053 | 0.61275 | 1.25812 | 187 |
| GO_NEGATIVE_REGULATION_OF_CELL_CYCLE_PROCESS | 0.00001 | 0.00042 | 0.61053 | 0.59737 | 1.23288 | 255 |
| GO_NEGATIVE_REGULATION_OF_MITOTIC_CELL_CYCLE | 0.00013 | 0.00658 | 0.51885 | 0.58192 | 1.20034 | 238 |
| GO_NEGATIVE_REGULATION_OF_MRNA_METABOLIC_PROCESS | 0.00054 | 0.02159 | 0.47727 | 0.66353 | 1.31769 | 61 |
| GO_NEGATIVE_REGULATION_OF_RNA_SPLICING | 0.00133 | 0.03980 | 0.45506 | 0.75784 | 1.42485 | 22 |
| GO_NEGATIVE_REGULATION_OF_TRANSCRIPTION_BY_RNA_POLYME | 0.00040 | 0.01718 | 0.49849 | 0.54323 | 1.13215 | 477 |
| GO_NUCLEAR_CHROMOSOME_SEGREGATION | 0.00011 | 0.00578 | 0.53843 | 0.58743 | 1.20856 | 212 |
| GO_NUCLEAR_ENVELOPE | 0.00007 | 0.00419 | 0.53843 | 0.56710 | 1.17543 | 326 |
| GO_NUCLEAR_EXPORT | 0.00000 | 0.00004 | 0.67496 | 0.63444 | 1.30183 | 174 |
| GO_NUCLEAR_MEMBRANE | 0.00063 | 0.02286 | 0.47727 | 0.57397 | 1.18079 | 211 |
| GO_NUCLEAR_SPECK | 0.00000 | 0.00000 | 0.76146 | 0.61649 | 1.27639 | 299 |
| GO_NUCLEAR_TRANSCRIBED_MRNA_CATABOLIC_PROCESS | 0.00000 | 0.00000 | | 0.76107 | 1.56303 | 181 |
| GO_NUCLEAR_TRANSCRIBED_MRNA_CATABOLIC_PROCESS_NONSEN | 0.00000 | 0.00000 | | 0.85489 | 1.73479 | 109 |
| GO_NUCLEAR_TRANSPORT | 0.00000 | 0.00005 | 0.67496 | 0.60613 | 1.25291 | 275 |
| GO_NUCLEOBASE_CONTAINING_COMPOUND_TRANSPORT | 0.00000 | 0.00013 | 0.64355 | 0.61544 | 1.26598 | 205 |
| GO_ORGANIC_CYCLIC_COMPOUND_CATABOLIC_PROCESS | 0.00000 | 0.00000 | | 0.67248 | 1.40138 | 452 |
| GO_POLYSOMAL_RIBOSOME | 0.00000 | 0.00000 | 0.77494 | 0.86740 | 1.66563 | 30 |
| GO_POLYSOME | 0.00000 | 0.00000 | 0.81404 | 0.79308 | 1.57114 | 59 |
| GO_POLY_PURINE_TRACT_BINDING | 0.00058 | 0.02207 | 0.47727 | 0.78304 | 1.46619 | 20 |
| GO_POSITIVE_REGULATION_OF_CELLULAR_AMIDE_METABOLIC_PRO | 0.00179 | 0.04952 | 0.45506 | 0.59942 | 1.21915 | 117 |
| GO_POSITIVE_REGULATION_OF_CELLULAR_PROTEIN_LOCALIZATION | 0.00054 | 0.02137 | 0.47727 | 0.56953 | 1.17476 | 236 |
| GO_POSITIVE_REGULATION_OF_CELL_CYCLE | 0.00183 | 0.04983 | 0.45506 | 0.55881 | 1.15313 | 254 |
| GO_POSITIVE_REGULATION_OF_CELL_CYCLE_G1_S_PHASE_TRANSIT | 0.00043 | 0.01810 | 0.49849 | 0.72033 | 1.39706 | 35 |
| GO_POSITIVE_REGULATION_OF_CELL_CYCLE_PHASE_TRANSITION | 0.00003 | 0.00199 | 0.57561 | 0.67840 | 1.35589 | 74 |
| GO_POSITIVE_REGULATION_OF_CELL_POPULATION_PROLIFERATION | 0.00004 | 0.00234 | 0.55733 | 0.55815 | 1.16182 | 418 |
| GO_POSITIVE_REGULATION_OF_G1_S_TRANSITION_OF_MITOTIC_CE | 0.00058 | 0.02207 | 0.47727 | 0.74296 | 1.41367 | 27 |
| GO_POSITIVE_REGULATION_OF_MITOTIC_CELL_CYCLE | 0.00010 | 0.00538 | 0.53843 | 0.63677 | 1.29150 | 108 |
| GO_POSITIVE_REGULATION_OF_NUCLEOCYTOPLASMIC_TRANSPORT | 0.00052 | 0.02107 | 0.47727 | 0.69064 | 1.35888 | 47 |
| GO_POSITIVE_REGULATION_OF_PROTEIN_LOCALIZATION_TO_NUCLE | 0.00083 | 0.02796 | 0.47727 | 0.65826 | 1.30722 | 61 |
| GO_POSITIVE_REGULATION_OF_TRANSLATION | 0.00059 | 0.02207 | 0.47727 | 0.62138 | 1.25758 | 100 |
| GO_POSITIVE_REGULATION_OF_T_CELL_PROLIFERATION | 0.00133 | 0.03980 | 0.45506 | 0.74627 | 1.40715 | 23 |
| GO_POSITIVE_REGULATION_OF_VIRAL_GENOME_REPLICATION | 0.00133 | 0.03980 | 0.45506 | 0.73039 | 1.39443 | 28 |
| GO_POSITIVE_REGULATION_OF_VIRAL_PROCESS | 0.00004 | 0.00266 | 0.55733 | 0.66704 | 1.33774 | 80 |
| GO_POSTTRANSCRIPTIONAL_REGULATION_OF_GENE_EXPRESSION | 0.00000 | 0.00000 | | 0.61142 | 1.27425 | 467 |
| GO_PRERIBOSOME | 0.00001 | 0.00065 | 0.59333 | 0.69918 | 1.39315 | 69 |
| GO_PROTEASOMAL_PROTEIN_CATABOLIC_PROCESS | 0.00135 | 0.03994 | 0.45506 | 0.54489 | 1.13301 | 381 |
| GO_PROTEASOME_CORE_COMPLEX | 0.00115 | 0.03594 | 0.45506 | 0.80780 | 1.48500 | 15 |

| | | | | | | |
|---|---|---|---|---|---|---|
| GO_PROTEIN_CONTAINING_COMPLEX_LOCALIZATION | 0.00115 | 0.03594 | 0.45506 | 0.57475 | 1.18216 | 204 |
| GO_PROTEIN_DNA_COMPLEX_SUBUNIT_ORGANIZATION | 0.00009 | 0.00490 | 0.53843 | 0.60427 | 1.23931 | 169 |
| GO_PROTEIN_FOLDING | 0.00124 | 0.03781 | 0.45506 | 0.58347 | 1.19554 | 164 |
| GO_PROTEIN_FOLDING_CHAPERONE | 0.00053 | 0.02115 | 0.47727 | 0.79830 | 1.48993 | 19 |
| GO_PROTEIN_LOCALIZATION_TO_ENDOPLASMIC_RETICULUM | 0.00000 | 0.00000 | | 0.83407 | 1.69928 | 122 |
| GO_PROTEIN_LOCALIZATION_TO_MEMBRANE | 0.00000 | 0.00000 | 0.73376 | 0.58824 | 1.22469 | 426 |
| GO_PROTEIN_LOCALIZATION_TO_NUCLEUS | 0.00003 | 0.00189 | 0.57561 | 0.60119 | 1.23565 | 197 |
| GO_PROTEIN_MODIFICATION_BY_SMALL_PROTEIN_REMOVAL | 0.00027 | 0.01183 | 0.49849 | 0.58067 | 1.19676 | 224 |
| GO_PROTEIN_POLYUBIQUITINATION | 0.00143 | 0.04109 | 0.45506 | 0.56076 | 1.15732 | 255 |
| GO_PROTEIN_STABILIZATION | 0.00098 | 0.03267 | 0.45506 | 0.59259 | 1.21082 | 139 |
| GO_PROTEIN_TARGETING | 0.00000 | 0.00000 | | 0.66178 | 1.37180 | 323 |
| GO_PROTEIN_TARGETING_TO_MEMBRANE | 0.00000 | 0.00000 | | 0.79135 | 1.61801 | 150 |
| GO_PROTEOGLYCAN_BINDING | 0.00009 | 0.00498 | 0.53843 | 0.84593 | 1.56948 | 17 |
| GO_REGULATION_OF_CELLULAR_AMIDE_METABOLIC_PROCESS | 0.00000 | 0.00000 | 0.74774 | 0.60698 | 1.25705 | 314 |
| GO_REGULATION_OF_CELLULAR_PROTEIN_LOCALIZATION | 0.00071 | 0.02509 | 0.47727 | 0.54836 | 1.14003 | 379 |
| GO_REGULATION_OF_CELL_CYCLE_G2_M_PHASE_TRANSITION | 0.00106 | 0.03434 | 0.45506 | 0.58593 | 1.20239 | 173 |
| GO_REGULATION_OF_CELL_CYCLE_PHASE_TRANSITION | 0.00000 | 0.00018 | 0.62726 | 0.58180 | 1.20675 | 343 |
| GO_REGULATION_OF_CHROMOSOME_ORGANIZATION | 0.00011 | 0.00598 | 0.53843 | 0.57187 | 1.18193 | 274 |
| GO_REGULATION_OF_CHROMOSOME_SEGREGATION | 0.00002 | 0.00168 | 0.57561 | 0.66119 | 1.33288 | 91 |
| GO_REGULATION_OF_CHROMOSOME_SEPARATION | 0.00008 | 0.00469 | 0.53843 | 0.69358 | 1.37253 | 56 |
| GO_REGULATION_OF_DNA_METABOLIC_PROCESS | 0.00128 | 0.03874 | 0.45506 | 0.56273 | 1.16123 | 253 |
| GO_REGULATION_OF_DNA_REPLICATION | 0.00076 | 0.02609 | 0.47727 | 0.62526 | 1.26043 | 90 |
| GO_REGULATION_OF_GENE_EXPRESSION_EPIGENETIC | 0.00115 | 0.03594 | 0.45506 | 0.60009 | 1.22270 | 123 |
| GO_REGULATION_OF_INTRINSIC_APOPTOTIC_SIGNALING_PATHWAY | 0.00064 | 0.02306 | 0.47727 | 0.61449 | 1.24610 | 107 |
| GO_REGULATION_OF_INTRINSIC_APOPTOTIC_SIGNALING_PATHWAY_ | 0.00023 | 0.01032 | 0.51885 | 0.81801 | 1.52366 | 18 |
| GO_REGULATION_OF_INTRINSIC_APOPTOTIC_SIGNALING_PATHWAY_ | 0.00150 | 0.04300 | 0.45506 | 0.74788 | 1.41905 | 25 |
| GO_REGULATION_OF_MITOTIC_CELL_CYCLE | 0.00002 | 0.00168 | 0.57561 | 0.55538 | 1.15746 | 467 |
| GO_REGULATION_OF_MRNA_CATABOLIC_PROCESS | 0.00000 | 0.00010 | 0.64355 | 0.63160 | 1.29486 | 167 |
| GO_REGULATION_OF_MRNA_METABOLIC_PROCESS | 0.00000 | 0.00000 | | 0.65374 | 1.35026 | 268 |
| GO_REGULATION_OF_MRNA_PROCESSING | 0.00000 | 0.00001 | 0.70498 | 0.67511 | 1.37304 | 114 |
| GO_REGULATION_OF_MRNA_SPLICING_VIA_SPLICEOSOME | 0.00000 | 0.00014 | 0.64355 | 0.69225 | 1.38874 | 81 |
| GO_REGULATION_OF_NUCLEOCYTOPLASMIC_TRANSPORT | 0.00056 | 0.02202 | 0.47727 | 0.64616 | 1.29403 | 77 |
| GO_REGULATION_OF_PROTEASOMAL_UBIQUITIN_DEPENDENT_PRO | 0.00102 | 0.03351 | 0.45506 | 0.62119 | 1.25619 | 98 |
| GO_REGULATION_OF_PROTEIN_LOCALIZATION_TO_NUCLEUS | 0.00058 | 0.02207 | 0.47727 | 0.62964 | 1.26846 | 89 |
| GO_REGULATION_OF_PROTEIN_MODIFICATION_BY_SMALL_PROTEIN | 0.00006 | 0.00340 | 0.55733 | 0.60993 | 1.25118 | 171 |
| GO_REGULATION_OF_PROTEIN_STABILITY | 0.00113 | 0.03594 | 0.45506 | 0.56894 | 1.17085 | 215 |
| GO_REGULATION_OF_RESPONSE_TO_DNA_DAMAGE_STIMULUS | 0.00024 | 0.01078 | 0.51885 | 0.60127 | 1.23170 | 161 |
| GO_REGULATION_OF_RNA_SPLICING | 0.00000 | 0.00001 | 0.70498 | 0.67747 | 1.37862 | 116 |
| GO_REGULATION_OF_SIGNAL_TRANSDUCTION_BY_P53_CLASS_MED | 0.00060 | 0.02214 | 0.47727 | 0.59905 | 1.22312 | 135 |
| GO_REGULATION_OF_SISTER_CHROMATID_SEGREGATION | 0.00007 | 0.00428 | 0.53843 | 0.66947 | 1.33575 | 72 |
| GO_REGULATION_OF_SYMBIOTIC_PROCESS | 0.00001 | 0.00101 | 0.59333 | 0.63040 | 1.28828 | 140 |
| GO_REGULATION_OF_TRANSLATIONAL_INITIATION | 0.00004 | 0.00236 | 0.55733 | 0.69119 | 1.37758 | 66 |
| GO_REGULATION_OF_UBIQUITIN_DEPENDENT_PROTEIN_CATABOLIC | 0.00012 | 0.00634 | 0.51885 | 0.63115 | 1.28407 | 115 |
| GO_REGULATION_OF_UBIQUITIN_PROTEIN_TRANSFERASE_ACTIVITY | 0.00055 | 0.02181 | 0.47727 | 0.69607 | 1.36626 | 45 |
| GO_REGULATION_OF_VIRAL_GENOME_REPLICATION | 0.00121 | 0.03747 | 0.45506 | 0.65793 | 1.30107 | 55 |
| GO_REGULATION_OF_VIRAL_LIFE_CYCLE | 0.00061 | 0.02233 | 0.47727 | 0.62988 | 1.26752 | 86 |
| GO_RESPONSE_TO_INTERLEUKIN_1 | 0.00111 | 0.03593 | 0.45506 | 0.61238 | 1.24094 | 105 |
| GO_RESPONSE_TO_LEUKEMIA_INHIBITORY_FACTOR | 0.00014 | 0.00664 | 0.51885 | 0.66608 | 1.33502 | 78 |
| GO_RIBONUCLEOPROTEIN_COMPLEX_BINDING | 0.00152 | 0.04311 | 0.45506 | 0.59985 | 1.22066 | 116 |
| GO_RIBONUCLEOPROTEIN_COMPLEX_BIOGENESIS | 0.00000 | 0.00000 | | 0.67819 | 1.41037 | 389 |
| GO_RIBONUCLEOPROTEIN_COMPLEX_SUBUNIT_ORGANIZATION | 0.00000 | 0.00000 | | 0.67736 | 1.38995 | 176 |
| GO_RIBONUCLEOPROTEIN_GRANULE | 0.00002 | 0.00116 | 0.57561 | 0.61283 | 1.25784 | 179 |
| GO_RIBOSOMAL_LARGE_SUBUNIT_ASSEMBLY | 0.00010 | 0.00531 | 0.53843 | 0.77651 | 1.48249 | 28 |
| GO_RIBOSOMAL_LARGE_SUBUNIT_BIOGENESIS | 0.00000 | 0.00000 | 0.76146 | 0.76213 | 1.51896 | 66 |
| GO_RIBOSOMAL_SMALL_SUBUNIT_BIOGENESIS | 0.00000 | 0.00009 | 0.65944 | 0.71416 | 1.42331 | 68 |
| GO_RIBOSOMAL_SUBUNIT | 0.00000 | 0.00000 | | 0.78198 | 1.60356 | 168 |
| GO_RIBOSOME | 0.00000 | 0.00000 | | 0.74887 | 1.54013 | 201 |
| GO_RIBOSOME_ASSEMBLY | 0.00000 | 0.00005 | 0.67496 | 0.73710 | 1.46381 | 61 |
| GO_RIBOSOME_BIOGENESIS | 0.00000 | 0.00000 | | 0.67843 | 1.40215 | 274 |
| GO_RNA_3_END_PROCESSING | 0.00060 | 0.02214 | 0.47727 | 0.60918 | 1.24142 | 124 |
| GO_RNA_CATABOLIC_PROCESS | 0.00000 | 0.00000 | | 0.69811 | 1.44763 | 336 |
| GO_RNA_EXPORT_FROM_NUCLEUS | 0.00000 | 0.00010 | 0.64355 | 0.65225 | 1.32986 | 126 |
| GO_RNA_LOCALIZATION | 0.00000 | 0.00000 | 0.74774 | 0.64068 | 1.31681 | 197 |
| GO_RNA_SPLICING | 0.00000 | 0.00000 | | 0.64155 | 1.33276 | 365 |
| GO_RNA_SPLICING_VIA_TRANSESTERIFICATION_REACTIONS | 0.00000 | 0.00000 | | 0.65490 | 1.35608 | 295 |
| GO_RRNA_BINDING | 0.00000 | 0.00000 | 0.73376 | 0.77487 | 1.53158 | 54 |
| GO_RRNA_METABOLIC_PROCESS | 0.00000 | 0.00000 | | 0.66961 | 1.37724 | 208 |
| GO_SIGNAL_TRANSDUCTION_BY_P53_CLASS_MEDIATOR | 0.00074 | 0.02576 | 0.47727 | 0.57743 | 1.18707 | 198 |
| GO_SINGLE_STRANDED_RNA_BINDING | 0.00006 | 0.00357 | 0.55733 | 0.67622 | 1.34431 | 62 |
| GO_SISTER_CHROMATID_SEGREGATION | 0.00001 | 0.00076 | 0.59333 | 0.62360 | 1.27747 | 160 |
| GO_SMALL_RIBOSOMAL_SUBUNIT | 0.00000 | 0.00000 | | 0.81226 | 1.61887 | 66 |
| GO_SNRNA_TRANSCRIPTION | 0.00044 | 0.01822 | 0.49849 | 0.65310 | 1.30161 | 68 |
| GO_SPINDLE | 0.00019 | 0.00880 | 0.51885 | 0.57029 | 1.17981 | 284 |
| GO_SPLICEOSOMAL_COMPLEX | 0.00000 | 0.00002 | 0.69013 | 0.64726 | 1.32631 | 165 |
| GO_SPLICEOSOMAL_COMPLEX_ASSEMBLY | 0.00084 | 0.02811 | 0.47727 | 0.68197 | 1.34183 | 47 |
| GO_STRUCTURAL_CONSTITUENT_OF_RIBOSOME | 0.00000 | 0.00000 | | 0.79567 | 1.62613 | 146 |

| | | | | | | |
|---|---|---|---|---|---|---|
| GO_STRUCTURAL_MOLECULE_ACTIVITY | 0.00000 | 0.00000 | | 0.66466 | 1.37847 | 337 |
| GO_TRANSCRIPTION_BY_RNA_POLYMERASE_I | 0.00176 | 0.04873 | 0.45506 | 0.66269 | 1.31195 | 57 |
| GO_TRANSCRIPTION_COREGULATOR_ACTIVITY | 0.00075 | 0.02592 | 0.47727 | 0.55146 | 1.14440 | 350 |
| GO_TRANSCRIPTION_ELONGATION_FROM_RNA_POLYMERASE_II_PR | 0.00044 | 0.01822 | 0.49849 | 0.65699 | 1.30880 | 70 |
| GO_TRANSCRIPTION_FACTOR_BINDING | 0.00011 | 0.00598 | 0.53843 | 0.55485 | 1.15498 | 415 |
| GO_TRANSCRIPTION_REGULATOR_COMPLEX | 0.00183 | 0.04983 | 0.45506 | 0.55699 | 1.15000 | 264 |
| GO_TRANSFERASE_COMPLEX_TRANSFERRING_PHOSPHORUS_CONT/ | 0.00080 | 0.02719 | 0.47727 | 0.58061 | 1.19306 | 193 |
| GO_TRANSLATIONAL_INITIATION | 0.00000 | 0.00000 | | 0.80559 | 1.65197 | 168 |
| GO_TRANSLATION_FACTOR_ACTIVITY_RNA_BINDING | 0.00014 | 0.00664 | 0.51885 | 0.67104 | 1.33709 | 69 |
| GO_TRANSLATION_INITIATION_FACTOR_ACTIVITY | 0.00049 | 0.02037 | 0.47727 | 0.69293 | 1.36217 | 46 |
| GO_TRANSLATION_INITIATION_FACTOR_BINDING | 0.00027 | 0.01195 | 0.49849 | 0.75826 | 1.44764 | 28 |
| GO_TRANSLATION_REGULATOR_ACTIVITY | 0.00000 | 0.00009 | 0.65944 | 0.66302 | 1.34602 | 110 |
| GO_TRANSLATION_REGULATOR_ACTIVITY_NUCLEIC_ACID_BINDING | 0.00000 | 0.00023 | 0.62726 | 0.68152 | 1.36957 | 84 |
| GO_U2_TYPE_SPLICEOSOMAL_COMPLEX | 0.00059 | 0.02207 | 0.47727 | 0.63385 | 1.27218 | 82 |
| GO_UBIQUITIN_LIKE_PROTEIN_LIGASE_BINDING | 0.00006 | 0.00367 | 0.53843 | 0.58652 | 1.20762 | 218 |
| GO_UNFOLDED_PROTEIN_BINDING | 0.00004 | 0.00248 | 0.55733 | 0.66156 | 1.32971 | 85 |
| GO_VESICLE_LUMEN | 0.00001 | 0.00093 | 0.59333 | 0.61108 | 1.25417 | 178 |
| GO_VIRAL_GENE_EXPRESSION | 0.00000 | 0.00000 | | 0.77970 | 1.59963 | 172 |
| GO_VIRAL_GENOME_REPLICATION | 0.00115 | 0.03594 | 0.45506 | 0.63968 | 1.27569 | 71 |
| GO_VIRAL_LIFE_CYCLE | 0.00001 | 0.00101 | 0.59333 | 0.60247 | 1.23947 | 210 |
| HP_ABNORMALITY_OF_LOWER_LIP | 0.00059 | 0.02207 | 0.47727 | 0.59559 | 1.21826 | 155 |
| HP_ABNORMALITY_OF_THE_NASAL_DORSUM | 0.00066 | 0.02376 | 0.47727 | 0.59566 | 1.21709 | 139 |
| HP_ABNORMAL_RETICULOCYTE_MORPHOLOGY | 0.00057 | 0.02207 | 0.47727 | 0.73378 | 1.40507 | 29 |
| HP_ACUTE_LEUKEMIA | 0.00000 | 0.00010 | 0.64355 | 0.71225 | 1.41890 | 70 |
| HP_ANEMIA_OF_INADEQUATE_PRODUCTION | 0.00002 | 0.00170 | 0.57561 | 0.65219 | 1.32023 | 101 |
| HP_DELAYED_PUBERTY | 0.00000 | 0.00008 | 0.65944 | 0.66437 | 1.35125 | 117 |
| HP_DEPRESSED_NASAL_RIDGE | 0.00002 | 0.00168 | 0.57561 | 0.68199 | 1.36692 | 78 |
| HP_LEUKEMIA | 0.00002 | 0.00117 | 0.57561 | 0.65722 | 1.32985 | 99 |
| HP_MACROCYTIC_ANEMIA | 0.00000 | 0.00002 | 0.69013 | 0.79882 | 1.55733 | 40 |
| HP_MIGRAINE | 0.00000 | 0.00000 | 0.76146 | 0.75303 | 1.50078 | 68 |
| HP_PALLOR | 0.00001 | 0.00042 | 0.61053 | 0.67418 | 1.35819 | 89 |
| HP_THICK_LOWER_LIP_VERMILION | 0.00019 | 0.00880 | 0.51885 | 0.62951 | 1.27623 | 106 |
| HP_THICK_VERMILION_BORDER | 0.00183 | 0.04983 | 0.45506 | 0.58182 | 1.19169 | 162 |
| **Cluster 10** | | | | | | |
| pathway | pval | padj | log2err | ES | NES | size |
| GO_90S_PRERIBOSOME | 0.00194 | 0.04151 | 0.45506 | 0.74255 | 1.38382 | 26 |
| GO_ACTIVATION_OF_IMMUNE_RESPONSE | 0.00027 | 0.00827 | 0.49849 | 0.58608 | 1.19057 | 243 |
| GO_ACTIVATION_OF_INNATE_IMMUNE_RESPONSE | 0.00000 | 0.00019 | 0.62726 | 0.68619 | 1.35975 | 95 |
| GO_AMIDE_TRANSMEMBRANE_TRANSPORTER_ACTIVITY | 0.00150 | 0.03396 | 0.45506 | 0.73742 | 1.38545 | 30 |
| GO_AMINE_METABOLIC_PROCESS | 0.00000 | 0.00001 | 0.71951 | 0.72729 | 1.43932 | 88 |
| GO_ANAPHASE_PROMOTING_COMPLEX_DEPENDENT_CATABOLIC_P | 0.00000 | 0.00000 | 0.78819 | 0.76574 | 1.50923 | 78 |
| GO_ANTIGEN_PROCESSING_AND_PRESENTATION | 0.00004 | 0.00165 | 0.55733 | 0.62764 | 1.26338 | 159 |
| GO_ANTIGEN_PROCESSING_AND_PRESENTATION_OF_EXOGENOUS_ | 0.00000 | 0.00001 | 0.70498 | 0.76130 | 1.48856 | 63 |
| GO_ANTIGEN_PROCESSING_AND_PRESENTATION_OF_PEPTIDE_ANTI | 0.00003 | 0.00135 | 0.55733 | 0.63812 | 1.27857 | 137 |
| GO_ANTIGEN_PROCESSING_AND_PRESENTATION_OF_PEPTIDE_ANTI | 0.00000 | 0.00009 | 0.64355 | 0.72721 | 1.43153 | 75 |
| GO_ANTIGEN_RECEPTOR_MEDIATED_SIGNALING_PATHWAY | 0.00003 | 0.00119 | 0.57561 | 0.63116 | 1.26778 | 145 |
| GO_APOPTOTIC_SIGNALING_PATHWAY | 0.00047 | 0.01373 | 0.49849 | 0.56151 | 1.15010 | 393 |
| GO_ATPASE_ACTIVITY | 0.00172 | 0.03730 | 0.45506 | 0.56069 | 1.14377 | 302 |
| GO_ATPASE_COMPLEX | 0.00117 | 0.02904 | 0.45506 | 0.65501 | 1.28418 | 67 |
| GO_BINDING_OF_SPERM_TO_ZONA_PELLUCIDA | 0.00098 | 0.02540 | 0.45506 | 0.81179 | 1.47237 | 17 |
| GO_CADHERIN_BINDING | 0.00000 | 0.00001 | 0.69013 | 0.62814 | 1.27653 | 255 |
| GO_CANONICAL_WNT_SIGNALING_PATHWAY | 0.00113 | 0.02836 | 0.45506 | 0.57910 | 1.17413 | 219 |
| GO_CATALYTIC_STEP_2_SPLICEOSOME | 0.00002 | 0.00091 | 0.57561 | 0.68953 | 1.36146 | 83 |
| GO_CELLULAR_AMINO_ACID_METABOLIC_PROCESS | 0.00002 | 0.00077 | 0.57561 | 0.60640 | 1.23022 | 226 |
| GO_CELLULAR_CARBOHYDRATE_CATABOLIC_PROCESS | 0.00007 | 0.00260 | 0.53843 | 0.78425 | 1.47343 | 30 |
| GO_CELLULAR_COMPONENT_DISASSEMBLY | 0.00008 | 0.00301 | 0.53843 | 0.56795 | 1.16419 | 423 |
| GO_CELLULAR_KETONE_METABOLIC_PROCESS | 0.00152 | 0.03416 | 0.45506 | 0.59128 | 1.19089 | 169 |
| GO_CELLULAR_NITROGEN_COMPOUND_CATABOLIC_PROCESS | 0.00000 | 0.00000 | | 0.68548 | 1.40675 | 461 |
| GO_CELLULAR_RESPONSE_TO_OXYGEN_LEVELS | 0.00001 | 0.00031 | 0.61053 | 0.64109 | 1.29049 | 160 |
| GO_CELL_ADHESION_MOLECULE_BINDING | 0.00000 | 0.00005 | 0.65944 | 0.59900 | 1.22577 | 348 |
| GO_CELL_CELL_RECOGNITION | 0.00010 | 0.00370 | 0.53843 | 0.78327 | 1.46756 | 29 |
| GO_CELL_CELL_SIGNALING_BY_WNT | 0.00016 | 0.00550 | 0.51885 | 0.57035 | 1.16702 | 347 |
| GO_CELL_CYCLE_G1_S_PHASE_TRANSITION | 0.00009 | 0.00321 | 0.53843 | 0.59850 | 1.21298 | 213 |
| GO_CELL_CYCLE_G2_M_PHASE_TRANSITION | 0.00008 | 0.00299 | 0.53843 | 0.59774 | 1.21277 | 228 |
| GO_CELL_CYCLE_PHASE_TRANSITION | 0.00000 | 0.00000 | 0.81404 | 0.60319 | 1.23841 | 493 |
| GO_CELL_DIVISION | 0.00005 | 0.00193 | 0.55733 | 0.56799 | 1.16476 | 439 |
| GO_CELL_SUBSTRATE_JUNCTION | 0.00000 | 0.00000 | 0.80122 | 0.63358 | 1.29273 | 307 |
| GO_CELL_SURFACE_RECEPTOR_SIGNALING_PATHWAY_INVOLVED_IN | 0.00128 | 0.03079 | 0.45506 | 0.55368 | 1.13394 | 385 |
| GO_CHAPERONE_COMPLEX | 0.00001 | 0.00039 | 0.61053 | 0.85683 | 1.58022 | 22 |
| GO_CHROMATIN_BINDING | 0.00004 | 0.00187 | 0.55733 | 0.57082 | 1.16961 | 403 |
| GO_CHROMOSOMAL_REGION | 0.00027 | 0.00827 | 0.49849 | 0.57431 | 1.17088 | 284 |
| GO_CHROMOSOME_SEGREGATION | 0.00003 | 0.00128 | 0.57561 | 0.59241 | 1.20483 | 267 |
| GO_CHROMOSOME_SEPARATION | 0.00111 | 0.02813 | 0.45506 | 0.63923 | 1.26161 | 82 |
| GO_CONDENSED_CHROMOSOME | 0.00132 | 0.03155 | 0.45506 | 0.59454 | 1.19727 | 167 |
| GO_COTRANSLATIONAL_PROTEIN_TARGETING_TO_MEMBRANE | 0.00000 | 0.00000 | | 0.84526 | 1.67449 | 92 |
| GO_COVALENT_CHROMATIN_MODIFICATION | 0.00036 | 0.01080 | 0.49849 | 0.56479 | 1.15614 | 364 |

| | | | | | | |
|---|---|---|---|---|---|---|
| GO_CYTOKINE_MEDIATED_SIGNALING_PATHWAY | 0.00073 | 0.01969 | 0.47727 | 0.56043 | 1.14828 | 398 |
| GO_CYTOPLASMIC_STRESS_GRANULE | 0.00075 | 0.02001 | 0.47727 | 0.66782 | 1.30579 | 63 |
| GO_CYTOPLASMIC_TRANSLATION | 0.00000 | 0.00000 | | 0.82738 | 1.63747 | 89 |
| GO_CYTOPLASMIC_TRANSLATIONAL_INITIATION | 0.00000 | 0.00006 | 0.65944 | 0.83059 | 1.56246 | 31 |
| GO_CYTOSOLIC_LARGE_RIBOSOMAL_SUBUNIT | 0.00000 | 0.00000 | | 0.87141 | 1.68912 | 54 |
| GO_CYTOSOLIC_RIBOSOME | 0.00000 | 0.00000 | | 0.86003 | 1.70400 | 96 |
| GO_CYTOSOLIC_SMALL_RIBOSOMAL_SUBUNIT | 0.00000 | 0.00000 | 0.80122 | 0.85336 | 1.62908 | 41 |
| GO_DNA_BINDING_TRANSCRIPTION_FACTOR_BINDING | 0.00037 | 0.01102 | 0.49849 | 0.58026 | 1.17880 | 246 |
| GO_DNA_BIOSYNTHETIC_PROCESS | 0.00005 | 0.00194 | 0.55733 | 0.62676 | 1.25894 | 145 |
| GO_DNA_CONFORMATION_CHANGE | 0.00059 | 0.01619 | 0.47727 | 0.58689 | 1.19053 | 223 |
| GO_DNA_PACKAGING | 0.00048 | 0.01386 | 0.49849 | 0.61949 | 1.23823 | 122 |
| GO_DNA_POLYMERASE_BINDING | 0.00209 | 0.04406 | 0.43171 | 0.78542 | 1.43061 | 18 |
| GO_DNA_REPAIR | 0.00119 | 0.02938 | 0.45506 | 0.55052 | 1.12875 | 435 |
| GO_DNA_REPLICATION | 0.00024 | 0.00766 | 0.51885 | 0.58688 | 1.19067 | 225 |
| GO_DNA_TEMPLATED_TRANSCRIPTION_ELONGATION | 0.00069 | 0.01878 | 0.47727 | 0.63330 | 1.25591 | 99 |
| GO_ENDOPEPTIDASE_COMPLEX | 0.00000 | 0.00001 | 0.70498 | 0.76053 | 1.48707 | 63 |
| GO_ESTABLISHMENT_OF_PROTEIN_LOCALIZATION_TO_ENDOPLASM | 0.00000 | 0.00000 | | 0.83553 | 1.65980 | 103 |
| GO_ESTABLISHMENT_OF_PROTEIN_LOCALIZATION_TO_MEMBRANE | 0.00000 | 0.00000 | | 0.70738 | 1.43797 | 257 |
| GO_ESTABLISHMENT_OF_PROTEIN_LOCALIZATION_TO_ORGANELLE | 0.00000 | 0.00000 | | 0.67249 | 1.37889 | 432 |
| GO_ESTABLISHMENT_OF_PROTEIN_LOCALIZATION_TO_TELOMERE | 0.00095 | 0.02455 | 0.47727 | 0.82499 | 1.49186 | 16 |
| GO_ESTABLISHMENT_OF_RNA_LOCALIZATION | 0.00004 | 0.00175 | 0.55733 | 0.62104 | 1.25082 | 169 |
| GO_ESTABLISHMENT_OF_TISSUE_POLARITY | 0.00001 | 0.00034 | 0.61053 | 0.68014 | 1.34777 | 95 |
| GO_EUKARYOTIC_TRANSLATION_INITIATION_FACTOR_3_COMPLEX | 0.00018 | 0.00606 | 0.51885 | 0.85324 | 1.54219 | 15 |
| GO_FC_EPSILON_RECEPTOR_SIGNALING_PATHWAY | 0.00001 | 0.00056 | 0.59333 | 0.68614 | 1.35788 | 88 |
| GO_FC_RECEPTOR_SIGNALING_PATHWAY | 0.00001 | 0.00054 | 0.59333 | 0.64718 | 1.29598 | 134 |
| GO_FICOLIN_1_RICH_GRANULE | 0.00000 | 0.00003 | 0.67496 | 0.70255 | 1.39324 | 99 |
| GO_GENE_SILENCING | 0.00102 | 0.02613 | 0.45506 | 0.59208 | 1.19390 | 177 |
| GO_GLYCOSYL_COMPOUND_METABOLIC_PROCESS | 0.00056 | 0.01565 | 0.47727 | 0.65983 | 1.29938 | 76 |
| GO_GTP_METABOLIC_PROCESS | 0.00224 | 0.04653 | 0.43171 | 0.76131 | 1.39776 | 20 |
| GO_GUANOSINE_CONTAINING_COMPOUND_METABOLIC_PROCESS | 0.00181 | 0.03889 | 0.45506 | 0.73412 | 1.37925 | 30 |
| GO_HEMATOPOIETIC_PROGENITOR_CELL_DIFFERENTIATION | 0.00000 | 0.00000 | 0.71951 | 0.69253 | 1.38288 | 120 |
| GO_HEMATOPOIETIC_STEM_CELL_DIFFERENTIATION | 0.00000 | 0.00000 | 0.74774 | 0.76038 | 1.49531 | 72 |
| GO_IMMUNE_RESPONSE_REGULATING_SIGNALING_PATHWAY | 0.00032 | 0.00958 | 0.49849 | 0.59137 | 1.19963 | 223 |
| GO_INNATE_IMMUNE_RESPONSE_ACTIVATING_SIGNAL_TRANSDUCT | 0.00001 | 0.00032 | 0.61053 | 0.70763 | 1.39470 | 78 |
| GO_INTERLEUKIN_1_MEDIATED_SIGNALING_PATHWAY | 0.00000 | 0.00001 | 0.70498 | 0.72510 | 1.43090 | 81 |
| GO_INTRINSIC_APOPTOTIC_SIGNALING_PATHWAY | 0.00002 | 0.00116 | 0.57561 | 0.60760 | 1.23057 | 204 |
| GO_LARGE_RIBOSOMAL_SUBUNIT | 0.00000 | 0.00000 | | 0.74189 | 1.47607 | 109 |
| GO_MATURATION_OF_LSU_RRNA | 0.00026 | 0.00826 | 0.49849 | 0.77922 | 1.45584 | 27 |
| GO_MATURATION_OF_LSU_RRNA_FROM_TRICISTRONIC_RRNA_TRA | 0.00141 | 0.03273 | 0.45506 | 0.81750 | 1.47760 | 15 |
| GO_MATURATION_OF_SSU_RRNA | 0.00133 | 0.03174 | 0.45506 | 0.68359 | 1.31352 | 47 |
| GO_METAPHASE_ANAPHASE_TRANSITION_OF_CELL_CYCLE | 0.00209 | 0.04406 | 0.43171 | 0.67420 | 1.31024 | 56 |
| GO_MITOTIC_NUCLEAR_DIVISION | 0.00012 | 0.00439 | 0.51885 | 0.59661 | 1.20943 | 216 |
| GO_MITOTIC_SISTER_CHROMATID_SEGREGATION | 0.00053 | 0.01477 | 0.47727 | 0.61086 | 1.22515 | 140 |
| GO_MORPHOGENESIS_OF_AN_EPITHELIUM | 0.00122 | 0.03017 | 0.45506 | 0.56106 | 1.14751 | 332 |
| GO_MORPHOGENESIS_OF_A_POLARIZED_EPITHELIUM | 0.00000 | 0.00019 | 0.62726 | 0.67090 | 1.33543 | 110 |
| GO_MRNA_3_END_PROCESSING | 0.00228 | 0.04714 | 0.43171 | 0.62846 | 1.24417 | 90 |
| GO_MRNA_3_UTR_BINDING | 0.00085 | 0.02242 | 0.47727 | 0.66272 | 1.30142 | 69 |
| GO_MRNA_5_UTR_BINDING | 0.00020 | 0.00648 | 0.51885 | 0.81262 | 1.49869 | 22 |
| GO_MRNA_BINDING | 0.00000 | 0.00000 | 0.81404 | 0.65784 | 1.33537 | 231 |
| GO_MRNA_EXPORT_FROM_NUCLEUS | 0.00011 | 0.00390 | 0.53843 | 0.64624 | 1.28240 | 101 |
| GO_MRNA_PROCESSING | 0.00000 | 0.00000 | | 0.61501 | 1.26093 | 431 |
| GO_MRNA_TRANSPORT | 0.00100 | 0.02576 | 0.45506 | 0.61866 | 1.23709 | 128 |
| GO_MYD88_INDEPENDENT_TOLL_LIKE_RECEPTOR_SIGNALING_PATH | 0.00051 | 0.01452 | 0.47727 | 0.79065 | 1.46357 | 23 |
| GO_MYELOID_LEUKOCYTE_ACTIVATION | 0.00137 | 0.03212 | 0.45506 | 0.55525 | 1.13726 | 379 |
| GO_MYELOID_LEUKOCYTE_MEDIATED_IMMUNITY | 0.00022 | 0.00708 | 0.51885 | 0.57306 | 1.17227 | 342 |
| GO_NCRNA_METABOLIC_PROCESS | 0.00000 | 0.00022 | 0.62726 | 0.58715 | 1.20262 | 393 |
| GO_NCRNA_PROCESSING | 0.00000 | 0.00004 | 0.67496 | 0.60494 | 1.23750 | 335 |
| GO_NEGATIVE_REGULATION_OF_CANONICAL_WNT_SIGNALING_PAT | 0.00022 | 0.00708 | 0.51885 | 0.62694 | 1.25192 | 120 |
| GO_NEGATIVE_REGULATION_OF_CELL_CYCLE | 0.00000 | 0.00001 | 0.70498 | 0.59083 | 1.21244 | 462 |
| GO_NEGATIVE_REGULATION_OF_CELL_CYCLE_G2_M_PHASE_TRANS | 0.00000 | 0.00004 | 0.65944 | 0.71377 | 1.41263 | 89 |
| GO_NEGATIVE_REGULATION_OF_CELL_CYCLE_PHASE_TRANSITION | 0.00000 | 0.00003 | 0.67496 | 0.63912 | 1.29336 | 199 |
| GO_NEGATIVE_REGULATION_OF_CELL_CYCLE_PROCESS | 0.00000 | 0.00000 | 0.71951 | 0.62867 | 1.27876 | 269 |
| GO_NEGATIVE_REGULATION_OF_MITOTIC_CELL_CYCLE | 0.00000 | 0.00020 | 0.62726 | 0.60994 | 1.23978 | 251 |
| GO_NEGATIVE_REGULATION_OF_UBIQUITIN_DEPENDENT_PROTEIN_ | 0.00224 | 0.04653 | 0.43171 | 0.67879 | 1.29407 | 40 |
| GO_NEGATIVE_REGULATION_OF_WNT_SIGNALING_PATHWAY | 0.00012 | 0.00439 | 0.51885 | 0.62561 | 1.25461 | 139 |
| GO_NIK_NF_KAPPAB_SIGNALING | 0.00000 | 0.00001 | 0.70498 | 0.69604 | 1.38825 | 116 |
| GO_NON_CANONICAL_WNT_SIGNALING_PATHWAY | 0.00000 | 0.00025 | 0.61053 | 0.67622 | 1.34758 | 113 |
| GO_NUCLEAR_CHROMOSOME_SEGREGATION | 0.00030 | 0.00921 | 0.49849 | 0.58920 | 1.19422 | 214 |
| GO_NUCLEAR_ENVELOPE | 0.00139 | 0.03242 | 0.45506 | 0.56062 | 1.14679 | 338 |
| GO_NUCLEAR_EXPORT | 0.00005 | 0.00204 | 0.55733 | 0.62052 | 1.25208 | 181 |
| GO_NUCLEAR_SPECK | 0.00000 | 0.00023 | 0.62726 | 0.59694 | 1.21847 | 312 |
| GO_NUCLEAR_TRANSCRIBED_MRNA_CATABOLIC_PROCESS | 0.00000 | 0.00000 | | 0.74978 | 1.51562 | 190 |
| GO_NUCLEAR_TRANSCRIBED_MRNA_CATABOLIC_PROCESS_NONSEN | 0.00000 | 0.00000 | | 0.84459 | 1.68310 | 113 |
| GO_NUCLEAR_TRANSPORT | 0.00000 | 0.00011 | 0.64355 | 0.60577 | 1.23495 | 283 |
| GO_NUCLEOBASE_CONTAINING_COMPOUND_TRANSPORT | 0.00027 | 0.00827 | 0.49849 | 0.59459 | 1.20449 | 210 |
| GO_NUCLEOSIDE_METABOLIC_PROCESS | 0.00015 | 0.00525 | 0.51885 | 0.69407 | 1.35707 | 64 |

| | | | | | | |
|---|---|---|---|---|---|---|
| GO_NUCLEOSIDE_PHOSPHATE_BIOSYNTHETIC_PROCESS | 0.00145 | 0.03320 | 0.45506 | 0.59595 | 1.19976 | 158 |
| GO_ORGANIC_CYCLIC_COMPOUND_CATABOLIC_PROCESS | 0.00000 | 0.00000 | | 0.68207 | 1.39960 | 476 |
| GO_PEPTIDASE_COMPLEX | 0.00000 | 0.00007 | 0.65944 | 0.71681 | 1.41333 | 79 |
| GO_PEPTIDE_TRANSMEMBRANE_TRANSPORTER_ACTIVITY | 0.00115 | 0.02870 | 0.45506 | 0.78005 | 1.43614 | 21 |
| GO_PEPTIDYL_LYSINE_MODIFICATION | 0.00137 | 0.03212 | 0.45506 | 0.56339 | 1.14913 | 300 |
| GO_POLYSOMAL_RIBOSOME | 0.00001 | 0.00045 | 0.59333 | 0.81296 | 1.52738 | 30 |
| GO_POLYSOME | 0.00000 | 0.00002 | 0.69013 | 0.75727 | 1.47909 | 62 |
| GO_POLY_PYRIMIDINE_TRACT_BINDING | 0.00022 | 0.00708 | 0.51885 | 0.82778 | 1.51469 | 19 |
| GO_POSITIVE_REGULATION_OF_CANONICAL_WNT_SIGNALING_PATH | 0.00000 | 0.00004 | 0.67496 | 0.68526 | 1.36402 | 110 |
| GO_POSITIVE_REGULATION_OF_CELLULAR_AMIDE_METABOLIC_PRO | 0.00066 | 0.01812 | 0.47727 | 0.62100 | 1.23959 | 118 |
| GO_POSITIVE_REGULATION_OF_CELLULAR_PROTEIN_LOCALIZATION | 0.00034 | 0.01036 | 0.49849 | 0.58071 | 1.17972 | 246 |
| GO_POSITIVE_REGULATION_OF_CELL_CYCLE | 0.00000 | 0.00010 | 0.64355 | 0.60966 | 1.23933 | 260 |
| GO_POSITIVE_REGULATION_OF_CELL_CYCLE_PHASE_TRANSITION | 0.00154 | 0.03416 | 0.45506 | 0.64603 | 1.27221 | 76 |
| GO_POSITIVE_REGULATION_OF_CELL_CYCLE_PROCESS | 0.00005 | 0.00207 | 0.55733 | 0.61550 | 1.24629 | 202 |
| GO_POSITIVE_REGULATION_OF_CELL_DEATH | 0.00028 | 0.00861 | 0.49849 | 0.56068 | 1.14902 | 416 |
| GO_POSITIVE_REGULATION_OF_CHROMOSOME_ORGANIZATION | 0.00053 | 0.01477 | 0.47727 | 0.61540 | 1.23518 | 142 |
| GO_POSITIVE_REGULATION_OF_DEFENSE_RESPONSE | 0.00058 | 0.01601 | 0.47727 | 0.59541 | 1.20371 | 192 |
| GO_POSITIVE_REGULATION_OF_IMMUNE_RESPONSE | 0.00086 | 0.02256 | 0.47727 | 0.56456 | 1.15336 | 323 |
| GO_POSITIVE_REGULATION_OF_INNATE_IMMUNE_RESPONSE | 0.00006 | 0.00246 | 0.53843 | 0.64705 | 1.29339 | 123 |
| GO_POSITIVE_REGULATION_OF_MITOTIC_CELL_CYCLE | 0.00011 | 0.00388 | 0.53843 | 0.64286 | 1.28366 | 119 |
| GO_POSITIVE_REGULATION_OF_MRNA_PROCESSING | 0.00006 | 0.00246 | 0.53843 | 0.79930 | 1.49337 | 27 |
| GO_POSITIVE_REGULATION_OF_MRNA_SPLICING_VIA_SPLICEOSOM | 0.00009 | 0.00321 | 0.53843 | 0.85291 | 1.54695 | 17 |
| GO_POSITIVE_REGULATION_OF_PROTEIN_LOCALIZATION_TO_NUCLE | 0.00000 | 0.00023 | 0.62726 | 0.72607 | 1.41968 | 63 |
| GO_POSITIVE_REGULATION_OF_RESPONSE_TO_BIOTIC_STIMULUS | 0.00004 | 0.00172 | 0.55733 | 0.63674 | 1.27801 | 142 |
| GO_POSITIVE_REGULATION_OF_RNA_SPLICING | 0.00050 | 0.01432 | 0.47727 | 0.75876 | 1.42165 | 29 |
| GO_POSITIVE_REGULATION_OF_TELOMERASE_RNA_LOCALIZATION_ | 0.00000 | 0.00011 | 0.64355 | 0.91435 | 1.65265 | 15 |
| GO_POSITIVE_REGULATION_OF_TRANSLATION | 0.00048 | 0.01386 | 0.49849 | 0.64033 | 1.27048 | 100 |
| GO_POSITIVE_REGULATION_OF_WNT_SIGNALING_PATHWAY | 0.00001 | 0.00038 | 0.61053 | 0.65414 | 1.30885 | 130 |
| GO_POSTTRANSCRIPTIONAL_REGULATION_OF_GENE_EXPRESSION | 0.00000 | 0.00000 | | 0.63952 | 1.31244 | 482 |
| GO_POST_TRANSLATIONAL_PROTEIN_MODIFICATION | 0.00008 | 0.00312 | 0.53843 | 0.58754 | 1.19438 | 259 |
| GO_POTASSIUM_ION_IMPORT_ACROSS_PLASMA_MEMBRANE | 0.00163 | 0.03582 | 0.45506 | 0.76586 | 1.40611 | 20 |
| GO_PRERIBOSOME | 0.00001 | 0.00052 | 0.59333 | 0.71525 | 1.40456 | 69 |
| GO_PROTEASOMAL_PROTEIN_CATABOLIC_PROCESS | 0.00002 | 0.00091 | 0.57561 | 0.57528 | 1.17876 | 396 |
| GO_PROTEASOMAL_UBIQUITIN_INDEPENDENT_PROTEIN_CATABOLI( | 0.00000 | 0.00028 | 0.61053 | 0.86280 | 1.58410 | 20 |
| GO_PROTEASOME_ACCESSORY_COMPLEX | 0.00002 | 0.00093 | 0.57561 | 0.82814 | 1.53565 | 24 |
| GO_PROTEASOME_CORE_COMPLEX | 0.00001 | 0.00056 | 0.59333 | 0.87159 | 1.58757 | 18 |
| GO_PROTEIN_CONTAINING_COMPLEX_DISASSEMBLY | 0.00028 | 0.00861 | 0.49849 | 0.57753 | 1.17452 | 265 |
| GO_PROTEIN_CONTAINING_COMPLEX_LOCALIZATION | 0.00014 | 0.00506 | 0.51885 | 0.59289 | 1.20198 | 218 |
| GO_PROTEIN_DNA_COMPLEX_SUBUNIT_ORGANIZATION | 0.00015 | 0.00516 | 0.51885 | 0.61208 | 1.23278 | 169 |
| GO_PROTEIN_FOLDING | 0.00043 | 0.01259 | 0.49849 | 0.59965 | 1.20841 | 175 |
| GO_PROTEIN_IMPORT | 0.00053 | 0.01477 | 0.47727 | 0.61085 | 1.22579 | 141 |
| GO_PROTEIN_IMPORT_INTO_MITOCHONDRIAL_MATRIX | 0.00176 | 0.03797 | 0.45506 | 0.79054 | 1.43994 | 18 |
| GO_PROTEIN_LOCALIZATION_TO_CHROMOSOME | 0.00003 | 0.00135 | 0.55733 | 0.69109 | 1.35905 | 72 |
| GO_PROTEIN_LOCALIZATION_TO_ENDOPLASMIC_RETICULUM | 0.00000 | 0.00000 | | 0.81103 | 1.62148 | 127 |
| GO_PROTEIN_LOCALIZATION_TO_MEMBRANE | 0.00000 | 0.00000 | | 0.61937 | 1.27100 | 462 |
| GO_PROTEIN_LOCALIZATION_TO_NUCLEUS | 0.00001 | 0.00056 | 0.59333 | 0.61794 | 1.24950 | 194 |
| GO_PROTEIN_MODIFICATION_BY_SMALL_PROTEIN_REMOVAL | 0.00001 | 0.00051 | 0.59333 | 0.61355 | 1.24532 | 230 |
| GO_PROTEIN_POLYUBIQUITINATION | 0.00001 | 0.00056 | 0.59333 | 0.60263 | 1.22585 | 268 |
| GO_PROTEIN_STABILIZATION | 0.00006 | 0.00257 | 0.53843 | 0.63076 | 1.26575 | 141 |
| GO_PROTEIN_TARGETING | 0.00000 | 0.00000 | | 0.67584 | 1.38259 | 341 |
| GO_PROTEIN_TARGETING_TO_MEMBRANE | 0.00000 | 0.00000 | | 0.77831 | 1.56642 | 157 |
| GO_PURINE_NUCLEOSIDE_BIOSYNTHETIC_PROCESS | 0.00235 | 0.04819 | 0.43171 | 0.80780 | 1.46006 | 15 |
| GO_PURINE_NUCLEOSIDE_METABOLIC_PROCESS | 0.00093 | 0.02418 | 0.47727 | 0.69687 | 1.32853 | 40 |
| GO_PURINE_NUCLEOSIDE_MONOPHOSPHATE_BIOSYNTHETIC_PROC | 0.00178 | 0.03823 | 0.45506 | 0.79853 | 1.44833 | 17 |
| GO_REGENERATION | 0.00165 | 0.03609 | 0.45506 | 0.61809 | 1.22921 | 107 |
| GO_REGULATION_OF_ANIMAL_ORGAN_MORPHOGENESIS | 0.00001 | 0.00035 | 0.61053 | 0.64232 | 1.29038 | 146 |
| GO_REGULATION_OF_APOPTOTIC_SIGNALING_PATHWAY | 0.00148 | 0.03380 | 0.45506 | 0.56803 | 1.15489 | 264 |
| GO_REGULATION_OF_BINDING | 0.00143 | 0.03290 | 0.45506 | 0.57489 | 1.16760 | 237 |
| GO_REGULATION_OF_CELLULAR_AMIDE_METABOLIC_PROCESS | 0.00000 | 0.00001 | 0.71951 | 0.61294 | 1.25288 | 329 |
| GO_REGULATION_OF_CELLULAR_AMINE_METABOLIC_PROCESS | 0.00000 | 0.00000 | 0.73376 | 0.76705 | 1.49623 | 61 |
| GO_REGULATION_OF_CELLULAR_AMINO_ACID_METABOLIC_PROCES | 0.00000 | 0.00000 | 0.71951 | 0.78343 | 1.52063 | 55 |
| GO_REGULATION_OF_CELLULAR_KETONE_METABOLIC_PROCESS | 0.00006 | 0.00235 | 0.55733 | 0.63384 | 1.26860 | 133 |
| GO_REGULATION_OF_CELLULAR_PROTEIN_LOCALIZATION | 0.00004 | 0.00186 | 0.55733 | 0.57101 | 1.17002 | 396 |
| GO_REGULATION_OF_CELL_CYCLE_G2_M_PHASE_TRANSITION | 0.00000 | 0.00011 | 0.64355 | 0.63615 | 1.28407 | 183 |
| GO_REGULATION_OF_CELL_CYCLE_PHASE_TRANSITION | 0.00000 | 0.00000 | 0.73376 | 0.60777 | 1.24412 | 363 |
| GO_REGULATION_OF_CHROMOSOME_ORGANIZATION | 0.00000 | 0.00011 | 0.64355 | 0.60716 | 1.23644 | 277 |
| GO_REGULATION_OF_CHROMOSOME_SEGREGATION | 0.00052 | 0.01471 | 0.47727 | 0.64504 | 1.27788 | 93 |
| GO_REGULATION_OF_CHROMOSOME_SEPARATION | 0.00205 | 0.04358 | 0.43171 | 0.66425 | 1.29416 | 60 |
| GO_REGULATION_OF_DNA_BIOSYNTHETIC_PROCESS | 0.00135 | 0.03205 | 0.45506 | 0.63666 | 1.25637 | 81 |
| GO_REGULATION_OF_DNA_METABOLIC_PROCESS | 0.00003 | 0.00135 | 0.55733 | 0.59698 | 1.21345 | 251 |
| GO_REGULATION_OF_DNA_METHYLATION | 0.00167 | 0.03637 | 0.45506 | 0.76560 | 1.41719 | 23 |
| GO_REGULATION_OF_DNA_TEMPLATED_TRANSCRIPTION_IN_RESPO | 0.00001 | 0.00051 | 0.59333 | 0.68032 | 1.34800 | 94 |
| GO_REGULATION_OF_GENE_EXPRESSION_EPIGENETIC | 0.00074 | 0.01985 | 0.47727 | 0.61816 | 1.23589 | 126 |
| GO_REGULATION_OF_HEMATOPOIETIC_PROGENITOR_CELL_DIFFERE | 0.00000 | 0.00000 | 0.73376 | 0.76056 | 1.49717 | 75 |
| GO_REGULATION_OF_HEMATOPOIETIC_STEM_CELL_DIFFERENTIATIC | 0.00000 | 0.00000 | 0.73376 | 0.76796 | 1.50386 | 65 |
| GO_REGULATION_OF_INNATE_IMMUNE_RESPONSE | 0.00124 | 0.03038 | 0.45506 | 0.59577 | 1.20085 | 176 |

| | | | | | | |
|---|---|---|---|---|---|---|
| GO_REGULATION_OF_INTRINSIC_APOPTOTIC_SIGNALING_PATHWAY | 0.00038 | 0.01123 | 0.49849 | 0.62930 | 1.25659 | 119 |
| GO_REGULATION_OF_MITOTIC_CELL_CYCLE | 0.00000 | 0.00001 | 0.70498 | 0.58879 | 1.20883 | 494 |
| GO_REGULATION_OF_MORPHOGENESIS_OF_AN_EPITHELIUM | 0.00000 | 0.00023 | 0.62726 | 0.65676 | 1.31272 | 122 |
| GO_REGULATION_OF_MRNA_CATABOLIC_PROCESS | 0.00000 | 0.00000 | 0.77494 | 0.67108 | 1.35215 | 173 |
| GO_REGULATION_OF_MRNA_METABOLIC_PROCESS | 0.00000 | 0.00000 | | 0.67102 | 1.36649 | 277 |
| GO_REGULATION_OF_MRNA_PROCESSING | 0.00001 | 0.00070 | 0.59333 | 0.65811 | 1.31412 | 119 |
| GO_REGULATION_OF_MRNA_SPLICING_VIA_SPLICEOSOME | 0.00001 | 0.00034 | 0.61053 | 0.69475 | 1.37322 | 85 |
| GO_REGULATION_OF_NUCLEAR_DIVISION | 0.00104 | 0.02649 | 0.45506 | 0.60460 | 1.21249 | 139 |
| GO_REGULATION_OF_PROTEIN_CATABOLIC_PROCESS | 0.00198 | 0.04215 | 0.43171 | 0.56499 | 1.15178 | 294 |
| GO_REGULATION_OF_PROTEIN_IMPORT | 0.00231 | 0.04775 | 0.43171 | 0.68719 | 1.31321 | 42 |
| GO_REGULATION_OF_PROTEIN_LOCALIZATION_TO_NUCLEUS | 0.00001 | 0.00056 | 0.59333 | 0.67933 | 1.34604 | 94 |
| GO_REGULATION_OF_PROTEIN_MODIFICATION_BY_SMALL_PROTEIN | 0.00017 | 0.00567 | 0.51885 | 0.60547 | 1.22185 | 182 |
| GO_REGULATION_OF_PROTEIN_STABILITY | 0.00016 | 0.00546 | 0.51885 | 0.58924 | 1.19547 | 225 |
| GO_REGULATION_OF_RESPONSE_TO_BIOTIC_STIMULUS | 0.00025 | 0.00786 | 0.49849 | 0.59035 | 1.19838 | 231 |
| GO_REGULATION_OF_RNA_SPLICING | 0.00000 | 0.00011 | 0.64355 | 0.66838 | 1.33416 | 118 |
| GO_REGULATION_OF_SIGNAL_TRANSDUCTION_BY_P53_CLASS_MED | 0.00128 | 0.03079 | 0.45506 | 0.60739 | 1.21910 | 142 |
| GO_REGULATION_OF_SISTER_CHROMATID_SEGREGATION | 0.00022 | 0.00718 | 0.51885 | 0.66854 | 1.31579 | 74 |
| GO_REGULATION_OF_SMALL_MOLECULE_METABOLIC_PROCESS | 0.00003 | 0.00122 | 0.57561 | 0.59257 | 1.20852 | 299 |
| GO_REGULATION_OF_STEM_CELL_DIFFERENTIATION | 0.00000 | 0.00000 | 0.76146 | 0.73443 | 1.45493 | 92 |
| GO_REGULATION_OF_TRANSCRIPTION_FROM_RNA_POLYMERASE_II | 0.00000 | 0.00004 | 0.67496 | 0.73884 | 1.44853 | 67 |
| GO_REGULATION_OF_TRANSLATIONAL_INITIATION | 0.00001 | 0.00052 | 0.59333 | 0.71538 | 1.39879 | 63 |
| GO_REGULATION_OF_UBIQUITIN_DEPENDENT_PROTEIN_CATABOLIC | 0.00150 | 0.03396 | 0.45506 | 0.61037 | 1.22038 | 124 |
| GO_REGULATION_OF_UBIQUITIN_PROTEIN_LIGASE_ACTIVITY | 0.00076 | 0.02017 | 0.47727 | 0.79023 | 1.45740 | 22 |
| GO_REGULATION_OF_UBIQUITIN_PROTEIN_TRANSFERASE_ACTIVITY | 0.00113 | 0.02836 | 0.45506 | 0.69300 | 1.32907 | 45 |
| GO_REGULATION_OF_WNT_SIGNALING_PATHWAY | 0.00010 | 0.00368 | 0.53843 | 0.58590 | 1.18987 | 241 |
| GO_RESPONSE_TO_HEAT | 0.00080 | 0.02130 | 0.47727 | 0.61508 | 1.22941 | 122 |
| GO_RESPONSE_TO_INTERLEUKIN_1 | 0.00000 | 0.00001 | 0.69013 | 0.68785 | 1.37495 | 123 |
| GO_RESPONSE_TO_INTERLEUKIN_12 | 0.00220 | 0.04608 | 0.43171 | 0.70727 | 1.34482 | 37 |
| GO_RESPONSE_TO_INTERLEUKIN_4 | 0.00057 | 0.01583 | 0.47727 | 0.77721 | 1.44121 | 24 |
| GO_RESPONSE_TO_LEUKEMIA_INHIBITORY_FACTOR | 0.00137 | 0.03212 | 0.45506 | 0.64896 | 1.27907 | 78 |
| GO_RESPONSE_TO_OXYGEN_LEVELS | 0.00009 | 0.00324 | 0.53843 | 0.59143 | 1.20202 | 254 |
| GO_RESPONSE_TO_TUMOR_NECROSIS_FACTOR | 0.00087 | 0.02270 | 0.47727 | 0.59445 | 1.19896 | 179 |
| GO_RIBONUCLEOPROTEIN_COMPLEX_BINDING | 0.00000 | 0.00009 | 0.64355 | 0.67201 | 1.34126 | 117 |
| GO_RIBONUCLEOPROTEIN_COMPLEX_BIOGENESIS | 0.00000 | 0.00000 | | 0.69112 | 1.41590 | 395 |
| GO_RIBONUCLEOPROTEIN_COMPLEX_SUBUNIT_ORGANIZATION | 0.00000 | 0.00000 | | 0.71411 | 1.44025 | 178 |
| GO_RIBONUCLEOPROTEIN_GRANULE | 0.00001 | 0.00050 | 0.59333 | 0.62179 | 1.25547 | 185 |
| GO_RIBONUCLEOSIDE_METABOLIC_PROCESS | 0.00143 | 0.03290 | 0.45506 | 0.69587 | 1.33261 | 43 |
| GO_RIBONUCLEOSIDE_TRIPHOSPHATE_METABOLIC_PROCESS | 0.00235 | 0.04819 | 0.43171 | 0.67672 | 1.30177 | 48 |
| GO_RIBOSOMAL_LARGE_SUBUNIT_ASSEMBLY | 0.00003 | 0.00122 | 0.57561 | 0.80735 | 1.51203 | 28 |
| GO_RIBOSOMAL_LARGE_SUBUNIT_BIOGENESIS | 0.00000 | 0.00000 | 0.77494 | 0.78124 | 1.53166 | 67 |
| GO_RIBOSOMAL_SMALL_SUBUNIT_ASSEMBLY | 0.00216 | 0.04546 | 0.43171 | 0.78469 | 1.42929 | 18 |
| GO_RIBOSOMAL_SMALL_SUBUNIT_BIOGENESIS | 0.00002 | 0.00104 | 0.57561 | 0.70622 | 1.38683 | 69 |
| GO_RIBOSOMAL_SUBUNIT | 0.00000 | 0.00000 | | 0.74815 | 1.50771 | 174 |
| GO_RIBOSOME | 0.00000 | 0.00000 | | 0.73378 | 1.48599 | 205 |
| GO_RIBOSOME_ASSEMBLY | 0.00000 | 0.00000 | 0.73376 | 0.78280 | 1.52512 | 60 |
| GO_RIBOSOME_BIOGENESIS | 0.00000 | 0.00000 | | 0.68607 | 1.39718 | 278 |
| GO_RNA_3_END_PROCESSING | 0.00242 | 0.04954 | 0.43171 | 0.60696 | 1.21350 | 126 |
| GO_RNA_CATABOLIC_PROCESS | 0.00000 | 0.00000 | | 0.72228 | 1.47789 | 347 |
| GO_RNA_DEPENDENT_DNA_BIOSYNTHETIC_PROCESS | 0.00148 | 0.03380 | 0.45506 | 0.65932 | 1.28609 | 61 |
| GO_RNA_EXPORT_FROM_NUCLEUS | 0.00007 | 0.00289 | 0.53843 | 0.64324 | 1.28624 | 128 |
| GO_RNA_LOCALIZATION | 0.00000 | 0.00006 | 0.65944 | 0.63312 | 1.28146 | 200 |
| GO_RNA_SPLICING | 0.00000 | 0.00000 | | 0.63359 | 1.29786 | 378 |
| GO_RNA_SPLICING_VIA_TRANSESTERIFICATION_REACTIONS | 0.00000 | 0.00000 | 0.83909 | 0.64393 | 1.31387 | 308 |
| GO_RRNA_BINDING | 0.00000 | 0.00002 | 0.69013 | 0.77159 | 1.49765 | 55 |
| GO_RRNA_METABOLIC_PROCESS | 0.00000 | 0.00000 | 0.82666 | 0.67081 | 1.35931 | 211 |
| GO_SCF_DEPENDENT_PROTEASOMAL_UBIQUITIN_DEPENDENT_PRO | 0.00000 | 0.00011 | 0.64355 | 0.70481 | 1.39084 | 81 |
| GO_SIGNAL_TRANSDUCTION_BY_P53_CLASS_MEDIATOR | 0.00056 | 0.01565 | 0.47727 | 0.58706 | 1.18984 | 215 |
| GO_SINGLE_FERTILIZATION | 0.00014 | 0.00493 | 0.51885 | 0.68819 | 1.34239 | 61 |
| GO_SINGLE_STRANDED_RNA_BINDING | 0.00015 | 0.00511 | 0.51885 | 0.68717 | 1.34040 | 61 |
| GO_SISTER_CHROMATID_COHESION | 0.00036 | 0.01072 | 0.49849 | 0.70667 | 1.35586 | 46 |
| GO_SISTER_CHROMATID_SEGREGATION | 0.00015 | 0.00525 | 0.51885 | 0.61201 | 1.23243 | 164 |
| GO_SMALL_RIBOSOMAL_SUBUNIT | 0.00000 | 0.00000 | 0.71951 | 0.75582 | 1.48372 | 68 |
| GO_SMALL_SUBUNIT_PROCESSOME | 0.00039 | 0.01158 | 0.49849 | 0.74733 | 1.40974 | 32 |
| GO_SPERM_EGG_RECOGNITION | 0.00011 | 0.00391 | 0.53843 | 0.80297 | 1.48897 | 24 |
| GO_SPINDLE_MIDZONE | 0.00128 | 0.03079 | 0.45506 | 0.72879 | 1.37097 | 31 |
| GO_SPLICEOSOMAL_COMPLEX | 0.00000 | 0.00001 | 0.69013 | 0.65652 | 1.32278 | 172 |
| GO_STEM_CELL_DIFFERENTIATION | 0.00000 | 0.00000 | 0.77494 | 0.67801 | 1.36562 | 163 |
| GO_STRUCTURAL_CONSTITUENT_OF_RIBOSOME | 0.00000 | 0.00000 | | 0.75622 | 1.52104 | 152 |
| GO_STRUCTURAL_MOLECULE_ACTIVITY | 0.00000 | 0.00000 | | 0.66355 | 1.35916 | 381 |
| GO_TELOMERASE_RNA_LOCALIZATION | 0.00018 | 0.00606 | 0.51885 | 0.83148 | 1.52144 | 19 |
| GO_THREONINE_TYPE_ENDOPEPTIDASE_ACTIVITY | 0.00001 | 0.00031 | 0.61053 | 0.88055 | 1.60389 | 18 |
| GO_THREONINE_TYPE_PEPTIDASE_ACTIVITY | 0.00001 | 0.00037 | 0.61053 | 0.85926 | 1.58197 | 21 |
| GO_TRANSCRIPTION_COACTIVATOR_ACTIVITY | 0.00159 | 0.03513 | 0.45506 | 0.57883 | 1.17241 | 209 |
| GO_TRANSCRIPTION_COREGULATOR_ACTIVITY | 0.00030 | 0.00921 | 0.49849 | 0.56254 | 1.15173 | 367 |
| GO_TRANSCRIPTION_COUPLED_NUCLEOTIDE_EXCISION_REPAIR | 0.00133 | 0.03174 | 0.45506 | 0.65632 | 1.28885 | 69 |
| GO_TRANSCRIPTION_FACTOR_BINDING | 0.00000 | 0.00029 | 0.61053 | 0.57520 | 1.17987 | 453 |

| | | | | | | |
|---|---|---|---|---|---|---|
| GO_TRANSLATIONAL_ELONGATION | 0.00126 | 0.03071 | 0.45506 | 0.61363 | 1.22489 | 118 |
| GO_TRANSLATIONAL_INITIATION | 0.00000 | 0.00000 | | 0.82136 | 1.65404 | 167 |
| GO_TRANSLATION_FACTOR_ACTIVITY_RNA_BINDING | 0.00000 | 0.00000 | | 0.81785 | 1.60307 | 66 |
| GO_TRANSLATION_INITIATION_FACTOR_ACTIVITY | 0.00000 | 0.00000 | 0.76146 | 0.82116 | 1.57551 | 46 |
| GO_TRANSLATION_INITIATION_FACTOR_BINDING | 0.00009 | 0.00334 | 0.53843 | 0.79111 | 1.48163 | 28 |
| GO_TRANSLATION_PREINITIATION_COMPLEX | 0.00003 | 0.00128 | 0.57561 | 0.88229 | 1.59549 | 16 |
| GO_TRANSLATION_REGULATOR_ACTIVITY | 0.00000 | 0.00000 | | 0.77836 | 1.54644 | 104 |
| GO_TRANSLATION_REGULATOR_ACTIVITY_NUCLEIC_ACID_BINDING | 0.00000 | 0.00000 | | 0.80538 | 1.58931 | 81 |
| GO_TUMOR_NECROSIS_FACTOR_MEDIATED_SIGNALING_PATHWAY | 0.00001 | 0.00051 | 0.59333 | 0.66557 | 1.32616 | 112 |
| GO_T_CELL_RECEPTOR_SIGNALING_PATHWAY | 0.00001 | 0.00033 | 0.61053 | 0.66472 | 1.32872 | 123 |
| GO_U2_TYPE_SPLICEOSOMAL_COMPLEX | 0.00017 | 0.00567 | 0.51885 | 0.65560 | 1.29621 | 86 |
| GO_UBIQUITIN_LIKE_PROTEIN_LIGASE_BINDING | 0.00002 | 0.00109 | 0.57561 | 0.60521 | 1.22914 | 236 |
| GO_UBIQUITIN_PROTEIN_TRANSFERASE_REGULATOR_ACTIVITY | 0.00088 | 0.02309 | 0.47727 | 0.82617 | 1.49400 | 16 |
| GO_UNFOLDED_PROTEIN_BINDING | 0.00000 | 0.00022 | 0.62726 | 0.69709 | 1.37961 | 89 |
| GO_VESICLE_LUMEN | 0.00000 | 0.00000 | 0.73376 | 0.65283 | 1.32024 | 195 |
| GO_VIRAL_GENE_EXPRESSION | 0.00000 | 0.00000 | | 0.76922 | 1.54989 | 173 |
| GO_VIRAL_LIFE_CYCLE | 0.00021 | 0.00697 | 0.51885 | 0.59155 | 1.19967 | 222 |
| HP_ABNORMALITY_OF_MOUTH_SIZE | 0.00168 | 0.03664 | 0.45506 | 0.57709 | 1.16960 | 213 |
| HP_ABNORMALITY_OF_THE_AUDITORY_CANAL | 0.00124 | 0.03038 | 0.45506 | 0.69318 | 1.32150 | 40 |
| HP_ABNORMALITY_OF_THE_NASAL_DORSUM | 0.00010 | 0.00361 | 0.53843 | 0.62127 | 1.24792 | 145 |
| HP_ABNORMAL_CIRCULATING_SERINE_FAMILY_AMINO_ACID_CONC | 0.00154 | 0.03416 | 0.45506 | 0.81582 | 1.47455 | 15 |
| HP_ACUTE_LEUKEMIA | 0.00003 | 0.00120 | 0.57561 | 0.69321 | 1.36331 | 73 |
| HP_ANEMIA_OF_INADEQUATE_PRODUCTION | 0.00110 | 0.02778 | 0.45506 | 0.62349 | 1.24049 | 109 |
| HP_ATRESIA_OF_THE_EXTERNAL_AUDITORY_CANAL | 0.00154 | 0.03416 | 0.45506 | 0.74697 | 1.39559 | 27 |
| HP_DELAYED_PUBERTY | 0.00024 | 0.00776 | 0.51885 | 0.63426 | 1.26424 | 114 |
| HP_DEPRESSED_NASAL_RIDGE | 0.00001 | 0.00056 | 0.59333 | 0.70709 | 1.39192 | 75 |
| HP_LEUKEMIA | 0.00154 | 0.03416 | 0.45506 | 0.62365 | 1.23922 | 105 |
| HP_MACROCYTIC_ANEMIA | 0.00000 | 0.00023 | 0.62726 | 0.77979 | 1.48862 | 41 |
| HP_MIGRAINE | 0.00008 | 0.00299 | 0.53843 | 0.68921 | 1.35457 | 71 |
| HP_THICK_VERMILION_BORDER | 0.00159 | 0.03513 | 0.45506 | 0.59373 | 1.19673 | 176 |
| **Cluster 11** | | | | | | |
| pathway | pval | padj | log2err | ES | NES | size |
| GO_ACTIVATION_OF_INNATE_IMMUNE_RESPONSE | 0.00005 | 0.00222 | 0.55733 | 0.66951 | 1.35525 | 85 |
| GO_AEROBIC_RESPIRATION | 0.00029 | 0.00923 | 0.49849 | 0.68239 | 1.36569 | 63 |
| GO_ALTERNATIVE_MRNA_SPLICING_VIA_SPLICEOSOME | 0.00065 | 0.01798 | 0.47727 | 0.67089 | 1.34268 | 63 |
| GO_AMINE_METABOLIC_PROCESS | 0.00000 | 0.00002 | 0.69013 | 0.72770 | 1.46740 | 77 |
| GO_ANAPHASE_PROMOTING_COMPLEX_DEPENDENT_CATABOLIC_P | 0.00000 | 0.00000 | 0.76146 | 0.75797 | 1.52776 | 74 |
| GO_ANCHORING_JUNCTION | 0.00000 | 0.00000 | 0.80122 | 0.60171 | 1.25530 | 483 |
| GO_ANIMAL_ORGAN_REGENERATION | 0.00004 | 0.00198 | 0.55733 | 0.75127 | 1.47937 | 40 |
| GO_ANTIGEN_PROCESSING_AND_PRESENTATION | 0.00034 | 0.01073 | 0.49849 | 0.61013 | 1.25035 | 144 |
| GO_ANTIGEN_PROCESSING_AND_PRESENTATION_OF_EXOGENOUS_ | 0.00000 | 0.00004 | 0.65944 | 0.75490 | 1.50599 | 59 |
| GO_ANTIGEN_PROCESSING_AND_PRESENTATION_OF_PEPTIDE_ANTI | 0.00004 | 0.00198 | 0.55733 | 0.64082 | 1.30939 | 124 |
| GO_ANTIGEN_PROCESSING_AND_PRESENTATION_OF_PEPTIDE_ANTI | 0.00000 | 0.00012 | 0.62726 | 0.71831 | 1.44443 | 71 |
| GO_ANTIGEN_RECEPTOR_MEDIATED_SIGNALING_PATHWAY | 0.00037 | 0.01120 | 0.49849 | 0.62043 | 1.27010 | 131 |
| GO_ATP_METABOLIC_PROCESS | 0.00067 | 0.01865 | 0.47727 | 0.58852 | 1.20944 | 199 |
| GO_ATP_SYNTHESIS_COUPLED_ELECTRON_TRANSPORT | 0.00023 | 0.00743 | 0.51885 | 0.66252 | 1.33597 | 77 |
| GO_CADHERIN_BINDING | 0.00000 | 0.00007 | 0.64355 | 0.61494 | 1.26897 | 248 |
| GO_CANONICAL_WNT_SIGNALING_PATHWAY | 0.00015 | 0.00518 | 0.51885 | 0.59729 | 1.22697 | 183 |
| GO_CARBOHYDRATE_BINDING | 0.00106 | 0.02673 | 0.45506 | 0.62951 | 1.27574 | 90 |
| GO_CATALYTIC_STEP_2_SPLICEOSOME | 0.00014 | 0.00514 | 0.51885 | 0.66304 | 1.34109 | 82 |
| GO_CELLULAR_AMINO_ACID_METABOLIC_PROCESS | 0.00005 | 0.00214 | 0.55733 | 0.59948 | 1.23317 | 210 |
| GO_CELLULAR_COMPONENT_DISASSEMBLY | 0.00152 | 0.03440 | 0.45506 | 0.54626 | 1.13565 | 391 |
| GO_CELLULAR_KETONE_METABOLIC_PROCESS | 0.00019 | 0.00647 | 0.51885 | 0.61830 | 1.26730 | 146 |
| GO_CELLULAR_NITROGEN_COMPOUND_CATABOLIC_PROCESS | 0.00000 | 0.00000 | | 0.72980 | 1.52023 | 439 |
| GO_CELLULAR_RESPIRATION | 0.00008 | 0.00320 | 0.53843 | 0.62937 | 1.28875 | 137 |
| GO_CELLULAR_RESPONSE_TO_OXYGEN_LEVELS | 0.00000 | 0.00001 | 0.69013 | 0.66200 | 1.35826 | 150 |
| GO_CELL_ACTIVATION_INVOLVED_IN_IMMUNE_RESPONSE | 0.00004 | 0.00180 | 0.55733 | 0.56852 | 1.17976 | 365 |
| GO_CELL_ADHESION_MOLECULE_BINDING | 0.00000 | 0.00005 | 0.65944 | 0.59976 | 1.24302 | 320 |
| GO_CELL_CELL_RECOGNITION | 0.00021 | 0.00708 | 0.51885 | 0.79431 | 1.51507 | 24 |
| GO_CELL_CELL_SIGNALING_BY_WNT | 0.00022 | 0.00725 | 0.51885 | 0.57278 | 1.18515 | 291 |
| GO_CELL_CYCLE_G2_M_PHASE_TRANSITION | 0.00002 | 0.00121 | 0.57561 | 0.60708 | 1.24874 | 212 |
| GO_CELL_CYCLE_PHASE_TRANSITION | 0.00015 | 0.00539 | 0.51885 | 0.55636 | 1.15941 | 447 |
| GO_CELL_SUBSTRATE_JUNCTION | 0.00000 | 0.00000 | | 0.69222 | 1.43229 | 291 |
| GO_CELL_SURFACE_RECEPTOR_SIGNALING_PATHWAY_INVOLVED_IN | 0.00121 | 0.02868 | 0.45506 | 0.55888 | 1.15815 | 318 |
| GO_CHAPERONE_COMPLEX | 0.00008 | 0.00329 | 0.53843 | 0.82724 | 1.55809 | 20 |
| GO_COTRANSLATIONAL_PROTEIN_TARGETING_TO_MEMBRANE | 0.00000 | 0.00000 | | 0.93365 | 1.89327 | 92 |
| GO_CYTOKINE_MEDIATED_SIGNALING_PATHWAY | 0.00000 | 0.00005 | 0.65944 | 0.59509 | 1.23296 | 335 |
| GO_CYTOPLASMIC_TRANSLATION | 0.00000 | 0.00000 | | 0.86454 | 1.75198 | 89 |
| GO_CYTOPLASMIC_TRANSLATIONAL_INITIATION | 0.00000 | 0.00000 | 0.71951 | 0.86154 | 1.66969 | 30 |
| GO_CYTOSOLIC_LARGE_RIBOSOMAL_SUBUNIT | 0.00000 | 0.00000 | | 0.94277 | 1.87706 | 54 |
| GO_CYTOSOLIC_RIBOSOME | 0.00000 | 0.00000 | | 0.94452 | 1.91587 | 97 |
| GO_CYTOSOLIC_SMALL_RIBOSOMAL_SUBUNIT | 0.00000 | 0.00000 | | 0.95404 | 1.87866 | 40 |
| GO_DEFENSE_RESPONSE_TO_OTHER_ORGANISM | 0.00053 | 0.01526 | 0.47727 | 0.55262 | 1.14980 | 415 |
| GO_DNA_TEMPLATED_TRANSCRIPTION_ELONGATION | 0.00061 | 0.01704 | 0.47727 | 0.63104 | 1.27986 | 96 |
| GO_ELECTRON_TRANSFER_ACTIVITY | 0.00174 | 0.03879 | 0.45506 | 0.63691 | 1.28432 | 77 |
| GO_ELECTRON_TRANSPORT_CHAIN | 0.00028 | 0.00889 | 0.49849 | 0.62402 | 1.27747 | 131 |

| | | | | | | |
|---|---|---|---|---|---|---|
| GO_ENDOPEPTIDASE_COMPLEX | 0.00000 | 0.00003 | 0.67496 | 0.75042 | 1.49907 | 61 |
| GO_ENERGY_DERIVATION_BY_OXIDATION_OF_ORGANIC_COMPOUN | 0.00011 | 0.00414 | 0.53843 | 0.60473 | 1.24146 | 179 |
| GO_ESTABLISHMENT_OF_PROTEIN_LOCALIZATION_TO_ENDOPLASM | 0.00000 | 0.00000 | | 0.91680 | 1.86149 | 103 |
| GO_ESTABLISHMENT_OF_PROTEIN_LOCALIZATION_TO_MEMBRANE | 0.00000 | 0.00000 | | 0.78222 | 1.61401 | 240 |
| GO_ESTABLISHMENT_OF_PROTEIN_LOCALIZATION_TO_ORGANELLE | 0.00000 | 0.00000 | | 0.71504 | 1.48745 | 407 |
| GO_ESTABLISHMENT_OF_RNA_LOCALIZATION | 0.00007 | 0.00295 | 0.53843 | 0.61674 | 1.26643 | 164 |
| GO_ESTABLISHMENT_OF_TISSUE_POLARITY | 0.00000 | 0.00002 | 0.67496 | 0.71007 | 1.43873 | 86 |
| GO_EUKARYOTIC_TRANSLATION_INITIATION_FACTOR_3_COMPLEX | 0.00001 | 0.00033 | 0.61053 | 0.90762 | 1.68482 | 15 |
| GO_FC_EPSILON_RECEPTOR_SIGNALING_PATHWAY | 0.00001 | 0.00050 | 0.59333 | 0.69167 | 1.39877 | 83 |
| GO_FC_RECEPTOR_SIGNALING_PATHWAY | 0.00004 | 0.00198 | 0.55733 | 0.63488 | 1.29761 | 125 |
| GO_FICOLIN_1_RICH_GRANULE | 0.00000 | 0.00000 | 0.77494 | 0.73236 | 1.48537 | 96 |
| GO_GENERATION_OF_PRECURSOR_METABOLITES_AND_ENERGY | 0.00110 | 0.02718 | 0.45506 | 0.55890 | 1.15800 | 327 |
| GO_GLYCOSYL_COMPOUND_METABOLIC_PROCESS | 0.00075 | 0.02058 | 0.47727 | 0.65808 | 1.32192 | 68 |
| GO_GTP_METABOLIC_PROCESS | 0.00181 | 0.04012 | 0.45506 | 0.77521 | 1.45368 | 19 |
| GO_HEMATOPOIETIC_PROGENITOR_CELL_DIFFERENTIATION | 0.00000 | 0.00001 | 0.71951 | 0.70235 | 1.42763 | 107 |
| GO_HEMATOPOIETIC_STEM_CELL_DIFFERENTIATION | 0.00000 | 0.00000 | 0.71951 | 0.76119 | 1.52843 | 67 |
| GO_INNATE_IMMUNE_RESPONSE | 0.00119 | 0.02848 | 0.45506 | 0.55771 | 1.15606 | 340 |
| GO_INNATE_IMMUNE_RESPONSE_ACTIVATING_SIGNAL_TRANSDUCT | 0.00001 | 0.00048 | 0.59333 | 0.70615 | 1.41849 | 68 |
| GO_INNER_MITOCHONDRIAL_MEMBRANE_PROTEIN_COMPLEX | 0.00111 | 0.02728 | 0.45506 | 0.62953 | 1.27573 | 89 |
| GO_INTERLEUKIN_1_MEDIATED_SIGNALING_PATHWAY | 0.00000 | 0.00000 | 0.71951 | 0.74488 | 1.50136 | 74 |
| GO_INTRINSIC_APOPTOTIC_SIGNALING_PATHWAY | 0.00086 | 0.02253 | 0.47727 | 0.57851 | 1.18857 | 185 |
| GO_LARGE_RIBOSOMAL_SUBUNIT | 0.00000 | 0.00000 | | 0.77706 | 1.58080 | 109 |
| GO_LEUKOCYTE_MEDIATED_IMMUNITY | 0.00008 | 0.00322 | 0.53843 | 0.56627 | 1.17666 | 385 |
| GO_LIVER_REGENERATION | 0.00183 | 0.04037 | 0.45506 | 0.79049 | 1.47238 | 17 |
| GO_LYSOSOMAL_LUMEN | 0.00165 | 0.03702 | 0.45506 | 0.67104 | 1.33604 | 54 |
| GO_MATURATION_OF_LSU_RRNA | 0.00058 | 0.01652 | 0.47727 | 0.75931 | 1.45931 | 27 |
| GO_MITOCHONDRIAL_PROTEIN_COMPLEX | 0.00009 | 0.00366 | 0.53843 | 0.59389 | 1.22126 | 207 |
| GO_MODIFICATION_DEPENDENT_MACROMOLECULE_CATABOLIC_PR | 0.00011 | 0.00414 | 0.53843 | 0.55180 | 1.15131 | 477 |
| GO_MONOCARBOXYLIC_ACID_TRANSMEMBRANE_TRANSPORTER_A | 0.00122 | 0.02899 | 0.45506 | 0.78468 | 1.47145 | 19 |
| GO_MORPHOGENESIS_OF_AN_EPITHELIUM | 0.00061 | 0.01704 | 0.47727 | 0.56482 | 1.16802 | 279 |
| GO_MORPHOGENESIS_OF_A_POLARIZED_EPITHELIUM | 0.00000 | 0.00002 | 0.69013 | 0.70011 | 1.42100 | 101 |
| GO_MRNA_5_UTR_BINDING | 0.00004 | 0.00171 | 0.55733 | 0.83325 | 1.57992 | 22 |
| GO_MRNA_BINDING | 0.00000 | 0.00000 | | 0.66114 | 1.36297 | 231 |
| GO_MRNA_EXPORT_FROM_NUCLEUS | 0.00079 | 0.02153 | 0.47727 | 0.62095 | 1.25953 | 97 |
| GO_MRNA_PROCESSING | 0.00000 | 0.00002 | 0.67496 | 0.58605 | 1.21918 | 410 |
| GO_MYD88_INDEPENDENT_TOLL_LIKE_RECEPTOR_SIGNALING_PATH | 0.00029 | 0.00923 | 0.49849 | 0.79582 | 1.50565 | 21 |
| GO_MYELOID_LEUKOCYTE_ACTIVATION | 0.00000 | 0.00008 | 0.64355 | 0.59185 | 1.22701 | 345 |
| GO_MYELOID_LEUKOCYTE_MEDIATED_IMMUNITY | 0.00000 | 0.00001 | 0.69013 | 0.61017 | 1.26432 | 314 |
| GO_NCRNA_METABOLIC_PROCESS | 0.00012 | 0.00447 | 0.53843 | 0.56449 | 1.17318 | 387 |
| GO_NCRNA_PROCESSING | 0.00002 | 0.00080 | 0.57561 | 0.58465 | 1.21121 | 328 |
| GO_NEGATIVE_REGULATION_OF_CANONICAL_WNT_SIGNALING_PAT | 0.00005 | 0.00222 | 0.55733 | 0.64747 | 1.31718 | 109 |
| GO_NEGATIVE_REGULATION_OF_CELL_CYCLE_G2_M_PHASE_TRANS | 0.00001 | 0.00072 | 0.59333 | 0.68092 | 1.37834 | 85 |
| GO_NEGATIVE_REGULATION_OF_CELL_CYCLE_PHASE_TRANSITION | 0.00087 | 0.02269 | 0.47727 | 0.58417 | 1.19924 | 176 |
| GO_NEGATIVE_REGULATION_OF_CELL_CYCLE_PROCESS | 0.00012 | 0.00444 | 0.53843 | 0.58551 | 1.20811 | 241 |
| GO_NEGATIVE_REGULATION_OF_CELL_DEATH | 0.00035 | 0.01081 | 0.49849 | 0.54646 | 1.14022 | 487 |
| GO_NEGATIVE_REGULATION_OF_MITOTIC_CELL_CYCLE | 0.00146 | 0.03328 | 0.45506 | 0.56971 | 1.17369 | 222 |
| GO_NEGATIVE_REGULATION_OF_MRNA_METABOLIC_PROCESS | 0.00113 | 0.02762 | 0.45506 | 0.66533 | 1.32993 | 62 |
| GO_NEGATIVE_REGULATION_OF_WNT_SIGNALING_PATHWAY | 0.00006 | 0.00242 | 0.55733 | 0.63875 | 1.30425 | 121 |
| GO_NEURON_TO_NEURON_SYNAPSE | 0.00036 | 0.01112 | 0.49849 | 0.60376 | 1.24020 | 161 |
| GO_NIK_NF_KAPPAB_SIGNALING | 0.00001 | 0.00029 | 0.61053 | 0.67022 | 1.36345 | 109 |
| GO_NON_CANONICAL_WNT_SIGNALING_PATHWAY | 0.00001 | 0.00050 | 0.59333 | 0.67605 | 1.37154 | 99 |
| GO_NUCLEAR_EXPORT | 0.00022 | 0.00719 | 0.51885 | 0.60859 | 1.24906 | 171 |
| GO_NUCLEAR_TRANSCRIBED_MRNA_CATABOLIC_PROCESS | 0.00000 | 0.00000 | | 0.82630 | 1.69774 | 186 |
| GO_NUCLEAR_TRANSCRIBED_MRNA_CATABOLIC_PROCESS_NONSEN | 0.00000 | 0.00000 | | 0.91126 | 1.85638 | 113 |
| GO_NUCLEAR_TRANSPORT | 0.00000 | 0.00015 | 0.62726 | 0.60097 | 1.24224 | 267 |
| GO_NUCLEOBASE_CONTAINING_COMPOUND_TRANSPORT | 0.00038 | 0.01135 | 0.49849 | 0.59355 | 1.21978 | 199 |
| GO_NUCLEOSIDE_METABOLIC_PROCESS | 0.00111 | 0.02728 | 0.45506 | 0.67338 | 1.34072 | 57 |
| GO_NUCLEOTIDE_EXCISION_REPAIR_DNA_DAMAGE_RECOGNITION | 0.00100 | 0.02568 | 0.45506 | 0.77576 | 1.47092 | 22 |
| GO_NUCLEOTIDE_EXCISION_REPAIR_DNA_DUPLEX_UNWINDING | 0.00172 | 0.03853 | 0.45506 | 0.77494 | 1.45960 | 20 |
| GO_ORGANIC_CYCLIC_COMPOUND_CATABOLIC_PROCESS | 0.00000 | 0.00000 | | 0.72248 | 1.50602 | 452 |
| GO_OXIDATIVE_PHOSPHORYLATION | 0.00143 | 0.03284 | 0.45506 | 0.61841 | 1.25518 | 101 |
| GO_OXIDOREDUCTASE_COMPLEX | 0.00178 | 0.03945 | 0.45506 | 0.62654 | 1.26729 | 81 |
| GO_PEPTIDASE_COMPLEX | 0.00000 | 0.00009 | 0.64355 | 0.70984 | 1.43139 | 77 |
| GO_PIGMENT_GRANULE | 0.00082 | 0.02218 | 0.47727 | 0.64518 | 1.30039 | 75 |
| GO_POLYSOMAL_RIBOSOME | 0.00000 | 0.00000 | | 0.93399 | 1.81394 | 31 |
| GO_POLYSOME | 0.00000 | 0.00000 | | 0.83497 | 1.66902 | 62 |
| GO_POLY_PURINE_TRACT_BINDING | 0.00060 | 0.01695 | 0.47727 | 0.79463 | 1.49668 | 20 |
| GO_POLY_PYRIMIDINE_TRACT_BINDING | 0.00018 | 0.00623 | 0.51885 | 0.81906 | 1.53591 | 19 |
| GO_POSITIVE_REGULATION_OF_CANONICAL_WNT_SIGNALING_PATH | 0.00000 | 0.00000 | 0.71951 | 0.71889 | 1.45778 | 92 |
| GO_POSITIVE_REGULATION_OF_CELLULAR_AMIDE_METABOLIC_PRC | 0.00005 | 0.00217 | 0.55733 | 0.63921 | 1.30520 | 119 |
| GO_POSITIVE_REGULATION_OF_CELLULAR_PROTEIN_LOCALIZATION | 0.00145 | 0.03313 | 0.45506 | 0.56884 | 1.17255 | 230 |
| GO_POSITIVE_REGULATION_OF_CELL_DEATH | 0.00078 | 0.02139 | 0.47727 | 0.55609 | 1.15381 | 358 |
| GO_POSITIVE_REGULATION_OF_CELL_POPULATION_PROLIFERATION | 0.00141 | 0.03254 | 0.45506 | 0.55091 | 1.14510 | 389 |
| GO_POSITIVE_REGULATION_OF_DEFENSE_RESPONSE | 0.00121 | 0.02868 | 0.45506 | 0.59180 | 1.21565 | 161 |
| GO_POSITIVE_REGULATION_OF_ESTABLISHMENT_OF_PROTEIN_LOCA | 0.00092 | 0.02377 | 0.47727 | 0.57312 | 1.18072 | 222 |
| GO_POSITIVE_REGULATION_OF_INNATE_IMMUNE_RESPONSE | 0.00040 | 0.01199 | 0.49849 | 0.62459 | 1.27064 | 109 |

| | | | | | | |
|---|---|---|---|---|---|---|
| GO_POSITIVE_REGULATION_OF_INTRINSIC_APOPTOTIC_SIGNALING_ | 0.00108 | 0.02696 | 0.45506 | 0.71760 | 1.40328 | 35 |
| GO_POSITIVE_REGULATION_OF_NUCLEOCYTOPLASMIC_TRANSPORT | 0.00102 | 0.02592 | 0.45506 | 0.68460 | 1.35881 | 49 |
| GO_POSITIVE_REGULATION_OF_PROTEIN_LOCALIZATION_TO_NUCLE | 0.00111 | 0.02728 | 0.45506 | 0.67415 | 1.34391 | 58 |
| GO_POSITIVE_REGULATION_OF_RESPONSE_TO_BIOTIC_STIMULUS | 0.00029 | 0.00932 | 0.49849 | 0.61709 | 1.26213 | 127 |
| GO_POSITIVE_REGULATION_OF_TRANSLATION | 0.00004 | 0.00181 | 0.55733 | 0.65766 | 1.33485 | 101 |
| GO_POSITIVE_REGULATION_OF_WNT_SIGNALING_PATHWAY | 0.00000 | 0.00003 | 0.67496 | 0.68474 | 1.39338 | 110 |
| GO_POSTSYNAPSE | 0.00007 | 0.00280 | 0.53843 | 0.58784 | 1.21554 | 270 |
| GO_POSTTRANSCRIPTIONAL_REGULATION_OF_GENE_EXPRESSION | 0.00000 | 0.00000 | | 0.64024 | 1.33566 | 461 |
| GO_POST_TRANSLATIONAL_PROTEIN_MODIFICATION | 0.00021 | 0.00708 | 0.51885 | 0.58553 | 1.20648 | 224 |
| GO_PRERIBOSOME | 0.00039 | 0.01184 | 0.49849 | 0.66304 | 1.33189 | 68 |
| GO_PROTEASOMAL_PROTEIN_CATABOLIC_PROCESS | 0.00085 | 0.02239 | 0.47727 | 0.55495 | 1.15190 | 370 |
| GO_PROTEASOMAL_UBIQUITIN_INDEPENDENT_PROTEIN_CATABOLIC | 0.00019 | 0.00647 | 0.51885 | 0.81743 | 1.53285 | 19 |
| GO_PROTEASOME_ACCESSORY_COMPLEX | 0.00011 | 0.00430 | 0.53843 | 0.81498 | 1.55118 | 23 |
| GO_PROTEASOME_CORE_COMPLEX | 0.00052 | 0.01507 | 0.47727 | 0.81829 | 1.52415 | 17 |
| GO_PROTEIN_DNA_COMPLEX_SUBUNIT_ORGANIZATION | 0.00102 | 0.02592 | 0.45506 | 0.58889 | 1.20873 | 166 |
| GO_PROTEIN_FOLDING | 0.00004 | 0.00194 | 0.55733 | 0.61506 | 1.26274 | 163 |
| GO_PROTEIN_IMPORT | 0.00159 | 0.03592 | 0.45506 | 0.59778 | 1.22386 | 134 |
| GO_PROTEIN_LOCALIZATION_TO_ENDOPLASMIC_RETICULUM | 0.00000 | 0.00000 | | 0.89104 | 1.82246 | 127 |
| GO_PROTEIN_LOCALIZATION_TO_MEMBRANE | 0.00000 | 0.00000 | | 0.68271 | 1.42056 | 416 |
| GO_PROTEIN_LOCALIZATION_TO_MITOCHONDRION | 0.00135 | 0.03139 | 0.45506 | 0.61490 | 1.24914 | 105 |
| GO_PROTEIN_LOCALIZATION_TO_NUCLEUS | 0.00097 | 0.02485 | 0.47727 | 0.58950 | 1.21057 | 181 |
| GO_PROTEIN_MODIFICATION_BY_SMALL_PROTEIN_REMOVAL | 0.00009 | 0.00359 | 0.53843 | 0.59677 | 1.22844 | 217 |
| GO_PROTEIN_POLYUBIQUITINATION | 0.00013 | 0.00476 | 0.51885 | 0.58974 | 1.21670 | 243 |
| GO_PROTEIN_STABILIZATION | 0.00043 | 0.01260 | 0.49849 | 0.61822 | 1.26558 | 131 |
| GO_PROTEIN_TARGETING | 0.00000 | 0.00000 | | 0.73772 | 1.52895 | 320 |
| GO_PROTEIN_TARGETING_TO_MEMBRANE | 0.00000 | 0.00000 | | 0.87382 | 1.79154 | 147 |
| GO_PROTEIN_TARGETING_TO_MITOCHONDRION | 0.00191 | 0.04167 | 0.45506 | 0.63137 | 1.27298 | 76 |
| GO_REGENERATION | 0.00001 | 0.00038 | 0.61053 | 0.68074 | 1.38042 | 92 |
| GO_REGULATION_OF_ANIMAL_ORGAN_MORPHOGENESIS | 0.00000 | 0.00019 | 0.62726 | 0.66385 | 1.35549 | 121 |
| GO_REGULATION_OF_APOPTOTIC_SIGNALING_PATHWAY | 0.00084 | 0.02235 | 0.47727 | 0.57313 | 1.18209 | 235 |
| GO_REGULATION_OF_CELLULAR_AMIDE_METABOLIC_PROCESS | 0.00000 | 0.00008 | 0.64355 | 0.59473 | 1.23191 | 312 |
| GO_REGULATION_OF_CELLULAR_AMINE_METABOLIC_PROCESS | 0.00000 | 0.00001 | 0.69013 | 0.76980 | 1.53089 | 55 |
| GO_REGULATION_OF_CELLULAR_AMINO_ACID_METABOLIC_PROCES | 0.00000 | 0.00002 | 0.69013 | 0.78050 | 1.55084 | 52 |
| GO_REGULATION_OF_CELLULAR_KETONE_METABOLIC_PROCESS | 0.00005 | 0.00214 | 0.55733 | 0.64656 | 1.31878 | 115 |
| GO_REGULATION_OF_CELLULAR_PROTEIN_CATABOLIC_PROCESS | 0.00205 | 0.04457 | 0.43171 | 0.57260 | 1.17624 | 183 |
| GO_REGULATION_OF_CELLULAR_PROTEIN_LOCALIZATION | 0.00119 | 0.02848 | 0.45506 | 0.54754 | 1.13624 | 365 |
| GO_REGULATION_OF_CELL_CYCLE_G2_M_PHASE_TRANSITION | 0.00002 | 0.00092 | 0.57561 | 0.62406 | 1.28092 | 172 |
| GO_REGULATION_OF_CELL_CYCLE_PHASE_TRANSITION | 0.00126 | 0.02974 | 0.45506 | 0.55506 | 1.14990 | 328 |
| GO_REGULATION_OF_DNA_TEMPLATED_TRANSCRIPTION_IN_RESPO | 0.00000 | 0.00000 | 0.74774 | 0.73200 | 1.48340 | 89 |
| GO_REGULATION_OF_ESTABLISHMENT_OF_PROTEIN_LOCALIZATION_ | 0.00087 | 0.02269 | 0.47727 | 0.68043 | 1.35190 | 50 |
| GO_REGULATION_OF_HEMATOPOIETIC_PROGENITOR_CELL_DIFFERE | 0.00000 | 0.00000 | 0.74774 | 0.75643 | 1.52131 | 72 |
| GO_REGULATION_OF_HEMATOPOIETIC_STEM_CELL_DIFFERENTIATIC | 0.00000 | 0.00000 | 0.71951 | 0.77098 | 1.54013 | 61 |
| GO_REGULATION_OF_INNATE_IMMUNE_RESPONSE | 0.00132 | 0.03066 | 0.45506 | 0.58724 | 1.20630 | 160 |
| GO_REGULATION_OF_INTRINSIC_APOPTOTIC_SIGNALING_PATHWAY | 0.00006 | 0.00245 | 0.55733 | 0.65173 | 1.32369 | 104 |
| GO_REGULATION_OF_MORPHOGENESIS_OF_AN_EPITHELIUM | 0.00001 | 0.00067 | 0.59333 | 0.67311 | 1.36556 | 99 |
| GO_REGULATION_OF_MRNA_CATABOLIC_PROCESS | 0.00000 | 0.00000 | 0.76146 | 0.67222 | 1.38024 | 169 |
| GO_REGULATION_OF_MRNA_METABOLIC_PROCESS | 0.00000 | 0.00000 | | 0.66074 | 1.36628 | 270 |
| GO_REGULATION_OF_MRNA_PROCESSING | 0.00003 | 0.00157 | 0.55733 | 0.64631 | 1.31775 | 114 |
| GO_REGULATION_OF_MRNA_SPLICING_VIA_SPLICEOSOME | 0.00002 | 0.00100 | 0.57561 | 0.68285 | 1.38119 | 81 |
| GO_REGULATION_OF_NUCLEOCYTOPLASMIC_TRANSPORT | 0.00128 | 0.03005 | 0.45506 | 0.63409 | 1.28129 | 79 |
| GO_REGULATION_OF_PROTEIN_CATABOLIC_PROCESS | 0.00013 | 0.00457 | 0.51885 | 0.58163 | 1.20237 | 266 |
| GO_REGULATION_OF_PROTEIN_IMPORT | 0.00085 | 0.02239 | 0.47727 | 0.71019 | 1.39849 | 40 |
| GO_REGULATION_OF_PROTEIN_LOCALIZATION_TO_NUCLEUS | 0.00087 | 0.02266 | 0.47727 | 0.64102 | 1.29758 | 85 |
| GO_REGULATION_OF_PROTEIN_MODIFICATION_BY_SMALL_PROTEIN | 0.00084 | 0.02235 | 0.47727 | 0.59271 | 1.21678 | 165 |
| GO_REGULATION_OF_RNA_SPLICING | 0.00000 | 0.00018 | 0.62726 | 0.67387 | 1.37277 | 113 |
| GO_REGULATION_OF_SMALL_MOLECULE_METABOLIC_PROCESS | 0.00119 | 0.02848 | 0.45506 | 0.56548 | 1.16842 | 261 |
| GO_REGULATION_OF_STEM_CELL_DIFFERENTIATION | 0.00000 | 0.00000 | 0.76146 | 0.75068 | 1.51811 | 83 |
| GO_REGULATION_OF_TRANSCRIPTION_FROM_RNA_POLYMERASE_II | 0.00000 | 0.00000 | 0.74774 | 0.78190 | 1.56195 | 61 |
| GO_REGULATION_OF_TRANSLATIONAL_INITIATION | 0.00027 | 0.00865 | 0.49849 | 0.68045 | 1.36291 | 64 |
| GO_REGULATION_OF_WNT_SIGNALING_PATHWAY | 0.00012 | 0.00452 | 0.51885 | 0.59646 | 1.22563 | 202 |
| GO_RESPIRASOME | 0.00130 | 0.03036 | 0.45506 | 0.64369 | 1.29639 | 73 |
| GO_RESPIRATORY_CHAIN_COMPLEX | 0.00110 | 0.02718 | 0.45506 | 0.66700 | 1.33326 | 62 |
| GO_RESPIRATORY_ELECTRON_TRANSPORT_CHAIN | 0.00009 | 0.00357 | 0.53843 | 0.65554 | 1.32843 | 89 |
| GO_RESPONSE_TO_INTERLEUKIN_1 | 0.00000 | 0.00000 | 0.71951 | 0.70392 | 1.43083 | 107 |
| GO_RESPONSE_TO_INTERLEUKIN_12 | 0.00010 | 0.00394 | 0.53843 | 0.75146 | 1.46949 | 35 |
| GO_RESPONSE_TO_INTERLEUKIN_4 | 0.00187 | 0.04102 | 0.45506 | 0.77408 | 1.45798 | 20 |
| GO_RESPONSE_TO_OXYGEN_LEVELS | 0.00020 | 0.00685 | 0.51885 | 0.58575 | 1.20693 | 224 |
| GO_RESPONSE_TO_TUMOR_NECROSIS_FACTOR | 0.00000 | 0.00015 | 0.62726 | 0.63579 | 1.30530 | 163 |
| GO_RIBONUCLEOPROTEIN_COMPLEX_BINDING | 0.00011 | 0.00429 | 0.53843 | 0.63080 | 1.28803 | 119 |
| GO_RIBONUCLEOPROTEIN_COMPLEX_BIOGENESIS | 0.00000 | 0.00000 | | 0.67547 | 1.40389 | 388 |
| GO_RIBONUCLEOPROTEIN_COMPLEX_SUBUNIT_ORGANIZATION | 0.00000 | 0.00000 | | 0.70704 | 1.45146 | 176 |
| GO_RIBONUCLEOPROTEIN_GRANULE | 0.00022 | 0.00722 | 0.51885 | 0.59829 | 1.22831 | 173 |
| GO_RIBONUCLEOSIDE_TRIPHOSPHATE_METABOLIC_PROCESS | 0.00108 | 0.02696 | 0.45506 | 0.69449 | 1.37083 | 43 |
| GO_RIBOSOMAL_LARGE_SUBUNIT_ASSEMBLY | 0.00003 | 0.00164 | 0.55733 | 0.79863 | 1.53756 | 28 |
| GO_RIBOSOMAL_LARGE_SUBUNIT_BIOGENESIS | 0.00000 | 0.00000 | 0.74774 | 0.76466 | 1.53439 | 66 |
| GO_RIBOSOMAL_SMALL_SUBUNIT_ASSEMBLY | 0.00000 | 0.00004 | 0.65944 | 0.90634 | 1.69450 | 18 |

| | | | | | | |
|---|---|---|---|---|---|---|
| GO_RIBOSOMAL_SMALL_SUBUNIT_BINDING | 0.00054 | 0.01563 | 0.47727 | 0.81746 | 1.52261 | 17 |
| GO_RIBOSOMAL_SMALL_SUBUNIT_BIOGENESIS | 0.00001 | 0.00051 | 0.59333 | 0.70528 | 1.41675 | 68 |
| GO_RIBOSOMAL_SUBUNIT | 0.00000 | 0.00000 | | 0.79484 | 1.63184 | 173 |
| GO_RIBOSOME | 0.00000 | 0.00000 | | 0.76978 | 1.58295 | 207 |
| GO_RIBOSOME_ASSEMBLY | 0.00000 | 0.00000 | 0.76146 | 0.78437 | 1.56688 | 61 |
| GO_RIBOSOME_BIOGENESIS | 0.00000 | 0.00000 | | 0.66875 | 1.38288 | 273 |
| GO_RNA_CATABOLIC_PROCESS | 0.00000 | 0.00000 | | 0.75787 | 1.57105 | 342 |
| GO_RNA_EXPORT_FROM_NUCLEUS | 0.00013 | 0.00476 | 0.51885 | 0.62833 | 1.28379 | 123 |
| GO_RNA_LOCALIZATION | 0.00000 | 0.00003 | 0.67496 | 0.63655 | 1.30822 | 195 |
| GO_RNA_SPLICING | 0.00000 | 0.00000 | 0.77494 | 0.61172 | 1.26905 | 362 |
| GO_RNA_SPLICING_VIA_TRANSESTERIFICATION_REACTIONS | 0.00000 | 0.00000 | 0.77494 | 0.62533 | 1.29488 | 297 |
| GO_ROUGH_ENDOPLASMIC_RETICULUM | 0.00019 | 0.00663 | 0.51885 | 0.71379 | 1.41750 | 51 |
| GO_RRNA_BINDING | 0.00000 | 0.00003 | 0.67496 | 0.77348 | 1.53840 | 53 |
| GO_RRNA_METABOLIC_PROCESS | 0.00000 | 0.00000 | 0.74774 | 0.64704 | 1.33056 | 207 |
| GO_SCF_DEPENDENT_PROTEASOMAL_UBIQUITIN_DEPENDENT_PRO | 0.00000 | 0.00015 | 0.62726 | 0.71191 | 1.43178 | 72 |
| GO_SECRETORY_GRANULE | 0.00000 | 0.00008 | 0.64355 | 0.57940 | 1.20624 | 423 |
| GO_SECRETORY_VESICLE | 0.00003 | 0.00148 | 0.57561 | 0.55505 | 1.15828 | 499 |
| GO_SINGLE_STRANDED_RNA_BINDING | 0.00035 | 0.01090 | 0.49849 | 0.68665 | 1.36978 | 60 |
| GO_SMALL_RIBOSOMAL_SUBUNIT | 0.00000 | 0.00000 | | 0.83077 | 1.66814 | 67 |
| GO_SPERM_EGG_RECOGNITION | 0.00019 | 0.00651 | 0.51885 | 0.81714 | 1.53231 | 19 |
| GO_SPLICEOSOMAL_COMPLEX | 0.00005 | 0.00214 | 0.55733 | 0.62212 | 1.27683 | 171 |
| GO_SPLICEOSOMAL_COMPLEX_ASSEMBLY | 0.00146 | 0.03328 | 0.45506 | 0.68797 | 1.36147 | 46 |
| GO_STEM_CELL_DIFFERENTIATION | 0.00000 | 0.00000 | 0.71951 | 0.67177 | 1.37607 | 141 |
| GO_STRUCTURAL_CONSTITUENT_OF_RIBOSOME | 0.00000 | 0.00000 | | 0.81518 | 1.67263 | 152 |
| GO_STRUCTURAL_MOLECULE_ACTIVITY | 0.00000 | 0.00000 | | 0.71051 | 1.47187 | 333 |
| GO_THREONINE_TYPE_ENDOPEPTIDASE_ACTIVITY | 0.00046 | 0.01347 | 0.49849 | 0.82024 | 1.52779 | 17 |
| GO_THREONINE_TYPE_PEPTIDASE_ACTIVITY | 0.00040 | 0.01192 | 0.49849 | 0.79138 | 1.49724 | 21 |
| GO_TRANSCRIPTION_ELONGATION_FROM_RNA_POLYMERASE_II_PR | 0.00054 | 0.01563 | 0.47727 | 0.65716 | 1.32145 | 71 |
| GO_TRANSCRIPTION_FACTOR_BINDING | 0.00076 | 0.02099 | 0.47727 | 0.54647 | 1.13683 | 410 |
| GO_TRANSLATIONAL_ELONGATION | 0.00056 | 0.01608 | 0.47727 | 0.61501 | 1.25578 | 121 |
| GO_TRANSLATIONAL_INITIATION | 0.00000 | 0.00000 | | 0.86191 | 1.76997 | 168 |
| GO_TRANSLATION_ELONGATION_FACTOR_ACTIVITY | 0.00005 | 0.00222 | 0.55733 | 0.88286 | 1.63886 | 15 |
| GO_TRANSLATION_FACTOR_ACTIVITY_RNA_BINDING | 0.00000 | 0.00000 | 0.78819 | 0.77198 | 1.55234 | 71 |
| GO_TRANSLATION_INITIATION_FACTOR_ACTIVITY | 0.00000 | 0.00007 | 0.64355 | 0.77610 | 1.53586 | 46 |
| GO_TRANSLATION_INITIATION_FACTOR_BINDING | 0.00004 | 0.00198 | 0.55733 | 0.79483 | 1.53024 | 28 |
| GO_TRANSLATION_PREINITIATION_COMPLEX | 0.00000 | 0.00004 | 0.65944 | 0.91478 | 1.69896 | 16 |
| GO_TRANSLATION_REGULATOR_ACTIVITY | 0.00000 | 0.00000 | 0.81404 | 0.73460 | 1.49444 | 109 |
| GO_TRANSLATION_REGULATOR_ACTIVITY_NUCLEIC_ACID_BINDING | 0.00000 | 0.00000 | 0.81404 | 0.75464 | 1.52905 | 86 |
| GO_TUMOR_NECROSIS_FACTOR_MEDIATED_SIGNALING_PATHWAY | 0.00000 | 0.00022 | 0.61053 | 0.67586 | 1.37270 | 104 |
| GO_T_CELL_RECEPTOR_SIGNALING_PATHWAY | 0.00004 | 0.00194 | 0.55733 | 0.64700 | 1.32020 | 116 |
| GO_U12_TYPE_SPLICEOSOMAL_COMPLEX | 0.00119 | 0.02848 | 0.45506 | 0.74871 | 1.43895 | 27 |
| GO_UBIQUITIN_LIKE_PROTEIN_LIGASE_BINDING | 0.00007 | 0.00287 | 0.53843 | 0.59826 | 1.23150 | 217 |
| GO_UNFOLDED_PROTEIN_BINDING | 0.00000 | 0.00013 | 0.62726 | 0.70254 | 1.41962 | 79 |
| GO_VACUOLAR_LUMEN | 0.00034 | 0.01073 | 0.49849 | 0.63709 | 1.29074 | 94 |
| GO_VESICLE_LUMEN | 0.00000 | 0.00000 | 0.77494 | 0.67437 | 1.38485 | 181 |
| GO_VIRAL_GENE_EXPRESSION | 0.00000 | 0.00000 | | 0.81283 | 1.66824 | 171 |
| GO_VIRAL_LIFE_CYCLE | 0.00038 | 0.01135 | 0.49849 | 0.58832 | 1.21015 | 212 |
| HP_ABNORMALITY_OF_CARDIOVASCULAR_SYSTEM_ELECTROPHYSIC | 0.00012 | 0.00447 | 0.53843 | 0.58906 | 1.21492 | 235 |
| HP_ABNORMALITY_OF_LOWER_LIP | 0.00020 | 0.00682 | 0.51885 | 0.62009 | 1.27074 | 143 |
| HP_ABNORMALITY_OF_THE_NASAL_DORSUM | 0.00000 | 0.00002 | 0.69013 | 0.67037 | 1.37271 | 138 |
| HP_ABNORMAL_ERYTHROCYTE_MORPHOLOGY | 0.00084 | 0.02235 | 0.47727 | 0.55714 | 1.15457 | 336 |
| HP_ABNORMAL_RETICULOCYTE_MORPHOLOGY | 0.00030 | 0.00941 | 0.49849 | 0.76984 | 1.47956 | 27 |
| HP_ACUTE_LEUKEMIA | 0.00000 | 0.00001 | 0.69013 | 0.75477 | 1.51319 | 65 |
| HP_ANEMIA_OF_INADEQUATE_PRODUCTION | 0.00000 | 0.00002 | 0.67496 | 0.69340 | 1.40790 | 103 |
| HP_DELAYED_PUBERTY | 0.00000 | 0.00009 | 0.64355 | 0.68120 | 1.38580 | 109 |
| HP_DEPRESSED_NASAL_RIDGE | 0.00000 | 0.00000 | 0.78819 | 0.76609 | 1.54481 | 77 |
| HP_FATIGUE | 0.00002 | 0.00092 | 0.57561 | 0.61835 | 1.26982 | 180 |
| HP_HEADACHE | 0.00000 | 0.00012 | 0.62726 | 0.65495 | 1.34115 | 138 |
| HP_HEMATOLOGICAL_NEOPLASM | 0.00048 | 0.01408 | 0.49849 | 0.60328 | 1.23783 | 152 |
| HP_LEUKEMIA | 0.00000 | 0.00003 | 0.67496 | 0.70610 | 1.43148 | 91 |
| HP_MACROCYTIC_ANEMIA | 0.00000 | 0.00000 | 0.78819 | 0.84774 | 1.66933 | 40 |
| HP_MIGRAINE | 0.00000 | 0.00000 | | 0.81298 | 1.62705 | 63 |
| HP_ORAL_CLEFT | 0.00194 | 0.04232 | 0.45506 | 0.55426 | 1.14872 | 320 |
| HP_PALLOR | 0.00000 | 0.00007 | 0.64355 | 0.69784 | 1.41473 | 91 |
| HP_SHORT_NOSE | 0.00005 | 0.00214 | 0.55733 | 0.60868 | 1.24964 | 173 |
| HP_THICK_LOWER_LIP_VERMILION | 0.00001 | 0.00050 | 0.59333 | 0.67664 | 1.37235 | 96 |
| HP_THICK_VERMILION_BORDER | 0.00005 | 0.00227 | 0.55733 | 0.62062 | 1.27337 | 153 |
| **Cluster 12** | | | | | | |
| pathway | pval | padj | log2err | ES | NES | size |
| GO_ANCHORING_JUNCTION | 0.00063 | 0.04902 | 0.47727 | 0.57257 | 1.12271 | 364 |
| GO_BONE_RESORPTION | 0.00063 | 0.04902 | 0.47727 | 0.83888 | 1.48177 | 15 |
| GO_CADHERIN_BINDING | 0.00002 | 0.00532 | 0.57561 | 0.61413 | 1.19339 | 211 |
| GO_CELLULAR_NITROGEN_COMPOUND_CATABOLIC_PROCESS | 0.00000 | 0.00007 | 0.71951 | 0.60911 | 1.19524 | 385 |
| GO_CELLULAR_PROTEIN_CATABOLIC_PROCESS | 0.00002 | 0.00601 | 0.57561 | 0.57828 | 1.13757 | 493 |
| GO_CELL_ADHESION_MOLECULE_BINDING | 0.00018 | 0.02630 | 0.51885 | 0.59263 | 1.15694 | 269 |
| GO_CELL_SUBSTRATE_JUNCTION | 0.00001 | 0.00421 | 0.59333 | 0.61692 | 1.20085 | 231 |

| | | | | | | |
|---|---|---|---|---|---|---|
| GO_CHAPERONE_COMPLEX | 0.00015 | 0.02251 | 0.51885 | 0.84075 | 1.50370 | 19 |
| GO_CYTOPLASMIC_TRANSLATION | 0.00004 | 0.00901 | 0.55733 | 0.67631 | 1.28768 | 87 |
| GO_CYTOSOLIC_RIBOSOME | 0.00002 | 0.00488 | 0.57561 | 0.67615 | 1.28966 | 91 |
| GO_DNA_BINDING_TRANSCRIPTION_FACTOR_BINDING | 0.00014 | 0.02251 | 0.51885 | 0.61379 | 1.18911 | 178 |
| GO_ESTABLISHMENT_OF_PROTEIN_LOCALIZATION_TO_ENDOPLASM | 0.00062 | 0.04902 | 0.47727 | 0.63809 | 1.22097 | 99 |
| GO_ESTABLISHMENT_OF_PROTEIN_LOCALIZATION_TO_MEMBRANE | 0.00063 | 0.04902 | 0.47727 | 0.59234 | 1.15164 | 217 |
| GO_ESTABLISHMENT_OF_PROTEIN_LOCALIZATION_TO_ORGANELLE | 0.00000 | 0.00171 | 0.61053 | 0.59471 | 1.16689 | 377 |
| GO_ESTABLISHMENT_OF_RNA_LOCALIZATION | 0.00021 | 0.02742 | 0.51885 | 0.61903 | 1.19583 | 154 |
| GO_FICOLIN_1_RICH_GRANULE | 0.00031 | 0.03303 | 0.49849 | 0.65731 | 1.25015 | 84 |
| GO_HYDROLASE_ACTIVITY_ACTING_ON_ACID_ANHYDRIDES | 0.00055 | 0.04852 | 0.47727 | 0.56995 | 1.11899 | 397 |
| GO_LEUKOCYTE_MEDIATED_IMMUNITY | 0.00028 | 0.03094 | 0.49849 | 0.58338 | 1.13971 | 294 |
| GO_MODIFICATION_DEPENDENT_MACROMOLECULE_CATABOLIC_PR | 0.00013 | 0.02210 | 0.51885 | 0.57283 | 1.12509 | 418 |
| GO_MRNA_BINDING | 0.00000 | 0.00073 | 0.64355 | 0.62944 | 1.22223 | 206 |
| GO_MULTI_ORGANISM_LOCALIZATION | 0.00054 | 0.04852 | 0.47727 | 0.69613 | 1.30026 | 48 |
| GO_MULTI_ORGANISM_PROCESS | 0.00019 | 0.02630 | 0.51885 | 0.57445 | 1.12789 | 394 |
| GO_MYELOID_LEUKOCYTE_ACTIVATION | 0.00019 | 0.02630 | 0.51885 | 0.59174 | 1.15426 | 260 |
| GO_MYELOID_LEUKOCYTE_MEDIATED_IMMUNITY | 0.00059 | 0.04902 | 0.47727 | 0.59096 | 1.15157 | 239 |
| GO_NUCLEAR_BODY | 0.00055 | 0.04852 | 0.47727 | 0.56126 | 1.10409 | 492 |
| GO_NUCLEAR_REPLICATION_FORK | 0.00055 | 0.04852 | 0.47727 | 0.73956 | 1.35988 | 35 |
| GO_NUCLEAR_SPECK | 0.00044 | 0.04251 | 0.49849 | 0.58666 | 1.14436 | 260 |
| GO_NUCLEAR_TRANSCRIBED_MRNA_CATABOLIC_PROCESS | 0.00001 | 0.00263 | 0.61053 | 0.63199 | 1.22422 | 175 |
| GO_NUCLEAR_TRANSCRIBED_MRNA_CATABOLIC_PROCESS_NONSEN | 0.00001 | 0.00488 | 0.59333 | 0.66730 | 1.28001 | 107 |
| GO_ORGANIC_CYCLIC_COMPOUND_CATABOLIC_PROCESS | 0.00000 | 0.00007 | 0.71951 | 0.60904 | 1.19574 | 392 |
| GO_PEPTIDE_BIOSYNTHETIC_PROCESS | 0.00007 | 0.01432 | 0.53843 | 0.57440 | 1.12937 | 478 |
| GO_PERINUCLEAR_REGION_OF_CYTOPLASM | 0.00038 | 0.03865 | 0.49849 | 0.57891 | 1.13293 | 330 |
| GO_PIGMENT_GRANULE | 0.00004 | 0.00901 | 0.55733 | 0.70979 | 1.33962 | 61 |
| GO_POSITIVE_REGULATION_OF_DNA_BIOSYNTHETIC_PROCESS | 0.00018 | 0.02630 | 0.51885 | 0.71398 | 1.33140 | 47 |
| GO_POSITIVE_REGULATION_OF_PROTEIN_LOCALIZATION_TO_NUCLE | 0.00007 | 0.01432 | 0.53843 | 0.71669 | 1.34488 | 53 |
| GO_POSTTRANSCRIPTIONAL_REGULATION_OF_GENE_EXPRESSION | 0.00000 | 0.00073 | 0.64355 | 0.59324 | 1.16536 | 422 |
| GO_PROTEIN_LOCALIZATION_TO_ENDOPLASMIC_RETICULUM | 0.00028 | 0.03094 | 0.49849 | 0.63394 | 1.21804 | 118 |
| GO_PROTEIN_STABILIZATION | 0.00030 | 0.03225 | 0.49849 | 0.63655 | 1.22156 | 108 |
| GO_PROTEIN_TARGETING | 0.00012 | 0.02054 | 0.53843 | 0.59028 | 1.15281 | 287 |
| GO_REGULATION_OF_CELLULAR_AMIDE_METABOLIC_PROCESS | 0.00002 | 0.00488 | 0.57561 | 0.60097 | 1.17349 | 275 |
| GO_REGULATION_OF_CELLULAR_RESPONSE_TO_STRESS | 0.00045 | 0.04301 | 0.49849 | 0.57258 | 1.12343 | 383 |
| GO_REGULATION_OF_CELL_CYCLE_PROCESS | 0.00041 | 0.04107 | 0.49849 | 0.56736 | 1.11526 | 456 |
| GO_REGULATION_OF_DNA_METABOLIC_PROCESS | 0.00035 | 0.03613 | 0.49849 | 0.59968 | 1.16466 | 207 |
| GO_REGULATION_OF_MRNA_METABOLIC_PROCESS | 0.00003 | 0.00684 | 0.57561 | 0.60614 | 1.18135 | 246 |
| GO_REGULATION_OF_PROTEIN_CONTAINING_COMPLEX_ASSEMBLY | 0.00007 | 0.01432 | 0.53843 | 0.60918 | 1.18277 | 202 |
| GO_REGULATION_OF_RNA_SPLICING | 0.00059 | 0.04902 | 0.47727 | 0.63860 | 1.22196 | 99 |
| GO_REGULATION_OF_TELOMERASE_ACTIVITY | 0.00061 | 0.04902 | 0.47727 | 0.73049 | 1.34185 | 34 |
| GO_RESPONSE_TO_CYTOKINE | 0.00000 | 0.00044 | 0.65944 | 0.59155 | 1.16286 | 460 |
| GO_RESPONSE_TO_LEUKEMIA_INHIBITORY_FACTOR | 0.00011 | 0.01995 | 0.53843 | 0.69311 | 1.30940 | 63 |
| GO_RESPONSE_TO_OSMOTIC_STRESS | 0.00047 | 0.04436 | 0.49849 | 0.73406 | 1.35227 | 36 |
| GO_RIBONUCLEOPROTEIN_COMPLEX_BIOGENESIS | 0.00000 | 0.00006 | 0.73376 | 0.61246 | 1.20131 | 372 |
| GO_RIBOSOME_BIOGENESIS | 0.00000 | 0.00127 | 0.62726 | 0.61107 | 1.19233 | 262 |
| GO_RNA_CATABOLIC_PROCESS | 0.00000 | 0.00006 | 0.73376 | 0.62341 | 1.21945 | 313 |
| GO_RNA_EXPORT_FROM_NUCLEUS | 0.00009 | 0.01602 | 0.53843 | 0.64295 | 1.23571 | 119 |
| GO_RNA_LOCALIZATION | 0.00000 | 0.00171 | 0.61053 | 0.63461 | 1.23037 | 184 |
| GO_RRNA_METABOLIC_PROCESS | 0.00025 | 0.03058 | 0.49849 | 0.60174 | 1.16813 | 199 |
| GO_TELOMERE_ORGANIZATION | 0.00026 | 0.03077 | 0.49849 | 0.63410 | 1.21797 | 114 |
| GO_TRANSLATIONAL_INITIATION | 0.00000 | 0.00015 | 0.69013 | 0.66389 | 1.28322 | 160 |
| GO_TRANSLATION_FACTOR_ACTIVITY_RNA_BINDING | 0.00023 | 0.02851 | 0.51885 | 0.68596 | 1.29588 | 63 |
| GO_TRANSLATION_REGULATOR_ACTIVITY | 0.00012 | 0.02045 | 0.53843 | 0.65552 | 1.25432 | 99 |
| GO_TRANSLATION_REGULATOR_ACTIVITY_NUCLEIC_ACID_BINDING | 0.00008 | 0.01509 | 0.53843 | 0.67347 | 1.27913 | 77 |
| GO_UNFOLDED_PROTEIN_BINDING | 0.00000 | 0.00014 | 0.69013 | 0.73387 | 1.39060 | 73 |
| GO_VESICLE_LUMEN | 0.00023 | 0.02851 | 0.51885 | 0.61825 | 1.19398 | 146 |
| GO_VIRAL_GENE_EXPRESSION | 0.00027 | 0.03077 | 0.49849 | 0.61553 | 1.19019 | 161 |
| GO_VIRAL_LIFE_CYCLE | 0.00022 | 0.02851 | 0.51885 | 0.60918 | 1.18028 | 181 |
| **Cluster 13** | | | | | | |
| pathway | pval | padj | log2err | ES | NES | size |
| GO_ACTIVATION_OF_IMMUNE_RESPONSE | 0.00000 | 0.00001 | 0.69013 | 0.63131 | 1.32006 | 198 |
| GO_ACTIVATION_OF_INNATE_IMMUNE_RESPONSE | 0.00000 | 0.00000 | 0.74774 | 0.74217 | 1.51711 | 82 |
| GO_AEROBIC_RESPIRATION | 0.00024 | 0.00672 | 0.51885 | 0.67786 | 1.37326 | 65 |
| GO_AMIDE_TRANSMEMBRANE_TRANSPORTER_ACTIVITY | 0.00053 | 0.01296 | 0.47727 | 0.73903 | 1.45082 | 32 |
| GO_AMINE_METABOLIC_PROCESS | 0.00000 | 0.00000 | 0.81404 | 0.76613 | 1.56033 | 74 |
| GO_ANAPHASE_PROMOTING_COMPLEX_DEPENDENT_CATABOLIC_P | 0.00000 | 0.00000 | 0.77494 | 0.75863 | 1.54599 | 76 |
| GO_ANCHORING_JUNCTION | 0.00000 | 0.00000 | | 0.63069 | 1.33795 | 453 |
| GO_ANTIGEN_PROCESSING_AND_PRESENTATION | 0.00000 | 0.00002 | 0.67496 | 0.66500 | 1.38073 | 136 |
| GO_ANTIGEN_PROCESSING_AND_PRESENTATION_OF_EXOGENOUS_ | 0.00000 | 0.00000 | 0.78819 | 0.80011 | 1.60426 | 55 |
| GO_ANTIGEN_PROCESSING_AND_PRESENTATION_OF_PEPTIDE_ANTI | 0.00000 | 0.00001 | 0.69013 | 0.68859 | 1.42398 | 118 |
| GO_ANTIGEN_PROCESSING_AND_PRESENTATION_OF_PEPTIDE_ANTI | 0.00000 | 0.00000 | 0.77494 | 0.77289 | 1.56692 | 66 |
| GO_ANTIGEN_RECEPTOR_MEDIATED_SIGNALING_PATHWAY | 0.00000 | 0.00016 | 0.62726 | 0.65708 | 1.36002 | 120 |
| GO_APOPTOTIC_SIGNALING_PATHWAY | 0.00170 | 0.03319 | 0.45506 | 0.54432 | 1.15194 | 336 |
| GO_AXONAL_GROWTH_CONE | 0.00038 | 0.01015 | 0.49849 | 0.82766 | 1.55109 | 15 |
| GO_AZUROPHIL_GRANULE_LUMEN | 0.00261 | 0.04750 | 0.43171 | 0.65808 | 1.31801 | 52 |
| GO_BINDING_OF_SPERM_TO_ZONA_PELLUCIDA | 0.00000 | 0.00003 | 0.65944 | 0.91957 | 1.72334 | 15 |

| | | | | | | |
|---|---|---|---|---|---|---|
| GO_BRUSH_BORDER | 0.00161 | 0.03185 | 0.45506 | 0.66074 | 1.32980 | 58 |
| GO_CADHERIN_BINDING | 0.00000 | 0.00000 | 0.81404 | 0.65058 | 1.36742 | 247 |
| GO_CANONICAL_WNT_SIGNALING_PATHWAY | 0.00000 | 0.00018 | 0.61053 | 0.62577 | 1.30754 | 181 |
| GO_CATALYTIC_STEP_2_SPLICEOSOME | 0.00011 | 0.00332 | 0.53843 | 0.66525 | 1.35900 | 81 |
| GO_CELLULAR_AMINO_ACID_METABOLIC_PROCESS | 0.00000 | 0.00001 | 0.67496 | 0.63315 | 1.32438 | 205 |
| GO_CELLULAR_CARBOHYDRATE_CATABOLIC_PROCESS | 0.00242 | 0.04532 | 0.43171 | 0.73681 | 1.43617 | 27 |
| GO_CELLULAR_COMPONENT_DISASSEMBLY | 0.00044 | 0.01128 | 0.49849 | 0.54955 | 1.16429 | 385 |
| GO_CELLULAR_KETONE_METABOLIC_PROCESS | 0.00000 | 0.00001 | 0.69013 | 0.65687 | 1.36740 | 148 |
| GO_CELLULAR_NITROGEN_COMPOUND_CATABOLIC_PROCESS | 0.00000 | 0.00000 | | 0.72954 | 1.54568 | 406 |
| GO_CELLULAR_RESPIRATION | 0.00010 | 0.00318 | 0.53843 | 0.61616 | 1.27864 | 135 |
| GO_CELLULAR_RESPONSE_TO_OXYGEN_LEVELS | 0.00000 | 0.00000 | 0.71951 | 0.66742 | 1.38998 | 153 |
| GO_CELL_ACTIVATION_INVOLVED_IN_IMMUNE_RESPONSE | 0.00000 | 0.00004 | 0.65944 | 0.58897 | 1.24624 | 340 |
| GO_CELL_ADHESION_MOLECULE_BINDING | 0.00000 | 0.00000 | 0.80122 | 0.62535 | 1.32085 | 319 |
| GO_CELL_BODY | 0.00017 | 0.00504 | 0.51885 | 0.57724 | 1.21338 | 249 |
| GO_CELL_CELL_RECOGNITION | 0.00000 | 0.00000 | 0.78819 | 0.90962 | 1.75555 | 25 |
| GO_CELL_CELL_SIGNALING_BY_WNT | 0.00000 | 0.00017 | 0.61053 | 0.59576 | 1.25537 | 281 |
| GO_CELL_CYCLE_G2_M_PHASE_TRANSITION | 0.00017 | 0.00493 | 0.51885 | 0.59041 | 1.23481 | 204 |
| GO_CELL_CYCLE_PHASE_TRANSITION | 0.00012 | 0.00361 | 0.53843 | 0.55201 | 1.17031 | 437 |
| GO_CELL_DEATH_IN_RESPONSE_TO_OXIDATIVE_STRESS | 0.00268 | 0.04853 | 0.43171 | 0.66253 | 1.32445 | 49 |
| GO_CELL_RECOGNITION | 0.00000 | 0.00000 | 0.73376 | 0.80294 | 1.60277 | 47 |
| GO_CELL_SUBSTRATE_JUNCTION | 0.00000 | 0.00000 | | 0.70832 | 1.49182 | 271 |
| GO_CELL_SURFACE_RECEPTOR_SIGNALING_PATHWAY_INVOLVED_IN | 0.00001 | 0.00055 | 0.59333 | 0.58599 | 1.23628 | 302 |
| GO_CHAPERONE_BINDING | 0.00060 | 0.01450 | 0.47727 | 0.66072 | 1.34347 | 70 |
| GO_CHAPERONE_COFACTOR_DEPENDENT_PROTEIN_REFOLDING | 0.00250 | 0.04639 | 0.43171 | 0.80490 | 1.51780 | 16 |
| GO_CHAPERONE_COMPLEX | 0.00001 | 0.00033 | 0.59333 | 0.85547 | 1.64226 | 21 |
| GO_COTRANSLATIONAL_PROTEIN_TARGETING_TO_MEMBRANE | 0.00000 | 0.00000 | | 0.87548 | 1.79705 | 92 |
| GO_CYTOKINE_MEDIATED_SIGNALING_PATHWAY | 0.00000 | 0.00000 | | 0.64850 | 1.36811 | 301 |
| GO_CYTOPLASMIC_TRANSLATION | 0.00000 | 0.00000 | | 0.82695 | 1.69712 | 88 |
| GO_CYTOPLASMIC_TRANSLATIONAL_INITIATION | 0.00000 | 0.00003 | 0.65944 | 0.83640 | 1.63413 | 30 |
| GO_CYTOSOLIC_LARGE_RIBOSOMAL_SUBUNIT | 0.00000 | 0.00000 | | 0.87837 | 1.75871 | 53 |
| GO_CYTOSOLIC_RIBOSOME | 0.00000 | 0.00000 | | 0.87188 | 1.79273 | 95 |
| GO_CYTOSOLIC_SMALL_RIBOSOMAL_SUBUNIT | 0.00000 | 0.00000 | | 0.86469 | 1.71220 | 40 |
| GO_DEFENSE_RESPONSE_TO_OTHER_ORGANISM | 0.00001 | 0.00022 | 0.61053 | 0.57372 | 1.21533 | 383 |
| GO_DE_NOVO_PROTEIN_FOLDING | 0.00042 | 0.01090 | 0.49849 | 0.78033 | 1.50542 | 24 |
| GO_DNA_BIOSYNTHETIC_PROCESS | 0.00066 | 0.01551 | 0.47727 | 0.60547 | 1.25713 | 136 |
| GO_ENDOPEPTIDASE_COMPLEX | 0.00000 | 0.00000 | 0.73376 | 0.77670 | 1.57042 | 63 |
| GO_ENERGY_DERIVATION_BY_OXIDATION_OF_ORGANIC_COMPOUN | 0.00168 | 0.03295 | 0.45506 | 0.58108 | 1.21294 | 176 |
| GO_ESTABLISHMENT_OF_PROTEIN_LOCALIZATION_TO_CHROMOSON | 0.00044 | 0.01110 | 0.49849 | 0.79173 | 1.52460 | 23 |
| GO_ESTABLISHMENT_OF_PROTEIN_LOCALIZATION_TO_ENDOPLASM | 0.00000 | 0.00000 | | 0.86515 | 1.78329 | 101 |
| GO_ESTABLISHMENT_OF_PROTEIN_LOCALIZATION_TO_MEMBRANE | 0.00000 | 0.00000 | | 0.75098 | 1.57701 | 239 |
| GO_ESTABLISHMENT_OF_PROTEIN_LOCALIZATION_TO_ORGANELLE | 0.00000 | 0.00000 | | 0.70380 | 1.49125 | 400 |
| GO_ESTABLISHMENT_OF_PROTEIN_LOCALIZATION_TO_TELOMERE | 0.00004 | 0.00128 | 0.55733 | 0.86143 | 1.62963 | 17 |
| GO_ESTABLISHMENT_OF_RNA_LOCALIZATION | 0.00159 | 0.03160 | 0.45506 | 0.57858 | 1.20591 | 163 |
| GO_ESTABLISHMENT_OF_TISSUE_POLARITY | 0.00000 | 0.00000 | 0.82666 | 0.76206 | 1.55938 | 83 |
| GO_EUKARYOTIC_TRANSLATION_INITIATION_FACTOR_3_COMPLEX | 0.00004 | 0.00149 | 0.55733 | 0.87076 | 1.63188 | 15 |
| GO_EXOCYTOSIS | 0.00000 | 0.00016 | 0.61053 | 0.56919 | 1.20762 | 462 |
| GO_FC_EPSILON_RECEPTOR_SIGNALING_PATHWAY | 0.00000 | 0.00000 | 0.80122 | 0.76454 | 1.56183 | 81 |
| GO_FC_RECEPTOR_SIGNALING_PATHWAY | 0.00000 | 0.00000 | | 0.74534 | 1.54208 | 121 |
| GO_FERTILIZATION | 0.00010 | 0.00318 | 0.53843 | 0.69758 | 1.41044 | 63 |
| GO_FICOLIN_1_RICH_GRANULE | 0.00000 | 0.00000 | 0.80122 | 0.74401 | 1.52822 | 93 |
| GO_GENERATION_OF_PRECURSOR_METABOLITES_AND_ENERGY | 0.00152 | 0.03046 | 0.45506 | 0.54735 | 1.15617 | 320 |
| GO_GLUTAMATERGIC_SYNAPSE | 0.00261 | 0.04750 | 0.43171 | 0.58914 | 1.22389 | 138 |
| GO_GLYCOSYL_COMPOUND_METABOLIC_PROCESS | 0.00027 | 0.00738 | 0.49849 | 0.66541 | 1.35008 | 67 |
| GO_HEAT_SHOCK_PROTEIN_BINDING | 0.00019 | 0.00552 | 0.51885 | 0.65378 | 1.33781 | 83 |
| GO_HEMATOPOIETIC_PROGENITOR_CELL_DIFFERENTIATION | 0.00000 | 0.00000 | 0.74774 | 0.71994 | 1.48382 | 100 |
| GO_HEMATOPOIETIC_STEM_CELL_DIFFERENTIATION | 0.00000 | 0.00000 | | 0.80267 | 1.62893 | 68 |
| GO_HSP70_PROTEIN_BINDING | 0.00235 | 0.04424 | 0.43171 | 0.73184 | 1.41932 | 26 |
| GO_IMMUNE_RESPONSE_REGULATING_SIGNALING_PATHWAY | 0.00000 | 0.00001 | 0.69013 | 0.64380 | 1.34608 | 190 |
| GO_INNATE_IMMUNE_RESPONSE | 0.00000 | 0.00009 | 0.62726 | 0.58658 | 1.23859 | 313 |
| GO_INNATE_IMMUNE_RESPONSE_ACTIVATING_SIGNAL_TRANSDUCT | 0.00000 | 0.00000 | | 0.79341 | 1.61014 | 68 |
| GO_INTERACTION_WITH_SYMBIONT | 0.00052 | 0.01285 | 0.47727 | 0.70108 | 1.39364 | 44 |
| GO_INTERLEUKIN_1_MEDIATED_SIGNALING_PATHWAY | 0.00000 | 0.00000 | | 0.80824 | 1.64202 | 69 |
| GO_ION_CHANNEL_BINDING | 0.00097 | 0.02140 | 0.47727 | 0.66862 | 1.34136 | 56 |
| GO_LARGE_RIBOSOMAL_SUBUNIT | 0.00000 | 0.00000 | | 0.75509 | 1.56037 | 108 |
| GO_LEUKOCYTE_MEDIATED_IMMUNITY | 0.00000 | 0.00002 | 0.67496 | 0.59174 | 1.25240 | 364 |
| GO_LYSOSOMAL_LUMEN | 0.00216 | 0.04103 | 0.43171 | 0.67800 | 1.34776 | 44 |
| GO_MACROMOLECULE_TRANSMEMBRANE_TRANSPORTER_ACTIVITY | 0.00063 | 0.01497 | 0.47727 | 0.83425 | 1.57315 | 16 |
| GO_MATURATION_OF_LSU_RRNA | 0.00072 | 0.01652 | 0.47727 | 0.75765 | 1.47679 | 27 |
| GO_MITOCHONDRIAL_PROTEIN_COMPLEX | 0.00002 | 0.00064 | 0.57561 | 0.60690 | 1.26948 | 205 |
| GO_MODIFICATION_DEPENDENT_MACROMOLECULE_CATABOLIC_PR | 0.00004 | 0.00148 | 0.55733 | 0.55099 | 1.16893 | 454 |
| GO_MODIFIED_AMINO_ACID_TRANSPORT | 0.00119 | 0.02509 | 0.45506 | 0.78110 | 1.48183 | 18 |
| GO_MONOSACCHARIDE_BIOSYNTHETIC_PROCESS | 0.00078 | 0.01769 | 0.47727 | 0.66541 | 1.33713 | 57 |
| GO_MORPHOGENESIS_OF_AN_EPITHELIUM | 0.00002 | 0.00068 | 0.57561 | 0.58482 | 1.23127 | 267 |
| GO_MORPHOGENESIS_OF_A_POLARIZED_EPITHELIUM | 0.00000 | 0.00000 | 0.82666 | 0.74550 | 1.53397 | 96 |
| GO_MRNA_3_UTR_BINDING | 0.00040 | 0.01046 | 0.49849 | 0.67615 | 1.36533 | 62 |
| GO_MRNA_5_UTR_BINDING | 0.00053 | 0.01296 | 0.47727 | 0.78779 | 1.51549 | 22 |

| | | | | | | |
|---|---|---|---|---|---|---|
| GO_MRNA_BINDING | 0.00000 | 0.00000 | 0.77494 | 0.65017 | 1.36251 | 218 |
| GO_MRNA_PROCESSING | 0.00000 | 0.00008 | 0.64355 | 0.57705 | 1.22255 | 407 |
| GO_MYD88_DEPENDENT_TOLL_LIKE_RECEPTOR_SIGNALING_PATHW | 0.00014 | 0.00420 | 0.51885 | 0.82801 | 1.57081 | 18 |
| GO_MYD88_INDEPENDENT_TOLL_LIKE_RECEPTOR_SIGNALING_PATH | 0.00007 | 0.00220 | 0.53843 | 0.86188 | 1.61523 | 15 |
| GO_MYELOID_LEUKOCYTE_ACTIVATION | 0.00000 | 0.00000 | 0.73376 | 0.60734 | 1.28289 | 320 |
| GO_MYELOID_LEUKOCYTE_MEDIATED_IMMUNITY | 0.00000 | 0.00000 | 0.74774 | 0.61791 | 1.30293 | 295 |
| GO_NADH_METABOLIC_PROCESS | 0.00119 | 0.02509 | 0.45506 | 0.76396 | 1.47386 | 24 |
| GO_NCRNA_METABOLIC_PROCESS | 0.00043 | 0.01104 | 0.49849 | 0.55485 | 1.17506 | 377 |
| GO_NCRNA_PROCESSING | 0.00010 | 0.00318 | 0.53843 | 0.56718 | 1.19819 | 322 |
| GO_NEGATIVE_REGULATION_OF_CANONICAL_WNT_SIGNALING_PAT | 0.00000 | 0.00000 | | 0.74625 | 1.53804 | 100 |
| GO_NEGATIVE_REGULATION_OF_CELLULAR_AMIDE_METABOLIC_PR | 0.00040 | 0.01046 | 0.49849 | 0.62561 | 1.29270 | 111 |
| GO_NEGATIVE_REGULATION_OF_CELLULAR_RESPONSE_TO_TRANSF | 0.00272 | 0.04888 | 0.43171 | 0.70633 | 1.39322 | 35 |
| GO_NEGATIVE_REGULATION_OF_CELL_CYCLE_G2_M_PHASE_TRANS | 0.00000 | 0.00003 | 0.65944 | 0.70534 | 1.44331 | 83 |
| GO_NEGATIVE_REGULATION_OF_CELL_CYCLE_PHASE_TRANSITION | 0.00004 | 0.00141 | 0.55733 | 0.60141 | 1.25516 | 174 |
| GO_NEGATIVE_REGULATION_OF_CELL_CYCLE_PROCESS | 0.00035 | 0.00958 | 0.49849 | 0.57566 | 1.20878 | 237 |
| GO_NEGATIVE_REGULATION_OF_CELL_DEATH | 0.00000 | 0.00006 | 0.64355 | 0.56792 | 1.20519 | 465 |
| GO_NEGATIVE_REGULATION_OF_MITOTIC_CELL_CYCLE | 0.00183 | 0.03558 | 0.45506 | 0.56641 | 1.18663 | 217 |
| GO_NEGATIVE_REGULATION_OF_MRNA_METABOLIC_PROCESS | 0.00168 | 0.03295 | 0.45506 | 0.65310 | 1.31556 | 59 |
| GO_NEGATIVE_REGULATION_OF_WNT_SIGNALING_PATHWAY | 0.00000 | 0.00000 | | 0.74396 | 1.53734 | 112 |
| GO_NEURAL_NUCLEUS_DEVELOPMENT | 0.00077 | 0.01763 | 0.47727 | 0.74933 | 1.46325 | 29 |
| GO_NEURON_TO_NEURON_SYNAPSE | 0.00017 | 0.00508 | 0.51885 | 0.60919 | 1.26868 | 149 |
| GO_NIK_NF_KAPPAB_SIGNALING | 0.00000 | 0.00000 | | 0.77059 | 1.58335 | 94 |
| GO_NON_CANONICAL_WNT_SIGNALING_PATHWAY | 0.00000 | 0.00000 | | 0.75391 | 1.55127 | 96 |
| GO_NUCLEAR_TRANSCRIBED_MRNA_CATABOLIC_PROCESS | 0.00000 | 0.00000 | | 0.79170 | 1.65357 | 178 |
| GO_NUCLEAR_TRANSCRIBED_MRNA_CATABOLIC_PROCESS_NONSEN | 0.00000 | 0.00000 | | 0.87139 | 1.80022 | 110 |
| GO_NUCLEAR_TRANSPORT | 0.00009 | 0.00276 | 0.53843 | 0.58107 | 1.22229 | 260 |
| GO_NUCLEOBASE_CONTAINING_COMPOUND_TRANSPORT | 0.00283 | 0.04990 | 0.43171 | 0.56912 | 1.18978 | 194 |
| GO_NUCLEOBASE_CONTAINING_SMALL_MOLECULE_BIOSYNTHETIC_ | 0.00283 | 0.04990 | 0.43171 | 0.64398 | 1.30036 | 61 |
| GO_NUCLEOSIDE_METABOLIC_PROCESS | 0.00021 | 0.00605 | 0.51885 | 0.69276 | 1.38707 | 53 |
| GO_NUCLEOSIDE_MONOPHOSPHATE_BIOSYNTHETIC_PROCESS | 0.00128 | 0.02623 | 0.45506 | 0.72556 | 1.42437 | 32 |
| GO_NUCLEOSIDE_MONOPHOSPHATE_METABOLIC_PROCESS | 0.00265 | 0.04802 | 0.43171 | 0.67096 | 1.33549 | 45 |
| GO_NUCLEOSIDE_TRIPHOSPHATE_BIOSYNTHETIC_PROCESS | 0.00089 | 0.01985 | 0.47727 | 0.69930 | 1.38389 | 39 |
| GO_ORGANELLE_INNER_MEMBRANE | 0.00071 | 0.01644 | 0.47727 | 0.55058 | 1.16521 | 361 |
| GO_ORGANIC_CYCLIC_COMPOUND_CATABOLIC_PROCESS | 0.00000 | 0.00000 | | 0.72361 | 1.53370 | 418 |
| GO_OXIDOREDUCTASE_ACTIVITY | 0.00253 | 0.04661 | 0.43171 | 0.54230 | 1.14844 | 378 |
| GO_PEPTIDASE_COMPLEX | 0.00000 | 0.00006 | 0.64355 | 0.70773 | 1.44418 | 78 |
| GO_PEPTIDE_BINDING | 0.00133 | 0.02696 | 0.45506 | 0.61664 | 1.27271 | 104 |
| GO_PEPTIDE_TRANSMEMBRANE_TRANSPORTER_ACTIVITY | 0.00045 | 0.01135 | 0.49849 | 0.79454 | 1.52530 | 21 |
| GO_PIGMENT_GRANULE | 0.00018 | 0.00525 | 0.51885 | 0.66217 | 1.34990 | 77 |
| GO_POLYSOMAL_RIBOSOME | 0.00000 | 0.00001 | 0.69013 | 0.85035 | 1.66611 | 31 |
| GO_POLYSOME | 0.00000 | 0.00000 | 0.76146 | 0.78140 | 1.57783 | 61 |
| GO_POLY_PYRIMIDINE_TRACT_BINDING | 0.00104 | 0.02267 | 0.45506 | 0.82773 | 1.56085 | 16 |
| GO_POSITIVE_REGULATION_OF_CANONICAL_WNT_SIGNALING_PATH | 0.00000 | 0.00000 | 0.76146 | 0.72988 | 1.50260 | 97 |
| GO_POSITIVE_REGULATION_OF_CELLULAR_AMIDE_METABOLIC_PRO | 0.00008 | 0.00274 | 0.53843 | 0.63746 | 1.31752 | 113 |
| GO_POSITIVE_REGULATION_OF_CELLULAR_PROTEIN_LOCALIZATION | 0.00078 | 0.01769 | 0.47727 | 0.57103 | 1.19782 | 224 |
| GO_POSITIVE_REGULATION_OF_CELL_DEATH | 0.00042 | 0.01084 | 0.49849 | 0.55314 | 1.17036 | 343 |
| GO_POSITIVE_REGULATION_OF_DEFENSE_RESPONSE | 0.00002 | 0.00086 | 0.57561 | 0.62831 | 1.30853 | 153 |
| GO_POSITIVE_REGULATION_OF_DNA_BIOSYNTHETIC_PROCESS | 0.00065 | 0.01528 | 0.47727 | 0.69102 | 1.38118 | 48 |
| GO_POSITIVE_REGULATION_OF_ESTABLISHMENT_OF_PROTEIN_LOC | 0.00086 | 0.01919 | 0.47727 | 0.57319 | 1.19928 | 209 |
| GO_POSITIVE_REGULATION_OF_IMMUNE_RESPONSE | 0.00002 | 0.00066 | 0.57561 | 0.58889 | 1.23930 | 263 |
| GO_POSITIVE_REGULATION_OF_IMMUNE_SYSTEM_PROCESS | 0.00005 | 0.00175 | 0.55733 | 0.56473 | 1.19613 | 376 |
| GO_POSITIVE_REGULATION_OF_INNATE_IMMUNE_RESPONSE | 0.00000 | 0.00003 | 0.65944 | 0.68246 | 1.40968 | 107 |
| GO_POSITIVE_REGULATION_OF_MRNA_PROCESSING | 0.00191 | 0.03688 | 0.45506 | 0.73706 | 1.42944 | 26 |
| GO_POSITIVE_REGULATION_OF_MRNA_SPLICING_VIA_SPLICEOSOM | 0.00124 | 0.02586 | 0.45506 | 0.80351 | 1.52006 | 17 |
| GO_POSITIVE_REGULATION_OF_ORGANELLE_ORGANIZATION | 0.00272 | 0.04888 | 0.43171 | 0.53970 | 1.14344 | 384 |
| GO_POSITIVE_REGULATION_OF_PROTEIN_CONTAINING_COMPLEX_ | 0.00110 | 0.02369 | 0.45506 | 0.59101 | 1.23081 | 149 |
| GO_POSITIVE_REGULATION_OF_PROTEIN_LOCALIZATION_TO_NUCLE | 0.00132 | 0.02679 | 0.45506 | 0.66492 | 1.33318 | 55 |
| GO_POSITIVE_REGULATION_OF_RESPONSE_TO_BIOTIC_STIMULUS | 0.00000 | 0.00004 | 0.65944 | 0.66699 | 1.38015 | 122 |
| GO_POSITIVE_REGULATION_OF_RESPONSE_TO_EXTERNAL_STIMULU | 0.00034 | 0.00943 | 0.49849 | 0.58436 | 1.22191 | 191 |
| GO_POSITIVE_REGULATION_OF_TELOMERASE_RNA_LOCALIZATION_ | 0.00002 | 0.00071 | 0.57561 | 0.88178 | 1.65252 | 15 |
| GO_POSITIVE_REGULATION_OF_TRANSLATION | 0.00018 | 0.00529 | 0.51885 | 0.64484 | 1.32686 | 96 |
| GO_POSITIVE_REGULATION_OF_VIRAL_GENOME_REPLICATION | 0.00126 | 0.02595 | 0.45506 | 0.74050 | 1.44599 | 29 |
| GO_POSITIVE_REGULATION_OF_WNT_SIGNALING_PATHWAY | 0.00000 | 0.00000 | 0.73376 | 0.70143 | 1.44973 | 113 |
| GO_POSTSYNAPSE | 0.00001 | 0.00039 | 0.59333 | 0.59611 | 1.25305 | 249 |
| GO_POSTTRANSCRIPTIONAL_REGULATION_OF_GENE_EXPRESSION | 0.00000 | 0.00000 | | 0.65172 | 1.38232 | 443 |
| GO_POST_TRANSLATIONAL_PROTEIN_MODIFICATION | 0.00002 | 0.00058 | 0.59333 | 0.59896 | 1.25569 | 220 |
| GO_PRERIBOSOME | 0.00087 | 0.01932 | 0.47727 | 0.65090 | 1.32094 | 68 |
| GO_PROTEASOMAL_PROTEIN_CATABOLIC_PROCESS | 0.00003 | 0.00115 | 0.55733 | 0.57178 | 1.21006 | 344 |
| GO_PROTEASOMAL_UBIQUITIN_INDEPENDENT_PROTEIN_CATABOLI | 0.00000 | 0.00000 | 0.71951 | 0.92000 | 1.74532 | 18 |
| GO_PROTEASOME_ACCESSORY_COMPLEX | 0.00001 | 0.00022 | 0.61053 | 0.84297 | 1.62627 | 24 |
| GO_PROTEASOME_CORE_COMPLEX | 0.00000 | 0.00003 | 0.65944 | 0.90864 | 1.71893 | 17 |
| GO_PROTEIN_CONTAINING_COMPLEX_DISASSEMBLY | 0.00102 | 0.02236 | 0.45506 | 0.56806 | 1.19385 | 246 |
| GO_PROTEIN_DNA_COMPLEX_SUBUNIT_ORGANIZATION | 0.00202 | 0.03876 | 0.43171 | 0.58497 | 1.21920 | 159 |
| GO_PROTEIN_FOLDING | 0.00000 | 0.00000 | 0.71951 | 0.66110 | 1.37745 | 155 |
| GO_PROTEIN_FOLDING_CHAPERONE | 0.00156 | 0.03098 | 0.45506 | 0.79855 | 1.49655 | 15 |
| GO_PROTEIN_IMPORT | 0.00071 | 0.01644 | 0.47727 | 0.60502 | 1.25240 | 124 |

| | | | | | | |
|---|---|---|---|---|---|---|
| GO_PROTEIN_LOCALIZATION_TO_CHROMOSOME_TELOMERIC_REGIO | 0.00061 | 0.01466 | 0.47727 | 0.76062 | 1.48257 | 27 |
| GO_PROTEIN_LOCALIZATION_TO_ENDOPLASMIC_RETICULUM | 0.00000 | 0.00000 | | 0.84317 | 1.74449 | 121 |
| GO_PROTEIN_LOCALIZATION_TO_MEMBRANE | 0.00000 | 0.00000 | | 0.66321 | 1.40542 | 399 |
| GO_PROTEIN_LOCALIZATION_TO_MITOCHONDRION | 0.00276 | 0.04907 | 0.43171 | 0.60705 | 1.25392 | 107 |
| GO_PROTEIN_LOCALIZATION_TO_NUCLEUS | 0.00039 | 0.01044 | 0.49849 | 0.58995 | 1.23114 | 173 |
| GO_PROTEIN_MODIFICATION_BY_SMALL_PROTEIN_REMOVAL | 0.00000 | 0.00000 | 0.71951 | 0.63580 | 1.33049 | 208 |
| GO_PROTEIN_REFOLDING | 0.00126 | 0.02595 | 0.45506 | 0.80239 | 1.51794 | 17 |
| GO_PROTEIN_STABILIZATION | 0.00077 | 0.01763 | 0.47727 | 0.60739 | 1.25875 | 129 |
| GO_PROTEIN_TARGETING | 0.00000 | 0.00000 | | 0.71415 | 1.50819 | 316 |
| GO_PROTEIN_TARGETING_TO_MEMBRANE | 0.00000 | 0.00000 | | 0.81632 | 1.70024 | 150 |
| GO_PROTEIN_TRANSMEMBRANE_TRANSPORT | 0.00080 | 0.01803 | 0.47727 | 0.68855 | 1.37625 | 48 |
| GO_PURINE_NUCLEOSIDE_TRIPHOSPHATE_METABOLIC_PROCESS | 0.00242 | 0.04532 | 0.43171 | 0.67126 | 1.33289 | 43 |
| GO_REGULATION_OF_ANIMAL_ORGAN_MORPHOGENESIS | 0.00000 | 0.00000 | 0.78819 | 0.71602 | 1.48045 | 117 |
| GO_REGULATION_OF_APOPTOTIC_SIGNALING_PATHWAY | 0.00029 | 0.00810 | 0.49849 | 0.57570 | 1.20774 | 226 |
| GO_REGULATION_OF_CELLULAR_AMIDE_METABOLIC_PROCESS | 0.00000 | 0.00001 | 0.67496 | 0.60795 | 1.28224 | 298 |
| GO_REGULATION_OF_CELLULAR_AMINE_METABOLIC_PROCESS | 0.00000 | 0.00000 | | 0.82215 | 1.64844 | 55 |
| GO_REGULATION_OF_CELLULAR_AMINO_ACID_METABOLIC_PROCES | 0.00000 | 0.00000 | | 0.83307 | 1.66800 | 53 |
| GO_REGULATION_OF_CELLULAR_KETONE_METABOLIC_PROCESS | 0.00000 | 0.00000 | 0.74774 | 0.70697 | 1.46086 | 114 |
| GO_REGULATION_OF_CELLULAR_PROTEIN_CATABOLIC_PROCESS | 0.00209 | 0.04004 | 0.43171 | 0.57664 | 1.20488 | 181 |
| GO_REGULATION_OF_CELLULAR_PROTEIN_LOCALIZATION | 0.00005 | 0.00157 | 0.55733 | 0.56670 | 1.19920 | 353 |
| GO_REGULATION_OF_CELLULAR_RESPONSE_TO_TRANSFORMING_G | 0.00015 | 0.00444 | 0.51885 | 0.68835 | 1.38323 | 57 |
| GO_REGULATION_OF_CELL_CYCLE_G2_M_PHASE_TRANSITION | 0.00000 | 0.00007 | 0.64355 | 0.63798 | 1.32971 | 163 |
| GO_REGULATION_OF_CELL_CYCLE_PHASE_TRANSITION | 0.00012 | 0.00376 | 0.51885 | 0.57130 | 1.20667 | 317 |
| GO_REGULATION_OF_DEFENSE_RESPONSE | 0.00121 | 0.02538 | 0.45506 | 0.55541 | 1.16983 | 270 |
| GO_REGULATION_OF_DNA_BIOSYNTHETIC_PROCESS | 0.00154 | 0.03072 | 0.45506 | 0.64315 | 1.30971 | 73 |
| GO_REGULATION_OF_DNA_TEMPLATED_TRANSCRIPTION_IN_RESPO | 0.00000 | 0.00000 | 0.80122 | 0.74028 | 1.51983 | 91 |
| GO_REGULATION_OF_HEMATOPOIETIC_PROGENITOR_CELL_DIFFERE | 0.00000 | 0.00000 | 0.80122 | 0.78078 | 1.58452 | 68 |
| GO_REGULATION_OF_HEMATOPOIETIC_STEM_CELL_DIFFERENTIATIO | 0.00000 | 0.00000 | | 0.80991 | 1.63541 | 61 |
| GO_REGULATION_OF_HEMOPOIESIS | 0.00000 | 0.00016 | 0.62726 | 0.60684 | 1.27221 | 220 |
| GO_REGULATION_OF_IMMUNE_RESPONSE | 0.00024 | 0.00682 | 0.51885 | 0.55893 | 1.18333 | 348 |
| GO_REGULATION_OF_INNATE_IMMUNE_RESPONSE | 0.00000 | 0.00006 | 0.64355 | 0.64536 | 1.34389 | 151 |
| GO_REGULATION_OF_INTRINSIC_APOPTOTIC_SIGNALING_PATHWAY | 0.00132 | 0.02679 | 0.45506 | 0.61708 | 1.27360 | 104 |
| GO_REGULATION_OF_MITOTIC_CELL_CYCLE | 0.00010 | 0.00326 | 0.53843 | 0.55346 | 1.17336 | 428 |
| GO_REGULATION_OF_MORPHOGENESIS_OF_AN_EPITHELIUM | 0.00000 | 0.00000 | 0.80122 | 0.73126 | 1.50732 | 101 |
| GO_REGULATION_OF_MRNA_CATABOLIC_PROCESS | 0.00000 | 0.00000 | 0.83909 | 0.69534 | 1.44860 | 154 |
| GO_REGULATION_OF_MRNA_METABOLIC_PROCESS | 0.00000 | 0.00000 | | 0.67978 | 1.42945 | 254 |
| GO_REGULATION_OF_MRNA_PROCESSING | 0.00002 | 0.00061 | 0.57561 | 0.65547 | 1.35444 | 114 |
| GO_REGULATION_OF_MRNA_SPLICING_VIA_SPLICEOSOME | 0.00005 | 0.00156 | 0.55733 | 0.67435 | 1.37758 | 81 |
| GO_REGULATION_OF_PLASMA_LIPOPROTEIN_PARTICLE_LEVELS | 0.00060 | 0.01450 | 0.47727 | 0.69540 | 1.38082 | 43 |
| GO_REGULATION_OF_PROTEIN_CONTAINING_COMPLEX_ASSEMBLY | 0.00004 | 0.00152 | 0.55733 | 0.58760 | 1.23500 | 250 |
| GO_REGULATION_OF_PROTEIN_LOCALIZATION_TO_NUCLEUS | 0.00058 | 0.01418 | 0.47727 | 0.64923 | 1.32351 | 77 |
| GO_REGULATION_OF_PROTEIN_MODIFICATION_BY_SMALL_PROTEIN | 0.00253 | 0.04661 | 0.43171 | 0.58330 | 1.21573 | 159 |
| GO_REGULATION_OF_PROTEIN_STABILITY | 0.00052 | 0.01285 | 0.47727 | 0.58046 | 1.21402 | 204 |
| GO_REGULATION_OF_PROTEOLYSIS | 0.00122 | 0.02567 | 0.45506 | 0.54894 | 1.16165 | 359 |
| GO_REGULATION_OF_RESPONSE_TO_BIOTIC_STIMULUS | 0.00001 | 0.00033 | 0.59333 | 0.61879 | 1.29378 | 189 |
| GO_REGULATION_OF_RNA_SPLICING | 0.00001 | 0.00045 | 0.59333 | 0.65558 | 1.35502 | 109 |
| GO_REGULATION_OF_SMALL_MOLECULE_METABOLIC_PROCESS | 0.00005 | 0.00170 | 0.55733 | 0.58535 | 1.23235 | 265 |
| GO_REGULATION_OF_STEM_CELL_DIFFERENTIATION | 0.00000 | 0.00000 | 0.77494 | 0.75587 | 1.54413 | 81 |
| GO_REGULATION_OF_SYMBIOTIC_PROCESS | 0.00100 | 0.02204 | 0.45506 | 0.60233 | 1.24796 | 126 |
| GO_REGULATION_OF_TRANSCRIPTION_FROM_RNA_POLYMERASE_II | 0.00000 | 0.00000 | 0.82666 | 0.80072 | 1.62043 | 64 |
| GO_REGULATION_OF_TRANSLATIONAL_INITIATION | 0.00013 | 0.00381 | 0.51885 | 0.68858 | 1.39042 | 62 |
| GO_REGULATION_OF_VIRAL_GENOME_REPLICATION | 0.00023 | 0.00663 | 0.51885 | 0.70900 | 1.41120 | 45 |
| GO_REGULATION_OF_WNT_SIGNALING_PATHWAY | 0.00000 | 0.00000 | 0.73376 | 0.64392 | 1.34634 | 200 |
| GO_RESPONSE_TO_DRUG | 0.00004 | 0.00141 | 0.55733 | 0.60151 | 1.25535 | 174 |
| GO_RESPONSE_TO_ETHANOL | 0.00115 | 0.02478 | 0.45506 | 0.68721 | 1.36915 | 46 |
| GO_RESPONSE_TO_INTERFERON_GAMMA | 0.00194 | 0.03747 | 0.45506 | 0.64030 | 1.29912 | 67 |
| GO_RESPONSE_TO_INTERLEUKIN_1 | 0.00000 | 0.00000 | 0.81404 | 0.73975 | 1.52372 | 99 |
| GO_RESPONSE_TO_INTERLEUKIN_12 | 0.00009 | 0.00287 | 0.53843 | 0.76348 | 1.50235 | 34 |
| GO_RESPONSE_TO_OXYGEN_LEVELS | 0.00000 | 0.00001 | 0.69013 | 0.62220 | 1.30529 | 226 |
| GO_RESPONSE_TO_TOPOLOGICALLY_INCORRECT_PROTEIN | 0.00045 | 0.01141 | 0.49849 | 0.60490 | 1.25662 | 138 |
| GO_RESPONSE_TO_TUMOR_NECROSIS_FACTOR | 0.00000 | 0.00001 | 0.67496 | 0.66125 | 1.37537 | 142 |
| GO_RESPONSE_TO_VIRUS | 0.00066 | 0.01551 | 0.47727 | 0.60043 | 1.24601 | 135 |
| GO_RESPONSE_TO_XENOBIOTIC_STIMULUS | 0.00283 | 0.04990 | 0.43171 | 0.69199 | 1.36817 | 38 |
| GO_RIBONUCLEOPROTEIN_COMPLEX_BINDING | 0.00000 | 0.00003 | 0.65944 | 0.68028 | 1.40570 | 114 |
| GO_RIBONUCLEOPROTEIN_COMPLEX_BIOGENESIS | 0.00000 | 0.00000 | | 0.66075 | 1.39989 | 384 |
| GO_RIBONUCLEOPROTEIN_COMPLEX_SUBUNIT_ORGANIZATION | 0.00000 | 0.00000 | | 0.70465 | 1.47062 | 174 |
| GO_RIBONUCLEOPROTEIN_GRANULE | 0.00004 | 0.00126 | 0.55733 | 0.61893 | 1.29020 | 168 |
| GO_RIBONUCLEOSIDE_MONOPHOSPHATE_METABOLIC_PROCESS | 0.00231 | 0.04369 | 0.43171 | 0.71431 | 1.39956 | 31 |
| GO_RIBONUCLEOSIDE_TRIPHOSPHATE_BIOSYNTHETIC_PROCESS | 0.00133 | 0.02696 | 0.45506 | 0.71694 | 1.40929 | 33 |
| GO_RIBONUCLEOSIDE_TRIPHOSPHATE_METABOLIC_PROCESS | 0.00119 | 0.02509 | 0.45506 | 0.69626 | 1.38190 | 42 |
| GO_RIBOSOMAL_LARGE_SUBUNIT_ASSEMBLY | 0.00066 | 0.01551 | 0.47727 | 0.75920 | 1.47981 | 27 |
| GO_RIBOSOMAL_LARGE_SUBUNIT_BIOGENESIS | 0.00000 | 0.00000 | 0.70498 | 0.74640 | 1.51323 | 66 |
| GO_RIBOSOMAL_SMALL_SUBUNIT_BIOGENESIS | 0.00023 | 0.00640 | 0.51885 | 0.67110 | 1.36057 | 66 |
| GO_RIBOSOMAL_SUBUNIT | 0.00000 | 0.00000 | | 0.75725 | 1.57994 | 172 |
| GO_RIBOSOME | 0.00000 | 0.00000 | | 0.74222 | 1.55252 | 205 |
| GO_RIBOSOME_ASSEMBLY | 0.00000 | 0.00004 | 0.65944 | 0.74856 | 1.50654 | 58 |

| | | | | | | |
|---|---|---|---|---|---|---|
| GO_RIBOSOME_BINDING | 0.00261 | 0.04750 | 0.43171 | 0.67093 | 1.33928 | 47 |
| GO_RIBOSOME_BIOGENESIS | 0.00000 | 0.00000 | 0.81404 | 0.64204 | 1.35197 | 268 |
| GO_RNA_CATABOLIC_PROCESS | 0.00000 | 0.00000 | | 0.75715 | 1.59930 | 318 |
| GO_RNA_DEPENDENT_DNA_BIOSYNTHETIC_PROCESS | 0.00012 | 0.00376 | 0.51885 | 0.69251 | 1.39494 | 59 |
| GO_RNA_LOCALIZATION | 0.00005 | 0.00170 | 0.55733 | 0.60196 | 1.25843 | 194 |
| GO_RNA_SPLICING | 0.00000 | 0.00006 | 0.64355 | 0.58702 | 1.24239 | 358 |
| GO_RNA_SPLICING_VIA_TRANSESTERIFICATION_REACTIONS | 0.00000 | 0.00005 | 0.64355 | 0.60166 | 1.26896 | 298 |
| GO_ROUGH_ENDOPLASMIC_RETICULUM | 0.00009 | 0.00278 | 0.53843 | 0.71973 | 1.44064 | 50 |
| GO_RRNA_BINDING | 0.00000 | 0.00000 | 0.71951 | 0.77222 | 1.54832 | 55 |
| GO_RRNA_METABOLIC_PROCESS | 0.00000 | 0.00003 | 0.65944 | 0.62662 | 1.31035 | 203 |
| GO_SCF_DEPENDENT_PROTEASOMAL_UBIQUITIN_DEPENDENT_PRO | 0.00000 | 0.00000 | 0.81404 | 0.77664 | 1.58041 | 72 |
| GO_SECRETORY_GRANULE | 0.00000 | 0.00000 | 0.78819 | 0.60486 | 1.28161 | 400 |
| GO_SECRETORY_VESICLE | 0.00000 | 0.00000 | 0.74774 | 0.58680 | 1.24526 | 465 |
| GO_SIGNAL_TRANSDUCTION_BY_PROTEIN_PHOSPHORYLATION | 0.00035 | 0.00951 | 0.49849 | 0.54934 | 1.16406 | 395 |
| GO_SINGLE_FERTILIZATION | 0.00001 | 0.00027 | 0.61053 | 0.74968 | 1.49841 | 48 |
| GO_SMALL_RIBOSOMAL_SUBUNIT | 0.00000 | 0.00000 | 0.73376 | 0.75164 | 1.52502 | 67 |
| GO_SPERM_EGG_RECOGNITION | 0.00000 | 0.00000 | 0.76146 | 0.91995 | 1.76604 | 21 |
| GO_SPLICEOSOMAL_COMPLEX | 0.00004 | 0.00141 | 0.55733 | 0.61267 | 1.27747 | 169 |
| GO_STEM_CELL_DIFFERENTIATION | 0.00000 | 0.00000 | | 0.71297 | 1.47954 | 135 |
| GO_STRUCTURAL_CONSTITUENT_OF_CYTOSKELETON | 0.00276 | 0.04907 | 0.43171 | 0.66713 | 1.33535 | 50 |
| GO_STRUCTURAL_CONSTITUENT_OF_RIBOSOME | 0.00000 | 0.00000 | | 0.77837 | 1.62087 | 151 |
| GO_STRUCTURAL_MOLECULE_ACTIVITY | 0.00000 | 0.00000 | | 0.71012 | 1.50152 | 326 |
| GO_SUBSTANTIA_NIGRA_DEVELOPMENT | 0.00038 | 0.01023 | 0.49849 | 0.77738 | 1.50032 | 25 |
| GO_TELOMERASE_RNA_LOCALIZATION | 0.00003 | 0.00114 | 0.55733 | 0.85476 | 1.62659 | 19 |
| GO_TELOMERE_MAINTENANCE_VIA_TELOMERE_LENGTHENING | 0.00148 | 0.02983 | 0.45506 | 0.64532 | 1.30830 | 66 |
| GO_THREONINE_TYPE_ENDOPEPTIDASE_ACTIVITY | 0.00000 | 0.00000 | 0.70498 | 0.92714 | 1.75394 | 17 |
| GO_THREONINE_TYPE_PEPTIDASE_ACTIVITY | 0.00000 | 0.00000 | 0.73376 | 0.91605 | 1.75224 | 20 |
| GO_TISSUE_MORPHOGENESIS | 0.00050 | 0.01250 | 0.47727 | 0.56178 | 1.18516 | 301 |
| GO_TOXIN_TRANSPORT | 0.00015 | 0.00449 | 0.51885 | 0.79478 | 1.53390 | 25 |
| GO_TRANSCRIPTION_COUPLED_NUCLEOTIDE_EXCISION_REPAIR | 0.00213 | 0.04061 | 0.43171 | 0.64578 | 1.31309 | 70 |
| GO_TRANSFORMING_GROWTH_FACTOR_BETA_RECEPTOR_SIGNALIN | 0.00276 | 0.04907 | 0.43171 | 0.61983 | 1.27316 | 93 |
| GO_TRANSLATIONAL_ELONGATION | 0.00039 | 0.01025 | 0.49849 | 0.61505 | 1.27206 | 119 |
| GO_TRANSLATIONAL_INITIATION | 0.00000 | 0.00000 | | 0.84322 | 1.75752 | 165 |
| GO_TRANSLATION_FACTOR_ACTIVITY_RNA_BINDING | 0.00000 | 0.00000 | | 0.81289 | 1.64967 | 68 |
| GO_TRANSLATION_INITIATION_FACTOR_ACTIVITY | 0.00000 | 0.00000 | 0.80122 | 0.82747 | 1.64700 | 45 |
| GO_TRANSLATION_INITIATION_FACTOR_BINDING | 0.00167 | 0.03282 | 0.45506 | 0.73572 | 1.43668 | 29 |
| GO_TRANSLATION_PREINITIATION_COMPLEX | 0.00001 | 0.00039 | 0.59333 | 0.88865 | 1.67573 | 16 |
| GO_TRANSLATION_REGULATOR_ACTIVITY | 0.00000 | 0.00000 | | 0.76724 | 1.58258 | 103 |
| GO_TRANSLATION_REGULATOR_ACTIVITY_NUCLEIC_ACID_BINDING | 0.00000 | 0.00000 | | 0.78305 | 1.59964 | 81 |
| GO_TRICARBOXYLIC_ACID_CYCLE | 0.00124 | 0.02586 | 0.45506 | 0.73726 | 1.43755 | 28 |
| GO_TUMOR_NECROSIS_FACTOR_MEDIATED_SIGNALING_PATHWAY | 0.00000 | 0.00000 | 0.82666 | 0.75799 | 1.55562 | 89 |
| GO_T_CELL_RECEPTOR_SIGNALING_PATHWAY | 0.00000 | 0.00003 | 0.65944 | 0.68018 | 1.40557 | 108 |
| GO_U2_TYPE_SPLICEOSOMAL_COMPLEX | 0.00250 | 0.04639 | 0.43171 | 0.62812 | 1.28587 | 84 |
| GO_UBIQUITIN_LIKE_PROTEIN_LIGASE_BINDING | 0.00000 | 0.00000 | 0.71951 | 0.64271 | 1.34438 | 205 |
| GO_UNFOLDED_PROTEIN_BINDING | 0.00000 | 0.00001 | 0.70498 | 0.73262 | 1.49351 | 77 |
| GO_VACUOLAR_LUMEN | 0.00012 | 0.00366 | 0.53843 | 0.66208 | 1.35681 | 85 |
| GO_VACUOLAR_MEMBRANE | 0.00119 | 0.02509 | 0.45506 | 0.55846 | 1.17719 | 284 |
| GO_VACUOLE | 0.00020 | 0.00582 | 0.51885 | 0.54597 | 1.15835 | 456 |
| GO_VESICLE_LUMEN | 0.00000 | 0.00000 | | 0.70821 | 1.47848 | 175 |
| GO_VIRAL_GENE_EXPRESSION | 0.00000 | 0.00000 | | 0.78299 | 1.63295 | 170 |
| GO_VIRAL_GENOME_REPLICATION | 0.00098 | 0.02172 | 0.45506 | 0.65926 | 1.33559 | 65 |
| GO_VIRAL_LIFE_CYCLE | 0.00027 | 0.00738 | 0.49849 | 0.58734 | 1.22813 | 191 |
| HP_ABNORMALITY_OF_CARDIOVASCULAR_SYSTEM_ELECTROPHYSIO | 0.00216 | 0.04103 | 0.43171 | 0.56322 | 1.18144 | 224 |
| HP_ABNORMALITY_OF_LOWER_LIP | 0.00065 | 0.01528 | 0.47727 | 0.60168 | 1.24811 | 133 |
| HP_ABNORMALITY_OF_THE_NASAL_DORSUM | 0.00039 | 0.01025 | 0.49849 | 0.61322 | 1.27161 | 131 |
| HP_ABNORMAL_ERYTHROCYTE_MORPHOLOGY | 0.00072 | 0.01652 | 0.47727 | 0.55217 | 1.16756 | 326 |
| HP_ACUTE_LEUKEMIA | 0.00000 | 0.00004 | 0.65944 | 0.72737 | 1.47611 | 68 |
| HP_ANEMIA_OF_INADEQUATE_PRODUCTION | 0.00002 | 0.00076 | 0.57561 | 0.66584 | 1.37104 | 98 |
| HP_DELAYED_PUBERTY | 0.00062 | 0.01482 | 0.47727 | 0.61972 | 1.28030 | 110 |
| HP_DEPRESSED_NASAL_RIDGE | 0.00002 | 0.00086 | 0.57561 | 0.69366 | 1.41154 | 72 |
| HP_FATIGUE | 0.00126 | 0.02595 | 0.45506 | 0.57291 | 1.19568 | 174 |
| HP_HEADACHE | 0.00106 | 0.02298 | 0.45506 | 0.59658 | 1.23802 | 135 |
| HP_HEMATOLOGICAL_NEOPLASM | 0.00043 | 0.01104 | 0.49849 | 0.59882 | 1.24559 | 145 |
| HP_LEUKEMIA | 0.00001 | 0.00048 | 0.59333 | 0.67554 | 1.38692 | 91 |
| HP_MACROCYTIC_ANEMIA | 0.00000 | 0.00001 | 0.70498 | 0.80975 | 1.60341 | 40 |
| HP_MIGRAINE | 0.00000 | 0.00002 | 0.67496 | 0.74292 | 1.50014 | 61 |
| HP_ORAL_CLEFT | 0.00119 | 0.02509 | 0.45506 | 0.55226 | 1.16622 | 314 |
| HP_PALLOR | 0.00054 | 0.01312 | 0.47727 | 0.64304 | 1.31859 | 87 |
| HP_SHORT_NOSE | 0.00253 | 0.04661 | 0.43171 | 0.58067 | 1.21030 | 165 |
| HP_THICK_LOWER_LIP_VERMILION | 0.00042 | 0.01084 | 0.49849 | 0.63169 | 1.29886 | 95 |
| **Cluster 14** | | | | | | |
| pathway | pval | padj | log2err | ES | NES | size |
| GO_ACTION_POTENTIAL | 0.00087 | 0.02538 | 0.47727 | 0.71846 | 1.34786 | 36 |
| GO_ALTERNATIVE_MRNA_SPLICING_VIA_SPLICEOSOME | 0.00062 | 0.02059 | 0.47727 | 0.67798 | 1.30027 | 59 |
| GO_ANAPHASE_PROMOTING_COMPLEX_DEPENDENT_CATABOLIC_P | 0.00122 | 0.03197 | 0.45506 | 0.65695 | 1.26811 | 72 |
| GO_ANATOMICAL_STRUCTURE_HOMEOSTASIS | 0.00141 | 0.03529 | 0.45506 | 0.58944 | 1.16618 | 195 |

| | | | | | | |
|---|---|---|---|---|---|---|
| GO_ANCHORING_JUNCTION | 0.00216 | 0.04731 | 0.43171 | 0.56471 | 1.12626 | 349 |
| GO_APOPTOTIC_SIGNALING_PATHWAY | 0.00065 | 0.02094 | 0.47727 | 0.58418 | 1.16100 | 263 |
| GO_ATP_DEPENDENT_CHROMATIN_REMODELING | 0.00187 | 0.04292 | 0.45506 | 0.66739 | 1.27348 | 54 |
| GO_AUTOPHAGOSOME_MEMBRANE | 0.00133 | 0.03396 | 0.45506 | 0.77965 | 1.41839 | 20 |
| GO_CADHERIN_BINDING | 0.00002 | 0.00183 | 0.57561 | 0.62036 | 1.22733 | 196 |
| GO_CARDIAC_MUSCLE_CELL_ACTION_POTENTIAL | 0.00161 | 0.03880 | 0.45506 | 0.77700 | 1.42117 | 22 |
| GO_CELLULAR_NITROGEN_COMPOUND_CATABOLIC_PROCESS | 0.00000 | 0.00011 | 0.69013 | 0.60621 | 1.20955 | 361 |
| GO_CELLULAR_PROTEIN_CATABOLIC_PROCESS | 0.00001 | 0.00117 | 0.59333 | 0.58192 | 1.16424 | 458 |
| GO_CELLULAR_RESPONSE_TO_ORGANIC_CYCLIC_COMPOUND | 0.00056 | 0.01888 | 0.47727 | 0.59337 | 1.17582 | 217 |
| GO_CELL_ACTIVATION_INVOLVED_IN_IMMUNE_RESPONSE | 0.00095 | 0.02704 | 0.47727 | 0.58178 | 1.15634 | 265 |
| GO_CELL_ADHESION_MOLECULE_BINDING | 0.00001 | 0.00145 | 0.59333 | 0.60906 | 1.21001 | 245 |
| GO_CELL_CELL_RECOGNITION | 0.00006 | 0.00385 | 0.55733 | 0.83820 | 1.52875 | 21 |
| GO_CELL_CYCLE_PHASE_TRANSITION | 0.00008 | 0.00534 | 0.53843 | 0.58224 | 1.16301 | 382 |
| GO_CELL_DIVISION | 0.00012 | 0.00622 | 0.53843 | 0.58335 | 1.16322 | 338 |
| GO_CELL_SUBSTRATE_JUNCTION | 0.00032 | 0.01310 | 0.49849 | 0.59729 | 1.18313 | 212 |
| GO_CHANNEL_REGULATOR_ACTIVITY | 0.00088 | 0.02550 | 0.47727 | 0.71010 | 1.33834 | 39 |
| GO_CHAPERONE_COMPLEX | 0.00001 | 0.00109 | 0.59333 | 0.87751 | 1.59063 | 19 |
| GO_CHROMATIN_BINDING | 0.00002 | 0.00204 | 0.57561 | 0.59423 | 1.18304 | 307 |
| GO_CHROMATIN_ORGANIZATION | 0.00000 | 0.00061 | 0.62726 | 0.59283 | 1.18486 | 425 |
| GO_CHROMATIN_REMODELING | 0.00012 | 0.00622 | 0.53843 | 0.64301 | 1.26065 | 121 |
| GO_CHROMOSOMAL_REGION | 0.00167 | 0.03936 | 0.45506 | 0.57571 | 1.14375 | 245 |
| GO_CHROMOSOME_SEGREGATION | 0.00117 | 0.03170 | 0.45506 | 0.58770 | 1.16523 | 224 |
| GO_COTRANSLATIONAL_PROTEIN_TARGETING_TO_MEMBRANE | 0.00001 | 0.00129 | 0.59333 | 0.69340 | 1.34632 | 84 |
| GO_COVALENT_CHROMATIN_MODIFICATION | 0.00012 | 0.00622 | 0.53843 | 0.59030 | 1.17424 | 286 |
| GO_CYSTEINE_TYPE_ENDOPEPTIDASE_ACTIVITY | 0.00003 | 0.00246 | 0.57561 | 0.74244 | 1.40760 | 44 |
| GO_CYSTEINE_TYPE_PEPTIDASE_ACTIVITY | 0.00035 | 0.01384 | 0.49849 | 0.68943 | 1.31924 | 57 |
| GO_CYTOKINE_MEDIATED_SIGNALING_PATHWAY | 0.00034 | 0.01377 | 0.49849 | 0.58838 | 1.16899 | 253 |
| GO_CYTOPLASMIC_STRESS_GRANULE | 0.00033 | 0.01325 | 0.49849 | 0.69315 | 1.32264 | 54 |
| GO_CYTOPLASMIC_TRANSLATION | 0.00012 | 0.00622 | 0.53843 | 0.67266 | 1.30510 | 82 |
| GO_CYTOSOLIC_LARGE_RIBOSOMAL_SUBUNIT | 0.00008 | 0.00535 | 0.53843 | 0.73103 | 1.38901 | 50 |
| GO_CYTOSOLIC_RIBOSOME | 0.00000 | 0.00025 | 0.65944 | 0.70274 | 1.36710 | 90 |
| GO_DEFENSE_RESPONSE | 0.00011 | 0.00622 | 0.53843 | 0.57058 | 1.14185 | 464 |
| GO_DEFENSE_RESPONSE_TO_OTHER_ORGANISM | 0.00001 | 0.00094 | 0.61053 | 0.59514 | 1.18690 | 332 |
| GO_DNA_BINDING_TRANSCRIPTION_FACTOR_BINDING | 0.00034 | 0.01377 | 0.49849 | 0.61294 | 1.20910 | 161 |
| GO_DNA_PACKAGING | 0.00016 | 0.00759 | 0.51885 | 0.64992 | 1.27018 | 103 |
| GO_DNA_POLYMERASE_BINDING | 0.00124 | 0.03211 | 0.45506 | 0.81875 | 1.47467 | 16 |
| GO_DNA_REPAIR | 0.00052 | 0.01771 | 0.47727 | 0.56967 | 1.13648 | 357 |
| GO_ENDOPEPTIDASE_ACTIVITY | 0.00071 | 0.02201 | 0.47727 | 0.62012 | 1.21793 | 127 |
| GO_ESTABLISHMENT_OF_PROTEIN_LOCALIZATION_TO_CHROMOSOM | 0.00119 | 0.03197 | 0.45506 | 0.80249 | 1.45428 | 18 |
| GO_ESTABLISHMENT_OF_PROTEIN_LOCALIZATION_TO_ENDOPLASM | 0.00001 | 0.00093 | 0.61053 | 0.67991 | 1.32569 | 96 |
| GO_ESTABLISHMENT_OF_PROTEIN_LOCALIZATION_TO_MEMBRANE | 0.00000 | 0.00025 | 0.65944 | 0.64074 | 1.26711 | 192 |
| GO_ESTABLISHMENT_OF_PROTEIN_LOCALIZATION_TO_ORGANELLE | 0.00000 | 0.00000 | 0.78819 | 0.63188 | 1.26018 | 337 |
| GO_ESTABLISHMENT_OF_RNA_LOCALIZATION | 0.00004 | 0.00294 | 0.55733 | 0.63716 | 1.25662 | 148 |
| GO_FICOLIN_1_RICH_GRANULE | 0.00005 | 0.00367 | 0.55733 | 0.69028 | 1.33529 | 75 |
| GO_HEMATOPOIETIC_PROGENITOR_CELL_DIFFERENTIATION | 0.00044 | 0.01575 | 0.49849 | 0.64452 | 1.25518 | 92 |
| GO_HEMATOPOIETIC_STEM_CELL_DIFFERENTIATION | 0.00231 | 0.04989 | 0.43171 | 0.66095 | 1.26984 | 61 |
| GO_HETEROCHROMATIN | 0.00006 | 0.00387 | 0.55733 | 0.73396 | 1.39172 | 45 |
| GO_HISTONE_BINDING | 0.00005 | 0.00355 | 0.55733 | 0.62917 | 1.24142 | 152 |
| GO_IMMUNE_EFFECTOR_PROCESS | 0.00041 | 0.01510 | 0.49849 | 0.56915 | 1.13754 | 425 |
| GO_INNATE_IMMUNE_RESPONSE | 0.00003 | 0.00271 | 0.55733 | 0.59683 | 1.18685 | 282 |
| GO_INTERACTION_WITH_SYMBIONT | 0.00069 | 0.02158 | 0.47727 | 0.72904 | 1.36946 | 37 |
| GO_INTRINSIC_APOPTOTIC_SIGNALING_PATHWAY | 0.00007 | 0.00489 | 0.53843 | 0.63422 | 1.24861 | 139 |
| GO_KINASE_BINDING | 0.00121 | 0.03197 | 0.45506 | 0.56359 | 1.12402 | 345 |
| GO_LEUKOCYTE_MEDIATED_IMMUNITY | 0.00056 | 0.01888 | 0.47727 | 0.57981 | 1.15334 | 291 |
| GO_METHYLTRANSFERASE_COMPLEX | 0.00065 | 0.02096 | 0.47727 | 0.66353 | 1.28566 | 77 |
| GO_MITOTIC_NUCLEAR_DIVISION | 0.00115 | 0.03137 | 0.45506 | 0.59904 | 1.18413 | 179 |
| GO_MITOTIC_SISTER_CHROMATID_SEGREGATION | 0.00122 | 0.03197 | 0.45506 | 0.61535 | 1.20637 | 118 |
| GO_MITOTIC_SPINDLE | 0.00040 | 0.01488 | 0.49849 | 0.65631 | 1.27420 | 83 |
| GO_MODIFICATION_DEPENDENT_MACROMOLECULE_CATABOLIC_PR | 0.00012 | 0.00622 | 0.53843 | 0.57728 | 1.15336 | 391 |
| GO_MODULATION_BY_HOST_OF_SYMBIONT_PROCESS | 0.00205 | 0.04610 | 0.43171 | 0.72224 | 1.34547 | 31 |
| GO_MRNA_BINDING | 0.00000 | 0.00059 | 0.62726 | 0.63310 | 1.25300 | 203 |
| GO_MRNA_CIS_SPLICING_VIA_SPLICEOSOME | 0.00047 | 0.01637 | 0.49849 | 0.78305 | 1.43792 | 24 |
| GO_MRNA_EXPORT_FROM_NUCLEUS | 0.00076 | 0.02357 | 0.47727 | 0.65380 | 1.27074 | 86 |
| GO_MRNA_PROCESSING | 0.00000 | 0.00036 | 0.64355 | 0.59908 | 1.19557 | 367 |
| GO_MRNA_TRANSPORT | 0.00029 | 0.01217 | 0.49849 | 0.64231 | 1.25678 | 109 |
| GO_MULTI_ORGANISM_PROCESS | 0.00121 | 0.03197 | 0.45506 | 0.55835 | 1.11583 | 419 |
| GO_MYELOID_LEUKOCYTE_ACTIVATION | 0.00066 | 0.02099 | 0.47727 | 0.58463 | 1.16133 | 251 |
| GO_MYELOID_LEUKOCYTE_MEDIATED_IMMUNITY | 0.00064 | 0.02078 | 0.47727 | 0.59092 | 1.17301 | 233 |
| GO_NEGATIVE_REGULATION_OF_CELLULAR_COMPONENT_ORGANIZ | 0.00163 | 0.03886 | 0.45506 | 0.56875 | 1.13307 | 320 |
| GO_NEGATIVE_REGULATION_OF_CELLULAR_PROTEIN_CATABOLIC_PI | 0.00106 | 0.02902 | 0.45506 | 0.69334 | 1.31739 | 50 |
| GO_NEGATIVE_REGULATION_OF_CELL_CYCLE | 0.00010 | 0.00605 | 0.53843 | 0.58419 | 1.16490 | 338 |
| GO_NEGATIVE_REGULATION_OF_CELL_CYCLE_PHASE_TRANSITION | 0.00228 | 0.04930 | 0.43171 | 0.60658 | 1.19621 | 147 |
| GO_NEGATIVE_REGULATION_OF_CELL_CYCLE_PROCESS | 0.00002 | 0.00216 | 0.57561 | 0.61669 | 1.22062 | 204 |
| GO_NEGATIVE_REGULATION_OF_CHROMOSOME_ORGANIZATION | 0.00097 | 0.02725 | 0.47727 | 0.63740 | 1.23999 | 90 |
| GO_NEGATIVE_REGULATION_OF_DNA_METABOLIC_PROCESS | 0.00178 | 0.04138 | 0.45506 | 0.66513 | 1.27561 | 59 |
| GO_NEGATIVE_REGULATION_OF_DNA_REPAIR | 0.00018 | 0.00841 | 0.51885 | 0.84254 | 1.52201 | 17 |

| | | | | | | |
|---|---|---|---|---|---|---|
| GO_NEGATIVE_REGULATION_OF_MITOTIC_CELL_CYCLE | 0.00135 | 0.03425 | 0.45506 | 0.59403 | 1.17489 | 190 |
| GO_NEGATIVE_REGULATION_OF_MOLECULAR_FUNCTION | 0.00194 | 0.04401 | 0.45506 | 0.55815 | 1.11535 | 396 |
| GO_NEGATIVE_REGULATION_OF_PROTEIN_CATABOLIC_PROCESS | 0.00040 | 0.01488 | 0.49849 | 0.66971 | 1.29368 | 73 |
| GO_NEGATIVE_REGULATION_OF_PROTEIN_METABOLIC_PROCESS | 0.00016 | 0.00756 | 0.51885 | 0.56824 | 1.13659 | 454 |
| GO_NEGATIVE_REGULATION_OF_PROTEIN_MODIFICATION_BY_SMA | 0.00038 | 0.01457 | 0.49849 | 0.71555 | 1.35741 | 47 |
| GO_NEGATIVE_REGULATION_OF_PROTEIN_MODIFICATION_PROCESS | 0.00004 | 0.00306 | 0.55733 | 0.60423 | 1.20077 | 262 |
| GO_NEGATIVE_REGULATION_OF_RESPONSE_TO_DNA_DAMAGE_STI | 0.00041 | 0.01516 | 0.49849 | 0.71809 | 1.35701 | 41 |
| GO_NEGATIVE_REGULATION_OF_TRANSCRIPTION_BY_RNA_POLYME | 0.00008 | 0.00534 | 0.53843 | 0.58312 | 1.16300 | 346 |
| GO_NUCLEAR_BODY | 0.00012 | 0.00627 | 0.53843 | 0.56943 | 1.13967 | 466 |
| GO_NUCLEAR_ENVELOPE | 0.00001 | 0.00086 | 0.61053 | 0.61230 | 1.21649 | 253 |
| GO_NUCLEAR_EXPORT | 0.00002 | 0.00188 | 0.57561 | 0.63875 | 1.26058 | 150 |
| GO_NUCLEAR_HETEROCHROMATIN | 0.00033 | 0.01325 | 0.49849 | 0.80778 | 1.46955 | 20 |
| GO_NUCLEAR_MEMBRANE | 0.00011 | 0.00622 | 0.53843 | 0.62545 | 1.23423 | 159 |
| GO_NUCLEAR_SPECK | 0.00132 | 0.03366 | 0.45506 | 0.58463 | 1.16123 | 248 |
| GO_NUCLEAR_TRANSCRIBED_MRNA_CATABOLIC_PROCESS | 0.00003 | 0.00276 | 0.55733 | 0.63479 | 1.25272 | 163 |
| GO_NUCLEAR_TRANSCRIBED_MRNA_CATABOLIC_PROCESS_NONSEN | 0.00000 | 0.00052 | 0.62726 | 0.68780 | 1.34400 | 102 |
| GO_NUCLEAR_TRANSPORT | 0.00000 | 0.00025 | 0.65944 | 0.62972 | 1.25027 | 234 |
| GO_NUCLEOBASE_CONTAINING_COMPOUND_TRANSPORT | 0.00010 | 0.00618 | 0.53843 | 0.61997 | 1.22487 | 174 |
| GO_ORGANIC_CYCLIC_COMPOUND_CATABOLIC_PROCESS | 0.00000 | 0.00017 | 0.67496 | 0.60571 | 1.20901 | 369 |
| GO_PATTERN_RECOGNITION_RECEPTOR_SIGNALING_PATHWAY | 0.00213 | 0.04713 | 0.43171 | 0.63663 | 1.23598 | 83 |
| GO_PEPTIDE_BIOSYNTHETIC_PROCESS | 0.00000 | 0.00035 | 0.64355 | 0.59230 | 1.18445 | 440 |
| GO_PEPTIDE_METABOLIC_PROCESS | 0.00000 | 0.00035 | 0.64355 | 0.58432 | 1.17070 | 500 |
| GO_PEPTIDYL_LYSINE_MODIFICATION | 0.00104 | 0.02883 | 0.45506 | 0.58964 | 1.16966 | 229 |
| GO_PIGMENT_GRANULE | 0.00087 | 0.02526 | 0.47727 | 0.66915 | 1.28718 | 66 |
| GO_POLYSOME | 0.00084 | 0.02490 | 0.47727 | 0.68512 | 1.31005 | 56 |
| GO_POSITIVE_REGULATION_OF_CANONICAL_WNT_SIGNALING_PATH | 0.00213 | 0.04713 | 0.43171 | 0.64865 | 1.25752 | 81 |
| GO_POSITIVE_REGULATION_OF_CELLULAR_AMIDE_METABOLIC_PRO | 0.00216 | 0.04731 | 0.43171 | 0.62542 | 1.21897 | 95 |
| GO_POSITIVE_REGULATION_OF_CELLULAR_PROTEIN_LOCALIZATION | 0.00004 | 0.00306 | 0.55733 | 0.62475 | 1.23504 | 181 |
| GO_POSITIVE_REGULATION_OF_CELL_CYCLE | 0.00006 | 0.00412 | 0.53843 | 0.60996 | 1.20728 | 204 |
| GO_POSITIVE_REGULATION_OF_CELL_CYCLE_PROCESS | 0.00026 | 0.01131 | 0.49849 | 0.62119 | 1.22566 | 156 |
| GO_POSITIVE_REGULATION_OF_CELL_DEATH | 0.00143 | 0.03539 | 0.45506 | 0.57997 | 1.15318 | 271 |
| GO_POSITIVE_REGULATION_OF_CHROMOSOME_ORGANIZATION | 0.00013 | 0.00637 | 0.51885 | 0.64055 | 1.25543 | 117 |
| GO_POSITIVE_REGULATION_OF_CHROMOSOME_SEGREGATION | 0.00163 | 0.03886 | 0.45506 | 0.73345 | 1.35679 | 28 |
| GO_POSITIVE_REGULATION_OF_DNA_BIOSYNTHETIC_PROCESS | 0.00051 | 0.01752 | 0.47727 | 0.70795 | 1.34242 | 45 |
| GO_POSITIVE_REGULATION_OF_DNA_METABOLIC_PROCESS | 0.00092 | 0.02640 | 0.47727 | 0.62779 | 1.23067 | 116 |
| GO_POSITIVE_REGULATION_OF_ESTABLISHMENT_OF_PROTEIN_LOC/ | 0.00124 | 0.03211 | 0.45506 | 0.59552 | 1.17631 | 173 |
| GO_POSITIVE_REGULATION_OF_IMMUNE_RESPONSE | 0.00067 | 0.02114 | 0.47727 | 0.58918 | 1.16877 | 228 |
| GO_POSITIVE_REGULATION_OF_IMMUNE_SYSTEM_PROCESS | 0.00046 | 0.01600 | 0.49849 | 0.57555 | 1.14612 | 312 |
| GO_POSITIVE_REGULATION_OF_INTRACELLULAR_PROTEIN_TRANSP( | 0.00128 | 0.03289 | 0.45506 | 0.62670 | 1.22480 | 103 |
| GO_POSITIVE_REGULATION_OF_INTRACELLULAR_SIGNAL_TRANSDU( | 0.00043 | 0.01543 | 0.49849 | 0.57296 | 1.14259 | 335 |
| GO_POSITIVE_REGULATION_OF_INTRINSIC_APOPTOTIC_SIGNALING_ | 0.00066 | 0.02099 | 0.47727 | 0.78062 | 1.43562 | 25 |
| GO_POSITIVE_REGULATION_OF_MITOTIC_CELL_CYCLE | 0.00086 | 0.02514 | 0.47727 | 0.63467 | 1.23839 | 97 |
| GO_POSITIVE_REGULATION_OF_PROTEIN_LOCALIZATION_TO_NUCLE | 0.00005 | 0.00369 | 0.55733 | 0.74321 | 1.40988 | 47 |
| GO_POSITIVE_REGULATION_OF_TELOMERASE_ACTIVITY | 0.00178 | 0.04138 | 0.45506 | 0.75066 | 1.38253 | 26 |
| GO_POSITIVE_REGULATION_OF_TELOMERE_MAINTENANCE | 0.00137 | 0.03454 | 0.45506 | 0.71298 | 1.33757 | 36 |
| GO_POSITIVE_REGULATION_OF_TRANSFERASE_ACTIVITY | 0.00145 | 0.03567 | 0.45506 | 0.57398 | 1.14136 | 278 |
| GO_POSITIVE_REGULATION_OF_WNT_SIGNALING_PATHWAY | 0.00017 | 0.00828 | 0.51885 | 0.65110 | 1.26952 | 96 |
| GO_POSTTRANSCRIPTIONAL_REGULATION_OF_GENE_EXPRESSION | 0.00000 | 0.00003 | 0.71951 | 0.60734 | 1.21365 | 396 |
| GO_POST_TRANSLATIONAL_PROTEIN_MODIFICATION | 0.00037 | 0.01433 | 0.49849 | 0.60895 | 1.20372 | 179 |
| GO_POTASSIUM_ION_TRANSPORT | 0.00082 | 0.02450 | 0.47727 | 0.68997 | 1.31392 | 52 |
| GO_PROTEIN_CONTAINING_COMPLEX_LOCALIZATION | 0.00026 | 0.01131 | 0.49849 | 0.60672 | 1.19940 | 178 |
| GO_PROTEIN_DOMAIN_SPECIFIC_BINDING | 0.00005 | 0.00372 | 0.55733 | 0.58922 | 1.17332 | 308 |
| GO_PROTEIN_FOLDING | 0.00044 | 0.01575 | 0.49849 | 0.62512 | 1.22837 | 130 |
| GO_PROTEIN_IMPORT | 0.00027 | 0.01140 | 0.49849 | 0.64264 | 1.25783 | 108 |
| GO_PROTEIN_LOCALIZATION_TO_CHROMOSOME | 0.00011 | 0.00622 | 0.53843 | 0.70110 | 1.34543 | 60 |
| GO_PROTEIN_LOCALIZATION_TO_CHROMOSOME_TELOMERIC_REGI( | 0.00010 | 0.00605 | 0.53843 | 0.81100 | 1.48656 | 23 |
| GO_PROTEIN_LOCALIZATION_TO_ENDOPLASMIC_RETICULUM | 0.00000 | 0.00063 | 0.61053 | 0.68104 | 1.33266 | 111 |
| GO_PROTEIN_LOCALIZATION_TO_MEMBRANE | 0.00001 | 0.00148 | 0.59333 | 0.59476 | 1.18467 | 319 |
| GO_PROTEIN_LOCALIZATION_TO_NUCLEUS | 0.00000 | 0.00025 | 0.65944 | 0.65674 | 1.29616 | 154 |
| GO_PROTEIN_POLYUBIQUITINATION | 0.00028 | 0.01208 | 0.49849 | 0.60384 | 1.19503 | 198 |
| GO_PROTEIN_STABILIZATION | 0.00039 | 0.01470 | 0.49849 | 0.63951 | 1.25129 | 109 |
| GO_PROTEIN_TARGETING | 0.00000 | 0.00059 | 0.62726 | 0.61787 | 1.22769 | 257 |
| GO_PROTEIN_TARGETING_TO_MEMBRANE | 0.00000 | 0.00063 | 0.61053 | 0.66852 | 1.31066 | 121 |
| GO_REGULATION_OF_APOPTOTIC_SIGNALING_PATHWAY | 0.00036 | 0.01427 | 0.49849 | 0.60670 | 1.19928 | 176 |
| GO_REGULATION_OF_CELLULAR_AMIDE_METABOLIC_PROCESS | 0.00003 | 0.00266 | 0.57561 | 0.60735 | 1.20674 | 259 |
| GO_REGULATION_OF_CELLULAR_CATABOLIC_PROCESS | 0.00024 | 0.01080 | 0.51885 | 0.56744 | 1.13646 | 476 |
| GO_REGULATION_OF_CELLULAR_LOCALIZATION | 0.00039 | 0.01470 | 0.49849 | 0.56570 | 1.13112 | 433 |
| GO_REGULATION_OF_CELLULAR_PROTEIN_CATABOLIC_PROCESS | 0.00081 | 0.02438 | 0.47727 | 0.60924 | 1.19980 | 142 |
| GO_REGULATION_OF_CELLULAR_PROTEIN_LOCALIZATION | 0.00002 | 0.00231 | 0.57561 | 0.59908 | 1.19174 | 290 |
| GO_REGULATION_OF_CELLULAR_RESPONSE_TO_STRESS | 0.00000 | 0.00036 | 0.64355 | 0.60050 | 1.19755 | 353 |
| GO_REGULATION_OF_CELL_CYCLE_PHASE_TRANSITION | 0.00026 | 0.01131 | 0.49849 | 0.59226 | 1.17741 | 274 |
| GO_REGULATION_OF_CELL_CYCLE_PROCESS | 0.00001 | 0.00149 | 0.59333 | 0.58573 | 1.17142 | 437 |
| GO_REGULATION_OF_CHROMATIN_ORGANIZATION | 0.00122 | 0.03197 | 0.45506 | 0.62882 | 1.22786 | 100 |
| GO_REGULATION_OF_CHROMOSOME_ORGANIZATION | 0.00001 | 0.00149 | 0.59333 | 0.61422 | 1.21783 | 223 |
| GO_REGULATION_OF_CHROMOSOME_SEGREGATION | 0.00202 | 0.04548 | 0.43171 | 0.63380 | 1.23160 | 85 |
| GO_REGULATION_OF_DNA_BIOSYNTHETIC_PROCESS | 0.00053 | 0.01790 | 0.47727 | 0.67925 | 1.30846 | 68 |

| | | | | | | |
|---|---|---|---|---|---|---|
| GO_REGULATION_OF_DNA_METABOLIC_PROCESS | 0.00000 | 0.00001 | 0.76146 | 0.65534 | 1.29772 | 208 |
| GO_REGULATION_OF_DNA_REPAIR | 0.00085 | 0.02502 | 0.47727 | 0.65499 | 1.26973 | 78 |
| GO_REGULATION_OF_HEMATOPOIETIC_STEM_CELL_DIFFERENTIATIO | 0.00157 | 0.03810 | 0.45506 | 0.66703 | 1.27638 | 57 |
| GO_REGULATION_OF_IMMUNE_RESPONSE | 0.00106 | 0.02902 | 0.45506 | 0.57477 | 1.14403 | 302 |
| GO_REGULATION_OF_INTRACELLULAR_PROTEIN_TRANSPORT | 0.00065 | 0.02096 | 0.47727 | 0.61681 | 1.21398 | 138 |
| GO_REGULATION_OF_INTRINSIC_APOPTOTIC_SIGNALING_PATHWAY | 0.00001 | 0.00097 | 0.61053 | 0.70570 | 1.36630 | 76 |
| GO_REGULATION_OF_MITOTIC_CELL_CYCLE | 0.00020 | 0.00924 | 0.51885 | 0.57415 | 1.14670 | 378 |
| GO_REGULATION_OF_MRNA_METABOLIC_PROCESS | 0.00000 | 0.00011 | 0.69013 | 0.63630 | 1.26333 | 234 |
| GO_REGULATION_OF_MRNA_PROCESSING | 0.00000 | 0.00052 | 0.62726 | 0.68214 | 1.33199 | 100 |
| GO_REGULATION_OF_MRNA_SPLICING_VIA_SPLICEOSOME | 0.00001 | 0.00121 | 0.59333 | 0.70201 | 1.35915 | 76 |
| GO_REGULATION_OF_NUCLEASE_ACTIVITY | 0.00122 | 0.03197 | 0.45506 | 0.81937 | 1.47577 | 16 |
| GO_REGULATION_OF_PROTEIN_CATABOLIC_PROCESS | 0.00024 | 0.01080 | 0.51885 | 0.59838 | 1.18493 | 208 |
| GO_REGULATION_OF_PROTEIN_IMPORT | 0.00165 | 0.03912 | 0.45506 | 0.71852 | 1.34712 | 34 |
| GO_REGULATION_OF_PROTEIN_LOCALIZATION | 0.00044 | 0.01575 | 0.49849 | 0.56444 | 1.12785 | 417 |
| GO_REGULATION_OF_PROTEIN_LOCALIZATION_TO_NUCLEUS | 0.00011 | 0.00622 | 0.53843 | 0.69179 | 1.33307 | 69 |
| GO_REGULATION_OF_PROTEIN_MODIFICATION_BY_SMALL_PROTEIN | 0.00003 | 0.00242 | 0.57561 | 0.64365 | 1.26744 | 141 |
| GO_REGULATION_OF_PROTEIN_STABILITY | 0.00152 | 0.03712 | 0.45506 | 0.60471 | 1.19345 | 162 |
| GO_REGULATION_OF_RESPONSE_TO_BIOTIC_STIMULUS | 0.00150 | 0.03685 | 0.45506 | 0.59426 | 1.17447 | 175 |
| GO_REGULATION_OF_RESPONSE_TO_DNA_DAMAGE_STIMULUS | 0.00020 | 0.00924 | 0.51885 | 0.63400 | 1.24581 | 130 |
| GO_REGULATION_OF_RNA_SPLICING | 0.00000 | 0.00072 | 0.61053 | 0.67864 | 1.32631 | 103 |
| GO_REGULATION_OF_TELOMERE_MAINTENANCE | 0.00039 | 0.01488 | 0.49849 | 0.68398 | 1.31178 | 59 |
| GO_REGULATION_OF_TRANSFERASE_ACTIVITY | 0.00004 | 0.00306 | 0.55733 | 0.58160 | 1.16241 | 425 |
| GO_REGULATION_OF_TRANSLATIONAL_INITIATION | 0.00021 | 0.00966 | 0.51885 | 0.69560 | 1.33280 | 58 |
| GO_REGULATION_OF_TRANSLATION_IN_RESPONSE_TO_STRESS | 0.00046 | 0.01600 | 0.49849 | 0.84578 | 1.52223 | 15 |
| GO_REGULATION_OF_VIRAL_GENOME_REPLICATION | 0.00157 | 0.03810 | 0.45506 | 0.68748 | 1.30359 | 45 |
| GO_RESPONSE_TO_BIOTIC_STIMULUS | 0.00000 | 0.00036 | 0.64355 | 0.58759 | 1.17564 | 459 |
| GO_RESPONSE_TO_CYTOKINE | 0.00000 | 0.00002 | 0.73376 | 0.61012 | 1.22019 | 437 |
| GO_RESPONSE_TO_HEAT | 0.00097 | 0.02725 | 0.47727 | 0.64394 | 1.25194 | 88 |
| GO_RESPONSE_TO_INTERLEUKIN_4 | 0.00191 | 0.04337 | 0.45506 | 0.79229 | 1.43580 | 18 |
| GO_RESPONSE_TO_LEUKEMIA_INHIBITORY_FACTOR | 0.00004 | 0.00306 | 0.55733 | 0.71663 | 1.37682 | 61 |
| GO_RESPONSE_TO_TEMPERATURE_STIMULUS | 0.00081 | 0.02438 | 0.47727 | 0.63376 | 1.23860 | 103 |
| GO_RESPONSE_TO_VIRUS | 0.00042 | 0.01538 | 0.49849 | 0.62466 | 1.22686 | 127 |
| GO_RIBONUCLEOPROTEIN_COMPLEX_BIOGENESIS | 0.00000 | 0.00000 | 0.81404 | 0.63027 | 1.25750 | 359 |
| GO_RIBONUCLEOPROTEIN_COMPLEX_SUBUNIT_ORGANIZATION | 0.00012 | 0.00632 | 0.51885 | 0.62746 | 1.23836 | 162 |
| GO_RIBONUCLEOPROTEIN_GRANULE | 0.00064 | 0.02078 | 0.47727 | 0.61371 | 1.21091 | 156 |
| GO_RIBOSOMAL_LARGE_SUBUNIT_BIOGENESIS | 0.00001 | 0.00093 | 0.61053 | 0.71638 | 1.37759 | 63 |
| GO_RIBOSOMAL_SUBUNIT | 0.00011 | 0.00622 | 0.53843 | 0.62260 | 1.22845 | 152 |
| GO_RIBOSOME | 0.00011 | 0.00622 | 0.53843 | 0.61464 | 1.21511 | 183 |
| GO_RIBOSOME_ASSEMBLY | 0.00181 | 0.04185 | 0.45506 | 0.67164 | 1.28349 | 55 |
| GO_RIBOSOME_BIOGENESIS | 0.00000 | 0.00002 | 0.71951 | 0.64313 | 1.27744 | 248 |
| GO_RNA_CATABOLIC_PROCESS | 0.00000 | 0.00032 | 0.64355 | 0.61566 | 1.22551 | 297 |
| GO_RNA_DEPENDENT_DNA_BIOSYNTHETIC_PROCESS | 0.00002 | 0.00231 | 0.57561 | 0.73556 | 1.40193 | 53 |
| GO_RNA_EXPORT_FROM_NUCLEUS | 0.00006 | 0.00385 | 0.55733 | 0.65075 | 1.27367 | 112 |
| GO_RNA_LOCALIZATION | 0.00000 | 0.00036 | 0.64355 | 0.64183 | 1.26870 | 176 |
| GO_RNA_POLYMERASE_II_SPECIFIC_DNA_BINDING_TRANSCRIPTION | 0.00220 | 0.04791 | 0.43171 | 0.60962 | 1.19817 | 133 |
| GO_RNA_SPLICING | 0.00000 | 0.00011 | 0.67496 | 0.61385 | 1.22300 | 322 |
| GO_RNA_SPLICING_VIA_TRANSESTERIFICATION_REACTIONS | 0.00000 | 0.00061 | 0.62726 | 0.61185 | 1.21633 | 273 |
| GO_RRNA_METABOLIC_PROCESS | 0.00014 | 0.00710 | 0.51885 | 0.61284 | 1.21160 | 186 |
| GO_SERINE_FAMILY_AMINO_ACID_METABOLIC_PROCESS | 0.00174 | 0.04091 | 0.45506 | 0.79377 | 1.43883 | 19 |
| GO_SISTER_CHROMATID_SEGREGATION | 0.00216 | 0.04731 | 0.43171 | 0.60450 | 1.19035 | 141 |
| GO_SPERM_EGG_RECOGNITION | 0.00014 | 0.00714 | 0.51885 | 0.84545 | 1.53214 | 18 |
| GO_SPINDLE | 0.00080 | 0.02438 | 0.47727 | 0.58374 | 1.15751 | 222 |
| GO_SPINDLE_MIDZONE | 0.00011 | 0.00622 | 0.53843 | 0.80971 | 1.48421 | 23 |
| GO_SPLICEOSOMAL_COMPLEX | 0.00191 | 0.04337 | 0.45506 | 0.60668 | 1.19722 | 155 |
| GO_STRUCTURAL_CONSTITUENT_OF_RIBOSOME | 0.00064 | 0.02078 | 0.47727 | 0.61960 | 1.21907 | 135 |
| GO_TELOMERASE_RNA_LOCALIZATION | 0.00010 | 0.00605 | 0.53843 | 0.85067 | 1.54160 | 18 |
| GO_TELOMERE_MAINTENANCE_VIA_TELOMERE_LENGTHENING | 0.00003 | 0.00271 | 0.55733 | 0.71008 | 1.36183 | 59 |
| GO_TELOMERE_ORGANIZATION | 0.00181 | 0.04185 | 0.45506 | 0.62541 | 1.22381 | 111 |
| GO_TRANSCRIPTION_FACTOR_BINDING | 0.00003 | 0.00276 | 0.55733 | 0.59604 | 1.18649 | 303 |
| GO_TRANSLATIONAL_INITIATION | 0.00000 | 0.00000 | 0.82666 | 0.70258 | 1.38625 | 152 |
| GO_TRANSLATION_FACTOR_ACTIVITY_RNA_BINDING | 0.00019 | 0.00867 | 0.51885 | 0.69133 | 1.32587 | 59 |
| GO_TRANSLATION_INITIATION_FACTOR_ACTIVITY | 0.00078 | 0.02399 | 0.47727 | 0.72097 | 1.36151 | 40 |
| GO_TRANSLATION_REGULATOR_ACTIVITY | 0.00015 | 0.00718 | 0.51885 | 0.65896 | 1.28331 | 92 |
| GO_TRANSLATION_REGULATOR_ACTIVITY_NUCLEIC_ACID_BINDING | 0.00031 | 0.01283 | 0.49849 | 0.67428 | 1.30156 | 72 |
| GO_UBIQUITIN_LIKE_PROTEIN_LIGASE_BINDING | 0.00005 | 0.00355 | 0.55733 | 0.62082 | 1.22697 | 175 |
| GO_UNFOLDED_PROTEIN_BINDING | 0.00000 | 0.00006 | 0.70498 | 0.74860 | 1.44319 | 70 |
| GO_VESICLE_LUMEN | 0.00018 | 0.00854 | 0.51885 | 0.63241 | 1.24297 | 133 |
| GO_VIRAL_GENE_EXPRESSION | 0.00000 | 0.00052 | 0.62726 | 0.65239 | 1.28751 | 150 |
| GO_VIRAL_LIFE_CYCLE | 0.00001 | 0.00100 | 0.59333 | 0.63632 | 1.25571 | 164 |
| HP_ABNORMAL_LIP_MORPHOLOGY | 0.00102 | 0.02864 | 0.45506 | 0.56398 | 1.12708 | 409 |
| HP_DEPRESSED_NASAL_RIDGE | 0.00213 | 0.04713 | 0.43171 | 0.65684 | 1.26351 | 66 |
| HP_GONOSOMAL_INHERITANCE | 0.00012 | 0.00622 | 0.53843 | 0.64471 | 1.26184 | 112 |
| HP_LEUKEMIA | 0.00104 | 0.02883 | 0.45506 | 0.64457 | 1.25150 | 84 |
| HP_X_LINKED_RECESSIVE_INHERITANCE | 0.00143 | 0.03539 | 0.45506 | 0.63954 | 1.24174 | 84 |
| **Cluster 15** | | | | | | |
| pathway | pval | padj | log2err | ES | NES | size |

| | | | | | | |
|---|---|---|---|---|---|---|
| GO_ANAPHASE_PROMOTING_COMPLEX_DEPENDENT_CATABOLIC_P | 0.00001 | 0.00070 | 0.59333 | 0.68776 | 1.34974 | 73 |
| GO_ATPASE_ACTIVITY | 0.00004 | 0.00217 | 0.55733 | 0.58102 | 1.18156 | 295 |
| GO_BINDING_OF_SPERM_TO_ZONA_PELLUCIDA | 0.00224 | 0.04188 | 0.43171 | 0.78209 | 1.42983 | 18 |
| GO_BRUSH_BORDER_MEMBRANE | 0.00143 | 0.03126 | 0.45506 | 0.73067 | 1.37533 | 30 |
| GO_CADHERIN_BINDING | 0.00043 | 0.01299 | 0.49849 | 0.58256 | 1.17820 | 221 |
| GO_CANONICAL_WNT_SIGNALING_PATHWAY | 0.00014 | 0.00586 | 0.51885 | 0.59423 | 1.19957 | 204 |
| GO_CARDIAC_MUSCLE_CELL_CONTRACTION | 0.00104 | 0.02487 | 0.45506 | 0.76683 | 1.42083 | 23 |
| GO_CATALYTIC_STEP_2_SPLICEOSOME | 0.00036 | 0.01179 | 0.49849 | 0.65858 | 1.29425 | 76 |
| GO_CELLULAR_NITROGEN_COMPOUND_CATABOLIC_PROCESS | 0.00000 | 0.00000 | 0.81404 | 0.60960 | 1.24671 | 420 |
| GO_CELLULAR_RESPONSE_TO_HEAT | 0.00030 | 0.01049 | 0.49849 | 0.64217 | 1.26620 | 88 |
| GO_CELLULAR_RESPONSE_TO_ORGANIC_CYCLIC_COMPOUND | 0.00095 | 0.02334 | 0.47727 | 0.56652 | 1.15134 | 286 |
| GO_CELLULAR_RESPONSE_TO_STEROID_HORMONE_STIMULUS | 0.00257 | 0.04674 | 0.43171 | 0.60030 | 1.19571 | 118 |
| GO_CELL_ADHESION_MOLECULE_BINDING | 0.00113 | 0.02620 | 0.45506 | 0.56483 | 1.14791 | 286 |
| GO_CELL_CELL_RECOGNITION | 0.00018 | 0.00680 | 0.51885 | 0.76501 | 1.43995 | 30 |
| GO_CELL_CELL_SIGNALING_BY_WNT | 0.00025 | 0.00905 | 0.49849 | 0.56965 | 1.15999 | 306 |
| GO_CELL_CYCLE_CHECKPOINT | 0.00008 | 0.00404 | 0.53843 | 0.60863 | 1.22466 | 168 |
| GO_CELL_CYCLE_G1_S_PHASE_TRANSITION | 0.00003 | 0.00173 | 0.57561 | 0.60897 | 1.22902 | 200 |
| GO_CELL_CYCLE_G2_M_PHASE_TRANSITION | 0.00020 | 0.00737 | 0.51885 | 0.59261 | 1.19731 | 212 |
| GO_CELL_CYCLE_PHASE_TRANSITION | 0.00000 | 0.00000 | 0.81404 | 0.60459 | 1.23817 | 456 |
| GO_CELL_DIVISION | 0.00000 | 0.00021 | 0.62726 | 0.57883 | 1.18368 | 412 |
| GO_CELL_SUBSTRATE_JUNCTION | 0.00005 | 0.00285 | 0.55733 | 0.58428 | 1.18432 | 255 |
| GO_CELL_SURFACE_RECEPTOR_SIGNALING_PATHWAY_INVOLVED_IN | 0.00145 | 0.03154 | 0.45506 | 0.55345 | 1.12928 | 333 |
| GO_CHAPERONE_COMPLEX | 0.00016 | 0.00639 | 0.51885 | 0.81760 | 1.49796 | 19 |
| GO_CHROMATIN_BINDING | 0.00000 | 0.00001 | 0.71951 | 0.59972 | 1.22519 | 370 |
| GO_CHROMATIN_ORGANIZATION_INVOLVED_IN_REGULATION_OF_T | 0.00020 | 0.00737 | 0.51885 | 0.67130 | 1.31320 | 68 |
| GO_CHROMATIN_REMODELING | 0.00017 | 0.00666 | 0.51885 | 0.62061 | 1.24160 | 135 |
| GO_CHROMATIN_SILENCING | 0.00135 | 0.02987 | 0.45506 | 0.71063 | 1.33947 | 31 |
| GO_CHROMOSOMAL_REGION | 0.00006 | 0.00310 | 0.55733 | 0.58326 | 1.18452 | 276 |
| GO_CHROMOSOME_CENTROMERIC_REGION | 0.00037 | 0.01187 | 0.49849 | 0.59970 | 1.20426 | 156 |
| GO_CHROMOSOME_SEGREGATION | 0.00000 | 0.00029 | 0.62726 | 0.60038 | 1.21695 | 255 |
| GO_CHROMOSOME_SEPARATION | 0.00113 | 0.02620 | 0.45506 | 0.64627 | 1.26896 | 74 |
| GO_CONDENSED_CHROMOSOME | 0.00039 | 0.01220 | 0.49849 | 0.59924 | 1.20446 | 164 |
| GO_COTRANSLATIONAL_PROTEIN_TARGETING_TO_MEMBRANE | 0.00000 | 0.00000 | 0.74774 | 0.73158 | 1.44054 | 84 |
| GO_COVALENT_CHROMATIN_MODIFICATION | 0.00000 | 0.00004 | 0.67496 | 0.60191 | 1.22814 | 336 |
| GO_CYTOPLASMIC_STRESS_GRANULE | 0.00073 | 0.01961 | 0.47727 | 0.65863 | 1.28504 | 62 |
| GO_CYTOPLASMIC_TRANSLATION | 0.00005 | 0.00284 | 0.55733 | 0.66621 | 1.31257 | 85 |
| GO_CYTOPLASMIC_TRANSLATIONAL_INITIATION | 0.00261 | 0.04697 | 0.43171 | 0.71493 | 1.34193 | 28 |
| GO_CYTOSOLIC_LARGE_RIBOSOMAL_SUBUNIT | 0.00000 | 0.00017 | 0.64355 | 0.75582 | 1.46097 | 49 |
| GO_CYTOSOLIC_RIBOSOME | 0.00000 | 0.00000 | 0.80122 | 0.74169 | 1.46243 | 88 |
| GO_CYTOSOLIC_SMALL_RIBOSOMAL_SUBUNIT | 0.00001 | 0.00098 | 0.59333 | 0.75743 | 1.44319 | 37 |
| GO_DEFENSE_RESPONSE_TO_OTHER_ORGANISM | 0.00005 | 0.00285 | 0.55733 | 0.56458 | 1.15454 | 417 |
| GO_DEFENSE_RESPONSE_TO_VIRUS | 0.00178 | 0.03579 | 0.45506 | 0.60759 | 1.20769 | 111 |
| GO_DEMETHYLATION | 0.00148 | 0.03222 | 0.45506 | 0.69497 | 1.32046 | 35 |
| GO_DNA_BINDING_TRANSCRIPTION_FACTOR_BINDING | 0.00001 | 0.00052 | 0.61053 | 0.61205 | 1.23740 | 216 |
| GO_DNA_CONFORMATION_CHANGE | 0.00005 | 0.00295 | 0.55733 | 0.59805 | 1.20870 | 214 |
| GO_DNA_INTEGRITY_CHECKPOINT | 0.00257 | 0.04674 | 0.43171 | 0.60143 | 1.19825 | 120 |
| GO_DNA_PACKAGING | 0.00018 | 0.00696 | 0.51885 | 0.62340 | 1.24198 | 119 |
| GO_DNA_REPAIR | 0.00005 | 0.00295 | 0.55733 | 0.56697 | 1.15951 | 419 |
| GO_DNA_REPLICATION | 0.00085 | 0.02199 | 0.47727 | 0.58001 | 1.17232 | 215 |
| GO_DNA_TEMPLATED_TRANSCRIPTION_ELONGATION | 0.00011 | 0.00474 | 0.53843 | 0.65032 | 1.28316 | 91 |
| GO_DNA_TEMPLATED_TRANSCRIPTION_INITIATION | 0.00001 | 0.00108 | 0.59333 | 0.62013 | 1.24771 | 167 |
| GO_DNA_TEMPLATED_TRANSCRIPTION_TERMINATION | 0.00168 | 0.03491 | 0.45506 | 0.65081 | 1.27076 | 64 |
| GO_ESTABLISHMENT_OF_PROTEIN_LOCALIZATION_TO_CHROMOSOM | 0.00065 | 0.01797 | 0.47727 | 0.77797 | 1.43892 | 22 |
| GO_ESTABLISHMENT_OF_PROTEIN_LOCALIZATION_TO_ENDOPLASM | 0.00000 | 0.00001 | 0.71951 | 0.71375 | 1.41103 | 95 |
| GO_ESTABLISHMENT_OF_PROTEIN_LOCALIZATION_TO_MEMBRANE | 0.00000 | 0.00002 | 0.69013 | 0.62260 | 1.25969 | 234 |
| GO_ESTABLISHMENT_OF_PROTEIN_LOCALIZATION_TO_ORGANELLE | 0.00000 | 0.00000 | | 0.64263 | 1.31425 | 396 |
| GO_ESTABLISHMENT_OF_PROTEIN_LOCALIZATION_TO_TELOMERE | 0.00011 | 0.00490 | 0.53843 | 0.84764 | 1.53542 | 16 |
| GO_ESTABLISHMENT_OF_RNA_LOCALIZATION | 0.00000 | 0.00005 | 0.67496 | 0.64059 | 1.28798 | 165 |
| GO_FICOLIN_1_RICH_GRANULE | 0.00004 | 0.00212 | 0.55733 | 0.66357 | 1.30794 | 86 |
| GO_GENERAL_TRANSCRIPTION_INITIATION_FACTOR_BINDING | 0.00016 | 0.00644 | 0.51885 | 0.73725 | 1.40287 | 36 |
| GO_GENE_SILENCING | 0.00037 | 0.01196 | 0.49849 | 0.59681 | 1.20045 | 166 |
| GO_HEAT_SHOCK_PROTEIN_BINDING | 0.00130 | 0.02876 | 0.45506 | 0.62569 | 1.23326 | 86 |
| GO_HETEROCHROMATIN | 0.00113 | 0.02620 | 0.45506 | 0.66021 | 1.28547 | 58 |
| GO_HISTONE_BINDING | 0.00001 | 0.00048 | 0.61053 | 0.62190 | 1.25218 | 171 |
| GO_HISTONE_METHYLTRANSFERASE_COMPLEX | 0.00163 | 0.03438 | 0.45506 | 0.64697 | 1.26331 | 65 |
| GO_HISTONE_PHOSPHORYLATION | 0.00261 | 0.04697 | 0.43171 | 0.72793 | 1.36146 | 27 |
| GO_HORMONE_MEDIATED_SIGNALING_PATHWAY | 0.00168 | 0.03491 | 0.45506 | 0.61435 | 1.21818 | 101 |
| GO_HORMONE_RECEPTOR_BINDING | 0.00059 | 0.01660 | 0.47727 | 0.62737 | 1.24475 | 104 |
| GO_HOST_CELLULAR_COMPONENT | 0.00108 | 0.02564 | 0.45506 | 0.67286 | 1.30060 | 49 |
| GO_IMPORT_INTO_NUCLEUS | 0.00002 | 0.00136 | 0.57561 | 0.64835 | 1.28870 | 111 |
| GO_INNATE_IMMUNE_RESPONSE | 0.00009 | 0.00413 | 0.53843 | 0.56637 | 1.15599 | 347 |
| GO_INTERACTION_WITH_HOST | 0.00007 | 0.00348 | 0.53843 | 0.63241 | 1.26105 | 123 |
| GO_INTERLEUKIN_1_MEDIATED_SIGNALING_PATHWAY | 0.00268 | 0.04800 | 0.43171 | 0.63068 | 1.23835 | 74 |
| GO_INTRINSIC_APOPTOTIC_SIGNALING_PATHWAY | 0.00025 | 0.00900 | 0.49849 | 0.59720 | 1.20351 | 180 |
| GO_INTRINSIC_APOPTOTIC_SIGNALING_PATHWAY_BY_P53_CLASS_M | 0.00152 | 0.03277 | 0.45506 | 0.68063 | 1.31326 | 46 |
| GO_KINASE_BINDING | 0.00253 | 0.04636 | 0.43171 | 0.54404 | 1.11408 | 455 |

| | | | | | | |
|---|---|---|---|---|---|---|
| GO_LARGE_RIBOSOMAL_SUBUNIT | 0.00033 | 0.01122 | 0.49849 | 0.63199 | 1.24818 | 94 |
| GO_METAPHASE_ANAPHASE_TRANSITION_OF_CELL_CYCLE | 0.00119 | 0.02683 | 0.45506 | 0.67045 | 1.30131 | 53 |
| GO_METHYLTRANSFERASE_COMPLEX | 0.00209 | 0.04028 | 0.43171 | 0.62135 | 1.22594 | 89 |
| GO_MITOTIC_CELL_CYCLE_CHECKPOINT | 0.00035 | 0.01156 | 0.49849 | 0.60630 | 1.21012 | 126 |
| GO_MITOTIC_NUCLEAR_DIVISION | 0.00002 | 0.00131 | 0.57561 | 0.60587 | 1.22331 | 205 |
| GO_MITOTIC_SISTER_CHROMATID_SEGREGATION | 0.00009 | 0.00406 | 0.53843 | 0.62323 | 1.24592 | 131 |
| GO_MITOTIC_SPINDLE_ASSEMBLY | 0.00122 | 0.02745 | 0.45506 | 0.68232 | 1.31980 | 50 |
| GO_MODIFICATION_DEPENDENT_MACROMOLECULE_CATABOLIC_PR | 0.00000 | 0.00000 | 0.77494 | 0.59643 | 1.22209 | 473 |
| GO_MODIFIED_AMINO_ACID_TRANSPORT | 0.00231 | 0.04298 | 0.43171 | 0.76236 | 1.39677 | 19 |
| GO_MODULATION_BY_VIRUS_OF_HOST_PROCESS | 0.00213 | 0.04072 | 0.43171 | 0.78351 | 1.43243 | 18 |
| GO_MORPHOGENESIS_OF_A_POLARIZED_EPITHELIUM | 0.00176 | 0.03579 | 0.45506 | 0.61609 | 1.21938 | 97 |
| GO_MOVEMENT_IN_HOST_ENVIRONMENT | 0.00087 | 0.02216 | 0.47727 | 0.63130 | 1.24632 | 92 |
| GO_MRNA_3_END_PROCESSING | 0.00012 | 0.00506 | 0.53843 | 0.65208 | 1.28529 | 86 |
| GO_MRNA_5_UTR_BINDING | 0.00034 | 0.01146 | 0.49849 | 0.79879 | 1.46788 | 20 |
| GO_MRNA_BINDING | 0.00000 | 0.00000 | 0.78819 | 0.64904 | 1.31256 | 227 |
| GO_MRNA_CIS_SPLICING_VIA_SPLICEOSOME | 0.00261 | 0.04697 | 0.43171 | 0.74674 | 1.38360 | 23 |
| GO_MRNA_EXPORT_FROM_NUCLEUS | 0.00000 | 0.00010 | 0.65944 | 0.68051 | 1.34766 | 98 |
| GO_MRNA_PROCESSING | 0.00000 | 0.00000 | | 0.61585 | 1.25929 | 401 |
| GO_MRNA_TRANSPORT | 0.00000 | 0.00031 | 0.62726 | 0.64853 | 1.29441 | 126 |
| GO_MULTI_ORGANISM_LOCALIZATION | 0.00010 | 0.00444 | 0.53843 | 0.70775 | 1.36806 | 49 |
| GO_MYELOID_LEUKOCYTE_MEDIATED_IMMUNITY | 0.00268 | 0.04800 | 0.43171 | 0.55620 | 1.13123 | 296 |
| GO_NCRNA_EXPORT_FROM_NUCLEUS | 0.00091 | 0.02273 | 0.47727 | 0.71928 | 1.36372 | 34 |
| GO_NCRNA_METABOLIC_PROCESS | 0.00001 | 0.00098 | 0.59333 | 0.57500 | 1.17492 | 375 |
| GO_NCRNA_PROCESSING | 0.00000 | 0.00017 | 0.64355 | 0.59560 | 1.21378 | 315 |
| GO_NCRNA_TRANSCRIPTION | 0.00161 | 0.03425 | 0.45506 | 0.62357 | 1.23032 | 89 |
| GO_NEGATIVE_REGULATION_OF_CANONICAL_WNT_SIGNALING_PAT | 0.00046 | 0.01395 | 0.49849 | 0.61810 | 1.22989 | 115 |
| GO_NEGATIVE_REGULATION_OF_CELLULAR_RESPONSE_TO_GROWTI | 0.00242 | 0.04461 | 0.43171 | 0.63145 | 1.23986 | 74 |
| GO_NEGATIVE_REGULATION_OF_CELLULAR_RESPONSE_TO_TRANSF( | 0.00042 | 0.01292 | 0.49849 | 0.69303 | 1.33596 | 45 |
| GO_NEGATIVE_REGULATION_OF_CELL_CYCLE | 0.00000 | 0.00001 | 0.73376 | 0.59526 | 1.21736 | 422 |
| GO_NEGATIVE_REGULATION_OF_CELL_CYCLE_G2_M_PHASE_TRANS | 0.00038 | 0.01203 | 0.49849 | 0.64336 | 1.26631 | 83 |
| GO_NEGATIVE_REGULATION_OF_CELL_CYCLE_PHASE_TRANSITION | 0.00002 | 0.00120 | 0.57561 | 0.61765 | 1.24417 | 178 |
| GO_NEGATIVE_REGULATION_OF_CELL_CYCLE_PROCESS | 0.00000 | 0.00004 | 0.67496 | 0.61781 | 1.25086 | 244 |
| GO_NEGATIVE_REGULATION_OF_CHROMATIN_ORGANIZATION | 0.00220 | 0.04145 | 0.43171 | 0.70311 | 1.33306 | 34 |
| GO_NEGATIVE_REGULATION_OF_CHROMOSOME_ORGANIZATION | 0.00100 | 0.02419 | 0.45506 | 0.62106 | 1.23221 | 104 |
| GO_NEGATIVE_REGULATION_OF_MITOTIC_CELL_CYCLE | 0.00001 | 0.00063 | 0.59333 | 0.60568 | 1.22486 | 228 |
| GO_NEGATIVE_REGULATION_OF_PROTEIN_MODIFICATION_BY_SMA | 0.00159 | 0.03398 | 0.45506 | 0.64505 | 1.25854 | 62 |
| GO_NEGATIVE_REGULATION_OF_TRANSCRIPTION_BY_RNA_POLYME | 0.00006 | 0.00322 | 0.53843 | 0.56323 | 1.15327 | 453 |
| GO_NEGATIVE_REGULATION_OF_TRANSMEMBRANE_RECEPTOR_PR( | 0.00006 | 0.00313 | 0.55733 | 0.69879 | 1.36059 | 58 |
| GO_NEGATIVE_REGULATION_OF_WNT_SIGNALING_PATHWAY | 0.00027 | 0.00958 | 0.49849 | 0.61170 | 1.22176 | 128 |
| GO_NIK_NF_KAPPAB_SIGNALING | 0.00117 | 0.02662 | 0.45506 | 0.61867 | 1.22828 | 106 |
| GO_NUCLEAR_CHROMOSOME_SEGREGATION | 0.00003 | 0.00171 | 0.57561 | 0.60421 | 1.21971 | 204 |
| GO_NUCLEAR_ENVELOPE | 0.00000 | 0.00008 | 0.65944 | 0.59925 | 1.22025 | 309 |
| GO_NUCLEAR_EXPORT | 0.00000 | 0.00034 | 0.61053 | 0.62709 | 1.26085 | 165 |
| GO_NUCLEAR_HORMONE_RECEPTOR_BINDING | 0.00039 | 0.01228 | 0.49849 | 0.63524 | 1.25582 | 95 |
| GO_NUCLEAR_MATRIX | 0.00124 | 0.02764 | 0.45506 | 0.64192 | 1.26152 | 76 |
| GO_NUCLEAR_MEMBRANE | 0.00000 | 0.00013 | 0.64355 | 0.62750 | 1.26523 | 191 |
| GO_NUCLEAR_PERIPHERY | 0.00003 | 0.00183 | 0.57561 | 0.65655 | 1.29946 | 97 |
| GO_NUCLEAR_PORE | 0.00087 | 0.02216 | 0.47727 | 0.66087 | 1.29041 | 64 |
| GO_NUCLEAR_SPECK | 0.00000 | 0.00000 | 0.74774 | 0.62313 | 1.26667 | 289 |
| GO_NUCLEAR_TRANSCRIBED_MRNA_CATABOLIC_PROCESS | 0.00000 | 0.00005 | 0.67496 | 0.64286 | 1.29480 | 174 |
| GO_NUCLEAR_TRANSCRIBED_MRNA_CATABOLIC_PROCESS_NONSEN | 0.00000 | 0.00000 | 0.76146 | 0.71315 | 1.41480 | 103 |
| GO_NUCLEAR_TRANSPORT | 0.00000 | 0.00001 | 0.71951 | 0.62345 | 1.26433 | 260 |
| GO_NUCLEOBASE_CONTAINING_COMPOUND_TRANSPORT | 0.00001 | 0.00048 | 0.61053 | 0.61239 | 1.23623 | 204 |
| GO_NUCLEUS_ORGANIZATION | 0.00009 | 0.00413 | 0.53843 | 0.64348 | 1.27311 | 96 |
| GO_OOCYTE_DIFFERENTIATION | 0.00154 | 0.03305 | 0.45506 | 0.70793 | 1.33439 | 31 |
| GO_ORGANELLE_FISSION | 0.00004 | 0.00217 | 0.55733 | 0.57750 | 1.17791 | 329 |
| GO_ORGANIC_CYCLIC_COMPOUND_CATABOLIC_PROCESS | 0.00000 | 0.00000 | 0.80122 | 0.60670 | 1.24158 | 433 |
| GO_PEPTIDE_BIOSYNTHETIC_PROCESS | 0.00000 | 0.00000 | 0.81404 | 0.59978 | 1.22898 | 492 |
| GO_PEPTIDYL_LYSINE_MODIFICATION | 0.00000 | 0.00009 | 0.65944 | 0.60667 | 1.23203 | 274 |
| GO_PHAGOPHORE_ASSEMBLY_SITE | 0.00152 | 0.03277 | 0.45506 | 0.73727 | 1.37892 | 27 |
| GO_POLYSOME | 0.00065 | 0.01789 | 0.47727 | 0.66715 | 1.30012 | 59 |
| GO_POSITIVE_REGULATION_OF_CANONICAL_WNT_SIGNALING_PATH | 0.00005 | 0.00283 | 0.55733 | 0.64136 | 1.27328 | 105 |
| GO_POSITIVE_REGULATION_OF_CATABOLIC_PROCESS | 0.00250 | 0.04583 | 0.43171 | 0.55734 | 1.13400 | 300 |
| GO_POSITIVE_REGULATION_OF_CELLULAR_AMIDE_METABOLIC_PRC | 0.00224 | 0.04188 | 0.43171 | 0.60512 | 1.20321 | 112 |
| GO_POSITIVE_REGULATION_OF_CELLULAR_PROTEIN_CATABOLIC_PR | 0.00205 | 0.03997 | 0.43171 | 0.61182 | 1.21047 | 96 |
| GO_POSITIVE_REGULATION_OF_CELLULAR_PROTEIN_LOCALIZATION | 0.00024 | 0.00889 | 0.51885 | 0.58218 | 1.17729 | 220 |
| GO_POSITIVE_REGULATION_OF_CELL_CYCLE | 0.00002 | 0.00130 | 0.57561 | 0.59833 | 1.21208 | 248 |
| GO_POSITIVE_REGULATION_OF_CELL_CYCLE_G1_S_PHASE_TRANSIT | 0.00025 | 0.00905 | 0.49849 | 0.74824 | 1.41634 | 33 |
| GO_POSITIVE_REGULATION_OF_CELL_CYCLE_PHASE_TRANSITION | 0.00069 | 0.01871 | 0.47727 | 0.66233 | 1.29855 | 71 |
| GO_POSITIVE_REGULATION_OF_CELL_CYCLE_PROCESS | 0.00001 | 0.00098 | 0.59333 | 0.61367 | 1.23734 | 191 |
| GO_POSITIVE_REGULATION_OF_CELL_POPULATION_PROLIFERATION | 0.00047 | 0.01401 | 0.49849 | 0.55699 | 1.13899 | 397 |
| GO_POSITIVE_REGULATION_OF_CHROMATIN_ORGANIZATION | 0.00095 | 0.02334 | 0.47727 | 0.64849 | 1.27222 | 72 |
| GO_POSITIVE_REGULATION_OF_CHROMOSOME_ORGANIZATION | 0.00008 | 0.00401 | 0.53843 | 0.62401 | 1.24836 | 136 |
| GO_POSITIVE_REGULATION_OF_DEFENSE_RESPONSE | 0.00023 | 0.00861 | 0.51885 | 0.60367 | 1.21535 | 170 |
| GO_POSITIVE_REGULATION_OF_DNA_BIOSYNTHETIC_PROCESS | 0.00042 | 0.01292 | 0.49849 | 0.68548 | 1.32933 | 52 |
| GO_POSITIVE_REGULATION_OF_DNA_TEMPLATED_TRANSCRIPTION_ | 0.00098 | 0.02405 | 0.45506 | 0.74719 | 1.39748 | 27 |

| | | | | | | |
|---|---|---|---|---|---|---|
| GO_POSITIVE_REGULATION_OF_ESTABLISHMENT_OF_PROTEIN_LOC/ | 0.00026 | 0.00932 | 0.49849 | 0.58350 | 1.17888 | 211 |
| GO_POSITIVE_REGULATION_OF_G1_S_TRANSITION_OF_MITOTIC_CE | 0.00090 | 0.02270 | 0.47727 | 0.74926 | 1.40135 | 27 |
| GO_POSITIVE_REGULATION_OF_GENE_EXPRESSION_EPIGENETIC | 0.00163 | 0.03438 | 0.45506 | 0.68377 | 1.31529 | 43 |
| GO_POSITIVE_REGULATION_OF_IMMUNE_RESPONSE | 0.00157 | 0.03371 | 0.45506 | 0.56020 | 1.13875 | 289 |
| GO_POSITIVE_REGULATION_OF_IMMUNE_SYSTEM_PROCESS | 0.00178 | 0.03579 | 0.45506 | 0.54573 | 1.11598 | 412 |
| GO_POSITIVE_REGULATION_OF_INNATE_IMMUNE_RESPONSE | 0.00017 | 0.00666 | 0.51885 | 0.63068 | 1.25359 | 111 |
| GO_POSITIVE_REGULATION_OF_INTRACELLULAR_PROTEIN_TRANSP( | 0.00117 | 0.02662 | 0.45506 | 0.60938 | 1.21380 | 118 |
| GO_POSITIVE_REGULATION_OF_INTRINSIC_APOPTOTIC_SIGNALING_ | 0.00003 | 0.00186 | 0.55733 | 0.75057 | 1.42609 | 35 |
| GO_POSITIVE_REGULATION_OF_MITOTIC_CELL_CYCLE | 0.00068 | 0.01863 | 0.47727 | 0.62096 | 1.23526 | 114 |
| GO_POSITIVE_REGULATION_OF_NUCLEOCYTOPLASMIC_TRANSPORT | 0.00040 | 0.01249 | 0.49849 | 0.69416 | 1.33815 | 45 |
| GO_POSITIVE_REGULATION_OF_ORGANELLE_ORGANIZATION | 0.00035 | 0.01156 | 0.49849 | 0.55788 | 1.14103 | 393 |
| GO_POSITIVE_REGULATION_OF_PROTEIN_CATABOLIC_PROCESS | 0.00100 | 0.02419 | 0.45506 | 0.59879 | 1.20077 | 146 |
| GO_POSITIVE_REGULATION_OF_PROTEIN_IMPORT | 0.00057 | 0.01633 | 0.47727 | 0.75762 | 1.41698 | 27 |
| GO_POSITIVE_REGULATION_OF_PROTEIN_LOCALIZATION_TO_NUCLE | 0.00000 | 0.00028 | 0.62726 | 0.72695 | 1.41667 | 59 |
| GO_POSITIVE_REGULATION_OF_RESPONSE_TO_BIOTIC_STIMULUS | 0.00009 | 0.00413 | 0.53843 | 0.62253 | 1.24340 | 128 |
| GO_POSITIVE_REGULATION_OF_RESPONSE_TO_EXTERNAL_STIMULU | 0.00054 | 0.01570 | 0.47727 | 0.57888 | 1.17057 | 224 |
| GO_POSITIVE_REGULATION_OF_RNA_SPLICING | 0.00220 | 0.04145 | 0.43171 | 0.71170 | 1.33842 | 29 |
| GO_POSITIVE_REGULATION_OF_WNT_SIGNALING_PATHWAY | 0.00007 | 0.00350 | 0.53843 | 0.63008 | 1.25596 | 122 |
| GO_POSTTRANSCRIPTIONAL_REGULATION_OF_GENE_EXPRESSION | 0.00000 | 0.00000 | | 0.61360 | 1.25673 | 457 |
| GO_POST_TRANSLATIONAL_PROTEIN_MODIFICATION | 0.00032 | 0.01089 | 0.49849 | 0.58126 | 1.17548 | 222 |
| GO_PRERIBOSOME | 0.00019 | 0.00709 | 0.51885 | 0.67708 | 1.32301 | 66 |
| GO_PROTEASOMAL_PROTEIN_CATABOLIC_PROCESS | 0.00000 | 0.00014 | 0.64355 | 0.58559 | 1.19568 | 360 |
| GO_PROTEIN_CONTAINING_COMPLEX_LOCALIZATION | 0.00006 | 0.00322 | 0.53843 | 0.59726 | 1.20711 | 214 |
| GO_PROTEIN_C_TERMINUS_BINDING | 0.00167 | 0.03478 | 0.45506 | 0.60710 | 1.20715 | 112 |
| GO_PROTEIN_DNA_COMPLEX_SUBUNIT_ORGANIZATION | 0.00001 | 0.00058 | 0.61053 | 0.62550 | 1.25850 | 167 |
| GO_PROTEIN_DOMAIN_SPECIFIC_BINDING | 0.00115 | 0.02641 | 0.45506 | 0.55040 | 1.12556 | 405 |
| GO_PROTEIN_FOLDING | 0.00083 | 0.02161 | 0.47727 | 0.59699 | 1.19891 | 160 |
| GO_PROTEIN_IMPORT | 0.00000 | 0.00034 | 0.61053 | 0.65037 | 1.29851 | 127 |
| GO_PROTEIN_LOCALIZATION_TO_CHROMOSOME | 0.00092 | 0.02286 | 0.47727 | 0.65160 | 1.27578 | 69 |
| GO_PROTEIN_LOCALIZATION_TO_CHROMOSOME_TELOMERIC_REGI( | 0.00047 | 0.01401 | 0.49849 | 0.76785 | 1.43290 | 26 |
| GO_PROTEIN_LOCALIZATION_TO_ENDOPLASMIC_RETICULUM | 0.00000 | 0.00000 | 0.76146 | 0.70467 | 1.40308 | 116 |
| GO_PROTEIN_LOCALIZATION_TO_MEMBRANE | 0.00121 | 0.02714 | 0.45506 | 0.55071 | 1.12613 | 398 |
| GO_PROTEIN_LOCALIZATION_TO_NUCLEUS | 0.00000 | 0.00000 | 0.77494 | 0.65804 | 1.32700 | 184 |
| GO_PROTEIN_MODIFICATION_BY_SMALL_PROTEIN_REMOVAL | 0.00003 | 0.00187 | 0.55733 | 0.59880 | 1.21086 | 224 |
| GO_PROTEIN_POLYUBIQUITINATION | 0.00006 | 0.00325 | 0.53843 | 0.59051 | 1.19606 | 247 |
| GO_PROTEIN_STABILIZATION | 0.00001 | 0.00069 | 0.59333 | 0.64333 | 1.28554 | 129 |
| GO_PROTEIN_SUMOYLATION | 0.00031 | 0.01084 | 0.49849 | 0.67848 | 1.32104 | 58 |
| GO_PROTEIN_TARGETING | 0.00000 | 0.00008 | 0.65944 | 0.59813 | 1.21773 | 305 |
| GO_PROTEIN_TARGETING_TO_MEMBRANE | 0.00000 | 0.00000 | 0.77494 | 0.68358 | 1.37033 | 143 |
| GO_REGULATION_OF_APOPTOTIC_SIGNALING_PATHWAY | 0.00239 | 0.04421 | 0.43171 | 0.56590 | 1.14517 | 237 |
| GO_REGULATION_OF_CELLULAR_AMIDE_METABOLIC_PROCESS | 0.00000 | 0.00005 | 0.67496 | 0.60012 | 1.22178 | 305 |
| GO_REGULATION_OF_CELLULAR_AMINE_METABOLIC_PROCESS | 0.00194 | 0.03820 | 0.45506 | 0.64725 | 1.26135 | 59 |
| GO_REGULATION_OF_CELLULAR_AMINO_ACID_METABOLIC_PROCES | 0.00209 | 0.04028 | 0.43171 | 0.65275 | 1.26831 | 54 |
| GO_REGULATION_OF_CELLULAR_PROTEIN_LOCALIZATION | 0.00115 | 0.02641 | 0.45506 | 0.55507 | 1.13348 | 361 |
| GO_REGULATION_OF_CELLULAR_RESPONSE_TO_HEAT | 0.00172 | 0.03529 | 0.45506 | 0.64960 | 1.26841 | 64 |
| GO_REGULATION_OF_CELLULAR_RESPONSE_TO_STRESS | 0.00031 | 0.01084 | 0.49849 | 0.55388 | 1.13492 | 473 |
| GO_REGULATION_OF_CELLULAR_RESPONSE_TO_TRANSFORMING_G | 0.00242 | 0.04461 | 0.43171 | 0.64385 | 1.25808 | 66 |
| GO_REGULATION_OF_CELL_CYCLE_CHECKPOINT | 0.00032 | 0.01086 | 0.49849 | 0.74472 | 1.40050 | 29 |
| GO_REGULATION_OF_CELL_CYCLE_G2_M_PHASE_TRANSITION | 0.00082 | 0.02161 | 0.47727 | 0.59568 | 1.19926 | 170 |
| GO_REGULATION_OF_CELL_CYCLE_PHASE_TRANSITION | 0.00000 | 0.00005 | 0.67496 | 0.60010 | 1.22428 | 332 |
| GO_REGULATION_OF_CHROMATIN_ORGANIZATION | 0.00000 | 0.00009 | 0.65944 | 0.66074 | 1.32033 | 129 |
| GO_REGULATION_OF_CHROMOSOME_ORGANIZATION | 0.00000 | 0.00000 | 0.76146 | 0.62812 | 1.27393 | 263 |
| GO_REGULATION_OF_CHROMOSOME_SEGREGATION | 0.00003 | 0.00174 | 0.57561 | 0.66358 | 1.31052 | 93 |
| GO_REGULATION_OF_CHROMOSOME_SEPARATION | 0.00057 | 0.01633 | 0.47727 | 0.66874 | 1.30149 | 57 |
| GO_REGULATION_OF_DNA_BIOSYNTHETIC_PROCESS | 0.00272 | 0.04836 | 0.43171 | 0.62716 | 1.23325 | 78 |
| GO_REGULATION_OF_DNA_METABOLIC_PROCESS | 0.00003 | 0.00184 | 0.57561 | 0.59543 | 1.20496 | 238 |
| GO_REGULATION_OF_DNA_TEMPLATED_TRANSCRIPTION_INITIATIO! | 0.00047 | 0.01401 | 0.49849 | 0.74663 | 1.41102 | 32 |
| GO_REGULATION_OF_DNA_TEMPLATED_TRANSCRIPTION_IN_RESPO | 0.00276 | 0.04857 | 0.43171 | 0.61863 | 1.21938 | 87 |
| GO_REGULATION_OF_GENE_EXPRESSION_EPIGENETIC | 0.00049 | 0.01433 | 0.47727 | 0.62472 | 1.24275 | 114 |
| GO_REGULATION_OF_GENE_SILENCING | 0.00043 | 0.01306 | 0.49849 | 0.63773 | 1.25478 | 81 |
| GO_REGULATION_OF_HEART_RATE | 0.00048 | 0.01414 | 0.49849 | 0.76740 | 1.43206 | 26 |
| GO_REGULATION_OF_HEMATOPOIETIC_STEM_CELL_DIFFERENTIATIC | 0.00220 | 0.04145 | 0.43171 | 0.64946 | 1.26395 | 57 |
| GO_REGULATION_OF_HISTONE_MODIFICATION | 0.00003 | 0.00173 | 0.57561 | 0.65614 | 1.30111 | 102 |
| GO_REGULATION_OF_INNATE_IMMUNE_RESPONSE | 0.00039 | 0.01220 | 0.49849 | 0.59968 | 1.20495 | 162 |
| GO_REGULATION_OF_INTRACELLULAR_PROTEIN_TRANSPORT | 0.00039 | 0.01220 | 0.49849 | 0.59155 | 1.19107 | 171 |
| GO_REGULATION_OF_INTRINSIC_APOPTOTIC_SIGNALING_PATHWAY | 0.00065 | 0.01789 | 0.47727 | 0.62671 | 1.24277 | 102 |
| GO_REGULATION_OF_MITOTIC_CELL_CYCLE | 0.00000 | 0.00002 | 0.69013 | 0.58588 | 1.19976 | 454 |
| GO_REGULATION_OF_MRNA_CATABOLIC_PROCESS | 0.00016 | 0.00658 | 0.51885 | 0.60727 | 1.22020 | 162 |
| GO_REGULATION_OF_MRNA_METABOLIC_PROCESS | 0.00000 | 0.00001 | 0.71951 | 0.62181 | 1.26113 | 262 |
| GO_REGULATION_OF_MRNA_PROCESSING | 0.00008 | 0.00377 | 0.53843 | 0.64020 | 1.27299 | 113 |
| GO_REGULATION_OF_MRNA_SPLICING_VIA_SPLICEOSOME | 0.00003 | 0.00184 | 0.57561 | 0.66831 | 1.31483 | 82 |
| GO_REGULATION_OF_NUCLEAR_DIVISION | 0.00022 | 0.00808 | 0.51885 | 0.61610 | 1.23214 | 134 |
| GO_REGULATION_OF_NUCLEASE_ACTIVITY | 0.00007 | 0.00345 | 0.53843 | 0.87108 | 1.57485 | 15 |
| GO_REGULATION_OF_NUCLEOCYTOPLASMIC_TRANSPORT | 0.00009 | 0.00413 | 0.53843 | 0.67598 | 1.32729 | 74 |
| GO_REGULATION_OF_PROTEASOMAL_UBIQUITIN_DEPENDENT_PRO | 0.00091 | 0.02273 | 0.47727 | 0.62770 | 1.23832 | 90 |
| GO_REGULATION_OF_PROTEIN_CATABOLIC_PROCESS | 0.00017 | 0.00666 | 0.51885 | 0.57988 | 1.17715 | 272 |

| | | | | | | |
|---|---|---|---|---|---|---|
| GO_REGULATION_OF_PROTEIN_IMPORT | 0.00005 | 0.00280 | 0.55733 | 0.74225 | 1.41428 | 37 |
| GO_REGULATION_OF_PROTEIN_LOCALIZATION_TO_NUCLEUS | 0.00000 | 0.00029 | 0.62726 | 0.68967 | 1.35801 | 84 |
| GO_REGULATION_OF_PROTEIN_MODIFICATION_BY_SMALL_PROTEIN | 0.00002 | 0.00130 | 0.57561 | 0.61876 | 1.24546 | 169 |
| GO_REGULATION_OF_PROTEIN_STABILITY | 0.00013 | 0.00572 | 0.51885 | 0.59522 | 1.20102 | 198 |
| GO_REGULATION_OF_RESPONSE_TO_BIOTIC_STIMULUS | 0.00014 | 0.00601 | 0.51885 | 0.59221 | 1.19588 | 208 |
| GO_REGULATION_OF_RESPONSE_TO_DNA_DAMAGE_STIMULUS | 0.00032 | 0.01089 | 0.49849 | 0.60161 | 1.20882 | 162 |
| GO_REGULATION_OF_RNA_SPLICING | 0.00003 | 0.00184 | 0.57561 | 0.64492 | 1.28327 | 115 |
| GO_REGULATION_OF_SIGNAL_TRANSDUCTION_BY_P53_CLASS_MED | 0.00015 | 0.00616 | 0.51885 | 0.61970 | 1.23932 | 132 |
| GO_REGULATION_OF_SISTER_CHROMATID_SEGREGATION | 0.00008 | 0.00377 | 0.53843 | 0.66565 | 1.30634 | 73 |
| GO_REGULATION_OF_STEM_CELL_DIFFERENTIATION | 0.00079 | 0.02104 | 0.47727 | 0.63622 | 1.25170 | 82 |
| GO_REGULATION_OF_SYMBIOTIC_PROCESS | 0.00194 | 0.03820 | 0.45506 | 0.59393 | 1.18704 | 130 |
| GO_REGULATION_OF_TRANSCRIPTION_INITIATION_FROM_RNA_POL | 0.00276 | 0.04857 | 0.43171 | 0.72550 | 1.35691 | 27 |
| GO_REGULATION_OF_TRANSLATIONAL_INITIATION | 0.00087 | 0.02230 | 0.47727 | 0.66053 | 1.28974 | 64 |
| GO_REGULATION_OF_UBIQUITIN_DEPENDENT_PROTEIN_CATABOLIC | 0.00102 | 0.02453 | 0.45506 | 0.61475 | 1.22133 | 109 |
| GO_REGULATION_OF_VIRAL_LIFE_CYCLE | 0.00124 | 0.02764 | 0.45506 | 0.62699 | 1.23365 | 81 |
| GO_REGULATION_OF_WNT_SIGNALING_PATHWAY | 0.00010 | 0.00461 | 0.53843 | 0.58972 | 1.19259 | 222 |
| GO_REPRESSING_TRANSCRIPTION_FACTOR_BINDING | 0.00097 | 0.02370 | 0.47727 | 0.67492 | 1.30459 | 49 |
| GO_RESPONSE_TO_HEAT | 0.00005 | 0.00295 | 0.55733 | 0.64038 | 1.27165 | 107 |
| GO_RESPONSE_TO_LEUKEMIA_INHIBITORY_FACTOR | 0.00110 | 0.02587 | 0.45506 | 0.63943 | 1.25737 | 78 |
| GO_RESPONSE_TO_TEMPERATURE_STIMULUS | 0.00013 | 0.00566 | 0.51885 | 0.62036 | 1.24064 | 132 |
| GO_RIBONUCLEOPROTEIN_COMPLEX_BINDING | 0.00111 | 0.02619 | 0.45506 | 0.61274 | 1.21792 | 111 |
| GO_RIBONUCLEOPROTEIN_COMPLEX_BIOGENESIS | 0.00000 | 0.00000 | | 0.66879 | 1.36643 | 368 |
| GO_RIBONUCLEOPROTEIN_COMPLEX_SUBUNIT_ORGANIZATION | 0.00000 | 0.00001 | 0.70498 | 0.65045 | 1.30781 | 165 |
| GO_RIBONUCLEOPROTEIN_GRANULE | 0.00110 | 0.02587 | 0.45506 | 0.58760 | 1.18426 | 182 |
| GO_RIBOSOMAL_LARGE_SUBUNIT_ASSEMBLY | 0.00202 | 0.03939 | 0.43171 | 0.74077 | 1.38237 | 26 |
| GO_RIBOSOMAL_LARGE_SUBUNIT_BIOGENESIS | 0.00000 | 0.00028 | 0.62726 | 0.71693 | 1.39877 | 62 |
| GO_RIBOSOMAL_SMALL_SUBUNIT_BIOGENESIS | 0.00088 | 0.02234 | 0.47727 | 0.65523 | 1.27944 | 65 |
| GO_RIBOSOMAL_SUBUNIT | 0.00000 | 0.00016 | 0.64355 | 0.64195 | 1.28792 | 150 |
| GO_RIBOSOME | 0.00000 | 0.00005 | 0.67496 | 0.64115 | 1.29253 | 183 |
| GO_RIBOSOME_ASSEMBLY | 0.00069 | 0.01871 | 0.47727 | 0.67719 | 1.31770 | 55 |
| GO_RIBOSOME_BIOGENESIS | 0.00000 | 0.00000 | | 0.67039 | 1.35922 | 259 |
| GO_RNA_3_END_PROCESSING | 0.00033 | 0.01098 | 0.49849 | 0.62560 | 1.24642 | 120 |
| GO_RNA_CAP_BINDING | 0.00191 | 0.03773 | 0.45506 | 0.80379 | 1.45601 | 16 |
| GO_RNA_CATABOLIC_PROCESS | 0.00000 | 0.00000 | 0.82666 | 0.62456 | 1.27361 | 325 |
| GO_RNA_EXPORT_FROM_NUCLEUS | 0.00000 | 0.00016 | 0.64355 | 0.66074 | 1.31755 | 123 |
| GO_RNA_LOCALIZATION | 0.00000 | 0.00000 | 0.74774 | 0.65320 | 1.31800 | 196 |
| GO_RNA_POLYMERASE_COMPLEX | 0.00119 | 0.02683 | 0.45506 | 0.62661 | 1.23551 | 88 |
| GO_RNA_POLYMERASE_II_SPECIFIC_DNA_BINDING_TRANSCRIPTION | 0.00017 | 0.00662 | 0.51885 | 0.60402 | 1.21666 | 175 |
| GO_RNA_SPLICING | 0.00000 | 0.00000 | | 0.62844 | 1.28255 | 350 |
| GO_RNA_SPLICING_VIA_TRANSESTERIFICATION_REACTIONS | 0.00000 | 0.00000 | 0.80122 | 0.62993 | 1.28014 | 285 |
| GO_RRNA_BINDING | 0.00191 | 0.03773 | 0.45506 | 0.67467 | 1.30585 | 51 |
| GO_RRNA_METABOLIC_PROCESS | 0.00000 | 0.00000 | 0.78819 | 0.65550 | 1.32299 | 199 |
| GO_SIGNAL_SEQUENCE_BINDING | 0.00088 | 0.02234 | 0.47727 | 0.72751 | 1.37709 | 33 |
| GO_SIGNAL_TRANSDUCTION_BY_P53_CLASS_MEDIATOR | 0.00012 | 0.00531 | 0.51885 | 0.59942 | 1.20861 | 191 |
| GO_SINGLE_STRANDED_RNA_BINDING | 0.00143 | 0.03126 | 0.45506 | 0.64665 | 1.26165 | 62 |
| GO_SISTER_CHROMATID_COHESION | 0.00202 | 0.03939 | 0.43171 | 0.67844 | 1.30903 | 46 |
| GO_SISTER_CHROMATID_SEGREGATION | 0.00001 | 0.00058 | 0.61053 | 0.63134 | 1.26803 | 157 |
| GO_SMALL_RIBOSOMAL_SUBUNIT | 0.00016 | 0.00644 | 0.51885 | 0.68358 | 1.33215 | 59 |
| GO_SMALL_SUBUNIT_PROCESSOME | 0.00020 | 0.00737 | 0.51885 | 0.73926 | 1.39345 | 31 |
| GO_SPERM_EGG_RECOGNITION | 0.00032 | 0.01086 | 0.49849 | 0.77913 | 1.45153 | 25 |
| GO_SPINDLE | 0.00035 | 0.01156 | 0.49849 | 0.56973 | 1.15667 | 273 |
| GO_SPINDLE_ASSEMBLY | 0.00075 | 0.01991 | 0.47727 | 0.63730 | 1.25561 | 85 |
| GO_SPINDLE_MIDZONE | 0.00170 | 0.03517 | 0.45506 | 0.72614 | 1.36297 | 28 |
| GO_SPLICEOSOMAL_COMPLEX | 0.00000 | 0.00014 | 0.64355 | 0.64477 | 1.29459 | 155 |
| GO_STEM_CELL_DIFFERENTIATION | 0.00059 | 0.01660 | 0.47727 | 0.60622 | 1.21498 | 141 |
| GO_STEROID_HORMONE_MEDIATED_SIGNALING_PATHWAY | 0.00172 | 0.03529 | 0.45506 | 0.62712 | 1.23381 | 82 |
| GO_STRUCTURAL_CONSTITUENT_OF_RIBOSOME | 0.00000 | 0.00014 | 0.64355 | 0.65432 | 1.30858 | 134 |
| GO_STRUCTURAL_MOLECULE_ACTIVITY | 0.00006 | 0.00316 | 0.55733 | 0.57793 | 1.17779 | 315 |
| GO_TBP_CLASS_PROTEIN_BINDING | 0.00216 | 0.04130 | 0.43171 | 0.73729 | 1.37196 | 24 |
| GO_TRANSCRIPTION_BY_RNA_POLYMERASE_I | 0.00052 | 0.01506 | 0.47727 | 0.67394 | 1.30948 | 54 |
| GO_TRANSCRIPTION_COACTIVATOR_ACTIVITY | 0.00048 | 0.01414 | 0.49849 | 0.58984 | 1.18973 | 187 |
| GO_TRANSCRIPTION_COREGULATOR_ACTIVITY | 0.00010 | 0.00444 | 0.53843 | 0.57553 | 1.17378 | 327 |
| GO_TRANSCRIPTION_COREPRESSOR_ACTIVITY | 0.00061 | 0.01704 | 0.47727 | 0.61225 | 1.21976 | 119 |
| GO_TRANSCRIPTION_FACTOR_BINDING | 0.00000 | 0.00001 | 0.71951 | 0.59934 | 1.22554 | 389 |
| GO_TRANSCRIPTION_INITIATION_FROM_RNA_POLYMERASE_II_PROI | 0.00074 | 0.01976 | 0.47727 | 0.60456 | 1.20830 | 130 |
| GO_TRANSCRIPTION_PREINITIATION_COMPLEX_ASSEMBLY | 0.00065 | 0.01797 | 0.47727 | 0.72296 | 1.36273 | 31 |
| GO_TRANSCRIPTION_REGULATOR_COMPLEX | 0.00039 | 0.01220 | 0.49849 | 0.58013 | 1.17576 | 253 |
| GO_TRANSFERASE_COMPLEX_TRANSFERRING_PHOSPHORUS_CONT/ | 0.00100 | 0.02419 | 0.45506 | 0.58346 | 1.17728 | 196 |
| GO_TRANSLATIONAL_INITIATION | 0.00000 | 0.00000 | | 0.71806 | 1.44212 | 158 |
| GO_TRANSLATION_FACTOR_ACTIVITY_RNA_BINDING | 0.00002 | 0.00135 | 0.57561 | 0.70804 | 1.38252 | 64 |
| GO_TRANSLATION_INITIATION_FACTOR_ACTIVITY | 0.00018 | 0.00684 | 0.51885 | 0.71294 | 1.36995 | 42 |
| GO_TRANSLATION_INITIATION_FACTOR_BINDING | 0.00165 | 0.03465 | 0.45506 | 0.72672 | 1.36406 | 28 |
| GO_TRANSLATION_REGULATOR_ACTIVITY | 0.00000 | 0.00003 | 0.69013 | 0.69140 | 1.37104 | 102 |
| GO_TRANSLATION_REGULATOR_ACTIVITY_NUCLEIC_ACID_BINDING | 0.00000 | 0.00006 | 0.67496 | 0.71107 | 1.39826 | 78 |
| GO_TRNA_TRANSPORT | 0.00187 | 0.03725 | 0.45506 | 0.71854 | 1.35794 | 32 |
| GO_UBIQUITIN_LIKE_PROTEIN_LIGASE_ACTIVITY | 0.00272 | 0.04836 | 0.43171 | 0.57406 | 1.15679 | 181 |

| | | | | | | |
|---|---|---|---|---|---|---|
| GO_UBIQUITIN_LIKE_PROTEIN_LIGASE_BINDING | 0.00276 | 0.04857 | 0.43171 | 0.56849 | 1.14932 | 217 |
| GO_UBIQUITIN_LIKE_PROTEIN_TRANSFERASE_ACTIVITY | 0.00083 | 0.02161 | 0.47727 | 0.56488 | 1.14843 | 292 |
| GO_UNFOLDED_PROTEIN_BINDING | 0.00001 | 0.00109 | 0.59333 | 0.67583 | 1.33022 | 83 |
| GO_VESICLE_LUMEN | 0.00002 | 0.00136 | 0.57561 | 0.61830 | 1.24401 | 167 |
| GO_VIRAL_GENE_EXPRESSION | 0.00000 | 0.00000 | 0.81404 | 0.68352 | 1.37309 | 161 |
| GO_VIRAL_GENOME_REPLICATION | 0.00228 | 0.04243 | 0.43171 | 0.62761 | 1.23169 | 73 |
| GO_VIRAL_LIFE_CYCLE | 0.00000 | 0.00012 | 0.64355 | 0.62358 | 1.25908 | 205 |
| HP_ABNORMALITY_OF_UPPER_LIP | 0.00179 | 0.03591 | 0.45506 | 0.54625 | 1.11777 | 430 |
| HP_ABNORMALITY_OF_UPPER_LIP_VERMILLION | 0.00167 | 0.03478 | 0.45506 | 0.57433 | 1.15940 | 204 |
| HP_ACUTE_LEUKEMIA | 0.00174 | 0.03554 | 0.45506 | 0.63830 | 1.25223 | 72 |
| HP_DELAYED_PUBERTY | 0.00113 | 0.02620 | 0.45506 | 0.61432 | 1.21960 | 105 |
| HP_DEVIATION_OF_THE_THUMB | 0.00209 | 0.04028 | 0.43171 | 0.73968 | 1.37803 | 25 |
| HP_DOWNSLANTED_PALPEBRAL_FISSURES | 0.00009 | 0.00420 | 0.53843 | 0.58938 | 1.19422 | 251 |
| HP_FACIAL_HYPERTRICHOSIS | 0.00178 | 0.03579 | 0.45506 | 0.62417 | 1.22903 | 84 |
| HP_GONOSOMAL_INHERITANCE | 0.00016 | 0.00644 | 0.51885 | 0.62177 | 1.24141 | 127 |
| HP_LONG_FINGERS | 0.00058 | 0.01651 | 0.47727 | 0.61365 | 1.22374 | 124 |
| HP_SANDAL_GAP | 0.00054 | 0.01570 | 0.47727 | 0.68304 | 1.32028 | 49 |
| HP_SEVERE_POSTNATAL_GROWTH_RETARDATION | 0.00213 | 0.04072 | 0.43171 | 0.76427 | 1.40026 | 19 |
| HP_SHORT_NOSE | 0.00179 | 0.03591 | 0.45506 | 0.57761 | 1.16300 | 171 |
| HP_TAPERED_FINGER | 0.00083 | 0.02161 | 0.47727 | 0.67448 | 1.31244 | 55 |
| HP_TETRAPLEGIA | 0.00220 | 0.04145 | 0.43171 | 0.78213 | 1.42312 | 17 |
| HP_THICK_LOWER_LIP_VERMILION | 0.00279 | 0.04907 | 0.43171 | 0.60952 | 1.20637 | 97 |
| HP_THICK_VERMILION_BORDER | 0.00011 | 0.00467 | 0.53843 | 0.61323 | 1.23030 | 150 |
| HP_X_LINKED_RECESSIVE_INHERITANCE | 0.00086 | 0.02213 | 0.47727 | 0.62269 | 1.23244 | 97 |
| **Cluster 16** | | | | | | |
| pathway | pval | padj | log2err | ES | NES | size |
| GO_ACTIN_FILAMENT_BINDING | 0.00124 | 0.02400 | 0.45506 | 0.57135 | 1.36330 | 87 |
| GO_ACTIVATION_OF_IMMUNE_RESPONSE | 0.00000 | 0.00000 | 0.77494 | 0.60795 | 1.48683 | 168 |
| GO_ACTIVATION_OF_INNATE_IMMUNE_RESPONSE | 0.00000 | 0.00002 | 0.67496 | 0.66498 | 1.57655 | 76 |
| GO_AEROBIC_RESPIRATION | 0.00002 | 0.00078 | 0.57561 | 0.64612 | 1.51574 | 62 |
| GO_AMINE_METABOLIC_PROCESS | 0.00000 | 0.00000 | 0.71951 | 0.69867 | 1.64463 | 67 |
| GO_ANAPHASE_PROMOTING_COMPLEX_DEPENDENT_CATABOLIC_P | 0.00000 | 0.00001 | 0.70498 | 0.68730 | 1.62370 | 73 |
| GO_ANATOMICAL_STRUCTURE_HOMEOSTASIS | 0.00176 | 0.03108 | 0.45506 | 0.50065 | 1.22792 | 193 |
| GO_ANCHORING_JUNCTION | 0.00000 | 0.00000 | | 0.59618 | 1.48001 | 385 |
| GO_ANIMAL_ORGAN_REGENERATION | 0.00309 | 0.04773 | 0.43171 | 0.66325 | 1.48012 | 27 |
| GO_ANTIGEN_PROCESSING_AND_PRESENTATION | 0.00000 | 0.00009 | 0.64355 | 0.59832 | 1.44515 | 126 |
| GO_ANTIGEN_PROCESSING_AND_PRESENTATION_OF_EXOGENOUS_ | 0.00000 | 0.00001 | 0.69013 | 0.72133 | 1.68108 | 53 |
| GO_ANTIGEN_PROCESSING_AND_PRESENTATION_OF_PEPTIDE_ANTI | 0.00000 | 0.00005 | 0.64355 | 0.61727 | 1.48528 | 114 |
| GO_ANTIGEN_PROCESSING_AND_PRESENTATION_OF_PEPTIDE_ANTI | 0.00000 | 0.00000 | 0.73376 | 0.70884 | 1.66611 | 65 |
| GO_ANTIGEN_RECEPTOR_MEDIATED_SIGNALING_PATHWAY | 0.00000 | 0.00007 | 0.64355 | 0.62824 | 1.50481 | 98 |
| GO_ANTIOXIDANT_ACTIVITY | 0.00030 | 0.00708 | 0.49849 | 0.69783 | 1.57746 | 32 |
| GO_APOPTOTIC_SIGNALING_PATHWAY | 0.00030 | 0.00708 | 0.49849 | 0.49910 | 1.23372 | 291 |
| GO_ATP_METABOLIC_PROCESS | 0.00004 | 0.00130 | 0.55733 | 0.54276 | 1.33018 | 179 |
| GO_BLOOD_MICROPARTICLE | 0.00065 | 0.01357 | 0.47727 | 0.75364 | 1.63776 | 20 |
| GO_CADHERIN_BINDING | 0.00000 | 0.00000 | | 0.60250 | 1.48327 | 223 |
| GO_CANONICAL_WNT_SIGNALING_PATHWAY | 0.00010 | 0.00268 | 0.53843 | 0.54913 | 1.33746 | 153 |
| GO_CATALYTIC_STEP_2_SPLICEOSOME | 0.00124 | 0.02400 | 0.45506 | 0.57583 | 1.36702 | 80 |
| GO_CELLULAR_AMINO_ACID_METABOLIC_PROCESS | 0.00002 | 0.00071 | 0.57561 | 0.54963 | 1.34614 | 173 |
| GO_CELLULAR_KETONE_METABOLIC_PROCESS | 0.00001 | 0.00036 | 0.59333 | 0.58232 | 1.40730 | 127 |
| GO_CELLULAR_NITROGEN_COMPOUND_CATABOLIC_PROCESS | 0.00000 | 0.00000 | | 0.67646 | 1.67877 | 380 |
| GO_CELLULAR_OXIDANT_DETOXIFICATION | 0.00139 | 0.02623 | 0.45506 | 0.64217 | 1.46814 | 39 |
| GO_CELLULAR_PROTEIN_CATABOLIC_PROCESS | 0.00009 | 0.00242 | 0.53843 | 0.48248 | 1.20083 | 469 |
| GO_CELLULAR_RESPIRATION | 0.00011 | 0.00298 | 0.53843 | 0.56290 | 1.36242 | 132 |
| GO_CELLULAR_RESPONSE_TO_OXYGEN_LEVELS | 0.00000 | 0.00000 | 0.71951 | 0.61654 | 1.49119 | 136 |
| GO_CELLULAR_RESPONSE_TO_TOXIC_SUBSTANCE | 0.00272 | 0.04294 | 0.43171 | 0.60835 | 1.40861 | 48 |
| GO_CELL_ACTIVATION | 0.00000 | 0.00014 | 0.62726 | 0.49962 | 1.24372 | 472 |
| GO_CELL_ACTIVATION_INVOLVED_IN_IMMUNE_RESPONSE | 0.00000 | 0.00010 | 0.62726 | 0.53411 | 1.31822 | 269 |
| GO_CELL_ADHESION_MOLECULE_BINDING | 0.00000 | 0.00000 | | 0.59497 | 1.46856 | 276 |
| GO_CELL_CELL_RECOGNITION | 0.00000 | 0.00007 | 0.64355 | 0.84646 | 1.81916 | 18 |
| GO_CELL_CELL_SIGNALING_BY_WNT | 0.00009 | 0.00255 | 0.53843 | 0.51834 | 1.27613 | 235 |
| GO_CELL_CYCLE_G2_M_PHASE_TRANSITION | 0.00090 | 0.01834 | 0.47727 | 0.51606 | 1.26487 | 181 |
| GO_CELL_CYCLE_PHASE_TRANSITION | 0.00150 | 0.02790 | 0.45506 | 0.47271 | 1.17349 | 385 |
| GO_CELL_DEATH_IN_RESPONSE_TO_OXIDATIVE_STRESS | 0.00272 | 0.04294 | 0.43171 | 0.61801 | 1.41748 | 41 |
| GO_CELL_RECOGNITION | 0.00000 | 0.00011 | 0.62726 | 0.76192 | 1.72504 | 33 |
| GO_CELL_REDOX_HOMEOSTASIS | 0.00145 | 0.02707 | 0.45506 | 0.63752 | 1.45312 | 36 |
| GO_CELL_SUBSTRATE_JUNCTION | 0.00000 | 0.00000 | | 0.67919 | 1.67318 | 245 |
| GO_CELL_SURFACE_RECEPTOR_SIGNALING_PATHWAY_INVOLVED_IN | 0.00010 | 0.00266 | 0.53843 | 0.51294 | 1.26448 | 251 |
| GO_CHAPERONE_COMPLEX | 0.00001 | 0.00046 | 0.59333 | 0.82332 | 1.77877 | 19 |
| GO_COTRANSLATIONAL_PROTEIN_TARGETING_TO_MEMBRANE | 0.00000 | 0.00000 | | 0.86697 | 2.07381 | 90 |
| GO_CYTOKINE_MEDIATED_SIGNALING_PATHWAY | 0.00000 | 0.00000 | 0.76146 | 0.56811 | 1.40052 | 257 |
| GO_CYTOPLASMIC_TRANSLATION | 0.00000 | 0.00000 | | 0.81303 | 1.93791 | 85 |
| GO_CYTOPLASMIC_TRANSLATIONAL_INITIATION | 0.00000 | 0.00003 | 0.65944 | 0.78948 | 1.77517 | 30 |
| GO_CYTOSOLIC_LARGE_RIBOSOMAL_SUBUNIT | 0.00000 | 0.00000 | | 0.88134 | 2.05398 | 53 |
| GO_CYTOSOLIC_RIBOSOME | 0.00000 | 0.00000 | | 0.86720 | 2.07617 | 94 |
| GO_CYTOSOLIC_SMALL_RIBOSOMAL_SUBUNIT | 0.00000 | 0.00000 | | 0.87026 | 1.98961 | 39 |
| GO_DEFENSE_RESPONSE_TO_OTHER_ORGANISM | 0.00060 | 0.01276 | 0.47727 | 0.48923 | 1.21077 | 313 |

| | | | | | | |
|---|---|---|---|---|---|---|
| GO_DETOXIFICATION | 0.00159 | 0.02918 | 0.45506 | 0.61136 | 1.41783 | 50 |
| GO_DNA_BIOSYNTHETIC_PROCESS | 0.00000 | 0.00019 | 0.61053 | 0.59501 | 1.43448 | 120 |
| GO_DNA_POLYMERASE_BINDING | 0.00261 | 0.04198 | 0.43171 | 0.73656 | 1.57563 | 17 |
| GO_ELECTRON_TRANSFER_ACTIVITY | 0.00019 | 0.00480 | 0.51885 | 0.60583 | 1.42761 | 70 |
| GO_ELECTRON_TRANSPORT_CHAIN | 0.00135 | 0.02583 | 0.45506 | 0.54290 | 1.30918 | 121 |
| GO_ENDOPEPTIDASE_COMPLEX | 0.00000 | 0.00000 | 0.70498 | 0.71706 | 1.67807 | 57 |
| GO_ENERGY_DERIVATION_BY_OXIDATION_OF_ORGANIC_COMPOUN | 0.00007 | 0.00203 | 0.53843 | 0.54153 | 1.32333 | 164 |
| GO_EPITHELIUM_DEVELOPMENT | 0.00038 | 0.00846 | 0.49849 | 0.47798 | 1.18950 | 444 |
| GO_ESTABLISHMENT_OF_PROTEIN_LOCALIZATION_TO_CHROMOSOM | 0.00023 | 0.00558 | 0.51885 | 0.74745 | 1.64160 | 22 |
| GO_ESTABLISHMENT_OF_PROTEIN_LOCALIZATION_TO_ENDOPLASM | 0.00000 | 0.00000 | | 0.85855 | 2.05826 | 99 |
| GO_ESTABLISHMENT_OF_PROTEIN_LOCALIZATION_TO_MEMBRANE | 0.00000 | 0.00000 | | 0.73426 | 1.80572 | 216 |
| GO_ESTABLISHMENT_OF_PROTEIN_LOCALIZATION_TO_ORGANELLE | 0.00000 | 0.00000 | | 0.63886 | 1.58536 | 375 |
| GO_ESTABLISHMENT_OF_PROTEIN_LOCALIZATION_TO_TELOMERE | 0.00002 | 0.00053 | 0.57561 | 0.84251 | 1.79556 | 16 |
| GO_ESTABLISHMENT_OF_TISSUE_POLARITY | 0.00000 | 0.00001 | 0.69013 | 0.68205 | 1.61131 | 73 |
| GO_EUKARYOTIC_TRANSLATION_INITIATION_FACTOR_3_COMPLEX | 0.00002 | 0.00069 | 0.57561 | 0.84759 | 1.79551 | 15 |
| GO_EXOCYTOSIS | 0.00000 | 0.00009 | 0.64355 | 0.51841 | 1.28544 | 358 |
| GO_FC_EPSILON_RECEPTOR_SIGNALING_PATHWAY | 0.00001 | 0.00032 | 0.59333 | 0.64353 | 1.52029 | 73 |
| GO_FC_RECEPTOR_SIGNALING_PATHWAY | 0.00000 | 0.00000 | 0.74774 | 0.64431 | 1.54968 | 112 |
| GO_FERTILIZATION | 0.00194 | 0.03363 | 0.45506 | 0.62980 | 1.44761 | 43 |
| GO_FICOLIN_1_RICH_GRANULE | 0.00000 | 0.00000 | 0.71951 | 0.67349 | 1.60318 | 83 |
| GO_GENERATION_OF_PRECURSOR_METABOLITES_AND_ENERGY | 0.00000 | 0.00003 | 0.65944 | 0.53885 | 1.33093 | 284 |
| GO_GLAND_DEVELOPMENT | 0.00113 | 0.02229 | 0.45506 | 0.51964 | 1.27070 | 169 |
| GO_GLYCOLYTIC_PROCESS_THROUGH_FRUCTOSE_6_PHOSPHATE | 0.00139 | 0.02623 | 0.45506 | 0.76270 | 1.62547 | 16 |
| GO_HEMATOPOIETIC_PROGENITOR_CELL_DIFFERENTIATION | 0.00000 | 0.00002 | 0.67496 | 0.64094 | 1.53316 | 91 |
| GO_HEMATOPOIETIC_STEM_CELL_DIFFERENTIATION | 0.00000 | 0.00001 | 0.70498 | 0.69612 | 1.63831 | 66 |
| GO_IMMUNE_EFFECTOR_PROCESS | 0.00006 | 0.00184 | 0.53843 | 0.49253 | 1.22449 | 421 |
| GO_IMMUNE_RESPONSE_REGULATING_SIGNALING_PATHWAY | 0.00000 | 0.00000 | 0.74774 | 0.60483 | 1.47367 | 157 |
| GO_INNATE_IMMUNE_RESPONSE | 0.00000 | 0.00002 | 0.67496 | 0.54947 | 1.35448 | 255 |
| GO_INNATE_IMMUNE_RESPONSE_ACTIVATING_SIGNAL_TRANSDUCT | 0.00000 | 0.00001 | 0.67496 | 0.69618 | 1.63276 | 61 |
| GO_INNER_MITOCHONDRIAL_MEMBRANE_PROTEIN_COMPLEX | 0.00224 | 0.03746 | 0.43171 | 0.55885 | 1.33136 | 84 |
| GO_INTERACTION_WITH_SYMBIONT | 0.00051 | 0.01100 | 0.47727 | 0.66829 | 1.52291 | 37 |
| GO_INTERLEUKIN_1_MEDIATED_SIGNALING_PATHWAY | 0.00000 | 0.00000 | 0.77494 | 0.72254 | 1.69832 | 65 |
| GO_INTRINSIC_APOPTOTIC_SIGNALING_PATHWAY | 0.00011 | 0.00294 | 0.53843 | 0.54854 | 1.33603 | 153 |
| GO_LARGE_RIBOSOMAL_SUBUNIT | 0.00000 | 0.00000 | | 0.73821 | 1.77429 | 108 |
| GO_LEUKOCYTE_MEDIATED_IMMUNITY | 0.00000 | 0.00018 | 0.61053 | 0.52898 | 1.30635 | 282 |
| GO_MATURATION_OF_LSU_RRNA | 0.00028 | 0.00681 | 0.49849 | 0.72359 | 1.61117 | 26 |
| GO_MEMBRANE_PROTEIN_COMPLEX | 0.00108 | 0.02146 | 0.45506 | 0.46846 | 1.16592 | 469 |
| GO_MIDBRAIN_DEVELOPMENT | 0.00283 | 0.04427 | 0.43171 | 0.63965 | 1.45842 | 38 |
| GO_MITOCHONDRIAL_MATRIX | 0.00065 | 0.01370 | 0.47727 | 0.48897 | 1.20964 | 300 |
| GO_MITOCHONDRIAL_PROTEIN_COMPLEX | 0.00037 | 0.00833 | 0.49849 | 0.52155 | 1.27997 | 199 |
| GO_MODIFICATION_DEPENDENT_MACROMOLECULE_CATABOLIC_PR | 0.00029 | 0.00699 | 0.49849 | 0.48109 | 1.19474 | 401 |
| GO_MONOSACCHARIDE_BIOSYNTHETIC_PROCESS | 0.00102 | 0.02052 | 0.45506 | 0.61403 | 1.43101 | 53 |
| GO_MORPHOGENESIS_OF_AN_EPITHELIUM | 0.00018 | 0.00463 | 0.51885 | 0.51710 | 1.27315 | 237 |
| GO_MORPHOGENESIS_OF_A_POLARIZED_EPITHELIUM | 0.00000 | 0.00000 | 0.77494 | 0.68684 | 1.63713 | 85 |
| GO_MRNA_3_UTR_BINDING | 0.00009 | 0.00252 | 0.53843 | 0.63250 | 1.48287 | 60 |
| GO_MRNA_5_UTR_BINDING | 0.00038 | 0.00846 | 0.49849 | 0.73498 | 1.61423 | 22 |
| GO_MRNA_BINDING | 0.00000 | 0.00000 | 0.73376 | 0.57923 | 1.42306 | 209 |
| GO_MUSCLE_CONTRACTION | 0.00183 | 0.03216 | 0.45506 | 0.55422 | 1.32465 | 89 |
| GO_MYD88_INDEPENDENT_TOLL_LIKE_RECEPTOR_SIGNALING_PATH | 0.00163 | 0.02942 | 0.45506 | 0.77005 | 1.63125 | 15 |
| GO_MYELOID_LEUKOCYTE_ACTIVATION | 0.00000 | 0.00019 | 0.61053 | 0.53211 | 1.31172 | 254 |
| GO_MYELOID_LEUKOCYTE_MEDIATED_IMMUNITY | 0.00000 | 0.00004 | 0.65944 | 0.55166 | 1.35783 | 234 |
| GO_NADH_METABOLIC_PROCESS | 0.00035 | 0.00803 | 0.49849 | 0.73787 | 1.62057 | 22 |
| GO_NAD_BINDING | 0.00092 | 0.01864 | 0.47727 | 0.64815 | 1.47735 | 36 |
| GO_NAD_METABOLIC_PROCESS | 0.00165 | 0.02944 | 0.45506 | 0.69120 | 1.53185 | 24 |
| GO_NCRNA_PROCESSING | 0.00163 | 0.02942 | 0.45506 | 0.48448 | 1.19701 | 285 |
| GO_NEGATIVE_REGULATION_OF_CANONICAL_WNT_SIGNALING_PAT | 0.00000 | 0.00013 | 0.62726 | 0.62812 | 1.50319 | 93 |
| GO_NEGATIVE_REGULATION_OF_CELL_CYCLE_G2_M_PHASE_TRANS | 0.00001 | 0.00027 | 0.61053 | 0.63555 | 1.50626 | 75 |
| GO_NEGATIVE_REGULATION_OF_CELL_CYCLE_PHASE_TRANSITION | 0.00250 | 0.04058 | 0.43171 | 0.51427 | 1.25211 | 152 |
| GO_NEGATIVE_REGULATION_OF_CELL_CYCLE_PROCESS | 0.00064 | 0.01343 | 0.47727 | 0.50724 | 1.24616 | 208 |
| GO_NEGATIVE_REGULATION_OF_CELL_DEATH | 0.00000 | 0.00010 | 0.62726 | 0.50779 | 1.26263 | 425 |
| GO_NEGATIVE_REGULATION_OF_MITOTIC_CELL_CYCLE | 0.00187 | 0.03257 | 0.45506 | 0.50757 | 1.24488 | 186 |
| GO_NEGATIVE_REGULATION_OF_RESPONSE_TO_OXIDATIVE_STRESS | 0.00108 | 0.02146 | 0.45506 | 0.72834 | 1.59156 | 21 |
| GO_NEGATIVE_REGULATION_OF_TRANSLATIONAL_INITIATION | 0.00067 | 0.01403 | 0.47727 | 0.77750 | 1.65703 | 16 |
| GO_NEGATIVE_REGULATION_OF_WNT_SIGNALING_PATHWAY | 0.00000 | 0.00018 | 0.61053 | 0.61024 | 1.46520 | 103 |
| GO_NEURAL_NUCLEUS_DEVELOPMENT | 0.00018 | 0.00472 | 0.51885 | 0.74946 | 1.65640 | 23 |
| GO_NEURON_DEATH | 0.00316 | 0.04872 | 0.43171 | 0.51413 | 1.24746 | 146 |
| GO_NEURON_TO_NEURON_SYNAPSE | 0.00038 | 0.00853 | 0.49849 | 0.55413 | 1.34093 | 133 |
| GO_NIK_NF_KAPPAB_SIGNALING | 0.00000 | 0.00001 | 0.70498 | 0.66127 | 1.57535 | 84 |
| GO_NON_CANONICAL_WNT_SIGNALING_PATHWAY | 0.00000 | 0.00001 | 0.70498 | 0.66835 | 1.59095 | 83 |
| GO_NUCLEAR_TRANSCRIBED_MRNA_CATABOLIC_PROCESS | 0.00000 | 0.00000 | | 0.77955 | 1.90486 | 166 |
| GO_NUCLEAR_TRANSCRIBED_MRNA_CATABOLIC_PROCESS_NONSEN | 0.00000 | 0.00000 | | 0.85929 | 2.06572 | 110 |
| GO_NUCLEOSIDE_DIPHOSPHATE_METABOLIC_PROCESS | 0.00235 | 0.03893 | 0.43171 | 0.55810 | 1.32957 | 84 |
| GO_NUCLEOTIDE_BINDING_DOMAIN_LEUCINE_RICH_REPEAT_CONT/ | 0.00072 | 0.01486 | 0.47727 | 0.74687 | 1.60513 | 18 |
| GO_NUCLEOTIDE_EXCISION_REPAIR | 0.00150 | 0.02790 | 0.45506 | 0.56461 | 1.34507 | 84 |
| GO_NUCLEOTIDE_PHOSPHORYLATION | 0.00087 | 0.01793 | 0.47727 | 0.57840 | 1.37128 | 76 |
| GO_ORGANELLE_ASSEMBLY | 0.00250 | 0.04058 | 0.43171 | 0.46970 | 1.16769 | 417 |

| | | | | | | |
|---|---|---|---|---|---|---|
| GO_ORGANIC_ACID_METABOLIC_PROCESS | 0.00001 | 0.00026 | 0.61053 | 0.49319 | 1.22906 | 499 |
| GO_ORGANIC_CYCLIC_COMPOUND_CATABOLIC_PROCESS | 0.00000 | 0.00000 | | 0.67131 | 1.66656 | 388 |
| GO_OXIDATION_REDUCTION_PROCESS | 0.00010 | 0.00265 | 0.53843 | 0.48262 | 1.20130 | 467 |
| GO_OXIDATIVE_PHOSPHORYLATION | 0.00097 | 0.01949 | 0.47727 | 0.56320 | 1.34718 | 90 |
| GO_OXIDOREDUCTASE_ACTIVITY | 0.00028 | 0.00681 | 0.49849 | 0.49300 | 1.22132 | 327 |
| GO_OXIDOREDUCTASE_COMPLEX | 0.00130 | 0.02497 | 0.45506 | 0.56803 | 1.34824 | 79 |
| GO_PEPTIDASE_COMPLEX | 0.00000 | 0.00012 | 0.62726 | 0.65740 | 1.55307 | 73 |
| GO_PEPTIDE_BIOSYNTHETIC_PROCESS | 0.00000 | 0.00000 | | 0.63964 | 1.59275 | 476 |
| GO_PIGMENT_GRANULE | 0.00000 | 0.00014 | 0.62726 | 0.66110 | 1.55619 | 69 |
| GO_POLYSOMAL_RIBOSOME | 0.00000 | 0.00000 | 0.82666 | 0.85716 | 1.92734 | 30 |
| GO_POLYSOME | 0.00000 | 0.00000 | | 0.80041 | 1.87311 | 57 |
| GO_POSITIVE_REGULATION_OF_APOPTOTIC_SIGNALING_PATHWAY | 0.00046 | 0.01009 | 0.49849 | 0.58968 | 1.39991 | 80 |
| GO_POSITIVE_REGULATION_OF_BINDING | 0.00139 | 0.02623 | 0.45506 | 0.57450 | 1.36386 | 80 |
| GO_POSITIVE_REGULATION_OF_CANONICAL_WNT_SIGNALING_PATH | 0.00001 | 0.00022 | 0.61053 | 0.63130 | 1.49841 | 79 |
| GO_POSITIVE_REGULATION_OF_CELLULAR_AMIDE_METABOLIC_PRC | 0.00037 | 0.00833 | 0.49849 | 0.57526 | 1.37756 | 96 |
| GO_POSITIVE_REGULATION_OF_CELLULAR_PROTEIN_LOCALIZATION | 0.00213 | 0.03584 | 0.43171 | 0.50228 | 1.23263 | 192 |
| GO_POSITIVE_REGULATION_OF_CELL_DEATH | 0.00004 | 0.00126 | 0.55733 | 0.51225 | 1.26555 | 288 |
| GO_POSITIVE_REGULATION_OF_DEFENSE_RESPONSE | 0.00002 | 0.00071 | 0.57561 | 0.57790 | 1.39820 | 129 |
| GO_POSITIVE_REGULATION_OF_DNA_BIOSYNTHETIC_PROCESS | 0.00004 | 0.00112 | 0.55733 | 0.66398 | 1.53988 | 50 |
| GO_POSITIVE_REGULATION_OF_DNA_METABOLIC_PROCESS | 0.00268 | 0.04276 | 0.43171 | 0.54425 | 1.30957 | 114 |
| GO_POSITIVE_REGULATION_OF_ESTABLISHMENT_OF_PROTEIN_LOC/ | 0.00048 | 0.01054 | 0.49849 | 0.52108 | 1.27705 | 179 |
| GO_POSITIVE_REGULATION_OF_IMMUNE_RESPONSE | 0.00000 | 0.00000 | 0.71951 | 0.57231 | 1.40746 | 216 |
| GO_POSITIVE_REGULATION_OF_IMMUNE_SYSTEM_PROCESS | 0.00021 | 0.00530 | 0.51885 | 0.50301 | 1.24431 | 298 |
| GO_POSITIVE_REGULATION_OF_INNATE_IMMUNE_RESPONSE | 0.00000 | 0.00004 | 0.65944 | 0.63304 | 1.51674 | 97 |
| GO_POSITIVE_REGULATION_OF_INTRINSIC_APOPTOTIC_SIGNALING_ | 0.00052 | 0.01115 | 0.47727 | 0.68028 | 1.53360 | 31 |
| GO_POSITIVE_REGULATION_OF_NEURON_APOPTOTIC_PROCESS | 0.00235 | 0.03893 | 0.43171 | 0.66961 | 1.49797 | 28 |
| GO_POSITIVE_REGULATION_OF_NEURON_DEATH | 0.00216 | 0.03635 | 0.43171 | 0.61367 | 1.41621 | 46 |
| GO_POSITIVE_REGULATION_OF_PROTEIN_CONTAINING_COMPLEX_[ | 0.00168 | 0.02988 | 0.45506 | 0.75012 | 1.60463 | 17 |
| GO_POSITIVE_REGULATION_OF_PROTEIN_LOCALIZATION_TO_NUCLE | 0.00040 | 0.00900 | 0.49849 | 0.64285 | 1.49569 | 52 |
| GO_POSITIVE_REGULATION_OF_RESPONSE_TO_BIOTIC_STIMULUS | 0.00000 | 0.00012 | 0.62726 | 0.60762 | 1.46041 | 108 |
| GO_POSITIVE_REGULATION_OF_RESPONSE_TO_EXTERNAL_STIMULU | 0.00324 | 0.04971 | 0.43171 | 0.51477 | 1.25753 | 162 |
| GO_POSITIVE_REGULATION_OF_TELOMERASE_ACTIVITY | 0.00265 | 0.04244 | 0.43171 | 0.66796 | 1.49428 | 28 |
| GO_POSITIVE_REGULATION_OF_TELOMERE_MAINTENANCE | 0.00001 | 0.00023 | 0.61053 | 0.71011 | 1.62872 | 41 |
| GO_POSITIVE_REGULATION_OF_TELOMERE_MAINTENANCE_VIA_TE | 0.00013 | 0.00342 | 0.51885 | 0.72609 | 1.63263 | 30 |
| GO_POSITIVE_REGULATION_OF_TRANSFERASE_ACTIVITY | 0.00253 | 0.04092 | 0.43171 | 0.48185 | 1.19083 | 290 |
| GO_POSITIVE_REGULATION_OF_TRANSLATION | 0.00022 | 0.00538 | 0.51885 | 0.58953 | 1.40445 | 84 |
| GO_POSITIVE_REGULATION_OF_WNT_SIGNALING_PATHWAY | 0.00001 | 0.00050 | 0.59333 | 0.60732 | 1.45327 | 92 |
| GO_POSTREPLICATION_REPAIR | 0.00110 | 0.02174 | 0.45506 | 0.65340 | 1.48897 | 37 |
| GO_POSTSYNAPSE | 0.00010 | 0.00265 | 0.53843 | 0.52156 | 1.28302 | 218 |
| GO_POSTTRANSCRIPTIONAL_REGULATION_OF_GENE_EXPRESSION | 0.00000 | 0.00000 | | 0.58480 | 1.45311 | 407 |
| GO_POST_TRANSLATIONAL_PROTEIN_MODIFICATION | 0.00001 | 0.00027 | 0.61053 | 0.55184 | 1.35425 | 192 |
| GO_PROTEASOMAL_PROTEIN_CATABOLIC_PROCESS | 0.00113 | 0.02229 | 0.45506 | 0.48878 | 1.20997 | 314 |
| GO_PROTEASOMAL_UBIQUITIN_INDEPENDENT_PROTEIN_CATABOLI( | 0.00007 | 0.00207 | 0.53843 | 0.81257 | 1.73823 | 17 |
| GO_PROTEASOME_ACCESSORY_COMPLEX | 0.00002 | 0.00074 | 0.57561 | 0.77061 | 1.70784 | 24 |
| GO_PROTEASOME_CORE_COMPLEX | 0.00001 | 0.00048 | 0.59333 | 0.85370 | 1.80845 | 15 |
| GO_PROTEIN_DNA_COMPLEX_SUBUNIT_ORGANIZATION | 0.00119 | 0.02311 | 0.45506 | 0.53055 | 1.28693 | 144 |
| GO_PROTEIN_DOMAIN_SPECIFIC_BINDING | 0.00165 | 0.02944 | 0.45506 | 0.47793 | 1.18529 | 342 |
| GO_PROTEIN_FOLDING | 0.00000 | 0.00002 | 0.67496 | 0.60130 | 1.45631 | 139 |
| GO_PROTEIN_FOLDING_CHAPERONE | 0.00145 | 0.02707 | 0.45506 | 0.76184 | 1.62363 | 16 |
| GO_PROTEIN_LOCALIZATION_TO_CHROMOSOME_TELOMERIC_REGI( | 0.00003 | 0.00092 | 0.57561 | 0.77204 | 1.71783 | 25 |
| GO_PROTEIN_LOCALIZATION_TO_ENDOPLASMIC_RETICULUM | 0.00000 | 0.00000 | | 0.83319 | 2.00562 | 117 |
| GO_PROTEIN_LOCALIZATION_TO_MEMBRANE | 0.00000 | 0.00000 | | 0.64333 | 1.59490 | 337 |
| GO_PROTEIN_LOCALIZATION_TO_NUCLEUS | 0.00004 | 0.00115 | 0.55733 | 0.55369 | 1.35087 | 160 |
| GO_PROTEIN_MODIFICATION_BY_SMALL_PROTEIN_REMOVAL | 0.00018 | 0.00476 | 0.51885 | 0.52832 | 1.29544 | 183 |
| GO_PROTEIN_STABILIZATION | 0.00003 | 0.00082 | 0.57561 | 0.58215 | 1.39958 | 109 |
| GO_PROTEIN_TARGETING | 0.00000 | 0.00000 | | 0.67810 | 1.67551 | 289 |
| GO_PROTEIN_TARGETING_TO_MEMBRANE | 0.00000 | 0.00000 | | 0.81324 | 1.96960 | 139 |
| GO_REGULATION_OF_ANIMAL_ORGAN_MORPHOGENESIS | 0.00000 | 0.00003 | 0.65944 | 0.62315 | 1.49643 | 104 |
| GO_REGULATION_OF_APOPTOTIC_SIGNALING_PATHWAY | 0.00030 | 0.00707 | 0.49849 | 0.52435 | 1.28654 | 198 |
| GO_REGULATION_OF_BINDING | 0.00050 | 0.01085 | 0.47727 | 0.53254 | 1.29792 | 158 |
| GO_REGULATION_OF_CELLULAR_AMIDE_METABOLIC_PROCESS | 0.00000 | 0.00004 | 0.65944 | 0.53813 | 1.32778 | 268 |
| GO_REGULATION_OF_CELLULAR_AMINE_METABOLIC_PROCESS | 0.00000 | 0.00000 | 0.73376 | 0.73639 | 1.71617 | 53 |
| GO_REGULATION_OF_CELLULAR_AMINO_ACID_METABOLIC_PROCES | 0.00000 | 0.00000 | 0.74774 | 0.75257 | 1.74391 | 49 |
| GO_REGULATION_OF_CELLULAR_KETONE_METABOLIC_PROCESS | 0.00000 | 0.00013 | 0.62726 | 0.62115 | 1.48824 | 97 |
| GO_REGULATION_OF_CELL_CYCLE_G2_M_PHASE_TRANSITION | 0.00008 | 0.00233 | 0.53843 | 0.55379 | 1.34367 | 146 |
| GO_REGULATION_OF_CYTOPLASMIC_TRANSLATION | 0.00023 | 0.00564 | 0.51885 | 0.76918 | 1.67154 | 20 |
| GO_REGULATION_OF_DNA_BIOSYNTHETIC_PROCESS | 0.00017 | 0.00438 | 0.51885 | 0.60761 | 1.43181 | 70 |
| GO_REGULATION_OF_DNA_TEMPLATED_TRANSCRIPTION_IN_RESPO | 0.00000 | 0.00000 | 0.74774 | 0.67570 | 1.60730 | 82 |
| GO_REGULATION_OF_HEMATOPOIETIC_PROGENITOR_CELL_DIFFERE | 0.00000 | 0.00000 | 0.70498 | 0.71014 | 1.66755 | 64 |
| GO_REGULATION_OF_HEMATOPOIETIC_STEM_CELL_DIFFERENTIATIC | 0.00000 | 0.00000 | 0.71951 | 0.71876 | 1.68289 | 58 |
| GO_REGULATION_OF_HEMOPOIESIS | 0.00061 | 0.01290 | 0.47727 | 0.52017 | 1.27445 | 174 |
| GO_REGULATION_OF_IMMUNE_RESPONSE | 0.00000 | 0.00019 | 0.61053 | 0.53035 | 1.30924 | 279 |
| GO_REGULATION_OF_INNATE_IMMUNE_RESPONSE | 0.00000 | 0.00012 | 0.62726 | 0.59666 | 1.44385 | 133 |
| GO_REGULATION_OF_INTRINSIC_APOPTOTIC_SIGNALING_PATHWAY | 0.00005 | 0.00138 | 0.55733 | 0.59657 | 1.42757 | 92 |
| GO_REGULATION_OF_MITOTIC_CELL_CYCLE | 0.00115 | 0.02248 | 0.45506 | 0.47619 | 1.18136 | 369 |
| GO_REGULATION_OF_MORPHOGENESIS_OF_AN_EPITHELIUM | 0.00000 | 0.00001 | 0.70498 | 0.65751 | 1.57151 | 89 |

| | | | | | | |
|---|---|---|---|---|---|---|
| GO_REGULATION_OF_MRNA_CATABOLIC_PROCESS | 0.00000 | 0.00000 | 0.78819 | 0.63252 | 1.53483 | 145 |
| GO_REGULATION_OF_MRNA_METABOLIC_PROCESS | 0.00000 | 0.00000 | 0.78819 | 0.58396 | 1.43775 | 237 |
| GO_REGULATION_OF_OXIDATIVE_STRESS_INDUCED_CELL_DEATH | 0.00026 | 0.00625 | 0.49849 | 0.70096 | 1.58453 | 32 |
| GO_REGULATION_OF_OXIDATIVE_STRESS_INDUCED_INTRINSIC_APO | 0.00213 | 0.03584 | 0.43171 | 0.71064 | 1.55288 | 21 |
| GO_REGULATION_OF_PROTEIN_CATABOLIC_PROCESS | 0.00178 | 0.03130 | 0.45506 | 0.49677 | 1.22280 | 226 |
| GO_REGULATION_OF_PROTEIN_LOCALIZATION_TO_NUCLEUS | 0.00003 | 0.00094 | 0.57561 | 0.61989 | 1.46914 | 75 |
| GO_REGULATION_OF_PROTEIN_STABILITY | 0.00031 | 0.00712 | 0.49849 | 0.53161 | 1.30084 | 171 |
| GO_REGULATION_OF_PROTEOLYSIS | 0.00154 | 0.02827 | 0.45506 | 0.48313 | 1.19599 | 315 |
| GO_REGULATION_OF_RESPONSE_TO_BIOTIC_STIMULUS | 0.00021 | 0.00515 | 0.51885 | 0.53915 | 1.31738 | 163 |
| GO_REGULATION_OF_RESPONSE_TO_OXIDATIVE_STRESS | 0.00023 | 0.00572 | 0.51885 | 0.65950 | 1.51264 | 41 |
| GO_REGULATION_OF_SMALL_MOLECULE_METABOLIC_PROCESS | 0.00058 | 0.01248 | 0.47727 | 0.51098 | 1.25814 | 222 |
| GO_REGULATION_OF_STEM_CELL_DIFFERENTIATION | 0.00000 | 0.00001 | 0.69013 | 0.66752 | 1.58357 | 78 |
| GO_REGULATION_OF_TELOMERE_MAINTENANCE | 0.00154 | 0.02827 | 0.45506 | 0.58572 | 1.37672 | 65 |
| GO_REGULATION_OF_TELOMERE_MAINTENANCE_VIA_TELOMERE_L | 0.00031 | 0.00712 | 0.49849 | 0.63453 | 1.47157 | 50 |
| GO_REGULATION_OF_TRANSCRIPTION_FROM_RNA_POLYMERASE_II | 0.00000 | 0.00000 | 0.77494 | 0.73609 | 1.72361 | 59 |
| GO_REGULATION_OF_TRANSFERASE_ACTIVITY | 0.00115 | 0.02248 | 0.45506 | 0.47202 | 1.17427 | 438 |
| GO_REGULATION_OF_TRANSLATIONAL_INITIATION | 0.00002 | 0.00061 | 0.57561 | 0.66631 | 1.55930 | 57 |
| GO_REGULATION_OF_VASCULAR_ASSOCIATED_SMOOTH_MUSCLE_C | 0.00161 | 0.02930 | 0.45506 | 0.72622 | 1.56075 | 18 |
| GO_REGULATION_OF_WNT_SIGNALING_PATHWAY | 0.00018 | 0.00463 | 0.51885 | 0.53714 | 1.31303 | 167 |
| GO_RESPIRASOME | 0.00213 | 0.03584 | 0.43171 | 0.57394 | 1.35473 | 72 |
| GO_RESPIRATORY_CHAIN_COMPLEX | 0.00084 | 0.01724 | 0.47727 | 0.60299 | 1.41370 | 60 |
| GO_RESPONSE_TO_BIOTIC_STIMULUS | 0.00000 | 0.00004 | 0.65944 | 0.50907 | 1.26685 | 444 |
| GO_RESPONSE_TO_CYTOKINE | 0.00000 | 0.00000 | | 0.56362 | 1.40226 | 439 |
| GO_RESPONSE_TO_DRUG | 0.00019 | 0.00483 | 0.51885 | 0.53852 | 1.31207 | 154 |
| GO_RESPONSE_TO_HEAT | 0.00089 | 0.01823 | 0.47727 | 0.56360 | 1.35036 | 97 |
| GO_RESPONSE_TO_HORMONE | 0.00202 | 0.03467 | 0.43171 | 0.47108 | 1.16867 | 362 |
| GO_RESPONSE_TO_INORGANIC_SUBSTANCE | 0.00042 | 0.00936 | 0.49849 | 0.51516 | 1.26654 | 214 |
| GO_RESPONSE_TO_INTERFERON_GAMMA | 0.00035 | 0.00810 | 0.49849 | 0.62576 | 1.46167 | 55 |
| GO_RESPONSE_TO_INTERLEUKIN_1 | 0.00000 | 0.00000 | 0.73376 | 0.67080 | 1.60327 | 89 |
| GO_RESPONSE_TO_INTERLEUKIN_12 | 0.00005 | 0.00141 | 0.55733 | 0.71922 | 1.63215 | 34 |
| GO_RESPONSE_TO_INTERLEUKIN_4 | 0.00165 | 0.02944 | 0.45506 | 0.75937 | 1.61837 | 16 |
| GO_RESPONSE_TO_NITROGEN_COMPOUND | 0.00152 | 0.02814 | 0.45506 | 0.46953 | 1.16821 | 441 |
| GO_RESPONSE_TO_ORGANIC_CYCLIC_COMPOUND | 0.00002 | 0.00053 | 0.57561 | 0.50635 | 1.25551 | 345 |
| GO_RESPONSE_TO_ORGANOPHOSPHORUS | 0.00161 | 0.02930 | 0.45506 | 0.65034 | 1.48279 | 38 |
| GO_RESPONSE_TO_OXIDATIVE_STRESS | 0.00253 | 0.04092 | 0.43171 | 0.50030 | 1.22849 | 201 |
| GO_RESPONSE_TO_OXYGEN_LEVELS | 0.00000 | 0.00000 | 0.73376 | 0.58036 | 1.42502 | 202 |
| GO_RESPONSE_TO_TOPOLOGICALLY_INCORRECT_PROTEIN | 0.00309 | 0.04773 | 0.43171 | 0.52471 | 1.26638 | 124 |
| GO_RESPONSE_TO_TOXIC_SUBSTANCE | 0.00191 | 0.03310 | 0.45506 | 0.55368 | 1.32737 | 99 |
| GO_RESPONSE_TO_TUMOR_NECROSIS_FACTOR | 0.00000 | 0.00010 | 0.62726 | 0.59493 | 1.43510 | 122 |
| GO_RIBONUCLEOPROTEIN_COMPLEX_BINDING | 0.00000 | 0.00000 | 0.71951 | 0.64955 | 1.55690 | 100 |
| GO_RIBONUCLEOPROTEIN_COMPLEX_BIOGENESIS | 0.00000 | 0.00000 | | 0.57214 | 1.41915 | 365 |
| GO_RIBONUCLEOPROTEIN_COMPLEX_SUBUNIT_ORGANIZATION | 0.00000 | 0.00000 | | 0.63347 | 1.54791 | 166 |
| GO_RIBONUCLEOPROTEIN_GRANULE | 0.00003 | 0.00089 | 0.57561 | 0.56665 | 1.37413 | 142 |
| GO_RIBOSOMAL_LARGE_SUBUNIT_ASSEMBLY | 0.00001 | 0.00038 | 0.59333 | 0.77116 | 1.71709 | 26 |
| GO_RIBOSOMAL_LARGE_SUBUNIT_BIOGENESIS | 0.00000 | 0.00000 | | 0.75227 | 1.76611 | 63 |
| GO_RIBOSOMAL_SMALL_SUBUNIT_BIOGENESIS | 0.00001 | 0.00047 | 0.59333 | 0.64491 | 1.51585 | 65 |
| GO_RIBOSOMAL_SUBUNIT | 0.00000 | 0.00000 | | 0.74152 | 1.81369 | 170 |
| GO_RIBOSOME | 0.00000 | 0.00000 | | 0.72231 | 1.77364 | 201 |
| GO_RIBOSOME_ASSEMBLY | 0.00000 | 0.00004 | 0.65944 | 0.70205 | 1.64262 | 56 |
| GO_RIBOSOME_BINDING | 0.00004 | 0.00118 | 0.55733 | 0.70336 | 1.60319 | 36 |
| GO_RIBOSOME_BIOGENESIS | 0.00000 | 0.00000 | 0.80122 | 0.57502 | 1.41749 | 254 |
| GO_RNA_CATABOLIC_PROCESS | 0.00000 | 0.00000 | | 0.71973 | 1.78092 | 305 |
| GO_RNA_DEPENDENT_DNA_BIOSYNTHETIC_PROCESS | 0.00003 | 0.00091 | 0.57561 | 0.66015 | 1.54489 | 57 |
| GO_RNA_LOCALIZATION | 0.00072 | 0.01486 | 0.47727 | 0.51538 | 1.26371 | 183 |
| GO_RNA_SPLICING | 0.00133 | 0.02558 | 0.45506 | 0.48405 | 1.20026 | 339 |
| GO_RNA_SPLICING_VIA_TRANSESTERIFICATION_REACTIONS | 0.00060 | 0.01276 | 0.47727 | 0.49587 | 1.22440 | 281 |
| GO_ROUGH_ENDOPLASMIC_RETICULUM | 0.00001 | 0.00020 | 0.61053 | 0.70293 | 1.62218 | 46 |
| GO_RRNA_BINDING | 0.00000 | 0.00000 | 0.78819 | 0.77516 | 1.79487 | 48 |
| GO_RRNA_METABOLIC_PROCESS | 0.00001 | 0.00022 | 0.61053 | 0.55235 | 1.35573 | 191 |
| GO_SCF_DEPENDENT_PROTEASOMAL_UBIQUITIN_DEPENDENT_PRO | 0.00000 | 0.00001 | 0.69013 | 0.68934 | 1.62234 | 66 |
| GO_SECRETORY_GRANULE | 0.00000 | 0.00009 | 0.62726 | 0.52872 | 1.30856 | 312 |
| GO_SECRETORY_VESICLE | 0.00000 | 0.00004 | 0.65944 | 0.52111 | 1.29245 | 361 |
| GO_SIGNAL_TRANSDUCTION_BY_PROTEIN_PHOSPHORYLATION | 0.00002 | 0.00061 | 0.57561 | 0.51272 | 1.27014 | 331 |
| GO_SINGLE_FERTILIZATION | 0.00009 | 0.00243 | 0.53843 | 0.70997 | 1.61117 | 34 |
| GO_SINGLE_STRANDED_RNA_BINDING | 0.00242 | 0.03989 | 0.43171 | 0.60819 | 1.41338 | 51 |
| GO_SMALL_RIBOSOMAL_SUBUNIT | 0.00000 | 0.00000 | | 0.75156 | 1.76653 | 65 |
| GO_SPERM_EGG_RECOGNITION | 0.00000 | 0.00016 | 0.61053 | 0.86000 | 1.83284 | 16 |
| GO_SPLICEOSOMAL_COMPLEX | 0.00019 | 0.00480 | 0.51885 | 0.53671 | 1.30945 | 160 |
| GO_STEM_CELL_DIFFERENTIATION | 0.00000 | 0.00000 | 0.73376 | 0.62463 | 1.50840 | 125 |
| GO_STRUCTURAL_CONSTITUENT_OF_RIBOSOME | 0.00000 | 0.00000 | | 0.77080 | 1.87528 | 150 |
| GO_STRUCTURAL_MOLECULE_ACTIVITY | 0.00000 | 0.00000 | | 0.69807 | 1.72519 | 290 |
| GO_SUBSTANTIA_NIGRA_DEVELOPMENT | 0.00015 | 0.00401 | 0.51885 | 0.76581 | 1.67343 | 21 |
| GO_SYNAPSE | 0.00000 | 0.00001 | 0.69013 | 0.51215 | 1.27459 | 449 |
| GO_TBP_CLASS_PROTEIN_BINDING | 0.00246 | 0.04024 | 0.43171 | 0.71475 | 1.53611 | 18 |
| GO_TELOMERASE_RNA_LOCALIZATION | 0.00009 | 0.00253 | 0.53843 | 0.79180 | 1.70169 | 18 |
| GO_TELOMERE_MAINTENANCE_VIA_TELOMERE_LENGTHENING | 0.00004 | 0.00131 | 0.55733 | 0.63127 | 1.48378 | 65 |

| | | | | | | |
|---|---|---|---|---|---|---|
| GO_TELOMERE_ORGANIZATION | 0.00198 | 0.03415 | 0.43171 | 0.53780 | 1.29325 | 111 |
| GO_THREONINE_TYPE_ENDOPEPTIDASE_ACTIVITY | 0.00000 | 0.00010 | 0.62726 | 0.87691 | 1.85764 | 15 |
| GO_THREONINE_TYPE_PEPTIDASE_ACTIVITY | 0.00000 | 0.00011 | 0.62726 | 0.85885 | 1.83723 | 17 |
| GO_TISSUE_MORPHOGENESIS | 0.00167 | 0.02966 | 0.45506 | 0.48936 | 1.20725 | 264 |
| GO_TOXIN_TRANSPORT | 0.00005 | 0.00147 | 0.55733 | 0.77547 | 1.70316 | 22 |
| GO_TRANSCRIPTION_COUPLED_NUCLEOTIDE_EXCISION_REPAIR | 0.00268 | 0.04276 | 0.43171 | 0.58507 | 1.37386 | 64 |
| GO_TRANSLATIONAL_ELONGATION | 0.00228 | 0.03796 | 0.43171 | 0.54323 | 1.30774 | 116 |
| GO_TRANSLATIONAL_INITIATION | 0.00000 | 0.00000 | | 0.83612 | 2.03890 | 159 |
| GO_TRANSLATION_FACTOR_ACTIVITY_RNA_BINDING | 0.00000 | 0.00000 | 0.78819 | 0.73817 | 1.73338 | 64 |
| GO_TRANSLATION_INITIATION_FACTOR_ACTIVITY | 0.00000 | 0.00002 | 0.67496 | 0.72890 | 1.68179 | 45 |
| GO_TRANSLATION_PREINITIATION_COMPLEX | 0.00001 | 0.00021 | 0.61053 | 0.85581 | 1.82390 | 16 |
| GO_TRANSLATION_REGULATOR_ACTIVITY | 0.00000 | 0.00000 | 0.82666 | 0.68525 | 1.64246 | 100 |
| GO_TRANSLATION_REGULATOR_ACTIVITY_NUCLEIC_ACID_BINDING | 0.00000 | 0.00000 | 0.76146 | 0.69649 | 1.65229 | 78 |
| GO_TRANSMEMBRANE_TRANSPORT | 0.00008 | 0.00225 | 0.53843 | 0.48079 | 1.19789 | 487 |
| GO_TRICARBOXYLIC_ACID_CYCLE | 0.00302 | 0.04687 | 0.43171 | 0.66431 | 1.48613 | 28 |
| GO_TUMOR_NECROSIS_FACTOR_MEDIATED_SIGNALING_PATHWAY | 0.00000 | 0.00000 | 0.71951 | 0.68299 | 1.62132 | 77 |
| GO_T_CELL_RECEPTOR_SIGNALING_PATHWAY | 0.00000 | 0.00003 | 0.65944 | 0.64651 | 1.54364 | 88 |
| GO_UBIQUITIN_LIKE_PROTEIN_LIGASE_BINDING | 0.00000 | 0.00000 | 0.76146 | 0.59793 | 1.46540 | 179 |
| GO_UNFOLDED_PROTEIN_BINDING | 0.00000 | 0.00000 | 0.74774 | 0.70217 | 1.65884 | 73 |
| GO_VACUOLAR_LUMEN | 0.00209 | 0.03582 | 0.43171 | 0.57481 | 1.35656 | 71 |
| GO_VESICLE_COAT | 0.00298 | 0.04644 | 0.43171 | 0.61545 | 1.41161 | 41 |
| GO_VESICLE_LUMEN | 0.00000 | 0.00001 | 0.69013 | 0.60610 | 1.47017 | 144 |
| GO_VIRAL_GENE_EXPRESSION | 0.00000 | 0.00000 | | 0.79049 | 1.93109 | 162 |
| GO_VIRAL_LIFE_CYCLE | 0.00213 | 0.03584 | 0.43171 | 0.50763 | 1.24398 | 178 |
| HP_ABNORMALITY_OF_CARDIOVASCULAR_SYSTEM_ELECTROPHYSIC | 0.00007 | 0.00205 | 0.53843 | 0.53594 | 1.31448 | 186 |
| HP_ABNORMALITY_OF_LOWER_LIP | 0.00023 | 0.00564 | 0.51885 | 0.56181 | 1.35338 | 118 |
| HP_ABNORMALITY_OF_THE_NASAL_DORSUM | 0.00019 | 0.00480 | 0.51885 | 0.57009 | 1.36884 | 106 |
| HP_ABNORMAL_ERYTHROCYTE_MORPHOLOGY | 0.00016 | 0.00416 | 0.51885 | 0.50989 | 1.25822 | 267 |
| HP_ABNORMAL_MEAN_CORPUSCULAR_VOLUME | 0.00187 | 0.03257 | 0.45506 | 0.76634 | 1.62339 | 15 |
| HP_ABNORMAL_RETICULOCYTE_MORPHOLOGY | 0.00213 | 0.03584 | 0.43171 | 0.69394 | 1.53371 | 23 |
| HP_ACUTE_LEUKEMIA | 0.00000 | 0.00000 | 0.74774 | 0.73105 | 1.71167 | 58 |
| HP_ANEMIA_OF_INADEQUATE_PRODUCTION | 0.00000 | 0.00001 | 0.69013 | 0.66693 | 1.58329 | 80 |
| HP_DELAYED_PUBERTY | 0.00001 | 0.00028 | 0.59333 | 0.62728 | 1.49675 | 87 |
| HP_DEPRESSED_NASAL_RIDGE | 0.00001 | 0.00047 | 0.59333 | 0.67356 | 1.57597 | 56 |
| HP_FATIGUE | 0.00027 | 0.00644 | 0.49849 | 0.54863 | 1.32694 | 136 |
| HP_HEADACHE | 0.00001 | 0.00040 | 0.59333 | 0.59659 | 1.43390 | 108 |
| HP_HEMATOLOGICAL_NEOPLASM | 0.00030 | 0.00708 | 0.49849 | 0.55545 | 1.34161 | 126 |
| HP_LETHARGY | 0.00272 | 0.04294 | 0.43171 | 0.55317 | 1.31853 | 85 |
| HP_LEUKEMIA | 0.00000 | 0.00004 | 0.65944 | 0.65770 | 1.56027 | 78 |
| HP_LEUKOENCEPHALOPATHY | 0.00242 | 0.03989 | 0.43171 | 0.53264 | 1.28055 | 109 |
| HP_LIMB_GIRDLE_MUSCLE_ATROPHY | 0.00276 | 0.04325 | 0.43171 | 0.75652 | 1.60261 | 15 |
| HP_MACROCYTIC_ANEMIA | 0.00000 | 0.00000 | 0.74774 | 0.81756 | 1.85532 | 34 |
| HP_MIGRAINE | 0.00000 | 0.00001 | 0.70498 | 0.73026 | 1.69905 | 52 |
| HP_PALLOR | 0.00000 | 0.00018 | 0.61053 | 0.63953 | 1.51621 | 76 |
| HP_SHORT_NOSE | 0.00276 | 0.04325 | 0.43171 | 0.52134 | 1.26458 | 144 |
| HP_THICK_LOWER_LIP_VERMILION | 0.00002 | 0.00053 | 0.57561 | 0.61605 | 1.46540 | 82 |
| HP_THICK_VERMILION_BORDER | 0.00046 | 0.01009 | 0.49849 | 0.54568 | 1.31939 | 128 |
| HP_THROMBOCYTOSIS | 0.00049 | 0.01070 | 0.47727 | 0.75595 | 1.62464 | 18 |
| HP_URINARY_BLADDER_SPHINCTER_DYSFUNCTION | 0.00246 | 0.04024 | 0.43171 | 0.75173 | 1.60209 | 16 |

### Supplementary Table S5

#### Mouse female GSEA of NMF clusters

| **Cluster 1** | | | | | | |
|---|---|---|---|---|---|---|
| pathway | pval | padj | log2err | ES | NES | size |
| GO_ACTIVATION_OF_IMMUNE_RESPONSE | 0.00009 | 0.00323 | 0.53843 | 0.60032 | 1.22317 | 206 |
| GO_ACTIVATION_OF_INNATE_IMMUNE_RESPONSE | 0.00018 | 0.00591 | 0.51885 | 0.66371 | 1.31881 | 81 |
| GO_ALTERNATIVE_MRNA_SPLICING_VIA_SPLICEOSOME | 0.00181 | 0.03845 | 0.45506 | 0.66912 | 1.31366 | 60 |
| GO_AMINE_METABOLIC_PROCESS | 0.00000 | 0.00000 | 0.76146 | 0.76012 | 1.50510 | 75 |
| GO_ANAPHASE_PROMOTING_COMPLEX_DEPENDENT_CATA | 0.00000 | 0.00002 | 0.69013 | 0.73178 | 1.45241 | 77 |
| GO_ANCHORING_JUNCTION | 0.00000 | 0.00017 | 0.62726 | 0.57965 | 1.19128 | 440 |
| GO_ANIMAL_ORGAN_REGENERATION | 0.00004 | 0.00178 | 0.55733 | 0.76226 | 1.45678 | 35 |
| GO_ANTIGEN_PROCESSING_AND_PRESENTATION | 0.00016 | 0.00552 | 0.51885 | 0.61793 | 1.24643 | 139 |
| GO_ANTIGEN_PROCESSING_AND_PRESENTATION_OF_EXOG | 0.00017 | 0.00561 | 0.51885 | 0.69703 | 1.36544 | 56 |
| GO_ANTIGEN_PROCESSING_AND_PRESENTATION_OF_PEPT | 0.00023 | 0.00727 | 0.51885 | 0.62928 | 1.26287 | 120 |
| GO_ANTIGEN_PROCESSING_AND_PRESENTATION_OF_PEPT | 0.00012 | 0.00423 | 0.53843 | 0.67866 | 1.33856 | 68 |
| GO_ANTIGEN_RECEPTOR_MEDIATED_SIGNALING_PATHWAY | 0.00003 | 0.00143 | 0.55733 | 0.63947 | 1.28602 | 129 |
| GO_ATPASE_ACTIVITY | 0.00081 | 0.02083 | 0.47727 | 0.57157 | 1.16746 | 257 |
| GO_AZUROPHIL_GRANULE_LUMEN | 0.00239 | 0.04753 | 0.43171 | 0.65936 | 1.28730 | 53 |
| GO_BINDING_OF_SPERM_TO_ZONA_PELLUCIDA | 0.00062 | 0.01690 | 0.47727 | 0.80422 | 1.48715 | 19 |
| GO_CADHERIN_BINDING | 0.00000 | 0.00001 | 0.70498 | 0.63485 | 1.29581 | 230 |
| GO_CANONICAL_WNT_SIGNALING_PATHWAY | 0.00000 | 0.00002 | 0.69013 | 0.65427 | 1.32690 | 169 |
| GO_CATALYTIC_STEP_2_SPLICEOSOME | 0.00003 | 0.00141 | 0.55733 | 0.68726 | 1.36405 | 77 |
| GO_CELLULAR_AMINO_ACID_METABOLIC_PROCESS | 0.00001 | 0.00045 | 0.61053 | 0.62011 | 1.26177 | 200 |
| GO_CELLULAR_BIOGENIC_AMINE_METABOLIC_PROCESS | 0.00033 | 0.00977 | 0.49849 | 0.79073 | 1.47816 | 23 |
| GO_CELLULAR_KETONE_METABOLIC_PROCESS | 0.00018 | 0.00603 | 0.51885 | 0.62670 | 1.26373 | 137 |
| GO_CELLULAR_NITROGEN_COMPOUND_CATABOLIC_PROCE | 0.00000 | 0.00000 | | 0.69326 | 1.42359 | 413 |
| GO_CELLULAR_RESPONSE_TO_HEAT | 0.00194 | 0.04027 | 0.45506 | 0.62864 | 1.25101 | 86 |
| GO_CELLULAR_RESPONSE_TO_HYDROGEN_PEROXIDE | 0.00220 | 0.04432 | 0.43171 | 0.66089 | 1.29029 | 53 |
| GO_CELLULAR_RESPONSE_TO_OXYGEN_LEVELS | 0.00020 | 0.00632 | 0.51885 | 0.61531 | 1.24453 | 153 |
| GO_CELL_ACTIVATION_INVOLVED_IN_IMMUNE_RESPONSE | 0.00005 | 0.00211 | 0.55733 | 0.57587 | 1.18126 | 340 |
| GO_CELL_ADHESION_MOLECULE_BINDING | 0.00000 | 0.00006 | 0.65944 | 0.60514 | 1.23947 | 303 |
| GO_CELL_BODY | 0.00100 | 0.02443 | 0.45506 | 0.57316 | 1.16944 | 236 |
| GO_CELL_CELL_RECOGNITION | 0.00002 | 0.00093 | 0.57561 | 0.78799 | 1.49975 | 31 |
| GO_CELL_CELL_SIGNALING_BY_WNT | 0.00000 | 0.00001 | 0.71951 | 0.62397 | 1.27585 | 268 |
| GO_CELL_CYCLE_G2_M_PHASE_TRANSITION | 0.00003 | 0.00133 | 0.57561 | 0.60590 | 1.23456 | 209 |
| GO_CELL_CYCLE_PHASE_TRANSITION | 0.00000 | 0.00005 | 0.67496 | 0.58664 | 1.20584 | 447 |
| GO_CELL_DIVISION | 0.00021 | 0.00661 | 0.51885 | 0.56150 | 1.15307 | 418 |
| GO_CELL_KILLING | 0.00132 | 0.02984 | 0.45506 | 0.69978 | 1.35063 | 42 |
| GO_CELL_RECOGNITION | 0.00110 | 0.02622 | 0.45506 | 0.67487 | 1.31758 | 53 |
| GO_CELL_SUBSTRATE_JUNCTION | 0.00000 | 0.00000 | | 0.65108 | 1.33117 | 274 |
| GO_CELL_SURFACE_RECEPTOR_SIGNALING_PATHWAY_INVO | 0.00000 | 0.00001 | 0.69013 | 0.61586 | 1.26056 | 293 |
| GO_CHAPERONE_COMPLEX | 0.00000 | 0.00022 | 0.62726 | 0.88049 | 1.63117 | 20 |
| GO_CIRCADIAN_RHYTHM | 0.00058 | 0.01614 | 0.47727 | 0.62484 | 1.25211 | 110 |
| GO_COTRANSLATIONAL_PROTEIN_TARGETING_TO_MEMBR | 0.00000 | 0.00000 | | 0.84468 | 1.68461 | 92 |
| GO_CYTOKINE_MEDIATED_SIGNALING_PATHWAY | 0.00003 | 0.00137 | 0.57561 | 0.57955 | 1.18850 | 332 |
| GO_CYTOPLASMIC_STRESS_GRANULE | 0.00115 | 0.02674 | 0.45506 | 0.67722 | 1.32357 | 54 |
| GO_CYTOPLASMIC_TRANSLATION | 0.00000 | 0.00000 | | 0.81889 | 1.62961 | 86 |
| GO_CYTOPLASMIC_TRANSLATIONAL_INITIATION | 0.00001 | 0.00053 | 0.59333 | 0.81519 | 1.54479 | 29 |
| GO_CYTOSOLIC_LARGE_RIBOSOMAL_SUBUNIT | 0.00000 | 0.00000 | | 0.83786 | 1.63579 | 53 |
| GO_CYTOSOLIC_RIBOSOME | 0.00000 | 0.00000 | | 0.84135 | 1.67823 | 93 |
| GO_CYTOSOLIC_SMALL_RIBOSOMAL_SUBUNIT | 0.00000 | 0.00000 | 0.76146 | 0.84417 | 1.62414 | 39 |
| GO_DEFENSE_RESPONSE_TO_OTHER_ORGANISM | 0.00001 | 0.00080 | 0.59333 | 0.57915 | 1.18952 | 406 |
| GO_DNA_BINDING_TRANSCRIPTION_FACTOR_BINDING | 0.00216 | 0.04405 | 0.43171 | 0.57690 | 1.17561 | 207 |
| GO_DNA_BIOSYNTHETIC_PROCESS | 0.00021 | 0.00661 | 0.51885 | 0.62427 | 1.25576 | 131 |
| GO_DNA_CONFORMATION_CHANGE | 0.00128 | 0.02924 | 0.45506 | 0.57744 | 1.17680 | 210 |
| GO_DNA_GEOMETRIC_CHANGE | 0.00191 | 0.03995 | 0.45506 | 0.62452 | 1.24572 | 93 |
| GO_DNA_REPAIR | 0.00159 | 0.03496 | 0.45506 | 0.54597 | 1.12118 | 399 |
| GO_DNA_REPLICATION | 0.00161 | 0.03523 | 0.45506 | 0.57561 | 1.17293 | 208 |
| GO_EMBRYO_DEVELOPMENT | 0.00115 | 0.02674 | 0.45506 | 0.54467 | 1.12095 | 494 |
| GO_EMBRYO_DEVELOPMENT_ENDING_IN_BIRTH_OR_EGG_ | 0.00095 | 0.02329 | 0.47727 | 0.55950 | 1.14743 | 335 |
| GO_ENDOPEPTIDASE_ACTIVITY | 0.00079 | 0.02065 | 0.47727 | 0.59933 | 1.21315 | 161 |
| GO_ENDOPEPTIDASE_COMPLEX | 0.00000 | 0.00020 | 0.62726 | 0.73828 | 1.45094 | 61 |
| GO_ESTABLISHMENT_OF_PROTEIN_LOCALIZATION_TO_CHR | 0.00194 | 0.04027 | 0.45506 | 0.75727 | 1.41561 | 23 |
| GO_ESTABLISHMENT_OF_PROTEIN_LOCALIZATION_TO_END | 0.00000 | 0.00000 | | 0.82999 | 1.66200 | 102 |
| GO_ESTABLISHMENT_OF_PROTEIN_LOCALIZATION_TO_MEM | 0.00000 | 0.00000 | | 0.69991 | 1.42835 | 239 |
| GO_ESTABLISHMENT_OF_PROTEIN_LOCALIZATION_TO_ORG | 0.00000 | 0.00000 | | 0.67130 | 1.37877 | 406 |
| GO_ESTABLISHMENT_OF_PROTEIN_LOCALIZATION_TO_TELO | 0.00089 | 0.02253 | 0.47727 | 0.81554 | 1.49964 | 17 |
| GO_ESTABLISHMENT_OF_RNA_LOCALIZATION | 0.00015 | 0.00518 | 0.51885 | 0.61403 | 1.24295 | 160 |
| GO_ESTABLISHMENT_OF_TISSUE_POLARITY | 0.00007 | 0.00257 | 0.53843 | 0.67896 | 1.34790 | 78 |
| GO_EUKARYOTIC_TRANSLATION_INITIATION_FACTOR_3_CO | 0.00000 | 0.00002 | 0.69013 | 0.93334 | 1.70155 | 15 |
| GO_FC_EPSILON_RECEPTOR_SIGNALING_PATHWAY | 0.00016 | 0.00548 | 0.51885 | 0.66779 | 1.32659 | 80 |
| GO_FC_RECEPTOR_SIGNALING_PATHWAY | 0.00004 | 0.00183 | 0.55733 | 0.64066 | 1.28591 | 121 |

**Cluster 1**

| pathway | pval | padj | log2err | ES | NES | size |
|---|---|---|---|---|---|---|
| GO_FERTILIZATION | 0.00090 | 0.02266 | 0.47727 | 0.65222 | 1.29145 | 75 |
| GO_FICOLIN_1_RICH_GRANULE | 0.00000 | 0.00010 | 0.64355 | 0.69440 | 1.38490 | 92 |
| GO_GLYCOSYL_COMPOUND_METABOLIC_PROCESS | 0.00178 | 0.03781 | 0.45506 | 0.65258 | 1.28628 | 66 |
| GO_GUANYL_NUCLEOTIDE_BINDING | 0.00172 | 0.03735 | 0.45506 | 0.57948 | 1.17694 | 188 |
| GO_HEAT_SHOCK_PROTEIN_BINDING | 0.00113 | 0.02674 | 0.45506 | 0.63803 | 1.26873 | 84 |
| GO_HEMATOPOIETIC_PROGENITOR_CELL_DIFFERENTIATION | 0.00002 | 0.00087 | 0.57561 | 0.67144 | 1.34247 | 99 |
| GO_HEMATOPOIETIC_STEM_CELL_DIFFERENTIATION | 0.00001 | 0.00077 | 0.59333 | 0.70449 | 1.38753 | 64 |
| GO_HEPATICOBILIARY_SYSTEM_DEVELOPMENT | 0.00178 | 0.03781 | 0.45506 | 0.63798 | 1.26489 | 76 |
| GO_HYDROLASE_ACTIVITY_ACTING_ON_ACID_ANHYDRIDES | 0.00078 | 0.02060 | 0.47727 | 0.54872 | 1.12882 | 463 |
| GO_IMMUNE_RESPONSE_REGULATING_SIGNALING_PATHW | 0.00006 | 0.00252 | 0.53843 | 0.60539 | 1.23065 | 196 |
| GO_INNATE_IMMUNE_RESPONSE | 0.00002 | 0.00089 | 0.57561 | 0.58312 | 1.19587 | 331 |
| GO_INNATE_IMMUNE_RESPONSE_ACTIVATING_SIGNAL_TR | 0.00005 | 0.00198 | 0.55733 | 0.69433 | 1.36771 | 65 |
| GO_INTERLEUKIN_1_MEDIATED_SIGNALING_PATHWAY | 0.00000 | 0.00024 | 0.62726 | 0.70822 | 1.40124 | 73 |
| GO_INTRINSIC_APOPTOTIC_SIGNALING_PATHWAY | 0.00003 | 0.00120 | 0.57561 | 0.61579 | 1.25035 | 182 |
| GO_LARGE_RIBOSOMAL_SUBUNIT | 0.00000 | 0.00000 | 0.73376 | 0.70707 | 1.41595 | 106 |
| GO_LEUKOCYTE_MEDIATED_IMMUNITY | 0.00001 | 0.00039 | 0.61053 | 0.58445 | 1.19975 | 358 |
| GO_LIVER_REGENERATION | 0.00089 | 0.02253 | 0.47727 | 0.81552 | 1.49960 | 17 |
| GO_MEMBRANE_REPOLARIZATION | 0.00017 | 0.00565 | 0.51885 | 0.85648 | 1.56144 | 15 |
| GO_MODIFICATION_DEPENDENT_MACROMOLECULE_CATAI | 0.00001 | 0.00040 | 0.61053 | 0.57101 | 1.17495 | 466 |
| GO_MORPHOGENESIS_OF_AN_EPITHELIUM | 0.00010 | 0.00363 | 0.53843 | 0.58518 | 1.19638 | 267 |
| GO_MORPHOGENESIS_OF_A_POLARIZED_EPITHELIUM | 0.00011 | 0.00404 | 0.53843 | 0.66524 | 1.32446 | 88 |
| GO_MRNA_3_UTR_BINDING | 0.00001 | 0.00072 | 0.59333 | 0.70775 | 1.39414 | 65 |
| GO_MRNA_5_UTR_BINDING | 0.00021 | 0.00671 | 0.51885 | 0.81275 | 1.51186 | 22 |
| GO_MRNA_BINDING | 0.00000 | 0.00000 | | 0.68682 | 1.40034 | 222 |
| GO_MRNA_EXPORT_FROM_NUCLEUS | 0.00205 | 0.04210 | 0.43171 | 0.61791 | 1.23308 | 94 |
| GO_MRNA_PROCESSING | 0.00000 | 0.00000 | 0.83909 | 0.62061 | 1.27420 | 393 |
| GO_MYELOID_LEUKOCYTE_ACTIVATION | 0.00002 | 0.00108 | 0.57561 | 0.58247 | 1.19403 | 317 |
| GO_MYELOID_LEUKOCYTE_MEDIATED_IMMUNITY | 0.00000 | 0.00023 | 0.62726 | 0.59846 | 1.22466 | 289 |
| GO_NCRNA_METABOLIC_PROCESS | 0.00001 | 0.00068 | 0.59333 | 0.57925 | 1.18918 | 378 |
| GO_NCRNA_PROCESSING | 0.00000 | 0.00026 | 0.61053 | 0.59493 | 1.21989 | 325 |
| GO_NEGATIVE_REGULATION_OF_AXON_EXTENSION | 0.00113 | 0.02674 | 0.45506 | 0.82673 | 1.50719 | 15 |
| GO_NEGATIVE_REGULATION_OF_CANONICAL_WNT_SIGNAI | 0.00000 | 0.00027 | 0.61053 | 0.67884 | 1.35468 | 94 |
| GO_NEGATIVE_REGULATION_OF_CELLULAR_AMIDE_METAB | 0.00071 | 0.01894 | 0.47727 | 0.62407 | 1.25007 | 108 |
| GO_NEGATIVE_REGULATION_OF_CELL_CYCLE | 0.00000 | 0.00022 | 0.62726 | 0.58489 | 1.20117 | 404 |
| GO_NEGATIVE_REGULATION_OF_CELL_CYCLE_G2_M_PHASI | 0.00001 | 0.00033 | 0.61053 | 0.69048 | 1.37379 | 85 |
| GO_NEGATIVE_REGULATION_OF_CELL_CYCLE_PHASE_TRAN | 0.00000 | 0.00007 | 0.65944 | 0.63452 | 1.28829 | 181 |
| GO_NEGATIVE_REGULATION_OF_CELL_CYCLE_PROCESS | 0.00000 | 0.00002 | 0.69013 | 0.62237 | 1.26968 | 243 |
| GO_NEGATIVE_REGULATION_OF_MITOTIC_CELL_CYCLE | 0.00000 | 0.00018 | 0.62726 | 0.61614 | 1.25645 | 223 |
| GO_NEGATIVE_REGULATION_OF_MRNA_METABOLIC_PROC | 0.00250 | 0.04904 | 0.43171 | 0.66071 | 1.29245 | 55 |
| GO_NEGATIVE_REGULATION_OF_WNT_SIGNALING_PATHW | 0.00000 | 0.00011 | 0.64355 | 0.67596 | 1.35365 | 106 |
| GO_NEURON_TO_NEURON_SYNAPSE | 0.00002 | 0.00094 | 0.57561 | 0.62837 | 1.27121 | 155 |
| GO_NIK_NF_KAPPAB_SIGNALING | 0.00002 | 0.00115 | 0.57561 | 0.66415 | 1.32974 | 101 |
| GO_NON_CANONICAL_WNT_SIGNALING_PATHWAY | 0.00000 | 0.00018 | 0.62726 | 0.68791 | 1.37114 | 91 |
| GO_NUCLEAR_EXPORT | 0.00050 | 0.01399 | 0.47727 | 0.59820 | 1.21216 | 165 |
| GO_NUCLEAR_SPECK | 0.00011 | 0.00399 | 0.53843 | 0.58304 | 1.19205 | 274 |
| GO_NUCLEAR_TRANSCRIBED_MRNA_CATABOLIC_PROCESS | 0.00000 | 0.00000 | | 0.75612 | 1.53440 | 174 |
| GO_NUCLEAR_TRANSCRIBED_MRNA_CATABOLIC_PROCESS_ | 0.00000 | 0.00000 | | 0.83647 | 1.67503 | 105 |
| GO_NUCLEAR_TRANSPORT | 0.00002 | 0.00091 | 0.57561 | 0.59693 | 1.21937 | 256 |
| GO_NUCLEOBASE_CONTAINING_COMPOUND_TRANSPORT | 0.00065 | 0.01740 | 0.47727 | 0.58295 | 1.18544 | 197 |
| GO_NUCLEOSIDE_METABOLIC_PROCESS | 0.00110 | 0.02622 | 0.45506 | 0.67473 | 1.31731 | 53 |
| GO_NUCLEOSIDE_PHOSPHATE_BIOSYNTHETIC_PROCESS | 0.00209 | 0.04270 | 0.43171 | 0.60077 | 1.21145 | 137 |
| GO_NUCLEOTIDE_EXCISION_REPAIR_DNA_DAMAGE_RECOG | 0.00220 | 0.04432 | 0.43171 | 0.76471 | 1.42250 | 22 |
| GO_ORGANIC_CYCLIC_COMPOUND_CATABOLIC_PROCESS | 0.00000 | 0.00000 | | 0.69191 | 1.42129 | 422 |
| GO_PEPTIDASE_ACTIVITY | 0.00035 | 0.01029 | 0.49849 | 0.57753 | 1.17985 | 259 |
| GO_PEPTIDASE_COMPLEX | 0.00002 | 0.00089 | 0.57561 | 0.69227 | 1.37432 | 78 |
| GO_PIGMENT_GRANULE | 0.00093 | 0.02304 | 0.47727 | 0.65692 | 1.29955 | 72 |
| GO_POLYSOMAL_RIBOSOME | 0.00000 | 0.00005 | 0.65944 | 0.83108 | 1.58176 | 31 |
| GO_POLYSOME | 0.00000 | 0.00000 | 0.74774 | 0.78046 | 1.53258 | 59 |
| GO_POSITIVE_REGULATION_OF_CANONICAL_WNT_SIGNAL | 0.00000 | 0.00001 | 0.71951 | 0.71308 | 1.42467 | 96 |
| GO_POSITIVE_REGULATION_OF_CELLULAR_AMIDE_METABO | 0.00006 | 0.00247 | 0.53843 | 0.64923 | 1.30075 | 109 |
| GO_POSITIVE_REGULATION_OF_CELLULAR_PROTEIN_LOCAI | 0.00015 | 0.00520 | 0.51885 | 0.59188 | 1.20671 | 220 |
| GO_POSITIVE_REGULATION_OF_CELL_CYCLE | 0.00080 | 0.02079 | 0.47727 | 0.57327 | 1.16990 | 239 |
| GO_POSITIVE_REGULATION_OF_CELL_CYCLE_PROCESS | 0.00097 | 0.02364 | 0.47727 | 0.58683 | 1.19118 | 184 |
| GO_POSITIVE_REGULATION_OF_DEFENSE_RESPONSE | 0.00048 | 0.01354 | 0.49849 | 0.60466 | 1.22302 | 154 |
| GO_POSITIVE_REGULATION_OF_DNA_BIOSYNTHETIC_PROC | 0.00026 | 0.00789 | 0.49849 | 0.70884 | 1.37952 | 49 |
| GO_POSITIVE_REGULATION_OF_ESTABLISHMENT_OF_PROT | 0.00128 | 0.02924 | 0.45506 | 0.58144 | 1.18446 | 205 |
| GO_POSITIVE_REGULATION_OF_IMMUNE_RESPONSE | 0.00006 | 0.00255 | 0.53843 | 0.59096 | 1.20819 | 266 |
| GO_POSITIVE_REGULATION_OF_IMMUNE_SYSTEM_PROCES | 0.00032 | 0.00968 | 0.49849 | 0.56091 | 1.15147 | 390 |
| GO_POSITIVE_REGULATION_OF_INNATE_IMMUNE_RESPON | 0.00052 | 0.01443 | 0.47727 | 0.62801 | 1.25763 | 106 |
| GO_POSITIVE_REGULATION_OF_INTRINSIC_APOPTOTIC_SIG | 0.00095 | 0.02329 | 0.47727 | 0.71090 | 1.36521 | 38 |
| GO_POSITIVE_REGULATION_OF_MRNA_METABOLIC_PROCE | 0.00185 | 0.03908 | 0.45506 | 0.66535 | 1.30762 | 61 |

**Cluster 1**

| pathway | pval | padj | log2err | ES | NES | size |
|---|---|---|---|---|---|---|
| GO_POSITIVE_REGULATION_OF_PROTEIN_LOCALIZATION_T | 0.00006 | 0.00249 | 0.53843 | 0.71570 | 1.40287 | 57 |
| GO_POSITIVE_REGULATION_OF_RESPONSE_TO_BIOTIC_STI | 0.00013 | 0.00468 | 0.51885 | 0.63296 | 1.27186 | 125 |
| GO_POSITIVE_REGULATION_OF_RNA_SPLICING | 0.00198 | 0.04089 | 0.43171 | 0.75024 | 1.40807 | 25 |
| GO_POSITIVE_REGULATION_OF_TELOMERASE_RNA_LOCALI | 0.00036 | 0.01058 | 0.49849 | 0.84340 | 1.53758 | 15 |
| GO_POSITIVE_REGULATION_OF_TELOMERE_MAINTENANCE | 0.00191 | 0.03995 | 0.45506 | 0.73119 | 1.38561 | 29 |
| GO_POSITIVE_REGULATION_OF_TRANSLATION | 0.00002 | 0.00082 | 0.59333 | 0.66856 | 1.33417 | 94 |
| GO_POSITIVE_REGULATION_OF_WNT_SIGNALING_PATHWA | 0.00000 | 0.00006 | 0.65944 | 0.68391 | 1.37049 | 110 |
| GO_POSTSYNAPSE | 0.00029 | 0.00891 | 0.49849 | 0.57867 | 1.18280 | 262 |
| GO_POSTTRANSCRIPTIONAL_REGULATION_OF_GENE_EXPR | 0.00000 | 0.00000 | | 0.63784 | 1.31174 | 454 |
| GO_POST_TRANSLATIONAL_PROTEIN_MODIFICATION | 0.00007 | 0.00264 | 0.53843 | 0.59444 | 1.21200 | 222 |
| GO_POTASSIUM_ION_TRANSPORT | 0.00033 | 0.00999 | 0.49849 | 0.68023 | 1.33576 | 59 |
| GO_PRERIBOSOME | 0.00102 | 0.02477 | 0.45506 | 0.66039 | 1.30243 | 67 |
| GO_PRE_MRNA_BINDING | 0.00115 | 0.02674 | 0.45506 | 0.73059 | 1.38078 | 28 |
| GO_PROTEASOMAL_PROTEIN_CATABOLIC_PROCESS | 0.00044 | 0.01264 | 0.49849 | 0.56332 | 1.15658 | 361 |
| GO_PROTEASOMAL_UBIQUITIN_INDEPENDENT_PROTEIN_C | 0.00046 | 0.01309 | 0.49849 | 0.81064 | 1.49902 | 19 |
| GO_PROTEASOME_ACCESSORY_COMPLEX | 0.00128 | 0.02924 | 0.45506 | 0.75871 | 1.42087 | 24 |
| GO_PROTEASOME_CORE_COMPLEX | 0.00063 | 0.01707 | 0.47727 | 0.82357 | 1.51440 | 17 |
| GO_PROTEIN_CONTAINING_COMPLEX_LOCALIZATION | 0.00081 | 0.02083 | 0.47727 | 0.59086 | 1.20166 | 198 |
| GO_PROTEIN_DNA_COMPLEX_SUBUNIT_ORGANIZATION | 0.00220 | 0.04432 | 0.43171 | 0.58867 | 1.19231 | 163 |
| GO_PROTEIN_FOLDING | 0.00003 | 0.00118 | 0.57561 | 0.63272 | 1.27958 | 151 |
| GO_PROTEIN_LOCALIZATION_TO_CHROMOSOME | 0.00012 | 0.00445 | 0.51885 | 0.68286 | 1.34742 | 69 |
| GO_PROTEIN_LOCALIZATION_TO_CHROMOSOME_TELOMEF | 0.00250 | 0.04904 | 0.43171 | 0.74461 | 1.39751 | 25 |
| GO_PROTEIN_LOCALIZATION_TO_ENDOPLASMIC_RETICULU | 0.00000 | 0.00000 | | 0.80785 | 1.62329 | 125 |
| GO_PROTEIN_LOCALIZATION_TO_MEMBRANE | 0.00000 | 0.00000 | 0.81404 | 0.61946 | 1.27230 | 406 |
| GO_PROTEIN_LOCALIZATION_TO_NUCLEUS | 0.00087 | 0.02227 | 0.47727 | 0.59308 | 1.20355 | 174 |
| GO_PROTEIN_MODIFICATION_BY_SMALL_PROTEIN_REMOV | 0.00002 | 0.00098 | 0.57561 | 0.60503 | 1.23340 | 216 |
| GO_PROTEIN_POLYUBIQUITINATION | 0.00011 | 0.00409 | 0.53843 | 0.58766 | 1.19922 | 241 |
| GO_PROTEIN_STABILIZATION | 0.00001 | 0.00033 | 0.61053 | 0.66396 | 1.33305 | 119 |
| GO_PROTEIN_TARGETING | 0.00000 | 0.00000 | | 0.67444 | 1.38289 | 320 |
| GO_PROTEIN_TARGETING_TO_MEMBRANE | 0.00000 | 0.00000 | | 0.76817 | 1.55352 | 151 |
| GO_PURINE_NUCLEOSIDE_METABOLIC_PROCESS | 0.00137 | 0.03097 | 0.45506 | 0.72369 | 1.38178 | 34 |
| GO_REGENERATION | 0.00000 | 0.00024 | 0.62726 | 0.69447 | 1.38200 | 86 |
| GO_REGULATION_OF_ANIMAL_ORGAN_MORPHOGENESIS | 0.00004 | 0.00160 | 0.55733 | 0.64325 | 1.29111 | 121 |
| GO_REGULATION_OF_APOPTOTIC_SIGNALING_PATHWAY | 0.00104 | 0.02511 | 0.45506 | 0.57290 | 1.16891 | 236 |
| GO_REGULATION_OF_CELLULAR_AMIDE_METABOLIC_PROC | 0.00000 | 0.00000 | 0.73376 | 0.62025 | 1.26962 | 299 |
| GO_REGULATION_OF_CELLULAR_AMINE_METABOLIC_PROC | 0.00000 | 0.00003 | 0.67496 | 0.76569 | 1.49781 | 55 |
| GO_REGULATION_OF_CELLULAR_AMINO_ACID_METABOLIC | 0.00000 | 0.00007 | 0.65944 | 0.76475 | 1.49075 | 51 |
| GO_REGULATION_OF_CELLULAR_KETONE_METABOLIC_PRO | 0.00001 | 0.00079 | 0.59333 | 0.65946 | 1.32096 | 108 |
| GO_REGULATION_OF_CELLULAR_PROTEIN_LOCALIZATION | 0.00035 | 0.01037 | 0.49849 | 0.56368 | 1.15663 | 350 |
| GO_REGULATION_OF_CELL_CYCLE_G2_M_PHASE_TRANSITI | 0.00003 | 0.00146 | 0.55733 | 0.61702 | 1.25149 | 171 |
| GO_REGULATION_OF_CELL_CYCLE_PHASE_TRANSITION | 0.00000 | 0.00018 | 0.62726 | 0.59557 | 1.22119 | 327 |
| GO_REGULATION_OF_CHROMOSOME_ORGANIZATION | 0.00079 | 0.02065 | 0.47727 | 0.57383 | 1.17193 | 253 |
| GO_REGULATION_OF_CHROMOSOME_SEGREGATION | 0.00205 | 0.04210 | 0.43171 | 0.63248 | 1.25923 | 88 |
| GO_REGULATION_OF_DNA_BIOSYNTHETIC_PROCESS | 0.00022 | 0.00675 | 0.51885 | 0.66494 | 1.31615 | 74 |
| GO_REGULATION_OF_DNA_METABOLIC_PROCESS | 0.00042 | 0.01199 | 0.49849 | 0.58792 | 1.19968 | 232 |
| GO_REGULATION_OF_DNA_TEMPLATED_TRANSCRIPTION_I | 0.00159 | 0.03496 | 0.45506 | 0.72082 | 1.37628 | 34 |
| GO_REGULATION_OF_DNA_TEMPLATED_TRANSCRIPTION_I | 0.00008 | 0.00315 | 0.53843 | 0.66706 | 1.32808 | 88 |
| GO_REGULATION_OF_HEMATOPOIETIC_PROGENITOR_CELL | 0.00002 | 0.00102 | 0.57561 | 0.70005 | 1.38133 | 69 |
| GO_REGULATION_OF_HEMATOPOIETIC_STEM_CELL_DIFFER | 0.00004 | 0.00168 | 0.55733 | 0.70792 | 1.39013 | 59 |
| GO_REGULATION_OF_IMMUNE_RESPONSE | 0.00042 | 0.01199 | 0.49849 | 0.56285 | 1.15561 | 357 |
| GO_REGULATION_OF_INNATE_IMMUNE_RESPONSE | 0.00130 | 0.02954 | 0.45506 | 0.59405 | 1.20179 | 155 |
| GO_REGULATION_OF_INTRINSIC_APOPTOTIC_SIGNALING_P | 0.00002 | 0.00096 | 0.57561 | 0.65865 | 1.31931 | 107 |
| GO_REGULATION_OF_KERATINOCYTE_DIFFERENTIATION | 0.00235 | 0.04696 | 0.43171 | 0.79308 | 1.45834 | 17 |
| GO_REGULATION_OF_MITOTIC_CELL_CYCLE | 0.00001 | 0.00038 | 0.61053 | 0.57562 | 1.18298 | 440 |
| GO_REGULATION_OF_MORPHOGENESIS_OF_AN_EPITHELIU | 0.00002 | 0.00089 | 0.57561 | 0.66748 | 1.33573 | 100 |
| GO_REGULATION_OF_MRNA_CATABOLIC_PROCESS | 0.00000 | 0.00001 | 0.69013 | 0.65904 | 1.33403 | 159 |
| GO_REGULATION_OF_MRNA_METABOLIC_PROCESS | 0.00000 | 0.00000 | | 0.66954 | 1.36780 | 254 |
| GO_REGULATION_OF_MRNA_PROCESSING | 0.00000 | 0.00022 | 0.62726 | 0.67387 | 1.34979 | 107 |
| GO_REGULATION_OF_MRNA_SPLICING_VIA_SPLICEOSOME | 0.00000 | 0.00023 | 0.62726 | 0.70479 | 1.39736 | 76 |
| GO_REGULATION_OF_POTASSIUM_ION_TRANSPORT | 0.00026 | 0.00799 | 0.49849 | 0.75060 | 1.43315 | 34 |
| GO_REGULATION_OF_PROTEIN_CATABOLIC_PROCESS | 0.00044 | 0.01245 | 0.49849 | 0.57450 | 1.17469 | 264 |
| GO_REGULATION_OF_PROTEIN_CONTAINING_COMPLEX_AS | 0.00115 | 0.02674 | 0.45506 | 0.56612 | 1.15652 | 255 |
| GO_REGULATION_OF_PROTEIN_LOCALIZATION_TO_NUCLEU | 0.00242 | 0.04810 | 0.43171 | 0.63362 | 1.25902 | 81 |
| GO_REGULATION_OF_PROTEIN_STABILITY | 0.00011 | 0.00401 | 0.53843 | 0.60236 | 1.22281 | 186 |
| GO_REGULATION_OF_RESPONSE_TO_BIOTIC_STIMULUS | 0.00060 | 0.01656 | 0.47727 | 0.59339 | 1.20681 | 198 |
| GO_REGULATION_OF_RNA_SPLICING | 0.00000 | 0.00001 | 0.70498 | 0.70158 | 1.40530 | 107 |
| GO_REGULATION_OF_SMALL_MOLECULE_METABOLIC_PRO | 0.00111 | 0.02654 | 0.45506 | 0.57145 | 1.16651 | 249 |
| GO_REGULATION_OF_STEM_CELL_DIFFERENTIATION | 0.00002 | 0.00089 | 0.57561 | 0.68668 | 1.36445 | 81 |
| GO_REGULATION_OF_TRANSCRIPTION_FROM_RNA_POLYM | 0.00000 | 0.00026 | 0.61053 | 0.72463 | 1.42601 | 63 |
| GO_REGULATION_OF_TRANSLATIONAL_INITIATION | 0.00001 | 0.00079 | 0.59333 | 0.71124 | 1.39965 | 63 |

| Cluster 1 | | | | | | |
|---|---|---|---|---|---|---|
| pathway | pval | padj | log2err | ES | NES | size |
| GO_REGULATION_OF_VIRAL_LIFE_CYCLE | 0.00156 | 0.03456 | 0.45506 | 0.63796 | 1.26799 | 82 |
| GO_REGULATION_OF_WNT_SIGNALING_PATHWAY | 0.00000 | 0.00001 | 0.70498 | 0.64592 | 1.31198 | 189 |
| GO_RESPONSE_TO_HEAT | 0.00065 | 0.01740 | 0.47727 | 0.62580 | 1.25321 | 106 |
| GO_RESPONSE_TO_INTERLEUKIN_1 | 0.00001 | 0.00045 | 0.61053 | 0.66425 | 1.33056 | 108 |
| GO_RESPONSE_TO_INTERLEUKIN_12 | 0.00082 | 0.02097 | 0.47727 | 0.71761 | 1.37145 | 35 |
| GO_RESPONSE_TO_INTERLEUKIN_4 | 0.00178 | 0.03781 | 0.45506 | 0.77846 | 1.44215 | 20 |
| GO_RESPONSE_TO_LEUKEMIA_INHIBITORY_FACTOR | 0.00001 | 0.00070 | 0.59333 | 0.71051 | 1.40047 | 66 |
| GO_RESPONSE_TO_TEMPERATURE_STIMULUS | 0.00157 | 0.03483 | 0.45506 | 0.61078 | 1.22696 | 123 |
| GO_RESPONSE_TO_TUMOR_NECROSIS_FACTOR | 0.00070 | 0.01879 | 0.47727 | 0.60062 | 1.21465 | 150 |
| GO_RHYTHMIC_PROCESS | 0.00093 | 0.02304 | 0.47727 | 0.59951 | 1.21257 | 153 |
| GO_RIBONUCLEOPROTEIN_COMPLEX_BINDING | 0.00004 | 0.00185 | 0.55733 | 0.64277 | 1.28989 | 114 |
| GO_RIBONUCLEOPROTEIN_COMPLEX_BIOGENESIS | 0.00000 | 0.00000 | | 0.68138 | 1.39880 | 382 |
| GO_RIBONUCLEOPROTEIN_COMPLEX_SUBUNIT_ORGANIZAT | 0.00000 | 0.00000 | | 0.69359 | 1.40665 | 169 |
| GO_RIBONUCLEOPROTEIN_GRANULE | 0.00005 | 0.00221 | 0.55733 | 0.61943 | 1.25427 | 162 |
| GO_RIBONUCLEOSIDE_METABOLIC_PROCESS | 0.00174 | 0.03760 | 0.45506 | 0.71929 | 1.37337 | 34 |
| GO_RIBOSOMAL_LARGE_SUBUNIT_ASSEMBLY | 0.00017 | 0.00557 | 0.51885 | 0.77738 | 1.46515 | 27 |
| GO_RIBOSOMAL_LARGE_SUBUNIT_BIOGENESIS | 0.00000 | 0.00002 | 0.69013 | 0.74687 | 1.47214 | 66 |
| GO_RIBOSOMAL_SMALL_SUBUNIT_BIOGENESIS | 0.00005 | 0.00190 | 0.55733 | 0.69780 | 1.37542 | 66 |
| GO_RIBOSOMAL_SUBUNIT | 0.00000 | 0.00000 | | 0.71785 | 1.45584 | 169 |
| GO_RIBOSOME | 0.00000 | 0.00000 | | 0.70563 | 1.43579 | 200 |
| GO_RIBOSOME_ASSEMBLY | 0.00000 | 0.00007 | 0.65944 | 0.74398 | 1.45924 | 58 |
| GO_RIBOSOME_BIOGENESIS | 0.00000 | 0.00000 | | 0.67182 | 1.37377 | 270 |
| GO_RNA_CATABOLIC_PROCESS | 0.00000 | 0.00000 | | 0.72523 | 1.48720 | 324 |
| GO_RNA_DEPENDENT_DNA_BIOSYNTHETIC_PROCESS | 0.00041 | 0.01185 | 0.49849 | 0.68203 | 1.33606 | 56 |
| GO_RNA_EXPORT_FROM_NUCLEUS | 0.00013 | 0.00447 | 0.51885 | 0.63520 | 1.27475 | 120 |
| GO_RNA_LOCALIZATION | 0.00000 | 0.00006 | 0.65944 | 0.63727 | 1.29492 | 191 |
| GO_RNA_SPLICING | 0.00000 | 0.00000 | 0.80122 | 0.62539 | 1.28287 | 343 |
| GO_RNA_SPLICING_VIA_TRANSESTERIFICATION_REACTIONS | 0.00000 | 0.00000 | 0.77494 | 0.63247 | 1.29369 | 282 |
| GO_ROUGH_ENDOPLASMIC_RETICULUM | 0.00246 | 0.04866 | 0.43171 | 0.67283 | 1.31262 | 52 |
| GO_RRNA_BINDING | 0.00004 | 0.00185 | 0.55733 | 0.71550 | 1.39691 | 53 |
| GO_RRNA_METABOLIC_PROCESS | 0.00000 | 0.00000 | 0.74774 | 0.65233 | 1.32914 | 206 |
| GO_SCF_DEPENDENT_PROTEASOMAL_UBIQUITIN_DEPEND | 0.00002 | 0.00082 | 0.59333 | 0.70653 | 1.39571 | 70 |
| GO_SECRETORY_GRANULE | 0.00001 | 0.00047 | 0.59333 | 0.57569 | 1.18192 | 392 |
| GO_SECRETORY_VESICLE | 0.00020 | 0.00636 | 0.51885 | 0.55558 | 1.14278 | 458 |
| GO_SINGLE_FERTILIZATION | 0.00035 | 0.01029 | 0.49849 | 0.67983 | 1.33497 | 59 |
| GO_SINGLE_STRANDED_RNA_BINDING | 0.00019 | 0.00614 | 0.51885 | 0.70124 | 1.37452 | 57 |
| GO_SMALL_RIBOSOMAL_SUBUNIT | 0.00000 | 0.00001 | 0.70498 | 0.75274 | 1.48371 | 66 |
| GO_SODIUM_CHANNEL_REGULATOR_ACTIVITY | 0.00178 | 0.03781 | 0.45506 | 0.79857 | 1.46071 | 16 |
| GO_SPERM_EGG_RECOGNITION | 0.00005 | 0.00211 | 0.55733 | 0.80426 | 1.50946 | 25 |
| GO_SPLICEOSOMAL_COMPLEX | 0.00000 | 0.00027 | 0.61053 | 0.64001 | 1.29507 | 157 |
| GO_STEM_CELL_DIFFERENTIATION | 0.00002 | 0.00110 | 0.57561 | 0.64793 | 1.30484 | 134 |
| GO_STRUCTURAL_CONSTITUENT_OF_RIBOSOME | 0.00000 | 0.00000 | | 0.72494 | 1.46437 | 147 |
| GO_STRUCTURAL_MOLECULE_ACTIVITY | 0.00000 | 0.00000 | | 0.63604 | 1.30448 | 337 |
| GO_TELOMERASE_RNA_LOCALIZATION | 0.00172 | 0.03735 | 0.45506 | 0.78455 | 1.45078 | 19 |
| GO_TELOMERE_MAINTENANCE_VIA_TELOMERE_LENGTHEN | 0.00076 | 0.02021 | 0.47727 | 0.66431 | 1.30730 | 63 |
| GO_THREONINE_TYPE_ENDOPEPTIDASE_ACTIVITY | 0.00024 | 0.00751 | 0.51885 | 0.83884 | 1.54248 | 17 |
| GO_THREONINE_TYPE_PEPTIDASE_ACTIVITY | 0.00020 | 0.00647 | 0.51885 | 0.82440 | 1.52447 | 19 |
| GO_TISSUE_MORPHOGENESIS | 0.00076 | 0.02007 | 0.47727 | 0.56371 | 1.15379 | 297 |
| GO_TOXIN_TRANSPORT | 0.00092 | 0.02302 | 0.47727 | 0.73800 | 1.39479 | 28 |
| GO_TRANSCRIPTION_FACTOR_BINDING | 0.00001 | 0.00045 | 0.61053 | 0.57807 | 1.18670 | 377 |
| GO_TRANSLATIONAL_INITIATION | 0.00000 | 0.00000 | | 0.81349 | 1.64840 | 165 |
| GO_TRANSLATION_ELONGATION_FACTOR_ACTIVITY | 0.00013 | 0.00447 | 0.51885 | 0.86420 | 1.57550 | 15 |
| GO_TRANSLATION_FACTOR_ACTIVITY_RNA_BINDING | 0.00000 | 0.00000 | 0.80122 | 0.77758 | 1.53367 | 68 |
| GO_TRANSLATION_INITIATION_FACTOR_ACTIVITY | 0.00000 | 0.00005 | 0.65944 | 0.79396 | 1.53728 | 44 |
| GO_TRANSLATION_INITIATION_FACTOR_BINDING | 0.00061 | 0.01673 | 0.47727 | 0.77311 | 1.44783 | 24 |
| GO_TRANSLATION_PREINITIATION_COMPLEX | 0.00000 | 0.00013 | 0.64355 | 0.90706 | 1.65916 | 16 |
| GO_TRANSLATION_REGULATOR_ACTIVITY | 0.00000 | 0.00000 | | 0.74478 | 1.49149 | 103 |
| GO_TRANSLATION_REGULATOR_ACTIVITY_NUCLEIC_ACID_E | 0.00000 | 0.00000 | 0.82666 | 0.76407 | 1.51850 | 83 |
| GO_TUMOR_NECROSIS_FACTOR_MEDIATED_SIGNALING_PA | 0.00015 | 0.00520 | 0.51885 | 0.65079 | 1.30021 | 96 |
| GO_T_CELL_RECEPTOR_SIGNALING_PATHWAY | 0.00000 | 0.00014 | 0.64355 | 0.66907 | 1.34266 | 114 |
| GO_U2_TYPE_SPLICEOSOMAL_COMPLEX | 0.00119 | 0.02748 | 0.45506 | 0.64738 | 1.28489 | 77 |
| GO_UBIQUITIN_LIKE_PROTEIN_LIGASE_BINDING | 0.00008 | 0.00321 | 0.53843 | 0.60262 | 1.22618 | 200 |
| GO_UNFOLDED_PROTEIN_BINDING | 0.00004 | 0.00183 | 0.55733 | 0.68401 | 1.35881 | 80 |
| GO_VESICLE_LUMEN | 0.00000 | 0.00001 | 0.70498 | 0.65912 | 1.33673 | 169 |
| GO_VIRAL_GENE_EXPRESSION | 0.00000 | 0.00000 | | 0.75549 | 1.53090 | 165 |
| GO_VIRAL_LIFE_CYCLE | 0.00015 | 0.00520 | 0.51885 | 0.59577 | 1.21390 | 206 |
| HP_ABNORMALITY_OF_THE_NASAL_DORSUM | 0.00228 | 0.04565 | 0.43171 | 0.60297 | 1.21382 | 133 |
| HP_ACUTE_LEUKEMIA | 0.00013 | 0.00473 | 0.51885 | 0.68109 | 1.34391 | 69 |
| HP_ANEMIA_OF_INADEQUATE_PRODUCTION | 0.00194 | 0.04027 | 0.45506 | 0.62708 | 1.24940 | 90 |
| HP_DEPRESSED_NASAL_RIDGE | 0.00004 | 0.00168 | 0.55733 | 0.68680 | 1.35887 | 73 |

| Cluster 1 | | | | | | |
|---|---|---|---|---|---|---|
| pathway | pval | padj | log2err | ES | NES | size |
| HP_LEUKEMIA | 0.00150 | 0.03360 | 0.45506 | 0.62694 | 1.25181 | 95 |
| HP_MACROCYTIC_ANEMIA | 0.00015 | 0.00520 | 0.51885 | 0.74566 | 1.43460 | 39 |
| HP_MIGRAINE | 0.00011 | 0.00404 | 0.53843 | 0.67905 | 1.33932 | 68 |
| HP_SHORT_THUMB | 0.00152 | 0.03388 | 0.45506 | 0.69580 | 1.34474 | 43 |
| HP_THICK_VERMILION_BORDER | 0.00139 | 0.03125 | 0.45506 | 0.59388 | 1.20036 | 149 |
| **Cluster 2** | | | | | | |
| pathway | pval | padj | log2err | ES | NES | size |
| GO_ACTIN_FILAMENT_BUNDLE | 0.00012 | 0.00643 | 0.51885 | 0.76245 | 1.53426 | 22 |
| GO_ACTOMYOSIN | 0.00009 | 0.00499 | 0.53843 | 0.76688 | 1.54319 | 22 |
| GO_AMIDE_BIOSYNTHETIC_PROCESS | 0.00000 | 0.00000 | | 0.56850 | 1.28754 | 366 |
| GO_ANAPHASE_PROMOTING_COMPLEX_DEPENDENT_CATA | 0.00143 | 0.04431 | 0.45506 | 0.59839 | 1.28596 | 62 |
| GO_ANCHORING_JUNCTION | 0.00000 | 0.00000 | | 0.61248 | 1.37696 | 241 |
| GO_CADHERIN_BINDING | 0.00000 | 0.00018 | 0.62726 | 0.57771 | 1.28705 | 166 |
| GO_CELLULAR_AMIDE_METABOLIC_PROCESS | 0.00000 | 0.00000 | | 0.55758 | 1.26538 | 444 |
| GO_CELLULAR_MACROMOLECULE_CATABOLIC_PROCESS | 0.00000 | 0.00000 | 0.77494 | 0.54291 | 1.23272 | 455 |
| GO_CELLULAR_NITROGEN_COMPOUND_CATABOLIC_PROCE | 0.00000 | 0.00000 | | 0.62134 | 1.39827 | 248 |
| GO_CELL_ACTIVATION | 0.00001 | 0.00068 | 0.59333 | 0.53523 | 1.20915 | 309 |
| GO_CELL_ACTIVATION_INVOLVED_IN_IMMUNE_RESPONSE | 0.00000 | 0.00018 | 0.62726 | 0.56498 | 1.26318 | 190 |
| GO_CELL_ADHESION_MOLECULE_BINDING | 0.00000 | 0.00004 | 0.67496 | 0.57404 | 1.28394 | 192 |
| GO_CELL_RECOGNITION | 0.00037 | 0.01584 | 0.49849 | 0.73943 | 1.48795 | 22 |
| GO_CELL_SUBSTRATE_JUNCTION | 0.00000 | 0.00000 | | 0.67120 | 1.49570 | 169 |
| GO_COTRANSLATIONAL_PROTEIN_TARGETING_TO_MEMBR | 0.00000 | 0.00000 | | 0.78989 | 1.72107 | 87 |
| GO_CYTOKINE_MEDIATED_SIGNALING_PATHWAY | 0.00007 | 0.00390 | 0.53843 | 0.55314 | 1.23323 | 174 |
| GO_CYTOPLASMIC_TRANSLATION | 0.00000 | 0.00007 | 0.65944 | 0.66579 | 1.43723 | 70 |
| GO_CYTOSOLIC_LARGE_RIBOSOMAL_SUBUNIT | 0.00000 | 0.00000 | 0.83909 | 0.78678 | 1.66606 | 50 |
| GO_CYTOSOLIC_RIBOSOME | 0.00000 | 0.00000 | | 0.78526 | 1.71036 | 86 |
| GO_CYTOSOLIC_SMALL_RIBOSOMAL_SUBUNIT | 0.00000 | 0.00000 | 0.71951 | 0.78693 | 1.64257 | 37 |
| GO_DENDRITIC_TREE | 0.00113 | 0.03678 | 0.45506 | 0.55229 | 1.21755 | 121 |
| GO_DNA_DAMAGE_RESPONSE_DETECTION_OF_DNA_DAM | 0.00089 | 0.03163 | 0.47727 | 0.72085 | 1.46307 | 24 |
| GO_ESTABLISHMENT_OF_PROTEIN_LOCALIZATION_TO_END | 0.00000 | 0.00000 | | 0.77162 | 1.68718 | 94 |
| GO_ESTABLISHMENT_OF_PROTEIN_LOCALIZATION_TO_MEN | 0.00000 | 0.00000 | | 0.71620 | 1.59386 | 163 |
| GO_ESTABLISHMENT_OF_PROTEIN_LOCALIZATION_TO_ORG | 0.00000 | 0.00000 | | 0.65747 | 1.48157 | 261 |
| GO_EXOCYTOSIS | 0.00001 | 0.00069 | 0.59333 | 0.53987 | 1.21477 | 247 |
| GO_FC_RECEPTOR_MEDIATED_STIMULATORY_SIGNALING_F | 0.00065 | 0.02485 | 0.47727 | 0.73343 | 1.48283 | 23 |
| GO_FC_RECEPTOR_SIGNALING_PATHWAY | 0.00045 | 0.01867 | 0.49849 | 0.60206 | 1.29872 | 69 |
| GO_FICOLIN_1_RICH_GRANULE | 0.00015 | 0.00751 | 0.51885 | 0.62047 | 1.34257 | 73 |
| GO_IMMUNE_EFFECTOR_PROCESS | 0.00001 | 0.00065 | 0.59333 | 0.54560 | 1.22928 | 259 |
| GO_INTERACTION_WITH_HOST | 0.00036 | 0.01564 | 0.49849 | 0.61924 | 1.33099 | 63 |
| GO_INTRACELLULAR_PROTEIN_TRANSPORT | 0.00000 | 0.00000 | | 0.55766 | 1.26704 | 468 |
| GO_LARGE_RIBOSOMAL_SUBUNIT | 0.00000 | 0.00000 | 0.73376 | 0.66187 | 1.44798 | 96 |
| GO_LEUKOCYTE_MEDIATED_IMMUNITY | 0.00000 | 0.00008 | 0.64355 | 0.56926 | 1.27277 | 190 |
| GO_MATURATION_OF_LSU_RRNA | 0.00059 | 0.02324 | 0.47727 | 0.74980 | 1.49173 | 19 |
| GO_MRNA_METABOLIC_PROCESS | 0.00000 | 0.00000 | 0.77494 | 0.54681 | 1.24062 | 423 |
| GO_MUSCLE_CELL_DEVELOPMENT | 0.00167 | 0.04957 | 0.45506 | 0.67210 | 1.38535 | 31 |
| GO_MYELOID_LEUKOCYTE_ACTIVATION | 0.00000 | 0.00006 | 0.65944 | 0.57962 | 1.29425 | 182 |
| GO_MYELOID_LEUKOCYTE_MEDIATED_IMMUNITY | 0.00000 | 0.00006 | 0.65944 | 0.58224 | 1.29750 | 170 |
| GO_NEURON_TO_NEURON_SYNAPSE | 0.00020 | 0.00960 | 0.51885 | 0.61784 | 1.33688 | 73 |
| GO_NUCLEAR_TRANSCRIBED_MRNA_CATABOLIC_PROCESS | 0.00000 | 0.00000 | | 0.71854 | 1.58267 | 119 |
| GO_NUCLEAR_TRANSCRIBED_MRNA_CATABOLIC_PROCESS_ | 0.00000 | 0.00000 | | 0.76979 | 1.68155 | 91 |
| GO_NUCLEOTIDE_EXCISION_REPAIR_DNA_DAMAGE_RECOG | 0.00130 | 0.04103 | 0.45506 | 0.77395 | 1.51411 | 16 |
| GO_ORGANELLE_ASSEMBLY | 0.00010 | 0.00546 | 0.53843 | 0.53193 | 1.19641 | 245 |
| GO_ORGANIC_CYCLIC_COMPOUND_CATABOLIC_PROCESS | 0.00000 | 0.00000 | | 0.62261 | 1.40222 | 254 |
| GO_PEPTIDE_BIOSYNTHETIC_PROCESS | 0.00000 | 0.00000 | | 0.59182 | 1.33887 | 327 |
| GO_PEPTIDE_METABOLIC_PROCESS | 0.00000 | 0.00000 | | 0.58406 | 1.32279 | 366 |
| GO_POLYSOMAL_RIBOSOME | 0.00041 | 0.01740 | 0.49849 | 0.71089 | 1.45331 | 27 |
| GO_POLYSOME | 0.00065 | 0.02493 | 0.47727 | 0.64988 | 1.36694 | 43 |
| GO_POSITIVE_REGULATION_OF_CELLULAR_AMIDE_METABO | 0.00079 | 0.02892 | 0.47727 | 0.60853 | 1.29863 | 56 |
| GO_POSITIVE_REGULATION_OF_DNA_BIOSYNTHETIC_PROC | 0.00042 | 0.01777 | 0.49849 | 0.70203 | 1.43950 | 29 |
| GO_POSITIVE_REGULATION_OF_NF_KAPPAB_TRANSCRIPTIO | 0.00027 | 0.01215 | 0.49849 | 0.70861 | 1.45299 | 29 |
| GO_POSITIVE_REGULATION_OF_TELOMERASE_RNA_LOCALI | 0.00007 | 0.00383 | 0.53843 | 0.84766 | 1.64524 | 15 |
| GO_POSTSYNAPSE | 0.00001 | 0.00089 | 0.59333 | 0.58674 | 1.29541 | 125 |
| GO_POSTTRANSCRIPTIONAL_REGULATION_OF_GENE_EXPR | 0.00030 | 0.01321 | 0.49849 | 0.52146 | 1.17483 | 257 |
| GO_PROTEIN_CONTAINING_COMPLEX_BINDING | 0.00000 | 0.00006 | 0.65944 | 0.53501 | 1.21166 | 365 |
| GO_PROTEIN_LOCALIZATION_TO_CHROMOSOME_TELOMEF | 0.00068 | 0.02543 | 0.47727 | 0.79844 | 1.54970 | 15 |
| GO_PROTEIN_LOCALIZATION_TO_ENDOPLASMIC_RETICULU | 0.00000 | 0.00000 | | 0.76822 | 1.68662 | 106 |
| GO_PROTEIN_LOCALIZATION_TO_MEMBRANE | 0.00000 | 0.00000 | | 0.64195 | 1.44277 | 237 |
| GO_PROTEIN_LOCALIZATION_TO_ORGANELLE | 0.00000 | 0.00000 | | 0.59955 | 1.35880 | 394 |
| GO_PROTEIN_STABILIZATION | 0.00053 | 0.02119 | 0.47727 | 0.60335 | 1.30245 | 70 |
| GO_PROTEIN_TARGETING | 0.00000 | 0.00000 | | 0.65990 | 1.47940 | 212 |

| Cluster 1 | | | | | | |
|---|---|---|---|---|---|---|
| pathway | pval | padj | log2err | ES | NES | size |
| GO_PROTEIN_TARGETING_TO_MEMBRANE | 0.00000 | 0.00000 | | 0.75578 | 1.66323 | 115 |
| GO_REGULATION_OF_CELLULAR_AMIDE_METABOLIC_PROC | 0.00069 | 0.02551 | 0.47727 | 0.53820 | 1.19724 | 157 |
| GO_REGULATION_OF_DNA_BIOSYNTHETIC_PROCESS | 0.00156 | 0.04708 | 0.45506 | 0.64651 | 1.35102 | 38 |
| GO_REGULATION_OF_DNA_TEMPLATED_TRANSCRIPTION_I | 0.00113 | 0.03678 | 0.45506 | 0.60725 | 1.29833 | 57 |
| GO_REGULATION_OF_PROTEIN_STABILITY | 0.00110 | 0.03625 | 0.45506 | 0.55064 | 1.21079 | 113 |
| GO_REGULATION_OF_SIGNAL_TRANSDUCTION_BY_P53_CL | 0.00159 | 0.04778 | 0.45506 | 0.59497 | 1.28349 | 68 |
| GO_REGULATION_OF_UBIQUITIN_PROTEIN_LIGASE_ACTIVIT | 0.00110 | 0.03625 | 0.45506 | 0.77853 | 1.52308 | 16 |
| GO_RESPONSE_TO_CYTOKINE | 0.00000 | 0.00012 | 0.64355 | 0.54295 | 1.22511 | 293 |
| GO_RESPONSE_TO_OXYGEN_CONTAINING_COMPOUND | 0.00133 | 0.04182 | 0.45506 | 0.50003 | 1.13277 | 370 |
| GO_RIBONUCLEOPROTEIN_COMPLEX | 0.00000 | 0.00000 | 0.81404 | 0.55650 | 1.26181 | 407 |
| GO_RIBONUCLEOPROTEIN_COMPLEX_BINDING | 0.00090 | 0.03163 | 0.47727 | 0.59719 | 1.29220 | 73 |
| GO_RIBONUCLEOPROTEIN_COMPLEX_BIOGENESIS | 0.00009 | 0.00471 | 0.53843 | 0.53354 | 1.19940 | 240 |
| GO_RIBONUCLEOPROTEIN_COMPLEX_SUBUNIT_ORGANIZAT | 0.00015 | 0.00752 | 0.51885 | 0.57130 | 1.25761 | 117 |
| GO_RIBOSOMAL_LARGE_SUBUNIT_ASSEMBLY | 0.00018 | 0.00898 | 0.51885 | 0.75675 | 1.52998 | 23 |
| GO_RIBOSOMAL_LARGE_SUBUNIT_BIOGENESIS | 0.00000 | 0.00001 | 0.71951 | 0.73205 | 1.55016 | 50 |
| GO_RIBOSOMAL_SUBUNIT | 0.00000 | 0.00000 | | 0.66803 | 1.48438 | 151 |
| GO_RIBOSOME | 0.00000 | 0.00000 | | 0.65916 | 1.46949 | 172 |
| GO_RIBOSOME_ASSEMBLY | 0.00148 | 0.04563 | 0.45506 | 0.63693 | 1.33881 | 42 |
| GO_RIBOSOME_BIOGENESIS | 0.00021 | 0.00979 | 0.51885 | 0.54870 | 1.22241 | 166 |
| GO_RNA_CATABOLIC_PROCESS | 0.00000 | 0.00000 | | 0.63911 | 1.43267 | 214 |
| GO_RRNA_BINDING | 0.00000 | 0.00025 | 0.62726 | 0.73048 | 1.53117 | 40 |
| GO_SECRETION | 0.00087 | 0.03127 | 0.47727 | 0.50358 | 1.14056 | 349 |
| GO_SECRETORY_GRANULE | 0.00000 | 0.00004 | 0.65944 | 0.56959 | 1.27683 | 214 |
| GO_SECRETORY_VESICLE | 0.00001 | 0.00050 | 0.61053 | 0.54787 | 1.23317 | 249 |
| GO_SMALL_RIBOSOMAL_SUBUNIT | 0.00001 | 0.00069 | 0.59333 | 0.67700 | 1.44877 | 58 |
| GO_STEM_CELL_DIFFERENTIATION | 0.00095 | 0.03259 | 0.47727 | 0.58415 | 1.26861 | 79 |
| GO_STRUCTURAL_CONSTITUENT_OF_RIBOSOME | 0.00000 | 0.00000 | | 0.69140 | 1.53216 | 137 |
| GO_STRUCTURAL_MOLECULE_ACTIVITY | 0.00000 | 0.00000 | | 0.61599 | 1.38476 | 240 |
| GO_SUPRAMOLECULAR_POLYMER | 0.00095 | 0.03259 | 0.47727 | 0.52993 | 1.18563 | 198 |
| GO_SYMBIOTIC_PROCESS | 0.00000 | 0.00000 | | 0.62617 | 1.41947 | 399 |
| GO_SYNAPSE | 0.00000 | 0.00025 | 0.62726 | 0.54650 | 1.23200 | 266 |
| GO_TELOMERASE_RNA_LOCALIZATION | 0.00003 | 0.00200 | 0.55733 | 0.84629 | 1.65563 | 16 |
| GO_TRANSLATIONAL_INITIATION | 0.00000 | 0.00000 | | 0.72964 | 1.61668 | 138 |
| GO_TRANSLATION_FACTOR_ACTIVITY_RNA_BINDING | 0.00014 | 0.00716 | 0.51885 | 0.64740 | 1.37273 | 51 |
| GO_TRANSLATION_REGULATOR_ACTIVITY | 0.00002 | 0.00095 | 0.57561 | 0.62810 | 1.36205 | 77 |
| GO_TRANSLATION_REGULATOR_ACTIVITY_NUCLEIC_ACID_E | 0.00026 | 0.01169 | 0.49849 | 0.62160 | 1.33400 | 60 |
| GO_UBIQUITIN_LIKE_PROTEIN_LIGASE_BINDING | 0.00000 | 0.00004 | 0.67496 | 0.61768 | 1.35922 | 114 |
| GO_UNFOLDED_PROTEIN_BINDING | 0.00019 | 0.00940 | 0.51885 | 0.63038 | 1.34777 | 57 |
| GO_VACUOLAR_LUMEN | 0.00090 | 0.03163 | 0.47727 | 0.63246 | 1.33861 | 49 |
| GO_VACUOLE | 0.00058 | 0.02314 | 0.47727 | 0.52703 | 1.18282 | 224 |
| GO_VESICLE_LUMEN | 0.00001 | 0.00071 | 0.59333 | 0.59526 | 1.30853 | 112 |
| GO_VIRAL_GENE_EXPRESSION | 0.00000 | 0.00000 | | 0.70755 | 1.56707 | 135 |
| GO_WHOLE_MEMBRANE | 0.00124 | 0.04000 | 0.45506 | 0.49458 | 1.12224 | 430 |
| HP_ABNORMALITY_OF_CARDIOVASCULAR_SYSTEM_ELECTR | 0.00044 | 0.01832 | 0.49849 | 0.56734 | 1.24658 | 111 |
| HP_ABNORMALITY_OF_THE_NASAL_DORSUM | 0.00009 | 0.00476 | 0.53843 | 0.63579 | 1.36446 | 60 |
| HP_ANEMIA_OF_INADEQUATE_PRODUCTION | 0.00008 | 0.00434 | 0.53843 | 0.66278 | 1.40109 | 48 |
| HP_DELAYED_CRANIAL_SUTURE_CLOSURE | 0.00104 | 0.03508 | 0.45506 | 0.74045 | 1.48583 | 21 |
| HP_DELAYED_PUBERTY | 0.00007 | 0.00383 | 0.53843 | 0.67582 | 1.42741 | 46 |
| HP_DEPRESSED_NASAL_RIDGE | 0.00066 | 0.02501 | 0.47727 | 0.65892 | 1.37720 | 39 |
| HP_HEADACHE | 0.00025 | 0.01163 | 0.49849 | 0.61477 | 1.32215 | 65 |
| HP_JUVENILE_ONSET | 0.00102 | 0.03479 | 0.45506 | 0.73479 | 1.46187 | 19 |
| HP_MACROCYTIC_ANEMIA | 0.00060 | 0.02334 | 0.47727 | 0.69308 | 1.42114 | 29 |
| HP_MIGRAINE | 0.00001 | 0.00050 | 0.61053 | 0.72448 | 1.51394 | 38 |
| HP_PALLOR | 0.00048 | 0.01957 | 0.49849 | 0.63093 | 1.33603 | 50 |
| HP_RETROGNATHIA | 0.00150 | 0.04580 | 0.45506 | 0.58786 | 1.26809 | 69 |
| HP_TELANGIECTASIA_OF_THE_SKIN | 0.00128 | 0.04081 | 0.45506 | 0.75506 | 1.48503 | 17 |
| HP_TRIANGULAR_SHAPED_PHALANGES_OF_THE_HAND | 0.00023 | 0.01072 | 0.51885 | 0.67280 | 1.41026 | 40 |
| **Cluster 3** | | | | | | |
| pathway | pval | padj | log2err | ES | NES | size |
| GO_ACTIVATION_OF_IMMUNE_RESPONSE | 0.00000 | 0.00001 | 0.71951 | 0.64261 | 1.28810 | 231 |
| GO_ACTIVATION_OF_INNATE_IMMUNE_RESPONSE | 0.00000 | 0.00000 | 0.73376 | 0.73678 | 1.44140 | 88 |
| GO_AEROBIC_RESPIRATION | 0.00097 | 0.02209 | 0.47727 | 0.66858 | 1.29103 | 55 |
| GO_AMIDE_TRANSMEMBRANE_TRANSPORTER_ACTIVITY | 0.00006 | 0.00214 | 0.53843 | 0.78702 | 1.47958 | 30 |
| GO_AMINE_METABOLIC_PROCESS | 0.00000 | 0.00000 | | 0.78226 | 1.52920 | 86 |
| GO_ANAPHASE_PROMOTING_COMPLEX_DEPENDENT_CATA | 0.00000 | 0.00001 | 0.70498 | 0.74079 | 1.44489 | 78 |
| GO_ANIMAL_ORGAN_REGENERATION | 0.00257 | 0.04648 | 0.43171 | 0.69420 | 1.32882 | 42 |
| GO_ANTIGEN_PROCESSING_AND_PRESENTATION | 0.00000 | 0.00002 | 0.67496 | 0.66983 | 1.33112 | 152 |
| GO_ANTIGEN_PROCESSING_AND_PRESENTATION_OF_EXOG | 0.00000 | 0.00000 | 0.73376 | 0.79444 | 1.53659 | 59 |
| GO_ANTIGEN_PROCESSING_AND_PRESENTATION_OF_PEPT | 0.00000 | 0.00000 | 0.74774 | 0.69541 | 1.37873 | 131 |
| GO_ANTIGEN_PROCESSING_AND_PRESENTATION_OF_PEPT | 0.00000 | 0.00000 | 0.71951 | 0.75313 | 1.46492 | 72 |

| Cluster 1 | | | | | | |
|---|---|---|---|---|---|---|
| pathway | pval | padj | log2err | ES | NES | size |
| GO_ANTIGEN_RECEPTOR_MEDIATED_SIGNALING_PATHWAY | 0.00000 | 0.00013 | 0.62726 | 0.65842 | 1.30660 | 136 |
| GO_APOPTOTIC_SIGNALING_PATHWAY | 0.00239 | 0.04381 | 0.43171 | 0.55615 | 1.12255 | 362 |
| GO_ATPASE_ACTIVITY | 0.00006 | 0.00192 | 0.55733 | 0.58998 | 1.18798 | 303 |
| GO_ATP_METABOLIC_PROCESS | 0.00097 | 0.02209 | 0.47727 | 0.59442 | 1.18646 | 182 |
| GO_AZUROPHIL_GRANULE_LUMEN | 0.00005 | 0.00176 | 0.55733 | 0.72263 | 1.39316 | 52 |
| GO_BINDING_OF_SPERM_TO_ZONA_PELLUCIDA | 0.00002 | 0.00090 | 0.57561 | 0.86188 | 1.56970 | 19 |
| GO_BLOOD_MICROPARTICLE | 0.00054 | 0.01418 | 0.47727 | 0.71795 | 1.36611 | 38 |
| GO_CADHERIN_BINDING | 0.00000 | 0.00000 | 0.71951 | 0.64031 | 1.28498 | 246 |
| GO_CANONICAL_WNT_SIGNALING_PATHWAY | 0.00000 | 0.00006 | 0.64355 | 0.63759 | 1.27502 | 198 |
| GO_CARBOHYDRATE_CATABOLIC_PROCESS | 0.00272 | 0.04841 | 0.43171 | 0.60391 | 1.19745 | 132 |
| GO_CATALYTIC_STEP_2_SPLICEOSOME | 0.00003 | 0.00129 | 0.55733 | 0.68818 | 1.34214 | 77 |
| GO_CELLULAR_AMINO_ACID_METABOLIC_PROCESS | 0.00000 | 0.00000 | 0.74774 | 0.65298 | 1.30713 | 220 |
| GO_CELLULAR_BIOGENIC_AMINE_METABOLIC_PROCESS | 0.00121 | 0.02595 | 0.45506 | 0.74179 | 1.39662 | 31 |
| GO_CELLULAR_CARBOHYDRATE_CATABOLIC_PROCESS | 0.00098 | 0.02243 | 0.45506 | 0.74711 | 1.40298 | 29 |
| GO_CELLULAR_KETONE_METABOLIC_PROCESS | 0.00000 | 0.00012 | 0.62726 | 0.65000 | 1.29376 | 159 |
| GO_CELLULAR_METABOLIC_COMPOUND_SALVAGE | 0.00058 | 0.01519 | 0.47727 | 0.78929 | 1.46368 | 23 |
| GO_CELLULAR_NITROGEN_COMPOUND_CATABOLIC_PROCE | 0.00000 | 0.00000 | | 0.65793 | 1.33058 | 434 |
| GO_CELLULAR_RESPONSE_TO_DRUG | 0.00213 | 0.04084 | 0.43171 | 0.70335 | 1.33212 | 35 |
| GO_CELLULAR_RESPONSE_TO_FATTY_ACID | 0.00216 | 0.04129 | 0.43171 | 0.76654 | 1.42589 | 25 |
| GO_CELLULAR_RESPONSE_TO_HEAT | 0.00150 | 0.03039 | 0.45506 | 0.63721 | 1.24836 | 92 |
| GO_CELLULAR_RESPONSE_TO_OXYGEN_LEVELS | 0.00000 | 0.00011 | 0.62726 | 0.65082 | 1.29387 | 154 |
| GO_CELL_ACTIVATION_INVOLVED_IN_IMMUNE_RESPONSE | 0.00000 | 0.00015 | 0.62726 | 0.59033 | 1.19202 | 378 |
| GO_CELL_ADHESION_MOLECULE_BINDING | 0.00000 | 0.00000 | 0.74774 | 0.62511 | 1.26001 | 326 |
| GO_CELL_BODY | 0.00049 | 0.01321 | 0.47727 | 0.58370 | 1.17290 | 268 |
| GO_CELL_CELL_RECOGNITION | 0.00000 | 0.00005 | 0.65944 | 0.85660 | 1.60537 | 28 |
| GO_CELL_CELL_SIGNALING_BY_WNT | 0.00001 | 0.00029 | 0.61053 | 0.60175 | 1.21202 | 309 |
| GO_CELL_CYCLE_DNA_REPLICATION | 0.00163 | 0.03267 | 0.45506 | 0.65872 | 1.27266 | 57 |
| GO_CELL_CYCLE_G2_M_PHASE_TRANSITION | 0.00001 | 0.00033 | 0.61053 | 0.61540 | 1.23230 | 224 |
| GO_CELL_CYCLE_PHASE_TRANSITION | 0.00000 | 0.00001 | 0.69013 | 0.59126 | 1.19749 | 478 |
| GO_CELL_DIVISION | 0.00156 | 0.03130 | 0.45506 | 0.55316 | 1.11897 | 438 |
| GO_CELL_RECOGNITION | 0.00005 | 0.00167 | 0.55733 | 0.72443 | 1.39758 | 53 |
| GO_CELL_SUBSTRATE_JUNCTION | 0.00000 | 0.00003 | 0.67496 | 0.61344 | 1.23521 | 303 |
| GO_CELL_SURFACE_RECEPTOR_SIGNALING_PATHWAY_INVO | 0.00002 | 0.00080 | 0.57561 | 0.59005 | 1.18999 | 342 |
| GO_CHAPERONE_COFACTOR_DEPENDENT_PROTEIN_REFOL | 0.00124 | 0.02655 | 0.45506 | 0.76830 | 1.40754 | 21 |
| GO_CHAPERONE_COMPLEX | 0.00001 | 0.00024 | 0.61053 | 0.86187 | 1.57897 | 21 |
| GO_CHROMATIN_BINDING | 0.00002 | 0.00072 | 0.57561 | 0.58570 | 1.18252 | 377 |
| GO_COTRANSLATIONAL_PROTEIN_TARGETING_TO_MEMBR | 0.00000 | 0.00001 | 0.69013 | 0.72189 | 1.41372 | 91 |
| GO_COVALENT_CHROMATIN_MODIFICATION | 0.00205 | 0.03955 | 0.43171 | 0.56286 | 1.13527 | 346 |
| GO_CYTOKINE_MEDIATED_SIGNALING_PATHWAY | 0.00000 | 0.00000 | 0.76146 | 0.61883 | 1.24800 | 344 |
| GO_CYTOPLASMIC_TRANSLATION | 0.00000 | 0.00000 | | 0.78554 | 1.53592 | 87 |
| GO_CYTOPLASMIC_TRANSLATIONAL_INITIATION | 0.00000 | 0.00001 | 0.70498 | 0.86355 | 1.62586 | 31 |
| GO_CYTOSOLIC_LARGE_RIBOSOMAL_SUBUNIT | 0.00000 | 0.00016 | 0.62726 | 0.75329 | 1.45327 | 53 |
| GO_CYTOSOLIC_RIBOSOME | 0.00000 | 0.00000 | 0.74774 | 0.74128 | 1.45426 | 95 |
| GO_CYTOSOLIC_SMALL_RIBOSOMAL_SUBUNIT | 0.00054 | 0.01429 | 0.47727 | 0.72148 | 1.37693 | 40 |
| GO_DEFENSE_RESPONSE_TO_OTHER_ORGANISM | 0.00000 | 0.00004 | 0.65944 | 0.59253 | 1.19876 | 447 |
| GO_DE_NOVO_PROTEIN_FOLDING | 0.00104 | 0.02341 | 0.45506 | 0.75836 | 1.42127 | 28 |
| GO_DNA_BINDING_TRANSCRIPTION_FACTOR_BINDING | 0.00071 | 0.01745 | 0.47727 | 0.58733 | 1.17782 | 234 |
| GO_DNA_BIOSYNTHETIC_PROCESS | 0.00008 | 0.00251 | 0.53843 | 0.63285 | 1.25741 | 143 |
| GO_DNA_REPAIR | 0.00202 | 0.03923 | 0.43171 | 0.54895 | 1.11013 | 432 |
| GO_DOUBLE_STRANDED_RNA_BINDING | 0.00126 | 0.02674 | 0.45506 | 0.68260 | 1.31925 | 56 |
| GO_DRUG_BINDING | 0.00272 | 0.04841 | 0.43171 | 0.66450 | 1.28197 | 53 |
| GO_ENDOPEPTIDASE_COMPLEX | 0.00000 | 0.00001 | 0.70498 | 0.76400 | 1.47971 | 64 |
| GO_ESTABLISHMENT_OF_PROTEIN_LOCALIZATION_TO_END | 0.00000 | 0.00000 | 0.73376 | 0.71756 | 1.41052 | 101 |
| GO_ESTABLISHMENT_OF_PROTEIN_LOCALIZATION_TO_MEM | 0.00000 | 0.00000 | 0.76146 | 0.64607 | 1.29654 | 247 |
| GO_ESTABLISHMENT_OF_PROTEIN_LOCALIZATION_TO_MIT | 0.00187 | 0.03659 | 0.45506 | 0.71866 | 1.35886 | 34 |
| GO_ESTABLISHMENT_OF_PROTEIN_LOCALIZATION_TO_ORG | 0.00000 | 0.00000 | | 0.63449 | 1.28219 | 417 |
| GO_ESTABLISHMENT_OF_PROTEIN_LOCALIZATION_TO_TEL | 0.00111 | 0.02451 | 0.45506 | 0.81128 | 1.46428 | 16 |
| GO_ESTABLISHMENT_OF_RNA_LOCALIZATION | 0.00000 | 0.00017 | 0.62726 | 0.64309 | 1.28030 | 164 |
| GO_ESTABLISHMENT_OF_TISSUE_POLARITY | 0.00000 | 0.00000 | 0.74774 | 0.74336 | 1.45344 | 87 |
| GO_EUKARYOTIC_TRANSLATION_INITIATION_FACTOR_3_CO | 0.00001 | 0.00045 | 0.59333 | 0.90138 | 1.61984 | 15 |
| GO_FC_EPSILON_RECEPTOR_SIGNALING_PATHWAY | 0.00000 | 0.00006 | 0.64355 | 0.71457 | 1.39848 | 89 |
| GO_FC_RECEPTOR_SIGNALING_PATHWAY | 0.00000 | 0.00000 | 0.74774 | 0.69559 | 1.38022 | 137 |
| GO_FERTILIZATION | 0.00037 | 0.01039 | 0.49849 | 0.66463 | 1.29634 | 78 |
| GO_FICOLIN_1_RICH_GRANULE | 0.00000 | 0.00000 | 0.81404 | 0.75467 | 1.48129 | 96 |
| GO_GENERATION_OF_PRECURSOR_METABOLITES_AND_EN | 0.00042 | 0.01155 | 0.49849 | 0.57758 | 1.16334 | 309 |
| GO_GLYCOSYL_COMPOUND_METABOLIC_PROCESS | 0.00080 | 0.01930 | 0.47727 | 0.66493 | 1.29256 | 71 |
| GO_GTP_METABOLIC_PROCESS | 0.00086 | 0.02012 | 0.47727 | 0.81830 | 1.47695 | 16 |
| GO_HEAT_SHOCK_PROTEIN_BINDING | 0.00047 | 0.01277 | 0.49849 | 0.64908 | 1.27056 | 90 |
| GO_HEMATOPOIETIC_PROGENITOR_CELL_DIFFERENTIATION | 0.00000 | 0.00013 | 0.62726 | 0.67801 | 1.33914 | 111 |
| GO_HEMATOPOIETIC_STEM_CELL_DIFFERENTIATION | 0.00000 | 0.00000 | 0.74774 | 0.77285 | 1.50051 | 69 |
| GO_IMMUNE_RESPONSE_REGULATING_SIGNALING_PATHW | 0.00000 | 0.00005 | 0.65944 | 0.63365 | 1.26812 | 214 |

| Cluster 1 | | | | | | |
|---|---|---|---|---|---|---|
| pathway | pval | padj | log2err | ES | NES | size |
| GO_INNATE_IMMUNE_RESPONSE | 0.00000 | 0.00003 | 0.67496 | 0.60436 | 1.21981 | 365 |
| GO_INNATE_IMMUNE_RESPONSE_ACTIVATING_SIGNAL_TR/ | 0.00000 | 0.00000 | 0.73376 | 0.75634 | 1.47116 | 72 |
| GO_INTERACTION_WITH_SYMBIONT | 0.00242 | 0.04435 | 0.43171 | 0.68676 | 1.31850 | 47 |
| GO_INTERLEUKIN_1_MEDIATED_SIGNALING_PATHWAY | 0.00000 | 0.00000 | 0.76146 | 0.75993 | 1.48118 | 76 |
| GO_ION_CHANNEL_BINDING | 0.00001 | 0.00024 | 0.61053 | 0.72962 | 1.41146 | 60 |
| GO_LARGE_RIBOSOMAL_SUBUNIT | 0.00004 | 0.00145 | 0.55733 | 0.66585 | 1.30887 | 101 |
| GO_LEUKOCYTE_MEDIATED_IMMUNITY | 0.00001 | 0.00027 | 0.61053 | 0.58907 | 1.19020 | 403 |
| GO_LIVER_REGENERATION | 0.00022 | 0.00673 | 0.51885 | 0.82386 | 1.50045 | 19 |
| GO_LYSOSOMAL_LUMEN | 0.00246 | 0.04475 | 0.43171 | 0.67531 | 1.30248 | 51 |
| GO_MATURATION_OF_LSU_RRNA | 0.00172 | 0.03416 | 0.45506 | 0.75523 | 1.41143 | 27 |
| GO_MODIFICATION_DEPENDENT_MACROMOLECULE_CATAI | 0.00000 | 0.00003 | 0.65944 | 0.58395 | 1.18375 | 498 |
| GO_MODULATION_BY_HOST_OF_SYMBIONT_PROCESS | 0.00115 | 0.02503 | 0.45506 | 0.70365 | 1.33891 | 38 |
| GO_MORPHOGENESIS_OF_AN_EPITHELIUM | 0.00137 | 0.02827 | 0.45506 | 0.57028 | 1.14856 | 306 |
| GO_MORPHOGENESIS_OF_A_POLARIZED_EPITHELIUM | 0.00000 | 0.00000 | 0.73376 | 0.71677 | 1.41002 | 102 |
| GO_MRNA_3_END_PROCESSING | 0.00124 | 0.02655 | 0.45506 | 0.63849 | 1.24911 | 88 |
| GO_MRNA_3_UTR_BINDING | 0.00183 | 0.03599 | 0.45506 | 0.65619 | 1.27401 | 69 |
| GO_MRNA_BINDING | 0.00000 | 0.00000 | 0.78819 | 0.65825 | 1.31945 | 231 |
| GO_MRNA_EXPORT_FROM_NUCLEUS | 0.00065 | 0.01635 | 0.47727 | 0.63899 | 1.25534 | 98 |
| GO_MRNA_PROCESSING | 0.00000 | 0.00000 | | 0.62833 | 1.26962 | 414 |
| GO_MRNA_TRANSPORT | 0.00050 | 0.01339 | 0.47727 | 0.62333 | 1.23507 | 123 |
| GO_MYELOID_LEUKOCYTE_ACTIVATION | 0.00000 | 0.00002 | 0.67496 | 0.60760 | 1.22612 | 359 |
| GO_MYELOID_LEUKOCYTE_MEDIATED_IMMUNITY | 0.00000 | 0.00000 | 0.71951 | 0.61777 | 1.24444 | 321 |
| GO_NCRNA_METABOLIC_PROCESS | 0.00000 | 0.00020 | 0.61053 | 0.59165 | 1.19466 | 390 |
| GO_NCRNA_PROCESSING | 0.00000 | 0.00004 | 0.65944 | 0.60653 | 1.22263 | 330 |
| GO_NEGATIVE_REGULATION_OF_CALCIUM_MEDIATED_SIGI | 0.00104 | 0.02341 | 0.45506 | 0.80508 | 1.46307 | 18 |
| GO_NEGATIVE_REGULATION_OF_CANONICAL_WNT_SIGNAI | 0.00000 | 0.00001 | 0.70498 | 0.69921 | 1.37979 | 110 |
| GO_NEGATIVE_REGULATION_OF_CELLULAR_AMIDE_METAB | 0.00104 | 0.02341 | 0.45506 | 0.61630 | 1.22114 | 123 |
| GO_NEGATIVE_REGULATION_OF_CELL_CYCLE | 0.00008 | 0.00251 | 0.53843 | 0.56994 | 1.15290 | 438 |
| GO_NEGATIVE_REGULATION_OF_CELL_CYCLE_G2_M_PHASI | 0.00000 | 0.00010 | 0.62726 | 0.70368 | 1.37664 | 88 |
| GO_NEGATIVE_REGULATION_OF_CELL_CYCLE_PHASE_TRAN | 0.00002 | 0.00087 | 0.57561 | 0.62130 | 1.24126 | 191 |
| GO_NEGATIVE_REGULATION_OF_CELL_CYCLE_PROCESS | 0.00002 | 0.00078 | 0.57561 | 0.60102 | 1.20696 | 259 |
| GO_NEGATIVE_REGULATION_OF_MITOTIC_CELL_CYCLE | 0.00036 | 0.01028 | 0.49849 | 0.58943 | 1.18207 | 238 |
| GO_NEGATIVE_REGULATION_OF_NEURON_DEATH | 0.00110 | 0.02437 | 0.45506 | 0.62736 | 1.23415 | 102 |
| GO_NEGATIVE_REGULATION_OF_RNA_CATABOLIC_PROCESS | 0.00239 | 0.04381 | 0.43171 | 0.69668 | 1.33355 | 42 |
| GO_NEGATIVE_REGULATION_OF_WNT_SIGNALING_PATHW/ | 0.00000 | 0.00000 | 0.74774 | 0.70209 | 1.39034 | 121 |
| GO_NEURON_DEATH_IN_RESPONSE_TO_OXIDATIVE_STRESS | 0.00095 | 0.02193 | 0.47727 | 0.79874 | 1.45470 | 19 |
| GO_NIK_NF_KAPPAB_SIGNALING | 0.00000 | 0.00000 | 0.77494 | 0.72553 | 1.43123 | 109 |
| GO_NON_CANONICAL_WNT_SIGNALING_PATHWAY | 0.00000 | 0.00000 | 0.74774 | 0.72514 | 1.42649 | 102 |
| GO_NUCLEAR_EXPORT | 0.00010 | 0.00304 | 0.53843 | 0.62047 | 1.23703 | 174 |
| GO_NUCLEAR_SPECK | 0.00003 | 0.00118 | 0.55733 | 0.59344 | 1.19511 | 307 |
| GO_NUCLEAR_TRANSCRIBED_MRNA_CATABOLIC_PROCESS | 0.00000 | 0.00001 | 0.70498 | 0.66218 | 1.32090 | 180 |
| GO_NUCLEAR_TRANSCRIBED_MRNA_CATABOLIC_PROCESS_ | 0.00000 | 0.00000 | 0.76146 | 0.71725 | 1.41665 | 111 |
| GO_NUCLEAR_TRANSPORT | 0.00000 | 0.00010 | 0.64355 | 0.61347 | 1.23357 | 275 |
| GO_NUCLEOBASE_CONTAINING_COMPOUND_TRANSPORT | 0.00002 | 0.00091 | 0.57561 | 0.61661 | 1.23362 | 208 |
| GO_NUCLEOBASE_CONTAINING_SMALL_MOLECULE_METAE | 0.00052 | 0.01376 | 0.47727 | 0.56464 | 1.13945 | 358 |
| GO_NUCLEOSIDE_METABOLIC_PROCESS | 0.00062 | 0.01600 | 0.47727 | 0.68076 | 1.31605 | 58 |
| GO_NUCLEOSIDE_MONOPHOSPHATE_BIOSYNTHETIC_PROC | 0.00082 | 0.01950 | 0.47727 | 0.74911 | 1.41039 | 31 |
| GO_NUCLEOSIDE_PHOSPHATE_BIOSYNTHETIC_PROCESS | 0.00070 | 0.01730 | 0.47727 | 0.61492 | 1.22178 | 143 |
| GO_ORGANIC_CYCLIC_COMPOUND_CATABOLIC_PROCESS | 0.00000 | 0.00000 | | 0.65287 | 1.32085 | 446 |
| GO_PEPTIDASE_COMPLEX | 0.00000 | 0.00002 | 0.67496 | 0.72864 | 1.42135 | 79 |
| GO_PEPTIDE_TRANSMEMBRANE_TRANSPORTER_ACTIVITY | 0.00039 | 0.01098 | 0.49849 | 0.81514 | 1.48456 | 19 |
| GO_PEPTIDYL_LYSINE_MODIFICATION | 0.00111 | 0.02451 | 0.45506 | 0.57211 | 1.15032 | 280 |
| GO_PIGMENT_GRANULE | 0.00108 | 0.02406 | 0.45506 | 0.64966 | 1.26714 | 78 |
| GO_POLYSOMAL_RIBOSOME | 0.00224 | 0.04228 | 0.43171 | 0.73171 | 1.37405 | 29 |
| GO_POLYSOME | 0.00003 | 0.00115 | 0.57561 | 0.71903 | 1.39074 | 59 |
| GO_POLY_PYRIMIDINE_TRACT_BINDING | 0.00083 | 0.01956 | 0.47727 | 0.81817 | 1.48329 | 17 |
| GO_POSITIVE_REGULATION_OF_CANONICAL_WNT_SIGNAL | 0.00000 | 0.00001 | 0.70498 | 0.71470 | 1.40718 | 105 |
| GO_POSITIVE_REGULATION_OF_CELLULAR_AMIDE_METABO | 0.00009 | 0.00275 | 0.53843 | 0.64739 | 1.28069 | 116 |
| GO_POSITIVE_REGULATION_OF_CELLULAR_PROTEIN_LOCAI | 0.00001 | 0.00033 | 0.61053 | 0.61294 | 1.22870 | 232 |
| GO_POSITIVE_REGULATION_OF_CELL_CYCLE | 0.00202 | 0.03923 | 0.43171 | 0.57158 | 1.14811 | 261 |
| GO_POSITIVE_REGULATION_OF_CELL_CYCLE_PROCESS | 0.00235 | 0.04368 | 0.43171 | 0.58251 | 1.16502 | 200 |
| GO_POSITIVE_REGULATION_OF_CHROMOSOME_ORGANIZA | 0.00081 | 0.01936 | 0.47727 | 0.60897 | 1.20865 | 138 |
| GO_POSITIVE_REGULATION_OF_DEFENSE_RESPONSE | 0.00000 | 0.00013 | 0.62726 | 0.64057 | 1.27660 | 170 |
| GO_POSITIVE_REGULATION_OF_DNA_BIOSYNTHETIC_PROC | 0.00009 | 0.00288 | 0.53843 | 0.71532 | 1.37906 | 52 |
| GO_POSITIVE_REGULATION_OF_DNA_METABOLIC_PROCESS | 0.00065 | 0.01635 | 0.47727 | 0.62088 | 1.23109 | 132 |
| GO_POSITIVE_REGULATION_OF_ESTABLISHMENT_OF_PROT | 0.00001 | 0.00059 | 0.59333 | 0.61832 | 1.23780 | 221 |
| GO_POSITIVE_REGULATION_OF_ESTABLISHMENT_OF_PROT | 0.00235 | 0.04368 | 0.43171 | 0.69343 | 1.31946 | 38 |
| GO_POSITIVE_REGULATION_OF_IMMUNE_RESPONSE | 0.00000 | 0.00020 | 0.61053 | 0.60424 | 1.21636 | 297 |
| GO_POSITIVE_REGULATION_OF_IMMUNE_SYSTEM_PROCES | 0.00013 | 0.00397 | 0.51885 | 0.57295 | 1.15785 | 424 |
| GO_POSITIVE_REGULATION_OF_INNATE_IMMUNE_RESPON | 0.00000 | 0.00010 | 0.64355 | 0.68424 | 1.35219 | 113 |
| GO_POSITIVE_REGULATION_OF_INTRACELLULAR_PROTEIN_ | 0.00146 | 0.02985 | 0.45506 | 0.61795 | 1.22466 | 128 |

| **Cluster 1** | | | | | | |
|---|---|---|---|---|---|---|
| pathway | pval | padj | log2err | ES | NES | size |
| GO_POSITIVE_REGULATION_OF_INTRACELLULAR_TRANSPO | 0.00080 | 0.01930 | 0.47727 | 0.60690 | 1.20684 | 155 |
| GO_POSITIVE_REGULATION_OF_MRNA_METABOLIC_PROCE | 0.00205 | 0.03955 | 0.43171 | 0.65160 | 1.26398 | 67 |
| GO_POSITIVE_REGULATION_OF_MRNA_PROCESSING | 0.00150 | 0.03039 | 0.45506 | 0.75894 | 1.41836 | 27 |
| GO_POSITIVE_REGULATION_OF_MRNA_SPLICING_VIA_SPLI | 0.00018 | 0.00549 | 0.51885 | 0.84859 | 1.53844 | 17 |
| GO_POSITIVE_REGULATION_OF_ORGANELLE_ORGANIZATIO | 0.00054 | 0.01429 | 0.47727 | 0.56093 | 1.13356 | 420 |
| GO_POSITIVE_REGULATION_OF_PROTEIN_LOCALIZATION_T | 0.00001 | 0.00031 | 0.61053 | 0.72104 | 1.39651 | 64 |
| GO_POSITIVE_REGULATION_OF_RESPONSE_TO_BIOTIC_STI | 0.00000 | 0.00006 | 0.64355 | 0.66877 | 1.32721 | 134 |
| GO_POSITIVE_REGULATION_OF_RESPONSE_TO_EXTERNAL_ | 0.00047 | 0.01277 | 0.49849 | 0.59234 | 1.18576 | 217 |
| GO_POSITIVE_REGULATION_OF_RNA_SPLICING | 0.00097 | 0.02209 | 0.47727 | 0.75944 | 1.42328 | 28 |
| GO_POSITIVE_REGULATION_OF_TRANSLATION | 0.00006 | 0.00216 | 0.53843 | 0.66851 | 1.31218 | 96 |
| GO_POSITIVE_REGULATION_OF_VIRAL_PROCESS | 0.00137 | 0.02827 | 0.45506 | 0.64873 | 1.26631 | 81 |
| GO_POSITIVE_REGULATION_OF_WNT_SIGNALING_PATHWA | 0.00000 | 0.00000 | 0.71951 | 0.69846 | 1.38380 | 122 |
| GO_POSTSYNAPTIC_SPECIALIZATION_MEMBRANE | 0.00027 | 0.00800 | 0.49849 | 0.76520 | 1.44069 | 31 |
| GO_POSTTRANSCRIPTIONAL_REGULATION_OF_GENE_EXPR | 0.00000 | 0.00000 | | 0.66322 | 1.34232 | 469 |
| GO_POST_TRANSLATIONAL_PROTEIN_MODIFICATION | 0.00001 | 0.00046 | 0.59333 | 0.61196 | 1.22810 | 246 |
| GO_PRERIBOSOME | 0.00000 | 0.00006 | 0.64355 | 0.73360 | 1.42371 | 68 |
| GO_PRE_MRNA_BINDING | 0.00119 | 0.02565 | 0.45506 | 0.75466 | 1.41433 | 28 |
| GO_PROTEASOMAL_PROTEIN_CATABOLIC_PROCESS | 0.00000 | 0.00004 | 0.65944 | 0.60087 | 1.21333 | 385 |
| GO_PROTEASOMAL_UBIQUITIN_INDEPENDENT_PROTEIN_C | 0.00000 | 0.00021 | 0.61053 | 0.86844 | 1.58882 | 20 |
| GO_PROTEASOME_ACCESSORY_COMPLEX | 0.00001 | 0.00055 | 0.59333 | 0.84367 | 1.56543 | 24 |
| GO_PROTEASOME_CORE_COMPLEX | 0.00002 | 0.00085 | 0.57561 | 0.86976 | 1.58061 | 18 |
| GO_PROTEIN_CONTAINING_COMPLEX_LOCALIZATION | 0.00031 | 0.00904 | 0.49849 | 0.59941 | 1.19895 | 206 |
| GO_PROTEIN_DNA_COMPLEX_SUBUNIT_ORGANIZATION | 0.00014 | 0.00433 | 0.51885 | 0.61787 | 1.23167 | 173 |
| GO_PROTEIN_DOMAIN_SPECIFIC_BINDING | 0.00156 | 0.03130 | 0.45506 | 0.55469 | 1.12181 | 435 |
| GO_PROTEIN_FOLDING | 0.00000 | 0.00005 | 0.65944 | 0.64920 | 1.29300 | 166 |
| GO_PROTEIN_HETERODIMERIZATION_ACTIVITY | 0.00261 | 0.04701 | 0.43171 | 0.59709 | 1.18606 | 142 |
| GO_PROTEIN_IMPORT | 0.00089 | 0.02090 | 0.47727 | 0.61885 | 1.22816 | 139 |
| GO_PROTEIN_INSERTION_INTO_MEMBRANE | 0.00065 | 0.01635 | 0.47727 | 0.71705 | 1.36603 | 39 |
| GO_PROTEIN_LOCALIZATION_TO_CHROMOSOME | 0.00052 | 0.01376 | 0.47727 | 0.67222 | 1.30567 | 70 |
| GO_PROTEIN_LOCALIZATION_TO_ENDOPLASMIC_RETICULU | 0.00000 | 0.00001 | 0.70498 | 0.69150 | 1.36990 | 124 |
| GO_PROTEIN_LOCALIZATION_TO_MEMBRANE | 0.00008 | 0.00254 | 0.53843 | 0.57356 | 1.15901 | 419 |
| GO_PROTEIN_LOCALIZATION_TO_MITOCHONDRION | 0.00005 | 0.00167 | 0.55733 | 0.66343 | 1.30412 | 101 |
| GO_PROTEIN_LOCALIZATION_TO_NUCLEUS | 0.00002 | 0.00094 | 0.57561 | 0.62258 | 1.24368 | 190 |
| GO_PROTEIN_MODIFICATION_BY_SMALL_PROTEIN_REMOV | 0.00033 | 0.00951 | 0.49849 | 0.59687 | 1.19497 | 222 |
| GO_PROTEIN_POLYUBIQUITINATION | 0.00001 | 0.00035 | 0.59333 | 0.61007 | 1.22467 | 255 |
| GO_PROTEIN_STABILIZATION | 0.00092 | 0.02137 | 0.47727 | 0.61801 | 1.22650 | 139 |
| GO_PROTEIN_TARGETING | 0.00000 | 0.00000 | 0.76146 | 0.62633 | 1.26214 | 323 |
| GO_PROTEIN_TARGETING_TO_MEMBRANE | 0.00000 | 0.00001 | 0.69013 | 0.66459 | 1.32125 | 154 |
| GO_PROTEIN_TRANSMEMBRANE_TRANSPORT | 0.00113 | 0.02473 | 0.45506 | 0.69348 | 1.33123 | 46 |
| GO_PURINE_CONTAINING_COMPOUND_BIOSYNTHETIC_PR | 0.00128 | 0.02694 | 0.45506 | 0.62451 | 1.23239 | 110 |
| GO_PURINE_CONTAINING_COMPOUND_METABOLIC_PROC | 0.00013 | 0.00407 | 0.51885 | 0.59078 | 1.18669 | 261 |
| GO_PURINE_NUCLEOSIDE_METABOLIC_PROCESS | 0.00111 | 0.02451 | 0.45506 | 0.71754 | 1.36194 | 36 |
| GO_PURINE_NUCLEOSIDE_MONOPHOSPHATE_BIOSYNTHET | 0.00113 | 0.02473 | 0.45506 | 0.80982 | 1.46815 | 17 |
| GO_REGENERATION | 0.00003 | 0.00101 | 0.57561 | 0.67073 | 1.31714 | 97 |
| GO_REGULATION_OF_ANIMAL_ORGAN_MORPHOGENESIS | 0.00000 | 0.00000 | 0.74774 | 0.69661 | 1.38110 | 131 |
| GO_REGULATION_OF_APOPTOTIC_SIGNALING_PATHWAY | 0.00265 | 0.04753 | 0.43171 | 0.57481 | 1.15275 | 238 |
| GO_REGULATION_OF_BINDING | 0.00047 | 0.01277 | 0.49849 | 0.59238 | 1.18582 | 217 |
| GO_REGULATION_OF_CELLULAR_AMIDE_METABOLIC_PROC | 0.00000 | 0.00000 | 0.78819 | 0.63199 | 1.27310 | 321 |
| GO_REGULATION_OF_CELLULAR_AMINE_METABOLIC_PROC | 0.00000 | 0.00000 | 0.81404 | 0.81494 | 1.57545 | 58 |
| GO_REGULATION_OF_CELLULAR_AMINO_ACID_METABOLIC | 0.00000 | 0.00000 | 0.81404 | 0.82856 | 1.60023 | 54 |
| GO_REGULATION_OF_CELLULAR_KETONE_METABOLIC_PRO | 0.00000 | 0.00001 | 0.70498 | 0.69208 | 1.37114 | 122 |
| GO_REGULATION_OF_CELLULAR_PROTEIN_CATABOLIC_PRO | 0.00064 | 0.01635 | 0.47727 | 0.59675 | 1.19258 | 193 |
| GO_REGULATION_OF_CELLULAR_PROTEIN_LOCALIZATION | 0.00002 | 0.00096 | 0.57561 | 0.58593 | 1.18287 | 375 |
| GO_REGULATION_OF_CELLULAR_RESPONSE_TO_STRESS | 0.00015 | 0.00457 | 0.51885 | 0.56426 | 1.14215 | 471 |
| GO_REGULATION_OF_CELL_CYCLE_G2_M_PHASE_TRANSITI | 0.00000 | 0.00003 | 0.67496 | 0.64816 | 1.29372 | 182 |
| GO_REGULATION_OF_CELL_CYCLE_PHASE_TRANSITION | 0.00000 | 0.00005 | 0.65944 | 0.60012 | 1.21054 | 353 |
| GO_REGULATION_OF_CHROMOSOME_ORGANIZATION | 0.00027 | 0.00800 | 0.49849 | 0.58780 | 1.18118 | 266 |
| GO_REGULATION_OF_DEFENSE_RESPONSE | 0.00033 | 0.00948 | 0.49849 | 0.58101 | 1.17034 | 313 |
| GO_REGULATION_OF_DNA_BIOSYNTHETIC_PROCESS | 0.00007 | 0.00230 | 0.53843 | 0.68161 | 1.32961 | 79 |
| GO_REGULATION_OF_DNA_METABOLIC_PROCESS | 0.00010 | 0.00310 | 0.53843 | 0.59973 | 1.20273 | 238 |
| GO_REGULATION_OF_DNA_TEMPLATED_TRANSCRIPTION_I | 0.00000 | 0.00001 | 0.69013 | 0.71763 | 1.40693 | 93 |
| GO_REGULATION_OF_ESTABLISHMENT_OF_PROTEIN_LOCA | 0.00077 | 0.01887 | 0.47727 | 0.68889 | 1.32378 | 48 |
| GO_REGULATION_OF_HEMATOPOIETIC_PROGENITOR_CELL | 0.00000 | 0.00000 | 0.74774 | 0.76099 | 1.48020 | 72 |
| GO_REGULATION_OF_HEMATOPOIETIC_STEM_CELL_DIFFER | 0.00000 | 0.00000 | 0.74774 | 0.78009 | 1.51011 | 63 |
| GO_REGULATION_OF_HEMOPOIESIS | 0.00005 | 0.00168 | 0.55733 | 0.60424 | 1.21259 | 246 |
| GO_REGULATION_OF_IMMUNE_RESPONSE | 0.00143 | 0.02931 | 0.45506 | 0.56149 | 1.13420 | 395 |
| GO_REGULATION_OF_INNATE_IMMUNE_RESPONSE | 0.00000 | 0.00012 | 0.62726 | 0.64456 | 1.28381 | 167 |
| GO_REGULATION_OF_INTRINSIC_APOPTOTIC_SIGNALING_P | 0.00066 | 0.01667 | 0.47727 | 0.63316 | 1.24808 | 107 |
| GO_REGULATION_OF_MEMBRANE_DEPOLARIZATION | 0.00083 | 0.01956 | 0.47727 | 0.77677 | 1.42306 | 21 |
| GO_REGULATION_OF_MITOTIC_CELL_CYCLE | 0.00004 | 0.00133 | 0.55733 | 0.57226 | 1.15853 | 473 |
| GO_REGULATION_OF_MORPHOGENESIS_OF_AN_EPITHELIU | 0.00000 | 0.00000 | 0.73376 | 0.71600 | 1.41213 | 108 |

| Cluster 1 | | | | | | |
|---|---|---|---|---|---|---|
| pathway | pval | padj | log2err | ES | NES | size |
| GO_REGULATION_OF_MRNA_CATABOLIC_PROCESS | 0.00000 | 0.00000 | 0.78819 | 0.68893 | 1.37175 | 165 |
| GO_REGULATION_OF_MRNA_METABOLIC_PROCESS | 0.00000 | 0.00000 | | 0.66571 | 1.33768 | 268 |
| GO_REGULATION_OF_MRNA_PROCESSING | 0.00014 | 0.00438 | 0.51885 | 0.64799 | 1.28186 | 115 |
| GO_REGULATION_OF_MRNA_SPLICING_VIA_SPLICEOSOME | 0.00008 | 0.00256 | 0.53843 | 0.67512 | 1.31858 | 83 |
| GO_REGULATION_OF_NUCLEOTIDE_METABOLIC_PROCESS | 0.00176 | 0.03477 | 0.45506 | 0.64117 | 1.24913 | 75 |
| GO_REGULATION_OF_PROTEASOMAL_PROTEIN_CATABOLIC | 0.00078 | 0.01901 | 0.47727 | 0.61115 | 1.21471 | 148 |
| GO_REGULATION_OF_PROTEASOMAL_UBIQUITIN_DEPENDI | 0.00137 | 0.02827 | 0.45506 | 0.62545 | 1.22945 | 101 |
| GO_REGULATION_OF_PROTEIN_CATABOLIC_PROCESS | 0.00004 | 0.00136 | 0.55733 | 0.59781 | 1.20208 | 275 |
| GO_REGULATION_OF_PROTEIN_IMPORT | 0.00216 | 0.04129 | 0.43171 | 0.69929 | 1.33855 | 42 |
| GO_REGULATION_OF_PROTEIN_LOCALIZATION_TO_NUCLEU | 0.00001 | 0.00033 | 0.61053 | 0.69325 | 1.35701 | 90 |
| GO_REGULATION_OF_PROTEIN_MODIFICATION_BY_SMALL_ | 0.00126 | 0.02674 | 0.45506 | 0.59811 | 1.19246 | 174 |
| GO_REGULATION_OF_PROTEIN_STABILITY | 0.00091 | 0.02125 | 0.47727 | 0.58880 | 1.17861 | 216 |
| GO_REGULATION_OF_PROTEOLYSIS | 0.00106 | 0.02373 | 0.45506 | 0.56010 | 1.13168 | 405 |
| GO_REGULATION_OF_RESPONSE_TO_BIOTIC_STIMULUS | 0.00003 | 0.00124 | 0.55733 | 0.61118 | 1.22362 | 218 |
| GO_REGULATION_OF_RNA_SPLICING | 0.00001 | 0.00029 | 0.61053 | 0.66702 | 1.31952 | 116 |
| GO_REGULATION_OF_SMALL_MOLECULE_METABOLIC_PRO | 0.00000 | 0.00000 | 0.71951 | 0.63263 | 1.27200 | 276 |
| GO_REGULATION_OF_STEM_CELL_DIFFERENTIATION | 0.00000 | 0.00000 | 0.77494 | 0.74561 | 1.45866 | 88 |
| GO_REGULATION_OF_SYMBIOTIC_PROCESS | 0.00023 | 0.00676 | 0.51885 | 0.62862 | 1.24644 | 132 |
| GO_REGULATION_OF_TELOMERE_MAINTENANCE | 0.00235 | 0.04368 | 0.43171 | 0.64332 | 1.24851 | 68 |
| GO_REGULATION_OF_TRANSCRIPTION_FROM_RNA_POLYM | 0.00000 | 0.00000 | 0.71951 | 0.76721 | 1.48618 | 65 |
| GO_REGULATION_OF_TRANSLATIONAL_INITIATION | 0.00015 | 0.00461 | 0.51885 | 0.69482 | 1.34597 | 65 |
| GO_REGULATION_OF_UBIQUITIN_PROTEIN_LIGASE_ACTIVIT | 0.00246 | 0.04475 | 0.43171 | 0.76772 | 1.40455 | 20 |
| GO_REGULATION_OF_UBIQUITIN_PROTEIN_TRANSFERASE_ | 0.00272 | 0.04841 | 0.43171 | 0.67626 | 1.29758 | 45 |
| GO_REGULATION_OF_WNT_SIGNALING_PATHWAY | 0.00000 | 0.00000 | 0.74774 | 0.65137 | 1.30392 | 217 |
| GO_RESPONSE_TO_DRUG | 0.00220 | 0.04186 | 0.43171 | 0.58700 | 1.17309 | 193 |
| GO_RESPONSE_TO_FATTY_ACID | 0.00070 | 0.01730 | 0.47727 | 0.71371 | 1.35804 | 38 |
| GO_RESPONSE_TO_HEAT | 0.00065 | 0.01651 | 0.47727 | 0.62973 | 1.24379 | 111 |
| GO_RESPONSE_TO_INTERLEUKIN_1 | 0.00000 | 0.00000 | 0.74774 | 0.71829 | 1.41949 | 113 |
| GO_RESPONSE_TO_INTERLEUKIN_12 | 0.00001 | 0.00033 | 0.61053 | 0.80825 | 1.52565 | 33 |
| GO_RESPONSE_TO_INTERLEUKIN_4 | 0.00037 | 0.01039 | 0.49849 | 0.79418 | 1.46471 | 22 |
| GO_RESPONSE_TO_LEUKEMIA_INHIBITORY_FACTOR | 0.00205 | 0.03955 | 0.43171 | 0.64197 | 1.25125 | 76 |
| GO_RESPONSE_TO_OXYGEN_LEVELS | 0.00027 | 0.00800 | 0.49849 | 0.59179 | 1.18650 | 236 |
| GO_RESPONSE_TO_TEMPERATURE_STIMULUS | 0.00132 | 0.02752 | 0.45506 | 0.61402 | 1.21857 | 139 |
| GO_RESPONSE_TO_TUMOR_NECROSIS_FACTOR | 0.00001 | 0.00035 | 0.59333 | 0.63780 | 1.26964 | 163 |
| GO_RIBONUCLEOPROTEIN_COMPLEX_BINDING | 0.00000 | 0.00001 | 0.70498 | 0.69959 | 1.38556 | 119 |
| GO_RIBONUCLEOPROTEIN_COMPLEX_BIOGENESIS | 0.00000 | 0.00000 | | 0.68651 | 1.38632 | 383 |
| GO_RIBONUCLEOPROTEIN_COMPLEX_SUBUNIT_ORGANIZAT | 0.00000 | 0.00000 | | 0.70573 | 1.40672 | 172 |
| GO_RIBONUCLEOPROTEIN_GRANULE | 0.00006 | 0.00192 | 0.55733 | 0.62081 | 1.23722 | 170 |
| GO_RIBONUCLEOSIDE_METABOLIC_PROCESS | 0.00043 | 0.01175 | 0.49849 | 0.72264 | 1.37670 | 39 |
| GO_RIBONUCLEOSIDE_MONOPHOSPHATE_BIOSYNTHETIC_I | 0.00257 | 0.04648 | 0.43171 | 0.76234 | 1.41807 | 25 |
| GO_RIBOSE_PHOSPHATE_BIOSYNTHETIC_PROCESS | 0.00228 | 0.04285 | 0.43171 | 0.61778 | 1.21841 | 108 |
| GO_RIBOSE_PHOSPHATE_METABOLIC_PROCESS | 0.00009 | 0.00275 | 0.53843 | 0.59506 | 1.19420 | 248 |
| GO_RIBOSOMAL_LARGE_SUBUNIT_ASSEMBLY | 0.00046 | 0.01270 | 0.49849 | 0.77370 | 1.45001 | 28 |
| GO_RIBOSOMAL_LARGE_SUBUNIT_BIOGENESIS | 0.00000 | 0.00001 | 0.71951 | 0.75758 | 1.46976 | 66 |
| GO_RIBOSOMAL_SMALL_SUBUNIT_BIOGENESIS | 0.00069 | 0.01721 | 0.47727 | 0.66753 | 1.29603 | 69 |
| GO_RIBOSOMAL_SUBUNIT | 0.00000 | 0.00001 | 0.69013 | 0.66873 | 1.33144 | 160 |
| GO_RIBOSOME | 0.00000 | 0.00000 | 0.74774 | 0.66836 | 1.33512 | 190 |
| GO_RIBOSOME_ASSEMBLY | 0.00001 | 0.00052 | 0.59333 | 0.73154 | 1.41598 | 61 |
| GO_RIBOSOME_BINDING | 0.00020 | 0.00615 | 0.51885 | 0.70768 | 1.36373 | 50 |
| GO_RIBOSOME_BIOGENESIS | 0.00000 | 0.00000 | | 0.67204 | 1.35097 | 272 |
| GO_RNA_CATABOLIC_PROCESS | 0.00000 | 0.00000 | | 0.67697 | 1.36467 | 333 |
| GO_RNA_DEPENDENT_DNA_BIOSYNTHETIC_PROCESS | 0.00062 | 0.01600 | 0.47727 | 0.67359 | 1.30139 | 57 |
| GO_RNA_EXPORT_FROM_NUCLEUS | 0.00004 | 0.00152 | 0.55733 | 0.64967 | 1.28704 | 124 |
| GO_RNA_LOCALIZATION | 0.00000 | 0.00000 | 0.77494 | 0.66923 | 1.33767 | 194 |
| GO_RNA_SPLICING | 0.00000 | 0.00000 | | 0.63768 | 1.28683 | 359 |
| GO_RNA_SPLICING_VIA_TRANSESTERIFICATION_REACTIONS | 0.00000 | 0.00000 | 0.82666 | 0.64971 | 1.30757 | 291 |
| GO_RNA_STABILIZATION | 0.00067 | 0.01683 | 0.47727 | 0.73152 | 1.39075 | 37 |
| GO_RRNA_BINDING | 0.00007 | 0.00239 | 0.53843 | 0.71768 | 1.38456 | 53 |
| GO_RRNA_METABOLIC_PROCESS | 0.00000 | 0.00000 | 0.73376 | 0.65761 | 1.31539 | 205 |
| GO_SCF_DEPENDENT_PROTEASOMAL_UBIQUITIN_DEPEND | 0.00000 | 0.00000 | 0.76146 | 0.75796 | 1.47838 | 78 |
| GO_SECRETORY_GRANULE | 0.00000 | 0.00002 | 0.67496 | 0.59177 | 1.19707 | 439 |
| GO_SIGNAL_TRANSDUCTION_BY_PROTEIN_PHOSPHORYLAT | 0.00181 | 0.03575 | 0.45506 | 0.55235 | 1.11781 | 463 |
| GO_SINGLE_FERTILIZATION | 0.00004 | 0.00148 | 0.55733 | 0.70277 | 1.36047 | 62 |
| GO_SINGLE_STRANDED_RNA_BINDING | 0.00033 | 0.00951 | 0.49849 | 0.68972 | 1.33336 | 58 |
| GO_SMALL_MOLECULE_BIOSYNTHETIC_PROCESS | 0.00119 | 0.02565 | 0.45506 | 0.55826 | 1.12799 | 404 |
| GO_SMALL_NUCLEAR_RIBONUCLEOPROTEIN_COMPLEX | 0.00132 | 0.02752 | 0.45506 | 0.67701 | 1.30611 | 53 |
| GO_SMALL_RIBOSOMAL_SUBUNIT | 0.00037 | 0.01039 | 0.49849 | 0.67618 | 1.30899 | 62 |
| GO_SMALL_SUBUNIT_PROCESSOME | 0.00065 | 0.01635 | 0.47727 | 0.75148 | 1.41485 | 31 |
| GO_SM_LIKE_PROTEIN_FAMILY_COMPLEX | 0.00072 | 0.01760 | 0.47727 | 0.67164 | 1.30082 | 64 |
| GO_SNORNA_BINDING | 0.00034 | 0.00973 | 0.49849 | 0.78313 | 1.45798 | 26 |

| **Cluster 1** | | | | | | |
|---|---|---|---|---|---|---|
| pathway | pval | padj | log2err | ES | NES | size |
| GO_SODIUM_ION_TRANSMEMBRANE_TRANSPORT | 0.00235 | 0.04368 | 0.43171 | 0.64873 | 1.25585 | 62 |
| GO_SOLUTE_CATION_SYMPORTER_ACTIVITY | 0.00239 | 0.04381 | 0.43171 | 0.69663 | 1.32952 | 40 |
| GO_SPERM_EGG_RECOGNITION | 0.00000 | 0.00012 | 0.62726 | 0.86609 | 1.61106 | 25 |
| GO_SPINDLE_MIDZONE | 0.00224 | 0.04228 | 0.43171 | 0.73022 | 1.37483 | 31 |
| GO_SPLICEOSOMAL_COMPLEX | 0.00000 | 0.00007 | 0.64355 | 0.65143 | 1.29679 | 163 |
| GO_STEM_CELL_DIFFERENTIATION | 0.00000 | 0.00000 | 0.74774 | 0.68820 | 1.36772 | 145 |
| GO_STRUCTURAL_CONSTITUENT_OF_RIBOSOME | 0.00000 | 0.00001 | 0.69013 | 0.67250 | 1.33490 | 140 |
| GO_STRUCTURAL_MOLECULE_ACTIVITY | 0.00000 | 0.00000 | 0.78819 | 0.62899 | 1.26852 | 342 |
| GO_TELOMERASE_RNA_LOCALIZATION | 0.00004 | 0.00153 | 0.55733 | 0.85915 | 1.56133 | 18 |
| GO_TELOMERE_MAINTENANCE_VIA_TELOMERE_LENGTHEN | 0.00128 | 0.02694 | 0.45506 | 0.66601 | 1.29015 | 65 |
| GO_TELOMERE_ORGANIZATION | 0.00135 | 0.02819 | 0.45506 | 0.61918 | 1.22663 | 124 |
| GO_THREONINE_TYPE_ENDOPEPTIDASE_ACTIVITY | 0.00001 | 0.00035 | 0.59333 | 0.88293 | 1.60454 | 18 |
| GO_THREONINE_TYPE_PEPTIDASE_ACTIVITY | 0.00001 | 0.00041 | 0.59333 | 0.85058 | 1.56874 | 22 |
| GO_TOXIN_TRANSPORT | 0.00167 | 0.03318 | 0.45506 | 0.74754 | 1.40097 | 28 |
| GO_TRANSCRIPTION_COREGULATOR_ACTIVITY | 0.00035 | 0.01007 | 0.49849 | 0.57398 | 1.15725 | 338 |
| GO_TRANSCRIPTION_COUPLED_NUCLEOTIDE_EXCISION_RE | 0.00081 | 0.01936 | 0.47727 | 0.66191 | 1.28458 | 68 |
| GO_TRANSCRIPTION_FACTOR_BINDING | 0.00000 | 0.00004 | 0.65944 | 0.59048 | 1.19329 | 420 |
| GO_TRANSLATIONAL_INITIATION | 0.00000 | 0.00000 | | 0.75026 | 1.49435 | 167 |
| GO_TRANSLATION_FACTOR_ACTIVITY_RNA_BINDING | 0.00000 | 0.00000 | | 0.81841 | 1.58963 | 70 |
| GO_TRANSLATION_INITIATION_FACTOR_ACTIVITY | 0.00000 | 0.00000 | 0.74774 | 0.82833 | 1.58936 | 45 |
| GO_TRANSLATION_INITIATION_FACTOR_BINDING | 0.00005 | 0.00171 | 0.55733 | 0.79047 | 1.48607 | 30 |
| GO_TRANSLATION_PREINITIATION_COMPLEX | 0.00013 | 0.00407 | 0.51885 | 0.85614 | 1.54525 | 16 |
| GO_TRANSLATION_REGULATOR_ACTIVITY | 0.00000 | 0.00000 | | 0.77365 | 1.52584 | 108 |
| GO_TRANSLATION_REGULATOR_ACTIVITY_NUCLEIC_ACID_E | 0.00000 | 0.00000 | | 0.80242 | 1.56872 | 85 |
| GO_TUMOR_NECROSIS_FACTOR_MEDIATED_SIGNALING_PA | 0.00000 | 0.00004 | 0.65944 | 0.69502 | 1.36724 | 103 |
| GO_T_CELL_RECEPTOR_SIGNALING_PATHWAY | 0.00000 | 0.00009 | 0.64355 | 0.67859 | 1.34382 | 120 |
| GO_U2_TYPE_SPLICEOSOMAL_COMPLEX | 0.00033 | 0.00948 | 0.49849 | 0.66671 | 1.30041 | 78 |
| GO_UBIQUITIN_LIKE_PROTEIN_LIGASE_BINDING | 0.00007 | 0.00237 | 0.53843 | 0.60447 | 1.21032 | 219 |
| GO_UNFOLDED_PROTEIN_BINDING | 0.00000 | 0.00001 | 0.70498 | 0.73367 | 1.43412 | 84 |
| GO_VACUOLAR_LUMEN | 0.00000 | 0.00021 | 0.61053 | 0.69691 | 1.36480 | 91 |
| GO_VESICLE_LUMEN | 0.00000 | 0.00000 | | 0.71289 | 1.42263 | 181 |
| GO_VIRAL_GENE_EXPRESSION | 0.00000 | 0.00000 | | 0.70297 | 1.40096 | 170 |
| GO_VIRAL_GENOME_REPLICATION | 0.00145 | 0.02958 | 0.45506 | 0.64144 | 1.24847 | 74 |
| GO_VIRAL_LIFE_CYCLE | 0.00005 | 0.00186 | 0.55733 | 0.60907 | 1.21892 | 211 |
| HP_BULBAR_PALSY | 0.00239 | 0.04381 | 0.43171 | 0.75492 | 1.39232 | 22 |
| HP_MACROGYRIA | 0.00165 | 0.03292 | 0.45506 | 0.80389 | 1.45740 | 17 |
| **Cluster 4** | | | | | | |
| pathway | pval | padj | log2err | ES | NES | size |
| GO_ACTIVATION_OF_IMMUNE_RESPONSE | 0.00000 | 0.00000 | 0.74774 | 0.66942 | 1.31439 | 217 |
| GO_ACTIVATION_OF_INNATE_IMMUNE_RESPONSE | 0.00000 | 0.00000 | 0.76146 | 0.77702 | 1.49801 | 83 |
| GO_AMIDE_TRANSMEMBRANE_TRANSPORTER_ACTIVITY | 0.00079 | 0.01913 | 0.47727 | 0.77922 | 1.45181 | 29 |
| GO_AMINE_METABOLIC_PROCESS | 0.00000 | 0.00000 | 0.74774 | 0.77003 | 1.48454 | 83 |
| GO_ANAPHASE_PROMOTING_COMPLEX_DEPENDENT_CATA | 0.00000 | 0.00000 | 0.80122 | 0.78920 | 1.51882 | 76 |
| GO_ANCHORING_JUNCTION | 0.00000 | 0.00000 | | 0.64018 | 1.27252 | 482 |
| GO_ANTIGEN_PROCESSING_AND_PRESENTATION | 0.00000 | 0.00001 | 0.69013 | 0.69241 | 1.35122 | 146 |
| GO_ANTIGEN_PROCESSING_AND_PRESENTATION_OF_EXOG | 0.00000 | 0.00000 | 0.76146 | 0.81354 | 1.54729 | 57 |
| GO_ANTIGEN_PROCESSING_AND_PRESENTATION_OF_PEPT | 0.00000 | 0.00000 | 0.71951 | 0.71247 | 1.38698 | 126 |
| GO_ANTIGEN_PROCESSING_AND_PRESENTATION_OF_PEPT | 0.00000 | 0.00000 | 0.74774 | 0.78524 | 1.50676 | 70 |
| GO_ANTIGEN_RECEPTOR_MEDIATED_SIGNALING_PATHWAY | 0.00000 | 0.00001 | 0.69013 | 0.70480 | 1.37223 | 129 |
| GO_APOPTOTIC_SIGNALING_PATHWAY | 0.00108 | 0.02408 | 0.45506 | 0.57690 | 1.14275 | 351 |
| GO_AXONAL_GROWTH_CONE | 0.00016 | 0.00508 | 0.51885 | 0.82877 | 1.51114 | 20 |
| GO_BINDING_OF_SPERM_TO_ZONA_PELLUCIDA | 0.00000 | 0.00012 | 0.62726 | 0.91764 | 1.66033 | 17 |
| GO_CADHERIN_BINDING | 0.00000 | 0.00000 | 0.78819 | 0.67747 | 1.33226 | 234 |
| GO_CANONICAL_WNT_SIGNALING_PATHWAY | 0.00004 | 0.00153 | 0.55733 | 0.63762 | 1.24846 | 188 |
| GO_CATALYTIC_STEP_2_SPLICEOSOME | 0.00076 | 0.01870 | 0.47727 | 0.67000 | 1.29111 | 81 |
| GO_CELLULAR_AMINO_ACID_METABOLIC_PROCESS | 0.00000 | 0.00000 | 0.73376 | 0.67160 | 1.31805 | 207 |
| GO_CELLULAR_COMPONENT_DISASSEMBLY | 0.00010 | 0.00347 | 0.53843 | 0.58458 | 1.15991 | 396 |
| GO_CELLULAR_HOMEOSTASIS | 0.00246 | 0.04748 | 0.43171 | 0.56574 | 1.12348 | 432 |
| GO_CELLULAR_KETONE_METABOLIC_PROCESS | 0.00000 | 0.00014 | 0.62726 | 0.66755 | 1.30315 | 147 |
| GO_CELLULAR_NITROGEN_COMPOUND_CATABOLIC_PROCE | 0.00000 | 0.00000 | | 0.73663 | 1.46284 | 432 |
| GO_CELLULAR_PROTEIN_COMPLEX_DISASSEMBLY | 0.00108 | 0.02408 | 0.45506 | 0.61175 | 1.19667 | 174 |
| GO_CELLULAR_RESPIRATION | 0.00100 | 0.02288 | 0.45506 | 0.63343 | 1.23345 | 133 |
| GO_CELLULAR_RESPONSE_TO_OXYGEN_LEVELS | 0.00000 | 0.00000 | 0.74774 | 0.70280 | 1.37319 | 155 |
| GO_CELL_ACTIVATION_INVOLVED_IN_IMMUNE_RESPONSE | 0.00000 | 0.00009 | 0.62726 | 0.61649 | 1.22126 | 353 |
| GO_CELL_ADHESION_MOLECULE_BINDING | 0.00000 | 0.00000 | | 0.66313 | 1.31062 | 314 |
| GO_CELL_BODY | 0.00017 | 0.00524 | 0.51885 | 0.60676 | 1.19630 | 273 |
| GO_CELL_CELL_RECOGNITION | 0.00000 | 0.00000 | 0.76146 | 0.90957 | 1.68786 | 27 |
| GO_CELL_CELL_SIGNALING_BY_WNT | 0.00002 | 0.00069 | 0.57561 | 0.61172 | 1.20881 | 305 |
| GO_CELL_CYCLE_G2_M_PHASE_TRANSITION | 0.00001 | 0.00027 | 0.61053 | 0.63623 | 1.24922 | 217 |
| GO_CELL_CYCLE_PHASE_TRANSITION | 0.00001 | 0.00047 | 0.59333 | 0.59333 | 1.17902 | 454 |

| Cluster 1 | | | | | | |
|---|---|---|---|---|---|---|
| pathway | pval | padj | log2err | ES | NES | size |
| GO_CELL_RECOGNITION | 0.00000 | 0.00002 | 0.67496 | 0.79330 | 1.50665 | 52 |
| GO_CELL_SUBSTRATE_JUNCTION | 0.00000 | 0.00000 | | 0.70988 | 1.40030 | 283 |
| GO_CELL_SURFACE_RECEPTOR_SIGNALING_PATHWAY_INVO | 0.00003 | 0.00119 | 0.55733 | 0.60122 | 1.19065 | 339 |
| GO_CHAPERONE_BINDING | 0.00031 | 0.00874 | 0.49849 | 0.69247 | 1.33088 | 74 |
| GO_CHAPERONE_COMPLEX | 0.00001 | 0.00028 | 0.61053 | 0.87986 | 1.61407 | 22 |
| GO_COTRANSLATIONAL_PROTEIN_TARGETING_TO_MEMBR | 0.00000 | 0.00000 | | 0.87698 | 1.69643 | 93 |
| GO_CYTOKINE_MEDIATED_SIGNALING_PATHWAY | 0.00000 | 0.00000 | 0.80122 | 0.64616 | 1.27880 | 329 |
| GO_CYTOPLASMIC_TRANSLATION | 0.00000 | 0.00000 | | 0.83513 | 1.61513 | 89 |
| GO_CYTOPLASMIC_TRANSLATIONAL_INITIATION | 0.00000 | 0.00007 | 0.64355 | 0.85134 | 1.58845 | 30 |
| GO_CYTOSOLIC_LARGE_RIBOSOMAL_SUBUNIT | 0.00000 | 0.00000 | | 0.88176 | 1.67634 | 54 |
| GO_CYTOSOLIC_RIBOSOME | 0.00000 | 0.00000 | | 0.87602 | 1.69714 | 95 |
| GO_CYTOSOLIC_SMALL_RIBOSOMAL_SUBUNIT | 0.00000 | 0.00000 | 0.80122 | 0.87644 | 1.65057 | 40 |
| GO_DEFENSE_RESPONSE_TO_OTHER_ORGANISM | 0.00000 | 0.00004 | 0.65944 | 0.60750 | 1.20626 | 416 |
| GO_DNA_BIOSYNTHETIC_PROCESS | 0.00220 | 0.04365 | 0.43171 | 0.62746 | 1.22184 | 133 |
| GO_DNA_POLYMERASE_BINDING | 0.00083 | 0.01993 | 0.47727 | 0.84132 | 1.52225 | 17 |
| GO_ENDOPEPTIDASE_COMPLEX | 0.00000 | 0.00000 | 0.73376 | 0.79517 | 1.51558 | 61 |
| GO_ENERGY_DERIVATION_BY_OXIDATION_OF_ORGANIC_CO | 0.00113 | 0.02512 | 0.45506 | 0.61015 | 1.19396 | 177 |
| GO_ESTABLISHMENT_OF_PROTEIN_LOCALIZATION_TO_END | 0.00000 | 0.00000 | | 0.87148 | 1.69176 | 103 |
| GO_ESTABLISHMENT_OF_PROTEIN_LOCALIZATION_TO_MEM | 0.00000 | 0.00000 | | 0.76107 | 1.49802 | 240 |
| GO_ESTABLISHMENT_OF_PROTEIN_LOCALIZATION_TO_ORG | 0.00000 | 0.00000 | | 0.70943 | 1.40869 | 417 |
| GO_ESTABLISHMENT_OF_PROTEIN_LOCALIZATION_TO_TELO | 0.00161 | 0.03331 | 0.45506 | 0.82945 | 1.50077 | 17 |
| GO_ESTABLISHMENT_OF_TISSUE_POLARITY | 0.00000 | 0.00000 | 0.78819 | 0.77013 | 1.48943 | 89 |
| GO_EUKARYOTIC_TRANSLATION_INITIATION_FACTOR_3_CO | 0.00016 | 0.00498 | 0.51885 | 0.88003 | 1.58617 | 15 |
| GO_EXOCYTOSIS | 0.00000 | 0.00001 | 0.69013 | 0.60426 | 1.20092 | 478 |
| GO_FC_EPSILON_RECEPTOR_SIGNALING_PATHWAY | 0.00000 | 0.00000 | | 0.80079 | 1.54397 | 82 |
| GO_FC_RECEPTOR_SIGNALING_PATHWAY | 0.00000 | 0.00000 | | 0.75895 | 1.47619 | 121 |
| GO_FERTILIZATION | 0.00023 | 0.00674 | 0.51885 | 0.69631 | 1.33826 | 74 |
| GO_FICOLIN_1_RICH_GRANULE | 0.00000 | 0.00000 | 0.73376 | 0.75070 | 1.45271 | 92 |
| GO_GLYCOSYL_COMPOUND_METABOLIC_PROCESS | 0.00016 | 0.00513 | 0.51885 | 0.70086 | 1.34595 | 72 |
| GO_HEAT_SHOCK_PROTEIN_BINDING | 0.00100 | 0.02288 | 0.45506 | 0.66110 | 1.27572 | 85 |
| GO_HEMATOPOIETIC_PROGENITOR_CELL_DIFFERENTIATION | 0.00000 | 0.00000 | 0.81404 | 0.75398 | 1.46397 | 110 |
| GO_HEMATOPOIETIC_STEM_CELL_DIFFERENTIATION | 0.00000 | 0.00000 | | 0.82187 | 1.57311 | 67 |
| GO_IMMUNE_RESPONSE_REGULATING_SIGNALING_PATHW | 0.00000 | 0.00000 | 0.74774 | 0.67865 | 1.32967 | 197 |
| GO_INNATE_IMMUNE_RESPONSE | 0.00000 | 0.00004 | 0.65944 | 0.61712 | 1.22230 | 346 |
| GO_INNATE_IMMUNE_RESPONSE_ACTIVATING_SIGNAL_TR | 0.00000 | 0.00000 | | 0.82993 | 1.58606 | 65 |
| GO_INNER_MITOCHONDRIAL_MEMBRANE_PROTEIN_COMP | 0.00133 | 0.02891 | 0.45506 | 0.65750 | 1.26878 | 85 |
| GO_INTERACTION_WITH_SYMBIONT | 0.00008 | 0.00296 | 0.53843 | 0.77402 | 1.45770 | 40 |
| GO_INTERLEUKIN_1_MEDIATED_SIGNALING_PATHWAY | 0.00000 | 0.00000 | | 0.81595 | 1.56698 | 72 |
| GO_INTRINSIC_APOPTOTIC_SIGNALING_PATHWAY | 0.00098 | 0.02264 | 0.45506 | 0.60803 | 1.18999 | 185 |
| GO_ION_CHANNEL_BINDING | 0.00023 | 0.00678 | 0.51885 | 0.70274 | 1.34510 | 67 |
| GO_LARGE_RIBOSOMAL_SUBUNIT | 0.00000 | 0.00000 | | 0.79097 | 1.53597 | 109 |
| GO_LEUKOCYTE_MEDIATED_IMMUNITY | 0.00000 | 0.00001 | 0.67496 | 0.61827 | 1.22648 | 380 |
| GO_LYSOSOMAL_LUMEN | 0.00050 | 0.01315 | 0.47727 | 0.70860 | 1.34743 | 55 |
| GO_MACROAUTOPHAGY | 0.00137 | 0.02920 | 0.45506 | 0.59792 | 1.17699 | 241 |
| GO_MACROMOLECULE_TRANSMEMBRANE_TRANSPORTER_ | 0.00098 | 0.02264 | 0.45506 | 0.83602 | 1.51266 | 17 |
| GO_MATURATION_OF_LSU_RRNA | 0.00261 | 0.04984 | 0.43171 | 0.76442 | 1.41851 | 27 |
| GO_MITOCHONDRIAL_LARGE_RIBOSOMAL_SUBUNIT | 0.00156 | 0.03252 | 0.45506 | 0.69424 | 1.31984 | 54 |
| GO_MITOCHONDRIAL_MATRIX | 0.00005 | 0.00164 | 0.55733 | 0.59341 | 1.17562 | 359 |
| GO_MITOCHONDRIAL_PROTEIN_COMPLEX | 0.00000 | 0.00001 | 0.70498 | 0.66724 | 1.30871 | 202 |
| GO_MITOCHONDRIAL_TRANSLATION | 0.00187 | 0.03770 | 0.45506 | 0.62529 | 1.21726 | 126 |
| GO_MITOCHONDRIAL_TRANSLATIONAL_TERMINATION | 0.00091 | 0.02120 | 0.47727 | 0.66554 | 1.28543 | 87 |
| GO_MODIFICATION_DEPENDENT_MACROMOLECULE_CATAB | 0.00007 | 0.00250 | 0.53843 | 0.58544 | 1.16350 | 464 |
| GO_MODULATION_BY_HOST_OF_SYMBIONT_PROCESS | 0.00205 | 0.04114 | 0.43171 | 0.75186 | 1.40530 | 32 |
| GO_MODULATION_OF_PROCESS_OF_OTHER_ORGANISM | 0.00119 | 0.02605 | 0.45506 | 0.69420 | 1.32006 | 55 |
| GO_MORPHOGENESIS_OF_AN_EPITHELIUM | 0.00005 | 0.00165 | 0.55733 | 0.61040 | 1.20419 | 286 |
| GO_MORPHOGENESIS_OF_A_POLARIZED_EPITHELIUM | 0.00000 | 0.00000 | 0.76146 | 0.74728 | 1.45066 | 103 |
| GO_MRNA_3_UTR_BINDING | 0.00043 | 0.01161 | 0.49849 | 0.69322 | 1.32550 | 66 |
| GO_MRNA_BINDING | 0.00000 | 0.00001 | 0.69013 | 0.65665 | 1.29123 | 227 |
| GO_MRNA_PROCESSING | 0.00004 | 0.00143 | 0.55733 | 0.59057 | 1.17271 | 412 |
| GO_MYD88_DEPENDENT_TOLL_LIKE_RECEPTOR_SIGNALING | 0.00040 | 0.01090 | 0.49849 | 0.83664 | 1.52012 | 19 |
| GO_MYD88_INDEPENDENT_TOLL_LIKE_RECEPTOR_SIGNALI | 0.00018 | 0.00550 | 0.51885 | 0.87152 | 1.57097 | 16 |
| GO_MYELOID_LEUKOCYTE_ACTIVATION | 0.00000 | 0.00000 | 0.74774 | 0.64337 | 1.27421 | 341 |
| GO_MYELOID_LEUKOCYTE_MEDIATED_IMMUNITY | 0.00000 | 0.00000 | 0.77494 | 0.65093 | 1.28584 | 303 |
| GO_NADH_METABOLIC_PROCESS | 0.00152 | 0.03186 | 0.45506 | 0.75656 | 1.41128 | 31 |
| GO_NCRNA_METABOLIC_PROCESS | 0.00031 | 0.00879 | 0.49849 | 0.58543 | 1.16131 | 381 |
| GO_NCRNA_PROCESSING | 0.00010 | 0.00347 | 0.53843 | 0.60024 | 1.18767 | 325 |
| GO_NEGATIVE_REGULATION_OF_CANONICAL_WNT_SIGNA | 0.00000 | 0.00000 | 0.74774 | 0.74329 | 1.44339 | 107 |
| GO_NEGATIVE_REGULATION_OF_CELLULAR_AMIDE_METAB | 0.00087 | 0.02052 | 0.47727 | 0.64186 | 1.24629 | 112 |
| GO_NEGATIVE_REGULATION_OF_CELL_CYCLE | 0.00161 | 0.03331 | 0.45506 | 0.56780 | 1.12693 | 403 |

| Cluster 1 | | | | | | |
|---|---|---|---|---|---|---|
| pathway | pval | padj | log2err | ES | NES | size |
| GO_NEGATIVE_REGULATION_OF_CELL_CYCLE_G2_M_PHASI | 0.00000 | 0.00002 | 0.67496 | 0.73867 | 1.42540 | 85 |
| GO_NEGATIVE_REGULATION_OF_CELL_CYCLE_PHASE_TRAN | 0.00004 | 0.00158 | 0.55733 | 0.63370 | 1.23979 | 176 |
| GO_NEGATIVE_REGULATION_OF_CELL_CYCLE_PROCESS | 0.00015 | 0.00477 | 0.51885 | 0.61747 | 1.21548 | 241 |
| GO_NEGATIVE_REGULATION_OF_CELL_DEATH | 0.00000 | 0.00004 | 0.65944 | 0.59593 | 1.18437 | 487 |
| GO_NEGATIVE_REGULATION_OF_MITOTIC_CELL_CYCLE | 0.00026 | 0.00767 | 0.49849 | 0.61402 | 1.20604 | 219 |
| GO_NEGATIVE_REGULATION_OF_NEURON_DEATH | 0.00141 | 0.02987 | 0.45506 | 0.65097 | 1.25923 | 93 |
| GO_NEGATIVE_REGULATION_OF_WNT_SIGNALING_PATHW/ | 0.00000 | 0.00000 | 0.81404 | 0.74200 | 1.44385 | 124 |
| GO_NEURAL_NUCLEUS_DEVELOPMENT | 0.00152 | 0.03186 | 0.45506 | 0.75669 | 1.41153 | 31 |
| GO_NEURON_DEATH | 0.00108 | 0.02408 | 0.45506 | 0.60719 | 1.18791 | 176 |
| GO_NEURON_PROJECTION_TERMINUS | 0.00135 | 0.02891 | 0.45506 | 0.67947 | 1.29364 | 59 |
| GO_NEURON_TO_NEURON_SYNAPSE | 0.00025 | 0.00731 | 0.49849 | 0.63042 | 1.23279 | 167 |
| GO_NIK_NF_KAPPAB_SIGNALING | 0.00000 | 0.00000 | | 0.79314 | 1.53893 | 101 |
| GO_NON_CANONICAL_WNT_SIGNALING_PATHWAY | 0.00000 | 0.00000 | 0.80122 | 0.75654 | 1.46864 | 103 |
| GO_NUCLEAR_TRANSCRIBED_MRNA_CATABOLIC_PROCESS | 0.00000 | 0.00000 | | 0.79296 | 1.55199 | 186 |
| GO_NUCLEAR_TRANSCRIBED_MRNA_CATABOLIC_PROCESS_ | 0.00000 | 0.00000 | | 0.86832 | 1.68727 | 114 |
| GO_NUCLEAR_TRANSPORT | 0.00178 | 0.03622 | 0.45506 | 0.58640 | 1.15601 | 263 |
| GO_NUCLEOBASE_CONTAINING_SMALL_MOLECULE_BIOSYI | 0.00191 | 0.03831 | 0.45506 | 0.67248 | 1.28583 | 66 |
| GO_NUCLEOSIDE_METABOLIC_PROCESS | 0.00016 | 0.00504 | 0.51885 | 0.72047 | 1.37128 | 58 |
| GO_NUCLEOSIDE_TRIPHOSPHATE_METABOLIC_PROCESS | 0.00075 | 0.01833 | 0.47727 | 0.69339 | 1.32230 | 62 |
| GO_ORGANELLAR_RIBOSOME | 0.00041 | 0.01117 | 0.49849 | 0.67269 | 1.29743 | 84 |
| GO_ORGANELLE_INNER_MEMBRANE | 0.00016 | 0.00502 | 0.51885 | 0.59093 | 1.17202 | 373 |
| GO_ORGANIC_CYCLIC_COMPOUND_CATABOLIC_PROCESS | 0.00000 | 0.00000 | | 0.72965 | 1.44935 | 442 |
| GO_OXIDOREDUCTASE_COMPLEX | 0.00163 | 0.03345 | 0.45506 | 0.66340 | 1.27741 | 79 |
| GO_PEPTIDASE_COMPLEX | 0.00000 | 0.00005 | 0.64355 | 0.74248 | 1.42865 | 77 |
| GO_PEPTIDE_TRANSMEMBRANE_TRANSPORTER_ACTIVITY | 0.00056 | 0.01451 | 0.47727 | 0.81307 | 1.48251 | 20 |
| GO_PIGMENT_BIOSYNTHETIC_PROCESS | 0.00261 | 0.04984 | 0.43171 | 0.71751 | 1.35278 | 41 |
| GO_PIGMENT_GRANULE | 0.00014 | 0.00472 | 0.51885 | 0.69202 | 1.33353 | 81 |
| GO_PLATELET_DEGRANULATION | 0.00235 | 0.04580 | 0.43171 | 0.68347 | 1.29965 | 55 |
| GO_POLYSOMAL_RIBOSOME | 0.00000 | 0.00006 | 0.64355 | 0.85441 | 1.59381 | 31 |
| GO_POLYSOME | 0.00000 | 0.00001 | 0.70498 | 0.78946 | 1.50290 | 60 |
| GO_POSITIVE_REGULATION_OF_CANONICAL_WNT_SIGNAL | 0.00000 | 0.00000 | 0.73376 | 0.74358 | 1.44134 | 97 |
| GO_POSITIVE_REGULATION_OF_CELLULAR_AMIDE_METABC | 0.00033 | 0.00916 | 0.49849 | 0.65098 | 1.26400 | 112 |
| GO_POSITIVE_REGULATION_OF_CELLULAR_PROTEIN_LOCAI | 0.00179 | 0.03646 | 0.45506 | 0.59471 | 1.16935 | 226 |
| GO_POSITIVE_REGULATION_OF_CELL_DEATH | 0.00062 | 0.01578 | 0.47727 | 0.58548 | 1.16054 | 366 |
| GO_POSITIVE_REGULATION_OF_DEFENSE_RESPONSE | 0.00000 | 0.00010 | 0.62726 | 0.66294 | 1.29597 | 170 |
| GO_POSITIVE_REGULATION_OF_ESTABLISHMENT_OF_PROT | 0.00102 | 0.02312 | 0.45506 | 0.60409 | 1.18596 | 214 |
| GO_POSITIVE_REGULATION_OF_IMMUNE_RESPONSE | 0.00001 | 0.00041 | 0.59333 | 0.61895 | 1.22083 | 285 |
| GO_POSITIVE_REGULATION_OF_IMMUNE_SYSTEM_PROCES | 0.00038 | 0.01033 | 0.49849 | 0.57567 | 1.14257 | 405 |
| GO_POSITIVE_REGULATION_OF_INNATE_IMMUNE_RESPON | 0.00000 | 0.00002 | 0.67496 | 0.71292 | 1.38441 | 109 |
| GO_POSITIVE_REGULATION_OF_MRNA_METABOLIC_PROCE | 0.00246 | 0.04748 | 0.43171 | 0.67470 | 1.28820 | 63 |
| GO_POSITIVE_REGULATION_OF_ORGANELLE_ORGANIZATIO | 0.00065 | 0.01627 | 0.47727 | 0.57373 | 1.13870 | 403 |
| GO_POSITIVE_REGULATION_OF_PROTEIN_LOCALIZATION_T | 0.00133 | 0.02891 | 0.45506 | 0.70768 | 1.34594 | 56 |
| GO_POSITIVE_REGULATION_OF_RESPONSE_TO_BIOTIC_STII | 0.00000 | 0.00002 | 0.67496 | 0.70144 | 1.36553 | 128 |
| GO_POSITIVE_REGULATION_OF_RESPONSE_TO_EXTERNAL_ | 0.00001 | 0.00054 | 0.59333 | 0.63181 | 1.24054 | 217 |
| GO_POSITIVE_REGULATION_OF_TELOMERASE_RNA_LOCALI | 0.00019 | 0.00583 | 0.51885 | 0.87629 | 1.57945 | 15 |
| GO_POSITIVE_REGULATION_OF_TELOMERE_MAINTENANCE | 0.00135 | 0.02891 | 0.45506 | 0.71802 | 1.35411 | 43 |
| GO_POSITIVE_REGULATION_OF_TELOMERE_MAINTENANCE | 0.00135 | 0.02891 | 0.45506 | 0.75916 | 1.41895 | 32 |
| GO_POSITIVE_REGULATION_OF_TRANSLATION | 0.00033 | 0.00908 | 0.49849 | 0.66703 | 1.29281 | 96 |
| GO_POSITIVE_REGULATION_OF_WNT_SIGNALING_PATHWA | 0.00000 | 0.00001 | 0.69013 | 0.71233 | 1.38339 | 113 |
| GO_POSTSYNAPSE | 0.00004 | 0.00159 | 0.55733 | 0.60990 | 1.20298 | 285 |
| GO_POSTTRANSCRIPTIONAL_REGULATION_OF_GENE_EXPR | 0.00000 | 0.00000 | | 0.68446 | 1.36016 | 453 |
| GO_POST_TRANSLATIONAL_PROTEIN_MODIFICATION | 0.00002 | 0.00071 | 0.57561 | 0.62374 | 1.22649 | 231 |
| GO_PROCESS_UTILIZING_AUTOPHAGIC_MECHANISM | 0.00015 | 0.00481 | 0.51885 | 0.58828 | 1.16696 | 377 |
| GO_PROTEASOMAL_PROTEIN_CATABOLIC_PROCESS | 0.00001 | 0.00033 | 0.59333 | 0.60396 | 1.19673 | 361 |
| GO_PROTEASOMAL_UBIQUITIN_INDEPENDENT_PROTEIN_C | 0.00000 | 0.00005 | 0.64355 | 0.89888 | 1.63898 | 20 |
| GO_PROTEASOME_ACCESSORY_COMPLEX | 0.00002 | 0.00079 | 0.57561 | 0.85844 | 1.57922 | 23 |
| GO_PROTEASOME_CORE_COMPLEX | 0.00000 | 0.00013 | 0.62726 | 0.90676 | 1.64696 | 18 |
| GO_PROTEIN_CONTAINING_COMPLEX_DISASSEMBLY | 0.00033 | 0.00925 | 0.49849 | 0.60431 | 1.19057 | 254 |
| GO_PROTEIN_DNA_COMPLEX_SUBUNIT_ORGANIZATION | 0.00014 | 0.00470 | 0.51885 | 0.63884 | 1.24854 | 162 |
| GO_PROTEIN_FOLDING | 0.00000 | 0.00000 | 0.71951 | 0.68786 | 1.34443 | 163 |
| GO_PROTEIN_LOCALIZATION_TO_ENDOPLASMIC_RETICULU | 0.00000 | 0.00000 | | 0.84951 | 1.65217 | 123 |
| GO_PROTEIN_LOCALIZATION_TO_MEMBRANE | 0.00000 | 0.00000 | | 0.68643 | 1.36348 | 422 |
| GO_PROTEIN_LOCALIZATION_TO_MITOCHONDRION | 0.00231 | 0.04523 | 0.43171 | 0.63352 | 1.23023 | 107 |
| GO_PROTEIN_LOCALIZATION_TO_NUCLEUS | 0.00052 | 0.01351 | 0.47727 | 0.61684 | 1.20706 | 177 |
| GO_PROTEIN_MODIFICATION_BY_SMALL_PROTEIN_REMOV | 0.00000 | 0.00001 | 0.69013 | 0.66135 | 1.29793 | 207 |
| GO_PROTEIN_POLYUBIQUITINATION | 0.00068 | 0.01690 | 0.47727 | 0.59949 | 1.18006 | 242 |
| GO_PROTEIN_REFOLDING | 0.00231 | 0.04523 | 0.43171 | 0.78959 | 1.43971 | 20 |
| GO_PROTEIN_STABILIZATION | 0.00017 | 0.00528 | 0.51885 | 0.65238 | 1.27017 | 129 |
| GO_PROTEIN_TARGETING | 0.00000 | 0.00000 | | 0.73056 | 1.44640 | 332 |
| GO_PROTEIN_TARGETING_TO_MEMBRANE | 0.00000 | 0.00000 | | 0.82557 | 1.61280 | 151 |

| Cluster 1 | | | | | | |
|---|---|---|---|---|---|---|
| pathway | pval | padj | log2err | ES | NES | size |
| GO_PURINE_NUCLEOSIDE_METABOLIC_PROCESS | 0.00163 | 0.03345 | 0.45506 | 0.73340 | 1.37772 | 37 |
| GO_REGULATION_OF_ANIMAL_ORGAN_MORPHOGENESIS | 0.00000 | 0.00000 | 0.74774 | 0.72079 | 1.40311 | 127 |
| GO_REGULATION_OF_APOPTOTIC_SIGNALING_PATHWAY | 0.00066 | 0.01666 | 0.47727 | 0.60185 | 1.18426 | 238 |
| GO_REGULATION_OF_CELLULAR_AMIDE_METABOLIC_PROC | 0.00000 | 0.00000 | 0.73376 | 0.64434 | 1.27281 | 302 |
| GO_REGULATION_OF_CELLULAR_AMINE_METABOLIC_PROC | 0.00000 | 0.00000 | 0.78819 | 0.82215 | 1.56482 | 58 |
| GO_REGULATION_OF_CELLULAR_AMINO_ACID_METABOLIC | 0.00000 | 0.00000 | 0.81404 | 0.83698 | 1.59122 | 54 |
| GO_REGULATION_OF_CELLULAR_KETONE_METABOLIC_PRO | 0.00000 | 0.00001 | 0.69013 | 0.71500 | 1.38859 | 113 |
| GO_REGULATION_OF_CELLULAR_PROTEIN_CATABOLIC_PRO | 0.00063 | 0.01588 | 0.47727 | 0.61304 | 1.20033 | 188 |
| GO_REGULATION_OF_CELLULAR_PROTEIN_LOCALIZATION | 0.00050 | 0.01315 | 0.47727 | 0.58507 | 1.15960 | 364 |
| GO_REGULATION_OF_CELL_CYCLE_G2_M_PHASE_TRANSITI | 0.00000 | 0.00002 | 0.67496 | 0.66898 | 1.30887 | 175 |
| GO_REGULATION_OF_CELL_CYCLE_PHASE_TRANSITION | 0.00002 | 0.00078 | 0.57561 | 0.60682 | 1.20151 | 336 |
| GO_REGULATION_OF_DEFENSE_RESPONSE | 0.00002 | 0.00092 | 0.57561 | 0.60944 | 1.20391 | 301 |
| GO_REGULATION_OF_DNA_TEMPLATED_TRANSCRIPTION_I | 0.00000 | 0.00000 | 0.76146 | 0.75636 | 1.46593 | 96 |
| GO_REGULATION_OF_HEMATOPOIETIC_PROGENITOR_CELL_ | 0.00000 | 0.00000 | 0.80122 | 0.81076 | 1.55456 | 69 |
| GO_REGULATION_OF_HEMATOPOIETIC_STEM_CELL_DIFFER | 0.00000 | 0.00000 | 0.81404 | 0.82840 | 1.57703 | 60 |
| GO_REGULATION_OF_HEMOPOIESIS | 0.00016 | 0.00504 | 0.51885 | 0.61447 | 1.20810 | 232 |
| GO_REGULATION_OF_IMMUNE_RESPONSE | 0.00014 | 0.00457 | 0.51885 | 0.59226 | 1.17466 | 373 |
| GO_REGULATION_OF_INNATE_IMMUNE_RESPONSE | 0.00000 | 0.00004 | 0.65944 | 0.67526 | 1.31939 | 155 |
| GO_REGULATION_OF_INTRINSIC_APOPTOTIC_SIGNALING_P | 0.00003 | 0.00116 | 0.55733 | 0.67728 | 1.31506 | 110 |
| GO_REGULATION_OF_MITOTIC_CELL_CYCLE | 0.00006 | 0.00211 | 0.55733 | 0.58796 | 1.16820 | 445 |
| GO_REGULATION_OF_MORPHOGENESIS_OF_AN_EPITHELIU | 0.00000 | 0.00000 | 0.81404 | 0.75251 | 1.46113 | 110 |
| GO_REGULATION_OF_MRNA_CATABOLIC_PROCESS | 0.00000 | 0.00000 | | 0.72568 | 1.41825 | 162 |
| GO_REGULATION_OF_MRNA_METABOLIC_PROCESS | 0.00000 | 0.00000 | | 0.69969 | 1.37932 | 262 |
| GO_REGULATION_OF_MRNA_PROCESSING | 0.00012 | 0.00415 | 0.51885 | 0.66411 | 1.29046 | 114 |
| GO_REGULATION_OF_MRNA_SPLICING_VIA_SPLICEOSOME | 0.00024 | 0.00708 | 0.51885 | 0.68557 | 1.32181 | 82 |
| GO_REGULATION_OF_OXIDATIVE_STRESS_INDUCED_CELL_D | 0.00143 | 0.03015 | 0.45506 | 0.72963 | 1.37217 | 38 |
| GO_REGULATION_OF_PEPTIDASE_ACTIVITY | 0.00085 | 0.02012 | 0.47727 | 0.60688 | 1.18975 | 199 |
| GO_REGULATION_OF_PROTEASOMAL_PROTEIN_CATABOLIC | 0.00178 | 0.03622 | 0.45506 | 0.62284 | 1.21474 | 142 |
| GO_REGULATION_OF_PROTEIN_CATABOLIC_PROCESS | 0.00019 | 0.00565 | 0.51885 | 0.60687 | 1.19660 | 266 |
| GO_REGULATION_OF_PROTEIN_LOCALIZATION_TO_NUCLEU | 0.00041 | 0.01117 | 0.49849 | 0.68681 | 1.32315 | 80 |
| GO_REGULATION_OF_PROTEIN_MODIFICATION_BY_SMALL_ | 0.00049 | 0.01303 | 0.47727 | 0.62446 | 1.22113 | 167 |
| GO_REGULATION_OF_PROTEIN_STABILITY | 0.00086 | 0.02025 | 0.47727 | 0.61271 | 1.20135 | 200 |
| GO_REGULATION_OF_PROTEOLYSIS | 0.00001 | 0.00048 | 0.59333 | 0.59906 | 1.18837 | 382 |
| GO_REGULATION_OF_RESPONSE_TO_BIOTIC_STIMULUS | 0.00000 | 0.00007 | 0.64355 | 0.65046 | 1.27590 | 203 |
| GO_REGULATION_OF_RNA_SPLICING | 0.00019 | 0.00565 | 0.51885 | 0.65850 | 1.27973 | 116 |
| GO_REGULATION_OF_SMALL_MOLECULE_METABOLIC_PRO | 0.00003 | 0.00115 | 0.57561 | 0.61878 | 1.21971 | 264 |
| GO_REGULATION_OF_STEM_CELL_DIFFERENTIATION | 0.00000 | 0.00000 | 0.74774 | 0.77589 | 1.49515 | 81 |
| GO_REGULATION_OF_TRANSCRIPTION_FROM_RNA_POLYM | 0.00000 | 0.00000 | 0.78819 | 0.80801 | 1.54498 | 66 |
| GO_REGULATION_OF_TRANSLATIONAL_INITIATION | 0.00054 | 0.01410 | 0.47727 | 0.69639 | 1.32803 | 62 |
| GO_REGULATION_OF_WNT_SIGNALING_PATHWAY | 0.00000 | 0.00005 | 0.64355 | 0.65086 | 1.27772 | 210 |
| GO_RESPONSE_TO_DRUG | 0.00063 | 0.01588 | 0.47727 | 0.61763 | 1.20853 | 178 |
| GO_RESPONSE_TO_INORGANIC_SUBSTANCE | 0.00044 | 0.01168 | 0.49849 | 0.59653 | 1.17684 | 286 |
| GO_RESPONSE_TO_INTERLEUKIN_1 | 0.00000 | 0.00000 | 0.78819 | 0.74976 | 1.45569 | 105 |
| GO_RESPONSE_TO_INTERLEUKIN_12 | 0.00012 | 0.00394 | 0.53843 | 0.78991 | 1.47686 | 33 |
| GO_RESPONSE_TO_OXYGEN_LEVELS | 0.00000 | 0.00000 | 0.73376 | 0.66381 | 1.30587 | 237 |
| GO_RESPONSE_TO_TOPOLOGICALLY_INCORRECT_PROTEIN | 0.00183 | 0.03708 | 0.45506 | 0.61855 | 1.20685 | 145 |
| GO_RESPONSE_TO_TUMOR_NECROSIS_FACTOR | 0.00000 | 0.00001 | 0.69013 | 0.68093 | 1.33110 | 159 |
| GO_RIBONUCLEOPROTEIN_COMPLEX_BINDING | 0.00000 | 0.00001 | 0.67496 | 0.70235 | 1.36513 | 118 |
| GO_RIBONUCLEOPROTEIN_COMPLEX_BIOGENESIS | 0.00000 | 0.00000 | | 0.69754 | 1.38335 | 388 |
| GO_RIBONUCLEOPROTEIN_COMPLEX_SUBUNIT_ORGANIZAT | 0.00000 | 0.00000 | | 0.72147 | 1.41130 | 174 |
| GO_RIBONUCLEOPROTEIN_GRANULE | 0.00014 | 0.00472 | 0.51885 | 0.63036 | 1.23300 | 172 |
| GO_RIBONUCLEOSIDE_METABOLIC_PROCESS | 0.00231 | 0.04523 | 0.43171 | 0.72779 | 1.36717 | 37 |
| GO_RIBOSOMAL_LARGE_SUBUNIT_ASSEMBLY | 0.00091 | 0.02120 | 0.47727 | 0.78659 | 1.45964 | 27 |
| GO_RIBOSOMAL_LARGE_SUBUNIT_BIOGENESIS | 0.00000 | 0.00000 | 0.70498 | 0.78270 | 1.49659 | 66 |
| GO_RIBOSOMAL_SMALL_SUBUNIT_ASSEMBLY | 0.00085 | 0.02012 | 0.47727 | 0.82773 | 1.50342 | 18 |
| GO_RIBOSOMAL_SMALL_SUBUNIT_BIOGENESIS | 0.00075 | 0.01833 | 0.47727 | 0.69189 | 1.32662 | 69 |
| GO_RIBOSOMAL_SUBUNIT | 0.00000 | 0.00000 | | 0.78753 | 1.54022 | 173 |
| GO_RIBOSOME | 0.00000 | 0.00000 | | 0.77261 | 1.51589 | 206 |
| GO_RIBOSOME_ASSEMBLY | 0.00000 | 0.00008 | 0.64355 | 0.76486 | 1.45607 | 60 |
| GO_RIBOSOME_BIOGENESIS | 0.00000 | 0.00000 | | 0.68211 | 1.34498 | 275 |
| GO_RNA_CATABOLIC_PROCESS | 0.00000 | 0.00000 | | 0.77096 | 1.52659 | 335 |
| GO_RNA_DEPENDENT_DNA_BIOSYNTHETIC_PROCESS | 0.00068 | 0.01690 | 0.47727 | 0.69782 | 1.33004 | 61 |
| GO_RNA_LOCALIZATION | 0.00115 | 0.02543 | 0.45506 | 0.60912 | 1.19333 | 196 |
| GO_RNA_SPLICING | 0.00009 | 0.00327 | 0.53843 | 0.59506 | 1.17951 | 365 |
| GO_RNA_SPLICING_VIA_TRANSESTERIFICATION_REACTIONS | 0.00001 | 0.00026 | 0.61053 | 0.61994 | 1.22344 | 295 |
| GO_RNA_STABILIZATION | 0.00159 | 0.03317 | 0.45506 | 0.74525 | 1.39336 | 33 |
| GO_ROUGH_ENDOPLASMIC_RETICULUM | 0.00052 | 0.01351 | 0.47727 | 0.71710 | 1.36278 | 53 |
| GO_RRNA_BINDING | 0.00000 | 0.00011 | 0.62726 | 0.77830 | 1.48026 | 56 |
| GO_RRNA_METABOLIC_PROCESS | 0.00000 | 0.00002 | 0.67496 | 0.65901 | 1.29385 | 208 |
| GO_SCF_DEPENDENT_PROTEASOMAL_UBIQUITIN_DEPEND | 0.00000 | 0.00000 | 0.74774 | 0.78124 | 1.50149 | 73 |

| Cluster 1 | | | | | | |
|---|---|---|---|---|---|---|
| pathway | pval | padj | log2err | ES | NES | size |
| GO_SECRETORY_GRANULE | 0.00000 | 0.00000 | 0.77494 | 0.63346 | 1.25729 | 404 |
| GO_SECRETORY_VESICLE | 0.00000 | 0.00000 | 0.76146 | 0.61827 | 1.22877 | 479 |
| GO_SIGNAL_TRANSDUCTION_BY_PROTEIN_PHOSPHORYLAT | 0.00030 | 0.00866 | 0.49849 | 0.58112 | 1.15410 | 430 |
| GO_SINGLE_FERTILIZATION | 0.00001 | 0.00040 | 0.59333 | 0.75248 | 1.43115 | 57 |
| GO_SMALL_MOLECULE_BIOSYNTHETIC_PROCESS | 0.00085 | 0.02012 | 0.47727 | 0.57526 | 1.14164 | 400 |
| GO_SMALL_RIBOSOMAL_SUBUNIT | 0.00000 | 0.00000 | 0.77494 | 0.79650 | 1.52455 | 67 |
| GO_SPERM_EGG_RECOGNITION | 0.00000 | 0.00000 | 0.74774 | 0.92245 | 1.70125 | 24 |
| GO_SPLICEOSOMAL_COMPLEX | 0.00068 | 0.01690 | 0.47727 | 0.61967 | 1.21160 | 168 |
| GO_STEM_CELL_DIFFERENTIATION | 0.00000 | 0.00000 | 0.73376 | 0.71305 | 1.39045 | 141 |
| GO_STRIATED_MUSCLE_CONTRACTION | 0.00078 | 0.01898 | 0.47727 | 0.68223 | 1.30380 | 65 |
| GO_STRUCTURAL_CONSTITUENT_OF_RIBOSOME | 0.00000 | 0.00000 | | 0.80942 | 1.58177 | 152 |
| GO_STRUCTURAL_MOLECULE_ACTIVITY | 0.00000 | 0.00000 | | 0.72556 | 1.43713 | 348 |
| GO_SUBSTANTIA_NIGRA_DEVELOPMENT | 0.00122 | 0.02676 | 0.45506 | 0.78145 | 1.45011 | 27 |
| GO_TELOMERASE_RNA_LOCALIZATION | 0.00030 | 0.00857 | 0.49849 | 0.84097 | 1.52798 | 19 |
| GO_TELOMERE_MAINTENANCE_VIA_TELOMERE_LENGTHEN | 0.00102 | 0.02312 | 0.45506 | 0.68698 | 1.31722 | 69 |
| GO_THREONINE_TYPE_ENDOPEPTIDASE_ACTIVITY | 0.00000 | 0.00003 | 0.65944 | 0.92080 | 1.67246 | 18 |
| GO_THREONINE_TYPE_PEPTIDASE_ACTIVITY | 0.00000 | 0.00004 | 0.65944 | 0.90029 | 1.64622 | 21 |
| GO_TISSUE_MORPHOGENESIS | 0.00224 | 0.04423 | 0.43171 | 0.57861 | 1.14472 | 324 |
| GO_TOXIN_TRANSPORT | 0.00095 | 0.02197 | 0.47727 | 0.78166 | 1.45007 | 26 |
| GO_TRANSCRIPTION_COUPLED_NUCLEOTIDE_EXCISION_RE | 0.00047 | 0.01260 | 0.49849 | 0.69260 | 1.32569 | 67 |
| GO_TRANSCRIPTION_FACTOR_BINDING | 0.00135 | 0.02891 | 0.45506 | 0.57324 | 1.13739 | 395 |
| GO_TRANSLATIONAL_ELONGATION | 0.00000 | 0.00017 | 0.61053 | 0.68764 | 1.33659 | 119 |
| GO_TRANSLATIONAL_INITIATION | 0.00000 | 0.00000 | | 0.84572 | 1.65317 | 165 |
| GO_TRANSLATIONAL_TERMINATION | 0.00032 | 0.00887 | 0.49849 | 0.66704 | 1.29460 | 99 |
| GO_TRANSLATION_FACTOR_ACTIVITY_RNA_BINDING | 0.00000 | 0.00000 | 0.81404 | 0.81653 | 1.56408 | 68 |
| GO_TRANSLATION_INITIATION_FACTOR_ACTIVITY | 0.00000 | 0.00001 | 0.69013 | 0.81717 | 1.54058 | 45 |
| GO_TRANSLATION_PREINITIATION_COMPLEX | 0.00018 | 0.00550 | 0.51885 | 0.87138 | 1.57073 | 16 |
| GO_TRANSLATION_REGULATOR_ACTIVITY | 0.00000 | 0.00000 | | 0.76765 | 1.49010 | 102 |
| GO_TRANSLATION_REGULATOR_ACTIVITY_NUCLEIC_ACID_B | 0.00000 | 0.00000 | 0.80122 | 0.78587 | 1.51519 | 82 |
| GO_TUMOR_NECROSIS_FACTOR_MEDIATED_SIGNALING_PA | 0.00000 | 0.00000 | 0.77494 | 0.75198 | 1.45871 | 98 |
| GO_T_CELL_RECEPTOR_SIGNALING_PATHWAY | 0.00000 | 0.00001 | 0.69013 | 0.71449 | 1.38759 | 113 |
| GO_UBIQUITIN_LIKE_PROTEIN_LIGASE_BINDING | 0.00000 | 0.00001 | 0.70498 | 0.65853 | 1.29301 | 217 |
| GO_UBIQUITIN_PROTEIN_TRANSFERASE_REGULATOR_ACTI | 0.00027 | 0.00777 | 0.49849 | 0.87210 | 1.57189 | 15 |
| GO_UNFOLDED_PROTEIN_BINDING | 0.00000 | 0.00006 | 0.64355 | 0.73845 | 1.42300 | 81 |
| GO_VACUOLAR_LUMEN | 0.00009 | 0.00325 | 0.53843 | 0.67789 | 1.31232 | 94 |
| GO_VACUOLAR_MEMBRANE | 0.00054 | 0.01410 | 0.47727 | 0.58811 | 1.16042 | 290 |
| GO_VACUOLE | 0.00015 | 0.00493 | 0.51885 | 0.57917 | 1.15101 | 484 |
| GO_VESICLE_LUMEN | 0.00000 | 0.00000 | | 0.72517 | 1.41845 | 172 |
| GO_VIRAL_GENE_EXPRESSION | 0.00000 | 0.00000 | | 0.80178 | 1.56767 | 168 |
| GO_VIRAL_GENOME_REPLICATION | 0.00058 | 0.01491 | 0.47727 | 0.69734 | 1.33708 | 69 |
| HP_ABNORMALITY_OF_CARDIOVASCULAR_SYSTEM_ELECTR | 0.00029 | 0.00848 | 0.49849 | 0.61002 | 1.19937 | 232 |
| HP_ABNORMALITY_OF_LOWER_LIP | 0.00119 | 0.02605 | 0.45506 | 0.62745 | 1.22374 | 142 |
| HP_ABNORMALITY_OF_THE_NASAL_DORSUM | 0.00006 | 0.00208 | 0.55733 | 0.65932 | 1.28533 | 140 |
| HP_ABNORMAL_ERYTHROCYTE_MORPHOLOGY | 0.00010 | 0.00333 | 0.53843 | 0.59343 | 1.17535 | 340 |
| HP_ACUTE_LEUKEMIA | 0.00000 | 0.00001 | 0.70498 | 0.77438 | 1.48222 | 67 |
| HP_ANEMIA_OF_INADEQUATE_PRODUCTION | 0.00003 | 0.00117 | 0.55733 | 0.68373 | 1.32764 | 104 |
| HP_DELAYED_PUBERTY | 0.00002 | 0.00071 | 0.57561 | 0.68638 | 1.33294 | 108 |
| HP_DEPRESSED_NASAL_RIDGE | 0.00000 | 0.00004 | 0.65944 | 0.74604 | 1.43726 | 80 |
| HP_EMOTIONAL_LABILITY | 0.00220 | 0.04365 | 0.43171 | 0.64929 | 1.25314 | 86 |
| HP_FATIGUE | 0.00028 | 0.00799 | 0.49849 | 0.61840 | 1.21027 | 185 |
| HP_HEADACHE | 0.00077 | 0.01884 | 0.47727 | 0.62722 | 1.22399 | 146 |
| HP_HEMATOLOGICAL_NEOPLASM | 0.00010 | 0.00347 | 0.53843 | 0.63902 | 1.24746 | 147 |
| HP_INAPPROPRIATE_BEHAVIOR | 0.00242 | 0.04708 | 0.43171 | 0.78768 | 1.43622 | 20 |
| HP_LEUKEMIA | 0.00000 | 0.00010 | 0.62726 | 0.70916 | 1.37180 | 93 |
| HP_MACROCYTIC_ANEMIA | 0.00000 | 0.00011 | 0.62726 | 0.80419 | 1.51456 | 42 |
| HP_MACROGYRIA | 0.00031 | 0.00879 | 0.49849 | 0.86349 | 1.55649 | 16 |
| HP_MIGRAINE | 0.00000 | 0.00013 | 0.62726 | 0.74765 | 1.42957 | 66 |
| HP_PALLOR | 0.00087 | 0.02052 | 0.47727 | 0.65732 | 1.27178 | 91 |
| HP_RETICULOCYTOPENIA | 0.00111 | 0.02481 | 0.45506 | 0.84740 | 1.52736 | 15 |
| HP_THICK_LOWER_LIP_VERMILION | 0.00216 | 0.04321 | 0.43171 | 0.64390 | 1.24969 | 99 |
| HP_THICK_VERMILION_BORDER | 0.00257 | 0.04946 | 0.43171 | 0.60771 | 1.18791 | 156 |
| **Cluster 5** | | | | | | |
| pathway | pval | padj | log2err | ES | NES | size |
| GO_ACTIVATION_OF_IMMUNE_RESPONSE | 0.00001 | 0.00069 | 0.59333 | 0.61577 | 1.20681 | 248 |
| GO_ACTIVATION_OF_INNATE_IMMUNE_RESPONSE | 0.00000 | 0.00029 | 0.61053 | 0.69513 | 1.33533 | 94 |
| GO_AMIDE_TRANSMEMBRANE_TRANSPORTER_ACTIVITY | 0.00047 | 0.01384 | 0.49849 | 0.76682 | 1.41314 | 29 |
| GO_AMINE_METABOLIC_PROCESS | 0.00000 | 0.00000 | 0.71951 | 0.74449 | 1.42700 | 87 |
| GO_ANAPHASE_PROMOTING_COMPLEX_DEPENDENT_CATA | 0.00000 | 0.00000 | 0.73376 | 0.75577 | 1.44312 | 78 |
| GO_ANCHORING_JUNCTION | 0.00000 | 0.00001 | 0.69013 | 0.60110 | 1.19026 | 499 |
| GO_ANTIGEN_PROCESSING_AND_PRESENTATION | 0.00006 | 0.00237 | 0.55733 | 0.63303 | 1.23207 | 161 |

| Cluster 1 | | | | | | |
|---|---|---|---|---|---|---|
| pathway | pval | padj | log2err | ES | NES | size |
| GO_ANTIGEN_PROCESSING_AND_PRESENTATION_OF_EXOG | 0.00000 | 0.00002 | 0.69013 | 0.77116 | 1.45926 | 61 |
| GO_ANTIGEN_PROCESSING_AND_PRESENTATION_OF_PEPT | 0.00004 | 0.00185 | 0.55733 | 0.64851 | 1.25709 | 138 |
| GO_ANTIGEN_PROCESSING_AND_PRESENTATION_OF_PEPT | 0.00000 | 0.00010 | 0.64355 | 0.73044 | 1.39107 | 74 |
| GO_ANTIGEN_RECEPTOR_MEDIATED_SIGNALING_PATHWAY | 0.00000 | 0.00020 | 0.62726 | 0.66602 | 1.29245 | 144 |
| GO_ATPASE_ACTIVITY | 0.00168 | 0.03842 | 0.45506 | 0.57594 | 1.13236 | 297 |
| GO_ATPASE_COMPLEX | 0.00148 | 0.03458 | 0.45506 | 0.67028 | 1.27383 | 68 |
| GO_CADHERIN_BINDING | 0.00000 | 0.00006 | 0.65944 | 0.63777 | 1.24994 | 251 |
| GO_CANONICAL_WNT_SIGNALING_PATHWAY | 0.00029 | 0.00935 | 0.49849 | 0.60479 | 1.18284 | 211 |
| GO_CATALYTIC_STEP_2_SPLICEOSOME | 0.00002 | 0.00109 | 0.57561 | 0.69768 | 1.33537 | 83 |
| GO_CELLULAR_AMINO_ACID_METABOLIC_PROCESS | 0.00008 | 0.00321 | 0.53843 | 0.61157 | 1.19725 | 229 |
| GO_CELLULAR_CARBOHYDRATE_CATABOLIC_PROCESS | 0.00004 | 0.00163 | 0.55733 | 0.79372 | 1.46642 | 32 |
| GO_CELLULAR_COMPONENT_DISASSEMBLY | 0.00030 | 0.00954 | 0.49849 | 0.57544 | 1.13661 | 410 |
| GO_CELLULAR_NITROGEN_COMPOUND_CATABOLIC_PROCE | 0.00000 | 0.00000 | | 0.70033 | 1.38504 | 450 |
| GO_CELLULAR_RESPONSE_TO_OXYGEN_LEVELS | 0.00009 | 0.00330 | 0.53843 | 0.63128 | 1.22821 | 160 |
| GO_CELL_ACTIVATION_INVOLVED_IN_IMMUNE_RESPONSE | 0.00026 | 0.00856 | 0.49849 | 0.57996 | 1.14503 | 392 |
| GO_CELL_ADHESION_MOLECULE_BINDING | 0.00000 | 0.00005 | 0.65944 | 0.61320 | 1.20719 | 336 |
| GO_CELL_CELL_RECOGNITION | 0.00104 | 0.02641 | 0.45506 | 0.74545 | 1.38061 | 33 |
| GO_CELL_CELL_SIGNALING_BY_WNT | 0.00015 | 0.00536 | 0.51885 | 0.58457 | 1.15094 | 335 |
| GO_CELL_CYCLE_G1_S_PHASE_TRANSITION | 0.00054 | 0.01549 | 0.47727 | 0.60381 | 1.18039 | 208 |
| GO_CELL_CYCLE_G2_M_PHASE_TRANSITION | 0.00009 | 0.00349 | 0.53843 | 0.61070 | 1.19499 | 223 |
| GO_CELL_CYCLE_PHASE_TRANSITION | 0.00000 | 0.00001 | 0.71951 | 0.60418 | 1.19590 | 478 |
| GO_CELL_DIVISION | 0.00095 | 0.02438 | 0.47727 | 0.56620 | 1.11881 | 431 |
| GO_CELL_SUBSTRATE_JUNCTION | 0.00000 | 0.00000 | | 0.67790 | 1.33269 | 294 |
| GO_CELL_SURFACE_RECEPTOR_SIGNALING_PATHWAY_INVC | 0.00068 | 0.01883 | 0.47727 | 0.57374 | 1.13123 | 366 |
| GO_CHAPERONE_COMPLEX | 0.00010 | 0.00358 | 0.53843 | 0.83408 | 1.51646 | 23 |
| GO_CHROMATIN_BINDING | 0.00007 | 0.00276 | 0.53843 | 0.58610 | 1.15678 | 387 |
| GO_COTRANSLATIONAL_PROTEIN_TARGETING_TO_MEMBR | 0.00000 | 0.00000 | | 0.86667 | 1.66528 | 92 |
| GO_COVALENT_CHROMATIN_MODIFICATION | 0.00090 | 0.02360 | 0.47727 | 0.57551 | 1.13426 | 356 |
| GO_CYTOKINE_MEDIATED_SIGNALING_PATHWAY | 0.00059 | 0.01676 | 0.47727 | 0.57448 | 1.13367 | 385 |
| GO_CYTOPLASMIC_TRANSLATION | 0.00000 | 0.00000 | | 0.81799 | 1.56888 | 89 |
| GO_CYTOPLASMIC_TRANSLATIONAL_INITIATION | 0.00001 | 0.00055 | 0.59333 | 0.82290 | 1.51744 | 30 |
| GO_CYTOSOLIC_LARGE_RIBOSOMAL_SUBUNIT | 0.00000 | 0.00000 | | 0.88350 | 1.66656 | 54 |
| GO_CYTOSOLIC_RIBOSOME | 0.00000 | 0.00000 | | 0.87563 | 1.68391 | 98 |
| GO_CYTOSOLIC_SMALL_RIBOSOMAL_SUBUNIT | 0.00000 | 0.00000 | 0.77494 | 0.86191 | 1.60897 | 41 |
| GO_DEFENSE_RESPONSE_TO_OTHER_ORGANISM | 0.00093 | 0.02411 | 0.47727 | 0.56206 | 1.11260 | 484 |
| GO_DNA_BINDING_TRANSCRIPTION_FACTOR_BINDING | 0.00024 | 0.00794 | 0.51885 | 0.60084 | 1.17682 | 237 |
| GO_DNA_BIOSYNTHETIC_PROCESS | 0.00085 | 0.02246 | 0.47727 | 0.61954 | 1.20239 | 147 |
| GO_DNA_CONFORMATION_CHANGE | 0.00176 | 0.03950 | 0.45506 | 0.59024 | 1.15497 | 224 |
| GO_DNA_REPAIR | 0.00165 | 0.03772 | 0.45506 | 0.56638 | 1.11924 | 434 |
| GO_DNA_TEMPLATED_TRANSCRIPTION_ELONGATION | 0.00042 | 0.01266 | 0.49849 | 0.65792 | 1.26612 | 99 |
| GO_DNA_TEMPLATED_TRANSCRIPTION_INITIATION | 0.00054 | 0.01568 | 0.47727 | 0.60906 | 1.18816 | 181 |
| GO_ENDOPEPTIDASE_COMPLEX | 0.00000 | 0.00000 | 0.73376 | 0.78132 | 1.48093 | 64 |
| GO_ESTABLISHMENT_OF_PROTEIN_LOCALIZATION_TO_END | 0.00000 | 0.00000 | | 0.85986 | 1.65497 | 101 |
| GO_ESTABLISHMENT_OF_PROTEIN_LOCALIZATION_TO_MEN | 0.00000 | 0.00000 | | 0.72831 | 1.42747 | 250 |
| GO_ESTABLISHMENT_OF_PROTEIN_LOCALIZATION_TO_ORG | 0.00000 | 0.00000 | | 0.68776 | 1.35838 | 420 |
| GO_ESTABLISHMENT_OF_RNA_LOCALIZATION | 0.00005 | 0.00200 | 0.55733 | 0.63449 | 1.23536 | 169 |
| GO_ESTABLISHMENT_OF_TISSUE_POLARITY | 0.00001 | 0.00049 | 0.59333 | 0.69077 | 1.32724 | 93 |
| GO_EUKARYOTIC_TRANSLATION_INITIATION_FACTOR_3_CO | 0.00001 | 0.00049 | 0.59333 | 0.90398 | 1.60253 | 15 |
| GO_FC_EPSILON_RECEPTOR_SIGNALING_PATHWAY | 0.00000 | 0.00019 | 0.62726 | 0.71300 | 1.36664 | 87 |
| GO_FC_RECEPTOR_SIGNALING_PATHWAY | 0.00003 | 0.00122 | 0.57561 | 0.65519 | 1.27005 | 137 |
| GO_FIBRILLAR_CENTER | 0.00108 | 0.02701 | 0.45506 | 0.65032 | 1.25061 | 98 |
| GO_FICOLIN_1_RICH_GRANULE | 0.00000 | 0.00001 | 0.71951 | 0.72692 | 1.39763 | 96 |
| GO_GLYCOSYL_COMPOUND_METABOLIC_PROCESS | 0.00023 | 0.00757 | 0.51885 | 0.68339 | 1.30271 | 75 |
| GO_GTP_METABOLIC_PROCESS | 0.00030 | 0.00954 | 0.49849 | 0.80617 | 1.45964 | 21 |
| GO_GUANOSINE_CONTAINING_COMPOUND_METABOLIC_P | 0.00111 | 0.02770 | 0.45506 | 0.75221 | 1.38622 | 29 |
| GO_HEMATOPOIETIC_PROGENITOR_CELL_DIFFERENTIATION | 0.00000 | 0.00008 | 0.65944 | 0.68406 | 1.32331 | 119 |
| GO_HEMATOPOIETIC_STEM_CELL_DIFFERENTIATION | 0.00000 | 0.00001 | 0.71951 | 0.76246 | 1.44945 | 70 |
| GO_IMMUNE_RESPONSE_REGULATING_SIGNALING_PATHW | 0.00009 | 0.00330 | 0.53843 | 0.61145 | 1.19646 | 223 |
| GO_INNATE_IMMUNE_RESPONSE | 0.00034 | 0.01066 | 0.49849 | 0.57674 | 1.13909 | 400 |
| GO_INNATE_IMMUNE_RESPONSE_ACTIVATING_SIGNAL_TR | 0.00000 | 0.00018 | 0.62726 | 0.72588 | 1.38464 | 76 |
| GO_INTERLEUKIN_1_MEDIATED_SIGNALING_PATHWAY | 0.00000 | 0.00001 | 0.70498 | 0.74718 | 1.42726 | 79 |
| GO_INTRINSIC_APOPTOTIC_SIGNALING_PATHWAY | 0.00132 | 0.03144 | 0.45506 | 0.59879 | 1.16936 | 199 |
| GO_LARGE_RIBOSOMAL_SUBUNIT | 0.00000 | 0.00000 | 0.81404 | 0.74343 | 1.43461 | 110 |
| GO_LEUKOCYTE_MEDIATED_IMMUNITY | 0.00037 | 0.01139 | 0.49849 | 0.57402 | 1.13427 | 430 |
| GO_MATURATION_OF_LSU_RRNA | 0.00019 | 0.00658 | 0.51885 | 0.80367 | 1.47086 | 27 |
| GO_MATURATION_OF_SSU_RRNA | 0.00133 | 0.03163 | 0.45506 | 0.70286 | 1.32151 | 47 |
| GO_MITOTIC_NUCLEAR_DIVISION | 0.00126 | 0.03036 | 0.45506 | 0.59218 | 1.15818 | 211 |
| GO_MODIFICATION_DEPENDENT_MACROMOLECULE_CATAI | 0.00000 | 0.00006 | 0.65944 | 0.59558 | 1.17931 | 496 |
| GO_MORPHOGENESIS_OF_AN_EPITHELIUM | 0.00054 | 0.01549 | 0.47727 | 0.58252 | 1.14568 | 301 |
| GO_MORPHOGENESIS_OF_A_POLARIZED_EPITHELIUM | 0.00000 | 0.00026 | 0.61053 | 0.68789 | 1.32547 | 107 |

| Cluster 1 | | | | | | |
|---|---|---|---|---|---|---|
| pathway | pval | padj | log2err | ES | NES | size |
| GO_MRNA_3_END_PROCESSING | 0.00067 | 0.01866 | 0.47727 | 0.64963 | 1.24671 | 90 |
| GO_MRNA_3_UTR_BINDING | 0.00115 | 0.02815 | 0.45506 | 0.67401 | 1.28092 | 68 |
| GO_MRNA_5_UTR_BINDING | 0.00010 | 0.00389 | 0.53843 | 0.82401 | 1.49354 | 22 |
| GO_MRNA_BINDING | 0.00000 | 0.00000 | 0.83909 | 0.67296 | 1.31818 | 233 |
| GO_MRNA_EXPORT_FROM_NUCLEUS | 0.00045 | 0.01350 | 0.49849 | 0.65390 | 1.25855 | 101 |
| GO_MRNA_PROCESSING | 0.00000 | 0.00000 | | 0.64526 | 1.27487 | 426 |
| GO_MRNA_TRANSPORT | 0.00133 | 0.03163 | 0.45506 | 0.62785 | 1.21587 | 126 |
| GO_MYD88_INDEPENDENT_TOLL_LIKE_RECEPTOR_SIGNALI | 0.00213 | 0.04676 | 0.43171 | 0.76591 | 1.38823 | 22 |
| GO_MYELOID_LEUKOCYTE_ACTIVATION | 0.00008 | 0.00294 | 0.53843 | 0.58860 | 1.16069 | 374 |
| GO_MYELOID_LEUKOCYTE_MEDIATED_IMMUNITY | 0.00002 | 0.00114 | 0.57561 | 0.59639 | 1.17421 | 335 |
| GO_NCRNA_METABOLIC_PROCESS | 0.00000 | 0.00005 | 0.65944 | 0.60764 | 1.20017 | 402 |
| GO_NCRNA_PROCESSING | 0.00000 | 0.00000 | 0.73376 | 0.62613 | 1.23298 | 339 |
| GO_NEGATIVE_REGULATION_OF_CANONICAL_WNT_SIGNAI | 0.00003 | 0.00150 | 0.55733 | 0.66211 | 1.28031 | 116 |
| GO_NEGATIVE_REGULATION_OF_CELLULAR_AMIDE_METAB | 0.00176 | 0.03950 | 0.45506 | 0.61659 | 1.19418 | 125 |
| GO_NEGATIVE_REGULATION_OF_CELL_CYCLE | 0.00001 | 0.00043 | 0.61053 | 0.59091 | 1.16771 | 434 |
| GO_NEGATIVE_REGULATION_OF_CELL_CYCLE_G2_M_PHASI | 0.00000 | 0.00010 | 0.64355 | 0.71358 | 1.36848 | 88 |
| GO_NEGATIVE_REGULATION_OF_CELL_CYCLE_PHASE_TRAN | 0.00001 | 0.00063 | 0.59333 | 0.63246 | 1.23416 | 190 |
| GO_NEGATIVE_REGULATION_OF_CELL_CYCLE_PROCESS | 0.00001 | 0.00035 | 0.61053 | 0.62718 | 1.22936 | 256 |
| GO_NEGATIVE_REGULATION_OF_INTERFERON_GAMMA_PR | 0.00026 | 0.00846 | 0.49849 | 0.86009 | 1.52474 | 15 |
| GO_NEGATIVE_REGULATION_OF_MITOTIC_CELL_CYCLE | 0.00015 | 0.00523 | 0.51885 | 0.60384 | 1.18279 | 233 |
| GO_NEGATIVE_REGULATION_OF_WNT_SIGNALING_PATHW/ | 0.00001 | 0.00055 | 0.59333 | 0.66257 | 1.28407 | 135 |
| GO_NIK_NF_KAPPAB_SIGNALING | 0.00000 | 0.00004 | 0.67496 | 0.69127 | 1.33700 | 118 |
| GO_NON_CANONICAL_WNT_SIGNALING_PATHWAY | 0.00001 | 0.00036 | 0.61053 | 0.68318 | 1.31947 | 111 |
| GO_NUCLEAR_ENVELOPE | 0.00224 | 0.04902 | 0.43171 | 0.56977 | 1.12126 | 327 |
| GO_NUCLEAR_EXPORT | 0.00002 | 0.00100 | 0.57561 | 0.63238 | 1.23279 | 177 |
| GO_NUCLEAR_PERIPHERY | 0.00113 | 0.02781 | 0.45506 | 0.63256 | 1.21861 | 106 |
| GO_NUCLEAR_SPECK | 0.00001 | 0.00069 | 0.59333 | 0.60502 | 1.18941 | 307 |
| GO_NUCLEAR_TRANSCRIBED_MRNA_CATABOLIC_PROCESS | 0.00000 | 0.00000 | | 0.76271 | 1.48764 | 188 |
| GO_NUCLEAR_TRANSCRIBED_MRNA_CATABOLIC_PROCESS_ | 0.00000 | 0.00000 | | 0.86147 | 1.66478 | 114 |
| GO_NUCLEAR_TRANSPORT | 0.00000 | 0.00006 | 0.65944 | 0.62563 | 1.22793 | 278 |
| GO_NUCLEOBASE_CONTAINING_COMPOUND_TRANSPORT | 0.00016 | 0.00549 | 0.51885 | 0.61245 | 1.19711 | 205 |
| GO_NUCLEOSIDE_METABOLIC_PROCESS | 0.00002 | 0.00105 | 0.57561 | 0.72182 | 1.36609 | 62 |
| GO_ORGANIC_CYCLIC_COMPOUND_CATABOLIC_PROCESS | 0.00000 | 0.00000 | | 0.69944 | 1.38355 | 462 |
| GO_PEPTIDASE_COMPLEX | 0.00000 | 0.00003 | 0.67496 | 0.73168 | 1.39835 | 80 |
| GO_PEPTIDE_TRANSMEMBRANE_TRANSPORTER_ACTIVITY | 0.00097 | 0.02475 | 0.47727 | 0.80469 | 1.45467 | 20 |
| GO_POLYSOMAL_RIBOSOME | 0.00001 | 0.00049 | 0.59333 | 0.82491 | 1.52113 | 30 |
| GO_POLYSOME | 0.00000 | 0.00001 | 0.70498 | 0.77335 | 1.46363 | 62 |
| GO_POLY_PYRIMIDINE_TRACT_BINDING | 0.00090 | 0.02360 | 0.47727 | 0.82022 | 1.47330 | 18 |
| GO_POSITIVE_REGULATION_OF_CANONICAL_WNT_SIGNAL | 0.00001 | 0.00032 | 0.61053 | 0.69161 | 1.33278 | 105 |
| GO_POSITIVE_REGULATION_OF_CELLULAR_AMIDE_METABC | 0.00012 | 0.00433 | 0.53843 | 0.64719 | 1.25164 | 117 |
| GO_POSITIVE_REGULATION_OF_CELLULAR_PROTEIN_LOCAI | 0.00111 | 0.02770 | 0.45506 | 0.59336 | 1.16193 | 236 |
| GO_POSITIVE_REGULATION_OF_CELL_CYCLE | 0.00020 | 0.00693 | 0.51885 | 0.59336 | 1.16323 | 258 |
| GO_POSITIVE_REGULATION_OF_CELL_CYCLE_PROCESS | 0.00070 | 0.01908 | 0.47727 | 0.60458 | 1.18065 | 199 |
| GO_POSITIVE_REGULATION_OF_DEFENSE_RESPONSE | 0.00032 | 0.00992 | 0.49849 | 0.61505 | 1.19991 | 182 |
| GO_POSITIVE_REGULATION_OF_ESTABLISHMENT_OF_PROT | 0.00205 | 0.04530 | 0.43171 | 0.58524 | 1.14626 | 237 |
| GO_POSITIVE_REGULATION_OF_IMMUNE_RESPONSE | 0.00035 | 0.01075 | 0.49849 | 0.58306 | 1.14753 | 328 |
| GO_POSITIVE_REGULATION_OF_INNATE_IMMUNE_RESPON | 0.00006 | 0.00237 | 0.55733 | 0.65539 | 1.26919 | 124 |
| GO_POSITIVE_REGULATION_OF_MRNA_PROCESSING | 0.00059 | 0.01676 | 0.47727 | 0.78411 | 1.43507 | 27 |
| GO_POSITIVE_REGULATION_OF_MRNA_SPLICING_VIA_SPLI | 0.00108 | 0.02701 | 0.45506 | 0.81963 | 1.46316 | 17 |
| GO_POSITIVE_REGULATION_OF_PROTEIN_LOCALIZATION_T | 0.00008 | 0.00297 | 0.53843 | 0.71485 | 1.35493 | 64 |
| GO_POSITIVE_REGULATION_OF_RESPONSE_TO_BIOTIC_STI | 0.00002 | 0.00103 | 0.57561 | 0.64531 | 1.25212 | 145 |
| GO_POSITIVE_REGULATION_OF_TELOMERASE_RNA_LOCALI | 0.00002 | 0.00099 | 0.57561 | 0.89563 | 1.58773 | 15 |
| GO_POSITIVE_REGULATION_OF_TRANSLATION | 0.00028 | 0.00886 | 0.49849 | 0.66164 | 1.27327 | 99 |
| GO_POSITIVE_REGULATION_OF_WNT_SIGNALING_PATHWA | 0.00004 | 0.00177 | 0.55733 | 0.65791 | 1.27408 | 124 |
| GO_POSTSYNAPSE | 0.00071 | 0.01924 | 0.47727 | 0.57828 | 1.13704 | 303 |
| GO_POSTTRANSCRIPTIONAL_REGULATION_OF_GENE_EXPR | 0.00000 | 0.00000 | | 0.66197 | 1.31037 | 481 |
| GO_POST_TRANSLATIONAL_PROTEIN_MODIFICATION | 0.00002 | 0.00088 | 0.57561 | 0.61860 | 1.21165 | 240 |
| GO_PRERIBOSOME | 0.00000 | 0.00010 | 0.64355 | 0.73433 | 1.39550 | 69 |
| GO_PRERIBOSOME_LARGE_SUBUNIT_PRECURSOR | 0.00156 | 0.03588 | 0.45506 | 0.77233 | 1.39988 | 22 |
| GO_PROTEASOMAL_PROTEIN_CATABOLIC_PROCESS | 0.00000 | 0.00024 | 0.62726 | 0.59747 | 1.17867 | 382 |
| GO_PROTEASOMAL_UBIQUITIN_INDEPENDENT_PROTEIN_C | 0.00002 | 0.00104 | 0.57561 | 0.86224 | 1.55871 | 20 |
| GO_PROTEASOME_ACCESSORY_COMPLEX | 0.00007 | 0.00287 | 0.53843 | 0.82663 | 1.50442 | 24 |
| GO_PROTEASOME_CORE_COMPLEX | 0.00003 | 0.00131 | 0.57561 | 0.87359 | 1.56915 | 18 |
| GO_PROTEIN_CONTAINING_COMPLEX_DISASSEMBLY | 0.00130 | 0.03113 | 0.45506 | 0.58189 | 1.14071 | 262 |
| GO_PROTEIN_CONTAINING_COMPLEX_LOCALIZATION | 0.00004 | 0.00170 | 0.55733 | 0.61631 | 1.20525 | 213 |
| GO_PROTEIN_DNA_COMPLEX_SUBUNIT_ORGANIZATION | 0.00010 | 0.00358 | 0.53843 | 0.61873 | 1.20619 | 177 |
| GO_PROTEIN_FOLDING | 0.00013 | 0.00466 | 0.51885 | 0.62500 | 1.21755 | 171 |
| GO_PROTEIN_IMPORT | 0.00093 | 0.02411 | 0.47727 | 0.62052 | 1.20376 | 142 |
| GO_PROTEIN_LOCALIZATION_TO_CHROMOSOME | 0.00104 | 0.02641 | 0.45506 | 0.67482 | 1.28178 | 67 |
| GO_PROTEIN_LOCALIZATION_TO_ENDOPLASMIC_RETICULU | 0.00000 | 0.00000 | | 0.83548 | 1.61795 | 123 |

| **Cluster 1** | | | | | | |
|---|---|---|---|---|---|---|
| pathway | pval | padj | log2err | ES | NES | size |
| GO_PROTEIN_LOCALIZATION_TO_MEMBRANE | 0.00000 | 0.00000 | | 0.65287 | 1.29042 | 438 |
| GO_PROTEIN_LOCALIZATION_TO_NUCLEUS | 0.00004 | 0.00168 | 0.55733 | 0.62532 | 1.22023 | 190 |
| GO_PROTEIN_MODIFICATION_BY_SMALL_PROTEIN_REMOV | 0.00002 | 0.00099 | 0.57561 | 0.61593 | 1.20618 | 231 |
| GO_PROTEIN_POLYUBIQUITINATION | 0.00003 | 0.00149 | 0.55733 | 0.60752 | 1.19114 | 259 |
| GO_PROTEIN_STABILIZATION | 0.00004 | 0.00163 | 0.55733 | 0.65189 | 1.26477 | 143 |
| GO_PROTEIN_TARGETING | 0.00000 | 0.00000 | | 0.70000 | 1.37781 | 330 |
| GO_PROTEIN_TARGETING_TO_MEMBRANE | 0.00000 | 0.00000 | | 0.80413 | 1.56285 | 154 |
| GO_PURINE_NUCLEOSIDE_METABOLIC_PROCESS | 0.00067 | 0.01866 | 0.47727 | 0.73466 | 1.37145 | 41 |
| GO_REGENERATION | 0.00065 | 0.01807 | 0.47727 | 0.65544 | 1.26045 | 98 |
| GO_REGULATION_OF_ANIMAL_ORGAN_MORPHOGENESIS | 0.00002 | 0.00085 | 0.57561 | 0.65506 | 1.26942 | 136 |
| GO_REGULATION_OF_CELLULAR_AMIDE_METABOLIC_PROC | 0.00000 | 0.00000 | 0.76146 | 0.63383 | 1.24718 | 326 |
| GO_REGULATION_OF_CELLULAR_AMINE_METABOLIC_PROC | 0.00000 | 0.00001 | 0.70498 | 0.77854 | 1.47322 | 61 |
| GO_REGULATION_OF_CELLULAR_AMINO_ACID_METABOLIC | 0.00000 | 0.00000 | 0.73376 | 0.79726 | 1.50469 | 56 |
| GO_REGULATION_OF_CELLULAR_KETONE_METABOLIC_PRO | 0.00013 | 0.00461 | 0.51885 | 0.64508 | 1.24935 | 125 |
| GO_REGULATION_OF_CELLULAR_PROTEIN_CATABOLIC_PRO | 0.00172 | 0.03911 | 0.45506 | 0.60035 | 1.17180 | 193 |
| GO_REGULATION_OF_CELLULAR_PROTEIN_LOCALIZATION | 0.00004 | 0.00161 | 0.55733 | 0.58777 | 1.15946 | 381 |
| GO_REGULATION_OF_CELL_CYCLE_G2_M_PHASE_TRANSITI | 0.00002 | 0.00088 | 0.57561 | 0.63257 | 1.23402 | 181 |
| GO_REGULATION_OF_CELL_CYCLE_PHASE_TRANSITION | 0.00000 | 0.00011 | 0.64355 | 0.60863 | 1.19939 | 350 |
| GO_REGULATION_OF_CHROMOSOME_ORGANIZATION | 0.00012 | 0.00455 | 0.51885 | 0.59771 | 1.17233 | 268 |
| GO_REGULATION_OF_CHROMOSOME_SEGREGATION | 0.00059 | 0.01676 | 0.47727 | 0.65823 | 1.26233 | 88 |
| GO_REGULATION_OF_DNA_METABOLIC_PROCESS | 0.00108 | 0.02701 | 0.45506 | 0.58641 | 1.14896 | 246 |
| GO_REGULATION_OF_DNA_TEMPLATED_TRANSCRIPTION_I | 0.00000 | 0.00017 | 0.62726 | 0.69996 | 1.34461 | 94 |
| GO_REGULATION_OF_GENE_EXPRESSION_EPIGENETIC | 0.00045 | 0.01343 | 0.49849 | 0.63105 | 1.22200 | 127 |
| GO_REGULATION_OF_HEMATOPOIETIC_PROGENITOR_CELL_ | 0.00000 | 0.00000 | 0.73376 | 0.75924 | 1.44592 | 74 |
| GO_REGULATION_OF_HEMATOPOIETIC_STEM_CELL_DIFFER | 0.00000 | 0.00003 | 0.67496 | 0.77151 | 1.46160 | 63 |
| GO_REGULATION_OF_INNATE_IMMUNE_RESPONSE | 0.00029 | 0.00925 | 0.49849 | 0.61484 | 1.19857 | 176 |
| GO_REGULATION_OF_INTRINSIC_APOPTOTIC_SIGNALING_P | 0.00053 | 0.01538 | 0.47727 | 0.63889 | 1.23394 | 111 |
| GO_REGULATION_OF_MITOTIC_CELL_CYCLE | 0.00001 | 0.00075 | 0.59333 | 0.58379 | 1.15568 | 473 |
| GO_REGULATION_OF_MORPHOGENESIS_OF_AN_EPITHELIU | 0.00000 | 0.00025 | 0.61053 | 0.67964 | 1.31338 | 114 |
| GO_REGULATION_OF_MRNA_CATABOLIC_PROCESS | 0.00000 | 0.00001 | 0.71951 | 0.67582 | 1.31704 | 174 |
| GO_REGULATION_OF_MRNA_METABOLIC_PROCESS | 0.00000 | 0.00000 | | 0.67654 | 1.32767 | 276 |
| GO_REGULATION_OF_MRNA_PROCESSING | 0.00003 | 0.00143 | 0.55733 | 0.66261 | 1.28130 | 116 |
| GO_REGULATION_OF_MRNA_SPLICING_VIA_SPLICEOSOME | 0.00005 | 0.00220 | 0.55733 | 0.68954 | 1.31979 | 83 |
| GO_REGULATION_OF_PROTEASOMAL_UBIQUITIN_DEPENDI | 0.00187 | 0.04167 | 0.45506 | 0.63071 | 1.21400 | 100 |
| GO_REGULATION_OF_PROTEIN_CATABOLIC_PROCESS | 0.00025 | 0.00835 | 0.49849 | 0.59230 | 1.16316 | 285 |
| GO_REGULATION_OF_PROTEIN_IMPORT | 0.00194 | 0.04300 | 0.45506 | 0.70856 | 1.31899 | 39 |
| GO_REGULATION_OF_PROTEIN_LOCALIZATION_TO_NUCLEU | 0.00000 | 0.00022 | 0.62726 | 0.69857 | 1.34064 | 90 |
| GO_REGULATION_OF_PROTEIN_MODIFICATION_BY_SMALL_ | 0.00021 | 0.00697 | 0.51885 | 0.61480 | 1.19917 | 180 |
| GO_REGULATION_OF_PROTEIN_STABILITY | 0.00073 | 0.01939 | 0.47727 | 0.59618 | 1.16678 | 225 |
| GO_REGULATION_OF_RESPONSE_TO_BIOTIC_STIMULUS | 0.00023 | 0.00757 | 0.51885 | 0.60481 | 1.18401 | 229 |
| GO_REGULATION_OF_RNA_SPLICING | 0.00000 | 0.00026 | 0.61053 | 0.68019 | 1.31528 | 116 |
| GO_REGULATION_OF_SIGNAL_TRANSDUCTION_BY_P53_CL | 0.00044 | 0.01336 | 0.49849 | 0.62917 | 1.21960 | 138 |
| GO_REGULATION_OF_SISTER_CHROMATID_SEGREGATION | 0.00157 | 0.03617 | 0.45506 | 0.66894 | 1.27127 | 68 |
| GO_REGULATION_OF_SMALL_MOLECULE_METABOLIC_PRO | 0.00007 | 0.00270 | 0.53843 | 0.59645 | 1.17258 | 294 |
| GO_REGULATION_OF_STEM_CELL_DIFFERENTIATION | 0.00000 | 0.00004 | 0.67496 | 0.71934 | 1.37968 | 89 |
| GO_REGULATION_OF_TRANSCRIPTION_FROM_RNA_POLYM | 0.00000 | 0.00010 | 0.64355 | 0.74909 | 1.42090 | 65 |
| GO_REGULATION_OF_TRANSLATIONAL_INITIATION | 0.00002 | 0.00114 | 0.57561 | 0.71260 | 1.35270 | 66 |
| GO_REGULATION_OF_UBIQUITIN_DEPENDENT_PROTEIN_C | 0.00154 | 0.03573 | 0.45506 | 0.62795 | 1.21454 | 118 |
| GO_REGULATION_OF_UBIQUITIN_PROTEIN_LIGASE_ACTIVIT | 0.00156 | 0.03588 | 0.45506 | 0.77553 | 1.40416 | 21 |
| GO_REGULATION_OF_UBIQUITIN_PROTEIN_TRANSFERASE_ | 0.00113 | 0.02781 | 0.45506 | 0.70389 | 1.32187 | 46 |
| GO_REGULATION_OF_WNT_SIGNALING_PATHWAY | 0.00004 | 0.00173 | 0.55733 | 0.61015 | 1.19515 | 233 |
| GO_RESPONSE_TO_INTERLEUKIN_1 | 0.00000 | 0.00002 | 0.69013 | 0.69622 | 1.34658 | 118 |
| GO_RESPONSE_TO_INTERLEUKIN_4 | 0.00016 | 0.00570 | 0.51885 | 0.80329 | 1.46604 | 25 |
| GO_RESPONSE_TO_LEUKEMIA_INHIBITORY_FACTOR | 0.00006 | 0.00240 | 0.55733 | 0.68924 | 1.31657 | 79 |
| GO_RIBONUCLEOPROTEIN_COMPLEX_BINDING | 0.00001 | 0.00039 | 0.61053 | 0.67298 | 1.30163 | 118 |
| GO_RIBONUCLEOPROTEIN_COMPLEX_BIOGENESIS | 0.00000 | 0.00000 | | 0.71313 | 1.40833 | 398 |
| GO_RIBONUCLEOPROTEIN_COMPLEX_SUBUNIT_ORGANIZAT | 0.00000 | 0.00000 | | 0.71382 | 1.39252 | 181 |
| GO_RIBONUCLEOPROTEIN_GRANULE | 0.00003 | 0.00123 | 0.57561 | 0.63064 | 1.23013 | 185 |
| GO_RIBONUCLEOSIDE_METABOLIC_PROCESS | 0.00018 | 0.00630 | 0.51885 | 0.74370 | 1.38945 | 42 |
| GO_RIBOSOMAL_LARGE_SUBUNIT_ASSEMBLY | 0.00002 | 0.00081 | 0.59333 | 0.82140 | 1.50832 | 28 |
| GO_RIBOSOMAL_LARGE_SUBUNIT_BIOGENESIS | 0.00000 | 0.00000 | 0.76146 | 0.78387 | 1.48891 | 67 |
| GO_RIBOSOMAL_SMALL_SUBUNIT_BIOGENESIS | 0.00001 | 0.00034 | 0.61053 | 0.72219 | 1.37243 | 69 |
| GO_RIBOSOMAL_SUBUNIT | 0.00000 | 0.00000 | | 0.74345 | 1.44893 | 175 |
| GO_RIBOSOME | 0.00000 | 0.00000 | | 0.73335 | 1.43363 | 208 |
| GO_RIBOSOME_ASSEMBLY | 0.00000 | 0.00000 | 0.77494 | 0.80169 | 1.51666 | 60 |
| GO_RIBOSOME_BIOGENESIS | 0.00000 | 0.00000 | | 0.70558 | 1.38510 | 279 |
| GO_RNA_CATABOLIC_PROCESS | 0.00000 | 0.00000 | | 0.72915 | 1.43688 | 349 |
| GO_RNA_DEPENDENT_DNA_BIOSYNTHETIC_PROCESS | 0.00095 | 0.02438 | 0.47727 | 0.68237 | 1.29093 | 60 |
| GO_RNA_EXPORT_FROM_NUCLEUS | 0.00002 | 0.00114 | 0.57561 | 0.65300 | 1.26450 | 127 |
| GO_RNA_LOCALIZATION | 0.00000 | 0.00011 | 0.64355 | 0.64463 | 1.25888 | 200 |

| Cluster 1 | | | | | | |
|---|---|---|---|---|---|---|
| pathway | pval | padj | log2err | ES | NES | size |
| GO_RNA_SPLICING | 0.00000 | 0.00000 | | 0.65675 | 1.29512 | 376 |
| GO_RNA_SPLICING_VIA_TRANSESTERIFICATION_REACTIONS | 0.00000 | 0.00000 | | 0.66603 | 1.30941 | 305 |
| GO_RRNA_BINDING | 0.00000 | 0.00011 | 0.64355 | 0.75580 | 1.42647 | 55 |
| GO_RRNA_METABOLIC_PROCESS | 0.00000 | 0.00000 | | 0.69630 | 1.36156 | 212 |
| GO_SCF_DEPENDENT_PROTEASOMAL_UBIQUITIN_DEPEND | 0.00000 | 0.00013 | 0.64355 | 0.72686 | 1.38558 | 75 |
| GO_SECONDARY_METABOLIC_PROCESS | 0.00176 | 0.03950 | 0.45506 | 0.77156 | 1.40420 | 24 |
| GO_SECRETORY_GRANULE | 0.00039 | 0.01203 | 0.49849 | 0.56984 | 1.12706 | 459 |
| GO_SIGNAL_TRANSDUCTION_BY_P53_CLASS_MEDIATOR | 0.00019 | 0.00650 | 0.51885 | 0.60760 | 1.18803 | 209 |
| GO_SINGLE_STRANDED_RNA_BINDING | 0.00063 | 0.01764 | 0.47727 | 0.68168 | 1.29012 | 62 |
| GO_SKELETAL_MUSCLE_TISSUE_REGENERATION | 0.00178 | 0.03976 | 0.45506 | 0.77069 | 1.39691 | 22 |
| GO_SMALL_NUCLEAR_RIBONUCLEOPROTEIN_COMPLEX | 0.00148 | 0.03458 | 0.45506 | 0.67608 | 1.27902 | 60 |
| GO_SMALL_RIBOSOMAL_SUBUNIT | 0.00000 | 0.00006 | 0.65944 | 0.74820 | 1.42191 | 68 |
| GO_SMALL_SUBUNIT_PROCESSOME | 0.00139 | 0.03269 | 0.45506 | 0.73925 | 1.36579 | 32 |
| GO_SM_LIKE_PROTEIN_FAMILY_COMPLEX | 0.00069 | 0.01900 | 0.47727 | 0.67314 | 1.28061 | 71 |
| GO_SNORNA_BINDING | 0.00073 | 0.01939 | 0.47727 | 0.76014 | 1.40082 | 29 |
| GO_SPINDLE_MIDZONE | 0.00087 | 0.02284 | 0.47727 | 0.76149 | 1.40419 | 30 |
| GO_SPLICEOSOMAL_COMPLEX | 0.00000 | 0.00000 | 0.74774 | 0.67625 | 1.31684 | 170 |
| GO_SPLICEOSOMAL_COMPLEX_ASSEMBLY | 0.00191 | 0.04234 | 0.45506 | 0.69560 | 1.30935 | 50 |
| GO_STEM_CELL_DIFFERENTIATION | 0.00000 | 0.00005 | 0.65944 | 0.67336 | 1.30887 | 155 |
| GO_STRUCTURAL_CONSTITUENT_OF_RIBOSOME | 0.00000 | 0.00000 | | 0.75828 | 1.47313 | 152 |
| GO_STRUCTURAL_MOLECULE_ACTIVITY | 0.00000 | 0.00000 | | 0.68250 | 1.34548 | 365 |
| GO_TELOMERASE_RNA_LOCALIZATION | 0.00027 | 0.00876 | 0.49849 | 0.82255 | 1.48263 | 19 |
| GO_THREONINE_TYPE_ENDOPEPTIDASE_ACTIVITY | 0.00003 | 0.00150 | 0.55733 | 0.87165 | 1.56568 | 18 |
| GO_THREONINE_TYPE_PEPTIDASE_ACTIVITY | 0.00002 | 0.00082 | 0.57561 | 0.84877 | 1.53843 | 22 |
| GO_TRANSCRIPTION_COACTIVATOR_ACTIVITY | 0.00117 | 0.02837 | 0.45506 | 0.59841 | 1.16914 | 202 |
| GO_TRANSCRIPTION_COREGULATOR_ACTIVITY | 0.00019 | 0.00654 | 0.51885 | 0.58467 | 1.15229 | 357 |
| GO_TRANSCRIPTION_COUPLED_NUCLEOTIDE_EXCISION_RE | 0.00072 | 0.01931 | 0.47727 | 0.67286 | 1.28008 | 71 |
| GO_TRANSCRIPTION_FACTOR_BINDING | 0.00001 | 0.00036 | 0.61053 | 0.59180 | 1.16976 | 439 |
| GO_TRANSLATIONAL_ELONGATION | 0.00121 | 0.02915 | 0.45506 | 0.62600 | 1.21183 | 121 |
| GO_TRANSLATIONAL_INITIATION | 0.00000 | 0.00000 | | 0.82956 | 1.61517 | 169 |
| GO_TRANSLATION_FACTOR_ACTIVITY_RNA_BINDING | 0.00000 | 0.00000 | | 0.81900 | 1.55642 | 69 |
| GO_TRANSLATION_INITIATION_FACTOR_ACTIVITY | 0.00000 | 0.00000 | 0.71951 | 0.82042 | 1.53829 | 45 |
| GO_TRANSLATION_INITIATION_FACTOR_BINDING | 0.00046 | 0.01364 | 0.49849 | 0.77465 | 1.42246 | 28 |
| GO_TRANSLATION_PREINITIATION_COMPLEX | 0.00001 | 0.00043 | 0.61053 | 0.91071 | 1.61939 | 16 |
| GO_TRANSLATION_REGULATOR_ACTIVITY | 0.00000 | 0.00000 | | 0.78326 | 1.51083 | 109 |
| GO_TRANSLATION_REGULATOR_ACTIVITY_NUCLEIC_ACID_E | 0.00000 | 0.00000 | | 0.80895 | 1.54951 | 84 |
| GO_TUMOR_NECROSIS_FACTOR_MEDIATED_SIGNALING_PA | 0.00002 | 0.00085 | 0.57561 | 0.66819 | 1.29122 | 113 |
| GO_T_CELL_RECEPTOR_SIGNALING_PATHWAY | 0.00000 | 0.00002 | 0.69013 | 0.69199 | 1.34009 | 126 |
| GO_U2_TYPE_SPLICEOSOMAL_COMPLEX | 0.00013 | 0.00480 | 0.51885 | 0.67295 | 1.28992 | 86 |
| GO_UBIQUITIN_LIKE_PROTEIN_LIGASE_BINDING | 0.00062 | 0.01746 | 0.47727 | 0.59775 | 1.17002 | 227 |
| GO_UBIQUITIN_PROTEIN_TRANSFERASE_REGULATOR_ACTI | 0.00113 | 0.02781 | 0.45506 | 0.82945 | 1.47041 | 15 |
| GO_UNFOLDED_PROTEIN_BINDING | 0.00000 | 0.00011 | 0.64355 | 0.71656 | 1.37336 | 85 |
| GO_VACUOLAR_LUMEN | 0.00117 | 0.02837 | 0.45506 | 0.63516 | 1.22345 | 104 |
| GO_VESICLE_LUMEN | 0.00000 | 0.00000 | 0.73376 | 0.66761 | 1.30272 | 192 |
| GO_VIRAL_GENE_EXPRESSION | 0.00000 | 0.00000 | | 0.78095 | 1.52168 | 173 |
| GO_VIRAL_LIFE_CYCLE | 0.00046 | 0.01364 | 0.49849 | 0.60084 | 1.17527 | 217 |
| HP_ABNORMALITY_OF_LOWER_LIP | 0.00137 | 0.03238 | 0.45506 | 0.60581 | 1.17908 | 163 |
| HP_ABNORMALITY_OF_THE_NASAL_DORSUM | 0.00023 | 0.00784 | 0.51885 | 0.62914 | 1.22104 | 148 |
| HP_ACUTE_LEUKEMIA | 0.00007 | 0.00273 | 0.53843 | 0.70226 | 1.33601 | 71 |
| HP_ANEMIA_OF_INADEQUATE_PRODUCTION | 0.00004 | 0.00167 | 0.55733 | 0.67482 | 1.30041 | 105 |
| HP_ATRESIA_OF_THE_EXTERNAL_AUDITORY_CANAL | 0.00014 | 0.00518 | 0.51885 | 0.79304 | 1.45623 | 28 |
| HP_DELAYED_PUBERTY | 0.00070 | 0.01908 | 0.47727 | 0.63039 | 1.21914 | 117 |
| HP_DEPRESSED_NASAL_RIDGE | 0.00000 | 0.00024 | 0.62726 | 0.71492 | 1.36767 | 82 |
| HP_LEUKEMIA | 0.00027 | 0.00866 | 0.49849 | 0.64879 | 1.24941 | 102 |
| HP_MACROCYTIC_ANEMIA | 0.00000 | 0.00022 | 0.62726 | 0.80017 | 1.49257 | 40 |
| HP_MIGRAINE | 0.00001 | 0.00064 | 0.59333 | 0.71868 | 1.36725 | 71 |
| HP_ORAL_CLEFT | 0.00072 | 0.01931 | 0.47727 | 0.57695 | 1.13704 | 355 |
| HP_THICK_LOWER_LIP_VERMILION | 0.00042 | 0.01274 | 0.49849 | 0.64400 | 1.24220 | 109 |
| HP_THICK_VERMILION_BORDER | 0.00034 | 0.01058 | 0.49849 | 0.62114 | 1.21015 | 172 |
| **Cluster 6** | | | | | | |
| pathway | pval | padj | log2err | ES | NES | size |
| GO_ACTIVATION_OF_IMMUNE_RESPONSE | 0.00000 | 0.00000 | | 0.65686 | 1.36705 | 224 |
| GO_ACTIVATION_OF_INNATE_IMMUNE_RESPONSE | 0.00000 | 0.00000 | | 0.75768 | 1.54184 | 89 |
| GO_AEROBIC_RESPIRATION | 0.00020 | 0.00485 | 0.51885 | 0.67533 | 1.35944 | 68 |
| GO_AMINE_METABOLIC_PROCESS | 0.00000 | 0.00000 | | 0.80547 | 1.63158 | 79 |
| GO_ANAPHASE_PROMOTING_COMPLEX_DEPENDENT_CATA | 0.00000 | 0.00000 | | 0.78206 | 1.58247 | 77 |
| GO_ANCHORING_JUNCTION | 0.00000 | 0.00000 | | 0.63415 | 1.33793 | 482 |
| GO_ANIMAL_ORGAN_MORPHOGENESIS | 0.00283 | 0.04583 | 0.43171 | 0.53392 | 1.12406 | 432 |
| GO_ANTIGEN_PROCESSING_AND_PRESENTATION | 0.00000 | 0.00000 | 0.78819 | 0.68827 | 1.42434 | 143 |

| Cluster 1 | | | | | | |
|---|---|---|---|---|---|---|
| pathway | pval | padj | log2err | ES | NES | size |
| GO_ANTIGEN_PROCESSING_AND_PRESENTATION_OF_EXOG | 0.00000 | 0.00000 | 0.83909 | 0.81145 | 1.62037 | 56 |
| GO_ANTIGEN_PROCESSING_AND_PRESENTATION_OF_PEPT | 0.00000 | 0.00000 | 0.81404 | 0.70435 | 1.45105 | 124 |
| GO_ANTIGEN_PROCESSING_AND_PRESENTATION_OF_PEPT | 0.00000 | 0.00000 | 0.82666 | 0.78860 | 1.58852 | 69 |
| GO_ANTIGEN_RECEPTOR_MEDIATED_SIGNALING_PATHWAY | 0.00000 | 0.00000 | | 0.71099 | 1.46924 | 134 |
| GO_ATP_METABOLIC_PROCESS | 0.00152 | 0.02744 | 0.45506 | 0.57183 | 1.18697 | 196 |
| GO_AUTOPHAGOSOME | 0.00309 | 0.04866 | 0.43171 | 0.62617 | 1.26240 | 71 |
| GO_AUTOPHAGOSOME_MEMBRANE | 0.00198 | 0.03436 | 0.43171 | 0.73779 | 1.41820 | 26 |
| GO_AXONAL_GROWTH_CONE | 0.00216 | 0.03702 | 0.43171 | 0.79675 | 1.48553 | 16 |
| GO_AZUROPHIL_GRANULE | 0.00012 | 0.00313 | 0.53843 | 0.64288 | 1.31571 | 100 |
| GO_AZUROPHIL_GRANULE_LUMEN | 0.00073 | 0.01525 | 0.47727 | 0.67556 | 1.34803 | 55 |
| GO_BLOOD_MICROPARTICLE | 0.00013 | 0.00328 | 0.51885 | 0.75052 | 1.47079 | 35 |
| GO_BRUSH_BORDER | 0.00011 | 0.00304 | 0.53843 | 0.68376 | 1.37034 | 63 |
| GO_CADHERIN_BINDING | 0.00000 | 0.00000 | | 0.66685 | 1.38951 | 245 |
| GO_CANONICAL_WNT_SIGNALING_PATHWAY | 0.00000 | 0.00002 | 0.67496 | 0.63197 | 1.31136 | 190 |
| GO_CATALYTIC_STEP_2_SPLICEOSOME | 0.00004 | 0.00130 | 0.55733 | 0.67104 | 1.36120 | 81 |
| GO_CELLULAR_AMINO_ACID_METABOLIC_PROCESS | 0.00000 | 0.00005 | 0.64355 | 0.61947 | 1.28919 | 222 |
| GO_CELLULAR_COMPONENT_DISASSEMBLY | 0.00111 | 0.02175 | 0.45506 | 0.54361 | 1.14326 | 397 |
| GO_CELLULAR_KETONE_METABOLIC_PROCESS | 0.00000 | 0.00001 | 0.69013 | 0.65743 | 1.36140 | 153 |
| GO_CELLULAR_NITROGEN_COMPOUND_CATABOLIC_PROCE | 0.00000 | 0.00000 | | 0.71601 | 1.50814 | 440 |
| GO_CELLULAR_RESPIRATION | 0.00128 | 0.02401 | 0.45506 | 0.59370 | 1.22814 | 140 |
| GO_CELLULAR_RESPONSE_TO_HEAT | 0.00004 | 0.00123 | 0.55733 | 0.65576 | 1.34092 | 98 |
| GO_CELLULAR_RESPONSE_TO_ORGANIC_CYCLIC_COMPOUI | 0.00128 | 0.02401 | 0.45506 | 0.55595 | 1.16181 | 278 |
| GO_CELLULAR_RESPONSE_TO_OXYGEN_LEVELS | 0.00000 | 0.00000 | 0.74774 | 0.67068 | 1.38871 | 156 |
| GO_CELL_ACTIVATION_INVOLVED_IN_IMMUNE_RESPONSE | 0.00000 | 0.00000 | | 0.62577 | 1.31443 | 363 |
| GO_CELL_ADHESION_MOLECULE_BINDING | 0.00000 | 0.00000 | | 0.63663 | 1.33336 | 317 |
| GO_CELL_BODY | 0.00003 | 0.00087 | 0.57561 | 0.58698 | 1.22555 | 268 |
| GO_CELL_CELL_RECOGNITION | 0.00000 | 0.00001 | 0.69013 | 0.88422 | 1.69040 | 23 |
| GO_CELL_CELL_SIGNALING_BY_WNT | 0.00000 | 0.00000 | 0.77494 | 0.62503 | 1.30750 | 294 |
| GO_CELL_CYCLE_G2_M_PHASE_TRANSITION | 0.00004 | 0.00123 | 0.55733 | 0.60322 | 1.25522 | 221 |
| GO_CELL_CYCLE_PHASE_TRANSITION | 0.00001 | 0.00046 | 0.59333 | 0.55937 | 1.17968 | 469 |
| GO_CELL_RECOGNITION | 0.00000 | 0.00004 | 0.65944 | 0.78226 | 1.54690 | 43 |
| GO_CELL_SUBSTRATE_JUNCTION | 0.00000 | 0.00000 | | 0.68333 | 1.42923 | 292 |
| GO_CELL_SURFACE_RECEPTOR_SIGNALING_PATHWAY_INVC | 0.00000 | 0.00000 | 0.74774 | 0.61379 | 1.28559 | 319 |
| GO_CHAPERONE_BINDING | 0.00015 | 0.00376 | 0.51885 | 0.66731 | 1.34862 | 75 |
| GO_CHAPERONE_COFACTOR_DEPENDENT_PROTEIN_REFOL | 0.00087 | 0.01762 | 0.47727 | 0.77871 | 1.47731 | 20 |
| GO_CHAPERONE_COMPLEX | 0.00000 | 0.00003 | 0.65944 | 0.87002 | 1.66327 | 23 |
| GO_CLATHRIN_COATED_VESICLE | 0.00316 | 0.04956 | 0.43171 | 0.60440 | 1.23646 | 99 |
| GO_CLATHRIN_COATED_VESICLE_MEMBRANE | 0.00117 | 0.02253 | 0.45506 | 0.68031 | 1.35525 | 51 |
| GO_CLUSTER_OF_ACTIN_BASED_CELL_PROJECTIONS | 0.00016 | 0.00400 | 0.51885 | 0.64878 | 1.31707 | 84 |
| GO_COATED_VESICLE | 0.00130 | 0.02420 | 0.45506 | 0.58628 | 1.21569 | 174 |
| GO_COATED_VESICLE_MEMBRANE | 0.00071 | 0.01498 | 0.47727 | 0.62521 | 1.27845 | 98 |
| GO_CONTRACTILE_FIBER | 0.00081 | 0.01668 | 0.47727 | 0.61451 | 1.26187 | 110 |
| GO_COP9_SIGNALOSOME | 0.00228 | 0.03846 | 0.43171 | 0.73824 | 1.41665 | 25 |
| GO_COTRANSLATIONAL_PROTEIN_TARGETING_TO_MEMBR | 0.00000 | 0.00000 | | 0.89237 | 1.81770 | 91 |
| GO_CYTOKINE_MEDIATED_SIGNALING_PATHWAY | 0.00000 | 0.00000 | | 0.65879 | 1.38159 | 338 |
| GO_CYTOPLASMIC_TRANSLATION | 0.00000 | 0.00000 | | 0.82933 | 1.68843 | 90 |
| GO_CYTOPLASMIC_TRANSLATIONAL_INITIATION | 0.00000 | 0.00008 | 0.62726 | 0.81642 | 1.58160 | 30 |
| GO_CYTOSOLIC_LARGE_RIBOSOMAL_SUBUNIT | 0.00000 | 0.00000 | | 0.88682 | 1.76823 | 53 |
| GO_CYTOSOLIC_RIBOSOME | 0.00000 | 0.00000 | | 0.88579 | 1.80786 | 94 |
| GO_CYTOSOLIC_SMALL_RIBOSOMAL_SUBUNIT | 0.00000 | 0.00000 | | 0.89114 | 1.75330 | 39 |
| GO_DEFENSE_RESPONSE_TO_OTHER_ORGANISM | 0.00000 | 0.00000 | 0.81404 | 0.60469 | 1.27291 | 427 |
| GO_DE_NOVO_PROTEIN_FOLDING | 0.00031 | 0.00717 | 0.49849 | 0.76547 | 1.47412 | 27 |
| GO_DISORDERED_DOMAIN_SPECIFIC_BINDING | 0.00042 | 0.00944 | 0.49849 | 0.76090 | 1.46532 | 27 |
| GO_DNA_BIOSYNTHETIC_PROCESS | 0.00098 | 0.01970 | 0.45506 | 0.59320 | 1.22787 | 145 |
| GO_DNA_DAMAGE_RESPONSE_DETECTION_OF_DNA_DAM | 0.00220 | 0.03743 | 0.43171 | 0.69757 | 1.36900 | 36 |
| GO_DRUG_BINDING | 0.00143 | 0.02602 | 0.45506 | 0.67835 | 1.34927 | 50 |
| GO_ENDOPEPTIDASE_COMPLEX | 0.00000 | 0.00000 | 0.74774 | 0.77771 | 1.55825 | 62 |
| GO_ENERGY_DERIVATION_BY_OXIDATION_OF_ORGANIC_C( | 0.00076 | 0.01590 | 0.47727 | 0.58252 | 1.20945 | 185 |
| GO_ERROR_FREE_TRANSLESION_SYNTHESIS | 0.00152 | 0.02744 | 0.45506 | 0.76547 | 1.45219 | 20 |
| GO_ERROR_PRONE_TRANSLESION_SYNTHESIS | 0.00290 | 0.04677 | 0.43171 | 0.75423 | 1.43087 | 20 |
| GO_ESTABLISHMENT_OF_PROTEIN_LOCALIZATION_TO_CHR | 0.00004 | 0.00123 | 0.55733 | 0.82317 | 1.57370 | 23 |
| GO_ESTABLISHMENT_OF_PROTEIN_LOCALIZATION_TO_END | 0.00000 | 0.00000 | | 0.86684 | 1.77420 | 101 |
| GO_ESTABLISHMENT_OF_PROTEIN_LOCALIZATION_TO_MEN | 0.00000 | 0.00000 | | 0.74875 | 1.56007 | 244 |
| GO_ESTABLISHMENT_OF_PROTEIN_LOCALIZATION_TO_ORG | 0.00000 | 0.00000 | | 0.69423 | 1.46140 | 426 |
| GO_ESTABLISHMENT_OF_PROTEIN_LOCALIZATION_TO_TEL( | 0.00001 | 0.00042 | 0.59333 | 0.88263 | 1.65132 | 17 |
| GO_ESTABLISHMENT_OF_RNA_LOCALIZATION | 0.00040 | 0.00900 | 0.49849 | 0.59594 | 1.23573 | 167 |
| GO_ESTABLISHMENT_OF_TISSUE_POLARITY | 0.00000 | 0.00000 | | 0.79044 | 1.60614 | 86 |
| GO_EUKARYOTIC_TRANSLATION_INITIATION_FACTOR_3_CO | 0.00004 | 0.00127 | 0.55733 | 0.87178 | 1.62096 | 15 |
| GO_EXOCYTOSIS | 0.00000 | 0.00000 | | 0.60795 | 1.28269 | 483 |

| Cluster 1 | | | | | | |
|---|---|---|---|---|---|---|
| pathway | pval | padj | log2err | ES | NES | size |
| GO_FC_EPSILON_RECEPTOR_SIGNALING_PATHWAY | 0.00000 | 0.00000 | 0.80122 | 0.75021 | 1.52439 | 86 |
| GO_FC_RECEPTOR_SIGNALING_PATHWAY | 0.00000 | 0.00000 | | 0.72449 | 1.49347 | 127 |
| GO_FERTILIZATION | 0.00016 | 0.00396 | 0.51885 | 0.69004 | 1.38204 | 61 |
| GO_FICOLIN_1_RICH_GRANULE | 0.00000 | 0.00000 | 0.80122 | 0.73420 | 1.50045 | 96 |
| GO_GENERATION_OF_PRECURSOR_METABOLITES_AND_EN | 0.00089 | 0.01811 | 0.47727 | 0.55246 | 1.15773 | 327 |
| GO_GLUTAMATERGIC_SYNAPSE | 0.00005 | 0.00143 | 0.55733 | 0.62341 | 1.29004 | 142 |
| GO_GLYCOSYL_COMPOUND_METABOLIC_PROCESS | 0.00073 | 0.01525 | 0.47727 | 0.65168 | 1.31394 | 70 |
| GO_GUANYL_NUCLEOTIDE_BINDING | 0.00005 | 0.00150 | 0.55733 | 0.59607 | 1.23965 | 217 |
| GO_HEAT_SHOCK_PROTEIN_BINDING | 0.00001 | 0.00039 | 0.59333 | 0.67719 | 1.37805 | 89 |
| GO_HEMATOPOIETIC_PROGENITOR_CELL_DIFFERENTIATION | 0.00000 | 0.00000 | 0.81404 | 0.72722 | 1.49322 | 109 |
| GO_HEMATOPOIETIC_STEM_CELL_DIFFERENTIATION | 0.00000 | 0.00000 | | 0.81143 | 1.63603 | 70 |
| GO_HSP70_PROTEIN_BINDING | 0.00055 | 0.01186 | 0.47727 | 0.74468 | 1.43573 | 28 |
| GO_HSP90_PROTEIN_BINDING | 0.00106 | 0.02081 | 0.45506 | 0.74218 | 1.43260 | 29 |
| GO_IMMUNE_RESPONSE_REGULATING_SIGNALING_PATHW | 0.00000 | 0.00000 | 0.82666 | 0.66192 | 1.37523 | 209 |
| GO_INFLAMMATORY_RESPONSE_TO_ANTIGENIC_STIMULUS | 0.00111 | 0.02175 | 0.45506 | 0.81401 | 1.51772 | 16 |
| GO_INNATE_IMMUNE_RESPONSE | 0.00000 | 0.00000 | | 0.63337 | 1.32933 | 354 |
| GO_INNATE_IMMUNE_RESPONSE_ACTIVATING_SIGNAL_TR | 0.00000 | 0.00000 | | 0.79000 | 1.59533 | 73 |
| GO_INTERACTION_WITH_SYMBIONT | 0.00126 | 0.02390 | 0.45506 | 0.69120 | 1.36683 | 43 |
| GO_INTERLEUKIN_1_MEDIATED_SIGNALING_PATHWAY | 0.00000 | 0.00000 | | 0.80579 | 1.62721 | 73 |
| GO_ION_CHANNEL_BINDING | 0.00006 | 0.00163 | 0.55733 | 0.69187 | 1.38659 | 63 |
| GO_LARGE_RIBOSOMAL_SUBUNIT | 0.00000 | 0.00000 | | 0.74201 | 1.52258 | 108 |
| GO_LEUKOCYTE_MEDIATED_IMMUNITY | 0.00000 | 0.00000 | | 0.62133 | 1.30617 | 389 |
| GO_LYSOSOMAL_LUMEN | 0.00017 | 0.00412 | 0.51885 | 0.70388 | 1.39943 | 49 |
| GO_MACROAUTOPHAGY | 0.00202 | 0.03490 | 0.43171 | 0.55631 | 1.15999 | 250 |
| GO_MATURATION_OF_LSU_RRNA | 0.00098 | 0.01970 | 0.45506 | 0.75090 | 1.44340 | 26 |
| GO_MODIFICATION_DEPENDENT_MACROMOLECULE_CATA | 0.00000 | 0.00007 | 0.64355 | 0.56827 | 1.19937 | 487 |
| GO_MODULATION_BY_HOST_OF_VIRAL_PROCESS | 0.00231 | 0.03886 | 0.43171 | 0.80305 | 1.49317 | 15 |
| GO_MONOSACCHARIDE_BIOSYNTHETIC_PROCESS | 0.00302 | 0.04776 | 0.43171 | 0.64458 | 1.29065 | 60 |
| GO_MORPHOGENESIS_OF_AN_EPITHELIUM | 0.00000 | 0.00001 | 0.69013 | 0.60639 | 1.26716 | 277 |
| GO_MORPHOGENESIS_OF_A_POLARIZED_EPITHELIUM | 0.00000 | 0.00000 | | 0.78359 | 1.60304 | 99 |
| GO_MRNA_3_UTR_BINDING | 0.00003 | 0.00103 | 0.55733 | 0.68850 | 1.38818 | 70 |
| GO_MRNA_5_UTR_BINDING | 0.00036 | 0.00831 | 0.49849 | 0.79987 | 1.52457 | 22 |
| GO_MRNA_BINDING | 0.00000 | 0.00000 | | 0.65728 | 1.36859 | 232 |
| GO_MRNA_EXPORT_FROM_NUCLEUS | 0.00298 | 0.04756 | 0.43171 | 0.60800 | 1.24325 | 98 |
| GO_MRNA_PROCESSING | 0.00000 | 0.00006 | 0.64355 | 0.57406 | 1.20767 | 417 |
| GO_MRNA_TRANSPORT | 0.00119 | 0.02265 | 0.45506 | 0.59944 | 1.23494 | 125 |
| GO_MULTI_ORGANISM_PROCESS | 0.00119 | 0.02265 | 0.45506 | 0.53378 | 1.12672 | 492 |
| GO_MYD88_DEPENDENT_TOLL_LIKE_RECEPTOR_SIGNALING | 0.00141 | 0.02593 | 0.45506 | 0.78769 | 1.48090 | 18 |
| GO_MYD88_INDEPENDENT_TOLL_LIKE_RECEPTOR_SIGNALI | 0.00002 | 0.00063 | 0.57561 | 0.83234 | 1.58517 | 21 |
| GO_MYELOID_LEUKOCYTE_ACTIVATION | 0.00000 | 0.00000 | | 0.64617 | 1.35555 | 344 |
| GO_MYELOID_LEUKOCYTE_MEDIATED_IMMUNITY | 0.00000 | 0.00000 | | 0.65462 | 1.37089 | 315 |
| GO_NADH_METABOLIC_PROCESS | 0.00018 | 0.00432 | 0.51885 | 0.77495 | 1.49237 | 27 |
| GO_NAD_BINDING | 0.00143 | 0.02602 | 0.45506 | 0.69885 | 1.37676 | 40 |
| GO_NAD_METABOLIC_PROCESS | 0.00250 | 0.04113 | 0.43171 | 0.71123 | 1.37783 | 30 |
| GO_NCRNA_PROCESSING | 0.00157 | 0.02817 | 0.45506 | 0.54875 | 1.15013 | 328 |
| GO_NEGATIVE_REGULATION_OF_CALCIUM_MEDIATED_SIG | 0.00132 | 0.02447 | 0.45506 | 0.78929 | 1.48392 | 18 |
| GO_NEGATIVE_REGULATION_OF_CANONICAL_WNT_SIGNA | 0.00000 | 0.00000 | 0.80122 | 0.72823 | 1.49430 | 106 |
| GO_NEGATIVE_REGULATION_OF_CELLULAR_AMIDE_METAB | 0.00011 | 0.00294 | 0.53843 | 0.63739 | 1.30967 | 112 |
| GO_NEGATIVE_REGULATION_OF_CELL_CYCLE | 0.00058 | 0.01241 | 0.47727 | 0.54392 | 1.14497 | 430 |
| GO_NEGATIVE_REGULATION_OF_CELL_CYCLE_G2_M_PHAS | 0.00000 | 0.00000 | 0.71951 | 0.72647 | 1.47615 | 86 |
| GO_NEGATIVE_REGULATION_OF_CELL_CYCLE_PHASE_TRAN | 0.00002 | 0.00059 | 0.57561 | 0.60974 | 1.26539 | 188 |
| GO_NEGATIVE_REGULATION_OF_CELL_CYCLE_PROCESS | 0.00001 | 0.00023 | 0.61053 | 0.59693 | 1.24455 | 251 |
| GO_NEGATIVE_REGULATION_OF_MITOTIC_CELL_CYCLE | 0.00016 | 0.00410 | 0.51885 | 0.58111 | 1.21012 | 234 |
| GO_NEGATIVE_REGULATION_OF_RNA_CATABOLIC_PROCESS | 0.00242 | 0.04026 | 0.43171 | 0.68066 | 1.34599 | 43 |
| GO_NEGATIVE_REGULATION_OF_WNT_SIGNALING_PATHW | 0.00000 | 0.00000 | | 0.71891 | 1.47985 | 118 |
| GO_NEURON_TO_NEURON_SYNAPSE | 0.00006 | 0.00168 | 0.55733 | 0.61284 | 1.26979 | 162 |
| GO_NIK_NF_KAPPAB_SIGNALING | 0.00000 | 0.00000 | | 0.76169 | 1.56003 | 102 |
| GO_NON_CANONICAL_WNT_SIGNALING_PATHWAY | 0.00000 | 0.00000 | | 0.78168 | 1.59913 | 99 |
| GO_NUCLEAR_EXPORT | 0.00043 | 0.00957 | 0.49849 | 0.59770 | 1.23918 | 172 |
| GO_NUCLEAR_PERIPHERY | 0.00130 | 0.02420 | 0.45506 | 0.61031 | 1.25106 | 104 |
| GO_NUCLEAR_TRANSCRIBED_MRNA_CATABOLIC_PROCESS | 0.00000 | 0.00000 | | 0.78049 | 1.62017 | 187 |
| GO_NUCLEAR_TRANSCRIBED_MRNA_CATABOLIC_PROCESS_ | 0.00000 | 0.00000 | | 0.87454 | 1.79815 | 114 |
| GO_NUCLEAR_TRANSPORT | 0.00007 | 0.00184 | 0.53843 | 0.58142 | 1.21470 | 272 |
| GO_NUCLEOBASE_CONTAINING_COMPOUND_TRANSPORT | 0.00013 | 0.00338 | 0.51885 | 0.59174 | 1.22972 | 203 |
| GO_NUCLEOBASE_CONTAINING_SMALL_MOLECULE_METAB | 0.00316 | 0.04956 | 0.43171 | 0.53870 | 1.13095 | 357 |
| GO_NUCLEOSIDE_METABOLIC_PROCESS | 0.00063 | 0.01333 | 0.47727 | 0.67429 | 1.34700 | 57 |
| GO_NUCLEOSIDE_PHOSPHATE_BIOSYNTHETIC_PROCESS | 0.00250 | 0.04113 | 0.43171 | 0.58347 | 1.20753 | 148 |
| GO_NUCLEOTIDE_EXCISION_REPAIR | 0.00133 | 0.02473 | 0.45506 | 0.61738 | 1.26243 | 98 |
| GO_NUCLEOTIDE_EXCISION_REPAIR_DNA_DAMAGE_RECOG | 0.00002 | 0.00068 | 0.57561 | 0.82965 | 1.58609 | 23 |

| Cluster 1 | | | | | | |
|---|---|---|---|---|---|---|
| pathway | pval | padj | log2err | ES | NES | size |
| GO_ORGANIC_CYCLIC_COMPOUND_CATABOLIC_PROCESS | 0.00000 | 0.00000 | | 0.70999 | 1.49611 | 451 |
| GO_PEPTIDASE_COMPLEX | 0.00000 | 0.00010 | 0.62726 | 0.71156 | 1.44064 | 78 |
| GO_PERINUCLEAR_REGION_OF_CYTOPLASM | 0.00011 | 0.00288 | 0.53843 | 0.55465 | 1.16709 | 423 |
| GO_PIGMENT_GRANULE | 0.00000 | 0.00005 | 0.64355 | 0.71405 | 1.44426 | 76 |
| GO_PLASMA_LIPOPROTEIN_PARTICLE_CLEARANCE | 0.00084 | 0.01718 | 0.47727 | 0.73360 | 1.42511 | 31 |
| GO_POLYSOMAL_RIBOSOME | 0.00000 | 0.00000 | 0.81404 | 0.88672 | 1.72258 | 31 |
| GO_POLYSOME | 0.00000 | 0.00000 | | 0.82116 | 1.64422 | 60 |
| GO_POLY_PYRIMIDINE_TRACT_BINDING | 0.00150 | 0.02727 | 0.45506 | 0.79045 | 1.49202 | 19 |
| GO_POSITIVE_REGULATION_OF_CANONICAL_WNT_SIGNAL | 0.00000 | 0.00000 | 0.80122 | 0.73528 | 1.50331 | 97 |
| GO_POSITIVE_REGULATION_OF_CELLULAR_AMIDE_METABO | 0.00076 | 0.01590 | 0.47727 | 0.61761 | 1.27124 | 119 |
| GO_POSITIVE_REGULATION_OF_CELLULAR_PROTEIN_LOCAL | 0.00005 | 0.00144 | 0.55733 | 0.58941 | 1.22708 | 238 |
| GO_POSITIVE_REGULATION_OF_CELL_DEATH | 0.00220 | 0.03743 | 0.43171 | 0.54138 | 1.13738 | 377 |
| GO_POSITIVE_REGULATION_OF_DEFENSE_RESPONSE | 0.00000 | 0.00000 | 0.74774 | 0.66393 | 1.37664 | 168 |
| GO_POSITIVE_REGULATION_OF_DNA_BIOSYNTHETIC_PROC | 0.00001 | 0.00020 | 0.61053 | 0.73340 | 1.46233 | 53 |
| GO_POSITIVE_REGULATION_OF_ESTABLISHMENT_OF_PROT | 0.00000 | 0.00004 | 0.64355 | 0.61510 | 1.28015 | 224 |
| GO_POSITIVE_REGULATION_OF_IMMUNE_RESPONSE | 0.00000 | 0.00000 | 0.82666 | 0.62976 | 1.31747 | 296 |
| GO_POSITIVE_REGULATION_OF_IMMUNE_SYSTEM_PROCES | 0.00000 | 0.00002 | 0.65944 | 0.57930 | 1.21868 | 417 |
| GO_POSITIVE_REGULATION_OF_INNATE_IMMUNE_RESPON | 0.00000 | 0.00000 | 0.73376 | 0.69826 | 1.43723 | 119 |
| GO_POSITIVE_REGULATION_OF_INTRINSIC_APOPTOTIC_SIG | 0.00054 | 0.01171 | 0.47727 | 0.72783 | 1.41951 | 34 |
| GO_POSITIVE_REGULATION_OF_MRNA_SPLICING_VIA_SPLI | 0.00159 | 0.02832 | 0.45506 | 0.80431 | 1.49962 | 16 |
| GO_POSITIVE_REGULATION_OF_PROTEIN_CONTAINING_CO | 0.00194 | 0.03382 | 0.45506 | 0.75769 | 1.44851 | 23 |
| GO_POSITIVE_REGULATION_OF_PROTEIN_IMPORT | 0.00287 | 0.04630 | 0.43171 | 0.72311 | 1.39578 | 29 |
| GO_POSITIVE_REGULATION_OF_PROTEIN_LOCALIZATION_T | 0.00001 | 0.00026 | 0.61053 | 0.72550 | 1.45364 | 62 |
| GO_POSITIVE_REGULATION_OF_REACTIVE_OXYGEN_SPECIE | 0.00224 | 0.03795 | 0.43171 | 0.79662 | 1.49040 | 17 |
| GO_POSITIVE_REGULATION_OF_RESPONSE_TO_BIOTIC_STI | 0.00000 | 0.00000 | 0.76146 | 0.68520 | 1.41595 | 134 |
| GO_POSITIVE_REGULATION_OF_RESPONSE_TO_EXTERNAL_ | 0.00000 | 0.00011 | 0.62726 | 0.61865 | 1.28593 | 215 |
| GO_POSITIVE_REGULATION_OF_TELOMERASE_ACTIVITY | 0.00011 | 0.00296 | 0.53843 | 0.76504 | 1.48207 | 30 |
| GO_POSITIVE_REGULATION_OF_TELOMERASE_RNA_LOCALI | 0.00000 | 0.00011 | 0.62726 | 0.90449 | 1.68178 | 15 |
| GO_POSITIVE_REGULATION_OF_TELOMERE_MAINTENANCE | 0.00046 | 0.01009 | 0.49849 | 0.70738 | 1.40108 | 44 |
| GO_POSITIVE_REGULATION_OF_TELOMERE_MAINTENANCE | 0.00124 | 0.02363 | 0.45506 | 0.73755 | 1.43595 | 32 |
| GO_POSITIVE_REGULATION_OF_TRANSLATION | 0.00023 | 0.00552 | 0.51885 | 0.62962 | 1.29065 | 104 |
| GO_POSITIVE_REGULATION_OF_TRANSPORT | 0.00003 | 0.00103 | 0.55733 | 0.55804 | 1.17504 | 437 |
| GO_POSITIVE_REGULATION_OF_VIRAL_GENOME_REPLICAT | 0.00250 | 0.04113 | 0.43171 | 0.72303 | 1.40769 | 32 |
| GO_POSITIVE_REGULATION_OF_VIRAL_LIFE_CYCLE | 0.00044 | 0.00964 | 0.49849 | 0.69170 | 1.37522 | 49 |
| GO_POSITIVE_REGULATION_OF_VIRAL_PROCESS | 0.00092 | 0.01861 | 0.47727 | 0.63822 | 1.29544 | 83 |
| GO_POSITIVE_REGULATION_OF_WNT_SIGNALING_PATHWA | 0.00000 | 0.00000 | 0.76146 | 0.71563 | 1.47106 | 113 |
| GO_POSTSYNAPSE | 0.00000 | 0.00000 | 0.70498 | 0.61593 | 1.28589 | 267 |
| GO_POSTTRANSCRIPTIONAL_REGULATION_OF_GENE_EXPR | 0.00000 | 0.00000 | | 0.65452 | 1.38027 | 468 |
| GO_POST_TRANSLATIONAL_PROTEIN_MODIFICATION | 0.00000 | 0.00008 | 0.62726 | 0.60872 | 1.26748 | 232 |
| GO_PROTEASOMAL_PROTEIN_CATABOLIC_PROCESS | 0.00000 | 0.00001 | 0.67496 | 0.58569 | 1.23050 | 372 |
| GO_PROTEASOMAL_UBIQUITIN_INDEPENDENT_PROTEIN_C | 0.00000 | 0.00001 | 0.70498 | 0.91188 | 1.72124 | 19 |
| GO_PROTEASOME_ACCESSORY_COMPLEX | 0.00000 | 0.00005 | 0.64355 | 0.85821 | 1.64462 | 24 |
| GO_PROTEASOME_CORE_COMPLEX | 0.00000 | 0.00000 | 0.70498 | 0.92688 | 1.73412 | 17 |
| GO_PROTEIN_CONTAINING_COMPLEX_DISASSEMBLY | 0.00100 | 0.02000 | 0.45506 | 0.56415 | 1.17637 | 253 |
| GO_PROTEIN_CONTAINING_COMPLEX_LOCALIZATION | 0.00017 | 0.00412 | 0.51885 | 0.58499 | 1.21535 | 200 |
| GO_PROTEIN_DNA_COMPLEX_SUBUNIT_ORGANIZATION | 0.00046 | 0.01003 | 0.49849 | 0.59565 | 1.23457 | 166 |
| GO_PROTEIN_DOMAIN_SPECIFIC_BINDING | 0.00159 | 0.02832 | 0.45506 | 0.53833 | 1.13330 | 429 |
| GO_PROTEIN_FOLDING | 0.00000 | 0.00000 | 0.77494 | 0.66845 | 1.38506 | 164 |
| GO_PROTEIN_FOLDING_CHAPERONE | 0.00298 | 0.04756 | 0.43171 | 0.76980 | 1.44727 | 18 |
| GO_PROTEIN_IMPORT | 0.00051 | 0.01101 | 0.47727 | 0.60250 | 1.24584 | 138 |
| GO_PROTEIN_LOCALIZATION_TO_CHROMOSOME_TELOMEI | 0.00006 | 0.00159 | 0.55733 | 0.77828 | 1.50051 | 28 |
| GO_PROTEIN_LOCALIZATION_TO_ENDOPLASMIC_RETICULU | 0.00000 | 0.00000 | | 0.84257 | 1.73545 | 123 |
| GO_PROTEIN_LOCALIZATION_TO_MEMBRANE | 0.00000 | 0.00000 | | 0.66393 | 1.39649 | 414 |
| GO_PROTEIN_LOCALIZATION_TO_NUCLEUS | 0.00000 | 0.00016 | 0.61053 | 0.61575 | 1.27809 | 189 |
| GO_PROTEIN_MODIFICATION_BY_SMALL_PROTEIN_REMOV | 0.00000 | 0.00009 | 0.62726 | 0.60913 | 1.26767 | 225 |
| GO_PROTEIN_POLYUBIQUITINATION | 0.00010 | 0.00274 | 0.53843 | 0.58168 | 1.21276 | 251 |
| GO_PROTEIN_REFOLDING | 0.00054 | 0.01171 | 0.47727 | 0.80659 | 1.51643 | 18 |
| GO_PROTEIN_STABILIZATION | 0.00001 | 0.00027 | 0.59333 | 0.64686 | 1.33508 | 130 |
| GO_PROTEIN_TARGETING | 0.00000 | 0.00000 | | 0.70418 | 1.47678 | 338 |
| GO_PROTEIN_TARGETING_TO_MEMBRANE | 0.00000 | 0.00000 | | 0.82072 | 1.69904 | 154 |
| GO_PURINE_CONTAINING_COMPOUND_METABOLIC_PROC | 0.00167 | 0.02954 | 0.45506 | 0.55419 | 1.15656 | 261 |
| GO_PURINE_NUCLEOSIDE_METABOLIC_PROCESS | 0.00143 | 0.02602 | 0.45506 | 0.70423 | 1.38207 | 36 |
| GO_RECEPTOR_MEDIATED_ENDOCYTOSIS | 0.00044 | 0.00970 | 0.49849 | 0.61573 | 1.26759 | 122 |
| GO_REGENERATION | 0.00283 | 0.04583 | 0.43171 | 0.61683 | 1.25422 | 87 |
| GO_REGULATION_OF_ANIMAL_ORGAN_MORPHOGENESIS | 0.00000 | 0.00000 | | 0.72174 | 1.48689 | 124 |
| GO_REGULATION_OF_CELLULAR_AMIDE_METABOLIC_PROC | 0.00000 | 0.00000 | | 0.63680 | 1.33356 | 310 |
| GO_REGULATION_OF_CELLULAR_AMINE_METABOLIC_PROC | 0.00000 | 0.00000 | | 0.84760 | 1.69633 | 59 |
| GO_REGULATION_OF_CELLULAR_AMINO_ACID_METABOLIC | 0.00000 | 0.00000 | | 0.85206 | 1.69980 | 54 |
| GO_REGULATION_OF_CELLULAR_KETONE_METABOLIC_PRO | 0.00000 | 0.00000 | 0.78819 | 0.70442 | 1.45003 | 118 |
| GO_REGULATION_OF_CELLULAR_PROTEIN_CATABOLIC_PRO | 0.00029 | 0.00686 | 0.49849 | 0.59090 | 1.22684 | 185 |

| Cluster 1 | | | | | | |
|---|---|---|---|---|---|---|
| pathway | pval | padj | log2err | ES | NES | size |
| GO_REGULATION_OF_CELLULAR_PROTEIN_LOCALIZATION | 0.00002 | 0.00063 | 0.57561 | 0.56879 | 1.19512 | 374 |
| GO_REGULATION_OF_CELLULAR_RESPONSE_TO_HEAT | 0.00068 | 0.01445 | 0.47727 | 0.65258 | 1.31575 | 70 |
| GO_REGULATION_OF_CELL_CYCLE_G2_M_PHASE_TRANSITI | 0.00000 | 0.00009 | 0.62726 | 0.63025 | 1.30734 | 178 |
| GO_REGULATION_OF_CELL_CYCLE_PHASE_TRANSITION | 0.00005 | 0.00151 | 0.55733 | 0.56680 | 1.18905 | 344 |
| GO_REGULATION_OF_DEFENSE_RESPONSE | 0.00000 | 0.00006 | 0.64355 | 0.59234 | 1.23998 | 305 |
| GO_REGULATION_OF_DNA_BIOSYNTHETIC_PROCESS | 0.00038 | 0.00866 | 0.49849 | 0.65632 | 1.33051 | 80 |
| GO_REGULATION_OF_DNA_TEMPLATED_TRANSCRIPTION_I | 0.00128 | 0.02401 | 0.45506 | 0.72565 | 1.41567 | 33 |
| GO_REGULATION_OF_DNA_TEMPLATED_TRANSCRIPTION_I | 0.00000 | 0.00000 | | 0.77987 | 1.58774 | 90 |
| GO_REGULATION_OF_HEMATOPOIETIC_PROGENITOR_CELL_ | 0.00000 | 0.00000 | | 0.79703 | 1.60688 | 71 |
| GO_REGULATION_OF_HEMATOPOIETIC_STEM_CELL_DIFFER | 0.00000 | 0.00000 | | 0.82239 | 1.64778 | 62 |
| GO_REGULATION_OF_HEMOPOIESIS | 0.00000 | 0.00001 | 0.67496 | 0.61859 | 1.28820 | 235 |
| GO_REGULATION_OF_IMMUNE_RESPONSE | 0.00000 | 0.00000 | 0.70498 | 0.59004 | 1.24058 | 390 |
| GO_REGULATION_OF_INNATE_IMMUNE_RESPONSE | 0.00000 | 0.00000 | 0.71951 | 0.65499 | 1.35812 | 169 |
| GO_REGULATION_OF_INTRINSIC_APOPTOTIC_SIGNALING_P | 0.00031 | 0.00717 | 0.49849 | 0.63142 | 1.29564 | 106 |
| GO_REGULATION_OF_MITOTIC_CELL_CYCLE | 0.00020 | 0.00489 | 0.51885 | 0.54797 | 1.15537 | 464 |
| GO_REGULATION_OF_MORPHOGENESIS_OF_AN_EPITHELIU | 0.00000 | 0.00000 | | 0.76729 | 1.57198 | 103 |
| GO_REGULATION_OF_MRNA_CATABOLIC_PROCESS | 0.00000 | 0.00000 | | 0.70709 | 1.46553 | 166 |
| GO_REGULATION_OF_MRNA_METABOLIC_PROCESS | 0.00000 | 0.00000 | | 0.66995 | 1.39867 | 267 |
| GO_REGULATION_OF_MRNA_PROCESSING | 0.00038 | 0.00859 | 0.49849 | 0.61982 | 1.27583 | 115 |
| GO_REGULATION_OF_MRNA_SPLICING_VIA_SPLICEOSOME | 0.00004 | 0.00126 | 0.55733 | 0.67198 | 1.36311 | 81 |
| GO_REGULATION_OF_NUCLEOCYTOPLASMIC_TRANSPORT | 0.00276 | 0.04489 | 0.43171 | 0.62973 | 1.27424 | 77 |
| GO_REGULATION_OF_PEPTIDE_TRANSPORT | 0.00156 | 0.02793 | 0.45506 | 0.54436 | 1.14259 | 356 |
| GO_REGULATION_OF_PLASMA_LIPOPROTEIN_PARTICLE_LE\ | 0.00113 | 0.02196 | 0.45506 | 0.68709 | 1.36539 | 47 |
| GO_REGULATION_OF_PROTEASOMAL_PROTEIN_CATABOLIC | 0.00119 | 0.02265 | 0.45506 | 0.59388 | 1.22801 | 138 |
| GO_REGULATION_OF_PROTEIN_CATABOLIC_PROCESS | 0.00005 | 0.00157 | 0.55733 | 0.57842 | 1.20800 | 269 |
| GO_REGULATION_OF_PROTEIN_CONTAINING_COMPLEX_AS | 0.00104 | 0.02052 | 0.45506 | 0.56207 | 1.17353 | 268 |
| GO_REGULATION_OF_PROTEIN_IMPORT | 0.00031 | 0.00717 | 0.49849 | 0.72522 | 1.42685 | 39 |
| GO_REGULATION_OF_PROTEIN_LOCALIZATION_TO_NUCLEU | 0.00000 | 0.00010 | 0.62726 | 0.69384 | 1.41115 | 88 |
| GO_REGULATION_OF_PROTEIN_MODIFICATION_BY_SMALL_ | 0.00015 | 0.00380 | 0.51885 | 0.60298 | 1.25010 | 171 |
| GO_REGULATION_OF_PROTEIN_STABILITY | 0.00000 | 0.00002 | 0.67496 | 0.62824 | 1.30548 | 207 |
| GO_REGULATION_OF_PROTEOLYSIS | 0.00014 | 0.00360 | 0.51885 | 0.55650 | 1.16975 | 387 |
| GO_REGULATION_OF_RESPONSE_TO_BIOTIC_STIMULUS | 0.00000 | 0.00002 | 0.67496 | 0.62658 | 1.30179 | 211 |
| GO_REGULATION_OF_RESPONSE_TO_EXTERNAL_STIMULUS | 0.00115 | 0.02224 | 0.45506 | 0.53910 | 1.13529 | 442 |
| GO_REGULATION_OF_RNA_SPLICING | 0.00001 | 0.00034 | 0.59333 | 0.65235 | 1.34279 | 115 |
| GO_REGULATION_OF_SMALL_MOLECULE_METABOLIC_PRO | 0.00000 | 0.00000 | 0.76146 | 0.62707 | 1.31008 | 273 |
| GO_REGULATION_OF_STEM_CELL_DIFFERENTIATION | 0.00000 | 0.00000 | | 0.77562 | 1.57435 | 83 |
| GO_REGULATION_OF_SYMBIOTIC_PROCESS | 0.00027 | 0.00645 | 0.49849 | 0.61540 | 1.27223 | 135 |
| GO_REGULATION_OF_TELOMERASE_ACTIVITY | 0.00035 | 0.00813 | 0.49849 | 0.72042 | 1.41645 | 38 |
| GO_REGULATION_OF_TELOMERE_MAINTENANCE_VIA_TELO | 0.00302 | 0.04776 | 0.43171 | 0.65291 | 1.30184 | 53 |
| GO_REGULATION_OF_TRANSCRIPTION_FROM_RNA_POLYM | 0.00000 | 0.00000 | | 0.81519 | 1.63473 | 64 |
| GO_REGULATION_OF_TRANSLATIONAL_INITIATION | 0.00018 | 0.00447 | 0.51885 | 0.67211 | 1.34781 | 64 |
| GO_REGULATION_OF_VIRAL_GENOME_REPLICATION | 0.00231 | 0.03886 | 0.43171 | 0.65990 | 1.31679 | 55 |
| GO_REGULATION_OF_VIRAL_LIFE_CYCLE | 0.00043 | 0.00957 | 0.49849 | 0.64294 | 1.30643 | 86 |
| GO_REGULATION_OF_WNT_SIGNALING_PATHWAY | 0.00000 | 0.00000 | 0.71951 | 0.63599 | 1.32130 | 208 |
| GO_RESPONSE_TO_ALCOHOL | 0.00035 | 0.00809 | 0.49849 | 0.62586 | 1.28424 | 108 |
| GO_RESPONSE_TO_DRUG | 0.00000 | 0.00014 | 0.62726 | 0.61729 | 1.28128 | 189 |
| GO_RESPONSE_TO_ETHANOL | 0.00276 | 0.04489 | 0.43171 | 0.66538 | 1.32347 | 50 |
| GO_RESPONSE_TO_HEAT | 0.00000 | 0.00011 | 0.62726 | 0.65842 | 1.35533 | 118 |
| GO_RESPONSE_TO_INORGANIC_SUBSTANCE | 0.00187 | 0.03283 | 0.45506 | 0.55333 | 1.15626 | 280 |
| GO_RESPONSE_TO_INTERFERON_GAMMA | 0.00003 | 0.00091 | 0.57561 | 0.68611 | 1.38661 | 75 |
| GO_RESPONSE_TO_INTERLEUKIN_1 | 0.00000 | 0.00000 | 0.81404 | 0.73586 | 1.50976 | 105 |
| GO_RESPONSE_TO_INTERLEUKIN_12 | 0.00002 | 0.00078 | 0.57561 | 0.77249 | 1.51384 | 35 |
| GO_RESPONSE_TO_INTERLEUKIN_4 | 0.00011 | 0.00293 | 0.53843 | 0.82118 | 1.56520 | 22 |
| GO_RESPONSE_TO_LEUKEMIA_INHIBITORY_FACTOR | 0.00194 | 0.03382 | 0.45506 | 0.64611 | 1.30063 | 68 |
| GO_RESPONSE_TO_ORGANIC_CYCLIC_COMPOUND | 0.00016 | 0.00394 | 0.51885 | 0.54973 | 1.15774 | 441 |
| GO_RESPONSE_TO_OXYGEN_LEVELS | 0.00000 | 0.00003 | 0.65944 | 0.61493 | 1.28021 | 238 |
| GO_RESPONSE_TO_TEMPERATURE_STIMULUS | 0.00003 | 0.00080 | 0.57561 | 0.62939 | 1.30271 | 144 |
| GO_RESPONSE_TO_TOPOLOGICALLY_INCORRECT_PROTEIN | 0.00077 | 0.01604 | 0.47727 | 0.59782 | 1.23744 | 147 |
| GO_RESPONSE_TO_TUMOR_NECROSIS_FACTOR | 0.00000 | 0.00000 | 0.82666 | 0.68999 | 1.42826 | 151 |
| GO_RETINA_HOMEOSTASIS | 0.00216 | 0.03702 | 0.43171 | 0.73961 | 1.41928 | 25 |
| GO_RIBONUCLEOPROTEIN_COMPLEX_BINDING | 0.00008 | 0.00231 | 0.53843 | 0.63949 | 1.31627 | 119 |
| GO_RIBONUCLEOPROTEIN_COMPLEX_BIOGENESIS | 0.00000 | 0.00000 | | 0.64700 | 1.35986 | 385 |
| GO_RIBONUCLEOPROTEIN_COMPLEX_SUBUNIT_ORGANIZAT | 0.00000 | 0.00000 | 0.77494 | 0.67023 | 1.38952 | 171 |
| GO_RIBONUCLEOPROTEIN_GRANULE | 0.00000 | 0.00017 | 0.61053 | 0.62775 | 1.30169 | 174 |
| GO_RIBONUCLEOSIDE_METABOLIC_PROCESS | 0.00235 | 0.03925 | 0.43171 | 0.69683 | 1.36756 | 36 |
| GO_RIBOSOMAL_LARGE_SUBUNIT_ASSEMBLY | 0.00006 | 0.00170 | 0.55733 | 0.78893 | 1.51929 | 27 |
| GO_RIBOSOMAL_LARGE_SUBUNIT_BIOGENESIS | 0.00000 | 0.00000 | 0.76146 | 0.76810 | 1.54237 | 65 |
| GO_RIBOSOMAL_SMALL_SUBUNIT_BIOGENESIS | 0.00081 | 0.01668 | 0.47727 | 0.65840 | 1.32459 | 67 |
| GO_RIBOSOMAL_SUBUNIT | 0.00000 | 0.00000 | | 0.74974 | 1.55438 | 171 |
| GO_RIBOSOME | 0.00000 | 0.00000 | | 0.73675 | 1.53107 | 203 |

| Cluster 1 | | | | | | |
|---|---|---|---|---|---|---|
| pathway | pval | padj | log2err | ES | NES | size |
| GO_RIBOSOME_ASSEMBLY | 0.00000 | 0.00006 | 0.64355 | 0.74606 | 1.49312 | 59 |
| GO_RIBOSOME_BINDING | 0.00104 | 0.02052 | 0.45506 | 0.68020 | 1.35236 | 49 |
| GO_RIBOSOME_BIOGENESIS | 0.00000 | 0.00000 | 0.81404 | 0.64062 | 1.33838 | 272 |
| GO_RNA_CATABOLIC_PROCESS | 0.00000 | 0.00000 | | 0.75284 | 1.57919 | 341 |
| GO_RNA_DEPENDENT_DNA_BIOSYNTHETIC_PROCESS | 0.00003 | 0.00080 | 0.57561 | 0.71369 | 1.42999 | 62 |
| GO_RNA_EXPORT_FROM_NUCLEUS | 0.00086 | 0.01749 | 0.47727 | 0.60296 | 1.24217 | 125 |
| GO_RNA_LOCALIZATION | 0.00001 | 0.00043 | 0.59333 | 0.61008 | 1.26695 | 198 |
| GO_RNA_SPLICING | 0.00000 | 0.00001 | 0.70498 | 0.59085 | 1.24146 | 368 |
| GO_RNA_SPLICING_VIA_TRANSESTERIFICATION_REACTIONS | 0.00000 | 0.00004 | 0.65944 | 0.60059 | 1.25673 | 299 |
| GO_RNA_STABILIZATION | 0.00154 | 0.02768 | 0.45506 | 0.70324 | 1.38014 | 36 |
| GO_ROUGH_ENDOPLASMIC_RETICULUM | 0.00000 | 0.00005 | 0.64355 | 0.75410 | 1.50251 | 52 |
| GO_RRNA_BINDING | 0.00001 | 0.00050 | 0.59333 | 0.72798 | 1.45263 | 55 |
| GO_RRNA_METABOLIC_PROCESS | 0.00000 | 0.00007 | 0.62726 | 0.61429 | 1.27656 | 206 |
| GO_SARCOPLASMIC_RETICULUM_CALCIUM_ION_TRANSPO | 0.00030 | 0.00715 | 0.49849 | 0.84322 | 1.56785 | 15 |
| GO_SCF_DEPENDENT_PROTEASOMAL_UBIQUITIN_DEPEND | 0.00000 | 0.00000 | 0.82666 | 0.78642 | 1.58825 | 74 |
| GO_SECRETORY_GRANULE | 0.00000 | 0.00000 | | 0.63862 | 1.34331 | 408 |
| GO_SECRETORY_VESICLE | 0.00000 | 0.00000 | | 0.61170 | 1.29057 | 481 |
| GO_SIGNAL_TRANSDUCTION_BY_PROTEIN_PHOSPHORYLAT | 0.00007 | 0.00201 | 0.53843 | 0.55781 | 1.17422 | 426 |
| GO_SINGLE_FERTILIZATION | 0.00016 | 0.00396 | 0.51885 | 0.71862 | 1.42621 | 46 |
| GO_SINGLE_STRANDED_RNA_BINDING | 0.00205 | 0.03544 | 0.43171 | 0.64596 | 1.29199 | 58 |
| GO_SMALL_MOLECULE_BIOSYNTHETIC_PROCESS | 0.00104 | 0.02052 | 0.45506 | 0.54395 | 1.14418 | 407 |
| GO_SMALL_RIBOSOMAL_SUBUNIT | 0.00000 | 0.00000 | | 0.79581 | 1.59963 | 66 |
| GO_SOMATODENDRITIC_COMPARTMENT | 0.00242 | 0.04026 | 0.43171 | 0.54040 | 1.13533 | 377 |
| GO_SPERM_EGG_RECOGNITION | 0.00000 | 0.00003 | 0.65944 | 0.89901 | 1.69694 | 19 |
| GO_SPLICEOSOMAL_COMPLEX | 0.00004 | 0.00110 | 0.55733 | 0.61646 | 1.27826 | 167 |
| GO_STEM_CELL_DIFFERENTIATION | 0.00000 | 0.00000 | | 0.73288 | 1.51448 | 134 |
| GO_STRUCTURAL_CONSTITUENT_OF_CYTOSKELETON | 0.00194 | 0.03382 | 0.45506 | 0.66098 | 1.31794 | 53 |
| GO_STRUCTURAL_CONSTITUENT_OF_RIBOSOME | 0.00000 | 0.00000 | | 0.77017 | 1.59399 | 149 |
| GO_STRUCTURAL_MOLECULE_ACTIVITY | 0.00000 | 0.00000 | | 0.69650 | 1.45973 | 329 |
| GO_TELOMERASE_RNA_LOCALIZATION | 0.00002 | 0.00051 | 0.57561 | 0.86396 | 1.63078 | 19 |
| GO_TELOMERE_MAINTENANCE_VIA_TELOMERE_LENGTHEN | 0.00009 | 0.00254 | 0.53843 | 0.67524 | 1.36144 | 70 |
| GO_THREONINE_TYPE_ENDOPEPTIDASE_ACTIVITY | 0.00000 | 0.00000 | 0.76146 | 0.94208 | 1.75650 | 16 |
| GO_THREONINE_TYPE_PEPTIDASE_ACTIVITY | 0.00000 | 0.00001 | 0.67496 | 0.90450 | 1.70730 | 19 |
| GO_TISSUE_MORPHOGENESIS | 0.00001 | 0.00046 | 0.59333 | 0.58022 | 1.21509 | 315 |
| GO_TOXIN_TRANSPORT | 0.00006 | 0.00165 | 0.55733 | 0.80395 | 1.54274 | 25 |
| GO_TRANSCRIPTION_COUPLED_NUCLEOTIDE_EXCISION_RE | 0.00009 | 0.00236 | 0.53843 | 0.67164 | 1.35407 | 71 |
| GO_TRANSCRIPTION_FACTOR_BINDING | 0.00039 | 0.00894 | 0.49849 | 0.54820 | 1.15299 | 400 |
| GO_TRANSLATIONAL_INITIATION | 0.00000 | 0.00000 | | 0.81603 | 1.69123 | 165 |
| GO_TRANSLATION_ELONGATION_FACTOR_ACTIVITY | 0.00012 | 0.00308 | 0.53843 | 0.85922 | 1.59759 | 15 |
| GO_TRANSLATION_FACTOR_ACTIVITY_RNA_BINDING | 0.00000 | 0.00000 | | 0.78126 | 1.57520 | 70 |
| GO_TRANSLATION_INITIATION_FACTOR_ACTIVITY | 0.00000 | 0.00003 | 0.65944 | 0.78298 | 1.55080 | 44 |
| GO_TRANSLATION_INITIATION_FACTOR_BINDING | 0.00294 | 0.04723 | 0.43171 | 0.72204 | 1.39372 | 29 |
| GO_TRANSLATION_PREINITIATION_COMPLEX | 0.00001 | 0.00023 | 0.61053 | 0.88728 | 1.65432 | 16 |
| GO_TRANSLATION_REGULATOR_ACTIVITY | 0.00000 | 0.00000 | | 0.74652 | 1.53182 | 108 |
| GO_TRANSLATION_REGULATOR_ACTIVITY_NUCLEIC_ACID_E | 0.00000 | 0.00000 | 0.82666 | 0.76179 | 1.54777 | 85 |
| GO_TRICARBOXYLIC_ACID_CYCLE | 0.00019 | 0.00462 | 0.51885 | 0.75542 | 1.46344 | 30 |
| GO_TUMOR_NECROSIS_FACTOR_MEDIATED_SIGNALING_PA | 0.00000 | 0.00000 | | 0.77651 | 1.58759 | 97 |
| GO_T_CELL_RECEPTOR_SIGNALING_PATHWAY | 0.00000 | 0.00000 | | 0.73288 | 1.50872 | 121 |
| GO_UBIQUITIN_LIKE_PROTEIN_LIGASE_BINDING | 0.00000 | 0.00000 | 0.76146 | 0.64148 | 1.33468 | 220 |
| GO_UNFOLDED_PROTEIN_BINDING | 0.00000 | 0.00000 | 0.73376 | 0.73001 | 1.48198 | 84 |
| GO_VACUOLAR_LUMEN | 0.00000 | 0.00004 | 0.65944 | 0.69328 | 1.41218 | 91 |
| GO_VACUOLAR_MEMBRANE | 0.00000 | 0.00006 | 0.64355 | 0.59403 | 1.24280 | 297 |
| GO_VACUOLE | 0.00000 | 0.00000 | 0.71951 | 0.58053 | 1.22529 | 488 |
| GO_VESICLE_LUMEN | 0.00000 | 0.00000 | | 0.68536 | 1.42170 | 179 |
| GO_VESICLE_MEMBRANE | 0.00302 | 0.04776 | 0.43171 | 0.53618 | 1.12783 | 405 |
| GO_VIRAL_GENE_EXPRESSION | 0.00000 | 0.00000 | | 0.76742 | 1.59125 | 169 |
| GO_VIRAL_GENOME_REPLICATION | 0.00235 | 0.03925 | 0.43171 | 0.63350 | 1.28028 | 75 |
| GO_VIRAL_LIFE_CYCLE | 0.00001 | 0.00021 | 0.61053 | 0.60741 | 1.26223 | 213 |
| HP_ABNORMALITY_OF_CARDIOVASCULAR_SYSTEM_ELECTR | 0.00016 | 0.00394 | 0.51885 | 0.58240 | 1.21297 | 241 |
| HP_ABNORMALITY_OF_THE_NASAL_DORSUM | 0.00025 | 0.00604 | 0.49849 | 0.61559 | 1.27261 | 135 |
| HP_ABNORMAL_ERYTHROCYTE_MORPHOLOGY | 0.00113 | 0.02196 | 0.45506 | 0.54643 | 1.14632 | 344 |
| HP_ACUTE_LEUKEMIA | 0.00000 | 0.00004 | 0.64355 | 0.72814 | 1.46575 | 68 |
| HP_ANEMIA_OF_INADEQUATE_PRODUCTION | 0.00006 | 0.00182 | 0.53843 | 0.64606 | 1.32322 | 102 |
| HP_BILATERAL_SENSORINEURAL_HEARING_IMPAIRMENT | 0.00187 | 0.03283 | 0.45506 | 0.71580 | 1.38669 | 30 |
| HP_DEGENERATION_OF_THE_LATERAL_CORTICOSPINAL_TR | 0.00041 | 0.00917 | 0.49849 | 0.83715 | 1.55657 | 15 |
| HP_DELAYED_PUBERTY | 0.00007 | 0.00193 | 0.53843 | 0.64249 | 1.31835 | 108 |
| HP_DEPRESSED_NASAL_RIDGE | 0.00012 | 0.00317 | 0.53843 | 0.67554 | 1.36226 | 72 |
| HP_ENTROPION | 0.00168 | 0.02978 | 0.45506 | 0.81158 | 1.50902 | 15 |
| HP_FATIGUE | 0.00047 | 0.01025 | 0.49849 | 0.58308 | 1.21038 | 187 |

| Cluster 1 | | | | | | |
|---|---|---|---|---|---|---|
| pathway | pval | padj | log2err | ES | NES | size |
| HP_HEADACHE | 0.00095 | 0.01910 | 0.47727 | 0.59150 | 1.22408 | 143 |
| HP_JUVENILE_ASEPTIC_NECROSIS | 0.00276 | 0.04489 | 0.43171 | 0.77159 | 1.45062 | 18 |
| HP_LEUKEMIA | 0.00002 | 0.00074 | 0.57561 | 0.65596 | 1.34055 | 96 |
| HP_MACROCYTIC_ANEMIA | 0.00000 | 0.00006 | 0.64355 | 0.78931 | 1.55757 | 41 |
| HP_MACROGYRIA | 0.00213 | 0.03660 | 0.43171 | 0.80019 | 1.49708 | 17 |
| HP_MIGRAINE | 0.00000 | 0.00004 | 0.65944 | 0.72650 | 1.45884 | 65 |
| HP_NEURODEGENERATION | 0.00305 | 0.04821 | 0.43171 | 0.66047 | 1.31595 | 52 |
| HP_PALLOR | 0.00046 | 0.01003 | 0.49849 | 0.64118 | 1.30477 | 89 |
| HP_RETICULOCYTOPENIA | 0.00003 | 0.00099 | 0.55733 | 0.87839 | 1.63324 | 15 |
| HP_SHORT_NOSE | 0.00005 | 0.00136 | 0.55733 | 0.60529 | 1.25562 | 176 |
| HP_THICK_LOWER_LIP_VERMILION | 0.00035 | 0.00813 | 0.49849 | 0.62306 | 1.27648 | 103 |
| HP_THICK_VERMILION_BORDER | 0.00019 | 0.00465 | 0.51885 | 0.60368 | 1.25033 | 159 |
| HP_THROMBOCYTOSIS | 0.00268 | 0.04405 | 0.43171 | 0.76189 | 1.45219 | 22 |
| HP_TRANSIENT_ISCHEMIC_ATTACK | 0.00139 | 0.02567 | 0.45506 | 0.74486 | 1.43180 | 26 |
| **Cluster 7** | | | | | | |
| pathway | pval | padj | log2err | ES | NES | size |
| GO_ACTIN_CYTOSKELETON | 0.00168 | 0.02511 | 0.45506 | 0.45949 | 1.26269 | 170 |
| GO_ACTIVATION_OF_IMMUNE_RESPONSE | 0.00000 | 0.00000 | 0.76146 | 0.58151 | 1.58353 | 139 |
| GO_ACTIVATION_OF_INNATE_IMMUNE_RESPONSE | 0.00000 | 0.00000 | | 0.70963 | 1.85323 | 65 |
| GO_AMIDE_BIOSYNTHETIC_PROCESS | 0.00000 | 0.00000 | | 0.56933 | 1.61176 | 469 |
| GO_AMINE_METABOLIC_PROCESS | 0.00000 | 0.00000 | 0.82666 | 0.70405 | 1.83190 | 62 |
| GO_ANAPHASE_PROMOTING_COMPLEX_DEPENDENT_CATA | 0.00000 | 0.00000 | | 0.71214 | 1.86814 | 70 |
| GO_ANATOMICAL_STRUCTURE_HOMEOSTASIS | 0.00102 | 0.01632 | 0.45506 | 0.47154 | 1.29253 | 158 |
| GO_ANCHORING_JUNCTION | 0.00000 | 0.00000 | | 0.56351 | 1.58316 | 336 |
| GO_ANIMAL_ORGAN_MORPHOGENESIS | 0.00010 | 0.00220 | 0.53843 | 0.45559 | 1.27266 | 272 |
| GO_ANTIGEN_PROCESSING_AND_PRESENTATION | 0.00000 | 0.00001 | 0.69013 | 0.58478 | 1.57251 | 108 |
| GO_ANTIGEN_PROCESSING_AND_PRESENTATION_OF_EXOG | 0.00000 | 0.00000 | | 0.75575 | 1.92382 | 47 |
| GO_ANTIGEN_PROCESSING_AND_PRESENTATION_OF_PEPT | 0.00000 | 0.00000 | 0.77494 | 0.62118 | 1.66075 | 98 |
| GO_ANTIGEN_PROCESSING_AND_PRESENTATION_OF_PEPT | 0.00000 | 0.00000 | | 0.71118 | 1.84374 | 59 |
| GO_ANTIGEN_RECEPTOR_MEDIATED_SIGNALING_PATHWAY | 0.00000 | 0.00000 | 0.74774 | 0.62373 | 1.65215 | 85 |
| GO_ANTIOXIDANT_ACTIVITY | 0.00294 | 0.03916 | 0.43171 | 0.62372 | 1.52113 | 26 |
| GO_AZUROPHIL_GRANULE_LUMEN | 0.00110 | 0.01737 | 0.45506 | 0.59693 | 1.49207 | 37 |
| GO_CADHERIN_BINDING | 0.00000 | 0.00000 | | 0.56465 | 1.56168 | 203 |
| GO_CANONICAL_WNT_SIGNALING_PATHWAY | 0.00000 | 0.00000 | 0.71951 | 0.56450 | 1.53559 | 134 |
| GO_CATALYTIC_STEP_2_SPLICEOSOME | 0.00054 | 0.00938 | 0.47727 | 0.55323 | 1.45221 | 72 |
| GO_CELLULAR_AMINO_ACID_METABOLIC_PROCESS | 0.00001 | 0.00032 | 0.59333 | 0.51466 | 1.40515 | 148 |
| GO_CELLULAR_KETONE_METABOLIC_PROCESS | 0.00000 | 0.00004 | 0.64355 | 0.57082 | 1.53267 | 104 |
| GO_CELLULAR_NITROGEN_COMPOUND_CATABOLIC_PROCE | 0.00000 | 0.00000 | | 0.63387 | 1.77820 | 323 |
| GO_CELLULAR_OXIDANT_DETOXIFICATION | 0.00279 | 0.03763 | 0.43171 | 0.60417 | 1.48900 | 29 |
| GO_CELLULAR_PROTEIN_CATABOLIC_PROCESS | 0.00013 | 0.00259 | 0.51885 | 0.43398 | 1.22483 | 421 |
| GO_CELLULAR_RESPIRATION | 0.00157 | 0.02355 | 0.45506 | 0.48451 | 1.30981 | 117 |
| GO_CELLULAR_RESPONSE_TO_CHEMICAL_STRESS | 0.00042 | 0.00736 | 0.49849 | 0.49067 | 1.33536 | 136 |
| GO_CELLULAR_RESPONSE_TO_OXYGEN_LEVELS | 0.00000 | 0.00000 | 0.77494 | 0.59168 | 1.60293 | 121 |
| GO_CELLULAR_RESPONSE_TO_TOPOLOGICALLY_INCORRECT | 0.00283 | 0.03802 | 0.43171 | 0.50285 | 1.33196 | 85 |
| GO_CELL_ACTIVATION | 0.00000 | 0.00002 | 0.65944 | 0.46703 | 1.31635 | 381 |
| GO_CELL_ACTIVATION_INVOLVED_IN_IMMUNE_RESPONSE | 0.00000 | 0.00000 | 0.73376 | 0.51750 | 1.43813 | 230 |
| GO_CELL_ADHESION_MOLECULE_BINDING | 0.00000 | 0.00000 | | 0.55449 | 1.54424 | 244 |
| GO_CELL_BODY | 0.00316 | 0.04149 | 0.43171 | 0.45554 | 1.25032 | 164 |
| GO_CELL_CELL_RECOGNITION | 0.00003 | 0.00075 | 0.55733 | 0.76608 | 1.78481 | 18 |
| GO_CELL_CELL_SIGNALING_BY_WNT | 0.00000 | 0.00001 | 0.69013 | 0.52159 | 1.44223 | 196 |
| GO_CELL_CYCLE_G2_M_PHASE_TRANSITION | 0.00000 | 0.00001 | 0.67496 | 0.53731 | 1.47282 | 158 |
| GO_CELL_CYCLE_PHASE_TRANSITION | 0.00022 | 0.00427 | 0.51885 | 0.44069 | 1.23879 | 342 |
| GO_CELL_KILLING | 0.00383 | 0.04785 | 0.43171 | 0.60579 | 1.48770 | 28 |
| GO_CELL_LEADING_EDGE | 0.00121 | 0.01886 | 0.45506 | 0.46987 | 1.28796 | 158 |
| GO_CELL_RECOGNITION | 0.00021 | 0.00409 | 0.51885 | 0.66783 | 1.64006 | 28 |
| GO_CELL_REDOX_HOMEOSTASIS | 0.00346 | 0.04421 | 0.43171 | 0.60268 | 1.49350 | 31 |
| GO_CELL_SUBSTRATE_JUNCTION | 0.00000 | 0.00000 | | 0.65219 | 1.80522 | 208 |
| GO_CELL_SURFACE | 0.00036 | 0.00648 | 0.49849 | 0.50430 | 1.36227 | 116 |
| GO_CELL_SURFACE_RECEPTOR_SIGNALING_PATHWAY_INVO | 0.00000 | 0.00002 | 0.65944 | 0.50655 | 1.40257 | 211 |
| GO_CHAPERONE_COMPLEX | 0.00003 | 0.00077 | 0.55733 | 0.77410 | 1.78916 | 17 |
| GO_CLUSTER_OF_ACTIN_BASED_CELL_PROJECTIONS | 0.00100 | 0.01608 | 0.45506 | 0.56847 | 1.45437 | 49 |
| GO_CONTRACTILE_FIBER | 0.00017 | 0.00345 | 0.51885 | 0.54444 | 1.43096 | 74 |
| GO_COTRANSLATIONAL_PROTEIN_TARGETING_TO_MEMBR | 0.00000 | 0.00000 | | 0.82211 | 2.18283 | 89 |
| GO_CYTOKINE_MEDIATED_SIGNALING_PATHWAY | 0.00000 | 0.00000 | | 0.60028 | 1.66199 | 212 |
| GO_CYTOPLASMIC_STRESS_GRANULE | 0.00178 | 0.02588 | 0.45506 | 0.58277 | 1.47574 | 43 |
| GO_CYTOPLASMIC_TRANSLATION | 0.00000 | 0.00000 | | 0.73305 | 1.93685 | 82 |
| GO_CYTOPLASMIC_TRANSLATIONAL_INITIATION | 0.00000 | 0.00004 | 0.64355 | 0.72883 | 1.80134 | 30 |
| GO_CYTOSOLIC_LARGE_RIBOSOMAL_SUBUNIT | 0.00000 | 0.00000 | | 0.84142 | 2.16438 | 53 |
| GO_CYTOSOLIC_RIBOSOME | 0.00000 | 0.00000 | | 0.81714 | 2.17905 | 94 |

| Cluster 1 | | | | | | |
|---|---|---|---|---|---|---|
| pathway | pval | padj | log2err | ES | NES | size |
| GO_CYTOSOLIC_SMALL_RIBOSOMAL_SUBUNIT | 0.00000 | 0.00000 | | 0.84025 | 2.10923 | 39 |
| GO_DEFENSE_RESPONSE | 0.00001 | 0.00015 | 0.61053 | 0.46291 | 1.30221 | 348 |
| GO_DEFENSE_RESPONSE_TO_OTHER_ORGANISM | 0.00000 | 0.00000 | 0.71951 | 0.51341 | 1.43012 | 248 |
| GO_DETOXIFICATION | 0.00026 | 0.00495 | 0.49849 | 0.62288 | 1.56057 | 38 |
| GO_DE_NOVO_PROTEIN_FOLDING | 0.00265 | 0.03597 | 0.43171 | 0.65405 | 1.55283 | 21 |
| GO_DNA_BIOSYNTHETIC_PROCESS | 0.00033 | 0.00601 | 0.49849 | 0.51499 | 1.37685 | 98 |
| GO_DOUBLE_STRANDED_RNA_BINDING | 0.00170 | 0.02522 | 0.45506 | 0.58374 | 1.47821 | 43 |
| GO_ENDOPEPTIDASE_COMPLEX | 0.00000 | 0.00000 | | 0.74463 | 1.91797 | 54 |
| GO_ENERGY_DERIVATION_BY_OXIDATION_OF_ORGANIC_CO | 0.00106 | 0.01685 | 0.45506 | 0.47464 | 1.29476 | 143 |
| GO_ENZYME_REGULATOR_ACTIVITY | 0.00335 | 0.04329 | 0.43171 | 0.41941 | 1.18120 | 372 |
| GO_EPITHELIUM_DEVELOPMENT | 0.00000 | 0.00013 | 0.61053 | 0.46102 | 1.29842 | 366 |
| GO_ESTABLISHMENT_OF_PROTEIN_LOCALIZATION_TO_END | 0.00000 | 0.00000 | | 0.81478 | 2.17836 | 98 |
| GO_ESTABLISHMENT_OF_PROTEIN_LOCALIZATION_TO_MEM | 0.00000 | 0.00000 | | 0.66801 | 1.84677 | 199 |
| GO_ESTABLISHMENT_OF_PROTEIN_LOCALIZATION_TO_ORG | 0.00000 | 0.00000 | | 0.57813 | 1.62329 | 331 |
| GO_ESTABLISHMENT_OF_TISSUE_POLARITY | 0.00000 | 0.00000 | | 0.69402 | 1.81279 | 66 |
| GO_EUKARYOTIC_TRANSLATION_INITIATION_FACTOR_3_CO | 0.00004 | 0.00098 | 0.55733 | 0.79616 | 1.81958 | 15 |
| GO_EXOCYTOSIS | 0.00000 | 0.00000 | 0.70498 | 0.48954 | 1.37299 | 316 |
| GO_FC_EPSILON_RECEPTOR_SIGNALING_PATHWAY | 0.00000 | 0.00000 | 0.82666 | 0.70125 | 1.82960 | 64 |
| GO_FC_RECEPTOR_SIGNALING_PATHWAY | 0.00000 | 0.00000 | | 0.66368 | 1.76810 | 93 |
| GO_FERTILIZATION | 0.00170 | 0.02522 | 0.45506 | 0.57593 | 1.45513 | 42 |
| GO_FICOLIN_1_RICH_GRANULE | 0.00000 | 0.00000 | 0.77494 | 0.64743 | 1.70961 | 81 |
| GO_GLYCOSYL_COMPOUND_METABOLIC_PROCESS | 0.00213 | 0.03002 | 0.43171 | 0.55100 | 1.41081 | 51 |
| GO_HEMATOPOIETIC_PROGENITOR_CELL_DIFFERENTIATION | 0.00000 | 0.00000 | | 0.68395 | 1.79664 | 73 |
| GO_HEMATOPOIETIC_STEM_CELL_DIFFERENTIATION | 0.00000 | 0.00000 | | 0.77465 | 1.98346 | 51 |
| GO_HORMONE_RECEPTOR_BINDING | 0.00126 | 0.01965 | 0.45506 | 0.52195 | 1.37184 | 74 |
| GO_IMMUNE_EFFECTOR_PROCESS | 0.00000 | 0.00001 | 0.67496 | 0.47983 | 1.34925 | 344 |
| GO_IMMUNE_RESPONSE_REGULATING_SIGNALING_PATHW | 0.00000 | 0.00000 | 0.76146 | 0.58286 | 1.58472 | 132 |
| GO_IMMUNE_SYSTEM_DEVELOPMENT | 0.00024 | 0.00462 | 0.51885 | 0.44818 | 1.25612 | 307 |
| GO_INNATE_IMMUNE_RESPONSE | 0.00000 | 0.00000 | 0.80122 | 0.54143 | 1.49910 | 210 |
| GO_INNATE_IMMUNE_RESPONSE_ACTIVATING_SIGNAL_TR | 0.00000 | 0.00000 | | 0.76432 | 1.96605 | 53 |
| GO_INTERACTION_WITH_SYMBIONT | 0.00148 | 0.02239 | 0.45506 | 0.60458 | 1.50342 | 34 |
| GO_INTERLEUKIN_1_MEDIATED_SIGNALING_PATHWAY | 0.00000 | 0.00000 | | 0.75908 | 1.96792 | 59 |
| GO_INTRINSIC_APOPTOTIC_SIGNALING_PATHWAY | 0.00130 | 0.02002 | 0.45506 | 0.47640 | 1.29393 | 127 |
| GO_IN_UTERO_EMBRYONIC_DEVELOPMENT | 0.00279 | 0.03763 | 0.43171 | 0.46370 | 1.27401 | 168 |
| GO_ION_CHANNEL_BINDING | 0.00143 | 0.02163 | 0.45506 | 0.58337 | 1.46876 | 40 |
| GO_LAMELLIPODIUM | 0.00191 | 0.02732 | 0.45506 | 0.51857 | 1.36900 | 80 |
| GO_LARGE_RIBOSOMAL_SUBUNIT | 0.00000 | 0.00000 | | 0.66524 | 1.78024 | 100 |
| GO_LEUKOCYTE_MEDIATED_IMMUNITY | 0.00000 | 0.00002 | 0.65944 | 0.49768 | 1.38646 | 247 |
| GO_LYSOSOMAL_LUMEN | 0.00156 | 0.02335 | 0.45506 | 0.60622 | 1.50931 | 35 |
| GO_MIDBRAIN_DEVELOPMENT | 0.00044 | 0.00757 | 0.49849 | 0.61372 | 1.53762 | 38 |
| GO_MODIFICATION_DEPENDENT_MACROMOLECULE_CATAI | 0.00061 | 0.01036 | 0.47727 | 0.43183 | 1.21581 | 363 |
| GO_MONOSACCHARIDE_BIOSYNTHETIC_PROCESS | 0.00191 | 0.02732 | 0.45506 | 0.58673 | 1.46999 | 38 |
| GO_MORPHOGENESIS_OF_AN_EPITHELIUM | 0.00000 | 0.00009 | 0.62726 | 0.50491 | 1.39576 | 201 |
| GO_MORPHOGENESIS_OF_A_POLARIZED_EPITHELIUM | 0.00000 | 0.00000 | | 0.66998 | 1.76501 | 78 |
| GO_MRNA_3_UTR_BINDING | 0.00035 | 0.00626 | 0.49849 | 0.58163 | 1.48923 | 51 |
| GO_MRNA_5_UTR_BINDING | 0.00006 | 0.00129 | 0.55733 | 0.73500 | 1.74501 | 21 |
| GO_MRNA_BINDING | 0.00000 | 0.00000 | | 0.56675 | 1.56005 | 179 |
| GO_MRNA_PROCESSING | 0.00011 | 0.00221 | 0.53843 | 0.44756 | 1.25756 | 337 |
| GO_MUSCLE_CELL_DIFFERENTIATION | 0.00313 | 0.04113 | 0.43171 | 0.49124 | 1.31901 | 104 |
| GO_MUSCLE_CONTRACTION | 0.00014 | 0.00287 | 0.51885 | 0.55941 | 1.47133 | 75 |
| GO_MUSCLE_SYSTEM_PROCESS | 0.00033 | 0.00601 | 0.49849 | 0.51868 | 1.38978 | 101 |
| GO_MYELOID_LEUKOCYTE_ACTIVATION | 0.00000 | 0.00000 | 0.76146 | 0.53175 | 1.47415 | 218 |
| GO_MYELOID_LEUKOCYTE_MEDIATED_IMMUNITY | 0.00000 | 0.00000 | 0.73376 | 0.53023 | 1.46764 | 208 |
| GO_NADH_METABOLIC_PROCESS | 0.00073 | 0.01215 | 0.47727 | 0.71499 | 1.65256 | 17 |
| GO_NAD_BINDING | 0.00261 | 0.03557 | 0.43171 | 0.60057 | 1.48819 | 32 |
| GO_NAD_METABOLIC_PROCESS | 0.00132 | 0.02016 | 0.45506 | 0.68027 | 1.59425 | 19 |
| GO_NEGATIVE_REGULATION_OF_CALCIUM_ION_TRANSPOF | 0.00087 | 0.01423 | 0.47727 | 0.72240 | 1.65941 | 16 |
| GO_NEGATIVE_REGULATION_OF_CANONICAL_WNT_SIGNAI | 0.00000 | 0.00000 | 0.82666 | 0.66433 | 1.75425 | 81 |
| GO_NEGATIVE_REGULATION_OF_CATABOLIC_PROCESS | 0.00342 | 0.04399 | 0.43171 | 0.46282 | 1.26233 | 144 |
| GO_NEGATIVE_REGULATION_OF_CELLULAR_AMIDE_METAB | 0.00213 | 0.03002 | 0.43171 | 0.51878 | 1.36810 | 79 |
| GO_NEGATIVE_REGULATION_OF_CELLULAR_CATABOLIC_PR | 0.00287 | 0.03829 | 0.43171 | 0.47220 | 1.28042 | 124 |
| GO_NEGATIVE_REGULATION_OF_CELL_CYCLE | 0.00055 | 0.00950 | 0.47727 | 0.44474 | 1.24657 | 311 |
| GO_NEGATIVE_REGULATION_OF_CELL_CYCLE_G2_M_PHASI | 0.00000 | 0.00000 | 0.78819 | 0.66480 | 1.74394 | 70 |
| GO_NEGATIVE_REGULATION_OF_CELL_CYCLE_PHASE_TRAN | 0.00001 | 0.00020 | 0.59333 | 0.53244 | 1.44913 | 138 |
| GO_NEGATIVE_REGULATION_OF_CELL_CYCLE_PROCESS | 0.00000 | 0.00013 | 0.61053 | 0.50579 | 1.39519 | 188 |
| GO_NEGATIVE_REGULATION_OF_CELL_DEATH | 0.00038 | 0.00670 | 0.49849 | 0.44022 | 1.23788 | 344 |
| GO_NEGATIVE_REGULATION_OF_MITOTIC_CELL_CYCLE | 0.00005 | 0.00105 | 0.55733 | 0.49252 | 1.35344 | 173 |
| GO_NEGATIVE_REGULATION_OF_MRNA_METABOLIC_PROC | 0.00001 | 0.00028 | 0.59333 | 0.63317 | 1.61989 | 49 |
| GO_NEGATIVE_REGULATION_OF_MRNA_SPLICING_VIA_SPL | 0.00070 | 0.01174 | 0.47727 | 0.71623 | 1.65541 | 17 |

| Cluster 1 | | | | | | |
|---|---|---|---|---|---|---|
| pathway | pval | padj | log2err | ES | NES | size |
| GO_NEGATIVE_REGULATION_OF_PROTEIN_BINDING | 0.00092 | 0.01508 | 0.47727 | 0.60118 | 1.49838 | 36 |
| GO_NEGATIVE_REGULATION_OF_RNA_SPLICING | 0.00001 | 0.00026 | 0.59333 | 0.76141 | 1.80772 | 21 |
| GO_NEGATIVE_REGULATION_OF_SIGNALING | 0.00020 | 0.00383 | 0.51885 | 0.42958 | 1.21543 | 455 |
| GO_NEGATIVE_REGULATION_OF_WNT_SIGNALING_PATHW/ | 0.00000 | 0.00000 | | 0.66841 | 1.77050 | 85 |
| GO_NEURAL_NUCLEUS_DEVELOPMENT | 0.00003 | 0.00059 | 0.57561 | 0.73344 | 1.77179 | 24 |
| GO_NEURON_TO_NEURON_SYNAPSE | 0.00019 | 0.00380 | 0.51885 | 0.52663 | 1.41291 | 103 |
| GO_NIK_NF_KAPPAB_SIGNALING | 0.00000 | 0.00000 | | 0.71331 | 1.87900 | 77 |
| GO_NON_CANONICAL_WNT_SIGNALING_PATHWAY | 0.00000 | 0.00000 | | 0.71505 | 1.87578 | 70 |
| GO_NUCLEAR_HORMONE_RECEPTOR_BINDING | 0.00302 | 0.03990 | 0.43171 | 0.52486 | 1.37685 | 70 |
| GO_NUCLEAR_TRANSCRIBED_MRNA_CATABOLIC_PROCESS | 0.00000 | 0.00000 | | 0.70415 | 1.92645 | 153 |
| GO_NUCLEAR_TRANSCRIBED_MRNA_CATABOLIC_PROCESS_ | 0.00000 | 0.00000 | | 0.80547 | 2.16271 | 104 |
| GO_NUCLEOSIDE_METABOLIC_PROCESS | 0.00095 | 0.01536 | 0.47727 | 0.59026 | 1.48824 | 41 |
| GO_ORGANELLE_ASSEMBLY | 0.00020 | 0.00386 | 0.51885 | 0.43681 | 1.22957 | 360 |
| GO_ORGANIC_ACID_METABOLIC_PROCESS | 0.00009 | 0.00189 | 0.53843 | 0.43800 | 1.23534 | 412 |
| GO_ORGANIC_CYCLIC_COMPOUND_CATABOLIC_PROCESS | 0.00000 | 0.00000 | | 0.62563 | 1.75623 | 327 |
| GO_OXIDATION_REDUCTION_PROCESS | 0.00178 | 0.02588 | 0.45506 | 0.42313 | 1.19272 | 390 |
| GO_OXIDOREDUCTASE_ACTIVITY_ACTING_ON_PEROXIDE_A | 0.00361 | 0.04584 | 0.43171 | 0.66375 | 1.54640 | 18 |
| GO_PEPTIDASE_COMPLEX | 0.00000 | 0.00000 | 0.80122 | 0.68209 | 1.78623 | 68 |
| GO_PEPTIDASE_REGULATOR_ACTIVITY | 0.00346 | 0.04421 | 0.43171 | 0.55491 | 1.41088 | 46 |
| GO_PEPTIDE_BIOSYNTHETIC_PROCESS | 0.00000 | 0.00000 | | 0.58664 | 1.65516 | 415 |
| GO_PEPTIDE_METABOLIC_PROCESS | 0.00000 | 0.00000 | | 0.58008 | 1.64192 | 464 |
| GO_PIGMENT_GRANULE | 0.00002 | 0.00044 | 0.57561 | 0.60758 | 1.57038 | 56 |
| GO_POLYMERIC_CYTOSKELETAL_FIBER | 0.00016 | 0.00316 | 0.51885 | 0.46044 | 1.27956 | 230 |
| GO_POLYSOMAL_RIBOSOME | 0.00000 | 0.00000 | 0.73376 | 0.76359 | 1.89224 | 31 |
| GO_POLYSOME | 0.00000 | 0.00000 | 0.80122 | 0.71148 | 1.83443 | 55 |
| GO_POLY_PYRIMIDINE_TRACT_BINDING | 0.00098 | 0.01585 | 0.45506 | 0.71986 | 1.65356 | 16 |
| GO_POSITIVE_REGULATION_OF_BINDING | 0.00132 | 0.02016 | 0.45506 | 0.52828 | 1.38771 | 73 |
| GO_POSITIVE_REGULATION_OF_CANONICAL_WNT_SIGNAL | 0.00000 | 0.00000 | 0.77494 | 0.66666 | 1.74997 | 72 |
| GO_POSITIVE_REGULATION_OF_CELLULAR_AMIDE_METABC | 0.00006 | 0.00130 | 0.55733 | 0.54976 | 1.45696 | 87 |
| GO_POSITIVE_REGULATION_OF_CELLULAR_COMPONENT_C | 0.00242 | 0.03346 | 0.43171 | 0.41418 | 1.17136 | 451 |
| GO_POSITIVE_REGULATION_OF_CELL_DEATH | 0.00097 | 0.01561 | 0.47727 | 0.44900 | 1.25072 | 248 |
| GO_POSITIVE_REGULATION_OF_CELL_POPULATION_PROLIF | 0.00324 | 0.04222 | 0.43171 | 0.44056 | 1.22551 | 235 |
| GO_POSITIVE_REGULATION_OF_DEFENSE_RESPONSE | 0.00000 | 0.00000 | | 0.63941 | 1.71741 | 105 |
| GO_POSITIVE_REGULATION_OF_DNA_BINDING | 0.00143 | 0.02163 | 0.45506 | 0.65421 | 1.56380 | 22 |
| GO_POSITIVE_REGULATION_OF_DNA_BIOSYNTHETIC_PROC | 0.00000 | 0.00011 | 0.61053 | 0.70377 | 1.75009 | 34 |
| GO_POSITIVE_REGULATION_OF_DNA_METABOLIC_PROCESS | 0.00027 | 0.00507 | 0.49849 | 0.54095 | 1.42844 | 81 |
| GO_POSITIVE_REGULATION_OF_IMMUNE_RESPONSE | 0.00000 | 0.00000 | 0.70498 | 0.53247 | 1.46716 | 182 |
| GO_POSITIVE_REGULATION_OF_IMMUNE_SYSTEM_PROCES | 0.00000 | 0.00013 | 0.61053 | 0.48724 | 1.35725 | 248 |
| GO_POSITIVE_REGULATION_OF_INNATE_IMMUNE_RESPON | 0.00000 | 0.00000 | | 0.68770 | 1.81547 | 80 |
| GO_POSITIVE_REGULATION_OF_INTRINSIC_APOPTOTIC_SIG | 0.00034 | 0.00623 | 0.49849 | 0.67054 | 1.64060 | 27 |
| GO_POSITIVE_REGULATION_OF_RESPONSE_TO_BIOTIC_STII | 0.00000 | 0.00000 | | 0.65887 | 1.75365 | 92 |
| GO_POSITIVE_REGULATION_OF_RESPONSE_TO_EXTERNAL_ | 0.00000 | 0.00000 | 0.82666 | 0.60182 | 1.63487 | 128 |
| GO_POSITIVE_REGULATION_OF_RESPONSE_TO_WOUNDING | 0.00313 | 0.04113 | 0.43171 | 0.68842 | 1.58135 | 16 |
| GO_POSITIVE_REGULATION_OF_TELOMERASE_ACTIVITY | 0.00014 | 0.00279 | 0.51885 | 0.72189 | 1.71391 | 21 |
| GO_POSITIVE_REGULATION_OF_TELOMERE_MAINTENANCE | 0.00000 | 0.00008 | 0.62726 | 0.73288 | 1.79982 | 28 |
| GO_POSITIVE_REGULATION_OF_TELOMERE_MAINTENANCE | 0.00000 | 0.00006 | 0.62726 | 0.78078 | 1.85372 | 21 |
| GO_POSITIVE_REGULATION_OF_TRANSLATION | 0.00018 | 0.00346 | 0.51885 | 0.55324 | 1.45897 | 79 |
| GO_POSITIVE_REGULATION_OF_TRANSPORTER_ACTIVITY | 0.00054 | 0.00925 | 0.47727 | 0.65281 | 1.60317 | 28 |
| GO_POSITIVE_REGULATION_OF_WNT_SIGNALING_PATHWA | 0.00000 | 0.00000 | 0.73376 | 0.62986 | 1.66421 | 82 |
| GO_POSTSYNAPSE | 0.00001 | 0.00015 | 0.61053 | 0.51105 | 1.40438 | 170 |
| GO_POSTTRANSCRIPTIONAL_REGULATION_OF_GENE_EXPR | 0.00000 | 0.00000 | | 0.54229 | 1.52615 | 354 |
| GO_POST_TRANSLATIONAL_PROTEIN_MODIFICATION | 0.00000 | 0.00005 | 0.64355 | 0.52223 | 1.43281 | 163 |
| GO_PRE_MRNA_BINDING | 0.00030 | 0.00560 | 0.49849 | 0.68400 | 1.63501 | 22 |
| GO_PROTEASOMAL_PROTEIN_CATABOLIC_PROCESS | 0.00000 | 0.00012 | 0.61053 | 0.47923 | 1.33924 | 275 |
| GO_PROTEASOMAL_UBIQUITIN_INDEPENDENT_PROTEIN_C | 0.00000 | 0.00000 | 0.71951 | 0.87135 | 2.00155 | 16 |
| GO_PROTEASOME_ACCESSORY_COMPLEX | 0.00000 | 0.00002 | 0.65944 | 0.78847 | 1.89422 | 23 |
| GO_PROTEASOME_CORE_COMPLEX | 0.00000 | 0.00000 | 0.76146 | 0.89613 | 2.04805 | 15 |
| GO_PROTEIN_CATABOLIC_PROCESS | 0.00003 | 0.00062 | 0.57561 | 0.43925 | 1.24385 | 478 |
| GO_PROTEIN_CONTAINING_COMPLEX_BINDING | 0.00000 | 0.00003 | 0.65944 | 0.45042 | 1.27609 | 487 |
| GO_PROTEIN_FOLDING | 0.00000 | 0.00004 | 0.64355 | 0.54585 | 1.48435 | 131 |
| GO_PROTEIN_LOCALIZATION_TO_CHROMOSOME_TELOMEF | 0.00001 | 0.00021 | 0.59333 | 0.79250 | 1.83170 | 17 |
| GO_PROTEIN_LOCALIZATION_TO_ENDOPLASMIC_RETICULU | 0.00000 | 0.00000 | | 0.79606 | 2.15054 | 114 |
| GO_PROTEIN_LOCALIZATION_TO_MEMBRANE | 0.00000 | 0.00000 | | 0.57783 | 1.61940 | 302 |
| GO_PROTEIN_MODIFICATION_BY_SMALL_PROTEIN_REMOV | 0.00003 | 0.00071 | 0.55733 | 0.50085 | 1.37414 | 163 |
| GO_PROTEIN_POLYUBIQUITINATION | 0.00000 | 0.00001 | 0.69013 | 0.53029 | 1.45870 | 176 |
| GO_PROTEIN_STABILIZATION | 0.00039 | 0.00697 | 0.49849 | 0.52150 | 1.38446 | 90 |
| GO_PROTEIN_TARGETING | 0.00000 | 0.00000 | | 0.59518 | 1.66259 | 272 |
| GO_PROTEIN_TARGETING_TO_MEMBRANE | 0.00000 | 0.00000 | | 0.74713 | 2.03302 | 135 |
| GO_PURINE_NUCLEOSIDE_TRIPHOSPHATE_METABOLIC_PR( | 0.00202 | 0.02872 | 0.43171 | 0.61648 | 1.52769 | 31 |

| Cluster 1 | | | | | | |
|---|---|---|---|---|---|---|
| pathway | pval | padj | log2err | ES | NES | size |
| GO_REGULATION_OF_ANATOMICAL_STRUCTURE_MORPHO | 0.00003 | 0.00072 | 0.55733 | 0.45617 | 1.27998 | 322 |
| GO_REGULATION_OF_ANIMAL_ORGAN_MORPHOGENESIS | 0.00000 | 0.00000 | | 0.65343 | 1.73471 | 90 |
| GO_REGULATION_OF_BINDING | 0.00024 | 0.00455 | 0.51885 | 0.49263 | 1.34372 | 142 |
| GO_REGULATION_OF_CATABOLIC_PROCESS | 0.00000 | 0.00002 | 0.65944 | 0.45509 | 1.28945 | 499 |
| GO_REGULATION_OF_CELLULAR_AMIDE_METABOLIC_PROC | 0.00000 | 0.00001 | 0.69013 | 0.50727 | 1.40825 | 226 |
| GO_REGULATION_OF_CELLULAR_AMINE_METABOLIC_PROC | 0.00000 | 0.00000 | | 0.75239 | 1.93045 | 52 |
| GO_REGULATION_OF_CELLULAR_AMINO_ACID_METABOLIC | 0.00000 | 0.00000 | | 0.76524 | 1.95778 | 49 |
| GO_REGULATION_OF_CELLULAR_CATABOLIC_PROCESS | 0.00000 | 0.00000 | 0.71951 | 0.47534 | 1.34167 | 424 |
| GO_REGULATION_OF_CELLULAR_KETONE_METABOLIC_PRO | 0.00000 | 0.00000 | 0.74774 | 0.63317 | 1.67588 | 84 |
| GO_REGULATION_OF_CELLULAR_PROTEIN_CATABOLIC_PRO | 0.00287 | 0.03829 | 0.43171 | 0.46957 | 1.27799 | 137 |
| GO_REGULATION_OF_CELL_CYCLE_G2_M_PHASE_TRANSITI | 0.00000 | 0.00000 | 0.74774 | 0.58423 | 1.58577 | 126 |
| GO_REGULATION_OF_CELL_CYCLE_PHASE_TRANSITION | 0.00002 | 0.00044 | 0.57561 | 0.47817 | 1.33187 | 245 |
| GO_REGULATION_OF_CELL_CYCLE_PROCESS | 0.00013 | 0.00276 | 0.51885 | 0.44020 | 1.24083 | 382 |
| GO_REGULATION_OF_CELL_POPULATION_PROLIFERATION | 0.00035 | 0.00626 | 0.49849 | 0.42647 | 1.20533 | 444 |
| GO_REGULATION_OF_CYTOPLASMIC_TRANSLATION | 0.00028 | 0.00520 | 0.49849 | 0.72935 | 1.69924 | 18 |
| GO_REGULATION_OF_DEFENSE_RESPONSE | 0.00000 | 0.00000 | 0.77494 | 0.54987 | 1.51399 | 180 |
| GO_REGULATION_OF_DNA_BIOSYNTHETIC_PROCESS | 0.00031 | 0.00572 | 0.49849 | 0.58219 | 1.49756 | 53 |
| GO_REGULATION_OF_DNA_METABOLIC_PROCESS | 0.00130 | 0.02002 | 0.45506 | 0.46846 | 1.28409 | 158 |
| GO_REGULATION_OF_DNA_TEMPLATED_TRANSCRIPTION_I | 0.00000 | 0.00000 | | 0.72274 | 1.89682 | 71 |
| GO_REGULATION_OF_HEMATOPOIETIC_PROGENITOR_CELL_ | 0.00000 | 0.00000 | | 0.75687 | 1.95626 | 56 |
| GO_REGULATION_OF_HEMATOPOIETIC_STEM_CELL_DIFFER | 0.00000 | 0.00000 | | 0.78208 | 2.00086 | 49 |
| GO_REGULATION_OF_HEMOPOIESIS | 0.00000 | 0.00005 | 0.64355 | 0.52897 | 1.44715 | 154 |
| GO_REGULATION_OF_IMMUNE_RESPONSE | 0.00000 | 0.00006 | 0.62726 | 0.49416 | 1.37227 | 227 |
| GO_REGULATION_OF_IMMUNE_SYSTEM_PROCESS | 0.00027 | 0.00501 | 0.49849 | 0.43683 | 1.23027 | 371 |
| GO_REGULATION_OF_INNATE_IMMUNE_RESPONSE | 0.00000 | 0.00000 | | 0.64563 | 1.73409 | 105 |
| GO_REGULATION_OF_INTRINSIC_APOPTOTIC_SIGNALING_P | 0.00012 | 0.00248 | 0.53843 | 0.55853 | 1.47293 | 79 |
| GO_REGULATION_OF_MITOTIC_CELL_CYCLE | 0.00022 | 0.00424 | 0.51885 | 0.44441 | 1.24699 | 322 |
| GO_REGULATION_OF_MORPHOGENESIS_OF_AN_EPITHELIU | 0.00000 | 0.00000 | | 0.67882 | 1.78830 | 78 |
| GO_REGULATION_OF_MRNA_CATABOLIC_PROCESS | 0.00000 | 0.00000 | | 0.61056 | 1.65684 | 125 |
| GO_REGULATION_OF_MRNA_METABOLIC_PROCESS | 0.00000 | 0.00000 | 0.82666 | 0.55112 | 1.52635 | 214 |
| GO_REGULATION_OF_MRNA_PROCESSING | 0.00130 | 0.02002 | 0.45506 | 0.50247 | 1.34633 | 101 |
| GO_REGULATION_OF_MRNA_SPLICING_VIA_SPLICEOSOME | 0.00040 | 0.00699 | 0.49849 | 0.54568 | 1.43523 | 75 |
| GO_REGULATION_OF_MUSCLE_CONTRACTION | 0.00324 | 0.04222 | 0.43171 | 0.59262 | 1.47101 | 33 |
| GO_REGULATION_OF_PROTEIN_CATABOLIC_PROCESS | 0.00049 | 0.00850 | 0.47727 | 0.46603 | 1.28865 | 198 |
| GO_REGULATION_OF_PROTEIN_MODIFICATION_BY_SMALL_ | 0.00339 | 0.04364 | 0.43171 | 0.46997 | 1.27437 | 124 |
| GO_REGULATION_OF_PROTEIN_STABILITY | 0.00061 | 0.01036 | 0.47727 | 0.47412 | 1.29457 | 147 |
| GO_REGULATION_OF_PROTEOLYSIS | 0.00216 | 0.03035 | 0.43171 | 0.43937 | 1.22651 | 260 |
| GO_REGULATION_OF_RESPONSE_TO_BIOTIC_STIMULUS | 0.00000 | 0.00000 | 0.82666 | 0.60308 | 1.63828 | 128 |
| GO_REGULATION_OF_RESPONSE_TO_EXTERNAL_STIMULUS | 0.00000 | 0.00001 | 0.67496 | 0.49836 | 1.38957 | 255 |
| GO_REGULATION_OF_RESPONSE_TO_STRESS | 0.00000 | 0.00002 | 0.65944 | 0.45270 | 1.28253 | 487 |
| GO_REGULATION_OF_RNA_SPLICING | 0.00001 | 0.00027 | 0.59333 | 0.55497 | 1.48255 | 97 |
| GO_REGULATION_OF_SMALL_MOLECULE_METABOLIC_PRO | 0.00002 | 0.00057 | 0.57561 | 0.49603 | 1.36463 | 177 |
| GO_REGULATION_OF_STEM_CELL_DIFFERENTIATION | 0.00000 | 0.00000 | | 0.71203 | 1.85948 | 65 |
| GO_REGULATION_OF_TELOMERASE_ACTIVITY | 0.00117 | 0.01841 | 0.45506 | 0.64867 | 1.58197 | 26 |
| GO_REGULATION_OF_TELOMERE_MAINTENANCE | 0.00006 | 0.00125 | 0.55733 | 0.62124 | 1.58142 | 47 |
| GO_REGULATION_OF_TELOMERE_MAINTENANCE_VIA_TELO | 0.00000 | 0.00008 | 0.62726 | 0.69185 | 1.72437 | 36 |
| GO_REGULATION_OF_TRANSCRIPTION_FROM_RNA_POLYM | 0.00000 | 0.00000 | | 0.75863 | 1.95599 | 55 |
| GO_REGULATION_OF_TRANSLATIONAL_INITIATION | 0.00364 | 0.04618 | 0.43171 | 0.55220 | 1.41680 | 52 |
| GO_REGULATION_OF_UBIQUITIN_PROTEIN_LIGASE_ACTIVIT | 0.00002 | 0.00043 | 0.57561 | 0.78252 | 1.80864 | 17 |
| GO_REGULATION_OF_UBIQUITIN_PROTEIN_TRANSFERASE_ | 0.00202 | 0.02872 | 0.43171 | 0.61640 | 1.52749 | 31 |
| GO_REGULATION_OF_WNT_SIGNALING_PATHWAY | 0.00000 | 0.00003 | 0.65944 | 0.54959 | 1.49663 | 139 |
| GO_REGULATION_OF_WOUND_HEALING | 0.00235 | 0.03254 | 0.43171 | 0.60416 | 1.49708 | 32 |
| GO_REPRODUCTIVE_SYSTEM_DEVELOPMENT | 0.00213 | 0.03002 | 0.43171 | 0.47020 | 1.28254 | 142 |
| GO_RESPONSE_TO_ABIOTIC_STIMULUS | 0.00004 | 0.00087 | 0.55733 | 0.43663 | 1.23535 | 455 |
| GO_RESPONSE_TO_ACID_CHEMICAL | 0.00331 | 0.04306 | 0.43171 | 0.57020 | 1.43560 | 40 |
| GO_RESPONSE_TO_ARSENIC_CONTAINING_SUBSTANCE | 0.00174 | 0.02560 | 0.45506 | 0.68674 | 1.59996 | 18 |
| GO_RESPONSE_TO_BIOTIC_STIMULUS | 0.00000 | 0.00000 | 0.80122 | 0.50060 | 1.40896 | 357 |
| GO_RESPONSE_TO_CYTOKINE | 0.00000 | 0.00000 | | 0.55075 | 1.55111 | 371 |
| GO_RESPONSE_TO_DRUG | 0.00031 | 0.00572 | 0.49849 | 0.49326 | 1.33974 | 127 |
| GO_RESPONSE_TO_INORGANIC_SUBSTANCE | 0.00002 | 0.00044 | 0.57561 | 0.49754 | 1.36991 | 180 |
| GO_RESPONSE_TO_INTERLEUKIN_1 | 0.00000 | 0.00000 | | 0.70248 | 1.84865 | 76 |
| GO_RESPONSE_TO_INTERLEUKIN_12 | 0.00000 | 0.00005 | 0.64355 | 0.71991 | 1.78390 | 32 |
| GO_RESPONSE_TO_INTERLEUKIN_4 | 0.00003 | 0.00071 | 0.55733 | 0.78557 | 1.80450 | 16 |
| GO_RESPONSE_TO_METAL_ION | 0.00235 | 0.03254 | 0.43171 | 0.48444 | 1.30616 | 112 |
| GO_RESPONSE_TO_OXIDATIVE_STRESS | 0.00095 | 0.01536 | 0.47727 | 0.46840 | 1.28568 | 165 |
| GO_RESPONSE_TO_OXYGEN_CONTAINING_COMPOUND | 0.00353 | 0.04503 | 0.43171 | 0.40834 | 1.15700 | 499 |
| GO_RESPONSE_TO_OXYGEN_LEVELS | 0.00000 | 0.00011 | 0.61053 | 0.51105 | 1.40436 | 173 |
| GO_RESPONSE_TO_TOPOLOGICALLY_INCORRECT_PROTEIN | 0.00003 | 0.00071 | 0.55733 | 0.54583 | 1.46251 | 101 |
| GO_RESPONSE_TO_TOXIC_SUBSTANCE | 0.00001 | 0.00026 | 0.59333 | 0.57634 | 1.51481 | 74 |

| Cluster 1 | | | | | | |
|---|---|---|---|---|---|---|
| pathway | pval | padj | log2err | ES | NES | size |
| GO_RESPONSE_TO_TUMOR_NECROSIS_FACTOR | 0.00000 | 0.00000 | | 0.64129 | 1.72672 | 110 |
| GO_RIBONUCLEOPROTEIN_COMPLEX | 0.00000 | 0.00000 | | 0.57555 | 1.63086 | 496 |
| GO_RIBONUCLEOPROTEIN_COMPLEX_BINDING | 0.00001 | 0.00030 | 0.59333 | 0.55627 | 1.47928 | 91 |
| GO_RIBONUCLEOPROTEIN_COMPLEX_BIOGENESIS | 0.00000 | 0.00000 | | 0.53108 | 1.48986 | 323 |
| GO_RIBONUCLEOPROTEIN_COMPLEX_SUBUNIT_ORGANIZAT | 0.00000 | 0.00000 | 0.80122 | 0.57860 | 1.58297 | 153 |
| GO_RIBONUCLEOPROTEIN_GRANULE | 0.00000 | 0.00008 | 0.62726 | 0.54062 | 1.46919 | 129 |
| GO_RIBONUCLEOSIDE_MONOPHOSPHATE_METABOLIC_PRO | 0.00176 | 0.02578 | 0.45506 | 0.63668 | 1.55776 | 27 |
| GO_RIBONUCLEOSIDE_TRIPHOSPHATE_METABOLIC_PROCES | 0.00119 | 0.01863 | 0.45506 | 0.61850 | 1.53261 | 32 |
| GO_RIBOSOMAL_LARGE_SUBUNIT_ASSEMBLY | 0.00006 | 0.00132 | 0.53843 | 0.70792 | 1.72646 | 26 |
| GO_RIBOSOMAL_LARGE_SUBUNIT_BIOGENESIS | 0.00000 | 0.00000 | 0.76146 | 0.68007 | 1.76308 | 59 |
| GO_RIBOSOMAL_SMALL_SUBUNIT_ASSEMBLY | 0.00376 | 0.04732 | 0.43171 | 0.69420 | 1.58656 | 15 |
| GO_RIBOSOMAL_SMALL_SUBUNIT_BIOGENESIS | 0.00003 | 0.00067 | 0.57561 | 0.61287 | 1.57860 | 54 |
| GO_RIBOSOMAL_SUBUNIT | 0.00000 | 0.00000 | | 0.68237 | 1.87046 | 159 |
| GO_RIBOSOME | 0.00000 | 0.00000 | | 0.64991 | 1.79332 | 189 |
| GO_RIBOSOME_ASSEMBLY | 0.00000 | 0.00000 | 0.71951 | 0.68323 | 1.74938 | 51 |
| GO_RIBOSOME_BIOGENESIS | 0.00000 | 0.00000 | 0.70498 | 0.51442 | 1.42740 | 225 |
| GO_RNA_CATABOLIC_PROCESS | 0.00000 | 0.00000 | | 0.67477 | 1.88493 | 267 |
| GO_RNA_DEPENDENT_DNA_BIOSYNTHETIC_PROCESS | 0.00000 | 0.00000 | 0.71951 | 0.71991 | 1.80716 | 39 |
| GO_RNA_POLYMERASE_II_PREINITIATION_COMPLEX_ASSEM | 0.00372 | 0.04698 | 0.43171 | 0.69571 | 1.59000 | 15 |
| GO_RNA_SPLICING | 0.00001 | 0.00019 | 0.61053 | 0.46854 | 1.31305 | 299 |
| GO_RNA_SPLICING_VIA_TRANSESTERIFICATION_REACTIONS | 0.00016 | 0.00328 | 0.51885 | 0.46237 | 1.28970 | 256 |
| GO_RNA_STABILIZATION | 0.00067 | 0.01136 | 0.47727 | 0.64902 | 1.59387 | 28 |
| GO_ROUGH_ENDOPLASMIC_RETICULUM | 0.00010 | 0.00212 | 0.53843 | 0.63268 | 1.59292 | 40 |
| GO_RRNA_BINDING | 0.00000 | 0.00001 | 0.69013 | 0.68712 | 1.75439 | 48 |
| GO_RRNA_METABOLIC_PROCESS | 0.00021 | 0.00406 | 0.51885 | 0.48973 | 1.34450 | 166 |
| GO_SCF_DEPENDENT_PROTEASOMAL_UBIQUITIN_DEPENDE | 0.00000 | 0.00000 | 0.80122 | 0.69648 | 1.81023 | 61 |
| GO_SECRETION | 0.00001 | 0.00026 | 0.59333 | 0.44713 | 1.26543 | 459 |
| GO_SECRETORY_GRANULE | 0.00000 | 0.00000 | 0.74774 | 0.51514 | 1.43828 | 261 |
| GO_SECRETORY_VESICLE | 0.00000 | 0.00003 | 0.64355 | 0.48138 | 1.34916 | 308 |
| GO_SIGNALING_RECEPTOR_BINDING | 0.00095 | 0.01536 | 0.47727 | 0.43568 | 1.22199 | 319 |
| GO_SIGNAL_TRANSDUCTION_BY_PROTEIN_PHOSPHORYLAT | 0.00231 | 0.03222 | 0.43171 | 0.43586 | 1.21685 | 262 |
| GO_SINGLE_FERTILIZATION | 0.00335 | 0.04329 | 0.43171 | 0.59090 | 1.46674 | 33 |
| GO_SINGLE_STRANDED_RNA_BINDING | 0.00065 | 0.01110 | 0.47727 | 0.58977 | 1.50583 | 48 |
| GO_SMALL_RIBOSOMAL_SUBUNIT | 0.00000 | 0.00000 | | 0.71379 | 1.85724 | 62 |
| GO_SPERM_EGG_RECOGNITION | 0.00002 | 0.00038 | 0.57561 | 0.81393 | 1.86020 | 15 |
| GO_SPLICEOSOMAL_COMPLEX | 0.00012 | 0.00244 | 0.53843 | 0.49981 | 1.36330 | 142 |
| GO_STEM_CELL_DIFFERENTIATION | 0.00000 | 0.00000 | 0.77494 | 0.61754 | 1.65536 | 102 |
| GO_STRUCTURAL_CONSTITUENT_OF_RIBOSOME | 0.00000 | 0.00000 | | 0.70707 | 1.92863 | 142 |
| GO_STRUCTURAL_MOLECULE_ACTIVITY | 0.00000 | 0.00000 | | 0.60697 | 1.69241 | 255 |
| GO_SUBSTANTIA_NIGRA_DEVELOPMENT | 0.00001 | 0.00031 | 0.59333 | 0.74638 | 1.78412 | 22 |
| GO_SUPRAMOLECULAR_COMPLEX | 0.00000 | 0.00000 | 0.71951 | 0.46453 | 1.31464 | 460 |
| GO_SUPRAMOLECULAR_POLYMER | 0.00003 | 0.00063 | 0.57561 | 0.46575 | 1.30346 | 288 |
| GO_SYNAPSE | 0.00000 | 0.00002 | 0.65944 | 0.47348 | 1.33264 | 357 |
| GO_TELOMERASE_RNA_LOCALIZATION | 0.00007 | 0.00148 | 0.53843 | 0.76304 | 1.76360 | 17 |
| GO_TELOMERE_MAINTENANCE_VIA_TELOMERE_LENGTHEN | 0.00000 | 0.00006 | 0.62726 | 0.66517 | 1.69326 | 47 |
| GO_TELOMERE_ORGANIZATION | 0.00074 | 0.01226 | 0.47727 | 0.52410 | 1.38825 | 85 |
| GO_THREONINE_TYPE_PEPTIDASE_ACTIVITY | 0.00000 | 0.00000 | 0.69013 | 0.86289 | 1.98210 | 16 |
| GO_TISSUE_MORPHOGENESIS | 0.00002 | 0.00039 | 0.57561 | 0.48224 | 1.33732 | 221 |
| GO_TOXIN_TRANSPORT | 0.00001 | 0.00031 | 0.59333 | 0.78355 | 1.82551 | 18 |
| GO_TRANSCRIPTION_FACTOR_BINDING | 0.00081 | 0.01338 | 0.47727 | 0.44461 | 1.24250 | 275 |
| GO_TRANSLATIONAL_INITIATION | 0.00000 | 0.00000 | | 0.75673 | 2.06929 | 151 |
| GO_TRANSLATION_FACTOR_ACTIVITY_RNA_BINDING | 0.00000 | 0.00000 | 0.81404 | 0.70432 | 1.82044 | 56 |
| GO_TRANSLATION_INITIATION_FACTOR_ACTIVITY | 0.00000 | 0.00000 | 0.70498 | 0.71379 | 1.79969 | 41 |
| GO_TRANSLATION_PREINITIATION_COMPLEX | 0.00001 | 0.00028 | 0.59333 | 0.80259 | 1.84361 | 16 |
| GO_TRANSLATION_REGULATOR_ACTIVITY | 0.00000 | 0.00000 | 0.82666 | 0.64981 | 1.72178 | 86 |
| GO_TRANSLATION_REGULATOR_ACTIVITY_NUCLEIC_ACID_E | 0.00000 | 0.00000 | 0.80122 | 0.67905 | 1.77826 | 68 |
| GO_TRANSMEMBRANE_TRANSPORT | 0.00006 | 0.00129 | 0.55733 | 0.43970 | 1.24058 | 415 |
| GO_TUMOR_NECROSIS_FACTOR_MEDIATED_SIGNALING_PA | 0.00000 | 0.00000 | | 0.72551 | 1.90445 | 72 |
| GO_T_CELL_RECEPTOR_SIGNALING_PATHWAY | 0.00000 | 0.00000 | 0.76146 | 0.64284 | 1.69750 | 81 |
| GO_U2_TYPE_CATALYTIC_STEP_2_SPLICEOSOME | 0.00141 | 0.02150 | 0.45506 | 0.61923 | 1.52611 | 29 |
| GO_UBIQUITIN_LIKE_PROTEIN_LIGASE_BINDING | 0.00000 | 0.00000 | 0.70498 | 0.55216 | 1.51060 | 154 |
| GO_UNFOLDED_PROTEIN_BINDING | 0.00000 | 0.00001 | 0.69013 | 0.63224 | 1.65768 | 69 |
| GO_VACUOLAR_LUMEN | 0.00013 | 0.00276 | 0.51885 | 0.58513 | 1.52441 | 63 |
| GO_VESICLE_LUMEN | 0.00000 | 0.00000 | | 0.63190 | 1.71366 | 123 |
| GO_VIRAL_GENE_EXPRESSION | 0.00000 | 0.00000 | | 0.72850 | 1.99308 | 153 |
| HP_ABNORMALITY_OF_BLOOD_AND_BLOOD_FORMING_TIS | 0.00117 | 0.01841 | 0.45506 | 0.41959 | 1.18428 | 428 |
| HP_ABNORMALITY_OF_CARDIOVASCULAR_SYSTEM_ELECTR | 0.00012 | 0.00251 | 0.53843 | 0.48906 | 1.34059 | 159 |
| HP_ABNORMALITY_OF_LOWER_LIP | 0.00220 | 0.03077 | 0.43171 | 0.49707 | 1.32187 | 91 |
| HP_ABNORMALITY_OF_THE_NASAL_DORSUM | 0.00003 | 0.00074 | 0.55733 | 0.54594 | 1.45183 | 91 |
| HP_ABNORMAL_BASAL_GANGLIA_MRI_SIGNAL_INTENSITY | 0.00383 | 0.04785 | 0.43171 | 0.64215 | 1.52459 | 21 |

| Cluster 1 | | | | | | |
|---|---|---|---|---|---|---|
| pathway | pval | padj | log2err | ES | NES | size |
| HP_ABNORMAL_CARDIOVASCULAR_SYSTEM_PHYSIOLOGY | 0.00172 | 0.02541 | 0.45506 | 0.42303 | 1.19260 | 384 |
| HP_ABNORMAL_CEREBRAL_ARTERY_MORPHOLOGY | 0.00298 | 0.03953 | 0.43171 | 0.59337 | 1.47733 | 35 |
| HP_ABNORMAL_ERYTHROCYTE_MORPHOLOGY | 0.00000 | 0.00005 | 0.64355 | 0.49537 | 1.37371 | 221 |
| HP_ABNORMAL_MYELOID_LEUKOCYTE_MORPHOLOGY | 0.00216 | 0.03035 | 0.43171 | 0.48290 | 1.30447 | 116 |
| HP_ACUTE_LEUKEMIA | 0.00000 | 0.00001 | 0.67496 | 0.67959 | 1.73518 | 48 |
| HP_ANEMIA_OF_INADEQUATE_PRODUCTION | 0.00000 | 0.00000 | 0.71951 | 0.64628 | 1.68936 | 67 |
| HP_ARTERIAL_STENOSIS | 0.00181 | 0.02616 | 0.45506 | 0.58768 | 1.46895 | 37 |
| HP_CEREBRAL_ISCHEMIA | 0.00068 | 0.01147 | 0.47727 | 0.67026 | 1.61917 | 24 |
| HP_CONSTITUTIONAL_SYMPTOM | 0.00261 | 0.03557 | 0.43171 | 0.43959 | 1.22454 | 252 |
| HP_DELAYED_PUBERTY | 0.00001 | 0.00032 | 0.59333 | 0.58877 | 1.54371 | 69 |
| HP_DEPRESSED_NASAL_RIDGE | 0.00001 | 0.00034 | 0.59333 | 0.61815 | 1.59851 | 57 |
| HP_FATIGUE | 0.00000 | 0.00005 | 0.64355 | 0.55368 | 1.49682 | 117 |
| HP_HEADACHE | 0.00018 | 0.00346 | 0.51885 | 0.53520 | 1.42721 | 94 |
| HP_HEMATOLOGICAL_NEOPLASM | 0.00039 | 0.00697 | 0.49849 | 0.51705 | 1.38720 | 103 |
| HP_HEPATITIS | 0.00076 | 0.01267 | 0.47727 | 0.69492 | 1.62859 | 19 |
| HP_LEUKEMIA | 0.00000 | 0.00013 | 0.61053 | 0.62203 | 1.62054 | 63 |
| HP_LEUKODYSTROPHY | 0.00250 | 0.03437 | 0.43171 | 0.53195 | 1.37907 | 59 |
| HP_MACROCYTIC_ANEMIA | 0.00000 | 0.00001 | 0.67496 | 0.73806 | 1.83535 | 34 |
| HP_MIGRAINE | 0.00000 | 0.00001 | 0.69013 | 0.70094 | 1.77499 | 43 |
| HP_ORAL_CLEFT | 0.00040 | 0.00699 | 0.49849 | 0.46182 | 1.27818 | 207 |
| HP_PALLOR | 0.00028 | 0.00520 | 0.49849 | 0.57438 | 1.49640 | 63 |
| HP_PROGRESSIVE_CEREBELLAR_ATAXIA | 0.00383 | 0.04785 | 0.43171 | 0.56770 | 1.42933 | 40 |
| HP_SUPERNUMERARY_NIPPLE | 0.00181 | 0.02616 | 0.45506 | 0.68591 | 1.59802 | 18 |
| HP_SUPRANUCLEAR_GAZE_PALSY | 0.00181 | 0.02616 | 0.45506 | 0.69267 | 1.60097 | 17 |
| HP_THICK_LOWER_LIP_VERMILION | 0.00268 | 0.03636 | 0.43171 | 0.52599 | 1.37983 | 70 |
| HP_THICK_VERMILION_BORDER | 0.00261 | 0.03557 | 0.43171 | 0.49260 | 1.32360 | 106 |
| HP_TRANSIENT_ISCHEMIC_ATTACK | 0.00015 | 0.00299 | 0.51885 | 0.77045 | 1.76081 | 15 |
| HP_TRIPHALANGEAL_THUMB | 0.00152 | 0.02287 | 0.45506 | 0.66814 | 1.58629 | 21 |
| **Cluster 8** | | | | | | |
| pathway | pval | padj | log2err | ES | NES | size |
| GO_ACTIN_BINDING | 0.00003 | 0.00097 | 0.55733 | 0.59714 | 1.27024 | 205 |
| GO_ACTIN_CYTOSKELETON | 0.00001 | 0.00019 | 0.61053 | 0.59023 | 1.26128 | 263 |
| GO_ACTIN_FILAMENT_BASED_PROCESS | 0.00104 | 0.01961 | 0.45506 | 0.53565 | 1.15210 | 387 |
| GO_ACTIN_FILAMENT_BINDING | 0.00010 | 0.00274 | 0.53843 | 0.63641 | 1.33430 | 105 |
| GO_ACTIN_FILAMENT_BUNDLE | 0.00066 | 0.01409 | 0.47727 | 0.69449 | 1.39990 | 44 |
| GO_ACTIN_FILAMENT_ORGANIZATION | 0.00031 | 0.00743 | 0.49849 | 0.57670 | 1.22856 | 214 |
| GO_ACTIVATION_OF_IMMUNE_RESPONSE | 0.00000 | 0.00000 | 0.71951 | 0.64689 | 1.37358 | 189 |
| GO_ACTIVATION_OF_INNATE_IMMUNE_RESPONSE | 0.00000 | 0.00002 | 0.67496 | 0.70759 | 1.46984 | 82 |
| GO_ACTOMYOSIN | 0.00130 | 0.02341 | 0.45506 | 0.67522 | 1.36903 | 49 |
| GO_AEROBIC_RESPIRATION | 0.00000 | 0.00011 | 0.62726 | 0.72180 | 1.48238 | 64 |
| GO_AMINE_METABOLIC_PROCESS | 0.00000 | 0.00000 | 0.73376 | 0.74714 | 1.54505 | 73 |
| GO_ANAPHASE_PROMOTING_COMPLEX_DEPENDENT_CATA | 0.00000 | 0.00000 | 0.74774 | 0.75771 | 1.56835 | 75 |
| GO_ANCHORING_JUNCTION | 0.00000 | 0.00000 | | 0.67041 | 1.44585 | 454 |
| GO_ANTIGEN_PROCESSING_AND_PRESENTATION | 0.00000 | 0.00000 | 0.74774 | 0.67741 | 1.43061 | 136 |
| GO_ANTIGEN_PROCESSING_AND_PRESENTATION_OF_EXOG | 0.00000 | 0.00000 | 0.76146 | 0.79197 | 1.61752 | 57 |
| GO_ANTIGEN_PROCESSING_AND_PRESENTATION_OF_PEPT | 0.00000 | 0.00000 | 0.76146 | 0.70430 | 1.48297 | 120 |
| GO_ANTIGEN_PROCESSING_AND_PRESENTATION_OF_PEPT | 0.00000 | 0.00000 | 0.71951 | 0.75370 | 1.54833 | 67 |
| GO_ANTIGEN_PROCESSING_AND_PRESENTATION_OF_PEPT | 0.00065 | 0.01392 | 0.47727 | 0.66596 | 1.36493 | 62 |
| GO_ANTIGEN_RECEPTOR_MEDIATED_SIGNALING_PATHWAY | 0.00001 | 0.00033 | 0.59333 | 0.65108 | 1.36835 | 112 |
| GO_ANTIOXIDANT_ACTIVITY | 0.00097 | 0.01841 | 0.47727 | 0.72640 | 1.44201 | 33 |
| GO_APOPTOTIC_SIGNALING_PATHWAY | 0.00008 | 0.00224 | 0.53843 | 0.55871 | 1.19897 | 320 |
| GO_ATP_METABOLIC_PROCESS | 0.00000 | 0.00000 | 0.71951 | 0.64837 | 1.37639 | 183 |
| GO_ATP_SYNTHESIS_COUPLED_ELECTRON_TRANSPORT | 0.00000 | 0.00002 | 0.67496 | 0.71765 | 1.48561 | 76 |
| GO_AZUROPHIL_GRANULE | 0.00216 | 0.03571 | 0.43171 | 0.60955 | 1.27430 | 95 |
| GO_BLOOD_MICROPARTICLE | 0.00045 | 0.01031 | 0.49849 | 0.79283 | 1.53542 | 21 |
| GO_BRUSH_BORDER | 0.00161 | 0.02785 | 0.45506 | 0.66820 | 1.35711 | 52 |
| GO_CADHERIN_BINDING | 0.00000 | 0.00000 | | 0.67026 | 1.43024 | 239 |
| GO_CADHERIN_BINDING_INVOLVED_IN_CELL_CELL_ADHESI | 0.00068 | 0.01437 | 0.47727 | 0.83564 | 1.58359 | 15 |
| GO_CANONICAL_WNT_SIGNALING_PATHWAY | 0.00081 | 0.01624 | 0.47727 | 0.57936 | 1.22981 | 174 |
| GO_CARBOHYDRATE_BIOSYNTHETIC_PROCESS | 0.00113 | 0.02127 | 0.45506 | 0.61693 | 1.29034 | 98 |
| GO_CARBOHYDRATE_CATABOLIC_PROCESS | 0.00126 | 0.02290 | 0.45506 | 0.60676 | 1.27461 | 109 |
| GO_CELLULAR_AMINO_ACID_METABOLIC_PROCESS | 0.00003 | 0.00106 | 0.55733 | 0.59717 | 1.26922 | 198 |
| GO_CELLULAR_COMPONENT_DISASSEMBLY | 0.00006 | 0.00172 | 0.55733 | 0.55949 | 1.20300 | 370 |
| GO_CELLULAR_HOMEOSTASIS | 0.00027 | 0.00667 | 0.49849 | 0.54631 | 1.17500 | 390 |
| GO_CELLULAR_KETONE_METABOLIC_PROCESS | 0.00000 | 0.00007 | 0.64355 | 0.64883 | 1.36901 | 134 |
| GO_CELLULAR_NITROGEN_COMPOUND_CATABOLIC_PROCE | 0.00000 | 0.00000 | | 0.71723 | 1.54395 | 417 |
| GO_CELLULAR_PROTEIN_COMPLEX_DISASSEMBLY | 0.00078 | 0.01581 | 0.47727 | 0.58876 | 1.24734 | 158 |
| GO_CELLULAR_RESPIRATION | 0.00000 | 0.00000 | 0.73376 | 0.67628 | 1.42760 | 135 |
| GO_CELLULAR_RESPONSE_TO_OXYGEN_LEVELS | 0.00000 | 0.00000 | 0.77494 | 0.68586 | 1.44847 | 136 |
| GO_CELL_ACTIVATION_INVOLVED_IN_IMMUNE_RESPONSE | 0.00000 | 0.00000 | 0.74774 | 0.60607 | 1.30128 | 330 |
| GO_CELL_ADHESION_MOLECULE_BINDING | 0.00000 | 0.00000 | | 0.65726 | 1.40966 | 312 |

| **Cluster 1** | | | | | | |
|---|---|---|---|---|---|---|
| pathway | pval | padj | log2err | ES | NES | size |
| GO_CELL_CELL_JUNCTION | 0.00018 | 0.00457 | 0.51885 | 0.57365 | 1.22312 | 235 |
| GO_CELL_CELL_RECOGNITION | 0.00000 | 0.00000 | 0.74774 | 0.90784 | 1.75907 | 22 |
| GO_CELL_CELL_SIGNALING_BY_WNT | 0.00001 | 0.00027 | 0.61053 | 0.58718 | 1.25578 | 268 |
| GO_CELL_CORTEX | 0.00065 | 0.01392 | 0.47727 | 0.58843 | 1.24670 | 159 |
| GO_CELL_CYCLE_G2_M_PHASE_TRANSITION | 0.00004 | 0.00115 | 0.55733 | 0.59209 | 1.25810 | 195 |
| GO_CELL_KILLING | 0.00154 | 0.02683 | 0.45506 | 0.69990 | 1.39827 | 36 |
| GO_CELL_RECOGNITION | 0.00000 | 0.00001 | 0.70498 | 0.80125 | 1.61361 | 42 |
| GO_CELL_SUBSTRATE_JUNCTION | 0.00000 | 0.00000 | | 0.71354 | 1.52802 | 278 |
| GO_CELL_SURFACE | 0.00017 | 0.00444 | 0.51885 | 0.58540 | 1.24407 | 196 |
| GO_CELL_SURFACE_RECEPTOR_SIGNALING_PATHWAY_INVC | 0.00001 | 0.00030 | 0.59333 | 0.57978 | 1.24232 | 290 |
| GO_CHAPERONE_BINDING | 0.00022 | 0.00567 | 0.51885 | 0.66505 | 1.36622 | 67 |
| GO_CHAPERONE_COMPLEX | 0.00002 | 0.00080 | 0.57561 | 0.83619 | 1.61938 | 21 |
| GO_CONTRACTILE_FIBER | 0.00001 | 0.00039 | 0.59333 | 0.65803 | 1.37866 | 104 |
| GO_COTRANSLATIONAL_PROTEIN_TARGETING_TO_MEMBR | 0.00000 | 0.00000 | | 0.90390 | 1.88565 | 92 |
| GO_CYTOKINE_MEDIATED_SIGNALING_PATHWAY | 0.00000 | 0.00000 | | 0.63323 | 1.35732 | 306 |
| GO_CYTOPLASMIC_TRANSLATION | 0.00000 | 0.00000 | | 0.85600 | 1.78270 | 87 |
| GO_CYTOPLASMIC_TRANSLATIONAL_INITIATION | 0.00000 | 0.00016 | 0.61053 | 0.81087 | 1.60391 | 31 |
| GO_CYTOSKELETAL_PROTEIN_BINDING | 0.00130 | 0.02341 | 0.45506 | 0.52903 | 1.14205 | 484 |
| GO_CYTOSOLIC_LARGE_RIBOSOMAL_SUBUNIT | 0.00000 | 0.00000 | | 0.92028 | 1.87171 | 53 |
| GO_CYTOSOLIC_RIBOSOME | 0.00000 | 0.00000 | | 0.91403 | 1.91082 | 95 |
| GO_CYTOSOLIC_SMALL_RIBOSOMAL_SUBUNIT | 0.00000 | 0.00000 | | 0.90818 | 1.82257 | 40 |
| GO_DETOXIFICATION | 0.00087 | 0.01715 | 0.47727 | 0.66553 | 1.35928 | 57 |
| GO_DISORDERED_DOMAIN_SPECIFIC_BINDING | 0.00035 | 0.00824 | 0.49849 | 0.79252 | 1.53561 | 22 |
| GO_ELECTRON_TRANSFER_ACTIVITY | 0.00001 | 0.00047 | 0.59333 | 0.69677 | 1.43590 | 70 |
| GO_ELECTRON_TRANSPORT_CHAIN | 0.00000 | 0.00002 | 0.67496 | 0.66996 | 1.41127 | 123 |
| GO_ENDOCYTIC_VESICLE | 0.00191 | 0.03192 | 0.45506 | 0.58006 | 1.22889 | 157 |
| GO_ENDOCYTIC_VESICLE_MEMBRANE | 0.00250 | 0.03975 | 0.43171 | 0.63072 | 1.29837 | 69 |
| GO_ENDOPEPTIDASE_COMPLEX | 0.00001 | 0.00027 | 0.61053 | 0.71563 | 1.46674 | 62 |
| GO_ENDOPEPTIDASE_REGULATOR_ACTIVITY | 0.00213 | 0.03531 | 0.43171 | 0.66618 | 1.34595 | 46 |
| GO_ENERGY_DERIVATION_BY_OXIDATION_OF_ORGANIC_CC | 0.00000 | 0.00000 | 0.76146 | 0.66586 | 1.41288 | 172 |
| GO_ENTRY_INTO_HOST | 0.00272 | 0.04204 | 0.43171 | 0.63130 | 1.30097 | 70 |
| GO_ENZYME_INHIBITOR_ACTIVITY | 0.00051 | 0.01137 | 0.47727 | 0.59620 | 1.26068 | 142 |
| GO_EPHRIN_RECEPTOR_SIGNALING_PATHWAY | 0.00143 | 0.02523 | 0.45506 | 0.65171 | 1.33217 | 58 |
| GO_EPITHELIUM_DEVELOPMENT | 0.00000 | 0.00004 | 0.65944 | 0.56473 | 1.21986 | 491 |
| GO_ESTABLISHMENT_OF_PROTEIN_LOCALIZATION_TO_END | 0.00000 | 0.00000 | | 0.88714 | 1.85703 | 101 |
| GO_ESTABLISHMENT_OF_PROTEIN_LOCALIZATION_TO_MEM | 0.00000 | 0.00000 | | 0.76447 | 1.62934 | 228 |
| GO_ESTABLISHMENT_OF_PROTEIN_LOCALIZATION_TO_ORG | 0.00000 | 0.00000 | | 0.68796 | 1.48083 | 403 |
| GO_ESTABLISHMENT_OF_PROTEIN_LOCALIZATION_TO_TELC | 0.00135 | 0.02416 | 0.45506 | 0.80234 | 1.52624 | 16 |
| GO_ESTABLISHMENT_OF_TISSUE_POLARITY | 0.00000 | 0.00000 | | 0.79394 | 1.64378 | 77 |
| GO_EUKARYOTIC_TRANSLATION_INITIATION_FACTOR_3_CO | 0.00029 | 0.00713 | 0.49849 | 0.85373 | 1.61785 | 15 |
| GO_EXOCYTOSIS | 0.00000 | 0.00000 | 0.78819 | 0.59533 | 1.28215 | 431 |
| GO_EXTRACELLULAR_MATRIX_BINDING | 0.00250 | 0.03975 | 0.43171 | 0.76262 | 1.46122 | 18 |
| GO_FC_EPSILON_RECEPTOR_SIGNALING_PATHWAY | 0.00000 | 0.00012 | 0.62726 | 0.69835 | 1.44770 | 79 |
| GO_FC_RECEPTOR_MEDIATED_STIMULATORY_SIGNALING_F | 0.00039 | 0.00905 | 0.49849 | 0.68795 | 1.39722 | 52 |
| GO_FC_RECEPTOR_SIGNALING_PATHWAY | 0.00000 | 0.00000 | 0.82666 | 0.71756 | 1.51094 | 119 |
| GO_FERTILIZATION | 0.00010 | 0.00287 | 0.53843 | 0.70271 | 1.43080 | 54 |
| GO_FICOLIN_1_RICH_GRANULE | 0.00000 | 0.00004 | 0.65944 | 0.69300 | 1.44415 | 90 |
| GO_GENERATION_OF_PRECURSOR_METABOLITES_AND_EN | 0.00000 | 0.00000 | 0.80122 | 0.62419 | 1.33768 | 304 |
| GO_GLUTAMATERGIC_SYNAPSE | 0.00228 | 0.03686 | 0.43171 | 0.58095 | 1.22726 | 138 |
| GO_GLYCOLYTIC_PROCESS_THROUGH_FRUCTOSE_6_PHOSP | 0.00181 | 0.03076 | 0.45506 | 0.79633 | 1.51481 | 16 |
| GO_HEMATOPOIETIC_PROGENITOR_CELL_DIFFERENTIATION | 0.00000 | 0.00010 | 0.62726 | 0.68678 | 1.43512 | 94 |
| GO_HEMATOPOIETIC_STEM_CELL_DIFFERENTIATION | 0.00000 | 0.00000 | 0.77494 | 0.77507 | 1.59177 | 64 |
| GO_HEXOSE_CATABOLIC_PROCESS | 0.00298 | 0.04540 | 0.43171 | 0.72084 | 1.41705 | 28 |
| GO_IMMUNE_EFFECTOR_PROCESS | 0.00000 | 0.00000 | 0.71951 | 0.57461 | 1.24137 | 497 |
| GO_IMMUNE_RESPONSE_REGULATING_SIGNALING_PATHW | 0.00000 | 0.00001 | 0.67496 | 0.63910 | 1.35695 | 181 |
| GO_INNATE_IMMUNE_RESPONSE_ACTIVATING_SIGNAL_TR/ | 0.00000 | 0.00000 | 0.71951 | 0.75134 | 1.54558 | 68 |
| GO_INNER_MITOCHONDRIAL_MEMBRANE_PROTEIN_COMF | 0.00001 | 0.00036 | 0.59333 | 0.68265 | 1.42168 | 87 |
| GO_INTERACTION_WITH_SYMBIONT | 0.00283 | 0.04351 | 0.43171 | 0.69067 | 1.38213 | 38 |
| GO_INTERLEUKIN_1_MEDIATED_SIGNALING_PATHWAY | 0.00000 | 0.00000 | 0.77494 | 0.77048 | 1.58931 | 71 |
| GO_INTRINSIC_APOPTOTIC_SIGNALING_PATHWAY | 0.00025 | 0.00628 | 0.49849 | 0.59397 | 1.26028 | 168 |
| GO_LARGE_RIBOSOMAL_SUBUNIT | 0.00000 | 0.00000 | | 0.77528 | 1.62721 | 107 |
| GO_LEUKOCYTE_MEDIATED_IMMUNITY | 0.00000 | 0.00000 | 0.74774 | 0.60355 | 1.29642 | 342 |
| GO_LYSOSOMAL_LUMEN | 0.00090 | 0.01750 | 0.47727 | 0.69876 | 1.40672 | 43 |
| GO_MEMBRANE_ORGANIZATION | 0.00001 | 0.00048 | 0.59333 | 0.55525 | 1.19792 | 460 |
| GO_MIDBRAIN_DEVELOPMENT | 0.00037 | 0.00856 | 0.49849 | 0.71909 | 1.44466 | 41 |
| GO_MITOCHONDRIAL_ELECTRON_TRANSPORT_NADH_TO_I | 0.00049 | 0.01100 | 0.47727 | 0.70964 | 1.42864 | 43 |
| GO_MITOCHONDRIAL_ENVELOPE | 0.00000 | 0.00000 | 0.78819 | 0.58577 | 1.26516 | 492 |
| GO_MITOCHONDRIAL_PROTEIN_COMPLEX | 0.00000 | 0.00000 | 0.73376 | 0.64009 | 1.36112 | 203 |
| GO_MITOCHONDRIAL_TRANSLATIONAL_TERMINATION | 0.00313 | 0.04725 | 0.43171 | 0.60784 | 1.26630 | 86 |
| GO_MITOCHONDRION_ORGANIZATION | 0.00071 | 0.01460 | 0.47727 | 0.54297 | 1.16646 | 362 |

| Cluster 1 | | | | | | |
|---|---|---|---|---|---|---|
| pathway | pval | padj | log2err | ES | NES | size |
| GO_MODIFICATION_DEPENDENT_MACROMOLECULE_CATAI | 0.00036 | 0.00849 | 0.49849 | 0.53934 | 1.16317 | 454 |
| GO_MODULATION_OF_PROCESS_OF_OTHER_ORGANISM | 0.00257 | 0.04021 | 0.43171 | 0.65983 | 1.34626 | 56 |
| GO_MONOSACCHARIDE_BIOSYNTHETIC_PROCESS | 0.00033 | 0.00780 | 0.49849 | 0.68599 | 1.39676 | 54 |
| GO_MONOSACCHARIDE_CATABOLIC_PROCESS | 0.00178 | 0.03032 | 0.45506 | 0.71094 | 1.41651 | 35 |
| GO_MONOSACCHARIDE_METABOLIC_PROCESS | 0.00156 | 0.02706 | 0.45506 | 0.58165 | 1.23150 | 152 |
| GO_MONOVALENT_INORGANIC_CATION_TRANSPORT | 0.00076 | 0.01550 | 0.47727 | 0.57742 | 1.22625 | 179 |
| GO_MORPHOGENESIS_OF_AN_EPITHELIUM | 0.00000 | 0.00000 | 0.73376 | 0.62591 | 1.33594 | 252 |
| GO_MORPHOGENESIS_OF_A_POLARIZED_EPITHELIUM | 0.00000 | 0.00000 | | 0.77979 | 1.62615 | 91 |
| GO_MRNA_5_UTR_BINDING | 0.00070 | 0.01453 | 0.47727 | 0.78026 | 1.51186 | 22 |
| GO_MRNA_BINDING | 0.00000 | 0.00018 | 0.61053 | 0.60482 | 1.28890 | 225 |
| GO_MUSCLE_CONTRACTION | 0.00035 | 0.00831 | 0.49849 | 0.61537 | 1.29330 | 112 |
| GO_MYD88_DEPENDENT_TOLL_LIKE_RECEPTOR_SIGNALINC | 0.00056 | 0.01239 | 0.47727 | 0.84012 | 1.59207 | 15 |
| GO_MYELOID_LEUKOCYTE_ACTIVATION | 0.00000 | 0.00000 | 0.76146 | 0.61110 | 1.31101 | 315 |
| GO_MYELOID_LEUKOCYTE_MEDIATED_IMMUNITY | 0.00000 | 0.00000 | 0.74774 | 0.61986 | 1.32825 | 288 |
| GO_NADH_DEHYDROGENASE_ACTIVITY | 0.00039 | 0.00911 | 0.49849 | 0.74112 | 1.47664 | 35 |
| GO_NADH_DEHYDROGENASE_COMPLEX | 0.00021 | 0.00547 | 0.51885 | 0.73808 | 1.47701 | 38 |
| GO_NADH_METABOLIC_PROCESS | 0.00139 | 0.02466 | 0.45506 | 0.75937 | 1.47631 | 23 |
| GO_NAD_BINDING | 0.00235 | 0.03772 | 0.43171 | 0.69029 | 1.37908 | 36 |
| GO_NEGATIVE_REGULATION_OF_CANONICAL_WNT_SIGNAI | 0.00000 | 0.00003 | 0.65944 | 0.68777 | 1.43886 | 97 |
| GO_NEGATIVE_REGULATION_OF_CELLULAR_PROTEIN_CATA | 0.00242 | 0.03880 | 0.43171 | 0.63823 | 1.30875 | 63 |
| GO_NEGATIVE_REGULATION_OF_CELL_CYCLE_G2_M_PHASI | 0.00000 | 0.00003 | 0.65944 | 0.71092 | 1.47375 | 79 |
| GO_NEGATIVE_REGULATION_OF_CELL_DEATH | 0.00000 | 0.00002 | 0.67496 | 0.57230 | 1.23427 | 454 |
| GO_NEGATIVE_REGULATION_OF_COAGULATION | 0.00224 | 0.03637 | 0.43171 | 0.79237 | 1.50729 | 16 |
| GO_NEGATIVE_REGULATION_OF_PEPTIDASE_ACTIVITY | 0.00070 | 0.01453 | 0.47727 | 0.64197 | 1.32914 | 77 |
| GO_NEGATIVE_REGULATION_OF_PROTEIN_BINDING | 0.00080 | 0.01612 | 0.47727 | 0.67689 | 1.37475 | 52 |
| GO_NEGATIVE_REGULATION_OF_PROTEOLYSIS | 0.00003 | 0.00083 | 0.57561 | 0.62313 | 1.31613 | 137 |
| GO_NEGATIVE_REGULATION_OF_WNT_SIGNALING_PATHW/ | 0.00000 | 0.00001 | 0.69013 | 0.69047 | 1.44842 | 106 |
| GO_NEURAL_NUCLEUS_DEVELOPMENT | 0.00043 | 0.00977 | 0.49849 | 0.78091 | 1.52160 | 24 |
| GO_NEURON_TO_NEURON_SYNAPSE | 0.00002 | 0.00058 | 0.57561 | 0.62825 | 1.32844 | 142 |
| GO_NIK_NF_KAPPAB_SIGNALING | 0.00000 | 0.00000 | | 0.77013 | 1.60601 | 91 |
| GO_NON_CANONICAL_WNT_SIGNALING_PATHWAY | 0.00000 | 0.00000 | | 0.75732 | 1.57820 | 90 |
| GO_NUCLEAR_TRANSCRIBED_MRNA_CATABOLIC_PROCESS | 0.00000 | 0.00000 | | 0.81406 | 1.72807 | 184 |
| GO_NUCLEAR_TRANSCRIBED_MRNA_CATABOLIC_PROCESS_ | 0.00000 | 0.00000 | | 0.88627 | 1.86210 | 110 |
| GO_NUCLEOTIDE_EXCISION_REPAIR_DNA_DAMAGE_RECOC | 0.00216 | 0.03571 | 0.43171 | 0.75856 | 1.46981 | 22 |
| GO_ORGANELLE_ASSEMBLY | 0.00002 | 0.00058 | 0.57561 | 0.55078 | 1.18958 | 489 |
| GO_ORGANELLE_INNER_MEMBRANE | 0.00000 | 0.00003 | 0.65944 | 0.58338 | 1.25319 | 361 |
| GO_ORGANIC_CYCLIC_COMPOUND_CATABOLIC_PROCESS | 0.00000 | 0.00000 | | 0.71007 | 1.52872 | 424 |
| GO_OXIDATIVE_PHOSPHORYLATION | 0.00000 | 0.00001 | 0.69013 | 0.69830 | 1.45983 | 95 |
| GO_OXIDOREDUCTASE_ACTIVITY | 0.00000 | 0.00001 | 0.67496 | 0.58412 | 1.25484 | 357 |
| GO_OXIDOREDUCTASE_ACTIVITY_ACTING_ON_CH_OH_GRC | 0.00253 | 0.04010 | 0.43171 | 0.64274 | 1.31383 | 58 |
| GO_OXIDOREDUCTASE_ACTIVITY_ACTING_ON_NAD_P_H | 0.00220 | 0.03588 | 0.43171 | 0.63643 | 1.30704 | 64 |
| GO_OXIDOREDUCTASE_ACTIVITY_ACTING_ON_NAD_P_H_C | 0.00086 | 0.01691 | 0.47727 | 0.70736 | 1.42110 | 41 |
| GO_OXIDOREDUCTASE_COMPLEX | 0.00000 | 0.00013 | 0.62726 | 0.69291 | 1.43746 | 80 |
| GO_PEPTIDASE_COMPLEX | 0.00002 | 0.00076 | 0.57561 | 0.68301 | 1.41565 | 78 |
| GO_PIGMENT_GRANULE | 0.00007 | 0.00212 | 0.53843 | 0.67566 | 1.39851 | 75 |
| GO_PLASMA_MEMBRANE_REGION | 0.00034 | 0.00809 | 0.49849 | 0.53499 | 1.15342 | 450 |
| GO_POLYMERIC_CYTOSKELETAL_FIBER | 0.00047 | 0.01063 | 0.49849 | 0.54594 | 1.17189 | 325 |
| GO_POLYSOMAL_RIBOSOME | 0.00000 | 0.00000 | | 0.89688 | 1.76963 | 30 |
| GO_POLYSOME | 0.00000 | 0.00000 | | 0.84079 | 1.72217 | 60 |
| GO_POSITIVE_REGULATION_OF_BINDING | 0.00093 | 0.01796 | 0.47727 | 0.62220 | 1.29751 | 91 |
| GO_POSITIVE_REGULATION_OF_CANONICAL_WNT_SIGNAL | 0.00002 | 0.00066 | 0.57561 | 0.66631 | 1.38950 | 91 |
| GO_POSITIVE_REGULATION_OF_CELLULAR_AMIDE_METABC | 0.00027 | 0.00659 | 0.49849 | 0.61899 | 1.30100 | 111 |
| GO_POSITIVE_REGULATION_OF_CELLULAR_PROTEIN_LOCAI | 0.00062 | 0.01354 | 0.47727 | 0.56828 | 1.21104 | 225 |
| GO_POSITIVE_REGULATION_OF_CELL_DEATH | 0.00021 | 0.00538 | 0.51885 | 0.54846 | 1.17802 | 340 |
| GO_POSITIVE_REGULATION_OF_CYSTEINE_TYPE_ENDOPEP1 | 0.00276 | 0.04249 | 0.43171 | 0.63110 | 1.30056 | 70 |
| GO_POSITIVE_REGULATION_OF_DEFENSE_RESPONSE | 0.00015 | 0.00404 | 0.51885 | 0.60080 | 1.27313 | 156 |
| GO_POSITIVE_REGULATION_OF_DNA_BIOSYNTHETIC_PROC | 0.00154 | 0.02683 | 0.45506 | 0.67418 | 1.36566 | 48 |
| GO_POSITIVE_REGULATION_OF_ESTABLISHMENT_OF_PROT | 0.00088 | 0.01726 | 0.47727 | 0.57317 | 1.21956 | 208 |
| GO_POSITIVE_REGULATION_OF_IMMUNE_RESPONSE | 0.00000 | 0.00015 | 0.62726 | 0.59547 | 1.27075 | 248 |
| GO_POSITIVE_REGULATION_OF_IMMUNE_SYSTEM_PROCES | 0.00008 | 0.00222 | 0.53843 | 0.55792 | 1.19844 | 353 |
| GO_POSITIVE_REGULATION_OF_INNATE_IMMUNE_RESPON | 0.00000 | 0.00012 | 0.62726 | 0.67001 | 1.40476 | 105 |
| GO_POSITIVE_REGULATION_OF_INTRINSIC_APOPTOTIC_SIG | 0.00054 | 0.01204 | 0.47727 | 0.73786 | 1.46775 | 34 |
| GO_POSITIVE_REGULATION_OF_ION_TRANSMEMBRANE_TI | 0.00331 | 0.04977 | 0.43171 | 0.66127 | 1.33950 | 48 |
| GO_POSITIVE_REGULATION_OF_ION_TRANSPORT | 0.00043 | 0.00977 | 0.49849 | 0.63941 | 1.32821 | 82 |
| GO_POSITIVE_REGULATION_OF_NITRIC_OXIDE_METABOLIC | 0.00220 | 0.03588 | 0.43171 | 0.79293 | 1.50834 | 16 |
| GO_POSITIVE_REGULATION_OF_ORGANELLE_ORGANIZATIO | 0.00126 | 0.02290 | 0.45506 | 0.53541 | 1.15155 | 390 |
| GO_POSITIVE_REGULATION_OF_PROTEIN_CONTAINING_CO | 0.00082 | 0.01631 | 0.47727 | 0.59298 | 1.25459 | 148 |
| GO_POSITIVE_REGULATION_OF_PROTEIN_CONTAINING_CO | 0.00122 | 0.02245 | 0.45506 | 0.77246 | 1.49597 | 21 |
| GO_POSITIVE_REGULATION_OF_PROTEIN_LOCALIZATION_T | 0.00066 | 0.01409 | 0.47727 | 0.66153 | 1.35507 | 61 |
| GO_POSITIVE_REGULATION_OF_PROTEIN_POLYMERIZATION | 0.00294 | 0.04496 | 0.43171 | 0.62644 | 1.29663 | 75 |

| **Cluster 1** | | | | | | |
|---|---|---|---|---|---|---|
| pathway | pval | padj | log2err | ES | NES | size |
| GO_POSITIVE_REGULATION_OF_REACTIVE_OXYGEN_SPECIE | 0.00074 | 0.01505 | 0.47727 | 0.79398 | 1.53167 | 20 |
| GO_POSITIVE_REGULATION_OF_RESPONSE_TO_BIOTIC_STI | 0.00004 | 0.00126 | 0.55733 | 0.63643 | 1.34010 | 119 |
| GO_POSITIVE_REGULATION_OF_RESPONSE_TO_EXTERNAL_ | 0.00235 | 0.03772 | 0.43171 | 0.56008 | 1.19009 | 195 |
| GO_POSITIVE_REGULATION_OF_SUPRAMOLECULAR_FIBER_ | 0.00066 | 0.01409 | 0.47727 | 0.60892 | 1.27974 | 112 |
| GO_POSITIVE_REGULATION_OF_TRANSLATION | 0.00028 | 0.00697 | 0.49849 | 0.63482 | 1.32713 | 95 |
| GO_POSITIVE_REGULATION_OF_TRANSPORT | 0.00001 | 0.00029 | 0.59333 | 0.56432 | 1.21384 | 392 |
| GO_POSITIVE_REGULATION_OF_VIRAL_PROCESS | 0.00220 | 0.03588 | 0.43171 | 0.62021 | 1.28662 | 80 |
| GO_POSITIVE_REGULATION_OF_WNT_SIGNALING_PATHWA | 0.00007 | 0.00197 | 0.53843 | 0.64068 | 1.34231 | 104 |
| GO_POSTSYNAPSE | 0.00000 | 0.00004 | 0.65944 | 0.60510 | 1.29132 | 243 |
| GO_POSTTRANSCRIPTIONAL_REGULATION_OF_GENE_EXPR | 0.00000 | 0.00000 | | 0.60783 | 1.31021 | 441 |
| GO_PROTEASOMAL_PROTEIN_CATABOLIC_PROCESS | 0.00009 | 0.00242 | 0.53843 | 0.55822 | 1.19916 | 352 |
| GO_PROTEASOMAL_UBIQUITIN_INDEPENDENT_PROTEIN_C | 0.00001 | 0.00041 | 0.59333 | 0.88486 | 1.68904 | 17 |
| GO_PROTEASOME_ACCESSORY_COMPLEX | 0.00025 | 0.00628 | 0.49849 | 0.79011 | 1.53954 | 24 |
| GO_PROTEASOME_CORE_COMPLEX | 0.00000 | 0.00011 | 0.62726 | 0.89422 | 1.70102 | 16 |
| GO_PROTEIN_CONTAINING_COMPLEX_DISASSEMBLY | 0.00011 | 0.00291 | 0.53843 | 0.57665 | 1.23027 | 238 |
| GO_PROTEIN_DOMAIN_SPECIFIC_BINDING | 0.00004 | 0.00109 | 0.55733 | 0.55598 | 1.19526 | 382 |
| GO_PROTEIN_FOLDING | 0.00000 | 0.00013 | 0.62726 | 0.63507 | 1.34546 | 158 |
| GO_PROTEIN_LOCALIZATION_TO_ENDOPLASMIC_RETICULU | 0.00000 | 0.00000 | | 0.86267 | 1.81647 | 119 |
| GO_PROTEIN_LOCALIZATION_TO_MEMBRANE | 0.00000 | 0.00000 | | 0.68756 | 1.47848 | 378 |
| GO_PROTEIN_LOCALIZATION_TO_NUCLEUS | 0.00290 | 0.04452 | 0.43171 | 0.56479 | 1.19890 | 174 |
| GO_PROTEIN_MODIFICATION_BY_SMALL_PROTEIN_REMOV | 0.00002 | 0.00069 | 0.57561 | 0.60187 | 1.27984 | 203 |
| GO_PROTEIN_STABILIZATION | 0.00002 | 0.00066 | 0.57561 | 0.64494 | 1.35826 | 121 |
| GO_PROTEIN_TARGETING | 0.00000 | 0.00000 | | 0.70620 | 1.51557 | 321 |
| GO_PROTEIN_TARGETING_TO_MEMBRANE | 0.00000 | 0.00000 | | 0.83148 | 1.75808 | 146 |
| GO_PROTON_TRANSMEMBRANE_TRANSPORT | 0.00083 | 0.01643 | 0.47727 | 0.65354 | 1.34809 | 71 |
| GO_RECEPTOR_MEDIATED_ENDOCYTOSIS | 0.00102 | 0.01933 | 0.45506 | 0.60620 | 1.27319 | 108 |
| GO_REGULATION_OF_ACTIN_FILAMENT_ORGANIZATION | 0.00073 | 0.01492 | 0.47727 | 0.59140 | 1.25069 | 144 |
| GO_REGULATION_OF_ANATOMICAL_STRUCTURE_MORPHO | 0.00159 | 0.02762 | 0.45506 | 0.52977 | 1.14268 | 458 |
| GO_REGULATION_OF_ANIMAL_ORGAN_MORPHOGENESIS | 0.00000 | 0.00000 | 0.78819 | 0.71444 | 1.50082 | 109 |
| GO_REGULATION_OF_APOPTOTIC_SIGNALING_PATHWAY | 0.00023 | 0.00570 | 0.51885 | 0.57847 | 1.23274 | 219 |
| GO_REGULATION_OF_BINDING | 0.00097 | 0.01841 | 0.47727 | 0.57107 | 1.21342 | 193 |
| GO_REGULATION_OF_CELLULAR_AMIDE_METABOLIC_PROC | 0.00002 | 0.00059 | 0.57561 | 0.57833 | 1.23926 | 300 |
| GO_REGULATION_OF_CELLULAR_AMINE_METABOLIC_PROC | 0.00000 | 0.00000 | 0.71951 | 0.78404 | 1.59716 | 55 |
| GO_REGULATION_OF_CELLULAR_AMINO_ACID_METABOLIC | 0.00000 | 0.00000 | 0.77494 | 0.80842 | 1.63978 | 50 |
| GO_REGULATION_OF_CELLULAR_KETONE_METABOLIC_PRO | 0.00000 | 0.00003 | 0.65944 | 0.68056 | 1.42686 | 105 |
| GO_REGULATION_OF_CELLULAR_PROTEIN_CATABOLIC_PRO | 0.00024 | 0.00599 | 0.51885 | 0.58317 | 1.23797 | 183 |
| GO_REGULATION_OF_CELLULAR_PROTEIN_LOCALIZATION | 0.00024 | 0.00599 | 0.51885 | 0.54953 | 1.18039 | 345 |
| GO_REGULATION_OF_CELLULAR_RESPONSE_TO_GROWTH_ | 0.00065 | 0.01392 | 0.47727 | 0.62084 | 1.29959 | 101 |
| GO_REGULATION_OF_CELL_CYCLE_G2_M_PHASE_TRANSITI | 0.00000 | 0.00015 | 0.62726 | 0.62885 | 1.33234 | 159 |
| GO_REGULATION_OF_CYSTEINE_TYPE_ENDOPEPTIDASE_AC | 0.00004 | 0.00115 | 0.55733 | 0.63924 | 1.34355 | 111 |
| GO_REGULATION_OF_CYTOPLASMIC_TRANSLATION | 0.00020 | 0.00505 | 0.51885 | 0.80081 | 1.55167 | 22 |
| GO_REGULATION_OF_DNA_TEMPLATED_TRANSCRIPTION_I | 0.00000 | 0.00000 | 0.76146 | 0.74030 | 1.54160 | 85 |
| GO_REGULATION_OF_HEMATOPOIETIC_PROGENITOR_CELL_ | 0.00000 | 0.00000 | 0.73376 | 0.76150 | 1.56390 | 65 |
| GO_REGULATION_OF_HEMATOPOIETIC_STEM_CELL_DIFFER | 0.00000 | 0.00000 | 0.74774 | 0.78521 | 1.60680 | 59 |
| GO_REGULATION_OF_IMMUNE_RESPONSE | 0.00015 | 0.00408 | 0.51885 | 0.56022 | 1.20259 | 326 |
| GO_REGULATION_OF_INNATE_IMMUNE_RESPONSE | 0.00008 | 0.00217 | 0.53843 | 0.61119 | 1.29264 | 145 |
| GO_REGULATION_OF_INTRINSIC_APOPTOTIC_SIGNALING_P | 0.00005 | 0.00145 | 0.55733 | 0.65712 | 1.37539 | 99 |
| GO_REGULATION_OF_ION_TRANSPORT | 0.00133 | 0.02391 | 0.45506 | 0.56814 | 1.20745 | 194 |
| GO_REGULATION_OF_MITOTIC_CELL_CYCLE | 0.00095 | 0.01819 | 0.47727 | 0.53097 | 1.14317 | 420 |
| GO_REGULATION_OF_MORPHOGENESIS_OF_AN_EPITHELIU | 0.00000 | 0.00000 | | 0.75664 | 1.57846 | 92 |
| GO_REGULATION_OF_MRNA_CATABOLIC_PROCESS | 0.00000 | 0.00000 | 0.70498 | 0.65747 | 1.39369 | 160 |
| GO_REGULATION_OF_MRNA_METABOLIC_PROCESS | 0.00000 | 0.00000 | 0.73376 | 0.62119 | 1.32743 | 263 |
| GO_REGULATION_OF_PEPTIDASE_ACTIVITY | 0.00001 | 0.00035 | 0.59333 | 0.61326 | 1.30221 | 175 |
| GO_REGULATION_OF_PROTEASOMAL_PROTEIN_CATABOLIC | 0.00179 | 0.03054 | 0.45506 | 0.58229 | 1.22973 | 136 |
| GO_REGULATION_OF_PROTEIN_BINDING | 0.00117 | 0.02174 | 0.45506 | 0.61149 | 1.28274 | 106 |
| GO_REGULATION_OF_PROTEIN_CATABOLIC_PROCESS | 0.00005 | 0.00145 | 0.55733 | 0.57660 | 1.23213 | 262 |
| GO_REGULATION_OF_PROTEIN_CONTAINING_COMPLEX_AS | 0.00082 | 0.01631 | 0.47727 | 0.55932 | 1.19366 | 251 |
| GO_REGULATION_OF_PROTEIN_LOCALIZATION | 0.00087 | 0.01703 | 0.47727 | 0.52919 | 1.14311 | 494 |
| GO_REGULATION_OF_PROTEIN_LOCALIZATION_TO_NUCLEU | 0.00065 | 0.01406 | 0.47727 | 0.63405 | 1.31707 | 82 |
| GO_REGULATION_OF_PROTEIN_MODIFICATION_BY_SMALL_ | 0.00178 | 0.03032 | 0.45506 | 0.58008 | 1.22929 | 155 |
| GO_REGULATION_OF_PROTEIN_POLYMERIZATION | 0.00253 | 0.04010 | 0.43171 | 0.58957 | 1.24146 | 122 |
| GO_REGULATION_OF_PROTEIN_STABILITY | 0.00007 | 0.00212 | 0.53843 | 0.59296 | 1.26018 | 194 |
| GO_REGULATION_OF_PROTEOLYSIS | 0.00003 | 0.00095 | 0.57561 | 0.56099 | 1.20516 | 357 |
| GO_REGULATION_OF_SMALL_MOLECULE_METABOLIC_PRO | 0.00016 | 0.00428 | 0.51885 | 0.57314 | 1.22304 | 242 |
| GO_REGULATION_OF_STEM_CELL_DIFFERENTIATION | 0.00000 | 0.00002 | 0.67496 | 0.71709 | 1.48627 | 78 |
| GO_REGULATION_OF_SUPRAMOLECULAR_FIBER_ORGANIZA | 0.00095 | 0.01819 | 0.47727 | 0.57399 | 1.21933 | 191 |
| GO_REGULATION_OF_SYMBIOTIC_PROCESS | 0.00122 | 0.02245 | 0.45506 | 0.59609 | 1.25717 | 128 |
| GO_REGULATION_OF_TRANSCRIPTION_FROM_RNA_POLYM | 0.00000 | 0.00000 | 0.74774 | 0.78614 | 1.60870 | 59 |
| GO_REGULATION_OF_UBIQUITIN_PROTEIN_TRANSFERASE_ | 0.00231 | 0.03735 | 0.43171 | 0.66970 | 1.34994 | 44 |
| GO_REGULATION_OF_WNT_SIGNALING_PATHWAY | 0.00067 | 0.01423 | 0.47727 | 0.57680 | 1.22505 | 187 |

| Cluster 1 | | | | | | |
|---|---|---|---|---|---|---|
| pathway | pval | padj | log2err | ES | NES | size |
| GO_RESPIRASOME | 0.00000 | 0.00007 | 0.64355 | 0.71999 | 1.48515 | 71 |
| GO_RESPIRATORY_CHAIN_COMPLEX | 0.00000 | 0.00016 | 0.61053 | 0.73196 | 1.49926 | 60 |
| GO_RESPIRATORY_ELECTRON_TRANSPORT_CHAIN | 0.00000 | 0.00003 | 0.65944 | 0.70304 | 1.46414 | 87 |
| GO_RESPONSE_TO_ALCOHOL | 0.00090 | 0.01750 | 0.47727 | 0.61736 | 1.28924 | 93 |
| GO_RESPONSE_TO_DRUG | 0.00018 | 0.00457 | 0.51885 | 0.59596 | 1.26456 | 172 |
| GO_RESPONSE_TO_GROWTH_FACTOR | 0.00146 | 0.02571 | 0.45506 | 0.54406 | 1.16595 | 304 |
| GO_RESPONSE_TO_INORGANIC_SUBSTANCE | 0.00012 | 0.00330 | 0.53843 | 0.57148 | 1.22073 | 259 |
| GO_RESPONSE_TO_INTERFERON_GAMMA | 0.00174 | 0.02988 | 0.45506 | 0.64375 | 1.32206 | 65 |
| GO_RESPONSE_TO_INTERLEUKIN_1 | 0.00000 | 0.00000 | 0.74774 | 0.72120 | 1.50951 | 99 |
| GO_RESPONSE_TO_INTERLEUKIN_12 | 0.00001 | 0.00046 | 0.59333 | 0.77547 | 1.54962 | 37 |
| GO_RESPONSE_TO_ORGANIC_CYCLIC_COMPOUND | 0.00145 | 0.02547 | 0.45506 | 0.53095 | 1.14278 | 401 |
| GO_RESPONSE_TO_OXYGEN_LEVELS | 0.00000 | 0.00001 | 0.67496 | 0.62496 | 1.33094 | 212 |
| GO_RESPONSE_TO_TOPOLOGICALLY_INCORRECT_PROTEIN | 0.00305 | 0.04639 | 0.43171 | 0.57755 | 1.22008 | 138 |
| GO_RESPONSE_TO_TOXIC_SUBSTANCE | 0.00272 | 0.04204 | 0.43171 | 0.59250 | 1.24523 | 112 |
| GO_RESPONSE_TO_TUMOR_NECROSIS_FACTOR | 0.00000 | 0.00000 | 0.78819 | 0.68754 | 1.45414 | 145 |
| GO_RETINA_HOMEOSTASIS | 0.00070 | 0.01453 | 0.47727 | 0.79459 | 1.52773 | 19 |
| GO_RIBONUCLEOPROTEIN_COMPLEX_BINDING | 0.00000 | 0.00014 | 0.62726 | 0.67046 | 1.40721 | 107 |
| GO_RIBONUCLEOPROTEIN_COMPLEX_BIOGENESIS | 0.00000 | 0.00000 | | 0.61523 | 1.32277 | 372 |
| GO_RIBONUCLEOPROTEIN_COMPLEX_SUBUNIT_ORGANIZAT | 0.00000 | 0.00000 | 0.83909 | 0.68110 | 1.44526 | 171 |
| GO_RIBONUCLEOPROTEIN_GRANULE | 0.00324 | 0.04879 | 0.43171 | 0.57766 | 1.22471 | 163 |
| GO_RIBOSOMAL_LARGE_SUBUNIT_ASSEMBLY | 0.00006 | 0.00190 | 0.53843 | 0.80601 | 1.57467 | 25 |
| GO_RIBOSOMAL_LARGE_SUBUNIT_BIOGENESIS | 0.00000 | 0.00012 | 0.62726 | 0.72262 | 1.48181 | 63 |
| GO_RIBOSOMAL_SMALL_SUBUNIT_ASSEMBLY | 0.00032 | 0.00762 | 0.49849 | 0.83916 | 1.60181 | 17 |
| GO_RIBOSOMAL_SMALL_SUBUNIT_BIOGENESIS | 0.00007 | 0.00195 | 0.53843 | 0.68804 | 1.41303 | 65 |
| GO_RIBOSOMAL_SUBUNIT | 0.00000 | 0.00000 | | 0.78206 | 1.65951 | 171 |
| GO_RIBOSOME | 0.00000 | 0.00000 | | 0.76523 | 1.62723 | 203 |
| GO_RIBOSOME_ASSEMBLY | 0.00000 | 0.00002 | 0.67496 | 0.76363 | 1.55804 | 56 |
| GO_RIBOSOME_BINDING | 0.00220 | 0.03588 | 0.43171 | 0.69163 | 1.38799 | 40 |
| GO_RIBOSOME_BIOGENESIS | 0.00000 | 0.00000 | 0.71951 | 0.61660 | 1.31682 | 257 |
| GO_RNA_CATABOLIC_PROCESS | 0.00000 | 0.00000 | | 0.76022 | 1.63242 | 328 |
| GO_RNA_DEPENDENT_DNA_BIOSYNTHETIC_PROCESS | 0.00202 | 0.03357 | 0.43171 | 0.66339 | 1.35353 | 56 |
| GO_RNA_SPLICING_VIA_TRANSESTERIFICATION_REACTIONS | 0.00075 | 0.01518 | 0.47727 | 0.55433 | 1.18745 | 294 |
| GO_ROUGH_ENDOPLASMIC_RETICULUM | 0.00010 | 0.00282 | 0.53843 | 0.71357 | 1.44171 | 46 |
| GO_RRNA_BINDING | 0.00000 | 0.00000 | 0.78819 | 0.80858 | 1.64220 | 52 |
| GO_RRNA_METABOLIC_PROCESS | 0.00016 | 0.00424 | 0.51885 | 0.58572 | 1.24488 | 198 |
| GO_SCF_DEPENDENT_PROTEASOMAL_UBIQUITIN_DEPEND | 0.00000 | 0.00002 | 0.67496 | 0.73173 | 1.50794 | 70 |
| GO_SECRETORY_GRANULE | 0.00000 | 0.00000 | 0.77494 | 0.60308 | 1.29676 | 373 |
| GO_SECRETORY_VESICLE | 0.00000 | 0.00000 | 0.76146 | 0.58831 | 1.26764 | 436 |
| GO_SIDE_OF_MEMBRANE | 0.00003 | 0.00100 | 0.55733 | 0.61115 | 1.29670 | 170 |
| GO_SIGNAL_TRANSDUCTION_BY_PROTEIN_PHOSPHORYLAT | 0.00119 | 0.02200 | 0.45506 | 0.53840 | 1.15760 | 379 |
| GO_SINGLE_FERTILIZATION | 0.00018 | 0.00478 | 0.51885 | 0.70476 | 1.42391 | 46 |
| GO_SMALL_RIBOSOMAL_SUBUNIT | 0.00000 | 0.00000 | | 0.79672 | 1.63671 | 67 |
| GO_SPERM_EGG_RECOGNITION | 0.00000 | 0.00001 | 0.70498 | 0.91116 | 1.74582 | 18 |
| GO_SPLICEOSOMAL_COMPLEX | 0.00044 | 0.01014 | 0.49849 | 0.58786 | 1.24596 | 162 |
| GO_STEM_CELL_DIFFERENTIATION | 0.00000 | 0.00003 | 0.65944 | 0.66716 | 1.40684 | 127 |
| GO_STRUCTURAL_CONSTITUENT_OF_CYTOSKELETON | 0.00014 | 0.00388 | 0.51885 | 0.70882 | 1.43210 | 46 |
| GO_STRUCTURAL_CONSTITUENT_OF_MUSCLE | 0.00124 | 0.02271 | 0.45506 | 0.81595 | 1.55750 | 17 |
| GO_STRUCTURAL_CONSTITUENT_OF_RIBOSOME | 0.00000 | 0.00000 | | 0.80104 | 1.69543 | 151 |
| GO_STRUCTURAL_MOLECULE_ACTIVITY | 0.00000 | 0.00000 | | 0.74172 | 1.59081 | 313 |
| GO_SUBSTANTIA_NIGRA_DEVELOPMENT | 0.00029 | 0.00713 | 0.49849 | 0.78853 | 1.53300 | 23 |
| GO_SUPRAMOLECULAR_FIBER_ORGANIZATION | 0.00010 | 0.00274 | 0.53843 | 0.55571 | 1.19341 | 334 |
| GO_SUPRAMOLECULAR_POLYMER | 0.00001 | 0.00048 | 0.59333 | 0.55718 | 1.19941 | 412 |
| GO_TELOMERASE_RNA_LOCALIZATION | 0.00046 | 0.01047 | 0.49849 | 0.80279 | 1.53818 | 18 |
| GO_THREONINE_TYPE_ENDOPEPTIDASE_ACTIVITY | 0.00000 | 0.00002 | 0.67496 | 0.91167 | 1.73421 | 16 |
| GO_THREONINE_TYPE_PEPTIDASE_ACTIVITY | 0.00000 | 0.00005 | 0.64355 | 0.88385 | 1.69934 | 19 |
| GO_TISSUE_MORPHOGENESIS | 0.00000 | 0.00002 | 0.67496 | 0.59675 | 1.27882 | 285 |
| GO_TRANSLATIONAL_ELONGATION | 0.00005 | 0.00163 | 0.55733 | 0.63485 | 1.33656 | 118 |
| GO_TRANSLATIONAL_INITIATION | 0.00000 | 0.00000 | | 0.84554 | 1.79368 | 167 |
| GO_TRANSLATIONAL_TERMINATION | 0.00187 | 0.03150 | 0.45506 | 0.61145 | 1.27919 | 97 |
| GO_TRANSLATION_FACTOR_ACTIVITY_RNA_BINDING | 0.00000 | 0.00003 | 0.65944 | 0.73875 | 1.51490 | 63 |
| GO_TRANSLATION_INITIATION_FACTOR_ACTIVITY | 0.00004 | 0.00114 | 0.55733 | 0.73558 | 1.48274 | 44 |
| GO_TRANSLATION_PREINITIATION_COMPLEX | 0.00003 | 0.00097 | 0.55733 | 0.86415 | 1.64382 | 16 |
| GO_TRANSLATION_REGULATOR_ACTIVITY | 0.00000 | 0.00000 | 0.73376 | 0.70479 | 1.47532 | 101 |
| GO_TRANSLATION_REGULATOR_ACTIVITY_NUCLEIC_ACID_E | 0.00000 | 0.00010 | 0.62726 | 0.70652 | 1.46280 | 77 |
| GO_TRICARBOXYLIC_ACID_CYCLE | 0.00117 | 0.02174 | 0.45506 | 0.73851 | 1.45179 | 28 |
| GO_TUMOR_NECROSIS_FACTOR_MEDIATED_SIGNALING_PA | 0.00000 | 0.00000 | | 0.76255 | 1.59244 | 93 |
| GO_T_CELL_RECEPTOR_SIGNALING_PATHWAY | 0.00000 | 0.00011 | 0.62726 | 0.66740 | 1.39793 | 103 |
| GO_UBIQUITIN_LIKE_PROTEIN_LIGASE_BINDING | 0.00000 | 0.00001 | 0.69013 | 0.62937 | 1.33868 | 204 |
| GO_UNFOLDED_PROTEIN_BINDING | 0.00001 | 0.00047 | 0.59333 | 0.68599 | 1.42207 | 79 |
| GO_VACUOLAR_LUMEN | 0.00053 | 0.01173 | 0.47727 | 0.64588 | 1.34368 | 84 |

| **Cluster 1** | | | | | | |
|---|---|---|---|---|---|---|
| pathway | pval | padj | log2err | ES | NES | size |
| GO_VACUOLE | 0.00065 | 0.01392 | 0.47727 | 0.53135 | 1.14550 | 443 |
| GO_VESICLE_LUMEN | 0.00000 | 0.00000 | 0.70498 | 0.64750 | 1.37398 | 171 |
| GO_VIRAL_GENE_EXPRESSION | 0.00000 | 0.00000 | | 0.81080 | 1.72042 | 169 |
| GO_VIRAL_GENOME_REPLICATION | 0.00257 | 0.04021 | 0.43171 | 0.63051 | 1.29526 | 67 |
| GO_VIRAL_LIFE_CYCLE | 0.00002 | 0.00058 | 0.57561 | 0.60075 | 1.27669 | 196 |
| HP_ABNORMALITY_OF_CARDIOVASCULAR_SYSTEM_ELECTR | 0.00001 | 0.00021 | 0.61053 | 0.59937 | 1.27718 | 216 |
| HP_ABNORMALITY_OF_LOWER_LIP | 0.00054 | 0.01204 | 0.47727 | 0.60911 | 1.28445 | 127 |
| HP_ABNORMALITY_OF_THE_MITOCHONDRION | 0.00100 | 0.01905 | 0.45506 | 0.60995 | 1.28153 | 110 |
| HP_ABNORMALITY_OF_THE_NASAL_DORSUM | 0.00004 | 0.00115 | 0.55733 | 0.63789 | 1.34296 | 118 |
| HP_ABNORMAL_ERYTHROCYTE_MORPHOLOGY | 0.00000 | 0.00000 | 0.70498 | 0.60226 | 1.29094 | 306 |
| HP_ABNORMAL_IMMUNE_SYSTEM_MORPHOLOGY | 0.00191 | 0.03192 | 0.45506 | 0.54981 | 1.17706 | 275 |
| HP_ABNORMAL_RETICULOCYTE_MORPHOLOGY | 0.00183 | 0.03097 | 0.45506 | 0.73015 | 1.43535 | 28 |
| HP_ABNORMAL_VENTRICULAR_SEPTUM_MORPHOLOGY | 0.00202 | 0.03357 | 0.43171 | 0.55906 | 1.19166 | 223 |
| HP_ACUTE_LEUKEMIA | 0.00000 | 0.00000 | 0.74774 | 0.76494 | 1.57095 | 65 |
| HP_ANEMIA_OF_INADEQUATE_PRODUCTION | 0.00000 | 0.00001 | 0.69013 | 0.70518 | 1.47110 | 92 |
| HP_DELAYED_PUBERTY | 0.00000 | 0.00001 | 0.67496 | 0.69381 | 1.45150 | 97 |
| HP_DEPRESSED_NASAL_RIDGE | 0.00000 | 0.00016 | 0.61053 | 0.71149 | 1.46163 | 67 |
| HP_ECHOLALIA | 0.00257 | 0.04021 | 0.43171 | 0.74533 | 1.45611 | 25 |
| HP_FATIGUE | 0.00001 | 0.00047 | 0.59333 | 0.62255 | 1.31974 | 161 |
| HP_HEADACHE | 0.00000 | 0.00010 | 0.62726 | 0.65694 | 1.38531 | 127 |
| HP_HEMATOLOGICAL_NEOPLASM | 0.00070 | 0.01453 | 0.47727 | 0.59735 | 1.26175 | 140 |
| HP_HYPERTROPHIC_CARDIOMYOPATHY | 0.00121 | 0.02227 | 0.45506 | 0.57908 | 1.22869 | 168 |
| HP_LETHARGY | 0.00137 | 0.02441 | 0.45506 | 0.61717 | 1.29022 | 95 |
| HP_LEUKEMIA | 0.00000 | 0.00003 | 0.65944 | 0.69689 | 1.45227 | 90 |
| HP_LEUKODYSTROPHY | 0.00071 | 0.01460 | 0.47727 | 0.64697 | 1.33569 | 72 |
| HP_LONG_NOSE | 0.00272 | 0.04204 | 0.43171 | 0.76055 | 1.45725 | 18 |
| HP_MACROCYTIC_ANEMIA | 0.00000 | 0.00000 | 0.80122 | 0.85438 | 1.71293 | 39 |
| HP_MIGRAINE | 0.00000 | 0.00000 | 0.78819 | 0.79333 | 1.62498 | 60 |
| HP_MUTISM | 0.00132 | 0.02366 | 0.45506 | 0.73670 | 1.44823 | 28 |
| HP_NAUSEA_AND_VOMITING | 0.00313 | 0.04725 | 0.43171 | 0.54988 | 1.17173 | 216 |
| HP_NEOPLASM | 0.00163 | 0.02807 | 0.45506 | 0.53502 | 1.15118 | 397 |
| HP_ORAL_CLEFT | 0.00034 | 0.00817 | 0.49849 | 0.55648 | 1.19245 | 282 |
| HP_PALLOR | 0.00000 | 0.00003 | 0.65944 | 0.69599 | 1.45020 | 89 |
| HP_POOR_HEAD_CONTROL | 0.00261 | 0.04067 | 0.43171 | 0.60465 | 1.26269 | 93 |
| HP_RETICULOCYTOPENIA | 0.00006 | 0.00181 | 0.53843 | 0.87300 | 1.65439 | 15 |
| HP_SHORT_NOSE | 0.00012 | 0.00334 | 0.51885 | 0.60646 | 1.28483 | 158 |
| HP_THICK_LOWER_LIP_VERMILION | 0.00003 | 0.00106 | 0.55733 | 0.65931 | 1.37377 | 89 |
| HP_THICK_VERMILION_BORDER | 0.00117 | 0.02174 | 0.45506 | 0.59277 | 1.25290 | 141 |
| HP_WEBBED_NECK | 0.00257 | 0.04021 | 0.43171 | 0.66434 | 1.34752 | 50 |
| **Cluster 9** | | | | | | |
| pathway | pval | padj | log2err | ES | NES | size |
| GO_ACTION_POTENTIAL | 0.00283 | 0.04728 | 0.43171 | 0.70574 | 1.31017 | 38 |
| GO_ACTIVATION_OF_IMMUNE_RESPONSE | 0.00067 | 0.01776 | 0.47727 | 0.60413 | 1.17934 | 183 |
| GO_AMINE_METABOLIC_PROCESS | 0.00122 | 0.02613 | 0.45506 | 0.65350 | 1.25305 | 77 |
| GO_ANAPHASE_PROMOTING_COMPLEX_DEPENDENT_CATA | 0.00098 | 0.02242 | 0.45506 | 0.66838 | 1.27687 | 70 |
| GO_ANATOMICAL_STRUCTURE_HOMEOSTASIS | 0.00018 | 0.00741 | 0.51885 | 0.61224 | 1.19917 | 206 |
| GO_ANCHORING_JUNCTION | 0.00039 | 0.01261 | 0.49849 | 0.57715 | 1.13986 | 387 |
| GO_ANTIGEN_PROCESSING_AND_PRESENTATION | 0.00024 | 0.00890 | 0.51885 | 0.63425 | 1.22905 | 129 |
| GO_ANTIGEN_PROCESSING_AND_PRESENTATION_OF_PEPT | 0.00059 | 0.01643 | 0.47727 | 0.64367 | 1.24265 | 111 |
| GO_APOPTOTIC_SIGNALING_PATHWAY | 0.00287 | 0.04744 | 0.43171 | 0.57525 | 1.13292 | 281 |
| GO_ATPASE_ACTIVITY | 0.00213 | 0.03827 | 0.43171 | 0.57816 | 1.13687 | 249 |
| GO_AUTOPHAGOSOME_MEMBRANE | 0.00110 | 0.02419 | 0.45506 | 0.76633 | 1.39571 | 25 |
| GO_BINDING_OF_SPERM_TO_ZONA_PELLUCIDA | 0.00121 | 0.02585 | 0.45506 | 0.82500 | 1.45682 | 16 |
| GO_CADHERIN_BINDING | 0.00002 | 0.00145 | 0.57561 | 0.62629 | 1.22567 | 201 |
| GO_CARDIAC_MUSCLE_CELL_ACTION_POTENTIAL | 0.00020 | 0.00788 | 0.51885 | 0.82044 | 1.48615 | 23 |
| GO_CARDIAC_MUSCLE_CELL_CONTRACTION | 0.00024 | 0.00886 | 0.51885 | 0.81377 | 1.46036 | 20 |
| GO_CELLULAR_NITROGEN_COMPOUND_CATABOLIC_PROCE | 0.00000 | 0.00000 | 0.76146 | 0.62600 | 1.23565 | 375 |
| GO_CELLULAR_PROTEIN_CATABOLIC_PROCESS | 0.00001 | 0.00048 | 0.61053 | 0.58714 | 1.16185 | 484 |
| GO_CELLULAR_RESPONSE_TO_HEAT | 0.00036 | 0.01172 | 0.49849 | 0.66040 | 1.26922 | 84 |
| GO_CELLULAR_RESPONSE_TO_ORGANIC_CYCLIC_COMPOUI | 0.00043 | 0.01311 | 0.49849 | 0.59596 | 1.16974 | 231 |
| GO_CELL_ACTIVATION_INVOLVED_IN_IMMUNE_RESPONSE | 0.00000 | 0.00027 | 0.62726 | 0.61988 | 1.22063 | 284 |
| GO_CELL_ADHESION_MOLECULE_BINDING | 0.00004 | 0.00229 | 0.55733 | 0.60853 | 1.19766 | 258 |
| GO_CELL_CELL_RECOGNITION | 0.00076 | 0.01938 | 0.47727 | 0.79548 | 1.44092 | 23 |
| GO_CELL_CYCLE_ARREST | 0.00194 | 0.03621 | 0.45506 | 0.62681 | 1.21201 | 116 |
| GO_CELL_CYCLE_G1_S_PHASE_TRANSITION | 0.00004 | 0.00258 | 0.55733 | 0.62742 | 1.22236 | 172 |
| GO_CELL_CYCLE_G2_M_PHASE_TRANSITION | 0.00013 | 0.00605 | 0.51885 | 0.61707 | 1.20689 | 193 |
| GO_CELL_CYCLE_PHASE_TRANSITION | 0.00000 | 0.00028 | 0.62726 | 0.59937 | 1.18424 | 399 |
| GO_CELL_DIVISION | 0.00024 | 0.00886 | 0.51885 | 0.58354 | 1.15195 | 362 |
| GO_CELL_SUBSTRATE_JUNCTION | 0.00001 | 0.00089 | 0.59333 | 0.61942 | 1.21578 | 231 |
| GO_CHAPERONE_COMPLEX | 0.00004 | 0.00228 | 0.55733 | 0.87341 | 1.54918 | 18 |
| GO_CHAPERONE_MEDIATED_PROTEIN_COMPLEX_ASSEMBL | 0.00115 | 0.02519 | 0.45506 | 0.82505 | 1.44768 | 15 |

| Cluster 1 | | | | | | |
|---|---|---|---|---|---|---|
| pathway | pval | padj | log2err | ES | NES | size |
| GO_CHROMATIN | 0.00088 | 0.02115 | 0.47727 | 0.56419 | 1.11645 | 484 |
| GO_CHROMATIN_BINDING | 0.00000 | 0.00016 | 0.65944 | 0.61336 | 1.20904 | 316 |
| GO_CHROMATIN_ORGANIZATION | 0.00000 | 0.00001 | 0.74774 | 0.61884 | 1.22305 | 443 |
| GO_CHROMATIN_REMODELING | 0.00007 | 0.00368 | 0.53843 | 0.65089 | 1.26021 | 124 |
| GO_CHROMOSOMAL_REGION | 0.00023 | 0.00866 | 0.51885 | 0.59926 | 1.17821 | 247 |
| GO_CHROMOSOME_SEGREGATION | 0.00023 | 0.00877 | 0.51885 | 0.60289 | 1.18343 | 230 |
| GO_CHROMOSOME_TELOMERIC_REGION | 0.00176 | 0.03388 | 0.45506 | 0.62655 | 1.20794 | 106 |
| GO_CONDENSED_CHROMOSOME | 0.00294 | 0.04820 | 0.43171 | 0.60227 | 1.17086 | 149 |
| GO_COTRANSLATIONAL_PROTEIN_TARGETING_TO_MEMBR | 0.00000 | 0.00038 | 0.62726 | 0.71079 | 1.36508 | 82 |
| GO_COVALENT_CHROMATIN_MODIFICATION | 0.00000 | 0.00016 | 0.65944 | 0.61923 | 1.21955 | 294 |
| GO_CYTOKINE_MEDIATED_SIGNALING_PATHWAY | 0.00042 | 0.01306 | 0.49849 | 0.58670 | 1.15548 | 283 |
| GO_CYTOPLASMIC_STRESS_GRANULE | 0.00001 | 0.00077 | 0.59333 | 0.74381 | 1.40402 | 55 |
| GO_CYTOPLASMIC_TRANSLATION | 0.00004 | 0.00231 | 0.55733 | 0.68637 | 1.31757 | 81 |
| GO_CYTOSOLIC_LARGE_RIBOSOMAL_SUBUNIT | 0.00004 | 0.00229 | 0.55733 | 0.74894 | 1.40767 | 47 |
| GO_CYTOSOLIC_RIBOSOME | 0.00000 | 0.00003 | 0.70498 | 0.73236 | 1.40822 | 85 |
| GO_CYTOSOLIC_SMALL_RIBOSOMAL_SUBUNIT | 0.00082 | 0.02028 | 0.47727 | 0.73920 | 1.37171 | 37 |
| GO_DEFENSE_RESPONSE_TO_OTHER_ORGANISM | 0.00015 | 0.00664 | 0.51885 | 0.58736 | 1.15954 | 360 |
| GO_DEMETHYLATION | 0.00198 | 0.03677 | 0.43171 | 0.72836 | 1.34268 | 33 |
| GO_DNA_BINDING_TRANSCRIPTION_FACTOR_BINDING | 0.00018 | 0.00741 | 0.51885 | 0.61860 | 1.20572 | 174 |
| GO_DNA_BIOSYNTHETIC_PROCESS | 0.00041 | 0.01302 | 0.49849 | 0.63761 | 1.23398 | 120 |
| GO_DNA_CONFORMATION_CHANGE | 0.00041 | 0.01295 | 0.49849 | 0.60777 | 1.18813 | 191 |
| GO_DNA_DAMAGE_RESPONSE_SIGNAL_TRANSDUCTION_BY | 0.00030 | 0.01038 | 0.49849 | 0.69280 | 1.31647 | 63 |
| GO_DNA_PACKAGING | 0.00075 | 0.01911 | 0.47727 | 0.63902 | 1.23253 | 108 |
| GO_DNA_POLYMERASE_BINDING | 0.00052 | 0.01487 | 0.47727 | 0.83635 | 1.48049 | 17 |
| GO_DNA_REPLICATION | 0.00033 | 0.01096 | 0.49849 | 0.60824 | 1.18870 | 189 |
| GO_ENDOPEPTIDASE_ACTIVITY | 0.00176 | 0.03388 | 0.45506 | 0.61659 | 1.19527 | 132 |
| GO_ENDOPEPTIDASE_COMPLEX | 0.00276 | 0.04650 | 0.43171 | 0.67499 | 1.27572 | 57 |
| GO_ENTRY_INTO_HOST | 0.00213 | 0.03827 | 0.43171 | 0.65703 | 1.24790 | 62 |
| GO_ESTABLISHMENT_OF_PROTEIN_LOCALIZATION_TO_CHR | 0.00043 | 0.01311 | 0.49849 | 0.83003 | 1.47222 | 18 |
| GO_ESTABLISHMENT_OF_PROTEIN_LOCALIZATION_TO_END | 0.00000 | 0.00027 | 0.64355 | 0.70151 | 1.35046 | 92 |
| GO_ESTABLISHMENT_OF_PROTEIN_LOCALIZATION_TO_MEM | 0.00000 | 0.00043 | 0.61053 | 0.63320 | 1.23929 | 200 |
| GO_ESTABLISHMENT_OF_PROTEIN_LOCALIZATION_TO_ORG | 0.00000 | 0.00000 | | 0.65533 | 1.29247 | 342 |
| GO_ESTABLISHMENT_OF_PROTEIN_LOCALIZATION_TO_TELO | 0.00052 | 0.01487 | 0.47727 | 0.84399 | 1.48092 | 15 |
| GO_ESTABLISHMENT_OF_RNA_LOCALIZATION | 0.00000 | 0.00027 | 0.64355 | 0.65830 | 1.28055 | 154 |
| GO_FICOLIN_1_RICH_GRANULE | 0.00000 | 0.00040 | 0.62726 | 0.72388 | 1.38439 | 72 |
| GO_GLYCOSAMINOGLYCAN_BINDING | 0.00290 | 0.04789 | 0.43171 | 0.70533 | 1.30583 | 36 |
| GO_HETEROCHROMATIN | 0.00032 | 0.01086 | 0.49849 | 0.70940 | 1.33525 | 50 |
| GO_HISTONE_BINDING | 0.00012 | 0.00578 | 0.53843 | 0.63192 | 1.22960 | 156 |
| GO_HISTONE_METHYLATION | 0.00209 | 0.03800 | 0.43171 | 0.63936 | 1.22878 | 84 |
| GO_HISTONE_METHYLTRANSFERASE_COMPLEX | 0.00065 | 0.01764 | 0.47727 | 0.70174 | 1.32173 | 53 |
| GO_HISTONE_UBIQUITINATION | 0.00187 | 0.03521 | 0.45506 | 0.71224 | 1.32223 | 38 |
| GO_HOST_CELLULAR_COMPONENT | 0.00046 | 0.01362 | 0.49849 | 0.71880 | 1.34761 | 45 |
| GO_HYDROLASE_ACTIVITY_ACTING_ON_ACID_ANHYDRIDES | 0.00298 | 0.04865 | 0.43171 | 0.56143 | 1.10931 | 431 |
| GO_ICOSANOID_BIOSYNTHETIC_PROCESS | 0.00124 | 0.02642 | 0.45506 | 0.82375 | 1.45461 | 16 |
| GO_ICOSANOID_METABOLIC_PROCESS | 0.00097 | 0.02220 | 0.47727 | 0.74131 | 1.36656 | 33 |
| GO_IMMUNE_EFFECTOR_PROCESS | 0.00001 | 0.00049 | 0.61053 | 0.58981 | 1.16629 | 458 |
| GO_IMMUNE_RESPONSE_REGULATING_SIGNALING_PATHW | 0.00231 | 0.04088 | 0.43171 | 0.59382 | 1.15770 | 175 |
| GO_IMPORT_INTO_NUCLEUS | 0.00021 | 0.00841 | 0.51885 | 0.65699 | 1.26508 | 97 |
| GO_INNATE_IMMUNE_RESPONSE | 0.00024 | 0.00886 | 0.51885 | 0.58971 | 1.16196 | 302 |
| GO_INTERACTION_WITH_HOST | 0.00079 | 0.01979 | 0.47727 | 0.63456 | 1.22313 | 103 |
| GO_INTRINSIC_APOPTOTIC_SIGNALING_PATHWAY | 0.00008 | 0.00424 | 0.53843 | 0.63456 | 1.23328 | 146 |
| GO_ION_CHANNEL_BINDING | 0.00220 | 0.03919 | 0.43171 | 0.69464 | 1.29934 | 43 |
| GO_KINASE_BINDING | 0.00268 | 0.04570 | 0.43171 | 0.56616 | 1.11764 | 358 |
| GO_LARGE_RIBOSOMAL_SUBUNIT | 0.00088 | 0.02115 | 0.47727 | 0.65241 | 1.25503 | 91 |
| GO_LEUKOCYTE_MEDIATED_IMMUNITY | 0.00000 | 0.00020 | 0.64355 | 0.61534 | 1.21298 | 313 |
| GO_LEUKOCYTE_PROLIFERATION | 0.00209 | 0.03800 | 0.43171 | 0.63676 | 1.22462 | 88 |
| GO_METHYLTRANSFERASE_COMPLEX | 0.00055 | 0.01566 | 0.47727 | 0.66726 | 1.27922 | 76 |
| GO_MITOTIC_NUCLEAR_DIVISION | 0.00044 | 0.01345 | 0.49849 | 0.60713 | 1.18521 | 183 |
| GO_MITOTIC_SPINDLE | 0.00121 | 0.02585 | 0.45506 | 0.64469 | 1.23971 | 87 |
| GO_MODIFICATION_DEPENDENT_MACROMOLECULE_CATAI | 0.00003 | 0.00217 | 0.55733 | 0.58764 | 1.16137 | 412 |
| GO_MODIFICATION_DEPENDENT_PROTEIN_BINDING | 0.00178 | 0.03410 | 0.45506 | 0.63392 | 1.22066 | 97 |
| GO_MOVEMENT_IN_HOST_ENVIRONMENT | 0.00220 | 0.03919 | 0.43171 | 0.64335 | 1.23359 | 77 |
| GO_MRNA_BINDING | 0.00000 | 0.00021 | 0.64355 | 0.64354 | 1.26091 | 210 |
| GO_MRNA_CIS_SPLICING_VIA_SPLICEOSOME | 0.00246 | 0.04306 | 0.43171 | 0.77250 | 1.39930 | 23 |
| GO_MRNA_EXPORT_FROM_NUCLEUS | 0.00017 | 0.00734 | 0.51885 | 0.67081 | 1.29019 | 90 |
| GO_MRNA_PROCESSING | 0.00000 | 0.00015 | 0.65944 | 0.60874 | 1.20189 | 366 |
| GO_MRNA_TRANSPORT | 0.00014 | 0.00656 | 0.51885 | 0.64957 | 1.25601 | 116 |
| GO_MULTI_ORGANISM_LOCALIZATION | 0.00005 | 0.00303 | 0.55733 | 0.73692 | 1.38294 | 46 |
| GO_MYELOID_LEUKOCYTE_ACTIVATION | 0.00001 | 0.00050 | 0.61053 | 0.61588 | 1.21287 | 275 |
| GO_MYELOID_LEUKOCYTE_MEDIATED_IMMUNITY | 0.00001 | 0.00048 | 0.61053 | 0.62007 | 1.21950 | 251 |
| GO_NEGATIVE_REGULATION_OF_CELLULAR_COMPONENT_ | 0.00202 | 0.03718 | 0.43171 | 0.57121 | 1.12623 | 330 |

| Cluster 1 | | | | | | |
|---|---|---|---|---|---|---|
| pathway | pval | padj | log2err | ES | NES | size |
| GO_NEGATIVE_REGULATION_OF_CELLULAR_PROTEIN_CATA | 0.00181 | 0.03468 | 0.45506 | 0.67744 | 1.27841 | 56 |
| GO_NEGATIVE_REGULATION_OF_CELL_CYCLE | 0.00000 | 0.00029 | 0.62726 | 0.60776 | 1.19976 | 352 |
| GO_NEGATIVE_REGULATION_OF_CELL_CYCLE_G2_M_PHASI | 0.00113 | 0.02489 | 0.45506 | 0.65504 | 1.25602 | 77 |
| GO_NEGATIVE_REGULATION_OF_CELL_CYCLE_PHASE_TRAN | 0.00033 | 0.01104 | 0.49849 | 0.62128 | 1.20896 | 158 |
| GO_NEGATIVE_REGULATION_OF_CELL_CYCLE_PROCESS | 0.00000 | 0.00029 | 0.62726 | 0.63482 | 1.24452 | 215 |
| GO_NEGATIVE_REGULATION_OF_CELL_DEATH | 0.00091 | 0.02151 | 0.47727 | 0.57069 | 1.12787 | 412 |
| GO_NEGATIVE_REGULATION_OF_CHROMOSOME_ORGANIZ | 0.00016 | 0.00707 | 0.51885 | 0.66313 | 1.27657 | 92 |
| GO_NEGATIVE_REGULATION_OF_DNA_METABOLIC_PROCES | 0.00022 | 0.00853 | 0.51885 | 0.69536 | 1.32296 | 65 |
| GO_NEGATIVE_REGULATION_OF_DNA_REPAIR | 0.00016 | 0.00707 | 0.51885 | 0.84570 | 1.50002 | 18 |
| GO_NEGATIVE_REGULATION_OF_GENE_EXPRESSION_EPIGE | 0.00152 | 0.03076 | 0.45506 | 0.70567 | 1.31820 | 42 |
| GO_NEGATIVE_REGULATION_OF_INTRINSIC_APOPTOTIC_SI | 0.00287 | 0.04744 | 0.43171 | 0.68032 | 1.27954 | 49 |
| GO_NEGATIVE_REGULATION_OF_MITOTIC_CELL_CYCLE | 0.00042 | 0.01306 | 0.49849 | 0.60645 | 1.18661 | 197 |
| GO_NEGATIVE_REGULATION_OF_PROTEIN_CATABOLIC_PRO | 0.00128 | 0.02709 | 0.45506 | 0.65761 | 1.25979 | 75 |
| GO_NEGATIVE_REGULATION_OF_PROTEIN_METABOLIC_PR | 0.00007 | 0.00384 | 0.53843 | 0.57461 | 1.13731 | 490 |
| GO_NEGATIVE_REGULATION_OF_PROTEIN_MODIFICATION_ | 0.00089 | 0.02127 | 0.47727 | 0.71030 | 1.33505 | 47 |
| GO_NEGATIVE_REGULATION_OF_PROTEIN_MODIFICATION_ | 0.00034 | 0.01135 | 0.49849 | 0.59160 | 1.16499 | 280 |
| GO_NEGATIVE_REGULATION_OF_RESPONSE_TO_DNA_DAM | 0.00066 | 0.01770 | 0.47727 | 0.71442 | 1.34279 | 47 |
| GO_NEGATIVE_REGULATION_OF_RESPONSE_TO_WOUNDIN | 0.00265 | 0.04522 | 0.43171 | 0.74117 | 1.35937 | 28 |
| GO_NEGATIVE_REGULATION_OF_TRANSCRIPTION_BY_RNA_ | 0.00026 | 0.00943 | 0.49849 | 0.58403 | 1.15297 | 360 |
| GO_NEGATIVE_REGULATION_OF_UBIQUITIN_DEPENDENT_ | 0.00202 | 0.03718 | 0.43171 | 0.72678 | 1.33691 | 32 |
| GO_NEGATIVE_REGULATION_OF_WOUND_HEALING | 0.00239 | 0.04205 | 0.43171 | 0.75137 | 1.36847 | 25 |
| GO_NUCLEAR_BODY | 0.00001 | 0.00068 | 0.59333 | 0.58531 | 1.15813 | 483 |
| GO_NUCLEAR_CHROMOSOME_SEGREGATION | 0.00194 | 0.03621 | 0.45506 | 0.59999 | 1.17150 | 184 |
| GO_NUCLEAR_ENVELOPE | 0.00004 | 0.00235 | 0.55733 | 0.60467 | 1.19094 | 276 |
| GO_NUCLEAR_EXPORT | 0.00000 | 0.00014 | 0.65944 | 0.66328 | 1.28961 | 151 |
| GO_NUCLEAR_HETEROCHROMATIN | 0.00294 | 0.04820 | 0.43171 | 0.75804 | 1.36506 | 21 |
| GO_NUCLEAR_MEMBRANE | 0.00167 | 0.03297 | 0.45506 | 0.60497 | 1.17838 | 171 |
| GO_NUCLEAR_PERIPHERY | 0.00279 | 0.04681 | 0.43171 | 0.63312 | 1.21761 | 88 |
| GO_NUCLEAR_PORE | 0.00054 | 0.01523 | 0.47727 | 0.68196 | 1.29525 | 62 |
| GO_NUCLEAR_SPECK | 0.00008 | 0.00403 | 0.53843 | 0.60757 | 1.19507 | 253 |
| GO_NUCLEAR_TRANSCRIBED_MRNA_CATABOLIC_PROCESS | 0.00000 | 0.00038 | 0.62726 | 0.65378 | 1.27255 | 161 |
| GO_NUCLEAR_TRANSCRIBED_MRNA_CATABOLIC_PROCESS_ | 0.00000 | 0.00008 | 0.67496 | 0.70888 | 1.36638 | 101 |
| GO_NUCLEAR_TRANSPORT | 0.00000 | 0.00000 | 0.76146 | 0.65873 | 1.29334 | 233 |
| GO_NUCLEOBASE_CONTAINING_COMPOUND_TRANSPORT | 0.00000 | 0.00038 | 0.62726 | 0.64154 | 1.25356 | 188 |
| GO_NUCLEUS_ORGANIZATION | 0.00185 | 0.03499 | 0.45506 | 0.64166 | 1.23413 | 90 |
| GO_ORGANIC_CYCLIC_COMPOUND_CATABOLIC_PROCESS | 0.00000 | 0.00001 | 0.73376 | 0.62336 | 1.23080 | 379 |
| GO_PEPTIDASE_ACTIVITY | 0.00137 | 0.02880 | 0.45506 | 0.59231 | 1.16142 | 217 |
| GO_PEPTIDASE_COMPLEX | 0.00141 | 0.02922 | 0.45506 | 0.66630 | 1.27160 | 69 |
| GO_PEPTIDE_BIOSYNTHETIC_PROCESS | 0.00000 | 0.00001 | 0.71951 | 0.60779 | 1.20132 | 451 |
| GO_PEPTIDYL_LYSINE_METHYLATION | 0.00287 | 0.04744 | 0.43171 | 0.64139 | 1.22778 | 74 |
| GO_PEPTIDYL_LYSINE_MODIFICATION | 0.00019 | 0.00785 | 0.51885 | 0.60348 | 1.18624 | 241 |
| GO_PEPTIDYL_SERINE_MODIFICATION | 0.00100 | 0.02254 | 0.45506 | 0.61049 | 1.18705 | 153 |
| GO_PERINUCLEAR_REGION_OF_CYTOPLASM | 0.00117 | 0.02527 | 0.45506 | 0.57339 | 1.13060 | 333 |
| GO_PIGMENT_GRANULE | 0.00036 | 0.01180 | 0.49849 | 0.68169 | 1.29989 | 68 |
| GO_PML_BODY | 0.00093 | 0.02164 | 0.47727 | 0.66828 | 1.27431 | 68 |
| GO_POLYSOME | 0.00011 | 0.00522 | 0.53843 | 0.71072 | 1.34589 | 58 |
| GO_POSITIVE_REGULATION_OF_CANONICAL_WNT_SIGNAL | 0.00174 | 0.03377 | 0.45506 | 0.65224 | 1.25114 | 80 |
| GO_POSITIVE_REGULATION_OF_CELLULAR_PROTEIN_LOCAI | 0.00000 | 0.00009 | 0.67496 | 0.65326 | 1.27551 | 185 |
| GO_POSITIVE_REGULATION_OF_CELL_CYCLE | 0.00023 | 0.00866 | 0.51885 | 0.60954 | 1.19392 | 207 |
| GO_POSITIVE_REGULATION_OF_CELL_CYCLE_PROCESS | 0.00104 | 0.02317 | 0.45506 | 0.61613 | 1.19913 | 159 |
| GO_POSITIVE_REGULATION_OF_CELL_DEATH | 0.00046 | 0.01368 | 0.49849 | 0.58814 | 1.15830 | 281 |
| GO_POSITIVE_REGULATION_OF_CELL_POPULATION_PROLIF | 0.00062 | 0.01701 | 0.47727 | 0.58125 | 1.14598 | 320 |
| GO_POSITIVE_REGULATION_OF_CHROMATIN_ORGANIZATIC | 0.00076 | 0.01938 | 0.47727 | 0.68417 | 1.29765 | 60 |
| GO_POSITIVE_REGULATION_OF_CHROMOSOME_ORGANIZA | 0.00001 | 0.00048 | 0.61053 | 0.67612 | 1.30785 | 119 |
| GO_POSITIVE_REGULATION_OF_DNA_BIOSYNTHETIC_PROC | 0.00095 | 0.02187 | 0.47727 | 0.69903 | 1.31378 | 48 |
| GO_POSITIVE_REGULATION_OF_DNA_METABOLIC_PROCESS | 0.00020 | 0.00800 | 0.51885 | 0.64674 | 1.25054 | 116 |
| GO_POSITIVE_REGULATION_OF_ESTABLISHMENT_OF_PROT | 0.00002 | 0.00125 | 0.57561 | 0.63151 | 1.23257 | 179 |
| GO_POSITIVE_REGULATION_OF_IMMUNE_RESPONSE | 0.00010 | 0.00483 | 0.53843 | 0.60335 | 1.18604 | 243 |
| GO_POSITIVE_REGULATION_OF_IMMUNE_SYSTEM_PROCES | 0.00040 | 0.01289 | 0.49849 | 0.58103 | 1.14560 | 332 |
| GO_POSITIVE_REGULATION_OF_INTRACELLULAR_PROTEIN_ | 0.00046 | 0.01362 | 0.49849 | 0.64113 | 1.23604 | 106 |
| GO_POSITIVE_REGULATION_OF_INTRACELLULAR_TRANSPO | 0.00272 | 0.04602 | 0.43171 | 0.61266 | 1.18754 | 130 |
| GO_POSITIVE_REGULATION_OF_INTRINSIC_APOPTOTIC_SIG | 0.00141 | 0.02922 | 0.45506 | 0.75271 | 1.38053 | 28 |
| GO_POSITIVE_REGULATION_OF_ORGANELLE_ORGANIZATIO | 0.00034 | 0.01135 | 0.49849 | 0.58201 | 1.14753 | 330 |
| GO_POSITIVE_REGULATION_OF_PROTEIN_LOCALIZATION_T | 0.00018 | 0.00761 | 0.51885 | 0.72959 | 1.37130 | 47 |
| GO_POSITIVE_REGULATION_OF_PROTEIN_MODIFICATION_I | 0.00152 | 0.03076 | 0.45506 | 0.55886 | 1.10579 | 482 |
| GO_POSITIVE_REGULATION_OF_TELOMERASE_ACTIVITY | 0.00161 | 0.03225 | 0.45506 | 0.74870 | 1.37318 | 28 |
| GO_POSITIVE_REGULATION_OF_TELOMERE_MAINTENANCE | 0.00066 | 0.01770 | 0.47727 | 0.72488 | 1.34984 | 39 |
| GO_POSITIVE_REGULATION_OF_TRANSCRIPTION_BY_RNA_ | 0.00007 | 0.00360 | 0.53843 | 0.57726 | 1.14215 | 475 |
| GO_POSITIVE_REGULATION_OF_TRANSPORT | 0.00146 | 0.03000 | 0.45506 | 0.57086 | 1.12584 | 339 |
| GO_POSITIVE_REGULATION_OF_WNT_SIGNALING_PATHWA | 0.00069 | 0.01806 | 0.47727 | 0.64720 | 1.24591 | 92 |
| GO_POSTTRANSCRIPTIONAL_REGULATION_OF_GENE_EXPR | 0.00000 | 0.00000 | 0.78819 | 0.62383 | 1.23265 | 410 |

| Cluster 1 | | | | | | |
|---|---|---|---|---|---|---|
| pathway | pval | padj | log2err | ES | NES | size |
| GO_POST_TRANSLATIONAL_PROTEIN_MODIFICATION | 0.00170 | 0.03357 | 0.45506 | 0.59582 | 1.16505 | 192 |
| GO_PROSTANOID_METABOLIC_PROCESS | 0.00050 | 0.01451 | 0.47727 | 0.82632 | 1.46564 | 18 |
| GO_PROTEASOMAL_PROTEIN_CATABOLIC_PROCESS | 0.00092 | 0.02162 | 0.47727 | 0.58200 | 1.14727 | 313 |
| GO_PROTEIN_CONTAINING_COMPLEX_LOCALIZATION | 0.00002 | 0.00167 | 0.57561 | 0.62886 | 1.22763 | 183 |
| GO_PROTEIN_DOMAIN_SPECIFIC_BINDING | 0.00001 | 0.00089 | 0.59333 | 0.60085 | 1.18469 | 332 |
| GO_PROTEIN_FOLDING | 0.00172 | 0.03367 | 0.45506 | 0.61992 | 1.20121 | 128 |
| GO_PROTEIN_IMPORT | 0.00004 | 0.00231 | 0.55733 | 0.66582 | 1.28521 | 110 |
| GO_PROTEIN_LOCALIZATION_TO_CHROMOSOME | 0.00014 | 0.00651 | 0.51885 | 0.71355 | 1.35272 | 59 |
| GO_PROTEIN_LOCALIZATION_TO_CHROMOSOME_TELOMEF | 0.00002 | 0.00111 | 0.57561 | 0.84055 | 1.52633 | 24 |
| GO_PROTEIN_LOCALIZATION_TO_ENDOPLASMIC_RETICULU | 0.00000 | 0.00010 | 0.67496 | 0.70098 | 1.35142 | 106 |
| GO_PROTEIN_LOCALIZATION_TO_MEMBRANE | 0.00000 | 0.00017 | 0.64355 | 0.61077 | 1.20451 | 335 |
| GO_PROTEIN_LOCALIZATION_TO_NUCLEUS | 0.00000 | 0.00010 | 0.67496 | 0.66827 | 1.29941 | 152 |
| GO_PROTEIN_MODIFICATION_BY_SMALL_PROTEIN_REMOV | 0.00213 | 0.03827 | 0.43171 | 0.59253 | 1.16019 | 205 |
| GO_PROTEIN_POLYUBIQUITINATION | 0.00026 | 0.00943 | 0.49849 | 0.60633 | 1.18906 | 219 |
| GO_PROTEIN_STABILIZATION | 0.00001 | 0.00061 | 0.61053 | 0.67670 | 1.30513 | 107 |
| GO_PROTEIN_TARGETING | 0.00000 | 0.00015 | 0.65944 | 0.62770 | 1.23559 | 259 |
| GO_PROTEIN_TARGETING_TO_MEMBRANE | 0.00000 | 0.00039 | 0.62726 | 0.67575 | 1.30825 | 122 |
| GO_REGULATION_OF_APOPTOTIC_SIGNALING_PATHWAY | 0.00098 | 0.02242 | 0.45506 | 0.59750 | 1.16726 | 187 |
| GO_REGULATION_OF_CELLULAR_AMIDE_METABOLIC_PROC | 0.00000 | 0.00041 | 0.61053 | 0.61810 | 1.21725 | 275 |
| GO_REGULATION_OF_CELLULAR_AMINE_METABOLIC_PROC | 0.00093 | 0.02164 | 0.47727 | 0.69759 | 1.31392 | 53 |
| GO_REGULATION_OF_CELLULAR_AMINO_ACID_METABOLIC | 0.00059 | 0.01643 | 0.47727 | 0.70047 | 1.31844 | 50 |
| GO_REGULATION_OF_CELLULAR_LOCALIZATION | 0.00001 | 0.00078 | 0.59333 | 0.58927 | 1.16502 | 457 |
| GO_REGULATION_OF_CELLULAR_PROTEIN_CATABOLIC_PRO | 0.00071 | 0.01835 | 0.47727 | 0.61195 | 1.19038 | 154 |
| GO_REGULATION_OF_CELLULAR_PROTEIN_LOCALIZATION | 0.00000 | 0.00002 | 0.70498 | 0.62771 | 1.23687 | 298 |
| GO_REGULATION_OF_CELLULAR_RESPONSE_TO_HEAT | 0.00231 | 0.04088 | 0.43171 | 0.66115 | 1.25453 | 61 |
| GO_REGULATION_OF_CELLULAR_RESPONSE_TO_STRESS | 0.00000 | 0.00044 | 0.61053 | 0.60145 | 1.18712 | 374 |
| GO_REGULATION_OF_CELL_CYCLE_G2_M_PHASE_TRANSITI | 0.00011 | 0.00523 | 0.53843 | 0.62887 | 1.22371 | 158 |
| GO_REGULATION_OF_CELL_CYCLE_PHASE_TRANSITION | 0.00003 | 0.00198 | 0.55733 | 0.60430 | 1.19027 | 287 |
| GO_REGULATION_OF_CELL_CYCLE_PROCESS | 0.00000 | 0.00001 | 0.74774 | 0.61166 | 1.20938 | 455 |
| GO_REGULATION_OF_CHROMATIN_ORGANIZATION | 0.00007 | 0.00384 | 0.53843 | 0.66489 | 1.28220 | 105 |
| GO_REGULATION_OF_CHROMOSOME_ORGANIZATION | 0.00000 | 0.00034 | 0.62726 | 0.63319 | 1.24269 | 227 |
| GO_REGULATION_OF_DNA_BIOSYNTHETIC_PROCESS | 0.00018 | 0.00761 | 0.51885 | 0.69138 | 1.32081 | 70 |
| GO_REGULATION_OF_DNA_METABOLIC_PROCESS | 0.00000 | 0.00001 | 0.74774 | 0.66701 | 1.30646 | 206 |
| GO_REGULATION_OF_DNA_REPLICATION | 0.00078 | 0.01975 | 0.47727 | 0.66215 | 1.26943 | 76 |
| GO_REGULATION_OF_HEMATOPOIETIC_STEM_CELL_DIFFER | 0.00183 | 0.03490 | 0.45506 | 0.67447 | 1.27313 | 55 |
| GO_REGULATION_OF_HISTONE_MODIFICATION | 0.00013 | 0.00610 | 0.51885 | 0.67711 | 1.30041 | 82 |
| GO_REGULATION_OF_IMMUNE_RESPONSE | 0.00059 | 0.01643 | 0.47727 | 0.58167 | 1.14681 | 320 |
| GO_REGULATION_OF_INTRACELLULAR_PROTEIN_TRANSPOI | 0.00017 | 0.00720 | 0.51885 | 0.62836 | 1.22122 | 146 |
| GO_REGULATION_OF_INTRACELLULAR_TRANSPORT | 0.00069 | 0.01806 | 0.47727 | 0.60142 | 1.17645 | 194 |
| GO_REGULATION_OF_INTRINSIC_APOPTOTIC_SIGNALING_P | 0.00005 | 0.00297 | 0.55733 | 0.68301 | 1.31113 | 81 |
| GO_REGULATION_OF_ION_TRANSMEMBRANE_TRANSPORT | 0.00265 | 0.04522 | 0.43171 | 0.62029 | 1.19633 | 107 |
| GO_REGULATION_OF_MITOTIC_CELL_CYCLE | 0.00001 | 0.00049 | 0.61053 | 0.59446 | 1.17409 | 388 |
| GO_REGULATION_OF_MRNA_CATABOLIC_PROCESS | 0.00067 | 0.01776 | 0.47727 | 0.61552 | 1.19663 | 149 |
| GO_REGULATION_OF_MRNA_METABOLIC_PROCESS | 0.00000 | 0.00003 | 0.70498 | 0.64472 | 1.26721 | 240 |
| GO_REGULATION_OF_MRNA_PROCESSING | 0.00000 | 0.00043 | 0.61053 | 0.68183 | 1.31423 | 103 |
| GO_REGULATION_OF_MRNA_SPLICING_VIA_SPLICEOSOME | 0.00009 | 0.00465 | 0.53843 | 0.68627 | 1.31567 | 76 |
| GO_REGULATION_OF_NUCLEOCYTOPLASMIC_TRANSPORT | 0.00065 | 0.01758 | 0.47727 | 0.67775 | 1.29059 | 66 |
| GO_REGULATION_OF_POTASSIUM_ION_TRANSPORT | 0.00100 | 0.02254 | 0.45506 | 0.75091 | 1.37643 | 30 |
| GO_REGULATION_OF_PROTEASOMAL_PROTEIN_CATABOLIC | 0.00279 | 0.04681 | 0.43171 | 0.61771 | 1.19317 | 114 |
| GO_REGULATION_OF_PROTEASOMAL_UBIQUITIN_DEPENDI | 0.00261 | 0.04488 | 0.43171 | 0.64285 | 1.23263 | 79 |
| GO_REGULATION_OF_PROTEIN_BINDING | 0.00257 | 0.04454 | 0.43171 | 0.63167 | 1.21426 | 86 |
| GO_REGULATION_OF_PROTEIN_CATABOLIC_PROCESS | 0.00006 | 0.00321 | 0.55733 | 0.61122 | 1.19924 | 222 |
| GO_REGULATION_OF_PROTEIN_LOCALIZATION | 0.00011 | 0.00529 | 0.53843 | 0.58152 | 1.14911 | 432 |
| GO_REGULATION_OF_PROTEIN_LOCALIZATION_TO_NUCLEU | 0.00095 | 0.02187 | 0.47727 | 0.66803 | 1.27384 | 68 |
| GO_REGULATION_OF_PROTEIN_MODIFICATION_BY_SMALL_ | 0.00019 | 0.00785 | 0.51885 | 0.63150 | 1.22615 | 142 |
| GO_REGULATION_OF_PROTEIN_STABILITY | 0.00074 | 0.01897 | 0.47727 | 0.60903 | 1.18515 | 165 |
| GO_REGULATION_OF_PROTEOLYSIS | 0.00032 | 0.01087 | 0.49849 | 0.58428 | 1.15136 | 301 |
| GO_REGULATION_OF_RESPONSE_TO_BIOTIC_STIMULUS | 0.00220 | 0.03919 | 0.43171 | 0.58907 | 1.15018 | 185 |
| GO_REGULATION_OF_RESPONSE_TO_DNA_DAMAGE_STIMU | 0.00028 | 0.00978 | 0.49849 | 0.63280 | 1.22707 | 134 |
| GO_REGULATION_OF_RNA_SPLICING | 0.00014 | 0.00646 | 0.51885 | 0.66980 | 1.29059 | 100 |
| GO_REGULATION_OF_SMALL_MOLECULE_METABOLIC_PRO | 0.00159 | 0.03213 | 0.45506 | 0.59332 | 1.16299 | 213 |
| GO_REGULATION_OF_SODIUM_ION_TRANSMEMBRANE_TR | 0.00191 | 0.03578 | 0.45506 | 0.75885 | 1.37796 | 24 |
| GO_REGULATION_OF_SODIUM_ION_TRANSMEMBRANE_TR | 0.00261 | 0.04488 | 0.43171 | 0.76653 | 1.37559 | 20 |
| GO_REGULATION_OF_SODIUM_ION_TRANSPORT | 0.00242 | 0.04255 | 0.43171 | 0.72791 | 1.33641 | 31 |
| GO_REGULATION_OF_STEM_CELL_DIFFERENTIATION | 0.00091 | 0.02151 | 0.47727 | 0.65987 | 1.26505 | 76 |
| GO_REGULATION_OF_TELOMERE_MAINTENANCE | 0.00018 | 0.00744 | 0.51885 | 0.69178 | 1.31264 | 61 |
| GO_REGULATION_OF_TELOMERE_MAINTENANCE_VIA_TELO | 0.00172 | 0.03367 | 0.45506 | 0.68874 | 1.29444 | 48 |
| GO_REGULATION_OF_TRANSFERASE_ACTIVITY | 0.00042 | 0.01306 | 0.49849 | 0.56921 | 1.12495 | 449 |
| GO_REGULATION_OF_UBIQUITIN_DEPENDENT_PROTEIN_C/ | 0.00148 | 0.03025 | 0.45506 | 0.64099 | 1.23421 | 94 |
| GO_REGULATORY_REGION_NUCLEIC_ACID_BINDING | 0.00165 | 0.03285 | 0.45506 | 0.56956 | 1.12420 | 374 |
| GO_RESPONSE_TO_BIOTIC_STIMULUS | 0.00000 | 0.00043 | 0.61053 | 0.58795 | 1.16367 | 495 |

| Cluster 1 | | | | | | |
|---|---|---|---|---|---|---|
| pathway | pval | padj | log2err | ES | NES | size |
| GO_RESPONSE_TO_CYTOKINE | 0.00000 | 0.00000 | 0.80122 | 0.61645 | 1.21997 | 487 |
| GO_RESPONSE_TO_HEAT | 0.00030 | 0.01038 | 0.49849 | 0.66070 | 1.27305 | 100 |
| GO_RESPONSE_TO_INTERLEUKIN_1 | 0.00100 | 0.02254 | 0.45506 | 0.64193 | 1.23569 | 93 |
| GO_RESPONSE_TO_LEUKEMIA_INHIBITORY_FACTOR | 0.00006 | 0.00324 | 0.55733 | 0.70539 | 1.34407 | 67 |
| GO_RESPONSE_TO_ORGANIC_CYCLIC_COMPOUND | 0.00070 | 0.01821 | 0.47727 | 0.57737 | 1.13955 | 354 |
| GO_RESPONSE_TO_TEMPERATURE_STIMULUS | 0.00043 | 0.01311 | 0.49849 | 0.63754 | 1.23149 | 114 |
| GO_RESPONSE_TO_TOPOLOGICALLY_INCORRECT_PROTEIN | 0.00079 | 0.01979 | 0.47727 | 0.63541 | 1.22501 | 106 |
| GO_RESPONSE_TO_VIRUS | 0.00054 | 0.01523 | 0.47727 | 0.62581 | 1.21313 | 132 |
| GO_RHYTHMIC_PROCESS | 0.00065 | 0.01758 | 0.47727 | 0.62435 | 1.21149 | 137 |
| GO_RIBONUCLEOPROTEIN_COMPLEX_BINDING | 0.00143 | 0.02948 | 0.45506 | 0.62955 | 1.21372 | 106 |
| GO_RIBONUCLEOPROTEIN_COMPLEX_BIOGENESIS | 0.00000 | 0.00000 | | 0.64779 | 1.27878 | 352 |
| GO_RIBONUCLEOPROTEIN_COMPLEX_SUBUNIT_ORGANIZAT | 0.00006 | 0.00327 | 0.55733 | 0.63575 | 1.23728 | 160 |
| GO_RIBONUCLEOPROTEIN_GRANULE | 0.00003 | 0.00170 | 0.57561 | 0.64597 | 1.25720 | 159 |
| GO_RIBOSOMAL_LARGE_SUBUNIT_ASSEMBLY | 0.00085 | 0.02056 | 0.47727 | 0.77025 | 1.40285 | 25 |
| GO_RIBOSOMAL_LARGE_SUBUNIT_BIOGENESIS | 0.00003 | 0.00190 | 0.57561 | 0.71373 | 1.35559 | 62 |
| GO_RIBOSOMAL_SUBUNIT | 0.00001 | 0.00050 | 0.61053 | 0.65540 | 1.27377 | 146 |
| GO_RIBOSOME | 0.00000 | 0.00038 | 0.62726 | 0.64622 | 1.26021 | 176 |
| GO_RIBOSOME_ASSEMBLY | 0.00102 | 0.02285 | 0.45506 | 0.68819 | 1.29603 | 52 |
| GO_RIBOSOME_BIOGENESIS | 0.00000 | 0.00001 | 0.71951 | 0.64891 | 1.27549 | 242 |
| GO_RNA_CATABOLIC_PROCESS | 0.00000 | 0.00001 | 0.73376 | 0.63440 | 1.24989 | 296 |
| GO_RNA_DEPENDENT_DNA_BIOSYNTHETIC_PROCESS | 0.00000 | 0.00031 | 0.62726 | 0.75911 | 1.43131 | 54 |
| GO_RNA_EXPORT_FROM_NUCLEUS | 0.00001 | 0.00061 | 0.61053 | 0.67338 | 1.30161 | 115 |
| GO_RNA_LOCALIZATION | 0.00000 | 0.00002 | 0.70498 | 0.66668 | 1.30154 | 181 |
| GO_RNA_SPLICING | 0.00000 | 0.00009 | 0.67496 | 0.61803 | 1.21825 | 316 |
| GO_RNA_SPLICING_VIA_TRANSESTERIFICATION_REACTIONS | 0.00000 | 0.00038 | 0.62726 | 0.61855 | 1.21794 | 268 |
| GO_RRNA_METABOLIC_PROCESS | 0.00005 | 0.00278 | 0.55733 | 0.62271 | 1.21586 | 185 |
| GO_SECRETORY_GRANULE | 0.00020 | 0.00803 | 0.51885 | 0.58622 | 1.15554 | 316 |
| GO_SEQUENCE_SPECIFIC_DNA_BINDING | 0.00027 | 0.00952 | 0.49849 | 0.57038 | 1.12830 | 469 |
| GO_SERINE_FAMILY_AMINO_ACID_METABOLIC_PROCESS | 0.00030 | 0.01038 | 0.49849 | 0.80236 | 1.44488 | 21 |
| GO_SIGNAL_TRANSDUCTION_BY_P53_CLASS_MEDIATOR | 0.00117 | 0.02527 | 0.45506 | 0.60907 | 1.18509 | 164 |
| GO_SIGNAL_TRANSDUCTION_IN_RESPONSE_TO_DNA_DAM | 0.00040 | 0.01289 | 0.49849 | 0.66995 | 1.28460 | 77 |
| GO_SISTER_CHROMATID_SEGREGATION | 0.00108 | 0.02388 | 0.45506 | 0.61712 | 1.19899 | 144 |
| GO_SMALL_RIBOSOMAL_SUBUNIT | 0.00048 | 0.01414 | 0.49849 | 0.69073 | 1.30804 | 58 |
| GO_SODIUM_ION_TRANSMEMBRANE_TRANSPORT | 0.00046 | 0.01362 | 0.49849 | 0.71215 | 1.33941 | 49 |
| GO_SODIUM_ION_TRANSPORT | 0.00209 | 0.03800 | 0.43171 | 0.65426 | 1.24990 | 71 |
| GO_SPERM_EGG_RECOGNITION | 0.00060 | 0.01659 | 0.47727 | 0.80756 | 1.44070 | 19 |
| GO_SPINDLE | 0.00253 | 0.04405 | 0.43171 | 0.58235 | 1.14303 | 231 |
| GO_SPINDLE_MIDZONE | 0.00083 | 0.02031 | 0.47727 | 0.79356 | 1.43745 | 23 |
| GO_SPLICEOSOMAL_COMPLEX | 0.00065 | 0.01764 | 0.47727 | 0.61186 | 1.18965 | 151 |
| GO_STEM_CELL_DIFFERENTIATION | 0.00080 | 0.01992 | 0.47727 | 0.63108 | 1.22073 | 119 |
| GO_STRUCTURAL_CONSTITUENT_OF_RIBOSOME | 0.00002 | 0.00115 | 0.57561 | 0.65931 | 1.27796 | 130 |
| GO_STRUCTURAL_MOLECULE_ACTIVITY | 0.00016 | 0.00689 | 0.51885 | 0.59585 | 1.17316 | 272 |
| GO_TELOMERE_MAINTENANCE_VIA_TELOMERE_LENGTHEN | 0.00001 | 0.00089 | 0.59333 | 0.72177 | 1.36955 | 61 |
| GO_TELOMERE_ORGANIZATION | 0.00032 | 0.01087 | 0.49849 | 0.64823 | 1.25127 | 110 |
| GO_TRANSCRIPTION_COACTIVATOR_ACTIVITY | 0.00130 | 0.02737 | 0.45506 | 0.60592 | 1.17866 | 154 |
| GO_TRANSCRIPTION_COREGULATOR_ACTIVITY | 0.00023 | 0.00886 | 0.51885 | 0.59326 | 1.16804 | 270 |
| GO_TRANSCRIPTION_FACTOR_BINDING | 0.00000 | 0.00015 | 0.65944 | 0.61785 | 1.21793 | 313 |
| GO_TRANSLATIONAL_INITIATION | 0.00000 | 0.00000 | 0.81404 | 0.70502 | 1.37084 | 153 |
| GO_TRANSLATION_FACTOR_ACTIVITY_RNA_BINDING | 0.00085 | 0.02056 | 0.47727 | 0.67357 | 1.27931 | 62 |
| GO_TRANSLATION_INITIATION_FACTOR_ACTIVITY | 0.00117 | 0.02527 | 0.45506 | 0.70805 | 1.32443 | 43 |
| GO_TRANSLATION_REGULATOR_ACTIVITY | 0.00008 | 0.00422 | 0.53843 | 0.66678 | 1.28435 | 96 |
| GO_TRANSLATION_REGULATOR_ACTIVITY_NUCLEIC_ACID_E | 0.00031 | 0.01078 | 0.49849 | 0.67523 | 1.29354 | 75 |
| GO_UBIQUITIN_LIKE_PROTEIN_LIGASE_BINDING | 0.00139 | 0.02907 | 0.45506 | 0.59798 | 1.16926 | 192 |
| GO_UNFOLDED_PROTEIN_BINDING | 0.00000 | 0.00043 | 0.61053 | 0.72835 | 1.39144 | 70 |
| GO_UNSATURATED_FATTY_ACID_BIOSYNTHETIC_PROCESS | 0.00087 | 0.02091 | 0.47727 | 0.78198 | 1.40817 | 21 |
| GO_VACUOLAR_LUMEN | 0.00083 | 0.02031 | 0.47727 | 0.65644 | 1.25659 | 74 |
| GO_VESICLE_LUMEN | 0.00001 | 0.00091 | 0.59333 | 0.65521 | 1.27059 | 135 |
| GO_VIRAL_GENE_EXPRESSION | 0.00002 | 0.00162 | 0.57561 | 0.64334 | 1.25092 | 153 |
| GO_VIRAL_GENOME_REPLICATION | 0.00272 | 0.04602 | 0.43171 | 0.65777 | 1.25053 | 64 |
| GO_VIRAL_LIFE_CYCLE | 0.00000 | 0.00015 | 0.65944 | 0.65087 | 1.27035 | 179 |
| HP_ABNORMALITY_OF_UPPER_LIP | 0.00250 | 0.04356 | 0.43171 | 0.56698 | 1.11935 | 364 |
| HP_ABNORMALITY_OF_UPPER_LIP_VERMILLION | 0.00185 | 0.03499 | 0.45506 | 0.60384 | 1.17619 | 171 |
| HP_ABNORMAL_LIP_MORPHOLOGY | 0.00015 | 0.00660 | 0.51885 | 0.57833 | 1.14300 | 441 |
| HP_ABNORMAL_TRABECULAR_BONE_MORPHOLOGY | 0.00050 | 0.01451 | 0.47727 | 0.84469 | 1.48215 | 15 |
| HP_ACUTE_LEUKEMIA | 0.00024 | 0.00890 | 0.51885 | 0.69462 | 1.31992 | 63 |
| HP_GONOSOMAL_INHERITANCE | 0.00001 | 0.00048 | 0.61053 | 0.68059 | 1.31475 | 113 |
| HP_HIGH_PALATE | 0.00167 | 0.03297 | 0.45506 | 0.57779 | 1.13848 | 302 |
| HP_LEUKEMIA | 0.00146 | 0.03000 | 0.45506 | 0.64287 | 1.23638 | 88 |
| HP_SEVERE_POSTNATAL_GROWTH_RETARDATION | 0.00161 | 0.03225 | 0.45506 | 0.81697 | 1.43352 | 15 |
| HP_SMALL_HAND | 0.00205 | 0.03773 | 0.43171 | 0.71066 | 1.31932 | 38 |
| HP_THIN_VERMILION_BORDER | 0.00174 | 0.03377 | 0.45506 | 0.60210 | 1.17145 | 163 |

| **Cluster 1** | | | | | | |
|---|---|---|---|---|---|---|
| pathway | pval | padj | log2err | ES | NES | size |
| HP_X_LINKED_RECESSIVE_INHERITANCE | 0.00003 | 0.00190 | 0.57561 | 0.69096 | 1.32701 | 82 |
| **Cluster 10** | | | | | | |
| pathway | pval | padj | log2err | ES | NES | size |
| GO_AMIDE_BIOSYNTHETIC_PROCESS | 0.00000 | 0.00000 | | 0.69265 | 1.25818 | 363 |
| GO_ANCHORING_JUNCTION | 0.00000 | 0.00003 | 0.67496 | 0.64769 | 1.17313 | 249 |
| GO_CELLULAR_AMIDE_METABOLIC_PROCESS | 0.00000 | 0.00000 | | 0.66411 | 1.20998 | 439 |
| GO_CELLULAR_MACROMOLECULE_CATABOLIC_PROCESS | 0.00000 | 0.00000 | | 0.67448 | 1.22886 | 436 |
| GO_CELLULAR_NITROGEN_COMPOUND_CATABOLIC_PROCE | 0.00000 | 0.00000 | | 0.74217 | 1.34421 | 248 |
| GO_CELL_SUBSTRATE_JUNCTION | 0.00000 | 0.00000 | 0.82666 | 0.70690 | 1.27409 | 166 |
| GO_COTRANSLATIONAL_PROTEIN_TARGETING_TO_MEMBR | 0.00000 | 0.00000 | | 0.92537 | 1.64301 | 86 |
| GO_CYTOKINE_MEDIATED_SIGNALING_PATHWAY | 0.00012 | 0.00655 | 0.53843 | 0.63908 | 1.15178 | 165 |
| GO_CYTOPLASMIC_TRANSLATION | 0.00000 | 0.00000 | 0.80122 | 0.79786 | 1.40999 | 70 |
| GO_CYTOSOLIC_LARGE_RIBOSOMAL_SUBUNIT | 0.00000 | 0.00000 | | 0.89155 | 1.55852 | 50 |
| GO_CYTOSOLIC_RIBOSOME | 0.00000 | 0.00000 | | 0.91625 | 1.62663 | 85 |
| GO_CYTOSOLIC_SMALL_RIBOSOMAL_SUBUNIT | 0.00000 | 0.00000 | | 0.93185 | 1.60701 | 36 |
| GO_ESTABLISHMENT_OF_PROTEIN_LOCALIZATION_TO_END | 0.00000 | 0.00000 | | 0.92545 | 1.64579 | 93 |
| GO_ESTABLISHMENT_OF_PROTEIN_LOCALIZATION_TO_MEM | 0.00000 | 0.00000 | | 0.80074 | 1.44252 | 157 |
| GO_ESTABLISHMENT_OF_PROTEIN_LOCALIZATION_TO_ORG | 0.00000 | 0.00000 | | 0.71261 | 1.29019 | 243 |
| GO_FC_RECEPTOR_SIGNALING_PATHWAY | 0.00035 | 0.01829 | 0.49849 | 0.69384 | 1.22488 | 67 |
| GO_INTRACELLULAR_PROTEIN_TRANSPORT | 0.00000 | 0.00000 | | 0.65129 | 1.18665 | 438 |
| GO_LARGE_RIBOSOMAL_SUBUNIT | 0.00000 | 0.00000 | | 0.77880 | 1.38530 | 95 |
| GO_MACROMOLECULE_CATABOLIC_PROCESS | 0.00000 | 0.00000 | | 0.66308 | 1.20905 | 490 |
| GO_MRNA_METABOLIC_PROCESS | 0.00000 | 0.00000 | | 0.65210 | 1.18702 | 418 |
| GO_NUCLEAR_TRANSCRIBED_MRNA_CATABOLIC_PROCESS | 0.00000 | 0.00000 | | 0.86862 | 1.55534 | 123 |
| GO_NUCLEAR_TRANSCRIBED_MRNA_CATABOLIC_PROCESS_ | 0.00000 | 0.00000 | | 0.90976 | 1.61790 | 93 |
| GO_ORGANIC_CYCLIC_COMPOUND_CATABOLIC_PROCESS | 0.00000 | 0.00000 | | 0.73979 | 1.33988 | 250 |
| GO_OXIDATION_REDUCTION_PROCESS | 0.00100 | 0.04820 | 0.45506 | 0.60788 | 1.10197 | 277 |
| GO_PEPTIDE_BIOSYNTHETIC_PROCESS | 0.00000 | 0.00000 | | 0.69531 | 1.26203 | 334 |
| GO_PEPTIDE_METABOLIC_PROCESS | 0.00000 | 0.00000 | | 0.68534 | 1.24513 | 369 |
| GO_POLYSOMAL_RIBOSOME | 0.00000 | 0.00000 | 0.78819 | 0.93377 | 1.58073 | 27 |
| GO_POLYSOME | 0.00000 | 0.00000 | 0.76146 | 0.85571 | 1.48888 | 45 |
| GO_PROTEIN_LOCALIZATION_TO_ENDOPLASMIC_RETICULU | 0.00000 | 0.00000 | | 0.90972 | 1.62250 | 102 |
| GO_PROTEIN_LOCALIZATION_TO_MEMBRANE | 0.00000 | 0.00000 | | 0.72415 | 1.31018 | 233 |
| GO_PROTEIN_LOCALIZATION_TO_ORGANELLE | 0.00000 | 0.00000 | 0.80122 | 0.65005 | 1.18079 | 363 |
| GO_PROTEIN_TARGETING | 0.00000 | 0.00000 | | 0.75504 | 1.36425 | 202 |
| GO_PROTEIN_TARGETING_TO_MEMBRANE | 0.00000 | 0.00000 | | 0.87228 | 1.55994 | 115 |
| GO_RIBONUCLEOPROTEIN_COMPLEX | 0.00000 | 0.00000 | | 0.64947 | 1.18219 | 415 |
| GO_RIBONUCLEOPROTEIN_COMPLEX_BIOGENESIS | 0.00004 | 0.00245 | 0.55733 | 0.62920 | 1.13915 | 244 |
| GO_RIBOSOMAL_LARGE_SUBUNIT_ASSEMBLY | 0.00007 | 0.00401 | 0.53843 | 0.87165 | 1.44674 | 19 |
| GO_RIBOSOMAL_LARGE_SUBUNIT_BIOGENESIS | 0.00000 | 0.00019 | 0.62726 | 0.77637 | 1.35639 | 49 |
| GO_RIBOSOMAL_SMALL_SUBUNIT_BIOGENESIS | 0.00000 | 0.00006 | 0.65944 | 0.81940 | 1.42189 | 42 |
| GO_RIBOSOMAL_SUBUNIT | 0.00000 | 0.00000 | | 0.78487 | 1.41284 | 150 |
| GO_RIBOSOME | 0.00000 | 0.00000 | | 0.76041 | 1.37103 | 174 |
| GO_RIBOSOME_ASSEMBLY | 0.00000 | 0.00006 | 0.65944 | 0.82805 | 1.42939 | 37 |
| GO_RIBOSOME_BIOGENESIS | 0.00000 | 0.00000 | 0.73376 | 0.68689 | 1.23794 | 165 |
| GO_RNA_CATABOLIC_PROCESS | 0.00000 | 0.00000 | | 0.76892 | 1.39032 | 215 |
| GO_RRNA_BINDING | 0.00000 | 0.00015 | 0.62726 | 0.82616 | 1.42796 | 38 |
| GO_RRNA_METABOLIC_PROCESS | 0.00055 | 0.02780 | 0.47727 | 0.64695 | 1.15848 | 121 |
| GO_SMALL_RIBOSOMAL_SUBUNIT | 0.00000 | 0.00000 | 0.76146 | 0.81651 | 1.43465 | 58 |
| GO_STRUCTURAL_CONSTITUENT_OF_RIBOSOME | 0.00000 | 0.00000 | | 0.78804 | 1.41562 | 138 |
| GO_STRUCTURAL_MOLECULE_ACTIVITY | 0.00000 | 0.00000 | | 0.73044 | 1.32144 | 231 |
| GO_SYMBIOTIC_PROCESS | 0.00000 | 0.00000 | | 0.67608 | 1.22963 | 391 |
| GO_TRANSLATIONAL_INITIATION | 0.00000 | 0.00000 | | 0.83878 | 1.50624 | 136 |
| GO_TRANSMEMBRANE_TRANSPORT | 0.00080 | 0.03905 | 0.47727 | 0.60547 | 1.09806 | 296 |
| GO_VIRAL_GENE_EXPRESSION | 0.00000 | 0.00000 | | 0.85044 | 1.52595 | 132 |
| HP_ABNORMALITY_OF_LOWER_LIP | 0.00000 | 0.00018 | 0.62726 | 0.76034 | 1.33618 | 59 |
| HP_ABNORMALITY_OF_THE_EXTERNAL_NOSE | 0.00051 | 0.02588 | 0.47727 | 0.62835 | 1.13280 | 173 |
| HP_ABNORMALITY_OF_THE_NASAL_DORSUM | 0.00004 | 0.00245 | 0.55733 | 0.73100 | 1.28441 | 58 |
| HP_ABNORMAL_HAIR_PATTERN | 0.00060 | 0.02966 | 0.47727 | 0.71105 | 1.23816 | 46 |
| HP_ABNORMAL_RETICULOCYTE_MORPHOLOGY | 0.00004 | 0.00227 | 0.55733 | 0.90234 | 1.49097 | 17 |
| HP_ACUTE_LEUKEMIA | 0.00001 | 0.00085 | 0.59333 | 0.78422 | 1.35793 | 40 |
| HP_ANEMIA_OF_INADEQUATE_PRODUCTION | 0.00000 | 0.00003 | 0.67496 | 0.77268 | 1.35669 | 57 |
| HP_DELAYED_PUBERTY | 0.00000 | 0.00012 | 0.64355 | 0.77743 | 1.35939 | 51 |
| HP_DEPRESSED_NASAL_RIDGE | 0.00000 | 0.00014 | 0.62726 | 0.81509 | 1.40940 | 39 |
| HP_HEADACHE | 0.00048 | 0.02485 | 0.49849 | 0.68759 | 1.21246 | 65 |
| HP_LEUKEMIA | 0.00004 | 0.00245 | 0.55733 | 0.74744 | 1.30660 | 50 |
| HP_MACROCYTIC_ANEMIA | 0.00000 | 0.00001 | 0.70498 | 0.88894 | 1.51990 | 31 |

| Cluster 1 | | | | | | |
|---|---|---|---|---|---|---|
| pathway | pval | padj | log2err | ES | NES | size |
| HP_MIGRAINE | 0.00000 | 0.00012 | 0.64355 | 0.84359 | 1.44454 | 32 |
| HP_ORAL_CLEFT | 0.00025 | 0.01345 | 0.49849 | 0.65139 | 1.16821 | 128 |
| HP_PALLOR | 0.00027 | 0.01421 | 0.49849 | 0.72488 | 1.26643 | 49 |
| HP_SHORT_NOSE | 0.00002 | 0.00133 | 0.57561 | 0.72073 | 1.27370 | 70 |
| HP_SHORT_THUMB | 0.00003 | 0.00171 | 0.57561 | 0.90840 | 1.49621 | 16 |
| HP_THICK_LOWER_LIP_VERMILION | 0.00000 | 0.00017 | 0.62726 | 0.78159 | 1.36417 | 48 |
| HP_THICK_VERMILION_BORDER | 0.00005 | 0.00270 | 0.55733 | 0.71408 | 1.25916 | 65 |
| **Cluster 11** | | | | | | |
| pathway | pval | padj | log2err | ES | NES | size |
| GO_ACTIN_CYTOSKELETON | 0.00111 | 0.01928 | 0.45506 | 0.52272 | 1.22467 | 203 |
| GO_ACTIN_FILAMENT_BINDING | 0.00183 | 0.02907 | 0.45506 | 0.56704 | 1.29385 | 85 |
| GO_ACTIVATION_OF_IMMUNE_RESPONSE | 0.00000 | 0.00000 | 0.76146 | 0.60395 | 1.41479 | 192 |
| GO_ACTIVATION_OF_INNATE_IMMUNE_RESPONSE | 0.00000 | 0.00000 | | 0.73329 | 1.66857 | 81 |
| GO_AEROBIC_RESPIRATION | 0.00035 | 0.00732 | 0.49849 | 0.62937 | 1.40889 | 59 |
| GO_AMIDE_BINDING | 0.00004 | 0.00116 | 0.55733 | 0.57599 | 1.33363 | 129 |
| GO_AMINE_METABOLIC_PROCESS | 0.00000 | 0.00000 | | 0.75244 | 1.70666 | 76 |
| GO_ANAPHASE_PROMOTING_COMPLEX_DEPENDENT_CATA | 0.00000 | 0.00000 | 0.73376 | 0.69073 | 1.56708 | 77 |
| GO_ANATOMICAL_STRUCTURE_HOMEOSTASIS | 0.00364 | 0.04732 | 0.43171 | 0.50511 | 1.18430 | 212 |
| GO_ANCHORING_JUNCTION | 0.00000 | 0.00000 | 0.74774 | 0.54580 | 1.29509 | 386 |
| GO_ANIMAL_ORGAN_REGENERATION | 0.00320 | 0.04353 | 0.43171 | 0.66440 | 1.42974 | 29 |
| GO_ANTIGEN_PROCESSING_AND_PRESENTATION | 0.00000 | 0.00000 | 0.83909 | 0.66305 | 1.53623 | 130 |
| GO_ANTIGEN_PROCESSING_AND_PRESENTATION_OF_EXOG | 0.00000 | 0.00000 | 0.77494 | 0.75536 | 1.68652 | 54 |
| GO_ANTIGEN_PROCESSING_AND_PRESENTATION_OF_PEPT | 0.00000 | 0.00000 | 0.81404 | 0.67431 | 1.55776 | 116 |
| GO_ANTIGEN_PROCESSING_AND_PRESENTATION_OF_PEPT | 0.00000 | 0.00000 | 0.77494 | 0.73202 | 1.64886 | 64 |
| GO_ANTIGEN_PROCESSING_AND_PRESENTATION_OF_PEPT | 0.00361 | 0.04708 | 0.43171 | 0.58873 | 1.32102 | 61 |
| GO_ANTIGEN_RECEPTOR_MEDIATED_SIGNALING_PATHWAY | 0.00000 | 0.00000 | 0.76146 | 0.65340 | 1.50971 | 115 |
| GO_APOPTOTIC_SIGNALING_PATHWAY | 0.00011 | 0.00253 | 0.53843 | 0.51662 | 1.21915 | 301 |
| GO_ATPASE_ACTIVITY | 0.00080 | 0.01461 | 0.47727 | 0.51198 | 1.20395 | 250 |
| GO_ATP_METABOLIC_PROCESS | 0.00003 | 0.00083 | 0.55733 | 0.56294 | 1.31355 | 164 |
| GO_BRUSH_BORDER | 0.00115 | 0.01965 | 0.45506 | 0.64239 | 1.41664 | 45 |
| GO_CADHERIN_BINDING | 0.00000 | 0.00000 | | 0.63133 | 1.48251 | 228 |
| GO_CANONICAL_WNT_SIGNALING_PATHWAY | 0.00000 | 0.00000 | 0.71951 | 0.60529 | 1.41262 | 165 |
| GO_CARBOHYDRATE_BIOSYNTHETIC_PROCESS | 0.00179 | 0.02866 | 0.45506 | 0.57048 | 1.30270 | 86 |
| GO_CARBOHYDRATE_CATABOLIC_PROCESS | 0.00069 | 0.01298 | 0.47727 | 0.57046 | 1.31203 | 102 |
| GO_CATALYTIC_STEP_2_SPLICEOSOME | 0.00004 | 0.00106 | 0.55733 | 0.62395 | 1.41844 | 79 |
| GO_CELLULAR_AMINO_ACID_METABOLIC_PROCESS | 0.00000 | 0.00000 | 0.73376 | 0.59539 | 1.39486 | 201 |
| GO_CELLULAR_CARBOHYDRATE_CATABOLIC_PROCESS | 0.00390 | 0.04920 | 0.43171 | 0.71579 | 1.49584 | 20 |
| GO_CELLULAR_COMPONENT_DISASSEMBLY | 0.00126 | 0.02119 | 0.45506 | 0.49414 | 1.17029 | 349 |
| GO_CELLULAR_KETONE_METABOLIC_PROCESS | 0.00000 | 0.00000 | 0.77494 | 0.63877 | 1.48442 | 137 |
| GO_CELLULAR_NITROGEN_COMPOUND_CATABOLIC_PROCE | 0.00000 | 0.00000 | | 0.61379 | 1.45643 | 386 |
| GO_CELLULAR_RESPIRATION | 0.00398 | 0.04977 | 0.40702 | 0.54876 | 1.26771 | 116 |
| GO_CELLULAR_RESPONSE_TO_HEAT | 0.00011 | 0.00254 | 0.53843 | 0.60692 | 1.38288 | 83 |
| GO_CELLULAR_RESPONSE_TO_OXYGEN_LEVELS | 0.00000 | 0.00000 | 0.82666 | 0.65942 | 1.53136 | 136 |
| GO_CELL_ACTIVATION_INVOLVED_IN_IMMUNE_RESPONSE | 0.00000 | 0.00000 | 0.74774 | 0.56255 | 1.32784 | 302 |
| GO_CELL_ADHESION_MOLECULE_BINDING | 0.00000 | 0.00000 | | 0.60956 | 1.43643 | 282 |
| GO_CELL_BODY | 0.00001 | 0.00032 | 0.59333 | 0.55212 | 1.29368 | 202 |
| GO_CELL_CELL_RECOGNITION | 0.00000 | 0.00000 | 0.80122 | 0.90419 | 1.89882 | 21 |
| GO_CELL_CELL_SIGNALING_BY_WNT | 0.00000 | 0.00000 | 0.73376 | 0.57387 | 1.34965 | 258 |
| GO_CELL_CYCLE_ARREST | 0.00235 | 0.03481 | 0.43171 | 0.53341 | 1.23996 | 138 |
| GO_CELL_CYCLE_G1_S_PHASE_TRANSITION | 0.00198 | 0.03103 | 0.43171 | 0.52347 | 1.22384 | 179 |
| GO_CELL_CYCLE_G2_M_PHASE_TRANSITION | 0.00000 | 0.00001 | 0.69013 | 0.57842 | 1.35520 | 205 |
| GO_CELL_CYCLE_PHASE_TRANSITION | 0.00000 | 0.00000 | 0.80122 | 0.54851 | 1.30362 | 420 |
| GO_CELL_DIVISION | 0.00042 | 0.00841 | 0.49849 | 0.50264 | 1.19164 | 371 |
| GO_CELL_RECOGNITION | 0.00001 | 0.00032 | 0.59333 | 0.71453 | 1.56918 | 41 |
| GO_CELL_SUBSTRATE_JUNCTION | 0.00000 | 0.00000 | | 0.61237 | 1.43873 | 238 |
| GO_CELL_SURFACE_RECEPTOR_SIGNALING_PATHWAY_INVO | 0.00000 | 0.00000 | 0.70498 | 0.56386 | 1.32859 | 274 |
| GO_CHAPERONE_COFACTOR_DEPENDENT_PROTEIN_REFOL | 0.00246 | 0.03604 | 0.43171 | 0.72455 | 1.50555 | 19 |
| GO_CHAPERONE_COMPLEX | 0.00000 | 0.00010 | 0.62726 | 0.84414 | 1.75404 | 19 |
| GO_CHROMATIN_BINDING | 0.00265 | 0.03821 | 0.43171 | 0.49266 | 1.16402 | 315 |
| GO_CHROMOSOMAL_REGION | 0.00353 | 0.04635 | 0.43171 | 0.49864 | 1.17259 | 255 |
| GO_CHROMOSOME_SEGREGATION | 0.00143 | 0.02374 | 0.45506 | 0.50600 | 1.18856 | 233 |
| GO_COTRANSLATIONAL_PROTEIN_TARGETING_TO_MEMBR | 0.00000 | 0.00000 | 0.82666 | 0.70394 | 1.61185 | 91 |
| GO_CYTOKINE_MEDIATED_SIGNALING_PATHWAY | 0.00000 | 0.00000 | | 0.60196 | 1.41764 | 271 |
| GO_CYTOPLASMIC_STRESS_GRANULE | 0.00202 | 0.03142 | 0.43171 | 0.61326 | 1.36824 | 53 |
| GO_CYTOPLASMIC_TRANSLATION | 0.00000 | 0.00000 | | 0.77190 | 1.75878 | 83 |
| GO_CYTOPLASMIC_TRANSLATIONAL_INITIATION | 0.00000 | 0.00000 | 0.74774 | 0.82332 | 1.77970 | 31 |
| GO_CYTOSOLIC_LARGE_RIBOSOMAL_SUBUNIT | 0.00000 | 0.00004 | 0.64355 | 0.71300 | 1.58708 | 52 |
| GO_CYTOSOLIC_RIBOSOME | 0.00000 | 0.00000 | 0.77494 | 0.68633 | 1.57193 | 93 |
| GO_CYTOSOLIC_SMALL_RIBOSOMAL_SUBUNIT | 0.00047 | 0.00936 | 0.49849 | 0.67437 | 1.47677 | 39 |
| GO_DEFENSE_RESPONSE | 0.00000 | 0.00000 | 0.70498 | 0.52840 | 1.25858 | 482 |

| **Cluster 1** | | | | | | |
|---|---|---|---|---|---|---|
| pathway | pval | padj | log2err | ES | NES | size |
| GO_DEFENSE_RESPONSE_TO_OTHER_ORGANISM | 0.00000 | 0.00000 | 0.74774 | 0.55644 | 1.31705 | 342 |
| GO_DETECTION_OF_STIMULUS | 0.00031 | 0.00656 | 0.49849 | 0.60929 | 1.38197 | 76 |
| GO_DISORDERED_DOMAIN_SPECIFIC_BINDING | 0.00228 | 0.03391 | 0.43171 | 0.69870 | 1.48069 | 24 |
| GO_DISTAL_AXON | 0.00283 | 0.03998 | 0.43171 | 0.54836 | 1.26443 | 110 |
| GO_DNA_BINDING_TRANSCRIPTION_FACTOR_BINDING | 0.00152 | 0.02487 | 0.45506 | 0.52551 | 1.22868 | 182 |
| GO_DNA_BIOSYNTHETIC_PROCESS | 0.00005 | 0.00135 | 0.55733 | 0.57764 | 1.33937 | 132 |
| GO_DNA_DAMAGE_RESPONSE_DETECTION_OF_DNA_DAM | 0.00035 | 0.00726 | 0.49849 | 0.69362 | 1.50718 | 34 |
| GO_DNA_REPAIR | 0.00198 | 0.03103 | 0.43171 | 0.48932 | 1.16044 | 379 |
| GO_DNA_REPLICATION | 0.00320 | 0.04353 | 0.43171 | 0.51803 | 1.21350 | 195 |
| GO_DOUBLE_STRANDED_RNA_BINDING | 0.00294 | 0.04088 | 0.43171 | 0.61572 | 1.36465 | 49 |
| GO_DRUG_BINDING | 0.00202 | 0.03142 | 0.43171 | 0.64180 | 1.41197 | 42 |
| GO_EMBRYO_IMPLANTATION | 0.00331 | 0.04423 | 0.43171 | 0.68594 | 1.44758 | 22 |
| GO_ENDOPEPTIDASE_ACTIVITY | 0.00003 | 0.00071 | 0.57561 | 0.57100 | 1.32703 | 141 |
| GO_ENDOPEPTIDASE_COMPLEX | 0.00000 | 0.00000 | | 0.76960 | 1.73068 | 63 |
| GO_ENERGY_DERIVATION_BY_OXIDATION_OF_ORGANIC_CO | 0.00302 | 0.04168 | 0.43171 | 0.53145 | 1.23718 | 147 |
| GO_EPITHELIUM_DEVELOPMENT | 0.00000 | 0.00004 | 0.64355 | 0.52064 | 1.23896 | 457 |
| GO_ESTABLISHMENT_OF_PROTEIN_LOCALIZATION_TO_CHR | 0.00006 | 0.00140 | 0.55733 | 0.79243 | 1.65600 | 20 |
| GO_ESTABLISHMENT_OF_PROTEIN_LOCALIZATION_TO_END | 0.00000 | 0.00000 | 0.80122 | 0.67863 | 1.55918 | 100 |
| GO_ESTABLISHMENT_OF_PROTEIN_LOCALIZATION_TO_MEM | 0.00000 | 0.00001 | 0.69013 | 0.57019 | 1.33914 | 231 |
| GO_ESTABLISHMENT_OF_PROTEIN_LOCALIZATION_TO_ORG | 0.00000 | 0.00000 | | 0.57113 | 1.35550 | 398 |
| GO_ESTABLISHMENT_OF_PROTEIN_LOCALIZATION_TO_TELO | 0.00005 | 0.00131 | 0.55733 | 0.83759 | 1.70562 | 15 |
| GO_ESTABLISHMENT_OF_RNA_LOCALIZATION | 0.00000 | 0.00004 | 0.64355 | 0.59144 | 1.37870 | 157 |
| GO_ESTABLISHMENT_OF_TISSUE_POLARITY | 0.00000 | 0.00000 | 0.81404 | 0.72403 | 1.64222 | 76 |
| GO_EUKARYOTIC_TRANSLATION_INITIATION_FACTOR_3_CO | 0.00003 | 0.00076 | 0.57561 | 0.84649 | 1.72375 | 15 |
| GO_EXOCYTOSIS | 0.00000 | 0.00000 | 0.74774 | 0.54435 | 1.29192 | 395 |
| GO_FC_EPSILON_RECEPTOR_SIGNALING_PATHWAY | 0.00000 | 0.00000 | 0.78819 | 0.70861 | 1.61089 | 79 |
| GO_FC_RECEPTOR_SIGNALING_PATHWAY | 0.00000 | 0.00000 | | 0.69310 | 1.60069 | 114 |
| GO_FERTILIZATION | 0.00028 | 0.00613 | 0.49849 | 0.64751 | 1.44129 | 52 |
| GO_FICOLIN_1_RICH_GRANULE | 0.00000 | 0.00000 | 0.82666 | 0.70599 | 1.61546 | 90 |
| GO_GENERATION_OF_PRECURSOR_METABOLITES_AND_EN | 0.00004 | 0.00113 | 0.55733 | 0.53050 | 1.24892 | 268 |
| GO_GLUTAMATERGIC_SYNAPSE | 0.00074 | 0.01364 | 0.47727 | 0.55644 | 1.28711 | 125 |
| GO_GLYCOSYL_COMPOUND_METABOLIC_PROCESS | 0.00121 | 0.02039 | 0.45506 | 0.61075 | 1.37872 | 67 |
| GO_GUANYL_NUCLEOTIDE_BINDING | 0.00039 | 0.00795 | 0.49849 | 0.53350 | 1.24600 | 175 |
| GO_HEAT_SHOCK_PROTEIN_BINDING | 0.00113 | 0.01947 | 0.45506 | 0.59680 | 1.35282 | 75 |
| GO_HEMATOPOIETIC_PROGENITOR_CELL_DIFFERENTIATION | 0.00000 | 0.00000 | 0.78819 | 0.68887 | 1.57775 | 93 |
| GO_HEMATOPOIETIC_STEM_CELL_DIFFERENTIATION | 0.00000 | 0.00000 | 0.78819 | 0.74562 | 1.67305 | 61 |
| GO_HYDROLASE_ACTIVITY_ACTING_ON_ACID_ANHYDRIDES | 0.00062 | 0.01177 | 0.47727 | 0.49032 | 1.16583 | 433 |
| GO_IMMUNE_EFFECTOR_PROCESS | 0.00000 | 0.00001 | 0.67496 | 0.52426 | 1.24797 | 470 |
| GO_IMMUNE_RESPONSE_REGULATING_SIGNALING_PATHW | 0.00000 | 0.00000 | 0.74774 | 0.61037 | 1.42700 | 179 |
| GO_IMMUNE_SYSTEM_DEVELOPMENT | 0.00020 | 0.00449 | 0.51885 | 0.50049 | 1.18786 | 398 |
| GO_IMPORT_INTO_NUCLEUS | 0.00074 | 0.01364 | 0.47727 | 0.57479 | 1.32371 | 105 |
| GO_INNATE_IMMUNE_RESPONSE | 0.00000 | 0.00000 | 0.80122 | 0.57615 | 1.35764 | 284 |
| GO_INNATE_IMMUNE_RESPONSE_ACTIVATING_SIGNAL_TR | 0.00000 | 0.00000 | | 0.76665 | 1.72806 | 66 |
| GO_INTERACTION_WITH_HOST | 0.00148 | 0.02442 | 0.45506 | 0.55652 | 1.28250 | 107 |
| GO_INTERACTION_WITH_SYMBIONT | 0.00082 | 0.01479 | 0.47727 | 0.65425 | 1.43519 | 40 |
| GO_INTERLEUKIN_1_MEDIATED_SIGNALING_PATHWAY | 0.00000 | 0.00000 | | 0.75576 | 1.71075 | 71 |
| GO_INTRINSIC_APOPTOTIC_SIGNALING_PATHWAY | 0.00043 | 0.00865 | 0.49849 | 0.54396 | 1.26843 | 159 |
| GO_ION_CHANNEL_BINDING | 0.00020 | 0.00449 | 0.51885 | 0.64823 | 1.44019 | 51 |
| GO_LAMELLIPODIUM | 0.00339 | 0.04486 | 0.43171 | 0.55891 | 1.28009 | 93 |
| GO_LARGE_RIBOSOMAL_SUBUNIT | 0.00044 | 0.00876 | 0.49849 | 0.57153 | 1.31544 | 103 |
| GO_LEUKOCYTE_MEDIATED_IMMUNITY | 0.00000 | 0.00000 | 0.73376 | 0.55495 | 1.31119 | 315 |
| GO_MALE_SEX_DIFFERENTIATION | 0.00220 | 0.03310 | 0.43171 | 0.60234 | 1.35280 | 62 |
| GO_MATURATION_OF_LSU_RRNA | 0.00209 | 0.03228 | 0.43171 | 0.68301 | 1.46566 | 27 |
| GO_MEMBRANE_ORGANIZATION | 0.00213 | 0.03265 | 0.43171 | 0.48371 | 1.15001 | 432 |
| GO_MICROTUBULE_CYTOSKELETON_ORGANIZATION_INVOL | 0.00159 | 0.02583 | 0.45506 | 0.55517 | 1.27780 | 103 |
| GO_MITOTIC_NUCLEAR_DIVISION | 0.00090 | 0.01600 | 0.47727 | 0.52371 | 1.22582 | 187 |
| GO_MITOTIC_SPINDLE_ORGANIZATION | 0.00117 | 0.01990 | 0.45506 | 0.57242 | 1.30613 | 85 |
| GO_MODIFICATION_DEPENDENT_MACROMOLECULE_CATAL | 0.00000 | 0.00000 | 0.82666 | 0.55221 | 1.31367 | 438 |
| GO_MONOSACCHARIDE_BIOSYNTHETIC_PROCESS | 0.00390 | 0.04920 | 0.43171 | 0.61000 | 1.35781 | 52 |
| GO_MORPHOGENESIS_OF_AN_EPITHELIUM | 0.00000 | 0.00006 | 0.64355 | 0.55439 | 1.30304 | 243 |
| GO_MORPHOGENESIS_OF_A_POLARIZED_EPITHELIUM | 0.00000 | 0.00000 | | 0.71406 | 1.63096 | 88 |
| GO_MRNA_3_UTR_BINDING | 0.00157 | 0.02569 | 0.45506 | 0.60156 | 1.34981 | 61 |
| GO_MRNA_5_UTR_BINDING | 0.00224 | 0.03346 | 0.43171 | 0.69374 | 1.46405 | 22 |
| GO_MRNA_BINDING | 0.00000 | 0.00000 | 0.81404 | 0.60974 | 1.42987 | 210 |
| GO_MRNA_EXPORT_FROM_NUCLEUS | 0.00018 | 0.00419 | 0.51885 | 0.59473 | 1.36323 | 94 |
| GO_MRNA_PROCESSING | 0.00000 | 0.00000 | 0.74774 | 0.54949 | 1.30376 | 389 |
| GO_MRNA_TRANSPORT | 0.00022 | 0.00485 | 0.51885 | 0.57409 | 1.32631 | 117 |
| GO_MULTI_ORGANISM_LOCALIZATION | 0.00390 | 0.04920 | 0.43171 | 0.61069 | 1.35639 | 50 |
| GO_MULTI_ORGANISM_PROCESS | 0.00209 | 0.03228 | 0.43171 | 0.48527 | 1.15181 | 402 |
| GO_MYELOID_LEUKOCYTE_ACTIVATION | 0.00000 | 0.00000 | 0.78819 | 0.58081 | 1.36862 | 276 |

| Cluster 1 | | | | | | |
|---|---|---|---|---|---|---|
| pathway | pval | padj | log2err | ES | NES | size |
| GO_MYELOID_LEUKOCYTE_MEDIATED_IMMUNITY | 0.00000 | 0.00000 | 0.80122 | 0.58612 | 1.37840 | 261 |
| GO_NADH_METABOLIC_PROCESS | 0.00257 | 0.03725 | 0.43171 | 0.71646 | 1.50458 | 21 |
| GO_NAD_BINDING | 0.00041 | 0.00829 | 0.49849 | 0.66807 | 1.45898 | 37 |
| GO_NAD_METABOLIC_PROCESS | 0.00220 | 0.03310 | 0.43171 | 0.69925 | 1.48187 | 24 |
| GO_NCRNA_EXPORT_FROM_NUCLEUS | 0.00398 | 0.04977 | 0.40702 | 0.64780 | 1.40763 | 34 |
| GO_NCRNA_METABOLIC_PROCESS | 0.00002 | 0.00069 | 0.57561 | 0.51636 | 1.22310 | 354 |
| GO_NCRNA_PROCESSING | 0.00000 | 0.00015 | 0.61053 | 0.53619 | 1.26574 | 304 |
| GO_NEGATIVE_REGULATION_OF_CANONICAL_WNT_SIGNAI | 0.00000 | 0.00000 | 0.78819 | 0.68546 | 1.56986 | 92 |
| GO_NEGATIVE_REGULATION_OF_CELLULAR_AMIDE_METAB | 0.00002 | 0.00049 | 0.57561 | 0.60762 | 1.39462 | 97 |
| GO_NEGATIVE_REGULATION_OF_CELLULAR_CATABOLIC_PR | 0.00294 | 0.04088 | 0.43171 | 0.52805 | 1.23006 | 152 |
| GO_NEGATIVE_REGULATION_OF_CELL_CYCLE | 0.00000 | 0.00001 | 0.69013 | 0.53772 | 1.27570 | 387 |
| GO_NEGATIVE_REGULATION_OF_CELL_CYCLE_G2_M_PHASI | 0.00000 | 0.00000 | 0.77494 | 0.69832 | 1.59210 | 84 |
| GO_NEGATIVE_REGULATION_OF_CELL_CYCLE_PHASE_TRAN | 0.00000 | 0.00000 | 0.74774 | 0.60879 | 1.42093 | 166 |
| GO_NEGATIVE_REGULATION_OF_CELL_CYCLE_PROCESS | 0.00000 | 0.00001 | 0.67496 | 0.57014 | 1.33845 | 227 |
| GO_NEGATIVE_REGULATION_OF_CELL_DEATH | 0.00000 | 0.00013 | 0.61053 | 0.51707 | 1.22889 | 423 |
| GO_NEGATIVE_REGULATION_OF_MAP_KINASE_ACTIVITY | 0.00272 | 0.03895 | 0.43171 | 0.66608 | 1.44160 | 32 |
| GO_NEGATIVE_REGULATION_OF_MITOTIC_CELL_CYCLE | 0.00000 | 0.00001 | 0.69013 | 0.57451 | 1.34683 | 211 |
| GO_NEGATIVE_REGULATION_OF_VIRAL_TRANSCRIPTION | 0.00372 | 0.04792 | 0.43171 | 0.75117 | 1.52963 | 15 |
| GO_NEGATIVE_REGULATION_OF_WNT_SIGNALING_PATHW/ | 0.00000 | 0.00000 | 0.76146 | 0.66653 | 1.53488 | 104 |
| GO_NEURAL_NUCLEUS_DEVELOPMENT | 0.00283 | 0.03998 | 0.43171 | 0.69329 | 1.46923 | 24 |
| GO_NEURON_PROJECTION | 0.00137 | 0.02290 | 0.45506 | 0.48217 | 1.14733 | 440 |
| GO_NIK_NF_KAPPAB_SIGNALING | 0.00000 | 0.00000 | | 0.71455 | 1.63953 | 96 |
| GO_NON_CANONICAL_WNT_SIGNALING_PATHWAY | 0.00000 | 0.00000 | | 0.71964 | 1.64206 | 85 |
| GO_NUCLEAR_BODY | 0.00148 | 0.02442 | 0.45506 | 0.48152 | 1.14753 | 493 |
| GO_NUCLEAR_CHROMOSOME_SEGREGATION | 0.00390 | 0.04920 | 0.43171 | 0.50973 | 1.19255 | 185 |
| GO_NUCLEAR_ENVELOPE | 0.00076 | 0.01383 | 0.47727 | 0.50999 | 1.20062 | 268 |
| GO_NUCLEAR_EXPORT | 0.00001 | 0.00028 | 0.59333 | 0.57494 | 1.34054 | 158 |
| GO_NUCLEAR_SPECK | 0.00063 | 0.01190 | 0.47727 | 0.51297 | 1.20672 | 265 |
| GO_NUCLEAR_TRANSCRIBED_MRNA_CATABOLIC_PROCESS | 0.00000 | 0.00000 | 0.80122 | 0.62375 | 1.45572 | 169 |
| GO_NUCLEAR_TRANSCRIBED_MRNA_CATABOLIC_PROCESS_ | 0.00000 | 0.00000 | 0.81404 | 0.67524 | 1.55699 | 110 |
| GO_NUCLEAR_TRANSPORT | 0.00000 | 0.00001 | 0.69013 | 0.56437 | 1.32716 | 250 |
| GO_NUCLEOBASE_CONTAINING_COMPOUND_TRANSPORT | 0.00002 | 0.00047 | 0.57561 | 0.56640 | 1.32611 | 189 |
| GO_NUCLEOBASE_CONTAINING_SMALL_MOLECULE_METAE | 0.00016 | 0.00362 | 0.51885 | 0.51504 | 1.21603 | 305 |
| GO_NUCLEOSIDE_DIPHOSPHATE_METABOLIC_PROCESS | 0.00224 | 0.03346 | 0.43171 | 0.56843 | 1.29802 | 86 |
| GO_NUCLEOSIDE_METABOLIC_PROCESS | 0.00028 | 0.00605 | 0.49849 | 0.64019 | 1.43129 | 55 |
| GO_NUCLEOSIDE_MONOPHOSPHATE_BIOSYNTHETIC_PROC | 0.00390 | 0.04920 | 0.43171 | 0.65819 | 1.42275 | 31 |
| GO_NUCLEOSIDE_PHOSPHATE_BIOSYNTHETIC_PROCESS | 0.00056 | 0.01093 | 0.47727 | 0.55346 | 1.28128 | 128 |
| GO_NUCLEOTIDE_EXCISION_REPAIR | 0.00183 | 0.02907 | 0.45506 | 0.57191 | 1.30954 | 91 |
| GO_NUCLEOTIDE_EXCISION_REPAIR_DNA_DAMAGE_RECOG | 0.00253 | 0.03692 | 0.43171 | 0.71704 | 1.50580 | 21 |
| GO_NUCLEOTIDE_PHOSPHORYLATION | 0.00108 | 0.01877 | 0.45506 | 0.57950 | 1.31472 | 77 |
| GO_NUCLEUS_ORGANIZATION | 0.00132 | 0.02205 | 0.45506 | 0.57551 | 1.31450 | 88 |
| GO_ORGANELLE_ASSEMBLY | 0.00113 | 0.01947 | 0.45506 | 0.48340 | 1.15139 | 482 |
| GO_ORGANIC_CYCLIC_COMPOUND_CATABOLIC_PROCESS | 0.00000 | 0.00000 | | 0.60854 | 1.44428 | 394 |
| GO_PEPTIDASE_ACTIVITY | 0.00008 | 0.00188 | 0.53843 | 0.53531 | 1.25624 | 226 |
| GO_PEPTIDASE_COMPLEX | 0.00000 | 0.00000 | 0.77494 | 0.70826 | 1.60911 | 78 |
| GO_PEPTIDE_BINDING | 0.00087 | 0.01556 | 0.47727 | 0.57916 | 1.32426 | 89 |
| GO_PEPTIDE_BIOSYNTHETIC_PROCESS | 0.00000 | 0.00000 | | 0.57600 | 1.37143 | 477 |
| GO_PIGMENT_GRANULE | 0.00001 | 0.00042 | 0.59333 | 0.64738 | 1.46416 | 70 |
| GO_POLYSOMAL_RIBOSOME | 0.00003 | 0.00070 | 0.57561 | 0.74911 | 1.61203 | 29 |
| GO_POLYSOME | 0.00000 | 0.00008 | 0.62726 | 0.69351 | 1.55045 | 57 |
| GO_POSITIVE_REGULATION_OF_CANONICAL_WNT_SIGNAL | 0.00000 | 0.00000 | 0.82666 | 0.70186 | 1.60750 | 93 |
| GO_POSITIVE_REGULATION_OF_CELLULAR_AMIDE_METABC | 0.00006 | 0.00151 | 0.55733 | 0.59026 | 1.35923 | 104 |
| GO_POSITIVE_REGULATION_OF_CELLULAR_PROTEIN_CATAE | 0.00179 | 0.02866 | 0.45506 | 0.56996 | 1.30420 | 90 |
| GO_POSITIVE_REGULATION_OF_CELLULAR_PROTEIN_LOCAI | 0.00000 | 0.00012 | 0.62726 | 0.55471 | 1.30085 | 214 |
| GO_POSITIVE_REGULATION_OF_CELL_CYCLE | 0.00235 | 0.03481 | 0.43171 | 0.51387 | 1.20500 | 213 |
| GO_POSITIVE_REGULATION_OF_CELL_CYCLE_PROCESS | 0.00246 | 0.03604 | 0.43171 | 0.53070 | 1.23867 | 167 |
| GO_POSITIVE_REGULATION_OF_CELL_DEATH | 0.00003 | 0.00072 | 0.57561 | 0.52409 | 1.23665 | 300 |
| GO_POSITIVE_REGULATION_OF_CELL_POPULATION_PROLIF | 0.00179 | 0.02866 | 0.45506 | 0.49470 | 1.16831 | 311 |
| GO_POSITIVE_REGULATION_OF_CHROMOSOME_ORGANIZA | 0.00073 | 0.01361 | 0.47727 | 0.55026 | 1.27387 | 128 |
| GO_POSITIVE_REGULATION_OF_DEFENSE_RESPONSE | 0.00000 | 0.00000 | 0.77494 | 0.64075 | 1.48779 | 135 |
| GO_POSITIVE_REGULATION_OF_DNA_BIOSYNTHETIC_PROC | 0.00006 | 0.00155 | 0.53843 | 0.67566 | 1.49428 | 48 |
| GO_POSITIVE_REGULATION_OF_DNA_METABOLIC_PROCESS | 0.00290 | 0.04058 | 0.43171 | 0.55185 | 1.27400 | 113 |
| GO_POSITIVE_REGULATION_OF_DNA_TEMPLATED_TRANSC | 0.00376 | 0.04792 | 0.43171 | 0.68032 | 1.44966 | 25 |
| GO_POSITIVE_REGULATION_OF_ESTABLISHMENT_OF_PROT | 0.00000 | 0.00015 | 0.61053 | 0.56286 | 1.31889 | 204 |
| GO_POSITIVE_REGULATION_OF_IMMUNE_RESPONSE | 0.00000 | 0.00000 | 0.71951 | 0.57251 | 1.34563 | 243 |
| GO_POSITIVE_REGULATION_OF_IMMUNE_SYSTEM_PROCES | 0.00000 | 0.00001 | 0.69013 | 0.54781 | 1.29558 | 329 |
| GO_POSITIVE_REGULATION_OF_INNATE_IMMUNE_RESPON | 0.00000 | 0.00000 | 0.82666 | 0.68145 | 1.56845 | 103 |
| GO_POSITIVE_REGULATION_OF_INTRACELLULAR_TRANSPO | 0.00287 | 0.04028 | 0.43171 | 0.53606 | 1.24852 | 149 |
| GO_POSITIVE_REGULATION_OF_INTRINSIC_APOPTOTIC_SIG | 0.00279 | 0.03968 | 0.43171 | 0.65110 | 1.41090 | 33 |
| GO_POSITIVE_REGULATION_OF_MRNA_PROCESSING | 0.00061 | 0.01169 | 0.47727 | 0.73073 | 1.54491 | 23 |
| GO_POSITIVE_REGULATION_OF_MRNA_SPLICING_VIA_SPLI | 0.00017 | 0.00381 | 0.51885 | 0.81835 | 1.66643 | 15 |

| Cluster 1 | | | | | | |
|---|---|---|---|---|---|---|
| pathway | pval | padj | log2err | ES | NES | size |
| GO_POSITIVE_REGULATION_OF_NUCLEOCYTOPLASMIC_TRA | 0.00279 | 0.03968 | 0.43171 | 0.62936 | 1.38791 | 45 |
| GO_POSITIVE_REGULATION_OF_ORGANELLE_ORGANIZATIO | 0.00191 | 0.03005 | 0.45506 | 0.48897 | 1.15865 | 363 |
| GO_POSITIVE_REGULATION_OF_PEPTIDE_SECRETION | 0.00313 | 0.04287 | 0.43171 | 0.59762 | 1.34907 | 67 |
| GO_POSITIVE_REGULATION_OF_PROTEIN_CATABOLIC_PROC | 0.00095 | 0.01675 | 0.47727 | 0.55471 | 1.28522 | 130 |
| GO_POSITIVE_REGULATION_OF_PROTEIN_LOCALIZATION_T | 0.00001 | 0.00022 | 0.61053 | 0.67958 | 1.51937 | 55 |
| GO_POSITIVE_REGULATION_OF_RESPONSE_TO_BIOTIC_STI | 0.00000 | 0.00000 | 0.80122 | 0.66573 | 1.53820 | 115 |
| GO_POSITIVE_REGULATION_OF_RESPONSE_TO_EXTERNAL_ | 0.00000 | 0.00001 | 0.67496 | 0.58738 | 1.37095 | 166 |
| GO_POSITIVE_REGULATION_OF_RNA_SPLICING | 0.00290 | 0.04058 | 0.43171 | 0.68388 | 1.46147 | 26 |
| GO_POSITIVE_REGULATION_OF_TRANSLATION | 0.00009 | 0.00222 | 0.53843 | 0.60414 | 1.37988 | 88 |
| GO_POSITIVE_REGULATION_OF_TRANSPORT | 0.00324 | 0.04381 | 0.43171 | 0.48467 | 1.14912 | 376 |
| GO_POSITIVE_REGULATION_OF_VIRAL_GENOME_REPLICAT | 0.00084 | 0.01501 | 0.47727 | 0.70005 | 1.50448 | 28 |
| GO_POSITIVE_REGULATION_OF_VIRAL_LIFE_CYCLE | 0.00111 | 0.01928 | 0.45506 | 0.64907 | 1.42542 | 41 |
| GO_POSITIVE_REGULATION_OF_VIRAL_PROCESS | 0.00104 | 0.01825 | 0.45506 | 0.59835 | 1.35633 | 75 |
| GO_POSITIVE_REGULATION_OF_WNT_SIGNALING_PATHWA | 0.00000 | 0.00000 | 0.80122 | 0.67373 | 1.55266 | 108 |
| GO_POSTSYNAPSE | 0.00009 | 0.00209 | 0.53843 | 0.53451 | 1.25340 | 218 |
| GO_POSTSYNAPSE_ORGANIZATION | 0.00287 | 0.04028 | 0.43171 | 0.58345 | 1.31512 | 66 |
| GO_POSTTRANSCRIPTIONAL_REGULATION_OF_GENE_EXPR | 0.00000 | 0.00000 | | 0.62133 | 1.47600 | 412 |
| GO_POST_TRANSLATIONAL_PROTEIN_MODIFICATION | 0.00000 | 0.00000 | 0.70498 | 0.59441 | 1.39243 | 195 |
| GO_PRECATALYTIC_SPLICEOSOME | 0.00313 | 0.04287 | 0.43171 | 0.61730 | 1.36316 | 46 |
| GO_PRERIBOSOME | 0.00213 | 0.03265 | 0.43171 | 0.59106 | 1.33523 | 68 |
| GO_PROCESS_UTILIZING_AUTOPHAGIC_MECHANISM | 0.00272 | 0.03895 | 0.43171 | 0.48962 | 1.15921 | 348 |
| GO_PROTEASOMAL_PROTEIN_CATABOLIC_PROCESS | 0.00000 | 0.00000 | 0.78819 | 0.56041 | 1.32625 | 336 |
| GO_PROTEASOMAL_UBIQUITIN_INDEPENDENT_PROTEIN_C | 0.00000 | 0.00008 | 0.62726 | 0.83326 | 1.74134 | 20 |
| GO_PROTEASOME_ACCESSORY_COMPLEX | 0.00000 | 0.00007 | 0.62726 | 0.81454 | 1.72620 | 24 |
| GO_PROTEASOME_CORE_COMPLEX | 0.00000 | 0.00009 | 0.62726 | 0.85804 | 1.77153 | 18 |
| GO_PROTEIN_CONTAINING_COMPLEX_DISASSEMBLY | 0.00327 | 0.04407 | 0.43171 | 0.50613 | 1.18673 | 220 |
| GO_PROTEIN_CONTAINING_COMPLEX_LOCALIZATION | 0.00002 | 0.00065 | 0.57561 | 0.55488 | 1.29923 | 188 |
| GO_PROTEIN_DNA_COMPLEX | 0.00376 | 0.04792 | 0.43171 | 0.56852 | 1.29164 | 78 |
| GO_PROTEIN_DNA_COMPLEX_SUBUNIT_ORGANIZATION | 0.00007 | 0.00165 | 0.53843 | 0.55814 | 1.30015 | 154 |
| GO_PROTEIN_DOMAIN_SPECIFIC_BINDING | 0.00044 | 0.00876 | 0.49849 | 0.50085 | 1.18618 | 352 |
| GO_PROTEIN_FOLDING | 0.00000 | 0.00001 | 0.69013 | 0.60517 | 1.40864 | 146 |
| GO_PROTEIN_IMPORT | 0.00064 | 0.01203 | 0.47727 | 0.55607 | 1.28663 | 126 |
| GO_PROTEIN_LOCALIZATION_TO_CHROMOSOME_TELOMEF | 0.00023 | 0.00508 | 0.51885 | 0.72608 | 1.55164 | 26 |
| GO_PROTEIN_LOCALIZATION_TO_ENDOPLASMIC_RETICULU | 0.00000 | 0.00000 | 0.81404 | 0.65978 | 1.52553 | 121 |
| GO_PROTEIN_LOCALIZATION_TO_MEMBRANE | 0.00007 | 0.00173 | 0.53843 | 0.51115 | 1.21137 | 366 |
| GO_PROTEIN_LOCALIZATION_TO_NUCLEUS | 0.00000 | 0.00001 | 0.67496 | 0.58790 | 1.37218 | 166 |
| GO_PROTEIN_MODIFICATION_BY_SMALL_PROTEIN_REMOV | 0.00000 | 0.00001 | 0.67496 | 0.57208 | 1.34156 | 210 |
| GO_PROTEIN_POLYUBIQUITINATION | 0.00000 | 0.00000 | 0.71951 | 0.57874 | 1.35681 | 223 |
| GO_PROTEIN_REFOLDING | 0.00012 | 0.00273 | 0.53843 | 0.82186 | 1.67924 | 16 |
| GO_PROTEIN_STABILIZATION | 0.00011 | 0.00261 | 0.53843 | 0.58914 | 1.36131 | 118 |
| GO_PROTEIN_SUMOYLATION | 0.00086 | 0.01529 | 0.47727 | 0.63064 | 1.41023 | 56 |
| GO_PROTEIN_TARGETING | 0.00000 | 0.00000 | 0.77494 | 0.56519 | 1.33538 | 315 |
| GO_PROTEIN_TARGETING_TO_MEMBRANE | 0.00000 | 0.00000 | 0.74774 | 0.62412 | 1.45338 | 148 |
| GO_PURINE_CONTAINING_COMPOUND_BIOSYNTHETIC_PR | 0.00372 | 0.04792 | 0.43171 | 0.55100 | 1.26467 | 97 |
| GO_PURINE_CONTAINING_COMPOUND_METABOLIC_PROC | 0.00039 | 0.00795 | 0.49849 | 0.52304 | 1.22673 | 224 |
| GO_PURINE_NUCLEOSIDE_TRIPHOSPHATE_METABOLIC_PR | 0.00353 | 0.04635 | 0.43171 | 0.62658 | 1.37449 | 40 |
| GO_PYRUVATE_METABOLIC_PROCESS | 0.00010 | 0.00237 | 0.53843 | 0.60165 | 1.37283 | 85 |
| GO_RECEPTOR_MEDIATED_ENDOCYTOSIS | 0.00339 | 0.04486 | 0.43171 | 0.54467 | 1.25363 | 103 |
| GO_REGULATION_OF_ANATOMICAL_STRUCTURE_MORPHO | 0.00010 | 0.00241 | 0.53843 | 0.50074 | 1.18926 | 410 |
| GO_REGULATION_OF_ANIMAL_ORGAN_MORPHOGENESIS | 0.00000 | 0.00000 | 0.80122 | 0.67395 | 1.55402 | 110 |
| GO_REGULATION_OF_APOPTOTIC_SIGNALING_PATHWAY | 0.00013 | 0.00312 | 0.51885 | 0.53587 | 1.25551 | 205 |
| GO_REGULATION_OF_ATP_METABOLIC_PROCESS | 0.00006 | 0.00144 | 0.55733 | 0.63003 | 1.42491 | 70 |
| GO_REGULATION_OF_BINDING | 0.00041 | 0.00835 | 0.49849 | 0.53879 | 1.25864 | 176 |
| GO_REGULATION_OF_CELLULAR_AMIDE_METABOLIC_PROC | 0.00000 | 0.00000 | 0.80122 | 0.58229 | 1.37212 | 276 |
| GO_REGULATION_OF_CELLULAR_AMINE_METABOLIC_PROC | 0.00000 | 0.00000 | | 0.80584 | 1.80199 | 56 |
| GO_REGULATION_OF_CELLULAR_AMINO_ACID_METABOLIC | 0.00000 | 0.00000 | | 0.80525 | 1.79657 | 53 |
| GO_REGULATION_OF_CELLULAR_KETONE_METABOLIC_PRO | 0.00000 | 0.00000 | 0.77494 | 0.67158 | 1.54661 | 105 |
| GO_REGULATION_OF_CELLULAR_LOCALIZATION | 0.00074 | 0.01364 | 0.47727 | 0.48735 | 1.16112 | 491 |
| GO_REGULATION_OF_CELLULAR_PROTEIN_CATABOLIC_PRO | 0.00002 | 0.00057 | 0.57561 | 0.56783 | 1.32534 | 167 |
| GO_REGULATION_OF_CELLULAR_PROTEIN_LOCALIZATION | 0.00004 | 0.00096 | 0.55733 | 0.51915 | 1.22777 | 329 |
| GO_REGULATION_OF_CELLULAR_RESPONSE_TO_HEAT | 0.00081 | 0.01473 | 0.47727 | 0.60613 | 1.36586 | 65 |
| GO_REGULATION_OF_CELLULAR_RESPONSE_TO_STRESS | 0.00018 | 0.00405 | 0.51885 | 0.50019 | 1.18731 | 403 |
| GO_REGULATION_OF_CELL_CYCLE_ARREST | 0.00316 | 0.04326 | 0.43171 | 0.58731 | 1.32289 | 64 |
| GO_REGULATION_OF_CELL_CYCLE_G2_M_PHASE_TRANSITI | 0.00000 | 0.00000 | 0.77494 | 0.61458 | 1.43444 | 166 |
| GO_REGULATION_OF_CELL_CYCLE_PHASE_TRANSITION | 0.00000 | 0.00000 | 0.71951 | 0.55643 | 1.31407 | 309 |
| GO_REGULATION_OF_CELL_CYCLE_PROCESS | 0.00000 | 0.00000 | 0.76146 | 0.53488 | 1.27402 | 484 |
| GO_REGULATION_OF_CHROMOSOME_ORGANIZATION | 0.00076 | 0.01383 | 0.47727 | 0.51426 | 1.20837 | 241 |
| GO_REGULATION_OF_DEFENSE_RESPONSE | 0.00000 | 0.00005 | 0.64355 | 0.55600 | 1.30681 | 243 |
| GO_REGULATION_OF_DNA_BIOSYNTHETIC_PROCESS | 0.00024 | 0.00526 | 0.51885 | 0.61780 | 1.39922 | 72 |
| GO_REGULATION_OF_DNA_METABOLIC_PROCESS | 0.00168 | 0.02724 | 0.45506 | 0.51447 | 1.20612 | 208 |
| GO_REGULATION_OF_DNA_TEMPLATED_TRANSCRIPTION_I | 0.00191 | 0.03005 | 0.45506 | 0.67819 | 1.45942 | 29 |

| **Cluster 1** | | | | | | |
|---|---|---|---|---|---|---|
| pathway | pval | padj | log2err | ES | NES | size |
| GO_REGULATION_OF_DNA_TEMPLATED_TRANSCRIPTION_I | 0.00000 | 0.00000 | 0.82666 | 0.71133 | 1.62079 | 83 |
| GO_REGULATION_OF_GENERATION_OF_PRECURSOR_META | 0.00049 | 0.00965 | 0.47727 | 0.58865 | 1.34208 | 84 |
| GO_REGULATION_OF_HEMATOPOIETIC_PROGENITOR_CELL_ | 0.00000 | 0.00000 | 0.78819 | 0.74474 | 1.67478 | 63 |
| GO_REGULATION_OF_HEMATOPOIETIC_STEM_CELL_DIFFER | 0.00000 | 0.00000 | 0.77494 | 0.75615 | 1.69088 | 56 |
| GO_REGULATION_OF_HEMOPOIESIS | 0.00000 | 0.00002 | 0.65944 | 0.57950 | 1.35753 | 192 |
| GO_REGULATION_OF_IMMUNE_RESPONSE | 0.00001 | 0.00016 | 0.61053 | 0.53414 | 1.26169 | 312 |
| GO_REGULATION_OF_IMMUNE_SYSTEM_PROCESS | 0.00001 | 0.00021 | 0.61053 | 0.50784 | 1.21057 | 499 |
| GO_REGULATION_OF_INNATE_IMMUNE_RESPONSE | 0.00000 | 0.00000 | 0.77494 | 0.63362 | 1.47359 | 144 |
| GO_REGULATION_OF_INTRACELLULAR_TRANSPORT | 0.00335 | 0.04460 | 0.43171 | 0.50546 | 1.18514 | 220 |
| GO_REGULATION_OF_INTRINSIC_APOPTOTIC_SIGNALING_P | 0.00002 | 0.00053 | 0.57561 | 0.61118 | 1.40143 | 95 |
| GO_REGULATION_OF_MITOTIC_CELL_CYCLE | 0.00000 | 0.00000 | 0.76146 | 0.54058 | 1.28414 | 411 |
| GO_REGULATION_OF_MORPHOGENESIS_OF_AN_EPITHELIU | 0.00000 | 0.00000 | 0.77494 | 0.68527 | 1.56950 | 93 |
| GO_REGULATION_OF_MRNA_CATABOLIC_PROCESS | 0.00000 | 0.00000 | 0.82666 | 0.64559 | 1.50363 | 149 |
| GO_REGULATION_OF_MRNA_METABOLIC_PROCESS | 0.00000 | 0.00000 | | 0.63386 | 1.48940 | 241 |
| GO_REGULATION_OF_MRNA_PROCESSING | 0.00001 | 0.00020 | 0.61053 | 0.61229 | 1.40996 | 104 |
| GO_REGULATION_OF_MRNA_SPLICING_VIA_SPLICEOSOME | 0.00001 | 0.00042 | 0.59333 | 0.64289 | 1.45817 | 76 |
| GO_REGULATION_OF_NUCLEAR_DIVISION | 0.00145 | 0.02397 | 0.45506 | 0.55384 | 1.27966 | 115 |
| GO_REGULATION_OF_NUCLEASE_ACTIVITY | 0.00062 | 0.01177 | 0.47727 | 0.77769 | 1.59977 | 17 |
| GO_REGULATION_OF_NUCLEOCYTOPLASMIC_TRANSPORT | 0.00082 | 0.01479 | 0.47727 | 0.59981 | 1.35774 | 71 |
| GO_REGULATION_OF_NUCLEOTIDE_METABOLIC_PROCESS | 0.00250 | 0.03648 | 0.43171 | 0.60069 | 1.34910 | 62 |
| GO_REGULATION_OF_PEPTIDE_TRANSPORT | 0.00152 | 0.02487 | 0.45506 | 0.50044 | 1.18174 | 310 |
| GO_REGULATION_OF_PROTEASOMAL_PROTEIN_CATABOLIC | 0.00005 | 0.00136 | 0.55733 | 0.57839 | 1.33828 | 126 |
| GO_REGULATION_OF_PROTEASOMAL_UBIQUITIN_DEPENDI | 0.00033 | 0.00703 | 0.49849 | 0.59098 | 1.35129 | 89 |
| GO_REGULATION_OF_PROTEIN_CATABOLIC_PROCESS | 0.00000 | 0.00000 | 0.71951 | 0.57487 | 1.35062 | 238 |
| GO_REGULATION_OF_PROTEIN_CONTAINING_COMPLEX_AS | 0.00038 | 0.00786 | 0.49849 | 0.52095 | 1.22254 | 226 |
| GO_REGULATION_OF_PROTEIN_IMPORT | 0.00076 | 0.01383 | 0.47727 | 0.67730 | 1.47425 | 35 |
| GO_REGULATION_OF_PROTEIN_LOCALIZATION | 0.00006 | 0.00140 | 0.55733 | 0.50066 | 1.19207 | 477 |
| GO_REGULATION_OF_PROTEIN_LOCALIZATION_TO_NUCLEU | 0.00001 | 0.00027 | 0.59333 | 0.64716 | 1.46786 | 76 |
| GO_REGULATION_OF_PROTEIN_MODIFICATION_BY_SMALL_ | 0.00014 | 0.00327 | 0.51885 | 0.56139 | 1.30838 | 156 |
| GO_REGULATION_OF_PROTEIN_STABILITY | 0.00010 | 0.00246 | 0.53843 | 0.55157 | 1.29139 | 189 |
| GO_REGULATION_OF_PROTEOLYSIS | 0.00031 | 0.00663 | 0.49849 | 0.50425 | 1.19295 | 334 |
| GO_REGULATION_OF_RESPONSE_TO_BIOTIC_STIMULUS | 0.00000 | 0.00000 | 0.76146 | 0.60656 | 1.41797 | 180 |
| GO_REGULATION_OF_RESPONSE_TO_EXTERNAL_STIMULUS | 0.00002 | 0.00055 | 0.57561 | 0.52162 | 1.23489 | 344 |
| GO_REGULATION_OF_RNA_SPLICING | 0.00021 | 0.00472 | 0.51885 | 0.58452 | 1.34712 | 106 |
| GO_REGULATION_OF_SMALL_MOLECULE_METABOLIC_PRO | 0.00000 | 0.00000 | 0.80122 | 0.60027 | 1.40787 | 224 |
| GO_REGULATION_OF_STEM_CELL_DIFFERENTIATION | 0.00000 | 0.00000 | 0.82666 | 0.72666 | 1.64859 | 77 |
| GO_REGULATION_OF_SYMBIOTIC_PROCESS | 0.00098 | 0.01734 | 0.45506 | 0.55957 | 1.29311 | 120 |
| GO_REGULATION_OF_TELOMERE_MAINTENANCE | 0.00361 | 0.04708 | 0.43171 | 0.58149 | 1.31033 | 65 |
| GO_REGULATION_OF_TRANSCRIPTION_FROM_RNA_POLYM | 0.00000 | 0.00000 | 0.80122 | 0.75210 | 1.68637 | 60 |
| GO_REGULATION_OF_TRANSCRIPTION_INITIATION_FROM_ | 0.00220 | 0.03310 | 0.43171 | 0.69946 | 1.48230 | 24 |
| GO_REGULATION_OF_TRANSFERASE_ACTIVITY | 0.00376 | 0.04792 | 0.43171 | 0.47376 | 1.12760 | 462 |
| GO_REGULATION_OF_TRANSLATIONAL_INITIATION | 0.00034 | 0.00710 | 0.49849 | 0.62959 | 1.40939 | 59 |
| GO_REGULATION_OF_UBIQUITIN_DEPENDENT_PROTEIN_CA | 0.00052 | 0.01016 | 0.47727 | 0.57461 | 1.32430 | 106 |
| GO_REGULATION_OF_VIRAL_GENOME_REPLICATION | 0.00220 | 0.03310 | 0.43171 | 0.62179 | 1.37811 | 49 |
| GO_REGULATION_OF_WNT_SIGNALING_PATHWAY | 0.00000 | 0.00000 | 0.76146 | 0.60421 | 1.41391 | 186 |
| GO_REPLISOME | 0.00313 | 0.04287 | 0.43171 | 0.68700 | 1.44982 | 22 |
| GO_RESPONSE_TO_BIOTIC_STIMULUS | 0.00000 | 0.00000 | 0.74774 | 0.53572 | 1.27573 | 480 |
| GO_RESPONSE_TO_CYTOKINE | 0.00000 | 0.00000 | | 0.58952 | 1.40315 | 466 |
| GO_RESPONSE_TO_DRUG | 0.00013 | 0.00293 | 0.51885 | 0.55161 | 1.28469 | 151 |
| GO_RESPONSE_TO_HEAT | 0.00006 | 0.00146 | 0.55733 | 0.60001 | 1.37955 | 101 |
| GO_RESPONSE_TO_INTERFERON_GAMMA | 0.00276 | 0.03937 | 0.43171 | 0.61625 | 1.37593 | 54 |
| GO_RESPONSE_TO_INTERLEUKIN_1 | 0.00000 | 0.00000 | | 0.69788 | 1.60180 | 97 |
| GO_RESPONSE_TO_INTERLEUKIN_12 | 0.00032 | 0.00670 | 0.49849 | 0.69653 | 1.51351 | 34 |
| GO_RESPONSE_TO_ORGANIC_CYCLIC_COMPOUND | 0.00033 | 0.00696 | 0.49849 | 0.49929 | 1.18298 | 358 |
| GO_RESPONSE_TO_OXYGEN_LEVELS | 0.00000 | 0.00000 | 0.81404 | 0.61084 | 1.43144 | 200 |
| GO_RESPONSE_TO_TEMPERATURE_STIMULUS | 0.00065 | 0.01233 | 0.47727 | 0.56389 | 1.30317 | 119 |
| GO_RESPONSE_TO_TUMOR_NECROSIS_FACTOR | 0.00000 | 0.00000 | | 0.65589 | 1.52294 | 135 |
| GO_RESPONSE_TO_VIRUS | 0.00353 | 0.04635 | 0.43171 | 0.53215 | 1.23350 | 131 |
| GO_RIBONUCLEOPROTEIN_COMPLEX_BINDING | 0.00000 | 0.00002 | 0.65944 | 0.62744 | 1.44646 | 109 |
| GO_RIBONUCLEOPROTEIN_COMPLEX_BIOGENESIS | 0.00000 | 0.00000 | | 0.61162 | 1.44931 | 365 |
| GO_RIBONUCLEOPROTEIN_COMPLEX_SUBUNIT_ORGANIZAT | 0.00000 | 0.00000 | 0.78819 | 0.62828 | 1.46473 | 161 |
| GO_RIBONUCLEOPROTEIN_GRANULE | 0.00000 | 0.00001 | 0.67496 | 0.60986 | 1.41759 | 139 |
| GO_RIBONUCLEOSIDE_TRIPHOSPHATE_METABOLIC_PROCES | 0.00220 | 0.03310 | 0.43171 | 0.63546 | 1.39399 | 40 |
| GO_RIBOSE_PHOSPHATE_METABOLIC_PROCESS | 0.00025 | 0.00543 | 0.49849 | 0.53043 | 1.24367 | 212 |
| GO_RIBOSOMAL_LARGE_SUBUNIT_ASSEMBLY | 0.00272 | 0.03895 | 0.43171 | 0.67661 | 1.45194 | 27 |
| GO_RIBOSOMAL_LARGE_SUBUNIT_BIOGENESIS | 0.00000 | 0.00008 | 0.62726 | 0.68340 | 1.53683 | 63 |
| GO_RIBOSOMAL_SMALL_SUBUNIT_BINDING | 0.00126 | 0.02119 | 0.45506 | 0.75892 | 1.56115 | 17 |
| GO_RIBOSOMAL_SUBUNIT | 0.00000 | 0.00002 | 0.65944 | 0.59655 | 1.39088 | 160 |
| GO_RIBOSOME | 0.00000 | 0.00000 | 0.70498 | 0.58783 | 1.37704 | 192 |
| GO_RIBOSOME_ASSEMBLY | 0.00220 | 0.03310 | 0.43171 | 0.61374 | 1.37230 | 58 |

| Cluster 1 | | | | | | |
|---|---|---|---|---|---|---|
| pathway | pval | padj | log2err | ES | NES | size |
| GO_RIBOSOME_BINDING | 0.00057 | 0.01107 | 0.47727 | 0.64860 | 1.43227 | 46 |
| GO_RIBOSOME_BIOGENESIS | 0.00000 | 0.00000 | 0.78819 | 0.58620 | 1.37864 | 259 |
| GO_RNA_CATABOLIC_PROCESS | 0.00000 | 0.00000 | | 0.63787 | 1.50614 | 307 |
| GO_RNA_DEPENDENT_DNA_BIOSYNTHETIC_PROCESS | 0.00019 | 0.00427 | 0.51885 | 0.64482 | 1.44159 | 57 |
| GO_RNA_EXPORT_FROM_NUCLEUS | 0.00002 | 0.00051 | 0.57561 | 0.59654 | 1.37862 | 119 |
| GO_RNA_LOCALIZATION | 0.00000 | 0.00000 | 0.76146 | 0.61080 | 1.42897 | 184 |
| GO_RNA_POLYMERASE_II_PREINITIATION_COMPLEX_ASSEM | 0.00205 | 0.03190 | 0.43171 | 0.74727 | 1.54282 | 18 |
| GO_RNA_SPLICING | 0.00000 | 0.00000 | 0.73376 | 0.55268 | 1.30843 | 344 |
| GO_RNA_SPLICING_VIA_TRANSESTERIFICATION_REACTIONS | 0.00000 | 0.00000 | 0.77494 | 0.57000 | 1.34311 | 281 |
| GO_ROUGH_ENDOPLASMIC_RETICULUM | 0.00327 | 0.04407 | 0.43171 | 0.62611 | 1.38074 | 45 |
| GO_RRNA_BINDING | 0.00006 | 0.00153 | 0.53843 | 0.66665 | 1.48844 | 54 |
| GO_RRNA_METABOLIC_PROCESS | 0.00000 | 0.00001 | 0.69013 | 0.58556 | 1.37159 | 194 |
| GO_SCF_DEPENDENT_PROTEASOMAL_UBIQUITIN_DEPEND | 0.00000 | 0.00000 | | 0.74127 | 1.67990 | 73 |
| GO_SECRETORY_GRANULE | 0.00000 | 0.00000 | 0.80122 | 0.56487 | 1.33681 | 336 |
| GO_SECRETORY_VESICLE | 0.00000 | 0.00000 | 0.74774 | 0.55029 | 1.30566 | 389 |
| GO_SIGNAL_SEQUENCE_BINDING | 0.00242 | 0.03581 | 0.43171 | 0.67146 | 1.44494 | 29 |
| GO_SIGNAL_TRANSDUCTION_BY_PROTEIN_PHOSPHORYLAT | 0.00000 | 0.00015 | 0.61053 | 0.52683 | 1.24767 | 353 |
| GO_SINGLE_FERTILIZATION | 0.00039 | 0.00795 | 0.49849 | 0.66746 | 1.46418 | 40 |
| GO_SINGLE_STRANDED_DNA_BINDING | 0.00172 | 0.02774 | 0.45506 | 0.57111 | 1.30413 | 86 |
| GO_SINGLE_STRANDED_RNA_BINDING | 0.00054 | 0.01065 | 0.47727 | 0.63385 | 1.40824 | 51 |
| GO_SISTER_CHROMATID_SEGREGATION | 0.00246 | 0.03604 | 0.43171 | 0.53728 | 1.25137 | 149 |
| GO_SMALL_MOLECULE_BIOSYNTHETIC_PROCESS | 0.00060 | 0.01155 | 0.47727 | 0.49530 | 1.17202 | 335 |
| GO_SMALL_RIBOSOMAL_SUBUNIT | 0.00056 | 0.01093 | 0.47727 | 0.62299 | 1.39687 | 60 |
| GO_SNORNA_BINDING | 0.00364 | 0.04732 | 0.43171 | 0.67119 | 1.44030 | 27 |
| GO_SOMATODENDRITIC_COMPARTMENT | 0.00029 | 0.00630 | 0.49849 | 0.51059 | 1.20430 | 296 |
| GO_SPERM_EGG_RECOGNITION | 0.00000 | 0.00000 | 0.78819 | 0.91711 | 1.89348 | 18 |
| GO_SPLICEOSOMAL_COMPLEX | 0.00001 | 0.00022 | 0.61053 | 0.57693 | 1.34529 | 159 |
| GO_SPLICEOSOMAL_COMPLEX_ASSEMBLY | 0.00331 | 0.04423 | 0.43171 | 0.62086 | 1.36799 | 43 |
| GO_STEM_CELL_DIFFERENTIATION | 0.00000 | 0.00000 | | 0.69975 | 1.61813 | 122 |
| GO_STRUCTURAL_CONSTITUENT_OF_RIBOSOME | 0.00000 | 0.00009 | 0.62726 | 0.58808 | 1.36671 | 141 |
| GO_STRUCTURAL_MOLECULE_ACTIVITY | 0.00000 | 0.00001 | 0.69013 | 0.55256 | 1.30278 | 290 |
| GO_SUBSTANTIA_NIGRA_DEVELOPMENT | 0.00216 | 0.03310 | 0.43171 | 0.69478 | 1.46624 | 22 |
| GO_SUPRAMOLECULAR_POLYMER | 0.00298 | 0.04128 | 0.43171 | 0.48518 | 1.14969 | 365 |
| GO_SYNAPSE | 0.00000 | 0.00000 | 0.76146 | 0.53936 | 1.28337 | 455 |
| GO_TBP_CLASS_PROTEIN_BINDING | 0.00376 | 0.04792 | 0.43171 | 0.71000 | 1.49101 | 21 |
| GO_TELOMERASE_RNA_LOCALIZATION | 0.00005 | 0.00130 | 0.55733 | 0.82494 | 1.69695 | 17 |
| GO_TELOMERE_MAINTENANCE_VIA_TELOMERE_LENGTHEN | 0.00026 | 0.00568 | 0.49849 | 0.62218 | 1.40144 | 64 |
| GO_TELOMERE_ORGANIZATION | 0.00108 | 0.01877 | 0.45506 | 0.56319 | 1.30135 | 118 |
| GO_THREONINE_TYPE_ENDOPEPTIDASE_ACTIVITY | 0.00000 | 0.00002 | 0.65944 | 0.87608 | 1.80877 | 18 |
| GO_THREONINE_TYPE_PEPTIDASE_ACTIVITY | 0.00000 | 0.00001 | 0.67496 | 0.85676 | 1.79044 | 20 |
| GO_TISSUE_MORPHOGENESIS | 0.00001 | 0.00026 | 0.59333 | 0.53899 | 1.26990 | 277 |
| GO_TOXIN_TRANSPORT | 0.00004 | 0.00097 | 0.55733 | 0.77177 | 1.62872 | 22 |
| GO_TRANSCRIPTION_COUPLED_NUCLEOTIDE_EXCISION_RE | 0.00083 | 0.01490 | 0.47727 | 0.61476 | 1.38776 | 67 |
| GO_TRANSCRIPTION_FACTOR_BINDING | 0.00000 | 0.00001 | 0.67496 | 0.53977 | 1.27642 | 327 |
| GO_TRANSLATIONAL_INITIATION | 0.00000 | 0.00000 | | 0.68768 | 1.60355 | 159 |
| GO_TRANSLATION_FACTOR_ACTIVITY_RNA_BINDING | 0.00000 | 0.00000 | | 0.75751 | 1.70699 | 65 |
| GO_TRANSLATION_INITIATION_FACTOR_ACTIVITY | 0.00000 | 0.00003 | 0.65944 | 0.73528 | 1.61973 | 44 |
| GO_TRANSLATION_PREINITIATION_COMPLEX | 0.00003 | 0.00075 | 0.57561 | 0.84699 | 1.73057 | 16 |
| GO_TRANSLATION_REGULATOR_ACTIVITY | 0.00000 | 0.00000 | 0.80122 | 0.68852 | 1.58141 | 99 |
| GO_TRANSLATION_REGULATOR_ACTIVITY_NUCLEIC_ACID_E | 0.00000 | 0.00000 | 0.80122 | 0.71844 | 1.63223 | 78 |
| GO_TRICARBOXYLIC_ACID_CYCLE | 0.00257 | 0.03725 | 0.43171 | 0.68078 | 1.46306 | 28 |
| GO_TUMOR_NECROSIS_FACTOR_MEDIATED_SIGNALING_PA | 0.00000 | 0.00000 | | 0.73532 | 1.67955 | 87 |
| GO_T_CELL_RECEPTOR_SIGNALING_PATHWAY | 0.00000 | 0.00000 | 0.77494 | 0.67249 | 1.54872 | 105 |
| GO_U2_SNRNP | 0.00324 | 0.04381 | 0.43171 | 0.74079 | 1.52386 | 17 |
| GO_U2_TYPE_SPLICEOSOMAL_COMPLEX | 0.00048 | 0.00951 | 0.49849 | 0.59540 | 1.35476 | 80 |
| GO_UBIQUITIN_LIKE_PROTEIN_LIGASE_BINDING | 0.00000 | 0.00003 | 0.65944 | 0.57714 | 1.35184 | 190 |
| GO_UNFOLDED_PROTEIN_BINDING | 0.00000 | 0.00000 | 0.70498 | 0.68853 | 1.56170 | 76 |
| GO_VACUOLAR_LUMEN | 0.00121 | 0.02039 | 0.45506 | 0.58748 | 1.33553 | 79 |
| GO_VACUOLAR_MEMBRANE | 0.00346 | 0.04573 | 0.43171 | 0.49796 | 1.17096 | 251 |
| GO_VACUOLE | 0.00026 | 0.00568 | 0.49849 | 0.49456 | 1.17497 | 414 |
| GO_VESICLE_LUMEN | 0.00000 | 0.00000 | 0.80122 | 0.63650 | 1.48240 | 151 |
| GO_VIRAL_GENE_EXPRESSION | 0.00000 | 0.00000 | 0.78819 | 0.61943 | 1.44577 | 166 |
| GO_VIRAL_GENOME_REPLICATION | 0.00372 | 0.04792 | 0.43171 | 0.58046 | 1.31127 | 68 |
| GO_VIRAL_LIFE_CYCLE | 0.00000 | 0.00014 | 0.61053 | 0.57243 | 1.33876 | 183 |
| HP_ABNORMAL_CNS_MYELINATION | 0.00331 | 0.04423 | 0.43171 | 0.53071 | 1.23367 | 138 |
| HP_BULBAR_PALSY | 0.00159 | 0.02583 | 0.45506 | 0.75399 | 1.55671 | 18 |
| HP_MACROGYRIA | 0.00115 | 0.01965 | 0.45506 | 0.77342 | 1.58025 | 16 |
| HP_PTERYGIUM | 0.00398 | 0.04977 | 0.40702 | 0.74849 | 1.52418 | 15 |
| **Cluster 12** | | | | | | |
| pathway | pval | padj | log2err | ES | NES | size |

| Cluster 1 | | | | | | |
|---|---|---|---|---|---|---|
| pathway | pval | padj | log2err | ES | NES | size |
| GO_ACTIVATION_OF_IMMUNE_RESPONSE | 0.00000 | 0.00000 | 0.77494 | 0.61761 | 1.38423 | 207 |
| GO_ACTIVATION_OF_INNATE_IMMUNE_RESPONSE | 0.00000 | 0.00000 | 0.78819 | 0.71645 | 1.56871 | 93 |
| GO_AEROBIC_RESPIRATION | 0.00000 | 0.00001 | 0.69013 | 0.70799 | 1.53052 | 65 |
| GO_ALTERNATIVE_MRNA_SPLICING_VIA_SPLICEOSOME | 0.00268 | 0.03951 | 0.43171 | 0.61510 | 1.32789 | 63 |
| GO_AMINE_METABOLIC_PROCESS | 0.00000 | 0.00000 | | 0.78559 | 1.70588 | 75 |
| GO_ANAPHASE_PROMOTING_COMPLEX_DEPENDENT_CATA | 0.00000 | 0.00000 | | 0.75263 | 1.63803 | 78 |
| GO_ANATOMICAL_STRUCTURE_HOMEOSTASIS | 0.00113 | 0.01980 | 0.45506 | 0.53656 | 1.20778 | 237 |
| GO_ANCHORING_JUNCTION | 0.00000 | 0.00000 | | 0.60905 | 1.38887 | 474 |
| GO_ANIMAL_ORGAN_REGENERATION | 0.00174 | 0.02842 | 0.45506 | 0.67535 | 1.41606 | 36 |
| GO_ANTIGEN_PROCESSING_AND_PRESENTATION | 0.00000 | 0.00000 | 0.76146 | 0.65428 | 1.45549 | 142 |
| GO_ANTIGEN_PROCESSING_AND_PRESENTATION_OF_EXOG | 0.00000 | 0.00000 | | 0.81848 | 1.76213 | 58 |
| GO_ANTIGEN_PROCESSING_AND_PRESENTATION_OF_PEPT | 0.00000 | 0.00000 | 0.74774 | 0.67073 | 1.48579 | 124 |
| GO_ANTIGEN_PROCESSING_AND_PRESENTATION_OF_PEPT | 0.00000 | 0.00000 | | 0.79688 | 1.72891 | 70 |
| GO_ANTIGEN_RECEPTOR_MEDIATED_SIGNALING_PATHWAY | 0.00000 | 0.00002 | 0.67496 | 0.64508 | 1.42907 | 123 |
| GO_ANTIOXIDANT_ACTIVITY | 0.00250 | 0.03757 | 0.43171 | 0.67110 | 1.41313 | 38 |
| GO_APOPTOTIC_SIGNALING_PATHWAY | 0.00044 | 0.00892 | 0.49849 | 0.51581 | 1.17106 | 357 |
| GO_ATP_BIOSYNTHETIC_PROCESS | 0.00143 | 0.02421 | 0.45506 | 0.76071 | 1.53623 | 21 |
| GO_ATP_METABOLIC_PROCESS | 0.00000 | 0.00005 | 0.64355 | 0.59043 | 1.32135 | 201 |
| GO_ATP_SYNTHESIS_COUPLED_ELECTRON_TRANSPORT | 0.00005 | 0.00142 | 0.55733 | 0.65407 | 1.42029 | 75 |
| GO_BLOOD_MICROPARTICLE | 0.00261 | 0.03893 | 0.43171 | 0.70368 | 1.45343 | 27 |
| GO_CADHERIN_BINDING | 0.00000 | 0.00000 | | 0.63897 | 1.43786 | 236 |
| GO_CANONICAL_WNT_SIGNALING_PATHWAY | 0.00001 | 0.00019 | 0.61053 | 0.58597 | 1.31018 | 190 |
| GO_CATALYTIC_STEP_2_SPLICEOSOME | 0.00000 | 0.00003 | 0.65944 | 0.67781 | 1.48017 | 83 |
| GO_CELLULAR_AMINO_ACID_METABOLIC_PROCESS | 0.00002 | 0.00053 | 0.57561 | 0.57016 | 1.27864 | 214 |
| GO_CELLULAR_COMPONENT_DISASSEMBLY | 0.00025 | 0.00551 | 0.49849 | 0.52142 | 1.18557 | 397 |
| GO_CELLULAR_HOMEOSTASIS | 0.00152 | 0.02539 | 0.45506 | 0.50409 | 1.14657 | 410 |
| GO_CELLULAR_KETONE_METABOLIC_PROCESS | 0.00000 | 0.00000 | 0.78819 | 0.65344 | 1.45483 | 149 |
| GO_CELLULAR_NITROGEN_COMPOUND_CATABOLIC_PROCE | 0.00000 | 0.00000 | | 0.71962 | 1.63806 | 427 |
| GO_CELLULAR_PROTEIN_COMPLEX_DISASSEMBLY | 0.00143 | 0.02421 | 0.45506 | 0.54557 | 1.21763 | 175 |
| GO_CELLULAR_RESPIRATION | 0.00000 | 0.00000 | 0.74774 | 0.65883 | 1.46365 | 137 |
| GO_CELLULAR_RESPONSE_TO_CHEMICAL_STRESS | 0.00305 | 0.04361 | 0.43171 | 0.52489 | 1.17684 | 211 |
| GO_CELLULAR_RESPONSE_TO_OXYGEN_LEVELS | 0.00000 | 0.00000 | | 0.66973 | 1.49354 | 154 |
| GO_CELL_ACTIVATION_INVOLVED_IN_IMMUNE_RESPONSE | 0.00000 | 0.00000 | 0.73376 | 0.57051 | 1.29392 | 339 |
| GO_CELL_ADHESION_MOLECULE_BINDING | 0.00000 | 0.00000 | | 0.62455 | 1.41245 | 306 |
| GO_CELL_BODY | 0.00080 | 0.01455 | 0.47727 | 0.53097 | 1.19748 | 255 |
| GO_CELL_CELL_RECOGNITION | 0.00000 | 0.00000 | 0.71951 | 0.90812 | 1.81828 | 18 |
| GO_CELL_CELL_SIGNALING_BY_WNT | 0.00001 | 0.00019 | 0.61053 | 0.55506 | 1.25474 | 298 |
| GO_CELL_CYCLE_G2_M_PHASE_TRANSITION | 0.00004 | 0.00098 | 0.55733 | 0.56835 | 1.27490 | 215 |
| GO_CELL_CYCLE_PHASE_TRANSITION | 0.00000 | 0.00004 | 0.64355 | 0.53785 | 1.22563 | 460 |
| GO_CELL_RECOGNITION | 0.00001 | 0.00037 | 0.59333 | 0.72897 | 1.54587 | 43 |
| GO_CELL_REDOX_HOMEOSTASIS | 0.00309 | 0.04392 | 0.43171 | 0.66557 | 1.39944 | 37 |
| GO_CELL_SUBSTRATE_JUNCTION | 0.00000 | 0.00000 | | 0.67981 | 1.53607 | 292 |
| GO_CELL_SURFACE_RECEPTOR_SIGNALING_PATHWAY_INVC | 0.00001 | 0.00037 | 0.59333 | 0.54504 | 1.23416 | 318 |
| GO_CHAPERONE_BINDING | 0.00012 | 0.00289 | 0.51885 | 0.63654 | 1.38375 | 76 |
| GO_CHAPERONE_COMPLEX | 0.00000 | 0.00006 | 0.64355 | 0.84970 | 1.72685 | 22 |
| GO_COLLAGEN_CONTAINING_EXTRACELLULAR_MATRIX | 0.00022 | 0.00486 | 0.51885 | 0.61099 | 1.34011 | 98 |
| GO_COP9_SIGNALOSOME | 0.00065 | 0.01228 | 0.47727 | 0.74620 | 1.52420 | 24 |
| GO_COTRANSLATIONAL_PROTEIN_TARGETING_TO_MEMBR | 0.00000 | 0.00000 | | 0.89774 | 1.96556 | 91 |
| GO_CYTOKINE_MEDIATED_SIGNALING_PATHWAY | 0.00000 | 0.00000 | | 0.61683 | 1.39781 | 324 |
| GO_CYTOPLASMIC_STRESS_GRANULE | 0.00159 | 0.02639 | 0.45506 | 0.63015 | 1.35667 | 58 |
| GO_CYTOPLASMIC_TRANSLATION | 0.00000 | 0.00000 | | 0.84890 | 1.85444 | 85 |
| GO_CYTOPLASMIC_TRANSLATIONAL_INITIATION | 0.00000 | 0.00000 | 0.71951 | 0.85042 | 1.76196 | 29 |
| GO_CYTOSOLIC_LARGE_RIBOSOMAL_SUBUNIT | 0.00000 | 0.00000 | | 0.89401 | 1.91777 | 54 |
| GO_CYTOSOLIC_RIBOSOME | 0.00000 | 0.00000 | | 0.89680 | 1.96611 | 97 |
| GO_CYTOSOLIC_SMALL_RIBOSOMAL_SUBUNIT | 0.00000 | 0.00000 | | 0.90512 | 1.90950 | 39 |
| GO_DEFENSE_RESPONSE_TO_OTHER_ORGANISM | 0.00004 | 0.00112 | 0.55733 | 0.53056 | 1.20636 | 397 |
| GO_DE_NOVO_PROTEIN_FOLDING | 0.00265 | 0.03927 | 0.43171 | 0.69555 | 1.42992 | 26 |
| GO_DISORDERED_DOMAIN_SPECIFIC_BINDING | 0.00075 | 0.01382 | 0.47727 | 0.74375 | 1.51918 | 24 |
| GO_DISULFIDE_OXIDOREDUCTASE_ACTIVITY | 0.00268 | 0.03951 | 0.43171 | 0.68902 | 1.42571 | 28 |
| GO_DNA_BIOSYNTHETIC_PROCESS | 0.00001 | 0.00034 | 0.59333 | 0.60756 | 1.34994 | 138 |
| GO_DNA_CONFORMATION_CHANGE | 0.00209 | 0.03310 | 0.43171 | 0.53257 | 1.19513 | 218 |
| GO_DNA_DAMAGE_RESPONSE_DETECTION_OF_DNA_DAM | 0.00076 | 0.01390 | 0.47727 | 0.69192 | 1.45082 | 36 |
| GO_DNA_GEOMETRIC_CHANGE | 0.00145 | 0.02445 | 0.45506 | 0.59094 | 1.29613 | 98 |
| GO_DNA_SYNTHESIS_INVOLVED_IN_DNA_REPAIR | 0.00093 | 0.01651 | 0.47727 | 0.68255 | 1.44351 | 41 |
| GO_DOUBLE_STRANDED_RNA_BINDING | 0.00272 | 0.03985 | 0.43171 | 0.62680 | 1.34406 | 53 |
| GO_ELECTRON_TRANSFER_ACTIVITY | 0.00000 | 0.00018 | 0.61053 | 0.68139 | 1.47880 | 72 |
| GO_ELECTRON_TRANSPORT_CHAIN | 0.00000 | 0.00002 | 0.67496 | 0.64875 | 1.43710 | 124 |
| GO_ENDOPEPTIDASE_COMPLEX | 0.00000 | 0.00000 | | 0.78715 | 1.70035 | 64 |
| GO_ENERGY_DERIVATION_BY_OXIDATION_OF_ORGANIC_CO | 0.00000 | 0.00000 | 0.74774 | 0.63021 | 1.40658 | 176 |

| Cluster 1 | | | | | | |
|---|---|---|---|---|---|---|
| pathway | pval | padj | log2err | ES | NES | size |
| GO_ENZYME_INHIBITOR_ACTIVITY | 0.00287 | 0.04128 | 0.43171 | 0.54590 | 1.21502 | 146 |
| GO_ERROR_FREE_TRANSLESION_SYNTHESIS | 0.00261 | 0.03893 | 0.43171 | 0.76060 | 1.53039 | 19 |
| GO_ESTABLISHMENT_OF_PROTEIN_LOCALIZATION_TO_CHR | 0.00002 | 0.00063 | 0.57561 | 0.81488 | 1.65784 | 23 |
| GO_ESTABLISHMENT_OF_PROTEIN_LOCALIZATION_TO_END | 0.00000 | 0.00000 | | 0.87408 | 1.92066 | 102 |
| GO_ESTABLISHMENT_OF_PROTEIN_LOCALIZATION_TO_MEM | 0.00000 | 0.00000 | | 0.73761 | 1.66168 | 249 |
| GO_ESTABLISHMENT_OF_PROTEIN_LOCALIZATION_TO_ORG | 0.00000 | 0.00000 | | 0.67525 | 1.53705 | 427 |
| GO_ESTABLISHMENT_OF_PROTEIN_LOCALIZATION_TO_TEL | 0.00002 | 0.00048 | 0.57561 | 0.85998 | 1.71055 | 17 |
| GO_ESTABLISHMENT_OF_RNA_LOCALIZATION | 0.00246 | 0.03721 | 0.43171 | 0.54523 | 1.21719 | 168 |
| GO_ESTABLISHMENT_OF_TISSUE_POLARITY | 0.00000 | 0.00000 | 0.78819 | 0.72062 | 1.57421 | 85 |
| GO_EUKARYOTIC_TRANSLATION_INITIATION_FACTOR_3_CO | 0.00002 | 0.00052 | 0.57561 | 0.87052 | 1.71734 | 15 |
| GO_EXOCYTOSIS | 0.00000 | 0.00003 | 0.65944 | 0.53853 | 1.22736 | 462 |
| GO_EXTRACELLULAR_MATRIX | 0.00027 | 0.00580 | 0.49849 | 0.58899 | 1.30354 | 119 |
| GO_EXTRACELLULAR_MATRIX_STRUCTURAL_CONSTITUENT | 0.00172 | 0.02828 | 0.45506 | 0.71113 | 1.46881 | 27 |
| GO_FC_EPSILON_RECEPTOR_SIGNALING_PATHWAY | 0.00000 | 0.00000 | | 0.74155 | 1.62088 | 87 |
| GO_FC_RECEPTOR_MEDIATED_STIMULATORY_SIGNALING_P | 0.00106 | 0.01862 | 0.45506 | 0.64020 | 1.37231 | 52 |
| GO_FC_RECEPTOR_SIGNALING_PATHWAY | 0.00000 | 0.00000 | 0.81404 | 0.69013 | 1.52918 | 126 |
| GO_FERTILIZATION | 0.00065 | 0.01215 | 0.47727 | 0.63534 | 1.36919 | 61 |
| GO_FICOLIN_1_RICH_GRANULE | 0.00000 | 0.00000 | 0.80122 | 0.72218 | 1.58119 | 91 |
| GO_GENERATION_OF_PRECURSOR_METABOLITES_AND_EN | 0.00000 | 0.00000 | 0.71951 | 0.57246 | 1.29715 | 326 |
| GO_GLYCOSAMINOGLYCAN_BINDING | 0.00077 | 0.01414 | 0.47727 | 0.66613 | 1.41261 | 43 |
| GO_GUANYL_NUCLEOTIDE_BINDING | 0.00235 | 0.03630 | 0.43171 | 0.53479 | 1.19961 | 215 |
| GO_HEAT_SHOCK_PROTEIN_BINDING | 0.00010 | 0.00249 | 0.53843 | 0.62487 | 1.36504 | 85 |
| GO_HEMATOPOIETIC_PROGENITOR_CELL_DIFFERENTIATION | 0.00000 | 0.00000 | 0.83909 | 0.70397 | 1.55392 | 112 |
| GO_HEMATOPOIETIC_STEM_CELL_DIFFERENTIATION | 0.00000 | 0.00000 | | 0.78651 | 1.70640 | 70 |
| GO_HEPARIN_BINDING | 0.00108 | 0.01889 | 0.45506 | 0.69551 | 1.44513 | 30 |
| GO_IMMUNE_RESPONSE_REGULATING_SIGNALING_PATHW | 0.00000 | 0.00000 | 0.71951 | 0.60903 | 1.36214 | 196 |
| GO_INFLAMMATORY_RESPONSE_TO_ANTIGENIC_STIMULUS | 0.00005 | 0.00144 | 0.55733 | 0.85355 | 1.69247 | 16 |
| GO_INNATE_IMMUNE_RESPONSE | 0.00000 | 0.00004 | 0.64355 | 0.55859 | 1.26593 | 328 |
| GO_INNATE_IMMUNE_RESPONSE_ACTIVATING_SIGNAL_TR | 0.00000 | 0.00000 | | 0.77132 | 1.67674 | 76 |
| GO_INNER_MITOCHONDRIAL_MEMBRANE_PROTEIN_COMP | 0.00004 | 0.00120 | 0.55733 | 0.63442 | 1.38800 | 89 |
| GO_INTERACTION_WITH_SYMBIONT | 0.00001 | 0.00021 | 0.61053 | 0.73292 | 1.55708 | 45 |
| GO_INTERLEUKIN_1_MEDIATED_SIGNALING_PATHWAY | 0.00000 | 0.00000 | | 0.80838 | 1.75437 | 73 |
| GO_INTRINSIC_APOPTOTIC_SIGNALING_PATHWAY | 0.00115 | 0.01994 | 0.45506 | 0.54758 | 1.22379 | 187 |
| GO_LARGE_RIBOSOMAL_SUBUNIT | 0.00000 | 0.00000 | | 0.76525 | 1.68769 | 110 |
| GO_LEUKOCYTE_MEDIATED_IMMUNITY | 0.00000 | 0.00000 | 0.70498 | 0.56420 | 1.28130 | 360 |
| GO_MATURATION_OF_LSU_RRNA | 0.00027 | 0.00588 | 0.49849 | 0.74855 | 1.54609 | 27 |
| GO_MATURATION_OF_SSU_RRNA_FROM_TRICISTRONIC_R | 0.00239 | 0.03657 | 0.43171 | 0.67417 | 1.41319 | 35 |
| GO_MIDBRAIN_DEVELOPMENT | 0.00124 | 0.02127 | 0.45506 | 0.66271 | 1.40701 | 44 |
| GO_MITOCHONDRIAL_ELECTRON_TRANSPORT_NADH_TO_ | 0.00331 | 0.04604 | 0.43171 | 0.64232 | 1.36211 | 43 |
| GO_MITOCHONDRIAL_LARGE_RIBOSOMAL_SUBUNIT | 0.00025 | 0.00559 | 0.49849 | 0.65799 | 1.41379 | 55 |
| GO_MITOCHONDRIAL_MATRIX | 0.00010 | 0.00251 | 0.53843 | 0.52713 | 1.19729 | 367 |
| GO_MITOCHONDRIAL_PROTEIN_COMPLEX | 0.00000 | 0.00000 | 0.77494 | 0.61645 | 1.38221 | 210 |
| GO_MITOCHONDRIAL_TRANSLATIONAL_TERMINATION | 0.00030 | 0.00649 | 0.49849 | 0.61850 | 1.35277 | 88 |
| GO_MITOCHONDRION_ORGANIZATION | 0.00209 | 0.03310 | 0.43171 | 0.50740 | 1.15375 | 395 |
| GO_MODIFICATION_DEPENDENT_MACROMOLECULE_CATA | 0.00002 | 0.00051 | 0.57561 | 0.52442 | 1.19674 | 491 |
| GO_MODULATION_BY_HOST_OF_SYMBIONT_PROCESS | 0.00053 | 0.01025 | 0.47727 | 0.69890 | 1.46545 | 36 |
| GO_MODULATION_BY_HOST_OF_VIRAL_PROCESS | 0.00316 | 0.04464 | 0.43171 | 0.77584 | 1.53056 | 15 |
| GO_MODULATION_OF_PROCESS_OF_OTHER_ORGANISM | 0.00037 | 0.00773 | 0.49849 | 0.64360 | 1.38943 | 63 |
| GO_MORPHOGENESIS_OF_AN_EPITHELIUM | 0.00002 | 0.00068 | 0.57561 | 0.55184 | 1.24567 | 284 |
| GO_MORPHOGENESIS_OF_A_POLARIZED_EPITHELIUM | 0.00000 | 0.00000 | 0.82666 | 0.70951 | 1.55747 | 100 |
| GO_MRNA_3_UTR_BINDING | 0.00001 | 0.00018 | 0.61053 | 0.69494 | 1.49763 | 61 |
| GO_MRNA_5_UTR_BINDING | 0.00055 | 0.01074 | 0.47727 | 0.76665 | 1.55807 | 22 |
| GO_MRNA_BINDING | 0.00000 | 0.00000 | | 0.65249 | 1.46603 | 222 |
| GO_MRNA_PROCESSING | 0.00000 | 0.00003 | 0.65944 | 0.54373 | 1.23755 | 420 |
| GO_MULTI_ORGANISM_PROCESS | 0.00090 | 0.01612 | 0.47727 | 0.50391 | 1.14901 | 473 |
| GO_MYD88_DEPENDENT_TOLL_LIKE_RECEPTOR_SIGNALING | 0.00007 | 0.00174 | 0.53843 | 0.82159 | 1.64501 | 18 |
| GO_MYD88_INDEPENDENT_TOLL_LIKE_RECEPTOR_SIGNALI | 0.00012 | 0.00289 | 0.51885 | 0.83108 | 1.65307 | 17 |
| GO_MYELOID_LEUKOCYTE_ACTIVATION | 0.00000 | 0.00000 | 0.74774 | 0.57926 | 1.31185 | 320 |
| GO_MYELOID_LEUKOCYTE_MEDIATED_IMMUNITY | 0.00000 | 0.00000 | 0.76146 | 0.58830 | 1.32943 | 294 |
| GO_NADH_DEHYDROGENASE_ACTIVITY | 0.00357 | 0.04928 | 0.43171 | 0.66499 | 1.39396 | 35 |
| GO_NADH_DEHYDROGENASE_COMPLEX | 0.00327 | 0.04586 | 0.43171 | 0.66406 | 1.39832 | 38 |
| GO_NAD_BINDING | 0.00059 | 0.01138 | 0.47727 | 0.69241 | 1.45801 | 38 |
| GO_NCRNA_METABOLIC_PROCESS | 0.00002 | 0.00051 | 0.57561 | 0.53671 | 1.21942 | 382 |
| GO_NCRNA_PROCESSING | 0.00001 | 0.00020 | 0.61053 | 0.54888 | 1.24378 | 331 |
| GO_NEGATIVE_REGULATION_OF_CANONICAL_WNT_SIGNA | 0.00000 | 0.00000 | 0.74774 | 0.67599 | 1.49148 | 111 |
| GO_NEGATIVE_REGULATION_OF_CELLULAR_AMIDE_METAB | 0.00003 | 0.00084 | 0.57561 | 0.61893 | 1.36614 | 114 |
| GO_NEGATIVE_REGULATION_OF_CELL_CYCLE | 0.00038 | 0.00789 | 0.49849 | 0.51488 | 1.17188 | 422 |
| GO_NEGATIVE_REGULATION_OF_CELL_CYCLE_G2_M_PHAS | 0.00000 | 0.00001 | 0.69013 | 0.68435 | 1.49597 | 86 |
| GO_NEGATIVE_REGULATION_OF_CELL_CYCLE_PHASE_TRAN | 0.00019 | 0.00439 | 0.51885 | 0.56806 | 1.26885 | 183 |
| GO_NEGATIVE_REGULATION_OF_CELL_CYCLE_PROCESS | 0.00001 | 0.00020 | 0.61053 | 0.56427 | 1.27120 | 249 |

| Cluster 1 | | | | | | |
|---|---|---|---|---|---|---|
| pathway | pval | padj | log2err | ES | NES | size |
| GO_NEGATIVE_REGULATION_OF_MITOTIC_CELL_CYCLE | 0.00010 | 0.00243 | 0.53843 | 0.55113 | 1.23928 | 228 |
| GO_NEGATIVE_REGULATION_OF_MRNA_METABOLIC_PROC | 0.00137 | 0.02341 | 0.45506 | 0.62528 | 1.34750 | 61 |
| GO_NEGATIVE_REGULATION_OF_MRNA_SPLICING_VIA_SPL | 0.00276 | 0.03998 | 0.43171 | 0.75769 | 1.52453 | 19 |
| GO_NEGATIVE_REGULATION_OF_PROTEIN_CATABOLIC_PRO | 0.00246 | 0.03721 | 0.43171 | 0.58727 | 1.28366 | 87 |
| GO_NEGATIVE_REGULATION_OF_PROTEOLYSIS | 0.00063 | 0.01192 | 0.47727 | 0.56512 | 1.25736 | 145 |
| GO_NEGATIVE_REGULATION_OF_RNA_SPLICING | 0.00012 | 0.00289 | 0.51885 | 0.79020 | 1.60763 | 23 |
| GO_NEGATIVE_REGULATION_OF_WNT_SIGNALING_PATHW/ | 0.00000 | 0.00000 | 0.74774 | 0.66118 | 1.46553 | 128 |
| GO_NEURAL_NUCLEUS_DEVELOPMENT | 0.00006 | 0.00153 | 0.55733 | 0.77460 | 1.59991 | 27 |
| GO_NEURON_TO_NEURON_SYNAPSE | 0.00005 | 0.00132 | 0.55733 | 0.58674 | 1.30907 | 159 |
| GO_NIK_NF_KAPPAB_SIGNALING | 0.00000 | 0.00000 | | 0.72888 | 1.60515 | 106 |
| GO_NON_CANONICAL_WNT_SIGNALING_PATHWAY | 0.00000 | 0.00000 | 0.81404 | 0.71504 | 1.56941 | 99 |
| GO_NUCLEAR_EXPORT | 0.00052 | 0.01010 | 0.47727 | 0.55522 | 1.23918 | 175 |
| GO_NUCLEAR_TRANSCRIBED_MRNA_CATABOLIC_PROCESS | 0.00000 | 0.00000 | | 0.79452 | 1.77499 | 184 |
| GO_NUCLEAR_TRANSCRIBED_MRNA_CATABOLIC_PROCESS_ | 0.00000 | 0.00000 | | 0.88803 | 1.96019 | 112 |
| GO_NUCLEAR_TRANSPORT | 0.00016 | 0.00357 | 0.51885 | 0.54483 | 1.22945 | 274 |
| GO_NUCLEOBASE_CONTAINING_COMPOUND_TRANSPORT | 0.00194 | 0.03119 | 0.45506 | 0.53794 | 1.20430 | 202 |
| GO_NUCLEOBASE_CONTAINING_SMALL_MOLECULE_METAE | 0.00327 | 0.04586 | 0.43171 | 0.50690 | 1.15117 | 360 |
| GO_NUCLEOID | 0.00239 | 0.03657 | 0.43171 | 0.66838 | 1.40145 | 36 |
| GO_NUCLEOSIDE_METABOLIC_PROCESS | 0.00150 | 0.02516 | 0.45506 | 0.61838 | 1.33189 | 60 |
| GO_NUCLEOSIDE_PHOSPHATE_BIOSYNTHETIC_PROCESS | 0.00041 | 0.00837 | 0.49849 | 0.57696 | 1.28266 | 140 |
| GO_NUCLEOSIDE_TRIPHOSPHATE_BIOSYNTHETIC_PROCESS | 0.00250 | 0.03757 | 0.43171 | 0.66134 | 1.39668 | 40 |
| GO_NUCLEOSIDE_TRIPHOSPHATE_METABOLIC_PROCESS | 0.00010 | 0.00243 | 0.53843 | 0.65500 | 1.41076 | 60 |
| GO_NUCLEOSOMAL_DNA_BINDING | 0.00121 | 0.02083 | 0.45506 | 0.76526 | 1.54542 | 21 |
| GO_NUCLEOSOME_BINDING | 0.00115 | 0.01994 | 0.45506 | 0.66187 | 1.40881 | 46 |
| GO_NUCLEOTIDE_EXCISION_REPAIR | 0.00010 | 0.00245 | 0.53843 | 0.61704 | 1.35431 | 99 |
| GO_NUCLEOTIDE_EXCISION_REPAIR_DNA_DAMAGE_RECOG | 0.00002 | 0.00061 | 0.57561 | 0.81550 | 1.65910 | 23 |
| GO_NUCLEOTIDE_EXCISION_REPAIR_DNA_GAP_FILLING | 0.00070 | 0.01301 | 0.47727 | 0.76073 | 1.54605 | 22 |
| GO_NUCLEOTIDE_EXCISION_REPAIR_DNA_INCISION | 0.00064 | 0.01202 | 0.47727 | 0.69145 | 1.45599 | 38 |
| GO_ORGANELLAR_RIBOSOME | 0.00037 | 0.00773 | 0.49849 | 0.61133 | 1.33546 | 85 |
| GO_ORGANELLE_INNER_MEMBRANE | 0.00000 | 0.00004 | 0.64355 | 0.54925 | 1.24790 | 382 |
| GO_ORGANIC_CYCLIC_COMPOUND_CATABOLIC_PROCESS | 0.00000 | 0.00000 | | 0.71290 | 1.62380 | 440 |
| GO_OXIDATIVE_PHOSPHORYLATION | 0.00001 | 0.00031 | 0.59333 | 0.64605 | 1.41563 | 95 |
| GO_OXIDOREDUCTASE_ACTIVITY | 0.00003 | 0.00091 | 0.55733 | 0.53096 | 1.20761 | 402 |
| GO_OXIDOREDUCTASE_ACTIVITY_ACTING_ON_CH_OH_GRC | 0.00068 | 0.01271 | 0.47727 | 0.62408 | 1.35050 | 66 |
| GO_OXIDOREDUCTASE_ACTIVITY_ACTING_ON_NAD_P_H_C | 0.00242 | 0.03694 | 0.43171 | 0.64672 | 1.37145 | 43 |
| GO_OXIDOREDUCTASE_ACTIVITY_ACTING_ON_PEROXIDE_A | 0.00242 | 0.03694 | 0.43171 | 0.70717 | 1.44897 | 25 |
| GO_OXIDOREDUCTASE_COMPLEX | 0.00001 | 0.00020 | 0.61053 | 0.66488 | 1.45073 | 82 |
| GO_PEPTIDASE_COMPLEX | 0.00000 | 0.00000 | 0.78819 | 0.73152 | 1.59530 | 80 |
| GO_PEPTIDASE_REGULATOR_ACTIVITY | 0.00178 | 0.02877 | 0.45506 | 0.61054 | 1.32120 | 66 |
| GO_PHAGOCYTOSIS | 0.00353 | 0.04889 | 0.43171 | 0.56453 | 1.25055 | 124 |
| GO_PIGMENT_GRANULE | 0.00000 | 0.00002 | 0.65944 | 0.68244 | 1.48855 | 81 |
| GO_POLYSOMAL_RIBOSOME | 0.00000 | 0.00000 | | 0.89913 | 1.87436 | 31 |
| GO_POLYSOME | 0.00000 | 0.00000 | | 0.82099 | 1.77053 | 62 |
| GO_POLY_PYRIMIDINE_TRACT_BINDING | 0.00014 | 0.00317 | 0.51885 | 0.80806 | 1.61793 | 18 |
| GO_POSITIVE_REGULATION_OF_APOPTOTIC_SIGNALING_PA | 0.00298 | 0.04277 | 0.43171 | 0.56400 | 1.24090 | 104 |
| GO_POSITIVE_REGULATION_OF_CANONICAL_WNT_SIGNAL | 0.00000 | 0.00000 | 0.73376 | 0.69154 | 1.51623 | 96 |
| GO_POSITIVE_REGULATION_OF_CELLULAR_AMIDE_METABO | 0.00004 | 0.00108 | 0.55733 | 0.60818 | 1.34587 | 118 |
| GO_POSITIVE_REGULATION_OF_CELLULAR_PROTEIN_LOCAL | 0.00011 | 0.00255 | 0.53843 | 0.54978 | 1.23647 | 231 |
| GO_POSITIVE_REGULATION_OF_CELL_DEATH | 0.00048 | 0.00952 | 0.49849 | 0.51786 | 1.17605 | 359 |
| GO_POSITIVE_REGULATION_OF_DEFENSE_RESPONSE | 0.00001 | 0.00020 | 0.61053 | 0.59947 | 1.33782 | 163 |
| GO_POSITIVE_REGULATION_OF_DNA_BIOSYNTHETIC_PROC | 0.00000 | 0.00014 | 0.61053 | 0.71595 | 1.53524 | 53 |
| GO_POSITIVE_REGULATION_OF_DNA_METABOLIC_PROCESS | 0.00064 | 0.01202 | 0.47727 | 0.57744 | 1.28059 | 130 |
| GO_POSITIVE_REGULATION_OF_DNA_TEMPLATED_TRANSC | 0.00327 | 0.04586 | 0.43171 | 0.69044 | 1.41941 | 26 |
| GO_POSITIVE_REGULATION_OF_ESTABLISHMENT_OF_PROT | 0.00004 | 0.00114 | 0.55733 | 0.55858 | 1.25503 | 222 |
| GO_POSITIVE_REGULATION_OF_IMMUNE_RESPONSE | 0.00000 | 0.00000 | 0.71951 | 0.58822 | 1.32723 | 275 |
| GO_POSITIVE_REGULATION_OF_IMMUNE_SYSTEM_PROCES | 0.00001 | 0.00024 | 0.61053 | 0.53809 | 1.22272 | 383 |
| GO_POSITIVE_REGULATION_OF_INNATE_IMMUNE_RESPON | 0.00000 | 0.00000 | 0.74774 | 0.66787 | 1.47812 | 119 |
| GO_POSITIVE_REGULATION_OF_INTRINSIC_APOPTOTIC_SIG | 0.00148 | 0.02492 | 0.45506 | 0.68551 | 1.43696 | 35 |
| GO_POSITIVE_REGULATION_OF_PROTEIN_LOCALIZATION_T | 0.00001 | 0.00026 | 0.59333 | 0.69062 | 1.48832 | 61 |
| GO_POSITIVE_REGULATION_OF_RESPONSE_TO_BIOTIC_STI | 0.00000 | 0.00001 | 0.69013 | 0.64349 | 1.42913 | 135 |
| GO_POSITIVE_REGULATION_OF_RESPONSE_TO_EXTERNAL_ | 0.00000 | 0.00011 | 0.62726 | 0.58011 | 1.30072 | 210 |
| GO_POSITIVE_REGULATION_OF_SIGNAL_TRANSDUCTION_B | 0.00050 | 0.00981 | 0.47727 | 0.81733 | 1.61241 | 15 |
| GO_POSITIVE_REGULATION_OF_TELOMERASE_ACTIVITY | 0.00268 | 0.03951 | 0.43171 | 0.70390 | 1.45839 | 29 |
| GO_POSITIVE_REGULATION_OF_TELOMERASE_RNA_LOCAL | 0.00000 | 0.00002 | 0.67496 | 0.90994 | 1.79509 | 15 |
| GO_POSITIVE_REGULATION_OF_TELOMERE_MAINTENANCE | 0.00010 | 0.00239 | 0.53843 | 0.71125 | 1.50722 | 42 |
| GO_POSITIVE_REGULATION_OF_TELOMERE_MAINTENANCE | 0.00023 | 0.00523 | 0.51885 | 0.73501 | 1.53223 | 31 |
| GO_POSITIVE_REGULATION_OF_TRANSLATION | 0.00007 | 0.00172 | 0.53843 | 0.62546 | 1.37184 | 98 |
| GO_POSITIVE_REGULATION_OF_TRANSPORT | 0.00239 | 0.03657 | 0.43171 | 0.49727 | 1.13239 | 434 |
| GO_POSITIVE_REGULATION_OF_WNT_SIGNALING_PATHWA | 0.00000 | 0.00003 | 0.65944 | 0.64151 | 1.41632 | 113 |
| GO_POSTREPLICATION_REPAIR | 0.00008 | 0.00213 | 0.53843 | 0.69897 | 1.48495 | 45 |

| Cluster 1 | | | | | | |
|---|---|---|---|---|---|---|
| pathway | pval | padj | log2err | ES | NES | size |
| GO_POSTSYNAPSE | 0.00000 | 0.00007 | 0.62726 | 0.56727 | 1.27946 | 266 |
| GO_POSTTRANSCRIPTIONAL_REGULATION_OF_GENE_EXPR | 0.00000 | 0.00000 | | 0.62791 | 1.43118 | 465 |
| GO_POST_TRANSLATIONAL_PROTEIN_MODIFICATION | 0.00000 | 0.00002 | 0.67496 | 0.59005 | 1.32817 | 237 |
| GO_PRECATALYTIC_SPLICEOSOME | 0.00305 | 0.04361 | 0.43171 | 0.63103 | 1.34635 | 49 |
| GO_PRERIBOSOME | 0.00040 | 0.00815 | 0.49849 | 0.63108 | 1.36859 | 69 |
| GO_PRE_MRNA_BINDING | 0.00220 | 0.03438 | 0.43171 | 0.70871 | 1.46836 | 29 |
| GO_PROTEASOMAL_PROTEIN_CATABOLIC_PROCESS | 0.00001 | 0.00046 | 0.59333 | 0.53549 | 1.21645 | 378 |
| GO_PROTEASOMAL_UBIQUITIN_INDEPENDENT_PROTEIN_C | 0.00000 | 0.00000 | 0.80122 | 0.92289 | 1.85840 | 20 |
| GO_PROTEASOME_ACCESSORY_COMPLEX | 0.00000 | 0.00003 | 0.65944 | 0.84521 | 1.72642 | 24 |
| GO_PROTEASOME_CORE_COMPLEX | 0.00000 | 0.00000 | 0.82666 | 0.94081 | 1.88372 | 18 |
| GO_PROTEIN_CONTAINING_COMPLEX_DISASSEMBLY | 0.00007 | 0.00188 | 0.53843 | 0.55075 | 1.24209 | 255 |
| GO_PROTEIN_CONTAINING_COMPLEX_LOCALIZATION | 0.00276 | 0.03998 | 0.43171 | 0.53198 | 1.19095 | 202 |
| GO_PROTEIN_DNA_COMPLEX_SUBUNIT_ORGANIZATION | 0.00007 | 0.00181 | 0.53843 | 0.57540 | 1.28454 | 168 |
| GO_PROTEIN_DOMAIN_SPECIFIC_BINDING | 0.00231 | 0.03592 | 0.43171 | 0.50285 | 1.14374 | 406 |
| GO_PROTEIN_FOLDING | 0.00000 | 0.00000 | 0.73376 | 0.63929 | 1.42674 | 161 |
| GO_PROTEIN_LOCALIZATION_TO_CHROMOSOME_TELOMEF | 0.00020 | 0.00447 | 0.51885 | 0.74228 | 1.53591 | 28 |
| GO_PROTEIN_LOCALIZATION_TO_ENDOPLASMIC_RETICULU | 0.00000 | 0.00000 | | 0.84762 | 1.87799 | 125 |
| GO_PROTEIN_LOCALIZATION_TO_MEMBRANE | 0.00000 | 0.00000 | | 0.64976 | 1.47816 | 413 |
| GO_PROTEIN_LOCALIZATION_TO_MITOCHONDRION | 0.00046 | 0.00910 | 0.49849 | 0.58735 | 1.29649 | 112 |
| GO_PROTEIN_LOCALIZATION_TO_NUCLEUS | 0.00010 | 0.00244 | 0.53843 | 0.56819 | 1.26988 | 188 |
| GO_PROTEIN_MODIFICATION_BY_SMALL_PROTEIN_REMOV | 0.00000 | 0.00002 | 0.65944 | 0.58370 | 1.31088 | 221 |
| GO_PROTEIN_POLYUBIQUITINATION | 0.00122 | 0.02108 | 0.45506 | 0.52961 | 1.19471 | 265 |
| GO_PROTEIN_REFOLDING | 0.00104 | 0.01836 | 0.45506 | 0.79925 | 1.58480 | 16 |
| GO_PROTEIN_STABILIZATION | 0.00004 | 0.00099 | 0.55733 | 0.60366 | 1.33851 | 131 |
| GO_PROTEIN_TARGETING | 0.00000 | 0.00000 | | 0.70951 | 1.60919 | 338 |
| GO_PROTEIN_TARGETING_TO_MEMBRANE | 0.00000 | 0.00000 | | 0.82304 | 1.83533 | 155 |
| GO_PROTEIN_TRANSMEMBRANE_TRANSPORT | 0.00361 | 0.04967 | 0.43171 | 0.62551 | 1.33457 | 49 |
| GO_PURINE_CONTAINING_COMPOUND_BIOSYNTHETIC_PR | 0.00272 | 0.03985 | 0.43171 | 0.56859 | 1.25286 | 107 |
| GO_PURINE_NUCLEOSIDE_TRIPHOSPHATE_METABOLIC_PR | 0.00027 | 0.00580 | 0.49849 | 0.68212 | 1.44916 | 45 |
| GO_REGENERATION | 0.00194 | 0.03119 | 0.45506 | 0.59082 | 1.29141 | 87 |
| GO_REGULATION_OF_ANIMAL_ORGAN_MORPHOGENESIS | 0.00000 | 0.00000 | 0.81404 | 0.68391 | 1.51488 | 127 |
| GO_REGULATION_OF_APOPTOTIC_SIGNALING_PATHWAY | 0.00009 | 0.00234 | 0.53843 | 0.55160 | 1.24215 | 239 |
| GO_REGULATION_OF_CELLULAR_AMIDE_METABOLIC_PROC | 0.00000 | 0.00000 | | 0.61031 | 1.38158 | 315 |
| GO_REGULATION_OF_CELLULAR_AMINE_METABOLIC_PROC | 0.00000 | 0.00000 | | 0.81678 | 1.75847 | 58 |
| GO_REGULATION_OF_CELLULAR_AMINO_ACID_METABOLIC | 0.00000 | 0.00000 | | 0.82716 | 1.77437 | 54 |
| GO_REGULATION_OF_CELLULAR_KETONE_METABOLIC_PRO | 0.00000 | 0.00000 | | 0.70601 | 1.56115 | 117 |
| GO_REGULATION_OF_CELLULAR_PROTEIN_CATABOLIC_PRO | 0.00235 | 0.03630 | 0.43171 | 0.53880 | 1.20377 | 182 |
| GO_REGULATION_OF_CELLULAR_PROTEIN_LOCALIZATION | 0.00033 | 0.00685 | 0.49849 | 0.51860 | 1.17781 | 365 |
| GO_REGULATION_OF_CELL_CYCLE_G2_M_PHASE_TRANSITI | 0.00000 | 0.00007 | 0.64355 | 0.60230 | 1.34443 | 170 |
| GO_REGULATION_OF_CELL_CYCLE_PHASE_TRANSITION | 0.00001 | 0.00037 | 0.59333 | 0.54278 | 1.23037 | 334 |
| GO_REGULATION_OF_DEFENSE_RESPONSE | 0.00076 | 0.01402 | 0.47727 | 0.52502 | 1.18644 | 294 |
| GO_REGULATION_OF_DNA_BIOSYNTHETIC_PROCESS | 0.00063 | 0.01192 | 0.47727 | 0.62067 | 1.34959 | 77 |
| GO_REGULATION_OF_DNA_METABOLIC_PROCESS | 0.00279 | 0.04032 | 0.43171 | 0.52196 | 1.17391 | 231 |
| GO_REGULATION_OF_DNA_TEMPLATED_TRANSCRIPTION_I | 0.00313 | 0.04423 | 0.43171 | 0.68145 | 1.42057 | 31 |
| GO_REGULATION_OF_DNA_TEMPLATED_TRANSCRIPTION_I | 0.00000 | 0.00000 | | 0.75260 | 1.64505 | 87 |
| GO_REGULATION_OF_HEMATOPOIETIC_PROGENITOR_CELL_ | 0.00000 | 0.00000 | | 0.76985 | 1.67128 | 74 |
| GO_REGULATION_OF_HEMATOPOIETIC_STEM_CELL_DIFFER | 0.00000 | 0.00000 | | 0.80347 | 1.73455 | 63 |
| GO_REGULATION_OF_HEMOPOIESIS | 0.00002 | 0.00046 | 0.59333 | 0.57048 | 1.28282 | 225 |
| GO_REGULATION_OF_IMMUNE_RESPONSE | 0.00000 | 0.00007 | 0.64355 | 0.54902 | 1.24669 | 358 |
| GO_REGULATION_OF_INNATE_IMMUNE_RESPONSE | 0.00000 | 0.00001 | 0.69013 | 0.61918 | 1.38214 | 165 |
| GO_REGULATION_OF_INTRINSIC_APOPTOTIC_SIGNALING_P | 0.00001 | 0.00038 | 0.59333 | 0.62454 | 1.37498 | 105 |
| GO_REGULATION_OF_MITOTIC_CELL_CYCLE | 0.00000 | 0.00013 | 0.62726 | 0.53309 | 1.21450 | 450 |
| GO_REGULATION_OF_MORPHOGENESIS_OF_AN_EPITHELIU | 0.00000 | 0.00000 | 0.82666 | 0.70787 | 1.55617 | 103 |
| GO_REGULATION_OF_MRNA_CATABOLIC_PROCESS | 0.00000 | 0.00000 | | 0.67161 | 1.49882 | 163 |
| GO_REGULATION_OF_MRNA_METABOLIC_PROCESS | 0.00000 | 0.00000 | | 0.64097 | 1.44599 | 264 |
| GO_REGULATION_OF_MRNA_PROCESSING | 0.00011 | 0.00264 | 0.53843 | 0.60295 | 1.33087 | 114 |
| GO_REGULATION_OF_MRNA_SPLICING_VIA_SPLICEOSOME | 0.00001 | 0.00038 | 0.59333 | 0.65699 | 1.43350 | 82 |
| GO_REGULATION_OF_NUCLEASE_ACTIVITY | 0.00115 | 0.01994 | 0.45506 | 0.79480 | 1.57598 | 16 |
| GO_REGULATION_OF_OXIDATIVE_STRESS_INDUCED_CELL_D | 0.00039 | 0.00799 | 0.49849 | 0.69762 | 1.47539 | 41 |
| GO_REGULATION_OF_OXIDATIVE_STRESS_INDUCED_INTRIN | 0.00261 | 0.03893 | 0.43171 | 0.73224 | 1.48813 | 22 |
| GO_REGULATION_OF_PEPTIDASE_ACTIVITY | 0.00028 | 0.00595 | 0.49849 | 0.56110 | 1.25424 | 191 |
| GO_REGULATION_OF_PROTEIN_CATABOLIC_PROCESS | 0.00076 | 0.01390 | 0.47727 | 0.52949 | 1.19397 | 269 |
| GO_REGULATION_OF_PROTEIN_LOCALIZATION_TO_NUCLEU | 0.00001 | 0.00038 | 0.59333 | 0.64237 | 1.40410 | 87 |
| GO_REGULATION_OF_PROTEIN_STABILITY | 0.00011 | 0.00257 | 0.53843 | 0.55670 | 1.24771 | 207 |
| GO_REGULATION_OF_PROTEOLYSIS | 0.00014 | 0.00317 | 0.51885 | 0.52463 | 1.19157 | 375 |
| GO_REGULATION_OF_RESPONSE_TO_BIOTIC_STIMULUS | 0.00001 | 0.00023 | 0.61053 | 0.57711 | 1.29327 | 206 |
| GO_REGULATION_OF_RESPONSE_TO_EXTERNAL_STIMULUS | 0.00061 | 0.01165 | 0.47727 | 0.50949 | 1.15975 | 428 |
| GO_REGULATION_OF_RESPONSE_TO_OXIDATIVE_STRESS | 0.00246 | 0.03721 | 0.43171 | 0.62940 | 1.34682 | 51 |
| GO_REGULATION_OF_RNA_SPLICING | 0.00000 | 0.00002 | 0.67496 | 0.64136 | 1.41731 | 116 |

| **Cluster 1** | | | | | | |
|---|---|---|---|---|---|---|
| pathway | pval | padj | log2err | ES | NES | size |
| GO_REGULATION_OF_SMALL_MOLECULE_METABOLIC_PRO | 0.00002 | 0.00061 | 0.57561 | 0.55996 | 1.26319 | 272 |
| GO_REGULATION_OF_STEM_CELL_DIFFERENTIATION | 0.00000 | 0.00000 | | 0.74612 | 1.63190 | 88 |
| GO_REGULATION_OF_TELOMERE_MAINTENANCE | 0.00093 | 0.01651 | 0.47727 | 0.62711 | 1.35796 | 68 |
| GO_REGULATION_OF_TELOMERE_MAINTENANCE_VIA_TELO | 0.00031 | 0.00657 | 0.49849 | 0.66171 | 1.41841 | 52 |
| GO_REGULATION_OF_TRANSCRIPTION_FROM_RNA_POLYM | 0.00000 | 0.00000 | | 0.80954 | 1.74871 | 64 |
| GO_REGULATION_OF_TRANSLATIONAL_INITIATION | 0.00039 | 0.00795 | 0.49849 | 0.64311 | 1.38836 | 63 |
| GO_REGULATION_OF_VIRAL_GENOME_REPLICATION | 0.00309 | 0.04392 | 0.43171 | 0.62380 | 1.33764 | 53 |
| GO_REGULATION_OF_WNT_SIGNALING_PATHWAY | 0.00002 | 0.00061 | 0.57561 | 0.57159 | 1.28218 | 215 |
| GO_RESPIRASOME | 0.00001 | 0.00037 | 0.59333 | 0.66933 | 1.45218 | 70 |
| GO_RESPIRATORY_CHAIN_COMPLEX | 0.00001 | 0.00026 | 0.59333 | 0.68361 | 1.47239 | 60 |
| GO_RESPIRATORY_ELECTRON_TRANSPORT_CHAIN | 0.00000 | 0.00011 | 0.62726 | 0.66261 | 1.44845 | 86 |
| GO_RESPONSE_TO_CADMIUM_ION | 0.00276 | 0.03998 | 0.43171 | 0.66864 | 1.40588 | 37 |
| GO_RESPONSE_TO_DRUG | 0.00313 | 0.04423 | 0.43171 | 0.53748 | 1.20122 | 187 |
| GO_RESPONSE_TO_HORMONE | 0.00159 | 0.02639 | 0.45506 | 0.50181 | 1.14343 | 457 |
| GO_RESPONSE_TO_INORGANIC_SUBSTANCE | 0.00060 | 0.01152 | 0.47727 | 0.53571 | 1.20889 | 274 |
| GO_RESPONSE_TO_INTERLEUKIN_1 | 0.00000 | 0.00000 | | 0.73612 | 1.62109 | 106 |
| GO_RESPONSE_TO_INTERLEUKIN_12 | 0.00007 | 0.00179 | 0.53843 | 0.74424 | 1.55324 | 32 |
| GO_RESPONSE_TO_INTERLEUKIN_4 | 0.00039 | 0.00799 | 0.49849 | 0.78181 | 1.57431 | 20 |
| GO_RESPONSE_TO_INTERLEUKIN_7 | 0.00265 | 0.03927 | 0.43171 | 0.74719 | 1.50893 | 21 |
| GO_RESPONSE_TO_METAL_ION | 0.00174 | 0.02842 | 0.45506 | 0.54365 | 1.21393 | 169 |
| GO_RESPONSE_TO_ORGANIC_CYCLIC_COMPOUND | 0.00009 | 0.00220 | 0.53843 | 0.52120 | 1.18720 | 443 |
| GO_RESPONSE_TO_ORGANOPHOSPHORUS | 0.00051 | 0.00996 | 0.47727 | 0.65187 | 1.39835 | 54 |
| GO_RESPONSE_TO_OXYGEN_LEVELS | 0.00000 | 0.00001 | 0.69013 | 0.59259 | 1.33297 | 233 |
| GO_RESPONSE_TO_TOPOLOGICALLY_INCORRECT_PROTEIN | 0.00209 | 0.03310 | 0.43171 | 0.55452 | 1.23313 | 141 |
| GO_RESPONSE_TO_TUMOR_NECROSIS_FACTOR | 0.00000 | 0.00000 | 0.78819 | 0.64839 | 1.44595 | 154 |
| GO_RETINA_HOMEOSTASIS | 0.00124 | 0.02127 | 0.45506 | 0.75120 | 1.52667 | 22 |
| GO_RIBONUCLEOPROTEIN_COMPLEX_BINDING | 0.00000 | 0.00001 | 0.69013 | 0.65737 | 1.45099 | 114 |
| GO_RIBONUCLEOPROTEIN_COMPLEX_BIOGENESIS | 0.00000 | 0.00000 | | 0.63654 | 1.44682 | 389 |
| GO_RIBONUCLEOPROTEIN_COMPLEX_SUBUNIT_ORGANIZAT | 0.00000 | 0.00000 | | 0.68224 | 1.52345 | 177 |
| GO_RIBONUCLEOPROTEIN_GRANULE | 0.00000 | 0.00003 | 0.65944 | 0.60885 | 1.35922 | 168 |
| GO_RIBONUCLEOSIDE_TRIPHOSPHATE_BIOSYNTHETIC_PRO | 0.00100 | 0.01776 | 0.45506 | 0.69954 | 1.46422 | 34 |
| GO_RIBONUCLEOSIDE_TRIPHOSPHATE_METABOLIC_PROCES | 0.00009 | 0.00233 | 0.53843 | 0.70204 | 1.49050 | 44 |
| GO_RIBOSE_PHOSPHATE_BIOSYNTHETIC_PROCESS | 0.00046 | 0.00910 | 0.49849 | 0.58295 | 1.28342 | 105 |
| GO_RIBOSE_PHOSPHATE_METABOLIC_PROCESS | 0.00216 | 0.03399 | 0.43171 | 0.52592 | 1.18484 | 250 |
| GO_RIBOSOMAL_LARGE_SUBUNIT_ASSEMBLY | 0.00004 | 0.00099 | 0.55733 | 0.76942 | 1.59207 | 28 |
| GO_RIBOSOMAL_LARGE_SUBUNIT_BIOGENESIS | 0.00000 | 0.00000 | 0.82666 | 0.76428 | 1.65390 | 66 |
| GO_RIBOSOMAL_SMALL_SUBUNIT_ASSEMBLY | 0.00042 | 0.00843 | 0.49849 | 0.81397 | 1.61904 | 17 |
| GO_RIBOSOMAL_SMALL_SUBUNIT_BIOGENESIS | 0.00001 | 0.00030 | 0.59333 | 0.68300 | 1.47896 | 68 |
| GO_RIBOSOMAL_SUBUNIT | 0.00000 | 0.00000 | | 0.75841 | 1.69284 | 173 |
| GO_RIBOSOME | 0.00000 | 0.00000 | | 0.74768 | 1.67574 | 207 |
| GO_RIBOSOME_ASSEMBLY | 0.00000 | 0.00000 | 0.74774 | 0.75218 | 1.62008 | 60 |
| GO_RIBOSOME_BINDING | 0.00031 | 0.00664 | 0.49849 | 0.68069 | 1.44612 | 45 |
| GO_RIBOSOME_BIOGENESIS | 0.00000 | 0.00000 | | 0.63313 | 1.42826 | 272 |
| GO_RNA_CATABOLIC_PROCESS | 0.00000 | 0.00000 | | 0.75375 | 1.70861 | 334 |
| GO_RNA_DEPENDENT_DNA_BIOSYNTHETIC_PROCESS | 0.00003 | 0.00090 | 0.55733 | 0.66789 | 1.43854 | 60 |
| GO_RNA_EXPORT_FROM_NUCLEUS | 0.00090 | 0.01612 | 0.47727 | 0.56887 | 1.26091 | 128 |
| GO_RNA_LOCALIZATION | 0.00009 | 0.00215 | 0.53843 | 0.56815 | 1.27098 | 198 |
| GO_RNA_SPLICING | 0.00000 | 0.00001 | 0.67496 | 0.55687 | 1.26490 | 374 |
| GO_RNA_SPLICING_VIA_TRANSESTERIFICATION_REACTIONS | 0.00000 | 0.00000 | 0.71951 | 0.57623 | 1.30317 | 306 |
| GO_ROUGH_ENDOPLASMIC_RETICULUM | 0.00004 | 0.00102 | 0.55733 | 0.69091 | 1.48099 | 52 |
| GO_RRNA_BINDING | 0.00000 | 0.00000 | 0.70498 | 0.74945 | 1.60766 | 54 |
| GO_RRNA_METABOLIC_PROCESS | 0.00000 | 0.00001 | 0.69013 | 0.60274 | 1.35013 | 204 |
| GO_SCF_DEPENDENT_PROTEASOMAL_UBIQUITIN_DEPEND | 0.00000 | 0.00000 | 0.80122 | 0.73887 | 1.60808 | 78 |
| GO_SECRETORY_GRANULE | 0.00000 | 0.00000 | 0.82666 | 0.58176 | 1.32283 | 395 |
| GO_SECRETORY_VESICLE | 0.00000 | 0.00000 | 0.71951 | 0.54852 | 1.25012 | 462 |
| GO_SIGNAL_TRANSDUCTION_BY_PROTEIN_PHOSPHORYLAT | 0.00000 | 0.00004 | 0.64355 | 0.54338 | 1.23662 | 419 |
| GO_SINGLE_FERTILIZATION | 0.00006 | 0.00155 | 0.55733 | 0.70884 | 1.50495 | 44 |
| GO_SINGLE_STRANDED_DNA_BINDING | 0.00090 | 0.01612 | 0.47727 | 0.59891 | 1.31129 | 91 |
| GO_SINGLE_STRANDED_RNA_BINDING | 0.00018 | 0.00416 | 0.51885 | 0.64652 | 1.39249 | 60 |
| GO_SMALL_RIBOSOMAL_SUBUNIT | 0.00000 | 0.00000 | | 0.78543 | 1.69967 | 66 |
| GO_SPERM_EGG_RECOGNITION | 0.00000 | 0.00001 | 0.69013 | 0.91357 | 1.81147 | 16 |
| GO_SPLICEOSOMAL_COMPLEX | 0.00002 | 0.00053 | 0.57561 | 0.58610 | 1.30831 | 172 |
| GO_STEM_CELL_DIFFERENTIATION | 0.00000 | 0.00000 | 0.81404 | 0.65841 | 1.46590 | 149 |
| GO_STRUCTURAL_CONSTITUENT_OF_CYTOSKELETON | 0.00163 | 0.02692 | 0.45506 | 0.63877 | 1.36163 | 48 |
| GO_STRUCTURAL_CONSTITUENT_OF_RIBOSOME | 0.00000 | 0.00000 | | 0.77202 | 1.72007 | 151 |
| GO_STRUCTURAL_MOLECULE_ACTIVITY | 0.00000 | 0.00000 | | 0.71853 | 1.62751 | 321 |
| GO_SUBSTANTIA_NIGRA_DEVELOPMENT | 0.00002 | 0.00049 | 0.57561 | 0.80627 | 1.64689 | 24 |
| GO_TELOMERASE_RNA_LOCALIZATION | 0.00001 | 0.00032 | 0.59333 | 0.85022 | 1.71070 | 19 |
| GO_TELOMERE_MAINTENANCE_VIA_TELOMERE_LENGTHEN | 0.00057 | 0.01106 | 0.47727 | 0.63519 | 1.37545 | 68 |
| GO_TELOMERE_ORGANIZATION | 0.00045 | 0.00898 | 0.49849 | 0.58630 | 1.29902 | 125 |

| Cluster 1 | | | | | | |
|---|---|---|---|---|---|---|
| pathway | pval | padj | log2err | ES | NES | size |
| GO_THREONINE_TYPE_ENDOPEPTIDASE_ACTIVITY | 0.00000 | 0.00000 | | 0.95352 | 1.90918 | 18 |
| GO_THREONINE_TYPE_PEPTIDASE_ACTIVITY | 0.00000 | 0.00000 | 0.78819 | 0.91786 | 1.85359 | 21 |
| GO_TISSUE_MORPHOGENESIS | 0.00026 | 0.00565 | 0.49849 | 0.52891 | 1.19771 | 317 |
| GO_TOXIN_TRANSPORT | 0.00010 | 0.00245 | 0.53843 | 0.77094 | 1.57964 | 25 |
| GO_TRANSCRIPTION_COUPLED_NUCLEOTIDE_EXCISION_RE | 0.00003 | 0.00097 | 0.55733 | 0.66012 | 1.43240 | 71 |
| GO_TRANSCRIPTION_FACTOR_BINDING | 0.00331 | 0.04604 | 0.43171 | 0.50251 | 1.14292 | 402 |
| GO_TRANSLATIONAL_ELONGATION | 0.00001 | 0.00025 | 0.61053 | 0.62139 | 1.37523 | 120 |
| GO_TRANSLATIONAL_INITIATION | 0.00000 | 0.00000 | | 0.83714 | 1.86866 | 165 |
| GO_TRANSLATIONAL_TERMINATION | 0.00007 | 0.00183 | 0.53843 | 0.62087 | 1.36272 | 99 |
| GO_TRANSLATION_FACTOR_ACTIVITY_RNA_BINDING | 0.00000 | 0.00000 | | 0.77412 | 1.67518 | 66 |
| GO_TRANSLATION_INITIATION_FACTOR_ACTIVITY | 0.00000 | 0.00000 | 0.73376 | 0.78258 | 1.66258 | 45 |
| GO_TRANSLATION_INITIATION_FACTOR_BINDING | 0.00050 | 0.00981 | 0.47727 | 0.72434 | 1.49880 | 28 |
| GO_TRANSLATION_PREINITIATION_COMPLEX | 0.00000 | 0.00013 | 0.62726 | 0.88753 | 1.75985 | 16 |
| GO_TRANSLATION_REGULATOR_ACTIVITY | 0.00000 | 0.00000 | | 0.74248 | 1.63465 | 105 |
| GO_TRANSLATION_REGULATOR_ACTIVITY_NUCLEIC_ACID_E | 0.00000 | 0.00000 | | 0.75200 | 1.63996 | 80 |
| GO_TRANSLESION_SYNTHESIS | 0.00032 | 0.00671 | 0.49849 | 0.71480 | 1.49836 | 35 |
| GO_TRICARBOXYLIC_ACID_CYCLE | 0.00018 | 0.00407 | 0.51885 | 0.74348 | 1.53839 | 28 |
| GO_TUMOR_NECROSIS_FACTOR_MEDIATED_SIGNALING_PA | 0.00000 | 0.00000 | | 0.73757 | 1.61774 | 98 |
| GO_T_CELL_RECEPTOR_SIGNALING_PATHWAY | 0.00000 | 0.00000 | 0.71951 | 0.66924 | 1.47596 | 110 |
| GO_U12_TYPE_SPLICEOSOMAL_COMPLEX | 0.00084 | 0.01517 | 0.47727 | 0.72654 | 1.50063 | 27 |
| GO_U2_SNRNP | 0.00013 | 0.00298 | 0.51885 | 0.80933 | 1.62048 | 18 |
| GO_U2_TYPE_CATALYTIC_STEP_2_SPLICEOSOME | 0.00067 | 0.01258 | 0.47727 | 0.70771 | 1.47047 | 30 |
| GO_U2_TYPE_SPLICEOSOMAL_COMPLEX | 0.00178 | 0.02877 | 0.45506 | 0.59371 | 1.29699 | 85 |
| GO_UBIQUITIN_LIKE_PROTEIN_LIGASE_BINDING | 0.00000 | 0.00000 | 0.77494 | 0.61263 | 1.37457 | 216 |
| GO_UBIQUITIN_PROTEIN_TRANSFERASE_REGULATOR_ACTI' | 0.00165 | 0.02715 | 0.45506 | 0.79175 | 1.56194 | 15 |
| GO_UNFOLDED_PROTEIN_BINDING | 0.00000 | 0.00000 | 0.77494 | 0.72346 | 1.57855 | 82 |
| GO_VACUOLAR_LUMEN | 0.00019 | 0.00430 | 0.51885 | 0.61995 | 1.35520 | 86 |
| GO_VESICLE_LUMEN | 0.00000 | 0.00000 | | 0.66802 | 1.49129 | 166 |
| GO_VIRAL_GENE_EXPRESSION | 0.00000 | 0.00000 | | 0.78978 | 1.76297 | 172 |
| GO_VIRAL_GENOME_REPLICATION | 0.00279 | 0.04032 | 0.43171 | 0.60561 | 1.31412 | 71 |
| GO_VIRAL_LIFE_CYCLE | 0.00041 | 0.00830 | 0.49849 | 0.54699 | 1.22595 | 207 |
| HP_ABNORMALITY_OF_ACID_BASE_HOMEOSTASIS | 0.00087 | 0.01561 | 0.47727 | 0.53355 | 1.19997 | 231 |
| HP_ABNORMALITY_OF_CARDIOVASCULAR_SYSTEM_ELECTR | 0.00005 | 0.00127 | 0.55733 | 0.55670 | 1.25181 | 228 |
| HP_ABNORMALITY_OF_LOWER_LIP | 0.00146 | 0.02468 | 0.45506 | 0.55650 | 1.23894 | 148 |
| HP_ABNORMALITY_OF_THE_MITOCHONDRION | 0.00035 | 0.00731 | 0.49849 | 0.59011 | 1.30253 | 114 |
| HP_ABNORMALITY_OF_THE_NASAL_DORSUM | 0.00033 | 0.00685 | 0.49849 | 0.57966 | 1.28688 | 134 |
| HP_ABNORMALITY_OF_VITAMIN_METABOLISM | 0.00154 | 0.02562 | 0.45506 | 0.70891 | 1.45739 | 26 |
| HP_ABNORMAL_ACTIVITY_OF_MITOCHONDRIAL_RESPIRATO | 0.00331 | 0.04604 | 0.43171 | 0.61926 | 1.33321 | 58 |
| HP_ABNORMAL_CSF_METABOLITE_LEVEL | 0.00080 | 0.01455 | 0.47727 | 0.61720 | 1.34203 | 77 |
| HP_ABNORMAL_ENZYME_COENZYME_ACTIVITY | 0.00194 | 0.03119 | 0.45506 | 0.57147 | 1.26145 | 112 |
| HP_ABNORMAL_ERYTHROCYTE_MORPHOLOGY | 0.00000 | 0.00002 | 0.67496 | 0.56523 | 1.28089 | 324 |
| HP_ABNORMAL_RETICULOCYTE_MORPHOLOGY | 0.00176 | 0.02863 | 0.45506 | 0.71088 | 1.46830 | 27 |
| HP_ACUTE_LEUKEMIA | 0.00000 | 0.00000 | 0.77494 | 0.74501 | 1.61219 | 66 |
| HP_ANEMIA_OF_INADEQUATE_PRODUCTION | 0.00001 | 0.00037 | 0.59333 | 0.64579 | 1.41393 | 91 |
| HP_DELAYED_PUBERTY | 0.00000 | 0.00003 | 0.65944 | 0.64984 | 1.43069 | 105 |
| HP_DEPRESSED_NASAL_RIDGE | 0.00000 | 0.00016 | 0.61053 | 0.67144 | 1.45764 | 74 |
| HP_ECHOLALIA | 0.00224 | 0.03487 | 0.43171 | 0.70064 | 1.44038 | 26 |
| HP_FATIGUE | 0.00000 | 0.00016 | 0.61053 | 0.59373 | 1.32626 | 180 |
| HP_HEADACHE | 0.00001 | 0.00019 | 0.61053 | 0.61275 | 1.36224 | 140 |
| HP_HEMATOLOGICAL_NEOPLASM | 0.00004 | 0.00100 | 0.55733 | 0.59216 | 1.31865 | 150 |
| HP_INCREASED_CSF_LACTATE | 0.00023 | 0.00509 | 0.51885 | 0.63729 | 1.37663 | 64 |
| HP_LEUKEMIA | 0.00000 | 0.00000 | 0.77494 | 0.70705 | 1.54851 | 94 |
| HP_LEUKODYSTROPHY | 0.00029 | 0.00623 | 0.49849 | 0.62039 | 1.35115 | 79 |
| HP_LEUKOENCEPHALOPATHY | 0.00213 | 0.03360 | 0.43171 | 0.56989 | 1.26243 | 124 |
| HP_MACROCYTIC_ANEMIA | 0.00000 | 0.00000 | 0.76146 | 0.81354 | 1.71629 | 39 |
| HP_MIGRAINE | 0.00000 | 0.00000 | | 0.76916 | 1.66275 | 65 |
| HP_NEOPLASM | 0.00350 | 0.04850 | 0.43171 | 0.49697 | 1.13226 | 445 |
| HP_ORAL_CLEFT | 0.00209 | 0.03310 | 0.43171 | 0.51234 | 1.16191 | 337 |
| HP_PALLOR | 0.00000 | 0.00009 | 0.62726 | 0.65471 | 1.43106 | 87 |
| HP_PAROXYSMAL_INVOLUNTARY_EYE_MOVEMENTS | 0.00216 | 0.03399 | 0.43171 | 0.69511 | 1.43832 | 28 |
| HP_POOR_HEAD_CONTROL | 0.00209 | 0.03310 | 0.43171 | 0.57813 | 1.26892 | 99 |
| HP_PROXIMAL_TUBULOPATHY | 0.00276 | 0.03998 | 0.43171 | 0.68407 | 1.42605 | 31 |
| HP_SHORT_NOSE | 0.00012 | 0.00287 | 0.53843 | 0.56601 | 1.26425 | 179 |
| HP_THICK_LOWER_LIP_VERMILION | 0.00026 | 0.00565 | 0.49849 | 0.60976 | 1.33741 | 98 |
| HP_THICK_VERMILION_BORDER | 0.00043 | 0.00867 | 0.49849 | 0.56573 | 1.26160 | 154 |
| HP_TRIANGULAR_SHAPED_PHALANGES_OF_THE_HAND | 0.00220 | 0.03438 | 0.43171 | 0.57781 | 1.26821 | 99 |
| **Cluster 13** | | | | | | |
| pathway | pval | padj | log2err | ES | NES | size |
| GO_ALTERNATIVE_MRNA_SPLICING_VIA_SPLICEOSOME | 0.00011 | 0.00585 | 0.53843 | 0.67580 | 1.38093 | 61 |
| GO_ANAPHASE_PROMOTING_COMPLEX_DEPENDENT_CATA | 0.00115 | 0.03441 | 0.45506 | 0.61761 | 1.27432 | 74 |

| **Cluster 1** | | | | | | |
|---|---|---|---|---|---|---|
| pathway | pval | padj | log2err | ES | NES | size |
| GO_CATALYTIC_STEP_2_SPLICEOSOME | 0.00001 | 0.00062 | 0.61053 | 0.67366 | 1.39266 | 77 |
| GO_CELLULAR_NITROGEN_COMPOUND_CATABOLIC_PROCE | 0.00000 | 0.00000 | | 0.64563 | 1.38684 | 414 |
| GO_CELLULAR_RESPONSE_TO_HEAT | 0.00043 | 0.01673 | 0.49849 | 0.62596 | 1.29472 | 83 |
| GO_CELL_CYCLE_G1_S_PHASE_TRANSITION | 0.00016 | 0.00722 | 0.51885 | 0.57593 | 1.22164 | 200 |
| GO_CELL_CYCLE_PHASE_TRANSITION | 0.00001 | 0.00074 | 0.59333 | 0.54721 | 1.17673 | 461 |
| GO_CELL_DIVISION | 0.00000 | 0.00000 | 0.73376 | 0.57640 | 1.23835 | 418 |
| GO_CELL_SUBSTRATE_JUNCTION | 0.00000 | 0.00000 | 0.73376 | 0.60329 | 1.28822 | 281 |
| GO_CHROMATIN_ASSEMBLY_OR_DISASSEMBLY | 0.00035 | 0.01436 | 0.49849 | 0.60373 | 1.25935 | 109 |
| GO_CHROMATIN_BINDING | 0.00000 | 0.00005 | 0.65944 | 0.57200 | 1.22731 | 376 |
| GO_CHROMATIN_ORGANIZATION_INVOLVED_IN_REGULATI | 0.00146 | 0.04121 | 0.45506 | 0.61644 | 1.27086 | 72 |
| GO_CHROMOSOMAL_REGION | 0.00015 | 0.00697 | 0.51885 | 0.56102 | 1.19754 | 278 |
| GO_CHROMOSOME_CENTROMERIC_REGION | 0.00005 | 0.00283 | 0.55733 | 0.59471 | 1.25346 | 159 |
| GO_CHROMOSOME_SEGREGATION | 0.00004 | 0.00277 | 0.55733 | 0.57004 | 1.21609 | 252 |
| GO_CONDENSED_CHROMOSOME | 0.00028 | 0.01172 | 0.49849 | 0.58209 | 1.22736 | 162 |
| GO_COTRANSLATIONAL_PROTEIN_TARGETING_TO_MEMBR | 0.00000 | 0.00000 | | 0.87364 | 1.81450 | 91 |
| GO_COVALENT_CHROMATIN_MODIFICATION | 0.00001 | 0.00045 | 0.61053 | 0.56634 | 1.21373 | 336 |
| GO_CYTOKINESIS | 0.00073 | 0.02473 | 0.47727 | 0.59565 | 1.24486 | 113 |
| GO_CYTOPLASMIC_STRESS_GRANULE | 0.00003 | 0.00168 | 0.57561 | 0.68969 | 1.40487 | 59 |
| GO_CYTOPLASMIC_TRANSLATION | 0.00000 | 0.00000 | 0.80122 | 0.72803 | 1.51081 | 88 |
| GO_CYTOSOLIC_LARGE_RIBOSOMAL_SUBUNIT | 0.00000 | 0.00000 | | 0.85040 | 1.72499 | 52 |
| GO_CYTOSOLIC_RIBOSOME | 0.00000 | 0.00000 | | 0.87398 | 1.81735 | 95 |
| GO_CYTOSOLIC_SMALL_RIBOSOMAL_SUBUNIT | 0.00000 | 0.00000 | | 0.91569 | 1.82991 | 41 |
| GO_DNA_BINDING_TRANSCRIPTION_FACTOR_BINDING | 0.00126 | 0.03677 | 0.45506 | 0.55282 | 1.17628 | 224 |
| GO_DNA_CONFORMATION_CHANGE | 0.00020 | 0.00884 | 0.51885 | 0.56852 | 1.20769 | 214 |
| GO_DNA_DAMAGE_RESPONSE_DETECTION_OF_DNA_DAM | 0.00054 | 0.01945 | 0.47727 | 0.71595 | 1.41998 | 33 |
| GO_DNA_PACKAGING | 0.00056 | 0.02032 | 0.47727 | 0.59587 | 1.24755 | 120 |
| GO_DNA_REPAIR | 0.00028 | 0.01162 | 0.49849 | 0.53686 | 1.15305 | 408 |
| GO_DNA_REPLICATION | 0.00008 | 0.00457 | 0.53843 | 0.57646 | 1.22417 | 208 |
| GO_DNA_TEMPLATED_TRANSCRIPTION_ELONGATION | 0.00003 | 0.00195 | 0.57561 | 0.63741 | 1.32542 | 95 |
| GO_DNA_TEMPLATED_TRANSCRIPTION_INITIATION | 0.00001 | 0.00106 | 0.59333 | 0.60139 | 1.27035 | 173 |
| GO_DNA_TEMPLATED_TRANSCRIPTION_TERMINATION | 0.00010 | 0.00518 | 0.53843 | 0.65714 | 1.34842 | 66 |
| GO_ESTABLISHMENT_OF_PROTEIN_LOCALIZATION_TO_END | 0.00000 | 0.00000 | | 0.85894 | 1.78891 | 101 |
| GO_ESTABLISHMENT_OF_PROTEIN_LOCALIZATION_TO_MEM | 0.00000 | 0.00000 | | 0.68308 | 1.45595 | 245 |
| GO_ESTABLISHMENT_OF_PROTEIN_LOCALIZATION_TO_ORG | 0.00000 | 0.00000 | | 0.63671 | 1.36744 | 398 |
| GO_ESTABLISHMENT_OF_RNA_LOCALIZATION | 0.00000 | 0.00000 | 0.74774 | 0.65309 | 1.37690 | 160 |
| GO_FICOLIN_1_RICH_GRANULE | 0.00013 | 0.00627 | 0.51885 | 0.62942 | 1.30753 | 92 |
| GO_HEAT_SHOCK_PROTEIN_BINDING | 0.00013 | 0.00627 | 0.51885 | 0.62741 | 1.30245 | 89 |
| GO_HETEROCHROMATIN | 0.00084 | 0.02703 | 0.47727 | 0.64744 | 1.31719 | 57 |
| GO_HISTONE_BINDING | 0.00087 | 0.02805 | 0.47727 | 0.56819 | 1.20049 | 177 |
| GO_HISTONE_H2A_UBIQUITINATION | 0.00089 | 0.02848 | 0.47727 | 0.74263 | 1.44418 | 23 |
| GO_HISTONE_UBIQUITINATION | 0.00135 | 0.03867 | 0.45506 | 0.67770 | 1.35593 | 42 |
| GO_HOST_CELLULAR_COMPONENT | 0.00156 | 0.04338 | 0.45506 | 0.66204 | 1.33153 | 46 |
| GO_IMPORT_INTO_NUCLEUS | 0.00018 | 0.00811 | 0.51885 | 0.61120 | 1.27683 | 112 |
| GO_KINETOCHORE | 0.00132 | 0.03818 | 0.45506 | 0.59447 | 1.24081 | 110 |
| GO_LARGE_RIBOSOMAL_SUBUNIT | 0.00000 | 0.00000 | | 0.72923 | 1.51852 | 100 |
| GO_LIVER_REGENERATION | 0.00183 | 0.04873 | 0.45506 | 0.76926 | 1.46019 | 17 |
| GO_MATURATION_OF_LSU_RRNA | 0.00001 | 0.00107 | 0.59333 | 0.79889 | 1.56671 | 26 |
| GO_METHYLTRANSFERASE_COMPLEX | 0.00051 | 0.01881 | 0.47727 | 0.61355 | 1.27394 | 90 |
| GO_MITOTIC_NUCLEAR_DIVISION | 0.00047 | 0.01780 | 0.49849 | 0.56592 | 1.20108 | 203 |
| GO_MITOTIC_SISTER_CHROMATID_SEGREGATION | 0.00061 | 0.02144 | 0.47727 | 0.58864 | 1.23388 | 128 |
| GO_MITOTIC_SPINDLE | 0.00074 | 0.02475 | 0.47727 | 0.60362 | 1.25679 | 99 |
| GO_MODIFICATION_DEPENDENT_MACROMOLECULE_CATAI | 0.00005 | 0.00283 | 0.55733 | 0.53864 | 1.15888 | 470 |
| GO_MRNA_3_END_PROCESSING | 0.00001 | 0.00085 | 0.59333 | 0.65141 | 1.35227 | 89 |
| GO_MRNA_3_UTR_BINDING | 0.00011 | 0.00584 | 0.53843 | 0.65547 | 1.34500 | 66 |
| GO_MRNA_5_UTR_BINDING | 0.00004 | 0.00276 | 0.55733 | 0.81305 | 1.56714 | 21 |
| GO_MRNA_BINDING | 0.00000 | 0.00000 | | 0.66329 | 1.41210 | 231 |
| GO_MRNA_CIS_SPLICING_VIA_SPLICEOSOME | 0.00178 | 0.04819 | 0.45506 | 0.72190 | 1.41573 | 26 |
| GO_MRNA_EXPORT_FROM_NUCLEUS | 0.00000 | 0.00008 | 0.65944 | 0.66689 | 1.38777 | 97 |
| GO_MRNA_PROCESSING | 0.00000 | 0.00000 | | 0.61940 | 1.33024 | 409 |
| GO_MRNA_TRANSPORT | 0.00000 | 0.00000 | 0.71951 | 0.66108 | 1.38538 | 122 |
| GO_MULTI_ORGANISM_LOCALIZATION | 0.00065 | 0.02232 | 0.47727 | 0.66910 | 1.35143 | 49 |
| GO_NCRNA_METABOLIC_PROCESS | 0.00000 | 0.00007 | 0.65944 | 0.57051 | 1.22411 | 358 |
| GO_NCRNA_PROCESSING | 0.00000 | 0.00002 | 0.69013 | 0.58874 | 1.25929 | 302 |
| GO_NCRNA_TRANSCRIPTION | 0.00002 | 0.00151 | 0.57561 | 0.64508 | 1.34017 | 93 |
| GO_NEGATIVE_REGULATION_OF_CELLULAR_AMIDE_METAB | 0.00156 | 0.04338 | 0.45506 | 0.58700 | 1.22679 | 113 |
| GO_NEGATIVE_REGULATION_OF_CELL_CYCLE | 0.00000 | 0.00004 | 0.67496 | 0.56540 | 1.21434 | 412 |
| GO_NEGATIVE_REGULATION_OF_CELL_CYCLE_PHASE_TRAN | 0.00179 | 0.04819 | 0.45506 | 0.56206 | 1.18739 | 176 |
| GO_NEGATIVE_REGULATION_OF_CELL_CYCLE_PROCESS | 0.00002 | 0.00139 | 0.57561 | 0.57532 | 1.22593 | 242 |
| GO_NEGATIVE_REGULATION_OF_CHROMOSOME_ORGANIZ | 0.00121 | 0.03551 | 0.45506 | 0.59607 | 1.24198 | 103 |
| GO_NEGATIVE_REGULATION_OF_INTRINSIC_APOPTOTIC_SI | 0.00187 | 0.04903 | 0.45506 | 0.77841 | 1.46873 | 15 |

| Cluster 1 | | | | | | |
|---|---|---|---|---|---|---|
| pathway | pval | padj | log2err | ES | NES | size |
| GO_NEGATIVE_REGULATION_OF_MITOTIC_CELL_CYCLE | 0.00168 | 0.04630 | 0.45506 | 0.55008 | 1.17045 | 224 |
| GO_NEGATIVE_REGULATION_OF_MRNA_METABOLIC_PROC | 0.00075 | 0.02477 | 0.47727 | 0.65519 | 1.33882 | 61 |
| GO_NEGATIVE_REGULATION_OF_TRANSCRIPTION_BY_RNA_ | 0.00012 | 0.00620 | 0.51885 | 0.53303 | 1.14693 | 475 |
| GO_NUCLEAR_CHROMOSOME_SEGREGATION | 0.00052 | 0.01903 | 0.47727 | 0.56528 | 1.19887 | 199 |
| GO_NUCLEAR_ENVELOPE | 0.00000 | 0.00001 | 0.71951 | 0.59219 | 1.26801 | 315 |
| GO_NUCLEAR_EXPORT | 0.00000 | 0.00000 | 0.82666 | 0.66571 | 1.40477 | 165 |
| GO_NUCLEAR_MEMBRANE | 0.00001 | 0.00052 | 0.61053 | 0.59419 | 1.25983 | 198 |
| GO_NUCLEAR_PORE | 0.00095 | 0.02974 | 0.47727 | 0.64346 | 1.31790 | 64 |
| GO_NUCLEAR_SPECK | 0.00000 | 0.00000 | 0.74774 | 0.60332 | 1.29040 | 296 |
| GO_NUCLEAR_TRANSCRIBED_MRNA_CATABOLIC_PROCESS | 0.00000 | 0.00000 | | 0.74151 | 1.56841 | 181 |
| GO_NUCLEAR_TRANSCRIBED_MRNA_CATABOLIC_PROCESS_ | 0.00000 | 0.00000 | | 0.83901 | 1.75203 | 111 |
| GO_NUCLEAR_TRANSPORT | 0.00000 | 0.00000 | 0.80122 | 0.61817 | 1.31874 | 263 |
| GO_NUCLEOBASE_CONTAINING_COMPOUND_TRANSPORT | 0.00000 | 0.00000 | 0.73376 | 0.62330 | 1.32125 | 196 |
| GO_NUCLEOTIDE_EXCISION_REPAIR | 0.00017 | 0.00772 | 0.51885 | 0.62511 | 1.29795 | 90 |
| GO_NUCLEOTIDE_EXCISION_REPAIR_DNA_DAMAGE_RECOG | 0.00108 | 0.03272 | 0.45506 | 0.75223 | 1.45549 | 22 |
| GO_NUCLEUS_ORGANIZATION | 0.00012 | 0.00591 | 0.53843 | 0.62147 | 1.29378 | 98 |
| GO_ORGANIC_CYCLIC_COMPOUND_CATABOLIC_PROCESS | 0.00000 | 0.00000 | | 0.64324 | 1.38220 | 434 |
| GO_PEPTIDYL_LYSINE_MODIFICATION | 0.00015 | 0.00707 | 0.51885 | 0.55412 | 1.18332 | 283 |
| GO_POLYSOMAL_RIBOSOME | 0.00000 | 0.00000 | 0.77494 | 0.86407 | 1.70669 | 30 |
| GO_POLYSOME | 0.00000 | 0.00000 | 0.77494 | 0.76692 | 1.56218 | 59 |
| GO_POLY_A_BINDING | 0.00013 | 0.00626 | 0.51885 | 0.83487 | 1.57526 | 15 |
| GO_POLY_PURINE_TRACT_BINDING | 0.00191 | 0.04955 | 0.45506 | 0.75775 | 1.45176 | 19 |
| GO_POSITIVE_REGULATION_OF_CELLULAR_AMIDE_METABO | 0.00143 | 0.04057 | 0.45506 | 0.58902 | 1.23180 | 117 |
| GO_POSITIVE_REGULATION_OF_CELL_CYCLE | 0.00179 | 0.04819 | 0.45506 | 0.54661 | 1.16508 | 245 |
| GO_POSITIVE_REGULATION_OF_CELL_POPULATION_PROLIF | 0.00081 | 0.02644 | 0.47727 | 0.53211 | 1.14272 | 400 |
| GO_POSITIVE_REGULATION_OF_CYTOKINESIS | 0.00191 | 0.04955 | 0.45506 | 0.72555 | 1.41097 | 23 |
| GO_POSITIVE_REGULATION_OF_G1_S_TRANSITION_OF_MI | 0.00069 | 0.02362 | 0.47727 | 0.73139 | 1.43814 | 28 |
| GO_POSITIVE_REGULATION_OF_GENE_EXPRESSION_EPIGEN | 0.00185 | 0.04877 | 0.45506 | 0.65921 | 1.32583 | 46 |
| GO_POSITIVE_REGULATION_OF_MITOTIC_CELL_CYCLE | 0.00185 | 0.04877 | 0.45506 | 0.58824 | 1.22673 | 108 |
| GO_POSITIVE_REGULATION_OF_NUCLEOCYTOPLASMIC_TRA | 0.00005 | 0.00310 | 0.55733 | 0.70707 | 1.41888 | 44 |
| GO_POSITIVE_REGULATION_OF_PROTEIN_IMPORT | 0.00063 | 0.02195 | 0.47727 | 0.71985 | 1.41252 | 27 |
| GO_POSITIVE_REGULATION_OF_PROTEIN_LOCALIZATION_T | 0.00008 | 0.00425 | 0.53843 | 0.67438 | 1.37544 | 60 |
| GO_POSITIVE_REGULATION_OF_RNA_SPLICING | 0.00135 | 0.03867 | 0.45506 | 0.71750 | 1.41566 | 29 |
| GO_POSITIVE_REGULATION_OF_TRANSLATION | 0.00044 | 0.01698 | 0.49849 | 0.60804 | 1.26636 | 101 |
| GO_POSITIVE_REGULATION_OF_T_CELL_PROLIFERATION | 0.00009 | 0.00457 | 0.53843 | 0.78304 | 1.52423 | 24 |
| GO_POSITIVE_REGULATION_OF_VIRAL_GENOME_REPLICAT | 0.00043 | 0.01673 | 0.49849 | 0.74048 | 1.46100 | 29 |
| GO_POSITIVE_REGULATION_OF_VIRAL_LIFE_CYCLE | 0.00110 | 0.03311 | 0.45506 | 0.67274 | 1.34440 | 41 |
| GO_POSITIVE_REGULATION_OF_VIRAL_PROCESS | 0.00005 | 0.00283 | 0.55733 | 0.65061 | 1.34453 | 78 |
| GO_POSTTRANSCRIPTIONAL_REGULATION_OF_GENE_EXPR | 0.00000 | 0.00000 | | 0.58868 | 1.26555 | 458 |
| GO_PRERIBOSOME | 0.00002 | 0.00131 | 0.57561 | 0.67304 | 1.38329 | 67 |
| GO_PRE_MRNA_BINDING | 0.00117 | 0.03461 | 0.45506 | 0.72072 | 1.41716 | 28 |
| GO_PROTEASOMAL_PROTEIN_CATABOLIC_PROCESS | 0.00057 | 0.02040 | 0.47727 | 0.53526 | 1.14812 | 367 |
| GO_PROTEIN_CONTAINING_COMPLEX_LOCALIZATION | 0.00013 | 0.00627 | 0.51885 | 0.57530 | 1.22189 | 211 |
| GO_PROTEIN_DNA_COMPLEX_SUBUNIT_ORGANIZATION | 0.00001 | 0.00108 | 0.59333 | 0.59967 | 1.26516 | 163 |
| GO_PROTEIN_IMPORT | 0.00006 | 0.00364 | 0.53843 | 0.61002 | 1.27871 | 128 |
| GO_PROTEIN_LOCALIZATION_TO_ENDOPLASMIC_RETICULU | 0.00000 | 0.00000 | | 0.83538 | 1.74802 | 119 |
| GO_PROTEIN_LOCALIZATION_TO_MEMBRANE | 0.00000 | 0.00001 | 0.70498 | 0.57191 | 1.22887 | 421 |
| GO_PROTEIN_LOCALIZATION_TO_NUCLEUS | 0.00000 | 0.00003 | 0.67496 | 0.61788 | 1.30795 | 188 |
| GO_PROTEIN_MODIFICATION_BY_SMALL_PROTEIN_REMOV | 0.00181 | 0.04846 | 0.45506 | 0.55044 | 1.16996 | 217 |
| GO_PROTEIN_STABILIZATION | 0.00081 | 0.02644 | 0.47727 | 0.58502 | 1.22705 | 133 |
| GO_PROTEIN_TARGETING | 0.00000 | 0.00000 | | 0.64730 | 1.38490 | 310 |
| GO_PROTEIN_TARGETING_TO_MEMBRANE | 0.00000 | 0.00000 | | 0.78772 | 1.65697 | 148 |
| GO_PROTEOGLYCAN_BINDING | 0.00005 | 0.00283 | 0.55733 | 0.85083 | 1.60537 | 15 |
| GO_REGENERATION | 0.00035 | 0.01436 | 0.49849 | 0.61735 | 1.28184 | 90 |
| GO_REGULATION_OF_CELLULAR_AMIDE_METABOLIC_PROC | 0.00000 | 0.00003 | 0.67496 | 0.58354 | 1.24850 | 310 |
| GO_REGULATION_OF_CELLULAR_RESPONSE_TO_HEAT | 0.00167 | 0.04602 | 0.45506 | 0.62443 | 1.27949 | 65 |
| GO_REGULATION_OF_CELL_CYCLE_PHASE_TRANSITION | 0.00057 | 0.02040 | 0.47727 | 0.54003 | 1.15709 | 334 |
| GO_REGULATION_OF_CHROMOSOME_ORGANIZATION | 0.00005 | 0.00283 | 0.55733 | 0.56376 | 1.20292 | 264 |
| GO_REGULATION_OF_CHROMOSOME_SEGREGATION | 0.00014 | 0.00671 | 0.51885 | 0.63094 | 1.30775 | 86 |
| GO_REGULATION_OF_DNA_REPLICATION | 0.00002 | 0.00126 | 0.57561 | 0.65022 | 1.34773 | 86 |
| GO_REGULATION_OF_GENE_EXPRESSION_EPIGENETIC | 0.00036 | 0.01444 | 0.49849 | 0.59686 | 1.24892 | 119 |
| GO_REGULATION_OF_INTRINSIC_APOPTOTIC_SIGNALING_P | 0.00049 | 0.01825 | 0.47727 | 0.60018 | 1.25195 | 109 |
| GO_REGULATION_OF_INTRINSIC_APOPTOTIC_SIGNALING_P | 0.00003 | 0.00196 | 0.55733 | 0.82744 | 1.58529 | 19 |
| GO_REGULATION_OF_KERATINOCYTE_DIFFERENTIATION | 0.00146 | 0.04121 | 0.45506 | 0.74275 | 1.43715 | 22 |
| GO_REGULATION_OF_MITOTIC_CELL_CYCLE | 0.00067 | 0.02313 | 0.47727 | 0.52946 | 1.13843 | 455 |
| GO_REGULATION_OF_MRNA_CATABOLIC_PROCESS | 0.00000 | 0.00017 | 0.64355 | 0.61224 | 1.29158 | 164 |
| GO_REGULATION_OF_MRNA_METABOLIC_PROCESS | 0.00000 | 0.00000 | | 0.63459 | 1.35410 | 265 |
| GO_REGULATION_OF_MRNA_PROCESSING | 0.00000 | 0.00002 | 0.69013 | 0.66372 | 1.38798 | 114 |
| GO_REGULATION_OF_MRNA_SPLICING_VIA_SPLICEOSOME | 0.00000 | 0.00013 | 0.64355 | 0.68188 | 1.40947 | 81 |
| GO_REGULATION_OF_NUCLEOCYTOPLASMIC_TRANSPORT | 0.00003 | 0.00193 | 0.57561 | 0.66482 | 1.37060 | 72 |

| Cluster 1 | | | | | | |
|---|---|---|---|---|---|---|
| pathway | pval | padj | log2err | ES | NES | size |
| GO_REGULATION_OF_PROTEASOMAL_UBIQUITIN_DEPENDI | 0.00133 | 0.03852 | 0.45506 | 0.60576 | 1.25875 | 94 |
| GO_REGULATION_OF_PROTEIN_IMPORT | 0.00113 | 0.03404 | 0.45506 | 0.69183 | 1.38117 | 37 |
| GO_REGULATION_OF_PROTEIN_MODIFICATION_BY_SMALL_ | 0.00001 | 0.00077 | 0.59333 | 0.60072 | 1.26802 | 166 |
| GO_REGULATION_OF_PROTEIN_STABILITY | 0.00179 | 0.04819 | 0.45506 | 0.55429 | 1.17556 | 199 |
| GO_REGULATION_OF_RESPONSE_TO_DNA_DAMAGE_STIMU | 0.00093 | 0.02949 | 0.47727 | 0.57265 | 1.20630 | 157 |
| GO_REGULATION_OF_RNA_SPLICING | 0.00000 | 0.00004 | 0.67496 | 0.65737 | 1.37386 | 113 |
| GO_REGULATION_OF_SIGNAL_TRANSDUCTION_BY_P53_CL | 0.00007 | 0.00410 | 0.53843 | 0.60632 | 1.27140 | 132 |
| GO_REGULATION_OF_SYMBIOTIC_PROCESS | 0.00002 | 0.00131 | 0.57561 | 0.61747 | 1.29512 | 133 |
| GO_REGULATION_OF_TRANSLATIONAL_INITIATION | 0.00106 | 0.03250 | 0.45506 | 0.62876 | 1.29020 | 66 |
| GO_REGULATION_OF_UBIQUITIN_DEPENDENT_PROTEIN_C | 0.00102 | 0.03154 | 0.45506 | 0.59633 | 1.24471 | 110 |
| GO_REGULATION_OF_UBIQUITIN_PROTEIN_TRANSFERASE_ | 0.00054 | 0.01945 | 0.47727 | 0.68628 | 1.37633 | 43 |
| GO_REGULATION_OF_VIRAL_GENOME_REPLICATION | 0.00017 | 0.00776 | 0.51885 | 0.67201 | 1.36686 | 56 |
| GO_REGULATION_OF_VIRAL_LIFE_CYCLE | 0.00010 | 0.00525 | 0.53843 | 0.64277 | 1.32788 | 80 |
| GO_RESPONSE_TO_HEAT | 0.00013 | 0.00644 | 0.51885 | 0.62450 | 1.30045 | 100 |
| GO_RESPONSE_TO_LEUKEMIA_INHIBITORY_FACTOR | 0.00045 | 0.01757 | 0.49849 | 0.63665 | 1.31346 | 73 |
| GO_RESPONSE_TO_TEMPERATURE_STIMULUS | 0.00108 | 0.03272 | 0.45506 | 0.58250 | 1.22029 | 126 |
| GO_RIBONUCLEOPROTEIN_COMPLEX_BIOGENESIS | 0.00000 | 0.00000 | | 0.65304 | 1.40145 | 380 |
| GO_RIBONUCLEOPROTEIN_COMPLEX_SUBUNIT_ORGANIZAT | 0.00000 | 0.00000 | 0.82666 | 0.65775 | 1.38971 | 177 |
| GO_RIBONUCLEOPROTEIN_GRANULE | 0.00000 | 0.00003 | 0.67496 | 0.62025 | 1.31000 | 172 |
| GO_RIBOSOMAL_LARGE_SUBUNIT_ASSEMBLY | 0.00021 | 0.00927 | 0.51885 | 0.74926 | 1.47327 | 28 |
| GO_RIBOSOMAL_LARGE_SUBUNIT_BIOGENESIS | 0.00000 | 0.00000 | 0.78819 | 0.75350 | 1.54397 | 65 |
| GO_RIBOSOMAL_SMALL_SUBUNIT_BIOGENESIS | 0.00001 | 0.00078 | 0.59333 | 0.68163 | 1.39867 | 66 |
| GO_RIBOSOMAL_SUBUNIT | 0.00000 | 0.00000 | | 0.76864 | 1.62049 | 161 |
| GO_RIBOSOME | 0.00000 | 0.00000 | | 0.72770 | 1.54226 | 193 |
| GO_RIBOSOME_ASSEMBLY | 0.00000 | 0.00027 | 0.62726 | 0.71231 | 1.44917 | 57 |
| GO_RIBOSOME_BIOGENESIS | 0.00000 | 0.00000 | | 0.66089 | 1.40988 | 263 |
| GO_RNA_3_END_PROCESSING | 0.00002 | 0.00131 | 0.57561 | 0.62081 | 1.30097 | 122 |
| GO_RNA_CATABOLIC_PROCESS | 0.00000 | 0.00000 | | 0.67406 | 1.44432 | 328 |
| GO_RNA_EXPORT_FROM_NUCLEUS | 0.00000 | 0.00000 | 0.77494 | 0.67942 | 1.42345 | 121 |
| GO_RNA_LOCALIZATION | 0.00000 | 0.00000 | | 0.65354 | 1.38459 | 190 |
| GO_RNA_SPLICING | 0.00000 | 0.00000 | | 0.62728 | 1.34589 | 357 |
| GO_RNA_SPLICING_VIA_TRANSESTERIFICATION_REACTIONS | 0.00000 | 0.00000 | | 0.63703 | 1.36219 | 293 |
| GO_RRNA_BINDING | 0.00000 | 0.00021 | 0.62726 | 0.73326 | 1.48481 | 51 |
| GO_RRNA_METABOLIC_PROCESS | 0.00000 | 0.00000 | | 0.65488 | 1.38837 | 197 |
| GO_SIGNAL_TRANSDUCTION_BY_P53_CLASS_MEDIATOR | 0.00001 | 0.00080 | 0.59333 | 0.59521 | 1.26108 | 192 |
| GO_SINGLE_STRANDED_RNA_BINDING | 0.00009 | 0.00457 | 0.53843 | 0.67268 | 1.37198 | 60 |
| GO_SISTER_CHROMATID_SEGREGATION | 0.00022 | 0.00969 | 0.51885 | 0.59151 | 1.24496 | 153 |
| GO_SMALL_RIBOSOMAL_SUBUNIT | 0.00000 | 0.00000 | | 0.83394 | 1.70721 | 63 |
| GO_SNRNA_TRANSCRIPTION | 0.00049 | 0.01825 | 0.47727 | 0.64371 | 1.32411 | 68 |
| GO_SPINDLE | 0.00016 | 0.00727 | 0.51885 | 0.55957 | 1.19435 | 272 |
| GO_SPLICEOSOMAL_COMPLEX | 0.00000 | 0.00001 | 0.71951 | 0.63456 | 1.33865 | 164 |
| GO_SPLICEOSOMAL_COMPLEX_ASSEMBLY | 0.00117 | 0.03461 | 0.45506 | 0.66932 | 1.35051 | 48 |
| GO_STRUCTURAL_CONSTITUENT_OF_RIBOSOME | 0.00000 | 0.00000 | | 0.78544 | 1.65058 | 141 |
| GO_STRUCTURAL_MOLECULE_ACTIVITY | 0.00000 | 0.00000 | | 0.64226 | 1.37600 | 330 |
| GO_TRANSCRIPTION_BY_RNA_POLYMERASE_I | 0.00098 | 0.03073 | 0.45506 | 0.64889 | 1.31742 | 55 |
| GO_TRANSCRIPTION_BY_RNA_POLYMERASE_III | 0.00075 | 0.02477 | 0.47727 | 0.68375 | 1.36606 | 39 |
| GO_TRANSCRIPTION_COACTIVATOR_ACTIVITY | 0.00095 | 0.02974 | 0.47727 | 0.56003 | 1.18658 | 191 |
| GO_TRANSCRIPTION_COREGULATOR_ACTIVITY | 0.00033 | 0.01353 | 0.49849 | 0.54655 | 1.17151 | 338 |
| GO_TRANSCRIPTION_COUPLED_NUCLEOTIDE_EXCISION_RE | 0.00058 | 0.02060 | 0.47727 | 0.63664 | 1.30635 | 66 |
| GO_TRANSCRIPTION_ELONGATION_FROM_RNA_POLYMERA | 0.00040 | 0.01606 | 0.49849 | 0.63901 | 1.31516 | 70 |
| GO_TRANSCRIPTION_FACTOR_BINDING | 0.00082 | 0.02659 | 0.47727 | 0.53068 | 1.13970 | 409 |
| GO_TRANSCRIPTION_REGULATOR_COMPLEX | 0.00074 | 0.02475 | 0.47727 | 0.54643 | 1.16593 | 264 |
| GO_TRANSFERASE_COMPLEX_TRANSFERRING_PHOSPHORU | 0.00024 | 0.01043 | 0.51885 | 0.57074 | 1.20922 | 192 |
| GO_TRANSLATIONAL_INITIATION | 0.00000 | 0.00000 | | 0.78257 | 1.65211 | 169 |
| GO_TRANSLATION_FACTOR_ACTIVITY_RNA_BINDING | 0.00005 | 0.00293 | 0.55733 | 0.66708 | 1.37217 | 68 |
| GO_TRANSLATION_INITIATION_FACTOR_ACTIVITY | 0.00038 | 0.01507 | 0.49849 | 0.68210 | 1.37188 | 46 |
| GO_TRANSLATION_REGULATOR_ACTIVITY | 0.00001 | 0.00083 | 0.59333 | 0.63981 | 1.33365 | 106 |
| GO_TRANSLATION_REGULATOR_ACTIVITY_NUCLEIC_ACID_E | 0.00001 | 0.00072 | 0.59333 | 0.66391 | 1.37322 | 83 |
| GO_U2_TYPE_SPLICEOSOMAL_COMPLEX | 0.00161 | 0.04471 | 0.45506 | 0.61876 | 1.27964 | 82 |
| GO_UBIQUITIN_LIGASE_COMPLEX | 0.00000 | 0.00037 | 0.61053 | 0.59357 | 1.26089 | 214 |
| GO_UBIQUITIN_LIKE_PROTEIN_LIGASE_BINDING | 0.00024 | 0.01043 | 0.51885 | 0.56463 | 1.20088 | 220 |
| GO_UBIQUITIN_LIKE_PROTEIN_TRANSFERASE_ACTIVITY | 0.00076 | 0.02493 | 0.47727 | 0.54326 | 1.16066 | 287 |
| GO_UNFOLDED_PROTEIN_BINDING | 0.00006 | 0.00354 | 0.53843 | 0.64596 | 1.33523 | 81 |
| GO_VIRAL_GENE_EXPRESSION | 0.00000 | 0.00000 | | 0.74080 | 1.56405 | 171 |
| GO_VIRAL_GENOME_REPLICATION | 0.00033 | 0.01362 | 0.49849 | 0.64344 | 1.32524 | 71 |
| GO_VIRAL_LIFE_CYCLE | 0.00004 | 0.00244 | 0.55733 | 0.58242 | 1.23487 | 198 |
| HP_ABNORMALITY_OF_LOWER_LIP | 0.00122 | 0.03587 | 0.45506 | 0.57671 | 1.21290 | 147 |
| HP_ABNORMALITY_OF_THE_NASAL_DORSUM | 0.00065 | 0.02232 | 0.47727 | 0.57928 | 1.21654 | 137 |
| HP_ACUTE_LEUKEMIA | 0.00015 | 0.00707 | 0.51885 | 0.65031 | 1.33840 | 70 |

| Cluster 1 | | | | | | |
|---|---|---|---|---|---|---|
| pathway | pval | padj | log2err | ES | NES | size |
| HP_ANEMIA_OF_INADEQUATE_PRODUCTION | 0.00047 | 0.01780 | 0.49849 | 0.61499 | 1.27766 | 93 |
| HP_DELAYED_PUBERTY | 0.00013 | 0.00627 | 0.51885 | 0.61587 | 1.28607 | 111 |
| HP_DEPRESSED_NASAL_RIDGE | 0.00005 | 0.00300 | 0.55733 | 0.66045 | 1.36318 | 75 |
| HP_DOWNSLANTED_PALPEBRAL_FISSURES | 0.00179 | 0.04819 | 0.45506 | 0.54410 | 1.16002 | 248 |
| HP_HEMATOLOGICAL_NEOPLASM | 0.00102 | 0.03154 | 0.45506 | 0.57182 | 1.20455 | 157 |
| HP_LEUKEMIA | 0.00026 | 0.01112 | 0.49849 | 0.61484 | 1.27945 | 97 |
| HP_MACROCYTIC_ANEMIA | 0.00000 | 0.00021 | 0.62726 | 0.78385 | 1.56128 | 35 |
| HP_MIGRAINE | 0.00000 | 0.00000 | 0.76146 | 0.74844 | 1.53293 | 64 |
| HP_PALLOR | 0.00009 | 0.00457 | 0.53843 | 0.64348 | 1.33095 | 83 |
| HP_THICK_LOWER_LIP_VERMILION | 0.00047 | 0.01780 | 0.49849 | 0.60924 | 1.26781 | 97 |
| HP_THICK_VERMILION_BORDER | 0.00047 | 0.01780 | 0.49849 | 0.58026 | 1.22074 | 152 |
| **Cluster 14** | | | | | | |
| pathway | pval | padj | log2err | ES | NES | size |
| GO_ACTION_POTENTIAL | 0.00183 | 0.03528 | 0.45506 | 0.68073 | 1.31855 | 43 |
| GO_ALTERNATIVE_MRNA_SPLICING_VIA_SPLICEOSOME | 0.00178 | 0.03456 | 0.45506 | 0.64875 | 1.27763 | 67 |
| GO_AMINE_METABOLIC_PROCESS | 0.00002 | 0.00119 | 0.57561 | 0.67463 | 1.33748 | 87 |
| GO_ANAPHASE_PROMOTING_COMPLEX_DEPENDENT_CATA | 0.00000 | 0.00003 | 0.69013 | 0.72175 | 1.42654 | 74 |
| GO_ANCHORED_COMPONENT_OF_PLASMA_MEMBRANE | 0.00018 | 0.00625 | 0.51885 | 0.83360 | 1.53050 | 17 |
| GO_ANCHORING_JUNCTION | 0.00047 | 0.01365 | 0.49849 | 0.55056 | 1.12845 | 456 |
| GO_ANTIGEN_PROCESSING_AND_PRESENTATION | 0.00220 | 0.04063 | 0.43171 | 0.59326 | 1.19554 | 145 |
| GO_ANTIGEN_PROCESSING_AND_PRESENTATION_OF_EXOG | 0.00163 | 0.03258 | 0.45506 | 0.65727 | 1.28653 | 59 |
| GO_ANTIGEN_PROCESSING_AND_PRESENTATION_OF_PEPT | 0.00198 | 0.03750 | 0.43171 | 0.60382 | 1.21163 | 124 |
| GO_ANTIGEN_PROCESSING_AND_PRESENTATION_OF_PEPT | 0.00106 | 0.02498 | 0.45506 | 0.64922 | 1.28176 | 71 |
| GO_ATPASE_ACTIVITY | 0.00028 | 0.00902 | 0.49849 | 0.57470 | 1.17135 | 299 |
| GO_AUTOPHAGOSOME_MEMBRANE | 0.00034 | 0.01046 | 0.49849 | 0.78149 | 1.46188 | 24 |
| GO_AZUROPHIL_GRANULE_LUMEN | 0.00034 | 0.01054 | 0.49849 | 0.69041 | 1.34855 | 55 |
| GO_BINDING_OF_SPERM_TO_ZONA_PELLUCIDA | 0.00216 | 0.04008 | 0.43171 | 0.76788 | 1.41475 | 18 |
| GO_BLASTOCYST_DEVELOPMENT | 0.00124 | 0.02784 | 0.45506 | 0.64385 | 1.26909 | 69 |
| GO_CADHERIN_BINDING | 0.00003 | 0.00154 | 0.57561 | 0.59894 | 1.21596 | 228 |
| GO_CARDIAC_MUSCLE_CELL_ACTION_POTENTIAL | 0.00276 | 0.04840 | 0.43171 | 0.74267 | 1.39262 | 25 |
| GO_CARDIAC_MUSCLE_CELL_CONTRACTION | 0.00239 | 0.04322 | 0.43171 | 0.74717 | 1.39377 | 23 |
| GO_CATALYTIC_STEP_2_SPLICEOSOME | 0.00012 | 0.00487 | 0.51885 | 0.66455 | 1.31538 | 79 |
| GO_CELLULAR_NITROGEN_COMPOUND_CATABOLIC_PROCE | 0.00000 | 0.00000 | | 0.65333 | 1.33781 | 426 |
| GO_CELLULAR_RESPONSE_TO_HEAT | 0.00257 | 0.04571 | 0.43171 | 0.62546 | 1.23912 | 85 |
| GO_CELLULAR_RESPONSE_TO_ORGANIC_CYCLIC_COMPOUI | 0.00064 | 0.01713 | 0.47727 | 0.56603 | 1.15391 | 288 |
| GO_CELLULAR_RESPONSE_TO_OXYGEN_LEVELS | 0.00080 | 0.02033 | 0.47727 | 0.60112 | 1.21141 | 152 |
| GO_CELL_ACTIVATION_INVOLVED_IN_IMMUNE_RESPONSE | 0.00052 | 0.01490 | 0.47727 | 0.55650 | 1.13702 | 357 |
| GO_CELL_ADHESION_MOLECULE_BINDING | 0.00002 | 0.00112 | 0.57561 | 0.58597 | 1.19463 | 303 |
| GO_CELL_CELL_RECOGNITION | 0.00015 | 0.00558 | 0.51885 | 0.78394 | 1.47927 | 27 |
| GO_CELL_CELL_SIGNALING_BY_WNT | 0.00172 | 0.03360 | 0.45506 | 0.55915 | 1.14009 | 295 |
| GO_CELL_CYCLE_G1_S_PHASE_TRANSITION | 0.00011 | 0.00466 | 0.53843 | 0.59332 | 1.20189 | 199 |
| GO_CELL_CYCLE_G2_M_PHASE_TRANSITION | 0.00143 | 0.03010 | 0.45506 | 0.57682 | 1.17016 | 212 |
| GO_CELL_CYCLE_PHASE_TRANSITION | 0.00000 | 0.00002 | 0.69013 | 0.58591 | 1.20091 | 456 |
| GO_CELL_DIVISION | 0.00001 | 0.00090 | 0.59333 | 0.57222 | 1.17153 | 413 |
| GO_CELL_SUBSTRATE_JUNCTION | 0.00000 | 0.00008 | 0.65944 | 0.60729 | 1.23682 | 274 |
| GO_CERAMIDE_METABOLIC_PROCESS | 0.00168 | 0.03299 | 0.45506 | 0.66814 | 1.30504 | 55 |
| GO_CHANNEL_REGULATOR_ACTIVITY | 0.00181 | 0.03504 | 0.45506 | 0.65861 | 1.28755 | 56 |
| GO_CHAPERONE_COMPLEX | 0.00008 | 0.00355 | 0.53843 | 0.82430 | 1.52874 | 21 |
| GO_CHROMATIN_ASSEMBLY_OR_DISASSEMBLY | 0.00074 | 0.01939 | 0.47727 | 0.62961 | 1.25205 | 97 |
| GO_CHROMATIN_BINDING | 0.00000 | 0.00020 | 0.62726 | 0.58878 | 1.20317 | 362 |
| GO_CHROMATIN_ORGANIZATION | 0.00000 | 0.00000 | 0.78819 | 0.59497 | 1.22032 | 496 |
| GO_CHROMATIN_REMODELING | 0.00253 | 0.04533 | 0.43171 | 0.59692 | 1.19961 | 130 |
| GO_CHROMOSOMAL_REGION | 0.00106 | 0.02498 | 0.45506 | 0.56945 | 1.15925 | 272 |
| GO_CHROMOSOME_SEGREGATION | 0.00008 | 0.00357 | 0.53843 | 0.58467 | 1.18838 | 250 |
| GO_CHROMOSOME_SEPARATION | 0.00015 | 0.00568 | 0.51885 | 0.66133 | 1.30712 | 74 |
| GO_CONDENSED_CHROMOSOME | 0.00163 | 0.03258 | 0.45506 | 0.59185 | 1.19332 | 157 |
| GO_COTRANSLATIONAL_PROTEIN_TARGETING_TO_MEMBR | 0.00000 | 0.00000 | | 0.78297 | 1.55257 | 89 |
| GO_COVALENT_CHROMATIN_MODIFICATION | 0.00001 | 0.00046 | 0.61053 | 0.58748 | 1.20004 | 334 |
| GO_CYTOPLASMIC_TRANSLATION | 0.00000 | 0.00000 | | 0.76292 | 1.51209 | 88 |
| GO_CYTOPLASMIC_TRANSLATIONAL_INITIATION | 0.00001 | 0.00045 | 0.61053 | 0.81021 | 1.53852 | 30 |
| GO_CYTOSOLIC_LARGE_RIBOSOMAL_SUBUNIT | 0.00000 | 0.00000 | 0.76146 | 0.79911 | 1.55851 | 54 |
| GO_CYTOSOLIC_RIBOSOME | 0.00000 | 0.00000 | | 0.79797 | 1.58600 | 94 |
| GO_CYTOSOLIC_SMALL_RIBOSOMAL_SUBUNIT | 0.00000 | 0.00013 | 0.64355 | 0.80008 | 1.53978 | 38 |
| GO_DEFENSE_RESPONSE_TO_OTHER_ORGANISM | 0.00005 | 0.00245 | 0.55733 | 0.56395 | 1.15488 | 429 |
| GO_DNA_BINDING_TRANSCRIPTION_FACTOR_BINDING | 0.00002 | 0.00134 | 0.57561 | 0.60538 | 1.22810 | 212 |
| GO_DNA_CONFORMATION_CHANGE | 0.00012 | 0.00487 | 0.51885 | 0.59540 | 1.20785 | 212 |
| GO_DNA_DAMAGE_RESPONSE_DETECTION_OF_DNA_DAM | 0.00092 | 0.02265 | 0.47727 | 0.70650 | 1.35380 | 35 |
| GO_DNA_DAMAGE_RESPONSE_SIGNAL_TRANSDUCTION_BY | 0.00287 | 0.04989 | 0.43171 | 0.63685 | 1.25773 | 73 |
| GO_DNA_PACKAGING | 0.00159 | 0.03218 | 0.45506 | 0.61390 | 1.22730 | 111 |
| GO_DNA_REPAIR | 0.00012 | 0.00487 | 0.51885 | 0.56540 | 1.15786 | 418 |
| GO_DNA_REPLICATION | 0.00035 | 0.01057 | 0.49849 | 0.58558 | 1.18819 | 217 |

| Cluster 1 | | | | | | |
|---|---|---|---|---|---|---|
| pathway | pval | padj | log2err | ES | NES | size |
| GO_DNA_TEMPLATED_TRANSCRIPTION_ELONGATION | 0.00002 | 0.00134 | 0.57561 | 0.66363 | 1.32026 | 98 |
| GO_DNA_TEMPLATED_TRANSCRIPTION_INITIATION | 0.00005 | 0.00252 | 0.55733 | 0.61503 | 1.24083 | 166 |
| GO_DNA_TEMPLATED_TRANSCRIPTION_TERMINATION | 0.00085 | 0.02121 | 0.47727 | 0.65786 | 1.29557 | 67 |
| GO_EMBRYO_DEVELOPMENT_ENDING_IN_BIRTH_OR_EGG_ | 0.00135 | 0.02907 | 0.45506 | 0.55195 | 1.12772 | 357 |
| GO_EMBRYO_IMPLANTATION | 0.00276 | 0.04840 | 0.43171 | 0.72852 | 1.37125 | 26 |
| GO_ENDOPEPTIDASE_COMPLEX | 0.00007 | 0.00340 | 0.53843 | 0.68902 | 1.34965 | 61 |
| GO_ESTABLISHMENT_OF_PROTEIN_LOCALIZATION_TO_END | 0.00000 | 0.00000 | | 0.77082 | 1.53382 | 100 |
| GO_ESTABLISHMENT_OF_PROTEIN_LOCALIZATION_TO_MEN | 0.00000 | 0.00000 | 0.77494 | 0.64574 | 1.31097 | 228 |
| GO_ESTABLISHMENT_OF_PROTEIN_LOCALIZATION_TO_ORG | 0.00000 | 0.00000 | | 0.64728 | 1.32509 | 410 |
| GO_ESTABLISHMENT_OF_PROTEIN_LOCALIZATION_TO_TEL( | 0.00028 | 0.00911 | 0.49849 | 0.84184 | 1.53955 | 16 |
| GO_ESTABLISHMENT_OF_RNA_LOCALIZATION | 0.00000 | 0.00017 | 0.64355 | 0.64048 | 1.29318 | 170 |
| GO_ESTABLISHMENT_OF_TISSUE_POLARITY | 0.00012 | 0.00487 | 0.53843 | 0.65649 | 1.30059 | 85 |
| GO_EUKARYOTIC_TRANSLATION_INITIATION_FACTOR_3_CO | 0.00021 | 0.00707 | 0.51885 | 0.85251 | 1.55166 | 15 |
| GO_FIBRILLAR_CENTER | 0.00119 | 0.02713 | 0.45506 | 0.62699 | 1.24616 | 94 |
| GO_FICOLIN_1_RICH_GRANULE | 0.00000 | 0.00001 | 0.70498 | 0.72435 | 1.43565 | 88 |
| GO_GENE_SILENCING | 0.00036 | 0.01073 | 0.49849 | 0.59960 | 1.20982 | 167 |
| GO_GLYCOSYL_COMPOUND_METABOLIC_PROCESS | 0.00224 | 0.04119 | 0.43171 | 0.63617 | 1.25326 | 68 |
| GO_HEMATOPOIETIC_PROGENITOR_CELL_DIFFERENTIATION | 0.00021 | 0.00703 | 0.51885 | 0.63069 | 1.26017 | 110 |
| GO_HEMATOPOIETIC_STEM_CELL_DIFFERENTIATION | 0.00003 | 0.00158 | 0.57561 | 0.69804 | 1.37310 | 66 |
| GO_HISTONE_BINDING | 0.00016 | 0.00577 | 0.51885 | 0.60674 | 1.22538 | 171 |
| GO_HISTONE_DEACETYLASE_BINDING | 0.00066 | 0.01770 | 0.47727 | 0.64402 | 1.27291 | 74 |
| GO_HISTONE_PHOSPHORYLATION | 0.00078 | 0.02004 | 0.47727 | 0.75421 | 1.42316 | 27 |
| GO_HORMONE_RECEPTOR_BINDING | 0.00063 | 0.01696 | 0.47727 | 0.62551 | 1.24569 | 102 |
| GO_IMPORT_INTO_NUCLEUS | 0.00152 | 0.03157 | 0.45506 | 0.60283 | 1.20752 | 118 |
| GO_INNATE_IMMUNE_RESPONSE | 0.00029 | 0.00921 | 0.49849 | 0.56339 | 1.15094 | 354 |
| GO_INTERACTION_WITH_HOST | 0.00213 | 0.03978 | 0.43171 | 0.60520 | 1.21394 | 123 |
| GO_INTERLEUKIN_1_MEDIATED_SIGNALING_PATHWAY | 0.00035 | 0.01065 | 0.49849 | 0.65889 | 1.30337 | 77 |
| GO_INTRINSIC_APOPTOTIC_SIGNALING_PATHWAY | 0.00016 | 0.00590 | 0.51885 | 0.60166 | 1.21552 | 176 |
| GO_INTRINSIC_APOPTOTIC_SIGNALING_PATHWAY_BY_P53_ | 0.00128 | 0.02834 | 0.45506 | 0.68237 | 1.32467 | 46 |
| GO_IN_UTERO_EMBRYONIC_DEVELOPMENT | 0.00202 | 0.03807 | 0.43171 | 0.56665 | 1.15114 | 237 |
| GO_KINASE_BINDING | 0.00106 | 0.02498 | 0.45506 | 0.54525 | 1.11760 | 457 |
| GO_LARGE_RIBOSOMAL_SUBUNIT | 0.00001 | 0.00058 | 0.61053 | 0.66202 | 1.31955 | 105 |
| GO_LEUKOCYTE_MEDIATED_IMMUNITY | 0.00119 | 0.02713 | 0.45506 | 0.55043 | 1.12546 | 380 |
| GO_LYSOSOMAL_LUMEN | 0.00168 | 0.03299 | 0.45506 | 0.67537 | 1.31474 | 48 |
| GO_MAINTENANCE_OF_CELL_NUMBER | 0.00062 | 0.01687 | 0.47727 | 0.62180 | 1.24094 | 108 |
| GO_MATURATION_OF_LSU_RRNA | 0.00017 | 0.00608 | 0.51885 | 0.78101 | 1.47373 | 27 |
| GO_MATURATION_OF_LSU_RRNA_FROM_TRICISTRONIC_RF | 0.00042 | 0.01251 | 0.49849 | 0.83991 | 1.52871 | 15 |
| GO_MATURATION_OF_SSU_RRNA | 0.00030 | 0.00959 | 0.49849 | 0.69789 | 1.35679 | 47 |
| GO_MATURATION_OF_SSU_RRNA_FROM_TRICISTRONIC_RF | 0.00111 | 0.02597 | 0.45506 | 0.70365 | 1.34833 | 35 |
| GO_MEMBRANE_REPOLARIZATION | 0.00137 | 0.02936 | 0.45506 | 0.79399 | 1.45778 | 17 |
| GO_METAPHASE_ANAPHASE_TRANSITION_OF_CELL_CYCLE | 0.00019 | 0.00653 | 0.51885 | 0.70224 | 1.36932 | 51 |
| GO_MITOTIC_NUCLEAR_DIVISION | 0.00006 | 0.00309 | 0.53843 | 0.59997 | 1.21616 | 207 |
| GO_MITOTIC_SISTER_CHROMATID_SEGREGATION | 0.00053 | 0.01509 | 0.47727 | 0.61293 | 1.23179 | 130 |
| GO_MODIFICATION_DEPENDENT_MACROMOLECULE_CATAI | 0.00001 | 0.00046 | 0.61053 | 0.56953 | 1.16765 | 470 |
| GO_MODIFIED_AMINO_ACID_TRANSPORT | 0.00110 | 0.02564 | 0.45506 | 0.81278 | 1.48640 | 16 |
| GO_MORPHOGENESIS_OF_A_POLARIZED_EPITHELIUM | 0.00031 | 0.00977 | 0.49849 | 0.63748 | 1.26701 | 95 |
| GO_MRNA_3_END_PROCESSING | 0.00007 | 0.00312 | 0.53843 | 0.66130 | 1.31106 | 87 |
| GO_MRNA_5_UTR_BINDING | 0.00019 | 0.00653 | 0.51885 | 0.80547 | 1.49988 | 22 |
| GO_MRNA_BINDING | 0.00000 | 0.00000 | 0.77494 | 0.64679 | 1.31276 | 225 |
| GO_MRNA_EXPORT_FROM_NUCLEUS | 0.00001 | 0.00059 | 0.59333 | 0.67054 | 1.33395 | 99 |
| GO_MRNA_PROCESSING | 0.00000 | 0.00000 | | 0.64199 | 1.31469 | 418 |
| GO_MRNA_TRANSPORT | 0.00002 | 0.00145 | 0.57561 | 0.63644 | 1.27906 | 129 |
| GO_MYELOID_LEUKOCYTE_ACTIVATION | 0.00003 | 0.00168 | 0.55733 | 0.57662 | 1.17810 | 338 |
| GO_MYELOID_LEUKOCYTE_MEDIATED_IMMUNITY | 0.00006 | 0.00283 | 0.55733 | 0.58126 | 1.18519 | 305 |
| GO_NCRNA_EXPORT_FROM_NUCLEUS | 0.00198 | 0.03750 | 0.43171 | 0.69392 | 1.32969 | 35 |
| GO_NCRNA_METABOLIC_PROCESS | 0.00000 | 0.00000 | 0.73376 | 0.60283 | 1.23254 | 387 |
| GO_NCRNA_PROCESSING | 0.00000 | 0.00000 | 0.81404 | 0.62697 | 1.28012 | 324 |
| GO_NCRNA_TRANSCRIPTION | 0.00074 | 0.01939 | 0.47727 | 0.63051 | 1.25256 | 91 |
| GO_NEGATIVE_REGULATION_OF_CANONICAL_WNT_SIGNAI | 0.00191 | 0.03658 | 0.45506 | 0.61409 | 1.22337 | 103 |
| GO_NEGATIVE_REGULATION_OF_CELLULAR_RESPONSE_TO_ | 0.00257 | 0.04571 | 0.43171 | 0.65362 | 1.28131 | 62 |
| GO_NEGATIVE_REGULATION_OF_CELLULAR_RESPONSE_TO_ | 0.00128 | 0.02834 | 0.45506 | 0.70100 | 1.34860 | 37 |
| GO_NEGATIVE_REGULATION_OF_CELL_CYCLE | 0.00001 | 0.00058 | 0.61053 | 0.57543 | 1.17818 | 414 |
| GO_NEGATIVE_REGULATION_OF_CELL_CYCLE_G2_M_PHASI | 0.00181 | 0.03504 | 0.45506 | 0.63252 | 1.25283 | 82 |
| GO_NEGATIVE_REGULATION_OF_CELL_CYCLE_PHASE_TRAN | 0.00009 | 0.00393 | 0.53843 | 0.60712 | 1.22741 | 179 |
| GO_NEGATIVE_REGULATION_OF_CELL_CYCLE_PROCESS | 0.00004 | 0.00212 | 0.55733 | 0.59597 | 1.21099 | 243 |
| GO_NEGATIVE_REGULATION_OF_CHROMOSOME_ORGANIZ | 0.00102 | 0.02441 | 0.45506 | 0.61940 | 1.23413 | 104 |
| GO_NEGATIVE_REGULATION_OF_CHROMOSOME_SEGREGA | 0.00216 | 0.04008 | 0.43171 | 0.69199 | 1.33127 | 37 |
| GO_NEGATIVE_REGULATION_OF_MITOTIC_CELL_CYCLE | 0.00037 | 0.01095 | 0.49849 | 0.58443 | 1.18651 | 227 |
| GO_NEGATIVE_REGULATION_OF_MRNA_METABOLIC_PROC | 0.00159 | 0.03218 | 0.45506 | 0.66246 | 1.29491 | 58 |
| GO_NEGATIVE_REGULATION_OF_PROTEIN_MODIFICATION_ | 0.00235 | 0.04282 | 0.43171 | 0.64425 | 1.26197 | 61 |
| GO_NEGATIVE_REGULATION_OF_TRANSMEMBRANE_RECEF | 0.00133 | 0.02900 | 0.45506 | 0.68090 | 1.32550 | 48 |

| **Cluster 1** | | | | | | |
|---|---|---|---|---|---|---|
| pathway | pval | padj | log2err | ES | NES | size |
| GO_NEGATIVE_REGULATION_OF_WNT_SIGNALING_PATHWA | 0.00239 | 0.04322 | 0.43171 | 0.60239 | 1.20608 | 116 |
| GO_NIK_NF_KAPPAB_SIGNALING | 0.00006 | 0.00292 | 0.55733 | 0.64749 | 1.29196 | 107 |
| GO_NUCLEAR_CHROMOSOME_SEGREGATION | 0.00046 | 0.01332 | 0.49849 | 0.58975 | 1.19466 | 200 |
| GO_NUCLEAR_ENVELOPE | 0.00000 | 0.00018 | 0.64355 | 0.59193 | 1.20724 | 313 |
| GO_NUCLEAR_EXPORT | 0.00001 | 0.00096 | 0.59333 | 0.62293 | 1.25869 | 174 |
| GO_NUCLEAR_HORMONE_RECEPTOR_BINDING | 0.00078 | 0.02004 | 0.47727 | 0.62936 | 1.25028 | 91 |
| GO_NUCLEAR_MEMBRANE | 0.00019 | 0.00653 | 0.51885 | 0.59544 | 1.20520 | 194 |
| GO_NUCLEAR_PORE | 0.00121 | 0.02734 | 0.45506 | 0.64658 | 1.27377 | 68 |
| GO_NUCLEAR_SPECK | 0.00000 | 0.00000 | 0.73376 | 0.62038 | 1.26469 | 288 |
| GO_NUCLEAR_TRANSCRIBED_MRNA_CATABOLIC_PROCESS | 0.00000 | 0.00000 | | 0.70345 | 1.42258 | 180 |
| GO_NUCLEAR_TRANSCRIBED_MRNA_CATABOLIC_PROCESS_ | 0.00000 | 0.00000 | | 0.77400 | 1.54651 | 110 |
| GO_NUCLEAR_TRANSPORT | 0.00000 | 0.00002 | 0.69013 | 0.61454 | 1.25197 | 277 |
| GO_NUCLEOBASE_CONTAINING_COMPOUND_TRANSPORT | 0.00001 | 0.00046 | 0.61053 | 0.61571 | 1.24806 | 207 |
| GO_NUCLEOSIDE_METABOLIC_PROCESS | 0.00168 | 0.03299 | 0.45506 | 0.66179 | 1.29360 | 58 |
| GO_NUCLEUS_ORGANIZATION | 0.00008 | 0.00353 | 0.53843 | 0.65212 | 1.29612 | 95 |
| GO_ORGANELLE_FISSION | 0.00272 | 0.04804 | 0.43171 | 0.55088 | 1.12519 | 333 |
| GO_ORGANIC_CYCLIC_COMPOUND_CATABOLIC_PROCESS | 0.00000 | 0.00000 | | 0.64588 | 1.32305 | 438 |
| GO_PEPTIDASE_COMPLEX | 0.00093 | 0.02269 | 0.47727 | 0.64614 | 1.27815 | 77 |
| GO_PEPTIDYL_LYSINE_MODIFICATION | 0.00000 | 0.00017 | 0.64355 | 0.60400 | 1.23019 | 275 |
| GO_POLYSOMAL_RIBOSOME | 0.00004 | 0.00213 | 0.55733 | 0.77402 | 1.47415 | 31 |
| GO_POLYSOME | 0.00001 | 0.00045 | 0.61053 | 0.71843 | 1.40727 | 61 |
| GO_POLY_PYRIMIDINE_TRACT_BINDING | 0.00279 | 0.04875 | 0.43171 | 0.76161 | 1.40320 | 18 |
| GO_POSITIVE_REGULATION_OF_CANONICAL_WNT_SIGNAL | 0.00000 | 0.00012 | 0.64355 | 0.68255 | 1.35827 | 101 |
| GO_POSITIVE_REGULATION_OF_CELLULAR_AMIDE_METABO | 0.00209 | 0.03921 | 0.43171 | 0.60926 | 1.21624 | 109 |
| GO_POSITIVE_REGULATION_OF_CELLULAR_PROTEIN_LOCAL | 0.00272 | 0.04804 | 0.43171 | 0.56647 | 1.14975 | 225 |
| GO_POSITIVE_REGULATION_OF_CELL_CYCLE | 0.00002 | 0.00135 | 0.57561 | 0.59527 | 1.21097 | 255 |
| GO_POSITIVE_REGULATION_OF_CELL_CYCLE_ARREST | 0.00279 | 0.04875 | 0.43171 | 0.65878 | 1.28428 | 53 |
| GO_POSITIVE_REGULATION_OF_CELL_CYCLE_PHASE_TRANS | 0.00095 | 0.02304 | 0.47727 | 0.64407 | 1.27185 | 72 |
| GO_POSITIVE_REGULATION_OF_CELL_CYCLE_PROCESS | 0.00007 | 0.00340 | 0.53843 | 0.60984 | 1.23439 | 195 |
| GO_POSITIVE_REGULATION_OF_CELL_DEATH | 0.00028 | 0.00902 | 0.49849 | 0.56268 | 1.15000 | 367 |
| GO_POSITIVE_REGULATION_OF_CHROMATIN_ORGANIZATIO | 0.00163 | 0.03258 | 0.45506 | 0.64736 | 1.27673 | 70 |
| GO_POSITIVE_REGULATION_OF_CHROMOSOME_ORGANIZA | 0.00001 | 0.00058 | 0.59333 | 0.64025 | 1.28782 | 134 |
| GO_POSITIVE_REGULATION_OF_DNA_BIOSYNTHETIC_PROC | 0.00156 | 0.03199 | 0.45506 | 0.66708 | 1.30102 | 54 |
| GO_POSITIVE_REGULATION_OF_ESTABLISHMENT_OF_PROT | 0.00010 | 0.00415 | 0.53843 | 0.59482 | 1.20678 | 216 |
| GO_POSITIVE_REGULATION_OF_INTRINSIC_APOPTOTIC_SIG | 0.00066 | 0.01770 | 0.47727 | 0.72588 | 1.38734 | 34 |
| GO_POSITIVE_REGULATION_OF_MITOTIC_CELL_CYCLE | 0.00043 | 0.01272 | 0.49849 | 0.62095 | 1.24322 | 116 |
| GO_POSITIVE_REGULATION_OF_NUCLEOCYTOPLASMIC_TRA | 0.00141 | 0.03004 | 0.45506 | 0.67884 | 1.32150 | 48 |
| GO_POSITIVE_REGULATION_OF_ORGANELLE_ORGANIZATIO | 0.00011 | 0.00456 | 0.53843 | 0.56331 | 1.15290 | 407 |
| GO_POSITIVE_REGULATION_OF_PROTEIN_IMPORT | 0.00098 | 0.02373 | 0.45506 | 0.73179 | 1.38392 | 28 |
| GO_POSITIVE_REGULATION_OF_PROTEIN_LOCALIZATION_T | 0.00001 | 0.00059 | 0.59333 | 0.72171 | 1.41480 | 62 |
| GO_POSITIVE_REGULATION_OF_RESPONSE_TO_BIOTIC_STI | 0.00092 | 0.02265 | 0.47727 | 0.60172 | 1.21019 | 133 |
| GO_POSITIVE_REGULATION_OF_TELOMERASE_RNA_LOCALI | 0.00121 | 0.02734 | 0.45506 | 0.81805 | 1.48893 | 15 |
| GO_POSITIVE_REGULATION_OF_TELOMERE_MAINTENANCE | 0.00124 | 0.02784 | 0.45506 | 0.71958 | 1.37046 | 31 |
| GO_POSITIVE_REGULATION_OF_VIRAL_PROCESS | 0.00028 | 0.00897 | 0.49849 | 0.65548 | 1.29828 | 81 |
| GO_POSITIVE_REGULATION_OF_WNT_SIGNALING_PATHWA | 0.00005 | 0.00232 | 0.55733 | 0.63665 | 1.27528 | 118 |
| GO_POSTTRANSCRIPTIONAL_REGULATION_OF_GENE_EXPR | 0.00000 | 0.00000 | | 0.62166 | 1.27413 | 454 |
| GO_POST_TRANSLATIONAL_PROTEIN_MODIFICATION | 0.00198 | 0.03750 | 0.43171 | 0.57461 | 1.16586 | 222 |
| GO_PRERIBOSOME | 0.00000 | 0.00025 | 0.62726 | 0.71122 | 1.40189 | 69 |
| GO_PROTEASOMAL_PROTEIN_CATABOLIC_PROCESS | 0.00021 | 0.00726 | 0.51885 | 0.56848 | 1.16155 | 360 |
| GO_PROTEASOMAL_UBIQUITIN_INDEPENDENT_PROTEIN_C | 0.00198 | 0.03750 | 0.43171 | 0.76074 | 1.40731 | 19 |
| GO_PROTEASOME_ACCESSORY_COMPLEX | 0.00154 | 0.03173 | 0.45506 | 0.75637 | 1.41490 | 24 |
| GO_PROTEASOME_CORE_COMPLEX | 0.00133 | 0.02900 | 0.45506 | 0.79537 | 1.46031 | 17 |
| GO_PROTEIN_CONTAINING_COMPLEX_LOCALIZATION | 0.00013 | 0.00493 | 0.51885 | 0.59840 | 1.21279 | 206 |
| GO_PROTEIN_DNA_COMPLEX_SUBUNIT_ORGANIZATION | 0.00001 | 0.00069 | 0.59333 | 0.62854 | 1.26750 | 162 |
| GO_PROTEIN_FOLDING | 0.00033 | 0.01037 | 0.49849 | 0.59849 | 1.20690 | 162 |
| GO_PROTEIN_IMPORT | 0.00006 | 0.00306 | 0.53843 | 0.62219 | 1.25150 | 134 |
| GO_PROTEIN_LOCALIZATION_TO_CHROMOSOME | 0.00023 | 0.00763 | 0.51885 | 0.67387 | 1.32709 | 67 |
| GO_PROTEIN_LOCALIZATION_TO_CHROMOSOME_TELOMER | 0.00033 | 0.01037 | 0.49849 | 0.76530 | 1.44049 | 26 |
| GO_PROTEIN_LOCALIZATION_TO_ENDOPLASMIC_RETICULU | 0.00000 | 0.00000 | | 0.74492 | 1.49366 | 121 |
| GO_PROTEIN_LOCALIZATION_TO_MEMBRANE | 0.00001 | 0.00047 | 0.61053 | 0.57817 | 1.18284 | 398 |
| GO_PROTEIN_LOCALIZATION_TO_NUCLEUS | 0.00000 | 0.00030 | 0.62726 | 0.63042 | 1.27521 | 190 |
| GO_PROTEIN_MODIFICATION_BY_SMALL_PROTEIN_REMOV | 0.00008 | 0.00353 | 0.53843 | 0.59536 | 1.20804 | 217 |
| GO_PROTEIN_POLYUBIQUITINATION | 0.00017 | 0.00617 | 0.51885 | 0.58293 | 1.18489 | 248 |
| GO_PROTEIN_STABILIZATION | 0.00012 | 0.00483 | 0.53843 | 0.61788 | 1.24190 | 131 |
| GO_PROTEIN_SUMOYLATION | 0.00013 | 0.00509 | 0.51885 | 0.68957 | 1.34976 | 59 |
| GO_PROTEIN_TARGETING | 0.00000 | 0.00000 | 0.80122 | 0.62571 | 1.27648 | 316 |
| GO_PROTEIN_TARGETING_TO_MEMBRANE | 0.00000 | 0.00000 | | 0.72949 | 1.47007 | 145 |
| GO_PROTEOGLYCAN_BINDING | 0.00010 | 0.00415 | 0.53843 | 0.86615 | 1.57648 | 15 |
| GO_REGULATION_OF_ANIMAL_ORGAN_MORPHOGENESIS | 0.00209 | 0.03921 | 0.43171 | 0.60457 | 1.21259 | 122 |
| GO_REGULATION_OF_APOPTOTIC_SIGNALING_PATHWAY | 0.00231 | 0.04241 | 0.43171 | 0.57078 | 1.15909 | 232 |

| Cluster 1 | | | | | | |
|---|---|---|---|---|---|---|
| pathway | pval | padj | log2err | ES | NES | size |
| GO_REGULATION_OF_CELLULAR_AMIDE_METABOLIC_PROC | 0.00000 | 0.00008 | 0.65944 | 0.60140 | 1.22610 | 303 |
| GO_REGULATION_OF_CELLULAR_AMINE_METABOLIC_PROC | 0.00003 | 0.00157 | 0.57561 | 0.70951 | 1.39088 | 62 |
| GO_REGULATION_OF_CELLULAR_AMINO_ACID_METABOLIC | 0.00001 | 0.00058 | 0.61053 | 0.72328 | 1.41399 | 56 |
| GO_REGULATION_OF_CELLULAR_PROTEIN_LOCALIZATION | 0.00143 | 0.03010 | 0.45506 | 0.55051 | 1.12493 | 361 |
| GO_REGULATION_OF_CELLULAR_RESPONSE_TO_STRESS | 0.00055 | 0.01557 | 0.47727 | 0.55376 | 1.13487 | 452 |
| GO_REGULATION_OF_CELLULAR_RESPONSE_TO_TRANSFOR | 0.00058 | 0.01603 | 0.47727 | 0.66945 | 1.31037 | 59 |
| GO_REGULATION_OF_CELL_CYCLE_G2_M_PHASE_TRANSITI | 0.00130 | 0.02842 | 0.45506 | 0.58634 | 1.18463 | 173 |
| GO_REGULATION_OF_CELL_CYCLE_PHASE_TRANSITION | 0.00000 | 0.00034 | 0.62726 | 0.58818 | 1.20162 | 336 |
| GO_REGULATION_OF_CHROMATIN_ORGANIZATION | 0.00054 | 0.01538 | 0.47727 | 0.61731 | 1.23814 | 122 |
| GO_REGULATION_OF_CHROMOSOME_ORGANIZATION | 0.00000 | 0.00001 | 0.71951 | 0.62221 | 1.26521 | 253 |
| GO_REGULATION_OF_CHROMOSOME_SEGREGATION | 0.00001 | 0.00074 | 0.59333 | 0.67819 | 1.34479 | 89 |
| GO_REGULATION_OF_CHROMOSOME_SEPARATION | 0.00017 | 0.00604 | 0.51885 | 0.69906 | 1.36543 | 55 |
| GO_REGULATION_OF_DNA_BIOSYNTHETIC_PROCESS | 0.00157 | 0.03214 | 0.45506 | 0.63878 | 1.26359 | 77 |
| GO_REGULATION_OF_DNA_METABOLIC_PROCESS | 0.00057 | 0.01585 | 0.47727 | 0.58174 | 1.18133 | 232 |
| GO_REGULATION_OF_DNA_TEMPLATED_TRANSCRIPTION_I | 0.00113 | 0.02629 | 0.45506 | 0.63041 | 1.24934 | 86 |
| GO_REGULATION_OF_GENE_EXPRESSION_EPIGENETIC | 0.00010 | 0.00426 | 0.53843 | 0.63036 | 1.26267 | 118 |
| GO_REGULATION_OF_GENE_SILENCING | 0.00069 | 0.01835 | 0.47727 | 0.64405 | 1.27566 | 82 |
| GO_REGULATION_OF_HEMATOPOIETIC_PROGENITOR_CELL_ | 0.00001 | 0.00092 | 0.59333 | 0.69463 | 1.36919 | 69 |
| GO_REGULATION_OF_HEMATOPOIETIC_STEM_CELL_DIFFER | 0.00004 | 0.00210 | 0.55733 | 0.71249 | 1.39269 | 58 |
| GO_REGULATION_OF_HISTONE_MODIFICATION | 0.00165 | 0.03283 | 0.45506 | 0.61961 | 1.23216 | 97 |
| GO_REGULATION_OF_INTRINSIC_APOPTOTIC_SIGNALING_P | 0.00017 | 0.00604 | 0.51885 | 0.64389 | 1.28094 | 99 |
| GO_REGULATION_OF_INTRINSIC_APOPTOTIC_SIGNALING_P | 0.00167 | 0.03297 | 0.45506 | 0.80126 | 1.46533 | 16 |
| GO_REGULATION_OF_MITOTIC_CELL_CYCLE | 0.00000 | 0.00015 | 0.64355 | 0.57767 | 1.18427 | 464 |
| GO_REGULATION_OF_MORPHOGENESIS_OF_AN_EPITHELIU | 0.00246 | 0.04428 | 0.43171 | 0.61017 | 1.21556 | 103 |
| GO_REGULATION_OF_MRNA_CATABOLIC_PROCESS | 0.00010 | 0.00415 | 0.53843 | 0.61878 | 1.24748 | 161 |
| GO_REGULATION_OF_MRNA_METABOLIC_PROCESS | 0.00000 | 0.00000 | 0.76146 | 0.63165 | 1.28569 | 263 |
| GO_REGULATION_OF_MRNA_PROCESSING | 0.00000 | 0.00034 | 0.62726 | 0.66199 | 1.32552 | 115 |
| GO_REGULATION_OF_MRNA_SPLICING_VIA_SPLICEOSOME | 0.00003 | 0.00166 | 0.57561 | 0.67776 | 1.34258 | 83 |
| GO_REGULATION_OF_NUCLEAR_DIVISION | 0.00009 | 0.00393 | 0.53843 | 0.61907 | 1.24577 | 136 |
| GO_REGULATION_OF_NUCLEASE_ACTIVITY | 0.00167 | 0.03297 | 0.45506 | 0.80160 | 1.46595 | 16 |
| GO_REGULATION_OF_NUCLEOCYTOPLASMIC_TRANSPORT | 0.00119 | 0.02713 | 0.45506 | 0.64044 | 1.26766 | 79 |
| GO_REGULATION_OF_POSTTRANSCRIPTIONAL_GENE_SILEN | 0.00087 | 0.02181 | 0.47727 | 0.64034 | 1.26563 | 74 |
| GO_REGULATION_OF_PROTEIN_CATABOLIC_PROCESS | 0.00135 | 0.02907 | 0.45506 | 0.56858 | 1.15708 | 261 |
| GO_REGULATION_OF_PROTEIN_IMPORT | 0.00076 | 0.01978 | 0.47727 | 0.70334 | 1.35649 | 39 |
| GO_REGULATION_OF_PROTEIN_LOCALIZATION_TO_NUCLEU | 0.00002 | 0.00104 | 0.57561 | 0.67515 | 1.33877 | 89 |
| GO_REGULATION_OF_PROTEIN_MODIFICATION_BY_SMALL_ | 0.00001 | 0.00092 | 0.59333 | 0.62337 | 1.25778 | 167 |
| GO_REGULATION_OF_PROTEIN_STABILITY | 0.00027 | 0.00887 | 0.49849 | 0.59291 | 1.20100 | 201 |
| GO_REGULATION_OF_RNA_SPLICING | 0.00000 | 0.00030 | 0.62726 | 0.66185 | 1.32511 | 116 |
| GO_REGULATION_OF_SIGNAL_TRANSDUCTION_BY_P53_CL | 0.00018 | 0.00621 | 0.51885 | 0.62278 | 1.25078 | 126 |
| GO_REGULATION_OF_SISTER_CHROMATID_SEGREGATION | 0.00001 | 0.00072 | 0.59333 | 0.69764 | 1.37511 | 69 |
| GO_REGULATION_OF_SMALL_MOLECULE_METABOLIC_PRO | 0.00015 | 0.00572 | 0.51885 | 0.58121 | 1.18265 | 260 |
| GO_REGULATION_OF_STEM_CELL_DIFFERENTIATION | 0.00004 | 0.00211 | 0.55733 | 0.67207 | 1.33116 | 82 |
| GO_REGULATION_OF_SYMBIOTIC_PROCESS | 0.00006 | 0.00300 | 0.53843 | 0.62482 | 1.25587 | 131 |
| GO_REGULATION_OF_TRANSCRIPTION_FROM_RNA_POLYM | 0.00003 | 0.00154 | 0.57561 | 0.69829 | 1.37201 | 64 |
| GO_REGULATION_OF_TRANSLATIONAL_INITIATION | 0.00005 | 0.00232 | 0.55733 | 0.69481 | 1.36352 | 63 |
| GO_REGULATION_OF_UBIQUITIN_PROTEIN_TRANSFERASE_ | 0.00143 | 0.03010 | 0.45506 | 0.68004 | 1.31812 | 44 |
| GO_REGULATION_OF_WNT_SIGNALING_PATHWAY | 0.00145 | 0.03037 | 0.45506 | 0.57842 | 1.17208 | 205 |
| GO_RESPONSE_TO_HEAT | 0.00056 | 0.01575 | 0.47727 | 0.62320 | 1.24217 | 105 |
| GO_RESPONSE_TO_INTERLEUKIN_1 | 0.00023 | 0.00766 | 0.51885 | 0.63270 | 1.26304 | 109 |
| GO_RESPONSE_TO_INTERLEUKIN_12 | 0.00130 | 0.02842 | 0.45506 | 0.71619 | 1.36667 | 33 |
| GO_RESPONSE_TO_LEUKEMIA_INHIBITORY_FACTOR | 0.00126 | 0.02815 | 0.45506 | 0.63958 | 1.26298 | 72 |
| GO_RESPONSE_TO_ORGANIC_CYCLIC_COMPOUND | 0.00108 | 0.02531 | 0.45506 | 0.54902 | 1.12461 | 445 |
| GO_RESPONSE_TO_TEMPERATURE_STIMULUS | 0.00057 | 0.01585 | 0.47727 | 0.61042 | 1.22677 | 129 |
| GO_RIBONUCLEOPROTEIN_COMPLEX_BINDING | 0.00000 | 0.00038 | 0.61053 | 0.66074 | 1.32302 | 115 |
| GO_RIBONUCLEOPROTEIN_COMPLEX_BIOGENESIS | 0.00000 | 0.00000 | | 0.68769 | 1.40606 | 382 |
| GO_RIBONUCLEOPROTEIN_COMPLEX_SUBUNIT_ORGANIZAT | 0.00000 | 0.00000 | 0.76146 | 0.66744 | 1.34849 | 173 |
| GO_RIBONUCLEOPROTEIN_GRANULE | 0.00159 | 0.03218 | 0.45506 | 0.57796 | 1.16870 | 184 |
| GO_RIBOSOMAL_LARGE_SUBUNIT_ASSEMBLY | 0.00005 | 0.00242 | 0.55733 | 0.77821 | 1.47170 | 28 |
| GO_RIBOSOMAL_LARGE_SUBUNIT_BIOGENESIS | 0.00000 | 0.00000 | 0.76146 | 0.76996 | 1.51458 | 66 |
| GO_RIBOSOMAL_SMALL_SUBUNIT_BIOGENESIS | 0.00003 | 0.00157 | 0.57561 | 0.69651 | 1.37168 | 67 |
| GO_RIBOSOMAL_SUBUNIT | 0.00000 | 0.00000 | | 0.68775 | 1.38713 | 164 |
| GO_RIBOSOME | 0.00000 | 0.00000 | | 0.68394 | 1.38437 | 196 |
| GO_RIBOSOME_ASSEMBLY | 0.00000 | 0.00040 | 0.61053 | 0.72649 | 1.42202 | 59 |
| GO_RIBOSOME_BIOGENESIS | 0.00000 | 0.00000 | | 0.68884 | 1.40176 | 268 |
| GO_RNA_3_END_PROCESSING | 0.00003 | 0.00184 | 0.55733 | 0.64091 | 1.28548 | 122 |
| GO_RNA_CAP_BINDING | 0.00085 | 0.02121 | 0.47727 | 0.81853 | 1.49691 | 16 |
| GO_RNA_CATABOLIC_PROCESS | 0.00000 | 0.00000 | | 0.67065 | 1.36969 | 330 |
| GO_RNA_EXPORT_FROM_NUCLEUS | 0.00001 | 0.00059 | 0.59333 | 0.65565 | 1.31643 | 125 |
| GO_RNA_LOCALIZATION | 0.00000 | 0.00000 | 0.76146 | 0.65345 | 1.32363 | 201 |
| GO_RNA_POLYMERASE_COMPLEX | 0.00093 | 0.02269 | 0.47727 | 0.63234 | 1.25633 | 92 |

| Cluster 1 | | | | | | |
|---|---|---|---|---|---|---|
| pathway | pval | padj | log2err | ES | NES | size |
| GO_RNA_POLYMERASE_II_SPECIFIC_DNA_BINDING_TRANSC | 0.00098 | 0.02373 | 0.45506 | 0.59114 | 1.19409 | 172 |
| GO_RNA_SPLICING | 0.00000 | 0.00000 | | 0.65363 | 1.33581 | 365 |
| GO_RNA_SPLICING_VIA_TRANSESTERIFICATION_REACTIONS | 0.00000 | 0.00000 | | 0.65452 | 1.33411 | 297 |
| GO_ROUGH_ENDOPLASMIC_RETICULUM | 0.00250 | 0.04481 | 0.43171 | 0.66078 | 1.28820 | 53 |
| GO_RRNA_BINDING | 0.00001 | 0.00046 | 0.61053 | 0.73898 | 1.44063 | 53 |
| GO_RRNA_METABOLIC_PROCESS | 0.00000 | 0.00000 | | 0.69129 | 1.40012 | 203 |
| GO_SCF_DEPENDENT_PROTEASOMAL_UBIQUITIN_DEPEND | 0.00015 | 0.00568 | 0.51885 | 0.67261 | 1.32794 | 71 |
| GO_SECRETORY_GRANULE | 0.00031 | 0.00968 | 0.49849 | 0.55552 | 1.13743 | 415 |
| GO_SIGNAL_TRANSDUCTION_BY_P53_CLASS_MEDIATOR | 0.00063 | 0.01696 | 0.47727 | 0.59566 | 1.20489 | 190 |
| GO_SIGNAL_TRANSDUCTION_INVOLVED_IN_CELL_CYCLE_C | 0.00146 | 0.03054 | 0.45506 | 0.66800 | 1.30226 | 53 |
| GO_SIGNAL_TRANSDUCTION_IN_RESPONSE_TO_DNA_DAM | 0.00054 | 0.01520 | 0.47727 | 0.63390 | 1.25930 | 91 |
| GO_SINGLE_STRANDED_RNA_BINDING | 0.00102 | 0.02441 | 0.45506 | 0.67054 | 1.31070 | 58 |
| GO_SISTER_CHROMATID_SEGREGATION | 0.00013 | 0.00498 | 0.51885 | 0.61156 | 1.23253 | 151 |
| GO_SMALL_RIBOSOMAL_SUBUNIT | 0.00001 | 0.00060 | 0.59333 | 0.72140 | 1.41418 | 62 |
| GO_SMALL_SUBUNIT_PROCESSOME | 0.00077 | 0.01999 | 0.47727 | 0.73059 | 1.39245 | 32 |
| GO_SPERM_EGG_RECOGNITION | 0.00022 | 0.00729 | 0.51885 | 0.78928 | 1.47646 | 24 |
| GO_SPHINGOLIPID_METABOLIC_PROCESS | 0.00122 | 0.02765 | 0.45506 | 0.63046 | 1.25135 | 90 |
| GO_SPINDLE | 0.00157 | 0.03214 | 0.45506 | 0.55974 | 1.14022 | 276 |
| GO_SPINDLE_MIDZONE | 0.00017 | 0.00612 | 0.51885 | 0.76017 | 1.43759 | 28 |
| GO_SPLICEOSOMAL_COMPLEX | 0.00000 | 0.00006 | 0.65944 | 0.64738 | 1.30543 | 163 |
| GO_STEM_CELL_DIFFERENTIATION | 0.00013 | 0.00504 | 0.51885 | 0.61360 | 1.23509 | 139 |
| GO_STRUCTURAL_CONSTITUENT_OF_RIBOSOME | 0.00000 | 0.00000 | 0.80122 | 0.69872 | 1.40806 | 145 |
| GO_STRUCTURAL_MOLECULE_ACTIVITY | 0.00000 | 0.00000 | 0.73376 | 0.61282 | 1.25159 | 331 |
| GO_TRANSCRIPTION_BY_RNA_POLYMERASE_I | 0.00079 | 0.02019 | 0.47727 | 0.67795 | 1.32421 | 55 |
| GO_TRANSCRIPTION_COACTIVATOR_ACTIVITY | 0.00010 | 0.00415 | 0.53843 | 0.60260 | 1.21826 | 182 |
| GO_TRANSCRIPTION_COREGULATOR_ACTIVITY | 0.00003 | 0.00157 | 0.57561 | 0.57887 | 1.18193 | 326 |
| GO_TRANSCRIPTION_COUPLED_NUCLEOTIDE_EXCISION_RE | 0.00088 | 0.02194 | 0.47727 | 0.65152 | 1.28351 | 68 |
| GO_TRANSCRIPTION_ELONGATION_FROM_RNA_POLYMERA | 0.00059 | 0.01620 | 0.47727 | 0.65185 | 1.28721 | 72 |
| GO_TRANSCRIPTION_FACTOR_BINDING | 0.00000 | 0.00001 | 0.70498 | 0.59480 | 1.21676 | 397 |
| GO_TRANSCRIPTION_INITIATION_FROM_RNA_POLYMERASE | 0.00044 | 0.01299 | 0.49849 | 0.60525 | 1.21653 | 131 |
| GO_TRANSFERASE_COMPLEX_TRANSFERRING_PHOSPHORU | 0.00008 | 0.00353 | 0.53843 | 0.59580 | 1.20690 | 199 |
| GO_TRANSLATIONAL_INITIATION | 0.00000 | 0.00000 | | 0.77397 | 1.56104 | 164 |
| GO_TRANSLATION_FACTOR_ACTIVITY_RNA_BINDING | 0.00000 | 0.00000 | 0.81404 | 0.78133 | 1.53872 | 67 |
| GO_TRANSLATION_INITIATION_FACTOR_ACTIVITY | 0.00000 | 0.00003 | 0.67496 | 0.79488 | 1.54149 | 45 |
| GO_TRANSLATION_INITIATION_FACTOR_BINDING | 0.00002 | 0.00134 | 0.57561 | 0.79027 | 1.49451 | 28 |
| GO_TRANSLATION_PREINITIATION_COMPLEX | 0.00054 | 0.01520 | 0.47727 | 0.82999 | 1.51788 | 16 |
| GO_TRANSLATION_REGULATOR_ACTIVITY | 0.00000 | 0.00000 | | 0.74204 | 1.47665 | 101 |
| GO_TRANSLATION_REGULATOR_ACTIVITY_NUCLEIC_ACID_E | 0.00000 | 0.00000 | 0.83909 | 0.76973 | 1.52457 | 81 |
| GO_TUMOR_NECROSIS_FACTOR_MEDIATED_SIGNALING_PA | 0.00044 | 0.01292 | 0.49849 | 0.63665 | 1.26537 | 94 |
| GO_T_CELL_RECEPTOR_SIGNALING_PATHWAY | 0.00084 | 0.02117 | 0.47727 | 0.61483 | 1.23247 | 120 |
| GO_U2_TYPE_SPLICEOSOMAL_COMPLEX | 0.00154 | 0.03173 | 0.45506 | 0.63043 | 1.24852 | 80 |
| GO_UBIQUITIN_LIKE_PROTEIN_LIGASE_BINDING | 0.00027 | 0.00877 | 0.49849 | 0.58642 | 1.18986 | 221 |
| GO_UNFOLDED_PROTEIN_BINDING | 0.00007 | 0.00330 | 0.53843 | 0.66784 | 1.32293 | 83 |
| GO_VACUOLAR_LUMEN | 0.00016 | 0.00595 | 0.51885 | 0.64714 | 1.28560 | 91 |
| GO_VESICLE_LUMEN | 0.00000 | 0.00000 | 0.81404 | 0.67645 | 1.36643 | 172 |
| GO_VIRAL_GENE_EXPRESSION | 0.00000 | 0.00000 | | 0.71162 | 1.43679 | 169 |
| GO_VIRAL_LIFE_CYCLE | 0.00001 | 0.00089 | 0.59333 | 0.61235 | 1.24061 | 204 |
| HP_ABNORMALITY_OF_LOWER_LIP | 0.00077 | 0.01999 | 0.47727 | 0.59830 | 1.20539 | 144 |
| HP_ABNORMALITY_OF_THE_AUDITORY_CANAL | 0.00135 | 0.02907 | 0.45506 | 0.69961 | 1.34060 | 35 |
| HP_ABNORMALITY_OF_THE_NASAL_DORSUM | 0.00006 | 0.00303 | 0.53843 | 0.61864 | 1.24523 | 139 |
| HP_ACUTE_LEUKEMIA | 0.00130 | 0.02842 | 0.45506 | 0.64278 | 1.26699 | 69 |
| HP_ATRESIA_OF_THE_EXTERNAL_AUDITORY_CANAL | 0.00216 | 0.04008 | 0.43171 | 0.74883 | 1.40416 | 25 |
| HP_DELAYED_PUBERTY | 0.00035 | 0.01057 | 0.49849 | 0.62667 | 1.25067 | 108 |
| HP_DEPRESSED_NASAL_RIDGE | 0.00002 | 0.00145 | 0.57561 | 0.68152 | 1.34898 | 79 |
| HP_LEUKEMIA | 0.00235 | 0.04282 | 0.43171 | 0.60969 | 1.21319 | 100 |
| HP_MACROCYTIC_ANEMIA | 0.00009 | 0.00379 | 0.53843 | 0.74253 | 1.42850 | 37 |
| HP_ORAL_CLEFT | 0.00246 | 0.04428 | 0.43171 | 0.55266 | 1.12870 | 330 |
| HP_OVOID_VERTEBRAL_BODIES | 0.00077 | 0.01999 | 0.47727 | 0.78080 | 1.44440 | 19 |
| HP_RETICULOCYTOPENIA | 0.00117 | 0.02704 | 0.45506 | 0.81896 | 1.49059 | 15 |
| HP_SHORT_NOSE | 0.00146 | 0.03054 | 0.45506 | 0.58704 | 1.18582 | 172 |
| HP_THICK_LOWER_LIP_VERMILION | 0.00024 | 0.00807 | 0.51885 | 0.64127 | 1.27522 | 96 |
| HP_THICK_VERMILION_BORDER | 0.00012 | 0.00477 | 0.53843 | 0.61558 | 1.23991 | 149 |

### Supplementary Table S6

#### Human male GSEA of NMF clusters

| Cluster 1 | | | | | | |
|---|---|---|---|---|---|---|
| pathway | pval | padj | log2err | ES | NES | size |
| GO_ACTIN_CYTOSKELETON | 0.00020 | 0.01269 | 0.51885 | 0.54374 | 1.19316 | 340 |
| GO_ACTIN_FILAMENT_ORGANIZATION | 0.00071 | 0.02727 | 0.47727 | 0.54407 | 1.18716 | 278 |
| GO_ACTIN_POLYMERIZATION_OR_DEPOLYMERIZATION | 0.00150 | 0.04445 | 0.45506 | 0.56802 | 1.22112 | 148 |
| GO_ACTIVATION_OF_IMMUNE_RESPONSE | 0.00163 | 0.04638 | 0.45506 | 0.54259 | 1.17921 | 230 |
| GO_ANAPHASE_PROMOTING_COMPLEX_DEPENDENT_CATA | 0.00006 | 0.00530 | 0.53843 | 0.66094 | 1.38943 | 70 |
| GO_ANATOMICAL_STRUCTURE_MATURATION | 0.00010 | 0.00797 | 0.53843 | 0.60303 | 1.29023 | 129 |
| GO_ANTERIOR_POSTERIOR_AXIS_SPECIFICATION | 0.00003 | 0.00331 | 0.55733 | 0.75447 | 1.53244 | 33 |
| GO_APOPTOTIC_SIGNALING_PATHWAY | 0.00056 | 0.02400 | 0.47727 | 0.53205 | 1.16971 | 393 |
| GO_AXIS_SPECIFICATION | 0.00052 | 0.02251 | 0.47727 | 0.66174 | 1.37433 | 54 |
| GO_AXONAL_GROWTH_CONE | 0.00141 | 0.04281 | 0.45506 | 0.76455 | 1.52084 | 21 |
| GO_BINDING_OF_SPERM_TO_ZONA_PELLUCIDA | 0.00095 | 0.03280 | 0.47727 | 0.75766 | 1.50900 | 22 |
| GO_CADHERIN_BINDING | 0.00000 | 0.00045 | 0.62726 | 0.57950 | 1.26268 | 260 |
| GO_CATALYTIC_STEP_2_SPLICEOSOME | 0.00078 | 0.02885 | 0.47727 | 0.61778 | 1.30522 | 82 |
| GO_CELLULAR_NITROGEN_COMPOUND_CATABOLIC_PROCE | 0.00000 | 0.00001 | 0.73376 | 0.57054 | 1.25566 | 441 |
| GO_CELLULAR_PROCESS_INVOLVED_IN_REPRODUCTION_IN | 0.00001 | 0.00092 | 0.61053 | 0.58828 | 1.27774 | 223 |
| GO_CELLULAR_RESPONSE_TO_ORGANIC_CYCLIC_COMPOUN | 0.00183 | 0.04966 | 0.45506 | 0.52738 | 1.15725 | 341 |
| GO_CELLULAR_RESPONSE_TO_OXYGEN_LEVELS | 0.00002 | 0.00250 | 0.57561 | 0.60568 | 1.30293 | 153 |
| GO_CELL_ADHESION_MOLECULE_BINDING | 0.00002 | 0.00238 | 0.57561 | 0.55013 | 1.20914 | 369 |
| GO_CELL_CORTEX | 0.00024 | 0.01387 | 0.51885 | 0.56744 | 1.23187 | 215 |
| GO_CELL_CYCLE_G1_S_PHASE_TRANSITION | 0.00017 | 0.01110 | 0.51885 | 0.56832 | 1.23236 | 206 |
| GO_CELL_CYCLE_G2_M_PHASE_TRANSITION | 0.00012 | 0.00904 | 0.53843 | 0.56730 | 1.23262 | 227 |
| GO_CELL_CYCLE_PHASE_TRANSITION | 0.00000 | 0.00002 | 0.70498 | 0.56486 | 1.24481 | 490 |
| GO_CELL_DIVISION | 0.00004 | 0.00365 | 0.55733 | 0.53758 | 1.18380 | 456 |
| GO_CELL_JUNCTION_ORGANIZATION | 0.00163 | 0.04638 | 0.45506 | 0.51782 | 1.13950 | 428 |
| GO_CELL_LEADING_EDGE | 0.00152 | 0.04445 | 0.45506 | 0.53068 | 1.15983 | 302 |
| GO_CELL_MATURATION | 0.00095 | 0.03280 | 0.47727 | 0.60315 | 1.28004 | 96 |
| GO_CELL_SUBSTRATE_JUNCTION | 0.00000 | 0.00000 | | 0.62443 | 1.36788 | 320 |
| GO_CHAPERONE_COMPLEX | 0.00008 | 0.00610 | 0.53843 | 0.82177 | 1.62066 | 19 |
| GO_CHROMATIN_BINDING | 0.00058 | 0.02445 | 0.47727 | 0.52646 | 1.15854 | 430 |
| GO_CHROMATIN_REMODELING | 0.00139 | 0.04267 | 0.45506 | 0.57029 | 1.22629 | 150 |
| GO_CHROMOSOMAL_REGION | 0.00008 | 0.00620 | 0.53843 | 0.56134 | 1.22467 | 277 |
| GO_CHROMOSOME_CENTROMERIC_REGION | 0.00009 | 0.00720 | 0.53843 | 0.58818 | 1.26918 | 166 |
| GO_CHROMOSOME_SEGREGATION | 0.00005 | 0.00471 | 0.55733 | 0.56490 | 1.23109 | 264 |
| GO_CORTICAL_ACTIN_CYTOSKELETON | 0.00058 | 0.02445 | 0.47727 | 0.65801 | 1.37293 | 59 |
| GO_COTRANSLATIONAL_PROTEIN_TARGETING_TO_MEMBRA | 0.00000 | 0.00000 | 0.81404 | 0.73713 | 1.56189 | 89 |
| GO_COVALENT_CHROMATIN_MODIFICATION | 0.00000 | 0.00003 | 0.69013 | 0.57681 | 1.26795 | 376 |
| GO_CYCLIN_DEPENDENT_PROTEIN_KINASE_HOLOENZYME_ | 0.00063 | 0.02583 | 0.47727 | 0.70545 | 1.44276 | 37 |
| GO_CYTOPLASMIC_TRANSLATION | 0.00002 | 0.00238 | 0.57561 | 0.65444 | 1.38656 | 86 |
| GO_CYTOSOLIC_LARGE_RIBOSOMAL_SUBUNIT | 0.00000 | 0.00012 | 0.65944 | 0.75421 | 1.56067 | 50 |
| GO_CYTOSOLIC_RIBOSOME | 0.00000 | 0.00000 | 0.82666 | 0.73599 | 1.56071 | 93 |
| GO_CYTOSOLIC_SMALL_RIBOSOMAL_SUBUNIT | 0.00001 | 0.00083 | 0.61053 | 0.76724 | 1.57264 | 39 |
| GO_DEVELOPMENTAL_MATURATION | 0.00020 | 0.01269 | 0.51885 | 0.58678 | 1.26404 | 159 |
| GO_DEVELOPMENT_OF_PRIMARY_SEXUAL_CHARACTERISTI | 0.00077 | 0.02868 | 0.47727 | 0.59331 | 1.26858 | 126 |
| GO_DNA_CONFORMATION_CHANGE | 0.00005 | 0.00465 | 0.55733 | 0.57273 | 1.24372 | 218 |
| GO_DNA_DAMAGE_RESPONSE_DETECTION_OF_DNA_DAMA | 0.00023 | 0.01359 | 0.51885 | 0.74041 | 1.49888 | 31 |
| GO_DNA_GEOMETRIC_CHANGE | 0.00097 | 0.03326 | 0.47727 | 0.59954 | 1.27042 | 92 |
| GO_DNA_METHYLATION | 0.00089 | 0.03178 | 0.47727 | 0.65297 | 1.35612 | 54 |
| GO_DNA_METHYLATION_OR_DEMETHYLATION | 0.00031 | 0.01619 | 0.49849 | 0.64665 | 1.35871 | 67 |
| GO_DNA_MODIFICATION | 0.00007 | 0.00560 | 0.53843 | 0.65062 | 1.37626 | 83 |
| GO_DNA_PACKAGING | 0.00121 | 0.03853 | 0.45506 | 0.58383 | 1.24910 | 127 |
| GO_DNA_REPAIR | 0.00161 | 0.04627 | 0.45506 | 0.52122 | 1.14681 | 414 |
| GO_EMBRYONIC_PATTERN_SPECIFICATION | 0.00132 | 0.04081 | 0.45506 | 0.67169 | 1.38116 | 42 |
| GO_ENZYME_ACTIVATOR_ACTIVITY | 0.00032 | 0.01623 | 0.49849 | 0.53741 | 1.17903 | 346 |
| GO_ESTABLISHMENT_OF_PROTEIN_LOCALIZATION_TO_END | 0.00000 | 0.00000 | 0.76146 | 0.71062 | 1.51092 | 100 |
| GO_ESTABLISHMENT_OF_PROTEIN_LOCALIZATION_TO_MEM | 0.00000 | 0.00000 | 0.77494 | 0.62114 | 1.35228 | 251 |
| GO_ESTABLISHMENT_OF_PROTEIN_LOCALIZATION_TO_ORG | 0.00000 | 0.00000 | | 0.59092 | 1.30066 | 436 |
| GO_ESTABLISHMENT_OF_TISSUE_POLARITY | 0.00159 | 0.04596 | 0.45506 | 0.59253 | 1.25555 | 92 |
| GO_FC_RECEPTOR_SIGNALING_PATHWAY | 0.00108 | 0.03626 | 0.45506 | 0.58261 | 1.24704 | 130 |
| GO_FEMALE_GAMETE_GENERATION | 0.00000 | 0.00008 | 0.67496 | 0.69111 | 1.46470 | 87 |
| GO_FEMALE_MEIOTIC_NUCLEAR_DIVISION | 0.00043 | 0.01977 | 0.49849 | 0.76921 | 1.53245 | 23 |
| GO_FICOLIN_1_RICH_GRANULE | 0.00050 | 0.02202 | 0.47727 | 0.62144 | 1.31780 | 93 |
| GO_GAMETE_GENERATION | 0.00000 | 0.00000 | 0.77494 | 0.58387 | 1.28417 | 409 |
| GO_GENE_SILENCING | 0.00001 | 0.00091 | 0.61053 | 0.60890 | 1.31539 | 175 |
| GO_GERM_CELL_DEVELOPMENT | 0.00002 | 0.00229 | 0.57561 | 0.60473 | 1.30464 | 164 |
| GO_HEMATOPOIETIC_PROGENITOR_CELL_DIFFERENTIATION | 0.00179 | 0.04909 | 0.45506 | 0.58727 | 1.25339 | 115 |
| GO_HISTONE_BINDING | 0.00003 | 0.00343 | 0.55733 | 0.59021 | 1.27670 | 187 |
| GO_HISTONE_H3_K4_METHYLATION | 0.00170 | 0.04782 | 0.45506 | 0.67122 | 1.38446 | 44 |
| GO_HISTONE_METHYLATION | 0.00004 | 0.00380 | 0.55733 | 0.62451 | 1.33125 | 108 |
| GO_IMMUNE_RESPONSE_REGULATING_SIGNALING_PATHW | 0.00040 | 0.01936 | 0.49849 | 0.55860 | 1.21346 | 224 |
| GO_INTRINSIC_APOPTOTIC_SIGNALING_PATHWAY | 0.00002 | 0.00234 | 0.57561 | 0.58668 | 1.27175 | 201 |

| Cluster 1 | | | | | | |
|---|---|---|---|---|---|---|
| GO_KINASE_ACTIVATOR_ACTIVITY | 0.00054 | 0.02338 | 0.47727 | 0.63957 | 1.34385 | 67 |
| GO_KINASE_REGULATOR_ACTIVITY | 0.00001 | 0.00091 | 0.61053 | 0.61290 | 1.31964 | 158 |
| GO_LARGE_RIBOSOMAL_SUBUNIT | 0.00005 | 0.00416 | 0.55733 | 0.63639 | 1.35059 | 96 |
| GO_LEUKOCYTE_MEDIATED_IMMUNITY | 0.00172 | 0.04812 | 0.45506 | 0.52276 | 1.14927 | 395 |
| GO_LIVER_REGENERATION | 0.00066 | 0.02583 | 0.47727 | 0.78493 | 1.54799 | 19 |
| GO_MALE_GAMETE_GENERATION | 0.00013 | 0.00949 | 0.51885 | 0.54662 | 1.19688 | 318 |
| GO_MALE_SEX_DIFFERENTIATION | 0.00133 | 0.04118 | 0.45506 | 0.60474 | 1.28076 | 91 |
| GO_MEIOTIC_CELL_CYCLE | 0.00066 | 0.02583 | 0.47727 | 0.56888 | 1.23058 | 184 |
| GO_MEIOTIC_CELL_CYCLE_PROCESS | 0.00027 | 0.01518 | 0.49849 | 0.58612 | 1.25802 | 138 |
| GO_METHYLATION | 0.00029 | 0.01562 | 0.49849 | 0.55865 | 1.21516 | 243 |
| GO_MICROTUBULE | 0.00042 | 0.01970 | 0.49849 | 0.53831 | 1.18104 | 339 |
| GO_MICROTUBULE_CYTOSKELETON_ORGANIZATION | 0.00009 | 0.00720 | 0.53843 | 0.53523 | 1.17797 | 438 |
| GO_MITOTIC_SISTER_CHROMATID_SEGREGATION | 0.00045 | 0.02008 | 0.49849 | 0.58767 | 1.26281 | 142 |
| GO_MODIFICATION_DEPENDENT_MACROMOLECULE_CATAI | 0.00003 | 0.00297 | 0.57561 | 0.53894 | 1.18788 | 498 |
| GO_MORPHOGENESIS_OF_A_POLARIZED_EPITHELIUM | 0.00148 | 0.04441 | 0.45506 | 0.59611 | 1.27023 | 107 |
| GO_MRNA_BINDING | 0.00002 | 0.00229 | 0.57561 | 0.57580 | 1.25190 | 240 |
| GO_MRNA_EXPORT_FROM_NUCLEUS | 0.00121 | 0.03853 | 0.45506 | 0.59924 | 1.27271 | 99 |
| GO_MRNA_PROCESSING | 0.00000 | 0.00068 | 0.61053 | 0.54755 | 1.20494 | 425 |
| GO_MULTICELLULAR_ORGANISM_REPRODUCTION | 0.00000 | 0.00000 | 0.76146 | 0.56986 | 1.25601 | 494 |
| GO_MYELOID_LEUKOCYTE_MEDIATED_IMMUNITY | 0.00076 | 0.02851 | 0.47727 | 0.53584 | 1.17328 | 318 |
| GO_NEGATIVE_REGULATION_OF_CATALYTIC_ACTIVITY | 0.00119 | 0.03833 | 0.45506 | 0.51616 | 1.13644 | 453 |
| GO_NEGATIVE_REGULATION_OF_CELLULAR_COMPONENT_( | 0.00001 | 0.00124 | 0.59333 | 0.54135 | 1.19275 | 487 |
| GO_NEGATIVE_REGULATION_OF_CELL_CYCLE | 0.00018 | 0.01164 | 0.51885 | 0.53354 | 1.17527 | 460 |
| GO_NEGATIVE_REGULATION_OF_CELL_CYCLE_PROCESS | 0.00095 | 0.03280 | 0.47727 | 0.54739 | 1.19263 | 256 |
| GO_NEGATIVE_REGULATION_OF_CHROMATIN_ORGANIZATI | 0.00033 | 0.01632 | 0.49849 | 0.71300 | 1.46032 | 38 |
| GO_NEGATIVE_REGULATION_OF_CHROMOSOME_ORGANIZ | 0.00001 | 0.00124 | 0.59333 | 0.64817 | 1.38087 | 106 |
| GO_NEGATIVE_REGULATION_OF_HISTONE_MODIFICATION | 0.00065 | 0.02583 | 0.47727 | 0.71354 | 1.44666 | 32 |
| GO_NEGATIVE_REGULATION_OF_MITOTIC_CELL_CYCLE | 0.00018 | 0.01146 | 0.51885 | 0.56050 | 1.22027 | 251 |
| GO_NEGATIVE_REGULATION_OF_ORGANELLE_ORGANIZATIO | 0.00000 | 0.00080 | 0.61053 | 0.57466 | 1.25511 | 291 |
| GO_NEGATIVE_REGULATION_OF_PHOSPHORUS_METABOLI( | 0.00075 | 0.02799 | 0.47727 | 0.52840 | 1.16151 | 372 |
| GO_NEGATIVE_REGULATION_OF_PROTEIN_MODIFICATION_ | 0.00004 | 0.00380 | 0.55733 | 0.54313 | 1.19468 | 412 |
| GO_NUCLEAR_CHROMOSOME_SEGREGATION | 0.00022 | 0.01358 | 0.51885 | 0.56840 | 1.23256 | 210 |
| GO_NUCLEAR_ENVELOPE | 0.00042 | 0.01970 | 0.49849 | 0.53614 | 1.17547 | 330 |
| GO_NUCLEAR_EXPORT | 0.00065 | 0.02583 | 0.47727 | 0.57013 | 1.23204 | 177 |
| GO_NUCLEAR_SPECK | 0.00001 | 0.00191 | 0.59333 | 0.56062 | 1.22646 | 308 |
| GO_NUCLEAR_TRANSCRIBED_MRNA_CATABOLIC_PROCESS | 0.00000 | 0.00000 | 0.81404 | 0.66020 | 1.42814 | 181 |
| GO_NUCLEAR_TRANSCRIBED_MRNA_CATABOLIC_PROCESS_ | 0.00000 | 0.00000 | | 0.73194 | 1.56074 | 109 |
| GO_NUCLEAR_TRANSPORT | 0.00001 | 0.00191 | 0.59333 | 0.56867 | 1.24131 | 284 |
| GO_NUCLEUS_ORGANIZATION | 0.00030 | 0.01598 | 0.49849 | 0.61673 | 1.31274 | 102 |
| GO_OOCYTE_DIFFERENTIATION | 0.00004 | 0.00380 | 0.55733 | 0.75122 | 1.53637 | 37 |
| GO_OOCYTE_MATURATION | 0.00152 | 0.04445 | 0.45506 | 0.76458 | 1.50786 | 19 |
| GO_OOGENESIS | 0.00004 | 0.00380 | 0.55733 | 0.69523 | 1.45092 | 60 |
| GO_ORGANIC_CYCLIC_COMPOUND_CATABOLIC_PROCESS | 0.00000 | 0.00001 | 0.73376 | 0.56460 | 1.24393 | 466 |
| GO_PATHWAY_RESTRICTED_SMAD_PROTEIN_PHOSPHORYL/ | 0.00148 | 0.04441 | 0.45506 | 0.70073 | 1.41856 | 31 |
| GO_PEPTIDYL_LYSINE_METHYLATION | 0.00029 | 0.01562 | 0.49849 | 0.61563 | 1.30450 | 92 |
| GO_PEPTIDYL_LYSINE_MODIFICATION | 0.00023 | 0.01360 | 0.51885 | 0.55341 | 1.20980 | 303 |
| GO_PHOSPHATIDYLINOSITOL_3_4_5_TRISPHOSPHATE_BIND | 0.00150 | 0.04445 | 0.45506 | 0.71538 | 1.44206 | 28 |
| GO_POLYMERIC_CYTOSKELETAL_FIBER | 0.00002 | 0.00258 | 0.57561 | 0.54047 | 1.18989 | 450 |
| GO_POLYSOMAL_RIBOSOME | 0.00001 | 0.00181 | 0.59333 | 0.78309 | 1.58062 | 29 |
| GO_POLYSOME | 0.00021 | 0.01286 | 0.51885 | 0.67277 | 1.40556 | 61 |
| GO_POSITIVE_REGULATION_OF_CELLULAR_COMPONENT_B | 0.00048 | 0.02136 | 0.49849 | 0.53497 | 1.17480 | 359 |
| GO_POSITIVE_REGULATION_OF_CELLULAR_PROTEIN_LOCAI | 0.00130 | 0.04064 | 0.45506 | 0.53910 | 1.17376 | 252 |
| GO_POSITIVE_REGULATION_OF_CELL_CYCLE | 0.00016 | 0.01073 | 0.51885 | 0.55658 | 1.21430 | 277 |
| GO_POSITIVE_REGULATION_OF_CELL_CYCLE_PROCESS | 0.00041 | 0.01957 | 0.49849 | 0.56054 | 1.21589 | 212 |
| GO_POSITIVE_REGULATION_OF_CELL_POPULATION_PROLIF | 0.00026 | 0.01506 | 0.49849 | 0.52521 | 1.15717 | 486 |
| GO_POSITIVE_REGULATION_OF_CHROMATIN_ORGANIZATIO | 0.00030 | 0.01608 | 0.49849 | 0.64563 | 1.36238 | 79 |
| GO_POSITIVE_REGULATION_OF_CHROMOSOME_ORGANIZA | 0.00039 | 0.01884 | 0.49849 | 0.58955 | 1.26594 | 139 |
| GO_POSITIVE_REGULATION_OF_HISTONE_METHYLATION | 0.00043 | 0.01983 | 0.49849 | 0.72315 | 1.46615 | 32 |
| GO_POSITIVE_REGULATION_OF_HYDROLASE_ACTIVITY | 0.00128 | 0.04047 | 0.45506 | 0.51777 | 1.14060 | 461 |
| GO_POSITIVE_REGULATION_OF_IMMUNE_SYSTEM_PROCES | 0.00152 | 0.04445 | 0.45506 | 0.51541 | 1.13556 | 466 |
| GO_POSITIVE_REGULATION_OF_MITOTIC_CELL_CYCLE | 0.00014 | 0.01036 | 0.51885 | 0.60717 | 1.29631 | 118 |
| GO_POSITIVE_REGULATION_OF_MUSCLE_CONTRACTION | 0.00028 | 0.01553 | 0.49849 | 0.79929 | 1.58478 | 20 |
| GO_POSITIVE_REGULATION_OF_ORGANELLE_ORGANIZATIO | 0.00000 | 0.00007 | 0.67496 | 0.55966 | 1.23241 | 456 |
| GO_POSITIVE_REGULATION_OF_PATHWAY_RESTRICTED_SM | 0.00113 | 0.03752 | 0.45506 | 0.75470 | 1.50310 | 22 |
| GO_POSITIVE_REGULATION_OF_PROTEIN_KINASE_ACTIVITY | 0.00051 | 0.02226 | 0.47727 | 0.53386 | 1.17079 | 329 |
| GO_POSITIVE_REGULATION_OF_TRANSFERASE_ACTIVITY | 0.00023 | 0.01374 | 0.51885 | 0.52682 | 1.15957 | 436 |
| GO_POSITIVE_REGULATION_OF_TRANSMEMBRANE_RECEPT | 0.00064 | 0.02583 | 0.47727 | 0.63650 | 1.33756 | 69 |
| GO_POSITIVE_REGULATION_OF_VIRAL_LIFE_CYCLE | 0.00113 | 0.03752 | 0.45506 | 0.66976 | 1.38407 | 46 |
| GO_POSITIVE_REGULATION_OF_VIRAL_PROCESS | 0.00007 | 0.00572 | 0.53843 | 0.64276 | 1.36134 | 85 |
| GO_POSTTRANSCRIPTIONAL_REGULATION_OF_GENE_EXPRI | 0.00000 | 0.00000 | | 0.59540 | 1.31189 | 479 |
| GO_POST_TRANSLATIONAL_PROTEIN_MODIFICATION | 0.00064 | 0.02583 | 0.47727 | 0.54358 | 1.18314 | 250 |
| GO_PROTEASOMAL_PROTEIN_CATABOLIC_PROCESS | 0.00015 | 0.01045 | 0.51885 | 0.53937 | 1.18564 | 376 |
| GO_PROTEIN_CONTAINING_COMPLEX_LOCALIZATION | 0.00095 | 0.03280 | 0.47727 | 0.54950 | 1.19496 | 238 |

| Cluster 1 | | | | | | |
|---|---|---|---|---|---|---|
| GO_PROTEIN_DEPHOSPHORYLATION | 0.00178 | 0.04880 | 0.45506 | 0.54135 | 1.17651 | 230 |
| GO_PROTEIN_DNA_COMPLEX_SUBUNIT_ORGANIZATION | 0.00066 | 0.02583 | 0.47727 | 0.57498 | 1.24146 | 172 |
| GO_PROTEIN_DOMAIN_SPECIFIC_BINDING | 0.00021 | 0.01316 | 0.51885 | 0.52654 | 1.16024 | 489 |
| GO_PROTEIN_LOCALIZATION_TO_ENDOPLASMIC_RETICULU | 0.00000 | 0.00000 | 0.76146 | 0.68504 | 1.46311 | 124 |
| GO_PROTEIN_LOCALIZATION_TO_MEMBRANE | 0.00000 | 0.00000 | 0.74774 | 0.57004 | 1.25620 | 469 |
| GO_PROTEIN_LOCALIZATION_TO_NUCLEUS | 0.00159 | 0.04596 | 0.45506 | 0.55117 | 1.19462 | 199 |
| GO_PROTEIN_METHYLATION | 0.00006 | 0.00498 | 0.55733 | 0.61824 | 1.32253 | 128 |
| GO_PROTEIN_MODIFICATION_BY_SMALL_PROTEIN_REMOV | 0.00167 | 0.04721 | 0.45506 | 0.54652 | 1.18750 | 228 |
| GO_PROTEIN_POLYMERIZATION | 0.00141 | 0.04281 | 0.45506 | 0.55178 | 1.19647 | 207 |
| GO_PROTEIN_POLYUBIQUITINATION | 0.00033 | 0.01642 | 0.49849 | 0.55181 | 1.20347 | 268 |
| GO_PROTEIN_SERINE_THREONINE_KINASE_ACTIVATOR_ACT | 0.00011 | 0.00838 | 0.53843 | 0.74411 | 1.51365 | 34 |
| GO_PROTEIN_TARGETING | 0.00000 | 0.00000 | 0.81404 | 0.60393 | 1.32504 | 337 |
| GO_PROTEIN_TARGETING_TO_MEMBRANE | 0.00000 | 0.00000 | 0.74774 | 0.65655 | 1.41331 | 156 |
| GO_RECEPTOR_SERINE_THREONINE_KINASE_BINDING | 0.00067 | 0.02603 | 0.47727 | 0.83157 | 1.61129 | 15 |
| GO_REGULATION_OF_ACTIN_FILAMENT_BASED_PROCESS | 0.00054 | 0.02314 | 0.47727 | 0.54524 | 1.18851 | 265 |
| GO_REGULATION_OF_ACTIN_FILAMENT_ORGANIZATION | 0.00104 | 0.03521 | 0.45506 | 0.56436 | 1.22081 | 181 |
| GO_REGULATION_OF_ANATOMICAL_STRUCTURE_SIZE | 0.00104 | 0.03521 | 0.45506 | 0.54229 | 1.18550 | 303 |
| GO_REGULATION_OF_CELLULAR_AMIDE_METABOLIC_PROC | 0.00000 | 0.00038 | 0.62726 | 0.56785 | 1.24539 | 331 |
| GO_REGULATION_OF_CELLULAR_COMPONENT_SIZE | 0.00079 | 0.02885 | 0.47727 | 0.54556 | 1.18818 | 254 |
| GO_REGULATION_OF_CELLULAR_PROTEIN_LOCALIZATION | 0.00041 | 0.01957 | 0.49849 | 0.52903 | 1.16368 | 412 |
| GO_REGULATION_OF_CELL_CYCLE_G2_M_PHASE_TRANSITI | 0.00017 | 0.01143 | 0.51885 | 0.58652 | 1.26808 | 180 |
| GO_REGULATION_OF_CELL_CYCLE_PHASE_TRANSITION | 0.00000 | 0.00062 | 0.62726 | 0.56592 | 1.24316 | 361 |
| GO_REGULATION_OF_CELL_MATURATION | 0.00174 | 0.04842 | 0.45506 | 0.81133 | 1.57207 | 15 |
| GO_REGULATION_OF_CHROMATIN_ORGANIZATION | 0.00002 | 0.00275 | 0.57561 | 0.60754 | 1.30522 | 141 |
| GO_REGULATION_OF_CHROMOSOME_ORGANIZATION | 0.00000 | 0.00034 | 0.64355 | 0.57705 | 1.25869 | 273 |
| GO_REGULATION_OF_CHROMOSOME_SEGREGATION | 0.00045 | 0.02008 | 0.49849 | 0.61046 | 1.29356 | 92 |
| GO_REGULATION_OF_CYTOSKELETON_ORGANIZATION | 0.00028 | 0.01553 | 0.49849 | 0.53673 | 1.18007 | 385 |
| GO_REGULATION_OF_GENE_EXPRESSION_EPIGENETIC | 0.00032 | 0.01623 | 0.49849 | 0.59980 | 1.28058 | 118 |
| GO_REGULATION_OF_GENE_SILENCING | 0.00126 | 0.04009 | 0.45506 | 0.61229 | 1.29680 | 85 |
| GO_REGULATION_OF_HISTONE_METHYLATION | 0.00032 | 0.01623 | 0.49849 | 0.67398 | 1.39779 | 53 |
| GO_REGULATION_OF_HISTONE_MODIFICATION | 0.00004 | 0.00365 | 0.55733 | 0.63111 | 1.34653 | 112 |
| GO_REGULATION_OF_INTRINSIC_APOPTOTIC_SIGNALING_P | 0.00183 | 0.04966 | 0.45506 | 0.58884 | 1.25447 | 106 |
| GO_REGULATION_OF_MACROAUTOPHAGY | 0.00115 | 0.03753 | 0.45506 | 0.57185 | 1.22738 | 138 |
| GO_REGULATION_OF_MRNA_CATABOLIC_PROCESS | 0.00013 | 0.00949 | 0.51885 | 0.58496 | 1.26180 | 163 |
| GO_REGULATION_OF_MRNA_METABOLIC_PROCESS | 0.00000 | 0.00019 | 0.65944 | 0.58356 | 1.27320 | 276 |
| GO_REGULATION_OF_PEPTIDE_TRANSPORT | 0.00157 | 0.04585 | 0.45506 | 0.52189 | 1.14800 | 411 |
| GO_REGULATION_OF_PROTEIN_CONTAINING_COMPLEX_AS | 0.00015 | 0.01049 | 0.51885 | 0.54985 | 1.20164 | 298 |
| GO_REGULATION_OF_PROTEIN_MODIFICATION_BY_SMALL_ | 0.00092 | 0.03257 | 0.47727 | 0.56136 | 1.21408 | 185 |
| GO_REGULATION_OF_PROTEIN_POLYMERIZATION | 0.00045 | 0.02008 | 0.49849 | 0.57537 | 1.23883 | 158 |
| GO_REGULATION_OF_PROTEIN_SERINE_THREONINE_KINAS | 0.00004 | 0.00365 | 0.55733 | 0.55471 | 1.21702 | 339 |
| GO_REGULATION_OF_REPRODUCTIVE_PROCESS | 0.00005 | 0.00471 | 0.55733 | 0.64341 | 1.36440 | 93 |
| GO_REGULATION_OF_RNA_SPLICING | 0.00100 | 0.03434 | 0.45506 | 0.58345 | 1.24682 | 125 |
| GO_REGULATION_OF_SMOOTH_MUSCLE_CONTRACTION | 0.00020 | 0.01271 | 0.51885 | 0.74585 | 1.50546 | 29 |
| GO_REGULATION_OF_SUPRAMOLECULAR_FIBER_ORGANIZA | 0.00015 | 0.01041 | 0.51885 | 0.56016 | 1.21915 | 249 |
| GO_REGULATION_OF_TRANSLATIONAL_INITIATION | 0.00086 | 0.03082 | 0.47727 | 0.64615 | 1.35445 | 64 |
| GO_REPRODUCTIVE_SYSTEM_DEVELOPMENT | 0.00073 | 0.02764 | 0.47727 | 0.54935 | 1.19691 | 256 |
| GO_RESPONSE_TO_OXYGEN_LEVELS | 0.00037 | 0.01832 | 0.49849 | 0.55532 | 1.20807 | 246 |
| GO_RESPONSE_TO_TEMPERATURE_STIMULUS | 0.00115 | 0.03753 | 0.45506 | 0.57431 | 1.23546 | 152 |
| GO_RIBONUCLEOPROTEIN_GRANULE | 0.00072 | 0.02746 | 0.47727 | 0.56160 | 1.21624 | 191 |
| GO_RIBOSOMAL_SUBUNIT | 0.00000 | 0.00003 | 0.69013 | 0.63691 | 1.37140 | 157 |
| GO_RIBOSOME | 0.00000 | 0.00019 | 0.65944 | 0.60936 | 1.32006 | 192 |
| GO_RNA_CATABOLIC_PROCESS | 0.00000 | 0.00000 | 0.76146 | 0.59494 | 1.30505 | 334 |
| GO_RNA_EXPORT_FROM_NUCLEUS | 0.00074 | 0.02782 | 0.47727 | 0.58991 | 1.25992 | 124 |
| GO_RNA_LOCALIZATION | 0.00145 | 0.04372 | 0.45506 | 0.55579 | 1.20478 | 198 |
| GO_RNA_SPLICING | 0.00002 | 0.00234 | 0.57561 | 0.55150 | 1.21227 | 373 |
| GO_RNA_SPLICING_VIA_TRANSESTERIFICATION_REACTIONS | 0.00005 | 0.00436 | 0.55733 | 0.55557 | 1.21421 | 302 |
| GO_SECRETORY_GRANULE | 0.00065 | 0.02583 | 0.47727 | 0.52150 | 1.14835 | 454 |
| GO_SEXUAL_REPRODUCTION | 0.00000 | 0.00001 | 0.73376 | 0.56539 | 1.24578 | 482 |
| GO_SEX_DIFFERENTIATION | 0.00016 | 0.01073 | 0.51885 | 0.58954 | 1.26822 | 153 |
| GO_SISTER_CHROMATID_SEGREGATION | 0.00010 | 0.00772 | 0.53843 | 0.58367 | 1.25948 | 168 |
| GO_SMALL_RIBOSOMAL_SUBUNIT | 0.00028 | 0.01562 | 0.49849 | 0.66203 | 1.38663 | 63 |
| GO_SPERM_EGG_RECOGNITION | 0.00032 | 0.01623 | 0.49849 | 0.73745 | 1.48848 | 29 |
| GO_SPINDLE | 0.00001 | 0.00123 | 0.59333 | 0.56554 | 1.23602 | 302 |
| GO_SPINDLE_POLE | 0.00016 | 0.01107 | 0.51885 | 0.59838 | 1.28350 | 136 |
| GO_SPLICEOSOMAL_COMPLEX | 0.00066 | 0.02583 | 0.47727 | 0.57245 | 1.23561 | 167 |
| GO_STRUCTURAL_CONSTITUENT_OF_CYTOSKELETON | 0.00115 | 0.03753 | 0.45506 | 0.63297 | 1.32820 | 66 |
| GO_STRUCTURAL_CONSTITUENT_OF_RIBOSOME | 0.00000 | 0.00012 | 0.65944 | 0.64198 | 1.37852 | 139 |
| GO_STRUCTURAL_MOLECULE_ACTIVITY | 0.00000 | 0.00000 | 0.81404 | 0.59123 | 1.30000 | 403 |
| GO_SUPRAMOLECULAR_FIBER_ORGANIZATION | 0.00087 | 0.03130 | 0.47727 | 0.52303 | 1.15232 | 463 |
| GO_SYNAPSE_ORGANIZATION | 0.00113 | 0.03752 | 0.45506 | 0.54092 | 1.17773 | 252 |
| GO_TRANSCRIPTION_COACTIVATOR_ACTIVITY | 0.00176 | 0.04850 | 0.45506 | 0.54852 | 1.18962 | 211 |
| GO_TRANSCRIPTION_COREGULATOR_ACTIVITY | 0.00044 | 0.02008 | 0.49849 | 0.53138 | 1.16832 | 383 |
| GO_TRANSLATIONAL_INITIATION | 0.00000 | 0.00000 | | 0.68350 | 1.47434 | 163 |
| GO_TRANSLATION_REGULATOR_ACTIVITY | 0.00063 | 0.02583 | 0.47727 | 0.59682 | 1.27223 | 108 |

| **Cluster 1** | | | | | | |
|---|---|---|---|---|---|---|
| GO_TRANSLATION_REGULATOR_ACTIVITY_NUCLEIC_ACID_E | 0.00081 | 0.02934 | 0.47727 | 0.61710 | 1.30699 | 85 |
| GO_TRANSMEMBRANE_RECEPTOR_PROTEIN_TYROSINE_KIN | 0.00117 | 0.03793 | 0.45506 | 0.51928 | 1.14284 | 445 |
| GO_UBIQUITIN_LIKE_PROTEIN_LIGASE_BINDING | 0.00041 | 0.01957 | 0.49849 | 0.55541 | 1.20863 | 248 |
| GO_VESICLE_LUMEN | 0.00027 | 0.01530 | 0.49849 | 0.57678 | 1.24640 | 177 |
| GO_VIRAL_GENE_EXPRESSION | 0.00000 | 0.00000 | 0.76146 | 0.65182 | 1.40692 | 167 |
| GO_VIRAL_LIFE_CYCLE | 0.00065 | 0.02583 | 0.47727 | 0.55134 | 1.19659 | 214 |
| HP_ABNORMALITY_OF_ESOPHAGUS_PHYSIOLOGY | 0.00023 | 0.01359 | 0.51885 | 0.53759 | 1.18160 | 369 |
| HP_ABNORMALITY_OF_MOUTH_SIZE | 0.00130 | 0.04064 | 0.45506 | 0.54962 | 1.19359 | 220 |
| HP_ABNORMALITY_OF_UPPER_LIP | 0.00168 | 0.04752 | 0.45506 | 0.51553 | 1.13593 | 482 |
| HP_ABNORMAL_ASTROCYTE_MORPHOLOGY | 0.00132 | 0.04081 | 0.45506 | 0.71163 | 1.43637 | 29 |
| HP_CONGENITAL_ABNORMAL_HAIR_PATTERN | 0.00176 | 0.04850 | 0.45506 | 0.57334 | 1.23112 | 139 |
| HP_FATIGUE | 0.00031 | 0.01623 | 0.49849 | 0.56268 | 1.22190 | 219 |
| HP_INFERTILITY | 0.00001 | 0.00125 | 0.59333 | 0.66103 | 1.40002 | 85 |
| HP_SLEEP_DISTURBANCE | 0.00079 | 0.02885 | 0.47727 | 0.54569 | 1.18846 | 254 |
| **Cluster 2** | | | | | | |
| pathway | pval | padj | log2err | ES | NES | size |
| GO_ACTIVATION_OF_IMMUNE_RESPONSE | 0.00000 | 0.00004 | 0.65944 | 0.61656 | 1.23895 | 255 |
| GO_ACTIVATION_OF_INNATE_IMMUNE_RESPONSE | 0.00000 | 0.00000 | 0.73376 | 0.71834 | 1.40612 | 100 |
| GO_ACTIVE_ION_TRANSMEMBRANE_TRANSPORTER_ACTIV | 0.00302 | 0.03933 | 0.43171 | 0.60872 | 1.19578 | 114 |
| GO_ACTIVE_TRANSMEMBRANE_TRANSPORTER_ACTIVITY | 0.00046 | 0.00919 | 0.49849 | 0.59372 | 1.18528 | 186 |
| GO_AEROBIC_RESPIRATION | 0.00000 | 0.00012 | 0.62726 | 0.72155 | 1.39807 | 70 |
| GO_AMIDE_BINDING | 0.00050 | 0.00974 | 0.47727 | 0.59006 | 1.18107 | 205 |
| GO_AMINE_METABOLIC_PROCESS | 0.00000 | 0.00000 | 0.82666 | 0.75939 | 1.48046 | 88 |
| GO_ANAPHASE_PROMOTING_COMPLEX_DEPENDENT_CATA | 0.00000 | 0.00000 | 0.81404 | 0.76923 | 1.49427 | 77 |
| GO_ANATOMICAL_STRUCTURE_HOMEOSTASIS | 0.00383 | 0.04728 | 0.43171 | 0.56007 | 1.12797 | 285 |
| GO_ANIMAL_ORGAN_REGENERATION | 0.00357 | 0.04506 | 0.43171 | 0.68679 | 1.30635 | 41 |
| GO_ANTIGEN_PROCESSING_AND_PRESENTATION | 0.00000 | 0.00000 | 0.71951 | 0.66563 | 1.32481 | 164 |
| GO_ANTIGEN_PROCESSING_AND_PRESENTATION_OF_EXOG | 0.00000 | 0.00000 | 0.74774 | 0.78576 | 1.51685 | 62 |
| GO_ANTIGEN_PROCESSING_AND_PRESENTATION_OF_PEPT | 0.00000 | 0.00000 | 0.71951 | 0.67151 | 1.33085 | 145 |
| GO_ANTIGEN_PROCESSING_AND_PRESENTATION_OF_PEPT | 0.00000 | 0.00000 | 0.76146 | 0.75522 | 1.46594 | 76 |
| GO_ANTIGEN_RECEPTOR_MEDIATED_SIGNALING_PATHWAY | 0.00004 | 0.00135 | 0.55733 | 0.62597 | 1.24237 | 150 |
| GO_ANTIOXIDANT_ACTIVITY | 0.00163 | 0.02413 | 0.45506 | 0.70641 | 1.34208 | 40 |
| GO_APOPTOTIC_SIGNALING_PATHWAY | 0.00001 | 0.00050 | 0.59333 | 0.57369 | 1.16111 | 427 |
| GO_ATPASE_ACTIVITY | 0.00178 | 0.02564 | 0.45506 | 0.56245 | 1.13375 | 312 |
| GO_ATP_BIOSYNTHETIC_PROCESS | 0.00012 | 0.00291 | 0.53843 | 0.79924 | 1.47936 | 23 |
| GO_ATP_METABOLIC_PROCESS | 0.00000 | 0.00000 | | 0.69164 | 1.38631 | 218 |
| GO_ATP_SYNTHESIS_COUPLED_ELECTRON_TRANSPORT | 0.00000 | 0.00000 | 0.80122 | 0.77658 | 1.50883 | 78 |
| GO_AZUROPHIL_GRANULE | 0.00036 | 0.00762 | 0.49849 | 0.62912 | 1.23427 | 111 |
| GO_AZUROPHIL_GRANULE_LUMEN | 0.00224 | 0.03081 | 0.43171 | 0.65394 | 1.26129 | 61 |
| GO_BINDING_OF_SPERM_TO_ZONA_PELLUCIDA | 0.00005 | 0.00158 | 0.55733 | 0.83818 | 1.53080 | 20 |
| GO_BLOOD_MICROPARTICLE | 0.00001 | 0.00045 | 0.59333 | 0.75279 | 1.43594 | 43 |
| GO_CADHERIN_BINDING | 0.00000 | 0.00000 | | 0.66042 | 1.32829 | 265 |
| GO_CADHERIN_BINDING_INVOLVED_IN_CELL_CELL_ADHESI | 0.00176 | 0.02556 | 0.45506 | 0.80273 | 1.44293 | 15 |
| GO_CALCIUM_DEPENDENT_PROTEIN_BINDING | 0.00018 | 0.00408 | 0.51885 | 0.72085 | 1.37415 | 42 |
| GO_CANONICAL_WNT_SIGNALING_PATHWAY | 0.00000 | 0.00019 | 0.61053 | 0.61118 | 1.22595 | 226 |
| GO_CARBOHYDRATE_BIOSYNTHETIC_PROCESS | 0.00154 | 0.02305 | 0.45506 | 0.60320 | 1.19143 | 132 |
| GO_CARBOHYDRATE_CATABOLIC_PROCESS | 0.00017 | 0.00404 | 0.51885 | 0.62253 | 1.23296 | 142 |
| GO_CARBOHYDRATE_METABOLIC_PROCESS | 0.00089 | 0.01526 | 0.47727 | 0.55774 | 1.12828 | 403 |
| GO_CARBOHYDRATE_TRANSMEMBRANE_TRANSPORTER_AC | 0.00001 | 0.00034 | 0.59333 | 0.86411 | 1.57814 | 20 |
| GO_CATALYTIC_ACTIVITY_ACTING_ON_RNA | 0.00085 | 0.01464 | 0.47727 | 0.56498 | 1.13909 | 315 |
| GO_CATALYTIC_STEP_2_SPLICEOSOME | 0.00000 | 0.00006 | 0.64355 | 0.70800 | 1.37801 | 82 |
| GO_CATION_CATION_ANTIPORTER_ACTIVITY | 0.00076 | 0.01371 | 0.47727 | 0.81901 | 1.47220 | 15 |
| GO_CELLULAR_AMINO_ACID_METABOLIC_PROCESS | 0.00000 | 0.00003 | 0.65944 | 0.63186 | 1.26680 | 220 |
| GO_CELLULAR_BIOGENIC_AMINE_METABOLIC_PROCESS | 0.00014 | 0.00344 | 0.51885 | 0.76696 | 1.43640 | 29 |
| GO_CELLULAR_COMPONENT_DISASSEMBLY | 0.00000 | 0.00000 | | 0.61592 | 1.24710 | 442 |
| GO_CELLULAR_IRON_ION_HOMEOSTASIS | 0.00405 | 0.04964 | 0.40702 | 0.66979 | 1.27761 | 43 |
| GO_CELLULAR_KETONE_METABOLIC_PROCESS | 0.00000 | 0.00005 | 0.64355 | 0.65008 | 1.29239 | 157 |
| GO_CELLULAR_MONOVALENT_INORGANIC_CATION_HOME( | 0.00390 | 0.04801 | 0.43171 | 0.63482 | 1.23225 | 76 |
| GO_CELLULAR_NITROGEN_COMPOUND_CATABOLIC_PROCE | 0.00000 | 0.00000 | | 0.67248 | 1.36278 | 463 |
| GO_CELLULAR_OXIDANT_DETOXIFICATION | 0.00181 | 0.02599 | 0.45506 | 0.68376 | 1.30767 | 48 |
| GO_CELLULAR_PROTEIN_COMPLEX_DISASSEMBLY | 0.00000 | 0.00000 | | 0.68899 | 1.37632 | 190 |
| GO_CELLULAR_RESPIRATION | 0.00000 | 0.00000 | | 0.72006 | 1.42788 | 147 |
| GO_CELLULAR_RESPONSE_TO_HEAT | 0.00009 | 0.00248 | 0.53843 | 0.64890 | 1.26856 | 97 |
| GO_CELLULAR_RESPONSE_TO_ORGANIC_CYCLIC_COMPOUI | 0.00364 | 0.04581 | 0.43171 | 0.55839 | 1.12590 | 316 |
| GO_CELLULAR_RESPONSE_TO_OXYGEN_LEVELS | 0.00000 | 0.00000 | 0.74774 | 0.67417 | 1.34210 | 165 |
| GO_CELLULAR_RESPONSE_TO_TOXIC_SUBSTANCE | 0.00191 | 0.02701 | 0.45506 | 0.65949 | 1.26803 | 57 |
| GO_CELL_ACTIVATION_INVOLVED_IN_IMMUNE_RESPONSE | 0.00000 | 0.00000 | | 0.62526 | 1.26575 | 431 |
| GO_CELL_ADHESION_MOLECULE_BINDING | 0.00000 | 0.00000 | 0.80122 | 0.62061 | 1.25321 | 364 |
| GO_CELL_BODY | 0.00128 | 0.02068 | 0.45506 | 0.56680 | 1.14225 | 309 |
| GO_CELL_CELL_RECOGNITION | 0.00010 | 0.00269 | 0.53843 | 0.74772 | 1.41756 | 37 |
| GO_CELL_CELL_SIGNALING_BY_WNT | 0.00011 | 0.00278 | 0.53843 | 0.57581 | 1.16230 | 353 |
| GO_CELL_CYCLE_G2_M_PHASE_TRANSITION | 0.00004 | 0.00115 | 0.55733 | 0.59872 | 1.20101 | 234 |
| GO_CELL_DIVISION | 0.00016 | 0.00384 | 0.51885 | 0.55663 | 1.12801 | 463 |

| Cluster 1 | | | | | | |
|---|---|---|---|---|---|---|
| GO_CELL_FATE_DETERMINATION | 0.00235 | 0.03199 | 0.43171 | 0.79236 | 1.43209 | 16 |
| GO_CELL_RECOGNITION | 0.00001 | 0.00050 | 0.59333 | 0.70238 | 1.36094 | 72 |
| GO_CELL_REDOX_HOMEOSTASIS | 0.00088 | 0.01515 | 0.47727 | 0.71177 | 1.35182 | 39 |
| GO_CELL_SUBSTRATE_JUNCTION | 0.00000 | 0.00000 | | 0.63606 | 1.28259 | 321 |
| GO_CELL_SURFACE_RECEPTOR_SIGNALING_PATHWAY_INVC | 0.00119 | 0.01944 | 0.45506 | 0.55622 | 1.12496 | 395 |
| GO_CHAPERONE_COMPLEX | 0.00006 | 0.00167 | 0.55733 | 0.83472 | 1.53287 | 21 |
| GO_CHAPERONE_MEDIATED_PROTEIN_COMPLEX_ASSEMBL | 0.00145 | 0.02226 | 0.45506 | 0.79100 | 1.43364 | 18 |
| GO_CHROMOSOME_SEGREGATION | 0.00121 | 0.01964 | 0.45506 | 0.57049 | 1.14793 | 272 |
| GO_COLLAGEN_CATABOLIC_PROCESS | 0.00342 | 0.04390 | 0.43171 | 0.76144 | 1.39064 | 20 |
| GO_CONTRACTILE_FIBER | 0.00152 | 0.02288 | 0.45506 | 0.59964 | 1.18685 | 140 |
| GO_CORTICAL_CYTOSKELETON | 0.00209 | 0.02929 | 0.43171 | 0.63417 | 1.23191 | 77 |
| GO_COTRANSLATIONAL_PROTEIN_TARGETING_TO_MEMBR/ | 0.00000 | 0.00000 | | 0.81340 | 1.58996 | 95 |
| GO_CYCLIN_DEPENDENT_PROTEIN_KINASE_HOLOENZYME_ | 0.00383 | 0.04728 | 0.43171 | 0.68618 | 1.30322 | 39 |
| GO_CYTOCHROME_COMPLEX | 0.00133 | 0.02140 | 0.45506 | 0.77449 | 1.42225 | 21 |
| GO_CYTOKINE_MEDIATED_SIGNALING_PATHWAY | 0.00000 | 0.00003 | 0.65944 | 0.59154 | 1.19706 | 411 |
| GO_CYTOPLASMIC_TRANSLATION | 0.00000 | 0.00000 | 0.73376 | 0.72322 | 1.41267 | 93 |
| GO_CYTOPLASMIC_TRANSLATIONAL_INITIATION | 0.00055 | 0.01048 | 0.47727 | 0.74388 | 1.39723 | 31 |
| GO_CYTOSKELETON_DEPENDENT_CYTOKINESIS | 0.00276 | 0.03649 | 0.43171 | 0.63913 | 1.23951 | 74 |
| GO_CYTOSOLIC_LARGE_RIBOSOMAL_SUBUNIT | 0.00000 | 0.00000 | 0.81404 | 0.82458 | 1.58116 | 53 |
| GO_CYTOSOLIC_RIBOSOME | 0.00000 | 0.00000 | | 0.80224 | 1.56866 | 98 |
| GO_CYTOSOLIC_SMALL_RIBOSOMAL_SUBUNIT | 0.00000 | 0.00002 | 0.67496 | 0.79329 | 1.51319 | 43 |
| GO_DEFENSE_RESPONSE_TO_GRAM_NEGATIVE_BACTERIUN | 0.00015 | 0.00358 | 0.51885 | 0.79678 | 1.47480 | 23 |
| GO_DETOXIFICATION | 0.00246 | 0.03321 | 0.43171 | 0.64245 | 1.24317 | 68 |
| GO_DISORDERED_DOMAIN_SPECIFIC_BINDING | 0.00220 | 0.03044 | 0.43171 | 0.74657 | 1.39660 | 28 |
| GO_DNA_BIOSYNTHETIC_PROCESS | 0.00024 | 0.00535 | 0.51885 | 0.61144 | 1.21462 | 153 |
| GO_DNA_DAMAGE_RESPONSE_DETECTION_OF_DNA_DAM/ | 0.00216 | 0.03019 | 0.43171 | 0.70429 | 1.33494 | 36 |
| GO_DNA_GEOMETRIC_CHANGE | 0.00228 | 0.03125 | 0.43171 | 0.61549 | 1.20532 | 102 |
| GO_DOUBLE_STRANDED_RNA_BINDING | 0.00026 | 0.00572 | 0.49849 | 0.68676 | 1.32045 | 57 |
| GO_ELECTRON_TRANSFER_ACTIVITY | 0.00000 | 0.00014 | 0.62726 | 0.70499 | 1.37062 | 79 |
| GO_ELECTRON_TRANSPORT_CHAIN | 0.00000 | 0.00000 | 0.82666 | 0.71096 | 1.40651 | 136 |
| GO_EMBRYO_IMPLANTATION | 0.00327 | 0.04227 | 0.43171 | 0.70839 | 1.33056 | 31 |
| GO_ENDOPEPTIDASE_COMPLEX | 0.00000 | 0.00004 | 0.65944 | 0.73894 | 1.42644 | 63 |
| GO_ENDOPEPTIDASE_REGULATOR_ACTIVITY | 0.00062 | 0.01145 | 0.47727 | 0.66593 | 1.29028 | 70 |
| GO_ENDOPLASMIC_RETICULUM_GOLGI_INTERMEDIATE_CC | 0.00143 | 0.02208 | 0.45506 | 0.61908 | 1.21069 | 99 |
| GO_ENERGY_DERIVATION_BY_OXIDATION_OF_ORGANIC_CC | 0.00000 | 0.00000 | 0.82666 | 0.67009 | 1.34063 | 201 |
| GO_EPHRIN_RECEPTOR_SIGNALING_PATHWAY | 0.00298 | 0.03893 | 0.43171 | 0.63985 | 1.23856 | 69 |
| GO_ESTABLISHMENT_OF_PROTEIN_LOCALIZATION_TO_CHR | 0.00016 | 0.00384 | 0.51885 | 0.81027 | 1.49404 | 22 |
| GO_ESTABLISHMENT_OF_PROTEIN_LOCALIZATION_TO_END | 0.00000 | 0.00000 | | 0.81295 | 1.59381 | 107 |
| GO_ESTABLISHMENT_OF_PROTEIN_LOCALIZATION_TO_MEN | 0.00000 | 0.00000 | | 0.68819 | 1.38435 | 270 |
| GO_ESTABLISHMENT_OF_PROTEIN_LOCALIZATION_TO_ORG | 0.00000 | 0.00000 | | 0.64777 | 1.31276 | 461 |
| GO_ESTABLISHMENT_OF_PROTEIN_LOCALIZATION_TO_TELC | 0.00005 | 0.00150 | 0.55733 | 0.85250 | 1.54335 | 17 |
| GO_ESTABLISHMENT_OF_RNA_LOCALIZATION | 0.00003 | 0.00096 | 0.55733 | 0.62091 | 1.23810 | 174 |
| GO_ESTABLISHMENT_OF_TISSUE_POLARITY | 0.00000 | 0.00002 | 0.67496 | 0.71086 | 1.38848 | 92 |
| GO_EUKARYOTIC_TRANSLATION_INITIATION_FACTOR_3_CO | 0.00091 | 0.01549 | 0.47727 | 0.81588 | 1.46658 | 15 |
| GO_FC_EPSILON_RECEPTOR_SIGNALING_PATHWAY | 0.00000 | 0.00002 | 0.67496 | 0.70710 | 1.38115 | 92 |
| GO_FC_RECEPTOR_SIGNALING_PATHWAY | 0.00000 | 0.00004 | 0.65944 | 0.66312 | 1.31336 | 142 |
| GO_FERTILIZATION | 0.00013 | 0.00314 | 0.51885 | 0.66806 | 1.30028 | 82 |
| GO_FICOLIN_1_RICH_GRANULE | 0.00000 | 0.00000 | | 0.74783 | 1.46520 | 105 |
| GO_GENERATION_OF_PRECURSOR_METABOLITES_AND_EN | 0.00000 | 0.00000 | | 0.65313 | 1.31895 | 366 |
| GO_GENE_SILENCING | 0.00075 | 0.01342 | 0.47727 | 0.59868 | 1.19462 | 184 |
| GO_GLYCOLYTIC_PROCESS_THROUGH_FRUCTOSE_6_PHOSP | 0.00028 | 0.00612 | 0.49849 | 0.78484 | 1.45270 | 23 |
| GO_GTPASE_ACTIVITY | 0.00000 | 0.00011 | 0.62726 | 0.62388 | 1.24996 | 211 |
| GO_GTP_METABOLIC_PROCESS | 0.00185 | 0.02634 | 0.45506 | 0.78658 | 1.42562 | 18 |
| GO_GUANYL_NUCLEOTIDE_BINDING | 0.00001 | 0.00021 | 0.61053 | 0.60847 | 1.22214 | 247 |
| GO_HEMATOPOIETIC_PROGENITOR_CELL_DIFFERENTIATION | 0.00003 | 0.00094 | 0.57561 | 0.65222 | 1.28366 | 121 |
| GO_HEMATOPOIETIC_STEM_CELL_DIFFERENTIATION | 0.00000 | 0.00001 | 0.67496 | 0.74051 | 1.43342 | 69 |
| GO_HEXOSE_CATABOLIC_PROCESS | 0.00372 | 0.04646 | 0.43171 | 0.67349 | 1.28386 | 42 |
| GO_HOST_CELLULAR_COMPONENT | 0.00117 | 0.01924 | 0.45506 | 0.67381 | 1.29382 | 55 |
| GO_HYDROGEN_PEROXIDE_METABOLIC_PROCESS | 0.00046 | 0.00919 | 0.49849 | 0.76882 | 1.42872 | 25 |
| GO_IMMUNE_RESPONSE_REGULATING_SIGNALING_PATHW | 0.00001 | 0.00036 | 0.59333 | 0.60617 | 1.21703 | 242 |
| GO_IMPORT_INTO_NUCLEUS | 0.00064 | 0.01168 | 0.47727 | 0.62375 | 1.23013 | 126 |
| GO_INNATE_IMMUNE_RESPONSE | 0.00000 | 0.00001 | 0.67496 | 0.59271 | 1.19923 | 408 |
| GO_INNATE_IMMUNE_RESPONSE_ACTIVATING_SIGNAL_TR/ | 0.00000 | 0.00000 | 0.70498 | 0.73350 | 1.42663 | 81 |
| GO_INNER_MITOCHONDRIAL_MEMBRANE_PROTEIN_COMF | 0.00000 | 0.00000 | 0.82666 | 0.75433 | 1.47343 | 93 |
| GO_INORGANIC_ION_TRANSMEMBRANE_TRANSPORT | 0.00257 | 0.03433 | 0.43171 | 0.55237 | 1.11742 | 403 |
| GO_INTERACTION_WITH_SYMBIONT | 0.00342 | 0.04390 | 0.43171 | 0.66622 | 1.27716 | 52 |
| GO_INTERLEUKIN_1_MEDIATED_SIGNALING_PATHWAY | 0.00000 | 0.00000 | 0.76146 | 0.74401 | 1.44913 | 83 |
| GO_INTRINSIC_APOPTOTIC_SIGNALING_PATHWAY | 0.00090 | 0.01538 | 0.47727 | 0.57852 | 1.16044 | 226 |
| GO_LARGE_RIBOSOMAL_SUBUNIT | 0.00000 | 0.00000 | | 0.78087 | 1.53175 | 108 |
| GO_LEUKOCYTE_MEDIATED_IMMUNITY | 0.00000 | 0.00000 | | 0.62547 | 1.26758 | 461 |
| GO_LYSOSOMAL_LUMEN | 0.00009 | 0.00239 | 0.53843 | 0.69899 | 1.34727 | 60 |
| GO_MATERNAL_PLACENTA_DEVELOPMENT | 0.00064 | 0.01168 | 0.47727 | 0.79237 | 1.45509 | 21 |
| GO_MATURATION_OF_SSU_RRNA | 0.00152 | 0.02288 | 0.45506 | 0.67480 | 1.28983 | 47 |

| Cluster 1 | | | | | | |
|---|---|---|---|---|---|---|
| GO_MEMBRANE_REGION | 0.00024 | 0.00544 | 0.51885 | 0.60068 | 1.20105 | 197 |
| GO_MITOCHONDRIAL_ELECTRON_TRANSPORT_NADH_TO_U | 0.00000 | 0.00012 | 0.62726 | 0.76629 | 1.46364 | 45 |
| GO_MITOCHONDRIAL_GENE_EXPRESSION | 0.00000 | 0.00000 | | 0.71661 | 1.42387 | 154 |
| GO_MITOCHONDRIAL_LARGE_RIBOSOMAL_SUBUNIT | 0.00000 | 0.00002 | 0.67496 | 0.77024 | 1.47862 | 54 |
| GO_MITOCHONDRIAL_MATRIX | 0.00000 | 0.00000 | | 0.65125 | 1.31667 | 389 |
| GO_MITOCHONDRIAL_MEMBRANE_ORGANIZATION | 0.00001 | 0.00031 | 0.59333 | 0.67542 | 1.32068 | 98 |
| GO_MITOCHONDRIAL_PROTEIN_COMPLEX | 0.00000 | 0.00000 | | 0.75039 | 1.50359 | 214 |
| GO_MITOCHONDRIAL_RESPIRATORY_CHAIN_COMPLEX_ASS | 0.00021 | 0.00471 | 0.51885 | 0.66024 | 1.28677 | 87 |
| GO_MITOCHONDRIAL_SMALL_RIBOSOMAL_SUBUNIT | 0.00287 | 0.03764 | 0.43171 | 0.74087 | 1.38594 | 28 |
| GO_MITOCHONDRIAL_TRANSLATION | 0.00000 | 0.00000 | | 0.73058 | 1.44277 | 130 |
| GO_MITOCHONDRIAL_TRANSLATIONAL_TERMINATION | 0.00000 | 0.00000 | 0.82666 | 0.76114 | 1.48388 | 88 |
| GO_MITOCHONDRIAL_TRANSPORT | 0.00008 | 0.00224 | 0.53843 | 0.60356 | 1.20803 | 204 |
| GO_MITOCHONDRION_ORGANIZATION | 0.00000 | 0.00000 | | 0.62176 | 1.25862 | 430 |
| GO_MITOTIC_NUCLEAR_DIVISION | 0.00079 | 0.01404 | 0.47727 | 0.58277 | 1.16907 | 230 |
| GO_MITOTIC_SISTER_CHROMATID_SEGREGATION | 0.00070 | 0.01275 | 0.47727 | 0.61113 | 1.21056 | 143 |
| GO_MITOTIC_SPINDLE_ORGANIZATION | 0.00261 | 0.03468 | 0.43171 | 0.61135 | 1.19764 | 104 |
| GO_MODULATION_BY_HOST_OF_VIRAL_PROCESS | 0.00239 | 0.03242 | 0.43171 | 0.76985 | 1.40601 | 20 |
| GO_MONOCARBOXYLIC_ACID_METABOLIC_PROCESS | 0.00019 | 0.00447 | 0.51885 | 0.56969 | 1.15106 | 377 |
| GO_MONOSACCHARIDE_BIOSYNTHETIC_PROCESS | 0.00053 | 0.01014 | 0.47727 | 0.67004 | 1.29550 | 66 |
| GO_MONOSACCHARIDE_METABOLIC_PROCESS | 0.00305 | 0.03973 | 0.43171 | 0.57324 | 1.14606 | 193 |
| GO_MONOVALENT_INORGANIC_CATION_TRANSMEMBRAN | 0.00002 | 0.00064 | 0.57561 | 0.63370 | 1.25777 | 151 |
| GO_MONOVALENT_INORGANIC_CATION_TRANSPORT | 0.00059 | 0.01104 | 0.47727 | 0.58075 | 1.16650 | 253 |
| GO_MORPHOGENESIS_OF_AN_EPITHELIUM | 0.00012 | 0.00291 | 0.53843 | 0.57826 | 1.16656 | 337 |
| GO_MORPHOGENESIS_OF_A_POLARIZED_EPITHELIUM | 0.00000 | 0.00000 | 0.76146 | 0.72100 | 1.41354 | 107 |
| GO_MRNA_3_END_PROCESSING | 0.00054 | 0.01037 | 0.47727 | 0.64030 | 1.25161 | 95 |
| GO_MRNA_5_UTR_BINDING | 0.00346 | 0.04402 | 0.43171 | 0.73635 | 1.36627 | 24 |
| GO_MRNA_BINDING | 0.00000 | 0.00000 | 0.78819 | 0.64648 | 1.29841 | 246 |
| GO_MRNA_EXPORT_FROM_NUCLEUS | 0.00024 | 0.00535 | 0.51885 | 0.63351 | 1.24121 | 105 |
| GO_MRNA_PROCESSING | 0.00000 | 0.00000 | | 0.62787 | 1.27130 | 442 |
| GO_MRNA_TRANSPORT | 0.00056 | 0.01062 | 0.47727 | 0.61672 | 1.21930 | 134 |
| GO_MULTI_ORGANISM_LOCALIZATION | 0.00137 | 0.02160 | 0.45506 | 0.67615 | 1.29827 | 56 |
| GO_MYELOID_LEUKOCYTE_ACTIVATION | 0.00000 | 0.00000 | | 0.63328 | 1.28110 | 400 |
| GO_MYELOID_LEUKOCYTE_MEDIATED_IMMUNITY | 0.00000 | 0.00000 | | 0.64665 | 1.30589 | 365 |
| GO_NADH_DEHYDROGENASE_ACTIVITY | 0.00002 | 0.00063 | 0.57561 | 0.77154 | 1.46317 | 38 |
| GO_NADH_DEHYDROGENASE_COMPLEX | 0.00001 | 0.00031 | 0.59333 | 0.77056 | 1.46569 | 41 |
| GO_NADH_DEHYDROGENASE_COMPLEX_ASSEMBLY | 0.00002 | 0.00065 | 0.57561 | 0.72828 | 1.39837 | 56 |
| GO_NADH_METABOLIC_PROCESS | 0.00028 | 0.00617 | 0.49849 | 0.75360 | 1.41550 | 31 |
| GO_NAD_METABOLIC_PROCESS | 0.00048 | 0.00947 | 0.49849 | 0.72249 | 1.37219 | 39 |
| GO_NCRNA_EXPORT_FROM_NUCLEUS | 0.00405 | 0.04964 | 0.40702 | 0.69320 | 1.31390 | 36 |
| GO_NCRNA_METABOLIC_PROCESS | 0.00000 | 0.00000 | 0.77494 | 0.61235 | 1.23878 | 404 |
| GO_NCRNA_PROCESSING | 0.00000 | 0.00000 | 0.76146 | 0.61631 | 1.24338 | 347 |
| GO_NEGATIVE_REGULATION_OF_APOPTOTIC_SIGNALING_P | 0.00165 | 0.02429 | 0.45506 | 0.59401 | 1.18065 | 156 |
| GO_NEGATIVE_REGULATION_OF_CANONICAL_WNT_SIGNAI | 0.00000 | 0.00000 | 0.76146 | 0.69785 | 1.37555 | 125 |
| GO_NEGATIVE_REGULATION_OF_CATALYTIC_ACTIVITY | 0.00135 | 0.02159 | 0.45506 | 0.55262 | 1.12021 | 466 |
| GO_NEGATIVE_REGULATION_OF_CELL_CYCLE | 0.00001 | 0.00035 | 0.59333 | 0.57118 | 1.15775 | 480 |
| GO_NEGATIVE_REGULATION_OF_CELL_CYCLE_G2_M_PHASI | 0.00000 | 0.00000 | 0.71951 | 0.72188 | 1.40732 | 88 |
| GO_NEGATIVE_REGULATION_OF_CELL_CYCLE_PHASE_TRAN | 0.00001 | 0.00022 | 0.61053 | 0.62619 | 1.25327 | 203 |
| GO_NEGATIVE_REGULATION_OF_CELL_CYCLE_PROCESS | 0.00000 | 0.00013 | 0.62726 | 0.60622 | 1.21983 | 272 |
| GO_NEGATIVE_REGULATION_OF_HYDROLASE_ACTIVITY | 0.00052 | 0.01000 | 0.47727 | 0.58814 | 1.17963 | 236 |
| GO_NEGATIVE_REGULATION_OF_MITOTIC_CELL_CYCLE | 0.00002 | 0.00060 | 0.57561 | 0.60023 | 1.20684 | 262 |
| GO_NEGATIVE_REGULATION_OF_MRNA_METABOLIC_PROC | 0.00122 | 0.01989 | 0.45506 | 0.65738 | 1.27129 | 67 |
| GO_NEGATIVE_REGULATION_OF_MRNA_SPLICING_VIA_SPL | 0.00117 | 0.01924 | 0.45506 | 0.77740 | 1.42761 | 21 |
| GO_NEGATIVE_REGULATION_OF_PEPTIDASE_ACTIVITY | 0.00028 | 0.00606 | 0.49849 | 0.63548 | 1.24670 | 112 |
| GO_NEGATIVE_REGULATION_OF_PROTEOLYSIS | 0.00071 | 0.01284 | 0.47727 | 0.59013 | 1.17765 | 183 |
| GO_NEGATIVE_REGULATION_OF_REACTIVE_OXYGEN_SPECI | 0.00346 | 0.04402 | 0.43171 | 0.77170 | 1.39865 | 18 |
| GO_NEGATIVE_REGULATION_OF_RNA_SPLICING | 0.00021 | 0.00475 | 0.51885 | 0.78891 | 1.46380 | 24 |
| GO_NEGATIVE_REGULATION_OF_WNT_SIGNALING_PATHWA | 0.00000 | 0.00001 | 0.70498 | 0.66549 | 1.31891 | 145 |
| GO_NIK_NF_KAPPAB_SIGNALING | 0.00000 | 0.00000 | 0.78819 | 0.71773 | 1.40992 | 114 |
| GO_NON_CANONICAL_WNT_SIGNALING_PATHWAY | 0.00000 | 0.00000 | 0.70498 | 0.69633 | 1.36604 | 109 |
| GO_NUCLEAR_ENVELOPE | 0.00088 | 0.01515 | 0.47727 | 0.56428 | 1.13824 | 344 |
| GO_NUCLEAR_EXPORT | 0.00004 | 0.00112 | 0.55733 | 0.61765 | 1.23246 | 184 |
| GO_NUCLEAR_PERIPHERY | 0.00152 | 0.02288 | 0.45506 | 0.61447 | 1.20728 | 116 |
| GO_NUCLEAR_PORE | 0.00137 | 0.02160 | 0.45506 | 0.65555 | 1.26749 | 66 |
| GO_NUCLEAR_TRANSCRIBED_MRNA_CATABOLIC_PROCESS | 0.00000 | 0.00000 | | 0.71216 | 1.42182 | 187 |
| GO_NUCLEAR_TRANSCRIBED_MRNA_CATABOLIC_PROCESS_ | 0.00000 | 0.00000 | | 0.80614 | 1.58396 | 115 |
| GO_NUCLEAR_TRANSPORT | 0.00000 | 0.00005 | 0.65944 | 0.60831 | 1.22558 | 294 |
| GO_NUCLEOBASE_CONTAINING_COMPOUND_TRANSPORT | 0.00010 | 0.00252 | 0.53843 | 0.60025 | 1.20381 | 221 |
| GO_NUCLEOBASE_CONTAINING_SMALL_MOLECULE_METAE | 0.00004 | 0.00133 | 0.55733 | 0.57606 | 1.16516 | 396 |
| GO_NUCLEOSIDE_DIPHOSPHATE_METABOLIC_PROCESS | 0.00008 | 0.00226 | 0.53843 | 0.64300 | 1.26150 | 111 |
| GO_NUCLEOSIDE_PHOSPHATE_BIOSYNTHETIC_PROCESS | 0.00100 | 0.01683 | 0.45506 | 0.59705 | 1.18815 | 163 |
| GO_NUCLEOSIDE_TRIPHOSPHATE_BIOSYNTHETIC_PROCESS | 0.00383 | 0.04728 | 0.43171 | 0.67488 | 1.28880 | 44 |
| GO_NUCLEOSIDE_TRIPHOSPHATE_METABOLIC_PROCESS | 0.00178 | 0.02564 | 0.45506 | 0.65091 | 1.25826 | 65 |
| GO_NUCLEOTIDE_EXCISION_REPAIR | 0.00376 | 0.04673 | 0.43171 | 0.60849 | 1.19203 | 104 |

| Cluster 1 | | | | | | |
|---|---|---|---|---|---|---|
| GO_NUCLEOTIDE_PHOSPHORYLATION | 0.00005 | 0.00150 | 0.55733 | 0.65531 | 1.28154 | 99 |
| GO_ORGANELLAR_RIBOSOME | 0.00000 | 0.00000 | 0.81404 | 0.75614 | 1.47318 | 84 |
| GO_ORGANELLE_ENVELOPE_LUMEN | 0.00121 | 0.01964 | 0.45506 | 0.64113 | 1.24542 | 77 |
| GO_ORGANELLE_FISSION | 0.00390 | 0.04801 | 0.43171 | 0.55420 | 1.11849 | 351 |
| GO_ORGANELLE_INNER_MEMBRANE | 0.00000 | 0.00000 | | 0.67166 | 1.35945 | 419 |
| GO_ORGANIC_CYCLIC_COMPOUND_CATABOLIC_PROCESS | 0.00000 | 0.00000 | | 0.67148 | 1.36140 | 484 |
| GO_ORGANIC_HYDROXY_COMPOUND_TRANSMEMBRANE_ | 0.00000 | 0.00012 | 0.62726 | 0.90338 | 1.63275 | 16 |
| GO_OUTER_MEMBRANE | 0.00083 | 0.01449 | 0.47727 | 0.60131 | 1.19319 | 149 |
| GO_OUTER_MITOCHONDRIAL_MEMBRANE_PROTEIN_COM | 0.00198 | 0.02793 | 0.43171 | 0.76574 | 1.40620 | 21 |
| GO_OXIDATIVE_PHOSPHORYLATION | 0.00000 | 0.00000 | | 0.75561 | 1.47919 | 101 |
| GO_OXIDOREDUCTASE_ACTIVITY | 0.00000 | 0.00000 | 0.77494 | 0.60116 | 1.21860 | 468 |
| GO_OXIDOREDUCTASE_ACTIVITY_ACTING_ON_A_HEME_GR | 0.00113 | 0.01878 | 0.45506 | 0.78671 | 1.43023 | 19 |
| GO_OXIDOREDUCTASE_ACTIVITY_ACTING_ON_NAD_P_H | 0.00001 | 0.00031 | 0.59333 | 0.69957 | 1.35792 | 76 |
| GO_OXIDOREDUCTASE_ACTIVITY_ACTING_ON_NAD_P_H_Q | 0.00006 | 0.00167 | 0.55733 | 0.73127 | 1.39777 | 47 |
| GO_OXIDOREDUCTASE_ACTIVITY_ACTING_ON_PEROXIDE_A | 0.00085 | 0.01464 | 0.47727 | 0.76052 | 1.41611 | 26 |
| GO_OXIDOREDUCTASE_COMPLEX | 0.00000 | 0.00000 | 0.76146 | 0.74367 | 1.44888 | 84 |
| GO_PEPTIDASE_COMPLEX | 0.00000 | 0.00010 | 0.62726 | 0.70838 | 1.37712 | 80 |
| GO_PEPTIDASE_REGULATOR_ACTIVITY | 0.00035 | 0.00740 | 0.49849 | 0.64001 | 1.25145 | 98 |
| GO_PEPTIDE_BINDING | 0.00011 | 0.00276 | 0.53843 | 0.62244 | 1.23543 | 151 |
| GO_PH_REDUCTION | 0.00253 | 0.03398 | 0.43171 | 0.67559 | 1.29257 | 49 |
| GO_PIGMENT_GRANULE | 0.00000 | 0.00000 | 0.70498 | 0.72283 | 1.41059 | 90 |
| GO_PLASMA_LIPOPROTEIN_PARTICLE_CLEARANCE | 0.00139 | 0.02173 | 0.45506 | 0.72654 | 1.36740 | 32 |
| GO_PLATELET_DEGRANULATION | 0.00044 | 0.00892 | 0.49849 | 0.67072 | 1.29656 | 65 |
| GO_POLYMERIC_CYTOSKELETAL_FIBER | 0.00167 | 0.02445 | 0.45506 | 0.55405 | 1.12127 | 424 |
| GO_POLYSOMAL_RIBOSOME | 0.00000 | 0.00007 | 0.64355 | 0.83458 | 1.56432 | 30 |
| GO_POLYSOME | 0.00000 | 0.00001 | 0.70498 | 0.75843 | 1.46611 | 65 |
| GO_POSITIVE_REGULATION_OF_ATPASE_ACTIVITY | 0.00209 | 0.02929 | 0.43171 | 0.67523 | 1.28983 | 46 |
| GO_POSITIVE_REGULATION_OF_CANONICAL_WNT_SIGNALI | 0.00000 | 0.00002 | 0.67496 | 0.68668 | 1.34712 | 109 |
| GO_POSITIVE_REGULATION_OF_CELLULAR_AMIDE_METABO | 0.00003 | 0.00096 | 0.55733 | 0.64892 | 1.27976 | 126 |
| GO_POSITIVE_REGULATION_OF_CELLULAR_PROTEIN_LOCAL | 0.00040 | 0.00828 | 0.49849 | 0.58024 | 1.16569 | 250 |
| GO_POSITIVE_REGULATION_OF_DEFENSE_RESPONSE | 0.00000 | 0.00015 | 0.62726 | 0.62841 | 1.25693 | 199 |
| GO_POSITIVE_REGULATION_OF_DNA_BIOSYNTHETIC_PROC | 0.00054 | 0.01037 | 0.47727 | 0.67843 | 1.30765 | 60 |
| GO_POSITIVE_REGULATION_OF_DNA_METABOLIC_PROCESS | 0.00078 | 0.01396 | 0.47727 | 0.60732 | 1.20432 | 147 |
| GO_POSITIVE_REGULATION_OF_ESTABLISHMENT_OF_PROT | 0.00028 | 0.00612 | 0.49849 | 0.58655 | 1.17922 | 260 |
| GO_POSITIVE_REGULATION_OF_IMMUNE_RESPONSE | 0.00000 | 0.00002 | 0.67496 | 0.60341 | 1.21720 | 342 |
| GO_POSITIVE_REGULATION_OF_IMMUNE_SYSTEM_PROCES | 0.00001 | 0.00049 | 0.59333 | 0.56726 | 1.15061 | 498 |
| GO_POSITIVE_REGULATION_OF_INNATE_IMMUNE_RESPON | 0.00000 | 0.00001 | 0.67496 | 0.67720 | 1.33738 | 131 |
| GO_POSITIVE_REGULATION_OF_INTRACELLULAR_TRANSPO | 0.00157 | 0.02343 | 0.45506 | 0.58753 | 1.17134 | 173 |
| GO_POSITIVE_REGULATION_OF_MRNA_PROCESSING | 0.00327 | 0.04227 | 0.43171 | 0.72676 | 1.35324 | 26 |
| GO_POSITIVE_REGULATION_OF_MULTI_ORGANISM_PROCE | 0.00085 | 0.01464 | 0.47727 | 0.78464 | 1.44090 | 21 |
| GO_POSITIVE_REGULATION_OF_ORGANELLE_ORGANIZATIO | 0.00157 | 0.02343 | 0.45506 | 0.55155 | 1.11806 | 466 |
| GO_POSITIVE_REGULATION_OF_PEPTIDASE_ACTIVITY | 0.00137 | 0.02160 | 0.45506 | 0.60159 | 1.18970 | 135 |
| GO_POSITIVE_REGULATION_OF_PROTEIN_LOCALIZATION_T | 0.00009 | 0.00244 | 0.53843 | 0.68709 | 1.32874 | 67 |
| GO_POSITIVE_REGULATION_OF_RESPONSE_TO_BIOTIC_STI | 0.00000 | 0.00002 | 0.67496 | 0.65852 | 1.30787 | 152 |
| GO_POSITIVE_REGULATION_OF_RESPONSE_TO_EXTERNAL_ | 0.00016 | 0.00373 | 0.51885 | 0.58459 | 1.17545 | 263 |
| GO_POSITIVE_REGULATION_OF_TELOMERASE_RNA_LOCALI | 0.00002 | 0.00061 | 0.57561 | 0.88559 | 1.59188 | 15 |
| GO_POSITIVE_REGULATION_OF_TELOMERE_MAINTENANCE | 0.00040 | 0.00828 | 0.49849 | 0.70634 | 1.34913 | 45 |
| GO_POSITIVE_REGULATION_OF_TELOMERE_MAINTENANCE | 0.00012 | 0.00301 | 0.53843 | 0.76558 | 1.44642 | 34 |
| GO_POSITIVE_REGULATION_OF_TRANSLATION | 0.00000 | 0.00006 | 0.64355 | 0.68372 | 1.34118 | 108 |
| GO_POSITIVE_REGULATION_OF_VIRAL_LIFE_CYCLE | 0.00074 | 0.01330 | 0.47727 | 0.68434 | 1.31020 | 50 |
| GO_POSITIVE_REGULATION_OF_VIRAL_PROCESS | 0.00000 | 0.00013 | 0.62726 | 0.69679 | 1.36011 | 91 |
| GO_POSITIVE_REGULATION_OF_VIRAL_TRANSCRIPTION | 0.00143 | 0.02208 | 0.45506 | 0.70619 | 1.33884 | 37 |
| GO_POSITIVE_REGULATION_OF_WNT_SIGNALING_PATHWA | 0.00001 | 0.00021 | 0.61053 | 0.65251 | 1.28860 | 130 |
| GO_POST_TRANSLATIONAL_PROTEIN_MODIFICATION | 0.00108 | 0.01792 | 0.45506 | 0.57180 | 1.14985 | 264 |
| GO_PRECATALYTIC_SPLICEOSOME | 0.00139 | 0.02173 | 0.45506 | 0.67847 | 1.29939 | 51 |
| GO_PRERIBOSOME | 0.00002 | 0.00072 | 0.57561 | 0.69734 | 1.35122 | 71 |
| GO_PROTEASOMAL_UBIQUITIN_INDEPENDENT_PROTEIN_C | 0.00028 | 0.00617 | 0.49849 | 0.82226 | 1.49029 | 18 |
| GO_PROTEASOME_ACCESSORY_COMPLEX | 0.00001 | 0.00040 | 0.59333 | 0.83313 | 1.54586 | 24 |
| GO_PROTEASOME_CORE_COMPLEX | 0.00013 | 0.00329 | 0.51885 | 0.83898 | 1.51888 | 17 |
| GO_PROTEIN_CONTAINING_COMPLEX_DISASSEMBLY | 0.00000 | 0.00000 | | 0.65685 | 1.32219 | 275 |
| GO_PROTEIN_CONTAINING_COMPLEX_LOCALIZATION | 0.00146 | 0.02237 | 0.45506 | 0.57755 | 1.15869 | 227 |
| GO_PROTEIN_DNA_COMPLEX_SUBUNIT_ORGANIZATION | 0.00059 | 0.01104 | 0.47727 | 0.59591 | 1.18946 | 185 |
| GO_PROTEIN_FOLDING | 0.00000 | 0.00000 | 0.70498 | 0.65548 | 1.30720 | 179 |
| GO_PROTEIN_FOLDING_CHAPERONE | 0.00036 | 0.00754 | 0.49849 | 0.80413 | 1.47669 | 21 |
| GO_PROTEIN_IMPORT | 0.00002 | 0.00062 | 0.57561 | 0.63109 | 1.25364 | 153 |
| GO_PROTEIN_IMPORT_INTO_MITOCHONDRIAL_MATRIX | 0.00283 | 0.03723 | 0.43171 | 0.77649 | 1.40733 | 18 |
| GO_PROTEIN_LIPID_COMPLEX_ASSEMBLY | 0.00011 | 0.00276 | 0.53843 | 0.83531 | 1.51858 | 19 |
| GO_PROTEIN_LIPID_COMPLEX_SUBUNIT_ORGANIZATION | 0.00061 | 0.01135 | 0.47727 | 0.79031 | 1.45724 | 22 |
| GO_PROTEIN_LOCALIZATION_TO_CHROMOSOME | 0.00202 | 0.02838 | 0.43171 | 0.64841 | 1.25635 | 70 |
| GO_PROTEIN_LOCALIZATION_TO_CHROMOSOME_TELOMER | 0.00002 | 0.00066 | 0.57561 | 0.81176 | 1.51857 | 28 |
| GO_PROTEIN_LOCALIZATION_TO_ENDOPLASMIC_RETICULU | 0.00000 | 0.00000 | | 0.78815 | 1.55599 | 129 |
| GO_PROTEIN_LOCALIZATION_TO_MEMBRANE | 0.00000 | 0.00000 | | 0.63179 | 1.28056 | 479 |
| GO_PROTEIN_LOCALIZATION_TO_MITOCHONDRION | 0.00001 | 0.00041 | 0.59333 | 0.65009 | 1.28023 | 122 |

| Cluster 1 | | | | | | |
|---|---|---|---|---|---|---|
| GO_PROTEIN_LOCALIZATION_TO_NUCLEUS | 0.00000 | 0.00002 | 0.67496 | 0.63186 | 1.26513 | 207 |
| GO_PROTEIN_MODIFICATION_BY_SMALL_PROTEIN_REMOV | 0.00055 | 0.01048 | 0.47727 | 0.58145 | 1.16636 | 234 |
| GO_PROTEIN_REFOLDING | 0.00324 | 0.04196 | 0.43171 | 0.76250 | 1.38622 | 19 |
| GO_PROTEIN_STABILIZATION | 0.00141 | 0.02190 | 0.45506 | 0.60300 | 1.19307 | 139 |
| GO_PROTEIN_TARGETING | 0.00000 | 0.00000 | | 0.66376 | 1.34001 | 358 |
| GO_PROTEIN_TARGETING_TO_MEMBRANE | 0.00000 | 0.00000 | | 0.74138 | 1.47590 | 166 |
| GO_PROTEIN_TARGETING_TO_MITOCHONDRION | 0.00004 | 0.00135 | 0.55733 | 0.67397 | 1.31309 | 84 |
| GO_PROTEIN_TRANSMEMBRANE_TRANSPORT | 0.00081 | 0.01425 | 0.47727 | 0.67993 | 1.30558 | 55 |
| GO_PROTON_TRANSMEMBRANE_TRANSPORT | 0.00000 | 0.00012 | 0.62726 | 0.69013 | 1.34544 | 88 |
| GO_PROTON_TRANSMEMBRANE_TRANSPORTER_ACTIVITY | 0.00052 | 0.01000 | 0.47727 | 0.66851 | 1.29528 | 70 |
| GO_PURINE_CONTAINING_COMPOUND_BIOSYNTHETIC_PR | 0.00092 | 0.01561 | 0.47727 | 0.61194 | 1.20574 | 124 |
| GO_PURINE_CONTAINING_COMPOUND_METABOLIC_PROC | 0.00009 | 0.00232 | 0.53843 | 0.58538 | 1.17875 | 288 |
| GO_PURINE_NUCLEOSIDE_TRIPHOSPHATE_METABOLIC_PR | 0.00019 | 0.00443 | 0.51885 | 0.71508 | 1.36758 | 48 |
| GO_PYRUVATE_METABOLIC_PROCESS | 0.00000 | 0.00011 | 0.62726 | 0.67814 | 1.32935 | 106 |
| GO_REACTIVE_OXYGEN_SPECIES_BIOSYNTHETIC_PROCESS | 0.00376 | 0.04673 | 0.43171 | 0.65286 | 1.25328 | 54 |
| GO_REACTIVE_OXYGEN_SPECIES_METABOLIC_PROCESS | 0.00032 | 0.00682 | 0.49849 | 0.61847 | 1.22493 | 142 |
| GO_REGENERATION | 0.00353 | 0.04487 | 0.43171 | 0.60747 | 1.19082 | 106 |
| GO_REGULATION_OF_ANIMAL_ORGAN_MORPHOGENESIS | 0.00000 | 0.00000 | 0.71951 | 0.68068 | 1.34668 | 138 |
| GO_REGULATION_OF_APOPTOTIC_SIGNALING_PATHWAY | 0.00036 | 0.00754 | 0.49849 | 0.57879 | 1.16554 | 282 |
| GO_REGULATION_OF_ATPASE_ACTIVITY | 0.00141 | 0.02190 | 0.45506 | 0.66859 | 1.29066 | 62 |
| GO_REGULATION_OF_ATP_METABOLIC_PROCESS | 0.00077 | 0.01384 | 0.47727 | 0.64582 | 1.25866 | 87 |
| GO_REGULATION_OF_CELLULAR_AMIDE_METABOLIC_PROC | 0.00000 | 0.00000 | 0.74774 | 0.61662 | 1.24385 | 342 |
| GO_REGULATION_OF_CELLULAR_AMINE_METABOLIC_PROC | 0.00000 | 0.00000 | 0.74774 | 0.78509 | 1.51556 | 62 |
| GO_REGULATION_OF_CELLULAR_AMINO_ACID_METABOLIC | 0.00000 | 0.00000 | 0.74774 | 0.79570 | 1.52750 | 54 |
| GO_REGULATION_OF_CELLULAR_KETONE_METABOLIC_PRO | 0.00000 | 0.00003 | 0.65944 | 0.68240 | 1.34306 | 121 |
| GO_REGULATION_OF_CELLULAR_PROTEIN_LOCALIZATION | 0.00009 | 0.00237 | 0.53843 | 0.56649 | 1.14617 | 408 |
| GO_REGULATION_OF_CELLULAR_RESPIRATION | 0.00316 | 0.04109 | 0.43171 | 0.76250 | 1.39257 | 20 |
| GO_REGULATION_OF_CELLULAR_RESPONSE_TO_HEAT | 0.00181 | 0.02599 | 0.45506 | 0.65149 | 1.25990 | 67 |
| GO_REGULATION_OF_CELL_CYCLE_G2_M_PHASE_TRANSITI | 0.00002 | 0.00063 | 0.57561 | 0.61563 | 1.22902 | 186 |
| GO_REGULATION_OF_CELL_CYCLE_PHASE_TRANSITION | 0.00005 | 0.00159 | 0.55733 | 0.57548 | 1.16275 | 377 |
| GO_REGULATION_OF_CHROMOSOME_ORGANIZATION | 0.00265 | 0.03509 | 0.43171 | 0.56333 | 1.13454 | 285 |
| GO_REGULATION_OF_CHROMOSOME_SEGREGATION | 0.00346 | 0.04402 | 0.43171 | 0.61835 | 1.20782 | 93 |
| GO_REGULATION_OF_CYSTEINE_TYPE_ENDOPEPTIDASE_AC | 0.00041 | 0.00835 | 0.49849 | 0.60820 | 1.20884 | 156 |
| GO_REGULATION_OF_DEFENSE_RESPONSE | 0.00007 | 0.00192 | 0.53843 | 0.57678 | 1.16452 | 361 |
| GO_REGULATION_OF_DNA_BINDING | 0.00224 | 0.03081 | 0.43171 | 0.63553 | 1.23301 | 75 |
| GO_REGULATION_OF_DNA_BIOSYNTHETIC_PROCESS | 0.00133 | 0.02140 | 0.45506 | 0.63610 | 1.23958 | 86 |
| GO_REGULATION_OF_DNA_TEMPLATED_TRANSCRIPTION_I | 0.00000 | 0.00005 | 0.64355 | 0.69155 | 1.35221 | 98 |
| GO_REGULATION_OF_GENERATION_OF_PRECURSOR_META | 0.00037 | 0.00775 | 0.49849 | 0.64037 | 1.25614 | 108 |
| GO_REGULATION_OF_GENE_SILENCING | 0.00021 | 0.00488 | 0.51885 | 0.65497 | 1.27847 | 91 |
| GO_REGULATION_OF_HEMATOPOIETIC_PROGENITOR_CELL_ | 0.00000 | 0.00000 | 0.71951 | 0.75447 | 1.46184 | 70 |
| GO_REGULATION_OF_HEMATOPOIETIC_STEM_CELL_DIFFER | 0.00000 | 0.00000 | 0.71951 | 0.76694 | 1.47923 | 61 |
| GO_REGULATION_OF_HEMOPOIESIS | 0.00016 | 0.00370 | 0.51885 | 0.58536 | 1.17724 | 268 |
| GO_REGULATION_OF_IMMUNE_RESPONSE | 0.00011 | 0.00282 | 0.53843 | 0.56608 | 1.14695 | 455 |
| GO_REGULATION_OF_INNATE_IMMUNE_RESPONSE | 0.00001 | 0.00022 | 0.61053 | 0.62818 | 1.25389 | 185 |
| GO_REGULATION_OF_INTRACELLULAR_PROTEIN_TRANSPOI | 0.00108 | 0.01792 | 0.45506 | 0.58263 | 1.16545 | 200 |
| GO_REGULATION_OF_INTRACELLULAR_TRANSPORT | 0.00039 | 0.00815 | 0.49849 | 0.57589 | 1.15844 | 270 |
| GO_REGULATION_OF_INTRINSIC_APOPTOTIC_SIGNALING_P | 0.00135 | 0.02159 | 0.45506 | 0.61197 | 1.20574 | 123 |
| GO_REGULATION_OF_MORPHOGENESIS_OF_AN_EPITHELIU | 0.00000 | 0.00002 | 0.67496 | 0.68635 | 1.34890 | 117 |
| GO_REGULATION_OF_MRNA_CATABOLIC_PROCESS | 0.00000 | 0.00000 | 0.76146 | 0.67132 | 1.33689 | 169 |
| GO_REGULATION_OF_MRNA_METABOLIC_PROCESS | 0.00000 | 0.00000 | | 0.65795 | 1.32463 | 280 |
| GO_REGULATION_OF_MRNA_PROCESSING | 0.00009 | 0.00237 | 0.53843 | 0.64161 | 1.26535 | 126 |
| GO_REGULATION_OF_MRNA_SPLICING_VIA_SPLICEOSOME | 0.00044 | 0.00888 | 0.49849 | 0.64768 | 1.26394 | 90 |
| GO_REGULATION_OF_MULTI_ORGANISM_PROCESS | 0.00039 | 0.00815 | 0.49849 | 0.71014 | 1.35372 | 42 |
| GO_REGULATION_OF_NUCLEAR_DIVISION | 0.00335 | 0.04313 | 0.43171 | 0.59443 | 1.17747 | 143 |
| GO_REGULATION_OF_NUCLEOCYTOPLASMIC_TRANSPORT | 0.00257 | 0.03433 | 0.43171 | 0.62172 | 1.21206 | 88 |
| GO_REGULATION_OF_OXIDATIVE_PHOSPHORYLATION | 0.00052 | 0.01000 | 0.47727 | 0.79199 | 1.46033 | 22 |
| GO_REGULATION_OF_PEPTIDASE_ACTIVITY | 0.00000 | 0.00007 | 0.64355 | 0.61553 | 1.23625 | 246 |
| GO_REGULATION_OF_PEPTIDE_TRANSPORT | 0.00137 | 0.02160 | 0.45506 | 0.55083 | 1.11501 | 429 |
| GO_REGULATION_OF_PLASMA_LIPOPROTEIN_PARTICLE_LEV | 0.00005 | 0.00147 | 0.55733 | 0.73414 | 1.40325 | 47 |
| GO_REGULATION_OF_POSTTRANSCRIPTIONAL_GENE_SILEN | 0.00035 | 0.00740 | 0.49849 | 0.65734 | 1.27850 | 81 |
| GO_REGULATION_OF_PROTEIN_CONTAINING_COMPLEX_AS | 0.00044 | 0.00888 | 0.49849 | 0.56950 | 1.14823 | 314 |
| GO_REGULATION_OF_PROTEIN_IMPORT | 0.00368 | 0.04609 | 0.43171 | 0.67133 | 1.28055 | 43 |
| GO_REGULATION_OF_PROTEIN_LOCALIZATION_TO_NUCLEU | 0.00000 | 0.00015 | 0.62726 | 0.68439 | 1.33778 | 95 |
| GO_REGULATION_OF_PROTEIN_POLYMERIZATION | 0.00095 | 0.01594 | 0.47727 | 0.59359 | 1.18167 | 165 |
| GO_REGULATION_OF_PROTEIN_STABILITY | 0.00176 | 0.02556 | 0.45506 | 0.57510 | 1.15378 | 227 |
| GO_REGULATION_OF_PROTEOLYSIS | 0.00084 | 0.01461 | 0.47727 | 0.55388 | 1.12220 | 457 |
| GO_REGULATION_OF_RESPONSE_TO_BIOTIC_STIMULUS | 0.00016 | 0.00370 | 0.51885 | 0.59317 | 1.19119 | 243 |
| GO_REGULATION_OF_RNA_SPLICING | 0.00000 | 0.00019 | 0.61053 | 0.65721 | 1.29544 | 125 |
| GO_REGULATION_OF_SMALL_MOLECULE_METABOLIC_PRO | 0.00001 | 0.00027 | 0.61053 | 0.60035 | 1.20877 | 281 |
| GO_REGULATION_OF_STEM_CELL_DIFFERENTIATION | 0.00000 | 0.00013 | 0.62726 | 0.69840 | 1.36104 | 85 |
| GO_REGULATION_OF_SUPRAMOLECULAR_FIBER_ORGANIZA | 0.00064 | 0.01168 | 0.47727 | 0.57933 | 1.16366 | 251 |
| GO_REGULATION_OF_SYMBIOTIC_PROCESS | 0.00000 | 0.00002 | 0.67496 | 0.65463 | 1.30143 | 157 |
| GO_REGULATION_OF_TELOMERE_MAINTENANCE_VIA_TEL( | 0.00080 | 0.01417 | 0.47727 | 0.68242 | 1.30856 | 53 |

| Cluster 1 | | | | | | |
|---|---|---|---|---|---|---|
| GO_REGULATION_OF_TRANSCRIPTION_FROM_RNA_POLYM | 0.00000 | 0.00001 | 0.69013 | 0.74846 | 1.44742 | 67 |
| GO_REGULATION_OF_TRANSLATIONAL_INITIATION | 0.00039 | 0.00801 | 0.49849 | 0.66907 | 1.29468 | 68 |
| GO_REGULATION_OF_VIRAL_GENOME_REPLICATION | 0.00150 | 0.02288 | 0.45506 | 0.66835 | 1.29021 | 62 |
| GO_REGULATION_OF_VIRAL_LIFE_CYCLE | 0.00174 | 0.02541 | 0.45506 | 0.62100 | 1.21403 | 97 |
| GO_REGULATION_OF_VIRAL_TRANSCRIPTION | 0.00145 | 0.02226 | 0.45506 | 0.67171 | 1.28981 | 55 |
| GO_REGULATION_OF_WNT_SIGNALING_PATHWAY | 0.00015 | 0.00362 | 0.51885 | 0.59036 | 1.18585 | 252 |
| GO_REPLACEMENT_OSSIFICATION | 0.00216 | 0.03019 | 0.43171 | 0.79856 | 1.43544 | 15 |
| GO_RESPIRASOME | 0.00000 | 0.00000 | 0.78819 | 0.77519 | 1.50338 | 74 |
| GO_RESPIRATORY_CHAIN_COMPLEX | 0.00000 | 0.00000 | 0.76146 | 0.77893 | 1.50364 | 63 |
| GO_RESPIRATORY_ELECTRON_TRANSPORT_CHAIN | 0.00000 | 0.00000 | 0.81404 | 0.75389 | 1.47254 | 92 |
| GO_RESPONSE_TO_ALCOHOL | 0.00124 | 0.02013 | 0.45506 | 0.60911 | 1.20016 | 124 |
| GO_RESPONSE_TO_CALCIUM_ION | 0.00006 | 0.00182 | 0.53843 | 0.66795 | 1.30098 | 83 |
| GO_RESPONSE_TO_CORTICOSTEROID | 0.00152 | 0.02288 | 0.45506 | 0.63465 | 1.23676 | 86 |
| GO_RESPONSE_TO_DRUG | 0.00062 | 0.01145 | 0.47727 | 0.58069 | 1.16479 | 226 |
| GO_RESPONSE_TO_HEAT | 0.00016 | 0.00384 | 0.51885 | 0.63890 | 1.25647 | 119 |
| GO_RESPONSE_TO_INORGANIC_SUBSTANCE | 0.00000 | 0.00000 | 0.71951 | 0.61356 | 1.23758 | 329 |
| GO_RESPONSE_TO_INTERLEUKIN_1 | 0.00000 | 0.00000 | 0.78819 | 0.70707 | 1.39372 | 125 |
| GO_RESPONSE_TO_INTERLEUKIN_12 | 0.00132 | 0.02122 | 0.45506 | 0.72716 | 1.36858 | 32 |
| GO_RESPONSE_TO_INTERLEUKIN_4 | 0.00220 | 0.03044 | 0.43171 | 0.74587 | 1.38393 | 24 |
| GO_RESPONSE_TO_METAL_ION | 0.00004 | 0.00135 | 0.55733 | 0.60694 | 1.21524 | 207 |
| GO_RESPONSE_TO_NUTRIENT | 0.00001 | 0.00029 | 0.61053 | 0.66955 | 1.30892 | 97 |
| GO_RESPONSE_TO_OXYGEN_LEVELS | 0.00000 | 0.00000 | 0.77494 | 0.63628 | 1.27952 | 264 |
| GO_RESPONSE_TO_TEMPERATURE_STIMULUS | 0.00050 | 0.00974 | 0.47727 | 0.60452 | 1.20088 | 153 |
| GO_RESPONSE_TO_TOXIC_SUBSTANCE | 0.00002 | 0.00063 | 0.57561 | 0.64323 | 1.27049 | 132 |
| GO_RESPONSE_TO_TUMOR_NECROSIS_FACTOR | 0.00002 | 0.00070 | 0.57561 | 0.61474 | 1.22724 | 186 |
| GO_RESPONSE_TO_VITAMIN | 0.00008 | 0.00231 | 0.53843 | 0.72961 | 1.39173 | 43 |
| GO_RETINA_HOMEOSTASIS | 0.00191 | 0.02701 | 0.45506 | 0.70727 | 1.34057 | 36 |
| GO_RIBONUCLEOPROTEIN_COMPLEX_BINDING | 0.00000 | 0.00001 | 0.70498 | 0.69318 | 1.36322 | 119 |
| GO_RIBONUCLEOPROTEIN_COMPLEX_BIOGENESIS | 0.00000 | 0.00000 | | 0.68523 | 1.38613 | 397 |
| GO_RIBONUCLEOPROTEIN_COMPLEX_SUBUNIT_ORGANIZAT | 0.00000 | 0.00000 | | 0.70532 | 1.40678 | 177 |
| GO_RIBONUCLEOPROTEIN_GRANULE | 0.00001 | 0.00032 | 0.59333 | 0.62051 | 1.23980 | 191 |
| GO_RIBONUCLEOSIDE_METABOLIC_PROCESS | 0.00279 | 0.03682 | 0.43171 | 0.68080 | 1.30011 | 44 |
| GO_RIBONUCLEOSIDE_TRIPHOSPHATE_BIOSYNTHETIC_PRO | 0.00115 | 0.01904 | 0.45506 | 0.71757 | 1.36009 | 36 |
| GO_RIBONUCLEOSIDE_TRIPHOSPHATE_METABOLIC_PROCES | 0.00010 | 0.00250 | 0.53843 | 0.72462 | 1.38418 | 46 |
| GO_RIBOSE_PHOSPHATE_BIOSYNTHETIC_PROCESS | 0.00137 | 0.02160 | 0.45506 | 0.61181 | 1.20544 | 123 |
| GO_RIBOSE_PHOSPHATE_METABOLIC_PROCESS | 0.00007 | 0.00190 | 0.53843 | 0.59353 | 1.19455 | 274 |
| GO_RIBOSOMAL_LARGE_SUBUNIT_ASSEMBLY | 0.00167 | 0.02445 | 0.45506 | 0.74882 | 1.39717 | 27 |
| GO_RIBOSOMAL_LARGE_SUBUNIT_BIOGENESIS | 0.00000 | 0.00013 | 0.62726 | 0.71871 | 1.39121 | 69 |
| GO_RIBOSOMAL_SMALL_SUBUNIT_ASSEMBLY | 0.00009 | 0.00248 | 0.53843 | 0.84437 | 1.53036 | 18 |
| GO_RIBOSOMAL_SMALL_SUBUNIT_BIOGENESIS | 0.00001 | 0.00037 | 0.59333 | 0.70855 | 1.37106 | 68 |
| GO_RIBOSOMAL_SUBUNIT | 0.00000 | 0.00000 | | 0.76902 | 1.53346 | 175 |
| GO_RIBOSOME | 0.00000 | 0.00000 | | 0.75904 | 1.52057 | 209 |
| GO_RIBOSOME_ASSEMBLY | 0.00000 | 0.00001 | 0.69013 | 0.76076 | 1.46274 | 57 |
| GO_RIBOSOME_BINDING | 0.00368 | 0.04609 | 0.43171 | 0.66255 | 1.26641 | 47 |
| GO_RIBOSOME_BIOGENESIS | 0.00000 | 0.00000 | | 0.67993 | 1.36890 | 280 |
| GO_RNA_3_END_PROCESSING | 0.00005 | 0.00157 | 0.55733 | 0.63445 | 1.25316 | 132 |
| GO_RNA_CATABOLIC_PROCESS | 0.00000 | 0.00000 | | 0.69240 | 1.39714 | 349 |
| GO_RNA_DEPENDENT_DNA_BIOSYNTHETIC_PROCESS | 0.00054 | 0.01037 | 0.47727 | 0.67837 | 1.31077 | 64 |
| GO_RNA_EXPORT_FROM_NUCLEUS | 0.00003 | 0.00097 | 0.55733 | 0.63770 | 1.25934 | 130 |
| GO_RNA_HELICASE_ACTIVITY | 0.00163 | 0.02413 | 0.45506 | 0.64844 | 1.25174 | 63 |
| GO_RNA_LOCALIZATION | 0.00000 | 0.00002 | 0.65944 | 0.63633 | 1.27369 | 205 |
| GO_RNA_POLYMERASE_II_CORE_COMPLEX | 0.00023 | 0.00516 | 0.51885 | 0.84296 | 1.51524 | 15 |
| GO_RNA_SPLICING | 0.00000 | 0.00000 | | 0.63204 | 1.27773 | 387 |
| GO_RNA_SPLICING_VIA_TRANSESTERIFICATION_REACTIONS | 0.00000 | 0.00000 | | 0.64865 | 1.30750 | 312 |
| GO_ROUGH_ENDOPLASMIC_RETICULUM | 0.00009 | 0.00232 | 0.53843 | 0.70155 | 1.34889 | 57 |
| GO_RRNA_BINDING | 0.00000 | 0.00001 | 0.70498 | 0.76787 | 1.47641 | 57 |
| GO_RRNA_METABOLIC_PROCESS | 0.00000 | 0.00000 | 0.81404 | 0.66459 | 1.33167 | 214 |
| GO_SARCOLEMMA | 0.00242 | 0.03285 | 0.43171 | 0.64385 | 1.24754 | 72 |
| GO_SARCOMERE_ORGANIZATION | 0.00152 | 0.02288 | 0.45506 | 0.77119 | 1.41619 | 21 |
| GO_SCF_DEPENDENT_PROTEASOMAL_UBIQUITIN_DEPENDI | 0.00000 | 0.00000 | 0.73376 | 0.74017 | 1.43782 | 77 |
| GO_SECRETORY_GRANULE_MEMBRANE | 0.00357 | 0.04506 | 0.43171 | 0.58186 | 1.16167 | 187 |
| GO_SIGNAL_SEQUENCE_BINDING | 0.00235 | 0.03199 | 0.43171 | 0.69423 | 1.31852 | 39 |
| GO_SINGLE_FERTILIZATION | 0.00001 | 0.00024 | 0.61053 | 0.71218 | 1.37859 | 69 |
| GO_SINGLE_STRANDED_RNA_BINDING | 0.00048 | 0.00947 | 0.49849 | 0.66916 | 1.29407 | 67 |
| GO_SISTER_CHROMATID_SEGREGATION | 0.00119 | 0.01944 | 0.45506 | 0.59541 | 1.18573 | 169 |
| GO_SMALL_MOLECULE_BIOSYNTHETIC_PROCESS | 0.00013 | 0.00309 | 0.51885 | 0.56616 | 1.14572 | 426 |
| GO_SMALL_NUCLEAR_RIBONUCLEOPROTEIN_COMPLEX | 0.00005 | 0.00141 | 0.55733 | 0.70683 | 1.36090 | 59 |
| GO_SMALL_RIBOSOMAL_SUBUNIT | 0.00000 | 0.00001 | 0.70498 | 0.74854 | 1.45036 | 70 |
| GO_SM_LIKE_PROTEIN_FAMILY_COMPLEX | 0.00002 | 0.00058 | 0.57561 | 0.70328 | 1.36135 | 69 |
| GO_SNORNA_BINDING | 0.00083 | 0.01449 | 0.47727 | 0.74119 | 1.38813 | 29 |
| GO_SOMATIC_STEM_CELL_POPULATION_MAINTENANCE | 0.00139 | 0.02173 | 0.45506 | 0.67563 | 1.29352 | 50 |
| GO_SPERM_EGG_RECOGNITION | 0.00010 | 0.00260 | 0.53843 | 0.79697 | 1.48701 | 27 |
| GO_SPLICEOSOMAL_COMPLEX | 0.00000 | 0.00000 | 0.73376 | 0.66037 | 1.31622 | 172 |

| Cluster 1 | | | | | | |
|---|---|---|---|---|---|---|
| GO_SPLICEOSOMAL_SNRNP_ASSEMBLY | 0.00047 | 0.00935 | 0.49849 | 0.74250 | 1.40531 | 35 |
| GO_SPLICEOSOMAL_TRI_SNRNP_COMPLEX | 0.00108 | 0.01792 | 0.45506 | 0.74335 | 1.39331 | 30 |
| GO_STEM_CELL_DIFFERENTIATION | 0.00000 | 0.00019 | 0.61053 | 0.63919 | 1.27166 | 161 |
| GO_STRUCTURAL_CONSTITUENT_OF_CYTOSKELETON | 0.00079 | 0.01404 | 0.47727 | 0.67315 | 1.30068 | 64 |
| GO_STRUCTURAL_CONSTITUENT_OF_RIBOSOME | 0.00000 | 0.00000 | | 0.78406 | 1.55720 | 152 |
| GO_STRUCTURAL_MOLECULE_ACTIVITY | 0.00000 | 0.00000 | | 0.70593 | 1.42777 | 394 |
| GO_SUPEROXIDE_ANION_GENERATION | 0.00364 | 0.04581 | 0.43171 | 0.78604 | 1.41293 | 15 |
| GO_TAU_PROTEIN_BINDING | 0.00383 | 0.04728 | 0.43171 | 0.70209 | 1.32647 | 34 |
| GO_TELOMERASE_RNA_LOCALIZATION | 0.00009 | 0.00234 | 0.53843 | 0.83831 | 1.52403 | 19 |
| GO_TELOMERE_MAINTENANCE_VIA_TELOMERE_LENGTHEN | 0.00027 | 0.00588 | 0.49849 | 0.67638 | 1.31054 | 70 |
| GO_TELOMERE_ORGANIZATION | 0.00185 | 0.02634 | 0.45506 | 0.59971 | 1.18453 | 132 |
| GO_THREONINE_TYPE_ENDOPEPTIDASE_ACTIVITY | 0.00006 | 0.00165 | 0.55733 | 0.85082 | 1.54205 | 18 |
| GO_THREONINE_TYPE_PEPTIDASE_ACTIVITY | 0.00001 | 0.00036 | 0.59333 | 0.85216 | 1.57127 | 22 |
| GO_TISSUE_MORPHOGENESIS | 0.00027 | 0.00588 | 0.49849 | 0.56327 | 1.13861 | 386 |
| GO_TOXIN_TRANSPORT | 0.00250 | 0.03363 | 0.43171 | 0.71629 | 1.34150 | 29 |
| GO_TRANSCRIPTION_COUPLED_NUCLEOTIDE_EXCISION_RE | 0.00065 | 0.01198 | 0.47727 | 0.66138 | 1.28151 | 72 |
| GO_TRANSLATIONAL_ELONGATION | 0.00000 | 0.00000 | | 0.73780 | 1.45506 | 126 |
| GO_TRANSLATIONAL_INITIATION | 0.00000 | 0.00000 | | 0.75043 | 1.49572 | 172 |
| GO_TRANSLATIONAL_TERMINATION | 0.00000 | 0.00000 | 0.81404 | 0.73745 | 1.44415 | 102 |
| GO_TRANSLATION_FACTOR_ACTIVITY_RNA_BINDING | 0.00000 | 0.00013 | 0.62726 | 0.71489 | 1.38644 | 74 |
| GO_TRANSLATION_INITIATION_FACTOR_ACTIVITY | 0.00005 | 0.00156 | 0.55733 | 0.73122 | 1.39679 | 46 |
| GO_TRANSLATION_PREINITIATION_COMPLEX | 0.00049 | 0.00962 | 0.47727 | 0.82865 | 1.49769 | 16 |
| GO_TRANSLATION_REGULATOR_ACTIVITY | 0.00000 | 0.00000 | 0.71951 | 0.69624 | 1.36794 | 116 |
| GO_TRANSLATION_REGULATOR_ACTIVITY_NUCLEIC_ACID_E | 0.00000 | 0.00003 | 0.65944 | 0.70561 | 1.37823 | 92 |
| GO_TRICARBOXYLIC_ACID_CYCLE | 0.00165 | 0.02429 | 0.45506 | 0.72358 | 1.35911 | 31 |
| GO_TUMOR_NECROSIS_FACTOR_MEDIATED_SIGNALING_PA | 0.00000 | 0.00004 | 0.65944 | 0.68001 | 1.33835 | 121 |
| GO_T_CELL_RECEPTOR_SIGNALING_PATHWAY | 0.00004 | 0.00111 | 0.55733 | 0.64509 | 1.27356 | 129 |
| GO_U12_TYPE_SPLICEOSOMAL_COMPLEX | 0.00220 | 0.03044 | 0.43171 | 0.74607 | 1.38431 | 24 |
| GO_U2_TYPE_CATALYTIC_STEP_2_SPLICEOSOME | 0.00046 | 0.00919 | 0.49849 | 0.75075 | 1.40604 | 29 |
| GO_U2_TYPE_SPLICEOSOMAL_COMPLEX | 0.00010 | 0.00262 | 0.53843 | 0.66762 | 1.30115 | 87 |
| GO_UBIQUITIN_LIKE_PROTEIN_LIGASE_BINDING | 0.00000 | 0.00005 | 0.64355 | 0.61291 | 1.23181 | 257 |
| GO_UNFOLDED_PROTEIN_BINDING | 0.00000 | 0.00004 | 0.65944 | 0.70180 | 1.37079 | 92 |
| GO_VACUOLAR_LUMEN | 0.00001 | 0.00029 | 0.59333 | 0.67075 | 1.31574 | 108 |
| GO_VACUOLAR_MEMBRANE | 0.00357 | 0.04506 | 0.43171 | 0.55476 | 1.11862 | 323 |
| GO_VESICLE_LUMEN | 0.00000 | 0.00000 | | 0.68833 | 1.37612 | 194 |
| GO_VESICLE_MEMBRANE | 0.00009 | 0.00237 | 0.53843 | 0.56152 | 1.13859 | 485 |
| GO_VIRAL_GENE_EXPRESSION | 0.00000 | 0.00000 | | 0.73673 | 1.46922 | 179 |
| GO_VIRAL_GENOME_REPLICATION | 0.00007 | 0.00194 | 0.53843 | 0.67550 | 1.31384 | 81 |
| GO_VIRAL_LIFE_CYCLE | 0.00000 | 0.00001 | 0.69013 | 0.63153 | 1.26694 | 238 |
| HP_ABNORMALITY_OF_ACID_BASE_HOMEOSTASIS | 0.00001 | 0.00047 | 0.59333 | 0.59995 | 1.20633 | 263 |
| HP_ABNORMALITY_OF_CARDIOVASCULAR_SYSTEM_ELECTR | 0.00146 | 0.02237 | 0.45506 | 0.56742 | 1.14165 | 271 |
| HP_ABNORMALITY_OF_THE_GALLBLADDER | 0.00054 | 0.01029 | 0.47727 | 0.68835 | 1.31789 | 50 |
| HP_ABNORMALITY_OF_THE_LARYNX | 0.00154 | 0.02305 | 0.45506 | 0.61901 | 1.21344 | 106 |
| HP_ABNORMALITY_OF_THE_OPTIC_DISC | 0.00024 | 0.00535 | 0.51885 | 0.55993 | 1.13369 | 441 |
| HP_ABNORMALITY_OF_THE_OPTIC_NERVE | 0.00011 | 0.00287 | 0.53843 | 0.55814 | 1.13211 | 498 |
| HP_ABNORMAL_ACTIVITY_OF_MITOCHONDRIAL_RESPIRATO | 0.00156 | 0.02327 | 0.45506 | 0.66791 | 1.28934 | 62 |
| HP_ABNORMAL_CIRCULATING_CARBOHYDRATE_CONCENTR | 0.00022 | 0.00502 | 0.51885 | 0.70869 | 1.35460 | 47 |
| HP_ABNORMAL_CNS_MYELINATION | 0.00003 | 0.00094 | 0.57561 | 0.61674 | 1.22995 | 178 |
| HP_ABNORMAL_CSF_METABOLITE_LEVEL | 0.00012 | 0.00305 | 0.51885 | 0.66878 | 1.30168 | 82 |
| HP_ABNORMAL_ERYTHROCYTE_MORPHOLOGY | 0.00002 | 0.00062 | 0.57561 | 0.57684 | 1.16604 | 386 |
| HP_ABNORMAL_MITOCHONDRIA_IN_MUSCLE_TISSUE | 0.00044 | 0.00888 | 0.49849 | 0.75148 | 1.40740 | 29 |
| HP_ABNORMAL_MYELINATION | 0.00044 | 0.00888 | 0.49849 | 0.56902 | 1.14784 | 342 |
| HP_ACUTE_LEUKEMIA | 0.00037 | 0.00775 | 0.49849 | 0.66088 | 1.28282 | 76 |
| HP_ANEMIA_OF_INADEQUATE_PRODUCTION | 0.00011 | 0.00288 | 0.53843 | 0.64679 | 1.26705 | 104 |
| HP_APHASIA | 0.00279 | 0.03682 | 0.43171 | 0.71344 | 1.33616 | 29 |
| HP_BLINDNESS | 0.00194 | 0.02747 | 0.45506 | 0.58798 | 1.17246 | 175 |
| HP_DECREASED_ACTIVITY_OF_MITOCHONDRIAL_COMPLEX_ | 0.00058 | 0.01093 | 0.47727 | 0.74138 | 1.40070 | 34 |
| HP_DEPRESSED_NASAL_RIDGE | 0.00168 | 0.02466 | 0.45506 | 0.63542 | 1.23676 | 82 |
| HP_DYSPHASIA | 0.00179 | 0.02585 | 0.45506 | 0.70866 | 1.34322 | 36 |
| HP_EMOTIONAL_LABILITY | 0.00095 | 0.01594 | 0.47727 | 0.63513 | 1.24150 | 95 |
| HP_FETAL_DISTRESS | 0.00033 | 0.00699 | 0.49849 | 0.75062 | 1.41274 | 32 |
| HP_HYPERTROPHIC_CARDIOMYOPATHY | 0.00095 | 0.01594 | 0.47727 | 0.58554 | 1.17076 | 195 |
| HP_INAPPROPRIATE_BEHAVIOR | 0.00253 | 0.03398 | 0.43171 | 0.73374 | 1.36353 | 25 |
| HP_INCREASED_CSF_LACTATE | 0.00006 | 0.00184 | 0.53843 | 0.68746 | 1.33074 | 69 |
| HP_INCREASED_SERUM_LACTATE | 0.00007 | 0.00206 | 0.53843 | 0.63197 | 1.25025 | 136 |
| HP_INCREASED_SERUM_PYRUVATE | 0.00033 | 0.00716 | 0.49849 | 0.73183 | 1.38786 | 38 |
| HP_LACTIC_ACIDOSIS | 0.00029 | 0.00625 | 0.49849 | 0.61894 | 1.22315 | 133 |
| HP_LETHARGY | 0.00087 | 0.01492 | 0.47727 | 0.62515 | 1.22466 | 104 |
| HP_LEUKODYSTROPHY | 0.00044 | 0.00892 | 0.49849 | 0.64979 | 1.26626 | 86 |
| HP_LEUKOENCEPHALOPATHY | 0.00081 | 0.01425 | 0.47727 | 0.60899 | 1.20615 | 142 |
| HP_MACROCYTIC_ANEMIA | 0.00016 | 0.00373 | 0.51885 | 0.74192 | 1.40699 | 38 |
| HP_METABOLIC_ACIDOSIS | 0.00346 | 0.04402 | 0.43171 | 0.61148 | 1.19752 | 103 |
| HP_MIGRAINE | 0.00067 | 0.01228 | 0.47727 | 0.65711 | 1.27328 | 71 |

| Cluster 1 | | | | | | |
|---|---|---|---|---|---|---|
| HP_OPTIC_NERVE_COLOBOMA | 0.00220 | 0.03044 | 0.43171 | 0.77715 | 1.40695 | 17 |
| HP_OPTIC_NEUROPATHY | 0.00002 | 0.00077 | 0.57561 | 0.77729 | 1.47329 | 36 |
| HP_PAROXYSMAL_INVOLUNTARY_EYE_MOVEMENTS | 0.00015 | 0.00354 | 0.51885 | 0.76244 | 1.43210 | 31 |
| HP_POOR_EYE_CONTACT | 0.00183 | 0.02620 | 0.45506 | 0.66484 | 1.27629 | 54 |
| HP_POOR_HEAD_CONTROL | 0.00071 | 0.01284 | 0.47727 | 0.62385 | 1.22782 | 121 |
| HP_PROXIMAL_TUBULOPATHY | 0.00261 | 0.03468 | 0.43171 | 0.71535 | 1.34931 | 33 |
| HP_REDUCED_CONSCIOUSNESS_CONFUSION | 0.00246 | 0.03321 | 0.43171 | 0.57873 | 1.15784 | 201 |
| HP_RENAL_TUBULAR_DYSFUNCTION | 0.00231 | 0.03169 | 0.43171 | 0.63938 | 1.24227 | 78 |
| HP_RESPIRATORY_FAILURE | 0.00146 | 0.02237 | 0.45506 | 0.61061 | 1.20305 | 123 |
| HP_RESPIRATORY_INSUFFICIENCY | 0.00012 | 0.00291 | 0.53843 | 0.57794 | 1.16597 | 339 |
| HP_SENSORY_NEUROPATHY | 0.00235 | 0.03199 | 0.43171 | 0.60766 | 1.19217 | 111 |
| HP_SUPERNUMERARY_BONES_OF_THE_AXIAL_SKELETON | 0.00178 | 0.02564 | 0.45506 | 0.80179 | 1.44914 | 16 |
| HP_VAGINAL_FISTULA | 0.00290 | 0.03804 | 0.43171 | 0.77198 | 1.39759 | 17 |
| **Cluster 3** | | | | | | |
| pathway | pval | padj | log2err | ES | NES | size |
| GO_ACTIN_CYTOSKELETON | 0.00003 | 0.00109 | 0.55733 | 0.57329 | 1.17717 | 347 |
| GO_ACTIN_FILAMENT_BUNDLE_ORGANIZATION | 0.00309 | 0.03938 | 0.43171 | 0.60412 | 1.21427 | 112 |
| GO_ACTIN_FILAMENT_ORGANIZATION | 0.00124 | 0.02105 | 0.45506 | 0.56061 | 1.14701 | 297 |
| GO_ACTIN_POLYMERIZATION_OR_DEPOLYMERIZATION | 0.00405 | 0.04880 | 0.40702 | 0.58329 | 1.18116 | 152 |
| GO_ACTIVATION_OF_IMMUNE_RESPONSE | 0.00000 | 0.00000 | | 0.64570 | 1.31907 | 273 |
| GO_ACTIVATION_OF_INNATE_IMMUNE_RESPONSE | 0.00000 | 0.00000 | | 0.73250 | 1.46912 | 105 |
| GO_ACTIVE_ION_TRANSMEMBRANE_TRANSPORTER_ACTIV | 0.00253 | 0.03467 | 0.43171 | 0.60748 | 1.22095 | 115 |
| GO_ACTIVE_TRANSMEMBRANE_TRANSPORTER_ACTIVITY | 0.00121 | 0.02064 | 0.45506 | 0.58296 | 1.18437 | 185 |
| GO_AEROBIC_RESPIRATION | 0.00067 | 0.01350 | 0.47727 | 0.65597 | 1.29897 | 70 |
| GO_ALTERNATIVE_MRNA_SPLICING_VIA_SPLICEOSOME | 0.00133 | 0.02182 | 0.45506 | 0.64582 | 1.28082 | 74 |
| GO_AMIDE_BINDING | 0.00133 | 0.02182 | 0.45506 | 0.57680 | 1.17519 | 210 |
| GO_AMINE_METABOLIC_PROCESS | 0.00000 | 0.00000 | 0.81404 | 0.73916 | 1.47792 | 97 |
| GO_ANAPHASE_PROMOTING_COMPLEX_DEPENDENT_CATA | 0.00000 | 0.00000 | 0.81404 | 0.76568 | 1.52040 | 77 |
| GO_ANATOMICAL_STRUCTURE_HOMEOSTASIS | 0.00191 | 0.02819 | 0.45506 | 0.55720 | 1.14038 | 303 |
| GO_ANIMAL_ORGAN_REGENERATION | 0.00290 | 0.03761 | 0.43171 | 0.67149 | 1.31217 | 52 |
| GO_ANTIGEN_PROCESSING_AND_PRESENTATION | 0.00000 | 0.00000 | 0.81404 | 0.68231 | 1.38328 | 169 |
| GO_ANTIGEN_PROCESSING_AND_PRESENTATION_OF_EXOG | 0.00000 | 0.00000 | 0.81404 | 0.79321 | 1.56649 | 65 |
| GO_ANTIGEN_PROCESSING_AND_PRESENTATION_OF_PEPT | 0.00000 | 0.00000 | 0.80122 | 0.68958 | 1.39532 | 146 |
| GO_ANTIGEN_PROCESSING_AND_PRESENTATION_OF_PEPT | 0.00000 | 0.00000 | 0.77494 | 0.75596 | 1.50110 | 77 |
| GO_ANTIGEN_PROCESSING_AND_PRESENTATION_OF_PEPT | 0.00072 | 0.01391 | 0.47727 | 0.64334 | 1.27945 | 80 |
| GO_ANTIGEN_RECEPTOR_MEDIATED_SIGNALING_PATHWAY | 0.00000 | 0.00000 | 0.73376 | 0.66137 | 1.34066 | 161 |
| GO_ANTIMICROBIAL_HUMORAL_RESPONSE | 0.00133 | 0.02182 | 0.45506 | 0.72505 | 1.38022 | 30 |
| GO_APICAL_PART_OF_CELL | 0.00092 | 0.01695 | 0.47727 | 0.57249 | 1.16749 | 227 |
| GO_APOPTOTIC_SIGNALING_PATHWAY | 0.00000 | 0.00009 | 0.62726 | 0.57395 | 1.18294 | 439 |
| GO_ATPASE_ACTIVITY_COUPLED_TO_TRANSMEMBRANE_M | 0.00216 | 0.03050 | 0.43171 | 0.78322 | 1.43573 | 18 |
| GO_ATP_BIOSYNTHETIC_PROCESS | 0.00035 | 0.00813 | 0.49849 | 0.78511 | 1.47268 | 24 |
| GO_ATP_METABOLIC_PROCESS | 0.00000 | 0.00000 | | 0.65821 | 1.34237 | 223 |
| GO_ATP_SYNTHESIS_COUPLED_ELECTRON_TRANSPORT | 0.00000 | 0.00000 | 0.71951 | 0.73656 | 1.46258 | 77 |
| GO_AZUROPHIL_GRANULE | 0.00045 | 0.00978 | 0.49849 | 0.62534 | 1.25565 | 109 |
| GO_AZUROPHIL_GRANULE_LUMEN | 0.00261 | 0.03481 | 0.43171 | 0.64523 | 1.27120 | 62 |
| GO_BASAL_PART_OF_CELL | 0.00216 | 0.03050 | 0.43171 | 0.67029 | 1.29906 | 44 |
| GO_BASAL_PLASMA_MEMBRANE | 0.00054 | 0.01138 | 0.47727 | 0.74197 | 1.41450 | 31 |
| GO_BASOLATERAL_PLASMA_MEMBRANE | 0.00035 | 0.00813 | 0.49849 | 0.60607 | 1.22516 | 142 |
| GO_BINDING_OF_SPERM_TO_ZONA_PELLUCIDA | 0.00001 | 0.00041 | 0.59333 | 0.85869 | 1.58913 | 21 |
| GO_BLOOD_MICROPARTICLE | 0.00000 | 0.00001 | 0.70498 | 0.78421 | 1.53243 | 52 |
| GO_BRUSH_BORDER | 0.00011 | 0.00310 | 0.53843 | 0.68687 | 1.35323 | 62 |
| GO_BRUSH_BORDER_MEMBRANE | 0.00170 | 0.02614 | 0.45506 | 0.71917 | 1.37104 | 31 |
| GO_CADHERIN_BINDING | 0.00000 | 0.00000 | | 0.66818 | 1.36513 | 277 |
| GO_CALCIUM_ION_BINDING | 0.00294 | 0.03786 | 0.43171 | 0.54579 | 1.11912 | 321 |
| GO_CANONICAL_WNT_SIGNALING_PATHWAY | 0.00000 | 0.00004 | 0.65944 | 0.61188 | 1.24999 | 260 |
| GO_CARBOHYDRATE_CATABOLIC_PROCESS | 0.00257 | 0.03481 | 0.43171 | 0.58927 | 1.19236 | 146 |
| GO_CARBOHYDRATE_DERIVATIVE_BIOSYNTHETIC_PROCESS | 0.00124 | 0.02105 | 0.45506 | 0.54292 | 1.11910 | 454 |
| GO_CARDIAC_CHAMBER_MORPHOGENESIS | 0.00276 | 0.03612 | 0.43171 | 0.62238 | 1.23778 | 80 |
| GO_CATALYTIC_STEP_2_SPLICEOSOME | 0.00001 | 0.00057 | 0.59333 | 0.68112 | 1.35575 | 83 |
| GO_CATENIN_COMPLEX | 0.00235 | 0.03255 | 0.43171 | 0.81283 | 1.47138 | 15 |
| GO_CATION_TRANSMEMBRANE_TRANSPORT | 0.00045 | 0.00978 | 0.49849 | 0.55083 | 1.13482 | 426 |
| GO_CATION_TRANSMEMBRANE_TRANSPORTER_ACTIVITY | 0.00005 | 0.00169 | 0.55733 | 0.58200 | 1.18905 | 277 |
| GO_CELLULAR_AMINO_ACID_METABOLIC_PROCESS | 0.00015 | 0.00405 | 0.51885 | 0.58255 | 1.18839 | 235 |
| GO_CELLULAR_COMPONENT_DISASSEMBLY | 0.00000 | 0.00000 | 0.73376 | 0.59142 | 1.21891 | 438 |
| GO_CELLULAR_ION_HOMEOSTASIS | 0.00000 | 0.00013 | 0.62726 | 0.58736 | 1.20596 | 343 |
| GO_CELLULAR_IRON_ION_HOMEOSTASIS | 0.00018 | 0.00481 | 0.51885 | 0.70268 | 1.36846 | 49 |
| GO_CELLULAR_KETONE_METABOLIC_PROCESS | 0.00000 | 0.00000 | 0.73376 | 0.66009 | 1.33816 | 173 |
| GO_CELLULAR_NITROGEN_COMPOUND_CATABOLIC_PROCE | 0.00000 | 0.00000 | | 0.65939 | 1.36049 | 476 |
| GO_CELLULAR_PROTEIN_COMPLEX_DISASSEMBLY | 0.00000 | 0.00002 | 0.67496 | 0.64013 | 1.29952 | 181 |
| GO_CELLULAR_RESPIRATION | 0.00000 | 0.00001 | 0.69013 | 0.66194 | 1.33941 | 146 |
| GO_CELLULAR_RESPONSE_TO_CHEMICAL_STRESS | 0.00202 | 0.02904 | 0.43171 | 0.56405 | 1.15242 | 258 |
| GO_CELLULAR_RESPONSE_TO_HEAT | 0.00011 | 0.00311 | 0.53843 | 0.64579 | 1.29511 | 104 |
| GO_CELLULAR_RESPONSE_TO_HORMONE_STIMULUS | 0.00191 | 0.02819 | 0.45506 | 0.54274 | 1.11801 | 419 |

| Cluster 1 | | | | | | |
|---|---|---|---|---|---|---|
| GO_CELLULAR_RESPONSE_TO_NITROGEN_COMPOUND | 0.00083 | 0.01566 | 0.47727 | 0.54579 | 1.12496 | 443 |
| GO_CELLULAR_RESPONSE_TO_ORGANIC_CYCLIC_COMPOUI | 0.00030 | 0.00715 | 0.49849 | 0.55799 | 1.14685 | 377 |
| GO_CELLULAR_RESPONSE_TO_OXYGEN_LEVELS | 0.00000 | 0.00000 | | 0.69902 | 1.41713 | 171 |
| GO_CELLULAR_RESPONSE_TO_REACTIVE_OXYGEN_SPECIES | 0.00335 | 0.04202 | 0.43171 | 0.60144 | 1.20892 | 114 |
| GO_CELLULAR_RESPONSE_TO_TOPOLOGICALLY_INCORRECT | 0.00086 | 0.01595 | 0.47727 | 0.60065 | 1.21407 | 140 |
| GO_CELLULAR_TRANSITION_METAL_ION_HOMEOSTASIS | 0.00139 | 0.02233 | 0.45506 | 0.63925 | 1.26828 | 75 |
| GO_CELL_ACTIVATION_INVOLVED_IN_IMMUNE_RESPONSE | 0.00000 | 0.00000 | 0.81404 | 0.60415 | 1.24544 | 446 |
| GO_CELL_ADHESION_MEDIATOR_ACTIVITY | 0.00187 | 0.02796 | 0.45506 | 0.70508 | 1.35925 | 38 |
| GO_CELL_ADHESION_MOLECULE_BINDING | 0.00000 | 0.00000 | | 0.65695 | 1.35095 | 389 |
| GO_CELL_CELL_ADHESION | 0.00413 | 0.04907 | 0.40702 | 0.53686 | 1.10666 | 455 |
| GO_CELL_CELL_RECOGNITION | 0.00001 | 0.00046 | 0.59333 | 0.78076 | 1.49625 | 35 |
| GO_CELL_CELL_SIGNALING_BY_WNT | 0.00000 | 0.00003 | 0.65944 | 0.58445 | 1.20249 | 400 |
| GO_CELL_CORTEX | 0.00228 | 0.03186 | 0.43171 | 0.57553 | 1.17256 | 208 |
| GO_CELL_CYCLE_G2_M_PHASE_TRANSITION | 0.00010 | 0.00290 | 0.53843 | 0.58583 | 1.19525 | 236 |
| GO_CELL_DIVISION | 0.00083 | 0.01566 | 0.47727 | 0.54712 | 1.12831 | 471 |
| GO_CELL_FATE_DETERMINATION | 0.00156 | 0.02428 | 0.45506 | 0.76204 | 1.41643 | 22 |
| GO_CELL_KILLING | 0.00063 | 0.01286 | 0.47727 | 0.67160 | 1.32411 | 63 |
| GO_CELL_PROJECTION_MEMBRANE | 0.00093 | 0.01708 | 0.47727 | 0.58286 | 1.18754 | 209 |
| GO_CELL_RECOGNITION | 0.00007 | 0.00213 | 0.53843 | 0.68108 | 1.34868 | 70 |
| GO_CELL_REDOX_HOMEOSTASIS | 0.00145 | 0.02316 | 0.45506 | 0.69936 | 1.34961 | 39 |
| GO_CELL_SUBSTRATE_JUNCTION | 0.00000 | 0.00000 | | 0.65205 | 1.33890 | 345 |
| GO_CELL_SURFACE | 0.00202 | 0.02904 | 0.43171 | 0.54868 | 1.12831 | 388 |
| GO_CELL_SURFACE_RECEPTOR_SIGNALING_PATHWAY_INVO | 0.00000 | 0.00002 | 0.67496 | 0.58132 | 1.19820 | 443 |
| GO_CHAPERONE_BINDING | 0.00261 | 0.03481 | 0.43171 | 0.61174 | 1.22095 | 90 |
| GO_CHAPERONE_COMPLEX | 0.00001 | 0.00043 | 0.59333 | 0.83540 | 1.55990 | 23 |
| GO_CHAPERONE_MEDIATED_PROTEIN_FOLDING | 0.00231 | 0.03224 | 0.43171 | 0.67527 | 1.31955 | 52 |
| GO_CHROMOSOME_SEGREGATION | 0.00327 | 0.04125 | 0.43171 | 0.55715 | 1.13824 | 263 |
| GO_CLUSTER_OF_ACTIN_BASED_CELL_PROJECTIONS | 0.00023 | 0.00583 | 0.51885 | 0.64543 | 1.28704 | 88 |
| GO_COLLAGEN_CONTAINING_EXTRACELLULAR_MATRIX | 0.00092 | 0.01695 | 0.47727 | 0.58304 | 1.18792 | 209 |
| GO_CONTRACTILE_FIBER | 0.00003 | 0.00094 | 0.57561 | 0.62726 | 1.26897 | 145 |
| GO_COPI_COATED_VESICLE | 0.00113 | 0.01964 | 0.45506 | 0.76202 | 1.42936 | 24 |
| GO_COPI_COATED_VESICLE_MEMBRANE | 0.00276 | 0.03612 | 0.43171 | 0.78951 | 1.43466 | 16 |
| GO_CORNIFICATION | 0.00040 | 0.00906 | 0.49849 | 0.73261 | 1.40217 | 34 |
| GO_CORTICAL_CYTOSKELETON | 0.00150 | 0.02375 | 0.45506 | 0.64437 | 1.27795 | 74 |
| GO_COTRANSLATIONAL_PROTEIN_TARGETING_TO_MEMBR/ | 0.00000 | 0.00000 | | 0.81048 | 1.62058 | 96 |
| GO_CYTOCHROME_COMPLEX | 0.00035 | 0.00813 | 0.49849 | 0.78262 | 1.46134 | 23 |
| GO_CYTOKINE_MEDIATED_SIGNALING_PATHWAY | 0.00000 | 0.00000 | | 0.60612 | 1.24970 | 461 |
| GO_CYTOPLASMIC_TRANSLATION | 0.00000 | 0.00000 | 0.73376 | 0.71611 | 1.43081 | 93 |
| GO_CYTOPLASMIC_TRANSLATIONAL_INITIATION | 0.00287 | 0.03728 | 0.43171 | 0.71013 | 1.35380 | 31 |
| GO_CYTOSOLIC_LARGE_RIBOSOMAL_SUBUNIT | 0.00000 | 0.00000 | 0.74774 | 0.80091 | 1.56507 | 52 |
| GO_CYTOSOLIC_RIBOSOME | 0.00000 | 0.00000 | | 0.79999 | 1.59984 | 98 |
| GO_CYTOSOLIC_SMALL_RIBOSOMAL_SUBUNIT | 0.00000 | 0.00000 | 0.71951 | 0.81934 | 1.58412 | 42 |
| GO_DIENCEPHALON_DEVELOPMENT | 0.00022 | 0.00569 | 0.51885 | 0.74071 | 1.42048 | 36 |
| GO_DISORDERED_DOMAIN_SPECIFIC_BINDING | 0.00148 | 0.02352 | 0.45506 | 0.72163 | 1.37571 | 31 |
| GO_DOUBLE_STRANDED_RNA_BINDING | 0.00062 | 0.01279 | 0.47727 | 0.66149 | 1.30157 | 61 |
| GO_DRUG_BINDING | 0.00057 | 0.01204 | 0.47727 | 0.66781 | 1.31368 | 60 |
| GO_ELECTRON_TRANSFER_ACTIVITY | 0.00000 | 0.00013 | 0.62726 | 0.68868 | 1.37140 | 84 |
| GO_ELECTRON_TRANSPORT_CHAIN | 0.00000 | 0.00000 | 0.77494 | 0.68601 | 1.38582 | 138 |
| GO_EMBRYONIC_PLACENTA_DEVELOPMENT | 0.00405 | 0.04880 | 0.40702 | 0.63530 | 1.25579 | 68 |
| GO_EMBRYONIC_PLACENTA_MORPHOGENESIS | 0.00063 | 0.01286 | 0.47727 | 0.79632 | 1.46296 | 19 |
| GO_ENDOCYTIC_VESICLE | 0.00346 | 0.04317 | 0.43171 | 0.57284 | 1.16708 | 208 |
| GO_ENDOCYTIC_VESICLE_MEMBRANE | 0.00353 | 0.04376 | 0.43171 | 0.60568 | 1.21383 | 103 |
| GO_ENDOPEPTIDASE_ACTIVITY | 0.00191 | 0.02819 | 0.45506 | 0.57072 | 1.16285 | 214 |
| GO_ENDOPEPTIDASE_COMPLEX | 0.00000 | 0.00000 | 0.77494 | 0.77699 | 1.53447 | 65 |
| GO_ENDOPLASMIC_RETICULUM_GOLGI_INTERMEDIATE_CC | 0.00097 | 0.01755 | 0.47727 | 0.66403 | 1.30169 | 56 |
| GO_ENDOPLASMIC_RETICULUM_LUMEN | 0.00335 | 0.04202 | 0.43171 | 0.57884 | 1.17503 | 178 |
| GO_ENDOPLASMIC_RETICULUM_TO_GOLGI_VESICLE_MEDI/ | 0.00098 | 0.01784 | 0.45506 | 0.58969 | 1.19535 | 161 |
| GO_ENDOSOME_LUMEN | 0.00383 | 0.04672 | 0.43171 | 0.75782 | 1.39223 | 19 |
| GO_ENERGY_DERIVATION_BY_OXIDATION_OF_ORGANIC_CC | 0.00000 | 0.00002 | 0.67496 | 0.63013 | 1.28295 | 203 |
| GO_EPHRIN_RECEPTOR_SIGNALING_PATHWAY | 0.00069 | 0.01362 | 0.47727 | 0.65251 | 1.28951 | 67 |
| GO_EPIDERMAL_CELL_DIFFERENTIATION | 0.00305 | 0.03898 | 0.43171 | 0.59515 | 1.20121 | 131 |
| GO_EPITHELIAL_CELL_DIFFERENTIATION | 0.00027 | 0.00641 | 0.49849 | 0.55753 | 1.14729 | 401 |
| GO_EPITHELIAL_CELL_DIFFERENTIATION_INVOLVED_IN_KID | 0.00346 | 0.04317 | 0.43171 | 0.72268 | 1.36788 | 28 |
| GO_EPITHELIAL_TO_MESENCHYMAL_TRANSITION | 0.00178 | 0.02689 | 0.45506 | 0.61523 | 1.23281 | 102 |
| GO_ERAD_PATHWAY | 0.00194 | 0.02835 | 0.45506 | 0.61916 | 1.23466 | 88 |
| GO_ESTABLISHMENT_OF_ORGANELLE_LOCALIZATION | 0.00122 | 0.02090 | 0.45506 | 0.55568 | 1.14046 | 335 |
| GO_ESTABLISHMENT_OF_PROTEIN_LOCALIZATION_TO_CHR | 0.00350 | 0.04355 | 0.43171 | 0.75409 | 1.39555 | 21 |
| GO_ESTABLISHMENT_OF_PROTEIN_LOCALIZATION_TO_END | 0.00000 | 0.00000 | | 0.80160 | 1.60886 | 108 |
| GO_ESTABLISHMENT_OF_PROTEIN_LOCALIZATION_TO_MEN | 0.00000 | 0.00000 | | 0.68126 | 1.39197 | 275 |
| GO_ESTABLISHMENT_OF_PROTEIN_LOCALIZATION_TO_ORG | 0.00000 | 0.00000 | | 0.64216 | 1.32465 | 474 |
| GO_ESTABLISHMENT_OF_PROTEIN_LOCALIZATION_TO_TELC | 0.00031 | 0.00721 | 0.49849 | 0.83600 | 1.51912 | 16 |
| GO_ESTABLISHMENT_OF_RNA_LOCALIZATION | 0.00054 | 0.01138 | 0.47727 | 0.59855 | 1.21472 | 177 |
| GO_ESTABLISHMENT_OF_TISSUE_POLARITY | 0.00000 | 0.00000 | | 0.74791 | 1.50002 | 105 |

| Cluster 1 | | | | | | |
|---|---|---|---|---|---|---|
| GO_EUKARYOTIC_TRANSLATION_INITIATION_FACTOR_3_CO | 0.00009 | 0.00265 | 0.53843 | 0.87451 | 1.58303 | 15 |
| GO_FC_EPSILON_RECEPTOR_SIGNALING_PATHWAY | 0.00000 | 0.00000 | 0.83909 | 0.75060 | 1.50085 | 96 |
| GO_FC_RECEPTOR_MEDIATED_STIMULATORY_SIGNALING_F | 0.00049 | 0.01054 | 0.47727 | 0.66305 | 1.30951 | 66 |
| GO_FC_RECEPTOR_SIGNALING_PATHWAY | 0.00000 | 0.00000 | 0.78819 | 0.68756 | 1.39094 | 145 |
| GO_FICOLIN_1_RICH_GRANULE | 0.00000 | 0.00000 | 0.71951 | 0.69715 | 1.39923 | 108 |
| GO_FOREBRAIN_GENERATION_OF_NEURONS | 0.00036 | 0.00831 | 0.49849 | 0.73548 | 1.40659 | 33 |
| GO_FOREBRAIN_NEURON_DIFFERENTIATION | 0.00002 | 0.00071 | 0.57561 | 0.80711 | 1.52769 | 28 |
| GO_GENERATION_OF_PRECURSOR_METABOLITES_AND_EN | 0.00000 | 0.00000 | 0.77494 | 0.60609 | 1.24580 | 378 |
| GO_GLUCOSE_CATABOLIC_PROCESS | 0.00031 | 0.00721 | 0.49849 | 0.76477 | 1.45144 | 29 |
| GO_GLUTAMATERGIC_SYNAPSE | 0.00084 | 0.01579 | 0.47727 | 0.58338 | 1.18777 | 205 |
| GO_GLYCOLYTIC_PROCESS_THROUGH_FRUCTOSE_6_PHOSP | 0.00040 | 0.00899 | 0.49849 | 0.78413 | 1.47412 | 25 |
| GO_GOLGI_LUMEN | 0.00172 | 0.02636 | 0.45506 | 0.67971 | 1.32176 | 47 |
| GO_GRANULOCYTE_CHEMOTAXIS | 0.00205 | 0.02938 | 0.43171 | 0.67728 | 1.31705 | 47 |
| GO_GRANULOCYTE_MIGRATION | 0.00041 | 0.00920 | 0.49849 | 0.67663 | 1.32638 | 56 |
| GO_GTPASE_ACTIVITY | 0.00022 | 0.00570 | 0.51885 | 0.58652 | 1.19632 | 221 |
| GO_GUANYL_NUCLEOTIDE_BINDING | 0.00003 | 0.00095 | 0.57561 | 0.58703 | 1.19952 | 267 |
| GO_HEAT_SHOCK_PROTEIN_BINDING | 0.00017 | 0.00452 | 0.51885 | 0.63555 | 1.27466 | 105 |
| GO_HEMATOPOIETIC_PROGENITOR_CELL_DIFFERENTIATION | 0.00000 | 0.00004 | 0.65944 | 0.66279 | 1.33779 | 130 |
| GO_HEMATOPOIETIC_STEM_CELL_DIFFERENTIATION | 0.00000 | 0.00000 | 0.82666 | 0.77303 | 1.53371 | 75 |
| GO_HEXOSE_CATABOLIC_PROCESS | 0.00327 | 0.04125 | 0.43171 | 0.68841 | 1.33097 | 42 |
| GO_HINDBRAIN_DEVELOPMENT | 0.00324 | 0.04103 | 0.43171 | 0.61212 | 1.22053 | 87 |
| GO_HORMONE_BINDING | 0.00025 | 0.00611 | 0.49849 | 0.73637 | 1.41958 | 38 |
| GO_HUMORAL_IMMUNE_RESPONSE | 0.00003 | 0.00114 | 0.55733 | 0.68233 | 1.35490 | 77 |
| GO_HYPEROSMOTIC_RESPONSE | 0.00261 | 0.03481 | 0.43171 | 0.77846 | 1.42701 | 18 |
| GO_IMMUNE_RESPONSE_REGULATING_SIGNALING_PATHW | 0.00000 | 0.00000 | 0.78819 | 0.63592 | 1.29907 | 257 |
| GO_IMPORT_INTO_NUCLEUS | 0.00079 | 0.01510 | 0.47727 | 0.59938 | 1.21068 | 135 |
| GO_INNATE_IMMUNE_RESPONSE | 0.00000 | 0.00000 | 0.71951 | 0.58535 | 1.20699 | 465 |
| GO_INNATE_IMMUNE_RESPONSE_ACTIVATING_SIGNAL_TRA | 0.00000 | 0.00000 | 0.82666 | 0.76239 | 1.51951 | 86 |
| GO_INNER_MITOCHONDRIAL_MEMBRANE_PROTEIN_COMP | 0.00000 | 0.00003 | 0.65944 | 0.69554 | 1.38970 | 93 |
| GO_INORGANIC_ION_TRANSMEMBRANE_TRANSPORT | 0.00146 | 0.02340 | 0.45506 | 0.54447 | 1.12079 | 411 |
| GO_INTERACTION_WITH_HOST | 0.00119 | 0.02043 | 0.45506 | 0.59090 | 1.19677 | 153 |
| GO_INTERACTION_WITH_SYMBIONT | 0.00135 | 0.02201 | 0.45506 | 0.66818 | 1.30835 | 55 |
| GO_INTERLEUKIN_1_MEDIATED_SIGNALING_PATHWAY | 0.00000 | 0.00000 | | 0.77164 | 1.53861 | 87 |
| GO_INTERMEDIATE_FILAMENT_CYTOSKELETON | 0.00368 | 0.04525 | 0.43171 | 0.62100 | 1.23481 | 81 |
| GO_INTRINSIC_APOPTOTIC_SIGNALING_PATHWAY | 0.00047 | 0.01018 | 0.49849 | 0.58027 | 1.18356 | 231 |
| GO_INTRINSIC_COMPONENT_OF_ENDOPLASMIC_RETICULU | 0.00154 | 0.02422 | 0.45506 | 0.60864 | 1.22737 | 125 |
| GO_ION_HOMEOSTASIS | 0.00023 | 0.00583 | 0.51885 | 0.55835 | 1.15017 | 423 |
| GO_ION_TRANSMEMBRANE_TRANSPORTER_ACTIVITY | 0.00018 | 0.00473 | 0.51885 | 0.55791 | 1.14818 | 403 |
| GO_IRON_ION_HOMEOSTASIS | 0.00046 | 0.00992 | 0.49849 | 0.66676 | 1.31195 | 61 |
| GO_IRON_ION_TRANSPORT | 0.00023 | 0.00571 | 0.51885 | 0.68635 | 1.34543 | 56 |
| GO_KERATINIZATION | 0.00064 | 0.01301 | 0.47727 | 0.69974 | 1.35177 | 41 |
| GO_LABYRINTHINE_LAYER_MORPHOGENESIS | 0.00137 | 0.02214 | 0.45506 | 0.80443 | 1.46176 | 16 |
| GO_LARGE_RIBOSOMAL_SUBUNIT | 0.00000 | 0.00000 | | 0.75409 | 1.51315 | 107 |
| GO_LEUKOCYTE_CHEMOTAXIS | 0.00191 | 0.02819 | 0.45506 | 0.62261 | 1.24674 | 101 |
| GO_LEUKOCYTE_MEDIATED_IMMUNITY | 0.00000 | 0.00000 | | 0.60922 | 1.25671 | 474 |
| GO_LEUKOCYTE_MIGRATION | 0.00014 | 0.00382 | 0.51885 | 0.58273 | 1.18876 | 235 |
| GO_LUMENAL_SIDE_OF_ENDOPLASMIC_RETICULUM_MEM | 0.00272 | 0.03578 | 0.43171 | 0.79095 | 1.44352 | 17 |
| GO_LUMENAL_SIDE_OF_MEMBRANE | 0.00405 | 0.04880 | 0.40702 | 0.74223 | 1.37960 | 22 |
| GO_LYSOSOMAL_LUMEN | 0.00065 | 0.01330 | 0.47727 | 0.66316 | 1.31085 | 68 |
| GO_L_AMINO_ACID_TRANSPORT | 0.00413 | 0.04907 | 0.40702 | 0.65878 | 1.27675 | 44 |
| GO_MACROAUTOPHAGY | 0.00235 | 0.03255 | 0.43171 | 0.55876 | 1.14153 | 263 |
| GO_MAINTENANCE_OF_CELL_NUMBER | 0.00413 | 0.04907 | 0.40702 | 0.59029 | 1.19192 | 134 |
| GO_MEMBRANE_REGION | 0.00019 | 0.00499 | 0.51885 | 0.58366 | 1.19027 | 227 |
| GO_MESENCHYMAL_CELL_DIFFERENTIATION | 0.00202 | 0.02904 | 0.43171 | 0.59305 | 1.19789 | 137 |
| GO_METAL_ION_HOMEOSTASIS | 0.00040 | 0.00899 | 0.49849 | 0.56323 | 1.15486 | 320 |
| GO_METAL_ION_TRANSPORT | 0.00148 | 0.02352 | 0.45506 | 0.54367 | 1.12057 | 441 |
| GO_MIDBRAIN_DEVELOPMENT | 0.00178 | 0.02689 | 0.45506 | 0.65266 | 1.28154 | 58 |
| GO_MITOCHONDRIAL_ELECTRON_TRANSPORT_NADH_TO_U | 0.00015 | 0.00416 | 0.51885 | 0.71175 | 1.37940 | 44 |
| GO_MITOCHONDRIAL_GENE_EXPRESSION | 0.00010 | 0.00290 | 0.53843 | 0.61273 | 1.24098 | 153 |
| GO_MITOCHONDRIAL_LARGE_RIBOSOMAL_SUBUNIT | 0.00005 | 0.00150 | 0.55733 | 0.71829 | 1.40510 | 54 |
| GO_MITOCHONDRIAL_MATRIX | 0.00008 | 0.00252 | 0.53843 | 0.56269 | 1.15712 | 389 |
| GO_MITOCHONDRIAL_PROTEIN_COMPLEX | 0.00000 | 0.00000 | | 0.67839 | 1.38238 | 212 |
| GO_MITOCHONDRIAL_SMALL_RIBOSOMAL_SUBUNIT | 0.00161 | 0.02496 | 0.45506 | 0.73818 | 1.39723 | 28 |
| GO_MITOCHONDRIAL_TRANSLATION | 0.00002 | 0.00059 | 0.57561 | 0.64038 | 1.29275 | 129 |
| GO_MITOCHONDRIAL_TRANSLATIONAL_TERMINATION | 0.00000 | 0.00001 | 0.69013 | 0.71439 | 1.42447 | 87 |
| GO_MITOCHONDRION_ORGANIZATION | 0.00004 | 0.00126 | 0.55733 | 0.56301 | 1.16025 | 430 |
| GO_MITOTIC_SPINDLE_ASSEMBLY | 0.00261 | 0.03481 | 0.43171 | 0.64692 | 1.27026 | 58 |
| GO_MONOAMINE_TRANSPORT | 0.00185 | 0.02781 | 0.45506 | 0.71851 | 1.36776 | 30 |
| GO_MONOCARBOXYLIC_ACID_BINDING | 0.00294 | 0.03786 | 0.43171 | 0.68779 | 1.31808 | 35 |
| GO_MONOSACCHARIDE_CATABOLIC_PROCESS | 0.00113 | 0.01964 | 0.45506 | 0.67822 | 1.32379 | 51 |
| GO_MONOVALENT_INORGANIC_CATION_HOMEOSTASIS | 0.00102 | 0.01830 | 0.45506 | 0.62592 | 1.24814 | 88 |
| GO_MONOVALENT_INORGANIC_CATION_TRANSMEMBRAN | 0.00000 | 0.00003 | 0.65944 | 0.65127 | 1.31846 | 149 |
| GO_MONOVALENT_INORGANIC_CATION_TRANSPORT | 0.00003 | 0.00091 | 0.57561 | 0.58922 | 1.20351 | 255 |

| Cluster 1 | | | | | | |
|---|---|---|---|---|---|---|
| GO_MORPHOGENESIS_OF_AN_EPITHELIUM | 0.00000 | 0.00001 | 0.69013 | 0.59107 | 1.21514 | 384 |
| GO_MORPHOGENESIS_OF_A_POLARIZED_EPITHELIUM | 0.00000 | 0.00000 | | 0.75909 | 1.52917 | 120 |
| GO_MOVEMENT_IN_HOST_ENVIRONMENT | 0.00095 | 0.01732 | 0.47727 | 0.60863 | 1.22471 | 118 |
| GO_MRNA_3_END_PROCESSING | 0.00170 | 0.02614 | 0.45506 | 0.61739 | 1.23356 | 93 |
| GO_MRNA_5_UTR_BINDING | 0.00069 | 0.01362 | 0.47727 | 0.77360 | 1.45432 | 25 |
| GO_MRNA_BINDING | 0.00000 | 0.00000 | 0.76146 | 0.63467 | 1.29622 | 252 |
| GO_MRNA_EXPORT_FROM_NUCLEUS | 0.00026 | 0.00635 | 0.49849 | 0.63804 | 1.27958 | 104 |
| GO_MRNA_PROCESSING | 0.00000 | 0.00000 | | 0.62242 | 1.28303 | 455 |
| GO_MRNA_TRANSPORT | 0.00042 | 0.00934 | 0.49849 | 0.61260 | 1.23719 | 136 |
| GO_MULTI_MULTICELLULAR_ORGANISM_PROCESS | 0.00320 | 0.04064 | 0.43171 | 0.58830 | 1.18829 | 137 |
| GO_MULTI_ORGANISM_LOCALIZATION | 0.00156 | 0.02428 | 0.45506 | 0.65400 | 1.28203 | 56 |
| GO_MYELOID_LEUKOCYTE_ACTIVATION | 0.00000 | 0.00000 | | 0.61976 | 1.27658 | 418 |
| GO_MYELOID_LEUKOCYTE_MEDIATED_IMMUNITY | 0.00000 | 0.00000 | | 0.63077 | 1.29625 | 373 |
| GO_MYELOID_LEUKOCYTE_MIGRATION | 0.00139 | 0.02233 | 0.45506 | 0.61881 | 1.23734 | 96 |
| GO_NADH_DEHYDROGENASE_ACTIVITY | 0.00154 | 0.02422 | 0.45506 | 0.70921 | 1.36351 | 37 |
| GO_NADH_DEHYDROGENASE_COMPLEX | 0.00110 | 0.01915 | 0.45506 | 0.69835 | 1.34803 | 40 |
| GO_NADH_METABOLIC_PROCESS | 0.00001 | 0.00043 | 0.59333 | 0.79757 | 1.52212 | 32 |
| GO_NAD_METABOLIC_PROCESS | 0.00024 | 0.00590 | 0.51885 | 0.73877 | 1.42035 | 37 |
| GO_NCRNA_METABOLIC_PROCESS | 0.00008 | 0.00250 | 0.53843 | 0.56259 | 1.15781 | 403 |
| GO_NCRNA_PROCESSING | 0.00008 | 0.00235 | 0.53843 | 0.57055 | 1.17160 | 346 |
| GO_NEGATIVE_REGULATION_OF_APOPTOTIC_SIGNALING_P | 0.00007 | 0.00218 | 0.53843 | 0.61149 | 1.23968 | 171 |
| GO_NEGATIVE_REGULATION_OF_CANONICAL_WNT_SIGNAI | 0.00000 | 0.00000 | | 0.70656 | 1.42852 | 143 |
| GO_NEGATIVE_REGULATION_OF_CELL_CYCLE | 0.00045 | 0.00985 | 0.49849 | 0.54840 | 1.13195 | 497 |
| GO_NEGATIVE_REGULATION_OF_CELL_CYCLE_G2_M_PHASI | 0.00000 | 0.00007 | 0.64355 | 0.69453 | 1.38691 | 91 |
| GO_NEGATIVE_REGULATION_OF_CELL_CYCLE_PHASE_TRAN | 0.00003 | 0.00106 | 0.55733 | 0.60196 | 1.22648 | 211 |
| GO_NEGATIVE_REGULATION_OF_CELL_CYCLE_PROCESS | 0.00003 | 0.00098 | 0.57561 | 0.58516 | 1.19622 | 283 |
| GO_NEGATIVE_REGULATION_OF_INTRINSIC_APOPTOTIC_SI( | 0.00242 | 0.03322 | 0.43171 | 0.63005 | 1.25259 | 79 |
| GO_NEGATIVE_REGULATION_OF_MITOTIC_CELL_CYCLE | 0.00023 | 0.00570 | 0.51885 | 0.57520 | 1.17542 | 268 |
| GO_NEGATIVE_REGULATION_OF_MRNA_METABOLIC_PROC | 0.00261 | 0.03481 | 0.43171 | 0.63991 | 1.26380 | 66 |
| GO_NEGATIVE_REGULATION_OF_PROTEOLYSIS | 0.00176 | 0.02673 | 0.45506 | 0.57166 | 1.16338 | 195 |
| GO_NEGATIVE_REGULATION_OF_RNA_SPLICING | 0.00413 | 0.04907 | 0.40702 | 0.72951 | 1.36838 | 24 |
| GO_NEGATIVE_REGULATION_OF_WNT_SIGNALING_PATHW/ | 0.00000 | 0.00000 | 0.81404 | 0.68096 | 1.38028 | 163 |
| GO_NEURAL_NUCLEUS_DEVELOPMENT | 0.00111 | 0.01942 | 0.45506 | 0.68759 | 1.32830 | 41 |
| GO_NEURON_PROJECTION_TERMINUS | 0.00198 | 0.02876 | 0.43171 | 0.63907 | 1.26454 | 69 |
| GO_NEUTROPHIL_CHEMOTAXIS | 0.00342 | 0.04287 | 0.43171 | 0.69029 | 1.32379 | 36 |
| GO_NEUTROPHIL_MIGRATION | 0.00170 | 0.02614 | 0.45506 | 0.68741 | 1.33399 | 45 |
| GO_NIK_NF_KAPPAB_SIGNALING | 0.00000 | 0.00000 | | 0.73053 | 1.47349 | 126 |
| GO_NON_CANONICAL_WNT_SIGNALING_PATHWAY | 0.00000 | 0.00000 | | 0.73776 | 1.48776 | 125 |
| GO_NUCLEAR_ENVELOPE | 0.00005 | 0.00162 | 0.55733 | 0.57291 | 1.17632 | 349 |
| GO_NUCLEAR_EXPORT | 0.00001 | 0.00031 | 0.59333 | 0.61925 | 1.25809 | 185 |
| GO_NUCLEAR_SPECK | 0.00024 | 0.00590 | 0.51885 | 0.56832 | 1.16591 | 330 |
| GO_NUCLEAR_TRANSCRIBED_MRNA_CATABOLIC_PROCESS | 0.00000 | 0.00000 | | 0.70228 | 1.42796 | 189 |
| GO_NUCLEAR_TRANSCRIBED_MRNA_CATABOLIC_PROCESS_ | 0.00000 | 0.00000 | | 0.77878 | 1.56538 | 114 |
| GO_NUCLEAR_TRANSPORT | 0.00000 | 0.00003 | 0.65944 | 0.59888 | 1.22611 | 306 |
| GO_NUCLEOBASE_CONTAINING_COMPOUND_TRANSPORT | 0.00068 | 0.01360 | 0.47727 | 0.57815 | 1.17913 | 219 |
| GO_NUCLEOBASE_CONTAINING_SMALL_MOLECULE_METAB | 0.00005 | 0.00150 | 0.55733 | 0.56190 | 1.15745 | 421 |
| GO_NUCLEOSIDE_DIPHOSPHATE_METABOLIC_PROCESS | 0.00265 | 0.03509 | 0.43171 | 0.59938 | 1.20697 | 119 |
| GO_NUCLEOSIDE_PHOSPHATE_BIOSYNTHETIC_PROCESS | 0.00023 | 0.00570 | 0.51885 | 0.59776 | 1.21181 | 168 |
| GO_NUCLEOSIDE_TRIPHOSPHATE_BIOSYNTHETIC_PROCESS | 0.00239 | 0.03285 | 0.43171 | 0.67588 | 1.31432 | 47 |
| GO_NUCLEOSIDE_TRIPHOSPHATE_METABOLIC_PROCESS | 0.00032 | 0.00740 | 0.49849 | 0.66105 | 1.30804 | 69 |
| GO_NUCLEOTIDE_PHOSPHORYLATION | 0.00209 | 0.02978 | 0.43171 | 0.61208 | 1.22665 | 103 |
| GO_OLIGODENDROCYTE_DIFFERENTIATION | 0.00110 | 0.01915 | 0.45506 | 0.65668 | 1.29804 | 68 |
| GO_ORGANELLAR_RIBOSOME | 0.00000 | 0.00004 | 0.65944 | 0.70023 | 1.39440 | 84 |
| GO_ORGANELLE_INNER_MEMBRANE | 0.00000 | 0.00000 | | 0.62911 | 1.29487 | 405 |
| GO_ORGANIC_ACID_TRANSMEMBRANE_TRANSPORTER_AC | 0.00191 | 0.02819 | 0.45506 | 0.62340 | 1.23992 | 82 |
| GO_ORGANIC_CATION_TRANSPORT | 0.00067 | 0.01350 | 0.47727 | 0.75283 | 1.42171 | 27 |
| GO_OUTER_MEMBRANE | 0.00170 | 0.02614 | 0.45506 | 0.59091 | 1.19655 | 155 |
| GO_OXIDATIVE_PHOSPHORYLATION | 0.00000 | 0.00000 | 0.76146 | 0.71446 | 1.42961 | 99 |
| GO_OXIDOREDUCTASE_ACTIVITY | 0.00000 | 0.00007 | 0.64355 | 0.57399 | 1.18458 | 494 |
| GO_OXIDOREDUCTASE_ACTIVITY_ACTING_ON_A_HEME_GR | 0.00126 | 0.02120 | 0.45506 | 0.77489 | 1.42949 | 20 |
| GO_OXIDOREDUCTASE_ACTIVITY_ACTING_ON_NAD_P_H | 0.00100 | 0.01807 | 0.45506 | 0.64236 | 1.27553 | 77 |
| GO_OXIDOREDUCTASE_ACTIVITY_ACTING_ON_NAD_P_H_Q | 0.00265 | 0.03509 | 0.43171 | 0.67371 | 1.31009 | 47 |
| GO_OXIDOREDUCTASE_ACTIVITY_ACTING_ON_PEROXIDE_A | 0.00272 | 0.03578 | 0.43171 | 0.71175 | 1.35489 | 30 |
| GO_OXIDOREDUCTASE_COMPLEX | 0.00000 | 0.00019 | 0.61053 | 0.69177 | 1.37554 | 81 |
| GO_PEPTIDASE_COMPLEX | 0.00000 | 0.00000 | 0.76146 | 0.73916 | 1.47016 | 82 |
| GO_PEPTIDE_BINDING | 0.00097 | 0.01755 | 0.47727 | 0.59436 | 1.20442 | 158 |
| GO_PEPTIDYL_ASPARAGINE_MODIFICATION | 0.00235 | 0.03255 | 0.43171 | 0.72522 | 1.37638 | 29 |
| GO_PHAGOCYTOSIS | 0.00011 | 0.00313 | 0.53843 | 0.61058 | 1.23783 | 172 |
| GO_PHAGOSOME_ACIDIFICATION | 0.00028 | 0.00669 | 0.49849 | 0.78584 | 1.46736 | 23 |
| GO_PHAGOSOME_MATURATION | 0.00071 | 0.01377 | 0.47727 | 0.72036 | 1.38144 | 36 |
| GO_PIGMENT_GRANULE | 0.00000 | 0.00000 | 0.78819 | 0.74145 | 1.48060 | 91 |
| GO_PLACENTA_DEVELOPMENT | 0.00110 | 0.01915 | 0.45506 | 0.61420 | 1.23498 | 116 |
| GO_PLASMA_LIPOPROTEIN_PARTICLE_CLEARANCE | 0.00298 | 0.03826 | 0.43171 | 0.68763 | 1.31778 | 35 |

| Cluster 1 | | | | | | |
|---|---|---|---|---|---|---|
| GO_POLYMERIC_CYTOSKELETAL_FIBER | 0.00261 | 0.03481 | 0.43171 | 0.53882 | 1.11068 | 451 |
| GO_POLYSOMAL_RIBOSOME | 0.00004 | 0.00135 | 0.55733 | 0.78293 | 1.49258 | 31 |
| GO_POLYSOME | 0.00000 | 0.00001 | 0.69013 | 0.73733 | 1.45621 | 66 |
| GO_POSITIVE_CHEMOTAXIS | 0.00364 | 0.04496 | 0.43171 | 0.71732 | 1.36140 | 29 |
| GO_POSITIVE_REGULATION_OF_ACTIN_FILAMENT_POLYME | 0.00228 | 0.03186 | 0.43171 | 0.63633 | 1.25911 | 69 |
| GO_POSITIVE_REGULATION_OF_ATPASE_ACTIVITY | 0.00085 | 0.01587 | 0.47727 | 0.69826 | 1.35504 | 45 |
| GO_POSITIVE_REGULATION_OF_BINDING | 0.00027 | 0.00641 | 0.49849 | 0.61840 | 1.24777 | 127 |
| GO_POSITIVE_REGULATION_OF_CANONICAL_WNT_SIGNALI | 0.00000 | 0.00001 | 0.69013 | 0.66611 | 1.34355 | 126 |
| GO_POSITIVE_REGULATION_OF_CELLULAR_AMIDE_METABO | 0.00019 | 0.00507 | 0.51885 | 0.62187 | 1.25514 | 131 |
| GO_POSITIVE_REGULATION_OF_CELLULAR_COMPONENT_B | 0.00137 | 0.02214 | 0.45506 | 0.54674 | 1.12384 | 376 |
| GO_POSITIVE_REGULATION_OF_CELLULAR_COMPONENT_M | 0.00035 | 0.00813 | 0.49849 | 0.55813 | 1.14654 | 362 |
| GO_POSITIVE_REGULATION_OF_CELLULAR_PROTEIN_LOCAL | 0.00132 | 0.02182 | 0.45506 | 0.56185 | 1.14807 | 265 |
| GO_POSITIVE_REGULATION_OF_CELL_DEATH | 0.00038 | 0.00857 | 0.49849 | 0.54647 | 1.12792 | 495 |
| GO_POSITIVE_REGULATION_OF_CELL_DEVELOPMENT | 0.00194 | 0.02835 | 0.45506 | 0.54766 | 1.12509 | 361 |
| GO_POSITIVE_REGULATION_OF_CELL_SUBSTRATE_ADHESIO | 0.00063 | 0.01286 | 0.47727 | 0.63887 | 1.27222 | 84 |
| GO_POSITIVE_REGULATION_OF_CHROMOSOME_ORGANIZA | 0.00133 | 0.02182 | 0.45506 | 0.59280 | 1.20008 | 149 |
| GO_POSITIVE_REGULATION_OF_CYTOSKELETON_ORGANIZA | 0.00242 | 0.03322 | 0.43171 | 0.58272 | 1.18093 | 166 |
| GO_POSITIVE_REGULATION_OF_DEFENSE_RESPONSE | 0.00000 | 0.00007 | 0.64355 | 0.61893 | 1.26137 | 215 |
| GO_POSITIVE_REGULATION_OF_DNA_BIOSYNTHETIC_PROC | 0.00126 | 0.02120 | 0.45506 | 0.65791 | 1.29185 | 58 |
| GO_POSITIVE_REGULATION_OF_ESTABLISHMENT_OF_PROT | 0.00016 | 0.00423 | 0.51885 | 0.57551 | 1.17650 | 283 |
| GO_POSITIVE_REGULATION_OF_IMMUNE_RESPONSE | 0.00000 | 0.00000 | | 0.62146 | 1.27668 | 367 |
| GO_POSITIVE_REGULATION_OF_INNATE_IMMUNE_RESPON | 0.00000 | 0.00000 | 0.71951 | 0.67466 | 1.36273 | 137 |
| GO_POSITIVE_REGULATION_OF_MUSCLE_CONTRACTION | 0.00128 | 0.02140 | 0.45506 | 0.75851 | 1.42277 | 24 |
| GO_POSITIVE_REGULATION_OF_ORGANELLE_ORGANIZATIO | 0.00000 | 0.00001 | 0.69013 | 0.57791 | 1.19240 | 484 |
| GO_POSITIVE_REGULATION_OF_PROTEIN_CONTAINING_CO | 0.00104 | 0.01858 | 0.45506 | 0.58832 | 1.19550 | 186 |
| GO_POSITIVE_REGULATION_OF_PROTEIN_KINASE_B_SIGNA | 0.00405 | 0.04880 | 0.40702 | 0.60817 | 1.21266 | 87 |
| GO_POSITIVE_REGULATION_OF_PROTEIN_LOCALIZATION_T | 0.00002 | 0.00090 | 0.57561 | 0.69664 | 1.37672 | 67 |
| GO_POSITIVE_REGULATION_OF_RESPONSE_TO_BIOTIC_STI | 0.00000 | 0.00002 | 0.67496 | 0.64903 | 1.31521 | 158 |
| GO_POSITIVE_REGULATION_OF_RESPONSE_TO_EXTERNAL_ | 0.00001 | 0.00023 | 0.61053 | 0.59373 | 1.21429 | 289 |
| GO_POSITIVE_REGULATION_OF_SIGNAL_TRANSDUCTION_B | 0.00028 | 0.00677 | 0.49849 | 0.85383 | 1.54560 | 15 |
| GO_POSITIVE_REGULATION_OF_SUPRAMOLECULAR_FIBER_ | 0.00327 | 0.04125 | 0.43171 | 0.58287 | 1.18027 | 151 |
| GO_POSITIVE_REGULATION_OF_TELOMERASE_ACTIVITY | 0.00148 | 0.02352 | 0.45506 | 0.71950 | 1.37313 | 32 |
| GO_POSITIVE_REGULATION_OF_TELOMERASE_RNA_LOCALI | 0.00001 | 0.00039 | 0.59333 | 0.90044 | 1.62997 | 15 |
| GO_POSITIVE_REGULATION_OF_TELOMERE_MAINTENANCE | 0.00020 | 0.00517 | 0.51885 | 0.71880 | 1.39490 | 45 |
| GO_POSITIVE_REGULATION_OF_TELOMERE_MAINTENANCE | 0.00006 | 0.00196 | 0.53843 | 0.76170 | 1.45784 | 34 |
| GO_POSITIVE_REGULATION_OF_TRANSLATION | 0.00005 | 0.00170 | 0.55733 | 0.64338 | 1.29240 | 110 |
| GO_POSITIVE_REGULATION_OF_VIRAL_PROCESS | 0.00005 | 0.00161 | 0.55733 | 0.65694 | 1.31258 | 93 |
| GO_POSITIVE_REGULATION_OF_VIRAL_TRANSCRIPTION | 0.00205 | 0.02938 | 0.43171 | 0.70338 | 1.35597 | 38 |
| GO_POSITIVE_REGULATION_OF_WNT_SIGNALING_PATHWA | 0.00000 | 0.00001 | 0.69013 | 0.65488 | 1.32608 | 151 |
| GO_POSTSYNAPSE | 0.00008 | 0.00237 | 0.53843 | 0.56561 | 1.16200 | 368 |
| GO_POSTSYNAPSE_ORGANIZATION | 0.00119 | 0.02043 | 0.45506 | 0.62077 | 1.24494 | 104 |
| GO_POST_TRANSLATIONAL_PROTEIN_MODIFICATION | 0.00009 | 0.00260 | 0.53843 | 0.58215 | 1.18933 | 271 |
| GO_PRECATALYTIC_SPLICEOSOME | 0.00102 | 0.01830 | 0.45506 | 0.67961 | 1.32651 | 51 |
| GO_PRERIBOSOME | 0.00005 | 0.00152 | 0.55733 | 0.68264 | 1.35241 | 71 |
| GO_PRE_MRNA_BINDING | 0.00353 | 0.04376 | 0.43171 | 0.69639 | 1.33888 | 37 |
| GO_PRIMARY_ACTIVE_TRANSMEMBRANE_TRANSPORTER_A | 0.00231 | 0.03224 | 0.43171 | 0.64813 | 1.27997 | 65 |
| GO_PROCESS_UTILIZING_AUTOPHAGIC_MECHANISM | 0.00239 | 0.03285 | 0.43171 | 0.53962 | 1.11213 | 431 |
| GO_PROTEASOMAL_PROTEIN_CATABOLIC_PROCESS | 0.00000 | 0.00002 | 0.67496 | 0.58313 | 1.20089 | 416 |
| GO_PROTEASOMAL_UBIQUITIN_INDEPENDENT_PROTEIN_C | 0.00000 | 0.00002 | 0.67496 | 0.90484 | 1.66231 | 19 |
| GO_PROTEASOME_ACCESSORY_COMPLEX | 0.00001 | 0.00023 | 0.61053 | 0.84860 | 1.59176 | 24 |
| GO_PROTEASOME_CORE_COMPLEX | 0.00000 | 0.00001 | 0.69013 | 0.92794 | 1.69352 | 17 |
| GO_PROTEIN_CONTAINING_COMPLEX_DISASSEMBLY | 0.00000 | 0.00000 | 0.77494 | 0.63104 | 1.28935 | 270 |
| GO_PROTEIN_CONTAINING_COMPLEX_LOCALIZATION | 0.00001 | 0.00029 | 0.61053 | 0.60046 | 1.22458 | 228 |
| GO_PROTEIN_DNA_COMPLEX_SUBUNIT_ORGANIZATION | 0.00133 | 0.02182 | 0.45506 | 0.58874 | 1.19676 | 188 |
| GO_PROTEIN_FOLDING | 0.00000 | 0.00003 | 0.65944 | 0.63799 | 1.29617 | 185 |
| GO_PROTEIN_FOLDING_CHAPERONE | 0.00073 | 0.01404 | 0.47727 | 0.76991 | 1.43761 | 23 |
| GO_PROTEIN_IMPORT | 0.00087 | 0.01620 | 0.47727 | 0.59325 | 1.20238 | 162 |
| GO_PROTEIN_INSERTION_INTO_MEMBRANE | 0.00405 | 0.04880 | 0.40702 | 0.65560 | 1.27964 | 51 |
| GO_PROTEIN_LOCALIZATION_TO_CHROMOSOME_TELOMER | 0.00110 | 0.01915 | 0.45506 | 0.74562 | 1.40810 | 27 |
| GO_PROTEIN_LOCALIZATION_TO_ENDOPLASMIC_RETICULU | 0.00000 | 0.00000 | | 0.77225 | 1.55866 | 131 |
| GO_PROTEIN_LOCALIZATION_TO_MEMBRANE | 0.00000 | 0.00000 | | 0.63285 | 1.30602 | 486 |
| GO_PROTEIN_LOCALIZATION_TO_MITOCHONDRION | 0.00038 | 0.00857 | 0.49849 | 0.61605 | 1.24231 | 124 |
| GO_PROTEIN_LOCALIZATION_TO_NUCLEUS | 0.00007 | 0.00224 | 0.53843 | 0.59014 | 1.20367 | 225 |
| GO_PROTEIN_MODIFICATION_BY_SMALL_PROTEIN_REMOV | 0.00002 | 0.00063 | 0.57561 | 0.59625 | 1.21685 | 240 |
| GO_PROTEIN_POLYUBIQUITINATION | 0.00085 | 0.01587 | 0.47727 | 0.56356 | 1.15220 | 286 |
| GO_PROTEIN_STABILIZATION | 0.00279 | 0.03653 | 0.43171 | 0.58620 | 1.18673 | 149 |
| GO_PROTEIN_TARGETING | 0.00000 | 0.00000 | | 0.65691 | 1.34959 | 366 |
| GO_PROTEIN_TARGETING_TO_MEMBRANE | 0.00000 | 0.00000 | | 0.72738 | 1.47458 | 168 |
| GO_PROTEIN_TARGETING_TO_MITOCHONDRION | 0.00106 | 0.01886 | 0.45506 | 0.63203 | 1.25860 | 84 |
| GO_PROTON_TRANSMEMBRANE_TRANSPORT | 0.00000 | 0.00009 | 0.62726 | 0.69080 | 1.37751 | 88 |
| GO_PROTON_TRANSMEMBRANE_TRANSPORTER_ACTIVITY | 0.00003 | 0.00103 | 0.57561 | 0.69385 | 1.37121 | 67 |
| GO_PURINE_CONTAINING_COMPOUND_BIOSYNTHETIC_PR | 0.00019 | 0.00503 | 0.51885 | 0.62599 | 1.26236 | 125 |
| GO_PURINE_CONTAINING_COMPOUND_METABOLIC_PROC | 0.00045 | 0.00978 | 0.49849 | 0.56519 | 1.15713 | 306 |

| Cluster 1 | | | | | | |
|---|---|---|---|---|---|---|
| GO_PURINE_NUCLEOSIDE_TRIPHOSPHATE_METABOLIC_PRO | 0.00067 | 0.01350 | 0.47727 | 0.68736 | 1.34040 | 50 |
| GO_PYRUVATE_METABOLIC_PROCESS | 0.00108 | 0.01903 | 0.45506 | 0.61558 | 1.23724 | 115 |
| GO_RECEPTOR_MEDIATED_ENDOCYTOSIS | 0.00066 | 0.01344 | 0.47727 | 0.59349 | 1.20267 | 165 |
| GO_REGENERATION | 0.00030 | 0.00707 | 0.49849 | 0.61185 | 1.23412 | 126 |
| GO_REGULATION_OF_ACTIN_FILAMENT_BASED_PROCESS | 0.00124 | 0.02105 | 0.45506 | 0.56225 | 1.14877 | 276 |
| GO_REGULATION_OF_ACTIN_FILAMENT_LENGTH | 0.00413 | 0.04907 | 0.40702 | 0.59237 | 1.19560 | 131 |
| GO_REGULATION_OF_ACTIN_FILAMENT_ORGANIZATION | 0.00055 | 0.01169 | 0.47727 | 0.58725 | 1.19440 | 190 |
| GO_REGULATION_OF_ANATOMICAL_STRUCTURE_SIZE | 0.00053 | 0.01125 | 0.47727 | 0.56755 | 1.16241 | 309 |
| GO_REGULATION_OF_ANIMAL_ORGAN_MORPHOGENESIS | 0.00000 | 0.00000 | | 0.69842 | 1.41592 | 171 |
| GO_REGULATION_OF_APOPTOTIC_SIGNALING_PATHWAY | 0.00002 | 0.00058 | 0.59333 | 0.58574 | 1.19871 | 301 |
| GO_REGULATION_OF_ATPASE_ACTIVITY | 0.00268 | 0.03543 | 0.43171 | 0.64303 | 1.26494 | 60 |
| GO_REGULATION_OF_ATP_METABOLIC_PROCESS | 0.00216 | 0.03050 | 0.43171 | 0.61379 | 1.22505 | 90 |
| GO_REGULATION_OF_BINDING | 0.00018 | 0.00491 | 0.51885 | 0.58139 | 1.18766 | 259 |
| GO_REGULATION_OF_CELLULAR_AMIDE_METABOLIC_PROC | 0.00000 | 0.00003 | 0.65944 | 0.59296 | 1.21794 | 357 |
| GO_REGULATION_OF_CELLULAR_AMINE_METABOLIC_PROC | 0.00000 | 0.00000 | 0.82666 | 0.80095 | 1.58178 | 65 |
| GO_REGULATION_OF_CELLULAR_AMINO_ACID_METABOLIC | 0.00000 | 0.00000 | 0.83909 | 0.81680 | 1.59936 | 55 |
| GO_REGULATION_OF_CELLULAR_COMPONENT_SIZE | 0.00055 | 0.01169 | 0.47727 | 0.57122 | 1.16664 | 252 |
| GO_REGULATION_OF_CELLULAR_KETONE_METABOLIC_PRO | 0.00000 | 0.00000 | 0.78819 | 0.69357 | 1.39994 | 133 |
| GO_REGULATION_OF_CELLULAR_PROTEIN_LOCALIZATION | 0.00025 | 0.00619 | 0.49849 | 0.55283 | 1.13941 | 435 |
| GO_REGULATION_OF_CELLULAR_RESPONSE_TO_HEAT | 0.00086 | 0.01595 | 0.47727 | 0.64866 | 1.28509 | 71 |
| GO_REGULATION_OF_CELL_CYCLE_G2_M_PHASE_TRANSITI | 0.00012 | 0.00331 | 0.53843 | 0.59891 | 1.21812 | 190 |
| GO_REGULATION_OF_CELL_CYCLE_PHASE_TRANSITION | 0.00002 | 0.00074 | 0.57561 | 0.56951 | 1.17081 | 384 |
| GO_REGULATION_OF_CELL_MIGRATION_INVOLVED_IN_SPR | 0.00130 | 0.02165 | 0.45506 | 0.74249 | 1.39989 | 26 |
| GO_REGULATION_OF_CELL_MORPHOGENESIS | 0.00021 | 0.00538 | 0.51885 | 0.56593 | 1.16090 | 325 |
| GO_REGULATION_OF_CELL_SHAPE | 0.00095 | 0.01732 | 0.47727 | 0.61838 | 1.24111 | 108 |
| GO_REGULATION_OF_CHROMOSOME_ORGANIZATION | 0.00239 | 0.03285 | 0.43171 | 0.55658 | 1.13836 | 290 |
| GO_REGULATION_OF_CYTOSKELETON_ORGANIZATION | 0.00071 | 0.01377 | 0.47727 | 0.54979 | 1.13120 | 397 |
| GO_REGULATION_OF_DEFENSE_RESPONSE | 0.00002 | 0.00069 | 0.57561 | 0.56834 | 1.17007 | 412 |
| GO_REGULATION_OF_DNA_BIOSYNTHETIC_PROCESS | 0.00213 | 0.03018 | 0.43171 | 0.61854 | 1.23343 | 88 |
| GO_REGULATION_OF_DNA_TEMPLATED_TRANSCRIPTION_I | 0.00000 | 0.00000 | 0.82666 | 0.73812 | 1.47726 | 100 |
| GO_REGULATION_OF_GENERATION_OF_PRECURSOR_META | 0.00305 | 0.03898 | 0.43171 | 0.60244 | 1.21093 | 114 |
| GO_REGULATION_OF_HEMATOPOIETIC_PROGENITOR_CELL_ | 0.00000 | 0.00000 | 0.78819 | 0.76094 | 1.50913 | 74 |
| GO_REGULATION_OF_HEMATOPOIETIC_STEM_CELL_DIFFER | 0.00000 | 0.00000 | 0.81404 | 0.78965 | 1.55821 | 64 |
| GO_REGULATION_OF_HEMOPOIESIS | 0.00003 | 0.00117 | 0.55733 | 0.58132 | 1.19032 | 307 |
| GO_REGULATION_OF_IMMUNE_RESPONSE | 0.00000 | 0.00000 | 0.80122 | 0.59608 | 1.23021 | 492 |
| GO_REGULATION_OF_INNATE_IMMUNE_RESPONSE | 0.00000 | 0.00004 | 0.65944 | 0.62842 | 1.27968 | 201 |
| GO_REGULATION_OF_INTRINSIC_APOPTOTIC_SIGNALING_P | 0.00010 | 0.00295 | 0.53843 | 0.62535 | 1.26240 | 129 |
| GO_REGULATION_OF_INTRINSIC_APOPTOTIC_SIGNALING_P | 0.00062 | 0.01279 | 0.47727 | 0.77295 | 1.44986 | 24 |
| GO_REGULATION_OF_MORPHOGENESIS_OF_AN_EPITHELIU | 0.00000 | 0.00000 | | 0.71599 | 1.44599 | 136 |
| GO_REGULATION_OF_MRNA_CATABOLIC_PROCESS | 0.00000 | 0.00000 | 0.78819 | 0.67025 | 1.35893 | 174 |
| GO_REGULATION_OF_MRNA_METABOLIC_PROCESS | 0.00000 | 0.00000 | | 0.65362 | 1.33684 | 290 |
| GO_REGULATION_OF_MRNA_PROCESSING | 0.00000 | 0.00003 | 0.65944 | 0.66414 | 1.34046 | 131 |
| GO_REGULATION_OF_MRNA_SPLICING_VIA_SPLICEOSOME | 0.00002 | 0.00069 | 0.57561 | 0.66220 | 1.32406 | 95 |
| GO_REGULATION_OF_NEURAL_PRECURSOR_CELL_PROLIFEF | 0.00108 | 0.01903 | 0.45506 | 0.67727 | 1.31898 | 49 |
| GO_REGULATION_OF_NUCLEOCYTOPLASMIC_TRANSPORT | 0.00135 | 0.02201 | 0.45506 | 0.61922 | 1.23816 | 96 |
| GO_REGULATION_OF_OLIGODENDROCYTE_DIFFERENTIATIO | 0.00077 | 0.01484 | 0.47727 | 0.75191 | 1.41766 | 26 |
| GO_REGULATION_OF_OXIDATIVE_PHOSPHORYLATION | 0.00353 | 0.04376 | 0.43171 | 0.75300 | 1.38911 | 20 |
| GO_REGULATION_OF_PEPTIDASE_ACTIVITY | 0.00017 | 0.00456 | 0.51885 | 0.57793 | 1.18065 | 262 |
| GO_REGULATION_OF_PEPTIDE_TRANSPORT | 0.00012 | 0.00335 | 0.53843 | 0.55642 | 1.14732 | 464 |
| GO_REGULATION_OF_PH | 0.00156 | 0.02428 | 0.45506 | 0.64250 | 1.27132 | 69 |
| GO_REGULATION_OF_PLASMA_LIPOPROTEIN_PARTICLE_LEV | 0.00305 | 0.03898 | 0.43171 | 0.65904 | 1.29329 | 57 |
| GO_REGULATION_OF_PROTEIN_BINDING | 0.00368 | 0.04525 | 0.43171 | 0.57988 | 1.17239 | 143 |
| GO_REGULATION_OF_PROTEIN_CONTAINING_COMPLEX_AS | 0.00044 | 0.00978 | 0.49849 | 0.55858 | 1.14535 | 321 |
| GO_REGULATION_OF_PROTEIN_IMPORT | 0.00202 | 0.02904 | 0.43171 | 0.66759 | 1.30306 | 51 |
| GO_REGULATION_OF_PROTEIN_LOCALIZATION_TO_NUCLEU | 0.00015 | 0.00420 | 0.51885 | 0.64285 | 1.28830 | 103 |
| GO_REGULATION_OF_PROTEIN_MODIFICATION_BY_SMALL_ | 0.00313 | 0.03977 | 0.43171 | 0.57182 | 1.16321 | 192 |
| GO_REGULATION_OF_PROTEIN_STABILITY | 0.00070 | 0.01372 | 0.47727 | 0.57287 | 1.16881 | 236 |
| GO_REGULATION_OF_PROTEOLYSIS | 0.00023 | 0.00570 | 0.51885 | 0.55181 | 1.13878 | 486 |
| GO_REGULATION_OF_RESPONSE_TO_BIOTIC_STIMULUS | 0.00000 | 0.00009 | 0.62726 | 0.60318 | 1.23233 | 261 |
| GO_REGULATION_OF_RNA_SPLICING | 0.00001 | 0.00022 | 0.61053 | 0.64281 | 1.29748 | 133 |
| GO_REGULATION_OF_SMALL_MOLECULE_METABOLIC_PRO | 0.00000 | 0.00001 | 0.69013 | 0.60304 | 1.23617 | 317 |
| GO_REGULATION_OF_SPROUTING_ANGIOGENESIS | 0.00026 | 0.00626 | 0.49849 | 0.70283 | 1.36212 | 44 |
| GO_REGULATION_OF_STEM_CELL_DIFFERENTIATION | 0.00000 | 0.00000 | 0.80122 | 0.74182 | 1.48218 | 93 |
| GO_REGULATION_OF_SUPRAMOLECULAR_FIBER_ORGANIZA | 0.00294 | 0.03786 | 0.43171 | 0.56031 | 1.14477 | 258 |
| GO_REGULATION_OF_SYMBIOTIC_PROCESS | 0.00002 | 0.00061 | 0.57561 | 0.62198 | 1.26050 | 166 |
| GO_REGULATION_OF_SYNAPSE_STRUCTURE_OR_ACTIVITY | 0.00283 | 0.03687 | 0.43171 | 0.59379 | 1.19849 | 132 |
| GO_REGULATION_OF_TELOMERE_MAINTENANCE_VIA_TELO | 0.00413 | 0.04907 | 0.40702 | 0.65065 | 1.27236 | 53 |
| GO_REGULATION_OF_TRANSCRIPTION_FROM_RNA_POLYM | 0.00000 | 0.00000 | 0.81404 | 0.78472 | 1.55080 | 67 |
| GO_REGULATION_OF_TRANSLATIONAL_INITIATION | 0.00194 | 0.02835 | 0.45506 | 0.64201 | 1.27131 | 70 |
| GO_REGULATION_OF_VIRAL_TRANSCRIPTION | 0.00026 | 0.00626 | 0.49849 | 0.68336 | 1.33957 | 56 |
| GO_REGULATION_OF_WNT_SIGNALING_PATHWAY | 0.00000 | 0.00006 | 0.64355 | 0.59907 | 1.22459 | 284 |
| GO_REPRODUCTIVE_SYSTEM_DEVELOPMENT | 0.00156 | 0.02428 | 0.45506 | 0.55813 | 1.14186 | 296 |
| GO_RESPIRASOME | 0.00000 | 0.00001 | 0.70498 | 0.72875 | 1.44529 | 74 |

| Cluster 1 | | | | | | |
|---|---|---|---|---|---|---|
| GO_RESPIRATORY_CHAIN_COMPLEX | 0.00000 | 0.00003 | 0.65944 | 0.73152 | 1.44350 | 64 |
| GO_RESPIRATORY_ELECTRON_TRANSPORT_CHAIN | 0.00000 | 0.00000 | 0.73376 | 0.72170 | 1.44041 | 90 |
| GO_RESPONSE_TO_ATP | 0.00078 | 0.01497 | 0.47727 | 0.80657 | 1.47855 | 18 |
| GO_RESPONSE_TO_DRUG | 0.00002 | 0.00059 | 0.57561 | 0.59418 | 1.21306 | 248 |
| GO_RESPONSE_TO_ENDOPLASMIC_RETICULUM_STRESS | 0.00003 | 0.00117 | 0.55733 | 0.59133 | 1.20680 | 240 |
| GO_RESPONSE_TO_ESTRADIOL | 0.00133 | 0.02182 | 0.45506 | 0.62692 | 1.25076 | 89 |
| GO_RESPONSE_TO_ESTROGEN | 0.00023 | 0.00571 | 0.51885 | 0.69407 | 1.35905 | 55 |
| GO_RESPONSE_TO_GROWTH_FACTOR | 0.00108 | 0.01903 | 0.45506 | 0.54207 | 1.11785 | 469 |
| GO_RESPONSE_TO_HEAT | 0.00003 | 0.00108 | 0.55733 | 0.63736 | 1.28640 | 131 |
| GO_RESPONSE_TO_INORGANIC_SUBSTANCE | 0.00009 | 0.00273 | 0.53843 | 0.56761 | 1.16545 | 348 |
| GO_RESPONSE_TO_INTERLEUKIN_1 | 0.00000 | 0.00000 | | 0.70748 | 1.42999 | 140 |
| GO_RESPONSE_TO_INTERLEUKIN_12 | 0.00021 | 0.00542 | 0.51885 | 0.73891 | 1.42447 | 38 |
| GO_RESPONSE_TO_INTERLEUKIN_4 | 0.00028 | 0.00660 | 0.49849 | 0.80397 | 1.48786 | 21 |
| GO_RESPONSE_TO_METAL_ION | 0.00257 | 0.03481 | 0.43171 | 0.56413 | 1.15052 | 219 |
| GO_RESPONSE_TO_OXIDATIVE_STRESS | 0.00398 | 0.04835 | 0.40702 | 0.54938 | 1.12542 | 313 |
| GO_RESPONSE_TO_OXYGEN_LEVELS | 0.00000 | 0.00000 | | 0.65196 | 1.33272 | 284 |
| GO_RESPONSE_TO_PEPTIDE_HORMONE | 0.00187 | 0.02796 | 0.45506 | 0.55729 | 1.14057 | 303 |
| GO_RESPONSE_TO_REACTIVE_OXYGEN_SPECIES | 0.00132 | 0.02182 | 0.45506 | 0.59099 | 1.19759 | 158 |
| GO_RESPONSE_TO_STEROID_HORMONE | 0.00364 | 0.04496 | 0.43171 | 0.56177 | 1.14557 | 224 |
| GO_RESPONSE_TO_TEMPERATURE_STIMULUS | 0.00012 | 0.00339 | 0.53843 | 0.60907 | 1.23442 | 167 |
| GO_RESPONSE_TO_TOPOLOGICALLY_INCORRECT_PROTEIN | 0.00005 | 0.00153 | 0.55733 | 0.61445 | 1.24568 | 171 |
| GO_RESPONSE_TO_TOXIC_SUBSTANCE | 0.00194 | 0.02835 | 0.45506 | 0.59065 | 1.19399 | 142 |
| GO_RESPONSE_TO_TUMOR_NECROSIS_FACTOR | 0.00000 | 0.00000 | 0.74774 | 0.64698 | 1.31701 | 196 |
| GO_RESPONSE_TO_VIRUS | 0.00100 | 0.01807 | 0.45506 | 0.58359 | 1.18868 | 207 |
| GO_RESPONSE_TO_WOUNDING | 0.00008 | 0.00234 | 0.53843 | 0.56196 | 1.15529 | 382 |
| GO_RETINA_HOMEOSTASIS | 0.00000 | 0.00007 | 0.64355 | 0.79604 | 1.53459 | 38 |
| GO_RETROGRADE_VESICLE_MEDIATED_TRANSPORT_GOLGI | 0.00194 | 0.02835 | 0.45506 | 0.63265 | 1.25776 | 79 |
| GO_RIBONUCLEOPROTEIN_COMPLEX_BINDING | 0.00001 | 0.00028 | 0.61053 | 0.65306 | 1.31508 | 119 |
| GO_RIBONUCLEOPROTEIN_COMPLEX_BIOGENESIS | 0.00000 | 0.00000 | | 0.65678 | 1.35169 | 402 |
| GO_RIBONUCLEOPROTEIN_COMPLEX_SUBUNIT_ORGANIZAT | 0.00000 | 0.00000 | 0.80122 | 0.67439 | 1.36983 | 184 |
| GO_RIBONUCLEOPROTEIN_GRANULE | 0.00001 | 0.00051 | 0.59333 | 0.61141 | 1.24451 | 197 |
| GO_RIBONUCLEOSIDE_TRIPHOSPHATE_METABOLIC_PROCES | 0.00070 | 0.01372 | 0.47727 | 0.69125 | 1.34538 | 48 |
| GO_RIBOSE_PHOSPHATE_BIOSYNTHETIC_PROCESS | 0.00023 | 0.00570 | 0.51885 | 0.61968 | 1.24962 | 124 |
| GO_RIBOSE_PHOSPHATE_METABOLIC_PROCESS | 0.00041 | 0.00920 | 0.49849 | 0.56770 | 1.16121 | 291 |
| GO_RIBOSOMAL_LARGE_SUBUNIT_ASSEMBLY | 0.00174 | 0.02658 | 0.45506 | 0.73778 | 1.39329 | 27 |
| GO_RIBOSOMAL_LARGE_SUBUNIT_BIOGENESIS | 0.00002 | 0.00063 | 0.57561 | 0.70480 | 1.39317 | 68 |
| GO_RIBOSOMAL_SMALL_SUBUNIT_ASSEMBLY | 0.00060 | 0.01249 | 0.47727 | 0.81355 | 1.49133 | 18 |
| GO_RIBOSOMAL_SMALL_SUBUNIT_BIOGENESIS | 0.00009 | 0.00260 | 0.53843 | 0.67893 | 1.34172 | 67 |
| GO_RIBOSOMAL_SUBUNIT | 0.00000 | 0.00000 | | 0.74702 | 1.51440 | 173 |
| GO_RIBOSOME | 0.00000 | 0.00000 | | 0.74278 | 1.51337 | 210 |
| GO_RIBOSOME_ASSEMBLY | 0.00001 | 0.00023 | 0.61053 | 0.73100 | 1.43450 | 57 |
| GO_RIBOSOME_BIOGENESIS | 0.00000 | 0.00000 | | 0.65672 | 1.34217 | 279 |
| GO_RNA_3_END_PROCESSING | 0.00071 | 0.01377 | 0.47727 | 0.60800 | 1.22738 | 129 |
| GO_RNA_CATABOLIC_PROCESS | 0.00000 | 0.00000 | | 0.69504 | 1.42765 | 356 |
| GO_RNA_DEPENDENT_DNA_BIOSYNTHETIC_PROCESS | 0.00183 | 0.02766 | 0.45506 | 0.65408 | 1.28957 | 63 |
| GO_RNA_EXPORT_FROM_NUCLEUS | 0.00005 | 0.00152 | 0.55733 | 0.63688 | 1.28550 | 130 |
| GO_RNA_LOCALIZATION | 0.00000 | 0.00020 | 0.61053 | 0.61351 | 1.24999 | 210 |
| GO_RNA_SPLICING | 0.00000 | 0.00000 | | 0.62846 | 1.29337 | 403 |
| GO_RNA_SPLICING_VIA_TRANSESTERIFICATION_REACTIONS | 0.00000 | 0.00000 | | 0.64537 | 1.32387 | 325 |
| GO_ROUGH_ENDOPLASMIC_RETICULUM | 0.00000 | 0.00006 | 0.64355 | 0.72195 | 1.42583 | 66 |
| GO_RRNA_BINDING | 0.00000 | 0.00010 | 0.62726 | 0.73682 | 1.44679 | 58 |
| GO_RRNA_METABOLIC_PROCESS | 0.00000 | 0.00000 | 0.76146 | 0.65155 | 1.32768 | 212 |
| GO_SCF_DEPENDENT_PROTEASOMAL_UBIQUITIN_DEPENDI | 0.00000 | 0.00000 | 0.80122 | 0.75746 | 1.50616 | 81 |
| GO_SECRETORY_GRANULE_MEMBRANE | 0.00261 | 0.03481 | 0.43171 | 0.58213 | 1.18149 | 179 |
| GO_SERINE_TYPE_ENDOPEPTIDASE_INHIBITOR_ACTIVITY | 0.00059 | 0.01234 | 0.47727 | 0.72367 | 1.38779 | 36 |
| GO_SINGLE_FERTILIZATION | 0.00209 | 0.02978 | 0.43171 | 0.65012 | 1.27655 | 58 |
| GO_SINGLE_STRANDED_RNA_BINDING | 0.00042 | 0.00927 | 0.49849 | 0.66029 | 1.30900 | 72 |
| GO_SMALL_MOLECULE_BIOSYNTHETIC_PROCESS | 0.00005 | 0.00166 | 0.55733 | 0.56109 | 1.15711 | 471 |
| GO_SMALL_NUCLEAR_RIBONUCLEOPROTEIN_COMPLEX | 0.00058 | 0.01219 | 0.47727 | 0.67339 | 1.32765 | 63 |
| GO_SMALL_RIBOSOMAL_SUBUNIT | 0.00000 | 0.00000 | 0.71951 | 0.74563 | 1.47539 | 69 |
| GO_SMOOTH_MUSCLE_CELL_PROLIFERATION | 0.00228 | 0.03186 | 0.43171 | 0.61700 | 1.23239 | 92 |
| GO_SM_LIKE_PROTEIN_FAMILY_COMPLEX | 0.00065 | 0.01315 | 0.47727 | 0.65544 | 1.29922 | 73 |
| GO_SNORNA_BINDING | 0.00261 | 0.03481 | 0.43171 | 0.72278 | 1.37176 | 29 |
| GO_SOMATIC_STEM_CELL_POPULATION_MAINTENANCE | 0.00265 | 0.03509 | 0.43171 | 0.64680 | 1.27002 | 58 |
| GO_SPERM_EGG_RECOGNITION | 0.00009 | 0.00271 | 0.53843 | 0.79469 | 1.49832 | 26 |
| GO_SPLICEOSOMAL_COMPLEX | 0.00000 | 0.00004 | 0.65944 | 0.63861 | 1.29555 | 175 |
| GO_SPROUTING_ANGIOGENESIS | 0.00372 | 0.04562 | 0.43171 | 0.62500 | 1.24256 | 79 |
| GO_STEM_CELL_DIFFERENTIATION | 0.00000 | 0.00000 | 0.76146 | 0.66341 | 1.34621 | 176 |
| GO_STEROID_METABOLIC_PROCESS | 0.00268 | 0.03543 | 0.43171 | 0.57057 | 1.16146 | 196 |
| GO_STRUCTURAL_CONSTITUENT_OF_CYTOSKELETON | 0.00005 | 0.00156 | 0.55733 | 0.68486 | 1.35616 | 70 |
| GO_STRUCTURAL_CONSTITUENT_OF_RIBOSOME | 0.00000 | 0.00000 | | 0.75841 | 1.53604 | 153 |
| GO_STRUCTURAL_MOLECULE_ACTIVITY | 0.00000 | 0.00000 | | 0.69309 | 1.42861 | 444 |

| Cluster 1 | | | | | | |
|---|---|---|---|---|---|---|
| GO_SUBSTANTIA_NIGRA_DEVELOPMENT | 0.00283 | 0.03687 | 0.43171 | 0.70288 | 1.34424 | 33 |
| GO_SUBSTRATE_ADHESION_DEPENDENT_CELL_SPREADING | 0.00290 | 0.03761 | 0.43171 | 0.62793 | 1.24838 | 79 |
| GO_SYNAPSE_ORGANIZATION | 0.00005 | 0.00163 | 0.55733 | 0.58945 | 1.20221 | 232 |
| GO_TELOMERASE_RNA_LOCALIZATION | 0.00011 | 0.00311 | 0.53843 | 0.82646 | 1.51832 | 19 |
| GO_TELOMERE_MAINTENANCE_VIA_TELOMERE_LENGTHEN | 0.00082 | 0.01558 | 0.47727 | 0.65324 | 1.29355 | 70 |
| GO_THREONINE_TYPE_ENDOPEPTIDASE_ACTIVITY | 0.00000 | 0.00000 | 0.74774 | 0.94530 | 1.72520 | 17 |
| GO_THREONINE_TYPE_PEPTIDASE_ACTIVITY | 0.00000 | 0.00000 | 0.76146 | 0.93037 | 1.72177 | 21 |
| GO_TISSUE_MORPHOGENESIS | 0.00000 | 0.00010 | 0.62726 | 0.57531 | 1.18590 | 445 |
| GO_TOXIN_TRANSPORT | 0.00068 | 0.01360 | 0.47727 | 0.72555 | 1.38760 | 33 |
| GO_TRANSCRIPTION_ELONGATION_FROM_RNA_POLYMERA | 0.00383 | 0.04672 | 0.43171 | 0.62246 | 1.23601 | 77 |
| GO_TRANSITION_METAL_ION_TRANSPORT | 0.00159 | 0.02473 | 0.45506 | 0.62918 | 1.25236 | 83 |
| GO_TRANSLATIONAL_ELONGATION | 0.00000 | 0.00000 | 0.78819 | 0.69972 | 1.41103 | 124 |
| GO_TRANSLATIONAL_INITIATION | 0.00000 | 0.00000 | | 0.74352 | 1.50751 | 174 |
| GO_TRANSLATIONAL_TERMINATION | 0.00000 | 0.00001 | 0.69013 | 0.70207 | 1.40587 | 101 |
| GO_TRANSLATION_FACTOR_ACTIVITY_RNA_BINDING | 0.00000 | 0.00021 | 0.61053 | 0.70482 | 1.39728 | 72 |
| GO_TRANSLATION_INITIATION_FACTOR_ACTIVITY | 0.00016 | 0.00423 | 0.51885 | 0.72024 | 1.39861 | 46 |
| GO_TRANSLATION_PREINITIATION_COMPLEX | 0.00018 | 0.00481 | 0.51885 | 0.84429 | 1.53419 | 16 |
| GO_TRANSLATION_REGULATOR_ACTIVITY | 0.00000 | 0.00002 | 0.67496 | 0.67405 | 1.35635 | 118 |
| GO_TRANSLATION_REGULATOR_ACTIVITY_NUCLEIC_ACID_E | 0.00000 | 0.00003 | 0.65944 | 0.70236 | 1.40254 | 91 |
| GO_TRANSMEMBRANE_RECEPTOR_PROTEIN_TYROSINE_KIN | 0.00198 | 0.02876 | 0.43171 | 0.54157 | 1.11731 | 475 |
| GO_TUMOR_NECROSIS_FACTOR_MEDIATED_SIGNALING_PA | 0.00000 | 0.00000 | 0.82666 | 0.72034 | 1.45137 | 121 |
| GO_T_CELL_RECEPTOR_SIGNALING_PATHWAY | 0.00000 | 0.00000 | 0.78819 | 0.68968 | 1.39401 | 140 |
| GO_U12_TYPE_SPLICEOSOMAL_COMPLEX | 0.00008 | 0.00235 | 0.53843 | 0.79034 | 1.49254 | 27 |
| GO_U1_SNRNP | 0.00420 | 0.04986 | 0.40702 | 0.76755 | 1.40702 | 18 |
| GO_U2_SNRNP | 0.00213 | 0.03018 | 0.43171 | 0.78375 | 1.43671 | 18 |
| GO_U2_TYPE_SPLICEOSOMAL_COMPLEX | 0.00019 | 0.00499 | 0.51885 | 0.64763 | 1.29143 | 88 |
| GO_UBIQUITIN_LIKE_PROTEIN_LIGASE_BINDING | 0.00000 | 0.00002 | 0.67496 | 0.61137 | 1.24925 | 264 |
| GO_UNFOLDED_PROTEIN_BINDING | 0.00000 | 0.00000 | 0.71951 | 0.70675 | 1.41312 | 97 |
| GO_VACUOLAR_LUMEN | 0.00000 | 0.00012 | 0.62726 | 0.66529 | 1.33770 | 116 |
| GO_VACUOLAR_MEMBRANE | 0.00390 | 0.04754 | 0.43171 | 0.54598 | 1.11998 | 325 |
| GO_VASCULAR_ENDOTHELIAL_GROWTH_FACTOR_SIGNALIN | 0.00194 | 0.02835 | 0.45506 | 0.75153 | 1.40329 | 23 |
| GO_VESICLE_BUDDING_FROM_MEMBRANE | 0.00261 | 0.03481 | 0.43171 | 0.62325 | 1.24457 | 91 |
| GO_VESICLE_LUMEN | 0.00000 | 0.00000 | 0.77494 | 0.64836 | 1.32105 | 214 |
| GO_VESICLE_MEMBRANE | 0.00052 | 0.01109 | 0.47727 | 0.54778 | 1.13068 | 497 |
| GO_VIRAL_GENE_EXPRESSION | 0.00000 | 0.00000 | | 0.72575 | 1.47328 | 180 |
| GO_VIRAL_LIFE_CYCLE | 0.00000 | 0.00019 | 0.61053 | 0.60074 | 1.22647 | 248 |
| GO_WOUND_HEALING | 0.00039 | 0.00895 | 0.49849 | 0.56379 | 1.15405 | 304 |
| HP_ABNORMALITY_OF_ACID_BASE_HOMEOSTASIS | 0.00137 | 0.02214 | 0.45506 | 0.56291 | 1.14976 | 255 |
| HP_ABNORMAL_CSF_METABOLITE_LEVEL | 0.00157 | 0.02450 | 0.45506 | 0.62998 | 1.25561 | 86 |
| HP_ABNORMAL_ERYTHROCYTE_MORPHOLOGY | 0.00128 | 0.02140 | 0.45506 | 0.54661 | 1.12520 | 410 |
| HP_ABNORMAL_RETICULOCYTE_MORPHOLOGY | 0.00376 | 0.04598 | 0.43171 | 0.68820 | 1.31978 | 36 |
| HP_ACUTE_LEUKEMIA | 0.00069 | 0.01362 | 0.47727 | 0.64869 | 1.28812 | 76 |
| HP_ANEMIA_OF_INADEQUATE_PRODUCTION | 0.00074 | 0.01418 | 0.47727 | 0.62062 | 1.24562 | 108 |
| HP_BULBAR_PALSY | 0.00043 | 0.00951 | 0.49849 | 0.76078 | 1.43672 | 27 |
| HP_CEREBRAL_ISCHEMIA | 0.00121 | 0.02064 | 0.45506 | 0.66063 | 1.30248 | 63 |
| HP_FATIGUE | 0.00126 | 0.02120 | 0.45506 | 0.56948 | 1.16134 | 229 |
| HP_INCREASED_CSF_LACTATE | 0.00205 | 0.02938 | 0.43171 | 0.63403 | 1.25610 | 71 |
| HP_MACROCYTIC_ANEMIA | 0.00005 | 0.00169 | 0.55733 | 0.74705 | 1.44435 | 42 |
| HP_MIGRAINE | 0.00012 | 0.00339 | 0.53843 | 0.66146 | 1.31527 | 81 |
| HP_MOTOR_POLYNEUROPATHY | 0.00115 | 0.01990 | 0.45506 | 0.82793 | 1.49872 | 15 |
| HP_MUTISM | 0.00176 | 0.02673 | 0.45506 | 0.70735 | 1.36363 | 38 |
| HP_PALLOR | 0.00261 | 0.03481 | 0.43171 | 0.60822 | 1.21890 | 103 |
| HP_POOR_HEAD_CONTROL | 0.00069 | 0.01362 | 0.47727 | 0.61337 | 1.23514 | 119 |
| HP_RESPIRATORY_INSUFFICIENCY | 0.00185 | 0.02781 | 0.45506 | 0.55129 | 1.13209 | 352 |
| HP_THROMBOCYTOSIS | 0.00413 | 0.04907 | 0.40702 | 0.73102 | 1.37122 | 24 |
| HP_TRANSIENT_ISCHEMIC_ATTACK | 0.00087 | 0.01620 | 0.47727 | 0.71620 | 1.37348 | 36 |
| **Cluster 4** | | | | | | |
| pathway | pval | padj | log2err | ES | NES | size |
| GO_ACTIN_BINDING | 0.00428 | 0.04497 | 0.40702 | 0.55470 | 1.13554 | 281 |
| GO_ACTIN_CYTOSKELETON | 0.00178 | 0.02448 | 0.45506 | 0.55660 | 1.14190 | 333 |
| GO_ACTIN_FILAMENT_BINDING | 0.00012 | 0.00300 | 0.53843 | 0.61481 | 1.24373 | 138 |
| GO_ACTIN_FILAMENT_ORGANIZATION | 0.00036 | 0.00736 | 0.49849 | 0.57085 | 1.16870 | 284 |
| GO_ACTIN_POLYMERIZATION_OR_DEPOLYMERIZATION | 0.00110 | 0.01734 | 0.45506 | 0.59200 | 1.19845 | 147 |
| GO_ACTIVATION_OF_IMMUNE_RESPONSE | 0.00022 | 0.00494 | 0.51885 | 0.58161 | 1.19010 | 264 |
| GO_ACTIVATION_OF_INNATE_IMMUNE_RESPONSE | 0.00001 | 0.00024 | 0.61053 | 0.67182 | 1.35178 | 106 |
| GO_ADP_BINDING | 0.00009 | 0.00233 | 0.53843 | 0.78631 | 1.49047 | 27 |
| GO_AEROBIC_RESPIRATION | 0.00000 | 0.00000 | 0.73376 | 0.74748 | 1.48359 | 72 |
| GO_ALCOHOL_BIOSYNTHETIC_PROCESS | 0.00413 | 0.04406 | 0.40702 | 0.60004 | 1.20735 | 106 |
| GO_ALTERNATIVE_MRNA_SPLICING_VIA_SPLICEOSOME | 0.00178 | 0.02448 | 0.45506 | 0.63875 | 1.26594 | 70 |
| GO_AMIDE_BINDING | 0.00220 | 0.02861 | 0.43171 | 0.57241 | 1.16669 | 208 |
| GO_AMIDE_TRANSMEMBRANE_TRANSPORTER_ACTIVITY | 0.00294 | 0.03542 | 0.43171 | 0.71837 | 1.36920 | 30 |
| GO_AMINE_METABOLIC_PROCESS | 0.00003 | 0.00108 | 0.55733 | 0.66713 | 1.33377 | 93 |
| GO_ANAPHASE_PROMOTING_COMPLEX_DEPENDENT_CATA | 0.00000 | 0.00001 | 0.70498 | 0.72875 | 1.45006 | 76 |

| Cluster 1 | | | | | | |
|---|---|---|---|---|---|---|
| GO_ANTIGEN_PROCESSING_AND_PRESENTATION | 0.00000 | 0.00000 | 0.76146 | 0.67739 | 1.37553 | 169 |
| GO_ANTIGEN_PROCESSING_AND_PRESENTATION_OF_EXOG | 0.00000 | 0.00009 | 0.64355 | 0.72224 | 1.42805 | 66 |
| GO_ANTIGEN_PROCESSING_AND_PRESENTATION_OF_PEPT | 0.00000 | 0.00000 | 0.77494 | 0.68699 | 1.39126 | 149 |
| GO_ANTIGEN_PROCESSING_AND_PRESENTATION_OF_PEPT | 0.00000 | 0.00004 | 0.65944 | 0.71666 | 1.42760 | 80 |
| GO_ANTIGEN_PROCESSING_AND_PRESENTATION_OF_PEPT | 0.00061 | 0.01103 | 0.47727 | 0.64723 | 1.28874 | 79 |
| GO_ANTIGEN_RECEPTOR_MEDIATED_SIGNALING_PATHWAY | 0.00124 | 0.01901 | 0.45506 | 0.58785 | 1.19098 | 155 |
| GO_ANTIMICROBIAL_HUMORAL_IMMUNE_RESPONSE_MED | 0.00185 | 0.02507 | 0.45506 | 0.82607 | 1.50857 | 15 |
| GO_ANTIOXIDANT_ACTIVITY | 0.00018 | 0.00418 | 0.51885 | 0.73340 | 1.41957 | 40 |
| GO_APOPTOTIC_SIGNALING_PATHWAY | 0.00003 | 0.00115 | 0.55733 | 0.56538 | 1.16413 | 423 |
| GO_ATP_BIOSYNTHETIC_PROCESS | 0.00000 | 0.00002 | 0.67496 | 0.89256 | 1.67690 | 22 |
| GO_ATP_METABOLIC_PROCESS | 0.00000 | 0.00000 | | 0.70163 | 1.43215 | 219 |
| GO_ATP_SYNTHESIS_COUPLED_ELECTRON_TRANSPORT | 0.00000 | 0.00000 | | 0.79398 | 1.58093 | 79 |
| GO_AZUROPHIL_GRANULE | 0.00004 | 0.00134 | 0.55733 | 0.65134 | 1.31130 | 107 |
| GO_AZUROPHIL_GRANULE_LUMEN | 0.00012 | 0.00306 | 0.51885 | 0.68846 | 1.35353 | 59 |
| GO_BINDING_OF_SPERM_TO_ZONA_PELLUCIDA | 0.00004 | 0.00128 | 0.55733 | 0.83385 | 1.56224 | 21 |
| GO_BLOOD_MICROPARTICLE | 0.00000 | 0.00004 | 0.65944 | 0.77100 | 1.50763 | 53 |
| GO_CADHERIN_BINDING | 0.00000 | 0.00000 | | 0.67362 | 1.37852 | 269 |
| GO_CANONICAL_WNT_SIGNALING_PATHWAY | 0.00003 | 0.00111 | 0.55733 | 0.59534 | 1.21502 | 227 |
| GO_CARBOHYDRATE_BINDING | 0.00216 | 0.02825 | 0.43171 | 0.59958 | 1.20920 | 115 |
| GO_CARBOHYDRATE_BIOSYNTHETIC_PROCESS | 0.00007 | 0.00192 | 0.53843 | 0.63150 | 1.27625 | 134 |
| GO_CARBOHYDRATE_CATABOLIC_PROCESS | 0.00001 | 0.00055 | 0.59333 | 0.63733 | 1.28925 | 142 |
| GO_CARBOHYDRATE_METABOLIC_PROCESS | 0.00059 | 0.01076 | 0.47727 | 0.55167 | 1.13579 | 407 |
| GO_CATALYTIC_STEP_2_SPLICEOSOME | 0.00010 | 0.00256 | 0.53843 | 0.66217 | 1.32001 | 83 |
| GO_CELLULAR_AMINO_ACID_METABOLIC_PROCESS | 0.00005 | 0.00153 | 0.55733 | 0.59408 | 1.21244 | 230 |
| GO_CELLULAR_COMPONENT_DISASSEMBLY | 0.00000 | 0.00000 | 0.82666 | 0.60831 | 1.25283 | 435 |
| GO_CELLULAR_KETONE_METABOLIC_PROCESS | 0.00005 | 0.00153 | 0.55733 | 0.61848 | 1.25471 | 162 |
| GO_CELLULAR_NITROGEN_COMPOUND_CATABOLIC_PROCE | 0.00000 | 0.00000 | | 0.64729 | 1.33348 | 462 |
| GO_CELLULAR_OXIDANT_DETOXIFICATION | 0.00012 | 0.00295 | 0.53843 | 0.71772 | 1.39768 | 48 |
| GO_CELLULAR_PROTEIN_COMPLEX_DISASSEMBLY | 0.00000 | 0.00000 | 0.76146 | 0.66015 | 1.34288 | 184 |
| GO_CELLULAR_RESPIRATION | 0.00000 | 0.00000 | | 0.72247 | 1.46342 | 150 |
| GO_CELLULAR_RESPONSE_TO_CHEMICAL_STRESS | 0.00115 | 0.01808 | 0.45506 | 0.56638 | 1.15755 | 252 |
| GO_CELLULAR_RESPONSE_TO_HEAT | 0.00298 | 0.03567 | 0.43171 | 0.60416 | 1.21255 | 100 |
| GO_CELLULAR_RESPONSE_TO_ORGANIC_CYCLIC_COMPOUN | 0.00056 | 0.01044 | 0.47727 | 0.55708 | 1.14295 | 336 |
| GO_CELLULAR_RESPONSE_TO_OXYGEN_LEVELS | 0.00000 | 0.00000 | 0.74774 | 0.67109 | 1.36162 | 161 |
| GO_CELLULAR_RESPONSE_TO_TOXIC_SUBSTANCE | 0.00009 | 0.00248 | 0.53843 | 0.69997 | 1.37506 | 57 |
| GO_CELL_ACTIVATION_INVOLVED_IN_IMMUNE_RESPONSE | 0.00000 | 0.00000 | | 0.62192 | 1.28055 | 426 |
| GO_CELL_ADHESION_MOLECULE_BINDING | 0.00000 | 0.00000 | | 0.64818 | 1.33285 | 378 |
| GO_CELL_BODY | 0.00246 | 0.03110 | 0.43171 | 0.55486 | 1.13834 | 333 |
| GO_CELL_CELL_JUNCTION | 0.00457 | 0.04748 | 0.40702 | 0.55013 | 1.12686 | 298 |
| GO_CELL_CELL_RECOGNITION | 0.00000 | 0.00022 | 0.61053 | 0.79259 | 1.52467 | 37 |
| GO_CELL_CELL_SIGNALING_BY_WNT | 0.00038 | 0.00763 | 0.49849 | 0.56199 | 1.15406 | 356 |
| GO_CELL_CORTEX | 0.00008 | 0.00226 | 0.53843 | 0.59687 | 1.21650 | 206 |
| GO_CELL_CYCLE_G2_M_PHASE_TRANSITION | 0.00002 | 0.00064 | 0.57561 | 0.60225 | 1.22912 | 230 |
| GO_CELL_DIVISION | 0.00006 | 0.00177 | 0.55733 | 0.56050 | 1.15453 | 451 |
| GO_CELL_KILLING | 0.00005 | 0.00146 | 0.55733 | 0.69996 | 1.38355 | 65 |
| GO_CELL_LEADING_EDGE | 0.00084 | 0.01396 | 0.47727 | 0.56293 | 1.15312 | 294 |
| GO_CELL_RECOGNITION | 0.00002 | 0.00075 | 0.57561 | 0.69721 | 1.38514 | 73 |
| GO_CELL_REDOX_HOMEOSTASIS | 0.00047 | 0.00931 | 0.49849 | 0.71879 | 1.38610 | 38 |
| GO_CELL_SUBSTRATE_JUNCTION | 0.00000 | 0.00000 | | 0.65735 | 1.34852 | 328 |
| GO_CELL_SURFACE | 0.00309 | 0.03612 | 0.43171 | 0.54970 | 1.12873 | 349 |
| GO_CELL_SURFACE_RECEPTOR_SIGNALING_PATHWAY_INVC | 0.00052 | 0.00988 | 0.47727 | 0.55367 | 1.13938 | 399 |
| GO_CHAPERONE_BINDING | 0.00037 | 0.00756 | 0.49849 | 0.65193 | 1.29878 | 82 |
| GO_CHAPERONE_COMPLEX | 0.00065 | 0.01147 | 0.47727 | 0.78394 | 1.46873 | 21 |
| GO_CHROMOSOMAL_REGION | 0.00261 | 0.03245 | 0.43171 | 0.55545 | 1.13716 | 284 |
| GO_CLATHRIN_COATED_ENDOCYTIC_VESICLE_MEMBRANE | 0.00019 | 0.00423 | 0.51885 | 0.80548 | 1.50908 | 21 |
| GO_COLLAGEN_CONTAINING_EXTRACELLULAR_MATRIX | 0.00004 | 0.00121 | 0.55733 | 0.61411 | 1.24766 | 177 |
| GO_CONTRACTILE_FIBER | 0.00002 | 0.00063 | 0.57561 | 0.64130 | 1.29646 | 135 |
| GO_CORTICAL_CYTOSKELETON | 0.00185 | 0.02507 | 0.45506 | 0.63687 | 1.26593 | 74 |
| GO_COTRANSLATIONAL_PROTEIN_TARGETING_TO_MEMBR/ | 0.00000 | 0.00000 | | 0.80991 | 1.62107 | 95 |
| GO_CYTOCHROME_COMPLEX | 0.00001 | 0.00037 | 0.59333 | 0.85248 | 1.60159 | 22 |
| GO_CYTOKINE_MEDIATED_SIGNALING_PATHWAY | 0.00000 | 0.00000 | 0.74774 | 0.59454 | 1.22437 | 437 |
| GO_CYTOKINE_PRODUCTION | 0.00055 | 0.01033 | 0.47727 | 0.55005 | 1.13266 | 419 |
| GO_CYTOPLASMIC_TRANSLATION | 0.00000 | 0.00000 | 0.80122 | 0.74906 | 1.49554 | 89 |
| GO_CYTOPLASMIC_TRANSLATIONAL_INITIATION | 0.00009 | 0.00246 | 0.53843 | 0.77268 | 1.47511 | 31 |
| GO_CYTOSOLIC_LARGE_RIBOSOMAL_SUBUNIT | 0.00000 | 0.00000 | 0.81404 | 0.82792 | 1.61891 | 53 |
| GO_CYTOSOLIC_RIBOSOME | 0.00000 | 0.00000 | | 0.81981 | 1.64469 | 99 |
| GO_CYTOSOLIC_SMALL_RIBOSOMAL_SUBUNIT | 0.00000 | 0.00000 | 0.73376 | 0.82908 | 1.60871 | 42 |
| GO_DEFENSE_RESPONSE_TO_GRAM_NEGATIVE_BACTERIUN | 0.00191 | 0.02555 | 0.45506 | 0.76351 | 1.43444 | 22 |
| GO_DETOXIFICATION | 0.00010 | 0.00258 | 0.53843 | 0.68199 | 1.34999 | 68 |
| GO_DISORDERED_DOMAIN_SPECIFIC_BINDING | 0.00213 | 0.02782 | 0.43171 | 0.72366 | 1.37929 | 30 |
| GO_DNA_CONFORMATION_CHANGE | 0.00052 | 0.00988 | 0.47727 | 0.57716 | 1.17813 | 234 |
| GO_DNA_PACKAGING | 0.00350 | 0.03950 | 0.43171 | 0.59395 | 1.20003 | 132 |
| GO_DOUBLE_STRANDED_RNA_BINDING | 0.00309 | 0.03612 | 0.43171 | 0.64411 | 1.26463 | 56 |

| Cluster 1 | | | | | | |
|---|---|---|---|---|---|---|
| GO_ELECTRON_TRANSFER_ACTIVITY | 0.00000 | 0.00000 | 0.73376 | 0.73136 | 1.45704 | 82 |
| GO_ELECTRON_TRANSPORT_CHAIN | 0.00000 | 0.00000 | | 0.71781 | 1.45180 | 140 |
| GO_ENDOLYSOSOME | 0.00265 | 0.03285 | 0.43171 | 0.77958 | 1.43140 | 16 |
| GO_ENDOPEPTIDASE_COMPLEX | 0.00009 | 0.00237 | 0.53843 | 0.69880 | 1.37714 | 62 |
| GO_ENDOPEPTIDASE_REGULATOR_ACTIVITY | 0.00014 | 0.00329 | 0.51885 | 0.67329 | 1.33633 | 72 |
| GO_ENDOPLASMIC_RETICULUM_GOLGI_INTERMEDIATE_CO | 0.00242 | 0.03069 | 0.43171 | 0.61812 | 1.23917 | 98 |
| GO_ENDOPLASMIC_RETICULUM_LUMEN | 0.00054 | 0.01014 | 0.47727 | 0.60187 | 1.22075 | 158 |
| GO_ENDOSOME_LUMEN | 0.00316 | 0.03678 | 0.43171 | 0.76070 | 1.41448 | 19 |
| GO_ENERGY_DERIVATION_BY_OXIDATION_OF_ORGANIC_CO | 0.00000 | 0.00000 | | 0.69820 | 1.42357 | 210 |
| GO_ENZYME_INHIBITOR_ACTIVITY | 0.00005 | 0.00162 | 0.55733 | 0.60077 | 1.22381 | 204 |
| GO_ESTABLISHMENT_OF_PROTEIN_LOCALIZATION_TO_CHR | 0.00111 | 0.01759 | 0.45506 | 0.77460 | 1.45527 | 22 |
| GO_ESTABLISHMENT_OF_PROTEIN_LOCALIZATION_TO_END | 0.00000 | 0.00000 | | 0.79200 | 1.59448 | 107 |
| GO_ESTABLISHMENT_OF_PROTEIN_LOCALIZATION_TO_MEM | 0.00000 | 0.00000 | | 0.67381 | 1.37918 | 273 |
| GO_ESTABLISHMENT_OF_PROTEIN_LOCALIZATION_TO_MITO | 0.00253 | 0.03184 | 0.43171 | 0.68724 | 1.33447 | 43 |
| GO_ESTABLISHMENT_OF_PROTEIN_LOCALIZATION_TO_ORG | 0.00000 | 0.00000 | | 0.65321 | 1.34634 | 473 |
| GO_ESTABLISHMENT_OF_PROTEIN_LOCALIZATION_TO_TELO | 0.00062 | 0.01117 | 0.47727 | 0.80509 | 1.48428 | 17 |
| GO_ESTABLISHMENT_OF_RNA_LOCALIZATION | 0.00167 | 0.02347 | 0.45506 | 0.58134 | 1.18218 | 181 |
| GO_ESTABLISHMENT_OF_TISSUE_POLARITY | 0.00000 | 0.00017 | 0.62726 | 0.67576 | 1.35569 | 99 |
| GO_EUKARYOTIC_TRANSLATION_INITIATION_FACTOR_3_CO | 0.00167 | 0.02347 | 0.45506 | 0.82912 | 1.51413 | 15 |
| GO_EXTRACELLULAR_MATRIX | 0.00093 | 0.01517 | 0.47727 | 0.57571 | 1.17531 | 218 |
| GO_FC_EPSILON_RECEPTOR_SIGNALING_PATHWAY | 0.00018 | 0.00414 | 0.51885 | 0.64822 | 1.29483 | 90 |
| GO_FC_RECEPTOR_SIGNALING_PATHWAY | 0.00009 | 0.00241 | 0.53843 | 0.62070 | 1.25562 | 142 |
| GO_FERTILIZATION | 0.00420 | 0.04456 | 0.40702 | 0.61286 | 1.22234 | 84 |
| GO_FICOLIN_1_RICH_GRANULE | 0.00000 | 0.00000 | 0.73376 | 0.71610 | 1.43793 | 102 |
| GO_GENERATION_OF_PRECURSOR_METABOLITES_AND_EN | 0.00000 | 0.00000 | | 0.67130 | 1.38013 | 372 |
| GO_GENE_SILENCING | 0.00428 | 0.04497 | 0.40702 | 0.57193 | 1.16162 | 175 |
| GO_GLUCOSE_CATABOLIC_PROCESS | 0.00069 | 0.01189 | 0.47727 | 0.75337 | 1.42804 | 27 |
| GO_GLUCOSE_METABOLIC_PROCESS | 0.00327 | 0.03773 | 0.43171 | 0.58949 | 1.19250 | 143 |
| GO_GLUTAMATERGIC_SYNAPSE | 0.00376 | 0.04133 | 0.43171 | 0.56330 | 1.14856 | 212 |
| GO_GLYCOLYTIC_PROCESS_THROUGH_FRUCTOSE_6_PHOSP | 0.00014 | 0.00340 | 0.51885 | 0.80214 | 1.51157 | 24 |
| GO_GTPASE_ACTIVITY | 0.00004 | 0.00122 | 0.55733 | 0.59643 | 1.21612 | 211 |
| GO_GUANYL_NUCLEOTIDE_BINDING | 0.00016 | 0.00372 | 0.51885 | 0.58421 | 1.19440 | 256 |
| GO_HEMATOPOIETIC_PROGENITOR_CELL_DIFFERENTIATION | 0.00003 | 0.00099 | 0.57561 | 0.64532 | 1.30265 | 122 |
| GO_HEMATOPOIETIC_STEM_CELL_DIFFERENTIATION | 0.00002 | 0.00072 | 0.57561 | 0.69766 | 1.38416 | 71 |
| GO_HEXOSE_CATABOLIC_PROCESS | 0.00161 | 0.02294 | 0.45506 | 0.69650 | 1.35026 | 41 |
| GO_HOMOTYPIC_CELL_CELL_ADHESION | 0.00010 | 0.00263 | 0.53843 | 0.69842 | 1.37224 | 58 |
| GO_HYDROGEN_PEROXIDE_METABOLIC_PROCESS | 0.00213 | 0.02782 | 0.43171 | 0.73363 | 1.38749 | 26 |
| GO_IMMUNE_RESPONSE_REGULATING_SIGNALING_PATHW | 0.00121 | 0.01872 | 0.45506 | 0.56994 | 1.16412 | 246 |
| GO_INNATE_IMMUNE_RESPONSE | 0.00000 | 0.00000 | 0.71951 | 0.59088 | 1.21678 | 443 |
| GO_INNATE_IMMUNE_RESPONSE_ACTIVATING_SIGNAL_TRA | 0.00008 | 0.00218 | 0.53843 | 0.66834 | 1.33292 | 86 |
| GO_INNER_MITOCHONDRIAL_MEMBRANE_ORGANIZATION | 0.00213 | 0.02782 | 0.43171 | 0.72840 | 1.38071 | 27 |
| GO_INNER_MITOCHONDRIAL_MEMBRANE_PROTEIN_COMP | 0.00000 | 0.00000 | | 0.78056 | 1.55936 | 92 |
| GO_INORGANIC_ION_TRANSMEMBRANE_TRANSPORT | 0.00283 | 0.03454 | 0.43171 | 0.54055 | 1.11305 | 413 |
| GO_INTERACTION_WITH_SYMBIONT | 0.00001 | 0.00047 | 0.59333 | 0.71853 | 1.41076 | 56 |
| GO_INTERLEUKIN_1_MEDIATED_SIGNALING_PATHWAY | 0.00000 | 0.00005 | 0.65944 | 0.70536 | 1.40611 | 83 |
| GO_INTRINSIC_APOPTOTIC_SIGNALING_PATHWAY | 0.00072 | 0.01226 | 0.47727 | 0.57654 | 1.17677 | 222 |
| GO_IRON_ION_BINDING | 0.00346 | 0.03922 | 0.43171 | 0.62907 | 1.24808 | 71 |
| GO_KILLING_OF_CELLS_OF_OTHER_ORGANISM | 0.00172 | 0.02403 | 0.45506 | 0.77423 | 1.43963 | 19 |
| GO_LARGE_RIBOSOMAL_SUBUNIT | 0.00000 | 0.00000 | | 0.74670 | 1.50345 | 108 |
| GO_LEUKOCYTE_MEDIATED_CYTOTOXICITY | 0.00086 | 0.01419 | 0.47727 | 0.68920 | 1.33986 | 47 |
| GO_LEUKOCYTE_MEDIATED_IMMUNITY | 0.00000 | 0.00000 | | 0.61874 | 1.27456 | 460 |
| GO_LEUKOCYTE_MIGRATION | 0.00413 | 0.04406 | 0.40702 | 0.56837 | 1.15765 | 201 |
| GO_LOW_DENSITY_LIPOPROTEIN_PARTICLE_RECEPTOR_BIN | 0.00398 | 0.04290 | 0.40702 | 0.80210 | 1.46479 | 15 |
| GO_LOW_DENSITY_LIPOPROTEIN_RECEPTOR_PARTICLE_ME | 0.00064 | 0.01137 | 0.47727 | 0.79737 | 1.48265 | 19 |
| GO_LUMENAL_SIDE_OF_MEMBRANE | 0.00054 | 0.01022 | 0.47727 | 0.78787 | 1.48232 | 23 |
| GO_MACROMOLECULE_TRANSMEMBRANE_TRANSPORTER_ | 0.00093 | 0.01517 | 0.47727 | 0.79052 | 1.46992 | 19 |
| GO_MAINTENANCE_OF_LOCATION | 0.00309 | 0.03612 | 0.43171 | 0.56996 | 1.16171 | 208 |
| GO_MAINTENANCE_OF_LOCATION_IN_CELL | 0.00031 | 0.00647 | 0.49849 | 0.61380 | 1.24085 | 135 |
| GO_MAINTENANCE_OF_PROTEIN_LOCATION | 0.00309 | 0.03612 | 0.43171 | 0.62795 | 1.24948 | 76 |
| GO_MAINTENANCE_OF_PROTEIN_LOCATION_IN_CELL | 0.00148 | 0.02164 | 0.45506 | 0.66935 | 1.30792 | 52 |
| GO_MALE_SEX_DIFFERENTIATION | 0.00383 | 0.04186 | 0.43171 | 0.60651 | 1.21474 | 96 |
| GO_MEMBRANE_REGION | 0.00178 | 0.02448 | 0.45506 | 0.57724 | 1.17567 | 203 |
| GO_MIDBRAIN_DEVELOPMENT | 0.00053 | 0.01002 | 0.47727 | 0.67515 | 1.32231 | 54 |
| GO_MITOCHONDRIAL_ELECTRON_TRANSPORT_NADH_TO_U | 0.00000 | 0.00001 | 0.70498 | 0.79061 | 1.53685 | 46 |
| GO_MITOCHONDRIAL_GENE_EXPRESSION | 0.00000 | 0.00002 | 0.67496 | 0.65408 | 1.32490 | 153 |
| GO_MITOCHONDRIAL_LARGE_RIBOSOMAL_SUBUNIT | 0.00002 | 0.00064 | 0.57561 | 0.72224 | 1.41454 | 54 |
| GO_MITOCHONDRIAL_MATRIX | 0.00000 | 0.00000 | 0.76146 | 0.60443 | 1.24338 | 390 |
| GO_MITOCHONDRIAL_MEMBRANE_ORGANIZATION | 0.00018 | 0.00411 | 0.51885 | 0.64466 | 1.29330 | 99 |
| GO_MITOCHONDRIAL_PROTEIN_COMPLEX | 0.00000 | 0.00000 | | 0.71770 | 1.46316 | 209 |
| GO_MITOCHONDRIAL_RESPIRATORY_CHAIN_COMPLEX_ASS | 0.00007 | 0.00210 | 0.53843 | 0.66875 | 1.33373 | 86 |
| GO_MITOCHONDRIAL_TRANSLATION | 0.00000 | 0.00002 | 0.67496 | 0.67021 | 1.35391 | 130 |
| GO_MITOCHONDRIAL_TRANSLATIONAL_TERMINATION | 0.00000 | 0.00006 | 0.64355 | 0.69861 | 1.39469 | 88 |
| GO_MITOCHONDRIAL_TRANSPORT | 0.00005 | 0.00156 | 0.55733 | 0.60257 | 1.22817 | 208 |

| Cluster 1 | | | | | | |
|---|---|---|---|---|---|---|
| GO_MITOCHONDRION_ORGANIZATION | 0.00000 | 0.00000 | 0.78819 | 0.60491 | 1.24555 | 426 |
| GO_MODULATION_BY_HOST_OF_SYMBIONT_PROCESS | 0.00066 | 0.01157 | 0.47727 | 0.70337 | 1.36698 | 44 |
| GO_MODULATION_BY_HOST_OF_VIRAL_PROCESS | 0.00364 | 0.04067 | 0.43171 | 0.76308 | 1.42353 | 20 |
| GO_MODULATION_OF_PROCESS_OF_OTHER_ORGANISM | 0.00137 | 0.02054 | 0.45506 | 0.63489 | 1.26195 | 75 |
| GO_MONOAMINE_TRANSPORT | 0.00294 | 0.03542 | 0.43171 | 0.71831 | 1.36909 | 30 |
| GO_MONOCARBOXYLIC_ACID_BINDING | 0.00480 | 0.04900 | 0.40702 | 0.70167 | 1.33636 | 29 |
| GO_MONOCARBOXYLIC_ACID_METABOLIC_PROCESS | 0.00003 | 0.00107 | 0.57561 | 0.56832 | 1.16932 | 397 |
| GO_MONOSACCHARIDE_BINDING | 0.00302 | 0.03584 | 0.43171 | 0.68509 | 1.32932 | 42 |
| GO_MONOSACCHARIDE_BIOSYNTHETIC_PROCESS | 0.00023 | 0.00507 | 0.51885 | 0.68112 | 1.34631 | 65 |
| GO_MONOSACCHARIDE_CATABOLIC_PROCESS | 0.00042 | 0.00848 | 0.49849 | 0.69606 | 1.35636 | 49 |
| GO_MONOSACCHARIDE_METABOLIC_PROCESS | 0.00198 | 0.02632 | 0.43171 | 0.57635 | 1.17364 | 194 |
| GO_MONOVALENT_INORGANIC_CATION_TRANSMEMBRAN | 0.00133 | 0.02008 | 0.45506 | 0.59808 | 1.20989 | 143 |
| GO_MONOVALENT_INORGANIC_CATION_TRANSPORT | 0.00088 | 0.01457 | 0.47727 | 0.57231 | 1.16877 | 244 |
| GO_MORPHOGENESIS_OF_AN_EPITHELIUM | 0.00015 | 0.00348 | 0.51885 | 0.56842 | 1.16778 | 363 |
| GO_MORPHOGENESIS_OF_A_POLARIZED_EPITHELIUM | 0.00000 | 0.00004 | 0.65944 | 0.68002 | 1.37138 | 116 |
| GO_MRNA_5_UTR_BINDING | 0.00057 | 0.01058 | 0.47727 | 0.76595 | 1.44492 | 25 |
| GO_MRNA_BINDING | 0.00000 | 0.00000 | 0.74774 | 0.63853 | 1.30380 | 238 |
| GO_MRNA_EXPORT_FROM_NUCLEUS | 0.00198 | 0.02632 | 0.43171 | 0.61068 | 1.22804 | 104 |
| GO_MRNA_PROCESSING | 0.00000 | 0.00000 | 0.76146 | 0.59682 | 1.22902 | 443 |
| GO_MRNA_TRANSPORT | 0.00305 | 0.03608 | 0.43171 | 0.58852 | 1.19043 | 137 |
| GO_MUSCLE_CELL_PROLIFERATION | 0.00021 | 0.00457 | 0.51885 | 0.62355 | 1.25755 | 115 |
| GO_MYELOID_LEUKOCYTE_ACTIVATION | 0.00000 | 0.00000 | | 0.62540 | 1.28677 | 397 |
| GO_MYELOID_LEUKOCYTE_MEDIATED_IMMUNITY | 0.00000 | 0.00000 | | 0.64272 | 1.32049 | 362 |
| GO_MYOFIBRIL_ASSEMBLY | 0.00287 | 0.03479 | 0.43171 | 0.71044 | 1.35857 | 32 |
| GO_NADH_DEHYDROGENASE_ACTIVITY | 0.00000 | 0.00001 | 0.69013 | 0.81551 | 1.56911 | 36 |
| GO_NADH_DEHYDROGENASE_COMPLEX | 0.00000 | 0.00001 | 0.70498 | 0.81514 | 1.57779 | 40 |
| GO_NADH_DEHYDROGENASE_COMPLEX_ASSEMBLY | 0.00000 | 0.00007 | 0.64355 | 0.74169 | 1.45447 | 55 |
| GO_NADH_METABOLIC_PROCESS | 0.00000 | 0.00022 | 0.61053 | 0.79980 | 1.53110 | 33 |
| GO_NAD_BINDING | 0.00335 | 0.03837 | 0.43171 | 0.66728 | 1.29596 | 45 |
| GO_NAD_METABOLIC_PROCESS | 0.00010 | 0.00254 | 0.53843 | 0.74225 | 1.43669 | 40 |
| GO_NATURAL_KILLER_CELL_MEDIATED_IMMUNITY | 0.00156 | 0.02245 | 0.45506 | 0.72092 | 1.37859 | 32 |
| GO_NCRNA_METABOLIC_PROCESS | 0.00163 | 0.02310 | 0.45506 | 0.54546 | 1.12209 | 395 |
| GO_NCRNA_PROCESSING | 0.00231 | 0.02975 | 0.43171 | 0.55169 | 1.13222 | 340 |
| GO_NEGATIVE_REGULATION_OF_APOPTOTIC_SIGNALING_P | 0.00110 | 0.01734 | 0.45506 | 0.59019 | 1.19690 | 159 |
| GO_NEGATIVE_REGULATION_OF_ATP_METABOLIC_PROCESS | 0.00364 | 0.04067 | 0.43171 | 0.75696 | 1.40752 | 19 |
| GO_NEGATIVE_REGULATION_OF_CALCIUM_MEDIATED_SIGI | 0.00257 | 0.03212 | 0.43171 | 0.77111 | 1.43850 | 20 |
| GO_NEGATIVE_REGULATION_OF_CANONICAL_WNT_SIGNAI | 0.00001 | 0.00044 | 0.59333 | 0.64917 | 1.31082 | 127 |
| GO_NEGATIVE_REGULATION_OF_CATALYTIC_ACTIVITY | 0.00025 | 0.00552 | 0.49849 | 0.55005 | 1.13372 | 481 |
| GO_NEGATIVE_REGULATION_OF_CELLULAR_AMIDE_METAB | 0.00146 | 0.02142 | 0.45506 | 0.60320 | 1.21750 | 125 |
| GO_NEGATIVE_REGULATION_OF_CELLULAR_COMPONENT_( | 0.00074 | 0.01251 | 0.47727 | 0.54583 | 1.12518 | 498 |
| GO_NEGATIVE_REGULATION_OF_CELL_CYCLE | 0.00132 | 0.01995 | 0.45506 | 0.54229 | 1.11767 | 472 |
| GO_NEGATIVE_REGULATION_OF_CELL_CYCLE_G2_M_PHASI | 0.00001 | 0.00029 | 0.61053 | 0.68062 | 1.35955 | 90 |
| GO_NEGATIVE_REGULATION_OF_CELL_CYCLE_PHASE_TRAN | 0.00024 | 0.00526 | 0.51885 | 0.59448 | 1.21083 | 202 |
| GO_NEGATIVE_REGULATION_OF_CELL_CYCLE_PROCESS | 0.00010 | 0.00256 | 0.53843 | 0.58137 | 1.18997 | 273 |
| GO_NEGATIVE_REGULATION_OF_DEFENSE_RESPONSE | 0.00054 | 0.01014 | 0.47727 | 0.61762 | 1.24634 | 120 |
| GO_NEGATIVE_REGULATION_OF_HYDROLASE_ACTIVITY | 0.00009 | 0.00237 | 0.53843 | 0.58291 | 1.19099 | 249 |
| GO_NEGATIVE_REGULATION_OF_MITOTIC_CELL_CYCLE | 0.00157 | 0.02257 | 0.45506 | 0.56815 | 1.16155 | 256 |
| GO_NEGATIVE_REGULATION_OF_MRNA_METABOLIC_PROC | 0.00161 | 0.02294 | 0.45506 | 0.65108 | 1.28538 | 64 |
| GO_NEGATIVE_REGULATION_OF_ORGANELLE_ORGANIZATI( | 0.00368 | 0.04087 | 0.43171 | 0.55416 | 1.13588 | 303 |
| GO_NEGATIVE_REGULATION_OF_PEPTIDASE_ACTIVITY | 0.00018 | 0.00414 | 0.51885 | 0.63414 | 1.27877 | 114 |
| GO_NEGATIVE_REGULATION_OF_PROTEIN_CONTAINING_C( | 0.00013 | 0.00308 | 0.51885 | 0.64413 | 1.29008 | 96 |
| GO_NEGATIVE_REGULATION_OF_PROTEIN_POLYMERIZATIO | 0.00372 | 0.04100 | 0.43171 | 0.66125 | 1.29159 | 51 |
| GO_NEGATIVE_REGULATION_OF_PROTEOLYSIS | 0.00008 | 0.00229 | 0.53843 | 0.60358 | 1.22777 | 183 |
| GO_NEGATIVE_REGULATION_OF_RESPONSE_TO_BIOTIC_ST | 0.00390 | 0.04246 | 0.43171 | 0.64178 | 1.26365 | 61 |
| GO_NEGATIVE_REGULATION_OF_RESPONSE_TO_EXTERNAL_ | 0.00383 | 0.04186 | 0.43171 | 0.56964 | 1.15998 | 194 |
| GO_NEGATIVE_REGULATION_OF_TRANSPORT | 0.00472 | 0.04863 | 0.40702 | 0.55899 | 1.14227 | 251 |
| GO_NEGATIVE_REGULATION_OF_VIRAL_GENOME_REPLICAT | 0.00178 | 0.02448 | 0.45506 | 0.70635 | 1.35878 | 37 |
| GO_NEGATIVE_REGULATION_OF_VIRAL_PROCESS | 0.00150 | 0.02185 | 0.45506 | 0.65166 | 1.28652 | 64 |
| GO_NEGATIVE_REGULATION_OF_WNT_SIGNALING_PATHW/ | 0.00013 | 0.00308 | 0.51885 | 0.62014 | 1.25468 | 145 |
| GO_NEURAL_NUCLEUS_DEVELOPMENT | 0.00034 | 0.00710 | 0.49849 | 0.72025 | 1.39178 | 39 |
| GO_NEURON_TO_NEURON_SYNAPSE | 0.00316 | 0.03678 | 0.43171 | 0.56408 | 1.15071 | 214 |
| GO_NIK_NF_KAPPAB_SIGNALING | 0.00000 | 0.00000 | 0.74774 | 0.69700 | 1.40592 | 119 |
| GO_NON_CANONICAL_WNT_SIGNALING_PATHWAY | 0.00002 | 0.00062 | 0.57561 | 0.65526 | 1.32144 | 116 |
| GO_NUCLEAR_CHROMOSOME_TELOMERIC_REGION | 0.00316 | 0.03678 | 0.43171 | 0.61114 | 1.22056 | 91 |
| GO_NUCLEAR_EXPORT | 0.00133 | 0.02008 | 0.45506 | 0.58485 | 1.18956 | 182 |
| GO_NUCLEAR_SPECK | 0.00320 | 0.03714 | 0.43171 | 0.54965 | 1.12760 | 329 |
| GO_NUCLEAR_TRANSCRIBED_MRNA_CATABOLIC_PROCESS | 0.00000 | 0.00000 | | 0.68941 | 1.40299 | 190 |
| GO_NUCLEAR_TRANSCRIBED_MRNA_CATABOLIC_PROCESS_ | 0.00000 | 0.00000 | | 0.78539 | 1.58394 | 115 |
| GO_NUCLEAR_TRANSPORT | 0.00007 | 0.00193 | 0.53843 | 0.57691 | 1.18187 | 299 |
| GO_NUCLEOBASE_CONTAINING_COMPOUND_TRANSPORT | 0.00205 | 0.02719 | 0.43171 | 0.56817 | 1.15992 | 218 |
| GO_NUCLEOBASE_CONTAINING_SMALL_MOLECULE_METAE | 0.00001 | 0.00052 | 0.59333 | 0.57435 | 1.18218 | 402 |
| GO_NUCLEOSIDE_DIPHOSPHATE_METABOLIC_PROCESS | 0.00003 | 0.00095 | 0.57561 | 0.64874 | 1.30663 | 110 |
| GO_NUCLEOSIDE_PHOSPHATE_BIOSYNTHETIC_PROCESS | 0.00331 | 0.03809 | 0.43171 | 0.58164 | 1.17997 | 162 |

| **Cluster 1** | | | | | | |
|---|---|---|---|---|---|---|
| GO_NUCLEOSIDE_TRIPHOSPHATE_BIOSYNTHETIC_PROCESS | 0.00005 | 0.00161 | 0.55733 | 0.72686 | 1.41167 | 45 |
| GO_NUCLEOSIDE_TRIPHOSPHATE_METABOLIC_PROCESS | 0.00025 | 0.00544 | 0.49849 | 0.66857 | 1.32307 | 67 |
| GO_NUCLEOSOME_ORGANIZATION | 0.00046 | 0.00912 | 0.49849 | 0.63655 | 1.27151 | 90 |
| GO_NUCLEOTIDE_PHOSPHORYLATION | 0.00005 | 0.00147 | 0.55733 | 0.65844 | 1.31933 | 97 |
| GO_ORGANELLAR_RIBOSOME | 0.00000 | 0.00002 | 0.67496 | 0.70650 | 1.40910 | 84 |
| GO_ORGANELLE_ENVELOPE_LUMEN | 0.00224 | 0.02898 | 0.43171 | 0.62887 | 1.24999 | 75 |
| GO_ORGANELLE_INNER_MEMBRANE | 0.00000 | 0.00000 | | 0.65010 | 1.33831 | 406 |
| GO_ORGANIC_ACID_BINDING | 0.00457 | 0.04748 | 0.40702 | 0.61330 | 1.22521 | 92 |
| GO_ORGANIC_CYCLIC_COMPOUND_CATABOLIC_PROCESS | 0.00000 | 0.00000 | | 0.64770 | 1.33491 | 482 |
| GO_OSTEOBLAST_DIFFERENTIATION | 0.00014 | 0.00329 | 0.51885 | 0.62263 | 1.25870 | 135 |
| GO_OUTER_MEMBRANE | 0.00049 | 0.00955 | 0.47727 | 0.60439 | 1.22440 | 154 |
| GO_OXIDATIVE_PHOSPHORYLATION | 0.00000 | 0.00000 | | 0.78667 | 1.58094 | 103 |
| GO_OXIDOREDUCTASE_ACTIVITY | 0.00000 | 0.00000 | | 0.63219 | 1.30291 | 469 |
| GO_OXIDOREDUCTASE_ACTIVITY_ACTING_ON_A_HEME_GR | 0.00010 | 0.00256 | 0.53843 | 0.83414 | 1.55609 | 20 |
| GO_OXIDOREDUCTASE_ACTIVITY_ACTING_ON_NAD_P_H | 0.00000 | 0.00003 | 0.65944 | 0.73777 | 1.46040 | 68 |
| GO_OXIDOREDUCTASE_ACTIVITY_ACTING_ON_NAD_P_H_Q | 0.00000 | 0.00009 | 0.64355 | 0.77812 | 1.51223 | 44 |
| GO_OXIDOREDUCTASE_ACTIVITY_ACTING_ON_PEROXIDE_A | 0.00033 | 0.00690 | 0.49849 | 0.76831 | 1.45309 | 26 |
| GO_OXIDOREDUCTASE_COMPLEX | 0.00000 | 0.00000 | 0.81404 | 0.75766 | 1.51037 | 83 |
| GO_PEPTIDASE_COMPLEX | 0.00004 | 0.00128 | 0.55733 | 0.67541 | 1.34450 | 78 |
| GO_PEPTIDASE_REGULATOR_ACTIVITY | 0.00016 | 0.00372 | 0.51885 | 0.64712 | 1.29665 | 97 |
| GO_PEPTIDE_BINDING | 0.00095 | 0.01539 | 0.47727 | 0.59571 | 1.20785 | 157 |
| GO_PEPTIDE_TRANSMEMBRANE_TRANSPORTER_ACTIVITY | 0.00058 | 0.01069 | 0.47727 | 0.78721 | 1.48106 | 23 |
| GO_PERINUCLEAR_REGION_OF_CYTOPLASM | 0.00183 | 0.02507 | 0.45506 | 0.53759 | 1.10795 | 489 |
| GO_PHOSPHATE_ION_TRANSPORT | 0.00198 | 0.02632 | 0.43171 | 0.82569 | 1.50787 | 15 |
| GO_PIGMENT_GRANULE | 0.00000 | 0.00021 | 0.61053 | 0.69385 | 1.38458 | 87 |
| GO_PLASMA_LIPOPROTEIN_PARTICLE_CLEARANCE | 0.00012 | 0.00306 | 0.51885 | 0.73264 | 1.42032 | 41 |
| GO_PLATELET_ACTIVATION | 0.00028 | 0.00599 | 0.49849 | 0.64164 | 1.28426 | 95 |
| GO_PLATELET_AGGREGATION | 0.00014 | 0.00329 | 0.51885 | 0.73443 | 1.42505 | 42 |
| GO_PLATELET_DEGRANULATION | 0.00179 | 0.02468 | 0.45506 | 0.64222 | 1.27151 | 69 |
| GO_POLYMERIC_CYTOSKELETAL_FIBER | 0.00052 | 0.00988 | 0.47727 | 0.55368 | 1.14038 | 432 |
| GO_POLYSACCHARIDE_CATABOLIC_PROCESS | 0.00372 | 0.04100 | 0.43171 | 0.75681 | 1.40723 | 19 |
| GO_POLYSOMAL_RIBOSOME | 0.00000 | 0.00005 | 0.65944 | 0.83475 | 1.59103 | 30 |
| GO_POLYSOME | 0.00000 | 0.00000 | 0.74774 | 0.77622 | 1.53166 | 63 |
| GO_POSITIVE_REGULATION_OF_ACTIN_FILAMENT_POLYME | 0.00261 | 0.03245 | 0.43171 | 0.63176 | 1.25208 | 70 |
| GO_POSITIVE_REGULATION_OF_BINDING | 0.00302 | 0.03584 | 0.43171 | 0.59606 | 1.20414 | 129 |
| GO_POSITIVE_REGULATION_OF_CANONICAL_WNT_SIGNALI | 0.00000 | 0.00015 | 0.62726 | 0.66687 | 1.34476 | 114 |
| GO_POSITIVE_REGULATION_OF_CELLULAR_AMIDE_METABO | 0.00002 | 0.00080 | 0.57561 | 0.64106 | 1.29439 | 126 |
| GO_POSITIVE_REGULATION_OF_CELLULAR_COMPONENT_B | 0.00250 | 0.03144 | 0.43171 | 0.54959 | 1.12979 | 370 |
| GO_POSITIVE_REGULATION_OF_CELLULAR_PROTEIN_LOCAL | 0.00095 | 0.01539 | 0.47727 | 0.57281 | 1.17052 | 251 |
| GO_POSITIVE_REGULATION_OF_CELL_DEATH | 0.00004 | 0.00139 | 0.55733 | 0.56213 | 1.15862 | 473 |
| GO_POSITIVE_REGULATION_OF_CHROMOSOME_ORGANIZA | 0.00239 | 0.03046 | 0.43171 | 0.58452 | 1.18332 | 147 |
| GO_POSITIVE_REGULATION_OF_CYSTEINE_TYPE_ENDOPEPT | 0.00005 | 0.00160 | 0.55733 | 0.65543 | 1.31491 | 99 |
| GO_POSITIVE_REGULATION_OF_CYTOKINE_PRODUCTION | 0.00079 | 0.01323 | 0.47727 | 0.57069 | 1.16520 | 237 |
| GO_POSITIVE_REGULATION_OF_CYTOKINE_SECRETION | 0.00119 | 0.01856 | 0.45506 | 0.73287 | 1.39910 | 31 |
| GO_POSITIVE_REGULATION_OF_CYTOSKELETON_ORGANIZA | 0.00102 | 0.01642 | 0.45506 | 0.59290 | 1.20297 | 161 |
| GO_POSITIVE_REGULATION_OF_DEFENSE_RESPONSE | 0.00000 | 0.00000 | 0.76146 | 0.64887 | 1.32302 | 212 |
| GO_POSITIVE_REGULATION_OF_ESTABLISHMENT_OF_PROT | 0.00034 | 0.00703 | 0.49849 | 0.57558 | 1.17805 | 271 |
| GO_POSITIVE_REGULATION_OF_HYDROLASE_ACTIVITY | 0.00298 | 0.03567 | 0.43171 | 0.53421 | 1.10102 | 477 |
| GO_POSITIVE_REGULATION_OF_IMMUNE_RESPONSE | 0.00004 | 0.00128 | 0.55733 | 0.57795 | 1.18664 | 346 |
| GO_POSITIVE_REGULATION_OF_INFLAMMATORY_RESPONS | 0.00014 | 0.00338 | 0.51885 | 0.68345 | 1.34928 | 64 |
| GO_POSITIVE_REGULATION_OF_INNATE_IMMUNE_RESPON | 0.00000 | 0.00000 | 0.71951 | 0.67539 | 1.36588 | 141 |
| GO_POSITIVE_REGULATION_OF_LIPASE_ACTIVITY | 0.00287 | 0.03479 | 0.43171 | 0.68531 | 1.32426 | 39 |
| GO_POSITIVE_REGULATION_OF_MRNA_METABOLIC_PROCE | 0.00033 | 0.00698 | 0.49849 | 0.66617 | 1.31718 | 66 |
| GO_POSITIVE_REGULATION_OF_MRNA_PROCESSING | 0.00324 | 0.03751 | 0.43171 | 0.72293 | 1.36726 | 26 |
| GO_POSITIVE_REGULATION_OF_MRNA_SPLICING_VIA_SPLI | 0.00361 | 0.04040 | 0.43171 | 0.77768 | 1.44166 | 18 |
| GO_POSITIVE_REGULATION_OF_ORGANELLE_ORGANIZATIO | 0.00000 | 0.00005 | 0.65944 | 0.58005 | 1.19549 | 470 |
| GO_POSITIVE_REGULATION_OF_PEPTIDASE_ACTIVITY | 0.00001 | 0.00042 | 0.59333 | 0.64682 | 1.30668 | 129 |
| GO_POSITIVE_REGULATION_OF_PROTEIN_CONTAINING_CO | 0.00044 | 0.00870 | 0.49849 | 0.59364 | 1.20721 | 181 |
| GO_POSITIVE_REGULATION_OF_PROTEIN_KINASE_B_SIGNA | 0.00009 | 0.00239 | 0.53843 | 0.66518 | 1.32519 | 82 |
| GO_POSITIVE_REGULATION_OF_PROTEIN_LOCALIZATION_T | 0.00078 | 0.01311 | 0.47727 | 0.66045 | 1.30735 | 68 |
| GO_POSITIVE_REGULATION_OF_PROTEIN_POLYMERIZATION | 0.00268 | 0.03318 | 0.43171 | 0.61154 | 1.22685 | 99 |
| GO_POSITIVE_REGULATION_OF_PROTEOLYSIS | 0.00004 | 0.00137 | 0.55733 | 0.58765 | 1.20028 | 247 |
| GO_POSITIVE_REGULATION_OF_REACTIVE_OXYGEN_SPECIE | 0.00480 | 0.04900 | 0.40702 | 0.65498 | 1.27934 | 51 |
| GO_POSITIVE_REGULATION_OF_RESPONSE_TO_BIOTIC_STI | 0.00000 | 0.00001 | 0.69013 | 0.65683 | 1.33259 | 160 |
| GO_POSITIVE_REGULATION_OF_RESPONSE_TO_EXTERNAL_ | 0.00000 | 0.00002 | 0.67496 | 0.61043 | 1.24963 | 282 |
| GO_POSITIVE_REGULATION_OF_SUPRAMOLECULAR_FIBER_ | 0.00037 | 0.00756 | 0.49849 | 0.60816 | 1.23043 | 144 |
| GO_POSITIVE_REGULATION_OF_TELOMERASE_RNA_LOCALI | 0.00019 | 0.00420 | 0.51885 | 0.86929 | 1.58750 | 15 |
| GO_POSITIVE_REGULATION_OF_TELOMERE_MAINTENANCE | 0.00390 | 0.04246 | 0.43171 | 0.70086 | 1.34170 | 33 |
| GO_POSITIVE_REGULATION_OF_TRANSLATION | 0.00002 | 0.00061 | 0.59333 | 0.66175 | 1.33240 | 108 |
| GO_POSITIVE_REGULATION_OF_VIRAL_LIFE_CYCLE | 0.00357 | 0.04020 | 0.43171 | 0.65897 | 1.28555 | 50 |
| GO_POSITIVE_REGULATION_OF_VIRAL_PROCESS | 0.00008 | 0.00226 | 0.53843 | 0.65562 | 1.30939 | 91 |
| GO_POSITIVE_REGULATION_OF_WNT_SIGNALING_PATHWA | 0.00004 | 0.00128 | 0.55733 | 0.63580 | 1.28443 | 133 |
| GO_POSTSYNAPSE | 0.00220 | 0.02861 | 0.43171 | 0.54736 | 1.12440 | 361 |

| Cluster 1 | | | | | | |
|---|---|---|---|---|---|---|
| GO_POSTTRANSCRIPTIONAL_REGULATION_OF_GENE_EXPRI | 0.00000 | 0.00000 | | 0.63737 | 1.31397 | 499 |
| GO_POST_TRANSLATIONAL_PROTEIN_MODIFICATION | 0.00012 | 0.00295 | 0.53843 | 0.58397 | 1.19414 | 258 |
| GO_PROTEASE_BINDING | 0.00383 | 0.04186 | 0.43171 | 0.62064 | 1.23849 | 87 |
| GO_PROTEASOMAL_PROTEIN_CATABOLIC_PROCESS | 0.00368 | 0.04087 | 0.43171 | 0.54309 | 1.11733 | 393 |
| GO_PROTEASOMAL_UBIQUITIN_INDEPENDENT_PROTEIN_C | 0.00002 | 0.00088 | 0.57561 | 0.86722 | 1.59881 | 17 |
| GO_PROTEASOME_ACCESSORY_COMPLEX | 0.00121 | 0.01872 | 0.45506 | 0.76141 | 1.43482 | 24 |
| GO_PROTEASOME_CORE_COMPLEX | 0.00022 | 0.00481 | 0.51885 | 0.84091 | 1.54400 | 16 |
| GO_PROTEIN_CONTAINING_COMPLEX_DISASSEMBLY | 0.00000 | 0.00000 | 0.78819 | 0.63557 | 1.30066 | 269 |
| GO_PROTEIN_CONTAINING_COMPLEX_LOCALIZATION | 0.00276 | 0.03396 | 0.43171 | 0.56668 | 1.15662 | 231 |
| GO_PROTEIN_DNA_COMPLEX_SUBUNIT_ORGANIZATION | 0.00026 | 0.00560 | 0.49849 | 0.59912 | 1.21719 | 177 |
| GO_PROTEIN_DOMAIN_SPECIFIC_BINDING | 0.00398 | 0.04290 | 0.40702 | 0.53571 | 1.10414 | 485 |
| GO_PROTEIN_FOLDING | 0.00000 | 0.00001 | 0.69013 | 0.64581 | 1.31098 | 167 |
| GO_PROTEIN_FOLDING_CHAPERONE | 0.00135 | 0.02031 | 0.45506 | 0.76969 | 1.44202 | 21 |
| GO_PROTEIN_HOMODIMERIZATION_ACTIVITY | 0.00420 | 0.04456 | 0.40702 | 0.53832 | 1.10859 | 417 |
| GO_PROTEIN_IMPORT | 0.00054 | 0.01022 | 0.47727 | 0.60165 | 1.22029 | 158 |
| GO_PROTEIN_LIPID_COMPLEX | 0.00287 | 0.03479 | 0.43171 | 0.75186 | 1.41256 | 22 |
| GO_PROTEIN_LIPID_COMPLEX_ASSEMBLY | 0.00139 | 0.02077 | 0.45506 | 0.74313 | 1.40546 | 26 |
| GO_PROTEIN_LIPID_COMPLEX_BINDING | 0.00350 | 0.03950 | 0.43171 | 0.76572 | 1.41168 | 17 |
| GO_PROTEIN_LIPID_COMPLEX_SUBUNIT_ORGANIZATION | 0.00115 | 0.01808 | 0.45506 | 0.72678 | 1.38980 | 32 |
| GO_PROTEIN_LOCALIZATION_TO_CHROMOSOME_TELOMEF | 0.00428 | 0.04497 | 0.40702 | 0.71483 | 1.35763 | 28 |
| GO_PROTEIN_LOCALIZATION_TO_ENDOPLASMIC_RETICULU | 0.00000 | 0.00000 | | 0.77168 | 1.55892 | 129 |
| GO_PROTEIN_LOCALIZATION_TO_MEMBRANE | 0.00000 | 0.00000 | | 0.62355 | 1.28513 | 474 |
| GO_PROTEIN_LOCALIZATION_TO_MITOCHONDRION | 0.00001 | 0.00036 | 0.59333 | 0.64816 | 1.30791 | 124 |
| GO_PROTEIN_LOCALIZATION_TO_NUCLEUS | 0.00006 | 0.00180 | 0.53843 | 0.59925 | 1.22316 | 219 |
| GO_PROTEIN_MODIFICATION_BY_SMALL_PROTEIN_REMOV | 0.00028 | 0.00599 | 0.49849 | 0.57953 | 1.18275 | 227 |
| GO_PROTEIN_POLYMERIZATION | 0.00066 | 0.01157 | 0.47727 | 0.57826 | 1.18011 | 216 |
| GO_PROTEIN_REFOLDING | 0.00167 | 0.02347 | 0.45506 | 0.78116 | 1.45726 | 20 |
| GO_PROTEIN_STABILIZATION | 0.00085 | 0.01407 | 0.47727 | 0.59789 | 1.21032 | 146 |
| GO_PROTEIN_TARGETING | 0.00000 | 0.00000 | | 0.66669 | 1.36972 | 364 |
| GO_PROTEIN_TARGETING_TO_MEMBRANE | 0.00000 | 0.00000 | | 0.73107 | 1.48406 | 167 |
| GO_PROTEIN_TARGETING_TO_MITOCHONDRION | 0.00008 | 0.00231 | 0.53843 | 0.66766 | 1.33156 | 86 |
| GO_PROTON_TRANSMEMBRANE_TRANSPORT | 0.00000 | 0.00007 | 0.64355 | 0.70270 | 1.40079 | 83 |
| GO_PROTON_TRANSMEMBRANE_TRANSPORTER_ACTIVITY | 0.00104 | 0.01667 | 0.45506 | 0.66247 | 1.30720 | 63 |
| GO_PURINE_CONTAINING_COMPOUND_BIOSYNTHETIC_PR | 0.00146 | 0.02142 | 0.45506 | 0.60324 | 1.21760 | 125 |
| GO_PURINE_CONTAINING_COMPOUND_METABOLIC_PROC | 0.00001 | 0.00023 | 0.61053 | 0.59588 | 1.22053 | 289 |
| GO_PURINE_NUCLEOSIDE_TRIPHOSPHATE_METABOLIC_PR | 0.00004 | 0.00123 | 0.55733 | 0.72294 | 1.41035 | 50 |
| GO_PYRUVATE_METABOLIC_PROCESS | 0.00001 | 0.00059 | 0.59333 | 0.66223 | 1.33248 | 106 |
| GO_REACTIVE_OXYGEN_SPECIES_METABOLIC_PROCESS | 0.00038 | 0.00770 | 0.49849 | 0.60741 | 1.22890 | 144 |
| GO_RECEPTOR_MEDIATED_ENDOCYTOSIS | 0.00015 | 0.00348 | 0.51885 | 0.60731 | 1.23211 | 160 |
| GO_REGULATION_OF_ACTIN_FILAMENT_BASED_PROCESS | 0.00027 | 0.00574 | 0.49849 | 0.57301 | 1.17259 | 270 |
| GO_REGULATION_OF_ACTIN_FILAMENT_LENGTH | 0.00039 | 0.00795 | 0.49849 | 0.61167 | 1.23565 | 130 |
| GO_REGULATION_OF_ACTIN_FILAMENT_ORGANIZATION | 0.00031 | 0.00647 | 0.49849 | 0.59166 | 1.20382 | 186 |
| GO_REGULATION_OF_ANATOMICAL_STRUCTURE_SIZE | 0.00044 | 0.00870 | 0.49849 | 0.56173 | 1.15129 | 310 |
| GO_REGULATION_OF_ANIMAL_ORGAN_MORPHOGENESIS | 0.00000 | 0.00014 | 0.62726 | 0.64027 | 1.29693 | 153 |
| GO_REGULATION_OF_APOPTOTIC_SIGNALING_PATHWAY | 0.00013 | 0.00308 | 0.51885 | 0.57285 | 1.17286 | 286 |
| GO_REGULATION_OF_ATPASE_ACTIVITY | 0.00209 | 0.02756 | 0.43171 | 0.65084 | 1.27955 | 59 |
| GO_REGULATION_OF_ATP_METABOLIC_PROCESS | 0.00076 | 0.01278 | 0.47727 | 0.62981 | 1.25805 | 90 |
| GO_REGULATION_OF_BINDING | 0.00110 | 0.01734 | 0.45506 | 0.57156 | 1.16854 | 256 |
| GO_REGULATION_OF_CARDIAC_MUSCLE_CONTRACTION_BY | 0.00048 | 0.00943 | 0.49849 | 0.80935 | 1.49212 | 17 |
| GO_REGULATION_OF_CELLULAR_AMIDE_METABOLIC_PROC | 0.00000 | 0.00001 | 0.69013 | 0.60307 | 1.23767 | 340 |
| GO_REGULATION_OF_CELLULAR_AMINE_METABOLIC_PROC | 0.00000 | 0.00018 | 0.62726 | 0.73535 | 1.44481 | 58 |
| GO_REGULATION_OF_CELLULAR_AMINO_ACID_METABOLIC | 0.00001 | 0.00027 | 0.61053 | 0.74947 | 1.46389 | 51 |
| GO_REGULATION_OF_CELLULAR_COMPONENT_SIZE | 0.00059 | 0.01076 | 0.47727 | 0.57393 | 1.17299 | 253 |
| GO_REGULATION_OF_CELLULAR_KETONE_METABOLIC_PRO | 0.00000 | 0.00004 | 0.65944 | 0.66689 | 1.34571 | 124 |
| GO_REGULATION_OF_CELLULAR_PROTEIN_LOCALIZATION | 0.00005 | 0.00157 | 0.55733 | 0.56451 | 1.16239 | 409 |
| GO_REGULATION_OF_CELL_CYCLE_G2_M_PHASE_TRANSITI | 0.00002 | 0.00069 | 0.57561 | 0.61884 | 1.25878 | 185 |
| GO_REGULATION_OF_CELL_CYCLE_PHASE_TRANSITION | 0.00013 | 0.00313 | 0.51885 | 0.56769 | 1.16699 | 370 |
| GO_REGULATION_OF_CELL_KILLING | 0.00361 | 0.04040 | 0.43171 | 0.68241 | 1.31594 | 38 |
| GO_REGULATION_OF_CHROMOSOME_ORGANIZATION | 0.00327 | 0.03773 | 0.43171 | 0.55380 | 1.13379 | 284 |
| GO_REGULATION_OF_CYSTEINE_TYPE_ENDOPEPTIDASE_AC | 0.00008 | 0.00218 | 0.53843 | 0.62282 | 1.26157 | 150 |
| GO_REGULATION_OF_CYTOPLASMIC_TRANSLATION | 0.00157 | 0.02257 | 0.45506 | 0.76767 | 1.44225 | 22 |
| GO_REGULATION_OF_CYTOSKELETON_ORGANIZATION | 0.00058 | 0.01069 | 0.47727 | 0.55568 | 1.14306 | 389 |
| GO_REGULATION_OF_DEFENSE_RESPONSE | 0.00000 | 0.00001 | 0.70498 | 0.59426 | 1.22231 | 386 |
| GO_REGULATION_OF_DNA_TEMPLATED_TRANSCRIPTION_I | 0.00000 | 0.00008 | 0.64355 | 0.69035 | 1.38176 | 95 |
| GO_REGULATION_OF_GENERATION_OF_PRECURSOR_META | 0.00239 | 0.03046 | 0.43171 | 0.59791 | 1.20585 | 115 |
| GO_REGULATION_OF_GENE_SILENCING | 0.00335 | 0.03837 | 0.43171 | 0.62238 | 1.24105 | 85 |
| GO_REGULATION_OF_HEMATOPOIETIC_PROGENITOR_CELL_ | 0.00000 | 0.00022 | 0.61053 | 0.70668 | 1.40056 | 70 |
| GO_REGULATION_OF_HEMATOPOIETIC_STEM_CELL_DIFFER | 0.00000 | 0.00015 | 0.62726 | 0.72915 | 1.43381 | 60 |
| GO_REGULATION_OF_HEMOPOIESIS | 0.00185 | 0.02507 | 0.45506 | 0.55819 | 1.14269 | 282 |
| GO_REGULATION_OF_IMMUNE_RESPONSE | 0.00004 | 0.00134 | 0.55733 | 0.56205 | 1.15805 | 463 |
| GO_REGULATION_OF_INNATE_IMMUNE_RESPONSE | 0.00000 | 0.00001 | 0.70498 | 0.64319 | 1.31005 | 201 |
| GO_REGULATION_OF_INTRACELLULAR_PROTEIN_TRANSPOI | 0.00442 | 0.04616 | 0.40702 | 0.56820 | 1.15730 | 202 |
| GO_REGULATION_OF_INTRACELLULAR_TRANSPORT | 0.00145 | 0.02134 | 0.45506 | 0.56115 | 1.14817 | 263 |

| Cluster 1 | | | | | | |
|---|---|---|---|---|---|---|
| GO_REGULATION_OF_INTRINSIC_APOPTOTIC_SIGNALING_P | 0.00050 | 0.00964 | 0.47727 | 0.61238 | 1.23604 | 125 |
| GO_REGULATION_OF_LIPASE_ACTIVITY | 0.00372 | 0.04100 | 0.43171 | 0.64780 | 1.27034 | 55 |
| GO_REGULATION_OF_LIPID_BIOSYNTHETIC_PROCESS | 0.00353 | 0.03985 | 0.43171 | 0.59157 | 1.19373 | 124 |
| GO_REGULATION_OF_MORPHOGENESIS_OF_AN_EPITHELIU | 0.00000 | 0.00011 | 0.62726 | 0.66135 | 1.33543 | 127 |
| GO_REGULATION_OF_MRNA_CATABOLIC_PROCESS | 0.00000 | 0.00000 | 0.74774 | 0.66330 | 1.34663 | 168 |
| GO_REGULATION_OF_MRNA_METABOLIC_PROCESS | 0.00000 | 0.00000 | | 0.65949 | 1.34986 | 280 |
| GO_REGULATION_OF_MRNA_PROCESSING | 0.00001 | 0.00025 | 0.61053 | 0.65310 | 1.31870 | 126 |
| GO_REGULATION_OF_MRNA_SPLICING_VIA_SPLICEOSOME | 0.00001 | 0.00054 | 0.59333 | 0.67151 | 1.34112 | 91 |
| GO_REGULATION_OF_MULTI_ORGANISM_PROCESS | 0.00156 | 0.02245 | 0.45506 | 0.69718 | 1.35378 | 43 |
| GO_REGULATION_OF_NIK_NF_KAPPAB_SIGNALING | 0.00213 | 0.02782 | 0.43171 | 0.64949 | 1.28565 | 68 |
| GO_REGULATION_OF_NUCLEOTIDE_BIOSYNTHETIC_PROCES | 0.00242 | 0.03069 | 0.43171 | 0.75979 | 1.42949 | 23 |
| GO_REGULATION_OF_OXIDATIVE_PHOSPHORYLATION | 0.00071 | 0.01214 | 0.47727 | 0.77279 | 1.45627 | 24 |
| GO_REGULATION_OF_OXIDATIVE_STRESS_INDUCED_CELL_D | 0.00398 | 0.04290 | 0.40702 | 0.67038 | 1.30286 | 44 |
| GO_REGULATION_OF_PEPTIDASE_ACTIVITY | 0.00000 | 0.00001 | 0.69013 | 0.62517 | 1.27652 | 238 |
| GO_REGULATION_OF_PEPTIDE_TRANSPORT | 0.00008 | 0.00226 | 0.53843 | 0.56307 | 1.15963 | 429 |
| GO_REGULATION_OF_PHOSPHOLIPID_METABOLIC_PROCESS | 0.00465 | 0.04794 | 0.40702 | 0.63643 | 1.24955 | 56 |
| GO_REGULATION_OF_PLASMA_LIPOPROTEIN_PARTICLE_LEV | 0.00001 | 0.00051 | 0.59333 | 0.71436 | 1.40472 | 60 |
| GO_REGULATION_OF_PLATELET_ACTIVATION | 0.00279 | 0.03429 | 0.43171 | 0.74604 | 1.40586 | 24 |
| GO_REGULATION_OF_POSTTRANSCRIPTIONAL_GENE_SILEN | 0.00141 | 0.02099 | 0.45506 | 0.63748 | 1.26844 | 76 |
| GO_REGULATION_OF_PROTEIN_BINDING | 0.00145 | 0.02134 | 0.45506 | 0.59485 | 1.20299 | 141 |
| GO_REGULATION_OF_PROTEIN_CONTAINING_COMPLEX_AS | 0.00000 | 0.00021 | 0.61053 | 0.59153 | 1.21257 | 311 |
| GO_REGULATION_OF_PROTEIN_LOCALIZATION_TO_NUCLEU | 0.00002 | 0.00074 | 0.57561 | 0.65949 | 1.32378 | 101 |
| GO_REGULATION_OF_PROTEIN_POLYMERIZATION | 0.00027 | 0.00574 | 0.49849 | 0.60400 | 1.22534 | 162 |
| GO_REGULATION_OF_PROTEIN_STABILITY | 0.00010 | 0.00256 | 0.53843 | 0.58897 | 1.20203 | 233 |
| GO_REGULATION_OF_PROTEOLYSIS | 0.00000 | 0.00022 | 0.61053 | 0.57739 | 1.18921 | 439 |
| GO_REGULATION_OF_RESPONSE_TO_BIOTIC_STIMULUS | 0.00000 | 0.00005 | 0.65944 | 0.60802 | 1.24337 | 259 |
| GO_REGULATION_OF_RNA_SPLICING | 0.00000 | 0.00019 | 0.61053 | 0.65227 | 1.31769 | 129 |
| GO_REGULATION_OF_RYANODINE_SENSITIVE_CALCIUM_RE | 0.00156 | 0.02245 | 0.45506 | 0.83055 | 1.51676 | 15 |
| GO_REGULATION_OF_SMALL_MOLECULE_METABOLIC_PRO | 0.00000 | 0.00022 | 0.61053 | 0.59575 | 1.22082 | 301 |
| GO_REGULATION_OF_SPINDLE_ORGANIZATION | 0.00390 | 0.04246 | 0.43171 | 0.70096 | 1.34189 | 33 |
| GO_REGULATION_OF_STEM_CELL_DIFFERENTIATION | 0.00005 | 0.00150 | 0.55733 | 0.66669 | 1.33097 | 88 |
| GO_REGULATION_OF_STEROID_METABOLIC_PROCESS | 0.00242 | 0.03069 | 0.43171 | 0.62674 | 1.24975 | 85 |
| GO_REGULATION_OF_SUPRAMOLECULAR_FIBER_ORGANIZA | 0.00004 | 0.00128 | 0.55733 | 0.59248 | 1.21089 | 252 |
| GO_REGULATION_OF_SYMBIOTIC_PROCESS | 0.00000 | 0.00006 | 0.64355 | 0.64376 | 1.30666 | 166 |
| GO_REGULATION_OF_TRANSCRIPTION_FROM_RNA_POLYM | 0.00000 | 0.00005 | 0.65944 | 0.74300 | 1.46610 | 63 |
| GO_REGULATION_OF_TRANSLATIONAL_INITIATION | 0.00017 | 0.00386 | 0.51885 | 0.68559 | 1.35515 | 65 |
| GO_REGULATION_OF_VIRAL_GENOME_REPLICATION | 0.00031 | 0.00647 | 0.49849 | 0.66755 | 1.32167 | 69 |
| GO_REGULATION_OF_VIRAL_LIFE_CYCLE | 0.00012 | 0.00306 | 0.51885 | 0.63966 | 1.28630 | 104 |
| GO_REGULATION_OF_VIRAL_TRANSCRIPTION | 0.00250 | 0.03144 | 0.43171 | 0.65568 | 1.28804 | 57 |
| GO_REGULATION_OF_WNT_SIGNALING_PATHWAY | 0.00029 | 0.00627 | 0.49849 | 0.58214 | 1.18959 | 251 |
| GO_RELEASE_OF_SEQUESTERED_CALCIUM_ION_INTO_CYTO | 0.00146 | 0.02142 | 0.45506 | 0.80068 | 1.48429 | 18 |
| GO_RESPIRASOME | 0.00000 | 0.00000 | | 0.80925 | 1.60853 | 75 |
| GO_RESPIRATORY_CHAIN_COMPLEX | 0.00000 | 0.00000 | | 0.82144 | 1.62088 | 63 |
| GO_RESPIRATORY_ELECTRON_TRANSPORT_CHAIN | 0.00000 | 0.00000 | | 0.75589 | 1.51123 | 93 |
| GO_RESPONSE_TO_ALCOHOL | 0.00047 | 0.00931 | 0.49849 | 0.61349 | 1.23872 | 126 |
| GO_RESPONSE_TO_COCAINE | 0.00435 | 0.04553 | 0.40702 | 0.70884 | 1.35104 | 30 |
| GO_RESPONSE_TO_CORTICOSTEROID | 0.00128 | 0.01948 | 0.45506 | 0.63483 | 1.26472 | 82 |
| GO_RESPONSE_TO_DRUG | 0.00228 | 0.02940 | 0.43171 | 0.56808 | 1.15949 | 231 |
| GO_RESPONSE_TO_ENDOPLASMIC_RETICULUM_STRESS | 0.00279 | 0.03429 | 0.43171 | 0.56011 | 1.14290 | 229 |
| GO_RESPONSE_TO_ETHANOL | 0.00018 | 0.00414 | 0.51885 | 0.68746 | 1.35651 | 63 |
| GO_RESPONSE_TO_GROWTH_FACTOR | 0.00157 | 0.02257 | 0.45506 | 0.54025 | 1.11285 | 454 |
| GO_RESPONSE_TO_INORGANIC_SUBSTANCE | 0.00065 | 0.01147 | 0.47727 | 0.55924 | 1.14725 | 328 |
| GO_RESPONSE_TO_INTERLEUKIN_1 | 0.00000 | 0.00005 | 0.65944 | 0.66311 | 1.33956 | 131 |
| GO_RESPONSE_TO_INTERLEUKIN_12 | 0.00051 | 0.00979 | 0.47727 | 0.72859 | 1.40156 | 37 |
| GO_RESPONSE_TO_INTERLEUKIN_4 | 0.00339 | 0.03859 | 0.43171 | 0.74848 | 1.40620 | 22 |
| GO_RESPONSE_TO_ISCHEMIA | 0.00046 | 0.00919 | 0.49849 | 0.72902 | 1.39737 | 34 |
| GO_RESPONSE_TO_MONOSACCHARIDE | 0.00257 | 0.03212 | 0.43171 | 0.59480 | 1.20011 | 121 |
| GO_RESPONSE_TO_OXIDATIVE_STRESS | 0.00067 | 0.01167 | 0.47727 | 0.56551 | 1.15835 | 298 |
| GO_RESPONSE_TO_OXYGEN_LEVELS | 0.00000 | 0.00000 | 0.77494 | 0.63916 | 1.30787 | 264 |
| GO_RESPONSE_TO_STEROID_HORMONE | 0.00170 | 0.02383 | 0.45506 | 0.57626 | 1.17450 | 205 |
| GO_RESPONSE_TO_TOPOLOGICALLY_INCORRECT_PROTEIN | 0.00050 | 0.00964 | 0.47727 | 0.59693 | 1.21106 | 160 |
| GO_RESPONSE_TO_TOXIC_SUBSTANCE | 0.00122 | 0.01882 | 0.45506 | 0.60498 | 1.22218 | 133 |
| GO_RESPONSE_TO_TUMOR_NECROSIS_FACTOR | 0.00007 | 0.00202 | 0.53843 | 0.60482 | 1.23109 | 191 |
| GO_RESPONSE_TO_VIRUS | 0.00001 | 0.00036 | 0.59333 | 0.61635 | 1.25479 | 192 |
| GO_RETINA_HOMEOSTASIS | 0.00077 | 0.01302 | 0.47727 | 0.71269 | 1.37433 | 38 |
| GO_RIBONUCLEOPROTEIN_COMPLEX_BINDING | 0.00004 | 0.00128 | 0.55733 | 0.64828 | 1.30678 | 113 |
| GO_RIBONUCLEOPROTEIN_COMPLEX_BIOGENESIS | 0.00000 | 0.00000 | | 0.62692 | 1.28967 | 395 |
| GO_RIBONUCLEOPROTEIN_COMPLEX_SUBUNIT_ORGANIZAT | 0.00000 | 0.00000 | | 0.68597 | 1.39365 | 177 |
| GO_RIBONUCLEOPROTEIN_GRANULE | 0.00006 | 0.00171 | 0.55733 | 0.60293 | 1.22716 | 187 |
| GO_RIBONUCLEOSIDE_TRIPHOSPHATE_BIOSYNTHETIC_PRO | 0.00001 | 0.00055 | 0.59333 | 0.77143 | 1.48430 | 36 |
| GO_RIBONUCLEOSIDE_TRIPHOSPHATE_METABOLIC_PROCES | 0.00001 | 0.00051 | 0.59333 | 0.74521 | 1.45121 | 48 |
| GO_RIBOSE_PHOSPHATE_BIOSYNTHETIC_PROCESS | 0.00087 | 0.01445 | 0.47727 | 0.61153 | 1.23403 | 120 |
| GO_RIBOSE_PHOSPHATE_METABOLIC_PROCESS | 0.00001 | 0.00051 | 0.59333 | 0.59390 | 1.21563 | 273 |

| Cluster 1 | | | | | | |
|---|---|---|---|---|---|---|
| GO_RIBOSOMAL_LARGE_SUBUNIT_ASSEMBLY | 0.00183 | 0.02507 | 0.45506 | 0.73650 | 1.39293 | 26 |
| GO_RIBOSOMAL_LARGE_SUBUNIT_BIOGENESIS | 0.00007 | 0.00193 | 0.53843 | 0.68722 | 1.36033 | 68 |
| GO_RIBOSOMAL_SMALL_SUBUNIT_ASSEMBLY | 0.00005 | 0.00151 | 0.55733 | 0.86188 | 1.59774 | 18 |
| GO_RIBOSOMAL_SMALL_SUBUNIT_BIOGENESIS | 0.00242 | 0.03069 | 0.43171 | 0.64854 | 1.28377 | 68 |
| GO_RIBOSOMAL_SUBUNIT | 0.00000 | 0.00000 | | 0.74369 | 1.51064 | 174 |
| GO_RIBOSOME | 0.00000 | 0.00000 | | 0.74111 | 1.51111 | 212 |
| GO_RIBOSOME_ASSEMBLY | 0.00000 | 0.00005 | 0.65944 | 0.75327 | 1.47976 | 57 |
| GO_RIBOSOME_BIOGENESIS | 0.00000 | 0.00006 | 0.64355 | 0.60481 | 1.23802 | 276 |
| GO_RNA_3_END_PROCESSING | 0.00465 | 0.04794 | 0.40702 | 0.59074 | 1.19293 | 128 |
| GO_RNA_CATABOLIC_PROCESS | 0.00000 | 0.00000 | | 0.66961 | 1.37489 | 348 |
| GO_RNA_DEPENDENT_DNA_BIOSYNTHETIC_PROCESS | 0.00231 | 0.02975 | 0.43171 | 0.64970 | 1.27926 | 61 |
| GO_RNA_EXPORT_FROM_NUCLEUS | 0.00122 | 0.01882 | 0.45506 | 0.59880 | 1.20965 | 130 |
| GO_RNA_LOCALIZATION | 0.00002 | 0.00064 | 0.57561 | 0.60686 | 1.23738 | 212 |
| GO_RNA_POLYMERASE_II_SPECIFIC_DNA_BINDING_TRANS( | 0.00428 | 0.04497 | 0.40702 | 0.56880 | 1.15852 | 202 |
| GO_RNA_SPLICING | 0.00000 | 0.00000 | 0.74774 | 0.60281 | 1.24008 | 395 |
| GO_RNA_SPLICING_VIA_TRANSESTERIFICATION_REACTIONS | 0.00000 | 0.00001 | 0.70498 | 0.61236 | 1.25511 | 316 |
| GO_ROUGH_ENDOPLASMIC_RETICULUM | 0.00002 | 0.00064 | 0.57561 | 0.70823 | 1.39240 | 59 |
| GO_RRNA_BINDING | 0.00000 | 0.00000 | 0.71951 | 0.77688 | 1.52614 | 57 |
| GO_RRNA_METABOLIC_PROCESS | 0.00071 | 0.01214 | 0.47727 | 0.58257 | 1.18740 | 208 |
| GO_SARCOMERE_ORGANIZATION | 0.00302 | 0.03584 | 0.43171 | 0.76133 | 1.41565 | 19 |
| GO_SARCOPLASMIC_RETICULUM_CALCIUM_ION_TRANSPO | 0.00122 | 0.01882 | 0.45506 | 0.77177 | 1.44593 | 21 |
| GO_SCF_DEPENDENT_PROTEASOMAL_UBIQUITIN_DEPENDI | 0.00000 | 0.00004 | 0.65944 | 0.71857 | 1.42828 | 75 |
| GO_SECRETORY_GRANULE | 0.00000 | 0.00000 | | 0.62761 | 1.29351 | 493 |
| GO_SECRETORY_GRANULE_MEMBRANE | 0.00283 | 0.03454 | 0.43171 | 0.57513 | 1.16922 | 180 |
| GO_SINGLE_FERTILIZATION | 0.00059 | 0.01076 | 0.47727 | 0.65955 | 1.30408 | 66 |
| GO_SINGLE_STRANDED_RNA_BINDING | 0.00205 | 0.02719 | 0.43171 | 0.64995 | 1.28656 | 68 |
| GO_SKELETAL_MUSCLE_CONTRACTION | 0.00031 | 0.00647 | 0.49849 | 0.81867 | 1.52723 | 20 |
| GO_SMALL_MOLECULE_BIOSYNTHETIC_PROCESS | 0.00000 | 0.00009 | 0.64355 | 0.57658 | 1.18766 | 447 |
| GO_SMALL_NUCLEAR_RIBONUCLEOPROTEIN_COMPLEX | 0.00019 | 0.00420 | 0.51885 | 0.69054 | 1.35677 | 58 |
| GO_SMALL_RIBOSOMAL_SUBUNIT | 0.00000 | 0.00000 | 0.76146 | 0.76556 | 1.51571 | 69 |
| GO_SMOOTH_MUSCLE_CELL_PROLIFERATION | 0.00420 | 0.04456 | 0.40702 | 0.61699 | 1.22914 | 81 |
| GO_SM_LIKE_PROTEIN_FAMILY_COMPLEX | 0.00037 | 0.00756 | 0.49849 | 0.66844 | 1.32317 | 68 |
| GO_SNORNA_BINDING | 0.00239 | 0.03046 | 0.43171 | 0.71615 | 1.36394 | 29 |
| GO_SOMATIC_STEM_CELL_POPULATION_MAINTENANCE | 0.00442 | 0.04616 | 0.40702 | 0.64508 | 1.26721 | 57 |
| GO_SPERM_EGG_RECOGNITION | 0.00001 | 0.00027 | 0.61053 | 0.82615 | 1.56906 | 28 |
| GO_SPINDLE_ORGANIZATION | 0.00405 | 0.04341 | 0.40702 | 0.57971 | 1.17426 | 153 |
| GO_SPLICEOSOMAL_COMPLEX | 0.00010 | 0.00255 | 0.53843 | 0.60721 | 1.23340 | 174 |
| GO_STEM_CELL_DIFFERENTIATION | 0.00007 | 0.00208 | 0.53843 | 0.60942 | 1.23790 | 174 |
| GO_STEROL_BIOSYNTHETIC_PROCESS | 0.00014 | 0.00340 | 0.51885 | 0.69487 | 1.36528 | 58 |
| GO_STEROL_METABOLIC_PROCESS | 0.00090 | 0.01479 | 0.47727 | 0.61785 | 1.24201 | 105 |
| GO_STRUCTURAL_CONSTITUENT_OF_CYTOSKELETON | 0.00008 | 0.00226 | 0.53843 | 0.68942 | 1.36108 | 64 |
| GO_STRUCTURAL_CONSTITUENT_OF_RIBOSOME | 0.00000 | 0.00000 | | 0.77599 | 1.57265 | 156 |
| GO_STRUCTURAL_MOLECULE_ACTIVITY | 0.00000 | 0.00000 | | 0.70510 | 1.45182 | 423 |
| GO_SUBSTANTIA_NIGRA_DEVELOPMENT | 0.00119 | 0.01856 | 0.45506 | 0.73598 | 1.40277 | 30 |
| GO_SUPRAMOLECULAR_FIBER_ORGANIZATION | 0.00002 | 0.00067 | 0.57561 | 0.56896 | 1.17207 | 458 |
| GO_TELOMERASE_RNA_LOCALIZATION | 0.00010 | 0.00254 | 0.53843 | 0.82985 | 1.54306 | 19 |
| GO_TELOMERE_MAINTENANCE_VIA_TELOMERE_LENGTHEN | 0.00076 | 0.01290 | 0.47727 | 0.66084 | 1.30812 | 68 |
| GO_THREONINE_TYPE_ENDOPEPTIDASE_ACTIVITY | 0.00005 | 0.00149 | 0.55733 | 0.86519 | 1.58859 | 16 |
| GO_THREONINE_TYPE_PEPTIDASE_ACTIVITY | 0.00003 | 0.00115 | 0.55733 | 0.85001 | 1.58569 | 20 |
| GO_TISSUE_MORPHOGENESIS | 0.00064 | 0.01137 | 0.47727 | 0.55067 | 1.13404 | 415 |
| GO_TRANSLATIONAL_ELONGATION | 0.00000 | 0.00001 | 0.69013 | 0.67630 | 1.36471 | 124 |
| GO_TRANSLATIONAL_INITIATION | 0.00000 | 0.00000 | | 0.77255 | 1.56876 | 169 |
| GO_TRANSLATIONAL_TERMINATION | 0.00000 | 0.00003 | 0.67496 | 0.69251 | 1.39056 | 102 |
| GO_TRANSLATION_FACTOR_ACTIVITY_RNA_BINDING | 0.00010 | 0.00254 | 0.53843 | 0.67527 | 1.33832 | 70 |
| GO_TRANSLATION_INITIATION_FACTOR_ACTIVITY | 0.00078 | 0.01311 | 0.47727 | 0.69314 | 1.34739 | 46 |
| GO_TRANSLATION_INITIATION_FACTOR_BINDING | 0.00100 | 0.01620 | 0.45506 | 0.74660 | 1.41520 | 27 |
| GO_TRANSLATION_PREINITIATION_COMPLEX | 0.00032 | 0.00664 | 0.49849 | 0.83439 | 1.53203 | 16 |
| GO_TRANSLATION_REGULATOR_ACTIVITY | 0.00000 | 0.00001 | 0.70498 | 0.69754 | 1.40607 | 113 |
| GO_TRANSLATION_REGULATOR_ACTIVITY_NUCLEIC_ACID_E | 0.00002 | 0.00076 | 0.57561 | 0.68095 | 1.35806 | 86 |
| GO_TRICARBOXYLIC_ACID_CYCLE | 0.00012 | 0.00306 | 0.51885 | 0.75840 | 1.45028 | 32 |
| GO_TUMOR_NECROSIS_FACTOR_MEDIATED_SIGNALING_PA | 0.00000 | 0.00002 | 0.67496 | 0.67914 | 1.37048 | 120 |
| GO_T_CELL_RECEPTOR_SIGNALING_PATHWAY | 0.00102 | 0.01642 | 0.45506 | 0.60202 | 1.21710 | 136 |
| GO_U12_TYPE_SPLICEOSOMAL_COMPLEX | 0.00170 | 0.02383 | 0.45506 | 0.74257 | 1.40082 | 25 |
| GO_U2_TYPE_SPLICEOSOMAL_COMPLEX | 0.00209 | 0.02756 | 0.43171 | 0.61618 | 1.23013 | 88 |
| GO_UBIQUITIN_LIKE_PROTEIN_LIGASE_BINDING | 0.00000 | 0.00003 | 0.67496 | 0.61149 | 1.25047 | 259 |
| GO_UBIQUITIN_PROTEIN_TRANSFERASE_REGULATOR_ACTI\ | 0.00405 | 0.04341 | 0.40702 | 0.76134 | 1.40361 | 17 |
| GO_UNFOLDED_PROTEIN_BINDING | 0.00000 | 0.00004 | 0.65944 | 0.69381 | 1.38774 | 94 |
| GO_VACUOLAR_LUMEN | 0.00001 | 0.00024 | 0.61053 | 0.66866 | 1.34617 | 107 |
| GO_VESICLE_LUMEN | 0.00000 | 0.00000 | | 0.68398 | 1.39313 | 201 |
| GO_VESICLE_MEMBRANE | 0.00294 | 0.03542 | 0.43171 | 0.53478 | 1.10216 | 489 |
| GO_VIRAL_GENE_EXPRESSION | 0.00000 | 0.00000 | | 0.73242 | 1.48900 | 180 |
| GO_VIRAL_GENOME_REPLICATION | 0.00018 | 0.00414 | 0.51885 | 0.64872 | 1.29582 | 90 |
| GO_VIRAL_LIFE_CYCLE | 0.00005 | 0.00158 | 0.55733 | 0.59599 | 1.21708 | 242 |

| Cluster 1 | | | | | | |
|---|---|---|---|---|---|---|
| HP_ABNORMALITY_OF_ACID_BASE_HOMEOSTASIS | 0.00002 | 0.00081 | 0.57561 | 0.59318 | 1.21328 | 260 |
| HP_ABNORMALITY_OF_BLOOD_CIRCULATION | 0.00398 | 0.04290 | 0.40702 | 0.57345 | 1.16484 | 174 |
| HP_ABNORMALITY_OF_BRAINSTEM_MORPHOLOGY | 0.00327 | 0.03773 | 0.43171 | 0.58954 | 1.19260 | 143 |
| HP_ABNORMALITY_OF_CARDIOVASCULAR_SYSTEM_ELECTR | 0.00069 | 0.01189 | 0.47727 | 0.56735 | 1.16152 | 275 |
| HP_ABNORMALITY_OF_THE_MITOCHONDRION | 0.00013 | 0.00316 | 0.51885 | 0.63272 | 1.27703 | 123 |
| HP_ABNORMALITY_OF_THE_NASAL_DORSUM | 0.00372 | 0.04100 | 0.43171 | 0.57957 | 1.17594 | 161 |
| HP_ABNORMALITY_OF_THE_OPTIC_NERVE | 0.00191 | 0.02555 | 0.45506 | 0.53828 | 1.10960 | 496 |
| HP_ABNORMAL_ABDOMEN_MORPHOLOGY | 0.00145 | 0.02134 | 0.45506 | 0.54309 | 1.11838 | 443 |
| HP_ABNORMAL_ACTIVITY_OF_MITOCHONDRIAL_RESPIRATO | 0.00010 | 0.00255 | 0.53843 | 0.69505 | 1.36854 | 61 |
| HP_ABNORMAL_ASTROCYTE_MORPHOLOGY | 0.00128 | 0.01948 | 0.45506 | 0.73505 | 1.40099 | 30 |
| HP_ABNORMAL_BASAL_GANGLIA_MRI_SIGNAL_INTENSITY | 0.00064 | 0.01137 | 0.47727 | 0.72372 | 1.38721 | 34 |
| HP_ABNORMAL_BLOOD_GLUCOSE_CONCENTRATION | 0.00294 | 0.03542 | 0.43171 | 0.58183 | 1.18028 | 163 |
| HP_ABNORMAL_CELLULAR_PHENOTYPE | 0.00194 | 0.02599 | 0.45506 | 0.55104 | 1.13059 | 337 |
| HP_ABNORMAL_CIRCULATING_CARBOHYDRATE_CONCENTR | 0.00298 | 0.03567 | 0.43171 | 0.66976 | 1.30077 | 45 |
| HP_ABNORMAL_CNS_MYELINATION | 0.00014 | 0.00340 | 0.51885 | 0.59968 | 1.21733 | 167 |
| HP_ABNORMAL_CSF_METABOLITE_LEVEL | 0.00000 | 0.00018 | 0.62726 | 0.69942 | 1.39230 | 78 |
| HP_ABNORMAL_ERYTHROCYTE_MORPHOLOGY | 0.00000 | 0.00001 | 0.69013 | 0.59261 | 1.21920 | 391 |
| HP_ABNORMAL_FOVEAL_MORPHOLOGY | 0.00178 | 0.02448 | 0.45506 | 0.77339 | 1.43806 | 19 |
| HP_ABNORMAL_GLUCOSE_HOMEOSTASIS | 0.00090 | 0.01479 | 0.47727 | 0.55267 | 1.13623 | 372 |
| HP_ABNORMAL_MITOCHONDRIA_IN_MUSCLE_TISSUE | 0.00106 | 0.01692 | 0.45506 | 0.74230 | 1.40982 | 28 |
| HP_ABNORMAL_MYELINATION | 0.00065 | 0.01147 | 0.47727 | 0.55898 | 1.14666 | 332 |
| HP_ABNORMAL_MYOCARDIUM_MORPHOLOGY | 0.00097 | 0.01565 | 0.47727 | 0.55869 | 1.14710 | 346 |
| HP_ABNORMAL_SOCIAL_BEHAVIOR | 0.00305 | 0.03608 | 0.43171 | 0.60177 | 1.21230 | 111 |
| HP_ABNORMAL_URINE_CYTOLOGY | 0.00398 | 0.04290 | 0.40702 | 0.61029 | 1.22151 | 95 |
| HP_ABNORMAL_VASCULAR_MORPHOLOGY | 0.00428 | 0.04497 | 0.40702 | 0.53119 | 1.09478 | 482 |
| HP_ACUTE_LEUKEMIA | 0.00048 | 0.00943 | 0.49849 | 0.65898 | 1.30918 | 73 |
| HP_ANEMIA_OF_INADEQUATE_PRODUCTION | 0.00004 | 0.00128 | 0.55733 | 0.65462 | 1.31448 | 102 |
| HP_APHASIA | 0.00055 | 0.01033 | 0.47727 | 0.74456 | 1.42143 | 31 |
| HP_APLASIA_HYPOPLASIA_INVOLVING_FOREARM_BONES | 0.00342 | 0.03887 | 0.43171 | 0.63428 | 1.26010 | 73 |
| HP_APLASIA_HYPOPLASIA_OF_THE_RADIUS | 0.00480 | 0.04900 | 0.40702 | 0.63875 | 1.25879 | 62 |
| HP_BLINDNESS | 0.00124 | 0.01901 | 0.45506 | 0.58463 | 1.18738 | 171 |
| HP_BULBAR_PALSY | 0.00339 | 0.03859 | 0.43171 | 0.71774 | 1.36049 | 27 |
| HP_CONGESTIVE_HEART_FAILURE | 0.00450 | 0.04686 | 0.40702 | 0.59580 | 1.20252 | 123 |
| HP_DECREASED_ACTIVITY_OF_MITOCHONDRIAL_COMPLEX | 0.00185 | 0.02507 | 0.45506 | 0.70632 | 1.35384 | 34 |
| HP_DECREASED_ACTIVITY_OF_THE_PYRUVATE_DEHYDROGE | 0.00049 | 0.00955 | 0.47727 | 0.75035 | 1.43015 | 30 |
| HP_DEPRESSED_NASAL_RIDGE | 0.00108 | 0.01717 | 0.45506 | 0.62706 | 1.25256 | 90 |
| HP_DYSPHAGIA | 0.00163 | 0.02310 | 0.45506 | 0.56679 | 1.15717 | 235 |
| HP_DYSPHASIA | 0.00133 | 0.02008 | 0.45506 | 0.69764 | 1.34810 | 39 |
| HP_DYSPNEA | 0.00065 | 0.01147 | 0.47727 | 0.57074 | 1.16612 | 249 |
| HP_ECHOLALIA | 0.00309 | 0.03612 | 0.43171 | 0.72269 | 1.37257 | 28 |
| HP_EMOTIONAL_LABILITY | 0.00034 | 0.00703 | 0.49849 | 0.64284 | 1.28423 | 92 |
| HP_ENCEPHALOPATHY | 0.00257 | 0.03212 | 0.43171 | 0.57475 | 1.16672 | 167 |
| HP_FATIGUE | 0.00004 | 0.00134 | 0.55733 | 0.59874 | 1.22209 | 222 |
| HP_FETAL_DISTRESS | 0.00170 | 0.02383 | 0.45506 | 0.73280 | 1.39177 | 28 |
| HP_FOCAL_T2_HYPERINTENSE_BRAINSTEM_LESION | 0.00480 | 0.04900 | 0.40702 | 0.71135 | 1.34839 | 27 |
| HP_FREQUENT_FALLS | 0.00465 | 0.04794 | 0.40702 | 0.62870 | 1.24417 | 67 |
| HP_GENERALIZED_MUSCLE_WEAKNESS | 0.00178 | 0.02448 | 0.45506 | 0.63433 | 1.26219 | 76 |
| HP_GLUCOSE_INTOLERANCE | 0.00191 | 0.02555 | 0.45506 | 0.56305 | 1.15046 | 250 |
| HP_HEADACHE | 0.00054 | 0.01022 | 0.47727 | 0.59840 | 1.21532 | 176 |
| HP_HEMATOLOGICAL_NEOPLASM | 0.00074 | 0.01251 | 0.47727 | 0.58958 | 1.19782 | 177 |
| HP_HYPERTRICHOSIS | 0.00309 | 0.03612 | 0.43171 | 0.58548 | 1.18568 | 149 |
| HP_HYPERTROPHIC_CARDIOMYOPATHY | 0.00002 | 0.00064 | 0.57561 | 0.61342 | 1.24860 | 191 |
| HP_IMPAIRED_SOCIAL_INTERACTIONS | 0.00405 | 0.04341 | 0.40702 | 0.63128 | 1.25296 | 72 |
| HP_IMPAIRMENT_IN_PERSONALITY_FUNCTIONING | 0.00145 | 0.02134 | 0.45506 | 0.54922 | 1.12943 | 383 |
| HP_INCOORDINATION | 0.00159 | 0.02278 | 0.45506 | 0.59620 | 1.20441 | 130 |
| HP_INCREASED_CSF_LACTATE | 0.00001 | 0.00025 | 0.61053 | 0.70613 | 1.39742 | 67 |
| HP_INCREASED_SERUM_LACTATE | 0.00018 | 0.00414 | 0.51885 | 0.62106 | 1.25479 | 132 |
| HP_INCREASED_SERUM_PYRUVATE | 0.00383 | 0.04186 | 0.43171 | 0.69213 | 1.33142 | 37 |
| HP_INTRAUTERINE_GROWTH_RETARDATION | 0.00480 | 0.04900 | 0.40702 | 0.54363 | 1.11638 | 353 |
| HP_IRREGULAR_HYPERPIGMENTATION | 0.00283 | 0.03454 | 0.43171 | 0.60685 | 1.22032 | 104 |
| HP_IRRITABILITY | 0.00239 | 0.03046 | 0.43171 | 0.60821 | 1.22084 | 101 |
| HP_LACTIC_ACIDOSIS | 0.00004 | 0.00134 | 0.55733 | 0.63278 | 1.27924 | 135 |
| HP_LETHARGY | 0.00342 | 0.03887 | 0.43171 | 0.60432 | 1.21710 | 109 |
| HP_LEUKEMIA | 0.00060 | 0.01090 | 0.47727 | 0.62409 | 1.25499 | 104 |
| HP_LEUKODYSTROPHY | 0.00005 | 0.00148 | 0.55733 | 0.67507 | 1.34485 | 81 |
| HP_LEUKOENCEPHALOPATHY | 0.00368 | 0.04087 | 0.43171 | 0.59332 | 1.19909 | 134 |
| HP_LIMB_UNDERGROWTH | 0.00435 | 0.04553 | 0.40702 | 0.56797 | 1.15308 | 168 |
| HP_LOWER_LIMB_SPASTICITY | 0.00224 | 0.02898 | 0.43171 | 0.59943 | 1.21143 | 134 |
| HP_MACROCYTIC_ANEMIA | 0.00004 | 0.00133 | 0.55733 | 0.75356 | 1.45860 | 40 |
| HP_MICROMELIA | 0.00420 | 0.04456 | 0.40702 | 0.61754 | 1.23022 | 81 |
| HP_MIGRAINE | 0.00013 | 0.00316 | 0.51885 | 0.66661 | 1.32799 | 81 |
| HP_MUTISM | 0.00001 | 0.00026 | 0.61053 | 0.78713 | 1.50875 | 34 |
| HP_MYOPATHY | 0.00185 | 0.02507 | 0.45506 | 0.57489 | 1.16993 | 190 |
| HP_NEOPLASM_OF_THE_NERVOUS_SYSTEM | 0.00361 | 0.04040 | 0.43171 | 0.59140 | 1.19412 | 126 |

| Cluster 1 | | | | | | |
|---|---|---|---|---|---|---|
| HP_OPHTHALMOPARESIS | 0.00121 | 0.01872 | 0.45506 | 0.58423 | 1.18660 | 175 |
| HP_OPTIC_NERVE_COLOBOMA | 0.00435 | 0.04553 | 0.40702 | 0.76009 | 1.40131 | 17 |
| HP_OPTIC_NEUROPATHY | 0.00056 | 0.01044 | 0.47727 | 0.72569 | 1.39598 | 37 |
| HP_ORAL_CLEFT | 0.00405 | 0.04341 | 0.40702 | 0.53888 | 1.10866 | 391 |
| HP_PALLOR | 0.00065 | 0.01147 | 0.47727 | 0.62731 | 1.25919 | 101 |
| HP_PARAPLEGIA_PARAPARESIS | 0.00268 | 0.03318 | 0.43171 | 0.58313 | 1.18141 | 151 |
| HP_PAROXYSMAL_INVOLUNTARY_EYE_MOVEMENTS | 0.00146 | 0.02142 | 0.45506 | 0.73210 | 1.39538 | 30 |
| HP_PERIPHERAL_NEUROPATHY | 0.00122 | 0.01882 | 0.45506 | 0.54640 | 1.12414 | 391 |
| HP_POOR_EYE_CONTACT | 0.00065 | 0.01147 | 0.47727 | 0.68156 | 1.33177 | 52 |
| HP_POOR_HEAD_CONTROL | 0.00191 | 0.02555 | 0.45506 | 0.61149 | 1.23155 | 109 |
| HP_PROGRESSIVE_SPASTIC_PARAPLEGIA | 0.00050 | 0.00964 | 0.47727 | 0.69758 | 1.35614 | 47 |
| HP_PROXIMAL_TUBULOPATHY | 0.00156 | 0.02245 | 0.45506 | 0.72095 | 1.37866 | 32 |
| HP_PTOSIS | 0.00191 | 0.02555 | 0.45506 | 0.54299 | 1.11802 | 424 |
| HP_REDUCED_CONSCIOUSNESS_CONFUSION | 0.00069 | 0.01189 | 0.47727 | 0.58330 | 1.18822 | 204 |
| HP_REDUCED_VISUAL_ACUITY | 0.00465 | 0.04794 | 0.40702 | 0.54908 | 1.12468 | 289 |
| HP_RENAL_TUBULAR_DYSFUNCTION | 0.00067 | 0.01167 | 0.47727 | 0.65460 | 1.29924 | 72 |
| HP_RESPIRATORY_FAILURE | 0.00305 | 0.03608 | 0.43171 | 0.59551 | 1.20121 | 119 |
| HP_RESPIRATORY_INSUFFICIENCY | 0.00002 | 0.00064 | 0.57561 | 0.58138 | 1.19274 | 333 |
| HP_SENSORY_NEUROPATHY | 0.00276 | 0.03396 | 0.43171 | 0.59683 | 1.20387 | 119 |
| HP_TACHYCARDIA | 0.00132 | 0.01995 | 0.45506 | 0.65830 | 1.29422 | 59 |
| HP_TETRAPARESIS | 0.00065 | 0.01147 | 0.47727 | 0.77485 | 1.46014 | 24 |
| HP_TETRAPLEGIA_TETRAPARESIS | 0.00003 | 0.00113 | 0.55733 | 0.73573 | 1.42890 | 45 |
| HP_THICK_LOWER_LIP_VERMILION | 0.00487 | 0.04968 | 0.40702 | 0.59865 | 1.20719 | 114 |
| HP_TRIGONOCEPHALY | 0.00302 | 0.03584 | 0.43171 | 0.71976 | 1.36433 | 27 |
| HP_VERTIGO | 0.00064 | 0.01137 | 0.47727 | 0.66691 | 1.31142 | 60 |
| HP_WEAKNESS_DUE_TO_UPPER_MOTOR_NEURON_DYSFU | 0.00026 | 0.00568 | 0.49849 | 0.57317 | 1.17314 | 279 |
| HP_WEIGHT_LOSS | 0.00339 | 0.03859 | 0.43171 | 0.57539 | 1.16894 | 172 |
| **Cluster 5** | | | | | | |
| pathway | pval | padj | log2err | ES | NES | size |
| GO_ACTIN_BINDING | 0.00098 | 0.01672 | 0.45506 | 0.58073 | 1.15592 | 266 |
| GO_ACTIN_CYTOSKELETON | 0.00146 | 0.02289 | 0.45506 | 0.57264 | 1.14346 | 318 |
| GO_ACTIN_FILAMENT_BASED_PROCESS | 0.00187 | 0.02717 | 0.45506 | 0.55611 | 1.11601 | 468 |
| GO_ACTIN_FILAMENT_BUNDLE_ORGANIZATION | 0.00272 | 0.03487 | 0.43171 | 0.63005 | 1.22648 | 97 |
| GO_ACTIN_FILAMENT_ORGANIZATION | 0.00062 | 0.01162 | 0.47727 | 0.58660 | 1.16724 | 262 |
| GO_ACTIVATION_OF_IMMUNE_RESPONSE | 0.00010 | 0.00291 | 0.53843 | 0.60496 | 1.20176 | 228 |
| GO_ACTIVATION_OF_INNATE_IMMUNE_RESPONSE | 0.00009 | 0.00255 | 0.53843 | 0.66288 | 1.29084 | 100 |
| GO_ACTIVE_ION_TRANSMEMBRANE_TRANSPORTER_ACTIV | 0.00007 | 0.00216 | 0.53843 | 0.66015 | 1.28918 | 108 |
| GO_ACTIVE_TRANSMEMBRANE_TRANSPORTER_ACTIVITY | 0.00008 | 0.00248 | 0.53843 | 0.62527 | 1.23284 | 167 |
| GO_ACUTE_INFLAMMATORY_RESPONSE | 0.00092 | 0.01581 | 0.47727 | 0.74950 | 1.39018 | 31 |
| GO_ADP_BINDING | 0.00287 | 0.03638 | 0.43171 | 0.75690 | 1.39058 | 26 |
| GO_AEROBIC_RESPIRATION | 0.00002 | 0.00091 | 0.57561 | 0.71308 | 1.37072 | 64 |
| GO_ALCOHOL_BIOSYNTHETIC_PROCESS | 0.00001 | 0.00058 | 0.59333 | 0.67858 | 1.32141 | 100 |
| GO_ALCOHOL_METABOLIC_PROCESS | 0.00003 | 0.00105 | 0.57561 | 0.62551 | 1.24068 | 209 |
| GO_AMIDE_BINDING | 0.00000 | 0.00022 | 0.61053 | 0.64316 | 1.27114 | 178 |
| GO_AMINE_METABOLIC_PROCESS | 0.00000 | 0.00005 | 0.65944 | 0.72043 | 1.39762 | 85 |
| GO_ANAPHASE_PROMOTING_COMPLEX_DEPENDENT_CATA | 0.00000 | 0.00004 | 0.65944 | 0.74014 | 1.43038 | 74 |
| GO_ANATOMICAL_STRUCTURE_HOMEOSTASIS | 0.00294 | 0.03701 | 0.43171 | 0.57782 | 1.14973 | 260 |
| GO_ANGIOGENESIS_INVOLVED_IN_WOUND_HEALING | 0.00302 | 0.03762 | 0.43171 | 0.79691 | 1.42897 | 16 |
| GO_ANTIGEN_BINDING | 0.00231 | 0.03141 | 0.43171 | 0.81795 | 1.45992 | 15 |
| GO_ANTIGEN_PROCESSING_AND_PRESENTATION | 0.00000 | 0.00013 | 0.62726 | 0.65441 | 1.28647 | 150 |
| GO_ANTIGEN_PROCESSING_AND_PRESENTATION_OF_EXOG | 0.00002 | 0.00081 | 0.57561 | 0.72106 | 1.38396 | 60 |
| GO_ANTIGEN_PROCESSING_AND_PRESENTATION_OF_PEPT | 0.00001 | 0.00045 | 0.59333 | 0.65929 | 1.29170 | 133 |
| GO_ANTIGEN_PROCESSING_AND_PRESENTATION_OF_PEPT | 0.00005 | 0.00165 | 0.55733 | 0.69624 | 1.34579 | 73 |
| GO_ANTIGEN_RECEPTOR_MEDIATED_SIGNALING_PATHWAY | 0.00364 | 0.04221 | 0.43171 | 0.60904 | 1.19306 | 131 |
| GO_ANTIMICROBIAL_HUMORAL_RESPONSE | 0.00065 | 0.01212 | 0.47727 | 0.78323 | 1.43896 | 26 |
| GO_ANTIOXIDANT_ACTIVITY | 0.00390 | 0.04461 | 0.43171 | 0.68673 | 1.29325 | 40 |
| GO_APICAL_JUNCTION_COMPLEX | 0.00272 | 0.03487 | 0.43171 | 0.63279 | 1.23124 | 95 |
| GO_APICAL_PART_OF_CELL | 0.00000 | 0.00005 | 0.65944 | 0.64577 | 1.27963 | 201 |
| GO_APICAL_PLASMA_MEMBRANE | 0.00000 | 0.00000 | 0.73376 | 0.67791 | 1.33600 | 164 |
| GO_ARACHIDONIC_ACID_METABOLIC_PROCESS | 0.00324 | 0.03973 | 0.43171 | 0.79737 | 1.43879 | 18 |
| GO_ATP_BIOSYNTHETIC_PROCESS | 0.00043 | 0.00903 | 0.49849 | 0.80907 | 1.47158 | 21 |
| GO_ATP_METABOLIC_PROCESS | 0.00000 | 0.00000 | 0.81404 | 0.68476 | 1.35765 | 207 |
| GO_ATP_SYNTHESIS_COUPLED_ELECTRON_TRANSPORT | 0.00000 | 0.00000 | 0.77494 | 0.76490 | 1.47853 | 76 |
| GO_AZUROPHIL_GRANULE | 0.00048 | 0.00994 | 0.49849 | 0.64170 | 1.25039 | 101 |
| GO_AZUROPHIL_GRANULE_LUMEN | 0.00187 | 0.02717 | 0.45506 | 0.67484 | 1.29283 | 56 |
| GO_BASAL_PART_OF_CELL | 0.00076 | 0.01381 | 0.47727 | 0.73778 | 1.37260 | 33 |
| GO_BASAL_PLASMA_MEMBRANE | 0.00051 | 0.01019 | 0.47727 | 0.77808 | 1.43432 | 27 |
| GO_BASOLATERAL_PLASMA_MEMBRANE | 0.00000 | 0.00001 | 0.69013 | 0.69679 | 1.36183 | 121 |
| GO_BINDING_OF_SPERM_TO_ZONA_PELLUCIDA | 0.00202 | 0.02829 | 0.43171 | 0.80892 | 1.45962 | 18 |
| GO_BLOOD_MICROPARTICLE | 0.00032 | 0.00724 | 0.49849 | 0.74750 | 1.39964 | 36 |
| GO_BRUSH_BORDER | 0.00000 | 0.00001 | 0.70498 | 0.76471 | 1.46996 | 64 |
| GO_BRUSH_BORDER_MEMBRANE | 0.00002 | 0.00071 | 0.57561 | 0.81044 | 1.50322 | 31 |
| GO_CADHERIN_BINDING | 0.00000 | 0.00000 | | 0.68042 | 1.35428 | 265 |
| GO_CALCIUM_DEPENDENT_PROTEIN_BINDING | 0.00272 | 0.03487 | 0.43171 | 0.69287 | 1.30481 | 40 |

| Cluster 1 | | | | | | |
|---|---|---|---|---|---|---|
| GO_CALCIUM_ION_BINDING | 0.00000 | 0.00018 | 0.62726 | 0.61778 | 1.22968 | 266 |
| GO_CARBOHYDRATE_BIOSYNTHETIC_PROCESS | 0.00265 | 0.03452 | 0.43171 | 0.61238 | 1.19797 | 125 |
| GO_CARDIAC_CONDUCTION | 0.00435 | 0.04823 | 0.40702 | 0.66286 | 1.27202 | 59 |
| GO_CATALYTIC_STEP_2_SPLICEOSOME | 0.00047 | 0.00979 | 0.49849 | 0.66313 | 1.28428 | 81 |
| GO_CATION_TRANSMEMBRANE_TRANSPORT | 0.00000 | 0.00010 | 0.64355 | 0.60027 | 1.20172 | 379 |
| GO_CATION_TRANSMEMBRANE_TRANSPORTER_ACTIVITY | 0.00000 | 0.00022 | 0.61053 | 0.62162 | 1.23576 | 250 |
| GO_CATION_TRANSPORT | 0.00000 | 0.00002 | 0.69013 | 0.59363 | 1.19219 | 497 |
| GO_CAVEOLA | 0.00413 | 0.04636 | 0.40702 | 0.68564 | 1.29120 | 40 |
| GO_CELLULAR_COMPONENT_DISASSEMBLY | 0.00000 | 0.00000 | 0.74774 | 0.61830 | 1.23976 | 410 |
| GO_CELLULAR_HOMEOSTASIS | 0.00036 | 0.00790 | 0.49849 | 0.56639 | 1.13635 | 462 |
| GO_CELLULAR_ION_HOMEOSTASIS | 0.00074 | 0.01339 | 0.47727 | 0.58140 | 1.16020 | 295 |
| GO_CELLULAR_KETONE_METABOLIC_PROCESS | 0.00018 | 0.00464 | 0.51885 | 0.62681 | 1.23326 | 154 |
| GO_CELLULAR_NITROGEN_COMPOUND_CATABOLIC_PROCE | 0.00000 | 0.00000 | | 0.67534 | 1.35457 | 422 |
| GO_CELLULAR_PROTEIN_COMPLEX_DISASSEMBLY | 0.00000 | 0.00002 | 0.67496 | 0.66164 | 1.30624 | 173 |
| GO_CELLULAR_RESPIRATION | 0.00000 | 0.00000 | 0.77494 | 0.70729 | 1.38751 | 137 |
| GO_CELLULAR_RESPONSE_TO_ALCOHOL | 0.00165 | 0.02486 | 0.45506 | 0.70111 | 1.32490 | 43 |
| GO_CELLULAR_RESPONSE_TO_ORGANIC_CYCLIC_COMPOUN | 0.00023 | 0.00568 | 0.51885 | 0.58836 | 1.17432 | 298 |
| GO_CELLULAR_RESPONSE_TO_OXYGEN_LEVELS | 0.00000 | 0.00002 | 0.69013 | 0.67056 | 1.31824 | 149 |
| GO_CELL_ACTIVATION_INVOLVED_IN_IMMUNE_RESPONSE | 0.00000 | 0.00000 | 0.80122 | 0.62435 | 1.25082 | 397 |
| GO_CELL_ADHESION_MOLECULE_BINDING | 0.00000 | 0.00000 | | 0.66861 | 1.33722 | 350 |
| GO_CELL_BODY | 0.00133 | 0.02144 | 0.45506 | 0.57363 | 1.14442 | 293 |
| GO_CELL_CELL_JUNCTION | 0.00007 | 0.00224 | 0.53843 | 0.59666 | 1.19054 | 294 |
| GO_CELL_CELL_JUNCTION_ASSEMBLY | 0.00163 | 0.02465 | 0.45506 | 0.65527 | 1.27184 | 86 |
| GO_CELL_CELL_JUNCTION_ORGANIZATION | 0.00250 | 0.03324 | 0.43171 | 0.61225 | 1.19719 | 124 |
| GO_CELL_CELL_RECOGNITION | 0.00002 | 0.00097 | 0.57561 | 0.79618 | 1.47989 | 32 |
| GO_CELL_CELL_SIGNALING_BY_WNT | 0.00157 | 0.02406 | 0.45506 | 0.56954 | 1.13747 | 321 |
| GO_CELL_COMMUNICATION_BY_ELECTRICAL_COUPLING | 0.00428 | 0.04758 | 0.40702 | 0.80311 | 1.43345 | 15 |
| GO_CELL_CORTEX | 0.00027 | 0.00625 | 0.49849 | 0.60705 | 1.20164 | 193 |
| GO_CELL_CYCLE_G2_M_PHASE_TRANSITION | 0.00141 | 0.02244 | 0.45506 | 0.58941 | 1.16899 | 210 |
| GO_CELL_CYCLE_PHASE_TRANSITION | 0.00209 | 0.02892 | 0.43171 | 0.55283 | 1.10909 | 463 |
| GO_CELL_JUNCTION_ORGANIZATION | 0.00209 | 0.02892 | 0.43171 | 0.56172 | 1.12443 | 373 |
| GO_CELL_LEADING_EDGE | 0.00025 | 0.00605 | 0.49849 | 0.59189 | 1.17815 | 266 |
| GO_CELL_PROJECTION_MEMBRANE | 0.00000 | 0.00002 | 0.67496 | 0.65654 | 1.29810 | 184 |
| GO_CELL_RECOGNITION | 0.00015 | 0.00399 | 0.51885 | 0.70550 | 1.35385 | 59 |
| GO_CELL_SUBSTRATE_ADHESION | 0.00420 | 0.04693 | 0.40702 | 0.58162 | 1.15251 | 199 |
| GO_CELL_SUBSTRATE_JUNCTION | 0.00000 | 0.00000 | | 0.67922 | 1.35585 | 309 |
| GO_CELL_SURFACE | 0.00001 | 0.00036 | 0.61053 | 0.60846 | 1.21344 | 288 |
| GO_CELL_SURFACE_RECEPTOR_SIGNALING_PATHWAY_INVC | 0.00265 | 0.03452 | 0.43171 | 0.56238 | 1.12488 | 356 |
| GO_CHAPERONE_BINDING | 0.00042 | 0.00887 | 0.49849 | 0.67394 | 1.30237 | 75 |
| GO_CHAPERONE_COMPLEX | 0.00239 | 0.03212 | 0.43171 | 0.77563 | 1.41076 | 21 |
| GO_CLUSTER_OF_ACTIN_BASED_CELL_PROJECTIONS | 0.00000 | 0.00001 | 0.70498 | 0.73659 | 1.43035 | 88 |
| GO_COLLAGEN_CONTAINING_EXTRACELLULAR_MATRIX | 0.00026 | 0.00613 | 0.49849 | 0.62978 | 1.23983 | 156 |
| GO_CONTRACTILE_FIBER | 0.00000 | 0.00002 | 0.67496 | 0.68373 | 1.33923 | 132 |
| GO_CORNIFICATION | 0.00420 | 0.04693 | 0.40702 | 0.72231 | 1.33547 | 29 |
| GO_CORTICAL_ACTIN_CYTOSKELETON | 0.00181 | 0.02682 | 0.45506 | 0.68114 | 1.30638 | 58 |
| GO_CORTICAL_CYTOSKELETON | 0.00008 | 0.00237 | 0.53843 | 0.69674 | 1.34619 | 72 |
| GO_COTRANSLATIONAL_PROTEIN_TARGETING_TO_MEMBR/ | 0.00000 | 0.00000 | | 0.82613 | 1.60628 | 92 |
| GO_CYTOCHROME_COMPLEX | 0.00041 | 0.00877 | 0.49849 | 0.81031 | 1.47383 | 21 |
| GO_CYTOKINE_MEDIATED_SIGNALING_PATHWAY | 0.00000 | 0.00001 | 0.70498 | 0.60846 | 1.21777 | 375 |
| GO_CYTOPLASMIC_STRESS_GRANULE | 0.00364 | 0.04221 | 0.43171 | 0.67009 | 1.28517 | 58 |
| GO_CYTOPLASMIC_TRANSLATION | 0.00000 | 0.00002 | 0.67496 | 0.72705 | 1.41182 | 88 |
| GO_CYTOPLASMIC_TRANSLATIONAL_INITIATION | 0.00353 | 0.04181 | 0.43171 | 0.73050 | 1.35309 | 30 |
| GO_CYTOSOLIC_LARGE_RIBOSOMAL_SUBUNIT | 0.00000 | 0.00000 | 0.80122 | 0.83213 | 1.58560 | 51 |
| GO_CYTOSOLIC_RIBOSOME | 0.00000 | 0.00000 | | 0.83278 | 1.62078 | 96 |
| GO_CYTOSOLIC_SMALL_RIBOSOMAL_SUBUNIT | 0.00000 | 0.00000 | | 0.87444 | 1.65246 | 43 |
| GO_DEFENSE_RESPONSE_TO_GRAM_NEGATIVE_BACTERIUI | 0.00398 | 0.04503 | 0.40702 | 0.80474 | 1.43636 | 15 |
| GO_DEFENSE_RESPONSE_TO_OTHER_ORGANISM | 0.00021 | 0.00518 | 0.51885 | 0.56578 | 1.13515 | 464 |
| GO_DISORDERED_DOMAIN_SPECIFIC_BINDING | 0.00457 | 0.04987 | 0.40702 | 0.74369 | 1.36631 | 26 |
| GO_ELECTRON_TRANSFER_ACTIVITY | 0.00000 | 0.00014 | 0.62726 | 0.72690 | 1.40469 | 75 |
| GO_ELECTRON_TRANSPORT_CHAIN | 0.00000 | 0.00000 | 0.78819 | 0.72262 | 1.41455 | 127 |
| GO_ENDOCYTIC_VESICLE | 0.00037 | 0.00804 | 0.49849 | 0.61404 | 1.21357 | 180 |
| GO_ENDOCYTIC_VESICLE_MEMBRANE | 0.00239 | 0.03212 | 0.43171 | 0.64195 | 1.24621 | 87 |
| GO_ENDOLYSOSOME | 0.00087 | 0.01528 | 0.47727 | 0.82554 | 1.48032 | 16 |
| GO_ENDOPEPTIDASE_COMPLEX | 0.00016 | 0.00411 | 0.51885 | 0.70138 | 1.34866 | 66 |
| GO_ENDOPEPTIDASE_REGULATOR_ACTIVITY | 0.00013 | 0.00358 | 0.51885 | 0.69852 | 1.34671 | 68 |
| GO_ENERGY_DERIVATION_BY_OXIDATION_OF_ORGANIC_C( | 0.00000 | 0.00000 | 0.76146 | 0.67493 | 1.33489 | 185 |
| GO_ENTRY_INTO_HOST | 0.00085 | 0.01502 | 0.47727 | 0.66404 | 1.28886 | 86 |
| GO_ENZYME_INHIBITOR_ACTIVITY | 0.00069 | 0.01272 | 0.47727 | 0.60434 | 1.19536 | 186 |
| GO_EPITHELIAL_CELL_DIFFERENTIATION | 0.00364 | 0.04221 | 0.43171 | 0.56567 | 1.13012 | 332 |
| GO_EPITHELIAL_TO_MESENCHYMAL_TRANSITION | 0.00350 | 0.04154 | 0.43171 | 0.64064 | 1.23962 | 79 |
| GO_ESTABLISHMENT_OF_PROTEIN_LOCALIZATION_TO_END | 0.00000 | 0.00000 | | 0.81090 | 1.58145 | 102 |
| GO_ESTABLISHMENT_OF_PROTEIN_LOCALIZATION_TO_MEN | 0.00000 | 0.00000 | | 0.69801 | 1.38773 | 243 |
| GO_ESTABLISHMENT_OF_PROTEIN_LOCALIZATION_TO_MIT( | 0.00178 | 0.02647 | 0.45506 | 0.71430 | 1.34033 | 38 |

| Cluster 1 | | | | | | |
|---|---|---|---|---|---|---|
| GO_ESTABLISHMENT_OF_PROTEIN_LOCALIZATION_TO_ORG | 0.00000 | 0.00000 | | 0.67644 | 1.35660 | 420 |
| GO_ESTABLISHMENT_OF_TISSUE_POLARITY | 0.00000 | 0.00021 | 0.61053 | 0.69762 | 1.35738 | 95 |
| GO_EUKARYOTIC_TRANSLATION_INITIATION_FACTOR_3_CO | 0.00305 | 0.03792 | 0.43171 | 0.81303 | 1.45115 | 15 |
| GO_EXTRACELLULAR_MATRIX | 0.00049 | 0.01006 | 0.47727 | 0.60106 | 1.19056 | 197 |
| GO_FATTY_ACID_DERIVATIVE_METABOLIC_PROCESS | 0.00006 | 0.00198 | 0.55733 | 0.70059 | 1.35113 | 69 |
| GO_FC_EPSILON_RECEPTOR_SIGNALING_PATHWAY | 0.00037 | 0.00804 | 0.49849 | 0.66022 | 1.28314 | 91 |
| GO_FC_RECEPTOR_SIGNALING_PATHWAY | 0.00008 | 0.00237 | 0.53843 | 0.64197 | 1.25955 | 138 |
| GO_FERTILIZATION | 0.00027 | 0.00625 | 0.49849 | 0.67947 | 1.31305 | 75 |
| GO_FIBROBLAST_GROWTH_FACTOR_RECEPTOR_SIGNALING | 0.00044 | 0.00919 | 0.49849 | 0.68696 | 1.31851 | 60 |
| GO_FICOLIN_1_RICH_GRANULE | 0.00000 | 0.00010 | 0.64355 | 0.70229 | 1.36765 | 98 |
| GO_GENERATION_OF_PRECURSOR_METABOLITES_AND_EN | 0.00000 | 0.00000 | | 0.65086 | 1.30023 | 335 |
| GO_GLUCOSE_CATABOLIC_PROCESS | 0.00051 | 0.01019 | 0.47727 | 0.78160 | 1.43255 | 24 |
| GO_GLYCOLYTIC_PROCESS_THROUGH_FRUCTOSE_6_PHOSP | 0.00021 | 0.00521 | 0.51885 | 0.81587 | 1.47817 | 20 |
| GO_GTPASE_ACTIVITY | 0.00089 | 0.01551 | 0.47727 | 0.60253 | 1.19111 | 181 |
| GO_GUANYL_NUCLEOTIDE_BINDING | 0.00257 | 0.03385 | 0.43171 | 0.58417 | 1.15899 | 217 |
| GO_HEMATOPOIETIC_PROGENITOR_CELL_DIFFERENTIATION | 0.00019 | 0.00468 | 0.51885 | 0.64352 | 1.25713 | 116 |
| GO_HEMATOPOIETIC_STEM_CELL_DIFFERENTIATION | 0.00003 | 0.00114 | 0.57561 | 0.70006 | 1.35123 | 71 |
| GO_HOMOTYPIC_CELL_CELL_ADHESION | 0.00117 | 0.01903 | 0.45506 | 0.70582 | 1.34261 | 49 |
| GO_HUMORAL_IMMUNE_RESPONSE | 0.00088 | 0.01540 | 0.47727 | 0.67376 | 1.29498 | 63 |
| GO_ICOSANOID_METABOLIC_PROCESS | 0.00055 | 0.01079 | 0.47727 | 0.72225 | 1.36014 | 40 |
| GO_IMMUNE_RESPONSE_REGULATING_SIGNALING_PATHW | 0.00086 | 0.01510 | 0.47727 | 0.59793 | 1.18627 | 216 |
| GO_IMPORT_INTO_CELL | 0.00045 | 0.00941 | 0.49849 | 0.65255 | 1.27028 | 97 |
| GO_INNATE_IMMUNE_RESPONSE | 0.00198 | 0.02803 | 0.43171 | 0.56191 | 1.12469 | 374 |
| GO_INNATE_IMMUNE_RESPONSE_ACTIVATING_SIGNAL_TR/ | 0.00030 | 0.00702 | 0.49849 | 0.67029 | 1.29950 | 83 |
| GO_INNER_MITOCHONDRIAL_MEMBRANE_PROTEIN_COMP | 0.00000 | 0.00000 | 0.81404 | 0.76944 | 1.49476 | 90 |
| GO_INORGANIC_ION_TRANSMEMBRANE_TRANSPORT | 0.00000 | 0.00006 | 0.65944 | 0.60430 | 1.20899 | 362 |
| GO_INTERACTION_WITH_HOST | 0.00027 | 0.00625 | 0.49849 | 0.62608 | 1.22817 | 139 |
| GO_INTERACTION_WITH_SYMBIONT | 0.00242 | 0.03254 | 0.43171 | 0.69368 | 1.31952 | 49 |
| GO_INTERLEUKIN_1_MEDIATED_SIGNALING_PATHWAY | 0.00000 | 0.00018 | 0.62726 | 0.71350 | 1.38183 | 81 |
| GO_INTERLEUKIN_2_PRODUCTION | 0.00013 | 0.00372 | 0.51885 | 0.81158 | 1.49102 | 26 |
| GO_INTERLEUKIN_4_PRODUCTION | 0.00246 | 0.03289 | 0.43171 | 0.80157 | 1.43733 | 16 |
| GO_INTRAMOLECULAR_OXIDOREDUCTASE_ACTIVITY | 0.00205 | 0.02860 | 0.43171 | 0.74027 | 1.37119 | 30 |
| GO_ION_CHANNEL_BINDING | 0.00268 | 0.03470 | 0.43171 | 0.64854 | 1.25328 | 75 |
| GO_ION_TRANSMEMBRANE_TRANSPORT | 0.00000 | 0.00008 | 0.64355 | 0.58911 | 1.18327 | 498 |
| GO_ION_TRANSMEMBRANE_TRANSPORTER_ACTIVITY | 0.00004 | 0.00146 | 0.55733 | 0.58917 | 1.17848 | 355 |
| GO_KILLING_OF_CELLS_OF_OTHER_ORGANISM | 0.00335 | 0.04034 | 0.43171 | 0.81021 | 1.44611 | 15 |
| GO_LARGE_RIBOSOMAL_SUBUNIT | 0.00000 | 0.00000 | | 0.76393 | 1.49061 | 105 |
| GO_LATERAL_PLASMA_MEMBRANE | 0.00018 | 0.00468 | 0.51885 | 0.73710 | 1.39112 | 42 |
| GO_LEUKOCYTE_MEDIATED_IMMUNITY | 0.00000 | 0.00000 | 0.80122 | 0.62273 | 1.24904 | 422 |
| GO_LEUKOCYTE_MIGRATION | 0.00064 | 0.01185 | 0.47727 | 0.60547 | 1.19824 | 191 |
| GO_LOW_DENSITY_LIPOPROTEIN_PARTICLE_CLEARANCE | 0.00106 | 0.01767 | 0.45506 | 0.77550 | 1.42475 | 26 |
| GO_LYSOSOMAL_LUMEN | 0.00272 | 0.03487 | 0.43171 | 0.66871 | 1.27968 | 55 |
| GO_MACROAUTOPHAGY | 0.00353 | 0.04181 | 0.43171 | 0.57584 | 1.14455 | 246 |
| GO_MAINTENANCE_OF_LOCATION | 0.00450 | 0.04951 | 0.40702 | 0.59201 | 1.17002 | 180 |
| GO_MAINTENANCE_OF_LOCATION_IN_CELL | 0.00205 | 0.02860 | 0.43171 | 0.61872 | 1.20868 | 116 |
| GO_MAIN_AXON | 0.00257 | 0.03385 | 0.43171 | 0.73457 | 1.35813 | 29 |
| GO_MEMBRANE_BIOGENESIS | 0.00054 | 0.01071 | 0.47727 | 0.79245 | 1.44586 | 22 |
| GO_MEMBRANE_REGION | 0.00000 | 0.00004 | 0.65944 | 0.65203 | 1.28918 | 184 |
| GO_MEMBRANE_REPOLARIZATION | 0.00268 | 0.03470 | 0.43171 | 0.79972 | 1.43402 | 16 |
| GO_MESENCHYMAL_CELL_DIFFERENTIATION | 0.00068 | 0.01259 | 0.47727 | 0.64132 | 1.25176 | 107 |
| GO_MESENCHYME_DEVELOPMENT | 0.00157 | 0.02406 | 0.45506 | 0.61635 | 1.20861 | 136 |
| GO_MESONEPHROS_DEVELOPMENT | 0.00390 | 0.04461 | 0.43171 | 0.66370 | 1.26462 | 50 |
| GO_METAL_ION_TRANSPORT | 0.00015 | 0.00399 | 0.51885 | 0.57759 | 1.15633 | 384 |
| GO_MICROVILLUS | 0.00290 | 0.03669 | 0.43171 | 0.69389 | 1.31485 | 46 |
| GO_MICROVILLUS_MEMBRANE | 0.00298 | 0.03731 | 0.43171 | 0.81367 | 1.45228 | 15 |
| GO_MIDBRAIN_DEVELOPMENT | 0.00194 | 0.02784 | 0.45506 | 0.67169 | 1.28383 | 53 |
| GO_MITOCHONDRIAL_ELECTRON_TRANSPORT_NADH_TO_U | 0.00002 | 0.00087 | 0.57561 | 0.76085 | 1.43955 | 44 |
| GO_MITOCHONDRIAL_GENE_EXPRESSION | 0.00000 | 0.00019 | 0.62726 | 0.65439 | 1.28608 | 148 |
| GO_MITOCHONDRIAL_LARGE_RIBOSOMAL_SUBUNIT | 0.00007 | 0.00229 | 0.53843 | 0.72013 | 1.37643 | 53 |
| GO_MITOCHONDRIAL_MATRIX | 0.00000 | 0.00007 | 0.64355 | 0.60843 | 1.21699 | 354 |
| GO_MITOCHONDRIAL_MEMBRANE_ORGANIZATION | 0.00051 | 0.01019 | 0.47727 | 0.65736 | 1.27758 | 91 |
| GO_MITOCHONDRIAL_PROTEIN_COMPLEX | 0.00000 | 0.00000 | | 0.71369 | 1.41446 | 204 |
| GO_MITOCHONDRIAL_RESPIRATORY_CHAIN_COMPLEX_ASS | 0.00087 | 0.01517 | 0.47727 | 0.65565 | 1.27153 | 84 |
| GO_MITOCHONDRIAL_TRANSLATION | 0.00000 | 0.00016 | 0.62726 | 0.67147 | 1.31513 | 128 |
| GO_MITOCHONDRIAL_TRANSLATIONAL_TERMINATION | 0.00000 | 0.00022 | 0.61053 | 0.70797 | 1.37477 | 88 |
| GO_MITOCHONDRIAL_TRANSPORT | 0.00051 | 0.01019 | 0.47727 | 0.60134 | 1.18974 | 189 |
| GO_MITOCHONDRION_ORGANIZATION | 0.00000 | 0.00000 | 0.71951 | 0.60926 | 1.22057 | 397 |
| GO_MODIFIED_AMINO_ACID_BINDING | 0.00110 | 0.01818 | 0.45506 | 0.71157 | 1.34295 | 42 |
| GO_MONOCARBOXYLIC_ACID_METABOLIC_PROCESS | 0.00019 | 0.00468 | 0.51885 | 0.58193 | 1.16284 | 340 |
| GO_MONOSACCHARIDE_BIOSYNTHETIC_PROCESS | 0.00050 | 0.01019 | 0.47727 | 0.69040 | 1.32607 | 62 |
| GO_MONOVALENT_INORGANIC_CATION_TRANSMEMBRAN | 0.00000 | 0.00000 | 0.78819 | 0.70691 | 1.38673 | 139 |
| GO_MONOVALENT_INORGANIC_CATION_TRANSPORT | 0.00000 | 0.00001 | 0.70498 | 0.65322 | 1.29750 | 226 |
| GO_MORPHOGENESIS_OF_AN_EPITHELIUM | 0.00000 | 0.00001 | 0.69013 | 0.62121 | 1.24059 | 324 |

| Cluster 1 | | | | | | |
|---|---|---|---|---|---|---|
| GO_MORPHOGENESIS_OF_A_POLARIZED_EPITHELIUM | 0.00000 | 0.00005 | 0.65944 | 0.69972 | 1.36647 | 111 |
| GO_MOVEMENT_IN_HOST_ENVIRONMENT | 0.00013 | 0.00355 | 0.51885 | 0.65985 | 1.28787 | 106 |
| GO_MRNA_BINDING | 0.00000 | 0.00001 | 0.70498 | 0.64429 | 1.28070 | 237 |
| GO_MRNA_PROCESSING | 0.00000 | 0.00001 | 0.71951 | 0.60622 | 1.21571 | 419 |
| GO_MUSCLE_CONTRACTION | 0.00117 | 0.01903 | 0.45506 | 0.60952 | 1.20075 | 159 |
| GO_MYD88_DEPENDENT_TOLL_LIKE_RECEPTOR_SIGNALING | 0.00150 | 0.02329 | 0.45506 | 0.81411 | 1.46899 | 18 |
| GO_MYELIN_SHEATH | 0.00450 | 0.04951 | 0.40702 | 0.71183 | 1.32311 | 32 |
| GO_MYELOID_LEUKOCYTE_ACTIVATION | 0.00000 | 0.00000 | 0.82666 | 0.63326 | 1.26749 | 370 |
| GO_MYELOID_LEUKOCYTE_MEDIATED_IMMUNITY | 0.00000 | 0.00000 | 0.81404 | 0.64042 | 1.27969 | 338 |
| GO_M_BAND | 0.00023 | 0.00568 | 0.51885 | 0.86146 | 1.53758 | 15 |
| GO_NADH_DEHYDROGENASE_ACTIVITY | 0.00006 | 0.00190 | 0.55733 | 0.77300 | 1.44738 | 36 |
| GO_NADH_DEHYDROGENASE_COMPLEX | 0.00003 | 0.00099 | 0.57561 | 0.77561 | 1.45736 | 39 |
| GO_NADH_DEHYDROGENASE_COMPLEX_ASSEMBLY | 0.00007 | 0.00216 | 0.53843 | 0.72865 | 1.39276 | 54 |
| GO_NADH_METABOLIC_PROCESS | 0.00001 | 0.00048 | 0.59333 | 0.83187 | 1.52469 | 24 |
| GO_NAD_METABOLIC_PROCESS | 0.00054 | 0.01060 | 0.47727 | 0.76265 | 1.41265 | 30 |
| GO_NCRNA_PROCESSING | 0.00405 | 0.04578 | 0.40702 | 0.56501 | 1.12836 | 324 |
| GO_NEGATIVE_REGULATION_OF_CALCIUM_ION_TRANSPOR | 0.00457 | 0.04987 | 0.40702 | 0.73835 | 1.35328 | 24 |
| GO_NEGATIVE_REGULATION_OF_CANONICAL_WNT_SIGNAI | 0.00018 | 0.00462 | 0.51885 | 0.64557 | 1.26137 | 118 |
| GO_NEGATIVE_REGULATION_OF_CELLULAR_COMPONENT_( | 0.00327 | 0.03985 | 0.43171 | 0.55675 | 1.11665 | 446 |
| GO_NEGATIVE_REGULATION_OF_CELL_CYCLE_G2_M_PHASI | 0.00027 | 0.00625 | 0.49849 | 0.66592 | 1.29273 | 87 |
| GO_NEGATIVE_REGULATION_OF_CELL_CYCLE_PROCESS | 0.00198 | 0.02803 | 0.43171 | 0.58013 | 1.15327 | 250 |
| GO_NEGATIVE_REGULATION_OF_PEPTIDASE_ACTIVITY | 0.00283 | 0.03614 | 0.43171 | 0.62629 | 1.22141 | 103 |
| GO_NEGATIVE_REGULATION_OF_PROTEOLYSIS | 0.00327 | 0.03985 | 0.43171 | 0.59651 | 1.17604 | 166 |
| GO_NEGATIVE_REGULATION_OF_WNT_SIGNALING_PATHW/ | 0.00033 | 0.00741 | 0.49849 | 0.63317 | 1.24020 | 132 |
| GO_NEURAL_CREST_CELL_DIFFERENTIATION | 0.00156 | 0.02396 | 0.45506 | 0.71798 | 1.34722 | 38 |
| GO_NEURAL_CREST_CELL_MIGRATION | 0.00346 | 0.04126 | 0.43171 | 0.75332 | 1.38257 | 25 |
| GO_NEURAL_NUCLEUS_DEVELOPMENT | 0.00168 | 0.02529 | 0.45506 | 0.72321 | 1.34550 | 33 |
| GO_NIK_NF_KAPPAB_SIGNALING | 0.00002 | 0.00067 | 0.57561 | 0.66985 | 1.30814 | 113 |
| GO_NON_CANONICAL_WNT_SIGNALING_PATHWAY | 0.00003 | 0.00112 | 0.57561 | 0.67311 | 1.31300 | 104 |
| GO_NUCLEAR_PERIPHERY | 0.00265 | 0.03452 | 0.43171 | 0.61825 | 1.20723 | 110 |
| GO_NUCLEAR_TRANSCRIBED_MRNA_CATABOLIC_PROCESS | 0.00000 | 0.00000 | | 0.71081 | 1.40485 | 178 |
| GO_NUCLEAR_TRANSCRIBED_MRNA_CATABOLIC_PROCESS_ | 0.00000 | 0.00000 | | 0.79512 | 1.55272 | 112 |
| GO_NUCLEOBASE_CONTAINING_SMALL_MOLECULE_METAE | 0.00091 | 0.01574 | 0.47727 | 0.57271 | 1.14514 | 344 |
| GO_NUCLEOSIDE_BISPHOSPHATE_METABOLIC_PROCESS | 0.00257 | 0.03385 | 0.43171 | 0.64735 | 1.25216 | 77 |
| GO_NUCLEOSIDE_PHOSPHATE_BIOSYNTHETIC_PROCESS | 0.00032 | 0.00720 | 0.49849 | 0.62559 | 1.22637 | 135 |
| GO_NUCLEOSIDE_TRIPHOSPHATE_BIOSYNTHETIC_PROCESS | 0.00132 | 0.02126 | 0.45506 | 0.71711 | 1.34745 | 39 |
| GO_NUCLEOSIDE_TRIPHOSPHATE_METABOLIC_PROCESS | 0.00033 | 0.00727 | 0.49849 | 0.69868 | 1.33850 | 56 |
| GO_NUCLEOSOMAL_DNA_BINDING | 0.00413 | 0.04636 | 0.40702 | 0.75972 | 1.38182 | 21 |
| GO_ORGANELLAR_RIBOSOME | 0.00000 | 0.00019 | 0.62726 | 0.70908 | 1.37471 | 83 |
| GO_ORGANELLE_ENVELOPE_LUMEN | 0.00305 | 0.03792 | 0.43171 | 0.65855 | 1.27078 | 70 |
| GO_ORGANELLE_INNER_MEMBRANE | 0.00000 | 0.00000 | | 0.65713 | 1.31564 | 385 |
| GO_ORGANIC_CYCLIC_COMPOUND_CATABOLIC_PROCESS | 0.00000 | 0.00000 | | 0.66812 | 1.34003 | 441 |
| GO_ORGANIC_HYDROXY_COMPOUND_BIOSYNTHETIC_PRO | 0.00013 | 0.00355 | 0.51885 | 0.63870 | 1.25315 | 138 |
| GO_ORGANIC_HYDROXY_COMPOUND_METABOLIC_PROCES | 0.00038 | 0.00811 | 0.49849 | 0.59090 | 1.17826 | 280 |
| GO_OSTEOBLAST_DIFFERENTIATION | 0.00152 | 0.02351 | 0.45506 | 0.62257 | 1.21672 | 123 |
| GO_OXIDATIVE_PHOSPHORYLATION | 0.00000 | 0.00000 | 0.82666 | 0.75832 | 1.47676 | 98 |
| GO_OXIDOREDUCTASE_ACTIVITY | 0.00000 | 0.00000 | | 0.63724 | 1.27716 | 404 |
| GO_OXIDOREDUCTASE_ACTIVITY_ACTING_ON_A_HEME_GR | 0.00011 | 0.00332 | 0.53843 | 0.85735 | 1.54703 | 18 |
| GO_OXIDOREDUCTASE_ACTIVITY_ACTING_ON_CH_OH_GRO | 0.00044 | 0.00935 | 0.49849 | 0.67794 | 1.30322 | 65 |
| GO_OXIDOREDUCTASE_ACTIVITY_ACTING_ON_NAD_P_H | 0.00000 | 0.00014 | 0.62726 | 0.73083 | 1.40946 | 69 |
| GO_OXIDOREDUCTASE_ACTIVITY_ACTING_ON_NAD_P_H_O | 0.00001 | 0.00048 | 0.59333 | 0.77028 | 1.45961 | 46 |
| GO_OXIDOREDUCTASE_ACTIVITY_ACTING_ON_PEROXIDE_A | 0.00202 | 0.02829 | 0.43171 | 0.75780 | 1.38892 | 24 |
| GO_OXIDOREDUCTASE_COMPLEX | 0.00000 | 0.00002 | 0.69013 | 0.74673 | 1.44303 | 75 |
| GO_PEPTIDASE_COMPLEX | 0.00058 | 0.01118 | 0.47727 | 0.66161 | 1.28134 | 81 |
| GO_PEPTIDASE_REGULATOR_ACTIVITY | 0.00100 | 0.01694 | 0.45506 | 0.65108 | 1.26309 | 85 |
| GO_PEPTIDE_BINDING | 0.00002 | 0.00083 | 0.57561 | 0.66192 | 1.29648 | 129 |
| GO_PERINUCLEAR_REGION_OF_CYTOPLASM | 0.00235 | 0.03176 | 0.43171 | 0.55632 | 1.11607 | 452 |
| GO_PHAGOSOME_MATURATION | 0.00357 | 0.04184 | 0.43171 | 0.70894 | 1.32482 | 35 |
| GO_PIGMENT_GRANULE | 0.00000 | 0.00016 | 0.62726 | 0.71793 | 1.38774 | 76 |
| GO_PLASMA_LIPOPROTEIN_PARTICLE_CLEARANCE | 0.00005 | 0.00175 | 0.55733 | 0.77598 | 1.45008 | 35 |
| GO_PLASMA_MEMBRANE_PROTEIN_COMPLEX | 0.00025 | 0.00596 | 0.49849 | 0.61001 | 1.20864 | 202 |
| GO_PLASMA_MEMBRANE_RAFT | 0.00294 | 0.03701 | 0.43171 | 0.66423 | 1.26958 | 53 |
| GO_PLATELET_ALPHA_GRANULE | 0.00198 | 0.02803 | 0.43171 | 0.71978 | 1.34506 | 35 |
| GO_PLATELET_DEGRANULATION | 0.00008 | 0.00237 | 0.53843 | 0.71872 | 1.38037 | 61 |
| GO_POLYMERIC_CYTOSKELETAL_FIBER | 0.00342 | 0.04099 | 0.43171 | 0.55678 | 1.11590 | 404 |
| GO_POLYSOMAL_RIBOSOME | 0.00001 | 0.00048 | 0.59333 | 0.82198 | 1.51975 | 29 |
| GO_POLYSOME | 0.00000 | 0.00001 | 0.69013 | 0.77615 | 1.49066 | 61 |
| GO_POSITIVE_REGULATION_OF_CANONICAL_WNT_SIGNALI | 0.00179 | 0.02661 | 0.45506 | 0.62994 | 1.22854 | 102 |
| GO_POSITIVE_REGULATION_OF_CD4_POSITIVE_ALPHA_BET | 0.00058 | 0.01118 | 0.47727 | 0.84404 | 1.50649 | 15 |
| GO_POSITIVE_REGULATION_OF_CELLULAR_AMIDE_METABO | 0.00146 | 0.02289 | 0.45506 | 0.61961 | 1.21053 | 122 |
| GO_POSITIVE_REGULATION_OF_CELLULAR_COMPONENT_M | 0.00346 | 0.04126 | 0.43171 | 0.56464 | 1.12689 | 296 |
| GO_POSITIVE_REGULATION_OF_CELLULAR_PROTEIN_LOCAL | 0.00018 | 0.00459 | 0.51885 | 0.60039 | 1.19267 | 228 |
| GO_POSITIVE_REGULATION_OF_CELL_POPULATION_PROLIF | 0.00020 | 0.00505 | 0.51885 | 0.57111 | 1.14562 | 434 |

| Cluster 1 | | | | | | |
|---|---|---|---|---|---|---|
| GO_POSITIVE_REGULATION_OF_CELL_SUBSTRATE_ADHESIO | 0.00157 | 0.02406 | 0.45506 | 0.65642 | 1.26882 | 73 |
| GO_POSITIVE_REGULATION_OF_CYTOSKELETON_ORGANIZA | 0.00309 | 0.03830 | 0.43171 | 0.60289 | 1.18397 | 145 |
| GO_POSITIVE_REGULATION_OF_DEFENSE_RESPONSE | 0.00015 | 0.00402 | 0.51885 | 0.61708 | 1.22121 | 191 |
| GO_POSITIVE_REGULATION_OF_DNA_BIOSYNTHETIC_PROC | 0.00261 | 0.03426 | 0.43171 | 0.68546 | 1.30612 | 51 |
| GO_POSITIVE_REGULATION_OF_ESTABLISHMENT_OF_PROT | 0.00146 | 0.02289 | 0.45506 | 0.58629 | 1.16525 | 236 |
| GO_POSITIVE_REGULATION_OF_IMMUNE_RESPONSE | 0.00002 | 0.00093 | 0.57561 | 0.59871 | 1.19485 | 307 |
| GO_POSITIVE_REGULATION_OF_IMMUNE_SYSTEM_PROCES | 0.00007 | 0.00222 | 0.53843 | 0.57386 | 1.15137 | 456 |
| GO_POSITIVE_REGULATION_OF_INNATE_IMMUNE_RESPON | 0.00031 | 0.00710 | 0.49849 | 0.63672 | 1.24640 | 127 |
| GO_POSITIVE_REGULATION_OF_ORGANELLE_ORGANIZATIO | 0.00004 | 0.00149 | 0.55733 | 0.58149 | 1.16635 | 424 |
| GO_POSITIVE_REGULATION_OF_PEPTIDASE_ACTIVITY | 0.00342 | 0.04099 | 0.43171 | 0.61162 | 1.19482 | 116 |
| GO_POSITIVE_REGULATION_OF_PROTEIN_KINASE_B_SIGNA | 0.00111 | 0.01833 | 0.45506 | 0.66142 | 1.27816 | 75 |
| GO_POSITIVE_REGULATION_OF_PROTEIN_LOCALIZATION_TO | 0.00161 | 0.02456 | 0.45506 | 0.67826 | 1.30266 | 61 |
| GO_POSITIVE_REGULATION_OF_RESPONSE_TO_BIOTIC_STIM | 0.00013 | 0.00358 | 0.51885 | 0.63209 | 1.24131 | 145 |
| GO_POSITIVE_REGULATION_OF_RESPONSE_TO_EXTERNAL_ | 0.00041 | 0.00883 | 0.49849 | 0.58918 | 1.17217 | 256 |
| GO_POSITIVE_REGULATION_OF_SUPRAMOLECULAR_FIBER_ | 0.00250 | 0.03324 | 0.43171 | 0.61268 | 1.20005 | 132 |
| GO_POSITIVE_REGULATION_OF_TELOMERASE_ACTIVITY | 0.00302 | 0.03762 | 0.43171 | 0.72005 | 1.33838 | 32 |
| GO_POSITIVE_REGULATION_OF_TELOMERASE_RNA_LOCALI | 0.00183 | 0.02703 | 0.45506 | 0.82552 | 1.47343 | 15 |
| GO_POSITIVE_REGULATION_OF_TELOMERE_MAINTENANCE | 0.00457 | 0.04987 | 0.40702 | 0.68263 | 1.28555 | 41 |
| GO_POSITIVE_REGULATION_OF_TELOMERE_MAINTENANCE | 0.00163 | 0.02465 | 0.45506 | 0.73639 | 1.36587 | 31 |
| GO_POSITIVE_REGULATION_OF_TRANSLATION | 0.00032 | 0.00720 | 0.49849 | 0.64890 | 1.26650 | 106 |
| GO_POSITIVE_REGULATION_OF_TRANSPORT | 0.00012 | 0.00344 | 0.53843 | 0.57004 | 1.14442 | 478 |
| GO_POSITIVE_REGULATION_OF_VIRAL_PROCESS | 0.00016 | 0.00415 | 0.51885 | 0.67143 | 1.30343 | 87 |
| GO_POSITIVE_REGULATION_OF_WNT_SIGNALING_PATHWA | 0.00047 | 0.00979 | 0.49849 | 0.63016 | 1.23153 | 119 |
| GO_POSTTRANSCRIPTIONAL_REGULATION_OF_GENE_EXPRE | 0.00000 | 0.00000 | | 0.61938 | 1.24351 | 479 |
| GO_PRIMARY_ACTIVE_TRANSMEMBRANE_TRANSPORTER_A | 0.00090 | 0.01563 | 0.47727 | 0.67355 | 1.29459 | 63 |
| GO_PROSTANOID_METABOLIC_PROCESS | 0.00372 | 0.04290 | 0.43171 | 0.75448 | 1.38049 | 23 |
| GO_PROSTATE_GLAND_DEVELOPMENT | 0.00032 | 0.00724 | 0.49849 | 0.80860 | 1.46500 | 20 |
| GO_PROTEASOMAL_UBIQUITIN_INDEPENDENT_PROTEIN_C | 0.00062 | 0.01162 | 0.47727 | 0.79777 | 1.44538 | 20 |
| GO_PROTEASOME_ACCESSORY_COMPLEX | 0.00361 | 0.04202 | 0.43171 | 0.75500 | 1.38145 | 23 |
| GO_PROTEASOME_CORE_COMPLEX | 0.00062 | 0.01162 | 0.47727 | 0.81382 | 1.46989 | 19 |
| GO_PROTEIN_CONTAINING_COMPLEX_DISASSEMBLY | 0.00000 | 0.00000 | 0.71951 | 0.64423 | 1.28122 | 254 |
| GO_PROTEIN_C_TERMINUS_BINDING | 0.00342 | 0.04099 | 0.43171 | 0.60912 | 1.19445 | 136 |
| GO_PROTEIN_DNA_COMPLEX_DISASSEMBLY | 0.00287 | 0.03638 | 0.43171 | 0.81449 | 1.45376 | 15 |
| GO_PROTEIN_FOLDING | 0.00084 | 0.01491 | 0.47727 | 0.61015 | 1.20278 | 163 |
| GO_PROTEIN_FOLDING_CHAPERONE | 0.00145 | 0.02289 | 0.45506 | 0.82295 | 1.48198 | 17 |
| GO_PROTEIN_IMPORT | 0.00121 | 0.01953 | 0.45506 | 0.61656 | 1.20998 | 142 |
| GO_PROTEIN_LOCALIZATION_TO_CELL_PERIPHERY | 0.00117 | 0.01903 | 0.45506 | 0.59375 | 1.17730 | 206 |
| GO_PROTEIN_LOCALIZATION_TO_ENDOPLASMIC_RETICULU | 0.00000 | 0.00000 | | 0.77615 | 1.51652 | 120 |
| GO_PROTEIN_LOCALIZATION_TO_MEMBRANE | 0.00000 | 0.00000 | | 0.66710 | 1.33826 | 427 |
| GO_PROTEIN_LOCALIZATION_TO_MITOCHONDRION | 0.00005 | 0.00179 | 0.55733 | 0.65720 | 1.28355 | 114 |
| GO_PROTEIN_LOCALIZATION_TO_NUCLEUS | 0.00231 | 0.03141 | 0.43171 | 0.59081 | 1.16948 | 193 |
| GO_PROTEIN_LOCALIZATION_TO_PLASMA_MEMBRANE | 0.00031 | 0.00717 | 0.49849 | 0.61138 | 1.20859 | 181 |
| GO_PROTEIN_POLYMERIZATION | 0.00111 | 0.01833 | 0.45506 | 0.59730 | 1.18239 | 194 |
| GO_PROTEIN_STABILIZATION | 0.00253 | 0.03358 | 0.43171 | 0.61176 | 1.19686 | 126 |
| GO_PROTEIN_TARGETING | 0.00000 | 0.00000 | | 0.68761 | 1.37330 | 319 |
| GO_PROTEIN_TARGETING_TO_MEMBRANE | 0.00000 | 0.00000 | | 0.74614 | 1.46731 | 151 |
| GO_PROTEIN_TARGETING_TO_MITOCHONDRION | 0.00011 | 0.00324 | 0.53843 | 0.67773 | 1.31186 | 80 |
| GO_PROTON_TRANSMEMBRANE_TRANSPORT | 0.00000 | 0.00000 | 0.73376 | 0.74641 | 1.44756 | 84 |
| GO_PROTON_TRANSMEMBRANE_TRANSPORTER_ACTIVITY | 0.00000 | 0.00018 | 0.62726 | 0.73166 | 1.40649 | 65 |
| GO_PURINE_CONTAINING_COMPOUND_BIOSYNTHETIC_PRO | 0.00006 | 0.00199 | 0.53843 | 0.66128 | 1.29139 | 108 |
| GO_PURINE_CONTAINING_COMPOUND_METABOLIC_PROC | 0.00002 | 0.00067 | 0.57561 | 0.61168 | 1.21713 | 259 |
| GO_PURINE_NUCLEOSIDE_TRIPHOSPHATE_METABOLIC_PRO | 0.00021 | 0.00514 | 0.51885 | 0.73921 | 1.39208 | 40 |
| GO_REACTIVE_OXYGEN_SPECIES_METABOLIC_PROCESS | 0.00372 | 0.04290 | 0.43171 | 0.60842 | 1.19185 | 131 |
| GO_RECEPTOR_INTERNALIZATION | 0.00357 | 0.04184 | 0.43171 | 0.66473 | 1.26659 | 50 |
| GO_RECEPTOR_MEDIATED_ENDOCYTOSIS | 0.00042 | 0.00887 | 0.49849 | 0.62579 | 1.22787 | 141 |
| GO_REGENERATION | 0.00298 | 0.03731 | 0.43171 | 0.62653 | 1.22189 | 102 |
| GO_REGULATION_OF_ACTIN_FILAMENT_BASED_PROCESS | 0.00058 | 0.01118 | 0.47727 | 0.59263 | 1.17750 | 229 |
| GO_REGULATION_OF_ACTIN_FILAMENT_LENGTH | 0.00246 | 0.03289 | 0.43171 | 0.61779 | 1.20715 | 117 |
| GO_REGULATION_OF_ACTIN_FILAMENT_ORGANIZATION | 0.00072 | 0.01314 | 0.47727 | 0.60965 | 1.20148 | 164 |
| GO_REGULATION_OF_ALCOHOL_BIOSYNTHETIC_PROCESS | 0.00004 | 0.00147 | 0.55733 | 0.73297 | 1.39661 | 50 |
| GO_REGULATION_OF_ANIMAL_ORGAN_MORPHOGENESIS | 0.00000 | 0.00008 | 0.64355 | 0.66790 | 1.31209 | 146 |
| GO_REGULATION_OF_CELLULAR_AMIDE_METABOLIC_PROC | 0.00001 | 0.00058 | 0.59333 | 0.60233 | 1.20223 | 314 |
| GO_REGULATION_OF_CELLULAR_AMINE_METABOLIC_PROC | 0.00001 | 0.00041 | 0.59333 | 0.73310 | 1.40808 | 62 |
| GO_REGULATION_OF_CELLULAR_AMINO_ACID_METABOLIC | 0.00000 | 0.00017 | 0.62726 | 0.75416 | 1.44147 | 53 |
| GO_REGULATION_OF_CELLULAR_KETONE_METABOLIC_PRO | 0.00003 | 0.00126 | 0.55733 | 0.65791 | 1.28554 | 117 |
| GO_REGULATION_OF_CELLULAR_PROTEIN_LOCALIZATION | 0.00016 | 0.00411 | 0.51885 | 0.57947 | 1.15931 | 362 |
| GO_REGULATION_OF_CELL_CYCLE_G2_M_PHASE_TRANSITI | 0.00071 | 0.01301 | 0.47727 | 0.60952 | 1.20360 | 174 |
| GO_REGULATION_OF_CELL_CYCLE_PHASE_TRANSITION | 0.00398 | 0.04503 | 0.40702 | 0.56076 | 1.12081 | 342 |
| GO_REGULATION_OF_CELL_SUBSTRATE_ADHESION | 0.00167 | 0.02508 | 0.45506 | 0.61762 | 1.20822 | 125 |
| GO_REGULATION_OF_CHOLESTEROL_BIOSYNTHETIC_PROCE | 0.00015 | 0.00402 | 0.51885 | 0.75262 | 1.41077 | 37 |
| GO_REGULATION_OF_CHOLESTEROL_METABOLIC_PROCESS | 0.00010 | 0.00299 | 0.53843 | 0.74337 | 1.40648 | 44 |
| GO_REGULATION_OF_CYSTEINE_TYPE_ENDOPEPTIDASE_AC | 0.00383 | 0.04401 | 0.43171 | 0.60643 | 1.18781 | 132 |
| GO_REGULATION_OF_CYTOSKELETON_ORGANIZATION | 0.00025 | 0.00596 | 0.49849 | 0.58092 | 1.16089 | 339 |

| Cluster 1 | | | | | | |
|---|---|---|---|---|---|---|
| GO_REGULATION_OF_DEFENSE_RESPONSE | 0.00033 | 0.00727 | 0.49849 | 0.57939 | 1.15751 | 334 |
| GO_REGULATION_OF_DNA_BINDING | 0.00398 | 0.04503 | 0.40702 | 0.63549 | 1.23011 | 80 |
| GO_REGULATION_OF_DNA_TEMPLATED_TRANSCRIPTION_I | 0.00016 | 0.00419 | 0.51885 | 0.66871 | 1.29966 | 91 |
| GO_REGULATION_OF_HEMATOPOIETIC_PROGENITOR_CELL_ | 0.00006 | 0.00190 | 0.55733 | 0.70903 | 1.36818 | 70 |
| GO_REGULATION_OF_HEMATOPOIETIC_STEM_CELL_DIFFER | 0.00001 | 0.00056 | 0.59333 | 0.72864 | 1.39951 | 62 |
| GO_REGULATION_OF_HEMOPOIESIS | 0.00086 | 0.01510 | 0.47727 | 0.58718 | 1.16835 | 258 |
| GO_REGULATION_OF_IMMUNE_RESPONSE | 0.00054 | 0.01060 | 0.47727 | 0.57114 | 1.14521 | 410 |
| GO_REGULATION_OF_INNATE_IMMUNE_RESPONSE | 0.00081 | 0.01450 | 0.47727 | 0.60437 | 1.19317 | 173 |
| GO_REGULATION_OF_LIPID_BIOSYNTHETIC_PROCESS | 0.00015 | 0.00406 | 0.51885 | 0.64555 | 1.26110 | 116 |
| GO_REGULATION_OF_LIPID_METABOLIC_PROCESS | 0.00008 | 0.00250 | 0.53843 | 0.60609 | 1.20453 | 230 |
| GO_REGULATION_OF_MACROAUTOPHAGY | 0.00320 | 0.03935 | 0.43171 | 0.60402 | 1.18490 | 139 |
| GO_REGULATION_OF_MITOTIC_CELL_CYCLE | 0.00172 | 0.02571 | 0.45506 | 0.55371 | 1.11084 | 463 |
| GO_REGULATION_OF_MORPHOGENESIS_OF_AN_EPITHELIU | 0.00000 | 0.00002 | 0.69013 | 0.69517 | 1.35834 | 117 |
| GO_REGULATION_OF_MRNA_CATABOLIC_PROCESS | 0.00001 | 0.00027 | 0.61053 | 0.64967 | 1.27984 | 159 |
| GO_REGULATION_OF_MRNA_METABOLIC_PROCESS | 0.00000 | 0.00000 | 0.77494 | 0.64818 | 1.29012 | 263 |
| GO_REGULATION_OF_MRNA_PROCESSING | 0.00013 | 0.00355 | 0.51885 | 0.64282 | 1.25626 | 119 |
| GO_REGULATION_OF_MRNA_SPLICING_VIA_SPLICEOSOME | 0.00034 | 0.00749 | 0.49849 | 0.66402 | 1.28819 | 85 |
| GO_REGULATION_OF_MULTI_ORGANISM_PROCESS | 0.00051 | 0.01019 | 0.47727 | 0.73491 | 1.38090 | 39 |
| GO_REGULATION_OF_MUSCLE_CONTRACTION | 0.00156 | 0.02396 | 0.45506 | 0.66428 | 1.28347 | 72 |
| GO_REGULATION_OF_OXIDATIVE_PHOSPHORYLATION | 0.00428 | 0.04758 | 0.40702 | 0.75083 | 1.36993 | 22 |
| GO_REGULATION_OF_PEPTIDASE_ACTIVITY | 0.00002 | 0.00071 | 0.57561 | 0.61978 | 1.22948 | 218 |
| GO_REGULATION_OF_PEPTIDE_TRANSPORT | 0.00290 | 0.03669 | 0.43171 | 0.55702 | 1.11516 | 378 |
| GO_REGULATION_OF_PLASMA_LIPOPROTEIN_PARTICLE_LE\ | 0.00000 | 0.00018 | 0.62726 | 0.75295 | 1.43915 | 53 |
| GO_REGULATION_OF_PROTEIN_CONTAINING_COMPLEX_AS | 0.00061 | 0.01155 | 0.47727 | 0.57967 | 1.15596 | 283 |
| GO_REGULATION_OF_PROTEIN_LOCALIZATION_TO_NUCLEU | 0.00022 | 0.00539 | 0.51885 | 0.66748 | 1.29577 | 87 |
| GO_REGULATION_OF_PROTEIN_MATURATION | 0.00198 | 0.02803 | 0.43171 | 0.72042 | 1.34273 | 34 |
| GO_REGULATION_OF_PROTEIN_POLYMERIZATION | 0.00034 | 0.00756 | 0.49849 | 0.62403 | 1.22630 | 147 |
| GO_REGULATION_OF_PROTEIN_STABILITY | 0.00121 | 0.01953 | 0.45506 | 0.59331 | 1.17642 | 206 |
| GO_REGULATION_OF_PROTEOLYSIS | 0.00036 | 0.00783 | 0.49849 | 0.57401 | 1.15096 | 410 |
| GO_REGULATION_OF_RESPONSE_TO_BIOTIC_STIMULUS | 0.00185 | 0.02710 | 0.45506 | 0.58752 | 1.16616 | 222 |
| GO_REGULATION_OF_RESPONSE_TO_EXTERNAL_STIMULUS | 0.00054 | 0.01071 | 0.47727 | 0.55995 | 1.12458 | 495 |
| GO_REGULATION_OF_RNA_SPLICING | 0.00004 | 0.00146 | 0.55733 | 0.65321 | 1.27657 | 119 |
| GO_REGULATION_OF_SMALL_MOLECULE_METABOLIC_PRO | 0.00000 | 0.00002 | 0.67496 | 0.62536 | 1.24716 | 281 |
| GO_REGULATION_OF_STEM_CELL_DIFFERENTIATION | 0.00002 | 0.00070 | 0.57561 | 0.69408 | 1.34651 | 85 |
| GO_REGULATION_OF_STEROID_BIOSYNTHETIC_PROCESS | 0.00005 | 0.00179 | 0.55733 | 0.72871 | 1.39641 | 57 |
| GO_REGULATION_OF_STEROID_METABOLIC_PROCESS | 0.00000 | 0.00012 | 0.62726 | 0.72885 | 1.40823 | 72 |
| GO_REGULATION_OF_SUPRAMOLECULAR_FIBER_ORGANIZA | 0.00003 | 0.00117 | 0.55733 | 0.61233 | 1.21593 | 224 |
| GO_REGULATION_OF_SYMBIOTIC_PROCESS | 0.00004 | 0.00154 | 0.55733 | 0.64050 | 1.25783 | 145 |
| GO_REGULATION_OF_TRANSCRIPTION_FROM_RNA_POLYM | 0.00001 | 0.00048 | 0.59333 | 0.72164 | 1.38717 | 64 |
| GO_REGULATION_OF_VIRAL_GENOME_REPLICATION | 0.00253 | 0.03358 | 0.43171 | 0.67000 | 1.28356 | 56 |
| GO_REGULATION_OF_VIRAL_LIFE_CYCLE | 0.00413 | 0.04636 | 0.40702 | 0.63369 | 1.23093 | 89 |
| GO_REGULATION_OF_VIRAL_TRANSCRIPTION | 0.00420 | 0.04693 | 0.40702 | 0.65968 | 1.26089 | 53 |
| GO_REGULATION_OF_WNT_SIGNALING_PATHWAY | 0.00398 | 0.04503 | 0.40702 | 0.58307 | 1.15815 | 226 |
| GO_REPRODUCTIVE_SYSTEM_DEVELOPMENT | 0.00008 | 0.00246 | 0.53843 | 0.59877 | 1.19021 | 251 |
| GO_RESPIRASOME | 0.00000 | 0.00000 | 0.77494 | 0.77873 | 1.50461 | 72 |
| GO_RESPIRATORY_CHAIN_COMPLEX | 0.00000 | 0.00000 | 0.73376 | 0.78987 | 1.51701 | 61 |
| GO_RESPIRATORY_ELECTRON_TRANSPORT_CHAIN | 0.00000 | 0.00000 | 0.76146 | 0.75165 | 1.45961 | 88 |
| GO_RESPONSE_TO_ALCOHOL | 0.00000 | 0.00001 | 0.70498 | 0.70320 | 1.37403 | 117 |
| GO_RESPONSE_TO_CORTICOSTEROID | 0.00191 | 0.02737 | 0.45506 | 0.64648 | 1.25093 | 79 |
| GO_RESPONSE_TO_DRUG | 0.00000 | 0.00020 | 0.62726 | 0.63606 | 1.26160 | 209 |
| GO_RESPONSE_TO_ETHANOL | 0.00004 | 0.00149 | 0.55733 | 0.72507 | 1.38907 | 56 |
| GO_RESPONSE_TO_EXTRACELLULAR_STIMULUS | 0.00061 | 0.01155 | 0.47727 | 0.58264 | 1.16205 | 287 |
| GO_RESPONSE_TO_FIBROBLAST_GROWTH_FACTOR | 0.00137 | 0.02191 | 0.45506 | 0.65418 | 1.26451 | 76 |
| GO_RESPONSE_TO_GROWTH_FACTOR | 0.00100 | 0.01694 | 0.45506 | 0.56567 | 1.13308 | 393 |
| GO_RESPONSE_TO_INORGANIC_SUBSTANCE | 0.00080 | 0.01438 | 0.47727 | 0.57812 | 1.15375 | 307 |
| GO_RESPONSE_TO_INTERLEUKIN_1 | 0.00001 | 0.00049 | 0.59333 | 0.67180 | 1.31207 | 114 |
| GO_RESPONSE_TO_INTERLEUKIN_4 | 0.00148 | 0.02306 | 0.45506 | 0.77519 | 1.41840 | 23 |
| GO_RESPONSE_TO_KETONE | 0.00383 | 0.04401 | 0.43171 | 0.61354 | 1.19551 | 101 |
| GO_RESPONSE_TO_LIPID | 0.00012 | 0.00349 | 0.53843 | 0.57579 | 1.15508 | 426 |
| GO_RESPONSE_TO_NUTRIENT | 0.00013 | 0.00355 | 0.51885 | 0.66729 | 1.29744 | 92 |
| GO_RESPONSE_TO_ORGANIC_CYCLIC_COMPOUND | 0.00000 | 0.00002 | 0.67496 | 0.59311 | 1.19058 | 474 |
| GO_RESPONSE_TO_OXYGEN_LEVELS | 0.00000 | 0.00000 | 0.77494 | 0.65451 | 1.30090 | 245 |
| GO_RESPONSE_TO_TOXIC_SUBSTANCE | 0.00063 | 0.01175 | 0.47727 | 0.62780 | 1.22814 | 125 |
| GO_RESPONSE_TO_TUMOR_NECROSIS_FACTOR | 0.00001 | 0.00036 | 0.61053 | 0.64296 | 1.26835 | 170 |
| GO_RESPONSE_TO_VIRUS | 0.00033 | 0.00727 | 0.49849 | 0.61799 | 1.21773 | 165 |
| GO_RESPONSE_TO_VITAMIN | 0.00003 | 0.00119 | 0.55733 | 0.75726 | 1.43102 | 43 |
| GO_RESPONSE_TO_WOUNDING | 0.00082 | 0.01462 | 0.47727 | 0.57543 | 1.14826 | 305 |
| GO_RESPONSE_TO_XENOBIOTIC_STIMULUS | 0.00052 | 0.01034 | 0.47727 | 0.71416 | 1.35638 | 48 |
| GO_RETINA_HOMEOSTASIS | 0.00021 | 0.00525 | 0.51885 | 0.79396 | 1.45520 | 24 |
| GO_RIBONUCLEOPROTEIN_COMPLEX_BINDING | 0.00018 | 0.00468 | 0.51885 | 0.65004 | 1.26907 | 109 |
| GO_RIBONUCLEOPROTEIN_COMPLEX_BIOGENESIS | 0.00000 | 0.00000 | | 0.63829 | 1.27756 | 376 |
| GO_RIBONUCLEOPROTEIN_COMPLEX_SUBUNIT_ORGANIZAT | 0.00000 | 0.00001 | 0.70498 | 0.67579 | 1.33294 | 169 |
| GO_RIBONUCLEOPROTEIN_GRANULE | 0.00019 | 0.00468 | 0.51885 | 0.62013 | 1.22534 | 177 |
| GO_RIBONUCLEOSIDE_TRIPHOSPHATE_BIOSYNTHETIC_PRO | 0.00111 | 0.01833 | 0.45506 | 0.74484 | 1.38155 | 31 |

| Cluster 1 | | | | | | |
|---|---|---|---|---|---|---|
| GO_RIBONUCLEOSIDE_TRIPHOSPHATE_METABOLIC_PROCES | 0.00074 | 0.01339 | 0.47727 | 0.73139 | 1.37239 | 38 |
| GO_RIBOSE_PHOSPHATE_BIOSYNTHETIC_PROCESS | 0.00006 | 0.00199 | 0.53843 | 0.66130 | 1.29142 | 108 |
| GO_RIBOSE_PHOSPHATE_METABOLIC_PROCESS | 0.00002 | 0.00083 | 0.57561 | 0.61174 | 1.21592 | 248 |
| GO_RIBOSOMAL_LARGE_SUBUNIT_BIOGENESIS | 0.00064 | 0.01185 | 0.47727 | 0.68758 | 1.32389 | 67 |
| GO_RIBOSOMAL_SMALL_SUBUNIT_ASSEMBLY | 0.00095 | 0.01623 | 0.47727 | 0.82235 | 1.48387 | 18 |
| GO_RIBOSOMAL_SMALL_SUBUNIT_BIOGENESIS | 0.00185 | 0.02710 | 0.45506 | 0.65822 | 1.26531 | 65 |
| GO_RIBOSOMAL_SUBUNIT | 0.00000 | 0.00000 | | 0.76162 | 1.50329 | 172 |
| GO_RIBOSOME | 0.00000 | 0.00000 | | 0.74475 | 1.47671 | 206 |
| GO_RIBOSOME_ASSEMBLY | 0.00004 | 0.00152 | 0.55733 | 0.73030 | 1.39945 | 57 |
| GO_RIBOSOME_BIOGENESIS | 0.00000 | 0.00006 | 0.65944 | 0.62366 | 1.24132 | 265 |
| GO_RNA_CATABOLIC_PROCESS | 0.00000 | 0.00000 | | 0.69257 | 1.38323 | 326 |
| GO_RNA_DEPENDENT_DNA_BIOSYNTHETIC_PROCESS | 0.00143 | 0.02267 | 0.45506 | 0.66881 | 1.28368 | 60 |
| GO_RNA_SPLICING | 0.00000 | 0.00000 | 0.73376 | 0.61572 | 1.23239 | 374 |
| GO_RNA_SPLICING_VIA_TRANSESTERIFICATION_REACTIONS | 0.00000 | 0.00001 | 0.70498 | 0.62683 | 1.25093 | 304 |
| GO_ROUGH_ENDOPLASMIC_RETICULUM | 0.00018 | 0.00464 | 0.51885 | 0.70989 | 1.35849 | 55 |
| GO_RRNA_BINDING | 0.00000 | 0.00003 | 0.67496 | 0.77511 | 1.48493 | 56 |
| GO_RRNA_METABOLIC_PROCESS | 0.00007 | 0.00216 | 0.53843 | 0.61521 | 1.21905 | 199 |
| GO_RUFFLE | 0.00202 | 0.02829 | 0.43171 | 0.62174 | 1.21514 | 121 |
| GO_SARCOLEMMA | 0.00003 | 0.00120 | 0.55733 | 0.73390 | 1.40635 | 57 |
| GO_SARCOMERE_ORGANIZATION | 0.00146 | 0.02289 | 0.45506 | 0.77479 | 1.41365 | 22 |
| GO_SCAFFOLD_PROTEIN_BINDING | 0.00335 | 0.04034 | 0.43171 | 0.72007 | 1.33560 | 31 |
| GO_SCF_DEPENDENT_PROTEASOMAL_UBIQUITIN_DEPENDI | 0.00032 | 0.00720 | 0.49849 | 0.68041 | 1.31222 | 69 |
| GO_SECRETORY_GRANULE | 0.00000 | 0.00000 | | 0.62734 | 1.25824 | 446 |
| GO_SECRETORY_GRANULE_MEMBRANE | 0.00146 | 0.02289 | 0.45506 | 0.61033 | 1.20095 | 155 |
| GO_SENSORY_PERCEPTION_OF_MECHANICAL_STIMULUS | 0.00019 | 0.00468 | 0.51885 | 0.67782 | 1.31021 | 76 |
| GO_SERINE_TYPE_ENDOPEPTIDASE_INHIBITOR_ACTIVITY | 0.00148 | 0.02306 | 0.45506 | 0.74974 | 1.38337 | 28 |
| GO_SINGLE_FERTILIZATION | 0.00004 | 0.00146 | 0.55733 | 0.73089 | 1.40060 | 57 |
| GO_SINGLE_STRANDED_RNA_BINDING | 0.00316 | 0.03898 | 0.43171 | 0.66768 | 1.28235 | 61 |
| GO_SMALL_MOLECULE_BIOSYNTHETIC_PROCESS | 0.00000 | 0.00021 | 0.61053 | 0.59179 | 1.18586 | 402 |
| GO_SMALL_NUCLEAR_RIBONUCLEOPROTEIN_COMPLEX | 0.00065 | 0.01198 | 0.47727 | 0.68975 | 1.32362 | 59 |
| GO_SMALL_RIBOSOMAL_SUBUNIT | 0.00000 | 0.00000 | 0.73376 | 0.77647 | 1.49833 | 70 |
| GO_SM_LIKE_PROTEIN_FAMILY_COMPLEX | 0.00049 | 0.01006 | 0.47727 | 0.67471 | 1.30123 | 69 |
| GO_SNORNA_BINDING | 0.00335 | 0.04034 | 0.43171 | 0.72701 | 1.34417 | 29 |
| GO_SODIUM_ION_TRANSMEMBRANE_TRANSPORT | 0.00268 | 0.03470 | 0.43171 | 0.64914 | 1.25452 | 74 |
| GO_SODIUM_ION_TRANSMEMBRANE_TRANSPORTER_ACTI | 0.00001 | 0.00059 | 0.59333 | 0.73977 | 1.41760 | 57 |
| GO_SODIUM_ION_TRANSPORT | 0.00087 | 0.01517 | 0.47727 | 0.63644 | 1.24146 | 104 |
| GO_SOLUTE_SODIUM_SYMPORTER_ACTIVITY | 0.00135 | 0.02168 | 0.45506 | 0.74651 | 1.38023 | 29 |
| GO_SPERM_EGG_RECOGNITION | 0.00006 | 0.00199 | 0.55733 | 0.81248 | 1.48915 | 24 |
| GO_SPINDLE_ORGANIZATION | 0.00191 | 0.02737 | 0.45506 | 0.61214 | 1.20129 | 142 |
| GO_SPLICEOSOMAL_COMPLEX | 0.00001 | 0.00033 | 0.61053 | 0.64146 | 1.26476 | 167 |
| GO_STEM_CELL_DIFFERENTIATION | 0.00000 | 0.00015 | 0.62726 | 0.65810 | 1.29482 | 153 |
| GO_STEROID_BIOSYNTHETIC_PROCESS | 0.00006 | 0.00199 | 0.53843 | 0.66133 | 1.29148 | 108 |
| GO_STEROID_METABOLIC_PROCESS | 0.00004 | 0.00141 | 0.55733 | 0.63582 | 1.25287 | 165 |
| GO_STEROL_BIOSYNTHETIC_PROCESS | 0.00000 | 0.00023 | 0.61053 | 0.75074 | 1.43862 | 57 |
| GO_STEROL_METABOLIC_PROCESS | 0.00000 | 0.00005 | 0.65944 | 0.70263 | 1.36777 | 97 |
| GO_STRUCTURAL_CONSTITUENT_OF_CYTOSKELETON | 0.00007 | 0.00212 | 0.53843 | 0.71936 | 1.37812 | 56 |
| GO_STRUCTURAL_CONSTITUENT_OF_MUSCLE | 0.00287 | 0.03638 | 0.43171 | 0.76785 | 1.39115 | 20 |
| GO_STRUCTURAL_CONSTITUENT_OF_RIBOSOME | 0.00000 | 0.00000 | | 0.78086 | 1.53561 | 151 |
| GO_STRUCTURAL_MOLECULE_ACTIVITY | 0.00000 | 0.00000 | | 0.72608 | 1.45357 | 379 |
| GO_SUBSTANTIA_NIGRA_DEVELOPMENT | 0.00191 | 0.02737 | 0.45506 | 0.74012 | 1.36841 | 29 |
| GO_SUPRAMOLECULAR_FIBER_ORGANIZATION | 0.00015 | 0.00408 | 0.51885 | 0.57261 | 1.14837 | 420 |
| GO_TELOMERE_MAINTENANCE_VIA_TELOMERE_LENGTHEN | 0.00398 | 0.04503 | 0.40702 | 0.66452 | 1.27949 | 67 |
| GO_THIOESTER_METABOLIC_PROCESS | 0.00133 | 0.02144 | 0.45506 | 0.68429 | 1.31130 | 57 |
| GO_THREONINE_TYPE_ENDOPEPTIDASE_ACTIVITY | 0.00015 | 0.00408 | 0.51885 | 0.82192 | 1.48913 | 20 |
| GO_THREONINE_TYPE_PEPTIDASE_ACTIVITY | 0.00013 | 0.00372 | 0.51885 | 0.82260 | 1.50514 | 23 |
| GO_TISSUE_MORPHOGENESIS | 0.00000 | 0.00002 | 0.67496 | 0.60711 | 1.21516 | 374 |
| GO_TRANSLATIONAL_ELONGATION | 0.00000 | 0.00004 | 0.65944 | 0.68860 | 1.34532 | 122 |
| GO_TRANSLATIONAL_INITIATION | 0.00000 | 0.00000 | | 0.76534 | 1.50978 | 170 |
| GO_TRANSLATIONAL_TERMINATION | 0.00000 | 0.00004 | 0.65944 | 0.70140 | 1.36585 | 100 |
| GO_TRANSLATION_ELONGATION_FACTOR_ACTIVITY | 0.00435 | 0.04823 | 0.40702 | 0.80147 | 1.43051 | 15 |
| GO_TRANSLATION_FACTOR_ACTIVITY_RNA_BINDING | 0.00061 | 0.01155 | 0.47727 | 0.68789 | 1.32449 | 67 |
| GO_TRANSLATION_PREINITIATION_COMPLEX | 0.00187 | 0.02717 | 0.45506 | 0.80838 | 1.44955 | 16 |
| GO_TRANSLATION_REGULATOR_ACTIVITY | 0.00001 | 0.00055 | 0.59333 | 0.67525 | 1.31866 | 108 |
| GO_TRANSLATION_REGULATOR_ACTIVITY_NUCLEIC_ACID_E | 0.00007 | 0.00226 | 0.53843 | 0.68415 | 1.32637 | 83 |
| GO_TUMOR_NECROSIS_FACTOR_MEDIATED_SIGNALING_PA | 0.00000 | 0.00004 | 0.65944 | 0.70209 | 1.37111 | 111 |
| GO_T_CELL_RECEPTOR_SIGNALING_PATHWAY | 0.00075 | 0.01352 | 0.47727 | 0.62922 | 1.22919 | 116 |
| GO_T_TUBULE | 0.00056 | 0.01093 | 0.47727 | 0.80511 | 1.46437 | 21 |
| GO_U2_TYPE_SPLICEOSOMAL_COMPLEX | 0.00191 | 0.02737 | 0.45506 | 0.64210 | 1.24566 | 85 |
| GO_UBIQUITIN_LIKE_PROTEIN_LIGASE_BINDING | 0.00006 | 0.00199 | 0.53843 | 0.60833 | 1.20798 | 224 |
| GO_UNFOLDED_PROTEIN_BINDING | 0.00001 | 0.00038 | 0.59333 | 0.70849 | 1.37266 | 82 |
| GO_UNSATURATED_FATTY_ACID_BIOSYNTHETIC_PROCESS | 0.00313 | 0.03860 | 0.43171 | 0.75819 | 1.38729 | 23 |
| GO_UNSATURATED_FATTY_ACID_METABOLIC_PROCESS | 0.00106 | 0.01767 | 0.45506 | 0.70839 | 1.34241 | 45 |
| GO_VACUOLAR_LUMEN | 0.00001 | 0.00026 | 0.61053 | 0.68589 | 1.33565 | 100 |

| Cluster 1 | | | | | | |
|---|---|---|---|---|---|---|
| GO_VACUOLAR_MEMBRANE | 0.00327 | 0.03985 | 0.43171 | 0.56984 | 1.13737 | 298 |
| GO_VESICLE_LUMEN | 0.00000 | 0.00000 | 0.82666 | 0.69064 | 1.36678 | 191 |
| GO_VESICLE_MEMBRANE | 0.00006 | 0.00199 | 0.53843 | 0.57896 | 1.16135 | 431 |
| GO_VIRAL_GENE_EXPRESSION | 0.00000 | 0.00000 | | 0.73473 | 1.45138 | 176 |
| GO_VIRAL_GENOME_REPLICATION | 0.00224 | 0.03075 | 0.43171 | 0.64862 | 1.25377 | 76 |
| GO_VIRAL_LIFE_CYCLE | 0.00003 | 0.00127 | 0.55733 | 0.61196 | 1.21521 | 224 |
| GO_ZYMOGEN_ACTIVATION | 0.00335 | 0.04034 | 0.43171 | 0.75259 | 1.38265 | 26 |
| HP_11_PAIRS_OF_RIBS | 0.00231 | 0.03141 | 0.43171 | 0.80369 | 1.44114 | 16 |
| HP_ABNORMALITY_OF_ACID_BASE_HOMEOSTASIS | 0.00106 | 0.01767 | 0.45506 | 0.59192 | 1.17646 | 241 |
| HP_ABNORMALITY_OF_CARDIOVASCULAR_SYSTEM_ELECTR | 0.00029 | 0.00686 | 0.49849 | 0.59882 | 1.19022 | 239 |
| HP_ABNORMALITY_OF_THE_INNER_EAR | 0.00102 | 0.01720 | 0.45506 | 0.55831 | 1.12124 | 496 |
| HP_ABNORMALITY_OF_THE_LARYNX | 0.00357 | 0.04184 | 0.43171 | 0.62681 | 1.22018 | 97 |
| HP_ABNORMALITY_OF_THE_MITOCHONDRION | 0.00198 | 0.02803 | 0.43171 | 0.62131 | 1.21330 | 112 |
| HP_ABNORMALITY_OF_THE_OPTIC_DISC | 0.00457 | 0.04987 | 0.40702 | 0.55343 | 1.10899 | 402 |
| HP_ABNORMALITY_OF_THE_OPTIC_NERVE | 0.00097 | 0.01650 | 0.47727 | 0.56079 | 1.12495 | 450 |
| HP_ABNORMAL_ACTIVITY_OF_MITOCHONDRIAL_RESPIRAT( | 0.00450 | 0.04951 | 0.40702 | 0.66200 | 1.27038 | 59 |
| HP_ABNORMAL_BLOOD_POTASSIUM_CONCENTRATION | 0.00268 | 0.03470 | 0.43171 | 0.71465 | 1.33548 | 35 |
| HP_ABNORMAL_CELLULAR_PHENOTYPE | 0.00220 | 0.03031 | 0.43171 | 0.56828 | 1.13499 | 320 |
| HP_ABNORMAL_CIRCULATING_CARBOHYDRATE_CONCENTR | 0.00163 | 0.02465 | 0.45506 | 0.70111 | 1.32653 | 44 |
| HP_ABNORMAL_CNS_MYELINATION | 0.00055 | 0.01079 | 0.47727 | 0.61324 | 1.20723 | 157 |
| HP_ABNORMAL_CSF_METABOLITE_LEVEL | 0.00008 | 0.00237 | 0.53843 | 0.68104 | 1.31780 | 79 |
| HP_ABNORMAL_ERYTHROCYTE_MORPHOLOGY | 0.00001 | 0.00048 | 0.59333 | 0.59216 | 1.18485 | 364 |
| HP_ABNORMAL_EYELID_PHYSIOLOGY | 0.00053 | 0.01049 | 0.47727 | 0.79704 | 1.45837 | 23 |
| HP_ABNORMAL_IMMUNE_SYSTEM_MORPHOLOGY | 0.00108 | 0.01793 | 0.45506 | 0.57070 | 1.14000 | 330 |
| HP_ABNORMAL_MITOCHONDRIA_IN_MUSCLE_TISSUE | 0.00457 | 0.04987 | 0.40702 | 0.72666 | 1.34077 | 28 |
| HP_ABNORMAL_VAGINA_MORPHOLOGY | 0.00327 | 0.03985 | 0.43171 | 0.66680 | 1.27052 | 50 |
| HP_ACUTE_LEUKEMIA | 0.00051 | 0.01019 | 0.47727 | 0.67183 | 1.29838 | 74 |
| HP_ANEMIA_OF_INADEQUATE_PRODUCTION | 0.00202 | 0.02829 | 0.43171 | 0.63034 | 1.22930 | 103 |
| HP_APHASIA | 0.00033 | 0.00734 | 0.49849 | 0.77289 | 1.42900 | 29 |
| HP_DEMENTIA | 0.00313 | 0.03860 | 0.43171 | 0.61795 | 1.20576 | 105 |
| HP_DUPLICATION_OF_THUMB_PHALANX | 0.00357 | 0.04184 | 0.43171 | 0.76181 | 1.38021 | 20 |
| HP_EMOTIONAL_LABILITY | 0.00078 | 0.01410 | 0.47727 | 0.65673 | 1.27363 | 84 |
| HP_FATIGUE | 0.00061 | 0.01155 | 0.47727 | 0.60156 | 1.19203 | 201 |
| HP_FETAL_DISTRESS | 0.00185 | 0.02710 | 0.45506 | 0.73008 | 1.35702 | 32 |
| HP_FRONTOTEMPORAL_DEMENTIA | 0.00228 | 0.03112 | 0.43171 | 0.78210 | 1.41260 | 19 |
| HP_HEADACHE | 0.00004 | 0.00149 | 0.55733 | 0.63549 | 1.24927 | 150 |
| HP_HEMATOLOGICAL_NEOPLASM | 0.00055 | 0.01079 | 0.47727 | 0.61472 | 1.21180 | 163 |
| HP_HYPOKALEMIA | 0.00006 | 0.00195 | 0.55733 | 0.84213 | 1.53171 | 21 |
| HP_INCREASED_CIRCULATING_ANTIBODY_LEVEL | 0.00205 | 0.02860 | 0.43171 | 0.71240 | 1.33676 | 38 |
| HP_INCREASED_CSF_LACTATE | 0.00030 | 0.00702 | 0.49849 | 0.68893 | 1.32823 | 68 |
| HP_INCREASED_SERUM_LACTATE | 0.00179 | 0.02661 | 0.45506 | 0.62096 | 1.21625 | 129 |
| HP_INCREASED_SERUM_PYRUVATE | 0.00390 | 0.04461 | 0.43171 | 0.70722 | 1.32421 | 36 |
| HP_LETHARGY | 0.00361 | 0.04202 | 0.43171 | 0.62677 | 1.22010 | 97 |
| HP_LEUKEMIA | 0.00046 | 0.00966 | 0.49849 | 0.64279 | 1.25423 | 105 |
| HP_LEUKODYSTROPHY | 0.00004 | 0.00154 | 0.55733 | 0.69181 | 1.33835 | 78 |
| HP_MACROCYTIC_ANEMIA | 0.00008 | 0.00237 | 0.53843 | 0.74654 | 1.40592 | 41 |
| HP_MENTAL_DETERIORATION | 0.00170 | 0.02550 | 0.45506 | 0.59121 | 1.16930 | 185 |
| HP_MIGRAINE | 0.00000 | 0.00021 | 0.61053 | 0.72648 | 1.40108 | 69 |
| HP_MISSING_RIBS | 0.00283 | 0.03614 | 0.43171 | 0.76966 | 1.39989 | 21 |
| HP_MUTISM | 0.00013 | 0.00362 | 0.51885 | 0.78377 | 1.45176 | 30 |
| HP_NAUSEA_AND_VOMITING | 0.00228 | 0.03112 | 0.43171 | 0.58314 | 1.15905 | 239 |
| HP_OPTIC_NERVE_COLOBOMA | 0.00335 | 0.04034 | 0.43171 | 0.81032 | 1.44632 | 15 |
| HP_OPTIC_NEUROPATHY | 0.00191 | 0.02737 | 0.45506 | 0.73292 | 1.35943 | 31 |
| HP_PAROXYSMAL_INVOLUNTARY_EYE_MOVEMENTS | 0.00357 | 0.04184 | 0.43171 | 0.72984 | 1.34664 | 28 |
| HP_POOR_HEAD_CONTROL | 0.00450 | 0.04951 | 0.40702 | 0.61260 | 1.19598 | 109 |
| HP_PROGRESSIVE_CEREBELLAR_ATAXIA | 0.00115 | 0.01889 | 0.45506 | 0.67120 | 1.29022 | 64 |
| HP_PROXIMAL_TUBULOPATHY | 0.00361 | 0.04202 | 0.43171 | 0.71848 | 1.33264 | 31 |
| HP_RENAL_TUBULAR_DYSFUNCTION | 0.00220 | 0.03031 | 0.43171 | 0.66532 | 1.28272 | 68 |
| HP_RESPIRATORY_FAILURE | 0.00350 | 0.04154 | 0.43171 | 0.61846 | 1.20775 | 108 |
| HP_RESPIRATORY_INSUFFICIENCY | 0.00018 | 0.00468 | 0.51885 | 0.58821 | 1.17386 | 304 |
| HP_THROMBOCYTOSIS | 0.00092 | 0.01581 | 0.47727 | 0.78148 | 1.42585 | 22 |
| HP_TRANSIENT_ISCHEMIC_ATTACK | 0.00098 | 0.01672 | 0.45506 | 0.78303 | 1.43275 | 23 |
| HP_VERTIGO | 0.00235 | 0.03176 | 0.43171 | 0.67772 | 1.29981 | 58 |
| HP_XEROSTOMIA | 0.00209 | 0.02892 | 0.43171 | 0.72835 | 1.35382 | 32 |
| **Cluster 6** | | | | | | |
| pathway | pval | padj | log2err | ES | NES | size |
| GO_ACTIVATION_OF_IMMUNE_RESPONSE | 0.00000 | 0.00000 | 0.76146 | 0.64380 | 1.30880 | 201 |
| GO_ACTIVATION_OF_INNATE_IMMUNE_RESPONSE | 0.00000 | 0.00003 | 0.65944 | 0.69608 | 1.37777 | 86 |
| GO_AEROBIC_RESPIRATION | 0.00005 | 0.00142 | 0.55733 | 0.68568 | 1.34429 | 66 |
| GO_AMIDE_BINDING | 0.00324 | 0.04309 | 0.43171 | 0.57855 | 1.16687 | 155 |
| GO_AMINE_METABOLIC_PROCESS | 0.00000 | 0.00000 | 0.76146 | 0.74300 | 1.46532 | 78 |
| GO_ANAPHASE_PROMOTING_COMPLEX_DEPENDENT_CATA | 0.00000 | 0.00012 | 0.62726 | 0.69788 | 1.37363 | 71 |
| GO_ANATOMICAL_STRUCTURE_HOMEOSTASIS | 0.00001 | 0.00022 | 0.61053 | 0.60371 | 1.22979 | 224 |
| GO_ANCHORING_JUNCTION | 0.00000 | 0.00000 | 0.71951 | 0.58355 | 1.20182 | 436 |

| Cluster 1 | | | | | | |
|---|---|---|---|---|---|---|
| GO_ANTIGEN_PROCESSING_AND_PRESENTATION | 0.00000 | 0.00001 | 0.70498 | 0.65892 | 1.32568 | 139 |
| GO_ANTIGEN_PROCESSING_AND_PRESENTATION_OF_EXOG | 0.00000 | 0.00002 | 0.69013 | 0.75436 | 1.47085 | 56 |
| GO_ANTIGEN_PROCESSING_AND_PRESENTATION_OF_PEPT | 0.00000 | 0.00009 | 0.62726 | 0.65435 | 1.31196 | 123 |
| GO_ANTIGEN_PROCESSING_AND_PRESENTATION_OF_PEPT | 0.00000 | 0.00006 | 0.64355 | 0.71653 | 1.40915 | 69 |
| GO_ANTIGEN_RECEPTOR_MEDIATED_SIGNALING_PATHWAY | 0.00003 | 0.00098 | 0.55733 | 0.63546 | 1.27269 | 119 |
| GO_APOPTOTIC_MITOCHONDRIAL_CHANGES | 0.00205 | 0.03026 | 0.43171 | 0.63487 | 1.24737 | 68 |
| GO_APOPTOTIC_SIGNALING_PATHWAY | 0.00007 | 0.00169 | 0.53843 | 0.56684 | 1.16397 | 330 |
| GO_ATPASE_ACTIVITY | 0.00257 | 0.03634 | 0.43171 | 0.55355 | 1.13081 | 263 |
| GO_ATPASE_ACTIVITY_COUPLED_TO_TRANSMEMBRANE_M | 0.00268 | 0.03733 | 0.43171 | 0.79532 | 1.43666 | 15 |
| GO_ATPASE_COUPLED_ION_TRANSMEMBRANE_TRANSPOR | 0.00014 | 0.00328 | 0.51885 | 0.77119 | 1.45465 | 30 |
| GO_ATP_METABOLIC_PROCESS | 0.00000 | 0.00000 | | 0.67627 | 1.37206 | 190 |
| GO_ATP_SYNTHESIS_COUPLED_ELECTRON_TRANSPORT | 0.00000 | 0.00000 | 0.74774 | 0.74111 | 1.46056 | 75 |
| GO_AZUROPHIL_GRANULE | 0.00022 | 0.00481 | 0.51885 | 0.64342 | 1.27246 | 84 |
| GO_AZUROPHIL_GRANULE_LUMEN | 0.00024 | 0.00532 | 0.51885 | 0.69838 | 1.35077 | 47 |
| GO_BINDING_OF_SPERM_TO_ZONA_PELLUCIDA | 0.00004 | 0.00123 | 0.55733 | 0.84846 | 1.54933 | 17 |
| GO_BLOOD_MICROPARTICLE | 0.00000 | 0.00003 | 0.65944 | 0.80407 | 1.53315 | 36 |
| GO_CADHERIN_BINDING | 0.00000 | 0.00000 | | 0.66299 | 1.34807 | 206 |
| GO_CATALYTIC_STEP_2_SPLICEOSOME | 0.00004 | 0.00107 | 0.55733 | 0.67808 | 1.33577 | 74 |
| GO_CATION_TRANSMEMBRANE_TRANSPORT | 0.00062 | 0.01168 | 0.47727 | 0.55753 | 1.14245 | 303 |
| GO_CATION_TRANSPORT | 0.00008 | 0.00202 | 0.53843 | 0.55953 | 1.15120 | 404 |
| GO_CELLULAR_AMINO_ACID_METABOLIC_PROCESS | 0.00000 | 0.00005 | 0.65944 | 0.62417 | 1.26550 | 183 |
| GO_CELLULAR_COMPONENT_DISASSEMBLY | 0.00000 | 0.00000 | 0.74774 | 0.60602 | 1.24482 | 334 |
| GO_CELLULAR_HOMEOSTASIS | 0.00000 | 0.00001 | 0.69013 | 0.58749 | 1.20867 | 391 |
| GO_CELLULAR_ION_HOMEOSTASIS | 0.00001 | 0.00023 | 0.61053 | 0.59690 | 1.21806 | 246 |
| GO_CELLULAR_KETONE_METABOLIC_PROCESS | 0.00005 | 0.00124 | 0.55733 | 0.62830 | 1.26275 | 133 |
| GO_CELLULAR_MONOVALENT_INORGANIC_CATION_HOME( | 0.00150 | 0.02377 | 0.45506 | 0.64889 | 1.26754 | 60 |
| GO_CELLULAR_NITROGEN_COMPOUND_CATABOLIC_PROCE | 0.00000 | 0.00000 | | 0.63994 | 1.31669 | 406 |
| GO_CELLULAR_PROTEIN_COMPLEX_DISASSEMBLY | 0.00000 | 0.00000 | 0.78819 | 0.67448 | 1.36020 | 154 |
| GO_CELLULAR_RESPIRATION | 0.00000 | 0.00000 | 0.78819 | 0.69186 | 1.39049 | 133 |
| GO_CELLULAR_RESPONSE_TO_CHEMICAL_STRESS | 0.00290 | 0.03971 | 0.43171 | 0.56682 | 1.15065 | 196 |
| GO_CELLULAR_RESPONSE_TO_GAMMA_RADIATION | 0.00261 | 0.03676 | 0.43171 | 0.75979 | 1.40446 | 20 |
| GO_CELLULAR_RESPONSE_TO_HEAT | 0.00004 | 0.00115 | 0.55733 | 0.65996 | 1.30550 | 85 |
| GO_CELLULAR_RESPONSE_TO_ORGANIC_CYCLIC_COMPOUI | 0.00342 | 0.04490 | 0.43171 | 0.55213 | 1.12803 | 266 |
| GO_CELLULAR_RESPONSE_TO_OXYGEN_LEVELS | 0.00000 | 0.00009 | 0.64355 | 0.64247 | 1.29349 | 143 |
| GO_CELL_ACTIVATION_INVOLVED_IN_IMMUNE_RESPONSE | 0.00000 | 0.00000 | 0.76146 | 0.60769 | 1.24654 | 323 |
| GO_CELL_ADHESION_MOLECULE_BINDING | 0.00000 | 0.00000 | | 0.64225 | 1.31263 | 273 |
| GO_CELL_CELL_RECOGNITION | 0.00022 | 0.00478 | 0.51885 | 0.76421 | 1.43873 | 29 |
| GO_CELL_CELL_SIGNALING_BY_WNT | 0.00124 | 0.02057 | 0.45506 | 0.55558 | 1.13569 | 272 |
| GO_CELL_CYCLE_G2_M_PHASE_TRANSITION | 0.00009 | 0.00211 | 0.53843 | 0.58967 | 1.19919 | 203 |
| GO_CELL_CYCLE_PHASE_TRANSITION | 0.00000 | 0.00003 | 0.65944 | 0.57277 | 1.17963 | 440 |
| GO_CELL_DIVISION | 0.00002 | 0.00060 | 0.57561 | 0.56394 | 1.16025 | 403 |
| GO_CELL_KILLING | 0.00128 | 0.02081 | 0.45506 | 0.68098 | 1.31552 | 46 |
| GO_CELL_RECOGNITION | 0.00006 | 0.00164 | 0.53843 | 0.72474 | 1.40006 | 46 |
| GO_CELL_SUBSTRATE_JUNCTION | 0.00000 | 0.00000 | 0.80122 | 0.63238 | 1.29111 | 258 |
| GO_CELL_SURFACE_RECEPTOR_SIGNALING_PATHWAY_INVC | 0.00220 | 0.03193 | 0.43171 | 0.54732 | 1.12078 | 297 |
| GO_CENTRAL_NERVOUS_SYSTEM_DEVELOPMENT | 0.00183 | 0.02766 | 0.45506 | 0.53872 | 1.10956 | 444 |
| GO_CHAPERONE_COMPLEX | 0.00001 | 0.00035 | 0.59333 | 0.87047 | 1.58953 | 17 |
| GO_CHEMICAL_HOMEOSTASIS | 0.00242 | 0.03451 | 0.43171 | 0.53768 | 1.10806 | 453 |
| GO_CHROMOSOMAL_REGION | 0.00001 | 0.00021 | 0.61053 | 0.59438 | 1.21339 | 253 |
| GO_CHROMOSOME_CENTROMERIC_REGION | 0.00020 | 0.00450 | 0.51885 | 0.60129 | 1.21306 | 156 |
| GO_CHROMOSOME_SEGREGATION | 0.00006 | 0.00154 | 0.55733 | 0.58823 | 1.19956 | 238 |
| GO_CONDENSED_CHROMOSOME | 0.00018 | 0.00416 | 0.51885 | 0.60483 | 1.22130 | 161 |
| GO_CONDENSED_CHROMOSOME_CENTROMERIC_REGION | 0.00072 | 0.01313 | 0.47727 | 0.62663 | 1.24465 | 93 |
| GO_CONTRACTILE_FIBER | 0.00364 | 0.04703 | 0.43171 | 0.59401 | 1.18510 | 108 |
| GO_COTRANSLATIONAL_PROTEIN_TARGETING_TO_MEMBR/ | 0.00000 | 0.00000 | | 0.75391 | 1.49672 | 91 |
| GO_CYTOCHROME_COMPLEX | 0.00065 | 0.01207 | 0.47727 | 0.77718 | 1.44390 | 22 |
| GO_CYTOKINE_MEDIATED_SIGNALING_PATHWAY | 0.00000 | 0.00001 | 0.69013 | 0.59557 | 1.22186 | 322 |
| GO_CYTOPLASMIC_TRANSLATION | 0.00002 | 0.00050 | 0.57561 | 0.65610 | 1.30284 | 92 |
| GO_CYTOSOLIC_LARGE_RIBOSOMAL_SUBUNIT | 0.00003 | 0.00088 | 0.57561 | 0.71600 | 1.39216 | 52 |
| GO_CYTOSOLIC_RIBOSOME | 0.00000 | 0.00000 | 0.77494 | 0.72878 | 1.44837 | 95 |
| GO_CYTOSOLIC_SMALL_RIBOSOMAL_SUBUNIT | 0.00000 | 0.00005 | 0.64355 | 0.78242 | 1.50240 | 40 |
| GO_DEFENSE_RESPONSE_TO_OTHER_ORGANISM | 0.00000 | 0.00000 | | 0.61321 | 1.26166 | 404 |
| GO_DETECTION_OF_STIMULUS | 0.00316 | 0.04241 | 0.43171 | 0.60753 | 1.20740 | 95 |
| GO_DISORDERED_DOMAIN_SPECIFIC_BINDING | 0.00002 | 0.00065 | 0.57561 | 0.82370 | 1.53552 | 24 |
| GO_DIVALENT_INORGANIC_CATION_TRANSPORT | 0.00302 | 0.04103 | 0.43171 | 0.57865 | 1.16837 | 159 |
| GO_DNA_BIOSYNTHETIC_PROCESS | 0.00030 | 0.00638 | 0.49849 | 0.61878 | 1.23978 | 120 |
| GO_DNA_CONFORMATION_CHANGE | 0.00100 | 0.01740 | 0.45506 | 0.57344 | 1.16546 | 200 |
| GO_DNA_DAMAGE_RESPONSE_DETECTION_OF_DNA_DAM/ | 0.00353 | 0.04592 | 0.43171 | 0.71010 | 1.34513 | 32 |
| GO_DNA_REPLICATION | 0.00157 | 0.02451 | 0.45506 | 0.57386 | 1.16478 | 192 |
| GO_ELECTRON_TRANSFER_ACTIVITY | 0.00004 | 0.00109 | 0.55733 | 0.67770 | 1.33430 | 73 |
| GO_ELECTRON_TRANSPORT_CHAIN | 0.00000 | 0.00000 | 0.76146 | 0.69197 | 1.38739 | 123 |
| GO_ENDOPEPTIDASE_ACTIVITY | 0.00031 | 0.00652 | 0.49849 | 0.60067 | 1.21105 | 152 |
| GO_ENDOPEPTIDASE_COMPLEX | 0.00001 | 0.00021 | 0.61053 | 0.71632 | 1.39674 | 58 |

| Cluster 1 | | | | | | |
|---|---|---|---|---|---|---|
| GO_ENERGY_DERIVATION_BY_OXIDATION_OF_ORGANIC_CO | 0.00000 | 0.00000 | 0.77494 | 0.66646 | 1.34944 | 175 |
| GO_ENZYME_LINKED_RECEPTOR_PROTEIN_SIGNALING_PAT | 0.00372 | 0.04776 | 0.43171 | 0.53049 | 1.09387 | 474 |
| GO_ESTABLISHMENT_OF_ORGANELLE_LOCALIZATION | 0.00037 | 0.00736 | 0.49849 | 0.56825 | 1.16082 | 263 |
| GO_ESTABLISHMENT_OF_PROTEIN_LOCALIZATION_TO_END | 0.00000 | 0.00000 | | 0.74109 | 1.47563 | 101 |
| GO_ESTABLISHMENT_OF_PROTEIN_LOCALIZATION_TO_MEM | 0.00000 | 0.00000 | | 0.67317 | 1.37056 | 221 |
| GO_ESTABLISHMENT_OF_PROTEIN_LOCALIZATION_TO_ORG | 0.00000 | 0.00000 | | 0.63160 | 1.29941 | 397 |
| GO_ESTABLISHMENT_OF_PROTEIN_LOCALIZATION_TO_TELO | 0.00305 | 0.04133 | 0.43171 | 0.79313 | 1.43270 | 15 |
| GO_ESTABLISHMENT_OF_RNA_LOCALIZATION | 0.00003 | 0.00092 | 0.55733 | 0.61733 | 1.24654 | 161 |
| GO_ESTABLISHMENT_OF_TISSUE_POLARITY | 0.00000 | 0.00010 | 0.62726 | 0.69412 | 1.36874 | 79 |
| GO_EUKARYOTIC_TRANSLATION_INITIATION_FACTOR_3_CO | 0.00030 | 0.00631 | 0.49849 | 0.83844 | 1.51456 | 15 |
| GO_EXOCYTOSIS | 0.00000 | 0.00005 | 0.64355 | 0.57722 | 1.18830 | 419 |
| GO_FC_EPSILON_RECEPTOR_SIGNALING_PATHWAY | 0.00051 | 0.00991 | 0.47727 | 0.64264 | 1.26943 | 82 |
| GO_FC_RECEPTOR_SIGNALING_PATHWAY | 0.00000 | 0.00015 | 0.62726 | 0.64976 | 1.30277 | 123 |
| GO_FERTILIZATION | 0.00135 | 0.02181 | 0.45506 | 0.64924 | 1.26910 | 61 |
| GO_FIBROBLAST_GROWTH_FACTOR_RECEPTOR_SIGNALING | 0.00194 | 0.02886 | 0.45506 | 0.64793 | 1.26463 | 59 |
| GO_FICOLIN_1_RICH_GRANULE | 0.00000 | 0.00007 | 0.64355 | 0.69072 | 1.36853 | 88 |
| GO_GENERATION_OF_PRECURSOR_METABOLITES_AND_EN | 0.00000 | 0.00000 | | 0.63561 | 1.30308 | 313 |
| GO_GLUTAMATERGIC_SYNAPSE | 0.00032 | 0.00659 | 0.49849 | 0.59662 | 1.20365 | 156 |
| GO_GTP_METABOLIC_PROCESS | 0.00305 | 0.04133 | 0.43171 | 0.79295 | 1.43237 | 15 |
| GO_HEAD_DEVELOPMENT | 0.00174 | 0.02671 | 0.45506 | 0.54714 | 1.12488 | 364 |
| GO_HEMATOPOIETIC_PROGENITOR_CELL_DIFFERENTIATION | 0.00002 | 0.00075 | 0.57561 | 0.64397 | 1.28408 | 107 |
| GO_HEMATOPOIETIC_STEM_CELL_DIFFERENTIATION | 0.00000 | 0.00016 | 0.61053 | 0.71265 | 1.39386 | 62 |
| GO_HISTONE_MRNA_METABOLIC_PROCESS | 0.00390 | 0.04945 | 0.43171 | 0.75954 | 1.40900 | 21 |
| GO_HOST_CELLULAR_COMPONENT | 0.00128 | 0.02081 | 0.45506 | 0.68109 | 1.31573 | 46 |
| GO_HYDROLASE_ACTIVITY_ACTING_ON_ACID_ANHYDRIDES | 0.00092 | 0.01611 | 0.47727 | 0.54081 | 1.11559 | 499 |
| GO_IMMUNE_RESPONSE_REGULATING_SIGNALING_PATHW | 0.00000 | 0.00016 | 0.61053 | 0.61547 | 1.24924 | 192 |
| GO_IMMUNE_SYSTEM_DEVELOPMENT | 0.00082 | 0.01468 | 0.47727 | 0.54042 | 1.11383 | 458 |
| GO_IMPORT_INTO_NUCLEUS | 0.00049 | 0.00962 | 0.47727 | 0.61772 | 1.23517 | 112 |
| GO_INNATE_IMMUNE_RESPONSE | 0.00000 | 0.00000 | 0.81404 | 0.62151 | 1.27550 | 327 |
| GO_INNATE_IMMUNE_RESPONSE_ACTIVATING_SIGNAL_TRA | 0.00002 | 0.00059 | 0.57561 | 0.68754 | 1.35251 | 70 |
| GO_INNER_MITOCHONDRIAL_MEMBRANE_PROTEIN_COMP | 0.00001 | 0.00042 | 0.59333 | 0.67319 | 1.33536 | 90 |
| GO_INORGANIC_ION_TRANSMEMBRANE_TRANSPORT | 0.00089 | 0.01568 | 0.47727 | 0.55625 | 1.13924 | 299 |
| GO_INTERACTION_WITH_HOST | 0.00075 | 0.01354 | 0.47727 | 0.60836 | 1.21796 | 117 |
| GO_INTERLEUKIN_1_BETA_PRODUCTION | 0.00342 | 0.04490 | 0.43171 | 0.71414 | 1.33549 | 27 |
| GO_INTERLEUKIN_1_MEDIATED_SIGNALING_PATHWAY | 0.00002 | 0.00065 | 0.57561 | 0.68406 | 1.34661 | 72 |
| GO_INTERLEUKIN_1_PRODUCTION | 0.00088 | 0.01562 | 0.47727 | 0.72362 | 1.37377 | 33 |
| GO_INTERLEUKIN_8_PRODUCTION | 0.00220 | 0.03193 | 0.43171 | 0.72293 | 1.35192 | 27 |
| GO_INTRINSIC_APOPTOTIC_SIGNALING_PATHWAY | 0.00025 | 0.00535 | 0.49849 | 0.59433 | 1.20339 | 175 |
| GO_ION_HOMEOSTASIS | 0.00028 | 0.00607 | 0.49849 | 0.56573 | 1.15825 | 294 |
| GO_IRON_ION_TRANSPORT | 0.00014 | 0.00328 | 0.51885 | 0.72054 | 1.39014 | 45 |
| GO_KINETOCHORE | 0.00042 | 0.00827 | 0.49849 | 0.61617 | 1.23170 | 111 |
| GO_LARGE_RIBOSOMAL_SUBUNIT | 0.00000 | 0.00000 | 0.81404 | 0.72483 | 1.44369 | 103 |
| GO_LEUKOCYTE_MEDIATED_IMMUNITY | 0.00000 | 0.00000 | 0.82666 | 0.61733 | 1.26839 | 348 |
| GO_MACROAUTOPHAGY | 0.00272 | 0.03775 | 0.43171 | 0.56535 | 1.14988 | 204 |
| GO_MACROMOLECULE_TRANSMEMBRANE_TRANSPORTER_ | 0.00265 | 0.03700 | 0.43171 | 0.77979 | 1.42395 | 17 |
| GO_MEMBRANE_ORGANIZATION | 0.00003 | 0.00087 | 0.57561 | 0.55791 | 1.14913 | 443 |
| GO_MEMBRANE_REGION | 0.00011 | 0.00257 | 0.53843 | 0.60917 | 1.22842 | 153 |
| GO_METAL_ION_HOMEOSTASIS | 0.00083 | 0.01475 | 0.47727 | 0.56963 | 1.16068 | 225 |
| GO_METAL_ION_TRANSPORT | 0.00176 | 0.02684 | 0.45506 | 0.55119 | 1.13017 | 315 |
| GO_MICROTUBULE_CYTOSKELETON_ORGANIZATION_INVOL | 0.00126 | 0.02069 | 0.45506 | 0.61344 | 1.22243 | 106 |
| GO_MICROVILLUS | 0.00062 | 0.01168 | 0.47727 | 0.70216 | 1.34829 | 40 |
| GO_MITOCHONDRIAL_ELECTRON_TRANSPORT_NADH_TO_U | 0.00018 | 0.00397 | 0.51885 | 0.71838 | 1.38500 | 44 |
| GO_MITOCHONDRIAL_ENVELOPE | 0.00000 | 0.00000 | | 0.61260 | 1.26364 | 490 |
| GO_MITOCHONDRIAL_GENE_EXPRESSION | 0.00000 | 0.00012 | 0.62726 | 0.63730 | 1.28218 | 139 |
| GO_MITOCHONDRIAL_LARGE_RIBOSOMAL_SUBUNIT | 0.00001 | 0.00042 | 0.59333 | 0.73627 | 1.42928 | 50 |
| GO_MITOCHONDRIAL_MATRIX | 0.00001 | 0.00023 | 0.61053 | 0.58251 | 1.19498 | 324 |
| GO_MITOCHONDRIAL_PROTEIN_COMPLEX | 0.00000 | 0.00000 | | 0.69576 | 1.41327 | 198 |
| GO_MITOCHONDRIAL_SMALL_RIBOSOMAL_SUBUNIT | 0.00076 | 0.01366 | 0.47727 | 0.75897 | 1.41633 | 25 |
| GO_MITOCHONDRIAL_TRANSLATION | 0.00000 | 0.00003 | 0.65944 | 0.67086 | 1.34317 | 118 |
| GO_MITOCHONDRIAL_TRANSLATIONAL_TERMINATION | 0.00000 | 0.00000 | 0.77494 | 0.73502 | 1.45173 | 81 |
| GO_MITOCHONDRION_ORGANIZATION | 0.00003 | 0.00090 | 0.55733 | 0.57075 | 1.17299 | 351 |
| GO_MITOTIC_CELL_CYCLE_CHECKPOINT | 0.00331 | 0.04386 | 0.43171 | 0.59366 | 1.18861 | 118 |
| GO_MITOTIC_NUCLEAR_DIVISION | 0.00154 | 0.02400 | 0.45506 | 0.57280 | 1.16214 | 190 |
| GO_MITOTIC_SISTER_CHROMATID_SEGREGATION | 0.00067 | 0.01246 | 0.47727 | 0.60396 | 1.21215 | 126 |
| GO_MITOTIC_SPINDLE | 0.00170 | 0.02622 | 0.45506 | 0.61496 | 1.22370 | 99 |
| GO_MODIFICATION_DEPENDENT_MACROMOLECULE_CATAB | 0.00000 | 0.00012 | 0.62726 | 0.56691 | 1.16757 | 440 |
| GO_MONOVALENT_INORGANIC_CATION_HOMEOSTASIS | 0.00119 | 0.01978 | 0.45506 | 0.64915 | 1.27267 | 66 |
| GO_MONOVALENT_INORGANIC_CATION_TRANSMEMBRAN | 0.00001 | 0.00046 | 0.59333 | 0.64896 | 1.29873 | 114 |
| GO_MORPHOGENESIS_OF_AN_EPITHELIUM | 0.00115 | 0.01934 | 0.45506 | 0.55635 | 1.13726 | 272 |
| GO_MORPHOGENESIS_OF_A_POLARIZED_EPITHELIUM | 0.00000 | 0.00004 | 0.65944 | 0.68673 | 1.36335 | 91 |
| GO_MRNA_3_END_PROCESSING | 0.00191 | 0.02846 | 0.45506 | 0.62604 | 1.23555 | 80 |
| GO_MRNA_BINDING | 0.00000 | 0.00001 | 0.69013 | 0.62049 | 1.26398 | 224 |
| GO_MRNA_EXPORT_FROM_NUCLEUS | 0.00025 | 0.00535 | 0.49849 | 0.64061 | 1.27180 | 91 |

| Cluster 1 | | | | | | |
|---|---|---|---|---|---|---|
| GO_MRNA_PROCESSING | 0.00000 | 0.00000 | 0.71951 | 0.59039 | 1.21439 | 393 |
| GO_MRNA_TRANSPORT | 0.00053 | 0.01019 | 0.47727 | 0.61306 | 1.22744 | 118 |
| GO_MULTIVESICULAR_BODY | 0.00015 | 0.00356 | 0.51885 | 0.77822 | 1.46113 | 28 |
| GO_MULTI_ORGANISM_LOCALIZATION | 0.00055 | 0.01065 | 0.47727 | 0.69521 | 1.34302 | 46 |
| GO_MULTI_ORGANISM_PROCESS | 0.00089 | 0.01568 | 0.47727 | 0.54104 | 1.11608 | 499 |
| GO_MYELOID_LEUKOCYTE_ACTIVATION | 0.00000 | 0.00000 | 0.78819 | 0.61743 | 1.26508 | 302 |
| GO_MYELOID_LEUKOCYTE_MEDIATED_IMMUNITY | 0.00000 | 0.00000 | 0.80122 | 0.63037 | 1.28924 | 278 |
| GO_NADH_DEHYDROGENASE_ACTIVITY | 0.00050 | 0.00976 | 0.47727 | 0.72921 | 1.39262 | 37 |
| GO_NADH_DEHYDROGENASE_COMPLEX | 0.00065 | 0.01207 | 0.47727 | 0.70125 | 1.34655 | 40 |
| GO_NADH_DEHYDROGENASE_COMPLEX_ASSEMBLY | 0.00117 | 0.01953 | 0.45506 | 0.66117 | 1.28804 | 55 |
| GO_NCRNA_METABOLIC_PROCESS | 0.00128 | 0.02081 | 0.45506 | 0.54811 | 1.12601 | 337 |
| GO_NCRNA_PROCESSING | 0.00178 | 0.02697 | 0.45506 | 0.55500 | 1.13608 | 288 |
| GO_NEGATIVE_REGULATION_OF_BLOOD_VESSEL_DIAMETE | 0.00246 | 0.03495 | 0.43171 | 0.72147 | 1.35458 | 28 |
| GO_NEGATIVE_REGULATION_OF_CANONICAL_WNT_SIGNAI | 0.00152 | 0.02399 | 0.45506 | 0.61185 | 1.21647 | 96 |
| GO_NEGATIVE_REGULATION_OF_CATABOLIC_PROCESS | 0.00068 | 0.01259 | 0.47727 | 0.58480 | 1.18398 | 174 |
| GO_NEGATIVE_REGULATION_OF_CATALYTIC_ACTIVITY | 0.00383 | 0.04873 | 0.43171 | 0.54357 | 1.11786 | 372 |
| GO_NEGATIVE_REGULATION_OF_CELLULAR_CATABOLIC_PR | 0.00353 | 0.04592 | 0.43171 | 0.58039 | 1.16957 | 149 |
| GO_NEGATIVE_REGULATION_OF_CELLULAR_COMPONENT_ | 0.00025 | 0.00535 | 0.49849 | 0.55229 | 1.13609 | 389 |
| GO_NEGATIVE_REGULATION_OF_CELLULAR_PROTEIN_CATA | 0.00185 | 0.02779 | 0.45506 | 0.65986 | 1.28457 | 54 |
| GO_NEGATIVE_REGULATION_OF_CELL_CYCLE | 0.00007 | 0.00171 | 0.53843 | 0.56231 | 1.15684 | 399 |
| GO_NEGATIVE_REGULATION_OF_CELL_CYCLE_G2_M_PHASI | 0.00000 | 0.00009 | 0.62726 | 0.68856 | 1.35996 | 81 |
| GO_NEGATIVE_REGULATION_OF_CELL_CYCLE_PHASE_TRAN | 0.00000 | 0.00003 | 0.65944 | 0.63043 | 1.27636 | 174 |
| GO_NEGATIVE_REGULATION_OF_CELL_CYCLE_PROCESS | 0.00000 | 0.00011 | 0.62726 | 0.60624 | 1.23579 | 229 |
| GO_NEGATIVE_REGULATION_OF_CELL_DEATH | 0.00072 | 0.01313 | 0.47727 | 0.54271 | 1.11910 | 481 |
| GO_NEGATIVE_REGULATION_OF_ESTABLISHMENT_OF_PRO | 0.00298 | 0.04063 | 0.43171 | 0.63352 | 1.24097 | 65 |
| GO_NEGATIVE_REGULATION_OF_MITOTIC_CELL_CYCLE | 0.00000 | 0.00006 | 0.64355 | 0.61092 | 1.24426 | 223 |
| GO_NEGATIVE_REGULATION_OF_ORGANELLE_ORGANIZATI | 0.00239 | 0.03407 | 0.43171 | 0.55941 | 1.14143 | 243 |
| GO_NEGATIVE_REGULATION_OF_PEPTIDASE_ACTIVITY | 0.00097 | 0.01681 | 0.47727 | 0.62099 | 1.23416 | 95 |
| GO_NEGATIVE_REGULATION_OF_PROTEIN_CATABOLIC_PRO | 0.00066 | 0.01233 | 0.47727 | 0.64605 | 1.27412 | 78 |
| GO_NEGATIVE_REGULATION_OF_PROTEOLYSIS | 0.00005 | 0.00130 | 0.55733 | 0.61281 | 1.23517 | 151 |
| GO_NEGATIVE_REGULATION_OF_TRANSPORT | 0.00010 | 0.00234 | 0.53843 | 0.59614 | 1.20983 | 191 |
| GO_NEGATIVE_REGULATION_OF_UBIQUITIN_DEPENDENT_ | 0.00346 | 0.04528 | 0.43171 | 0.70601 | 1.34281 | 34 |
| GO_NIK_NF_KAPPAB_SIGNALING | 0.00004 | 0.00116 | 0.55733 | 0.64733 | 1.28755 | 98 |
| GO_NON_CANONICAL_WNT_SIGNALING_PATHWAY | 0.00000 | 0.00012 | 0.62726 | 0.67724 | 1.34453 | 91 |
| GO_NUCLEAR_CHROMOSOME_SEGREGATION | 0.00035 | 0.00711 | 0.49849 | 0.58532 | 1.18754 | 190 |
| GO_NUCLEAR_ENVELOPE | 0.00146 | 0.02325 | 0.45506 | 0.55754 | 1.14008 | 277 |
| GO_NUCLEAR_EXPORT | 0.00000 | 0.00004 | 0.65944 | 0.63677 | 1.28586 | 160 |
| GO_NUCLEAR_MATRIX | 0.00071 | 0.01305 | 0.47727 | 0.63723 | 1.25656 | 79 |
| GO_NUCLEAR_PERIPHERY | 0.00033 | 0.00676 | 0.49849 | 0.63339 | 1.25921 | 97 |
| GO_NUCLEAR_SPECK | 0.00005 | 0.00128 | 0.55733 | 0.57986 | 1.18491 | 270 |
| GO_NUCLEAR_TRANSCRIBED_MRNA_CATABOLIC_PROCESS | 0.00000 | 0.00000 | 0.76146 | 0.65672 | 1.32907 | 173 |
| GO_NUCLEAR_TRANSCRIBED_MRNA_CATABOLIC_PROCESS_ | 0.00000 | 0.00000 | 0.74774 | 0.69911 | 1.39402 | 107 |
| GO_NUCLEAR_TRANSPORT | 0.00000 | 0.00001 | 0.69013 | 0.61135 | 1.24833 | 260 |
| GO_NUCLEOBASE_CONTAINING_COMPOUND_TRANSPORT | 0.00006 | 0.00149 | 0.55733 | 0.59855 | 1.21473 | 191 |
| GO_ORGANELLAR_RIBOSOME | 0.00000 | 0.00000 | 0.73376 | 0.74233 | 1.46372 | 76 |
| GO_ORGANELLE_FISSION | 0.00170 | 0.02622 | 0.45506 | 0.55029 | 1.12696 | 298 |
| GO_ORGANELLE_INNER_MEMBRANE | 0.00000 | 0.00000 | | 0.62946 | 1.29398 | 354 |
| GO_ORGANIC_CYCLIC_COMPOUND_CATABOLIC_PROCESS | 0.00000 | 0.00000 | | 0.63794 | 1.31349 | 424 |
| GO_OXIDATIVE_PHOSPHORYLATION | 0.00000 | 0.00000 | 0.76146 | 0.72559 | 1.44205 | 95 |
| GO_OXIDOREDUCTASE_ACTIVITY | 0.00016 | 0.00378 | 0.51885 | 0.55664 | 1.14500 | 379 |
| GO_OXIDOREDUCTASE_ACTIVITY_ACTING_ON_A_HEME_GR | 0.00097 | 0.01681 | 0.47727 | 0.80112 | 1.46978 | 18 |
| GO_OXIDOREDUCTASE_ACTIVITY_ACTING_ON_NAD_P_H | 0.00059 | 0.01128 | 0.47727 | 0.65731 | 1.28654 | 64 |
| GO_OXIDOREDUCTASE_ACTIVITY_ACTING_ON_NAD_P_H_Q | 0.00126 | 0.02069 | 0.45506 | 0.67649 | 1.30272 | 43 |
| GO_OXIDOREDUCTASE_COMPLEX | 0.00000 | 0.00016 | 0.61053 | 0.70258 | 1.38533 | 76 |
| GO_PEPTIDASE_COMPLEX | 0.00001 | 0.00021 | 0.61053 | 0.69646 | 1.37123 | 73 |
| GO_PEPTIDE_BINDING | 0.00209 | 0.03072 | 0.43171 | 0.59994 | 1.20028 | 113 |
| GO_PEPTIDE_BIOSYNTHETIC_PROCESS | 0.00000 | 0.00000 | | 0.64821 | 1.33716 | 500 |
| GO_PEPTIDE_TRANSMEMBRANE_TRANSPORTER_ACTIVITY | 0.00021 | 0.00472 | 0.51885 | 0.79410 | 1.47535 | 22 |
| GO_PHAGOCYTOSIS | 0.00265 | 0.03700 | 0.43171 | 0.59714 | 1.19469 | 113 |
| GO_PHAGOSOME_ACIDIFICATION | 0.00013 | 0.00320 | 0.51885 | 0.80030 | 1.48685 | 22 |
| GO_PHAGOSOME_MATURATION | 0.00038 | 0.00751 | 0.49849 | 0.75257 | 1.41954 | 30 |
| GO_PIGMENT_GRANULE | 0.00000 | 0.00000 | 0.74774 | 0.73915 | 1.45487 | 71 |
| GO_PLASMA_LIPOPROTEIN_PARTICLE_CLEARANCE | 0.00342 | 0.04490 | 0.43171 | 0.73464 | 1.36949 | 24 |
| GO_POLYSOMAL_RIBOSOME | 0.00031 | 0.00645 | 0.49849 | 0.75826 | 1.43026 | 30 |
| GO_POLYSOME | 0.00006 | 0.00154 | 0.55733 | 0.68598 | 1.34091 | 61 |
| GO_POSITIVE_REGULATION_OF_BINDING | 0.00121 | 0.02002 | 0.45506 | 0.61345 | 1.22015 | 98 |
| GO_POSITIVE_REGULATION_OF_CANONICAL_WNT_SIGNALI | 0.00000 | 0.00003 | 0.65944 | 0.69031 | 1.36875 | 89 |
| GO_POSITIVE_REGULATION_OF_CELLULAR_AMIDE_METABO | 0.00016 | 0.00360 | 0.51885 | 0.63374 | 1.26272 | 104 |
| GO_POSITIVE_REGULATION_OF_CELLULAR_PROTEIN_LOCAL | 0.00000 | 0.00014 | 0.62726 | 0.61588 | 1.25008 | 192 |
| GO_POSITIVE_REGULATION_OF_CELL_POPULATION_PROLIF | 0.00383 | 0.04873 | 0.43171 | 0.54013 | 1.11088 | 375 |
| GO_POSITIVE_REGULATION_OF_CHROMOSOME_ORGANIZA | 0.00357 | 0.04629 | 0.43171 | 0.59780 | 1.19633 | 114 |
| GO_POSITIVE_REGULATION_OF_DEFENSE_RESPONSE | 0.00000 | 0.00000 | 0.73376 | 0.65707 | 1.32680 | 163 |
| GO_POSITIVE_REGULATION_OF_DNA_BIOSYNTHETIC_PROC | 0.00030 | 0.00631 | 0.49849 | 0.69858 | 1.35300 | 48 |

| Cluster 1 | | | | | | |
|---|---|---|---|---|---|---|
| GO_POSITIVE_REGULATION_OF_DNA_METABOLIC_PROCESS | 0.00012 | 0.00294 | 0.51885 | 0.62305 | 1.24783 | 119 |
| GO_POSITIVE_REGULATION_OF_ESTABLISHMENT_OF_PROT | 0.00000 | 0.00001 | 0.69013 | 0.63308 | 1.28463 | 189 |
| GO_POSITIVE_REGULATION_OF_IMMUNE_RESPONSE | 0.00000 | 0.00000 | 0.81404 | 0.63458 | 1.29672 | 270 |
| GO_POSITIVE_REGULATION_OF_IMMUNE_SYSTEM_PROCES | 0.00000 | 0.00002 | 0.67496 | 0.58514 | 1.20366 | 383 |
| GO_POSITIVE_REGULATION_OF_INNATE_IMMUNE_RESPON | 0.00000 | 0.00003 | 0.67496 | 0.67107 | 1.34297 | 114 |
| GO_POSITIVE_REGULATION_OF_INTRACELLULAR_PROTEIN_ | 0.00023 | 0.00494 | 0.51885 | 0.62359 | 1.24410 | 108 |
| GO_POSITIVE_REGULATION_OF_INTRACELLULAR_TRANSPO | 0.00003 | 0.00082 | 0.57561 | 0.62719 | 1.26016 | 131 |
| GO_POSITIVE_REGULATION_OF_MRNA_METABOLIC_PROCE | 0.00191 | 0.02846 | 0.45506 | 0.64889 | 1.26649 | 59 |
| GO_POSITIVE_REGULATION_OF_MRNA_PROCESSING | 0.00239 | 0.03407 | 0.43171 | 0.74239 | 1.38223 | 23 |
| GO_POSITIVE_REGULATION_OF_MRNA_SPLICING_VIA_SPLI | 0.00253 | 0.03591 | 0.43171 | 0.79634 | 1.43850 | 15 |
| GO_POSITIVE_REGULATION_OF_ORGANELLE_ORGANIZATIO | 0.00006 | 0.00156 | 0.55733 | 0.56856 | 1.16861 | 358 |
| GO_POSITIVE_REGULATION_OF_PROTEIN_LOCALIZATION_T | 0.00001 | 0.00040 | 0.59333 | 0.72206 | 1.40527 | 53 |
| GO_POSITIVE_REGULATION_OF_RESPONSE_TO_BIOTIC_STI | 0.00000 | 0.00002 | 0.67496 | 0.67300 | 1.35176 | 129 |
| GO_POSITIVE_REGULATION_OF_RESPONSE_TO_EXTERNAL_ | 0.00000 | 0.00002 | 0.67496 | 0.62181 | 1.26435 | 206 |
| GO_POSITIVE_REGULATION_OF_SECRETION | 0.00145 | 0.02302 | 0.45506 | 0.59251 | 1.19094 | 136 |
| GO_POSITIVE_REGULATION_OF_TRANSLATION | 0.00012 | 0.00294 | 0.51885 | 0.64505 | 1.28125 | 93 |
| GO_POSITIVE_REGULATION_OF_TRANSPORT | 0.00000 | 0.00001 | 0.69013 | 0.58529 | 1.20395 | 389 |
| GO_POSITIVE_REGULATION_OF_WNT_SIGNALING_PATHWA | 0.00000 | 0.00004 | 0.65944 | 0.67657 | 1.34822 | 105 |
| GO_POSTSYNAPSE | 0.00034 | 0.00696 | 0.49849 | 0.57376 | 1.17087 | 249 |
| GO_POSTTRANSCRIPTIONAL_REGULATION_OF_GENE_EXPR | 0.00000 | 0.00000 | | 0.63558 | 1.30932 | 447 |
| GO_POST_TRANSLATIONAL_PROTEIN_MODIFICATION | 0.00012 | 0.00291 | 0.53843 | 0.58725 | 1.19482 | 214 |
| GO_PRIMARY_ACTIVE_TRANSMEMBRANE_TRANSPORTER_A | 0.00002 | 0.00074 | 0.57561 | 0.72851 | 1.41423 | 50 |
| GO_PROTEASOMAL_PROTEIN_CATABOLIC_PROCESS | 0.00000 | 0.00014 | 0.62726 | 0.58197 | 1.19557 | 336 |
| GO_PROTEASOMAL_UBIQUITIN_INDEPENDENT_PROTEIN_C | 0.00002 | 0.00075 | 0.57561 | 0.86495 | 1.58690 | 18 |
| GO_PROTEASOME_CORE_COMPLEX | 0.00001 | 0.00030 | 0.59333 | 0.87296 | 1.59408 | 17 |
| GO_PROTEIN_CONTAINING_COMPLEX_DISASSEMBLY | 0.00000 | 0.00000 | 0.76146 | 0.63467 | 1.29287 | 224 |
| GO_PROTEIN_CONTAINING_COMPLEX_LOCALIZATION | 0.00010 | 0.00247 | 0.53843 | 0.59140 | 1.20197 | 200 |
| GO_PROTEIN_DNA_COMPLEX_SUBUNIT_ORGANIZATION | 0.00026 | 0.00563 | 0.49849 | 0.60201 | 1.21560 | 161 |
| GO_PROTEIN_DOMAIN_SPECIFIC_BINDING | 0.00006 | 0.00158 | 0.53843 | 0.56215 | 1.15630 | 393 |
| GO_PROTEIN_FOLDING | 0.00002 | 0.00070 | 0.57561 | 0.62400 | 1.25752 | 150 |
| GO_PROTEIN_FOLDING_CHAPERONE | 0.00368 | 0.04739 | 0.43171 | 0.75334 | 1.39254 | 20 |
| GO_PROTEIN_IMPORT | 0.00003 | 0.00089 | 0.57561 | 0.63336 | 1.27252 | 130 |
| GO_PROTEIN_LOCALIZATION_TO_CHROMOSOME_TELOMER | 0.00309 | 0.04173 | 0.43171 | 0.73242 | 1.36678 | 25 |
| GO_PROTEIN_LOCALIZATION_TO_ENDOPLASMIC_RETICULU | 0.00000 | 0.00000 | 0.80122 | 0.70968 | 1.42089 | 118 |
| GO_PROTEIN_LOCALIZATION_TO_MEMBRANE | 0.00000 | 0.00000 | 0.82666 | 0.61305 | 1.26037 | 368 |
| GO_PROTEIN_LOCALIZATION_TO_MITOCHONDRION | 0.00276 | 0.03817 | 0.43171 | 0.59747 | 1.18966 | 101 |
| GO_PROTEIN_LOCALIZATION_TO_NUCLEUS | 0.00000 | 0.00016 | 0.61053 | 0.62656 | 1.26910 | 177 |
| GO_PROTEIN_MODIFICATION_BY_SMALL_PROTEIN_REMOV | 0.00056 | 0.01079 | 0.47727 | 0.57727 | 1.17378 | 206 |
| GO_PROTEIN_N_TERMINUS_BINDING | 0.00228 | 0.03292 | 0.43171 | 0.62523 | 1.23064 | 71 |
| GO_PROTEIN_STABILIZATION | 0.00008 | 0.00203 | 0.53843 | 0.62917 | 1.25973 | 116 |
| GO_PROTEIN_TARGETING | 0.00000 | 0.00000 | | 0.62845 | 1.28768 | 302 |
| GO_PROTEIN_TARGETING_TO_MEMBRANE | 0.00000 | 0.00000 | | 0.70969 | 1.42782 | 139 |
| GO_PROTEIN_TRANSMEMBRANE_TRANSPORT | 0.00113 | 0.01915 | 0.45506 | 0.67863 | 1.30685 | 43 |
| GO_PROTON_TRANSMEMBRANE_TRANSPORT | 0.00001 | 0.00028 | 0.59333 | 0.69375 | 1.36569 | 72 |
| GO_PROTON_TRANSMEMBRANE_TRANSPORTER_ACTIVITY | 0.00004 | 0.00117 | 0.55733 | 0.70504 | 1.37469 | 56 |
| GO_PROTON_TRANSPORTING_TWO_SECTOR_ATPASE_COM | 0.00108 | 0.01856 | 0.45506 | 0.76163 | 1.41804 | 23 |
| GO_PYROPHOSPHATE_HYDROLYSIS_DRIVEN_PROTON_TRAN | 0.00239 | 0.03407 | 0.43171 | 0.76166 | 1.40791 | 20 |
| GO_PYRUVATE_METABOLIC_PROCESS | 0.00048 | 0.00947 | 0.49849 | 0.63515 | 1.25991 | 90 |
| GO_REGULATION_OF_ANIMAL_ORGAN_MORPHOGENESIS | 0.00001 | 0.00038 | 0.59333 | 0.63825 | 1.28027 | 124 |
| GO_REGULATION_OF_APOPTOTIC_SIGNALING_PATHWAY | 0.00135 | 0.02181 | 0.45506 | 0.57089 | 1.16185 | 217 |
| GO_REGULATION_OF_ATP_METABOLIC_PROCESS | 0.00074 | 0.01342 | 0.47727 | 0.64483 | 1.26958 | 73 |
| GO_REGULATION_OF_BINDING | 0.00113 | 0.01915 | 0.45506 | 0.57531 | 1.16737 | 193 |
| GO_REGULATION_OF_CELLULAR_AMIDE_METABOLIC_PROC | 0.00000 | 0.00001 | 0.69013 | 0.60505 | 1.23861 | 292 |
| GO_REGULATION_OF_CELLULAR_AMINE_METABOLIC_PROC | 0.00000 | 0.00000 | 0.73376 | 0.77412 | 1.50808 | 55 |
| GO_REGULATION_OF_CELLULAR_AMINO_ACID_METABOLIC | 0.00000 | 0.00002 | 0.67496 | 0.77129 | 1.49382 | 48 |
| GO_REGULATION_OF_CELLULAR_COMPONENT_BIOGENESIS | 0.00316 | 0.04241 | 0.43171 | 0.53053 | 1.09421 | 483 |
| GO_REGULATION_OF_CELLULAR_KETONE_METABOLIC_PRO | 0.00001 | 0.00041 | 0.59333 | 0.66136 | 1.31667 | 100 |
| GO_REGULATION_OF_CELLULAR_LOCALIZATION | 0.00000 | 0.00002 | 0.67496 | 0.57099 | 1.17786 | 496 |
| GO_REGULATION_OF_CELLULAR_PROTEIN_LOCALIZATION | 0.00000 | 0.00008 | 0.64355 | 0.58922 | 1.20861 | 320 |
| GO_REGULATION_OF_CELLULAR_RESPONSE_TO_HEAT | 0.00176 | 0.02684 | 0.45506 | 0.65024 | 1.26789 | 58 |
| GO_REGULATION_OF_CELLULAR_RESPONSE_TO_STRESS | 0.00313 | 0.04212 | 0.43171 | 0.53603 | 1.10405 | 431 |
| GO_REGULATION_OF_CELL_CYCLE_G2_M_PHASE_TRANSITI | 0.00022 | 0.00478 | 0.51885 | 0.60017 | 1.21248 | 165 |
| GO_REGULATION_OF_CELL_CYCLE_PHASE_TRANSITION | 0.00001 | 0.00030 | 0.59333 | 0.57935 | 1.18840 | 323 |
| GO_REGULATION_OF_CELL_CYCLE_PROCESS | 0.00000 | 0.00006 | 0.64355 | 0.56892 | 1.17356 | 489 |
| GO_REGULATION_OF_CHROMOSOME_ORGANIZATION | 0.00034 | 0.00703 | 0.49849 | 0.57397 | 1.16994 | 230 |
| GO_REGULATION_OF_CHROMOSOME_SEGREGATION | 0.00145 | 0.02302 | 0.45506 | 0.62302 | 1.23315 | 86 |
| GO_REGULATION_OF_CHROMOSOME_SEPARATION | 0.00390 | 0.04945 | 0.43171 | 0.64294 | 1.25252 | 55 |
| GO_REGULATION_OF_CYSTEINE_TYPE_ENDOPEPTIDASE_AC | 0.00185 | 0.02779 | 0.45506 | 0.59704 | 1.19813 | 127 |
| GO_REGULATION_OF_DEFENSE_RESPONSE | 0.00000 | 0.00001 | 0.69013 | 0.60517 | 1.23865 | 287 |
| GO_REGULATION_OF_DNA_BIOSYNTHETIC_PROCESS | 0.00009 | 0.00224 | 0.53843 | 0.67711 | 1.32956 | 67 |
| GO_REGULATION_OF_DNA_METABOLIC_PROCESS | 0.00005 | 0.00139 | 0.55733 | 0.59492 | 1.21005 | 210 |
| GO_REGULATION_OF_DNA_TEMPLATED_TRANSCRIPTION_I | 0.00000 | 0.00003 | 0.67496 | 0.70150 | 1.38551 | 81 |
| GO_REGULATION_OF_GENERATION_OF_PRECURSOR_META | 0.00039 | 0.00776 | 0.49849 | 0.63596 | 1.26256 | 91 |

| Cluster 1 | | | | | | |
|---|---|---|---|---|---|---|
| GO_REGULATION_OF_GLYCOPROTEIN_METABOLIC_PROCES | 0.00115 | 0.01934 | 0.45506 | 0.76078 | 1.41647 | 23 |
| GO_REGULATION_OF_HEMATOPOIETIC_PROGENITOR_CELL_ | 0.00004 | 0.00108 | 0.55733 | 0.68683 | 1.34541 | 65 |
| GO_REGULATION_OF_HEMATOPOIETIC_STEM_CELL_DIFFER | 0.00001 | 0.00023 | 0.61053 | 0.72329 | 1.40906 | 55 |
| GO_REGULATION_OF_IMMUNE_RESPONSE | 0.00000 | 0.00000 | 0.76146 | 0.60479 | 1.24246 | 345 |
| GO_REGULATION_OF_INNATE_IMMUNE_RESPONSE | 0.00000 | 0.00005 | 0.64355 | 0.63902 | 1.28869 | 154 |
| GO_REGULATION_OF_INTRACELLULAR_PROTEIN_TRANSPOI | 0.00021 | 0.00472 | 0.51885 | 0.60410 | 1.21954 | 158 |
| GO_REGULATION_OF_INTRACELLULAR_TRANSPORT | 0.00006 | 0.00159 | 0.53843 | 0.59398 | 1.20814 | 210 |
| GO_REGULATION_OF_INTRINSIC_APOPTOTIC_SIGNALING_P | 0.00283 | 0.03899 | 0.43171 | 0.60834 | 1.20902 | 95 |
| GO_REGULATION_OF_MITOTIC_CELL_CYCLE | 0.00000 | 0.00006 | 0.64355 | 0.57313 | 1.18030 | 430 |
| GO_REGULATION_OF_MORPHOGENESIS_OF_AN_EPITHELIU | 0.00001 | 0.00022 | 0.61053 | 0.66853 | 1.33030 | 99 |
| GO_REGULATION_OF_MRNA_CATABOLIC_PROCESS | 0.00000 | 0.00000 | 0.74774 | 0.67123 | 1.35357 | 153 |
| GO_REGULATION_OF_MRNA_METABOLIC_PROCESS | 0.00000 | 0.00000 | | 0.66846 | 1.36440 | 252 |
| GO_REGULATION_OF_MRNA_PROCESSING | 0.00000 | 0.00014 | 0.62726 | 0.65810 | 1.31592 | 112 |
| GO_REGULATION_OF_MRNA_SPLICING_VIA_SPLICEOSOME | 0.00001 | 0.00028 | 0.61053 | 0.67470 | 1.33545 | 86 |
| GO_REGULATION_OF_MULTI_ORGANISM_PROCESS | 0.00110 | 0.01876 | 0.45506 | 0.72774 | 1.37514 | 31 |
| GO_REGULATION_OF_NUCLEAR_DIVISION | 0.00202 | 0.02980 | 0.43171 | 0.60054 | 1.20274 | 119 |
| GO_REGULATION_OF_NUCLEOCYTOPLASMIC_TRANSPORT | 0.00003 | 0.00097 | 0.55733 | 0.66222 | 1.30696 | 80 |
| GO_REGULATION_OF_OXIDATIVE_PHOSPHORYLATION | 0.00017 | 0.00385 | 0.51885 | 0.81517 | 1.50683 | 20 |
| GO_REGULATION_OF_PEPTIDASE_ACTIVITY | 0.00154 | 0.02400 | 0.45506 | 0.57322 | 1.16364 | 196 |
| GO_REGULATION_OF_PEPTIDE_TRANSPORT | 0.00000 | 0.00009 | 0.62726 | 0.59061 | 1.21091 | 314 |
| GO_REGULATION_OF_PLASMA_LIPOPROTEIN_PARTICLE_LEV | 0.00331 | 0.04386 | 0.43171 | 0.69769 | 1.32453 | 33 |
| GO_REGULATION_OF_PROTEIN_CATABOLIC_PROCESS | 0.00159 | 0.02466 | 0.45506 | 0.56296 | 1.14808 | 239 |
| GO_REGULATION_OF_PROTEIN_IMPORT | 0.00033 | 0.00690 | 0.49849 | 0.71053 | 1.36436 | 40 |
| GO_REGULATION_OF_PROTEIN_LOCALIZATION | 0.00000 | 0.00002 | 0.67496 | 0.57391 | 1.18378 | 485 |
| GO_REGULATION_OF_PROTEIN_LOCALIZATION_TO_MEMBR | 0.00220 | 0.03193 | 0.43171 | 0.60030 | 1.19530 | 101 |
| GO_REGULATION_OF_PROTEIN_LOCALIZATION_TO_NUCLEU | 0.00001 | 0.00030 | 0.59333 | 0.67346 | 1.32914 | 80 |
| GO_REGULATION_OF_PROTEIN_MODIFICATION_BY_SMALL_ | 0.00376 | 0.04812 | 0.43171 | 0.57393 | 1.15826 | 157 |
| GO_REGULATION_OF_PROTEIN_STABILITY | 0.00054 | 0.01033 | 0.47727 | 0.58212 | 1.18056 | 185 |
| GO_REGULATION_OF_PROTEOLYSIS | 0.00035 | 0.00711 | 0.49849 | 0.55850 | 1.14829 | 365 |
| GO_REGULATION_OF_RESPONSE_TO_BIOTIC_STIMULUS | 0.00000 | 0.00001 | 0.69013 | 0.63548 | 1.29124 | 199 |
| GO_REGULATION_OF_RESPONSE_TO_EXTERNAL_STIMULUS | 0.00011 | 0.00257 | 0.53843 | 0.56063 | 1.15388 | 416 |
| GO_REGULATION_OF_RNA_SPLICING | 0.00004 | 0.00107 | 0.55733 | 0.63856 | 1.27850 | 118 |
| GO_REGULATION_OF_SISTER_CHROMATID_SEGREGATION | 0.00265 | 0.03700 | 0.43171 | 0.63525 | 1.24736 | 67 |
| GO_REGULATION_OF_SMALL_MOLECULE_METABOLIC_PRO | 0.00001 | 0.00025 | 0.61053 | 0.60309 | 1.22832 | 223 |
| GO_REGULATION_OF_STEM_CELL_DIFFERENTIATION | 0.00001 | 0.00035 | 0.59333 | 0.68144 | 1.34607 | 82 |
| GO_REGULATION_OF_SYMBIOTIC_PROCESS | 0.00239 | 0.03407 | 0.43171 | 0.59156 | 1.18726 | 126 |
| GO_REGULATION_OF_TELOMERE_MAINTENANCE | 0.00290 | 0.03971 | 0.43171 | 0.64732 | 1.26107 | 55 |
| GO_REGULATION_OF_TELOMERE_MAINTENANCE_VIA_TELO | 0.00268 | 0.03733 | 0.43171 | 0.66217 | 1.27516 | 43 |
| GO_REGULATION_OF_TRANSCRIPTION_FROM_RNA_POLYM | 0.00000 | 0.00001 | 0.69013 | 0.73918 | 1.44391 | 60 |
| GO_REGULATION_OF_TRANSLATIONAL_INITIATION | 0.00000 | 0.00015 | 0.62726 | 0.71173 | 1.39256 | 63 |
| GO_RESPIRASOME | 0.00000 | 0.00002 | 0.67496 | 0.72038 | 1.41911 | 74 |
| GO_RESPIRATORY_CHAIN_COMPLEX | 0.00000 | 0.00004 | 0.65944 | 0.72489 | 1.41831 | 63 |
| GO_RESPIRATORY_ELECTRON_TRANSPORT_CHAIN | 0.00000 | 0.00000 | 0.77494 | 0.73617 | 1.45588 | 84 |
| GO_RESPONSE_TO_COCAINE | 0.00060 | 0.01141 | 0.47727 | 0.79175 | 1.46353 | 20 |
| GO_RESPONSE_TO_DRUG | 0.00003 | 0.00099 | 0.55733 | 0.60598 | 1.22771 | 178 |
| GO_RESPONSE_TO_FIBROBLAST_GROWTH_FACTOR | 0.00220 | 0.03193 | 0.43171 | 0.63327 | 1.24576 | 70 |
| GO_RESPONSE_TO_HEAT | 0.00028 | 0.00590 | 0.49849 | 0.62495 | 1.24537 | 105 |
| GO_RESPONSE_TO_HYDROGEN_PEROXIDE | 0.00290 | 0.03971 | 0.43171 | 0.62551 | 1.23360 | 78 |
| GO_RESPONSE_TO_INORGANIC_SUBSTANCE | 0.00194 | 0.02886 | 0.45506 | 0.55944 | 1.14226 | 259 |
| GO_RESPONSE_TO_INTERLEUKIN_1 | 0.00005 | 0.00139 | 0.55733 | 0.64613 | 1.28693 | 103 |
| GO_RESPONSE_TO_LEUKEMIA_INHIBITORY_FACTOR | 0.00350 | 0.04565 | 0.43171 | 0.62623 | 1.23041 | 68 |
| GO_RESPONSE_TO_ORGANIC_CYCLIC_COMPOUND | 0.00003 | 0.00097 | 0.55733 | 0.56280 | 1.15861 | 419 |
| GO_RESPONSE_TO_OXIDATIVE_STRESS | 0.00154 | 0.02400 | 0.45506 | 0.56324 | 1.14865 | 239 |
| GO_RESPONSE_TO_OXYGEN_LEVELS | 0.00000 | 0.00006 | 0.64355 | 0.61218 | 1.24543 | 213 |
| GO_RESPONSE_TO_STEROID_HORMONE | 0.00077 | 0.01395 | 0.47727 | 0.59177 | 1.19464 | 158 |
| GO_RESPONSE_TO_TEMPERATURE_STIMULUS | 0.00113 | 0.01915 | 0.45506 | 0.59463 | 1.19512 | 132 |
| GO_RESPONSE_TO_TUMOR_NECROSIS_FACTOR | 0.00001 | 0.00028 | 0.59333 | 0.63589 | 1.27889 | 137 |
| GO_RETINA_HOMEOSTASIS | 0.00065 | 0.01207 | 0.47727 | 0.72865 | 1.38332 | 33 |
| GO_RIBONUCLEOPROTEIN_COMPLEX_BINDING | 0.00000 | 0.00009 | 0.62726 | 0.66885 | 1.33283 | 105 |
| GO_RIBONUCLEOPROTEIN_COMPLEX_BIOGENESIS | 0.00000 | 0.00000 | | 0.63409 | 1.30300 | 349 |
| GO_RIBONUCLEOPROTEIN_COMPLEX_SUBUNIT_ORGANIZAT | 0.00000 | 0.00008 | 0.64355 | 0.63513 | 1.28309 | 165 |
| GO_RIBONUCLEOPROTEIN_GRANULE | 0.00033 | 0.00683 | 0.49849 | 0.59164 | 1.19657 | 169 |
| GO_RIBONUCLEOSIDE_TRIPHOSPHATE_METABOLIC_PROCES | 0.00178 | 0.02697 | 0.45506 | 0.69197 | 1.32733 | 39 |
| GO_RIBOSOMAL_LARGE_SUBUNIT_ASSEMBLY | 0.00383 | 0.04873 | 0.43171 | 0.74311 | 1.38061 | 22 |
| GO_RIBOSOMAL_LARGE_SUBUNIT_BIOGENESIS | 0.00001 | 0.00045 | 0.59333 | 0.70070 | 1.36968 | 61 |
| GO_RIBOSOMAL_SMALL_SUBUNIT_BIOGENESIS | 0.00053 | 0.01019 | 0.47727 | 0.66832 | 1.30269 | 57 |
| GO_RIBOSOMAL_SUBUNIT | 0.00000 | 0.00000 | | 0.74285 | 1.50027 | 164 |
| GO_RIBOSOME | 0.00000 | 0.00000 | | 0.73035 | 1.48352 | 198 |
| GO_RIBOSOME_ASSEMBLY | 0.00001 | 0.00038 | 0.59333 | 0.73766 | 1.43198 | 50 |
| GO_RIBOSOME_BIOGENESIS | 0.00000 | 0.00000 | 0.78819 | 0.64109 | 1.30724 | 237 |
| GO_RNA_3_END_PROCESSING | 0.00066 | 0.01233 | 0.47727 | 0.61483 | 1.23043 | 114 |
| GO_RNA_CATABOLIC_PROCESS | 0.00000 | 0.00000 | | 0.66506 | 1.36402 | 318 |
| GO_RNA_DEPENDENT_DNA_BIOSYNTHETIC_PROCESS | 0.00086 | 0.01519 | 0.47727 | 0.67877 | 1.31653 | 49 |

| Cluster 1 | | | | | | |
|---|---|---|---|---|---|---|
| GO_RNA_EXPORT_FROM_NUCLEUS | 0.00001 | 0.00042 | 0.59333 | 0.64726 | 1.29592 | 118 |
| GO_RNA_LOCALIZATION | 0.00000 | 0.00001 | 0.69013 | 0.63320 | 1.28469 | 190 |
| GO_RNA_SPLICING | 0.00000 | 0.00002 | 0.67496 | 0.59100 | 1.21492 | 353 |
| GO_RNA_SPLICING_VIA_TRANSESTERIFICATION_REACTIONS | 0.00000 | 0.00000 | 0.71951 | 0.60692 | 1.24235 | 288 |
| GO_ROUGH_ENDOPLASMIC_RETICULUM | 0.00005 | 0.00125 | 0.55733 | 0.73384 | 1.41579 | 45 |
| GO_RRNA_BINDING | 0.00002 | 0.00060 | 0.57561 | 0.71022 | 1.38361 | 55 |
| GO_RRNA_METABOLIC_PROCESS | 0.00001 | 0.00036 | 0.59333 | 0.61369 | 1.24352 | 179 |
| GO_SCF_DEPENDENT_PROTEASOMAL_UBIQUITIN_DEPENDI | 0.00003 | 0.00094 | 0.55733 | 0.67950 | 1.33784 | 73 |
| GO_SECRETORY_GRANULE | 0.00000 | 0.00000 | 0.81404 | 0.61223 | 1.25873 | 369 |
| GO_SECRETORY_GRANULE_MEMBRANE | 0.00137 | 0.02204 | 0.45506 | 0.59527 | 1.19636 | 133 |
| GO_SECRETORY_VESICLE | 0.00000 | 0.00000 | | 0.60236 | 1.24055 | 436 |
| GO_SIGNAL_SEQUENCE_BINDING | 0.00320 | 0.04280 | 0.43171 | 0.70136 | 1.33478 | 35 |
| GO_SIGNAL_TRANSDUCTION_BY_P53_CLASS_MEDIATOR | 0.00106 | 0.01830 | 0.45506 | 0.57564 | 1.16741 | 185 |
| GO_SINGLE_FERTILIZATION | 0.00132 | 0.02134 | 0.45506 | 0.67500 | 1.31035 | 50 |
| GO_SINGLE_STRANDED_RNA_BINDING | 0.00017 | 0.00382 | 0.51885 | 0.67734 | 1.32312 | 60 |
| GO_SISTER_CHROMATID_SEGREGATION | 0.00035 | 0.00711 | 0.49849 | 0.60598 | 1.22041 | 148 |
| GO_SMALL_RIBOSOMAL_SUBUNIT | 0.00000 | 0.00000 | 0.76146 | 0.76684 | 1.50092 | 64 |
| GO_SPERM_EGG_RECOGNITION | 0.00006 | 0.00159 | 0.53843 | 0.83049 | 1.54062 | 21 |
| GO_SPINDLE_ORGANIZATION | 0.00279 | 0.03858 | 0.43171 | 0.58476 | 1.17490 | 131 |
| GO_SPLICEOSOMAL_COMPLEX | 0.00016 | 0.00369 | 0.51885 | 0.60604 | 1.22346 | 158 |
| GO_STEM_CELL_DIFFERENTIATION | 0.00000 | 0.00005 | 0.65944 | 0.64364 | 1.29455 | 138 |
| GO_STRUCTURAL_CONSTITUENT_OF_RIBOSOME | 0.00000 | 0.00000 | | 0.73817 | 1.48751 | 149 |
| GO_STRUCTURAL_MOLECULE_ACTIVITY | 0.00000 | 0.00000 | | 0.66509 | 1.36615 | 334 |
| GO_SUPRAMOLECULAR_POLYMER | 0.00159 | 0.02466 | 0.45506 | 0.54130 | 1.11466 | 429 |
| GO_TELOMERE_MAINTENANCE_VIA_TELOMERE_LENGTHEN | 0.00143 | 0.02286 | 0.45506 | 0.66436 | 1.29333 | 54 |
| GO_TELOMERE_ORGANIZATION | 0.00002 | 0.00073 | 0.57561 | 0.65366 | 1.30192 | 103 |
| GO_THREONINE_TYPE_ENDOPEPTIDASE_ACTIVITY | 0.00000 | 0.00012 | 0.62726 | 0.88928 | 1.62389 | 17 |
| GO_THREONINE_TYPE_PEPTIDASE_ACTIVITY | 0.00002 | 0.00071 | 0.57561 | 0.85515 | 1.57356 | 19 |
| GO_TISSUE_MORPHOGENESIS | 0.00183 | 0.02766 | 0.45506 | 0.55021 | 1.12800 | 310 |
| GO_TOXIN_TRANSPORT | 0.00154 | 0.02400 | 0.45506 | 0.75293 | 1.40184 | 23 |
| GO_TRANSFERRIN_TRANSPORT | 0.00014 | 0.00331 | 0.51885 | 0.79944 | 1.48525 | 22 |
| GO_TRANSLATIONAL_ELONGATION | 0.00000 | 0.00000 | 0.80122 | 0.71278 | 1.42526 | 112 |
| GO_TRANSLATIONAL_INITIATION | 0.00000 | 0.00000 | | 0.71132 | 1.43719 | 166 |
| GO_TRANSLATIONAL_TERMINATION | 0.00000 | 0.00000 | 0.78819 | 0.72500 | 1.44005 | 93 |
| GO_TRANSLATION_FACTOR_ACTIVITY_RNA_BINDING | 0.00005 | 0.00125 | 0.55733 | 0.68752 | 1.35082 | 68 |
| GO_TRANSLATION_INITIATION_FACTOR_ACTIVITY | 0.00010 | 0.00254 | 0.53843 | 0.71952 | 1.38998 | 46 |
| GO_TRANSLATION_PREINITIATION_COMPLEX | 0.00111 | 0.01901 | 0.45506 | 0.81634 | 1.48210 | 16 |
| GO_TRANSLATION_REGULATOR_ACTIVITY | 0.00000 | 0.00018 | 0.61053 | 0.65779 | 1.31163 | 107 |
| GO_TRANSLATION_REGULATOR_ACTIVITY_NUCLEIC_ACID_E | 0.00000 | 0.00009 | 0.64355 | 0.68989 | 1.36437 | 84 |
| GO_TRANSPORT_VESICLE | 0.00342 | 0.04490 | 0.43171 | 0.56681 | 1.15015 | 189 |
| GO_TUMOR_NECROSIS_FACTOR_MEDIATED_SIGNALING_PA | 0.00000 | 0.00003 | 0.65944 | 0.69668 | 1.38196 | 90 |
| GO_T_CELL_RECEPTOR_SIGNALING_PATHWAY | 0.00006 | 0.00147 | 0.55733 | 0.64179 | 1.27878 | 104 |
| GO_UBIQUITIN_LIKE_PROTEIN_LIGASE_BINDING | 0.00002 | 0.00063 | 0.57561 | 0.60064 | 1.22131 | 207 |
| GO_UNFOLDED_PROTEIN_BINDING | 0.00001 | 0.00046 | 0.59333 | 0.67859 | 1.34044 | 82 |
| GO_VACUOLAR_LUMEN | 0.00001 | 0.00023 | 0.61053 | 0.68472 | 1.35021 | 79 |
| GO_VACUOLE | 0.00035 | 0.00711 | 0.49849 | 0.55002 | 1.13175 | 408 |
| GO_VESICLE_LUMEN | 0.00000 | 0.00001 | 0.70498 | 0.64818 | 1.30875 | 159 |
| GO_VESICLE_MEMBRANE | 0.00202 | 0.02980 | 0.43171 | 0.54225 | 1.11481 | 368 |
| GO_VIRAL_GENE_EXPRESSION | 0.00000 | 0.00000 | 0.81404 | 0.67819 | 1.36936 | 159 |
| GO_VIRAL_LIFE_CYCLE | 0.00000 | 0.00003 | 0.65944 | 0.62604 | 1.26997 | 188 |
| GO_VIRION_ASSEMBLY | 0.00082 | 0.01468 | 0.47727 | 0.76355 | 1.42338 | 24 |
| HP_ABNORMALITY_OF_ACID_BASE_HOMEOSTASIS | 0.00083 | 0.01475 | 0.47727 | 0.56999 | 1.16111 | 224 |
| HP_ABNORMALITY_OF_CARDIOVASCULAR_SYSTEM_ELECTR | 0.00220 | 0.03193 | 0.43171 | 0.56725 | 1.15377 | 209 |
| HP_ABNORMAL_CIRCULATING_METABOLITE_CONCENTRATI | 0.00117 | 0.01953 | 0.45506 | 0.53556 | 1.10478 | 497 |
| HP_ABNORMAL_RETICULOCYTE_MORPHOLOGY | 0.00324 | 0.04309 | 0.43171 | 0.75595 | 1.39735 | 20 |
| HP_ACUTE_LEUKEMIA | 0.00045 | 0.00897 | 0.49849 | 0.65782 | 1.28858 | 65 |
| HP_ANEMIA_OF_INADEQUATE_PRODUCTION | 0.00126 | 0.02069 | 0.45506 | 0.62409 | 1.23796 | 90 |
| HP_DEPRESSED_NASAL_RIDGE | 0.00235 | 0.03390 | 0.43171 | 0.63008 | 1.24054 | 73 |
| HP_MACROCYTIC_ANEMIA | 0.00017 | 0.00394 | 0.51885 | 0.74986 | 1.42708 | 35 |
| HP_MIGRAINE | 0.00110 | 0.01876 | 0.45506 | 0.65158 | 1.27533 | 64 |
| HP_MUTISM | 0.00361 | 0.04666 | 0.43171 | 0.71208 | 1.33162 | 27 |
| **Cluster 7** | | | | | | |
| pathway | pval | padj | log2err | ES | NES | size |
| GO_ACTIVATION_OF_IMMUNE_RESPONSE | 0.00000 | 0.00000 | 0.78819 | 0.65054 | 1.33715 | 225 |
| GO_ACTIVATION_OF_INNATE_IMMUNE_RESPONSE | 0.00000 | 0.00000 | 0.70498 | 0.70896 | 1.42735 | 94 |
| GO_AEROBIC_RESPIRATION | 0.00004 | 0.00113 | 0.55733 | 0.69243 | 1.37979 | 67 |
| GO_ALTERNATIVE_MRNA_SPLICING_VIA_SPLICEOSOME | 0.00001 | 0.00032 | 0.59333 | 0.70322 | 1.40465 | 71 |
| GO_AMIDE_BINDING | 0.00213 | 0.02938 | 0.43171 | 0.57265 | 1.17145 | 174 |
| GO_AMINE_METABOLIC_PROCESS | 0.00000 | 0.00000 | 0.73376 | 0.72414 | 1.45456 | 87 |
| GO_ANAPHASE_PROMOTING_COMPLEX_DEPENDENT_CATA | 0.00000 | 0.00000 | 0.77494 | 0.75848 | 1.51732 | 75 |
| GO_ANATOMICAL_STRUCTURE_HOMEOSTASIS | 0.00246 | 0.03304 | 0.43171 | 0.56109 | 1.15536 | 244 |
| GO_ANTIGEN_PROCESSING_AND_PRESENTATION | 0.00000 | 0.00000 | 0.71951 | 0.65728 | 1.34117 | 154 |
| GO_ANTIGEN_PROCESSING_AND_PRESENTATION_OF_EXOG | 0.00000 | 0.00000 | 0.73376 | 0.77041 | 1.52906 | 60 |

| Cluster 1 | | | | | | |
|---|---|---|---|---|---|---|
| GO_ANTIGEN_PROCESSING_AND_PRESENTATION_OF_PEPT | 0.00000 | 0.00001 | 0.67496 | 0.66238 | 1.34658 | 135 |
| GO_ANTIGEN_PROCESSING_AND_PRESENTATION_OF_PEPT | 0.00000 | 0.00002 | 0.65944 | 0.73238 | 1.46290 | 71 |
| GO_ANTIGEN_RECEPTOR_MEDIATED_SIGNALING_PATHWAY | 0.00000 | 0.00005 | 0.64355 | 0.64679 | 1.31579 | 137 |
| GO_ANTIMICROBIAL_HUMORAL_RESPONSE | 0.00012 | 0.00282 | 0.53843 | 0.81984 | 1.54140 | 21 |
| GO_ANTIOXIDANT_ACTIVITY | 0.00044 | 0.00853 | 0.49849 | 0.71267 | 1.39256 | 42 |
| GO_APOPTOTIC_SIGNALING_PATHWAY | 0.00049 | 0.00919 | 0.47727 | 0.55093 | 1.14289 | 382 |
| GO_ATPASE_ACTIVITY | 0.00376 | 0.04627 | 0.43171 | 0.54900 | 1.13262 | 275 |
| GO_ATPASE_BINDING | 0.00316 | 0.04043 | 0.43171 | 0.66355 | 1.30711 | 50 |
| GO_ATP_BIOSYNTHETIC_PROCESS | 0.00020 | 0.00435 | 0.51885 | 0.82644 | 1.54197 | 19 |
| GO_ATP_METABOLIC_PROCESS | 0.00000 | 0.00000 | | 0.69098 | 1.41790 | 206 |
| GO_ATP_SYNTHESIS_COUPLED_ELECTRON_TRANSPORT | 0.00000 | 0.00000 | 0.83909 | 0.77824 | 1.55685 | 75 |
| GO_AZUROPHIL_GRANULE_LUMEN | 0.00041 | 0.00810 | 0.49849 | 0.69890 | 1.37389 | 47 |
| GO_BINDING_OF_SPERM_TO_ZONA_PELLUCIDA | 0.00067 | 0.01162 | 0.47727 | 0.82944 | 1.52346 | 15 |
| GO_BLOOD_MICROPARTICLE | 0.00013 | 0.00315 | 0.51885 | 0.76330 | 1.46511 | 31 |
| GO_CADHERIN_BINDING | 0.00000 | 0.00000 | | 0.67157 | 1.38342 | 256 |
| GO_CALCIUM_DEPENDENT_PROTEIN_BINDING | 0.00017 | 0.00390 | 0.51885 | 0.72509 | 1.41683 | 42 |
| GO_CALCIUM_ION_BINDING | 0.00272 | 0.03619 | 0.43171 | 0.56265 | 1.15678 | 227 |
| GO_CANONICAL_WNT_SIGNALING_PATHWAY | 0.00004 | 0.00106 | 0.55733 | 0.60313 | 1.23793 | 212 |
| GO_CARBOHYDRATE_BIOSYNTHETIC_PROCESS | 0.00075 | 0.01273 | 0.47727 | 0.61168 | 1.23952 | 119 |
| GO_CATALYTIC_STEP_2_SPLICEOSOME | 0.00000 | 0.00003 | 0.65944 | 0.69924 | 1.40254 | 83 |
| GO_CATION_TRANSMEMBRANE_TRANSPORTER_ACTIVITY | 0.00339 | 0.04252 | 0.43171 | 0.56848 | 1.16462 | 190 |
| GO_CATION_TRANSPORT | 0.00132 | 0.02014 | 0.45506 | 0.54175 | 1.12471 | 417 |
| GO_CELLULAR_AMINO_ACID_METABOLIC_PROCESS | 0.00002 | 0.00056 | 0.57561 | 0.60371 | 1.23728 | 198 |
| GO_CELLULAR_COMPONENT_DISASSEMBLY | 0.00000 | 0.00000 | 0.80122 | 0.60917 | 1.26445 | 394 |
| GO_CELLULAR_HOMEOSTASIS | 0.00030 | 0.00629 | 0.49849 | 0.55235 | 1.14681 | 415 |
| GO_CELLULAR_ION_HOMEOSTASIS | 0.00092 | 0.01499 | 0.47727 | 0.56868 | 1.17149 | 250 |
| GO_CELLULAR_KETONE_METABOLIC_PROCESS | 0.00000 | 0.00000 | 0.71951 | 0.66502 | 1.35545 | 148 |
| GO_CELLULAR_NITROGEN_COMPOUND_CATABOLIC_PROCE | 0.00000 | 0.00000 | | 0.68644 | 1.42689 | 444 |
| GO_CELLULAR_OXIDANT_DETOXIFICATION | 0.00054 | 0.00989 | 0.47727 | 0.69342 | 1.36594 | 50 |
| GO_CELLULAR_PROTEIN_COMPLEX_DISASSEMBLY | 0.00000 | 0.00000 | 0.70498 | 0.64810 | 1.32581 | 174 |
| GO_CELLULAR_RESPIRATION | 0.00000 | 0.00000 | | 0.70727 | 1.43829 | 136 |
| GO_CELLULAR_RESPONSE_TO_CHEMICAL_STRESS | 0.00405 | 0.04900 | 0.40702 | 0.55729 | 1.14509 | 221 |
| GO_CELLULAR_RESPONSE_TO_HEAT | 0.00040 | 0.00800 | 0.49849 | 0.63918 | 1.28687 | 94 |
| GO_CELLULAR_RESPONSE_TO_OXYGEN_LEVELS | 0.00000 | 0.00000 | 0.77494 | 0.67354 | 1.37410 | 152 |
| GO_CELLULAR_RESPONSE_TO_TOXIC_SUBSTANCE | 0.00065 | 0.01124 | 0.47727 | 0.66801 | 1.32582 | 60 |
| GO_CELL_ACTIVATION_INVOLVED_IN_IMMUNE_RESPONSE | 0.00000 | 0.00000 | 0.76146 | 0.61009 | 1.26328 | 344 |
| GO_CELL_ADHESION_MOLECULE_BINDING | 0.00000 | 0.00000 | | 0.65561 | 1.35685 | 328 |
| GO_CELL_BODY | 0.00039 | 0.00788 | 0.49849 | 0.56531 | 1.16507 | 267 |
| GO_CELL_CELL_RECOGNITION | 0.00057 | 0.01024 | 0.47727 | 0.76500 | 1.44663 | 25 |
| GO_CELL_CELL_SIGNALING_BY_WNT | 0.00000 | 0.00012 | 0.62726 | 0.58826 | 1.21745 | 330 |
| GO_CELL_CORTEX | 0.00283 | 0.03715 | 0.43171 | 0.57251 | 1.17284 | 183 |
| GO_CELL_CYCLE_G2_M_PHASE_TRANSITION | 0.00000 | 0.00008 | 0.62726 | 0.61208 | 1.25826 | 226 |
| GO_CELL_CYCLE_PHASE_TRANSITION | 0.00000 | 0.00002 | 0.67496 | 0.57152 | 1.18920 | 491 |
| GO_CELL_DEATH_IN_RESPONSE_TO_OXIDATIVE_STRESS | 0.00191 | 0.02689 | 0.45506 | 0.66229 | 1.31019 | 56 |
| GO_CELL_DIVISION | 0.00005 | 0.00142 | 0.55733 | 0.55850 | 1.16013 | 433 |
| GO_CELL_KILLING | 0.00087 | 0.01462 | 0.47727 | 0.69707 | 1.36550 | 44 |
| GO_CELL_LEADING_EDGE | 0.00272 | 0.03619 | 0.43171 | 0.55475 | 1.14326 | 262 |
| GO_CELL_RECOGNITION | 0.00006 | 0.00144 | 0.55733 | 0.71495 | 1.41124 | 52 |
| GO_CELL_SUBSTRATE_JUNCTION | 0.00000 | 0.00000 | | 0.65112 | 1.34395 | 293 |
| GO_CELL_SURFACE_RECEPTOR_SIGNALING_PATHWAY_INVO | 0.00002 | 0.00054 | 0.57561 | 0.57614 | 1.19390 | 359 |
| GO_CHAPERONE_COMPLEX | 0.00002 | 0.00065 | 0.57561 | 0.83145 | 1.56455 | 22 |
| GO_CHAPERONE_MEDIATED_PROTEIN_FOLDING | 0.00335 | 0.04242 | 0.43171 | 0.67726 | 1.32337 | 42 |
| GO_CHROMOSOMAL_REGION | 0.00145 | 0.02155 | 0.45506 | 0.56059 | 1.15581 | 270 |
| GO_CHROMOSOME_SEGREGATION | 0.00055 | 0.01013 | 0.47727 | 0.56797 | 1.17015 | 253 |
| GO_CONTRACTILE_FIBER | 0.00000 | 0.00017 | 0.61053 | 0.65632 | 1.32838 | 114 |
| GO_CORTICAL_CYTOSKELETON | 0.00413 | 0.04958 | 0.40702 | 0.63620 | 1.26360 | 61 |
| GO_COTRANSLATIONAL_PROTEIN_TARGETING_TO_MEMBR | 0.00000 | 0.00000 | | 0.83574 | 1.67987 | 89 |
| GO_CYTOCHROME_COMPLEX | 0.00005 | 0.00140 | 0.55733 | 0.83195 | 1.56417 | 21 |
| GO_CYTOKINE_MEDIATED_SIGNALING_PATHWAY | 0.00000 | 0.00000 | 0.78819 | 0.61332 | 1.27144 | 365 |
| GO_CYTOPLASMIC_STRESS_GRANULE | 0.00018 | 0.00401 | 0.51885 | 0.68558 | 1.36167 | 61 |
| GO_CYTOPLASMIC_TRANSLATION | 0.00000 | 0.00000 | | 0.76759 | 1.54290 | 89 |
| GO_CYTOPLASMIC_TRANSLATIONAL_INITIATION | 0.00002 | 0.00059 | 0.57561 | 0.79644 | 1.52558 | 30 |
| GO_CYTOSOLIC_LARGE_RIBOSOMAL_SUBUNIT | 0.00000 | 0.00000 | | 0.82838 | 1.63515 | 52 |
| GO_CYTOSOLIC_RIBOSOME | 0.00000 | 0.00000 | | 0.82901 | 1.66929 | 95 |
| GO_CYTOSOLIC_SMALL_RIBOSOMAL_SUBUNIT | 0.00000 | 0.00000 | 0.82666 | 0.85328 | 1.66557 | 41 |
| GO_DEFENSE_RESPONSE_TO_OTHER_ORGANISM | 0.00000 | 0.00000 | 0.73376 | 0.59033 | 1.22699 | 447 |
| GO_DETECTION_OF_STIMULUS | 0.00041 | 0.00807 | 0.49849 | 0.64155 | 1.28950 | 91 |
| GO_DETOXIFICATION | 0.00081 | 0.01366 | 0.47727 | 0.67650 | 1.34142 | 59 |
| GO_DIENCEPHALON_DEVELOPMENT | 0.00087 | 0.01450 | 0.47727 | 0.74927 | 1.42408 | 27 |
| GO_DISORDERED_DOMAIN_SPECIFIC_BINDING | 0.00126 | 0.01950 | 0.45506 | 0.74833 | 1.41925 | 26 |
| GO_DNA_BIOSYNTHETIC_PROCESS | 0.00065 | 0.01124 | 0.47727 | 0.60272 | 1.22785 | 143 |
| GO_DNA_CONFORMATION_CHANGE | 0.00016 | 0.00374 | 0.51885 | 0.58119 | 1.19441 | 224 |
| GO_DOUBLE_STRANDED_RNA_BINDING | 0.00055 | 0.01013 | 0.47727 | 0.67796 | 1.34093 | 55 |

| Cluster 1 | | | | | | |
|---|---|---|---|---|---|---|
| GO_ELECTRON_TRANSFER_ACTIVITY | 0.00001 | 0.00019 | 0.61053 | 0.69933 | 1.39836 | 74 |
| GO_ELECTRON_TRANSPORT_CHAIN | 0.00000 | 0.00000 | | 0.72706 | 1.47620 | 123 |
| GO_ENDOPEPTIDASE_COMPLEX | 0.00000 | 0.00000 | 0.71951 | 0.76430 | 1.51802 | 61 |
| GO_ENERGY_DERIVATION_BY_OXIDATION_OF_ORGANIC_CO | 0.00000 | 0.00000 | 0.80122 | 0.66415 | 1.36060 | 190 |
| GO_ENZYME_INHIBITOR_ACTIVITY | 0.00119 | 0.01861 | 0.45506 | 0.58162 | 1.18960 | 173 |
| GO_EPHRIN_RECEPTOR_SIGNALING_PATHWAY | 0.00224 | 0.03070 | 0.43171 | 0.66325 | 1.31372 | 57 |
| GO_ESTABLISHMENT_OF_PROTEIN_LOCALIZATION_TO_CHR | 0.00080 | 0.01355 | 0.47727 | 0.78318 | 1.47248 | 21 |
| GO_ESTABLISHMENT_OF_PROTEIN_LOCALIZATION_TO_END | 0.00000 | 0.00000 | | 0.81451 | 1.64179 | 102 |
| GO_ESTABLISHMENT_OF_PROTEIN_LOCALIZATION_TO_MEM | 0.00000 | 0.00000 | | 0.68946 | 1.42035 | 252 |
| GO_ESTABLISHMENT_OF_PROTEIN_LOCALIZATION_TO_ORG | 0.00000 | 0.00000 | | 0.65159 | 1.35312 | 428 |
| GO_ESTABLISHMENT_OF_PROTEIN_LOCALIZATION_TO_TELO | 0.00001 | 0.00032 | 0.59333 | 0.89114 | 1.64236 | 16 |
| GO_ESTABLISHMENT_OF_RNA_LOCALIZATION | 0.00000 | 0.00005 | 0.64355 | 0.63713 | 1.30208 | 169 |
| GO_ESTABLISHMENT_OF_TISSUE_POLARITY | 0.00000 | 0.00000 | | 0.75113 | 1.51225 | 94 |
| GO_EUKARYOTIC_TRANSLATION_INITIATION_FACTOR_3_CO | 0.00016 | 0.00365 | 0.51885 | 0.85554 | 1.57140 | 15 |
| GO_EXOCYTOSIS | 0.00000 | 0.00000 | 0.70498 | 0.58395 | 1.21386 | 455 |
| GO_FC_EPSILON_RECEPTOR_SIGNALING_PATHWAY | 0.00000 | 0.00000 | 0.70498 | 0.71226 | 1.43164 | 91 |
| GO_FC_RECEPTOR_SIGNALING_PATHWAY | 0.00000 | 0.00000 | 0.70498 | 0.67266 | 1.36674 | 133 |
| GO_FERTILIZATION | 0.00011 | 0.00269 | 0.53843 | 0.69578 | 1.37815 | 57 |
| GO_FIBROBLAST_GROWTH_FACTOR_RECEPTOR_SIGNALING | 0.00185 | 0.02630 | 0.45506 | 0.64700 | 1.28528 | 62 |
| GO_FICOLIN_1_RICH_GRANULE | 0.00000 | 0.00000 | 0.70498 | 0.70531 | 1.42054 | 96 |
| GO_FOREBRAIN_GENERATION_OF_NEURONS | 0.00168 | 0.02443 | 0.45506 | 0.76408 | 1.43030 | 20 |
| GO_GENERATION_OF_PRECURSOR_METABOLITES_AND_EN | 0.00000 | 0.00000 | | 0.63388 | 1.31231 | 341 |
| GO_GLUCOSE_CATABOLIC_PROCESS | 0.00250 | 0.03346 | 0.43171 | 0.75103 | 1.41321 | 22 |
| GO_GLUTAMATERGIC_SYNAPSE | 0.00179 | 0.02583 | 0.45506 | 0.57997 | 1.18461 | 167 |
| GO_GLYCOLYTIC_PROCESS_THROUGH_FRUCTOSE_6_PHOSP | 0.00126 | 0.01950 | 0.45506 | 0.77162 | 1.44442 | 20 |
| GO_GLYCOSAMINOGLYCAN_BINDING | 0.00324 | 0.04128 | 0.43171 | 0.64111 | 1.27335 | 61 |
| GO_GUANYL_NUCLEOTIDE_BINDING | 0.00205 | 0.02857 | 0.43171 | 0.56179 | 1.15632 | 240 |
| GO_HEAT_SHOCK_PROTEIN_BINDING | 0.00346 | 0.04335 | 0.43171 | 0.61485 | 1.23588 | 89 |
| GO_HEMATOPOIETIC_PROGENITOR_CELL_DIFFERENTIATION | 0.00000 | 0.00000 | 0.77494 | 0.70165 | 1.42270 | 120 |
| GO_HEMATOPOIETIC_STEM_CELL_DIFFERENTIATION | 0.00000 | 0.00000 | 0.77494 | 0.76026 | 1.51843 | 72 |
| GO_HEXOSE_CATABOLIC_PROCESS | 0.00231 | 0.03156 | 0.43171 | 0.69763 | 1.34933 | 35 |
| GO_HOMOTYPIC_CELL_CELL_ADHESION | 0.00128 | 0.01968 | 0.45506 | 0.68863 | 1.34895 | 44 |
| GO_HUMORAL_IMMUNE_RESPONSE | 0.00035 | 0.00726 | 0.49849 | 0.69782 | 1.37461 | 50 |
| GO_HYDROGEN_PEROXIDE_METABOLIC_PROCESS | 0.00000 | 0.00012 | 0.62726 | 0.84614 | 1.60473 | 26 |
| GO_IMMUNE_RESPONSE_REGULATING_SIGNALING_PATHW | 0.00000 | 0.00000 | 0.76146 | 0.64265 | 1.31999 | 219 |
| GO_INFLAMMATORY_RESPONSE_TO_ANTIGENIC_STIMULUS | 0.00181 | 0.02591 | 0.45506 | 0.75755 | 1.42549 | 22 |
| GO_INNATE_IMMUNE_RESPONSE | 0.00000 | 0.00001 | 0.70498 | 0.59642 | 1.23601 | 362 |
| GO_INNATE_IMMUNE_RESPONSE_ACTIVATING_SIGNAL_TRA | 0.00000 | 0.00002 | 0.67496 | 0.71175 | 1.42535 | 78 |
| GO_INNER_MITOCHONDRIAL_MEMBRANE_PROTEIN_COMP | 0.00000 | 0.00000 | 0.74774 | 0.72583 | 1.45902 | 90 |
| GO_INORGANIC_ION_TRANSMEMBRANE_TRANSPORT | 0.00242 | 0.03270 | 0.43171 | 0.55038 | 1.13801 | 308 |
| GO_INTERACTION_WITH_SYMBIONT | 0.00089 | 0.01475 | 0.47727 | 0.70277 | 1.37322 | 42 |
| GO_INTERLEUKIN_1_BETA_PRODUCTION | 0.00339 | 0.04252 | 0.43171 | 0.70526 | 1.35094 | 30 |
| GO_INTERLEUKIN_1_MEDIATED_SIGNALING_PATHWAY | 0.00000 | 0.00000 | 0.77494 | 0.74833 | 1.49895 | 80 |
| GO_INTERLEUKIN_1_PRODUCTION | 0.00033 | 0.00674 | 0.49849 | 0.73326 | 1.42511 | 38 |
| GO_INTERMEDIATE_FILAMENT_CYTOSKELETON | 0.00287 | 0.03755 | 0.43171 | 0.64944 | 1.28640 | 58 |
| GO_LARGE_RIBOSOMAL_SUBUNIT | 0.00000 | 0.00000 | | 0.76718 | 1.54966 | 107 |
| GO_LEUKOCYTE_MEDIATED_IMMUNITY | 0.00000 | 0.00000 | | 0.63023 | 1.30599 | 359 |
| GO_MAINTENANCE_OF_CELL_NUMBER | 0.00420 | 0.04996 | 0.40702 | 0.59382 | 1.20054 | 111 |
| GO_MAINTENANCE_OF_LOCATION_IN_CELL | 0.00132 | 0.02014 | 0.45506 | 0.61512 | 1.23924 | 100 |
| GO_MEMBRANE_REGION | 0.00002 | 0.00064 | 0.57561 | 0.61906 | 1.26318 | 154 |
| GO_MESENCHYMAL_CELL_DIFFERENTIATION | 0.00305 | 0.03927 | 0.43171 | 0.59442 | 1.20127 | 110 |
| GO_MESENCHYME_DEVELOPMENT | 0.00220 | 0.03026 | 0.43171 | 0.59122 | 1.20191 | 135 |
| GO_METAL_ION_HOMEOSTASIS | 0.00413 | 0.04958 | 0.40702 | 0.55707 | 1.14584 | 235 |
| GO_MICROVILLUS | 0.00361 | 0.04473 | 0.43171 | 0.65793 | 1.29334 | 47 |
| GO_MIDBRAIN_DEVELOPMENT | 0.00018 | 0.00401 | 0.51885 | 0.69141 | 1.36949 | 57 |
| GO_MITOCHONDRIAL_ELECTRON_TRANSPORT_NADH_TO_U | 0.00000 | 0.00012 | 0.62726 | 0.76593 | 1.49823 | 43 |
| GO_MITOCHONDRIAL_GENE_EXPRESSION | 0.00002 | 0.00059 | 0.57561 | 0.62541 | 1.27528 | 151 |
| GO_MITOCHONDRIAL_LARGE_RIBOSOMAL_SUBUNIT | 0.00006 | 0.00158 | 0.53843 | 0.71600 | 1.41510 | 54 |
| GO_MITOCHONDRIAL_MATRIX | 0.00000 | 0.00005 | 0.64355 | 0.58952 | 1.22157 | 357 |
| GO_MITOCHONDRIAL_PROTEIN_COMPLEX | 0.00000 | 0.00000 | | 0.71022 | 1.45746 | 205 |
| GO_MITOCHONDRIAL_RESPIRATORY_CHAIN_COMPLEX_ASS | 0.00056 | 0.01024 | 0.47727 | 0.63860 | 1.28092 | 83 |
| GO_MITOCHONDRIAL_SMALL_RIBOSOMAL_SUBUNIT | 0.00368 | 0.04555 | 0.43171 | 0.71644 | 1.36567 | 28 |
| GO_MITOCHONDRIAL_TRANSLATION | 0.00002 | 0.00054 | 0.57561 | 0.63952 | 1.29911 | 128 |
| GO_MITOCHONDRIAL_TRANSLATIONAL_TERMINATION | 0.00000 | 0.00004 | 0.65944 | 0.69536 | 1.39772 | 88 |
| GO_MITOCHONDRION_ORGANIZATION | 0.00000 | 0.00001 | 0.67496 | 0.58640 | 1.21739 | 398 |
| GO_MITOTIC_SPINDLE_ORGANIZATION | 0.00383 | 0.04678 | 0.43171 | 0.60684 | 1.22117 | 93 |
| GO_MODIFICATION_DEPENDENT_MACROMOLECULE_CATAB | 0.00017 | 0.00390 | 0.51885 | 0.54932 | 1.14293 | 481 |
| GO_MODULATION_OF_PROCESS_OF_OTHER_ORGANISM | 0.00361 | 0.04473 | 0.43171 | 0.64072 | 1.27440 | 65 |
| GO_MONOAMINE_TRANSPORT | 0.00089 | 0.01475 | 0.47727 | 0.78022 | 1.46691 | 21 |
| GO_MONOSACCHARIDE_BIOSYNTHETIC_PROCESS | 0.00001 | 0.00029 | 0.59333 | 0.70629 | 1.40377 | 63 |
| GO_MONOSACCHARIDE_CATABOLIC_PROCESS | 0.00191 | 0.02689 | 0.45506 | 0.68888 | 1.34468 | 41 |
| GO_MONOVALENT_INORGANIC_CATION_TRANSMEMBRAN | 0.00042 | 0.00823 | 0.49849 | 0.63002 | 1.27088 | 104 |
| GO_MONOVALENT_INORGANIC_CATION_TRANSPORT | 0.00042 | 0.00816 | 0.49849 | 0.58584 | 1.20013 | 189 |

| Cluster 1 | | | | | | |
|---|---|---|---|---|---|---|
| GO_MORPHOGENESIS_OF_AN_EPITHELIUM | 0.00000 | 0.00006 | 0.64355 | 0.59298 | 1.22686 | 320 |
| GO_MORPHOGENESIS_OF_A_POLARIZED_EPITHELIUM | 0.00000 | 0.00000 | | 0.75286 | 1.52147 | 110 |
| GO_MRNA_3_UTR_BINDING | 0.00152 | 0.02253 | 0.45506 | 0.63470 | 1.26765 | 72 |
| GO_MRNA_5_UTR_BINDING | 0.00018 | 0.00401 | 0.51885 | 0.78462 | 1.48374 | 25 |
| GO_MRNA_BINDING | 0.00000 | 0.00000 | | 0.69978 | 1.43974 | 236 |
| GO_MRNA_EXPORT_FROM_NUCLEUS | 0.00008 | 0.00205 | 0.53843 | 0.64428 | 1.29800 | 100 |
| GO_MRNA_PROCESSING | 0.00000 | 0.00000 | | 0.63370 | 1.31633 | 433 |
| GO_MRNA_TRANSPORT | 0.00001 | 0.00040 | 0.59333 | 0.63162 | 1.28294 | 130 |
| GO_MUSCLE_CONTRACTION | 0.00170 | 0.02463 | 0.45506 | 0.59480 | 1.20957 | 136 |
| GO_MYD88_DEPENDENT_TOLL_LIKE_RECEPTOR_SIGNALING | 0.00065 | 0.01124 | 0.47727 | 0.78828 | 1.48206 | 21 |
| GO_MYELOID_LEUKOCYTE_ACTIVATION | 0.00000 | 0.00000 | | 0.63341 | 1.31050 | 320 |
| GO_MYELOID_LEUKOCYTE_MEDIATED_IMMUNITY | 0.00000 | 0.00000 | | 0.64324 | 1.32778 | 286 |
| GO_NADH_DEHYDROGENASE_ACTIVITY | 0.00003 | 0.00081 | 0.57561 | 0.76040 | 1.47614 | 37 |
| GO_NADH_DEHYDROGENASE_COMPLEX | 0.00002 | 0.00066 | 0.57561 | 0.76469 | 1.48773 | 39 |
| GO_NADH_DEHYDROGENASE_COMPLEX_ASSEMBLY | 0.00060 | 0.01060 | 0.47727 | 0.68476 | 1.35335 | 54 |
| GO_NADH_METABOLIC_PROCESS | 0.00021 | 0.00455 | 0.51885 | 0.78980 | 1.49063 | 23 |
| GO_NAD_METABOLIC_PROCESS | 0.00056 | 0.01024 | 0.47727 | 0.73842 | 1.41735 | 31 |
| GO_NCRNA_METABOLIC_PROCESS | 0.00009 | 0.00216 | 0.53843 | 0.56413 | 1.17043 | 386 |
| GO_NCRNA_PROCESSING | 0.00001 | 0.00018 | 0.61053 | 0.58241 | 1.20543 | 336 |
| GO_NEGATIVE_REGULATION_OF_APOPTOTIC_SIGNALING_P | 0.00420 | 0.04996 | 0.40702 | 0.58600 | 1.19168 | 136 |
| GO_NEGATIVE_REGULATION_OF_CANONICAL_WNT_SIGNAI | 0.00000 | 0.00000 | 0.74774 | 0.69718 | 1.41436 | 122 |
| GO_NEGATIVE_REGULATION_OF_CATALYTIC_ACTIVITY | 0.00420 | 0.04996 | 0.40702 | 0.53628 | 1.11345 | 415 |
| GO_NEGATIVE_REGULATION_OF_CELLULAR_AMIDE_METAB | 0.00050 | 0.00933 | 0.47727 | 0.61666 | 1.24921 | 118 |
| GO_NEGATIVE_REGULATION_OF_CELLULAR_PROTEIN_CATA | 0.00413 | 0.04958 | 0.40702 | 0.63621 | 1.26361 | 61 |
| GO_NEGATIVE_REGULATION_OF_CELL_CYCLE | 0.00013 | 0.00305 | 0.51885 | 0.55275 | 1.14885 | 449 |
| GO_NEGATIVE_REGULATION_OF_CELL_CYCLE_G2_M_PHASI | 0.00000 | 0.00006 | 0.64355 | 0.69161 | 1.39017 | 88 |
| GO_NEGATIVE_REGULATION_OF_CELL_CYCLE_PHASE_TRAN | 0.00001 | 0.00030 | 0.59333 | 0.61252 | 1.25510 | 197 |
| GO_NEGATIVE_REGULATION_OF_CELL_CYCLE_PROCESS | 0.00002 | 0.00063 | 0.57561 | 0.58655 | 1.20870 | 264 |
| GO_NEGATIVE_REGULATION_OF_INTERFERON_GAMMA_PR | 0.00091 | 0.01493 | 0.47727 | 0.81463 | 1.50135 | 16 |
| GO_NEGATIVE_REGULATION_OF_MITOTIC_CELL_CYCLE | 0.00001 | 0.00025 | 0.59333 | 0.59638 | 1.22877 | 251 |
| GO_NEGATIVE_REGULATION_OF_MRNA_METABOLIC_PROC | 0.00032 | 0.00660 | 0.49849 | 0.66671 | 1.32511 | 63 |
| GO_NEGATIVE_REGULATION_OF_PEPTIDASE_ACTIVITY | 0.00276 | 0.03635 | 0.43171 | 0.61815 | 1.24165 | 87 |
| GO_NEGATIVE_REGULATION_OF_PROTEOLYSIS | 0.00111 | 0.01769 | 0.45506 | 0.58831 | 1.20023 | 152 |
| GO_NEGATIVE_REGULATION_OF_WNT_SIGNALING_PATHW/ | 0.00000 | 0.00000 | 0.73376 | 0.67847 | 1.37928 | 135 |
| GO_NEURAL_NUCLEUS_DEVELOPMENT | 0.00090 | 0.01486 | 0.47727 | 0.71742 | 1.39001 | 36 |
| GO_NIK_NF_KAPPAB_SIGNALING | 0.00000 | 0.00000 | 0.74774 | 0.69754 | 1.41182 | 114 |
| GO_NON_CANONICAL_WNT_SIGNALING_PATHWAY | 0.00000 | 0.00000 | 0.82666 | 0.73318 | 1.48131 | 109 |
| GO_NUCLEAR_ENVELOPE | 0.00006 | 0.00151 | 0.55733 | 0.57593 | 1.18908 | 296 |
| GO_NUCLEAR_EXPORT | 0.00000 | 0.00002 | 0.67496 | 0.63820 | 1.30559 | 175 |
| GO_NUCLEAR_PERIPHERY | 0.00022 | 0.00475 | 0.51885 | 0.62543 | 1.26333 | 107 |
| GO_NUCLEAR_SPECK | 0.00072 | 0.01233 | 0.47727 | 0.55902 | 1.15617 | 312 |
| GO_NUCLEAR_TRANSCRIBED_MRNA_CATABOLIC_PROCESS | 0.00000 | 0.00000 | | 0.72269 | 1.48048 | 189 |
| GO_NUCLEAR_TRANSCRIBED_MRNA_CATABOLIC_PROCESS_ | 0.00000 | 0.00000 | | 0.79621 | 1.61152 | 114 |
| GO_NUCLEAR_TRANSPORT | 0.00000 | 0.00000 | 0.71951 | 0.61507 | 1.26965 | 287 |
| GO_NUCLEOBASE_CONTAINING_COMPOUND_TRANSPORT | 0.00001 | 0.00020 | 0.61053 | 0.60674 | 1.24496 | 204 |
| GO_NUCLEOBASE_CONTAINING_SMALL_MOLECULE_METAE | 0.00005 | 0.00141 | 0.55733 | 0.57076 | 1.18183 | 344 |
| GO_NUCLEOSIDE_DIPHOSPHATE_METABOLIC_PROCESS | 0.00194 | 0.02722 | 0.45506 | 0.61083 | 1.23218 | 104 |
| GO_NUCLEOSIDE_PHOSPHATE_BIOSYNTHETIC_PROCESS | 0.00052 | 0.00961 | 0.47727 | 0.61004 | 1.24224 | 142 |
| GO_NUCLEOSIDE_TRIPHOSPHATE_METABOLIC_PROCESS | 0.00126 | 0.01950 | 0.45506 | 0.66487 | 1.31696 | 58 |
| GO_NUCLEOSOMAL_DNA_BINDING | 0.00157 | 0.02317 | 0.45506 | 0.75397 | 1.42299 | 23 |
| GO_NUCLEOTIDE_PHOSPHORYLATION | 0.00350 | 0.04372 | 0.43171 | 0.61777 | 1.24170 | 91 |
| GO_ORGANELLAR_RIBOSOME | 0.00000 | 0.00005 | 0.64355 | 0.69603 | 1.39638 | 84 |
| GO_ORGANELLE_INNER_MEMBRANE | 0.00000 | 0.00000 | | 0.64779 | 1.34348 | 375 |
| GO_ORGANIC_CYCLIC_COMPOUND_CATABOLIC_PROCESS | 0.00000 | 0.00000 | | 0.67670 | 1.40732 | 465 |
| GO_OSTEOBLAST_DIFFERENTIATION | 0.00006 | 0.00153 | 0.55733 | 0.63680 | 1.28630 | 107 |
| GO_OXIDATIVE_PHOSPHORYLATION | 0.00000 | 0.00000 | | 0.75743 | 1.52541 | 97 |
| GO_OXIDOREDUCTASE_ACTIVITY | 0.00000 | 0.00001 | 0.69013 | 0.58770 | 1.22002 | 400 |
| GO_OXIDOREDUCTASE_ACTIVITY_ACTING_ON_A_HEME_GR | 0.00034 | 0.00709 | 0.49849 | 0.81258 | 1.50968 | 18 |
| GO_OXIDOREDUCTASE_ACTIVITY_ACTING_ON_NAD_P_H | 0.00001 | 0.00017 | 0.61053 | 0.70532 | 1.40760 | 69 |
| GO_OXIDOREDUCTASE_ACTIVITY_ACTING_ON_NAD_P_H_Q | 0.00003 | 0.00083 | 0.57561 | 0.74919 | 1.46759 | 44 |
| GO_OXIDOREDUCTASE_ACTIVITY_ACTING_ON_PEROXIDE_A | 0.00023 | 0.00497 | 0.51885 | 0.76995 | 1.46768 | 28 |
| GO_OXIDOREDUCTASE_COMPLEX | 0.00000 | 0.00002 | 0.67496 | 0.70960 | 1.42133 | 79 |
| GO_PEPTIDASE_COMPLEX | 0.00000 | 0.00000 | 0.73376 | 0.73719 | 1.47603 | 77 |
| GO_PEPTIDE_BINDING | 0.00117 | 0.01837 | 0.45506 | 0.59423 | 1.20699 | 130 |
| GO_PHOSPHATIDYLSERINE_BINDING | 0.00165 | 0.02395 | 0.45506 | 0.75962 | 1.42938 | 22 |
| GO_PIGMENT_GRANULE | 0.00031 | 0.00653 | 0.49849 | 0.64824 | 1.29794 | 77 |
| GO_POLYMERIC_CYTOSKELETAL_FIBER | 0.00010 | 0.00254 | 0.53843 | 0.56265 | 1.16737 | 384 |
| GO_POLYSOMAL_RIBOSOME | 0.00000 | 0.00013 | 0.62726 | 0.81730 | 1.56170 | 29 |
| GO_POLYSOME | 0.00000 | 0.00000 | 0.70498 | 0.76121 | 1.51189 | 61 |
| GO_POSITIVE_REGULATION_OF_ATP_METABOLIC_PROCESS | 0.00106 | 0.01691 | 0.45506 | 0.76287 | 1.43980 | 23 |
| GO_POSITIVE_REGULATION_OF_BINDING | 0.00060 | 0.01060 | 0.47727 | 0.62521 | 1.26118 | 104 |
| GO_POSITIVE_REGULATION_OF_CANONICAL_WNT_SIGNALI | 0.00000 | 0.00006 | 0.64355 | 0.67388 | 1.35833 | 102 |
| GO_POSITIVE_REGULATION_OF_CELLULAR_AMIDE_METABC | 0.00000 | 0.00000 | 0.70498 | 0.67403 | 1.36570 | 117 |

| Cluster 1 | | | | | | |
|---|---|---|---|---|---|---|
| GO_POSITIVE_REGULATION_OF_CELLULAR_COMPONENT_B | 0.00141 | 0.02121 | 0.45506 | 0.55348 | 1.14478 | 314 |
| GO_POSITIVE_REGULATION_OF_CELLULAR_PROTEIN_LOCAL | 0.00012 | 0.00290 | 0.53843 | 0.58343 | 1.19946 | 228 |
| GO_POSITIVE_REGULATION_OF_CELL_POPULATION_PROLIF | 0.00022 | 0.00472 | 0.51885 | 0.55236 | 1.14663 | 420 |
| GO_POSITIVE_REGULATION_OF_CYTOSKELETON_ORGANIZA | 0.00093 | 0.01510 | 0.47727 | 0.59618 | 1.21283 | 137 |
| GO_POSITIVE_REGULATION_OF_DEFENSE_RESPONSE | 0.00000 | 0.00000 | 0.74774 | 0.65564 | 1.34150 | 177 |
| GO_POSITIVE_REGULATION_OF_DNA_BINDING | 0.00339 | 0.04252 | 0.43171 | 0.69778 | 1.34651 | 34 |
| GO_POSITIVE_REGULATION_OF_DNA_BIOSYNTHETIC_PROC | 0.00010 | 0.00241 | 0.53843 | 0.69903 | 1.38261 | 55 |
| GO_POSITIVE_REGULATION_OF_DNA_METABOLIC_PROCESS | 0.00220 | 0.03026 | 0.43171 | 0.59090 | 1.20165 | 136 |
| GO_POSITIVE_REGULATION_OF_ESTABLISHMENT_OF_PROT | 0.00001 | 0.00030 | 0.59333 | 0.59805 | 1.22978 | 232 |
| GO_POSITIVE_REGULATION_OF_IMMUNE_RESPONSE | 0.00000 | 0.00000 | 0.82666 | 0.63026 | 1.30182 | 301 |
| GO_POSITIVE_REGULATION_OF_IMMUNE_SYSTEM_PROCES | 0.00000 | 0.00000 | 0.77494 | 0.59700 | 1.24064 | 439 |
| GO_POSITIVE_REGULATION_OF_INNATE_IMMUNE_RESPON | 0.00000 | 0.00000 | 0.71951 | 0.68298 | 1.38689 | 124 |
| GO_POSITIVE_REGULATION_OF_INTRACELLULAR_TRANSPO | 0.00194 | 0.02722 | 0.45506 | 0.57912 | 1.18162 | 160 |
| GO_POSITIVE_REGULATION_OF_MRNA_METABOLIC_PROCE | 0.00176 | 0.02537 | 0.45506 | 0.63777 | 1.27091 | 68 |
| GO_POSITIVE_REGULATION_OF_MUSCLE_CONTRACTION | 0.00065 | 0.01137 | 0.47727 | 0.82157 | 1.51414 | 16 |
| GO_POSITIVE_REGULATION_OF_NUCLEOTIDE_METABOLIC_ | 0.00181 | 0.02591 | 0.45506 | 0.80205 | 1.47817 | 16 |
| GO_POSITIVE_REGULATION_OF_ORGANELLE_ORGANIZATIO | 0.00000 | 0.00004 | 0.65944 | 0.57843 | 1.20100 | 424 |
| GO_POSITIVE_REGULATION_OF_PEPTIDE_SECRETION | 0.00026 | 0.00558 | 0.49849 | 0.64659 | 1.29964 | 91 |
| GO_POSITIVE_REGULATION_OF_PROTEIN_CONTAINING_CO | 0.00121 | 0.01885 | 0.45506 | 0.58938 | 1.20277 | 159 |
| GO_POSITIVE_REGULATION_OF_PROTEIN_LOCALIZATION_T | 0.00037 | 0.00750 | 0.49849 | 0.67573 | 1.34210 | 61 |
| GO_POSITIVE_REGULATION_OF_REACTIVE_OXYGEN_SPECIE | 0.00198 | 0.02767 | 0.43171 | 0.77775 | 1.45111 | 19 |
| GO_POSITIVE_REGULATION_OF_REACTIVE_OXYGEN_SPECIE | 0.00128 | 0.01968 | 0.45506 | 0.69657 | 1.35969 | 41 |
| GO_POSITIVE_REGULATION_OF_RESPONSE_TO_BIOTIC_STI | 0.00000 | 0.00000 | 0.71951 | 0.67196 | 1.36833 | 142 |
| GO_POSITIVE_REGULATION_OF_RESPONSE_TO_EXTERNAL_ | 0.00000 | 0.00000 | 0.74774 | 0.63481 | 1.30508 | 228 |
| GO_POSITIVE_REGULATION_OF_TELOMERASE_ACTIVITY | 0.00017 | 0.00390 | 0.51885 | 0.76462 | 1.46465 | 30 |
| GO_POSITIVE_REGULATION_OF_TELOMERASE_RNA_LOCALI | 0.00008 | 0.00200 | 0.53843 | 0.86638 | 1.59131 | 15 |
| GO_POSITIVE_REGULATION_OF_TELOMERE_MAINTENANCE | 0.00049 | 0.00919 | 0.47727 | 0.71208 | 1.38996 | 41 |
| GO_POSITIVE_REGULATION_OF_TELOMERE_MAINTENANCE | 0.00014 | 0.00323 | 0.51885 | 0.76293 | 1.46440 | 31 |
| GO_POSITIVE_REGULATION_OF_TRANSLATION | 0.00000 | 0.00000 | 0.71951 | 0.70430 | 1.42051 | 103 |
| GO_POSITIVE_REGULATION_OF_VIRAL_LIFE_CYCLE | 0.00361 | 0.04473 | 0.43171 | 0.65790 | 1.29329 | 47 |
| GO_POSITIVE_REGULATION_OF_VIRAL_PROCESS | 0.00001 | 0.00020 | 0.61053 | 0.68000 | 1.36589 | 87 |
| GO_POSITIVE_REGULATION_OF_VIRAL_TRANSCRIPTION | 0.00104 | 0.01671 | 0.45506 | 0.70923 | 1.37982 | 39 |
| GO_POSITIVE_REGULATION_OF_WNT_SIGNALING_PATHWA | 0.00000 | 0.00015 | 0.61053 | 0.65690 | 1.33115 | 119 |
| GO_POSTSYNAPSE | 0.00145 | 0.02155 | 0.45506 | 0.55353 | 1.14270 | 295 |
| GO_POSTTRANSCRIPTIONAL_REGULATION_OF_GENE_EXPR | 0.00000 | 0.00000 | | 0.66886 | 1.39158 | 484 |
| GO_POST_TRANSLATIONAL_PROTEIN_MODIFICATION | 0.00133 | 0.02032 | 0.45506 | 0.56705 | 1.16569 | 226 |
| GO_PRECATALYTIC_SPLICEOSOME | 0.00040 | 0.00794 | 0.49849 | 0.70150 | 1.38320 | 51 |
| GO_PRERIBOSOME | 0.00033 | 0.00674 | 0.49849 | 0.66574 | 1.32979 | 71 |
| GO_PRE_MRNA_BINDING | 0.00037 | 0.00748 | 0.49849 | 0.74575 | 1.43142 | 31 |
| GO_PROTEASOMAL_PROTEIN_CATABOLIC_PROCESS | 0.00004 | 0.00098 | 0.55733 | 0.56691 | 1.17503 | 363 |
| GO_PROTEASOMAL_UBIQUITIN_INDEPENDENT_PROTEIN_C | 0.00000 | 0.00014 | 0.61053 | 0.87971 | 1.63439 | 18 |
| GO_PROTEASOME_ACCESSORY_COMPLEX | 0.00004 | 0.00110 | 0.55733 | 0.80924 | 1.52917 | 24 |
| GO_PROTEASOME_CORE_COMPLEX | 0.00000 | 0.00008 | 0.62726 | 0.90782 | 1.67311 | 16 |
| GO_PROTEIN_CONTAINING_COMPLEX_DISASSEMBLY | 0.00000 | 0.00000 | 0.78819 | 0.63588 | 1.30998 | 249 |
| GO_PROTEIN_CONTAINING_COMPLEX_LOCALIZATION | 0.00002 | 0.00059 | 0.57561 | 0.60376 | 1.23958 | 215 |
| GO_PROTEIN_DNA_COMPLEX_DISASSEMBLY | 0.00235 | 0.03185 | 0.43171 | 0.79973 | 1.47866 | 17 |
| GO_PROTEIN_DNA_COMPLEX_SUBUNIT_ORGANIZATION | 0.00088 | 0.01473 | 0.47727 | 0.58584 | 1.19848 | 175 |
| GO_PROTEIN_DOMAIN_SPECIFIC_BINDING | 0.00076 | 0.01300 | 0.47727 | 0.54308 | 1.12890 | 454 |
| GO_PROTEIN_FOLDING | 0.00000 | 0.00000 | 0.70498 | 0.66066 | 1.34700 | 149 |
| GO_PROTEIN_FOLDING_CHAPERONE | 0.00294 | 0.03809 | 0.43171 | 0.79536 | 1.47059 | 17 |
| GO_PROTEIN_IMPORT | 0.00097 | 0.01561 | 0.47727 | 0.59819 | 1.21879 | 145 |
| GO_PROTEIN_INSERTION_INTO_MEMBRANE | 0.00095 | 0.01536 | 0.47727 | 0.68375 | 1.34512 | 48 |
| GO_PROTEIN_LOCALIZATION_TO_CHROMOSOME_TELOMEF | 0.00004 | 0.00098 | 0.55733 | 0.80274 | 1.52571 | 27 |
| GO_PROTEIN_LOCALIZATION_TO_ENDOPLASMIC_RETICULU | 0.00000 | 0.00000 | | 0.79097 | 1.60231 | 118 |
| GO_PROTEIN_LOCALIZATION_TO_MEMBRANE | 0.00000 | 0.00000 | | 0.61610 | 1.28002 | 434 |
| GO_PROTEIN_LOCALIZATION_TO_MITOCHONDRION | 0.00012 | 0.00286 | 0.53843 | 0.63221 | 1.28017 | 115 |
| GO_PROTEIN_LOCALIZATION_TO_NUCLEUS | 0.00004 | 0.00105 | 0.55733 | 0.59942 | 1.22880 | 200 |
| GO_PROTEIN_MODIFICATION_BY_SMALL_PROTEIN_REMOV | 0.00000 | 0.00012 | 0.62726 | 0.60216 | 1.23839 | 231 |
| GO_PROTEIN_POLYUBIQUITINATION | 0.00036 | 0.00742 | 0.49849 | 0.57449 | 1.18379 | 265 |
| GO_PROTEIN_TARGETING | 0.00000 | 0.00000 | | 0.66872 | 1.38397 | 327 |
| GO_PROTEIN_TARGETING_TO_MEMBRANE | 0.00000 | 0.00000 | | 0.74462 | 1.51933 | 153 |
| GO_PROTEIN_TARGETING_TO_MITOCHONDRION | 0.00057 | 0.01024 | 0.47727 | 0.64210 | 1.28614 | 79 |
| GO_PROTON_TRANSMEMBRANE_TRANSPORT | 0.00003 | 0.00083 | 0.57561 | 0.68894 | 1.37490 | 69 |
| GO_PROTON_TRANSMEMBRANE_TRANSPORTER_ACTIVITY | 0.00058 | 0.01037 | 0.47727 | 0.69155 | 1.36226 | 50 |
| GO_PURINE_CONTAINING_COMPOUND_BIOSYNTHETIC_PR | 0.00015 | 0.00343 | 0.51885 | 0.62925 | 1.27105 | 107 |
| GO_PURINE_CONTAINING_COMPOUND_METABOLIC_PROC | 0.00005 | 0.00133 | 0.55733 | 0.58969 | 1.21476 | 248 |
| GO_PURINE_NUCLEOSIDE_TRIPHOSPHATE_METABOLIC_PR | 0.00044 | 0.00844 | 0.49849 | 0.71261 | 1.39392 | 43 |
| GO_PYRUVATE_METABOLIC_PROCESS | 0.00194 | 0.02722 | 0.45506 | 0.61036 | 1.23029 | 102 |
| GO_REACTIVE_OXYGEN_SPECIES_BIOSYNTHETIC_PROCESS | 0.00145 | 0.02155 | 0.45506 | 0.67806 | 1.33699 | 51 |
| GO_REACTIVE_OXYGEN_SPECIES_METABOLIC_PROCESS | 0.00025 | 0.00545 | 0.49849 | 0.61551 | 1.25041 | 129 |
| GO_REGULATION_OF_ANIMAL_ORGAN_MORPHOGENESIS | 0.00000 | 0.00000 | 0.81404 | 0.69247 | 1.40991 | 141 |
| GO_REGULATION_OF_APOPTOTIC_SIGNALING_PATHWAY | 0.00191 | 0.02689 | 0.45506 | 0.55702 | 1.14746 | 259 |
| GO_REGULATION_OF_ATP_METABOLIC_PROCESS | 0.00135 | 0.02049 | 0.45506 | 0.63339 | 1.26991 | 82 |

| Cluster 1 | | | | | | |
|---|---|---|---|---|---|---|
| GO_REGULATION_OF_BINDING | 0.00115 | 0.01813 | 0.45506 | 0.56724 | 1.16616 | 228 |
| GO_REGULATION_OF_CELLULAR_AMIDE_METABOLIC_PROC | 0.00000 | 0.00000 | | 0.62948 | 1.30250 | 321 |
| GO_REGULATION_OF_CELLULAR_AMINE_METABOLIC_PROC | 0.00000 | 0.00000 | 0.71951 | 0.76519 | 1.51978 | 61 |
| GO_REGULATION_OF_CELLULAR_AMINO_ACID_METABOLIC | 0.00000 | 0.00000 | 0.73376 | 0.78449 | 1.55047 | 54 |
| GO_REGULATION_OF_CELLULAR_KETONE_METABOLIC_PRO | 0.00000 | 0.00000 | 0.70498 | 0.67571 | 1.36910 | 117 |
| GO_REGULATION_OF_CELLULAR_PROTEIN_LOCALIZATION | 0.00015 | 0.00356 | 0.51885 | 0.55985 | 1.16076 | 370 |
| GO_REGULATION_OF_CELL_CYCLE_G2_M_PHASE_TRANSITI | 0.00001 | 0.00017 | 0.61053 | 0.62272 | 1.27555 | 184 |
| GO_REGULATION_OF_CELL_CYCLE_PHASE_TRANSITION | 0.00002 | 0.00051 | 0.57561 | 0.57184 | 1.18544 | 365 |
| GO_REGULATION_OF_COAGULATION | 0.00420 | 0.04996 | 0.40702 | 0.70065 | 1.33880 | 29 |
| GO_REGULATION_OF_CYTOSKELETON_ORGANIZATION | 0.00163 | 0.02374 | 0.45506 | 0.55023 | 1.13895 | 337 |
| GO_REGULATION_OF_DEFENSE_RESPONSE | 0.00000 | 0.00005 | 0.64355 | 0.59013 | 1.22059 | 314 |
| GO_REGULATION_OF_DNA_BINDING | 0.00037 | 0.00750 | 0.49849 | 0.64772 | 1.29738 | 79 |
| GO_REGULATION_OF_DNA_BIOSYNTHETIC_PROCESS | 0.00253 | 0.03388 | 0.43171 | 0.62313 | 1.24816 | 80 |
| GO_REGULATION_OF_DNA_METABOLIC_PROCESS | 0.00246 | 0.03304 | 0.43171 | 0.56100 | 1.15503 | 243 |
| GO_REGULATION_OF_DNA_TEMPLATED_TRANSCRIPTION_I | 0.00000 | 0.00001 | 0.69013 | 0.70255 | 1.41444 | 94 |
| GO_REGULATION_OF_HEART_RATE | 0.00163 | 0.02374 | 0.45506 | 0.71402 | 1.37783 | 34 |
| GO_REGULATION_OF_HEMATOPOIETIC_PROGENITOR_CELL_ | 0.00000 | 0.00000 | 0.73376 | 0.74044 | 1.47940 | 73 |
| GO_REGULATION_OF_HEMATOPOIETIC_STEM_CELL_DIFFER | 0.00000 | 0.00000 | 0.74774 | 0.77376 | 1.53709 | 62 |
| GO_REGULATION_OF_HEMOPOIESIS | 0.00007 | 0.00181 | 0.53843 | 0.58040 | 1.19633 | 268 |
| GO_REGULATION_OF_IMMUNE_RESPONSE | 0.00000 | 0.00000 | 0.76146 | 0.60340 | 1.25243 | 395 |
| GO_REGULATION_OF_INNATE_IMMUNE_RESPONSE | 0.00000 | 0.00000 | 0.71951 | 0.64844 | 1.32650 | 174 |
| GO_REGULATION_OF_MITOTIC_CELL_CYCLE | 0.00000 | 0.00004 | 0.65944 | 0.57083 | 1.18772 | 493 |
| GO_REGULATION_OF_MORPHOGENESIS_OF_AN_EPITHELIU | 0.00000 | 0.00000 | | 0.72863 | 1.47650 | 119 |
| GO_REGULATION_OF_MRNA_CATABOLIC_PROCESS | 0.00000 | 0.00000 | | 0.68599 | 1.40151 | 168 |
| GO_REGULATION_OF_MRNA_METABOLIC_PROCESS | 0.00000 | 0.00000 | | 0.67617 | 1.39528 | 280 |
| GO_REGULATION_OF_MRNA_PROCESSING | 0.00000 | 0.00002 | 0.67496 | 0.66634 | 1.35376 | 127 |
| GO_REGULATION_OF_MRNA_SPLICING_VIA_SPLICEOSOME | 0.00000 | 0.00004 | 0.65944 | 0.69004 | 1.38860 | 93 |
| GO_REGULATION_OF_MUSCLE_CONTRACTION | 0.00061 | 0.01070 | 0.47727 | 0.65765 | 1.30710 | 63 |
| GO_REGULATION_OF_OXIDATIVE_PHOSPHORYLATION | 0.00048 | 0.00914 | 0.49849 | 0.79686 | 1.49819 | 21 |
| GO_REGULATION_OF_OXIDATIVE_STRESS_INDUCED_CELL_D | 0.00205 | 0.02857 | 0.43171 | 0.68747 | 1.34191 | 41 |
| GO_REGULATION_OF_PEPTIDASE_ACTIVITY | 0.00049 | 0.00919 | 0.47727 | 0.58025 | 1.18919 | 198 |
| GO_REGULATION_OF_POSTTRANSCRIPTIONAL_GENE_SILEN | 0.00115 | 0.01813 | 0.45506 | 0.63202 | 1.26570 | 78 |
| GO_REGULATION_OF_PROTEIN_CONTAINING_COMPLEX_AS | 0.00110 | 0.01745 | 0.45506 | 0.56276 | 1.16087 | 274 |
| GO_REGULATION_OF_PROTEIN_LOCALIZATION_TO_NUCLEU | 0.00038 | 0.00766 | 0.49849 | 0.63357 | 1.27497 | 93 |
| GO_REGULATION_OF_PROTEIN_STABILITY | 0.00327 | 0.04166 | 0.43171 | 0.56410 | 1.15763 | 209 |
| GO_REGULATION_OF_PROTEOLYSIS | 0.00059 | 0.01051 | 0.47727 | 0.55120 | 1.14362 | 385 |
| GO_REGULATION_OF_REACTIVE_OXYGEN_SPECIES_BIOSYNT | 0.00106 | 0.01691 | 0.45506 | 0.69926 | 1.36493 | 41 |
| GO_REGULATION_OF_REACTIVE_OXYGEN_SPECIES_METABO | 0.00235 | 0.03185 | 0.43171 | 0.62189 | 1.25004 | 88 |
| GO_REGULATION_OF_RESPONSE_TO_BIOTIC_STIMULUS | 0.00000 | 0.00001 | 0.69013 | 0.62630 | 1.28670 | 220 |
| GO_REGULATION_OF_RESPONSE_TO_EXTERNAL_STIMULUS | 0.00001 | 0.00032 | 0.59333 | 0.56621 | 1.17697 | 444 |
| GO_REGULATION_OF_RESPONSE_TO_OXIDATIVE_STRESS | 0.00298 | 0.03849 | 0.43171 | 0.66060 | 1.30396 | 52 |
| GO_REGULATION_OF_RNA_SPLICING | 0.00000 | 0.00000 | 0.70498 | 0.67606 | 1.37366 | 126 |
| GO_REGULATION_OF_SMALL_MOLECULE_METABOLIC_PRO | 0.00001 | 0.00017 | 0.61053 | 0.59600 | 1.22827 | 262 |
| GO_REGULATION_OF_STEM_CELL_DIFFERENTIATION | 0.00000 | 0.00000 | 0.70498 | 0.71431 | 1.43481 | 87 |
| GO_REGULATION_OF_SYMBIOTIC_PROCESS | 0.00002 | 0.00048 | 0.57561 | 0.63144 | 1.28654 | 145 |
| GO_REGULATION_OF_TELOMERASE_ACTIVITY | 0.00085 | 0.01424 | 0.47727 | 0.71885 | 1.39710 | 38 |
| GO_REGULATION_OF_TELOMERE_MAINTENANCE | 0.00335 | 0.04242 | 0.43171 | 0.64002 | 1.27391 | 66 |
| GO_REGULATION_OF_TELOMERE_MAINTENANCE_VIA_TEL | 0.00054 | 0.01003 | 0.47727 | 0.69284 | 1.36406 | 49 |
| GO_REGULATION_OF_TRANSCRIPTION_FROM_RNA_POLYM | 0.00000 | 0.00000 | 0.71951 | 0.75701 | 1.50677 | 66 |
| GO_REGULATION_OF_TRANSLATIONAL_INITIATION | 0.00005 | 0.00131 | 0.55733 | 0.68870 | 1.36880 | 63 |
| GO_REGULATION_OF_VIRAL_TRANSCRIPTION | 0.00089 | 0.01475 | 0.47727 | 0.67280 | 1.33099 | 56 |
| GO_REGULATION_OF_WNT_SIGNALING_PATHWAY | 0.00001 | 0.00041 | 0.59333 | 0.59860 | 1.23064 | 228 |
| GO_RESPIRASOME | 0.00000 | 0.00000 | 0.77494 | 0.76758 | 1.53319 | 71 |
| GO_RESPIRATORY_CHAIN_COMPLEX | 0.00000 | 0.00000 | 0.73376 | 0.77088 | 1.53109 | 61 |
| GO_RESPIRATORY_ELECTRON_TRANSPORT_CHAIN | 0.00000 | 0.00000 | | 0.76599 | 1.53708 | 85 |
| GO_RESPONSE_TO_CALCIUM_ION | 0.00405 | 0.04900 | 0.40702 | 0.62819 | 1.25390 | 70 |
| GO_RESPONSE_TO_CORTICOSTEROID | 0.00185 | 0.02630 | 0.45506 | 0.63417 | 1.26708 | 73 |
| GO_RESPONSE_TO_DRUG | 0.00091 | 0.01493 | 0.47727 | 0.57784 | 1.18581 | 205 |
| GO_RESPONSE_TO_ESTROGEN | 0.00057 | 0.01024 | 0.47727 | 0.69264 | 1.36157 | 47 |
| GO_RESPONSE_TO_FIBROBLAST_GROWTH_FACTOR | 0.00122 | 0.01908 | 0.45506 | 0.63845 | 1.27799 | 76 |
| GO_RESPONSE_TO_HEAT | 0.00020 | 0.00435 | 0.51885 | 0.62263 | 1.25938 | 112 |
| GO_RESPONSE_TO_HORMONE | 0.00161 | 0.02365 | 0.45506 | 0.53600 | 1.11532 | 496 |
| GO_RESPONSE_TO_INORGANIC_SUBSTANCE | 0.00003 | 0.00081 | 0.57561 | 0.58320 | 1.20351 | 283 |
| GO_RESPONSE_TO_INTERLEUKIN_1 | 0.00000 | 0.00000 | | 0.71724 | 1.45325 | 117 |
| GO_RESPONSE_TO_INTERLEUKIN_12 | 0.00135 | 0.02049 | 0.45506 | 0.70760 | 1.36862 | 35 |
| GO_RESPONSE_TO_INTERLEUKIN_4 | 0.00316 | 0.04043 | 0.43171 | 0.75186 | 1.41358 | 21 |
| GO_RESPONSE_TO_ORGANIC_CYCLIC_COMPOUND | 0.00020 | 0.00444 | 0.51885 | 0.55044 | 1.14498 | 469 |
| GO_RESPONSE_TO_OXIDATIVE_STRESS | 0.00044 | 0.00844 | 0.49849 | 0.57223 | 1.17898 | 258 |
| GO_RESPONSE_TO_OXYGEN_LEVELS | 0.00000 | 0.00000 | 0.76146 | 0.63454 | 1.30490 | 234 |
| GO_RESPONSE_TO_REACTIVE_OXYGEN_SPECIES | 0.00398 | 0.04830 | 0.40702 | 0.58662 | 1.19173 | 129 |
| GO_RESPONSE_TO_STEROID_HORMONE | 0.00294 | 0.03809 | 0.43171 | 0.56605 | 1.15946 | 188 |
| GO_RESPONSE_TO_TEMPERATURE_STIMULUS | 0.00102 | 0.01646 | 0.45506 | 0.59547 | 1.21138 | 137 |
| GO_RESPONSE_TO_TOXIC_SUBSTANCE | 0.00010 | 0.00240 | 0.53843 | 0.63335 | 1.28246 | 115 |

| Cluster 1 | | | | | | |
|---|---|---|---|---|---|---|
| GO_RESPONSE_TO_TUMOR_NECROSIS_FACTOR | 0.00000 | 0.00001 | 0.69013 | 0.64897 | 1.32421 | 154 |
| GO_RESPONSE_TO_VIRUS | 0.00339 | 0.04252 | 0.43171 | 0.57830 | 1.18015 | 159 |
| GO_RETINA_HOMEOSTASIS | 0.00008 | 0.00207 | 0.53843 | 0.79915 | 1.51122 | 25 |
| GO_RIBONUCLEOPROTEIN_COMPLEX_BINDING | 0.00000 | 0.00014 | 0.61053 | 0.65914 | 1.33318 | 113 |
| GO_RIBONUCLEOPROTEIN_COMPLEX_BIOGENESIS | 0.00000 | 0.00000 | | 0.67093 | 1.39207 | 389 |
| GO_RIBONUCLEOPROTEIN_COMPLEX_SUBUNIT_ORGANIZAT | 0.00000 | 0.00000 | | 0.68906 | 1.40868 | 171 |
| GO_RIBONUCLEOPROTEIN_GRANULE | 0.00000 | 0.00000 | 0.77494 | 0.65704 | 1.34486 | 180 |
| GO_RIBONUCLEOSIDE_TRIPHOSPHATE_BIOSYNTHETIC_PRO | 0.00054 | 0.00989 | 0.47727 | 0.72343 | 1.39271 | 33 |
| GO_RIBONUCLEOSIDE_TRIPHOSPHATE_METABOLIC_PROCES | 0.00011 | 0.00266 | 0.53843 | 0.72947 | 1.42539 | 42 |
| GO_RIBOSE_PHOSPHATE_BIOSYNTHETIC_PROCESS | 0.00044 | 0.00847 | 0.49849 | 0.61887 | 1.25069 | 110 |
| GO_RIBOSE_PHOSPHATE_METABOLIC_PROCESS | 0.00009 | 0.00224 | 0.53843 | 0.58659 | 1.20736 | 240 |
| GO_RIBOSOMAL_LARGE_SUBUNIT_ASSEMBLY | 0.00148 | 0.02204 | 0.45506 | 0.74868 | 1.41578 | 25 |
| GO_RIBOSOMAL_LARGE_SUBUNIT_BIOGENESIS | 0.00000 | 0.00009 | 0.62726 | 0.72403 | 1.44112 | 66 |
| GO_RIBOSOMAL_SMALL_SUBUNIT_ASSEMBLY | 0.00044 | 0.00847 | 0.49849 | 0.80809 | 1.50134 | 18 |
| GO_RIBOSOMAL_SMALL_SUBUNIT_BIOGENESIS | 0.00133 | 0.02032 | 0.45506 | 0.64174 | 1.27883 | 68 |
| GO_RIBOSOMAL_SUBUNIT | 0.00000 | 0.00000 | | 0.76429 | 1.56307 | 172 |
| GO_RIBOSOME | 0.00000 | 0.00000 | | 0.75467 | 1.54868 | 205 |
| GO_RIBOSOME_ASSEMBLY | 0.00002 | 0.00050 | 0.57561 | 0.71921 | 1.42251 | 55 |
| GO_RIBOSOME_BIOGENESIS | 0.00000 | 0.00000 | | 0.65934 | 1.36027 | 275 |
| GO_RNA_3_END_PROCESSING | 0.00031 | 0.00653 | 0.49849 | 0.61620 | 1.25185 | 125 |
| GO_RNA_CATABOLIC_PROCESS | 0.00000 | 0.00000 | | 0.71173 | 1.47401 | 347 |
| GO_RNA_DEPENDENT_DNA_BIOSYNTHETIC_PROCESS | 0.00037 | 0.00750 | 0.49849 | 0.68750 | 1.36322 | 59 |
| GO_RNA_EXPORT_FROM_NUCLEUS | 0.00000 | 0.00005 | 0.64355 | 0.65814 | 1.33724 | 126 |
| GO_RNA_LOCALIZATION | 0.00000 | 0.00000 | 0.77494 | 0.65445 | 1.34196 | 201 |
| GO_RNA_SPLICING | 0.00000 | 0.00000 | | 0.63503 | 1.31758 | 389 |
| GO_RNA_SPLICING_VIA_TRANSESTERIFICATION_REACTIONS | 0.00000 | 0.00000 | | 0.64850 | 1.34130 | 316 |
| GO_RNA_STABILIZATION | 0.00383 | 0.04678 | 0.43171 | 0.68413 | 1.32553 | 36 |
| GO_ROUGH_ENDOPLASMIC_RETICULUM | 0.00239 | 0.03227 | 0.43171 | 0.67394 | 1.32088 | 45 |
| GO_RRNA_BINDING | 0.00000 | 0.00002 | 0.65944 | 0.75334 | 1.49030 | 56 |
| GO_RRNA_METABOLIC_PROCESS | 0.00000 | 0.00000 | 0.80122 | 0.65560 | 1.34539 | 209 |
| GO_SARCOMERE_ORGANIZATION | 0.00209 | 0.02901 | 0.43171 | 0.77704 | 1.44978 | 19 |
| GO_SCF_DEPENDENT_PROTEASOMAL_UBIQUITIN_DEPENDI | 0.00000 | 0.00001 | 0.69013 | 0.72696 | 1.45516 | 76 |
| GO_SECRETORY_GRANULE | 0.00000 | 0.00000 | 0.81404 | 0.61499 | 1.27560 | 376 |
| GO_SECRETORY_VESICLE | 0.00000 | 0.00000 | 0.77494 | 0.59324 | 1.23316 | 454 |
| GO_SIGNAL_TRANSDUCTION_BY_PROTEIN_PHOSPHORYLAT | 0.00276 | 0.03635 | 0.43171 | 0.53726 | 1.11727 | 462 |
| GO_SINGLE_FERTILIZATION | 0.00004 | 0.00109 | 0.55733 | 0.73055 | 1.43382 | 46 |
| GO_SINGLE_STRANDED_DNA_BINDING | 0.00069 | 0.01190 | 0.47727 | 0.63510 | 1.27570 | 87 |
| GO_SINGLE_STRANDED_RNA_BINDING | 0.00003 | 0.00083 | 0.57561 | 0.69662 | 1.38658 | 66 |
| GO_SMALL_MOLECULE_BIOSYNTHETIC_PROCESS | 0.00030 | 0.00629 | 0.49849 | 0.55581 | 1.15278 | 378 |
| GO_SMALL_NUCLEAR_RIBONUCLEOPROTEIN_COMPLEX | 0.00005 | 0.00126 | 0.55733 | 0.70734 | 1.40105 | 57 |
| GO_SMALL_RIBOSOMAL_SUBUNIT | 0.00000 | 0.00000 | 0.76146 | 0.76060 | 1.51569 | 68 |
| GO_SM_LIKE_PROTEIN_FAMILY_COMPLEX | 0.00014 | 0.00323 | 0.51885 | 0.68099 | 1.35700 | 67 |
| GO_SOMATIC_STEM_CELL_POPULATION_MAINTENANCE | 0.00235 | 0.03185 | 0.43171 | 0.67201 | 1.31892 | 46 |
| GO_SPERM_EGG_RECOGNITION | 0.00041 | 0.00810 | 0.49849 | 0.80986 | 1.50463 | 18 |
| GO_SPLICEOSOMAL_COMPLEX | 0.00000 | 0.00000 | 0.76146 | 0.66446 | 1.35839 | 171 |
| GO_SPLICEOSOMAL_COMPLEX_ASSEMBLY | 0.00039 | 0.00781 | 0.49849 | 0.70044 | 1.37691 | 47 |
| GO_SPLICEOSOMAL_TRI_SNRNP_COMPLEX | 0.00066 | 0.01149 | 0.47727 | 0.73978 | 1.41705 | 30 |
| GO_STEM_CELL_DIFFERENTIATION | 0.00000 | 0.00000 | 0.76146 | 0.67362 | 1.37446 | 153 |
| GO_STEROID_HORMONE_RECEPTOR_BINDING | 0.00228 | 0.03113 | 0.43171 | 0.65196 | 1.29396 | 60 |
| GO_STRUCTURAL_CONSTITUENT_OF_CYTOSKELETON | 0.00075 | 0.01273 | 0.47727 | 0.68215 | 1.34820 | 54 |
| GO_STRUCTURAL_CONSTITUENT_OF_MUSCLE | 0.00021 | 0.00463 | 0.51885 | 0.80893 | 1.51425 | 20 |
| GO_STRUCTURAL_CONSTITUENT_OF_RIBOSOME | 0.00000 | 0.00000 | | 0.77595 | 1.58226 | 151 |
| GO_STRUCTURAL_MOLECULE_ACTIVITY | 0.00000 | 0.00000 | | 0.71193 | 1.47548 | 361 |
| GO_SUBSTANTIA_NIGRA_DEVELOPMENT | 0.00092 | 0.01499 | 0.47727 | 0.72798 | 1.39984 | 32 |
| GO_SUPEROXIDE_METABOLIC_PROCESS | 0.00113 | 0.01794 | 0.45506 | 0.72620 | 1.39104 | 30 |
| GO_SUPRAMOLECULAR_FIBER_ORGANIZATION | 0.00357 | 0.04455 | 0.43171 | 0.53775 | 1.11638 | 401 |
| GO_SUPRAMOLECULAR_POLYMER | 0.00000 | 0.00011 | 0.62726 | 0.56448 | 1.17458 | 489 |
| GO_SYNAPSE_ORGANIZATION | 0.00290 | 0.03795 | 0.43171 | 0.57054 | 1.16868 | 184 |
| GO_TELOMERASE_RNA_LOCALIZATION | 0.00156 | 0.02296 | 0.45506 | 0.78139 | 1.45791 | 19 |
| GO_TELOMERE_MAINTENANCE_VIA_TELOMERE_LENGTHEN | 0.00008 | 0.00212 | 0.53843 | 0.68653 | 1.36648 | 66 |
| GO_TERTIARY_GRANULE | 0.00213 | 0.02938 | 0.43171 | 0.64107 | 1.27414 | 63 |
| GO_THREONINE_TYPE_ENDOPEPTIDASE_ACTIVITY | 0.00000 | 0.00001 | 0.69013 | 0.92886 | 1.70607 | 15 |
| GO_THREONINE_TYPE_PEPTIDASE_ACTIVITY | 0.00000 | 0.00002 | 0.65944 | 0.90394 | 1.67942 | 18 |
| GO_TISSUE_MORPHOGENESIS | 0.00000 | 0.00003 | 0.65944 | 0.58798 | 1.21841 | 360 |
| GO_TOXIN_TRANSPORT | 0.00021 | 0.00455 | 0.51885 | 0.77181 | 1.47121 | 28 |
| GO_TRANSCRIPTION_COUPLED_NUCLEOTIDE_EXCISION_RE | 0.00294 | 0.03809 | 0.43171 | 0.63074 | 1.25876 | 69 |
| GO_TRANSCRIPTION_ELONGATION_FROM_RNA_POLYMERA | 0.00181 | 0.02591 | 0.45506 | 0.63477 | 1.26827 | 73 |
| GO_TRANSLATIONAL_ELONGATION | 0.00000 | 0.00000 | 0.71951 | 0.68581 | 1.39123 | 121 |
| GO_TRANSLATIONAL_INITIATION | 0.00000 | 0.00000 | | 0.77621 | 1.58516 | 166 |
| GO_TRANSLATIONAL_TERMINATION | 0.00000 | 0.00001 | 0.67496 | 0.68789 | 1.38658 | 102 |
| GO_TRANSLATION_FACTOR_ACTIVITY_RNA_BINDING | 0.00000 | 0.00002 | 0.65944 | 0.72463 | 1.44641 | 70 |
| GO_TRANSLATION_INITIATION_FACTOR_ACTIVITY | 0.00004 | 0.00099 | 0.55733 | 0.73277 | 1.43617 | 45 |

| Cluster 1 | | | | | | |
|---|---|---|---|---|---|---|
| GO_TRANSLATION_INITIATION_FACTOR_BINDING | 0.00011 | 0.00266 | 0.53843 | 0.79236 | 1.49727 | 24 |
| GO_TRANSLATION_PREINITIATION_COMPLEX | 0.00026 | 0.00558 | 0.49849 | 0.84201 | 1.55182 | 16 |
| GO_TRANSLATION_REGULATOR_ACTIVITY | 0.00000 | 0.00000 | 0.74774 | 0.69934 | 1.41387 | 111 |
| GO_TRANSLATION_REGULATOR_ACTIVITY_NUCLEIC_ACID_E | 0.00000 | 0.00001 | 0.69013 | 0.72036 | 1.44551 | 85 |
| GO_TRANSMEMBRANE_RECEPTOR_PROTEIN_TYROSINE_KIN | 0.00302 | 0.03888 | 0.43171 | 0.54259 | 1.12574 | 386 |
| GO_TUMOR_NECROSIS_FACTOR_MEDIATED_SIGNALING_PA | 0.00000 | 0.00000 | 0.76146 | 0.72046 | 1.45106 | 98 |
| GO_T_CELL_RECEPTOR_SIGNALING_PATHWAY | 0.00000 | 0.00006 | 0.64355 | 0.66032 | 1.33889 | 120 |
| GO_U12_TYPE_SPLICEOSOMAL_COMPLEX | 0.00027 | 0.00566 | 0.49849 | 0.77922 | 1.47782 | 26 |
| GO_U1_SNRNP | 0.00015 | 0.00356 | 0.51885 | 0.85613 | 1.57248 | 15 |
| GO_U2_SNRNP | 0.00049 | 0.00919 | 0.47727 | 0.80594 | 1.49735 | 18 |
| GO_U2_TYPE_SPLICEOSOMAL_COMPLEX | 0.00011 | 0.00269 | 0.53843 | 0.65232 | 1.30971 | 86 |
| GO_U5_SNRNP | 0.00143 | 0.02144 | 0.45506 | 0.81282 | 1.49293 | 15 |
| GO_UBIQUITIN_LIKE_PROTEIN_LIGASE_BINDING | 0.00000 | 0.00011 | 0.62726 | 0.60937 | 1.25269 | 226 |
| GO_UNFOLDED_PROTEIN_BINDING | 0.00000 | 0.00000 | 0.76146 | 0.74375 | 1.48918 | 77 |
| GO_VACUOLAR_LUMEN | 0.00026 | 0.00558 | 0.49849 | 0.66171 | 1.31861 | 68 |
| GO_VESICLE_LUMEN | 0.00000 | 0.00000 | | 0.70462 | 1.44103 | 172 |
| GO_VIRAL_GENE_EXPRESSION | 0.00000 | 0.00000 | | 0.75146 | 1.53731 | 176 |
| GO_VIRAL_GENOME_REPLICATION | 0.00048 | 0.00914 | 0.49849 | 0.64935 | 1.29740 | 73 |
| GO_VIRAL_LIFE_CYCLE | 0.00061 | 0.01070 | 0.47727 | 0.58004 | 1.19101 | 213 |
| GO_VOLTAGE_GATED_ION_CHANNEL_ACTIVITY | 0.00383 | 0.04678 | 0.43171 | 0.68471 | 1.32665 | 36 |
| HP_ABNORMALITY_OF_ACID_BASE_HOMEOSTASIS | 0.00079 | 0.01343 | 0.47727 | 0.57263 | 1.17766 | 233 |
| HP_ABNORMAL_ACTIVITY_OF_MITOCHONDRIAL_RESPIRATO | 0.00390 | 0.04759 | 0.43171 | 0.64271 | 1.27560 | 60 |
| HP_ABNORMAL_CSF_METABOLITE_LEVEL | 0.00048 | 0.00914 | 0.49849 | 0.64315 | 1.28775 | 77 |
| HP_ABNORMAL_ERYTHROCYTE_MORPHOLOGY | 0.00398 | 0.04830 | 0.40702 | 0.54241 | 1.12320 | 346 |
| HP_ACUTE_LEUKEMIA | 0.00309 | 0.03966 | 0.43171 | 0.63453 | 1.26745 | 71 |
| HP_ANEMIA_OF_INADEQUATE_PRODUCTION | 0.00018 | 0.00405 | 0.51885 | 0.63404 | 1.27802 | 102 |
| HP_APHASIA | 0.00372 | 0.04591 | 0.43171 | 0.79159 | 1.46361 | 17 |
| HP_DEPRESSED_NASAL_RIDGE | 0.00004 | 0.00104 | 0.55733 | 0.68612 | 1.36727 | 68 |
| HP_ECHOLALIA | 0.00156 | 0.02296 | 0.45506 | 0.76609 | 1.43407 | 20 |
| HP_EMOTIONAL_LABILITY | 0.00276 | 0.03635 | 0.43171 | 0.61861 | 1.24203 | 86 |
| HP_FATIGUE | 0.00057 | 0.01024 | 0.47727 | 0.58198 | 1.19252 | 197 |
| HP_FETAL_DISTRESS | 0.00383 | 0.04678 | 0.43171 | 0.70232 | 1.34531 | 30 |
| HP_HEMATOLOGICAL_NEOPLASM | 0.00294 | 0.03809 | 0.43171 | 0.57574 | 1.17550 | 165 |
| HP_HYPERTROPHIC_CARDIOMYOPATHY | 0.00279 | 0.03675 | 0.43171 | 0.57627 | 1.17890 | 175 |
| HP_INCREASED_CSF_LACTATE | 0.00052 | 0.00961 | 0.47727 | 0.66574 | 1.32511 | 66 |
| HP_INCREASED_SERUM_LACTATE | 0.00163 | 0.02374 | 0.45506 | 0.59933 | 1.21747 | 128 |
| HP_MACROCYTIC_ANEMIA | 0.00002 | 0.00056 | 0.57561 | 0.76702 | 1.49227 | 39 |
| HP_MIGRAINE | 0.00019 | 0.00428 | 0.51885 | 0.67230 | 1.34195 | 70 |
| HP_MUTISM | 0.00057 | 0.01024 | 0.47727 | 0.76537 | 1.45155 | 26 |
| HP_OPTIC_NEUROPATHY | 0.00420 | 0.04996 | 0.40702 | 0.68771 | 1.32393 | 33 |
| HP_POOR_HEAD_CONTROL | 0.00276 | 0.03635 | 0.43171 | 0.60636 | 1.22297 | 103 |
| HP_RIMMED_VACUOLES | 0.00139 | 0.02099 | 0.45506 | 0.76990 | 1.44751 | 21 |
| **Cluster 8** | | | | | | |
| pathway | pval | padj | log2err | ES | NES | size |
| GO_ACTIVATION_OF_IMMUNE_RESPONSE | 0.00062 | 0.02068 | 0.47727 | 0.61765 | 1.15658 | 240 |
| GO_ACTIVATION_OF_INNATE_IMMUNE_RESPONSE | 0.00001 | 0.00095 | 0.59333 | 0.70925 | 1.30626 | 94 |
| GO_ALTERNATIVE_MRNA_SPLICING_VIA_SPLICEOSOME | 0.00000 | 0.00042 | 0.62726 | 0.74698 | 1.36671 | 69 |
| GO_AMINE_METABOLIC_PROCESS | 0.00038 | 0.01439 | 0.49849 | 0.69092 | 1.26929 | 87 |
| GO_ANAPHASE_PROMOTING_COMPLEX_DEPENDENT_CATA | 0.00000 | 0.00025 | 0.62726 | 0.74311 | 1.36297 | 76 |
| GO_ANTIGEN_PROCESSING_AND_PRESENTATION | 0.00154 | 0.03948 | 0.45506 | 0.63103 | 1.17247 | 161 |
| GO_ANTIGEN_PROCESSING_AND_PRESENTATION_OF_EXOG | 0.00082 | 0.02507 | 0.47727 | 0.70381 | 1.28517 | 63 |
| GO_APOPTOTIC_SIGNALING_PATHWAY | 0.00008 | 0.00498 | 0.53843 | 0.61213 | 1.15273 | 370 |
| GO_ATP_BIOSYNTHETIC_PROCESS | 0.00130 | 0.03527 | 0.45506 | 0.78995 | 1.39645 | 23 |
| GO_ATP_METABOLIC_PROCESS | 0.00027 | 0.01130 | 0.49849 | 0.63690 | 1.18944 | 203 |
| GO_CADHERIN_BINDING | 0.00003 | 0.00254 | 0.57561 | 0.63865 | 1.19610 | 244 |
| GO_CATALYTIC_STEP_2_SPLICEOSOME | 0.00000 | 0.00003 | 0.69013 | 0.76392 | 1.40251 | 82 |
| GO_CELLULAR_COMPONENT_DISASSEMBLY | 0.00032 | 0.01268 | 0.49849 | 0.60128 | 1.13319 | 404 |
| GO_CELLULAR_KETONE_METABOLIC_PROCESS | 0.00008 | 0.00500 | 0.53843 | 0.65144 | 1.20986 | 156 |
| GO_CELLULAR_NITROGEN_COMPOUND_CATABOLIC_PROCE | 0.00000 | 0.00000 | | 0.65535 | 1.23549 | 440 |
| GO_CELLULAR_RESPONSE_TO_HEAT | 0.00014 | 0.00749 | 0.51885 | 0.68017 | 1.25334 | 98 |
| GO_CELLULAR_RESPONSE_TO_OXYGEN_LEVELS | 0.00004 | 0.00323 | 0.55733 | 0.66269 | 1.23100 | 157 |
| GO_CELLULAR_RESPONSE_TO_STARVATION | 0.00018 | 0.00872 | 0.51885 | 0.67372 | 1.24369 | 109 |
| GO_CELL_ADHESION_MOLECULE_BINDING | 0.00019 | 0.00901 | 0.51885 | 0.61030 | 1.14864 | 338 |
| GO_CELL_CELL_RECOGNITION | 0.00152 | 0.03948 | 0.45506 | 0.73812 | 1.33335 | 40 |
| GO_CELL_CYCLE_G2_M_PHASE_TRANSITION | 0.00167 | 0.04225 | 0.45506 | 0.61438 | 1.14938 | 224 |
| GO_CELL_CYCLE_PHASE_TRANSITION | 0.00000 | 0.00032 | 0.62726 | 0.61416 | 1.15875 | 489 |
| GO_CELL_DIVISION | 0.00004 | 0.00310 | 0.55733 | 0.60475 | 1.14000 | 451 |
| GO_CELL_RECOGNITION | 0.00065 | 0.02143 | 0.47727 | 0.69205 | 1.26754 | 71 |
| GO_CELL_SUBSTRATE_JUNCTION | 0.00005 | 0.00356 | 0.55733 | 0.62561 | 1.17607 | 302 |
| GO_CHROMOSOMAL_REGION | 0.00016 | 0.00781 | 0.51885 | 0.62036 | 1.16440 | 282 |
| GO_CHROMOSOME_CENTROMERIC_REGION | 0.00031 | 0.01258 | 0.49849 | 0.63799 | 1.18582 | 166 |
| GO_CHROMOSOME_SEGREGATION | 0.00000 | 0.00018 | 0.64355 | 0.64701 | 1.21311 | 262 |
| GO_CONDENSED_CHROMOSOME_CENTROMERIC_REGION | 0.00064 | 0.02106 | 0.47727 | 0.66258 | 1.22093 | 98 |

| Cluster 1 | | | | | | |
|---|---|---|---|---|---|---|
| GO_COTRANSLATIONAL_PROTEIN_TARGETING_TO_MEMBR/ | 0.00000 | 0.00005 | 0.67496 | 0.74126 | 1.36534 | 95 |
| GO_COVALENT_CHROMATIN_MODIFICATION | 0.00059 | 0.01998 | 0.47727 | 0.59966 | 1.12924 | 361 |
| GO_CYTOKINE_MEDIATED_SIGNALING_PATHWAY | 0.00025 | 0.01088 | 0.49849 | 0.60532 | 1.14014 | 374 |
| GO_CYTOPLASMIC_TRANSLATION | 0.00000 | 0.00005 | 0.67496 | 0.74603 | 1.37252 | 90 |
| GO_CYTOPLASMIC_TRANSLATIONAL_INITIATION | 0.00048 | 0.01712 | 0.49849 | 0.78590 | 1.40342 | 30 |
| GO_CYTOSOLIC_LARGE_RIBOSOMAL_SUBUNIT | 0.00009 | 0.00538 | 0.53843 | 0.75218 | 1.36900 | 53 |
| GO_CYTOSOLIC_RIBOSOME | 0.00000 | 0.00000 | 0.74774 | 0.74695 | 1.37732 | 100 |
| GO_CYTOSOLIC_SMALL_RIBOSOMAL_SUBUNIT | 0.00036 | 0.01387 | 0.49849 | 0.74904 | 1.35351 | 43 |
| GO_DEFENSE_RESPONSE_TO_OTHER_ORGANISM | 0.00021 | 0.00932 | 0.51885 | 0.59484 | 1.12206 | 480 |
| GO_DEVELOPMENT_OF_PRIMARY_SEXUAL_CHARACTERISTI | 0.00040 | 0.01515 | 0.49849 | 0.65437 | 1.21097 | 125 |
| GO_DNA_BINDING_TRANSCRIPTION_FACTOR_BINDING | 0.00073 | 0.02320 | 0.47727 | 0.61549 | 1.15213 | 239 |
| GO_DNA_BIOSYNTHETIC_PROCESS | 0.00095 | 0.02774 | 0.47727 | 0.64475 | 1.19632 | 143 |
| GO_DNA_CONFORMATION_CHANGE | 0.00163 | 0.04149 | 0.45506 | 0.61193 | 1.14485 | 226 |
| GO_DNA_TEMPLATED_TRANSCRIPTION_ELONGATION | 0.00008 | 0.00519 | 0.53843 | 0.69138 | 1.27442 | 99 |
| GO_DNA_TEMPLATED_TRANSCRIPTION_INITIATION | 0.00088 | 0.02663 | 0.47727 | 0.63507 | 1.18262 | 179 |
| GO_ENDOPEPTIDASE_COMPLEX | 0.00028 | 0.01142 | 0.49849 | 0.71446 | 1.30325 | 60 |
| GO_ENDOTHELIAL_CELL_CHEMOTAXIS | 0.00010 | 0.00584 | 0.53843 | 0.89544 | 1.55007 | 15 |
| GO_ESTABLISHMENT_OF_PROTEIN_LOCALIZATION_TO_END | 0.00000 | 0.00002 | 0.70498 | 0.73659 | 1.35940 | 105 |
| GO_ESTABLISHMENT_OF_PROTEIN_LOCALIZATION_TO_MEN | 0.00000 | 0.00001 | 0.71951 | 0.66716 | 1.25004 | 252 |
| GO_ESTABLISHMENT_OF_PROTEIN_LOCALIZATION_TO_ORG | 0.00000 | 0.00000 | | 0.65675 | 1.23803 | 443 |
| GO_ESTABLISHMENT_OF_RNA_LOCALIZATION | 0.00004 | 0.00305 | 0.55733 | 0.65666 | 1.22243 | 172 |
| GO_FIBRILLAR_CENTER | 0.00020 | 0.00904 | 0.51885 | 0.68506 | 1.26183 | 95 |
| GO_FIBROBLAST_GROWTH_FACTOR_RECEPTOR_SIGNALING | 0.00064 | 0.02106 | 0.47727 | 0.69900 | 1.28104 | 72 |
| GO_FICOLIN_1_RICH_GRANULE | 0.00035 | 0.01360 | 0.49849 | 0.68264 | 1.25663 | 92 |
| GO_HEAT_SHOCK_PROTEIN_BINDING | 0.00006 | 0.00403 | 0.55733 | 0.70882 | 1.30332 | 89 |
| GO_HEMATOPOIETIC_PROGENITOR_CELL_DIFFERENTIATION | 0.00071 | 0.02310 | 0.47727 | 0.65033 | 1.20309 | 123 |
| GO_HEMATOPOIETIC_STEM_CELL_DIFFERENTIATION | 0.00047 | 0.01699 | 0.49849 | 0.69825 | 1.27755 | 69 |
| GO_INNATE_IMMUNE_RESPONSE | 0.00045 | 0.01673 | 0.49849 | 0.59843 | 1.12769 | 392 |
| GO_INNATE_IMMUNE_RESPONSE_ACTIVATING_SIGNAL_TR/ | 0.00024 | 0.01055 | 0.51885 | 0.69515 | 1.27550 | 78 |
| GO_INTERLEUKIN_1_MEDIATED_SIGNALING_PATHWAY | 0.00009 | 0.00546 | 0.53843 | 0.70152 | 1.28729 | 79 |
| GO_INTRINSIC_APOPTOTIC_SIGNALING_PATHWAY | 0.00003 | 0.00253 | 0.57561 | 0.65161 | 1.21675 | 199 |
| GO_INTRINSIC_APOPTOTIC_SIGNALING_PATHWAY_BY_P53_ | 0.00073 | 0.02320 | 0.47727 | 0.71167 | 1.29600 | 56 |
| GO_IN_UTERO_EMBRYONIC_DEVELOPMENT | 0.00185 | 0.04615 | 0.45506 | 0.60981 | 1.14250 | 255 |
| GO_KINASE_REGULATOR_ACTIVITY | 0.00027 | 0.01130 | 0.49849 | 0.64902 | 1.20568 | 152 |
| GO_LARGE_RIBOSOMAL_SUBUNIT | 0.00095 | 0.02774 | 0.47727 | 0.66139 | 1.22087 | 108 |
| GO_MALE_SEX_DIFFERENTIATION | 0.00198 | 0.04839 | 0.43171 | 0.66131 | 1.21795 | 94 |
| GO_MATURATION_OF_LSU_RRNA | 0.00205 | 0.04960 | 0.43171 | 0.77858 | 1.38411 | 26 |
| GO_MITOCHONDRION_ORGANIZATION | 0.00061 | 0.02048 | 0.47727 | 0.59777 | 1.12627 | 378 |
| GO_MITOTIC_NUCLEAR_DIVISION | 0.00011 | 0.00613 | 0.53843 | 0.63543 | 1.18825 | 221 |
| GO_MITOTIC_SISTER_CHROMATID_SEGREGATION | 0.00006 | 0.00433 | 0.53843 | 0.66504 | 1.23397 | 143 |
| GO_MORPHOGENESIS_OF_A_POLARIZED_EPITHELIUM | 0.00143 | 0.03792 | 0.45506 | 0.65596 | 1.21029 | 104 |
| GO_MRNA_3_END_PROCESSING | 0.00010 | 0.00598 | 0.53843 | 0.69492 | 1.27924 | 92 |
| GO_MRNA_BINDING | 0.00000 | 0.00000 | | 0.69528 | 1.30194 | 240 |
| GO_MRNA_EXPORT_FROM_NUCLEUS | 0.00003 | 0.00272 | 0.55733 | 0.69053 | 1.27329 | 100 |
| GO_MRNA_PROCESSING | 0.00000 | 0.00000 | | 0.70344 | 1.32592 | 432 |
| GO_MRNA_TRANSPORT | 0.00018 | 0.00853 | 0.51885 | 0.66189 | 1.22623 | 129 |
| GO_NCRNA_METABOLIC_PROCESS | 0.00000 | 0.00055 | 0.61053 | 0.62093 | 1.17010 | 381 |
| GO_NCRNA_PROCESSING | 0.00000 | 0.00030 | 0.62726 | 0.63704 | 1.19952 | 323 |
| GO_NEGATIVE_REGULATION_OF_CATALYTIC_ACTIVITY | 0.00014 | 0.00745 | 0.51885 | 0.60161 | 1.13388 | 433 |
| GO_NEGATIVE_REGULATION_OF_CELL_CYCLE | 0.00001 | 0.00117 | 0.59333 | 0.61222 | 1.15412 | 453 |
| GO_NEGATIVE_REGULATION_OF_CELL_CYCLE_G2_M_PHASI | 0.00047 | 0.01699 | 0.49849 | 0.68889 | 1.26556 | 87 |
| GO_NEGATIVE_REGULATION_OF_CELL_CYCLE_PHASE_TRAN | 0.00001 | 0.00135 | 0.59333 | 0.65654 | 1.22389 | 190 |
| GO_NEGATIVE_REGULATION_OF_CELL_CYCLE_PROCESS | 0.00000 | 0.00020 | 0.64355 | 0.65083 | 1.21955 | 256 |
| GO_NEGATIVE_REGULATION_OF_MITOTIC_CELL_CYCLE | 0.00003 | 0.00272 | 0.55733 | 0.63557 | 1.19072 | 251 |
| GO_NEGATIVE_REGULATION_OF_MRNA_PROCESSING | 0.00191 | 0.04734 | 0.45506 | 0.75994 | 1.35491 | 29 |
| GO_NEGATIVE_REGULATION_OF_PROTEIN_MODIFICATION_ | 0.00198 | 0.04839 | 0.43171 | 0.59046 | 1.11270 | 395 |
| GO_NEGATIVE_REGULATION_OF_RNA_SPLICING | 0.00113 | 0.03205 | 0.45506 | 0.79331 | 1.40634 | 24 |
| GO_NIK_NF_KAPPAB_SIGNALING | 0.00004 | 0.00291 | 0.55733 | 0.68849 | 1.27097 | 109 |
| GO_NUCLEAR_CHROMOSOME_SEGREGATION | 0.00004 | 0.00314 | 0.55733 | 0.64721 | 1.20964 | 209 |
| GO_NUCLEAR_EXPORT | 0.00001 | 0.00106 | 0.59333 | 0.66438 | 1.23679 | 173 |
| GO_NUCLEAR_MATRIX | 0.00119 | 0.03315 | 0.45506 | 0.67682 | 1.24402 | 86 |
| GO_NUCLEAR_PERIPHERY | 0.00015 | 0.00754 | 0.51885 | 0.68297 | 1.26045 | 105 |
| GO_NUCLEAR_SPECK | 0.00000 | 0.00000 | 0.76146 | 0.66475 | 1.24864 | 298 |
| GO_NUCLEAR_TRANSCRIBED_MRNA_CATABOLIC_PROCESS | 0.00000 | 0.00001 | 0.71951 | 0.69058 | 1.28599 | 180 |
| GO_NUCLEAR_TRANSCRIBED_MRNA_CATABOLIC_PROCESS_ | 0.00000 | 0.00000 | 0.76146 | 0.73997 | 1.36663 | 114 |
| GO_NUCLEAR_TRANSPORT | 0.00000 | 0.00002 | 0.70498 | 0.65410 | 1.22825 | 289 |
| GO_NUCLEOBASE_CONTAINING_COMPOUND_TRANSPORT | 0.00005 | 0.00364 | 0.55733 | 0.64487 | 1.20517 | 208 |
| GO_NUCLEUS_ORGANIZATION | 0.00038 | 0.01448 | 0.49849 | 0.67498 | 1.24526 | 103 |
| GO_ORGANELLE_FISSION | 0.00009 | 0.00536 | 0.53843 | 0.61328 | 1.15428 | 340 |
| GO_ORGANIC_CYCLIC_COMPOUND_CATABOLIC_PROCESS | 0.00000 | 0.00000 | | 0.65219 | 1.22950 | 452 |
| GO_OXIDATIVE_PHOSPHORYLATION | 0.00148 | 0.03905 | 0.45506 | 0.66444 | 1.22384 | 95 |
| GO_PEPTIDASE_COMPLEX | 0.00027 | 0.01130 | 0.49849 | 0.69603 | 1.27652 | 75 |
| GO_POLYSOMAL_RIBOSOME | 0.00079 | 0.02473 | 0.47727 | 0.77714 | 1.38778 | 30 |

| Cluster 1 | | | | | | |
|---|---|---|---|---|---|---|
| GO_POLYSOME | 0.00089 | 0.02665 | 0.47727 | 0.70209 | 1.28211 | 64 |
| GO_POSITIVE_REGULATION_OF_ATPASE_ACTIVITY | 0.00150 | 0.03936 | 0.45506 | 0.73886 | 1.33469 | 40 |
| GO_POSITIVE_REGULATION_OF_CELLULAR_AMIDE_METABO | 0.00050 | 0.01756 | 0.47727 | 0.66284 | 1.22657 | 126 |
| GO_POSITIVE_REGULATION_OF_CELL_CYCLE | 0.00124 | 0.03422 | 0.45506 | 0.60949 | 1.14286 | 268 |
| GO_POSITIVE_REGULATION_OF_CELL_CYCLE_PROCESS | 0.00031 | 0.01248 | 0.49849 | 0.63374 | 1.18367 | 202 |
| GO_POSITIVE_REGULATION_OF_CELL_DEATH | 0.00052 | 0.01800 | 0.47727 | 0.59769 | 1.12658 | 419 |
| GO_POSITIVE_REGULATION_OF_CELL_POPULATION_PROLIF | 0.00012 | 0.00672 | 0.51885 | 0.59795 | 1.12792 | 479 |
| GO_POSITIVE_REGULATION_OF_CYSTEINE_TYPE_ENDOPEPT | 0.00018 | 0.00853 | 0.51885 | 0.69218 | 1.27173 | 85 |
| GO_POSITIVE_REGULATION_OF_DEFENSE_RESPONSE | 0.00003 | 0.00260 | 0.55733 | 0.65341 | 1.21676 | 180 |
| GO_POSITIVE_REGULATION_OF_IMMUNE_RESPONSE | 0.00022 | 0.01000 | 0.51885 | 0.61301 | 1.15427 | 323 |
| GO_POSITIVE_REGULATION_OF_INNATE_IMMUNE_RESPON | 0.00000 | 0.00056 | 0.61053 | 0.69460 | 1.28478 | 121 |
| GO_POSITIVE_REGULATION_OF_ORGANELLE_ORGANIZATIO | 0.00025 | 0.01100 | 0.49849 | 0.59672 | 1.12491 | 444 |
| GO_POSITIVE_REGULATION_OF_PEPTIDASE_ACTIVITY | 0.00135 | 0.03660 | 0.45506 | 0.65382 | 1.20743 | 113 |
| GO_POSITIVE_REGULATION_OF_RESPONSE_TO_BIOTIC_STI | 0.00011 | 0.00614 | 0.53843 | 0.66584 | 1.23467 | 140 |
| GO_POSITIVE_REGULATION_OF_RESPONSE_TO_EXTERNAL_ | 0.00008 | 0.00498 | 0.53843 | 0.63250 | 1.18439 | 240 |
| GO_POSITIVE_REGULATION_OF_RNA_SPLICING | 0.00172 | 0.04347 | 0.45506 | 0.76170 | 1.36021 | 30 |
| GO_POSITIVE_REGULATION_OF_TRANSLATION | 0.00017 | 0.00845 | 0.51885 | 0.67724 | 1.24985 | 107 |
| GO_POSITIVE_REGULATION_OF_VIRAL_PROCESS | 0.00090 | 0.02666 | 0.47727 | 0.68072 | 1.25165 | 89 |
| GO_POSTTRANSCRIPTIONAL_REGULATION_OF_GENE_EXPRI | 0.00000 | 0.00000 | | 0.67140 | 1.26669 | 484 |
| GO_PRECATALYTIC_SPLICEOSOME | 0.00065 | 0.02125 | 0.47727 | 0.72600 | 1.31747 | 49 |
| GO_PRERIBOSOME | 0.00033 | 0.01296 | 0.49849 | 0.70547 | 1.29153 | 70 |
| GO_PRE_MRNA_BINDING | 0.00009 | 0.00554 | 0.53843 | 0.80519 | 1.44396 | 32 |
| GO_PROTEASOMAL_PROTEIN_CATABOLIC_PROCESS | 0.00046 | 0.01686 | 0.49849 | 0.60337 | 1.13680 | 388 |
| GO_PROTEIN_CONTAINING_COMPLEX_DISASSEMBLY | 0.00082 | 0.02507 | 0.47727 | 0.61050 | 1.14467 | 263 |
| GO_PROTEIN_CONTAINING_COMPLEX_LOCALIZATION | 0.00137 | 0.03677 | 0.45506 | 0.61533 | 1.15114 | 224 |
| GO_PROTEIN_C_TERMINUS_BINDING | 0.00102 | 0.02947 | 0.45506 | 0.64578 | 1.19747 | 140 |
| GO_PROTEIN_DNA_COMPLEX_SUBUNIT_ORGANIZATION | 0.00046 | 0.01679 | 0.49849 | 0.63770 | 1.18712 | 173 |
| GO_PROTEIN_DOMAIN_SPECIFIC_BINDING | 0.00034 | 0.01360 | 0.49849 | 0.59436 | 1.12052 | 449 |
| GO_PROTEIN_FOLDING | 0.00014 | 0.00749 | 0.51885 | 0.64380 | 1.19873 | 175 |
| GO_PROTEIN_FOLDING_CHAPERONE | 0.00007 | 0.00454 | 0.53843 | 0.85413 | 1.50575 | 21 |
| GO_PROTEIN_LOCALIZATION_TO_ENDOPLASMIC_RETICULU | 0.00000 | 0.00001 | 0.71951 | 0.72328 | 1.33826 | 124 |
| GO_PROTEIN_LOCALIZATION_TO_MEMBRANE | 0.00000 | 0.00011 | 0.65944 | 0.62184 | 1.17225 | 453 |
| GO_PROTEIN_LOCALIZATION_TO_MITOCHONDRION | 0.00079 | 0.02473 | 0.47727 | 0.66280 | 1.22346 | 108 |
| GO_PROTEIN_LOCALIZATION_TO_NUCLEUS | 0.00000 | 0.00020 | 0.64355 | 0.66539 | 1.24279 | 201 |
| GO_PROTEIN_MODIFICATION_BY_SMALL_PROTEIN_REMOV | 0.00037 | 0.01431 | 0.49849 | 0.62801 | 1.17470 | 222 |
| GO_PROTEIN_REFOLDING | 0.00079 | 0.02473 | 0.47727 | 0.81791 | 1.44189 | 21 |
| GO_PROTEIN_STABILIZATION | 0.00183 | 0.04607 | 0.45506 | 0.63419 | 1.17555 | 136 |
| GO_PROTEIN_TARGETING | 0.00000 | 0.00000 | 0.80122 | 0.66024 | 1.24272 | 342 |
| GO_PROTEIN_TARGETING_TO_MEMBRANE | 0.00000 | 0.00001 | 0.73376 | 0.70109 | 1.30287 | 162 |
| GO_PURINE_NUCLEOSIDE_TRIPHOSPHATE_METABOLIC_PRO | 0.00035 | 0.01360 | 0.49849 | 0.74136 | 1.34428 | 48 |
| GO_REGULATION_OF_ANIMAL_ORGAN_MORPHOGENESIS | 0.00072 | 0.02320 | 0.47727 | 0.64300 | 1.19355 | 145 |
| GO_REGULATION_OF_APOPTOTIC_SIGNALING_PATHWAY | 0.00185 | 0.04615 | 0.45506 | 0.61151 | 1.14574 | 253 |
| GO_REGULATION_OF_ATP_METABOLIC_PROCESS | 0.00198 | 0.04839 | 0.43171 | 0.66898 | 1.22759 | 79 |
| GO_REGULATION_OF_CELLULAR_AMIDE_METABOLIC_PROC | 0.00000 | 0.00004 | 0.69013 | 0.64109 | 1.20690 | 334 |
| GO_REGULATION_OF_CELLULAR_AMINE_METABOLIC_PROC | 0.00018 | 0.00857 | 0.51885 | 0.72539 | 1.32418 | 62 |
| GO_REGULATION_OF_CELLULAR_AMINO_ACID_METABOLIC | 0.00013 | 0.00714 | 0.51885 | 0.74865 | 1.36192 | 52 |
| GO_REGULATION_OF_CELLULAR_KETONE_METABOLIC_PRO | 0.00022 | 0.01000 | 0.51885 | 0.67165 | 1.24131 | 116 |
| GO_REGULATION_OF_CELLULAR_RESPONSE_TO_STRESS | 0.00011 | 0.00606 | 0.53843 | 0.59607 | 1.12465 | 499 |
| GO_REGULATION_OF_CELL_CYCLE_PHASE_TRANSITION | 0.00027 | 0.01130 | 0.49849 | 0.60682 | 1.14249 | 357 |
| GO_REGULATION_OF_CHROMOSOME_ORGANIZATION | 0.00002 | 0.00213 | 0.57561 | 0.63597 | 1.19261 | 267 |
| GO_REGULATION_OF_CHROMOSOME_SEGREGATION | 0.00087 | 0.02649 | 0.47727 | 0.67154 | 1.23621 | 92 |
| GO_REGULATION_OF_CYSTEINE_TYPE_ENDOPEPTIDASE_AC | 0.00012 | 0.00640 | 0.53843 | 0.65202 | 1.20895 | 139 |
| GO_REGULATION_OF_DNA_TEMPLATED_TRANSCRIPTION_I | 0.00000 | 0.00051 | 0.61053 | 0.72465 | 1.33436 | 91 |
| GO_REGULATION_OF_FERTILIZATION | 0.00010 | 0.00584 | 0.53843 | 0.88921 | 1.54631 | 16 |
| GO_REGULATION_OF_GENE_EXPRESSION_EPIGENETIC | 0.00122 | 0.03387 | 0.45506 | 0.65393 | 1.20791 | 115 |
| GO_REGULATION_OF_HEMATOPOIETIC_STEM_CELL_DIFFER | 0.00113 | 0.03205 | 0.45506 | 0.70030 | 1.27839 | 62 |
| GO_REGULATION_OF_INNATE_IMMUNE_RESPONSE | 0.00001 | 0.00059 | 0.61053 | 0.66727 | 1.24146 | 170 |
| GO_REGULATION_OF_INTRINSIC_APOPTOTIC_SIGNALING_P | 0.00027 | 0.01130 | 0.49849 | 0.66337 | 1.22516 | 114 |
| GO_REGULATION_OF_MITOTIC_CELL_CYCLE | 0.00005 | 0.00356 | 0.55733 | 0.60081 | 1.13345 | 495 |
| GO_REGULATION_OF_MORPHOGENESIS_OF_AN_EPITHELIU | 0.00073 | 0.02320 | 0.47727 | 0.66051 | 1.21979 | 112 |
| GO_REGULATION_OF_MRNA_CATABOLIC_PROCESS | 0.00004 | 0.00304 | 0.55733 | 0.65851 | 1.22381 | 163 |
| GO_REGULATION_OF_MRNA_METABOLIC_PROCESS | 0.00000 | 0.00000 | | 0.68939 | 1.29305 | 273 |
| GO_REGULATION_OF_MRNA_PROCESSING | 0.00000 | 0.00002 | 0.70498 | 0.71499 | 1.32315 | 125 |
| GO_REGULATION_OF_MRNA_SPLICING_VIA_SPLICEOSOME | 0.00000 | 0.00007 | 0.67496 | 0.74232 | 1.36568 | 90 |
| GO_REGULATION_OF_MULTI_ORGANISM_PROCESS | 0.00043 | 0.01581 | 0.49849 | 0.75829 | 1.36980 | 40 |
| GO_REGULATION_OF_NUCLEAR_DIVISION | 0.00083 | 0.02523 | 0.47727 | 0.64896 | 1.20337 | 140 |
| GO_REGULATION_OF_REPRODUCTIVE_PROCESS | 0.00054 | 0.01843 | 0.47727 | 0.68840 | 1.26562 | 88 |
| GO_REGULATION_OF_RESPONSE_TO_BIOTIC_STIMULUS | 0.00039 | 0.01473 | 0.49849 | 0.62757 | 1.17387 | 222 |
| GO_REGULATION_OF_RNA_SPLICING | 0.00000 | 0.00000 | 0.74774 | 0.72559 | 1.34333 | 127 |
| GO_REGULATION_OF_SIGNAL_TRANSDUCTION_BY_P53_CL | 0.00006 | 0.00430 | 0.53843 | 0.65902 | 1.22192 | 139 |
| GO_REGULATION_OF_SMALL_MOLECULE_METABOLIC_PRO | 0.00006 | 0.00436 | 0.53843 | 0.62770 | 1.17728 | 271 |
| GO_REGULATION_OF_STEM_CELL_DIFFERENTIATION | 0.00053 | 0.01821 | 0.47727 | 0.66960 | 1.23322 | 96 |
| GO_REGULATION_OF_SYMBIOTIC_PROCESS | 0.00003 | 0.00246 | 0.57561 | 0.67075 | 1.24457 | 144 |

| Cluster 1 | | | | | | |
|---|---|---|---|---|---|---|
| GO_REGULATION_OF_TRANSCRIPTION_FROM_RNA_POLYM | 0.00015 | 0.00765 | 0.51885 | 0.72524 | 1.32431 | 63 |
| GO_REGULATION_OF_TRANSLATIONAL_INITIATION | 0.00054 | 0.01864 | 0.47727 | 0.70670 | 1.29250 | 67 |
| GO_REGULATION_OF_UBIQUITIN_PROTEIN_LIGASE_ACTIVIT | 0.00163 | 0.04149 | 0.45506 | 0.78643 | 1.39022 | 23 |
| GO_REGULATION_OF_VIRAL_TRANSCRIPTION | 0.00050 | 0.01756 | 0.47727 | 0.72863 | 1.32437 | 51 |
| GO_REPRODUCTIVE_SYSTEM_DEVELOPMENT | 0.00078 | 0.02473 | 0.47727 | 0.61442 | 1.15142 | 258 |
| GO_RESPONSE_TO_FIBROBLAST_GROWTH_FACTOR | 0.00020 | 0.00908 | 0.51885 | 0.70072 | 1.28712 | 84 |
| GO_RESPONSE_TO_HEAT | 0.00011 | 0.00614 | 0.53843 | 0.66650 | 1.23301 | 123 |
| GO_RESPONSE_TO_INTERLEUKIN_1 | 0.00028 | 0.01153 | 0.49849 | 0.66982 | 1.23793 | 116 |
| GO_RESPONSE_TO_OXYGEN_LEVELS | 0.00008 | 0.00498 | 0.53843 | 0.62874 | 1.17785 | 247 |
| GO_RESPONSE_TO_STARVATION | 0.00137 | 0.03677 | 0.45506 | 0.64170 | 1.18885 | 133 |
| GO_RESPONSE_TO_TEMPERATURE_STIMULUS | 0.00014 | 0.00749 | 0.51885 | 0.65642 | 1.21967 | 160 |
| GO_RESPONSE_TO_TUMOR_NECROSIS_FACTOR | 0.00081 | 0.02505 | 0.47727 | 0.64047 | 1.19002 | 160 |
| GO_RIBONUCLEOPROTEIN_COMPLEX_BINDING | 0.00003 | 0.00235 | 0.57561 | 0.68659 | 1.26950 | 117 |
| GO_RIBONUCLEOPROTEIN_COMPLEX_BIOGENESIS | 0.00000 | 0.00000 | | 0.70521 | 1.32871 | 384 |
| GO_RIBONUCLEOPROTEIN_COMPLEX_SUBUNIT_ORGANIZAT | 0.00000 | 0.00000 | 0.81404 | 0.71241 | 1.32648 | 175 |
| GO_RIBONUCLEOPROTEIN_GRANULE | 0.00009 | 0.00555 | 0.53843 | 0.64780 | 1.20670 | 181 |
| GO_RIBONUCLEOSIDE_TRIPHOSPHATE_METABOLIC_PROCES | 0.00048 | 0.01712 | 0.49849 | 0.73825 | 1.33552 | 44 |
| GO_RIBOSOMAL_LARGE_SUBUNIT_BIOGENESIS | 0.00000 | 0.00020 | 0.64355 | 0.75895 | 1.38848 | 68 |
| GO_RIBOSOMAL_SUBUNIT | 0.00000 | 0.00020 | 0.64355 | 0.67627 | 1.25914 | 174 |
| GO_RIBOSOME | 0.00000 | 0.00002 | 0.70498 | 0.67712 | 1.26585 | 211 |
| GO_RIBOSOME_ASSEMBLY | 0.00008 | 0.00517 | 0.53843 | 0.73373 | 1.33622 | 57 |
| GO_RIBOSOME_BIOGENESIS | 0.00000 | 0.00000 | | 0.70039 | 1.31341 | 267 |
| GO_RNA_3_END_PROCESSING | 0.00003 | 0.00245 | 0.57561 | 0.67813 | 1.25545 | 127 |
| GO_RNA_CATABOLIC_PROCESS | 0.00000 | 0.00000 | | 0.67758 | 1.27527 | 338 |
| GO_RNA_EXPORT_FROM_NUCLEUS | 0.00003 | 0.00245 | 0.57561 | 0.67831 | 1.25579 | 127 |
| GO_RNA_LOCALIZATION | 0.00000 | 0.00004 | 0.69013 | 0.67691 | 1.26416 | 203 |
| GO_RNA_POLYMERASE_BINDING | 0.00026 | 0.01111 | 0.49849 | 0.72171 | 1.31318 | 55 |
| GO_RNA_POLYMERASE_COMPLEX | 0.00097 | 0.02815 | 0.47727 | 0.67058 | 1.23444 | 92 |
| GO_RNA_POLYMERASE_II_HOLOENZYME | 0.00102 | 0.02947 | 0.45506 | 0.69755 | 1.27575 | 67 |
| GO_RNA_POLYMERASE_II_TRANSCRIPTION_REGULATOR_CO | 0.00119 | 0.03315 | 0.45506 | 0.65453 | 1.21081 | 119 |
| GO_RNA_SPLICING | 0.00000 | 0.00000 | | 0.70036 | 1.31957 | 384 |
| GO_RNA_SPLICING_VIA_TRANSESTERIFICATION_REACTIONS | 0.00000 | 0.00000 | | 0.70874 | 1.33336 | 309 |
| GO_RRNA_BINDING | 0.00000 | 0.00013 | 0.65944 | 0.78077 | 1.42065 | 55 |
| GO_RRNA_METABOLIC_PROCESS | 0.00000 | 0.00000 | 0.76146 | 0.69522 | 1.29818 | 199 |
| GO_SCF_DEPENDENT_PROTEASOMAL_UBIQUITIN_DEPENDE | 0.00024 | 0.01055 | 0.51885 | 0.70151 | 1.28668 | 76 |
| GO_SEX_DIFFERENTIATION | 0.00126 | 0.03442 | 0.45506 | 0.63469 | 1.17910 | 153 |
| GO_SIGNAL_TRANSDUCTION_BY_P53_CLASS_MEDIATOR | 0.00001 | 0.00095 | 0.59333 | 0.65573 | 1.22474 | 201 |
| GO_SINGLE_FERTILIZATION | 0.00090 | 0.02666 | 0.47727 | 0.68460 | 1.25565 | 76 |
| GO_SISTER_CHROMATID_SEGREGATION | 0.00002 | 0.00166 | 0.57561 | 0.66187 | 1.23007 | 165 |
| GO_SMALL_NUCLEAR_RIBONUCLEOPROTEIN_COMPLEX | 0.00052 | 0.01800 | 0.47727 | 0.71257 | 1.29767 | 57 |
| GO_SMALL_RIBOSOMAL_SUBUNIT | 0.00080 | 0.02489 | 0.47727 | 0.69323 | 1.26837 | 69 |
| GO_SMALL_SUBUNIT_PROCESSOME | 0.00205 | 0.04960 | 0.43171 | 0.74674 | 1.34273 | 33 |
| GO_SM_LIKE_PROTEIN_FAMILY_COMPLEX | 0.00056 | 0.01915 | 0.47727 | 0.70624 | 1.29166 | 67 |
| GO_SNORNA_BINDING | 0.00043 | 0.01581 | 0.49849 | 0.78721 | 1.40092 | 28 |
| GO_SPERM_EGG_RECOGNITION | 0.00152 | 0.03948 | 0.45506 | 0.75201 | 1.35221 | 33 |
| GO_SPLICEOSOMAL_COMPLEX | 0.00000 | 0.00000 | 0.81404 | 0.72415 | 1.34728 | 169 |
| GO_SPLICEOSOMAL_COMPLEX_ASSEMBLY | 0.00194 | 0.04806 | 0.45506 | 0.70984 | 1.28816 | 49 |
| GO_STEM_CELL_DIFFERENTIATION | 0.00141 | 0.03760 | 0.45506 | 0.62468 | 1.16347 | 177 |
| GO_STRUCTURAL_CONSTITUENT_OF_RIBOSOME | 0.00000 | 0.00018 | 0.64355 | 0.68838 | 1.27885 | 153 |
| GO_STRUCTURAL_MOLECULE_ACTIVITY | 0.00001 | 0.00132 | 0.59333 | 0.61669 | 1.16210 | 392 |
| GO_TELOMERE_MAINTENANCE_VIA_TELOMERE_LENGTHEN | 0.00017 | 0.00829 | 0.51885 | 0.70821 | 1.29578 | 69 |
| GO_TELOMERE_ORGANIZATION | 0.00015 | 0.00765 | 0.51885 | 0.66266 | 1.22631 | 125 |
| GO_TRANSCRIPTION_COREGULATOR_ACTIVITY | 0.00001 | 0.00073 | 0.61053 | 0.62021 | 1.16792 | 359 |
| GO_TRANSCRIPTION_COUPLED_NUCLEOTIDE_EXCISION_RE | 0.00119 | 0.03315 | 0.45506 | 0.69527 | 1.27159 | 67 |
| GO_TRANSCRIPTION_ELONGATION_FROM_RNA_POLYMERA | 0.00019 | 0.00897 | 0.51885 | 0.70827 | 1.29786 | 73 |
| GO_TRANSCRIPTION_FACTOR_BINDING | 0.00002 | 0.00231 | 0.57561 | 0.60830 | 1.14670 | 437 |
| GO_TRANSCRIPTION_INITIATION_FROM_RNA_POLYMERASE | 0.00113 | 0.03205 | 0.45506 | 0.64488 | 1.19581 | 140 |
| GO_TRANSCRIPTION_REGULATOR_COMPLEX | 0.00121 | 0.03351 | 0.45506 | 0.60624 | 1.13831 | 286 |
| GO_TRANSFERASE_COMPLEX_TRANSFERRING_PHOSPHORU | 0.00008 | 0.00527 | 0.53843 | 0.64204 | 1.19996 | 209 |
| GO_TRANSLATIONAL_INITIATION | 0.00000 | 0.00000 | 0.82666 | 0.72787 | 1.35419 | 169 |
| GO_TRANSLATION_FACTOR_ACTIVITY_RNA_BINDING | 0.00001 | 0.00060 | 0.61053 | 0.74124 | 1.35828 | 73 |
| GO_TRANSLATION_INITIATION_FACTOR_ACTIVITY | 0.00016 | 0.00809 | 0.51885 | 0.75121 | 1.36064 | 45 |
| GO_TRANSLATION_REGULATOR_ACTIVITY | 0.00000 | 0.00005 | 0.67496 | 0.72707 | 1.34224 | 110 |
| GO_TRANSLATION_REGULATOR_ACTIVITY_NUCLEIC_ACID_B | 0.00000 | 0.00040 | 0.62726 | 0.73202 | 1.34583 | 88 |
| GO_TUMOR_NECROSIS_FACTOR_MEDIATED_SIGNALING_PA | 0.00011 | 0.00621 | 0.53843 | 0.68063 | 1.25504 | 100 |
| GO_T_CELL_RECEPTOR_SIGNALING_PATHWAY | 0.00202 | 0.04910 | 0.43171 | 0.64586 | 1.19463 | 121 |
| GO_U12_TYPE_SPLICEOSOMAL_COMPLEX | 0.00154 | 0.03948 | 0.45506 | 0.78558 | 1.39655 | 26 |
| GO_U2_TYPE_CATALYTIC_STEP_2_SPLICEOSOME | 0.00146 | 0.03873 | 0.45506 | 0.76657 | 1.36891 | 30 |
| GO_U2_TYPE_SPLICEOSOMAL_COMPLEX | 0.00000 | 0.00018 | 0.64355 | 0.74295 | 1.36471 | 84 |
| GO_UBIQUITIN_LIKE_PROTEIN_LIGASE_BINDING | 0.00003 | 0.00272 | 0.55733 | 0.63557 | 1.18972 | 239 |
| GO_UNFOLDED_PROTEIN_BINDING | 0.00016 | 0.00813 | 0.51885 | 0.68287 | 1.25767 | 94 |
| GO_VESICLE_LUMEN | 0.00154 | 0.03948 | 0.45506 | 0.62392 | 1.16205 | 177 |
| GO_VIRAL_GENE_EXPRESSION | 0.00000 | 0.00000 | | 0.72308 | 1.34577 | 171 |

| Cluster 1 | | | | | | |
|---|---|---|---|---|---|---|
| GO_VIRAL_LIFE_CYCLE | 0.00029 | 0.01201 | 0.49849 | 0.63191 | 1.18155 | 217 |
| HP_INCREASED_LACTATE_DEHYDROGENASE_LEVEL | 0.00089 | 0.02665 | 0.47727 | 0.85731 | 1.48405 | 15 |
| HP_PTERYGIUM | 0.00126 | 0.03442 | 0.45506 | 0.82911 | 1.45094 | 18 |
| HP_SPARSE_OR_ABSENT_EYELASHES | 0.00111 | 0.03198 | 0.45506 | 0.73390 | 1.32641 | 42 |
| **Cluster 9** | | | | | | |
| pathway | pval | padj | log2err | ES | NES | size |
| GO_ACTIN_BINDING | 0.00335 | 0.03382 | 0.43171 | 0.56433 | 1.14057 | 248 |
| GO_ACTIN_CYTOSKELETON | 0.00024 | 0.00430 | 0.51885 | 0.57528 | 1.16778 | 307 |
| GO_ACTIN_FILAMENT_BINDING | 0.00335 | 0.03382 | 0.43171 | 0.60068 | 1.19484 | 127 |
| GO_ACTIVATION_OF_IMMUNE_RESPONSE | 0.00000 | 0.00015 | 0.61053 | 0.61479 | 1.24055 | 230 |
| GO_ACTIVATION_OF_INNATE_IMMUNE_RESPONSE | 0.00000 | 0.00004 | 0.65944 | 0.70474 | 1.39179 | 89 |
| GO_ACTIVE_ION_TRANSMEMBRANE_TRANSPORTER_ACTIV | 0.00242 | 0.02619 | 0.43171 | 0.61140 | 1.21385 | 109 |
| GO_ACTIVE_TRANSMEMBRANE_TRANSPORTER_ACTIVITY | 0.00033 | 0.00555 | 0.49849 | 0.60176 | 1.20698 | 175 |
| GO_ADP_BINDING | 0.00398 | 0.03856 | 0.40702 | 0.72320 | 1.35963 | 29 |
| GO_AEROBIC_RESPIRATION | 0.00001 | 0.00022 | 0.61053 | 0.71535 | 1.40030 | 67 |
| GO_ALCOHOL_BIOSYNTHETIC_PROCESS | 0.00000 | 0.00009 | 0.62726 | 0.68050 | 1.35038 | 103 |
| GO_ALCOHOL_METABOLIC_PROCESS | 0.00036 | 0.00599 | 0.49849 | 0.58967 | 1.18838 | 218 |
| GO_AMEBOIDAL_TYPE_CELL_MIGRATION | 0.00547 | 0.04780 | 0.40702 | 0.56225 | 1.13443 | 232 |
| GO_AMIDE_BINDING | 0.00001 | 0.00017 | 0.61053 | 0.62636 | 1.25714 | 181 |
| GO_AMINE_METABOLIC_PROCESS | 0.00000 | 0.00004 | 0.65944 | 0.70958 | 1.39925 | 86 |
| GO_AMINO_ACID_IMPORT | 0.00040 | 0.00647 | 0.49849 | 0.79795 | 1.47388 | 22 |
| GO_AMINO_ACID_TRANSMEMBRANE_TRANSPORT | 0.00019 | 0.00345 | 0.51885 | 0.71680 | 1.37900 | 44 |
| GO_AMINO_ACID_TRANSMEMBRANE_TRANSPORTER_ACTI | 0.00015 | 0.00286 | 0.51885 | 0.73455 | 1.40610 | 40 |
| GO_AMINO_ACID_TRANSPORT | 0.00004 | 0.00098 | 0.55733 | 0.68559 | 1.34468 | 72 |
| GO_AMYLOID_PRECURSOR_PROTEIN_METABOLIC_PROCESS | 0.00390 | 0.03790 | 0.43171 | 0.67071 | 1.29297 | 46 |
| GO_ANAPHASE_PROMOTING_COMPLEX_DEPENDENT_CATA | 0.00000 | 0.00000 | 0.76146 | 0.77088 | 1.51279 | 73 |
| GO_ANATOMICAL_STRUCTURE_HOMEOSTASIS | 0.00428 | 0.04059 | 0.40702 | 0.55921 | 1.13039 | 259 |
| GO_ANION_TRANSMEMBRANE_TRANSPORT | 0.00001 | 0.00032 | 0.59333 | 0.63616 | 1.27254 | 157 |
| GO_ANION_TRANSMEMBRANE_TRANSPORTER_ACTIVITY | 0.00002 | 0.00045 | 0.57561 | 0.62646 | 1.25680 | 174 |
| GO_ANION_TRANSPORT | 0.00000 | 0.00007 | 0.64355 | 0.60169 | 1.22216 | 319 |
| GO_ANTIGEN_BINDING | 0.00037 | 0.00603 | 0.49849 | 0.82000 | 1.49549 | 18 |
| GO_ANTIGEN_PROCESSING_AND_PRESENTATION | 0.00000 | 0.00001 | 0.70498 | 0.66682 | 1.33240 | 151 |
| GO_ANTIGEN_PROCESSING_AND_PRESENTATION_OF_EXOG | 0.00000 | 0.00000 | 0.74774 | 0.79287 | 1.54623 | 60 |
| GO_ANTIGEN_PROCESSING_AND_PRESENTATION_OF_PEPT | 0.00000 | 0.00001 | 0.69013 | 0.68158 | 1.35631 | 132 |
| GO_ANTIGEN_PROCESSING_AND_PRESENTATION_OF_PEPT | 0.00000 | 0.00000 | 0.76146 | 0.75776 | 1.48623 | 72 |
| GO_ANTIGEN_RECEPTOR_MEDIATED_SIGNALING_PATHWAY | 0.00004 | 0.00100 | 0.55733 | 0.63652 | 1.26708 | 134 |
| GO_ANTIOXIDANT_ACTIVITY | 0.00072 | 0.01027 | 0.47727 | 0.71039 | 1.35855 | 39 |
| GO_APICAL_PART_OF_CELL | 0.00000 | 0.00002 | 0.67496 | 0.62722 | 1.26453 | 221 |
| GO_APICAL_PLASMA_MEMBRANE | 0.00000 | 0.00006 | 0.64355 | 0.64119 | 1.28660 | 178 |
| GO_APOPTOTIC_SIGNALING_PATHWAY | 0.00001 | 0.00042 | 0.59333 | 0.58167 | 1.18397 | 373 |
| GO_ATPASE_ACTIVITY | 0.00561 | 0.04881 | 0.40702 | 0.55000 | 1.11637 | 305 |
| GO_ATP_BIOSYNTHETIC_PROCESS | 0.00020 | 0.00365 | 0.51885 | 0.81615 | 1.50404 | 21 |
| GO_ATP_METABOLIC_PROCESS | 0.00000 | 0.00000 | | 0.68083 | 1.37066 | 209 |
| GO_ATP_SYNTHESIS_COUPLED_ELECTRON_TRANSPORT | 0.00000 | 0.00000 | 0.73376 | 0.74259 | 1.46058 | 78 |
| GO_AZUROPHIL_GRANULE | 0.00000 | 0.00000 | 0.71951 | 0.71167 | 1.41236 | 104 |
| GO_AZUROPHIL_GRANULE_LUMEN | 0.00000 | 0.00008 | 0.62726 | 0.73677 | 1.43618 | 59 |
| GO_AZUROPHIL_GRANULE_MEMBRANE | 0.00041 | 0.00654 | 0.49849 | 0.71894 | 1.37491 | 39 |
| GO_BASAL_PART_OF_CELL | 0.00000 | 0.00000 | 0.74774 | 0.82770 | 1.58584 | 41 |
| GO_BASAL_PLASMA_MEMBRANE | 0.00000 | 0.00000 | 0.82666 | 0.88767 | 1.68564 | 34 |
| GO_BASEMENT_MEMBRANE | 0.00309 | 0.03190 | 0.43171 | 0.67250 | 1.29377 | 44 |
| GO_BASOLATERAL_PLASMA_MEMBRANE | 0.00000 | 0.00000 | 0.71951 | 0.68153 | 1.35624 | 133 |
| GO_BINDING_OF_SPERM_TO_ZONA_PELLUCIDA | 0.00000 | 0.00005 | 0.64355 | 0.88991 | 1.63832 | 20 |
| GO_BLOOD_MICROPARTICLE | 0.00000 | 0.00012 | 0.62726 | 0.80521 | 1.53470 | 36 |
| GO_BRUSH_BORDER | 0.00000 | 0.00015 | 0.61053 | 0.72002 | 1.40944 | 67 |
| GO_BRUSH_BORDER_MEMBRANE | 0.00041 | 0.00651 | 0.49849 | 0.74202 | 1.41217 | 35 |
| GO_CADHERIN_BINDING | 0.00000 | 0.00000 | | 0.66267 | 1.34024 | 265 |
| GO_CALCIUM_DEPENDENT_PROTEIN_BINDING | 0.00272 | 0.02857 | 0.43171 | 0.67450 | 1.29762 | 44 |
| GO_CALCIUM_ION_BINDING | 0.00014 | 0.00273 | 0.51885 | 0.58686 | 1.18781 | 273 |
| GO_CARBOHYDRATE_BIOSYNTHETIC_PROCESS | 0.00090 | 0.01227 | 0.47727 | 0.61684 | 1.22698 | 127 |
| GO_CARBOHYDRATE_CATABOLIC_PROCESS | 0.00115 | 0.01465 | 0.45506 | 0.61190 | 1.21795 | 129 |
| GO_CARBOHYDRATE_DERIVATIVE_BIOSYNTHETIC_PROCESS | 0.00004 | 0.00100 | 0.55733 | 0.56626 | 1.15296 | 416 |
| GO_CARBOHYDRATE_METABOLIC_PROCESS | 0.00026 | 0.00456 | 0.49849 | 0.56348 | 1.14714 | 377 |
| GO_CARBOHYDRATE_TRANSMEMBRANE_TRANSPORTER_A( | 0.00111 | 0.01440 | 0.45506 | 0.80956 | 1.46675 | 15 |
| GO_CATALYTIC_STEP_2_SPLICEOSOME | 0.00097 | 0.01283 | 0.47727 | 0.64542 | 1.27072 | 81 |
| GO_CATION_TRANSMEMBRANE_TRANSPORT | 0.00001 | 0.00027 | 0.59333 | 0.58127 | 1.18313 | 388 |
| GO_CATION_TRANSMEMBRANE_TRANSPORTER_ACTIVITY | 0.00000 | 0.00005 | 0.64355 | 0.61253 | 1.23797 | 257 |
| GO_CELLULAR_AMINO_ACID_METABOLIC_PROCESS | 0.00004 | 0.00100 | 0.55733 | 0.60502 | 1.21904 | 216 |
| GO_CELLULAR_COMPONENT_DISASSEMBLY | 0.00000 | 0.00002 | 0.67496 | 0.58905 | 1.19951 | 419 |
| GO_CELLULAR_HOMEOSTASIS | 0.00000 | 0.00002 | 0.67496 | 0.58119 | 1.18557 | 485 |
| GO_CELLULAR_ION_HOMEOSTASIS | 0.00000 | 0.00016 | 0.61053 | 0.59731 | 1.21249 | 309 |
| GO_CELLULAR_IRON_ION_HOMEOSTASIS | 0.00061 | 0.00907 | 0.47727 | 0.70008 | 1.35001 | 47 |
| GO_CELLULAR_KETONE_METABOLIC_PROCESS | 0.00000 | 0.00000 | 0.73376 | 0.67664 | 1.35153 | 149 |
| GO_CELLULAR_MODIFIED_AMINO_ACID_METABOLIC_PROC | 0.00085 | 0.01158 | 0.47727 | 0.63449 | 1.25919 | 104 |

| Cluster 1 | | | | | | |
|---|---|---|---|---|---|---|
| GO_CELLULAR_NITROGEN_COMPOUND_CATABOLIC_PROCE | 0.00000 | 0.00000 | | 0.66931 | 1.36465 | 442 |
| GO_CELLULAR_OXIDANT_DETOXIFICATION | 0.00191 | 0.02177 | 0.45506 | 0.67834 | 1.30808 | 47 |
| GO_CELLULAR_PROTEIN_COMPLEX_DISASSEMBLY | 0.00002 | 0.00050 | 0.57561 | 0.61642 | 1.23725 | 180 |
| GO_CELLULAR_RESPIRATION | 0.00000 | 0.00000 | 0.80122 | 0.69745 | 1.39133 | 142 |
| GO_CELLULAR_RESPONSE_TO_CALCIUM_ION | 0.00034 | 0.00566 | 0.49849 | 0.73524 | 1.40363 | 37 |
| GO_CELLULAR_RESPONSE_TO_HEAT | 0.00015 | 0.00286 | 0.51885 | 0.64967 | 1.28459 | 93 |
| GO_CELLULAR_RESPONSE_TO_ORGANIC_CYCLIC_COMPOUN | 0.00235 | 0.02558 | 0.43171 | 0.56117 | 1.13751 | 289 |
| GO_CELLULAR_RESPONSE_TO_OXYGEN_LEVELS | 0.00000 | 0.00000 | 0.77494 | 0.69270 | 1.38326 | 146 |
| GO_CELLULAR_RESPONSE_TO_TOPOLOGICALLY_INCORRECT | 0.00079 | 0.01105 | 0.47727 | 0.61523 | 1.22457 | 129 |
| GO_CELLULAR_RESPONSE_TO_TOXIC_SUBSTANCE | 0.00133 | 0.01647 | 0.45506 | 0.67111 | 1.30800 | 58 |
| GO_CELLULAR_TRANSITION_METAL_ION_HOMEOSTASIS | 0.00509 | 0.04561 | 0.40702 | 0.63279 | 1.23944 | 68 |
| GO_CELL_ACTIVATION_INVOLVED_IN_IMMUNE_RESPONSE | 0.00000 | 0.00000 | | 0.63729 | 1.29730 | 404 |
| GO_CELL_ADHESION_MEDIATOR_ACTIVITY | 0.00017 | 0.00318 | 0.51885 | 0.74421 | 1.42129 | 38 |
| GO_CELL_ADHESION_MOLECULE_BINDING | 0.00000 | 0.00000 | | 0.65668 | 1.33611 | 353 |
| GO_CELL_BODY | 0.00006 | 0.00129 | 0.55733 | 0.58875 | 1.19293 | 285 |
| GO_CELL_CELL_ADHESION | 0.00084 | 0.01148 | 0.47727 | 0.55947 | 1.13845 | 360 |
| GO_CELL_CELL_JUNCTION | 0.00022 | 0.00389 | 0.51885 | 0.57748 | 1.17023 | 286 |
| GO_CELL_CELL_RECOGNITION | 0.00000 | 0.00015 | 0.61053 | 0.80710 | 1.52979 | 33 |
| GO_CELL_COMMUNICATION_INVOLVED_IN_CARDIAC_CONI | 0.00372 | 0.03651 | 0.43171 | 0.71227 | 1.33360 | 27 |
| GO_CELL_CORTEX | 0.00000 | 0.00014 | 0.61053 | 0.62570 | 1.25702 | 189 |
| GO_CELL_KILLING | 0.00002 | 0.00046 | 0.57561 | 0.71788 | 1.39936 | 59 |
| GO_CELL_LEADING_EDGE | 0.00228 | 0.02505 | 0.43171 | 0.56289 | 1.13923 | 272 |
| GO_CELL_PROJECTION_MEMBRANE | 0.00002 | 0.00047 | 0.57561 | 0.61965 | 1.24458 | 188 |
| GO_CELL_RECOGNITION | 0.00002 | 0.00053 | 0.57561 | 0.71388 | 1.39372 | 61 |
| GO_CELL_REDOX_HOMEOSTASIS | 0.00002 | 0.00054 | 0.57561 | 0.78403 | 1.49433 | 36 |
| GO_CELL_SUBSTRATE_JUNCTION | 0.00000 | 0.00000 | | 0.66195 | 1.34493 | 325 |
| GO_CELL_SURFACE | 0.00000 | 0.00000 | | 0.66517 | 1.35049 | 311 |
| GO_CERAMIDE_METABOLIC_PROCESS | 0.00176 | 0.02036 | 0.45506 | 0.66714 | 1.29747 | 56 |
| GO_CHAPERONE_BINDING | 0.00011 | 0.00213 | 0.53843 | 0.67156 | 1.31922 | 75 |
| GO_CHAPERONE_COFACTOR_DEPENDENT_PROTEIN_REFOL | 0.00472 | 0.04318 | 0.40702 | 0.73051 | 1.35497 | 23 |
| GO_CHAPERONE_COMPLEX | 0.00009 | 0.00190 | 0.53843 | 0.82598 | 1.52216 | 21 |
| GO_CHAPERONE_MEDIATED_PROTEIN_FOLDING | 0.00137 | 0.01676 | 0.45506 | 0.68484 | 1.31751 | 44 |
| GO_CLUSTER_OF_ACTIN_BASED_CELL_PROJECTIONS | 0.00004 | 0.00087 | 0.55733 | 0.67714 | 1.33825 | 91 |
| GO_COATED_VESICLE | 0.00015 | 0.00289 | 0.51885 | 0.60515 | 1.21521 | 186 |
| GO_COATED_VESICLE_MEMBRANE | 0.00068 | 0.00986 | 0.47727 | 0.62469 | 1.24023 | 109 |
| GO_COLLAGEN_BINDING | 0.00119 | 0.01506 | 0.45506 | 0.74393 | 1.39862 | 29 |
| GO_COLLAGEN_CATABOLIC_PROCESS | 0.00126 | 0.01579 | 0.45506 | 0.80103 | 1.45924 | 17 |
| GO_COLLAGEN_CONTAINING_EXTRACELLULAR_MATRIX | 0.00000 | 0.00000 | 0.76146 | 0.68622 | 1.37039 | 148 |
| GO_COLLAGEN_METABOLIC_PROCESS | 0.00465 | 0.04283 | 0.40702 | 0.67279 | 1.28789 | 40 |
| GO_CONTRACTILE_FIBER | 0.00000 | 0.00017 | 0.61053 | 0.66013 | 1.31308 | 127 |
| GO_CORNIFICATION | 0.00010 | 0.00211 | 0.53843 | 0.79704 | 1.49012 | 26 |
| GO_CORTICAL_ACTIN_CYTOSKELETON | 0.00106 | 0.01383 | 0.45506 | 0.66534 | 1.29894 | 61 |
| GO_CORTICAL_CYTOSKELETON | 0.00008 | 0.00161 | 0.53843 | 0.67950 | 1.33482 | 76 |
| GO_COTRANSLATIONAL_PROTEIN_TARGETING_TO_MEMBR/ | 0.00000 | 0.00000 | | 0.82012 | 1.62210 | 95 |
| GO_CYTOCHROME_COMPLEX | 0.00035 | 0.00578 | 0.49849 | 0.80041 | 1.47843 | 22 |
| GO_CYTOKINE_MEDIATED_SIGNALING_PATHWAY | 0.00000 | 0.00008 | 0.64355 | 0.59233 | 1.20561 | 372 |
| GO_CYTOKINE_PRODUCTION | 0.00065 | 0.00940 | 0.47727 | 0.56109 | 1.14213 | 362 |
| GO_CYTOKINE_PRODUCTION_INVOLVED_IN_IMMUNE_RESI | 0.00198 | 0.02228 | 0.43171 | 0.67853 | 1.30538 | 44 |
| GO_CYTOPLASMIC_SIDE_OF_MEMBRANE | 0.00257 | 0.02760 | 0.43171 | 0.61790 | 1.22574 | 102 |
| GO_CYTOPLASMIC_TRANSLATION | 0.00000 | 0.00000 | 0.80122 | 0.75359 | 1.48741 | 88 |
| GO_CYTOPLASMIC_TRANSLATIONAL_INITIATION | 0.00539 | 0.04742 | 0.40702 | 0.70688 | 1.33342 | 31 |
| GO_CYTOSOLIC_LARGE_RIBOSOMAL_SUBUNIT | 0.00000 | 0.00000 | 0.82666 | 0.82990 | 1.60654 | 52 |
| GO_CYTOSOLIC_RIBOSOME | 0.00000 | 0.00000 | | 0.81815 | 1.61956 | 97 |
| GO_CYTOSOLIC_SMALL_RIBOSOMAL_SUBUNIT | 0.00000 | 0.00000 | 0.71951 | 0.82072 | 1.57574 | 42 |
| GO_DECIDUALIZATION | 0.00008 | 0.00159 | 0.53843 | 0.86405 | 1.56802 | 16 |
| GO_DEFENSE_RESPONSE_TO_OTHER_ORGANISM | 0.00004 | 0.00101 | 0.55733 | 0.56372 | 1.14993 | 472 |
| GO_DENDRITIC_CELL_MIGRATION | 0.00569 | 0.04917 | 0.40702 | 0.77699 | 1.41003 | 16 |
| GO_DESMOSOME | 0.00335 | 0.03382 | 0.43171 | 0.77559 | 1.41449 | 18 |
| GO_DETOXIFICATION | 0.00084 | 0.01148 | 0.47727 | 0.67581 | 1.31796 | 60 |
| GO_DE_NOVO_PROTEIN_FOLDING | 0.00145 | 0.01741 | 0.45506 | 0.73730 | 1.38807 | 30 |
| GO_DICARBOXYLIC_ACID_METABOLIC_PROCESS | 0.00547 | 0.04780 | 0.40702 | 0.64973 | 1.26362 | 56 |
| GO_DIENCEPHALON_DEVELOPMENT | 0.00457 | 0.04247 | 0.40702 | 0.75373 | 1.39221 | 22 |
| GO_DISORDERED_DOMAIN_SPECIFIC_BINDING | 0.00480 | 0.04360 | 0.40702 | 0.72646 | 1.36154 | 28 |
| GO_DISULFIDE_OXIDOREDUCTASE_ACTIVITY | 0.00174 | 0.02022 | 0.45506 | 0.74991 | 1.39864 | 25 |
| GO_DIVALENT_INORGANIC_CATION_HOMEOSTASIS | 0.00191 | 0.02177 | 0.45506 | 0.57621 | 1.15938 | 204 |
| GO_DIVALENT_INORGANIC_CATION_TRANSPORT | 0.00063 | 0.00927 | 0.47727 | 0.58460 | 1.17616 | 203 |
| GO_DNA_BIOSYNTHETIC_PROCESS | 0.00495 | 0.04468 | 0.40702 | 0.58334 | 1.16489 | 147 |
| GO_DOUBLE_STRANDED_RNA_BINDING | 0.00055 | 0.00831 | 0.47727 | 0.69469 | 1.34777 | 54 |
| GO_ELECTRON_TRANSFER_ACTIVITY | 0.00000 | 0.00011 | 0.62726 | 0.70980 | 1.39321 | 74 |
| GO_ELECTRON_TRANSPORT_CHAIN | 0.00000 | 0.00000 | 0.77494 | 0.70174 | 1.39676 | 129 |
| GO_EMBRYO_IMPLANTATION | 0.00019 | 0.00348 | 0.51885 | 0.77588 | 1.45867 | 29 |
| GO_ENDOPEPTIDASE_ACTIVITY | 0.00000 | 0.00016 | 0.61053 | 0.62474 | 1.25656 | 198 |
| GO_ENDOPEPTIDASE_COMPLEX | 0.00000 | 0.00000 | 0.74774 | 0.78051 | 1.52144 | 59 |

| Cluster 1 | | | | | | |
|---|---|---|---|---|---|---|
| GO_ENDOPEPTIDASE_REGULATOR_ACTIVITY | 0.00012 | 0.00243 | 0.53843 | 0.69689 | 1.36116 | 62 |
| GO_ENDOPLASMIC_RETICULUM_GOLGI_INTERMEDIATE_CO | 0.00002 | 0.00055 | 0.57561 | 0.67335 | 1.33291 | 97 |
| GO_ENDOPLASMIC_RETICULUM_GOLGI_INTERMEDIATE_CO | 0.00009 | 0.00186 | 0.53843 | 0.71737 | 1.38869 | 52 |
| GO_ENDOPLASMIC_RETICULUM_LUMEN | 0.00000 | 0.00000 | 0.76146 | 0.67909 | 1.35763 | 155 |
| GO_ENDOPLASMIC_RETICULUM_ORGANIZATION | 0.00265 | 0.02799 | 0.43171 | 0.66444 | 1.28624 | 52 |
| GO_ENDOPLASMIC_RETICULUM_TO_CYTOSOL_TRANSPORT | 0.00279 | 0.02919 | 0.43171 | 0.73449 | 1.36735 | 24 |
| GO_ENDOPLASMIC_RETICULUM_TO_GOLGI_VESICLE_MEDI | 0.00335 | 0.03382 | 0.43171 | 0.58199 | 1.16464 | 158 |
| GO_ENDOPLASMIC_RETICULUM_UNFOLDED_PROTEIN_RES | 0.00517 | 0.04607 | 0.40702 | 0.61224 | 1.21193 | 97 |
| GO_ENDOSOME_LUMEN | 0.00027 | 0.00460 | 0.49849 | 0.83152 | 1.51478 | 17 |
| GO_ENERGY_DERIVATION_BY_OXIDATION_OF_ORGANIC_CO | 0.00000 | 0.00001 | 0.70498 | 0.64338 | 1.29388 | 197 |
| GO_ENTRY_INTO_HOST | 0.00102 | 0.01347 | 0.45506 | 0.62894 | 1.24360 | 93 |
| GO_ENZYME_INHIBITOR_ACTIVITY | 0.00465 | 0.04283 | 0.40702 | 0.57542 | 1.15433 | 172 |
| GO_EPIDERMAL_CELL_DIFFERENTIATION | 0.00013 | 0.00263 | 0.51885 | 0.64273 | 1.27601 | 112 |
| GO_EPIDERMIS_DEVELOPMENT | 0.00174 | 0.02022 | 0.45506 | 0.58418 | 1.17189 | 172 |
| GO_EPITHELIAL_CELL_DIFFERENTIATION | 0.00003 | 0.00070 | 0.57561 | 0.58496 | 1.18830 | 321 |
| GO_EPITHELIAL_TO_MESENCHYMAL_TRANSITION | 0.00316 | 0.03238 | 0.43171 | 0.62422 | 1.22776 | 78 |
| GO_ESTABLISHMENT_OF_EPITHELIAL_CELL_POLARITY | 0.00569 | 0.04917 | 0.40702 | 0.71649 | 1.33385 | 24 |
| GO_ESTABLISHMENT_OF_PROTEIN_LOCALIZATION_TO_CHR | 0.00272 | 0.02857 | 0.43171 | 0.74330 | 1.37870 | 23 |
| GO_ESTABLISHMENT_OF_PROTEIN_LOCALIZATION_TO_END | 0.00000 | 0.00000 | | 0.80860 | 1.60544 | 106 |
| GO_ESTABLISHMENT_OF_PROTEIN_LOCALIZATION_TO_MEM | 0.00000 | 0.00000 | | 0.67938 | 1.37306 | 257 |
| GO_ESTABLISHMENT_OF_PROTEIN_LOCALIZATION_TO_ORG | 0.00000 | 0.00000 | | 0.64770 | 1.32066 | 447 |
| GO_ESTABLISHMENT_OF_PROTEIN_LOCALIZATION_TO_TELO | 0.00076 | 0.01062 | 0.47727 | 0.80680 | 1.47141 | 18 |
| GO_ESTABLISHMENT_OF_RNA_LOCALIZATION | 0.00435 | 0.04103 | 0.40702 | 0.57480 | 1.15253 | 169 |
| GO_ESTABLISHMENT_OF_TISSUE_POLARITY | 0.00000 | 0.00001 | 0.69013 | 0.70917 | 1.40241 | 94 |
| GO_EUKARYOTIC_TRANSLATION_INITIATION_FACTOR_3_CO | 0.00168 | 0.01981 | 0.45506 | 0.79881 | 1.44727 | 15 |
| GO_EXCRETION | 0.00176 | 0.02036 | 0.45506 | 0.77107 | 1.42423 | 22 |
| GO_EXOGENOUS_PROTEIN_BINDING | 0.00063 | 0.00927 | 0.47727 | 0.70942 | 1.36206 | 42 |
| GO_EXPORT_ACROSS_PLASMA_MEMBRANE | 0.00517 | 0.04607 | 0.40702 | 0.72934 | 1.35280 | 23 |
| GO_EXTERNAL_SIDE_OF_PLASMA_MEMBRANE | 0.00000 | 0.00006 | 0.64355 | 0.68095 | 1.35190 | 112 |
| GO_EXTRACELLULAR_MATRIX | 0.00000 | 0.00001 | 0.69013 | 0.64190 | 1.28957 | 189 |
| GO_EXTRACELLULAR_MATRIX_DISASSEMBLY | 0.00209 | 0.02331 | 0.43171 | 0.70587 | 1.34806 | 38 |
| GO_EXTRACELLULAR_STRUCTURE_ORGANIZATION | 0.00015 | 0.00292 | 0.51885 | 0.61681 | 1.23266 | 152 |
| GO_EXTRINSIC_COMPONENT_OF_PLASMA_MEMBRANE | 0.00265 | 0.02799 | 0.43171 | 0.63399 | 1.24907 | 82 |
| GO_FATTY_ACID_BIOSYNTHETIC_PROCESS | 0.00139 | 0.01691 | 0.45506 | 0.63837 | 1.25860 | 85 |
| GO_FATTY_ACID_DERIVATIVE_BIOSYNTHETIC_PROCESS | 0.00146 | 0.01752 | 0.45506 | 0.68412 | 1.31924 | 47 |
| GO_FATTY_ACID_DERIVATIVE_METABOLIC_PROCESS | 0.00202 | 0.02257 | 0.43171 | 0.64052 | 1.25627 | 72 |
| GO_FATTY_ACYL_COA_BIOSYNTHETIC_PROCESS | 0.00045 | 0.00703 | 0.49849 | 0.79592 | 1.47013 | 22 |
| GO_FATTY_ACYL_COA_METABOLIC_PROCESS | 0.00465 | 0.04283 | 0.40702 | 0.72719 | 1.36291 | 28 |
| GO_FC_EPSILON_RECEPTOR_SIGNALING_PATHWAY | 0.00000 | 0.00007 | 0.64355 | 0.70396 | 1.38792 | 85 |
| GO_FC_RECEPTOR_SIGNALING_PATHWAY | 0.00000 | 0.00007 | 0.64355 | 0.66619 | 1.32514 | 127 |
| GO_FERTILIZATION | 0.00000 | 0.00012 | 0.62726 | 0.72012 | 1.41051 | 68 |
| GO_FICOLIN_1_RICH_GRANULE | 0.00000 | 0.00000 | 0.76146 | 0.72181 | 1.42838 | 96 |
| GO_GABA_ERGIC_SYNAPSE | 0.00001 | 0.00019 | 0.61053 | 0.83846 | 1.56378 | 25 |
| GO_GENERATION_OF_PRECURSOR_METABOLITES_AND_EN | 0.00000 | 0.00000 | | 0.62920 | 1.28002 | 351 |
| GO_GLOMERULUS_DEVELOPMENT | 0.00353 | 0.03498 | 0.43171 | 0.72834 | 1.36168 | 26 |
| GO_GLUCOSE_CATABOLIC_PROCESS | 0.00004 | 0.00092 | 0.55733 | 0.82151 | 1.52376 | 23 |
| GO_GLUCOSE_METABOLIC_PROCESS | 0.00413 | 0.03955 | 0.40702 | 0.59745 | 1.18890 | 132 |
| GO_GLUTAMATERGIC_SYNAPSE | 0.00110 | 0.01422 | 0.45506 | 0.58819 | 1.17993 | 172 |
| GO_GLYCOLYTIC_PROCESS_THROUGH_FRUCTOSE_6_PHOSP | 0.00010 | 0.00197 | 0.53843 | 0.83395 | 1.53531 | 20 |
| GO_GLYCOSAMINOGLYCAN_BINDING | 0.00002 | 0.00061 | 0.57561 | 0.69169 | 1.35637 | 71 |
| GO_GLYCOSPHINGOLIPID_METABOLIC_PROCESS | 0.00084 | 0.01148 | 0.47727 | 0.70794 | 1.35387 | 39 |
| GO_GOLGI_ASSOCIATED_VESICLE | 0.00043 | 0.00673 | 0.49849 | 0.61899 | 1.23048 | 120 |
| GO_GUANYL_NUCLEOTIDE_BINDING | 0.00239 | 0.02589 | 0.43171 | 0.57073 | 1.15062 | 223 |
| GO_HEAT_SHOCK_PROTEIN_BINDING | 0.00067 | 0.00975 | 0.47727 | 0.64293 | 1.27064 | 91 |
| GO_HEMATOPOIETIC_PROGENITOR_CELL_DIFFERENTIATION | 0.00000 | 0.00002 | 0.67496 | 0.68094 | 1.35214 | 113 |
| GO_HEMATOPOIETIC_STEM_CELL_DIFFERENTIATION | 0.00000 | 0.00002 | 0.67496 | 0.74434 | 1.45704 | 67 |
| GO_HEPARIN_BINDING | 0.00002 | 0.00046 | 0.57561 | 0.73748 | 1.42989 | 53 |
| GO_HETEROTYPIC_CELL_CELL_ADHESION | 0.00261 | 0.02779 | 0.43171 | 0.73337 | 1.37108 | 26 |
| GO_HEXOSE_CATABOLIC_PROCESS | 0.00011 | 0.00224 | 0.53843 | 0.74862 | 1.42916 | 37 |
| GO_HUMORAL_IMMUNE_RESPONSE | 0.00324 | 0.03303 | 0.43171 | 0.64394 | 1.25973 | 64 |
| GO_HYDROGEN_PEROXIDE_METABOLIC_PROCESS | 0.00040 | 0.00647 | 0.49849 | 0.77109 | 1.44160 | 26 |
| GO_IMMUNE_RESPONSE_REGULATING_SIGNALING_PATHW | 0.00004 | 0.00085 | 0.55733 | 0.60255 | 1.21403 | 215 |
| GO_IMPORT_ACROSS_PLASMA_MEMBRANE | 0.00000 | 0.00005 | 0.64355 | 0.72294 | 1.41872 | 73 |
| GO_IMPORT_INTO_CELL | 0.00000 | 0.00000 | 0.73376 | 0.70576 | 1.40122 | 107 |
| GO_IMPORT_INTO_NUCLEUS | 0.00113 | 0.01448 | 0.45506 | 0.60577 | 1.20533 | 125 |
| GO_INNATE_IMMUNE_RESPONSE | 0.00004 | 0.00100 | 0.55733 | 0.57229 | 1.16488 | 382 |
| GO_INNATE_IMMUNE_RESPONSE_ACTIVATING_SIGNAL_TRA | 0.00000 | 0.00003 | 0.65944 | 0.72927 | 1.43114 | 73 |
| GO_INNER_EAR_RECEPTOR_CELL_STEREOCILIUM_ORGANIZ | 0.00539 | 0.04742 | 0.40702 | 0.77511 | 1.40433 | 15 |
| GO_INNER_MITOCHONDRIAL_MEMBRANE_PROTEIN_COMP | 0.00000 | 0.00004 | 0.65944 | 0.69928 | 1.38181 | 92 |
| GO_INORGANIC_ION_TRANSMEMBRANE_TRANSPORT | 0.00008 | 0.00177 | 0.53843 | 0.57444 | 1.16948 | 375 |
| GO_INTEGRIN_BINDING | 0.00261 | 0.02779 | 0.43171 | 0.64420 | 1.26102 | 67 |
| GO_INTERACTION_WITH_HOST | 0.00080 | 0.01108 | 0.47727 | 0.60069 | 1.19875 | 144 |
| GO_INTERACTION_WITH_SYMBIONT | 0.00126 | 0.01579 | 0.45506 | 0.69246 | 1.33088 | 43 |

| Cluster 1 | | | | | | |
|---|---|---|---|---|---|---|
| GO_INTERCALATED_DISC | 0.00154 | 0.01833 | 0.45506 | 0.73842 | 1.38826 | 29 |
| GO_INTERLEUKIN_1_MEDIATED_SIGNALING_PATHWAY | 0.00000 | 0.00001 | 0.69013 | 0.72831 | 1.43294 | 79 |
| GO_INTERMEDIATE_FILAMENT_CYTOSKELETON | 0.00268 | 0.02828 | 0.43171 | 0.65389 | 1.27463 | 59 |
| GO_INTRAMOLECULAR_OXIDOREDUCTASE_ACTIVITY | 0.00026 | 0.00450 | 0.49849 | 0.75054 | 1.42524 | 34 |
| GO_INTRINSIC_APOPTOTIC_SIGNALING_PATHWAY | 0.00039 | 0.00639 | 0.49849 | 0.59761 | 1.20194 | 200 |
| GO_INTRINSIC_APOPTOTIC_SIGNALING_PATHWAY_BY_P53_ | 0.00135 | 0.01663 | 0.45506 | 0.67720 | 1.31094 | 52 |
| GO_INTRINSIC_COMPONENT_OF_ENDOPLASMIC_RETICULU | 0.00000 | 0.00002 | 0.67496 | 0.68508 | 1.36164 | 119 |
| GO_INTRINSIC_COMPONENT_OF_ORGANELLE_MEMBRANE | 0.00004 | 0.00088 | 0.55733 | 0.59098 | 1.19744 | 285 |
| GO_ION_HOMEOSTASIS | 0.00000 | 0.00005 | 0.65944 | 0.58889 | 1.19867 | 381 |
| GO_ION_TRANSMEMBRANE_TRANSPORTER_ACTIVITY | 0.00000 | 0.00000 | 0.76146 | 0.60995 | 1.24174 | 378 |
| GO_IRON_ION_HOMEOSTASIS | 0.00235 | 0.02558 | 0.43171 | 0.66338 | 1.29017 | 56 |
| GO_IRON_ION_TRANSPORT | 0.00442 | 0.04154 | 0.40702 | 0.65491 | 1.27060 | 54 |
| GO_I_BAND | 0.00009 | 0.00186 | 0.53843 | 0.68515 | 1.34201 | 68 |
| GO_KERATINIZATION | 0.00015 | 0.00289 | 0.51885 | 0.76700 | 1.44682 | 31 |
| GO_KERATINOCYTE_DIFFERENTIATION | 0.00029 | 0.00501 | 0.49849 | 0.65920 | 1.29698 | 79 |
| GO_LARGE_RIBOSOMAL_SUBUNIT | 0.00000 | 0.00000 | | 0.76205 | 1.51296 | 107 |
| GO_LATERAL_PLASMA_MEMBRANE | 0.00023 | 0.00415 | 0.51885 | 0.71499 | 1.37876 | 47 |
| GO_LEUKOCYTE_MEDIATED_IMMUNITY | 0.00000 | 0.00000 | | 0.64651 | 1.31743 | 429 |
| GO_LEUKOCYTE_MIGRATION | 0.00000 | 0.00017 | 0.61053 | 0.62677 | 1.25960 | 192 |
| GO_LIVER_REGENERATION | 0.00145 | 0.01741 | 0.45506 | 0.77511 | 1.43170 | 22 |
| GO_LONG_CHAIN_FATTY_ACYL_COA_METABOLIC_PROCESS | 0.00350 | 0.03467 | 0.43171 | 0.78923 | 1.43225 | 16 |
| GO_LOW_DENSITY_LIPOPROTEIN_PARTICLE_CLEARANCE | 0.00045 | 0.00703 | 0.49849 | 0.77671 | 1.44596 | 24 |
| GO_LUMENAL_SIDE_OF_MEMBRANE | 0.00130 | 0.01609 | 0.45506 | 0.79948 | 1.45641 | 17 |
| GO_LYASE_ACTIVITY | 0.00290 | 0.03020 | 0.43171 | 0.60525 | 1.20245 | 115 |
| GO_LYSOSOMAL_LUMEN | 0.00000 | 0.00005 | 0.64355 | 0.75604 | 1.47441 | 60 |
| GO_L_ALPHA_AMINO_ACID_TRANSMEMBRANE_TRANSPOR | 0.00051 | 0.00772 | 0.47727 | 0.74888 | 1.41265 | 31 |
| GO_L_AMINO_ACID_TRANSMEMBRANE_TRANSPORTER_AC | 0.00018 | 0.00326 | 0.51885 | 0.76338 | 1.43999 | 31 |
| GO_L_AMINO_ACID_TRANSPORT | 0.00041 | 0.00651 | 0.49849 | 0.74228 | 1.41476 | 36 |
| GO_MATERNAL_PLACENTA_DEVELOPMENT | 0.00001 | 0.00034 | 0.59333 | 0.82843 | 1.54508 | 25 |
| GO_MATERNAL_PROCESS_INVOLVED_IN_FEMALE_PREGNAI | 0.00037 | 0.00603 | 0.49849 | 0.72065 | 1.37817 | 39 |
| GO_MATURATION_OF_SSU_RRNA_FROM_TRICISTRONIC_RF | 0.00235 | 0.02558 | 0.43171 | 0.70809 | 1.33997 | 32 |
| GO_MEMBRANE_LIPID_CATABOLIC_PROCESS | 0.00194 | 0.02199 | 0.45506 | 0.74580 | 1.38840 | 24 |
| GO_MEMBRANE_PROTEIN_PROTEOLYSIS | 0.00539 | 0.04742 | 0.40702 | 0.67659 | 1.29392 | 39 |
| GO_MEMBRANE_REGION | 0.00000 | 0.00002 | 0.67496 | 0.64641 | 1.29692 | 177 |
| GO_MESENCHYMAL_CELL_DIFFERENTIATION | 0.00047 | 0.00724 | 0.49849 | 0.62930 | 1.24941 | 107 |
| GO_METAL_ION_HOMEOSTASIS | 0.00001 | 0.00023 | 0.61053 | 0.59675 | 1.20943 | 287 |
| GO_METAL_ION_TRANSPORT | 0.00001 | 0.00037 | 0.59333 | 0.57837 | 1.17704 | 399 |
| GO_MITOCHONDRIAL_ELECTRON_TRANSPORT_NADH_TO_U | 0.00038 | 0.00620 | 0.49849 | 0.71067 | 1.36810 | 45 |
| GO_MITOCHONDRIAL_GENE_EXPRESSION | 0.00001 | 0.00018 | 0.61053 | 0.64190 | 1.28280 | 152 |
| GO_MITOCHONDRIAL_LARGE_RIBOSOMAL_SUBUNIT | 0.00018 | 0.00326 | 0.51885 | 0.71090 | 1.37921 | 54 |
| GO_MITOCHONDRIAL_MATRIX | 0.00000 | 0.00001 | 0.69013 | 0.60273 | 1.22700 | 369 |
| GO_MITOCHONDRIAL_PROTEIN_COMPLEX | 0.00000 | 0.00000 | | 0.68857 | 1.38699 | 211 |
| GO_MITOCHONDRIAL_RESPIRATORY_CHAIN_COMPLEX_ASS | 0.00202 | 0.02257 | 0.43171 | 0.63304 | 1.24811 | 85 |
| GO_MITOCHONDRIAL_TRANSLATION | 0.00000 | 0.00009 | 0.62726 | 0.66128 | 1.31623 | 129 |
| GO_MITOCHONDRIAL_TRANSLATIONAL_TERMINATION | 0.00000 | 0.00016 | 0.61053 | 0.69129 | 1.36446 | 88 |
| GO_MITOCHONDRION_ORGANIZATION | 0.00001 | 0.00035 | 0.59333 | 0.57134 | 1.16324 | 411 |
| GO_MITOTIC_CYTOKINESIS | 0.00350 | 0.03467 | 0.43171 | 0.64632 | 1.26181 | 61 |
| GO_MODIFIED_AMINO_ACID_BINDING | 0.00003 | 0.00066 | 0.57561 | 0.75938 | 1.45365 | 40 |
| GO_MODIFIED_AMINO_ACID_TRANSPORT | 0.00198 | 0.02228 | 0.43171 | 0.79021 | 1.43953 | 17 |
| GO_MODULATION_BY_HOST_OF_VIRAL_PROCESS | 0.00376 | 0.03675 | 0.43171 | 0.78782 | 1.42968 | 16 |
| GO_MONOAMINE_TRANSPORT | 0.00039 | 0.00626 | 0.49849 | 0.77850 | 1.44398 | 23 |
| GO_MONOCARBOXYLIC_ACID_BIOSYNTHETIC_PROCESS | 0.00065 | 0.00940 | 0.47727 | 0.61604 | 1.22502 | 121 |
| GO_MONOCARBOXYLIC_ACID_METABOLIC_PROCESS | 0.00002 | 0.00046 | 0.57561 | 0.58121 | 1.18326 | 368 |
| GO_MONOSACCHARIDE_BIOSYNTHETIC_PROCESS | 0.00006 | 0.00125 | 0.55733 | 0.70089 | 1.37025 | 63 |
| GO_MONOSACCHARIDE_CATABOLIC_PROCESS | 0.00015 | 0.00284 | 0.51885 | 0.72094 | 1.38697 | 44 |
| GO_MONOSACCHARIDE_METABOLIC_PROCESS | 0.00026 | 0.00450 | 0.49849 | 0.60161 | 1.20699 | 179 |
| GO_MONOVALENT_INORGANIC_CATION_TRANSMEMBRAN | 0.00007 | 0.00149 | 0.53843 | 0.62080 | 1.23889 | 144 |
| GO_MONOVALENT_INORGANIC_CATION_TRANSPORT | 0.00000 | 0.00015 | 0.61053 | 0.61627 | 1.24391 | 235 |
| GO_MORPHOGENESIS_OF_AN_EPITHELIUM | 0.00000 | 0.00010 | 0.62726 | 0.59457 | 1.20768 | 316 |
| GO_MORPHOGENESIS_OF_A_POLARIZED_EPITHELIUM | 0.00000 | 0.00000 | 0.81404 | 0.72957 | 1.44818 | 110 |
| GO_MOVEMENT_IN_HOST_ENVIRONMENT | 0.00145 | 0.01741 | 0.45506 | 0.60997 | 1.21121 | 113 |
| GO_MRNA_5_UTR_BINDING | 0.00111 | 0.01440 | 0.45506 | 0.75743 | 1.41006 | 24 |
| GO_MRNA_BINDING | 0.00000 | 0.00000 | 0.76146 | 0.64891 | 1.30822 | 223 |
| GO_MRNA_PROCESSING | 0.00000 | 0.00002 | 0.67496 | 0.58820 | 1.19794 | 423 |
| GO_MULTI_MULTICELLULAR_ORGANISM_PROCESS | 0.00000 | 0.00011 | 0.62726 | 0.67504 | 1.34005 | 111 |
| GO_MUSCLE_SYSTEM_PROCESS | 0.00054 | 0.00808 | 0.47727 | 0.59656 | 1.19781 | 185 |
| GO_MYD88_DEPENDENT_TOLL_LIKE_RECEPTOR_SIGNALINC | 0.00287 | 0.02986 | 0.43171 | 0.76640 | 1.40280 | 19 |
| GO_MYELOID_LEUKOCYTE_ACTIVATION | 0.00000 | 0.00000 | | 0.65532 | 1.33396 | 374 |
| GO_MYELOID_LEUKOCYTE_MEDIATED_IMMUNITY | 0.00000 | 0.00000 | | 0.66724 | 1.35695 | 339 |
| GO_NADH_DEHYDROGENASE_ACTIVITY | 0.00170 | 0.01995 | 0.45506 | 0.70641 | 1.34858 | 37 |
| GO_NADH_DEHYDROGENASE_COMPLEX | 0.00073 | 0.01035 | 0.47727 | 0.70571 | 1.35090 | 40 |
| GO_NADH_DEHYDROGENASE_COMPLEX_ASSEMBLY | 0.00346 | 0.03459 | 0.43171 | 0.65718 | 1.27810 | 56 |
| GO_NADH_METABOLIC_PROCESS | 0.00000 | 0.00006 | 0.64355 | 0.84348 | 1.57928 | 27 |

| Cluster 1 | | | | | | |
|---|---|---|---|---|---|---|
| GO_NAD_METABOLIC_PROCESS | 0.00002 | 0.00059 | 0.57561 | 0.81731 | 1.53181 | 28 |
| GO_NCRNA_METABOLIC_PROCESS | 0.00000 | 0.00004 | 0.65944 | 0.59075 | 1.20238 | 386 |
| GO_NCRNA_PROCESSING | 0.00000 | 0.00006 | 0.64355 | 0.59779 | 1.21552 | 338 |
| GO_NEGATIVE_REGULATION_OF_APOPTOTIC_SIGNALING_P | 0.00026 | 0.00456 | 0.49849 | 0.61240 | 1.22207 | 143 |
| GO_NEGATIVE_REGULATION_OF_CANONICAL_WNT_SIGNAI | 0.00002 | 0.00056 | 0.57561 | 0.64574 | 1.28365 | 120 |
| GO_NEGATIVE_REGULATION_OF_CATALYTIC_ACTIVITY | 0.00442 | 0.04154 | 0.40702 | 0.54261 | 1.10481 | 414 |
| GO_NEGATIVE_REGULATION_OF_CELLULAR_AMIDE_METAB | 0.00383 | 0.03730 | 0.43171 | 0.60094 | 1.19328 | 113 |
| GO_NEGATIVE_REGULATION_OF_CELLULAR_RESPONSE_TO_ | 0.00216 | 0.02404 | 0.43171 | 0.63921 | 1.25371 | 72 |
| GO_NEGATIVE_REGULATION_OF_CELLULAR_RESPONSE_TO_ | 0.00198 | 0.02228 | 0.43171 | 0.68352 | 1.31369 | 43 |
| GO_NEGATIVE_REGULATION_OF_CELL_CYCLE_G2_M_PHASI | 0.00001 | 0.00023 | 0.61053 | 0.69007 | 1.36014 | 84 |
| GO_NEGATIVE_REGULATION_OF_CELL_CYCLE_PHASE_TRAN | 0.00316 | 0.03238 | 0.43171 | 0.57936 | 1.16421 | 193 |
| GO_NEGATIVE_REGULATION_OF_CELL_CYCLE_PROCESS | 0.00115 | 0.01465 | 0.45506 | 0.57258 | 1.15734 | 255 |
| GO_NEGATIVE_REGULATION_OF_HYDROLASE_ACTIVITY | 0.00368 | 0.03626 | 0.43171 | 0.56936 | 1.14739 | 217 |
| GO_NEGATIVE_REGULATION_OF_INTRINSIC_APOPTOTIC_SI( | 0.00057 | 0.00856 | 0.47727 | 0.65635 | 1.28708 | 71 |
| GO_NEGATIVE_REGULATION_OF_NEURAL_PRECURSOR_CEL | 0.00139 | 0.01691 | 0.45506 | 0.80392 | 1.45653 | 15 |
| GO_NEGATIVE_REGULATION_OF_PEPTIDASE_ACTIVITY | 0.00007 | 0.00142 | 0.53843 | 0.66333 | 1.31448 | 99 |
| GO_NEGATIVE_REGULATION_OF_PROTEIN_CATABOLIC_PRO | 0.00502 | 0.04514 | 0.40702 | 0.61302 | 1.21065 | 89 |
| GO_NEGATIVE_REGULATION_OF_PROTEOLYSIS | 0.00002 | 0.00046 | 0.57561 | 0.62663 | 1.25521 | 163 |
| GO_NEGATIVE_REGULATION_OF_REACTIVE_OXYGEN_SPECI | 0.00279 | 0.02919 | 0.43171 | 0.79332 | 1.43967 | 16 |
| GO_NEGATIVE_REGULATION_OF_RNA_SPLICING | 0.00547 | 0.04780 | 0.40702 | 0.74886 | 1.38321 | 22 |
| GO_NEGATIVE_REGULATION_OF_TRANSMEMBRANE_RECEP | 0.00253 | 0.02725 | 0.43171 | 0.65451 | 1.27583 | 59 |
| GO_NEGATIVE_REGULATION_OF_WNT_SIGNALING_PATHW/ | 0.00030 | 0.00506 | 0.49849 | 0.61942 | 1.23303 | 134 |
| GO_NEPHRON_DEVELOPMENT | 0.00250 | 0.02690 | 0.43171 | 0.64679 | 1.26673 | 66 |
| GO_NEURAL_NUCLEUS_DEVELOPMENT | 0.00224 | 0.02473 | 0.43171 | 0.71800 | 1.36091 | 33 |
| GO_NEUTRAL_AMINO_ACID_TRANSMEMBRANE_TRANSPOI | 0.00342 | 0.03445 | 0.43171 | 0.73931 | 1.37130 | 23 |
| GO_NEUTRAL_AMINO_ACID_TRANSPORT | 0.00242 | 0.02619 | 0.43171 | 0.73492 | 1.37397 | 26 |
| GO_NIK_NF_KAPPAB_SIGNALING | 0.00000 | 0.00000 | 0.78819 | 0.72347 | 1.43608 | 110 |
| GO_NON_CANONICAL_WNT_SIGNALING_PATHWAY | 0.00000 | 0.00003 | 0.65944 | 0.68800 | 1.36599 | 106 |
| GO_NUCLEAR_ENVELOPE | 0.00005 | 0.00115 | 0.55733 | 0.57707 | 1.17330 | 336 |
| GO_NUCLEAR_EXPORT | 0.00228 | 0.02505 | 0.43171 | 0.58564 | 1.17468 | 173 |
| GO_NUCLEAR_TRANSCRIBED_MRNA_CATABOLIC_PROCESS | 0.00000 | 0.00000 | | 0.69750 | 1.40023 | 182 |
| GO_NUCLEAR_TRANSCRIBED_MRNA_CATABOLIC_PROCESS_ | 0.00000 | 0.00000 | | 0.79084 | 1.57037 | 113 |
| GO_NUCLEAR_TRANSPORT | 0.00020 | 0.00368 | 0.51885 | 0.58009 | 1.17506 | 282 |
| GO_NUCLEOBASE_CONTAINING_COMPOUND_TRANSPORT | 0.00157 | 0.01870 | 0.45506 | 0.57569 | 1.15932 | 210 |
| GO_NUCLEOBASE_CONTAINING_SMALL_MOLECULE_METAE | 0.00000 | 0.00003 | 0.65944 | 0.59179 | 1.20478 | 377 |
| GO_NUCLEOSIDE_BISPHOSPHATE_BIOSYNTHETIC_PROCESS | 0.00074 | 0.01043 | 0.47727 | 0.69470 | 1.33921 | 46 |
| GO_NUCLEOSIDE_DIPHOSPHATE_METABOLIC_PROCESS | 0.00167 | 0.01963 | 0.45506 | 0.62722 | 1.24476 | 104 |
| GO_NUCLEOSIDE_PHOSPHATE_BIOSYNTHETIC_PROCESS | 0.00001 | 0.00037 | 0.59333 | 0.63764 | 1.27411 | 151 |
| GO_NUCLEOSIDE_TRIPHOSPHATE_BIOSYNTHETIC_PROCESS | 0.00106 | 0.01383 | 0.45506 | 0.69992 | 1.34382 | 42 |
| GO_NUCLEOSIDE_TRIPHOSPHATE_METABOLIC_PROCESS | 0.00088 | 0.01205 | 0.47727 | 0.66728 | 1.30274 | 61 |
| GO_NUCLEOTIDE_PHOSPHORYLATION | 0.00113 | 0.01448 | 0.45506 | 0.63694 | 1.25881 | 91 |
| GO_ORGANELLAR_RIBOSOME | 0.00001 | 0.00018 | 0.61053 | 0.69312 | 1.36616 | 84 |
| GO_ORGANELLE_ENVELOPE_LUMEN | 0.00071 | 0.01016 | 0.47727 | 0.65300 | 1.28277 | 75 |
| GO_ORGANELLE_INNER_MEMBRANE | 0.00000 | 0.00000 | | 0.64087 | 1.30455 | 402 |
| GO_ORGANIC_ACID_BIOSYNTHETIC_PROCESS | 0.00006 | 0.00140 | 0.53843 | 0.60409 | 1.21440 | 194 |
| GO_ORGANIC_ACID_TRANSMEMBRANE_TRANSPORT | 0.00001 | 0.00027 | 0.59333 | 0.69340 | 1.36384 | 78 |
| GO_ORGANIC_ACID_TRANSMEMBRANE_TRANSPORTER_AC | 0.00002 | 0.00053 | 0.57561 | 0.69703 | 1.36787 | 73 |
| GO_ORGANIC_ACID_TRANSPORT | 0.00000 | 0.00006 | 0.64355 | 0.64844 | 1.29710 | 157 |
| GO_ORGANIC_ANION_TRANSMEMBRANE_TRANSPORTER_/ | 0.00000 | 0.00010 | 0.62726 | 0.66852 | 1.32815 | 115 |
| GO_ORGANIC_ANION_TRANSPORT | 0.00000 | 0.00005 | 0.64355 | 0.61424 | 1.24125 | 252 |
| GO_ORGANIC_CYCLIC_COMPOUND_CATABOLIC_PROCESS | 0.00000 | 0.00000 | | 0.66011 | 1.34687 | 466 |
| GO_ORGANIC_HYDROXY_COMPOUND_BIOSYNTHETIC_PRO | 0.00001 | 0.00018 | 0.61053 | 0.63857 | 1.27586 | 150 |
| GO_ORGANIC_HYDROXY_COMPOUND_METABOLIC_PROCES | 0.00092 | 0.01241 | 0.47727 | 0.56879 | 1.15352 | 296 |
| GO_ORGANOPHOSPHATE_BIOSYNTHETIC_PROCESS | 0.00081 | 0.01118 | 0.47727 | 0.55776 | 1.13548 | 367 |
| GO_OXIDATIVE_PHOSPHORYLATION | 0.00000 | 0.00000 | 0.74774 | 0.73048 | 1.44670 | 98 |
| GO_OXIDOREDUCTASE_ACTIVITY | 0.00000 | 0.00000 | | 0.62551 | 1.27513 | 435 |
| GO_OXIDOREDUCTASE_ACTIVITY_ACTING_ON_A_HEME_GR | 0.00009 | 0.00179 | 0.53843 | 0.84548 | 1.54195 | 18 |
| GO_OXIDOREDUCTASE_ACTIVITY_ACTING_ON_A_SULFUR_( | 0.00163 | 0.01928 | 0.45506 | 0.69174 | 1.32416 | 40 |
| GO_OXIDOREDUCTASE_ACTIVITY_ACTING_ON_CH_OH_GRC | 0.00128 | 0.01599 | 0.45506 | 0.65760 | 1.28790 | 66 |
| GO_OXIDOREDUCTASE_ACTIVITY_ACTING_ON_NAD_P_H | 0.00113 | 0.01448 | 0.45506 | 0.65963 | 1.29187 | 66 |
| GO_OXIDOREDUCTASE_ACTIVITY_ACTING_ON_NAD_P_H_Q | 0.00130 | 0.01609 | 0.45506 | 0.68673 | 1.32116 | 44 |
| GO_OXIDOREDUCTASE_ACTIVITY_ACTING_ON_PEROXIDE_A | 0.00016 | 0.00302 | 0.51885 | 0.77535 | 1.45171 | 27 |
| GO_OXIDOREDUCTASE_COMPLEX | 0.00002 | 0.00056 | 0.57561 | 0.68686 | 1.35139 | 79 |
| GO_PEPTIDASE_ACTIVITY | 0.00034 | 0.00572 | 0.49849 | 0.57433 | 1.16490 | 297 |
| GO_PEPTIDASE_COMPLEX | 0.00000 | 0.00007 | 0.64355 | 0.71784 | 1.41014 | 76 |
| GO_PEPTIDASE_REGULATOR_ACTIVITY | 0.00095 | 0.01264 | 0.47727 | 0.64444 | 1.26981 | 83 |
| GO_PEPTIDE_BINDING | 0.00019 | 0.00345 | 0.51885 | 0.62477 | 1.24480 | 136 |
| GO_PEPTIDE_SECRETION | 0.00135 | 0.01663 | 0.45506 | 0.56948 | 1.15084 | 250 |
| GO_PEPTIDYL_ASPARAGINE_MODIFICATION | 0.00428 | 0.04059 | 0.40702 | 0.72240 | 1.35813 | 29 |
| GO_PERINUCLEAR_REGION_OF_CYTOPLASM | 0.00040 | 0.00647 | 0.49849 | 0.55597 | 1.13393 | 452 |
| GO_PHOSPHATIDYLSERINE_BINDING | 0.00413 | 0.03955 | 0.40702 | 0.73525 | 1.36377 | 23 |
| GO_PHOSPHOLIPASE_C_ACTIVATING_G_PROTEIN_COUPLED | 0.00364 | 0.03596 | 0.43171 | 0.71268 | 1.33438 | 27 |
| GO_PIGMENT_GRANULE | 0.00000 | 0.00000 | | 0.79892 | 1.57515 | 85 |

| Cluster 1 | | | | | | |
|---|---|---|---|---|---|---|
| GO_PLACENTA_DEVELOPMENT | 0.00346 | 0.03459 | 0.43171 | 0.62005 | 1.22798 | 98 |
| GO_PLASMA_LIPOPROTEIN_PARTICLE_CLEARANCE | 0.00017 | 0.00325 | 0.51885 | 0.75493 | 1.43673 | 35 |
| GO_PLASMA_MEMBRANE_PROTEIN_COMPLEX | 0.00033 | 0.00560 | 0.49849 | 0.58978 | 1.18853 | 219 |
| GO_POLYMERIC_CYTOSKELETAL_FIBER | 0.00075 | 0.01051 | 0.47727 | 0.55775 | 1.13525 | 388 |
| GO_POLYSOMAL_RIBOSOME | 0.00001 | 0.00039 | 0.59333 | 0.81695 | 1.53590 | 29 |
| GO_POLYSOME | 0.00000 | 0.00000 | 0.77494 | 0.79088 | 1.54166 | 59 |
| GO_POSITIVE_REGULATION_OF_ATPASE_ACTIVITY | 0.00509 | 0.04561 | 0.40702 | 0.66868 | 1.28517 | 43 |
| GO_POSITIVE_REGULATION_OF_ATP_METABOLIC_PROCESS | 0.00113 | 0.01448 | 0.45506 | 0.74075 | 1.38694 | 27 |
| GO_POSITIVE_REGULATION_OF_CANONICAL_WNT_SIGNALI | 0.00004 | 0.00092 | 0.55733 | 0.66103 | 1.31185 | 104 |
| GO_POSITIVE_REGULATION_OF_CELLULAR_AMIDE_METABO | 0.00002 | 0.00056 | 0.57561 | 0.65020 | 1.29173 | 116 |
| GO_POSITIVE_REGULATION_OF_CELLULAR_PROTEIN_LOCAL | 0.00045 | 0.00703 | 0.49849 | 0.57884 | 1.16815 | 231 |
| GO_POSITIVE_REGULATION_OF_CELL_DEATH | 0.00346 | 0.03459 | 0.43171 | 0.54567 | 1.11102 | 409 |
| GO_POSITIVE_REGULATION_OF_CELL_POPULATION_PROLIF | 0.00009 | 0.00186 | 0.53843 | 0.56356 | 1.14899 | 438 |
| GO_POSITIVE_REGULATION_OF_CYTOKINE_PRODUCTION | 0.00216 | 0.02404 | 0.43171 | 0.57522 | 1.15739 | 204 |
| GO_POSITIVE_REGULATION_OF_CYTOSKELETON_ORGANIZA | 0.00092 | 0.01241 | 0.47727 | 0.60116 | 1.19965 | 143 |
| GO_POSITIVE_REGULATION_OF_DEFENSE_RESPONSE | 0.00000 | 0.00008 | 0.64355 | 0.63085 | 1.26736 | 189 |
| GO_POSITIVE_REGULATION_OF_DNA_BIOSYNTHETIC_PROC | 0.00095 | 0.01264 | 0.47727 | 0.68358 | 1.32846 | 55 |
| GO_POSITIVE_REGULATION_OF_ESTABLISHMENT_OF_PROT | 0.00004 | 0.00100 | 0.55733 | 0.59246 | 1.19623 | 237 |
| GO_POSITIVE_REGULATION_OF_IMMUNE_RESPONSE | 0.00001 | 0.00022 | 0.61053 | 0.59335 | 1.20484 | 314 |
| GO_POSITIVE_REGULATION_OF_IMMUNE_SYSTEM_PROCES | 0.00003 | 0.00078 | 0.55733 | 0.56669 | 1.15620 | 464 |
| GO_POSITIVE_REGULATION_OF_INFLAMMATORY_RESPONS | 0.00261 | 0.02779 | 0.43171 | 0.65220 | 1.26964 | 57 |
| GO_POSITIVE_REGULATION_OF_INNATE_IMMUNE_RESPON | 0.00002 | 0.00056 | 0.57561 | 0.64963 | 1.29229 | 122 |
| GO_POSITIVE_REGULATION_OF_ION_TRANSMEMBRANE_TI | 0.00428 | 0.04059 | 0.40702 | 0.63844 | 1.24897 | 64 |
| GO_POSITIVE_REGULATION_OF_ION_TRANSPORT | 0.00064 | 0.00931 | 0.47727 | 0.62515 | 1.24114 | 109 |
| GO_POSITIVE_REGULATION_OF_ORGANELLE_ORGANIZATIO | 0.00069 | 0.00995 | 0.47727 | 0.55562 | 1.13241 | 432 |
| GO_POSITIVE_REGULATION_OF_PEPTIDASE_ACTIVITY | 0.00472 | 0.04318 | 0.40702 | 0.60207 | 1.19649 | 118 |
| GO_POSITIVE_REGULATION_OF_PEPTIDE_SECRETION | 0.00554 | 0.04824 | 0.40702 | 0.61151 | 1.21179 | 99 |
| GO_POSITIVE_REGULATION_OF_PROTEIN_CONTAINING_CO | 0.00350 | 0.03467 | 0.43171 | 0.58213 | 1.16616 | 164 |
| GO_POSITIVE_REGULATION_OF_PROTEIN_LOCALIZATION_T | 0.00001 | 0.00038 | 0.59333 | 0.72016 | 1.40380 | 59 |
| GO_POSITIVE_REGULATION_OF_PROTEIN_MATURATION | 0.00016 | 0.00302 | 0.51885 | 0.84306 | 1.53581 | 17 |
| GO_POSITIVE_REGULATION_OF_PROTEOLYSIS | 0.00121 | 0.01523 | 0.45506 | 0.57660 | 1.16368 | 234 |
| GO_POSITIVE_REGULATION_OF_RESPONSE_TO_BIOTIC_STII | 0.00002 | 0.00057 | 0.57561 | 0.63614 | 1.26800 | 139 |
| GO_POSITIVE_REGULATION_OF_RESPONSE_TO_EXTERNAL_ | 0.00005 | 0.00103 | 0.55733 | 0.59430 | 1.20084 | 245 |
| GO_POSITIVE_REGULATION_OF_SUPRAMOLECULAR_FIBER_ | 0.00110 | 0.01422 | 0.45506 | 0.60950 | 1.21317 | 130 |
| GO_POSITIVE_REGULATION_OF_TELOMERASE_ACTIVITY | 0.00146 | 0.01752 | 0.45506 | 0.73991 | 1.39105 | 29 |
| GO_POSITIVE_REGULATION_OF_TELOMERASE_RNA_LOCALI | 0.00003 | 0.00078 | 0.55733 | 0.87702 | 1.58898 | 15 |
| GO_POSITIVE_REGULATION_OF_TELOMERE_MAINTENANCE | 0.00037 | 0.00607 | 0.49849 | 0.71349 | 1.37129 | 43 |
| GO_POSITIVE_REGULATION_OF_TELOMERE_MAINTENANCE | 0.00004 | 0.00095 | 0.55733 | 0.77832 | 1.47524 | 33 |
| GO_POSITIVE_REGULATION_OF_TRANSLATION | 0.00003 | 0.00079 | 0.55733 | 0.66783 | 1.32261 | 98 |
| GO_POSITIVE_REGULATION_OF_TRANSPORT | 0.00165 | 0.01946 | 0.45506 | 0.54596 | 1.11368 | 476 |
| GO_POSITIVE_REGULATION_OF_VIRAL_PROCESS | 0.00005 | 0.00118 | 0.55733 | 0.67395 | 1.32796 | 83 |
| GO_POSITIVE_REGULATION_OF_VIRAL_TRANSCRIPTION | 0.00194 | 0.02199 | 0.45506 | 0.71285 | 1.34897 | 32 |
| GO_POSITIVE_REGULATION_OF_WNT_SIGNALING_PATHWA | 0.00062 | 0.00918 | 0.47727 | 0.61537 | 1.22418 | 123 |
| GO_POSTSYNAPSE | 0.00045 | 0.00701 | 0.49849 | 0.56836 | 1.15365 | 305 |
| GO_POSTSYNAPSE_ORGANIZATION | 0.00472 | 0.04318 | 0.40702 | 0.62174 | 1.22583 | 85 |
| GO_POSTTRANSCRIPTIONAL_REGULATION_OF_GENE_EXPRI | 0.00000 | 0.00000 | | 0.64084 | 1.30735 | 470 |
| GO_POST_TRANSLATIONAL_PROTEIN_MODIFICATION | 0.00030 | 0.00506 | 0.49849 | 0.58015 | 1.17261 | 247 |
| GO_PRECATALYTIC_SPLICEOSOME | 0.00547 | 0.04780 | 0.40702 | 0.65463 | 1.26482 | 50 |
| GO_PRERIBOSOME | 0.00054 | 0.00819 | 0.47727 | 0.66701 | 1.30568 | 67 |
| GO_PRIMARY_ACTIVE_TRANSMEMBRANE_TRANSPORTER_A | 0.00472 | 0.04318 | 0.40702 | 0.63688 | 1.24511 | 63 |
| GO_PROTEASOMAL_PROTEIN_CATABOLIC_PROCESS | 0.00265 | 0.02799 | 0.43171 | 0.55469 | 1.12900 | 372 |
| GO_PROTEASOMAL_UBIQUITIN_INDEPENDENT_PROTEIN_C | 0.00003 | 0.00074 | 0.57561 | 0.87721 | 1.59191 | 16 |
| GO_PROTEASOME_ACCESSORY_COMPLEX | 0.00006 | 0.00136 | 0.53843 | 0.81121 | 1.51018 | 24 |
| GO_PROTEASOME_CORE_COMPLEX | 0.00001 | 0.00039 | 0.59333 | 0.88773 | 1.61100 | 16 |
| GO_PROTEIN_CONTAINING_COMPLEX_DISASSEMBLY | 0.00000 | 0.00015 | 0.61053 | 0.60738 | 1.22821 | 262 |
| GO_PROTEIN_CONTAINING_COMPLEX_LOCALIZATION | 0.00376 | 0.03675 | 0.43171 | 0.56915 | 1.14697 | 217 |
| GO_PROTEIN_EXIT_FROM_ENDOPLASMIC_RETICULUM | 0.00346 | 0.03459 | 0.43171 | 0.69541 | 1.32809 | 38 |
| GO_PROTEIN_FOLDING | 0.00000 | 0.00000 | 0.77494 | 0.67814 | 1.35886 | 166 |
| GO_PROTEIN_FOLDING_CHAPERONE | 0.00029 | 0.00495 | 0.49849 | 0.82278 | 1.50056 | 18 |
| GO_PROTEIN_IMPORT | 0.00020 | 0.00363 | 0.51885 | 0.61373 | 1.22587 | 149 |
| GO_PROTEIN_LIPID_COMPLEX_SUBUNIT_ORGANIZATION | 0.00324 | 0.03303 | 0.43171 | 0.73512 | 1.37777 | 28 |
| GO_PROTEIN_LOCALIZATION_TO_CHROMOSOME_TELOMER | 0.00110 | 0.01422 | 0.45506 | 0.75705 | 1.41887 | 28 |
| GO_PROTEIN_LOCALIZATION_TO_ENDOPLASMIC_RETICULU | 0.00000 | 0.00000 | | 0.79320 | 1.57778 | 127 |
| GO_PROTEIN_LOCALIZATION_TO_MEMBRANE | 0.00000 | 0.00000 | | 0.63630 | 1.29740 | 446 |
| GO_PROTEIN_LOCALIZATION_TO_NUCLEUS | 0.00002 | 0.00043 | 0.59333 | 0.61354 | 1.23424 | 199 |
| GO_PROTEIN_MATURATION | 0.00001 | 0.00018 | 0.61053 | 0.62998 | 1.26361 | 176 |
| GO_PROTEIN_MODIFICATION_BY_SMALL_PROTEIN_REMOV | 0.00442 | 0.04154 | 0.40702 | 0.56634 | 1.14128 | 219 |
| GO_PROTEIN_N_LINKED_GLYCOSYLATION | 0.00450 | 0.04191 | 0.40702 | 0.63739 | 1.24612 | 63 |
| GO_PROTEIN_PROCESSING | 0.00001 | 0.00019 | 0.61053 | 0.65296 | 1.29924 | 125 |
| GO_PROTEIN_STABILIZATION | 0.00121 | 0.01523 | 0.45506 | 0.60532 | 1.20443 | 125 |
| GO_PROTEIN_TARGETING | 0.00000 | 0.00000 | | 0.65512 | 1.33231 | 339 |
| GO_PROTEIN_TARGETING_TO_MEMBRANE | 0.00000 | 0.00000 | | 0.73619 | 1.47219 | 156 |
| GO_PROTEIN_TARGETING_TO_MITOCHONDRION | 0.00078 | 0.01095 | 0.47727 | 0.64750 | 1.27483 | 81 |

| Cluster 1 | | | | | | |
|---|---|---|---|---|---|---|
| GO_PROTON_TRANSMEMBRANE_TRANSPORT | 0.00012 | 0.00239 | 0.53843 | 0.66795 | 1.31716 | 86 |
| GO_PURINE_CONTAINING_COMPOUND_BIOSYNTHETIC_PR | 0.00000 | 0.00003 | 0.65944 | 0.68140 | 1.35423 | 117 |
| GO_PURINE_CONTAINING_COMPOUND_METABOLIC_PROC | 0.00000 | 0.00001 | 0.69013 | 0.61805 | 1.25147 | 277 |
| GO_PURINE_NUCLEOSIDE_TRIPHOSPHATE_METABOLIC_PR | 0.00021 | 0.00378 | 0.51885 | 0.72047 | 1.38696 | 45 |
| GO_PYRUVATE_METABOLIC_PROCESS | 0.00002 | 0.00051 | 0.57561 | 0.66506 | 1.31823 | 100 |
| GO_REACTIVE_OXYGEN_SPECIES_BIOSYNTHETIC_PROCESS | 0.00137 | 0.01676 | 0.45506 | 0.67683 | 1.31023 | 52 |
| GO_REACTIVE_OXYGEN_SPECIES_METABOLIC_PROCESS | 0.00524 | 0.04659 | 0.40702 | 0.59546 | 1.18522 | 129 |
| GO_RECEPTOR_MEDIATED_ENDOCYTOSIS | 0.00172 | 0.02013 | 0.45506 | 0.59853 | 1.19358 | 141 |
| GO_REGULATION_OF_ACTIN_FILAMENT_ORGANIZATION | 0.00357 | 0.03529 | 0.43171 | 0.58421 | 1.17064 | 166 |
| GO_REGULATION_OF_ALCOHOL_BIOSYNTHETIC_PROCESS | 0.00027 | 0.00460 | 0.49849 | 0.71235 | 1.37367 | 47 |
| GO_REGULATION_OF_ANIMAL_ORGAN_MORPHOGENESIS | 0.00000 | 0.00000 | 0.74774 | 0.68913 | 1.37291 | 137 |
| GO_REGULATION_OF_APOPTOTIC_SIGNALING_PATHWAY | 0.00047 | 0.00724 | 0.49849 | 0.58005 | 1.17217 | 252 |
| GO_REGULATION_OF_ATPASE_ACTIVITY | 0.00405 | 0.03915 | 0.40702 | 0.64485 | 1.25532 | 57 |
| GO_REGULATION_OF_ATP_METABOLIC_PROCESS | 0.00073 | 0.01035 | 0.47727 | 0.64752 | 1.27588 | 83 |
| GO_REGULATION_OF_BINDING | 0.00480 | 0.04360 | 0.40702 | 0.56905 | 1.14625 | 211 |
| GO_REGULATION_OF_BLOOD_CIRCULATION | 0.00509 | 0.04561 | 0.40702 | 0.59963 | 1.19037 | 111 |
| GO_REGULATION_OF_CELLULAR_AMIDE_METABOLIC_PROC | 0.00000 | 0.00001 | 0.70498 | 0.61124 | 1.24163 | 317 |
| GO_REGULATION_OF_CELLULAR_AMINE_METABOLIC_PROC | 0.00000 | 0.00001 | 0.70498 | 0.76813 | 1.49710 | 58 |
| GO_REGULATION_OF_CELLULAR_AMINO_ACID_METABOLIC | 0.00000 | 0.00000 | 0.74774 | 0.80036 | 1.55182 | 53 |
| GO_REGULATION_OF_CELLULAR_KETONE_METABOLIC_PRO | 0.00000 | 0.00000 | 0.78819 | 0.71885 | 1.42746 | 114 |
| GO_REGULATION_OF_CELLULAR_PROTEIN_CATABOLIC_PRO | 0.00428 | 0.04059 | 0.40702 | 0.56941 | 1.14353 | 187 |
| GO_REGULATION_OF_CELLULAR_PROTEIN_LOCALIZATION | 0.00014 | 0.00277 | 0.51885 | 0.56645 | 1.15296 | 383 |
| GO_REGULATION_OF_CELLULAR_RESPONSE_TO_HEAT | 0.00261 | 0.02779 | 0.43171 | 0.64774 | 1.26829 | 65 |
| GO_REGULATION_OF_CELLULAR_RESPONSE_TO_STRESS | 0.00080 | 0.01108 | 0.47727 | 0.54562 | 1.11302 | 486 |
| GO_REGULATION_OF_CELLULAR_RESPONSE_TO_TRANSFOR | 0.00051 | 0.00772 | 0.47727 | 0.67027 | 1.31124 | 64 |
| GO_REGULATION_OF_CELL_CYCLE_G2_M_PHASE_TRANSITI | 0.00080 | 0.01108 | 0.47727 | 0.59256 | 1.18855 | 176 |
| GO_REGULATION_OF_CHOLESTEROL_BIOSYNTHETIC_PROCE | 0.00095 | 0.01264 | 0.47727 | 0.71783 | 1.37091 | 38 |
| GO_REGULATION_OF_CHOLESTEROL_METABOLIC_PROCESS | 0.00043 | 0.00678 | 0.49849 | 0.70335 | 1.35589 | 46 |
| GO_REGULATION_OF_CYTOSKELETON_ORGANIZATION | 0.00372 | 0.03651 | 0.43171 | 0.54811 | 1.11460 | 344 |
| GO_REGULATION_OF_DEFENSE_RESPONSE | 0.00239 | 0.02589 | 0.43171 | 0.55445 | 1.12749 | 342 |
| GO_REGULATION_OF_DNA_BIOSYNTHETIC_PROCESS | 0.00279 | 0.02919 | 0.43171 | 0.62951 | 1.23891 | 80 |
| GO_REGULATION_OF_DNA_TEMPLATED_TRANSCRIPTION_I | 0.00000 | 0.00005 | 0.64355 | 0.70493 | 1.39137 | 88 |
| GO_REGULATION_OF_GENERATION_OF_PRECURSOR_META | 0.00569 | 0.04917 | 0.40702 | 0.60490 | 1.20081 | 105 |
| GO_REGULATION_OF_HEMATOPOIETIC_PROGENITOR_CELL_ | 0.00000 | 0.00011 | 0.62726 | 0.72344 | 1.41676 | 69 |
| GO_REGULATION_OF_HEMATOPOIETIC_STEM_CELL_DIFFER | 0.00000 | 0.00001 | 0.67496 | 0.76762 | 1.49700 | 60 |
| GO_REGULATION_OF_IMMUNE_RESPONSE | 0.00003 | 0.00075 | 0.55733 | 0.56944 | 1.15946 | 415 |
| GO_REGULATION_OF_INNATE_IMMUNE_RESPONSE | 0.00041 | 0.00654 | 0.49849 | 0.59971 | 1.20287 | 175 |
| GO_REGULATION_OF_INTRINSIC_APOPTOTIC_SIGNALING_P | 0.00004 | 0.00084 | 0.55733 | 0.65241 | 1.29514 | 111 |
| GO_REGULATION_OF_ION_TRANSPORT | 0.00146 | 0.01752 | 0.45506 | 0.56807 | 1.14852 | 261 |
| GO_REGULATION_OF_LIPASE_ACTIVITY | 0.00117 | 0.01486 | 0.45506 | 0.68860 | 1.31935 | 41 |
| GO_REGULATION_OF_LIPID_BIOSYNTHETIC_PROCESS | 0.00010 | 0.00201 | 0.53843 | 0.64381 | 1.27962 | 119 |
| GO_REGULATION_OF_LIPID_METABOLIC_PROCESS | 0.00413 | 0.03955 | 0.40702 | 0.56598 | 1.14258 | 236 |
| GO_REGULATION_OF_MORPHOGENESIS_OF_AN_EPITHELIU | 0.00000 | 0.00001 | 0.70498 | 0.68948 | 1.36909 | 113 |
| GO_REGULATION_OF_MRNA_CATABOLIC_PROCESS | 0.00000 | 0.00006 | 0.64355 | 0.65120 | 1.30138 | 152 |
| GO_REGULATION_OF_MRNA_METABOLIC_PROCESS | 0.00000 | 0.00000 | 0.76146 | 0.63624 | 1.28601 | 255 |
| GO_REGULATION_OF_MRNA_PROCESSING | 0.00152 | 0.01815 | 0.45506 | 0.61501 | 1.22229 | 117 |
| GO_REGULATION_OF_MRNA_SPLICING_VIA_SPLICEOSOME | 0.00435 | 0.04103 | 0.40702 | 0.61914 | 1.22034 | 84 |
| GO_REGULATION_OF_MULTI_ORGANISM_PROCESS | 0.00001 | 0.00018 | 0.61053 | 0.79989 | 1.52457 | 36 |
| GO_REGULATION_OF_MUSCLE_CONTRACTION | 0.00020 | 0.00360 | 0.51885 | 0.67792 | 1.32813 | 70 |
| GO_REGULATION_OF_MUSCLE_SYSTEM_PROCESS | 0.00080 | 0.01108 | 0.47727 | 0.63615 | 1.25988 | 98 |
| GO_REGULATION_OF_NEURAL_PRECURSOR_CELL_PROLIFEF | 0.00524 | 0.04659 | 0.40702 | 0.67083 | 1.28413 | 40 |
| GO_REGULATION_OF_OXIDATIVE_PHOSPHORYLATION | 0.00023 | 0.00415 | 0.51885 | 0.81824 | 1.50638 | 20 |
| GO_REGULATION_OF_PEPTIDASE_ACTIVITY | 0.00002 | 0.00056 | 0.57561 | 0.60619 | 1.22161 | 217 |
| GO_REGULATION_OF_PEPTIDE_SECRETION | 0.00502 | 0.04514 | 0.40702 | 0.57536 | 1.15628 | 192 |
| GO_REGULATION_OF_PEPTIDE_TRANSPORT | 0.00005 | 0.00106 | 0.55733 | 0.56925 | 1.15882 | 393 |
| GO_REGULATION_OF_PLASMA_LIPOPROTEIN_PARTICLE_LEV | 0.00000 | 0.00006 | 0.64355 | 0.76076 | 1.47503 | 53 |
| GO_REGULATION_OF_PROTEIN_CATABOLIC_PROCESS | 0.00046 | 0.00706 | 0.49849 | 0.57268 | 1.15915 | 274 |
| GO_REGULATION_OF_PROTEIN_CONTAINING_COMPLEX_AS | 0.00191 | 0.02177 | 0.45506 | 0.56264 | 1.14016 | 286 |
| GO_REGULATION_OF_PROTEIN_LOCALIZATION_TO_NUCLEU | 0.00004 | 0.00097 | 0.55733 | 0.66664 | 1.31655 | 89 |
| GO_REGULATION_OF_PROTEIN_MATURATION | 0.00074 | 0.01043 | 0.47727 | 0.72286 | 1.38052 | 38 |
| GO_REGULATION_OF_PROTEIN_STABILITY | 0.00098 | 0.01301 | 0.45506 | 0.57841 | 1.16431 | 208 |
| GO_REGULATION_OF_PROTEOLYSIS | 0.00001 | 0.00025 | 0.59333 | 0.57870 | 1.17813 | 413 |
| GO_REGULATION_OF_REACTIVE_OXYGEN_SPECIES_BIOSYNT | 0.00075 | 0.01051 | 0.47727 | 0.70602 | 1.35553 | 42 |
| GO_REGULATION_OF_RESPONSE_TO_BIOTIC_STIMULUS | 0.00025 | 0.00437 | 0.49849 | 0.58580 | 1.18063 | 220 |
| GO_REGULATION_OF_RESPONSE_TO_EXTERNAL_STIMULUS | 0.00495 | 0.04468 | 0.40702 | 0.53921 | 1.10017 | 493 |
| GO_REGULATION_OF_RESPONSE_TO_OXIDATIVE_STRESS | 0.00309 | 0.03190 | 0.43171 | 0.66476 | 1.28538 | 51 |
| GO_REGULATION_OF_RNA_SPLICING | 0.00064 | 0.00931 | 0.47727 | 0.61601 | 1.22542 | 122 |
| GO_REGULATION_OF_SMALL_MOLECULE_METABOLIC_PRO | 0.00000 | 0.00000 | 0.81404 | 0.64003 | 1.29588 | 276 |
| GO_REGULATION_OF_SMOOTH_MUSCLE_CONTRACTION | 0.00472 | 0.04318 | 0.40702 | 0.75455 | 1.38112 | 19 |
| GO_REGULATION_OF_STEM_CELL_DIFFERENTIATION | 0.00000 | 0.00000 | 0.73376 | 0.74068 | 1.45927 | 82 |
| GO_REGULATION_OF_STEROID_BIOSYNTHETIC_PROCESS | 0.00010 | 0.00208 | 0.53843 | 0.70275 | 1.37049 | 60 |
| GO_REGULATION_OF_STEROID_METABOLIC_PROCESS | 0.00006 | 0.00126 | 0.55733 | 0.68516 | 1.34457 | 73 |
| GO_REGULATION_OF_SUPRAMOLECULAR_FIBER_ORGANIZA | 0.00033 | 0.00549 | 0.49849 | 0.59057 | 1.19091 | 224 |
| GO_REGULATION_OF_SYMBIOTIC_PROCESS | 0.00002 | 0.00059 | 0.57561 | 0.64038 | 1.27590 | 136 |

| Cluster 1 | | | | | | |
|---|---|---|---|---|---|---|
| GO_REGULATION_OF_SYSTEM_PROCESS | 0.00465 | 0.04283 | 0.40702 | 0.56686 | 1.14354 | 228 |
| GO_REGULATION_OF_TELOMERE_MAINTENANCE | 0.00302 | 0.03124 | 0.43171 | 0.64284 | 1.25836 | 67 |
| GO_REGULATION_OF_TELOMERE_MAINTENANCE_VIA_TELO | 0.00022 | 0.00396 | 0.51885 | 0.70252 | 1.35839 | 51 |
| GO_REGULATION_OF_TRANSCRIPTION_FROM_RNA_POLYM | 0.00000 | 0.00001 | 0.69013 | 0.75933 | 1.48312 | 62 |
| GO_REGULATION_OF_TRANSLATIONAL_INITIATION | 0.00405 | 0.03915 | 0.40702 | 0.64248 | 1.25800 | 65 |
| GO_REGULATION_OF_VIRAL_TRANSCRIPTION | 0.00224 | 0.02473 | 0.43171 | 0.67649 | 1.30453 | 47 |
| GO_RENAL_SYSTEM_PROCESS | 0.00170 | 0.01995 | 0.45506 | 0.66768 | 1.29852 | 56 |
| GO_REPRODUCTIVE_SYSTEM_DEVELOPMENT | 0.00265 | 0.02799 | 0.43171 | 0.57021 | 1.15144 | 239 |
| GO_RESPIRASOME | 0.00000 | 0.00001 | 0.70498 | 0.73871 | 1.44886 | 72 |
| GO_RESPIRATORY_CHAIN_COMPLEX | 0.00000 | 0.00013 | 0.62726 | 0.72822 | 1.42368 | 63 |
| GO_RESPIRATORY_ELECTRON_TRANSPORT_CHAIN | 0.00000 | 0.00000 | 0.74774 | 0.73624 | 1.45453 | 90 |
| GO_RESPONSE_TO_ALCOHOL | 0.00178 | 0.02053 | 0.45506 | 0.60726 | 1.20584 | 113 |
| GO_RESPONSE_TO_ATP | 0.00032 | 0.00535 | 0.49849 | 0.82721 | 1.50695 | 17 |
| GO_RESPONSE_TO_CALCIUM_ION | 0.00009 | 0.00180 | 0.53843 | 0.67581 | 1.32549 | 72 |
| GO_RESPONSE_TO_DRUG | 0.00002 | 0.00057 | 0.57561 | 0.60997 | 1.22800 | 209 |
| GO_RESPONSE_TO_ENDOPLASMIC_RETICULUM_STRESS | 0.00001 | 0.00034 | 0.59333 | 0.61394 | 1.23729 | 218 |
| GO_RESPONSE_TO_HEAT | 0.00014 | 0.00270 | 0.51885 | 0.63526 | 1.26207 | 115 |
| GO_RESPONSE_TO_HORMONE | 0.00001 | 0.00020 | 0.61053 | 0.56985 | 1.16244 | 485 |
| GO_RESPONSE_TO_INORGANIC_SUBSTANCE | 0.00001 | 0.00027 | 0.59333 | 0.59092 | 1.19899 | 299 |
| GO_RESPONSE_TO_INTERLEUKIN_1 | 0.00000 | 0.00000 | 0.73376 | 0.70268 | 1.39599 | 116 |
| GO_RESPONSE_TO_INTERLEUKIN_12 | 0.00002 | 0.00059 | 0.57561 | 0.78071 | 1.48252 | 34 |
| GO_RESPONSE_TO_INTERLEUKIN_4 | 0.00040 | 0.00645 | 0.49849 | 0.80500 | 1.47346 | 19 |
| GO_RESPONSE_TO_METAL_ION | 0.00006 | 0.00129 | 0.55733 | 0.60495 | 1.21489 | 187 |
| GO_RESPONSE_TO_NUTRIENT | 0.00122 | 0.01540 | 0.45506 | 0.63059 | 1.24885 | 98 |
| GO_RESPONSE_TO_ORGANIC_CYCLIC_COMPOUND | 0.00001 | 0.00024 | 0.59333 | 0.57219 | 1.16725 | 478 |
| GO_RESPONSE_TO_OXIDATIVE_STRESS | 0.00235 | 0.02558 | 0.43171 | 0.56247 | 1.13909 | 279 |
| GO_RESPONSE_TO_OXYGEN_LEVELS | 0.00000 | 0.00000 | | 0.66619 | 1.34524 | 239 |
| GO_RESPONSE_TO_TEMPERATURE_STIMULUS | 0.00029 | 0.00495 | 0.49849 | 0.61500 | 1.22632 | 140 |
| GO_RESPONSE_TO_TOPOLOGICALLY_INCORRECT_PROTEIN | 0.00002 | 0.00055 | 0.57561 | 0.63101 | 1.26223 | 157 |
| GO_RESPONSE_TO_TOXIC_SUBSTANCE | 0.00235 | 0.02558 | 0.43171 | 0.60212 | 1.19778 | 122 |
| GO_RESPONSE_TO_TUMOR_NECROSIS_FACTOR | 0.00000 | 0.00000 | 0.71951 | 0.65965 | 1.32267 | 169 |
| GO_RETINA_HOMEOSTASIS | 0.00001 | 0.00041 | 0.59333 | 0.80329 | 1.51529 | 31 |
| GO_RETROGRADE_VESICLE_MEDIATED_TRANSPORT_GOLGI | 0.00450 | 0.04191 | 0.40702 | 0.62890 | 1.23442 | 74 |
| GO_RIBONUCLEOPROTEIN_COMPLEX_BINDING | 0.00001 | 0.00027 | 0.59333 | 0.66536 | 1.32073 | 110 |
| GO_RIBONUCLEOPROTEIN_COMPLEX_BIOGENESIS | 0.00000 | 0.00000 | | 0.66602 | 1.35562 | 388 |
| GO_RIBONUCLEOPROTEIN_COMPLEX_SUBUNIT_ORGANIZAT | 0.00000 | 0.00000 | | 0.69387 | 1.39173 | 175 |
| GO_RIBONUCLEOPROTEIN_GRANULE | 0.00006 | 0.00128 | 0.55733 | 0.61609 | 1.23574 | 176 |
| GO_RIBONUCLEOSIDE_TRIPHOSPHATE_BIOSYNTHETIC_PRO | 0.00095 | 0.01264 | 0.47727 | 0.72863 | 1.38362 | 34 |
| GO_RIBONUCLEOSIDE_TRIPHOSPHATE_METABOLIC_PROCES | 0.00033 | 0.00560 | 0.49849 | 0.72079 | 1.38389 | 42 |
| GO_RIBOSE_PHOSPHATE_BIOSYNTHETIC_PROCESS | 0.00000 | 0.00007 | 0.64355 | 0.67304 | 1.33709 | 116 |
| GO_RIBOSE_PHOSPHATE_METABOLIC_PROCESS | 0.00000 | 0.00001 | 0.70498 | 0.62407 | 1.26209 | 263 |
| GO_RIBOSOMAL_LARGE_SUBUNIT_ASSEMBLY | 0.00098 | 0.01301 | 0.45506 | 0.75899 | 1.41557 | 25 |
| GO_RIBOSOMAL_LARGE_SUBUNIT_BIOGENESIS | 0.00003 | 0.00079 | 0.55733 | 0.69921 | 1.36871 | 67 |
| GO_RIBOSOMAL_SMALL_SUBUNIT_ASSEMBLY | 0.00037 | 0.00607 | 0.49849 | 0.81971 | 1.49496 | 18 |
| GO_RIBOSOMAL_SMALL_SUBUNIT_BIOGENESIS | 0.00016 | 0.00296 | 0.51885 | 0.68217 | 1.33571 | 65 |
| GO_RIBOSOMAL_SUBUNIT | 0.00000 | 0.00000 | | 0.75170 | 1.50777 | 173 |
| GO_RIBOSOME | 0.00000 | 0.00000 | | 0.74354 | 1.49665 | 206 |
| GO_RIBOSOME_ASSEMBLY | 0.00000 | 0.00011 | 0.62726 | 0.74244 | 1.44391 | 56 |
| GO_RIBOSOME_BIOGENESIS | 0.00000 | 0.00000 | | 0.66013 | 1.33593 | 271 |
| GO_RNA_3_END_PROCESSING | 0.00205 | 0.02294 | 0.43171 | 0.60089 | 1.19562 | 125 |
| GO_RNA_CATABOLIC_PROCESS | 0.00000 | 0.00000 | | 0.68252 | 1.38715 | 330 |
| GO_RNA_DEPENDENT_DNA_BIOSYNTHETIC_PROCESS | 0.00018 | 0.00326 | 0.51885 | 0.69041 | 1.34790 | 61 |
| GO_RNA_EXPORT_FROM_NUCLEUS | 0.00313 | 0.03222 | 0.43171 | 0.59798 | 1.19006 | 126 |
| GO_RNA_LOCALIZATION | 0.00017 | 0.00318 | 0.51885 | 0.60430 | 1.21540 | 200 |
| GO_RNA_SPLICING | 0.00000 | 0.00001 | 0.70498 | 0.60265 | 1.22690 | 375 |
| GO_RNA_SPLICING_VIA_TRANSESTERIFICATION_REACTIONS | 0.00000 | 0.00002 | 0.67496 | 0.61159 | 1.24119 | 301 |
| GO_ROUGH_ENDOPLASMIC_RETICULUM | 0.00000 | 0.00007 | 0.64355 | 0.73726 | 1.43936 | 61 |
| GO_RRNA_BINDING | 0.00000 | 0.00008 | 0.64355 | 0.74785 | 1.45338 | 55 |
| GO_RRNA_METABOLIC_PROCESS | 0.00000 | 0.00000 | 0.74774 | 0.64991 | 1.30767 | 204 |
| GO_SARCOLEMMA | 0.00009 | 0.00194 | 0.53843 | 0.69889 | 1.36053 | 57 |
| GO_SARCOMERE_ORGANIZATION | 0.00457 | 0.04247 | 0.40702 | 0.72566 | 1.35341 | 25 |
| GO_SCAFFOLD_PROTEIN_BINDING | 0.00576 | 0.04960 | 0.40702 | 0.69544 | 1.32060 | 34 |
| GO_SCF_DEPENDENT_PROTEASOMAL_UBIQUITIN_DEPENDE | 0.00002 | 0.00047 | 0.57561 | 0.69305 | 1.35931 | 72 |
| GO_SECRETORY_GRANULE | 0.00000 | 0.00000 | | 0.64488 | 1.31500 | 444 |
| GO_SECRETORY_GRANULE_MEMBRANE | 0.00007 | 0.00155 | 0.53843 | 0.62061 | 1.24262 | 159 |
| GO_SERINE_TYPE_ENDOPEPTIDASE_INHIBITOR_ACTIVITY | 0.00000 | 0.00010 | 0.62726 | 0.84822 | 1.58198 | 25 |
| GO_SIDE_OF_MEMBRANE | 0.00000 | 0.00000 | 0.71951 | 0.63965 | 1.28946 | 222 |
| GO_SIGNAL_TRANSDUCTION_BY_PROTEIN_PHOSPHORYLAT | 0.00069 | 0.00995 | 0.47727 | 0.54734 | 1.11652 | 475 |
| GO_SINGLE_FERTILIZATION | 0.00000 | 0.00005 | 0.64355 | 0.75115 | 1.46227 | 57 |
| GO_SINGLE_STRANDED_RNA_BINDING | 0.00316 | 0.03238 | 0.43171 | 0.64773 | 1.26457 | 61 |
| GO_SKIN_DEVELOPMENT | 0.00013 | 0.00251 | 0.51885 | 0.61742 | 1.23323 | 149 |
| GO_SMALL_MOLECULE_BIOSYNTHETIC_PROCESS | 0.00000 | 0.00000 | 0.82666 | 0.61405 | 1.25151 | 432 |
| GO_SMALL_NUCLEAR_RIBONUCLEOPROTEIN_COMPLEX | 0.00042 | 0.00667 | 0.49849 | 0.68922 | 1.34041 | 56 |

| Cluster 1 | | | | | | |
|---|---|---|---|---|---|---|
| GO_SMALL_RIBOSOMAL_SUBUNIT | 0.00000 | 0.00005 | 0.64355 | 0.73319 | 1.43585 | 69 |
| GO_SMOOTH_ENDOPLASMIC_RETICULUM | 0.00093 | 0.01250 | 0.47727 | 0.76088 | 1.41131 | 23 |
| GO_SM_LIKE_PROTEIN_FAMILY_COMPLEX | 0.00044 | 0.00696 | 0.49849 | 0.66869 | 1.30931 | 65 |
| GO_SNORNA_BINDING | 0.00191 | 0.02177 | 0.45506 | 0.72998 | 1.36676 | 27 |
| GO_SODIUM_ION_TRANSMEMBRANE_TRANSPORTER_ACTI | 0.00420 | 0.04013 | 0.40702 | 0.66280 | 1.28509 | 53 |
| GO_SOLUTE_CATION_SYMPORTER_ACTIVITY | 0.00327 | 0.03335 | 0.43171 | 0.67914 | 1.30391 | 42 |
| GO_SPERM_EGG_RECOGNITION | 0.00000 | 0.00004 | 0.65944 | 0.85907 | 1.60609 | 26 |
| GO_SPHINGOLIPID_METABOLIC_PROCESS | 0.00013 | 0.00259 | 0.51885 | 0.65105 | 1.28649 | 92 |
| GO_SPLICEOSOMAL_COMPLEX | 0.00013 | 0.00266 | 0.51885 | 0.61160 | 1.22584 | 167 |
| GO_STEM_CELL_DIFFERENTIATION | 0.00000 | 0.00000 | 0.77494 | 0.68886 | 1.37566 | 148 |
| GO_STEROID_BIOSYNTHETIC_PROCESS | 0.00004 | 0.00092 | 0.55733 | 0.64594 | 1.28327 | 116 |
| GO_STEROID_METABOLIC_PROCESS | 0.00000 | 0.00007 | 0.64355 | 0.63517 | 1.27418 | 172 |
| GO_STEROL_BIOSYNTHETIC_PROCESS | 0.00000 | 0.00011 | 0.62726 | 0.73010 | 1.42538 | 61 |
| GO_STEROL_HOMEOSTASIS | 0.00435 | 0.04103 | 0.40702 | 0.66715 | 1.28348 | 44 |
| GO_STEROL_METABOLIC_PROCESS | 0.00001 | 0.00020 | 0.61053 | 0.67202 | 1.33420 | 108 |
| GO_STRUCTURAL_CONSTITUENT_OF_CYTOSKELETON | 0.00298 | 0.03091 | 0.43171 | 0.65717 | 1.28083 | 58 |
| GO_STRUCTURAL_CONSTITUENT_OF_RIBOSOME | 0.00000 | 0.00000 | | 0.76567 | 1.53014 | 152 |
| GO_STRUCTURAL_MOLECULE_ACTIVITY | 0.00000 | 0.00000 | | 0.71643 | 1.45851 | 378 |
| GO_SUBSTANTIA_NIGRA_DEVELOPMENT | 0.00224 | 0.02473 | 0.43171 | 0.73303 | 1.37812 | 29 |
| GO_SULFUR_COMPOUND_BINDING | 0.00268 | 0.02828 | 0.43171 | 0.60848 | 1.20803 | 112 |
| GO_SULFUR_COMPOUND_BIOSYNTHETIC_PROCESS | 0.00450 | 0.04191 | 0.40702 | 0.59587 | 1.18491 | 121 |
| GO_SULFUR_COMPOUND_METABOLIC_PROCESS | 0.00143 | 0.01729 | 0.45506 | 0.57090 | 1.15194 | 233 |
| GO_SUPRAMOLECULAR_FIBER_ORGANIZATION | 0.00050 | 0.00764 | 0.47727 | 0.55654 | 1.13332 | 419 |
| GO_SUPRAMOLECULAR_POLYMER | 0.00002 | 0.00046 | 0.57561 | 0.56448 | 1.15187 | 498 |
| GO_SYMPORTER_ACTIVITY | 0.00021 | 0.00378 | 0.51885 | 0.68802 | 1.34322 | 61 |
| GO_TELOMERASE_RNA_LOCALIZATION | 0.00014 | 0.00280 | 0.51885 | 0.82464 | 1.50941 | 19 |
| GO_TELOMERE_MAINTENANCE_VIA_TELOMERE_LENGTHEN | 0.00020 | 0.00365 | 0.51885 | 0.67612 | 1.32432 | 68 |
| GO_TERTIARY_GRANULE | 0.00383 | 0.03730 | 0.43171 | 0.62099 | 1.22728 | 91 |
| GO_THIOESTER_BIOSYNTHETIC_PROCESS | 0.00027 | 0.00460 | 0.49849 | 0.74872 | 1.42493 | 35 |
| GO_THREONINE_TYPE_ENDOPEPTIDASE_ACTIVITY | 0.00002 | 0.00047 | 0.57561 | 0.88535 | 1.60668 | 16 |
| GO_THREONINE_TYPE_PEPTIDASE_ACTIVITY | 0.00001 | 0.00031 | 0.59333 | 0.87473 | 1.59530 | 18 |
| GO_TISSUE_MORPHOGENESIS | 0.00001 | 0.00024 | 0.61053 | 0.58239 | 1.18507 | 356 |
| GO_TOXIN_TRANSPORT | 0.00339 | 0.03413 | 0.43171 | 0.73378 | 1.37527 | 28 |
| GO_TRANSCRIPTION_COUPLED_NUCLEOTIDE_EXCISION_RE | 0.00554 | 0.04824 | 0.40702 | 0.62469 | 1.22523 | 72 |
| GO_TRANSITION_METAL_ION_HOMEOSTASIS | 0.00539 | 0.04742 | 0.40702 | 0.61739 | 1.21471 | 79 |
| GO_TRANSITION_METAL_ION_TRANSPORT | 0.00202 | 0.02257 | 0.43171 | 0.63555 | 1.25129 | 81 |
| GO_TRANSLATIONAL_ELONGATION | 0.00000 | 0.00000 | 0.74774 | 0.69525 | 1.38308 | 123 |
| GO_TRANSLATIONAL_INITIATION | 0.00000 | 0.00000 | | 0.75208 | 1.50799 | 169 |
| GO_TRANSLATIONAL_TERMINATION | 0.00001 | 0.00034 | 0.59333 | 0.66734 | 1.32381 | 102 |
| GO_TRANSLATION_FACTOR_ACTIVITY_RNA_BINDING | 0.00008 | 0.00159 | 0.53843 | 0.69040 | 1.35258 | 70 |
| GO_TRANSLATION_INITIATION_FACTOR_ACTIVITY | 0.00122 | 0.01540 | 0.45506 | 0.68758 | 1.32549 | 46 |
| GO_TRANSLATION_PREINITIATION_COMPLEX | 0.00132 | 0.01628 | 0.45506 | 0.80835 | 1.46694 | 16 |
| GO_TRANSLATION_REGULATOR_ACTIVITY | 0.00000 | 0.00010 | 0.62726 | 0.67786 | 1.34580 | 108 |
| GO_TRANSLATION_REGULATOR_ACTIVITY_NUCLEIC_ACID_B | 0.00002 | 0.00046 | 0.57561 | 0.68776 | 1.35623 | 86 |
| GO_TRANSMEMBRANE_RECEPTOR_PROTEIN_TYROSINE_KIN | 0.00450 | 0.04191 | 0.40702 | 0.54338 | 1.10609 | 402 |
| GO_TRANSPORT_VESICLE | 0.00060 | 0.00895 | 0.47727 | 0.57887 | 1.16668 | 220 |
| GO_TUMOR_NECROSIS_FACTOR_MEDIATED_SIGNALING_PA | 0.00000 | 0.00000 | 0.77494 | 0.72220 | 1.43367 | 105 |
| GO_T_CELL_RECEPTOR_SIGNALING_PATHWAY | 0.00000 | 0.00011 | 0.62726 | 0.66700 | 1.32513 | 115 |
| GO_U12_TYPE_SPLICEOSOMAL_COMPLEX | 0.00191 | 0.02177 | 0.45506 | 0.74681 | 1.39029 | 24 |
| GO_UBIQUITIN_LIKE_PROTEIN_LIGASE_BINDING | 0.00001 | 0.00028 | 0.59333 | 0.60623 | 1.22383 | 236 |
| GO_UNFOLDED_PROTEIN_BINDING | 0.00000 | 0.00001 | 0.70498 | 0.71875 | 1.41998 | 90 |
| GO_UNSATURATED_FATTY_ACID_BIOSYNTHETIC_PROCESS | 0.00194 | 0.02199 | 0.45506 | 0.72863 | 1.36423 | 27 |
| GO_UROGENITAL_SYSTEM_DEVELOPMENT | 0.00576 | 0.04960 | 0.40702 | 0.57688 | 1.15564 | 164 |
| GO_VACUOLAR_LUMEN | 0.00000 | 0.00000 | 0.82666 | 0.73873 | 1.46648 | 105 |
| GO_VACUOLAR_MEMBRANE | 0.00044 | 0.00691 | 0.49849 | 0.57145 | 1.15974 | 301 |
| GO_VESICLE_LUMEN | 0.00000 | 0.00000 | | 0.69352 | 1.39348 | 190 |
| GO_VESICLE_MEMBRANE | 0.00000 | 0.00004 | 0.65944 | 0.58397 | 1.19100 | 451 |
| GO_VIRAL_GENE_EXPRESSION | 0.00000 | 0.00000 | | 0.74080 | 1.48536 | 169 |
| GO_VIRAL_LIFE_CYCLE | 0.00000 | 0.00005 | 0.65944 | 0.62229 | 1.25404 | 219 |
| GO_VITAMIN_METABOLIC_PROCESS | 0.00539 | 0.04742 | 0.40702 | 0.62026 | 1.21889 | 77 |
| GO_WATER_SOLUBLE_VITAMIN_METABOLIC_PROCESS | 0.00502 | 0.04514 | 0.40702 | 0.64230 | 1.25204 | 59 |
| GO_ZYMOGEN_ACTIVATION | 0.00053 | 0.00796 | 0.47727 | 0.77162 | 1.43123 | 23 |
| HP_11_PAIRS_OF_RIBS | 0.00194 | 0.02199 | 0.45506 | 0.79065 | 1.44034 | 17 |
| HP_ABNORMALITY_OF_ACID_BASE_HOMEOSTASIS | 0.00006 | 0.00136 | 0.53843 | 0.59265 | 1.19767 | 250 |
| HP_ABNORMALITY_OF_CARDIOVASCULAR_SYSTEM_ELECTR | 0.00413 | 0.03955 | 0.40702 | 0.56374 | 1.13853 | 242 |
| HP_ABNORMALITY_OF_CIRCULATING_ENZYME_LEVEL | 0.00420 | 0.04013 | 0.40702 | 0.57511 | 1.15315 | 169 |
| HP_ABNORMALITY_OF_HEAD_BLOOD_VESSEL | 0.00174 | 0.02022 | 0.45506 | 0.71445 | 1.35199 | 32 |
| HP_ABNORMALITY_OF_THE_ANKLES | 0.00383 | 0.03730 | 0.43171 | 0.65754 | 1.27571 | 54 |
| HP_ABNORMAL_ABDOMEN_MORPHOLOGY | 0.00335 | 0.03382 | 0.43171 | 0.54507 | 1.11001 | 420 |
| HP_ABNORMAL_CNS_MYELINATION | 0.00532 | 0.04718 | 0.40702 | 0.58005 | 1.16259 | 167 |
| HP_ABNORMAL_CORONARY_ARTERY_MORPHOLOGY | 0.00156 | 0.01851 | 0.45506 | 0.77361 | 1.42892 | 22 |
| HP_ABNORMAL_CSF_METABOLITE_LEVEL | 0.00023 | 0.00410 | 0.51885 | 0.65512 | 1.29126 | 84 |
| HP_ABNORMAL_ERYTHROCYTE_MORPHOLOGY | 0.00003 | 0.00065 | 0.57561 | 0.57614 | 1.17279 | 374 |

| Cluster 1 | | | | | | |
|---|---|---|---|---|---|---|
| HP_ABNORMAL_EYELID_PHYSIOLOGY | 0.00487 | 0.04414 | 0.40702 | 0.75674 | 1.39317 | 20 |
| HP_ABNORMAL_SYSTEMIC_ARTERIAL_MORPHOLOGY | 0.00350 | 0.03467 | 0.43171 | 0.56737 | 1.14506 | 234 |
| HP_ABNORMAL_URINE_PROTEIN_LEVEL | 0.00569 | 0.04917 | 0.40702 | 0.59825 | 1.18762 | 111 |
| HP_ABNORMAL_VASCULAR_MORPHOLOGY | 0.00076 | 0.01072 | 0.47727 | 0.55268 | 1.12684 | 437 |
| HP_ACUTE_LEUKEMIA | 0.00052 | 0.00784 | 0.47727 | 0.65587 | 1.28735 | 74 |
| HP_ANEMIA_OF_INADEQUATE_PRODUCTION | 0.00051 | 0.00772 | 0.47727 | 0.63180 | 1.25374 | 103 |
| HP_AORTIC_ROOT_ANEURYSM | 0.00413 | 0.03955 | 0.40702 | 0.75368 | 1.38890 | 21 |
| HP_CEREBRAL_ISCHEMIA | 0.00194 | 0.02199 | 0.45506 | 0.68046 | 1.30374 | 41 |
| HP_ECHOLALIA | 0.00159 | 0.01888 | 0.45506 | 0.74929 | 1.39490 | 24 |
| HP_EMG_CHRONIC_DENERVATION_SIGNS | 0.00465 | 0.04283 | 0.40702 | 0.76581 | 1.39666 | 18 |
| HP_EMOTIONAL_LABILITY | 0.00231 | 0.02541 | 0.43171 | 0.63417 | 1.25056 | 86 |
| HP_FASCICULATIONS | 0.00130 | 0.01609 | 0.45506 | 0.67778 | 1.31720 | 55 |
| HP_FATIGUE | 0.00028 | 0.00490 | 0.49849 | 0.58948 | 1.18656 | 207 |
| HP_FUNCTIONAL_ABNORMALITY_OF_THE_GASTROINTESTII | 0.00487 | 0.04414 | 0.40702 | 0.53619 | 1.09380 | 478 |
| HP_HEADACHE | 0.00067 | 0.00975 | 0.47727 | 0.60010 | 1.20002 | 156 |
| HP_HYPERTROPHIC_CARDIOMYOPATHY | 0.00091 | 0.01237 | 0.47727 | 0.58609 | 1.17630 | 181 |
| HP_INCOORDINATION | 0.00517 | 0.04607 | 0.40702 | 0.59697 | 1.18541 | 113 |
| HP_INCREASED_CSF_LACTATE | 0.00064 | 0.00931 | 0.47727 | 0.66462 | 1.30207 | 70 |
| HP_INCREASED_SERUM_LACTATE | 0.00064 | 0.00931 | 0.47727 | 0.61280 | 1.22094 | 136 |
| HP_LACTIC_ACIDOSIS | 0.00141 | 0.01710 | 0.45506 | 0.60769 | 1.20947 | 131 |
| HP_LETHARGY | 0.00250 | 0.02690 | 0.43171 | 0.61530 | 1.22098 | 103 |
| HP_LOWER_LIMB_ASYMMETRY | 0.00092 | 0.01241 | 0.47727 | 0.81594 | 1.47831 | 15 |
| HP_MACROCYTIC_ANEMIA | 0.00022 | 0.00399 | 0.51885 | 0.72795 | 1.39347 | 40 |
| HP_MACROGYRIA | 0.00287 | 0.02986 | 0.43171 | 0.78771 | 1.42716 | 15 |
| HP_MENTAL_DETERIORATION | 0.00350 | 0.03467 | 0.43171 | 0.57212 | 1.14897 | 187 |
| HP_METABOLIC_ACIDOSIS | 0.00113 | 0.01448 | 0.45506 | 0.63246 | 1.24903 | 89 |
| HP_MIGRAINE | 0.00004 | 0.00087 | 0.55733 | 0.69852 | 1.36796 | 69 |
| HP_MUSCLE_SPASM | 0.00185 | 0.02134 | 0.45506 | 0.62246 | 1.23094 | 94 |
| HP_MUTISM | 0.00104 | 0.01368 | 0.45506 | 0.72328 | 1.36871 | 32 |
| HP_NAUSEA_AND_VOMITING | 0.00435 | 0.04103 | 0.40702 | 0.56350 | 1.13862 | 251 |
| HP_OPTIC_NERVE_COLOBOMA | 0.00130 | 0.01609 | 0.45506 | 0.80964 | 1.46929 | 16 |
| HP_PACE_OF_PROGRESSION | 0.00316 | 0.03238 | 0.43171 | 0.55855 | 1.13275 | 296 |
| HP_PALLOR | 0.00071 | 0.01016 | 0.47727 | 0.63622 | 1.25837 | 95 |
| HP_PAROXYSMAL_INVOLUNTARY_EYE_MOVEMENTS | 0.00097 | 0.01283 | 0.47727 | 0.72448 | 1.37099 | 32 |
| HP_PEDIATRIC_ONSET | 0.00480 | 0.04360 | 0.40702 | 0.54748 | 1.11404 | 360 |
| HP_POOR_HEAD_CONTROL | 0.00106 | 0.01383 | 0.45506 | 0.62553 | 1.24177 | 105 |
| HP_PROPORTIONATE_SHORT_STATURE | 0.00554 | 0.04824 | 0.40702 | 0.61757 | 1.21759 | 85 |
| HP_REDUCED_CONSCIOUSNESS_CONFUSION | 0.00450 | 0.04191 | 0.40702 | 0.57692 | 1.15930 | 193 |
| HP_RESPIRATORY_INSUFFICIENCY | 0.00003 | 0.00073 | 0.57561 | 0.58861 | 1.19555 | 318 |
| HP_SENSORY_NEUROPATHY | 0.00137 | 0.01676 | 0.45506 | 0.62582 | 1.23758 | 94 |
| HP_SPARSE_EYELASHES | 0.00576 | 0.04960 | 0.40702 | 0.66906 | 1.28075 | 40 |
| HP_STROKE | 0.00539 | 0.04742 | 0.40702 | 0.63742 | 1.24808 | 65 |
| HP_THROMBOCYTOSIS | 0.00390 | 0.03790 | 0.43171 | 0.75549 | 1.39545 | 22 |
| HP_TRANSIENT_ISCHEMIC_ATTACK | 0.00046 | 0.00706 | 0.49849 | 0.77656 | 1.44567 | 24 |
| HP_VARIABLE_EXPRESSIVITY | 0.00480 | 0.04360 | 0.40702 | 0.57265 | 1.14996 | 186 |
| **Cluster 10** | | | | | | |
| pathway | pval | padj | log2err | ES | NES | size |
| GO_ACTIN_FILAMENT_BASED_MOVEMENT | 0.00001 | 0.00061 | 0.61053 | 0.70069 | 1.33868 | 86 |
| GO_ACTIN_FILAMENT_BASED_PROCESS | 0.00098 | 0.01882 | 0.45506 | 0.55973 | 1.11030 | 480 |
| GO_ACTIN_MEDIATED_CELL_CONTRACTION | 0.00065 | 0.01480 | 0.47727 | 0.68035 | 1.28605 | 62 |
| GO_ACTION_POTENTIAL | 0.00006 | 0.00297 | 0.55733 | 0.68487 | 1.30448 | 78 |
| GO_AGING | 0.00198 | 0.02979 | 0.43171 | 0.59762 | 1.16500 | 179 |
| GO_ALTERNATIVE_MRNA_SPLICING_VIA_SPLICEOSOME | 0.00000 | 0.00041 | 0.61053 | 0.72969 | 1.38370 | 68 |
| GO_AMIDE_BINDING | 0.00272 | 0.03578 | 0.43171 | 0.59075 | 1.15215 | 184 |
| GO_ANATOMICAL_STRUCTURE_HOMEOSTASIS | 0.00014 | 0.00529 | 0.51885 | 0.59631 | 1.17289 | 261 |
| GO_ANTIMICROBIAL_HUMORAL_RESPONSE | 0.00152 | 0.02533 | 0.45506 | 0.77396 | 1.39898 | 24 |
| GO_ATPASE_ACTIVITY | 0.00006 | 0.00305 | 0.53843 | 0.59875 | 1.17996 | 285 |
| GO_ATPASE_COMPLEX | 0.00008 | 0.00371 | 0.53843 | 0.71012 | 1.34064 | 60 |
| GO_ATP_DEPENDENT_CHROMATIN_REMODELING | 0.00100 | 0.01888 | 0.45506 | 0.68299 | 1.28440 | 54 |
| GO_ATP_SYNTHESIS_COUPLED_ELECTRON_TRANSPORT | 0.00168 | 0.02700 | 0.45506 | 0.67230 | 1.27584 | 69 |
| GO_AU_RICH_ELEMENT_BINDING | 0.00413 | 0.04673 | 0.40702 | 0.75519 | 1.36084 | 23 |
| GO_A_BAND | 0.00066 | 0.01490 | 0.47727 | 0.79840 | 1.43724 | 22 |
| GO_BASAL_TRANSCRIPTION_MACHINERY_BINDING | 0.00152 | 0.02533 | 0.45506 | 0.70611 | 1.32091 | 45 |
| GO_BHLH_TRANSCRIPTION_FACTOR_BINDING | 0.00398 | 0.04597 | 0.40702 | 0.77314 | 1.37618 | 18 |
| GO_CADHERIN_BINDING | 0.00000 | 0.00000 | | 0.66998 | 1.31567 | 244 |
| GO_CARDIAC_CHAMBER_DEVELOPMENT | 0.00442 | 0.04833 | 0.40702 | 0.61550 | 1.18398 | 106 |
| GO_CARDIAC_CONDUCTION | 0.00010 | 0.00403 | 0.53843 | 0.67993 | 1.29807 | 84 |
| GO_CARDIAC_MUSCLE_CELL_ACTION_POTENTIAL | 0.00005 | 0.00238 | 0.55733 | 0.74699 | 1.39776 | 44 |
| GO_CARDIAC_MUSCLE_CELL_CONTRACTION | 0.00022 | 0.00722 | 0.51885 | 0.74566 | 1.38529 | 39 |
| GO_CARDIAC_MUSCLE_CELL_MEMBRANE_REPOLARIZATIOI | 0.00030 | 0.00890 | 0.49849 | 0.82722 | 1.47760 | 19 |
| GO_CARDIAC_MUSCLE_CONTRACTION | 0.00004 | 0.00202 | 0.55733 | 0.69912 | 1.33121 | 76 |
| GO_CARDIAC_MUSCLE_TISSUE_DEVELOPMENT | 0.00065 | 0.01465 | 0.47727 | 0.62585 | 1.20978 | 125 |
| GO_CARDIOCYTE_DIFFERENTIATION | 0.00435 | 0.04794 | 0.40702 | 0.62892 | 1.20057 | 83 |
| GO_CATALYTIC_STEP_2_SPLICEOSOME | 0.00056 | 0.01342 | 0.47727 | 0.67196 | 1.27820 | 75 |

| Cluster 1 | | | | | | |
|---|---|---|---|---|---|---|
| GO_CELLULAR_NITROGEN_COMPOUND_CATABOLIC_PROCE | 0.00000 | 0.00000 | | 0.66717 | 1.32041 | 394 |
| GO_CELLULAR_RESPONSE_TO_ABIOTIC_STIMULUS | 0.00170 | 0.02715 | 0.45506 | 0.59392 | 1.16082 | 199 |
| GO_CELLULAR_RESPONSE_TO_HORMONE_STIMULUS | 0.00008 | 0.00380 | 0.53843 | 0.58287 | 1.15271 | 367 |
| GO_CELLULAR_RESPONSE_TO_LIPID | 0.00012 | 0.00465 | 0.53843 | 0.59099 | 1.16507 | 297 |
| GO_CELLULAR_RESPONSE_TO_ORGANIC_CYCLIC_COMPOUI | 0.00000 | 0.00002 | 0.69013 | 0.61819 | 1.22066 | 324 |
| GO_CELLULAR_RESPONSE_TO_STEROID_HORMONE_STIMU | 0.00002 | 0.00139 | 0.57561 | 0.65660 | 1.26923 | 125 |
| GO_CELL_ACTIVATION_INVOLVED_IN_IMMUNE_RESPONSE | 0.00039 | 0.01062 | 0.49849 | 0.57395 | 1.13528 | 370 |
| GO_CELL_ADHESION_MOLECULE_BINDING | 0.00000 | 0.00000 | | 0.65406 | 1.29272 | 351 |
| GO_CELL_BODY | 0.00068 | 0.01509 | 0.47727 | 0.58054 | 1.14554 | 314 |
| GO_CELL_CELL_JUNCTION | 0.00413 | 0.04673 | 0.40702 | 0.56957 | 1.12285 | 297 |
| GO_CELL_CELL_SIGNALING_BY_WNT | 0.00097 | 0.01852 | 0.47727 | 0.57238 | 1.13088 | 343 |
| GO_CELL_COMMUNICATION_BY_ELECTRICAL_COUPLING | 0.00056 | 0.01342 | 0.47727 | 0.81985 | 1.45934 | 18 |
| GO_CELL_COMMUNICATION_BY_ELECTRICAL_COUPLING_IN | 0.00055 | 0.01341 | 0.47727 | 0.84816 | 1.49917 | 16 |
| GO_CELL_COMMUNICATION_INVOLVED_IN_CARDIAC_CONI | 0.00022 | 0.00722 | 0.51885 | 0.74830 | 1.38396 | 35 |
| GO_CELL_CORTEX | 0.00033 | 0.00961 | 0.49849 | 0.61187 | 1.19325 | 183 |
| GO_CELL_CYCLE_G2_M_PHASE_TRANSITION | 0.00364 | 0.04308 | 0.43171 | 0.58890 | 1.14884 | 187 |
| GO_CELL_CYCLE_PHASE_TRANSITION | 0.00001 | 0.00099 | 0.59333 | 0.58375 | 1.15633 | 431 |
| GO_CELL_DIFFERENTIATION_INVOLVED_IN_KIDNEY_DEVELC | 0.00376 | 0.04414 | 0.43171 | 0.72604 | 1.32968 | 30 |
| GO_CELL_DIVISION | 0.00059 | 0.01402 | 0.47727 | 0.56936 | 1.12682 | 394 |
| GO_CELL_JUNCTION_ASSEMBLY | 0.00224 | 0.03168 | 0.43171 | 0.57831 | 1.13566 | 248 |
| GO_CELL_LEADING_EDGE | 0.00004 | 0.00203 | 0.55733 | 0.60390 | 1.18877 | 266 |
| GO_CELL_SUBSTRATE_JUNCTION | 0.00000 | 0.00000 | | 0.70365 | 1.38648 | 291 |
| GO_CENTRAL_NERVOUS_SYSTEM_NEURON_DIFFERENTIATI( | 0.00220 | 0.03145 | 0.43171 | 0.62341 | 1.19919 | 106 |
| GO_CHROMATIN_ASSEMBLY_OR_DISASSEMBLY | 0.00008 | 0.00380 | 0.53843 | 0.66592 | 1.27678 | 98 |
| GO_CHROMATIN_BINDING | 0.00000 | 0.00001 | 0.70498 | 0.61046 | 1.20899 | 413 |
| GO_CHROMATIN_ORGANIZATION_INVOLVED_IN_REGULATI( | 0.00163 | 0.02667 | 0.45506 | 0.66588 | 1.26218 | 66 |
| GO_CHROMATIN_REMODELING | 0.00000 | 0.00027 | 0.62726 | 0.66252 | 1.28387 | 137 |
| GO_CHROMOSOMAL_REGION | 0.00261 | 0.03454 | 0.43171 | 0.58173 | 1.14199 | 241 |
| GO_CHROMOSOME_TELOMERIC_REGION | 0.00435 | 0.04794 | 0.40702 | 0.61862 | 1.18839 | 103 |
| GO_CIRCADIAN_REGULATION_OF_GENE_EXPRESSION | 0.00085 | 0.01725 | 0.47727 | 0.71070 | 1.32593 | 42 |
| GO_CIRCADIAN_RHYTHM | 0.00194 | 0.02945 | 0.45506 | 0.61617 | 1.19118 | 126 |
| GO_CLATHRIN_DEPENDENT_ENDOCYTOSIS | 0.00235 | 0.03288 | 0.43171 | 0.71154 | 1.32061 | 37 |
| GO_CORTICAL_CYTOSKELETON | 0.00235 | 0.03288 | 0.43171 | 0.65253 | 1.23702 | 67 |
| GO_COTRANSLATIONAL_PROTEIN_TARGETING_TO_MEMBR/ | 0.00000 | 0.00000 | | 0.83671 | 1.60202 | 91 |
| GO_COVALENT_CHROMATIN_MODIFICATION | 0.00000 | 0.00003 | 0.67496 | 0.61640 | 1.21747 | 331 |
| GO_CYTOCHROME_COMPLEX | 0.00239 | 0.03302 | 0.43171 | 0.79075 | 1.41247 | 19 |
| GO_CYTOKINE_PRODUCTION | 0.00146 | 0.02495 | 0.45506 | 0.56525 | 1.11836 | 384 |
| GO_CYTOKINE_PRODUCTION_INVOLVED_IN_IMMUNE_RESI | 0.00398 | 0.04597 | 0.40702 | 0.67669 | 1.27106 | 51 |
| GO_CYTOPLASMIC_SIDE_OF_MEMBRANE | 0.00135 | 0.02392 | 0.45506 | 0.62958 | 1.21107 | 106 |
| GO_CYTOPLASMIC_STRESS_GRANULE | 0.00000 | 0.00015 | 0.64355 | 0.74672 | 1.41152 | 62 |
| GO_CYTOPLASMIC_TRANSLATION | 0.00000 | 0.00000 | 0.77494 | 0.76652 | 1.46215 | 81 |
| GO_CYTOPLASMIC_TRANSLATIONAL_INITIATION | 0.00250 | 0.03365 | 0.43171 | 0.74631 | 1.35175 | 25 |
| GO_CYTOSOLIC_LARGE_RIBOSOMAL_SUBUNIT | 0.00000 | 0.00000 | 0.81404 | 0.83889 | 1.57442 | 50 |
| GO_CYTOSOLIC_RIBOSOME | 0.00000 | 0.00000 | | 0.84069 | 1.60879 | 92 |
| GO_CYTOSOLIC_SMALL_RIBOSOMAL_SUBUNIT | 0.00000 | 0.00000 | 0.77494 | 0.86681 | 1.60905 | 38 |
| GO_DENDRITIC_TREE | 0.00008 | 0.00356 | 0.53843 | 0.58626 | 1.15936 | 366 |
| GO_DEVELOPMENTAL_GROWTH | 0.00001 | 0.00050 | 0.61053 | 0.59119 | 1.17014 | 401 |
| GO_DE_NOVO_PROTEIN_FOLDING | 0.00209 | 0.03067 | 0.43171 | 0.75115 | 1.36052 | 25 |
| GO_DISORDERED_DOMAIN_SPECIFIC_BINDING | 0.00413 | 0.04673 | 0.40702 | 0.73534 | 1.33533 | 26 |
| GO_DNA_BINDING_TRANSCRIPTION_FACTOR_BINDING | 0.00022 | 0.00722 | 0.51885 | 0.59654 | 1.17072 | 238 |
| GO_DNA_CONFORMATION_CHANGE | 0.00033 | 0.00948 | 0.49849 | 0.60466 | 1.18129 | 198 |
| GO_DNA_METHYLATION | 0.00168 | 0.02700 | 0.45506 | 0.70326 | 1.31559 | 45 |
| GO_DNA_METHYLATION_OR_DEMETHYLATION | 0.00011 | 0.00452 | 0.53843 | 0.71382 | 1.34321 | 55 |
| GO_DNA_MODIFICATION | 0.00069 | 0.01524 | 0.47727 | 0.68493 | 1.29981 | 69 |
| GO_DNA_PACKAGING | 0.00261 | 0.03454 | 0.43171 | 0.62109 | 1.19717 | 112 |
| GO_DNA_REPLICATION | 0.00398 | 0.04597 | 0.40702 | 0.58760 | 1.14846 | 199 |
| GO_DNA_TEMPLATED_TRANSCRIPTION_ELONGATION | 0.00106 | 0.01979 | 0.45506 | 0.65586 | 1.25210 | 84 |
| GO_DOPAMINE_RECEPTOR_SIGNALING_PATHWAY | 0.00027 | 0.00831 | 0.49849 | 0.82181 | 1.47194 | 20 |
| GO_DOUBLE_STRANDED_RNA_BINDING | 0.00013 | 0.00508 | 0.51885 | 0.71237 | 1.33965 | 54 |
| GO_EPITHELIAL_CELL_DIFFERENTIATION | 0.00023 | 0.00736 | 0.51885 | 0.58117 | 1.14825 | 343 |
| GO_EPITHELIAL_CELL_DIFFERENTIATION_INVOLVED_IN_KID | 0.00224 | 0.03168 | 0.43171 | 0.74849 | 1.35571 | 25 |
| GO_ERYTHROCYTE_HOMEOSTASIS | 0.00276 | 0.03596 | 0.43171 | 0.63998 | 1.22129 | 82 |
| GO_ESTABLISHMENT_OF_CELL_POLARITY | 0.00364 | 0.04308 | 0.43171 | 0.62216 | 1.19370 | 101 |
| GO_ESTABLISHMENT_OF_PROTEIN_LOCALIZATION_TO_END | 0.00000 | 0.00000 | | 0.80410 | 1.54471 | 103 |
| GO_ESTABLISHMENT_OF_PROTEIN_LOCALIZATION_TO_MEN | 0.00000 | 0.00000 | | 0.70125 | 1.37711 | 245 |
| GO_ESTABLISHMENT_OF_PROTEIN_LOCALIZATION_TO_ORG | 0.00000 | 0.00000 | | 0.66535 | 1.31724 | 405 |
| GO_ESTABLISHMENT_OF_RNA_LOCALIZATION | 0.00006 | 0.00305 | 0.53843 | 0.63142 | 1.22586 | 155 |
| GO_ESTABLISHMENT_OR_MAINTENANCE_OF_CELL_POLARI | 0.00029 | 0.00875 | 0.49849 | 0.61987 | 1.20296 | 147 |
| GO_EXPORT_ACROSS_PLASMA_MEMBRANE | 0.00031 | 0.00917 | 0.49849 | 0.78300 | 1.43078 | 29 |
| GO_EXTRINSIC_COMPONENT_OF_CYTOPLASMIC_SIDE_OF_ | 0.00185 | 0.02876 | 0.45506 | 0.69087 | 1.29574 | 49 |
| GO_EXTRINSIC_COMPONENT_OF_MEMBRANE | 0.00067 | 0.01500 | 0.47727 | 0.60725 | 1.18273 | 172 |
| GO_EXTRINSIC_COMPONENT_OF_PLASMA_MEMBRANE | 0.00011 | 0.00462 | 0.53843 | 0.67572 | 1.29252 | 89 |
| GO_FILOPODIUM | 0.00242 | 0.03323 | 0.43171 | 0.65766 | 1.24711 | 68 |

| Cluster 1 | | | | | | |
|---|---|---|---|---|---|---|
| GO_GENE_SILENCING | 0.00002 | 0.00135 | 0.57561 | 0.63379 | 1.23493 | 176 |
| GO_GLAND_DEVELOPMENT | 0.00001 | 0.00076 | 0.59333 | 0.61073 | 1.20237 | 274 |
| GO_GLAND_MORPHOGENESIS | 0.00209 | 0.03067 | 0.43171 | 0.64352 | 1.22804 | 82 |
| GO_GLOMERULAR_EPITHELIUM_DEVELOPMENT | 0.00074 | 0.01569 | 0.47727 | 0.84568 | 1.48843 | 15 |
| GO_G_PROTEIN_COUPLED_RECEPTOR_SIGNALING_PATHWA | 0.00163 | 0.02667 | 0.45506 | 0.66239 | 1.25742 | 71 |
| GO_HEAD_DEVELOPMENT | 0.00004 | 0.00209 | 0.55733 | 0.57492 | 1.14043 | 483 |
| GO_HEART_DEVELOPMENT | 0.00213 | 0.03067 | 0.43171 | 0.56664 | 1.11962 | 344 |
| GO_HEART_GROWTH | 0.00045 | 0.01184 | 0.49849 | 0.69241 | 1.30602 | 58 |
| GO_HEART_VALVE_DEVELOPMENT | 0.00135 | 0.02392 | 0.45506 | 0.71457 | 1.33152 | 41 |
| GO_HELICASE_ACTIVITY | 0.00128 | 0.02308 | 0.45506 | 0.62386 | 1.20348 | 116 |
| GO_HEMATOPOIETIC_PROGENITOR_CELL_DIFFERENTIATION | 0.00086 | 0.01738 | 0.47727 | 0.63217 | 1.21665 | 107 |
| GO_HEPATICOBILIARY_SYSTEM_DEVELOPMENT | 0.00063 | 0.01434 | 0.47727 | 0.65096 | 1.24460 | 88 |
| GO_HISTONE_BINDING | 0.00000 | 0.00011 | 0.64355 | 0.65316 | 1.27215 | 172 |
| GO_HISTONE_DEACETYLASE_COMPLEX | 0.00320 | 0.03926 | 0.43171 | 0.66176 | 1.24934 | 60 |
| GO_HISTONE_H3_K4_METHYLATION | 0.00239 | 0.03302 | 0.43171 | 0.70177 | 1.30666 | 40 |
| GO_HISTONE_H3_K4_TRIMETHYLATION | 0.00202 | 0.03020 | 0.43171 | 0.80041 | 1.42100 | 17 |
| GO_HISTONE_LYSINE_N_METHYLTRANSFERASE_ACTIVITY | 0.00141 | 0.02434 | 0.45506 | 0.74746 | 1.36536 | 28 |
| GO_HISTONE_METHYLATION | 0.00000 | 0.00029 | 0.62726 | 0.69234 | 1.32643 | 94 |
| GO_HISTONE_METHYLTRANSFERASE_ACTIVITY | 0.00036 | 0.00994 | 0.49849 | 0.75562 | 1.39140 | 33 |
| GO_HORMONE_MEDIATED_SIGNALING_PATHWAY | 0.00018 | 0.00640 | 0.51885 | 0.64203 | 1.23889 | 117 |
| GO_INTERLEUKIN_2_PRODUCTION | 0.00092 | 0.01803 | 0.47727 | 0.77219 | 1.39863 | 25 |
| GO_INTRACELLULAR_RECEPTOR_SIGNALING_PATHWAY | 0.00465 | 0.04997 | 0.40702 | 0.59374 | 1.15579 | 169 |
| GO_INTRACELLULAR_STEROID_HORMONE_RECEPTOR_SIGN | 0.00026 | 0.00821 | 0.49849 | 0.66967 | 1.27552 | 78 |
| GO_ION_CHANNEL_BINDING | 0.00161 | 0.02651 | 0.45506 | 0.64770 | 1.23367 | 78 |
| GO_LAMELLIPODIUM | 0.00287 | 0.03686 | 0.43171 | 0.61190 | 1.18486 | 135 |
| GO_LARGE_RIBOSOMAL_SUBUNIT | 0.00000 | 0.00032 | 0.62726 | 0.70037 | 1.34249 | 96 |
| GO_LEUKOCYTE_MEDIATED_IMMUNITY | 0.00008 | 0.00371 | 0.53843 | 0.58511 | 1.15780 | 387 |
| GO_LYSINE_ACETYLATED_HISTONE_BINDING | 0.00246 | 0.03359 | 0.43171 | 0.76964 | 1.38166 | 21 |
| GO_LYSINE_N_METHYLTRANSFERASE_ACTIVITY | 0.00174 | 0.02753 | 0.45506 | 0.72745 | 1.33952 | 33 |
| GO_MAINTENANCE_OF_CELL_NUMBER | 0.00077 | 0.01624 | 0.47727 | 0.62873 | 1.21321 | 117 |
| GO_MEMBRANE_REGION | 0.00006 | 0.00305 | 0.53843 | 0.61717 | 1.20423 | 188 |
| GO_MEMBRANE_REPOLARIZATION | 0.00012 | 0.00474 | 0.51885 | 0.81579 | 1.47004 | 23 |
| GO_METHYLATION | 0.00000 | 0.00029 | 0.62726 | 0.62939 | 1.23423 | 222 |
| GO_METHYLTRANSFERASE_COMPLEX | 0.00111 | 0.02045 | 0.45506 | 0.66242 | 1.26131 | 77 |
| GO_MIDBRAIN_DEVELOPMENT | 0.00021 | 0.00719 | 0.51885 | 0.71193 | 1.34085 | 56 |
| GO_MIRNA_MEDIATED_INHIBITION_OF_TRANSLATION | 0.00261 | 0.03454 | 0.43171 | 0.81528 | 1.43491 | 15 |
| GO_MITOTIC_CELL_CYCLE_CHECKPOINT | 0.00457 | 0.04942 | 0.40702 | 0.60058 | 1.16087 | 124 |
| GO_MODULATION_OF_PROCESS_OF_OTHER_ORGANISM | 0.00335 | 0.04051 | 0.43171 | 0.65648 | 1.24095 | 62 |
| GO_MORPHOGENESIS_OF_A_BRANCHING_STRUCTURE | 0.00316 | 0.03912 | 0.43171 | 0.61292 | 1.18271 | 117 |
| GO_MRNA_3_END_PROCESSING | 0.00413 | 0.04673 | 0.40702 | 0.63465 | 1.21111 | 82 |
| GO_MRNA_3_UTR_BINDING | 0.00008 | 0.00353 | 0.53843 | 0.68945 | 1.31279 | 76 |
| GO_MRNA_5_UTR_BINDING | 0.00405 | 0.04640 | 0.40702 | 0.75580 | 1.36194 | 23 |
| GO_MRNA_BINDING | 0.00000 | 0.00000 | | 0.70694 | 1.38829 | 245 |
| GO_MRNA_EXPORT_FROM_NUCLEUS | 0.00072 | 0.01560 | 0.47727 | 0.65162 | 1.24390 | 83 |
| GO_MRNA_PROCESSING | 0.00000 | 0.00000 | | 0.66341 | 1.31259 | 383 |
| GO_MRNA_TRANSPORT | 0.00020 | 0.00688 | 0.51885 | 0.65056 | 1.25337 | 111 |
| GO_MULTICELLULAR_ORGANISMAL_SIGNALING | 0.00066 | 0.01490 | 0.47727 | 0.63739 | 1.22799 | 111 |
| GO_MUSCLE_CELL_DIFFERENTIATION | 0.00420 | 0.04723 | 0.40702 | 0.58790 | 1.14849 | 193 |
| GO_MUSCLE_CONTRACTION | 0.00049 | 0.01265 | 0.47727 | 0.60411 | 1.17939 | 189 |
| GO_MUSCLE_ORGAN_DEVELOPMENT | 0.00047 | 0.01228 | 0.49849 | 0.59880 | 1.17316 | 214 |
| GO_MUSCLE_STRUCTURE_DEVELOPMENT | 0.00002 | 0.00116 | 0.57561 | 0.59229 | 1.17118 | 357 |
| GO_MUSCLE_SYSTEM_PROCESS | 0.00141 | 0.02434 | 0.45506 | 0.58824 | 1.15447 | 236 |
| GO_MUSCLE_TISSUE_DEVELOPMENT | 0.00011 | 0.00452 | 0.53843 | 0.60499 | 1.18597 | 219 |
| GO_MYELOID_CELL_DIFFERENTIATION | 0.00056 | 0.01342 | 0.47727 | 0.59296 | 1.16425 | 247 |
| GO_MYELOID_LEUKOCYTE_ACTIVATION | 0.00183 | 0.02854 | 0.45506 | 0.56859 | 1.12338 | 343 |
| GO_MYELOID_LEUKOCYTE_MEDIATED_IMMUNITY | 0.00036 | 0.00994 | 0.49849 | 0.58603 | 1.15588 | 301 |
| GO_M_BAND | 0.00035 | 0.00977 | 0.49849 | 0.83277 | 1.47846 | 17 |
| GO_NCRNA_METABOLIC_PROCESS | 0.00004 | 0.00209 | 0.55733 | 0.59612 | 1.17597 | 305 |
| GO_NCRNA_PROCESSING | 0.00004 | 0.00213 | 0.55733 | 0.60350 | 1.18758 | 263 |
| GO_NEGATIVE_REGULATION_OF_CATABOLIC_PROCESS | 0.00010 | 0.00403 | 0.53843 | 0.61396 | 1.20133 | 205 |
| GO_NEGATIVE_REGULATION_OF_CELLULAR_AMIDE_METAB | 0.00213 | 0.03067 | 0.43171 | 0.61580 | 1.19035 | 125 |
| GO_NEGATIVE_REGULATION_OF_CELLULAR_CATABOLIC_PR | 0.00003 | 0.00156 | 0.57561 | 0.62953 | 1.22487 | 167 |
| GO_NEGATIVE_REGULATION_OF_CELLULAR_COMPONENT_ | 0.00014 | 0.00527 | 0.51885 | 0.57495 | 1.13993 | 454 |
| GO_NEGATIVE_REGULATION_OF_CELL_CYCLE | 0.00055 | 0.01341 | 0.47727 | 0.56864 | 1.12605 | 412 |
| GO_NEGATIVE_REGULATION_OF_CELL_CYCLE_PHASE_TRAN | 0.00039 | 0.01062 | 0.49849 | 0.61197 | 1.19069 | 167 |
| GO_NEGATIVE_REGULATION_OF_CELL_CYCLE_PROCESS | 0.00053 | 0.01305 | 0.47727 | 0.59891 | 1.17488 | 227 |
| GO_NEGATIVE_REGULATION_OF_CELL_DEVELOPMENT | 0.00052 | 0.01303 | 0.47727 | 0.60301 | 1.17800 | 193 |
| GO_NEGATIVE_REGULATION_OF_CELL_DIFFERENTIATION | 0.00053 | 0.01305 | 0.47727 | 0.57138 | 1.13060 | 385 |
| GO_NEGATIVE_REGULATION_OF_CHROMOSOME_ORGANIZ | 0.00191 | 0.02918 | 0.45506 | 0.63488 | 1.21591 | 93 |
| GO_NEGATIVE_REGULATION_OF_GENE_EXPRESSION_EPIGE | 0.00239 | 0.03302 | 0.43171 | 0.68944 | 1.29006 | 44 |
| GO_NEGATIVE_REGULATION_OF_MITOTIC_CELL_CYCLE | 0.00043 | 0.01150 | 0.49849 | 0.59782 | 1.17161 | 216 |
| GO_NEGATIVE_REGULATION_OF_MRNA_METABOLIC_PROC | 0.00063 | 0.01434 | 0.47727 | 0.69077 | 1.30760 | 63 |
| GO_NEGATIVE_REGULATION_OF_NEURON_DIFFERENTIATIO | 0.00176 | 0.02767 | 0.45506 | 0.61108 | 1.18214 | 132 |

| Cluster 1 | | | | | | |
|---|---|---|---|---|---|---|
| GO_NEGATIVE_REGULATION_OF_ORGANELLE_ORGANIZATIO | 0.00130 | 0.02335 | 0.45506 | 0.58102 | 1.14335 | 263 |
| GO_NEGATIVE_REGULATION_OF_PROTEIN_CONTAINING_CO | 0.00364 | 0.04308 | 0.43171 | 0.62826 | 1.20120 | 88 |
| GO_NEGATIVE_REGULATION_OF_RNA_CATABOLIC_PROCESS | 0.00139 | 0.02430 | 0.45506 | 0.69837 | 1.30677 | 44 |
| GO_NEGATIVE_REGULATION_OF_RNA_SPLICING | 0.00372 | 0.04379 | 0.43171 | 0.75945 | 1.36852 | 23 |
| GO_NEURAL_NUCLEUS_DEVELOPMENT | 0.00035 | 0.00977 | 0.49849 | 0.74015 | 1.37505 | 39 |
| GO_NUCLEAR_ENVELOPE | 0.00045 | 0.01184 | 0.49849 | 0.58073 | 1.14560 | 305 |
| GO_NUCLEAR_EXPORT | 0.00003 | 0.00170 | 0.57561 | 0.63460 | 1.23292 | 158 |
| GO_NUCLEAR_HORMONE_RECEPTOR_BINDING | 0.00346 | 0.04161 | 0.43171 | 0.61754 | 1.18790 | 106 |
| GO_NUCLEAR_SPECK | 0.00000 | 0.00000 | 0.78819 | 0.64744 | 1.27599 | 284 |
| GO_NUCLEAR_TRANSCRIBED_MRNA_CATABOLIC_PROCESS | 0.00000 | 0.00000 | | 0.74102 | 1.44352 | 173 |
| GO_NUCLEAR_TRANSCRIBED_MRNA_CATABOLIC_PROCESS_ | 0.00000 | 0.00000 | | 0.81791 | 1.57434 | 108 |
| GO_NUCLEAR_TRANSPORT | 0.00000 | 0.00005 | 0.67496 | 0.62600 | 1.23228 | 266 |
| GO_NUCLEOBASE_CONTAINING_COMPOUND_TRANSPORT | 0.00012 | 0.00465 | 0.53843 | 0.61592 | 1.20135 | 186 |
| GO_NUCLEOSOME_ORGANIZATION | 0.00004 | 0.00233 | 0.55733 | 0.69637 | 1.32596 | 76 |
| GO_N_METHYLTRANSFERASE_ACTIVITY | 0.00167 | 0.02684 | 0.45506 | 0.69121 | 1.29835 | 51 |
| GO_ORGANIC_CYCLIC_COMPOUND_CATABOLIC_PROCESS | 0.00000 | 0.00000 | | 0.66534 | 1.31754 | 412 |
| GO_ORGAN_GROWTH | 0.00001 | 0.00098 | 0.59333 | 0.66908 | 1.28851 | 109 |
| GO_OSTEOBLAST_DIFFERENTIATION | 0.00298 | 0.03782 | 0.43171 | 0.60608 | 1.17209 | 128 |
| GO_OXIDATIVE_PHOSPHORYLATION | 0.00272 | 0.03578 | 0.43171 | 0.63280 | 1.20989 | 88 |
| GO_PEPTIDE_BIOSYNTHETIC_PROCESS | 0.00000 | 0.00000 | | 0.66634 | 1.32154 | 474 |
| GO_PEPTIDYL_LYSINE_ACETYLATION | 0.00442 | 0.04833 | 0.40702 | 0.60714 | 1.17371 | 126 |
| GO_PEPTIDYL_LYSINE_METHYLATION | 0.00009 | 0.00387 | 0.53843 | 0.68083 | 1.29925 | 82 |
| GO_PEPTIDYL_LYSINE_MODIFICATION | 0.00001 | 0.00100 | 0.59333 | 0.60849 | 1.19795 | 274 |
| GO_PEPTIDYL_LYSINE_TRIMETHYLATION | 0.00085 | 0.01725 | 0.47727 | 0.74791 | 1.37490 | 32 |
| GO_PERINUCLEAR_REGION_OF_CYTOPLASM | 0.00145 | 0.02477 | 0.45506 | 0.55958 | 1.10949 | 456 |
| GO_PHOSPHATASE_COMPLEX | 0.00283 | 0.03677 | 0.43171 | 0.70873 | 1.31293 | 36 |
| GO_PIGMENT_GRANULE | 0.00002 | 0.00116 | 0.57561 | 0.71647 | 1.36006 | 71 |
| GO_POLYMERIC_CYTOSKELETAL_FIBER | 0.00276 | 0.03596 | 0.43171 | 0.56338 | 1.11469 | 383 |
| GO_POLYSOMAL_RIBOSOME | 0.00003 | 0.00170 | 0.57561 | 0.81151 | 1.48287 | 29 |
| GO_POLYSOME | 0.00000 | 0.00012 | 0.64355 | 0.75008 | 1.41786 | 62 |
| GO_POLY_A_BINDING | 0.00001 | 0.00094 | 0.59333 | 0.86504 | 1.54936 | 20 |
| GO_POLY_PURINE_TRACT_BINDING | 0.00021 | 0.00715 | 0.51885 | 0.79729 | 1.44409 | 25 |
| GO_POLY_PYRIMIDINE_TRACT_BINDING | 0.00082 | 0.01686 | 0.47727 | 0.78378 | 1.41672 | 24 |
| GO_POSITIVE_REGULATION_OF_CELLULAR_AMIDE_METABO | 0.00001 | 0.00073 | 0.59333 | 0.67378 | 1.29872 | 112 |
| GO_POSITIVE_REGULATION_OF_CELLULAR_PROTEIN_LOCAL | 0.00034 | 0.00965 | 0.49849 | 0.59655 | 1.17211 | 252 |
| GO_POSITIVE_REGULATION_OF_CELL_ACTIVATION | 0.00150 | 0.02530 | 0.45506 | 0.60514 | 1.17473 | 154 |
| GO_POSITIVE_REGULATION_OF_CELL_CYCLE | 0.00093 | 0.01810 | 0.47727 | 0.58219 | 1.14614 | 267 |
| GO_POSITIVE_REGULATION_OF_CELL_CYCLE_PROCESS | 0.00055 | 0.01341 | 0.47727 | 0.59951 | 1.17334 | 207 |
| GO_POSITIVE_REGULATION_OF_CELL_DEVELOPMENT | 0.00004 | 0.00233 | 0.55733 | 0.59604 | 1.17675 | 322 |
| GO_POSITIVE_REGULATION_OF_CELL_POPULATION_PROLIF | 0.00000 | 0.00010 | 0.65944 | 0.59020 | 1.17074 | 480 |
| GO_POSITIVE_REGULATION_OF_CELL_SUBSTRATE_ADHESIO | 0.00305 | 0.03821 | 0.43171 | 0.66166 | 1.25565 | 69 |
| GO_POSITIVE_REGULATION_OF_CHROMOSOME_ORGANIZA | 0.00009 | 0.00387 | 0.53843 | 0.63885 | 1.23801 | 137 |
| GO_POSITIVE_REGULATION_OF_CYTOKINE_PRODUCTION | 0.00428 | 0.04789 | 0.40702 | 0.58066 | 1.13874 | 224 |
| GO_POSITIVE_REGULATION_OF_CYTOKINE_PRODUCTION_I | 0.00350 | 0.04198 | 0.43171 | 0.75308 | 1.37166 | 27 |
| GO_POSITIVE_REGULATION_OF_ESTABLISHMENT_OF_PROT | 0.00108 | 0.02002 | 0.45506 | 0.58442 | 1.14923 | 258 |
| GO_POSITIVE_REGULATION_OF_HEART_GROWTH | 0.00220 | 0.03145 | 0.43171 | 0.74883 | 1.35632 | 25 |
| GO_POSITIVE_REGULATION_OF_IMMUNE_EFFECTOR_PROC | 0.00205 | 0.03053 | 0.43171 | 0.63867 | 1.22284 | 91 |
| GO_POSITIVE_REGULATION_OF_IMMUNE_RESPONSE | 0.00305 | 0.03821 | 0.43171 | 0.56565 | 1.11691 | 324 |
| GO_POSITIVE_REGULATION_OF_IMMUNE_SYSTEM_PROCES | 0.00165 | 0.02668 | 0.45506 | 0.55675 | 1.10431 | 476 |
| GO_POSITIVE_REGULATION_OF_INTRACELLULAR_PROTEIN_ | 0.00242 | 0.03323 | 0.43171 | 0.60456 | 1.17289 | 142 |
| GO_POSITIVE_REGULATION_OF_INTRACELLULAR_TRANSPO | 0.00117 | 0.02140 | 0.45506 | 0.60768 | 1.18293 | 169 |
| GO_POSITIVE_REGULATION_OF_KIDNEY_DEVELOPMENT | 0.00465 | 0.04997 | 0.40702 | 0.76499 | 1.37016 | 20 |
| GO_POSITIVE_REGULATION_OF_LYMPHOCYTE_ACTIVATION | 0.00035 | 0.00977 | 0.49849 | 0.63582 | 1.22949 | 127 |
| GO_POSITIVE_REGULATION_OF_MRNA_CATABOLIC_PROCES | 0.00141 | 0.02434 | 0.45506 | 0.70938 | 1.32083 | 40 |
| GO_POSITIVE_REGULATION_OF_MRNA_METABOLIC_PROCE | 0.00035 | 0.00977 | 0.49849 | 0.68686 | 1.30106 | 64 |
| GO_POSITIVE_REGULATION_OF_NERVOUS_SYSTEM_DEVELO | 0.00119 | 0.02168 | 0.45506 | 0.57698 | 1.13848 | 308 |
| GO_POSITIVE_REGULATION_OF_ORGANELLE_ORGANIZATIO | 0.00001 | 0.00054 | 0.61053 | 0.58744 | 1.16400 | 438 |
| GO_POSITIVE_REGULATION_OF_ORGAN_GROWTH | 0.00152 | 0.02533 | 0.45506 | 0.74085 | 1.36191 | 32 |
| GO_POSITIVE_REGULATION_OF_POTASSIUM_ION_TRANSPO | 0.00091 | 0.01797 | 0.47727 | 0.78190 | 1.41334 | 24 |
| GO_POSITIVE_REGULATION_OF_PRODUCTION_OF_MOLECU | 0.00305 | 0.03821 | 0.43171 | 0.68972 | 1.28678 | 42 |
| GO_POSITIVE_REGULATION_OF_PROTEIN_DEPHOSPHORYLA | 0.00457 | 0.04942 | 0.40702 | 0.73289 | 1.33088 | 26 |
| GO_POSITIVE_REGULATION_OF_PROTEIN_KINASE_ACTIVITY | 0.00442 | 0.04833 | 0.40702 | 0.56425 | 1.11416 | 325 |
| GO_POSITIVE_REGULATION_OF_SODIUM_ION_TRANSMEM | 0.00181 | 0.02840 | 0.45506 | 0.82152 | 1.45207 | 16 |
| GO_POSITIVE_REGULATION_OF_SODIUM_ION_TRANSPORT | 0.00090 | 0.01796 | 0.47727 | 0.78239 | 1.41422 | 24 |
| GO_POSITIVE_REGULATION_OF_SUBSTRATE_ADHESION_DE | 0.00165 | 0.02668 | 0.45506 | 0.75261 | 1.36668 | 26 |
| GO_POSITIVE_REGULATION_OF_TRANSFERASE_ACTIVITY | 0.00316 | 0.03912 | 0.43171 | 0.55619 | 1.10181 | 426 |
| GO_POSITIVE_REGULATION_OF_TRANSLATION | 0.00001 | 0.00071 | 0.59333 | 0.68295 | 1.30844 | 94 |
| GO_POSITIVE_REGULATION_OF_VIRAL_PROCESS | 0.00111 | 0.02045 | 0.45506 | 0.65193 | 1.24173 | 78 |
| GO_POSITIVE_REGULATION_OF_WNT_SIGNALING_PATHWA | 0.00435 | 0.04794 | 0.40702 | 0.61130 | 1.17975 | 118 |
| GO_POSTSYNAPSE | 0.00017 | 0.00604 | 0.51885 | 0.57688 | 1.14146 | 389 |
| GO_POSTTRANSCRIPTIONAL_REGULATION_OF_GENE_EXPR | 0.00000 | 0.00000 | | 0.63218 | 1.25298 | 448 |
| GO_PRE_MRNA_BINDING | 0.00191 | 0.02918 | 0.45506 | 0.73686 | 1.35458 | 32 |
| GO_PRIMARY_ACTIVE_TRANSMEMBRANE_TRANSPORTER_A | 0.00405 | 0.04640 | 0.40702 | 0.64950 | 1.22774 | 62 |

| Cluster 1 | | | | | | |
|---|---|---|---|---|---|---|
| GO_PRODUCTION_OF_MOLECULAR_MEDIATOR_OF_IMMUI | 0.00148 | 0.02512 | 0.45506 | 0.62670 | 1.20214 | 100 |
| GO_PROTEIN_ACETYLATION | 0.00135 | 0.02392 | 0.45506 | 0.61167 | 1.18698 | 143 |
| GO_PROTEIN_ACYLATION | 0.00100 | 0.01888 | 0.45506 | 0.60554 | 1.17860 | 168 |
| GO_PROTEIN_CONTAINING_COMPLEX_LOCALIZATION | 0.00137 | 0.02411 | 0.45506 | 0.59248 | 1.15929 | 205 |
| GO_PROTEIN_C_TERMINUS_BINDING | 0.00313 | 0.03905 | 0.43171 | 0.61042 | 1.18200 | 135 |
| GO_PROTEIN_DNA_COMPLEX_SUBUNIT_ORGANIZATION | 0.00013 | 0.00497 | 0.51885 | 0.62897 | 1.22052 | 150 |
| GO_PROTEIN_DOMAIN_SPECIFIC_BINDING | 0.00000 | 0.00001 | 0.71951 | 0.60375 | 1.19748 | 468 |
| GO_PROTEIN_FOLDING | 0.00062 | 0.01434 | 0.47727 | 0.61899 | 1.20125 | 146 |
| GO_PROTEIN_IMPORT | 0.00139 | 0.02430 | 0.45506 | 0.60979 | 1.18303 | 142 |
| GO_PROTEIN_KINASE_A_BINDING | 0.00033 | 0.00961 | 0.49849 | 0.74258 | 1.37339 | 35 |
| GO_PROTEIN_KINASE_A_SIGNALING | 0.00049 | 0.01265 | 0.47727 | 0.81886 | 1.46267 | 19 |
| GO_PROTEIN_LOCALIZATION_TO_CELL_PERIPHERY | 0.00194 | 0.02945 | 0.45506 | 0.58792 | 1.15343 | 228 |
| GO_PROTEIN_LOCALIZATION_TO_ENDOPLASMIC_RETICULU | 0.00000 | 0.00000 | | 0.76549 | 1.48023 | 127 |
| GO_PROTEIN_LOCALIZATION_TO_MEMBRANE | 0.00000 | 0.00000 | | 0.64817 | 1.28498 | 451 |
| GO_PROTEIN_LOCALIZATION_TO_MITOCHONDRION | 0.00150 | 0.02530 | 0.45506 | 0.62804 | 1.20745 | 105 |
| GO_PROTEIN_LOCALIZATION_TO_NUCLEUS | 0.00067 | 0.01500 | 0.47727 | 0.60216 | 1.17605 | 192 |
| GO_PROTEIN_LOCALIZATION_TO_PLASMA_MEMBRANE | 0.00435 | 0.04794 | 0.40702 | 0.59034 | 1.15345 | 196 |
| GO_PROTEIN_METHYLATION | 0.00001 | 0.00054 | 0.61053 | 0.67066 | 1.29377 | 116 |
| GO_PROTEIN_STABILIZATION | 0.00261 | 0.03454 | 0.43171 | 0.61340 | 1.18777 | 135 |
| GO_PROTEIN_TARGETING | 0.00000 | 0.00000 | | 0.67596 | 1.33319 | 303 |
| GO_PROTEIN_TARGETING_TO_MEMBRANE | 0.00000 | 0.00000 | | 0.75078 | 1.45690 | 150 |
| GO_RECEPTOR_MEDIATED_ENDOCYTOSIS | 0.00011 | 0.00452 | 0.53843 | 0.62964 | 1.22235 | 151 |
| GO_REGULATION_OF_BINDING | 0.00015 | 0.00556 | 0.51885 | 0.60074 | 1.17940 | 242 |
| GO_REGULATION_OF_B_CELL_ACTIVATION | 0.00372 | 0.04379 | 0.43171 | 0.66825 | 1.25858 | 56 |
| GO_REGULATION_OF_CARDIAC_CONDUCTION | 0.00324 | 0.03948 | 0.43171 | 0.68171 | 1.27793 | 47 |
| GO_REGULATION_OF_CARDIAC_MUSCLE_CONTRACTION | 0.00074 | 0.01569 | 0.47727 | 0.70668 | 1.32539 | 49 |
| GO_REGULATION_OF_CARDIAC_MUSCLE_CONTRACTION_BY | 0.00152 | 0.02533 | 0.45506 | 0.80536 | 1.43065 | 17 |
| GO_REGULATION_OF_CELLULAR_AMIDE_METABOLIC_PROC | 0.00000 | 0.00000 | 0.76146 | 0.63577 | 1.25448 | 308 |
| GO_REGULATION_OF_CELLULAR_PROTEIN_LOCALIZATION | 0.00022 | 0.00722 | 0.51885 | 0.57813 | 1.14452 | 404 |
| GO_REGULATION_OF_CELLULAR_RESPONSE_TO_STRESS | 0.00001 | 0.00097 | 0.59333 | 0.57797 | 1.14702 | 498 |
| GO_REGULATION_OF_CELL_CYCLE_G2_M_PHASE_TRANSITI | 0.00145 | 0.02477 | 0.45506 | 0.60687 | 1.17764 | 150 |
| GO_REGULATION_OF_CELL_CYCLE_PHASE_TRANSITION | 0.00009 | 0.00380 | 0.53843 | 0.59198 | 1.16811 | 314 |
| GO_REGULATION_OF_CHROMATIN_ORGANIZATION | 0.00075 | 0.01572 | 0.47727 | 0.62720 | 1.21405 | 134 |
| GO_REGULATION_OF_CHROMOSOME_ORGANIZATION | 0.00002 | 0.00105 | 0.57561 | 0.61268 | 1.20316 | 248 |
| GO_REGULATION_OF_CYTOKINE_PRODUCTION_INVOLVED_ | 0.00167 | 0.02684 | 0.45506 | 0.71684 | 1.33046 | 37 |
| GO_REGULATION_OF_DEVELOPMENTAL_GROWTH | 0.00294 | 0.03742 | 0.43171 | 0.59012 | 1.15408 | 202 |
| GO_REGULATION_OF_DNA_METABOLIC_PROCESS | 0.00012 | 0.00474 | 0.51885 | 0.60517 | 1.18732 | 229 |
| GO_REGULATION_OF_GENE_EXPRESSION_EPIGENETIC | 0.00000 | 0.00027 | 0.62726 | 0.68775 | 1.32382 | 108 |
| GO_REGULATION_OF_GENE_SILENCING | 0.00316 | 0.03912 | 0.43171 | 0.63954 | 1.21813 | 78 |
| GO_REGULATION_OF_HEART_GROWTH | 0.00063 | 0.01434 | 0.47727 | 0.70806 | 1.32541 | 46 |
| GO_REGULATION_OF_HEART_RATE_BY_CARDIAC_CONDUCT | 0.00287 | 0.03686 | 0.43171 | 0.78613 | 1.40421 | 19 |
| GO_REGULATION_OF_HISTONE_METHYLATION | 0.00435 | 0.04794 | 0.40702 | 0.67501 | 1.26599 | 49 |
| GO_REGULATION_OF_HISTONE_MODIFICATION | 0.00305 | 0.03821 | 0.43171 | 0.62339 | 1.19992 | 108 |
| GO_REGULATION_OF_IMMUNE_EFFECTOR_PROCESS | 0.00450 | 0.04896 | 0.40702 | 0.59536 | 1.15919 | 171 |
| GO_REGULATION_OF_INTRACELLULAR_PROTEIN_TRANSPOI | 0.00261 | 0.03454 | 0.43171 | 0.58761 | 1.14928 | 203 |
| GO_REGULATION_OF_INTRACELLULAR_TRANSPORT | 0.00030 | 0.00895 | 0.49849 | 0.59123 | 1.16384 | 266 |
| GO_REGULATION_OF_LYMPHOCYTE_ACTIVATION | 0.00194 | 0.02945 | 0.45506 | 0.59055 | 1.15552 | 205 |
| GO_REGULATION_OF_MEMBRANE_REPOLARIZATION | 0.00435 | 0.04794 | 0.40702 | 0.78140 | 1.38725 | 17 |
| GO_REGULATION_OF_MITOTIC_CELL_CYCLE | 0.00001 | 0.00090 | 0.59333 | 0.58310 | 1.15543 | 437 |
| GO_REGULATION_OF_MRNA_CATABOLIC_PROCESS | 0.00000 | 0.00035 | 0.61053 | 0.65827 | 1.27748 | 146 |
| GO_REGULATION_OF_MRNA_METABOLIC_PROCESS | 0.00000 | 0.00000 | | 0.67154 | 1.31945 | 252 |
| GO_REGULATION_OF_MRNA_PROCESSING | 0.00000 | 0.00009 | 0.65944 | 0.69086 | 1.33463 | 121 |
| GO_REGULATION_OF_MRNA_SPLICING_VIA_SPLICEOSOME | 0.00000 | 0.00008 | 0.65944 | 0.71967 | 1.37495 | 86 |
| GO_REGULATION_OF_MUSCLE_SYSTEM_PROCESS | 0.00224 | 0.03168 | 0.43171 | 0.60995 | 1.17898 | 124 |
| GO_REGULATION_OF_NEURAL_PRECURSOR_CELL_PROLIFEF | 0.00239 | 0.03302 | 0.43171 | 0.67005 | 1.26007 | 54 |
| GO_REGULATION_OF_NEURON_DIFFERENTIATION | 0.00191 | 0.02918 | 0.45506 | 0.56129 | 1.11094 | 400 |
| GO_REGULATION_OF_NITRIC_OXIDE_BIOSYNTHETIC_PROCE | 0.00100 | 0.01888 | 0.45506 | 0.76030 | 1.38930 | 29 |
| GO_REGULATION_OF_NUCLEOCYTOPLASMIC_TRANSPORT | 0.00025 | 0.00800 | 0.49849 | 0.66388 | 1.27111 | 91 |
| GO_REGULATION_OF_ORGAN_GROWTH | 0.00024 | 0.00765 | 0.51885 | 0.68360 | 1.29716 | 70 |
| GO_REGULATION_OF_PEPTIDE_TRANSPORT | 0.00124 | 0.02255 | 0.45506 | 0.56302 | 1.11525 | 429 |
| GO_REGULATION_OF_PRODUCTION_OF_MOLECULAR_MED | 0.00100 | 0.01888 | 0.45506 | 0.67808 | 1.28181 | 61 |
| GO_REGULATION_OF_PROTEIN_STABILITY | 0.00061 | 0.01429 | 0.47727 | 0.59859 | 1.17154 | 207 |
| GO_REGULATION_OF_RNA_SPLICING | 0.00000 | 0.00003 | 0.67496 | 0.69841 | 1.34900 | 120 |
| GO_REGULATION_OF_SODIUM_ION_TRANSMEMBRANE_TR | 0.00024 | 0.00776 | 0.51885 | 0.73609 | 1.37057 | 40 |
| GO_REGULATION_OF_SODIUM_ION_TRANSMEMBRANE_TR | 0.00050 | 0.01273 | 0.47727 | 0.74958 | 1.38027 | 33 |
| GO_REGULATION_OF_SODIUM_ION_TRANSPORT | 0.00060 | 0.01413 | 0.47727 | 0.69059 | 1.29950 | 55 |
| GO_REGULATION_OF_STRIATED_MUSCLE_CONTRACTION | 0.00070 | 0.01533 | 0.47727 | 0.69077 | 1.29850 | 53 |
| GO_REGULATION_OF_SUBSTRATE_ADHESION_DEPENDENT_ | 0.00062 | 0.01434 | 0.47727 | 0.73152 | 1.35294 | 35 |
| GO_REGULATION_OF_SYMBIOTIC_PROCESS | 0.00054 | 0.01340 | 0.47727 | 0.62812 | 1.21522 | 131 |
| GO_REGULATION_OF_TRANSLATIONAL_INITIATION | 0.00044 | 0.01158 | 0.49849 | 0.68895 | 1.29834 | 57 |
| GO_REGULATION_OF_WNT_SIGNALING_PATHWAY | 0.00128 | 0.02308 | 0.45506 | 0.58473 | 1.14763 | 237 |
| GO_REGULATORY_RNA_BINDING | 0.00316 | 0.03912 | 0.43171 | 0.71402 | 1.31478 | 33 |
| GO_REPRODUCTIVE_SYSTEM_DEVELOPMENT | 0.00022 | 0.00722 | 0.51885 | 0.58912 | 1.15973 | 272 |

| Cluster 1 | | | | | | |
|---|---|---|---|---|---|---|
| GO_RESPIRASOME | 0.00450 | 0.04896 | 0.40702 | 0.64768 | 1.22818 | 68 |
| GO_RESPIRATORY_CHAIN_COMPLEX | 0.00442 | 0.04833 | 0.40702 | 0.65500 | 1.23436 | 57 |
| GO_RESPIRATORY_ELECTRON_TRANSPORT_CHAIN | 0.00205 | 0.03053 | 0.43171 | 0.64379 | 1.22856 | 82 |
| GO_RESPIRATORY_SYSTEM_DEVELOPMENT | 0.00287 | 0.03686 | 0.43171 | 0.60656 | 1.17302 | 128 |
| GO_RESPONSE_TO_CATECHOLAMINE | 0.00198 | 0.02979 | 0.43171 | 0.69209 | 1.29503 | 44 |
| GO_RESPONSE_TO_CORTICOSTEROID | 0.00009 | 0.00380 | 0.53843 | 0.68995 | 1.31046 | 73 |
| GO_RESPONSE_TO_DOPAMINE | 0.00209 | 0.03067 | 0.43171 | 0.71712 | 1.32326 | 34 |
| GO_RESPONSE_TO_ELECTRICAL_STIMULUS | 0.00250 | 0.03365 | 0.43171 | 0.76904 | 1.38439 | 22 |
| GO_RESPONSE_TO_FIBROBLAST_GROWTH_FACTOR | 0.00174 | 0.02753 | 0.45506 | 0.64286 | 1.22820 | 86 |
| GO_RESPONSE_TO_GROWTH_FACTOR | 0.00242 | 0.03323 | 0.43171 | 0.55917 | 1.10763 | 420 |
| GO_RESPONSE_TO_INTERLEUKIN_7 | 0.00165 | 0.02668 | 0.45506 | 0.77159 | 1.39470 | 24 |
| GO_RESPONSE_TO_KETONE | 0.00187 | 0.02897 | 0.45506 | 0.62742 | 1.20598 | 104 |
| GO_RESPONSE_TO_LEUKEMIA_INHIBITORY_FACTOR | 0.00390 | 0.04546 | 0.43171 | 0.65788 | 1.24849 | 69 |
| GO_RESPONSE_TO_LIPID | 0.00000 | 0.00001 | 0.70498 | 0.60087 | 1.19130 | 459 |
| GO_RESPONSE_TO_STEROID_HORMONE | 0.00000 | 0.00015 | 0.64355 | 0.64236 | 1.25487 | 193 |
| GO_RHYTHMIC_PROCESS | 0.00081 | 0.01679 | 0.47727 | 0.60331 | 1.17584 | 178 |
| GO_RIBONUCLEOPROTEIN_COMPLEX_BINDING | 0.00097 | 0.01852 | 0.47727 | 0.63884 | 1.22522 | 99 |
| GO_RIBONUCLEOPROTEIN_COMPLEX_BIOGENESIS | 0.00000 | 0.00000 | | 0.65109 | 1.28544 | 322 |
| GO_RIBONUCLEOPROTEIN_COMPLEX_SUBUNIT_ORGANIZAT | 0.00000 | 0.00004 | 0.67496 | 0.67535 | 1.31050 | 149 |
| GO_RIBONUCLEOPROTEIN_GRANULE | 0.00000 | 0.00008 | 0.65944 | 0.65445 | 1.27578 | 179 |
| GO_RIBOSOMAL_LARGE_SUBUNIT_BIOGENESIS | 0.00405 | 0.04640 | 0.40702 | 0.65922 | 1.24455 | 60 |
| GO_RIBOSOMAL_SMALL_SUBUNIT_BIOGENESIS | 0.00090 | 0.01796 | 0.47727 | 0.69853 | 1.31210 | 51 |
| GO_RIBOSOMAL_SUBUNIT | 0.00000 | 0.00000 | | 0.72639 | 1.41019 | 152 |
| GO_RIBOSOME | 0.00000 | 0.00000 | | 0.71525 | 1.39533 | 187 |
| GO_RIBOSOME_ASSEMBLY | 0.00087 | 0.01758 | 0.47727 | 0.70925 | 1.32478 | 43 |
| GO_RIBOSOME_BIOGENESIS | 0.00000 | 0.00004 | 0.67496 | 0.64383 | 1.26212 | 218 |
| GO_RNA_3_END_PROCESSING | 0.00250 | 0.03365 | 0.43171 | 0.63154 | 1.21223 | 102 |
| GO_RNA_CATABOLIC_PROCESS | 0.00000 | 0.00000 | | 0.69601 | 1.37340 | 309 |
| GO_RNA_EXPORT_FROM_NUCLEUS | 0.00060 | 0.01413 | 0.47727 | 0.63694 | 1.22662 | 109 |
| GO_RNA_LOCALIZATION | 0.00003 | 0.00170 | 0.57561 | 0.63215 | 1.23246 | 180 |
| GO_RNA_POLYMERASE_BINDING | 0.00022 | 0.00725 | 0.51885 | 0.71902 | 1.35059 | 51 |
| GO_RNA_POLYMERASE_CORE_ENZYME_BINDING | 0.00202 | 0.03020 | 0.43171 | 0.71381 | 1.32484 | 37 |
| GO_RNA_POLYMERASE_II_SPECIFIC_DNA_BINDING_TRANS( | 0.00027 | 0.00837 | 0.49849 | 0.60828 | 1.18689 | 188 |
| GO_RNA_SPLICING | 0.00000 | 0.00000 | | 0.67232 | 1.32794 | 337 |
| GO_RNA_SPLICING_VIA_TRANSESTERIFICATION_REACTIONS | 0.00000 | 0.00000 | | 0.67206 | 1.32329 | 275 |
| GO_RNA_STABILIZATION | 0.00063 | 0.01434 | 0.47727 | 0.73053 | 1.35111 | 35 |
| GO_ROUGH_ENDOPLASMIC_RETICULUM | 0.00068 | 0.01509 | 0.47727 | 0.68949 | 1.29662 | 54 |
| GO_RRNA_BINDING | 0.00102 | 0.01916 | 0.45506 | 0.70170 | 1.31539 | 47 |
| GO_RRNA_METABOLIC_PROCESS | 0.00000 | 0.00011 | 0.64355 | 0.65420 | 1.27348 | 169 |
| GO_RUFFLE | 0.00457 | 0.04942 | 0.40702 | 0.60885 | 1.17452 | 116 |
| GO_SALIVARY_GLAND_DEVELOPMENT | 0.00159 | 0.02627 | 0.45506 | 0.75194 | 1.37401 | 29 |
| GO_SARCOLEMMA | 0.00331 | 0.04022 | 0.43171 | 0.64512 | 1.22716 | 75 |
| GO_SIDE_OF_MEMBRANE | 0.00016 | 0.00560 | 0.51885 | 0.59548 | 1.16953 | 249 |
| GO_SINGLE_STRANDED_DNA_BINDING | 0.00239 | 0.03302 | 0.43171 | 0.64554 | 1.23138 | 81 |
| GO_SINGLE_STRANDED_RNA_BINDING | 0.00000 | 0.00034 | 0.62726 | 0.74488 | 1.40627 | 60 |
| GO_SISTER_CHROMATID_COHESION | 0.00327 | 0.03985 | 0.43171 | 0.69535 | 1.29472 | 40 |
| GO_SKIN_DEVELOPMENT | 0.00082 | 0.01686 | 0.47727 | 0.61759 | 1.19832 | 148 |
| GO_SMALL_RIBOSOMAL_SUBUNIT | 0.00000 | 0.00000 | 0.73376 | 0.79128 | 1.49323 | 59 |
| GO_SOMATODENDRITIC_COMPARTMENT | 0.00053 | 0.01305 | 0.47727 | 0.56510 | 1.12074 | 474 |
| GO_SPINAL_CORD_MOTOR_NEURON_DIFFERENTIATION | 0.00261 | 0.03454 | 0.43171 | 0.78719 | 1.40120 | 18 |
| GO_SPINDLE | 0.00224 | 0.03168 | 0.43171 | 0.57682 | 1.13328 | 251 |
| GO_SPLICEOSOMAL_COMPLEX | 0.00000 | 0.00030 | 0.62726 | 0.65399 | 1.26961 | 151 |
| GO_SPLICEOSOMAL_COMPLEX_ASSEMBLY | 0.00213 | 0.03067 | 0.43171 | 0.68676 | 1.28999 | 51 |
| GO_STEM_CELL_DIFFERENTIATION | 0.00073 | 0.01560 | 0.47727 | 0.60919 | 1.18570 | 168 |
| GO_STEROID_HORMONE_MEDIATED_SIGNALING_PATHWAY | 0.00050 | 0.01273 | 0.47727 | 0.64427 | 1.23449 | 95 |
| GO_STRESS_GRANULE_ASSEMBLY | 0.00231 | 0.03258 | 0.43171 | 0.77107 | 1.38423 | 21 |
| GO_STRIATED_MUSCLE_CONTRACTION | 0.00002 | 0.00104 | 0.59333 | 0.67683 | 1.29743 | 97 |
| GO_STRUCTURAL_CONSTITUENT_OF_RIBOSOME | 0.00000 | 0.00000 | | 0.74051 | 1.43500 | 138 |
| GO_STRUCTURAL_MOLECULE_ACTIVITY | 0.00000 | 0.00000 | | 0.66448 | 1.31479 | 389 |
| GO_SUBSTANTIA_NIGRA_DEVELOPMENT | 0.00071 | 0.01548 | 0.47727 | 0.75386 | 1.38064 | 30 |
| GO_SUPRAMOLECULAR_FIBER_ORGANIZATION | 0.00353 | 0.04217 | 0.43171 | 0.55743 | 1.10418 | 425 |
| GO_TAU_PROTEIN_BINDING | 0.00073 | 0.01560 | 0.47727 | 0.75047 | 1.37960 | 32 |
| GO_TELOMERE_ORGANIZATION | 0.00050 | 0.01273 | 0.47727 | 0.63918 | 1.23093 | 109 |
| GO_TRANSCRIPTION_COACTIVATOR_ACTIVITY | 0.00000 | 0.00015 | 0.64355 | 0.64448 | 1.25706 | 186 |
| GO_TRANSCRIPTION_COREGULATOR_ACTIVITY | 0.00000 | 0.00000 | 0.78819 | 0.63389 | 1.25249 | 348 |
| GO_TRANSCRIPTION_COREPRESSOR_ACTIVITY | 0.00002 | 0.00135 | 0.57561 | 0.64420 | 1.24979 | 142 |
| GO_TRANSCRIPTION_ELONGATION_FROM_RNA_POLYMERA | 0.00121 | 0.02195 | 0.45506 | 0.67544 | 1.27681 | 61 |
| GO_TRANSCRIPTION_FACTOR_BINDING | 0.00001 | 0.00061 | 0.59333 | 0.58751 | 1.16454 | 449 |
| GO_TRANSCRIPTION_REGULATOR_COMPLEX | 0.00004 | 0.00233 | 0.55733 | 0.60136 | 1.18422 | 276 |
| GO_TRANSCRIPTION_REPRESSOR_COMPLEX | 0.00032 | 0.00926 | 0.49849 | 0.69797 | 1.31650 | 58 |
| GO_TRANSLATIONAL_INITIATION | 0.00000 | 0.00000 | | 0.79217 | 1.53721 | 150 |
| GO_TRANSLATION_FACTOR_ACTIVITY_RNA_BINDING | 0.00051 | 0.01286 | 0.47727 | 0.68937 | 1.30146 | 60 |
| GO_TRANSLATION_INITIATION_FACTOR_ACTIVITY | 0.00242 | 0.03323 | 0.43171 | 0.70901 | 1.31615 | 38 |

| Cluster 1 | | | | | | |
|---|---|---|---|---|---|---|
| GO_TRANSLATION_REGULATOR_ACTIVITY | 0.00001 | 0.00090 | 0.59333 | 0.68055 | 1.30400 | 95 |
| GO_TRANSLATION_REGULATOR_ACTIVITY_NUCLEIC_ACID_B | 0.00006 | 0.00305 | 0.53843 | 0.69979 | 1.32854 | 72 |
| GO_TUMOR_NECROSIS_FACTOR_SECRETION | 0.00435 | 0.04794 | 0.40702 | 0.77141 | 1.37311 | 18 |
| GO_T_TUBULE | 0.00172 | 0.02737 | 0.45506 | 0.71503 | 1.32244 | 35 |
| GO_UNFOLDED_PROTEIN_BINDING | 0.00035 | 0.00977 | 0.49849 | 0.69438 | 1.31775 | 69 |
| GO_VESICLE_LUMEN | 0.00181 | 0.02840 | 0.45506 | 0.60304 | 1.17089 | 156 |
| GO_VIRAL_GENE_EXPRESSION | 0.00000 | 0.00000 | | 0.77178 | 1.49830 | 152 |
| GO_VIRAL_GENOME_REPLICATION | 0.00390 | 0.04546 | 0.43171 | 0.64333 | 1.22375 | 75 |
| GO_VIRAL_LIFE_CYCLE | 0.00012 | 0.00474 | 0.53843 | 0.61268 | 1.19882 | 205 |
| GO_ZINC_ION_BINDING | 0.00442 | 0.04833 | 0.40702 | 0.55244 | 1.09577 | 472 |
| GO_ZYMOGEN_ACTIVATION | 0.00093 | 0.01810 | 0.47727 | 0.77175 | 1.39784 | 25 |
| HP_ABNORMALITY_OF_CARDIOVASCULAR_SYSTEM_ELECTR | 0.00009 | 0.00380 | 0.53843 | 0.60662 | 1.18966 | 224 |
| HP_ABNORMALITY_OF_CORNEAL_SIZE | 0.00353 | 0.04217 | 0.43171 | 0.62260 | 1.19456 | 101 |
| HP_ABNORMALITY_OF_CRANIAL_SUTURES | 0.00045 | 0.01191 | 0.49849 | 0.60655 | 1.18296 | 184 |
| HP_ABNORMALITY_OF_DENTAL_ERUPTION | 0.00087 | 0.01751 | 0.47727 | 0.62596 | 1.21166 | 134 |
| HP_ABNORMALITY_OF_DENTAL_MORPHOLOGY | 0.00019 | 0.00653 | 0.51885 | 0.63269 | 1.22409 | 129 |
| HP_ABNORMALITY_OF_DENTAL_STRUCTURE | 0.00383 | 0.04493 | 0.43171 | 0.59119 | 1.15146 | 172 |
| HP_ABNORMALITY_OF_ESOPHAGUS_PHYSIOLOGY | 0.00316 | 0.03912 | 0.43171 | 0.56265 | 1.11292 | 369 |
| HP_ABNORMALITY_OF_LOWER_LIP | 0.00011 | 0.00462 | 0.53843 | 0.62440 | 1.21323 | 159 |
| HP_ABNORMALITY_OF_MOUTH_SHAPE | 0.00012 | 0.00474 | 0.53843 | 0.63633 | 1.23468 | 148 |
| HP_ABNORMALITY_OF_MOUTH_SIZE | 0.00005 | 0.00237 | 0.55733 | 0.61277 | 1.19918 | 206 |
| HP_ABNORMALITY_OF_ODONTOID_TISSUE | 0.00037 | 0.01014 | 0.49849 | 0.64645 | 1.24351 | 106 |
| HP_ABNORMALITY_OF_REFRACTION | 0.00019 | 0.00653 | 0.51885 | 0.58625 | 1.15670 | 313 |
| HP_ABNORMALITY_OF_THE_AUDITORY_CANAL | 0.00250 | 0.03365 | 0.43171 | 0.70518 | 1.31403 | 41 |
| HP_ABNORMALITY_OF_THE_CHEEK | 0.00002 | 0.00138 | 0.57561 | 0.65566 | 1.26751 | 126 |
| HP_ABNORMALITY_OF_THE_CHIN | 0.00324 | 0.03948 | 0.43171 | 0.59416 | 1.15826 | 179 |
| HP_ABNORMALITY_OF_THE_CHOANAE | 0.00257 | 0.03454 | 0.43171 | 0.64113 | 1.22349 | 82 |
| HP_ABNORMALITY_OF_THE_CLAVICLE | 0.00209 | 0.03067 | 0.43171 | 0.64456 | 1.22770 | 78 |
| HP_ABNORMALITY_OF_THE_COLUMELLA | 0.00420 | 0.04723 | 0.40702 | 0.70037 | 1.29746 | 36 |
| HP_ABNORMALITY_OF_THE_DENTITION | 0.00030 | 0.00895 | 0.49849 | 0.56303 | 1.11736 | 498 |
| HP_ABNORMALITY_OF_THE_DIAPHRAGM | 0.00100 | 0.01888 | 0.45506 | 0.68405 | 1.29487 | 63 |
| HP_ABNORMALITY_OF_THE_ELBOW | 0.00143 | 0.02459 | 0.45506 | 0.61098 | 1.18565 | 143 |
| HP_ABNORMALITY_OF_THE_FONTANELLES_OR_CRANIAL_S | 0.00012 | 0.00471 | 0.53843 | 0.60921 | 1.19505 | 226 |
| HP_ABNORMALITY_OF_THE_FOREARM | 0.00287 | 0.03686 | 0.43171 | 0.60242 | 1.16814 | 140 |
| HP_ABNORMALITY_OF_THE_FRONTAL_HAIRLINE | 0.00016 | 0.00562 | 0.51885 | 0.67209 | 1.28404 | 86 |
| HP_ABNORMALITY_OF_THE_HALLUX | 0.00080 | 0.01671 | 0.47727 | 0.64783 | 1.23689 | 85 |
| HP_ABNORMALITY_OF_THE_LOWER_URINARY_TRACT | 0.00346 | 0.04161 | 0.43171 | 0.55749 | 1.10408 | 411 |
| HP_ABNORMALITY_OF_THE_MIDDLE_EAR | 0.00027 | 0.00837 | 0.49849 | 0.59548 | 1.17241 | 268 |
| HP_ABNORMALITY_OF_THE_MIDFACE | 0.00002 | 0.00133 | 0.57561 | 0.60076 | 1.18653 | 332 |
| HP_ABNORMALITY_OF_THE_NAIL | 0.00294 | 0.03742 | 0.43171 | 0.58160 | 1.14194 | 247 |
| HP_ABNORMALITY_OF_THE_NASAL_DORSUM | 0.00000 | 0.00004 | 0.67496 | 0.67838 | 1.31583 | 141 |
| HP_ABNORMALITY_OF_THE_PALM | 0.00357 | 0.04253 | 0.43171 | 0.58557 | 1.14694 | 213 |
| HP_ABNORMALITY_OF_THE_PALPEBRAL_FISSURES | 0.00005 | 0.00262 | 0.55733 | 0.57797 | 1.14581 | 451 |
| HP_ABNORMALITY_OF_THE_PHILTRUM | 0.00133 | 0.02380 | 0.45506 | 0.57112 | 1.12904 | 353 |
| HP_ABNORMALITY_OF_THE_PINNA | 0.00004 | 0.00232 | 0.55733 | 0.58665 | 1.16159 | 407 |
| HP_ABNORMALITY_OF_THE_ULNA | 0.00095 | 0.01834 | 0.47727 | 0.63261 | 1.21347 | 100 |
| HP_ABNORMALITY_OF_THE_URETER | 0.00095 | 0.01834 | 0.47727 | 0.60349 | 1.17658 | 181 |
| HP_ABNORMALITY_OF_THE_URETHRA | 0.00420 | 0.04723 | 0.40702 | 0.57907 | 1.13554 | 222 |
| HP_ABNORMALITY_OF_THE_ZYGOMATIC_BONE | 0.00413 | 0.04673 | 0.40702 | 0.59167 | 1.14953 | 158 |
| HP_ABNORMALITY_OF_UPPER_LIP | 0.00002 | 0.00135 | 0.57561 | 0.58041 | 1.15072 | 459 |
| HP_ABNORMAL_5TH_FINGER_MORPHOLOGY | 0.00361 | 0.04289 | 0.43171 | 0.57794 | 1.13769 | 265 |
| HP_ABNORMAL_AORTIC_ARCH_MORPHOLOGY | 0.00191 | 0.02918 | 0.45506 | 0.75346 | 1.36470 | 25 |
| HP_ABNORMAL_CARDIAC_ATRIUM_MORPHOLOGY | 0.00016 | 0.00583 | 0.51885 | 0.60022 | 1.17852 | 247 |
| HP_ABNORMAL_CARDIAC_SEPTUM_MORPHOLOGY | 0.00111 | 0.02045 | 0.45506 | 0.56649 | 1.12063 | 377 |
| HP_ABNORMAL_CORPUS_CALLOSUM_MORPHOLOGY | 0.00324 | 0.03948 | 0.43171 | 0.55191 | 1.09459 | 469 |
| HP_ABNORMAL_DERMATOGLYPHICS | 0.00097 | 0.01852 | 0.47727 | 0.60817 | 1.18066 | 151 |
| HP_ABNORMAL_EYEBROW_MORPHOLOGY | 0.00029 | 0.00875 | 0.49849 | 0.58656 | 1.15669 | 299 |
| HP_ABNORMAL_EYELASH_MORPHOLOGY | 0.00003 | 0.00188 | 0.55733 | 0.65006 | 1.25921 | 136 |
| HP_ABNORMAL_FEAR_ANXIETY_RELATED_BEHAVIOR | 0.00014 | 0.00529 | 0.51885 | 0.61478 | 1.20062 | 191 |
| HP_ABNORMAL_FINGERNAIL_MORPHOLOGY | 0.00015 | 0.00534 | 0.51885 | 0.64826 | 1.24993 | 115 |
| HP_ABNORMAL_HAIR_PATTERN | 0.00012 | 0.00474 | 0.51885 | 0.62722 | 1.21785 | 156 |
| HP_ABNORMAL_IMMUNE_SYSTEM_MORPHOLOGY | 0.00051 | 0.01286 | 0.47727 | 0.58345 | 1.15154 | 316 |
| HP_ABNORMAL_INTESTINE_MORPHOLOGY | 0.00294 | 0.03742 | 0.43171 | 0.56137 | 1.11070 | 381 |
| HP_ABNORMAL_LOCATION_OF_EARS | 0.00009 | 0.00380 | 0.53843 | 0.57732 | 1.14388 | 436 |
| HP_ABNORMAL_MEAN_CORPUSCULAR_VOLUME | 0.00302 | 0.03813 | 0.43171 | 0.81281 | 1.43058 | 15 |
| HP_ABNORMAL_MORPHOLOGY_OF_THE_GREAT_VESSELS | 0.00283 | 0.03677 | 0.43171 | 0.57038 | 1.12539 | 313 |
| HP_ABNORMAL_NUMBER_OF_PERMANENT_TEETH | 0.00261 | 0.03454 | 0.43171 | 0.78735 | 1.40149 | 18 |
| HP_ABNORMAL_NUMBER_OF_TEETH | 0.00156 | 0.02587 | 0.45506 | 0.59211 | 1.15885 | 207 |
| HP_ABNORMAL_PALMAR_DERMATOGLYPHICS | 0.00092 | 0.01803 | 0.47727 | 0.62250 | 1.20330 | 125 |
| HP_ABNORMAL_PATELLA_MORPHOLOGY | 0.00091 | 0.01797 | 0.47727 | 0.67934 | 1.28024 | 57 |
| HP_ABNORMAL_RENAL_PELVIS_MORPHOLOGY | 0.00235 | 0.03288 | 0.43171 | 0.62170 | 1.20083 | 120 |
| HP_ABNORMAL_SCALP_MORPHOLOGY | 0.00033 | 0.00961 | 0.49849 | 0.60455 | 1.18107 | 198 |
| HP_ABNORMAL_SIZE_OF_THE_PALPEBRAL_FISSURES | 0.00009 | 0.00383 | 0.53843 | 0.62425 | 1.21502 | 168 |
| HP_ABNORMAL_SKIN_MORPHOLOGY_OF_THE_PALM | 0.00014 | 0.00527 | 0.51885 | 0.62462 | 1.21265 | 155 |

| Cluster 1 | | | | | | |
|---|---|---|---|---|---|---|
| HP_ABNORMAL_THUMB_MORPHOLOGY | 0.00047 | 0.01228 | 0.49849 | 0.60090 | 1.17605 | 207 |
| HP_ABNORMAL_URETER_PHYSIOLOGY | 0.00006 | 0.00305 | 0.53843 | 0.66982 | 1.28572 | 102 |
| HP_ABNORMAL_VASCULAR_MORPHOLOGY | 0.00320 | 0.03926 | 0.43171 | 0.55273 | 1.09505 | 435 |
| HP_ACUTE_LEUKEMIA | 0.00001 | 0.00046 | 0.61053 | 0.72384 | 1.37419 | 72 |
| HP_ACUTE_MYELOID_LEUKEMIA | 0.00390 | 0.04546 | 0.43171 | 0.71700 | 1.31523 | 31 |
| HP_ANEMIA_OF_INADEQUATE_PRODUCTION | 0.00250 | 0.03365 | 0.43171 | 0.63685 | 1.21872 | 92 |
| HP_APLASIA_HYPOPLASIA_INVOLVING_BONES_OF_THE_FEE | 0.00070 | 0.01533 | 0.47727 | 0.61008 | 1.18529 | 158 |
| HP_APLASIA_HYPOPLASIA_INVOLVING_BONES_OF_THE_LOV | 0.00246 | 0.03359 | 0.43171 | 0.59615 | 1.16609 | 204 |
| HP_APLASIA_HYPOPLASIA_INVOLVING_BONES_OF_THE_UP | 0.00021 | 0.00703 | 0.51885 | 0.58783 | 1.15990 | 308 |
| HP_APLASIA_HYPOPLASIA_OF_FINGERS | 0.00021 | 0.00700 | 0.51885 | 0.60195 | 1.17801 | 206 |
| HP_APLASIA_HYPOPLASIA_OF_THE_EXTREMITIES | 0.00146 | 0.02495 | 0.45506 | 0.56715 | 1.12226 | 387 |
| HP_APLASIA_HYPOPLASIA_OF_THE_NIPPLES | 0.00088 | 0.01771 | 0.47727 | 0.75098 | 1.37535 | 30 |
| HP_APLASIA_HYPOPLASIA_OF_THE_PHALANGES_OF_THE_H | 0.00205 | 0.03053 | 0.43171 | 0.60097 | 1.16664 | 154 |
| HP_APLASIA_HYPOPLASIA_OF_THE_THUMB | 0.00087 | 0.01758 | 0.47727 | 0.66179 | 1.26012 | 76 |
| HP_ATRESIA_OF_THE_EXTERNAL_AUDITORY_CANAL | 0.00194 | 0.02945 | 0.45506 | 0.74667 | 1.36439 | 29 |
| HP_BILATERAL_SINGLE_TRANSVERSE_PALMAR_CREASES | 0.00141 | 0.02434 | 0.45506 | 0.69891 | 1.30547 | 43 |
| HP_BLEPHAROPHIMOSIS | 0.00220 | 0.03145 | 0.43171 | 0.64989 | 1.23624 | 75 |
| HP_BRACHYDACTYLY | 0.00231 | 0.03258 | 0.43171 | 0.58556 | 1.14723 | 214 |
| HP_BRADYCARDIA | 0.00405 | 0.04640 | 0.40702 | 0.75588 | 1.36209 | 23 |
| HP_BROAD_FOREHEAD | 0.00276 | 0.03596 | 0.43171 | 0.63578 | 1.21667 | 92 |
| HP_BROAD_TOE | 0.00465 | 0.04997 | 0.40702 | 0.69840 | 1.29381 | 36 |
| HP_CARDIAC_ARREST | 0.00335 | 0.04051 | 0.43171 | 0.65606 | 1.24272 | 64 |
| HP_CHRONIC_DIARRHEA | 0.00250 | 0.03365 | 0.43171 | 0.70847 | 1.31031 | 35 |
| HP_CLEFT_LIP | 0.00176 | 0.02767 | 0.45506 | 0.62562 | 1.20405 | 107 |
| HP_CLINODACTYLY | 0.00024 | 0.00765 | 0.51885 | 0.58730 | 1.15780 | 297 |
| HP_COARSE_FACIAL_FEATURES | 0.00055 | 0.01341 | 0.47727 | 0.62242 | 1.20370 | 128 |
| HP_CONGENITAL_ABNORMAL_HAIR_PATTERN | 0.00001 | 0.00090 | 0.59333 | 0.64897 | 1.25904 | 142 |
| HP_CONGENITAL_DIAPHRAGMATIC_HERNIA | 0.00013 | 0.00502 | 0.51885 | 0.71122 | 1.33832 | 55 |
| HP_CONGENITAL_MALFORMATION_OF_THE_GREAT_ARTER | 0.00213 | 0.03067 | 0.43171 | 0.58835 | 1.15358 | 221 |
| HP_CRYPTORCHIDISM | 0.00062 | 0.01434 | 0.47727 | 0.57038 | 1.12961 | 413 |
| HP_DELAYED_SPEECH_AND_LANGUAGE_DEVELOPMENT | 0.00290 | 0.03718 | 0.43171 | 0.55617 | 1.10285 | 464 |
| HP_DEPRESSED_NASAL_RIDGE | 0.00000 | 0.00002 | 0.70498 | 0.75294 | 1.43156 | 74 |
| HP_DEVIATION_OF_THE_HAND_OR_OF_FINGERS_OF_THE_ | 0.00405 | 0.04640 | 0.40702 | 0.57168 | 1.12544 | 267 |
| HP_DISPLACEMENT_OF_THE_URETHRAL_MEATUS | 0.00198 | 0.02979 | 0.43171 | 0.59370 | 1.15992 | 194 |
| HP_DOWNSLANTED_PALPEBRAL_FISSURES | 0.00007 | 0.00335 | 0.53843 | 0.59885 | 1.17898 | 271 |
| HP_DOWNTURNED_CORNERS_OF_MOUTH | 0.00026 | 0.00821 | 0.49849 | 0.67431 | 1.28625 | 81 |
| HP_EPICANTHUS | 0.00006 | 0.00305 | 0.53843 | 0.59221 | 1.16824 | 305 |
| HP_FATIGUE | 0.00091 | 0.01797 | 0.47727 | 0.59157 | 1.15967 | 219 |
| HP_FULL_CHEEKS | 0.00042 | 0.01119 | 0.49849 | 0.67366 | 1.27952 | 73 |
| HP_FUNCTIONAL_ABNORMALITY_OF_THE_MIDDLE_EAR | 0.00001 | 0.00046 | 0.61053 | 0.63709 | 1.24265 | 186 |
| HP_GASTROESOPHAGEAL_REFLUX | 0.00075 | 0.01572 | 0.47727 | 0.60120 | 1.17417 | 192 |
| HP_GENERALIZED_HIRSUTISM | 0.00346 | 0.04161 | 0.43171 | 0.65880 | 1.24152 | 57 |
| HP_HEMATOLOGICAL_NEOPLASM | 0.00003 | 0.00158 | 0.57561 | 0.63982 | 1.24411 | 163 |
| HP_HIGHLY_ARCHED_EYEBROW | 0.00148 | 0.02512 | 0.45506 | 0.63246 | 1.21238 | 97 |
| HP_HIGH_FOREHEAD | 0.00004 | 0.00233 | 0.55733 | 0.63883 | 1.24192 | 161 |
| HP_HIP_DYSPLASIA | 0.00029 | 0.00880 | 0.49849 | 0.66782 | 1.27200 | 78 |
| HP_HIRSUTISM | 0.00034 | 0.00965 | 0.49849 | 0.64770 | 1.24272 | 101 |
| HP_HYPERACTIVITY | 0.00054 | 0.01323 | 0.47727 | 0.58952 | 1.16015 | 264 |
| HP_HYPERMETROPIA | 0.00056 | 0.01342 | 0.47727 | 0.64035 | 1.23112 | 105 |
| HP_HYPERTELORISM | 0.00320 | 0.03926 | 0.43171 | 0.55491 | 1.09905 | 419 |
| HP_HYPOPLASIA_OF_DENTAL_ENAMEL | 0.00287 | 0.03686 | 0.43171 | 0.70269 | 1.30546 | 39 |
| HP_HYPOPLASIA_OF_TEETH | 0.00165 | 0.02668 | 0.45506 | 0.68024 | 1.27870 | 53 |
| HP_HYPOPLASIA_OF_THE_CORPUS_CALLOSUM | 0.00320 | 0.03926 | 0.43171 | 0.57018 | 1.12244 | 272 |
| HP_HYPOPLASTIC_FEMALE_EXTERNAL_GENITALIA | 0.00428 | 0.04789 | 0.40702 | 0.70389 | 1.29885 | 34 |
| HP_HYPOPLASTIC_NIPPLES | 0.00276 | 0.03596 | 0.43171 | 0.76396 | 1.37664 | 23 |
| HP_INCREASED_NUCHAL_TRANSLUCENCY | 0.00111 | 0.02045 | 0.45506 | 0.78307 | 1.40964 | 22 |
| HP_INTESTINAL_FISTULA | 0.00305 | 0.03821 | 0.43171 | 0.79046 | 1.40335 | 17 |
| HP_INTESTINAL_MALROTATION | 0.00065 | 0.01465 | 0.47727 | 0.69623 | 1.31127 | 56 |
| HP_JOINT_HYPERMOBILITY | 0.00073 | 0.01560 | 0.47727 | 0.60077 | 1.17375 | 194 |
| HP_LEUKEMIA | 0.00000 | 0.00014 | 0.64355 | 0.69459 | 1.33509 | 104 |
| HP_LONG_EYELASHES | 0.00016 | 0.00562 | 0.51885 | 0.70229 | 1.32587 | 60 |
| HP_LOW_ANTERIOR_HAIRLINE | 0.00213 | 0.03067 | 0.43171 | 0.66679 | 1.25769 | 58 |
| HP_LOW_POSTERIOR_HAIRLINE | 0.00005 | 0.00274 | 0.55733 | 0.71082 | 1.34369 | 61 |
| HP_MACROCYTIC_ANEMIA | 0.00002 | 0.00135 | 0.57561 | 0.77992 | 1.44776 | 38 |
| HP_MICRODONTIA | 0.00290 | 0.03718 | 0.43171 | 0.65990 | 1.25085 | 66 |
| HP_MIDFACE_RETRUSION | 0.00028 | 0.00870 | 0.49849 | 0.61272 | 1.19376 | 174 |
| HP_MIGRAINE | 0.00011 | 0.00462 | 0.53843 | 0.69239 | 1.31707 | 75 |
| HP_MISALIGNMENT_OF_TEETH | 0.00081 | 0.01679 | 0.47727 | 0.59964 | 1.17003 | 188 |
| HP_MYOPIA | 0.00302 | 0.03813 | 0.43171 | 0.58557 | 1.14645 | 210 |
| HP_OPEN_BITE | 0.00132 | 0.02354 | 0.45506 | 0.73865 | 1.35494 | 31 |
| HP_ORAL_CLEFT | 0.00000 | 0.00029 | 0.62726 | 0.60370 | 1.19382 | 363 |
| HP_OTITIS_MEDIA | 0.00435 | 0.04794 | 0.40702 | 0.60871 | 1.17707 | 127 |
| HP_PATENT_DUCTUS_ARTERIOSUS | 0.00075 | 0.01572 | 0.47727 | 0.60999 | 1.18741 | 169 |
| HP_PLAGIOCEPHALY | 0.00039 | 0.01059 | 0.49849 | 0.69237 | 1.30713 | 60 |

| Cluster 1 | | | | | | |
|---|---|---|---|---|---|---|
| HP_POSTERIORLY_ROTATED_EARS | 0.00003 | 0.00158 | 0.57561 | 0.61467 | 1.20597 | 229 |
| HP_PREAURICULAR_SKIN_TAG | 0.00435 | 0.04794 | 0.40702 | 0.70359 | 1.29828 | 34 |
| HP_RADIOULNAR_SYNOSTOSIS | 0.00213 | 0.03067 | 0.43171 | 0.68792 | 1.29021 | 49 |
| HP_RECTAL_FISTULA | 0.00268 | 0.03545 | 0.43171 | 0.81417 | 1.43297 | 15 |
| HP_REGIONAL_ABNORMALITY_OF_SKIN | 0.00073 | 0.01560 | 0.47727 | 0.59466 | 1.16541 | 216 |
| HP_RIB_FUSION | 0.00213 | 0.03067 | 0.43171 | 0.78180 | 1.40027 | 20 |
| HP_SHORT_ATTENTION_SPAN | 0.00108 | 0.02002 | 0.45506 | 0.59160 | 1.15861 | 212 |
| HP_SHORT_DIGIT | 0.00079 | 0.01657 | 0.47727 | 0.57550 | 1.13657 | 326 |
| HP_SHORT_FOOT | 0.00216 | 0.03113 | 0.43171 | 0.65394 | 1.24517 | 77 |
| HP_SHORT_NOSE | 0.00000 | 0.00001 | 0.71951 | 0.66665 | 1.30052 | 187 |
| HP_SHORT_PALPEBRAL_FISSURE | 0.00137 | 0.02411 | 0.45506 | 0.65837 | 1.24811 | 67 |
| HP_SHORT_THUMB | 0.00157 | 0.02611 | 0.45506 | 0.68220 | 1.28484 | 56 |
| HP_SINGLE_TRANSVERSE_PALMAR_CREASE | 0.00420 | 0.04723 | 0.40702 | 0.62254 | 1.19360 | 98 |
| HP_SMALL_HAND | 0.00018 | 0.00644 | 0.51885 | 0.70528 | 1.33505 | 63 |
| HP_SMALL_NAIL | 0.00084 | 0.01718 | 0.47727 | 0.66205 | 1.26061 | 76 |
| HP_SMOOTH_PHILTRUM | 0.00457 | 0.04942 | 0.40702 | 0.63340 | 1.20873 | 82 |
| HP_SOMATIC_MUTATION | 0.00413 | 0.04673 | 0.40702 | 0.60224 | 1.16517 | 129 |
| HP_SPINA_BIFIDA_OCCULTA | 0.00398 | 0.04597 | 0.40702 | 0.68685 | 1.28488 | 45 |
| HP_SPORADIC | 0.00132 | 0.02354 | 0.45506 | 0.66026 | 1.25348 | 72 |
| HP_SYNOSTOSIS_INVOLVING_BONES_OF_THE_UPPER_LIMB | 0.00170 | 0.02715 | 0.45506 | 0.65554 | 1.24698 | 75 |
| HP_SYNOSTOSIS_OF_JOINTS | 0.00191 | 0.02918 | 0.45506 | 0.62670 | 1.20739 | 111 |
| HP_THICK_EYEBROW | 0.00029 | 0.00880 | 0.49849 | 0.67412 | 1.28361 | 76 |
| HP_THICK_LOWER_LIP_VERMILION | 0.00028 | 0.00846 | 0.49849 | 0.65408 | 1.25550 | 102 |
| HP_THICK_VERMILION_BORDER | 0.00009 | 0.00403 | 0.53843 | 0.63001 | 1.22327 | 156 |
| HP_THIN_UPPER_LIP_VERMILION | 0.00035 | 0.00977 | 0.49849 | 0.61460 | 1.19479 | 160 |
| HP_THIN_VERMILION_BORDER | 0.00003 | 0.00158 | 0.57561 | 0.61618 | 1.20586 | 206 |
| HP_TRUNCAL_OBESITY | 0.00213 | 0.03067 | 0.43171 | 0.71327 | 1.32383 | 37 |
| HP_UPPER_EXTREMITY_JOINT_DISLOCATION | 0.00053 | 0.01305 | 0.47727 | 0.68876 | 1.30031 | 60 |
| HP_VENTRICULAR_FIBRILLATION | 0.00390 | 0.04546 | 0.43171 | 0.75753 | 1.36506 | 23 |
| HP_VENTRICULOMEGALY | 0.00353 | 0.04217 | 0.43171 | 0.56193 | 1.11125 | 366 |
| HP_VERTEBRAL_SEGMENTATION_DEFECT | 0.00183 | 0.02854 | 0.45506 | 0.64586 | 1.23017 | 78 |
| HP_WIDELY_SPACED_TEETH | 0.00011 | 0.00452 | 0.53843 | 0.73869 | 1.38274 | 46 |
| HP_WIDE_MOUTH | 0.00209 | 0.03067 | 0.43171 | 0.62463 | 1.20091 | 105 |
| HP_WIDE_NASAL_BRIDGE | 0.00159 | 0.02627 | 0.45506 | 0.57581 | 1.13486 | 293 |
| **Cluster 11** | | | | | | |
| pathway | pval | padj | log2err | ES | NES | size |
| GO_ACTIN_BASED_CELL_PROJECTION | 0.00071 | 0.01016 | 0.47727 | 0.60779 | 1.23199 | 121 |
| GO_ACTIN_CYTOSKELETON | 0.00047 | 0.00727 | 0.49849 | 0.56269 | 1.16631 | 314 |
| GO_ACTIN_FILAMENT_BASED_PROCESS | 0.00283 | 0.02988 | 0.43171 | 0.53105 | 1.10608 | 467 |
| GO_ACTIN_FILAMENT_BINDING | 0.00279 | 0.02959 | 0.43171 | 0.58768 | 1.19633 | 134 |
| GO_ACTIN_FILAMENT_BUNDLE_ORGANIZATION | 0.00170 | 0.02040 | 0.45506 | 0.61667 | 1.24149 | 94 |
| GO_ACTIN_FILAMENT_ORGANIZATION | 0.00002 | 0.00045 | 0.57561 | 0.58586 | 1.21049 | 261 |
| GO_ACTIN_NUCLEATION | 0.00457 | 0.04261 | 0.40702 | 0.68523 | 1.32378 | 36 |
| GO_ACTIN_POLYMERIZATION_OR_DEPOLYMERIZATION | 0.00181 | 0.02122 | 0.45506 | 0.58676 | 1.19505 | 137 |
| GO_ACTIVATION_OF_IMMUNE_RESPONSE | 0.00000 | 0.00000 | 0.73376 | 0.62228 | 1.28270 | 229 |
| GO_ACTIVATION_OF_INNATE_IMMUNE_RESPONSE | 0.00000 | 0.00001 | 0.70498 | 0.70531 | 1.41995 | 94 |
| GO_ACTOMYOSIN_STRUCTURE_ORGANIZATION | 0.00076 | 0.01068 | 0.47727 | 0.60991 | 1.23554 | 116 |
| GO_ADHERENS_JUNCTION | 0.00093 | 0.01264 | 0.47727 | 0.61903 | 1.24964 | 99 |
| GO_ADP_BINDING | 0.00095 | 0.01283 | 0.47727 | 0.74927 | 1.42170 | 26 |
| GO_AEROBIC_RESPIRATION | 0.00000 | 0.00000 | 0.70498 | 0.73836 | 1.47493 | 72 |
| GO_ALCOHOL_BIOSYNTHETIC_PROCESS | 0.00034 | 0.00562 | 0.49849 | 0.63089 | 1.27532 | 103 |
| GO_ALCOHOL_METABOLIC_PROCESS | 0.00067 | 0.00977 | 0.47727 | 0.57565 | 1.18319 | 206 |
| GO_ALTERNATIVE_MRNA_SPLICING_VIA_SPLICEOSOME | 0.00235 | 0.02561 | 0.43171 | 0.63275 | 1.26329 | 71 |
| GO_AMIDE_BINDING | 0.00001 | 0.00040 | 0.59333 | 0.60565 | 1.24194 | 191 |
| GO_AMINE_METABOLIC_PROCESS | 0.00000 | 0.00000 | 0.76146 | 0.73625 | 1.47802 | 82 |
| GO_ANAPHASE_PROMOTING_COMPLEX_DEPENDENT_CATA | 0.00000 | 0.00000 | 0.74774 | 0.73616 | 1.47510 | 76 |
| GO_ANATOMICAL_STRUCTURE_HOMEOSTASIS | 0.00361 | 0.03559 | 0.43171 | 0.55188 | 1.14070 | 267 |
| GO_ANTERIOR_POSTERIOR_AXIS_SPECIFICATION | 0.00547 | 0.04911 | 0.40702 | 0.68885 | 1.32100 | 32 |
| GO_ANTIGEN_PROCESSING_AND_PRESENTATION | 0.00000 | 0.00000 | 0.74774 | 0.66030 | 1.34761 | 153 |
| GO_ANTIGEN_PROCESSING_AND_PRESENTATION_OF_EXOG | 0.00000 | 0.00000 | 0.82666 | 0.79901 | 1.58458 | 60 |
| GO_ANTIGEN_PROCESSING_AND_PRESENTATION_OF_PEPT | 0.00000 | 0.00000 | 0.74774 | 0.67840 | 1.38024 | 133 |
| GO_ANTIGEN_PROCESSING_AND_PRESENTATION_OF_PEPT | 0.00000 | 0.00000 | 0.76146 | 0.76114 | 1.51834 | 70 |
| GO_ANTIGEN_RECEPTOR_MEDIATED_SIGNALING_PATHWAY | 0.00000 | 0.00014 | 0.61053 | 0.64454 | 1.31134 | 133 |
| GO_ANTIMICROBIAL_HUMORAL_RESPONSE | 0.00383 | 0.03700 | 0.43171 | 0.76764 | 1.41690 | 18 |
| GO_ANTIOXIDANT_ACTIVITY | 0.00016 | 0.00307 | 0.51885 | 0.72338 | 1.41038 | 42 |
| GO_APICAL_PART_OF_CELL | 0.00000 | 0.00005 | 0.64355 | 0.61377 | 1.26192 | 209 |
| GO_APICAL_PLASMA_MEMBRANE | 0.00011 | 0.00222 | 0.53843 | 0.60329 | 1.23277 | 169 |
| GO_APOPTOTIC_SIGNALING_PATHWAY | 0.00003 | 0.00067 | 0.57561 | 0.56553 | 1.17475 | 379 |
| GO_ATP_BIOSYNTHETIC_PROCESS | 0.00000 | 0.00004 | 0.64355 | 0.88141 | 1.63191 | 19 |
| GO_ATP_METABOLIC_PROCESS | 0.00000 | 0.00000 | | 0.71045 | 1.46190 | 214 |
| GO_ATP_SYNTHESIS_COUPLED_ELECTRON_TRANSPORT | 0.00000 | 0.00000 | | 0.81398 | 1.63216 | 78 |
| GO_AUTONOMIC_NERVOUS_SYSTEM_DEVELOPMENT | 0.00294 | 0.03067 | 0.43171 | 0.75820 | 1.40378 | 19 |
| GO_AZUROPHIL_GRANULE | 0.00013 | 0.00262 | 0.51885 | 0.63095 | 1.27586 | 107 |

| Cluster 1 | | | | | | |
|---|---|---|---|---|---|---|
| GO_AZUROPHIL_GRANULE_LUMEN | 0.00480 | 0.04435 | 0.40702 | 0.63643 | 1.26323 | 61 |
| GO_BASAL_PART_OF_CELL | 0.00022 | 0.00396 | 0.51885 | 0.72545 | 1.40862 | 39 |
| GO_BASAL_PLASMA_MEMBRANE | 0.00078 | 0.01100 | 0.47727 | 0.73820 | 1.41299 | 30 |
| GO_BASEMENT_MEMBRANE | 0.00331 | 0.03338 | 0.43171 | 0.67725 | 1.31710 | 40 |
| GO_BASOLATERAL_PLASMA_MEMBRANE | 0.00084 | 0.01161 | 0.47727 | 0.60853 | 1.23347 | 120 |
| GO_BINDING_OF_SPERM_TO_ZONA_PELLUCIDA | 0.00003 | 0.00068 | 0.57561 | 0.87705 | 1.61027 | 16 |
| GO_BLOOD_MICROPARTICLE | 0.00000 | 0.00000 | 0.77494 | 0.83356 | 1.62108 | 40 |
| GO_BRUSH_BORDER | 0.00000 | 0.00010 | 0.62726 | 0.71060 | 1.41362 | 66 |
| GO_BRUSH_BORDER_MEMBRANE | 0.00007 | 0.00153 | 0.53843 | 0.76156 | 1.46044 | 32 |
| GO_CADHERIN_BINDING | 0.00000 | 0.00000 | | 0.66287 | 1.36969 | 265 |
| GO_CALCIUM_DEPENDENT_PHOSPHOLIPID_BINDING | 0.00324 | 0.03296 | 0.43171 | 0.76148 | 1.41623 | 20 |
| GO_CALCIUM_DEPENDENT_PROTEIN_BINDING | 0.00008 | 0.00180 | 0.53843 | 0.73058 | 1.42354 | 41 |
| GO_CALCIUM_ION_BINDING | 0.00008 | 0.00172 | 0.53843 | 0.57668 | 1.19160 | 262 |
| GO_CANONICAL_WNT_SIGNALING_PATHWAY | 0.00000 | 0.00002 | 0.65944 | 0.62579 | 1.28496 | 200 |
| GO_CARBOHYDRATE_BIOSYNTHETIC_PROCESS | 0.00150 | 0.01845 | 0.45506 | 0.60002 | 1.21622 | 121 |
| GO_CARBOHYDRATE_CATABOLIC_PROCESS | 0.00242 | 0.02622 | 0.43171 | 0.58363 | 1.18867 | 137 |
| GO_CARBOHYDRATE_DERIVATIVE_BIOSYNTHETIC_PROCESS | 0.00172 | 0.02050 | 0.45506 | 0.54117 | 1.12522 | 412 |
| GO_CATALYTIC_STEP_2_SPLICEOSOME | 0.00005 | 0.00111 | 0.55733 | 0.66680 | 1.33859 | 82 |
| GO_CATION_TRANSMEMBRANE_TRANSPORT | 0.00008 | 0.00175 | 0.53843 | 0.56233 | 1.16802 | 372 |
| GO_CATION_TRANSMEMBRANE_TRANSPORTER_ACTIVITY | 0.00041 | 0.00653 | 0.49849 | 0.57169 | 1.17892 | 235 |
| GO_CATION_TRANSPORT | 0.00000 | 0.00014 | 0.61053 | 0.56057 | 1.16824 | 491 |
| GO_CELLULAR_AMINO_ACID_METABOLIC_PROCESS | 0.00000 | 0.00011 | 0.62726 | 0.60570 | 1.24649 | 216 |
| GO_CELLULAR_COMPONENT_DISASSEMBLY | 0.00000 | 0.00000 | | 0.61195 | 1.27211 | 408 |
| GO_CELLULAR_HOMEOSTASIS | 0.00001 | 0.00029 | 0.59333 | 0.55741 | 1.16124 | 481 |
| GO_CELLULAR_ION_HOMEOSTASIS | 0.00011 | 0.00228 | 0.53843 | 0.56795 | 1.17630 | 295 |
| GO_CELLULAR_IRON_ION_HOMEOSTASIS | 0.00320 | 0.03275 | 0.43171 | 0.66861 | 1.30559 | 46 |
| GO_CELLULAR_KETONE_METABOLIC_PROCESS | 0.00001 | 0.00018 | 0.61053 | 0.63519 | 1.29594 | 146 |
| GO_CELLULAR_NITROGEN_COMPOUND_CATABOLIC_PROCE | 0.00000 | 0.00000 | | 0.65614 | 1.36628 | 451 |
| GO_CELLULAR_OXIDANT_DETOXIFICATION | 0.00066 | 0.00971 | 0.47727 | 0.67137 | 1.32697 | 54 |
| GO_CELLULAR_PROTEIN_COMPLEX_DISASSEMBLY | 0.00000 | 0.00000 | | 0.67954 | 1.39086 | 181 |
| GO_CELLULAR_RESPIRATION | 0.00000 | 0.00000 | | 0.74439 | 1.51881 | 145 |
| GO_CELLULAR_RESPONSE_TO_CHEMICAL_STRESS | 0.00420 | 0.03976 | 0.40702 | 0.55074 | 1.13540 | 230 |
| GO_CELLULAR_RESPONSE_TO_HEAT | 0.00235 | 0.02561 | 0.43171 | 0.60531 | 1.22270 | 100 |
| GO_CELLULAR_RESPONSE_TO_ORGANIC_CYCLIC_COMPOUI | 0.00228 | 0.02516 | 0.43171 | 0.54960 | 1.13867 | 307 |
| GO_CELLULAR_RESPONSE_TO_OXYGEN_LEVELS | 0.00000 | 0.00000 | | 0.69962 | 1.42793 | 151 |
| GO_CELLULAR_RESPONSE_TO_TOXIC_SUBSTANCE | 0.00050 | 0.00763 | 0.47727 | 0.66616 | 1.32354 | 65 |
| GO_CELLULAR_TRANSITION_METAL_ION_HOMEOSTASIS | 0.00290 | 0.03039 | 0.43171 | 0.63351 | 1.26027 | 66 |
| GO_CELL_ACTIVATION_INVOLVED_IN_IMMUNE_RESPONSE | 0.00000 | 0.00000 | | 0.61678 | 1.28215 | 408 |
| GO_CELL_ADHESION_MEDIATOR_ACTIVITY | 0.00141 | 0.01753 | 0.45506 | 0.72061 | 1.37935 | 31 |
| GO_CELL_ADHESION_MOLECULE_BINDING | 0.00000 | 0.00000 | | 0.64454 | 1.33783 | 343 |
| GO_CELL_BODY | 0.00011 | 0.00228 | 0.53843 | 0.57549 | 1.19103 | 283 |
| GO_CELL_CELL_JUNCTION | 0.00006 | 0.00133 | 0.55733 | 0.57948 | 1.19929 | 283 |
| GO_CELL_CELL_JUNCTION_ASSEMBLY | 0.00176 | 0.02077 | 0.45506 | 0.61992 | 1.24522 | 85 |
| GO_CELL_CELL_JUNCTION_ORGANIZATION | 0.00108 | 0.01426 | 0.45506 | 0.60677 | 1.22975 | 118 |
| GO_CELL_CELL_RECOGNITION | 0.00000 | 0.00002 | 0.65944 | 0.83362 | 1.59251 | 29 |
| GO_CELL_CELL_SIGNALING_BY_WNT | 0.00000 | 0.00003 | 0.65944 | 0.59200 | 1.22709 | 315 |
| GO_CELL_CORTEX | 0.00117 | 0.01508 | 0.45506 | 0.57010 | 1.17014 | 196 |
| GO_CELL_CORTEX_REGION | 0.00383 | 0.03700 | 0.43171 | 0.69768 | 1.33546 | 31 |
| GO_CELL_CYCLE_G2_M_PHASE_TRANSITION | 0.00163 | 0.01971 | 0.45506 | 0.56221 | 1.15888 | 229 |
| GO_CELL_CYCLE_PHASE_TRANSITION | 0.00044 | 0.00693 | 0.49849 | 0.53955 | 1.12468 | 498 |
| GO_CELL_DIVISION | 0.00509 | 0.04618 | 0.40702 | 0.53116 | 1.10589 | 444 |
| GO_CELL_KILLING | 0.00052 | 0.00785 | 0.47727 | 0.68128 | 1.34117 | 51 |
| GO_CELL_LEADING_EDGE | 0.00025 | 0.00431 | 0.49849 | 0.56642 | 1.17078 | 269 |
| GO_CELL_PROJECTION_MEMBRANE | 0.00004 | 0.00101 | 0.55733 | 0.60553 | 1.23870 | 175 |
| GO_CELL_RECOGNITION | 0.00000 | 0.00001 | 0.67496 | 0.74883 | 1.48207 | 57 |
| GO_CELL_REDOX_HOMEOSTASIS | 0.00130 | 0.01646 | 0.45506 | 0.70302 | 1.36356 | 38 |
| GO_CELL_SUBSTRATE_JUNCTION | 0.00000 | 0.00000 | | 0.64873 | 1.34399 | 304 |
| GO_CELL_SURFACE | 0.00000 | 0.00000 | 0.76146 | 0.61828 | 1.27985 | 286 |
| GO_CELL_SURFACE_RECEPTOR_SIGNALING_PATHWAY_INVC | 0.00001 | 0.00036 | 0.59333 | 0.57463 | 1.19302 | 348 |
| GO_CEREBELLAR_PURKINJE_CELL_LAYER_DEVELOPMENT | 0.00083 | 0.01153 | 0.47727 | 0.81566 | 1.49754 | 16 |
| GO_CHAPERONE_COMPLEX | 0.00000 | 0.00013 | 0.61053 | 0.85792 | 1.60776 | 22 |
| GO_CHAPERONE_MEDIATED_PROTEIN_FOLDING | 0.00350 | 0.03495 | 0.43171 | 0.66662 | 1.30198 | 44 |
| GO_CLUSTER_OF_ACTIN_BASED_CELL_PROJECTIONS | 0.00000 | 0.00004 | 0.64355 | 0.68755 | 1.38264 | 89 |
| GO_COLLAGEN_CONTAINING_EXTRACELLULAR_MATRIX | 0.00000 | 0.00003 | 0.65944 | 0.65577 | 1.33420 | 133 |
| GO_CONTRACTILE_FIBER | 0.00000 | 0.00006 | 0.64355 | 0.65305 | 1.32792 | 131 |
| GO_COPI_COATED_VESICLE | 0.00383 | 0.03700 | 0.43171 | 0.72313 | 1.36648 | 25 |
| GO_CORTICAL_CYTOSKELETON | 0.00020 | 0.00368 | 0.51885 | 0.65967 | 1.31704 | 71 |
| GO_COSTAMERE | 0.00150 | 0.01845 | 0.45506 | 0.79413 | 1.45802 | 16 |
| GO_COTRANSLATIONAL_PROTEIN_TARGETING_TO_MEMBR | 0.00000 | 0.00000 | | 0.77839 | 1.56774 | 95 |
| GO_CYTOCHROME_COMPLEX | 0.00003 | 0.00075 | 0.57561 | 0.82878 | 1.55315 | 22 |
| GO_CYTOKINE_MEDIATED_SIGNALING_PATHWAY | 0.00000 | 0.00000 | 0.74774 | 0.59961 | 1.24538 | 378 |
| GO_CYTOPLASMIC_STRESS_GRANULE | 0.00302 | 0.03128 | 0.43171 | 0.63903 | 1.26862 | 62 |
| GO_CYTOPLASMIC_TRANSLATION | 0.00000 | 0.00000 | 0.74774 | 0.72004 | 1.44799 | 89 |

| Cluster 1 | | | | | | |
|---|---|---|---|---|---|---|
| GO_CYTOPLASMIC_TRANSLATIONAL_INITIATION | 0.00035 | 0.00575 | 0.49849 | 0.74642 | 1.42875 | 31 |
| GO_CYTOSOLIC_LARGE_RIBOSOMAL_SUBUNIT | 0.00000 | 0.00000 | 0.73376 | 0.79029 | 1.55787 | 52 |
| GO_CYTOSOLIC_RIBOSOME | 0.00000 | 0.00000 | | 0.76381 | 1.53973 | 97 |
| GO_CYTOSOLIC_SMALL_RIBOSOMAL_SUBUNIT | 0.00001 | 0.00023 | 0.61053 | 0.76275 | 1.48622 | 41 |
| GO_DEFENSE_RESPONSE_TO_OTHER_ORGANISM | 0.00000 | 0.00010 | 0.62726 | 0.56531 | 1.17755 | 471 |
| GO_DETECTION_OF_STIMULUS | 0.00002 | 0.00051 | 0.57561 | 0.65091 | 1.31609 | 105 |
| GO_DETOXIFICATION | 0.00012 | 0.00247 | 0.51885 | 0.67306 | 1.34063 | 68 |
| GO_DEVELOPMENT_OF_PRIMARY_SEXUAL_CHARACTERISTI | 0.00070 | 0.01005 | 0.47727 | 0.61168 | 1.23912 | 116 |
| GO_DICARBOXYLIC_ACID_METABOLIC_PROCESS | 0.00012 | 0.00241 | 0.53843 | 0.68431 | 1.35524 | 59 |
| GO_DISORDERED_DOMAIN_SPECIFIC_BINDING | 0.00023 | 0.00403 | 0.51885 | 0.75752 | 1.44712 | 29 |
| GO_DNA_BIOSYNTHETIC_PROCESS | 0.00014 | 0.00278 | 0.51885 | 0.60767 | 1.24004 | 149 |
| GO_DNA_DAMAGE_RESPONSE_DETECTION_OF_DNA_DAM/ | 0.00045 | 0.00700 | 0.49849 | 0.72607 | 1.40266 | 36 |
| GO_DNA_GEOMETRIC_CHANGE | 0.00036 | 0.00587 | 0.49849 | 0.62107 | 1.25595 | 106 |
| GO_DOUBLE_STRANDED_RNA_BINDING | 0.00028 | 0.00472 | 0.49849 | 0.68326 | 1.35354 | 58 |
| GO_ELECTRON_TRANSFER_ACTIVITY | 0.00000 | 0.00000 | 0.70498 | 0.72229 | 1.44754 | 77 |
| GO_ELECTRON_TRANSPORT_CHAIN | 0.00000 | 0.00000 | | 0.73597 | 1.49653 | 131 |
| GO_EMBRYO_IMPLANTATION | 0.00141 | 0.01753 | 0.45506 | 0.72045 | 1.37904 | 31 |
| GO_ENDOCYTIC_VESICLE_MEMBRANE | 0.00342 | 0.03444 | 0.43171 | 0.60844 | 1.22493 | 94 |
| GO_ENDOPEPTIDASE_ACTIVITY | 0.00034 | 0.00562 | 0.49849 | 0.58111 | 1.19214 | 193 |
| GO_ENDOPEPTIDASE_COMPLEX | 0.00000 | 0.00001 | 0.69013 | 0.75233 | 1.49290 | 63 |
| GO_ENDOPEPTIDASE_REGULATOR_ACTIVITY | 0.00023 | 0.00413 | 0.51885 | 0.67522 | 1.33637 | 57 |
| GO_ENERGY_DERIVATION_BY_OXIDATION_OF_ORGANIC_CC | 0.00000 | 0.00000 | | 0.69792 | 1.43249 | 196 |
| GO_ENZYME_INHIBITOR_ACTIVITY | 0.00006 | 0.00140 | 0.55733 | 0.60619 | 1.23880 | 170 |
| GO_EPITHELIAL_CELL_DIFFERENTIATION | 0.00102 | 0.01371 | 0.45506 | 0.55532 | 1.15102 | 319 |
| GO_ESTABLISHMENT_OF_PROTEIN_LOCALIZATION_TO_CHR | 0.00013 | 0.00253 | 0.51885 | 0.79913 | 1.50264 | 23 |
| GO_ESTABLISHMENT_OF_PROTEIN_LOCALIZATION_TO_END | 0.00000 | 0.00000 | | 0.76433 | 1.54566 | 106 |
| GO_ESTABLISHMENT_OF_PROTEIN_LOCALIZATION_TO_MEN | 0.00000 | 0.00000 | | 0.65340 | 1.35029 | 263 |
| GO_ESTABLISHMENT_OF_PROTEIN_LOCALIZATION_TO_ORG | 0.00000 | 0.00000 | | 0.64219 | 1.33724 | 447 |
| GO_ESTABLISHMENT_OF_PROTEIN_LOCALIZATION_TO_TELC | 0.00002 | 0.00047 | 0.57561 | 0.86251 | 1.59201 | 18 |
| GO_ESTABLISHMENT_OF_RNA_LOCALIZATION | 0.00011 | 0.00224 | 0.53843 | 0.59862 | 1.22456 | 175 |
| GO_ESTABLISHMENT_OF_TISSUE_POLARITY | 0.00000 | 0.00000 | | 0.75409 | 1.51577 | 88 |
| GO_EUKARYOTIC_TRANSLATION_INITIATION_FACTOR_3_CO | 0.00350 | 0.03495 | 0.43171 | 0.78972 | 1.44312 | 15 |
| GO_EXCRETION | 0.00205 | 0.02309 | 0.43171 | 0.76446 | 1.43262 | 22 |
| GO_EXTERNAL_SIDE_OF_PLASMA_MEMBRANE | 0.00053 | 0.00797 | 0.47727 | 0.63754 | 1.28257 | 90 |
| GO_EXTRACELLULAR_MATRIX | 0.00076 | 0.01068 | 0.47727 | 0.58691 | 1.19949 | 171 |
| GO_FATTY_ACID_DERIVATIVE_METABOLIC_PROCESS | 0.00327 | 0.03317 | 0.43171 | 0.62525 | 1.25160 | 75 |
| GO_FC_EPSILON_RECEPTOR_SIGNALING_PATHWAY | 0.00000 | 0.00001 | 0.69013 | 0.69941 | 1.40651 | 89 |
| GO_FC_RECEPTOR_SIGNALING_PATHWAY | 0.00000 | 0.00000 | 0.70498 | 0.66395 | 1.35174 | 135 |
| GO_FERTILIZATION | 0.00008 | 0.00175 | 0.53843 | 0.68696 | 1.36367 | 64 |
| GO_FICOLIN_1_RICH_GRANULE | 0.00000 | 0.00000 | 0.77494 | 0.71432 | 1.44289 | 100 |
| GO_FOREBRAIN_GENERATION_OF_NEURONS | 0.00320 | 0.03275 | 0.43171 | 0.74213 | 1.39546 | 23 |
| GO_FOREBRAIN_NEURON_DIFFERENTIATION | 0.00390 | 0.03748 | 0.43171 | 0.77291 | 1.41906 | 16 |
| GO_GABA_ERGIC_SYNAPSE | 0.00532 | 0.04792 | 0.40702 | 0.71219 | 1.35134 | 26 |
| GO_GENERATION_OF_PRECURSOR_METABOLITES_AND_EN | 0.00000 | 0.00000 | | 0.65978 | 1.36996 | 355 |
| GO_GLUCOSE_CATABOLIC_PROCESS | 0.00067 | 0.00977 | 0.47727 | 0.73930 | 1.40977 | 28 |
| GO_GLUTAMATERGIC_SYNAPSE | 0.00117 | 0.01508 | 0.45506 | 0.57650 | 1.18008 | 183 |
| GO_GLYCOLYTIC_PROCESS_THROUGH_FRUCTOSE_6_PHOSP | 0.00420 | 0.03976 | 0.40702 | 0.74525 | 1.40532 | 24 |
| GO_GTPASE_ACTIVITY | 0.00025 | 0.00431 | 0.49849 | 0.58732 | 1.20596 | 200 |
| GO_GUANYL_NUCLEOTIDE_BINDING | 0.00015 | 0.00286 | 0.51885 | 0.57638 | 1.18962 | 246 |
| GO_HEAT_SHOCK_PROTEIN_BINDING | 0.00465 | 0.04324 | 0.40702 | 0.60896 | 1.22348 | 87 |
| GO_HEMATOPOIETIC_PROGENITOR_CELL_DIFFERENTIATION | 0.00000 | 0.00000 | 0.78819 | 0.69842 | 1.41761 | 126 |
| GO_HEMATOPOIETIC_STEM_CELL_DIFFERENTIATION | 0.00000 | 0.00000 | 0.74774 | 0.75889 | 1.51593 | 72 |
| GO_HUMORAL_IMMUNE_RESPONSE | 0.00106 | 0.01414 | 0.45506 | 0.66645 | 1.31561 | 53 |
| GO_HYDROGEN_PEROXIDE_METABOLIC_PROCESS | 0.00073 | 0.01037 | 0.47727 | 0.75377 | 1.42439 | 25 |
| GO_IMMUNE_RESPONSE_REGULATING_SIGNALING_PATHW | 0.00000 | 0.00005 | 0.64355 | 0.61527 | 1.26677 | 219 |
| GO_IMPORT_ACROSS_PLASMA_MEMBRANE | 0.00115 | 0.01497 | 0.45506 | 0.65437 | 1.30010 | 65 |
| GO_IMPORT_INTO_CELL | 0.00128 | 0.01630 | 0.45506 | 0.61976 | 1.24771 | 94 |
| GO_IMPORT_INTO_NUCLEUS | 0.00038 | 0.00614 | 0.49849 | 0.61135 | 1.24143 | 127 |
| GO_INFLAMMATORY_RESPONSE_TO_ANTIGENIC_STIMULUS | 0.00357 | 0.03540 | 0.43171 | 0.74895 | 1.41230 | 24 |
| GO_INNATE_IMMUNE_RESPONSE | 0.00007 | 0.00152 | 0.53843 | 0.55984 | 1.16298 | 385 |
| GO_INNATE_IMMUNE_RESPONSE_ACTIVATING_SIGNAL_TR/ | 0.00000 | 0.00001 | 0.67496 | 0.71953 | 1.44276 | 78 |
| GO_INNER_MITOCHONDRIAL_MEMBRANE_PROTEIN_COMP | 0.00000 | 0.00000 | | 0.77093 | 1.55145 | 93 |
| GO_INORGANIC_ION_TRANSMEMBRANE_TRANSPORT | 0.00005 | 0.00111 | 0.55733 | 0.56348 | 1.17033 | 359 |
| GO_INTERACTION_WITH_HOST | 0.00137 | 0.01718 | 0.45506 | 0.59642 | 1.21496 | 138 |
| GO_INTERACTION_WITH_SYMBIONT | 0.00198 | 0.02255 | 0.43171 | 0.66448 | 1.30986 | 52 |
| GO_INTERLEUKIN_1_MEDIATED_SIGNALING_PATHWAY | 0.00000 | 0.00000 | 0.80122 | 0.74754 | 1.50119 | 84 |
| GO_INTERMEDIATE_FILAMENT_CYTOSKELETON | 0.00405 | 0.03878 | 0.40702 | 0.64567 | 1.28124 | 63 |
| GO_INTRAMOLECULAR_OXIDOREDUCTASE_ACTIVITY | 0.00106 | 0.01414 | 0.45506 | 0.71951 | 1.37979 | 32 |
| GO_INTRINSIC_APOPTOTIC_SIGNALING_PATHWAY | 0.00152 | 0.01864 | 0.45506 | 0.57330 | 1.17644 | 194 |
| GO_ION_CHANNEL_BINDING | 0.00084 | 0.01161 | 0.47727 | 0.64940 | 1.29723 | 72 |
| GO_ION_HOMEOSTASIS | 0.00035 | 0.00581 | 0.49849 | 0.55451 | 1.15165 | 358 |
| GO_ION_TRANSMEMBRANE_TRANSPORT | 0.00043 | 0.00684 | 0.49849 | 0.53802 | 1.12147 | 499 |
| GO_ION_TRANSMEMBRANE_TRANSPORTER_ACTIVITY | 0.00253 | 0.02718 | 0.43171 | 0.54475 | 1.13088 | 349 |

| Cluster 1 | | | | | | |
|---|---|---|---|---|---|---|
| GO_IRON_ION_HOMEOSTASIS | 0.00235 | 0.02561 | 0.43171 | 0.65349 | 1.29004 | 53 |
| GO_IRON_ION_TRANSPORT | 0.00174 | 0.02064 | 0.45506 | 0.66129 | 1.30182 | 51 |
| GO_ISOMERASE_ACTIVITY | 0.00170 | 0.02040 | 0.45506 | 0.60073 | 1.21581 | 111 |
| GO_I_BAND | 0.00163 | 0.01971 | 0.45506 | 0.63842 | 1.27621 | 73 |
| GO_KIDNEY_EPITHELIUM_DEVELOPMENT | 0.00390 | 0.03748 | 0.43171 | 0.63613 | 1.25983 | 59 |
| GO_LARGE_RIBOSOMAL_SUBUNIT | 0.00000 | 0.00000 | | 0.78273 | 1.58276 | 107 |
| GO_LEUKOCYTE_MEDIATED_IMMUNITY | 0.00000 | 0.00000 | | 0.62267 | 1.29598 | 432 |
| GO_LIPOPROTEIN_PARTICLE_RECEPTOR_BINDING | 0.00003 | 0.00070 | 0.57561 | 0.87551 | 1.59988 | 15 |
| GO_LOW_DENSITY_LIPOPROTEIN_PARTICLE_CLEARANCE | 0.00268 | 0.02851 | 0.43171 | 0.72656 | 1.37861 | 26 |
| GO_LOW_DENSITY_LIPOPROTEIN_RECEPTOR_PARTICLE_ME | 0.00324 | 0.03296 | 0.43171 | 0.77112 | 1.42333 | 18 |
| GO_LYASE_ACTIVITY | 0.00082 | 0.01143 | 0.47727 | 0.60603 | 1.22841 | 121 |
| GO_LYSOSOMAL_LUMEN | 0.00191 | 0.02200 | 0.45506 | 0.65466 | 1.29688 | 58 |
| GO_MALE_SEX_DIFFERENTIATION | 0.00023 | 0.00405 | 0.51885 | 0.64635 | 1.29860 | 87 |
| GO_MATURATION_OF_LSU_RRNA | 0.00457 | 0.04261 | 0.40702 | 0.71395 | 1.35998 | 27 |
| GO_MEMBRANE_REGION | 0.00000 | 0.00002 | 0.67496 | 0.63263 | 1.29498 | 183 |
| GO_MESONEPHRIC_TUBULE_MORPHOGENESIS | 0.00539 | 0.04851 | 0.40702 | 0.70942 | 1.35135 | 27 |
| GO_MESONEPHROS_DEVELOPMENT | 0.00030 | 0.00510 | 0.49849 | 0.71261 | 1.38853 | 41 |
| GO_METALLOPEPTIDASE_ACTIVITY | 0.00111 | 0.01459 | 0.45506 | 0.62735 | 1.25948 | 83 |
| GO_METAL_ION_HOMEOSTASIS | 0.00045 | 0.00706 | 0.49849 | 0.56655 | 1.17066 | 265 |
| GO_METAL_ION_TRANSPORT | 0.00198 | 0.02255 | 0.43171 | 0.53940 | 1.12045 | 382 |
| GO_MICROVILLUS | 0.00062 | 0.00917 | 0.47727 | 0.68902 | 1.35045 | 48 |
| GO_MICROVILLUS_MEMBRANE | 0.00003 | 0.00067 | 0.57561 | 0.87660 | 1.60188 | 15 |
| GO_MIDBRAIN_DEVELOPMENT | 0.00183 | 0.02139 | 0.45506 | 0.65481 | 1.29423 | 54 |
| GO_MITOCHONDRIAL_ELECTRON_TRANSPORT_NADH_TO_U | 0.00000 | 0.00000 | 0.77494 | 0.82436 | 1.61160 | 45 |
| GO_MITOCHONDRIAL_GENE_EXPRESSION | 0.00000 | 0.00000 | 0.82666 | 0.68169 | 1.39127 | 153 |
| GO_MITOCHONDRIAL_LARGE_RIBOSOMAL_SUBUNIT | 0.00000 | 0.00000 | 0.71951 | 0.77359 | 1.52901 | 54 |
| GO_MITOCHONDRIAL_MATRIX | 0.00000 | 0.00000 | | 0.62396 | 1.29633 | 389 |
| GO_MITOCHONDRIAL_MEMBRANE_ORGANIZATION | 0.00242 | 0.02622 | 0.43171 | 0.61211 | 1.23232 | 94 |
| GO_MITOCHONDRIAL_PROTEIN_COMPLEX | 0.00000 | 0.00000 | | 0.74886 | 1.54060 | 213 |
| GO_MITOCHONDRIAL_RESPIRATORY_CHAIN_COMPLEX_ASS | 0.00001 | 0.00027 | 0.59333 | 0.67552 | 1.35721 | 87 |
| GO_MITOCHONDRIAL_SMALL_RIBOSOMAL_SUBUNIT | 0.00037 | 0.00595 | 0.49849 | 0.75010 | 1.43036 | 28 |
| GO_MITOCHONDRIAL_TRANSLATION | 0.00000 | 0.00000 | | 0.70692 | 1.43723 | 130 |
| GO_MITOCHONDRIAL_TRANSLATIONAL_TERMINATION | 0.00000 | 0.00000 | | 0.75353 | 1.51465 | 88 |
| GO_MITOCHONDRIAL_TRANSPORT | 0.00025 | 0.00431 | 0.49849 | 0.58573 | 1.20196 | 194 |
| GO_MITOCHONDRION_ORGANIZATION | 0.00000 | 0.00000 | 0.77494 | 0.59662 | 1.24062 | 414 |
| GO_MODIFIED_AMINO_ACID_BINDING | 0.00000 | 0.00009 | 0.62726 | 0.75471 | 1.48083 | 49 |
| GO_MODULATION_BY_HOST_OF_VIRAL_PROCESS | 0.00224 | 0.02484 | 0.43171 | 0.76489 | 1.41617 | 19 |
| GO_MONOCARBOXYLIC_ACID_METABOLIC_PROCESS | 0.00020 | 0.00365 | 0.51885 | 0.55245 | 1.14755 | 382 |
| GO_MONOSACCHARIDE_BIOSYNTHETIC_PROCESS | 0.00029 | 0.00497 | 0.49849 | 0.67188 | 1.33382 | 62 |
| GO_MONOVALENT_INORGANIC_CATION_TRANSMEMBRAN | 0.00002 | 0.00048 | 0.57561 | 0.64073 | 1.29873 | 120 |
| GO_MONOVALENT_INORGANIC_CATION_TRANSPORT | 0.00001 | 0.00024 | 0.59333 | 0.60198 | 1.23934 | 218 |
| GO_MORPHOGENESIS_OF_AN_EPITHELIUM | 0.00000 | 0.00000 | 0.82666 | 0.62762 | 1.30077 | 312 |
| GO_MORPHOGENESIS_OF_A_POLARIZED_EPITHELIUM | 0.00000 | 0.00000 | | 0.75605 | 1.52779 | 102 |
| GO_MRNA_3_END_PROCESSING | 0.00353 | 0.03526 | 0.43171 | 0.60932 | 1.22534 | 89 |
| GO_MRNA_5_UTR_BINDING | 0.00002 | 0.00067 | 0.57561 | 0.81490 | 1.53991 | 25 |
| GO_MRNA_BINDING | 0.00000 | 0.00000 | 0.83909 | 0.64531 | 1.33060 | 237 |
| GO_MRNA_EXPORT_FROM_NUCLEUS | 0.00187 | 0.02165 | 0.45506 | 0.61204 | 1.23689 | 101 |
| GO_MRNA_PROCESSING | 0.00000 | 0.00000 | 0.82666 | 0.60342 | 1.25635 | 438 |
| GO_MRNA_TRANSPORT | 0.00172 | 0.02050 | 0.45506 | 0.59884 | 1.21769 | 131 |
| GO_MULTI_MULTICELLULAR_ORGANISM_PROCESS | 0.00010 | 0.00219 | 0.53843 | 0.63874 | 1.29074 | 102 |
| GO_MULTI_ORGANISM_LOCALIZATION | 0.00428 | 0.04034 | 0.40702 | 0.63852 | 1.26191 | 55 |
| GO_MUSCLE_CELL_DEVELOPMENT | 0.00097 | 0.01303 | 0.47727 | 0.63091 | 1.26650 | 81 |
| GO_MUSCLE_CONTRACTION | 0.00457 | 0.04261 | 0.40702 | 0.56598 | 1.15634 | 167 |
| GO_MYELOID_LEUKOCYTE_ACTIVATION | 0.00000 | 0.00000 | | 0.63103 | 1.31072 | 380 |
| GO_MYELOID_LEUKOCYTE_MEDIATED_IMMUNITY | 0.00000 | 0.00000 | | 0.63662 | 1.32152 | 345 |
| GO_MYOSIN_BINDING | 0.00191 | 0.02200 | 0.45506 | 0.67551 | 1.31905 | 46 |
| GO_NADH_DEHYDROGENASE_ACTIVITY | 0.00000 | 0.00000 | 0.74774 | 0.83200 | 1.61049 | 37 |
| GO_NADH_DEHYDROGENASE_COMPLEX | 0.00000 | 0.00000 | 0.73376 | 0.81655 | 1.58799 | 40 |
| GO_NADH_DEHYDROGENASE_COMPLEX_ASSEMBLY | 0.00000 | 0.00007 | 0.64355 | 0.73372 | 1.45110 | 56 |
| GO_NADH_METABOLIC_PROCESS | 0.00001 | 0.00041 | 0.59333 | 0.78658 | 1.50842 | 32 |
| GO_NAD_METABOLIC_PROCESS | 0.00016 | 0.00307 | 0.51885 | 0.73338 | 1.42243 | 38 |
| GO_NATURAL_KILLER_CELL_MEDIATED_IMMUNITY | 0.00420 | 0.03976 | 0.40702 | 0.73492 | 1.38191 | 23 |
| GO_NCRNA_METABOLIC_PROCESS | 0.00000 | 0.00002 | 0.67496 | 0.57980 | 1.20497 | 397 |
| GO_NCRNA_PROCESSING | 0.00000 | 0.00002 | 0.67496 | 0.59077 | 1.22622 | 343 |
| GO_NEGATIVE_REGULATION_OF_APOPTOTIC_SIGNALING_P | 0.00509 | 0.04618 | 0.40702 | 0.58642 | 1.19649 | 145 |
| GO_NEGATIVE_REGULATION_OF_CANONICAL_WNT_SIGNAL | 0.00000 | 0.00000 | 0.71951 | 0.69589 | 1.40689 | 109 |
| GO_NEGATIVE_REGULATION_OF_CATALYTIC_ACTIVITY | 0.00002 | 0.00054 | 0.57561 | 0.56024 | 1.16591 | 428 |
| GO_NEGATIVE_REGULATION_OF_CELLULAR_COMPONENT_ | 0.00517 | 0.04664 | 0.40702 | 0.57149 | 1.16663 | 157 |
| GO_NEGATIVE_REGULATION_OF_CELL_CYCLE | 0.00161 | 0.01961 | 0.45506 | 0.53628 | 1.11671 | 451 |
| GO_NEGATIVE_REGULATION_OF_CELL_CYCLE_G2_M_PHASE | 0.00000 | 0.00005 | 0.64355 | 0.68578 | 1.37908 | 89 |
| GO_NEGATIVE_REGULATION_OF_CELL_CYCLE_PHASE_TRAN | 0.00037 | 0.00595 | 0.49849 | 0.58476 | 1.20070 | 200 |
| GO_NEGATIVE_REGULATION_OF_CELL_CYCLE_PROCESS | 0.00016 | 0.00307 | 0.51885 | 0.57083 | 1.17982 | 268 |
| GO_NEGATIVE_REGULATION_OF_COAGULATION | 0.00372 | 0.03639 | 0.43171 | 0.77891 | 1.43457 | 17 |

| Cluster 1 | | | | | | |
|---|---|---|---|---|---|---|
| GO_NEGATIVE_REGULATION_OF_HYDROLASE_ACTIVITY | 0.00002 | 0.00050 | 0.57561 | 0.59490 | 1.22511 | 222 |
| GO_NEGATIVE_REGULATION_OF_MITOTIC_CELL_CYCLE | 0.00080 | 0.01123 | 0.47727 | 0.56497 | 1.16635 | 251 |
| GO_NEGATIVE_REGULATION_OF_MRNA_METABOLIC_PROC | 0.00320 | 0.03275 | 0.43171 | 0.63752 | 1.26561 | 62 |
| GO_NEGATIVE_REGULATION_OF_PEPTIDASE_ACTIVITY | 0.00065 | 0.00960 | 0.47727 | 0.62182 | 1.25472 | 98 |
| GO_NEGATIVE_REGULATION_OF_PROTEOLYSIS | 0.00257 | 0.02748 | 0.43171 | 0.57362 | 1.17194 | 167 |
| GO_NEGATIVE_REGULATION_OF_WNT_SIGNALING_PATHWA | 0.00000 | 0.00001 | 0.69013 | 0.67618 | 1.37060 | 120 |
| GO_NEURAL_CREST_CELL_DIFFERENTIATION | 0.00202 | 0.02280 | 0.43171 | 0.67990 | 1.32559 | 42 |
| GO_NEURAL_CREST_CELL_MIGRATION | 0.00027 | 0.00459 | 0.49849 | 0.75531 | 1.44030 | 28 |
| GO_NEURAL_NUCLEUS_DEVELOPMENT | 0.00179 | 0.02108 | 0.45506 | 0.70071 | 1.35367 | 36 |
| GO_NIK_NF_KAPPAB_SIGNALING | 0.00000 | 0.00000 | 0.81404 | 0.72077 | 1.45718 | 109 |
| GO_NON_CANONICAL_WNT_SIGNALING_PATHWAY | 0.00000 | 0.00000 | | 0.74231 | 1.50055 | 103 |
| GO_NUCLEAR_ENVELOPE | 0.00029 | 0.00491 | 0.49849 | 0.55744 | 1.15670 | 337 |
| GO_NUCLEAR_EXPORT | 0.00003 | 0.00070 | 0.57561 | 0.60480 | 1.23772 | 177 |
| GO_NUCLEAR_PERIPHERY | 0.00181 | 0.02122 | 0.45506 | 0.60572 | 1.22474 | 105 |
| GO_NUCLEAR_SPECK | 0.00150 | 0.01845 | 0.45506 | 0.55301 | 1.14574 | 307 |
| GO_NUCLEAR_TRANSCRIBED_MRNA_CATABOLIC_PROCESS | 0.00000 | 0.00000 | | 0.69234 | 1.41861 | 188 |
| GO_NUCLEAR_TRANSCRIBED_MRNA_CATABOLIC_PROCESS_ | 0.00000 | 0.00000 | | 0.75240 | 1.52325 | 115 |
| GO_NUCLEAR_TRANSPORT | 0.00000 | 0.00006 | 0.64355 | 0.59516 | 1.23184 | 287 |
| GO_NUCLEOBASE_CONTAINING_COMPOUND_TRANSPORT | 0.00029 | 0.00491 | 0.49849 | 0.58241 | 1.19843 | 215 |
| GO_NUCLEOBASE_CONTAINING_SMALL_MOLECULE_INTERC | 0.00320 | 0.03275 | 0.43171 | 0.74191 | 1.39504 | 23 |
| GO_NUCLEOBASE_CONTAINING_SMALL_MOLECULE_METAB | 0.00000 | 0.00009 | 0.62726 | 0.57084 | 1.18618 | 394 |
| GO_NUCLEOSIDE_DIPHOSPHATE_METABOLIC_PROCESS | 0.00113 | 0.01476 | 0.45506 | 0.60470 | 1.22385 | 111 |
| GO_NUCLEOSIDE_MONOPHOSPHATE_METABOLIC_PROCESS | 0.00361 | 0.03559 | 0.43171 | 0.64133 | 1.27188 | 60 |
| GO_NUCLEOSIDE_PHOSPHATE_BIOSYNTHETIC_PROCESS | 0.00001 | 0.00040 | 0.59333 | 0.62397 | 1.27390 | 158 |
| GO_NUCLEOSIDE_TRIPHOSPHATE_BIOSYNTHETIC_PROCESS | 0.00013 | 0.00262 | 0.51885 | 0.72849 | 1.42034 | 42 |
| GO_NUCLEOSIDE_TRIPHOSPHATE_METABOLIC_PROCESS | 0.00174 | 0.02064 | 0.45506 | 0.64858 | 1.28748 | 64 |
| GO_NUCLEOTIDE_EXCISION_REPAIR_DNA_INCISION | 0.00194 | 0.02225 | 0.45506 | 0.68954 | 1.33888 | 39 |
| GO_NUCLEOTIDE_PHOSPHORYLATION | 0.00040 | 0.00648 | 0.49849 | 0.62117 | 1.25218 | 97 |
| GO_ORGANELLAR_RIBOSOME | 0.00000 | 0.00000 | 0.81404 | 0.75491 | 1.51598 | 84 |
| GO_ORGANELLE_ENVELOPE_LUMEN | 0.00001 | 0.00018 | 0.61053 | 0.68818 | 1.37917 | 77 |
| GO_ORGANELLE_INNER_MEMBRANE | 0.00000 | 0.00000 | | 0.65827 | 1.36855 | 411 |
| GO_ORGANIC_CYCLIC_COMPOUND_CATABOLIC_PROCESS | 0.00000 | 0.00000 | | 0.65266 | 1.35951 | 471 |
| GO_ORGANIC_HYDROXY_COMPOUND_BIOSYNTHETIC_PRO | 0.00095 | 0.01283 | 0.47727 | 0.58674 | 1.19600 | 139 |
| GO_ORGANIC_HYDROXY_COMPOUND_METABOLIC_PROCES | 0.00265 | 0.02822 | 0.43171 | 0.55092 | 1.13979 | 278 |
| GO_ORGANOPHOSPHATE_BIOSYNTHETIC_PROCESS | 0.00327 | 0.03317 | 0.43171 | 0.53682 | 1.11507 | 381 |
| GO_OSTEOBLAST_DIFFERENTIATION | 0.00298 | 0.03101 | 0.43171 | 0.60023 | 1.21374 | 107 |
| GO_OXIDATIVE_PHOSPHORYLATION | 0.00000 | 0.00000 | | 0.79494 | 1.60476 | 99 |
| GO_OXIDOREDUCTASE_ACTIVITY | 0.00000 | 0.00000 | | 0.62767 | 1.30717 | 455 |
| GO_OXIDOREDUCTASE_ACTIVITY_ACTING_ON_A_HEME_GR | 0.00082 | 0.01143 | 0.47727 | 0.79236 | 1.46703 | 19 |
| GO_OXIDOREDUCTASE_ACTIVITY_ACTING_ON_CH_OH_GRO | 0.00133 | 0.01682 | 0.45506 | 0.64426 | 1.28695 | 72 |
| GO_OXIDOREDUCTASE_ACTIVITY_ACTING_ON_NAD_P_H | 0.00000 | 0.00000 | 0.80122 | 0.76181 | 1.52440 | 74 |
| GO_OXIDOREDUCTASE_ACTIVITY_ACTING_ON_NAD_P_H_Q | 0.00000 | 0.00000 | 0.74774 | 0.80252 | 1.56707 | 46 |
| GO_OXIDOREDUCTASE_ACTIVITY_ACTING_ON_PEROXIDE_A | 0.00007 | 0.00152 | 0.53843 | 0.79846 | 1.50883 | 25 |
| GO_OXIDOREDUCTASE_COMPLEX | 0.00000 | 0.00000 | 0.82666 | 0.76373 | 1.53262 | 80 |
| GO_PEPTIDASE_ACTIVITY | 0.00013 | 0.00253 | 0.51885 | 0.56926 | 1.17929 | 300 |
| GO_PEPTIDASE_COMPLEX | 0.00000 | 0.00000 | 0.71951 | 0.72409 | 1.45306 | 80 |
| GO_PEPTIDASE_REGULATOR_ACTIVITY | 0.00170 | 0.02040 | 0.45506 | 0.62291 | 1.24970 | 79 |
| GO_PEPTIDE_BINDING | 0.00007 | 0.00153 | 0.53843 | 0.62142 | 1.26722 | 140 |
| GO_PHOSPHATIDYLSERINE_BINDING | 0.00001 | 0.00028 | 0.59333 | 0.80762 | 1.54005 | 28 |
| GO_PIGMENT_GRANULE | 0.00000 | 0.00000 | 0.70498 | 0.71147 | 1.43011 | 88 |
| GO_PLASMA_LIPOPROTEIN_PARTICLE_CLEARANCE | 0.00017 | 0.00311 | 0.51885 | 0.74346 | 1.43470 | 35 |
| GO_PLATELET_DEGRANULATION | 0.00428 | 0.04034 | 0.40702 | 0.64003 | 1.27037 | 61 |
| GO_POLYSOMAL_RIBOSOME | 0.00000 | 0.00015 | 0.61053 | 0.80582 | 1.54241 | 30 |
| GO_POLYSOME | 0.00000 | 0.00002 | 0.65944 | 0.74593 | 1.48019 | 63 |
| GO_POSITIVE_REGULATION_OF_ACTIN_FILAMENT_POLYME | 0.00110 | 0.01441 | 0.45506 | 0.65309 | 1.29652 | 62 |
| GO_POSITIVE_REGULATION_OF_ATP_METABOLIC_PROCESS | 0.00122 | 0.01569 | 0.45506 | 0.74552 | 1.41458 | 26 |
| GO_POSITIVE_REGULATION_OF_CANONICAL_WNT_SIGNALI | 0.00000 | 0.00000 | 0.76146 | 0.71398 | 1.44221 | 100 |
| GO_POSITIVE_REGULATION_OF_CELLULAR_AMIDE_METABO | 0.00001 | 0.00019 | 0.61053 | 0.64625 | 1.31064 | 123 |
| GO_POSITIVE_REGULATION_OF_CELLULAR_COMPONENT_B | 0.00250 | 0.02688 | 0.43171 | 0.54663 | 1.13391 | 330 |
| GO_POSITIVE_REGULATION_OF_CELLULAR_PROTEIN_LOCAL | 0.00119 | 0.01525 | 0.45506 | 0.56676 | 1.16869 | 236 |
| GO_POSITIVE_REGULATION_OF_CELL_POPULATION_PROLIF | 0.00383 | 0.03700 | 0.43171 | 0.53252 | 1.10874 | 438 |
| GO_POSITIVE_REGULATION_OF_CELL_SUBSTRATE_ADHESIO | 0.00202 | 0.02280 | 0.43171 | 0.63934 | 1.27467 | 69 |
| GO_POSITIVE_REGULATION_OF_CYTOSKELETON_ORGANIZA | 0.00011 | 0.00224 | 0.53843 | 0.61767 | 1.26026 | 145 |
| GO_POSITIVE_REGULATION_OF_DEFENSE_RESPONSE | 0.00004 | 0.00091 | 0.55733 | 0.59955 | 1.22715 | 181 |
| GO_POSITIVE_REGULATION_OF_DNA_BIOSYNTHETIC_PROC | 0.00002 | 0.00054 | 0.57561 | 0.71311 | 1.41266 | 58 |
| GO_POSITIVE_REGULATION_OF_DNA_METABOLIC_PROCESS | 0.00051 | 0.00773 | 0.47727 | 0.60186 | 1.22749 | 141 |
| GO_POSITIVE_REGULATION_OF_ESTABLISHMENT_OF_PROT | 0.00073 | 0.01037 | 0.47727 | 0.56707 | 1.16949 | 239 |
| GO_POSITIVE_REGULATION_OF_FIBROBLAST_PROLIFERATIO | 0.00075 | 0.01060 | 0.47727 | 0.75308 | 1.42309 | 25 |
| GO_POSITIVE_REGULATION_OF_IMMUNE_RESPONSE | 0.00000 | 0.00001 | 0.70498 | 0.60339 | 1.25039 | 309 |
| GO_POSITIVE_REGULATION_OF_IMMUNE_SYSTEM_PROCES | 0.00000 | 0.00009 | 0.62726 | 0.56896 | 1.18481 | 449 |
| GO_POSITIVE_REGULATION_OF_INNATE_IMMUNE_RESPON | 0.00003 | 0.00077 | 0.55733 | 0.63637 | 1.29089 | 124 |
| GO_POSITIVE_REGULATION_OF_INTRACELLULAR_TRANSPO | 0.00390 | 0.03748 | 0.43171 | 0.57668 | 1.17763 | 159 |
| GO_POSITIVE_REGULATION_OF_ION_TRANSPORT | 0.00143 | 0.01769 | 0.45506 | 0.60723 | 1.22807 | 110 |

| Cluster 1 | | | | | | |
|---|---|---|---|---|---|---|
| GO_POSITIVE_REGULATION_OF_KIDNEY_DEVELOPMENT | 0.00001 | 0.00020 | 0.61053 | 0.87332 | 1.61197 | 18 |
| GO_POSITIVE_REGULATION_OF_MRNA_PROCESSING | 0.00213 | 0.02383 | 0.43171 | 0.73469 | 1.38834 | 25 |
| GO_POSITIVE_REGULATION_OF_MRNA_SPLICING_VIA_SPLI | 0.00239 | 0.02592 | 0.43171 | 0.78854 | 1.45230 | 17 |
| GO_POSITIVE_REGULATION_OF_NITRIC_OXIDE_METABOLIC | 0.00324 | 0.03296 | 0.43171 | 0.77826 | 1.42889 | 16 |
| GO_POSITIVE_REGULATION_OF_ORGANELLE_ORGANIZATIO | 0.00008 | 0.00178 | 0.53843 | 0.55563 | 1.15693 | 439 |
| GO_POSITIVE_REGULATION_OF_PROTEIN_CONTAINING_CO | 0.00042 | 0.00664 | 0.49849 | 0.59113 | 1.20801 | 170 |
| GO_POSITIVE_REGULATION_OF_PROTEIN_LOCALIZATION_T | 0.00004 | 0.00100 | 0.55733 | 0.70570 | 1.39799 | 58 |
| GO_POSITIVE_REGULATION_OF_PROTEIN_POLYMERIZATION | 0.00018 | 0.00339 | 0.51885 | 0.64208 | 1.29121 | 89 |
| GO_POSITIVE_REGULATION_OF_REACTIVE_OXYGEN_SPECIE | 0.00067 | 0.00977 | 0.47727 | 0.78203 | 1.46555 | 22 |
| GO_POSITIVE_REGULATION_OF_RESPONSE_TO_BIOTIC_STI | 0.00008 | 0.00176 | 0.53843 | 0.61368 | 1.25204 | 143 |
| GO_POSITIVE_REGULATION_OF_RESPONSE_TO_EXTERNAL_ | 0.00006 | 0.00138 | 0.55733 | 0.58112 | 1.19815 | 238 |
| GO_POSITIVE_REGULATION_OF_RNA_SPLICING | 0.00021 | 0.00379 | 0.51885 | 0.77523 | 1.47672 | 27 |
| GO_POSITIVE_REGULATION_OF_SUBSTRATE_ADHESION_DE | 0.00128 | 0.01630 | 0.45506 | 0.72277 | 1.38074 | 29 |
| GO_POSITIVE_REGULATION_OF_SUPRAMOLECULAR_FIBER_ | 0.00007 | 0.00150 | 0.53843 | 0.62667 | 1.27427 | 131 |
| GO_POSITIVE_REGULATION_OF_TELOMERASE_ACTIVITY | 0.00050 | 0.00763 | 0.47727 | 0.74072 | 1.41784 | 31 |
| GO_POSITIVE_REGULATION_OF_TELOMERASE_RNA_LOCALI | 0.00002 | 0.00054 | 0.57561 | 0.87975 | 1.60763 | 15 |
| GO_POSITIVE_REGULATION_OF_TELOMERE_MAINTENANCE | 0.00003 | 0.00076 | 0.57561 | 0.72907 | 1.42365 | 46 |
| GO_POSITIVE_REGULATION_OF_TELOMERE_MAINTENANCE | 0.00000 | 0.00007 | 0.62726 | 0.80496 | 1.55340 | 35 |
| GO_POSITIVE_REGULATION_OF_TRANSLATION | 0.00003 | 0.00069 | 0.57561 | 0.65573 | 1.32553 | 103 |
| GO_POSITIVE_REGULATION_OF_TRANSPORT | 0.00051 | 0.00773 | 0.47727 | 0.54116 | 1.12752 | 485 |
| GO_POSITIVE_REGULATION_OF_VIRAL_PROCESS | 0.00036 | 0.00593 | 0.49849 | 0.63949 | 1.28697 | 92 |
| GO_POSITIVE_REGULATION_OF_WNT_SIGNALING_PATHWA | 0.00000 | 0.00003 | 0.65944 | 0.67027 | 1.35862 | 119 |
| GO_POSTSYNAPSE | 0.00022 | 0.00396 | 0.51885 | 0.56428 | 1.16933 | 309 |
| GO_POSTTRANSCRIPTIONAL_REGULATION_OF_GENE_EXPR | 0.00000 | 0.00000 | | 0.63974 | 1.33305 | 487 |
| GO_POST_TRANSLATIONAL_PROTEIN_MODIFICATION | 0.00006 | 0.00142 | 0.53843 | 0.58292 | 1.20349 | 253 |
| GO_PRECATALYTIC_SPLICEOSOME | 0.00062 | 0.00917 | 0.47727 | 0.67789 | 1.33450 | 51 |
| GO_PRERIBOSOME | 0.00024 | 0.00420 | 0.51885 | 0.66312 | 1.32281 | 70 |
| GO_PRERIBOSOME_LARGE_SUBUNIT_PRECURSOR | 0.00487 | 0.04490 | 0.40702 | 0.74329 | 1.39295 | 22 |
| GO_PROTEASOMAL_PROTEIN_CATABOLIC_PROCESS | 0.00130 | 0.01646 | 0.45506 | 0.54376 | 1.12965 | 386 |
| GO_PROTEASOMAL_UBIQUITIN_INDEPENDENT_PROTEIN_C | 0.00000 | 0.00010 | 0.62726 | 0.88292 | 1.62968 | 18 |
| GO_PROTEASOME_ACCESSORY_COMPLEX | 0.00004 | 0.00108 | 0.55733 | 0.82750 | 1.56043 | 24 |
| GO_PROTEASOME_CORE_COMPLEX | 0.00000 | 0.00008 | 0.62726 | 0.89154 | 1.64200 | 17 |
| GO_PROTEIN_CONTAINING_COMPLEX_DISASSEMBLY | 0.00000 | 0.00000 | | 0.64807 | 1.33910 | 265 |
| GO_PROTEIN_CONTAINING_COMPLEX_LOCALIZATION | 0.00049 | 0.00753 | 0.47727 | 0.57373 | 1.18160 | 223 |
| GO_PROTEIN_DOMAIN_SPECIFIC_BINDING | 0.00065 | 0.00960 | 0.47727 | 0.54058 | 1.12605 | 477 |
| GO_PROTEIN_FOLDING | 0.00001 | 0.00019 | 0.61053 | 0.62540 | 1.27774 | 166 |
| GO_PROTEIN_FOLDING_CHAPERONE | 0.00024 | 0.00427 | 0.51885 | 0.82287 | 1.51884 | 18 |
| GO_PROTEIN_IMPORT | 0.00000 | 0.00007 | 0.64355 | 0.63311 | 1.29194 | 155 |
| GO_PROTEIN_LIPID_COMPLEX_ASSEMBLY | 0.00176 | 0.02077 | 0.45506 | 0.77649 | 1.43764 | 19 |
| GO_PROTEIN_LIPID_COMPLEX_SUBUNIT_ORGANIZATION | 0.00390 | 0.03748 | 0.43171 | 0.73618 | 1.38427 | 23 |
| GO_PROTEIN_LOCALIZATION_TO_CHROMOSOME_TELOMER | 0.00002 | 0.00063 | 0.57561 | 0.79110 | 1.51127 | 29 |
| GO_PROTEIN_LOCALIZATION_TO_ENDOPLASMIC_RETICULU | 0.00000 | 0.00000 | | 0.73718 | 1.49747 | 128 |
| GO_PROTEIN_LOCALIZATION_TO_MEMBRANE | 0.00000 | 0.00000 | 0.80122 | 0.59508 | 1.23925 | 458 |
| GO_PROTEIN_LOCALIZATION_TO_MITOCHONDRION | 0.00002 | 0.00046 | 0.57561 | 0.64244 | 1.30049 | 112 |
| GO_PROTEIN_LOCALIZATION_TO_NUCLEUS | 0.00001 | 0.00021 | 0.61053 | 0.60952 | 1.25150 | 202 |
| GO_PROTEIN_MODIFICATION_BY_SMALL_PROTEIN_REMOV | 0.00003 | 0.00067 | 0.57561 | 0.58981 | 1.21601 | 232 |
| GO_PROTEIN_POLYMERIZATION | 0.00163 | 0.01971 | 0.45506 | 0.57085 | 1.17178 | 197 |
| GO_PROTEIN_PROCESSING | 0.00357 | 0.03540 | 0.43171 | 0.59201 | 1.19984 | 118 |
| GO_PROTEIN_TARGETING | 0.00000 | 0.00000 | | 0.63567 | 1.31982 | 354 |
| GO_PROTEIN_TARGETING_TO_MEMBRANE | 0.00000 | 0.00000 | | 0.69753 | 1.42472 | 163 |
| GO_PROTEIN_TARGETING_TO_MITOCHONDRION | 0.00004 | 0.00104 | 0.55733 | 0.66528 | 1.33551 | 81 |
| GO_PROTON_TRANSMEMBRANE_TRANSPORT | 0.00000 | 0.00004 | 0.64355 | 0.70366 | 1.41171 | 79 |
| GO_PROTON_TRANSMEMBRANE_TRANSPORTER_ACTIVITY | 0.00019 | 0.00349 | 0.51885 | 0.68647 | 1.36255 | 61 |
| GO_PURINE_CONTAINING_COMPOUND_BIOSYNTHETIC_PR | 0.00000 | 0.00007 | 0.62726 | 0.65311 | 1.32385 | 121 |
| GO_PURINE_CONTAINING_COMPOUND_METABOLIC_PROC | 0.00000 | 0.00007 | 0.62726 | 0.59385 | 1.22929 | 286 |
| GO_PURINE_NUCLEOSIDE_TRIPHOSPHATE_METABOLIC_PR | 0.00007 | 0.00165 | 0.53843 | 0.71555 | 1.39980 | 47 |
| GO_PYRUVATE_METABOLIC_PROCESS | 0.00003 | 0.00081 | 0.55733 | 0.64957 | 1.31323 | 109 |
| GO_REACTIVE_OXYGEN_SPECIES_BIOSYNTHETIC_PROCESS | 0.00290 | 0.03039 | 0.43171 | 0.65010 | 1.28334 | 53 |
| GO_REACTIVE_OXYGEN_SPECIES_METABOLIC_PROCESS | 0.00228 | 0.02516 | 0.43171 | 0.59185 | 1.20415 | 133 |
| GO_REGULATION_OF_ACTIN_FILAMENT_BASED_PROCESS | 0.00135 | 0.01698 | 0.45506 | 0.56234 | 1.15974 | 239 |
| GO_REGULATION_OF_ACTIN_FILAMENT_LENGTH | 0.00058 | 0.00874 | 0.47727 | 0.61148 | 1.23945 | 120 |
| GO_REGULATION_OF_ACTIN_FILAMENT_ORGANIZATION | 0.00001 | 0.00042 | 0.59333 | 0.61450 | 1.25530 | 165 |
| GO_REGULATION_OF_ANIMAL_ORGAN_MORPHOGENESIS | 0.00000 | 0.00000 | | 0.72211 | 1.46849 | 132 |
| GO_REGULATION_OF_APOPTOTIC_SIGNALING_PATHWAY | 0.00031 | 0.00516 | 0.49849 | 0.56666 | 1.17088 | 262 |
| GO_REGULATION_OF_ATP_METABOLIC_PROCESS | 0.00039 | 0.00627 | 0.49849 | 0.64034 | 1.28653 | 87 |
| GO_REGULATION_OF_BINDING | 0.00185 | 0.02152 | 0.45506 | 0.55859 | 1.15034 | 222 |
| GO_REGULATION_OF_BODY_FLUID_LEVELS | 0.00480 | 0.04435 | 0.40702 | 0.55323 | 1.14079 | 236 |
| GO_REGULATION_OF_CELLULAR_AMIDE_METABOLIC_PROC | 0.00000 | 0.00000 | 0.74774 | 0.60381 | 1.25235 | 329 |
| GO_REGULATION_OF_CELLULAR_AMINE_METABOLIC_PROC | 0.00000 | 0.00000 | 0.74774 | 0.78283 | 1.54822 | 56 |
| GO_REGULATION_OF_CELLULAR_AMINO_ACID_METABOLIC | 0.00000 | 0.00000 | 0.76146 | 0.80096 | 1.57890 | 52 |
| GO_REGULATION_OF_CELLULAR_KETONE_METABOLIC_PRO | 0.00000 | 0.00001 | 0.67496 | 0.68108 | 1.37889 | 113 |
| GO_REGULATION_OF_CELLULAR_PROTEIN_LOCALIZATION | 0.00044 | 0.00693 | 0.49849 | 0.54571 | 1.13371 | 391 |
| GO_REGULATION_OF_CELLULAR_RESPIRATION | 0.00205 | 0.02309 | 0.43171 | 0.76978 | 1.43168 | 20 |

| Cluster 1 | | | | | | |
|---|---|---|---|---|---|---|
| GO_REGULATION_OF_CELLULAR_RESPONSE_TO_TRANSFOR | 0.00361 | 0.03559 | 0.43171 | 0.63498 | 1.25581 | 56 |
| GO_REGULATION_OF_CELL_CYCLE_G2_M_PHASE_TRANSITI | 0.00034 | 0.00562 | 0.49849 | 0.58806 | 1.20455 | 187 |
| GO_REGULATION_OF_CELL_CYCLE_PHASE_TRANSITION | 0.00231 | 0.02539 | 0.43171 | 0.54172 | 1.12532 | 368 |
| GO_REGULATION_OF_CYSTEINE_TYPE_ENDOPEPTIDASE_AC | 0.00087 | 0.01206 | 0.47727 | 0.59809 | 1.21766 | 135 |
| GO_REGULATION_OF_CYTOSKELETON_ORGANIZATION | 0.00128 | 0.01630 | 0.45506 | 0.54457 | 1.13105 | 359 |
| GO_REGULATION_OF_DEFENSE_RESPONSE | 0.00179 | 0.02108 | 0.45506 | 0.54909 | 1.13933 | 336 |
| GO_REGULATION_OF_DNA_BIOSYNTHETIC_PROCESS | 0.00011 | 0.00224 | 0.53843 | 0.65550 | 1.31599 | 83 |
| GO_REGULATION_OF_DNA_METABOLIC_PROCESS | 0.00554 | 0.04963 | 0.40702 | 0.54882 | 1.13315 | 252 |
| GO_REGULATION_OF_DNA_TEMPLATED_TRANSCRIPTION_I | 0.00000 | 0.00000 | 0.77494 | 0.72694 | 1.46292 | 93 |
| GO_REGULATION_OF_EPITHELIAL_CELL_MIGRATION | 0.00320 | 0.03275 | 0.43171 | 0.58808 | 1.19647 | 133 |
| GO_REGULATION_OF_GENERATION_OF_PRECURSOR_META | 0.00383 | 0.03700 | 0.43171 | 0.60147 | 1.21589 | 108 |
| GO_REGULATION_OF_HEMATOPOIETIC_PROGENITOR_CELL_ | 0.00000 | 0.00000 | 0.76146 | 0.76049 | 1.51913 | 72 |
| GO_REGULATION_OF_HEMATOPOIETIC_STEM_CELL_DIFFER | 0.00000 | 0.00000 | 0.77494 | 0.77559 | 1.53971 | 62 |
| GO_REGULATION_OF_HEMOPOIESIS | 0.00133 | 0.01682 | 0.45506 | 0.56192 | 1.16011 | 254 |
| GO_REGULATION_OF_IMMUNE_RESPONSE | 0.00000 | 0.00012 | 0.62726 | 0.57164 | 1.18866 | 414 |
| GO_REGULATION_OF_INNATE_IMMUNE_RESPONSE | 0.00017 | 0.00321 | 0.51885 | 0.59388 | 1.21486 | 175 |
| GO_REGULATION_OF_INTRACELLULAR_TRANSPORT | 0.00398 | 0.03813 | 0.40702 | 0.54853 | 1.13293 | 258 |
| GO_REGULATION_OF_INTRINSIC_APOPTOTIC_SIGNALING_P | 0.00108 | 0.01426 | 0.45506 | 0.60794 | 1.23082 | 113 |
| GO_REGULATION_OF_KIDNEY_DEVELOPMENT | 0.00009 | 0.00191 | 0.53843 | 0.78517 | 1.48982 | 26 |
| GO_REGULATION_OF_LIPID_METABOLIC_PROCESS | 0.00167 | 0.02008 | 0.45506 | 0.56099 | 1.15675 | 237 |
| GO_REGULATION_OF_MITOTIC_CELL_CYCLE | 0.00198 | 0.02255 | 0.43171 | 0.53268 | 1.11031 | 497 |
| GO_REGULATION_OF_MORPHOGENESIS_OF_AN_EPITHELIU | 0.00000 | 0.00000 | | 0.74326 | 1.50456 | 112 |
| GO_REGULATION_OF_MRNA_CATABOLIC_PROCESS | 0.00000 | 0.00000 | 0.73376 | 0.66060 | 1.34930 | 163 |
| GO_REGULATION_OF_MRNA_METABOLIC_PROCESS | 0.00000 | 0.00000 | | 0.64581 | 1.33523 | 273 |
| GO_REGULATION_OF_MRNA_PROCESSING | 0.00004 | 0.00089 | 0.55733 | 0.63414 | 1.28637 | 124 |
| GO_REGULATION_OF_MRNA_SPLICING_VIA_SPLICEOSOME | 0.00003 | 0.00075 | 0.57561 | 0.66380 | 1.33550 | 91 |
| GO_REGULATION_OF_MULTI_ORGANISM_PROCESS | 0.00231 | 0.02539 | 0.43171 | 0.70502 | 1.35202 | 32 |
| GO_REGULATION_OF_MUSCLE_CONTRACTION | 0.00480 | 0.04435 | 0.40702 | 0.62360 | 1.24659 | 73 |
| GO_REGULATION_OF_NUCLEASE_ACTIVITY | 0.00194 | 0.02225 | 0.45506 | 0.79156 | 1.45787 | 17 |
| GO_REGULATION_OF_NUCLEOTIDE_BIOSYNTHETIC_PROCES | 0.00331 | 0.03338 | 0.43171 | 0.76114 | 1.41560 | 20 |
| GO_REGULATION_OF_OXIDATIVE_PHOSPHORYLATION | 0.00012 | 0.00244 | 0.53843 | 0.82141 | 1.52769 | 20 |
| GO_REGULATION_OF_PEPTIDASE_ACTIVITY | 0.00005 | 0.00111 | 0.55733 | 0.59384 | 1.22168 | 213 |
| GO_REGULATION_OF_PEPTIDE_TRANSPORT | 0.00346 | 0.03470 | 0.43171 | 0.53627 | 1.11449 | 397 |
| GO_REGULATION_OF_PHOSPHOLIPASE_ACTIVITY | 0.00509 | 0.04618 | 0.40702 | 0.69925 | 1.33340 | 28 |
| GO_REGULATION_OF_PHOSPHOLIPID_METABOLIC_PROCESS | 0.00044 | 0.00693 | 0.49849 | 0.67563 | 1.33525 | 55 |
| GO_REGULATION_OF_PLASMA_LIPOPROTEIN_PARTICLE_LEV | 0.00001 | 0.00023 | 0.61053 | 0.74607 | 1.46225 | 48 |
| GO_REGULATION_OF_PROTEIN_CATABOLIC_PROCESS | 0.00357 | 0.03540 | 0.43171 | 0.54832 | 1.13490 | 287 |
| GO_REGULATION_OF_PROTEIN_CONTAINING_COMPLEX_AS | 0.00037 | 0.00595 | 0.49849 | 0.56429 | 1.16836 | 292 |
| GO_REGULATION_OF_PROTEIN_IMPORT | 0.00235 | 0.02561 | 0.43171 | 0.67231 | 1.31308 | 44 |
| GO_REGULATION_OF_PROTEIN_LOCALIZATION_TO_NUCLEU | 0.00000 | 0.00009 | 0.62726 | 0.68104 | 1.36956 | 89 |
| GO_REGULATION_OF_PROTEIN_POLYMERIZATION | 0.00012 | 0.00244 | 0.53843 | 0.60977 | 1.24455 | 151 |
| GO_REGULATION_OF_PROTEOLYSIS | 0.00002 | 0.00063 | 0.57561 | 0.55937 | 1.16316 | 413 |
| GO_REGULATION_OF_REACTIVE_OXYGEN_SPECIES_BIOSYNT | 0.00220 | 0.02452 | 0.43171 | 0.67369 | 1.31579 | 44 |
| GO_REGULATION_OF_RESPONSE_TO_BIOTIC_STIMULUS | 0.00023 | 0.00413 | 0.51885 | 0.57583 | 1.18610 | 227 |
| GO_REGULATION_OF_RESPONSE_TO_EXTERNAL_STIMULUS | 0.00290 | 0.03039 | 0.43171 | 0.52891 | 1.10223 | 493 |
| GO_REGULATION_OF_RESPONSE_TO_REACTIVE_OXYGEN_S | 0.00495 | 0.04552 | 0.40702 | 0.72951 | 1.37174 | 23 |
| GO_REGULATION_OF_RNA_SPLICING | 0.00001 | 0.00017 | 0.61053 | 0.64601 | 1.31122 | 126 |
| GO_REGULATION_OF_SMALL_MOLECULE_METABOLIC_PRO | 0.00000 | 0.00001 | 0.67496 | 0.60893 | 1.25953 | 277 |
| GO_REGULATION_OF_SMOOTH_MUSCLE_CONTRACTION | 0.00283 | 0.02988 | 0.43171 | 0.75584 | 1.41054 | 21 |
| GO_REGULATION_OF_STEM_CELL_DIFFERENTIATION | 0.00000 | 0.00000 | 0.77494 | 0.73483 | 1.47583 | 86 |
| GO_REGULATION_OF_SUBSTRATE_ADHESION_DEPENDENT_ | 0.00372 | 0.03639 | 0.43171 | 0.67590 | 1.31448 | 40 |
| GO_REGULATION_OF_SUPRAMOLECULAR_FIBER_ORGANIZA | 0.00001 | 0.00027 | 0.59333 | 0.59951 | 1.23534 | 228 |
| GO_REGULATION_OF_SYMBIOTIC_PROCESS | 0.00010 | 0.00216 | 0.53843 | 0.61038 | 1.24556 | 149 |
| GO_REGULATION_OF_TELOMERASE_ACTIVITY | 0.00087 | 0.01206 | 0.47727 | 0.70727 | 1.37331 | 39 |
| GO_REGULATION_OF_TELOMERE_MAINTENANCE | 0.00216 | 0.02420 | 0.43171 | 0.63620 | 1.27085 | 72 |
| GO_REGULATION_OF_TELOMERE_MAINTENANCE_VIA_TELO | 0.00044 | 0.00695 | 0.49849 | 0.67796 | 1.33998 | 54 |
| GO_REGULATION_OF_TRANSCRIPTION_FROM_RNA_POLYM | 0.00000 | 0.00000 | 0.76146 | 0.76828 | 1.52643 | 65 |
| GO_REGULATION_OF_TRANSLATIONAL_INITIATION | 0.00108 | 0.01426 | 0.45506 | 0.64702 | 1.29000 | 69 |
| GO_REGULATION_OF_VIRAL_GENOME_REPLICATION | 0.00313 | 0.03232 | 0.43171 | 0.63710 | 1.26001 | 56 |
| GO_REGULATION_OF_WNT_SIGNALING_PATHWAY | 0.00001 | 0.00029 | 0.59333 | 0.60219 | 1.23914 | 214 |
| GO_REPRODUCTIVE_SYSTEM_DEVELOPMENT | 0.00294 | 0.03067 | 0.43171 | 0.56019 | 1.15557 | 241 |
| GO_RESPIRASOME | 0.00000 | 0.00000 | | 0.80604 | 1.61290 | 74 |
| GO_RESPIRATORY_CHAIN_COMPLEX | 0.00000 | 0.00000 | | 0.81535 | 1.61795 | 63 |
| GO_RESPIRATORY_ELECTRON_TRANSPORT_CHAIN | 0.00000 | 0.00000 | | 0.78801 | 1.58526 | 90 |
| GO_RESPONSE_TO_ALCOHOL | 0.00004 | 0.00105 | 0.55733 | 0.64528 | 1.30456 | 109 |
| GO_RESPONSE_TO_CORTICOSTEROID | 0.00346 | 0.03470 | 0.43171 | 0.61133 | 1.22497 | 76 |
| GO_RESPONSE_TO_DRUG | 0.00005 | 0.00119 | 0.55733 | 0.59649 | 1.22416 | 195 |
| GO_RESPONSE_TO_ESTRADIOL | 0.00368 | 0.03614 | 0.43171 | 0.63004 | 1.25337 | 66 |
| GO_RESPONSE_TO_ESTROGEN | 0.00060 | 0.00899 | 0.47727 | 0.69317 | 1.35382 | 44 |
| GO_RESPONSE_TO_ETHANOL | 0.00027 | 0.00465 | 0.49849 | 0.67158 | 1.33003 | 59 |
| GO_RESPONSE_TO_HEAT | 0.00117 | 0.01508 | 0.45506 | 0.61160 | 1.24008 | 122 |
| GO_RESPONSE_TO_HYDROGEN_PEROXIDE | 0.00450 | 0.04212 | 0.40702 | 0.61027 | 1.22771 | 90 |
| GO_RESPONSE_TO_INORGANIC_SUBSTANCE | 0.00001 | 0.00030 | 0.59333 | 0.58505 | 1.21194 | 299 |

| Cluster 1 | | | | | | |
|---|---|---|---|---|---|---|
| GO_RESPONSE_TO_INTERLEUKIN_1 | 0.00000 | 0.00000 | | 0.72620 | 1.47198 | 120 |
| GO_RESPONSE_TO_INTERLEUKIN_12 | 0.00023 | 0.00407 | 0.51885 | 0.74901 | 1.44071 | 33 |
| GO_RESPONSE_TO_INTERLEUKIN_4 | 0.00011 | 0.00220 | 0.53843 | 0.81435 | 1.51973 | 21 |
| GO_RESPONSE_TO_LEUKEMIA_INHIBITORY_FACTOR | 0.00554 | 0.04963 | 0.40702 | 0.63234 | 1.25524 | 64 |
| GO_RESPONSE_TO_METAL_ION | 0.00021 | 0.00386 | 0.51885 | 0.59113 | 1.20961 | 179 |
| GO_RESPONSE_TO_ORGANIC_CYCLIC_COMPOUND | 0.00000 | 0.00013 | 0.61053 | 0.56570 | 1.17842 | 479 |
| GO_RESPONSE_TO_OXIDATIVE_STRESS | 0.00133 | 0.01682 | 0.45506 | 0.55487 | 1.14796 | 278 |
| GO_RESPONSE_TO_OXYGEN_LEVELS | 0.00000 | 0.00000 | | 0.66699 | 1.37544 | 235 |
| GO_RESPONSE_TO_PEPTIDE | 0.00480 | 0.04435 | 0.40702 | 0.54423 | 1.12711 | 293 |
| GO_RESPONSE_TO_PEPTIDE_HORMONE | 0.00420 | 0.03976 | 0.40702 | 0.54821 | 1.13226 | 258 |
| GO_RESPONSE_TO_REACTIVE_OXYGEN_SPECIES | 0.00368 | 0.03614 | 0.43171 | 0.58906 | 1.20189 | 145 |
| GO_RESPONSE_TO_STEROID_HORMONE | 0.00316 | 0.03265 | 0.43171 | 0.56256 | 1.15359 | 191 |
| GO_RESPONSE_TO_TEMPERATURE_STIMULUS | 0.00202 | 0.02280 | 0.43171 | 0.58612 | 1.19611 | 147 |
| GO_RESPONSE_TO_TOXIC_SUBSTANCE | 0.00008 | 0.00176 | 0.53843 | 0.62712 | 1.27213 | 124 |
| GO_RESPONSE_TO_TUMOR_NECROSIS_FACTOR | 0.00000 | 0.00001 | 0.69013 | 0.63464 | 1.29878 | 177 |
| GO_RESPONSE_TO_VIRUS | 0.00287 | 0.03021 | 0.43171 | 0.57140 | 1.16742 | 167 |
| GO_RETINA_HOMEOSTASIS | 0.00100 | 0.01349 | 0.45506 | 0.72646 | 1.38779 | 29 |
| GO_RIBONUCLEOPROTEIN_COMPLEX_BINDING | 0.00000 | 0.00000 | 0.71951 | 0.68874 | 1.39394 | 111 |
| GO_RIBONUCLEOPROTEIN_COMPLEX_BIOGENESIS | 0.00000 | 0.00000 | | 0.66701 | 1.38571 | 391 |
| GO_RIBONUCLEOPROTEIN_COMPLEX_SUBUNIT_ORGANIZAT | 0.00000 | 0.00000 | | 0.67936 | 1.38914 | 173 |
| GO_RIBONUCLEOPROTEIN_GRANULE | 0.00000 | 0.00015 | 0.61053 | 0.61603 | 1.26187 | 185 |
| GO_RIBONUCLEOSIDE_TRIPHOSPHATE_BIOSYNTHETIC_PRO | 0.00010 | 0.00212 | 0.53843 | 0.75982 | 1.46150 | 33 |
| GO_RIBONUCLEOSIDE_TRIPHOSPHATE_METABOLIC_PROCES | 0.00010 | 0.00204 | 0.53843 | 0.72069 | 1.40758 | 44 |
| GO_RIBOSE_PHOSPHATE_BIOSYNTHETIC_PROCESS | 0.00001 | 0.00034 | 0.59333 | 0.65115 | 1.31986 | 119 |
| GO_RIBOSE_PHOSPHATE_METABOLIC_PROCESS | 0.00000 | 0.00009 | 0.62726 | 0.59471 | 1.22941 | 270 |
| GO_RIBOSOMAL_LARGE_SUBUNIT_ASSEMBLY | 0.00088 | 0.01213 | 0.47727 | 0.75029 | 1.41781 | 25 |
| GO_RIBOSOMAL_LARGE_SUBUNIT_BIOGENESIS | 0.00000 | 0.00009 | 0.62726 | 0.71678 | 1.42643 | 67 |
| GO_RIBOSOMAL_SMALL_SUBUNIT_ASSEMBLY | 0.00110 | 0.01441 | 0.45506 | 0.79025 | 1.45862 | 18 |
| GO_RIBOSOMAL_SMALL_SUBUNIT_BIOGENESIS | 0.00013 | 0.00262 | 0.51885 | 0.67216 | 1.33883 | 68 |
| GO_RIBOSOMAL_SUBUNIT | 0.00000 | 0.00000 | | 0.76258 | 1.55900 | 172 |
| GO_RIBOSOME | 0.00000 | 0.00000 | | 0.75264 | 1.54773 | 210 |
| GO_RIBOSOME_ASSEMBLY | 0.00000 | 0.00001 | 0.69013 | 0.76328 | 1.50846 | 55 |
| GO_RIBOSOME_BINDING | 0.00331 | 0.03338 | 0.43171 | 0.66898 | 1.30430 | 42 |
| GO_RIBOSOME_BIOGENESIS | 0.00000 | 0.00000 | | 0.66638 | 1.37778 | 275 |
| GO_RNA_3_END_PROCESSING | 0.00010 | 0.00206 | 0.53843 | 0.62394 | 1.26619 | 125 |
| GO_RNA_CATABOLIC_PROCESS | 0.00000 | 0.00000 | | 0.68655 | 1.42503 | 343 |
| GO_RNA_DEPENDENT_DNA_BIOSYNTHETIC_PROCESS | 0.00026 | 0.00444 | 0.49849 | 0.67408 | 1.33809 | 64 |
| GO_RNA_EXPORT_FROM_NUCLEUS | 0.00006 | 0.00148 | 0.53843 | 0.62107 | 1.26162 | 128 |
| GO_RNA_LOCALIZATION | 0.00000 | 0.00016 | 0.61053 | 0.61156 | 1.25708 | 207 |
| GO_RNA_SPLICING | 0.00000 | 0.00000 | 0.82666 | 0.60927 | 1.26586 | 387 |
| GO_RNA_SPLICING_VIA_TRANSESTERIFICATION_REACTIONS | 0.00000 | 0.00000 | 0.82666 | 0.62674 | 1.29895 | 312 |
| GO_ROUGH_ENDOPLASMIC_RETICULUM | 0.00009 | 0.00202 | 0.53843 | 0.70156 | 1.38663 | 54 |
| GO_RRNA_BINDING | 0.00000 | 0.00003 | 0.65944 | 0.74869 | 1.47979 | 54 |
| GO_RRNA_METABOLIC_PROCESS | 0.00000 | 0.00000 | 0.80122 | 0.65023 | 1.33689 | 209 |
| GO_RUFFLE | 0.00139 | 0.01737 | 0.45506 | 0.60152 | 1.21992 | 123 |
| GO_SARCOLEMMA | 0.00357 | 0.03540 | 0.43171 | 0.63531 | 1.25647 | 56 |
| GO_SCF_DEPENDENT_PROTEASOMAL_UBIQUITIN_DEPENDI | 0.00000 | 0.00001 | 0.70498 | 0.71908 | 1.44089 | 76 |
| GO_SECRETORY_GRANULE | 0.00000 | 0.00000 | | 0.62435 | 1.30004 | 446 |
| GO_SECRETORY_GRANULE_MEMBRANE | 0.00093 | 0.01264 | 0.47727 | 0.58740 | 1.19912 | 157 |
| GO_SERINE_TYPE_ENDOPEPTIDASE_INHIBITOR_ACTIVITY | 0.00088 | 0.01213 | 0.47727 | 0.76701 | 1.44224 | 23 |
| GO_SEX_DIFFERENTIATION | 0.00117 | 0.01508 | 0.45506 | 0.59625 | 1.21589 | 140 |
| GO_SIDE_OF_MEMBRANE | 0.00011 | 0.00228 | 0.53843 | 0.59012 | 1.21167 | 202 |
| GO_SIGNAL_TRANSDUCTION_BY_PROTEIN_PHOSPHORYLAT | 0.00013 | 0.00250 | 0.51885 | 0.54634 | 1.13831 | 485 |
| GO_SINGLE_FERTILIZATION | 0.00003 | 0.00070 | 0.57561 | 0.71571 | 1.41286 | 53 |
| GO_SINGLE_STRANDED_RNA_BINDING | 0.00018 | 0.00324 | 0.51885 | 0.68672 | 1.36271 | 63 |
| GO_SMALL_MOLECULE_BIOSYNTHETIC_PROCESS | 0.00000 | 0.00004 | 0.65944 | 0.57385 | 1.19398 | 426 |
| GO_SMALL_NUCLEAR_RIBONUCLEOPROTEIN_COMPLEX | 0.00013 | 0.00250 | 0.51885 | 0.68206 | 1.34991 | 57 |
| GO_SMALL_RIBOSOMAL_SUBUNIT | 0.00000 | 0.00000 | 0.74774 | 0.75478 | 1.50338 | 68 |
| GO_SM_LIKE_PROTEIN_FAMILY_COMPLEX | 0.00028 | 0.00486 | 0.49849 | 0.66640 | 1.32616 | 67 |
| GO_SNORNA_BINDING | 0.00037 | 0.00601 | 0.49849 | 0.74951 | 1.43182 | 29 |
| GO_SPERM_EGG_RECOGNITION | 0.00000 | 0.00004 | 0.64355 | 0.87135 | 1.62610 | 21 |
| GO_SPLICEOSOMAL_COMPLEX | 0.00000 | 0.00003 | 0.65944 | 0.62958 | 1.28735 | 173 |
| GO_SPLICEOSOMAL_SNRNP_ASSEMBLY | 0.00420 | 0.03976 | 0.40702 | 0.69004 | 1.33163 | 35 |
| GO_SPLICEOSOMAL_TRI_SNRNP_COMPLEX | 0.00220 | 0.02452 | 0.43171 | 0.71681 | 1.37204 | 30 |
| GO_STEM_CELL_DIFFERENTIATION | 0.00000 | 0.00000 | | 0.69579 | 1.42035 | 154 |
| GO_STEROID_BIOSYNTHETIC_PROCESS | 0.00205 | 0.02309 | 0.43171 | 0.59853 | 1.21035 | 106 |
| GO_STEROID_METABOLIC_PROCESS | 0.00146 | 0.01811 | 0.45506 | 0.58209 | 1.18884 | 162 |
| GO_STEROL_BIOSYNTHETIC_PROCESS | 0.00056 | 0.00850 | 0.47727 | 0.67227 | 1.33324 | 60 |
| GO_STEROL_METABOLIC_PROCESS | 0.00062 | 0.00917 | 0.47727 | 0.61973 | 1.25233 | 102 |
| GO_STRUCTURAL_CONSTITUENT_OF_CYTOSKELETON | 0.00058 | 0.00874 | 0.47727 | 0.66358 | 1.31735 | 62 |
| GO_STRUCTURAL_CONSTITUENT_OF_RIBOSOME | 0.00000 | 0.00000 | | 0.77552 | 1.58277 | 153 |
| GO_STRUCTURAL_MOLECULE_ACTIVITY | 0.00000 | 0.00000 | | 0.72925 | 1.51464 | 378 |

| Cluster 1 | | | | | | |
|---|---|---|---|---|---|---|
| GO_SUBSTANTIA_NIGRA_DEVELOPMENT | 0.00143 | 0.01769 | 0.45506 | 0.71307 | 1.37157 | 33 |
| GO_SUBSTRATE_ADHESION_DEPENDENT_CELL_SPREADING | 0.00383 | 0.03700 | 0.43171 | 0.62534 | 1.24849 | 71 |
| GO_SUPRAMOLECULAR_FIBER_ORGANIZATION | 0.00000 | 0.00013 | 0.61053 | 0.56838 | 1.18189 | 413 |
| GO_TELOMERASE_RNA_LOCALIZATION | 0.00032 | 0.00530 | 0.49849 | 0.80987 | 1.49945 | 19 |
| GO_TELOMERE_MAINTENANCE_VIA_TELOMERE_LENGTHEN | 0.00003 | 0.00067 | 0.57561 | 0.68170 | 1.36101 | 71 |
| GO_TELOMERE_ORGANIZATION | 0.00327 | 0.03317 | 0.43171 | 0.58605 | 1.19300 | 134 |
| GO_TERTIARY_GRANULE | 0.00450 | 0.04212 | 0.40702 | 0.60838 | 1.22204 | 85 |
| GO_THREONINE_TYPE_ENDOPEPTIDASE_ACTIVITY | 0.00000 | 0.00001 | 0.69013 | 0.91142 | 1.67862 | 17 |
| GO_THREONINE_TYPE_PEPTIDASE_ACTIVITY | 0.00000 | 0.00002 | 0.67496 | 0.88214 | 1.64624 | 21 |
| GO_TISSUE_MORPHOGENESIS | 0.00000 | 0.00000 | 0.78819 | 0.60638 | 1.25899 | 354 |
| GO_TOXIN_TRANSPORT | 0.00068 | 0.00988 | 0.47727 | 0.73598 | 1.40597 | 29 |
| GO_TRANSCRIPTION_COUPLED_NUCLEOTIDE_EXCISION_RE | 0.00165 | 0.01990 | 0.45506 | 0.64130 | 1.28104 | 72 |
| GO_TRANSFERRIN_TRANSPORT | 0.00097 | 0.01303 | 0.47727 | 0.72768 | 1.39012 | 29 |
| GO_TRANSITION_METAL_ION_HOMEOSTASIS | 0.00185 | 0.02152 | 0.45506 | 0.61956 | 1.24147 | 76 |
| GO_TRANSLATIONAL_ELONGATION | 0.00000 | 0.00000 | | 0.72507 | 1.47083 | 124 |
| GO_TRANSLATIONAL_INITIATION | 0.00000 | 0.00000 | | 0.74224 | 1.51741 | 172 |
| GO_TRANSLATIONAL_TERMINATION | 0.00000 | 0.00000 | | 0.74679 | 1.50910 | 102 |
| GO_TRANSLATION_ELONGATION_FACTOR_ACTIVITY | 0.00517 | 0.04664 | 0.40702 | 0.76585 | 1.40610 | 16 |
| GO_TRANSLATION_FACTOR_ACTIVITY_RNA_BINDING | 0.00000 | 0.00002 | 0.65944 | 0.72661 | 1.44728 | 68 |
| GO_TRANSLATION_INITIATION_FACTOR_ACTIVITY | 0.00005 | 0.00111 | 0.55733 | 0.73203 | 1.43110 | 45 |
| GO_TRANSLATION_PREINITIATION_COMPLEX | 0.00442 | 0.04168 | 0.40702 | 0.76975 | 1.41326 | 16 |
| GO_TRANSLATION_REGULATOR_ACTIVITY | 0.00000 | 0.00000 | 0.71951 | 0.69810 | 1.41123 | 108 |
| GO_TRANSLATION_REGULATOR_ACTIVITY_NUCLEIC_ACID_E | 0.00000 | 0.00000 | 0.70498 | 0.72064 | 1.44677 | 83 |
| GO_TRICARBOXYLIC_ACID_CYCLE | 0.00076 | 0.01079 | 0.47727 | 0.72842 | 1.40111 | 33 |
| GO_TUMOR_NECROSIS_FACTOR_MEDIATED_SIGNALING_PA | 0.00000 | 0.00000 | 0.78819 | 0.70914 | 1.43418 | 110 |
| GO_T_CELL_RECEPTOR_SIGNALING_PATHWAY | 0.00000 | 0.00001 | 0.67496 | 0.66977 | 1.35744 | 118 |
| GO_U12_TYPE_SPLICEOSOMAL_COMPLEX | 0.00006 | 0.00148 | 0.53843 | 0.78922 | 1.49749 | 26 |
| GO_U1_SNRNP | 0.00517 | 0.04664 | 0.40702 | 0.76656 | 1.40741 | 16 |
| GO_U2_SNRNP | 0.00420 | 0.03976 | 0.40702 | 0.76473 | 1.41152 | 18 |
| GO_U2_TYPE_SPLICEOSOMAL_COMPLEX | 0.00018 | 0.00336 | 0.51885 | 0.64746 | 1.30034 | 86 |
| GO_UBIQUITIN_LIKE_PROTEIN_LIGASE_BINDING | 0.00016 | 0.00306 | 0.51885 | 0.57596 | 1.18875 | 246 |
| GO_UNFOLDED_PROTEIN_BINDING | 0.00000 | 0.00004 | 0.64355 | 0.68741 | 1.38237 | 89 |
| GO_VACUOLAR_LUMEN | 0.00026 | 0.00444 | 0.49849 | 0.62706 | 1.26789 | 105 |
| GO_VESICLE_LUMEN | 0.00000 | 0.00000 | | 0.66525 | 1.36529 | 195 |
| GO_VESICLE_MEMBRANE | 0.00013 | 0.00256 | 0.51885 | 0.54953 | 1.14425 | 446 |
| GO_VIRAL_GENE_EXPRESSION | 0.00000 | 0.00000 | | 0.71306 | 1.45912 | 179 |
| GO_VIRAL_GENOME_REPLICATION | 0.00135 | 0.01698 | 0.45506 | 0.62477 | 1.25191 | 76 |
| GO_VIRAL_LIFE_CYCLE | 0.00000 | 0.00011 | 0.62726 | 0.60192 | 1.23976 | 226 |
| HP_ABNORMALITY_OF_ACID_BASE_HOMEOSTASIS | 0.00000 | 0.00000 | 0.70498 | 0.61540 | 1.27139 | 260 |
| HP_ABNORMALITY_OF_CARDIOVASCULAR_SYSTEM_ELECTR | 0.00063 | 0.00926 | 0.47727 | 0.56621 | 1.16891 | 251 |
| HP_ABNORMALITY_OF_PERIPHERAL_NERVOUS_SYSTEM_EL | 0.00231 | 0.02539 | 0.43171 | 0.61806 | 1.23846 | 76 |
| HP_ABNORMALITY_OF_THE_CEREBROSPINAL_FLUID | 0.00509 | 0.04618 | 0.40702 | 0.54264 | 1.12474 | 314 |
| HP_ABNORMALITY_OF_THE_MITOCHONDRION | 0.00090 | 0.01235 | 0.47727 | 0.60524 | 1.22775 | 124 |
| HP_ABNORMALITY_OF_THE_OPTIC_DISC | 0.00016 | 0.00301 | 0.51885 | 0.55033 | 1.14528 | 430 |
| HP_ABNORMALITY_OF_THE_OPTIC_NERVE | 0.00011 | 0.00228 | 0.53843 | 0.54795 | 1.14155 | 481 |
| HP_ABNORMALITY_OF_URINE_HOMEOSTASIS | 0.00290 | 0.03039 | 0.43171 | 0.53943 | 1.12065 | 361 |
| HP_ABNORMAL_ABDOMEN_MORPHOLOGY | 0.00157 | 0.01924 | 0.45506 | 0.53807 | 1.11952 | 425 |
| HP_ABNORMAL_ACTIVITY_OF_MITOCHONDRIAL_RESPIRATO | 0.00046 | 0.00716 | 0.49849 | 0.67524 | 1.33992 | 63 |
| HP_ABNORMAL_BASAL_GANGLIA_MRI_SIGNAL_INTENSITY | 0.00279 | 0.02959 | 0.43171 | 0.69295 | 1.33868 | 36 |
| HP_ABNORMAL_BLOOD_GLUCOSE_CONCENTRATION | 0.00502 | 0.04592 | 0.40702 | 0.57639 | 1.17663 | 154 |
| HP_ABNORMAL_BRAINSTEM_MRI_SIGNAL_INTENSITY | 0.00159 | 0.01942 | 0.45506 | 0.71126 | 1.36809 | 33 |
| HP_ABNORMAL_CELLULAR_PHENOTYPE | 0.00257 | 0.02748 | 0.43171 | 0.54424 | 1.12918 | 332 |
| HP_ABNORMAL_CIRCULATING_CARBOHYDRATE_CONCENTR | 0.00017 | 0.00321 | 0.51885 | 0.70692 | 1.38040 | 46 |
| HP_ABNORMAL_CNS_MYELINATION | 0.00002 | 0.00045 | 0.57561 | 0.61478 | 1.25644 | 171 |
| HP_ABNORMAL_CSF_METABOLITE_LEVEL | 0.00000 | 0.00014 | 0.61053 | 0.68470 | 1.37449 | 81 |
| HP_ABNORMAL_ELECTROPHYSIOLOGY_OF_SINOATRIAL_NO | 0.00202 | 0.02280 | 0.43171 | 0.69556 | 1.34909 | 38 |
| HP_ABNORMAL_ERYTHROCYTE_MORPHOLOGY | 0.00001 | 0.00042 | 0.59333 | 0.57136 | 1.18678 | 372 |
| HP_ABNORMAL_MITOCHONDRIA_IN_MUSCLE_TISSUE | 0.00023 | 0.00405 | 0.51885 | 0.75734 | 1.44678 | 29 |
| HP_ABNORMAL_MYELINATION | 0.00005 | 0.00111 | 0.55733 | 0.56715 | 1.17565 | 323 |
| HP_ABNORMAL_MYOCARDIUM_MORPHOLOGY | 0.00030 | 0.00503 | 0.49849 | 0.55688 | 1.15548 | 333 |
| HP_ABNORMAL_RENAL_PHYSIOLOGY | 0.00187 | 0.02165 | 0.45506 | 0.54471 | 1.13156 | 360 |
| HP_ABNORMAL_SOCIAL_BEHAVIOR | 0.00077 | 0.01089 | 0.47727 | 0.61665 | 1.24682 | 105 |
| HP_ABNORMAL_TRICUSPID_VALVE_PHYSIOLOGY | 0.00420 | 0.03976 | 0.40702 | 0.70262 | 1.33982 | 28 |
| HP_ACUTE_LEUKEMIA | 0.00016 | 0.00306 | 0.51885 | 0.66689 | 1.33496 | 75 |
| HP_ANEMIA_OF_INADEQUATE_PRODUCTION | 0.00194 | 0.02225 | 0.45506 | 0.60021 | 1.21375 | 106 |
| HP_BLINDNESS | 0.00119 | 0.01525 | 0.45506 | 0.57923 | 1.18342 | 167 |
| HP_BONE_MARROW_HYPERCELLULARITY | 0.00224 | 0.02484 | 0.43171 | 0.77849 | 1.43691 | 18 |
| HP_CHRONIC_INFECTION | 0.00509 | 0.04618 | 0.40702 | 0.76723 | 1.40864 | 16 |
| HP_CONGESTIVE_HEART_FAILURE | 0.00383 | 0.03700 | 0.43171 | 0.59330 | 1.20118 | 113 |
| HP_CRANIAL_NERVE_COMPRESSION | 0.00231 | 0.02539 | 0.43171 | 0.75787 | 1.42913 | 24 |
| HP_DECREASED_ACTIVITY_OF_MITOCHONDRIAL_COMPLEX | 0.00179 | 0.02108 | 0.45506 | 0.70627 | 1.36295 | 35 |
| HP_DECREASED_ACTIVITY_OF_THE_PYRUVATE_DEHYDROGE | 0.00305 | 0.03161 | 0.43171 | 0.69886 | 1.34018 | 32 |
| HP_DEPRESSED_NASAL_RIDGE | 0.00213 | 0.02383 | 0.43171 | 0.62641 | 1.25704 | 80 |

| **Cluster 1** | | | | | | |
|---|---|---|---|---|---|---|
| HP_DEPRESSED_NASAL_TIP | 0.00070 | 0.01005 | 0.47727 | 0.82514 | 1.50784 | 15 |
| HP_DYSTONIA | 0.00041 | 0.00659 | 0.49849 | 0.56540 | 1.16975 | 279 |
| HP_EMOTIONAL_LABILITY | 0.00013 | 0.00262 | 0.51885 | 0.65119 | 1.31002 | 90 |
| HP_ENCEPHALOPATHY | 0.00450 | 0.04212 | 0.40702 | 0.56803 | 1.16081 | 170 |
| HP_FATIGUE | 0.00502 | 0.04592 | 0.40702 | 0.55989 | 1.15088 | 207 |
| HP_FETAL_DISTRESS | 0.00044 | 0.00695 | 0.49849 | 0.73313 | 1.40592 | 32 |
| HP_FOCAL_T2_HYPERINTENSE_BRAINSTEM_LESION | 0.00108 | 0.01426 | 0.45506 | 0.73053 | 1.39304 | 28 |
| HP_HAMARTOMA | 0.00383 | 0.03700 | 0.43171 | 0.63423 | 1.25434 | 56 |
| HP_HEADACHE | 0.00046 | 0.00716 | 0.49849 | 0.59986 | 1.22433 | 151 |
| HP_HEPATOMEGALY | 0.00113 | 0.01476 | 0.45506 | 0.55183 | 1.14470 | 330 |
| HP_HYPERTROPHIC_CARDIOMYOPATHY | 0.00006 | 0.00130 | 0.55733 | 0.59837 | 1.22605 | 188 |
| HP_IMPAIRED_SOCIAL_INTERACTIONS | 0.00063 | 0.00926 | 0.47727 | 0.65238 | 1.30068 | 69 |
| HP_INCREASED_CSF_LACTATE | 0.00000 | 0.00014 | 0.61053 | 0.70759 | 1.40940 | 68 |
| HP_INCREASED_SERUM_LACTATE | 0.00000 | 0.00010 | 0.62726 | 0.64325 | 1.31035 | 138 |
| HP_INCREASED_SERUM_PYRUVATE | 0.00014 | 0.00274 | 0.51885 | 0.73448 | 1.42457 | 38 |
| HP_INFANTILE_ONSET | 0.00093 | 0.01264 | 0.47727 | 0.55624 | 1.15118 | 288 |
| HP_LACTIC_ACIDOSIS | 0.00002 | 0.00045 | 0.57561 | 0.63510 | 1.29321 | 136 |
| HP_LETHARGY | 0.00012 | 0.00247 | 0.51885 | 0.63309 | 1.27969 | 104 |
| HP_LEUKEMIA | 0.00268 | 0.02851 | 0.43171 | 0.59349 | 1.20016 | 106 |
| HP_LEUKODYSTROPHY | 0.00006 | 0.00142 | 0.53843 | 0.66035 | 1.32610 | 84 |
| HP_LEUKOENCEPHALOPATHY | 0.00047 | 0.00727 | 0.49849 | 0.60474 | 1.23119 | 135 |
| HP_LIMB_GIRDLE_MUSCLE_ATROPHY | 0.00246 | 0.02658 | 0.43171 | 0.73038 | 1.38018 | 25 |
| HP_LIPID_ACCUMULATION_IN_HEPATOCYTES | 0.00239 | 0.02592 | 0.43171 | 0.61976 | 1.24680 | 90 |
| HP_LOWER_LIMB_ASYMMETRY | 0.00364 | 0.03590 | 0.43171 | 0.78916 | 1.44209 | 15 |
| HP_MACROCYTIC_ANEMIA | 0.00015 | 0.00289 | 0.51885 | 0.72559 | 1.41469 | 42 |
| HP_METABOLIC_ACIDOSIS | 0.00016 | 0.00306 | 0.51885 | 0.64097 | 1.29394 | 99 |
| HP_MICRONODULAR_CIRRHOSIS | 0.00194 | 0.02225 | 0.45506 | 0.79113 | 1.45708 | 17 |
| HP_MIGRAINE | 0.00010 | 0.00214 | 0.53843 | 0.67353 | 1.34357 | 70 |
| HP_MITOCHONDRIAL_MYOPATHY | 0.00450 | 0.04212 | 0.40702 | 0.67225 | 1.30531 | 39 |
| HP_MUSCLE_ABNORMALITY_RELATED_TO_MITOCHONDRIA | 0.00172 | 0.02050 | 0.45506 | 0.66673 | 1.31429 | 52 |
| HP_MUTISM | 0.00487 | 0.04490 | 0.40702 | 0.69418 | 1.32874 | 31 |
| HP_NAUSEA_AND_VOMITING | 0.00065 | 0.00951 | 0.47727 | 0.56316 | 1.16315 | 258 |
| HP_OPTIC_DISC_PALLOR | 0.00111 | 0.01459 | 0.45506 | 0.62242 | 1.25110 | 88 |
| HP_OPTIC_NERVE_COLOBOMA | 0.00110 | 0.01441 | 0.45506 | 0.81495 | 1.48921 | 15 |
| HP_OPTIC_NEUROPATHY | 0.00016 | 0.00306 | 0.51885 | 0.74425 | 1.43623 | 35 |
| HP_PALLOR | 0.00019 | 0.00346 | 0.51885 | 0.63439 | 1.28009 | 98 |
| HP_PARALYSIS | 0.00253 | 0.02718 | 0.43171 | 0.61603 | 1.23977 | 92 |
| HP_PARAPLEGIA_PARAPARESIS | 0.00502 | 0.04592 | 0.40702 | 0.57543 | 1.17424 | 149 |
| HP_PAROXYSMAL_INVOLUNTARY_EYE_MOVEMENTS | 0.00010 | 0.00217 | 0.53843 | 0.76667 | 1.46751 | 31 |
| HP_PEDIATRIC_ONSET | 0.00044 | 0.00695 | 0.49849 | 0.55328 | 1.14908 | 358 |
| HP_PIGMENTARY_RETINOPATHY | 0.00157 | 0.01924 | 0.45506 | 0.60386 | 1.22238 | 112 |
| HP_POOR_EYE_CONTACT | 0.00061 | 0.00910 | 0.47727 | 0.68762 | 1.34920 | 49 |
| HP_POOR_HEAD_CONTROL | 0.00009 | 0.00184 | 0.53843 | 0.63541 | 1.28643 | 113 |
| HP_PROGRESSIVE_SPASTIC_PARAPLEGIA | 0.00302 | 0.03128 | 0.43171 | 0.66415 | 1.30170 | 48 |
| HP_PROXIMAL_TUBULOPATHY | 0.00070 | 0.01005 | 0.47727 | 0.72353 | 1.39559 | 34 |
| HP_REDUCED_CONSCIOUSNESS_CONFUSION | 0.00032 | 0.00536 | 0.49849 | 0.58423 | 1.19735 | 189 |
| HP_REDUCED_VISUAL_ACUITY | 0.00069 | 0.00999 | 0.47727 | 0.56223 | 1.16238 | 272 |
| HP_RENAL_TUBULAR_DYSFUNCTION | 0.00007 | 0.00158 | 0.53843 | 0.67238 | 1.34544 | 74 |
| HP_RESPIRATORY_INSUFFICIENCY | 0.00003 | 0.00070 | 0.57561 | 0.57228 | 1.18636 | 324 |
| HP_SENSORY_NEUROPATHY | 0.00509 | 0.04618 | 0.40702 | 0.59267 | 1.19834 | 105 |
| HP_TACHYCARDIA | 0.00502 | 0.04592 | 0.40702 | 0.65194 | 1.28105 | 50 |
| HP_VERTIGO | 0.00003 | 0.00083 | 0.55733 | 0.70035 | 1.38612 | 57 |
| HP_VESTIBULAR_DYSFUNCTION | 0.00005 | 0.00128 | 0.55733 | 0.68166 | 1.35978 | 70 |
| HP_VOMITING | 0.00250 | 0.02688 | 0.43171 | 0.58580 | 1.19517 | 146 |
| **Cluster 12** | | | | | | |
| pathway | pval | padj | log2err | ES | NES | size |
| GO_ACTIVATION_OF_ADENYLATE_CYCLASE_ACTIVITY | 0.00157 | 0.02863 | 0.45506 | 0.81209 | 1.43000 | 17 |
| GO_ACTIVATION_OF_IMMUNE_RESPONSE | 0.00000 | 0.00000 | 0.76146 | 0.64794 | 1.25487 | 270 |
| GO_ACTIVATION_OF_INNATE_IMMUNE_RESPONSE | 0.00000 | 0.00000 | 0.82666 | 0.75359 | 1.42681 | 103 |
| GO_AEROBIC_ELECTRON_TRANSPORT_CHAIN | 0.00030 | 0.00767 | 0.49849 | 0.85713 | 1.50283 | 15 |
| GO_ALTERNATIVE_MRNA_SPLICING_VIA_SPLICEOSOME | 0.00097 | 0.01922 | 0.47727 | 0.66786 | 1.25215 | 70 |
| GO_AMINE_METABOLIC_PROCESS | 0.00000 | 0.00000 | 0.76146 | 0.74403 | 1.40513 | 93 |
| GO_ANAPHASE_PROMOTING_COMPLEX_DEPENDENT_CATA | 0.00000 | 0.00000 | 0.80122 | 0.77091 | 1.45109 | 80 |
| GO_ANTIGEN_PROCESSING_AND_PRESENTATION | 0.00000 | 0.00001 | 0.69013 | 0.67308 | 1.29114 | 169 |
| GO_ANTIGEN_PROCESSING_AND_PRESENTATION_OF_EXOG | 0.00000 | 0.00001 | 0.70498 | 0.76449 | 1.43228 | 67 |
| GO_ANTIGEN_PROCESSING_AND_PRESENTATION_OF_PEPT | 0.00000 | 0.00001 | 0.70498 | 0.68043 | 1.30159 | 152 |
| GO_ANTIGEN_PROCESSING_AND_PRESENTATION_OF_PEPT | 0.00000 | 0.00004 | 0.65944 | 0.72590 | 1.36739 | 81 |
| GO_ANTIGEN_RECEPTOR_MEDIATED_SIGNALING_PATHWAY | 0.00000 | 0.00006 | 0.64355 | 0.65849 | 1.26176 | 163 |
| GO_ANTIMICROBIAL_HUMORAL_IMMUNE_RESPONSE_MED | 0.00298 | 0.04745 | 0.43171 | 0.80527 | 1.41559 | 16 |
| GO_APOPTOTIC_SIGNALING_PATHWAY | 0.00261 | 0.04294 | 0.43171 | 0.56714 | 1.10619 | 419 |
| GO_ATP_BIOSYNTHETIC_PROCESS | 0.00305 | 0.04791 | 0.43171 | 0.75731 | 1.35301 | 24 |
| GO_ATP_METABOLIC_PROCESS | 0.00000 | 0.00000 | | 0.68111 | 1.31424 | 225 |
| GO_ATP_SYNTHESIS_COUPLED_ELECTRON_TRANSPORT | 0.00000 | 0.00003 | 0.65944 | 0.73412 | 1.38046 | 78 |
| GO_AUTOPHAGY_OF_MITOCHONDRION | 0.00174 | 0.03112 | 0.45506 | 0.66050 | 1.24043 | 73 |

| Cluster 1 | | | | | | |
|---|---|---|---|---|---|---|
| GO_BINDING_OF_SPERM_TO_ZONA_PELLUCIDA | 0.00043 | 0.01014 | 0.49849 | 0.76349 | 1.37655 | 28 |
| GO_BLOOD_MICROPARTICLE | 0.00027 | 0.00697 | 0.49849 | 0.72580 | 1.34070 | 45 |
| GO_CADHERIN_BINDING | 0.00000 | 0.00001 | 0.70498 | 0.64491 | 1.24707 | 251 |
| GO_CAJAL_BODY | 0.00119 | 0.02270 | 0.45506 | 0.67545 | 1.25968 | 58 |
| GO_CARBOHYDRATE_CATABOLIC_PROCESS | 0.00030 | 0.00767 | 0.49849 | 0.62842 | 1.19998 | 143 |
| GO_CATALYTIC_ACTIVITY_ACTING_ON_RNA | 0.00003 | 0.00095 | 0.57561 | 0.60222 | 1.16950 | 319 |
| GO_CATALYTIC_STEP_2_SPLICEOSOME | 0.00000 | 0.00000 | 0.76146 | 0.74835 | 1.41081 | 83 |
| GO_CELLULAR_AMINO_ACID_METABOLIC_PROCESS | 0.00290 | 0.04697 | 0.43171 | 0.58804 | 1.13507 | 227 |
| GO_CELLULAR_COMPONENT_DISASSEMBLY | 0.00000 | 0.00000 | | 0.65545 | 1.28023 | 451 |
| GO_CELLULAR_KETONE_METABOLIC_PROCESS | 0.00000 | 0.00015 | 0.62726 | 0.65020 | 1.24753 | 171 |
| GO_CELLULAR_NITROGEN_COMPOUND_CATABOLIC_PROCE | 0.00000 | 0.00000 | | 0.68948 | 1.34787 | 479 |
| GO_CELLULAR_PROTEIN_COMPLEX_DISASSEMBLY | 0.00000 | 0.00000 | | 0.69751 | 1.34057 | 186 |
| GO_CELLULAR_RESPIRATION | 0.00000 | 0.00006 | 0.64355 | 0.66753 | 1.27466 | 143 |
| GO_CELLULAR_RESPONSE_TO_HEAT | 0.00001 | 0.00048 | 0.59333 | 0.68599 | 1.29748 | 97 |
| GO_CELLULAR_RESPONSE_TO_OXYGEN_LEVELS | 0.00000 | 0.00002 | 0.67496 | 0.66471 | 1.27548 | 173 |
| GO_CELL_ACTIVATION_INVOLVED_IN_IMMUNE_RESPONSE | 0.00002 | 0.00057 | 0.57561 | 0.59266 | 1.15614 | 422 |
| GO_CELL_ADHESION_MOLECULE_BINDING | 0.00000 | 0.00000 | 0.73376 | 0.62724 | 1.22019 | 347 |
| GO_CELL_CELL_RECOGNITION | 0.00019 | 0.00523 | 0.51885 | 0.73656 | 1.36252 | 47 |
| GO_CELL_CYCLE_G2_M_PHASE_TRANSITION | 0.00006 | 0.00194 | 0.55733 | 0.61138 | 1.18095 | 239 |
| GO_CELL_DIVISION | 0.00003 | 0.00114 | 0.55733 | 0.58263 | 1.13882 | 475 |
| GO_CELL_RECOGNITION | 0.00008 | 0.00250 | 0.53843 | 0.68645 | 1.29464 | 85 |
| GO_CELL_SUBSTRATE_JUNCTION | 0.00000 | 0.00000 | 0.76146 | 0.64106 | 1.24512 | 323 |
| GO_CHAPERONE_COMPLEX | 0.00016 | 0.00453 | 0.51885 | 0.83015 | 1.47765 | 22 |
| GO_CHROMOSOMAL_REGION | 0.00004 | 0.00128 | 0.55733 | 0.60762 | 1.17856 | 297 |
| GO_CHROMOSOME_CENTROMERIC_REGION | 0.00005 | 0.00179 | 0.55733 | 0.63315 | 1.21504 | 172 |
| GO_CHROMOSOME_SEGREGATION | 0.00013 | 0.00391 | 0.51885 | 0.60058 | 1.16404 | 281 |
| GO_CONDENSED_CHROMOSOME | 0.00185 | 0.03264 | 0.45506 | 0.60257 | 1.15771 | 183 |
| GO_COP9_SIGNALOSOME | 0.00220 | 0.03720 | 0.43171 | 0.73199 | 1.31976 | 28 |
| GO_COTRANSLATIONAL_PROTEIN_TARGETING_TO_MEMBR/ | 0.00000 | 0.00000 | | 0.82953 | 1.56756 | 95 |
| GO_CYTOCHROME_COMPLEX | 0.00156 | 0.02838 | 0.45506 | 0.77004 | 1.37178 | 23 |
| GO_CYTOKINE_MEDIATED_SIGNALING_PATHWAY | 0.00001 | 0.00025 | 0.61053 | 0.59486 | 1.16099 | 430 |
| GO_CYTOPLASMIC_TRANSLATION | 0.00000 | 0.00000 | 0.74774 | 0.74287 | 1.40380 | 95 |
| GO_CYTOPLASMIC_TRANSLATIONAL_INITIATION | 0.00055 | 0.01245 | 0.47727 | 0.76545 | 1.38879 | 31 |
| GO_CYTOSOLIC_LARGE_RIBOSOMAL_SUBUNIT | 0.00000 | 0.00000 | | 0.84380 | 1.56883 | 53 |
| GO_CYTOSOLIC_RIBOSOME | 0.00000 | 0.00000 | | 0.83315 | 1.57496 | 96 |
| GO_CYTOSOLIC_SMALL_RIBOSOMAL_SUBUNIT | 0.00000 | 0.00000 | 0.74774 | 0.83916 | 1.54122 | 42 |
| GO_DEFENSE_RESPONSE_TO_GRAM_NEGATIVE_BACTERIUM | 0.00028 | 0.00706 | 0.49849 | 0.79875 | 1.42292 | 23 |
| GO_DENDRITIC_CELL_MIGRATION | 0.00213 | 0.03614 | 0.43171 | 0.81241 | 1.42815 | 16 |
| GO_DISORDERED_DOMAIN_SPECIFIC_BINDING | 0.00060 | 0.01307 | 0.47727 | 0.75790 | 1.36649 | 28 |
| GO_DNA_BIOSYNTHETIC_PROCESS | 0.00001 | 0.00053 | 0.59333 | 0.64365 | 1.23200 | 158 |
| GO_DNA_CONFORMATION_CHANGE | 0.00095 | 0.01903 | 0.47727 | 0.59301 | 1.14547 | 239 |
| GO_DNA_REPAIR | 0.00021 | 0.00586 | 0.51885 | 0.57834 | 1.13040 | 464 |
| GO_DNA_TEMPLATED_TRANSCRIPTION_INITIATION | 0.00253 | 0.04183 | 0.43171 | 0.59625 | 1.14734 | 197 |
| GO_DNA_TEMPLATED_TRANSCRIPTION_TERMINATION | 0.00106 | 0.02060 | 0.45506 | 0.66601 | 1.24868 | 70 |
| GO_ELECTRON_TRANSFER_ACTIVITY | 0.00062 | 0.01338 | 0.47727 | 0.67251 | 1.26404 | 77 |
| GO_ELECTRON_TRANSPORT_CHAIN | 0.00000 | 0.00001 | 0.69013 | 0.68907 | 1.31305 | 132 |
| GO_EMBRYO_IMPLANTATION | 0.00191 | 0.03343 | 0.45506 | 0.73633 | 1.32759 | 28 |
| GO_ENDOPEPTIDASE_COMPLEX | 0.00001 | 0.00031 | 0.59333 | 0.72559 | 1.35940 | 67 |
| GO_ENERGY_DERIVATION_BY_OXIDATION_OF_ORGANIC_CC | 0.00000 | 0.00015 | 0.62726 | 0.64204 | 1.23600 | 201 |
| GO_ESTABLISHMENT_OF_ORGANELLE_LOCALIZATION | 0.00294 | 0.04709 | 0.43171 | 0.57327 | 1.11440 | 337 |
| GO_ESTABLISHMENT_OF_PROTEIN_LOCALIZATION_TO_END | 0.00000 | 0.00000 | | 0.82078 | 1.55473 | 108 |
| GO_ESTABLISHMENT_OF_PROTEIN_LOCALIZATION_TO_MEN | 0.00000 | 0.00000 | | 0.71541 | 1.38537 | 269 |
| GO_ESTABLISHMENT_OF_PROTEIN_LOCALIZATION_TO_MITC | 0.00294 | 0.04709 | 0.43171 | 0.69200 | 1.26988 | 41 |
| GO_ESTABLISHMENT_OF_PROTEIN_LOCALIZATION_TO_ORG | 0.00000 | 0.00000 | | 0.68469 | 1.33825 | 464 |
| GO_ESTABLISHMENT_OF_PROTEIN_LOCALIZATION_TO_TELC | 0.00087 | 0.01778 | 0.47727 | 0.81682 | 1.44009 | 18 |
| GO_ESTABLISHMENT_OF_RNA_LOCALIZATION | 0.00001 | 0.00032 | 0.59333 | 0.64295 | 1.23495 | 179 |
| GO_ESTABLISHMENT_OF_TISSUE_POLARITY | 0.00000 | 0.00000 | 0.74774 | 0.73813 | 1.39533 | 96 |
| GO_EUKARYOTIC_TRANSLATION_INITIATION_FACTOR_3_CO | 0.00170 | 0.03054 | 0.45506 | 0.82172 | 1.44075 | 15 |
| GO_FC_EPSILON_RECEPTOR_SIGNALING_PATHWAY | 0.00000 | 0.00002 | 0.67496 | 0.70934 | 1.34216 | 98 |
| GO_FC_RECEPTOR_SIGNALING_PATHWAY | 0.00000 | 0.00014 | 0.62726 | 0.66263 | 1.26675 | 147 |
| GO_FIBRILLAR_CENTER | 0.00041 | 0.00986 | 0.49849 | 0.65376 | 1.23828 | 101 |
| GO_FICOLIN_1_RICH_GRANULE | 0.00000 | 0.00007 | 0.64355 | 0.70487 | 1.33320 | 97 |
| GO_FOAM_CELL_DIFFERENTIATION | 0.00122 | 0.02326 | 0.45506 | 0.81807 | 1.44054 | 17 |
| GO_GENERATION_OF_PRECURSOR_METABOLITES_AND_EN | 0.00000 | 0.00000 | 0.77494 | 0.63218 | 1.23073 | 367 |
| GO_GENE_SILENCING | 0.00002 | 0.00076 | 0.57561 | 0.62993 | 1.21112 | 193 |
| GO_GTPASE_ACTIVITY | 0.00036 | 0.00896 | 0.49849 | 0.60700 | 1.17033 | 217 |
| GO_GUANYL_NUCLEOTIDE_BINDING | 0.00054 | 0.01208 | 0.47727 | 0.59749 | 1.15604 | 258 |
| GO_HEMATOPOIETIC_PROGENITOR_CELL_DIFFERENTIATION | 0.00000 | 0.00001 | 0.70498 | 0.70359 | 1.34005 | 127 |
| GO_HEMATOPOIETIC_STEM_CELL_DIFFERENTIATION | 0.00000 | 0.00000 | 0.71951 | 0.76594 | 1.43845 | 73 |
| GO_HOST_CELLULAR_COMPONENT | 0.00016 | 0.00457 | 0.51885 | 0.71791 | 1.33535 | 54 |
| GO_IMMUNE_RESPONSE_REGULATING_SIGNALING_PATHW | 0.00002 | 0.00064 | 0.57561 | 0.61917 | 1.19799 | 258 |
| GO_IMPORT_INTO_NUCLEUS | 0.00000 | 0.00019 | 0.61053 | 0.67654 | 1.28854 | 127 |
| GO_INFLAMMATORY_RESPONSE_TO_ANTIGENIC_STIMULUS | 0.00228 | 0.03815 | 0.43171 | 0.76382 | 1.36465 | 24 |

| Cluster 1 | | | | | | |
|---|---|---|---|---|---|---|
| GO_INNATE_IMMUNE_RESPONSE | 0.00000 | 0.00001 | 0.69013 | 0.60742 | 1.18633 | 445 |
| GO_INNATE_IMMUNE_RESPONSE_ACTIVATING_SIGNAL_TRA | 0.00000 | 0.00000 | 0.71951 | 0.74478 | 1.40465 | 85 |
| GO_INNER_MITOCHONDRIAL_MEMBRANE_PROTEIN_COMP | 0.00000 | 0.00004 | 0.65944 | 0.71060 | 1.34252 | 94 |
| GO_INTERACTION_WITH_SYMBIONT | 0.00042 | 0.00996 | 0.49849 | 0.70777 | 1.31520 | 52 |
| GO_INTERLEUKIN_1_MEDIATED_SIGNALING_PATHWAY | 0.00000 | 0.00000 | 0.73376 | 0.74750 | 1.41062 | 88 |
| GO_INTESTINAL_ABSORPTION | 0.00205 | 0.03507 | 0.43171 | 0.79737 | 1.40581 | 18 |
| GO_INTRINSIC_APOPTOTIC_SIGNALING_PATHWAY | 0.00070 | 0.01481 | 0.47727 | 0.60369 | 1.16313 | 212 |
| GO_KILLING_OF_CELLS_OF_OTHER_ORGANISM | 0.00294 | 0.04709 | 0.43171 | 0.81140 | 1.42265 | 15 |
| GO_KINETOCHORE | 0.00071 | 0.01495 | 0.47727 | 0.63695 | 1.21140 | 120 |
| GO_LARGE_RIBOSOMAL_SUBUNIT | 0.00000 | 0.00000 | | 0.80951 | 1.53337 | 108 |
| GO_LEUKOCYTE_MEDIATED_IMMUNITY | 0.00001 | 0.00031 | 0.59333 | 0.59572 | 1.16351 | 449 |
| GO_MAINTENANCE_OF_BLOOD_BRAIN_BARRIER | 0.00148 | 0.02735 | 0.45506 | 0.79068 | 1.40739 | 22 |
| GO_MALE_SEX_DIFFERENTIATION | 0.00191 | 0.03343 | 0.45506 | 0.63311 | 1.19915 | 101 |
| GO_MATERNAL_PLACENTA_DEVELOPMENT | 0.00235 | 0.03928 | 0.43171 | 0.79355 | 1.40330 | 19 |
| GO_MATERNAL_PROCESS_INVOLVED_IN_FEMALE_PREGNAI | 0.00058 | 0.01276 | 0.47727 | 0.74681 | 1.35757 | 33 |
| GO_MATURATION_OF_LSU_RRNA | 0.00110 | 0.02119 | 0.45506 | 0.77879 | 1.40012 | 27 |
| GO_MATURATION_OF_SSU_RRNA | 0.00077 | 0.01599 | 0.47727 | 0.71582 | 1.32414 | 47 |
| GO_MATURATION_OF_SSU_RRNA_FROM_TRICISTRONIC_RI | 0.00161 | 0.02922 | 0.45506 | 0.73144 | 1.33234 | 35 |
| GO_METHYLTRANSFERASE_COMPLEX | 0.00152 | 0.02795 | 0.45506 | 0.64030 | 1.21108 | 97 |
| GO_MICROTUBULE_CYTOSKELETON_ORGANIZATION_INVOL | 0.00042 | 0.00993 | 0.49849 | 0.63924 | 1.21688 | 123 |
| GO_MITOCHONDRIAL_ELECTRON_TRANSPORT_NADH_TO_U | 0.00021 | 0.00564 | 0.51885 | 0.72874 | 1.34613 | 45 |
| GO_MITOCHONDRIAL_GENE_EXPRESSION | 0.00000 | 0.00000 | | 0.71898 | 1.37526 | 154 |
| GO_MITOCHONDRIAL_LARGE_RIBOSOMAL_SUBUNIT | 0.00000 | 0.00000 | 0.74774 | 0.80789 | 1.50273 | 54 |
| GO_MITOCHONDRIAL_MATRIX | 0.00000 | 0.00000 | 0.76146 | 0.62866 | 1.22460 | 380 |
| GO_MITOCHONDRIAL_PROTEIN_COMPLEX | 0.00000 | 0.00000 | | 0.73556 | 1.41735 | 213 |
| GO_MITOCHONDRIAL_SMALL_RIBOSOMAL_SUBUNIT | 0.00025 | 0.00664 | 0.49849 | 0.77246 | 1.39273 | 28 |
| GO_MITOCHONDRIAL_TRANSLATION | 0.00000 | 0.00000 | | 0.73627 | 1.40233 | 130 |
| GO_MITOCHONDRIAL_TRANSLATIONAL_TERMINATION | 0.00000 | 0.00000 | | 0.78652 | 1.48425 | 88 |
| GO_MITOCHONDRIAL_TRANSPORT | 0.00205 | 0.03507 | 0.43171 | 0.59826 | 1.15120 | 197 |
| GO_MITOCHONDRION_ORGANIZATION | 0.00000 | 0.00001 | 0.69013 | 0.61110 | 1.19237 | 426 |
| GO_MITOTIC_NUCLEAR_DIVISION | 0.00012 | 0.00356 | 0.53843 | 0.60750 | 1.17359 | 240 |
| GO_MITOTIC_SISTER_CHROMATID_SEGREGATION | 0.00024 | 0.00633 | 0.51885 | 0.63218 | 1.20854 | 147 |
| GO_MITOTIC_SPINDLE | 0.00283 | 0.04600 | 0.43171 | 0.62408 | 1.18345 | 111 |
| GO_MITOTIC_SPINDLE_ASSEMBLY | 0.00042 | 0.00993 | 0.49849 | 0.68537 | 1.27696 | 57 |
| GO_MITOTIC_SPINDLE_ORGANIZATION | 0.00066 | 0.01413 | 0.47727 | 0.64766 | 1.22672 | 101 |
| GO_MORPHOGENESIS_OF_AN_EPITHELIUM | 0.00027 | 0.00691 | 0.49849 | 0.58528 | 1.13854 | 348 |
| GO_MORPHOGENESIS_OF_A_POLARIZED_EPITHELIUM | 0.00000 | 0.00000 | 0.76146 | 0.73045 | 1.38514 | 111 |
| GO_MRNA_3_END_PROCESSING | 0.00070 | 0.01481 | 0.47727 | 0.65216 | 1.23238 | 95 |
| GO_MRNA_5_UTR_BINDING | 0.00104 | 0.02043 | 0.45506 | 0.76938 | 1.38020 | 26 |
| GO_MRNA_BINDING | 0.00000 | 0.00000 | | 0.68840 | 1.33160 | 256 |
| GO_MRNA_EXPORT_FROM_NUCLEUS | 0.00027 | 0.00697 | 0.49849 | 0.65830 | 1.24611 | 104 |
| GO_MRNA_PROCESSING | 0.00000 | 0.00000 | | 0.67561 | 1.31959 | 447 |
| GO_MRNA_TRANSPORT | 0.00017 | 0.00468 | 0.51885 | 0.63817 | 1.21640 | 137 |
| GO_MULTI_ORGANISM_LOCALIZATION | 0.00016 | 0.00448 | 0.51885 | 0.72041 | 1.34038 | 55 |
| GO_MYELOID_LEUKOCYTE_ACTIVATION | 0.00001 | 0.00044 | 0.59333 | 0.59928 | 1.16766 | 387 |
| GO_MYELOID_LEUKOCYTE_MEDIATED_IMMUNITY | 0.00000 | 0.00003 | 0.65944 | 0.61375 | 1.19398 | 353 |
| GO_NADH_DEHYDROGENASE_ACTIVITY | 0.00048 | 0.01108 | 0.49849 | 0.72319 | 1.32159 | 38 |
| GO_NADH_DEHYDROGENASE_COMPLEX | 0.00059 | 0.01291 | 0.47727 | 0.72299 | 1.32674 | 41 |
| GO_NADH_DEHYDROGENASE_COMPLEX_ASSEMBLY | 0.00058 | 0.01276 | 0.47727 | 0.69711 | 1.29867 | 56 |
| GO_NCRNA_EXPORT_FROM_NUCLEUS | 0.00305 | 0.04791 | 0.43171 | 0.72046 | 1.31235 | 35 |
| GO_NCRNA_METABOLIC_PROCESS | 0.00000 | 0.00000 | | 0.64225 | 1.25259 | 418 |
| GO_NCRNA_PROCESSING | 0.00000 | 0.00000 | | 0.67078 | 1.30498 | 351 |
| GO_NEGATIVE_REGULATION_OF_CANONICAL_WNT_SIGNAI | 0.00000 | 0.00002 | 0.67496 | 0.69180 | 1.31792 | 125 |
| GO_NEGATIVE_REGULATION_OF_CELLULAR_AMIDE_METAB | 0.00130 | 0.02444 | 0.45506 | 0.62478 | 1.19024 | 125 |
| GO_NEGATIVE_REGULATION_OF_CELL_CYCLE | 0.00000 | 0.00002 | 0.67496 | 0.60041 | 1.17382 | 481 |
| GO_NEGATIVE_REGULATION_OF_CELL_CYCLE_G2_M_PHASI | 0.00000 | 0.00000 | 0.70498 | 0.73889 | 1.39436 | 88 |
| GO_NEGATIVE_REGULATION_OF_CELL_CYCLE_PHASE_TRAN | 0.00000 | 0.00002 | 0.67496 | 0.65330 | 1.25747 | 199 |
| GO_NEGATIVE_REGULATION_OF_CELL_CYCLE_PROCESS | 0.00000 | 0.00000 | 0.74774 | 0.64526 | 1.24968 | 270 |
| GO_NEGATIVE_REGULATION_OF_CHROMOSOME_ORGANIZ | 0.00133 | 0.02499 | 0.45506 | 0.63216 | 1.20100 | 116 |
| GO_NEGATIVE_REGULATION_OF_HYDROLASE_ACTIVITY | 0.00305 | 0.04791 | 0.43171 | 0.58480 | 1.13009 | 244 |
| GO_NEGATIVE_REGULATION_OF_MITOTIC_CELL_CYCLE | 0.00000 | 0.00006 | 0.64355 | 0.62662 | 1.21213 | 260 |
| GO_NEGATIVE_REGULATION_OF_ORGANELLE_ORGANIZATIO | 0.00265 | 0.04333 | 0.43171 | 0.57786 | 1.12198 | 309 |
| GO_NEGATIVE_REGULATION_OF_PEPTIDE_SECRETION | 0.00091 | 0.01841 | 0.47727 | 0.69584 | 1.29373 | 53 |
| GO_NEGATIVE_REGULATION_OF_WNT_SIGNALING_PATHWA | 0.00000 | 0.00006 | 0.64355 | 0.66764 | 1.27545 | 144 |
| GO_NIK_NF_KAPPAB_SIGNALING | 0.00000 | 0.00000 | 0.71951 | 0.70754 | 1.34730 | 124 |
| GO_NON_CANONICAL_WNT_SIGNALING_PATHWAY | 0.00000 | 0.00000 | 0.73376 | 0.71485 | 1.35839 | 117 |
| GO_NUCLEAR_CHROMOSOME_SEGREGATION | 0.00126 | 0.02382 | 0.45506 | 0.59335 | 1.14488 | 224 |
| GO_NUCLEAR_ENVELOPE | 0.00006 | 0.00204 | 0.53843 | 0.59222 | 1.15182 | 345 |
| GO_NUCLEAR_EXPORT | 0.00000 | 0.00005 | 0.64355 | 0.65575 | 1.25990 | 183 |
| GO_NUCLEAR_LOCALIZATION_SEQUENCE_BINDING | 0.00209 | 0.03560 | 0.43171 | 0.78496 | 1.39721 | 22 |
| GO_NUCLEAR_MATRIX | 0.00037 | 0.00904 | 0.49849 | 0.65939 | 1.24605 | 95 |
| GO_NUCLEAR_PERIPHERY | 0.00009 | 0.00283 | 0.53843 | 0.65715 | 1.24849 | 116 |
| GO_NUCLEAR_PORE | 0.00276 | 0.04492 | 0.43171 | 0.66070 | 1.23784 | 67 |

| Cluster 1 | | | | | | |
|---|---|---|---|---|---|---|
| GO_NUCLEAR_SPECK | 0.00000 | 0.00001 | 0.70498 | 0.62620 | 1.21669 | 332 |
| GO_NUCLEAR_TRANSCRIBED_MRNA_CATABOLIC_PROCESS | 0.00000 | 0.00000 | | 0.73108 | 1.40568 | 188 |
| GO_NUCLEAR_TRANSCRIBED_MRNA_CATABOLIC_PROCESS_ | 0.00000 | 0.00000 | | 0.80310 | 1.52409 | 114 |
| GO_NUCLEAR_TRANSPORT | 0.00000 | 0.00000 | 0.80122 | 0.65164 | 1.26363 | 294 |
| GO_NUCLEIC_ACID_PHOSPHODIESTER_BOND_HYDROLYSIS | 0.00145 | 0.02682 | 0.45506 | 0.58677 | 1.13502 | 256 |
| GO_NUCLEOBASE_CONTAINING_COMPOUND_TRANSPORT | 0.00001 | 0.00052 | 0.59333 | 0.63007 | 1.21537 | 222 |
| GO_NUCLEOCYTOPLASMIC_CARRIER_ACTIVITY | 0.00224 | 0.03772 | 0.43171 | 0.73210 | 1.32297 | 29 |
| GO_NUCLEOSIDE_DIPHOSPHATE_METABOLIC_PROCESS | 0.00022 | 0.00591 | 0.51885 | 0.65071 | 1.23437 | 113 |
| GO_NUCLEOSIDE_TRIPHOSPHATE_METABOLIC_PROCESS | 0.00074 | 0.01533 | 0.47727 | 0.67865 | 1.27204 | 68 |
| GO_NUCLEOTIDE_PHOSPHORYLATION | 0.00011 | 0.00324 | 0.53843 | 0.66120 | 1.25187 | 102 |
| GO_NUCLEUS_ORGANIZATION | 0.00082 | 0.01677 | 0.47727 | 0.63442 | 1.20353 | 112 |
| GO_ORGANELLAR_RIBOSOME | 0.00000 | 0.00000 | | 0.78971 | 1.48910 | 84 |
| GO_ORGANELLE_DISASSEMBLY | 0.00053 | 0.01196 | 0.47727 | 0.64857 | 1.22823 | 100 |
| GO_ORGANELLE_INNER_MEMBRANE | 0.00000 | 0.00000 | | 0.67273 | 1.31184 | 413 |
| GO_ORGANIC_CYCLIC_COMPOUND_CATABOLIC_PROCESS | 0.00000 | 0.00000 | | 0.68671 | 1.34251 | 494 |
| GO_OSTEOBLAST_DIFFERENTIATION | 0.00294 | 0.04709 | 0.43171 | 0.61849 | 1.17833 | 126 |
| GO_OXIDATIVE_PHOSPHORYLATION | 0.00000 | 0.00000 | 0.71951 | 0.72487 | 1.37297 | 101 |
| GO_OXIDOREDUCTASE_ACTIVITY | 0.00106 | 0.02060 | 0.45506 | 0.56840 | 1.11102 | 465 |
| GO_OXIDOREDUCTASE_ACTIVITY_ACTING_ON_A_HEME_GR | 0.00198 | 0.03462 | 0.43171 | 0.79076 | 1.40292 | 20 |
| GO_OXIDOREDUCTASE_ACTIVITY_ACTING_ON_NAD_P_H | 0.00097 | 0.01922 | 0.47727 | 0.66305 | 1.24665 | 75 |
| GO_OXIDOREDUCTASE_ACTIVITY_ACTING_ON_NAD_P_H_Q | 0.00185 | 0.03264 | 0.45506 | 0.70224 | 1.29332 | 44 |
| GO_OXIDOREDUCTASE_COMPLEX | 0.00001 | 0.00036 | 0.59333 | 0.70345 | 1.32617 | 82 |
| GO_PEPTIDASE_COMPLEX | 0.00001 | 0.00022 | 0.61053 | 0.70944 | 1.33800 | 85 |
| GO_PHAGOPHORE_ASSEMBLY_SITE | 0.00040 | 0.00978 | 0.49849 | 0.76147 | 1.37605 | 29 |
| GO_PHAGOPHORE_ASSEMBLY_SITE_MEMBRANE | 0.00018 | 0.00502 | 0.51885 | 0.86477 | 1.51623 | 15 |
| GO_PIGMENT_GRANULE | 0.00011 | 0.00324 | 0.53843 | 0.66921 | 1.26400 | 92 |
| GO_POLYSOMAL_RIBOSOME | 0.00001 | 0.00025 | 0.61053 | 0.82740 | 1.49723 | 30 |
| GO_POLYSOME | 0.00000 | 0.00000 | 0.73376 | 0.78849 | 1.47517 | 65 |
| GO_POSITIVE_REGULATION_OF_BLOOD_CIRCULATION | 0.00004 | 0.00145 | 0.55733 | 0.79283 | 1.43271 | 29 |
| GO_POSITIVE_REGULATION_OF_BLOOD_VESSEL_DIAMETER | 0.00022 | 0.00601 | 0.51885 | 0.77770 | 1.40731 | 30 |
| GO_POSITIVE_REGULATION_OF_CANONICAL_WNT_SIGNALI | 0.00027 | 0.00691 | 0.49849 | 0.64865 | 1.23233 | 116 |
| GO_POSITIVE_REGULATION_OF_CELLULAR_AMIDE_METABO | 0.00000 | 0.00004 | 0.65944 | 0.68893 | 1.31212 | 127 |
| GO_POSITIVE_REGULATION_OF_CELLULAR_PROTEIN_LOCAL | 0.00079 | 0.01631 | 0.47727 | 0.59482 | 1.14947 | 244 |
| GO_POSITIVE_REGULATION_OF_CELL_CYCLE | 0.00072 | 0.01510 | 0.47727 | 0.58944 | 1.14257 | 285 |
| GO_POSITIVE_REGULATION_OF_CELL_CYCLE_PROCESS | 0.00250 | 0.04133 | 0.43171 | 0.59427 | 1.14631 | 222 |
| GO_POSITIVE_REGULATION_OF_CELL_DEATH | 0.00057 | 0.01264 | 0.47727 | 0.57345 | 1.12084 | 464 |
| GO_POSITIVE_REGULATION_OF_DEFENSE_RESPONSE | 0.00000 | 0.00001 | 0.69013 | 0.65656 | 1.26456 | 206 |
| GO_POSITIVE_REGULATION_OF_DNA_BINDING | 0.00050 | 0.01138 | 0.47727 | 0.73809 | 1.34646 | 37 |
| GO_POSITIVE_REGULATION_OF_DNA_BIOSYNTHETIC_PROC | 0.00000 | 0.00018 | 0.61053 | 0.74578 | 1.39151 | 60 |
| GO_POSITIVE_REGULATION_OF_DNA_METABOLIC_PROCESS | 0.00082 | 0.01677 | 0.47727 | 0.62104 | 1.18766 | 149 |
| GO_POSITIVE_REGULATION_OF_FIBROBLAST_PROLIFERATIC | 0.00016 | 0.00444 | 0.51885 | 0.77902 | 1.40455 | 28 |
| GO_POSITIVE_REGULATION_OF_GENE_EXPRESSION_EPIGEN | 0.00202 | 0.03462 | 0.43171 | 0.69757 | 1.28928 | 46 |
| GO_POSITIVE_REGULATION_OF_IMMUNE_RESPONSE | 0.00000 | 0.00002 | 0.67496 | 0.61767 | 1.20208 | 361 |
| GO_POSITIVE_REGULATION_OF_INNATE_IMMUNE_RESPON | 0.00000 | 0.00000 | 0.76146 | 0.71077 | 1.35408 | 134 |
| GO_POSITIVE_REGULATION_OF_MITOTIC_CELL_CYCLE | 0.00163 | 0.02947 | 0.45506 | 0.62535 | 1.19104 | 127 |
| GO_POSITIVE_REGULATION_OF_MRNA_SPLICING_VIA_SPLIC | 0.00202 | 0.03462 | 0.43171 | 0.80572 | 1.41879 | 17 |
| GO_POSITIVE_REGULATION_OF_ORGANELLE_ORGANIZATIO | 0.00111 | 0.02142 | 0.45506 | 0.56716 | 1.10858 | 472 |
| GO_POSITIVE_REGULATION_OF_PEPTIDASE_ACTIVITY | 0.00165 | 0.02972 | 0.45506 | 0.61677 | 1.17560 | 137 |
| GO_POSITIVE_REGULATION_OF_PROTEIN_LOCALIZATION_T | 0.00057 | 0.01264 | 0.47727 | 0.69031 | 1.28974 | 62 |
| GO_POSITIVE_REGULATION_OF_RESPONSE_TO_BIOTIC_STI | 0.00000 | 0.00000 | 0.71951 | 0.68221 | 1.30493 | 154 |
| GO_POSITIVE_REGULATION_OF_RESPONSE_TO_EXTERNAL_ | 0.00000 | 0.00004 | 0.65944 | 0.63093 | 1.22233 | 274 |
| GO_POSITIVE_REGULATION_OF_TELOMERASE_ACTIVITY | 0.00033 | 0.00838 | 0.49849 | 0.75425 | 1.37110 | 33 |
| GO_POSITIVE_REGULATION_OF_TELOMERASE_RNA_LOCALI | 0.00031 | 0.00772 | 0.49849 | 0.85666 | 1.50202 | 15 |
| GO_POSITIVE_REGULATION_OF_TELOMERE_MAINTENANCE | 0.00106 | 0.02060 | 0.45506 | 0.72718 | 1.32493 | 36 |
| GO_POSITIVE_REGULATION_OF_TRANSLATION | 0.00000 | 0.00001 | 0.69013 | 0.70810 | 1.34128 | 108 |
| GO_POSITIVE_REGULATION_OF_VIRAL_PROCESS | 0.00004 | 0.00140 | 0.55733 | 0.68627 | 1.29567 | 90 |
| GO_POSITIVE_REGULATION_OF_VIRAL_TRANSCRIPTION | 0.00302 | 0.04780 | 0.43171 | 0.70919 | 1.29374 | 37 |
| GO_POSITIVE_REGULATION_OF_WNT_SIGNALING_PATHWA | 0.00154 | 0.02820 | 0.45506 | 0.62024 | 1.18268 | 139 |
| GO_PRECATALYTIC_SPLICEOSOME | 0.00019 | 0.00523 | 0.51885 | 0.71785 | 1.33300 | 51 |
| GO_PRERIBOSOME | 0.00000 | 0.00001 | 0.70498 | 0.75461 | 1.41544 | 71 |
| GO_PRERIBOSOME_LARGE_SUBUNIT_PRECURSOR | 0.00202 | 0.03462 | 0.43171 | 0.76626 | 1.36503 | 23 |
| GO_PRE_MRNA_BINDING | 0.00031 | 0.00772 | 0.49849 | 0.77371 | 1.40378 | 31 |
| GO_PROTEASOMAL_PROTEIN_CATABOLIC_PROCESS | 0.00010 | 0.00304 | 0.53843 | 0.58151 | 1.13398 | 412 |
| GO_PROTEASOMAL_UBIQUITIN_INDEPENDENT_PROTEIN_C | 0.00000 | 0.00016 | 0.62726 | 0.88871 | 1.57670 | 20 |
| GO_PROTEASOME_ACCESSORY_COMPLEX | 0.00042 | 0.00996 | 0.49849 | 0.79690 | 1.42375 | 24 |
| GO_PROTEASOME_CORE_COMPLEX | 0.00000 | 0.00006 | 0.64355 | 0.90934 | 1.60805 | 19 |
| GO_PROTEIN_CONTAINING_COMPLEX_DISASSEMBLY | 0.00000 | 0.00000 | | 0.67358 | 1.30495 | 274 |
| GO_PROTEIN_CONTAINING_COMPLEX_LOCALIZATION | 0.00006 | 0.00183 | 0.55733 | 0.61573 | 1.18906 | 237 |
| GO_PROTEIN_DNA_COMPLEX_SUBUNIT_ORGANIZATION | 0.00023 | 0.00624 | 0.51885 | 0.61471 | 1.18179 | 189 |
| GO_PROTEIN_FOLDING | 0.00002 | 0.00072 | 0.57561 | 0.63631 | 1.22304 | 187 |
| GO_PROTEIN_FOLDING_CHAPERONE | 0.00056 | 0.01257 | 0.47727 | 0.81048 | 1.44263 | 22 |
| GO_PROTEIN_IMPORT | 0.00001 | 0.00037 | 0.59333 | 0.65789 | 1.25852 | 153 |
| GO_PROTEIN_LIPID_COMPLEX_ASSEMBLY | 0.00002 | 0.00068 | 0.57561 | 0.88516 | 1.56059 | 18 |

| Cluster 1 | | | | | | |
|---|---|---|---|---|---|---|
| GO_PROTEIN_LIPID_COMPLEX_SUBUNIT_ORGANIZATION | 0.00002 | 0.00065 | 0.57561 | 0.87006 | 1.54360 | 20 |
| GO_PROTEIN_LOCALIZATION_TO_CHROMOSOME_TELOMER | 0.00087 | 0.01778 | 0.47727 | 0.75004 | 1.35231 | 28 |
| GO_PROTEIN_LOCALIZATION_TO_ENDOPLASMIC_RETICULU | 0.00000 | 0.00000 | | 0.79846 | 1.52151 | 132 |
| GO_PROTEIN_LOCALIZATION_TO_MEMBRANE | 0.00000 | 0.00000 | | 0.63355 | 1.23835 | 471 |
| GO_PROTEIN_LOCALIZATION_TO_MITOCHONDRION | 0.00030 | 0.00767 | 0.49849 | 0.64398 | 1.22441 | 118 |
| GO_PROTEIN_LOCALIZATION_TO_NUCLEUS | 0.00000 | 0.00000 | 0.73376 | 0.66259 | 1.27675 | 213 |
| GO_PROTEIN_MODIFICATION_BY_SMALL_PROTEIN_REMOV | 0.00047 | 0.01090 | 0.49849 | 0.59771 | 1.15510 | 247 |
| GO_PROTEIN_POLYUBIQUITINATION | 0.00141 | 0.02622 | 0.45506 | 0.58301 | 1.13054 | 294 |
| GO_PROTEIN_STABILIZATION | 0.00009 | 0.00277 | 0.53843 | 0.64133 | 1.22368 | 141 |
| GO_PROTEIN_TARGETING | 0.00000 | 0.00000 | | 0.68533 | 1.33414 | 364 |
| GO_PROTEIN_TARGETING_TO_MEMBRANE | 0.00000 | 0.00000 | | 0.76971 | 1.47687 | 170 |
| GO_PROTEIN_TARGETING_TO_MITOCHONDRION | 0.00064 | 0.01363 | 0.47727 | 0.65861 | 1.24163 | 82 |
| GO_PURINE_NUCLEOSIDE_TRIPHOSPHATE_METABOLIC_PRO | 0.00009 | 0.00289 | 0.53843 | 0.72847 | 1.35273 | 51 |
| GO_PYRUVATE_METABOLIC_PROCESS | 0.00006 | 0.00194 | 0.55733 | 0.66909 | 1.26812 | 109 |
| GO_RECEPTOR_INHIBITOR_ACTIVITY | 0.00093 | 0.01872 | 0.47727 | 0.83649 | 1.46665 | 15 |
| GO_REGULATION_OF_ANIMAL_ORGAN_MORPHOGENESIS | 0.00000 | 0.00000 | 0.76146 | 0.70157 | 1.34167 | 149 |
| GO_REGULATION_OF_APOPTOTIC_SIGNALING_PATHWAY | 0.00057 | 0.01264 | 0.47727 | 0.58847 | 1.14020 | 278 |
| GO_REGULATION_OF_ATP_METABOLIC_PROCESS | 0.00009 | 0.00283 | 0.53843 | 0.67692 | 1.27819 | 91 |
| GO_REGULATION_OF_CARBOHYDRATE_CATABOLIC_PROCES | 0.00242 | 0.04031 | 0.43171 | 0.66302 | 1.24219 | 67 |
| GO_REGULATION_OF_CELLULAR_AMIDE_METABOLIC_PROC | 0.00000 | 0.00000 | 0.80122 | 0.63916 | 1.24313 | 345 |
| GO_REGULATION_OF_CELLULAR_AMINE_METABOLIC_PROC | 0.00000 | 0.00001 | 0.69013 | 0.77567 | 1.45118 | 65 |
| GO_REGULATION_OF_CELLULAR_AMINO_ACID_METABOLIC | 0.00000 | 0.00000 | 0.73376 | 0.79615 | 1.48130 | 55 |
| GO_REGULATION_OF_CELLULAR_KETONE_METABOLIC_PRO | 0.00000 | 0.00001 | 0.70498 | 0.69480 | 1.32333 | 130 |
| GO_REGULATION_OF_CELLULAR_PROTEIN_LOCALIZATION | 0.00012 | 0.00345 | 0.53843 | 0.58406 | 1.13835 | 397 |
| GO_REGULATION_OF_CELLULAR_RESPONSE_TO_HEAT | 0.00006 | 0.00207 | 0.53843 | 0.70238 | 1.31653 | 69 |
| GO_REGULATION_OF_CELL_CYCLE_G2_M_PHASE_TRANSITI | 0.00000 | 0.00008 | 0.64355 | 0.64716 | 1.24445 | 191 |
| GO_REGULATION_OF_CELL_CYCLE_PHASE_TRANSITION | 0.00001 | 0.00029 | 0.61053 | 0.59922 | 1.16705 | 376 |
| GO_REGULATION_OF_CHROMOSOME_ORGANIZATION | 0.00002 | 0.00057 | 0.59333 | 0.61271 | 1.18784 | 287 |
| GO_REGULATION_OF_CHROMOSOME_SEGREGATION | 0.00156 | 0.02838 | 0.45506 | 0.64370 | 1.21640 | 95 |
| GO_REGULATION_OF_CHROMOSOME_SEPARATION | 0.00309 | 0.04837 | 0.43171 | 0.66496 | 1.24237 | 62 |
| GO_REGULATION_OF_CYTOPLASMIC_TRANSLATION | 0.00122 | 0.02326 | 0.45506 | 0.74413 | 1.34164 | 28 |
| GO_REGULATION_OF_DEFENSE_RESPONSE | 0.00004 | 0.00131 | 0.55733 | 0.59131 | 1.15176 | 378 |
| GO_REGULATION_OF_DNA_BIOSYNTHETIC_PROCESS | 0.00000 | 0.00004 | 0.65944 | 0.71614 | 1.35100 | 87 |
| GO_REGULATION_OF_DNA_METABOLIC_PROCESS | 0.00202 | 0.03462 | 0.43171 | 0.58123 | 1.12568 | 270 |
| GO_REGULATION_OF_DNA_TEMPLATED_TRANSCRIPTION_I | 0.00000 | 0.00000 | 0.71951 | 0.73000 | 1.37996 | 96 |
| GO_REGULATION_OF_GENERATION_OF_PRECURSOR_META | 0.00034 | 0.00857 | 0.49849 | 0.64634 | 1.22794 | 116 |
| GO_REGULATION_OF_GENE_SILENCING | 0.00011 | 0.00336 | 0.53843 | 0.66993 | 1.26568 | 94 |
| GO_REGULATION_OF_HEMATOPOIETIC_PROGENITOR_CELL_ | 0.00000 | 0.00000 | 0.73376 | 0.76413 | 1.43669 | 75 |
| GO_REGULATION_OF_HEMATOPOIETIC_STEM_CELL_DIFFER | 0.00000 | 0.00000 | 0.71951 | 0.77950 | 1.45736 | 63 |
| GO_REGULATION_OF_HEMOPOIESIS | 0.00242 | 0.04031 | 0.43171 | 0.57860 | 1.12203 | 293 |
| GO_REGULATION_OF_IMMUNE_RESPONSE | 0.00000 | 0.00013 | 0.62726 | 0.59184 | 1.15707 | 481 |
| GO_REGULATION_OF_INNATE_IMMUNE_RESPONSE | 0.00000 | 0.00000 | 0.77494 | 0.67623 | 1.30021 | 188 |
| GO_REGULATION_OF_INTRINSIC_APOPTOTIC_SIGNALING_P | 0.00045 | 0.01045 | 0.49849 | 0.64410 | 1.22369 | 116 |
| GO_REGULATION_OF_MORPHOGENESIS_OF_AN_EPITHELIU | 0.00000 | 0.00000 | 0.81404 | 0.73172 | 1.39204 | 121 |
| GO_REGULATION_OF_MRNA_CATABOLIC_PROCESS | 0.00000 | 0.00000 | 0.78819 | 0.69170 | 1.32727 | 173 |
| GO_REGULATION_OF_MRNA_METABOLIC_PROCESS | 0.00000 | 0.00000 | | 0.69095 | 1.33949 | 284 |
| GO_REGULATION_OF_MRNA_PROCESSING | 0.00000 | 0.00011 | 0.62726 | 0.68118 | 1.29737 | 127 |
| GO_REGULATION_OF_MRNA_SPLICING_VIA_SPLICEOSOME | 0.00002 | 0.00057 | 0.59333 | 0.69381 | 1.30989 | 90 |
| GO_REGULATION_OF_NUCLEAR_DIVISION | 0.00054 | 0.01208 | 0.47727 | 0.62237 | 1.19047 | 154 |
| GO_REGULATION_OF_NUCLEOCYTOPLASMIC_TRANSPORT | 0.00126 | 0.02382 | 0.45506 | 0.65565 | 1.23744 | 89 |
| GO_REGULATION_OF_PEPTIDASE_ACTIVITY | 0.00074 | 0.01533 | 0.47727 | 0.59619 | 1.15286 | 251 |
| GO_REGULATION_OF_PHAGOCYTOSIS | 0.00313 | 0.04883 | 0.43171 | 0.68889 | 1.26698 | 43 |
| GO_REGULATION_OF_PLASMA_LIPOPROTEIN_PARTICLE_LEV | 0.00011 | 0.00336 | 0.53843 | 0.72396 | 1.34119 | 49 |
| GO_REGULATION_OF_POSTTRANSCRIPTIONAL_GENE_SILEN | 0.00015 | 0.00439 | 0.51885 | 0.68005 | 1.28232 | 84 |
| GO_REGULATION_OF_PROTEIN_LOCALIZATION_TO_NUCLEU | 0.00005 | 0.00169 | 0.55733 | 0.67744 | 1.27938 | 93 |
| GO_REGULATION_OF_PROTEIN_STABILITY | 0.00023 | 0.00604 | 0.51885 | 0.60666 | 1.17102 | 231 |
| GO_REGULATION_OF_PROTEOLYSIS | 0.00066 | 0.01413 | 0.47727 | 0.57054 | 1.11502 | 461 |
| GO_REGULATION_OF_RESPONSE_TO_BIOTIC_STIMULUS | 0.00000 | 0.00020 | 0.61053 | 0.62335 | 1.20458 | 244 |
| GO_REGULATION_OF_RNA_SPLICING | 0.00000 | 0.00001 | 0.69013 | 0.69718 | 1.32796 | 128 |
| GO_REGULATION_OF_SISTER_CHROMATID_SEGREGATION | 0.00220 | 0.03720 | 0.43171 | 0.65440 | 1.23030 | 76 |
| GO_REGULATION_OF_SMALL_MOLECULE_METABOLIC_PRO | 0.00000 | 0.00020 | 0.61053 | 0.61418 | 1.19239 | 306 |
| GO_REGULATION_OF_STEM_CELL_DIFFERENTIATION | 0.00000 | 0.00000 | 0.71951 | 0.73132 | 1.38167 | 94 |
| GO_REGULATION_OF_SYMBIOTIC_PROCESS | 0.00000 | 0.00018 | 0.61053 | 0.65982 | 1.26217 | 152 |
| GO_REGULATION_OF_TELOMERASE_ACTIVITY | 0.00042 | 0.00993 | 0.49849 | 0.72914 | 1.33915 | 42 |
| GO_REGULATION_OF_TELOMERE_MAINTENANCE_VIA_TELO | 0.00090 | 0.01828 | 0.47727 | 0.69698 | 1.29680 | 55 |
| GO_REGULATION_OF_TRANSCRIPTION_FROM_RNA_POLYM | 0.00000 | 0.00000 | 0.74774 | 0.78337 | 1.46832 | 68 |
| GO_REGULATION_OF_TRANSLATIONAL_INITIATION | 0.00013 | 0.00375 | 0.51885 | 0.69763 | 1.30761 | 68 |
| GO_REGULATION_OF_UBIQUITIN_PROTEIN_LIGASE_ACTIVIT | 0.00038 | 0.00934 | 0.49849 | 0.79860 | 1.42678 | 24 |
| GO_REGULATION_OF_UBIQUITIN_PROTEIN_TRANSFERASE_ | 0.00176 | 0.03127 | 0.45506 | 0.69708 | 1.29007 | 48 |
| GO_REGULATION_OF_VIRAL_TRANSCRIPTION | 0.00056 | 0.01257 | 0.47727 | 0.70200 | 1.30520 | 53 |
| GO_RESPIRASOME | 0.00000 | 0.00006 | 0.64355 | 0.73409 | 1.38021 | 75 |
| GO_RESPIRATORY_CHAIN_COMPLEX | 0.00002 | 0.00068 | 0.57561 | 0.73011 | 1.36593 | 65 |
| GO_RESPIRATORY_ELECTRON_TRANSPORT_CHAIN | 0.00000 | 0.00002 | 0.67496 | 0.71908 | 1.35760 | 90 |

| Cluster 1 | | | | | | |
|---|---|---|---|---|---|---|
| GO_RESPONSE_TO_DRUG | 0.00064 | 0.01363 | 0.47727 | 0.60119 | 1.16048 | 230 |
| GO_RESPONSE_TO_FIBROBLAST_GROWTH_FACTOR | 0.00202 | 0.03462 | 0.43171 | 0.64504 | 1.21606 | 83 |
| GO_RESPONSE_TO_HEAT | 0.00002 | 0.00079 | 0.57561 | 0.66505 | 1.26600 | 123 |
| GO_RESPONSE_TO_INTERLEUKIN_1 | 0.00000 | 0.00000 | 0.76146 | 0.71214 | 1.35669 | 134 |
| GO_RESPONSE_TO_INTERLEUKIN_4 | 0.00305 | 0.04791 | 0.43171 | 0.77339 | 1.37661 | 22 |
| GO_RESPONSE_TO_OXYGEN_LEVELS | 0.00000 | 0.00003 | 0.65944 | 0.63096 | 1.22201 | 272 |
| GO_RESPONSE_TO_TEMPERATURE_STIMULUS | 0.00005 | 0.00152 | 0.55733 | 0.63876 | 1.22317 | 161 |
| GO_RESPONSE_TO_TUMOR_NECROSIS_FACTOR | 0.00007 | 0.00230 | 0.53843 | 0.62174 | 1.19537 | 193 |
| GO_RESPONSE_TO_VIRUS | 0.00202 | 0.03462 | 0.43171 | 0.60139 | 1.15544 | 182 |
| GO_RIBONUCLEOPROTEIN_COMPLEX_BINDING | 0.00000 | 0.00000 | 0.71951 | 0.71158 | 1.35292 | 119 |
| GO_RIBONUCLEOPROTEIN_COMPLEX_BIOGENESIS | 0.00000 | 0.00000 | | 0.73515 | 1.43285 | 400 |
| GO_RIBONUCLEOPROTEIN_COMPLEX_SUBUNIT_ORGANIZAT | 0.00000 | 0.00000 | | 0.73980 | 1.42058 | 178 |
| GO_RIBONUCLEOPROTEIN_GRANULE | 0.00007 | 0.00220 | 0.53843 | 0.62424 | 1.20172 | 201 |
| GO_RIBONUCLEOSIDE_TRIPHOSPHATE_METABOLIC_PROCES | 0.00017 | 0.00485 | 0.51885 | 0.72087 | 1.33546 | 49 |
| GO_RIBOSOMAL_LARGE_SUBUNIT_ASSEMBLY | 0.00063 | 0.01353 | 0.47727 | 0.78892 | 1.41832 | 27 |
| GO_RIBOSOMAL_LARGE_SUBUNIT_BIOGENESIS | 0.00000 | 0.00000 | 0.77494 | 0.78753 | 1.47612 | 69 |
| GO_RIBOSOMAL_SMALL_SUBUNIT_ASSEMBLY | 0.00044 | 0.01038 | 0.49849 | 0.83252 | 1.46778 | 18 |
| GO_RIBOSOMAL_SMALL_SUBUNIT_BIOGENESIS | 0.00000 | 0.00013 | 0.62726 | 0.73499 | 1.37763 | 68 |
| GO_RIBOSOMAL_SUBUNIT | 0.00000 | 0.00000 | | 0.78975 | 1.51590 | 174 |
| GO_RIBOSOME | 0.00000 | 0.00000 | | 0.78206 | 1.50656 | 209 |
| GO_RIBOSOME_ASSEMBLY | 0.00000 | 0.00001 | 0.70498 | 0.77739 | 1.44842 | 57 |
| GO_RIBOSOME_BINDING | 0.00202 | 0.03462 | 0.43171 | 0.69383 | 1.28406 | 48 |
| GO_RIBOSOME_BIOGENESIS | 0.00000 | 0.00000 | | 0.73053 | 1.41602 | 282 |
| GO_RNA_3_END_PROCESSING | 0.00000 | 0.00014 | 0.62726 | 0.66735 | 1.27166 | 132 |
| GO_RNA_CATABOLIC_PROCESS | 0.00000 | 0.00000 | | 0.71464 | 1.39051 | 355 |
| GO_RNA_DEPENDENT_DNA_BIOSYNTHETIC_PROCESS | 0.00011 | 0.00336 | 0.53843 | 0.70311 | 1.31627 | 66 |
| GO_RNA_EXPORT_FROM_NUCLEUS | 0.00001 | 0.00031 | 0.59333 | 0.66501 | 1.26659 | 130 |
| GO_RNA_HELICASE_ACTIVITY | 0.00228 | 0.03815 | 0.43171 | 0.65654 | 1.23092 | 70 |
| GO_RNA_LOCALIZATION | 0.00000 | 0.00001 | 0.69013 | 0.65657 | 1.26500 | 212 |
| GO_RNA_PHOSPHODIESTER_BOND_HYDROLYSIS | 0.00049 | 0.01121 | 0.47727 | 0.63000 | 1.20049 | 132 |
| GO_RNA_SPLICING | 0.00000 | 0.00000 | | 0.68609 | 1.33714 | 395 |
| GO_RNA_SPLICING_VIA_TRANSESTERIFICATION_REACTIONS | 0.00000 | 0.00000 | | 0.69579 | 1.35104 | 316 |
| GO_ROUGH_ENDOPLASMIC_RETICULUM | 0.00002 | 0.00057 | 0.59333 | 0.72990 | 1.36147 | 59 |
| GO_RRNA_BINDING | 0.00000 | 0.00000 | 0.74774 | 0.79344 | 1.47832 | 57 |
| GO_RRNA_METABOLIC_PROCESS | 0.00000 | 0.00000 | | 0.72546 | 1.39857 | 216 |
| GO_SCF_DEPENDENT_PROTEASOMAL_UBIQUITIN_DEPENDI | 0.00000 | 0.00001 | 0.69013 | 0.73706 | 1.38954 | 82 |
| GO_SIGNAL_SEQUENCE_BINDING | 0.00011 | 0.00324 | 0.53843 | 0.75989 | 1.39038 | 39 |
| GO_SINGLE_FERTILIZATION | 0.00040 | 0.00978 | 0.49849 | 0.66539 | 1.25443 | 82 |
| GO_SINGLE_STRANDED_RNA_BINDING | 0.00009 | 0.00282 | 0.53843 | 0.70024 | 1.31252 | 69 |
| GO_SISTER_CHROMATID_SEGREGATION | 0.00026 | 0.00685 | 0.49849 | 0.62190 | 1.19334 | 173 |
| GO_SMALL_NUCLEAR_RIBONUCLEOPROTEIN_COMPLEX | 0.00002 | 0.00066 | 0.57561 | 0.72907 | 1.36033 | 60 |
| GO_SMALL_NUCLEOLAR_RIBONUCLEOPROTEIN_COMPLEX | 0.00104 | 0.02043 | 0.45506 | 0.76958 | 1.38057 | 26 |
| GO_SMALL_RIBOSOMAL_SUBUNIT | 0.00000 | 0.00000 | 0.71951 | 0.76837 | 1.44020 | 69 |
| GO_SMALL_SUBUNIT_PROCESSOME | 0.00014 | 0.00409 | 0.51885 | 0.76665 | 1.39364 | 33 |
| GO_SM_LIKE_PROTEIN_FAMILY_COMPLEX | 0.00000 | 0.00018 | 0.61053 | 0.72297 | 1.35546 | 70 |
| GO_SNORNA_BINDING | 0.00004 | 0.00119 | 0.55733 | 0.80539 | 1.45741 | 30 |
| GO_SOMATIC_STEM_CELL_POPULATION_MAINTENANCE | 0.00302 | 0.04780 | 0.43171 | 0.67586 | 1.25909 | 56 |
| GO_SPERM_EGG_RECOGNITION | 0.00013 | 0.00379 | 0.51885 | 0.75742 | 1.38001 | 36 |
| GO_SPINDLE | 0.00265 | 0.04333 | 0.43171 | 0.57537 | 1.11730 | 312 |
| GO_SPINDLE_ORGANIZATION | 0.00298 | 0.04745 | 0.43171 | 0.60500 | 1.15851 | 161 |
| GO_SPLICEOSOMAL_COMPLEX | 0.00000 | 0.00000 | | 0.70420 | 1.35195 | 175 |
| GO_SPLICEOSOMAL_COMPLEX_ASSEMBLY | 0.00044 | 0.01032 | 0.49849 | 0.70918 | 1.31381 | 49 |
| GO_SPLICEOSOMAL_SNRNP_ASSEMBLY | 0.00023 | 0.00624 | 0.51885 | 0.75063 | 1.36766 | 36 |
| GO_SPLICEOSOMAL_TRI_SNRNP_COMPLEX | 0.00183 | 0.03249 | 0.45506 | 0.73205 | 1.32470 | 30 |
| GO_STEM_CELL_DIFFERENTIATION | 0.00000 | 0.00000 | 0.76146 | 0.68503 | 1.31439 | 170 |
| GO_STRUCTURAL_CONSTITUENT_OF_RIBOSOME | 0.00000 | 0.00000 | | 0.80839 | 1.54627 | 154 |
| GO_STRUCTURAL_MOLECULE_ACTIVITY | 0.00000 | 0.00000 | | 0.71676 | 1.39770 | 413 |
| GO_TELOMERASE_RNA_LOCALIZATION | 0.00111 | 0.02142 | 0.45506 | 0.80982 | 1.43206 | 19 |
| GO_TELOMERE_MAINTENANCE_VIA_TELOMERE_LENGTHEN | 0.00022 | 0.00591 | 0.51885 | 0.68393 | 1.28542 | 74 |
| GO_TELOMERE_ORGANIZATION | 0.00133 | 0.02499 | 0.45506 | 0.62435 | 1.19034 | 138 |
| GO_THREONINE_TYPE_ENDOPEPTIDASE_ACTIVITY | 0.00000 | 0.00002 | 0.67496 | 0.91008 | 1.61460 | 20 |
| GO_THREONINE_TYPE_PEPTIDASE_ACTIVITY | 0.00000 | 0.00000 | 0.73376 | 0.89443 | 1.60453 | 26 |
| GO_TONGUE_DEVELOPMENT | 0.00036 | 0.00888 | 0.49849 | 0.84710 | 1.48911 | 16 |
| GO_TRANSCRIPTION_COUPLED_NUCLEOTIDE_EXCISION_RE | 0.00046 | 0.01073 | 0.49849 | 0.67881 | 1.27481 | 73 |
| GO_TRANSCRIPTION_FACTOR_BINDING | 0.00076 | 0.01585 | 0.47727 | 0.56764 | 1.10983 | 486 |
| GO_TRANSLATIONAL_ELONGATION | 0.00000 | 0.00000 | | 0.75955 | 1.44699 | 125 |
| GO_TRANSLATIONAL_INITIATION | 0.00000 | 0.00000 | | 0.77322 | 1.48386 | 172 |
| GO_TRANSLATIONAL_TERMINATION | 0.00000 | 0.00000 | | 0.76456 | 1.44757 | 102 |
| GO_TRANSLATION_FACTOR_ACTIVITY_RNA_BINDING | 0.00000 | 0.00010 | 0.62726 | 0.73539 | 1.38107 | 73 |
| GO_TRANSLATION_INITIATION_FACTOR_ACTIVITY | 0.00002 | 0.00079 | 0.57561 | 0.75420 | 1.39396 | 46 |
| GO_TRANSLATION_PREINITIATION_COMPLEX | 0.00108 | 0.02090 | 0.45506 | 0.82835 | 1.45615 | 16 |
| GO_TRANSLATION_REGULATOR_ACTIVITY | 0.00000 | 0.00000 | 0.73376 | 0.71272 | 1.35337 | 115 |

| Cluster 1 | | | | | | |
|---|---|---|---|---|---|---|
| GO_TRANSLATION_REGULATOR_ACTIVITY_NUCLEIC_ACID_E | 0.00000 | 0.00001 | 0.70498 | 0.72939 | 1.37727 | 91 |
| GO_TRNA_TRANSPORT | 0.00148 | 0.02735 | 0.45506 | 0.72297 | 1.31702 | 34 |
| GO_TUMOR_NECROSIS_FACTOR_MEDIATED_SIGNALING_PA | 0.00002 | 0.00083 | 0.57561 | 0.66588 | 1.26603 | 119 |
| GO_T_CELL_RECEPTOR_SIGNALING_PATHWAY | 0.00000 | 0.00001 | 0.69013 | 0.67899 | 1.29542 | 140 |
| GO_U12_TYPE_SPLICEOSOMAL_COMPLEX | 0.00113 | 0.02171 | 0.45506 | 0.76689 | 1.37573 | 26 |
| GO_U1_SNRNP | 0.00022 | 0.00591 | 0.51885 | 0.85600 | 1.50477 | 16 |
| GO_U2_SNRNP | 0.00100 | 0.01989 | 0.45506 | 0.81374 | 1.43466 | 18 |
| GO_U2_TYPE_CATALYTIC_STEP_2_SPLICEOSOME | 0.00024 | 0.00643 | 0.51885 | 0.77513 | 1.40265 | 30 |
| GO_U2_TYPE_SPLICEOSOMAL_COMPLEX | 0.00000 | 0.00012 | 0.62726 | 0.71178 | 1.34320 | 88 |
| GO_U5_SNRNP | 0.00290 | 0.04697 | 0.43171 | 0.81169 | 1.42317 | 15 |
| GO_UBIQUITIN_LIKE_PROTEIN_LIGASE_BINDING | 0.00002 | 0.00073 | 0.57561 | 0.61618 | 1.19182 | 253 |
| GO_UNFOLDED_PROTEIN_BINDING | 0.00005 | 0.00168 | 0.55733 | 0.67695 | 1.27895 | 94 |
| GO_VACUOLAR_LUMEN | 0.00049 | 0.01121 | 0.47727 | 0.64587 | 1.22302 | 105 |
| GO_VASODILATION | 0.00104 | 0.02043 | 0.45506 | 0.82329 | 1.44973 | 17 |
| GO_VESICLE_LUMEN | 0.00002 | 0.00060 | 0.57561 | 0.63254 | 1.21812 | 203 |
| GO_VIRAL_GENE_EXPRESSION | 0.00000 | 0.00000 | | 0.76060 | 1.46022 | 175 |
| GO_VIRAL_GENOME_REPLICATION | 0.00039 | 0.00952 | 0.49849 | 0.67569 | 1.27143 | 79 |
| GO_VIRAL_LIFE_CYCLE | 0.00000 | 0.00004 | 0.65944 | 0.64091 | 1.23758 | 235 |
| HP_ACUTE_LEUKEMIA | 0.00074 | 0.01533 | 0.47727 | 0.67286 | 1.26461 | 74 |
| HP_ANEMIA_OF_INADEQUATE_PRODUCTION | 0.00042 | 0.00993 | 0.49849 | 0.64317 | 1.21934 | 110 |
| HP_HEMATOLOGICAL_NEOPLASM | 0.00176 | 0.03127 | 0.45506 | 0.60676 | 1.16517 | 176 |
| HP_INTRAUTERINE_GROWTH_RETARDATION | 0.00276 | 0.04492 | 0.43171 | 0.57170 | 1.11251 | 359 |
| HP_LEUKEMIA | 0.00061 | 0.01322 | 0.47727 | 0.64670 | 1.22453 | 106 |
| HP_MACROCYTIC_ANEMIA | 0.00000 | 0.00006 | 0.64355 | 0.79390 | 1.46012 | 43 |
| HP_MIGRAINE | 0.00097 | 0.01922 | 0.47727 | 0.66745 | 1.25139 | 70 |
| HP_MUTISM | 0.00141 | 0.02622 | 0.45506 | 0.72336 | 1.31797 | 36 |
| HP_RECTAL_FISTULA | 0.00246 | 0.04082 | 0.43171 | 0.81011 | 1.42410 | 16 |
| HP_SUPERNUMERARY_BONES_OF_THE_AXIAL_SKELETON | 0.00170 | 0.03054 | 0.45506 | 0.80101 | 1.41222 | 18 |

### Supplementary Table S7
#### Human female GSEA of NMF clusters

| Cluster 1 | | | | | | |
|---|---|---|---|---|---|---|
| pathway | pval | padj | log2err | ES | NES | size |
| GO_ACTIVATION_OF_IMMUNE_RESPONSE | 0.00000 | 0.00000 | 0.74774 | 0.65038 | 1.26999 | 235 |
| GO_ACTIVATION_OF_INNATE_IMMUNE_RESPONSE | 0.00000 | 0.00001 | 0.69013 | 0.71967 | 1.37861 | 96 |
| GO_AEROBIC_RESPIRATION | 0.00000 | 0.00000 | 0.73376 | 0.76280 | 1.44478 | 72 |
| GO_ALCOHOL_BIOSYNTHETIC_PROCESS | 0.00032 | 0.00572 | 0.49849 | 0.64679 | 1.24367 | 108 |
| GO_ALCOHOL_METABOLIC_PROCESS | 0.00364 | 0.03677 | 0.43171 | 0.58112 | 1.13424 | 228 |
| GO_AMEBOIDAL_TYPE_CELL_MIGRATION | 0.00457 | 0.04386 | 0.40702 | 0.57755 | 1.12854 | 247 |
| GO_AMIDE_BINDING | 0.00014 | 0.00297 | 0.51885 | 0.61533 | 1.19780 | 199 |
| GO_AMINE_METABOLIC_PROCESS | 0.00000 | 0.00000 | 0.76146 | 0.73732 | 1.41097 | 92 |
| GO_AMYLOID_BETA_CLEARANCE | 0.00561 | 0.04971 | 0.40702 | 0.74081 | 1.33976 | 24 |
| GO_ANAPHASE_PROMOTING_COMPLEX_DEPENDENT_CATABOL | 0.00000 | 0.00000 | 0.76146 | 0.77107 | 1.46112 | 73 |
| GO_ANATOMICAL_STRUCTURE_HOMEOSTASIS | 0.00020 | 0.00390 | 0.51885 | 0.59307 | 1.16021 | 271 |
| GO_ANIMAL_ORGAN_REGENERATION | 0.00012 | 0.00269 | 0.51885 | 0.71251 | 1.33510 | 52 |
| GO_ANTIGEN_PROCESSING_AND_PRESENTATION | 0.00000 | 0.00000 | 0.74774 | 0.68785 | 1.33064 | 149 |
| GO_ANTIGEN_PROCESSING_AND_PRESENTATION_OF_EXOGENO | 0.00000 | 0.00000 | 0.76146 | 0.79644 | 1.50237 | 62 |
| GO_ANTIGEN_PROCESSING_AND_PRESENTATION_OF_PEPTIDE_ | 0.00000 | 0.00000 | 0.74774 | 0.70107 | 1.35144 | 130 |
| GO_ANTIGEN_PROCESSING_AND_PRESENTATION_OF_PEPTIDE_ | 0.00000 | 0.00000 | 0.78819 | 0.77566 | 1.47012 | 74 |
| GO_ANTIGEN_RECEPTOR_MEDIATED_SIGNALING_PATHWAY | 0.00000 | 0.00009 | 0.62726 | 0.66362 | 1.28080 | 136 |
| GO_APICAL_PART_OF_CELL | 0.00135 | 0.01808 | 0.45506 | 0.58926 | 1.14966 | 220 |
| GO_APOPTOTIC_SIGNALING_PATHWAY | 0.00006 | 0.00152 | 0.53843 | 0.58638 | 1.15586 | 393 |
| GO_ATPASE_ACTIVITY | 0.00050 | 0.00802 | 0.47727 | 0.58216 | 1.14385 | 316 |
| GO_ATPASE_ACTIVITY_COUPLED_TO_TRANSMEMBRANE_MOVE | 0.00420 | 0.04132 | 0.40702 | 0.78005 | 1.38741 | 18 |
| GO_ATP_BIOSYNTHETIC_PROCESS | 0.00013 | 0.00276 | 0.51885 | 0.83897 | 1.50801 | 21 |
| GO_ATP_METABOLIC_PROCESS | 0.00000 | 0.00000 | | 0.72178 | 1.40814 | 219 |
| GO_ATP_SYNTHESIS_COUPLED_ELECTRON_TRANSPORT | 0.00000 | 0.00000 | | 0.80353 | 1.52425 | 76 |
| GO_AZUROPHIL_GRANULE | 0.00025 | 0.00466 | 0.49849 | 0.64781 | 1.24551 | 107 |
| GO_AZUROPHIL_GRANULE_LUMEN | 0.00405 | 0.04011 | 0.40702 | 0.65625 | 1.23793 | 62 |
| GO_BASAL_PART_OF_CELL | 0.00509 | 0.04656 | 0.40702 | 0.68515 | 1.27697 | 45 |
| GO_BASAL_PLASMA_MEMBRANE | 0.00346 | 0.03548 | 0.43171 | 0.71184 | 1.31104 | 35 |
| GO_BASEMENT_MEMBRANE | 0.00279 | 0.03056 | 0.43171 | 0.68935 | 1.28452 | 46 |
| GO_BASOLATERAL_PLASMA_MEMBRANE | 0.00011 | 0.00247 | 0.53843 | 0.63209 | 1.21993 | 136 |
| GO_BINDING_OF_SPERM_TO_ZONA_PELLUCIDA | 0.00000 | 0.00006 | 0.64355 | 0.90095 | 1.60966 | 19 |
| GO_BLOOD_MICROPARTICLE | 0.00000 | 0.00000 | 0.70498 | 0.81982 | 1.52763 | 46 |
| GO_BRUSH_BORDER | 0.00060 | 0.00923 | 0.47727 | 0.66897 | 1.26790 | 74 |
| GO_CADHERIN_BINDING | 0.00000 | 0.00000 | | 0.68759 | 1.34435 | 253 |
| GO_CALCIUM_DEPENDENT_PHOSPHOLIPID_BINDING | 0.00152 | 0.01958 | 0.45506 | 0.75021 | 1.37120 | 28 |
| GO_CALCIUM_DEPENDENT_PROTEIN_BINDING | 0.00009 | 0.00211 | 0.53843 | 0.74254 | 1.38394 | 45 |
| GO_CANONICAL_WNT_SIGNALING_PATHWAY | 0.00004 | 0.00109 | 0.55733 | 0.61740 | 1.20458 | 222 |
| GO_CARBOHYDRATE_CATABOLIC_PROCESS | 0.00135 | 0.01808 | 0.45506 | 0.61179 | 1.18186 | 144 |
| GO_CATALYTIC_STEP_2_SPLICEOSOME | 0.00001 | 0.00022 | 0.61053 | 0.70290 | 1.33765 | 80 |
| GO_CATION_TRANSMEMBRANE_TRANSPORT | 0.00327 | 0.03409 | 0.43171 | 0.56060 | 1.10531 | 406 |
| GO_CATION_TRANSMEMBRANE_TRANSPORTER_ACTIVITY | 0.00097 | 0.01387 | 0.47727 | 0.58196 | 1.13846 | 265 |
| GO_CELLULAR_AMINO_ACID_METABOLIC_PROCESS | 0.00002 | 0.00060 | 0.57561 | 0.61438 | 1.19930 | 231 |
| GO_CELLULAR_COMPONENT_DISASSEMBLY | 0.00000 | 0.00000 | | 0.64070 | 1.26342 | 408 |
| GO_CELLULAR_ION_HOMEOSTASIS | 0.00027 | 0.00493 | 0.49849 | 0.58658 | 1.15340 | 328 |
| GO_CELLULAR_IRON_ION_HOMEOSTASIS | 0.00327 | 0.03409 | 0.43171 | 0.68360 | 1.27374 | 47 |
| GO_CELLULAR_KETONE_METABOLIC_PROCESS | 0.00000 | 0.00010 | 0.62726 | 0.65358 | 1.26726 | 164 |
| GO_CELLULAR_MONOVALENT_INORGANIC_CATION_HOMEOSTA | 0.00179 | 0.02183 | 0.45506 | 0.64747 | 1.22821 | 76 |
| GO_CELLULAR_NITROGEN_COMPOUND_CATABOLIC_PROCESS | 0.00000 | 0.00000 | | 0.67915 | 1.34096 | 456 |
| GO_CELLULAR_PROTEIN_COMPLEX_DISASSEMBLY | 0.00000 | 0.00000 | | 0.69566 | 1.35184 | 179 |
| GO_CELLULAR_RESPIRATION | 0.00000 | 0.00000 | | 0.74205 | 1.43370 | 145 |
| GO_CELLULAR_RESPONSE_TO_HEAT | 0.00004 | 0.00109 | 0.55733 | 0.67186 | 1.28698 | 95 |
| GO_CELLULAR_RESPONSE_TO_HORMONE_STIMULUS | 0.00309 | 0.03299 | 0.43171 | 0.56453 | 1.11191 | 374 |
| GO_CELLULAR_RESPONSE_TO_ORGANIC_CYCLIC_COMPOUND | 0.00502 | 0.04608 | 0.40702 | 0.56512 | 1.11134 | 329 |
| GO_CELLULAR_RESPONSE_TO_OXYGEN_LEVELS | 0.00000 | 0.00000 | | 0.71461 | 1.38224 | 151 |
| GO_CELL_ACTIVATION_INVOLVED_IN_IMMUNE_RESPONSE | 0.00000 | 0.00000 | | 0.63924 | 1.26108 | 422 |
| GO_CELL_ADHESION_MEDIATOR_ACTIVITY | 0.00157 | 0.02009 | 0.45506 | 0.71837 | 1.32488 | 36 |
| GO_CELL_ADHESION_MOLECULE_BINDING | 0.00000 | 0.00000 | | 0.67109 | 1.32112 | 353 |
| GO_CELL_BODY | 0.00004 | 0.00109 | 0.55733 | 0.59973 | 1.17541 | 293 |
| GO_CELL_CELL_RECOGNITION | 0.00000 | 0.00006 | 0.64355 | 0.82001 | 1.51026 | 35 |
| GO_CELL_CELL_SIGNALING_BY_WNT | 0.00004 | 0.00109 | 0.55733 | 0.59042 | 1.16141 | 339 |
| GO_CELL_CYCLE_G1_S_PHASE_TRANSITION | 0.00376 | 0.03783 | 0.43171 | 0.58112 | 1.13378 | 220 |
| GO_CELL_CYCLE_G2_M_PHASE_TRANSITION | 0.00024 | 0.00459 | 0.51885 | 0.59807 | 1.16746 | 231 |
| GO_CELL_CYCLE_PHASE_TRANSITION | 0.00000 | 0.00001 | 0.69013 | 0.59417 | 1.17348 | 499 |
| GO_CELL_DIVISION | 0.00003 | 0.00071 | 0.57561 | 0.58314 | 1.15094 | 449 |
| GO_CELL_KILLING | 0.00205 | 0.02440 | 0.43171 | 0.66727 | 1.25696 | 60 |
| GO_CELL_RECOGNITION | 0.00000 | 0.00000 | 0.73376 | 0.75953 | 1.43954 | 74 |
| GO_CELL_REDOX_HOMEOSTASIS | 0.00163 | 0.02050 | 0.45506 | 0.71772 | 1.32665 | 38 |

| | | | | | | |
|---|---|---|---|---|---|---|
| GO_CELL_SUBSTRATE_JUNCTION | 0.00000 | 0.00000 | | 0.65742 | 1.29171 | 316 |
| GO_CELL_SURFACE | 0.00000 | 0.00000 | 0.73376 | 0.62572 | 1.23070 | 330 |
| GO_CELL_SURFACE_RECEPTOR_SIGNALING_PATHWAY_INVOLVEI | 0.00038 | 0.00640 | 0.49849 | 0.57762 | 1.13770 | 374 |
| GO_CHAPERONE_BINDING | 0.00495 | 0.04573 | 0.40702 | 0.62860 | 1.19935 | 86 |
| GO_CHAPERONE_COMPLEX | 0.00001 | 0.00028 | 0.59333 | 0.86337 | 1.55587 | 22 |
| GO_CHAPERONE_MEDIATED_PROTEIN_COMPLEX_ASSEMBLY | 0.00100 | 0.01430 | 0.45506 | 0.80787 | 1.44337 | 19 |
| GO_CHAPERONE_MEDIATED_PROTEIN_FOLDING | 0.00168 | 0.02098 | 0.45506 | 0.71110 | 1.32288 | 44 |
| GO_CHROMOSOMAL_REGION | 0.00018 | 0.00349 | 0.51885 | 0.59323 | 1.16180 | 286 |
| GO_CHROMOSOME_CENTROMERIC_REGION | 0.00172 | 0.02135 | 0.45506 | 0.60447 | 1.17248 | 168 |
| GO_CHROMOSOME_SEGREGATION | 0.00003 | 0.00087 | 0.55733 | 0.60463 | 1.18280 | 270 |
| GO_CLUSTER_OF_ACTIN_BASED_CELL_PROJECTIONS | 0.00074 | 0.01107 | 0.47727 | 0.64184 | 1.23044 | 99 |
| GO_COATED_VESICLE | 0.00213 | 0.02490 | 0.43171 | 0.59512 | 1.15846 | 199 |
| GO_COATED_VESICLE_MEMBRANE | 0.00060 | 0.00923 | 0.47727 | 0.63443 | 1.22207 | 115 |
| GO_COLLAGEN_BINDING | 0.00465 | 0.04430 | 0.40702 | 0.73471 | 1.33628 | 26 |
| GO_COLLAGEN_CONTAINING_EXTRACELLULAR_MATRIX | 0.00000 | 0.00006 | 0.64355 | 0.65358 | 1.26542 | 157 |
| GO_CONDENSED_CHROMOSOME | 0.00350 | 0.03580 | 0.43171 | 0.59394 | 1.15417 | 179 |
| GO_CONTRACTILE_FIBER | 0.00064 | 0.00972 | 0.47727 | 0.62217 | 1.19973 | 134 |
| GO_COPI_COATED_VESICLE | 0.00316 | 0.03344 | 0.43171 | 0.75688 | 1.37234 | 25 |
| GO_CORNIFICATION | 0.00027 | 0.00500 | 0.49849 | 0.77970 | 1.42512 | 28 |
| GO_CORTICAL_CYTOSKELETON | 0.00502 | 0.04608 | 0.40702 | 0.64388 | 1.21870 | 71 |
| GO_COTRANSLATIONAL_PROTEIN_TARGETING_TO_MEMBRANE | 0.00000 | 0.00000 | | 0.78223 | 1.49819 | 94 |
| GO_CYTOCHROME_COMPLEX | 0.00035 | 0.00610 | 0.49849 | 0.80796 | 1.45601 | 22 |
| GO_CYTOKINE_MEDIATED_SIGNALING_PATHWAY | 0.00000 | 0.00002 | 0.67496 | 0.60742 | 1.19735 | 392 |
| GO_CYTOPLASMIC_STRESS_GRANULE | 0.00014 | 0.00302 | 0.51885 | 0.69390 | 1.31009 | 63 |
| GO_CYTOPLASMIC_TRANSLATION | 0.00000 | 0.00001 | 0.70498 | 0.71773 | 1.37349 | 92 |
| GO_CYTOPLASMIC_TRANSLATIONAL_INITIATION | 0.00178 | 0.02182 | 0.45506 | 0.73517 | 1.35039 | 31 |
| GO_CYTOSOLIC_LARGE_RIBOSOMAL_SUBUNIT | 0.00000 | 0.00005 | 0.64355 | 0.77240 | 1.44935 | 53 |
| GO_CYTOSOLIC_RIBOSOME | 0.00000 | 0.00000 | | 0.77483 | 1.48437 | 97 |
| GO_CYTOSOLIC_SMALL_RIBOSOMAL_SUBUNIT | 0.00000 | 0.00010 | 0.62726 | 0.79384 | 1.47033 | 40 |
| GO_DETECTION_OF_STIMULUS | 0.00020 | 0.00393 | 0.51885 | 0.65208 | 1.25384 | 108 |
| GO_DETOXIFICATION | 0.00413 | 0.04065 | 0.40702 | 0.65399 | 1.23684 | 70 |
| GO_DEVELOPMENT_OF_PRIMARY_FEMALE_SEXUAL_CHARACTE | 0.00179 | 0.02183 | 0.45506 | 0.68410 | 1.28057 | 51 |
| GO_DICARBOXYLIC_ACID_METABOLIC_PROCESS | 0.00261 | 0.02910 | 0.43171 | 0.65541 | 1.23747 | 65 |
| GO_DISORDERED_DOMAIN_SPECIFIC_BINDING | 0.00051 | 0.00812 | 0.47727 | 0.76867 | 1.40343 | 27 |
| GO_DISULFIDE_OXIDOREDUCTASE_ACTIVITY | 0.00435 | 0.04238 | 0.40702 | 0.71445 | 1.31232 | 31 |
| GO_DNA_BIOSYNTHETIC_PROCESS | 0.00017 | 0.00334 | 0.51885 | 0.62540 | 1.20833 | 145 |
| GO_DNA_CONFORMATION_CHANGE | 0.00327 | 0.03409 | 0.43171 | 0.57979 | 1.13178 | 231 |
| GO_DNA_DAMAGE_RESPONSE_DETECTION_OF_DNA_DAMAGE | 0.00305 | 0.03271 | 0.43171 | 0.71625 | 1.31916 | 35 |
| GO_DNA_GEOMETRIC_CHANGE | 0.00141 | 0.01850 | 0.45506 | 0.62642 | 1.20356 | 105 |
| GO_DOUBLE_STRANDED_RNA_BINDING | 0.00039 | 0.00658 | 0.49849 | 0.69158 | 1.30365 | 61 |
| GO_DRUG_BINDING | 0.00450 | 0.04342 | 0.40702 | 0.66245 | 1.24130 | 52 |
| GO_ELECTRON_TRANSFER_ACTIVITY | 0.00000 | 0.00001 | 0.67496 | 0.72917 | 1.38765 | 80 |
| GO_ELECTRON_TRANSPORT_CHAIN | 0.00000 | 0.00000 | | 0.73336 | 1.41392 | 133 |
| GO_EMBRYO_IMPLANTATION | 0.00044 | 0.00706 | 0.49849 | 0.75968 | 1.39146 | 29 |
| GO_ENDOCYTIC_VESICLE_MEMBRANE | 0.00524 | 0.04770 | 0.40702 | 0.62563 | 1.19564 | 89 |
| GO_ENDOPEPTIDASE_ACTIVITY | 0.00065 | 0.00981 | 0.47727 | 0.60123 | 1.16998 | 196 |
| GO_ENDOPEPTIDASE_COMPLEX | 0.00000 | 0.00001 | 0.70498 | 0.76593 | 1.44609 | 63 |
| GO_ENDOPEPTIDASE_REGULATOR_ACTIVITY | 0.00006 | 0.00145 | 0.55733 | 0.70425 | 1.33001 | 68 |
| GO_ENDOPLASMIC_RETICULUM_GOLGI_INTERMEDIATE_COMPA | 0.00055 | 0.00863 | 0.47727 | 0.64758 | 1.24050 | 96 |
| GO_ENDOPLASMIC_RETICULUM_GOLGI_INTERMEDIATE_COMPA | 0.00148 | 0.01923 | 0.45506 | 0.68944 | 1.29056 | 51 |
| GO_ENDOPLASMIC_RETICULUM_TO_GOLGI_VESICLE_MEDIATED | 0.00532 | 0.04803 | 0.40702 | 0.59194 | 1.14697 | 162 |
| GO_ENERGY_DERIVATION_BY_OXIDATION_OF_ORGANIC_COMP | 0.00000 | 0.00000 | | 0.69995 | 1.36305 | 202 |
| GO_ENZYME_INHIBITOR_ACTIVITY | 0.00246 | 0.02780 | 0.43171 | 0.58913 | 1.14643 | 191 |
| GO_EPIDERMAL_CELL_DIFFERENTIATION | 0.00216 | 0.02519 | 0.43171 | 0.62667 | 1.20589 | 112 |
| GO_EPITHELIAL_CELL_DIFFERENTIATION | 0.00054 | 0.00857 | 0.47727 | 0.57819 | 1.13725 | 334 |
| GO_ESTABLISHMENT_OF_ORGANELLE_LOCALIZATION | 0.00122 | 0.01679 | 0.45506 | 0.57737 | 1.13489 | 323 |
| GO_ESTABLISHMENT_OF_PROTEIN_LOCALIZATION_TO_CHROMO | 0.00016 | 0.00321 | 0.51885 | 0.81363 | 1.46985 | 23 |
| GO_ESTABLISHMENT_OF_PROTEIN_LOCALIZATION_TO_ENDOPLA | 0.00000 | 0.00000 | | 0.77241 | 1.48479 | 106 |
| GO_ESTABLISHMENT_OF_PROTEIN_LOCALIZATION_TO_MEMBRA | 0.00000 | 0.00000 | | 0.67484 | 1.31991 | 260 |
| GO_ESTABLISHMENT_OF_PROTEIN_LOCALIZATION_TO_ORGANE | 0.00000 | 0.00000 | | 0.65856 | 1.30056 | 461 |
| GO_ESTABLISHMENT_OF_PROTEIN_LOCALIZATION_TO_TELOME | 0.00000 | 0.00006 | 0.64355 | 0.90447 | 1.60870 | 18 |
| GO_ESTABLISHMENT_OF_RNA_LOCALIZATION | 0.00000 | 0.00005 | 0.64355 | 0.64271 | 1.24830 | 176 |
| GO_ESTABLISHMENT_OF_TISSUE_POLARITY | 0.00000 | 0.00000 | 0.81404 | 0.75693 | 1.44999 | 96 |
| GO_EUKARYOTIC_TRANSLATION_INITIATION_FACTOR_3_COMPL | 0.00143 | 0.01863 | 0.45506 | 0.83263 | 1.46100 | 15 |
| GO_EXTERNAL_SIDE_OF_PLASMA_MEMBRANE | 0.00159 | 0.02016 | 0.45506 | 0.61775 | 1.19022 | 118 |
| GO_EXTRACELLULAR_MATRIX | 0.00042 | 0.00695 | 0.49849 | 0.61138 | 1.18938 | 189 |
| GO_FATTY_ACID_DERIVATIVE_METABOLIC_PROCESS | 0.00043 | 0.00695 | 0.49849 | 0.66266 | 1.26212 | 81 |
| GO_FATTY_ACYL_COA_METABOLIC_PROCESS | 0.00502 | 0.04608 | 0.40702 | 0.72264 | 1.32082 | 28 |
| GO_FC_EPSILON_RECEPTOR_SIGNALING_PATHWAY | 0.00000 | 0.00000 | 0.71951 | 0.72370 | 1.38610 | 94 |
| GO_FC_RECEPTOR_SIGNALING_PATHWAY | 0.00000 | 0.00000 | 0.70498 | 0.68815 | 1.32879 | 140 |

| | | | | | | |
|---|---|---|---|---|---|---|
| GO_FERTILIZATION | 0.00000 | 0.00005 | 0.64355 | 0.73257 | 1.38752 | 72 |
| GO_FIBRILLAR_CENTER | 0.00246 | 0.02780 | 0.43171 | 0.62638 | 1.19990 | 96 |
| GO_FIBROBLAST_GROWTH_FACTOR_RECEPTOR_SIGNALING_PAT | 0.00495 | 0.04573 | 0.40702 | 0.64888 | 1.22552 | 67 |
| GO_FIBROBLAST_PROLIFERATION | 0.00032 | 0.00575 | 0.49849 | 0.70676 | 1.32618 | 53 |
| GO_FICOLIN_1_RICH_GRANULE | 0.00000 | 0.00000 | 0.70498 | 0.71544 | 1.37138 | 98 |
| GO_GABA_ERGIC_SYNAPSE | 0.00022 | 0.00427 | 0.51885 | 0.76430 | 1.40406 | 32 |
| GO_GENERATION_OF_PRECURSOR_METABOLITES_AND_ENERGY | 0.00000 | 0.00000 | | 0.67527 | 1.32968 | 368 |
| GO_GENE_SILENCING | 0.00019 | 0.00376 | 0.51885 | 0.61625 | 1.19788 | 184 |
| GO_GLUCOSE_CATABOLIC_PROCESS | 0.00495 | 0.04573 | 0.40702 | 0.72314 | 1.32174 | 28 |
| GO_GLUTAMATERGIC_SYNAPSE | 0.00135 | 0.01808 | 0.45506 | 0.60052 | 1.16783 | 186 |
| GO_GLYCOLYTIC_PROCESS_THROUGH_FRUCTOSE_6_PHOSPHATE | 0.00554 | 0.04968 | 0.40702 | 0.74156 | 1.34111 | 24 |
| GO_GOLGI_ASSOCIATED_VESICLE | 0.00085 | 0.01239 | 0.47727 | 0.62294 | 1.20037 | 129 |
| GO_GOLGI_ASSOCIATED_VESICLE_MEMBRANE | 0.00313 | 0.03310 | 0.43171 | 0.64208 | 1.21753 | 75 |
| GO_GTPASE_ACTIVITY | 0.00004 | 0.00098 | 0.55733 | 0.61546 | 1.19974 | 210 |
| GO_GUANYL_NUCLEOTIDE_BINDING | 0.00005 | 0.00126 | 0.55733 | 0.60899 | 1.19065 | 252 |
| GO_HEAT_SHOCK_PROTEIN_BINDING | 0.00001 | 0.00034 | 0.59333 | 0.68258 | 1.30633 | 93 |
| GO_HEMATOPOIETIC_PROGENITOR_CELL_DIFFERENTIATION | 0.00000 | 0.00000 | 0.71951 | 0.70099 | 1.35150 | 121 |
| GO_HEMATOPOIETIC_STEM_CELL_DIFFERENTIATION | 0.00000 | 0.00000 | 0.76146 | 0.77246 | 1.46307 | 72 |
| GO_HISTONE_MRNA_METABOLIC_PROCESS | 0.00265 | 0.02936 | 0.43171 | 0.76832 | 1.38459 | 22 |
| GO_HOST_CELLULAR_COMPONENT | 0.00043 | 0.00695 | 0.49849 | 0.69729 | 1.31074 | 54 |
| GO_IMMUNE_RESPONSE_REGULATING_SIGNALING_PATHWAY | 0.00000 | 0.00003 | 0.65944 | 0.63620 | 1.24112 | 225 |
| GO_IMPORT_INTO_CELL | 0.00428 | 0.04185 | 0.40702 | 0.61709 | 1.18655 | 108 |
| GO_IMPORT_INTO_NUCLEUS | 0.00039 | 0.00652 | 0.49849 | 0.62707 | 1.20905 | 131 |
| GO_INNATE_IMMUNE_RESPONSE | 0.00000 | 0.00003 | 0.65944 | 0.59893 | 1.18089 | 406 |
| GO_INNATE_IMMUNE_RESPONSE_ACTIVATING_SIGNAL_TRANSD | 0.00000 | 0.00001 | 0.69013 | 0.73348 | 1.39585 | 80 |
| GO_INNER_MITOCHONDRIAL_MEMBRANE_PROTEIN_COMPLEX | 0.00000 | 0.00000 | | 0.76576 | 1.46666 | 94 |
| GO_INORGANIC_ION_TRANSMEMBRANE_TRANSPORT | 0.00141 | 0.01850 | 0.45506 | 0.56404 | 1.11185 | 401 |
| GO_INTERACTION_WITH_HOST | 0.00139 | 0.01850 | 0.45506 | 0.60855 | 1.17710 | 151 |
| GO_INTERLEUKIN_1_MEDIATED_SIGNALING_PATHWAY | 0.00000 | 0.00000 | 0.77494 | 0.75654 | 1.44176 | 83 |
| GO_INTRAMOLECULAR_OXIDOREDUCTASE_ACTIVITY | 0.00051 | 0.00812 | 0.47727 | 0.74506 | 1.37081 | 34 |
| GO_INTRINSIC_APOPTOTIC_SIGNALING_PATHWAY | 0.00156 | 0.01993 | 0.45506 | 0.59427 | 1.15817 | 207 |
| GO_ION_HOMEOSTASIS | 0.00170 | 0.02117 | 0.45506 | 0.56164 | 1.10717 | 403 |
| GO_ION_TRANSMEMBRANE_TRANSPORTER_ACTIVITY | 0.00106 | 0.01495 | 0.45506 | 0.56696 | 1.11734 | 390 |
| GO_IRON_ION_HOMEOSTASIS | 0.00361 | 0.03657 | 0.43171 | 0.65837 | 1.23983 | 59 |
| GO_IRON_ION_TRANSPORT | 0.00054 | 0.00857 | 0.47727 | 0.69556 | 1.30334 | 52 |
| GO_ISOMERASE_ACTIVITY | 0.00413 | 0.04065 | 0.40702 | 0.61065 | 1.17496 | 113 |
| GO_KERATINIZATION | 0.00038 | 0.00640 | 0.49849 | 0.75690 | 1.39013 | 33 |
| GO_KILLING_OF_CELLS_OF_OTHER_ORGANISM | 0.00495 | 0.04573 | 0.40702 | 0.78614 | 1.38643 | 16 |
| GO_LARGE_RIBOSOMAL_SUBUNIT | 0.00000 | 0.00000 | | 0.77656 | 1.49319 | 108 |
| GO_LATERAL_PLASMA_MEMBRANE | 0.00524 | 0.04770 | 0.40702 | 0.67385 | 1.25557 | 47 |
| GO_LEUKOCYTE_MEDIATED_IMMUNITY | 0.00000 | 0.00000 | | 0.64010 | 1.26331 | 447 |
| GO_LIPOPROTEIN_PARTICLE_RECEPTOR_BINDING | 0.00126 | 0.01715 | 0.45506 | 0.80503 | 1.42497 | 17 |
| GO_LIVER_REGENERATION | 0.00093 | 0.01337 | 0.47727 | 0.78442 | 1.41708 | 23 |
| GO_LOW_DENSITY_LIPOPROTEIN_RECEPTOR_PARTICLE_METABO | 0.00327 | 0.03409 | 0.43171 | 0.78520 | 1.39657 | 18 |
| GO_LYASE_ACTIVITY | 0.00152 | 0.01958 | 0.45506 | 0.61769 | 1.19079 | 122 |
| GO_LYSOSOMAL_LUMEN | 0.00143 | 0.01863 | 0.45506 | 0.67371 | 1.26872 | 59 |
| GO_MAINTENANCE_OF_CELL_NUMBER | 0.00435 | 0.04238 | 0.40702 | 0.60761 | 1.17073 | 117 |
| GO_MATURATION_OF_SSU_RRNA | 0.00126 | 0.01715 | 0.45506 | 0.69615 | 1.29711 | 47 |
| GO_MATURATION_OF_SSU_RRNA_FROM_TRICISTRONIC_RRNA_ | 0.00554 | 0.04968 | 0.40702 | 0.70205 | 1.29301 | 35 |
| GO_MEMBRANE_REGION | 0.00000 | 0.00003 | 0.65944 | 0.64496 | 1.25542 | 194 |
| GO_METALLOPEPTIDASE_ACTIVITY | 0.00487 | 0.04559 | 0.40702 | 0.62714 | 1.19577 | 85 |
| GO_METAL_ION_HOMEOSTASIS | 0.00032 | 0.00575 | 0.49849 | 0.58816 | 1.15376 | 301 |
| GO_METAL_ION_TRANSPORT | 0.00390 | 0.03895 | 0.43171 | 0.55877 | 1.10227 | 419 |
| GO_METAPHASE_ANAPHASE_TRANSITION_OF_CELL_CYCLE | 0.00320 | 0.03371 | 0.43171 | 0.66563 | 1.25122 | 54 |
| GO_MICROTUBULE_CYTOSKELETON_ORGANIZATION_INVOLVED_ | 0.00220 | 0.02548 | 0.43171 | 0.61295 | 1.18118 | 127 |
| GO_MICROVILLUS | 0.00168 | 0.02098 | 0.45506 | 0.68128 | 1.28184 | 56 |
| GO_MIDBODY | 0.00561 | 0.04971 | 0.40702 | 0.59866 | 1.15606 | 141 |
| GO_MITOCHONDRIAL_ELECTRON_TRANSPORT_NADH_TO_UBIQ | 0.00000 | 0.00006 | 0.64355 | 0.79447 | 1.47799 | 44 |
| GO_MITOCHONDRIAL_GENE_EXPRESSION | 0.00000 | 0.00000 | 0.78819 | 0.70230 | 1.35844 | 151 |
| GO_MITOCHONDRIAL_LARGE_RIBOSOMAL_SUBUNIT | 0.00000 | 0.00000 | 0.71951 | 0.79015 | 1.48529 | 54 |
| GO_MITOCHONDRIAL_MATRIX | 0.00000 | 0.00000 | 0.82666 | 0.63141 | 1.24412 | 387 |
| GO_MITOCHONDRIAL_MEMBRANE_ORGANIZATION | 0.00050 | 0.00802 | 0.47727 | 0.64578 | 1.23686 | 94 |
| GO_MITOCHONDRIAL_PROTEIN_COMPLEX | 0.00000 | 0.00000 | | 0.75011 | 1.46215 | 212 |
| GO_MITOCHONDRIAL_RESPIRATORY_CHAIN_COMPLEX_ASSEMB | 0.00001 | 0.00033 | 0.59333 | 0.69650 | 1.32804 | 85 |
| GO_MITOCHONDRIAL_SMALL_RIBOSOMAL_SUBUNIT | 0.00119 | 0.01640 | 0.45506 | 0.75384 | 1.37785 | 28 |
| GO_MITOCHONDRIAL_TRANSLATION | 0.00000 | 0.00000 | 0.80122 | 0.71435 | 1.37651 | 128 |
| GO_MITOCHONDRIAL_TRANSLATIONAL_TERMINATION | 0.00000 | 0.00000 | 0.81404 | 0.75760 | 1.44657 | 87 |
| GO_MITOCHONDRIAL_TRANSMEMBRANE_TRANSPORT | 0.00213 | 0.02490 | 0.43171 | 0.66300 | 1.25387 | 70 |
| GO_MITOCHONDRIAL_TRANSPORT | 0.00010 | 0.00227 | 0.53843 | 0.61119 | 1.19060 | 203 |
| GO_MITOCHONDRION_ORGANIZATION | 0.00000 | 0.00000 | 0.74774 | 0.61590 | 1.21471 | 416 |

| | | | | | | |
|---|---|---|---|---|---|---|
| GO_MITOTIC_NUCLEAR_DIVISION | 0.00008 | 0.00182 | 0.53843 | 0.60435 | 1.17901 | 223 |
| GO_MITOTIC_SISTER_CHROMATID_SEGREGATION | 0.00014 | 0.00302 | 0.51885 | 0.62652 | 1.21048 | 145 |
| GO_MITOTIC_SPINDLE_ORGANIZATION | 0.00078 | 0.01164 | 0.47727 | 0.63943 | 1.22821 | 104 |
| GO_MODIFIED_AMINO_ACID_BINDING | 0.00002 | 0.00048 | 0.57561 | 0.74200 | 1.38679 | 49 |
| GO_MODULATION_BY_HOST_OF_VIRAL_PROCESS | 0.00216 | 0.02519 | 0.43171 | 0.77232 | 1.38066 | 20 |
| GO_MONOCARBOXYLIC_ACID_METABOLIC_PROCESS | 0.00012 | 0.00265 | 0.53843 | 0.57846 | 1.14053 | 406 |
| GO_MONOSACCHARIDE_BIOSYNTHETIC_PROCESS | 0.00305 | 0.03271 | 0.43171 | 0.65908 | 1.24472 | 68 |
| GO_MONOVALENT_INORGANIC_CATION_HOMEOSTASIS | 0.00108 | 0.01507 | 0.45506 | 0.63643 | 1.21783 | 91 |
| GO_MONOVALENT_INORGANIC_CATION_TRANSMEMBRANE_TR | 0.00001 | 0.00026 | 0.59333 | 0.65186 | 1.25996 | 147 |
| GO_MONOVALENT_INORGANIC_CATION_TRANSPORT | 0.00008 | 0.00178 | 0.53843 | 0.60205 | 1.17659 | 248 |
| GO_MORPHOGENESIS_OF_AN_EPITHELIUM | 0.00000 | 0.00009 | 0.62726 | 0.60570 | 1.19137 | 334 |
| GO_MORPHOGENESIS_OF_A_POLARIZED_EPITHELIUM | 0.00000 | 0.00000 | 0.83909 | 0.74384 | 1.43057 | 110 |
| GO_MOVEMENT_IN_HOST_ENVIRONMENT | 0.00268 | 0.02961 | 0.43171 | 0.61282 | 1.18078 | 117 |
| GO_MRNA_3_END_PROCESSING | 0.00087 | 0.01270 | 0.47727 | 0.64010 | 1.22502 | 93 |
| GO_MRNA_5_UTR_BINDING | 0.00002 | 0.00063 | 0.57561 | 0.83759 | 1.51313 | 23 |
| GO_MRNA_BINDING | 0.00000 | 0.00000 | | 0.67496 | 1.31810 | 239 |
| GO_MRNA_EXPORT_FROM_NUCLEUS | 0.00006 | 0.00159 | 0.53843 | 0.66290 | 1.27265 | 103 |
| GO_MRNA_PROCESSING | 0.00000 | 0.00000 | | 0.62959 | 1.24255 | 437 |
| GO_MRNA_TRANSPORT | 0.00007 | 0.00171 | 0.53843 | 0.64103 | 1.23563 | 132 |
| GO_MULTI_MULTICELLULAR_ORGANISM_PROCESS | 0.00041 | 0.00687 | 0.49849 | 0.63087 | 1.21515 | 116 |
| GO_MULTI_ORGANISM_LOCALIZATION | 0.00017 | 0.00334 | 0.51885 | 0.71077 | 1.33608 | 54 |
| GO_MUSCLE_CONTRACTION | 0.00335 | 0.03468 | 0.43171 | 0.59925 | 1.16199 | 165 |
| GO_MUSCLE_FILAMENT_SLIDING | 0.00324 | 0.03393 | 0.43171 | 0.77980 | 1.39322 | 19 |
| GO_MYELOID_LEUKOCYTE_ACTIVATION | 0.00000 | 0.00000 | | 0.65288 | 1.28712 | 395 |
| GO_MYELOID_LEUKOCYTE_MEDIATED_IMMUNITY | 0.00000 | 0.00000 | | 0.65734 | 1.29408 | 356 |
| GO_NADH_DEHYDROGENASE_ACTIVITY | 0.00001 | 0.00019 | 0.61053 | 0.79360 | 1.46691 | 38 |
| GO_NADH_DEHYDROGENASE_COMPLEX | 0.00001 | 0.00028 | 0.59333 | 0.77749 | 1.44215 | 41 |
| GO_NADH_DEHYDROGENASE_COMPLEX_ASSEMBLY | 0.00001 | 0.00035 | 0.59333 | 0.74044 | 1.39281 | 55 |
| GO_NADH_METABOLIC_PROCESS | 0.00034 | 0.00599 | 0.49849 | 0.75773 | 1.39166 | 33 |
| GO_NAD_METABOLIC_PROCESS | 0.00435 | 0.04238 | 0.40702 | 0.70570 | 1.30442 | 37 |
| GO_NCRNA_EXPORT_FROM_NUCLEUS | 0.00205 | 0.02440 | 0.43171 | 0.71473 | 1.31817 | 36 |
| GO_NCRNA_METABOLIC_PROCESS | 0.00000 | 0.00000 | 0.70498 | 0.61139 | 1.20509 | 399 |
| GO_NCRNA_PROCESSING | 0.00000 | 0.00000 | 0.73376 | 0.62238 | 1.22425 | 341 |
| GO_NEGATIVE_REGULATION_OF_APOPTOTIC_SIGNALING_PATHW | 0.00364 | 0.03677 | 0.43171 | 0.59958 | 1.15998 | 150 |
| GO_NEGATIVE_REGULATION_OF_ATP_METABOLIC_PROCESS | 0.00302 | 0.03248 | 0.43171 | 0.78586 | 1.39103 | 17 |
| GO_NEGATIVE_REGULATION_OF_CANONICAL_WNT_SIGNALING | 0.00000 | 0.00000 | 0.71951 | 0.69629 | 1.34180 | 123 |
| GO_NEGATIVE_REGULATION_OF_CATALYTIC_ACTIVITY | 0.00145 | 0.01879 | 0.45506 | 0.56238 | 1.11005 | 450 |
| GO_NEGATIVE_REGULATION_OF_CELLULAR_AMIDE_METABOLIC | 0.00268 | 0.02961 | 0.43171 | 0.61074 | 1.17686 | 124 |
| GO_NEGATIVE_REGULATION_OF_CELLULAR_COMPONENT_ORG | 0.00320 | 0.03371 | 0.43171 | 0.55470 | 1.09534 | 470 |
| GO_NEGATIVE_REGULATION_OF_CELL_CYCLE | 0.00023 | 0.00445 | 0.51885 | 0.57348 | 1.13228 | 455 |
| GO_NEGATIVE_REGULATION_OF_CELL_CYCLE_G2_M_PHASE_TR | 0.00000 | 0.00007 | 0.62726 | 0.70549 | 1.34606 | 86 |
| GO_NEGATIVE_REGULATION_OF_CELL_CYCLE_PHASE_TRANSITIO | 0.00000 | 0.00000 | 0.70498 | 0.65532 | 1.27560 | 194 |
| GO_NEGATIVE_REGULATION_OF_CELL_CYCLE_PROCESS | 0.00000 | 0.00005 | 0.64355 | 0.62725 | 1.22670 | 259 |
| GO_NEGATIVE_REGULATION_OF_HYDROLASE_ACTIVITY | 0.00216 | 0.02519 | 0.43171 | 0.58306 | 1.13858 | 238 |
| GO_NEGATIVE_REGULATION_OF_INTERLEUKIN_1_PRODUCTION | 0.00205 | 0.02440 | 0.43171 | 0.82297 | 1.44404 | 15 |
| GO_NEGATIVE_REGULATION_OF_MITOTIC_CELL_CYCLE | 0.00000 | 0.00010 | 0.62726 | 0.62191 | 1.21541 | 248 |
| GO_NEGATIVE_REGULATION_OF_MRNA_METABOLIC_PROCESS | 0.00213 | 0.02490 | 0.43171 | 0.66731 | 1.25879 | 62 |
| GO_NEGATIVE_REGULATION_OF_MRNA_SPLICING_VIA_SPLICEO | 0.00339 | 0.03501 | 0.43171 | 0.76309 | 1.36417 | 20 |
| GO_NEGATIVE_REGULATION_OF_NUCLEAR_DIVISION | 0.00335 | 0.03468 | 0.43171 | 0.67724 | 1.26685 | 50 |
| GO_NEGATIVE_REGULATION_OF_ORGANELLE_ORGANIZATION | 0.00487 | 0.04559 | 0.40702 | 0.57038 | 1.11722 | 288 |
| GO_NEGATIVE_REGULATION_OF_PEPTIDASE_ACTIVITY | 0.00036 | 0.00621 | 0.49849 | 0.64044 | 1.23159 | 109 |
| GO_NEGATIVE_REGULATION_OF_PROTEIN_CATABOLIC_PROCESS | 0.00342 | 0.03521 | 0.43171 | 0.62320 | 1.19268 | 93 |
| GO_NEGATIVE_REGULATION_OF_PROTEOLYSIS | 0.00205 | 0.02440 | 0.43171 | 0.60172 | 1.16902 | 178 |
| GO_NEGATIVE_REGULATION_OF_REACTIVE_OXYGEN_SPECIES_B | 0.00036 | 0.00621 | 0.49849 | 0.83108 | 1.47818 | 18 |
| GO_NEGATIVE_REGULATION_OF_REPRODUCTIVE_PROCESS | 0.00457 | 0.04386 | 0.40702 | 0.74845 | 1.35356 | 24 |
| GO_NEGATIVE_REGULATION_OF_RNA_SPLICING | 0.00183 | 0.02219 | 0.45506 | 0.76824 | 1.38935 | 24 |
| GO_NEGATIVE_REGULATION_OF_WNT_SIGNALING_PATHWAY | 0.00000 | 0.00002 | 0.67496 | 0.67731 | 1.30721 | 136 |
| GO_NEURAL_CREST_CELL_MIGRATION | 0.00265 | 0.02936 | 0.43171 | 0.73967 | 1.35049 | 27 |
| GO_NEURON_RECOGNITION | 0.00390 | 0.03895 | 0.43171 | 0.71862 | 1.31998 | 31 |
| GO_NIK_NF_KAPPAB_SIGNALING | 0.00000 | 0.00000 | 0.77494 | 0.72329 | 1.39253 | 114 |
| GO_NON_CANONICAL_WNT_SIGNALING_PATHWAY | 0.00000 | 0.00000 | | 0.75756 | 1.45654 | 107 |
| GO_NUCLEAR_CHROMOSOME_SEGREGATION | 0.00098 | 0.01407 | 0.45506 | 0.59156 | 1.15357 | 215 |
| GO_NUCLEAR_ENVELOPE | 0.00005 | 0.00131 | 0.55733 | 0.59109 | 1.16270 | 341 |
| GO_NUCLEAR_EXPORT | 0.00000 | 0.00004 | 0.65944 | 0.64915 | 1.26157 | 181 |
| GO_NUCLEAR_MATRIX | 0.00502 | 0.04608 | 0.40702 | 0.62777 | 1.19868 | 87 |
| GO_NUCLEAR_MEMBRANE | 0.00185 | 0.02237 | 0.45506 | 0.58925 | 1.14853 | 211 |
| GO_NUCLEAR_PERIPHERY | 0.00043 | 0.00701 | 0.49849 | 0.64264 | 1.23558 | 107 |
| GO_NUCLEAR_PORE | 0.00561 | 0.04971 | 0.40702 | 0.64876 | 1.22522 | 68 |
| GO_NUCLEAR_SPECK | 0.00015 | 0.00312 | 0.51885 | 0.59159 | 1.16089 | 306 |
| GO_NUCLEAR_TRANSCRIBED_MRNA_CATABOLIC_PROCESS | 0.00000 | 0.00000 | | 0.70362 | 1.36820 | 185 |

| | | | | | | |
|---|---|---|---|---|---|---|
| GO_NUCLEAR_TRANSCRIBED_MRNA_CATABOLIC_PROCESS_NON | 0.00000 | 0.00000 | | 0.74239 | 1.42931 | 114 |
| GO_NUCLEAR_TRANSPORT | 0.00000 | 0.00000 | 0.71951 | 0.62920 | 1.23383 | 297 |
| GO_NUCLEOBASE_CONTAINING_COMPOUND_TRANSPORT | 0.00000 | 0.00010 | 0.62726 | 0.62975 | 1.22823 | 216 |
| GO_NUCLEOBASE_CONTAINING_SMALL_MOLECULE_METABOLI( | 0.00000 | 0.00003 | 0.65944 | 0.60153 | 1.18594 | 405 |
| GO_NUCLEOSIDE_DIPHOSPHATE_METABOLIC_PROCESS | 0.00016 | 0.00319 | 0.51885 | 0.64491 | 1.24260 | 117 |
| GO_NUCLEOSIDE_PHOSPHATE_BIOSYNTHETIC_PROCESS | 0.00002 | 0.00044 | 0.59333 | 0.64092 | 1.24175 | 161 |
| GO_NUCLEOSIDE_TRIPHOSPHATE_BIOSYNTHETIC_PROCESS | 0.00040 | 0.00676 | 0.49849 | 0.72727 | 1.35074 | 42 |
| GO_NUCLEOSIDE_TRIPHOSPHATE_METABOLIC_PROCESS | 0.00009 | 0.00196 | 0.53843 | 0.69306 | 1.30854 | 65 |
| GO_NUCLEOTIDE_PHOSPHORYLATION | 0.00016 | 0.00328 | 0.51885 | 0.65541 | 1.25767 | 102 |
| GO_OOCYTE_DIFFERENTIATION | 0.00450 | 0.04342 | 0.40702 | 0.73513 | 1.33705 | 26 |
| GO_ORGANELLAR_RIBOSOME | 0.00000 | 0.00000 | 0.76146 | 0.75712 | 1.44311 | 84 |
| GO_ORGANELLE_ENVELOPE_LUMEN | 0.00026 | 0.00482 | 0.49849 | 0.67265 | 1.27716 | 78 |
| GO_ORGANELLE_FISSION | 0.00154 | 0.01974 | 0.45506 | 0.57133 | 1.12383 | 342 |
| GO_ORGANELLE_INNER_MEMBRANE | 0.00000 | 0.00000 | | 0.68423 | 1.34915 | 412 |
| GO_ORGANIC_CYCLIC_COMPOUND_CATABOLIC_PROCESS | 0.00000 | 0.00000 | | 0.66975 | 1.32263 | 478 |
| GO_ORGANIC_HYDROXY_COMPOUND_BIOSYNTHETIC_PROCESS | 0.00141 | 0.01850 | 0.45506 | 0.60712 | 1.17627 | 161 |
| GO_OXIDATIVE_PHOSPHORYLATION | 0.00000 | 0.00000 | | 0.78902 | 1.51260 | 99 |
| GO_OXIDOREDUCTASE_ACTIVITY | 0.00000 | 0.00000 | | 0.63268 | 1.24935 | 468 |
| GO_OXIDOREDUCTASE_ACTIVITY_ACTING_ON_A_HEME_GROUP | 0.00176 | 0.02168 | 0.45506 | 0.79383 | 1.41828 | 19 |
| GO_OXIDOREDUCTASE_ACTIVITY_ACTING_ON_NAD_P_H | 0.00002 | 0.00056 | 0.57561 | 0.71468 | 1.35162 | 70 |
| GO_OXIDOREDUCTASE_ACTIVITY_ACTING_ON_NAD_P_H_QUIN( | 0.00004 | 0.00113 | 0.55733 | 0.74511 | 1.38835 | 47 |
| GO_OXIDOREDUCTASE_ACTIVITY_ACTING_ON_PEROXIDE_AS_A( | 0.00106 | 0.01495 | 0.45506 | 0.73897 | 1.35352 | 29 |
| GO_OXIDOREDUCTASE_COMPLEX | 0.00000 | 0.00000 | 0.73376 | 0.74898 | 1.42731 | 82 |
| GO_PEPTIDASE_ACTIVITY | 0.00133 | 0.01799 | 0.45506 | 0.57681 | 1.13188 | 306 |
| GO_PEPTIDASE_COMPLEX | 0.00000 | 0.00002 | 0.67496 | 0.73250 | 1.39240 | 79 |
| GO_PEPTIDASE_REGULATOR_ACTIVITY | 0.00313 | 0.03310 | 0.43171 | 0.63434 | 1.21189 | 88 |
| GO_PEPTIDE_BINDING | 0.00005 | 0.00124 | 0.55733 | 0.63708 | 1.23187 | 148 |
| GO_PERINUCLEAR_REGION_OF_CYTOPLASM | 0.00276 | 0.03021 | 0.43171 | 0.55814 | 1.10207 | 466 |
| GO_PHAGOCYTOSIS | 0.00554 | 0.04968 | 0.40702 | 0.59337 | 1.14837 | 155 |
| GO_PHAGOSOME_ACIDIFICATION | 0.00059 | 0.00916 | 0.47727 | 0.78902 | 1.42694 | 24 |
| GO_PHOSPHATIDYLSERINE_BINDING | 0.00035 | 0.00610 | 0.49849 | 0.76387 | 1.39912 | 29 |
| GO_PH_REDUCTION | 0.00302 | 0.03248 | 0.43171 | 0.68821 | 1.28238 | 46 |
| GO_PIGMENT_GRANULE | 0.00000 | 0.00000 | 0.76146 | 0.74333 | 1.41934 | 87 |
| GO_PLASMA_LIPOPROTEIN_PARTICLE_CLEARANCE | 0.00106 | 0.01495 | 0.45506 | 0.73201 | 1.35304 | 37 |
| GO_POLYSOMAL_RIBOSOME | 0.00008 | 0.00180 | 0.53843 | 0.78840 | 1.44683 | 30 |
| GO_POLYSOME | 0.00000 | 0.00003 | 0.65944 | 0.75964 | 1.43296 | 62 |
| GO_POSITIVE_REGULATION_OF_ATP_METABOLIC_PROCESS | 0.00174 | 0.02149 | 0.45506 | 0.73011 | 1.33729 | 29 |
| GO_POSITIVE_REGULATION_OF_BINDING | 0.00390 | 0.03895 | 0.43171 | 0.61042 | 1.17582 | 115 |
| GO_POSITIVE_REGULATION_OF_CANONICAL_WNT_SIGNALING_ | 0.00000 | 0.00000 | 0.73376 | 0.72036 | 1.38514 | 108 |
| GO_POSITIVE_REGULATION_OF_CELLULAR_AMIDE_METABOLIC_ | 0.00001 | 0.00036 | 0.59333 | 0.66485 | 1.28095 | 125 |
| GO_POSITIVE_REGULATION_OF_CELLULAR_PROTEIN_LOCALIZAT | 0.00242 | 0.02748 | 0.43171 | 0.58048 | 1.13378 | 244 |
| GO_POSITIVE_REGULATION_OF_CELL_POPULATION_PROLIFERAT | 0.00040 | 0.00670 | 0.49849 | 0.56967 | 1.12503 | 461 |
| GO_POSITIVE_REGULATION_OF_CYSTEINE_TYPE_ENDOPEPTIDA: | 0.00547 | 0.04923 | 0.40702 | 0.62225 | 1.19077 | 92 |
| GO_POSITIVE_REGULATION_OF_CYTOKINE_PRODUCTION | 0.00283 | 0.03081 | 0.43171 | 0.58672 | 1.14373 | 210 |
| GO_POSITIVE_REGULATION_OF_DEFENSE_RESPONSE | 0.00000 | 0.00000 | 0.73376 | 0.66200 | 1.28884 | 200 |
| GO_POSITIVE_REGULATION_OF_DNA_BIOSYNTHETIC_PROCESS | 0.00010 | 0.00228 | 0.53843 | 0.71224 | 1.33977 | 55 |
| GO_POSITIVE_REGULATION_OF_DNA_METABOLIC_PROCESS | 0.00020 | 0.00387 | 0.51885 | 0.62771 | 1.21205 | 139 |
| GO_POSITIVE_REGULATION_OF_ESTABLISHMENT_OF_PROTEIN_ | 0.00194 | 0.02331 | 0.45506 | 0.58158 | 1.13709 | 253 |
| GO_POSITIVE_REGULATION_OF_FIBROBLAST_PROLIFERATION | 0.00017 | 0.00338 | 0.51885 | 0.77423 | 1.42213 | 31 |
| GO_POSITIVE_REGULATION_OF_IMMUNE_RESPONSE | 0.00000 | 0.00000 | 0.71951 | 0.63064 | 1.23937 | 321 |
| GO_POSITIVE_REGULATION_OF_IMMUNE_SYSTEM_PROCESS | 0.00000 | 0.00002 | 0.67496 | 0.59748 | 1.17992 | 475 |
| GO_POSITIVE_REGULATION_OF_INNATE_IMMUNE_RESPONSE | 0.00000 | 0.00000 | 0.70498 | 0.68885 | 1.32737 | 128 |
| GO_POSITIVE_REGULATION_OF_MITOTIC_CELL_CYCLE | 0.00178 | 0.02182 | 0.45506 | 0.61563 | 1.18683 | 122 |
| GO_POSITIVE_REGULATION_OF_MRNA_METABOLIC_PROCESS | 0.00157 | 0.02009 | 0.45506 | 0.66330 | 1.25231 | 63 |
| GO_POSITIVE_REGULATION_OF_MRNA_PROCESSING | 0.00250 | 0.02807 | 0.43171 | 0.76194 | 1.38151 | 25 |
| GO_POSITIVE_REGULATION_OF_MRNA_SPLICING_VIA_SPLICEOS | 0.00132 | 0.01782 | 0.45506 | 0.81851 | 1.44353 | 16 |
| GO_POSITIVE_REGULATION_OF_NITRIC_OXIDE_METABOLIC_PR( | 0.00133 | 0.01799 | 0.45506 | 0.80807 | 1.43726 | 18 |
| GO_POSITIVE_REGULATION_OF_ORGANELLE_ORGANIZATION | 0.00010 | 0.00218 | 0.53843 | 0.57473 | 1.13449 | 452 |
| GO_POSITIVE_REGULATION_OF_PROTEIN_LOCALIZATION_TO_N | 0.00012 | 0.00265 | 0.53843 | 0.71018 | 1.33697 | 58 |
| GO_POSITIVE_REGULATION_OF_PROTEOLYSIS | 0.00313 | 0.03310 | 0.43171 | 0.58059 | 1.13381 | 237 |
| GO_POSITIVE_REGULATION_OF_REACTIVE_OXYGEN_SPECIES_BI | 0.00115 | 0.01593 | 0.45506 | 0.79865 | 1.43554 | 21 |
| GO_POSITIVE_REGULATION_OF_REACTIVE_OXYGEN_SPECIES_M | 0.00174 | 0.02149 | 0.45506 | 0.69785 | 1.30035 | 46 |
| GO_POSITIVE_REGULATION_OF_RESPONSE_TO_BIOTIC_STIMUL | 0.00000 | 0.00000 | 0.70498 | 0.67507 | 1.30603 | 150 |
| GO_POSITIVE_REGULATION_OF_RESPONSE_TO_EXTERNAL_STIM | 0.00001 | 0.00016 | 0.61053 | 0.62123 | 1.21505 | 260 |
| GO_POSITIVE_REGULATION_OF_RNA_SPLICING | 0.00265 | 0.02936 | 0.43171 | 0.74051 | 1.35202 | 27 |
| GO_POSITIVE_REGULATION_OF_SIGNAL_TRANSDUCTION_BY_P5 | 0.00390 | 0.03895 | 0.43171 | 0.80515 | 1.41278 | 15 |
| GO_POSITIVE_REGULATION_OF_TELOMERASE_ACTIVITY | 0.00069 | 0.01043 | 0.47727 | 0.75265 | 1.38122 | 30 |
| GO_POSITIVE_REGULATION_OF_TELOMERASE_RNA_LOCALIZATI | 0.00007 | 0.00169 | 0.53843 | 0.88616 | 1.55493 | 15 |
| GO_POSITIVE_REGULATION_OF_TELOMERE_MAINTENANCE | 0.00026 | 0.00486 | 0.49849 | 0.73697 | 1.37102 | 44 |
| GO_POSITIVE_REGULATION_OF_TELOMERE_MAINTENANCE_VIA | 0.00001 | 0.00040 | 0.59333 | 0.80196 | 1.47289 | 33 |

| | | | | | | |
|---|---|---|---|---|---|---|
| GO_POSITIVE_REGULATION_OF_TRANSFERASE_ACTIVITY | 0.00108 | 0.01507 | 0.45506 | 0.56501 | 1.11376 | 401 |
| GO_POSITIVE_REGULATION_OF_TRANSLATION | 0.00000 | 0.00016 | 0.61053 | 0.68408 | 1.31526 | 107 |
| GO_POSITIVE_REGULATION_OF_VIRAL_PROCESS | 0.00026 | 0.00486 | 0.49849 | 0.66195 | 1.26462 | 88 |
| GO_POSITIVE_REGULATION_OF_WNT_SIGNALING_PATHWAY | 0.00000 | 0.00000 | 0.73376 | 0.69924 | 1.34747 | 127 |
| GO_POSTSYNAPSE | 0.00239 | 0.02716 | 0.43171 | 0.57062 | 1.12116 | 316 |
| GO_POST_TRANSLATIONAL_PROTEIN_MODIFICATION | 0.00024 | 0.00452 | 0.51885 | 0.60046 | 1.17392 | 251 |
| GO_PRECATALYTIC_SPLICEOSOME | 0.00141 | 0.01850 | 0.45506 | 0.69251 | 1.29540 | 50 |
| GO_PRERIBOSOME | 0.00006 | 0.00154 | 0.53843 | 0.69866 | 1.32020 | 69 |
| GO_PROTEASOMAL_PROTEIN_CATABOLIC_PROCESS | 0.00007 | 0.00169 | 0.53843 | 0.58188 | 1.14652 | 387 |
| GO_PROTEASOMAL_UBIQUITIN_INDEPENDENT_PROTEIN_CATAB | 0.00000 | 0.00003 | 0.65944 | 0.91182 | 1.62178 | 18 |
| GO_PROTEASOME_ACCESSORY_COMPLEX | 0.00005 | 0.00135 | 0.55733 | 0.82638 | 1.49450 | 24 |
| GO_PROTEASOME_CORE_COMPLEX | 0.00000 | 0.00004 | 0.65944 | 0.91231 | 1.61486 | 17 |
| GO_PROTEIN_CONTAINING_COMPLEX_DISASSEMBLY | 0.00000 | 0.00000 | 0.81404 | 0.66285 | 1.29656 | 261 |
| GO_PROTEIN_CONTAINING_COMPLEX_LOCALIZATION | 0.00018 | 0.00352 | 0.51885 | 0.60451 | 1.17943 | 226 |
| GO_PROTEIN_DNA_COMPLEX_SUBUNIT_ORGANIZATION | 0.00052 | 0.00825 | 0.47727 | 0.60887 | 1.18342 | 183 |
| GO_PROTEIN_DOMAIN_SPECIFIC_BINDING | 0.00003 | 0.00088 | 0.55733 | 0.57828 | 1.14190 | 470 |
| GO_PROTEIN_FOLDING | 0.00000 | 0.00000 | 0.73376 | 0.67425 | 1.30867 | 171 |
| GO_PROTEIN_FOLDING_CHAPERONE | 0.00079 | 0.01170 | 0.47727 | 0.81632 | 1.45193 | 18 |
| GO_PROTEIN_HETEROOLIGOMERIZATION | 0.00442 | 0.04297 | 0.40702 | 0.78998 | 1.39322 | 16 |
| GO_PROTEIN_IMPORT | 0.00001 | 0.00040 | 0.59333 | 0.64197 | 1.24306 | 158 |
| GO_PROTEIN_IMPORT_INTO_MITOCHONDRIAL_MATRIX | 0.00231 | 0.02642 | 0.43171 | 0.79530 | 1.41455 | 18 |
| GO_PROTEIN_INSERTION_INTO_MEMBRANE | 0.00191 | 0.02299 | 0.45506 | 0.68322 | 1.27694 | 49 |
| GO_PROTEIN_LOCALIZATION_TO_CHROMOSOME_TELOMERIC_F | 0.00001 | 0.00031 | 0.59333 | 0.82810 | 1.51357 | 28 |
| GO_PROTEIN_LOCALIZATION_TO_ENDOPLASMIC_RETICULUM | 0.00000 | 0.00000 | | 0.73850 | 1.42303 | 129 |
| GO_PROTEIN_LOCALIZATION_TO_MEMBRANE | 0.00000 | 0.00000 | 0.77494 | 0.61216 | 1.20881 | 462 |
| GO_PROTEIN_LOCALIZATION_TO_MITOCHONDRION | 0.00015 | 0.00310 | 0.51885 | 0.64234 | 1.23759 | 118 |
| GO_PROTEIN_LOCALIZATION_TO_NUCLEUS | 0.00000 | 0.00014 | 0.61053 | 0.63315 | 1.23408 | 208 |
| GO_PROTEIN_MATURATION | 0.00357 | 0.03638 | 0.43171 | 0.59209 | 1.14965 | 174 |
| GO_PROTEIN_MODIFICATION_BY_SMALL_PROTEIN_REMOVAL | 0.00002 | 0.00052 | 0.57561 | 0.61535 | 1.20119 | 231 |
| GO_PROTEIN_REFOLDING | 0.00465 | 0.04430 | 0.40702 | 0.77561 | 1.37288 | 17 |
| GO_PROTEIN_STABILIZATION | 0.00081 | 0.01194 | 0.47727 | 0.61881 | 1.19577 | 146 |
| GO_PROTEIN_SUMOYLATION | 0.00450 | 0.04342 | 0.40702 | 0.65043 | 1.22846 | 67 |
| GO_PROTEIN_TARGETING | 0.00000 | 0.00000 | | 0.66223 | 1.30361 | 358 |
| GO_PROTEIN_TARGETING_TO_MEMBRANE | 0.00000 | 0.00000 | | 0.71675 | 1.38879 | 162 |
| GO_PROTEIN_TARGETING_TO_MITOCHONDRION | 0.00113 | 0.01571 | 0.45506 | 0.65407 | 1.24648 | 83 |
| GO_PROTEIN_TRANSMEMBRANE_TRANSPORT | 0.00110 | 0.01526 | 0.45506 | 0.68173 | 1.28238 | 55 |
| GO_PROTON_TRANSMEMBRANE_TRANSPORT | 0.00000 | 0.00001 | 0.69013 | 0.71234 | 1.36309 | 91 |
| GO_PROTON_TRANSMEMBRANE_TRANSPORTER_ACTIVITY | 0.00004 | 0.00115 | 0.55733 | 0.70138 | 1.32535 | 69 |
| GO_PROTON_TRANSPORTING_TWO_SECTOR_ATPASE_COMPLEX | 0.00083 | 0.01218 | 0.47727 | 0.77254 | 1.40508 | 26 |
| GO_PROTON_TRANSPORTING_V_TYPE_ATPASE_COMPLEX | 0.00239 | 0.02716 | 0.43171 | 0.76100 | 1.37626 | 24 |
| GO_PURINE_CONTAINING_COMPOUND_BIOSYNTHETIC_PROCES | 0.00001 | 0.00040 | 0.59333 | 0.65863 | 1.26901 | 126 |
| GO_PURINE_CONTAINING_COMPOUND_METABOLIC_PROCESS | 0.00000 | 0.00000 | 0.73376 | 0.63132 | 1.23831 | 300 |
| GO_PURINE_NUCLEOSIDE_TRIPHOSPHATE_METABOLIC_PROCES | 0.00033 | 0.00582 | 0.49849 | 0.70887 | 1.32487 | 49 |
| GO_PYROPHOSPHATE_HYDROLYSIS_DRIVEN_PROTON_TRANSMI | 0.00532 | 0.04803 | 0.40702 | 0.74470 | 1.34678 | 24 |
| GO_PYRUVATE_METABOLIC_PROCESS | 0.00001 | 0.00018 | 0.61053 | 0.67194 | 1.29289 | 113 |
| GO_P_BODY | 0.00472 | 0.04480 | 0.40702 | 0.65470 | 1.23501 | 62 |
| GO_REACTIVE_OXYGEN_SPECIES_BIOSYNTHETIC_PROCESS | 0.00040 | 0.00670 | 0.49849 | 0.70271 | 1.31857 | 53 |
| GO_REACTIVE_OXYGEN_SPECIES_METABOLIC_PROCESS | 0.00021 | 0.00409 | 0.51885 | 0.62881 | 1.21472 | 144 |
| GO_REGULATION_OF_ALCOHOL_BIOSYNTHETIC_PROCESS | 0.00228 | 0.02614 | 0.43171 | 0.68438 | 1.28020 | 50 |
| GO_REGULATION_OF_ANIMAL_ORGAN_MORPHOGENESIS | 0.00000 | 0.00000 | | 0.72330 | 1.39922 | 149 |
| GO_REGULATION_OF_APOPTOTIC_SIGNALING_PATHWAY | 0.00079 | 0.01170 | 0.47727 | 0.58599 | 1.14641 | 266 |
| GO_REGULATION_OF_ATP_METABOLIC_PROCESS | 0.00000 | 0.00006 | 0.64355 | 0.70523 | 1.34777 | 89 |
| GO_REGULATION_OF_BINDING | 0.00428 | 0.04185 | 0.40702 | 0.58264 | 1.13664 | 225 |
| GO_REGULATION_OF_BLOOD_PRESSURE | 0.00290 | 0.03156 | 0.43171 | 0.65319 | 1.23428 | 69 |
| GO_REGULATION_OF_CARBOHYDRATE_CATABOLIC_PROCESS | 0.00324 | 0.03393 | 0.43171 | 0.65541 | 1.23787 | 67 |
| GO_REGULATION_OF_CELLULAR_AMIDE_METABOLIC_PROCESS | 0.00000 | 0.00000 | 0.78819 | 0.63116 | 1.24143 | 337 |
| GO_REGULATION_OF_CELLULAR_AMINE_METABOLIC_PROCESS | 0.00000 | 0.00000 | 0.74774 | 0.78955 | 1.48730 | 60 |
| GO_REGULATION_OF_CELLULAR_AMINO_ACID_METABOLIC_PR( | 0.00000 | 0.00000 | 0.76146 | 0.80341 | 1.51022 | 54 |
| GO_REGULATION_OF_CELLULAR_KETONE_METABOLIC_PROCESS | 0.00000 | 0.00000 | 0.70498 | 0.69239 | 1.33405 | 126 |
| GO_REGULATION_OF_CELLULAR_PH | 0.00442 | 0.04297 | 0.40702 | 0.65360 | 1.23436 | 68 |
| GO_REGULATION_OF_CELLULAR_PROTEIN_LOCALIZATION | 0.00228 | 0.02614 | 0.43171 | 0.56216 | 1.10830 | 402 |
| GO_REGULATION_OF_CELLULAR_RESPIRATION | 0.00428 | 0.04185 | 0.40702 | 0.77183 | 1.37897 | 19 |
| GO_REGULATION_OF_CELLULAR_RESPONSE_TO_HEAT | 0.00091 | 0.01314 | 0.47727 | 0.67360 | 1.27221 | 67 |
| GO_REGULATION_OF_CELL_CYCLE_G2_M_PHASE_TRANSITION | 0.00016 | 0.00332 | 0.51885 | 0.61554 | 1.19738 | 187 |
| GO_REGULATION_OF_CELL_CYCLE_PHASE_TRANSITION | 0.00000 | 0.00014 | 0.61053 | 0.59968 | 1.18091 | 369 |
| GO_REGULATION_OF_CHOLESTEROL_BIOSYNTHETIC_PROCESS | 0.00209 | 0.02470 | 0.43171 | 0.71476 | 1.32117 | 38 |
| GO_REGULATION_OF_CHOLESTEROL_METABOLIC_PROCESS | 0.00113 | 0.01571 | 0.45506 | 0.69865 | 1.30177 | 47 |
| GO_REGULATION_OF_CHROMOSOME_ORGANIZATION | 0.00036 | 0.00622 | 0.49849 | 0.58896 | 1.15261 | 278 |
| GO_REGULATION_OF_CHROMOSOME_SEGREGATION | 0.00079 | 0.01170 | 0.47727 | 0.64131 | 1.22718 | 91 |
| GO_REGULATION_OF_CHROMOSOME_SEPARATION | 0.00253 | 0.02838 | 0.43171 | 0.66349 | 1.24947 | 59 |

| | | | | | | |
|---|---|---|---|---|---|---|
| GO_REGULATION_OF_CYSTEINE_TYPE_ENDOPEPTIDASE_ACTIVIT | 0.00044 | 0.00719 | 0.49849 | 0.62610 | 1.20905 | 141 |
| GO_REGULATION_OF_DEFENSE_RESPONSE | 0.00006 | 0.00150 | 0.55733 | 0.58834 | 1.15824 | 356 |
| GO_REGULATION_OF_DNA_BIOSYNTHETIC_PROCESS | 0.00043 | 0.00695 | 0.49849 | 0.66709 | 1.27130 | 83 |
| GO_REGULATION_OF_DNA_METABOLIC_PROCESS | 0.00055 | 0.00863 | 0.47727 | 0.58909 | 1.15134 | 249 |
| GO_REGULATION_OF_DNA_TEMPLATED_TRANSCRIPTION_IN_RI | 0.00000 | 0.00000 | 0.71951 | 0.72410 | 1.38687 | 94 |
| GO_REGULATION_OF_GENERATION_OF_PRECURSOR_METABOLI | 0.00004 | 0.00114 | 0.55733 | 0.65536 | 1.26115 | 111 |
| GO_REGULATION_OF_GENE_SILENCING | 0.00023 | 0.00434 | 0.51885 | 0.65464 | 1.25268 | 91 |
| GO_REGULATION_OF_HEMATOPOIETIC_PROGENITOR_CELL_DIF | 0.00000 | 0.00000 | 0.73376 | 0.76273 | 1.44465 | 72 |
| GO_REGULATION_OF_HEMATOPOIETIC_STEM_CELL_DIFFERENT | 0.00000 | 0.00000 | 0.74774 | 0.78891 | 1.48817 | 62 |
| GO_REGULATION_OF_HEMOPOIESIS | 0.00390 | 0.03895 | 0.43171 | 0.57739 | 1.12909 | 258 |
| GO_REGULATION_OF_IMMUNE_RESPONSE | 0.00000 | 0.00006 | 0.64355 | 0.60032 | 1.18447 | 433 |
| GO_REGULATION_OF_INNATE_IMMUNE_RESPONSE | 0.00000 | 0.00007 | 0.62726 | 0.64536 | 1.25422 | 181 |
| GO_REGULATION_OF_INTRINSIC_APOPTOTIC_SIGNALING_PATH\ | 0.00179 | 0.02183 | 0.45506 | 0.61548 | 1.18550 | 116 |
| GO_REGULATION_OF_MITOTIC_CELL_CYCLE | 0.00000 | 0.00003 | 0.65944 | 0.58992 | 1.16508 | 499 |
| GO_REGULATION_OF_MORPHOGENESIS_OF_AN_EPITHELIUM | 0.00000 | 0.00000 | | 0.74680 | 1.43884 | 125 |
| GO_REGULATION_OF_MRNA_CATABOLIC_PROCESS | 0.00000 | 0.00000 | | 0.71126 | 1.37919 | 165 |
| GO_REGULATION_OF_MRNA_METABOLIC_PROCESS | 0.00000 | 0.00000 | | 0.67178 | 1.31458 | 276 |
| GO_REGULATION_OF_MRNA_PROCESSING | 0.00008 | 0.00186 | 0.53843 | 0.65189 | 1.25598 | 125 |
| GO_REGULATION_OF_MRNA_SPLICING_VIA_SPLICEOSOME | 0.00012 | 0.00256 | 0.53843 | 0.66950 | 1.28033 | 90 |
| GO_REGULATION_OF_MULTI_ORGANISM_PROCESS | 0.00008 | 0.00190 | 0.53843 | 0.76905 | 1.42150 | 37 |
| GO_REGULATION_OF_MUSCLE_CONTRACTION | 0.00163 | 0.02050 | 0.45506 | 0.64879 | 1.23071 | 76 |
| GO_REGULATION_OF_NUCLEAR_DIVISION | 0.00060 | 0.00923 | 0.47727 | 0.62443 | 1.20552 | 137 |
| GO_REGULATION_OF_NUCLEASE_ACTIVITY | 0.00059 | 0.00916 | 0.47727 | 0.81786 | 1.44767 | 17 |
| GO_REGULATION_OF_NUCLEOTIDE_BIOSYNTHETIC_PROCESS | 0.00302 | 0.03248 | 0.43171 | 0.78902 | 1.40337 | 18 |
| GO_REGULATION_OF_NUCLEOTIDE_METABOLIC_PROCESS | 0.00033 | 0.00588 | 0.49849 | 0.66858 | 1.26942 | 78 |
| GO_REGULATION_OF_OXIDATIVE_PHOSPHORYLATION | 0.00005 | 0.00130 | 0.55733 | 0.85284 | 1.53294 | 21 |
| GO_REGULATION_OF_PEPTIDASE_ACTIVITY | 0.00006 | 0.00152 | 0.53843 | 0.61192 | 1.19388 | 226 |
| GO_REGULATION_OF_PEPTIDE_TRANSPORT | 0.00250 | 0.02807 | 0.43171 | 0.56252 | 1.10917 | 412 |
| GO_REGULATION_OF_PH | 0.00561 | 0.04971 | 0.40702 | 0.64191 | 1.21498 | 71 |
| GO_REGULATION_OF_PLASMA_LIPOPROTEIN_PARTICLE_LEVELS | 0.00006 | 0.00145 | 0.55733 | 0.72431 | 1.36154 | 54 |
| GO_REGULATION_OF_POSTTRANSCRIPTIONAL_GENE_SILENCING | 0.00026 | 0.00486 | 0.49849 | 0.66803 | 1.27236 | 81 |
| GO_REGULATION_OF_PROTEIN_CATABOLIC_PROCESS | 0.00309 | 0.03299 | 0.43171 | 0.57360 | 1.12399 | 291 |
| GO_REGULATION_OF_PROTEIN_IMPORT | 0.00487 | 0.04559 | 0.40702 | 0.67491 | 1.25754 | 47 |
| GO_REGULATION_OF_PROTEIN_LOCALIZATION_TO_NUCLEUS | 0.00000 | 0.00013 | 0.61053 | 0.69035 | 1.32101 | 91 |
| GO_REGULATION_OF_PROTEIN_STABILITY | 0.00066 | 0.01007 | 0.47727 | 0.59308 | 1.15811 | 235 |
| GO_REGULATION_OF_PROTEOLYSIS | 0.00007 | 0.00169 | 0.53843 | 0.57923 | 1.14299 | 431 |
| GO_REGULATION_OF_REACTIVE_OXYGEN_SPECIES_BIOSYNTHET | 0.00022 | 0.00427 | 0.51885 | 0.73154 | 1.36343 | 45 |
| GO_REGULATION_OF_REACTIVE_OXYGEN_SPECIES_METABOLIC_ | 0.00088 | 0.01280 | 0.47727 | 0.63948 | 1.22592 | 99 |
| GO_REGULATION_OF_RESPONSE_TO_BIOTIC_STIMULUS | 0.00001 | 0.00039 | 0.59333 | 0.61773 | 1.20625 | 235 |
| GO_REGULATION_OF_RNA_SPLICING | 0.00011 | 0.00241 | 0.53843 | 0.64982 | 1.25198 | 125 |
| GO_REGULATION_OF_SISTER_CHROMATID_SEGREGATION | 0.00405 | 0.04011 | 0.40702 | 0.64401 | 1.22034 | 73 |
| GO_REGULATION_OF_SMALL_MOLECULE_METABOLIC_PROCESS | 0.00000 | 0.00000 | 0.77494 | 0.64342 | 1.26057 | 290 |
| GO_REGULATION_OF_SMOOTH_MUSCLE_CONTRACTION | 0.00007 | 0.00171 | 0.53843 | 0.82642 | 1.49295 | 23 |
| GO_REGULATION_OF_STEM_CELL_DIFFERENTIATION | 0.00000 | 0.00000 | 0.73376 | 0.73434 | 1.40292 | 88 |
| GO_REGULATION_OF_STEROID_METABOLIC_PROCESS | 0.00524 | 0.04770 | 0.40702 | 0.64061 | 1.21567 | 77 |
| GO_REGULATION_OF_SUPEROXIDE_METABOLIC_PROCESS | 0.00346 | 0.03548 | 0.43171 | 0.77826 | 1.39046 | 19 |
| GO_REGULATION_OF_SUPRAMOLECULAR_FIBER_ORGANIZATIOI | 0.00509 | 0.04656 | 0.40702 | 0.57599 | 1.12367 | 223 |
| GO_REGULATION_OF_SYMBIOTIC_PROCESS | 0.00009 | 0.00211 | 0.53843 | 0.62950 | 1.21763 | 151 |
| GO_REGULATION_OF_TELOMERASE_ACTIVITY | 0.00087 | 0.01263 | 0.47727 | 0.72147 | 1.33426 | 39 |
| GO_REGULATION_OF_TELOMERE_MAINTENANCE | 0.00547 | 0.04923 | 0.40702 | 0.64959 | 1.22679 | 68 |
| GO_REGULATION_OF_TELOMERE_MAINTENANCE_VIA_TELOME | 0.00020 | 0.00387 | 0.51885 | 0.72043 | 1.34856 | 51 |
| GO_REGULATION_OF_TRANSCRIPTION_FROM_RNA_POLYMERA! | 0.00000 | 0.00000 | 0.70498 | 0.76931 | 1.45239 | 64 |
| GO_REGULATION_OF_TRANSLATIONAL_INITIATION | 0.00023 | 0.00437 | 0.51885 | 0.68479 | 1.29399 | 69 |
| GO_REGULATION_OF_VIRAL_TRANSCRIPTION | 0.00361 | 0.03657 | 0.43171 | 0.67497 | 1.26259 | 50 |
| GO_REGULATION_OF_WNT_SIGNALING_PATHWAY | 0.00015 | 0.00312 | 0.51885 | 0.60129 | 1.17419 | 238 |
| GO_RESPIRASOME | 0.00000 | 0.00000 | 0.81404 | 0.78326 | 1.48524 | 75 |
| GO_RESPIRATORY_CHAIN_COMPLEX | 0.00000 | 0.00000 | 0.77494 | 0.79183 | 1.49492 | 64 |
| GO_RESPIRATORY_ELECTRON_TRANSPORT_CHAIN | 0.00000 | 0.00000 | | 0.78275 | 1.49591 | 89 |
| GO_RESPONSE_TO_ALCOHOL | 0.00015 | 0.00310 | 0.51885 | 0.63794 | 1.22915 | 126 |
| GO_RESPONSE_TO_ATP | 0.00087 | 0.01270 | 0.47727 | 0.81086 | 1.43528 | 17 |
| GO_RESPONSE_TO_CALCIUM_ION | 0.00124 | 0.01701 | 0.45506 | 0.65196 | 1.24071 | 80 |
| GO_RESPONSE_TO_CORTICOSTEROID | 0.00010 | 0.00220 | 0.53843 | 0.66894 | 1.27841 | 89 |
| GO_RESPONSE_TO_DRUG | 0.00000 | 0.00007 | 0.62726 | 0.63080 | 1.23070 | 220 |
| GO_RESPONSE_TO_ETHANOL | 0.00168 | 0.02098 | 0.45506 | 0.66543 | 1.25679 | 67 |
| GO_RESPONSE_TO_FIBROBLAST_GROWTH_FACTOR | 0.00357 | 0.03638 | 0.43171 | 0.63469 | 1.20976 | 84 |
| GO_RESPONSE_TO_GROWTH_FACTOR | 0.00272 | 0.02986 | 0.43171 | 0.56235 | 1.10805 | 387 |
| GO_RESPONSE_TO_HEAT | 0.00000 | 0.00013 | 0.62726 | 0.66982 | 1.29069 | 119 |
| GO_RESPONSE_TO_HYDROGEN_PEROXIDE | 0.00532 | 0.04803 | 0.40702 | 0.62255 | 1.19054 | 90 |
| GO_RESPONSE_TO_INORGANIC_SUBSTANCE | 0.00016 | 0.00328 | 0.51885 | 0.58928 | 1.15810 | 321 |
| GO_RESPONSE_TO_INTERLEUKIN_1 | 0.00000 | 0.00000 | 0.78819 | 0.71590 | 1.37950 | 124 |

| | | | | | | |
|---|---|---|---|---|---|---|
| GO_RESPONSE_TO_INTERLEUKIN_12 | 0.00036 | 0.00622 | 0.49849 | 0.74158 | 1.36769 | 36 |
| GO_RESPONSE_TO_INTERLEUKIN_4 | 0.00224 | 0.02581 | 0.43171 | 0.77088 | 1.38921 | 22 |
| GO_RESPONSE_TO_LEUKEMIA_INHIBITORY_FACTOR | 0.00076 | 0.01129 | 0.47727 | 0.66148 | 1.25288 | 72 |
| GO_RESPONSE_TO_LIPID | 0.00043 | 0.00695 | 0.49849 | 0.56669 | 1.11902 | 470 |
| GO_RESPONSE_TO_METAL_ION | 0.00143 | 0.01863 | 0.45506 | 0.59895 | 1.16592 | 199 |
| GO_RESPONSE_TO_NUTRIENT | 0.00294 | 0.03185 | 0.43171 | 0.62596 | 1.19987 | 98 |
| GO_RESPONSE_TO_ORGANOPHOSPHORUS | 0.00294 | 0.03185 | 0.43171 | 0.64154 | 1.21511 | 72 |
| GO_RESPONSE_TO_OXIDATIVE_STRESS | 0.00495 | 0.04573 | 0.40702 | 0.57010 | 1.11758 | 294 |
| GO_RESPONSE_TO_OXYGEN_LEVELS | 0.00000 | 0.00000 | | 0.67661 | 1.32155 | 244 |
| GO_RESPONSE_TO_PURINE_CONTAINING_COMPOUND | 0.00509 | 0.04656 | 0.40702 | 0.63176 | 1.20394 | 82 |
| GO_RESPONSE_TO_STEROID_HORMONE | 0.00009 | 0.00200 | 0.53843 | 0.61312 | 1.19360 | 201 |
| GO_RESPONSE_TO_TEMPERATURE_STIMULUS | 0.00002 | 0.00047 | 0.57561 | 0.64468 | 1.24782 | 153 |
| GO_RESPONSE_TO_TOXIC_SUBSTANCE | 0.00011 | 0.00235 | 0.53843 | 0.63892 | 1.23203 | 134 |
| GO_RESPONSE_TO_TUMOR_NECROSIS_FACTOR | 0.00000 | 0.00000 | 0.70498 | 0.66331 | 1.28815 | 175 |
| GO_RESPONSE_TO_VIRUS | 0.00261 | 0.02910 | 0.43171 | 0.59428 | 1.15408 | 175 |
| GO_RETINA_HOMEOSTASIS | 0.00034 | 0.00606 | 0.49849 | 0.75202 | 1.38362 | 34 |
| GO_RIBONUCLEOPROTEIN_COMPLEX_BINDING | 0.00000 | 0.00001 | 0.69013 | 0.69839 | 1.34459 | 114 |
| GO_RIBONUCLEOPROTEIN_COMPLEX_BIOGENESIS | 0.00000 | 0.00000 | | 0.68714 | 1.35448 | 393 |
| GO_RIBONUCLEOPROTEIN_COMPLEX_SUBUNIT_ORGANIZATION | 0.00000 | 0.00000 | 0.81404 | 0.69112 | 1.34183 | 173 |
| GO_RIBONUCLEOPROTEIN_GRANULE | 0.00000 | 0.00000 | 0.71951 | 0.66271 | 1.28875 | 186 |
| GO_RIBONUCLEOSIDE_TRIPHOSPHATE_BIOSYNTHETIC_PROCESS | 0.00009 | 0.00210 | 0.53843 | 0.77194 | 1.42026 | 34 |
| GO_RIBONUCLEOSIDE_TRIPHOSPHATE_METABOLIC_PROCESS | 0.00013 | 0.00280 | 0.51885 | 0.73558 | 1.37065 | 46 |
| GO_RIBOSE_PHOSPHATE_BIOSYNTHETIC_PROCESS | 0.00002 | 0.00056 | 0.57561 | 0.65814 | 1.26821 | 124 |
| GO_RIBOSE_PHOSPHATE_METABOLIC_PROCESS | 0.00000 | 0.00000 | 0.74774 | 0.63837 | 1.25001 | 285 |
| GO_RIBOSOMAL_LARGE_SUBUNIT_BIOGENESIS | 0.00001 | 0.00031 | 0.59333 | 0.72435 | 1.36807 | 67 |
| GO_RIBOSOMAL_SMALL_SUBUNIT_ASSEMBLY | 0.00405 | 0.04011 | 0.40702 | 0.78074 | 1.38865 | 18 |
| GO_RIBOSOMAL_SMALL_SUBUNIT_BINDING | 0.00532 | 0.04803 | 0.40702 | 0.77341 | 1.36899 | 17 |
| GO_RIBOSOMAL_SMALL_SUBUNIT_BIOGENESIS | 0.00005 | 0.00138 | 0.55733 | 0.70510 | 1.33163 | 68 |
| GO_RIBOSOMAL_SUBUNIT | 0.00000 | 0.00000 | | 0.76292 | 1.48089 | 172 |
| GO_RIBOSOME | 0.00000 | 0.00000 | | 0.75411 | 1.46984 | 208 |
| GO_RIBOSOME_ASSEMBLY | 0.00001 | 0.00026 | 0.59333 | 0.74485 | 1.40144 | 56 |
| GO_RIBOSOME_BIOGENESIS | 0.00000 | 0.00000 | | 0.68899 | 1.34833 | 277 |
| GO_RNA_3_END_PROCESSING | 0.00002 | 0.00046 | 0.57561 | 0.65576 | 1.26361 | 128 |
| GO_RNA_CATABOLIC_PROCESS | 0.00000 | 0.00000 | | 0.70701 | 1.39073 | 341 |
| GO_RNA_DEPENDENT_DNA_BIOSYNTHETIC_PROCESS | 0.00003 | 0.00081 | 0.57561 | 0.71669 | 1.34965 | 59 |
| GO_RNA_EXPORT_FROM_NUCLEUS | 0.00000 | 0.00016 | 0.61053 | 0.66306 | 1.27817 | 130 |
| GO_RNA_LOCALIZATION | 0.00000 | 0.00000 | 0.76146 | 0.66638 | 1.29886 | 208 |
| GO_RNA_SPLICING | 0.00000 | 0.00000 | | 0.63512 | 1.25129 | 384 |
| GO_RNA_SPLICING_VIA_TRANSESTERIFICATION_REACTIONS | 0.00000 | 0.00000 | 0.80122 | 0.64478 | 1.26586 | 311 |
| GO_ROUGH_ENDOPLASMIC_RETICULUM | 0.00014 | 0.00293 | 0.51885 | 0.70363 | 1.32507 | 59 |
| GO_RRNA_BINDING | 0.00000 | 0.00001 | 0.69013 | 0.77638 | 1.45941 | 54 |
| GO_RRNA_METABOLIC_PROCESS | 0.00000 | 0.00000 | | 0.68430 | 1.33386 | 209 |
| GO_SCF_DEPENDENT_PROTEASOMAL_UBIQUITIN_DEPENDENT_ | 0.00000 | 0.00008 | 0.62726 | 0.71957 | 1.36498 | 76 |
| GO_SECRETORY_GRANULE | 0.00000 | 0.00000 | | 0.62978 | 1.24363 | 481 |
| GO_SECRETORY_GRANULE_MEMBRANE | 0.00032 | 0.00575 | 0.49849 | 0.61375 | 1.19134 | 172 |
| GO_SERINE_TYPE_ENDOPEPTIDASE_INHIBITOR_ACTIVITY | 0.00038 | 0.00640 | 0.49849 | 0.77388 | 1.41447 | 28 |
| GO_SEXUAL_REPRODUCTION | 0.00472 | 0.04480 | 0.40702 | 0.55748 | 1.10026 | 439 |
| GO_SIDE_OF_MEMBRANE | 0.00495 | 0.04573 | 0.40702 | 0.57741 | 1.12751 | 235 |
| GO_SIGNAL_TRANSDUCTION_BY_PROTEIN_PHOSPHORYLATION | 0.00224 | 0.02581 | 0.43171 | 0.55690 | 1.09975 | 500 |
| GO_SINGLE_FERTILIZATION | 0.00000 | 0.00002 | 0.67496 | 0.76307 | 1.43842 | 61 |
| GO_SINGLE_STRANDED_RNA_BINDING | 0.00001 | 0.00042 | 0.59333 | 0.71960 | 1.35864 | 66 |
| GO_SISTER_CHROMATID_SEGREGATION | 0.00042 | 0.00689 | 0.49849 | 0.60926 | 1.18209 | 170 |
| GO_SKIN_DEVELOPMENT | 0.00076 | 0.01140 | 0.47727 | 0.61452 | 1.18945 | 153 |
| GO_SMALL_MOLECULE_BIOSYNTHETIC_PROCESS | 0.00001 | 0.00021 | 0.61053 | 0.58728 | 1.15957 | 459 |
| GO_SMALL_NUCLEAR_RIBONUCLEOPROTEIN_COMPLEX | 0.00042 | 0.00689 | 0.49849 | 0.69542 | 1.30936 | 57 |
| GO_SMALL_RIBOSOMAL_SUBUNIT | 0.00000 | 0.00001 | 0.69013 | 0.75767 | 1.43100 | 67 |
| GO_SM_LIKE_PROTEIN_FAMILY_COMPLEX | 0.00145 | 0.01879 | 0.45506 | 0.66792 | 1.26149 | 67 |
| GO_SNORNA_BINDING | 0.00038 | 0.00646 | 0.49849 | 0.76224 | 1.39614 | 29 |
| GO_SPERM_EGG_RECOGNITION | 0.00000 | 0.00002 | 0.65944 | 0.87469 | 1.59087 | 26 |
| GO_SPINDLE_MIDZONE | 0.00361 | 0.03657 | 0.43171 | 0.73316 | 1.33859 | 27 |
| GO_SPINDLE_ORGANIZATION | 0.00272 | 0.02986 | 0.43171 | 0.60076 | 1.16362 | 159 |
| GO_SPLICEOSOMAL_COMPLEX | 0.00000 | 0.00008 | 0.62726 | 0.65256 | 1.26657 | 171 |
| GO_SPLICEOSOMAL_TRI_SNRNP_COMPLEX | 0.00168 | 0.02098 | 0.45506 | 0.73474 | 1.34835 | 30 |
| GO_STEM_CELL_DIFFERENTIATION | 0.00000 | 0.00000 | 0.82666 | 0.70006 | 1.35554 | 158 |
| GO_STEROID_BIOSYNTHETIC_PROCESS | 0.00465 | 0.04430 | 0.40702 | 0.61242 | 1.17992 | 125 |
| GO_STEROID_METABOLIC_PROCESS | 0.00313 | 0.03310 | 0.43171 | 0.59525 | 1.15748 | 185 |
| GO_STEROL_BIOSYNTHETIC_PROCESS | 0.00005 | 0.00120 | 0.55733 | 0.71876 | 1.35584 | 62 |
| GO_STEROL_METABOLIC_PROCESS | 0.00003 | 0.00073 | 0.57561 | 0.66485 | 1.27854 | 109 |
| GO_STRESS_GRANULE_ASSEMBLY | 0.00465 | 0.04430 | 0.40702 | 0.77288 | 1.38921 | 21 |
| GO_STRUCTURAL_CONSTITUENT_OF_RIBOSOME | 0.00000 | 0.00000 | | 0.76380 | 1.47839 | 153 |

| | | | | | | |
|---|---|---|---|---|---|---|
| GO_STRUCTURAL_MOLECULE_ACTIVITY | 0.00000 | 0.00000 | | 0.71760 | 1.41436 | 391 |
| GO_SUPEROXIDE_METABOLIC_PROCESS | 0.00272 | 0.02986 | 0.43171 | 0.72028 | 1.32659 | 35 |
| GO_TELOMERASE_RNA_LOCALIZATION | 0.00054 | 0.00857 | 0.47727 | 0.81801 | 1.46148 | 19 |
| GO_TELOMERE_MAINTENANCE_VIA_TELOMERE_LENGTHENING | 0.00005 | 0.00138 | 0.55733 | 0.70715 | 1.33512 | 66 |
| GO_TELOMERE_ORGANIZATION | 0.00159 | 0.02016 | 0.45506 | 0.61666 | 1.18834 | 127 |
| GO_TERTIARY_GRANULE | 0.00008 | 0.00186 | 0.53843 | 0.66372 | 1.27005 | 91 |
| GO_TERTIARY_GRANULE_MEMBRANE | 0.00102 | 0.01453 | 0.45506 | 0.71940 | 1.33244 | 40 |
| GO_THREONINE_TYPE_ENDOPEPTIDASE_ACTIVITY | 0.00000 | 0.00000 | 0.71951 | 0.93456 | 1.65424 | 17 |
| GO_THREONINE_TYPE_PEPTIDASE_ACTIVITY | 0.00000 | 0.00004 | 0.65944 | 0.89807 | 1.61424 | 21 |
| GO_TISSUE_MORPHOGENESIS | 0.00000 | 0.00006 | 0.64355 | 0.59467 | 1.17150 | 380 |
| GO_TOXIN_TRANSPORT | 0.00064 | 0.00972 | 0.47727 | 0.77713 | 1.41343 | 26 |
| GO_TRANSCRIPTION_COUPLED_NUCLEOTIDE_EXCISION_REPAIR | 0.00231 | 0.02642 | 0.43171 | 0.65429 | 1.23840 | 71 |
| GO_TRANSFERRIN_TRANSPORT | 0.00141 | 0.01850 | 0.45506 | 0.75134 | 1.37179 | 27 |
| GO_TRANSLATIONAL_ELONGATION | 0.00000 | 0.00000 | | 0.75263 | 1.45028 | 124 |
| GO_TRANSLATIONAL_INITIATION | 0.00000 | 0.00000 | | 0.75139 | 1.45839 | 171 |
| GO_TRANSLATIONAL_TERMINATION | 0.00000 | 0.00000 | 0.81404 | 0.75110 | 1.44066 | 101 |
| GO_TRANSLATION_ELONGATION_FACTOR_ACTIVITY | 0.00390 | 0.03895 | 0.43171 | 0.80505 | 1.41260 | 15 |
| GO_TRANSLATION_FACTOR_ACTIVITY_RNA_BINDING | 0.00000 | 0.00000 | 0.74774 | 0.77244 | 1.46085 | 70 |
| GO_TRANSLATION_INITIATION_FACTOR_ACTIVITY | 0.00000 | 0.00011 | 0.62726 | 0.77773 | 1.44952 | 45 |
| GO_TRANSLATION_PREINITIATION_COMPLEX | 0.00342 | 0.03521 | 0.43171 | 0.79777 | 1.40694 | 16 |
| GO_TRANSLATION_REGULATOR_ACTIVITY | 0.00000 | 0.00000 | 0.83909 | 0.74000 | 1.42403 | 111 |
| GO_TRANSLATION_REGULATOR_ACTIVITY_NUCLEIC_ACID_BIND | 0.00000 | 0.00000 | 0.83909 | 0.76441 | 1.45959 | 87 |
| GO_TRANSMEMBRANE_RECEPTOR_PROTEIN_TYROSINE_KINASE | 0.00168 | 0.02098 | 0.45506 | 0.56326 | 1.11076 | 411 |
| GO_TRICARBOXYLIC_ACID_CYCLE | 0.00159 | 0.02016 | 0.45506 | 0.73558 | 1.35097 | 33 |
| GO_TRNA_TRANSPORT | 0.00342 | 0.03521 | 0.43171 | 0.71247 | 1.31220 | 35 |
| GO_TUMOR_NECROSIS_FACTOR_MEDIATED_SIGNALING_PATHW | 0.00000 | 0.00000 | 0.80122 | 0.73921 | 1.42138 | 108 |
| GO_T_CELL_RECEPTOR_SIGNALING_PATHWAY | 0.00000 | 0.00001 | 0.70498 | 0.69185 | 1.33314 | 119 |
| GO_U12_TYPE_SPLICEOSOMAL_COMPLEX | 0.00487 | 0.04559 | 0.40702 | 0.73423 | 1.33541 | 26 |
| GO_U2_SNRNP | 0.00141 | 0.01850 | 0.45506 | 0.80259 | 1.42065 | 17 |
| GO_U2_TYPE_CATALYTIC_STEP_2_SPLICEOSOME | 0.00268 | 0.02961 | 0.43171 | 0.72225 | 1.32288 | 29 |
| GO_U2_TYPE_SPLICEOSOMAL_COMPLEX | 0.00033 | 0.00595 | 0.49849 | 0.66590 | 1.26925 | 84 |
| GO_UBIQUITIN_LIKE_PROTEIN_LIGASE_BINDING | 0.00000 | 0.00007 | 0.62726 | 0.62616 | 1.22416 | 251 |
| GO_UNFOLDED_PROTEIN_BINDING | 0.00000 | 0.00001 | 0.69013 | 0.72112 | 1.37766 | 88 |
| GO_VACUOLAR_LUMEN | 0.00016 | 0.00319 | 0.51885 | 0.65416 | 1.25772 | 107 |
| GO_VESICLE_BUDDING_FROM_MEMBRANE | 0.00480 | 0.04537 | 0.40702 | 0.62411 | 1.19353 | 90 |
| GO_VESICLE_LUMEN | 0.00000 | 0.00000 | 0.77494 | 0.67028 | 1.30631 | 207 |
| GO_VESICLE_MEMBRANE | 0.00001 | 0.00026 | 0.59333 | 0.58430 | 1.15387 | 476 |
| GO_VIRAL_GENE_EXPRESSION | 0.00000 | 0.00000 | | 0.70963 | 1.37745 | 172 |
| GO_VIRAL_LIFE_CYCLE | 0.00000 | 0.00001 | 0.69013 | 0.63985 | 1.24950 | 238 |
| GO_VIRION_ASSEMBLY | 0.00457 | 0.04386 | 0.40702 | 0.69689 | 1.28528 | 36 |
| HP_ABNORMALITY_OF_ACID_BASE_HOMEOSTASIS | 0.00000 | 0.00010 | 0.62726 | 0.62283 | 1.21827 | 261 |
| HP_ABNORMALITY_OF_BLOOD_CIRCULATION | 0.00502 | 0.04608 | 0.40702 | 0.59228 | 1.14883 | 167 |
| HP_ABNORMALITY_OF_CARDIOVASCULAR_SYSTEM_ELECTROPH | 0.00011 | 0.00237 | 0.53843 | 0.60202 | 1.17758 | 261 |
| HP_ABNORMALITY_OF_SKIN_ADNEXA_PHYSIOLOGY | 0.00242 | 0.02748 | 0.43171 | 0.61294 | 1.18094 | 118 |
| HP_ABNORMALITY_OF_THE_ANKLES | 0.00313 | 0.03310 | 0.43171 | 0.65369 | 1.23418 | 63 |
| HP_ABNORMALITY_OF_THE_CEREBRAL_VASCULATURE | 0.00357 | 0.03638 | 0.43171 | 0.59764 | 1.15722 | 158 |
| HP_ABNORMALITY_OF_THE_CEREBROSPINAL_FLUID | 0.00030 | 0.00549 | 0.49849 | 0.58221 | 1.14439 | 325 |
| HP_ABNORMALITY_OF_THE_MITOCHONDRION | 0.00126 | 0.01715 | 0.45506 | 0.62149 | 1.19764 | 120 |
| HP_ABNORMALITY_OF_THE_OPTIC_DISC | 0.00198 | 0.02370 | 0.43171 | 0.56624 | 1.11727 | 432 |
| HP_ABNORMALITY_OF_THE_OPTIC_NERVE | 0.00036 | 0.00622 | 0.49849 | 0.56723 | 1.12034 | 486 |
| HP_ABNORMAL_ABDOMEN_MORPHOLOGY | 0.00089 | 0.01291 | 0.47727 | 0.56424 | 1.11364 | 441 |
| HP_ABNORMAL_ACTIVITY_OF_MITOCHONDRIAL_RESPIRATORY_ | 0.00014 | 0.00300 | 0.51885 | 0.70303 | 1.32524 | 61 |
| HP_ABNORMAL_BASAL_GANGLIA_MRI_SIGNAL_INTENSITY | 0.00179 | 0.02183 | 0.45506 | 0.71597 | 1.32046 | 36 |
| HP_ABNORMAL_BRAINSTEM_MRI_SIGNAL_INTENSITY | 0.00398 | 0.03962 | 0.40702 | 0.71528 | 1.31370 | 33 |
| HP_ABNORMAL_CELLULAR_PHENOTYPE | 0.00209 | 0.02470 | 0.43171 | 0.56937 | 1.11992 | 334 |
| HP_ABNORMAL_CIRCULATING_CARBOHYDRATE_CONCENTRATIC | 0.00194 | 0.02331 | 0.45506 | 0.69518 | 1.29539 | 46 |
| HP_ABNORMAL_CNS_MYELINATION | 0.00014 | 0.00300 | 0.51885 | 0.62318 | 1.20954 | 171 |
| HP_ABNORMAL_CSF_METABOLITE_LEVEL | 0.00001 | 0.00027 | 0.59333 | 0.70531 | 1.34409 | 82 |
| HP_ABNORMAL_ELECTROPHYSIOLOGY_OF_SINOATRIAL_NODE_ | 0.00029 | 0.00536 | 0.49849 | 0.73172 | 1.35725 | 41 |
| HP_ABNORMAL_ENZYME_COENZYME_ACTIVITY | 0.00561 | 0.04971 | 0.40702 | 0.60490 | 1.16567 | 123 |
| HP_ABNORMAL_ERYTHROCYTE_MORPHOLOGY | 0.00001 | 0.00023 | 0.61053 | 0.58835 | 1.15904 | 380 |
| HP_ABNORMAL_HEART_VALVE_PHYSIOLOGY | 0.00405 | 0.04011 | 0.40702 | 0.59004 | 1.14474 | 169 |
| HP_ABNORMAL_MITOCHONDRIA_IN_MUSCLE_TISSUE | 0.00075 | 0.01118 | 0.47727 | 0.74633 | 1.36700 | 29 |
| HP_ABNORMAL_MYELINATION | 0.00007 | 0.00169 | 0.53843 | 0.59258 | 1.16479 | 324 |
| HP_ABNORMAL_MYOCARDIUM_MORPHOLOGY | 0.00008 | 0.00192 | 0.53843 | 0.58647 | 1.15365 | 340 |
| HP_ABNORMAL_RENAL_PHYSIOLOGY | 0.00450 | 0.04342 | 0.40702 | 0.55484 | 1.09317 | 385 |
| HP_ABNORMAL_TRICUSPID_VALVE_PHYSIOLOGY | 0.00231 | 0.02642 | 0.43171 | 0.72968 | 1.34029 | 31 |
| HP_ABNORMAL_VITREOUS_HUMOR_MORPHOLOGY | 0.00181 | 0.02201 | 0.45506 | 0.79632 | 1.40955 | 17 |
| HP_ACUTE_LEUKEMIA | 0.00013 | 0.00276 | 0.51885 | 0.68145 | 1.29219 | 75 |
| HP_ANEMIA_OF_INADEQUATE_PRODUCTION | 0.00561 | 0.04971 | 0.40702 | 0.61298 | 1.17855 | 107 |

| | | | | | | |
|---|---|---|---|---|---|---|
| HP_APHASIA | 0.00213 | 0.02490 | 0.43171 | 0.72858 | 1.33704 | 30 |
| HP_BLINDNESS | 0.00018 | 0.00355 | 0.51885 | 0.61672 | 1.19768 | 175 |
| HP_BONE_MARROW_HYPERCELLULARITY | 0.00045 | 0.00725 | 0.49849 | 0.82169 | 1.46806 | 19 |
| HP_CHOREOATHETOSIS | 0.00561 | 0.04971 | 0.40702 | 0.64956 | 1.22631 | 64 |
| HP_CHRONIC_INFECTION | 0.00121 | 0.01658 | 0.45506 | 0.81916 | 1.44468 | 16 |
| HP_CONGESTIVE_HEART_FAILURE | 0.00495 | 0.04573 | 0.40702 | 0.60734 | 1.16929 | 114 |
| HP_CRANIAL_NERVE_COMPRESSION | 0.00331 | 0.03441 | 0.43171 | 0.75830 | 1.36989 | 23 |
| HP_DECREASED_ACTIVITY_OF_MITOCHONDRIAL_COMPLEX_I | 0.00283 | 0.03081 | 0.43171 | 0.71940 | 1.32496 | 35 |
| HP_DECREASED_ACTIVITY_OF_THE_PYRUVATE_DEHYDROGENAS | 0.00062 | 0.00949 | 0.47727 | 0.75390 | 1.38478 | 31 |
| HP_DEPRESSED_NASAL_RIDGE | 0.00041 | 0.00681 | 0.49849 | 0.66218 | 1.26189 | 82 |
| HP_DYSPNEA | 0.00250 | 0.02807 | 0.43171 | 0.58107 | 1.13348 | 224 |
| HP_DYSTONIA | 0.00069 | 0.01043 | 0.47727 | 0.58101 | 1.13736 | 279 |
| HP_EMOTIONAL_LABILITY | 0.00016 | 0.00321 | 0.51885 | 0.66683 | 1.27522 | 90 |
| HP_EPISODIC_ATAXIA | 0.00487 | 0.04559 | 0.40702 | 0.79934 | 1.40259 | 15 |
| HP_FASCICULATIONS | 0.00130 | 0.01761 | 0.45506 | 0.68128 | 1.28064 | 54 |
| HP_FATIGUE | 0.00042 | 0.00689 | 0.49849 | 0.59902 | 1.16764 | 212 |
| HP_FETAL_DISTRESS | 0.00179 | 0.02183 | 0.45506 | 0.72928 | 1.33973 | 32 |
| HP_FLUSHING | 0.00561 | 0.04971 | 0.40702 | 0.74782 | 1.33686 | 20 |
| HP_FOCAL_T2_HYPERINTENSE_BRAINSTEM_LESION | 0.00159 | 0.02016 | 0.45506 | 0.74970 | 1.36879 | 27 |
| HP_HEADACHE | 0.00016 | 0.00319 | 0.51885 | 0.62537 | 1.21179 | 163 |
| HP_HEPATOMEGALY | 0.00108 | 0.01507 | 0.45506 | 0.57439 | 1.12990 | 335 |
| HP_HYPERTROPHIC_CARDIOMYOPATHY | 0.00002 | 0.00050 | 0.57561 | 0.62629 | 1.21908 | 194 |
| HP_IMPAIRED_SOCIAL_INTERACTIONS | 0.00055 | 0.00863 | 0.47727 | 0.66555 | 1.26058 | 72 |
| HP_INCREASED_CSF_LACTATE | 0.00001 | 0.00034 | 0.59333 | 0.71937 | 1.35857 | 68 |
| HP_INCREASED_SERUM_LACTATE | 0.00001 | 0.00028 | 0.59333 | 0.65837 | 1.26932 | 133 |
| HP_INCREASED_SERUM_PYRUVATE | 0.00110 | 0.01526 | 0.45506 | 0.73143 | 1.35198 | 37 |
| HP_INFANTILE_ONSET | 0.00487 | 0.04559 | 0.40702 | 0.56559 | 1.10939 | 300 |
| HP_INTRACRANIAL_CYSTIC_LESION | 0.00055 | 0.00863 | 0.47727 | 0.62760 | 1.20885 | 116 |
| HP_INTRAUTERINE_GROWTH_RETARDATION | 0.00305 | 0.03271 | 0.43171 | 0.56594 | 1.11393 | 350 |
| HP_LACTIC_ACIDOSIS | 0.00018 | 0.00355 | 0.51885 | 0.63312 | 1.22071 | 131 |
| HP_LETHARGY | 0.00084 | 0.01228 | 0.47727 | 0.63361 | 1.21845 | 109 |
| HP_LEUKODYSTROPHY | 0.00029 | 0.00529 | 0.49849 | 0.66598 | 1.26915 | 82 |
| HP_LEUKOENCEPHALOPATHY | 0.00008 | 0.00190 | 0.53843 | 0.64132 | 1.23667 | 134 |
| HP_LIPID_ACCUMULATION_IN_HEPATOCYTES | 0.00220 | 0.02548 | 0.43171 | 0.62794 | 1.20159 | 91 |
| HP_LOWER_LIMB_SPASTICITY | 0.00154 | 0.01974 | 0.45506 | 0.60929 | 1.17648 | 139 |
| HP_MACROCYTIC_ANEMIA | 0.00030 | 0.00542 | 0.49849 | 0.73108 | 1.35606 | 41 |
| HP_METABOLIC_ACIDOSIS | 0.00034 | 0.00599 | 0.49849 | 0.64831 | 1.24464 | 103 |
| HP_MICRONODULAR_CIRRHOSIS | 0.00324 | 0.03393 | 0.43171 | 0.78723 | 1.40018 | 18 |
| HP_MIGRAINE | 0.00028 | 0.00515 | 0.49849 | 0.67309 | 1.27635 | 75 |
| HP_MUSCLE_ABNORMALITY_RELATED_TO_MITOCHONDRIAL_DY | 0.00209 | 0.02470 | 0.43171 | 0.68258 | 1.27573 | 49 |
| HP_MUTISM | 0.00283 | 0.03081 | 0.43171 | 0.71937 | 1.32490 | 35 |
| HP_NAUSEA_AND_VOMITING | 0.00022 | 0.00417 | 0.51885 | 0.59383 | 1.16158 | 264 |
| HP_NEOPLASM_OF_THE_ENDOCRINE_SYSTEM | 0.00495 | 0.04573 | 0.40702 | 0.63166 | 1.20308 | 81 |
| HP_OPTIC_NEUROPATHY | 0.00135 | 0.01808 | 0.45506 | 0.73837 | 1.35610 | 33 |
| HP_PACE_OF_PROGRESSION | 0.00487 | 0.04559 | 0.40702 | 0.56814 | 1.11416 | 298 |
| HP_PALLOR | 0.00036 | 0.00622 | 0.49849 | 0.64861 | 1.24463 | 102 |
| HP_PARAPLEGIA | 0.00121 | 0.01658 | 0.45506 | 0.63584 | 1.22133 | 104 |
| HP_PARAPLEGIA_PARAPARESIS | 0.00036 | 0.00621 | 0.49849 | 0.62047 | 1.20040 | 150 |
| HP_PAROXYSMAL_INVOLUNTARY_EYE_MOVEMENTS | 0.00054 | 0.00849 | 0.47727 | 0.75650 | 1.38957 | 31 |
| HP_PEDIATRIC_ONSET | 0.00561 | 0.04971 | 0.40702 | 0.55915 | 1.10116 | 371 |
| HP_PIGMENTARY_RETINOPATHY | 0.00065 | 0.00981 | 0.47727 | 0.63260 | 1.21720 | 113 |
| HP_POOR_EYE_CONTACT | 0.00017 | 0.00341 | 0.51885 | 0.71279 | 1.33749 | 53 |
| HP_POOR_HEAD_CONTROL | 0.00000 | 0.00005 | 0.64355 | 0.67819 | 1.30665 | 118 |
| HP_PROGRESSIVE | 0.00194 | 0.02331 | 0.45506 | 0.59790 | 1.16273 | 186 |
| HP_PROGRESSIVE_CEREBELLAR_ATAXIA | 0.00364 | 0.03677 | 0.43171 | 0.64788 | 1.22325 | 65 |
| HP_PROGRESSIVE_SPASTIC_PARAPLEGIA | 0.00253 | 0.02838 | 0.43171 | 0.69655 | 1.29822 | 45 |
| HP_PROXIMAL_TUBULOPATHY | 0.00141 | 0.01850 | 0.45506 | 0.72579 | 1.33535 | 34 |
| HP_PTOSIS | 0.00472 | 0.04480 | 0.40702 | 0.55582 | 1.09637 | 418 |
| HP_REDUCED_CONSCIOUSNESS_CONFUSION | 0.00007 | 0.00164 | 0.53843 | 0.61227 | 1.19295 | 205 |
| HP_REDUCED_VISUAL_ACUITY | 0.00053 | 0.00837 | 0.47727 | 0.58618 | 1.14774 | 283 |
| HP_RENAL_TUBULAR_DYSFUNCTION | 0.00013 | 0.00284 | 0.51885 | 0.68108 | 1.29150 | 75 |
| HP_RESPIRATORY_DISTRESS | 0.00108 | 0.01507 | 0.45506 | 0.62693 | 1.20762 | 115 |
| HP_RESPIRATORY_INSUFFICIENCY | 0.00000 | 0.00010 | 0.62726 | 0.60536 | 1.18998 | 326 |
| HP_SINUS_TACHYCARDIA | 0.00213 | 0.02490 | 0.43171 | 0.77203 | 1.39128 | 22 |
| HP_SPARSE_EYELASHES | 0.00150 | 0.01943 | 0.45506 | 0.71355 | 1.32161 | 40 |
| HP_SPARSE_OR_ABSENT_EYELASHES | 0.00220 | 0.02548 | 0.43171 | 0.67331 | 1.26567 | 54 |
| HP_SPASTIC_PARAPLEGIA | 0.00098 | 0.01407 | 0.45506 | 0.64059 | 1.22712 | 96 |
| HP_THROMBOCYTOSIS | 0.00413 | 0.04065 | 0.40702 | 0.75210 | 1.35870 | 23 |
| HP_TONGUE_FASCICULATIONS | 0.00457 | 0.04386 | 0.40702 | 0.80056 | 1.40472 | 15 |
| HP_TRANSIENT_ISCHEMIC_ATTACK | 0.00480 | 0.04537 | 0.40702 | 0.74902 | 1.35809 | 25 |

| | | | | | | |
|---|---|---|---|---|---|---|
| HP_VERTIGO | 0.00002 | 0.00063 | 0.57561 | 0.72114 | 1.35844 | 60 |
| HP_VESTIBULAR_DYSFUNCTION | 0.00028 | 0.00522 | 0.49849 | 0.67279 | 1.27577 | 75 |
| HP_VOMITING | 0.00072 | 0.01082 | 0.47727 | 0.61459 | 1.18792 | 147 |
| HP_WEAKNESS_DUE_TO_UPPER_MOTOR_NEURON_DYSFUNCTI | 0.00532 | 0.04803 | 0.40702 | 0.56829 | 1.11186 | 272 |
| HP_WEAKNESS_OF_MUSCLES_OF_RESPIRATION | 0.00163 | 0.02050 | 0.45506 | 0.66272 | 1.25123 | 63 |
| **Cluster 1** | | | | | | |
| pathway | pval | padj | log2err | ES | NES | size |
| GO_ACTIVATION_OF_IMMUNE_RESPONSE | 0.00000 | 0.00000 | 0.74774 | 0.65038 | 1.26999 | 235 |
| GO_ACTIVATION_OF_INNATE_IMMUNE_RESPONSE | 0.00000 | 0.00001 | 0.69013 | 0.71967 | 1.37861 | 96 |
| GO_AEROBIC_RESPIRATION | 0.00000 | 0.00000 | 0.73376 | 0.76280 | 1.44478 | 72 |
| GO_ALCOHOL_BIOSYNTHETIC_PROCESS | 0.00032 | 0.00572 | 0.49849 | 0.64679 | 1.24367 | 108 |
| GO_ALCOHOL_METABOLIC_PROCESS | 0.00364 | 0.03677 | 0.43171 | 0.58112 | 1.13424 | 228 |
| GO_AMEBOIDAL_TYPE_CELL_MIGRATION | 0.00457 | 0.04386 | 0.40702 | 0.57755 | 1.12854 | 247 |
| GO_AMIDE_BINDING | 0.00014 | 0.00297 | 0.51885 | 0.61533 | 1.19780 | 199 |
| GO_AMINE_METABOLIC_PROCESS | 0.00000 | 0.00000 | 0.76146 | 0.73732 | 1.41097 | 92 |
| GO_AMYLOID_BETA_CLEARANCE | 0.00561 | 0.04971 | 0.40702 | 0.74081 | 1.33976 | 24 |
| GO_ANAPHASE_PROMOTING_COMPLEX_DEPENDENT_CATABOL | 0.00000 | 0.00000 | 0.76146 | 0.77107 | 1.46112 | 73 |
| GO_ANATOMICAL_STRUCTURE_HOMEOSTASIS | 0.00020 | 0.00390 | 0.51885 | 0.59307 | 1.16021 | 271 |
| GO_ANIMAL_ORGAN_REGENERATION | 0.00012 | 0.00269 | 0.51885 | 0.71251 | 1.33510 | 52 |
| GO_ANTIGEN_PROCESSING_AND_PRESENTATION | 0.00000 | 0.00000 | 0.74774 | 0.68785 | 1.33064 | 149 |
| GO_ANTIGEN_PROCESSING_AND_PRESENTATION_OF_EXOGENO | 0.00000 | 0.00000 | 0.76146 | 0.79644 | 1.50237 | 62 |
| GO_ANTIGEN_PROCESSING_AND_PRESENTATION_OF_PEPTIDE_ | 0.00000 | 0.00000 | 0.74774 | 0.70107 | 1.35144 | 130 |
| GO_ANTIGEN_PROCESSING_AND_PRESENTATION_OF_PEPTIDE_ | 0.00000 | 0.00000 | 0.78819 | 0.77566 | 1.47012 | 74 |
| GO_ANTIGEN_RECEPTOR_MEDIATED_SIGNALING_PATHWAY | 0.00000 | 0.00009 | 0.62726 | 0.66362 | 1.28080 | 136 |
| GO_APICAL_PART_OF_CELL | 0.00135 | 0.01808 | 0.45506 | 0.58926 | 1.14966 | 220 |
| GO_APOPTOTIC_SIGNALING_PATHWAY | 0.00006 | 0.00152 | 0.53843 | 0.58638 | 1.15586 | 393 |
| GO_ATPASE_ACTIVITY | 0.00050 | 0.00802 | 0.47727 | 0.58216 | 1.14385 | 316 |
| GO_ATPASE_ACTIVITY_COUPLED_TO_TRANSMEMBRANE_MOVE | 0.00420 | 0.04132 | 0.40702 | 0.78005 | 1.38741 | 18 |
| GO_ATP_BIOSYNTHETIC_PROCESS | 0.00013 | 0.00276 | 0.51885 | 0.83897 | 1.50801 | 21 |
| GO_ATP_METABOLIC_PROCESS | 0.00000 | 0.00000 | | 0.72178 | 1.40814 | 219 |
| GO_ATP_SYNTHESIS_COUPLED_ELECTRON_TRANSPORT | 0.00000 | 0.00000 | | 0.80353 | 1.52425 | 76 |
| GO_AZUROPHIL_GRANULE | 0.00025 | 0.00466 | 0.49849 | 0.64781 | 1.24551 | 107 |
| GO_AZUROPHIL_GRANULE_LUMEN | 0.00405 | 0.04011 | 0.40702 | 0.65625 | 1.23793 | 62 |
| GO_BASAL_PART_OF_CELL | 0.00509 | 0.04656 | 0.40702 | 0.68515 | 1.27697 | 45 |
| GO_BASAL_PLASMA_MEMBRANE | 0.00346 | 0.03548 | 0.43171 | 0.71184 | 1.31104 | 35 |
| GO_BASEMENT_MEMBRANE | 0.00279 | 0.03056 | 0.43171 | 0.68935 | 1.28452 | 46 |
| GO_BASOLATERAL_PLASMA_MEMBRANE | 0.00011 | 0.00247 | 0.53843 | 0.63209 | 1.21993 | 136 |
| GO_BINDING_OF_SPERM_TO_ZONA_PELLUCIDA | 0.00000 | 0.00006 | 0.64355 | 0.90095 | 1.60966 | 19 |
| GO_BLOOD_MICROPARTICLE | 0.00000 | 0.00000 | 0.70498 | 0.81982 | 1.52763 | 46 |
| GO_BRUSH_BORDER | 0.00060 | 0.00923 | 0.47727 | 0.66897 | 1.26790 | 74 |
| GO_CADHERIN_BINDING | 0.00000 | 0.00000 | | 0.68759 | 1.34435 | 253 |
| GO_CALCIUM_DEPENDENT_PHOSPHOLIPID_BINDING | 0.00152 | 0.01958 | 0.45506 | 0.75021 | 1.37120 | 28 |
| GO_CALCIUM_DEPENDENT_PROTEIN_BINDING | 0.00009 | 0.00211 | 0.53843 | 0.74254 | 1.38394 | 45 |
| GO_CANONICAL_WNT_SIGNALING_PATHWAY | 0.00004 | 0.00109 | 0.55733 | 0.61740 | 1.20458 | 222 |
| GO_CARBOHYDRATE_CATABOLIC_PROCESS | 0.00135 | 0.01808 | 0.45506 | 0.61179 | 1.18186 | 144 |
| GO_CATALYTIC_STEP_2_SPLICEOSOME | 0.00001 | 0.00022 | 0.61053 | 0.70290 | 1.33765 | 80 |
| GO_CATION_TRANSMEMBRANE_TRANSPORT | 0.00327 | 0.03409 | 0.43171 | 0.56060 | 1.10531 | 406 |
| GO_CATION_TRANSMEMBRANE_TRANSPORTER_ACTIVITY | 0.00097 | 0.01387 | 0.47727 | 0.58196 | 1.13846 | 265 |
| GO_CELLULAR_AMINO_ACID_METABOLIC_PROCESS | 0.00002 | 0.00060 | 0.57561 | 0.61438 | 1.19930 | 231 |
| GO_CELLULAR_COMPONENT_DISASSEMBLY | 0.00000 | 0.00000 | | 0.64070 | 1.26342 | 408 |
| GO_CELLULAR_ION_HOMEOSTASIS | 0.00027 | 0.00493 | 0.49849 | 0.58658 | 1.15340 | 328 |
| GO_CELLULAR_IRON_ION_HOMEOSTASIS | 0.00327 | 0.03409 | 0.43171 | 0.68360 | 1.27374 | 47 |
| GO_CELLULAR_KETONE_METABOLIC_PROCESS | 0.00000 | 0.00010 | 0.62726 | 0.65358 | 1.26726 | 164 |
| GO_CELLULAR_MONOVALENT_INORGANIC_CATION_HOMEOSTA | 0.00179 | 0.02183 | 0.45506 | 0.64747 | 1.22821 | 76 |
| GO_CELLULAR_NITROGEN_COMPOUND_CATABOLIC_PROCESS | 0.00000 | 0.00000 | | 0.67915 | 1.34096 | 456 |
| GO_CELLULAR_PROTEIN_COMPLEX_DISASSEMBLY | 0.00000 | 0.00000 | | 0.69566 | 1.35184 | 179 |
| GO_CELLULAR_RESPIRATION | 0.00000 | 0.00000 | | 0.74205 | 1.43370 | 145 |
| GO_CELLULAR_RESPONSE_TO_HEAT | 0.00004 | 0.00109 | 0.55733 | 0.67186 | 1.28698 | 95 |
| GO_CELLULAR_RESPONSE_TO_HORMONE_STIMULUS | 0.00309 | 0.03299 | 0.43171 | 0.56453 | 1.11191 | 374 |
| GO_CELLULAR_RESPONSE_TO_ORGANIC_CYCLIC_COMPOUND | 0.00502 | 0.04608 | 0.40702 | 0.56512 | 1.11134 | 329 |
| GO_CELLULAR_RESPONSE_TO_OXYGEN_LEVELS | 0.00000 | 0.00000 | | 0.71461 | 1.38224 | 151 |
| GO_CELL_ACTIVATION_INVOLVED_IN_IMMUNE_RESPONSE | 0.00000 | 0.00000 | | 0.63924 | 1.26108 | 422 |
| GO_CELL_ADHESION_MEDIATOR_ACTIVITY | 0.00157 | 0.02009 | 0.45506 | 0.71837 | 1.32488 | 36 |
| GO_CELL_ADHESION_MOLECULE_BINDING | 0.00000 | 0.00000 | | 0.67109 | 1.32112 | 353 |
| GO_CELL_BODY | 0.00004 | 0.00109 | 0.55733 | 0.59973 | 1.17541 | 293 |
| GO_CELL_CELL_RECOGNITION | 0.00000 | 0.00006 | 0.64355 | 0.82001 | 1.51026 | 35 |
| GO_CELL_CELL_SIGNALING_BY_WNT | 0.00004 | 0.00109 | 0.55733 | 0.59042 | 1.16141 | 339 |
| GO_CELL_CYCLE_G1_S_PHASE_TRANSITION | 0.00376 | 0.03783 | 0.43171 | 0.58112 | 1.13378 | 220 |
| GO_CELL_CYCLE_G2_M_PHASE_TRANSITION | 0.00024 | 0.00459 | 0.51885 | 0.59807 | 1.16746 | 231 |
| GO_CELL_CYCLE_PHASE_TRANSITION | 0.00000 | 0.00001 | 0.69013 | 0.59417 | 1.17348 | 499 |

| | | | | | | |
|---|---|---|---|---|---|---|
| GO_CELL_DIVISION | 0.00003 | 0.00071 | 0.57561 | 0.58314 | 1.15094 | 449 |
| GO_CELL_KILLING | 0.00205 | 0.02440 | 0.43171 | 0.66727 | 1.25696 | 60 |
| GO_CELL_RECOGNITION | 0.00000 | 0.00000 | 0.73376 | 0.75953 | 1.43954 | 74 |
| GO_CELL_REDOX_HOMEOSTASIS | 0.00163 | 0.02050 | 0.45506 | 0.71772 | 1.32665 | 38 |
| GO_CELL_SUBSTRATE_JUNCTION | 0.00000 | 0.00000 | | 0.65742 | 1.29171 | 316 |
| GO_CELL_SURFACE | 0.00000 | 0.00000 | 0.73376 | 0.62572 | 1.23070 | 330 |
| GO_CELL_SURFACE_RECEPTOR_SIGNALING_PATHWAY_INVOLVEI | 0.00038 | 0.00640 | 0.49849 | 0.57762 | 1.13770 | 374 |
| GO_CHAPERONE_BINDING | 0.00495 | 0.04573 | 0.40702 | 0.62860 | 1.19935 | 86 |
| GO_CHAPERONE_COMPLEX | 0.00001 | 0.00028 | 0.59333 | 0.86337 | 1.55587 | 22 |
| GO_CHAPERONE_MEDIATED_PROTEIN_COMPLEX_ASSEMBLY | 0.00100 | 0.01430 | 0.45506 | 0.80787 | 1.44337 | 19 |
| GO_CHAPERONE_MEDIATED_PROTEIN_FOLDING | 0.00168 | 0.02098 | 0.45506 | 0.71110 | 1.32288 | 44 |
| GO_CHROMOSOMAL_REGION | 0.00018 | 0.00349 | 0.51885 | 0.59323 | 1.16180 | 286 |
| GO_CHROMOSOME_CENTROMERIC_REGION | 0.00172 | 0.02135 | 0.45506 | 0.60447 | 1.17248 | 168 |
| GO_CHROMOSOME_SEGREGATION | 0.00003 | 0.00087 | 0.55733 | 0.60463 | 1.18280 | 270 |
| GO_CLUSTER_OF_ACTIN_BASED_CELL_PROJECTIONS | 0.00074 | 0.01107 | 0.47727 | 0.64184 | 1.23044 | 99 |
| GO_COATED_VESICLE | 0.00213 | 0.02490 | 0.43171 | 0.59512 | 1.15846 | 199 |
| GO_COATED_VESICLE_MEMBRANE | 0.00060 | 0.00923 | 0.47727 | 0.63443 | 1.22207 | 115 |
| GO_COLLAGEN_BINDING | 0.00465 | 0.04430 | 0.40702 | 0.73471 | 1.33628 | 26 |
| GO_COLLAGEN_CONTAINING_EXTRACELLULAR_MATRIX | 0.00000 | 0.00006 | 0.64355 | 0.65358 | 1.26542 | 157 |
| GO_CONDENSED_CHROMOSOME | 0.00350 | 0.03580 | 0.43171 | 0.59394 | 1.15417 | 179 |
| GO_CONTRACTILE_FIBER | 0.00064 | 0.00972 | 0.47727 | 0.62217 | 1.19973 | 134 |
| GO_COPI_COATED_VESICLE | 0.00316 | 0.03344 | 0.43171 | 0.75688 | 1.37234 | 25 |
| GO_CORNIFICATION | 0.00027 | 0.00500 | 0.49849 | 0.77970 | 1.42512 | 28 |
| GO_CORTICAL_CYTOSKELETON | 0.00502 | 0.04608 | 0.40702 | 0.64388 | 1.21870 | 71 |
| GO_COTRANSLATIONAL_PROTEIN_TARGETING_TO_MEMBRANE | 0.00000 | 0.00000 | | 0.78223 | 1.49819 | 94 |
| GO_CYTOCHROME_COMPLEX | 0.00035 | 0.00610 | 0.49849 | 0.80796 | 1.45601 | 22 |
| GO_CYTOKINE_MEDIATED_SIGNALING_PATHWAY | 0.00000 | 0.00002 | 0.67496 | 0.60742 | 1.19735 | 392 |
| GO_CYTOPLASMIC_STRESS_GRANULE | 0.00014 | 0.00302 | 0.51885 | 0.69390 | 1.31009 | 63 |
| GO_CYTOPLASMIC_TRANSLATION | 0.00000 | 0.00001 | 0.70498 | 0.71773 | 1.37349 | 92 |
| GO_CYTOPLASMIC_TRANSLATIONAL_INITIATION | 0.00178 | 0.02182 | 0.45506 | 0.73517 | 1.35039 | 31 |
| GO_CYTOSOLIC_LARGE_RIBOSOMAL_SUBUNIT | 0.00000 | 0.00005 | 0.64355 | 0.77240 | 1.44935 | 53 |
| GO_CYTOSOLIC_RIBOSOME | 0.00000 | 0.00000 | | 0.77483 | 1.48437 | 97 |
| GO_CYTOSOLIC_SMALL_RIBOSOMAL_SUBUNIT | 0.00000 | 0.00010 | 0.62726 | 0.79384 | 1.47033 | 40 |
| GO_DETECTION_OF_STIMULUS | 0.00020 | 0.00393 | 0.51885 | 0.65208 | 1.25384 | 108 |
| GO_DETOXIFICATION | 0.00413 | 0.04065 | 0.40702 | 0.65399 | 1.23684 | 70 |
| GO_DEVELOPMENT_OF_PRIMARY_FEMALE_SEXUAL_CHARACTE | 0.00179 | 0.02183 | 0.45506 | 0.68410 | 1.28057 | 51 |
| GO_DICARBOXYLIC_ACID_METABOLIC_PROCESS | 0.00261 | 0.02910 | 0.43171 | 0.65541 | 1.23747 | 65 |
| GO_DISORDERED_DOMAIN_SPECIFIC_BINDING | 0.00051 | 0.00812 | 0.47727 | 0.76867 | 1.40343 | 27 |
| GO_DISULFIDE_OXIDOREDUCTASE_ACTIVITY | 0.00435 | 0.04238 | 0.40702 | 0.71445 | 1.31232 | 31 |
| GO_DNA_BIOSYNTHETIC_PROCESS | 0.00017 | 0.00334 | 0.51885 | 0.62540 | 1.20833 | 145 |
| GO_DNA_CONFORMATION_CHANGE | 0.00327 | 0.03409 | 0.43171 | 0.57979 | 1.13178 | 231 |
| GO_DNA_DAMAGE_RESPONSE_DETECTION_OF_DNA_DAMAGE | 0.00305 | 0.03271 | 0.43171 | 0.71625 | 1.31916 | 35 |
| GO_DNA_GEOMETRIC_CHANGE | 0.00141 | 0.01850 | 0.45506 | 0.62642 | 1.20356 | 105 |
| GO_DOUBLE_STRANDED_RNA_BINDING | 0.00039 | 0.00658 | 0.49849 | 0.69158 | 1.30365 | 61 |
| GO_DRUG_BINDING | 0.00450 | 0.04342 | 0.40702 | 0.66245 | 1.24130 | 52 |
| GO_ELECTRON_TRANSFER_ACTIVITY | 0.00000 | 0.00001 | 0.67496 | 0.72917 | 1.38765 | 80 |
| GO_ELECTRON_TRANSPORT_CHAIN | 0.00000 | 0.00000 | | 0.73336 | 1.41392 | 133 |
| GO_EMBRYO_IMPLANTATION | 0.00044 | 0.00706 | 0.49849 | 0.75968 | 1.39146 | 29 |
| GO_ENDOCYTIC_VESICLE_MEMBRANE | 0.00524 | 0.04770 | 0.40702 | 0.62563 | 1.19564 | 89 |
| GO_ENDOPEPTIDASE_ACTIVITY | 0.00065 | 0.00981 | 0.47727 | 0.60123 | 1.16998 | 196 |
| GO_ENDOPEPTIDASE_COMPLEX | 0.00000 | 0.00001 | 0.70498 | 0.76593 | 1.44609 | 63 |
| GO_ENDOPEPTIDASE_REGULATOR_ACTIVITY | 0.00006 | 0.00145 | 0.55733 | 0.70425 | 1.33001 | 68 |
| GO_ENDOPLASMIC_RETICULUM_GOLGI_INTERMEDIATE_COMPA | 0.00055 | 0.00863 | 0.47727 | 0.64758 | 1.24050 | 96 |
| GO_ENDOPLASMIC_RETICULUM_GOLGI_INTERMEDIATE_COMPA | 0.00148 | 0.01923 | 0.45506 | 0.68944 | 1.29056 | 51 |
| GO_ENDOPLASMIC_RETICULUM_TO_GOLGI_VESICLE_MEDIATED | 0.00532 | 0.04803 | 0.40702 | 0.59194 | 1.14697 | 162 |
| GO_ENERGY_DERIVATION_BY_OXIDATION_OF_ORGANIC_COMP | 0.00000 | 0.00000 | | 0.69995 | 1.36305 | 202 |
| GO_ENZYME_INHIBITOR_ACTIVITY | 0.00246 | 0.02780 | 0.43171 | 0.58913 | 1.14643 | 191 |
| GO_EPIDERMAL_CELL_DIFFERENTIATION | 0.00216 | 0.02519 | 0.43171 | 0.62667 | 1.20589 | 112 |
| GO_EPITHELIAL_CELL_DIFFERENTIATION | 0.00054 | 0.00857 | 0.47727 | 0.57819 | 1.13725 | 334 |
| GO_ESTABLISHMENT_OF_ORGANELLE_LOCALIZATION | 0.00122 | 0.01679 | 0.45506 | 0.57737 | 1.13489 | 323 |
| GO_ESTABLISHMENT_OF_PROTEIN_LOCALIZATION_TO_CHROMO | 0.00016 | 0.00321 | 0.51885 | 0.81363 | 1.46985 | 23 |
| GO_ESTABLISHMENT_OF_PROTEIN_LOCALIZATION_TO_ENDOPLA | 0.00000 | 0.00000 | | 0.77241 | 1.48479 | 106 |
| GO_ESTABLISHMENT_OF_PROTEIN_LOCALIZATION_TO_MEMBRA | 0.00000 | 0.00000 | | 0.67484 | 1.31991 | 260 |
| GO_ESTABLISHMENT_OF_PROTEIN_LOCALIZATION_TO_ORGANE | 0.00000 | 0.00000 | | 0.65856 | 1.30056 | 461 |
| GO_ESTABLISHMENT_OF_PROTEIN_LOCALIZATION_TO_TELOME | 0.00000 | 0.00006 | 0.64355 | 0.90447 | 1.60870 | 18 |
| GO_ESTABLISHMENT_OF_RNA_LOCALIZATION | 0.00000 | 0.00005 | 0.64355 | 0.64271 | 1.24830 | 176 |
| GO_ESTABLISHMENT_OF_TISSUE_POLARITY | 0.00000 | 0.00000 | 0.81404 | 0.75693 | 1.44999 | 96 |
| GO_EUKARYOTIC_TRANSLATION_INITIATION_FACTOR_3_COMPL | 0.00143 | 0.01863 | 0.45506 | 0.83263 | 1.46100 | 15 |
| GO_EXTERNAL_SIDE_OF_PLASMA_MEMBRANE | 0.00159 | 0.02016 | 0.45506 | 0.61775 | 1.19022 | 118 |
| GO_EXTRACELLULAR_MATRIX | 0.00042 | 0.00695 | 0.49849 | 0.61138 | 1.18938 | 189 |

| | | | | | | |
|---|---|---|---|---|---|---|
| GO_FATTY_ACID_DERIVATIVE_METABOLIC_PROCESS | 0.00043 | 0.00695 | 0.49849 | 0.66266 | 1.26212 | 81 |
| GO_FATTY_ACYL_COA_METABOLIC_PROCESS | 0.00502 | 0.04608 | 0.40702 | 0.72264 | 1.32082 | 28 |
| GO_FC_EPSILON_RECEPTOR_SIGNALING_PATHWAY | 0.00000 | 0.00000 | 0.71951 | 0.72370 | 1.38610 | 94 |
| GO_FC_RECEPTOR_SIGNALING_PATHWAY | 0.00000 | 0.00000 | 0.70498 | 0.68815 | 1.32879 | 140 |
| GO_FERTILIZATION | 0.00000 | 0.00005 | 0.64355 | 0.73257 | 1.38752 | 72 |
| GO_FIBRILLAR_CENTER | 0.00246 | 0.02780 | 0.43171 | 0.62638 | 1.19990 | 96 |
| GO_FIBROBLAST_GROWTH_FACTOR_RECEPTOR_SIGNALING_PAT | 0.00495 | 0.04573 | 0.40702 | 0.64888 | 1.22552 | 67 |
| GO_FIBROBLAST_PROLIFERATION | 0.00032 | 0.00575 | 0.49849 | 0.70676 | 1.32618 | 53 |
| GO_FICOLIN_1_RICH_GRANULE | 0.00000 | 0.00000 | 0.70498 | 0.71544 | 1.37138 | 98 |
| GO_GABA_ERGIC_SYNAPSE | 0.00022 | 0.00427 | 0.51885 | 0.76430 | 1.40406 | 32 |
| GO_GENERATION_OF_PRECURSOR_METABOLITES_AND_ENERGY | 0.00000 | 0.00000 | | 0.67527 | 1.32968 | 368 |
| GO_GENE_SILENCING | 0.00019 | 0.00376 | 0.51885 | 0.61625 | 1.19788 | 184 |
| GO_GLUCOSE_CATABOLIC_PROCESS | 0.00495 | 0.04573 | 0.40702 | 0.72314 | 1.32174 | 28 |
| GO_GLUTAMATERGIC_SYNAPSE | 0.00135 | 0.01808 | 0.45506 | 0.60052 | 1.16783 | 186 |
| GO_GLYCOLYTIC_PROCESS_THROUGH_FRUCTOSE_6_PHOSPHATI | 0.00554 | 0.04968 | 0.40702 | 0.74156 | 1.34111 | 24 |
| GO_GOLGI_ASSOCIATED_VESICLE | 0.00085 | 0.01239 | 0.47727 | 0.62294 | 1.20037 | 129 |
| GO_GOLGI_ASSOCIATED_VESICLE_MEMBRANE | 0.00313 | 0.03310 | 0.43171 | 0.64208 | 1.21753 | 75 |
| GO_GTPASE_ACTIVITY | 0.00004 | 0.00098 | 0.55733 | 0.61546 | 1.19974 | 210 |
| GO_GUANYL_NUCLEOTIDE_BINDING | 0.00005 | 0.00126 | 0.55733 | 0.60899 | 1.19065 | 252 |
| GO_HEAT_SHOCK_PROTEIN_BINDING | 0.00001 | 0.00034 | 0.59333 | 0.68258 | 1.30633 | 93 |
| GO_HEMATOPOIETIC_PROGENITOR_CELL_DIFFERENTIATION | 0.00000 | 0.00000 | 0.71951 | 0.70099 | 1.35150 | 121 |
| GO_HEMATOPOIETIC_STEM_CELL_DIFFERENTIATION | 0.00000 | 0.00000 | 0.76146 | 0.77246 | 1.46307 | 72 |
| GO_HISTONE_MRNA_METABOLIC_PROCESS | 0.00265 | 0.02936 | 0.43171 | 0.76832 | 1.38459 | 22 |
| GO_HOST_CELLULAR_COMPONENT | 0.00043 | 0.00695 | 0.49849 | 0.69729 | 1.31074 | 54 |
| GO_IMMUNE_RESPONSE_REGULATING_SIGNALING_PATHWAY | 0.00000 | 0.00003 | 0.65944 | 0.63620 | 1.24112 | 225 |
| GO_IMPORT_INTO_CELL | 0.00428 | 0.04185 | 0.40702 | 0.61709 | 1.18655 | 108 |
| GO_IMPORT_INTO_NUCLEUS | 0.00039 | 0.00652 | 0.49849 | 0.62707 | 1.20905 | 131 |
| GO_INNATE_IMMUNE_RESPONSE | 0.00000 | 0.00003 | 0.65944 | 0.59893 | 1.18089 | 406 |
| GO_INNATE_IMMUNE_RESPONSE_ACTIVATING_SIGNAL_TRANSD | 0.00000 | 0.00001 | 0.69013 | 0.73348 | 1.39585 | 80 |
| GO_INNER_MITOCHONDRIAL_MEMBRANE_PROTEIN_COMPLEX | 0.00000 | 0.00000 | | 0.76576 | 1.46666 | 94 |
| GO_INORGANIC_ION_TRANSMEMBRANE_TRANSPORT | 0.00141 | 0.01850 | 0.45506 | 0.56404 | 1.11185 | 401 |
| GO_INTERACTION_WITH_HOST | 0.00139 | 0.01850 | 0.45506 | 0.60855 | 1.17710 | 151 |
| GO_INTERLEUKIN_1_MEDIATED_SIGNALING_PATHWAY | 0.00000 | 0.00000 | 0.77494 | 0.75654 | 1.44176 | 83 |
| GO_INTRAMOLECULAR_OXIDOREDUCTASE_ACTIVITY | 0.00051 | 0.00812 | 0.47727 | 0.74506 | 1.37081 | 34 |
| GO_INTRINSIC_APOPTOTIC_SIGNALING_PATHWAY | 0.00156 | 0.01993 | 0.45506 | 0.59427 | 1.15817 | 207 |
| GO_ION_HOMEOSTASIS | 0.00170 | 0.02117 | 0.45506 | 0.56164 | 1.10717 | 403 |
| GO_ION_TRANSMEMBRANE_TRANSPORTER_ACTIVITY | 0.00106 | 0.01495 | 0.45506 | 0.56696 | 1.11734 | 390 |
| GO_IRON_ION_HOMEOSTASIS | 0.00361 | 0.03657 | 0.43171 | 0.65837 | 1.23983 | 59 |
| GO_IRON_ION_TRANSPORT | 0.00054 | 0.00857 | 0.47727 | 0.69556 | 1.30334 | 52 |
| GO_ISOMERASE_ACTIVITY | 0.00413 | 0.04065 | 0.40702 | 0.61065 | 1.17496 | 113 |
| GO_KERATINIZATION | 0.00038 | 0.00640 | 0.49849 | 0.75690 | 1.39013 | 33 |
| GO_KILLING_OF_CELLS_OF_OTHER_ORGANISM | 0.00495 | 0.04573 | 0.40702 | 0.78614 | 1.38643 | 16 |
| GO_LARGE_RIBOSOMAL_SUBUNIT | 0.00000 | 0.00000 | | 0.77656 | 1.49319 | 108 |
| GO_LATERAL_PLASMA_MEMBRANE | 0.00524 | 0.04770 | 0.40702 | 0.67385 | 1.25557 | 47 |
| GO_LEUKOCYTE_MEDIATED_IMMUNITY | 0.00000 | 0.00000 | | 0.64010 | 1.26331 | 447 |
| GO_LIPOPROTEIN_PARTICLE_RECEPTOR_BINDING | 0.00126 | 0.01715 | 0.45506 | 0.80503 | 1.42497 | 17 |
| GO_LIVER_REGENERATION | 0.00093 | 0.01337 | 0.47727 | 0.78442 | 1.41708 | 23 |
| GO_LOW_DENSITY_LIPOPROTEIN_RECEPTOR_PARTICLE_METABO | 0.00327 | 0.03409 | 0.43171 | 0.78520 | 1.39657 | 18 |
| GO_LYASE_ACTIVITY | 0.00152 | 0.01958 | 0.45506 | 0.61769 | 1.19079 | 122 |
| GO_LYSOSOMAL_LUMEN | 0.00143 | 0.01863 | 0.45506 | 0.67371 | 1.26872 | 59 |
| GO_MAINTENANCE_OF_CELL_NUMBER | 0.00435 | 0.04238 | 0.40702 | 0.60761 | 1.17073 | 117 |
| GO_MATURATION_OF_SSU_RRNA | 0.00126 | 0.01715 | 0.45506 | 0.69615 | 1.29711 | 47 |
| GO_MATURATION_OF_SSU_RRNA_FROM_TRICISTRONIC_RRNA_ | 0.00554 | 0.04968 | 0.40702 | 0.70205 | 1.29301 | 35 |
| GO_MEMBRANE_REGION | 0.00000 | 0.00003 | 0.65944 | 0.64496 | 1.25542 | 194 |
| GO_METALLOPEPTIDASE_ACTIVITY | 0.00487 | 0.04559 | 0.40702 | 0.62714 | 1.19577 | 85 |
| GO_METAL_ION_HOMEOSTASIS | 0.00032 | 0.00575 | 0.49849 | 0.58816 | 1.15376 | 301 |
| GO_METAL_ION_TRANSPORT | 0.00390 | 0.03895 | 0.43171 | 0.55877 | 1.10227 | 419 |
| GO_METAPHASE_ANAPHASE_TRANSITION_OF_CELL_CYCLE | 0.00320 | 0.03371 | 0.43171 | 0.66563 | 1.25122 | 54 |
| GO_MICROTUBULE_CYTOSKELETON_ORGANIZATION_INVOLVED_ | 0.00220 | 0.02548 | 0.43171 | 0.61295 | 1.18118 | 127 |
| GO_MICROVILLUS | 0.00168 | 0.02098 | 0.45506 | 0.68128 | 1.28184 | 56 |
| GO_MIDBODY | 0.00561 | 0.04971 | 0.40702 | 0.59866 | 1.15606 | 141 |
| GO_MITOCHONDRIAL_ELECTRON_TRANSPORT_NADH_TO_UBIQ | 0.00000 | 0.00006 | 0.64355 | 0.79447 | 1.47799 | 44 |
| GO_MITOCHONDRIAL_GENE_EXPRESSION | 0.00000 | 0.00000 | 0.78819 | 0.70230 | 1.35844 | 151 |
| GO_MITOCHONDRIAL_LARGE_RIBOSOMAL_SUBUNIT | 0.00000 | 0.00000 | 0.71951 | 0.79015 | 1.48529 | 54 |
| GO_MITOCHONDRIAL_MATRIX | 0.00000 | 0.00000 | 0.82666 | 0.63141 | 1.24412 | 387 |
| GO_MITOCHONDRIAL_MEMBRANE_ORGANIZATION | 0.00050 | 0.00802 | 0.47727 | 0.64578 | 1.23686 | 94 |
| GO_MITOCHONDRIAL_PROTEIN_COMPLEX | 0.00000 | 0.00000 | | 0.75011 | 1.46215 | 212 |
| GO_MITOCHONDRIAL_RESPIRATORY_CHAIN_COMPLEX_ASSEMB | 0.00001 | 0.00033 | 0.59333 | 0.69650 | 1.32804 | 85 |
| GO_MITOCHONDRIAL_SMALL_RIBOSOMAL_SUBUNIT | 0.00119 | 0.01640 | 0.45506 | 0.75384 | 1.37785 | 28 |
| GO_MITOCHONDRIAL_TRANSLATION | 0.00000 | 0.00000 | 0.80122 | 0.71435 | 1.37651 | 128 |

| | | | | | | |
|---|---|---|---|---|---|---|
| GO_MITOCHONDRIAL_TRANSLATIONAL_TERMINATION | 0.00000 | 0.00000 | 0.81404 | 0.75760 | 1.44657 | 87 |
| GO_MITOCHONDRIAL_TRANSMEMBRANE_TRANSPORT | 0.00213 | 0.02490 | 0.43171 | 0.66300 | 1.25387 | 70 |
| GO_MITOCHONDRIAL_TRANSPORT | 0.00010 | 0.00227 | 0.53843 | 0.61119 | 1.19060 | 203 |
| GO_MITOCHONDRION_ORGANIZATION | 0.00000 | 0.00000 | 0.74774 | 0.61590 | 1.21471 | 416 |
| GO_MITOTIC_NUCLEAR_DIVISION | 0.00008 | 0.00182 | 0.53843 | 0.60435 | 1.17901 | 223 |
| GO_MITOTIC_SISTER_CHROMATID_SEGREGATION | 0.00014 | 0.00302 | 0.51885 | 0.62652 | 1.21048 | 145 |
| GO_MITOTIC_SPINDLE_ORGANIZATION | 0.00078 | 0.01164 | 0.47727 | 0.63943 | 1.22821 | 104 |
| GO_MODIFIED_AMINO_ACID_BINDING | 0.00002 | 0.00048 | 0.57561 | 0.74200 | 1.38679 | 49 |
| GO_MODULATION_BY_HOST_OF_VIRAL_PROCESS | 0.00216 | 0.02519 | 0.43171 | 0.77232 | 1.38066 | 20 |
| GO_MONOCARBOXYLIC_ACID_METABOLIC_PROCESS | 0.00012 | 0.00265 | 0.53843 | 0.57846 | 1.14053 | 406 |
| GO_MONOSACCHARIDE_BIOSYNTHETIC_PROCESS | 0.00305 | 0.03271 | 0.43171 | 0.65908 | 1.24472 | 68 |
| GO_MONOVALENT_INORGANIC_CATION_HOMEOSTASIS | 0.00108 | 0.01507 | 0.45506 | 0.63643 | 1.21783 | 91 |
| GO_MONOVALENT_INORGANIC_CATION_TRANSMEMBRANE_TR | 0.00001 | 0.00026 | 0.59333 | 0.65186 | 1.25996 | 147 |
| GO_MONOVALENT_INORGANIC_CATION_TRANSPORT | 0.00008 | 0.00178 | 0.53843 | 0.60205 | 1.17659 | 248 |
| GO_MORPHOGENESIS_OF_AN_EPITHELIUM | 0.00000 | 0.00009 | 0.62726 | 0.60570 | 1.19137 | 334 |
| GO_MORPHOGENESIS_OF_A_POLARIZED_EPITHELIUM | 0.00000 | 0.00000 | 0.83909 | 0.74384 | 1.43057 | 110 |
| GO_MOVEMENT_IN_HOST_ENVIRONMENT | 0.00268 | 0.02961 | 0.43171 | 0.61282 | 1.18078 | 117 |
| GO_MRNA_3_END_PROCESSING | 0.00087 | 0.01270 | 0.47727 | 0.64010 | 1.22502 | 93 |
| GO_MRNA_5_UTR_BINDING | 0.00002 | 0.00063 | 0.57561 | 0.83759 | 1.51313 | 23 |
| GO_MRNA_BINDING | 0.00000 | 0.00000 | | 0.67496 | 1.31810 | 239 |
| GO_MRNA_EXPORT_FROM_NUCLEUS | 0.00006 | 0.00159 | 0.53843 | 0.66290 | 1.27265 | 103 |
| GO_MRNA_PROCESSING | 0.00000 | 0.00000 | | 0.62959 | 1.24255 | 437 |
| GO_MRNA_TRANSPORT | 0.00007 | 0.00171 | 0.53843 | 0.64103 | 1.23563 | 132 |
| GO_MULTI_MULTICELLULAR_ORGANISM_PROCESS | 0.00041 | 0.00687 | 0.49849 | 0.63087 | 1.21515 | 116 |
| GO_MULTI_ORGANISM_LOCALIZATION | 0.00017 | 0.00334 | 0.51885 | 0.71077 | 1.33608 | 54 |
| GO_MUSCLE_CONTRACTION | 0.00335 | 0.03468 | 0.43171 | 0.59925 | 1.16199 | 165 |
| GO_MUSCLE_FILAMENT_SLIDING | 0.00324 | 0.03393 | 0.43171 | 0.77980 | 1.39322 | 19 |
| GO_MYELOID_LEUKOCYTE_ACTIVATION | 0.00000 | 0.00000 | | 0.65288 | 1.28712 | 395 |
| GO_MYELOID_LEUKOCYTE_MEDIATED_IMMUNITY | 0.00000 | 0.00000 | | 0.65734 | 1.29408 | 356 |
| GO_NADH_DEHYDROGENASE_ACTIVITY | 0.00001 | 0.00019 | 0.61053 | 0.79360 | 1.46691 | 38 |
| GO_NADH_DEHYDROGENASE_COMPLEX | 0.00001 | 0.00028 | 0.59333 | 0.77749 | 1.44215 | 41 |
| GO_NADH_DEHYDROGENASE_COMPLEX_ASSEMBLY | 0.00001 | 0.00035 | 0.59333 | 0.74044 | 1.39281 | 55 |
| GO_NADH_METABOLIC_PROCESS | 0.00034 | 0.00599 | 0.49849 | 0.75773 | 1.39166 | 33 |
| GO_NAD_METABOLIC_PROCESS | 0.00435 | 0.04238 | 0.40702 | 0.70570 | 1.30442 | 37 |
| GO_NCRNA_EXPORT_FROM_NUCLEUS | 0.00205 | 0.02440 | 0.43171 | 0.71473 | 1.31817 | 36 |
| GO_NCRNA_METABOLIC_PROCESS | 0.00000 | 0.00000 | 0.70498 | 0.61139 | 1.20509 | 399 |
| GO_NCRNA_PROCESSING | 0.00000 | 0.00000 | 0.73376 | 0.62238 | 1.22425 | 341 |
| GO_NEGATIVE_REGULATION_OF_APOPTOTIC_SIGNALING_PATHW | 0.00364 | 0.03677 | 0.43171 | 0.59958 | 1.15998 | 150 |
| GO_NEGATIVE_REGULATION_OF_ATP_METABOLIC_PROCESS | 0.00302 | 0.03248 | 0.43171 | 0.78586 | 1.39103 | 17 |
| GO_NEGATIVE_REGULATION_OF_CANONICAL_WNT_SIGNALING | 0.00000 | 0.00000 | 0.71951 | 0.69629 | 1.34180 | 123 |
| GO_NEGATIVE_REGULATION_OF_CATALYTIC_ACTIVITY | 0.00145 | 0.01879 | 0.45506 | 0.56238 | 1.11005 | 450 |
| GO_NEGATIVE_REGULATION_OF_CELLULAR_AMIDE_METABOLIC | 0.00268 | 0.02961 | 0.43171 | 0.61074 | 1.17686 | 124 |
| GO_NEGATIVE_REGULATION_OF_CELLULAR_COMPONENT_ORG | 0.00320 | 0.03371 | 0.43171 | 0.55470 | 1.09534 | 470 |
| GO_NEGATIVE_REGULATION_OF_CELL_CYCLE | 0.00023 | 0.00445 | 0.51885 | 0.57348 | 1.13228 | 455 |
| GO_NEGATIVE_REGULATION_OF_CELL_CYCLE_G2_M_PHASE_TR | 0.00000 | 0.00007 | 0.62726 | 0.70549 | 1.34606 | 86 |
| GO_NEGATIVE_REGULATION_OF_CELL_CYCLE_PHASE_TRANSITI | 0.00000 | 0.00000 | 0.70498 | 0.65532 | 1.27560 | 194 |
| GO_NEGATIVE_REGULATION_OF_CELL_CYCLE_PROCESS | 0.00000 | 0.00005 | 0.64355 | 0.62725 | 1.22670 | 259 |
| GO_NEGATIVE_REGULATION_OF_HYDROLASE_ACTIVITY | 0.00216 | 0.02519 | 0.43171 | 0.58306 | 1.13858 | 238 |
| GO_NEGATIVE_REGULATION_OF_INTERLEUKIN_1_PRODUCTION | 0.00205 | 0.02440 | 0.43171 | 0.82297 | 1.44404 | 15 |
| GO_NEGATIVE_REGULATION_OF_MITOTIC_CELL_CYCLE | 0.00000 | 0.00010 | 0.62726 | 0.62191 | 1.21541 | 248 |
| GO_NEGATIVE_REGULATION_OF_MRNA_METABOLIC_PROCESS | 0.00213 | 0.02490 | 0.43171 | 0.66731 | 1.25879 | 62 |
| GO_NEGATIVE_REGULATION_OF_MRNA_SPLICING_VIA_SPLICEO | 0.00339 | 0.03501 | 0.43171 | 0.76309 | 1.36417 | 20 |
| GO_NEGATIVE_REGULATION_OF_NUCLEAR_DIVISION | 0.00335 | 0.03468 | 0.43171 | 0.67724 | 1.26685 | 50 |
| GO_NEGATIVE_REGULATION_OF_ORGANELLE_ORGANIZATION | 0.00487 | 0.04559 | 0.40702 | 0.57038 | 1.11722 | 288 |
| GO_NEGATIVE_REGULATION_OF_PEPTIDASE_ACTIVITY | 0.00036 | 0.00621 | 0.49849 | 0.64044 | 1.23159 | 109 |
| GO_NEGATIVE_REGULATION_OF_PROTEIN_CATABOLIC_PROCESS | 0.00342 | 0.03521 | 0.43171 | 0.62320 | 1.19268 | 93 |
| GO_NEGATIVE_REGULATION_OF_PROTEOLYSIS | 0.00205 | 0.02440 | 0.43171 | 0.60172 | 1.16902 | 178 |
| GO_NEGATIVE_REGULATION_OF_REACTIVE_OXYGEN_SPECIES_B | 0.00036 | 0.00621 | 0.49849 | 0.83108 | 1.47818 | 18 |
| GO_NEGATIVE_REGULATION_OF_REPRODUCTIVE_PROCESS | 0.00457 | 0.04386 | 0.40702 | 0.74845 | 1.35356 | 24 |
| GO_NEGATIVE_REGULATION_OF_RNA_SPLICING | 0.00183 | 0.02219 | 0.45506 | 0.76824 | 1.38935 | 24 |
| GO_NEGATIVE_REGULATION_OF_WNT_SIGNALING_PATHWAY | 0.00000 | 0.00002 | 0.67496 | 0.67731 | 1.30721 | 136 |
| GO_NEURAL_CREST_CELL_MIGRATION | 0.00265 | 0.02936 | 0.43171 | 0.73967 | 1.35049 | 27 |
| GO_NEURON_RECOGNITION | 0.00390 | 0.03895 | 0.43171 | 0.71862 | 1.31998 | 31 |
| GO_NIK_NF_KAPPAB_SIGNALING | 0.00000 | 0.00000 | 0.77494 | 0.72329 | 1.39253 | 114 |
| GO_NON_CANONICAL_WNT_SIGNALING_PATHWAY | 0.00000 | 0.00000 | | 0.75756 | 1.45654 | 107 |
| GO_NUCLEAR_CHROMOSOME_SEGREGATION | 0.00098 | 0.01407 | 0.45506 | 0.59156 | 1.15357 | 215 |
| GO_NUCLEAR_ENVELOPE | 0.00005 | 0.00131 | 0.55733 | 0.59109 | 1.16270 | 341 |
| GO_NUCLEAR_EXPORT | 0.00000 | 0.00004 | 0.65944 | 0.64915 | 1.26157 | 181 |
| GO_NUCLEAR_MATRIX | 0.00502 | 0.04608 | 0.40702 | 0.62777 | 1.19868 | 87 |
| GO_NUCLEAR_MEMBRANE | 0.00185 | 0.02237 | 0.45506 | 0.58925 | 1.14853 | 211 |

| | | | | | | |
|---|---|---|---|---|---|---|
| GO_NUCLEAR_PERIPHERY | 0.00043 | 0.00701 | 0.49849 | 0.64264 | 1.23558 | 107 |
| GO_NUCLEAR_PORE | 0.00561 | 0.04971 | 0.40702 | 0.64876 | 1.22522 | 68 |
| GO_NUCLEAR_SPECK | 0.00015 | 0.00312 | 0.51885 | 0.59159 | 1.16089 | 306 |
| GO_NUCLEAR_TRANSCRIBED_MRNA_CATABOLIC_PROCESS | 0.00000 | 0.00000 | | 0.70362 | 1.36820 | 185 |
| GO_NUCLEAR_TRANSCRIBED_MRNA_CATABOLIC_PROCESS_NON | 0.00000 | 0.00000 | | 0.74239 | 1.42931 | 114 |
| GO_NUCLEAR_TRANSPORT | 0.00000 | 0.00000 | 0.71951 | 0.62920 | 1.23383 | 297 |
| GO_NUCLEOBASE_CONTAINING_COMPOUND_TRANSPORT | 0.00000 | 0.00010 | 0.62726 | 0.62975 | 1.22823 | 216 |
| GO_NUCLEOBASE_CONTAINING_SMALL_MOLECULE_METABOLI( | 0.00000 | 0.00003 | 0.65944 | 0.60153 | 1.18594 | 405 |
| GO_NUCLEOSIDE_DIPHOSPHATE_METABOLIC_PROCESS | 0.00016 | 0.00319 | 0.51885 | 0.64491 | 1.24260 | 117 |
| GO_NUCLEOSIDE_PHOSPHATE_BIOSYNTHETIC_PROCESS | 0.00002 | 0.00044 | 0.59333 | 0.64092 | 1.24175 | 161 |
| GO_NUCLEOSIDE_TRIPHOSPHATE_BIOSYNTHETIC_PROCESS | 0.00040 | 0.00676 | 0.49849 | 0.72727 | 1.35074 | 42 |
| GO_NUCLEOSIDE_TRIPHOSPHATE_METABOLIC_PROCESS | 0.00009 | 0.00196 | 0.53843 | 0.69306 | 1.30854 | 65 |
| GO_NUCLEOTIDE_PHOSPHORYLATION | 0.00016 | 0.00328 | 0.51885 | 0.65541 | 1.25767 | 102 |
| GO_OOCYTE_DIFFERENTIATION | 0.00450 | 0.04342 | 0.40702 | 0.73513 | 1.33705 | 26 |
| GO_ORGANELLAR_RIBOSOME | 0.00000 | 0.00000 | 0.76146 | 0.75712 | 1.44311 | 84 |
| GO_ORGANELLE_ENVELOPE_LUMEN | 0.00026 | 0.00482 | 0.49849 | 0.67265 | 1.27716 | 78 |
| GO_ORGANELLE_FISSION | 0.00154 | 0.01974 | 0.45506 | 0.57133 | 1.12383 | 342 |
| GO_ORGANELLE_INNER_MEMBRANE | 0.00000 | 0.00000 | | 0.68423 | 1.34915 | 412 |
| GO_ORGANIC_CYCLIC_COMPOUND_CATABOLIC_PROCESS | 0.00000 | 0.00000 | | 0.66975 | 1.32263 | 478 |
| GO_ORGANIC_HYDROXY_COMPOUND_BIOSYNTHETIC_PROCESS | 0.00141 | 0.01850 | 0.45506 | 0.60712 | 1.17627 | 161 |
| GO_OXIDATIVE_PHOSPHORYLATION | 0.00000 | 0.00000 | | 0.78902 | 1.51260 | 99 |
| GO_OXIDOREDUCTASE_ACTIVITY | 0.00000 | 0.00000 | | 0.63268 | 1.24935 | 468 |
| GO_OXIDOREDUCTASE_ACTIVITY_ACTING_ON_A_HEME_GROUP | 0.00176 | 0.02168 | 0.45506 | 0.79383 | 1.41828 | 19 |
| GO_OXIDOREDUCTASE_ACTIVITY_ACTING_ON_NAD_P_H | 0.00002 | 0.00056 | 0.57561 | 0.71468 | 1.35162 | 70 |
| GO_OXIDOREDUCTASE_ACTIVITY_ACTING_ON_NAD_P_H_QUIN( | 0.00004 | 0.00113 | 0.55733 | 0.74511 | 1.38835 | 47 |
| GO_OXIDOREDUCTASE_ACTIVITY_ACTING_ON_PEROXIDE_AS_A( | 0.00106 | 0.01495 | 0.45506 | 0.73897 | 1.35352 | 29 |
| GO_OXIDOREDUCTASE_COMPLEX | 0.00000 | 0.00000 | 0.73376 | 0.74898 | 1.42731 | 82 |
| GO_PEPTIDASE_ACTIVITY | 0.00133 | 0.01799 | 0.45506 | 0.57681 | 1.13188 | 306 |
| GO_PEPTIDASE_COMPLEX | 0.00000 | 0.00002 | 0.67496 | 0.73250 | 1.39240 | 79 |
| GO_PEPTIDASE_REGULATOR_ACTIVITY | 0.00313 | 0.03310 | 0.43171 | 0.63434 | 1.21189 | 88 |
| GO_PEPTIDE_BINDING | 0.00005 | 0.00124 | 0.55733 | 0.63708 | 1.23187 | 148 |
| GO_PERINUCLEAR_REGION_OF_CYTOPLASM | 0.00276 | 0.03021 | 0.43171 | 0.55814 | 1.10207 | 466 |
| GO_PHAGOCYTOSIS | 0.00554 | 0.04968 | 0.40702 | 0.59337 | 1.14837 | 155 |
| GO_PHAGOSOME_ACIDIFICATION | 0.00059 | 0.00916 | 0.47727 | 0.78902 | 1.42694 | 24 |
| GO_PHOSPHATIDYLSERINE_BINDING | 0.00035 | 0.00610 | 0.49849 | 0.76387 | 1.39912 | 29 |
| GO_PH_REDUCTION | 0.00302 | 0.03248 | 0.43171 | 0.68821 | 1.28238 | 46 |
| GO_PIGMENT_GRANULE | 0.00000 | 0.00000 | 0.76146 | 0.74333 | 1.41934 | 87 |
| GO_PLASMA_LIPOPROTEIN_PARTICLE_CLEARANCE | 0.00106 | 0.01495 | 0.45506 | 0.73201 | 1.35304 | 37 |
| GO_POLYSOMAL_RIBOSOME | 0.00008 | 0.00180 | 0.53843 | 0.78840 | 1.44683 | 30 |
| GO_POLYSOME | 0.00000 | 0.00003 | 0.65944 | 0.75964 | 1.43296 | 62 |
| GO_POSITIVE_REGULATION_OF_ATP_METABOLIC_PROCESS | 0.00174 | 0.02149 | 0.45506 | 0.73011 | 1.33729 | 29 |
| GO_POSITIVE_REGULATION_OF_BINDING | 0.00390 | 0.03895 | 0.43171 | 0.61042 | 1.17582 | 115 |
| GO_POSITIVE_REGULATION_OF_CANONICAL_WNT_SIGNALING_ | 0.00000 | 0.00000 | 0.73376 | 0.72036 | 1.38514 | 108 |
| GO_POSITIVE_REGULATION_OF_CELLULAR_AMIDE_METABOLIC_ | 0.00001 | 0.00036 | 0.59333 | 0.66485 | 1.28095 | 125 |
| GO_POSITIVE_REGULATION_OF_CELLULAR_PROTEIN_LOCALIZAT | 0.00242 | 0.02748 | 0.43171 | 0.58048 | 1.13378 | 244 |
| GO_POSITIVE_REGULATION_OF_CELL_POPULATION_PROLIFERAT | 0.00040 | 0.00670 | 0.49849 | 0.56967 | 1.12503 | 461 |
| GO_POSITIVE_REGULATION_OF_CYSTEINE_TYPE_ENDOPEPTIDA: | 0.00547 | 0.04923 | 0.40702 | 0.62225 | 1.19077 | 92 |
| GO_POSITIVE_REGULATION_OF_CYTOKINE_PRODUCTION | 0.00283 | 0.03081 | 0.43171 | 0.58672 | 1.14373 | 210 |
| GO_POSITIVE_REGULATION_OF_DEFENSE_RESPONSE | 0.00000 | 0.00000 | 0.73376 | 0.66200 | 1.28884 | 200 |
| GO_POSITIVE_REGULATION_OF_DNA_BIOSYNTHETIC_PROCESS | 0.00010 | 0.00228 | 0.53843 | 0.71224 | 1.33977 | 55 |
| GO_POSITIVE_REGULATION_OF_DNA_METABOLIC_PROCESS | 0.00020 | 0.00387 | 0.51885 | 0.62771 | 1.21205 | 139 |
| GO_POSITIVE_REGULATION_OF_ESTABLISHMENT_OF_PROTEIN_ | 0.00194 | 0.02331 | 0.45506 | 0.58158 | 1.13709 | 253 |
| GO_POSITIVE_REGULATION_OF_FIBROBLAST_PROLIFERATION | 0.00017 | 0.00338 | 0.51885 | 0.77423 | 1.42213 | 31 |
| GO_POSITIVE_REGULATION_OF_IMMUNE_RESPONSE | 0.00000 | 0.00000 | 0.71951 | 0.63064 | 1.23937 | 321 |
| GO_POSITIVE_REGULATION_OF_IMMUNE_SYSTEM_PROCESS | 0.00000 | 0.00002 | 0.67496 | 0.59748 | 1.17992 | 475 |
| GO_POSITIVE_REGULATION_OF_INNATE_IMMUNE_RESPONSE | 0.00000 | 0.00000 | 0.70498 | 0.68885 | 1.32737 | 128 |
| GO_POSITIVE_REGULATION_OF_MITOTIC_CELL_CYCLE | 0.00178 | 0.02182 | 0.45506 | 0.61563 | 1.18683 | 122 |
| GO_POSITIVE_REGULATION_OF_MRNA_METABOLIC_PROCESS | 0.00157 | 0.02009 | 0.45506 | 0.66330 | 1.25231 | 63 |
| GO_POSITIVE_REGULATION_OF_MRNA_PROCESSING | 0.00250 | 0.02807 | 0.43171 | 0.76194 | 1.38151 | 25 |
| GO_POSITIVE_REGULATION_OF_MRNA_SPLICING_VIA_SPLICEOS | 0.00132 | 0.01782 | 0.45506 | 0.81851 | 1.44353 | 16 |
| GO_POSITIVE_REGULATION_OF_NITRIC_OXIDE_METABOLIC_PR( | 0.00133 | 0.01799 | 0.45506 | 0.80807 | 1.43726 | 18 |
| GO_POSITIVE_REGULATION_OF_ORGANELLE_ORGANIZATION | 0.00010 | 0.00218 | 0.53843 | 0.57473 | 1.13449 | 452 |
| GO_POSITIVE_REGULATION_OF_PROTEIN_LOCALIZATION_TO_N | 0.00012 | 0.00265 | 0.53843 | 0.71018 | 1.33697 | 58 |
| GO_POSITIVE_REGULATION_OF_PROTEOLYSIS | 0.00313 | 0.03310 | 0.43171 | 0.58059 | 1.13381 | 237 |
| GO_POSITIVE_REGULATION_OF_REACTIVE_OXYGEN_SPECIES_BI | 0.00115 | 0.01593 | 0.45506 | 0.79865 | 1.43554 | 21 |
| GO_POSITIVE_REGULATION_OF_REACTIVE_OXYGEN_SPECIES_M | 0.00174 | 0.02149 | 0.45506 | 0.69785 | 1.30035 | 46 |
| GO_POSITIVE_REGULATION_OF_RESPONSE_TO_BIOTIC_STIMUL | 0.00000 | 0.00000 | 0.70498 | 0.67507 | 1.30603 | 150 |
| GO_POSITIVE_REGULATION_OF_RESPONSE_TO_EXTERNAL_STIM | 0.00001 | 0.00016 | 0.61053 | 0.62123 | 1.21505 | 260 |
| GO_POSITIVE_REGULATION_OF_RNA_SPLICING | 0.00265 | 0.02936 | 0.43171 | 0.74051 | 1.35202 | 27 |
| GO_POSITIVE_REGULATION_OF_SIGNAL_TRANSDUCTION_BY_P! | 0.00390 | 0.03895 | 0.43171 | 0.80515 | 1.41278 | 15 |

| | | | | | | |
|---|---|---|---|---|---|---|
| GO_POSITIVE_REGULATION_OF_TELOMERASE_ACTIVITY | 0.00069 | 0.01043 | 0.47727 | 0.75265 | 1.38122 | 30 |
| GO_POSITIVE_REGULATION_OF_TELOMERASE_RNA_LOCALIZATI | 0.00007 | 0.00169 | 0.53843 | 0.88616 | 1.55493 | 15 |
| GO_POSITIVE_REGULATION_OF_TELOMERE_MAINTENANCE | 0.00026 | 0.00486 | 0.49849 | 0.73697 | 1.37102 | 44 |
| GO_POSITIVE_REGULATION_OF_TELOMERE_MAINTENANCE_VIA | 0.00001 | 0.00040 | 0.59333 | 0.80196 | 1.47289 | 33 |
| GO_POSITIVE_REGULATION_OF_TRANSFERASE_ACTIVITY | 0.00108 | 0.01507 | 0.45506 | 0.56501 | 1.11376 | 401 |
| GO_POSITIVE_REGULATION_OF_TRANSLATION | 0.00000 | 0.00016 | 0.61053 | 0.68408 | 1.31526 | 107 |
| GO_POSITIVE_REGULATION_OF_VIRAL_PROCESS | 0.00026 | 0.00486 | 0.49849 | 0.66195 | 1.26462 | 88 |
| GO_POSITIVE_REGULATION_OF_WNT_SIGNALING_PATHWAY | 0.00000 | 0.00000 | 0.73376 | 0.69924 | 1.34747 | 127 |
| GO_POSTSYNAPSE | 0.00239 | 0.02716 | 0.43171 | 0.57062 | 1.12116 | 316 |
| GO_POST_TRANSLATIONAL_PROTEIN_MODIFICATION | 0.00024 | 0.00452 | 0.51885 | 0.60046 | 1.17392 | 251 |
| GO_PRECATALYTIC_SPLICEOSOME | 0.00141 | 0.01850 | 0.45506 | 0.69251 | 1.29540 | 50 |
| GO_PRERIBOSOME | 0.00006 | 0.00154 | 0.53843 | 0.69866 | 1.32020 | 69 |
| GO_PROTEASOMAL_PROTEIN_CATABOLIC_PROCESS | 0.00007 | 0.00169 | 0.53843 | 0.58188 | 1.14652 | 387 |
| GO_PROTEASOMAL_UBIQUITIN_INDEPENDENT_PROTEIN_CATAE | 0.00000 | 0.00003 | 0.65944 | 0.91182 | 1.62178 | 18 |
| GO_PROTEASOME_ACCESSORY_COMPLEX | 0.00005 | 0.00135 | 0.55733 | 0.82638 | 1.49450 | 24 |
| GO_PROTEASOME_CORE_COMPLEX | 0.00000 | 0.00004 | 0.65944 | 0.91231 | 1.61486 | 17 |
| GO_PROTEIN_CONTAINING_COMPLEX_DISASSEMBLY | 0.00000 | 0.00000 | 0.81404 | 0.66285 | 1.29656 | 261 |
| GO_PROTEIN_CONTAINING_COMPLEX_LOCALIZATION | 0.00018 | 0.00352 | 0.51885 | 0.60451 | 1.17943 | 226 |
| GO_PROTEIN_DNA_COMPLEX_SUBUNIT_ORGANIZATION | 0.00052 | 0.00825 | 0.47727 | 0.60887 | 1.18342 | 183 |
| GO_PROTEIN_DOMAIN_SPECIFIC_BINDING | 0.00003 | 0.00088 | 0.55733 | 0.57828 | 1.14190 | 470 |
| GO_PROTEIN_FOLDING | 0.00000 | 0.00000 | 0.73376 | 0.67425 | 1.30867 | 171 |
| GO_PROTEIN_FOLDING_CHAPERONE | 0.00079 | 0.01170 | 0.47727 | 0.81632 | 1.45193 | 18 |
| GO_PROTEIN_HETEROOLIGOMERIZATION | 0.00442 | 0.04297 | 0.40702 | 0.78998 | 1.39322 | 16 |
| GO_PROTEIN_IMPORT | 0.00001 | 0.00040 | 0.59333 | 0.64197 | 1.24306 | 158 |
| GO_PROTEIN_IMPORT_INTO_MITOCHONDRIAL_MATRIX | 0.00231 | 0.02642 | 0.43171 | 0.79530 | 1.41455 | 18 |
| GO_PROTEIN_INSERTION_INTO_MEMBRANE | 0.00191 | 0.02299 | 0.45506 | 0.68322 | 1.27694 | 49 |
| GO_PROTEIN_LOCALIZATION_TO_CHROMOSOME_TELOMERIC_I | 0.00001 | 0.00031 | 0.59333 | 0.82810 | 1.51357 | 28 |
| GO_PROTEIN_LOCALIZATION_TO_ENDOPLASMIC_RETICULUM | 0.00000 | 0.00000 | | 0.73850 | 1.42303 | 129 |
| GO_PROTEIN_LOCALIZATION_TO_MEMBRANE | 0.00000 | 0.00000 | 0.77494 | 0.61216 | 1.20881 | 462 |
| GO_PROTEIN_LOCALIZATION_TO_MITOCHONDRION | 0.00015 | 0.00310 | 0.51885 | 0.64234 | 1.23759 | 118 |
| GO_PROTEIN_LOCALIZATION_TO_NUCLEUS | 0.00000 | 0.00014 | 0.61053 | 0.63315 | 1.23408 | 208 |
| GO_PROTEIN_MATURATION | 0.00357 | 0.03638 | 0.43171 | 0.59209 | 1.14965 | 174 |
| GO_PROTEIN_MODIFICATION_BY_SMALL_PROTEIN_REMOVAL | 0.00002 | 0.00052 | 0.57561 | 0.61535 | 1.20119 | 231 |
| GO_PROTEIN_REFOLDING | 0.00465 | 0.04430 | 0.40702 | 0.77561 | 1.37288 | 17 |
| GO_PROTEIN_STABILIZATION | 0.00081 | 0.01194 | 0.47727 | 0.61881 | 1.19577 | 146 |
| GO_PROTEIN_SUMOYLATION | 0.00450 | 0.04342 | 0.40702 | 0.65043 | 1.22846 | 67 |
| GO_PROTEIN_TARGETING | 0.00000 | 0.00000 | | 0.66223 | 1.30361 | 358 |
| GO_PROTEIN_TARGETING_TO_MEMBRANE | 0.00000 | 0.00000 | | 0.71675 | 1.38879 | 162 |
| GO_PROTEIN_TARGETING_TO_MITOCHONDRION | 0.00113 | 0.01571 | 0.45506 | 0.65407 | 1.24648 | 83 |
| GO_PROTEIN_TRANSMEMBRANE_TRANSPORT | 0.00110 | 0.01526 | 0.45506 | 0.68173 | 1.28238 | 55 |
| GO_PROTON_TRANSMEMBRANE_TRANSPORT | 0.00000 | 0.00001 | 0.69013 | 0.71234 | 1.36309 | 91 |
| GO_PROTON_TRANSMEMBRANE_TRANSPORTER_ACTIVITY | 0.00004 | 0.00115 | 0.55733 | 0.70138 | 1.32535 | 69 |
| GO_PROTON_TRANSPORTING_TWO_SECTOR_ATPASE_COMPLEX | 0.00083 | 0.01218 | 0.47727 | 0.77254 | 1.40508 | 26 |
| GO_PROTON_TRANSPORTING_V_TYPE_ATPASE_COMPLEX | 0.00239 | 0.02716 | 0.43171 | 0.76100 | 1.37626 | 24 |
| GO_PURINE_CONTAINING_COMPOUND_BIOSYNTHETIC_PROCES | 0.00001 | 0.00040 | 0.59333 | 0.65863 | 1.26901 | 126 |
| GO_PURINE_CONTAINING_COMPOUND_METABOLIC_PROCESS | 0.00000 | 0.00000 | 0.73376 | 0.63132 | 1.23831 | 300 |
| GO_PURINE_NUCLEOSIDE_TRIPHOSPHATE_METABOLIC_PROCES | 0.00033 | 0.00582 | 0.49849 | 0.70887 | 1.32487 | 49 |
| GO_PYROPHOSPHATE_HYDROLYSIS_DRIVEN_PROTON_TRANSMI | 0.00532 | 0.04803 | 0.40702 | 0.74470 | 1.34678 | 24 |
| GO_PYRUVATE_METABOLIC_PROCESS | 0.00001 | 0.00018 | 0.61053 | 0.67194 | 1.29289 | 113 |
| GO_P_BODY | 0.00472 | 0.04480 | 0.40702 | 0.65470 | 1.23501 | 62 |
| GO_REACTIVE_OXYGEN_SPECIES_BIOSYNTHETIC_PROCESS | 0.00040 | 0.00670 | 0.49849 | 0.70271 | 1.31857 | 53 |
| GO_REACTIVE_OXYGEN_SPECIES_METABOLIC_PROCESS | 0.00021 | 0.00409 | 0.51885 | 0.62881 | 1.21472 | 144 |
| GO_REGULATION_OF_ALCOHOL_BIOSYNTHETIC_PROCESS | 0.00228 | 0.02614 | 0.43171 | 0.68438 | 1.28020 | 50 |
| GO_REGULATION_OF_ANIMAL_ORGAN_MORPHOGENESIS | 0.00000 | 0.00000 | | 0.72330 | 1.39922 | 149 |
| GO_REGULATION_OF_APOPTOTIC_SIGNALING_PATHWAY | 0.00079 | 0.01170 | 0.47727 | 0.58599 | 1.14641 | 266 |
| GO_REGULATION_OF_ATP_METABOLIC_PROCESS | 0.00000 | 0.00006 | 0.64355 | 0.70523 | 1.34777 | 89 |
| GO_REGULATION_OF_BINDING | 0.00428 | 0.04185 | 0.40702 | 0.58264 | 1.13664 | 225 |
| GO_REGULATION_OF_BLOOD_PRESSURE | 0.00290 | 0.03156 | 0.43171 | 0.65319 | 1.23428 | 69 |
| GO_REGULATION_OF_CARBOHYDRATE_CATABOLIC_PROCESS | 0.00324 | 0.03393 | 0.43171 | 0.65541 | 1.23787 | 67 |
| GO_REGULATION_OF_CELLULAR_AMIDE_METABOLIC_PROCESS | 0.00000 | 0.00000 | 0.78819 | 0.63116 | 1.24143 | 337 |
| GO_REGULATION_OF_CELLULAR_AMINE_METABOLIC_PROCESS | 0.00000 | 0.00000 | 0.74774 | 0.78955 | 1.48730 | 60 |
| GO_REGULATION_OF_CELLULAR_AMINO_ACID_METABOLIC_PR( | 0.00000 | 0.00000 | 0.76146 | 0.80341 | 1.51022 | 54 |
| GO_REGULATION_OF_CELLULAR_KETONE_METABOLIC_PROCESS | 0.00000 | 0.00000 | 0.70498 | 0.69239 | 1.33405 | 126 |
| GO_REGULATION_OF_CELLULAR_PH | 0.00442 | 0.04297 | 0.40702 | 0.65360 | 1.23436 | 68 |
| GO_REGULATION_OF_CELLULAR_PROTEIN_LOCALIZATION | 0.00228 | 0.02614 | 0.43171 | 0.56216 | 1.10830 | 402 |
| GO_REGULATION_OF_CELLULAR_RESPIRATION | 0.00428 | 0.04185 | 0.40702 | 0.77183 | 1.37897 | 19 |
| GO_REGULATION_OF_CELLULAR_RESPONSE_TO_HEAT | 0.00091 | 0.01314 | 0.47727 | 0.67360 | 1.27221 | 67 |
| GO_REGULATION_OF_CELL_CYCLE_G2_M_PHASE_TRANSITION | 0.00016 | 0.00332 | 0.51885 | 0.61554 | 1.19738 | 187 |
| GO_REGULATION_OF_CELL_CYCLE_PHASE_TRANSITION | 0.00000 | 0.00014 | 0.61053 | 0.59968 | 1.18091 | 369 |
| GO_REGULATION_OF_CHOLESTEROL_BIOSYNTHETIC_PROCESS | 0.00209 | 0.02470 | 0.43171 | 0.71476 | 1.32117 | 38 |

| | | | | | | |
|---|---|---|---|---|---|---|
| GO_REGULATION_OF_CHOLESTEROL_METABOLIC_PROCESS | 0.00113 | 0.01571 | 0.45506 | 0.69865 | 1.30177 | 47 |
| GO_REGULATION_OF_CHROMOSOME_ORGANIZATION | 0.00036 | 0.00622 | 0.49849 | 0.58896 | 1.15261 | 278 |
| GO_REGULATION_OF_CHROMOSOME_SEGREGATION | 0.00079 | 0.01170 | 0.47727 | 0.64131 | 1.22718 | 91 |
| GO_REGULATION_OF_CHROMOSOME_SEPARATION | 0.00253 | 0.02838 | 0.43171 | 0.66349 | 1.24947 | 59 |
| GO_REGULATION_OF_CYSTEINE_TYPE_ENDOPEPTIDASE_ACTIVIT | 0.00044 | 0.00719 | 0.49849 | 0.62610 | 1.20905 | 141 |
| GO_REGULATION_OF_DEFENSE_RESPONSE | 0.00006 | 0.00150 | 0.55733 | 0.58834 | 1.15824 | 356 |
| GO_REGULATION_OF_DNA_BIOSYNTHETIC_PROCESS | 0.00043 | 0.00695 | 0.49849 | 0.66709 | 1.27130 | 83 |
| GO_REGULATION_OF_DNA_METABOLIC_PROCESS | 0.00055 | 0.00863 | 0.47727 | 0.58909 | 1.15134 | 249 |
| GO_REGULATION_OF_DNA_TEMPLATED_TRANSCRIPTION_IN_RE | 0.00000 | 0.00000 | 0.71951 | 0.72410 | 1.38687 | 94 |
| GO_REGULATION_OF_GENERATION_OF_PRECURSOR_METABOLI | 0.00004 | 0.00114 | 0.55733 | 0.65536 | 1.26115 | 111 |
| GO_REGULATION_OF_GENE_SILENCING | 0.00023 | 0.00434 | 0.51885 | 0.65464 | 1.25268 | 91 |
| GO_REGULATION_OF_HEMATOPOIETIC_PROGENITOR_CELL_DIF | 0.00000 | 0.00000 | 0.73376 | 0.76273 | 1.44465 | 72 |
| GO_REGULATION_OF_HEMATOPOIETIC_STEM_CELL_DIFFERENT | 0.00000 | 0.00000 | 0.74774 | 0.78891 | 1.48817 | 62 |
| GO_REGULATION_OF_HEMOPOIESIS | 0.00390 | 0.03895 | 0.43171 | 0.57739 | 1.12909 | 258 |
| GO_REGULATION_OF_IMMUNE_RESPONSE | 0.00000 | 0.00006 | 0.64355 | 0.60032 | 1.18447 | 433 |
| GO_REGULATION_OF_INNATE_IMMUNE_RESPONSE | 0.00000 | 0.00007 | 0.62726 | 0.64536 | 1.25422 | 181 |
| GO_REGULATION_OF_INTRINSIC_APOPTOTIC_SIGNALING_PATH\ | 0.00179 | 0.02183 | 0.45506 | 0.61548 | 1.18550 | 116 |
| GO_REGULATION_OF_MITOTIC_CELL_CYCLE | 0.00000 | 0.00003 | 0.65944 | 0.58992 | 1.16508 | 499 |
| GO_REGULATION_OF_MORPHOGENESIS_OF_AN_EPITHELIUM | 0.00000 | 0.00000 | | 0.74680 | 1.43884 | 125 |
| GO_REGULATION_OF_MRNA_CATABOLIC_PROCESS | 0.00000 | 0.00000 | | 0.71126 | 1.37919 | 165 |
| GO_REGULATION_OF_MRNA_METABOLIC_PROCESS | 0.00000 | 0.00000 | | 0.67178 | 1.31458 | 276 |
| GO_REGULATION_OF_MRNA_PROCESSING | 0.00008 | 0.00186 | 0.53843 | 0.65189 | 1.25598 | 125 |
| GO_REGULATION_OF_MRNA_SPLICING_VIA_SPLICEOSOME | 0.00012 | 0.00256 | 0.53843 | 0.66950 | 1.28033 | 90 |
| GO_REGULATION_OF_MULTI_ORGANISM_PROCESS | 0.00008 | 0.00190 | 0.53843 | 0.76905 | 1.42150 | 37 |
| GO_REGULATION_OF_MUSCLE_CONTRACTION | 0.00163 | 0.02050 | 0.45506 | 0.64879 | 1.23071 | 76 |
| GO_REGULATION_OF_NUCLEAR_DIVISION | 0.00060 | 0.00923 | 0.47727 | 0.62443 | 1.20552 | 137 |
| GO_REGULATION_OF_NUCLEASE_ACTIVITY | 0.00059 | 0.00916 | 0.47727 | 0.81786 | 1.44767 | 17 |
| GO_REGULATION_OF_NUCLEOTIDE_BIOSYNTHETIC_PROCESS | 0.00302 | 0.03248 | 0.43171 | 0.78902 | 1.40337 | 18 |
| GO_REGULATION_OF_NUCLEOTIDE_METABOLIC_PROCESS | 0.00033 | 0.00588 | 0.49849 | 0.66858 | 1.26942 | 78 |
| GO_REGULATION_OF_OXIDATIVE_PHOSPHORYLATION | 0.00005 | 0.00130 | 0.55733 | 0.85284 | 1.53294 | 21 |
| GO_REGULATION_OF_PEPTIDASE_ACTIVITY | 0.00006 | 0.00152 | 0.53843 | 0.61192 | 1.19388 | 226 |
| GO_REGULATION_OF_PEPTIDE_TRANSPORT | 0.00250 | 0.02807 | 0.43171 | 0.56252 | 1.10917 | 412 |
| GO_REGULATION_OF_PH | 0.00561 | 0.04971 | 0.40702 | 0.64191 | 1.21498 | 71 |
| GO_REGULATION_OF_PLASMA_LIPOPROTEIN_PARTICLE_LEVELS | 0.00006 | 0.00145 | 0.55733 | 0.72431 | 1.36154 | 54 |
| GO_REGULATION_OF_POSTTRANSCRIPTIONAL_GENE_SILENCING | 0.00026 | 0.00486 | 0.49849 | 0.66803 | 1.27236 | 81 |
| GO_REGULATION_OF_PROTEIN_CATABOLIC_PROCESS | 0.00309 | 0.03299 | 0.43171 | 0.57360 | 1.12399 | 291 |
| GO_REGULATION_OF_PROTEIN_IMPORT | 0.00487 | 0.04559 | 0.40702 | 0.67491 | 1.25754 | 47 |
| GO_REGULATION_OF_PROTEIN_LOCALIZATION_TO_NUCLEUS | 0.00000 | 0.00013 | 0.61053 | 0.69035 | 1.32101 | 91 |
| GO_REGULATION_OF_PROTEIN_STABILITY | 0.00066 | 0.01007 | 0.47727 | 0.59308 | 1.15811 | 235 |
| GO_REGULATION_OF_PROTEOLYSIS | 0.00007 | 0.00169 | 0.53843 | 0.57923 | 1.14299 | 431 |
| GO_REGULATION_OF_REACTIVE_OXYGEN_SPECIES_BIOSYNTHET | 0.00022 | 0.00427 | 0.51885 | 0.73154 | 1.36343 | 45 |
| GO_REGULATION_OF_REACTIVE_OXYGEN_SPECIES_METABOLIC_ | 0.00088 | 0.01280 | 0.47727 | 0.63948 | 1.22592 | 99 |
| GO_REGULATION_OF_RESPONSE_TO_BIOTIC_STIMULUS | 0.00001 | 0.00039 | 0.59333 | 0.61773 | 1.20625 | 235 |
| GO_REGULATION_OF_RNA_SPLICING | 0.00011 | 0.00241 | 0.53843 | 0.64982 | 1.25198 | 125 |
| GO_REGULATION_OF_SISTER_CHROMATID_SEGREGATION | 0.00405 | 0.04011 | 0.40702 | 0.64401 | 1.22034 | 73 |
| GO_REGULATION_OF_SMALL_MOLECULE_METABOLIC_PROCESS | 0.00000 | 0.00000 | 0.77494 | 0.64342 | 1.26057 | 290 |
| GO_REGULATION_OF_SMOOTH_MUSCLE_CONTRACTION | 0.00007 | 0.00171 | 0.53843 | 0.82642 | 1.49295 | 23 |
| GO_REGULATION_OF_STEM_CELL_DIFFERENTIATION | 0.00000 | 0.00000 | 0.73376 | 0.73434 | 1.40292 | 88 |
| GO_REGULATION_OF_STEROID_METABOLIC_PROCESS | 0.00524 | 0.04770 | 0.40702 | 0.64061 | 1.21567 | 77 |
| GO_REGULATION_OF_SUPEROXIDE_METABOLIC_PROCESS | 0.00346 | 0.03548 | 0.43171 | 0.77826 | 1.39046 | 19 |
| GO_REGULATION_OF_SUPRAMOLECULAR_FIBER_ORGANIZATIO | 0.00509 | 0.04656 | 0.40702 | 0.57599 | 1.12367 | 223 |
| GO_REGULATION_OF_SYMBIOTIC_PROCESS | 0.00009 | 0.00211 | 0.53843 | 0.62950 | 1.21763 | 151 |
| GO_REGULATION_OF_TELOMERASE_ACTIVITY | 0.00087 | 0.01263 | 0.47727 | 0.72147 | 1.33426 | 39 |
| GO_REGULATION_OF_TELOMERE_MAINTENANCE | 0.00547 | 0.04923 | 0.40702 | 0.64959 | 1.22679 | 68 |
| GO_REGULATION_OF_TELOMERE_MAINTENANCE_VIA_TELOME | 0.00020 | 0.00387 | 0.51885 | 0.72043 | 1.34856 | 51 |
| GO_REGULATION_OF_TRANSCRIPTION_FROM_RNA_POLYMERA | 0.00000 | 0.00000 | 0.70498 | 0.76931 | 1.45239 | 64 |
| GO_REGULATION_OF_TRANSLATIONAL_INITIATION | 0.00023 | 0.00437 | 0.51885 | 0.68479 | 1.29399 | 69 |
| GO_REGULATION_OF_VIRAL_TRANSCRIPTION | 0.00361 | 0.03657 | 0.43171 | 0.67497 | 1.26259 | 50 |
| GO_REGULATION_OF_WNT_SIGNALING_PATHWAY | 0.00015 | 0.00312 | 0.51885 | 0.60129 | 1.17419 | 238 |
| GO_RESPIRASOME | 0.00000 | 0.00000 | 0.81404 | 0.78326 | 1.48524 | 75 |
| GO_RESPIRATORY_CHAIN_COMPLEX | 0.00000 | 0.00000 | 0.77494 | 0.79183 | 1.49492 | 64 |
| GO_RESPIRATORY_ELECTRON_TRANSPORT_CHAIN | 0.00000 | 0.00000 | | 0.78275 | 1.49591 | 89 |
| GO_RESPONSE_TO_ALCOHOL | 0.00015 | 0.00310 | 0.51885 | 0.63794 | 1.22915 | 126 |
| GO_RESPONSE_TO_ATP | 0.00087 | 0.01270 | 0.47727 | 0.81086 | 1.43528 | 17 |
| GO_RESPONSE_TO_CALCIUM_ION | 0.00124 | 0.01701 | 0.45506 | 0.65196 | 1.24071 | 80 |
| GO_RESPONSE_TO_CORTICOSTEROID | 0.00010 | 0.00220 | 0.53843 | 0.66894 | 1.27841 | 89 |
| GO_RESPONSE_TO_DRUG | 0.00000 | 0.00007 | 0.62726 | 0.63080 | 1.23070 | 220 |
| GO_RESPONSE_TO_ETHANOL | 0.00168 | 0.02098 | 0.45506 | 0.66543 | 1.25679 | 67 |
| GO_RESPONSE_TO_FIBROBLAST_GROWTH_FACTOR | 0.00357 | 0.03638 | 0.43171 | 0.63469 | 1.20976 | 84 |
| GO_RESPONSE_TO_GROWTH_FACTOR | 0.00272 | 0.02986 | 0.43171 | 0.56235 | 1.10805 | 387 |

| | | | | | | |
|---|---|---|---|---|---|---|
| GO_RESPONSE_TO_HEAT | 0.00000 | 0.00013 | 0.62726 | 0.66982 | 1.29069 | 119 |
| GO_RESPONSE_TO_HYDROGEN_PEROXIDE | 0.00532 | 0.04803 | 0.40702 | 0.62255 | 1.19054 | 90 |
| GO_RESPONSE_TO_INORGANIC_SUBSTANCE | 0.00016 | 0.00328 | 0.51885 | 0.58928 | 1.15810 | 321 |
| GO_RESPONSE_TO_INTERLEUKIN_1 | 0.00000 | 0.00000 | 0.78819 | 0.71590 | 1.37950 | 124 |
| GO_RESPONSE_TO_INTERLEUKIN_12 | 0.00036 | 0.00622 | 0.49849 | 0.74158 | 1.36769 | 36 |
| GO_RESPONSE_TO_INTERLEUKIN_4 | 0.00224 | 0.02581 | 0.43171 | 0.77088 | 1.38921 | 22 |
| GO_RESPONSE_TO_LEUKEMIA_INHIBITORY_FACTOR | 0.00076 | 0.01129 | 0.47727 | 0.66148 | 1.25288 | 72 |
| GO_RESPONSE_TO_LIPID | 0.00043 | 0.00695 | 0.49849 | 0.56669 | 1.11902 | 470 |
| GO_RESPONSE_TO_METAL_ION | 0.00143 | 0.01863 | 0.45506 | 0.59895 | 1.16592 | 199 |
| GO_RESPONSE_TO_NUTRIENT | 0.00294 | 0.03185 | 0.43171 | 0.62596 | 1.19987 | 98 |
| GO_RESPONSE_TO_ORGANOPHOSPHORUS | 0.00294 | 0.03185 | 0.43171 | 0.64154 | 1.21511 | 72 |
| GO_RESPONSE_TO_OXIDATIVE_STRESS | 0.00495 | 0.04573 | 0.40702 | 0.57010 | 1.11758 | 294 |
| GO_RESPONSE_TO_OXYGEN_LEVELS | 0.00000 | 0.00000 | | 0.67661 | 1.32155 | 244 |
| GO_RESPONSE_TO_PURINE_CONTAINING_COMPOUND | 0.00509 | 0.04656 | 0.40702 | 0.63176 | 1.20394 | 82 |
| GO_RESPONSE_TO_STEROID_HORMONE | 0.00009 | 0.00200 | 0.53843 | 0.61312 | 1.19360 | 201 |
| GO_RESPONSE_TO_TEMPERATURE_STIMULUS | 0.00002 | 0.00047 | 0.57561 | 0.64468 | 1.24782 | 153 |
| GO_RESPONSE_TO_TOXIC_SUBSTANCE | 0.00011 | 0.00235 | 0.53843 | 0.63892 | 1.23203 | 134 |
| GO_RESPONSE_TO_TUMOR_NECROSIS_FACTOR | 0.00000 | 0.00000 | 0.70498 | 0.66331 | 1.28815 | 175 |
| GO_RESPONSE_TO_VIRUS | 0.00261 | 0.02910 | 0.43171 | 0.59428 | 1.15408 | 175 |
| GO_RETINA_HOMEOSTASIS | 0.00034 | 0.00606 | 0.49849 | 0.75202 | 1.38362 | 34 |
| GO_RIBONUCLEOPROTEIN_COMPLEX_BINDING | 0.00000 | 0.00001 | 0.69013 | 0.69839 | 1.34459 | 114 |
| GO_RIBONUCLEOPROTEIN_COMPLEX_BIOGENESIS | 0.00000 | 0.00000 | | 0.68714 | 1.35448 | 393 |
| GO_RIBONUCLEOPROTEIN_COMPLEX_SUBUNIT_ORGANIZATION | 0.00000 | 0.00000 | 0.81404 | 0.69112 | 1.34183 | 173 |
| GO_RIBONUCLEOPROTEIN_GRANULE | 0.00000 | 0.00000 | 0.71951 | 0.66271 | 1.28875 | 186 |
| GO_RIBONUCLEOSIDE_TRIPHOSPHATE_BIOSYNTHETIC_PROCESS | 0.00009 | 0.00210 | 0.53843 | 0.77194 | 1.42026 | 34 |
| GO_RIBONUCLEOSIDE_TRIPHOSPHATE_METABOLIC_PROCESS | 0.00013 | 0.00280 | 0.51885 | 0.73558 | 1.37065 | 46 |
| GO_RIBOSE_PHOSPHATE_BIOSYNTHETIC_PROCESS | 0.00002 | 0.00056 | 0.57561 | 0.65814 | 1.26821 | 124 |
| GO_RIBOSE_PHOSPHATE_METABOLIC_PROCESS | 0.00000 | 0.00000 | 0.74774 | 0.63837 | 1.25001 | 285 |
| GO_RIBOSOMAL_LARGE_SUBUNIT_BIOGENESIS | 0.00001 | 0.00031 | 0.59333 | 0.72435 | 1.36807 | 67 |
| GO_RIBOSOMAL_SMALL_SUBUNIT_ASSEMBLY | 0.00405 | 0.04011 | 0.40702 | 0.78074 | 1.38865 | 18 |
| GO_RIBOSOMAL_SMALL_SUBUNIT_BINDING | 0.00532 | 0.04803 | 0.40702 | 0.77341 | 1.36899 | 17 |
| GO_RIBOSOMAL_SMALL_SUBUNIT_BIOGENESIS | 0.00005 | 0.00138 | 0.55733 | 0.70510 | 1.33163 | 68 |
| GO_RIBOSOMAL_SUBUNIT | 0.00000 | 0.00000 | | 0.76292 | 1.48089 | 172 |
| GO_RIBOSOME | 0.00000 | 0.00000 | | 0.75411 | 1.46984 | 208 |
| GO_RIBOSOME_ASSEMBLY | 0.00001 | 0.00026 | 0.59333 | 0.74485 | 1.40144 | 56 |
| GO_RIBOSOME_BIOGENESIS | 0.00000 | 0.00000 | | 0.68899 | 1.34833 | 277 |
| GO_RNA_3_END_PROCESSING | 0.00002 | 0.00046 | 0.57561 | 0.65576 | 1.26361 | 128 |
| GO_RNA_CATABOLIC_PROCESS | 0.00000 | 0.00000 | | 0.70701 | 1.39073 | 341 |
| GO_RNA_DEPENDENT_DNA_BIOSYNTHETIC_PROCESS | 0.00003 | 0.00081 | 0.57561 | 0.71669 | 1.34965 | 59 |
| GO_RNA_EXPORT_FROM_NUCLEUS | 0.00000 | 0.00016 | 0.61053 | 0.66306 | 1.27817 | 130 |
| GO_RNA_LOCALIZATION | 0.00000 | 0.00000 | 0.76146 | 0.66638 | 1.29886 | 208 |
| GO_RNA_SPLICING | 0.00000 | 0.00000 | | 0.63512 | 1.25129 | 384 |
| GO_RNA_SPLICING_VIA_TRANSESTERIFICATION_REACTIONS | 0.00000 | 0.00000 | 0.80122 | 0.64478 | 1.26586 | 311 |
| GO_ROUGH_ENDOPLASMIC_RETICULUM | 0.00014 | 0.00293 | 0.51885 | 0.70363 | 1.32507 | 59 |
| GO_RRNA_BINDING | 0.00000 | 0.00001 | 0.69013 | 0.77638 | 1.45941 | 54 |
| GO_RRNA_METABOLIC_PROCESS | 0.00000 | 0.00000 | | 0.68430 | 1.33386 | 209 |
| GO_SCF_DEPENDENT_PROTEASOMAL_UBIQUITIN_DEPENDENT_ | 0.00000 | 0.00008 | 0.62726 | 0.71957 | 1.36498 | 76 |
| GO_SECRETORY_GRANULE | 0.00000 | 0.00000 | | 0.62978 | 1.24363 | 481 |
| GO_SECRETORY_GRANULE_MEMBRANE | 0.00032 | 0.00575 | 0.49849 | 0.61375 | 1.19134 | 172 |
| GO_SERINE_TYPE_ENDOPEPTIDASE_INHIBITOR_ACTIVITY | 0.00038 | 0.00640 | 0.49849 | 0.77388 | 1.41447 | 28 |
| GO_SEXUAL_REPRODUCTION | 0.00472 | 0.04480 | 0.40702 | 0.55748 | 1.10026 | 439 |
| GO_SIDE_OF_MEMBRANE | 0.00495 | 0.04573 | 0.40702 | 0.57741 | 1.12751 | 235 |
| GO_SIGNAL_TRANSDUCTION_BY_PROTEIN_PHOSPHORYLATION | 0.00224 | 0.02581 | 0.43171 | 0.55690 | 1.09975 | 500 |
| GO_SINGLE_FERTILIZATION | 0.00000 | 0.00002 | 0.67496 | 0.76307 | 1.43842 | 61 |
| GO_SINGLE_STRANDED_RNA_BINDING | 0.00001 | 0.00042 | 0.59333 | 0.71960 | 1.35864 | 66 |
| GO_SISTER_CHROMATID_SEGREGATION | 0.00042 | 0.00689 | 0.49849 | 0.60926 | 1.18209 | 170 |
| GO_SKIN_DEVELOPMENT | 0.00076 | 0.01140 | 0.47727 | 0.61452 | 1.18945 | 153 |
| GO_SMALL_MOLECULE_BIOSYNTHETIC_PROCESS | 0.00001 | 0.00021 | 0.61053 | 0.58728 | 1.15957 | 459 |
| GO_SMALL_NUCLEAR_RIBONUCLEOPROTEIN_COMPLEX | 0.00042 | 0.00689 | 0.49849 | 0.69542 | 1.30936 | 57 |
| GO_SMALL_RIBOSOMAL_SUBUNIT | 0.00000 | 0.00001 | 0.69013 | 0.75767 | 1.43100 | 67 |
| GO_SM_LIKE_PROTEIN_FAMILY_COMPLEX | 0.00145 | 0.01879 | 0.45506 | 0.66792 | 1.26149 | 67 |
| GO_SNORNA_BINDING | 0.00038 | 0.00646 | 0.49849 | 0.76224 | 1.39614 | 29 |
| GO_SPERM_EGG_RECOGNITION | 0.00000 | 0.00002 | 0.65944 | 0.87469 | 1.59087 | 26 |
| GO_SPINDLE_MIDZONE | 0.00361 | 0.03657 | 0.43171 | 0.73316 | 1.33859 | 27 |
| GO_SPINDLE_ORGANIZATION | 0.00272 | 0.02986 | 0.43171 | 0.60076 | 1.16362 | 159 |
| GO_SPLICEOSOMAL_COMPLEX | 0.00000 | 0.00008 | 0.62726 | 0.65256 | 1.26657 | 171 |
| GO_SPLICEOSOMAL_TRI_SNRNP_COMPLEX | 0.00168 | 0.02098 | 0.45506 | 0.73474 | 1.34835 | 30 |
| GO_STEM_CELL_DIFFERENTIATION | 0.00000 | 0.00000 | 0.82666 | 0.70006 | 1.35554 | 158 |
| GO_STEROID_BIOSYNTHETIC_PROCESS | 0.00465 | 0.04430 | 0.40702 | 0.61242 | 1.17992 | 125 |
| GO_STEROID_METABOLIC_PROCESS | 0.00313 | 0.03310 | 0.43171 | 0.59525 | 1.15748 | 185 |

| | | | | | | |
|---|---|---|---|---|---|---|
| GO_STEROL_BIOSYNTHETIC_PROCESS | 0.00005 | 0.00120 | 0.55733 | 0.71876 | 1.35584 | 62 |
| GO_STEROL_METABOLIC_PROCESS | 0.00003 | 0.00073 | 0.57561 | 0.66485 | 1.27854 | 109 |
| GO_STRESS_GRANULE_ASSEMBLY | 0.00465 | 0.04430 | 0.40702 | 0.77288 | 1.38921 | 21 |
| GO_STRUCTURAL_CONSTITUENT_OF_RIBOSOME | 0.00000 | 0.00000 | | 0.76380 | 1.47839 | 153 |
| GO_STRUCTURAL_MOLECULE_ACTIVITY | 0.00000 | 0.00000 | | 0.71760 | 1.41436 | 391 |
| GO_SUPEROXIDE_METABOLIC_PROCESS | 0.00272 | 0.02986 | 0.43171 | 0.72028 | 1.32659 | 35 |
| GO_TELOMERASE_RNA_LOCALIZATION | 0.00054 | 0.00857 | 0.47727 | 0.81801 | 1.46148 | 19 |
| GO_TELOMERE_MAINTENANCE_VIA_TELOMERE_LENGTHENING | 0.00005 | 0.00138 | 0.55733 | 0.70715 | 1.33512 | 66 |
| GO_TELOMERE_ORGANIZATION | 0.00159 | 0.02016 | 0.45506 | 0.61666 | 1.18834 | 127 |
| GO_TERTIARY_GRANULE | 0.00008 | 0.00186 | 0.53843 | 0.66372 | 1.27005 | 91 |
| GO_TERTIARY_GRANULE_MEMBRANE | 0.00102 | 0.01453 | 0.45506 | 0.71940 | 1.33244 | 40 |
| GO_THREONINE_TYPE_ENDOPEPTIDASE_ACTIVITY | 0.00000 | 0.00000 | 0.71951 | 0.93456 | 1.65424 | 17 |
| GO_THREONINE_TYPE_PEPTIDASE_ACTIVITY | 0.00000 | 0.00004 | 0.65944 | 0.89807 | 1.61424 | 21 |
| GO_TISSUE_MORPHOGENESIS | 0.00000 | 0.00006 | 0.64355 | 0.59467 | 1.17150 | 380 |
| GO_TOXIN_TRANSPORT | 0.00064 | 0.00972 | 0.47727 | 0.77713 | 1.41343 | 26 |
| GO_TRANSCRIPTION_COUPLED_NUCLEOTIDE_EXCISION_REPAIR | 0.00231 | 0.02642 | 0.43171 | 0.65429 | 1.23840 | 71 |
| GO_TRANSFERRIN_TRANSPORT | 0.00141 | 0.01850 | 0.45506 | 0.75134 | 1.37179 | 27 |
| GO_TRANSLATIONAL_ELONGATION | 0.00000 | 0.00000 | | 0.75263 | 1.45028 | 124 |
| GO_TRANSLATIONAL_INITIATION | 0.00000 | 0.00000 | | 0.75139 | 1.45839 | 171 |
| GO_TRANSLATIONAL_TERMINATION | 0.00000 | 0.00000 | 0.81404 | 0.75110 | 1.44066 | 101 |
| GO_TRANSLATION_ELONGATION_FACTOR_ACTIVITY | 0.00390 | 0.03895 | 0.43171 | 0.80505 | 1.41260 | 15 |
| GO_TRANSLATION_FACTOR_ACTIVITY_RNA_BINDING | 0.00000 | 0.00000 | 0.74774 | 0.77244 | 1.46085 | 70 |
| GO_TRANSLATION_INITIATION_FACTOR_ACTIVITY | 0.00000 | 0.00011 | 0.62726 | 0.77773 | 1.44952 | 45 |
| GO_TRANSLATION_PREINITIATION_COMPLEX | 0.00342 | 0.03521 | 0.43171 | 0.79777 | 1.40694 | 16 |
| GO_TRANSLATION_REGULATOR_ACTIVITY | 0.00000 | 0.00000 | 0.83909 | 0.74000 | 1.42403 | 111 |
| GO_TRANSLATION_REGULATOR_ACTIVITY_NUCLEIC_ACID_BIND | 0.00000 | 0.00000 | 0.83909 | 0.76441 | 1.45959 | 87 |
| GO_TRANSMEMBRANE_RECEPTOR_PROTEIN_TYROSINE_KINASE | 0.00168 | 0.02098 | 0.45506 | 0.56326 | 1.11076 | 411 |
| GO_TRICARBOXYLIC_ACID_CYCLE | 0.00159 | 0.02016 | 0.45506 | 0.73558 | 1.35097 | 33 |
| GO_TRNA_TRANSPORT | 0.00342 | 0.03521 | 0.43171 | 0.71247 | 1.31220 | 35 |
| GO_TUMOR_NECROSIS_FACTOR_MEDIATED_SIGNALING_PATHW | 0.00000 | 0.00000 | 0.80122 | 0.73921 | 1.42138 | 108 |
| GO_T_CELL_RECEPTOR_SIGNALING_PATHWAY | 0.00000 | 0.00001 | 0.70498 | 0.69185 | 1.33314 | 119 |
| GO_U12_TYPE_SPLICEOSOMAL_COMPLEX | 0.00487 | 0.04559 | 0.40702 | 0.73423 | 1.33541 | 26 |
| GO_U2_SNRNP | 0.00141 | 0.01850 | 0.45506 | 0.80259 | 1.42065 | 17 |
| GO_U2_TYPE_CATALYTIC_STEP_2_SPLICEOSOME | 0.00268 | 0.02961 | 0.43171 | 0.72225 | 1.32288 | 29 |
| GO_U2_TYPE_SPLICEOSOMAL_COMPLEX | 0.00033 | 0.00595 | 0.49849 | 0.66590 | 1.26925 | 84 |
| GO_UBIQUITIN_LIKE_PROTEIN_LIGASE_BINDING | 0.00000 | 0.00007 | 0.62726 | 0.62616 | 1.22416 | 251 |
| GO_UNFOLDED_PROTEIN_BINDING | 0.00000 | 0.00001 | 0.69013 | 0.72112 | 1.37766 | 88 |
| GO_VACUOLAR_LUMEN | 0.00016 | 0.00319 | 0.51885 | 0.65416 | 1.25772 | 107 |
| GO_VESICLE_BUDDING_FROM_MEMBRANE | 0.00480 | 0.04537 | 0.40702 | 0.62411 | 1.19353 | 90 |
| GO_VESICLE_LUMEN | 0.00000 | 0.00000 | 0.77494 | 0.67028 | 1.30631 | 207 |
| GO_VESICLE_MEMBRANE | 0.00001 | 0.00026 | 0.59333 | 0.58430 | 1.15387 | 476 |
| GO_VIRAL_GENE_EXPRESSION | 0.00000 | 0.00000 | | 0.70963 | 1.37745 | 172 |
| GO_VIRAL_LIFE_CYCLE | 0.00000 | 0.00001 | 0.69013 | 0.63985 | 1.24950 | 238 |
| GO_VIRION_ASSEMBLY | 0.00457 | 0.04386 | 0.40702 | 0.69689 | 1.28528 | 36 |
| HP_ABNORMALITY_OF_ACID_BASE_HOMEOSTASIS | 0.00000 | 0.00010 | 0.62726 | 0.62283 | 1.21827 | 261 |
| HP_ABNORMALITY_OF_BLOOD_CIRCULATION | 0.00502 | 0.04608 | 0.40702 | 0.59228 | 1.14883 | 167 |
| HP_ABNORMALITY_OF_CARDIOVASCULAR_SYSTEM_ELECTROPH | 0.00011 | 0.00237 | 0.53843 | 0.60202 | 1.17758 | 261 |
| HP_ABNORMALITY_OF_SKIN_ADNEXA_PHYSIOLOGY | 0.00242 | 0.02748 | 0.43171 | 0.61294 | 1.18094 | 118 |
| HP_ABNORMALITY_OF_THE_ANKLES | 0.00313 | 0.03310 | 0.43171 | 0.65369 | 1.23418 | 63 |
| HP_ABNORMALITY_OF_THE_CEREBRAL_VASCULATURE | 0.00357 | 0.03638 | 0.43171 | 0.59764 | 1.15722 | 158 |
| HP_ABNORMALITY_OF_THE_CEREBROSPINAL_FLUID | 0.00030 | 0.00549 | 0.49849 | 0.58221 | 1.14439 | 325 |
| HP_ABNORMALITY_OF_THE_MITOCHONDRION | 0.00126 | 0.01715 | 0.45506 | 0.62149 | 1.19764 | 120 |
| HP_ABNORMALITY_OF_THE_OPTIC_DISC | 0.00198 | 0.02370 | 0.43171 | 0.56624 | 1.11727 | 432 |
| HP_ABNORMALITY_OF_THE_OPTIC_NERVE | 0.00036 | 0.00622 | 0.49849 | 0.56723 | 1.12034 | 486 |
| HP_ABNORMAL_ABDOMEN_MORPHOLOGY | 0.00089 | 0.01291 | 0.47727 | 0.56424 | 1.11364 | 441 |
| HP_ABNORMAL_ACTIVITY_OF_MITOCHONDRIAL_RESPIRATORY_ | 0.00014 | 0.00300 | 0.51885 | 0.70303 | 1.32524 | 61 |
| HP_ABNORMAL_BASAL_GANGLIA_MRI_SIGNAL_INTENSITY | 0.00179 | 0.02183 | 0.45506 | 0.71597 | 1.32046 | 36 |
| HP_ABNORMAL_BRAINSTEM_MRI_SIGNAL_INTENSITY | 0.00398 | 0.03962 | 0.40702 | 0.71528 | 1.31370 | 33 |
| HP_ABNORMAL_CELLULAR_PHENOTYPE | 0.00209 | 0.02470 | 0.43171 | 0.56937 | 1.11992 | 334 |
| HP_ABNORMAL_CIRCULATING_CARBOHYDRATE_CONCENTRATIC | 0.00194 | 0.02331 | 0.45506 | 0.69518 | 1.29539 | 46 |
| HP_ABNORMAL_CNS_MYELINATION | 0.00014 | 0.00300 | 0.51885 | 0.62318 | 1.20954 | 171 |
| HP_ABNORMAL_CSF_METABOLITE_LEVEL | 0.00001 | 0.00027 | 0.59333 | 0.70531 | 1.34409 | 82 |
| HP_ABNORMAL_ELECTROPHYSIOLOGY_OF_SINOATRIAL_NODE_ | 0.00029 | 0.00536 | 0.49849 | 0.73172 | 1.35725 | 41 |
| HP_ABNORMAL_ENZYME_COENZYME_ACTIVITY | 0.00561 | 0.04971 | 0.40702 | 0.60490 | 1.16567 | 123 |
| HP_ABNORMAL_ERYTHROCYTE_MORPHOLOGY | 0.00001 | 0.00023 | 0.61053 | 0.58835 | 1.15904 | 380 |
| HP_ABNORMAL_HEART_VALVE_PHYSIOLOGY | 0.00405 | 0.04011 | 0.40702 | 0.59004 | 1.14474 | 169 |
| HP_ABNORMAL_MITOCHONDRIA_IN_MUSCLE_TISSUE | 0.00075 | 0.01118 | 0.47727 | 0.74633 | 1.36700 | 29 |
| HP_ABNORMAL_MYELINATION | 0.00007 | 0.00169 | 0.53843 | 0.59258 | 1.16479 | 324 |
| HP_ABNORMAL_MYOCARDIUM_MORPHOLOGY | 0.00008 | 0.00192 | 0.53843 | 0.58647 | 1.15365 | 340 |
| HP_ABNORMAL_RENAL_PHYSIOLOGY | 0.00450 | 0.04342 | 0.40702 | 0.55484 | 1.09317 | 385 |

| | | | | | | |
|---|---|---|---|---|---|---|
| HP_ABNORMAL_TRICUSPID_VALVE_PHYSIOLOGY | 0.00231 | 0.02642 | 0.43171 | 0.72968 | 1.34029 | 31 |
| HP_ABNORMAL_VITREOUS_HUMOR_MORPHOLOGY | 0.00181 | 0.02201 | 0.45506 | 0.79632 | 1.40955 | 17 |
| HP_ACUTE_LEUKEMIA | 0.00013 | 0.00276 | 0.51885 | 0.68145 | 1.29219 | 75 |
| HP_ANEMIA_OF_INADEQUATE_PRODUCTION | 0.00561 | 0.04971 | 0.40702 | 0.61298 | 1.17855 | 107 |
| HP_APHASIA | 0.00213 | 0.02490 | 0.43171 | 0.72858 | 1.33704 | 30 |
| HP_BLINDNESS | 0.00018 | 0.00355 | 0.51885 | 0.61672 | 1.19768 | 175 |
| HP_BONE_MARROW_HYPERCELLULARITY | 0.00045 | 0.00725 | 0.49849 | 0.82169 | 1.46806 | 19 |
| HP_CHOREOATHETOSIS | 0.00561 | 0.04971 | 0.40702 | 0.64956 | 1.22631 | 64 |
| HP_CHRONIC_INFECTION | 0.00121 | 0.01658 | 0.45506 | 0.81916 | 1.44468 | 16 |
| HP_CONGESTIVE_HEART_FAILURE | 0.00495 | 0.04573 | 0.40702 | 0.60734 | 1.16929 | 114 |
| HP_CRANIAL_NERVE_COMPRESSION | 0.00331 | 0.03441 | 0.43171 | 0.75830 | 1.36989 | 23 |
| HP_DECREASED_ACTIVITY_OF_MITOCHONDRIAL_COMPLEX_I | 0.00283 | 0.03081 | 0.43171 | 0.71940 | 1.32496 | 35 |
| HP_DECREASED_ACTIVITY_OF_THE_PYRUVATE_DEHYDROGENAS | 0.00062 | 0.00949 | 0.47727 | 0.75390 | 1.38478 | 31 |
| HP_DEPRESSED_NASAL_RIDGE | 0.00041 | 0.00681 | 0.49849 | 0.66218 | 1.26189 | 82 |
| HP_DYSPNEA | 0.00250 | 0.02807 | 0.43171 | 0.58107 | 1.13348 | 224 |
| HP_DYSTONIA | 0.00069 | 0.01043 | 0.47727 | 0.58101 | 1.13736 | 279 |
| HP_EMOTIONAL_LABILITY | 0.00016 | 0.00321 | 0.51885 | 0.66683 | 1.27522 | 90 |
| HP_EPISODIC_ATAXIA | 0.00487 | 0.04559 | 0.40702 | 0.79934 | 1.40259 | 15 |
| HP_FASCICULATIONS | 0.00130 | 0.01761 | 0.45506 | 0.68128 | 1.28064 | 54 |
| HP_FATIGUE | 0.00042 | 0.00689 | 0.49849 | 0.59902 | 1.16764 | 212 |
| HP_FETAL_DISTRESS | 0.00179 | 0.02183 | 0.45506 | 0.72928 | 1.33973 | 32 |
| HP_FLUSHING | 0.00561 | 0.04971 | 0.40702 | 0.74782 | 1.33686 | 20 |
| HP_FOCAL_T2_HYPERINTENSE_BRAINSTEM_LESION | 0.00159 | 0.02016 | 0.45506 | 0.74970 | 1.36879 | 27 |
| HP_HEADACHE | 0.00016 | 0.00319 | 0.51885 | 0.62537 | 1.21179 | 163 |
| HP_HEPATOMEGALY | 0.00108 | 0.01507 | 0.45506 | 0.57439 | 1.12990 | 335 |
| HP_HYPERTROPHIC_CARDIOMYOPATHY | 0.00002 | 0.00050 | 0.57561 | 0.62629 | 1.21908 | 194 |
| HP_IMPAIRED_SOCIAL_INTERACTIONS | 0.00055 | 0.00863 | 0.47727 | 0.66555 | 1.26058 | 72 |
| HP_INCREASED_CSF_LACTATE | 0.00001 | 0.00034 | 0.59333 | 0.71937 | 1.35857 | 68 |
| HP_INCREASED_SERUM_LACTATE | 0.00001 | 0.00028 | 0.59333 | 0.65837 | 1.26932 | 133 |
| HP_INCREASED_SERUM_PYRUVATE | 0.00110 | 0.01526 | 0.45506 | 0.73143 | 1.35198 | 37 |
| HP_INFANTILE_ONSET | 0.00487 | 0.04559 | 0.40702 | 0.56559 | 1.10939 | 300 |
| HP_INTRACRANIAL_CYSTIC_LESION | 0.00055 | 0.00863 | 0.47727 | 0.62760 | 1.20885 | 116 |
| HP_INTRAUTERINE_GROWTH_RETARDATION | 0.00305 | 0.03271 | 0.43171 | 0.56594 | 1.11393 | 350 |
| HP_LACTIC_ACIDOSIS | 0.00018 | 0.00355 | 0.51885 | 0.63312 | 1.22071 | 131 |
| HP_LETHARGY | 0.00084 | 0.01228 | 0.47727 | 0.63361 | 1.21845 | 109 |
| HP_LEUKODYSTROPHY | 0.00029 | 0.00529 | 0.49849 | 0.66598 | 1.26915 | 82 |
| HP_LEUKOENCEPHALOPATHY | 0.00008 | 0.00190 | 0.53843 | 0.64132 | 1.23667 | 134 |
| HP_LIPID_ACCUMULATION_IN_HEPATOCYTES | 0.00220 | 0.02548 | 0.43171 | 0.62794 | 1.20159 | 91 |
| HP_LOWER_LIMB_SPASTICITY | 0.00154 | 0.01974 | 0.45506 | 0.60929 | 1.17648 | 139 |
| HP_MACROCYTIC_ANEMIA | 0.00030 | 0.00542 | 0.49849 | 0.73108 | 1.35606 | 41 |
| HP_METABOLIC_ACIDOSIS | 0.00034 | 0.00599 | 0.49849 | 0.64831 | 1.24464 | 103 |
| HP_MICRONODULAR_CIRRHOSIS | 0.00324 | 0.03393 | 0.43171 | 0.78723 | 1.40018 | 18 |
| HP_MIGRAINE | 0.00028 | 0.00515 | 0.49849 | 0.67309 | 1.27635 | 75 |
| HP_MUSCLE_ABNORMALITY_RELATED_TO_MITOCHONDRIAL_DY | 0.00209 | 0.02470 | 0.43171 | 0.68258 | 1.27573 | 49 |
| HP_MUTISM | 0.00283 | 0.03081 | 0.43171 | 0.71937 | 1.32490 | 35 |
| HP_NAUSEA_AND_VOMITING | 0.00022 | 0.00417 | 0.51885 | 0.59383 | 1.16158 | 264 |
| HP_NEOPLASM_OF_THE_ENDOCRINE_SYSTEM | 0.00495 | 0.04573 | 0.40702 | 0.63166 | 1.20308 | 81 |
| HP_OPTIC_NEUROPATHY | 0.00135 | 0.01808 | 0.45506 | 0.73837 | 1.35610 | 33 |
| HP_PACE_OF_PROGRESSION | 0.00487 | 0.04559 | 0.40702 | 0.56814 | 1.11416 | 298 |
| HP_PALLOR | 0.00036 | 0.00622 | 0.49849 | 0.64861 | 1.24463 | 102 |
| HP_PARAPLEGIA | 0.00121 | 0.01658 | 0.45506 | 0.63584 | 1.22133 | 104 |
| HP_PARAPLEGIA_PARAPARESIS | 0.00036 | 0.00621 | 0.49849 | 0.62047 | 1.20040 | 150 |
| HP_PAROXYSMAL_INVOLUNTARY_EYE_MOVEMENTS | 0.00054 | 0.00849 | 0.47727 | 0.75650 | 1.38957 | 31 |
| HP_PEDIATRIC_ONSET | 0.00561 | 0.04971 | 0.40702 | 0.55915 | 1.10116 | 371 |
| HP_PIGMENTARY_RETINOPATHY | 0.00065 | 0.00981 | 0.47727 | 0.63260 | 1.21720 | 113 |
| HP_POOR_EYE_CONTACT | 0.00017 | 0.00341 | 0.51885 | 0.71279 | 1.33749 | 53 |
| HP_POOR_HEAD_CONTROL | 0.00000 | 0.00005 | 0.64355 | 0.67819 | 1.30665 | 118 |
| HP_PROGRESSIVE | 0.00194 | 0.02331 | 0.45506 | 0.59790 | 1.16273 | 186 |
| HP_PROGRESSIVE_CEREBELLAR_ATAXIA | 0.00364 | 0.03677 | 0.43171 | 0.64788 | 1.22325 | 65 |
| HP_PROGRESSIVE_SPASTIC_PARAPLEGIA | 0.00253 | 0.02838 | 0.43171 | 0.69655 | 1.29822 | 45 |
| HP_PROXIMAL_TUBULOPATHY | 0.00141 | 0.01850 | 0.45506 | 0.72579 | 1.33535 | 34 |
| HP_PTOSIS | 0.00472 | 0.04480 | 0.40702 | 0.55582 | 1.09637 | 418 |
| HP_REDUCED_CONSCIOUSNESS_CONFUSION | 0.00007 | 0.00164 | 0.53843 | 0.61227 | 1.19295 | 205 |
| HP_REDUCED_VISUAL_ACUITY | 0.00053 | 0.00837 | 0.47727 | 0.58618 | 1.14774 | 283 |
| HP_RENAL_TUBULAR_DYSFUNCTION | 0.00013 | 0.00284 | 0.51885 | 0.68108 | 1.29150 | 75 |
| HP_RESPIRATORY_DISTRESS | 0.00108 | 0.01507 | 0.45506 | 0.62693 | 1.20762 | 115 |
| HP_RESPIRATORY_INSUFFICIENCY | 0.00000 | 0.00010 | 0.62726 | 0.60536 | 1.18998 | 326 |
| HP_SINUS_TACHYCARDIA | 0.00213 | 0.02490 | 0.43171 | 0.77203 | 1.39128 | 22 |
| HP_SPARSE_EYELASHES | 0.00150 | 0.01943 | 0.45506 | 0.71355 | 1.32161 | 40 |
| HP_SPARSE_OR_ABSENT_EYELASHES | 0.00220 | 0.02548 | 0.43171 | 0.67331 | 1.26567 | 54 |

| | | | | | | |
|---|---|---|---|---|---|---|
| HP_SPASTIC_PARAPLEGIA | 0.00098 | 0.01407 | 0.45506 | 0.64059 | 1.22712 | 96 |
| HP_THROMBOCYTOSIS | 0.00413 | 0.04065 | 0.40702 | 0.75210 | 1.35870 | 23 |
| HP_TONGUE_FASCICULATIONS | 0.00457 | 0.04386 | 0.40702 | 0.80056 | 1.40472 | 15 |
| HP_TRANSIENT_ISCHEMIC_ATTACK | 0.00480 | 0.04537 | 0.40702 | 0.74902 | 1.35809 | 25 |
| HP_VERTIGO | 0.00002 | 0.00063 | 0.57561 | 0.72114 | 1.35844 | 60 |
| HP_VESTIBULAR_DYSFUNCTION | 0.00028 | 0.00522 | 0.49849 | 0.67279 | 1.27577 | 75 |
| HP_VOMITING | 0.00072 | 0.01082 | 0.47727 | 0.61459 | 1.18792 | 147 |
| HP_WEAKNESS_DUE_TO_UPPER_MOTOR_NEURON_DYSFUNCTI | 0.00532 | 0.04803 | 0.40702 | 0.56829 | 1.11186 | 272 |
| HP_WEAKNESS_OF_MUSCLES_OF_RESPIRATION | 0.00163 | 0.02050 | 0.45506 | 0.66272 | 1.25123 | 63 |
| **Cluster 1** | | | | | | |
| pathway | pval | padj | log2err | ES | NES | size |
| GO_ACTIVATION_OF_IMMUNE_RESPONSE | 0.00000 | 0.00000 | 0.74774 | 0.65038 | 1.26999 | 235 |
| GO_ACTIVATION_OF_INNATE_IMMUNE_RESPONSE | 0.00000 | 0.00001 | 0.69013 | 0.71967 | 1.37861 | 96 |
| GO_AEROBIC_RESPIRATION | 0.00000 | 0.00000 | 0.73376 | 0.76280 | 1.44478 | 72 |
| GO_ALCOHOL_BIOSYNTHETIC_PROCESS | 0.00032 | 0.00572 | 0.49849 | 0.64679 | 1.24367 | 108 |
| GO_ALCOHOL_METABOLIC_PROCESS | 0.00364 | 0.03677 | 0.43171 | 0.58112 | 1.13424 | 228 |
| GO_AMEBOIDAL_TYPE_CELL_MIGRATION | 0.00457 | 0.04386 | 0.40702 | 0.57755 | 1.12854 | 247 |
| GO_AMIDE_BINDING | 0.00014 | 0.00297 | 0.51885 | 0.61533 | 1.19780 | 199 |
| GO_AMINE_METABOLIC_PROCESS | 0.00000 | 0.00000 | 0.76146 | 0.73732 | 1.41097 | 92 |
| GO_AMYLOID_BETA_CLEARANCE | 0.00561 | 0.04971 | 0.40702 | 0.74081 | 1.33976 | 24 |
| GO_ANAPHASE_PROMOTING_COMPLEX_DEPENDENT_CATABOL | 0.00000 | 0.00000 | 0.76146 | 0.77107 | 1.46112 | 73 |
| GO_ANATOMICAL_STRUCTURE_HOMEOSTASIS | 0.00020 | 0.00390 | 0.51885 | 0.59307 | 1.16021 | 271 |
| GO_ANIMAL_ORGAN_REGENERATION | 0.00012 | 0.00269 | 0.51885 | 0.71251 | 1.33510 | 52 |
| GO_ANTIGEN_PROCESSING_AND_PRESENTATION | 0.00000 | 0.00000 | 0.74774 | 0.68785 | 1.33064 | 149 |
| GO_ANTIGEN_PROCESSING_AND_PRESENTATION_OF_EXOGENC | 0.00000 | 0.00000 | 0.76146 | 0.79644 | 1.50237 | 62 |
| GO_ANTIGEN_PROCESSING_AND_PRESENTATION_OF_PEPTIDE_ | 0.00000 | 0.00000 | 0.74774 | 0.70107 | 1.35144 | 130 |
| GO_ANTIGEN_PROCESSING_AND_PRESENTATION_OF_PEPTIDE_ | 0.00000 | 0.00000 | 0.78819 | 0.77566 | 1.47012 | 74 |
| GO_ANTIGEN_RECEPTOR_MEDIATED_SIGNALING_PATHWAY | 0.00000 | 0.00009 | 0.62726 | 0.66362 | 1.28080 | 136 |
| GO_APICAL_PART_OF_CELL | 0.00135 | 0.01808 | 0.45506 | 0.58926 | 1.14966 | 220 |
| GO_APOPTOTIC_SIGNALING_PATHWAY | 0.00006 | 0.00152 | 0.53843 | 0.58638 | 1.15586 | 393 |
| GO_ATPASE_ACTIVITY | 0.00050 | 0.00802 | 0.47727 | 0.58216 | 1.14385 | 316 |
| GO_ATPASE_ACTIVITY_COUPLED_TO_TRANSMEMBRANE_MOVE | 0.00420 | 0.04132 | 0.40702 | 0.78005 | 1.38741 | 18 |
| GO_ATP_BIOSYNTHETIC_PROCESS | 0.00013 | 0.00276 | 0.51885 | 0.83897 | 1.50801 | 21 |
| GO_ATP_METABOLIC_PROCESS | 0.00000 | 0.00000 | | 0.72178 | 1.40814 | 219 |
| GO_ATP_SYNTHESIS_COUPLED_ELECTRON_TRANSPORT | 0.00000 | 0.00000 | | 0.80353 | 1.52425 | 76 |
| GO_AZUROPHIL_GRANULE | 0.00025 | 0.00466 | 0.49849 | 0.64781 | 1.24551 | 107 |
| GO_AZUROPHIL_GRANULE_LUMEN | 0.00405 | 0.04011 | 0.40702 | 0.65625 | 1.23793 | 62 |
| GO_BASAL_PART_OF_CELL | 0.00509 | 0.04656 | 0.40702 | 0.68515 | 1.27697 | 45 |
| GO_BASAL_PLASMA_MEMBRANE | 0.00346 | 0.03548 | 0.43171 | 0.71184 | 1.31104 | 35 |
| GO_BASEMENT_MEMBRANE | 0.00279 | 0.03056 | 0.43171 | 0.68935 | 1.28452 | 46 |
| GO_BASOLATERAL_PLASMA_MEMBRANE | 0.00011 | 0.00247 | 0.53843 | 0.63209 | 1.21993 | 136 |
| GO_BINDING_OF_SPERM_TO_ZONA_PELLUCIDA | 0.00000 | 0.00006 | 0.64355 | 0.90095 | 1.60966 | 19 |
| GO_BLOOD_MICROPARTICLE | 0.00000 | 0.00000 | 0.70498 | 0.81982 | 1.52763 | 46 |
| GO_BRUSH_BORDER | 0.00060 | 0.00923 | 0.47727 | 0.66897 | 1.26790 | 74 |
| GO_CADHERIN_BINDING | 0.00000 | 0.00000 | | 0.68759 | 1.34435 | 253 |
| GO_CALCIUM_DEPENDENT_PHOSPHOLIPID_BINDING | 0.00152 | 0.01958 | 0.45506 | 0.75021 | 1.37120 | 28 |
| GO_CALCIUM_DEPENDENT_PROTEIN_BINDING | 0.00009 | 0.00211 | 0.53843 | 0.74254 | 1.38394 | 45 |
| GO_CANONICAL_WNT_SIGNALING_PATHWAY | 0.00004 | 0.00109 | 0.55733 | 0.61740 | 1.20458 | 222 |
| GO_CARBOHYDRATE_CATABOLIC_PROCESS | 0.00135 | 0.01808 | 0.45506 | 0.61179 | 1.18186 | 144 |
| GO_CATALYTIC_STEP_2_SPLICEOSOME | 0.00001 | 0.00022 | 0.61053 | 0.70290 | 1.33765 | 80 |
| GO_CATION_TRANSMEMBRANE_TRANSPORT | 0.00327 | 0.03409 | 0.43171 | 0.56060 | 1.10531 | 406 |
| GO_CATION_TRANSMEMBRANE_TRANSPORTER_ACTIVITY | 0.00097 | 0.01387 | 0.47727 | 0.58196 | 1.13846 | 265 |
| GO_CELLULAR_AMINO_ACID_METABOLIC_PROCESS | 0.00002 | 0.00060 | 0.57561 | 0.61438 | 1.19930 | 231 |
| GO_CELLULAR_COMPONENT_DISASSEMBLY | 0.00000 | 0.00000 | | 0.64070 | 1.26342 | 408 |
| GO_CELLULAR_ION_HOMEOSTASIS | 0.00027 | 0.00493 | 0.49849 | 0.58658 | 1.15340 | 328 |
| GO_CELLULAR_IRON_ION_HOMEOSTASIS | 0.00327 | 0.03409 | 0.43171 | 0.68360 | 1.27374 | 47 |
| GO_CELLULAR_KETONE_METABOLIC_PROCESS | 0.00000 | 0.00010 | 0.62726 | 0.65358 | 1.26726 | 164 |
| GO_CELLULAR_MONOVALENT_INORGANIC_CATION_HOMEOSTA | 0.00179 | 0.02183 | 0.45506 | 0.64747 | 1.22821 | 76 |
| GO_CELLULAR_NITROGEN_COMPOUND_CATABOLIC_PROCESS | 0.00000 | 0.00000 | | 0.67915 | 1.34096 | 456 |
| GO_CELLULAR_PROTEIN_COMPLEX_DISASSEMBLY | 0.00000 | 0.00000 | | 0.69566 | 1.35184 | 179 |
| GO_CELLULAR_RESPIRATION | 0.00000 | 0.00000 | | 0.74205 | 1.43370 | 145 |
| GO_CELLULAR_RESPONSE_TO_HEAT | 0.00004 | 0.00109 | 0.55733 | 0.67186 | 1.28698 | 95 |
| GO_CELLULAR_RESPONSE_TO_HORMONE_STIMULUS | 0.00309 | 0.03299 | 0.43171 | 0.56453 | 1.11191 | 374 |
| GO_CELLULAR_RESPONSE_TO_ORGANIC_CYCLIC_COMPOUND | 0.00502 | 0.04608 | 0.40702 | 0.56512 | 1.11134 | 329 |
| GO_CELLULAR_RESPONSE_TO_OXYGEN_LEVELS | 0.00000 | 0.00000 | | 0.71461 | 1.38224 | 151 |
| GO_CELL_ACTIVATION_INVOLVED_IN_IMMUNE_RESPONSE | 0.00000 | 0.00000 | | 0.63924 | 1.26108 | 422 |
| GO_CELL_ADHESION_MEDIATOR_ACTIVITY | 0.00157 | 0.02009 | 0.45506 | 0.71837 | 1.32488 | 36 |
| GO_CELL_ADHESION_MOLECULE_BINDING | 0.00000 | 0.00000 | | 0.67109 | 1.32112 | 353 |
| GO_CELL_BODY | 0.00004 | 0.00109 | 0.55733 | 0.59973 | 1.17541 | 293 |
| GO_CELL_CELL_RECOGNITION | 0.00000 | 0.00006 | 0.64355 | 0.82001 | 1.51026 | 35 |

| | | | | | | |
|---|---|---|---|---|---|---|
| GO_CELL_CELL_SIGNALING_BY_WNT | 0.00004 | 0.00109 | 0.55733 | 0.59042 | 1.16141 | 339 |
| GO_CELL_CYCLE_G1_S_PHASE_TRANSITION | 0.00376 | 0.03783 | 0.43171 | 0.58112 | 1.13378 | 220 |
| GO_CELL_CYCLE_G2_M_PHASE_TRANSITION | 0.00024 | 0.00459 | 0.51885 | 0.59807 | 1.16746 | 231 |
| GO_CELL_CYCLE_PHASE_TRANSITION | 0.00000 | 0.00001 | 0.69013 | 0.59417 | 1.17348 | 499 |
| GO_CELL_DIVISION | 0.00003 | 0.00071 | 0.57561 | 0.58314 | 1.15094 | 449 |
| GO_CELL_KILLING | 0.00205 | 0.02440 | 0.43171 | 0.66727 | 1.25696 | 60 |
| GO_CELL_RECOGNITION | 0.00000 | 0.00000 | 0.73376 | 0.75953 | 1.43954 | 74 |
| GO_CELL_REDOX_HOMEOSTASIS | 0.00163 | 0.02050 | 0.45506 | 0.71772 | 1.32665 | 38 |
| GO_CELL_SUBSTRATE_JUNCTION | 0.00000 | 0.00000 | | 0.65742 | 1.29171 | 316 |
| GO_CELL_SURFACE | 0.00000 | 0.00000 | 0.73376 | 0.62572 | 1.23070 | 330 |
| GO_CELL_SURFACE_RECEPTOR_SIGNALING_PATHWAY_INVOLVEI | 0.00038 | 0.00640 | 0.49849 | 0.57762 | 1.13770 | 374 |
| GO_CHAPERONE_BINDING | 0.00495 | 0.04573 | 0.40702 | 0.62860 | 1.19935 | 86 |
| GO_CHAPERONE_COMPLEX | 0.00001 | 0.00028 | 0.59333 | 0.86337 | 1.55587 | 22 |
| GO_CHAPERONE_MEDIATED_PROTEIN_COMPLEX_ASSEMBLY | 0.00100 | 0.01430 | 0.45506 | 0.80787 | 1.44337 | 19 |
| GO_CHAPERONE_MEDIATED_PROTEIN_FOLDING | 0.00168 | 0.02098 | 0.45506 | 0.71110 | 1.32288 | 44 |
| GO_CHROMOSOMAL_REGION | 0.00018 | 0.00349 | 0.51885 | 0.59323 | 1.16180 | 286 |
| GO_CHROMOSOME_CENTROMERIC_REGION | 0.00172 | 0.02135 | 0.45506 | 0.60447 | 1.17248 | 168 |
| GO_CHROMOSOME_SEGREGATION | 0.00003 | 0.00087 | 0.55733 | 0.60463 | 1.18280 | 270 |
| GO_CLUSTER_OF_ACTIN_BASED_CELL_PROJECTIONS | 0.00074 | 0.01107 | 0.47727 | 0.64184 | 1.23044 | 99 |
| GO_COATED_VESICLE | 0.00213 | 0.02490 | 0.43171 | 0.59512 | 1.15846 | 199 |
| GO_COATED_VESICLE_MEMBRANE | 0.00060 | 0.00923 | 0.47727 | 0.63443 | 1.22207 | 115 |
| GO_COLLAGEN_BINDING | 0.00465 | 0.04430 | 0.40702 | 0.73471 | 1.33628 | 26 |
| GO_COLLAGEN_CONTAINING_EXTRACELLULAR_MATRIX | 0.00000 | 0.00006 | 0.64355 | 0.65358 | 1.26542 | 157 |
| GO_CONDENSED_CHROMOSOME | 0.00350 | 0.03580 | 0.43171 | 0.59394 | 1.15417 | 179 |
| GO_CONTRACTILE_FIBER | 0.00064 | 0.00972 | 0.47727 | 0.62217 | 1.19973 | 134 |
| GO_COPI_COATED_VESICLE | 0.00316 | 0.03344 | 0.43171 | 0.75688 | 1.37234 | 25 |
| GO_CORNIFICATION | 0.00027 | 0.00500 | 0.49849 | 0.77970 | 1.42512 | 28 |
| GO_CORTICAL_CYTOSKELETON | 0.00502 | 0.04608 | 0.40702 | 0.64388 | 1.21870 | 71 |
| GO_COTRANSLATIONAL_PROTEIN_TARGETING_TO_MEMBRANE | 0.00000 | 0.00000 | | 0.78223 | 1.49819 | 94 |
| GO_CYTOCHROME_COMPLEX | 0.00035 | 0.00610 | 0.49849 | 0.80796 | 1.45601 | 22 |
| GO_CYTOKINE_MEDIATED_SIGNALING_PATHWAY | 0.00000 | 0.00002 | 0.67496 | 0.60742 | 1.19735 | 392 |
| GO_CYTOPLASMIC_STRESS_GRANULE | 0.00014 | 0.00302 | 0.51885 | 0.69390 | 1.31009 | 63 |
| GO_CYTOPLASMIC_TRANSLATION | 0.00000 | 0.00001 | 0.70498 | 0.71773 | 1.37349 | 92 |
| GO_CYTOPLASMIC_TRANSLATIONAL_INITIATION | 0.00178 | 0.02182 | 0.45506 | 0.73517 | 1.35039 | 31 |
| GO_CYTOSOLIC_LARGE_RIBOSOMAL_SUBUNIT | 0.00000 | 0.00005 | 0.64355 | 0.77240 | 1.44935 | 53 |
| GO_CYTOSOLIC_RIBOSOME | 0.00000 | 0.00000 | | 0.77483 | 1.48437 | 97 |
| GO_CYTOSOLIC_SMALL_RIBOSOMAL_SUBUNIT | 0.00000 | 0.00010 | 0.62726 | 0.79384 | 1.47033 | 40 |
| GO_DETECTION_OF_STIMULUS | 0.00020 | 0.00393 | 0.51885 | 0.65208 | 1.25384 | 108 |
| GO_DETOXIFICATION | 0.00413 | 0.04065 | 0.40702 | 0.65399 | 1.23684 | 70 |
| GO_DEVELOPMENT_OF_PRIMARY_FEMALE_SEXUAL_CHARACTE | 0.00179 | 0.02183 | 0.45506 | 0.68410 | 1.28057 | 51 |
| GO_DICARBOXYLIC_ACID_METABOLIC_PROCESS | 0.00261 | 0.02910 | 0.43171 | 0.65541 | 1.23747 | 65 |
| GO_DISORDERED_DOMAIN_SPECIFIC_BINDING | 0.00051 | 0.00812 | 0.47727 | 0.76867 | 1.40343 | 27 |
| GO_DISULFIDE_OXIDOREDUCTASE_ACTIVITY | 0.00435 | 0.04238 | 0.40702 | 0.71445 | 1.31232 | 31 |
| GO_DNA_BIOSYNTHETIC_PROCESS | 0.00017 | 0.00334 | 0.51885 | 0.62540 | 1.20833 | 145 |
| GO_DNA_CONFORMATION_CHANGE | 0.00327 | 0.03409 | 0.43171 | 0.57979 | 1.13178 | 231 |
| GO_DNA_DAMAGE_RESPONSE_DETECTION_OF_DNA_DAMAGE | 0.00305 | 0.03271 | 0.43171 | 0.71625 | 1.31916 | 35 |
| GO_DNA_GEOMETRIC_CHANGE | 0.00141 | 0.01850 | 0.45506 | 0.62642 | 1.20356 | 105 |
| GO_DOUBLE_STRANDED_RNA_BINDING | 0.00039 | 0.00658 | 0.49849 | 0.69158 | 1.30365 | 61 |
| GO_DRUG_BINDING | 0.00450 | 0.04342 | 0.40702 | 0.66245 | 1.24130 | 52 |
| GO_ELECTRON_TRANSFER_ACTIVITY | 0.00000 | 0.00001 | 0.67496 | 0.72917 | 1.38765 | 80 |
| GO_ELECTRON_TRANSPORT_CHAIN | 0.00000 | 0.00000 | | 0.73336 | 1.41392 | 133 |
| GO_EMBRYO_IMPLANTATION | 0.00044 | 0.00706 | 0.49849 | 0.75968 | 1.39146 | 29 |
| GO_ENDOCYTIC_VESICLE_MEMBRANE | 0.00524 | 0.04770 | 0.40702 | 0.62563 | 1.19564 | 89 |
| GO_ENDOPEPTIDASE_ACTIVITY | 0.00065 | 0.00981 | 0.47727 | 0.60123 | 1.16998 | 196 |
| GO_ENDOPEPTIDASE_COMPLEX | 0.00000 | 0.00001 | 0.70498 | 0.76593 | 1.44609 | 63 |
| GO_ENDOPEPTIDASE_REGULATOR_ACTIVITY | 0.00006 | 0.00145 | 0.55733 | 0.70425 | 1.33001 | 68 |
| GO_ENDOPLASMIC_RETICULUM_GOLGI_INTERMEDIATE_COMPA | 0.00055 | 0.00863 | 0.47727 | 0.64758 | 1.24050 | 96 |
| GO_ENDOPLASMIC_RETICULUM_GOLGI_INTERMEDIATE_COMPA | 0.00148 | 0.01923 | 0.45506 | 0.68944 | 1.29056 | 51 |
| GO_ENDOPLASMIC_RETICULUM_TO_GOLGI_VESICLE_MEDIATED | 0.00532 | 0.04803 | 0.40702 | 0.59194 | 1.14697 | 162 |
| GO_ENERGY_DERIVATION_BY_OXIDATION_OF_ORGANIC_COMP | 0.00000 | 0.00000 | | 0.69995 | 1.36305 | 202 |
| GO_ENZYME_INHIBITOR_ACTIVITY | 0.00246 | 0.02780 | 0.43171 | 0.58913 | 1.14643 | 191 |
| GO_EPIDERMAL_CELL_DIFFERENTIATION | 0.00216 | 0.02519 | 0.43171 | 0.62667 | 1.20589 | 112 |
| GO_EPITHELIAL_CELL_DIFFERENTIATION | 0.00054 | 0.00857 | 0.47727 | 0.57819 | 1.13725 | 334 |
| GO_ESTABLISHMENT_OF_ORGANELLE_LOCALIZATION | 0.00122 | 0.01679 | 0.45506 | 0.57737 | 1.13489 | 323 |
| GO_ESTABLISHMENT_OF_PROTEIN_LOCALIZATION_TO_CHROMO | 0.00016 | 0.00321 | 0.51885 | 0.81363 | 1.46985 | 23 |
| GO_ESTABLISHMENT_OF_PROTEIN_LOCALIZATION_TO_ENDOPL/ | 0.00000 | 0.00000 | | 0.77241 | 1.48479 | 106 |
| GO_ESTABLISHMENT_OF_PROTEIN_LOCALIZATION_TO_MEMBR/ | 0.00000 | 0.00000 | | 0.67484 | 1.31991 | 260 |
| GO_ESTABLISHMENT_OF_PROTEIN_LOCALIZATION_TO_ORGANE | 0.00000 | 0.00000 | | 0.65856 | 1.30056 | 461 |
| GO_ESTABLISHMENT_OF_PROTEIN_LOCALIZATION_TO_TELOME | 0.00000 | 0.00006 | 0.64355 | 0.90447 | 1.60870 | 18 |
| GO_ESTABLISHMENT_OF_RNA_LOCALIZATION | 0.00000 | 0.00005 | 0.64355 | 0.64271 | 1.24830 | 176 |

| | | | | | | |
|---|---|---|---|---|---|---|
| GO_ESTABLISHMENT_OF_TISSUE_POLARITY | 0.00000 | 0.00000 | 0.81404 | 0.75693 | 1.44999 | 96 |
| GO_EUKARYOTIC_TRANSLATION_INITIATION_FACTOR_3_COMPL | 0.00143 | 0.01863 | 0.45506 | 0.83263 | 1.46100 | 15 |
| GO_EXTERNAL_SIDE_OF_PLASMA_MEMBRANE | 0.00159 | 0.02016 | 0.45506 | 0.61775 | 1.19022 | 118 |
| GO_EXTRACELLULAR_MATRIX | 0.00042 | 0.00695 | 0.49849 | 0.61138 | 1.18938 | 189 |
| GO_FATTY_ACID_DERIVATIVE_METABOLIC_PROCESS | 0.00043 | 0.00695 | 0.49849 | 0.66266 | 1.26212 | 81 |
| GO_FATTY_ACYL_COA_METABOLIC_PROCESS | 0.00502 | 0.04608 | 0.40702 | 0.72264 | 1.32082 | 28 |
| GO_FC_EPSILON_RECEPTOR_SIGNALING_PATHWAY | 0.00000 | 0.00000 | 0.71951 | 0.72370 | 1.38610 | 94 |
| GO_FC_RECEPTOR_SIGNALING_PATHWAY | 0.00000 | 0.00000 | 0.70498 | 0.68815 | 1.32879 | 140 |
| GO_FERTILIZATION | 0.00000 | 0.00005 | 0.64355 | 0.73257 | 1.38752 | 72 |
| GO_FIBRILLAR_CENTER | 0.00246 | 0.02780 | 0.43171 | 0.62638 | 1.19990 | 96 |
| GO_FIBROBLAST_GROWTH_FACTOR_RECEPTOR_SIGNALING_PAT | 0.00495 | 0.04573 | 0.40702 | 0.64888 | 1.22552 | 67 |
| GO_FIBROBLAST_PROLIFERATION | 0.00032 | 0.00575 | 0.49849 | 0.70676 | 1.32618 | 53 |
| GO_FICOLIN_1_RICH_GRANULE | 0.00000 | 0.00000 | 0.70498 | 0.71544 | 1.37138 | 98 |
| GO_GABA_ERGIC_SYNAPSE | 0.00022 | 0.00427 | 0.51885 | 0.76430 | 1.40406 | 32 |
| GO_GENERATION_OF_PRECURSOR_METABOLITES_AND_ENERGY | 0.00000 | 0.00000 | | 0.67527 | 1.32968 | 368 |
| GO_GENE_SILENCING | 0.00019 | 0.00376 | 0.51885 | 0.61625 | 1.19788 | 184 |
| GO_GLUCOSE_CATABOLIC_PROCESS | 0.00495 | 0.04573 | 0.40702 | 0.72314 | 1.32174 | 28 |
| GO_GLUTAMATERGIC_SYNAPSE | 0.00135 | 0.01808 | 0.45506 | 0.60052 | 1.16783 | 186 |
| GO_GLYCOLYTIC_PROCESS_THROUGH_FRUCTOSE_6_PHOSPHATE | 0.00554 | 0.04968 | 0.40702 | 0.74156 | 1.34111 | 24 |
| GO_GOLGI_ASSOCIATED_VESICLE | 0.00085 | 0.01239 | 0.47727 | 0.62294 | 1.20037 | 129 |
| GO_GOLGI_ASSOCIATED_VESICLE_MEMBRANE | 0.00313 | 0.03310 | 0.43171 | 0.64208 | 1.21753 | 75 |
| GO_GTPASE_ACTIVITY | 0.00004 | 0.00098 | 0.55733 | 0.61546 | 1.19974 | 210 |
| GO_GUANYL_NUCLEOTIDE_BINDING | 0.00005 | 0.00126 | 0.55733 | 0.60899 | 1.19065 | 252 |
| GO_HEAT_SHOCK_PROTEIN_BINDING | 0.00001 | 0.00034 | 0.59333 | 0.68258 | 1.30633 | 93 |
| GO_HEMATOPOIETIC_PROGENITOR_CELL_DIFFERENTIATION | 0.00000 | 0.00000 | 0.71951 | 0.70099 | 1.35150 | 121 |
| GO_HEMATOPOIETIC_STEM_CELL_DIFFERENTIATION | 0.00000 | 0.00000 | 0.76146 | 0.77246 | 1.46307 | 72 |
| GO_HISTONE_MRNA_METABOLIC_PROCESS | 0.00265 | 0.02936 | 0.43171 | 0.76832 | 1.38459 | 22 |
| GO_HOST_CELLULAR_COMPONENT | 0.00043 | 0.00695 | 0.49849 | 0.69729 | 1.31074 | 54 |
| GO_IMMUNE_RESPONSE_REGULATING_SIGNALING_PATHWAY | 0.00000 | 0.00003 | 0.65944 | 0.63620 | 1.24112 | 225 |
| GO_IMPORT_INTO_CELL | 0.00428 | 0.04185 | 0.40702 | 0.61709 | 1.18655 | 108 |
| GO_IMPORT_INTO_NUCLEUS | 0.00039 | 0.00652 | 0.49849 | 0.62707 | 1.20905 | 131 |
| GO_INNATE_IMMUNE_RESPONSE | 0.00000 | 0.00003 | 0.65944 | 0.59893 | 1.18089 | 406 |
| GO_INNATE_IMMUNE_RESPONSE_ACTIVATING_SIGNAL_TRANSD | 0.00000 | 0.00001 | 0.69013 | 0.73348 | 1.39585 | 80 |
| GO_INNER_MITOCHONDRIAL_MEMBRANE_PROTEIN_COMPLEX | 0.00000 | 0.00000 | | 0.76576 | 1.46666 | 94 |
| GO_INORGANIC_ION_TRANSMEMBRANE_TRANSPORT | 0.00141 | 0.01850 | 0.45506 | 0.56404 | 1.11185 | 401 |
| GO_INTERACTION_WITH_HOST | 0.00139 | 0.01850 | 0.45506 | 0.60855 | 1.17710 | 151 |
| GO_INTERLEUKIN_1_MEDIATED_SIGNALING_PATHWAY | 0.00000 | 0.00000 | 0.77494 | 0.75654 | 1.44176 | 83 |
| GO_INTRAMOLECULAR_OXIDOREDUCTASE_ACTIVITY | 0.00051 | 0.00812 | 0.47727 | 0.74506 | 1.37081 | 34 |
| GO_INTRINSIC_APOPTOTIC_SIGNALING_PATHWAY | 0.00156 | 0.01993 | 0.45506 | 0.59427 | 1.15817 | 207 |
| GO_ION_HOMEOSTASIS | 0.00170 | 0.02117 | 0.45506 | 0.56164 | 1.10717 | 403 |
| GO_ION_TRANSMEMBRANE_TRANSPORTER_ACTIVITY | 0.00106 | 0.01495 | 0.45506 | 0.56696 | 1.11734 | 390 |
| GO_IRON_ION_HOMEOSTASIS | 0.00361 | 0.03657 | 0.43171 | 0.65837 | 1.23983 | 59 |
| GO_IRON_ION_TRANSPORT | 0.00054 | 0.00857 | 0.47727 | 0.69556 | 1.30334 | 52 |
| GO_ISOMERASE_ACTIVITY | 0.00413 | 0.04065 | 0.40702 | 0.61065 | 1.17496 | 113 |
| GO_KERATINIZATION | 0.00038 | 0.00640 | 0.49849 | 0.75690 | 1.39013 | 33 |
| GO_KILLING_OF_CELLS_OF_OTHER_ORGANISM | 0.00495 | 0.04573 | 0.40702 | 0.78614 | 1.38643 | 16 |
| GO_LARGE_RIBOSOMAL_SUBUNIT | 0.00000 | 0.00000 | | 0.77656 | 1.49319 | 108 |
| GO_LATERAL_PLASMA_MEMBRANE | 0.00524 | 0.04770 | 0.40702 | 0.67385 | 1.25557 | 47 |
| GO_LEUKOCYTE_MEDIATED_IMMUNITY | 0.00000 | 0.00000 | | 0.64010 | 1.26331 | 447 |
| GO_LIPOPROTEIN_PARTICLE_RECEPTOR_BINDING | 0.00126 | 0.01715 | 0.45506 | 0.80503 | 1.42497 | 17 |
| GO_LIVER_REGENERATION | 0.00093 | 0.01337 | 0.47727 | 0.78442 | 1.41708 | 23 |
| GO_LOW_DENSITY_LIPOPROTEIN_RECEPTOR_PARTICLE_METABO | 0.00327 | 0.03409 | 0.43171 | 0.78520 | 1.39657 | 18 |
| GO_LYASE_ACTIVITY | 0.00152 | 0.01958 | 0.45506 | 0.61769 | 1.19079 | 122 |
| GO_LYSOSOMAL_LUMEN | 0.00143 | 0.01863 | 0.45506 | 0.67371 | 1.26872 | 59 |
| GO_MAINTENANCE_OF_CELL_NUMBER | 0.00435 | 0.04238 | 0.40702 | 0.60761 | 1.17073 | 117 |
| GO_MATURATION_OF_SSU_RRNA | 0.00126 | 0.01715 | 0.45506 | 0.69615 | 1.29711 | 47 |
| GO_MATURATION_OF_SSU_RRNA_FROM_TRICISTRONIC_RRNA_ | 0.00554 | 0.04968 | 0.40702 | 0.70205 | 1.29301 | 35 |
| GO_MEMBRANE_REGION | 0.00000 | 0.00003 | 0.65944 | 0.64496 | 1.25542 | 194 |
| GO_METALLOPEPTIDASE_ACTIVITY | 0.00487 | 0.04559 | 0.40702 | 0.62714 | 1.19577 | 85 |
| GO_METAL_ION_HOMEOSTASIS | 0.00032 | 0.00575 | 0.49849 | 0.58816 | 1.15376 | 301 |
| GO_METAL_ION_TRANSPORT | 0.00390 | 0.03895 | 0.43171 | 0.55877 | 1.10227 | 419 |
| GO_METAPHASE_ANAPHASE_TRANSITION_OF_CELL_CYCLE | 0.00320 | 0.03371 | 0.43171 | 0.66563 | 1.25122 | 54 |
| GO_MICROTUBULE_CYTOSKELETON_ORGANIZATION_INVOLVED_ | 0.00220 | 0.02548 | 0.43171 | 0.61295 | 1.18118 | 127 |
| GO_MICROVILLUS | 0.00168 | 0.02098 | 0.45506 | 0.68128 | 1.28184 | 56 |
| GO_MIDBODY | 0.00561 | 0.04971 | 0.40702 | 0.59866 | 1.15606 | 141 |
| GO_MITOCHONDRIAL_ELECTRON_TRANSPORT_NADH_TO_UBIQ | 0.00000 | 0.00006 | 0.64355 | 0.79447 | 1.47799 | 44 |
| GO_MITOCHONDRIAL_GENE_EXPRESSION | 0.00000 | 0.00000 | 0.78819 | 0.70230 | 1.35844 | 151 |
| GO_MITOCHONDRIAL_LARGE_RIBOSOMAL_SUBUNIT | 0.00000 | 0.00000 | 0.71951 | 0.79015 | 1.48529 | 54 |
| GO_MITOCHONDRIAL_MATRIX | 0.00000 | 0.00000 | 0.82666 | 0.63141 | 1.24412 | 387 |
| GO_MITOCHONDRIAL_MEMBRANE_ORGANIZATION | 0.00050 | 0.00802 | 0.47727 | 0.64578 | 1.23686 | 94 |

| | | | | | | |
|---|---|---|---|---|---|---|
| GO_MITOCHONDRIAL_PROTEIN_COMPLEX | 0.00000 | 0.00000 | | 0.75011 | 1.46215 | 212 |
| GO_MITOCHONDRIAL_RESPIRATORY_CHAIN_COMPLEX_ASSEMB | 0.00001 | 0.00033 | 0.59333 | 0.69650 | 1.32804 | 85 |
| GO_MITOCHONDRIAL_SMALL_RIBOSOMAL_SUBUNIT | 0.00119 | 0.01640 | 0.45506 | 0.75384 | 1.37785 | 28 |
| GO_MITOCHONDRIAL_TRANSLATION | 0.00000 | 0.00000 | 0.80122 | 0.71435 | 1.37651 | 128 |
| GO_MITOCHONDRIAL_TRANSLATIONAL_TERMINATION | 0.00000 | 0.00000 | 0.81404 | 0.75760 | 1.44657 | 87 |
| GO_MITOCHONDRIAL_TRANSMEMBRANE_TRANSPORT | 0.00213 | 0.02490 | 0.43171 | 0.66300 | 1.25387 | 70 |
| GO_MITOCHONDRIAL_TRANSPORT | 0.00010 | 0.00227 | 0.53843 | 0.61119 | 1.19060 | 203 |
| GO_MITOCHONDRION_ORGANIZATION | 0.00000 | 0.00000 | 0.74774 | 0.61590 | 1.21471 | 416 |
| GO_MITOTIC_NUCLEAR_DIVISION | 0.00008 | 0.00182 | 0.53843 | 0.60435 | 1.17901 | 223 |
| GO_MITOTIC_SISTER_CHROMATID_SEGREGATION | 0.00014 | 0.00302 | 0.51885 | 0.62652 | 1.21048 | 145 |
| GO_MITOTIC_SPINDLE_ORGANIZATION | 0.00078 | 0.01164 | 0.47727 | 0.63943 | 1.22821 | 104 |
| GO_MODIFIED_AMINO_ACID_BINDING | 0.00002 | 0.00048 | 0.57561 | 0.74200 | 1.38679 | 49 |
| GO_MODULATION_BY_HOST_OF_VIRAL_PROCESS | 0.00216 | 0.02519 | 0.43171 | 0.77232 | 1.38066 | 20 |
| GO_MONOCARBOXYLIC_ACID_METABOLIC_PROCESS | 0.00012 | 0.00265 | 0.53843 | 0.57846 | 1.14053 | 406 |
| GO_MONOSACCHARIDE_BIOSYNTHETIC_PROCESS | 0.00305 | 0.03271 | 0.43171 | 0.65908 | 1.24472 | 68 |
| GO_MONOVALENT_INORGANIC_CATION_HOMEOSTASIS | 0.00108 | 0.01507 | 0.45506 | 0.63643 | 1.21783 | 91 |
| GO_MONOVALENT_INORGANIC_CATION_TRANSMEMBRANE_TR | 0.00001 | 0.00026 | 0.59333 | 0.65186 | 1.25996 | 147 |
| GO_MONOVALENT_INORGANIC_CATION_TRANSPORT | 0.00008 | 0.00178 | 0.53843 | 0.60205 | 1.17659 | 248 |
| GO_MORPHOGENESIS_OF_AN_EPITHELIUM | 0.00000 | 0.00009 | 0.62726 | 0.60570 | 1.19137 | 334 |
| GO_MORPHOGENESIS_OF_A_POLARIZED_EPITHELIUM | 0.00000 | 0.00000 | 0.83909 | 0.74384 | 1.43057 | 110 |
| GO_MOVEMENT_IN_HOST_ENVIRONMENT | 0.00268 | 0.02961 | 0.43171 | 0.61282 | 1.18078 | 117 |
| GO_MRNA_3_END_PROCESSING | 0.00087 | 0.01270 | 0.47727 | 0.64010 | 1.22502 | 93 |
| GO_MRNA_5_UTR_BINDING | 0.00002 | 0.00063 | 0.57561 | 0.83759 | 1.51313 | 23 |
| GO_MRNA_BINDING | 0.00000 | 0.00000 | | 0.67496 | 1.31810 | 239 |
| GO_MRNA_EXPORT_FROM_NUCLEUS | 0.00006 | 0.00159 | 0.53843 | 0.66290 | 1.27265 | 103 |
| GO_MRNA_PROCESSING | 0.00000 | 0.00000 | | 0.62959 | 1.24255 | 437 |
| GO_MRNA_TRANSPORT | 0.00007 | 0.00171 | 0.53843 | 0.64103 | 1.23563 | 132 |
| GO_MULTI_MULTICELLULAR_ORGANISM_PROCESS | 0.00041 | 0.00687 | 0.49849 | 0.63087 | 1.21515 | 116 |
| GO_MULTI_ORGANISM_LOCALIZATION | 0.00017 | 0.00334 | 0.51885 | 0.71077 | 1.33608 | 54 |
| GO_MUSCLE_CONTRACTION | 0.00335 | 0.03468 | 0.43171 | 0.59925 | 1.16199 | 165 |
| GO_MUSCLE_FILAMENT_SLIDING | 0.00324 | 0.03393 | 0.43171 | 0.77980 | 1.39322 | 19 |
| GO_MYELOID_LEUKOCYTE_ACTIVATION | 0.00000 | 0.00000 | | 0.65288 | 1.28712 | 395 |
| GO_MYELOID_LEUKOCYTE_MEDIATED_IMMUNITY | 0.00000 | 0.00000 | | 0.65734 | 1.29408 | 356 |
| GO_NADH_DEHYDROGENASE_ACTIVITY | 0.00001 | 0.00019 | 0.61053 | 0.79360 | 1.46691 | 38 |
| GO_NADH_DEHYDROGENASE_COMPLEX | 0.00001 | 0.00028 | 0.59333 | 0.77749 | 1.44215 | 41 |
| GO_NADH_DEHYDROGENASE_COMPLEX_ASSEMBLY | 0.00001 | 0.00035 | 0.59333 | 0.74044 | 1.39281 | 55 |
| GO_NADH_METABOLIC_PROCESS | 0.00034 | 0.00599 | 0.49849 | 0.75773 | 1.39166 | 33 |
| GO_NAD_METABOLIC_PROCESS | 0.00435 | 0.04238 | 0.40702 | 0.70570 | 1.30442 | 37 |
| GO_NCRNA_EXPORT_FROM_NUCLEUS | 0.00205 | 0.02440 | 0.43171 | 0.71473 | 1.31817 | 36 |
| GO_NCRNA_METABOLIC_PROCESS | 0.00000 | 0.00000 | 0.70498 | 0.61139 | 1.20509 | 399 |
| GO_NCRNA_PROCESSING | 0.00000 | 0.00000 | 0.73376 | 0.62238 | 1.22425 | 341 |
| GO_NEGATIVE_REGULATION_OF_APOPTOTIC_SIGNALING_PATH\ | 0.00364 | 0.03677 | 0.43171 | 0.59958 | 1.15998 | 150 |
| GO_NEGATIVE_REGULATION_OF_ATP_METABOLIC_PROCESS | 0.00302 | 0.03248 | 0.43171 | 0.78586 | 1.39103 | 17 |
| GO_NEGATIVE_REGULATION_OF_CANONICAL_WNT_SIGNALING | 0.00000 | 0.00000 | 0.71951 | 0.69629 | 1.34180 | 123 |
| GO_NEGATIVE_REGULATION_OF_CATALYTIC_ACTIVITY | 0.00145 | 0.01879 | 0.45506 | 0.56238 | 1.11005 | 450 |
| GO_NEGATIVE_REGULATION_OF_CELLULAR_AMIDE_METABOLIC | 0.00268 | 0.02961 | 0.43171 | 0.61074 | 1.17686 | 124 |
| GO_NEGATIVE_REGULATION_OF_CELLULAR_COMPONENT_ORG | 0.00320 | 0.03371 | 0.43171 | 0.55470 | 1.09534 | 470 |
| GO_NEGATIVE_REGULATION_OF_CELL_CYCLE | 0.00023 | 0.00445 | 0.51885 | 0.57348 | 1.13228 | 455 |
| GO_NEGATIVE_REGULATION_OF_CELL_CYCLE_G2_M_PHASE_TR | 0.00000 | 0.00007 | 0.62726 | 0.70549 | 1.34606 | 86 |
| GO_NEGATIVE_REGULATION_OF_CELL_CYCLE_PHASE_TRANSITI( | 0.00000 | 0.00000 | 0.70498 | 0.65532 | 1.27560 | 194 |
| GO_NEGATIVE_REGULATION_OF_CELL_CYCLE_PROCESS | 0.00000 | 0.00005 | 0.64355 | 0.62725 | 1.22670 | 259 |
| GO_NEGATIVE_REGULATION_OF_HYDROLASE_ACTIVITY | 0.00216 | 0.02519 | 0.43171 | 0.58306 | 1.13858 | 238 |
| GO_NEGATIVE_REGULATION_OF_INTERLEUKIN_1_PRODUCTION | 0.00205 | 0.02440 | 0.43171 | 0.82297 | 1.44404 | 15 |
| GO_NEGATIVE_REGULATION_OF_MITOTIC_CELL_CYCLE | 0.00000 | 0.00010 | 0.62726 | 0.62191 | 1.21541 | 248 |
| GO_NEGATIVE_REGULATION_OF_MRNA_METABOLIC_PROCESS | 0.00213 | 0.02490 | 0.43171 | 0.66731 | 1.25879 | 62 |
| GO_NEGATIVE_REGULATION_OF_MRNA_SPLICING_VIA_SPLICEO | 0.00339 | 0.03501 | 0.43171 | 0.76309 | 1.36417 | 20 |
| GO_NEGATIVE_REGULATION_OF_NUCLEAR_DIVISION | 0.00335 | 0.03468 | 0.43171 | 0.67724 | 1.26685 | 50 |
| GO_NEGATIVE_REGULATION_OF_ORGANELLE_ORGANIZATION | 0.00487 | 0.04559 | 0.40702 | 0.57038 | 1.11722 | 288 |
| GO_NEGATIVE_REGULATION_OF_PEPTIDASE_ACTIVITY | 0.00036 | 0.00621 | 0.49849 | 0.64044 | 1.23159 | 109 |
| GO_NEGATIVE_REGULATION_OF_PROTEIN_CATABOLIC_PROCESS | 0.00342 | 0.03521 | 0.43171 | 0.62320 | 1.19268 | 93 |
| GO_NEGATIVE_REGULATION_OF_PROTEOLYSIS | 0.00205 | 0.02440 | 0.43171 | 0.60172 | 1.16902 | 178 |
| GO_NEGATIVE_REGULATION_OF_REACTIVE_OXYGEN_SPECIES_B | 0.00036 | 0.00621 | 0.49849 | 0.83108 | 1.47818 | 18 |
| GO_NEGATIVE_REGULATION_OF_REPRODUCTIVE_PROCESS | 0.00457 | 0.04386 | 0.40702 | 0.74845 | 1.35356 | 24 |
| GO_NEGATIVE_REGULATION_OF_RNA_SPLICING | 0.00183 | 0.02219 | 0.45506 | 0.76824 | 1.38935 | 24 |
| GO_NEGATIVE_REGULATION_OF_WNT_SIGNALING_PATHWAY | 0.00000 | 0.00002 | 0.67496 | 0.67731 | 1.30721 | 136 |
| GO_NEURAL_CREST_CELL_MIGRATION | 0.00265 | 0.02936 | 0.43171 | 0.73967 | 1.35049 | 27 |
| GO_NEURON_RECOGNITION | 0.00390 | 0.03895 | 0.43171 | 0.71862 | 1.31998 | 31 |
| GO_NIK_NF_KAPPAB_SIGNALING | 0.00000 | 0.00000 | 0.77494 | 0.72329 | 1.39253 | 114 |
| GO_NON_CANONICAL_WNT_SIGNALING_PATHWAY | 0.00000 | 0.00000 | | 0.75756 | 1.45654 | 107 |
| GO_NUCLEAR_CHROMOSOME_SEGREGATION | 0.00098 | 0.01407 | 0.45506 | 0.59156 | 1.15357 | 215 |

| | | | | | | |
|---|---|---|---|---|---|---|
| GO_NUCLEAR_ENVELOPE | 0.00005 | 0.00131 | 0.55733 | 0.59109 | 1.16270 | 341 |
| GO_NUCLEAR_EXPORT | 0.00000 | 0.00004 | 0.65944 | 0.64915 | 1.26157 | 181 |
| GO_NUCLEAR_MATRIX | 0.00502 | 0.04608 | 0.40702 | 0.62777 | 1.19868 | 87 |
| GO_NUCLEAR_MEMBRANE | 0.00185 | 0.02237 | 0.45506 | 0.58925 | 1.14853 | 211 |
| GO_NUCLEAR_PERIPHERY | 0.00043 | 0.00701 | 0.49849 | 0.64264 | 1.23558 | 107 |
| GO_NUCLEAR_PORE | 0.00561 | 0.04971 | 0.40702 | 0.64876 | 1.22522 | 68 |
| GO_NUCLEAR_SPECK | 0.00015 | 0.00312 | 0.51885 | 0.59159 | 1.16089 | 306 |
| GO_NUCLEAR_TRANSCRIBED_MRNA_CATABOLIC_PROCESS | 0.00000 | 0.00000 | | 0.70362 | 1.36820 | 185 |
| GO_NUCLEAR_TRANSCRIBED_MRNA_CATABOLIC_PROCESS_NON | 0.00000 | 0.00000 | | 0.74239 | 1.42931 | 114 |
| GO_NUCLEAR_TRANSPORT | 0.00000 | 0.00000 | 0.71951 | 0.62920 | 1.23383 | 297 |
| GO_NUCLEOBASE_CONTAINING_COMPOUND_TRANSPORT | 0.00000 | 0.00010 | 0.62726 | 0.62975 | 1.22823 | 216 |
| GO_NUCLEOBASE_CONTAINING_SMALL_MOLECULE_METABOLIC | 0.00000 | 0.00003 | 0.65944 | 0.60153 | 1.18594 | 405 |
| GO_NUCLEOSIDE_DIPHOSPHATE_METABOLIC_PROCESS | 0.00016 | 0.00319 | 0.51885 | 0.64491 | 1.24260 | 117 |
| GO_NUCLEOSIDE_PHOSPHATE_BIOSYNTHETIC_PROCESS | 0.00002 | 0.00044 | 0.59333 | 0.64092 | 1.24175 | 161 |
| GO_NUCLEOSIDE_TRIPHOSPHATE_BIOSYNTHETIC_PROCESS | 0.00040 | 0.00676 | 0.49849 | 0.72727 | 1.35074 | 42 |
| GO_NUCLEOSIDE_TRIPHOSPHATE_METABOLIC_PROCESS | 0.00009 | 0.00196 | 0.53843 | 0.69306 | 1.30854 | 65 |
| GO_NUCLEOTIDE_PHOSPHORYLATION | 0.00016 | 0.00328 | 0.51885 | 0.65541 | 1.25767 | 102 |
| GO_OOCYTE_DIFFERENTIATION | 0.00450 | 0.04342 | 0.40702 | 0.73513 | 1.33705 | 26 |
| GO_ORGANELLAR_RIBOSOME | 0.00000 | 0.00000 | 0.76146 | 0.75712 | 1.44311 | 84 |
| GO_ORGANELLE_ENVELOPE_LUMEN | 0.00026 | 0.00482 | 0.49849 | 0.67265 | 1.27716 | 78 |
| GO_ORGANELLE_FISSION | 0.00154 | 0.01974 | 0.45506 | 0.57133 | 1.12383 | 342 |
| GO_ORGANELLE_INNER_MEMBRANE | 0.00000 | 0.00000 | | 0.68423 | 1.34915 | 412 |
| GO_ORGANIC_CYCLIC_COMPOUND_CATABOLIC_PROCESS | 0.00000 | 0.00000 | | 0.66975 | 1.32263 | 478 |
| GO_ORGANIC_HYDROXY_COMPOUND_BIOSYNTHETIC_PROCESS | 0.00141 | 0.01850 | 0.45506 | 0.60712 | 1.17627 | 161 |
| GO_OXIDATIVE_PHOSPHORYLATION | 0.00000 | 0.00000 | | 0.78902 | 1.51260 | 99 |
| GO_OXIDOREDUCTASE_ACTIVITY | 0.00000 | 0.00000 | | 0.63268 | 1.24935 | 468 |
| GO_OXIDOREDUCTASE_ACTIVITY_ACTING_ON_A_HEME_GROUP | 0.00176 | 0.02168 | 0.45506 | 0.79383 | 1.41828 | 19 |
| GO_OXIDOREDUCTASE_ACTIVITY_ACTING_ON_NAD_P_H | 0.00002 | 0.00056 | 0.57561 | 0.71468 | 1.35162 | 70 |
| GO_OXIDOREDUCTASE_ACTIVITY_ACTING_ON_NAD_P_H_QUIN | 0.00004 | 0.00113 | 0.55733 | 0.74511 | 1.38835 | 47 |
| GO_OXIDOREDUCTASE_ACTIVITY_ACTING_ON_PEROXIDE_AS_A | 0.00106 | 0.01495 | 0.45506 | 0.73897 | 1.35352 | 29 |
| GO_OXIDOREDUCTASE_COMPLEX | 0.00000 | 0.00000 | 0.73376 | 0.74898 | 1.42731 | 82 |
| GO_PEPTIDASE_ACTIVITY | 0.00133 | 0.01799 | 0.45506 | 0.57681 | 1.13188 | 306 |
| GO_PEPTIDASE_COMPLEX | 0.00000 | 0.00002 | 0.67496 | 0.73250 | 1.39240 | 79 |
| GO_PEPTIDASE_REGULATOR_ACTIVITY | 0.00313 | 0.03310 | 0.43171 | 0.63434 | 1.21189 | 88 |
| GO_PEPTIDE_BINDING | 0.00005 | 0.00124 | 0.55733 | 0.63708 | 1.23187 | 148 |
| GO_PERINUCLEAR_REGION_OF_CYTOPLASM | 0.00276 | 0.03021 | 0.43171 | 0.55814 | 1.10207 | 466 |
| GO_PHAGOCYTOSIS | 0.00554 | 0.04968 | 0.40702 | 0.59337 | 1.14837 | 155 |
| GO_PHAGOSOME_ACIDIFICATION | 0.00059 | 0.00916 | 0.47727 | 0.78902 | 1.42694 | 24 |
| GO_PHOSPHATIDYLSERINE_BINDING | 0.00035 | 0.00610 | 0.49849 | 0.76387 | 1.39912 | 29 |
| GO_PH_REDUCTION | 0.00302 | 0.03248 | 0.43171 | 0.68821 | 1.28238 | 46 |
| GO_PIGMENT_GRANULE | 0.00000 | 0.00000 | 0.76146 | 0.74333 | 1.41934 | 87 |
| GO_PLASMA_LIPOPROTEIN_PARTICLE_CLEARANCE | 0.00106 | 0.01495 | 0.45506 | 0.73201 | 1.35304 | 37 |
| GO_POLYSOMAL_RIBOSOME | 0.00008 | 0.00180 | 0.53843 | 0.78840 | 1.44683 | 30 |
| GO_POLYSOME | 0.00000 | 0.00003 | 0.65944 | 0.75964 | 1.43296 | 62 |
| GO_POSITIVE_REGULATION_OF_ATP_METABOLIC_PROCESS | 0.00174 | 0.02149 | 0.45506 | 0.73011 | 1.33729 | 29 |
| GO_POSITIVE_REGULATION_OF_BINDING | 0.00390 | 0.03895 | 0.43171 | 0.61042 | 1.17582 | 115 |
| GO_POSITIVE_REGULATION_OF_CANONICAL_WNT_SIGNALING_ | 0.00000 | 0.00000 | 0.73376 | 0.72036 | 1.38514 | 108 |
| GO_POSITIVE_REGULATION_OF_CELLULAR_AMIDE_METABOLIC_ | 0.00001 | 0.00036 | 0.59333 | 0.66485 | 1.28095 | 125 |
| GO_POSITIVE_REGULATION_OF_CELLULAR_PROTEIN_LOCALIZAT | 0.00242 | 0.02748 | 0.43171 | 0.58048 | 1.13378 | 244 |
| GO_POSITIVE_REGULATION_OF_CELL_POPULATION_PROLIFERAT | 0.00040 | 0.00670 | 0.49849 | 0.56967 | 1.12503 | 461 |
| GO_POSITIVE_REGULATION_OF_CYSTEINE_TYPE_ENDOPEPTIDA | 0.00547 | 0.04923 | 0.40702 | 0.62225 | 1.19077 | 92 |
| GO_POSITIVE_REGULATION_OF_CYTOKINE_PRODUCTION | 0.00283 | 0.03081 | 0.43171 | 0.58672 | 1.14373 | 210 |
| GO_POSITIVE_REGULATION_OF_DEFENSE_RESPONSE | 0.00000 | 0.00000 | 0.73376 | 0.66200 | 1.28884 | 200 |
| GO_POSITIVE_REGULATION_OF_DNA_BIOSYNTHETIC_PROCESS | 0.00010 | 0.00228 | 0.53843 | 0.71224 | 1.33977 | 55 |
| GO_POSITIVE_REGULATION_OF_DNA_METABOLIC_PROCESS | 0.00020 | 0.00387 | 0.51885 | 0.62771 | 1.21205 | 139 |
| GO_POSITIVE_REGULATION_OF_ESTABLISHMENT_OF_PROTEIN_ | 0.00194 | 0.02331 | 0.45506 | 0.58158 | 1.13709 | 253 |
| GO_POSITIVE_REGULATION_OF_FIBROBLAST_PROLIFERATION | 0.00017 | 0.00338 | 0.51885 | 0.77423 | 1.42213 | 31 |
| GO_POSITIVE_REGULATION_OF_IMMUNE_RESPONSE | 0.00000 | 0.00000 | 0.71951 | 0.63064 | 1.23937 | 321 |
| GO_POSITIVE_REGULATION_OF_IMMUNE_SYSTEM_PROCESS | 0.00000 | 0.00002 | 0.67496 | 0.59748 | 1.17992 | 475 |
| GO_POSITIVE_REGULATION_OF_INNATE_IMMUNE_RESPONSE | 0.00000 | 0.00000 | 0.70498 | 0.68885 | 1.32737 | 128 |
| GO_POSITIVE_REGULATION_OF_MITOTIC_CELL_CYCLE | 0.00178 | 0.02182 | 0.45506 | 0.61563 | 1.18683 | 122 |
| GO_POSITIVE_REGULATION_OF_MRNA_METABOLIC_PROCESS | 0.00157 | 0.02009 | 0.45506 | 0.66330 | 1.25231 | 63 |
| GO_POSITIVE_REGULATION_OF_MRNA_PROCESSING | 0.00250 | 0.02807 | 0.43171 | 0.76194 | 1.38151 | 25 |
| GO_POSITIVE_REGULATION_OF_MRNA_SPLICING_VIA_SPLICEOS | 0.00132 | 0.01782 | 0.45506 | 0.81851 | 1.44353 | 16 |
| GO_POSITIVE_REGULATION_OF_NITRIC_OXIDE_METABOLIC_PR | 0.00133 | 0.01799 | 0.45506 | 0.80807 | 1.43726 | 18 |
| GO_POSITIVE_REGULATION_OF_ORGANELLE_ORGANIZATION | 0.00010 | 0.00218 | 0.53843 | 0.57473 | 1.13449 | 452 |
| GO_POSITIVE_REGULATION_OF_PROTEIN_LOCALIZATION_TO_N | 0.00012 | 0.00265 | 0.53843 | 0.71018 | 1.33697 | 58 |
| GO_POSITIVE_REGULATION_OF_PROTEOLYSIS | 0.00313 | 0.03310 | 0.43171 | 0.58059 | 1.13381 | 237 |
| GO_POSITIVE_REGULATION_OF_REACTIVE_OXYGEN_SPECIES_BI | 0.00115 | 0.01593 | 0.45506 | 0.79865 | 1.43554 | 21 |
| GO_POSITIVE_REGULATION_OF_REACTIVE_OXYGEN_SPECIES_M | 0.00174 | 0.02149 | 0.45506 | 0.69785 | 1.30035 | 46 |

| | | | | | | |
|---|---|---|---|---|---|---|
| GO_POSITIVE_REGULATION_OF_RESPONSE_TO_BIOTIC_STIMUL | 0.00000 | 0.00000 | 0.70498 | 0.67507 | 1.30603 | 150 |
| GO_POSITIVE_REGULATION_OF_RESPONSE_TO_EXTERNAL_STIM | 0.00001 | 0.00016 | 0.61053 | 0.62123 | 1.21505 | 260 |
| GO_POSITIVE_REGULATION_OF_RNA_SPLICING | 0.00265 | 0.02936 | 0.43171 | 0.74051 | 1.35202 | 27 |
| GO_POSITIVE_REGULATION_OF_SIGNAL_TRANSDUCTION_BY_P5 | 0.00390 | 0.03895 | 0.43171 | 0.80515 | 1.41278 | 15 |
| GO_POSITIVE_REGULATION_OF_TELOMERASE_ACTIVITY | 0.00069 | 0.01043 | 0.47727 | 0.75265 | 1.38122 | 30 |
| GO_POSITIVE_REGULATION_OF_TELOMERASE_RNA_LOCALIZATI | 0.00007 | 0.00169 | 0.53843 | 0.88616 | 1.55493 | 15 |
| GO_POSITIVE_REGULATION_OF_TELOMERE_MAINTENANCE | 0.00026 | 0.00486 | 0.49849 | 0.73697 | 1.37102 | 44 |
| GO_POSITIVE_REGULATION_OF_TELOMERE_MAINTENANCE_VIA | 0.00001 | 0.00040 | 0.59333 | 0.80196 | 1.47289 | 33 |
| GO_POSITIVE_REGULATION_OF_TRANSFERASE_ACTIVITY | 0.00108 | 0.01507 | 0.45506 | 0.56501 | 1.11376 | 401 |
| GO_POSITIVE_REGULATION_OF_TRANSLATION | 0.00000 | 0.00016 | 0.61053 | 0.68408 | 1.31526 | 107 |
| GO_POSITIVE_REGULATION_OF_VIRAL_PROCESS | 0.00026 | 0.00486 | 0.49849 | 0.66195 | 1.26462 | 88 |
| GO_POSITIVE_REGULATION_OF_WNT_SIGNALING_PATHWAY | 0.00000 | 0.00000 | 0.73376 | 0.69924 | 1.34747 | 127 |
| GO_POSTSYNAPSE | 0.00239 | 0.02716 | 0.43171 | 0.57062 | 1.12116 | 316 |
| GO_POST_TRANSLATIONAL_PROTEIN_MODIFICATION | 0.00024 | 0.00452 | 0.51885 | 0.60046 | 1.17392 | 251 |
| GO_PRECATALYTIC_SPLICEOSOME | 0.00141 | 0.01850 | 0.45506 | 0.69251 | 1.29540 | 50 |
| GO_PRERIBOSOME | 0.00006 | 0.00154 | 0.53843 | 0.69866 | 1.32020 | 69 |
| GO_PROTEASOMAL_PROTEIN_CATABOLIC_PROCESS | 0.00007 | 0.00169 | 0.53843 | 0.58188 | 1.14652 | 387 |
| GO_PROTEASOMAL_UBIQUITIN_INDEPENDENT_PROTEIN_CATAB | 0.00000 | 0.00003 | 0.65944 | 0.91182 | 1.62178 | 18 |
| GO_PROTEASOME_ACCESSORY_COMPLEX | 0.00005 | 0.00135 | 0.55733 | 0.82638 | 1.49450 | 24 |
| GO_PROTEASOME_CORE_COMPLEX | 0.00000 | 0.00004 | 0.65944 | 0.91231 | 1.61486 | 17 |
| GO_PROTEIN_CONTAINING_COMPLEX_DISASSEMBLY | 0.00000 | 0.00000 | 0.81404 | 0.66285 | 1.29656 | 261 |
| GO_PROTEIN_CONTAINING_COMPLEX_LOCALIZATION | 0.00018 | 0.00352 | 0.51885 | 0.60451 | 1.17943 | 226 |
| GO_PROTEIN_DNA_COMPLEX_SUBUNIT_ORGANIZATION | 0.00052 | 0.00825 | 0.47727 | 0.60887 | 1.18342 | 183 |
| GO_PROTEIN_DOMAIN_SPECIFIC_BINDING | 0.00003 | 0.00088 | 0.55733 | 0.57828 | 1.14190 | 470 |
| GO_PROTEIN_FOLDING | 0.00000 | 0.00000 | 0.73376 | 0.67425 | 1.30867 | 171 |
| GO_PROTEIN_FOLDING_CHAPERONE | 0.00079 | 0.01170 | 0.47727 | 0.81632 | 1.45193 | 18 |
| GO_PROTEIN_HETEROOLIGOMERIZATION | 0.00442 | 0.04297 | 0.40702 | 0.78998 | 1.39322 | 16 |
| GO_PROTEIN_IMPORT | 0.00001 | 0.00040 | 0.59333 | 0.64197 | 1.24306 | 158 |
| GO_PROTEIN_IMPORT_INTO_MITOCHONDRIAL_MATRIX | 0.00231 | 0.02642 | 0.43171 | 0.79530 | 1.41455 | 18 |
| GO_PROTEIN_INSERTION_INTO_MEMBRANE | 0.00191 | 0.02299 | 0.45506 | 0.68322 | 1.27694 | 49 |
| GO_PROTEIN_LOCALIZATION_TO_CHROMOSOME_TELOMERIC_F | 0.00001 | 0.00031 | 0.59333 | 0.82810 | 1.51357 | 28 |
| GO_PROTEIN_LOCALIZATION_TO_ENDOPLASMIC_RETICULUM | 0.00000 | 0.00000 | | 0.73850 | 1.42303 | 129 |
| GO_PROTEIN_LOCALIZATION_TO_MEMBRANE | 0.00000 | 0.00000 | 0.77494 | 0.61216 | 1.20881 | 462 |
| GO_PROTEIN_LOCALIZATION_TO_MITOCHONDRION | 0.00015 | 0.00310 | 0.51885 | 0.64234 | 1.23759 | 118 |
| GO_PROTEIN_LOCALIZATION_TO_NUCLEUS | 0.00000 | 0.00014 | 0.61053 | 0.63315 | 1.23408 | 208 |
| GO_PROTEIN_MATURATION | 0.00357 | 0.03638 | 0.43171 | 0.59209 | 1.14965 | 174 |
| GO_PROTEIN_MODIFICATION_BY_SMALL_PROTEIN_REMOVAL | 0.00002 | 0.00052 | 0.57561 | 0.61535 | 1.20119 | 231 |
| GO_PROTEIN_REFOLDING | 0.00465 | 0.04430 | 0.40702 | 0.77561 | 1.37288 | 17 |
| GO_PROTEIN_STABILIZATION | 0.00081 | 0.01194 | 0.47727 | 0.61881 | 1.19577 | 146 |
| GO_PROTEIN_SUMOYLATION | 0.00450 | 0.04342 | 0.40702 | 0.65043 | 1.22846 | 67 |
| GO_PROTEIN_TARGETING | 0.00000 | 0.00000 | | 0.66223 | 1.30361 | 358 |
| GO_PROTEIN_TARGETING_TO_MEMBRANE | 0.00000 | 0.00000 | | 0.71675 | 1.38879 | 162 |
| GO_PROTEIN_TARGETING_TO_MITOCHONDRION | 0.00113 | 0.01571 | 0.45506 | 0.65407 | 1.24648 | 83 |
| GO_PROTEIN_TRANSMEMBRANE_TRANSPORT | 0.00110 | 0.01526 | 0.45506 | 0.68173 | 1.28238 | 55 |
| GO_PROTON_TRANSMEMBRANE_TRANSPORT | 0.00000 | 0.00001 | 0.69013 | 0.71234 | 1.36309 | 91 |
| GO_PROTON_TRANSMEMBRANE_TRANSPORTER_ACTIVITY | 0.00004 | 0.00115 | 0.55733 | 0.70138 | 1.32535 | 69 |
| GO_PROTON_TRANSPORTING_TWO_SECTOR_ATPASE_COMPLEX | 0.00083 | 0.01218 | 0.47727 | 0.77254 | 1.40508 | 26 |
| GO_PROTON_TRANSPORTING_V_TYPE_ATPASE_COMPLEX | 0.00239 | 0.02716 | 0.43171 | 0.76100 | 1.37626 | 24 |
| GO_PURINE_CONTAINING_COMPOUND_BIOSYNTHETIC_PROCES | 0.00001 | 0.00040 | 0.59333 | 0.65863 | 1.26901 | 126 |
| GO_PURINE_CONTAINING_COMPOUND_METABOLIC_PROCESS | 0.00000 | 0.00000 | 0.73376 | 0.63132 | 1.23831 | 300 |
| GO_PURINE_NUCLEOSIDE_TRIPHOSPHATE_METABOLIC_PROCES | 0.00033 | 0.00582 | 0.49849 | 0.70887 | 1.32487 | 49 |
| GO_PYROPHOSPHATE_HYDROLYSIS_DRIVEN_PROTON_TRANSME | 0.00532 | 0.04803 | 0.40702 | 0.74470 | 1.34678 | 24 |
| GO_PYRUVATE_METABOLIC_PROCESS | 0.00001 | 0.00018 | 0.61053 | 0.67194 | 1.29289 | 113 |
| GO_P_BODY | 0.00472 | 0.04480 | 0.40702 | 0.65470 | 1.23501 | 62 |
| GO_REACTIVE_OXYGEN_SPECIES_BIOSYNTHETIC_PROCESS | 0.00040 | 0.00670 | 0.49849 | 0.70271 | 1.31857 | 53 |
| GO_REACTIVE_OXYGEN_SPECIES_METABOLIC_PROCESS | 0.00021 | 0.00409 | 0.51885 | 0.62881 | 1.21472 | 144 |
| GO_REGULATION_OF_ALCOHOL_BIOSYNTHETIC_PROCESS | 0.00228 | 0.02614 | 0.43171 | 0.68438 | 1.28020 | 50 |
| GO_REGULATION_OF_ANIMAL_ORGAN_MORPHOGENESIS | 0.00000 | 0.00000 | | 0.72330 | 1.39922 | 149 |
| GO_REGULATION_OF_APOPTOTIC_SIGNALING_PATHWAY | 0.00079 | 0.01170 | 0.47727 | 0.58599 | 1.14641 | 266 |
| GO_REGULATION_OF_ATP_METABOLIC_PROCESS | 0.00000 | 0.00006 | 0.64355 | 0.70523 | 1.34777 | 89 |
| GO_REGULATION_OF_BINDING | 0.00428 | 0.04185 | 0.40702 | 0.58264 | 1.13664 | 225 |
| GO_REGULATION_OF_BLOOD_PRESSURE | 0.00290 | 0.03156 | 0.43171 | 0.65319 | 1.23428 | 69 |
| GO_REGULATION_OF_CARBOHYDRATE_CATABOLIC_PROCESS | 0.00324 | 0.03393 | 0.43171 | 0.65541 | 1.23787 | 67 |
| GO_REGULATION_OF_CELLULAR_AMIDE_METABOLIC_PROCESS | 0.00000 | 0.00000 | 0.78819 | 0.63116 | 1.24143 | 337 |
| GO_REGULATION_OF_CELLULAR_AMINE_METABOLIC_PROCESS | 0.00000 | 0.00000 | 0.74774 | 0.78955 | 1.48730 | 60 |
| GO_REGULATION_OF_CELLULAR_AMINO_ACID_METABOLIC_PR( | 0.00000 | 0.00000 | 0.76146 | 0.80341 | 1.51022 | 54 |
| GO_REGULATION_OF_CELLULAR_KETONE_METABOLIC_PROCESS | 0.00000 | 0.00000 | 0.70498 | 0.69239 | 1.33405 | 126 |
| GO_REGULATION_OF_CELLULAR_PH | 0.00442 | 0.04297 | 0.40702 | 0.65360 | 1.23436 | 68 |
| GO_REGULATION_OF_CELLULAR_PROTEIN_LOCALIZATION | 0.00228 | 0.02614 | 0.43171 | 0.56216 | 1.10830 | 402 |
| GO_REGULATION_OF_CELLULAR_RESPIRATION | 0.00428 | 0.04185 | 0.40702 | 0.77183 | 1.37897 | 19 |

| | | | | | | |
|---|---|---|---|---|---|---|
| GO_REGULATION_OF_CELLULAR_RESPONSE_TO_HEAT | 0.00091 | 0.01314 | 0.47727 | 0.67360 | 1.27221 | 67 |
| GO_REGULATION_OF_CELL_CYCLE_G2_M_PHASE_TRANSITION | 0.00016 | 0.00332 | 0.51885 | 0.61554 | 1.19738 | 187 |
| GO_REGULATION_OF_CELL_CYCLE_PHASE_TRANSITION | 0.00000 | 0.00014 | 0.61053 | 0.59968 | 1.18091 | 369 |
| GO_REGULATION_OF_CHOLESTEROL_BIOSYNTHETIC_PROCESS | 0.00209 | 0.02470 | 0.43171 | 0.71476 | 1.32117 | 38 |
| GO_REGULATION_OF_CHOLESTEROL_METABOLIC_PROCESS | 0.00113 | 0.01571 | 0.45506 | 0.69865 | 1.30177 | 47 |
| GO_REGULATION_OF_CHROMOSOME_ORGANIZATION | 0.00036 | 0.00622 | 0.49849 | 0.58896 | 1.15261 | 278 |
| GO_REGULATION_OF_CHROMOSOME_SEGREGATION | 0.00079 | 0.01170 | 0.47727 | 0.64131 | 1.22718 | 91 |
| GO_REGULATION_OF_CHROMOSOME_SEPARATION | 0.00253 | 0.02838 | 0.43171 | 0.66349 | 1.24947 | 59 |
| GO_REGULATION_OF_CYSTEINE_TYPE_ENDOPEPTIDASE_ACTIVIT | 0.00044 | 0.00719 | 0.49849 | 0.62610 | 1.20905 | 141 |
| GO_REGULATION_OF_DEFENSE_RESPONSE | 0.00006 | 0.00150 | 0.55733 | 0.58834 | 1.15824 | 356 |
| GO_REGULATION_OF_DNA_BIOSYNTHETIC_PROCESS | 0.00043 | 0.00695 | 0.49849 | 0.66709 | 1.27130 | 83 |
| GO_REGULATION_OF_DNA_METABOLIC_PROCESS | 0.00055 | 0.00863 | 0.47727 | 0.58909 | 1.15134 | 249 |
| GO_REGULATION_OF_DNA_TEMPLATED_TRANSCRIPTION_IN_RE | 0.00000 | 0.00000 | 0.71951 | 0.72410 | 1.38687 | 94 |
| GO_REGULATION_OF_GENERATION_OF_PRECURSOR_METABOLI | 0.00004 | 0.00114 | 0.55733 | 0.65536 | 1.26115 | 111 |
| GO_REGULATION_OF_GENE_SILENCING | 0.00023 | 0.00434 | 0.51885 | 0.65464 | 1.25268 | 91 |
| GO_REGULATION_OF_HEMATOPOIETIC_PROGENITOR_CELL_DIF | 0.00000 | 0.00000 | 0.73376 | 0.76273 | 1.44465 | 72 |
| GO_REGULATION_OF_HEMATOPOIETIC_STEM_CELL_DIFFERENT | 0.00000 | 0.00000 | 0.74774 | 0.78891 | 1.48817 | 62 |
| GO_REGULATION_OF_HEMOPOIESIS | 0.00390 | 0.03895 | 0.43171 | 0.57739 | 1.12909 | 258 |
| GO_REGULATION_OF_IMMUNE_RESPONSE | 0.00000 | 0.00006 | 0.64355 | 0.60032 | 1.18447 | 433 |
| GO_REGULATION_OF_INNATE_IMMUNE_RESPONSE | 0.00000 | 0.00007 | 0.62726 | 0.64536 | 1.25422 | 181 |
| GO_REGULATION_OF_INTRINSIC_APOPTOTIC_SIGNALING_PATHV | 0.00179 | 0.02183 | 0.45506 | 0.61548 | 1.18550 | 116 |
| GO_REGULATION_OF_MITOTIC_CELL_CYCLE | 0.00000 | 0.00003 | 0.65944 | 0.58992 | 1.16508 | 499 |
| GO_REGULATION_OF_MORPHOGENESIS_OF_AN_EPITHELIUM | 0.00000 | 0.00000 | | 0.74680 | 1.43884 | 125 |
| GO_REGULATION_OF_MRNA_CATABOLIC_PROCESS | 0.00000 | 0.00000 | | 0.71126 | 1.37919 | 165 |
| GO_REGULATION_OF_MRNA_METABOLIC_PROCESS | 0.00000 | 0.00000 | | 0.67178 | 1.31458 | 276 |
| GO_REGULATION_OF_MRNA_PROCESSING | 0.00008 | 0.00186 | 0.53843 | 0.65189 | 1.25598 | 125 |
| GO_REGULATION_OF_MRNA_SPLICING_VIA_SPLICEOSOME | 0.00012 | 0.00256 | 0.53843 | 0.66950 | 1.28033 | 90 |
| GO_REGULATION_OF_MULTI_ORGANISM_PROCESS | 0.00008 | 0.00190 | 0.53843 | 0.76905 | 1.42150 | 37 |
| GO_REGULATION_OF_MUSCLE_CONTRACTION | 0.00163 | 0.02050 | 0.45506 | 0.64879 | 1.23071 | 76 |
| GO_REGULATION_OF_NUCLEAR_DIVISION | 0.00060 | 0.00923 | 0.47727 | 0.62443 | 1.20552 | 137 |
| GO_REGULATION_OF_NUCLEASE_ACTIVITY | 0.00059 | 0.00916 | 0.47727 | 0.81786 | 1.44767 | 17 |
| GO_REGULATION_OF_NUCLEOTIDE_BIOSYNTHETIC_PROCESS | 0.00302 | 0.03248 | 0.43171 | 0.78902 | 1.40337 | 18 |
| GO_REGULATION_OF_NUCLEOTIDE_METABOLIC_PROCESS | 0.00033 | 0.00588 | 0.49849 | 0.66858 | 1.26942 | 78 |
| GO_REGULATION_OF_OXIDATIVE_PHOSPHORYLATION | 0.00005 | 0.00130 | 0.55733 | 0.85284 | 1.53294 | 21 |
| GO_REGULATION_OF_PEPTIDASE_ACTIVITY | 0.00006 | 0.00152 | 0.53843 | 0.61192 | 1.19388 | 226 |
| GO_REGULATION_OF_PEPTIDE_TRANSPORT | 0.00250 | 0.02807 | 0.43171 | 0.56252 | 1.10917 | 412 |
| GO_REGULATION_OF_PH | 0.00561 | 0.04971 | 0.40702 | 0.64191 | 1.21498 | 71 |
| GO_REGULATION_OF_PLASMA_LIPOPROTEIN_PARTICLE_LEVELS | 0.00006 | 0.00145 | 0.55733 | 0.72431 | 1.36154 | 54 |
| GO_REGULATION_OF_POSTTRANSCRIPTIONAL_GENE_SILENCING | 0.00026 | 0.00486 | 0.49849 | 0.66803 | 1.27236 | 81 |
| GO_REGULATION_OF_PROTEIN_CATABOLIC_PROCESS | 0.00309 | 0.03299 | 0.43171 | 0.57360 | 1.12399 | 291 |
| GO_REGULATION_OF_PROTEIN_IMPORT | 0.00487 | 0.04559 | 0.40702 | 0.67491 | 1.25754 | 47 |
| GO_REGULATION_OF_PROTEIN_LOCALIZATION_TO_NUCLEUS | 0.00000 | 0.00013 | 0.61053 | 0.69035 | 1.32101 | 91 |
| GO_REGULATION_OF_PROTEIN_STABILITY | 0.00066 | 0.01007 | 0.47727 | 0.59308 | 1.15811 | 235 |
| GO_REGULATION_OF_PROTEOLYSIS | 0.00007 | 0.00169 | 0.53843 | 0.57923 | 1.14299 | 431 |
| GO_REGULATION_OF_REACTIVE_OXYGEN_SPECIES_BIOSYNTHET | 0.00022 | 0.00427 | 0.51885 | 0.73154 | 1.36343 | 45 |
| GO_REGULATION_OF_REACTIVE_OXYGEN_SPECIES_METABOLIC_ | 0.00088 | 0.01280 | 0.47727 | 0.63948 | 1.22592 | 99 |
| GO_REGULATION_OF_RESPONSE_TO_BIOTIC_STIMULUS | 0.00001 | 0.00039 | 0.59333 | 0.61773 | 1.20625 | 235 |
| GO_REGULATION_OF_RNA_SPLICING | 0.00011 | 0.00241 | 0.53843 | 0.64982 | 1.25198 | 125 |
| GO_REGULATION_OF_SISTER_CHROMATID_SEGREGATION | 0.00405 | 0.04011 | 0.40702 | 0.64401 | 1.22034 | 73 |
| GO_REGULATION_OF_SMALL_MOLECULE_METABOLIC_PROCESS | 0.00000 | 0.00000 | 0.77494 | 0.64342 | 1.26057 | 290 |
| GO_REGULATION_OF_SMOOTH_MUSCLE_CONTRACTION | 0.00007 | 0.00171 | 0.53843 | 0.82642 | 1.49295 | 23 |
| GO_REGULATION_OF_STEM_CELL_DIFFERENTIATION | 0.00000 | 0.00000 | 0.73376 | 0.73434 | 1.40292 | 88 |
| GO_REGULATION_OF_STEROID_METABOLIC_PROCESS | 0.00524 | 0.04770 | 0.40702 | 0.64061 | 1.21567 | 77 |
| GO_REGULATION_OF_SUPEROXIDE_METABOLIC_PROCESS | 0.00346 | 0.03548 | 0.43171 | 0.77826 | 1.39046 | 19 |
| GO_REGULATION_OF_SUPRAMOLECULAR_FIBER_ORGANIZATIO | 0.00509 | 0.04656 | 0.40702 | 0.57599 | 1.12367 | 223 |
| GO_REGULATION_OF_SYMBIOTIC_PROCESS | 0.00009 | 0.00211 | 0.53843 | 0.62950 | 1.21763 | 151 |
| GO_REGULATION_OF_TELOMERASE_ACTIVITY | 0.00087 | 0.01263 | 0.47727 | 0.72147 | 1.33426 | 39 |
| GO_REGULATION_OF_TELOMERE_MAINTENANCE | 0.00547 | 0.04923 | 0.40702 | 0.64959 | 1.22679 | 68 |
| GO_REGULATION_OF_TELOMERE_MAINTENANCE_VIA_TELOME | 0.00020 | 0.00387 | 0.51885 | 0.72043 | 1.34856 | 51 |
| GO_REGULATION_OF_TRANSCRIPTION_FROM_RNA_POLYMERA | 0.00000 | 0.00000 | 0.70498 | 0.76931 | 1.45239 | 64 |
| GO_REGULATION_OF_TRANSLATIONAL_INITIATION | 0.00023 | 0.00437 | 0.51885 | 0.68479 | 1.29399 | 69 |
| GO_REGULATION_OF_VIRAL_TRANSCRIPTION | 0.00361 | 0.03657 | 0.43171 | 0.67497 | 1.26259 | 50 |
| GO_REGULATION_OF_WNT_SIGNALING_PATHWAY | 0.00015 | 0.00312 | 0.51885 | 0.60129 | 1.17419 | 238 |
| GO_RESPIRASOME | 0.00000 | 0.00000 | 0.81404 | 0.78326 | 1.48524 | 75 |
| GO_RESPIRATORY_CHAIN_COMPLEX | 0.00000 | 0.00000 | 0.77494 | 0.79183 | 1.49492 | 64 |
| GO_RESPIRATORY_ELECTRON_TRANSPORT_CHAIN | 0.00000 | 0.00000 | | 0.78275 | 1.49591 | 89 |
| GO_RESPONSE_TO_ALCOHOL | 0.00015 | 0.00310 | 0.51885 | 0.63794 | 1.22915 | 126 |
| GO_RESPONSE_TO_ATP | 0.00087 | 0.01270 | 0.47727 | 0.81086 | 1.43528 | 17 |
| GO_RESPONSE_TO_CALCIUM_ION | 0.00124 | 0.01701 | 0.45506 | 0.65196 | 1.24071 | 80 |
| GO_RESPONSE_TO_CORTICOSTEROID | 0.00010 | 0.00220 | 0.53843 | 0.66894 | 1.27841 | 89 |

| | | | | | | |
|---|---|---|---|---|---|---|
| GO_RESPONSE_TO_DRUG | 0.00000 | 0.00007 | 0.62726 | 0.63080 | 1.23070 | 220 |
| GO_RESPONSE_TO_ETHANOL | 0.00168 | 0.02098 | 0.45506 | 0.66543 | 1.25679 | 67 |
| GO_RESPONSE_TO_FIBROBLAST_GROWTH_FACTOR | 0.00357 | 0.03638 | 0.43171 | 0.63469 | 1.20976 | 84 |
| GO_RESPONSE_TO_GROWTH_FACTOR | 0.00272 | 0.02986 | 0.43171 | 0.56235 | 1.10805 | 387 |
| GO_RESPONSE_TO_HEAT | 0.00000 | 0.00013 | 0.62726 | 0.66982 | 1.29069 | 119 |
| GO_RESPONSE_TO_HYDROGEN_PEROXIDE | 0.00532 | 0.04803 | 0.40702 | 0.62255 | 1.19054 | 90 |
| GO_RESPONSE_TO_INORGANIC_SUBSTANCE | 0.00016 | 0.00328 | 0.51885 | 0.58928 | 1.15810 | 321 |
| GO_RESPONSE_TO_INTERLEUKIN_1 | 0.00000 | 0.00000 | 0.78819 | 0.71590 | 1.37950 | 124 |
| GO_RESPONSE_TO_INTERLEUKIN_12 | 0.00036 | 0.00622 | 0.49849 | 0.74158 | 1.36769 | 36 |
| GO_RESPONSE_TO_INTERLEUKIN_4 | 0.00224 | 0.02581 | 0.43171 | 0.77088 | 1.38921 | 22 |
| GO_RESPONSE_TO_LEUKEMIA_INHIBITORY_FACTOR | 0.00076 | 0.01129 | 0.47727 | 0.66148 | 1.25288 | 72 |
| GO_RESPONSE_TO_LIPID | 0.00043 | 0.00695 | 0.49849 | 0.56669 | 1.11902 | 470 |
| GO_RESPONSE_TO_METAL_ION | 0.00143 | 0.01863 | 0.45506 | 0.59895 | 1.16592 | 199 |
| GO_RESPONSE_TO_NUTRIENT | 0.00294 | 0.03185 | 0.43171 | 0.62596 | 1.19987 | 98 |
| GO_RESPONSE_TO_ORGANOPHOSPHORUS | 0.00294 | 0.03185 | 0.43171 | 0.64154 | 1.21511 | 72 |
| GO_RESPONSE_TO_OXIDATIVE_STRESS | 0.00495 | 0.04573 | 0.40702 | 0.57010 | 1.11758 | 294 |
| GO_RESPONSE_TO_OXYGEN_LEVELS | 0.00000 | 0.00000 | | 0.67661 | 1.32155 | 244 |
| GO_RESPONSE_TO_PURINE_CONTAINING_COMPOUND | 0.00509 | 0.04656 | 0.40702 | 0.63176 | 1.20394 | 82 |
| GO_RESPONSE_TO_STEROID_HORMONE | 0.00009 | 0.00200 | 0.53843 | 0.61312 | 1.19360 | 201 |
| GO_RESPONSE_TO_TEMPERATURE_STIMULUS | 0.00002 | 0.00047 | 0.57561 | 0.64468 | 1.24782 | 153 |
| GO_RESPONSE_TO_TOXIC_SUBSTANCE | 0.00011 | 0.00235 | 0.53843 | 0.63892 | 1.23203 | 134 |
| GO_RESPONSE_TO_TUMOR_NECROSIS_FACTOR | 0.00000 | 0.00000 | 0.70498 | 0.66331 | 1.28815 | 175 |
| GO_RESPONSE_TO_VIRUS | 0.00261 | 0.02910 | 0.43171 | 0.59428 | 1.15408 | 175 |
| GO_RETINA_HOMEOSTASIS | 0.00034 | 0.00606 | 0.49849 | 0.75202 | 1.38362 | 34 |
| GO_RIBONUCLEOPROTEIN_COMPLEX_BINDING | 0.00000 | 0.00001 | 0.69013 | 0.69839 | 1.34459 | 114 |
| GO_RIBONUCLEOPROTEIN_COMPLEX_BIOGENESIS | 0.00000 | 0.00000 | | 0.68714 | 1.35448 | 393 |
| GO_RIBONUCLEOPROTEIN_COMPLEX_SUBUNIT_ORGANIZATION | 0.00000 | 0.00000 | 0.81404 | 0.69112 | 1.34183 | 173 |
| GO_RIBONUCLEOPROTEIN_GRANULE | 0.00000 | 0.00000 | 0.71951 | 0.66271 | 1.28875 | 186 |
| GO_RIBONUCLEOSIDE_TRIPHOSPHATE_BIOSYNTHETIC_PROCESS | 0.00009 | 0.00210 | 0.53843 | 0.77194 | 1.42026 | 34 |
| GO_RIBONUCLEOSIDE_TRIPHOSPHATE_METABOLIC_PROCESS | 0.00013 | 0.00280 | 0.51885 | 0.73558 | 1.37065 | 46 |
| GO_RIBOSE_PHOSPHATE_BIOSYNTHETIC_PROCESS | 0.00002 | 0.00056 | 0.57561 | 0.65814 | 1.26821 | 124 |
| GO_RIBOSE_PHOSPHATE_METABOLIC_PROCESS | 0.00000 | 0.00000 | 0.74774 | 0.63837 | 1.25001 | 285 |
| GO_RIBOSOMAL_LARGE_SUBUNIT_BIOGENESIS | 0.00001 | 0.00031 | 0.59333 | 0.72435 | 1.36807 | 67 |
| GO_RIBOSOMAL_SMALL_SUBUNIT_ASSEMBLY | 0.00405 | 0.04011 | 0.40702 | 0.78074 | 1.38865 | 18 |
| GO_RIBOSOMAL_SMALL_SUBUNIT_BINDING | 0.00532 | 0.04803 | 0.40702 | 0.77341 | 1.36899 | 17 |
| GO_RIBOSOMAL_SMALL_SUBUNIT_BIOGENESIS | 0.00005 | 0.00138 | 0.55733 | 0.70510 | 1.33163 | 68 |
| GO_RIBOSOMAL_SUBUNIT | 0.00000 | 0.00000 | | 0.76292 | 1.48089 | 172 |
| GO_RIBOSOME | 0.00000 | 0.00000 | | 0.75411 | 1.46984 | 208 |
| GO_RIBOSOME_ASSEMBLY | 0.00001 | 0.00026 | 0.59333 | 0.74485 | 1.40144 | 56 |
| GO_RIBOSOME_BIOGENESIS | 0.00000 | 0.00000 | | 0.68899 | 1.34833 | 277 |
| GO_RNA_3_END_PROCESSING | 0.00002 | 0.00046 | 0.57561 | 0.65576 | 1.26361 | 128 |
| GO_RNA_CATABOLIC_PROCESS | 0.00000 | 0.00000 | | 0.70701 | 1.39073 | 341 |
| GO_RNA_DEPENDENT_DNA_BIOSYNTHETIC_PROCESS | 0.00003 | 0.00081 | 0.57561 | 0.71669 | 1.34965 | 59 |
| GO_RNA_EXPORT_FROM_NUCLEUS | 0.00000 | 0.00016 | 0.61053 | 0.66306 | 1.27817 | 130 |
| GO_RNA_LOCALIZATION | 0.00000 | 0.00000 | 0.76146 | 0.66638 | 1.29886 | 208 |
| GO_RNA_SPLICING | 0.00000 | 0.00000 | | 0.63512 | 1.25129 | 384 |
| GO_RNA_SPLICING_VIA_TRANSESTERIFICATION_REACTIONS | 0.00000 | 0.00000 | 0.80122 | 0.64478 | 1.26586 | 311 |
| GO_ROUGH_ENDOPLASMIC_RETICULUM | 0.00014 | 0.00293 | 0.51885 | 0.70363 | 1.32507 | 59 |
| GO_RRNA_BINDING | 0.00000 | 0.00001 | 0.69013 | 0.77638 | 1.45941 | 54 |
| GO_RRNA_METABOLIC_PROCESS | 0.00000 | 0.00000 | | 0.68430 | 1.33386 | 209 |
| GO_SCF_DEPENDENT_PROTEASOMAL_UBIQUITIN_DEPENDENT_ | 0.00000 | 0.00008 | 0.62726 | 0.71957 | 1.36498 | 76 |
| GO_SECRETORY_GRANULE | 0.00000 | 0.00000 | | 0.62978 | 1.24363 | 481 |
| GO_SECRETORY_GRANULE_MEMBRANE | 0.00032 | 0.00575 | 0.49849 | 0.61375 | 1.19134 | 172 |
| GO_SERINE_TYPE_ENDOPEPTIDASE_INHIBITOR_ACTIVITY | 0.00038 | 0.00640 | 0.49849 | 0.77388 | 1.41447 | 28 |
| GO_SEXUAL_REPRODUCTION | 0.00472 | 0.04480 | 0.40702 | 0.55748 | 1.10026 | 439 |
| GO_SIDE_OF_MEMBRANE | 0.00495 | 0.04573 | 0.40702 | 0.57741 | 1.12751 | 235 |
| GO_SIGNAL_TRANSDUCTION_BY_PROTEIN_PHOSPHORYLATION | 0.00224 | 0.02581 | 0.43171 | 0.55690 | 1.09975 | 500 |
| GO_SINGLE_FERTILIZATION | 0.00000 | 0.00002 | 0.67496 | 0.76307 | 1.43842 | 61 |
| GO_SINGLE_STRANDED_RNA_BINDING | 0.00001 | 0.00042 | 0.59333 | 0.71960 | 1.35864 | 66 |
| GO_SISTER_CHROMATID_SEGREGATION | 0.00042 | 0.00689 | 0.49849 | 0.60926 | 1.18209 | 170 |
| GO_SKIN_DEVELOPMENT | 0.00076 | 0.01140 | 0.47727 | 0.61452 | 1.18945 | 153 |
| GO_SMALL_MOLECULE_BIOSYNTHETIC_PROCESS | 0.00001 | 0.00021 | 0.61053 | 0.58728 | 1.15957 | 459 |
| GO_SMALL_NUCLEAR_RIBONUCLEOPROTEIN_COMPLEX | 0.00042 | 0.00689 | 0.49849 | 0.69542 | 1.30936 | 57 |
| GO_SMALL_RIBOSOMAL_SUBUNIT | 0.00000 | 0.00001 | 0.69013 | 0.75767 | 1.43100 | 67 |
| GO_SM_LIKE_PROTEIN_FAMILY_COMPLEX | 0.00145 | 0.01879 | 0.45506 | 0.66792 | 1.26149 | 67 |
| GO_SNORNA_BINDING | 0.00038 | 0.00646 | 0.49849 | 0.76224 | 1.39614 | 29 |
| GO_SPERM_EGG_RECOGNITION | 0.00000 | 0.00002 | 0.65944 | 0.87469 | 1.59087 | 26 |
| GO_SPINDLE_MIDZONE | 0.00361 | 0.03657 | 0.43171 | 0.73316 | 1.33859 | 27 |
| GO_SPINDLE_ORGANIZATION | 0.00272 | 0.02986 | 0.43171 | 0.60076 | 1.16362 | 159 |
| GO_SPLICEOSOMAL_COMPLEX | 0.00000 | 0.00008 | 0.62726 | 0.65256 | 1.26657 | 171 |

| | | | | | | |
|---|---|---|---|---|---|---|
| GO_SPLICEOSOMAL_TRI_SNRNP_COMPLEX | 0.00168 | 0.02098 | 0.45506 | 0.73474 | 1.34835 | 30 |
| GO_STEM_CELL_DIFFERENTIATION | 0.00000 | 0.00000 | 0.82666 | 0.70006 | 1.35554 | 158 |
| GO_STEROID_BIOSYNTHETIC_PROCESS | 0.00465 | 0.04430 | 0.40702 | 0.61242 | 1.17992 | 125 |
| GO_STEROID_METABOLIC_PROCESS | 0.00313 | 0.03310 | 0.43171 | 0.59525 | 1.15748 | 185 |
| GO_STEROL_BIOSYNTHETIC_PROCESS | 0.00005 | 0.00120 | 0.55733 | 0.71876 | 1.35584 | 62 |
| GO_STEROL_METABOLIC_PROCESS | 0.00003 | 0.00073 | 0.57561 | 0.66485 | 1.27854 | 109 |
| GO_STRESS_GRANULE_ASSEMBLY | 0.00465 | 0.04430 | 0.40702 | 0.77288 | 1.38921 | 21 |
| GO_STRUCTURAL_CONSTITUENT_OF_RIBOSOME | 0.00000 | 0.00000 | | 0.76380 | 1.47839 | 153 |
| GO_STRUCTURAL_MOLECULE_ACTIVITY | 0.00000 | 0.00000 | | 0.71760 | 1.41436 | 391 |
| GO_SUPEROXIDE_METABOLIC_PROCESS | 0.00272 | 0.02986 | 0.43171 | 0.72028 | 1.32659 | 35 |
| GO_TELOMERASE_RNA_LOCALIZATION | 0.00054 | 0.00857 | 0.47727 | 0.81801 | 1.46148 | 19 |
| GO_TELOMERE_MAINTENANCE_VIA_TELOMERE_LENGTHENING | 0.00005 | 0.00138 | 0.55733 | 0.70715 | 1.33512 | 66 |
| GO_TELOMERE_ORGANIZATION | 0.00159 | 0.02016 | 0.45506 | 0.61666 | 1.18834 | 127 |
| GO_TERTIARY_GRANULE | 0.00008 | 0.00186 | 0.53843 | 0.66372 | 1.27005 | 91 |
| GO_TERTIARY_GRANULE_MEMBRANE | 0.00102 | 0.01453 | 0.45506 | 0.71940 | 1.33244 | 40 |
| GO_THREONINE_TYPE_ENDOPEPTIDASE_ACTIVITY | 0.00000 | 0.00000 | 0.71951 | 0.93456 | 1.65424 | 17 |
| GO_THREONINE_TYPE_PEPTIDASE_ACTIVITY | 0.00000 | 0.00004 | 0.65944 | 0.89807 | 1.61424 | 21 |
| GO_TISSUE_MORPHOGENESIS | 0.00000 | 0.00006 | 0.64355 | 0.59467 | 1.17150 | 380 |
| GO_TOXIN_TRANSPORT | 0.00064 | 0.00972 | 0.47727 | 0.77713 | 1.41343 | 26 |
| GO_TRANSCRIPTION_COUPLED_NUCLEOTIDE_EXCISION_REPAIR | 0.00231 | 0.02642 | 0.43171 | 0.65429 | 1.23840 | 71 |
| GO_TRANSFERRIN_TRANSPORT | 0.00141 | 0.01850 | 0.45506 | 0.75134 | 1.37179 | 27 |
| GO_TRANSLATIONAL_ELONGATION | 0.00000 | 0.00000 | | 0.75263 | 1.45028 | 124 |
| GO_TRANSLATIONAL_INITIATION | 0.00000 | 0.00000 | | 0.75139 | 1.45839 | 171 |
| GO_TRANSLATIONAL_TERMINATION | 0.00000 | 0.00000 | 0.81404 | 0.75110 | 1.44066 | 101 |
| GO_TRANSLATION_ELONGATION_FACTOR_ACTIVITY | 0.00390 | 0.03895 | 0.43171 | 0.80505 | 1.41260 | 15 |
| GO_TRANSLATION_FACTOR_ACTIVITY_RNA_BINDING | 0.00000 | 0.00000 | 0.74774 | 0.77244 | 1.46085 | 70 |
| GO_TRANSLATION_INITIATION_FACTOR_ACTIVITY | 0.00000 | 0.00011 | 0.62726 | 0.77773 | 1.44952 | 45 |
| GO_TRANSLATION_PREINITIATION_COMPLEX | 0.00342 | 0.03521 | 0.43171 | 0.79777 | 1.40694 | 16 |
| GO_TRANSLATION_REGULATOR_ACTIVITY | 0.00000 | 0.00000 | 0.83909 | 0.74000 | 1.42403 | 111 |
| GO_TRANSLATION_REGULATOR_ACTIVITY_NUCLEIC_ACID_BIND | 0.00000 | 0.00000 | 0.83909 | 0.76441 | 1.45959 | 87 |
| GO_TRANSMEMBRANE_RECEPTOR_PROTEIN_TYROSINE_KINASE | 0.00168 | 0.02098 | 0.45506 | 0.56326 | 1.11076 | 411 |
| GO_TRICARBOXYLIC_ACID_CYCLE | 0.00159 | 0.02016 | 0.45506 | 0.73558 | 1.35097 | 33 |
| GO_TRNA_TRANSPORT | 0.00342 | 0.03521 | 0.43171 | 0.71247 | 1.31220 | 35 |
| GO_TUMOR_NECROSIS_FACTOR_MEDIATED_SIGNALING_PATHW | 0.00000 | 0.00000 | 0.80122 | 0.73921 | 1.42138 | 108 |
| GO_T_CELL_RECEPTOR_SIGNALING_PATHWAY | 0.00000 | 0.00001 | 0.70498 | 0.69185 | 1.33314 | 119 |
| GO_U12_TYPE_SPLICEOSOMAL_COMPLEX | 0.00487 | 0.04559 | 0.40702 | 0.73423 | 1.33541 | 26 |
| GO_U2_SNRNP | 0.00141 | 0.01850 | 0.45506 | 0.80259 | 1.42065 | 17 |
| GO_U2_TYPE_CATALYTIC_STEP_2_SPLICEOSOME | 0.00268 | 0.02961 | 0.43171 | 0.72225 | 1.32288 | 29 |
| GO_U2_TYPE_SPLICEOSOMAL_COMPLEX | 0.00033 | 0.00595 | 0.49849 | 0.66590 | 1.26925 | 84 |
| GO_UBIQUITIN_LIKE_PROTEIN_LIGASE_BINDING | 0.00000 | 0.00007 | 0.62726 | 0.62616 | 1.22416 | 251 |
| GO_UNFOLDED_PROTEIN_BINDING | 0.00000 | 0.00001 | 0.69013 | 0.72112 | 1.37766 | 88 |
| GO_VACUOLAR_LUMEN | 0.00016 | 0.00319 | 0.51885 | 0.65416 | 1.25772 | 107 |
| GO_VESICLE_BUDDING_FROM_MEMBRANE | 0.00480 | 0.04537 | 0.40702 | 0.62411 | 1.19353 | 90 |
| GO_VESICLE_LUMEN | 0.00000 | 0.00000 | 0.77494 | 0.67028 | 1.30631 | 207 |
| GO_VESICLE_MEMBRANE | 0.00001 | 0.00026 | 0.59333 | 0.58430 | 1.15387 | 476 |
| GO_VIRAL_GENE_EXPRESSION | 0.00000 | 0.00000 | | 0.70963 | 1.37745 | 172 |
| GO_VIRAL_LIFE_CYCLE | 0.00000 | 0.00001 | 0.69013 | 0.63985 | 1.24950 | 238 |
| GO_VIRION_ASSEMBLY | 0.00457 | 0.04386 | 0.40702 | 0.69689 | 1.28528 | 36 |
| HP_ABNORMALITY_OF_ACID_BASE_HOMEOSTASIS | 0.00000 | 0.00010 | 0.62726 | 0.62283 | 1.21827 | 261 |
| HP_ABNORMALITY_OF_BLOOD_CIRCULATION | 0.00502 | 0.04608 | 0.40702 | 0.59228 | 1.14883 | 167 |
| HP_ABNORMALITY_OF_CARDIOVASCULAR_SYSTEM_ELECTROPH | 0.00011 | 0.00237 | 0.53843 | 0.60202 | 1.17758 | 261 |
| HP_ABNORMALITY_OF_SKIN_ADNEXA_PHYSIOLOGY | 0.00242 | 0.02748 | 0.43171 | 0.61294 | 1.18094 | 118 |
| HP_ABNORMALITY_OF_THE_ANKLES | 0.00313 | 0.03310 | 0.43171 | 0.65369 | 1.23418 | 63 |
| HP_ABNORMALITY_OF_THE_CEREBRAL_VASCULATURE | 0.00357 | 0.03638 | 0.43171 | 0.59764 | 1.15722 | 158 |
| HP_ABNORMALITY_OF_THE_CEREBROSPINAL_FLUID | 0.00030 | 0.00549 | 0.49849 | 0.58221 | 1.14439 | 325 |
| HP_ABNORMALITY_OF_THE_MITOCHONDRION | 0.00126 | 0.01715 | 0.45506 | 0.62149 | 1.19764 | 120 |
| HP_ABNORMALITY_OF_THE_OPTIC_DISC | 0.00198 | 0.02370 | 0.43171 | 0.56624 | 1.11727 | 432 |
| HP_ABNORMALITY_OF_THE_OPTIC_NERVE | 0.00036 | 0.00622 | 0.49849 | 0.56723 | 1.12034 | 486 |
| HP_ABNORMAL_ABDOMEN_MORPHOLOGY | 0.00089 | 0.01291 | 0.47727 | 0.56424 | 1.11364 | 441 |
| HP_ABNORMAL_ACTIVITY_OF_MITOCHONDRIAL_RESPIRATORY_ | 0.00014 | 0.00300 | 0.51885 | 0.70303 | 1.32524 | 61 |
| HP_ABNORMAL_BASAL_GANGLIA_MRI_SIGNAL_INTENSITY | 0.00179 | 0.02183 | 0.45506 | 0.71597 | 1.32046 | 36 |
| HP_ABNORMAL_BRAINSTEM_MRI_SIGNAL_INTENSITY | 0.00398 | 0.03962 | 0.40702 | 0.71528 | 1.31370 | 33 |
| HP_ABNORMAL_CELLULAR_PHENOTYPE | 0.00209 | 0.02470 | 0.43171 | 0.56937 | 1.11992 | 334 |
| HP_ABNORMAL_CIRCULATING_CARBOHYDRATE_CONCENTRATIC | 0.00194 | 0.02331 | 0.45506 | 0.69518 | 1.29539 | 46 |
| HP_ABNORMAL_CNS_MYELINATION | 0.00014 | 0.00300 | 0.51885 | 0.62318 | 1.20954 | 171 |
| HP_ABNORMAL_CSF_METABOLITE_LEVEL | 0.00001 | 0.00027 | 0.59333 | 0.70531 | 1.34409 | 82 |
| HP_ABNORMAL_ELECTROPHYSIOLOGY_OF_SINOATRIAL_NODE_ | 0.00029 | 0.00536 | 0.49849 | 0.73172 | 1.35725 | 41 |
| HP_ABNORMAL_ENZYME_COENZYME_ACTIVITY | 0.00561 | 0.04971 | 0.40702 | 0.60490 | 1.16567 | 123 |
| HP_ABNORMAL_ERYTHROCYTE_MORPHOLOGY | 0.00001 | 0.00023 | 0.61053 | 0.58835 | 1.15904 | 380 |
| HP_ABNORMAL_HEART_VALVE_PHYSIOLOGY | 0.00405 | 0.04011 | 0.40702 | 0.59004 | 1.14474 | 169 |

| | | | | | | |
|---|---|---|---|---|---|---|
| HP_ABNORMAL_MITOCHONDRIA_IN_MUSCLE_TISSUE | 0.00075 | 0.01118 | 0.47727 | 0.74633 | 1.36700 | 29 |
| HP_ABNORMAL_MYELINATION | 0.00007 | 0.00169 | 0.53843 | 0.59258 | 1.16479 | 324 |
| HP_ABNORMAL_MYOCARDIUM_MORPHOLOGY | 0.00008 | 0.00192 | 0.53843 | 0.58647 | 1.15365 | 340 |
| HP_ABNORMAL_RENAL_PHYSIOLOGY | 0.00450 | 0.04342 | 0.40702 | 0.55484 | 1.09317 | 385 |
| HP_ABNORMAL_TRICUSPID_VALVE_PHYSIOLOGY | 0.00231 | 0.02642 | 0.43171 | 0.72968 | 1.34029 | 31 |
| HP_ABNORMAL_VITREOUS_HUMOR_MORPHOLOGY | 0.00181 | 0.02201 | 0.45506 | 0.79632 | 1.40955 | 17 |
| HP_ACUTE_LEUKEMIA | 0.00013 | 0.00276 | 0.51885 | 0.68145 | 1.29219 | 75 |
| HP_ANEMIA_OF_INADEQUATE_PRODUCTION | 0.00561 | 0.04971 | 0.40702 | 0.61298 | 1.17855 | 107 |
| HP_APHASIA | 0.00213 | 0.02490 | 0.43171 | 0.72858 | 1.33704 | 30 |
| HP_BLINDNESS | 0.00018 | 0.00355 | 0.51885 | 0.61672 | 1.19768 | 175 |
| HP_BONE_MARROW_HYPERCELLULARITY | 0.00045 | 0.00725 | 0.49849 | 0.82169 | 1.46806 | 19 |
| HP_CHOREOATHETOSIS | 0.00561 | 0.04971 | 0.40702 | 0.64956 | 1.22631 | 64 |
| HP_CHRONIC_INFECTION | 0.00121 | 0.01658 | 0.45506 | 0.81916 | 1.44468 | 16 |
| HP_CONGESTIVE_HEART_FAILURE | 0.00495 | 0.04573 | 0.40702 | 0.60734 | 1.16929 | 114 |
| HP_CRANIAL_NERVE_COMPRESSION | 0.00331 | 0.03441 | 0.43171 | 0.75830 | 1.36989 | 23 |
| HP_DECREASED_ACTIVITY_OF_MITOCHONDRIAL_COMPLEX_I | 0.00283 | 0.03081 | 0.43171 | 0.71940 | 1.32496 | 35 |
| HP_DECREASED_ACTIVITY_OF_THE_PYRUVATE_DEHYDROGENAS | 0.00062 | 0.00949 | 0.47727 | 0.75390 | 1.38478 | 31 |
| HP_DEPRESSED_NASAL_RIDGE | 0.00041 | 0.00681 | 0.49849 | 0.66218 | 1.26189 | 82 |
| HP_DYSPNEA | 0.00250 | 0.02807 | 0.43171 | 0.58107 | 1.13348 | 224 |
| HP_DYSTONIA | 0.00069 | 0.01043 | 0.47727 | 0.58101 | 1.13736 | 279 |
| HP_EMOTIONAL_LABILITY | 0.00016 | 0.00321 | 0.51885 | 0.66683 | 1.27522 | 90 |
| HP_EPISODIC_ATAXIA | 0.00487 | 0.04559 | 0.40702 | 0.79934 | 1.40259 | 15 |
| HP_FASCICULATIONS | 0.00130 | 0.01761 | 0.45506 | 0.68128 | 1.28064 | 54 |
| HP_FATIGUE | 0.00042 | 0.00689 | 0.49849 | 0.59902 | 1.16764 | 212 |
| HP_FETAL_DISTRESS | 0.00179 | 0.02183 | 0.45506 | 0.72928 | 1.33973 | 32 |
| HP_FLUSHING | 0.00561 | 0.04971 | 0.40702 | 0.74782 | 1.33686 | 20 |
| HP_FOCAL_T2_HYPERINTENSE_BRAINSTEM_LESION | 0.00159 | 0.02016 | 0.45506 | 0.74970 | 1.36879 | 27 |
| HP_HEADACHE | 0.00016 | 0.00319 | 0.51885 | 0.62537 | 1.21179 | 163 |
| HP_HEPATOMEGALY | 0.00108 | 0.01507 | 0.45506 | 0.57439 | 1.12990 | 335 |
| HP_HYPERTROPHIC_CARDIOMYOPATHY | 0.00002 | 0.00050 | 0.57561 | 0.62629 | 1.21908 | 194 |
| HP_IMPAIRED_SOCIAL_INTERACTIONS | 0.00055 | 0.00863 | 0.47727 | 0.66555 | 1.26058 | 72 |
| HP_INCREASED_CSF_LACTATE | 0.00001 | 0.00034 | 0.59333 | 0.71937 | 1.35857 | 68 |
| HP_INCREASED_SERUM_LACTATE | 0.00001 | 0.00028 | 0.59333 | 0.65837 | 1.26932 | 133 |
| HP_INCREASED_SERUM_PYRUVATE | 0.00110 | 0.01526 | 0.45506 | 0.73143 | 1.35198 | 37 |
| HP_INFANTILE_ONSET | 0.00487 | 0.04559 | 0.40702 | 0.56559 | 1.10939 | 300 |
| HP_INTRACRANIAL_CYSTIC_LESION | 0.00055 | 0.00863 | 0.47727 | 0.62760 | 1.20885 | 116 |
| HP_INTRAUTERINE_GROWTH_RETARDATION | 0.00305 | 0.03271 | 0.43171 | 0.56594 | 1.11393 | 350 |
| HP_LACTIC_ACIDOSIS | 0.00018 | 0.00355 | 0.51885 | 0.63312 | 1.22071 | 131 |
| HP_LETHARGY | 0.00084 | 0.01228 | 0.47727 | 0.63361 | 1.21845 | 109 |
| HP_LEUKODYSTROPHY | 0.00029 | 0.00529 | 0.49849 | 0.66598 | 1.26915 | 82 |
| HP_LEUKOENCEPHALOPATHY | 0.00008 | 0.00190 | 0.53843 | 0.64132 | 1.23667 | 134 |
| HP_LIPID_ACCUMULATION_IN_HEPATOCYTES | 0.00220 | 0.02548 | 0.43171 | 0.62794 | 1.20159 | 91 |
| HP_LOWER_LIMB_SPASTICITY | 0.00154 | 0.01974 | 0.45506 | 0.60929 | 1.17648 | 139 |
| HP_MACROCYTIC_ANEMIA | 0.00030 | 0.00542 | 0.49849 | 0.73108 | 1.35606 | 41 |
| HP_METABOLIC_ACIDOSIS | 0.00034 | 0.00599 | 0.49849 | 0.64831 | 1.24464 | 103 |
| HP_MICRONODULAR_CIRRHOSIS | 0.00324 | 0.03393 | 0.43171 | 0.78723 | 1.40018 | 18 |
| HP_MIGRAINE | 0.00028 | 0.00515 | 0.49849 | 0.67309 | 1.27635 | 75 |
| HP_MUSCLE_ABNORMALITY_RELATED_TO_MITOCHONDRIAL_DY | 0.00209 | 0.02470 | 0.43171 | 0.68258 | 1.27573 | 49 |
| HP_MUTISM | 0.00283 | 0.03081 | 0.43171 | 0.71937 | 1.32490 | 35 |
| HP_NAUSEA_AND_VOMITING | 0.00022 | 0.00417 | 0.51885 | 0.59383 | 1.16158 | 264 |
| HP_NEOPLASM_OF_THE_ENDOCRINE_SYSTEM | 0.00495 | 0.04573 | 0.40702 | 0.63166 | 1.20308 | 81 |
| HP_OPTIC_NEUROPATHY | 0.00135 | 0.01808 | 0.45506 | 0.73837 | 1.35610 | 33 |
| HP_PACE_OF_PROGRESSION | 0.00487 | 0.04559 | 0.40702 | 0.56814 | 1.11416 | 298 |
| HP_PALLOR | 0.00036 | 0.00622 | 0.49849 | 0.64861 | 1.24463 | 102 |
| HP_PARAPLEGIA | 0.00121 | 0.01658 | 0.45506 | 0.63584 | 1.22133 | 104 |
| HP_PARAPLEGIA_PARAPARESIS | 0.00036 | 0.00621 | 0.49849 | 0.62047 | 1.20040 | 150 |
| HP_PAROXYSMAL_INVOLUNTARY_EYE_MOVEMENTS | 0.00054 | 0.00849 | 0.47727 | 0.75650 | 1.38957 | 31 |
| HP_PEDIATRIC_ONSET | 0.00561 | 0.04971 | 0.40702 | 0.55915 | 1.10116 | 371 |
| HP_PIGMENTARY_RETINOPATHY | 0.00065 | 0.00981 | 0.47727 | 0.63260 | 1.21720 | 113 |
| HP_POOR_EYE_CONTACT | 0.00017 | 0.00341 | 0.51885 | 0.71279 | 1.33749 | 53 |
| HP_POOR_HEAD_CONTROL | 0.00000 | 0.00005 | 0.64355 | 0.67819 | 1.30665 | 118 |
| HP_PROGRESSIVE | 0.00194 | 0.02331 | 0.45506 | 0.59790 | 1.16273 | 186 |
| HP_PROGRESSIVE_CEREBELLAR_ATAXIA | 0.00364 | 0.03677 | 0.43171 | 0.64788 | 1.22325 | 65 |
| HP_PROGRESSIVE_SPASTIC_PARAPLEGIA | 0.00253 | 0.02838 | 0.43171 | 0.69655 | 1.29822 | 45 |
| HP_PROXIMAL_TUBULOPATHY | 0.00141 | 0.01850 | 0.45506 | 0.72579 | 1.33535 | 34 |
| HP_PTOSIS | 0.00472 | 0.04480 | 0.40702 | 0.55582 | 1.09637 | 418 |
| HP_REDUCED_CONSCIOUSNESS_CONFUSION | 0.00007 | 0.00164 | 0.53843 | 0.61227 | 1.19295 | 205 |
| HP_REDUCED_VISUAL_ACUITY | 0.00053 | 0.00837 | 0.47727 | 0.58618 | 1.14774 | 283 |
| HP_RENAL_TUBULAR_DYSFUNCTION | 0.00013 | 0.00284 | 0.51885 | 0.68108 | 1.29150 | 75 |
| HP_RESPIRATORY_DISTRESS | 0.00108 | 0.01507 | 0.45506 | 0.62693 | 1.20762 | 115 |

| | | | | | | |
|---|---|---|---|---|---|---|
| HP_RESPIRATORY_INSUFFICIENCY | 0.00000 | 0.00010 | 0.62726 | 0.60536 | 1.18998 | 326 |
| HP_SINUS_TACHYCARDIA | 0.00213 | 0.02490 | 0.43171 | 0.77203 | 1.39128 | 22 |
| HP_SPARSE_EYELASHES | 0.00150 | 0.01943 | 0.45506 | 0.71355 | 1.32161 | 40 |
| HP_SPARSE_OR_ABSENT_EYELASHES | 0.00220 | 0.02548 | 0.43171 | 0.67331 | 1.26567 | 54 |
| HP_SPASTIC_PARAPLEGIA | 0.00098 | 0.01407 | 0.45506 | 0.64059 | 1.22712 | 96 |
| HP_THROMBOCYTOSIS | 0.00413 | 0.04065 | 0.40702 | 0.75210 | 1.35870 | 23 |
| HP_TONGUE_FASCICULATIONS | 0.00457 | 0.04386 | 0.40702 | 0.80056 | 1.40472 | 15 |
| HP_TRANSIENT_ISCHEMIC_ATTACK | 0.00480 | 0.04537 | 0.40702 | 0.74902 | 1.35809 | 25 |
| HP_VERTIGO | 0.00002 | 0.00063 | 0.57561 | 0.72114 | 1.35844 | 60 |
| HP_VESTIBULAR_DYSFUNCTION | 0.00028 | 0.00522 | 0.49849 | 0.67279 | 1.27577 | 75 |
| HP_VOMITING | 0.00072 | 0.01082 | 0.47727 | 0.61459 | 1.18792 | 147 |
| HP_WEAKNESS_DUE_TO_UPPER_MOTOR_NEURON_DYSFUNCTI | 0.00532 | 0.04803 | 0.40702 | 0.56829 | 1.11186 | 272 |
| HP_WEAKNESS_OF_MUSCLES_OF_RESPIRATION | 0.00163 | 0.02050 | 0.45506 | 0.66272 | 1.25123 | 63 |
| **Cluster 1** | | | | | | |
| pathway | pval | padj | log2err | ES | NES | size |
| GO_ACTIVATION_OF_IMMUNE_RESPONSE | 0.00000 | 0.00000 | 0.74774 | 0.65038 | 1.26999 | 235 |
| GO_ACTIVATION_OF_INNATE_IMMUNE_RESPONSE | 0.00000 | 0.00001 | 0.69013 | 0.71967 | 1.37861 | 96 |
| GO_AEROBIC_RESPIRATION | 0.00000 | 0.00000 | 0.73376 | 0.76280 | 1.44478 | 72 |
| GO_ALCOHOL_BIOSYNTHETIC_PROCESS | 0.00032 | 0.00572 | 0.49849 | 0.64679 | 1.24367 | 108 |
| GO_ALCOHOL_METABOLIC_PROCESS | 0.00364 | 0.03677 | 0.43171 | 0.58112 | 1.13424 | 228 |
| GO_AMEBOIDAL_TYPE_CELL_MIGRATION | 0.00457 | 0.04386 | 0.40702 | 0.57755 | 1.12854 | 247 |
| GO_AMIDE_BINDING | 0.00014 | 0.00297 | 0.51885 | 0.61533 | 1.19780 | 199 |
| GO_AMINE_METABOLIC_PROCESS | 0.00000 | 0.00000 | 0.76146 | 0.73732 | 1.41097 | 92 |
| GO_AMYLOID_BETA_CLEARANCE | 0.00561 | 0.04971 | 0.40702 | 0.74081 | 1.33976 | 24 |
| GO_ANAPHASE_PROMOTING_COMPLEX_DEPENDENT_CATABOL | 0.00000 | 0.00000 | 0.76146 | 0.77107 | 1.46112 | 73 |
| GO_ANATOMICAL_STRUCTURE_HOMEOSTASIS | 0.00020 | 0.00390 | 0.51885 | 0.59307 | 1.16021 | 271 |
| GO_ANIMAL_ORGAN_REGENERATION | 0.00012 | 0.00269 | 0.51885 | 0.71251 | 1.33510 | 52 |
| GO_ANTIGEN_PROCESSING_AND_PRESENTATION | 0.00000 | 0.00000 | 0.74774 | 0.68785 | 1.33064 | 149 |
| GO_ANTIGEN_PROCESSING_AND_PRESENTATION_OF_EXOGENC | 0.00000 | 0.00000 | 0.76146 | 0.79644 | 1.50237 | 62 |
| GO_ANTIGEN_PROCESSING_AND_PRESENTATION_OF_PEPTIDE_ | 0.00000 | 0.00000 | 0.74774 | 0.70107 | 1.35144 | 130 |
| GO_ANTIGEN_PROCESSING_AND_PRESENTATION_OF_PEPTIDE_ | 0.00000 | 0.00000 | 0.78819 | 0.77566 | 1.47012 | 74 |
| GO_ANTIGEN_RECEPTOR_MEDIATED_SIGNALING_PATHWAY | 0.00000 | 0.00009 | 0.62726 | 0.66362 | 1.28080 | 136 |
| GO_APICAL_PART_OF_CELL | 0.00135 | 0.01808 | 0.45506 | 0.58926 | 1.14966 | 220 |
| GO_APOPTOTIC_SIGNALING_PATHWAY | 0.00006 | 0.00152 | 0.53843 | 0.58638 | 1.15586 | 393 |
| GO_ATPASE_ACTIVITY | 0.00050 | 0.00802 | 0.47727 | 0.58216 | 1.14385 | 316 |
| GO_ATPASE_ACTIVITY_COUPLED_TO_TRANSMEMBRANE_MOVE | 0.00420 | 0.04132 | 0.40702 | 0.78005 | 1.38741 | 18 |
| GO_ATP_BIOSYNTHETIC_PROCESS | 0.00013 | 0.00276 | 0.51885 | 0.83897 | 1.50801 | 21 |
| GO_ATP_METABOLIC_PROCESS | 0.00000 | 0.00000 | | 0.72178 | 1.40814 | 219 |
| GO_ATP_SYNTHESIS_COUPLED_ELECTRON_TRANSPORT | 0.00000 | 0.00000 | | 0.80353 | 1.52425 | 76 |
| GO_AZUROPHIL_GRANULE | 0.00025 | 0.00466 | 0.49849 | 0.64781 | 1.24551 | 107 |
| GO_AZUROPHIL_GRANULE_LUMEN | 0.00405 | 0.04011 | 0.40702 | 0.65625 | 1.23793 | 62 |
| GO_BASAL_PART_OF_CELL | 0.00509 | 0.04656 | 0.40702 | 0.68515 | 1.27697 | 45 |
| GO_BASAL_PLASMA_MEMBRANE | 0.00346 | 0.03548 | 0.43171 | 0.71184 | 1.31104 | 35 |
| GO_BASEMENT_MEMBRANE | 0.00279 | 0.03056 | 0.43171 | 0.68935 | 1.28452 | 46 |
| GO_BASOLATERAL_PLASMA_MEMBRANE | 0.00011 | 0.00247 | 0.53843 | 0.63209 | 1.21993 | 136 |
| GO_BINDING_OF_SPERM_TO_ZONA_PELLUCIDA | 0.00000 | 0.00006 | 0.64355 | 0.90095 | 1.60966 | 19 |
| GO_BLOOD_MICROPARTICLE | 0.00000 | 0.00000 | 0.70498 | 0.81982 | 1.52763 | 46 |
| GO_BRUSH_BORDER | 0.00060 | 0.00923 | 0.47727 | 0.66897 | 1.26790 | 74 |
| GO_CADHERIN_BINDING | 0.00000 | 0.00000 | | 0.68759 | 1.34435 | 253 |
| GO_CALCIUM_DEPENDENT_PHOSPHOLIPID_BINDING | 0.00152 | 0.01958 | 0.45506 | 0.75021 | 1.37120 | 28 |
| GO_CALCIUM_DEPENDENT_PROTEIN_BINDING | 0.00009 | 0.00211 | 0.53843 | 0.74254 | 1.38394 | 45 |
| GO_CANONICAL_WNT_SIGNALING_PATHWAY | 0.00004 | 0.00109 | 0.55733 | 0.61740 | 1.20458 | 222 |
| GO_CARBOHYDRATE_CATABOLIC_PROCESS | 0.00135 | 0.01808 | 0.45506 | 0.61179 | 1.18186 | 144 |
| GO_CATALYTIC_STEP_2_SPLICEOSOME | 0.00001 | 0.00022 | 0.61053 | 0.70290 | 1.33765 | 80 |
| GO_CATION_TRANSMEMBRANE_TRANSPORT | 0.00327 | 0.03409 | 0.43171 | 0.56060 | 1.10531 | 406 |
| GO_CATION_TRANSMEMBRANE_TRANSPORTER_ACTIVITY | 0.00097 | 0.01387 | 0.47727 | 0.58196 | 1.13846 | 265 |
| GO_CELLULAR_AMINO_ACID_METABOLIC_PROCESS | 0.00002 | 0.00060 | 0.57561 | 0.61438 | 1.19930 | 231 |
| GO_CELLULAR_COMPONENT_DISASSEMBLY | 0.00000 | 0.00000 | | 0.64070 | 1.26342 | 408 |
| GO_CELLULAR_ION_HOMEOSTASIS | 0.00027 | 0.00493 | 0.49849 | 0.58658 | 1.15340 | 328 |
| GO_CELLULAR_IRON_ION_HOMEOSTASIS | 0.00327 | 0.03409 | 0.43171 | 0.68360 | 1.27374 | 47 |
| GO_CELLULAR_KETONE_METABOLIC_PROCESS | 0.00000 | 0.00010 | 0.62726 | 0.65358 | 1.26726 | 164 |
| GO_CELLULAR_MONOVALENT_INORGANIC_CATION_HOMEOSTA | 0.00179 | 0.02183 | 0.45506 | 0.64747 | 1.22821 | 76 |
| GO_CELLULAR_NITROGEN_COMPOUND_CATABOLIC_PROCESS | 0.00000 | 0.00000 | | 0.67915 | 1.34096 | 456 |
| GO_CELLULAR_PROTEIN_COMPLEX_DISASSEMBLY | 0.00000 | 0.00000 | | 0.69566 | 1.35184 | 179 |
| GO_CELLULAR_RESPIRATION | 0.00000 | 0.00000 | | 0.74205 | 1.43370 | 145 |
| GO_CELLULAR_RESPONSE_TO_HEAT | 0.00004 | 0.00109 | 0.55733 | 0.67186 | 1.28698 | 95 |
| GO_CELLULAR_RESPONSE_TO_HORMONE_STIMULUS | 0.00309 | 0.03299 | 0.43171 | 0.56453 | 1.11191 | 374 |
| GO_CELLULAR_RESPONSE_TO_ORGANIC_CYCLIC_COMPOUND | 0.00502 | 0.04608 | 0.40702 | 0.56512 | 1.11134 | 329 |
| GO_CELLULAR_RESPONSE_TO_OXYGEN_LEVELS | 0.00000 | 0.00000 | | 0.71461 | 1.38224 | 151 |
| GO_CELL_ACTIVATION_INVOLVED_IN_IMMUNE_RESPONSE | 0.00000 | 0.00000 | | 0.63924 | 1.26108 | 422 |

| | | | | | | |
|---|---|---|---|---|---|---|
| GO_CELL_ADHESION_MEDIATOR_ACTIVITY | 0.00157 | 0.02009 | 0.45506 | 0.71837 | 1.32488 | 36 |
| GO_CELL_ADHESION_MOLECULE_BINDING | 0.00000 | 0.00000 | | 0.67109 | 1.32112 | 353 |
| GO_CELL_BODY | 0.00004 | 0.00109 | 0.55733 | 0.59973 | 1.17541 | 293 |
| GO_CELL_CELL_RECOGNITION | 0.00000 | 0.00006 | 0.64355 | 0.82001 | 1.51026 | 35 |
| GO_CELL_CELL_SIGNALING_BY_WNT | 0.00004 | 0.00109 | 0.55733 | 0.59042 | 1.16141 | 339 |
| GO_CELL_CYCLE_G1_S_PHASE_TRANSITION | 0.00376 | 0.03783 | 0.43171 | 0.58112 | 1.13378 | 220 |
| GO_CELL_CYCLE_G2_M_PHASE_TRANSITION | 0.00024 | 0.00459 | 0.51885 | 0.59807 | 1.16746 | 231 |
| GO_CELL_CYCLE_PHASE_TRANSITION | 0.00000 | 0.00001 | 0.69013 | 0.59417 | 1.17348 | 499 |
| GO_CELL_DIVISION | 0.00003 | 0.00071 | 0.57561 | 0.58314 | 1.15094 | 449 |
| GO_CELL_KILLING | 0.00205 | 0.02440 | 0.43171 | 0.66727 | 1.25696 | 60 |
| GO_CELL_RECOGNITION | 0.00000 | 0.00000 | 0.73376 | 0.75953 | 1.43954 | 74 |
| GO_CELL_REDOX_HOMEOSTASIS | 0.00163 | 0.02050 | 0.45506 | 0.71772 | 1.32665 | 38 |
| GO_CELL_SUBSTRATE_JUNCTION | 0.00000 | 0.00000 | | 0.65742 | 1.29171 | 316 |
| GO_CELL_SURFACE | 0.00000 | 0.00000 | 0.73376 | 0.62572 | 1.23070 | 330 |
| GO_CELL_SURFACE_RECEPTOR_SIGNALING_PATHWAY_INVOLVEI | 0.00038 | 0.00640 | 0.49849 | 0.57762 | 1.13770 | 374 |
| GO_CHAPERONE_BINDING | 0.00495 | 0.04573 | 0.40702 | 0.62860 | 1.19935 | 86 |
| GO_CHAPERONE_COMPLEX | 0.00001 | 0.00028 | 0.59333 | 0.86337 | 1.55587 | 22 |
| GO_CHAPERONE_MEDIATED_PROTEIN_COMPLEX_ASSEMBLY | 0.00100 | 0.01430 | 0.45506 | 0.80787 | 1.44337 | 19 |
| GO_CHAPERONE_MEDIATED_PROTEIN_FOLDING | 0.00168 | 0.02098 | 0.45506 | 0.71110 | 1.32288 | 44 |
| GO_CHROMOSOMAL_REGION | 0.00018 | 0.00349 | 0.51885 | 0.59323 | 1.16180 | 286 |
| GO_CHROMOSOME_CENTROMERIC_REGION | 0.00172 | 0.02135 | 0.45506 | 0.60447 | 1.17248 | 168 |
| GO_CHROMOSOME_SEGREGATION | 0.00003 | 0.00087 | 0.55733 | 0.60463 | 1.18280 | 270 |
| GO_CLUSTER_OF_ACTIN_BASED_CELL_PROJECTIONS | 0.00074 | 0.01107 | 0.47727 | 0.64184 | 1.23044 | 99 |
| GO_COATED_VESICLE | 0.00213 | 0.02490 | 0.43171 | 0.59512 | 1.15846 | 199 |
| GO_COATED_VESICLE_MEMBRANE | 0.00060 | 0.00923 | 0.47727 | 0.63443 | 1.22207 | 115 |
| GO_COLLAGEN_BINDING | 0.00465 | 0.04430 | 0.40702 | 0.73471 | 1.33628 | 26 |
| GO_COLLAGEN_CONTAINING_EXTRACELLULAR_MATRIX | 0.00000 | 0.00006 | 0.64355 | 0.65358 | 1.26542 | 157 |
| GO_CONDENSED_CHROMOSOME | 0.00350 | 0.03580 | 0.43171 | 0.59394 | 1.15417 | 179 |
| GO_CONTRACTILE_FIBER | 0.00064 | 0.00972 | 0.47727 | 0.62217 | 1.19973 | 134 |
| GO_COPI_COATED_VESICLE | 0.00316 | 0.03344 | 0.43171 | 0.75688 | 1.37234 | 25 |
| GO_CORNIFICATION | 0.00027 | 0.00500 | 0.49849 | 0.77970 | 1.42512 | 28 |
| GO_CORTICAL_CYTOSKELETON | 0.00502 | 0.04608 | 0.40702 | 0.64388 | 1.21870 | 71 |
| GO_COTRANSLATIONAL_PROTEIN_TARGETING_TO_MEMBRANE | 0.00000 | 0.00000 | | 0.78223 | 1.49819 | 94 |
| GO_CYTOCHROME_COMPLEX | 0.00035 | 0.00610 | 0.49849 | 0.80796 | 1.45601 | 22 |
| GO_CYTOKINE_MEDIATED_SIGNALING_PATHWAY | 0.00000 | 0.00002 | 0.67496 | 0.60742 | 1.19735 | 392 |
| GO_CYTOPLASMIC_STRESS_GRANULE | 0.00014 | 0.00302 | 0.51885 | 0.69390 | 1.31009 | 63 |
| GO_CYTOPLASMIC_TRANSLATION | 0.00000 | 0.00001 | 0.70498 | 0.71773 | 1.37349 | 92 |
| GO_CYTOPLASMIC_TRANSLATIONAL_INITIATION | 0.00178 | 0.02182 | 0.45506 | 0.73517 | 1.35039 | 31 |
| GO_CYTOSOLIC_LARGE_RIBOSOMAL_SUBUNIT | 0.00000 | 0.00005 | 0.64355 | 0.77240 | 1.44935 | 53 |
| GO_CYTOSOLIC_RIBOSOME | 0.00000 | 0.00000 | | 0.77483 | 1.48437 | 97 |
| GO_CYTOSOLIC_SMALL_RIBOSOMAL_SUBUNIT | 0.00000 | 0.00010 | 0.62726 | 0.79384 | 1.47033 | 40 |
| GO_DETECTION_OF_STIMULUS | 0.00020 | 0.00393 | 0.51885 | 0.65208 | 1.25384 | 108 |
| GO_DETOXIFICATION | 0.00413 | 0.04065 | 0.40702 | 0.65399 | 1.23684 | 70 |
| GO_DEVELOPMENT_OF_PRIMARY_FEMALE_SEXUAL_CHARACTE | 0.00179 | 0.02183 | 0.45506 | 0.68410 | 1.28057 | 51 |
| GO_DICARBOXYLIC_ACID_METABOLIC_PROCESS | 0.00261 | 0.02910 | 0.43171 | 0.65541 | 1.23747 | 65 |
| GO_DISORDERED_DOMAIN_SPECIFIC_BINDING | 0.00051 | 0.00812 | 0.47727 | 0.76867 | 1.40343 | 27 |
| GO_DISULFIDE_OXIDOREDUCTASE_ACTIVITY | 0.00435 | 0.04238 | 0.40702 | 0.71445 | 1.31232 | 31 |
| GO_DNA_BIOSYNTHETIC_PROCESS | 0.00017 | 0.00334 | 0.51885 | 0.62540 | 1.20833 | 145 |
| GO_DNA_CONFORMATION_CHANGE | 0.00327 | 0.03409 | 0.43171 | 0.57979 | 1.13178 | 231 |
| GO_DNA_DAMAGE_RESPONSE_DETECTION_OF_DNA_DAMAGE | 0.00305 | 0.03271 | 0.43171 | 0.71625 | 1.31916 | 35 |
| GO_DNA_GEOMETRIC_CHANGE | 0.00141 | 0.01850 | 0.45506 | 0.62642 | 1.20356 | 105 |
| GO_DOUBLE_STRANDED_RNA_BINDING | 0.00039 | 0.00658 | 0.49849 | 0.69158 | 1.30365 | 61 |
| GO_DRUG_BINDING | 0.00450 | 0.04342 | 0.40702 | 0.66245 | 1.24130 | 52 |
| GO_ELECTRON_TRANSFER_ACTIVITY | 0.00000 | 0.00001 | 0.67496 | 0.72917 | 1.38765 | 80 |
| GO_ELECTRON_TRANSPORT_CHAIN | 0.00000 | 0.00000 | | 0.73336 | 1.41392 | 133 |
| GO_EMBRYO_IMPLANTATION | 0.00044 | 0.00706 | 0.49849 | 0.75968 | 1.39146 | 29 |
| GO_ENDOCYTIC_VESICLE_MEMBRANE | 0.00524 | 0.04770 | 0.40702 | 0.62563 | 1.19564 | 89 |
| GO_ENDOPEPTIDASE_ACTIVITY | 0.00065 | 0.00981 | 0.47727 | 0.60123 | 1.16998 | 196 |
| GO_ENDOPEPTIDASE_COMPLEX | 0.00000 | 0.00001 | 0.70498 | 0.76593 | 1.44609 | 63 |
| GO_ENDOPEPTIDASE_REGULATOR_ACTIVITY | 0.00006 | 0.00145 | 0.55733 | 0.70425 | 1.33001 | 68 |
| GO_ENDOPLASMIC_RETICULUM_GOLGI_INTERMEDIATE_COMPA | 0.00055 | 0.00863 | 0.47727 | 0.64758 | 1.24050 | 96 |
| GO_ENDOPLASMIC_RETICULUM_GOLGI_INTERMEDIATE_COMPA | 0.00148 | 0.01923 | 0.45506 | 0.68944 | 1.29056 | 51 |
| GO_ENDOPLASMIC_RETICULUM_TO_GOLGI_VESICLE_MEDIATED | 0.00532 | 0.04803 | 0.40702 | 0.59194 | 1.14697 | 162 |
| GO_ENERGY_DERIVATION_BY_OXIDATION_OF_ORGANIC_COMP | 0.00000 | 0.00000 | | 0.69995 | 1.36305 | 202 |
| GO_ENZYME_INHIBITOR_ACTIVITY | 0.00246 | 0.02780 | 0.43171 | 0.58913 | 1.14643 | 191 |
| GO_EPIDERMAL_CELL_DIFFERENTIATION | 0.00216 | 0.02519 | 0.43171 | 0.62667 | 1.20589 | 112 |
| GO_EPITHELIAL_CELL_DIFFERENTIATION | 0.00054 | 0.00857 | 0.47727 | 0.57819 | 1.13725 | 334 |
| GO_ESTABLISHMENT_OF_ORGANELLE_LOCALIZATION | 0.00122 | 0.01679 | 0.45506 | 0.57737 | 1.13489 | 323 |
| GO_ESTABLISHMENT_OF_PROTEIN_LOCALIZATION_TO_CHROMO | 0.00016 | 0.00321 | 0.51885 | 0.81363 | 1.46985 | 23 |
| GO_ESTABLISHMENT_OF_PROTEIN_LOCALIZATION_TO_ENDOPL | 0.00000 | 0.00000 | | 0.77241 | 1.48479 | 106 |

| | | | | | | |
|---|---|---|---|---|---|---|
| GO_ESTABLISHMENT_OF_PROTEIN_LOCALIZATION_TO_MEMBRA | 0.00000 | 0.00000 | | 0.67484 | 1.31991 | 260 |
| GO_ESTABLISHMENT_OF_PROTEIN_LOCALIZATION_TO_ORGANE | 0.00000 | 0.00000 | | 0.65856 | 1.30056 | 461 |
| GO_ESTABLISHMENT_OF_PROTEIN_LOCALIZATION_TO_TELOME | 0.00000 | 0.00006 | 0.64355 | 0.90447 | 1.60870 | 18 |
| GO_ESTABLISHMENT_OF_RNA_LOCALIZATION | 0.00000 | 0.00005 | 0.64355 | 0.64271 | 1.24830 | 176 |
| GO_ESTABLISHMENT_OF_TISSUE_POLARITY | 0.00000 | 0.00000 | 0.81404 | 0.75693 | 1.44999 | 96 |
| GO_EUKARYOTIC_TRANSLATION_INITIATION_FACTOR_3_COMPL | 0.00143 | 0.01863 | 0.45506 | 0.83263 | 1.46100 | 15 |
| GO_EXTERNAL_SIDE_OF_PLASMA_MEMBRANE | 0.00159 | 0.02016 | 0.45506 | 0.61775 | 1.19022 | 118 |
| GO_EXTRACELLULAR_MATRIX | 0.00042 | 0.00695 | 0.49849 | 0.61138 | 1.18938 | 189 |
| GO_FATTY_ACID_DERIVATIVE_METABOLIC_PROCESS | 0.00043 | 0.00695 | 0.49849 | 0.66266 | 1.26212 | 81 |
| GO_FATTY_ACYL_COA_METABOLIC_PROCESS | 0.00502 | 0.04608 | 0.40702 | 0.72264 | 1.32082 | 28 |
| GO_FC_EPSILON_RECEPTOR_SIGNALING_PATHWAY | 0.00000 | 0.00000 | 0.71951 | 0.72370 | 1.38610 | 94 |
| GO_FC_RECEPTOR_SIGNALING_PATHWAY | 0.00000 | 0.00000 | 0.70498 | 0.68815 | 1.32879 | 140 |
| GO_FERTILIZATION | 0.00000 | 0.00005 | 0.64355 | 0.73257 | 1.38752 | 72 |
| GO_FIBRILLAR_CENTER | 0.00246 | 0.02780 | 0.43171 | 0.62638 | 1.19990 | 96 |
| GO_FIBROBLAST_GROWTH_FACTOR_RECEPTOR_SIGNALING_PAT | 0.00495 | 0.04573 | 0.40702 | 0.64888 | 1.22552 | 67 |
| GO_FIBROBLAST_PROLIFERATION | 0.00032 | 0.00575 | 0.49849 | 0.70676 | 1.32618 | 53 |
| GO_FICOLIN_1_RICH_GRANULE | 0.00000 | 0.00000 | 0.70498 | 0.71544 | 1.37138 | 98 |
| GO_GABA_ERGIC_SYNAPSE | 0.00022 | 0.00427 | 0.51885 | 0.76430 | 1.40406 | 32 |
| GO_GENERATION_OF_PRECURSOR_METABOLITES_AND_ENERGY | 0.00000 | 0.00000 | | 0.67527 | 1.32968 | 368 |
| GO_GENE_SILENCING | 0.00019 | 0.00376 | 0.51885 | 0.61625 | 1.19788 | 184 |
| GO_GLUCOSE_CATABOLIC_PROCESS | 0.00495 | 0.04573 | 0.40702 | 0.72314 | 1.32174 | 28 |
| GO_GLUTAMATERGIC_SYNAPSE | 0.00135 | 0.01808 | 0.45506 | 0.60052 | 1.16783 | 186 |
| GO_GLYCOLYTIC_PROCESS_THROUGH_FRUCTOSE_6_PHOSPHATI | 0.00554 | 0.04968 | 0.40702 | 0.74156 | 1.34111 | 24 |
| GO_GOLGI_ASSOCIATED_VESICLE | 0.00085 | 0.01239 | 0.47727 | 0.62294 | 1.20037 | 129 |
| GO_GOLGI_ASSOCIATED_VESICLE_MEMBRANE | 0.00313 | 0.03310 | 0.43171 | 0.64208 | 1.21753 | 75 |
| GO_GTPASE_ACTIVITY | 0.00004 | 0.00098 | 0.55733 | 0.61546 | 1.19974 | 210 |
| GO_GUANYL_NUCLEOTIDE_BINDING | 0.00005 | 0.00126 | 0.55733 | 0.60899 | 1.19065 | 252 |
| GO_HEAT_SHOCK_PROTEIN_BINDING | 0.00001 | 0.00034 | 0.59333 | 0.68258 | 1.30633 | 93 |
| GO_HEMATOPOIETIC_PROGENITOR_CELL_DIFFERENTIATION | 0.00000 | 0.00000 | 0.71951 | 0.70099 | 1.35150 | 121 |
| GO_HEMATOPOIETIC_STEM_CELL_DIFFERENTIATION | 0.00000 | 0.00000 | 0.76146 | 0.77246 | 1.46307 | 72 |
| GO_HISTONE_MRNA_METABOLIC_PROCESS | 0.00265 | 0.02936 | 0.43171 | 0.76832 | 1.38459 | 22 |
| GO_HOST_CELLULAR_COMPONENT | 0.00043 | 0.00695 | 0.49849 | 0.69729 | 1.31074 | 54 |
| GO_IMMUNE_RESPONSE_REGULATING_SIGNALING_PATHWAY | 0.00000 | 0.00003 | 0.65944 | 0.63620 | 1.24112 | 225 |
| GO_IMPORT_INTO_CELL | 0.00428 | 0.04185 | 0.40702 | 0.61709 | 1.18655 | 108 |
| GO_IMPORT_INTO_NUCLEUS | 0.00039 | 0.00652 | 0.49849 | 0.62707 | 1.20905 | 131 |
| GO_INNATE_IMMUNE_RESPONSE | 0.00000 | 0.00003 | 0.65944 | 0.59893 | 1.18089 | 406 |
| GO_INNATE_IMMUNE_RESPONSE_ACTIVATING_SIGNAL_TRANSI | 0.00000 | 0.00001 | 0.69013 | 0.73348 | 1.39585 | 80 |
| GO_INNER_MITOCHONDRIAL_MEMBRANE_PROTEIN_COMPLEX | 0.00000 | 0.00000 | | 0.76576 | 1.46666 | 94 |
| GO_INORGANIC_ION_TRANSMEMBRANE_TRANSPORT | 0.00141 | 0.01850 | 0.45506 | 0.56404 | 1.11185 | 401 |
| GO_INTERACTION_WITH_HOST | 0.00139 | 0.01850 | 0.45506 | 0.60855 | 1.17710 | 151 |
| GO_INTERLEUKIN_1_MEDIATED_SIGNALING_PATHWAY | 0.00000 | 0.00000 | 0.77494 | 0.75654 | 1.44176 | 83 |
| GO_INTRAMOLECULAR_OXIDOREDUCTASE_ACTIVITY | 0.00051 | 0.00812 | 0.47727 | 0.74506 | 1.37081 | 34 |
| GO_INTRINSIC_APOPTOTIC_SIGNALING_PATHWAY | 0.00156 | 0.01993 | 0.45506 | 0.59427 | 1.15817 | 207 |
| GO_ION_HOMEOSTASIS | 0.00170 | 0.02117 | 0.45506 | 0.56164 | 1.10717 | 403 |
| GO_ION_TRANSMEMBRANE_TRANSPORTER_ACTIVITY | 0.00106 | 0.01495 | 0.45506 | 0.56696 | 1.11734 | 390 |
| GO_IRON_ION_HOMEOSTASIS | 0.00361 | 0.03657 | 0.43171 | 0.65837 | 1.23983 | 59 |
| GO_IRON_ION_TRANSPORT | 0.00054 | 0.00857 | 0.47727 | 0.69556 | 1.30334 | 52 |
| GO_ISOMERASE_ACTIVITY | 0.00413 | 0.04065 | 0.40702 | 0.61065 | 1.17496 | 113 |
| GO_KERATINIZATION | 0.00038 | 0.00640 | 0.49849 | 0.75690 | 1.39013 | 33 |
| GO_KILLING_OF_CELLS_OF_OTHER_ORGANISM | 0.00495 | 0.04573 | 0.40702 | 0.78614 | 1.38643 | 16 |
| GO_LARGE_RIBOSOMAL_SUBUNIT | 0.00000 | 0.00000 | | 0.77656 | 1.49319 | 108 |
| GO_LATERAL_PLASMA_MEMBRANE | 0.00524 | 0.04770 | 0.40702 | 0.67385 | 1.25557 | 47 |
| GO_LEUKOCYTE_MEDIATED_IMMUNITY | 0.00000 | 0.00000 | | 0.64010 | 1.26331 | 447 |
| GO_LIPOPROTEIN_PARTICLE_RECEPTOR_BINDING | 0.00126 | 0.01715 | 0.45506 | 0.80503 | 1.42497 | 17 |
| GO_LIVER_REGENERATION | 0.00093 | 0.01337 | 0.47727 | 0.78442 | 1.41708 | 23 |
| GO_LOW_DENSITY_LIPOPROTEIN_RECEPTOR_PARTICLE_METABO | 0.00327 | 0.03409 | 0.43171 | 0.78520 | 1.39657 | 18 |
| GO_LYASE_ACTIVITY | 0.00152 | 0.01958 | 0.45506 | 0.61769 | 1.19079 | 122 |
| GO_LYSOSOMAL_LUMEN | 0.00143 | 0.01863 | 0.45506 | 0.67371 | 1.26872 | 59 |
| GO_MAINTENANCE_OF_CELL_NUMBER | 0.00435 | 0.04238 | 0.40702 | 0.60761 | 1.17073 | 117 |
| GO_MATURATION_OF_SSU_RRNA | 0.00126 | 0.01715 | 0.45506 | 0.69615 | 1.29711 | 47 |
| GO_MATURATION_OF_SSU_RRNA_FROM_TRICISTRONIC_RRNA_ | 0.00554 | 0.04968 | 0.40702 | 0.70205 | 1.29301 | 35 |
| GO_MEMBRANE_REGION | 0.00000 | 0.00003 | 0.65944 | 0.64496 | 1.25542 | 194 |
| GO_METALLOPEPTIDASE_ACTIVITY | 0.00487 | 0.04559 | 0.40702 | 0.62714 | 1.19577 | 85 |
| GO_METAL_ION_HOMEOSTASIS | 0.00032 | 0.00575 | 0.49849 | 0.58816 | 1.15376 | 301 |
| GO_METAL_ION_TRANSPORT | 0.00390 | 0.03895 | 0.43171 | 0.55877 | 1.10227 | 419 |
| GO_METAPHASE_ANAPHASE_TRANSITION_OF_CELL_CYCLE | 0.00320 | 0.03371 | 0.43171 | 0.66563 | 1.25122 | 54 |
| GO_MICROTUBULE_CYTOSKELETON_ORGANIZATION_INVOLVED_ | 0.00220 | 0.02548 | 0.43171 | 0.61295 | 1.18118 | 127 |
| GO_MICROVILLUS | 0.00168 | 0.02098 | 0.45506 | 0.68128 | 1.28184 | 56 |
| GO_MIDBODY | 0.00561 | 0.04971 | 0.40702 | 0.59866 | 1.15606 | 141 |
| GO_MITOCHONDRIAL_ELECTRON_TRANSPORT_NADH_TO_UBIQ | 0.00000 | 0.00006 | 0.64355 | 0.79447 | 1.47799 | 44 |

| | | | | | | |
|---|---|---|---|---|---|---|
| GO_MITOCHONDRIAL_GENE_EXPRESSION | 0.00000 | 0.00000 | 0.78819 | 0.70230 | 1.35844 | 151 |
| GO_MITOCHONDRIAL_LARGE_RIBOSOMAL_SUBUNIT | 0.00000 | 0.00000 | 0.71951 | 0.79015 | 1.48529 | 54 |
| GO_MITOCHONDRIAL_MATRIX | 0.00000 | 0.00000 | 0.82666 | 0.63141 | 1.24412 | 387 |
| GO_MITOCHONDRIAL_MEMBRANE_ORGANIZATION | 0.00050 | 0.00802 | 0.47727 | 0.64578 | 1.23686 | 94 |
| GO_MITOCHONDRIAL_PROTEIN_COMPLEX | 0.00000 | 0.00000 | | 0.75011 | 1.46215 | 212 |
| GO_MITOCHONDRIAL_RESPIRATORY_CHAIN_COMPLEX_ASSEMB | 0.00001 | 0.00033 | 0.59333 | 0.69650 | 1.32804 | 85 |
| GO_MITOCHONDRIAL_SMALL_RIBOSOMAL_SUBUNIT | 0.00119 | 0.01640 | 0.45506 | 0.75384 | 1.37785 | 28 |
| GO_MITOCHONDRIAL_TRANSLATION | 0.00000 | 0.00000 | 0.80122 | 0.71435 | 1.37651 | 128 |
| GO_MITOCHONDRIAL_TRANSLATIONAL_TERMINATION | 0.00000 | 0.00000 | 0.81404 | 0.75760 | 1.44657 | 87 |
| GO_MITOCHONDRIAL_TRANSMEMBRANE_TRANSPORT | 0.00213 | 0.02490 | 0.43171 | 0.66300 | 1.25387 | 70 |
| GO_MITOCHONDRIAL_TRANSPORT | 0.00010 | 0.00227 | 0.53843 | 0.61119 | 1.19060 | 203 |
| GO_MITOCHONDRION_ORGANIZATION | 0.00000 | 0.00000 | 0.74774 | 0.61590 | 1.21471 | 416 |
| GO_MITOTIC_NUCLEAR_DIVISION | 0.00008 | 0.00182 | 0.53843 | 0.60435 | 1.17901 | 223 |
| GO_MITOTIC_SISTER_CHROMATID_SEGREGATION | 0.00014 | 0.00302 | 0.51885 | 0.62652 | 1.21048 | 145 |
| GO_MITOTIC_SPINDLE_ORGANIZATION | 0.00078 | 0.01164 | 0.47727 | 0.63943 | 1.22821 | 104 |
| GO_MODIFIED_AMINO_ACID_BINDING | 0.00002 | 0.00048 | 0.57561 | 0.74200 | 1.38679 | 49 |
| GO_MODULATION_BY_HOST_OF_VIRAL_PROCESS | 0.00216 | 0.02519 | 0.43171 | 0.77232 | 1.38066 | 20 |
| GO_MONOCARBOXYLIC_ACID_METABOLIC_PROCESS | 0.00012 | 0.00265 | 0.53843 | 0.57846 | 1.14053 | 406 |
| GO_MONOSACCHARIDE_BIOSYNTHETIC_PROCESS | 0.00305 | 0.03271 | 0.43171 | 0.65908 | 1.24472 | 68 |
| GO_MONOVALENT_INORGANIC_CATION_HOMEOSTASIS | 0.00108 | 0.01507 | 0.45506 | 0.63643 | 1.21783 | 91 |
| GO_MONOVALENT_INORGANIC_CATION_TRANSMEMBRANE_TR | 0.00001 | 0.00026 | 0.59333 | 0.65186 | 1.25996 | 147 |
| GO_MONOVALENT_INORGANIC_CATION_TRANSPORT | 0.00008 | 0.00178 | 0.53843 | 0.60205 | 1.17659 | 248 |
| GO_MORPHOGENESIS_OF_AN_EPITHELIUM | 0.00000 | 0.00009 | 0.62726 | 0.60570 | 1.19137 | 334 |
| GO_MORPHOGENESIS_OF_A_POLARIZED_EPITHELIUM | 0.00000 | 0.00000 | 0.83909 | 0.74384 | 1.43057 | 110 |
| GO_MOVEMENT_IN_HOST_ENVIRONMENT | 0.00268 | 0.02961 | 0.43171 | 0.61282 | 1.18078 | 117 |
| GO_MRNA_3_END_PROCESSING | 0.00087 | 0.01270 | 0.47727 | 0.64010 | 1.22502 | 93 |
| GO_MRNA_5_UTR_BINDING | 0.00002 | 0.00063 | 0.57561 | 0.83759 | 1.51313 | 23 |
| GO_MRNA_BINDING | 0.00000 | 0.00000 | | 0.67496 | 1.31810 | 239 |
| GO_MRNA_EXPORT_FROM_NUCLEUS | 0.00006 | 0.00159 | 0.53843 | 0.66290 | 1.27265 | 103 |
| GO_MRNA_PROCESSING | 0.00000 | 0.00000 | | 0.62959 | 1.24255 | 437 |
| GO_MRNA_TRANSPORT | 0.00007 | 0.00171 | 0.53843 | 0.64103 | 1.23563 | 132 |
| GO_MULTI_MULTICELLULAR_ORGANISM_PROCESS | 0.00041 | 0.00687 | 0.49849 | 0.63087 | 1.21515 | 116 |
| GO_MULTI_ORGANISM_LOCALIZATION | 0.00017 | 0.00334 | 0.51885 | 0.71077 | 1.33608 | 54 |
| GO_MUSCLE_CONTRACTION | 0.00335 | 0.03468 | 0.43171 | 0.59925 | 1.16199 | 165 |
| GO_MUSCLE_FILAMENT_SLIDING | 0.00324 | 0.03393 | 0.43171 | 0.77980 | 1.39322 | 19 |
| GO_MYELOID_LEUKOCYTE_ACTIVATION | 0.00000 | 0.00000 | | 0.65288 | 1.28712 | 395 |
| GO_MYELOID_LEUKOCYTE_MEDIATED_IMMUNITY | 0.00000 | 0.00000 | | 0.65734 | 1.29408 | 356 |
| GO_NADH_DEHYDROGENASE_ACTIVITY | 0.00001 | 0.00019 | 0.61053 | 0.79360 | 1.46691 | 38 |
| GO_NADH_DEHYDROGENASE_COMPLEX | 0.00001 | 0.00028 | 0.59333 | 0.77749 | 1.44215 | 41 |
| GO_NADH_DEHYDROGENASE_COMPLEX_ASSEMBLY | 0.00001 | 0.00035 | 0.59333 | 0.74044 | 1.39281 | 55 |
| GO_NADH_METABOLIC_PROCESS | 0.00034 | 0.00599 | 0.49849 | 0.75773 | 1.39166 | 33 |
| GO_NAD_METABOLIC_PROCESS | 0.00435 | 0.04238 | 0.40702 | 0.70570 | 1.30442 | 37 |
| GO_NCRNA_EXPORT_FROM_NUCLEUS | 0.00205 | 0.02440 | 0.43171 | 0.71473 | 1.31817 | 36 |
| GO_NCRNA_METABOLIC_PROCESS | 0.00000 | 0.00000 | 0.70498 | 0.61139 | 1.20509 | 399 |
| GO_NCRNA_PROCESSING | 0.00000 | 0.00000 | 0.73376 | 0.62238 | 1.22425 | 341 |
| GO_NEGATIVE_REGULATION_OF_APOPTOTIC_SIGNALING_PATHW | 0.00364 | 0.03677 | 0.43171 | 0.59958 | 1.15998 | 150 |
| GO_NEGATIVE_REGULATION_OF_ATP_METABOLIC_PROCESS | 0.00302 | 0.03248 | 0.43171 | 0.78586 | 1.39103 | 17 |
| GO_NEGATIVE_REGULATION_OF_CANONICAL_WNT_SIGNALING | 0.00000 | 0.00000 | 0.71951 | 0.69629 | 1.34180 | 123 |
| GO_NEGATIVE_REGULATION_OF_CATALYTIC_ACTIVITY | 0.00145 | 0.01879 | 0.45506 | 0.56238 | 1.11005 | 450 |
| GO_NEGATIVE_REGULATION_OF_CELLULAR_AMIDE_METABOLIC | 0.00268 | 0.02961 | 0.43171 | 0.61074 | 1.17686 | 124 |
| GO_NEGATIVE_REGULATION_OF_CELLULAR_COMPONENT_ORGA | 0.00320 | 0.03371 | 0.43171 | 0.55470 | 1.09534 | 470 |
| GO_NEGATIVE_REGULATION_OF_CELL_CYCLE | 0.00023 | 0.00445 | 0.51885 | 0.57348 | 1.13228 | 455 |
| GO_NEGATIVE_REGULATION_OF_CELL_CYCLE_G2_M_PHASE_TR | 0.00000 | 0.00007 | 0.62726 | 0.70549 | 1.34606 | 86 |
| GO_NEGATIVE_REGULATION_OF_CELL_CYCLE_PHASE_TRANSITIO | 0.00000 | 0.00000 | 0.70498 | 0.65532 | 1.27560 | 194 |
| GO_NEGATIVE_REGULATION_OF_CELL_CYCLE_PROCESS | 0.00000 | 0.00005 | 0.64355 | 0.62725 | 1.22670 | 259 |
| GO_NEGATIVE_REGULATION_OF_HYDROLASE_ACTIVITY | 0.00216 | 0.02519 | 0.43171 | 0.58306 | 1.13858 | 238 |
| GO_NEGATIVE_REGULATION_OF_INTERLEUKIN_1_PRODUCTION | 0.00205 | 0.02440 | 0.43171 | 0.82297 | 1.44404 | 15 |
| GO_NEGATIVE_REGULATION_OF_MITOTIC_CELL_CYCLE | 0.00000 | 0.00010 | 0.62726 | 0.62191 | 1.21541 | 248 |
| GO_NEGATIVE_REGULATION_OF_MRNA_METABOLIC_PROCESS | 0.00213 | 0.02490 | 0.43171 | 0.66731 | 1.25879 | 62 |
| GO_NEGATIVE_REGULATION_OF_MRNA_SPLICING_VIA_SPLICEO | 0.00339 | 0.03501 | 0.43171 | 0.76309 | 1.36417 | 20 |
| GO_NEGATIVE_REGULATION_OF_NUCLEAR_DIVISION | 0.00335 | 0.03468 | 0.43171 | 0.67724 | 1.26685 | 50 |
| GO_NEGATIVE_REGULATION_OF_ORGANELLE_ORGANIZATION | 0.00487 | 0.04559 | 0.40702 | 0.57038 | 1.11722 | 288 |
| GO_NEGATIVE_REGULATION_OF_PEPTIDASE_ACTIVITY | 0.00036 | 0.00621 | 0.49849 | 0.64044 | 1.23159 | 109 |
| GO_NEGATIVE_REGULATION_OF_PROTEIN_CATABOLIC_PROCESS | 0.00342 | 0.03521 | 0.43171 | 0.62320 | 1.19268 | 93 |
| GO_NEGATIVE_REGULATION_OF_PROTEOLYSIS | 0.00205 | 0.02440 | 0.43171 | 0.60172 | 1.16902 | 178 |
| GO_NEGATIVE_REGULATION_OF_REACTIVE_OXYGEN_SPECIES_B | 0.00036 | 0.00621 | 0.49849 | 0.83108 | 1.47818 | 18 |
| GO_NEGATIVE_REGULATION_OF_REPRODUCTIVE_PROCESS | 0.00457 | 0.04386 | 0.40702 | 0.74845 | 1.35356 | 24 |
| GO_NEGATIVE_REGULATION_OF_RNA_SPLICING | 0.00183 | 0.02219 | 0.45506 | 0.76824 | 1.38935 | 24 |
| GO_NEGATIVE_REGULATION_OF_WNT_SIGNALING_PATHWAY | 0.00000 | 0.00002 | 0.67496 | 0.67731 | 1.30721 | 136 |
| GO_NEURAL_CREST_CELL_MIGRATION | 0.00265 | 0.02936 | 0.43171 | 0.73967 | 1.35049 | 27 |

| | | | | | | |
|---|---|---|---|---|---|---|
| GO_NEURON_RECOGNITION | 0.00390 | 0.03895 | 0.43171 | 0.71862 | 1.31998 | 31 |
| GO_NIK_NF_KAPPAB_SIGNALING | 0.00000 | 0.00000 | 0.77494 | 0.72329 | 1.39253 | 114 |
| GO_NON_CANONICAL_WNT_SIGNALING_PATHWAY | 0.00000 | 0.00000 | | 0.75756 | 1.45654 | 107 |
| GO_NUCLEAR_CHROMOSOME_SEGREGATION | 0.00098 | 0.01407 | 0.45506 | 0.59156 | 1.15357 | 215 |
| GO_NUCLEAR_ENVELOPE | 0.00005 | 0.00131 | 0.55733 | 0.59109 | 1.16270 | 341 |
| GO_NUCLEAR_EXPORT | 0.00000 | 0.00004 | 0.65944 | 0.64915 | 1.26157 | 181 |
| GO_NUCLEAR_MATRIX | 0.00502 | 0.04608 | 0.40702 | 0.62777 | 1.19868 | 87 |
| GO_NUCLEAR_MEMBRANE | 0.00185 | 0.02237 | 0.45506 | 0.58925 | 1.14853 | 211 |
| GO_NUCLEAR_PERIPHERY | 0.00043 | 0.00701 | 0.49849 | 0.64264 | 1.23558 | 107 |
| GO_NUCLEAR_PORE | 0.00561 | 0.04971 | 0.40702 | 0.64876 | 1.22522 | 68 |
| GO_NUCLEAR_SPECK | 0.00015 | 0.00312 | 0.51885 | 0.59159 | 1.16089 | 306 |
| GO_NUCLEAR_TRANSCRIBED_MRNA_CATABOLIC_PROCESS | 0.00000 | 0.00000 | | 0.70362 | 1.36820 | 185 |
| GO_NUCLEAR_TRANSCRIBED_MRNA_CATABOLIC_PROCESS_NON | 0.00000 | 0.00000 | | 0.74239 | 1.42931 | 114 |
| GO_NUCLEAR_TRANSPORT | 0.00000 | 0.00000 | 0.71951 | 0.62920 | 1.23383 | 297 |
| GO_NUCLEOBASE_CONTAINING_COMPOUND_TRANSPORT | 0.00000 | 0.00010 | 0.62726 | 0.62975 | 1.22823 | 216 |
| GO_NUCLEOBASE_CONTAINING_SMALL_MOLECULE_METABOLIC | 0.00000 | 0.00003 | 0.65944 | 0.60153 | 1.18594 | 405 |
| GO_NUCLEOSIDE_DIPHOSPHATE_METABOLIC_PROCESS | 0.00016 | 0.00319 | 0.51885 | 0.64491 | 1.24260 | 117 |
| GO_NUCLEOSIDE_PHOSPHATE_BIOSYNTHETIC_PROCESS | 0.00002 | 0.00044 | 0.59333 | 0.64092 | 1.24175 | 161 |
| GO_NUCLEOSIDE_TRIPHOSPHATE_BIOSYNTHETIC_PROCESS | 0.00040 | 0.00676 | 0.49849 | 0.72727 | 1.35074 | 42 |
| GO_NUCLEOSIDE_TRIPHOSPHATE_METABOLIC_PROCESS | 0.00009 | 0.00196 | 0.53843 | 0.69306 | 1.30854 | 65 |
| GO_NUCLEOTIDE_PHOSPHORYLATION | 0.00016 | 0.00328 | 0.51885 | 0.65541 | 1.25767 | 102 |
| GO_OOCYTE_DIFFERENTIATION | 0.00450 | 0.04342 | 0.40702 | 0.73513 | 1.33705 | 26 |
| GO_ORGANELLAR_RIBOSOME | 0.00000 | 0.00000 | 0.76146 | 0.75712 | 1.44311 | 84 |
| GO_ORGANELLE_ENVELOPE_LUMEN | 0.00026 | 0.00482 | 0.49849 | 0.67265 | 1.27716 | 78 |
| GO_ORGANELLE_FISSION | 0.00154 | 0.01974 | 0.45506 | 0.57133 | 1.12383 | 342 |
| GO_ORGANELLE_INNER_MEMBRANE | 0.00000 | 0.00000 | | 0.68423 | 1.34915 | 412 |
| GO_ORGANIC_CYCLIC_COMPOUND_CATABOLIC_PROCESS | 0.00000 | 0.00000 | | 0.66975 | 1.32263 | 478 |
| GO_ORGANIC_HYDROXY_COMPOUND_BIOSYNTHETIC_PROCESS | 0.00141 | 0.01850 | 0.45506 | 0.60712 | 1.17627 | 161 |
| GO_OXIDATIVE_PHOSPHORYLATION | 0.00000 | 0.00000 | | 0.78902 | 1.51260 | 99 |
| GO_OXIDOREDUCTASE_ACTIVITY | 0.00000 | 0.00000 | | 0.63268 | 1.24935 | 468 |
| GO_OXIDOREDUCTASE_ACTIVITY_ACTING_ON_A_HEME_GROUP | 0.00176 | 0.02168 | 0.45506 | 0.79383 | 1.41828 | 19 |
| GO_OXIDOREDUCTASE_ACTIVITY_ACTING_ON_NAD_P_H | 0.00002 | 0.00056 | 0.57561 | 0.71468 | 1.35162 | 70 |
| GO_OXIDOREDUCTASE_ACTIVITY_ACTING_ON_NAD_P_H_QUIN | 0.00004 | 0.00113 | 0.55733 | 0.74511 | 1.38835 | 47 |
| GO_OXIDOREDUCTASE_ACTIVITY_ACTING_ON_PEROXIDE_AS_A | 0.00106 | 0.01495 | 0.45506 | 0.73897 | 1.35352 | 29 |
| GO_OXIDOREDUCTASE_COMPLEX | 0.00000 | 0.00000 | 0.73376 | 0.74898 | 1.42731 | 82 |
| GO_PEPTIDASE_ACTIVITY | 0.00133 | 0.01799 | 0.45506 | 0.57681 | 1.13188 | 306 |
| GO_PEPTIDASE_COMPLEX | 0.00000 | 0.00002 | 0.67496 | 0.73250 | 1.39240 | 79 |
| GO_PEPTIDASE_REGULATOR_ACTIVITY | 0.00313 | 0.03310 | 0.43171 | 0.63434 | 1.21189 | 88 |
| GO_PEPTIDE_BINDING | 0.00005 | 0.00124 | 0.55733 | 0.63708 | 1.23187 | 148 |
| GO_PERINUCLEAR_REGION_OF_CYTOPLASM | 0.00276 | 0.03021 | 0.43171 | 0.55814 | 1.10207 | 466 |
| GO_PHAGOCYTOSIS | 0.00554 | 0.04968 | 0.40702 | 0.59337 | 1.14837 | 155 |
| GO_PHAGOSOME_ACIDIFICATION | 0.00059 | 0.00916 | 0.47727 | 0.78902 | 1.42694 | 24 |
| GO_PHOSPHATIDYLSERINE_BINDING | 0.00035 | 0.00610 | 0.49849 | 0.76387 | 1.39912 | 29 |
| GO_PH_REDUCTION | 0.00302 | 0.03248 | 0.43171 | 0.68821 | 1.28238 | 46 |
| GO_PIGMENT_GRANULE | 0.00000 | 0.00000 | 0.76146 | 0.74333 | 1.41934 | 87 |
| GO_PLASMA_LIPOPROTEIN_PARTICLE_CLEARANCE | 0.00106 | 0.01495 | 0.45506 | 0.73201 | 1.35304 | 37 |
| GO_POLYSOMAL_RIBOSOME | 0.00008 | 0.00180 | 0.53843 | 0.78840 | 1.44683 | 30 |
| GO_POLYSOME | 0.00000 | 0.00003 | 0.65944 | 0.75964 | 1.43296 | 62 |
| GO_POSITIVE_REGULATION_OF_ATP_METABOLIC_PROCESS | 0.00174 | 0.02149 | 0.45506 | 0.73011 | 1.33729 | 29 |
| GO_POSITIVE_REGULATION_OF_BINDING | 0.00390 | 0.03895 | 0.43171 | 0.61042 | 1.17582 | 115 |
| GO_POSITIVE_REGULATION_OF_CANONICAL_WNT_SIGNALING_ | 0.00000 | 0.00000 | 0.73376 | 0.72036 | 1.38514 | 108 |
| GO_POSITIVE_REGULATION_OF_CELLULAR_AMIDE_METABOLIC_ | 0.00001 | 0.00036 | 0.59333 | 0.66485 | 1.28095 | 125 |
| GO_POSITIVE_REGULATION_OF_CELLULAR_PROTEIN_LOCALIZAT | 0.00242 | 0.02748 | 0.43171 | 0.58048 | 1.13378 | 244 |
| GO_POSITIVE_REGULATION_OF_CELL_POPULATION_PROLIFERAT | 0.00040 | 0.00670 | 0.49849 | 0.56967 | 1.12503 | 461 |
| GO_POSITIVE_REGULATION_OF_CYSTEINE_TYPE_ENDOPEPTIDA | 0.00547 | 0.04923 | 0.40702 | 0.62225 | 1.19077 | 92 |
| GO_POSITIVE_REGULATION_OF_CYTOKINE_PRODUCTION | 0.00283 | 0.03081 | 0.43171 | 0.58672 | 1.14373 | 210 |
| GO_POSITIVE_REGULATION_OF_DEFENSE_RESPONSE | 0.00000 | 0.00000 | 0.73376 | 0.66200 | 1.28884 | 200 |
| GO_POSITIVE_REGULATION_OF_DNA_BIOSYNTHETIC_PROCESS | 0.00010 | 0.00228 | 0.53843 | 0.71224 | 1.33977 | 55 |
| GO_POSITIVE_REGULATION_OF_DNA_METABOLIC_PROCESS | 0.00020 | 0.00387 | 0.51885 | 0.62771 | 1.21205 | 139 |
| GO_POSITIVE_REGULATION_OF_ESTABLISHMENT_OF_PROTEIN_ | 0.00194 | 0.02331 | 0.45506 | 0.58158 | 1.13709 | 253 |
| GO_POSITIVE_REGULATION_OF_FIBROBLAST_PROLIFERATION | 0.00017 | 0.00338 | 0.51885 | 0.77423 | 1.42213 | 31 |
| GO_POSITIVE_REGULATION_OF_IMMUNE_RESPONSE | 0.00000 | 0.00000 | 0.71951 | 0.63064 | 1.23937 | 321 |
| GO_POSITIVE_REGULATION_OF_IMMUNE_SYSTEM_PROCESS | 0.00000 | 0.00002 | 0.67496 | 0.59748 | 1.17992 | 475 |
| GO_POSITIVE_REGULATION_OF_INNATE_IMMUNE_RESPONSE | 0.00000 | 0.00000 | 0.70498 | 0.68885 | 1.32737 | 128 |
| GO_POSITIVE_REGULATION_OF_MITOTIC_CELL_CYCLE | 0.00178 | 0.02182 | 0.45506 | 0.61563 | 1.18683 | 122 |
| GO_POSITIVE_REGULATION_OF_MRNA_METABOLIC_PROCESS | 0.00157 | 0.02009 | 0.45506 | 0.66330 | 1.25231 | 63 |
| GO_POSITIVE_REGULATION_OF_MRNA_PROCESSING | 0.00250 | 0.02807 | 0.43171 | 0.76194 | 1.38151 | 25 |
| GO_POSITIVE_REGULATION_OF_MRNA_SPLICING_VIA_SPLICEOS | 0.00132 | 0.01782 | 0.45506 | 0.81851 | 1.44353 | 16 |
| GO_POSITIVE_REGULATION_OF_NITRIC_OXIDE_METABOLIC_PR | 0.00133 | 0.01799 | 0.45506 | 0.80807 | 1.43726 | 18 |
| GO_POSITIVE_REGULATION_OF_ORGANELLE_ORGANIZATION | 0.00010 | 0.00218 | 0.53843 | 0.57473 | 1.13449 | 452 |

| | | | | | | |
|---|---|---|---|---|---|---|
| GO_POSITIVE_REGULATION_OF_PROTEIN_LOCALIZATION_TO_N | 0.00012 | 0.00265 | 0.53843 | 0.71018 | 1.33697 | 58 |
| GO_POSITIVE_REGULATION_OF_PROTEOLYSIS | 0.00313 | 0.03310 | 0.43171 | 0.58059 | 1.13381 | 237 |
| GO_POSITIVE_REGULATION_OF_REACTIVE_OXYGEN_SPECIES_BI | 0.00115 | 0.01593 | 0.45506 | 0.79865 | 1.43554 | 21 |
| GO_POSITIVE_REGULATION_OF_REACTIVE_OXYGEN_SPECIES_M | 0.00174 | 0.02149 | 0.45506 | 0.69785 | 1.30035 | 46 |
| GO_POSITIVE_REGULATION_OF_RESPONSE_TO_BIOTIC_STIMULI | 0.00000 | 0.00000 | 0.70498 | 0.67507 | 1.30603 | 150 |
| GO_POSITIVE_REGULATION_OF_RESPONSE_TO_EXTERNAL_STIM | 0.00001 | 0.00016 | 0.61053 | 0.62123 | 1.21505 | 260 |
| GO_POSITIVE_REGULATION_OF_RNA_SPLICING | 0.00265 | 0.02936 | 0.43171 | 0.74051 | 1.35202 | 27 |
| GO_POSITIVE_REGULATION_OF_SIGNAL_TRANSDUCTION_BY_P5 | 0.00390 | 0.03895 | 0.43171 | 0.80515 | 1.41278 | 15 |
| GO_POSITIVE_REGULATION_OF_TELOMERASE_ACTIVITY | 0.00069 | 0.01043 | 0.47727 | 0.75265 | 1.38122 | 30 |
| GO_POSITIVE_REGULATION_OF_TELOMERASE_RNA_LOCALIZATI | 0.00007 | 0.00169 | 0.53843 | 0.88616 | 1.55493 | 15 |
| GO_POSITIVE_REGULATION_OF_TELOMERE_MAINTENANCE | 0.00026 | 0.00486 | 0.49849 | 0.73697 | 1.37102 | 44 |
| GO_POSITIVE_REGULATION_OF_TELOMERE_MAINTENANCE_VIA | 0.00001 | 0.00040 | 0.59333 | 0.80196 | 1.47289 | 33 |
| GO_POSITIVE_REGULATION_OF_TRANSFERASE_ACTIVITY | 0.00108 | 0.01507 | 0.45506 | 0.56501 | 1.11376 | 401 |
| GO_POSITIVE_REGULATION_OF_TRANSLATION | 0.00000 | 0.00016 | 0.61053 | 0.68408 | 1.31526 | 107 |
| GO_POSITIVE_REGULATION_OF_VIRAL_PROCESS | 0.00026 | 0.00486 | 0.49849 | 0.66195 | 1.26462 | 88 |
| GO_POSITIVE_REGULATION_OF_WNT_SIGNALING_PATHWAY | 0.00000 | 0.00000 | 0.73376 | 0.69924 | 1.34747 | 127 |
| GO_POSTSYNAPSE | 0.00239 | 0.02716 | 0.43171 | 0.57062 | 1.12116 | 316 |
| GO_POST_TRANSLATIONAL_PROTEIN_MODIFICATION | 0.00024 | 0.00452 | 0.51885 | 0.60046 | 1.17392 | 251 |
| GO_PRECATALYTIC_SPLICEOSOME | 0.00141 | 0.01850 | 0.45506 | 0.69251 | 1.29540 | 50 |
| GO_PRERIBOSOME | 0.00006 | 0.00154 | 0.53843 | 0.69866 | 1.32020 | 69 |
| GO_PROTEASOMAL_PROTEIN_CATABOLIC_PROCESS | 0.00007 | 0.00169 | 0.53843 | 0.58188 | 1.14652 | 387 |
| GO_PROTEASOMAL_UBIQUITIN_INDEPENDENT_PROTEIN_CATAB | 0.00000 | 0.00003 | 0.65944 | 0.91182 | 1.62178 | 18 |
| GO_PROTEASOME_ACCESSORY_COMPLEX | 0.00005 | 0.00135 | 0.55733 | 0.82638 | 1.49450 | 24 |
| GO_PROTEASOME_CORE_COMPLEX | 0.00000 | 0.00004 | 0.65944 | 0.91231 | 1.61486 | 17 |
| GO_PROTEIN_CONTAINING_COMPLEX_DISASSEMBLY | 0.00000 | 0.00000 | 0.81404 | 0.66285 | 1.29656 | 261 |
| GO_PROTEIN_CONTAINING_COMPLEX_LOCALIZATION | 0.00018 | 0.00352 | 0.51885 | 0.60451 | 1.17943 | 226 |
| GO_PROTEIN_DNA_COMPLEX_SUBUNIT_ORGANIZATION | 0.00052 | 0.00825 | 0.47727 | 0.60887 | 1.18342 | 183 |
| GO_PROTEIN_DOMAIN_SPECIFIC_BINDING | 0.00003 | 0.00088 | 0.55733 | 0.57828 | 1.14190 | 470 |
| GO_PROTEIN_FOLDING | 0.00000 | 0.00000 | 0.73376 | 0.67425 | 1.30867 | 171 |
| GO_PROTEIN_FOLDING_CHAPERONE | 0.00079 | 0.01170 | 0.47727 | 0.81632 | 1.45193 | 18 |
| GO_PROTEIN_HETEROOLIGOMERIZATION | 0.00442 | 0.04297 | 0.40702 | 0.78998 | 1.39322 | 16 |
| GO_PROTEIN_IMPORT | 0.00001 | 0.00040 | 0.59333 | 0.64197 | 1.24306 | 158 |
| GO_PROTEIN_IMPORT_INTO_MITOCHONDRIAL_MATRIX | 0.00231 | 0.02642 | 0.43171 | 0.79530 | 1.41455 | 18 |
| GO_PROTEIN_INSERTION_INTO_MEMBRANE | 0.00191 | 0.02299 | 0.45506 | 0.68322 | 1.27694 | 49 |
| GO_PROTEIN_LOCALIZATION_TO_CHROMOSOME_TELOMERIC_F | 0.00001 | 0.00031 | 0.59333 | 0.82810 | 1.51357 | 28 |
| GO_PROTEIN_LOCALIZATION_TO_ENDOPLASMIC_RETICULUM | 0.00000 | 0.00000 | | 0.73850 | 1.42303 | 129 |
| GO_PROTEIN_LOCALIZATION_TO_MEMBRANE | 0.00000 | 0.00000 | 0.77494 | 0.61216 | 1.20881 | 462 |
| GO_PROTEIN_LOCALIZATION_TO_MITOCHONDRION | 0.00015 | 0.00310 | 0.51885 | 0.64234 | 1.23759 | 118 |
| GO_PROTEIN_LOCALIZATION_TO_NUCLEUS | 0.00000 | 0.00014 | 0.61053 | 0.63315 | 1.23408 | 208 |
| GO_PROTEIN_MATURATION | 0.00357 | 0.03638 | 0.43171 | 0.59209 | 1.14965 | 174 |
| GO_PROTEIN_MODIFICATION_BY_SMALL_PROTEIN_REMOVAL | 0.00002 | 0.00052 | 0.57561 | 0.61535 | 1.20119 | 231 |
| GO_PROTEIN_REFOLDING | 0.00465 | 0.04430 | 0.40702 | 0.77561 | 1.37288 | 17 |
| GO_PROTEIN_STABILIZATION | 0.00081 | 0.01194 | 0.47727 | 0.61881 | 1.19577 | 146 |
| GO_PROTEIN_SUMOYLATION | 0.00450 | 0.04342 | 0.40702 | 0.65043 | 1.22846 | 67 |
| GO_PROTEIN_TARGETING | 0.00000 | 0.00000 | | 0.66223 | 1.30361 | 358 |
| GO_PROTEIN_TARGETING_TO_MEMBRANE | 0.00000 | 0.00000 | | 0.71675 | 1.38879 | 162 |
| GO_PROTEIN_TARGETING_TO_MITOCHONDRION | 0.00113 | 0.01571 | 0.45506 | 0.65407 | 1.24648 | 83 |
| GO_PROTEIN_TRANSMEMBRANE_TRANSPORT | 0.00110 | 0.01526 | 0.45506 | 0.68173 | 1.28238 | 55 |
| GO_PROTON_TRANSMEMBRANE_TRANSPORT | 0.00000 | 0.00001 | 0.69013 | 0.71234 | 1.36309 | 91 |
| GO_PROTON_TRANSMEMBRANE_TRANSPORTER_ACTIVITY | 0.00004 | 0.00115 | 0.55733 | 0.70138 | 1.32535 | 69 |
| GO_PROTON_TRANSPORTING_TWO_SECTOR_ATPASE_COMPLEX | 0.00083 | 0.01218 | 0.47727 | 0.77254 | 1.40508 | 26 |
| GO_PROTON_TRANSPORTING_V_TYPE_ATPASE_COMPLEX | 0.00239 | 0.02716 | 0.43171 | 0.76100 | 1.37626 | 24 |
| GO_PURINE_CONTAINING_COMPOUND_BIOSYNTHETIC_PROCES | 0.00001 | 0.00040 | 0.59333 | 0.65863 | 1.26901 | 126 |
| GO_PURINE_CONTAINING_COMPOUND_METABOLIC_PROCESS | 0.00000 | 0.00000 | 0.73376 | 0.63132 | 1.23831 | 300 |
| GO_PURINE_NUCLEOSIDE_TRIPHOSPHATE_METABOLIC_PROCES | 0.00033 | 0.00582 | 0.49849 | 0.70887 | 1.32487 | 49 |
| GO_PYROPHOSPHATE_HYDROLYSIS_DRIVEN_PROTON_TRANSME | 0.00532 | 0.04803 | 0.40702 | 0.74470 | 1.34678 | 24 |
| GO_PYRUVATE_METABOLIC_PROCESS | 0.00001 | 0.00018 | 0.61053 | 0.67194 | 1.29289 | 113 |
| GO_P_BODY | 0.00472 | 0.04480 | 0.40702 | 0.65470 | 1.23501 | 62 |
| GO_REACTIVE_OXYGEN_SPECIES_BIOSYNTHETIC_PROCESS | 0.00040 | 0.00670 | 0.49849 | 0.70271 | 1.31857 | 53 |
| GO_REACTIVE_OXYGEN_SPECIES_METABOLIC_PROCESS | 0.00021 | 0.00409 | 0.51885 | 0.62881 | 1.21472 | 144 |
| GO_REGULATION_OF_ALCOHOL_BIOSYNTHETIC_PROCESS | 0.00228 | 0.02614 | 0.43171 | 0.68438 | 1.28020 | 50 |
| GO_REGULATION_OF_ANIMAL_ORGAN_MORPHOGENESIS | 0.00000 | 0.00000 | | 0.72330 | 1.39922 | 149 |
| GO_REGULATION_OF_APOPTOTIC_SIGNALING_PATHWAY | 0.00079 | 0.01170 | 0.47727 | 0.58599 | 1.14641 | 266 |
| GO_REGULATION_OF_ATP_METABOLIC_PROCESS | 0.00000 | 0.00006 | 0.64355 | 0.70523 | 1.34777 | 89 |
| GO_REGULATION_OF_BINDING | 0.00428 | 0.04185 | 0.40702 | 0.58264 | 1.13664 | 225 |
| GO_REGULATION_OF_BLOOD_PRESSURE | 0.00290 | 0.03156 | 0.43171 | 0.65319 | 1.23428 | 69 |
| GO_REGULATION_OF_CARBOHYDRATE_CATABOLIC_PROCESS | 0.00324 | 0.03393 | 0.43171 | 0.65541 | 1.23787 | 67 |
| GO_REGULATION_OF_CELLULAR_AMIDE_METABOLIC_PROCESS | 0.00000 | 0.00000 | 0.78819 | 0.63116 | 1.24143 | 337 |
| GO_REGULATION_OF_CELLULAR_AMINE_METABOLIC_PROCESS | 0.00000 | 0.00000 | 0.74774 | 0.78955 | 1.48730 | 60 |
| GO_REGULATION_OF_CELLULAR_AMINO_ACID_METABOLIC_PRO | 0.00000 | 0.00000 | 0.76146 | 0.80341 | 1.51022 | 54 |

| | | | | | | |
|---|---|---|---|---|---|---|
| GO_REGULATION_OF_CELLULAR_KETONE_METABOLIC_PROCESS | 0.00000 | 0.00000 | 0.70498 | 0.69239 | 1.33405 | 126 |
| GO_REGULATION_OF_CELLULAR_PH | 0.00442 | 0.04297 | 0.40702 | 0.65360 | 1.23436 | 68 |
| GO_REGULATION_OF_CELLULAR_PROTEIN_LOCALIZATION | 0.00228 | 0.02614 | 0.43171 | 0.56216 | 1.10830 | 402 |
| GO_REGULATION_OF_CELLULAR_RESPIRATION | 0.00428 | 0.04185 | 0.40702 | 0.77183 | 1.37897 | 19 |
| GO_REGULATION_OF_CELLULAR_RESPONSE_TO_HEAT | 0.00091 | 0.01314 | 0.47727 | 0.67360 | 1.27221 | 67 |
| GO_REGULATION_OF_CELL_CYCLE_G2_M_PHASE_TRANSITION | 0.00016 | 0.00332 | 0.51885 | 0.61554 | 1.19738 | 187 |
| GO_REGULATION_OF_CELL_CYCLE_PHASE_TRANSITION | 0.00000 | 0.00014 | 0.61053 | 0.59968 | 1.18091 | 369 |
| GO_REGULATION_OF_CHOLESTEROL_BIOSYNTHETIC_PROCESS | 0.00209 | 0.02470 | 0.43171 | 0.71476 | 1.32117 | 38 |
| GO_REGULATION_OF_CHOLESTEROL_METABOLIC_PROCESS | 0.00113 | 0.01571 | 0.45506 | 0.69865 | 1.30177 | 47 |
| GO_REGULATION_OF_CHROMOSOME_ORGANIZATION | 0.00036 | 0.00622 | 0.49849 | 0.58896 | 1.15261 | 278 |
| GO_REGULATION_OF_CHROMOSOME_SEGREGATION | 0.00079 | 0.01170 | 0.47727 | 0.64131 | 1.22718 | 91 |
| GO_REGULATION_OF_CHROMOSOME_SEPARATION | 0.00253 | 0.02838 | 0.43171 | 0.66349 | 1.24947 | 59 |
| GO_REGULATION_OF_CYSTEINE_TYPE_ENDOPEPTIDASE_ACTIVIT | 0.00044 | 0.00719 | 0.49849 | 0.62610 | 1.20905 | 141 |
| GO_REGULATION_OF_DEFENSE_RESPONSE | 0.00006 | 0.00150 | 0.55733 | 0.58834 | 1.15824 | 356 |
| GO_REGULATION_OF_DNA_BIOSYNTHETIC_PROCESS | 0.00043 | 0.00695 | 0.49849 | 0.66709 | 1.27130 | 83 |
| GO_REGULATION_OF_DNA_METABOLIC_PROCESS | 0.00055 | 0.00863 | 0.47727 | 0.58909 | 1.15134 | 249 |
| GO_REGULATION_OF_DNA_TEMPLATED_TRANSCRIPTION_IN_RE | 0.00000 | 0.00000 | 0.71951 | 0.72410 | 1.38687 | 94 |
| GO_REGULATION_OF_GENERATION_OF_PRECURSOR_METABOLI | 0.00004 | 0.00114 | 0.55733 | 0.65536 | 1.26115 | 111 |
| GO_REGULATION_OF_GENE_SILENCING | 0.00023 | 0.00434 | 0.51885 | 0.65464 | 1.25268 | 91 |
| GO_REGULATION_OF_HEMATOPOIETIC_PROGENITOR_CELL_DIF | 0.00000 | 0.00000 | 0.73376 | 0.76273 | 1.44465 | 72 |
| GO_REGULATION_OF_HEMATOPOIETIC_STEM_CELL_DIFFERENT | 0.00000 | 0.00000 | 0.74774 | 0.78891 | 1.48817 | 62 |
| GO_REGULATION_OF_HEMOPOIESIS | 0.00390 | 0.03895 | 0.43171 | 0.57739 | 1.12909 | 258 |
| GO_REGULATION_OF_IMMUNE_RESPONSE | 0.00000 | 0.00006 | 0.64355 | 0.60032 | 1.18447 | 433 |
| GO_REGULATION_OF_INNATE_IMMUNE_RESPONSE | 0.00000 | 0.00007 | 0.62726 | 0.64536 | 1.25422 | 181 |
| GO_REGULATION_OF_INTRINSIC_APOPTOTIC_SIGNALING_PATHW | 0.00179 | 0.02183 | 0.45506 | 0.61548 | 1.18550 | 116 |
| GO_REGULATION_OF_MITOTIC_CELL_CYCLE | 0.00000 | 0.00003 | 0.65944 | 0.58992 | 1.16508 | 499 |
| GO_REGULATION_OF_MORPHOGENESIS_OF_AN_EPITHELIUM | 0.00000 | 0.00000 | | 0.74680 | 1.43884 | 125 |
| GO_REGULATION_OF_MRNA_CATABOLIC_PROCESS | 0.00000 | 0.00000 | | 0.71126 | 1.37919 | 165 |
| GO_REGULATION_OF_MRNA_METABOLIC_PROCESS | 0.00000 | 0.00000 | | 0.67178 | 1.31458 | 276 |
| GO_REGULATION_OF_MRNA_PROCESSING | 0.00008 | 0.00186 | 0.53843 | 0.65189 | 1.25598 | 125 |
| GO_REGULATION_OF_MRNA_SPLICING_VIA_SPLICEOSOME | 0.00012 | 0.00256 | 0.53843 | 0.66950 | 1.28033 | 90 |
| GO_REGULATION_OF_MULTI_ORGANISM_PROCESS | 0.00008 | 0.00190 | 0.53843 | 0.76905 | 1.42150 | 37 |
| GO_REGULATION_OF_MUSCLE_CONTRACTION | 0.00163 | 0.02050 | 0.45506 | 0.64879 | 1.23071 | 76 |
| GO_REGULATION_OF_NUCLEAR_DIVISION | 0.00060 | 0.00923 | 0.47727 | 0.62443 | 1.20552 | 137 |
| GO_REGULATION_OF_NUCLEASE_ACTIVITY | 0.00059 | 0.00916 | 0.47727 | 0.81786 | 1.44767 | 17 |
| GO_REGULATION_OF_NUCLEOTIDE_BIOSYNTHETIC_PROCESS | 0.00302 | 0.03248 | 0.43171 | 0.78902 | 1.40337 | 18 |
| GO_REGULATION_OF_NUCLEOTIDE_METABOLIC_PROCESS | 0.00033 | 0.00588 | 0.49849 | 0.66858 | 1.26942 | 78 |
| GO_REGULATION_OF_OXIDATIVE_PHOSPHORYLATION | 0.00005 | 0.00130 | 0.55733 | 0.85284 | 1.53294 | 21 |
| GO_REGULATION_OF_PEPTIDASE_ACTIVITY | 0.00006 | 0.00152 | 0.53843 | 0.61192 | 1.19388 | 226 |
| GO_REGULATION_OF_PEPTIDE_TRANSPORT | 0.00250 | 0.02807 | 0.43171 | 0.56252 | 1.10917 | 412 |
| GO_REGULATION_OF_PH | 0.00561 | 0.04971 | 0.40702 | 0.64191 | 1.21498 | 71 |
| GO_REGULATION_OF_PLASMA_LIPOPROTEIN_PARTICLE_LEVELS | 0.00006 | 0.00145 | 0.55733 | 0.72431 | 1.36154 | 54 |
| GO_REGULATION_OF_POSTTRANSCRIPTIONAL_GENE_SILENCING | 0.00026 | 0.00486 | 0.49849 | 0.66803 | 1.27236 | 81 |
| GO_REGULATION_OF_PROTEIN_CATABOLIC_PROCESS | 0.00309 | 0.03299 | 0.43171 | 0.57360 | 1.12399 | 291 |
| GO_REGULATION_OF_PROTEIN_IMPORT | 0.00487 | 0.04559 | 0.40702 | 0.67491 | 1.25754 | 47 |
| GO_REGULATION_OF_PROTEIN_LOCALIZATION_TO_NUCLEUS | 0.00000 | 0.00013 | 0.61053 | 0.69035 | 1.32101 | 91 |
| GO_REGULATION_OF_PROTEIN_STABILITY | 0.00066 | 0.01007 | 0.47727 | 0.59308 | 1.15811 | 235 |
| GO_REGULATION_OF_PROTEOLYSIS | 0.00007 | 0.00169 | 0.53843 | 0.57923 | 1.14299 | 431 |
| GO_REGULATION_OF_REACTIVE_OXYGEN_SPECIES_BIOSYNTHET | 0.00022 | 0.00427 | 0.51885 | 0.73154 | 1.36343 | 45 |
| GO_REGULATION_OF_REACTIVE_OXYGEN_SPECIES_METABOLIC_ | 0.00088 | 0.01280 | 0.47727 | 0.63948 | 1.22592 | 99 |
| GO_REGULATION_OF_RESPONSE_TO_BIOTIC_STIMULUS | 0.00001 | 0.00039 | 0.59333 | 0.61773 | 1.20625 | 235 |
| GO_REGULATION_OF_RNA_SPLICING | 0.00011 | 0.00241 | 0.53843 | 0.64982 | 1.25198 | 125 |
| GO_REGULATION_OF_SISTER_CHROMATID_SEGREGATION | 0.00405 | 0.04011 | 0.40702 | 0.64401 | 1.22034 | 73 |
| GO_REGULATION_OF_SMALL_MOLECULE_METABOLIC_PROCESS | 0.00000 | 0.00000 | 0.77494 | 0.64342 | 1.26057 | 290 |
| GO_REGULATION_OF_SMOOTH_MUSCLE_CONTRACTION | 0.00007 | 0.00171 | 0.53843 | 0.82642 | 1.49295 | 23 |
| GO_REGULATION_OF_STEM_CELL_DIFFERENTIATION | 0.00000 | 0.00000 | 0.73376 | 0.73434 | 1.40292 | 88 |
| GO_REGULATION_OF_STEROID_METABOLIC_PROCESS | 0.00524 | 0.04770 | 0.40702 | 0.64061 | 1.21567 | 77 |
| GO_REGULATION_OF_SUPEROXIDE_METABOLIC_PROCESS | 0.00346 | 0.03548 | 0.43171 | 0.77826 | 1.39046 | 19 |
| GO_REGULATION_OF_SUPRAMOLECULAR_FIBER_ORGANIZATIO | 0.00509 | 0.04656 | 0.40702 | 0.57599 | 1.12367 | 223 |
| GO_REGULATION_OF_SYMBIOTIC_PROCESS | 0.00009 | 0.00211 | 0.53843 | 0.62950 | 1.21763 | 151 |
| GO_REGULATION_OF_TELOMERASE_ACTIVITY | 0.00087 | 0.01263 | 0.47727 | 0.72147 | 1.33426 | 39 |
| GO_REGULATION_OF_TELOMERE_MAINTENANCE | 0.00547 | 0.04923 | 0.40702 | 0.64959 | 1.22679 | 68 |
| GO_REGULATION_OF_TELOMERE_MAINTENANCE_VIA_TELOME | 0.00020 | 0.00387 | 0.51885 | 0.72043 | 1.34856 | 51 |
| GO_REGULATION_OF_TRANSCRIPTION_FROM_RNA_POLYMERAS | 0.00000 | 0.00000 | 0.70498 | 0.76931 | 1.45239 | 64 |
| GO_REGULATION_OF_TRANSLATIONAL_INITIATION | 0.00023 | 0.00437 | 0.51885 | 0.68479 | 1.29399 | 69 |
| GO_REGULATION_OF_VIRAL_TRANSCRIPTION | 0.00361 | 0.03657 | 0.43171 | 0.67497 | 1.26259 | 50 |
| GO_REGULATION_OF_WNT_SIGNALING_PATHWAY | 0.00015 | 0.00312 | 0.51885 | 0.60129 | 1.17419 | 238 |
| GO_RESPIRASOME | 0.00000 | 0.00000 | 0.81404 | 0.78326 | 1.48524 | 75 |
| GO_RESPIRATORY_CHAIN_COMPLEX | 0.00000 | 0.00000 | 0.77494 | 0.79183 | 1.49492 | 64 |
| GO_RESPIRATORY_ELECTRON_TRANSPORT_CHAIN | 0.00000 | 0.00000 | | 0.78275 | 1.49591 | 89 |

| | | | | | | |
|---|---|---|---|---|---|---|
| GO_RESPONSE_TO_ALCOHOL | 0.00015 | 0.00310 | 0.51885 | 0.63794 | 1.22915 | 126 |
| GO_RESPONSE_TO_ATP | 0.00087 | 0.01270 | 0.47727 | 0.81086 | 1.43528 | 17 |
| GO_RESPONSE_TO_CALCIUM_ION | 0.00124 | 0.01701 | 0.45506 | 0.65196 | 1.24071 | 80 |
| GO_RESPONSE_TO_CORTICOSTEROID | 0.00010 | 0.00220 | 0.53843 | 0.66894 | 1.27841 | 89 |
| GO_RESPONSE_TO_DRUG | 0.00000 | 0.00007 | 0.62726 | 0.63080 | 1.23070 | 220 |
| GO_RESPONSE_TO_ETHANOL | 0.00168 | 0.02098 | 0.45506 | 0.66543 | 1.25679 | 67 |
| GO_RESPONSE_TO_FIBROBLAST_GROWTH_FACTOR | 0.00357 | 0.03638 | 0.43171 | 0.63469 | 1.20976 | 84 |
| GO_RESPONSE_TO_GROWTH_FACTOR | 0.00272 | 0.02986 | 0.43171 | 0.56235 | 1.10805 | 387 |
| GO_RESPONSE_TO_HEAT | 0.00000 | 0.00013 | 0.62726 | 0.66982 | 1.29069 | 119 |
| GO_RESPONSE_TO_HYDROGEN_PEROXIDE | 0.00532 | 0.04803 | 0.40702 | 0.62255 | 1.19054 | 90 |
| GO_RESPONSE_TO_INORGANIC_SUBSTANCE | 0.00016 | 0.00328 | 0.51885 | 0.58928 | 1.15810 | 321 |
| GO_RESPONSE_TO_INTERLEUKIN_1 | 0.00000 | 0.00000 | 0.78819 | 0.71590 | 1.37950 | 124 |
| GO_RESPONSE_TO_INTERLEUKIN_12 | 0.00036 | 0.00622 | 0.49849 | 0.74158 | 1.36769 | 36 |
| GO_RESPONSE_TO_INTERLEUKIN_4 | 0.00224 | 0.02581 | 0.43171 | 0.77088 | 1.38921 | 22 |
| GO_RESPONSE_TO_LEUKEMIA_INHIBITORY_FACTOR | 0.00076 | 0.01129 | 0.47727 | 0.66148 | 1.25288 | 72 |
| GO_RESPONSE_TO_LIPID | 0.00043 | 0.00695 | 0.49849 | 0.56669 | 1.11902 | 470 |
| GO_RESPONSE_TO_METAL_ION | 0.00143 | 0.01863 | 0.45506 | 0.59895 | 1.16592 | 199 |
| GO_RESPONSE_TO_NUTRIENT | 0.00294 | 0.03185 | 0.43171 | 0.62596 | 1.19987 | 98 |
| GO_RESPONSE_TO_ORGANOPHOSPHORUS | 0.00294 | 0.03185 | 0.43171 | 0.64154 | 1.21511 | 72 |
| GO_RESPONSE_TO_OXIDATIVE_STRESS | 0.00495 | 0.04573 | 0.40702 | 0.57010 | 1.11758 | 294 |
| GO_RESPONSE_TO_OXYGEN_LEVELS | 0.00000 | 0.00000 | | 0.67661 | 1.32155 | 244 |
| GO_RESPONSE_TO_PURINE_CONTAINING_COMPOUND | 0.00509 | 0.04656 | 0.40702 | 0.63176 | 1.20394 | 82 |
| GO_RESPONSE_TO_STEROID_HORMONE | 0.00009 | 0.00200 | 0.53843 | 0.61312 | 1.19360 | 201 |
| GO_RESPONSE_TO_TEMPERATURE_STIMULUS | 0.00002 | 0.00047 | 0.57561 | 0.64468 | 1.24782 | 153 |
| GO_RESPONSE_TO_TOXIC_SUBSTANCE | 0.00011 | 0.00235 | 0.53843 | 0.63892 | 1.23203 | 134 |
| GO_RESPONSE_TO_TUMOR_NECROSIS_FACTOR | 0.00000 | 0.00000 | 0.70498 | 0.66331 | 1.28815 | 175 |
| GO_RESPONSE_TO_VIRUS | 0.00261 | 0.02910 | 0.43171 | 0.59428 | 1.15408 | 175 |
| GO_RETINA_HOMEOSTASIS | 0.00034 | 0.00606 | 0.49849 | 0.75202 | 1.38362 | 34 |
| GO_RIBONUCLEOPROTEIN_COMPLEX_BINDING | 0.00000 | 0.00001 | 0.69013 | 0.69839 | 1.34459 | 114 |
| GO_RIBONUCLEOPROTEIN_COMPLEX_BIOGENESIS | 0.00000 | 0.00000 | | 0.68714 | 1.35448 | 393 |
| GO_RIBONUCLEOPROTEIN_COMPLEX_SUBUNIT_ORGANIZATION | 0.00000 | 0.00000 | 0.81404 | 0.69112 | 1.34183 | 173 |
| GO_RIBONUCLEOPROTEIN_GRANULE | 0.00000 | 0.00000 | 0.71951 | 0.66271 | 1.28875 | 186 |
| GO_RIBONUCLEOSIDE_TRIPHOSPHATE_BIOSYNTHETIC_PROCESS | 0.00009 | 0.00210 | 0.53843 | 0.77194 | 1.42026 | 34 |
| GO_RIBONUCLEOSIDE_TRIPHOSPHATE_METABOLIC_PROCESS | 0.00013 | 0.00280 | 0.51885 | 0.73558 | 1.37065 | 46 |
| GO_RIBOSE_PHOSPHATE_BIOSYNTHETIC_PROCESS | 0.00002 | 0.00056 | 0.57561 | 0.65814 | 1.26821 | 124 |
| GO_RIBOSE_PHOSPHATE_METABOLIC_PROCESS | 0.00000 | 0.00000 | 0.74774 | 0.63837 | 1.25001 | 285 |
| GO_RIBOSOMAL_LARGE_SUBUNIT_BIOGENESIS | 0.00001 | 0.00031 | 0.59333 | 0.72435 | 1.36807 | 67 |
| GO_RIBOSOMAL_SMALL_SUBUNIT_ASSEMBLY | 0.00405 | 0.04011 | 0.40702 | 0.78074 | 1.38865 | 18 |
| GO_RIBOSOMAL_SMALL_SUBUNIT_BINDING | 0.00532 | 0.04803 | 0.40702 | 0.77341 | 1.36899 | 17 |
| GO_RIBOSOMAL_SMALL_SUBUNIT_BIOGENESIS | 0.00005 | 0.00138 | 0.55733 | 0.70510 | 1.33163 | 68 |
| GO_RIBOSOMAL_SUBUNIT | 0.00000 | 0.00000 | | 0.76292 | 1.48089 | 172 |
| GO_RIBOSOME | 0.00000 | 0.00000 | | 0.75411 | 1.46984 | 208 |
| GO_RIBOSOME_ASSEMBLY | 0.00001 | 0.00026 | 0.59333 | 0.74485 | 1.40144 | 56 |
| GO_RIBOSOME_BIOGENESIS | 0.00000 | 0.00000 | | 0.68899 | 1.34833 | 277 |
| GO_RNA_3_END_PROCESSING | 0.00002 | 0.00046 | 0.57561 | 0.65576 | 1.26361 | 128 |
| GO_RNA_CATABOLIC_PROCESS | 0.00000 | 0.00000 | | 0.70701 | 1.39073 | 341 |
| GO_RNA_DEPENDENT_DNA_BIOSYNTHETIC_PROCESS | 0.00003 | 0.00081 | 0.57561 | 0.71669 | 1.34965 | 59 |
| GO_RNA_EXPORT_FROM_NUCLEUS | 0.00000 | 0.00016 | 0.61053 | 0.66306 | 1.27817 | 130 |
| GO_RNA_LOCALIZATION | 0.00000 | 0.00000 | 0.76146 | 0.66638 | 1.29886 | 208 |
| GO_RNA_SPLICING | 0.00000 | 0.00000 | | 0.63512 | 1.25129 | 384 |
| GO_RNA_SPLICING_VIA_TRANSESTERIFICATION_REACTIONS | 0.00000 | 0.00000 | 0.80122 | 0.64478 | 1.26586 | 311 |
| GO_ROUGH_ENDOPLASMIC_RETICULUM | 0.00014 | 0.00293 | 0.51885 | 0.70363 | 1.32507 | 59 |
| GO_RRNA_BINDING | 0.00000 | 0.00001 | 0.69013 | 0.77638 | 1.45941 | 54 |
| GO_RRNA_METABOLIC_PROCESS | 0.00000 | 0.00000 | | 0.68430 | 1.33386 | 209 |
| GO_SCF_DEPENDENT_PROTEASOMAL_UBIQUITIN_DEPENDENT_ | 0.00000 | 0.00008 | 0.62726 | 0.71957 | 1.36498 | 76 |
| GO_SECRETORY_GRANULE | 0.00000 | 0.00000 | | 0.62978 | 1.24363 | 481 |
| GO_SECRETORY_GRANULE_MEMBRANE | 0.00032 | 0.00575 | 0.49849 | 0.61375 | 1.19134 | 172 |
| GO_SERINE_TYPE_ENDOPEPTIDASE_INHIBITOR_ACTIVITY | 0.00038 | 0.00640 | 0.49849 | 0.77388 | 1.41447 | 28 |
| GO_SEXUAL_REPRODUCTION | 0.00472 | 0.04480 | 0.40702 | 0.55748 | 1.10026 | 439 |
| GO_SIDE_OF_MEMBRANE | 0.00495 | 0.04573 | 0.40702 | 0.57741 | 1.12751 | 235 |
| GO_SIGNAL_TRANSDUCTION_BY_PROTEIN_PHOSPHORYLATION | 0.00224 | 0.02581 | 0.43171 | 0.55690 | 1.09975 | 500 |
| GO_SINGLE_FERTILIZATION | 0.00000 | 0.00002 | 0.67496 | 0.76307 | 1.43842 | 61 |
| GO_SINGLE_STRANDED_RNA_BINDING | 0.00001 | 0.00042 | 0.59333 | 0.71960 | 1.35864 | 66 |
| GO_SISTER_CHROMATID_SEGREGATION | 0.00042 | 0.00689 | 0.49849 | 0.60926 | 1.18209 | 170 |
| GO_SKIN_DEVELOPMENT | 0.00076 | 0.01140 | 0.47727 | 0.61452 | 1.18945 | 153 |
| GO_SMALL_MOLECULE_BIOSYNTHETIC_PROCESS | 0.00001 | 0.00021 | 0.61053 | 0.58728 | 1.15957 | 459 |
| GO_SMALL_NUCLEAR_RIBONUCLEOPROTEIN_COMPLEX | 0.00042 | 0.00689 | 0.49849 | 0.69542 | 1.30936 | 57 |
| GO_SMALL_RIBOSOMAL_SUBUNIT | 0.00000 | 0.00001 | 0.69013 | 0.75767 | 1.43100 | 67 |
| GO_SM_LIKE_PROTEIN_FAMILY_COMPLEX | 0.00145 | 0.01879 | 0.45506 | 0.66792 | 1.26149 | 67 |
| GO_SNORNA_BINDING | 0.00038 | 0.00646 | 0.49849 | 0.76224 | 1.39614 | 29 |

| | | | | | | |
|---|---|---|---|---|---|---|
| GO_SPERM_EGG_RECOGNITION | 0.00000 | 0.00002 | 0.65944 | 0.87469 | 1.59087 | 26 |
| GO_SPINDLE_MIDZONE | 0.00361 | 0.03657 | 0.43171 | 0.73316 | 1.33859 | 27 |
| GO_SPINDLE_ORGANIZATION | 0.00272 | 0.02986 | 0.43171 | 0.60076 | 1.16362 | 159 |
| GO_SPLICEOSOMAL_COMPLEX | 0.00000 | 0.00008 | 0.62726 | 0.65256 | 1.26657 | 171 |
| GO_SPLICEOSOMAL_TRI_SNRNP_COMPLEX | 0.00168 | 0.02098 | 0.45506 | 0.73474 | 1.34835 | 30 |
| GO_STEM_CELL_DIFFERENTIATION | 0.00000 | 0.00000 | 0.82666 | 0.70006 | 1.35554 | 158 |
| GO_STEROID_BIOSYNTHETIC_PROCESS | 0.00465 | 0.04430 | 0.40702 | 0.61242 | 1.17992 | 125 |
| GO_STEROID_METABOLIC_PROCESS | 0.00313 | 0.03310 | 0.43171 | 0.59525 | 1.15748 | 185 |
| GO_STEROL_BIOSYNTHETIC_PROCESS | 0.00005 | 0.00120 | 0.55733 | 0.71876 | 1.35584 | 62 |
| GO_STEROL_METABOLIC_PROCESS | 0.00003 | 0.00073 | 0.57561 | 0.66485 | 1.27854 | 109 |
| GO_STRESS_GRANULE_ASSEMBLY | 0.00465 | 0.04430 | 0.40702 | 0.77288 | 1.38921 | 21 |
| GO_STRUCTURAL_CONSTITUENT_OF_RIBOSOME | 0.00000 | 0.00000 | | 0.76380 | 1.47839 | 153 |
| GO_STRUCTURAL_MOLECULE_ACTIVITY | 0.00000 | 0.00000 | | 0.71760 | 1.41436 | 391 |
| GO_SUPEROXIDE_METABOLIC_PROCESS | 0.00272 | 0.02986 | 0.43171 | 0.72028 | 1.32659 | 35 |
| GO_TELOMERASE_RNA_LOCALIZATION | 0.00054 | 0.00857 | 0.47727 | 0.81801 | 1.46148 | 19 |
| GO_TELOMERE_MAINTENANCE_VIA_TELOMERE_LENGTHENING | 0.00005 | 0.00138 | 0.55733 | 0.70715 | 1.33512 | 66 |
| GO_TELOMERE_ORGANIZATION | 0.00159 | 0.02016 | 0.45506 | 0.61666 | 1.18834 | 127 |
| GO_TERTIARY_GRANULE | 0.00008 | 0.00186 | 0.53843 | 0.66372 | 1.27005 | 91 |
| GO_TERTIARY_GRANULE_MEMBRANE | 0.00102 | 0.01453 | 0.45506 | 0.71940 | 1.33244 | 40 |
| GO_THREONINE_TYPE_ENDOPEPTIDASE_ACTIVITY | 0.00000 | 0.00000 | 0.71951 | 0.93456 | 1.65424 | 17 |
| GO_THREONINE_TYPE_PEPTIDASE_ACTIVITY | 0.00000 | 0.00004 | 0.65944 | 0.89807 | 1.61424 | 21 |
| GO_TISSUE_MORPHOGENESIS | 0.00000 | 0.00006 | 0.64355 | 0.59467 | 1.17150 | 380 |
| GO_TOXIN_TRANSPORT | 0.00064 | 0.00972 | 0.47727 | 0.77713 | 1.41343 | 26 |
| GO_TRANSCRIPTION_COUPLED_NUCLEOTIDE_EXCISION_REPAIR | 0.00231 | 0.02642 | 0.43171 | 0.65429 | 1.23840 | 71 |
| GO_TRANSFERRIN_TRANSPORT | 0.00141 | 0.01850 | 0.45506 | 0.75134 | 1.37179 | 27 |
| GO_TRANSLATIONAL_ELONGATION | 0.00000 | 0.00000 | | 0.75263 | 1.45028 | 124 |
| GO_TRANSLATIONAL_INITIATION | 0.00000 | 0.00000 | | 0.75139 | 1.45839 | 171 |
| GO_TRANSLATIONAL_TERMINATION | 0.00000 | 0.00000 | 0.81404 | 0.75110 | 1.44066 | 101 |
| GO_TRANSLATION_ELONGATION_FACTOR_ACTIVITY | 0.00390 | 0.03895 | 0.43171 | 0.80505 | 1.41260 | 15 |
| GO_TRANSLATION_FACTOR_ACTIVITY_RNA_BINDING | 0.00000 | 0.00000 | 0.74774 | 0.77244 | 1.46085 | 70 |
| GO_TRANSLATION_INITIATION_FACTOR_ACTIVITY | 0.00000 | 0.00011 | 0.62726 | 0.77773 | 1.44952 | 45 |
| GO_TRANSLATION_PREINITIATION_COMPLEX | 0.00342 | 0.03521 | 0.43171 | 0.79777 | 1.40694 | 16 |
| GO_TRANSLATION_REGULATOR_ACTIVITY | 0.00000 | 0.00000 | 0.83909 | 0.74000 | 1.42403 | 111 |
| GO_TRANSLATION_REGULATOR_ACTIVITY_NUCLEIC_ACID_BIND | 0.00000 | 0.00000 | 0.83909 | 0.76441 | 1.45959 | 87 |
| GO_TRANSMEMBRANE_RECEPTOR_PROTEIN_TYROSINE_KINASE | 0.00168 | 0.02098 | 0.45506 | 0.56326 | 1.11076 | 411 |
| GO_TRICARBOXYLIC_ACID_CYCLE | 0.00159 | 0.02016 | 0.45506 | 0.73558 | 1.35097 | 33 |
| GO_TRNA_TRANSPORT | 0.00342 | 0.03521 | 0.43171 | 0.71247 | 1.31220 | 35 |
| GO_TUMOR_NECROSIS_FACTOR_MEDIATED_SIGNALING_PATHW | 0.00000 | 0.00000 | 0.80122 | 0.73921 | 1.42138 | 108 |
| GO_T_CELL_RECEPTOR_SIGNALING_PATHWAY | 0.00000 | 0.00001 | 0.70498 | 0.69185 | 1.33314 | 119 |
| GO_U12_TYPE_SPLICEOSOMAL_COMPLEX | 0.00487 | 0.04559 | 0.40702 | 0.73423 | 1.33541 | 26 |
| GO_U2_SNRNP | 0.00141 | 0.01850 | 0.45506 | 0.80259 | 1.42065 | 17 |
| GO_U2_TYPE_CATALYTIC_STEP_2_SPLICEOSOME | 0.00268 | 0.02961 | 0.43171 | 0.72225 | 1.32288 | 29 |
| GO_U2_TYPE_SPLICEOSOMAL_COMPLEX | 0.00033 | 0.00595 | 0.49849 | 0.66590 | 1.26925 | 84 |
| GO_UBIQUITIN_LIKE_PROTEIN_LIGASE_BINDING | 0.00000 | 0.00007 | 0.62726 | 0.62616 | 1.22416 | 251 |
| GO_UNFOLDED_PROTEIN_BINDING | 0.00000 | 0.00001 | 0.69013 | 0.72112 | 1.37766 | 88 |
| GO_VACUOLAR_LUMEN | 0.00016 | 0.00319 | 0.51885 | 0.65416 | 1.25772 | 107 |
| GO_VESICLE_BUDDING_FROM_MEMBRANE | 0.00480 | 0.04537 | 0.40702 | 0.62411 | 1.19353 | 90 |
| GO_VESICLE_LUMEN | 0.00000 | 0.00000 | 0.77494 | 0.67028 | 1.30631 | 207 |
| GO_VESICLE_MEMBRANE | 0.00001 | 0.00026 | 0.59333 | 0.58430 | 1.15387 | 476 |
| GO_VIRAL_GENE_EXPRESSION | 0.00000 | 0.00000 | | 0.70963 | 1.37745 | 172 |
| GO_VIRAL_LIFE_CYCLE | 0.00000 | 0.00001 | 0.69013 | 0.63985 | 1.24950 | 238 |
| GO_VIRION_ASSEMBLY | 0.00457 | 0.04386 | 0.40702 | 0.69689 | 1.28528 | 36 |
| HP_ABNORMALITY_OF_ACID_BASE_HOMEOSTASIS | 0.00000 | 0.00010 | 0.62726 | 0.62283 | 1.21827 | 261 |
| HP_ABNORMALITY_OF_BLOOD_CIRCULATION | 0.00502 | 0.04608 | 0.40702 | 0.59228 | 1.14883 | 167 |
| HP_ABNORMALITY_OF_CARDIOVASCULAR_SYSTEM_ELECTROPH | 0.00011 | 0.00237 | 0.53843 | 0.60202 | 1.17758 | 261 |
| HP_ABNORMALITY_OF_SKIN_ADNEXA_PHYSIOLOGY | 0.00242 | 0.02748 | 0.43171 | 0.61294 | 1.18094 | 118 |
| HP_ABNORMALITY_OF_THE_ANKLES | 0.00313 | 0.03310 | 0.43171 | 0.65369 | 1.23418 | 63 |
| HP_ABNORMALITY_OF_THE_CEREBRAL_VASCULATURE | 0.00357 | 0.03638 | 0.43171 | 0.59764 | 1.15722 | 158 |
| HP_ABNORMALITY_OF_THE_CEREBROSPINAL_FLUID | 0.00030 | 0.00549 | 0.49849 | 0.58221 | 1.14439 | 325 |
| HP_ABNORMALITY_OF_THE_MITOCHONDRION | 0.00126 | 0.01715 | 0.45506 | 0.62149 | 1.19764 | 120 |
| HP_ABNORMALITY_OF_THE_OPTIC_DISC | 0.00198 | 0.02370 | 0.43171 | 0.56624 | 1.11727 | 432 |
| HP_ABNORMALITY_OF_THE_OPTIC_NERVE | 0.00036 | 0.00622 | 0.49849 | 0.56723 | 1.12034 | 486 |
| HP_ABNORMAL_ABDOMEN_MORPHOLOGY | 0.00089 | 0.01291 | 0.47727 | 0.56424 | 1.11364 | 441 |
| HP_ABNORMAL_ACTIVITY_OF_MITOCHONDRIAL_RESPIRATORY_ | 0.00014 | 0.00300 | 0.51885 | 0.70303 | 1.32524 | 61 |
| HP_ABNORMAL_BASAL_GANGLIA_MRI_SIGNAL_INTENSITY | 0.00179 | 0.02183 | 0.45506 | 0.71597 | 1.32046 | 36 |
| HP_ABNORMAL_BRAINSTEM_MRI_SIGNAL_INTENSITY | 0.00398 | 0.03962 | 0.40702 | 0.71528 | 1.31370 | 33 |
| HP_ABNORMAL_CELLULAR_PHENOTYPE | 0.00209 | 0.02470 | 0.43171 | 0.56937 | 1.11992 | 334 |
| HP_ABNORMAL_CIRCULATING_CARBOHYDRATE_CONCENTRATIO | 0.00194 | 0.02331 | 0.45506 | 0.69518 | 1.29539 | 46 |
| HP_ABNORMAL_CNS_MYELINATION | 0.00014 | 0.00300 | 0.51885 | 0.62318 | 1.20954 | 171 |
| HP_ABNORMAL_CSF_METABOLITE_LEVEL | 0.00001 | 0.00027 | 0.59333 | 0.70531 | 1.34409 | 82 |

| | | | | | | |
|---|---|---|---|---|---|---|
| HP_ABNORMAL_ELECTROPHYSIOLOGY_OF_SINOATRIAL_NODE_( | 0.00029 | 0.00536 | 0.49849 | 0.73172 | 1.35725 | 41 |
| HP_ABNORMAL_ENZYME_COENZYME_ACTIVITY | 0.00561 | 0.04971 | 0.40702 | 0.60490 | 1.16567 | 123 |
| HP_ABNORMAL_ERYTHROCYTE_MORPHOLOGY | 0.00001 | 0.00023 | 0.61053 | 0.58835 | 1.15904 | 380 |
| HP_ABNORMAL_HEART_VALVE_PHYSIOLOGY | 0.00405 | 0.04011 | 0.40702 | 0.59004 | 1.14474 | 169 |
| HP_ABNORMAL_MITOCHONDRIA_IN_MUSCLE_TISSUE | 0.00075 | 0.01118 | 0.47727 | 0.74633 | 1.36700 | 29 |
| HP_ABNORMAL_MYELINATION | 0.00007 | 0.00169 | 0.53843 | 0.59258 | 1.16479 | 324 |
| HP_ABNORMAL_MYOCARDIUM_MORPHOLOGY | 0.00008 | 0.00192 | 0.53843 | 0.58647 | 1.15365 | 340 |
| HP_ABNORMAL_RENAL_PHYSIOLOGY | 0.00450 | 0.04342 | 0.40702 | 0.55484 | 1.09317 | 385 |
| HP_ABNORMAL_TRICUSPID_VALVE_PHYSIOLOGY | 0.00231 | 0.02642 | 0.43171 | 0.72968 | 1.34029 | 31 |
| HP_ABNORMAL_VITREOUS_HUMOR_MORPHOLOGY | 0.00181 | 0.02201 | 0.45506 | 0.79632 | 1.40955 | 17 |
| HP_ACUTE_LEUKEMIA | 0.00013 | 0.00276 | 0.51885 | 0.68145 | 1.29219 | 75 |
| HP_ANEMIA_OF_INADEQUATE_PRODUCTION | 0.00561 | 0.04971 | 0.40702 | 0.61298 | 1.17855 | 107 |
| HP_APHASIA | 0.00213 | 0.02490 | 0.43171 | 0.72858 | 1.33704 | 30 |
| HP_BLINDNESS | 0.00018 | 0.00355 | 0.51885 | 0.61672 | 1.19768 | 175 |
| HP_BONE_MARROW_HYPERCELLULARITY | 0.00045 | 0.00725 | 0.49849 | 0.82169 | 1.46806 | 19 |
| HP_CHOREOATHETOSIS | 0.00561 | 0.04971 | 0.40702 | 0.64956 | 1.22631 | 64 |
| HP_CHRONIC_INFECTION | 0.00121 | 0.01658 | 0.45506 | 0.81916 | 1.44468 | 16 |
| HP_CONGESTIVE_HEART_FAILURE | 0.00495 | 0.04573 | 0.40702 | 0.60734 | 1.16929 | 114 |
| HP_CRANIAL_NERVE_COMPRESSION | 0.00331 | 0.03441 | 0.43171 | 0.75830 | 1.36989 | 23 |
| HP_DECREASED_ACTIVITY_OF_MITOCHONDRIAL_COMPLEX_I | 0.00283 | 0.03081 | 0.43171 | 0.71940 | 1.32496 | 35 |
| HP_DECREASED_ACTIVITY_OF_THE_PYRUVATE_DEHYDROGENAS | 0.00062 | 0.00949 | 0.47727 | 0.75390 | 1.38478 | 31 |
| HP_DEPRESSED_NASAL_RIDGE | 0.00041 | 0.00681 | 0.49849 | 0.66218 | 1.26189 | 82 |
| HP_DYSPNEA | 0.00250 | 0.02807 | 0.43171 | 0.58107 | 1.13348 | 224 |
| HP_DYSTONIA | 0.00069 | 0.01043 | 0.47727 | 0.58101 | 1.13736 | 279 |
| HP_EMOTIONAL_LABILITY | 0.00016 | 0.00321 | 0.51885 | 0.66683 | 1.27522 | 90 |
| HP_EPISODIC_ATAXIA | 0.00487 | 0.04559 | 0.40702 | 0.79934 | 1.40259 | 15 |
| HP_FASCICULATIONS | 0.00130 | 0.01761 | 0.45506 | 0.68128 | 1.28064 | 54 |
| HP_FATIGUE | 0.00042 | 0.00689 | 0.49849 | 0.59902 | 1.16764 | 212 |
| HP_FETAL_DISTRESS | 0.00179 | 0.02183 | 0.45506 | 0.72928 | 1.33973 | 32 |
| HP_FLUSHING | 0.00561 | 0.04971 | 0.40702 | 0.74782 | 1.33686 | 20 |
| HP_FOCAL_T2_HYPERINTENSE_BRAINSTEM_LESION | 0.00159 | 0.02016 | 0.45506 | 0.74970 | 1.36879 | 27 |
| HP_HEADACHE | 0.00016 | 0.00319 | 0.51885 | 0.62537 | 1.21179 | 163 |
| HP_HEPATOMEGALY | 0.00108 | 0.01507 | 0.45506 | 0.57439 | 1.12990 | 335 |
| HP_HYPERTROPHIC_CARDIOMYOPATHY | 0.00002 | 0.00050 | 0.57561 | 0.62629 | 1.21908 | 194 |
| HP_IMPAIRED_SOCIAL_INTERACTIONS | 0.00055 | 0.00863 | 0.47727 | 0.66555 | 1.26058 | 72 |
| HP_INCREASED_CSF_LACTATE | 0.00001 | 0.00034 | 0.59333 | 0.71937 | 1.35857 | 68 |
| HP_INCREASED_SERUM_LACTATE | 0.00001 | 0.00028 | 0.59333 | 0.65837 | 1.26932 | 133 |
| HP_INCREASED_SERUM_PYRUVATE | 0.00110 | 0.01526 | 0.45506 | 0.73143 | 1.35198 | 37 |
| HP_INFANTILE_ONSET | 0.00487 | 0.04559 | 0.40702 | 0.56559 | 1.10939 | 300 |
| HP_INTRACRANIAL_CYSTIC_LESION | 0.00055 | 0.00863 | 0.47727 | 0.62760 | 1.20885 | 116 |
| HP_INTRAUTERINE_GROWTH_RETARDATION | 0.00305 | 0.03271 | 0.43171 | 0.56594 | 1.11393 | 350 |
| HP_LACTIC_ACIDOSIS | 0.00018 | 0.00355 | 0.51885 | 0.63312 | 1.22071 | 131 |
| HP_LETHARGY | 0.00084 | 0.01228 | 0.47727 | 0.63361 | 1.21845 | 109 |
| HP_LEUKODYSTROPHY | 0.00029 | 0.00529 | 0.49849 | 0.66598 | 1.26915 | 82 |
| HP_LEUKOENCEPHALOPATHY | 0.00008 | 0.00190 | 0.53843 | 0.64132 | 1.23667 | 134 |
| HP_LIPID_ACCUMULATION_IN_HEPATOCYTES | 0.00220 | 0.02548 | 0.43171 | 0.62794 | 1.20159 | 91 |
| HP_LOWER_LIMB_SPASTICITY | 0.00154 | 0.01974 | 0.45506 | 0.60929 | 1.17648 | 139 |
| HP_MACROCYTIC_ANEMIA | 0.00030 | 0.00542 | 0.49849 | 0.73108 | 1.35606 | 41 |
| HP_METABOLIC_ACIDOSIS | 0.00034 | 0.00599 | 0.49849 | 0.64831 | 1.24464 | 103 |
| HP_MICRONODULAR_CIRRHOSIS | 0.00324 | 0.03393 | 0.43171 | 0.78723 | 1.40018 | 18 |
| HP_MIGRAINE | 0.00028 | 0.00515 | 0.49849 | 0.67309 | 1.27635 | 75 |
| HP_MUSCLE_ABNORMALITY_RELATED_TO_MITOCHONDRIAL_DY | 0.00209 | 0.02470 | 0.43171 | 0.68258 | 1.27573 | 49 |
| HP_MUTISM | 0.00283 | 0.03081 | 0.43171 | 0.71937 | 1.32490 | 35 |
| HP_NAUSEA_AND_VOMITING | 0.00022 | 0.00417 | 0.51885 | 0.59383 | 1.16158 | 264 |
| HP_NEOPLASM_OF_THE_ENDOCRINE_SYSTEM | 0.00495 | 0.04573 | 0.40702 | 0.63166 | 1.20308 | 81 |
| HP_OPTIC_NEUROPATHY | 0.00135 | 0.01808 | 0.45506 | 0.73837 | 1.35610 | 33 |
| HP_PACE_OF_PROGRESSION | 0.00487 | 0.04559 | 0.40702 | 0.56814 | 1.11416 | 298 |
| HP_PALLOR | 0.00036 | 0.00622 | 0.49849 | 0.64861 | 1.24463 | 102 |
| HP_PARAPLEGIA | 0.00121 | 0.01658 | 0.45506 | 0.63584 | 1.22133 | 104 |
| HP_PARAPLEGIA_PARAPARESIS | 0.00036 | 0.00621 | 0.49849 | 0.62047 | 1.20040 | 150 |
| HP_PAROXYSMAL_INVOLUNTARY_EYE_MOVEMENTS | 0.00054 | 0.00849 | 0.47727 | 0.75650 | 1.38957 | 31 |
| HP_PEDIATRIC_ONSET | 0.00561 | 0.04971 | 0.40702 | 0.55915 | 1.10116 | 371 |
| HP_PIGMENTARY_RETINOPATHY | 0.00065 | 0.00981 | 0.47727 | 0.63260 | 1.21720 | 113 |
| HP_POOR_EYE_CONTACT | 0.00017 | 0.00341 | 0.51885 | 0.71279 | 1.33749 | 53 |
| HP_POOR_HEAD_CONTROL | 0.00000 | 0.00005 | 0.64355 | 0.67819 | 1.30665 | 118 |
| HP_PROGRESSIVE | 0.00194 | 0.02331 | 0.45506 | 0.59790 | 1.16273 | 186 |
| HP_PROGRESSIVE_CEREBELLAR_ATAXIA | 0.00364 | 0.03677 | 0.43171 | 0.64788 | 1.22325 | 65 |
| HP_PROGRESSIVE_SPASTIC_PARAPLEGIA | 0.00253 | 0.02838 | 0.43171 | 0.69655 | 1.29822 | 45 |
| HP_PROXIMAL_TUBULOPATHY | 0.00141 | 0.01850 | 0.45506 | 0.72579 | 1.33535 | 34 |
| HP_PTOSIS | 0.00472 | 0.04480 | 0.40702 | 0.55582 | 1.09637 | 418 |

| | | | | | | |
|---|---|---|---|---|---|---|
| HP_REDUCED_CONSCIOUSNESS_CONFUSION | 0.00007 | 0.00164 | 0.53843 | 0.61227 | 1.19295 | 205 |
| HP_REDUCED_VISUAL_ACUITY | 0.00053 | 0.00837 | 0.47727 | 0.58618 | 1.14774 | 283 |
| HP_RENAL_TUBULAR_DYSFUNCTION | 0.00013 | 0.00284 | 0.51885 | 0.68108 | 1.29150 | 75 |
| HP_RESPIRATORY_DISTRESS | 0.00108 | 0.01507 | 0.45506 | 0.62693 | 1.20762 | 115 |
| HP_RESPIRATORY_INSUFFICIENCY | 0.00000 | 0.00010 | 0.62726 | 0.60536 | 1.18998 | 326 |
| HP_SINUS_TACHYCARDIA | 0.00213 | 0.02490 | 0.43171 | 0.77203 | 1.39128 | 22 |
| HP_SPARSE_EYELASHES | 0.00150 | 0.01943 | 0.45506 | 0.71355 | 1.32161 | 40 |
| HP_SPARSE_OR_ABSENT_EYELASHES | 0.00220 | 0.02548 | 0.43171 | 0.67331 | 1.26567 | 54 |
| HP_SPASTIC_PARAPLEGIA | 0.00098 | 0.01407 | 0.45506 | 0.64059 | 1.22712 | 96 |
| HP_THROMBOCYTOSIS | 0.00413 | 0.04065 | 0.40702 | 0.75210 | 1.35870 | 23 |
| HP_TONGUE_FASCICULATIONS | 0.00457 | 0.04386 | 0.40702 | 0.80056 | 1.40472 | 15 |
| HP_TRANSIENT_ISCHEMIC_ATTACK | 0.00480 | 0.04537 | 0.40702 | 0.74902 | 1.35809 | 25 |
| HP_VERTIGO | 0.00002 | 0.00063 | 0.57561 | 0.72114 | 1.35844 | 60 |
| HP_VESTIBULAR_DYSFUNCTION | 0.00028 | 0.00522 | 0.49849 | 0.67279 | 1.27577 | 75 |
| HP_VOMITING | 0.00072 | 0.01082 | 0.47727 | 0.61459 | 1.18792 | 147 |
| HP_WEAKNESS_DUE_TO_UPPER_MOTOR_NEURON_DYSFUNCTI | 0.00532 | 0.04803 | 0.40702 | 0.56829 | 1.11186 | 272 |
| HP_WEAKNESS_OF_MUSCLES_OF_RESPIRATION | 0.00163 | 0.02050 | 0.45506 | 0.66272 | 1.25123 | 63 |
| **Cluster 1** | | | | | | |
| pathway | pval | padj | log2err | ES | NES | size |
| GO_ACTIVATION_OF_IMMUNE_RESPONSE | 0.00000 | 0.00000 | 0.74774 | 0.65038 | 1.26999 | 235 |
| GO_ACTIVATION_OF_INNATE_IMMUNE_RESPONSE | 0.00000 | 0.00001 | 0.69013 | 0.71967 | 1.37861 | 96 |
| GO_AEROBIC_RESPIRATION | 0.00000 | 0.00000 | 0.73376 | 0.76280 | 1.44478 | 72 |
| GO_ALCOHOL_BIOSYNTHETIC_PROCESS | 0.00032 | 0.00572 | 0.49849 | 0.64679 | 1.24367 | 108 |
| GO_ALCOHOL_METABOLIC_PROCESS | 0.00364 | 0.03677 | 0.43171 | 0.58112 | 1.13424 | 228 |
| GO_AMEBOIDAL_TYPE_CELL_MIGRATION | 0.00457 | 0.04386 | 0.40702 | 0.57755 | 1.12854 | 247 |
| GO_AMIDE_BINDING | 0.00014 | 0.00297 | 0.51885 | 0.61533 | 1.19780 | 199 |
| GO_AMINE_METABOLIC_PROCESS | 0.00000 | 0.00000 | 0.76146 | 0.73732 | 1.41097 | 92 |
| GO_AMYLOID_BETA_CLEARANCE | 0.00561 | 0.04971 | 0.40702 | 0.74081 | 1.33976 | 24 |
| GO_ANAPHASE_PROMOTING_COMPLEX_DEPENDENT_CATABOL | 0.00000 | 0.00000 | 0.76146 | 0.77107 | 1.46112 | 73 |
| GO_ANATOMICAL_STRUCTURE_HOMEOSTASIS | 0.00020 | 0.00390 | 0.51885 | 0.59307 | 1.16021 | 271 |
| GO_ANIMAL_ORGAN_REGENERATION | 0.00012 | 0.00269 | 0.51885 | 0.71251 | 1.33510 | 52 |
| GO_ANTIGEN_PROCESSING_AND_PRESENTATION | 0.00000 | 0.00000 | 0.74774 | 0.68785 | 1.33064 | 149 |
| GO_ANTIGEN_PROCESSING_AND_PRESENTATION_OF_EXOGENC | 0.00000 | 0.00000 | 0.76146 | 0.79644 | 1.50237 | 62 |
| GO_ANTIGEN_PROCESSING_AND_PRESENTATION_OF_PEPTIDE_ | 0.00000 | 0.00000 | 0.74774 | 0.70107 | 1.35144 | 130 |
| GO_ANTIGEN_PROCESSING_AND_PRESENTATION_OF_PEPTIDE_ | 0.00000 | 0.00000 | 0.78819 | 0.77566 | 1.47012 | 74 |
| GO_ANTIGEN_RECEPTOR_MEDIATED_SIGNALING_PATHWAY | 0.00000 | 0.00009 | 0.62726 | 0.66362 | 1.28080 | 136 |
| GO_APICAL_PART_OF_CELL | 0.00135 | 0.01808 | 0.45506 | 0.58926 | 1.14966 | 220 |
| GO_APOPTOTIC_SIGNALING_PATHWAY | 0.00006 | 0.00152 | 0.53843 | 0.58638 | 1.15586 | 393 |
| GO_ATPASE_ACTIVITY | 0.00050 | 0.00802 | 0.47727 | 0.58216 | 1.14385 | 316 |
| GO_ATPASE_ACTIVITY_COUPLED_TO_TRANSMEMBRANE_MOVE | 0.00420 | 0.04132 | 0.40702 | 0.78005 | 1.38741 | 18 |
| GO_ATP_BIOSYNTHETIC_PROCESS | 0.00013 | 0.00276 | 0.51885 | 0.83897 | 1.50801 | 21 |
| GO_ATP_METABOLIC_PROCESS | 0.00000 | 0.00000 | | 0.72178 | 1.40814 | 219 |
| GO_ATP_SYNTHESIS_COUPLED_ELECTRON_TRANSPORT | 0.00000 | 0.00000 | | 0.80353 | 1.52425 | 76 |
| GO_AZUROPHIL_GRANULE | 0.00025 | 0.00466 | 0.49849 | 0.64781 | 1.24551 | 107 |
| GO_AZUROPHIL_GRANULE_LUMEN | 0.00405 | 0.04011 | 0.40702 | 0.65625 | 1.23793 | 62 |
| GO_BASAL_PART_OF_CELL | 0.00509 | 0.04656 | 0.40702 | 0.68515 | 1.27697 | 45 |
| GO_BASAL_PLASMA_MEMBRANE | 0.00346 | 0.03548 | 0.43171 | 0.71184 | 1.31104 | 35 |
| GO_BASEMENT_MEMBRANE | 0.00279 | 0.03056 | 0.43171 | 0.68935 | 1.28452 | 46 |
| GO_BASOLATERAL_PLASMA_MEMBRANE | 0.00011 | 0.00247 | 0.53843 | 0.63209 | 1.21993 | 136 |
| GO_BINDING_OF_SPERM_TO_ZONA_PELLUCIDA | 0.00000 | 0.00006 | 0.64355 | 0.90095 | 1.60966 | 19 |
| GO_BLOOD_MICROPARTICLE | 0.00000 | 0.00000 | 0.70498 | 0.81982 | 1.52763 | 46 |
| GO_BRUSH_BORDER | 0.00060 | 0.00923 | 0.47727 | 0.66897 | 1.26790 | 74 |
| GO_CADHERIN_BINDING | 0.00000 | 0.00000 | | 0.68759 | 1.34435 | 253 |
| GO_CALCIUM_DEPENDENT_PHOSPHOLIPID_BINDING | 0.00152 | 0.01958 | 0.45506 | 0.75021 | 1.37120 | 28 |
| GO_CALCIUM_DEPENDENT_PROTEIN_BINDING | 0.00009 | 0.00211 | 0.53843 | 0.74254 | 1.38394 | 45 |
| GO_CANONICAL_WNT_SIGNALING_PATHWAY | 0.00004 | 0.00109 | 0.55733 | 0.61740 | 1.20458 | 222 |
| GO_CARBOHYDRATE_CATABOLIC_PROCESS | 0.00135 | 0.01808 | 0.45506 | 0.61179 | 1.18186 | 144 |
| GO_CATALYTIC_STEP_2_SPLICEOSOME | 0.00001 | 0.00022 | 0.61053 | 0.70290 | 1.33765 | 80 |
| GO_CATION_TRANSMEMBRANE_TRANSPORT | 0.00327 | 0.03409 | 0.43171 | 0.56060 | 1.10531 | 406 |
| GO_CATION_TRANSMEMBRANE_TRANSPORTER_ACTIVITY | 0.00097 | 0.01387 | 0.47727 | 0.58196 | 1.13846 | 265 |
| GO_CELLULAR_AMINO_ACID_METABOLIC_PROCESS | 0.00002 | 0.00060 | 0.57561 | 0.61438 | 1.19930 | 231 |
| GO_CELLULAR_COMPONENT_DISASSEMBLY | 0.00000 | 0.00000 | | 0.64070 | 1.26342 | 408 |
| GO_CELLULAR_ION_HOMEOSTASIS | 0.00027 | 0.00493 | 0.49849 | 0.58658 | 1.15340 | 328 |
| GO_CELLULAR_IRON_ION_HOMEOSTASIS | 0.00327 | 0.03409 | 0.43171 | 0.68360 | 1.27374 | 47 |
| GO_CELLULAR_KETONE_METABOLIC_PROCESS | 0.00000 | 0.00010 | 0.62726 | 0.65358 | 1.26726 | 164 |
| GO_CELLULAR_MONOVALENT_INORGANIC_CATION_HOMEOSTA | 0.00179 | 0.02183 | 0.45506 | 0.64747 | 1.22821 | 76 |
| GO_CELLULAR_NITROGEN_COMPOUND_CATABOLIC_PROCESS | 0.00000 | 0.00000 | | 0.67915 | 1.34096 | 456 |
| GO_CELLULAR_PROTEIN_COMPLEX_DISASSEMBLY | 0.00000 | 0.00000 | | 0.69566 | 1.35184 | 179 |
| GO_CELLULAR_RESPIRATION | 0.00000 | 0.00000 | | 0.74205 | 1.43370 | 145 |
| GO_CELLULAR_RESPONSE_TO_HEAT | 0.00004 | 0.00109 | 0.55733 | 0.67186 | 1.28698 | 95 |

| | | | | | | |
|---|---|---|---|---|---|---|
| GO_CELLULAR_RESPONSE_TO_HORMONE_STIMULUS | 0.00309 | 0.03299 | 0.43171 | 0.56453 | 1.11191 | 374 |
| GO_CELLULAR_RESPONSE_TO_ORGANIC_CYCLIC_COMPOUND | 0.00502 | 0.04608 | 0.40702 | 0.56512 | 1.11134 | 329 |
| GO_CELLULAR_RESPONSE_TO_OXYGEN_LEVELS | 0.00000 | 0.00000 | | 0.71461 | 1.38224 | 151 |
| GO_CELL_ACTIVATION_INVOLVED_IN_IMMUNE_RESPONSE | 0.00000 | 0.00000 | | 0.63924 | 1.26108 | 422 |
| GO_CELL_ADHESION_MEDIATOR_ACTIVITY | 0.00157 | 0.02009 | 0.45506 | 0.71837 | 1.32488 | 36 |
| GO_CELL_ADHESION_MOLECULE_BINDING | 0.00000 | 0.00000 | | 0.67109 | 1.32112 | 353 |
| GO_CELL_BODY | 0.00004 | 0.00109 | 0.55733 | 0.59973 | 1.17541 | 293 |
| GO_CELL_CELL_RECOGNITION | 0.00000 | 0.00006 | 0.64355 | 0.82001 | 1.51026 | 35 |
| GO_CELL_CELL_SIGNALING_BY_WNT | 0.00004 | 0.00109 | 0.55733 | 0.59042 | 1.16141 | 339 |
| GO_CELL_CYCLE_G1_S_PHASE_TRANSITION | 0.00376 | 0.03783 | 0.43171 | 0.58112 | 1.13378 | 220 |
| GO_CELL_CYCLE_G2_M_PHASE_TRANSITION | 0.00024 | 0.00459 | 0.51885 | 0.59807 | 1.16746 | 231 |
| GO_CELL_CYCLE_PHASE_TRANSITION | 0.00000 | 0.00001 | 0.69013 | 0.59417 | 1.17348 | 499 |
| GO_CELL_DIVISION | 0.00003 | 0.00071 | 0.57561 | 0.58314 | 1.15094 | 449 |
| GO_CELL_KILLING | 0.00205 | 0.02440 | 0.43171 | 0.66727 | 1.25696 | 60 |
| GO_CELL_RECOGNITION | 0.00000 | 0.00000 | 0.73376 | 0.75953 | 1.43954 | 74 |
| GO_CELL_REDOX_HOMEOSTASIS | 0.00163 | 0.02050 | 0.45506 | 0.71772 | 1.32665 | 38 |
| GO_CELL_SUBSTRATE_JUNCTION | 0.00000 | 0.00000 | | 0.65742 | 1.29171 | 316 |
| GO_CELL_SURFACE | 0.00000 | 0.00000 | 0.73376 | 0.62572 | 1.23070 | 330 |
| GO_CELL_SURFACE_RECEPTOR_SIGNALING_PATHWAY_INVOLVEI | 0.00038 | 0.00640 | 0.49849 | 0.57762 | 1.13770 | 374 |
| GO_CHAPERONE_BINDING | 0.00495 | 0.04573 | 0.40702 | 0.62860 | 1.19935 | 86 |
| GO_CHAPERONE_COMPLEX | 0.00001 | 0.00028 | 0.59333 | 0.86337 | 1.55587 | 22 |
| GO_CHAPERONE_MEDIATED_PROTEIN_COMPLEX_ASSEMBLY | 0.00100 | 0.01430 | 0.45506 | 0.80787 | 1.44337 | 19 |
| GO_CHAPERONE_MEDIATED_PROTEIN_FOLDING | 0.00168 | 0.02098 | 0.45506 | 0.71110 | 1.32288 | 44 |
| GO_CHROMOSOMAL_REGION | 0.00018 | 0.00349 | 0.51885 | 0.59323 | 1.16180 | 286 |
| GO_CHROMOSOME_CENTROMERIC_REGION | 0.00172 | 0.02135 | 0.45506 | 0.60447 | 1.17248 | 168 |
| GO_CHROMOSOME_SEGREGATION | 0.00003 | 0.00087 | 0.55733 | 0.60463 | 1.18280 | 270 |
| GO_CLUSTER_OF_ACTIN_BASED_CELL_PROJECTIONS | 0.00074 | 0.01107 | 0.47727 | 0.64184 | 1.23044 | 99 |
| GO_COATED_VESICLE | 0.00213 | 0.02490 | 0.43171 | 0.59512 | 1.15846 | 199 |
| GO_COATED_VESICLE_MEMBRANE | 0.00060 | 0.00923 | 0.47727 | 0.63443 | 1.22207 | 115 |
| GO_COLLAGEN_BINDING | 0.00465 | 0.04430 | 0.40702 | 0.73471 | 1.33628 | 26 |
| GO_COLLAGEN_CONTAINING_EXTRACELLULAR_MATRIX | 0.00000 | 0.00006 | 0.64355 | 0.65358 | 1.26542 | 157 |
| GO_CONDENSED_CHROMOSOME | 0.00350 | 0.03580 | 0.43171 | 0.59394 | 1.15417 | 179 |
| GO_CONTRACTILE_FIBER | 0.00064 | 0.00972 | 0.47727 | 0.62217 | 1.19973 | 134 |
| GO_COPI_COATED_VESICLE | 0.00316 | 0.03344 | 0.43171 | 0.75688 | 1.37234 | 25 |
| GO_CORNIFICATION | 0.00027 | 0.00500 | 0.49849 | 0.77970 | 1.42512 | 28 |
| GO_CORTICAL_CYTOSKELETON | 0.00502 | 0.04608 | 0.40702 | 0.64388 | 1.21870 | 71 |
| GO_COTRANSLATIONAL_PROTEIN_TARGETING_TO_MEMBRANE | 0.00000 | 0.00000 | | 0.78223 | 1.49819 | 94 |
| GO_CYTOCHROME_COMPLEX | 0.00035 | 0.00610 | 0.49849 | 0.80796 | 1.45601 | 22 |
| GO_CYTOKINE_MEDIATED_SIGNALING_PATHWAY | 0.00000 | 0.00002 | 0.67496 | 0.60742 | 1.19735 | 392 |
| GO_CYTOPLASMIC_STRESS_GRANULE | 0.00014 | 0.00302 | 0.51885 | 0.69390 | 1.31009 | 63 |
| GO_CYTOPLASMIC_TRANSLATION | 0.00000 | 0.00001 | 0.70498 | 0.71773 | 1.37349 | 92 |
| GO_CYTOPLASMIC_TRANSLATIONAL_INITIATION | 0.00178 | 0.02182 | 0.45506 | 0.73517 | 1.35039 | 31 |
| GO_CYTOSOLIC_LARGE_RIBOSOMAL_SUBUNIT | 0.00000 | 0.00005 | 0.64355 | 0.77240 | 1.44935 | 53 |
| GO_CYTOSOLIC_RIBOSOME | 0.00000 | 0.00000 | | 0.77483 | 1.48437 | 97 |
| GO_CYTOSOLIC_SMALL_RIBOSOMAL_SUBUNIT | 0.00000 | 0.00010 | 0.62726 | 0.79384 | 1.47033 | 40 |
| GO_DETECTION_OF_STIMULUS | 0.00020 | 0.00393 | 0.51885 | 0.65208 | 1.25384 | 108 |
| GO_DETOXIFICATION | 0.00413 | 0.04065 | 0.40702 | 0.65399 | 1.23684 | 70 |
| GO_DEVELOPMENT_OF_PRIMARY_FEMALE_SEXUAL_CHARACTE | 0.00179 | 0.02183 | 0.45506 | 0.68410 | 1.28057 | 51 |
| GO_DICARBOXYLIC_ACID_METABOLIC_PROCESS | 0.00261 | 0.02910 | 0.43171 | 0.65541 | 1.23747 | 65 |
| GO_DISORDERED_DOMAIN_SPECIFIC_BINDING | 0.00051 | 0.00812 | 0.47727 | 0.76867 | 1.40343 | 27 |
| GO_DISULFIDE_OXIDOREDUCTASE_ACTIVITY | 0.00435 | 0.04238 | 0.40702 | 0.71445 | 1.31232 | 31 |
| GO_DNA_BIOSYNTHETIC_PROCESS | 0.00017 | 0.00334 | 0.51885 | 0.62540 | 1.20833 | 145 |
| GO_DNA_CONFORMATION_CHANGE | 0.00327 | 0.03409 | 0.43171 | 0.57979 | 1.13178 | 231 |
| GO_DNA_DAMAGE_RESPONSE_DETECTION_OF_DNA_DAMAGE | 0.00305 | 0.03271 | 0.43171 | 0.71625 | 1.31916 | 35 |
| GO_DNA_GEOMETRIC_CHANGE | 0.00141 | 0.01850 | 0.45506 | 0.62642 | 1.20356 | 105 |
| GO_DOUBLE_STRANDED_RNA_BINDING | 0.00039 | 0.00658 | 0.49849 | 0.69158 | 1.30365 | 61 |
| GO_DRUG_BINDING | 0.00450 | 0.04342 | 0.40702 | 0.66245 | 1.24130 | 52 |
| GO_ELECTRON_TRANSFER_ACTIVITY | 0.00000 | 0.00001 | 0.67496 | 0.72917 | 1.38765 | 80 |
| GO_ELECTRON_TRANSPORT_CHAIN | 0.00000 | 0.00000 | | 0.73336 | 1.41392 | 133 |
| GO_EMBRYO_IMPLANTATION | 0.00044 | 0.00706 | 0.49849 | 0.75968 | 1.39146 | 29 |
| GO_ENDOCYTIC_VESICLE_MEMBRANE | 0.00524 | 0.04770 | 0.40702 | 0.62563 | 1.19564 | 89 |
| GO_ENDOPEPTIDASE_ACTIVITY | 0.00065 | 0.00981 | 0.47727 | 0.60123 | 1.16998 | 196 |
| GO_ENDOPEPTIDASE_COMPLEX | 0.00000 | 0.00001 | 0.70498 | 0.76593 | 1.44609 | 63 |
| GO_ENDOPEPTIDASE_REGULATOR_ACTIVITY | 0.00006 | 0.00145 | 0.55733 | 0.70425 | 1.33001 | 68 |
| GO_ENDOPLASMIC_RETICULUM_GOLGI_INTERMEDIATE_COMPA | 0.00055 | 0.00863 | 0.47727 | 0.64758 | 1.24050 | 96 |
| GO_ENDOPLASMIC_RETICULUM_GOLGI_INTERMEDIATE_COMPA | 0.00148 | 0.01923 | 0.45506 | 0.68944 | 1.29056 | 51 |
| GO_ENDOPLASMIC_RETICULUM_TO_GOLGI_VESICLE_MEDIATED | 0.00532 | 0.04803 | 0.40702 | 0.59194 | 1.14697 | 162 |
| GO_ENERGY_DERIVATION_BY_OXIDATION_OF_ORGANIC_COMP | 0.00000 | 0.00000 | | 0.69995 | 1.36305 | 202 |
| GO_ENZYME_INHIBITOR_ACTIVITY | 0.00246 | 0.02780 | 0.43171 | 0.58913 | 1.14643 | 191 |
| GO_EPIDERMAL_CELL_DIFFERENTIATION | 0.00216 | 0.02519 | 0.43171 | 0.62667 | 1.20589 | 112 |

| | | | | | | |
|---|---|---|---|---|---|---|
| GO_EPITHELIAL_CELL_DIFFERENTIATION | 0.00054 | 0.00857 | 0.47727 | 0.57819 | 1.13725 | 334 |
| GO_ESTABLISHMENT_OF_ORGANELLE_LOCALIZATION | 0.00122 | 0.01679 | 0.45506 | 0.57737 | 1.13489 | 323 |
| GO_ESTABLISHMENT_OF_PROTEIN_LOCALIZATION_TO_CHROMO | 0.00016 | 0.00321 | 0.51885 | 0.81363 | 1.46985 | 23 |
| GO_ESTABLISHMENT_OF_PROTEIN_LOCALIZATION_TO_ENDOPL | 0.00000 | 0.00000 | | 0.77241 | 1.48479 | 106 |
| GO_ESTABLISHMENT_OF_PROTEIN_LOCALIZATION_TO_MEMBRA | 0.00000 | 0.00000 | | 0.67484 | 1.31991 | 260 |
| GO_ESTABLISHMENT_OF_PROTEIN_LOCALIZATION_TO_ORGANE | 0.00000 | 0.00000 | | 0.65856 | 1.30056 | 461 |
| GO_ESTABLISHMENT_OF_PROTEIN_LOCALIZATION_TO_TELOME | 0.00000 | 0.00006 | 0.64355 | 0.90447 | 1.60870 | 18 |
| GO_ESTABLISHMENT_OF_RNA_LOCALIZATION | 0.00000 | 0.00005 | 0.64355 | 0.64271 | 1.24830 | 176 |
| GO_ESTABLISHMENT_OF_TISSUE_POLARITY | 0.00000 | 0.00000 | 0.81404 | 0.75693 | 1.44999 | 96 |
| GO_EUKARYOTIC_TRANSLATION_INITIATION_FACTOR_3_COMPL | 0.00143 | 0.01863 | 0.45506 | 0.83263 | 1.46100 | 15 |
| GO_EXTERNAL_SIDE_OF_PLASMA_MEMBRANE | 0.00159 | 0.02016 | 0.45506 | 0.61775 | 1.19022 | 118 |
| GO_EXTRACELLULAR_MATRIX | 0.00042 | 0.00695 | 0.49849 | 0.61138 | 1.18938 | 189 |
| GO_FATTY_ACID_DERIVATIVE_METABOLIC_PROCESS | 0.00043 | 0.00695 | 0.49849 | 0.66266 | 1.26212 | 81 |
| GO_FATTY_ACYL_COA_METABOLIC_PROCESS | 0.00502 | 0.04608 | 0.40702 | 0.72264 | 1.32082 | 28 |
| GO_FC_EPSILON_RECEPTOR_SIGNALING_PATHWAY | 0.00000 | 0.00000 | 0.71951 | 0.72370 | 1.38610 | 94 |
| GO_FC_RECEPTOR_SIGNALING_PATHWAY | 0.00000 | 0.00000 | 0.70498 | 0.68815 | 1.32879 | 140 |
| GO_FERTILIZATION | 0.00000 | 0.00005 | 0.64355 | 0.73257 | 1.38752 | 72 |
| GO_FIBRILLAR_CENTER | 0.00246 | 0.02780 | 0.43171 | 0.62638 | 1.19990 | 96 |
| GO_FIBROBLAST_GROWTH_FACTOR_RECEPTOR_SIGNALING_PAT | 0.00495 | 0.04573 | 0.40702 | 0.64888 | 1.22552 | 67 |
| GO_FIBROBLAST_PROLIFERATION | 0.00032 | 0.00575 | 0.49849 | 0.70676 | 1.32618 | 53 |
| GO_FICOLIN_1_RICH_GRANULE | 0.00000 | 0.00000 | 0.70498 | 0.71544 | 1.37138 | 98 |
| GO_GABA_ERGIC_SYNAPSE | 0.00022 | 0.00427 | 0.51885 | 0.76430 | 1.40406 | 32 |
| GO_GENERATION_OF_PRECURSOR_METABOLITES_AND_ENERGY | 0.00000 | 0.00000 | | 0.67527 | 1.32968 | 368 |
| GO_GENE_SILENCING | 0.00019 | 0.00376 | 0.51885 | 0.61625 | 1.19788 | 184 |
| GO_GLUCOSE_CATABOLIC_PROCESS | 0.00495 | 0.04573 | 0.40702 | 0.72314 | 1.32174 | 28 |
| GO_GLUTAMATERGIC_SYNAPSE | 0.00135 | 0.01808 | 0.45506 | 0.60052 | 1.16783 | 186 |
| GO_GLYCOLYTIC_PROCESS_THROUGH_FRUCTOSE_6_PHOSPHATE | 0.00554 | 0.04968 | 0.40702 | 0.74156 | 1.34111 | 24 |
| GO_GOLGI_ASSOCIATED_VESICLE | 0.00085 | 0.01239 | 0.47727 | 0.62294 | 1.20037 | 129 |
| GO_GOLGI_ASSOCIATED_VESICLE_MEMBRANE | 0.00313 | 0.03310 | 0.43171 | 0.64208 | 1.21753 | 75 |
| GO_GTPASE_ACTIVITY | 0.00004 | 0.00098 | 0.55733 | 0.61546 | 1.19974 | 210 |
| GO_GUANYL_NUCLEOTIDE_BINDING | 0.00005 | 0.00126 | 0.55733 | 0.60899 | 1.19065 | 252 |
| GO_HEAT_SHOCK_PROTEIN_BINDING | 0.00001 | 0.00034 | 0.59333 | 0.68258 | 1.30633 | 93 |
| GO_HEMATOPOIETIC_PROGENITOR_CELL_DIFFERENTIATION | 0.00000 | 0.00000 | 0.71951 | 0.70099 | 1.35150 | 121 |
| GO_HEMATOPOIETIC_STEM_CELL_DIFFERENTIATION | 0.00000 | 0.00000 | 0.76146 | 0.77246 | 1.46307 | 72 |
| GO_HISTONE_MRNA_METABOLIC_PROCESS | 0.00265 | 0.02936 | 0.43171 | 0.76832 | 1.38459 | 22 |
| GO_HOST_CELLULAR_COMPONENT | 0.00043 | 0.00695 | 0.49849 | 0.69729 | 1.31074 | 54 |
| GO_IMMUNE_RESPONSE_REGULATING_SIGNALING_PATHWAY | 0.00000 | 0.00003 | 0.65944 | 0.63620 | 1.24112 | 225 |
| GO_IMPORT_INTO_CELL | 0.00428 | 0.04185 | 0.40702 | 0.61709 | 1.18655 | 108 |
| GO_IMPORT_INTO_NUCLEUS | 0.00039 | 0.00652 | 0.49849 | 0.62707 | 1.20905 | 131 |
| GO_INNATE_IMMUNE_RESPONSE | 0.00000 | 0.00003 | 0.65944 | 0.59893 | 1.18089 | 406 |
| GO_INNATE_IMMUNE_RESPONSE_ACTIVATING_SIGNAL_TRANSD | 0.00000 | 0.00001 | 0.69013 | 0.73348 | 1.39585 | 80 |
| GO_INNER_MITOCHONDRIAL_MEMBRANE_PROTEIN_COMPLEX | 0.00000 | 0.00000 | | 0.76576 | 1.46666 | 94 |
| GO_INORGANIC_ION_TRANSMEMBRANE_TRANSPORT | 0.00141 | 0.01850 | 0.45506 | 0.56404 | 1.11185 | 401 |
| GO_INTERACTION_WITH_HOST | 0.00139 | 0.01850 | 0.45506 | 0.60855 | 1.17710 | 151 |
| GO_INTERLEUKIN_1_MEDIATED_SIGNALING_PATHWAY | 0.00000 | 0.00000 | 0.77494 | 0.75654 | 1.44176 | 83 |
| GO_INTRAMOLECULAR_OXIDOREDUCTASE_ACTIVITY | 0.00051 | 0.00812 | 0.47727 | 0.74506 | 1.37081 | 34 |
| GO_INTRINSIC_APOPTOTIC_SIGNALING_PATHWAY | 0.00156 | 0.01993 | 0.45506 | 0.59427 | 1.15817 | 207 |
| GO_ION_HOMEOSTASIS | 0.00170 | 0.02117 | 0.45506 | 0.56164 | 1.10717 | 403 |
| GO_ION_TRANSMEMBRANE_TRANSPORTER_ACTIVITY | 0.00106 | 0.01495 | 0.45506 | 0.56696 | 1.11734 | 390 |
| GO_IRON_ION_HOMEOSTASIS | 0.00361 | 0.03657 | 0.43171 | 0.65837 | 1.23983 | 59 |
| GO_IRON_ION_TRANSPORT | 0.00054 | 0.00857 | 0.47727 | 0.69556 | 1.30334 | 52 |
| GO_ISOMERASE_ACTIVITY | 0.00413 | 0.04065 | 0.40702 | 0.61065 | 1.17496 | 113 |
| GO_KERATINIZATION | 0.00038 | 0.00640 | 0.49849 | 0.75690 | 1.39013 | 33 |
| GO_KILLING_OF_CELLS_OF_OTHER_ORGANISM | 0.00495 | 0.04573 | 0.40702 | 0.78614 | 1.38643 | 16 |
| GO_LARGE_RIBOSOMAL_SUBUNIT | 0.00000 | 0.00000 | | 0.77656 | 1.49319 | 108 |
| GO_LATERAL_PLASMA_MEMBRANE | 0.00524 | 0.04770 | 0.40702 | 0.67385 | 1.25557 | 47 |
| GO_LEUKOCYTE_MEDIATED_IMMUNITY | 0.00000 | 0.00000 | | 0.64010 | 1.26331 | 447 |
| GO_LIPOPROTEIN_PARTICLE_RECEPTOR_BINDING | 0.00126 | 0.01715 | 0.45506 | 0.80503 | 1.42497 | 17 |
| GO_LIVER_REGENERATION | 0.00093 | 0.01337 | 0.47727 | 0.78442 | 1.41708 | 23 |
| GO_LOW_DENSITY_LIPOPROTEIN_RECEPTOR_PARTICLE_METABO | 0.00327 | 0.03409 | 0.43171 | 0.78520 | 1.39657 | 18 |
| GO_LYASE_ACTIVITY | 0.00152 | 0.01958 | 0.45506 | 0.61769 | 1.19079 | 122 |
| GO_LYSOSOMAL_LUMEN | 0.00143 | 0.01863 | 0.45506 | 0.67371 | 1.26872 | 59 |
| GO_MAINTENANCE_OF_CELL_NUMBER | 0.00435 | 0.04238 | 0.40702 | 0.60761 | 1.17073 | 117 |
| GO_MATURATION_OF_SSU_RRNA | 0.00126 | 0.01715 | 0.45506 | 0.69615 | 1.29711 | 47 |
| GO_MATURATION_OF_SSU_RRNA_FROM_TRICISTRONIC_RRNA_ | 0.00554 | 0.04968 | 0.40702 | 0.70205 | 1.29301 | 35 |
| GO_MEMBRANE_REGION | 0.00000 | 0.00003 | 0.65944 | 0.64496 | 1.25542 | 194 |
| GO_METALLOPEPTIDASE_ACTIVITY | 0.00487 | 0.04559 | 0.40702 | 0.62714 | 1.19577 | 85 |
| GO_METAL_ION_HOMEOSTASIS | 0.00032 | 0.00575 | 0.49849 | 0.58816 | 1.15376 | 301 |
| GO_METAL_ION_TRANSPORT | 0.00390 | 0.03895 | 0.43171 | 0.55877 | 1.10227 | 419 |
| GO_METAPHASE_ANAPHASE_TRANSITION_OF_CELL_CYCLE | 0.00320 | 0.03371 | 0.43171 | 0.66563 | 1.25122 | 54 |

| | | | | | | |
|---|---|---|---|---|---|---|
| GO_MICROTUBULE_CYTOSKELETON_ORGANIZATION_INVOLVED_ | 0.00220 | 0.02548 | 0.43171 | 0.61295 | 1.18118 | 127 |
| GO_MICROVILLUS | 0.00168 | 0.02098 | 0.45506 | 0.68128 | 1.28184 | 56 |
| GO_MIDBODY | 0.00561 | 0.04971 | 0.40702 | 0.59866 | 1.15606 | 141 |
| GO_MITOCHONDRIAL_ELECTRON_TRANSPORT_NADH_TO_UBIQ | 0.00000 | 0.00006 | 0.64355 | 0.79447 | 1.47799 | 44 |
| GO_MITOCHONDRIAL_GENE_EXPRESSION | 0.00000 | 0.00000 | 0.78819 | 0.70230 | 1.35844 | 151 |
| GO_MITOCHONDRIAL_LARGE_RIBOSOMAL_SUBUNIT | 0.00000 | 0.00000 | 0.71951 | 0.79015 | 1.48529 | 54 |
| GO_MITOCHONDRIAL_MATRIX | 0.00000 | 0.00000 | 0.82666 | 0.63141 | 1.24412 | 387 |
| GO_MITOCHONDRIAL_MEMBRANE_ORGANIZATION | 0.00050 | 0.00802 | 0.47727 | 0.64578 | 1.23686 | 94 |
| GO_MITOCHONDRIAL_PROTEIN_COMPLEX | 0.00000 | 0.00000 | | 0.75011 | 1.46215 | 212 |
| GO_MITOCHONDRIAL_RESPIRATORY_CHAIN_COMPLEX_ASSEMB | 0.00001 | 0.00033 | 0.59333 | 0.69650 | 1.32804 | 85 |
| GO_MITOCHONDRIAL_SMALL_RIBOSOMAL_SUBUNIT | 0.00119 | 0.01640 | 0.45506 | 0.75384 | 1.37785 | 28 |
| GO_MITOCHONDRIAL_TRANSLATION | 0.00000 | 0.00000 | 0.80122 | 0.71435 | 1.37651 | 128 |
| GO_MITOCHONDRIAL_TRANSLATIONAL_TERMINATION | 0.00000 | 0.00000 | 0.81404 | 0.75760 | 1.44657 | 87 |
| GO_MITOCHONDRIAL_TRANSMEMBRANE_TRANSPORT | 0.00213 | 0.02490 | 0.43171 | 0.66300 | 1.25387 | 70 |
| GO_MITOCHONDRIAL_TRANSPORT | 0.00010 | 0.00227 | 0.53843 | 0.61119 | 1.19060 | 203 |
| GO_MITOCHONDRION_ORGANIZATION | 0.00000 | 0.00000 | 0.74774 | 0.61590 | 1.21471 | 416 |
| GO_MITOTIC_NUCLEAR_DIVISION | 0.00008 | 0.00182 | 0.53843 | 0.60435 | 1.17901 | 223 |
| GO_MITOTIC_SISTER_CHROMATID_SEGREGATION | 0.00014 | 0.00302 | 0.51885 | 0.62652 | 1.21048 | 145 |
| GO_MITOTIC_SPINDLE_ORGANIZATION | 0.00078 | 0.01164 | 0.47727 | 0.63943 | 1.22821 | 104 |
| GO_MODIFIED_AMINO_ACID_BINDING | 0.00002 | 0.00048 | 0.57561 | 0.74200 | 1.38679 | 49 |
| GO_MODULATION_BY_HOST_OF_VIRAL_PROCESS | 0.00216 | 0.02519 | 0.43171 | 0.77232 | 1.38066 | 20 |
| GO_MONOCARBOXYLIC_ACID_METABOLIC_PROCESS | 0.00012 | 0.00265 | 0.53843 | 0.57846 | 1.14053 | 406 |
| GO_MONOSACCHARIDE_BIOSYNTHETIC_PROCESS | 0.00305 | 0.03271 | 0.43171 | 0.65908 | 1.24472 | 68 |
| GO_MONOVALENT_INORGANIC_CATION_HOMEOSTASIS | 0.00108 | 0.01507 | 0.45506 | 0.63643 | 1.21783 | 91 |
| GO_MONOVALENT_INORGANIC_CATION_TRANSMEMBRANE_TR | 0.00001 | 0.00026 | 0.59333 | 0.65186 | 1.25996 | 147 |
| GO_MONOVALENT_INORGANIC_CATION_TRANSPORT | 0.00008 | 0.00178 | 0.53843 | 0.60205 | 1.17659 | 248 |
| GO_MORPHOGENESIS_OF_AN_EPITHELIUM | 0.00000 | 0.00009 | 0.62726 | 0.60570 | 1.19137 | 334 |
| GO_MORPHOGENESIS_OF_A_POLARIZED_EPITHELIUM | 0.00000 | 0.00000 | 0.83909 | 0.74384 | 1.43057 | 110 |
| GO_MOVEMENT_IN_HOST_ENVIRONMENT | 0.00268 | 0.02961 | 0.43171 | 0.61282 | 1.18078 | 117 |
| GO_MRNA_3_END_PROCESSING | 0.00087 | 0.01270 | 0.47727 | 0.64010 | 1.22502 | 93 |
| GO_MRNA_5_UTR_BINDING | 0.00002 | 0.00063 | 0.57561 | 0.83759 | 1.51313 | 23 |
| GO_MRNA_BINDING | 0.00000 | 0.00000 | | 0.67496 | 1.31810 | 239 |
| GO_MRNA_EXPORT_FROM_NUCLEUS | 0.00006 | 0.00159 | 0.53843 | 0.66290 | 1.27265 | 103 |
| GO_MRNA_PROCESSING | 0.00000 | 0.00000 | | 0.62959 | 1.24255 | 437 |
| GO_MRNA_TRANSPORT | 0.00007 | 0.00171 | 0.53843 | 0.64103 | 1.23563 | 132 |
| GO_MULTI_MULTICELLULAR_ORGANISM_PROCESS | 0.00041 | 0.00687 | 0.49849 | 0.63087 | 1.21515 | 116 |
| GO_MULTI_ORGANISM_LOCALIZATION | 0.00017 | 0.00334 | 0.51885 | 0.71077 | 1.33608 | 54 |
| GO_MUSCLE_CONTRACTION | 0.00335 | 0.03468 | 0.43171 | 0.59925 | 1.16199 | 165 |
| GO_MUSCLE_FILAMENT_SLIDING | 0.00324 | 0.03393 | 0.43171 | 0.77980 | 1.39322 | 19 |
| GO_MYELOID_LEUKOCYTE_ACTIVATION | 0.00000 | 0.00000 | | 0.65288 | 1.28712 | 395 |
| GO_MYELOID_LEUKOCYTE_MEDIATED_IMMUNITY | 0.00000 | 0.00000 | | 0.65734 | 1.29408 | 356 |
| GO_NADH_DEHYDROGENASE_ACTIVITY | 0.00001 | 0.00019 | 0.61053 | 0.79360 | 1.46691 | 38 |
| GO_NADH_DEHYDROGENASE_COMPLEX | 0.00001 | 0.00028 | 0.59333 | 0.77749 | 1.44215 | 41 |
| GO_NADH_DEHYDROGENASE_COMPLEX_ASSEMBLY | 0.00001 | 0.00035 | 0.59333 | 0.74044 | 1.39281 | 55 |
| GO_NADH_METABOLIC_PROCESS | 0.00034 | 0.00599 | 0.49849 | 0.75773 | 1.39166 | 33 |
| GO_NAD_METABOLIC_PROCESS | 0.00435 | 0.04238 | 0.40702 | 0.70570 | 1.30442 | 37 |
| GO_NCRNA_EXPORT_FROM_NUCLEUS | 0.00205 | 0.02440 | 0.43171 | 0.71473 | 1.31817 | 36 |
| GO_NCRNA_METABOLIC_PROCESS | 0.00000 | 0.00000 | 0.70498 | 0.61139 | 1.20509 | 399 |
| GO_NCRNA_PROCESSING | 0.00000 | 0.00000 | 0.73376 | 0.62238 | 1.22425 | 341 |
| GO_NEGATIVE_REGULATION_OF_APOPTOTIC_SIGNALING_PATHW | 0.00364 | 0.03677 | 0.43171 | 0.59958 | 1.15998 | 150 |
| GO_NEGATIVE_REGULATION_OF_ATP_METABOLIC_PROCESS | 0.00302 | 0.03248 | 0.43171 | 0.78586 | 1.39103 | 17 |
| GO_NEGATIVE_REGULATION_OF_CANONICAL_WNT_SIGNALING | 0.00000 | 0.00000 | 0.71951 | 0.69629 | 1.34180 | 123 |
| GO_NEGATIVE_REGULATION_OF_CATALYTIC_ACTIVITY | 0.00145 | 0.01879 | 0.45506 | 0.56238 | 1.11005 | 450 |
| GO_NEGATIVE_REGULATION_OF_CELLULAR_AMIDE_METABOLIC | 0.00268 | 0.02961 | 0.43171 | 0.61074 | 1.17686 | 124 |
| GO_NEGATIVE_REGULATION_OF_CELLULAR_COMPONENT_ORG | 0.00320 | 0.03371 | 0.43171 | 0.55470 | 1.09534 | 470 |
| GO_NEGATIVE_REGULATION_OF_CELL_CYCLE | 0.00023 | 0.00445 | 0.51885 | 0.57348 | 1.13228 | 455 |
| GO_NEGATIVE_REGULATION_OF_CELL_CYCLE_G2_M_PHASE_TR | 0.00000 | 0.00007 | 0.62726 | 0.70549 | 1.34606 | 86 |
| GO_NEGATIVE_REGULATION_OF_CELL_CYCLE_PHASE_TRANSITIO | 0.00000 | 0.00000 | 0.70498 | 0.65532 | 1.27560 | 194 |
| GO_NEGATIVE_REGULATION_OF_CELL_CYCLE_PROCESS | 0.00000 | 0.00005 | 0.64355 | 0.62725 | 1.22670 | 259 |
| GO_NEGATIVE_REGULATION_OF_HYDROLASE_ACTIVITY | 0.00216 | 0.02519 | 0.43171 | 0.58306 | 1.13858 | 238 |
| GO_NEGATIVE_REGULATION_OF_INTERLEUKIN_1_PRODUCTION | 0.00205 | 0.02440 | 0.43171 | 0.82297 | 1.44404 | 15 |
| GO_NEGATIVE_REGULATION_OF_MITOTIC_CELL_CYCLE | 0.00000 | 0.00010 | 0.62726 | 0.62191 | 1.21541 | 248 |
| GO_NEGATIVE_REGULATION_OF_MRNA_METABOLIC_PROCESS | 0.00213 | 0.02490 | 0.43171 | 0.66731 | 1.25879 | 62 |
| GO_NEGATIVE_REGULATION_OF_MRNA_SPLICING_VIA_SPLICEO | 0.00339 | 0.03501 | 0.43171 | 0.76309 | 1.36417 | 20 |
| GO_NEGATIVE_REGULATION_OF_NUCLEAR_DIVISION | 0.00335 | 0.03468 | 0.43171 | 0.67724 | 1.26685 | 50 |
| GO_NEGATIVE_REGULATION_OF_ORGANELLE_ORGANIZATION | 0.00487 | 0.04559 | 0.40702 | 0.57038 | 1.11722 | 288 |
| GO_NEGATIVE_REGULATION_OF_PEPTIDASE_ACTIVITY | 0.00036 | 0.00621 | 0.49849 | 0.64044 | 1.23159 | 109 |
| GO_NEGATIVE_REGULATION_OF_PROTEIN_CATABOLIC_PROCESS | 0.00342 | 0.03521 | 0.43171 | 0.62320 | 1.19268 | 93 |
| GO_NEGATIVE_REGULATION_OF_PROTEOLYSIS | 0.00205 | 0.02440 | 0.43171 | 0.60172 | 1.16902 | 178 |
| GO_NEGATIVE_REGULATION_OF_REACTIVE_OXYGEN_SPECIES_B | 0.00036 | 0.00621 | 0.49849 | 0.83108 | 1.47818 | 18 |

| | | | | | | |
|---|---|---|---|---|---|---|
| GO_NEGATIVE_REGULATION_OF_REPRODUCTIVE_PROCESS | 0.00457 | 0.04386 | 0.40702 | 0.74845 | 1.35356 | 24 |
| GO_NEGATIVE_REGULATION_OF_RNA_SPLICING | 0.00183 | 0.02219 | 0.45506 | 0.76824 | 1.38935 | 24 |
| GO_NEGATIVE_REGULATION_OF_WNT_SIGNALING_PATHWAY | 0.00000 | 0.00002 | 0.67496 | 0.67731 | 1.30721 | 136 |
| GO_NEURAL_CREST_CELL_MIGRATION | 0.00265 | 0.02936 | 0.43171 | 0.73967 | 1.35049 | 27 |
| GO_NEURON_RECOGNITION | 0.00390 | 0.03895 | 0.43171 | 0.71862 | 1.31998 | 31 |
| GO_NIK_NF_KAPPAB_SIGNALING | 0.00000 | 0.00000 | 0.77494 | 0.72329 | 1.39253 | 114 |
| GO_NON_CANONICAL_WNT_SIGNALING_PATHWAY | 0.00000 | 0.00000 | | 0.75756 | 1.45654 | 107 |
| GO_NUCLEAR_CHROMOSOME_SEGREGATION | 0.00098 | 0.01407 | 0.45506 | 0.59156 | 1.15357 | 215 |
| GO_NUCLEAR_ENVELOPE | 0.00005 | 0.00131 | 0.55733 | 0.59109 | 1.16270 | 341 |
| GO_NUCLEAR_EXPORT | 0.00000 | 0.00004 | 0.65944 | 0.64915 | 1.26157 | 181 |
| GO_NUCLEAR_MATRIX | 0.00502 | 0.04608 | 0.40702 | 0.62777 | 1.19868 | 87 |
| GO_NUCLEAR_MEMBRANE | 0.00185 | 0.02237 | 0.45506 | 0.58925 | 1.14853 | 211 |
| GO_NUCLEAR_PERIPHERY | 0.00043 | 0.00701 | 0.49849 | 0.64264 | 1.23558 | 107 |
| GO_NUCLEAR_PORE | 0.00561 | 0.04971 | 0.40702 | 0.64876 | 1.22522 | 68 |
| GO_NUCLEAR_SPECK | 0.00015 | 0.00312 | 0.51885 | 0.59159 | 1.16089 | 306 |
| GO_NUCLEAR_TRANSCRIBED_MRNA_CATABOLIC_PROCESS | 0.00000 | 0.00000 | | 0.70362 | 1.36820 | 185 |
| GO_NUCLEAR_TRANSCRIBED_MRNA_CATABOLIC_PROCESS_NON | 0.00000 | 0.00000 | | 0.74239 | 1.42931 | 114 |
| GO_NUCLEAR_TRANSPORT | 0.00000 | 0.00000 | 0.71951 | 0.62920 | 1.23383 | 297 |
| GO_NUCLEOBASE_CONTAINING_COMPOUND_TRANSPORT | 0.00000 | 0.00010 | 0.62726 | 0.62975 | 1.22823 | 216 |
| GO_NUCLEOBASE_CONTAINING_SMALL_MOLECULE_METABOLIC | 0.00000 | 0.00003 | 0.65944 | 0.60153 | 1.18594 | 405 |
| GO_NUCLEOSIDE_DIPHOSPHATE_METABOLIC_PROCESS | 0.00016 | 0.00319 | 0.51885 | 0.64491 | 1.24260 | 117 |
| GO_NUCLEOSIDE_PHOSPHATE_BIOSYNTHETIC_PROCESS | 0.00002 | 0.00044 | 0.59333 | 0.64092 | 1.24175 | 161 |
| GO_NUCLEOSIDE_TRIPHOSPHATE_BIOSYNTHETIC_PROCESS | 0.00040 | 0.00676 | 0.49849 | 0.72727 | 1.35074 | 42 |
| GO_NUCLEOSIDE_TRIPHOSPHATE_METABOLIC_PROCESS | 0.00009 | 0.00196 | 0.53843 | 0.69306 | 1.30854 | 65 |
| GO_NUCLEOTIDE_PHOSPHORYLATION | 0.00016 | 0.00328 | 0.51885 | 0.65541 | 1.25767 | 102 |
| GO_OOCYTE_DIFFERENTIATION | 0.00450 | 0.04342 | 0.40702 | 0.73513 | 1.33705 | 26 |
| GO_ORGANELLAR_RIBOSOME | 0.00000 | 0.00000 | 0.76146 | 0.75712 | 1.44311 | 84 |
| GO_ORGANELLE_ENVELOPE_LUMEN | 0.00026 | 0.00482 | 0.49849 | 0.67265 | 1.27716 | 78 |
| GO_ORGANELLE_FISSION | 0.00154 | 0.01974 | 0.45506 | 0.57133 | 1.12383 | 342 |
| GO_ORGANELLE_INNER_MEMBRANE | 0.00000 | 0.00000 | | 0.68423 | 1.34915 | 412 |
| GO_ORGANIC_CYCLIC_COMPOUND_CATABOLIC_PROCESS | 0.00000 | 0.00000 | | 0.66975 | 1.32263 | 478 |
| GO_ORGANIC_HYDROXY_COMPOUND_BIOSYNTHETIC_PROCESS | 0.00141 | 0.01850 | 0.45506 | 0.60712 | 1.17627 | 161 |
| GO_OXIDATIVE_PHOSPHORYLATION | 0.00000 | 0.00000 | | 0.78902 | 1.51260 | 99 |
| GO_OXIDOREDUCTASE_ACTIVITY | 0.00000 | 0.00000 | | 0.63268 | 1.24935 | 468 |
| GO_OXIDOREDUCTASE_ACTIVITY_ACTING_ON_A_HEME_GROUP | 0.00176 | 0.02168 | 0.45506 | 0.79383 | 1.41828 | 19 |
| GO_OXIDOREDUCTASE_ACTIVITY_ACTING_ON_NAD_P_H | 0.00002 | 0.00056 | 0.57561 | 0.71468 | 1.35162 | 70 |
| GO_OXIDOREDUCTASE_ACTIVITY_ACTING_ON_NAD_P_H_QUIN | 0.00004 | 0.00113 | 0.55733 | 0.74511 | 1.38835 | 47 |
| GO_OXIDOREDUCTASE_ACTIVITY_ACTING_ON_PEROXIDE_AS_A | 0.00106 | 0.01495 | 0.45506 | 0.73897 | 1.35352 | 29 |
| GO_OXIDOREDUCTASE_COMPLEX | 0.00000 | 0.00000 | 0.73376 | 0.74898 | 1.42731 | 82 |
| GO_PEPTIDASE_ACTIVITY | 0.00133 | 0.01799 | 0.45506 | 0.57681 | 1.13188 | 306 |
| GO_PEPTIDASE_COMPLEX | 0.00000 | 0.00002 | 0.67496 | 0.73250 | 1.39240 | 79 |
| GO_PEPTIDASE_REGULATOR_ACTIVITY | 0.00313 | 0.03310 | 0.43171 | 0.63434 | 1.21189 | 88 |
| GO_PEPTIDE_BINDING | 0.00005 | 0.00124 | 0.55733 | 0.63708 | 1.23187 | 148 |
| GO_PERINUCLEAR_REGION_OF_CYTOPLASM | 0.00276 | 0.03021 | 0.43171 | 0.55814 | 1.10207 | 466 |
| GO_PHAGOCYTOSIS | 0.00554 | 0.04968 | 0.40702 | 0.59337 | 1.14837 | 155 |
| GO_PHAGOSOME_ACIDIFICATION | 0.00059 | 0.00916 | 0.47727 | 0.78902 | 1.42694 | 24 |
| GO_PHOSPHATIDYLSERINE_BINDING | 0.00035 | 0.00610 | 0.49849 | 0.76387 | 1.39912 | 29 |
| GO_PH_REDUCTION | 0.00302 | 0.03248 | 0.43171 | 0.68821 | 1.28238 | 46 |
| GO_PIGMENT_GRANULE | 0.00000 | 0.00000 | 0.76146 | 0.74333 | 1.41934 | 87 |
| GO_PLASMA_LIPOPROTEIN_PARTICLE_CLEARANCE | 0.00106 | 0.01495 | 0.45506 | 0.73201 | 1.35304 | 37 |
| GO_POLYSOMAL_RIBOSOME | 0.00008 | 0.00180 | 0.53843 | 0.78840 | 1.44683 | 30 |
| GO_POLYSOME | 0.00000 | 0.00003 | 0.65944 | 0.75964 | 1.43296 | 62 |
| GO_POSITIVE_REGULATION_OF_ATP_METABOLIC_PROCESS | 0.00174 | 0.02149 | 0.45506 | 0.73011 | 1.33729 | 29 |
| GO_POSITIVE_REGULATION_OF_BINDING | 0.00390 | 0.03895 | 0.43171 | 0.61042 | 1.17582 | 115 |
| GO_POSITIVE_REGULATION_OF_CANONICAL_WNT_SIGNALING_ | 0.00000 | 0.00000 | 0.73376 | 0.72036 | 1.38514 | 108 |
| GO_POSITIVE_REGULATION_OF_CELLULAR_AMIDE_METABOLIC_ | 0.00001 | 0.00036 | 0.59333 | 0.66485 | 1.28095 | 125 |
| GO_POSITIVE_REGULATION_OF_CELLULAR_PROTEIN_LOCALIZAT | 0.00242 | 0.02748 | 0.43171 | 0.58048 | 1.13378 | 244 |
| GO_POSITIVE_REGULATION_OF_CELL_POPULATION_PROLIFERAT | 0.00040 | 0.00670 | 0.49849 | 0.56967 | 1.12503 | 461 |
| GO_POSITIVE_REGULATION_OF_CYSTEINE_TYPE_ENDOPEPTIDAS | 0.00547 | 0.04923 | 0.40702 | 0.62225 | 1.19077 | 92 |
| GO_POSITIVE_REGULATION_OF_CYTOKINE_PRODUCTION | 0.00283 | 0.03081 | 0.43171 | 0.58672 | 1.14373 | 210 |
| GO_POSITIVE_REGULATION_OF_DEFENSE_RESPONSE | 0.00000 | 0.00000 | 0.73376 | 0.66200 | 1.28884 | 200 |
| GO_POSITIVE_REGULATION_OF_DNA_BIOSYNTHETIC_PROCESS | 0.00010 | 0.00228 | 0.53843 | 0.71224 | 1.33977 | 55 |
| GO_POSITIVE_REGULATION_OF_DNA_METABOLIC_PROCESS | 0.00020 | 0.00387 | 0.51885 | 0.62771 | 1.21205 | 139 |
| GO_POSITIVE_REGULATION_OF_ESTABLISHMENT_OF_PROTEIN_ | 0.00194 | 0.02331 | 0.45506 | 0.58158 | 1.13709 | 253 |
| GO_POSITIVE_REGULATION_OF_FIBROBLAST_PROLIFERATION | 0.00017 | 0.00338 | 0.51885 | 0.77423 | 1.42213 | 31 |
| GO_POSITIVE_REGULATION_OF_IMMUNE_RESPONSE | 0.00000 | 0.00000 | 0.71951 | 0.63064 | 1.23937 | 321 |
| GO_POSITIVE_REGULATION_OF_IMMUNE_SYSTEM_PROCESS | 0.00000 | 0.00002 | 0.67496 | 0.59748 | 1.17992 | 475 |
| GO_POSITIVE_REGULATION_OF_INNATE_IMMUNE_RESPONSE | 0.00000 | 0.00000 | 0.70498 | 0.68885 | 1.32737 | 128 |
| GO_POSITIVE_REGULATION_OF_MITOTIC_CELL_CYCLE | 0.00178 | 0.02182 | 0.45506 | 0.61563 | 1.18683 | 122 |
| GO_POSITIVE_REGULATION_OF_MRNA_METABOLIC_PROCESS | 0.00157 | 0.02009 | 0.45506 | 0.66330 | 1.25231 | 63 |

| | | | | | | |
|---|---|---|---|---|---|---|
| GO_POSITIVE_REGULATION_OF_MRNA_PROCESSING | 0.00250 | 0.02807 | 0.43171 | 0.76194 | 1.38151 | 25 |
| GO_POSITIVE_REGULATION_OF_MRNA_SPLICING_VIA_SPLICEOS | 0.00132 | 0.01782 | 0.45506 | 0.81851 | 1.44353 | 16 |
| GO_POSITIVE_REGULATION_OF_NITRIC_OXIDE_METABOLIC_PRO | 0.00133 | 0.01799 | 0.45506 | 0.80807 | 1.43726 | 18 |
| GO_POSITIVE_REGULATION_OF_ORGANELLE_ORGANIZATION | 0.00010 | 0.00218 | 0.53843 | 0.57473 | 1.13449 | 452 |
| GO_POSITIVE_REGULATION_OF_PROTEIN_LOCALIZATION_TO_N | 0.00012 | 0.00265 | 0.53843 | 0.71018 | 1.33697 | 58 |
| GO_POSITIVE_REGULATION_OF_PROTEOLYSIS | 0.00313 | 0.03310 | 0.43171 | 0.58059 | 1.13381 | 237 |
| GO_POSITIVE_REGULATION_OF_REACTIVE_OXYGEN_SPECIES_BI | 0.00115 | 0.01593 | 0.45506 | 0.79865 | 1.43554 | 21 |
| GO_POSITIVE_REGULATION_OF_REACTIVE_OXYGEN_SPECIES_M | 0.00174 | 0.02149 | 0.45506 | 0.69785 | 1.30035 | 46 |
| GO_POSITIVE_REGULATION_OF_RESPONSE_TO_BIOTIC_STIMUL | 0.00000 | 0.00000 | 0.70498 | 0.67507 | 1.30603 | 150 |
| GO_POSITIVE_REGULATION_OF_RESPONSE_TO_EXTERNAL_STIM | 0.00001 | 0.00016 | 0.61053 | 0.62123 | 1.21505 | 260 |
| GO_POSITIVE_REGULATION_OF_RNA_SPLICING | 0.00265 | 0.02936 | 0.43171 | 0.74051 | 1.35202 | 27 |
| GO_POSITIVE_REGULATION_OF_SIGNAL_TRANSDUCTION_BY_P5 | 0.00390 | 0.03895 | 0.43171 | 0.80515 | 1.41278 | 15 |
| GO_POSITIVE_REGULATION_OF_TELOMERASE_ACTIVITY | 0.00069 | 0.01043 | 0.47727 | 0.75265 | 1.38122 | 30 |
| GO_POSITIVE_REGULATION_OF_TELOMERASE_RNA_LOCALIZATI | 0.00007 | 0.00169 | 0.53843 | 0.88616 | 1.55493 | 15 |
| GO_POSITIVE_REGULATION_OF_TELOMERE_MAINTENANCE | 0.00026 | 0.00486 | 0.49849 | 0.73697 | 1.37102 | 44 |
| GO_POSITIVE_REGULATION_OF_TELOMERE_MAINTENANCE_VIA | 0.00001 | 0.00040 | 0.59333 | 0.80196 | 1.47289 | 33 |
| GO_POSITIVE_REGULATION_OF_TRANSFERASE_ACTIVITY | 0.00108 | 0.01507 | 0.45506 | 0.56501 | 1.11376 | 401 |
| GO_POSITIVE_REGULATION_OF_TRANSLATION | 0.00000 | 0.00016 | 0.61053 | 0.68408 | 1.31526 | 107 |
| GO_POSITIVE_REGULATION_OF_VIRAL_PROCESS | 0.00026 | 0.00486 | 0.49849 | 0.66195 | 1.26462 | 88 |
| GO_POSITIVE_REGULATION_OF_WNT_SIGNALING_PATHWAY | 0.00000 | 0.00000 | 0.73376 | 0.69924 | 1.34747 | 127 |
| GO_POSTSYNAPSE | 0.00239 | 0.02716 | 0.43171 | 0.57062 | 1.12116 | 316 |
| GO_POST_TRANSLATIONAL_PROTEIN_MODIFICATION | 0.00024 | 0.00452 | 0.51885 | 0.60046 | 1.17392 | 251 |
| GO_PRECATALYTIC_SPLICEOSOME | 0.00141 | 0.01850 | 0.45506 | 0.69251 | 1.29540 | 50 |
| GO_PRERIBOSOME | 0.00006 | 0.00154 | 0.53843 | 0.69866 | 1.32020 | 69 |
| GO_PROTEASOMAL_PROTEIN_CATABOLIC_PROCESS | 0.00007 | 0.00169 | 0.53843 | 0.58188 | 1.14652 | 387 |
| GO_PROTEASOMAL_UBIQUITIN_INDEPENDENT_PROTEIN_CATAB | 0.00000 | 0.00003 | 0.65944 | 0.91182 | 1.62178 | 18 |
| GO_PROTEASOME_ACCESSORY_COMPLEX | 0.00005 | 0.00135 | 0.55733 | 0.82638 | 1.49450 | 24 |
| GO_PROTEASOME_CORE_COMPLEX | 0.00000 | 0.00004 | 0.65944 | 0.91231 | 1.61486 | 17 |
| GO_PROTEIN_CONTAINING_COMPLEX_DISASSEMBLY | 0.00000 | 0.00000 | 0.81404 | 0.66285 | 1.29656 | 261 |
| GO_PROTEIN_CONTAINING_COMPLEX_LOCALIZATION | 0.00018 | 0.00352 | 0.51885 | 0.60451 | 1.17943 | 226 |
| GO_PROTEIN_DNA_COMPLEX_SUBUNIT_ORGANIZATION | 0.00052 | 0.00825 | 0.47727 | 0.60887 | 1.18342 | 183 |
| GO_PROTEIN_DOMAIN_SPECIFIC_BINDING | 0.00003 | 0.00088 | 0.55733 | 0.57828 | 1.14190 | 470 |
| GO_PROTEIN_FOLDING | 0.00000 | 0.00000 | 0.73376 | 0.67425 | 1.30867 | 171 |
| GO_PROTEIN_FOLDING_CHAPERONE | 0.00079 | 0.01170 | 0.47727 | 0.81632 | 1.45193 | 18 |
| GO_PROTEIN_HETEROOLIGOMERIZATION | 0.00442 | 0.04297 | 0.40702 | 0.78998 | 1.39322 | 16 |
| GO_PROTEIN_IMPORT | 0.00001 | 0.00040 | 0.59333 | 0.64197 | 1.24306 | 158 |
| GO_PROTEIN_IMPORT_INTO_MITOCHONDRIAL_MATRIX | 0.00231 | 0.02642 | 0.43171 | 0.79530 | 1.41455 | 18 |
| GO_PROTEIN_INSERTION_INTO_MEMBRANE | 0.00191 | 0.02299 | 0.45506 | 0.68322 | 1.27694 | 49 |
| GO_PROTEIN_LOCALIZATION_TO_CHROMOSOME_TELOMERIC_F | 0.00001 | 0.00031 | 0.59333 | 0.82810 | 1.51357 | 28 |
| GO_PROTEIN_LOCALIZATION_TO_ENDOPLASMIC_RETICULUM | 0.00000 | 0.00000 | | 0.73850 | 1.42303 | 129 |
| GO_PROTEIN_LOCALIZATION_TO_MEMBRANE | 0.00000 | 0.00000 | 0.77494 | 0.61216 | 1.20881 | 462 |
| GO_PROTEIN_LOCALIZATION_TO_MITOCHONDRION | 0.00015 | 0.00310 | 0.51885 | 0.64234 | 1.23759 | 118 |
| GO_PROTEIN_LOCALIZATION_TO_NUCLEUS | 0.00000 | 0.00014 | 0.61053 | 0.63315 | 1.23408 | 208 |
| GO_PROTEIN_MATURATION | 0.00357 | 0.03638 | 0.43171 | 0.59209 | 1.14965 | 174 |
| GO_PROTEIN_MODIFICATION_BY_SMALL_PROTEIN_REMOVAL | 0.00002 | 0.00052 | 0.57561 | 0.61535 | 1.20119 | 231 |
| GO_PROTEIN_REFOLDING | 0.00465 | 0.04430 | 0.40702 | 0.77561 | 1.37288 | 17 |
| GO_PROTEIN_STABILIZATION | 0.00081 | 0.01194 | 0.47727 | 0.61881 | 1.19577 | 146 |
| GO_PROTEIN_SUMOYLATION | 0.00450 | 0.04342 | 0.40702 | 0.65043 | 1.22846 | 67 |
| GO_PROTEIN_TARGETING | 0.00000 | 0.00000 | | 0.66223 | 1.30361 | 358 |
| GO_PROTEIN_TARGETING_TO_MEMBRANE | 0.00000 | 0.00000 | | 0.71675 | 1.38879 | 162 |
| GO_PROTEIN_TARGETING_TO_MITOCHONDRION | 0.00113 | 0.01571 | 0.45506 | 0.65407 | 1.24648 | 83 |
| GO_PROTEIN_TRANSMEMBRANE_TRANSPORT | 0.00110 | 0.01526 | 0.45506 | 0.68173 | 1.28238 | 55 |
| GO_PROTON_TRANSMEMBRANE_TRANSPORT | 0.00000 | 0.00001 | 0.69013 | 0.71234 | 1.36309 | 91 |
| GO_PROTON_TRANSMEMBRANE_TRANSPORTER_ACTIVITY | 0.00004 | 0.00115 | 0.55733 | 0.70138 | 1.32535 | 69 |
| GO_PROTON_TRANSPORTING_TWO_SECTOR_ATPASE_COMPLEX | 0.00083 | 0.01218 | 0.47727 | 0.77254 | 1.40508 | 26 |
| GO_PROTON_TRANSPORTING_V_TYPE_ATPASE_COMPLEX | 0.00239 | 0.02716 | 0.43171 | 0.76100 | 1.37626 | 24 |
| GO_PURINE_CONTAINING_COMPOUND_BIOSYNTHETIC_PROCES | 0.00001 | 0.00040 | 0.59333 | 0.65863 | 1.26901 | 126 |
| GO_PURINE_CONTAINING_COMPOUND_METABOLIC_PROCESS | 0.00000 | 0.00000 | 0.73376 | 0.63132 | 1.23831 | 300 |
| GO_PURINE_NUCLEOSIDE_TRIPHOSPHATE_METABOLIC_PROCES | 0.00033 | 0.00582 | 0.49849 | 0.70887 | 1.32487 | 49 |
| GO_PYROPHOSPHATE_HYDROLYSIS_DRIVEN_PROTON_TRANSME | 0.00532 | 0.04803 | 0.40702 | 0.74470 | 1.34678 | 24 |
| GO_PYRUVATE_METABOLIC_PROCESS | 0.00001 | 0.00018 | 0.61053 | 0.67194 | 1.29289 | 113 |
| GO_P_BODY | 0.00472 | 0.04480 | 0.40702 | 0.65470 | 1.23501 | 62 |
| GO_REACTIVE_OXYGEN_SPECIES_BIOSYNTHETIC_PROCESS | 0.00040 | 0.00670 | 0.49849 | 0.70271 | 1.31857 | 53 |
| GO_REACTIVE_OXYGEN_SPECIES_METABOLIC_PROCESS | 0.00021 | 0.00409 | 0.51885 | 0.62881 | 1.21472 | 144 |
| GO_REGULATION_OF_ALCOHOL_BIOSYNTHETIC_PROCESS | 0.00228 | 0.02614 | 0.43171 | 0.68438 | 1.28020 | 50 |
| GO_REGULATION_OF_ANIMAL_ORGAN_MORPHOGENESIS | 0.00000 | 0.00000 | | 0.72330 | 1.39922 | 149 |
| GO_REGULATION_OF_APOPTOTIC_SIGNALING_PATHWAY | 0.00079 | 0.01170 | 0.47727 | 0.58599 | 1.14641 | 266 |
| GO_REGULATION_OF_ATP_METABOLIC_PROCESS | 0.00000 | 0.00006 | 0.64355 | 0.70523 | 1.34777 | 89 |
| GO_REGULATION_OF_BINDING | 0.00428 | 0.04185 | 0.40702 | 0.58264 | 1.13664 | 225 |
| GO_REGULATION_OF_BLOOD_PRESSURE | 0.00290 | 0.03156 | 0.43171 | 0.65319 | 1.23428 | 69 |

| | | | | | | |
|---|---|---|---|---|---|---|
| GO_REGULATION_OF_CARBOHYDRATE_CATABOLIC_PROCESS | 0.00324 | 0.03393 | 0.43171 | 0.65541 | 1.23787 | 67 |
| GO_REGULATION_OF_CELLULAR_AMIDE_METABOLIC_PROCESS | 0.00000 | 0.00000 | 0.78819 | 0.63116 | 1.24143 | 337 |
| GO_REGULATION_OF_CELLULAR_AMINE_METABOLIC_PROCESS | 0.00000 | 0.00000 | 0.74774 | 0.78955 | 1.48730 | 60 |
| GO_REGULATION_OF_CELLULAR_AMINO_ACID_METABOLIC_PR( | 0.00000 | 0.00000 | 0.76146 | 0.80341 | 1.51022 | 54 |
| GO_REGULATION_OF_CELLULAR_KETONE_METABOLIC_PROCESS | 0.00000 | 0.00000 | 0.70498 | 0.69239 | 1.33405 | 126 |
| GO_REGULATION_OF_CELLULAR_PH | 0.00442 | 0.04297 | 0.40702 | 0.65360 | 1.23436 | 68 |
| GO_REGULATION_OF_CELLULAR_PROTEIN_LOCALIZATION | 0.00228 | 0.02614 | 0.43171 | 0.56216 | 1.10830 | 402 |
| GO_REGULATION_OF_CELLULAR_RESPIRATION | 0.00428 | 0.04185 | 0.40702 | 0.77183 | 1.37897 | 19 |
| GO_REGULATION_OF_CELLULAR_RESPONSE_TO_HEAT | 0.00091 | 0.01314 | 0.47727 | 0.67360 | 1.27221 | 67 |
| GO_REGULATION_OF_CELL_CYCLE_G2_M_PHASE_TRANSITION | 0.00016 | 0.00332 | 0.51885 | 0.61554 | 1.19738 | 187 |
| GO_REGULATION_OF_CELL_CYCLE_PHASE_TRANSITION | 0.00000 | 0.00014 | 0.61053 | 0.59968 | 1.18091 | 369 |
| GO_REGULATION_OF_CHOLESTEROL_BIOSYNTHETIC_PROCESS | 0.00209 | 0.02470 | 0.43171 | 0.71476 | 1.32117 | 38 |
| GO_REGULATION_OF_CHOLESTEROL_METABOLIC_PROCESS | 0.00113 | 0.01571 | 0.45506 | 0.69865 | 1.30177 | 47 |
| GO_REGULATION_OF_CHROMOSOME_ORGANIZATION | 0.00036 | 0.00622 | 0.49849 | 0.58896 | 1.15261 | 278 |
| GO_REGULATION_OF_CHROMOSOME_SEGREGATION | 0.00079 | 0.01170 | 0.47727 | 0.64131 | 1.22718 | 91 |
| GO_REGULATION_OF_CHROMOSOME_SEPARATION | 0.00253 | 0.02838 | 0.43171 | 0.66349 | 1.24947 | 59 |
| GO_REGULATION_OF_CYSTEINE_TYPE_ENDOPEPTIDASE_ACTIVIT | 0.00044 | 0.00719 | 0.49849 | 0.62610 | 1.20905 | 141 |
| GO_REGULATION_OF_DEFENSE_RESPONSE | 0.00006 | 0.00150 | 0.55733 | 0.58834 | 1.15824 | 356 |
| GO_REGULATION_OF_DNA_BIOSYNTHETIC_PROCESS | 0.00043 | 0.00695 | 0.49849 | 0.66709 | 1.27130 | 83 |
| GO_REGULATION_OF_DNA_METABOLIC_PROCESS | 0.00055 | 0.00863 | 0.47727 | 0.58909 | 1.15134 | 249 |
| GO_REGULATION_OF_DNA_TEMPLATED_TRANSCRIPTION_IN_RE | 0.00000 | 0.00000 | 0.71951 | 0.72410 | 1.38687 | 94 |
| GO_REGULATION_OF_GENERATION_OF_PRECURSOR_METABOLI | 0.00004 | 0.00114 | 0.55733 | 0.65536 | 1.26115 | 111 |
| GO_REGULATION_OF_GENE_SILENCING | 0.00023 | 0.00434 | 0.51885 | 0.65464 | 1.25268 | 91 |
| GO_REGULATION_OF_HEMATOPOIETIC_PROGENITOR_CELL_DIF | 0.00000 | 0.00000 | 0.73376 | 0.76273 | 1.44465 | 72 |
| GO_REGULATION_OF_HEMATOPOIETIC_STEM_CELL_DIFFERENT | 0.00000 | 0.00000 | 0.74774 | 0.78891 | 1.48817 | 62 |
| GO_REGULATION_OF_HEMOPOIESIS | 0.00390 | 0.03895 | 0.43171 | 0.57739 | 1.12909 | 258 |
| GO_REGULATION_OF_IMMUNE_RESPONSE | 0.00000 | 0.00006 | 0.64355 | 0.60032 | 1.18447 | 433 |
| GO_REGULATION_OF_INNATE_IMMUNE_RESPONSE | 0.00000 | 0.00007 | 0.62726 | 0.64536 | 1.25422 | 181 |
| GO_REGULATION_OF_INTRINSIC_APOPTOTIC_SIGNALING_PATH\ | 0.00179 | 0.02183 | 0.45506 | 0.61548 | 1.18550 | 116 |
| GO_REGULATION_OF_MITOTIC_CELL_CYCLE | 0.00000 | 0.00003 | 0.65944 | 0.58992 | 1.16508 | 499 |
| GO_REGULATION_OF_MORPHOGENESIS_OF_AN_EPITHELIUM | 0.00000 | 0.00000 | | 0.74680 | 1.43884 | 125 |
| GO_REGULATION_OF_MRNA_CATABOLIC_PROCESS | 0.00000 | 0.00000 | | 0.71126 | 1.37919 | 165 |
| GO_REGULATION_OF_MRNA_METABOLIC_PROCESS | 0.00000 | 0.00000 | | 0.67178 | 1.31458 | 276 |
| GO_REGULATION_OF_MRNA_PROCESSING | 0.00008 | 0.00186 | 0.53843 | 0.65189 | 1.25598 | 125 |
| GO_REGULATION_OF_MRNA_SPLICING_VIA_SPLICEOSOME | 0.00012 | 0.00256 | 0.53843 | 0.66950 | 1.28033 | 90 |
| GO_REGULATION_OF_MULTI_ORGANISM_PROCESS | 0.00008 | 0.00190 | 0.53843 | 0.76905 | 1.42150 | 37 |
| GO_REGULATION_OF_MUSCLE_CONTRACTION | 0.00163 | 0.02050 | 0.45506 | 0.64879 | 1.23071 | 76 |
| GO_REGULATION_OF_NUCLEAR_DIVISION | 0.00060 | 0.00923 | 0.47727 | 0.62443 | 1.20552 | 137 |
| GO_REGULATION_OF_NUCLEASE_ACTIVITY | 0.00059 | 0.00916 | 0.47727 | 0.81786 | 1.44767 | 17 |
| GO_REGULATION_OF_NUCLEOTIDE_BIOSYNTHETIC_PROCESS | 0.00302 | 0.03248 | 0.43171 | 0.78902 | 1.40337 | 18 |
| GO_REGULATION_OF_NUCLEOTIDE_METABOLIC_PROCESS | 0.00033 | 0.00588 | 0.49849 | 0.66858 | 1.26942 | 78 |
| GO_REGULATION_OF_OXIDATIVE_PHOSPHORYLATION | 0.00005 | 0.00130 | 0.55733 | 0.85284 | 1.53294 | 21 |
| GO_REGULATION_OF_PEPTIDASE_ACTIVITY | 0.00006 | 0.00152 | 0.53843 | 0.61192 | 1.19388 | 226 |
| GO_REGULATION_OF_PEPTIDE_TRANSPORT | 0.00250 | 0.02807 | 0.43171 | 0.56252 | 1.10917 | 412 |
| GO_REGULATION_OF_PH | 0.00561 | 0.04971 | 0.40702 | 0.64191 | 1.21498 | 71 |
| GO_REGULATION_OF_PLASMA_LIPOPROTEIN_PARTICLE_LEVELS | 0.00006 | 0.00145 | 0.55733 | 0.72431 | 1.36154 | 54 |
| GO_REGULATION_OF_POSTTRANSCRIPTIONAL_GENE_SILENCIN( | 0.00026 | 0.00486 | 0.49849 | 0.66803 | 1.27236 | 81 |
| GO_REGULATION_OF_PROTEIN_CATABOLIC_PROCESS | 0.00309 | 0.03299 | 0.43171 | 0.57360 | 1.12399 | 291 |
| GO_REGULATION_OF_PROTEIN_IMPORT | 0.00487 | 0.04559 | 0.40702 | 0.67491 | 1.25754 | 47 |
| GO_REGULATION_OF_PROTEIN_LOCALIZATION_TO_NUCLEUS | 0.00000 | 0.00013 | 0.61053 | 0.69035 | 1.32101 | 91 |
| GO_REGULATION_OF_PROTEIN_STABILITY | 0.00066 | 0.01007 | 0.47727 | 0.59308 | 1.15811 | 235 |
| GO_REGULATION_OF_PROTEOLYSIS | 0.00007 | 0.00169 | 0.53843 | 0.57923 | 1.14299 | 431 |
| GO_REGULATION_OF_REACTIVE_OXYGEN_SPECIES_BIOSYNTHET | 0.00022 | 0.00427 | 0.51885 | 0.73154 | 1.36343 | 45 |
| GO_REGULATION_OF_REACTIVE_OXYGEN_SPECIES_METABOLIC_ | 0.00088 | 0.01280 | 0.47727 | 0.63948 | 1.22592 | 99 |
| GO_REGULATION_OF_RESPONSE_TO_BIOTIC_STIMULUS | 0.00001 | 0.00039 | 0.59333 | 0.61773 | 1.20625 | 235 |
| GO_REGULATION_OF_RNA_SPLICING | 0.00011 | 0.00241 | 0.53843 | 0.64982 | 1.25198 | 125 |
| GO_REGULATION_OF_SISTER_CHROMATID_SEGREGATION | 0.00405 | 0.04011 | 0.40702 | 0.64401 | 1.22034 | 73 |
| GO_REGULATION_OF_SMALL_MOLECULE_METABOLIC_PROCESS | 0.00000 | 0.00000 | 0.77494 | 0.64342 | 1.26057 | 290 |
| GO_REGULATION_OF_SMOOTH_MUSCLE_CONTRACTION | 0.00007 | 0.00171 | 0.53843 | 0.82642 | 1.49295 | 23 |
| GO_REGULATION_OF_STEM_CELL_DIFFERENTIATION | 0.00000 | 0.00000 | 0.73376 | 0.73434 | 1.40292 | 88 |
| GO_REGULATION_OF_STEROID_METABOLIC_PROCESS | 0.00524 | 0.04770 | 0.40702 | 0.64061 | 1.21567 | 77 |
| GO_REGULATION_OF_SUPEROXIDE_METABOLIC_PROCESS | 0.00346 | 0.03548 | 0.43171 | 0.77826 | 1.39046 | 19 |
| GO_REGULATION_OF_SUPRAMOLECULAR_FIBER_ORGANIZATIO | 0.00509 | 0.04656 | 0.40702 | 0.57599 | 1.12367 | 223 |
| GO_REGULATION_OF_SYMBIOTIC_PROCESS | 0.00009 | 0.00211 | 0.53843 | 0.62950 | 1.21763 | 151 |
| GO_REGULATION_OF_TELOMERASE_ACTIVITY | 0.00087 | 0.01263 | 0.47727 | 0.72147 | 1.33426 | 39 |
| GO_REGULATION_OF_TELOMERE_MAINTENANCE | 0.00547 | 0.04923 | 0.40702 | 0.64959 | 1.22679 | 68 |
| GO_REGULATION_OF_TELOMERE_MAINTENANCE_VIA_TELOME | 0.00020 | 0.00387 | 0.51885 | 0.72043 | 1.34856 | 51 |
| GO_REGULATION_OF_TRANSCRIPTION_FROM_RNA_POLYMERA | 0.00000 | 0.00000 | 0.70498 | 0.76931 | 1.45239 | 64 |
| GO_REGULATION_OF_TRANSLATIONAL_INITIATION | 0.00023 | 0.00437 | 0.51885 | 0.68479 | 1.29399 | 69 |
| GO_REGULATION_OF_VIRAL_TRANSCRIPTION | 0.00361 | 0.03657 | 0.43171 | 0.67497 | 1.26259 | 50 |

| | | | | | | |
|---|---|---|---|---|---|---|
| GO_REGULATION_OF_WNT_SIGNALING_PATHWAY | 0.00015 | 0.00312 | 0.51885 | 0.60129 | 1.17419 | 238 |
| GO_RESPIRASOME | 0.00000 | 0.00000 | 0.81404 | 0.78326 | 1.48524 | 75 |
| GO_RESPIRATORY_CHAIN_COMPLEX | 0.00000 | 0.00000 | 0.77494 | 0.79183 | 1.49492 | 64 |
| GO_RESPIRATORY_ELECTRON_TRANSPORT_CHAIN | 0.00000 | 0.00000 | | 0.78275 | 1.49591 | 89 |
| GO_RESPONSE_TO_ALCOHOL | 0.00015 | 0.00310 | 0.51885 | 0.63794 | 1.22915 | 126 |
| GO_RESPONSE_TO_ATP | 0.00087 | 0.01270 | 0.47727 | 0.81086 | 1.43528 | 17 |
| GO_RESPONSE_TO_CALCIUM_ION | 0.00124 | 0.01701 | 0.45506 | 0.65196 | 1.24071 | 80 |
| GO_RESPONSE_TO_CORTICOSTEROID | 0.00010 | 0.00220 | 0.53843 | 0.66894 | 1.27841 | 89 |
| GO_RESPONSE_TO_DRUG | 0.00000 | 0.00007 | 0.62726 | 0.63080 | 1.23070 | 220 |
| GO_RESPONSE_TO_ETHANOL | 0.00168 | 0.02098 | 0.45506 | 0.66543 | 1.25679 | 67 |
| GO_RESPONSE_TO_FIBROBLAST_GROWTH_FACTOR | 0.00357 | 0.03638 | 0.43171 | 0.63469 | 1.20976 | 84 |
| GO_RESPONSE_TO_GROWTH_FACTOR | 0.00272 | 0.02986 | 0.43171 | 0.56235 | 1.10805 | 387 |
| GO_RESPONSE_TO_HEAT | 0.00000 | 0.00013 | 0.62726 | 0.66982 | 1.29069 | 119 |
| GO_RESPONSE_TO_HYDROGEN_PEROXIDE | 0.00532 | 0.04803 | 0.40702 | 0.62255 | 1.19054 | 90 |
| GO_RESPONSE_TO_INORGANIC_SUBSTANCE | 0.00016 | 0.00328 | 0.51885 | 0.58928 | 1.15810 | 321 |
| GO_RESPONSE_TO_INTERLEUKIN_1 | 0.00000 | 0.00000 | 0.78819 | 0.71590 | 1.37950 | 124 |
| GO_RESPONSE_TO_INTERLEUKIN_12 | 0.00036 | 0.00622 | 0.49849 | 0.74158 | 1.36769 | 36 |
| GO_RESPONSE_TO_INTERLEUKIN_4 | 0.00224 | 0.02581 | 0.43171 | 0.77088 | 1.38921 | 22 |
| GO_RESPONSE_TO_LEUKEMIA_INHIBITORY_FACTOR | 0.00076 | 0.01129 | 0.47727 | 0.66148 | 1.25288 | 72 |
| GO_RESPONSE_TO_LIPID | 0.00043 | 0.00695 | 0.49849 | 0.56669 | 1.11902 | 470 |
| GO_RESPONSE_TO_METAL_ION | 0.00143 | 0.01863 | 0.45506 | 0.59895 | 1.16592 | 199 |
| GO_RESPONSE_TO_NUTRIENT | 0.00294 | 0.03185 | 0.43171 | 0.62596 | 1.19987 | 98 |
| GO_RESPONSE_TO_ORGANOPHOSPHORUS | 0.00294 | 0.03185 | 0.43171 | 0.64154 | 1.21511 | 72 |
| GO_RESPONSE_TO_OXIDATIVE_STRESS | 0.00495 | 0.04573 | 0.40702 | 0.57010 | 1.11758 | 294 |
| GO_RESPONSE_TO_OXYGEN_LEVELS | 0.00000 | 0.00000 | | 0.67661 | 1.32155 | 244 |
| GO_RESPONSE_TO_PURINE_CONTAINING_COMPOUND | 0.00509 | 0.04656 | 0.40702 | 0.63176 | 1.20394 | 82 |
| GO_RESPONSE_TO_STEROID_HORMONE | 0.00009 | 0.00200 | 0.53843 | 0.61312 | 1.19360 | 201 |
| GO_RESPONSE_TO_TEMPERATURE_STIMULUS | 0.00002 | 0.00047 | 0.57561 | 0.64468 | 1.24782 | 153 |
| GO_RESPONSE_TO_TOXIC_SUBSTANCE | 0.00011 | 0.00235 | 0.53843 | 0.63892 | 1.23203 | 134 |
| GO_RESPONSE_TO_TUMOR_NECROSIS_FACTOR | 0.00000 | 0.00000 | 0.70498 | 0.66331 | 1.28815 | 175 |
| GO_RESPONSE_TO_VIRUS | 0.00261 | 0.02910 | 0.43171 | 0.59428 | 1.15408 | 175 |
| GO_RETINA_HOMEOSTASIS | 0.00034 | 0.00606 | 0.49849 | 0.75202 | 1.38362 | 34 |
| GO_RIBONUCLEOPROTEIN_COMPLEX_BINDING | 0.00000 | 0.00001 | 0.69013 | 0.69839 | 1.34459 | 114 |
| GO_RIBONUCLEOPROTEIN_COMPLEX_BIOGENESIS | 0.00000 | 0.00000 | | 0.68714 | 1.35448 | 393 |
| GO_RIBONUCLEOPROTEIN_COMPLEX_SUBUNIT_ORGANIZATION | 0.00000 | 0.00000 | 0.81404 | 0.69112 | 1.34183 | 173 |
| GO_RIBONUCLEOPROTEIN_GRANULE | 0.00000 | 0.00000 | 0.71951 | 0.66271 | 1.28875 | 186 |
| GO_RIBONUCLEOSIDE_TRIPHOSPHATE_BIOSYNTHETIC_PROCESS | 0.00009 | 0.00210 | 0.53843 | 0.77194 | 1.42026 | 34 |
| GO_RIBONUCLEOSIDE_TRIPHOSPHATE_METABOLIC_PROCESS | 0.00013 | 0.00280 | 0.51885 | 0.73558 | 1.37065 | 46 |
| GO_RIBOSE_PHOSPHATE_BIOSYNTHETIC_PROCESS | 0.00002 | 0.00056 | 0.57561 | 0.65814 | 1.26821 | 124 |
| GO_RIBOSE_PHOSPHATE_METABOLIC_PROCESS | 0.00000 | 0.00000 | 0.74774 | 0.63837 | 1.25001 | 285 |
| GO_RIBOSOMAL_LARGE_SUBUNIT_BIOGENESIS | 0.00001 | 0.00031 | 0.59333 | 0.72435 | 1.36807 | 67 |
| GO_RIBOSOMAL_SMALL_SUBUNIT_ASSEMBLY | 0.00405 | 0.04011 | 0.40702 | 0.78074 | 1.38865 | 18 |
| GO_RIBOSOMAL_SMALL_SUBUNIT_BINDING | 0.00532 | 0.04803 | 0.40702 | 0.77341 | 1.36899 | 17 |
| GO_RIBOSOMAL_SMALL_SUBUNIT_BIOGENESIS | 0.00005 | 0.00138 | 0.55733 | 0.70510 | 1.33163 | 68 |
| GO_RIBOSOMAL_SUBUNIT | 0.00000 | 0.00000 | | 0.76292 | 1.48089 | 172 |
| GO_RIBOSOME | 0.00000 | 0.00000 | | 0.75411 | 1.46984 | 208 |
| GO_RIBOSOME_ASSEMBLY | 0.00001 | 0.00026 | 0.59333 | 0.74485 | 1.40144 | 56 |
| GO_RIBOSOME_BIOGENESIS | 0.00000 | 0.00000 | | 0.68899 | 1.34833 | 277 |
| GO_RNA_3_END_PROCESSING | 0.00002 | 0.00046 | 0.57561 | 0.65576 | 1.26361 | 128 |
| GO_RNA_CATABOLIC_PROCESS | 0.00000 | 0.00000 | | 0.70701 | 1.39073 | 341 |
| GO_RNA_DEPENDENT_DNA_BIOSYNTHETIC_PROCESS | 0.00003 | 0.00081 | 0.57561 | 0.71669 | 1.34965 | 59 |
| GO_RNA_EXPORT_FROM_NUCLEUS | 0.00000 | 0.00016 | 0.61053 | 0.66306 | 1.27817 | 130 |
| GO_RNA_LOCALIZATION | 0.00000 | 0.00000 | 0.76146 | 0.66638 | 1.29886 | 208 |
| GO_RNA_SPLICING | 0.00000 | 0.00000 | | 0.63512 | 1.25129 | 384 |
| GO_RNA_SPLICING_VIA_TRANSESTERIFICATION_REACTIONS | 0.00000 | 0.00000 | 0.80122 | 0.64478 | 1.26586 | 311 |
| GO_ROUGH_ENDOPLASMIC_RETICULUM | 0.00014 | 0.00293 | 0.51885 | 0.70363 | 1.32507 | 59 |
| GO_RRNA_BINDING | 0.00000 | 0.00001 | 0.69013 | 0.77638 | 1.45941 | 54 |
| GO_RRNA_METABOLIC_PROCESS | 0.00000 | 0.00000 | | 0.68430 | 1.33386 | 209 |
| GO_SCF_DEPENDENT_PROTEASOMAL_UBIQUITIN_DEPENDENT_ | 0.00000 | 0.00008 | 0.62726 | 0.71957 | 1.36498 | 76 |
| GO_SECRETORY_GRANULE | 0.00000 | 0.00000 | | 0.62978 | 1.24363 | 481 |
| GO_SECRETORY_GRANULE_MEMBRANE | 0.00032 | 0.00575 | 0.49849 | 0.61375 | 1.19134 | 172 |
| GO_SERINE_TYPE_ENDOPEPTIDASE_INHIBITOR_ACTIVITY | 0.00038 | 0.00640 | 0.49849 | 0.77388 | 1.41447 | 28 |
| GO_SEXUAL_REPRODUCTION | 0.00472 | 0.04480 | 0.40702 | 0.55748 | 1.10026 | 439 |
| GO_SIDE_OF_MEMBRANE | 0.00495 | 0.04573 | 0.40702 | 0.57741 | 1.12751 | 235 |
| GO_SIGNAL_TRANSDUCTION_BY_PROTEIN_PHOSPHORYLATION | 0.00224 | 0.02581 | 0.43171 | 0.55690 | 1.09975 | 500 |
| GO_SINGLE_FERTILIZATION | 0.00000 | 0.00002 | 0.67496 | 0.76307 | 1.43842 | 61 |
| GO_SINGLE_STRANDED_RNA_BINDING | 0.00001 | 0.00042 | 0.59333 | 0.71960 | 1.35864 | 66 |
| GO_SISTER_CHROMATID_SEGREGATION | 0.00042 | 0.00689 | 0.49849 | 0.60926 | 1.18209 | 170 |
| GO_SKIN_DEVELOPMENT | 0.00076 | 0.01140 | 0.47727 | 0.61452 | 1.18945 | 153 |
| GO_SMALL_MOLECULE_BIOSYNTHETIC_PROCESS | 0.00001 | 0.00021 | 0.61053 | 0.58728 | 1.15957 | 459 |

| | | | | | | |
|---|---|---|---|---|---|---|
| GO_SMALL_NUCLEAR_RIBONUCLEOPROTEIN_COMPLEX | 0.00042 | 0.00689 | 0.49849 | 0.69542 | 1.30936 | 57 |
| GO_SMALL_RIBOSOMAL_SUBUNIT | 0.00000 | 0.00001 | 0.69013 | 0.75767 | 1.43100 | 67 |
| GO_SM_LIKE_PROTEIN_FAMILY_COMPLEX | 0.00145 | 0.01879 | 0.45506 | 0.66792 | 1.26149 | 67 |
| GO_SNORNA_BINDING | 0.00038 | 0.00646 | 0.49849 | 0.76224 | 1.39614 | 29 |
| GO_SPERM_EGG_RECOGNITION | 0.00000 | 0.00002 | 0.65944 | 0.87469 | 1.59087 | 26 |
| GO_SPINDLE_MIDZONE | 0.00361 | 0.03657 | 0.43171 | 0.73316 | 1.33859 | 27 |
| GO_SPINDLE_ORGANIZATION | 0.00272 | 0.02986 | 0.43171 | 0.60076 | 1.16362 | 159 |
| GO_SPLICEOSOMAL_COMPLEX | 0.00000 | 0.00008 | 0.62726 | 0.65256 | 1.26657 | 171 |
| GO_SPLICEOSOMAL_TRI_SNRNP_COMPLEX | 0.00168 | 0.02098 | 0.45506 | 0.73474 | 1.34835 | 30 |
| GO_STEM_CELL_DIFFERENTIATION | 0.00000 | 0.00000 | 0.82666 | 0.70006 | 1.35554 | 158 |
| GO_STEROID_BIOSYNTHETIC_PROCESS | 0.00465 | 0.04430 | 0.40702 | 0.61242 | 1.17992 | 125 |
| GO_STEROID_METABOLIC_PROCESS | 0.00313 | 0.03310 | 0.43171 | 0.59525 | 1.15748 | 185 |
| GO_STEROL_BIOSYNTHETIC_PROCESS | 0.00005 | 0.00120 | 0.55733 | 0.71876 | 1.35584 | 62 |
| GO_STEROL_METABOLIC_PROCESS | 0.00003 | 0.00073 | 0.57561 | 0.66485 | 1.27854 | 109 |
| GO_STRESS_GRANULE_ASSEMBLY | 0.00465 | 0.04430 | 0.40702 | 0.77288 | 1.38921 | 21 |
| GO_STRUCTURAL_CONSTITUENT_OF_RIBOSOME | 0.00000 | 0.00000 | | 0.76380 | 1.47839 | 153 |
| GO_STRUCTURAL_MOLECULE_ACTIVITY | 0.00000 | 0.00000 | | 0.71760 | 1.41436 | 391 |
| GO_SUPEROXIDE_METABOLIC_PROCESS | 0.00272 | 0.02986 | 0.43171 | 0.72028 | 1.32659 | 35 |
| GO_TELOMERASE_RNA_LOCALIZATION | 0.00054 | 0.00857 | 0.47727 | 0.81801 | 1.46148 | 19 |
| GO_TELOMERE_MAINTENANCE_VIA_TELOMERE_LENGTHENING | 0.00005 | 0.00138 | 0.55733 | 0.70715 | 1.33512 | 66 |
| GO_TELOMERE_ORGANIZATION | 0.00159 | 0.02016 | 0.45506 | 0.61666 | 1.18834 | 127 |
| GO_TERTIARY_GRANULE | 0.00008 | 0.00186 | 0.53843 | 0.66372 | 1.27005 | 91 |
| GO_TERTIARY_GRANULE_MEMBRANE | 0.00102 | 0.01453 | 0.45506 | 0.71940 | 1.33244 | 40 |
| GO_THREONINE_TYPE_ENDOPEPTIDASE_ACTIVITY | 0.00000 | 0.00000 | 0.71951 | 0.93456 | 1.65424 | 17 |
| GO_THREONINE_TYPE_PEPTIDASE_ACTIVITY | 0.00000 | 0.00004 | 0.65944 | 0.89807 | 1.61424 | 21 |
| GO_TISSUE_MORPHOGENESIS | 0.00000 | 0.00006 | 0.64355 | 0.59467 | 1.17150 | 380 |
| GO_TOXIN_TRANSPORT | 0.00064 | 0.00972 | 0.47727 | 0.77713 | 1.41343 | 26 |
| GO_TRANSCRIPTION_COUPLED_NUCLEOTIDE_EXCISION_REPAIR | 0.00231 | 0.02642 | 0.43171 | 0.65429 | 1.23840 | 71 |
| GO_TRANSFERRIN_TRANSPORT | 0.00141 | 0.01850 | 0.45506 | 0.75134 | 1.37179 | 27 |
| GO_TRANSLATIONAL_ELONGATION | 0.00000 | 0.00000 | | 0.75263 | 1.45028 | 124 |
| GO_TRANSLATIONAL_INITIATION | 0.00000 | 0.00000 | | 0.75139 | 1.45839 | 171 |
| GO_TRANSLATIONAL_TERMINATION | 0.00000 | 0.00000 | 0.81404 | 0.75110 | 1.44066 | 101 |
| GO_TRANSLATION_ELONGATION_FACTOR_ACTIVITY | 0.00390 | 0.03895 | 0.43171 | 0.80505 | 1.41260 | 15 |
| GO_TRANSLATION_FACTOR_ACTIVITY_RNA_BINDING | 0.00000 | 0.00000 | 0.74774 | 0.77244 | 1.46085 | 70 |
| GO_TRANSLATION_INITIATION_FACTOR_ACTIVITY | 0.00000 | 0.00011 | 0.62726 | 0.77773 | 1.44952 | 45 |
| GO_TRANSLATION_PREINITIATION_COMPLEX | 0.00342 | 0.03521 | 0.43171 | 0.79777 | 1.40694 | 16 |
| GO_TRANSLATION_REGULATOR_ACTIVITY | 0.00000 | 0.00000 | 0.83909 | 0.74000 | 1.42403 | 111 |
| GO_TRANSLATION_REGULATOR_ACTIVITY_NUCLEIC_ACID_BIND | 0.00000 | 0.00000 | 0.83909 | 0.76441 | 1.45959 | 87 |
| GO_TRANSMEMBRANE_RECEPTOR_PROTEIN_TYROSINE_KINASE | 0.00168 | 0.02098 | 0.45506 | 0.56326 | 1.11076 | 411 |
| GO_TRICARBOXYLIC_ACID_CYCLE | 0.00159 | 0.02016 | 0.45506 | 0.73558 | 1.35097 | 33 |
| GO_TRNA_TRANSPORT | 0.00342 | 0.03521 | 0.43171 | 0.71247 | 1.31220 | 35 |
| GO_TUMOR_NECROSIS_FACTOR_MEDIATED_SIGNALING_PATHW | 0.00000 | 0.00000 | 0.80122 | 0.73921 | 1.42138 | 108 |
| GO_T_CELL_RECEPTOR_SIGNALING_PATHWAY | 0.00000 | 0.00001 | 0.70498 | 0.69185 | 1.33314 | 119 |
| GO_U12_TYPE_SPLICEOSOMAL_COMPLEX | 0.00487 | 0.04559 | 0.40702 | 0.73423 | 1.33541 | 26 |
| GO_U2_SNRNP | 0.00141 | 0.01850 | 0.45506 | 0.80259 | 1.42065 | 17 |
| GO_U2_TYPE_CATALYTIC_STEP_2_SPLICEOSOME | 0.00268 | 0.02961 | 0.43171 | 0.72225 | 1.32288 | 29 |
| GO_U2_TYPE_SPLICEOSOMAL_COMPLEX | 0.00033 | 0.00595 | 0.49849 | 0.66590 | 1.26925 | 84 |
| GO_UBIQUITIN_LIKE_PROTEIN_LIGASE_BINDING | 0.00000 | 0.00007 | 0.62726 | 0.62616 | 1.22416 | 251 |
| GO_UNFOLDED_PROTEIN_BINDING | 0.00000 | 0.00001 | 0.69013 | 0.72112 | 1.37766 | 88 |
| GO_VACUOLAR_LUMEN | 0.00016 | 0.00319 | 0.51885 | 0.65416 | 1.25772 | 107 |
| GO_VESICLE_BUDDING_FROM_MEMBRANE | 0.00480 | 0.04537 | 0.40702 | 0.62411 | 1.19353 | 90 |
| GO_VESICLE_LUMEN | 0.00000 | 0.00000 | 0.77494 | 0.67028 | 1.30631 | 207 |
| GO_VESICLE_MEMBRANE | 0.00001 | 0.00026 | 0.59333 | 0.58430 | 1.15387 | 476 |
| GO_VIRAL_GENE_EXPRESSION | 0.00000 | 0.00000 | | 0.70963 | 1.37745 | 172 |
| GO_VIRAL_LIFE_CYCLE | 0.00000 | 0.00001 | 0.69013 | 0.63985 | 1.24950 | 238 |
| GO_VIRION_ASSEMBLY | 0.00457 | 0.04386 | 0.40702 | 0.69689 | 1.28528 | 36 |
| HP_ABNORMALITY_OF_ACID_BASE_HOMEOSTASIS | 0.00000 | 0.00010 | 0.62726 | 0.62283 | 1.21827 | 261 |
| HP_ABNORMALITY_OF_BLOOD_CIRCULATION | 0.00502 | 0.04608 | 0.40702 | 0.59228 | 1.14883 | 167 |
| HP_ABNORMALITY_OF_CARDIOVASCULAR_SYSTEM_ELECTROPH | 0.00011 | 0.00237 | 0.53843 | 0.60202 | 1.17758 | 261 |
| HP_ABNORMALITY_OF_SKIN_ADNEXA_PHYSIOLOGY | 0.00242 | 0.02748 | 0.43171 | 0.61294 | 1.18094 | 118 |
| HP_ABNORMALITY_OF_THE_ANKLES | 0.00313 | 0.03310 | 0.43171 | 0.65369 | 1.23418 | 63 |
| HP_ABNORMALITY_OF_THE_CEREBRAL_VASCULATURE | 0.00357 | 0.03638 | 0.43171 | 0.59764 | 1.15722 | 158 |
| HP_ABNORMALITY_OF_THE_CEREBROSPINAL_FLUID | 0.00030 | 0.00549 | 0.49849 | 0.58221 | 1.14439 | 325 |
| HP_ABNORMALITY_OF_THE_MITOCHONDRION | 0.00126 | 0.01715 | 0.45506 | 0.62149 | 1.19764 | 120 |
| HP_ABNORMALITY_OF_THE_OPTIC_DISC | 0.00198 | 0.02370 | 0.43171 | 0.56624 | 1.11727 | 432 |
| HP_ABNORMALITY_OF_THE_OPTIC_NERVE | 0.00036 | 0.00622 | 0.49849 | 0.56723 | 1.12034 | 486 |
| HP_ABNORMAL_ABDOMEN_MORPHOLOGY | 0.00089 | 0.01291 | 0.47727 | 0.56424 | 1.11364 | 441 |
| HP_ABNORMAL_ACTIVITY_OF_MITOCHONDRIAL_RESPIRATORY_ | 0.00014 | 0.00300 | 0.51885 | 0.70303 | 1.32524 | 61 |
| HP_ABNORMAL_BASAL_GANGLIA_MRI_SIGNAL_INTENSITY | 0.00179 | 0.02183 | 0.45506 | 0.71597 | 1.32046 | 36 |
| HP_ABNORMAL_BRAINSTEM_MRI_SIGNAL_INTENSITY | 0.00398 | 0.03962 | 0.40702 | 0.71528 | 1.31370 | 33 |

| | | | | | | |
|---|---|---|---|---|---|---|
| HP_ABNORMAL_CELLULAR_PHENOTYPE | 0.00209 | 0.02470 | 0.43171 | 0.56937 | 1.11992 | 334 |
| HP_ABNORMAL_CIRCULATING_CARBOHYDRATE_CONCENTRATIC | 0.00194 | 0.02331 | 0.45506 | 0.69518 | 1.29539 | 46 |
| HP_ABNORMAL_CNS_MYELINATION | 0.00014 | 0.00300 | 0.51885 | 0.62318 | 1.20954 | 171 |
| HP_ABNORMAL_CSF_METABOLITE_LEVEL | 0.00001 | 0.00027 | 0.59333 | 0.70531 | 1.34409 | 82 |
| HP_ABNORMAL_ELECTROPHYSIOLOGY_OF_SINOATRIAL_NODE_ | 0.00029 | 0.00536 | 0.49849 | 0.73172 | 1.35725 | 41 |
| HP_ABNORMAL_ENZYME_COENZYME_ACTIVITY | 0.00561 | 0.04971 | 0.40702 | 0.60490 | 1.16567 | 123 |
| HP_ABNORMAL_ERYTHROCYTE_MORPHOLOGY | 0.00001 | 0.00023 | 0.61053 | 0.58835 | 1.15904 | 380 |
| HP_ABNORMAL_HEART_VALVE_PHYSIOLOGY | 0.00405 | 0.04011 | 0.40702 | 0.59004 | 1.14474 | 169 |
| HP_ABNORMAL_MITOCHONDRIA_IN_MUSCLE_TISSUE | 0.00075 | 0.01118 | 0.47727 | 0.74633 | 1.36700 | 29 |
| HP_ABNORMAL_MYELINATION | 0.00007 | 0.00169 | 0.53843 | 0.59258 | 1.16479 | 324 |
| HP_ABNORMAL_MYOCARDIUM_MORPHOLOGY | 0.00008 | 0.00192 | 0.53843 | 0.58647 | 1.15365 | 340 |
| HP_ABNORMAL_RENAL_PHYSIOLOGY | 0.00450 | 0.04342 | 0.40702 | 0.55484 | 1.09317 | 385 |
| HP_ABNORMAL_TRICUSPID_VALVE_PHYSIOLOGY | 0.00231 | 0.02642 | 0.43171 | 0.72968 | 1.34029 | 31 |
| HP_ABNORMAL_VITREOUS_HUMOR_MORPHOLOGY | 0.00181 | 0.02201 | 0.45506 | 0.79632 | 1.40955 | 17 |
| HP_ACUTE_LEUKEMIA | 0.00013 | 0.00276 | 0.51885 | 0.68145 | 1.29219 | 75 |
| HP_ANEMIA_OF_INADEQUATE_PRODUCTION | 0.00561 | 0.04971 | 0.40702 | 0.61298 | 1.17855 | 107 |
| HP_APHASIA | 0.00213 | 0.02490 | 0.43171 | 0.72858 | 1.33704 | 30 |
| HP_BLINDNESS | 0.00018 | 0.00355 | 0.51885 | 0.61672 | 1.19768 | 175 |
| HP_BONE_MARROW_HYPERCELLULARITY | 0.00045 | 0.00725 | 0.49849 | 0.82169 | 1.46806 | 19 |
| HP_CHOREOATHETOSIS | 0.00561 | 0.04971 | 0.40702 | 0.64956 | 1.22631 | 64 |
| HP_CHRONIC_INFECTION | 0.00121 | 0.01658 | 0.45506 | 0.81916 | 1.44468 | 16 |
| HP_CONGESTIVE_HEART_FAILURE | 0.00495 | 0.04573 | 0.40702 | 0.60734 | 1.16929 | 114 |
| HP_CRANIAL_NERVE_COMPRESSION | 0.00331 | 0.03441 | 0.43171 | 0.75830 | 1.36989 | 23 |
| HP_DECREASED_ACTIVITY_OF_MITOCHONDRIAL_COMPLEX_I | 0.00283 | 0.03081 | 0.43171 | 0.71940 | 1.32496 | 35 |
| HP_DECREASED_ACTIVITY_OF_THE_PYRUVATE_DEHYDROGENAS | 0.00062 | 0.00949 | 0.47727 | 0.75390 | 1.38478 | 31 |
| HP_DEPRESSED_NASAL_RIDGE | 0.00041 | 0.00681 | 0.49849 | 0.66218 | 1.26189 | 82 |
| HP_DYSPNEA | 0.00250 | 0.02807 | 0.43171 | 0.58107 | 1.13348 | 224 |
| HP_DYSTONIA | 0.00069 | 0.01043 | 0.47727 | 0.58101 | 1.13736 | 279 |
| HP_EMOTIONAL_LABILITY | 0.00016 | 0.00321 | 0.51885 | 0.66683 | 1.27522 | 90 |
| HP_EPISODIC_ATAXIA | 0.00487 | 0.04559 | 0.40702 | 0.79934 | 1.40259 | 15 |
| HP_FASCICULATIONS | 0.00130 | 0.01761 | 0.45506 | 0.68128 | 1.28064 | 54 |
| HP_FATIGUE | 0.00042 | 0.00689 | 0.49849 | 0.59902 | 1.16764 | 212 |
| HP_FETAL_DISTRESS | 0.00179 | 0.02183 | 0.45506 | 0.72928 | 1.33973 | 32 |
| HP_FLUSHING | 0.00561 | 0.04971 | 0.40702 | 0.74782 | 1.33686 | 20 |
| HP_FOCAL_T2_HYPERINTENSE_BRAINSTEM_LESION | 0.00159 | 0.02016 | 0.45506 | 0.74970 | 1.36879 | 27 |
| HP_HEADACHE | 0.00016 | 0.00319 | 0.51885 | 0.62537 | 1.21179 | 163 |
| HP_HEPATOMEGALY | 0.00108 | 0.01507 | 0.45506 | 0.57439 | 1.12990 | 335 |
| HP_HYPERTROPHIC_CARDIOMYOPATHY | 0.00002 | 0.00050 | 0.57561 | 0.62629 | 1.21908 | 194 |
| HP_IMPAIRED_SOCIAL_INTERACTIONS | 0.00055 | 0.00863 | 0.47727 | 0.66555 | 1.26058 | 72 |
| HP_INCREASED_CSF_LACTATE | 0.00001 | 0.00034 | 0.59333 | 0.71937 | 1.35857 | 68 |
| HP_INCREASED_SERUM_LACTATE | 0.00001 | 0.00028 | 0.59333 | 0.65837 | 1.26932 | 133 |
| HP_INCREASED_SERUM_PYRUVATE | 0.00110 | 0.01526 | 0.45506 | 0.73143 | 1.35198 | 37 |
| HP_INFANTILE_ONSET | 0.00487 | 0.04559 | 0.40702 | 0.56559 | 1.10939 | 300 |
| HP_INTRACRANIAL_CYSTIC_LESION | 0.00055 | 0.00863 | 0.47727 | 0.62760 | 1.20885 | 116 |
| HP_INTRAUTERINE_GROWTH_RETARDATION | 0.00305 | 0.03271 | 0.43171 | 0.56594 | 1.11393 | 350 |
| HP_LACTIC_ACIDOSIS | 0.00018 | 0.00355 | 0.51885 | 0.63312 | 1.22071 | 131 |
| HP_LETHARGY | 0.00084 | 0.01228 | 0.47727 | 0.63361 | 1.21845 | 109 |
| HP_LEUKODYSTROPHY | 0.00029 | 0.00529 | 0.49849 | 0.66598 | 1.26915 | 82 |
| HP_LEUKOENCEPHALOPATHY | 0.00008 | 0.00190 | 0.53843 | 0.64132 | 1.23667 | 134 |
| HP_LIPID_ACCUMULATION_IN_HEPATOCYTES | 0.00220 | 0.02548 | 0.43171 | 0.62794 | 1.20159 | 91 |
| HP_LOWER_LIMB_SPASTICITY | 0.00154 | 0.01974 | 0.45506 | 0.60929 | 1.17648 | 139 |
| HP_MACROCYTIC_ANEMIA | 0.00030 | 0.00542 | 0.49849 | 0.73108 | 1.35606 | 41 |
| HP_METABOLIC_ACIDOSIS | 0.00034 | 0.00599 | 0.49849 | 0.64831 | 1.24464 | 103 |
| HP_MICRONODULAR_CIRRHOSIS | 0.00324 | 0.03393 | 0.43171 | 0.78723 | 1.40018 | 18 |
| HP_MIGRAINE | 0.00028 | 0.00515 | 0.49849 | 0.67309 | 1.27635 | 75 |
| HP_MUSCLE_ABNORMALITY_RELATED_TO_MITOCHONDRIAL_DY | 0.00209 | 0.02470 | 0.43171 | 0.68258 | 1.27573 | 49 |
| HP_MUTISM | 0.00283 | 0.03081 | 0.43171 | 0.71937 | 1.32490 | 35 |
| HP_NAUSEA_AND_VOMITING | 0.00022 | 0.00417 | 0.51885 | 0.59383 | 1.16158 | 264 |
| HP_NEOPLASM_OF_THE_ENDOCRINE_SYSTEM | 0.00495 | 0.04573 | 0.40702 | 0.63166 | 1.20308 | 81 |
| HP_OPTIC_NEUROPATHY | 0.00135 | 0.01808 | 0.45506 | 0.73837 | 1.35610 | 33 |
| HP_PACE_OF_PROGRESSION | 0.00487 | 0.04559 | 0.40702 | 0.56814 | 1.11416 | 298 |
| HP_PALLOR | 0.00036 | 0.00622 | 0.49849 | 0.64861 | 1.24463 | 102 |
| HP_PARAPLEGIA | 0.00121 | 0.01658 | 0.45506 | 0.63584 | 1.22133 | 104 |
| HP_PARAPLEGIA_PARAPARESIS | 0.00036 | 0.00621 | 0.49849 | 0.62047 | 1.20040 | 150 |
| HP_PAROXYSMAL_INVOLUNTARY_EYE_MOVEMENTS | 0.00054 | 0.00849 | 0.47727 | 0.75650 | 1.38957 | 31 |
| HP_PEDIATRIC_ONSET | 0.00561 | 0.04971 | 0.40702 | 0.55915 | 1.10116 | 371 |
| HP_PIGMENTARY_RETINOPATHY | 0.00065 | 0.00981 | 0.47727 | 0.63260 | 1.21720 | 113 |
| HP_POOR_EYE_CONTACT | 0.00017 | 0.00341 | 0.51885 | 0.71279 | 1.33749 | 53 |
| HP_POOR_HEAD_CONTROL | 0.00000 | 0.00005 | 0.64355 | 0.67819 | 1.30665 | 118 |
| HP_PROGRESSIVE | 0.00194 | 0.02331 | 0.45506 | 0.59790 | 1.16273 | 186 |

| | | | | | | |
|---|---|---|---|---|---|---|
| HP_PROGRESSIVE_CEREBELLAR_ATAXIA | 0.00364 | 0.03677 | 0.43171 | 0.64788 | 1.22325 | 65 |
| HP_PROGRESSIVE_SPASTIC_PARAPLEGIA | 0.00253 | 0.02838 | 0.43171 | 0.69655 | 1.29822 | 45 |
| HP_PROXIMAL_TUBULOPATHY | 0.00141 | 0.01850 | 0.45506 | 0.72579 | 1.33535 | 34 |
| HP_PTOSIS | 0.00472 | 0.04480 | 0.40702 | 0.55582 | 1.09637 | 418 |
| HP_REDUCED_CONSCIOUSNESS_CONFUSION | 0.00007 | 0.00164 | 0.53843 | 0.61227 | 1.19295 | 205 |
| HP_REDUCED_VISUAL_ACUITY | 0.00053 | 0.00837 | 0.47727 | 0.58618 | 1.14774 | 283 |
| HP_RENAL_TUBULAR_DYSFUNCTION | 0.00013 | 0.00284 | 0.51885 | 0.68108 | 1.29150 | 75 |
| HP_RESPIRATORY_DISTRESS | 0.00108 | 0.01507 | 0.45506 | 0.62693 | 1.20762 | 115 |
| HP_RESPIRATORY_INSUFFICIENCY | 0.00000 | 0.00010 | 0.62726 | 0.60536 | 1.18998 | 326 |
| HP_SINUS_TACHYCARDIA | 0.00213 | 0.02490 | 0.43171 | 0.77203 | 1.39128 | 22 |
| HP_SPARSE_EYELASHES | 0.00150 | 0.01943 | 0.45506 | 0.71355 | 1.32161 | 40 |
| HP_SPARSE_OR_ABSENT_EYELASHES | 0.00220 | 0.02548 | 0.43171 | 0.67331 | 1.26567 | 54 |
| HP_SPASTIC_PARAPLEGIA | 0.00098 | 0.01407 | 0.45506 | 0.64059 | 1.22712 | 96 |
| HP_THROMBOCYTOSIS | 0.00413 | 0.04065 | 0.40702 | 0.75210 | 1.35870 | 23 |
| HP_TONGUE_FASCICULATIONS | 0.00457 | 0.04386 | 0.40702 | 0.80056 | 1.40472 | 15 |
| HP_TRANSIENT_ISCHEMIC_ATTACK | 0.00480 | 0.04537 | 0.40702 | 0.74902 | 1.35809 | 25 |
| HP_VERTIGO | 0.00002 | 0.00063 | 0.57561 | 0.72114 | 1.35844 | 60 |
| HP_VESTIBULAR_DYSFUNCTION | 0.00028 | 0.00522 | 0.49849 | 0.67279 | 1.27577 | 75 |
| HP_VOMITING | 0.00072 | 0.01082 | 0.47727 | 0.61459 | 1.18792 | 147 |
| HP_WEAKNESS_DUE_TO_UPPER_MOTOR_NEURON_DYSFUNCTI | 0.00532 | 0.04803 | 0.40702 | 0.56829 | 1.11186 | 272 |
| HP_WEAKNESS_OF_MUSCLES_OF_RESPIRATION | 0.00163 | 0.02050 | 0.45506 | 0.66272 | 1.25123 | 63 |
| **Cluster 1** | | | | | | |
| pathway | pval | padj | log2err | ES | NES | size |
| GO_ACTIVATION_OF_IMMUNE_RESPONSE | 0.00000 | 0.00000 | 0.74774 | 0.65038 | 1.26999 | 235 |
| GO_ACTIVATION_OF_INNATE_IMMUNE_RESPONSE | 0.00000 | 0.00001 | 0.69013 | 0.71967 | 1.37861 | 96 |
| GO_AEROBIC_RESPIRATION | 0.00000 | 0.00000 | 0.73376 | 0.76280 | 1.44478 | 72 |
| GO_ALCOHOL_BIOSYNTHETIC_PROCESS | 0.00032 | 0.00572 | 0.49849 | 0.64679 | 1.24367 | 108 |
| GO_ALCOHOL_METABOLIC_PROCESS | 0.00364 | 0.03677 | 0.43171 | 0.58112 | 1.13424 | 228 |
| GO_AMEBOIDAL_TYPE_CELL_MIGRATION | 0.00457 | 0.04386 | 0.40702 | 0.57755 | 1.12854 | 247 |
| GO_AMIDE_BINDING | 0.00014 | 0.00297 | 0.51885 | 0.61533 | 1.19780 | 199 |
| GO_AMINE_METABOLIC_PROCESS | 0.00000 | 0.00000 | 0.76146 | 0.73732 | 1.41097 | 92 |
| GO_AMYLOID_BETA_CLEARANCE | 0.00561 | 0.04971 | 0.40702 | 0.74081 | 1.33976 | 24 |
| GO_ANAPHASE_PROMOTING_COMPLEX_DEPENDENT_CATABOL | 0.00000 | 0.00000 | 0.76146 | 0.77107 | 1.46112 | 73 |
| GO_ANATOMICAL_STRUCTURE_HOMEOSTASIS | 0.00020 | 0.00390 | 0.51885 | 0.59307 | 1.16021 | 271 |
| GO_ANIMAL_ORGAN_REGENERATION | 0.00012 | 0.00269 | 0.51885 | 0.71251 | 1.33510 | 52 |
| GO_ANTIGEN_PROCESSING_AND_PRESENTATION | 0.00000 | 0.00000 | 0.74774 | 0.68785 | 1.33064 | 149 |
| GO_ANTIGEN_PROCESSING_AND_PRESENTATION_OF_EXOGENC | 0.00000 | 0.00000 | 0.76146 | 0.79644 | 1.50237 | 62 |
| GO_ANTIGEN_PROCESSING_AND_PRESENTATION_OF_PEPTIDE_ | 0.00000 | 0.00000 | 0.74774 | 0.70107 | 1.35144 | 130 |
| GO_ANTIGEN_PROCESSING_AND_PRESENTATION_OF_PEPTIDE_ | 0.00000 | 0.00000 | 0.78819 | 0.77566 | 1.47012 | 74 |
| GO_ANTIGEN_RECEPTOR_MEDIATED_SIGNALING_PATHWAY | 0.00000 | 0.00009 | 0.62726 | 0.66362 | 1.28080 | 136 |
| GO_APICAL_PART_OF_CELL | 0.00135 | 0.01808 | 0.45506 | 0.58926 | 1.14966 | 220 |
| GO_APOPTOTIC_SIGNALING_PATHWAY | 0.00006 | 0.00152 | 0.53843 | 0.58638 | 1.15586 | 393 |
| GO_ATPASE_ACTIVITY | 0.00050 | 0.00802 | 0.47727 | 0.58216 | 1.14385 | 316 |
| GO_ATPASE_ACTIVITY_COUPLED_TO_TRANSMEMBRANE_MOVE | 0.00420 | 0.04132 | 0.40702 | 0.78005 | 1.38741 | 18 |
| GO_ATP_BIOSYNTHETIC_PROCESS | 0.00013 | 0.00276 | 0.51885 | 0.83897 | 1.50801 | 21 |
| GO_ATP_METABOLIC_PROCESS | 0.00000 | 0.00000 | | 0.72178 | 1.40814 | 219 |
| GO_ATP_SYNTHESIS_COUPLED_ELECTRON_TRANSPORT | 0.00000 | 0.00000 | | 0.80353 | 1.52425 | 76 |
| GO_AZUROPHIL_GRANULE | 0.00025 | 0.00466 | 0.49849 | 0.64781 | 1.24551 | 107 |
| GO_AZUROPHIL_GRANULE_LUMEN | 0.00405 | 0.04011 | 0.40702 | 0.65625 | 1.23793 | 62 |
| GO_BASAL_PART_OF_CELL | 0.00509 | 0.04656 | 0.40702 | 0.68515 | 1.27697 | 45 |
| GO_BASAL_PLASMA_MEMBRANE | 0.00346 | 0.03548 | 0.43171 | 0.71184 | 1.31104 | 35 |
| GO_BASEMENT_MEMBRANE | 0.00279 | 0.03056 | 0.43171 | 0.68935 | 1.28452 | 46 |
| GO_BASOLATERAL_PLASMA_MEMBRANE | 0.00011 | 0.00247 | 0.53843 | 0.63209 | 1.21993 | 136 |
| GO_BINDING_OF_SPERM_TO_ZONA_PELLUCIDA | 0.00000 | 0.00006 | 0.64355 | 0.90095 | 1.60966 | 19 |
| GO_BLOOD_MICROPARTICLE | 0.00000 | 0.00000 | 0.70498 | 0.81982 | 1.52763 | 46 |
| GO_BRUSH_BORDER | 0.00060 | 0.00923 | 0.47727 | 0.66897 | 1.26790 | 74 |
| GO_CADHERIN_BINDING | 0.00000 | 0.00000 | | 0.68759 | 1.34435 | 253 |
| GO_CALCIUM_DEPENDENT_PHOSPHOLIPID_BINDING | 0.00152 | 0.01958 | 0.45506 | 0.75021 | 1.37120 | 28 |
| GO_CALCIUM_DEPENDENT_PROTEIN_BINDING | 0.00009 | 0.00211 | 0.53843 | 0.74254 | 1.38394 | 45 |
| GO_CANONICAL_WNT_SIGNALING_PATHWAY | 0.00004 | 0.00109 | 0.55733 | 0.61740 | 1.20458 | 222 |
| GO_CARBOHYDRATE_CATABOLIC_PROCESS | 0.00135 | 0.01808 | 0.45506 | 0.61179 | 1.18186 | 144 |
| GO_CATALYTIC_STEP_2_SPLICEOSOME | 0.00001 | 0.00022 | 0.61053 | 0.70290 | 1.33765 | 80 |
| GO_CATION_TRANSMEMBRANE_TRANSPORT | 0.00327 | 0.03409 | 0.43171 | 0.56060 | 1.10531 | 406 |
| GO_CATION_TRANSMEMBRANE_TRANSPORTER_ACTIVITY | 0.00097 | 0.01387 | 0.47727 | 0.58196 | 1.13846 | 265 |
| GO_CELLULAR_AMINO_ACID_METABOLIC_PROCESS | 0.00002 | 0.00060 | 0.57561 | 0.61438 | 1.19930 | 231 |
| GO_CELLULAR_COMPONENT_DISASSEMBLY | 0.00000 | 0.00000 | | 0.64070 | 1.26342 | 408 |
| GO_CELLULAR_ION_HOMEOSTASIS | 0.00027 | 0.00493 | 0.49849 | 0.58658 | 1.15340 | 328 |
| GO_CELLULAR_IRON_ION_HOMEOSTASIS | 0.00327 | 0.03409 | 0.43171 | 0.68360 | 1.27374 | 47 |
| GO_CELLULAR_KETONE_METABOLIC_PROCESS | 0.00000 | 0.00010 | 0.62726 | 0.65358 | 1.26726 | 164 |
| GO_CELLULAR_MONOVALENT_INORGANIC_CATION_HOMEOSTA | 0.00179 | 0.02183 | 0.45506 | 0.64747 | 1.22821 | 76 |

| | | | | | | |
|---|---|---|---|---|---|---|
| GO_CELLULAR_NITROGEN_COMPOUND_CATABOLIC_PROCESS | 0.00000 | 0.00000 | | 0.67915 | 1.34096 | 456 |
| GO_CELLULAR_PROTEIN_COMPLEX_DISASSEMBLY | 0.00000 | 0.00000 | | 0.69566 | 1.35184 | 179 |
| GO_CELLULAR_RESPIRATION | 0.00000 | 0.00000 | | 0.74205 | 1.43370 | 145 |
| GO_CELLULAR_RESPONSE_TO_HEAT | 0.00004 | 0.00109 | 0.55733 | 0.67186 | 1.28698 | 95 |
| GO_CELLULAR_RESPONSE_TO_HORMONE_STIMULUS | 0.00309 | 0.03299 | 0.43171 | 0.56453 | 1.11191 | 374 |
| GO_CELLULAR_RESPONSE_TO_ORGANIC_CYCLIC_COMPOUND | 0.00502 | 0.04608 | 0.40702 | 0.56512 | 1.11134 | 329 |
| GO_CELLULAR_RESPONSE_TO_OXYGEN_LEVELS | 0.00000 | 0.00000 | | 0.71461 | 1.38224 | 151 |
| GO_CELL_ACTIVATION_INVOLVED_IN_IMMUNE_RESPONSE | 0.00000 | 0.00000 | | 0.63924 | 1.26108 | 422 |
| GO_CELL_ADHESION_MEDIATOR_ACTIVITY | 0.00157 | 0.02009 | 0.45506 | 0.71837 | 1.32488 | 36 |
| GO_CELL_ADHESION_MOLECULE_BINDING | 0.00000 | 0.00000 | | 0.67109 | 1.32112 | 353 |
| GO_CELL_BODY | 0.00004 | 0.00109 | 0.55733 | 0.59973 | 1.17541 | 293 |
| GO_CELL_CELL_RECOGNITION | 0.00000 | 0.00006 | 0.64355 | 0.82001 | 1.51026 | 35 |
| GO_CELL_CELL_SIGNALING_BY_WNT | 0.00004 | 0.00109 | 0.55733 | 0.59042 | 1.16141 | 339 |
| GO_CELL_CYCLE_G1_S_PHASE_TRANSITION | 0.00376 | 0.03783 | 0.43171 | 0.58112 | 1.13378 | 220 |
| GO_CELL_CYCLE_G2_M_PHASE_TRANSITION | 0.00024 | 0.00459 | 0.51885 | 0.59807 | 1.16746 | 231 |
| GO_CELL_CYCLE_PHASE_TRANSITION | 0.00000 | 0.00001 | 0.69013 | 0.59417 | 1.17348 | 499 |
| GO_CELL_DIVISION | 0.00003 | 0.00071 | 0.57561 | 0.58314 | 1.15094 | 449 |
| GO_CELL_KILLING | 0.00205 | 0.02440 | 0.43171 | 0.66727 | 1.25696 | 60 |
| GO_CELL_RECOGNITION | 0.00000 | 0.00000 | 0.73376 | 0.75953 | 1.43954 | 74 |
| GO_CELL_REDOX_HOMEOSTASIS | 0.00163 | 0.02050 | 0.45506 | 0.71772 | 1.32665 | 38 |
| GO_CELL_SUBSTRATE_JUNCTION | 0.00000 | 0.00000 | | 0.65742 | 1.29171 | 316 |
| GO_CELL_SURFACE | 0.00000 | 0.00000 | 0.73376 | 0.62572 | 1.23070 | 330 |
| GO_CELL_SURFACE_RECEPTOR_SIGNALING_PATHWAY_INVOLVEI | 0.00038 | 0.00640 | 0.49849 | 0.57762 | 1.13770 | 374 |
| GO_CHAPERONE_BINDING | 0.00495 | 0.04573 | 0.40702 | 0.62860 | 1.19935 | 86 |
| GO_CHAPERONE_COMPLEX | 0.00001 | 0.00028 | 0.59333 | 0.86337 | 1.55587 | 22 |
| GO_CHAPERONE_MEDIATED_PROTEIN_COMPLEX_ASSEMBLY | 0.00100 | 0.01430 | 0.45506 | 0.80787 | 1.44337 | 19 |
| GO_CHAPERONE_MEDIATED_PROTEIN_FOLDING | 0.00168 | 0.02098 | 0.45506 | 0.71110 | 1.32288 | 44 |
| GO_CHROMOSOMAL_REGION | 0.00018 | 0.00349 | 0.51885 | 0.59323 | 1.16180 | 286 |
| GO_CHROMOSOME_CENTROMERIC_REGION | 0.00172 | 0.02135 | 0.45506 | 0.60447 | 1.17248 | 168 |
| GO_CHROMOSOME_SEGREGATION | 0.00003 | 0.00087 | 0.55733 | 0.60463 | 1.18280 | 270 |
| GO_CLUSTER_OF_ACTIN_BASED_CELL_PROJECTIONS | 0.00074 | 0.01107 | 0.47727 | 0.64184 | 1.23044 | 99 |
| GO_COATED_VESICLE | 0.00213 | 0.02490 | 0.43171 | 0.59512 | 1.15846 | 199 |
| GO_COATED_VESICLE_MEMBRANE | 0.00060 | 0.00923 | 0.47727 | 0.63443 | 1.22207 | 115 |
| GO_COLLAGEN_BINDING | 0.00465 | 0.04430 | 0.40702 | 0.73471 | 1.33628 | 26 |
| GO_COLLAGEN_CONTAINING_EXTRACELLULAR_MATRIX | 0.00000 | 0.00006 | 0.64355 | 0.65358 | 1.26542 | 157 |
| GO_CONDENSED_CHROMOSOME | 0.00350 | 0.03580 | 0.43171 | 0.59394 | 1.15417 | 179 |
| GO_CONTRACTILE_FIBER | 0.00064 | 0.00972 | 0.47727 | 0.62217 | 1.19973 | 134 |
| GO_COPI_COATED_VESICLE | 0.00316 | 0.03344 | 0.43171 | 0.75688 | 1.37234 | 25 |
| GO_CORNIFICATION | 0.00027 | 0.00500 | 0.49849 | 0.77970 | 1.42512 | 28 |
| GO_CORTICAL_CYTOSKELETON | 0.00502 | 0.04608 | 0.40702 | 0.64388 | 1.21870 | 71 |
| GO_COTRANSLATIONAL_PROTEIN_TARGETING_TO_MEMBRANE | 0.00000 | 0.00000 | | 0.78223 | 1.49819 | 94 |
| GO_CYTOCHROME_COMPLEX | 0.00035 | 0.00610 | 0.49849 | 0.80796 | 1.45601 | 22 |
| GO_CYTOKINE_MEDIATED_SIGNALING_PATHWAY | 0.00000 | 0.00002 | 0.67496 | 0.60742 | 1.19735 | 392 |
| GO_CYTOPLASMIC_STRESS_GRANULE | 0.00014 | 0.00302 | 0.51885 | 0.69390 | 1.31009 | 63 |
| GO_CYTOPLASMIC_TRANSLATION | 0.00000 | 0.00001 | 0.70498 | 0.71773 | 1.37349 | 92 |
| GO_CYTOPLASMIC_TRANSLATIONAL_INITIATION | 0.00178 | 0.02182 | 0.45506 | 0.73517 | 1.35039 | 31 |
| GO_CYTOSOLIC_LARGE_RIBOSOMAL_SUBUNIT | 0.00000 | 0.00005 | 0.64355 | 0.77240 | 1.44935 | 53 |
| GO_CYTOSOLIC_RIBOSOME | 0.00000 | 0.00000 | | 0.77483 | 1.48437 | 97 |
| GO_CYTOSOLIC_SMALL_RIBOSOMAL_SUBUNIT | 0.00000 | 0.00010 | 0.62726 | 0.79384 | 1.47033 | 40 |
| GO_DETECTION_OF_STIMULUS | 0.00020 | 0.00393 | 0.51885 | 0.65208 | 1.25384 | 108 |
| GO_DETOXIFICATION | 0.00413 | 0.04065 | 0.40702 | 0.65399 | 1.23684 | 70 |
| GO_DEVELOPMENT_OF_PRIMARY_FEMALE_SEXUAL_CHARACTE | 0.00179 | 0.02183 | 0.45506 | 0.68410 | 1.28057 | 51 |
| GO_DICARBOXYLIC_ACID_METABOLIC_PROCESS | 0.00261 | 0.02910 | 0.43171 | 0.65541 | 1.23747 | 65 |
| GO_DISORDERED_DOMAIN_SPECIFIC_BINDING | 0.00051 | 0.00812 | 0.47727 | 0.76867 | 1.40343 | 27 |
| GO_DISULFIDE_OXIDOREDUCTASE_ACTIVITY | 0.00435 | 0.04238 | 0.40702 | 0.71445 | 1.31232 | 31 |
| GO_DNA_BIOSYNTHETIC_PROCESS | 0.00017 | 0.00334 | 0.51885 | 0.62540 | 1.20833 | 145 |
| GO_DNA_CONFORMATION_CHANGE | 0.00327 | 0.03409 | 0.43171 | 0.57979 | 1.13178 | 231 |
| GO_DNA_DAMAGE_RESPONSE_DETECTION_OF_DNA_DAMAGE | 0.00305 | 0.03271 | 0.43171 | 0.71625 | 1.31916 | 35 |
| GO_DNA_GEOMETRIC_CHANGE | 0.00141 | 0.01850 | 0.45506 | 0.62642 | 1.20356 | 105 |
| GO_DOUBLE_STRANDED_RNA_BINDING | 0.00039 | 0.00658 | 0.49849 | 0.69158 | 1.30365 | 61 |
| GO_DRUG_BINDING | 0.00450 | 0.04342 | 0.40702 | 0.66245 | 1.24130 | 52 |
| GO_ELECTRON_TRANSFER_ACTIVITY | 0.00000 | 0.00001 | 0.67496 | 0.72917 | 1.38765 | 80 |
| GO_ELECTRON_TRANSPORT_CHAIN | 0.00000 | 0.00000 | | 0.73336 | 1.41392 | 133 |
| GO_EMBRYO_IMPLANTATION | 0.00044 | 0.00706 | 0.49849 | 0.75968 | 1.39146 | 29 |
| GO_ENDOCYTIC_VESICLE_MEMBRANE | 0.00524 | 0.04770 | 0.40702 | 0.62563 | 1.19564 | 89 |
| GO_ENDOPEPTIDASE_ACTIVITY | 0.00065 | 0.00981 | 0.47727 | 0.60123 | 1.16998 | 196 |
| GO_ENDOPEPTIDASE_COMPLEX | 0.00000 | 0.00001 | 0.70498 | 0.76593 | 1.44609 | 63 |
| GO_ENDOPEPTIDASE_REGULATOR_ACTIVITY | 0.00006 | 0.00145 | 0.55733 | 0.70425 | 1.33001 | 68 |
| GO_ENDOPLASMIC_RETICULUM_GOLGI_INTERMEDIATE_COMPA | 0.00055 | 0.00863 | 0.47727 | 0.64758 | 1.24050 | 96 |
| GO_ENDOPLASMIC_RETICULUM_GOLGI_INTERMEDIATE_COMPA | 0.00148 | 0.01923 | 0.45506 | 0.68944 | 1.29056 | 51 |

| | | | | | | |
|---|---|---|---|---|---|---|
| GO_ENDOPLASMIC_RETICULUM_TO_GOLGI_VESICLE_MEDIATED | 0.00532 | 0.04803 | 0.40702 | 0.59194 | 1.14697 | 162 |
| GO_ENERGY_DERIVATION_BY_OXIDATION_OF_ORGANIC_COMP | 0.00000 | 0.00000 | | 0.69995 | 1.36305 | 202 |
| GO_ENZYME_INHIBITOR_ACTIVITY | 0.00246 | 0.02780 | 0.43171 | 0.58913 | 1.14643 | 191 |
| GO_EPIDERMAL_CELL_DIFFERENTIATION | 0.00216 | 0.02519 | 0.43171 | 0.62667 | 1.20589 | 112 |
| GO_EPITHELIAL_CELL_DIFFERENTIATION | 0.00054 | 0.00857 | 0.47727 | 0.57819 | 1.13725 | 334 |
| GO_ESTABLISHMENT_OF_ORGANELLE_LOCALIZATION | 0.00122 | 0.01679 | 0.45506 | 0.57737 | 1.13489 | 323 |
| GO_ESTABLISHMENT_OF_PROTEIN_LOCALIZATION_TO_CHROMO | 0.00016 | 0.00321 | 0.51885 | 0.81363 | 1.46985 | 23 |
| GO_ESTABLISHMENT_OF_PROTEIN_LOCALIZATION_TO_ENDOPLA | 0.00000 | 0.00000 | | 0.77241 | 1.48479 | 106 |
| GO_ESTABLISHMENT_OF_PROTEIN_LOCALIZATION_TO_MEMBRA | 0.00000 | 0.00000 | | 0.67484 | 1.31991 | 260 |
| GO_ESTABLISHMENT_OF_PROTEIN_LOCALIZATION_TO_ORGANE | 0.00000 | 0.00000 | | 0.65856 | 1.30056 | 461 |
| GO_ESTABLISHMENT_OF_PROTEIN_LOCALIZATION_TO_TELOME | 0.00000 | 0.00006 | 0.64355 | 0.90447 | 1.60870 | 18 |
| GO_ESTABLISHMENT_OF_RNA_LOCALIZATION | 0.00000 | 0.00005 | 0.64355 | 0.64271 | 1.24830 | 176 |
| GO_ESTABLISHMENT_OF_TISSUE_POLARITY | 0.00000 | 0.00000 | 0.81404 | 0.75693 | 1.44999 | 96 |
| GO_EUKARYOTIC_TRANSLATION_INITIATION_FACTOR_3_COMPL | 0.00143 | 0.01863 | 0.45506 | 0.83263 | 1.46100 | 15 |
| GO_EXTERNAL_SIDE_OF_PLASMA_MEMBRANE | 0.00159 | 0.02016 | 0.45506 | 0.61775 | 1.19022 | 118 |
| GO_EXTRACELLULAR_MATRIX | 0.00042 | 0.00695 | 0.49849 | 0.61138 | 1.18938 | 189 |
| GO_FATTY_ACID_DERIVATIVE_METABOLIC_PROCESS | 0.00043 | 0.00695 | 0.49849 | 0.66266 | 1.26212 | 81 |
| GO_FATTY_ACYL_COA_METABOLIC_PROCESS | 0.00502 | 0.04608 | 0.40702 | 0.72264 | 1.32082 | 28 |
| GO_FC_EPSILON_RECEPTOR_SIGNALING_PATHWAY | 0.00000 | 0.00000 | 0.71951 | 0.72370 | 1.38610 | 94 |
| GO_FC_RECEPTOR_SIGNALING_PATHWAY | 0.00000 | 0.00000 | 0.70498 | 0.68815 | 1.32879 | 140 |
| GO_FERTILIZATION | 0.00000 | 0.00005 | 0.64355 | 0.73257 | 1.38752 | 72 |
| GO_FIBRILLAR_CENTER | 0.00246 | 0.02780 | 0.43171 | 0.62638 | 1.19990 | 96 |
| GO_FIBROBLAST_GROWTH_FACTOR_RECEPTOR_SIGNALING_PAT | 0.00495 | 0.04573 | 0.40702 | 0.64888 | 1.22552 | 67 |
| GO_FIBROBLAST_PROLIFERATION | 0.00032 | 0.00575 | 0.49849 | 0.70676 | 1.32618 | 53 |
| GO_FICOLIN_1_RICH_GRANULE | 0.00000 | 0.00000 | 0.70498 | 0.71544 | 1.37138 | 98 |
| GO_GABA_ERGIC_SYNAPSE | 0.00022 | 0.00427 | 0.51885 | 0.76430 | 1.40406 | 32 |
| GO_GENERATION_OF_PRECURSOR_METABOLITES_AND_ENERGY | 0.00000 | 0.00000 | | 0.67527 | 1.32968 | 368 |
| GO_GENE_SILENCING | 0.00019 | 0.00376 | 0.51885 | 0.61625 | 1.19788 | 184 |
| GO_GLUCOSE_CATABOLIC_PROCESS | 0.00495 | 0.04573 | 0.40702 | 0.72314 | 1.32174 | 28 |
| GO_GLUTAMATERGIC_SYNAPSE | 0.00135 | 0.01808 | 0.45506 | 0.60052 | 1.16783 | 186 |
| GO_GLYCOLYTIC_PROCESS_THROUGH_FRUCTOSE_6_PHOSPHATE | 0.00554 | 0.04968 | 0.40702 | 0.74156 | 1.34111 | 24 |
| GO_GOLGI_ASSOCIATED_VESICLE | 0.00085 | 0.01239 | 0.47727 | 0.62294 | 1.20037 | 129 |
| GO_GOLGI_ASSOCIATED_VESICLE_MEMBRANE | 0.00313 | 0.03310 | 0.43171 | 0.64208 | 1.21753 | 75 |
| GO_GTPASE_ACTIVITY | 0.00004 | 0.00098 | 0.55733 | 0.61546 | 1.19974 | 210 |
| GO_GUANYL_NUCLEOTIDE_BINDING | 0.00005 | 0.00126 | 0.55733 | 0.60899 | 1.19065 | 252 |
| GO_HEAT_SHOCK_PROTEIN_BINDING | 0.00001 | 0.00034 | 0.59333 | 0.68258 | 1.30633 | 93 |
| GO_HEMATOPOIETIC_PROGENITOR_CELL_DIFFERENTIATION | 0.00000 | 0.00000 | 0.71951 | 0.70099 | 1.35150 | 121 |
| GO_HEMATOPOIETIC_STEM_CELL_DIFFERENTIATION | 0.00000 | 0.00000 | 0.76146 | 0.77246 | 1.46307 | 72 |
| GO_HISTONE_MRNA_METABOLIC_PROCESS | 0.00265 | 0.02936 | 0.43171 | 0.76832 | 1.38459 | 22 |
| GO_HOST_CELLULAR_COMPONENT | 0.00043 | 0.00695 | 0.49849 | 0.69729 | 1.31074 | 54 |
| GO_IMMUNE_RESPONSE_REGULATING_SIGNALING_PATHWAY | 0.00000 | 0.00003 | 0.65944 | 0.63620 | 1.24112 | 225 |
| GO_IMPORT_INTO_CELL | 0.00428 | 0.04185 | 0.40702 | 0.61709 | 1.18655 | 108 |
| GO_IMPORT_INTO_NUCLEUS | 0.00039 | 0.00652 | 0.49849 | 0.62707 | 1.20905 | 131 |
| GO_INNATE_IMMUNE_RESPONSE | 0.00000 | 0.00003 | 0.65944 | 0.59893 | 1.18089 | 406 |
| GO_INNATE_IMMUNE_RESPONSE_ACTIVATING_SIGNAL_TRANSD | 0.00000 | 0.00001 | 0.69013 | 0.73348 | 1.39585 | 80 |
| GO_INNER_MITOCHONDRIAL_MEMBRANE_PROTEIN_COMPLEX | 0.00000 | 0.00000 | | 0.76576 | 1.46666 | 94 |
| GO_INORGANIC_ION_TRANSMEMBRANE_TRANSPORT | 0.00141 | 0.01850 | 0.45506 | 0.56404 | 1.11185 | 401 |
| GO_INTERACTION_WITH_HOST | 0.00139 | 0.01850 | 0.45506 | 0.60855 | 1.17710 | 151 |
| GO_INTERLEUKIN_1_MEDIATED_SIGNALING_PATHWAY | 0.00000 | 0.00000 | 0.77494 | 0.75654 | 1.44176 | 83 |
| GO_INTRAMOLECULAR_OXIDOREDUCTASE_ACTIVITY | 0.00051 | 0.00812 | 0.47727 | 0.74506 | 1.37081 | 34 |
| GO_INTRINSIC_APOPTOTIC_SIGNALING_PATHWAY | 0.00156 | 0.01993 | 0.45506 | 0.59427 | 1.15817 | 207 |
| GO_ION_HOMEOSTASIS | 0.00170 | 0.02117 | 0.45506 | 0.56164 | 1.10717 | 403 |
| GO_ION_TRANSMEMBRANE_TRANSPORTER_ACTIVITY | 0.00106 | 0.01495 | 0.45506 | 0.56696 | 1.11734 | 390 |
| GO_IRON_ION_HOMEOSTASIS | 0.00361 | 0.03657 | 0.43171 | 0.65837 | 1.23983 | 59 |
| GO_IRON_ION_TRANSPORT | 0.00054 | 0.00857 | 0.47727 | 0.69556 | 1.30334 | 52 |
| GO_ISOMERASE_ACTIVITY | 0.00413 | 0.04065 | 0.40702 | 0.61065 | 1.17496 | 113 |
| GO_KERATINIZATION | 0.00038 | 0.00640 | 0.49849 | 0.75690 | 1.39013 | 33 |
| GO_KILLING_OF_CELLS_OF_OTHER_ORGANISM | 0.00495 | 0.04573 | 0.40702 | 0.78614 | 1.38643 | 16 |
| GO_LARGE_RIBOSOMAL_SUBUNIT | 0.00000 | 0.00000 | | 0.77656 | 1.49319 | 108 |
| GO_LATERAL_PLASMA_MEMBRANE | 0.00524 | 0.04770 | 0.40702 | 0.67385 | 1.25557 | 47 |
| GO_LEUKOCYTE_MEDIATED_IMMUNITY | 0.00000 | 0.00000 | | 0.64010 | 1.26331 | 447 |
| GO_LIPOPROTEIN_PARTICLE_RECEPTOR_BINDING | 0.00126 | 0.01715 | 0.45506 | 0.80503 | 1.42497 | 17 |
| GO_LIVER_REGENERATION | 0.00093 | 0.01337 | 0.47727 | 0.78442 | 1.41708 | 23 |
| GO_LOW_DENSITY_LIPOPROTEIN_RECEPTOR_PARTICLE_METABO | 0.00327 | 0.03409 | 0.43171 | 0.78520 | 1.39657 | 18 |
| GO_LYASE_ACTIVITY | 0.00152 | 0.01958 | 0.45506 | 0.61769 | 1.19079 | 122 |
| GO_LYSOSOMAL_LUMEN | 0.00143 | 0.01863 | 0.45506 | 0.67371 | 1.26872 | 59 |
| GO_MAINTENANCE_OF_CELL_NUMBER | 0.00435 | 0.04238 | 0.40702 | 0.60761 | 1.17073 | 117 |
| GO_MATURATION_OF_SSU_RRNA | 0.00126 | 0.01715 | 0.45506 | 0.69615 | 1.29711 | 47 |
| GO_MATURATION_OF_SSU_RRNA_FROM_TRICISTRONIC_RRNA_ | 0.00554 | 0.04968 | 0.40702 | 0.70205 | 1.29301 | 35 |
| GO_MEMBRANE_REGION | 0.00000 | 0.00003 | 0.65944 | 0.64496 | 1.25542 | 194 |

| | | | | | | |
|---|---|---|---|---|---|---|
| GO_METALLOPEPTIDASE_ACTIVITY | 0.00487 | 0.04559 | 0.40702 | 0.62714 | 1.19577 | 85 |
| GO_METAL_ION_HOMEOSTASIS | 0.00032 | 0.00575 | 0.49849 | 0.58816 | 1.15376 | 301 |
| GO_METAL_ION_TRANSPORT | 0.00390 | 0.03895 | 0.43171 | 0.55877 | 1.10227 | 419 |
| GO_METAPHASE_ANAPHASE_TRANSITION_OF_CELL_CYCLE | 0.00320 | 0.03371 | 0.43171 | 0.66563 | 1.25122 | 54 |
| GO_MICROTUBULE_CYTOSKELETON_ORGANIZATION_INVOLVED_ | 0.00220 | 0.02548 | 0.43171 | 0.61295 | 1.18118 | 127 |
| GO_MICROVILLUS | 0.00168 | 0.02098 | 0.45506 | 0.68128 | 1.28184 | 56 |
| GO_MIDBODY | 0.00561 | 0.04971 | 0.40702 | 0.59866 | 1.15606 | 141 |
| GO_MITOCHONDRIAL_ELECTRON_TRANSPORT_NADH_TO_UBIQ | 0.00000 | 0.00006 | 0.64355 | 0.79447 | 1.47799 | 44 |
| GO_MITOCHONDRIAL_GENE_EXPRESSION | 0.00000 | 0.00000 | 0.78819 | 0.70230 | 1.35844 | 151 |
| GO_MITOCHONDRIAL_LARGE_RIBOSOMAL_SUBUNIT | 0.00000 | 0.00000 | 0.71951 | 0.79015 | 1.48529 | 54 |
| GO_MITOCHONDRIAL_MATRIX | 0.00000 | 0.00000 | 0.82666 | 0.63141 | 1.24412 | 387 |
| GO_MITOCHONDRIAL_MEMBRANE_ORGANIZATION | 0.00050 | 0.00802 | 0.47727 | 0.64578 | 1.23686 | 94 |
| GO_MITOCHONDRIAL_PROTEIN_COMPLEX | 0.00000 | 0.00000 | | 0.75011 | 1.46215 | 212 |
| GO_MITOCHONDRIAL_RESPIRATORY_CHAIN_COMPLEX_ASSEMB | 0.00001 | 0.00033 | 0.59333 | 0.69650 | 1.32804 | 85 |
| GO_MITOCHONDRIAL_SMALL_RIBOSOMAL_SUBUNIT | 0.00119 | 0.01640 | 0.45506 | 0.75384 | 1.37785 | 28 |
| GO_MITOCHONDRIAL_TRANSLATION | 0.00000 | 0.00000 | 0.80122 | 0.71435 | 1.37651 | 128 |
| GO_MITOCHONDRIAL_TRANSLATIONAL_TERMINATION | 0.00000 | 0.00000 | 0.81404 | 0.75760 | 1.44657 | 87 |
| GO_MITOCHONDRIAL_TRANSMEMBRANE_TRANSPORT | 0.00213 | 0.02490 | 0.43171 | 0.66300 | 1.25387 | 70 |
| GO_MITOCHONDRIAL_TRANSPORT | 0.00010 | 0.00227 | 0.53843 | 0.61119 | 1.19060 | 203 |
| GO_MITOCHONDRION_ORGANIZATION | 0.00000 | 0.00000 | 0.74774 | 0.61590 | 1.21471 | 416 |
| GO_MITOTIC_NUCLEAR_DIVISION | 0.00008 | 0.00182 | 0.53843 | 0.60435 | 1.17901 | 223 |
| GO_MITOTIC_SISTER_CHROMATID_SEGREGATION | 0.00014 | 0.00302 | 0.51885 | 0.62652 | 1.21048 | 145 |
| GO_MITOTIC_SPINDLE_ORGANIZATION | 0.00078 | 0.01164 | 0.47727 | 0.63943 | 1.22821 | 104 |
| GO_MODIFIED_AMINO_ACID_BINDING | 0.00002 | 0.00048 | 0.57561 | 0.74200 | 1.38679 | 49 |
| GO_MODULATION_BY_HOST_OF_VIRAL_PROCESS | 0.00216 | 0.02519 | 0.43171 | 0.77232 | 1.38066 | 20 |
| GO_MONOCARBOXYLIC_ACID_METABOLIC_PROCESS | 0.00012 | 0.00265 | 0.53843 | 0.57846 | 1.14053 | 406 |
| GO_MONOSACCHARIDE_BIOSYNTHETIC_PROCESS | 0.00305 | 0.03271 | 0.43171 | 0.65908 | 1.24472 | 68 |
| GO_MONOVALENT_INORGANIC_CATION_HOMEOSTASIS | 0.00108 | 0.01507 | 0.45506 | 0.63643 | 1.21783 | 91 |
| GO_MONOVALENT_INORGANIC_CATION_TRANSMEMBRANE_TR | 0.00001 | 0.00026 | 0.59333 | 0.65186 | 1.25996 | 147 |
| GO_MONOVALENT_INORGANIC_CATION_TRANSPORT | 0.00008 | 0.00178 | 0.53843 | 0.60205 | 1.17659 | 248 |
| GO_MORPHOGENESIS_OF_AN_EPITHELIUM | 0.00000 | 0.00009 | 0.62726 | 0.60570 | 1.19137 | 334 |
| GO_MORPHOGENESIS_OF_A_POLARIZED_EPITHELIUM | 0.00000 | 0.00000 | 0.83909 | 0.74384 | 1.43057 | 110 |
| GO_MOVEMENT_IN_HOST_ENVIRONMENT | 0.00268 | 0.02961 | 0.43171 | 0.61282 | 1.18078 | 117 |
| GO_MRNA_3_END_PROCESSING | 0.00087 | 0.01270 | 0.47727 | 0.64010 | 1.22502 | 93 |
| GO_MRNA_5_UTR_BINDING | 0.00002 | 0.00063 | 0.57561 | 0.83759 | 1.51313 | 23 |
| GO_MRNA_BINDING | 0.00000 | 0.00000 | | 0.67496 | 1.31810 | 239 |
| GO_MRNA_EXPORT_FROM_NUCLEUS | 0.00006 | 0.00159 | 0.53843 | 0.66290 | 1.27265 | 103 |
| GO_MRNA_PROCESSING | 0.00000 | 0.00000 | | 0.62959 | 1.24255 | 437 |
| GO_MRNA_TRANSPORT | 0.00007 | 0.00171 | 0.53843 | 0.64103 | 1.23563 | 132 |
| GO_MULTI_MULTICELLULAR_ORGANISM_PROCESS | 0.00041 | 0.00687 | 0.49849 | 0.63087 | 1.21515 | 116 |
| GO_MULTI_ORGANISM_LOCALIZATION | 0.00017 | 0.00334 | 0.51885 | 0.71077 | 1.33608 | 54 |
| GO_MUSCLE_CONTRACTION | 0.00335 | 0.03468 | 0.43171 | 0.59925 | 1.16199 | 165 |
| GO_MUSCLE_FILAMENT_SLIDING | 0.00324 | 0.03393 | 0.43171 | 0.77980 | 1.39322 | 19 |
| GO_MYELOID_LEUKOCYTE_ACTIVATION | 0.00000 | 0.00000 | | 0.65288 | 1.28712 | 395 |
| GO_MYELOID_LEUKOCYTE_MEDIATED_IMMUNITY | 0.00000 | 0.00000 | | 0.65734 | 1.29408 | 356 |
| GO_NADH_DEHYDROGENASE_ACTIVITY | 0.00001 | 0.00019 | 0.61053 | 0.79360 | 1.46691 | 38 |
| GO_NADH_DEHYDROGENASE_COMPLEX | 0.00001 | 0.00028 | 0.59333 | 0.77749 | 1.44215 | 41 |
| GO_NADH_DEHYDROGENASE_COMPLEX_ASSEMBLY | 0.00001 | 0.00035 | 0.59333 | 0.74044 | 1.39281 | 55 |
| GO_NADH_METABOLIC_PROCESS | 0.00034 | 0.00599 | 0.49849 | 0.75773 | 1.39166 | 33 |
| GO_NAD_METABOLIC_PROCESS | 0.00435 | 0.04238 | 0.40702 | 0.70570 | 1.30442 | 37 |
| GO_NCRNA_EXPORT_FROM_NUCLEUS | 0.00205 | 0.02440 | 0.43171 | 0.71473 | 1.31817 | 36 |
| GO_NCRNA_METABOLIC_PROCESS | 0.00000 | 0.00000 | 0.70498 | 0.61139 | 1.20509 | 399 |
| GO_NCRNA_PROCESSING | 0.00000 | 0.00000 | 0.73376 | 0.62238 | 1.22425 | 341 |
| GO_NEGATIVE_REGULATION_OF_APOPTOTIC_SIGNALING_PATHW | 0.00364 | 0.03677 | 0.43171 | 0.59958 | 1.15998 | 150 |
| GO_NEGATIVE_REGULATION_OF_ATP_METABOLIC_PROCESS | 0.00302 | 0.03248 | 0.43171 | 0.78586 | 1.39103 | 17 |
| GO_NEGATIVE_REGULATION_OF_CANONICAL_WNT_SIGNALING | 0.00000 | 0.00000 | 0.71951 | 0.69629 | 1.34180 | 123 |
| GO_NEGATIVE_REGULATION_OF_CATALYTIC_ACTIVITY | 0.00145 | 0.01879 | 0.45506 | 0.56238 | 1.11005 | 450 |
| GO_NEGATIVE_REGULATION_OF_CELLULAR_AMIDE_METABOLIC | 0.00268 | 0.02961 | 0.43171 | 0.61074 | 1.17686 | 124 |
| GO_NEGATIVE_REGULATION_OF_CELLULAR_COMPONENT_ORGA | 0.00320 | 0.03371 | 0.43171 | 0.55470 | 1.09534 | 470 |
| GO_NEGATIVE_REGULATION_OF_CELL_CYCLE | 0.00023 | 0.00445 | 0.51885 | 0.57348 | 1.13228 | 455 |
| GO_NEGATIVE_REGULATION_OF_CELL_CYCLE_G2_M_PHASE_TR | 0.00000 | 0.00007 | 0.62726 | 0.70549 | 1.34606 | 86 |
| GO_NEGATIVE_REGULATION_OF_CELL_CYCLE_PHASE_TRANSITIO | 0.00000 | 0.00000 | 0.70498 | 0.65532 | 1.27560 | 194 |
| GO_NEGATIVE_REGULATION_OF_CELL_CYCLE_PROCESS | 0.00000 | 0.00005 | 0.64355 | 0.62725 | 1.22670 | 259 |
| GO_NEGATIVE_REGULATION_OF_HYDROLASE_ACTIVITY | 0.00216 | 0.02519 | 0.43171 | 0.58306 | 1.13858 | 238 |
| GO_NEGATIVE_REGULATION_OF_INTERLEUKIN_1_PRODUCTION | 0.00205 | 0.02440 | 0.43171 | 0.82297 | 1.44404 | 15 |
| GO_NEGATIVE_REGULATION_OF_MITOTIC_CELL_CYCLE | 0.00000 | 0.00010 | 0.62726 | 0.62191 | 1.21541 | 248 |
| GO_NEGATIVE_REGULATION_OF_MRNA_METABOLIC_PROCESS | 0.00213 | 0.02490 | 0.43171 | 0.66731 | 1.25879 | 62 |
| GO_NEGATIVE_REGULATION_OF_MRNA_SPLICING_VIA_SPLICEO | 0.00339 | 0.03501 | 0.43171 | 0.76309 | 1.36417 | 20 |
| GO_NEGATIVE_REGULATION_OF_NUCLEAR_DIVISION | 0.00335 | 0.03468 | 0.43171 | 0.67724 | 1.26685 | 50 |
| GO_NEGATIVE_REGULATION_OF_ORGANELLE_ORGANIZATION | 0.00487 | 0.04559 | 0.40702 | 0.57038 | 1.11722 | 288 |

| | | | | | | |
|---|---|---|---|---|---|---|
| GO_NEGATIVE_REGULATION_OF_PEPTIDASE_ACTIVITY | 0.00036 | 0.00621 | 0.49849 | 0.64044 | 1.23159 | 109 |
| GO_NEGATIVE_REGULATION_OF_PROTEIN_CATABOLIC_PROCESS | 0.00342 | 0.03521 | 0.43171 | 0.62320 | 1.19268 | 93 |
| GO_NEGATIVE_REGULATION_OF_PROTEOLYSIS | 0.00205 | 0.02440 | 0.43171 | 0.60172 | 1.16902 | 178 |
| GO_NEGATIVE_REGULATION_OF_REACTIVE_OXYGEN_SPECIES_B | 0.00036 | 0.00621 | 0.49849 | 0.83108 | 1.47818 | 18 |
| GO_NEGATIVE_REGULATION_OF_REPRODUCTIVE_PROCESS | 0.00457 | 0.04386 | 0.40702 | 0.74845 | 1.35356 | 24 |
| GO_NEGATIVE_REGULATION_OF_RNA_SPLICING | 0.00183 | 0.02219 | 0.45506 | 0.76824 | 1.38935 | 24 |
| GO_NEGATIVE_REGULATION_OF_WNT_SIGNALING_PATHWAY | 0.00000 | 0.00002 | 0.67496 | 0.67731 | 1.30721 | 136 |
| GO_NEURAL_CREST_CELL_MIGRATION | 0.00265 | 0.02936 | 0.43171 | 0.73967 | 1.35049 | 27 |
| GO_NEURON_RECOGNITION | 0.00390 | 0.03895 | 0.43171 | 0.71862 | 1.31998 | 31 |
| GO_NIK_NF_KAPPAB_SIGNALING | 0.00000 | 0.00000 | 0.77494 | 0.72329 | 1.39253 | 114 |
| GO_NON_CANONICAL_WNT_SIGNALING_PATHWAY | 0.00000 | 0.00000 | | 0.75756 | 1.45654 | 107 |
| GO_NUCLEAR_CHROMOSOME_SEGREGATION | 0.00098 | 0.01407 | 0.45506 | 0.59156 | 1.15357 | 215 |
| GO_NUCLEAR_ENVELOPE | 0.00005 | 0.00131 | 0.55733 | 0.59109 | 1.16270 | 341 |
| GO_NUCLEAR_EXPORT | 0.00000 | 0.00004 | 0.65944 | 0.64915 | 1.26157 | 181 |
| GO_NUCLEAR_MATRIX | 0.00502 | 0.04608 | 0.40702 | 0.62777 | 1.19868 | 87 |
| GO_NUCLEAR_MEMBRANE | 0.00185 | 0.02237 | 0.45506 | 0.58925 | 1.14853 | 211 |
| GO_NUCLEAR_PERIPHERY | 0.00043 | 0.00701 | 0.49849 | 0.64264 | 1.23558 | 107 |
| GO_NUCLEAR_PORE | 0.00561 | 0.04971 | 0.40702 | 0.64876 | 1.22522 | 68 |
| GO_NUCLEAR_SPECK | 0.00015 | 0.00312 | 0.51885 | 0.59159 | 1.16089 | 306 |
| GO_NUCLEAR_TRANSCRIBED_MRNA_CATABOLIC_PROCESS | 0.00000 | 0.00000 | | 0.70362 | 1.36820 | 185 |
| GO_NUCLEAR_TRANSCRIBED_MRNA_CATABOLIC_PROCESS_NON | 0.00000 | 0.00000 | | 0.74239 | 1.42931 | 114 |
| GO_NUCLEAR_TRANSPORT | 0.00000 | 0.00000 | 0.71951 | 0.62920 | 1.23383 | 297 |
| GO_NUCLEOBASE_CONTAINING_COMPOUND_TRANSPORT | 0.00000 | 0.00010 | 0.62726 | 0.62975 | 1.22823 | 216 |
| GO_NUCLEOBASE_CONTAINING_SMALL_MOLECULE_METABOLIC | 0.00000 | 0.00003 | 0.65944 | 0.60153 | 1.18594 | 405 |
| GO_NUCLEOSIDE_DIPHOSPHATE_METABOLIC_PROCESS | 0.00016 | 0.00319 | 0.51885 | 0.64491 | 1.24260 | 117 |
| GO_NUCLEOSIDE_PHOSPHATE_BIOSYNTHETIC_PROCESS | 0.00002 | 0.00044 | 0.59333 | 0.64092 | 1.24175 | 161 |
| GO_NUCLEOSIDE_TRIPHOSPHATE_BIOSYNTHETIC_PROCESS | 0.00040 | 0.00676 | 0.49849 | 0.72727 | 1.35074 | 42 |
| GO_NUCLEOSIDE_TRIPHOSPHATE_METABOLIC_PROCESS | 0.00009 | 0.00196 | 0.53843 | 0.69306 | 1.30854 | 65 |
| GO_NUCLEOTIDE_PHOSPHORYLATION | 0.00016 | 0.00328 | 0.51885 | 0.65541 | 1.25767 | 102 |
| GO_OOCYTE_DIFFERENTIATION | 0.00450 | 0.04342 | 0.40702 | 0.73513 | 1.33705 | 26 |
| GO_ORGANELLAR_RIBOSOME | 0.00000 | 0.00000 | 0.76146 | 0.75712 | 1.44311 | 84 |
| GO_ORGANELLE_ENVELOPE_LUMEN | 0.00026 | 0.00482 | 0.49849 | 0.67265 | 1.27716 | 78 |
| GO_ORGANELLE_FISSION | 0.00154 | 0.01974 | 0.45506 | 0.57133 | 1.12383 | 342 |
| GO_ORGANELLE_INNER_MEMBRANE | 0.00000 | 0.00000 | | 0.68423 | 1.34915 | 412 |
| GO_ORGANIC_CYCLIC_COMPOUND_CATABOLIC_PROCESS | 0.00000 | 0.00000 | | 0.66975 | 1.32263 | 478 |
| GO_ORGANIC_HYDROXY_COMPOUND_BIOSYNTHETIC_PROCESS | 0.00141 | 0.01850 | 0.45506 | 0.60712 | 1.17627 | 161 |
| GO_OXIDATIVE_PHOSPHORYLATION | 0.00000 | 0.00000 | | 0.78902 | 1.51260 | 99 |
| GO_OXIDOREDUCTASE_ACTIVITY | 0.00000 | 0.00000 | | 0.63268 | 1.24935 | 468 |
| GO_OXIDOREDUCTASE_ACTIVITY_ACTING_ON_A_HEME_GROUP | 0.00176 | 0.02168 | 0.45506 | 0.79383 | 1.41828 | 19 |
| GO_OXIDOREDUCTASE_ACTIVITY_ACTING_ON_NAD_P_H | 0.00002 | 0.00056 | 0.57561 | 0.71468 | 1.35162 | 70 |
| GO_OXIDOREDUCTASE_ACTIVITY_ACTING_ON_NAD_P_H_QUIN | 0.00004 | 0.00113 | 0.55733 | 0.74511 | 1.38835 | 47 |
| GO_OXIDOREDUCTASE_ACTIVITY_ACTING_ON_PEROXIDE_AS_A | 0.00106 | 0.01495 | 0.45506 | 0.73897 | 1.35352 | 29 |
| GO_OXIDOREDUCTASE_COMPLEX | 0.00000 | 0.00000 | 0.73376 | 0.74898 | 1.42731 | 82 |
| GO_PEPTIDASE_ACTIVITY | 0.00133 | 0.01799 | 0.45506 | 0.57681 | 1.13188 | 306 |
| GO_PEPTIDASE_COMPLEX | 0.00000 | 0.00002 | 0.67496 | 0.73250 | 1.39240 | 79 |
| GO_PEPTIDASE_REGULATOR_ACTIVITY | 0.00313 | 0.03310 | 0.43171 | 0.63434 | 1.21189 | 88 |
| GO_PEPTIDE_BINDING | 0.00005 | 0.00124 | 0.55733 | 0.63708 | 1.23187 | 148 |
| GO_PERINUCLEAR_REGION_OF_CYTOPLASM | 0.00276 | 0.03021 | 0.43171 | 0.55814 | 1.10207 | 466 |
| GO_PHAGOCYTOSIS | 0.00554 | 0.04968 | 0.40702 | 0.59337 | 1.14837 | 155 |
| GO_PHAGOSOME_ACIDIFICATION | 0.00059 | 0.00916 | 0.47727 | 0.78902 | 1.42694 | 24 |
| GO_PHOSPHATIDYLSERINE_BINDING | 0.00035 | 0.00610 | 0.49849 | 0.76387 | 1.39912 | 29 |
| GO_PH_REDUCTION | 0.00302 | 0.03248 | 0.43171 | 0.68821 | 1.28238 | 46 |
| GO_PIGMENT_GRANULE | 0.00000 | 0.00000 | 0.76146 | 0.74333 | 1.41934 | 87 |
| GO_PLASMA_LIPOPROTEIN_PARTICLE_CLEARANCE | 0.00106 | 0.01495 | 0.45506 | 0.73201 | 1.35304 | 37 |
| GO_POLYSOMAL_RIBOSOME | 0.00008 | 0.00180 | 0.53843 | 0.78840 | 1.44683 | 30 |
| GO_POLYSOME | 0.00000 | 0.00003 | 0.65944 | 0.75964 | 1.43296 | 62 |
| GO_POSITIVE_REGULATION_OF_ATP_METABOLIC_PROCESS | 0.00174 | 0.02149 | 0.45506 | 0.73011 | 1.33729 | 29 |
| GO_POSITIVE_REGULATION_OF_BINDING | 0.00390 | 0.03895 | 0.43171 | 0.61042 | 1.17582 | 115 |
| GO_POSITIVE_REGULATION_OF_CANONICAL_WNT_SIGNALING_ | 0.00000 | 0.00000 | 0.73376 | 0.72036 | 1.38514 | 108 |
| GO_POSITIVE_REGULATION_OF_CELLULAR_AMIDE_METABOLIC_ | 0.00001 | 0.00036 | 0.59333 | 0.66485 | 1.28095 | 125 |
| GO_POSITIVE_REGULATION_OF_CELLULAR_PROTEIN_LOCALIZAT | 0.00242 | 0.02748 | 0.43171 | 0.58048 | 1.13378 | 244 |
| GO_POSITIVE_REGULATION_OF_CELL_POPULATION_PROLIFERAT | 0.00040 | 0.00670 | 0.49849 | 0.56967 | 1.12503 | 461 |
| GO_POSITIVE_REGULATION_OF_CYSTEINE_TYPE_ENDOPEPTIDAS | 0.00547 | 0.04923 | 0.40702 | 0.62225 | 1.19077 | 92 |
| GO_POSITIVE_REGULATION_OF_CYTOKINE_PRODUCTION | 0.00283 | 0.03081 | 0.43171 | 0.58672 | 1.14373 | 210 |
| GO_POSITIVE_REGULATION_OF_DEFENSE_RESPONSE | 0.00000 | 0.00000 | 0.73376 | 0.66200 | 1.28884 | 200 |
| GO_POSITIVE_REGULATION_OF_DNA_BIOSYNTHETIC_PROCESS | 0.00010 | 0.00228 | 0.53843 | 0.71224 | 1.33977 | 55 |
| GO_POSITIVE_REGULATION_OF_DNA_METABOLIC_PROCESS | 0.00020 | 0.00387 | 0.51885 | 0.62771 | 1.21205 | 139 |
| GO_POSITIVE_REGULATION_OF_ESTABLISHMENT_OF_PROTEIN_ | 0.00194 | 0.02331 | 0.45506 | 0.58158 | 1.13709 | 253 |
| GO_POSITIVE_REGULATION_OF_FIBROBLAST_PROLIFERATION | 0.00017 | 0.00338 | 0.51885 | 0.77423 | 1.42213 | 31 |
| GO_POSITIVE_REGULATION_OF_IMMUNE_RESPONSE | 0.00000 | 0.00000 | 0.71951 | 0.63064 | 1.23937 | 321 |

| | | | | | | |
|---|---|---|---|---|---|---|
| GO_POSITIVE_REGULATION_OF_IMMUNE_SYSTEM_PROCESS | 0.00000 | 0.00002 | 0.67496 | 0.59748 | 1.17992 | 475 |
| GO_POSITIVE_REGULATION_OF_INNATE_IMMUNE_RESPONSE | 0.00000 | 0.00000 | 0.70498 | 0.68885 | 1.32737 | 128 |
| GO_POSITIVE_REGULATION_OF_MITOTIC_CELL_CYCLE | 0.00178 | 0.02182 | 0.45506 | 0.61563 | 1.18683 | 122 |
| GO_POSITIVE_REGULATION_OF_MRNA_METABOLIC_PROCESS | 0.00157 | 0.02009 | 0.45506 | 0.66330 | 1.25231 | 63 |
| GO_POSITIVE_REGULATION_OF_MRNA_PROCESSING | 0.00250 | 0.02807 | 0.43171 | 0.76194 | 1.38151 | 25 |
| GO_POSITIVE_REGULATION_OF_MRNA_SPLICING_VIA_SPLICEOS | 0.00132 | 0.01782 | 0.45506 | 0.81851 | 1.44353 | 16 |
| GO_POSITIVE_REGULATION_OF_NITRIC_OXIDE_METABOLIC_PR( | 0.00133 | 0.01799 | 0.45506 | 0.80807 | 1.43726 | 18 |
| GO_POSITIVE_REGULATION_OF_ORGANELLE_ORGANIZATION | 0.00010 | 0.00218 | 0.53843 | 0.57473 | 1.13449 | 452 |
| GO_POSITIVE_REGULATION_OF_PROTEIN_LOCALIZATION_TO_N | 0.00012 | 0.00265 | 0.53843 | 0.71018 | 1.33697 | 58 |
| GO_POSITIVE_REGULATION_OF_PROTEOLYSIS | 0.00313 | 0.03310 | 0.43171 | 0.58059 | 1.13381 | 237 |
| GO_POSITIVE_REGULATION_OF_REACTIVE_OXYGEN_SPECIES_BI | 0.00115 | 0.01593 | 0.45506 | 0.79865 | 1.43554 | 21 |
| GO_POSITIVE_REGULATION_OF_REACTIVE_OXYGEN_SPECIES_M | 0.00174 | 0.02149 | 0.45506 | 0.69785 | 1.30035 | 46 |
| GO_POSITIVE_REGULATION_OF_RESPONSE_TO_BIOTIC_STIMUL | 0.00000 | 0.00000 | 0.70498 | 0.67507 | 1.30603 | 150 |
| GO_POSITIVE_REGULATION_OF_RESPONSE_TO_EXTERNAL_STIM | 0.00001 | 0.00016 | 0.61053 | 0.62123 | 1.21505 | 260 |
| GO_POSITIVE_REGULATION_OF_RNA_SPLICING | 0.00265 | 0.02936 | 0.43171 | 0.74051 | 1.35202 | 27 |
| GO_POSITIVE_REGULATION_OF_SIGNAL_TRANSDUCTION_BY_P5 | 0.00390 | 0.03895 | 0.43171 | 0.80515 | 1.41278 | 15 |
| GO_POSITIVE_REGULATION_OF_TELOMERASE_ACTIVITY | 0.00069 | 0.01043 | 0.47727 | 0.75265 | 1.38122 | 30 |
| GO_POSITIVE_REGULATION_OF_TELOMERASE_RNA_LOCALIZATI | 0.00007 | 0.00169 | 0.53843 | 0.88616 | 1.55493 | 15 |
| GO_POSITIVE_REGULATION_OF_TELOMERE_MAINTENANCE | 0.00026 | 0.00486 | 0.49849 | 0.73697 | 1.37102 | 44 |
| GO_POSITIVE_REGULATION_OF_TELOMERE_MAINTENANCE_VIA | 0.00001 | 0.00040 | 0.59333 | 0.80196 | 1.47289 | 33 |
| GO_POSITIVE_REGULATION_OF_TRANSFERASE_ACTIVITY | 0.00108 | 0.01507 | 0.45506 | 0.56501 | 1.11376 | 401 |
| GO_POSITIVE_REGULATION_OF_TRANSLATION | 0.00000 | 0.00016 | 0.61053 | 0.68408 | 1.31526 | 107 |
| GO_POSITIVE_REGULATION_OF_VIRAL_PROCESS | 0.00026 | 0.00486 | 0.49849 | 0.66195 | 1.26462 | 88 |
| GO_POSITIVE_REGULATION_OF_WNT_SIGNALING_PATHWAY | 0.00000 | 0.00000 | 0.73376 | 0.69924 | 1.34747 | 127 |
| GO_POSTSYNAPSE | 0.00239 | 0.02716 | 0.43171 | 0.57062 | 1.12116 | 316 |
| GO_POST_TRANSLATIONAL_PROTEIN_MODIFICATION | 0.00024 | 0.00452 | 0.51885 | 0.60046 | 1.17392 | 251 |
| GO_PRECATALYTIC_SPLICEOSOME | 0.00141 | 0.01850 | 0.45506 | 0.69251 | 1.29540 | 50 |
| GO_PRERIBOSOME | 0.00006 | 0.00154 | 0.53843 | 0.69866 | 1.32020 | 69 |
| GO_PROTEASOMAL_PROTEIN_CATABOLIC_PROCESS | 0.00007 | 0.00169 | 0.53843 | 0.58188 | 1.14652 | 387 |
| GO_PROTEASOMAL_UBIQUITIN_INDEPENDENT_PROTEIN_CATAE | 0.00000 | 0.00003 | 0.65944 | 0.91182 | 1.62178 | 18 |
| GO_PROTEASOME_ACCESSORY_COMPLEX | 0.00005 | 0.00135 | 0.55733 | 0.82638 | 1.49450 | 24 |
| GO_PROTEASOME_CORE_COMPLEX | 0.00000 | 0.00004 | 0.65944 | 0.91231 | 1.61486 | 17 |
| GO_PROTEIN_CONTAINING_COMPLEX_DISASSEMBLY | 0.00000 | 0.00000 | 0.81404 | 0.66285 | 1.29656 | 261 |
| GO_PROTEIN_CONTAINING_COMPLEX_LOCALIZATION | 0.00018 | 0.00352 | 0.51885 | 0.60451 | 1.17943 | 226 |
| GO_PROTEIN_DNA_COMPLEX_SUBUNIT_ORGANIZATION | 0.00052 | 0.00825 | 0.47727 | 0.60887 | 1.18342 | 183 |
| GO_PROTEIN_DOMAIN_SPECIFIC_BINDING | 0.00003 | 0.00088 | 0.55733 | 0.57828 | 1.14190 | 470 |
| GO_PROTEIN_FOLDING | 0.00000 | 0.00000 | 0.73376 | 0.67425 | 1.30867 | 171 |
| GO_PROTEIN_FOLDING_CHAPERONE | 0.00079 | 0.01170 | 0.47727 | 0.81632 | 1.45193 | 18 |
| GO_PROTEIN_HETEROOLIGOMERIZATION | 0.00442 | 0.04297 | 0.40702 | 0.78998 | 1.39322 | 16 |
| GO_PROTEIN_IMPORT | 0.00001 | 0.00040 | 0.59333 | 0.64197 | 1.24306 | 158 |
| GO_PROTEIN_IMPORT_INTO_MITOCHONDRIAL_MATRIX | 0.00231 | 0.02642 | 0.43171 | 0.79530 | 1.41455 | 18 |
| GO_PROTEIN_INSERTION_INTO_MEMBRANE | 0.00191 | 0.02299 | 0.45506 | 0.68322 | 1.27694 | 49 |
| GO_PROTEIN_LOCALIZATION_TO_CHROMOSOME_TELOMERIC_F | 0.00001 | 0.00031 | 0.59333 | 0.82810 | 1.51357 | 28 |
| GO_PROTEIN_LOCALIZATION_TO_ENDOPLASMIC_RETICULUM | 0.00000 | 0.00000 | | 0.73850 | 1.42303 | 129 |
| GO_PROTEIN_LOCALIZATION_TO_MEMBRANE | 0.00000 | 0.00000 | 0.77494 | 0.61216 | 1.20881 | 462 |
| GO_PROTEIN_LOCALIZATION_TO_MITOCHONDRION | 0.00015 | 0.00310 | 0.51885 | 0.64234 | 1.23759 | 118 |
| GO_PROTEIN_LOCALIZATION_TO_NUCLEUS | 0.00000 | 0.00014 | 0.61053 | 0.63315 | 1.23408 | 208 |
| GO_PROTEIN_MATURATION | 0.00357 | 0.03638 | 0.43171 | 0.59209 | 1.14965 | 174 |
| GO_PROTEIN_MODIFICATION_BY_SMALL_PROTEIN_REMOVAL | 0.00002 | 0.00052 | 0.57561 | 0.61535 | 1.20119 | 231 |
| GO_PROTEIN_REFOLDING | 0.00465 | 0.04430 | 0.40702 | 0.77561 | 1.37288 | 17 |
| GO_PROTEIN_STABILIZATION | 0.00081 | 0.01194 | 0.47727 | 0.61881 | 1.19577 | 146 |
| GO_PROTEIN_SUMOYLATION | 0.00450 | 0.04342 | 0.40702 | 0.65043 | 1.22846 | 67 |
| GO_PROTEIN_TARGETING | 0.00000 | 0.00000 | | 0.66223 | 1.30361 | 358 |
| GO_PROTEIN_TARGETING_TO_MEMBRANE | 0.00000 | 0.00000 | | 0.71675 | 1.38879 | 162 |
| GO_PROTEIN_TARGETING_TO_MITOCHONDRION | 0.00113 | 0.01571 | 0.45506 | 0.65407 | 1.24648 | 83 |
| GO_PROTEIN_TRANSMEMBRANE_TRANSPORT | 0.00110 | 0.01526 | 0.45506 | 0.68173 | 1.28238 | 55 |
| GO_PROTON_TRANSMEMBRANE_TRANSPORT | 0.00000 | 0.00001 | 0.69013 | 0.71234 | 1.36309 | 91 |
| GO_PROTON_TRANSMEMBRANE_TRANSPORTER_ACTIVITY | 0.00004 | 0.00115 | 0.55733 | 0.70138 | 1.32535 | 69 |
| GO_PROTON_TRANSPORTING_TWO_SECTOR_ATPASE_COMPLEX | 0.00083 | 0.01218 | 0.47727 | 0.77254 | 1.40508 | 26 |
| GO_PROTON_TRANSPORTING_V_TYPE_ATPASE_COMPLEX | 0.00239 | 0.02716 | 0.43171 | 0.76100 | 1.37626 | 24 |
| GO_PURINE_CONTAINING_COMPOUND_BIOSYNTHETIC_PROCES | 0.00001 | 0.00040 | 0.59333 | 0.65863 | 1.26901 | 126 |
| GO_PURINE_CONTAINING_COMPOUND_METABOLIC_PROCESS | 0.00000 | 0.00000 | 0.73376 | 0.63132 | 1.23831 | 300 |
| GO_PURINE_NUCLEOSIDE_TRIPHOSPHATE_METABOLIC_PROCES | 0.00033 | 0.00582 | 0.49849 | 0.70887 | 1.32487 | 49 |
| GO_PYROPHOSPHATE_HYDROLYSIS_DRIVEN_PROTON_TRANSMI | 0.00532 | 0.04803 | 0.40702 | 0.74470 | 1.34678 | 24 |
| GO_PYRUVATE_METABOLIC_PROCESS | 0.00001 | 0.00018 | 0.61053 | 0.67194 | 1.29289 | 113 |
| GO_P_BODY | 0.00472 | 0.04480 | 0.40702 | 0.65470 | 1.23501 | 62 |
| GO_REACTIVE_OXYGEN_SPECIES_BIOSYNTHETIC_PROCESS | 0.00040 | 0.00670 | 0.49849 | 0.70271 | 1.31857 | 53 |
| GO_REACTIVE_OXYGEN_SPECIES_METABOLIC_PROCESS | 0.00021 | 0.00409 | 0.51885 | 0.62881 | 1.21472 | 144 |
| GO_REGULATION_OF_ALCOHOL_BIOSYNTHETIC_PROCESS | 0.00228 | 0.02614 | 0.43171 | 0.68438 | 1.28020 | 50 |
| GO_REGULATION_OF_ANIMAL_ORGAN_MORPHOGENESIS | 0.00000 | 0.00000 | | 0.72330 | 1.39922 | 149 |

| | | | | | | |
|---|---|---|---|---|---|---|
| GO_REGULATION_OF_APOPTOTIC_SIGNALING_PATHWAY | 0.00079 | 0.01170 | 0.47727 | 0.58599 | 1.14641 | 266 |
| GO_REGULATION_OF_ATP_METABOLIC_PROCESS | 0.00000 | 0.00006 | 0.64355 | 0.70523 | 1.34777 | 89 |
| GO_REGULATION_OF_BINDING | 0.00428 | 0.04185 | 0.40702 | 0.58264 | 1.13664 | 225 |
| GO_REGULATION_OF_BLOOD_PRESSURE | 0.00290 | 0.03156 | 0.43171 | 0.65319 | 1.23428 | 69 |
| GO_REGULATION_OF_CARBOHYDRATE_CATABOLIC_PROCESS | 0.00324 | 0.03393 | 0.43171 | 0.65541 | 1.23787 | 67 |
| GO_REGULATION_OF_CELLULAR_AMIDE_METABOLIC_PROCESS | 0.00000 | 0.00000 | 0.78819 | 0.63116 | 1.24143 | 337 |
| GO_REGULATION_OF_CELLULAR_AMINE_METABOLIC_PROCESS | 0.00000 | 0.00000 | 0.74774 | 0.78955 | 1.48730 | 60 |
| GO_REGULATION_OF_CELLULAR_AMINO_ACID_METABOLIC_PR( | 0.00000 | 0.00000 | 0.76146 | 0.80341 | 1.51022 | 54 |
| GO_REGULATION_OF_CELLULAR_KETONE_METABOLIC_PROCESS | 0.00000 | 0.00000 | 0.70498 | 0.69239 | 1.33405 | 126 |
| GO_REGULATION_OF_CELLULAR_PH | 0.00442 | 0.04297 | 0.40702 | 0.65360 | 1.23436 | 68 |
| GO_REGULATION_OF_CELLULAR_PROTEIN_LOCALIZATION | 0.00228 | 0.02614 | 0.43171 | 0.56216 | 1.10830 | 402 |
| GO_REGULATION_OF_CELLULAR_RESPIRATION | 0.00428 | 0.04185 | 0.40702 | 0.77183 | 1.37897 | 19 |
| GO_REGULATION_OF_CELLULAR_RESPONSE_TO_HEAT | 0.00091 | 0.01314 | 0.47727 | 0.67360 | 1.27221 | 67 |
| GO_REGULATION_OF_CELL_CYCLE_G2_M_PHASE_TRANSITION | 0.00016 | 0.00332 | 0.51885 | 0.61554 | 1.19738 | 187 |
| GO_REGULATION_OF_CELL_CYCLE_PHASE_TRANSITION | 0.00000 | 0.00014 | 0.61053 | 0.59968 | 1.18091 | 369 |
| GO_REGULATION_OF_CHOLESTEROL_BIOSYNTHETIC_PROCESS | 0.00209 | 0.02470 | 0.43171 | 0.71476 | 1.32117 | 38 |
| GO_REGULATION_OF_CHOLESTEROL_METABOLIC_PROCESS | 0.00113 | 0.01571 | 0.45506 | 0.69865 | 1.30177 | 47 |
| GO_REGULATION_OF_CHROMOSOME_ORGANIZATION | 0.00036 | 0.00622 | 0.49849 | 0.58896 | 1.15261 | 278 |
| GO_REGULATION_OF_CHROMOSOME_SEGREGATION | 0.00079 | 0.01170 | 0.47727 | 0.64131 | 1.22718 | 91 |
| GO_REGULATION_OF_CHROMOSOME_SEPARATION | 0.00253 | 0.02838 | 0.43171 | 0.66349 | 1.24947 | 59 |
| GO_REGULATION_OF_CYSTEINE_TYPE_ENDOPEPTIDASE_ACTIVIT | 0.00044 | 0.00719 | 0.49849 | 0.62610 | 1.20905 | 141 |
| GO_REGULATION_OF_DEFENSE_RESPONSE | 0.00006 | 0.00150 | 0.55733 | 0.58834 | 1.15824 | 356 |
| GO_REGULATION_OF_DNA_BIOSYNTHETIC_PROCESS | 0.00043 | 0.00695 | 0.49849 | 0.66709 | 1.27130 | 83 |
| GO_REGULATION_OF_DNA_METABOLIC_PROCESS | 0.00055 | 0.00863 | 0.47727 | 0.58909 | 1.15134 | 249 |
| GO_REGULATION_OF_DNA_TEMPLATED_TRANSCRIPTION_IN_RI | 0.00000 | 0.00000 | 0.71951 | 0.72410 | 1.38687 | 94 |
| GO_REGULATION_OF_GENERATION_OF_PRECURSOR_METABOLI | 0.00004 | 0.00114 | 0.55733 | 0.65536 | 1.26115 | 111 |
| GO_REGULATION_OF_GENE_SILENCING | 0.00023 | 0.00434 | 0.51885 | 0.65464 | 1.25268 | 91 |
| GO_REGULATION_OF_HEMATOPOIETIC_PROGENITOR_CELL_DIF | 0.00000 | 0.00000 | 0.73376 | 0.76273 | 1.44465 | 72 |
| GO_REGULATION_OF_HEMATOPOIETIC_STEM_CELL_DIFFERENT | 0.00000 | 0.00000 | 0.74774 | 0.78891 | 1.48817 | 62 |
| GO_REGULATION_OF_HEMOPOIESIS | 0.00390 | 0.03895 | 0.43171 | 0.57739 | 1.12909 | 258 |
| GO_REGULATION_OF_IMMUNE_RESPONSE | 0.00000 | 0.00006 | 0.64355 | 0.60032 | 1.18447 | 433 |
| GO_REGULATION_OF_INNATE_IMMUNE_RESPONSE | 0.00000 | 0.00007 | 0.62726 | 0.64536 | 1.25422 | 181 |
| GO_REGULATION_OF_INTRINSIC_APOPTOTIC_SIGNALING_PATH\ | 0.00179 | 0.02183 | 0.45506 | 0.61548 | 1.18550 | 116 |
| GO_REGULATION_OF_MITOTIC_CELL_CYCLE | 0.00000 | 0.00003 | 0.65944 | 0.58992 | 1.16508 | 499 |
| GO_REGULATION_OF_MORPHOGENESIS_OF_AN_EPITHELIUM | 0.00000 | 0.00000 | | 0.74680 | 1.43884 | 125 |
| GO_REGULATION_OF_MRNA_CATABOLIC_PROCESS | 0.00000 | 0.00000 | | 0.71126 | 1.37919 | 165 |
| GO_REGULATION_OF_MRNA_METABOLIC_PROCESS | 0.00000 | 0.00000 | | 0.67178 | 1.31458 | 276 |
| GO_REGULATION_OF_MRNA_PROCESSING | 0.00008 | 0.00186 | 0.53843 | 0.65189 | 1.25598 | 125 |
| GO_REGULATION_OF_MRNA_SPLICING_VIA_SPLICEOSOME | 0.00012 | 0.00256 | 0.53843 | 0.66950 | 1.28033 | 90 |
| GO_REGULATION_OF_MULTI_ORGANISM_PROCESS | 0.00008 | 0.00190 | 0.53843 | 0.76905 | 1.42150 | 37 |
| GO_REGULATION_OF_MUSCLE_CONTRACTION | 0.00163 | 0.02050 | 0.45506 | 0.64879 | 1.23071 | 76 |
| GO_REGULATION_OF_NUCLEAR_DIVISION | 0.00060 | 0.00923 | 0.47727 | 0.62443 | 1.20552 | 137 |
| GO_REGULATION_OF_NUCLEASE_ACTIVITY | 0.00059 | 0.00916 | 0.47727 | 0.81786 | 1.44767 | 17 |
| GO_REGULATION_OF_NUCLEOTIDE_BIOSYNTHETIC_PROCESS | 0.00302 | 0.03248 | 0.43171 | 0.78902 | 1.40337 | 18 |
| GO_REGULATION_OF_NUCLEOTIDE_METABOLIC_PROCESS | 0.00033 | 0.00588 | 0.49849 | 0.66858 | 1.26942 | 78 |
| GO_REGULATION_OF_OXIDATIVE_PHOSPHORYLATION | 0.00005 | 0.00130 | 0.55733 | 0.85284 | 1.53294 | 21 |
| GO_REGULATION_OF_PEPTIDASE_ACTIVITY | 0.00006 | 0.00152 | 0.53843 | 0.61192 | 1.19388 | 226 |
| GO_REGULATION_OF_PEPTIDE_TRANSPORT | 0.00250 | 0.02807 | 0.43171 | 0.56252 | 1.10917 | 412 |
| GO_REGULATION_OF_PH | 0.00561 | 0.04971 | 0.40702 | 0.64191 | 1.21498 | 71 |
| GO_REGULATION_OF_PLASMA_LIPOPROTEIN_PARTICLE_LEVELS | 0.00006 | 0.00145 | 0.55733 | 0.72431 | 1.36154 | 54 |
| GO_REGULATION_OF_POSTTRANSCRIPTIONAL_GENE_SILENCING | 0.00026 | 0.00486 | 0.49849 | 0.66803 | 1.27236 | 81 |
| GO_REGULATION_OF_PROTEIN_CATABOLIC_PROCESS | 0.00309 | 0.03299 | 0.43171 | 0.57360 | 1.12399 | 291 |
| GO_REGULATION_OF_PROTEIN_IMPORT | 0.00487 | 0.04559 | 0.40702 | 0.67491 | 1.25754 | 47 |
| GO_REGULATION_OF_PROTEIN_LOCALIZATION_TO_NUCLEUS | 0.00000 | 0.00013 | 0.61053 | 0.69035 | 1.32101 | 91 |
| GO_REGULATION_OF_PROTEIN_STABILITY | 0.00066 | 0.01007 | 0.47727 | 0.59308 | 1.15811 | 235 |
| GO_REGULATION_OF_PROTEOLYSIS | 0.00007 | 0.00169 | 0.53843 | 0.57923 | 1.14299 | 431 |
| GO_REGULATION_OF_REACTIVE_OXYGEN_SPECIES_BIOSYNTHET | 0.00022 | 0.00427 | 0.51885 | 0.73154 | 1.36343 | 45 |
| GO_REGULATION_OF_REACTIVE_OXYGEN_SPECIES_METABOLIC_ | 0.00088 | 0.01280 | 0.47727 | 0.63948 | 1.22592 | 99 |
| GO_REGULATION_OF_RESPONSE_TO_BIOTIC_STIMULUS | 0.00001 | 0.00039 | 0.59333 | 0.61773 | 1.20625 | 235 |
| GO_REGULATION_OF_RNA_SPLICING | 0.00011 | 0.00241 | 0.53843 | 0.64982 | 1.25198 | 125 |
| GO_REGULATION_OF_SISTER_CHROMATID_SEGREGATION | 0.00405 | 0.04011 | 0.40702 | 0.64401 | 1.22034 | 73 |
| GO_REGULATION_OF_SMALL_MOLECULE_METABOLIC_PROCESS | 0.00000 | 0.00000 | 0.77494 | 0.64342 | 1.26057 | 290 |
| GO_REGULATION_OF_SMOOTH_MUSCLE_CONTRACTION | 0.00007 | 0.00171 | 0.53843 | 0.82642 | 1.49295 | 23 |
| GO_REGULATION_OF_STEM_CELL_DIFFERENTIATION | 0.00000 | 0.00000 | 0.73376 | 0.73434 | 1.40292 | 88 |
| GO_REGULATION_OF_STEROID_METABOLIC_PROCESS | 0.00524 | 0.04770 | 0.40702 | 0.64061 | 1.21567 | 77 |
| GO_REGULATION_OF_SUPEROXIDE_METABOLIC_PROCESS | 0.00346 | 0.03548 | 0.43171 | 0.77826 | 1.39046 | 19 |
| GO_REGULATION_OF_SUPRAMOLECULAR_FIBER_ORGANIZATIOI | 0.00509 | 0.04656 | 0.40702 | 0.57599 | 1.12367 | 223 |
| GO_REGULATION_OF_SYMBIOTIC_PROCESS | 0.00009 | 0.00211 | 0.53843 | 0.62950 | 1.21763 | 151 |
| GO_REGULATION_OF_TELOMERASE_ACTIVITY | 0.00087 | 0.01263 | 0.47727 | 0.72147 | 1.33426 | 39 |
| GO_REGULATION_OF_TELOMERE_MAINTENANCE | 0.00547 | 0.04923 | 0.40702 | 0.64959 | 1.22679 | 68 |

| | | | | | | |
|---|---|---|---|---|---|---|
| GO_REGULATION_OF_TELOMERE_MAINTENANCE_VIA_TELOME | 0.00020 | 0.00387 | 0.51885 | 0.72043 | 1.34856 | 51 |
| GO_REGULATION_OF_TRANSCRIPTION_FROM_RNA_POLYMERA: | 0.00000 | 0.00000 | 0.70498 | 0.76931 | 1.45239 | 64 |
| GO_REGULATION_OF_TRANSLATIONAL_INITIATION | 0.00023 | 0.00437 | 0.51885 | 0.68479 | 1.29399 | 69 |
| GO_REGULATION_OF_VIRAL_TRANSCRIPTION | 0.00361 | 0.03657 | 0.43171 | 0.67497 | 1.26259 | 50 |
| GO_REGULATION_OF_WNT_SIGNALING_PATHWAY | 0.00015 | 0.00312 | 0.51885 | 0.60129 | 1.17419 | 238 |
| GO_RESPIRASOME | 0.00000 | 0.00000 | 0.81404 | 0.78326 | 1.48524 | 75 |
| GO_RESPIRATORY_CHAIN_COMPLEX | 0.00000 | 0.00000 | 0.77494 | 0.79183 | 1.49492 | 64 |
| GO_RESPIRATORY_ELECTRON_TRANSPORT_CHAIN | 0.00000 | 0.00000 | | 0.78275 | 1.49591 | 89 |
| GO_RESPONSE_TO_ALCOHOL | 0.00015 | 0.00310 | 0.51885 | 0.63794 | 1.22915 | 126 |
| GO_RESPONSE_TO_ATP | 0.00087 | 0.01270 | 0.47727 | 0.81086 | 1.43528 | 17 |
| GO_RESPONSE_TO_CALCIUM_ION | 0.00124 | 0.01701 | 0.45506 | 0.65196 | 1.24071 | 80 |
| GO_RESPONSE_TO_CORTICOSTEROID | 0.00010 | 0.00220 | 0.53843 | 0.66894 | 1.27841 | 89 |
| GO_RESPONSE_TO_DRUG | 0.00000 | 0.00007 | 0.62726 | 0.63080 | 1.23070 | 220 |
| GO_RESPONSE_TO_ETHANOL | 0.00168 | 0.02098 | 0.45506 | 0.66543 | 1.25679 | 67 |
| GO_RESPONSE_TO_FIBROBLAST_GROWTH_FACTOR | 0.00357 | 0.03638 | 0.43171 | 0.63469 | 1.20976 | 84 |
| GO_RESPONSE_TO_GROWTH_FACTOR | 0.00272 | 0.02986 | 0.43171 | 0.56235 | 1.10805 | 387 |
| GO_RESPONSE_TO_HEAT | 0.00000 | 0.00013 | 0.62726 | 0.66982 | 1.29069 | 119 |
| GO_RESPONSE_TO_HYDROGEN_PEROXIDE | 0.00532 | 0.04803 | 0.40702 | 0.62255 | 1.19054 | 90 |
| GO_RESPONSE_TO_INORGANIC_SUBSTANCE | 0.00016 | 0.00328 | 0.51885 | 0.58928 | 1.15810 | 321 |
| GO_RESPONSE_TO_INTERLEUKIN_1 | 0.00000 | 0.00000 | 0.78819 | 0.71590 | 1.37950 | 124 |
| GO_RESPONSE_TO_INTERLEUKIN_12 | 0.00036 | 0.00622 | 0.49849 | 0.74158 | 1.36769 | 36 |
| GO_RESPONSE_TO_INTERLEUKIN_4 | 0.00224 | 0.02581 | 0.43171 | 0.77088 | 1.38921 | 22 |
| GO_RESPONSE_TO_LEUKEMIA_INHIBITORY_FACTOR | 0.00076 | 0.01129 | 0.47727 | 0.66148 | 1.25288 | 72 |
| GO_RESPONSE_TO_LIPID | 0.00043 | 0.00695 | 0.49849 | 0.56669 | 1.11902 | 470 |
| GO_RESPONSE_TO_METAL_ION | 0.00143 | 0.01863 | 0.45506 | 0.59895 | 1.16592 | 199 |
| GO_RESPONSE_TO_NUTRIENT | 0.00294 | 0.03185 | 0.43171 | 0.62596 | 1.19987 | 98 |
| GO_RESPONSE_TO_ORGANOPHOSPHORUS | 0.00294 | 0.03185 | 0.43171 | 0.64154 | 1.21511 | 72 |
| GO_RESPONSE_TO_OXIDATIVE_STRESS | 0.00495 | 0.04573 | 0.40702 | 0.57010 | 1.11758 | 294 |
| GO_RESPONSE_TO_OXYGEN_LEVELS | 0.00000 | 0.00000 | | 0.67661 | 1.32155 | 244 |
| GO_RESPONSE_TO_PURINE_CONTAINING_COMPOUND | 0.00509 | 0.04656 | 0.40702 | 0.63176 | 1.20394 | 82 |
| GO_RESPONSE_TO_STEROID_HORMONE | 0.00009 | 0.00200 | 0.53843 | 0.61312 | 1.19360 | 201 |
| GO_RESPONSE_TO_TEMPERATURE_STIMULUS | 0.00002 | 0.00047 | 0.57561 | 0.64468 | 1.24782 | 153 |
| GO_RESPONSE_TO_TOXIC_SUBSTANCE | 0.00011 | 0.00235 | 0.53843 | 0.63892 | 1.23203 | 134 |
| GO_RESPONSE_TO_TUMOR_NECROSIS_FACTOR | 0.00000 | 0.00000 | 0.70498 | 0.66331 | 1.28815 | 175 |
| GO_RESPONSE_TO_VIRUS | 0.00261 | 0.02910 | 0.43171 | 0.59428 | 1.15408 | 175 |
| GO_RETINA_HOMEOSTASIS | 0.00034 | 0.00606 | 0.49849 | 0.75202 | 1.38362 | 34 |
| GO_RIBONUCLEOPROTEIN_COMPLEX_BINDING | 0.00000 | 0.00001 | 0.69013 | 0.69839 | 1.34459 | 114 |
| GO_RIBONUCLEOPROTEIN_COMPLEX_BIOGENESIS | 0.00000 | 0.00000 | | 0.68714 | 1.35448 | 393 |
| GO_RIBONUCLEOPROTEIN_COMPLEX_SUBUNIT_ORGANIZATION | 0.00000 | 0.00000 | 0.81404 | 0.69112 | 1.34183 | 173 |
| GO_RIBONUCLEOPROTEIN_GRANULE | 0.00000 | 0.00000 | 0.71951 | 0.66271 | 1.28875 | 186 |
| GO_RIBONUCLEOSIDE_TRIPHOSPHATE_BIOSYNTHETIC_PROCESS | 0.00009 | 0.00210 | 0.53843 | 0.77194 | 1.42026 | 34 |
| GO_RIBONUCLEOSIDE_TRIPHOSPHATE_METABOLIC_PROCESS | 0.00013 | 0.00280 | 0.51885 | 0.73558 | 1.37065 | 46 |
| GO_RIBOSE_PHOSPHATE_BIOSYNTHETIC_PROCESS | 0.00002 | 0.00056 | 0.57561 | 0.65814 | 1.26821 | 124 |
| GO_RIBOSE_PHOSPHATE_METABOLIC_PROCESS | 0.00000 | 0.00000 | 0.74774 | 0.63837 | 1.25001 | 285 |
| GO_RIBOSOMAL_LARGE_SUBUNIT_BIOGENESIS | 0.00001 | 0.00031 | 0.59333 | 0.72435 | 1.36807 | 67 |
| GO_RIBOSOMAL_SMALL_SUBUNIT_ASSEMBLY | 0.00405 | 0.04011 | 0.40702 | 0.78074 | 1.38865 | 18 |
| GO_RIBOSOMAL_SMALL_SUBUNIT_BINDING | 0.00532 | 0.04803 | 0.40702 | 0.77341 | 1.36899 | 17 |
| GO_RIBOSOMAL_SMALL_SUBUNIT_BIOGENESIS | 0.00005 | 0.00138 | 0.55733 | 0.70510 | 1.33163 | 68 |
| GO_RIBOSOMAL_SUBUNIT | 0.00000 | 0.00000 | | 0.76292 | 1.48089 | 172 |
| GO_RIBOSOME | 0.00000 | 0.00000 | | 0.75411 | 1.46984 | 208 |
| GO_RIBOSOME_ASSEMBLY | 0.00001 | 0.00026 | 0.59333 | 0.74485 | 1.40144 | 56 |
| GO_RIBOSOME_BIOGENESIS | 0.00000 | 0.00000 | | 0.68899 | 1.34833 | 277 |
| GO_RNA_3_END_PROCESSING | 0.00002 | 0.00046 | 0.57561 | 0.65576 | 1.26361 | 128 |
| GO_RNA_CATABOLIC_PROCESS | 0.00000 | 0.00000 | | 0.70701 | 1.39073 | 341 |
| GO_RNA_DEPENDENT_DNA_BIOSYNTHETIC_PROCESS | 0.00003 | 0.00081 | 0.57561 | 0.71669 | 1.34965 | 59 |
| GO_RNA_EXPORT_FROM_NUCLEUS | 0.00000 | 0.00016 | 0.61053 | 0.66306 | 1.27817 | 130 |
| GO_RNA_LOCALIZATION | 0.00000 | 0.00000 | 0.76146 | 0.66638 | 1.29886 | 208 |
| GO_RNA_SPLICING | 0.00000 | 0.00000 | | 0.63512 | 1.25129 | 384 |
| GO_RNA_SPLICING_VIA_TRANSESTERIFICATION_REACTIONS | 0.00000 | 0.00000 | 0.80122 | 0.64478 | 1.26586 | 311 |
| GO_ROUGH_ENDOPLASMIC_RETICULUM | 0.00014 | 0.00293 | 0.51885 | 0.70363 | 1.32507 | 59 |
| GO_RRNA_BINDING | 0.00000 | 0.00001 | 0.69013 | 0.77638 | 1.45941 | 54 |
| GO_RRNA_METABOLIC_PROCESS | 0.00000 | 0.00000 | | 0.68430 | 1.33386 | 209 |
| GO_SCF_DEPENDENT_PROTEASOMAL_UBIQUITIN_DEPENDENT_ | 0.00000 | 0.00008 | 0.62726 | 0.71957 | 1.36498 | 76 |
| GO_SECRETORY_GRANULE | 0.00000 | 0.00000 | | 0.62978 | 1.24363 | 481 |
| GO_SECRETORY_GRANULE_MEMBRANE | 0.00032 | 0.00575 | 0.49849 | 0.61375 | 1.19134 | 172 |
| GO_SERINE_TYPE_ENDOPEPTIDASE_INHIBITOR_ACTIVITY | 0.00038 | 0.00640 | 0.49849 | 0.77388 | 1.41447 | 28 |
| GO_SEXUAL_REPRODUCTION | 0.00472 | 0.04480 | 0.40702 | 0.55748 | 1.10026 | 439 |
| GO_SIDE_OF_MEMBRANE | 0.00495 | 0.04573 | 0.40702 | 0.57741 | 1.12751 | 235 |
| GO_SIGNAL_TRANSDUCTION_BY_PROTEIN_PHOSPHORYLATION | 0.00224 | 0.02581 | 0.43171 | 0.55690 | 1.09975 | 500 |
| GO_SINGLE_FERTILIZATION | 0.00000 | 0.00002 | 0.67496 | 0.76307 | 1.43842 | 61 |

| | | | | | | |
|---|---|---|---|---|---|---|
| GO_SINGLE_STRANDED_RNA_BINDING | 0.00001 | 0.00042 | 0.59333 | 0.71960 | 1.35864 | 66 |
| GO_SISTER_CHROMATID_SEGREGATION | 0.00042 | 0.00689 | 0.49849 | 0.60926 | 1.18209 | 170 |
| GO_SKIN_DEVELOPMENT | 0.00076 | 0.01140 | 0.47727 | 0.61452 | 1.18945 | 153 |
| GO_SMALL_MOLECULE_BIOSYNTHETIC_PROCESS | 0.00001 | 0.00021 | 0.61053 | 0.58728 | 1.15957 | 459 |
| GO_SMALL_NUCLEAR_RIBONUCLEOPROTEIN_COMPLEX | 0.00042 | 0.00689 | 0.49849 | 0.69542 | 1.30936 | 57 |
| GO_SMALL_RIBOSOMAL_SUBUNIT | 0.00000 | 0.00001 | 0.69013 | 0.75767 | 1.43100 | 67 |
| GO_SM_LIKE_PROTEIN_FAMILY_COMPLEX | 0.00145 | 0.01879 | 0.45506 | 0.66792 | 1.26149 | 67 |
| GO_SNORNA_BINDING | 0.00038 | 0.00646 | 0.49849 | 0.76224 | 1.39614 | 29 |
| GO_SPERM_EGG_RECOGNITION | 0.00000 | 0.00002 | 0.65944 | 0.87469 | 1.59087 | 26 |
| GO_SPINDLE_MIDZONE | 0.00361 | 0.03657 | 0.43171 | 0.73316 | 1.33859 | 27 |
| GO_SPINDLE_ORGANIZATION | 0.00272 | 0.02986 | 0.43171 | 0.60076 | 1.16362 | 159 |
| GO_SPLICEOSOMAL_COMPLEX | 0.00000 | 0.00008 | 0.62726 | 0.65256 | 1.26657 | 171 |
| GO_SPLICEOSOMAL_TRI_SNRNP_COMPLEX | 0.00168 | 0.02098 | 0.45506 | 0.73474 | 1.34835 | 30 |
| GO_STEM_CELL_DIFFERENTIATION | 0.00000 | 0.00000 | 0.82666 | 0.70006 | 1.35554 | 158 |
| GO_STEROID_BIOSYNTHETIC_PROCESS | 0.00465 | 0.04430 | 0.40702 | 0.61242 | 1.17992 | 125 |
| GO_STEROID_METABOLIC_PROCESS | 0.00313 | 0.03310 | 0.43171 | 0.59525 | 1.15748 | 185 |
| GO_STEROL_BIOSYNTHETIC_PROCESS | 0.00005 | 0.00120 | 0.55733 | 0.71876 | 1.35584 | 62 |
| GO_STEROL_METABOLIC_PROCESS | 0.00003 | 0.00073 | 0.57561 | 0.66485 | 1.27854 | 109 |
| GO_STRESS_GRANULE_ASSEMBLY | 0.00465 | 0.04430 | 0.40702 | 0.77288 | 1.38921 | 21 |
| GO_STRUCTURAL_CONSTITUENT_OF_RIBOSOME | 0.00000 | 0.00000 | | 0.76380 | 1.47839 | 153 |
| GO_STRUCTURAL_MOLECULE_ACTIVITY | 0.00000 | 0.00000 | | 0.71760 | 1.41436 | 391 |
| GO_SUPEROXIDE_METABOLIC_PROCESS | 0.00272 | 0.02986 | 0.43171 | 0.72028 | 1.32659 | 35 |
| GO_TELOMERASE_RNA_LOCALIZATION | 0.00054 | 0.00857 | 0.47727 | 0.81801 | 1.46148 | 19 |
| GO_TELOMERE_MAINTENANCE_VIA_TELOMERE_LENGTHENING | 0.00005 | 0.00138 | 0.55733 | 0.70715 | 1.33512 | 66 |
| GO_TELOMERE_ORGANIZATION | 0.00159 | 0.02016 | 0.45506 | 0.61666 | 1.18834 | 127 |
| GO_TERTIARY_GRANULE | 0.00008 | 0.00186 | 0.53843 | 0.66372 | 1.27005 | 91 |
| GO_TERTIARY_GRANULE_MEMBRANE | 0.00102 | 0.01453 | 0.45506 | 0.71940 | 1.33244 | 40 |
| GO_THREONINE_TYPE_ENDOPEPTIDASE_ACTIVITY | 0.00000 | 0.00000 | 0.71951 | 0.93456 | 1.65424 | 17 |
| GO_THREONINE_TYPE_PEPTIDASE_ACTIVITY | 0.00000 | 0.00004 | 0.65944 | 0.89807 | 1.61424 | 21 |
| GO_TISSUE_MORPHOGENESIS | 0.00000 | 0.00006 | 0.64355 | 0.59467 | 1.17150 | 380 |
| GO_TOXIN_TRANSPORT | 0.00064 | 0.00972 | 0.47727 | 0.77713 | 1.41343 | 26 |
| GO_TRANSCRIPTION_COUPLED_NUCLEOTIDE_EXCISION_REPAIR | 0.00231 | 0.02642 | 0.43171 | 0.65429 | 1.23840 | 71 |
| GO_TRANSFERRIN_TRANSPORT | 0.00141 | 0.01850 | 0.45506 | 0.75134 | 1.37179 | 27 |
| GO_TRANSLATIONAL_ELONGATION | 0.00000 | 0.00000 | | 0.75263 | 1.45028 | 124 |
| GO_TRANSLATIONAL_INITIATION | 0.00000 | 0.00000 | | 0.75139 | 1.45839 | 171 |
| GO_TRANSLATIONAL_TERMINATION | 0.00000 | 0.00000 | 0.81404 | 0.75110 | 1.44066 | 101 |
| GO_TRANSLATION_ELONGATION_FACTOR_ACTIVITY | 0.00390 | 0.03895 | 0.43171 | 0.80505 | 1.41260 | 15 |
| GO_TRANSLATION_FACTOR_ACTIVITY_RNA_BINDING | 0.00000 | 0.00000 | 0.74774 | 0.77244 | 1.46085 | 70 |
| GO_TRANSLATION_INITIATION_FACTOR_ACTIVITY | 0.00000 | 0.00011 | 0.62726 | 0.77773 | 1.44952 | 45 |
| GO_TRANSLATION_PREINITIATION_COMPLEX | 0.00342 | 0.03521 | 0.43171 | 0.79777 | 1.40694 | 16 |
| GO_TRANSLATION_REGULATOR_ACTIVITY | 0.00000 | 0.00000 | 0.83909 | 0.74000 | 1.42403 | 111 |
| GO_TRANSLATION_REGULATOR_ACTIVITY_NUCLEIC_ACID_BIND | 0.00000 | 0.00000 | 0.83909 | 0.76441 | 1.45959 | 87 |
| GO_TRANSMEMBRANE_RECEPTOR_PROTEIN_TYROSINE_KINASE | 0.00168 | 0.02098 | 0.45506 | 0.56326 | 1.11076 | 411 |
| GO_TRICARBOXYLIC_ACID_CYCLE | 0.00159 | 0.02016 | 0.45506 | 0.73558 | 1.35097 | 33 |
| GO_TRNA_TRANSPORT | 0.00342 | 0.03521 | 0.43171 | 0.71247 | 1.31220 | 35 |
| GO_TUMOR_NECROSIS_FACTOR_MEDIATED_SIGNALING_PATHW | 0.00000 | 0.00000 | 0.80122 | 0.73921 | 1.42138 | 108 |
| GO_T_CELL_RECEPTOR_SIGNALING_PATHWAY | 0.00000 | 0.00001 | 0.70498 | 0.69185 | 1.33314 | 119 |
| GO_U12_TYPE_SPLICEOSOMAL_COMPLEX | 0.00487 | 0.04559 | 0.40702 | 0.73423 | 1.33541 | 26 |
| GO_U2_SNRNP | 0.00141 | 0.01850 | 0.45506 | 0.80259 | 1.42065 | 17 |
| GO_U2_TYPE_CATALYTIC_STEP_2_SPLICEOSOME | 0.00268 | 0.02961 | 0.43171 | 0.72225 | 1.32288 | 29 |
| GO_U2_TYPE_SPLICEOSOMAL_COMPLEX | 0.00033 | 0.00595 | 0.49849 | 0.66590 | 1.26925 | 84 |
| GO_UBIQUITIN_LIKE_PROTEIN_LIGASE_BINDING | 0.00000 | 0.00007 | 0.62726 | 0.62616 | 1.22416 | 251 |
| GO_UNFOLDED_PROTEIN_BINDING | 0.00000 | 0.00001 | 0.69013 | 0.72112 | 1.37766 | 88 |
| GO_VACUOLAR_LUMEN | 0.00016 | 0.00319 | 0.51885 | 0.65416 | 1.25772 | 107 |
| GO_VESICLE_BUDDING_FROM_MEMBRANE | 0.00480 | 0.04537 | 0.40702 | 0.62411 | 1.19353 | 90 |
| GO_VESICLE_LUMEN | 0.00000 | 0.00000 | 0.77494 | 0.67028 | 1.30631 | 207 |
| GO_VESICLE_MEMBRANE | 0.00001 | 0.00026 | 0.59333 | 0.58430 | 1.15387 | 476 |
| GO_VIRAL_GENE_EXPRESSION | 0.00000 | 0.00000 | | 0.70963 | 1.37745 | 172 |
| GO_VIRAL_LIFE_CYCLE | 0.00000 | 0.00001 | 0.69013 | 0.63985 | 1.24950 | 238 |
| GO_VIRION_ASSEMBLY | 0.00457 | 0.04386 | 0.40702 | 0.69689 | 1.28528 | 36 |
| HP_ABNORMALITY_OF_ACID_BASE_HOMEOSTASIS | 0.00000 | 0.00010 | 0.62726 | 0.62283 | 1.21827 | 261 |
| HP_ABNORMALITY_OF_BLOOD_CIRCULATION | 0.00502 | 0.04608 | 0.40702 | 0.59228 | 1.14883 | 167 |
| HP_ABNORMALITY_OF_CARDIOVASCULAR_SYSTEM_ELECTROPH | 0.00011 | 0.00237 | 0.53843 | 0.60202 | 1.17758 | 261 |
| HP_ABNORMALITY_OF_SKIN_ADNEXA_PHYSIOLOGY | 0.00242 | 0.02748 | 0.43171 | 0.61294 | 1.18094 | 118 |
| HP_ABNORMALITY_OF_THE_ANKLES | 0.00313 | 0.03310 | 0.43171 | 0.65369 | 1.23418 | 63 |
| HP_ABNORMALITY_OF_THE_CEREBRAL_VASCULATURE | 0.00357 | 0.03638 | 0.43171 | 0.59764 | 1.15722 | 158 |
| HP_ABNORMALITY_OF_THE_CEREBROSPINAL_FLUID | 0.00030 | 0.00549 | 0.49849 | 0.58221 | 1.14439 | 325 |
| HP_ABNORMALITY_OF_THE_MITOCHONDRION | 0.00126 | 0.01715 | 0.45506 | 0.62149 | 1.19764 | 120 |
| HP_ABNORMALITY_OF_THE_OPTIC_DISC | 0.00198 | 0.02370 | 0.43171 | 0.56624 | 1.11727 | 432 |
| HP_ABNORMALITY_OF_THE_OPTIC_NERVE | 0.00036 | 0.00622 | 0.49849 | 0.56723 | 1.12034 | 486 |

| | | | | | | |
|---|---|---|---|---|---|---|
| HP_ABNORMAL_ABDOMEN_MORPHOLOGY | 0.00089 | 0.01291 | 0.47727 | 0.56424 | 1.11364 | 441 |
| HP_ABNORMAL_ACTIVITY_OF_MITOCHONDRIAL_RESPIRATORY_ | 0.00014 | 0.00300 | 0.51885 | 0.70303 | 1.32524 | 61 |
| HP_ABNORMAL_BASAL_GANGLIA_MRI_SIGNAL_INTENSITY | 0.00179 | 0.02183 | 0.45506 | 0.71597 | 1.32046 | 36 |
| HP_ABNORMAL_BRAINSTEM_MRI_SIGNAL_INTENSITY | 0.00398 | 0.03962 | 0.40702 | 0.71528 | 1.31370 | 33 |
| HP_ABNORMAL_CELLULAR_PHENOTYPE | 0.00209 | 0.02470 | 0.43171 | 0.56937 | 1.11992 | 334 |
| HP_ABNORMAL_CIRCULATING_CARBOHYDRATE_CONCENTRATIC | 0.00194 | 0.02331 | 0.45506 | 0.69518 | 1.29539 | 46 |
| HP_ABNORMAL_CNS_MYELINATION | 0.00014 | 0.00300 | 0.51885 | 0.62318 | 1.20954 | 171 |
| HP_ABNORMAL_CSF_METABOLITE_LEVEL | 0.00001 | 0.00027 | 0.59333 | 0.70531 | 1.34409 | 82 |
| HP_ABNORMAL_ELECTROPHYSIOLOGY_OF_SINOATRIAL_NODE_ | 0.00029 | 0.00536 | 0.49849 | 0.73172 | 1.35725 | 41 |
| HP_ABNORMAL_ENZYME_COENZYME_ACTIVITY | 0.00561 | 0.04971 | 0.40702 | 0.60490 | 1.16567 | 123 |
| HP_ABNORMAL_ERYTHROCYTE_MORPHOLOGY | 0.00001 | 0.00023 | 0.61053 | 0.58835 | 1.15904 | 380 |
| HP_ABNORMAL_HEART_VALVE_PHYSIOLOGY | 0.00405 | 0.04011 | 0.40702 | 0.59004 | 1.14474 | 169 |
| HP_ABNORMAL_MITOCHONDRIA_IN_MUSCLE_TISSUE | 0.00075 | 0.01118 | 0.47727 | 0.74633 | 1.36700 | 29 |
| HP_ABNORMAL_MYELINATION | 0.00007 | 0.00169 | 0.53843 | 0.59258 | 1.16479 | 324 |
| HP_ABNORMAL_MYOCARDIUM_MORPHOLOGY | 0.00008 | 0.00192 | 0.53843 | 0.58647 | 1.15365 | 340 |
| HP_ABNORMAL_RENAL_PHYSIOLOGY | 0.00450 | 0.04342 | 0.40702 | 0.55484 | 1.09317 | 385 |
| HP_ABNORMAL_TRICUSPID_VALVE_PHYSIOLOGY | 0.00231 | 0.02642 | 0.43171 | 0.72968 | 1.34029 | 31 |
| HP_ABNORMAL_VITREOUS_HUMOR_MORPHOLOGY | 0.00181 | 0.02201 | 0.45506 | 0.79632 | 1.40955 | 17 |
| HP_ACUTE_LEUKEMIA | 0.00013 | 0.00276 | 0.51885 | 0.68145 | 1.29219 | 75 |
| HP_ANEMIA_OF_INADEQUATE_PRODUCTION | 0.00561 | 0.04971 | 0.40702 | 0.61298 | 1.17855 | 107 |
| HP_APHASIA | 0.00213 | 0.02490 | 0.43171 | 0.72858 | 1.33704 | 30 |
| HP_BLINDNESS | 0.00018 | 0.00355 | 0.51885 | 0.61672 | 1.19768 | 175 |
| HP_BONE_MARROW_HYPERCELLULARITY | 0.00045 | 0.00725 | 0.49849 | 0.82169 | 1.46806 | 19 |
| HP_CHOREOATHETOSIS | 0.00561 | 0.04971 | 0.40702 | 0.64956 | 1.22631 | 64 |
| HP_CHRONIC_INFECTION | 0.00121 | 0.01658 | 0.45506 | 0.81916 | 1.44468 | 16 |
| HP_CONGESTIVE_HEART_FAILURE | 0.00495 | 0.04573 | 0.40702 | 0.60734 | 1.16929 | 114 |
| HP_CRANIAL_NERVE_COMPRESSION | 0.00331 | 0.03441 | 0.43171 | 0.75830 | 1.36989 | 23 |
| HP_DECREASED_ACTIVITY_OF_MITOCHONDRIAL_COMPLEX_I | 0.00283 | 0.03081 | 0.43171 | 0.71940 | 1.32496 | 35 |
| HP_DECREASED_ACTIVITY_OF_THE_PYRUVATE_DEHYDROGENAS | 0.00062 | 0.00949 | 0.47727 | 0.75390 | 1.38478 | 31 |
| HP_DEPRESSED_NASAL_RIDGE | 0.00041 | 0.00681 | 0.49849 | 0.66218 | 1.26189 | 82 |
| HP_DYSPNEA | 0.00250 | 0.02807 | 0.43171 | 0.58107 | 1.13348 | 224 |
| HP_DYSTONIA | 0.00069 | 0.01043 | 0.47727 | 0.58101 | 1.13736 | 279 |
| HP_EMOTIONAL_LABILITY | 0.00016 | 0.00321 | 0.51885 | 0.66683 | 1.27522 | 90 |
| HP_EPISODIC_ATAXIA | 0.00487 | 0.04559 | 0.40702 | 0.79934 | 1.40259 | 15 |
| HP_FASCICULATIONS | 0.00130 | 0.01761 | 0.45506 | 0.68128 | 1.28064 | 54 |
| HP_FATIGUE | 0.00042 | 0.00689 | 0.49849 | 0.59902 | 1.16764 | 212 |
| HP_FETAL_DISTRESS | 0.00179 | 0.02183 | 0.45506 | 0.72928 | 1.33973 | 32 |
| HP_FLUSHING | 0.00561 | 0.04971 | 0.40702 | 0.74782 | 1.33686 | 20 |
| HP_FOCAL_T2_HYPERINTENSE_BRAINSTEM_LESION | 0.00159 | 0.02016 | 0.45506 | 0.74970 | 1.36879 | 27 |
| HP_HEADACHE | 0.00016 | 0.00319 | 0.51885 | 0.62537 | 1.21179 | 163 |
| HP_HEPATOMEGALY | 0.00108 | 0.01507 | 0.45506 | 0.57439 | 1.12990 | 335 |
| HP_HYPERTROPHIC_CARDIOMYOPATHY | 0.00002 | 0.00050 | 0.57561 | 0.62629 | 1.21908 | 194 |
| HP_IMPAIRED_SOCIAL_INTERACTIONS | 0.00055 | 0.00863 | 0.47727 | 0.66555 | 1.26058 | 72 |
| HP_INCREASED_CSF_LACTATE | 0.00001 | 0.00034 | 0.59333 | 0.71937 | 1.35857 | 68 |
| HP_INCREASED_SERUM_LACTATE | 0.00001 | 0.00028 | 0.59333 | 0.65837 | 1.26932 | 133 |
| HP_INCREASED_SERUM_PYRUVATE | 0.00110 | 0.01526 | 0.45506 | 0.73143 | 1.35198 | 37 |
| HP_INFANTILE_ONSET | 0.00487 | 0.04559 | 0.40702 | 0.56559 | 1.10939 | 300 |
| HP_INTRACRANIAL_CYSTIC_LESION | 0.00055 | 0.00863 | 0.47727 | 0.62760 | 1.20885 | 116 |
| HP_INTRAUTERINE_GROWTH_RETARDATION | 0.00305 | 0.03271 | 0.43171 | 0.56594 | 1.11393 | 350 |
| HP_LACTIC_ACIDOSIS | 0.00018 | 0.00355 | 0.51885 | 0.63312 | 1.22071 | 131 |
| HP_LETHARGY | 0.00084 | 0.01228 | 0.47727 | 0.63361 | 1.21845 | 109 |
| HP_LEUKODYSTROPHY | 0.00029 | 0.00529 | 0.49849 | 0.66598 | 1.26915 | 82 |
| HP_LEUKOENCEPHALOPATHY | 0.00008 | 0.00190 | 0.53843 | 0.64132 | 1.23667 | 134 |
| HP_LIPID_ACCUMULATION_IN_HEPATOCYTES | 0.00220 | 0.02548 | 0.43171 | 0.62794 | 1.20159 | 91 |
| HP_LOWER_LIMB_SPASTICITY | 0.00154 | 0.01974 | 0.45506 | 0.60929 | 1.17648 | 139 |
| HP_MACROCYTIC_ANEMIA | 0.00030 | 0.00542 | 0.49849 | 0.73108 | 1.35606 | 41 |
| HP_METABOLIC_ACIDOSIS | 0.00034 | 0.00599 | 0.49849 | 0.64831 | 1.24464 | 103 |
| HP_MICRONODULAR_CIRRHOSIS | 0.00324 | 0.03393 | 0.43171 | 0.78723 | 1.40018 | 18 |
| HP_MIGRAINE | 0.00028 | 0.00515 | 0.49849 | 0.67309 | 1.27635 | 75 |
| HP_MUSCLE_ABNORMALITY_RELATED_TO_MITOCHONDRIAL_DY | 0.00209 | 0.02470 | 0.43171 | 0.68258 | 1.27573 | 49 |
| HP_MUTISM | 0.00283 | 0.03081 | 0.43171 | 0.71937 | 1.32490 | 35 |
| HP_NAUSEA_AND_VOMITING | 0.00022 | 0.00417 | 0.51885 | 0.59383 | 1.16158 | 264 |
| HP_NEOPLASM_OF_THE_ENDOCRINE_SYSTEM | 0.00495 | 0.04573 | 0.40702 | 0.63166 | 1.20308 | 81 |
| HP_OPTIC_NEUROPATHY | 0.00135 | 0.01808 | 0.45506 | 0.73837 | 1.35610 | 33 |
| HP_PACE_OF_PROGRESSION | 0.00487 | 0.04559 | 0.40702 | 0.56814 | 1.11416 | 298 |
| HP_PALLOR | 0.00036 | 0.00622 | 0.49849 | 0.64861 | 1.24463 | 102 |
| HP_PARAPLEGIA | 0.00121 | 0.01658 | 0.45506 | 0.63584 | 1.22133 | 104 |
| HP_PARAPLEGIA_PARAPARESIS | 0.00036 | 0.00621 | 0.49849 | 0.62047 | 1.20040 | 150 |
| HP_PAROXYSMAL_INVOLUNTARY_EYE_MOVEMENTS | 0.00054 | 0.00849 | 0.47727 | 0.75650 | 1.38957 | 31 |
| HP_PEDIATRIC_ONSET | 0.00561 | 0.04971 | 0.40702 | 0.55915 | 1.10116 | 371 |

| | | | | | | |
|---|---|---|---|---|---|---|
| HP_PIGMENTARY_RETINOPATHY | 0.00065 | 0.00981 | 0.47727 | 0.63260 | 1.21720 | 113 |
| HP_POOR_EYE_CONTACT | 0.00017 | 0.00341 | 0.51885 | 0.71279 | 1.33749 | 53 |
| HP_POOR_HEAD_CONTROL | 0.00000 | 0.00005 | 0.64355 | 0.67819 | 1.30665 | 118 |
| HP_PROGRESSIVE | 0.00194 | 0.02331 | 0.45506 | 0.59790 | 1.16273 | 186 |
| HP_PROGRESSIVE_CEREBELLAR_ATAXIA | 0.00364 | 0.03677 | 0.43171 | 0.64788 | 1.22325 | 65 |
| HP_PROGRESSIVE_SPASTIC_PARAPLEGIA | 0.00253 | 0.02838 | 0.43171 | 0.69655 | 1.29822 | 45 |
| HP_PROXIMAL_TUBULOPATHY | 0.00141 | 0.01850 | 0.45506 | 0.72579 | 1.33535 | 34 |
| HP_PTOSIS | 0.00472 | 0.04480 | 0.40702 | 0.55582 | 1.09637 | 418 |
| HP_REDUCED_CONSCIOUSNESS_CONFUSION | 0.00007 | 0.00164 | 0.53843 | 0.61227 | 1.19295 | 205 |
| HP_REDUCED_VISUAL_ACUITY | 0.00053 | 0.00837 | 0.47727 | 0.58618 | 1.14774 | 283 |
| HP_RENAL_TUBULAR_DYSFUNCTION | 0.00013 | 0.00284 | 0.51885 | 0.68108 | 1.29150 | 75 |
| HP_RESPIRATORY_DISTRESS | 0.00108 | 0.01507 | 0.45506 | 0.62693 | 1.20762 | 115 |
| HP_RESPIRATORY_INSUFFICIENCY | 0.00000 | 0.00010 | 0.62726 | 0.60536 | 1.18998 | 326 |
| HP_SINUS_TACHYCARDIA | 0.00213 | 0.02490 | 0.43171 | 0.77203 | 1.39128 | 22 |
| HP_SPARSE_EYELASHES | 0.00150 | 0.01943 | 0.45506 | 0.71355 | 1.32161 | 40 |
| HP_SPARSE_OR_ABSENT_EYELASHES | 0.00220 | 0.02548 | 0.43171 | 0.67331 | 1.26567 | 54 |
| HP_SPASTIC_PARAPLEGIA | 0.00098 | 0.01407 | 0.45506 | 0.64059 | 1.22712 | 96 |
| HP_THROMBOCYTOSIS | 0.00413 | 0.04065 | 0.40702 | 0.75210 | 1.35870 | 23 |
| HP_TONGUE_FASCICULATIONS | 0.00457 | 0.04386 | 0.40702 | 0.80056 | 1.40472 | 15 |
| HP_TRANSIENT_ISCHEMIC_ATTACK | 0.00480 | 0.04537 | 0.40702 | 0.74902 | 1.35809 | 25 |
| HP_VERTIGO | 0.00002 | 0.00063 | 0.57561 | 0.72114 | 1.35844 | 60 |
| HP_VESTIBULAR_DYSFUNCTION | 0.00028 | 0.00522 | 0.49849 | 0.67279 | 1.27577 | 75 |
| HP_VOMITING | 0.00072 | 0.01082 | 0.47727 | 0.61459 | 1.18792 | 147 |
| HP_WEAKNESS_DUE_TO_UPPER_MOTOR_NEURON_DYSFUNCTI | 0.00532 | 0.04803 | 0.40702 | 0.56829 | 1.11186 | 272 |
| HP_WEAKNESS_OF_MUSCLES_OF_RESPIRATION | 0.00163 | 0.02050 | 0.45506 | 0.66272 | 1.25123 | 63 |
| **Cluster 1** | | | | | | |
| pathway | pval | padj | log2err | ES | NES | size |
| GO_ACTIVATION_OF_IMMUNE_RESPONSE | 0.00000 | 0.00000 | 0.74774 | 0.65038 | 1.26999 | 235 |
| GO_ACTIVATION_OF_INNATE_IMMUNE_RESPONSE | 0.00000 | 0.00001 | 0.69013 | 0.71967 | 1.37861 | 96 |
| GO_AEROBIC_RESPIRATION | 0.00000 | 0.00000 | 0.73376 | 0.76280 | 1.44478 | 72 |
| GO_ALCOHOL_BIOSYNTHETIC_PROCESS | 0.00032 | 0.00572 | 0.49849 | 0.64679 | 1.24367 | 108 |
| GO_ALCOHOL_METABOLIC_PROCESS | 0.00364 | 0.03677 | 0.43171 | 0.58112 | 1.13424 | 228 |
| GO_AMEBOIDAL_TYPE_CELL_MIGRATION | 0.00457 | 0.04386 | 0.40702 | 0.57755 | 1.12854 | 247 |
| GO_AMIDE_BINDING | 0.00014 | 0.00297 | 0.51885 | 0.61533 | 1.19780 | 199 |
| GO_AMINE_METABOLIC_PROCESS | 0.00000 | 0.00000 | 0.76146 | 0.73732 | 1.41097 | 92 |
| GO_AMYLOID_BETA_CLEARANCE | 0.00561 | 0.04971 | 0.40702 | 0.74081 | 1.33976 | 24 |
| GO_ANAPHASE_PROMOTING_COMPLEX_DEPENDENT_CATABOL | 0.00000 | 0.00000 | 0.76146 | 0.77107 | 1.46112 | 73 |
| GO_ANATOMICAL_STRUCTURE_HOMEOSTASIS | 0.00020 | 0.00390 | 0.51885 | 0.59307 | 1.16021 | 271 |
| GO_ANIMAL_ORGAN_REGENERATION | 0.00012 | 0.00269 | 0.51885 | 0.71251 | 1.33510 | 52 |
| GO_ANTIGEN_PROCESSING_AND_PRESENTATION | 0.00000 | 0.00000 | 0.74774 | 0.68785 | 1.33064 | 149 |
| GO_ANTIGEN_PROCESSING_AND_PRESENTATION_OF_EXOGENC | 0.00000 | 0.00000 | 0.76146 | 0.79644 | 1.50237 | 62 |
| GO_ANTIGEN_PROCESSING_AND_PRESENTATION_OF_PEPTIDE_ | 0.00000 | 0.00000 | 0.74774 | 0.70107 | 1.35144 | 130 |
| GO_ANTIGEN_PROCESSING_AND_PRESENTATION_OF_PEPTIDE_ | 0.00000 | 0.00000 | 0.78819 | 0.77566 | 1.47012 | 74 |
| GO_ANTIGEN_RECEPTOR_MEDIATED_SIGNALING_PATHWAY | 0.00000 | 0.00009 | 0.62726 | 0.66362 | 1.28080 | 136 |
| GO_APICAL_PART_OF_CELL | 0.00135 | 0.01808 | 0.45506 | 0.58926 | 1.14966 | 220 |
| GO_APOPTOTIC_SIGNALING_PATHWAY | 0.00006 | 0.00152 | 0.53843 | 0.58638 | 1.15586 | 393 |
| GO_ATPASE_ACTIVITY | 0.00050 | 0.00802 | 0.47727 | 0.58216 | 1.14385 | 316 |
| GO_ATPASE_ACTIVITY_COUPLED_TO_TRANSMEMBRANE_MOVE | 0.00420 | 0.04132 | 0.40702 | 0.78005 | 1.38741 | 18 |
| GO_ATP_BIOSYNTHETIC_PROCESS | 0.00013 | 0.00276 | 0.51885 | 0.83897 | 1.50801 | 21 |
| GO_ATP_METABOLIC_PROCESS | 0.00000 | 0.00000 | | 0.72178 | 1.40814 | 219 |
| GO_ATP_SYNTHESIS_COUPLED_ELECTRON_TRANSPORT | 0.00000 | 0.00000 | | 0.80353 | 1.52425 | 76 |
| GO_AZUROPHIL_GRANULE | 0.00025 | 0.00466 | 0.49849 | 0.64781 | 1.24551 | 107 |
| GO_AZUROPHIL_GRANULE_LUMEN | 0.00405 | 0.04011 | 0.40702 | 0.65625 | 1.23793 | 62 |
| GO_BASAL_PART_OF_CELL | 0.00509 | 0.04656 | 0.40702 | 0.68515 | 1.27697 | 45 |
| GO_BASAL_PLASMA_MEMBRANE | 0.00346 | 0.03548 | 0.43171 | 0.71184 | 1.31104 | 35 |
| GO_BASEMENT_MEMBRANE | 0.00279 | 0.03056 | 0.43171 | 0.68935 | 1.28452 | 46 |
| GO_BASOLATERAL_PLASMA_MEMBRANE | 0.00011 | 0.00247 | 0.53843 | 0.63209 | 1.21993 | 136 |
| GO_BINDING_OF_SPERM_TO_ZONA_PELLUCIDA | 0.00000 | 0.00006 | 0.64355 | 0.90095 | 1.60966 | 19 |
| GO_BLOOD_MICROPARTICLE | 0.00000 | 0.00000 | 0.70498 | 0.81982 | 1.52763 | 46 |
| GO_BRUSH_BORDER | 0.00060 | 0.00923 | 0.47727 | 0.66897 | 1.26790 | 74 |
| GO_CADHERIN_BINDING | 0.00000 | 0.00000 | | 0.68759 | 1.34435 | 253 |
| GO_CALCIUM_DEPENDENT_PHOSPHOLIPID_BINDING | 0.00152 | 0.01958 | 0.45506 | 0.75021 | 1.37120 | 28 |
| GO_CALCIUM_DEPENDENT_PROTEIN_BINDING | 0.00009 | 0.00211 | 0.53843 | 0.74254 | 1.38394 | 45 |
| GO_CANONICAL_WNT_SIGNALING_PATHWAY | 0.00004 | 0.00109 | 0.55733 | 0.61740 | 1.20458 | 222 |
| GO_CARBOHYDRATE_CATABOLIC_PROCESS | 0.00135 | 0.01808 | 0.45506 | 0.61179 | 1.18186 | 144 |
| GO_CATALYTIC_STEP_2_SPLICEOSOME | 0.00001 | 0.00022 | 0.61053 | 0.70290 | 1.33765 | 80 |
| GO_CATION_TRANSMEMBRANE_TRANSPORT | 0.00327 | 0.03409 | 0.43171 | 0.56060 | 1.10531 | 406 |
| GO_CATION_TRANSMEMBRANE_TRANSPORTER_ACTIVITY | 0.00097 | 0.01387 | 0.47727 | 0.58196 | 1.13846 | 265 |
| GO_CELLULAR_AMINO_ACID_METABOLIC_PROCESS | 0.00002 | 0.00060 | 0.57561 | 0.61438 | 1.19930 | 231 |
| GO_CELLULAR_COMPONENT_DISASSEMBLY | 0.00000 | 0.00000 | | 0.64070 | 1.26342 | 408 |

| | | | | | | |
|---|---|---|---|---|---|---|
| GO_CELLULAR_ION_HOMEOSTASIS | 0.00027 | 0.00493 | 0.49849 | 0.58658 | 1.15340 | 328 |
| GO_CELLULAR_IRON_ION_HOMEOSTASIS | 0.00327 | 0.03409 | 0.43171 | 0.68360 | 1.27374 | 47 |
| GO_CELLULAR_KETONE_METABOLIC_PROCESS | 0.00000 | 0.00010 | 0.62726 | 0.65358 | 1.26726 | 164 |
| GO_CELLULAR_MONOVALENT_INORGANIC_CATION_HOMEOSTA | 0.00179 | 0.02183 | 0.45506 | 0.64747 | 1.22821 | 76 |
| GO_CELLULAR_NITROGEN_COMPOUND_CATABOLIC_PROCESS | 0.00000 | 0.00000 | | 0.67915 | 1.34096 | 456 |
| GO_CELLULAR_PROTEIN_COMPLEX_DISASSEMBLY | 0.00000 | 0.00000 | | 0.69566 | 1.35184 | 179 |
| GO_CELLULAR_RESPIRATION | 0.00000 | 0.00000 | | 0.74205 | 1.43370 | 145 |
| GO_CELLULAR_RESPONSE_TO_HEAT | 0.00004 | 0.00109 | 0.55733 | 0.67186 | 1.28698 | 95 |
| GO_CELLULAR_RESPONSE_TO_HORMONE_STIMULUS | 0.00309 | 0.03299 | 0.43171 | 0.56453 | 1.11191 | 374 |
| GO_CELLULAR_RESPONSE_TO_ORGANIC_CYCLIC_COMPOUND | 0.00502 | 0.04608 | 0.40702 | 0.56512 | 1.11134 | 329 |
| GO_CELLULAR_RESPONSE_TO_OXYGEN_LEVELS | 0.00000 | 0.00000 | | 0.71461 | 1.38224 | 151 |
| GO_CELL_ACTIVATION_INVOLVED_IN_IMMUNE_RESPONSE | 0.00000 | 0.00000 | | 0.63924 | 1.26108 | 422 |
| GO_CELL_ADHESION_MEDIATOR_ACTIVITY | 0.00157 | 0.02009 | 0.45506 | 0.71837 | 1.32488 | 36 |
| GO_CELL_ADHESION_MOLECULE_BINDING | 0.00000 | 0.00000 | | 0.67109 | 1.32112 | 353 |
| GO_CELL_BODY | 0.00004 | 0.00109 | 0.55733 | 0.59973 | 1.17541 | 293 |
| GO_CELL_CELL_RECOGNITION | 0.00000 | 0.00006 | 0.64355 | 0.82001 | 1.51026 | 35 |
| GO_CELL_CELL_SIGNALING_BY_WNT | 0.00004 | 0.00109 | 0.55733 | 0.59042 | 1.16141 | 339 |
| GO_CELL_CYCLE_G1_S_PHASE_TRANSITION | 0.00376 | 0.03783 | 0.43171 | 0.58112 | 1.13378 | 220 |
| GO_CELL_CYCLE_G2_M_PHASE_TRANSITION | 0.00024 | 0.00459 | 0.51885 | 0.59807 | 1.16746 | 231 |
| GO_CELL_CYCLE_PHASE_TRANSITION | 0.00000 | 0.00001 | 0.69013 | 0.59417 | 1.17348 | 499 |
| GO_CELL_DIVISION | 0.00003 | 0.00071 | 0.57561 | 0.58314 | 1.15094 | 449 |
| GO_CELL_KILLING | 0.00205 | 0.02440 | 0.43171 | 0.66727 | 1.25696 | 60 |
| GO_CELL_RECOGNITION | 0.00000 | 0.00000 | 0.73376 | 0.75953 | 1.43954 | 74 |
| GO_CELL_REDOX_HOMEOSTASIS | 0.00163 | 0.02050 | 0.45506 | 0.71772 | 1.32665 | 38 |
| GO_CELL_SUBSTRATE_JUNCTION | 0.00000 | 0.00000 | | 0.65742 | 1.29171 | 316 |
| GO_CELL_SURFACE | 0.00000 | 0.00000 | 0.73376 | 0.62572 | 1.23070 | 330 |
| GO_CELL_SURFACE_RECEPTOR_SIGNALING_PATHWAY_INVOLVEI | 0.00038 | 0.00640 | 0.49849 | 0.57762 | 1.13770 | 374 |
| GO_CHAPERONE_BINDING | 0.00495 | 0.04573 | 0.40702 | 0.62860 | 1.19935 | 86 |
| GO_CHAPERONE_COMPLEX | 0.00001 | 0.00028 | 0.59333 | 0.86337 | 1.55587 | 22 |
| GO_CHAPERONE_MEDIATED_PROTEIN_COMPLEX_ASSEMBLY | 0.00100 | 0.01430 | 0.45506 | 0.80787 | 1.44337 | 19 |
| GO_CHAPERONE_MEDIATED_PROTEIN_FOLDING | 0.00168 | 0.02098 | 0.45506 | 0.71110 | 1.32288 | 44 |
| GO_CHROMOSOMAL_REGION | 0.00018 | 0.00349 | 0.51885 | 0.59323 | 1.16180 | 286 |
| GO_CHROMOSOME_CENTROMERIC_REGION | 0.00172 | 0.02135 | 0.45506 | 0.60447 | 1.17248 | 168 |
| GO_CHROMOSOME_SEGREGATION | 0.00003 | 0.00087 | 0.55733 | 0.60463 | 1.18280 | 270 |
| GO_CLUSTER_OF_ACTIN_BASED_CELL_PROJECTIONS | 0.00074 | 0.01107 | 0.47727 | 0.64184 | 1.23044 | 99 |
| GO_COATED_VESICLE | 0.00213 | 0.02490 | 0.43171 | 0.59512 | 1.15846 | 199 |
| GO_COATED_VESICLE_MEMBRANE | 0.00060 | 0.00923 | 0.47727 | 0.63443 | 1.22207 | 115 |
| GO_COLLAGEN_BINDING | 0.00465 | 0.04430 | 0.40702 | 0.73471 | 1.33628 | 26 |
| GO_COLLAGEN_CONTAINING_EXTRACELLULAR_MATRIX | 0.00000 | 0.00006 | 0.64355 | 0.65358 | 1.26542 | 157 |
| GO_CONDENSED_CHROMOSOME | 0.00350 | 0.03580 | 0.43171 | 0.59394 | 1.15417 | 179 |
| GO_CONTRACTILE_FIBER | 0.00064 | 0.00972 | 0.47727 | 0.62217 | 1.19973 | 134 |
| GO_COPI_COATED_VESICLE | 0.00316 | 0.03344 | 0.43171 | 0.75688 | 1.37234 | 25 |
| GO_CORNIFICATION | 0.00027 | 0.00500 | 0.49849 | 0.77970 | 1.42512 | 28 |
| GO_CORTICAL_CYTOSKELETON | 0.00502 | 0.04608 | 0.40702 | 0.64388 | 1.21870 | 71 |
| GO_COTRANSLATIONAL_PROTEIN_TARGETING_TO_MEMBRANE | 0.00000 | 0.00000 | | 0.78223 | 1.49819 | 94 |
| GO_CYTOCHROME_COMPLEX | 0.00035 | 0.00610 | 0.49849 | 0.80796 | 1.45601 | 22 |
| GO_CYTOKINE_MEDIATED_SIGNALING_PATHWAY | 0.00000 | 0.00002 | 0.67496 | 0.60742 | 1.19735 | 392 |
| GO_CYTOPLASMIC_STRESS_GRANULE | 0.00014 | 0.00302 | 0.51885 | 0.69390 | 1.31009 | 63 |
| GO_CYTOPLASMIC_TRANSLATION | 0.00000 | 0.00001 | 0.70498 | 0.71773 | 1.37349 | 92 |
| GO_CYTOPLASMIC_TRANSLATIONAL_INITIATION | 0.00178 | 0.02182 | 0.45506 | 0.73517 | 1.35039 | 31 |
| GO_CYTOSOLIC_LARGE_RIBOSOMAL_SUBUNIT | 0.00000 | 0.00005 | 0.64355 | 0.77240 | 1.44935 | 53 |
| GO_CYTOSOLIC_RIBOSOME | 0.00000 | 0.00000 | | 0.77483 | 1.48437 | 97 |
| GO_CYTOSOLIC_SMALL_RIBOSOMAL_SUBUNIT | 0.00000 | 0.00010 | 0.62726 | 0.79384 | 1.47033 | 40 |
| GO_DETECTION_OF_STIMULUS | 0.00020 | 0.00393 | 0.51885 | 0.65208 | 1.25384 | 108 |
| GO_DETOXIFICATION | 0.00413 | 0.04065 | 0.40702 | 0.65399 | 1.23684 | 70 |
| GO_DEVELOPMENT_OF_PRIMARY_FEMALE_SEXUAL_CHARACTE | 0.00179 | 0.02183 | 0.45506 | 0.68410 | 1.28057 | 51 |
| GO_DICARBOXYLIC_ACID_METABOLIC_PROCESS | 0.00261 | 0.02910 | 0.43171 | 0.65541 | 1.23747 | 65 |
| GO_DISORDERED_DOMAIN_SPECIFIC_BINDING | 0.00051 | 0.00812 | 0.47727 | 0.76867 | 1.40343 | 27 |
| GO_DISULFIDE_OXIDOREDUCTASE_ACTIVITY | 0.00435 | 0.04238 | 0.40702 | 0.71445 | 1.31232 | 31 |
| GO_DNA_BIOSYNTHETIC_PROCESS | 0.00017 | 0.00334 | 0.51885 | 0.62540 | 1.20833 | 145 |
| GO_DNA_CONFORMATION_CHANGE | 0.00327 | 0.03409 | 0.43171 | 0.57979 | 1.13178 | 231 |
| GO_DNA_DAMAGE_RESPONSE_DETECTION_OF_DNA_DAMAGE | 0.00305 | 0.03271 | 0.43171 | 0.71625 | 1.31916 | 35 |
| GO_DNA_GEOMETRIC_CHANGE | 0.00141 | 0.01850 | 0.45506 | 0.62642 | 1.20356 | 105 |
| GO_DOUBLE_STRANDED_RNA_BINDING | 0.00039 | 0.00658 | 0.49849 | 0.69158 | 1.30365 | 61 |
| GO_DRUG_BINDING | 0.00450 | 0.04342 | 0.40702 | 0.66245 | 1.24130 | 52 |
| GO_ELECTRON_TRANSFER_ACTIVITY | 0.00000 | 0.00001 | 0.67496 | 0.72917 | 1.38765 | 80 |
| GO_ELECTRON_TRANSPORT_CHAIN | 0.00000 | 0.00000 | | 0.73336 | 1.41392 | 133 |
| GO_EMBRYO_IMPLANTATION | 0.00044 | 0.00706 | 0.49849 | 0.75968 | 1.39146 | 29 |
| GO_ENDOCYTIC_VESICLE_MEMBRANE | 0.00524 | 0.04770 | 0.40702 | 0.62563 | 1.19564 | 89 |
| GO_ENDOPEPTIDASE_ACTIVITY | 0.00065 | 0.00981 | 0.47727 | 0.60123 | 1.16998 | 196 |

| | | | | | | |
|---|---|---|---|---|---|---|
| GO_ENDOPEPTIDASE_COMPLEX | 0.00000 | 0.00001 | 0.70498 | 0.76593 | 1.44609 | 63 |
| GO_ENDOPEPTIDASE_REGULATOR_ACTIVITY | 0.00006 | 0.00145 | 0.55733 | 0.70425 | 1.33001 | 68 |
| GO_ENDOPLASMIC_RETICULUM_GOLGI_INTERMEDIATE_COMPA | 0.00055 | 0.00863 | 0.47727 | 0.64758 | 1.24050 | 96 |
| GO_ENDOPLASMIC_RETICULUM_GOLGI_INTERMEDIATE_COMPA | 0.00148 | 0.01923 | 0.45506 | 0.68944 | 1.29056 | 51 |
| GO_ENDOPLASMIC_RETICULUM_TO_GOLGI_VESICLE_MEDIATED | 0.00532 | 0.04803 | 0.40702 | 0.59194 | 1.14697 | 162 |
| GO_ENERGY_DERIVATION_BY_OXIDATION_OF_ORGANIC_COMP | 0.00000 | 0.00000 | | 0.69995 | 1.36305 | 202 |
| GO_ENZYME_INHIBITOR_ACTIVITY | 0.00246 | 0.02780 | 0.43171 | 0.58913 | 1.14643 | 191 |
| GO_EPIDERMAL_CELL_DIFFERENTIATION | 0.00216 | 0.02519 | 0.43171 | 0.62667 | 1.20589 | 112 |
| GO_EPITHELIAL_CELL_DIFFERENTIATION | 0.00054 | 0.00857 | 0.47727 | 0.57819 | 1.13725 | 334 |
| GO_ESTABLISHMENT_OF_ORGANELLE_LOCALIZATION | 0.00122 | 0.01679 | 0.45506 | 0.57737 | 1.13489 | 323 |
| GO_ESTABLISHMENT_OF_PROTEIN_LOCALIZATION_TO_CHROM( | 0.00016 | 0.00321 | 0.51885 | 0.81363 | 1.46985 | 23 |
| GO_ESTABLISHMENT_OF_PROTEIN_LOCALIZATION_TO_ENDOPL | 0.00000 | 0.00000 | | 0.77241 | 1.48479 | 106 |
| GO_ESTABLISHMENT_OF_PROTEIN_LOCALIZATION_TO_MEMBR | 0.00000 | 0.00000 | | 0.67484 | 1.31991 | 260 |
| GO_ESTABLISHMENT_OF_PROTEIN_LOCALIZATION_TO_ORGANE | 0.00000 | 0.00000 | | 0.65856 | 1.30056 | 461 |
| GO_ESTABLISHMENT_OF_PROTEIN_LOCALIZATION_TO_TELOME | 0.00000 | 0.00006 | 0.64355 | 0.90447 | 1.60870 | 18 |
| GO_ESTABLISHMENT_OF_RNA_LOCALIZATION | 0.00000 | 0.00005 | 0.64355 | 0.64271 | 1.24830 | 176 |
| GO_ESTABLISHMENT_OF_TISSUE_POLARITY | 0.00000 | 0.00000 | 0.81404 | 0.75693 | 1.44999 | 96 |
| GO_EUKARYOTIC_TRANSLATION_INITIATION_FACTOR_3_COMPL | 0.00143 | 0.01863 | 0.45506 | 0.83263 | 1.46100 | 15 |
| GO_EXTERNAL_SIDE_OF_PLASMA_MEMBRANE | 0.00159 | 0.02016 | 0.45506 | 0.61775 | 1.19022 | 118 |
| GO_EXTRACELLULAR_MATRIX | 0.00042 | 0.00695 | 0.49849 | 0.61138 | 1.18938 | 189 |
| GO_FATTY_ACID_DERIVATIVE_METABOLIC_PROCESS | 0.00043 | 0.00695 | 0.49849 | 0.66266 | 1.26212 | 81 |
| GO_FATTY_ACYL_COA_METABOLIC_PROCESS | 0.00502 | 0.04608 | 0.40702 | 0.72264 | 1.32082 | 28 |
| GO_FC_EPSILON_RECEPTOR_SIGNALING_PATHWAY | 0.00000 | 0.00000 | 0.71951 | 0.72370 | 1.38610 | 94 |
| GO_FC_RECEPTOR_SIGNALING_PATHWAY | 0.00000 | 0.00000 | 0.70498 | 0.68815 | 1.32879 | 140 |
| GO_FERTILIZATION | 0.00000 | 0.00005 | 0.64355 | 0.73257 | 1.38752 | 72 |
| GO_FIBRILLAR_CENTER | 0.00246 | 0.02780 | 0.43171 | 0.62638 | 1.19990 | 96 |
| GO_FIBROBLAST_GROWTH_FACTOR_RECEPTOR_SIGNALING_PAT | 0.00495 | 0.04573 | 0.40702 | 0.64888 | 1.22552 | 67 |
| GO_FIBROBLAST_PROLIFERATION | 0.00032 | 0.00575 | 0.49849 | 0.70676 | 1.32618 | 53 |
| GO_FICOLIN_1_RICH_GRANULE | 0.00000 | 0.00000 | 0.70498 | 0.71544 | 1.37138 | 98 |
| GO_GABA_ERGIC_SYNAPSE | 0.00022 | 0.00427 | 0.51885 | 0.76430 | 1.40406 | 32 |
| GO_GENERATION_OF_PRECURSOR_METABOLITES_AND_ENERGY | 0.00000 | 0.00000 | | 0.67527 | 1.32968 | 368 |
| GO_GENE_SILENCING | 0.00019 | 0.00376 | 0.51885 | 0.61625 | 1.19788 | 184 |
| GO_GLUCOSE_CATABOLIC_PROCESS | 0.00495 | 0.04573 | 0.40702 | 0.72314 | 1.32174 | 28 |
| GO_GLUTAMATERGIC_SYNAPSE | 0.00135 | 0.01808 | 0.45506 | 0.60052 | 1.16783 | 186 |
| GO_GLYCOLYTIC_PROCESS_THROUGH_FRUCTOSE_6_PHOSPHAT | 0.00554 | 0.04968 | 0.40702 | 0.74156 | 1.34111 | 24 |
| GO_GOLGI_ASSOCIATED_VESICLE | 0.00085 | 0.01239 | 0.47727 | 0.62294 | 1.20037 | 129 |
| GO_GOLGI_ASSOCIATED_VESICLE_MEMBRANE | 0.00313 | 0.03310 | 0.43171 | 0.64208 | 1.21753 | 75 |
| GO_GTPASE_ACTIVITY | 0.00004 | 0.00098 | 0.55733 | 0.61546 | 1.19974 | 210 |
| GO_GUANYL_NUCLEOTIDE_BINDING | 0.00005 | 0.00126 | 0.55733 | 0.60899 | 1.19065 | 252 |
| GO_HEAT_SHOCK_PROTEIN_BINDING | 0.00001 | 0.00034 | 0.59333 | 0.68258 | 1.30633 | 93 |
| GO_HEMATOPOIETIC_PROGENITOR_CELL_DIFFERENTIATION | 0.00000 | 0.00000 | 0.71951 | 0.70099 | 1.35150 | 121 |
| GO_HEMATOPOIETIC_STEM_CELL_DIFFERENTIATION | 0.00000 | 0.00000 | 0.76146 | 0.77246 | 1.46307 | 72 |
| GO_HISTONE_MRNA_METABOLIC_PROCESS | 0.00265 | 0.02936 | 0.43171 | 0.76832 | 1.38459 | 22 |
| GO_HOST_CELLULAR_COMPONENT | 0.00043 | 0.00695 | 0.49849 | 0.69729 | 1.31074 | 54 |
| GO_IMMUNE_RESPONSE_REGULATING_SIGNALING_PATHWAY | 0.00000 | 0.00003 | 0.65944 | 0.63620 | 1.24112 | 225 |
| GO_IMPORT_INTO_CELL | 0.00428 | 0.04185 | 0.40702 | 0.61709 | 1.18655 | 108 |
| GO_IMPORT_INTO_NUCLEUS | 0.00039 | 0.00652 | 0.49849 | 0.62707 | 1.20905 | 131 |
| GO_INNATE_IMMUNE_RESPONSE | 0.00000 | 0.00003 | 0.65944 | 0.59893 | 1.18089 | 406 |
| GO_INNATE_IMMUNE_RESPONSE_ACTIVATING_SIGNAL_TRANS | 0.00000 | 0.00001 | 0.69013 | 0.73348 | 1.39585 | 80 |
| GO_INNER_MITOCHONDRIAL_MEMBRANE_PROTEIN_COMPLEX | 0.00000 | 0.00000 | | 0.76576 | 1.46666 | 94 |
| GO_INORGANIC_ION_TRANSMEMBRANE_TRANSPORT | 0.00141 | 0.01850 | 0.45506 | 0.56404 | 1.11185 | 401 |
| GO_INTERACTION_WITH_HOST | 0.00139 | 0.01850 | 0.45506 | 0.60855 | 1.17710 | 151 |
| GO_INTERLEUKIN_1_MEDIATED_SIGNALING_PATHWAY | 0.00000 | 0.00000 | 0.77494 | 0.75654 | 1.44176 | 83 |
| GO_INTRAMOLECULAR_OXIDOREDUCTASE_ACTIVITY | 0.00051 | 0.00812 | 0.47727 | 0.74506 | 1.37081 | 34 |
| GO_INTRINSIC_APOPTOTIC_SIGNALING_PATHWAY | 0.00156 | 0.01993 | 0.45506 | 0.59427 | 1.15817 | 207 |
| GO_ION_HOMEOSTASIS | 0.00170 | 0.02117 | 0.45506 | 0.56164 | 1.10717 | 403 |
| GO_ION_TRANSMEMBRANE_TRANSPORTER_ACTIVITY | 0.00106 | 0.01495 | 0.45506 | 0.56696 | 1.11734 | 390 |
| GO_IRON_ION_HOMEOSTASIS | 0.00361 | 0.03657 | 0.43171 | 0.65837 | 1.23983 | 59 |
| GO_IRON_ION_TRANSPORT | 0.00054 | 0.00857 | 0.47727 | 0.69556 | 1.30334 | 52 |
| GO_ISOMERASE_ACTIVITY | 0.00413 | 0.04065 | 0.40702 | 0.61065 | 1.17496 | 113 |
| GO_KERATINIZATION | 0.00038 | 0.00640 | 0.49849 | 0.75690 | 1.39013 | 33 |
| GO_KILLING_OF_CELLS_OF_OTHER_ORGANISM | 0.00495 | 0.04573 | 0.40702 | 0.78614 | 1.38643 | 16 |
| GO_LARGE_RIBOSOMAL_SUBUNIT | 0.00000 | 0.00000 | | 0.77656 | 1.49319 | 108 |
| GO_LATERAL_PLASMA_MEMBRANE | 0.00524 | 0.04770 | 0.40702 | 0.67385 | 1.25557 | 47 |
| GO_LEUKOCYTE_MEDIATED_IMMUNITY | 0.00000 | 0.00000 | | 0.64010 | 1.26331 | 447 |
| GO_LIPOPROTEIN_PARTICLE_RECEPTOR_BINDING | 0.00126 | 0.01715 | 0.45506 | 0.80503 | 1.42497 | 17 |
| GO_LIVER_REGENERATION | 0.00093 | 0.01337 | 0.47727 | 0.78442 | 1.41708 | 23 |
| GO_LOW_DENSITY_LIPOPROTEIN_RECEPTOR_PARTICLE_METAB( | 0.00327 | 0.03409 | 0.43171 | 0.78520 | 1.39657 | 18 |
| GO_LYASE_ACTIVITY | 0.00152 | 0.01958 | 0.45506 | 0.61769 | 1.19079 | 122 |
| GO_LYSOSOMAL_LUMEN | 0.00143 | 0.01863 | 0.45506 | 0.67371 | 1.26872 | 59 |

| | | | | | | |
|---|---|---|---|---|---|---|
| GO_MAINTENANCE_OF_CELL_NUMBER | 0.00435 | 0.04238 | 0.40702 | 0.60761 | 1.17073 | 117 |
| GO_MATURATION_OF_SSU_RRNA | 0.00126 | 0.01715 | 0.45506 | 0.69615 | 1.29711 | 47 |
| GO_MATURATION_OF_SSU_RRNA_FROM_TRICISTRONIC_RRNA_ | 0.00554 | 0.04968 | 0.40702 | 0.70205 | 1.29301 | 35 |
| GO_MEMBRANE_REGION | 0.00000 | 0.00003 | 0.65944 | 0.64496 | 1.25542 | 194 |
| GO_METALLOPEPTIDASE_ACTIVITY | 0.00487 | 0.04559 | 0.40702 | 0.62714 | 1.19577 | 85 |
| GO_METAL_ION_HOMEOSTASIS | 0.00032 | 0.00575 | 0.49849 | 0.58816 | 1.15376 | 301 |
| GO_METAL_ION_TRANSPORT | 0.00390 | 0.03895 | 0.43171 | 0.55877 | 1.10227 | 419 |
| GO_METAPHASE_ANAPHASE_TRANSITION_OF_CELL_CYCLE | 0.00320 | 0.03371 | 0.43171 | 0.66563 | 1.25122 | 54 |
| GO_MICROTUBULE_CYTOSKELETON_ORGANIZATION_INVOLVED_ | 0.00220 | 0.02548 | 0.43171 | 0.61295 | 1.18118 | 127 |
| GO_MICROVILLUS | 0.00168 | 0.02098 | 0.45506 | 0.68128 | 1.28184 | 56 |
| GO_MIDBODY | 0.00561 | 0.04971 | 0.40702 | 0.59866 | 1.15606 | 141 |
| GO_MITOCHONDRIAL_ELECTRON_TRANSPORT_NADH_TO_UBIQ | 0.00000 | 0.00006 | 0.64355 | 0.79447 | 1.47799 | 44 |
| GO_MITOCHONDRIAL_GENE_EXPRESSION | 0.00000 | 0.00000 | 0.78819 | 0.70230 | 1.35844 | 151 |
| GO_MITOCHONDRIAL_LARGE_RIBOSOMAL_SUBUNIT | 0.00000 | 0.00000 | 0.71951 | 0.79015 | 1.48529 | 54 |
| GO_MITOCHONDRIAL_MATRIX | 0.00000 | 0.00000 | 0.82666 | 0.63141 | 1.24412 | 387 |
| GO_MITOCHONDRIAL_MEMBRANE_ORGANIZATION | 0.00050 | 0.00802 | 0.47727 | 0.64578 | 1.23686 | 94 |
| GO_MITOCHONDRIAL_PROTEIN_COMPLEX | 0.00000 | 0.00000 | | 0.75011 | 1.46215 | 212 |
| GO_MITOCHONDRIAL_RESPIRATORY_CHAIN_COMPLEX_ASSEME | 0.00001 | 0.00033 | 0.59333 | 0.69650 | 1.32804 | 85 |
| GO_MITOCHONDRIAL_SMALL_RIBOSOMAL_SUBUNIT | 0.00119 | 0.01640 | 0.45506 | 0.75384 | 1.37785 | 28 |
| GO_MITOCHONDRIAL_TRANSLATION | 0.00000 | 0.00000 | 0.80122 | 0.71435 | 1.37651 | 128 |
| GO_MITOCHONDRIAL_TRANSLATIONAL_TERMINATION | 0.00000 | 0.00000 | 0.81404 | 0.75760 | 1.44657 | 87 |
| GO_MITOCHONDRIAL_TRANSMEMBRANE_TRANSPORT | 0.00213 | 0.02490 | 0.43171 | 0.66300 | 1.25387 | 70 |
| GO_MITOCHONDRIAL_TRANSPORT | 0.00010 | 0.00227 | 0.53843 | 0.61119 | 1.19060 | 203 |
| GO_MITOCHONDRION_ORGANIZATION | 0.00000 | 0.00000 | 0.74774 | 0.61590 | 1.21471 | 416 |
| GO_MITOTIC_NUCLEAR_DIVISION | 0.00008 | 0.00182 | 0.53843 | 0.60435 | 1.17901 | 223 |
| GO_MITOTIC_SISTER_CHROMATID_SEGREGATION | 0.00014 | 0.00302 | 0.51885 | 0.62652 | 1.21048 | 145 |
| GO_MITOTIC_SPINDLE_ORGANIZATION | 0.00078 | 0.01164 | 0.47727 | 0.63943 | 1.22821 | 104 |
| GO_MODIFIED_AMINO_ACID_BINDING | 0.00002 | 0.00048 | 0.57561 | 0.74200 | 1.38679 | 49 |
| GO_MODULATION_BY_HOST_OF_VIRAL_PROCESS | 0.00216 | 0.02519 | 0.43171 | 0.77232 | 1.38066 | 20 |
| GO_MONOCARBOXYLIC_ACID_METABOLIC_PROCESS | 0.00012 | 0.00265 | 0.53843 | 0.57846 | 1.14053 | 406 |
| GO_MONOSACCHARIDE_BIOSYNTHETIC_PROCESS | 0.00305 | 0.03271 | 0.43171 | 0.65908 | 1.24472 | 68 |
| GO_MONOVALENT_INORGANIC_CATION_HOMEOSTASIS | 0.00108 | 0.01507 | 0.45506 | 0.63643 | 1.21783 | 91 |
| GO_MONOVALENT_INORGANIC_CATION_TRANSMEMBRANE_TR | 0.00001 | 0.00026 | 0.59333 | 0.65186 | 1.25996 | 147 |
| GO_MONOVALENT_INORGANIC_CATION_TRANSPORT | 0.00008 | 0.00178 | 0.53843 | 0.60205 | 1.17659 | 248 |
| GO_MORPHOGENESIS_OF_AN_EPITHELIUM | 0.00000 | 0.00009 | 0.62726 | 0.60570 | 1.19137 | 334 |
| GO_MORPHOGENESIS_OF_A_POLARIZED_EPITHELIUM | 0.00000 | 0.00000 | 0.83909 | 0.74384 | 1.43057 | 110 |
| GO_MOVEMENT_IN_HOST_ENVIRONMENT | 0.00268 | 0.02961 | 0.43171 | 0.61282 | 1.18078 | 117 |
| GO_MRNA_3_END_PROCESSING | 0.00087 | 0.01270 | 0.47727 | 0.64010 | 1.22502 | 93 |
| GO_MRNA_5_UTR_BINDING | 0.00002 | 0.00063 | 0.57561 | 0.83759 | 1.51313 | 23 |
| GO_MRNA_BINDING | 0.00000 | 0.00000 | | 0.67496 | 1.31810 | 239 |
| GO_MRNA_EXPORT_FROM_NUCLEUS | 0.00006 | 0.00159 | 0.53843 | 0.66290 | 1.27265 | 103 |
| GO_MRNA_PROCESSING | 0.00000 | 0.00000 | | 0.62959 | 1.24255 | 437 |
| GO_MRNA_TRANSPORT | 0.00007 | 0.00171 | 0.53843 | 0.64103 | 1.23563 | 132 |
| GO_MULTI_MULTICELLULAR_ORGANISM_PROCESS | 0.00041 | 0.00687 | 0.49849 | 0.63087 | 1.21515 | 116 |
| GO_MULTI_ORGANISM_LOCALIZATION | 0.00017 | 0.00334 | 0.51885 | 0.71077 | 1.33608 | 54 |
| GO_MUSCLE_CONTRACTION | 0.00335 | 0.03468 | 0.43171 | 0.59925 | 1.16199 | 165 |
| GO_MUSCLE_FILAMENT_SLIDING | 0.00324 | 0.03393 | 0.43171 | 0.77980 | 1.39322 | 19 |
| GO_MYELOID_LEUKOCYTE_ACTIVATION | 0.00000 | 0.00000 | | 0.65288 | 1.28712 | 395 |
| GO_MYELOID_LEUKOCYTE_MEDIATED_IMMUNITY | 0.00000 | 0.00000 | | 0.65734 | 1.29408 | 356 |
| GO_NADH_DEHYDROGENASE_ACTIVITY | 0.00001 | 0.00019 | 0.61053 | 0.79360 | 1.46691 | 38 |
| GO_NADH_DEHYDROGENASE_COMPLEX | 0.00001 | 0.00028 | 0.59333 | 0.77749 | 1.44215 | 41 |
| GO_NADH_DEHYDROGENASE_COMPLEX_ASSEMBLY | 0.00001 | 0.00035 | 0.59333 | 0.74044 | 1.39281 | 55 |
| GO_NADH_METABOLIC_PROCESS | 0.00034 | 0.00599 | 0.49849 | 0.75773 | 1.39166 | 33 |
| GO_NAD_METABOLIC_PROCESS | 0.00435 | 0.04238 | 0.40702 | 0.70570 | 1.30442 | 37 |
| GO_NCRNA_EXPORT_FROM_NUCLEUS | 0.00205 | 0.02440 | 0.43171 | 0.71473 | 1.31817 | 36 |
| GO_NCRNA_METABOLIC_PROCESS | 0.00000 | 0.00000 | 0.70498 | 0.61139 | 1.20509 | 399 |
| GO_NCRNA_PROCESSING | 0.00000 | 0.00000 | 0.73376 | 0.62238 | 1.22425 | 341 |
| GO_NEGATIVE_REGULATION_OF_APOPTOTIC_SIGNALING_PATHV | 0.00364 | 0.03677 | 0.43171 | 0.59958 | 1.15998 | 150 |
| GO_NEGATIVE_REGULATION_OF_ATP_METABOLIC_PROCESS | 0.00302 | 0.03248 | 0.43171 | 0.78586 | 1.39103 | 17 |
| GO_NEGATIVE_REGULATION_OF_CANONICAL_WNT_SIGNALING | 0.00000 | 0.00000 | 0.71951 | 0.69629 | 1.34180 | 123 |
| GO_NEGATIVE_REGULATION_OF_CATALYTIC_ACTIVITY | 0.00145 | 0.01879 | 0.45506 | 0.56238 | 1.11005 | 450 |
| GO_NEGATIVE_REGULATION_OF_CELLULAR_AMIDE_METABOLIC | 0.00268 | 0.02961 | 0.43171 | 0.61074 | 1.17686 | 124 |
| GO_NEGATIVE_REGULATION_OF_CELLULAR_COMPONENT_ORG | 0.00320 | 0.03371 | 0.43171 | 0.55470 | 1.09534 | 470 |
| GO_NEGATIVE_REGULATION_OF_CELL_CYCLE | 0.00023 | 0.00445 | 0.51885 | 0.57348 | 1.13228 | 455 |
| GO_NEGATIVE_REGULATION_OF_CELL_CYCLE_G2_M_PHASE_TR | 0.00000 | 0.00007 | 0.62726 | 0.70549 | 1.34606 | 86 |
| GO_NEGATIVE_REGULATION_OF_CELL_CYCLE_PHASE_TRANSITI( | 0.00000 | 0.00000 | 0.70498 | 0.65532 | 1.27560 | 194 |
| GO_NEGATIVE_REGULATION_OF_CELL_CYCLE_PROCESS | 0.00000 | 0.00005 | 0.64355 | 0.62725 | 1.22670 | 259 |
| GO_NEGATIVE_REGULATION_OF_HYDROLASE_ACTIVITY | 0.00216 | 0.02519 | 0.43171 | 0.58306 | 1.13858 | 238 |
| GO_NEGATIVE_REGULATION_OF_INTERLEUKIN_1_PRODUCTION | 0.00205 | 0.02440 | 0.43171 | 0.82297 | 1.44404 | 15 |
| GO_NEGATIVE_REGULATION_OF_MITOTIC_CELL_CYCLE | 0.00000 | 0.00010 | 0.62726 | 0.62191 | 1.21541 | 248 |

| | | | | | | |
|---|---|---|---|---|---|---|
| GO_NEGATIVE_REGULATION_OF_MRNA_METABOLIC_PROCESS | 0.00213 | 0.02490 | 0.43171 | 0.66731 | 1.25879 | 62 |
| GO_NEGATIVE_REGULATION_OF_MRNA_SPLICING_VIA_SPLICEO | 0.00339 | 0.03501 | 0.43171 | 0.76309 | 1.36417 | 20 |
| GO_NEGATIVE_REGULATION_OF_NUCLEAR_DIVISION | 0.00335 | 0.03468 | 0.43171 | 0.67724 | 1.26685 | 50 |
| GO_NEGATIVE_REGULATION_OF_ORGANELLE_ORGANIZATION | 0.00487 | 0.04559 | 0.40702 | 0.57038 | 1.11722 | 288 |
| GO_NEGATIVE_REGULATION_OF_PEPTIDASE_ACTIVITY | 0.00036 | 0.00621 | 0.49849 | 0.64044 | 1.23159 | 109 |
| GO_NEGATIVE_REGULATION_OF_PROTEIN_CATABOLIC_PROCESS | 0.00342 | 0.03521 | 0.43171 | 0.62320 | 1.19268 | 93 |
| GO_NEGATIVE_REGULATION_OF_PROTEOLYSIS | 0.00205 | 0.02440 | 0.43171 | 0.60172 | 1.16902 | 178 |
| GO_NEGATIVE_REGULATION_OF_REACTIVE_OXYGEN_SPECIES_B | 0.00036 | 0.00621 | 0.49849 | 0.83108 | 1.47818 | 18 |
| GO_NEGATIVE_REGULATION_OF_REPRODUCTIVE_PROCESS | 0.00457 | 0.04386 | 0.40702 | 0.74845 | 1.35356 | 24 |
| GO_NEGATIVE_REGULATION_OF_RNA_SPLICING | 0.00183 | 0.02219 | 0.45506 | 0.76824 | 1.38935 | 24 |
| GO_NEGATIVE_REGULATION_OF_WNT_SIGNALING_PATHWAY | 0.00000 | 0.00002 | 0.67496 | 0.67731 | 1.30721 | 136 |
| GO_NEURAL_CREST_CELL_MIGRATION | 0.00265 | 0.02936 | 0.43171 | 0.73967 | 1.35049 | 27 |
| GO_NEURON_RECOGNITION | 0.00390 | 0.03895 | 0.43171 | 0.71862 | 1.31998 | 31 |
| GO_NIK_NF_KAPPAB_SIGNALING | 0.00000 | 0.00000 | 0.77494 | 0.72329 | 1.39253 | 114 |
| GO_NON_CANONICAL_WNT_SIGNALING_PATHWAY | 0.00000 | 0.00000 | | 0.75756 | 1.45654 | 107 |
| GO_NUCLEAR_CHROMOSOME_SEGREGATION | 0.00098 | 0.01407 | 0.45506 | 0.59156 | 1.15357 | 215 |
| GO_NUCLEAR_ENVELOPE | 0.00005 | 0.00131 | 0.55733 | 0.59109 | 1.16270 | 341 |
| GO_NUCLEAR_EXPORT | 0.00000 | 0.00004 | 0.65944 | 0.64915 | 1.26157 | 181 |
| GO_NUCLEAR_MATRIX | 0.00502 | 0.04608 | 0.40702 | 0.62777 | 1.19868 | 87 |
| GO_NUCLEAR_MEMBRANE | 0.00185 | 0.02237 | 0.45506 | 0.58925 | 1.14853 | 211 |
| GO_NUCLEAR_PERIPHERY | 0.00043 | 0.00701 | 0.49849 | 0.64264 | 1.23558 | 107 |
| GO_NUCLEAR_PORE | 0.00561 | 0.04971 | 0.40702 | 0.64876 | 1.22522 | 68 |
| GO_NUCLEAR_SPECK | 0.00015 | 0.00312 | 0.51885 | 0.59159 | 1.16089 | 306 |
| GO_NUCLEAR_TRANSCRIBED_MRNA_CATABOLIC_PROCESS | 0.00000 | 0.00000 | | 0.70362 | 1.36820 | 185 |
| GO_NUCLEAR_TRANSCRIBED_MRNA_CATABOLIC_PROCESS_NON | 0.00000 | 0.00000 | | 0.74239 | 1.42931 | 114 |
| GO_NUCLEAR_TRANSPORT | 0.00000 | 0.00000 | 0.71951 | 0.62920 | 1.23383 | 297 |
| GO_NUCLEOBASE_CONTAINING_COMPOUND_TRANSPORT | 0.00000 | 0.00010 | 0.62726 | 0.62975 | 1.22823 | 216 |
| GO_NUCLEOBASE_CONTAINING_SMALL_MOLECULE_METABOLIC | 0.00000 | 0.00003 | 0.65944 | 0.60153 | 1.18594 | 405 |
| GO_NUCLEOSIDE_DIPHOSPHATE_METABOLIC_PROCESS | 0.00016 | 0.00319 | 0.51885 | 0.64491 | 1.24260 | 117 |
| GO_NUCLEOSIDE_PHOSPHATE_BIOSYNTHETIC_PROCESS | 0.00002 | 0.00044 | 0.59333 | 0.64092 | 1.24175 | 161 |
| GO_NUCLEOSIDE_TRIPHOSPHATE_BIOSYNTHETIC_PROCESS | 0.00040 | 0.00676 | 0.49849 | 0.72727 | 1.35074 | 42 |
| GO_NUCLEOSIDE_TRIPHOSPHATE_METABOLIC_PROCESS | 0.00009 | 0.00196 | 0.53843 | 0.69306 | 1.30854 | 65 |
| GO_NUCLEOTIDE_PHOSPHORYLATION | 0.00016 | 0.00328 | 0.51885 | 0.65541 | 1.25767 | 102 |
| GO_OOCYTE_DIFFERENTIATION | 0.00450 | 0.04342 | 0.40702 | 0.73513 | 1.33705 | 26 |
| GO_ORGANELLAR_RIBOSOME | 0.00000 | 0.00000 | 0.76146 | 0.75712 | 1.44311 | 84 |
| GO_ORGANELLE_ENVELOPE_LUMEN | 0.00026 | 0.00482 | 0.49849 | 0.67265 | 1.27716 | 78 |
| GO_ORGANELLE_FISSION | 0.00154 | 0.01974 | 0.45506 | 0.57133 | 1.12383 | 342 |
| GO_ORGANELLE_INNER_MEMBRANE | 0.00000 | 0.00000 | | 0.68423 | 1.34915 | 412 |
| GO_ORGANIC_CYCLIC_COMPOUND_CATABOLIC_PROCESS | 0.00000 | 0.00000 | | 0.66975 | 1.32263 | 478 |
| GO_ORGANIC_HYDROXY_COMPOUND_BIOSYNTHETIC_PROCESS | 0.00141 | 0.01850 | 0.45506 | 0.60712 | 1.17627 | 161 |
| GO_OXIDATIVE_PHOSPHORYLATION | 0.00000 | 0.00000 | | 0.78902 | 1.51260 | 99 |
| GO_OXIDOREDUCTASE_ACTIVITY | 0.00000 | 0.00000 | | 0.63268 | 1.24935 | 468 |
| GO_OXIDOREDUCTASE_ACTIVITY_ACTING_ON_A_HEME_GROUP | 0.00176 | 0.02168 | 0.45506 | 0.79383 | 1.41828 | 19 |
| GO_OXIDOREDUCTASE_ACTIVITY_ACTING_ON_NAD_P_H | 0.00002 | 0.00056 | 0.57561 | 0.71468 | 1.35162 | 70 |
| GO_OXIDOREDUCTASE_ACTIVITY_ACTING_ON_NAD_P_H_QUIN | 0.00004 | 0.00113 | 0.55733 | 0.74511 | 1.38835 | 47 |
| GO_OXIDOREDUCTASE_ACTIVITY_ACTING_ON_PEROXIDE_AS_A | 0.00106 | 0.01495 | 0.45506 | 0.73897 | 1.35352 | 29 |
| GO_OXIDOREDUCTASE_COMPLEX | 0.00000 | 0.00000 | 0.73376 | 0.74898 | 1.42731 | 82 |
| GO_PEPTIDASE_ACTIVITY | 0.00133 | 0.01799 | 0.45506 | 0.57681 | 1.13188 | 306 |
| GO_PEPTIDASE_COMPLEX | 0.00000 | 0.00002 | 0.67496 | 0.73250 | 1.39240 | 79 |
| GO_PEPTIDASE_REGULATOR_ACTIVITY | 0.00313 | 0.03310 | 0.43171 | 0.63434 | 1.21189 | 88 |
| GO_PEPTIDE_BINDING | 0.00005 | 0.00124 | 0.55733 | 0.63708 | 1.23187 | 148 |
| GO_PERINUCLEAR_REGION_OF_CYTOPLASM | 0.00276 | 0.03021 | 0.43171 | 0.55814 | 1.10207 | 466 |
| GO_PHAGOCYTOSIS | 0.00554 | 0.04968 | 0.40702 | 0.59337 | 1.14837 | 155 |
| GO_PHAGOSOME_ACIDIFICATION | 0.00059 | 0.00916 | 0.47727 | 0.78902 | 1.42694 | 24 |
| GO_PHOSPHATIDYLSERINE_BINDING | 0.00035 | 0.00610 | 0.49849 | 0.76387 | 1.39912 | 29 |
| GO_PH_REDUCTION | 0.00302 | 0.03248 | 0.43171 | 0.68821 | 1.28238 | 46 |
| GO_PIGMENT_GRANULE | 0.00000 | 0.00000 | 0.76146 | 0.74333 | 1.41934 | 87 |
| GO_PLASMA_LIPOPROTEIN_PARTICLE_CLEARANCE | 0.00106 | 0.01495 | 0.45506 | 0.73201 | 1.35304 | 37 |
| GO_POLYSOMAL_RIBOSOME | 0.00008 | 0.00180 | 0.53843 | 0.78840 | 1.44683 | 30 |
| GO_POLYSOME | 0.00000 | 0.00003 | 0.65944 | 0.75964 | 1.43296 | 62 |
| GO_POSITIVE_REGULATION_OF_ATP_METABOLIC_PROCESS | 0.00174 | 0.02149 | 0.45506 | 0.73011 | 1.33729 | 29 |
| GO_POSITIVE_REGULATION_OF_BINDING | 0.00390 | 0.03895 | 0.43171 | 0.61042 | 1.17582 | 115 |
| GO_POSITIVE_REGULATION_OF_CANONICAL_WNT_SIGNALING_ | 0.00000 | 0.00000 | 0.73376 | 0.72036 | 1.38514 | 108 |
| GO_POSITIVE_REGULATION_OF_CELLULAR_AMIDE_METABOLIC_ | 0.00001 | 0.00036 | 0.59333 | 0.66485 | 1.28095 | 125 |
| GO_POSITIVE_REGULATION_OF_CELLULAR_PROTEIN_LOCALIZAT | 0.00242 | 0.02748 | 0.43171 | 0.58048 | 1.13378 | 244 |
| GO_POSITIVE_REGULATION_OF_CELL_POPULATION_PROLIFERAT | 0.00040 | 0.00670 | 0.49849 | 0.56967 | 1.12503 | 461 |
| GO_POSITIVE_REGULATION_OF_CYSTEINE_TYPE_ENDOPEPTIDAS | 0.00547 | 0.04923 | 0.40702 | 0.62225 | 1.19077 | 92 |
| GO_POSITIVE_REGULATION_OF_CYTOKINE_PRODUCTION | 0.00283 | 0.03081 | 0.43171 | 0.58672 | 1.14373 | 210 |
| GO_POSITIVE_REGULATION_OF_DEFENSE_RESPONSE | 0.00000 | 0.00000 | 0.73376 | 0.66200 | 1.28884 | 200 |
| GO_POSITIVE_REGULATION_OF_DNA_BIOSYNTHETIC_PROCESS | 0.00010 | 0.00228 | 0.53843 | 0.71224 | 1.33977 | 55 |

| | | | | | | |
|---|---|---|---|---|---|---|
| GO_POSITIVE_REGULATION_OF_DNA_METABOLIC_PROCESS | 0.00020 | 0.00387 | 0.51885 | 0.62771 | 1.21205 | 139 |
| GO_POSITIVE_REGULATION_OF_ESTABLISHMENT_OF_PROTEIN_ | 0.00194 | 0.02331 | 0.45506 | 0.58158 | 1.13709 | 253 |
| GO_POSITIVE_REGULATION_OF_FIBROBLAST_PROLIFERATION | 0.00017 | 0.00338 | 0.51885 | 0.77423 | 1.42213 | 31 |
| GO_POSITIVE_REGULATION_OF_IMMUNE_RESPONSE | 0.00000 | 0.00000 | 0.71951 | 0.63064 | 1.23937 | 321 |
| GO_POSITIVE_REGULATION_OF_IMMUNE_SYSTEM_PROCESS | 0.00000 | 0.00002 | 0.67496 | 0.59748 | 1.17992 | 475 |
| GO_POSITIVE_REGULATION_OF_INNATE_IMMUNE_RESPONSE | 0.00000 | 0.00000 | 0.70498 | 0.68885 | 1.32737 | 128 |
| GO_POSITIVE_REGULATION_OF_MITOTIC_CELL_CYCLE | 0.00178 | 0.02182 | 0.45506 | 0.61563 | 1.18683 | 122 |
| GO_POSITIVE_REGULATION_OF_MRNA_METABOLIC_PROCESS | 0.00157 | 0.02009 | 0.45506 | 0.66330 | 1.25231 | 63 |
| GO_POSITIVE_REGULATION_OF_MRNA_PROCESSING | 0.00250 | 0.02807 | 0.43171 | 0.76194 | 1.38151 | 25 |
| GO_POSITIVE_REGULATION_OF_MRNA_SPLICING_VIA_SPLICEOS | 0.00132 | 0.01782 | 0.45506 | 0.81851 | 1.44353 | 16 |
| GO_POSITIVE_REGULATION_OF_NITRIC_OXIDE_METABOLIC_PRO | 0.00133 | 0.01799 | 0.45506 | 0.80807 | 1.43726 | 18 |
| GO_POSITIVE_REGULATION_OF_ORGANELLE_ORGANIZATION | 0.00010 | 0.00218 | 0.53843 | 0.57473 | 1.13449 | 452 |
| GO_POSITIVE_REGULATION_OF_PROTEIN_LOCALIZATION_TO_N | 0.00012 | 0.00265 | 0.53843 | 0.71018 | 1.33697 | 58 |
| GO_POSITIVE_REGULATION_OF_PROTEOLYSIS | 0.00313 | 0.03310 | 0.43171 | 0.58059 | 1.13381 | 237 |
| GO_POSITIVE_REGULATION_OF_REACTIVE_OXYGEN_SPECIES_BI | 0.00115 | 0.01593 | 0.45506 | 0.79865 | 1.43554 | 21 |
| GO_POSITIVE_REGULATION_OF_REACTIVE_OXYGEN_SPECIES_M | 0.00174 | 0.02149 | 0.45506 | 0.69785 | 1.30035 | 46 |
| GO_POSITIVE_REGULATION_OF_RESPONSE_TO_BIOTIC_STIMUL | 0.00000 | 0.00000 | 0.70498 | 0.67507 | 1.30603 | 150 |
| GO_POSITIVE_REGULATION_OF_RESPONSE_TO_EXTERNAL_STIM | 0.00001 | 0.00016 | 0.61053 | 0.62123 | 1.21505 | 260 |
| GO_POSITIVE_REGULATION_OF_RNA_SPLICING | 0.00265 | 0.02936 | 0.43171 | 0.74051 | 1.35202 | 27 |
| GO_POSITIVE_REGULATION_OF_SIGNAL_TRANSDUCTION_BY_P5 | 0.00390 | 0.03895 | 0.43171 | 0.80515 | 1.41278 | 15 |
| GO_POSITIVE_REGULATION_OF_TELOMERASE_ACTIVITY | 0.00069 | 0.01043 | 0.47727 | 0.75265 | 1.38122 | 30 |
| GO_POSITIVE_REGULATION_OF_TELOMERASE_RNA_LOCALIZATI | 0.00007 | 0.00169 | 0.53843 | 0.88616 | 1.55493 | 15 |
| GO_POSITIVE_REGULATION_OF_TELOMERE_MAINTENANCE | 0.00026 | 0.00486 | 0.49849 | 0.73697 | 1.37102 | 44 |
| GO_POSITIVE_REGULATION_OF_TELOMERE_MAINTENANCE_VIA | 0.00001 | 0.00040 | 0.59333 | 0.80196 | 1.47289 | 33 |
| GO_POSITIVE_REGULATION_OF_TRANSFERASE_ACTIVITY | 0.00108 | 0.01507 | 0.45506 | 0.56501 | 1.11376 | 401 |
| GO_POSITIVE_REGULATION_OF_TRANSLATION | 0.00000 | 0.00016 | 0.61053 | 0.68408 | 1.31526 | 107 |
| GO_POSITIVE_REGULATION_OF_VIRAL_PROCESS | 0.00026 | 0.00486 | 0.49849 | 0.66195 | 1.26462 | 88 |
| GO_POSITIVE_REGULATION_OF_WNT_SIGNALING_PATHWAY | 0.00000 | 0.00000 | 0.73376 | 0.69924 | 1.34747 | 127 |
| GO_POSTSYNAPSE | 0.00239 | 0.02716 | 0.43171 | 0.57062 | 1.12116 | 316 |
| GO_POST_TRANSLATIONAL_PROTEIN_MODIFICATION | 0.00024 | 0.00452 | 0.51885 | 0.60046 | 1.17392 | 251 |
| GO_PRECATALYTIC_SPLICEOSOME | 0.00141 | 0.01850 | 0.45506 | 0.69251 | 1.29540 | 50 |
| GO_PRERIBOSOME | 0.00006 | 0.00154 | 0.53843 | 0.69866 | 1.32020 | 69 |
| GO_PROTEASOMAL_PROTEIN_CATABOLIC_PROCESS | 0.00007 | 0.00169 | 0.53843 | 0.58188 | 1.14652 | 387 |
| GO_PROTEASOMAL_UBIQUITIN_INDEPENDENT_PROTEIN_CATAB | 0.00000 | 0.00003 | 0.65944 | 0.91182 | 1.62178 | 18 |
| GO_PROTEASOME_ACCESSORY_COMPLEX | 0.00005 | 0.00135 | 0.55733 | 0.82638 | 1.49450 | 24 |
| GO_PROTEASOME_CORE_COMPLEX | 0.00000 | 0.00004 | 0.65944 | 0.91231 | 1.61486 | 17 |
| GO_PROTEIN_CONTAINING_COMPLEX_DISASSEMBLY | 0.00000 | 0.00000 | 0.81404 | 0.66285 | 1.29656 | 261 |
| GO_PROTEIN_CONTAINING_COMPLEX_LOCALIZATION | 0.00018 | 0.00352 | 0.51885 | 0.60451 | 1.17943 | 226 |
| GO_PROTEIN_DNA_COMPLEX_SUBUNIT_ORGANIZATION | 0.00052 | 0.00825 | 0.47727 | 0.60887 | 1.18342 | 183 |
| GO_PROTEIN_DOMAIN_SPECIFIC_BINDING | 0.00003 | 0.00088 | 0.55733 | 0.57828 | 1.14190 | 470 |
| GO_PROTEIN_FOLDING | 0.00000 | 0.00000 | 0.73376 | 0.67425 | 1.30867 | 171 |
| GO_PROTEIN_FOLDING_CHAPERONE | 0.00079 | 0.01170 | 0.47727 | 0.81632 | 1.45193 | 18 |
| GO_PROTEIN_HETEROOLIGOMERIZATION | 0.00442 | 0.04297 | 0.40702 | 0.78998 | 1.39322 | 16 |
| GO_PROTEIN_IMPORT | 0.00001 | 0.00040 | 0.59333 | 0.64197 | 1.24306 | 158 |
| GO_PROTEIN_IMPORT_INTO_MITOCHONDRIAL_MATRIX | 0.00231 | 0.02642 | 0.43171 | 0.79530 | 1.41455 | 18 |
| GO_PROTEIN_INSERTION_INTO_MEMBRANE | 0.00191 | 0.02299 | 0.45506 | 0.68322 | 1.27694 | 49 |
| GO_PROTEIN_LOCALIZATION_TO_CHROMOSOME_TELOMERIC_R | 0.00001 | 0.00031 | 0.59333 | 0.82810 | 1.51357 | 28 |
| GO_PROTEIN_LOCALIZATION_TO_ENDOPLASMIC_RETICULUM | 0.00000 | 0.00000 | | 0.73850 | 1.42303 | 129 |
| GO_PROTEIN_LOCALIZATION_TO_MEMBRANE | 0.00000 | 0.00000 | 0.77494 | 0.61216 | 1.20881 | 462 |
| GO_PROTEIN_LOCALIZATION_TO_MITOCHONDRION | 0.00015 | 0.00310 | 0.51885 | 0.64234 | 1.23759 | 118 |
| GO_PROTEIN_LOCALIZATION_TO_NUCLEUS | 0.00000 | 0.00014 | 0.61053 | 0.63315 | 1.23408 | 208 |
| GO_PROTEIN_MATURATION | 0.00357 | 0.03638 | 0.43171 | 0.59209 | 1.14965 | 174 |
| GO_PROTEIN_MODIFICATION_BY_SMALL_PROTEIN_REMOVAL | 0.00002 | 0.00052 | 0.57561 | 0.61535 | 1.20119 | 231 |
| GO_PROTEIN_REFOLDING | 0.00465 | 0.04430 | 0.40702 | 0.77561 | 1.37288 | 17 |
| GO_PROTEIN_STABILIZATION | 0.00081 | 0.01194 | 0.47727 | 0.61881 | 1.19577 | 146 |
| GO_PROTEIN_SUMOYLATION | 0.00450 | 0.04342 | 0.40702 | 0.65043 | 1.22846 | 67 |
| GO_PROTEIN_TARGETING | 0.00000 | 0.00000 | | 0.66223 | 1.30361 | 358 |
| GO_PROTEIN_TARGETING_TO_MEMBRANE | 0.00000 | 0.00000 | | 0.71675 | 1.38879 | 162 |
| GO_PROTEIN_TARGETING_TO_MITOCHONDRION | 0.00113 | 0.01571 | 0.45506 | 0.65407 | 1.24648 | 83 |
| GO_PROTEIN_TRANSMEMBRANE_TRANSPORT | 0.00110 | 0.01526 | 0.45506 | 0.68173 | 1.28238 | 55 |
| GO_PROTON_TRANSMEMBRANE_TRANSPORT | 0.00000 | 0.00001 | 0.69013 | 0.71234 | 1.36309 | 91 |
| GO_PROTON_TRANSMEMBRANE_TRANSPORTER_ACTIVITY | 0.00004 | 0.00115 | 0.55733 | 0.70138 | 1.32535 | 69 |
| GO_PROTON_TRANSPORTING_TWO_SECTOR_ATPASE_COMPLEX | 0.00083 | 0.01218 | 0.47727 | 0.77254 | 1.40508 | 26 |
| GO_PROTON_TRANSPORTING_V_TYPE_ATPASE_COMPLEX | 0.00239 | 0.02716 | 0.43171 | 0.76100 | 1.37626 | 24 |
| GO_PURINE_CONTAINING_COMPOUND_BIOSYNTHETIC_PROCES | 0.00001 | 0.00040 | 0.59333 | 0.65863 | 1.26901 | 126 |
| GO_PURINE_CONTAINING_COMPOUND_METABOLIC_PROCESS | 0.00000 | 0.00000 | 0.73376 | 0.63132 | 1.23831 | 300 |
| GO_PURINE_NUCLEOSIDE_TRIPHOSPHATE_METABOLIC_PROCES | 0.00033 | 0.00582 | 0.49849 | 0.70887 | 1.32487 | 49 |
| GO_PYROPHOSPHATE_HYDROLYSIS_DRIVEN_PROTON_TRANSME | 0.00532 | 0.04803 | 0.40702 | 0.74470 | 1.34678 | 24 |
| GO_PYRUVATE_METABOLIC_PROCESS | 0.00001 | 0.00018 | 0.61053 | 0.67194 | 1.29289 | 113 |
| GO_P_BODY | 0.00472 | 0.04480 | 0.40702 | 0.65470 | 1.23501 | 62 |

| | | | | | | |
|---|---|---|---|---|---|---|
| GO_REACTIVE_OXYGEN_SPECIES_BIOSYNTHETIC_PROCESS | 0.00040 | 0.00670 | 0.49849 | 0.70271 | 1.31857 | 53 |
| GO_REACTIVE_OXYGEN_SPECIES_METABOLIC_PROCESS | 0.00021 | 0.00409 | 0.51885 | 0.62881 | 1.21472 | 144 |
| GO_REGULATION_OF_ALCOHOL_BIOSYNTHETIC_PROCESS | 0.00228 | 0.02614 | 0.43171 | 0.68438 | 1.28020 | 50 |
| GO_REGULATION_OF_ANIMAL_ORGAN_MORPHOGENESIS | 0.00000 | 0.00000 | | 0.72330 | 1.39922 | 149 |
| GO_REGULATION_OF_APOPTOTIC_SIGNALING_PATHWAY | 0.00079 | 0.01170 | 0.47727 | 0.58599 | 1.14641 | 266 |
| GO_REGULATION_OF_ATP_METABOLIC_PROCESS | 0.00000 | 0.00006 | 0.64355 | 0.70523 | 1.34777 | 89 |
| GO_REGULATION_OF_BINDING | 0.00428 | 0.04185 | 0.40702 | 0.58264 | 1.13664 | 225 |
| GO_REGULATION_OF_BLOOD_PRESSURE | 0.00290 | 0.03156 | 0.43171 | 0.65319 | 1.23428 | 69 |
| GO_REGULATION_OF_CARBOHYDRATE_CATABOLIC_PROCESS | 0.00324 | 0.03393 | 0.43171 | 0.65541 | 1.23787 | 67 |
| GO_REGULATION_OF_CELLULAR_AMIDE_METABOLIC_PROCESS | 0.00000 | 0.00000 | 0.78819 | 0.63116 | 1.24143 | 337 |
| GO_REGULATION_OF_CELLULAR_AMINE_METABOLIC_PROCESS | 0.00000 | 0.00000 | 0.74774 | 0.78955 | 1.48730 | 60 |
| GO_REGULATION_OF_CELLULAR_AMINO_ACID_METABOLIC_PR( | 0.00000 | 0.00000 | 0.76146 | 0.80341 | 1.51022 | 54 |
| GO_REGULATION_OF_CELLULAR_KETONE_METABOLIC_PROCESS | 0.00000 | 0.00000 | 0.70498 | 0.69239 | 1.33405 | 126 |
| GO_REGULATION_OF_CELLULAR_PH | 0.00442 | 0.04297 | 0.40702 | 0.65360 | 1.23436 | 68 |
| GO_REGULATION_OF_CELLULAR_PROTEIN_LOCALIZATION | 0.00228 | 0.02614 | 0.43171 | 0.56216 | 1.10830 | 402 |
| GO_REGULATION_OF_CELLULAR_RESPIRATION | 0.00428 | 0.04185 | 0.40702 | 0.77183 | 1.37897 | 19 |
| GO_REGULATION_OF_CELLULAR_RESPONSE_TO_HEAT | 0.00091 | 0.01314 | 0.47727 | 0.67360 | 1.27221 | 67 |
| GO_REGULATION_OF_CELL_CYCLE_G2_M_PHASE_TRANSITION | 0.00016 | 0.00332 | 0.51885 | 0.61554 | 1.19738 | 187 |
| GO_REGULATION_OF_CELL_CYCLE_PHASE_TRANSITION | 0.00000 | 0.00014 | 0.61053 | 0.59968 | 1.18091 | 369 |
| GO_REGULATION_OF_CHOLESTEROL_BIOSYNTHETIC_PROCESS | 0.00209 | 0.02470 | 0.43171 | 0.71476 | 1.32117 | 38 |
| GO_REGULATION_OF_CHOLESTEROL_METABOLIC_PROCESS | 0.00113 | 0.01571 | 0.45506 | 0.69865 | 1.30177 | 47 |
| GO_REGULATION_OF_CHROMOSOME_ORGANIZATION | 0.00036 | 0.00622 | 0.49849 | 0.58896 | 1.15261 | 278 |
| GO_REGULATION_OF_CHROMOSOME_SEGREGATION | 0.00079 | 0.01170 | 0.47727 | 0.64131 | 1.22718 | 91 |
| GO_REGULATION_OF_CHROMOSOME_SEPARATION | 0.00253 | 0.02838 | 0.43171 | 0.66349 | 1.24947 | 59 |
| GO_REGULATION_OF_CYSTEINE_TYPE_ENDOPEPTIDASE_ACTIVIT | 0.00044 | 0.00719 | 0.49849 | 0.62610 | 1.20905 | 141 |
| GO_REGULATION_OF_DEFENSE_RESPONSE | 0.00006 | 0.00150 | 0.55733 | 0.58834 | 1.15824 | 356 |
| GO_REGULATION_OF_DNA_BIOSYNTHETIC_PROCESS | 0.00043 | 0.00695 | 0.49849 | 0.66709 | 1.27130 | 83 |
| GO_REGULATION_OF_DNA_METABOLIC_PROCESS | 0.00055 | 0.00863 | 0.47727 | 0.58909 | 1.15134 | 249 |
| GO_REGULATION_OF_DNA_TEMPLATED_TRANSCRIPTION_IN_RI | 0.00000 | 0.00000 | 0.71951 | 0.72410 | 1.38687 | 94 |
| GO_REGULATION_OF_GENERATION_OF_PRECURSOR_METABOLI | 0.00004 | 0.00114 | 0.55733 | 0.65536 | 1.26115 | 111 |
| GO_REGULATION_OF_GENE_SILENCING | 0.00023 | 0.00434 | 0.51885 | 0.65464 | 1.25268 | 91 |
| GO_REGULATION_OF_HEMATOPOIETIC_PROGENITOR_CELL_DIF | 0.00000 | 0.00000 | 0.73376 | 0.76273 | 1.44465 | 72 |
| GO_REGULATION_OF_HEMATOPOIETIC_STEM_CELL_DIFFERENT | 0.00000 | 0.00000 | 0.74774 | 0.78891 | 1.48817 | 62 |
| GO_REGULATION_OF_HEMOPOIESIS | 0.00390 | 0.03895 | 0.43171 | 0.57739 | 1.12909 | 258 |
| GO_REGULATION_OF_IMMUNE_RESPONSE | 0.00000 | 0.00006 | 0.64355 | 0.60032 | 1.18447 | 433 |
| GO_REGULATION_OF_INNATE_IMMUNE_RESPONSE | 0.00000 | 0.00007 | 0.62726 | 0.64536 | 1.25422 | 181 |
| GO_REGULATION_OF_INTRINSIC_APOPTOTIC_SIGNALING_PATH\ | 0.00179 | 0.02183 | 0.45506 | 0.61548 | 1.18550 | 116 |
| GO_REGULATION_OF_MITOTIC_CELL_CYCLE | 0.00000 | 0.00003 | 0.65944 | 0.58992 | 1.16508 | 499 |
| GO_REGULATION_OF_MORPHOGENESIS_OF_AN_EPITHELIUM | 0.00000 | 0.00000 | | 0.74680 | 1.43884 | 125 |
| GO_REGULATION_OF_MRNA_CATABOLIC_PROCESS | 0.00000 | 0.00000 | | 0.71126 | 1.37919 | 165 |
| GO_REGULATION_OF_MRNA_METABOLIC_PROCESS | 0.00000 | 0.00000 | | 0.67178 | 1.31458 | 276 |
| GO_REGULATION_OF_MRNA_PROCESSING | 0.00008 | 0.00186 | 0.53843 | 0.65189 | 1.25598 | 125 |
| GO_REGULATION_OF_MRNA_SPLICING_VIA_SPLICEOSOME | 0.00012 | 0.00256 | 0.53843 | 0.66950 | 1.28033 | 90 |
| GO_REGULATION_OF_MULTI_ORGANISM_PROCESS | 0.00008 | 0.00190 | 0.53843 | 0.76905 | 1.42150 | 37 |
| GO_REGULATION_OF_MUSCLE_CONTRACTION | 0.00163 | 0.02050 | 0.45506 | 0.64879 | 1.23071 | 76 |
| GO_REGULATION_OF_NUCLEAR_DIVISION | 0.00060 | 0.00923 | 0.47727 | 0.62443 | 1.20552 | 137 |
| GO_REGULATION_OF_NUCLEASE_ACTIVITY | 0.00059 | 0.00916 | 0.47727 | 0.81786 | 1.44767 | 17 |
| GO_REGULATION_OF_NUCLEOTIDE_BIOSYNTHETIC_PROCESS | 0.00302 | 0.03248 | 0.43171 | 0.78902 | 1.40337 | 18 |
| GO_REGULATION_OF_NUCLEOTIDE_METABOLIC_PROCESS | 0.00033 | 0.00588 | 0.49849 | 0.66858 | 1.26942 | 78 |
| GO_REGULATION_OF_OXIDATIVE_PHOSPHORYLATION | 0.00005 | 0.00130 | 0.55733 | 0.85284 | 1.53294 | 21 |
| GO_REGULATION_OF_PEPTIDASE_ACTIVITY | 0.00006 | 0.00152 | 0.53843 | 0.61192 | 1.19388 | 226 |
| GO_REGULATION_OF_PEPTIDE_TRANSPORT | 0.00250 | 0.02807 | 0.43171 | 0.56252 | 1.10917 | 412 |
| GO_REGULATION_OF_PH | 0.00561 | 0.04971 | 0.40702 | 0.64191 | 1.21498 | 71 |
| GO_REGULATION_OF_PLASMA_LIPOPROTEIN_PARTICLE_LEVELS | 0.00006 | 0.00145 | 0.55733 | 0.72431 | 1.36154 | 54 |
| GO_REGULATION_OF_POSTTRANSCRIPTIONAL_GENE_SILENCIN( | 0.00026 | 0.00486 | 0.49849 | 0.66803 | 1.27236 | 81 |
| GO_REGULATION_OF_PROTEIN_CATABOLIC_PROCESS | 0.00309 | 0.03299 | 0.43171 | 0.57360 | 1.12399 | 291 |
| GO_REGULATION_OF_PROTEIN_IMPORT | 0.00487 | 0.04559 | 0.40702 | 0.67491 | 1.25754 | 47 |
| GO_REGULATION_OF_PROTEIN_LOCALIZATION_TO_NUCLEUS | 0.00000 | 0.00013 | 0.61053 | 0.69035 | 1.32101 | 91 |
| GO_REGULATION_OF_PROTEIN_STABILITY | 0.00066 | 0.01007 | 0.47727 | 0.59308 | 1.15811 | 235 |
| GO_REGULATION_OF_PROTEOLYSIS | 0.00007 | 0.00169 | 0.53843 | 0.57923 | 1.14299 | 431 |
| GO_REGULATION_OF_REACTIVE_OXYGEN_SPECIES_BIOSYNTHET | 0.00022 | 0.00427 | 0.51885 | 0.73154 | 1.36343 | 45 |
| GO_REGULATION_OF_REACTIVE_OXYGEN_SPECIES_METABOLIC_ | 0.00088 | 0.01280 | 0.47727 | 0.63948 | 1.22592 | 99 |
| GO_REGULATION_OF_RESPONSE_TO_BIOTIC_STIMULUS | 0.00001 | 0.00039 | 0.59333 | 0.61773 | 1.20625 | 235 |
| GO_REGULATION_OF_RNA_SPLICING | 0.00011 | 0.00241 | 0.53843 | 0.64982 | 1.25198 | 125 |
| GO_REGULATION_OF_SISTER_CHROMATID_SEGREGATION | 0.00405 | 0.04011 | 0.40702 | 0.64401 | 1.22034 | 73 |
| GO_REGULATION_OF_SMALL_MOLECULE_METABOLIC_PROCESS | 0.00000 | 0.00000 | 0.77494 | 0.64342 | 1.26057 | 290 |
| GO_REGULATION_OF_SMOOTH_MUSCLE_CONTRACTION | 0.00007 | 0.00171 | 0.53843 | 0.82642 | 1.49295 | 23 |
| GO_REGULATION_OF_STEM_CELL_DIFFERENTIATION | 0.00000 | 0.00000 | 0.73376 | 0.73434 | 1.40292 | 88 |
| GO_REGULATION_OF_STEROID_METABOLIC_PROCESS | 0.00524 | 0.04770 | 0.40702 | 0.64061 | 1.21567 | 77 |
| GO_REGULATION_OF_SUPEROXIDE_METABOLIC_PROCESS | 0.00346 | 0.03548 | 0.43171 | 0.77826 | 1.39046 | 19 |

| | | | | | | |
|---|---|---|---|---|---|---|
| GO_REGULATION_OF_SUPRAMOLECULAR_FIBER_ORGANIZATIOI | 0.00509 | 0.04656 | 0.40702 | 0.57599 | 1.12367 | 223 |
| GO_REGULATION_OF_SYMBIOTIC_PROCESS | 0.00009 | 0.00211 | 0.53843 | 0.62950 | 1.21763 | 151 |
| GO_REGULATION_OF_TELOMERASE_ACTIVITY | 0.00087 | 0.01263 | 0.47727 | 0.72147 | 1.33426 | 39 |
| GO_REGULATION_OF_TELOMERE_MAINTENANCE | 0.00547 | 0.04923 | 0.40702 | 0.64959 | 1.22679 | 68 |
| GO_REGULATION_OF_TELOMERE_MAINTENANCE_VIA_TELOME | 0.00020 | 0.00387 | 0.51885 | 0.72043 | 1.34856 | 51 |
| GO_REGULATION_OF_TRANSCRIPTION_FROM_RNA_POLYMERA: | 0.00000 | 0.00000 | 0.70498 | 0.76931 | 1.45239 | 64 |
| GO_REGULATION_OF_TRANSLATIONAL_INITIATION | 0.00023 | 0.00437 | 0.51885 | 0.68479 | 1.29399 | 69 |
| GO_REGULATION_OF_VIRAL_TRANSCRIPTION | 0.00361 | 0.03657 | 0.43171 | 0.67497 | 1.26259 | 50 |
| GO_REGULATION_OF_WNT_SIGNALING_PATHWAY | 0.00015 | 0.00312 | 0.51885 | 0.60129 | 1.17419 | 238 |
| GO_RESPIRASOME | 0.00000 | 0.00000 | 0.81404 | 0.78326 | 1.48524 | 75 |
| GO_RESPIRATORY_CHAIN_COMPLEX | 0.00000 | 0.00000 | 0.77494 | 0.79183 | 1.49492 | 64 |
| GO_RESPIRATORY_ELECTRON_TRANSPORT_CHAIN | 0.00000 | 0.00000 | | 0.78275 | 1.49591 | 89 |
| GO_RESPONSE_TO_ALCOHOL | 0.00015 | 0.00310 | 0.51885 | 0.63794 | 1.22915 | 126 |
| GO_RESPONSE_TO_ATP | 0.00087 | 0.01270 | 0.47727 | 0.81086 | 1.43528 | 17 |
| GO_RESPONSE_TO_CALCIUM_ION | 0.00124 | 0.01701 | 0.45506 | 0.65196 | 1.24071 | 80 |
| GO_RESPONSE_TO_CORTICOSTEROID | 0.00010 | 0.00220 | 0.53843 | 0.66894 | 1.27841 | 89 |
| GO_RESPONSE_TO_DRUG | 0.00000 | 0.00007 | 0.62726 | 0.63080 | 1.23070 | 220 |
| GO_RESPONSE_TO_ETHANOL | 0.00168 | 0.02098 | 0.45506 | 0.66543 | 1.25679 | 67 |
| GO_RESPONSE_TO_FIBROBLAST_GROWTH_FACTOR | 0.00357 | 0.03638 | 0.43171 | 0.63469 | 1.20976 | 84 |
| GO_RESPONSE_TO_GROWTH_FACTOR | 0.00272 | 0.02986 | 0.43171 | 0.56235 | 1.10805 | 387 |
| GO_RESPONSE_TO_HEAT | 0.00000 | 0.00013 | 0.62726 | 0.66982 | 1.29069 | 119 |
| GO_RESPONSE_TO_HYDROGEN_PEROXIDE | 0.00532 | 0.04803 | 0.40702 | 0.62255 | 1.19054 | 90 |
| GO_RESPONSE_TO_INORGANIC_SUBSTANCE | 0.00016 | 0.00328 | 0.51885 | 0.58928 | 1.15810 | 321 |
| GO_RESPONSE_TO_INTERLEUKIN_1 | 0.00000 | 0.00000 | 0.78819 | 0.71590 | 1.37950 | 124 |
| GO_RESPONSE_TO_INTERLEUKIN_12 | 0.00036 | 0.00622 | 0.49849 | 0.74158 | 1.36769 | 36 |
| GO_RESPONSE_TO_INTERLEUKIN_4 | 0.00224 | 0.02581 | 0.43171 | 0.77088 | 1.38921 | 22 |
| GO_RESPONSE_TO_LEUKEMIA_INHIBITORY_FACTOR | 0.00076 | 0.01129 | 0.47727 | 0.66148 | 1.25288 | 72 |
| GO_RESPONSE_TO_LIPID | 0.00043 | 0.00695 | 0.49849 | 0.56669 | 1.11902 | 470 |
| GO_RESPONSE_TO_METAL_ION | 0.00143 | 0.01863 | 0.45506 | 0.59895 | 1.16592 | 199 |
| GO_RESPONSE_TO_NUTRIENT | 0.00294 | 0.03185 | 0.43171 | 0.62596 | 1.19987 | 98 |
| GO_RESPONSE_TO_ORGANOPHOSPHORUS | 0.00294 | 0.03185 | 0.43171 | 0.64154 | 1.21511 | 72 |
| GO_RESPONSE_TO_OXIDATIVE_STRESS | 0.00495 | 0.04573 | 0.40702 | 0.57010 | 1.11758 | 294 |
| GO_RESPONSE_TO_OXYGEN_LEVELS | 0.00000 | 0.00000 | | 0.67661 | 1.32155 | 244 |
| GO_RESPONSE_TO_PURINE_CONTAINING_COMPOUND | 0.00509 | 0.04656 | 0.40702 | 0.63176 | 1.20394 | 82 |
| GO_RESPONSE_TO_STEROID_HORMONE | 0.00009 | 0.00200 | 0.53843 | 0.61312 | 1.19360 | 201 |
| GO_RESPONSE_TO_TEMPERATURE_STIMULUS | 0.00002 | 0.00047 | 0.57561 | 0.64468 | 1.24782 | 153 |
| GO_RESPONSE_TO_TOXIC_SUBSTANCE | 0.00011 | 0.00235 | 0.53843 | 0.63892 | 1.23203 | 134 |
| GO_RESPONSE_TO_TUMOR_NECROSIS_FACTOR | 0.00000 | 0.00000 | 0.70498 | 0.66331 | 1.28815 | 175 |
| GO_RESPONSE_TO_VIRUS | 0.00261 | 0.02910 | 0.43171 | 0.59428 | 1.15408 | 175 |
| GO_RETINA_HOMEOSTASIS | 0.00034 | 0.00606 | 0.49849 | 0.75202 | 1.38362 | 34 |
| GO_RIBONUCLEOPROTEIN_COMPLEX_BINDING | 0.00000 | 0.00001 | 0.69013 | 0.69839 | 1.34459 | 114 |
| GO_RIBONUCLEOPROTEIN_COMPLEX_BIOGENESIS | 0.00000 | 0.00000 | | 0.68714 | 1.35448 | 393 |
| GO_RIBONUCLEOPROTEIN_COMPLEX_SUBUNIT_ORGANIZATION | 0.00000 | 0.00000 | 0.81404 | 0.69112 | 1.34183 | 173 |
| GO_RIBONUCLEOPROTEIN_GRANULE | 0.00000 | 0.00000 | 0.71951 | 0.66271 | 1.28875 | 186 |
| GO_RIBONUCLEOSIDE_TRIPHOSPHATE_BIOSYNTHETIC_PROCESS | 0.00009 | 0.00210 | 0.53843 | 0.77194 | 1.42026 | 34 |
| GO_RIBONUCLEOSIDE_TRIPHOSPHATE_METABOLIC_PROCESS | 0.00013 | 0.00280 | 0.51885 | 0.73558 | 1.37065 | 46 |
| GO_RIBOSE_PHOSPHATE_BIOSYNTHETIC_PROCESS | 0.00002 | 0.00056 | 0.57561 | 0.65814 | 1.26821 | 124 |
| GO_RIBOSE_PHOSPHATE_METABOLIC_PROCESS | 0.00000 | 0.00000 | 0.74774 | 0.63837 | 1.25001 | 285 |
| GO_RIBOSOMAL_LARGE_SUBUNIT_BIOGENESIS | 0.00001 | 0.00031 | 0.59333 | 0.72435 | 1.36807 | 67 |
| GO_RIBOSOMAL_SMALL_SUBUNIT_ASSEMBLY | 0.00405 | 0.04011 | 0.40702 | 0.78074 | 1.38865 | 18 |
| GO_RIBOSOMAL_SMALL_SUBUNIT_BINDING | 0.00532 | 0.04803 | 0.40702 | 0.77341 | 1.36899 | 17 |
| GO_RIBOSOMAL_SMALL_SUBUNIT_BIOGENESIS | 0.00005 | 0.00138 | 0.55733 | 0.70510 | 1.33163 | 68 |
| GO_RIBOSOMAL_SUBUNIT | 0.00000 | 0.00000 | | 0.76292 | 1.48089 | 172 |
| GO_RIBOSOME | 0.00000 | 0.00000 | | 0.75411 | 1.46984 | 208 |
| GO_RIBOSOME_ASSEMBLY | 0.00001 | 0.00026 | 0.59333 | 0.74485 | 1.40144 | 56 |
| GO_RIBOSOME_BIOGENESIS | 0.00000 | 0.00000 | | 0.68899 | 1.34833 | 277 |
| GO_RNA_3_END_PROCESSING | 0.00002 | 0.00046 | 0.57561 | 0.65576 | 1.26361 | 128 |
| GO_RNA_CATABOLIC_PROCESS | 0.00000 | 0.00000 | | 0.70701 | 1.39073 | 341 |
| GO_RNA_DEPENDENT_DNA_BIOSYNTHETIC_PROCESS | 0.00003 | 0.00081 | 0.57561 | 0.71669 | 1.34965 | 59 |
| GO_RNA_EXPORT_FROM_NUCLEUS | 0.00000 | 0.00016 | 0.61053 | 0.66306 | 1.27817 | 130 |
| GO_RNA_LOCALIZATION | 0.00000 | 0.00000 | 0.76146 | 0.66638 | 1.29886 | 208 |
| GO_RNA_SPLICING | 0.00000 | 0.00000 | | 0.63512 | 1.25129 | 384 |
| GO_RNA_SPLICING_VIA_TRANSESTERIFICATION_REACTIONS | 0.00000 | 0.00000 | 0.80122 | 0.64478 | 1.26586 | 311 |
| GO_ROUGH_ENDOPLASMIC_RETICULUM | 0.00014 | 0.00293 | 0.51885 | 0.70363 | 1.32507 | 59 |
| GO_RRNA_BINDING | 0.00000 | 0.00001 | 0.69013 | 0.77638 | 1.45941 | 54 |
| GO_RRNA_METABOLIC_PROCESS | 0.00000 | 0.00000 | | 0.68430 | 1.33386 | 209 |
| GO_SCF_DEPENDENT_PROTEASOMAL_UBIQUITIN_DEPENDENT_ | 0.00000 | 0.00008 | 0.62726 | 0.71957 | 1.36498 | 76 |
| GO_SECRETORY_GRANULE | 0.00000 | 0.00000 | | 0.62978 | 1.24363 | 481 |
| GO_SECRETORY_GRANULE_MEMBRANE | 0.00032 | 0.00575 | 0.49849 | 0.61375 | 1.19134 | 172 |
| GO_SERINE_TYPE_ENDOPEPTIDASE_INHIBITOR_ACTIVITY | 0.00038 | 0.00640 | 0.49849 | 0.77388 | 1.41447 | 28 |

| | | | | | | |
|---|---|---|---|---|---|---|
| GO_SEXUAL_REPRODUCTION | 0.00472 | 0.04480 | 0.40702 | 0.55748 | 1.10026 | 439 |
| GO_SIDE_OF_MEMBRANE | 0.00495 | 0.04573 | 0.40702 | 0.57741 | 1.12751 | 235 |
| GO_SIGNAL_TRANSDUCTION_BY_PROTEIN_PHOSPHORYLATION | 0.00224 | 0.02581 | 0.43171 | 0.55690 | 1.09975 | 500 |
| GO_SINGLE_FERTILIZATION | 0.00000 | 0.00002 | 0.67496 | 0.76307 | 1.43842 | 61 |
| GO_SINGLE_STRANDED_RNA_BINDING | 0.00001 | 0.00042 | 0.59333 | 0.71960 | 1.35864 | 66 |
| GO_SISTER_CHROMATID_SEGREGATION | 0.00042 | 0.00689 | 0.49849 | 0.60926 | 1.18209 | 170 |
| GO_SKIN_DEVELOPMENT | 0.00076 | 0.01140 | 0.47727 | 0.61452 | 1.18945 | 153 |
| GO_SMALL_MOLECULE_BIOSYNTHETIC_PROCESS | 0.00001 | 0.00021 | 0.61053 | 0.58728 | 1.15957 | 459 |
| GO_SMALL_NUCLEAR_RIBONUCLEOPROTEIN_COMPLEX | 0.00042 | 0.00689 | 0.49849 | 0.69542 | 1.30936 | 57 |
| GO_SMALL_RIBOSOMAL_SUBUNIT | 0.00000 | 0.00001 | 0.69013 | 0.75767 | 1.43100 | 67 |
| GO_SM_LIKE_PROTEIN_FAMILY_COMPLEX | 0.00145 | 0.01879 | 0.45506 | 0.66792 | 1.26149 | 67 |
| GO_SNORNA_BINDING | 0.00038 | 0.00646 | 0.49849 | 0.76224 | 1.39614 | 29 |
| GO_SPERM_EGG_RECOGNITION | 0.00000 | 0.00002 | 0.65944 | 0.87469 | 1.59087 | 26 |
| GO_SPINDLE_MIDZONE | 0.00361 | 0.03657 | 0.43171 | 0.73316 | 1.33859 | 27 |
| GO_SPINDLE_ORGANIZATION | 0.00272 | 0.02986 | 0.43171 | 0.60076 | 1.16362 | 159 |
| GO_SPLICEOSOMAL_COMPLEX | 0.00000 | 0.00008 | 0.62726 | 0.65256 | 1.26657 | 171 |
| GO_SPLICEOSOMAL_TRI_SNRNP_COMPLEX | 0.00168 | 0.02098 | 0.45506 | 0.73474 | 1.34835 | 30 |
| GO_STEM_CELL_DIFFERENTIATION | 0.00000 | 0.00000 | 0.82666 | 0.70006 | 1.35554 | 158 |
| GO_STEROID_BIOSYNTHETIC_PROCESS | 0.00465 | 0.04430 | 0.40702 | 0.61242 | 1.17992 | 125 |
| GO_STEROID_METABOLIC_PROCESS | 0.00313 | 0.03310 | 0.43171 | 0.59525 | 1.15748 | 185 |
| GO_STEROL_BIOSYNTHETIC_PROCESS | 0.00005 | 0.00120 | 0.55733 | 0.71876 | 1.35584 | 62 |
| GO_STEROL_METABOLIC_PROCESS | 0.00003 | 0.00073 | 0.57561 | 0.66485 | 1.27854 | 109 |
| GO_STRESS_GRANULE_ASSEMBLY | 0.00465 | 0.04430 | 0.40702 | 0.77288 | 1.38921 | 21 |
| GO_STRUCTURAL_CONSTITUENT_OF_RIBOSOME | 0.00000 | 0.00000 | | 0.76380 | 1.47839 | 153 |
| GO_STRUCTURAL_MOLECULE_ACTIVITY | 0.00000 | 0.00000 | | 0.71760 | 1.41436 | 391 |
| GO_SUPEROXIDE_METABOLIC_PROCESS | 0.00272 | 0.02986 | 0.43171 | 0.72028 | 1.32659 | 35 |
| GO_TELOMERASE_RNA_LOCALIZATION | 0.00054 | 0.00857 | 0.47727 | 0.81801 | 1.46148 | 19 |
| GO_TELOMERE_MAINTENANCE_VIA_TELOMERE_LENGTHENING | 0.00005 | 0.00138 | 0.55733 | 0.70715 | 1.33512 | 66 |
| GO_TELOMERE_ORGANIZATION | 0.00159 | 0.02016 | 0.45506 | 0.61666 | 1.18834 | 127 |
| GO_TERTIARY_GRANULE | 0.00008 | 0.00186 | 0.53843 | 0.66372 | 1.27005 | 91 |
| GO_TERTIARY_GRANULE_MEMBRANE | 0.00102 | 0.01453 | 0.45506 | 0.71940 | 1.33244 | 40 |
| GO_THREONINE_TYPE_ENDOPEPTIDASE_ACTIVITY | 0.00000 | 0.00000 | 0.71951 | 0.93456 | 1.65424 | 17 |
| GO_THREONINE_TYPE_PEPTIDASE_ACTIVITY | 0.00000 | 0.00004 | 0.65944 | 0.89807 | 1.61424 | 21 |
| GO_TISSUE_MORPHOGENESIS | 0.00000 | 0.00006 | 0.64355 | 0.59467 | 1.17150 | 380 |
| GO_TOXIN_TRANSPORT | 0.00064 | 0.00972 | 0.47727 | 0.77713 | 1.41343 | 26 |
| GO_TRANSCRIPTION_COUPLED_NUCLEOTIDE_EXCISION_REPAIR | 0.00231 | 0.02642 | 0.43171 | 0.65429 | 1.23840 | 71 |
| GO_TRANSFERRIN_TRANSPORT | 0.00141 | 0.01850 | 0.45506 | 0.75134 | 1.37179 | 27 |
| GO_TRANSLATIONAL_ELONGATION | 0.00000 | 0.00000 | | 0.75263 | 1.45028 | 124 |
| GO_TRANSLATIONAL_INITIATION | 0.00000 | 0.00000 | | 0.75139 | 1.45839 | 171 |
| GO_TRANSLATIONAL_TERMINATION | 0.00000 | 0.00000 | 0.81404 | 0.75110 | 1.44066 | 101 |
| GO_TRANSLATION_ELONGATION_FACTOR_ACTIVITY | 0.00390 | 0.03895 | 0.43171 | 0.80505 | 1.41260 | 15 |
| GO_TRANSLATION_FACTOR_ACTIVITY_RNA_BINDING | 0.00000 | 0.00000 | 0.74774 | 0.77244 | 1.46085 | 70 |
| GO_TRANSLATION_INITIATION_FACTOR_ACTIVITY | 0.00000 | 0.00011 | 0.62726 | 0.77773 | 1.44952 | 45 |
| GO_TRANSLATION_PREINITIATION_COMPLEX | 0.00342 | 0.03521 | 0.43171 | 0.79777 | 1.40694 | 16 |
| GO_TRANSLATION_REGULATOR_ACTIVITY | 0.00000 | 0.00000 | 0.83909 | 0.74000 | 1.42403 | 111 |
| GO_TRANSLATION_REGULATOR_ACTIVITY_NUCLEIC_ACID_BIND | 0.00000 | 0.00000 | 0.83909 | 0.76441 | 1.45959 | 87 |
| GO_TRANSMEMBRANE_RECEPTOR_PROTEIN_TYROSINE_KINASE | 0.00168 | 0.02098 | 0.45506 | 0.56326 | 1.11076 | 411 |
| GO_TRICARBOXYLIC_ACID_CYCLE | 0.00159 | 0.02016 | 0.45506 | 0.73558 | 1.35097 | 33 |
| GO_TRNA_TRANSPORT | 0.00342 | 0.03521 | 0.43171 | 0.71247 | 1.31220 | 35 |
| GO_TUMOR_NECROSIS_FACTOR_MEDIATED_SIGNALING_PATHW | 0.00000 | 0.00000 | 0.80122 | 0.73921 | 1.42138 | 108 |
| GO_T_CELL_RECEPTOR_SIGNALING_PATHWAY | 0.00000 | 0.00001 | 0.70498 | 0.69185 | 1.33314 | 119 |
| GO_U12_TYPE_SPLICEOSOMAL_COMPLEX | 0.00487 | 0.04559 | 0.40702 | 0.73423 | 1.33541 | 26 |
| GO_U2_SNRNP | 0.00141 | 0.01850 | 0.45506 | 0.80259 | 1.42065 | 17 |
| GO_U2_TYPE_CATALYTIC_STEP_2_SPLICEOSOME | 0.00268 | 0.02961 | 0.43171 | 0.72225 | 1.32288 | 29 |
| GO_U2_TYPE_SPLICEOSOMAL_COMPLEX | 0.00033 | 0.00595 | 0.49849 | 0.66590 | 1.26925 | 84 |
| GO_UBIQUITIN_LIKE_PROTEIN_LIGASE_BINDING | 0.00000 | 0.00007 | 0.62726 | 0.62616 | 1.22416 | 251 |
| GO_UNFOLDED_PROTEIN_BINDING | 0.00000 | 0.00001 | 0.69013 | 0.72112 | 1.37766 | 88 |
| GO_VACUOLAR_LUMEN | 0.00016 | 0.00319 | 0.51885 | 0.65416 | 1.25772 | 107 |
| GO_VESICLE_BUDDING_FROM_MEMBRANE | 0.00480 | 0.04537 | 0.40702 | 0.62411 | 1.19353 | 90 |
| GO_VESICLE_LUMEN | 0.00000 | 0.00000 | 0.77494 | 0.67028 | 1.30631 | 207 |
| GO_VESICLE_MEMBRANE | 0.00001 | 0.00026 | 0.59333 | 0.58430 | 1.15387 | 476 |
| GO_VIRAL_GENE_EXPRESSION | 0.00000 | 0.00000 | | 0.70963 | 1.37745 | 172 |
| GO_VIRAL_LIFE_CYCLE | 0.00000 | 0.00001 | 0.69013 | 0.63985 | 1.24950 | 238 |
| GO_VIRION_ASSEMBLY | 0.00457 | 0.04386 | 0.40702 | 0.69689 | 1.28528 | 36 |
| HP_ABNORMALITY_OF_ACID_BASE_HOMEOSTASIS | 0.00000 | 0.00010 | 0.62726 | 0.62283 | 1.21827 | 261 |
| HP_ABNORMALITY_OF_BLOOD_CIRCULATION | 0.00502 | 0.04608 | 0.40702 | 0.59228 | 1.14883 | 167 |
| HP_ABNORMALITY_OF_CARDIOVASCULAR_SYSTEM_ELECTROPH | 0.00011 | 0.00237 | 0.53843 | 0.60202 | 1.17758 | 261 |
| HP_ABNORMALITY_OF_SKIN_ADNEXA_PHYSIOLOGY | 0.00242 | 0.02748 | 0.43171 | 0.61294 | 1.18094 | 118 |
| HP_ABNORMALITY_OF_THE_ANKLES | 0.00313 | 0.03310 | 0.43171 | 0.65369 | 1.23418 | 63 |
| HP_ABNORMALITY_OF_THE_CEREBRAL_VASCULATURE | 0.00357 | 0.03638 | 0.43171 | 0.59764 | 1.15722 | 158 |

| | | | | | | |
|---|---|---|---|---|---|---|
| HP_ABNORMALITY_OF_THE_CEREBROSPINAL_FLUID | 0.00030 | 0.00549 | 0.49849 | 0.58221 | 1.14439 | 325 |
| HP_ABNORMALITY_OF_THE_MITOCHONDRION | 0.00126 | 0.01715 | 0.45506 | 0.62149 | 1.19764 | 120 |
| HP_ABNORMALITY_OF_THE_OPTIC_DISC | 0.00198 | 0.02370 | 0.43171 | 0.56624 | 1.11727 | 432 |
| HP_ABNORMALITY_OF_THE_OPTIC_NERVE | 0.00036 | 0.00622 | 0.49849 | 0.56723 | 1.12034 | 486 |
| HP_ABNORMAL_ABDOMEN_MORPHOLOGY | 0.00089 | 0.01291 | 0.47727 | 0.56424 | 1.11364 | 441 |
| HP_ABNORMAL_ACTIVITY_OF_MITOCHONDRIAL_RESPIRATORY_ | 0.00014 | 0.00300 | 0.51885 | 0.70303 | 1.32524 | 61 |
| HP_ABNORMAL_BASAL_GANGLIA_MRI_SIGNAL_INTENSITY | 0.00179 | 0.02183 | 0.45506 | 0.71597 | 1.32046 | 36 |
| HP_ABNORMAL_BRAINSTEM_MRI_SIGNAL_INTENSITY | 0.00398 | 0.03962 | 0.40702 | 0.71528 | 1.31370 | 33 |
| HP_ABNORMAL_CELLULAR_PHENOTYPE | 0.00209 | 0.02470 | 0.43171 | 0.56937 | 1.11992 | 334 |
| HP_ABNORMAL_CIRCULATING_CARBOHYDRATE_CONCENTRATIC | 0.00194 | 0.02331 | 0.45506 | 0.69518 | 1.29539 | 46 |
| HP_ABNORMAL_CNS_MYELINATION | 0.00014 | 0.00300 | 0.51885 | 0.62318 | 1.20954 | 171 |
| HP_ABNORMAL_CSF_METABOLITE_LEVEL | 0.00001 | 0.00027 | 0.59333 | 0.70531 | 1.34409 | 82 |
| HP_ABNORMAL_ELECTROPHYSIOLOGY_OF_SINOATRIAL_NODE_ | 0.00029 | 0.00536 | 0.49849 | 0.73172 | 1.35725 | 41 |
| HP_ABNORMAL_ENZYME_COENZYME_ACTIVITY | 0.00561 | 0.04971 | 0.40702 | 0.60490 | 1.16567 | 123 |
| HP_ABNORMAL_ERYTHROCYTE_MORPHOLOGY | 0.00001 | 0.00023 | 0.61053 | 0.58835 | 1.15904 | 380 |
| HP_ABNORMAL_HEART_VALVE_PHYSIOLOGY | 0.00405 | 0.04011 | 0.40702 | 0.59004 | 1.14474 | 169 |
| HP_ABNORMAL_MITOCHONDRIA_IN_MUSCLE_TISSUE | 0.00075 | 0.01118 | 0.47727 | 0.74633 | 1.36700 | 29 |
| HP_ABNORMAL_MYELINATION | 0.00007 | 0.00169 | 0.53843 | 0.59258 | 1.16479 | 324 |
| HP_ABNORMAL_MYOCARDIUM_MORPHOLOGY | 0.00008 | 0.00192 | 0.53843 | 0.58647 | 1.15365 | 340 |
| HP_ABNORMAL_RENAL_PHYSIOLOGY | 0.00450 | 0.04342 | 0.40702 | 0.55484 | 1.09317 | 385 |
| HP_ABNORMAL_TRICUSPID_VALVE_PHYSIOLOGY | 0.00231 | 0.02642 | 0.43171 | 0.72968 | 1.34029 | 31 |
| HP_ABNORMAL_VITREOUS_HUMOR_MORPHOLOGY | 0.00181 | 0.02201 | 0.45506 | 0.79632 | 1.40955 | 17 |
| HP_ACUTE_LEUKEMIA | 0.00013 | 0.00276 | 0.51885 | 0.68145 | 1.29219 | 75 |
| HP_ANEMIA_OF_INADEQUATE_PRODUCTION | 0.00561 | 0.04971 | 0.40702 | 0.61298 | 1.17855 | 107 |
| HP_APHASIA | 0.00213 | 0.02490 | 0.43171 | 0.72858 | 1.33704 | 30 |
| HP_BLINDNESS | 0.00018 | 0.00355 | 0.51885 | 0.61672 | 1.19768 | 175 |
| HP_BONE_MARROW_HYPERCELLULARITY | 0.00045 | 0.00725 | 0.49849 | 0.82169 | 1.46806 | 19 |
| HP_CHOREOATHETOSIS | 0.00561 | 0.04971 | 0.40702 | 0.64956 | 1.22631 | 64 |
| HP_CHRONIC_INFECTION | 0.00121 | 0.01658 | 0.45506 | 0.81916 | 1.44468 | 16 |
| HP_CONGESTIVE_HEART_FAILURE | 0.00495 | 0.04573 | 0.40702 | 0.60734 | 1.16929 | 114 |
| HP_CRANIAL_NERVE_COMPRESSION | 0.00331 | 0.03441 | 0.43171 | 0.75830 | 1.36989 | 23 |
| HP_DECREASED_ACTIVITY_OF_MITOCHONDRIAL_COMPLEX_I | 0.00283 | 0.03081 | 0.43171 | 0.71940 | 1.32496 | 35 |
| HP_DECREASED_ACTIVITY_OF_THE_PYRUVATE_DEHYDROGENAS | 0.00062 | 0.00949 | 0.47727 | 0.75390 | 1.38478 | 31 |
| HP_DEPRESSED_NASAL_RIDGE | 0.00041 | 0.00681 | 0.49849 | 0.66218 | 1.26189 | 82 |
| HP_DYSPNEA | 0.00250 | 0.02807 | 0.43171 | 0.58107 | 1.13348 | 224 |
| HP_DYSTONIA | 0.00069 | 0.01043 | 0.47727 | 0.58101 | 1.13736 | 279 |
| HP_EMOTIONAL_LABILITY | 0.00016 | 0.00321 | 0.51885 | 0.66683 | 1.27522 | 90 |
| HP_EPISODIC_ATAXIA | 0.00487 | 0.04559 | 0.40702 | 0.79934 | 1.40259 | 15 |
| HP_FASCICULATIONS | 0.00130 | 0.01761 | 0.45506 | 0.68128 | 1.28064 | 54 |
| HP_FATIGUE | 0.00042 | 0.00689 | 0.49849 | 0.59902 | 1.16764 | 212 |
| HP_FETAL_DISTRESS | 0.00179 | 0.02183 | 0.45506 | 0.72928 | 1.33973 | 32 |
| HP_FLUSHING | 0.00561 | 0.04971 | 0.40702 | 0.74782 | 1.33686 | 20 |
| HP_FOCAL_T2_HYPERINTENSE_BRAINSTEM_LESION | 0.00159 | 0.02016 | 0.45506 | 0.74970 | 1.36879 | 27 |
| HP_HEADACHE | 0.00016 | 0.00319 | 0.51885 | 0.62537 | 1.21179 | 163 |
| HP_HEPATOMEGALY | 0.00108 | 0.01507 | 0.45506 | 0.57439 | 1.12990 | 335 |
| HP_HYPERTROPHIC_CARDIOMYOPATHY | 0.00002 | 0.00050 | 0.57561 | 0.62629 | 1.21908 | 194 |
| HP_IMPAIRED_SOCIAL_INTERACTIONS | 0.00055 | 0.00863 | 0.47727 | 0.66555 | 1.26058 | 72 |
| HP_INCREASED_CSF_LACTATE | 0.00001 | 0.00034 | 0.59333 | 0.71937 | 1.35857 | 68 |
| HP_INCREASED_SERUM_LACTATE | 0.00001 | 0.00028 | 0.59333 | 0.65837 | 1.26932 | 133 |
| HP_INCREASED_SERUM_PYRUVATE | 0.00110 | 0.01526 | 0.45506 | 0.73143 | 1.35198 | 37 |
| HP_INFANTILE_ONSET | 0.00487 | 0.04559 | 0.40702 | 0.56559 | 1.10939 | 300 |
| HP_INTRACRANIAL_CYSTIC_LESION | 0.00055 | 0.00863 | 0.47727 | 0.62760 | 1.20885 | 116 |
| HP_INTRAUTERINE_GROWTH_RETARDATION | 0.00305 | 0.03271 | 0.43171 | 0.56594 | 1.11393 | 350 |
| HP_LACTIC_ACIDOSIS | 0.00018 | 0.00355 | 0.51885 | 0.63312 | 1.22071 | 131 |
| HP_LETHARGY | 0.00084 | 0.01228 | 0.47727 | 0.63361 | 1.21845 | 109 |
| HP_LEUKODYSTROPHY | 0.00029 | 0.00529 | 0.49849 | 0.66598 | 1.26915 | 82 |
| HP_LEUKOENCEPHALOPATHY | 0.00008 | 0.00190 | 0.53843 | 0.64132 | 1.23667 | 134 |
| HP_LIPID_ACCUMULATION_IN_HEPATOCYTES | 0.00220 | 0.02548 | 0.43171 | 0.62794 | 1.20159 | 91 |
| HP_LOWER_LIMB_SPASTICITY | 0.00154 | 0.01974 | 0.45506 | 0.60929 | 1.17648 | 139 |
| HP_MACROCYTIC_ANEMIA | 0.00030 | 0.00542 | 0.49849 | 0.73108 | 1.35606 | 41 |
| HP_METABOLIC_ACIDOSIS | 0.00034 | 0.00599 | 0.49849 | 0.64831 | 1.24464 | 103 |
| HP_MICRONODULAR_CIRRHOSIS | 0.00324 | 0.03393 | 0.43171 | 0.78723 | 1.40018 | 18 |
| HP_MIGRAINE | 0.00028 | 0.00515 | 0.49849 | 0.67309 | 1.27635 | 75 |
| HP_MUSCLE_ABNORMALITY_RELATED_TO_MITOCHONDRIAL_DY | 0.00209 | 0.02470 | 0.43171 | 0.68258 | 1.27573 | 49 |
| HP_MUTISM | 0.00283 | 0.03081 | 0.43171 | 0.71937 | 1.32490 | 35 |
| HP_NAUSEA_AND_VOMITING | 0.00022 | 0.00417 | 0.51885 | 0.59383 | 1.16158 | 264 |
| HP_NEOPLASM_OF_THE_ENDOCRINE_SYSTEM | 0.00495 | 0.04573 | 0.40702 | 0.63166 | 1.20308 | 81 |
| HP_OPTIC_NEUROPATHY | 0.00135 | 0.01808 | 0.45506 | 0.73837 | 1.35610 | 33 |
| HP_PACE_OF_PROGRESSION | 0.00487 | 0.04559 | 0.40702 | 0.56814 | 1.11416 | 298 |
| HP_PALLOR | 0.00036 | 0.00622 | 0.49849 | 0.64861 | 1.24463 | 102 |

| | | | | | | |
|---|---|---|---|---|---|---|
| HP_PARAPLEGIA | 0.00121 | 0.01658 | 0.45506 | 0.63584 | 1.22133 | 104 |
| HP_PARAPLEGIA_PARAPARESIS | 0.00036 | 0.00621 | 0.49849 | 0.62047 | 1.20040 | 150 |
| HP_PAROXYSMAL_INVOLUNTARY_EYE_MOVEMENTS | 0.00054 | 0.00849 | 0.47727 | 0.75650 | 1.38957 | 31 |
| HP_PEDIATRIC_ONSET | 0.00561 | 0.04971 | 0.40702 | 0.55915 | 1.10116 | 371 |
| HP_PIGMENTARY_RETINOPATHY | 0.00065 | 0.00981 | 0.47727 | 0.63260 | 1.21720 | 113 |
| HP_POOR_EYE_CONTACT | 0.00017 | 0.00341 | 0.51885 | 0.71279 | 1.33749 | 53 |
| HP_POOR_HEAD_CONTROL | 0.00000 | 0.00005 | 0.64355 | 0.67819 | 1.30665 | 118 |
| HP_PROGRESSIVE | 0.00194 | 0.02331 | 0.45506 | 0.59790 | 1.16273 | 186 |
| HP_PROGRESSIVE_CEREBELLAR_ATAXIA | 0.00364 | 0.03677 | 0.43171 | 0.64788 | 1.22325 | 65 |
| HP_PROGRESSIVE_SPASTIC_PARAPLEGIA | 0.00253 | 0.02838 | 0.43171 | 0.69655 | 1.29822 | 45 |
| HP_PROXIMAL_TUBULOPATHY | 0.00141 | 0.01850 | 0.45506 | 0.72579 | 1.33535 | 34 |
| HP_PTOSIS | 0.00472 | 0.04480 | 0.40702 | 0.55582 | 1.09637 | 418 |
| HP_REDUCED_CONSCIOUSNESS_CONFUSION | 0.00007 | 0.00164 | 0.53843 | 0.61227 | 1.19295 | 205 |
| HP_REDUCED_VISUAL_ACUITY | 0.00053 | 0.00837 | 0.47727 | 0.58618 | 1.14774 | 283 |
| HP_RENAL_TUBULAR_DYSFUNCTION | 0.00013 | 0.00284 | 0.51885 | 0.68108 | 1.29150 | 75 |
| HP_RESPIRATORY_DISTRESS | 0.00108 | 0.01507 | 0.45506 | 0.62693 | 1.20762 | 115 |
| HP_RESPIRATORY_INSUFFICIENCY | 0.00000 | 0.00010 | 0.62726 | 0.60536 | 1.18998 | 326 |
| HP_SINUS_TACHYCARDIA | 0.00213 | 0.02490 | 0.43171 | 0.77203 | 1.39128 | 22 |
| HP_SPARSE_EYELASHES | 0.00150 | 0.01943 | 0.45506 | 0.71355 | 1.32161 | 40 |
| HP_SPARSE_OR_ABSENT_EYELASHES | 0.00220 | 0.02548 | 0.43171 | 0.67331 | 1.26567 | 54 |
| HP_SPASTIC_PARAPLEGIA | 0.00098 | 0.01407 | 0.45506 | 0.64059 | 1.22712 | 96 |
| HP_THROMBOCYTOSIS | 0.00413 | 0.04065 | 0.40702 | 0.75210 | 1.35870 | 23 |
| HP_TONGUE_FASCICULATIONS | 0.00457 | 0.04386 | 0.40702 | 0.80056 | 1.40472 | 15 |
| HP_TRANSIENT_ISCHEMIC_ATTACK | 0.00480 | 0.04537 | 0.40702 | 0.74902 | 1.35809 | 25 |
| HP_VERTIGO | 0.00002 | 0.00063 | 0.57561 | 0.72114 | 1.35844 | 60 |
| HP_VESTIBULAR_DYSFUNCTION | 0.00028 | 0.00522 | 0.49849 | 0.67279 | 1.27577 | 75 |
| HP_VOMITING | 0.00072 | 0.01082 | 0.47727 | 0.61459 | 1.18792 | 147 |
| HP_WEAKNESS_DUE_TO_UPPER_MOTOR_NEURON_DYSFUNCTI | 0.00532 | 0.04803 | 0.40702 | 0.56829 | 1.11186 | 272 |
| HP_WEAKNESS_OF_MUSCLES_OF_RESPIRATION | 0.00163 | 0.02050 | 0.45506 | 0.66272 | 1.25123 | 63 |
| **Cluster 1** | | | | | | |
| pathway | pval | padj | log2err | ES | NES | size |
| GO_ACTIVATION_OF_IMMUNE_RESPONSE | 0.00000 | 0.00000 | 0.74774 | 0.65038 | 1.26999 | 235 |
| GO_ACTIVATION_OF_INNATE_IMMUNE_RESPONSE | 0.00000 | 0.00001 | 0.69013 | 0.71967 | 1.37861 | 96 |
| GO_AEROBIC_RESPIRATION | 0.00000 | 0.00000 | 0.73376 | 0.76280 | 1.44478 | 72 |
| GO_ALCOHOL_BIOSYNTHETIC_PROCESS | 0.00032 | 0.00572 | 0.49849 | 0.64679 | 1.24367 | 108 |
| GO_ALCOHOL_METABOLIC_PROCESS | 0.00364 | 0.03677 | 0.43171 | 0.58112 | 1.13424 | 228 |
| GO_AMEBOIDAL_TYPE_CELL_MIGRATION | 0.00457 | 0.04386 | 0.40702 | 0.57755 | 1.12854 | 247 |
| GO_AMIDE_BINDING | 0.00014 | 0.00297 | 0.51885 | 0.61533 | 1.19780 | 199 |
| GO_AMINE_METABOLIC_PROCESS | 0.00000 | 0.00000 | 0.76146 | 0.73732 | 1.41097 | 92 |
| GO_AMYLOID_BETA_CLEARANCE | 0.00561 | 0.04971 | 0.40702 | 0.74081 | 1.33976 | 24 |
| GO_ANAPHASE_PROMOTING_COMPLEX_DEPENDENT_CATABOL | 0.00000 | 0.00000 | 0.76146 | 0.77107 | 1.46112 | 73 |
| GO_ANATOMICAL_STRUCTURE_HOMEOSTASIS | 0.00020 | 0.00390 | 0.51885 | 0.59307 | 1.16021 | 271 |
| GO_ANIMAL_ORGAN_REGENERATION | 0.00012 | 0.00269 | 0.51885 | 0.71251 | 1.33510 | 52 |
| GO_ANTIGEN_PROCESSING_AND_PRESENTATION | 0.00000 | 0.00000 | 0.74774 | 0.68785 | 1.33064 | 149 |
| GO_ANTIGEN_PROCESSING_AND_PRESENTATION_OF_EXOGENO | 0.00000 | 0.00000 | 0.76146 | 0.79644 | 1.50237 | 62 |
| GO_ANTIGEN_PROCESSING_AND_PRESENTATION_OF_PEPTIDE_ | 0.00000 | 0.00000 | 0.74774 | 0.70107 | 1.35144 | 130 |
| GO_ANTIGEN_PROCESSING_AND_PRESENTATION_OF_PEPTIDE_ | 0.00000 | 0.00000 | 0.78819 | 0.77566 | 1.47012 | 74 |
| GO_ANTIGEN_RECEPTOR_MEDIATED_SIGNALING_PATHWAY | 0.00000 | 0.00009 | 0.62726 | 0.66362 | 1.28080 | 136 |
| GO_APICAL_PART_OF_CELL | 0.00135 | 0.01808 | 0.45506 | 0.58926 | 1.14966 | 220 |
| GO_APOPTOTIC_SIGNALING_PATHWAY | 0.00006 | 0.00152 | 0.53843 | 0.58638 | 1.15586 | 393 |
| GO_ATPASE_ACTIVITY | 0.00050 | 0.00802 | 0.47727 | 0.58216 | 1.14385 | 316 |
| GO_ATPASE_ACTIVITY_COUPLED_TO_TRANSMEMBRANE_MOVE | 0.00420 | 0.04132 | 0.40702 | 0.78005 | 1.38741 | 18 |
| GO_ATP_BIOSYNTHETIC_PROCESS | 0.00013 | 0.00276 | 0.51885 | 0.83897 | 1.50801 | 21 |
| GO_ATP_METABOLIC_PROCESS | 0.00000 | 0.00000 | | 0.72178 | 1.40814 | 219 |
| GO_ATP_SYNTHESIS_COUPLED_ELECTRON_TRANSPORT | 0.00000 | 0.00000 | | 0.80353 | 1.52425 | 76 |
| GO_AZUROPHIL_GRANULE | 0.00025 | 0.00466 | 0.49849 | 0.64781 | 1.24551 | 107 |
| GO_AZUROPHIL_GRANULE_LUMEN | 0.00405 | 0.04011 | 0.40702 | 0.65625 | 1.23793 | 62 |
| GO_BASAL_PART_OF_CELL | 0.00509 | 0.04656 | 0.40702 | 0.68515 | 1.27697 | 45 |
| GO_BASAL_PLASMA_MEMBRANE | 0.00346 | 0.03548 | 0.43171 | 0.71184 | 1.31104 | 35 |
| GO_BASEMENT_MEMBRANE | 0.00279 | 0.03056 | 0.43171 | 0.68935 | 1.28452 | 46 |
| GO_BASOLATERAL_PLASMA_MEMBRANE | 0.00011 | 0.00247 | 0.53843 | 0.63209 | 1.21993 | 136 |
| GO_BINDING_OF_SPERM_TO_ZONA_PELLUCIDA | 0.00000 | 0.00006 | 0.64355 | 0.90095 | 1.60966 | 19 |
| GO_BLOOD_MICROPARTICLE | 0.00000 | 0.00000 | 0.70498 | 0.81982 | 1.52763 | 46 |
| GO_BRUSH_BORDER | 0.00060 | 0.00923 | 0.47727 | 0.66897 | 1.26790 | 74 |
| GO_CADHERIN_BINDING | 0.00000 | 0.00000 | | 0.68759 | 1.34435 | 253 |
| GO_CALCIUM_DEPENDENT_PHOSPHOLIPID_BINDING | 0.00152 | 0.01958 | 0.45506 | 0.75021 | 1.37120 | 28 |
| GO_CALCIUM_DEPENDENT_PROTEIN_BINDING | 0.00009 | 0.00211 | 0.53843 | 0.74254 | 1.38394 | 45 |
| GO_CANONICAL_WNT_SIGNALING_PATHWAY | 0.00004 | 0.00109 | 0.55733 | 0.61740 | 1.20458 | 222 |
| GO_CARBOHYDRATE_CATABOLIC_PROCESS | 0.00135 | 0.01808 | 0.45506 | 0.61179 | 1.18186 | 144 |
| GO_CATALYTIC_STEP_2_SPLICEOSOME | 0.00001 | 0.00022 | 0.61053 | 0.70290 | 1.33765 | 80 |

| | | | | | | |
|---|---|---|---|---|---|---|
| GO_CATION_TRANSMEMBRANE_TRANSPORT | 0.00327 | 0.03409 | 0.43171 | 0.56060 | 1.10531 | 406 |
| GO_CATION_TRANSMEMBRANE_TRANSPORTER_ACTIVITY | 0.00097 | 0.01387 | 0.47727 | 0.58196 | 1.13846 | 265 |
| GO_CELLULAR_AMINO_ACID_METABOLIC_PROCESS | 0.00002 | 0.00060 | 0.57561 | 0.61438 | 1.19930 | 231 |
| GO_CELLULAR_COMPONENT_DISASSEMBLY | 0.00000 | 0.00000 | | 0.64070 | 1.26342 | 408 |
| GO_CELLULAR_ION_HOMEOSTASIS | 0.00027 | 0.00493 | 0.49849 | 0.58658 | 1.15340 | 328 |
| GO_CELLULAR_IRON_ION_HOMEOSTASIS | 0.00327 | 0.03409 | 0.43171 | 0.68360 | 1.27374 | 47 |
| GO_CELLULAR_KETONE_METABOLIC_PROCESS | 0.00000 | 0.00010 | 0.62726 | 0.65358 | 1.26726 | 164 |
| GO_CELLULAR_MONOVALENT_INORGANIC_CATION_HOMEOSTA | 0.00179 | 0.02183 | 0.45506 | 0.64747 | 1.22821 | 76 |
| GO_CELLULAR_NITROGEN_COMPOUND_CATABOLIC_PROCESS | 0.00000 | 0.00000 | | 0.67915 | 1.34096 | 456 |
| GO_CELLULAR_PROTEIN_COMPLEX_DISASSEMBLY | 0.00000 | 0.00000 | | 0.69566 | 1.35184 | 179 |
| GO_CELLULAR_RESPIRATION | 0.00000 | 0.00000 | | 0.74205 | 1.43370 | 145 |
| GO_CELLULAR_RESPONSE_TO_HEAT | 0.00004 | 0.00109 | 0.55733 | 0.67186 | 1.28698 | 95 |
| GO_CELLULAR_RESPONSE_TO_HORMONE_STIMULUS | 0.00309 | 0.03299 | 0.43171 | 0.56453 | 1.11191 | 374 |
| GO_CELLULAR_RESPONSE_TO_ORGANIC_CYCLIC_COMPOUND | 0.00502 | 0.04608 | 0.40702 | 0.56512 | 1.11134 | 329 |
| GO_CELLULAR_RESPONSE_TO_OXYGEN_LEVELS | 0.00000 | 0.00000 | | 0.71461 | 1.38224 | 151 |
| GO_CELL_ACTIVATION_INVOLVED_IN_IMMUNE_RESPONSE | 0.00000 | 0.00000 | | 0.63924 | 1.26108 | 422 |
| GO_CELL_ADHESION_MEDIATOR_ACTIVITY | 0.00157 | 0.02009 | 0.45506 | 0.71837 | 1.32488 | 36 |
| GO_CELL_ADHESION_MOLECULE_BINDING | 0.00000 | 0.00000 | | 0.67109 | 1.32112 | 353 |
| GO_CELL_BODY | 0.00004 | 0.00109 | 0.55733 | 0.59973 | 1.17541 | 293 |
| GO_CELL_CELL_RECOGNITION | 0.00000 | 0.00006 | 0.64355 | 0.82001 | 1.51026 | 35 |
| GO_CELL_CELL_SIGNALING_BY_WNT | 0.00004 | 0.00109 | 0.55733 | 0.59042 | 1.16141 | 339 |
| GO_CELL_CYCLE_G1_S_PHASE_TRANSITION | 0.00376 | 0.03783 | 0.43171 | 0.58112 | 1.13378 | 220 |
| GO_CELL_CYCLE_G2_M_PHASE_TRANSITION | 0.00024 | 0.00459 | 0.51885 | 0.59807 | 1.16746 | 231 |
| GO_CELL_CYCLE_PHASE_TRANSITION | 0.00000 | 0.00001 | 0.69013 | 0.59417 | 1.17348 | 499 |
| GO_CELL_DIVISION | 0.00003 | 0.00071 | 0.57561 | 0.58314 | 1.15094 | 449 |
| GO_CELL_KILLING | 0.00205 | 0.02440 | 0.43171 | 0.66727 | 1.25696 | 60 |
| GO_CELL_RECOGNITION | 0.00000 | 0.00000 | 0.73376 | 0.75953 | 1.43954 | 74 |
| GO_CELL_REDOX_HOMEOSTASIS | 0.00163 | 0.02050 | 0.45506 | 0.71772 | 1.32665 | 38 |
| GO_CELL_SUBSTRATE_JUNCTION | 0.00000 | 0.00000 | | 0.65742 | 1.29171 | 316 |
| GO_CELL_SURFACE | 0.00000 | 0.00000 | 0.73376 | 0.62572 | 1.23070 | 330 |
| GO_CELL_SURFACE_RECEPTOR_SIGNALING_PATHWAY_INVOLVEI | 0.00038 | 0.00640 | 0.49849 | 0.57762 | 1.13770 | 374 |
| GO_CHAPERONE_BINDING | 0.00495 | 0.04573 | 0.40702 | 0.62860 | 1.19935 | 86 |
| GO_CHAPERONE_COMPLEX | 0.00001 | 0.00028 | 0.59333 | 0.86337 | 1.55587 | 22 |
| GO_CHAPERONE_MEDIATED_PROTEIN_COMPLEX_ASSEMBLY | 0.00100 | 0.01430 | 0.45506 | 0.80787 | 1.44337 | 19 |
| GO_CHAPERONE_MEDIATED_PROTEIN_FOLDING | 0.00168 | 0.02098 | 0.45506 | 0.71110 | 1.32288 | 44 |
| GO_CHROMOSOMAL_REGION | 0.00018 | 0.00349 | 0.51885 | 0.59323 | 1.16180 | 286 |
| GO_CHROMOSOME_CENTROMERIC_REGION | 0.00172 | 0.02135 | 0.45506 | 0.60447 | 1.17248 | 168 |
| GO_CHROMOSOME_SEGREGATION | 0.00003 | 0.00087 | 0.55733 | 0.60463 | 1.18280 | 270 |
| GO_CLUSTER_OF_ACTIN_BASED_CELL_PROJECTIONS | 0.00074 | 0.01107 | 0.47727 | 0.64184 | 1.23044 | 99 |
| GO_COATED_VESICLE | 0.00213 | 0.02490 | 0.43171 | 0.59512 | 1.15846 | 199 |
| GO_COATED_VESICLE_MEMBRANE | 0.00060 | 0.00923 | 0.47727 | 0.63443 | 1.22207 | 115 |
| GO_COLLAGEN_BINDING | 0.00465 | 0.04430 | 0.40702 | 0.73471 | 1.33628 | 26 |
| GO_COLLAGEN_CONTAINING_EXTRACELLULAR_MATRIX | 0.00000 | 0.00006 | 0.64355 | 0.65358 | 1.26542 | 157 |
| GO_CONDENSED_CHROMOSOME | 0.00350 | 0.03580 | 0.43171 | 0.59394 | 1.15417 | 179 |
| GO_CONTRACTILE_FIBER | 0.00064 | 0.00972 | 0.47727 | 0.62217 | 1.19973 | 134 |
| GO_COPI_COATED_VESICLE | 0.00316 | 0.03344 | 0.43171 | 0.75688 | 1.37234 | 25 |
| GO_CORNIFICATION | 0.00027 | 0.00500 | 0.49849 | 0.77970 | 1.42512 | 28 |
| GO_CORTICAL_CYTOSKELETON | 0.00502 | 0.04608 | 0.40702 | 0.64388 | 1.21870 | 71 |
| GO_COTRANSLATIONAL_PROTEIN_TARGETING_TO_MEMBRANE | 0.00000 | 0.00000 | | 0.78223 | 1.49819 | 94 |
| GO_CYTOCHROME_COMPLEX | 0.00035 | 0.00610 | 0.49849 | 0.80796 | 1.45601 | 22 |
| GO_CYTOKINE_MEDIATED_SIGNALING_PATHWAY | 0.00000 | 0.00002 | 0.67496 | 0.60742 | 1.19735 | 392 |
| GO_CYTOPLASMIC_STRESS_GRANULE | 0.00014 | 0.00302 | 0.51885 | 0.69390 | 1.31009 | 63 |
| GO_CYTOPLASMIC_TRANSLATION | 0.00000 | 0.00001 | 0.70498 | 0.71773 | 1.37349 | 92 |
| GO_CYTOPLASMIC_TRANSLATIONAL_INITIATION | 0.00178 | 0.02182 | 0.45506 | 0.73517 | 1.35039 | 31 |
| GO_CYTOSOLIC_LARGE_RIBOSOMAL_SUBUNIT | 0.00000 | 0.00005 | 0.64355 | 0.77240 | 1.44935 | 53 |
| GO_CYTOSOLIC_RIBOSOME | 0.00000 | 0.00000 | | 0.77483 | 1.48437 | 97 |
| GO_CYTOSOLIC_SMALL_RIBOSOMAL_SUBUNIT | 0.00000 | 0.00010 | 0.62726 | 0.79384 | 1.47033 | 40 |
| GO_DETECTION_OF_STIMULUS | 0.00020 | 0.00393 | 0.51885 | 0.65208 | 1.25384 | 108 |
| GO_DETOXIFICATION | 0.00413 | 0.04065 | 0.40702 | 0.65399 | 1.23684 | 70 |
| GO_DEVELOPMENT_OF_PRIMARY_FEMALE_SEXUAL_CHARACTE | 0.00179 | 0.02183 | 0.45506 | 0.68410 | 1.28057 | 51 |
| GO_DICARBOXYLIC_ACID_METABOLIC_PROCESS | 0.00261 | 0.02910 | 0.43171 | 0.65541 | 1.23747 | 65 |
| GO_DISORDERED_DOMAIN_SPECIFIC_BINDING | 0.00051 | 0.00812 | 0.47727 | 0.76867 | 1.40343 | 27 |
| GO_DISULFIDE_OXIDOREDUCTASE_ACTIVITY | 0.00435 | 0.04238 | 0.40702 | 0.71445 | 1.31232 | 31 |
| GO_DNA_BIOSYNTHETIC_PROCESS | 0.00017 | 0.00334 | 0.51885 | 0.62540 | 1.20833 | 145 |
| GO_DNA_CONFORMATION_CHANGE | 0.00327 | 0.03409 | 0.43171 | 0.57979 | 1.13178 | 231 |
| GO_DNA_DAMAGE_RESPONSE_DETECTION_OF_DNA_DAMAGE | 0.00305 | 0.03271 | 0.43171 | 0.71625 | 1.31916 | 35 |
| GO_DNA_GEOMETRIC_CHANGE | 0.00141 | 0.01850 | 0.45506 | 0.62642 | 1.20356 | 105 |
| GO_DOUBLE_STRANDED_RNA_BINDING | 0.00039 | 0.00658 | 0.49849 | 0.69158 | 1.30365 | 61 |
| GO_DRUG_BINDING | 0.00450 | 0.04342 | 0.40702 | 0.66245 | 1.24130 | 52 |
| GO_ELECTRON_TRANSFER_ACTIVITY | 0.00000 | 0.00001 | 0.67496 | 0.72917 | 1.38765 | 80 |

| | | | | | | |
|---|---|---|---|---|---|---|
| GO_ELECTRON_TRANSPORT_CHAIN | 0.00000 | 0.00000 | | 0.73336 | 1.41392 | 133 |
| GO_EMBRYO_IMPLANTATION | 0.00044 | 0.00706 | 0.49849 | 0.75968 | 1.39146 | 29 |
| GO_ENDOCYTIC_VESICLE_MEMBRANE | 0.00524 | 0.04770 | 0.40702 | 0.62563 | 1.19564 | 89 |
| GO_ENDOPEPTIDASE_ACTIVITY | 0.00065 | 0.00981 | 0.47727 | 0.60123 | 1.16998 | 196 |
| GO_ENDOPEPTIDASE_COMPLEX | 0.00000 | 0.00001 | 0.70498 | 0.76593 | 1.44609 | 63 |
| GO_ENDOPEPTIDASE_REGULATOR_ACTIVITY | 0.00006 | 0.00145 | 0.55733 | 0.70425 | 1.33001 | 68 |
| GO_ENDOPLASMIC_RETICULUM_GOLGI_INTERMEDIATE_COMPA | 0.00055 | 0.00863 | 0.47727 | 0.64758 | 1.24050 | 96 |
| GO_ENDOPLASMIC_RETICULUM_GOLGI_INTERMEDIATE_COMPA | 0.00148 | 0.01923 | 0.45506 | 0.68944 | 1.29056 | 51 |
| GO_ENDOPLASMIC_RETICULUM_TO_GOLGI_VESICLE_MEDIATED | 0.00532 | 0.04803 | 0.40702 | 0.59194 | 1.14697 | 162 |
| GO_ENERGY_DERIVATION_BY_OXIDATION_OF_ORGANIC_COMP | 0.00000 | 0.00000 | | 0.69995 | 1.36305 | 202 |
| GO_ENZYME_INHIBITOR_ACTIVITY | 0.00246 | 0.02780 | 0.43171 | 0.58913 | 1.14643 | 191 |
| GO_EPIDERMAL_CELL_DIFFERENTIATION | 0.00216 | 0.02519 | 0.43171 | 0.62667 | 1.20589 | 112 |
| GO_EPITHELIAL_CELL_DIFFERENTIATION | 0.00054 | 0.00857 | 0.47727 | 0.57819 | 1.13725 | 334 |
| GO_ESTABLISHMENT_OF_ORGANELLE_LOCALIZATION | 0.00122 | 0.01679 | 0.45506 | 0.57737 | 1.13489 | 323 |
| GO_ESTABLISHMENT_OF_PROTEIN_LOCALIZATION_TO_CHROMO | 0.00016 | 0.00321 | 0.51885 | 0.81363 | 1.46985 | 23 |
| GO_ESTABLISHMENT_OF_PROTEIN_LOCALIZATION_TO_ENDOPL | 0.00000 | 0.00000 | | 0.77241 | 1.48479 | 106 |
| GO_ESTABLISHMENT_OF_PROTEIN_LOCALIZATION_TO_MEMBRA | 0.00000 | 0.00000 | | 0.67484 | 1.31991 | 260 |
| GO_ESTABLISHMENT_OF_PROTEIN_LOCALIZATION_TO_ORGANE | 0.00000 | 0.00000 | | 0.65856 | 1.30056 | 461 |
| GO_ESTABLISHMENT_OF_PROTEIN_LOCALIZATION_TO_TELOME | 0.00000 | 0.00006 | 0.64355 | 0.90447 | 1.60870 | 18 |
| GO_ESTABLISHMENT_OF_RNA_LOCALIZATION | 0.00000 | 0.00005 | 0.64355 | 0.64271 | 1.24830 | 176 |
| GO_ESTABLISHMENT_OF_TISSUE_POLARITY | 0.00000 | 0.00000 | 0.81404 | 0.75693 | 1.44999 | 96 |
| GO_EUKARYOTIC_TRANSLATION_INITIATION_FACTOR_3_COMPL | 0.00143 | 0.01863 | 0.45506 | 0.83263 | 1.46100 | 15 |
| GO_EXTERNAL_SIDE_OF_PLASMA_MEMBRANE | 0.00159 | 0.02016 | 0.45506 | 0.61775 | 1.19022 | 118 |
| GO_EXTRACELLULAR_MATRIX | 0.00042 | 0.00695 | 0.49849 | 0.61138 | 1.18938 | 189 |
| GO_FATTY_ACID_DERIVATIVE_METABOLIC_PROCESS | 0.00043 | 0.00695 | 0.49849 | 0.66266 | 1.26212 | 81 |
| GO_FATTY_ACYL_COA_METABOLIC_PROCESS | 0.00502 | 0.04608 | 0.40702 | 0.72264 | 1.32082 | 28 |
| GO_FC_EPSILON_RECEPTOR_SIGNALING_PATHWAY | 0.00000 | 0.00000 | 0.71951 | 0.72370 | 1.38610 | 94 |
| GO_FC_RECEPTOR_SIGNALING_PATHWAY | 0.00000 | 0.00000 | 0.70498 | 0.68815 | 1.32879 | 140 |
| GO_FERTILIZATION | 0.00000 | 0.00005 | 0.64355 | 0.73257 | 1.38752 | 72 |
| GO_FIBRILLAR_CENTER | 0.00246 | 0.02780 | 0.43171 | 0.62638 | 1.19990 | 96 |
| GO_FIBROBLAST_GROWTH_FACTOR_RECEPTOR_SIGNALING_PAT | 0.00495 | 0.04573 | 0.40702 | 0.64888 | 1.22552 | 67 |
| GO_FIBROBLAST_PROLIFERATION | 0.00032 | 0.00575 | 0.49849 | 0.70676 | 1.32618 | 53 |
| GO_FICOLIN_1_RICH_GRANULE | 0.00000 | 0.00000 | 0.70498 | 0.71544 | 1.37138 | 98 |
| GO_GABA_ERGIC_SYNAPSE | 0.00022 | 0.00427 | 0.51885 | 0.76430 | 1.40406 | 32 |
| GO_GENERATION_OF_PRECURSOR_METABOLITES_AND_ENERGY | 0.00000 | 0.00000 | | 0.67527 | 1.32968 | 368 |
| GO_GENE_SILENCING | 0.00019 | 0.00376 | 0.51885 | 0.61625 | 1.19788 | 184 |
| GO_GLUCOSE_CATABOLIC_PROCESS | 0.00495 | 0.04573 | 0.40702 | 0.72314 | 1.32174 | 28 |
| GO_GLUTAMATERGIC_SYNAPSE | 0.00135 | 0.01808 | 0.45506 | 0.60052 | 1.16783 | 186 |
| GO_GLYCOLYTIC_PROCESS_THROUGH_FRUCTOSE_6_PHOSPHATI | 0.00554 | 0.04968 | 0.40702 | 0.74156 | 1.34111 | 24 |
| GO_GOLGI_ASSOCIATED_VESICLE | 0.00085 | 0.01239 | 0.47727 | 0.62294 | 1.20037 | 129 |
| GO_GOLGI_ASSOCIATED_VESICLE_MEMBRANE | 0.00313 | 0.03310 | 0.43171 | 0.64208 | 1.21753 | 75 |
| GO_GTPASE_ACTIVITY | 0.00004 | 0.00098 | 0.55733 | 0.61546 | 1.19974 | 210 |
| GO_GUANYL_NUCLEOTIDE_BINDING | 0.00005 | 0.00126 | 0.55733 | 0.60899 | 1.19065 | 252 |
| GO_HEAT_SHOCK_PROTEIN_BINDING | 0.00001 | 0.00034 | 0.59333 | 0.68258 | 1.30633 | 93 |
| GO_HEMATOPOIETIC_PROGENITOR_CELL_DIFFERENTIATION | 0.00000 | 0.00000 | 0.71951 | 0.70099 | 1.35150 | 121 |
| GO_HEMATOPOIETIC_STEM_CELL_DIFFERENTIATION | 0.00000 | 0.00000 | 0.76146 | 0.77246 | 1.46307 | 72 |
| GO_HISTONE_MRNA_METABOLIC_PROCESS | 0.00265 | 0.02936 | 0.43171 | 0.76832 | 1.38459 | 22 |
| GO_HOST_CELLULAR_COMPONENT | 0.00043 | 0.00695 | 0.49849 | 0.69729 | 1.31074 | 54 |
| GO_IMMUNE_RESPONSE_REGULATING_SIGNALING_PATHWAY | 0.00000 | 0.00003 | 0.65944 | 0.63620 | 1.24112 | 225 |
| GO_IMPORT_INTO_CELL | 0.00428 | 0.04185 | 0.40702 | 0.61709 | 1.18655 | 108 |
| GO_IMPORT_INTO_NUCLEUS | 0.00039 | 0.00652 | 0.49849 | 0.62707 | 1.20905 | 131 |
| GO_INNATE_IMMUNE_RESPONSE | 0.00000 | 0.00003 | 0.65944 | 0.59893 | 1.18089 | 406 |
| GO_INNATE_IMMUNE_RESPONSE_ACTIVATING_SIGNAL_TRANSI | 0.00000 | 0.00001 | 0.69013 | 0.73348 | 1.39585 | 80 |
| GO_INNER_MITOCHONDRIAL_MEMBRANE_PROTEIN_COMPLEX | 0.00000 | 0.00000 | | 0.76576 | 1.46666 | 94 |
| GO_INORGANIC_ION_TRANSMEMBRANE_TRANSPORT | 0.00141 | 0.01850 | 0.45506 | 0.56404 | 1.11185 | 401 |
| GO_INTERACTION_WITH_HOST | 0.00139 | 0.01850 | 0.45506 | 0.60855 | 1.17710 | 151 |
| GO_INTERLEUKIN_1_MEDIATED_SIGNALING_PATHWAY | 0.00000 | 0.00000 | 0.77494 | 0.75654 | 1.44176 | 83 |
| GO_INTRAMOLECULAR_OXIDOREDUCTASE_ACTIVITY | 0.00051 | 0.00812 | 0.47727 | 0.74506 | 1.37081 | 34 |
| GO_INTRINSIC_APOPTOTIC_SIGNALING_PATHWAY | 0.00156 | 0.01993 | 0.45506 | 0.59427 | 1.15817 | 207 |
| GO_ION_HOMEOSTASIS | 0.00170 | 0.02117 | 0.45506 | 0.56164 | 1.10717 | 403 |
| GO_ION_TRANSMEMBRANE_TRANSPORTER_ACTIVITY | 0.00106 | 0.01495 | 0.45506 | 0.56696 | 1.11734 | 390 |
| GO_IRON_ION_HOMEOSTASIS | 0.00361 | 0.03657 | 0.43171 | 0.65837 | 1.23983 | 59 |
| GO_IRON_ION_TRANSPORT | 0.00054 | 0.00857 | 0.47727 | 0.69556 | 1.30334 | 52 |
| GO_ISOMERASE_ACTIVITY | 0.00413 | 0.04065 | 0.40702 | 0.61065 | 1.17496 | 113 |
| GO_KERATINIZATION | 0.00038 | 0.00640 | 0.49849 | 0.75690 | 1.39013 | 33 |
| GO_KILLING_OF_CELLS_OF_OTHER_ORGANISM | 0.00495 | 0.04573 | 0.40702 | 0.78614 | 1.38643 | 16 |
| GO_LARGE_RIBOSOMAL_SUBUNIT | 0.00000 | 0.00000 | | 0.77656 | 1.49319 | 108 |
| GO_LATERAL_PLASMA_MEMBRANE | 0.00524 | 0.04770 | 0.40702 | 0.67385 | 1.25557 | 47 |
| GO_LEUKOCYTE_MEDIATED_IMMUNITY | 0.00000 | 0.00000 | | 0.64010 | 1.26331 | 447 |
| GO_LIPOPROTEIN_PARTICLE_RECEPTOR_BINDING | 0.00126 | 0.01715 | 0.45506 | 0.80503 | 1.42497 | 17 |

| | | | | | | |
|---|---|---|---|---|---|---|
| GO_LIVER_REGENERATION | 0.00093 | 0.01337 | 0.47727 | 0.78442 | 1.41708 | 23 |
| GO_LOW_DENSITY_LIPOPROTEIN_RECEPTOR_PARTICLE_METABO | 0.00327 | 0.03409 | 0.43171 | 0.78520 | 1.39657 | 18 |
| GO_LYASE_ACTIVITY | 0.00152 | 0.01958 | 0.45506 | 0.61769 | 1.19079 | 122 |
| GO_LYSOSOMAL_LUMEN | 0.00143 | 0.01863 | 0.45506 | 0.67371 | 1.26872 | 59 |
| GO_MAINTENANCE_OF_CELL_NUMBER | 0.00435 | 0.04238 | 0.40702 | 0.60761 | 1.17073 | 117 |
| GO_MATURATION_OF_SSU_RRNA | 0.00126 | 0.01715 | 0.45506 | 0.69615 | 1.29711 | 47 |
| GO_MATURATION_OF_SSU_RRNA_FROM_TRICISTRONIC_RRNA_ | 0.00554 | 0.04968 | 0.40702 | 0.70205 | 1.29301 | 35 |
| GO_MEMBRANE_REGION | 0.00000 | 0.00003 | 0.65944 | 0.64496 | 1.25542 | 194 |
| GO_METALLOPEPTIDASE_ACTIVITY | 0.00487 | 0.04559 | 0.40702 | 0.62714 | 1.19577 | 85 |
| GO_METAL_ION_HOMEOSTASIS | 0.00032 | 0.00575 | 0.49849 | 0.58816 | 1.15376 | 301 |
| GO_METAL_ION_TRANSPORT | 0.00390 | 0.03895 | 0.43171 | 0.55877 | 1.10227 | 419 |
| GO_METAPHASE_ANAPHASE_TRANSITION_OF_CELL_CYCLE | 0.00320 | 0.03371 | 0.43171 | 0.66563 | 1.25122 | 54 |
| GO_MICROTUBULE_CYTOSKELETON_ORGANIZATION_INVOLVED_ | 0.00220 | 0.02548 | 0.43171 | 0.61295 | 1.18118 | 127 |
| GO_MICROVILLUS | 0.00168 | 0.02098 | 0.45506 | 0.68128 | 1.28184 | 56 |
| GO_MIDBODY | 0.00561 | 0.04971 | 0.40702 | 0.59866 | 1.15606 | 141 |
| GO_MITOCHONDRIAL_ELECTRON_TRANSPORT_NADH_TO_UBIQ | 0.00000 | 0.00006 | 0.64355 | 0.79447 | 1.47799 | 44 |
| GO_MITOCHONDRIAL_GENE_EXPRESSION | 0.00000 | 0.00000 | 0.78819 | 0.70230 | 1.35844 | 151 |
| GO_MITOCHONDRIAL_LARGE_RIBOSOMAL_SUBUNIT | 0.00000 | 0.00000 | 0.71951 | 0.79015 | 1.48529 | 54 |
| GO_MITOCHONDRIAL_MATRIX | 0.00000 | 0.00000 | 0.82666 | 0.63141 | 1.24412 | 387 |
| GO_MITOCHONDRIAL_MEMBRANE_ORGANIZATION | 0.00050 | 0.00802 | 0.47727 | 0.64578 | 1.23686 | 94 |
| GO_MITOCHONDRIAL_PROTEIN_COMPLEX | 0.00000 | 0.00000 | | 0.75011 | 1.46215 | 212 |
| GO_MITOCHONDRIAL_RESPIRATORY_CHAIN_COMPLEX_ASSEMB | 0.00001 | 0.00033 | 0.59333 | 0.69650 | 1.32804 | 85 |
| GO_MITOCHONDRIAL_SMALL_RIBOSOMAL_SUBUNIT | 0.00119 | 0.01640 | 0.45506 | 0.75384 | 1.37785 | 28 |
| GO_MITOCHONDRIAL_TRANSLATION | 0.00000 | 0.00000 | 0.80122 | 0.71435 | 1.37651 | 128 |
| GO_MITOCHONDRIAL_TRANSLATIONAL_TERMINATION | 0.00000 | 0.00000 | 0.81404 | 0.75760 | 1.44657 | 87 |
| GO_MITOCHONDRIAL_TRANSMEMBRANE_TRANSPORT | 0.00213 | 0.02490 | 0.43171 | 0.66300 | 1.25387 | 70 |
| GO_MITOCHONDRIAL_TRANSPORT | 0.00010 | 0.00227 | 0.53843 | 0.61119 | 1.19060 | 203 |
| GO_MITOCHONDRION_ORGANIZATION | 0.00000 | 0.00000 | 0.74774 | 0.61590 | 1.21471 | 416 |
| GO_MITOTIC_NUCLEAR_DIVISION | 0.00008 | 0.00182 | 0.53843 | 0.60435 | 1.17901 | 223 |
| GO_MITOTIC_SISTER_CHROMATID_SEGREGATION | 0.00014 | 0.00302 | 0.51885 | 0.62652 | 1.21048 | 145 |
| GO_MITOTIC_SPINDLE_ORGANIZATION | 0.00078 | 0.01164 | 0.47727 | 0.63943 | 1.22821 | 104 |
| GO_MODIFIED_AMINO_ACID_BINDING | 0.00002 | 0.00048 | 0.57561 | 0.74200 | 1.38679 | 49 |
| GO_MODULATION_BY_HOST_OF_VIRAL_PROCESS | 0.00216 | 0.02519 | 0.43171 | 0.77232 | 1.38066 | 20 |
| GO_MONOCARBOXYLIC_ACID_METABOLIC_PROCESS | 0.00012 | 0.00265 | 0.53843 | 0.57846 | 1.14053 | 406 |
| GO_MONOSACCHARIDE_BIOSYNTHETIC_PROCESS | 0.00305 | 0.03271 | 0.43171 | 0.65908 | 1.24472 | 68 |
| GO_MONOVALENT_INORGANIC_CATION_HOMEOSTASIS | 0.00108 | 0.01507 | 0.45506 | 0.63643 | 1.21783 | 91 |
| GO_MONOVALENT_INORGANIC_CATION_TRANSMEMBRANE_TR | 0.00001 | 0.00026 | 0.59333 | 0.65186 | 1.25996 | 147 |
| GO_MONOVALENT_INORGANIC_CATION_TRANSPORT | 0.00008 | 0.00178 | 0.53843 | 0.60205 | 1.17659 | 248 |
| GO_MORPHOGENESIS_OF_AN_EPITHELIUM | 0.00000 | 0.00009 | 0.62726 | 0.60570 | 1.19137 | 334 |
| GO_MORPHOGENESIS_OF_A_POLARIZED_EPITHELIUM | 0.00000 | 0.00000 | 0.83909 | 0.74384 | 1.43057 | 110 |
| GO_MOVEMENT_IN_HOST_ENVIRONMENT | 0.00268 | 0.02961 | 0.43171 | 0.61282 | 1.18078 | 117 |
| GO_MRNA_3_END_PROCESSING | 0.00087 | 0.01270 | 0.47727 | 0.64010 | 1.22502 | 93 |
| GO_MRNA_5_UTR_BINDING | 0.00002 | 0.00063 | 0.57561 | 0.83759 | 1.51313 | 23 |
| GO_MRNA_BINDING | 0.00000 | 0.00000 | | 0.67496 | 1.31810 | 239 |
| GO_MRNA_EXPORT_FROM_NUCLEUS | 0.00006 | 0.00159 | 0.53843 | 0.66290 | 1.27265 | 103 |
| GO_MRNA_PROCESSING | 0.00000 | 0.00000 | | 0.62959 | 1.24255 | 437 |
| GO_MRNA_TRANSPORT | 0.00007 | 0.00171 | 0.53843 | 0.64103 | 1.23563 | 132 |
| GO_MULTI_MULTICELLULAR_ORGANISM_PROCESS | 0.00041 | 0.00687 | 0.49849 | 0.63087 | 1.21515 | 116 |
| GO_MULTI_ORGANISM_LOCALIZATION | 0.00017 | 0.00334 | 0.51885 | 0.71077 | 1.33608 | 54 |
| GO_MUSCLE_CONTRACTION | 0.00335 | 0.03468 | 0.43171 | 0.59925 | 1.16199 | 165 |
| GO_MUSCLE_FILAMENT_SLIDING | 0.00324 | 0.03393 | 0.43171 | 0.77980 | 1.39322 | 19 |
| GO_MYELOID_LEUKOCYTE_ACTIVATION | 0.00000 | 0.00000 | | 0.65288 | 1.28712 | 395 |
| GO_MYELOID_LEUKOCYTE_MEDIATED_IMMUNITY | 0.00000 | 0.00000 | | 0.65734 | 1.29408 | 356 |
| GO_NADH_DEHYDROGENASE_ACTIVITY | 0.00001 | 0.00019 | 0.61053 | 0.79360 | 1.46691 | 38 |
| GO_NADH_DEHYDROGENASE_COMPLEX | 0.00001 | 0.00028 | 0.59333 | 0.77749 | 1.44215 | 41 |
| GO_NADH_DEHYDROGENASE_COMPLEX_ASSEMBLY | 0.00001 | 0.00035 | 0.59333 | 0.74044 | 1.39281 | 55 |
| GO_NADH_METABOLIC_PROCESS | 0.00034 | 0.00599 | 0.49849 | 0.75773 | 1.39166 | 33 |
| GO_NAD_METABOLIC_PROCESS | 0.00435 | 0.04238 | 0.40702 | 0.70570 | 1.30442 | 37 |
| GO_NCRNA_EXPORT_FROM_NUCLEUS | 0.00205 | 0.02440 | 0.43171 | 0.71473 | 1.31817 | 36 |
| GO_NCRNA_METABOLIC_PROCESS | 0.00000 | 0.00000 | 0.70498 | 0.61139 | 1.20509 | 399 |
| GO_NCRNA_PROCESSING | 0.00000 | 0.00000 | 0.73376 | 0.62238 | 1.22425 | 341 |
| GO_NEGATIVE_REGULATION_OF_APOPTOTIC_SIGNALING_PATHW | 0.00364 | 0.03677 | 0.43171 | 0.59958 | 1.15998 | 150 |
| GO_NEGATIVE_REGULATION_OF_ATP_METABOLIC_PROCESS | 0.00302 | 0.03248 | 0.43171 | 0.78586 | 1.39103 | 17 |
| GO_NEGATIVE_REGULATION_OF_CANONICAL_WNT_SIGNALING_ | 0.00000 | 0.00000 | 0.71951 | 0.69629 | 1.34180 | 123 |
| GO_NEGATIVE_REGULATION_OF_CATALYTIC_ACTIVITY | 0.00145 | 0.01879 | 0.45506 | 0.56238 | 1.11005 | 450 |
| GO_NEGATIVE_REGULATION_OF_CELLULAR_AMIDE_METABOLIC | 0.00268 | 0.02961 | 0.43171 | 0.61074 | 1.17686 | 124 |
| GO_NEGATIVE_REGULATION_OF_CELLULAR_COMPONENT_ORGA | 0.00320 | 0.03371 | 0.43171 | 0.55470 | 1.09534 | 470 |
| GO_NEGATIVE_REGULATION_OF_CELL_CYCLE | 0.00023 | 0.00445 | 0.51885 | 0.57348 | 1.13228 | 455 |
| GO_NEGATIVE_REGULATION_OF_CELL_CYCLE_G2_M_PHASE_TR | 0.00000 | 0.00007 | 0.62726 | 0.70549 | 1.34606 | 86 |
| GO_NEGATIVE_REGULATION_OF_CELL_CYCLE_PHASE_TRANSITIO | 0.00000 | 0.00000 | 0.70498 | 0.65532 | 1.27560 | 194 |

| | | | | | | |
|---|---|---|---|---|---|---|
| GO_NEGATIVE_REGULATION_OF_CELL_CYCLE_PROCESS | 0.00000 | 0.00005 | 0.64355 | 0.62725 | 1.22670 | 259 |
| GO_NEGATIVE_REGULATION_OF_HYDROLASE_ACTIVITY | 0.00216 | 0.02519 | 0.43171 | 0.58306 | 1.13858 | 238 |
| GO_NEGATIVE_REGULATION_OF_INTERLEUKIN_1_PRODUCTION | 0.00205 | 0.02440 | 0.43171 | 0.82297 | 1.44404 | 15 |
| GO_NEGATIVE_REGULATION_OF_MITOTIC_CELL_CYCLE | 0.00000 | 0.00010 | 0.62726 | 0.62191 | 1.21541 | 248 |
| GO_NEGATIVE_REGULATION_OF_MRNA_METABOLIC_PROCESS | 0.00213 | 0.02490 | 0.43171 | 0.66731 | 1.25879 | 62 |
| GO_NEGATIVE_REGULATION_OF_MRNA_SPLICING_VIA_SPLICEO | 0.00339 | 0.03501 | 0.43171 | 0.76309 | 1.36417 | 20 |
| GO_NEGATIVE_REGULATION_OF_NUCLEAR_DIVISION | 0.00335 | 0.03468 | 0.43171 | 0.67724 | 1.26685 | 50 |
| GO_NEGATIVE_REGULATION_OF_ORGANELLE_ORGANIZATION | 0.00487 | 0.04559 | 0.40702 | 0.57038 | 1.11722 | 288 |
| GO_NEGATIVE_REGULATION_OF_PEPTIDASE_ACTIVITY | 0.00036 | 0.00621 | 0.49849 | 0.64044 | 1.23159 | 109 |
| GO_NEGATIVE_REGULATION_OF_PROTEIN_CATABOLIC_PROCESS | 0.00342 | 0.03521 | 0.43171 | 0.62320 | 1.19268 | 93 |
| GO_NEGATIVE_REGULATION_OF_PROTEOLYSIS | 0.00205 | 0.02440 | 0.43171 | 0.60172 | 1.16902 | 178 |
| GO_NEGATIVE_REGULATION_OF_REACTIVE_OXYGEN_SPECIES_B | 0.00036 | 0.00621 | 0.49849 | 0.83108 | 1.47818 | 18 |
| GO_NEGATIVE_REGULATION_OF_REPRODUCTIVE_PROCESS | 0.00457 | 0.04386 | 0.40702 | 0.74845 | 1.35356 | 24 |
| GO_NEGATIVE_REGULATION_OF_RNA_SPLICING | 0.00183 | 0.02219 | 0.45506 | 0.76824 | 1.38935 | 24 |
| GO_NEGATIVE_REGULATION_OF_WNT_SIGNALING_PATHWAY | 0.00000 | 0.00002 | 0.67496 | 0.67731 | 1.30721 | 136 |
| GO_NEURAL_CREST_CELL_MIGRATION | 0.00265 | 0.02936 | 0.43171 | 0.73967 | 1.35049 | 27 |
| GO_NEURON_RECOGNITION | 0.00390 | 0.03895 | 0.43171 | 0.71862 | 1.31998 | 31 |
| GO_NIK_NF_KAPPAB_SIGNALING | 0.00000 | 0.00000 | 0.77494 | 0.72329 | 1.39253 | 114 |
| GO_NON_CANONICAL_WNT_SIGNALING_PATHWAY | 0.00000 | 0.00000 | | 0.75756 | 1.45654 | 107 |
| GO_NUCLEAR_CHROMOSOME_SEGREGATION | 0.00098 | 0.01407 | 0.45506 | 0.59156 | 1.15357 | 215 |
| GO_NUCLEAR_ENVELOPE | 0.00005 | 0.00131 | 0.55733 | 0.59109 | 1.16270 | 341 |
| GO_NUCLEAR_EXPORT | 0.00000 | 0.00004 | 0.65944 | 0.64915 | 1.26157 | 181 |
| GO_NUCLEAR_MATRIX | 0.00502 | 0.04608 | 0.40702 | 0.62777 | 1.19868 | 87 |
| GO_NUCLEAR_MEMBRANE | 0.00185 | 0.02237 | 0.45506 | 0.58925 | 1.14853 | 211 |
| GO_NUCLEAR_PERIPHERY | 0.00043 | 0.00701 | 0.49849 | 0.64264 | 1.23558 | 107 |
| GO_NUCLEAR_PORE | 0.00561 | 0.04971 | 0.40702 | 0.64876 | 1.22522 | 68 |
| GO_NUCLEAR_SPECK | 0.00015 | 0.00312 | 0.51885 | 0.59159 | 1.16089 | 306 |
| GO_NUCLEAR_TRANSCRIBED_MRNA_CATABOLIC_PROCESS | 0.00000 | 0.00000 | | 0.70362 | 1.36820 | 185 |
| GO_NUCLEAR_TRANSCRIBED_MRNA_CATABOLIC_PROCESS_NON | 0.00000 | 0.00000 | | 0.74239 | 1.42931 | 114 |
| GO_NUCLEAR_TRANSPORT | 0.00000 | 0.00000 | 0.71951 | 0.62920 | 1.23383 | 297 |
| GO_NUCLEOBASE_CONTAINING_COMPOUND_TRANSPORT | 0.00000 | 0.00010 | 0.62726 | 0.62975 | 1.22823 | 216 |
| GO_NUCLEOBASE_CONTAINING_SMALL_MOLECULE_METABOLIC | 0.00000 | 0.00003 | 0.65944 | 0.60153 | 1.18594 | 405 |
| GO_NUCLEOSIDE_DIPHOSPHATE_METABOLIC_PROCESS | 0.00016 | 0.00319 | 0.51885 | 0.64491 | 1.24260 | 117 |
| GO_NUCLEOSIDE_PHOSPHATE_BIOSYNTHETIC_PROCESS | 0.00002 | 0.00044 | 0.59333 | 0.64092 | 1.24175 | 161 |
| GO_NUCLEOSIDE_TRIPHOSPHATE_BIOSYNTHETIC_PROCESS | 0.00040 | 0.00676 | 0.49849 | 0.72727 | 1.35074 | 42 |
| GO_NUCLEOSIDE_TRIPHOSPHATE_METABOLIC_PROCESS | 0.00009 | 0.00196 | 0.53843 | 0.69306 | 1.30854 | 65 |
| GO_NUCLEOTIDE_PHOSPHORYLATION | 0.00016 | 0.00328 | 0.51885 | 0.65541 | 1.25767 | 102 |
| GO_OOCYTE_DIFFERENTIATION | 0.00450 | 0.04342 | 0.40702 | 0.73513 | 1.33705 | 26 |
| GO_ORGANELLAR_RIBOSOME | 0.00000 | 0.00000 | 0.76146 | 0.75712 | 1.44311 | 84 |
| GO_ORGANELLE_ENVELOPE_LUMEN | 0.00026 | 0.00482 | 0.49849 | 0.67265 | 1.27716 | 78 |
| GO_ORGANELLE_FISSION | 0.00154 | 0.01974 | 0.45506 | 0.57133 | 1.12383 | 342 |
| GO_ORGANELLE_INNER_MEMBRANE | 0.00000 | 0.00000 | | 0.68423 | 1.34915 | 412 |
| GO_ORGANIC_CYCLIC_COMPOUND_CATABOLIC_PROCESS | 0.00000 | 0.00000 | | 0.66975 | 1.32263 | 478 |
| GO_ORGANIC_HYDROXY_COMPOUND_BIOSYNTHETIC_PROCESS | 0.00141 | 0.01850 | 0.45506 | 0.60712 | 1.17627 | 161 |
| GO_OXIDATIVE_PHOSPHORYLATION | 0.00000 | 0.00000 | | 0.78902 | 1.51260 | 99 |
| GO_OXIDOREDUCTASE_ACTIVITY | 0.00000 | 0.00000 | | 0.63268 | 1.24935 | 468 |
| GO_OXIDOREDUCTASE_ACTIVITY_ACTING_ON_A_HEME_GROUP | 0.00176 | 0.02168 | 0.45506 | 0.79383 | 1.41828 | 19 |
| GO_OXIDOREDUCTASE_ACTIVITY_ACTING_ON_NAD_P_H | 0.00002 | 0.00056 | 0.57561 | 0.71468 | 1.35162 | 70 |
| GO_OXIDOREDUCTASE_ACTIVITY_ACTING_ON_NAD_P_H_QUIN | 0.00004 | 0.00113 | 0.55733 | 0.74511 | 1.38835 | 47 |
| GO_OXIDOREDUCTASE_ACTIVITY_ACTING_ON_PEROXIDE_AS_A | 0.00106 | 0.01495 | 0.45506 | 0.73897 | 1.35352 | 29 |
| GO_OXIDOREDUCTASE_COMPLEX | 0.00000 | 0.00000 | 0.73376 | 0.74898 | 1.42731 | 82 |
| GO_PEPTIDASE_ACTIVITY | 0.00133 | 0.01799 | 0.45506 | 0.57681 | 1.13188 | 306 |
| GO_PEPTIDASE_COMPLEX | 0.00000 | 0.00002 | 0.67496 | 0.73250 | 1.39240 | 79 |
| GO_PEPTIDASE_REGULATOR_ACTIVITY | 0.00313 | 0.03310 | 0.43171 | 0.63434 | 1.21189 | 88 |
| GO_PEPTIDE_BINDING | 0.00005 | 0.00124 | 0.55733 | 0.63708 | 1.23187 | 148 |
| GO_PERINUCLEAR_REGION_OF_CYTOPLASM | 0.00276 | 0.03021 | 0.43171 | 0.55814 | 1.10207 | 466 |
| GO_PHAGOCYTOSIS | 0.00554 | 0.04968 | 0.40702 | 0.59337 | 1.14837 | 155 |
| GO_PHAGOSOME_ACIDIFICATION | 0.00059 | 0.00916 | 0.47727 | 0.78902 | 1.42694 | 24 |
| GO_PHOSPHATIDYLSERINE_BINDING | 0.00035 | 0.00610 | 0.49849 | 0.76387 | 1.39912 | 29 |
| GO_PH_REDUCTION | 0.00302 | 0.03248 | 0.43171 | 0.68821 | 1.28238 | 46 |
| GO_PIGMENT_GRANULE | 0.00000 | 0.00000 | 0.76146 | 0.74333 | 1.41934 | 87 |
| GO_PLASMA_LIPOPROTEIN_PARTICLE_CLEARANCE | 0.00106 | 0.01495 | 0.45506 | 0.73201 | 1.35304 | 37 |
| GO_POLYSOMAL_RIBOSOME | 0.00008 | 0.00180 | 0.53843 | 0.78840 | 1.44683 | 30 |
| GO_POLYSOME | 0.00000 | 0.00003 | 0.65944 | 0.75964 | 1.43296 | 62 |
| GO_POSITIVE_REGULATION_OF_ATP_METABOLIC_PROCESS | 0.00174 | 0.02149 | 0.45506 | 0.73011 | 1.33729 | 29 |
| GO_POSITIVE_REGULATION_OF_BINDING | 0.00390 | 0.03895 | 0.43171 | 0.61042 | 1.17582 | 115 |
| GO_POSITIVE_REGULATION_OF_CANONICAL_WNT_SIGNALING_ | 0.00000 | 0.00000 | 0.73376 | 0.72036 | 1.38514 | 108 |
| GO_POSITIVE_REGULATION_OF_CELLULAR_AMIDE_METABOLIC_ | 0.00001 | 0.00036 | 0.59333 | 0.66485 | 1.28095 | 125 |
| GO_POSITIVE_REGULATION_OF_CELLULAR_PROTEIN_LOCALIZAT | 0.00242 | 0.02748 | 0.43171 | 0.58048 | 1.13378 | 244 |
| GO_POSITIVE_REGULATION_OF_CELL_POPULATION_PROLIFERAT | 0.00040 | 0.00670 | 0.49849 | 0.56967 | 1.12503 | 461 |

| | | | | | | |
|---|---|---|---|---|---|---|
| GO_POSITIVE_REGULATION_OF_CYSTEINE_TYPE_ENDOPEPTIDAS | 0.00547 | 0.04923 | 0.40702 | 0.62225 | 1.19077 | 92 |
| GO_POSITIVE_REGULATION_OF_CYTOKINE_PRODUCTION | 0.00283 | 0.03081 | 0.43171 | 0.58672 | 1.14373 | 210 |
| GO_POSITIVE_REGULATION_OF_DEFENSE_RESPONSE | 0.00000 | 0.00000 | 0.73376 | 0.66200 | 1.28884 | 200 |
| GO_POSITIVE_REGULATION_OF_DNA_BIOSYNTHETIC_PROCESS | 0.00010 | 0.00228 | 0.53843 | 0.71224 | 1.33977 | 55 |
| GO_POSITIVE_REGULATION_OF_DNA_METABOLIC_PROCESS | 0.00020 | 0.00387 | 0.51885 | 0.62771 | 1.21205 | 139 |
| GO_POSITIVE_REGULATION_OF_ESTABLISHMENT_OF_PROTEIN_ | 0.00194 | 0.02331 | 0.45506 | 0.58158 | 1.13709 | 253 |
| GO_POSITIVE_REGULATION_OF_FIBROBLAST_PROLIFERATION | 0.00017 | 0.00338 | 0.51885 | 0.77423 | 1.42213 | 31 |
| GO_POSITIVE_REGULATION_OF_IMMUNE_RESPONSE | 0.00000 | 0.00000 | 0.71951 | 0.63064 | 1.23937 | 321 |
| GO_POSITIVE_REGULATION_OF_IMMUNE_SYSTEM_PROCESS | 0.00000 | 0.00002 | 0.67496 | 0.59748 | 1.17992 | 475 |
| GO_POSITIVE_REGULATION_OF_INNATE_IMMUNE_RESPONSE | 0.00000 | 0.00000 | 0.70498 | 0.68885 | 1.32737 | 128 |
| GO_POSITIVE_REGULATION_OF_MITOTIC_CELL_CYCLE | 0.00178 | 0.02182 | 0.45506 | 0.61563 | 1.18683 | 122 |
| GO_POSITIVE_REGULATION_OF_MRNA_METABOLIC_PROCESS | 0.00157 | 0.02009 | 0.45506 | 0.66330 | 1.25231 | 63 |
| GO_POSITIVE_REGULATION_OF_MRNA_PROCESSING | 0.00250 | 0.02807 | 0.43171 | 0.76194 | 1.38151 | 25 |
| GO_POSITIVE_REGULATION_OF_MRNA_SPLICING_VIA_SPLICEOS | 0.00132 | 0.01782 | 0.45506 | 0.81851 | 1.44353 | 16 |
| GO_POSITIVE_REGULATION_OF_NITRIC_OXIDE_METABOLIC_PRO | 0.00133 | 0.01799 | 0.45506 | 0.80807 | 1.43726 | 18 |
| GO_POSITIVE_REGULATION_OF_ORGANELLE_ORGANIZATION | 0.00010 | 0.00218 | 0.53843 | 0.57473 | 1.13449 | 452 |
| GO_POSITIVE_REGULATION_OF_PROTEIN_LOCALIZATION_TO_N | 0.00012 | 0.00265 | 0.53843 | 0.71018 | 1.33697 | 58 |
| GO_POSITIVE_REGULATION_OF_PROTEOLYSIS | 0.00313 | 0.03310 | 0.43171 | 0.58059 | 1.13381 | 237 |
| GO_POSITIVE_REGULATION_OF_REACTIVE_OXYGEN_SPECIES_BI | 0.00115 | 0.01593 | 0.45506 | 0.79865 | 1.43554 | 21 |
| GO_POSITIVE_REGULATION_OF_REACTIVE_OXYGEN_SPECIES_M | 0.00174 | 0.02149 | 0.45506 | 0.69785 | 1.30035 | 46 |
| GO_POSITIVE_REGULATION_OF_RESPONSE_TO_BIOTIC_STIMUL | 0.00000 | 0.00000 | 0.70498 | 0.67507 | 1.30603 | 150 |
| GO_POSITIVE_REGULATION_OF_RESPONSE_TO_EXTERNAL_STIM | 0.00001 | 0.00016 | 0.61053 | 0.62123 | 1.21505 | 260 |
| GO_POSITIVE_REGULATION_OF_RNA_SPLICING | 0.00265 | 0.02936 | 0.43171 | 0.74051 | 1.35202 | 27 |
| GO_POSITIVE_REGULATION_OF_SIGNAL_TRANSDUCTION_BY_P5 | 0.00390 | 0.03895 | 0.43171 | 0.80515 | 1.41278 | 15 |
| GO_POSITIVE_REGULATION_OF_TELOMERASE_ACTIVITY | 0.00069 | 0.01043 | 0.47727 | 0.75265 | 1.38122 | 30 |
| GO_POSITIVE_REGULATION_OF_TELOMERASE_RNA_LOCALIZATI | 0.00007 | 0.00169 | 0.53843 | 0.88616 | 1.55493 | 15 |
| GO_POSITIVE_REGULATION_OF_TELOMERE_MAINTENANCE | 0.00026 | 0.00486 | 0.49849 | 0.73697 | 1.37102 | 44 |
| GO_POSITIVE_REGULATION_OF_TELOMERE_MAINTENANCE_VIA | 0.00001 | 0.00040 | 0.59333 | 0.80196 | 1.47289 | 33 |
| GO_POSITIVE_REGULATION_OF_TRANSFERASE_ACTIVITY | 0.00108 | 0.01507 | 0.45506 | 0.56501 | 1.11376 | 401 |
| GO_POSITIVE_REGULATION_OF_TRANSLATION | 0.00000 | 0.00016 | 0.61053 | 0.68408 | 1.31526 | 107 |
| GO_POSITIVE_REGULATION_OF_VIRAL_PROCESS | 0.00026 | 0.00486 | 0.49849 | 0.66195 | 1.26462 | 88 |
| GO_POSITIVE_REGULATION_OF_WNT_SIGNALING_PATHWAY | 0.00000 | 0.00000 | 0.73376 | 0.69924 | 1.34747 | 127 |
| GO_POSTSYNAPSE | 0.00239 | 0.02716 | 0.43171 | 0.57062 | 1.12116 | 316 |
| GO_POST_TRANSLATIONAL_PROTEIN_MODIFICATION | 0.00024 | 0.00452 | 0.51885 | 0.60046 | 1.17392 | 251 |
| GO_PRECATALYTIC_SPLICEOSOME | 0.00141 | 0.01850 | 0.45506 | 0.69251 | 1.29540 | 50 |
| GO_PRERIBOSOME | 0.00006 | 0.00154 | 0.53843 | 0.69866 | 1.32020 | 69 |
| GO_PROTEASOMAL_PROTEIN_CATABOLIC_PROCESS | 0.00007 | 0.00169 | 0.53843 | 0.58188 | 1.14652 | 387 |
| GO_PROTEASOMAL_UBIQUITIN_INDEPENDENT_PROTEIN_CATAB | 0.00000 | 0.00003 | 0.65944 | 0.91182 | 1.62178 | 18 |
| GO_PROTEASOME_ACCESSORY_COMPLEX | 0.00005 | 0.00135 | 0.55733 | 0.82638 | 1.49450 | 24 |
| GO_PROTEASOME_CORE_COMPLEX | 0.00000 | 0.00004 | 0.65944 | 0.91231 | 1.61486 | 17 |
| GO_PROTEIN_CONTAINING_COMPLEX_DISASSEMBLY | 0.00000 | 0.00000 | 0.81404 | 0.66285 | 1.29656 | 261 |
| GO_PROTEIN_CONTAINING_COMPLEX_LOCALIZATION | 0.00018 | 0.00352 | 0.51885 | 0.60451 | 1.17943 | 226 |
| GO_PROTEIN_DNA_COMPLEX_SUBUNIT_ORGANIZATION | 0.00052 | 0.00825 | 0.47727 | 0.60887 | 1.18342 | 183 |
| GO_PROTEIN_DOMAIN_SPECIFIC_BINDING | 0.00003 | 0.00088 | 0.55733 | 0.57828 | 1.14190 | 470 |
| GO_PROTEIN_FOLDING | 0.00000 | 0.00000 | 0.73376 | 0.67425 | 1.30867 | 171 |
| GO_PROTEIN_FOLDING_CHAPERONE | 0.00079 | 0.01170 | 0.47727 | 0.81632 | 1.45193 | 18 |
| GO_PROTEIN_HETEROOLIGOMERIZATION | 0.00442 | 0.04297 | 0.40702 | 0.78998 | 1.39322 | 16 |
| GO_PROTEIN_IMPORT | 0.00001 | 0.00040 | 0.59333 | 0.64197 | 1.24306 | 158 |
| GO_PROTEIN_IMPORT_INTO_MITOCHONDRIAL_MATRIX | 0.00231 | 0.02642 | 0.43171 | 0.79530 | 1.41455 | 18 |
| GO_PROTEIN_INSERTION_INTO_MEMBRANE | 0.00191 | 0.02299 | 0.45506 | 0.68322 | 1.27694 | 49 |
| GO_PROTEIN_LOCALIZATION_TO_CHROMOSOME_TELOMERIC_R | 0.00001 | 0.00031 | 0.59333 | 0.82810 | 1.51357 | 28 |
| GO_PROTEIN_LOCALIZATION_TO_ENDOPLASMIC_RETICULUM | 0.00000 | 0.00000 | | 0.73850 | 1.42303 | 129 |
| GO_PROTEIN_LOCALIZATION_TO_MEMBRANE | 0.00000 | 0.00000 | 0.77494 | 0.61216 | 1.20881 | 462 |
| GO_PROTEIN_LOCALIZATION_TO_MITOCHONDRION | 0.00015 | 0.00310 | 0.51885 | 0.64234 | 1.23759 | 118 |
| GO_PROTEIN_LOCALIZATION_TO_NUCLEUS | 0.00000 | 0.00014 | 0.61053 | 0.63315 | 1.23408 | 208 |
| GO_PROTEIN_MATURATION | 0.00357 | 0.03638 | 0.43171 | 0.59209 | 1.14965 | 174 |
| GO_PROTEIN_MODIFICATION_BY_SMALL_PROTEIN_REMOVAL | 0.00002 | 0.00052 | 0.57561 | 0.61535 | 1.20119 | 231 |
| GO_PROTEIN_REFOLDING | 0.00465 | 0.04430 | 0.40702 | 0.77561 | 1.37288 | 17 |
| GO_PROTEIN_STABILIZATION | 0.00081 | 0.01194 | 0.47727 | 0.61881 | 1.19577 | 146 |
| GO_PROTEIN_SUMOYLATION | 0.00450 | 0.04342 | 0.40702 | 0.65043 | 1.22846 | 67 |
| GO_PROTEIN_TARGETING | 0.00000 | 0.00000 | | 0.66223 | 1.30361 | 358 |
| GO_PROTEIN_TARGETING_TO_MEMBRANE | 0.00000 | 0.00000 | | 0.71675 | 1.38879 | 162 |
| GO_PROTEIN_TARGETING_TO_MITOCHONDRION | 0.00113 | 0.01571 | 0.45506 | 0.65407 | 1.24648 | 83 |
| GO_PROTEIN_TRANSMEMBRANE_TRANSPORT | 0.00110 | 0.01526 | 0.45506 | 0.68173 | 1.28238 | 55 |
| GO_PROTON_TRANSMEMBRANE_TRANSPORT | 0.00000 | 0.00001 | 0.69013 | 0.71234 | 1.36309 | 91 |
| GO_PROTON_TRANSMEMBRANE_TRANSPORTER_ACTIVITY | 0.00004 | 0.00115 | 0.55733 | 0.70138 | 1.32535 | 69 |
| GO_PROTON_TRANSPORTING_TWO_SECTOR_ATPASE_COMPLEX | 0.00083 | 0.01218 | 0.47727 | 0.77254 | 1.40508 | 26 |
| GO_PROTON_TRANSPORTING_V_TYPE_ATPASE_COMPLEX | 0.00239 | 0.02716 | 0.43171 | 0.76100 | 1.37626 | 24 |
| GO_PURINE_CONTAINING_COMPOUND_BIOSYNTHETIC_PROCES | 0.00001 | 0.00040 | 0.59333 | 0.65863 | 1.26901 | 126 |
| GO_PURINE_CONTAINING_COMPOUND_METABOLIC_PROCESS | 0.00000 | 0.00000 | 0.73376 | 0.63132 | 1.23831 | 300 |

| | | | | | | |
|---|---|---|---|---|---|---|
| GO_PURINE_NUCLEOSIDE_TRIPHOSPHATE_METABOLIC_PROCES | 0.00033 | 0.00582 | 0.49849 | 0.70887 | 1.32487 | 49 |
| GO_PYROPHOSPHATE_HYDROLYSIS_DRIVEN_PROTON_TRANSMI | 0.00532 | 0.04803 | 0.40702 | 0.74470 | 1.34678 | 24 |
| GO_PYRUVATE_METABOLIC_PROCESS | 0.00001 | 0.00018 | 0.61053 | 0.67194 | 1.29289 | 113 |
| GO_P_BODY | 0.00472 | 0.04480 | 0.40702 | 0.65470 | 1.23501 | 62 |
| GO_REACTIVE_OXYGEN_SPECIES_BIOSYNTHETIC_PROCESS | 0.00040 | 0.00670 | 0.49849 | 0.70271 | 1.31857 | 53 |
| GO_REACTIVE_OXYGEN_SPECIES_METABOLIC_PROCESS | 0.00021 | 0.00409 | 0.51885 | 0.62881 | 1.21472 | 144 |
| GO_REGULATION_OF_ALCOHOL_BIOSYNTHETIC_PROCESS | 0.00228 | 0.02614 | 0.43171 | 0.68438 | 1.28020 | 50 |
| GO_REGULATION_OF_ANIMAL_ORGAN_MORPHOGENESIS | 0.00000 | 0.00000 | | 0.72330 | 1.39922 | 149 |
| GO_REGULATION_OF_APOPTOTIC_SIGNALING_PATHWAY | 0.00079 | 0.01170 | 0.47727 | 0.58599 | 1.14641 | 266 |
| GO_REGULATION_OF_ATP_METABOLIC_PROCESS | 0.00000 | 0.00006 | 0.64355 | 0.70523 | 1.34777 | 89 |
| GO_REGULATION_OF_BINDING | 0.00428 | 0.04185 | 0.40702 | 0.58264 | 1.13664 | 225 |
| GO_REGULATION_OF_BLOOD_PRESSURE | 0.00290 | 0.03156 | 0.43171 | 0.65319 | 1.23428 | 69 |
| GO_REGULATION_OF_CARBOHYDRATE_CATABOLIC_PROCESS | 0.00324 | 0.03393 | 0.43171 | 0.65541 | 1.23787 | 67 |
| GO_REGULATION_OF_CELLULAR_AMIDE_METABOLIC_PROCESS | 0.00000 | 0.00000 | 0.78819 | 0.63116 | 1.24143 | 337 |
| GO_REGULATION_OF_CELLULAR_AMINE_METABOLIC_PROCESS | 0.00000 | 0.00000 | 0.74774 | 0.78955 | 1.48730 | 60 |
| GO_REGULATION_OF_CELLULAR_AMINO_ACID_METABOLIC_PR( | 0.00000 | 0.00000 | 0.76146 | 0.80341 | 1.51022 | 54 |
| GO_REGULATION_OF_CELLULAR_KETONE_METABOLIC_PROCESS | 0.00000 | 0.00000 | 0.70498 | 0.69239 | 1.33405 | 126 |
| GO_REGULATION_OF_CELLULAR_PH | 0.00442 | 0.04297 | 0.40702 | 0.65360 | 1.23436 | 68 |
| GO_REGULATION_OF_CELLULAR_PROTEIN_LOCALIZATION | 0.00228 | 0.02614 | 0.43171 | 0.56216 | 1.10830 | 402 |
| GO_REGULATION_OF_CELLULAR_RESPIRATION | 0.00428 | 0.04185 | 0.40702 | 0.77183 | 1.37897 | 19 |
| GO_REGULATION_OF_CELLULAR_RESPONSE_TO_HEAT | 0.00091 | 0.01314 | 0.47727 | 0.67360 | 1.27221 | 67 |
| GO_REGULATION_OF_CELL_CYCLE_G2_M_PHASE_TRANSITION | 0.00016 | 0.00332 | 0.51885 | 0.61554 | 1.19738 | 187 |
| GO_REGULATION_OF_CELL_CYCLE_PHASE_TRANSITION | 0.00000 | 0.00014 | 0.61053 | 0.59968 | 1.18091 | 369 |
| GO_REGULATION_OF_CHOLESTEROL_BIOSYNTHETIC_PROCESS | 0.00209 | 0.02470 | 0.43171 | 0.71476 | 1.32117 | 38 |
| GO_REGULATION_OF_CHOLESTEROL_METABOLIC_PROCESS | 0.00113 | 0.01571 | 0.45506 | 0.69865 | 1.30177 | 47 |
| GO_REGULATION_OF_CHROMOSOME_ORGANIZATION | 0.00036 | 0.00622 | 0.49849 | 0.58896 | 1.15261 | 278 |
| GO_REGULATION_OF_CHROMOSOME_SEGREGATION | 0.00079 | 0.01170 | 0.47727 | 0.64131 | 1.22718 | 91 |
| GO_REGULATION_OF_CHROMOSOME_SEPARATION | 0.00253 | 0.02838 | 0.43171 | 0.66349 | 1.24947 | 59 |
| GO_REGULATION_OF_CYSTEINE_TYPE_ENDOPEPTIDASE_ACTIVIT | 0.00044 | 0.00719 | 0.49849 | 0.62610 | 1.20905 | 141 |
| GO_REGULATION_OF_DEFENSE_RESPONSE | 0.00006 | 0.00150 | 0.55733 | 0.58834 | 1.15824 | 356 |
| GO_REGULATION_OF_DNA_BIOSYNTHETIC_PROCESS | 0.00043 | 0.00695 | 0.49849 | 0.66709 | 1.27130 | 83 |
| GO_REGULATION_OF_DNA_METABOLIC_PROCESS | 0.00055 | 0.00863 | 0.47727 | 0.58909 | 1.15134 | 249 |
| GO_REGULATION_OF_DNA_TEMPLATED_TRANSCRIPTION_IN_RI | 0.00000 | 0.00000 | 0.71951 | 0.72410 | 1.38687 | 94 |
| GO_REGULATION_OF_GENERATION_OF_PRECURSOR_METABOLI | 0.00004 | 0.00114 | 0.55733 | 0.65536 | 1.26115 | 111 |
| GO_REGULATION_OF_GENE_SILENCING | 0.00023 | 0.00434 | 0.51885 | 0.65464 | 1.25268 | 91 |
| GO_REGULATION_OF_HEMATOPOIETIC_PROGENITOR_CELL_DIF | 0.00000 | 0.00000 | 0.73376 | 0.76273 | 1.44465 | 72 |
| GO_REGULATION_OF_HEMATOPOIETIC_STEM_CELL_DIFFERENT | 0.00000 | 0.00000 | 0.74774 | 0.78891 | 1.48817 | 62 |
| GO_REGULATION_OF_HEMOPOIESIS | 0.00390 | 0.03895 | 0.43171 | 0.57739 | 1.12909 | 258 |
| GO_REGULATION_OF_IMMUNE_RESPONSE | 0.00000 | 0.00006 | 0.64355 | 0.60032 | 1.18447 | 433 |
| GO_REGULATION_OF_INNATE_IMMUNE_RESPONSE | 0.00000 | 0.00007 | 0.62726 | 0.64536 | 1.25422 | 181 |
| GO_REGULATION_OF_INTRINSIC_APOPTOTIC_SIGNALING_PATHV | 0.00179 | 0.02183 | 0.45506 | 0.61548 | 1.18550 | 116 |
| GO_REGULATION_OF_MITOTIC_CELL_CYCLE | 0.00000 | 0.00003 | 0.65944 | 0.58992 | 1.16508 | 499 |
| GO_REGULATION_OF_MORPHOGENESIS_OF_AN_EPITHELIUM | 0.00000 | 0.00000 | | 0.74680 | 1.43884 | 125 |
| GO_REGULATION_OF_MRNA_CATABOLIC_PROCESS | 0.00000 | 0.00000 | | 0.71126 | 1.37919 | 165 |
| GO_REGULATION_OF_MRNA_METABOLIC_PROCESS | 0.00000 | 0.00000 | | 0.67178 | 1.31458 | 276 |
| GO_REGULATION_OF_MRNA_PROCESSING | 0.00008 | 0.00186 | 0.53843 | 0.65189 | 1.25598 | 125 |
| GO_REGULATION_OF_MRNA_SPLICING_VIA_SPLICEOSOME | 0.00012 | 0.00256 | 0.53843 | 0.66950 | 1.28033 | 90 |
| GO_REGULATION_OF_MULTI_ORGANISM_PROCESS | 0.00008 | 0.00190 | 0.53843 | 0.76905 | 1.42150 | 37 |
| GO_REGULATION_OF_MUSCLE_CONTRACTION | 0.00163 | 0.02050 | 0.45506 | 0.64879 | 1.23071 | 76 |
| GO_REGULATION_OF_NUCLEAR_DIVISION | 0.00060 | 0.00923 | 0.47727 | 0.62443 | 1.20552 | 137 |
| GO_REGULATION_OF_NUCLEASE_ACTIVITY | 0.00059 | 0.00916 | 0.47727 | 0.81786 | 1.44767 | 17 |
| GO_REGULATION_OF_NUCLEOTIDE_BIOSYNTHETIC_PROCESS | 0.00302 | 0.03248 | 0.43171 | 0.78902 | 1.40337 | 18 |
| GO_REGULATION_OF_NUCLEOTIDE_METABOLIC_PROCESS | 0.00033 | 0.00588 | 0.49849 | 0.66858 | 1.26942 | 78 |
| GO_REGULATION_OF_OXIDATIVE_PHOSPHORYLATION | 0.00005 | 0.00130 | 0.55733 | 0.85284 | 1.53294 | 21 |
| GO_REGULATION_OF_PEPTIDASE_ACTIVITY | 0.00006 | 0.00152 | 0.53843 | 0.61192 | 1.19388 | 226 |
| GO_REGULATION_OF_PEPTIDE_TRANSPORT | 0.00250 | 0.02807 | 0.43171 | 0.56252 | 1.10917 | 412 |
| GO_REGULATION_OF_PH | 0.00561 | 0.04971 | 0.40702 | 0.64191 | 1.21498 | 71 |
| GO_REGULATION_OF_PLASMA_LIPOPROTEIN_PARTICLE_LEVELS | 0.00006 | 0.00145 | 0.55733 | 0.72431 | 1.36154 | 54 |
| GO_REGULATION_OF_POSTTRANSCRIPTIONAL_GENE_SILENCINC | 0.00026 | 0.00486 | 0.49849 | 0.66803 | 1.27236 | 81 |
| GO_REGULATION_OF_PROTEIN_CATABOLIC_PROCESS | 0.00309 | 0.03299 | 0.43171 | 0.57360 | 1.12399 | 291 |
| GO_REGULATION_OF_PROTEIN_IMPORT | 0.00487 | 0.04559 | 0.40702 | 0.67491 | 1.25754 | 47 |
| GO_REGULATION_OF_PROTEIN_LOCALIZATION_TO_NUCLEUS | 0.00000 | 0.00013 | 0.61053 | 0.69035 | 1.32101 | 91 |
| GO_REGULATION_OF_PROTEIN_STABILITY | 0.00066 | 0.01007 | 0.47727 | 0.59308 | 1.15811 | 235 |
| GO_REGULATION_OF_PROTEOLYSIS | 0.00007 | 0.00169 | 0.53843 | 0.57923 | 1.14299 | 431 |
| GO_REGULATION_OF_REACTIVE_OXYGEN_SPECIES_BIOSYNTHET | 0.00022 | 0.00427 | 0.51885 | 0.73154 | 1.36343 | 45 |
| GO_REGULATION_OF_REACTIVE_OXYGEN_SPECIES_METABOLIC_ | 0.00088 | 0.01280 | 0.47727 | 0.63948 | 1.22592 | 99 |
| GO_REGULATION_OF_RESPONSE_TO_BIOTIC_STIMULUS | 0.00001 | 0.00039 | 0.59333 | 0.61773 | 1.20625 | 235 |
| GO_REGULATION_OF_RNA_SPLICING | 0.00011 | 0.00241 | 0.53843 | 0.64982 | 1.25198 | 125 |
| GO_REGULATION_OF_SISTER_CHROMATID_SEGREGATION | 0.00405 | 0.04011 | 0.40702 | 0.64401 | 1.22034 | 73 |
| GO_REGULATION_OF_SMALL_MOLECULE_METABOLIC_PROCESS | 0.00000 | 0.00000 | 0.77494 | 0.64342 | 1.26057 | 290 |

| | | | | | | |
|---|---|---|---|---|---|---|
| GO_REGULATION_OF_SMOOTH_MUSCLE_CONTRACTION | 0.00007 | 0.00171 | 0.53843 | 0.82642 | 1.49295 | 23 |
| GO_REGULATION_OF_STEM_CELL_DIFFERENTIATION | 0.00000 | 0.00000 | 0.73376 | 0.73434 | 1.40292 | 88 |
| GO_REGULATION_OF_STEROID_METABOLIC_PROCESS | 0.00524 | 0.04770 | 0.40702 | 0.64061 | 1.21567 | 77 |
| GO_REGULATION_OF_SUPEROXIDE_METABOLIC_PROCESS | 0.00346 | 0.03548 | 0.43171 | 0.77826 | 1.39046 | 19 |
| GO_REGULATION_OF_SUPRAMOLECULAR_FIBER_ORGANIZATIO | 0.00509 | 0.04656 | 0.40702 | 0.57599 | 1.12367 | 223 |
| GO_REGULATION_OF_SYMBIOTIC_PROCESS | 0.00009 | 0.00211 | 0.53843 | 0.62950 | 1.21763 | 151 |
| GO_REGULATION_OF_TELOMERASE_ACTIVITY | 0.00087 | 0.01263 | 0.47727 | 0.72147 | 1.33426 | 39 |
| GO_REGULATION_OF_TELOMERE_MAINTENANCE | 0.00547 | 0.04923 | 0.40702 | 0.64959 | 1.22679 | 68 |
| GO_REGULATION_OF_TELOMERE_MAINTENANCE_VIA_TELOME | 0.00020 | 0.00387 | 0.51885 | 0.72043 | 1.34856 | 51 |
| GO_REGULATION_OF_TRANSCRIPTION_FROM_RNA_POLYMERA | 0.00000 | 0.00000 | 0.70498 | 0.76931 | 1.45239 | 64 |
| GO_REGULATION_OF_TRANSLATIONAL_INITIATION | 0.00023 | 0.00437 | 0.51885 | 0.68479 | 1.29399 | 69 |
| GO_REGULATION_OF_VIRAL_TRANSCRIPTION | 0.00361 | 0.03657 | 0.43171 | 0.67497 | 1.26259 | 50 |
| GO_REGULATION_OF_WNT_SIGNALING_PATHWAY | 0.00015 | 0.00312 | 0.51885 | 0.60129 | 1.17419 | 238 |
| GO_RESPIRASOME | 0.00000 | 0.00000 | 0.81404 | 0.78326 | 1.48524 | 75 |
| GO_RESPIRATORY_CHAIN_COMPLEX | 0.00000 | 0.00000 | 0.77494 | 0.79183 | 1.49492 | 64 |
| GO_RESPIRATORY_ELECTRON_TRANSPORT_CHAIN | 0.00000 | 0.00000 | | 0.78275 | 1.49591 | 89 |
| GO_RESPONSE_TO_ALCOHOL | 0.00015 | 0.00310 | 0.51885 | 0.63794 | 1.22915 | 126 |
| GO_RESPONSE_TO_ATP | 0.00087 | 0.01270 | 0.47727 | 0.81086 | 1.43528 | 17 |
| GO_RESPONSE_TO_CALCIUM_ION | 0.00124 | 0.01701 | 0.45506 | 0.65196 | 1.24071 | 80 |
| GO_RESPONSE_TO_CORTICOSTEROID | 0.00010 | 0.00220 | 0.53843 | 0.66894 | 1.27841 | 89 |
| GO_RESPONSE_TO_DRUG | 0.00000 | 0.00007 | 0.62726 | 0.63080 | 1.23070 | 220 |
| GO_RESPONSE_TO_ETHANOL | 0.00168 | 0.02098 | 0.45506 | 0.66543 | 1.25679 | 67 |
| GO_RESPONSE_TO_FIBROBLAST_GROWTH_FACTOR | 0.00357 | 0.03638 | 0.43171 | 0.63469 | 1.20976 | 84 |
| GO_RESPONSE_TO_GROWTH_FACTOR | 0.00272 | 0.02986 | 0.43171 | 0.56235 | 1.10805 | 387 |
| GO_RESPONSE_TO_HEAT | 0.00000 | 0.00013 | 0.62726 | 0.66982 | 1.29069 | 119 |
| GO_RESPONSE_TO_HYDROGEN_PEROXIDE | 0.00532 | 0.04803 | 0.40702 | 0.62255 | 1.19054 | 90 |
| GO_RESPONSE_TO_INORGANIC_SUBSTANCE | 0.00016 | 0.00328 | 0.51885 | 0.58928 | 1.15810 | 321 |
| GO_RESPONSE_TO_INTERLEUKIN_1 | 0.00000 | 0.00000 | 0.78819 | 0.71590 | 1.37950 | 124 |
| GO_RESPONSE_TO_INTERLEUKIN_12 | 0.00036 | 0.00622 | 0.49849 | 0.74158 | 1.36769 | 36 |
| GO_RESPONSE_TO_INTERLEUKIN_4 | 0.00224 | 0.02581 | 0.43171 | 0.77088 | 1.38921 | 22 |
| GO_RESPONSE_TO_LEUKEMIA_INHIBITORY_FACTOR | 0.00076 | 0.01129 | 0.47727 | 0.66148 | 1.25288 | 72 |
| GO_RESPONSE_TO_LIPID | 0.00043 | 0.00695 | 0.49849 | 0.56669 | 1.11902 | 470 |
| GO_RESPONSE_TO_METAL_ION | 0.00143 | 0.01863 | 0.45506 | 0.59895 | 1.16592 | 199 |
| GO_RESPONSE_TO_NUTRIENT | 0.00294 | 0.03185 | 0.43171 | 0.62596 | 1.19987 | 98 |
| GO_RESPONSE_TO_ORGANOPHOSPHORUS | 0.00294 | 0.03185 | 0.43171 | 0.64154 | 1.21511 | 72 |
| GO_RESPONSE_TO_OXIDATIVE_STRESS | 0.00495 | 0.04573 | 0.40702 | 0.57010 | 1.11758 | 294 |
| GO_RESPONSE_TO_OXYGEN_LEVELS | 0.00000 | 0.00000 | | 0.67661 | 1.32155 | 244 |
| GO_RESPONSE_TO_PURINE_CONTAINING_COMPOUND | 0.00509 | 0.04656 | 0.40702 | 0.63176 | 1.20394 | 82 |
| GO_RESPONSE_TO_STEROID_HORMONE | 0.00009 | 0.00200 | 0.53843 | 0.61312 | 1.19360 | 201 |
| GO_RESPONSE_TO_TEMPERATURE_STIMULUS | 0.00002 | 0.00047 | 0.57561 | 0.64468 | 1.24782 | 153 |
| GO_RESPONSE_TO_TOXIC_SUBSTANCE | 0.00011 | 0.00235 | 0.53843 | 0.63892 | 1.23203 | 134 |
| GO_RESPONSE_TO_TUMOR_NECROSIS_FACTOR | 0.00000 | 0.00000 | 0.70498 | 0.66331 | 1.28815 | 175 |
| GO_RESPONSE_TO_VIRUS | 0.00261 | 0.02910 | 0.43171 | 0.59428 | 1.15408 | 175 |
| GO_RETINA_HOMEOSTASIS | 0.00034 | 0.00606 | 0.49849 | 0.75202 | 1.38362 | 34 |
| GO_RIBONUCLEOPROTEIN_COMPLEX_BINDING | 0.00000 | 0.00001 | 0.69013 | 0.69839 | 1.34459 | 114 |
| GO_RIBONUCLEOPROTEIN_COMPLEX_BIOGENESIS | 0.00000 | 0.00000 | | 0.68714 | 1.35448 | 393 |
| GO_RIBONUCLEOPROTEIN_COMPLEX_SUBUNIT_ORGANIZATION | 0.00000 | 0.00000 | 0.81404 | 0.69112 | 1.34183 | 173 |
| GO_RIBONUCLEOPROTEIN_GRANULE | 0.00000 | 0.00000 | 0.71951 | 0.66271 | 1.28875 | 186 |
| GO_RIBONUCLEOSIDE_TRIPHOSPHATE_BIOSYNTHETIC_PROCESS | 0.00009 | 0.00210 | 0.53843 | 0.77194 | 1.42026 | 34 |
| GO_RIBONUCLEOSIDE_TRIPHOSPHATE_METABOLIC_PROCESS | 0.00013 | 0.00280 | 0.51885 | 0.73558 | 1.37065 | 46 |
| GO_RIBOSE_PHOSPHATE_BIOSYNTHETIC_PROCESS | 0.00002 | 0.00056 | 0.57561 | 0.65814 | 1.26821 | 124 |
| GO_RIBOSE_PHOSPHATE_METABOLIC_PROCESS | 0.00000 | 0.00000 | 0.74774 | 0.63837 | 1.25001 | 285 |
| GO_RIBOSOMAL_LARGE_SUBUNIT_BIOGENESIS | 0.00001 | 0.00031 | 0.59333 | 0.72435 | 1.36807 | 67 |
| GO_RIBOSOMAL_SMALL_SUBUNIT_ASSEMBLY | 0.00405 | 0.04011 | 0.40702 | 0.78074 | 1.38865 | 18 |
| GO_RIBOSOMAL_SMALL_SUBUNIT_BINDING | 0.00532 | 0.04803 | 0.40702 | 0.77341 | 1.36899 | 17 |
| GO_RIBOSOMAL_SMALL_SUBUNIT_BIOGENESIS | 0.00005 | 0.00138 | 0.55733 | 0.70510 | 1.33163 | 68 |
| GO_RIBOSOMAL_SUBUNIT | 0.00000 | 0.00000 | | 0.76292 | 1.48089 | 172 |
| GO_RIBOSOME | 0.00000 | 0.00000 | | 0.75411 | 1.46984 | 208 |
| GO_RIBOSOME_ASSEMBLY | 0.00001 | 0.00026 | 0.59333 | 0.74485 | 1.40144 | 56 |
| GO_RIBOSOME_BIOGENESIS | 0.00000 | 0.00000 | | 0.68899 | 1.34833 | 277 |
| GO_RNA_3_END_PROCESSING | 0.00002 | 0.00046 | 0.57561 | 0.65576 | 1.26361 | 128 |
| GO_RNA_CATABOLIC_PROCESS | 0.00000 | 0.00000 | | 0.70701 | 1.39073 | 341 |
| GO_RNA_DEPENDENT_DNA_BIOSYNTHETIC_PROCESS | 0.00003 | 0.00081 | 0.57561 | 0.71669 | 1.34965 | 59 |
| GO_RNA_EXPORT_FROM_NUCLEUS | 0.00000 | 0.00016 | 0.61053 | 0.66306 | 1.27817 | 130 |
| GO_RNA_LOCALIZATION | 0.00000 | 0.00000 | 0.76146 | 0.66638 | 1.29886 | 208 |
| GO_RNA_SPLICING | 0.00000 | 0.00000 | | 0.63512 | 1.25129 | 384 |
| GO_RNA_SPLICING_VIA_TRANSESTERIFICATION_REACTIONS | 0.00000 | 0.00000 | 0.80122 | 0.64478 | 1.26586 | 311 |
| GO_ROUGH_ENDOPLASMIC_RETICULUM | 0.00014 | 0.00293 | 0.51885 | 0.70363 | 1.32507 | 59 |
| GO_RRNA_BINDING | 0.00000 | 0.00001 | 0.69013 | 0.77638 | 1.45941 | 54 |
| GO_RRNA_METABOLIC_PROCESS | 0.00000 | 0.00000 | | 0.68430 | 1.33386 | 209 |

| | | | | | | |
|---|---|---|---|---|---|---|
| GO_SCF_DEPENDENT_PROTEASOMAL_UBIQUITIN_DEPENDENT_ | 0.00000 | 0.00008 | 0.62726 | 0.71957 | 1.36498 | 76 |
| GO_SECRETORY_GRANULE | 0.00000 | 0.00000 | | 0.62978 | 1.24363 | 481 |
| GO_SECRETORY_GRANULE_MEMBRANE | 0.00032 | 0.00575 | 0.49849 | 0.61375 | 1.19134 | 172 |
| GO_SERINE_TYPE_ENDOPEPTIDASE_INHIBITOR_ACTIVITY | 0.00038 | 0.00640 | 0.49849 | 0.77388 | 1.41447 | 28 |
| GO_SEXUAL_REPRODUCTION | 0.00472 | 0.04480 | 0.40702 | 0.55748 | 1.10026 | 439 |
| GO_SIDE_OF_MEMBRANE | 0.00495 | 0.04573 | 0.40702 | 0.57741 | 1.12751 | 235 |
| GO_SIGNAL_TRANSDUCTION_BY_PROTEIN_PHOSPHORYLATION | 0.00224 | 0.02581 | 0.43171 | 0.55690 | 1.09975 | 500 |
| GO_SINGLE_FERTILIZATION | 0.00000 | 0.00002 | 0.67496 | 0.76307 | 1.43842 | 61 |
| GO_SINGLE_STRANDED_RNA_BINDING | 0.00001 | 0.00042 | 0.59333 | 0.71960 | 1.35864 | 66 |
| GO_SISTER_CHROMATID_SEGREGATION | 0.00042 | 0.00689 | 0.49849 | 0.60926 | 1.18209 | 170 |
| GO_SKIN_DEVELOPMENT | 0.00076 | 0.01140 | 0.47727 | 0.61452 | 1.18945 | 153 |
| GO_SMALL_MOLECULE_BIOSYNTHETIC_PROCESS | 0.00001 | 0.00021 | 0.61053 | 0.58728 | 1.15957 | 459 |
| GO_SMALL_NUCLEAR_RIBONUCLEOPROTEIN_COMPLEX | 0.00042 | 0.00689 | 0.49849 | 0.69542 | 1.30936 | 57 |
| GO_SMALL_RIBOSOMAL_SUBUNIT | 0.00000 | 0.00001 | 0.69013 | 0.75767 | 1.43100 | 67 |
| GO_SM_LIKE_PROTEIN_FAMILY_COMPLEX | 0.00145 | 0.01879 | 0.45506 | 0.66792 | 1.26149 | 67 |
| GO_SNORNA_BINDING | 0.00038 | 0.00646 | 0.49849 | 0.76224 | 1.39614 | 29 |
| GO_SPERM_EGG_RECOGNITION | 0.00000 | 0.00002 | 0.65944 | 0.87469 | 1.59087 | 26 |
| GO_SPINDLE_MIDZONE | 0.00361 | 0.03657 | 0.43171 | 0.73316 | 1.33859 | 27 |
| GO_SPINDLE_ORGANIZATION | 0.00272 | 0.02986 | 0.43171 | 0.60076 | 1.16362 | 159 |
| GO_SPLICEOSOMAL_COMPLEX | 0.00000 | 0.00008 | 0.62726 | 0.65256 | 1.26657 | 171 |
| GO_SPLICEOSOMAL_TRI_SNRNP_COMPLEX | 0.00168 | 0.02098 | 0.45506 | 0.73474 | 1.34835 | 30 |
| GO_STEM_CELL_DIFFERENTIATION | 0.00000 | 0.00000 | 0.82666 | 0.70006 | 1.35554 | 158 |
| GO_STEROID_BIOSYNTHETIC_PROCESS | 0.00465 | 0.04430 | 0.40702 | 0.61242 | 1.17992 | 125 |
| GO_STEROID_METABOLIC_PROCESS | 0.00313 | 0.03310 | 0.43171 | 0.59525 | 1.15748 | 185 |
| GO_STEROL_BIOSYNTHETIC_PROCESS | 0.00005 | 0.00120 | 0.55733 | 0.71876 | 1.35584 | 62 |
| GO_STEROL_METABOLIC_PROCESS | 0.00003 | 0.00073 | 0.57561 | 0.66485 | 1.27854 | 109 |
| GO_STRESS_GRANULE_ASSEMBLY | 0.00465 | 0.04430 | 0.40702 | 0.77288 | 1.38921 | 21 |
| GO_STRUCTURAL_CONSTITUENT_OF_RIBOSOME | 0.00000 | 0.00000 | | 0.76380 | 1.47839 | 153 |
| GO_STRUCTURAL_MOLECULE_ACTIVITY | 0.00000 | 0.00000 | | 0.71760 | 1.41436 | 391 |
| GO_SUPEROXIDE_METABOLIC_PROCESS | 0.00272 | 0.02986 | 0.43171 | 0.72028 | 1.32659 | 35 |
| GO_TELOMERASE_RNA_LOCALIZATION | 0.00054 | 0.00857 | 0.47727 | 0.81801 | 1.46148 | 19 |
| GO_TELOMERE_MAINTENANCE_VIA_TELOMERE_LENGTHENING | 0.00005 | 0.00138 | 0.55733 | 0.70715 | 1.33512 | 66 |
| GO_TELOMERE_ORGANIZATION | 0.00159 | 0.02016 | 0.45506 | 0.61666 | 1.18834 | 127 |
| GO_TERTIARY_GRANULE | 0.00008 | 0.00186 | 0.53843 | 0.66372 | 1.27005 | 91 |
| GO_TERTIARY_GRANULE_MEMBRANE | 0.00102 | 0.01453 | 0.45506 | 0.71940 | 1.33244 | 40 |
| GO_THREONINE_TYPE_ENDOPEPTIDASE_ACTIVITY | 0.00000 | 0.00000 | 0.71951 | 0.93456 | 1.65424 | 17 |
| GO_THREONINE_TYPE_PEPTIDASE_ACTIVITY | 0.00000 | 0.00004 | 0.65944 | 0.89807 | 1.61424 | 21 |
| GO_TISSUE_MORPHOGENESIS | 0.00000 | 0.00006 | 0.64355 | 0.59467 | 1.17150 | 380 |
| GO_TOXIN_TRANSPORT | 0.00064 | 0.00972 | 0.47727 | 0.77713 | 1.41343 | 26 |
| GO_TRANSCRIPTION_COUPLED_NUCLEOTIDE_EXCISION_REPAIR | 0.00231 | 0.02642 | 0.43171 | 0.65429 | 1.23840 | 71 |
| GO_TRANSFERRIN_TRANSPORT | 0.00141 | 0.01850 | 0.45506 | 0.75134 | 1.37179 | 27 |
| GO_TRANSLATIONAL_ELONGATION | 0.00000 | 0.00000 | | 0.75263 | 1.45028 | 124 |
| GO_TRANSLATIONAL_INITIATION | 0.00000 | 0.00000 | | 0.75139 | 1.45839 | 171 |
| GO_TRANSLATIONAL_TERMINATION | 0.00000 | 0.00000 | 0.81404 | 0.75110 | 1.44066 | 101 |
| GO_TRANSLATION_ELONGATION_FACTOR_ACTIVITY | 0.00390 | 0.03895 | 0.43171 | 0.80505 | 1.41260 | 15 |
| GO_TRANSLATION_FACTOR_ACTIVITY_RNA_BINDING | 0.00000 | 0.00000 | 0.74774 | 0.77244 | 1.46085 | 70 |
| GO_TRANSLATION_INITIATION_FACTOR_ACTIVITY | 0.00000 | 0.00011 | 0.62726 | 0.77773 | 1.44952 | 45 |
| GO_TRANSLATION_PREINITIATION_COMPLEX | 0.00342 | 0.03521 | 0.43171 | 0.79777 | 1.40694 | 16 |
| GO_TRANSLATION_REGULATOR_ACTIVITY | 0.00000 | 0.00000 | 0.83909 | 0.74000 | 1.42403 | 111 |
| GO_TRANSLATION_REGULATOR_ACTIVITY_NUCLEIC_ACID_BIND | 0.00000 | 0.00000 | 0.83909 | 0.76441 | 1.45959 | 87 |
| GO_TRANSMEMBRANE_RECEPTOR_PROTEIN_TYROSINE_KINASE | 0.00168 | 0.02098 | 0.45506 | 0.56326 | 1.11076 | 411 |
| GO_TRICARBOXYLIC_ACID_CYCLE | 0.00159 | 0.02016 | 0.45506 | 0.73558 | 1.35097 | 33 |
| GO_TRNA_TRANSPORT | 0.00342 | 0.03521 | 0.43171 | 0.71247 | 1.31220 | 35 |
| GO_TUMOR_NECROSIS_FACTOR_MEDIATED_SIGNALING_PATHW | 0.00000 | 0.00000 | 0.80122 | 0.73921 | 1.42138 | 108 |
| GO_T_CELL_RECEPTOR_SIGNALING_PATHWAY | 0.00000 | 0.00001 | 0.70498 | 0.69185 | 1.33314 | 119 |
| GO_U12_TYPE_SPLICEOSOMAL_COMPLEX | 0.00487 | 0.04559 | 0.40702 | 0.73423 | 1.33541 | 26 |
| GO_U2_SNRNP | 0.00141 | 0.01850 | 0.45506 | 0.80259 | 1.42065 | 17 |
| GO_U2_TYPE_CATALYTIC_STEP_2_SPLICEOSOME | 0.00268 | 0.02961 | 0.43171 | 0.72225 | 1.32288 | 29 |
| GO_U2_TYPE_SPLICEOSOMAL_COMPLEX | 0.00033 | 0.00595 | 0.49849 | 0.66590 | 1.26925 | 84 |
| GO_UBIQUITIN_LIKE_PROTEIN_LIGASE_BINDING | 0.00000 | 0.00007 | 0.62726 | 0.62616 | 1.22416 | 251 |
| GO_UNFOLDED_PROTEIN_BINDING | 0.00000 | 0.00001 | 0.69013 | 0.72112 | 1.37766 | 88 |
| GO_VACUOLAR_LUMEN | 0.00016 | 0.00319 | 0.51885 | 0.65416 | 1.25772 | 107 |
| GO_VESICLE_BUDDING_FROM_MEMBRANE | 0.00480 | 0.04537 | 0.40702 | 0.62411 | 1.19353 | 90 |
| GO_VESICLE_LUMEN | 0.00000 | 0.00000 | 0.77494 | 0.67028 | 1.30631 | 207 |
| GO_VESICLE_MEMBRANE | 0.00001 | 0.00026 | 0.59333 | 0.58430 | 1.15387 | 476 |
| GO_VIRAL_GENE_EXPRESSION | 0.00000 | 0.00000 | | 0.70963 | 1.37745 | 172 |
| GO_VIRAL_LIFE_CYCLE | 0.00000 | 0.00001 | 0.69013 | 0.63985 | 1.24950 | 238 |
| GO_VIRION_ASSEMBLY | 0.00457 | 0.04386 | 0.40702 | 0.69689 | 1.28528 | 36 |
| HP_ABNORMALITY_OF_ACID_BASE_HOMEOSTASIS | 0.00000 | 0.00010 | 0.62726 | 0.62283 | 1.21827 | 261 |
| HP_ABNORMALITY_OF_BLOOD_CIRCULATION | 0.00502 | 0.04608 | 0.40702 | 0.59228 | 1.14883 | 167 |

| | | | | | | |
|---|---|---|---|---|---|---|
| HP_ABNORMALITY_OF_CARDIOVASCULAR_SYSTEM_ELECTROPH | 0.00011 | 0.00237 | 0.53843 | 0.60202 | 1.17758 | 261 |
| HP_ABNORMALITY_OF_SKIN_ADNEXA_PHYSIOLOGY | 0.00242 | 0.02748 | 0.43171 | 0.61294 | 1.18094 | 118 |
| HP_ABNORMALITY_OF_THE_ANKLES | 0.00313 | 0.03310 | 0.43171 | 0.65369 | 1.23418 | 63 |
| HP_ABNORMALITY_OF_THE_CEREBRAL_VASCULATURE | 0.00357 | 0.03638 | 0.43171 | 0.59764 | 1.15722 | 158 |
| HP_ABNORMALITY_OF_THE_CEREBROSPINAL_FLUID | 0.00030 | 0.00549 | 0.49849 | 0.58221 | 1.14439 | 325 |
| HP_ABNORMALITY_OF_THE_MITOCHONDRION | 0.00126 | 0.01715 | 0.45506 | 0.62149 | 1.19764 | 120 |
| HP_ABNORMALITY_OF_THE_OPTIC_DISC | 0.00198 | 0.02370 | 0.43171 | 0.56624 | 1.11727 | 432 |
| HP_ABNORMALITY_OF_THE_OPTIC_NERVE | 0.00036 | 0.00622 | 0.49849 | 0.56723 | 1.12034 | 486 |
| HP_ABNORMAL_ABDOMEN_MORPHOLOGY | 0.00089 | 0.01291 | 0.47727 | 0.56424 | 1.11364 | 441 |
| HP_ABNORMAL_ACTIVITY_OF_MITOCHONDRIAL_RESPIRATORY_ | 0.00014 | 0.00300 | 0.51885 | 0.70303 | 1.32524 | 61 |
| HP_ABNORMAL_BASAL_GANGLIA_MRI_SIGNAL_INTENSITY | 0.00179 | 0.02183 | 0.45506 | 0.71597 | 1.32046 | 36 |
| HP_ABNORMAL_BRAINSTEM_MRI_SIGNAL_INTENSITY | 0.00398 | 0.03962 | 0.40702 | 0.71528 | 1.31370 | 33 |
| HP_ABNORMAL_CELLULAR_PHENOTYPE | 0.00209 | 0.02470 | 0.43171 | 0.56937 | 1.11992 | 334 |
| HP_ABNORMAL_CIRCULATING_CARBOHYDRATE_CONCENTRATIC | 0.00194 | 0.02331 | 0.45506 | 0.69518 | 1.29539 | 46 |
| HP_ABNORMAL_CNS_MYELINATION | 0.00014 | 0.00300 | 0.51885 | 0.62318 | 1.20954 | 171 |
| HP_ABNORMAL_CSF_METABOLITE_LEVEL | 0.00001 | 0.00027 | 0.59333 | 0.70531 | 1.34409 | 82 |
| HP_ABNORMAL_ELECTROPHYSIOLOGY_OF_SINOATRIAL_NODE_ | 0.00029 | 0.00536 | 0.49849 | 0.73172 | 1.35725 | 41 |
| HP_ABNORMAL_ENZYME_COENZYME_ACTIVITY | 0.00561 | 0.04971 | 0.40702 | 0.60490 | 1.16567 | 123 |
| HP_ABNORMAL_ERYTHROCYTE_MORPHOLOGY | 0.00001 | 0.00023 | 0.61053 | 0.58835 | 1.15904 | 380 |
| HP_ABNORMAL_HEART_VALVE_PHYSIOLOGY | 0.00405 | 0.04011 | 0.40702 | 0.59004 | 1.14474 | 169 |
| HP_ABNORMAL_MITOCHONDRIA_IN_MUSCLE_TISSUE | 0.00075 | 0.01118 | 0.47727 | 0.74633 | 1.36700 | 29 |
| HP_ABNORMAL_MYELINATION | 0.00007 | 0.00169 | 0.53843 | 0.59258 | 1.16479 | 324 |
| HP_ABNORMAL_MYOCARDIUM_MORPHOLOGY | 0.00008 | 0.00192 | 0.53843 | 0.58647 | 1.15365 | 340 |
| HP_ABNORMAL_RENAL_PHYSIOLOGY | 0.00450 | 0.04342 | 0.40702 | 0.55484 | 1.09317 | 385 |
| HP_ABNORMAL_TRICUSPID_VALVE_PHYSIOLOGY | 0.00231 | 0.02642 | 0.43171 | 0.72968 | 1.34029 | 31 |
| HP_ABNORMAL_VITREOUS_HUMOR_MORPHOLOGY | 0.00181 | 0.02201 | 0.45506 | 0.79632 | 1.40955 | 17 |
| HP_ACUTE_LEUKEMIA | 0.00013 | 0.00276 | 0.51885 | 0.68145 | 1.29219 | 75 |
| HP_ANEMIA_OF_INADEQUATE_PRODUCTION | 0.00561 | 0.04971 | 0.40702 | 0.61298 | 1.17855 | 107 |
| HP_APHASIA | 0.00213 | 0.02490 | 0.43171 | 0.72858 | 1.33704 | 30 |
| HP_BLINDNESS | 0.00018 | 0.00355 | 0.51885 | 0.61672 | 1.19768 | 175 |
| HP_BONE_MARROW_HYPERCELLULARITY | 0.00045 | 0.00725 | 0.49849 | 0.82169 | 1.46806 | 19 |
| HP_CHOREOATHETOSIS | 0.00561 | 0.04971 | 0.40702 | 0.64956 | 1.22631 | 64 |
| HP_CHRONIC_INFECTION | 0.00121 | 0.01658 | 0.45506 | 0.81916 | 1.44468 | 16 |
| HP_CONGESTIVE_HEART_FAILURE | 0.00495 | 0.04573 | 0.40702 | 0.60734 | 1.16929 | 114 |
| HP_CRANIAL_NERVE_COMPRESSION | 0.00331 | 0.03441 | 0.43171 | 0.75830 | 1.36989 | 23 |
| HP_DECREASED_ACTIVITY_OF_MITOCHONDRIAL_COMPLEX_I | 0.00283 | 0.03081 | 0.43171 | 0.71940 | 1.32496 | 35 |
| HP_DECREASED_ACTIVITY_OF_THE_PYRUVATE_DEHYDROGENAS | 0.00062 | 0.00949 | 0.47727 | 0.75390 | 1.38478 | 31 |
| HP_DEPRESSED_NASAL_RIDGE | 0.00041 | 0.00681 | 0.49849 | 0.66218 | 1.26189 | 82 |
| HP_DYSPNEA | 0.00250 | 0.02807 | 0.43171 | 0.58107 | 1.13348 | 224 |
| HP_DYSTONIA | 0.00069 | 0.01043 | 0.47727 | 0.58101 | 1.13736 | 279 |
| HP_EMOTIONAL_LABILITY | 0.00016 | 0.00321 | 0.51885 | 0.66683 | 1.27522 | 90 |
| HP_EPISODIC_ATAXIA | 0.00487 | 0.04559 | 0.40702 | 0.79934 | 1.40259 | 15 |
| HP_FASCICULATIONS | 0.00130 | 0.01761 | 0.45506 | 0.68128 | 1.28064 | 54 |
| HP_FATIGUE | 0.00042 | 0.00689 | 0.49849 | 0.59902 | 1.16764 | 212 |
| HP_FETAL_DISTRESS | 0.00179 | 0.02183 | 0.45506 | 0.72928 | 1.33973 | 32 |
| HP_FLUSHING | 0.00561 | 0.04971 | 0.40702 | 0.74782 | 1.33686 | 20 |
| HP_FOCAL_T2_HYPERINTENSE_BRAINSTEM_LESION | 0.00159 | 0.02016 | 0.45506 | 0.74970 | 1.36879 | 27 |
| HP_HEADACHE | 0.00016 | 0.00319 | 0.51885 | 0.62537 | 1.21179 | 163 |
| HP_HEPATOMEGALY | 0.00108 | 0.01507 | 0.45506 | 0.57439 | 1.12990 | 335 |
| HP_HYPERTROPHIC_CARDIOMYOPATHY | 0.00002 | 0.00050 | 0.57561 | 0.62629 | 1.21908 | 194 |
| HP_IMPAIRED_SOCIAL_INTERACTIONS | 0.00055 | 0.00863 | 0.47727 | 0.66555 | 1.26058 | 72 |
| HP_INCREASED_CSF_LACTATE | 0.00001 | 0.00034 | 0.59333 | 0.71937 | 1.35857 | 68 |
| HP_INCREASED_SERUM_LACTATE | 0.00001 | 0.00028 | 0.59333 | 0.65837 | 1.26932 | 133 |
| HP_INCREASED_SERUM_PYRUVATE | 0.00110 | 0.01526 | 0.45506 | 0.73143 | 1.35198 | 37 |
| HP_INFANTILE_ONSET | 0.00487 | 0.04559 | 0.40702 | 0.56559 | 1.10939 | 300 |
| HP_INTRACRANIAL_CYSTIC_LESION | 0.00055 | 0.00863 | 0.47727 | 0.62760 | 1.20885 | 116 |
| HP_INTRAUTERINE_GROWTH_RETARDATION | 0.00305 | 0.03271 | 0.43171 | 0.56594 | 1.11393 | 350 |
| HP_LACTIC_ACIDOSIS | 0.00018 | 0.00355 | 0.51885 | 0.63312 | 1.22071 | 131 |
| HP_LETHARGY | 0.00084 | 0.01228 | 0.47727 | 0.63361 | 1.21845 | 109 |
| HP_LEUKODYSTROPHY | 0.00029 | 0.00529 | 0.49849 | 0.66598 | 1.26915 | 82 |
| HP_LEUKOENCEPHALOPATHY | 0.00008 | 0.00190 | 0.53843 | 0.64132 | 1.23667 | 134 |
| HP_LIPID_ACCUMULATION_IN_HEPATOCYTES | 0.00220 | 0.02548 | 0.43171 | 0.62794 | 1.20159 | 91 |
| HP_LOWER_LIMB_SPASTICITY | 0.00154 | 0.01974 | 0.45506 | 0.60929 | 1.17648 | 139 |
| HP_MACROCYTIC_ANEMIA | 0.00030 | 0.00542 | 0.49849 | 0.73108 | 1.35606 | 41 |
| HP_METABOLIC_ACIDOSIS | 0.00034 | 0.00599 | 0.49849 | 0.64831 | 1.24464 | 103 |
| HP_MICRONODULAR_CIRRHOSIS | 0.00324 | 0.03393 | 0.43171 | 0.78723 | 1.40018 | 18 |
| HP_MIGRAINE | 0.00028 | 0.00515 | 0.49849 | 0.67309 | 1.27635 | 75 |
| HP_MUSCLE_ABNORMALITY_RELATED_TO_MITOCHONDRIAL_DY | 0.00209 | 0.02470 | 0.43171 | 0.68258 | 1.27573 | 49 |
| HP_MUTISM | 0.00283 | 0.03081 | 0.43171 | 0.71937 | 1.32490 | 35 |
| HP_NAUSEA_AND_VOMITING | 0.00022 | 0.00417 | 0.51885 | 0.59383 | 1.16158 | 264 |

| | | | | | | |
|---|---|---|---|---|---|---|
| HP_NEOPLASM_OF_THE_ENDOCRINE_SYSTEM | 0.00495 | 0.04573 | 0.40702 | 0.63166 | 1.20308 | 81 |
| HP_OPTIC_NEUROPATHY | 0.00135 | 0.01808 | 0.45506 | 0.73837 | 1.35610 | 33 |
| HP_PACE_OF_PROGRESSION | 0.00487 | 0.04559 | 0.40702 | 0.56814 | 1.11416 | 298 |
| HP_PALLOR | 0.00036 | 0.00622 | 0.49849 | 0.64861 | 1.24463 | 102 |
| HP_PARAPLEGIA | 0.00121 | 0.01658 | 0.45506 | 0.63584 | 1.22133 | 104 |
| HP_PARAPLEGIA_PARAPARESIS | 0.00036 | 0.00621 | 0.49849 | 0.62047 | 1.20040 | 150 |
| HP_PAROXYSMAL_INVOLUNTARY_EYE_MOVEMENTS | 0.00054 | 0.00849 | 0.47727 | 0.75650 | 1.38957 | 31 |
| HP_PEDIATRIC_ONSET | 0.00561 | 0.04971 | 0.40702 | 0.55915 | 1.10116 | 371 |
| HP_PIGMENTARY_RETINOPATHY | 0.00065 | 0.00981 | 0.47727 | 0.63260 | 1.21720 | 113 |
| HP_POOR_EYE_CONTACT | 0.00017 | 0.00341 | 0.51885 | 0.71279 | 1.33749 | 53 |
| HP_POOR_HEAD_CONTROL | 0.00000 | 0.00005 | 0.64355 | 0.67819 | 1.30665 | 118 |
| HP_PROGRESSIVE | 0.00194 | 0.02331 | 0.45506 | 0.59790 | 1.16273 | 186 |
| HP_PROGRESSIVE_CEREBELLAR_ATAXIA | 0.00364 | 0.03677 | 0.43171 | 0.64788 | 1.22325 | 65 |
| HP_PROGRESSIVE_SPASTIC_PARAPLEGIA | 0.00253 | 0.02838 | 0.43171 | 0.69655 | 1.29822 | 45 |
| HP_PROXIMAL_TUBULOPATHY | 0.00141 | 0.01850 | 0.45506 | 0.72579 | 1.33535 | 34 |
| HP_PTOSIS | 0.00472 | 0.04480 | 0.40702 | 0.55582 | 1.09637 | 418 |
| HP_REDUCED_CONSCIOUSNESS_CONFUSION | 0.00007 | 0.00164 | 0.53843 | 0.61227 | 1.19295 | 205 |
| HP_REDUCED_VISUAL_ACUITY | 0.00053 | 0.00837 | 0.47727 | 0.58618 | 1.14774 | 283 |
| HP_RENAL_TUBULAR_DYSFUNCTION | 0.00013 | 0.00284 | 0.51885 | 0.68108 | 1.29150 | 75 |
| HP_RESPIRATORY_DISTRESS | 0.00108 | 0.01507 | 0.45506 | 0.62693 | 1.20762 | 115 |
| HP_RESPIRATORY_INSUFFICIENCY | 0.00000 | 0.00010 | 0.62726 | 0.60536 | 1.18998 | 326 |
| HP_SINUS_TACHYCARDIA | 0.00213 | 0.02490 | 0.43171 | 0.77203 | 1.39128 | 22 |
| HP_SPARSE_EYELASHES | 0.00150 | 0.01943 | 0.45506 | 0.71355 | 1.32161 | 40 |
| HP_SPARSE_OR_ABSENT_EYELASHES | 0.00220 | 0.02548 | 0.43171 | 0.67331 | 1.26567 | 54 |
| HP_SPASTIC_PARAPLEGIA | 0.00098 | 0.01407 | 0.45506 | 0.64059 | 1.22712 | 96 |
| HP_THROMBOCYTOSIS | 0.00413 | 0.04065 | 0.40702 | 0.75210 | 1.35870 | 23 |
| HP_TONGUE_FASCICULATIONS | 0.00457 | 0.04386 | 0.40702 | 0.80056 | 1.40472 | 15 |
| HP_TRANSIENT_ISCHEMIC_ATTACK | 0.00480 | 0.04537 | 0.40702 | 0.74902 | 1.35809 | 25 |
| HP_VERTIGO | 0.00002 | 0.00063 | 0.57561 | 0.72114 | 1.35844 | 60 |
| HP_VESTIBULAR_DYSFUNCTION | 0.00028 | 0.00522 | 0.49849 | 0.67279 | 1.27577 | 75 |
| HP_VOMITING | 0.00072 | 0.01082 | 0.47727 | 0.61459 | 1.18792 | 147 |
| HP_WEAKNESS_DUE_TO_UPPER_MOTOR_NEURON_DYSFUNCTI | 0.00532 | 0.04803 | 0.40702 | 0.56829 | 1.11186 | 272 |
| HP_WEAKNESS_OF_MUSCLES_OF_RESPIRATION | 0.00163 | 0.02050 | 0.45506 | 0.66272 | 1.25123 | 63 |
| **Cluster 1** | | | | | | |
| pathway | pval | padj | log2err | ES | NES | size |
| GO_ACTIVATION_OF_IMMUNE_RESPONSE | 0.00000 | 0.00000 | 0.74774 | 0.65038 | 1.26999 | 235 |
| GO_ACTIVATION_OF_INNATE_IMMUNE_RESPONSE | 0.00000 | 0.00001 | 0.69013 | 0.71967 | 1.37861 | 96 |
| GO_AEROBIC_RESPIRATION | 0.00000 | 0.00000 | 0.73376 | 0.76280 | 1.44478 | 72 |
| GO_ALCOHOL_BIOSYNTHETIC_PROCESS | 0.00032 | 0.00572 | 0.49849 | 0.64679 | 1.24367 | 108 |
| GO_ALCOHOL_METABOLIC_PROCESS | 0.00364 | 0.03677 | 0.43171 | 0.58112 | 1.13424 | 228 |
| GO_AMEBOIDAL_TYPE_CELL_MIGRATION | 0.00457 | 0.04386 | 0.40702 | 0.57755 | 1.12854 | 247 |
| GO_AMIDE_BINDING | 0.00014 | 0.00297 | 0.51885 | 0.61533 | 1.19780 | 199 |
| GO_AMINE_METABOLIC_PROCESS | 0.00000 | 0.00000 | 0.76146 | 0.73732 | 1.41097 | 92 |
| GO_AMYLOID_BETA_CLEARANCE | 0.00561 | 0.04971 | 0.40702 | 0.74081 | 1.33976 | 24 |
| GO_ANAPHASE_PROMOTING_COMPLEX_DEPENDENT_CATABOL | 0.00000 | 0.00000 | 0.76146 | 0.77107 | 1.46112 | 73 |
| GO_ANATOMICAL_STRUCTURE_HOMEOSTASIS | 0.00020 | 0.00390 | 0.51885 | 0.59307 | 1.16021 | 271 |
| GO_ANIMAL_ORGAN_REGENERATION | 0.00012 | 0.00269 | 0.51885 | 0.71251 | 1.33510 | 52 |
| GO_ANTIGEN_PROCESSING_AND_PRESENTATION | 0.00000 | 0.00000 | 0.74774 | 0.68785 | 1.33064 | 149 |
| GO_ANTIGEN_PROCESSING_AND_PRESENTATION_OF_EXOGENC | 0.00000 | 0.00000 | 0.76146 | 0.79644 | 1.50237 | 62 |
| GO_ANTIGEN_PROCESSING_AND_PRESENTATION_OF_PEPTIDE_ | 0.00000 | 0.00000 | 0.74774 | 0.70107 | 1.35144 | 130 |
| GO_ANTIGEN_PROCESSING_AND_PRESENTATION_OF_PEPTIDE_ | 0.00000 | 0.00000 | 0.78819 | 0.77566 | 1.47012 | 74 |
| GO_ANTIGEN_RECEPTOR_MEDIATED_SIGNALING_PATHWAY | 0.00000 | 0.00009 | 0.62726 | 0.66362 | 1.28080 | 136 |
| GO_APICAL_PART_OF_CELL | 0.00135 | 0.01808 | 0.45506 | 0.58926 | 1.14966 | 220 |
| GO_APOPTOTIC_SIGNALING_PATHWAY | 0.00006 | 0.00152 | 0.53843 | 0.58638 | 1.15586 | 393 |
| GO_ATPASE_ACTIVITY | 0.00050 | 0.00802 | 0.47727 | 0.58216 | 1.14385 | 316 |
| GO_ATPASE_ACTIVITY_COUPLED_TO_TRANSMEMBRANE_MOVE | 0.00420 | 0.04132 | 0.40702 | 0.78005 | 1.38741 | 18 |
| GO_ATP_BIOSYNTHETIC_PROCESS | 0.00013 | 0.00276 | 0.51885 | 0.83897 | 1.50801 | 21 |
| GO_ATP_METABOLIC_PROCESS | 0.00000 | 0.00000 | | 0.72178 | 1.40814 | 219 |
| GO_ATP_SYNTHESIS_COUPLED_ELECTRON_TRANSPORT | 0.00000 | 0.00000 | | 0.80353 | 1.52425 | 76 |
| GO_AZUROPHIL_GRANULE | 0.00025 | 0.00466 | 0.49849 | 0.64781 | 1.24551 | 107 |
| GO_AZUROPHIL_GRANULE_LUMEN | 0.00405 | 0.04011 | 0.40702 | 0.65625 | 1.23793 | 62 |
| GO_BASAL_PART_OF_CELL | 0.00509 | 0.04656 | 0.40702 | 0.68515 | 1.27697 | 45 |
| GO_BASAL_PLASMA_MEMBRANE | 0.00346 | 0.03548 | 0.43171 | 0.71184 | 1.31104 | 35 |
| GO_BASEMENT_MEMBRANE | 0.00279 | 0.03056 | 0.43171 | 0.68935 | 1.28452 | 46 |
| GO_BASOLATERAL_PLASMA_MEMBRANE | 0.00011 | 0.00247 | 0.53843 | 0.63209 | 1.21993 | 136 |
| GO_BINDING_OF_SPERM_TO_ZONA_PELLUCIDA | 0.00000 | 0.00006 | 0.64355 | 0.90095 | 1.60966 | 19 |
| GO_BLOOD_MICROPARTICLE | 0.00000 | 0.00000 | 0.70498 | 0.81982 | 1.52763 | 46 |
| GO_BRUSH_BORDER | 0.00060 | 0.00923 | 0.47727 | 0.66897 | 1.26790 | 74 |
| GO_CADHERIN_BINDING | 0.00000 | 0.00000 | | 0.68759 | 1.34435 | 253 |
| GO_CALCIUM_DEPENDENT_PHOSPHOLIPID_BINDING | 0.00152 | 0.01958 | 0.45506 | 0.75021 | 1.37120 | 28 |

| | | | | | | |
|---|---|---|---|---|---|---|
| GO_CALCIUM_DEPENDENT_PROTEIN_BINDING | 0.00009 | 0.00211 | 0.53843 | 0.74254 | 1.38394 | 45 |
| GO_CANONICAL_WNT_SIGNALING_PATHWAY | 0.00004 | 0.00109 | 0.55733 | 0.61740 | 1.20458 | 222 |
| GO_CARBOHYDRATE_CATABOLIC_PROCESS | 0.00135 | 0.01808 | 0.45506 | 0.61179 | 1.18186 | 144 |
| GO_CATALYTIC_STEP_2_SPLICEOSOME | 0.00001 | 0.00022 | 0.61053 | 0.70290 | 1.33765 | 80 |
| GO_CATION_TRANSMEMBRANE_TRANSPORT | 0.00327 | 0.03409 | 0.43171 | 0.56060 | 1.10531 | 406 |
| GO_CATION_TRANSMEMBRANE_TRANSPORTER_ACTIVITY | 0.00097 | 0.01387 | 0.47727 | 0.58196 | 1.13846 | 265 |
| GO_CELLULAR_AMINO_ACID_METABOLIC_PROCESS | 0.00002 | 0.00060 | 0.57561 | 0.61438 | 1.19930 | 231 |
| GO_CELLULAR_COMPONENT_DISASSEMBLY | 0.00000 | 0.00000 | | 0.64070 | 1.26342 | 408 |
| GO_CELLULAR_ION_HOMEOSTASIS | 0.00027 | 0.00493 | 0.49849 | 0.58658 | 1.15340 | 328 |
| GO_CELLULAR_IRON_ION_HOMEOSTASIS | 0.00327 | 0.03409 | 0.43171 | 0.68360 | 1.27374 | 47 |
| GO_CELLULAR_KETONE_METABOLIC_PROCESS | 0.00000 | 0.00010 | 0.62726 | 0.65358 | 1.26726 | 164 |
| GO_CELLULAR_MONOVALENT_INORGANIC_CATION_HOMEOSTA | 0.00179 | 0.02183 | 0.45506 | 0.64747 | 1.22821 | 76 |
| GO_CELLULAR_NITROGEN_COMPOUND_CATABOLIC_PROCESS | 0.00000 | 0.00000 | | 0.67915 | 1.34096 | 456 |
| GO_CELLULAR_PROTEIN_COMPLEX_DISASSEMBLY | 0.00000 | 0.00000 | | 0.69566 | 1.35184 | 179 |
| GO_CELLULAR_RESPIRATION | 0.00000 | 0.00000 | | 0.74205 | 1.43370 | 145 |
| GO_CELLULAR_RESPONSE_TO_HEAT | 0.00004 | 0.00109 | 0.55733 | 0.67186 | 1.28698 | 95 |
| GO_CELLULAR_RESPONSE_TO_HORMONE_STIMULUS | 0.00309 | 0.03299 | 0.43171 | 0.56453 | 1.11191 | 374 |
| GO_CELLULAR_RESPONSE_TO_ORGANIC_CYCLIC_COMPOUND | 0.00502 | 0.04608 | 0.40702 | 0.56512 | 1.11134 | 329 |
| GO_CELLULAR_RESPONSE_TO_OXYGEN_LEVELS | 0.00000 | 0.00000 | | 0.71461 | 1.38224 | 151 |
| GO_CELL_ACTIVATION_INVOLVED_IN_IMMUNE_RESPONSE | 0.00000 | 0.00000 | | 0.63924 | 1.26108 | 422 |
| GO_CELL_ADHESION_MEDIATOR_ACTIVITY | 0.00157 | 0.02009 | 0.45506 | 0.71837 | 1.32488 | 36 |
| GO_CELL_ADHESION_MOLECULE_BINDING | 0.00000 | 0.00000 | | 0.67109 | 1.32112 | 353 |
| GO_CELL_BODY | 0.00004 | 0.00109 | 0.55733 | 0.59973 | 1.17541 | 293 |
| GO_CELL_CELL_RECOGNITION | 0.00000 | 0.00006 | 0.64355 | 0.82001 | 1.51026 | 35 |
| GO_CELL_CELL_SIGNALING_BY_WNT | 0.00004 | 0.00109 | 0.55733 | 0.59042 | 1.16141 | 339 |
| GO_CELL_CYCLE_G1_S_PHASE_TRANSITION | 0.00376 | 0.03783 | 0.43171 | 0.58112 | 1.13378 | 220 |
| GO_CELL_CYCLE_G2_M_PHASE_TRANSITION | 0.00024 | 0.00459 | 0.51885 | 0.59807 | 1.16746 | 231 |
| GO_CELL_CYCLE_PHASE_TRANSITION | 0.00000 | 0.00001 | 0.69013 | 0.59417 | 1.17348 | 499 |
| GO_CELL_DIVISION | 0.00003 | 0.00071 | 0.57561 | 0.58314 | 1.15094 | 449 |
| GO_CELL_KILLING | 0.00205 | 0.02440 | 0.43171 | 0.66727 | 1.25696 | 60 |
| GO_CELL_RECOGNITION | 0.00000 | 0.00000 | 0.73376 | 0.75953 | 1.43954 | 74 |
| GO_CELL_REDOX_HOMEOSTASIS | 0.00163 | 0.02050 | 0.45506 | 0.71772 | 1.32665 | 38 |
| GO_CELL_SUBSTRATE_JUNCTION | 0.00000 | 0.00000 | | 0.65742 | 1.29171 | 316 |
| GO_CELL_SURFACE | 0.00000 | 0.00000 | 0.73376 | 0.62572 | 1.23070 | 330 |
| GO_CELL_SURFACE_RECEPTOR_SIGNALING_PATHWAY_INVOLVEI | 0.00038 | 0.00640 | 0.49849 | 0.57762 | 1.13770 | 374 |
| GO_CHAPERONE_BINDING | 0.00495 | 0.04573 | 0.40702 | 0.62860 | 1.19935 | 86 |
| GO_CHAPERONE_COMPLEX | 0.00001 | 0.00028 | 0.59333 | 0.86337 | 1.55587 | 22 |
| GO_CHAPERONE_MEDIATED_PROTEIN_COMPLEX_ASSEMBLY | 0.00100 | 0.01430 | 0.45506 | 0.80787 | 1.44337 | 19 |
| GO_CHAPERONE_MEDIATED_PROTEIN_FOLDING | 0.00168 | 0.02098 | 0.45506 | 0.71110 | 1.32288 | 44 |
| GO_CHROMOSOMAL_REGION | 0.00018 | 0.00349 | 0.51885 | 0.59323 | 1.16180 | 286 |
| GO_CHROMOSOME_CENTROMERIC_REGION | 0.00172 | 0.02135 | 0.45506 | 0.60447 | 1.17248 | 168 |
| GO_CHROMOSOME_SEGREGATION | 0.00003 | 0.00087 | 0.55733 | 0.60463 | 1.18280 | 270 |
| GO_CLUSTER_OF_ACTIN_BASED_CELL_PROJECTIONS | 0.00074 | 0.01107 | 0.47727 | 0.64184 | 1.23044 | 99 |
| GO_COATED_VESICLE | 0.00213 | 0.02490 | 0.43171 | 0.59512 | 1.15846 | 199 |
| GO_COATED_VESICLE_MEMBRANE | 0.00060 | 0.00923 | 0.47727 | 0.63443 | 1.22207 | 115 |
| GO_COLLAGEN_BINDING | 0.00465 | 0.04430 | 0.40702 | 0.73471 | 1.33628 | 26 |
| GO_COLLAGEN_CONTAINING_EXTRACELLULAR_MATRIX | 0.00000 | 0.00006 | 0.64355 | 0.65358 | 1.26542 | 157 |
| GO_CONDENSED_CHROMOSOME | 0.00350 | 0.03580 | 0.43171 | 0.59394 | 1.15417 | 179 |
| GO_CONTRACTILE_FIBER | 0.00064 | 0.00972 | 0.47727 | 0.62217 | 1.19973 | 134 |
| GO_COPI_COATED_VESICLE | 0.00316 | 0.03344 | 0.43171 | 0.75688 | 1.37234 | 25 |
| GO_CORNIFICATION | 0.00027 | 0.00500 | 0.49849 | 0.77970 | 1.42512 | 28 |
| GO_CORTICAL_CYTOSKELETON | 0.00502 | 0.04608 | 0.40702 | 0.64388 | 1.21870 | 71 |
| GO_COTRANSLATIONAL_PROTEIN_TARGETING_TO_MEMBRANE | 0.00000 | 0.00000 | | 0.78223 | 1.49819 | 94 |
| GO_CYTOCHROME_COMPLEX | 0.00035 | 0.00610 | 0.49849 | 0.80796 | 1.45601 | 22 |
| GO_CYTOKINE_MEDIATED_SIGNALING_PATHWAY | 0.00000 | 0.00002 | 0.67496 | 0.60742 | 1.19735 | 392 |
| GO_CYTOPLASMIC_STRESS_GRANULE | 0.00014 | 0.00302 | 0.51885 | 0.69390 | 1.31009 | 63 |
| GO_CYTOPLASMIC_TRANSLATION | 0.00000 | 0.00001 | 0.70498 | 0.71773 | 1.37349 | 92 |
| GO_CYTOPLASMIC_TRANSLATIONAL_INITIATION | 0.00178 | 0.02182 | 0.45506 | 0.73517 | 1.35039 | 31 |
| GO_CYTOSOLIC_LARGE_RIBOSOMAL_SUBUNIT | 0.00000 | 0.00005 | 0.64355 | 0.77240 | 1.44935 | 53 |
| GO_CYTOSOLIC_RIBOSOME | 0.00000 | 0.00000 | | 0.77483 | 1.48437 | 97 |
| GO_CYTOSOLIC_SMALL_RIBOSOMAL_SUBUNIT | 0.00000 | 0.00010 | 0.62726 | 0.79384 | 1.47033 | 40 |
| GO_DETECTION_OF_STIMULUS | 0.00020 | 0.00393 | 0.51885 | 0.65208 | 1.25384 | 108 |
| GO_DETOXIFICATION | 0.00413 | 0.04065 | 0.40702 | 0.65399 | 1.23684 | 70 |
| GO_DEVELOPMENT_OF_PRIMARY_FEMALE_SEXUAL_CHARACTE | 0.00179 | 0.02183 | 0.45506 | 0.68410 | 1.28057 | 51 |
| GO_DICARBOXYLIC_ACID_METABOLIC_PROCESS | 0.00261 | 0.02910 | 0.43171 | 0.65541 | 1.23747 | 65 |
| GO_DISORDERED_DOMAIN_SPECIFIC_BINDING | 0.00051 | 0.00812 | 0.47727 | 0.76867 | 1.40343 | 27 |
| GO_DISULFIDE_OXIDOREDUCTASE_ACTIVITY | 0.00435 | 0.04238 | 0.40702 | 0.71445 | 1.31232 | 31 |
| GO_DNA_BIOSYNTHETIC_PROCESS | 0.00017 | 0.00334 | 0.51885 | 0.62540 | 1.20833 | 145 |
| GO_DNA_CONFORMATION_CHANGE | 0.00327 | 0.03409 | 0.43171 | 0.57979 | 1.13178 | 231 |
| GO_DNA_DAMAGE_RESPONSE_DETECTION_OF_DNA_DAMAGE | 0.00305 | 0.03271 | 0.43171 | 0.71625 | 1.31916 | 35 |

| | | | | | | |
|---|---|---|---|---|---|---|
| GO_DNA_GEOMETRIC_CHANGE | 0.00141 | 0.01850 | 0.45506 | 0.62642 | 1.20356 | 105 |
| GO_DOUBLE_STRANDED_RNA_BINDING | 0.00039 | 0.00658 | 0.49849 | 0.69158 | 1.30365 | 61 |
| GO_DRUG_BINDING | 0.00450 | 0.04342 | 0.40702 | 0.66245 | 1.24130 | 52 |
| GO_ELECTRON_TRANSFER_ACTIVITY | 0.00000 | 0.00001 | 0.67496 | 0.72917 | 1.38765 | 80 |
| GO_ELECTRON_TRANSPORT_CHAIN | 0.00000 | 0.00000 | | 0.73336 | 1.41392 | 133 |
| GO_EMBRYO_IMPLANTATION | 0.00044 | 0.00706 | 0.49849 | 0.75968 | 1.39146 | 29 |
| GO_ENDOCYTIC_VESICLE_MEMBRANE | 0.00524 | 0.04770 | 0.40702 | 0.62563 | 1.19564 | 89 |
| GO_ENDOPEPTIDASE_ACTIVITY | 0.00065 | 0.00981 | 0.47727 | 0.60123 | 1.16998 | 196 |
| GO_ENDOPEPTIDASE_COMPLEX | 0.00000 | 0.00001 | 0.70498 | 0.76593 | 1.44609 | 63 |
| GO_ENDOPEPTIDASE_REGULATOR_ACTIVITY | 0.00006 | 0.00145 | 0.55733 | 0.70425 | 1.33001 | 68 |
| GO_ENDOPLASMIC_RETICULUM_GOLGI_INTERMEDIATE_COMPA | 0.00055 | 0.00863 | 0.47727 | 0.64758 | 1.24050 | 96 |
| GO_ENDOPLASMIC_RETICULUM_GOLGI_INTERMEDIATE_COMPA | 0.00148 | 0.01923 | 0.45506 | 0.68944 | 1.29056 | 51 |
| GO_ENDOPLASMIC_RETICULUM_TO_GOLGI_VESICLE_MEDIATED | 0.00532 | 0.04803 | 0.40702 | 0.59194 | 1.14697 | 162 |
| GO_ENERGY_DERIVATION_BY_OXIDATION_OF_ORGANIC_COMP | 0.00000 | 0.00000 | | 0.69995 | 1.36305 | 202 |
| GO_ENZYME_INHIBITOR_ACTIVITY | 0.00246 | 0.02780 | 0.43171 | 0.58913 | 1.14643 | 191 |
| GO_EPIDERMAL_CELL_DIFFERENTIATION | 0.00216 | 0.02519 | 0.43171 | 0.62667 | 1.20589 | 112 |
| GO_EPITHELIAL_CELL_DIFFERENTIATION | 0.00054 | 0.00857 | 0.47727 | 0.57819 | 1.13725 | 334 |
| GO_ESTABLISHMENT_OF_ORGANELLE_LOCALIZATION | 0.00122 | 0.01679 | 0.45506 | 0.57737 | 1.13489 | 323 |
| GO_ESTABLISHMENT_OF_PROTEIN_LOCALIZATION_TO_CHROMO | 0.00016 | 0.00321 | 0.51885 | 0.81363 | 1.46985 | 23 |
| GO_ESTABLISHMENT_OF_PROTEIN_LOCALIZATION_TO_ENDOPL | 0.00000 | 0.00000 | | 0.77241 | 1.48479 | 106 |
| GO_ESTABLISHMENT_OF_PROTEIN_LOCALIZATION_TO_MEMBRA | 0.00000 | 0.00000 | | 0.67484 | 1.31991 | 260 |
| GO_ESTABLISHMENT_OF_PROTEIN_LOCALIZATION_TO_ORGANE | 0.00000 | 0.00000 | | 0.65856 | 1.30056 | 461 |
| GO_ESTABLISHMENT_OF_PROTEIN_LOCALIZATION_TO_TELOME | 0.00000 | 0.00006 | 0.64355 | 0.90447 | 1.60870 | 18 |
| GO_ESTABLISHMENT_OF_RNA_LOCALIZATION | 0.00000 | 0.00005 | 0.64355 | 0.64271 | 1.24830 | 176 |
| GO_ESTABLISHMENT_OF_TISSUE_POLARITY | 0.00000 | 0.00000 | 0.81404 | 0.75693 | 1.44999 | 96 |
| GO_EUKARYOTIC_TRANSLATION_INITIATION_FACTOR_3_COMPL | 0.00143 | 0.01863 | 0.45506 | 0.83263 | 1.46100 | 15 |
| GO_EXTERNAL_SIDE_OF_PLASMA_MEMBRANE | 0.00159 | 0.02016 | 0.45506 | 0.61775 | 1.19022 | 118 |
| GO_EXTRACELLULAR_MATRIX | 0.00042 | 0.00695 | 0.49849 | 0.61138 | 1.18938 | 189 |
| GO_FATTY_ACID_DERIVATIVE_METABOLIC_PROCESS | 0.00043 | 0.00695 | 0.49849 | 0.66266 | 1.26212 | 81 |
| GO_FATTY_ACYL_COA_METABOLIC_PROCESS | 0.00502 | 0.04608 | 0.40702 | 0.72264 | 1.32082 | 28 |
| GO_FC_EPSILON_RECEPTOR_SIGNALING_PATHWAY | 0.00000 | 0.00000 | 0.71951 | 0.72370 | 1.38610 | 94 |
| GO_FC_RECEPTOR_SIGNALING_PATHWAY | 0.00000 | 0.00000 | 0.70498 | 0.68815 | 1.32879 | 140 |
| GO_FERTILIZATION | 0.00000 | 0.00005 | 0.64355 | 0.73257 | 1.38752 | 72 |
| GO_FIBRILLAR_CENTER | 0.00246 | 0.02780 | 0.43171 | 0.62638 | 1.19990 | 96 |
| GO_FIBROBLAST_GROWTH_FACTOR_RECEPTOR_SIGNALING_PAT | 0.00495 | 0.04573 | 0.40702 | 0.64888 | 1.22552 | 67 |
| GO_FIBROBLAST_PROLIFERATION | 0.00032 | 0.00575 | 0.49849 | 0.70676 | 1.32618 | 53 |
| GO_FICOLIN_1_RICH_GRANULE | 0.00000 | 0.00000 | 0.70498 | 0.71544 | 1.37138 | 98 |
| GO_GABA_ERGIC_SYNAPSE | 0.00022 | 0.00427 | 0.51885 | 0.76430 | 1.40406 | 32 |
| GO_GENERATION_OF_PRECURSOR_METABOLITES_AND_ENERGY | 0.00000 | 0.00000 | | 0.67527 | 1.32968 | 368 |
| GO_GENE_SILENCING | 0.00019 | 0.00376 | 0.51885 | 0.61625 | 1.19788 | 184 |
| GO_GLUCOSE_CATABOLIC_PROCESS | 0.00495 | 0.04573 | 0.40702 | 0.72314 | 1.32174 | 28 |
| GO_GLUTAMATERGIC_SYNAPSE | 0.00135 | 0.01808 | 0.45506 | 0.60052 | 1.16783 | 186 |
| GO_GLYCOLYTIC_PROCESS_THROUGH_FRUCTOSE_6_PHOSPHATE | 0.00554 | 0.04968 | 0.40702 | 0.74156 | 1.34111 | 24 |
| GO_GOLGI_ASSOCIATED_VESICLE | 0.00085 | 0.01239 | 0.47727 | 0.62294 | 1.20037 | 129 |
| GO_GOLGI_ASSOCIATED_VESICLE_MEMBRANE | 0.00313 | 0.03310 | 0.43171 | 0.64208 | 1.21753 | 75 |
| GO_GTPASE_ACTIVITY | 0.00004 | 0.00098 | 0.55733 | 0.61546 | 1.19974 | 210 |
| GO_GUANYL_NUCLEOTIDE_BINDING | 0.00005 | 0.00126 | 0.55733 | 0.60899 | 1.19065 | 252 |
| GO_HEAT_SHOCK_PROTEIN_BINDING | 0.00001 | 0.00034 | 0.59333 | 0.68258 | 1.30633 | 93 |
| GO_HEMATOPOIETIC_PROGENITOR_CELL_DIFFERENTIATION | 0.00000 | 0.00000 | 0.71951 | 0.70099 | 1.35150 | 121 |
| GO_HEMATOPOIETIC_STEM_CELL_DIFFERENTIATION | 0.00000 | 0.00000 | 0.76146 | 0.77246 | 1.46307 | 72 |
| GO_HISTONE_MRNA_METABOLIC_PROCESS | 0.00265 | 0.02936 | 0.43171 | 0.76832 | 1.38459 | 22 |
| GO_HOST_CELLULAR_COMPONENT | 0.00043 | 0.00695 | 0.49849 | 0.69729 | 1.31074 | 54 |
| GO_IMMUNE_RESPONSE_REGULATING_SIGNALING_PATHWAY | 0.00000 | 0.00003 | 0.65944 | 0.63620 | 1.24112 | 225 |
| GO_IMPORT_INTO_CELL | 0.00428 | 0.04185 | 0.40702 | 0.61709 | 1.18655 | 108 |
| GO_IMPORT_INTO_NUCLEUS | 0.00039 | 0.00652 | 0.49849 | 0.62707 | 1.20905 | 131 |
| GO_INNATE_IMMUNE_RESPONSE | 0.00000 | 0.00003 | 0.65944 | 0.59893 | 1.18089 | 406 |
| GO_INNATE_IMMUNE_RESPONSE_ACTIVATING_SIGNAL_TRANSD | 0.00000 | 0.00001 | 0.69013 | 0.73348 | 1.39585 | 80 |
| GO_INNER_MITOCHONDRIAL_MEMBRANE_PROTEIN_COMPLEX | 0.00000 | 0.00000 | | 0.76576 | 1.46666 | 94 |
| GO_INORGANIC_ION_TRANSMEMBRANE_TRANSPORT | 0.00141 | 0.01850 | 0.45506 | 0.56404 | 1.11185 | 401 |
| GO_INTERACTION_WITH_HOST | 0.00139 | 0.01850 | 0.45506 | 0.60855 | 1.17710 | 151 |
| GO_INTERLEUKIN_1_MEDIATED_SIGNALING_PATHWAY | 0.00000 | 0.00000 | 0.77494 | 0.75654 | 1.44176 | 83 |
| GO_INTRAMOLECULAR_OXIDOREDUCTASE_ACTIVITY | 0.00051 | 0.00812 | 0.47727 | 0.74506 | 1.37081 | 34 |
| GO_INTRINSIC_APOPTOTIC_SIGNALING_PATHWAY | 0.00156 | 0.01993 | 0.45506 | 0.59427 | 1.15817 | 207 |
| GO_ION_HOMEOSTASIS | 0.00170 | 0.02117 | 0.45506 | 0.56164 | 1.10717 | 403 |
| GO_ION_TRANSMEMBRANE_TRANSPORTER_ACTIVITY | 0.00106 | 0.01495 | 0.45506 | 0.56696 | 1.11734 | 390 |
| GO_IRON_ION_HOMEOSTASIS | 0.00361 | 0.03657 | 0.43171 | 0.65837 | 1.23983 | 59 |
| GO_IRON_ION_TRANSPORT | 0.00054 | 0.00857 | 0.47727 | 0.69556 | 1.30334 | 52 |
| GO_ISOMERASE_ACTIVITY | 0.00413 | 0.04065 | 0.40702 | 0.61065 | 1.17496 | 113 |
| GO_KERATINIZATION | 0.00038 | 0.00640 | 0.49849 | 0.75690 | 1.39013 | 33 |
| GO_KILLING_OF_CELLS_OF_OTHER_ORGANISM | 0.00495 | 0.04573 | 0.40702 | 0.78614 | 1.38643 | 16 |

| | | | | | | |
|---|---|---|---|---|---|---|
| GO_LARGE_RIBOSOMAL_SUBUNIT | 0.00000 | 0.00000 | | 0.77656 | 1.49319 | 108 |
| GO_LATERAL_PLASMA_MEMBRANE | 0.00524 | 0.04770 | 0.40702 | 0.67385 | 1.25557 | 47 |
| GO_LEUKOCYTE_MEDIATED_IMMUNITY | 0.00000 | 0.00000 | | 0.64010 | 1.26331 | 447 |
| GO_LIPOPROTEIN_PARTICLE_RECEPTOR_BINDING | 0.00126 | 0.01715 | 0.45506 | 0.80503 | 1.42497 | 17 |
| GO_LIVER_REGENERATION | 0.00093 | 0.01337 | 0.47727 | 0.78442 | 1.41708 | 23 |
| GO_LOW_DENSITY_LIPOPROTEIN_RECEPTOR_PARTICLE_METABO | 0.00327 | 0.03409 | 0.43171 | 0.78520 | 1.39657 | 18 |
| GO_LYASE_ACTIVITY | 0.00152 | 0.01958 | 0.45506 | 0.61769 | 1.19079 | 122 |
| GO_LYSOSOMAL_LUMEN | 0.00143 | 0.01863 | 0.45506 | 0.67371 | 1.26872 | 59 |
| GO_MAINTENANCE_OF_CELL_NUMBER | 0.00435 | 0.04238 | 0.40702 | 0.60761 | 1.17073 | 117 |
| GO_MATURATION_OF_SSU_RRNA | 0.00126 | 0.01715 | 0.45506 | 0.69615 | 1.29711 | 47 |
| GO_MATURATION_OF_SSU_RRNA_FROM_TRICISTRONIC_RRNA_ | 0.00554 | 0.04968 | 0.40702 | 0.70205 | 1.29301 | 35 |
| GO_MEMBRANE_REGION | 0.00000 | 0.00003 | 0.65944 | 0.64496 | 1.25542 | 194 |
| GO_METALLOPEPTIDASE_ACTIVITY | 0.00487 | 0.04559 | 0.40702 | 0.62714 | 1.19577 | 85 |
| GO_METAL_ION_HOMEOSTASIS | 0.00032 | 0.00575 | 0.49849 | 0.58816 | 1.15376 | 301 |
| GO_METAL_ION_TRANSPORT | 0.00390 | 0.03895 | 0.43171 | 0.55877 | 1.10227 | 419 |
| GO_METAPHASE_ANAPHASE_TRANSITION_OF_CELL_CYCLE | 0.00320 | 0.03371 | 0.43171 | 0.66563 | 1.25122 | 54 |
| GO_MICROTUBULE_CYTOSKELETON_ORGANIZATION_INVOLVED_ | 0.00220 | 0.02548 | 0.43171 | 0.61295 | 1.18118 | 127 |
| GO_MICROVILLUS | 0.00168 | 0.02098 | 0.45506 | 0.68128 | 1.28184 | 56 |
| GO_MIDBODY | 0.00561 | 0.04971 | 0.40702 | 0.59866 | 1.15606 | 141 |
| GO_MITOCHONDRIAL_ELECTRON_TRANSPORT_NADH_TO_UBIQ | 0.00000 | 0.00006 | 0.64355 | 0.79447 | 1.47799 | 44 |
| GO_MITOCHONDRIAL_GENE_EXPRESSION | 0.00000 | 0.00000 | 0.78819 | 0.70230 | 1.35844 | 151 |
| GO_MITOCHONDRIAL_LARGE_RIBOSOMAL_SUBUNIT | 0.00000 | 0.00000 | 0.71951 | 0.79015 | 1.48529 | 54 |
| GO_MITOCHONDRIAL_MATRIX | 0.00000 | 0.00000 | 0.82666 | 0.63141 | 1.24412 | 387 |
| GO_MITOCHONDRIAL_MEMBRANE_ORGANIZATION | 0.00050 | 0.00802 | 0.47727 | 0.64578 | 1.23686 | 94 |
| GO_MITOCHONDRIAL_PROTEIN_COMPLEX | 0.00000 | 0.00000 | | 0.75011 | 1.46215 | 212 |
| GO_MITOCHONDRIAL_RESPIRATORY_CHAIN_COMPLEX_ASSEMB | 0.00001 | 0.00033 | 0.59333 | 0.69650 | 1.32804 | 85 |
| GO_MITOCHONDRIAL_SMALL_RIBOSOMAL_SUBUNIT | 0.00119 | 0.01640 | 0.45506 | 0.75384 | 1.37785 | 28 |
| GO_MITOCHONDRIAL_TRANSLATION | 0.00000 | 0.00000 | 0.80122 | 0.71435 | 1.37651 | 128 |
| GO_MITOCHONDRIAL_TRANSLATIONAL_TERMINATION | 0.00000 | 0.00000 | 0.81404 | 0.75760 | 1.44657 | 87 |
| GO_MITOCHONDRIAL_TRANSMEMBRANE_TRANSPORT | 0.00213 | 0.02490 | 0.43171 | 0.66300 | 1.25387 | 70 |
| GO_MITOCHONDRIAL_TRANSPORT | 0.00010 | 0.00227 | 0.53843 | 0.61119 | 1.19060 | 203 |
| GO_MITOCHONDRION_ORGANIZATION | 0.00000 | 0.00000 | 0.74774 | 0.61590 | 1.21471 | 416 |
| GO_MITOTIC_NUCLEAR_DIVISION | 0.00008 | 0.00182 | 0.53843 | 0.60435 | 1.17901 | 223 |
| GO_MITOTIC_SISTER_CHROMATID_SEGREGATION | 0.00014 | 0.00302 | 0.51885 | 0.62652 | 1.21048 | 145 |
| GO_MITOTIC_SPINDLE_ORGANIZATION | 0.00078 | 0.01164 | 0.47727 | 0.63943 | 1.22821 | 104 |
| GO_MODIFIED_AMINO_ACID_BINDING | 0.00002 | 0.00048 | 0.57561 | 0.74200 | 1.38679 | 49 |
| GO_MODULATION_BY_HOST_OF_VIRAL_PROCESS | 0.00216 | 0.02519 | 0.43171 | 0.77232 | 1.38066 | 20 |
| GO_MONOCARBOXYLIC_ACID_METABOLIC_PROCESS | 0.00012 | 0.00265 | 0.53843 | 0.57846 | 1.14053 | 406 |
| GO_MONOSACCHARIDE_BIOSYNTHETIC_PROCESS | 0.00305 | 0.03271 | 0.43171 | 0.65908 | 1.24472 | 68 |
| GO_MONOVALENT_INORGANIC_CATION_HOMEOSTASIS | 0.00108 | 0.01507 | 0.45506 | 0.63643 | 1.21783 | 91 |
| GO_MONOVALENT_INORGANIC_CATION_TRANSMEMBRANE_TR | 0.00001 | 0.00026 | 0.59333 | 0.65186 | 1.25996 | 147 |
| GO_MONOVALENT_INORGANIC_CATION_TRANSPORT | 0.00008 | 0.00178 | 0.53843 | 0.60205 | 1.17659 | 248 |
| GO_MORPHOGENESIS_OF_AN_EPITHELIUM | 0.00000 | 0.00009 | 0.62726 | 0.60570 | 1.19137 | 334 |
| GO_MORPHOGENESIS_OF_A_POLARIZED_EPITHELIUM | 0.00000 | 0.00000 | 0.83909 | 0.74384 | 1.43057 | 110 |
| GO_MOVEMENT_IN_HOST_ENVIRONMENT | 0.00268 | 0.02961 | 0.43171 | 0.61282 | 1.18078 | 117 |
| GO_MRNA_3_END_PROCESSING | 0.00087 | 0.01270 | 0.47727 | 0.64010 | 1.22502 | 93 |
| GO_MRNA_5_UTR_BINDING | 0.00002 | 0.00063 | 0.57561 | 0.83759 | 1.51313 | 23 |
| GO_MRNA_BINDING | 0.00000 | 0.00000 | | 0.67496 | 1.31810 | 239 |
| GO_MRNA_EXPORT_FROM_NUCLEUS | 0.00006 | 0.00159 | 0.53843 | 0.66290 | 1.27265 | 103 |
| GO_MRNA_PROCESSING | 0.00000 | 0.00000 | | 0.62959 | 1.24255 | 437 |
| GO_MRNA_TRANSPORT | 0.00007 | 0.00171 | 0.53843 | 0.64103 | 1.23563 | 132 |
| GO_MULTI_MULTICELLULAR_ORGANISM_PROCESS | 0.00041 | 0.00687 | 0.49849 | 0.63087 | 1.21515 | 116 |
| GO_MULTI_ORGANISM_LOCALIZATION | 0.00017 | 0.00334 | 0.51885 | 0.71077 | 1.33608 | 54 |
| GO_MUSCLE_CONTRACTION | 0.00335 | 0.03468 | 0.43171 | 0.59925 | 1.16199 | 165 |
| GO_MUSCLE_FILAMENT_SLIDING | 0.00324 | 0.03393 | 0.43171 | 0.77980 | 1.39322 | 19 |
| GO_MYELOID_LEUKOCYTE_ACTIVATION | 0.00000 | 0.00000 | | 0.65288 | 1.28712 | 395 |
| GO_MYELOID_LEUKOCYTE_MEDIATED_IMMUNITY | 0.00000 | 0.00000 | | 0.65734 | 1.29408 | 356 |
| GO_NADH_DEHYDROGENASE_ACTIVITY | 0.00001 | 0.00019 | 0.61053 | 0.79360 | 1.46691 | 38 |
| GO_NADH_DEHYDROGENASE_COMPLEX | 0.00001 | 0.00028 | 0.59333 | 0.77749 | 1.44215 | 41 |
| GO_NADH_DEHYDROGENASE_COMPLEX_ASSEMBLY | 0.00001 | 0.00035 | 0.59333 | 0.74044 | 1.39281 | 55 |
| GO_NADH_METABOLIC_PROCESS | 0.00034 | 0.00599 | 0.49849 | 0.75773 | 1.39166 | 33 |
| GO_NAD_METABOLIC_PROCESS | 0.00435 | 0.04238 | 0.40702 | 0.70570 | 1.30442 | 37 |
| GO_NCRNA_EXPORT_FROM_NUCLEUS | 0.00205 | 0.02440 | 0.43171 | 0.71473 | 1.31817 | 36 |
| GO_NCRNA_METABOLIC_PROCESS | 0.00000 | 0.00000 | 0.70498 | 0.61139 | 1.20509 | 399 |
| GO_NCRNA_PROCESSING | 0.00000 | 0.00000 | 0.73376 | 0.62238 | 1.22425 | 341 |
| GO_NEGATIVE_REGULATION_OF_APOPTOTIC_SIGNALING_PATHW | 0.00364 | 0.03677 | 0.43171 | 0.59958 | 1.15998 | 150 |
| GO_NEGATIVE_REGULATION_OF_ATP_METABOLIC_PROCESS | 0.00302 | 0.03248 | 0.43171 | 0.78586 | 1.39103 | 17 |
| GO_NEGATIVE_REGULATION_OF_CANONICAL_WNT_SIGNALING | 0.00000 | 0.00000 | 0.71951 | 0.69629 | 1.34180 | 123 |
| GO_NEGATIVE_REGULATION_OF_CATALYTIC_ACTIVITY | 0.00145 | 0.01879 | 0.45506 | 0.56238 | 1.11005 | 450 |
| GO_NEGATIVE_REGULATION_OF_CELLULAR_AMIDE_METABOLIC | 0.00268 | 0.02961 | 0.43171 | 0.61074 | 1.17686 | 124 |

| | | | | | | |
|---|---|---|---|---|---|---|
| GO_NEGATIVE_REGULATION_OF_CELLULAR_COMPONENT_ORG | 0.00320 | 0.03371 | 0.43171 | 0.55470 | 1.09534 | 470 |
| GO_NEGATIVE_REGULATION_OF_CELL_CYCLE | 0.00023 | 0.00445 | 0.51885 | 0.57348 | 1.13228 | 455 |
| GO_NEGATIVE_REGULATION_OF_CELL_CYCLE_G2_M_PHASE_TR | 0.00000 | 0.00007 | 0.62726 | 0.70549 | 1.34606 | 86 |
| GO_NEGATIVE_REGULATION_OF_CELL_CYCLE_PHASE_TRANSITIO | 0.00000 | 0.00000 | 0.70498 | 0.65532 | 1.27560 | 194 |
| GO_NEGATIVE_REGULATION_OF_CELL_CYCLE_PROCESS | 0.00000 | 0.00005 | 0.64355 | 0.62725 | 1.22670 | 259 |
| GO_NEGATIVE_REGULATION_OF_HYDROLASE_ACTIVITY | 0.00216 | 0.02519 | 0.43171 | 0.58306 | 1.13858 | 238 |
| GO_NEGATIVE_REGULATION_OF_INTERLEUKIN_1_PRODUCTION | 0.00205 | 0.02440 | 0.43171 | 0.82297 | 1.44404 | 15 |
| GO_NEGATIVE_REGULATION_OF_MITOTIC_CELL_CYCLE | 0.00000 | 0.00010 | 0.62726 | 0.62191 | 1.21541 | 248 |
| GO_NEGATIVE_REGULATION_OF_MRNA_METABOLIC_PROCESS | 0.00213 | 0.02490 | 0.43171 | 0.66731 | 1.25879 | 62 |
| GO_NEGATIVE_REGULATION_OF_MRNA_SPLICING_VIA_SPLICEO | 0.00339 | 0.03501 | 0.43171 | 0.76309 | 1.36417 | 20 |
| GO_NEGATIVE_REGULATION_OF_NUCLEAR_DIVISION | 0.00335 | 0.03468 | 0.43171 | 0.67724 | 1.26685 | 50 |
| GO_NEGATIVE_REGULATION_OF_ORGANELLE_ORGANIZATION | 0.00487 | 0.04559 | 0.40702 | 0.57038 | 1.11722 | 288 |
| GO_NEGATIVE_REGULATION_OF_PEPTIDASE_ACTIVITY | 0.00036 | 0.00621 | 0.49849 | 0.64044 | 1.23159 | 109 |
| GO_NEGATIVE_REGULATION_OF_PROTEIN_CATABOLIC_PROCESS | 0.00342 | 0.03521 | 0.43171 | 0.62320 | 1.19268 | 93 |
| GO_NEGATIVE_REGULATION_OF_PROTEOLYSIS | 0.00205 | 0.02440 | 0.43171 | 0.60172 | 1.16902 | 178 |
| GO_NEGATIVE_REGULATION_OF_REACTIVE_OXYGEN_SPECIES_B | 0.00036 | 0.00621 | 0.49849 | 0.83108 | 1.47818 | 18 |
| GO_NEGATIVE_REGULATION_OF_REPRODUCTIVE_PROCESS | 0.00457 | 0.04386 | 0.40702 | 0.74845 | 1.35356 | 24 |
| GO_NEGATIVE_REGULATION_OF_RNA_SPLICING | 0.00183 | 0.02219 | 0.45506 | 0.76824 | 1.38935 | 24 |
| GO_NEGATIVE_REGULATION_OF_WNT_SIGNALING_PATHWAY | 0.00000 | 0.00002 | 0.67496 | 0.67731 | 1.30721 | 136 |
| GO_NEURAL_CREST_CELL_MIGRATION | 0.00265 | 0.02936 | 0.43171 | 0.73967 | 1.35049 | 27 |
| GO_NEURON_RECOGNITION | 0.00390 | 0.03895 | 0.43171 | 0.71862 | 1.31998 | 31 |
| GO_NIK_NF_KAPPAB_SIGNALING | 0.00000 | 0.00000 | 0.77494 | 0.72329 | 1.39253 | 114 |
| GO_NON_CANONICAL_WNT_SIGNALING_PATHWAY | 0.00000 | 0.00000 | | 0.75756 | 1.45654 | 107 |
| GO_NUCLEAR_CHROMOSOME_SEGREGATION | 0.00098 | 0.01407 | 0.45506 | 0.59156 | 1.15357 | 215 |
| GO_NUCLEAR_ENVELOPE | 0.00005 | 0.00131 | 0.55733 | 0.59109 | 1.16270 | 341 |
| GO_NUCLEAR_EXPORT | 0.00000 | 0.00004 | 0.65944 | 0.64915 | 1.26157 | 181 |
| GO_NUCLEAR_MATRIX | 0.00502 | 0.04608 | 0.40702 | 0.62777 | 1.19868 | 87 |
| GO_NUCLEAR_MEMBRANE | 0.00185 | 0.02237 | 0.45506 | 0.58925 | 1.14853 | 211 |
| GO_NUCLEAR_PERIPHERY | 0.00043 | 0.00701 | 0.49849 | 0.64264 | 1.23558 | 107 |
| GO_NUCLEAR_PORE | 0.00561 | 0.04971 | 0.40702 | 0.64876 | 1.22522 | 68 |
| GO_NUCLEAR_SPECK | 0.00015 | 0.00312 | 0.51885 | 0.59159 | 1.16089 | 306 |
| GO_NUCLEAR_TRANSCRIBED_MRNA_CATABOLIC_PROCESS | 0.00000 | 0.00000 | | 0.70362 | 1.36820 | 185 |
| GO_NUCLEAR_TRANSCRIBED_MRNA_CATABOLIC_PROCESS_NON | 0.00000 | 0.00000 | | 0.74239 | 1.42931 | 114 |
| GO_NUCLEAR_TRANSPORT | 0.00000 | 0.00000 | 0.71951 | 0.62920 | 1.23383 | 297 |
| GO_NUCLEOBASE_CONTAINING_COMPOUND_TRANSPORT | 0.00000 | 0.00010 | 0.62726 | 0.62975 | 1.22823 | 216 |
| GO_NUCLEOBASE_CONTAINING_SMALL_MOLECULE_METABOLIC | 0.00000 | 0.00003 | 0.65944 | 0.60153 | 1.18594 | 405 |
| GO_NUCLEOSIDE_DIPHOSPHATE_METABOLIC_PROCESS | 0.00016 | 0.00319 | 0.51885 | 0.64491 | 1.24260 | 117 |
| GO_NUCLEOSIDE_PHOSPHATE_BIOSYNTHETIC_PROCESS | 0.00002 | 0.00044 | 0.59333 | 0.64092 | 1.24175 | 161 |
| GO_NUCLEOSIDE_TRIPHOSPHATE_BIOSYNTHETIC_PROCESS | 0.00040 | 0.00676 | 0.49849 | 0.72727 | 1.35074 | 42 |
| GO_NUCLEOSIDE_TRIPHOSPHATE_METABOLIC_PROCESS | 0.00009 | 0.00196 | 0.53843 | 0.69306 | 1.30854 | 65 |
| GO_NUCLEOTIDE_PHOSPHORYLATION | 0.00016 | 0.00328 | 0.51885 | 0.65541 | 1.25767 | 102 |
| GO_OOCYTE_DIFFERENTIATION | 0.00450 | 0.04342 | 0.40702 | 0.73513 | 1.33705 | 26 |
| GO_ORGANELLAR_RIBOSOME | 0.00000 | 0.00000 | 0.76146 | 0.75712 | 1.44311 | 84 |
| GO_ORGANELLE_ENVELOPE_LUMEN | 0.00026 | 0.00482 | 0.49849 | 0.67265 | 1.27716 | 78 |
| GO_ORGANELLE_FISSION | 0.00154 | 0.01974 | 0.45506 | 0.57133 | 1.12383 | 342 |
| GO_ORGANELLE_INNER_MEMBRANE | 0.00000 | 0.00000 | | 0.68423 | 1.34915 | 412 |
| GO_ORGANIC_CYCLIC_COMPOUND_CATABOLIC_PROCESS | 0.00000 | 0.00000 | | 0.66975 | 1.32263 | 478 |
| GO_ORGANIC_HYDROXY_COMPOUND_BIOSYNTHETIC_PROCESS | 0.00141 | 0.01850 | 0.45506 | 0.60712 | 1.17627 | 161 |
| GO_OXIDATIVE_PHOSPHORYLATION | 0.00000 | 0.00000 | | 0.78902 | 1.51260 | 99 |
| GO_OXIDOREDUCTASE_ACTIVITY | 0.00000 | 0.00000 | | 0.63268 | 1.24935 | 468 |
| GO_OXIDOREDUCTASE_ACTIVITY_ACTING_ON_A_HEME_GROUP | 0.00176 | 0.02168 | 0.45506 | 0.79383 | 1.41828 | 19 |
| GO_OXIDOREDUCTASE_ACTIVITY_ACTING_ON_NAD_P_H | 0.00002 | 0.00056 | 0.57561 | 0.71468 | 1.35162 | 70 |
| GO_OXIDOREDUCTASE_ACTIVITY_ACTING_ON_NAD_P_H_QUINO | 0.00004 | 0.00113 | 0.55733 | 0.74511 | 1.38835 | 47 |
| GO_OXIDOREDUCTASE_ACTIVITY_ACTING_ON_PEROXIDE_AS_AC | 0.00106 | 0.01495 | 0.45506 | 0.73897 | 1.35352 | 29 |
| GO_OXIDOREDUCTASE_COMPLEX | 0.00000 | 0.00000 | 0.73376 | 0.74898 | 1.42731 | 82 |
| GO_PEPTIDASE_ACTIVITY | 0.00133 | 0.01799 | 0.45506 | 0.57681 | 1.13188 | 306 |
| GO_PEPTIDASE_COMPLEX | 0.00000 | 0.00002 | 0.67496 | 0.73250 | 1.39240 | 79 |
| GO_PEPTIDASE_REGULATOR_ACTIVITY | 0.00313 | 0.03310 | 0.43171 | 0.63434 | 1.21189 | 88 |
| GO_PEPTIDE_BINDING | 0.00005 | 0.00124 | 0.55733 | 0.63708 | 1.23187 | 148 |
| GO_PERINUCLEAR_REGION_OF_CYTOPLASM | 0.00276 | 0.03021 | 0.43171 | 0.55814 | 1.10207 | 466 |
| GO_PHAGOCYTOSIS | 0.00554 | 0.04968 | 0.40702 | 0.59337 | 1.14837 | 155 |
| GO_PHAGOSOME_ACIDIFICATION | 0.00059 | 0.00916 | 0.47727 | 0.78902 | 1.42694 | 24 |
| GO_PHOSPHATIDYLSERINE_BINDING | 0.00035 | 0.00610 | 0.49849 | 0.76387 | 1.39912 | 29 |
| GO_PH_REDUCTION | 0.00302 | 0.03248 | 0.43171 | 0.68821 | 1.28238 | 46 |
| GO_PIGMENT_GRANULE | 0.00000 | 0.00000 | 0.76146 | 0.74333 | 1.41934 | 87 |
| GO_PLASMA_LIPOPROTEIN_PARTICLE_CLEARANCE | 0.00106 | 0.01495 | 0.45506 | 0.73201 | 1.35304 | 37 |
| GO_POLYSOMAL_RIBOSOME | 0.00008 | 0.00180 | 0.53843 | 0.78840 | 1.44683 | 30 |
| GO_POLYSOME | 0.00000 | 0.00003 | 0.65944 | 0.75964 | 1.43296 | 62 |
| GO_POSITIVE_REGULATION_OF_ATP_METABOLIC_PROCESS | 0.00174 | 0.02149 | 0.45506 | 0.73011 | 1.33729 | 29 |
| GO_POSITIVE_REGULATION_OF_BINDING | 0.00390 | 0.03895 | 0.43171 | 0.61042 | 1.17582 | 115 |

| | | | | | | |
|---|---|---|---|---|---|---|
| GO_POSITIVE_REGULATION_OF_CANONICAL_WNT_SIGNALING_ | 0.00000 | 0.00000 | 0.73376 | 0.72036 | 1.38514 | 108 |
| GO_POSITIVE_REGULATION_OF_CELLULAR_AMIDE_METABOLIC_ | 0.00001 | 0.00036 | 0.59333 | 0.66485 | 1.28095 | 125 |
| GO_POSITIVE_REGULATION_OF_CELLULAR_PROTEIN_LOCALIZAT | 0.00242 | 0.02748 | 0.43171 | 0.58048 | 1.13378 | 244 |
| GO_POSITIVE_REGULATION_OF_CELL_POPULATION_PROLIFERAT | 0.00040 | 0.00670 | 0.49849 | 0.56967 | 1.12503 | 461 |
| GO_POSITIVE_REGULATION_OF_CYSTEINE_TYPE_ENDOPEPTIDAS | 0.00547 | 0.04923 | 0.40702 | 0.62225 | 1.19077 | 92 |
| GO_POSITIVE_REGULATION_OF_CYTOKINE_PRODUCTION | 0.00283 | 0.03081 | 0.43171 | 0.58672 | 1.14373 | 210 |
| GO_POSITIVE_REGULATION_OF_DEFENSE_RESPONSE | 0.00000 | 0.00000 | 0.73376 | 0.66200 | 1.28884 | 200 |
| GO_POSITIVE_REGULATION_OF_DNA_BIOSYNTHETIC_PROCESS | 0.00010 | 0.00228 | 0.53843 | 0.71224 | 1.33977 | 55 |
| GO_POSITIVE_REGULATION_OF_DNA_METABOLIC_PROCESS | 0.00020 | 0.00387 | 0.51885 | 0.62771 | 1.21205 | 139 |
| GO_POSITIVE_REGULATION_OF_ESTABLISHMENT_OF_PROTEIN_ | 0.00194 | 0.02331 | 0.45506 | 0.58158 | 1.13709 | 253 |
| GO_POSITIVE_REGULATION_OF_FIBROBLAST_PROLIFERATION | 0.00017 | 0.00338 | 0.51885 | 0.77423 | 1.42213 | 31 |
| GO_POSITIVE_REGULATION_OF_IMMUNE_RESPONSE | 0.00000 | 0.00000 | 0.71951 | 0.63064 | 1.23937 | 321 |
| GO_POSITIVE_REGULATION_OF_IMMUNE_SYSTEM_PROCESS | 0.00000 | 0.00002 | 0.67496 | 0.59748 | 1.17992 | 475 |
| GO_POSITIVE_REGULATION_OF_INNATE_IMMUNE_RESPONSE | 0.00000 | 0.00000 | 0.70498 | 0.68885 | 1.32737 | 128 |
| GO_POSITIVE_REGULATION_OF_MITOTIC_CELL_CYCLE | 0.00178 | 0.02182 | 0.45506 | 0.61563 | 1.18683 | 122 |
| GO_POSITIVE_REGULATION_OF_MRNA_METABOLIC_PROCESS | 0.00157 | 0.02009 | 0.45506 | 0.66330 | 1.25231 | 63 |
| GO_POSITIVE_REGULATION_OF_MRNA_PROCESSING | 0.00250 | 0.02807 | 0.43171 | 0.76194 | 1.38151 | 25 |
| GO_POSITIVE_REGULATION_OF_MRNA_SPLICING_VIA_SPLICEOS | 0.00132 | 0.01782 | 0.45506 | 0.81851 | 1.44353 | 16 |
| GO_POSITIVE_REGULATION_OF_NITRIC_OXIDE_METABOLIC_PR( | 0.00133 | 0.01799 | 0.45506 | 0.80807 | 1.43726 | 18 |
| GO_POSITIVE_REGULATION_OF_ORGANELLE_ORGANIZATION | 0.00010 | 0.00218 | 0.53843 | 0.57473 | 1.13449 | 452 |
| GO_POSITIVE_REGULATION_OF_PROTEIN_LOCALIZATION_TO_N | 0.00012 | 0.00265 | 0.53843 | 0.71018 | 1.33697 | 58 |
| GO_POSITIVE_REGULATION_OF_PROTEOLYSIS | 0.00313 | 0.03310 | 0.43171 | 0.58059 | 1.13381 | 237 |
| GO_POSITIVE_REGULATION_OF_REACTIVE_OXYGEN_SPECIES_BI | 0.00115 | 0.01593 | 0.45506 | 0.79865 | 1.43554 | 21 |
| GO_POSITIVE_REGULATION_OF_REACTIVE_OXYGEN_SPECIES_M | 0.00174 | 0.02149 | 0.45506 | 0.69785 | 1.30035 | 46 |
| GO_POSITIVE_REGULATION_OF_RESPONSE_TO_BIOTIC_STIMUL | 0.00000 | 0.00000 | 0.70498 | 0.67507 | 1.30603 | 150 |
| GO_POSITIVE_REGULATION_OF_RESPONSE_TO_EXTERNAL_STIM | 0.00001 | 0.00016 | 0.61053 | 0.62123 | 1.21505 | 260 |
| GO_POSITIVE_REGULATION_OF_RNA_SPLICING | 0.00265 | 0.02936 | 0.43171 | 0.74051 | 1.35202 | 27 |
| GO_POSITIVE_REGULATION_OF_SIGNAL_TRANSDUCTION_BY_P5 | 0.00390 | 0.03895 | 0.43171 | 0.80515 | 1.41278 | 15 |
| GO_POSITIVE_REGULATION_OF_TELOMERASE_ACTIVITY | 0.00069 | 0.01043 | 0.47727 | 0.75265 | 1.38122 | 30 |
| GO_POSITIVE_REGULATION_OF_TELOMERASE_RNA_LOCALIZATI | 0.00007 | 0.00169 | 0.53843 | 0.88616 | 1.55493 | 15 |
| GO_POSITIVE_REGULATION_OF_TELOMERE_MAINTENANCE | 0.00026 | 0.00486 | 0.49849 | 0.73697 | 1.37102 | 44 |
| GO_POSITIVE_REGULATION_OF_TELOMERE_MAINTENANCE_VIA | 0.00001 | 0.00040 | 0.59333 | 0.80196 | 1.47289 | 33 |
| GO_POSITIVE_REGULATION_OF_TRANSFERASE_ACTIVITY | 0.00108 | 0.01507 | 0.45506 | 0.56501 | 1.11376 | 401 |
| GO_POSITIVE_REGULATION_OF_TRANSLATION | 0.00000 | 0.00016 | 0.61053 | 0.68408 | 1.31526 | 107 |
| GO_POSITIVE_REGULATION_OF_VIRAL_PROCESS | 0.00026 | 0.00486 | 0.49849 | 0.66195 | 1.26462 | 88 |
| GO_POSITIVE_REGULATION_OF_WNT_SIGNALING_PATHWAY | 0.00000 | 0.00000 | 0.73376 | 0.69924 | 1.34747 | 127 |
| GO_POSTSYNAPSE | 0.00239 | 0.02716 | 0.43171 | 0.57062 | 1.12116 | 316 |
| GO_POST_TRANSLATIONAL_PROTEIN_MODIFICATION | 0.00024 | 0.00452 | 0.51885 | 0.60046 | 1.17392 | 251 |
| GO_PRECATALYTIC_SPLICEOSOME | 0.00141 | 0.01850 | 0.45506 | 0.69251 | 1.29540 | 50 |
| GO_PRERIBOSOME | 0.00006 | 0.00154 | 0.53843 | 0.69866 | 1.32020 | 69 |
| GO_PROTEASOMAL_PROTEIN_CATABOLIC_PROCESS | 0.00007 | 0.00169 | 0.53843 | 0.58188 | 1.14652 | 387 |
| GO_PROTEASOMAL_UBIQUITIN_INDEPENDENT_PROTEIN_CATAE | 0.00000 | 0.00003 | 0.65944 | 0.91182 | 1.62178 | 18 |
| GO_PROTEASOME_ACCESSORY_COMPLEX | 0.00005 | 0.00135 | 0.55733 | 0.82638 | 1.49450 | 24 |
| GO_PROTEASOME_CORE_COMPLEX | 0.00000 | 0.00004 | 0.65944 | 0.91231 | 1.61486 | 17 |
| GO_PROTEIN_CONTAINING_COMPLEX_DISASSEMBLY | 0.00000 | 0.00000 | 0.81404 | 0.66285 | 1.29656 | 261 |
| GO_PROTEIN_CONTAINING_COMPLEX_LOCALIZATION | 0.00018 | 0.00352 | 0.51885 | 0.60451 | 1.17943 | 226 |
| GO_PROTEIN_DNA_COMPLEX_SUBUNIT_ORGANIZATION | 0.00052 | 0.00825 | 0.47727 | 0.60887 | 1.18342 | 183 |
| GO_PROTEIN_DOMAIN_SPECIFIC_BINDING | 0.00003 | 0.00088 | 0.55733 | 0.57828 | 1.14190 | 470 |
| GO_PROTEIN_FOLDING | 0.00000 | 0.00000 | 0.73376 | 0.67425 | 1.30867 | 171 |
| GO_PROTEIN_FOLDING_CHAPERONE | 0.00079 | 0.01170 | 0.47727 | 0.81632 | 1.45193 | 18 |
| GO_PROTEIN_HETEROOLIGOMERIZATION | 0.00442 | 0.04297 | 0.40702 | 0.78998 | 1.39322 | 16 |
| GO_PROTEIN_IMPORT | 0.00001 | 0.00040 | 0.59333 | 0.64197 | 1.24306 | 158 |
| GO_PROTEIN_IMPORT_INTO_MITOCHONDRIAL_MATRIX | 0.00231 | 0.02642 | 0.43171 | 0.79530 | 1.41455 | 18 |
| GO_PROTEIN_INSERTION_INTO_MEMBRANE | 0.00191 | 0.02299 | 0.45506 | 0.68322 | 1.27694 | 49 |
| GO_PROTEIN_LOCALIZATION_TO_CHROMOSOME_TELOMERIC_F | 0.00001 | 0.00031 | 0.59333 | 0.82810 | 1.51357 | 28 |
| GO_PROTEIN_LOCALIZATION_TO_ENDOPLASMIC_RETICULUM | 0.00000 | 0.00000 | | 0.73850 | 1.42303 | 129 |
| GO_PROTEIN_LOCALIZATION_TO_MEMBRANE | 0.00000 | 0.00000 | 0.77494 | 0.61216 | 1.20881 | 462 |
| GO_PROTEIN_LOCALIZATION_TO_MITOCHONDRION | 0.00015 | 0.00310 | 0.51885 | 0.64234 | 1.23759 | 118 |
| GO_PROTEIN_LOCALIZATION_TO_NUCLEUS | 0.00000 | 0.00014 | 0.61053 | 0.63315 | 1.23408 | 208 |
| GO_PROTEIN_MATURATION | 0.00357 | 0.03638 | 0.43171 | 0.59209 | 1.14965 | 174 |
| GO_PROTEIN_MODIFICATION_BY_SMALL_PROTEIN_REMOVAL | 0.00002 | 0.00052 | 0.57561 | 0.61535 | 1.20119 | 231 |
| GO_PROTEIN_REFOLDING | 0.00465 | 0.04430 | 0.40702 | 0.77561 | 1.37288 | 17 |
| GO_PROTEIN_STABILIZATION | 0.00081 | 0.01194 | 0.47727 | 0.61881 | 1.19577 | 146 |
| GO_PROTEIN_SUMOYLATION | 0.00450 | 0.04342 | 0.40702 | 0.65043 | 1.22846 | 67 |
| GO_PROTEIN_TARGETING | 0.00000 | 0.00000 | | 0.66223 | 1.30361 | 358 |
| GO_PROTEIN_TARGETING_TO_MEMBRANE | 0.00000 | 0.00000 | | 0.71675 | 1.38879 | 162 |
| GO_PROTEIN_TARGETING_TO_MITOCHONDRION | 0.00113 | 0.01571 | 0.45506 | 0.65407 | 1.24648 | 83 |
| GO_PROTEIN_TRANSMEMBRANE_TRANSPORT | 0.00110 | 0.01526 | 0.45506 | 0.68173 | 1.28238 | 55 |
| GO_PROTON_TRANSMEMBRANE_TRANSPORT | 0.00000 | 0.00001 | 0.69013 | 0.71234 | 1.36309 | 91 |
| GO_PROTON_TRANSMEMBRANE_TRANSPORTER_ACTIVITY | 0.00004 | 0.00115 | 0.55733 | 0.70138 | 1.32535 | 69 |

| | | | | | | |
|---|---|---|---|---|---|---|
| GO_PROTON_TRANSPORTING_TWO_SECTOR_ATPASE_COMPLEX | 0.00083 | 0.01218 | 0.47727 | 0.77254 | 1.40508 | 26 |
| GO_PROTON_TRANSPORTING_V_TYPE_ATPASE_COMPLEX | 0.00239 | 0.02716 | 0.43171 | 0.76100 | 1.37626 | 24 |
| GO_PURINE_CONTAINING_COMPOUND_BIOSYNTHETIC_PROCES | 0.00001 | 0.00040 | 0.59333 | 0.65863 | 1.26901 | 126 |
| GO_PURINE_CONTAINING_COMPOUND_METABOLIC_PROCESS | 0.00000 | 0.00000 | 0.73376 | 0.63132 | 1.23831 | 300 |
| GO_PURINE_NUCLEOSIDE_TRIPHOSPHATE_METABOLIC_PROCES | 0.00033 | 0.00582 | 0.49849 | 0.70887 | 1.32487 | 49 |
| GO_PYROPHOSPHATE_HYDROLYSIS_DRIVEN_PROTON_TRANSMI | 0.00532 | 0.04803 | 0.40702 | 0.74470 | 1.34678 | 24 |
| GO_PYRUVATE_METABOLIC_PROCESS | 0.00001 | 0.00018 | 0.61053 | 0.67194 | 1.29289 | 113 |
| GO_P_BODY | 0.00472 | 0.04480 | 0.40702 | 0.65470 | 1.23501 | 62 |
| GO_REACTIVE_OXYGEN_SPECIES_BIOSYNTHETIC_PROCESS | 0.00040 | 0.00670 | 0.49849 | 0.70271 | 1.31857 | 53 |
| GO_REACTIVE_OXYGEN_SPECIES_METABOLIC_PROCESS | 0.00021 | 0.00409 | 0.51885 | 0.62881 | 1.21472 | 144 |
| GO_REGULATION_OF_ALCOHOL_BIOSYNTHETIC_PROCESS | 0.00228 | 0.02614 | 0.43171 | 0.68438 | 1.28020 | 50 |
| GO_REGULATION_OF_ANIMAL_ORGAN_MORPHOGENESIS | 0.00000 | 0.00000 | | 0.72330 | 1.39922 | 149 |
| GO_REGULATION_OF_APOPTOTIC_SIGNALING_PATHWAY | 0.00079 | 0.01170 | 0.47727 | 0.58599 | 1.14641 | 266 |
| GO_REGULATION_OF_ATP_METABOLIC_PROCESS | 0.00000 | 0.00006 | 0.64355 | 0.70523 | 1.34777 | 89 |
| GO_REGULATION_OF_BINDING | 0.00428 | 0.04185 | 0.40702 | 0.58264 | 1.13664 | 225 |
| GO_REGULATION_OF_BLOOD_PRESSURE | 0.00290 | 0.03156 | 0.43171 | 0.65319 | 1.23428 | 69 |
| GO_REGULATION_OF_CARBOHYDRATE_CATABOLIC_PROCESS | 0.00324 | 0.03393 | 0.43171 | 0.65541 | 1.23787 | 67 |
| GO_REGULATION_OF_CELLULAR_AMIDE_METABOLIC_PROCESS | 0.00000 | 0.00000 | 0.78819 | 0.63116 | 1.24143 | 337 |
| GO_REGULATION_OF_CELLULAR_AMINE_METABOLIC_PROCESS | 0.00000 | 0.00000 | 0.74774 | 0.78955 | 1.48730 | 60 |
| GO_REGULATION_OF_CELLULAR_AMINO_ACID_METABOLIC_PR( | 0.00000 | 0.00000 | 0.76146 | 0.80341 | 1.51022 | 54 |
| GO_REGULATION_OF_CELLULAR_KETONE_METABOLIC_PROCESS | 0.00000 | 0.00000 | 0.70498 | 0.69239 | 1.33405 | 126 |
| GO_REGULATION_OF_CELLULAR_PH | 0.00442 | 0.04297 | 0.40702 | 0.65360 | 1.23436 | 68 |
| GO_REGULATION_OF_CELLULAR_PROTEIN_LOCALIZATION | 0.00228 | 0.02614 | 0.43171 | 0.56216 | 1.10830 | 402 |
| GO_REGULATION_OF_CELLULAR_RESPIRATION | 0.00428 | 0.04185 | 0.40702 | 0.77183 | 1.37897 | 19 |
| GO_REGULATION_OF_CELLULAR_RESPONSE_TO_HEAT | 0.00091 | 0.01314 | 0.47727 | 0.67360 | 1.27221 | 67 |
| GO_REGULATION_OF_CELL_CYCLE_G2_M_PHASE_TRANSITION | 0.00016 | 0.00332 | 0.51885 | 0.61554 | 1.19738 | 187 |
| GO_REGULATION_OF_CELL_CYCLE_PHASE_TRANSITION | 0.00000 | 0.00014 | 0.61053 | 0.59968 | 1.18091 | 369 |
| GO_REGULATION_OF_CHOLESTEROL_BIOSYNTHETIC_PROCESS | 0.00209 | 0.02470 | 0.43171 | 0.71476 | 1.32117 | 38 |
| GO_REGULATION_OF_CHOLESTEROL_METABOLIC_PROCESS | 0.00113 | 0.01571 | 0.45506 | 0.69865 | 1.30177 | 47 |
| GO_REGULATION_OF_CHROMOSOME_ORGANIZATION | 0.00036 | 0.00622 | 0.49849 | 0.58896 | 1.15261 | 278 |
| GO_REGULATION_OF_CHROMOSOME_SEGREGATION | 0.00079 | 0.01170 | 0.47727 | 0.64131 | 1.22718 | 91 |
| GO_REGULATION_OF_CHROMOSOME_SEPARATION | 0.00253 | 0.02838 | 0.43171 | 0.66349 | 1.24947 | 59 |
| GO_REGULATION_OF_CYSTEINE_TYPE_ENDOPEPTIDASE_ACTIVIT | 0.00044 | 0.00719 | 0.49849 | 0.62610 | 1.20905 | 141 |
| GO_REGULATION_OF_DEFENSE_RESPONSE | 0.00006 | 0.00150 | 0.55733 | 0.58834 | 1.15824 | 356 |
| GO_REGULATION_OF_DNA_BIOSYNTHETIC_PROCESS | 0.00043 | 0.00695 | 0.49849 | 0.66709 | 1.27130 | 83 |
| GO_REGULATION_OF_DNA_METABOLIC_PROCESS | 0.00055 | 0.00863 | 0.47727 | 0.58909 | 1.15134 | 249 |
| GO_REGULATION_OF_DNA_TEMPLATED_TRANSCRIPTION_IN_RI | 0.00000 | 0.00000 | 0.71951 | 0.72410 | 1.38687 | 94 |
| GO_REGULATION_OF_GENERATION_OF_PRECURSOR_METABOLI | 0.00004 | 0.00114 | 0.55733 | 0.65536 | 1.26115 | 111 |
| GO_REGULATION_OF_GENE_SILENCING | 0.00023 | 0.00434 | 0.51885 | 0.65464 | 1.25268 | 91 |
| GO_REGULATION_OF_HEMATOPOIETIC_PROGENITOR_CELL_DIF | 0.00000 | 0.00000 | 0.73376 | 0.76273 | 1.44465 | 72 |
| GO_REGULATION_OF_HEMATOPOIETIC_STEM_CELL_DIFFERENT | 0.00000 | 0.00000 | 0.74774 | 0.78891 | 1.48817 | 62 |
| GO_REGULATION_OF_HEMOPOIESIS | 0.00390 | 0.03895 | 0.43171 | 0.57739 | 1.12909 | 258 |
| GO_REGULATION_OF_IMMUNE_RESPONSE | 0.00000 | 0.00006 | 0.64355 | 0.60032 | 1.18447 | 433 |
| GO_REGULATION_OF_INNATE_IMMUNE_RESPONSE | 0.00000 | 0.00007 | 0.62726 | 0.64536 | 1.25422 | 181 |
| GO_REGULATION_OF_INTRINSIC_APOPTOTIC_SIGNALING_PATH\ | 0.00179 | 0.02183 | 0.45506 | 0.61548 | 1.18550 | 116 |
| GO_REGULATION_OF_MITOTIC_CELL_CYCLE | 0.00000 | 0.00003 | 0.65944 | 0.58992 | 1.16508 | 499 |
| GO_REGULATION_OF_MORPHOGENESIS_OF_AN_EPITHELIUM | 0.00000 | 0.00000 | | 0.74680 | 1.43884 | 125 |
| GO_REGULATION_OF_MRNA_CATABOLIC_PROCESS | 0.00000 | 0.00000 | | 0.71126 | 1.37919 | 165 |
| GO_REGULATION_OF_MRNA_METABOLIC_PROCESS | 0.00000 | 0.00000 | | 0.67178 | 1.31458 | 276 |
| GO_REGULATION_OF_MRNA_PROCESSING | 0.00008 | 0.00186 | 0.53843 | 0.65189 | 1.25598 | 125 |
| GO_REGULATION_OF_MRNA_SPLICING_VIA_SPLICEOSOME | 0.00012 | 0.00256 | 0.53843 | 0.66950 | 1.28033 | 90 |
| GO_REGULATION_OF_MULTI_ORGANISM_PROCESS | 0.00008 | 0.00190 | 0.53843 | 0.76905 | 1.42150 | 37 |
| GO_REGULATION_OF_MUSCLE_CONTRACTION | 0.00163 | 0.02050 | 0.45506 | 0.64879 | 1.23071 | 76 |
| GO_REGULATION_OF_NUCLEAR_DIVISION | 0.00060 | 0.00923 | 0.47727 | 0.62443 | 1.20552 | 137 |
| GO_REGULATION_OF_NUCLEASE_ACTIVITY | 0.00059 | 0.00916 | 0.47727 | 0.81786 | 1.44767 | 17 |
| GO_REGULATION_OF_NUCLEOTIDE_BIOSYNTHETIC_PROCESS | 0.00302 | 0.03248 | 0.43171 | 0.78902 | 1.40337 | 18 |
| GO_REGULATION_OF_NUCLEOTIDE_METABOLIC_PROCESS | 0.00033 | 0.00588 | 0.49849 | 0.66858 | 1.26942 | 78 |
| GO_REGULATION_OF_OXIDATIVE_PHOSPHORYLATION | 0.00005 | 0.00130 | 0.55733 | 0.85284 | 1.53294 | 21 |
| GO_REGULATION_OF_PEPTIDASE_ACTIVITY | 0.00006 | 0.00152 | 0.53843 | 0.61192 | 1.19388 | 226 |
| GO_REGULATION_OF_PEPTIDE_TRANSPORT | 0.00250 | 0.02807 | 0.43171 | 0.56252 | 1.10917 | 412 |
| GO_REGULATION_OF_PH | 0.00561 | 0.04971 | 0.40702 | 0.64191 | 1.21498 | 71 |
| GO_REGULATION_OF_PLASMA_LIPOPROTEIN_PARTICLE_LEVELS | 0.00006 | 0.00145 | 0.55733 | 0.72431 | 1.36154 | 54 |
| GO_REGULATION_OF_POSTTRANSCRIPTIONAL_GENE_SILENCIN( | 0.00026 | 0.00486 | 0.49849 | 0.66803 | 1.27236 | 81 |
| GO_REGULATION_OF_PROTEIN_CATABOLIC_PROCESS | 0.00309 | 0.03299 | 0.43171 | 0.57360 | 1.12399 | 291 |
| GO_REGULATION_OF_PROTEIN_IMPORT | 0.00487 | 0.04559 | 0.40702 | 0.67491 | 1.25754 | 47 |
| GO_REGULATION_OF_PROTEIN_LOCALIZATION_TO_NUCLEUS | 0.00000 | 0.00013 | 0.61053 | 0.69035 | 1.32101 | 91 |
| GO_REGULATION_OF_PROTEIN_STABILITY | 0.00066 | 0.01007 | 0.47727 | 0.59308 | 1.15811 | 235 |
| GO_REGULATION_OF_PROTEOLYSIS | 0.00007 | 0.00169 | 0.53843 | 0.57923 | 1.14299 | 431 |
| GO_REGULATION_OF_REACTIVE_OXYGEN_SPECIES_BIOSYNTHET | 0.00022 | 0.00427 | 0.51885 | 0.73154 | 1.36343 | 45 |
| GO_REGULATION_OF_REACTIVE_OXYGEN_SPECIES_METABOLIC_ | 0.00088 | 0.01280 | 0.47727 | 0.63948 | 1.22592 | 99 |

| | | | | | | |
|---|---|---|---|---|---|---|
| GO_REGULATION_OF_RESPONSE_TO_BIOTIC_STIMULUS | 0.00001 | 0.00039 | 0.59333 | 0.61773 | 1.20625 | 235 |
| GO_REGULATION_OF_RNA_SPLICING | 0.00011 | 0.00241 | 0.53843 | 0.64982 | 1.25198 | 125 |
| GO_REGULATION_OF_SISTER_CHROMATID_SEGREGATION | 0.00405 | 0.04011 | 0.40702 | 0.64401 | 1.22034 | 73 |
| GO_REGULATION_OF_SMALL_MOLECULE_METABOLIC_PROCESS | 0.00000 | 0.00000 | 0.77494 | 0.64342 | 1.26057 | 290 |
| GO_REGULATION_OF_SMOOTH_MUSCLE_CONTRACTION | 0.00007 | 0.00171 | 0.53843 | 0.82642 | 1.49295 | 23 |
| GO_REGULATION_OF_STEM_CELL_DIFFERENTIATION | 0.00000 | 0.00000 | 0.73376 | 0.73434 | 1.40292 | 88 |
| GO_REGULATION_OF_STEROID_METABOLIC_PROCESS | 0.00524 | 0.04770 | 0.40702 | 0.64061 | 1.21567 | 77 |
| GO_REGULATION_OF_SUPEROXIDE_METABOLIC_PROCESS | 0.00346 | 0.03548 | 0.43171 | 0.77826 | 1.39046 | 19 |
| GO_REGULATION_OF_SUPRAMOLECULAR_FIBER_ORGANIZATIO | 0.00509 | 0.04656 | 0.40702 | 0.57599 | 1.12367 | 223 |
| GO_REGULATION_OF_SYMBIOTIC_PROCESS | 0.00009 | 0.00211 | 0.53843 | 0.62950 | 1.21763 | 151 |
| GO_REGULATION_OF_TELOMERASE_ACTIVITY | 0.00087 | 0.01263 | 0.47727 | 0.72147 | 1.33426 | 39 |
| GO_REGULATION_OF_TELOMERE_MAINTENANCE | 0.00547 | 0.04923 | 0.40702 | 0.64959 | 1.22679 | 68 |
| GO_REGULATION_OF_TELOMERE_MAINTENANCE_VIA_TELOME | 0.00020 | 0.00387 | 0.51885 | 0.72043 | 1.34856 | 51 |
| GO_REGULATION_OF_TRANSCRIPTION_FROM_RNA_POLYMERA | 0.00000 | 0.00000 | 0.70498 | 0.76931 | 1.45239 | 64 |
| GO_REGULATION_OF_TRANSLATIONAL_INITIATION | 0.00023 | 0.00437 | 0.51885 | 0.68479 | 1.29399 | 69 |
| GO_REGULATION_OF_VIRAL_TRANSCRIPTION | 0.00361 | 0.03657 | 0.43171 | 0.67497 | 1.26259 | 50 |
| GO_REGULATION_OF_WNT_SIGNALING_PATHWAY | 0.00015 | 0.00312 | 0.51885 | 0.60129 | 1.17419 | 238 |
| GO_RESPIRASOME | 0.00000 | 0.00000 | 0.81404 | 0.78326 | 1.48524 | 75 |
| GO_RESPIRATORY_CHAIN_COMPLEX | 0.00000 | 0.00000 | 0.77494 | 0.79183 | 1.49492 | 64 |
| GO_RESPIRATORY_ELECTRON_TRANSPORT_CHAIN | 0.00000 | 0.00000 | | 0.78275 | 1.49591 | 89 |
| GO_RESPONSE_TO_ALCOHOL | 0.00015 | 0.00310 | 0.51885 | 0.63794 | 1.22915 | 126 |
| GO_RESPONSE_TO_ATP | 0.00087 | 0.01270 | 0.47727 | 0.81086 | 1.43528 | 17 |
| GO_RESPONSE_TO_CALCIUM_ION | 0.00124 | 0.01701 | 0.45506 | 0.65196 | 1.24071 | 80 |
| GO_RESPONSE_TO_CORTICOSTEROID | 0.00010 | 0.00220 | 0.53843 | 0.66894 | 1.27841 | 89 |
| GO_RESPONSE_TO_DRUG | 0.00000 | 0.00007 | 0.62726 | 0.63080 | 1.23070 | 220 |
| GO_RESPONSE_TO_ETHANOL | 0.00168 | 0.02098 | 0.45506 | 0.66543 | 1.25679 | 67 |
| GO_RESPONSE_TO_FIBROBLAST_GROWTH_FACTOR | 0.00357 | 0.03638 | 0.43171 | 0.63469 | 1.20976 | 84 |
| GO_RESPONSE_TO_GROWTH_FACTOR | 0.00272 | 0.02986 | 0.43171 | 0.56235 | 1.10805 | 387 |
| GO_RESPONSE_TO_HEAT | 0.00000 | 0.00013 | 0.62726 | 0.66982 | 1.29069 | 119 |
| GO_RESPONSE_TO_HYDROGEN_PEROXIDE | 0.00532 | 0.04803 | 0.40702 | 0.62255 | 1.19054 | 90 |
| GO_RESPONSE_TO_INORGANIC_SUBSTANCE | 0.00016 | 0.00328 | 0.51885 | 0.58928 | 1.15810 | 321 |
| GO_RESPONSE_TO_INTERLEUKIN_1 | 0.00000 | 0.00000 | 0.78819 | 0.71590 | 1.37950 | 124 |
| GO_RESPONSE_TO_INTERLEUKIN_12 | 0.00036 | 0.00622 | 0.49849 | 0.74158 | 1.36769 | 36 |
| GO_RESPONSE_TO_INTERLEUKIN_4 | 0.00224 | 0.02581 | 0.43171 | 0.77088 | 1.38921 | 22 |
| GO_RESPONSE_TO_LEUKEMIA_INHIBITORY_FACTOR | 0.00076 | 0.01129 | 0.47727 | 0.66148 | 1.25288 | 72 |
| GO_RESPONSE_TO_LIPID | 0.00043 | 0.00695 | 0.49849 | 0.56669 | 1.11902 | 470 |
| GO_RESPONSE_TO_METAL_ION | 0.00143 | 0.01863 | 0.45506 | 0.59895 | 1.16592 | 199 |
| GO_RESPONSE_TO_NUTRIENT | 0.00294 | 0.03185 | 0.43171 | 0.62596 | 1.19987 | 98 |
| GO_RESPONSE_TO_ORGANOPHOSPHORUS | 0.00294 | 0.03185 | 0.43171 | 0.64154 | 1.21511 | 72 |
| GO_RESPONSE_TO_OXIDATIVE_STRESS | 0.00495 | 0.04573 | 0.40702 | 0.57010 | 1.11758 | 294 |
| GO_RESPONSE_TO_OXYGEN_LEVELS | 0.00000 | 0.00000 | | 0.67661 | 1.32155 | 244 |
| GO_RESPONSE_TO_PURINE_CONTAINING_COMPOUND | 0.00509 | 0.04656 | 0.40702 | 0.63176 | 1.20394 | 82 |
| GO_RESPONSE_TO_STEROID_HORMONE | 0.00009 | 0.00200 | 0.53843 | 0.61312 | 1.19360 | 201 |
| GO_RESPONSE_TO_TEMPERATURE_STIMULUS | 0.00002 | 0.00047 | 0.57561 | 0.64468 | 1.24782 | 153 |
| GO_RESPONSE_TO_TOXIC_SUBSTANCE | 0.00011 | 0.00235 | 0.53843 | 0.63892 | 1.23203 | 134 |
| GO_RESPONSE_TO_TUMOR_NECROSIS_FACTOR | 0.00000 | 0.00000 | 0.70498 | 0.66331 | 1.28815 | 175 |
| GO_RESPONSE_TO_VIRUS | 0.00261 | 0.02910 | 0.43171 | 0.59428 | 1.15408 | 175 |
| GO_RETINA_HOMEOSTASIS | 0.00034 | 0.00606 | 0.49849 | 0.75202 | 1.38362 | 34 |
| GO_RIBONUCLEOPROTEIN_COMPLEX_BINDING | 0.00000 | 0.00001 | 0.69013 | 0.69839 | 1.34459 | 114 |
| GO_RIBONUCLEOPROTEIN_COMPLEX_BIOGENESIS | 0.00000 | 0.00000 | | 0.68714 | 1.35448 | 393 |
| GO_RIBONUCLEOPROTEIN_COMPLEX_SUBUNIT_ORGANIZATION | 0.00000 | 0.00000 | 0.81404 | 0.69112 | 1.34183 | 173 |
| GO_RIBONUCLEOPROTEIN_GRANULE | 0.00000 | 0.00000 | 0.71951 | 0.66271 | 1.28875 | 186 |
| GO_RIBONUCLEOSIDE_TRIPHOSPHATE_BIOSYNTHETIC_PROCESS | 0.00009 | 0.00210 | 0.53843 | 0.77194 | 1.42026 | 34 |
| GO_RIBONUCLEOSIDE_TRIPHOSPHATE_METABOLIC_PROCESS | 0.00013 | 0.00280 | 0.51885 | 0.73558 | 1.37065 | 46 |
| GO_RIBOSE_PHOSPHATE_BIOSYNTHETIC_PROCESS | 0.00002 | 0.00056 | 0.57561 | 0.65814 | 1.26821 | 124 |
| GO_RIBOSE_PHOSPHATE_METABOLIC_PROCESS | 0.00000 | 0.00000 | 0.74774 | 0.63837 | 1.25001 | 285 |
| GO_RIBOSOMAL_LARGE_SUBUNIT_BIOGENESIS | 0.00001 | 0.00031 | 0.59333 | 0.72435 | 1.36807 | 67 |
| GO_RIBOSOMAL_SMALL_SUBUNIT_ASSEMBLY | 0.00405 | 0.04011 | 0.40702 | 0.78074 | 1.38865 | 18 |
| GO_RIBOSOMAL_SMALL_SUBUNIT_BINDING | 0.00532 | 0.04803 | 0.40702 | 0.77341 | 1.36899 | 17 |
| GO_RIBOSOMAL_SMALL_SUBUNIT_BIOGENESIS | 0.00005 | 0.00138 | 0.55733 | 0.70510 | 1.33163 | 68 |
| GO_RIBOSOMAL_SUBUNIT | 0.00000 | 0.00000 | | 0.76292 | 1.48089 | 172 |
| GO_RIBOSOME | 0.00000 | 0.00000 | | 0.75411 | 1.46984 | 208 |
| GO_RIBOSOME_ASSEMBLY | 0.00001 | 0.00026 | 0.59333 | 0.74485 | 1.40144 | 56 |
| GO_RIBOSOME_BIOGENESIS | 0.00000 | 0.00000 | | 0.68899 | 1.34833 | 277 |
| GO_RNA_3_END_PROCESSING | 0.00002 | 0.00046 | 0.57561 | 0.65576 | 1.26361 | 128 |
| GO_RNA_CATABOLIC_PROCESS | 0.00000 | 0.00000 | | 0.70701 | 1.39073 | 341 |
| GO_RNA_DEPENDENT_DNA_BIOSYNTHETIC_PROCESS | 0.00003 | 0.00081 | 0.57561 | 0.71669 | 1.34965 | 59 |
| GO_RNA_EXPORT_FROM_NUCLEUS | 0.00000 | 0.00016 | 0.61053 | 0.66306 | 1.27817 | 130 |
| GO_RNA_LOCALIZATION | 0.00000 | 0.00000 | 0.76146 | 0.66638 | 1.29886 | 208 |
| GO_RNA_SPLICING | 0.00000 | 0.00000 | | 0.63512 | 1.25129 | 384 |

| | | | | | | |
|---|---|---|---|---|---|---|
| GO_RNA_SPLICING_VIA_TRANSESTERIFICATION_REACTIONS | 0.00000 | 0.00000 | 0.80122 | 0.64478 | 1.26586 | 311 |
| GO_ROUGH_ENDOPLASMIC_RETICULUM | 0.00014 | 0.00293 | 0.51885 | 0.70363 | 1.32507 | 59 |
| GO_RRNA_BINDING | 0.00000 | 0.00001 | 0.69013 | 0.77638 | 1.45941 | 54 |
| GO_RRNA_METABOLIC_PROCESS | 0.00000 | 0.00000 | | 0.68430 | 1.33386 | 209 |
| GO_SCF_DEPENDENT_PROTEASOMAL_UBIQUITIN_DEPENDENT_ | 0.00000 | 0.00008 | 0.62726 | 0.71957 | 1.36498 | 76 |
| GO_SECRETORY_GRANULE | 0.00000 | 0.00000 | | 0.62978 | 1.24363 | 481 |
| GO_SECRETORY_GRANULE_MEMBRANE | 0.00032 | 0.00575 | 0.49849 | 0.61375 | 1.19134 | 172 |
| GO_SERINE_TYPE_ENDOPEPTIDASE_INHIBITOR_ACTIVITY | 0.00038 | 0.00640 | 0.49849 | 0.77388 | 1.41447 | 28 |
| GO_SEXUAL_REPRODUCTION | 0.00472 | 0.04480 | 0.40702 | 0.55748 | 1.10026 | 439 |
| GO_SIDE_OF_MEMBRANE | 0.00495 | 0.04573 | 0.40702 | 0.57741 | 1.12751 | 235 |
| GO_SIGNAL_TRANSDUCTION_BY_PROTEIN_PHOSPHORYLATION | 0.00224 | 0.02581 | 0.43171 | 0.55690 | 1.09975 | 500 |
| GO_SINGLE_FERTILIZATION | 0.00000 | 0.00002 | 0.67496 | 0.76307 | 1.43842 | 61 |
| GO_SINGLE_STRANDED_RNA_BINDING | 0.00001 | 0.00042 | 0.59333 | 0.71960 | 1.35864 | 66 |
| GO_SISTER_CHROMATID_SEGREGATION | 0.00042 | 0.00689 | 0.49849 | 0.60926 | 1.18209 | 170 |
| GO_SKIN_DEVELOPMENT | 0.00076 | 0.01140 | 0.47727 | 0.61452 | 1.18945 | 153 |
| GO_SMALL_MOLECULE_BIOSYNTHETIC_PROCESS | 0.00001 | 0.00021 | 0.61053 | 0.58728 | 1.15957 | 459 |
| GO_SMALL_NUCLEAR_RIBONUCLEOPROTEIN_COMPLEX | 0.00042 | 0.00689 | 0.49849 | 0.69542 | 1.30936 | 57 |
| GO_SMALL_RIBOSOMAL_SUBUNIT | 0.00000 | 0.00001 | 0.69013 | 0.75767 | 1.43100 | 67 |
| GO_SM_LIKE_PROTEIN_FAMILY_COMPLEX | 0.00145 | 0.01879 | 0.45506 | 0.66792 | 1.26149 | 67 |
| GO_SNORNA_BINDING | 0.00038 | 0.00646 | 0.49849 | 0.76224 | 1.39614 | 29 |
| GO_SPERM_EGG_RECOGNITION | 0.00000 | 0.00002 | 0.65944 | 0.87469 | 1.59087 | 26 |
| GO_SPINDLE_MIDZONE | 0.00361 | 0.03657 | 0.43171 | 0.73316 | 1.33859 | 27 |
| GO_SPINDLE_ORGANIZATION | 0.00272 | 0.02986 | 0.43171 | 0.60076 | 1.16362 | 159 |
| GO_SPLICEOSOMAL_COMPLEX | 0.00000 | 0.00008 | 0.62726 | 0.65256 | 1.26657 | 171 |
| GO_SPLICEOSOMAL_TRI_SNRNP_COMPLEX | 0.00168 | 0.02098 | 0.45506 | 0.73474 | 1.34835 | 30 |
| GO_STEM_CELL_DIFFERENTIATION | 0.00000 | 0.00000 | 0.82666 | 0.70006 | 1.35554 | 158 |
| GO_STEROID_BIOSYNTHETIC_PROCESS | 0.00465 | 0.04430 | 0.40702 | 0.61242 | 1.17992 | 125 |
| GO_STEROID_METABOLIC_PROCESS | 0.00313 | 0.03310 | 0.43171 | 0.59525 | 1.15748 | 185 |
| GO_STEROL_BIOSYNTHETIC_PROCESS | 0.00005 | 0.00120 | 0.55733 | 0.71876 | 1.35584 | 62 |
| GO_STEROL_METABOLIC_PROCESS | 0.00003 | 0.00073 | 0.57561 | 0.66485 | 1.27854 | 109 |
| GO_STRESS_GRANULE_ASSEMBLY | 0.00465 | 0.04430 | 0.40702 | 0.77288 | 1.38921 | 21 |
| GO_STRUCTURAL_CONSTITUENT_OF_RIBOSOME | 0.00000 | 0.00000 | | 0.76380 | 1.47839 | 153 |
| GO_STRUCTURAL_MOLECULE_ACTIVITY | 0.00000 | 0.00000 | | 0.71760 | 1.41436 | 391 |
| GO_SUPEROXIDE_METABOLIC_PROCESS | 0.00272 | 0.02986 | 0.43171 | 0.72028 | 1.32659 | 35 |
| GO_TELOMERASE_RNA_LOCALIZATION | 0.00054 | 0.00857 | 0.47727 | 0.81801 | 1.46148 | 19 |
| GO_TELOMERE_MAINTENANCE_VIA_TELOMERE_LENGTHENING | 0.00005 | 0.00138 | 0.55733 | 0.70715 | 1.33512 | 66 |
| GO_TELOMERE_ORGANIZATION | 0.00159 | 0.02016 | 0.45506 | 0.61666 | 1.18834 | 127 |
| GO_TERTIARY_GRANULE | 0.00008 | 0.00186 | 0.53843 | 0.66372 | 1.27005 | 91 |
| GO_TERTIARY_GRANULE_MEMBRANE | 0.00102 | 0.01453 | 0.45506 | 0.71940 | 1.33244 | 40 |
| GO_THREONINE_TYPE_ENDOPEPTIDASE_ACTIVITY | 0.00000 | 0.00000 | 0.71951 | 0.93456 | 1.65424 | 17 |
| GO_THREONINE_TYPE_PEPTIDASE_ACTIVITY | 0.00000 | 0.00004 | 0.65944 | 0.89807 | 1.61424 | 21 |
| GO_TISSUE_MORPHOGENESIS | 0.00000 | 0.00006 | 0.64355 | 0.59467 | 1.17150 | 380 |
| GO_TOXIN_TRANSPORT | 0.00064 | 0.00972 | 0.47727 | 0.77713 | 1.41343 | 26 |
| GO_TRANSCRIPTION_COUPLED_NUCLEOTIDE_EXCISION_REPAIR | 0.00231 | 0.02642 | 0.43171 | 0.65429 | 1.23840 | 71 |
| GO_TRANSFERRIN_TRANSPORT | 0.00141 | 0.01850 | 0.45506 | 0.75134 | 1.37179 | 27 |
| GO_TRANSLATIONAL_ELONGATION | 0.00000 | 0.00000 | | 0.75263 | 1.45028 | 124 |
| GO_TRANSLATIONAL_INITIATION | 0.00000 | 0.00000 | | 0.75139 | 1.45839 | 171 |
| GO_TRANSLATIONAL_TERMINATION | 0.00000 | 0.00000 | 0.81404 | 0.75110 | 1.44066 | 101 |
| GO_TRANSLATION_ELONGATION_FACTOR_ACTIVITY | 0.00390 | 0.03895 | 0.43171 | 0.80505 | 1.41260 | 15 |
| GO_TRANSLATION_FACTOR_ACTIVITY_RNA_BINDING | 0.00000 | 0.00000 | 0.74774 | 0.77244 | 1.46085 | 70 |
| GO_TRANSLATION_INITIATION_FACTOR_ACTIVITY | 0.00000 | 0.00011 | 0.62726 | 0.77773 | 1.44952 | 45 |
| GO_TRANSLATION_PREINITIATION_COMPLEX | 0.00342 | 0.03521 | 0.43171 | 0.79777 | 1.40694 | 16 |
| GO_TRANSLATION_REGULATOR_ACTIVITY | 0.00000 | 0.00000 | 0.83909 | 0.74000 | 1.42403 | 111 |
| GO_TRANSLATION_REGULATOR_ACTIVITY_NUCLEIC_ACID_BIND | 0.00000 | 0.00000 | 0.83909 | 0.76441 | 1.45959 | 87 |
| GO_TRANSMEMBRANE_RECEPTOR_PROTEIN_TYROSINE_KINASE | 0.00168 | 0.02098 | 0.45506 | 0.56326 | 1.11076 | 411 |
| GO_TRICARBOXYLIC_ACID_CYCLE | 0.00159 | 0.02016 | 0.45506 | 0.73558 | 1.35097 | 33 |
| GO_TRNA_TRANSPORT | 0.00342 | 0.03521 | 0.43171 | 0.71247 | 1.31220 | 35 |
| GO_TUMOR_NECROSIS_FACTOR_MEDIATED_SIGNALING_PATHW | 0.00000 | 0.00000 | 0.80122 | 0.73921 | 1.42138 | 108 |
| **Cluster 2** | | | | | | |
| pathway | pval | padj | log2err | ES | NES | size |
| GO_9PLUS2_MOTILE_CILIUM | 0.00174 | 0.03134 | 0.45506 | 0.70225 | 1.27654 | 49 |
| GO_ACTIVATION_OF_IMMUNE_RESPONSE | 0.00002 | 0.00065 | 0.59333 | 0.63663 | 1.21336 | 247 |
| GO_ACTIVATION_OF_INNATE_IMMUNE_RESPONSE | 0.00001 | 0.00063 | 0.59333 | 0.69571 | 1.29576 | 98 |
| GO_ACTIVE_ION_TRANSMEMBRANE_TRANSPORTER_ACTIVITY | 0.00178 | 0.03154 | 0.45506 | 0.65168 | 1.21532 | 102 |
| GO_ACTIVE_TRANSMEMBRANE_TRANSPORTER_ACTIVITY | 0.00191 | 0.03296 | 0.45506 | 0.61645 | 1.16494 | 162 |
| GO_ADP_BINDING | 0.00287 | 0.04513 | 0.43171 | 0.76611 | 1.35664 | 27 |
| GO_AEROBIC_RESPIRATION | 0.00006 | 0.00217 | 0.53843 | 0.71801 | 1.32306 | 70 |
| GO_AMIDE_BINDING | 0.00027 | 0.00749 | 0.49849 | 0.62300 | 1.18207 | 198 |
| GO_AMINE_METABOLIC_PROCESS | 0.00000 | 0.00000 | 0.77494 | 0.76681 | 1.42442 | 90 |
| GO_ANAPHASE_PROMOTING_COMPLEX_DEPENDENT_CATABOL | 0.00000 | 0.00000 | 0.73376 | 0.77782 | 1.43560 | 73 |

| | | | | | | |
|---|---|---|---|---|---|---|
| GO_ANTIGEN_BINDING | 0.00168 | 0.03061 | 0.45506 | 0.81490 | 1.41240 | 17 |
| GO_ANTIGEN_PROCESSING_AND_PRESENTATION | 0.00000 | 0.00000 | 0.71951 | 0.69725 | 1.31580 | 154 |
| GO_ANTIGEN_PROCESSING_AND_PRESENTATION_OF_EXOGENC | 0.00000 | 0.00005 | 0.65944 | 0.77796 | 1.42602 | 60 |
| GO_ANTIGEN_PROCESSING_AND_PRESENTATION_OF_PEPTIDE_ | 0.00000 | 0.00002 | 0.67496 | 0.69466 | 1.30630 | 138 |
| GO_ANTIGEN_PROCESSING_AND_PRESENTATION_OF_PEPTIDE_ | 0.00000 | 0.00005 | 0.65944 | 0.75531 | 1.39407 | 73 |
| GO_ANTIGEN_RECEPTOR_MEDIATED_SIGNALING_PATHWAY | 0.00001 | 0.00051 | 0.59333 | 0.66948 | 1.25963 | 140 |
| GO_ANTIMICROBIAL_HUMORAL_RESPONSE | 0.00044 | 0.01092 | 0.49849 | 0.76789 | 1.36649 | 32 |
| GO_ANTIOXIDANT_ACTIVITY | 0.00037 | 0.00953 | 0.49849 | 0.75598 | 1.36116 | 39 |
| GO_APOPTOTIC_SIGNALING_PATHWAY | 0.00108 | 0.02178 | 0.45506 | 0.58074 | 1.11422 | 402 |
| GO_ATP_BIOSYNTHETIC_PROCESS | 0.00010 | 0.00308 | 0.53843 | 0.83413 | 1.46663 | 23 |
| GO_ATP_METABOLIC_PROCESS | 0.00000 | 0.00000 | | 0.70572 | 1.34141 | 218 |
| GO_ATP_SYNTHESIS_COUPLED_ELECTRON_TRANSPORT | 0.00000 | 0.00000 | 0.77494 | 0.78343 | 1.45020 | 78 |
| GO_AZUROPHIL_GRANULE | 0.00100 | 0.02077 | 0.45506 | 0.65146 | 1.21651 | 107 |
| GO_BINDING_OF_SPERM_TO_ZONA_PELLUCIDA | 0.00004 | 0.00137 | 0.55733 | 0.87181 | 1.52029 | 19 |
| GO_BLOOD_MICROPARTICLE | 0.00018 | 0.00532 | 0.51885 | 0.75140 | 1.35858 | 43 |
| GO_CADHERIN_BINDING | 0.00000 | 0.00000 | 0.74774 | 0.66327 | 1.26494 | 257 |
| GO_CANONICAL_WNT_SIGNALING_PATHWAY | 0.00034 | 0.00890 | 0.49849 | 0.61893 | 1.17590 | 213 |
| GO_CARDIAC_MUSCLE_CELL_MEMBRANE_REPOLARIZATION | 0.00231 | 0.03796 | 0.43171 | 0.80925 | 1.40548 | 18 |
| GO_CATALYTIC_STEP_2_SPLICEOSOME | 0.00031 | 0.00820 | 0.49849 | 0.68678 | 1.27147 | 81 |
| GO_CATION_TRANSMEMBRANE_TRANSPORT | 0.00090 | 0.01933 | 0.47727 | 0.58158 | 1.11582 | 402 |
| GO_CAVEOLA | 0.00163 | 0.02997 | 0.45506 | 0.74182 | 1.33108 | 37 |
| GO_CELLULAR_AMINO_ACID_METABOLIC_PROCESS | 0.00143 | 0.02689 | 0.45506 | 0.60907 | 1.15694 | 212 |
| GO_CELLULAR_COMPONENT_DISASSEMBLY | 0.00000 | 0.00000 | 0.80122 | 0.63954 | 1.22788 | 429 |
| GO_CELLULAR_ION_HOMEOSTASIS | 0.00148 | 0.02776 | 0.45506 | 0.59025 | 1.13004 | 326 |
| GO_CELLULAR_KETONE_METABOLIC_PROCESS | 0.00001 | 0.00048 | 0.59333 | 0.66351 | 1.25249 | 156 |
| GO_CELLULAR_MONOVALENT_INORGANIC_CATION_HOMEOSTA | 0.00187 | 0.03269 | 0.45506 | 0.67172 | 1.23978 | 73 |
| GO_CELLULAR_NITROGEN_COMPOUND_CATABOLIC_PROCESS | 0.00000 | 0.00000 | | 0.67365 | 1.29466 | 451 |
| GO_CELLULAR_OXIDANT_DETOXIFICATION | 0.00037 | 0.00953 | 0.49849 | 0.72579 | 1.32062 | 50 |
| GO_CELLULAR_PROTEIN_COMPLEX_DISASSEMBLY | 0.00000 | 0.00000 | 0.73376 | 0.69191 | 1.30921 | 176 |
| GO_CELLULAR_RESPIRATION | 0.00000 | 0.00000 | 0.81404 | 0.72669 | 1.36974 | 147 |
| GO_CELLULAR_RESPONSE_TO_OXYGEN_LEVELS | 0.00000 | 0.00000 | 0.73376 | 0.69398 | 1.31106 | 160 |
| GO_CELLULAR_RESPONSE_TO_OXYGEN_RADICAL | 0.00168 | 0.03061 | 0.45506 | 0.81560 | 1.41650 | 18 |
| GO_CELL_ACTIVATION_INVOLVED_IN_IMMUNE_RESPONSE | 0.00000 | 0.00000 | 0.74774 | 0.63272 | 1.21455 | 420 |
| GO_CELL_ADHESION_MEDIATOR_ACTIVITY | 0.00183 | 0.03224 | 0.45506 | 0.72734 | 1.31283 | 41 |
| GO_CELL_ADHESION_MOLECULE_BINDING | 0.00000 | 0.00000 | 0.76146 | 0.64512 | 1.23563 | 346 |
| GO_CELL_BODY | 0.00163 | 0.02997 | 0.45506 | 0.59203 | 1.13277 | 299 |
| GO_CELL_CELL_JUNCTION | 0.00034 | 0.00886 | 0.49849 | 0.60639 | 1.15860 | 283 |
| GO_CELL_CELL_RECOGNITION | 0.00000 | 0.00011 | 0.64355 | 0.83451 | 1.49133 | 35 |
| GO_CELL_CELL_SIGNALING_BY_WNT | 0.00172 | 0.03110 | 0.45506 | 0.58652 | 1.12303 | 334 |
| GO_CELL_COMMUNICATION_INVOLVED_IN_CARDIAC_CONDUCT | 0.00331 | 0.04896 | 0.43171 | 0.73806 | 1.31198 | 30 |
| GO_CELL_CYCLE_G2_M_PHASE_TRANSITION | 0.00029 | 0.00782 | 0.49849 | 0.61541 | 1.17000 | 227 |
| GO_CELL_CYCLE_PHASE_TRANSITION | 0.00093 | 0.01964 | 0.47727 | 0.57548 | 1.10721 | 489 |
| GO_CELL_RECOGNITION | 0.00000 | 0.00008 | 0.64355 | 0.75538 | 1.38905 | 65 |
| GO_CELL_SUBSTRATE_JUNCTION | 0.00000 | 0.00000 | 0.81404 | 0.66567 | 1.27420 | 308 |
| GO_CELL_SURFACE | 0.00007 | 0.00238 | 0.53843 | 0.61276 | 1.17265 | 302 |
| GO_CHAPERONE_BINDING | 0.00024 | 0.00673 | 0.51885 | 0.68572 | 1.27012 | 82 |
| GO_CHAPERONE_COMPLEX | 0.00178 | 0.03154 | 0.45506 | 0.80216 | 1.40489 | 21 |
| GO_CONTRACTILE_FIBER | 0.00029 | 0.00782 | 0.49849 | 0.64798 | 1.22031 | 142 |
| GO_COTRANSLATIONAL_PROTEIN_TARGETING_TO_MEMBRANE | 0.00000 | 0.00000 | | 0.79318 | 1.47412 | 91 |
| GO_CYTOCHROME_COMPLEX | 0.00257 | 0.04100 | 0.43171 | 0.79261 | 1.38817 | 21 |
| GO_CYTOKINE_MEDIATED_SIGNALING_PATHWAY | 0.00000 | 0.00000 | 0.80122 | 0.64013 | 1.22810 | 392 |
| GO_CYTOPLASMIC_TRANSLATION | 0.00000 | 0.00015 | 0.62726 | 0.72163 | 1.33936 | 88 |
| GO_CYTOPLASMIC_TRANSLATIONAL_INITIATION | 0.00320 | 0.04826 | 0.43171 | 0.73910 | 1.31382 | 30 |
| GO_CYTOSOLIC_LARGE_RIBOSOMAL_SUBUNIT | 0.00000 | 0.00001 | 0.69013 | 0.80794 | 1.47035 | 51 |
| GO_CYTOSOLIC_RIBOSOME | 0.00000 | 0.00000 | | 0.79212 | 1.47482 | 96 |
| GO_CYTOSOLIC_SMALL_RIBOSOMAL_SUBUNIT | 0.00001 | 0.00054 | 0.59333 | 0.79124 | 1.42989 | 42 |
| GO_DETOXIFICATION | 0.00087 | 0.01887 | 0.47727 | 0.68579 | 1.26269 | 69 |
| GO_ELECTRON_TRANSFER_ACTIVITY | 0.00005 | 0.00177 | 0.55733 | 0.71672 | 1.32554 | 77 |
| GO_ELECTRON_TRANSPORT_CHAIN | 0.00000 | 0.00000 | 0.77494 | 0.72319 | 1.35865 | 133 |
| GO_EMBRYO_IMPLANTATION | 0.00313 | 0.04797 | 0.43171 | 0.73931 | 1.31421 | 30 |
| GO_ENDOPEPTIDASE_COMPLEX | 0.00000 | 0.00009 | 0.64355 | 0.75389 | 1.38631 | 65 |
| GO_ENDOPEPTIDASE_REGULATOR_ACTIVITY | 0.00019 | 0.00553 | 0.51885 | 0.70365 | 1.29558 | 69 |
| GO_ENERGY_DERIVATION_BY_OXIDATION_OF_ORGANIC_COMP | 0.00000 | 0.00000 | 0.78819 | 0.68963 | 1.30885 | 199 |
| GO_ENZYME_INHIBITOR_ACTIVITY | 0.00283 | 0.04466 | 0.43171 | 0.61053 | 1.15740 | 188 |
| GO_ESTABLISHMENT_OF_PROTEIN_LOCALIZATION_TO_CHROMC | 0.00283 | 0.04466 | 0.43171 | 0.79339 | 1.38728 | 20 |
| GO_ESTABLISHMENT_OF_PROTEIN_LOCALIZATION_TO_ENDOPL/ | 0.00000 | 0.00000 | | 0.77669 | 1.44855 | 103 |
| GO_ESTABLISHMENT_OF_PROTEIN_LOCALIZATION_TO_MEMBR/ | 0.00000 | 0.00000 | | 0.70341 | 1.34123 | 256 |
| GO_ESTABLISHMENT_OF_PROTEIN_LOCALIZATION_TO_MITOCH | 0.00143 | 0.02689 | 0.45506 | 0.73125 | 1.31989 | 41 |
| GO_ESTABLISHMENT_OF_PROTEIN_LOCALIZATION_TO_ORGANE | 0.00000 | 0.00000 | | 0.67237 | 1.29187 | 446 |
| GO_ESTABLISHMENT_OF_PROTEIN_LOCALIZATION_TO_TELOME | 0.00132 | 0.02536 | 0.45506 | 0.83719 | 1.44608 | 15 |

| | | | | | | |
|---|---|---|---|---|---|---|
| GO_ESTABLISHMENT_OF_TISSUE_POLARITY | 0.00001 | 0.00039 | 0.59333 | 0.71446 | 1.32902 | 93 |
| GO_EUKARYOTIC_TRANSLATION_INITIATION_FACTOR_3_COMPL | 0.00331 | 0.04896 | 0.43171 | 0.81491 | 1.40759 | 15 |
| GO_FC_EPSILON_RECEPTOR_SIGNALING_PATHWAY | 0.00001 | 0.00037 | 0.61053 | 0.70905 | 1.31855 | 92 |
| GO_FC_RECEPTOR_SIGNALING_PATHWAY | 0.00001 | 0.00048 | 0.59333 | 0.67453 | 1.26870 | 139 |
| GO_FERROUS_IRON_BINDING | 0.00313 | 0.04797 | 0.43171 | 0.80352 | 1.39552 | 18 |
| GO_FERTILIZATION | 0.00000 | 0.00024 | 0.61053 | 0.73591 | 1.36056 | 76 |
| GO_FICOLIN_1_RICH_GRANULE | 0.00000 | 0.00000 | 0.71951 | 0.73591 | 1.37239 | 102 |
| GO_GENERATION_OF_PRECURSOR_METABOLITES_AND_ENERGY | 0.00000 | 0.00000 | | 0.66570 | 1.27560 | 360 |
| GO_GLUCOSE_CATABOLIC_PROCESS | 0.00015 | 0.00447 | 0.51885 | 0.82104 | 1.44763 | 24 |
| GO_GLUCOSE_METABOLIC_PROCESS | 0.00231 | 0.03796 | 0.43171 | 0.62502 | 1.17423 | 133 |
| GO_GLYCOLYTIC_PROCESS_THROUGH_FRUCTOSE_6_PHOSPHATE | 0.00020 | 0.00570 | 0.51885 | 0.84024 | 1.46919 | 20 |
| GO_GTPASE_ACTIVITY | 0.00071 | 0.01617 | 0.47727 | 0.61635 | 1.16963 | 197 |
| GO_GUANYL_NUCLEOTIDE_BINDING | 0.00052 | 0.01239 | 0.47727 | 0.61185 | 1.16436 | 232 |
| GO_HEMATOPOIETIC_PROGENITOR_CELL_DIFFERENTIATION | 0.00003 | 0.00114 | 0.57561 | 0.67692 | 1.26847 | 122 |
| GO_HEMATOPOIETIC_STEM_CELL_DIFFERENTIATION | 0.00000 | 0.00012 | 0.62726 | 0.74304 | 1.36797 | 68 |
| GO_HEXOSE_CATABOLIC_PROCESS | 0.00050 | 0.01200 | 0.47727 | 0.75646 | 1.35541 | 36 |
| GO_HUMORAL_IMMUNE_RESPONSE | 0.00320 | 0.04826 | 0.43171 | 0.65714 | 1.21636 | 80 |
| GO_IMMUNE_RESPONSE_REGULATING_SIGNALING_PATHWAY | 0.00002 | 0.00078 | 0.57561 | 0.63533 | 1.20866 | 229 |
| GO_INNATE_IMMUNE_RESPONSE | 0.00023 | 0.00644 | 0.51885 | 0.59250 | 1.13650 | 400 |
| GO_INNATE_IMMUNE_RESPONSE_ACTIVATING_SIGNAL_TRANSD | 0.00003 | 0.00106 | 0.57561 | 0.70891 | 1.31226 | 78 |
| GO_INNER_MITOCHONDRIAL_MEMBRANE_PROTEIN_COMPLEX | 0.00000 | 0.00000 | | 0.78696 | 1.46185 | 90 |
| GO_INORGANIC_ION_TRANSMEMBRANE_TRANSPORT | 0.00007 | 0.00240 | 0.53843 | 0.60097 | 1.15300 | 387 |
| GO_INTERACTION_WITH_SYMBIONT | 0.00242 | 0.03926 | 0.43171 | 0.71324 | 1.29426 | 47 |
| GO_INTERLEUKIN_1_MEDIATED_SIGNALING_PATHWAY | 0.00000 | 0.00012 | 0.64355 | 0.73623 | 1.36278 | 80 |
| GO_LARGE_RIBOSOMAL_SUBUNIT | 0.00000 | 0.00000 | | 0.78427 | 1.46366 | 105 |
| GO_LEUKOCYTE_MEDIATED_IMMUNITY | 0.00000 | 0.00000 | 0.78819 | 0.63347 | 1.21768 | 458 |
| GO_LIPID_DROPLET | 0.00216 | 0.03631 | 0.43171 | 0.69151 | 1.26355 | 57 |
| GO_LUMENAL_SIDE_OF_MEMBRANE | 0.00198 | 0.03367 | 0.43171 | 0.81255 | 1.41122 | 18 |
| GO_LYSOSOMAL_LUMEN | 0.00010 | 0.00315 | 0.53843 | 0.73068 | 1.33351 | 55 |
| GO_MEMBRANE_REGION | 0.00124 | 0.02448 | 0.45506 | 0.61812 | 1.17067 | 183 |
| GO_MEMBRANE_REPOLARIZATION | 0.00076 | 0.01690 | 0.47727 | 0.79438 | 1.40062 | 24 |
| GO_MIDBRAIN_DEVELOPMENT | 0.00316 | 0.04818 | 0.43171 | 0.68060 | 1.24214 | 54 |
| GO_MITOCHONDRIAL_ELECTRON_TRANSPORT_NADH_TO_UBIQ | 0.00001 | 0.00030 | 0.61053 | 0.79069 | 1.43187 | 45 |
| GO_MITOCHONDRIAL_GENE_EXPRESSION | 0.00000 | 0.00001 | 0.70498 | 0.68953 | 1.30054 | 151 |
| GO_MITOCHONDRIAL_LARGE_RIBOSOMAL_SUBUNIT | 0.00000 | 0.00024 | 0.61053 | 0.76888 | 1.40182 | 53 |
| GO_MITOCHONDRIAL_MATRIX | 0.00000 | 0.00000 | 0.77494 | 0.64107 | 1.22922 | 378 |
| GO_MITOCHONDRIAL_MEMBRANE_ORGANIZATION | 0.00007 | 0.00238 | 0.53843 | 0.68438 | 1.27423 | 96 |
| GO_MITOCHONDRIAL_PROTEIN_COMPLEX | 0.00000 | 0.00000 | | 0.75346 | 1.43125 | 206 |
| GO_MITOCHONDRIAL_RESPIRATORY_CHAIN_COMPLEX_ASSEMB | 0.00011 | 0.00334 | 0.53843 | 0.69420 | 1.28685 | 84 |
| GO_MITOCHONDRIAL_TRANSLATION | 0.00000 | 0.00002 | 0.67496 | 0.70430 | 1.32263 | 129 |
| GO_MITOCHONDRIAL_TRANSLATIONAL_TERMINATION | 0.00000 | 0.00001 | 0.70498 | 0.74532 | 1.38332 | 88 |
| GO_MITOCHONDRIAL_TRANSMEMBRANE_TRANSPORT | 0.00011 | 0.00331 | 0.53843 | 0.71602 | 1.31732 | 66 |
| GO_MITOCHONDRIAL_TRANSPORT | 0.00008 | 0.00257 | 0.53843 | 0.63435 | 1.20415 | 200 |
| GO_MITOCHONDRION_ORGANIZATION | 0.00000 | 0.00000 | 0.80122 | 0.64288 | 1.23393 | 413 |
| GO_MONOCARBOXYLIC_ACID_METABOLIC_PROCESS | 0.00069 | 0.01588 | 0.47727 | 0.59090 | 1.13288 | 372 |
| GO_MONOSACCHARIDE_CATABOLIC_PROCESS | 0.00030 | 0.00799 | 0.49849 | 0.73576 | 1.33239 | 45 |
| GO_MONOSACCHARIDE_METABOLIC_PROCESS | 0.00168 | 0.03061 | 0.45506 | 0.61479 | 1.16409 | 182 |
| GO_MONOVALENT_INORGANIC_CATION_HOMEOSTASIS | 0.00178 | 0.03154 | 0.45506 | 0.65858 | 1.22256 | 89 |
| GO_MONOVALENT_INORGANIC_CATION_TRANSMEMBRANE_TR | 0.00000 | 0.00021 | 0.62726 | 0.68050 | 1.28155 | 142 |
| GO_MONOVALENT_INORGANIC_CATION_TRANSPORT | 0.00002 | 0.00068 | 0.57561 | 0.62977 | 1.19951 | 238 |
| GO_MORPHOGENESIS_OF_AN_EPITHELIUM | 0.00000 | 0.00021 | 0.62726 | 0.62322 | 1.19318 | 328 |
| GO_MORPHOGENESIS_OF_A_POLARIZED_EPITHELIUM | 0.00000 | 0.00002 | 0.69013 | 0.72354 | 1.35136 | 108 |
| GO_MRNA_BINDING | 0.00000 | 0.00010 | 0.64355 | 0.64596 | 1.23060 | 239 |
| GO_MRNA_PROCESSING | 0.00000 | 0.00000 | 0.73376 | 0.62953 | 1.20848 | 424 |
| GO_MYELOID_LEUKOCYTE_ACTIVATION | 0.00000 | 0.00000 | 0.74774 | 0.63675 | 1.22153 | 399 |
| GO_MYELOID_LEUKOCYTE_MEDIATED_IMMUNITY | 0.00000 | 0.00000 | 0.81404 | 0.65174 | 1.24860 | 358 |
| GO_M_BAND | 0.00081 | 0.01787 | 0.47727 | 0.84574 | 1.46084 | 15 |
| GO_NADH_DEHYDROGENASE_ACTIVITY | 0.00003 | 0.00129 | 0.55733 | 0.79834 | 1.43045 | 36 |
| GO_NADH_DEHYDROGENASE_COMPLEX | 0.00001 | 0.00048 | 0.59333 | 0.80113 | 1.44244 | 39 |
| GO_NADH_DEHYDROGENASE_COMPLEX_ASSEMBLY | 0.00002 | 0.00069 | 0.57561 | 0.75194 | 1.37233 | 54 |
| GO_NADH_METABOLIC_PROCESS | 0.00004 | 0.00143 | 0.55733 | 0.84258 | 1.49027 | 26 |
| GO_NAD_METABOLIC_PROCESS | 0.00007 | 0.00232 | 0.53843 | 0.79709 | 1.42011 | 33 |
| GO_NCRNA_METABOLIC_PROCESS | 0.00004 | 0.00140 | 0.55733 | 0.60406 | 1.15893 | 387 |
| GO_NCRNA_PROCESSING | 0.00000 | 0.00011 | 0.64355 | 0.62521 | 1.19699 | 331 |
| GO_NEGATIVE_REGULATION_OF_CANONICAL_WNT_SIGNALING | 0.00000 | 0.00021 | 0.62726 | 0.69493 | 1.30147 | 119 |
| GO_NEGATIVE_REGULATION_OF_CELL_CYCLE_G2_M_PHASE_TR | 0.00006 | 0.00203 | 0.55733 | 0.69717 | 1.29396 | 88 |
| GO_NEGATIVE_REGULATION_OF_CELL_CYCLE_PHASE_TRANSITI | 0.00031 | 0.00825 | 0.49849 | 0.62224 | 1.18095 | 199 |
| GO_NEGATIVE_REGULATION_OF_CELL_CYCLE_PROCESS | 0.00011 | 0.00341 | 0.53843 | 0.61170 | 1.16676 | 260 |
| GO_NEGATIVE_REGULATION_OF_HYDROLASE_ACTIVITY | 0.00002 | 0.00090 | 0.57561 | 0.63033 | 1.19980 | 233 |
| GO_NEGATIVE_REGULATION_OF_MITOTIC_CELL_CYCLE | 0.00070 | 0.01603 | 0.47727 | 0.60330 | 1.15042 | 255 |

| | | | | | | |
|---|---|---|---|---|---|---|
| GO_NEGATIVE_REGULATION_OF_MRNA_METABOLIC_PROCESS | 0.00327 | 0.04877 | 0.43171 | 0.67102 | 1.23100 | 61 |
| GO_NEGATIVE_REGULATION_OF_PEPTIDASE_ACTIVITY | 0.00007 | 0.00238 | 0.53843 | 0.67749 | 1.26565 | 109 |
| GO_NEGATIVE_REGULATION_OF_PROTEIN_CONTAINING_COMP | 0.00076 | 0.01690 | 0.47727 | 0.66856 | 1.24400 | 94 |
| GO_NEGATIVE_REGULATION_OF_PROTEOLYSIS | 0.00213 | 0.03589 | 0.43171 | 0.61886 | 1.17147 | 178 |
| GO_NEGATIVE_REGULATION_OF_WNT_SIGNALING_PATHWAY | 0.00001 | 0.00055 | 0.59333 | 0.67296 | 1.26532 | 137 |
| GO_NEURAL_NUCLEUS_DEVELOPMENT | 0.00161 | 0.02990 | 0.45506 | 0.73649 | 1.31616 | 35 |
| GO_NIK_NF_KAPPAB_SIGNALING | 0.00000 | 0.00001 | 0.69013 | 0.72029 | 1.34603 | 110 |
| GO_NON_CANONICAL_WNT_SIGNALING_PATHWAY | 0.00001 | 0.00039 | 0.59333 | 0.69555 | 1.29938 | 109 |
| GO_NUCLEAR_PERIPHERY | 0.00187 | 0.03269 | 0.45506 | 0.63950 | 1.19603 | 113 |
| GO_NUCLEAR_TRANSCRIBED_MRNA_CATABOLIC_PROCESS | 0.00000 | 0.00000 | 0.78819 | 0.69595 | 1.31934 | 188 |
| GO_NUCLEAR_TRANSCRIBED_MRNA_CATABOLIC_PROCESS_NON | 0.00000 | 0.00000 | | 0.76459 | 1.43028 | 114 |
| GO_NUCLEAR_TRANSPORT | 0.00132 | 0.02536 | 0.45506 | 0.59751 | 1.14155 | 279 |
| GO_NUCLEOBASE_CONTAINING_SMALL_MOLECULE_METABOLI( | 0.00092 | 0.01952 | 0.47727 | 0.58965 | 1.13054 | 376 |
| GO_NUCLEOSIDE_DIPHOSPHATE_METABOLIC_PROCESS | 0.00191 | 0.03296 | 0.45506 | 0.64494 | 1.20363 | 105 |
| GO_NUCLEOSIDE_PHOSPHATE_BIOSYNTHETIC_PROCESS | 0.00130 | 0.02532 | 0.45506 | 0.62740 | 1.18336 | 151 |
| GO_ORGANELLAR_RIBOSOME | 0.00000 | 0.00003 | 0.67496 | 0.74595 | 1.38287 | 83 |
| GO_ORGANELLE_ENVELOPE_LUMEN | 0.00170 | 0.03086 | 0.45506 | 0.67208 | 1.24155 | 75 |
| GO_ORGANELLE_INNER_MEMBRANE | 0.00000 | 0.00000 | | 0.68472 | 1.31390 | 405 |
| GO_ORGANIC_CYCLIC_COMPOUND_CATABOLIC_PROCESS | 0.00000 | 0.00000 | | 0.67278 | 1.29349 | 467 |
| GO_OXIDATIVE_PHOSPHORYLATION | 0.00000 | 0.00000 | | 0.77229 | 1.44023 | 102 |
| GO_OXIDOREDUCTASE_ACTIVITY | 0.00000 | 0.00000 | | 0.64717 | 1.24319 | 444 |
| GO_OXIDOREDUCTASE_ACTIVITY_ACTING_ON_A_HEME_GROUP | 0.00139 | 0.02652 | 0.45506 | 0.81066 | 1.41365 | 19 |
| GO_OXIDOREDUCTASE_ACTIVITY_ACTING_ON_NAD_P_H | 0.00001 | 0.00056 | 0.59333 | 0.73317 | 1.34993 | 69 |
| GO_OXIDOREDUCTASE_ACTIVITY_ACTING_ON_NAD_P_H_QUIN( | 0.00010 | 0.00308 | 0.53843 | 0.76431 | 1.38166 | 44 |
| GO_OXIDOREDUCTASE_ACTIVITY_ACTING_ON_PEROXIDE_AS_A( | 0.00146 | 0.02750 | 0.45506 | 0.78205 | 1.38320 | 26 |
| GO_OXIDOREDUCTASE_COMPLEX | 0.00000 | 0.00002 | 0.67496 | 0.75372 | 1.39514 | 80 |
| GO_PEPTIDASE_COMPLEX | 0.00006 | 0.00212 | 0.53843 | 0.70107 | 1.29855 | 82 |
| GO_PEPTIDASE_REGULATOR_ACTIVITY | 0.00049 | 0.01187 | 0.47727 | 0.66920 | 1.24310 | 90 |
| GO_PEPTIDE_BINDING | 0.00015 | 0.00443 | 0.51885 | 0.65367 | 1.23102 | 142 |
| GO_PERINUCLEAR_REGION_OF_CYTOPLASM | 0.00239 | 0.03886 | 0.43171 | 0.57054 | 1.09708 | 469 |
| GO_PIGMENT_GRANULE | 0.00050 | 0.01200 | 0.47727 | 0.68309 | 1.26482 | 79 |
| GO_PLASMA_LIPOPROTEIN_PARTICLE_CLEARANCE | 0.00220 | 0.03673 | 0.43171 | 0.74543 | 1.32564 | 31 |
| GO_PLASMA_MEMBRANE_RAFT | 0.00090 | 0.01933 | 0.47727 | 0.70577 | 1.28676 | 53 |
| GO_PLATELET_DEGRANULATION | 0.00216 | 0.03631 | 0.43171 | 0.67494 | 1.24112 | 65 |
| GO_POLYSOMAL_RIBOSOME | 0.00001 | 0.00064 | 0.59333 | 0.83569 | 1.48329 | 29 |
| GO_POLYSOME | 0.00000 | 0.00001 | 0.70498 | 0.77957 | 1.43221 | 63 |
| GO_POSITIVE_REGULATION_OF_CANONICAL_WNT_SIGNALING_ | 0.00002 | 0.00079 | 0.57561 | 0.69458 | 1.29531 | 102 |
| GO_POSITIVE_REGULATION_OF_CELLULAR_AMIDE_METABOLIC_ | 0.00017 | 0.00498 | 0.51885 | 0.65474 | 1.22870 | 127 |
| GO_POSITIVE_REGULATION_OF_CELLULAR_PROTEIN_LOCALIZAT | 0.00095 | 0.01990 | 0.47727 | 0.60612 | 1.15372 | 233 |
| GO_POSITIVE_REGULATION_OF_DEFENSE_RESPONSE | 0.00000 | 0.00009 | 0.64355 | 0.66258 | 1.25626 | 190 |
| GO_POSITIVE_REGULATION_OF_ESTABLISHMENT_OF_PROTEIN_ | 0.00006 | 0.00217 | 0.53843 | 0.61932 | 1.18052 | 250 |
| GO_POSITIVE_REGULATION_OF_IMMUNE_RESPONSE | 0.00000 | 0.00003 | 0.67496 | 0.63205 | 1.21007 | 326 |
| GO_POSITIVE_REGULATION_OF_IMMUNE_SYSTEM_PROCESS | 0.00001 | 0.00051 | 0.59333 | 0.59708 | 1.14869 | 479 |
| GO_POSITIVE_REGULATION_OF_INNATE_IMMUNE_RESPONSE | 0.00000 | 0.00012 | 0.62726 | 0.69499 | 1.30476 | 128 |
| GO_POSITIVE_REGULATION_OF_ORGANELLE_ORGANIZATION | 0.00042 | 0.01044 | 0.49849 | 0.58771 | 1.12852 | 433 |
| GO_POSITIVE_REGULATION_OF_PROTEIN_LOCALIZATION_TO_N | 0.00179 | 0.03168 | 0.45506 | 0.68920 | 1.26504 | 62 |
| GO_POSITIVE_REGULATION_OF_RESPONSE_TO_BIOTIC_STIMULI | 0.00000 | 0.00012 | 0.62726 | 0.67712 | 1.27630 | 147 |
| GO_POSITIVE_REGULATION_OF_RESPONSE_TO_EXTERNAL_STIM | 0.00019 | 0.00549 | 0.51885 | 0.61397 | 1.17069 | 256 |
| GO_POSITIVE_REGULATION_OF_TELOMERASE_RNA_LOCALIZATI | 0.00028 | 0.00780 | 0.49849 | 0.86612 | 1.49605 | 15 |
| GO_POSITIVE_REGULATION_OF_TELOMERE_MAINTENANCE_VIA | 0.00239 | 0.03886 | 0.43171 | 0.76419 | 1.35770 | 28 |
| GO_POSITIVE_REGULATION_OF_TRANSLATION | 0.00003 | 0.00111 | 0.57561 | 0.68469 | 1.27829 | 106 |
| GO_POSITIVE_REGULATION_OF_VIRAL_LIFE_CYCLE | 0.00294 | 0.04582 | 0.43171 | 0.69231 | 1.25847 | 49 |
| GO_POSITIVE_REGULATION_OF_VIRAL_PROCESS | 0.00028 | 0.00758 | 0.49849 | 0.68428 | 1.26896 | 87 |
| GO_POSITIVE_REGULATION_OF_WNT_SIGNALING_PATHWAY | 0.00001 | 0.00052 | 0.59333 | 0.68009 | 1.27418 | 121 |
| GO_POSTTRANSCRIPTIONAL_REGULATION_OF_GENE_EXPRESSI( | 0.00000 | 0.00000 | | 0.64843 | 1.24809 | 499 |
| GO_PRERIBOSOME | 0.00069 | 0.01588 | 0.47727 | 0.69044 | 1.27124 | 69 |
| GO_PROTEASOME_ACCESSORY_COMPLEX | 0.00002 | 0.00092 | 0.57561 | 0.85729 | 1.50735 | 23 |
| GO_PROTEASOME_CORE_COMPLEX | 0.00121 | 0.02383 | 0.45506 | 0.82293 | 1.42923 | 18 |
| GO_PROTEIN_CONTAINING_COMPLEX_DISASSEMBLY | 0.00000 | 0.00000 | 0.74774 | 0.66184 | 1.26253 | 261 |
| GO_PROTEIN_FOLDING | 0.00001 | 0.00037 | 0.61053 | 0.65782 | 1.24471 | 176 |
| GO_PROTEIN_FOLDING_CHAPERONE | 0.00108 | 0.02178 | 0.45506 | 0.81160 | 1.42143 | 21 |
| GO_PROTEIN_IMPORT | 0.00008 | 0.00248 | 0.53843 | 0.64473 | 1.21586 | 149 |
| GO_PROTEIN_IMPORT_INTO_MITOCHONDRIAL_MATRIX | 0.00095 | 0.01990 | 0.47727 | 0.82920 | 1.44013 | 18 |
| GO_PROTEIN_INSERTION_INTO_MEMBRANE | 0.00294 | 0.04582 | 0.43171 | 0.69229 | 1.25843 | 49 |
| GO_PROTEIN_LOCALIZATION_TO_ENDOPLASMIC_RETICULUM | 0.00000 | 0.00000 | | 0.76025 | 1.42464 | 122 |
| GO_PROTEIN_LOCALIZATION_TO_MEMBRANE | 0.00000 | 0.00000 | | 0.66406 | 1.27517 | 434 |
| GO_PROTEIN_LOCALIZATION_TO_MITOCHONDRION | 0.00005 | 0.00184 | 0.55733 | 0.67294 | 1.26029 | 119 |
| GO_PROTEIN_LOCALIZATION_TO_NUCLEUS | 0.00025 | 0.00709 | 0.49849 | 0.62698 | 1.19025 | 201 |
| GO_PROTEIN_MODIFICATION_BY_SMALL_PROTEIN_REMOVAL | 0.00194 | 0.03332 | 0.45506 | 0.60585 | 1.15125 | 214 |
| GO_PROTEIN_TARGETING | 0.00000 | 0.00000 | | 0.68521 | 1.31230 | 342 |

| | | | | | | |
|---|---|---|---|---|---|---|
| GO_PROTEIN_TARGETING_TO_MEMBRANE | 0.00000 | 0.00000 | 0.82666 | 0.72593 | 1.36992 | 154 |
| GO_PROTEIN_TARGETING_TO_MITOCHONDRION | 0.00027 | 0.00749 | 0.49849 | 0.68610 | 1.27191 | 83 |
| GO_PROTEIN_TRANSMEMBRANE_IMPORT_INTO_INTRACELLULA | 0.00009 | 0.00294 | 0.53843 | 0.79134 | 1.40988 | 33 |
| GO_PROTEIN_TRANSMEMBRANE_TRANSPORT | 0.00026 | 0.00718 | 0.49849 | 0.72109 | 1.31603 | 54 |
| GO_PROTON_TRANSMEMBRANE_TRANSPORT | 0.00001 | 0.00036 | 0.61053 | 0.71385 | 1.32384 | 86 |
| GO_PROTON_TRANSMEMBRANE_TRANSPORTER_ACTIVITY | 0.00017 | 0.00502 | 0.51885 | 0.71098 | 1.30805 | 66 |
| GO_PURINE_CONTAINING_COMPOUND_BIOSYNTHETIC_PROCES | 0.00110 | 0.02200 | 0.45506 | 0.64117 | 1.20025 | 117 |
| GO_PURINE_CONTAINING_COMPOUND_METABOLIC_PROCESS | 0.00119 | 0.02362 | 0.45506 | 0.59839 | 1.14322 | 279 |
| GO_PURINE_NUCLEOSIDE_TRIPHOSPHATE_METABOLIC_PROCES | 0.00055 | 0.01322 | 0.47727 | 0.73862 | 1.33523 | 44 |
| GO_PYRUVATE_METABOLIC_PROCESS | 0.00058 | 0.01377 | 0.47727 | 0.65883 | 1.22874 | 103 |
| GO_REGULATION_OF_ACTIN_FILAMENT_ORGANIZATION | 0.00132 | 0.02536 | 0.45506 | 0.61541 | 1.16373 | 166 |
| GO_REGULATION_OF_ANIMAL_ORGAN_MORPHOGENESIS | 0.00000 | 0.00021 | 0.62726 | 0.67338 | 1.27012 | 150 |
| GO_REGULATION_OF_CELLULAR_AMIDE_METABOLIC_PROCESS | 0.00001 | 0.00056 | 0.59333 | 0.61227 | 1.17260 | 338 |
| GO_REGULATION_OF_CELLULAR_AMINE_METABOLIC_PROCESS | 0.00000 | 0.00002 | 0.69013 | 0.78473 | 1.44038 | 62 |
| GO_REGULATION_OF_CELLULAR_AMINO_ACID_METABOLIC_PR( | 0.00000 | 0.00002 | 0.69013 | 0.79578 | 1.45235 | 54 |
| GO_REGULATION_OF_CELLULAR_KETONE_METABOLIC_PROCESS | 0.00001 | 0.00052 | 0.59333 | 0.68283 | 1.27868 | 118 |
| GO_REGULATION_OF_CELLULAR_PROTEIN_LOCALIZATION | 0.00045 | 0.01110 | 0.49849 | 0.59136 | 1.13390 | 378 |
| GO_REGULATION_OF_CELL_CYCLE_G2_M_PHASE_TRANSITION | 0.00011 | 0.00341 | 0.53843 | 0.63750 | 1.20666 | 181 |
| GO_REGULATION_OF_CELL_CYCLE_PHASE_TRANSITION | 0.00086 | 0.01855 | 0.47727 | 0.58641 | 1.12386 | 363 |
| GO_REGULATION_OF_CYSTEINE_TYPE_ENDOPEPTIDASE_ACTIVIT | 0.00038 | 0.00972 | 0.49849 | 0.63968 | 1.20568 | 146 |
| GO_REGULATION_OF_CYTOSKELETON_ORGANIZATION | 0.00309 | 0.04776 | 0.43171 | 0.58340 | 1.11737 | 349 |
| GO_REGULATION_OF_DEFENSE_RESPONSE | 0.00012 | 0.00350 | 0.53843 | 0.60233 | 1.15362 | 349 |
| GO_REGULATION_OF_DNA_TEMPLATED_TRANSCRIPTION_IN_RE | 0.00005 | 0.00168 | 0.55733 | 0.69623 | 1.29549 | 94 |
| GO_REGULATION_OF_HEMATOPOIETIC_PROGENITOR_CELL_DIF | 0.00000 | 0.00010 | 0.64355 | 0.75398 | 1.38715 | 66 |
| GO_REGULATION_OF_HEMATOPOIETIC_STEM_CELL_DIFFERENT | 0.00000 | 0.00021 | 0.62726 | 0.76740 | 1.40545 | 59 |
| GO_REGULATION_OF_HEMOPOIESIS | 0.00141 | 0.02679 | 0.45506 | 0.59995 | 1.14474 | 265 |
| GO_REGULATION_OF_IMMUNE_RESPONSE | 0.00003 | 0.00112 | 0.57561 | 0.60159 | 1.15501 | 429 |
| GO_REGULATION_OF_INNATE_IMMUNE_RESPONSE | 0.00004 | 0.00136 | 0.55733 | 0.65323 | 1.23654 | 178 |
| GO_REGULATION_OF_INTRINSIC_APOPTOTIC_SIGNALING_PATH\ | 0.00198 | 0.03367 | 0.43171 | 0.63885 | 1.19482 | 113 |
| GO_REGULATION_OF_MITOTIC_CELL_CYCLE | 0.00046 | 0.01130 | 0.49849 | 0.57976 | 1.11574 | 493 |
| GO_REGULATION_OF_MORPHOGENESIS_OF_AN_EPITHELIUM | 0.00001 | 0.00036 | 0.61053 | 0.68991 | 1.29208 | 119 |
| GO_REGULATION_OF_MRNA_CATABOLIC_PROCESS | 0.00000 | 0.00019 | 0.62726 | 0.67021 | 1.26697 | 163 |
| GO_REGULATION_OF_MRNA_METABOLIC_PROCESS | 0.00000 | 0.00001 | 0.70498 | 0.65292 | 1.24687 | 270 |
| GO_REGULATION_OF_MULTI_ORGANISM_PROCESS | 0.00042 | 0.01044 | 0.49849 | 0.75477 | 1.35697 | 38 |
| GO_REGULATION_OF_OXIDATIVE_PHOSPHORYLATION | 0.00049 | 0.01187 | 0.47727 | 0.80421 | 1.41795 | 24 |
| GO_REGULATION_OF_PEPTIDASE_ACTIVITY | 0.00000 | 0.00009 | 0.64355 | 0.64678 | 1.23127 | 234 |
| GO_REGULATION_OF_PEPTIDE_TRANSPORT | 0.00309 | 0.04776 | 0.43171 | 0.57551 | 1.10443 | 408 |
| GO_REGULATION_OF_PLASMA_LIPOPROTEIN_PARTICLE_LEVELS | 0.00008 | 0.00251 | 0.53843 | 0.75660 | 1.37014 | 45 |
| GO_REGULATION_OF_PROTEIN_CONTAINING_COMPLEX_ASSEM | 0.00316 | 0.04818 | 0.43171 | 0.59211 | 1.13216 | 287 |
| GO_REGULATION_OF_PROTEIN_LOCALIZATION_TO_NUCLEUS | 0.00011 | 0.00334 | 0.53843 | 0.69012 | 1.28110 | 89 |
| GO_REGULATION_OF_PROTEOLYSIS | 0.00005 | 0.00174 | 0.55733 | 0.59864 | 1.14974 | 439 |
| GO_REGULATION_OF_RESPONSE_TO_BIOTIC_STIMULUS | 0.00008 | 0.00244 | 0.53843 | 0.62488 | 1.18915 | 232 |
| GO_REGULATION_OF_RNA_SPLICING | 0.00075 | 0.01682 | 0.47727 | 0.64601 | 1.20986 | 119 |
| GO_REGULATION_OF_STEM_CELL_DIFFERENTIATION | 0.00001 | 0.00030 | 0.61053 | 0.71658 | 1.32890 | 86 |
| GO_REGULATION_OF_SUPRAMOLECULAR_FIBER_ORGANIZATIOI | 0.00097 | 0.02021 | 0.47727 | 0.60693 | 1.15389 | 227 |
| GO_REGULATION_OF_SYMBIOTIC_PROCESS | 0.00003 | 0.00124 | 0.55733 | 0.65988 | 1.24528 | 154 |
| GO_REGULATION_OF_TELOMERE_MAINTENANCE_VIA_TELOME | 0.00167 | 0.03055 | 0.45506 | 0.71793 | 1.30119 | 46 |
| GO_REGULATION_OF_TRANSCRIPTION_FROM_RNA_POLYMERA: | 0.00001 | 0.00031 | 0.61053 | 0.74345 | 1.36778 | 66 |
| GO_REGULATION_OF_VIRAL_GENOME_REPLICATION | 0.00098 | 0.02046 | 0.45506 | 0.68754 | 1.26314 | 63 |
| GO_REGULATION_OF_VIRAL_LIFE_CYCLE | 0.00198 | 0.03367 | 0.43171 | 0.64928 | 1.21003 | 99 |
| GO_REGULATION_OF_WNT_SIGNALING_PATHWAY | 0.00324 | 0.04870 | 0.43171 | 0.59825 | 1.13888 | 234 |
| GO_RESPIRASOME | 0.00000 | 0.00000 | 0.76146 | 0.78771 | 1.45386 | 73 |
| GO_RESPIRATORY_CHAIN_COMPLEX | 0.00000 | 0.00000 | 0.73376 | 0.80017 | 1.46794 | 61 |
| GO_RESPIRATORY_ELECTRON_TRANSPORT_CHAIN | 0.00000 | 0.00000 | 0.78819 | 0.76278 | 1.41846 | 92 |
| GO_RESPONSE_TO_ALCOHOL | 0.00056 | 0.01338 | 0.47727 | 0.65163 | 1.21897 | 114 |
| GO_RESPONSE_TO_CALCIUM_ION | 0.00228 | 0.03776 | 0.43171 | 0.66813 | 1.23314 | 73 |
| GO_RESPONSE_TO_INORGANIC_SUBSTANCE | 0.00001 | 0.00032 | 0.61053 | 0.62111 | 1.18902 | 321 |
| GO_RESPONSE_TO_INTERLEUKIN_1 | 0.00000 | 0.00006 | 0.65944 | 0.70359 | 1.31769 | 119 |
| GO_RESPONSE_TO_ISCHEMIA | 0.00224 | 0.03725 | 0.43171 | 0.74511 | 1.32507 | 31 |
| GO_RESPONSE_TO_METAL_ION | 0.00030 | 0.00799 | 0.49849 | 0.61968 | 1.17664 | 203 |
| GO_RESPONSE_TO_NUTRIENT | 0.00034 | 0.00890 | 0.49849 | 0.67010 | 1.24764 | 96 |
| GO_RESPONSE_TO_OXYGEN_LEVELS | 0.00000 | 0.00000 | 0.73376 | 0.66271 | 1.26352 | 253 |
| GO_RESPONSE_TO_OXYGEN_RADICAL | 0.00327 | 0.04877 | 0.43171 | 0.78900 | 1.37959 | 20 |
| GO_RESPONSE_TO_TOXIC_SUBSTANCE | 0.00194 | 0.03332 | 0.45506 | 0.63370 | 1.19026 | 132 |
| GO_RESPONSE_TO_TUMOR_NECROSIS_FACTOR | 0.00005 | 0.00169 | 0.55733 | 0.64378 | 1.21798 | 175 |
| GO_RESPONSE_TO_VIRUS | 0.00294 | 0.04582 | 0.43171 | 0.60789 | 1.15062 | 181 |
| GO_RESPONSE_TO_VITAMIN | 0.00078 | 0.01739 | 0.47727 | 0.72665 | 1.31383 | 43 |
| GO_RETINA_HOMEOSTASIS | 0.00012 | 0.00366 | 0.51885 | 0.78719 | 1.40083 | 32 |
| GO_RIBONUCLEOPROTEIN_COMPLEX_BINDING | 0.00009 | 0.00297 | 0.53843 | 0.67615 | 1.26283 | 108 |
| GO_RIBONUCLEOPROTEIN_COMPLEX_BIOGENESIS | 0.00000 | 0.00000 | | 0.67099 | 1.28658 | 381 |

| | | | | | | |
|---|---|---|---|---|---|---|
| GO_RIBONUCLEOPROTEIN_COMPLEX_SUBUNIT_ORGANIZATION | 0.00000 | 0.00002 | 0.69013 | 0.68131 | 1.28870 | 169 |
| GO_RIBONUCLEOPROTEIN_GRANULE | 0.00194 | 0.03332 | 0.45506 | 0.61462 | 1.16403 | 183 |
| GO_RIBONUCLEOSIDE_TRIPHOSPHATE_METABOLIC_PROCESS | 0.00220 | 0.03673 | 0.43171 | 0.72408 | 1.30694 | 41 |
| GO_RIBOSE_PHOSPHATE_BIOSYNTHETIC_PROCESS | 0.00213 | 0.03589 | 0.43171 | 0.63756 | 1.19361 | 116 |
| GO_RIBOSE_PHOSPHATE_METABOLIC_PROCESS | 0.00130 | 0.02532 | 0.45506 | 0.59797 | 1.14098 | 264 |
| GO_RIBOSOMAL_LARGE_SUBUNIT_BIOGENESIS | 0.00010 | 0.00309 | 0.53843 | 0.71756 | 1.32014 | 66 |
| GO_RIBOSOMAL_SMALL_SUBUNIT_ASSEMBLY | 0.00135 | 0.02590 | 0.45506 | 0.82133 | 1.42645 | 18 |
| GO_RIBOSOMAL_SMALL_SUBUNIT_BIOGENESIS | 0.00132 | 0.02536 | 0.45506 | 0.68696 | 1.26385 | 66 |
| GO_RIBOSOMAL_SUBUNIT | 0.00000 | 0.00000 | | 0.77285 | 1.46205 | 171 |
| GO_RIBOSOME | 0.00000 | 0.00000 | | 0.76556 | 1.45409 | 204 |
| GO_RIBOSOME_ASSEMBLY | 0.00002 | 0.00075 | 0.57561 | 0.74828 | 1.36729 | 57 |
| GO_RIBOSOME_BIOGENESIS | 0.00000 | 0.00000 | 0.77494 | 0.66867 | 1.27681 | 269 |
| GO_RNA_CATABOLIC_PROCESS | 0.00000 | 0.00000 | | 0.68929 | 1.32008 | 341 |
| GO_RNA_DEPENDENT_DNA_BIOSYNTHETIC_PROCESS | 0.00191 | 0.03296 | 0.45506 | 0.69321 | 1.26785 | 58 |
| GO_RNA_LOCALIZATION | 0.00075 | 0.01682 | 0.47727 | 0.61524 | 1.16817 | 202 |
| GO_RNA_SPLICING | 0.00000 | 0.00001 | 0.69013 | 0.63085 | 1.20949 | 371 |
| GO_RNA_SPLICING_VIA_TRANSESTERIFICATION_REACTIONS | 0.00000 | 0.00002 | 0.67496 | 0.64231 | 1.22913 | 303 |
| GO_ROUGH_ENDOPLASMIC_RETICULUM | 0.00150 | 0.02802 | 0.45506 | 0.70089 | 1.28364 | 59 |
| GO_RRNA_BINDING | 0.00001 | 0.00039 | 0.59333 | 0.75941 | 1.38598 | 54 |
| GO_RRNA_METABOLIC_PROCESS | 0.00000 | 0.00003 | 0.67496 | 0.65867 | 1.25107 | 204 |
| GO_SARCOLEMMA | 0.00012 | 0.00355 | 0.53843 | 0.71539 | 1.31617 | 66 |
| GO_SARCOMERE_ORGANIZATION | 0.00082 | 0.01801 | 0.47727 | 0.81696 | 1.43081 | 21 |
| GO_SCAFFOLD_PROTEIN_BINDING | 0.00316 | 0.04818 | 0.43171 | 0.72404 | 1.29390 | 35 |
| GO_SCF_DEPENDENT_PROTEASOMAL_UBIQUITIN_DEPENDENT_ | 0.00000 | 0.00009 | 0.64355 | 0.74732 | 1.38165 | 76 |
| GO_SECRETORY_GRANULE | 0.00000 | 0.00000 | | 0.64132 | 1.23389 | 481 |
| GO_SECRETORY_GRANULE_MEMBRANE | 0.00179 | 0.03168 | 0.45506 | 0.61470 | 1.16277 | 172 |
| GO_SIGNAL_TRANSDUCTION_BY_PROTEIN_PHOSPHORYLATION | 0.00007 | 0.00243 | 0.53843 | 0.58745 | 1.13024 | 489 |
| GO_SINGLE_FERTILIZATION | 0.00000 | 0.00000 | 0.73376 | 0.78722 | 1.44626 | 63 |
| GO_SINGLE_STRANDED_RNA_BINDING | 0.00257 | 0.04100 | 0.43171 | 0.67235 | 1.23522 | 63 |
| GO_SMALL_RIBOSOMAL_SUBUNIT | 0.00000 | 0.00015 | 0.62726 | 0.74818 | 1.37756 | 69 |
| GO_SNORNA_BINDING | 0.00102 | 0.02101 | 0.45506 | 0.76208 | 1.35467 | 30 |
| GO_SODIUM_ION_TRANSMEMBRANE_TRANSPORT | 0.00046 | 0.01123 | 0.49849 | 0.68410 | 1.26668 | 79 |
| GO_SODIUM_ION_TRANSMEMBRANE_TRANSPORTER_ACTIVITY | 0.00143 | 0.02689 | 0.45506 | 0.69423 | 1.26702 | 54 |
| GO_SPERM_EGG_RECOGNITION | 0.00001 | 0.00031 | 0.61053 | 0.86544 | 1.53069 | 26 |
| GO_SPINDLE_ORGANIZATION | 0.00080 | 0.01773 | 0.47727 | 0.63835 | 1.20217 | 142 |
| GO_SPLICEOSOMAL_COMPLEX | 0.00002 | 0.00090 | 0.57561 | 0.65344 | 1.23523 | 165 |
| GO_STEM_CELL_DIFFERENTIATION | 0.00000 | 0.00012 | 0.62726 | 0.67138 | 1.26789 | 159 |
| GO_STRUCTURAL_CONSTITUENT_OF_CYTOSKELETON | 0.00008 | 0.00248 | 0.53843 | 0.72088 | 1.32247 | 61 |
| GO_STRUCTURAL_CONSTITUENT_OF_RIBOSOME | 0.00000 | 0.00000 | | 0.79707 | 1.50271 | 148 |
| GO_STRUCTURAL_MOLECULE_ACTIVITY | 0.00000 | 0.00000 | | 0.73627 | 1.41266 | 389 |
| GO_SUBSTANTIA_NIGRA_DEVELOPMENT | 0.00279 | 0.04431 | 0.43171 | 0.74166 | 1.31837 | 30 |
| GO_SUPEROXIDE_METABOLIC_PROCESS | 0.00062 | 0.01447 | 0.47727 | 0.75195 | 1.34378 | 35 |
| GO_SUPRAMOLECULAR_FIBER_ORGANIZATION | 0.00327 | 0.04877 | 0.43171 | 0.57744 | 1.10842 | 423 |
| GO_TELOMERASE_RNA_LOCALIZATION | 0.00108 | 0.02178 | 0.45506 | 0.81491 | 1.42107 | 19 |
| GO_TELOMERE_MAINTENANCE_VIA_TELOMERE_LENGTHENING | 0.00076 | 0.01705 | 0.47727 | 0.69308 | 1.27347 | 64 |
| GO_THREONINE_TYPE_ENDOPEPTIDASE_ACTIVITY | 0.00121 | 0.02383 | 0.45506 | 0.81277 | 1.41734 | 19 |
| GO_THREONINE_TYPE_PEPTIDASE_ACTIVITY | 0.00029 | 0.00782 | 0.49849 | 0.81174 | 1.43123 | 24 |
| GO_TISSUE_MORPHOGENESIS | 0.00002 | 0.00086 | 0.57561 | 0.60988 | 1.16933 | 376 |
| GO_TRANSLATIONAL_ELONGATION | 0.00000 | 0.00000 | 0.76146 | 0.73191 | 1.37087 | 120 |
| GO_TRANSLATIONAL_INITIATION | 0.00000 | 0.00000 | | 0.76532 | 1.44760 | 169 |
| GO_TRANSLATIONAL_TERMINATION | 0.00000 | 0.00000 | 0.73376 | 0.74004 | 1.37944 | 101 |
| GO_TRANSLATION_FACTOR_ACTIVITY_RNA_BINDING | 0.00001 | 0.00043 | 0.59333 | 0.72995 | 1.34388 | 68 |
| GO_TRANSLATION_INITIATION_FACTOR_ACTIVITY | 0.00061 | 0.01431 | 0.47727 | 0.73655 | 1.33148 | 44 |
| GO_TRANSLATION_PREINITIATION_COMPLEX | 0.00242 | 0.03926 | 0.43171 | 0.81289 | 1.40745 | 16 |
| GO_TRANSLATION_REGULATOR_ACTIVITY | 0.00001 | 0.00055 | 0.59333 | 0.69148 | 1.29178 | 109 |
| GO_TRANSLATION_REGULATOR_ACTIVITY_NUCLEIC_ACID_BIND | 0.00001 | 0.00058 | 0.59333 | 0.71355 | 1.32299 | 85 |
| GO_TUMOR_NECROSIS_FACTOR_MEDIATED_SIGNALING_PATHW | 0.00000 | 0.00007 | 0.65944 | 0.70657 | 1.32148 | 113 |
| GO_T_CELL_RECEPTOR_SIGNALING_PATHWAY | 0.00000 | 0.00013 | 0.62726 | 0.69241 | 1.29726 | 121 |
| GO_U2_TYPE_SPLICEOSOMAL_COMPLEX | 0.00294 | 0.04582 | 0.43171 | 0.65470 | 1.21388 | 85 |
| GO_UBIQUITIN_LIKE_PROTEIN_LIGASE_BINDING | 0.00001 | 0.00062 | 0.59333 | 0.62839 | 1.19740 | 242 |
| GO_UNFOLDED_PROTEIN_BINDING | 0.00003 | 0.00125 | 0.55733 | 0.70289 | 1.30750 | 93 |
| GO_VACUOLAR_LUMEN | 0.00000 | 0.00007 | 0.65944 | 0.71505 | 1.33285 | 101 |
| GO_VESICLE_LUMEN | 0.00000 | 0.00000 | 0.81404 | 0.69445 | 1.31800 | 199 |
| GO_VIRAL_GENE_EXPRESSION | 0.00000 | 0.00000 | | 0.71915 | 1.36062 | 174 |
| GO_VIRAL_GENOME_REPLICATION | 0.00033 | 0.00858 | 0.49849 | 0.68607 | 1.27017 | 81 |
| GO_VIRAL_LIFE_CYCLE | 0.00001 | 0.00054 | 0.59333 | 0.63764 | 1.21242 | 224 |
| HP_11_PAIRS_OF_RIBS | 0.00008 | 0.00244 | 0.53843 | 0.86238 | 1.50384 | 19 |
| HP_ABNORMALITY_OF_ACID_BASE_HOMEOSTASIS | 0.00031 | 0.00825 | 0.49849 | 0.61011 | 1.16313 | 254 |
| HP_ABNORMALITY_OF_CARDIOVASCULAR_SYSTEM_ELECTROPH | 0.00231 | 0.03796 | 0.43171 | 0.59414 | 1.13296 | 255 |
| HP_ABNORMALITY_OF_THE_MITOCHONDRION | 0.00086 | 0.01855 | 0.47727 | 0.64320 | 1.20404 | 117 |

| | | | | | | |
|---|---|---|---|---|---|---|
| HP_ABNORMALITY_OF_THE_OPTIC_DISC | 0.00013 | 0.00371 | 0.51885 | 0.59276 | 1.13796 | 422 |
| HP_ABNORMALITY_OF_THE_OPTIC_NERVE | 0.00007 | 0.00238 | 0.53843 | 0.59015 | 1.13525 | 478 |
| HP_ABNORMAL_ABDOMEN_MORPHOLOGY | 0.00040 | 0.01018 | 0.49849 | 0.58596 | 1.12502 | 427 |
| HP_ABNORMAL_ACTIVITY_OF_MITOCHONDRIAL_RESPIRATORY_ | 0.00102 | 0.02101 | 0.45506 | 0.68859 | 1.26324 | 61 |
| HP_ABNORMAL_CELLULAR_PHENOTYPE | 0.00298 | 0.04628 | 0.43171 | 0.58760 | 1.12476 | 320 |
| HP_ABNORMAL_CIRCULATING_CARBOHYDRATE_CONCENTRATIC | 0.00239 | 0.03886 | 0.43171 | 0.71172 | 1.28993 | 46 |
| HP_ABNORMAL_CNS_MYELINATION | 0.00002 | 0.00086 | 0.57561 | 0.65337 | 1.23579 | 168 |
| HP_ABNORMAL_CSF_METABOLITE_LEVEL | 0.00006 | 0.00212 | 0.53843 | 0.70732 | 1.30967 | 79 |
| HP_ABNORMAL_ERYTHROCYTE_MORPHOLOGY | 0.00003 | 0.00100 | 0.57561 | 0.60931 | 1.16754 | 360 |
| HP_ABNORMAL_IMMUNE_SYSTEM_MORPHOLOGY | 0.00106 | 0.02169 | 0.45506 | 0.58845 | 1.12652 | 330 |
| HP_ABNORMAL_MYELINATION | 0.00044 | 0.01085 | 0.49849 | 0.59637 | 1.14186 | 324 |
| HP_ACUTE_LEUKEMIA | 0.00339 | 0.04994 | 0.43171 | 0.66652 | 1.22611 | 67 |
| HP_ANEMIA_OF_INADEQUATE_PRODUCTION | 0.00110 | 0.02200 | 0.45506 | 0.65653 | 1.22354 | 99 |
| HP_APHASIA | 0.00003 | 0.00105 | 0.57561 | 0.82834 | 1.47024 | 29 |
| HP_BLINDNESS | 0.00135 | 0.02590 | 0.45506 | 0.62341 | 1.17910 | 167 |
| HP_CACHEXIA | 0.00176 | 0.03154 | 0.45506 | 0.73179 | 1.31564 | 38 |
| HP_CEREBRAL_ISCHEMIA | 0.00231 | 0.03796 | 0.43171 | 0.71452 | 1.29166 | 44 |
| HP_DECREASED_ACTIVITY_OF_MITOCHONDRIAL_COMPLEX_I | 0.00257 | 0.04100 | 0.43171 | 0.73839 | 1.31554 | 33 |
| HP_DYSPHASIA | 0.00074 | 0.01674 | 0.47727 | 0.74961 | 1.34314 | 36 |
| HP_DYSTONIA | 0.00083 | 0.01814 | 0.47727 | 0.59849 | 1.14319 | 278 |
| HP_ECHOLALIA | 0.00111 | 0.02230 | 0.45506 | 0.78904 | 1.39557 | 26 |
| HP_EMOTIONAL_LABILITY | 0.00065 | 0.01499 | 0.47727 | 0.67273 | 1.25141 | 93 |
| HP_ENCEPHALOPATHY | 0.00257 | 0.04100 | 0.43171 | 0.61623 | 1.16559 | 169 |
| HP_ESOPHAGEAL_VARIX | 0.00092 | 0.01952 | 0.47727 | 0.84342 | 1.45684 | 15 |
| HP_FETAL_DISTRESS | 0.00320 | 0.04826 | 0.43171 | 0.73913 | 1.31387 | 30 |
| HP_GONOSOMAL_INHERITANCE | 0.00065 | 0.01515 | 0.47727 | 0.64424 | 1.21007 | 132 |
| HP_HEADACHE | 0.00178 | 0.03154 | 0.45506 | 0.62045 | 1.17288 | 165 |
| HP_HEPATOMEGALY | 0.00033 | 0.00878 | 0.49849 | 0.59841 | 1.14567 | 323 |
| HP_HYPERTROPHIC_CARDIOMYOPATHY | 0.00108 | 0.02178 | 0.45506 | 0.61673 | 1.16905 | 187 |
| HP_INCREASED_CSF_LACTATE | 0.00004 | 0.00139 | 0.55733 | 0.71607 | 1.31831 | 68 |
| HP_INCREASED_SERUM_LACTATE | 0.00117 | 0.02333 | 0.45506 | 0.63874 | 1.19973 | 132 |
| HP_INTRAUTERINE_GROWTH_RETARDATION | 0.00250 | 0.04035 | 0.43171 | 0.58379 | 1.11768 | 331 |
| HP_LACTIC_ACIDOSIS | 0.00046 | 0.01123 | 0.49849 | 0.64508 | 1.21154 | 130 |
| HP_LETHARGY | 0.00065 | 0.01499 | 0.47727 | 0.65717 | 1.22564 | 103 |
| HP_LEUKODYSTROPHY | 0.00002 | 0.00068 | 0.57561 | 0.71453 | 1.32286 | 81 |
| HP_MACROCYTIC_ANEMIA | 0.00202 | 0.03421 | 0.43171 | 0.72744 | 1.30783 | 38 |
| HP_MIGRAINE | 0.00040 | 0.01018 | 0.49849 | 0.69197 | 1.27715 | 73 |
| HP_MISSING_RIBS | 0.00061 | 0.01431 | 0.47727 | 0.80723 | 1.42333 | 25 |
| HP_MUSCLE_ABNORMALITY_RELATED_TO_MITOCHONDRIAL_DY | 0.00272 | 0.04325 | 0.43171 | 0.69440 | 1.26350 | 50 |
| HP_MUTISM | 0.00001 | 0.00051 | 0.59333 | 0.81730 | 1.45441 | 32 |
| HP_NAUSEA_AND_VOMITING | 0.00327 | 0.04877 | 0.43171 | 0.59278 | 1.13017 | 253 |
| HP_OPTIC_DISC_PALLOR | 0.00191 | 0.03296 | 0.45506 | 0.66153 | 1.22636 | 83 |
| HP_OPTIC_NERVE_COLOBOMA | 0.00040 | 0.01018 | 0.49849 | 0.85010 | 1.47188 | 16 |
| HP_OPTIC_NEUROPATHY | 0.00128 | 0.02512 | 0.45506 | 0.74790 | 1.33093 | 32 |
| HP_PROGRESSIVE_CEREBELLAR_ATAXIA | 0.00320 | 0.04826 | 0.43171 | 0.66762 | 1.22924 | 69 |
| HP_PROXIMAL_TUBULOPATHY | 0.00331 | 0.04896 | 0.43171 | 0.73249 | 1.30349 | 32 |
| HP_PTOSIS | 0.00313 | 0.04797 | 0.43171 | 0.57766 | 1.10826 | 396 |
| HP_REDUCED_CONSCIOUSNESS_CONFUSION | 0.00253 | 0.04084 | 0.43171 | 0.60744 | 1.15214 | 195 |
| HP_REDUCED_VISUAL_ACUITY | 0.00091 | 0.01946 | 0.47727 | 0.59868 | 1.14334 | 276 |
| HP_RENAL_TUBULAR_DYSFUNCTION | 0.00085 | 0.01848 | 0.47727 | 0.68290 | 1.26154 | 75 |
| HP_RESPIRATORY_FAILURE | 0.00159 | 0.02965 | 0.45506 | 0.64138 | 1.19955 | 113 |
| HP_RESPIRATORY_INSUFFICIENCY | 0.00001 | 0.00051 | 0.59333 | 0.61917 | 1.18496 | 313 |
| HP_SENSORY_NEUROPATHY | 0.00098 | 0.02046 | 0.45506 | 0.65935 | 1.22961 | 102 |
| HP_X_LINKED_RECESSIVE_INHERITANCE | 0.00163 | 0.02997 | 0.45506 | 0.65410 | 1.21769 | 95 |
| **Cluster 3** | | | | | | |
| pathway | pval | padj | log2err | ES | NES | size |
| GO_ACTIVATION_OF_IMMUNE_RESPONSE | 0.00000 | 0.00000 | 0.77494 | 0.63038 | 1.27979 | 267 |
| GO_ACTIVATION_OF_INNATE_IMMUNE_RESPONSE | 0.00000 | 0.00000 | | 0.74540 | 1.47786 | 104 |
| GO_AEROBIC_ELECTRON_TRANSPORT_CHAIN | 0.00152 | 0.02557 | 0.45506 | 0.80649 | 1.45095 | 15 |
| GO_ALTERNATIVE_MRNA_SPLICING_VIA_SPLICEOSOME | 0.00141 | 0.02397 | 0.45506 | 0.63953 | 1.25089 | 71 |
| GO_AMINE_METABOLIC_PROCESS | 0.00000 | 0.00000 | | 0.78221 | 1.54345 | 89 |
| GO_ANAPHASE_PROMOTING_COMPLEX_DEPENDENT_CATABOL | 0.00000 | 0.00000 | | 0.79004 | 1.55117 | 80 |
| GO_ANATOMICAL_STRUCTURE_HOMEOSTASIS | 0.00013 | 0.00344 | 0.51885 | 0.57262 | 1.16464 | 298 |
| GO_ANTIGEN_PROCESSING_AND_PRESENTATION | 0.00000 | 0.00000 | 0.73376 | 0.65696 | 1.32141 | 167 |
| GO_ANTIGEN_PROCESSING_AND_PRESENTATION_OF_EXOGENC | 0.00000 | 0.00000 | 0.80122 | 0.78244 | 1.52543 | 66 |
| GO_ANTIGEN_PROCESSING_AND_PRESENTATION_OF_PEPTIDE_ | 0.00000 | 0.00000 | 0.70498 | 0.66294 | 1.32948 | 151 |
| GO_ANTIGEN_PROCESSING_AND_PRESENTATION_OF_PEPTIDE_ | 0.00000 | 0.00000 | 0.71951 | 0.72798 | 1.42934 | 80 |
| GO_ANTIGEN_RECEPTOR_MEDIATED_SIGNALING_PATHWAY | 0.00000 | 0.00003 | 0.65944 | 0.64358 | 1.29292 | 158 |
| GO_APOPTOTIC_SIGNALING_PATHWAY | 0.00172 | 0.02830 | 0.45506 | 0.54793 | 1.11885 | 409 |
| GO_ATPASE_ACTIVITY | 0.00053 | 0.01054 | 0.47727 | 0.56352 | 1.14705 | 323 |

| | | | | | | |
|---|---|---|---|---|---|---|
| GO_ATP_BIOSYNTHETIC_PROCESS | 0.00324 | 0.04749 | 0.43171 | 0.74862 | 1.38602 | 23 |
| GO_ATP_METABOLIC_PROCESS | 0.00000 | 0.00000 | | 0.67543 | 1.36617 | 218 |
| GO_ATP_SYNTHESIS_COUPLED_ELECTRON_TRANSPORT | 0.00000 | 0.00002 | 0.67496 | 0.70841 | 1.38973 | 78 |
| GO_AZUROPHIL_GRANULE | 0.00039 | 0.00843 | 0.49849 | 0.62704 | 1.24489 | 108 |
| GO_BINDING_OF_SPERM_TO_ZONA_PELLUCIDA | 0.00032 | 0.00701 | 0.49849 | 0.78250 | 1.46249 | 26 |
| GO_BLOOD_MICROPARTICLE | 0.00050 | 0.01002 | 0.47727 | 0.71075 | 1.36621 | 42 |
| GO_CADHERIN_BINDING | 0.00000 | 0.00000 | 0.73376 | 0.62804 | 1.27298 | 246 |
| GO_CARBOHYDRATE_CATABOLIC_PROCESS | 0.00002 | 0.00058 | 0.57561 | 0.63824 | 1.27564 | 138 |
| GO_CATALYTIC_ACTIVITY_ACTING_ON_RNA | 0.00047 | 0.00966 | 0.49849 | 0.56760 | 1.15532 | 319 |
| GO_CATALYTIC_STEP_2_SPLICEOSOME | 0.00000 | 0.00000 | 0.71951 | 0.72455 | 1.42403 | 83 |
| GO_CELLULAR_AMINO_ACID_METABOLIC_PROCESS | 0.00030 | 0.00678 | 0.49849 | 0.58252 | 1.17924 | 230 |
| GO_CELLULAR_COMPONENT_DISASSEMBLY | 0.00000 | 0.00000 | | 0.61921 | 1.26607 | 439 |
| GO_CELLULAR_KETONE_METABOLIC_PROCESS | 0.00001 | 0.00035 | 0.59333 | 0.63473 | 1.27358 | 154 |
| GO_CELLULAR_NITROGEN_COMPOUND_CATABOLIC_PROCESS | 0.00000 | 0.00000 | | 0.66426 | 1.35925 | 471 |
| GO_CELLULAR_PROTEIN_COMPLEX_DISASSEMBLY | 0.00000 | 0.00000 | 0.77494 | 0.66245 | 1.33432 | 184 |
| GO_CELLULAR_RESPIRATION | 0.00001 | 0.00035 | 0.59333 | 0.63710 | 1.27647 | 145 |
| GO_CELLULAR_RESPONSE_TO_HEAT | 0.00000 | 0.00015 | 0.61053 | 0.67619 | 1.33841 | 98 |
| GO_CELLULAR_RESPONSE_TO_OXYGEN_LEVELS | 0.00000 | 0.00000 | 0.71951 | 0.65726 | 1.32273 | 172 |
| GO_CELL_ACTIVATION_INVOLVED_IN_IMMUNE_RESPONSE | 0.00000 | 0.00001 | 0.69013 | 0.58560 | 1.19651 | 423 |
| GO_CELL_ADHESION_MOLECULE_BINDING | 0.00000 | 0.00001 | 0.67496 | 0.59731 | 1.21725 | 341 |
| GO_CELL_CELL_RECOGNITION | 0.00048 | 0.00981 | 0.49849 | 0.72094 | 1.38451 | 41 |
| GO_CELL_CYCLE_G2_M_PHASE_TRANSITION | 0.00001 | 0.00021 | 0.61053 | 0.60465 | 1.22437 | 234 |
| GO_CELL_DIVISION | 0.00000 | 0.00008 | 0.62726 | 0.57799 | 1.18272 | 473 |
| GO_CELL_FATE_DETERMINATION | 0.00071 | 0.01347 | 0.47727 | 0.79295 | 1.44736 | 19 |
| GO_CELL_RECOGNITION | 0.00038 | 0.00812 | 0.49849 | 0.65841 | 1.28929 | 74 |
| GO_CELL_SUBSTRATE_JUNCTION | 0.00000 | 0.00000 | 0.77494 | 0.62391 | 1.26994 | 319 |
| GO_CHAPERONE_COMPLEX | 0.00001 | 0.00041 | 0.59333 | 0.84820 | 1.56751 | 22 |
| GO_CHROMOSOMAL_REGION | 0.00000 | 0.00001 | 0.69013 | 0.61076 | 1.24220 | 299 |
| GO_CHROMOSOME_CENTROMERIC_REGION | 0.00000 | 0.00011 | 0.62726 | 0.63225 | 1.27319 | 175 |
| GO_CHROMOSOME_SEGREGATION | 0.00001 | 0.00048 | 0.59333 | 0.59229 | 1.20346 | 282 |
| GO_CHROMOSOME_TELOMERIC_REGION | 0.00339 | 0.04943 | 0.43171 | 0.59780 | 1.19236 | 127 |
| GO_CONDENSED_CHROMOSOME | 0.00005 | 0.00141 | 0.55733 | 0.60746 | 1.22406 | 187 |
| GO_CONDENSED_CHROMOSOME_CENTROMERIC_REGION | 0.00023 | 0.00544 | 0.51885 | 0.63837 | 1.26631 | 105 |
| GO_COTRANSLATIONAL_PROTEIN_TARGETING_TO_MEMBRANE | 0.00000 | 0.00000 | | 0.84111 | 1.66456 | 95 |
| GO_CYTOCHROME_COMPLEX | 0.00224 | 0.03488 | 0.43171 | 0.75699 | 1.40151 | 23 |
| GO_CYTOKINESIS | 0.00113 | 0.01981 | 0.45506 | 0.60814 | 1.21298 | 127 |
| GO_CYTOKINE_MEDIATED_SIGNALING_PATHWAY | 0.00000 | 0.00010 | 0.62726 | 0.58036 | 1.18510 | 407 |
| GO_CYTOPLASMIC_TRANSLATION | 0.00000 | 0.00000 | 0.73376 | 0.71701 | 1.41827 | 94 |
| GO_CYTOPLASMIC_TRANSLATIONAL_INITIATION | 0.00003 | 0.00096 | 0.55733 | 0.77341 | 1.46330 | 31 |
| GO_CYTOSKELETON_DEPENDENT_CYTOKINESIS | 0.00216 | 0.03406 | 0.43171 | 0.63758 | 1.24924 | 75 |
| GO_CYTOSOLIC_LARGE_RIBOSOMAL_SUBUNIT | 0.00000 | 0.00000 | | 0.84029 | 1.62784 | 53 |
| GO_CYTOSOLIC_RIBOSOME | 0.00000 | 0.00000 | | 0.83299 | 1.64826 | 96 |
| GO_CYTOSOLIC_SMALL_RIBOSOMAL_SUBUNIT | 0.00000 | 0.00000 | 0.77494 | 0.83760 | 1.61005 | 42 |
| GO_DISORDERED_DOMAIN_SPECIFIC_BINDING | 0.00145 | 0.02440 | 0.45506 | 0.73464 | 1.38108 | 28 |
| GO_DNA_BIOSYNTHETIC_PROCESS | 0.00013 | 0.00339 | 0.51885 | 0.61524 | 1.23481 | 155 |
| GO_DNA_CONFORMATION_CHANGE | 0.00038 | 0.00812 | 0.49849 | 0.58187 | 1.17906 | 241 |
| GO_DNA_PACKAGING | 0.00224 | 0.03488 | 0.43171 | 0.59366 | 1.18635 | 136 |
| GO_DNA_REPAIR | 0.00003 | 0.00100 | 0.55733 | 0.56417 | 1.15408 | 459 |
| GO_DNA_REPLICATION | 0.00054 | 0.01069 | 0.47727 | 0.58164 | 1.17812 | 236 |
| GO_DNA_TEMPLATED_TRANSCRIPTION_ELONGATION | 0.00167 | 0.02783 | 0.45506 | 0.61803 | 1.22533 | 104 |
| GO_DNA_TEMPLATED_TRANSCRIPTION_TERMINATION | 0.00037 | 0.00811 | 0.49849 | 0.66748 | 1.30389 | 70 |
| GO_ELECTRON_TRANSFER_ACTIVITY | 0.00057 | 0.01131 | 0.47727 | 0.64970 | 1.27429 | 77 |
| GO_ELECTRON_TRANSPORT_CHAIN | 0.00000 | 0.00004 | 0.65944 | 0.65861 | 1.31473 | 132 |
| GO_ENDOPEPTIDASE_COMPLEX | 0.00000 | 0.00002 | 0.67496 | 0.73843 | 1.44057 | 67 |
| GO_ENERGY_DERIVATION_BY_OXIDATION_OF_ORGANIC_COMP | 0.00001 | 0.00022 | 0.61053 | 0.61553 | 1.24204 | 198 |
| GO_ESTABLISHMENT_OF_PROTEIN_LOCALIZATION_TO_ENDOPL/ | 0.00000 | 0.00000 | | 0.82384 | 1.63561 | 108 |
| GO_ESTABLISHMENT_OF_PROTEIN_LOCALIZATION_TO_MEMBR/ | 0.00000 | 0.00000 | | 0.68886 | 1.39835 | 268 |
| GO_ESTABLISHMENT_OF_PROTEIN_LOCALIZATION_TO_ORGANE | 0.00000 | 0.00000 | | 0.65698 | 1.34418 | 467 |
| GO_ESTABLISHMENT_OF_PROTEIN_LOCALIZATION_TO_TELOME | 0.00027 | 0.00607 | 0.49849 | 0.82169 | 1.49647 | 18 |
| GO_ESTABLISHMENT_OF_RNA_LOCALIZATION | 0.00000 | 0.00001 | 0.69013 | 0.64374 | 1.29679 | 180 |
| GO_ESTABLISHMENT_OF_TISSUE_POLARITY | 0.00000 | 0.00000 | 0.74774 | 0.72513 | 1.43503 | 95 |
| GO_EUKARYOTIC_TRANSLATION_INITIATION_FACTOR_3_COMPL | 0.00042 | 0.00871 | 0.49849 | 0.83556 | 1.50326 | 15 |
| GO_EXORIBONUCLEASE_COMPLEX | 0.00313 | 0.04608 | 0.43171 | 0.73160 | 1.36233 | 25 |
| GO_FC_EPSILON_RECEPTOR_SIGNALING_PATHWAY | 0.00000 | 0.00000 | 0.71951 | 0.70853 | 1.40242 | 98 |
| GO_FC_RECEPTOR_SIGNALING_PATHWAY | 0.00000 | 0.00010 | 0.62726 | 0.64603 | 1.29453 | 146 |
| GO_FIBRILLAR_CENTER | 0.00006 | 0.00161 | 0.55733 | 0.64880 | 1.28561 | 102 |
| GO_FIBROBLAST_GROWTH_FACTOR_RECEPTOR_SIGNALING_PAT | 0.00128 | 0.02208 | 0.45506 | 0.64771 | 1.26469 | 69 |
| GO_FICOLIN_1_RICH_GRANULE | 0.00000 | 0.00001 | 0.67496 | 0.69034 | 1.36665 | 99 |
| GO_GENERATION_OF_PRECURSOR_METABOLITES_AND_ENERGY | 0.00000 | 0.00000 | | 0.62069 | 1.26475 | 355 |
| GO_GENE_SILENCING | 0.00001 | 0.00022 | 0.61053 | 0.62044 | 1.25080 | 191 |

| | | | | | | |
|---|---|---|---|---|---|---|
| GO_GTPASE_ACTIVITY | 0.00111 | 0.01960 | 0.45506 | 0.58030 | 1.17265 | 208 |
| GO_GTP_METABOLIC_PROCESS | 0.00246 | 0.03787 | 0.43171 | 0.76714 | 1.40025 | 19 |
| GO_GUANYL_NUCLEOTIDE_BINDING | 0.00159 | 0.02667 | 0.45506 | 0.57145 | 1.15875 | 250 |
| GO_HEAT_SHOCK_PROTEIN_BINDING | 0.00128 | 0.02208 | 0.45506 | 0.61737 | 1.22220 | 99 |
| GO_HEMATOPOIETIC_PROGENITOR_CELL_DIFFERENTIATION | 0.00000 | 0.00000 | 0.70498 | 0.67411 | 1.34568 | 132 |
| GO_HEMATOPOIETIC_STEM_CELL_DIFFERENTIATION | 0.00000 | 0.00000 | 0.82666 | 0.77320 | 1.51549 | 76 |
| GO_HOST_CELLULAR_COMPONENT | 0.00008 | 0.00211 | 0.53843 | 0.71353 | 1.38339 | 54 |
| GO_HSP90_PROTEIN_BINDING | 0.00242 | 0.03739 | 0.43171 | 0.69766 | 1.32918 | 35 |
| GO_IMMUNE_RESPONSE_REGULATING_SIGNALING_PATHWAY | 0.00001 | 0.00046 | 0.59333 | 0.59807 | 1.21316 | 255 |
| GO_IMPORT_INTO_NUCLEUS | 0.00000 | 0.00003 | 0.65944 | 0.66172 | 1.32011 | 128 |
| GO_INNATE_IMMUNE_RESPONSE | 0.00000 | 0.00000 | 0.73376 | 0.59529 | 1.21676 | 430 |
| GO_INNATE_IMMUNE_RESPONSE_ACTIVATING_SIGNAL_TRANSI | 0.00000 | 0.00000 | 0.77494 | 0.73527 | 1.44754 | 86 |
| GO_INNER_MITOCHONDRIAL_MEMBRANE_PROTEIN_COMPLEX | 0.00000 | 0.00006 | 0.64355 | 0.68685 | 1.35802 | 93 |
| GO_INTERACTION_WITH_HOST | 0.00209 | 0.03323 | 0.43171 | 0.59569 | 1.19156 | 140 |
| GO_INTERLEUKIN_1_MEDIATED_SIGNALING_PATHWAY | 0.00000 | 0.00000 | | 0.77143 | 1.52087 | 88 |
| GO_INTRINSIC_APOPTOTIC_SIGNALING_PATHWAY | 0.00054 | 0.01084 | 0.47727 | 0.58692 | 1.18524 | 203 |
| GO_KINETOCHORE | 0.00010 | 0.00271 | 0.53843 | 0.63519 | 1.26638 | 123 |
| GO_LARGE_RIBOSOMAL_SUBUNIT | 0.00000 | 0.00000 | | 0.78157 | 1.55170 | 108 |
| GO_LEUKOCYTE_MEDIATED_IMMUNITY | 0.00000 | 0.00001 | 0.67496 | 0.58519 | 1.19675 | 451 |
| GO_MAINTENANCE_OF_CELL_NUMBER | 0.00089 | 0.01626 | 0.47727 | 0.61028 | 1.21724 | 127 |
| GO_MATERNAL_PLACENTA_DEVELOPMENT | 0.00034 | 0.00756 | 0.49849 | 0.80482 | 1.47364 | 20 |
| GO_MATERNAL_PROCESS_INVOLVED_IN_FEMALE_PREGNANCY | 0.00018 | 0.00438 | 0.51885 | 0.73599 | 1.40219 | 35 |
| GO_MATURATION_OF_5_8S_RRNA | 0.00170 | 0.02814 | 0.45506 | 0.71640 | 1.36269 | 34 |
| GO_MATURATION_OF_LSU_RRNA | 0.00043 | 0.00884 | 0.49849 | 0.77558 | 1.44956 | 26 |
| GO_MATURATION_OF_SSU_RRNA | 0.00030 | 0.00673 | 0.49849 | 0.70311 | 1.35564 | 47 |
| GO_MATURATION_OF_SSU_RRNA_FROM_TRICISTRONIC_RRNA_ | 0.00066 | 0.01276 | 0.47727 | 0.72088 | 1.37340 | 35 |
| GO_METAPHASE_ANAPHASE_TRANSITION_OF_CELL_CYCLE | 0.00133 | 0.02284 | 0.45506 | 0.66314 | 1.28787 | 56 |
| GO_MICROTUBULE_CYTOSKELETON_ORGANIZATION_INVOLVED | 0.00001 | 0.00023 | 0.61053 | 0.65637 | 1.30860 | 123 |
| GO_MITOCHONDRIAL_ELECTRON_TRANSPORT_NADH_TO_UBIC | 0.00077 | 0.01447 | 0.47727 | 0.69704 | 1.34343 | 45 |
| GO_MITOCHONDRIAL_GENE_EXPRESSION | 0.00000 | 0.00000 | | 0.70463 | 1.41360 | 153 |
| GO_MITOCHONDRIAL_LARGE_RIBOSOMAL_SUBUNIT | 0.00000 | 0.00000 | 0.73376 | 0.78666 | 1.52518 | 54 |
| GO_MITOCHONDRIAL_MATRIX | 0.00000 | 0.00000 | 0.71951 | 0.60199 | 1.22780 | 380 |
| GO_MITOCHONDRIAL_PROTEIN_COMPLEX | 0.00000 | 0.00000 | | 0.70652 | 1.42777 | 212 |
| GO_MITOCHONDRIAL_SMALL_RIBOSOMAL_SUBUNIT | 0.00062 | 0.01210 | 0.47727 | 0.74869 | 1.40749 | 28 |
| GO_MITOCHONDRIAL_TRANSLATION | 0.00000 | 0.00000 | | 0.72251 | 1.44226 | 130 |
| GO_MITOCHONDRIAL_TRANSLATIONAL_TERMINATION | 0.00000 | 0.00000 | | 0.77366 | 1.52529 | 88 |
| GO_MITOCHONDRIAL_TRANSPORT | 0.00063 | 0.01217 | 0.47727 | 0.58582 | 1.18137 | 193 |
| GO_MITOCHONDRION_ORGANIZATION | 0.00000 | 0.00000 | 0.74774 | 0.59715 | 1.21987 | 421 |
| GO_MITOTIC_CYTOKINESIS | 0.00170 | 0.02814 | 0.45506 | 0.64431 | 1.25418 | 64 |
| GO_MITOTIC_NUCLEAR_DIVISION | 0.00000 | 0.00011 | 0.62726 | 0.60866 | 1.23293 | 238 |
| GO_MITOTIC_SISTER_CHROMATID_SEGREGATION | 0.00021 | 0.00501 | 0.51885 | 0.61958 | 1.24218 | 148 |
| GO_MITOTIC_SPINDLE | 0.00064 | 0.01231 | 0.47727 | 0.61912 | 1.23015 | 111 |
| GO_MITOTIC_SPINDLE_ASSEMBLY | 0.00012 | 0.00301 | 0.53843 | 0.69947 | 1.35843 | 56 |
| GO_MITOTIC_SPINDLE_ORGANIZATION | 0.00003 | 0.00080 | 0.57561 | 0.65599 | 1.29986 | 102 |
| GO_MODULATION_BY_SYMBIONT_OF_HOST_CELLULAR_PROCE: | 0.00283 | 0.04262 | 0.43171 | 0.74408 | 1.36973 | 21 |
| GO_MORPHOGENESIS_OF_AN_EPITHELIUM | 0.00030 | 0.00678 | 0.49849 | 0.56470 | 1.15035 | 344 |
| GO_MORPHOGENESIS_OF_A_POLARIZED_EPITHELIUM | 0.00000 | 0.00000 | 0.78819 | 0.71425 | 1.41899 | 110 |
| GO_MRNA_3_END_PROCESSING | 0.00009 | 0.00235 | 0.53843 | 0.65418 | 1.29461 | 95 |
| GO_MRNA_5_UTR_BINDING | 0.00027 | 0.00607 | 0.49849 | 0.78159 | 1.45542 | 25 |
| GO_MRNA_BINDING | 0.00000 | 0.00000 | | 0.66658 | 1.35204 | 252 |
| GO_MRNA_EXPORT_FROM_NUCLEUS | 0.00002 | 0.00070 | 0.57561 | 0.65949 | 1.30695 | 103 |
| GO_MRNA_PROCESSING | 0.00000 | 0.00000 | | 0.64854 | 1.32639 | 452 |
| GO_MRNA_TRANSPORT | 0.00002 | 0.00072 | 0.57561 | 0.63414 | 1.26725 | 136 |
| GO_MULTI_ORGANISM_LOCALIZATION | 0.00009 | 0.00241 | 0.53843 | 0.71045 | 1.37742 | 54 |
| GO_MYELOID_LEUKOCYTE_ACTIVATION | 0.00000 | 0.00002 | 0.67496 | 0.59352 | 1.21094 | 386 |
| GO_MYELOID_LEUKOCYTE_MEDIATED_IMMUNITY | 0.00000 | 0.00000 | 0.74774 | 0.60477 | 1.23231 | 356 |
| GO_NADH_DEHYDROGENASE_ACTIVITY | 0.00224 | 0.03488 | 0.43171 | 0.70030 | 1.34185 | 38 |
| GO_NADH_DEHYDROGENASE_COMPLEX | 0.00185 | 0.03002 | 0.45506 | 0.69828 | 1.34099 | 41 |
| GO_NADH_DEHYDROGENASE_COMPLEX_ASSEMBLY | 0.00135 | 0.02309 | 0.45506 | 0.66274 | 1.28710 | 56 |
| GO_NCRNA_3_END_PROCESSING | 0.00290 | 0.04352 | 0.43171 | 0.69533 | 1.32474 | 35 |
| GO_NCRNA_EXPORT_FROM_NUCLEUS | 0.00003 | 0.00100 | 0.55733 | 0.75739 | 1.44297 | 35 |
| GO_NCRNA_METABOLIC_PROCESS | 0.00000 | 0.00000 | | 0.62040 | 1.26677 | 411 |
| GO_NCRNA_PROCESSING | 0.00000 | 0.00000 | | 0.64534 | 1.31481 | 345 |
| GO_NEGATIVE_REGULATION_OF_CANONICAL_WNT_SIGNALING | 0.00000 | 0.00001 | 0.70498 | 0.67967 | 1.35401 | 120 |
| GO_NEGATIVE_REGULATION_OF_CATALYTIC_ACTIVITY | 0.00320 | 0.04706 | 0.43171 | 0.54088 | 1.10618 | 450 |
| GO_NEGATIVE_REGULATION_OF_CELL_CYCLE | 0.00000 | 0.00003 | 0.65944 | 0.57447 | 1.17592 | 476 |
| GO_NEGATIVE_REGULATION_OF_CELL_CYCLE_G2_M_PHASE_TR | 0.00000 | 0.00001 | 0.67496 | 0.70662 | 1.39185 | 87 |
| GO_NEGATIVE_REGULATION_OF_CELL_CYCLE_PHASE_TRANSITI( | 0.00000 | 0.00001 | 0.70498 | 0.64011 | 1.29211 | 200 |
| GO_NEGATIVE_REGULATION_OF_CELL_CYCLE_PROCESS | 0.00000 | 0.00000 | 0.76146 | 0.63012 | 1.27907 | 270 |
| GO_NEGATIVE_REGULATION_OF_CHROMOSOME_ORGANIZATIO | 0.00124 | 0.02162 | 0.45506 | 0.61672 | 1.22771 | 118 |

| | | | | | | |
|---|---|---|---|---|---|---|
| GO_NEGATIVE_REGULATION_OF_MITOTIC_CELL_CYCLE | 0.00000 | 0.00001 | 0.69013 | 0.62060 | 1.25956 | 258 |
| GO_NEGATIVE_REGULATION_OF_ORGANELLE_ORGANIZATION | 0.00213 | 0.03374 | 0.43171 | 0.55803 | 1.13508 | 303 |
| GO_NEGATIVE_REGULATION_OF_RNA_SPLICING | 0.00239 | 0.03691 | 0.43171 | 0.74493 | 1.38269 | 24 |
| GO_NEGATIVE_REGULATION_OF_WNT_SIGNALING_PATHWAY | 0.00000 | 0.00003 | 0.65944 | 0.65546 | 1.31004 | 137 |
| GO_NIK_NF_KAPPAB_SIGNALING | 0.00000 | 0.00000 | 0.76146 | 0.69817 | 1.39088 | 120 |
| GO_NON_CANONICAL_WNT_SIGNALING_PATHWAY | 0.00000 | 0.00000 | 0.80122 | 0.71541 | 1.42149 | 111 |
| GO_NUCLEAR_CHROMOSOME_SEGREGATION | 0.00087 | 0.01607 | 0.47727 | 0.57769 | 1.16966 | 226 |
| GO_NUCLEAR_CHROMOSOME_TELOMERIC_REGION | 0.00268 | 0.04078 | 0.43171 | 0.61459 | 1.21627 | 95 |
| GO_NUCLEAR_ENVELOPE | 0.00001 | 0.00024 | 0.61053 | 0.58266 | 1.18666 | 336 |
| GO_NUCLEAR_EXPORT | 0.00000 | 0.00000 | 0.70498 | 0.64896 | 1.30731 | 180 |
| GO_NUCLEAR_LOCALIZATION_SEQUENCE_BINDING | 0.00257 | 0.03929 | 0.43171 | 0.75895 | 1.40258 | 22 |
| GO_NUCLEAR_MATRIX | 0.00041 | 0.00860 | 0.49849 | 0.63656 | 1.25859 | 93 |
| GO_NUCLEAR_MEMBRANE | 0.00154 | 0.02581 | 0.45506 | 0.57985 | 1.17178 | 212 |
| GO_NUCLEAR_PERIPHERY | 0.00003 | 0.00092 | 0.55733 | 0.64232 | 1.27723 | 114 |
| GO_NUCLEAR_PORE | 0.00024 | 0.00561 | 0.51885 | 0.67067 | 1.30851 | 68 |
| GO_NUCLEAR_SPECK | 0.00000 | 0.00000 | 0.73376 | 0.61337 | 1.24856 | 329 |
| GO_NUCLEAR_TRANSCRIBED_MRNA_CATABOLIC_PROCESS | 0.00000 | 0.00000 | | 0.71510 | 1.44111 | 188 |
| GO_NUCLEAR_TRANSCRIBED_MRNA_CATABOLIC_PROCESS_NON | 0.00000 | 0.00000 | | 0.78100 | 1.55299 | 114 |
| GO_NUCLEAR_TRANSPORT | 0.00000 | 0.00000 | | 0.64327 | 1.30805 | 293 |
| GO_NUCLEIC_ACID_PHOSPHODIESTER_BOND_HYDROLYSIS | 0.00342 | 0.04962 | 0.43171 | 0.56339 | 1.14284 | 253 |
| GO_NUCLEOBASE_CONTAINING_COMPOUND_TRANSPORT | 0.00000 | 0.00003 | 0.65944 | 0.62346 | 1.26176 | 221 |
| GO_NUCLEOBASE_CONTAINING_SMALL_MOLECULE_METABOLIC | 0.00279 | 0.04226 | 0.43171 | 0.54452 | 1.11186 | 404 |
| GO_NUCLEOSIDE_DIPHOSPHATE_METABOLIC_PROCESS | 0.00000 | 0.00004 | 0.65944 | 0.67617 | 1.34244 | 108 |
| GO_NUCLEOSIDE_TRIPHOSPHATE_BIOSYNTHETIC_PROCESS | 0.00335 | 0.04900 | 0.43171 | 0.66790 | 1.28776 | 47 |
| GO_NUCLEOSIDE_TRIPHOSPHATE_METABOLIC_PROCESS | 0.00039 | 0.00836 | 0.49849 | 0.66544 | 1.29818 | 67 |
| GO_NUCLEOTIDE_PHOSPHORYLATION | 0.00000 | 0.00002 | 0.65944 | 0.69191 | 1.36952 | 98 |
| GO_NUCLEUS_ORGANIZATION | 0.00092 | 0.01662 | 0.47727 | 0.61454 | 1.22224 | 113 |
| GO_ORGANELLAR_RIBOSOME | 0.00000 | 0.00000 | | 0.76282 | 1.50075 | 84 |
| GO_ORGANELLE_DISASSEMBLY | 0.00170 | 0.02814 | 0.45506 | 0.61373 | 1.21498 | 99 |
| GO_ORGANELLE_FISSION | 0.00023 | 0.00540 | 0.51885 | 0.56240 | 1.14704 | 373 |
| GO_ORGANELLE_INNER_MEMBRANE | 0.00000 | 0.00000 | | 0.64013 | 1.30706 | 408 |
| GO_ORGANIC_CYCLIC_COMPOUND_CATABOLIC_PROCESS | 0.00000 | 0.00000 | | 0.66111 | 1.35380 | 489 |
| GO_OXIDATIVE_PHOSPHORYLATION | 0.00000 | 0.00001 | 0.67496 | 0.69492 | 1.37606 | 100 |
| GO_OXIDOREDUCTASE_COMPLEX | 0.00020 | 0.00480 | 0.51885 | 0.65322 | 1.28383 | 83 |
| GO_PEPTIDASE_COMPLEX | 0.00000 | 0.00004 | 0.64355 | 0.69708 | 1.37141 | 84 |
| GO_PEPTIDE_BINDING | 0.00102 | 0.01823 | 0.45506 | 0.59675 | 1.19829 | 156 |
| GO_PHAGOPHORE_ASSEMBLY_SITE | 0.00309 | 0.04564 | 0.43171 | 0.71122 | 1.33993 | 29 |
| GO_PIGMENT_GRANULE | 0.00000 | 0.00003 | 0.65944 | 0.69543 | 1.36910 | 86 |
| GO_POLYSOMAL_RIBOSOME | 0.00000 | 0.00003 | 0.65944 | 0.82134 | 1.55398 | 31 |
| GO_POLYSOME | 0.00000 | 0.00000 | 0.71951 | 0.74419 | 1.44860 | 64 |
| GO_POSITIVE_REGULATION_OF_BINDING | 0.00049 | 0.00993 | 0.47727 | 0.61581 | 1.22707 | 121 |
| GO_POSITIVE_REGULATION_OF_BLOOD_CIRCULATION | 0.00074 | 0.01391 | 0.47727 | 0.74537 | 1.40126 | 28 |
| GO_POSITIVE_REGULATION_OF_BLOOD_VESSEL_DIAMETER | 0.00009 | 0.00233 | 0.53843 | 0.77050 | 1.45390 | 30 |
| GO_POSITIVE_REGULATION_OF_CANONICAL_WNT_SIGNALING_ | 0.00009 | 0.00236 | 0.53843 | 0.63688 | 1.26605 | 112 |
| GO_POSITIVE_REGULATION_OF_CELLULAR_AMIDE_METABOLIC_ | 0.00000 | 0.00004 | 0.64355 | 0.67066 | 1.33733 | 125 |
| GO_POSITIVE_REGULATION_OF_CELLULAR_PROTEIN_LOCALIZAT | 0.00011 | 0.00296 | 0.53843 | 0.58522 | 1.18605 | 245 |
| GO_POSITIVE_REGULATION_OF_CELL_CYCLE | 0.00111 | 0.01960 | 0.45506 | 0.56867 | 1.15478 | 276 |
| GO_POSITIVE_REGULATION_OF_CELL_CYCLE_PROCESS | 0.00097 | 0.01730 | 0.47727 | 0.58048 | 1.17328 | 214 |
| GO_POSITIVE_REGULATION_OF_CELL_POPULATION_PROLIFERAT | 0.00014 | 0.00358 | 0.51885 | 0.55576 | 1.13752 | 475 |
| GO_POSITIVE_REGULATION_OF_DEFENSE_RESPONSE | 0.00000 | 0.00000 | 0.73376 | 0.64745 | 1.30746 | 203 |
| GO_POSITIVE_REGULATION_OF_DNA_BINDING | 0.00023 | 0.00544 | 0.51885 | 0.73624 | 1.40858 | 37 |
| GO_POSITIVE_REGULATION_OF_DNA_BIOSYNTHETIC_PROCESS | 0.00001 | 0.00033 | 0.59333 | 0.72539 | 1.40974 | 57 |
| GO_POSITIVE_REGULATION_OF_DNA_METABOLIC_PROCESS | 0.00092 | 0.01662 | 0.47727 | 0.59670 | 1.19476 | 144 |
| GO_POSITIVE_REGULATION_OF_ESTABLISHMENT_OF_PROTEIN_ | 0.00216 | 0.03406 | 0.43171 | 0.56675 | 1.15029 | 258 |
| GO_POSITIVE_REGULATION_OF_FIBROBLAST_PROLIFERATION | 0.00181 | 0.02958 | 0.45506 | 0.75069 | 1.40303 | 26 |
| GO_POSITIVE_REGULATION_OF_GENE_EXPRESSION_EPIGENETIC | 0.00309 | 0.04564 | 0.43171 | 0.67475 | 1.30059 | 46 |
| GO_POSITIVE_REGULATION_OF_IMMUNE_RESPONSE | 0.00000 | 0.00003 | 0.65944 | 0.59084 | 1.20377 | 353 |
| GO_POSITIVE_REGULATION_OF_INNATE_IMMUNE_RESPONSE | 0.00000 | 0.00000 | | 0.70841 | 1.41476 | 134 |
| GO_POSITIVE_REGULATION_OF_MITOTIC_CELL_CYCLE | 0.00047 | 0.00966 | 0.49849 | 0.61879 | 1.23354 | 122 |
| GO_POSITIVE_REGULATION_OF_MRNA_METABOLIC_PROCESS | 0.00071 | 0.01347 | 0.47727 | 0.65902 | 1.28579 | 68 |
| GO_POSITIVE_REGULATION_OF_MUSCLE_CONTRACTION | 0.00305 | 0.04542 | 0.43171 | 0.74945 | 1.38756 | 23 |
| GO_POSITIVE_REGULATION_OF_ORGANELLE_ORGANIZATION | 0.00067 | 0.01290 | 0.47727 | 0.54853 | 1.12211 | 461 |
| GO_POSITIVE_REGULATION_OF_PROTEIN_LOCALIZATION_TO_N | 0.00011 | 0.00295 | 0.53843 | 0.68981 | 1.34204 | 61 |
| GO_POSITIVE_REGULATION_OF_RESPONSE_TO_BIOTIC_STIMUL | 0.00000 | 0.00000 | 0.73376 | 0.67310 | 1.35043 | 152 |
| GO_POSITIVE_REGULATION_OF_RESPONSE_TO_EXTERNAL_STIM | 0.00000 | 0.00006 | 0.64355 | 0.60556 | 1.22943 | 266 |
| GO_POSITIVE_REGULATION_OF_RNA_SPLICING | 0.00178 | 0.02913 | 0.45506 | 0.72149 | 1.35928 | 29 |
| GO_POSITIVE_REGULATION_OF_TELOMERASE_ACTIVITY | 0.00008 | 0.00226 | 0.53843 | 0.76031 | 1.44190 | 32 |
| GO_POSITIVE_REGULATION_OF_TELOMERASE_RNA_LOCALIZATI | 0.00018 | 0.00447 | 0.51885 | 0.84651 | 1.52295 | 15 |
| GO_POSITIVE_REGULATION_OF_TELOMERE_MAINTENANCE | 0.00106 | 0.01878 | 0.45506 | 0.69350 | 1.33672 | 46 |
| GO_POSITIVE_REGULATION_OF_TELOMERE_MAINTENANCE_VIA | 0.00010 | 0.00261 | 0.53843 | 0.74420 | 1.41783 | 35 |

| | | | | | | |
|---|---|---|---|---|---|---|
| GO_POSITIVE_REGULATION_OF_TRANSCRIPTION_REGULATORY_ | 0.00309 | 0.04564 | 0.43171 | 0.79298 | 1.42664 | 15 |
| GO_POSITIVE_REGULATION_OF_TRANSLATION | 0.00000 | 0.00001 | 0.69013 | 0.69688 | 1.38237 | 105 |
| GO_POSITIVE_REGULATION_OF_VIRAL_PROCESS | 0.00019 | 0.00456 | 0.51885 | 0.65221 | 1.28584 | 88 |
| GO_POSITIVE_REGULATION_OF_WNT_SIGNALING_PATHWAY | 0.00027 | 0.00615 | 0.49849 | 0.61243 | 1.22308 | 134 |
| GO_PRECATALYTIC_SPLICEOSOME | 0.00041 | 0.00867 | 0.49849 | 0.68976 | 1.33312 | 51 |
| GO_PRERIBOSOME | 0.00000 | 0.00001 | 0.69013 | 0.73667 | 1.43906 | 70 |
| GO_PRERIBOSOME_LARGE_SUBUNIT_PRECURSOR | 0.00305 | 0.04542 | 0.43171 | 0.75348 | 1.39247 | 22 |
| GO_PRE_MRNA_BINDING | 0.00044 | 0.00909 | 0.49849 | 0.73850 | 1.40053 | 32 |
| GO_PROTEASOMAL_PROTEIN_CATABOLIC_PROCESS | 0.00000 | 0.00011 | 0.62726 | 0.57718 | 1.17852 | 405 |
| GO_PROTEASOMAL_UBIQUITIN_INDEPENDENT_PROTEIN_CATAE | 0.00000 | 0.00008 | 0.62726 | 0.87414 | 1.60056 | 20 |
| GO_PROTEASOME_ACCESSORY_COMPLEX | 0.00001 | 0.00035 | 0.59333 | 0.83351 | 1.54710 | 24 |
| GO_PROTEASOME_CORE_COMPLEX | 0.00000 | 0.00005 | 0.64355 | 0.89029 | 1.62502 | 19 |
| GO_PROTEIN_CONTAINING_COMPLEX_DISASSEMBLY | 0.00000 | 0.00000 | 0.78819 | 0.63646 | 1.29194 | 270 |
| GO_PROTEIN_CONTAINING_COMPLEX_LOCALIZATION | 0.00000 | 0.00006 | 0.64355 | 0.61055 | 1.23605 | 232 |
| GO_PROTEIN_DNA_COMPLEX_SUBUNIT_ORGANIZATION | 0.00002 | 0.00072 | 0.57561 | 0.61317 | 1.23555 | 187 |
| GO_PROTEIN_FOLDING | 0.00003 | 0.00085 | 0.57561 | 0.61332 | 1.23536 | 184 |
| GO_PROTEIN_FOLDING_CHAPERONE | 0.00018 | 0.00428 | 0.51885 | 0.80961 | 1.49034 | 21 |
| GO_PROTEIN_IMPORT | 0.00000 | 0.00003 | 0.65944 | 0.64733 | 1.29985 | 156 |
| GO_PROTEIN_LIPID_COMPLEX_ASSEMBLY | 0.00095 | 0.01702 | 0.47727 | 0.81366 | 1.47193 | 16 |
| GO_PROTEIN_LIPID_COMPLEX_SUBUNIT_ORGANIZATION | 0.00038 | 0.00819 | 0.49849 | 0.80252 | 1.46942 | 20 |
| GO_PROTEIN_LOCALIZATION_TO_CHROMOSOME | 0.00198 | 0.03196 | 0.43171 | 0.64293 | 1.25854 | 73 |
| GO_PROTEIN_LOCALIZATION_TO_CHROMOSOME_TELOMERIC_F | 0.00086 | 0.01582 | 0.47727 | 0.73695 | 1.38841 | 29 |
| GO_PROTEIN_LOCALIZATION_TO_ENDOPLASMIC_RETICULUM | 0.00000 | 0.00000 | | 0.80411 | 1.60515 | 130 |
| GO_PROTEIN_LOCALIZATION_TO_MEMBRANE | 0.00000 | 0.00000 | | 0.60657 | 1.24097 | 464 |
| GO_PROTEIN_LOCALIZATION_TO_MITOCHONDRION | 0.00065 | 0.01262 | 0.47727 | 0.61662 | 1.22630 | 116 |
| GO_PROTEIN_LOCALIZATION_TO_NUCLEUS | 0.00000 | 0.00000 | 0.77494 | 0.64963 | 1.31256 | 207 |
| GO_PROTEIN_MODIFICATION_BY_SMALL_PROTEIN_REMOVAL | 0.00036 | 0.00793 | 0.49849 | 0.57993 | 1.17546 | 246 |
| GO_PROTEIN_POLYUBIQUITINATION | 0.00076 | 0.01417 | 0.47727 | 0.56895 | 1.15605 | 282 |
| GO_PROTEIN_STABILIZATION | 0.00087 | 0.01607 | 0.47727 | 0.60729 | 1.21441 | 139 |
| GO_PROTEIN_SUMOYLATION | 0.00006 | 0.00176 | 0.53843 | 0.68238 | 1.33239 | 69 |
| GO_PROTEIN_TARGETING | 0.00000 | 0.00000 | | 0.65096 | 1.32678 | 360 |
| GO_PROTEIN_TARGETING_TO_MEMBRANE | 0.00000 | 0.00000 | | 0.74931 | 1.50716 | 167 |
| GO_PROTEIN_TARGETING_TO_MITOCHONDRION | 0.00058 | 0.01146 | 0.47727 | 0.64633 | 1.26902 | 80 |
| GO_PROTEIN_TRANSMEMBRANE_TRANSPORT | 0.00068 | 0.01303 | 0.47727 | 0.67480 | 1.31052 | 56 |
| GO_PROTON_TRANSMEMBRANE_TRANSPORT | 0.00342 | 0.04962 | 0.43171 | 0.62306 | 1.22457 | 83 |
| GO_PURINE_CONTAINING_COMPOUND_METABOLIC_PROCESS | 0.00143 | 0.02422 | 0.45506 | 0.56364 | 1.14626 | 296 |
| GO_PURINE_NUCLEOSIDE_TRIPHOSPHATE_METABOLIC_PROCES | 0.00104 | 0.01851 | 0.45506 | 0.68719 | 1.32637 | 49 |
| GO_PYRUVATE_METABOLIC_PROCESS | 0.00000 | 0.00006 | 0.64355 | 0.68457 | 1.35794 | 105 |
| GO_REGENERATION | 0.00202 | 0.03239 | 0.43171 | 0.60404 | 1.20110 | 114 |
| GO_REGULATION_OF_ANIMAL_ORGAN_MORPHOGENESIS | 0.00000 | 0.00000 | 0.80122 | 0.69376 | 1.38795 | 143 |
| GO_REGULATION_OF_ATP_METABOLIC_PROCESS | 0.00001 | 0.00033 | 0.59333 | 0.68097 | 1.34132 | 87 |
| GO_REGULATION_OF_BINDING | 0.00027 | 0.00607 | 0.49849 | 0.58036 | 1.17641 | 243 |
| GO_REGULATION_OF_CARBOHYDRATE_CATABOLIC_PROCESS | 0.00015 | 0.00365 | 0.51885 | 0.68794 | 1.33873 | 63 |
| GO_REGULATION_OF_CELLULAR_AMIDE_METABOLIC_PROCESS | 0.00000 | 0.00000 | 0.76146 | 0.61386 | 1.25098 | 341 |
| GO_REGULATION_OF_CELLULAR_AMINE_METABOLIC_PROCESS | 0.00000 | 0.00000 | | 0.82273 | 1.60064 | 61 |
| GO_REGULATION_OF_CELLULAR_AMINO_ACID_METABOLIC_PR( | 0.00000 | 0.00000 | | 0.84000 | 1.62859 | 54 |
| GO_REGULATION_OF_CELLULAR_KETONE_METABOLIC_PROCESS | 0.00000 | 0.00000 | 0.73376 | 0.69809 | 1.38813 | 114 |
| GO_REGULATION_OF_CELLULAR_PROTEIN_LOCALIZATION | 0.00001 | 0.00048 | 0.59333 | 0.57248 | 1.16869 | 400 |
| GO_REGULATION_OF_CELLULAR_RESPONSE_TO_HEAT | 0.00002 | 0.00053 | 0.57561 | 0.69487 | 1.35574 | 68 |
| GO_REGULATION_OF_CELL_CYCLE_G2_M_PHASE_TRANSITION | 0.00001 | 0.00039 | 0.59333 | 0.61711 | 1.24352 | 186 |
| GO_REGULATION_OF_CELL_CYCLE_PHASE_TRANSITION | 0.00000 | 0.00006 | 0.64355 | 0.58544 | 1.19395 | 372 |
| GO_REGULATION_OF_CHROMOSOME_ORGANIZATION | 0.00002 | 0.00050 | 0.57561 | 0.58646 | 1.19228 | 288 |
| GO_REGULATION_OF_CHROMOSOME_SEGREGATION | 0.00016 | 0.00383 | 0.51885 | 0.64616 | 1.27875 | 95 |
| GO_REGULATION_OF_CHROMOSOME_SEPARATION | 0.00172 | 0.02830 | 0.45506 | 0.65139 | 1.26737 | 62 |
| GO_REGULATION_OF_DEFENSE_RESPONSE | 0.00000 | 0.00001 | 0.67496 | 0.59062 | 1.20440 | 368 |
| GO_REGULATION_OF_DNA_BINDING | 0.00017 | 0.00415 | 0.51885 | 0.65691 | 1.28999 | 81 |
| GO_REGULATION_OF_DNA_BIOSYNTHETIC_PROCESS | 0.00000 | 0.00007 | 0.64355 | 0.69361 | 1.36458 | 84 |
| GO_REGULATION_OF_DNA_METABOLIC_PROCESS | 0.00034 | 0.00756 | 0.49849 | 0.57773 | 1.17271 | 264 |
| GO_REGULATION_OF_DNA_TEMPLATED_TRANSCRIPTION_IN_RE | 0.00000 | 0.00000 | 0.76146 | 0.72679 | 1.43832 | 95 |
| GO_REGULATION_OF_GENERATION_OF_PRECURSOR_METABOLI | 0.00008 | 0.00231 | 0.53843 | 0.64470 | 1.28049 | 109 |
| GO_REGULATION_OF_GENE_SILENCING | 0.00012 | 0.00306 | 0.53843 | 0.65830 | 1.30118 | 92 |
| GO_REGULATION_OF_HEMATOPOIETIC_PROGENITOR_CELL_DIF | 0.00000 | 0.00000 | 0.78819 | 0.75794 | 1.48658 | 77 |
| GO_REGULATION_OF_HEMATOPOIETIC_STEM_CELL_DIFFERENT | 0.00000 | 0.00000 | 0.82666 | 0.79079 | 1.54172 | 66 |
| GO_REGULATION_OF_HEMOPOIESIS | 0.00021 | 0.00489 | 0.51885 | 0.57251 | 1.16386 | 287 |
| GO_REGULATION_OF_IMMUNE_RESPONSE | 0.00001 | 0.00043 | 0.59333 | 0.56602 | 1.15821 | 472 |
| GO_REGULATION_OF_INNATE_IMMUNE_RESPONSE | 0.00000 | 0.00000 | 0.78819 | 0.66449 | 1.33899 | 186 |
| GO_REGULATION_OF_INTRACELLULAR_PROTEIN_TRANSPORT | 0.00050 | 0.01002 | 0.47727 | 0.58782 | 1.18612 | 198 |
| GO_REGULATION_OF_INTRACELLULAR_TRANSPORT | 0.00084 | 0.01553 | 0.47727 | 0.57154 | 1.16016 | 264 |
| GO_REGULATION_OF_MORPHOGENESIS_OF_AN_EPITHELIUM | 0.00000 | 0.00000 | | 0.73212 | 1.45795 | 119 |
| GO_REGULATION_OF_MRNA_CATABOLIC_PROCESS | 0.00000 | 0.00000 | 0.78819 | 0.67368 | 1.35577 | 172 |

| | | | | | | |
|---|---|---|---|---|---|---|
| GO_REGULATION_OF_MRNA_METABOLIC_PROCESS | 0.00000 | 0.00000 | | 0.66102 | 1.34353 | 285 |
| GO_REGULATION_OF_MRNA_PROCESSING | 0.00000 | 0.00002 | 0.67496 | 0.66285 | 1.32256 | 129 |
| GO_REGULATION_OF_MRNA_SPLICING_VIA_SPLICEOSOME | 0.00002 | 0.00063 | 0.57561 | 0.67416 | 1.33253 | 92 |
| GO_REGULATION_OF_NUCLEAR_DIVISION | 0.00004 | 0.00110 | 0.55733 | 0.62386 | 1.25163 | 152 |
| GO_REGULATION_OF_NUCLEOCYTOPLASMIC_TRANSPORT | 0.00009 | 0.00236 | 0.53843 | 0.65908 | 1.29938 | 88 |
| GO_REGULATION_OF_NUCLEOTIDE_METABOLIC_PROCESS | 0.00023 | 0.00544 | 0.51885 | 0.65304 | 1.28238 | 81 |
| GO_REGULATION_OF_OXIDATIVE_PHOSPHORYLATION | 0.00010 | 0.00257 | 0.53843 | 0.82226 | 1.51364 | 21 |
| GO_REGULATION_OF_PLASMA_LIPOPROTEIN_PARTICLE_LEVELS | 0.00031 | 0.00686 | 0.49849 | 0.71028 | 1.36896 | 45 |
| GO_REGULATION_OF_POSTTRANSCRIPTIONAL_GENE_SILENCINC | 0.00004 | 0.00106 | 0.55733 | 0.66992 | 1.31617 | 82 |
| GO_REGULATION_OF_PROTEIN_IMPORT | 0.00126 | 0.02188 | 0.45506 | 0.68996 | 1.32978 | 45 |
| GO_REGULATION_OF_PROTEIN_LOCALIZATION_TO_NUCLEUS | 0.00001 | 0.00035 | 0.59333 | 0.68126 | 1.34504 | 90 |
| GO_REGULATION_OF_PROTEIN_STABILITY | 0.00113 | 0.01981 | 0.45506 | 0.57561 | 1.16544 | 226 |
| GO_REGULATION_OF_PROTEOLYSIS | 0.00342 | 0.04962 | 0.43171 | 0.54112 | 1.10650 | 449 |
| GO_REGULATION_OF_RESPONSE_TO_BIOTIC_STIMULUS | 0.00000 | 0.00000 | 0.70498 | 0.62562 | 1.26753 | 239 |
| GO_REGULATION_OF_RNA_SPLICING | 0.00000 | 0.00001 | 0.69013 | 0.66594 | 1.32872 | 129 |
| GO_REGULATION_OF_SISTER_CHROMATID_SEGREGATION | 0.00040 | 0.00853 | 0.49849 | 0.65059 | 1.27518 | 76 |
| GO_REGULATION_OF_SMALL_MOLECULE_METABOLIC_PROCESS | 0.00000 | 0.00001 | 0.69013 | 0.61443 | 1.24846 | 282 |
| GO_REGULATION_OF_STEM_CELL_DIFFERENTIATION | 0.00000 | 0.00000 | 0.80122 | 0.73911 | 1.46270 | 95 |
| GO_REGULATION_OF_SYMBIOTIC_PROCESS | 0.00002 | 0.00076 | 0.57561 | 0.62990 | 1.26222 | 146 |
| GO_REGULATION_OF_TELOMERASE_ACTIVITY | 0.00038 | 0.00812 | 0.49849 | 0.71460 | 1.37362 | 42 |
| GO_REGULATION_OF_TELOMERE_MAINTENANCE | 0.00055 | 0.01098 | 0.47727 | 0.65802 | 1.28736 | 72 |
| GO_REGULATION_OF_TELOMERE_MAINTENANCE_VIA_TELOME | 0.00040 | 0.00853 | 0.49849 | 0.69104 | 1.33979 | 54 |
| GO_REGULATION_OF_TRANSCRIPTION_FROM_RNA_POLYMERAS | 0.00000 | 0.00000 | 0.82666 | 0.78192 | 1.52558 | 68 |
| GO_REGULATION_OF_TRANSLATIONAL_INITIATION | 0.00004 | 0.00121 | 0.55733 | 0.69666 | 1.35821 | 66 |
| GO_REGULATION_OF_UBIQUITIN_PROTEIN_LIGASE_ACTIVITY | 0.00006 | 0.00170 | 0.55733 | 0.80562 | 1.49533 | 24 |
| GO_REGULATION_OF_VIRAL_TRANSCRIPTION | 0.00276 | 0.04181 | 0.43171 | 0.66154 | 1.28078 | 52 |
| GO_RESPIRASOME | 0.00000 | 0.00012 | 0.62726 | 0.70439 | 1.38015 | 75 |
| GO_RESPIRATORY_CHAIN_COMPLEX | 0.00007 | 0.00197 | 0.53843 | 0.69491 | 1.35413 | 65 |
| GO_RESPIRATORY_ELECTRON_TRANSPORT_CHAIN | 0.00000 | 0.00013 | 0.62726 | 0.69071 | 1.36369 | 90 |
| GO_RESPONSE_TO_DRUG | 0.00216 | 0.03406 | 0.43171 | 0.57397 | 1.16139 | 219 |
| GO_RESPONSE_TO_HEAT | 0.00001 | 0.00030 | 0.59333 | 0.65404 | 1.30417 | 125 |
| GO_RESPONSE_TO_INTERLEUKIN_1 | 0.00000 | 0.00000 | | 0.73609 | 1.46869 | 129 |
| GO_RESPONSE_TO_INTERLEUKIN_4 | 0.00015 | 0.00380 | 0.51885 | 0.80585 | 1.49197 | 23 |
| GO_RESPONSE_TO_LEUKEMIA_INHIBITORY_FACTOR | 0.00168 | 0.02806 | 0.45506 | 0.63488 | 1.24438 | 76 |
| GO_RESPONSE_TO_OXYGEN_LEVELS | 0.00000 | 0.00001 | 0.70498 | 0.61867 | 1.25604 | 266 |
| GO_RESPONSE_TO_TEMPERATURE_STIMULUS | 0.00000 | 0.00014 | 0.61053 | 0.63636 | 1.27781 | 156 |
| GO_RESPONSE_TO_TUMOR_NECROSIS_FACTOR | 0.00000 | 0.00007 | 0.62726 | 0.63102 | 1.27133 | 185 |
| GO_RIBONUCLEOPROTEIN_COMPLEX_BINDING | 0.00000 | 0.00000 | 0.77494 | 0.70522 | 1.40437 | 119 |
| GO_RIBONUCLEOPROTEIN_COMPLEX_BIOGENESIS | 0.00000 | 0.00000 | | 0.71879 | 1.46713 | 395 |
| GO_RIBONUCLEOPROTEIN_COMPLEX_SUBUNIT_ORGANIZATION | 0.00000 | 0.00000 | | 0.71902 | 1.44811 | 177 |
| GO_RIBONUCLEOPROTEIN_GRANULE | 0.00001 | 0.00023 | 0.61053 | 0.61550 | 1.24295 | 203 |
| GO_RIBONUCLEOSIDE_TRIPHOSPHATE_METABOLIC_PROCESS | 0.00011 | 0.00290 | 0.53843 | 0.71353 | 1.37582 | 48 |
| GO_RIBOSE_PHOSPHATE_METABOLIC_PROCESS | 0.00014 | 0.00348 | 0.51885 | 0.57893 | 1.17605 | 280 |
| GO_RIBOSOMAL_LARGE_SUBUNIT_ASSEMBLY | 0.00014 | 0.00362 | 0.51885 | 0.79375 | 1.48351 | 26 |
| GO_RIBOSOMAL_LARGE_SUBUNIT_BIOGENESIS | 0.00000 | 0.00000 | 0.77494 | 0.76722 | 1.49689 | 68 |
| GO_RIBOSOMAL_SMALL_SUBUNIT_ASSEMBLY | 0.00111 | 0.01960 | 0.45506 | 0.79197 | 1.44235 | 18 |
| GO_RIBOSOMAL_SMALL_SUBUNIT_BINDING | 0.00239 | 0.03691 | 0.43171 | 0.79176 | 1.43582 | 17 |
| GO_RIBOSOMAL_SMALL_SUBUNIT_BIOGENESIS | 0.00000 | 0.00010 | 0.62726 | 0.71356 | 1.39219 | 68 |
| GO_RIBOSOMAL_SUBUNIT | 0.00000 | 0.00000 | | 0.77320 | 1.55686 | 174 |
| GO_RIBOSOME | 0.00000 | 0.00000 | | 0.76517 | 1.54614 | 210 |
| GO_RIBOSOME_ASSEMBLY | 0.00000 | 0.00001 | 0.69013 | 0.76389 | 1.48353 | 56 |
| GO_RIBOSOME_BINDING | 0.00290 | 0.04352 | 0.43171 | 0.66499 | 1.28222 | 48 |
| GO_RIBOSOME_BIOGENESIS | 0.00000 | 0.00000 | | 0.71577 | 1.45377 | 277 |
| GO_RNA_3_END_PROCESSING | 0.00000 | 0.00001 | 0.69013 | 0.66861 | 1.33471 | 132 |
| GO_RNA_CATABOLIC_PROCESS | 0.00000 | 0.00000 | | 0.69520 | 1.41627 | 352 |
| GO_RNA_DEPENDENT_DNA_BIOSYNTHETIC_PROCESS | 0.00017 | 0.00424 | 0.51885 | 0.68671 | 1.33815 | 65 |
| GO_RNA_EXPORT_FROM_NUCLEUS | 0.00000 | 0.00001 | 0.70498 | 0.67060 | 1.33803 | 129 |
| GO_RNA_LOCALIZATION | 0.00000 | 0.00000 | 0.81404 | 0.66502 | 1.34390 | 212 |
| GO_RNA_PHOSPHODIESTER_BOND_HYDROLYSIS | 0.00079 | 0.01477 | 0.47727 | 0.60620 | 1.20953 | 129 |
| GO_RNA_SPLICING | 0.00000 | 0.00000 | | 0.65400 | 1.33514 | 399 |
| GO_RNA_SPLICING_VIA_TRANSESTERIFICATION_REACTIONS | 0.00000 | 0.00000 | | 0.66603 | 1.35580 | 320 |
| GO_ROUGH_ENDOPLASMIC_RETICULUM | 0.00000 | 0.00012 | 0.62726 | 0.72115 | 1.40300 | 61 |
| GO_RRNA_BINDING | 0.00000 | 0.00000 | 0.73376 | 0.77910 | 1.51413 | 57 |
| GO_RRNA_CONTAINING_RIBONUCLEOPROTEIN_COMPLEX_EXPC | 0.00063 | 0.01217 | 0.47727 | 0.82509 | 1.48443 | 15 |
| GO_RRNA_METABOLIC_PROCESS | 0.00000 | 0.00000 | | 0.71279 | 1.44058 | 211 |
| GO_SCF_DEPENDENT_PROTEASOMAL_UBIQUITIN_DEPENDENT_ | 0.00000 | 0.00000 | 0.74774 | 0.74148 | 1.45605 | 81 |
| GO_SECRETORY_GRANULE | 0.00000 | 0.00000 | 0.71951 | 0.58442 | 1.19681 | 500 |
| GO_SIGNAL_SEQUENCE_BINDING | 0.00020 | 0.00471 | 0.51885 | 0.73408 | 1.40976 | 41 |
| GO_SINGLE_FERTILIZATION | 0.00088 | 0.01614 | 0.47727 | 0.64186 | 1.25806 | 76 |
| GO_SINGLE_STRANDED_RNA_BINDING | 0.00042 | 0.00871 | 0.49849 | 0.66611 | 1.30123 | 70 |

| | | | | | | |
|---|---|---|---|---|---|---|
| GO_SISTER_CHROMATID_SEGREGATION | 0.00009 | 0.00252 | 0.53843 | 0.60810 | 1.22443 | 174 |
| GO_SMALL_NUCLEAR_RIBONUCLEOPROTEIN_COMPLEX | 0.00010 | 0.00263 | 0.53843 | 0.69135 | 1.34452 | 60 |
| GO_SMALL_NUCLEOLAR_RIBONUCLEOPROTEIN_COMPLEX | 0.00076 | 0.01417 | 0.47727 | 0.75791 | 1.41132 | 25 |
| GO_SMALL_RIBOSOMAL_SUBUNIT | 0.00000 | 0.00000 | 0.74774 | 0.75769 | 1.47942 | 69 |
| GO_SMALL_SUBUNIT_PROCESSOME | 0.00028 | 0.00623 | 0.49849 | 0.74257 | 1.40991 | 33 |
| GO_SM_LIKE_PROTEIN_FAMILY_COMPLEX | 0.00008 | 0.00209 | 0.53843 | 0.68608 | 1.34023 | 70 |
| GO_SNORNA_BINDING | 0.00006 | 0.00170 | 0.55733 | 0.78346 | 1.47286 | 28 |
| GO_SOMATIC_STEM_CELL_POPULATION_MAINTENANCE | 0.00261 | 0.03975 | 0.43171 | 0.67102 | 1.29991 | 53 |
| GO_SPERM_EGG_RECOGNITION | 0.00004 | 0.00111 | 0.55733 | 0.76961 | 1.45953 | 32 |
| GO_SPINDLE | 0.00021 | 0.00501 | 0.51885 | 0.57099 | 1.16202 | 311 |
| GO_SPINDLE_ASSEMBLY | 0.00220 | 0.03456 | 0.43171 | 0.61535 | 1.21850 | 100 |
| GO_SPINDLE_ORGANIZATION | 0.00016 | 0.00383 | 0.51885 | 0.60854 | 1.22300 | 160 |
| GO_SPLICEOSOMAL_COMPLEX | 0.00000 | 0.00000 | 0.76146 | 0.66276 | 1.33480 | 177 |
| GO_SPLICEOSOMAL_COMPLEX_ASSEMBLY | 0.00083 | 0.01541 | 0.47727 | 0.68561 | 1.32197 | 48 |
| GO_SPLICEOSOMAL_SNRNP_ASSEMBLY | 0.00061 | 0.01196 | 0.47727 | 0.71875 | 1.37512 | 37 |
| GO_STEM_CELL_DIFFERENTIATION | 0.00000 | 0.00000 | 0.77494 | 0.66991 | 1.34820 | 172 |
| GO_STRUCTURAL_CONSTITUENT_OF_RIBOSOME | 0.00000 | 0.00000 | | 0.78265 | 1.57039 | 154 |
| GO_STRUCTURAL_MOLECULE_ACTIVITY | 0.00000 | 0.00000 | | 0.69725 | 1.42343 | 399 |
| GO_TELOMERASE_RNA_LOCALIZATION | 0.00181 | 0.02958 | 0.45506 | 0.77352 | 1.41188 | 19 |
| GO_TELOMERE_MAINTENANCE_VIA_TELOMERE_LENGTHENING | 0.00015 | 0.00365 | 0.51885 | 0.67268 | 1.31678 | 73 |
| GO_TELOMERE_ORGANIZATION | 0.00010 | 0.00261 | 0.53843 | 0.62464 | 1.24845 | 138 |
| GO_TERMINATION_OF_RNA_POLYMERASE_II_TRANSCRIPTION | 0.00205 | 0.03290 | 0.43171 | 0.70065 | 1.33486 | 35 |
| GO_THREONINE_TYPE_ENDOPEPTIDASE_ACTIVITY | 0.00000 | 0.00001 | 0.70498 | 0.90580 | 1.65852 | 20 |
| GO_THREONINE_TYPE_PEPTIDASE_ACTIVITY | 0.00000 | 0.00000 | 0.73376 | 0.89355 | 1.67003 | 26 |
| GO_TISSUE_MORPHOGENESIS | 0.00202 | 0.03239 | 0.43171 | 0.54980 | 1.12220 | 392 |
| GO_TRANSCRIPTION_COUPLED_NUCLEOTIDE_EXCISION_REPAIR | 0.00050 | 0.01002 | 0.47727 | 0.66075 | 1.29341 | 73 |
| GO_TRANSCRIPTION_ELONGATION_FROM_RNA_POLYMERASE_I | 0.00183 | 0.02980 | 0.45506 | 0.63550 | 1.24644 | 77 |
| GO_TRANSLATIONAL_ELONGATION | 0.00000 | 0.00000 | | 0.75457 | 1.50463 | 125 |
| GO_TRANSLATIONAL_INITIATION | 0.00000 | 0.00000 | | 0.75742 | 1.52368 | 169 |
| GO_TRANSLATIONAL_TERMINATION | 0.00000 | 0.00000 | | 0.75005 | 1.48624 | 102 |
| GO_TRANSLATION_ELONGATION_FACTOR_ACTIVITY | 0.00209 | 0.03323 | 0.43171 | 0.79434 | 1.43699 | 16 |
| GO_TRANSLATION_FACTOR_ACTIVITY_RNA_BINDING | 0.00000 | 0.00000 | 0.78819 | 0.76857 | 1.50364 | 72 |
| GO_TRANSLATION_INITIATION_FACTOR_ACTIVITY | 0.00000 | 0.00001 | 0.69013 | 0.78979 | 1.52220 | 45 |
| GO_TRANSLATION_PREINITIATION_COMPLEX | 0.00046 | 0.00944 | 0.49849 | 0.82850 | 1.49878 | 16 |
| GO_TRANSLATION_REGULATOR_ACTIVITY | 0.00000 | 0.00000 | 0.78819 | 0.71144 | 1.41496 | 113 |
| GO_TRANSLATION_REGULATOR_ACTIVITY_NUCLEIC_ACID_BIND | 0.00000 | 0.00000 | 0.81404 | 0.74854 | 1.47701 | 89 |
| GO_TRNA_TRANSPORT | 0.00023 | 0.00537 | 0.51885 | 0.74517 | 1.41741 | 34 |
| GO_TUMOR_NECROSIS_FACTOR_MEDIATED_SIGNALING_PATHW | 0.00000 | 0.00001 | 0.69013 | 0.67548 | 1.34367 | 117 |
| GO_T_CELL_RECEPTOR_SIGNALING_PATHWAY | 0.00000 | 0.00000 | 0.71951 | 0.67090 | 1.34089 | 137 |
| GO_T_CELL_SELECTION | 0.00298 | 0.04453 | 0.43171 | 0.77028 | 1.40283 | 18 |
| GO_U12_TYPE_SPLICEOSOMAL_COMPLEX | 0.00072 | 0.01361 | 0.47727 | 0.74572 | 1.40192 | 28 |
| GO_U1_SNRNP | 0.00032 | 0.00701 | 0.49849 | 0.83692 | 1.51400 | 16 |
| GO_U2_SNRNP | 0.00228 | 0.03537 | 0.43171 | 0.77909 | 1.41889 | 18 |
| GO_U2_TYPE_CATALYTIC_STEP_2_SPLICEOSOME | 0.00024 | 0.00561 | 0.51885 | 0.75389 | 1.42255 | 30 |
| GO_U2_TYPE_SPLICEOSOMAL_COMPLEX | 0.00003 | 0.00077 | 0.57561 | 0.66921 | 1.31935 | 88 |
| GO_UBIQUITIN_LIKE_PROTEIN_LIGASE_BINDING | 0.00003 | 0.00096 | 0.55733 | 0.59180 | 1.19993 | 249 |
| GO_UNFOLDED_PROTEIN_BINDING | 0.00003 | 0.00082 | 0.57561 | 0.66084 | 1.30659 | 93 |
| GO_VACUOLAR_LUMEN | 0.00063 | 0.01217 | 0.47727 | 0.62461 | 1.23950 | 106 |
| GO_VASODILATION | 0.00093 | 0.01673 | 0.47727 | 0.81722 | 1.47025 | 15 |
| GO_VESICLE_LUMEN | 0.00001 | 0.00022 | 0.61053 | 0.61671 | 1.24439 | 197 |
| GO_VIRAL_GENE_EXPRESSION | 0.00000 | 0.00000 | | 0.74275 | 1.49556 | 174 |
| GO_VIRAL_GENOME_REPLICATION | 0.00209 | 0.03323 | 0.43171 | 0.63279 | 1.24028 | 76 |
| GO_VIRAL_LIFE_CYCLE | 0.00000 | 0.00000 | 0.70498 | 0.63198 | 1.27957 | 226 |
| HP_ACUTE_LEUKEMIA | 0.00040 | 0.00853 | 0.49849 | 0.65070 | 1.27540 | 76 |
| HP_ANEMIA_OF_INADEQUATE_PRODUCTION | 0.00049 | 0.00993 | 0.47727 | 0.62622 | 1.24270 | 106 |
| HP_ATRESIA_OF_THE_EXTERNAL_AUDITORY_CANAL | 0.00145 | 0.02440 | 0.45506 | 0.71315 | 1.35405 | 33 |
| HP_DELAYED_PUBERTY | 0.00133 | 0.02284 | 0.45506 | 0.60928 | 1.21459 | 122 |
| HP_ECHOLALIA | 0.00253 | 0.03882 | 0.43171 | 0.71971 | 1.35302 | 28 |
| HP_GONOSOMAL_INHERITANCE | 0.00043 | 0.00884 | 0.49849 | 0.60320 | 1.21143 | 157 |
| HP_HEMATOLOGICAL_NEOPLASM | 0.00133 | 0.02284 | 0.45506 | 0.58721 | 1.18263 | 178 |
| HP_INCREASED_CSF_LACTATE | 0.00283 | 0.04262 | 0.43171 | 0.63769 | 1.24511 | 69 |
| HP_LEUKEMIA | 0.00088 | 0.01614 | 0.47727 | 0.62619 | 1.24236 | 107 |
| HP_MACROCYTIC_ANEMIA | 0.00000 | 0.00005 | 0.64355 | 0.78956 | 1.51456 | 39 |
| HP_MIGRAINE | 0.00005 | 0.00142 | 0.55733 | 0.68392 | 1.33877 | 73 |
| HP_SPARSE_OR_ABSENT_EYELASHES | 0.00090 | 0.01638 | 0.47727 | 0.69549 | 1.34044 | 45 |
| HP_SUPERNUMERARY_BONES_OF_THE_AXIAL_SKELETON | 0.00187 | 0.03024 | 0.45506 | 0.77279 | 1.41056 | 19 |
| HP_THICK_LOWER_LIP_VERMILION | 0.00253 | 0.03882 | 0.43171 | 0.60336 | 1.19993 | 116 |
| HP_X_LINKED_RECESSIVE_INHERITANCE | 0.00121 | 0.02104 | 0.45506 | 0.61181 | 1.21656 | 114 |
| **Cluster 4** | | | | | | |
| pathway | pval | padj | log2err | ES | NES | size |

| | | | | | | |
|---|---|---|---|---|---|---|
| GO_ACTIN_CYTOSKELETON | 0.00165 | 0.02630 | 0.45506 | 0.56232 | 1.13377 | 355 |
| GO_ACTIN_FILAMENT_BINDING | 0.00383 | 0.04936 | 0.43171 | 0.59551 | 1.17950 | 142 |
| GO_ACTIVATION_OF_IMMUNE_RESPONSE | 0.00000 | 0.00000 | 0.78819 | 0.64619 | 1.29763 | 267 |
| GO_ACTIVATION_OF_INNATE_IMMUNE_RESPONSE | 0.00000 | 0.00000 | 0.71951 | 0.71098 | 1.39477 | 100 |
| GO_ADP_BINDING | 0.00302 | 0.04189 | 0.43171 | 0.71939 | 1.35164 | 31 |
| GO_AEROBIC_ELECTRON_TRANSPORT_CHAIN | 0.00014 | 0.00361 | 0.51885 | 0.87807 | 1.56814 | 15 |
| GO_AEROBIC_RESPIRATION | 0.00003 | 0.00089 | 0.57561 | 0.69285 | 1.35060 | 73 |
| GO_ALTERNATIVE_MRNA_SPLICING_VIA_SPLICEOSOME | 0.00095 | 0.01733 | 0.47727 | 0.65057 | 1.26818 | 73 |
| GO_AMIDE_BINDING | 0.00045 | 0.00951 | 0.49849 | 0.59195 | 1.18274 | 211 |
| GO_AMINE_METABOLIC_PROCESS | 0.00000 | 0.00000 | 0.76146 | 0.73755 | 1.44483 | 93 |
| GO_AMINO_ACID_TRANSMEMBRANE_TRANSPORTER_ACTIVITY | 0.00165 | 0.02630 | 0.45506 | 0.69030 | 1.32643 | 47 |
| GO_ANAPHASE_PROMOTING_COMPLEX_DEPENDENT_CATABOL | 0.00000 | 0.00000 | | 0.79661 | 1.55485 | 77 |
| GO_ANTIGEN_BINDING | 0.00305 | 0.04213 | 0.43171 | 0.75513 | 1.39876 | 23 |
| GO_ANTIGEN_PROCESSING_AND_PRESENTATION | 0.00000 | 0.00000 | 0.77494 | 0.68002 | 1.35266 | 168 |
| GO_ANTIGEN_PROCESSING_AND_PRESENTATION_OF_EXOGENO | 0.00000 | 0.00000 | | 0.81431 | 1.58106 | 64 |
| GO_ANTIGEN_PROCESSING_AND_PRESENTATION_OF_PEPTIDE_ | 0.00000 | 0.00000 | 0.78819 | 0.69996 | 1.38668 | 143 |
| GO_ANTIGEN_PROCESSING_AND_PRESENTATION_OF_PEPTIDE_ | 0.00000 | 0.00000 | 0.82666 | 0.77971 | 1.52034 | 76 |
| GO_ANTIGEN_RECEPTOR_MEDIATED_SIGNALING_PATHWAY | 0.00000 | 0.00000 | 0.73376 | 0.66891 | 1.32931 | 160 |
| GO_ANTIMICROBIAL_HUMORAL_RESPONSE | 0.00130 | 0.02203 | 0.45506 | 0.75642 | 1.40466 | 25 |
| GO_ANTIOXIDANT_ACTIVITY | 0.00342 | 0.04557 | 0.43171 | 0.68906 | 1.32063 | 45 |
| GO_APICAL_PART_OF_CELL | 0.00143 | 0.02360 | 0.45506 | 0.57917 | 1.15883 | 228 |
| GO_APOPTOTIC_SIGNALING_PATHWAY | 0.00000 | 0.00004 | 0.65944 | 0.58554 | 1.18262 | 434 |
| GO_ATP_BIOSYNTHETIC_PROCESS | 0.00268 | 0.03886 | 0.43171 | 0.77792 | 1.42947 | 20 |
| GO_ATP_METABOLIC_PROCESS | 0.00000 | 0.00000 | | 0.67863 | 1.35675 | 221 |
| GO_ATP_SYNTHESIS_COUPLED_ELECTRON_TRANSPORT | 0.00000 | 0.00000 | 0.78819 | 0.76890 | 1.50077 | 77 |
| GO_AZUROPHIL_GRANULE | 0.00004 | 0.00110 | 0.55733 | 0.65056 | 1.28030 | 108 |
| GO_AZUROPHIL_GRANULE_LUMEN | 0.00012 | 0.00314 | 0.53843 | 0.69901 | 1.35563 | 62 |
| GO_BASAL_PART_OF_CELL | 0.00040 | 0.00877 | 0.49849 | 0.71135 | 1.36720 | 48 |
| GO_BASAL_PLASMA_MEMBRANE | 0.00001 | 0.00041 | 0.59333 | 0.77497 | 1.46921 | 38 |
| GO_BASOLATERAL_PLASMA_MEMBRANE | 0.00002 | 0.00074 | 0.57561 | 0.64831 | 1.28189 | 137 |
| GO_BINDING_OF_SPERM_TO_ZONA_PELLUCIDA | 0.00001 | 0.00020 | 0.61053 | 0.88447 | 1.61747 | 19 |
| GO_BLOOD_MICROPARTICLE | 0.00000 | 0.00003 | 0.65944 | 0.76490 | 1.47656 | 54 |
| GO_BRUSH_BORDER | 0.00102 | 0.01837 | 0.45506 | 0.66468 | 1.28946 | 65 |
| GO_CADHERIN_BINDING | 0.00000 | 0.00000 | | 0.66857 | 1.34310 | 274 |
| GO_CALCIUM_DEPENDENT_PROTEIN_BINDING | 0.00146 | 0.02408 | 0.45506 | 0.69226 | 1.33019 | 47 |
| GO_CALCIUM_ION_BINDING | 0.00342 | 0.04557 | 0.43171 | 0.56283 | 1.13222 | 299 |
| GO_CANONICAL_WNT_SIGNALING_PATHWAY | 0.00003 | 0.00085 | 0.57561 | 0.60265 | 1.20790 | 245 |
| GO_CARBOHYDRATE_BIOSYNTHETIC_PROCESS | 0.00213 | 0.03220 | 0.43171 | 0.59791 | 1.18275 | 139 |
| GO_CARBOHYDRATE_CATABOLIC_PROCESS | 0.00076 | 0.01452 | 0.47727 | 0.60805 | 1.20538 | 145 |
| GO_CARBOHYDRATE_TRANSMEMBRANE_TRANSPORTER_ACTIVI | 0.00320 | 0.04352 | 0.43171 | 0.76045 | 1.39912 | 21 |
| GO_CATALYTIC_STEP_2_SPLICEOSOME | 0.00000 | 0.00004 | 0.65944 | 0.71636 | 1.40026 | 83 |
| GO_CATENIN_COMPLEX | 0.00000 | 0.00017 | 0.61053 | 0.89795 | 1.61626 | 16 |
| GO_CATION_TRANSMEMBRANE_TRANSPORT | 0.00276 | 0.03940 | 0.43171 | 0.54984 | 1.11018 | 422 |
| GO_CATION_TRANSMEMBRANE_TRANSPORTER_ACTIVITY | 0.00122 | 0.02106 | 0.45506 | 0.57069 | 1.14659 | 280 |
| GO_CELLULAR_AMINO_ACID_METABOLIC_PROCESS | 0.00002 | 0.00073 | 0.57561 | 0.60996 | 1.22061 | 229 |
| GO_CELLULAR_COMPONENT_DISASSEMBLY | 0.00000 | 0.00000 | 0.74774 | 0.60333 | 1.21863 | 441 |
| GO_CELLULAR_ION_HOMEOSTASIS | 0.00040 | 0.00870 | 0.49849 | 0.57377 | 1.15587 | 338 |
| GO_CELLULAR_IRON_ION_HOMEOSTASIS | 0.00028 | 0.00647 | 0.49849 | 0.71389 | 1.37175 | 47 |
| GO_CELLULAR_KETONE_METABOLIC_PROCESS | 0.00000 | 0.00000 | 0.73376 | 0.65993 | 1.31295 | 172 |
| GO_CELLULAR_NITROGEN_COMPOUND_CATABOLIC_PROCESS | 0.00000 | 0.00000 | | 0.67680 | 1.36796 | 467 |
| GO_CELLULAR_OXIDANT_DETOXIFICATION | 0.00213 | 0.03220 | 0.43171 | 0.67003 | 1.29567 | 57 |
| GO_CELLULAR_PROTEIN_COMPLEX_DISASSEMBLY | 0.00000 | 0.00001 | 0.69013 | 0.65192 | 1.29979 | 183 |
| GO_CELLULAR_RESPIRATION | 0.00000 | 0.00000 | 0.77494 | 0.69020 | 1.36948 | 149 |
| GO_CELLULAR_RESPONSE_TO_HEAT | 0.00012 | 0.00319 | 0.53843 | 0.64560 | 1.26899 | 105 |
| GO_CELLULAR_RESPONSE_TO_OXYGEN_LEVELS | 0.00000 | 0.00000 | | 0.70413 | 1.40134 | 174 |
| GO_CELLULAR_RESPONSE_TO_TOPOLOGICALLY_INCORRECT_PR | 0.00167 | 0.02647 | 0.45506 | 0.60255 | 1.19238 | 140 |
| GO_CELLULAR_RESPONSE_TO_TOXIC_SUBSTANCE | 0.00053 | 0.01087 | 0.47727 | 0.67177 | 1.30546 | 68 |
| GO_CELLULAR_TRANSITION_METAL_ION_HOMEOSTASIS | 0.00152 | 0.02461 | 0.45506 | 0.65417 | 1.27256 | 70 |
| GO_CELL_ACTIVATION_INVOLVED_IN_IMMUNE_RESPONSE | 0.00000 | 0.00000 | | 0.63332 | 1.27943 | 446 |
| GO_CELL_ADHESION_MEDIATOR_ACTIVITY | 0.00161 | 0.02584 | 0.45506 | 0.70594 | 1.33987 | 39 |
| GO_CELL_ADHESION_MOLECULE_BINDING | 0.00000 | 0.00000 | | 0.65145 | 1.31429 | 371 |
| GO_CELL_BODY | 0.00253 | 0.03714 | 0.43171 | 0.55833 | 1.12395 | 329 |
| GO_CELL_CELL_RECOGNITION | 0.00000 | 0.00003 | 0.65944 | 0.84226 | 1.58250 | 31 |
| GO_CELL_CELL_SIGNALING_BY_WNT | 0.00014 | 0.00361 | 0.51885 | 0.57029 | 1.15120 | 379 |
| GO_CELL_CYCLE_G2_M_PHASE_TRANSITION | 0.00002 | 0.00074 | 0.57561 | 0.60790 | 1.21774 | 238 |
| GO_CELL_DIVISION | 0.00006 | 0.00175 | 0.55733 | 0.56443 | 1.14089 | 473 |
| GO_CELL_KILLING | 0.00242 | 0.03592 | 0.43171 | 0.66322 | 1.28313 | 58 |
| GO_CELL_RECOGNITION | 0.00001 | 0.00037 | 0.59333 | 0.71237 | 1.38312 | 64 |
| GO_CELL_REDOX_HOMEOSTASIS | 0.00287 | 0.04045 | 0.43171 | 0.71092 | 1.34397 | 36 |
| GO_CELL_SUBSTRATE_JUNCTION | 0.00000 | 0.00000 | | 0.64335 | 1.29633 | 345 |

| | | | | | | |
|---|---|---|---|---|---|---|
| GO_CELL_SURFACE | 0.00298 | 0.04146 | 0.43171 | 0.55319 | 1.11663 | 376 |
| GO_CELL_SURFACE_RECEPTOR_SIGNALING_PATHWAY_INVOLVEI | 0.00040 | 0.00870 | 0.49849 | 0.56204 | 1.13480 | 422 |
| GO_CHAPERONE_COMPLEX | 0.00046 | 0.00976 | 0.49849 | 0.80469 | 1.48618 | 22 |
| GO_CHAPERONE_MEDIATED_PROTEIN_FOLDING | 0.00063 | 0.01247 | 0.47727 | 0.69076 | 1.33097 | 51 |
| GO_CHROMOSOME_SEGREGATION | 0.00130 | 0.02203 | 0.45506 | 0.56745 | 1.13954 | 272 |
| GO_COLLAGEN_CONTAINING_EXTRACELLULAR_MATRIX | 0.00011 | 0.00292 | 0.53843 | 0.60972 | 1.21580 | 189 |
| GO_CONTRACTILE_FIBER | 0.00015 | 0.00380 | 0.51885 | 0.62054 | 1.23183 | 150 |
| GO_CORNIFICATION | 0.00009 | 0.00250 | 0.53843 | 0.76178 | 1.44338 | 37 |
| GO_CORTICAL_CYTOSKELETON | 0.00383 | 0.04936 | 0.43171 | 0.63105 | 1.23297 | 80 |
| GO_COTRANSLATIONAL_PROTEIN_TARGETING_TO_MEMBRANE | 0.00000 | 0.00000 | | 0.82195 | 1.61066 | 94 |
| GO_CYTOCHROME_COMPLEX | 0.00014 | 0.00365 | 0.51885 | 0.80902 | 1.49858 | 23 |
| GO_CYTOKINE_MEDIATED_SIGNALING_PATHWAY | 0.00000 | 0.00000 | 0.78819 | 0.60864 | 1.22942 | 445 |
| GO_CYTOPLASMIC_TRANSLATION | 0.00000 | 0.00000 | 0.81404 | 0.75305 | 1.47428 | 90 |
| GO_CYTOPLASMIC_TRANSLATIONAL_INITIATION | 0.00044 | 0.00931 | 0.49849 | 0.76054 | 1.42624 | 30 |
| GO_CYTOSOLIC_LARGE_RIBOSOMAL_SUBUNIT | 0.00000 | 0.00000 | 0.82666 | 0.83456 | 1.60892 | 52 |
| GO_CYTOSOLIC_RIBOSOME | 0.00000 | 0.00000 | | 0.82222 | 1.61121 | 96 |
| GO_CYTOSOLIC_SMALL_RIBOSOMAL_SUBUNIT | 0.00000 | 0.00000 | 0.70498 | 0.82908 | 1.57485 | 40 |
| GO_DETOXIFICATION | 0.00016 | 0.00405 | 0.51885 | 0.67151 | 1.30841 | 74 |
| GO_DIENCEPHALON_DEVELOPMENT | 0.00024 | 0.00569 | 0.51885 | 0.76319 | 1.43394 | 31 |
| GO_DISORDERED_DOMAIN_SPECIFIC_BINDING | 0.00070 | 0.01354 | 0.47727 | 0.74536 | 1.40043 | 31 |
| GO_DISULFIDE_OXIDOREDUCTASE_ACTIVITY | 0.00213 | 0.03220 | 0.43171 | 0.74007 | 1.38230 | 28 |
| GO_DNA_BIOSYNTHETIC_PROCESS | 0.00026 | 0.00599 | 0.49849 | 0.61118 | 1.21463 | 158 |
| GO_DOUBLE_STRANDED_RNA_BINDING | 0.00086 | 0.01593 | 0.47727 | 0.67817 | 1.31206 | 58 |
| GO_DRUG_BINDING | 0.00176 | 0.02759 | 0.45506 | 0.67225 | 1.30045 | 56 |
| GO_ELECTRON_TRANSFER_ACTIVITY | 0.00001 | 0.00046 | 0.59333 | 0.68875 | 1.34628 | 83 |
| GO_ELECTRON_TRANSPORT_CHAIN | 0.00000 | 0.00000 | 0.82666 | 0.71221 | 1.40773 | 135 |
| GO_EMBRYONIC_PLACENTA_MORPHOGENESIS | 0.00048 | 0.01009 | 0.49849 | 0.81516 | 1.49071 | 19 |
| GO_ENDOCYTIC_VESICLE_MEMBRANE | 0.00368 | 0.04822 | 0.43171 | 0.60670 | 1.19253 | 105 |
| GO_ENDOPEPTIDASE_ACTIVITY | 0.00019 | 0.00462 | 0.51885 | 0.59992 | 1.19935 | 217 |
| GO_ENDOPEPTIDASE_COMPLEX | 0.00000 | 0.00000 | 0.81404 | 0.79758 | 1.54730 | 65 |
| GO_ENDOPEPTIDASE_REGULATOR_ACTIVITY | 0.00003 | 0.00088 | 0.57561 | 0.69180 | 1.34794 | 74 |
| GO_ENDOPLASMIC_RETICULUM_GOLGI_INTERMEDIATE_COMPA | 0.00067 | 0.01321 | 0.47727 | 0.62697 | 1.23158 | 103 |
| GO_ENDOPLASMIC_RETICULUM_GOLGI_INTERMEDIATE_COMPA | 0.00224 | 0.03349 | 0.43171 | 0.66948 | 1.29461 | 57 |
| GO_ENERGY_DERIVATION_BY_OXIDATION_OF_ORGANIC_COMP | 0.00000 | 0.00000 | 0.74774 | 0.65579 | 1.30966 | 206 |
| GO_EPIDERMAL_CELL_DIFFERENTIATION | 0.00006 | 0.00179 | 0.53843 | 0.63582 | 1.25574 | 132 |
| GO_EPITHELIAL_CELL_DIFFERENTIATION | 0.00026 | 0.00599 | 0.49849 | 0.56975 | 1.15006 | 376 |
| GO_ESTABLISHMENT_OF_ORGANELLE_LOCALIZATION | 0.00115 | 0.02007 | 0.45506 | 0.56246 | 1.13338 | 341 |
| GO_ESTABLISHMENT_OF_PROTEIN_LOCALIZATION_TO_CHROMO | 0.00294 | 0.04113 | 0.43171 | 0.76642 | 1.41551 | 22 |
| GO_ESTABLISHMENT_OF_PROTEIN_LOCALIZATION_TO_ENDOPL | 0.00000 | 0.00000 | | 0.81846 | 1.60878 | 105 |
| GO_ESTABLISHMENT_OF_PROTEIN_LOCALIZATION_TO_MEMBRA | 0.00000 | 0.00000 | | 0.67440 | 1.35442 | 271 |
| GO_ESTABLISHMENT_OF_PROTEIN_LOCALIZATION_TO_ORGANE | 0.00000 | 0.00000 | | 0.65482 | 1.32359 | 472 |
| GO_ESTABLISHMENT_OF_PROTEIN_LOCALIZATION_TO_TELOME | 0.00036 | 0.00797 | 0.49849 | 0.84093 | 1.52181 | 17 |
| GO_ESTABLISHMENT_OF_RNA_LOCALIZATION | 0.00051 | 0.01052 | 0.47727 | 0.59800 | 1.19175 | 180 |
| GO_ESTABLISHMENT_OF_TISSUE_POLARITY | 0.00000 | 0.00000 | | 0.75165 | 1.47649 | 103 |
| GO_EUKARYOTIC_TRANSLATION_INITIATION_FACTOR_3_COMPL | 0.00272 | 0.03904 | 0.43171 | 0.82217 | 1.46831 | 15 |
| GO_EXPORT_ACROSS_PLASMA_MEMBRANE | 0.00062 | 0.01232 | 0.47727 | 0.75413 | 1.40694 | 27 |
| GO_FATTY_ACID_DERIVATIVE_METABOLIC_PROCESS | 0.00220 | 0.03301 | 0.43171 | 0.62775 | 1.22897 | 90 |
| GO_FC_EPSILON_RECEPTOR_SIGNALING_PATHWAY | 0.00000 | 0.00000 | | 0.75608 | 1.48161 | 96 |
| GO_FC_RECEPTOR_SIGNALING_PATHWAY | 0.00000 | 0.00000 | 0.73376 | 0.68601 | 1.35874 | 142 |
| GO_FERTILIZATION | 0.00029 | 0.00663 | 0.49849 | 0.66580 | 1.29787 | 73 |
| GO_FICOLIN_1_RICH_GRANULE | 0.00000 | 0.00000 | 0.76146 | 0.72230 | 1.41976 | 105 |
| GO_GENERATION_OF_PRECURSOR_METABOLITES_AND_ENERGY | 0.00000 | 0.00000 | | 0.63273 | 1.27720 | 376 |
| GO_GENE_SILENCING | 0.00156 | 0.02508 | 0.45506 | 0.58726 | 1.17103 | 189 |
| GO_GLUCOSE_CATABOLIC_PROCESS | 0.00028 | 0.00647 | 0.49849 | 0.78427 | 1.45991 | 26 |
| GO_GLYCOLYTIC_PROCESS_THROUGH_FRUCTOSE_6_PHOSPHAT | 0.00010 | 0.00275 | 0.53843 | 0.80687 | 1.49721 | 24 |
| GO_GOLGI_LUMEN | 0.00342 | 0.04557 | 0.43171 | 0.67651 | 1.29992 | 47 |
| GO_GUANYL_NUCLEOTIDE_BINDING | 0.00198 | 0.03047 | 0.43171 | 0.56792 | 1.14048 | 268 |
| GO_HEAT_SHOCK_PROTEIN_BINDING | 0.00011 | 0.00300 | 0.53843 | 0.64698 | 1.27088 | 103 |
| GO_HEMATOPOIETIC_PROGENITOR_CELL_DIFFERENTIATION | 0.00000 | 0.00001 | 0.70498 | 0.68370 | 1.34976 | 131 |
| GO_HEMATOPOIETIC_STEM_CELL_DIFFERENTIATION | 0.00000 | 0.00000 | | 0.79105 | 1.54244 | 76 |
| GO_HEXOSE_CATABOLIC_PROCESS | 0.00156 | 0.02508 | 0.45506 | 0.70628 | 1.33899 | 38 |
| GO_HORMONE_BINDING | 0.00148 | 0.02426 | 0.45506 | 0.72372 | 1.36817 | 36 |
| GO_HUMORAL_IMMUNE_RESPONSE | 0.00087 | 0.01605 | 0.47727 | 0.65604 | 1.27682 | 71 |
| GO_HYDROGEN_PEROXIDE_CATABOLIC_PROCESS | 0.00051 | 0.01052 | 0.47727 | 0.83463 | 1.51042 | 17 |
| GO_HYDROGEN_PEROXIDE_METABOLIC_PROCESS | 0.00029 | 0.00663 | 0.49849 | 0.76645 | 1.43734 | 30 |
| GO_IMMUNE_RESPONSE_REGULATING_SIGNALING_PATHWAY | 0.00000 | 0.00000 | 0.74774 | 0.63907 | 1.28164 | 254 |
| GO_IMPORT_ACROSS_PLASMA_MEMBRANE | 0.00287 | 0.04045 | 0.43171 | 0.63479 | 1.24029 | 80 |
| GO_IMPORT_INTO_NUCLEUS | 0.00108 | 0.01920 | 0.45506 | 0.60846 | 1.20265 | 135 |
| GO_INNATE_IMMUNE_RESPONSE | 0.00000 | 0.00005 | 0.64355 | 0.58374 | 1.17898 | 443 |
| GO_INNATE_IMMUNE_RESPONSE_ACTIVATING_SIGNAL_TRANSI | 0.00000 | 0.00000 | 0.76146 | 0.75359 | 1.47312 | 84 |

| | | | | | | |
|---|---|---|---|---|---|---|
| GO_INNER_MITOCHONDRIAL_MEMBRANE_PROTEIN_COMPLEX | 0.00000 | 0.00000 | 0.73376 | 0.72135 | 1.41352 | 94 |
| GO_INORGANIC_ION_TRANSMEMBRANE_TRANSPORT | 0.00361 | 0.04735 | 0.43171 | 0.55146 | 1.11390 | 407 |
| GO_INTERACTION_WITH_HOST | 0.00276 | 0.03940 | 0.43171 | 0.59666 | 1.18388 | 149 |
| GO_INTERACTION_WITH_SYMBIONT | 0.00097 | 0.01752 | 0.47727 | 0.67939 | 1.31043 | 53 |
| GO_INTERLEUKIN_1_MEDIATED_SIGNALING_PATHWAY | 0.00000 | 0.00000 | | 0.77514 | 1.51690 | 89 |
| GO_INTRINSIC_APOPTOTIC_SIGNALING_PATHWAY | 0.00068 | 0.01327 | 0.47727 | 0.58278 | 1.16662 | 230 |
| GO_IN_UTERO_EMBRYONIC_DEVELOPMENT | 0.00279 | 0.03966 | 0.43171 | 0.55920 | 1.12427 | 290 |
| GO_IRON_ION_HOMEOSTASIS | 0.00106 | 0.01892 | 0.45506 | 0.66860 | 1.29362 | 59 |
| GO_I_BAND | 0.00331 | 0.04474 | 0.43171 | 0.62625 | 1.22552 | 89 |
| GO_KERATINIZATION | 0.00065 | 0.01293 | 0.47727 | 0.71509 | 1.36319 | 42 |
| GO_KERATINOCYTE_DIFFERENTIATION | 0.00071 | 0.01368 | 0.47727 | 0.63908 | 1.25231 | 94 |
| GO_LABYRINTHINE_LAYER_MORPHOGENESIS | 0.00035 | 0.00780 | 0.49849 | 0.83753 | 1.50750 | 16 |
| GO_LARGE_RIBOSOMAL_SUBUNIT | 0.00000 | 0.00000 | | 0.78431 | 1.54256 | 107 |
| GO_LATERAL_PLASMA_MEMBRANE | 0.00113 | 0.01991 | 0.45506 | 0.68640 | 1.32013 | 49 |
| GO_LEUKOCYTE_MEDIATED_IMMUNITY | 0.00000 | 0.00000 | | 0.64222 | 1.29810 | 466 |
| GO_LEUKOCYTE_MIGRATION | 0.00049 | 0.01024 | 0.47727 | 0.58851 | 1.17657 | 221 |
| GO_LOW_DENSITY_LIPOPROTEIN_PARTICLE_CLEARANCE | 0.00261 | 0.03787 | 0.43171 | 0.74324 | 1.38354 | 26 |
| GO_LYASE_ACTIVITY | 0.00097 | 0.01752 | 0.47727 | 0.61664 | 1.21529 | 118 |
| GO_LYSOSOMAL_LUMEN | 0.00012 | 0.00314 | 0.53843 | 0.69133 | 1.34117 | 65 |
| GO_L_AMINO_ACID_TRANSMEMBRANE_TRANSPORTER_ACTIVIT | 0.00261 | 0.03787 | 0.43171 | 0.72095 | 1.36229 | 35 |
| GO_L_AMINO_ACID_TRANSPORT | 0.00117 | 0.02028 | 0.45506 | 0.70544 | 1.34200 | 41 |
| GO_MACROAUTOPHAGY | 0.00202 | 0.03089 | 0.43171 | 0.56865 | 1.14136 | 263 |
| GO_MAINTENANCE_OF_LOCATION_IN_CELL | 0.00316 | 0.04320 | 0.43171 | 0.60308 | 1.19000 | 128 |
| GO_MAST_CELL_MEDIATED_IMMUNITY | 0.00279 | 0.03966 | 0.43171 | 0.71568 | 1.34950 | 33 |
| GO_MEMBRANE_REGION | 0.00001 | 0.00020 | 0.61053 | 0.61899 | 1.23662 | 208 |
| GO_METAL_ION_TRANSPORT | 0.00331 | 0.04474 | 0.43171 | 0.54751 | 1.10590 | 439 |
| GO_MIDBRAIN_DEVELOPMENT | 0.00209 | 0.03187 | 0.43171 | 0.66913 | 1.29442 | 56 |
| GO_MITOCHONDRIAL_ELECTRON_TRANSPORT_NADH_TO_UBIC | 0.00002 | 0.00055 | 0.57561 | 0.76012 | 1.45506 | 44 |
| GO_MITOCHONDRIAL_GENE_EXPRESSION | 0.00000 | 0.00007 | 0.64355 | 0.64764 | 1.28580 | 153 |
| GO_MITOCHONDRIAL_LARGE_RIBOSOMAL_SUBUNIT | 0.00001 | 0.00024 | 0.61053 | 0.74018 | 1.42884 | 54 |
| GO_MITOCHONDRIAL_MATRIX | 0.00000 | 0.00004 | 0.64355 | 0.59321 | 1.19777 | 387 |
| GO_MITOCHONDRIAL_PROTEIN_COMPLEX | 0.00000 | 0.00000 | | 0.71752 | 1.43394 | 214 |
| GO_MITOCHONDRIAL_RESPIRATORY_CHAIN_COMPLEX_ASSEME | 0.00030 | 0.00679 | 0.49849 | 0.65162 | 1.27454 | 86 |
| GO_MITOCHONDRIAL_SMALL_RIBOSOMAL_SUBUNIT | 0.00081 | 0.01517 | 0.47727 | 0.75885 | 1.41737 | 28 |
| GO_MITOCHONDRIAL_TRANSLATION | 0.00000 | 0.00005 | 0.64355 | 0.67091 | 1.32391 | 129 |
| GO_MITOCHONDRIAL_TRANSLATIONAL_TERMINATION | 0.00000 | 0.00000 | 0.76146 | 0.74239 | 1.45244 | 87 |
| GO_MITOCHONDRION_ORGANIZATION | 0.00001 | 0.00031 | 0.59333 | 0.57897 | 1.16912 | 431 |
| GO_MITOTIC_SPINDLE_ORGANIZATION | 0.00361 | 0.04735 | 0.43171 | 0.60701 | 1.19315 | 105 |
| GO_MODULATION_BY_HOST_OF_VIRAL_PROCESS | 0.00187 | 0.02890 | 0.45506 | 0.77488 | 1.43113 | 22 |
| GO_MONOAMINE_TRANSPORT | 0.00178 | 0.02781 | 0.45506 | 0.74958 | 1.39534 | 26 |
| GO_MONOCARBOXYLIC_ACID_METABOLIC_PROCESS | 0.00081 | 0.01517 | 0.47727 | 0.56054 | 1.13205 | 416 |
| GO_MONOVALENT_INORGANIC_CATION_TRANSMEMBRANE_TR | 0.00002 | 0.00073 | 0.57561 | 0.63597 | 1.26225 | 151 |
| GO_MONOVALENT_INORGANIC_CATION_TRANSPORT | 0.00002 | 0.00062 | 0.57561 | 0.60294 | 1.20917 | 254 |
| GO_MORPHOGENESIS_OF_AN_EPITHELIUM | 0.00000 | 0.00001 | 0.69013 | 0.60334 | 1.21656 | 361 |
| GO_MORPHOGENESIS_OF_A_POLARIZED_EPITHELIUM | 0.00000 | 0.00000 | | 0.76352 | 1.50467 | 119 |
| GO_MRNA_3_END_PROCESSING | 0.00327 | 0.04443 | 0.43171 | 0.62160 | 1.21805 | 94 |
| GO_MRNA_5_UTR_BINDING | 0.00024 | 0.00575 | 0.51885 | 0.78835 | 1.46396 | 25 |
| GO_MRNA_BINDING | 0.00000 | 0.00000 | 0.80122 | 0.65015 | 1.30398 | 252 |
| GO_MRNA_EXPORT_FROM_NUCLEUS | 0.00050 | 0.01040 | 0.47727 | 0.63028 | 1.23809 | 103 |
| GO_MRNA_PROCESSING | 0.00000 | 0.00000 | | 0.62831 | 1.26958 | 452 |
| GO_MRNA_TRANSPORT | 0.00115 | 0.02007 | 0.45506 | 0.60682 | 1.19960 | 136 |
| GO_MULTI_MULTICELLULAR_ORGANISM_PROCESS | 0.00001 | 0.00050 | 0.59333 | 0.64717 | 1.27634 | 125 |
| GO_MYD88_DEPENDENT_TOLL_LIKE_RECEPTOR_SIGNALING_PA | 0.00342 | 0.04557 | 0.43171 | 0.76063 | 1.40481 | 22 |
| GO_MYELOID_LEUKOCYTE_ACTIVATION | 0.00000 | 0.00000 | | 0.64219 | 1.29696 | 418 |
| GO_MYELOID_LEUKOCYTE_MEDIATED_IMMUNITY | 0.00000 | 0.00000 | | 0.66338 | 1.33834 | 372 |
| GO_NADH_DEHYDROGENASE_ACTIVITY | 0.00019 | 0.00462 | 0.51885 | 0.75022 | 1.42149 | 37 |
| GO_NADH_DEHYDROGENASE_COMPLEX | 0.00006 | 0.00171 | 0.55733 | 0.75863 | 1.44104 | 40 |
| GO_NADH_DEHYDROGENASE_COMPLEX_ASSEMBLY | 0.00075 | 0.01422 | 0.47727 | 0.68202 | 1.31826 | 55 |
| GO_NADH_METABOLIC_PROCESS | 0.00006 | 0.00179 | 0.53843 | 0.79449 | 1.48992 | 30 |
| GO_NAD_METABOLIC_PROCESS | 0.00036 | 0.00805 | 0.49849 | 0.75426 | 1.42523 | 35 |
| GO_NCRNA_METABOLIC_PROCESS | 0.00000 | 0.00018 | 0.61053 | 0.58365 | 1.17865 | 398 |
| GO_NCRNA_PROCESSING | 0.00000 | 0.00002 | 0.65944 | 0.60089 | 1.21076 | 343 |
| GO_NEGATIVE_REGULATION_OF_APOPTOTIC_SIGNALING_PATH\ | 0.00008 | 0.00216 | 0.53843 | 0.61752 | 1.22781 | 166 |
| GO_NEGATIVE_REGULATION_OF_CANONICAL_WNT_SIGNALING | 0.00000 | 0.00000 | | 0.72939 | 1.44101 | 133 |
| GO_NEGATIVE_REGULATION_OF_CATALYTIC_ACTIVITY | 0.00183 | 0.02839 | 0.45506 | 0.54673 | 1.10541 | 482 |
| GO_NEGATIVE_REGULATION_OF_CELL_CYCLE | 0.00007 | 0.00211 | 0.53843 | 0.56455 | 1.14167 | 491 |
| GO_NEGATIVE_REGULATION_OF_CELL_CYCLE_G2_M_PHASE_TR | 0.00000 | 0.00002 | 0.65944 | 0.70677 | 1.38366 | 90 |
| GO_NEGATIVE_REGULATION_OF_CELL_CYCLE_PHASE_TRANSITI( | 0.00000 | 0.00010 | 0.62726 | 0.62702 | 1.25243 | 207 |
| GO_NEGATIVE_REGULATION_OF_CELL_CYCLE_PROCESS | 0.00000 | 0.00016 | 0.61053 | 0.60575 | 1.21699 | 277 |
| GO_NEGATIVE_REGULATION_OF_INTRINSIC_APOPTOTIC_SIGNAI | 0.00346 | 0.04597 | 0.43171 | 0.63921 | 1.24607 | 75 |

| | | | | | | |
|---|---|---|---|---|---|---|
| GO_NEGATIVE_REGULATION_OF_MITOTIC_CELL_CYCLE | 0.00001 | 0.00029 | 0.59333 | 0.60523 | 1.21538 | 267 |
| GO_NEGATIVE_REGULATION_OF_PEPTIDASE_ACTIVITY | 0.00010 | 0.00273 | 0.53843 | 0.63961 | 1.26057 | 118 |
| GO_NEGATIVE_REGULATION_OF_PROTEOLYSIS | 0.00372 | 0.04850 | 0.43171 | 0.57726 | 1.15146 | 190 |
| GO_NEGATIVE_REGULATION_OF_RNA_SPLICING | 0.00141 | 0.02360 | 0.45506 | 0.76188 | 1.41372 | 24 |
| GO_NEGATIVE_REGULATION_OF_WNT_SIGNALING_PATHWAY | 0.00000 | 0.00000 | | 0.70654 | 1.40273 | 153 |
| GO_NEURAL_NUCLEUS_DEVELOPMENT | 0.00235 | 0.03498 | 0.43171 | 0.70732 | 1.33640 | 34 |
| GO_NEURON_PROJECTION_TERMINUS | 0.00246 | 0.03638 | 0.43171 | 0.64950 | 1.26349 | 70 |
| GO_NIK_NF_KAPPAB_SIGNALING | 0.00000 | 0.00000 | | 0.73302 | 1.44467 | 120 |
| GO_NON_CANONICAL_WNT_SIGNALING_PATHWAY | 0.00000 | 0.00000 | | 0.73533 | 1.44944 | 122 |
| GO_NUCLEAR_CHROMOSOME_TELOMERIC_REGION | 0.00202 | 0.03089 | 0.43171 | 0.62525 | 1.22485 | 93 |
| GO_NUCLEAR_ENVELOPE | 0.00111 | 0.01975 | 0.45506 | 0.56314 | 1.13509 | 353 |
| GO_NUCLEAR_EXPORT | 0.00003 | 0.00093 | 0.57561 | 0.61905 | 1.23426 | 183 |
| GO_NUCLEAR_PERIPHERY | 0.00033 | 0.00734 | 0.49849 | 0.63109 | 1.24340 | 115 |
| GO_NUCLEAR_SPECK | 0.00011 | 0.00292 | 0.53843 | 0.57774 | 1.16298 | 331 |
| GO_NUCLEAR_TRANSCRIBED_MRNA_CATABOLIC_PROCESS | 0.00000 | 0.00000 | | 0.72137 | 1.43808 | 188 |
| GO_NUCLEAR_TRANSCRIBED_MRNA_CATABOLIC_PROCESS_NON | 0.00000 | 0.00000 | | 0.79649 | 1.56803 | 112 |
| GO_NUCLEAR_TRANSPORT | 0.00000 | 0.00005 | 0.64355 | 0.60551 | 1.21833 | 303 |
| GO_NUCLEOBASE_CONTAINING_COMPOUND_TRANSPORT | 0.00078 | 0.01483 | 0.47727 | 0.58014 | 1.15977 | 220 |
| GO_NUCLEOBASE_CONTAINING_SMALL_MOLECULE_METABOLI( | 0.00009 | 0.00241 | 0.53843 | 0.56895 | 1.14908 | 413 |
| GO_NUCLEOSIDE_DIPHOSPHATE_METABOLIC_PROCESS | 0.00313 | 0.04306 | 0.43171 | 0.60868 | 1.19926 | 117 |
| GO_NUCLEOSIDE_PHOSPHATE_BIOSYNTHETIC_PROCESS | 0.00018 | 0.00441 | 0.51885 | 0.61062 | 1.21444 | 169 |
| GO_NUCLEOSIDE_TRIPHOSPHATE_BIOSYNTHETIC_PROCESS | 0.00287 | 0.04045 | 0.43171 | 0.68842 | 1.30960 | 41 |
| GO_NUCLEOSIDE_TRIPHOSPHATE_METABOLIC_PROCESS | 0.00220 | 0.03301 | 0.43171 | 0.65817 | 1.27642 | 62 |
| GO_NUCLEOTIDE_PHOSPHORYLATION | 0.00161 | 0.02584 | 0.45506 | 0.62341 | 1.22389 | 102 |
| GO_ORGANELLAR_RIBOSOME | 0.00000 | 0.00001 | 0.69013 | 0.72476 | 1.41677 | 84 |
| GO_ORGANELLE_ENVELOPE_LUMEN | 0.00383 | 0.04936 | 0.43171 | 0.63759 | 1.24290 | 75 |
| GO_ORGANELLE_INNER_MEMBRANE | 0.00000 | 0.00000 | | 0.64567 | 1.30408 | 410 |
| GO_ORGANIC_ACID_BIOSYNTHETIC_PROCESS | 0.00372 | 0.04850 | 0.43171 | 0.57372 | 1.14656 | 214 |
| GO_ORGANIC_ACID_TRANSMEMBRANE_TRANSPORTER_ACTIVIT | 0.00167 | 0.02647 | 0.45506 | 0.62952 | 1.23131 | 86 |
| GO_ORGANIC_CYCLIC_COMPOUND_CATABOLIC_PROCESS | 0.00000 | 0.00000 | | 0.66618 | 1.34739 | 495 |
| GO_OXIDATIVE_PHOSPHORYLATION | 0.00000 | 0.00000 | 0.82666 | 0.75288 | 1.47600 | 97 |
| GO_OXIDOREDUCTASE_ACTIVITY | 0.00000 | 0.00000 | 0.77494 | 0.60009 | 1.21323 | 480 |
| GO_OXIDOREDUCTASE_ACTIVITY_ACTING_ON_A_HEME_GROUF | 0.00013 | 0.00342 | 0.51885 | 0.83401 | 1.53255 | 20 |
| GO_OXIDOREDUCTASE_ACTIVITY_ACTING_ON_A_SULFUR_GROU | 0.00257 | 0.03751 | 0.43171 | 0.68876 | 1.31847 | 44 |
| GO_OXIDOREDUCTASE_ACTIVITY_ACTING_ON_NAD_P_H | 0.00001 | 0.00023 | 0.61053 | 0.70567 | 1.37734 | 77 |
| GO_OXIDOREDUCTASE_ACTIVITY_ACTING_ON_NAD_P_H_QUIN( | 0.00010 | 0.00276 | 0.53843 | 0.73105 | 1.40243 | 46 |
| GO_OXIDOREDUCTASE_ACTIVITY_ACTING_ON_PEROXIDE_AS_A( | 0.00350 | 0.04617 | 0.43171 | 0.71899 | 1.34833 | 30 |
| GO_OXIDOREDUCTASE_COMPLEX | 0.00000 | 0.00001 | 0.67496 | 0.72428 | 1.41561 | 81 |
| GO_PEPTIDASE_COMPLEX | 0.00000 | 0.00000 | 0.74774 | 0.74651 | 1.45914 | 82 |
| GO_PEPTIDASE_REGULATOR_ACTIVITY | 0.00130 | 0.02203 | 0.45506 | 0.62758 | 1.23079 | 98 |
| GO_PEPTIDE_BINDING | 0.00041 | 0.00884 | 0.49849 | 0.60053 | 1.19340 | 159 |
| GO_PHAGOCYTOSIS | 0.00316 | 0.04320 | 0.43171 | 0.58866 | 1.17010 | 162 |
| GO_PIGMENT_GRANULE | 0.00000 | 0.00000 | 0.76146 | 0.74388 | 1.45571 | 89 |
| GO_PLACENTA_DEVELOPMENT | 0.00069 | 0.01341 | 0.47727 | 0.62200 | 1.22452 | 112 |
| GO_PLASMA_LIPOPROTEIN_PARTICLE_CLEARANCE | 0.00350 | 0.04617 | 0.43171 | 0.69181 | 1.31155 | 38 |
| GO_POLYMERIC_CYTOSKELETAL_FIBER | 0.00139 | 0.02335 | 0.45506 | 0.55094 | 1.11311 | 448 |
| GO_POLYSOMAL_RIBOSOME | 0.00001 | 0.00038 | 0.59333 | 0.81007 | 1.51520 | 29 |
| GO_POLYSOME | 0.00000 | 0.00000 | 0.73376 | 0.77797 | 1.51045 | 63 |
| GO_POSITIVE_REGULATION_OF_CANONICAL_WNT_SIGNALING_ | 0.00000 | 0.00004 | 0.64355 | 0.67627 | 1.33282 | 118 |
| GO_POSITIVE_REGULATION_OF_CATION_CHANNEL_ACTIVITY | 0.00172 | 0.02714 | 0.45506 | 0.73352 | 1.36848 | 27 |
| GO_POSITIVE_REGULATION_OF_CELLULAR_AMIDE_METABOLIC_ | 0.00001 | 0.00019 | 0.61053 | 0.65130 | 1.28674 | 133 |
| GO_POSITIVE_REGULATION_OF_CELLULAR_COMPONENT_BIOGI | 0.00383 | 0.04936 | 0.43171 | 0.55204 | 1.11358 | 366 |
| GO_POSITIVE_REGULATION_OF_CELLULAR_PROTEIN_LOCALIZAT | 0.00115 | 0.02007 | 0.45506 | 0.57247 | 1.14956 | 266 |
| GO_POSITIVE_REGULATION_OF_DEFENSE_RESPONSE | 0.00000 | 0.00001 | 0.69013 | 0.63659 | 1.27179 | 208 |
| GO_POSITIVE_REGULATION_OF_DNA_BIOSYNTHETIC_PROCESS | 0.00026 | 0.00599 | 0.49849 | 0.68857 | 1.33537 | 62 |
| GO_POSITIVE_REGULATION_OF_ESTABLISHMENT_OF_PROTEIN_ | 0.00014 | 0.00370 | 0.51885 | 0.58679 | 1.17889 | 277 |
| GO_POSITIVE_REGULATION_OF_IMMUNE_RESPONSE | 0.00000 | 0.00000 | 0.82666 | 0.62431 | 1.25863 | 356 |
| GO_POSITIVE_REGULATION_OF_INNATE_IMMUNE_RESPONSE | 0.00000 | 0.00002 | 0.65944 | 0.67515 | 1.33289 | 131 |
| GO_POSITIVE_REGULATION_OF_INTRACELLULAR_TRANSPORT | 0.00294 | 0.04113 | 0.43171 | 0.58708 | 1.17023 | 181 |
| GO_POSITIVE_REGULATION_OF_MUSCLE_CONTRACTION | 0.00068 | 0.01327 | 0.47727 | 0.80781 | 1.48441 | 20 |
| GO_POSITIVE_REGULATION_OF_ORGANELLE_ORGANIZATION | 0.00000 | 0.00016 | 0.61053 | 0.57807 | 1.16873 | 481 |
| GO_POSITIVE_REGULATION_OF_PROTEIN_CONTAINING_COMPL | 0.00316 | 0.04320 | 0.43171 | 0.58287 | 1.16205 | 186 |
| GO_POSITIVE_REGULATION_OF_PROTEIN_LOCALIZATION_TO_N | 0.00034 | 0.00772 | 0.49849 | 0.67679 | 1.31553 | 67 |
| GO_POSITIVE_REGULATION_OF_RESPONSE_TO_BIOTIC_STIMUL | 0.00000 | 0.00002 | 0.65944 | 0.65642 | 1.30324 | 153 |
| GO_POSITIVE_REGULATION_OF_RESPONSE_TO_EXTERNAL_STIM | 0.00000 | 0.00001 | 0.67496 | 0.61755 | 1.24066 | 278 |
| GO_POSITIVE_REGULATION_OF_SIGNAL_TRANSDUCTION_BY_P5 | 0.00272 | 0.03904 | 0.43171 | 0.82170 | 1.46747 | 15 |
| GO_POSITIVE_REGULATION_OF_TELOMERASE_ACTIVITY | 0.00216 | 0.03261 | 0.43171 | 0.71973 | 1.35715 | 33 |
| GO_POSITIVE_REGULATION_OF_TELOMERASE_RNA_LOCALIZATI | 0.00001 | 0.00039 | 0.59333 | 0.91013 | 1.62540 | 15 |
| GO_POSITIVE_REGULATION_OF_TELOMERE_MAINTENANCE | 0.00025 | 0.00584 | 0.49849 | 0.71578 | 1.37538 | 47 |
| GO_POSITIVE_REGULATION_OF_TELOMERE_MAINTENANCE_VIA | 0.00001 | 0.00030 | 0.59333 | 0.79643 | 1.50562 | 36 |

| | | | | | | |
|---|---|---|---|---|---|---|
| GO_POSITIVE_REGULATION_OF_TRANSLATION | 0.00001 | 0.00021 | 0.61053 | 0.66818 | 1.31532 | 111 |
| GO_POSITIVE_REGULATION_OF_VIRAL_PROCESS | 0.00002 | 0.00078 | 0.57561 | 0.67043 | 1.31373 | 94 |
| GO_POSITIVE_REGULATION_OF_VIRAL_TRANSCRIPTION | 0.00135 | 0.02285 | 0.45506 | 0.70942 | 1.34494 | 38 |
| GO_POSITIVE_REGULATION_OF_WNT_SIGNALING_PATHWAY | 0.00003 | 0.00085 | 0.57561 | 0.63986 | 1.26621 | 140 |
| GO_POSTSYNAPSE | 0.00320 | 0.04352 | 0.43171 | 0.55941 | 1.12734 | 349 |
| GO_POST_TRANSLATIONAL_PROTEIN_MODIFICATION | 0.00001 | 0.00026 | 0.61053 | 0.60620 | 1.21732 | 267 |
| GO_PRECATALYTIC_SPLICEOSOME | 0.00004 | 0.00127 | 0.55733 | 0.72616 | 1.39918 | 51 |
| GO_PRERIBOSOME | 0.00016 | 0.00408 | 0.51885 | 0.68159 | 1.32591 | 70 |
| GO_PROTEASOMAL_PROTEIN_CATABOLIC_PROCESS | 0.00001 | 0.00027 | 0.61053 | 0.58249 | 1.17634 | 415 |
| GO_PROTEASOMAL_UBIQUITIN_INDEPENDENT_PROTEIN_CATAB | 0.00000 | 0.00000 | 0.70498 | 0.92005 | 1.68254 | 19 |
| GO_PROTEASOME_ACCESSORY_COMPLEX | 0.00000 | 0.00008 | 0.62726 | 0.85984 | 1.59550 | 24 |
| GO_PROTEASOME_CORE_COMPLEX | 0.00000 | 0.00001 | 0.70498 | 0.93086 | 1.68457 | 17 |
| GO_PROTEIN_CONTAINING_COMPLEX_DISASSEMBLY | 0.00000 | 0.00000 | 0.76146 | 0.63869 | 1.28271 | 271 |
| GO_PROTEIN_CONTAINING_COMPLEX_LOCALIZATION | 0.00033 | 0.00734 | 0.49849 | 0.58817 | 1.17740 | 230 |
| GO_PROTEIN_DNA_COMPLEX_SUBUNIT_ORGANIZATION | 0.00113 | 0.01991 | 0.45506 | 0.59683 | 1.18966 | 181 |
| GO_PROTEIN_FOLDING | 0.00000 | 0.00002 | 0.67496 | 0.64644 | 1.28879 | 186 |
| GO_PROTEIN_FOLDING_CHAPERONE | 0.00010 | 0.00268 | 0.53843 | 0.80704 | 1.49752 | 24 |
| GO_PROTEIN_IMPORT | 0.00117 | 0.02028 | 0.45506 | 0.60056 | 1.19394 | 164 |
| GO_PROTEIN_LOCALIZATION_TO_CHROMOSOME_TELOMERIC_F | 0.00024 | 0.00575 | 0.51885 | 0.77825 | 1.45360 | 28 |
| GO_PROTEIN_LOCALIZATION_TO_ENDOPLASMIC_RETICULUM | 0.00000 | 0.00000 | | 0.79236 | 1.56354 | 127 |
| GO_PROTEIN_LOCALIZATION_TO_MEMBRANE | 0.00000 | 0.00000 | | 0.62844 | 1.27056 | 481 |
| GO_PROTEIN_LOCALIZATION_TO_MITOCHONDRION | 0.00148 | 0.02426 | 0.45506 | 0.61437 | 1.21129 | 124 |
| GO_PROTEIN_LOCALIZATION_TO_NUCLEUS | 0.00003 | 0.00101 | 0.55733 | 0.60571 | 1.21111 | 219 |
| GO_PROTEIN_MODIFICATION_BY_SMALL_PROTEIN_REMOVAL | 0.00016 | 0.00408 | 0.51885 | 0.59387 | 1.19022 | 242 |
| GO_PROTEIN_POLYUBIQUITINATION | 0.00060 | 0.01211 | 0.47727 | 0.57578 | 1.15679 | 279 |
| GO_PROTEIN_TARGETING | 0.00000 | 0.00000 | | 0.66563 | 1.34215 | 361 |
| GO_PROTEIN_TARGETING_TO_MEMBRANE | 0.00000 | 0.00000 | | 0.73383 | 1.45865 | 162 |
| GO_PROTEIN_TARGETING_TO_MITOCHONDRION | 0.00092 | 0.01698 | 0.47727 | 0.63997 | 1.25113 | 85 |
| GO_PROTON_TRANSMEMBRANE_TRANSPORT | 0.00000 | 0.00011 | 0.62726 | 0.70196 | 1.37220 | 84 |
| GO_PROTON_TRANSMEMBRANE_TRANSPORTER_ACTIVITY | 0.00048 | 0.01009 | 0.49849 | 0.67178 | 1.30579 | 67 |
| GO_PURINE_CONTAINING_COMPOUND_BIOSYNTHETIC_PROCES | 0.00006 | 0.00175 | 0.55733 | 0.64280 | 1.26845 | 129 |
| GO_PURINE_CONTAINING_COMPOUND_METABOLIC_PROCESS | 0.00004 | 0.00108 | 0.55733 | 0.58797 | 1.18311 | 307 |
| GO_PURINE_NUCLEOSIDE_TRIPHOSPHATE_METABOLIC_PROCES | 0.00135 | 0.02285 | 0.45506 | 0.69215 | 1.32781 | 46 |
| GO_PYRUVATE_METABOLIC_PROCESS | 0.00041 | 0.00891 | 0.49849 | 0.62911 | 1.23938 | 116 |
| GO_REGULATION_OF_ANIMAL_ORGAN_MORPHOGENESIS | 0.00000 | 0.00000 | | 0.70642 | 1.40435 | 163 |
| GO_REGULATION_OF_APOPTOTIC_SIGNALING_PATHWAY | 0.00004 | 0.00126 | 0.55733 | 0.58791 | 1.18236 | 295 |
| GO_REGULATION_OF_ATPASE_ACTIVITY | 0.00253 | 0.03714 | 0.43171 | 0.65150 | 1.26150 | 60 |
| GO_REGULATION_OF_ATP_METABOLIC_PROCESS | 0.00257 | 0.03751 | 0.43171 | 0.63010 | 1.23306 | 89 |
| GO_REGULATION_OF_CELLULAR_AMIDE_METABOLIC_PROCESS | 0.00000 | 0.00001 | 0.69013 | 0.60858 | 1.22694 | 358 |
| GO_REGULATION_OF_CELLULAR_AMINE_METABOLIC_PROCESS | 0.00000 | 0.00000 | 0.74774 | 0.78979 | 1.52927 | 60 |
| GO_REGULATION_OF_CELLULAR_AMINO_ACID_METABOLIC_PR( | 0.00000 | 0.00000 | 0.78819 | 0.80754 | 1.56087 | 55 |
| GO_REGULATION_OF_CELLULAR_KETONE_METABOLIC_PROCESS | 0.00000 | 0.00001 | 0.69013 | 0.67542 | 1.33396 | 132 |
| GO_REGULATION_OF_CELLULAR_PROTEIN_LOCALIZATION | 0.00018 | 0.00443 | 0.51885 | 0.56445 | 1.13981 | 431 |
| GO_REGULATION_OF_CELLULAR_RESPONSE_TO_HEAT | 0.00102 | 0.01837 | 0.45506 | 0.65390 | 1.27339 | 72 |
| GO_REGULATION_OF_CELL_CYCLE_G2_M_PHASE_TRANSITION | 0.00001 | 0.00046 | 0.59333 | 0.62056 | 1.23782 | 190 |
| GO_REGULATION_OF_CELL_CYCLE_PHASE_TRANSITION | 0.00004 | 0.00123 | 0.55733 | 0.57778 | 1.16630 | 383 |
| GO_REGULATION_OF_CHROMOSOME_ORGANIZATION | 0.00342 | 0.04557 | 0.43171 | 0.56255 | 1.13116 | 293 |
| GO_REGULATION_OF_DEFENSE_RESPONSE | 0.00032 | 0.00718 | 0.49849 | 0.56421 | 1.13936 | 394 |
| GO_REGULATION_OF_DNA_BIOSYNTHETIC_PROCESS | 0.00024 | 0.00569 | 0.51885 | 0.65198 | 1.27720 | 93 |
| GO_REGULATION_OF_DNA_TEMPLATED_TRANSCRIPTION_IN_RE | 0.00000 | 0.00000 | 0.76146 | 0.72623 | 1.42446 | 99 |
| GO_REGULATION_OF_GENERATION_OF_PRECURSOR_METABOLI | 0.00150 | 0.02450 | 0.45506 | 0.61368 | 1.20813 | 112 |
| GO_REGULATION_OF_HEMATOPOIETIC_PROGENITOR_CELL_DIF | 0.00000 | 0.00000 | 0.78819 | 0.77007 | 1.50154 | 76 |
| GO_REGULATION_OF_HEMATOPOIETIC_STEM_CELL_DIFFERENT | 0.00000 | 0.00000 | | 0.80698 | 1.56709 | 66 |
| GO_REGULATION_OF_HEMOPOIESIS | 0.00074 | 0.01408 | 0.47727 | 0.56980 | 1.14613 | 296 |
| GO_REGULATION_OF_IMMUNE_RESPONSE | 0.00000 | 0.00001 | 0.70498 | 0.59161 | 1.19579 | 476 |
| GO_REGULATION_OF_INNATE_IMMUNE_RESPONSE | 0.00000 | 0.00010 | 0.62726 | 0.63251 | 1.26178 | 192 |
| GO_REGULATION_OF_INTRINSIC_APOPTOTIC_SIGNALING_PATH\ | 0.00079 | 0.01491 | 0.47727 | 0.61839 | 1.22001 | 126 |
| GO_REGULATION_OF_MACROAUTOPHAGY | 0.00170 | 0.02698 | 0.45506 | 0.59850 | 1.18691 | 146 |
| GO_REGULATION_OF_MORPHOGENESIS_OF_AN_EPITHELIUM | 0.00000 | 0.00000 | | 0.73057 | 1.44157 | 128 |
| GO_REGULATION_OF_MRNA_CATABOLIC_PROCESS | 0.00000 | 0.00000 | 0.74774 | 0.66354 | 1.32013 | 172 |
| GO_REGULATION_OF_MRNA_METABOLIC_PROCESS | 0.00000 | 0.00000 | | 0.66114 | 1.32907 | 287 |
| GO_REGULATION_OF_MRNA_PROCESSING | 0.00000 | 0.00007 | 0.64355 | 0.66939 | 1.32130 | 130 |
| GO_REGULATION_OF_MRNA_SPLICING_VIA_SPLICEOSOME | 0.00004 | 0.00109 | 0.55733 | 0.66766 | 1.30831 | 94 |
| GO_REGULATION_OF_MULTI_ORGANISM_PROCESS | 0.00253 | 0.03714 | 0.43171 | 0.69903 | 1.32781 | 40 |
| GO_REGULATION_OF_MUSCLE_CONTRACTION | 0.00143 | 0.02360 | 0.45506 | 0.63725 | 1.24558 | 82 |
| GO_REGULATION_OF_OXIDATIVE_PHOSPHORYLATION | 0.00056 | 0.01151 | 0.47727 | 0.81586 | 1.48418 | 18 |
| GO_REGULATION_OF_PEPTIDASE_ACTIVITY | 0.00130 | 0.02203 | 0.45506 | 0.57450 | 1.15224 | 252 |
| GO_REGULATION_OF_PEPTIDE_TRANSPORT | 0.00054 | 0.01117 | 0.47727 | 0.55875 | 1.12917 | 453 |
| GO_REGULATION_OF_PLASMA_LIPOPROTEIN_PARTICLE_LEVELS | 0.00179 | 0.02803 | 0.45506 | 0.66123 | 1.28234 | 62 |
| GO_REGULATION_OF_PROTEIN_CONTAINING_COMPLEX_ASSEM | 0.00072 | 0.01382 | 0.47727 | 0.56873 | 1.14466 | 318 |

| | | | | | | |
|---|---|---|---|---|---|---|
| GO_REGULATION_OF_PROTEIN_LOCALIZATION_TO_NUCLEUS | 0.00024 | 0.00569 | 0.51885 | 0.64091 | 1.25779 | 101 |
| GO_REGULATION_OF_PROTEOLYSIS | 0.00137 | 0.02310 | 0.45506 | 0.55126 | 1.11442 | 478 |
| GO_REGULATION_OF_RESPONSE_TO_BIOTIC_STIMULUS | 0.00000 | 0.00008 | 0.62726 | 0.61251 | 1.22782 | 247 |
| GO_REGULATION_OF_RNA_SPLICING | 0.00000 | 0.00016 | 0.61053 | 0.65623 | 1.29606 | 132 |
| GO_REGULATION_OF_SMALL_MOLECULE_METABOLIC_PROCESS | 0.00000 | 0.00002 | 0.67496 | 0.60792 | 1.22367 | 312 |
| GO_REGULATION_OF_STEM_CELL_DIFFERENTIATION | 0.00000 | 0.00000 | 0.78819 | 0.73294 | 1.43611 | 95 |
| GO_REGULATION_OF_SYMBIOTIC_PROCESS | 0.00002 | 0.00068 | 0.57561 | 0.62942 | 1.25087 | 158 |
| GO_REGULATION_OF_TELOMERE_MAINTENANCE_VIA_TELOME | 0.00087 | 0.01618 | 0.47727 | 0.67943 | 1.31324 | 55 |
| GO_REGULATION_OF_TRANSCRIPTION_FROM_RNA_POLYMERA | 0.00000 | 0.00000 | 0.74774 | 0.77265 | 1.50151 | 68 |
| GO_REGULATION_OF_TRANSLATIONAL_INITIATION | 0.00058 | 0.01185 | 0.47727 | 0.66831 | 1.30008 | 70 |
| GO_REGULATION_OF_VIRAL_TRANSCRIPTION | 0.00066 | 0.01307 | 0.47727 | 0.68463 | 1.32330 | 55 |
| GO_REGULATION_OF_WNT_SIGNALING_PATHWAY | 0.00172 | 0.02714 | 0.45506 | 0.56895 | 1.14253 | 268 |
| GO_REPRODUCTIVE_SYSTEM_DEVELOPMENT | 0.00143 | 0.02360 | 0.45506 | 0.56878 | 1.14315 | 284 |
| GO_RESPIRASOME | 0.00000 | 0.00000 | 0.77494 | 0.76805 | 1.49721 | 75 |
| GO_RESPIRATORY_CHAIN_COMPLEX | 0.00000 | 0.00000 | 0.71951 | 0.76445 | 1.48426 | 64 |
| GO_RESPIRATORY_ELECTRON_TRANSPORT_CHAIN | 0.00000 | 0.00000 | 0.80122 | 0.75907 | 1.48544 | 89 |
| GO_RESPONSE_TO_DRUG | 0.00008 | 0.00221 | 0.53843 | 0.59471 | 1.19167 | 240 |
| GO_RESPONSE_TO_ENDOPLASMIC_RETICULUM_STRESS | 0.00037 | 0.00812 | 0.49849 | 0.58874 | 1.17993 | 242 |
| GO_RESPONSE_TO_ESTROGEN | 0.00059 | 0.01200 | 0.47727 | 0.69096 | 1.33664 | 56 |
| GO_RESPONSE_TO_FIBROBLAST_GROWTH_FACTOR | 0.00174 | 0.02736 | 0.45506 | 0.62899 | 1.23058 | 87 |
| GO_RESPONSE_TO_GROWTH_FACTOR | 0.00054 | 0.01102 | 0.47727 | 0.55400 | 1.11923 | 447 |
| GO_RESPONSE_TO_HEAT | 0.00005 | 0.00150 | 0.55733 | 0.63714 | 1.25836 | 132 |
| GO_RESPONSE_TO_INORGANIC_SUBSTANCE | 0.00068 | 0.01327 | 0.47727 | 0.56691 | 1.14244 | 351 |
| GO_RESPONSE_TO_INTERLEUKIN_1 | 0.00000 | 0.00000 | | 0.71403 | 1.41133 | 135 |
| GO_RESPONSE_TO_INTERLEUKIN_12 | 0.00014 | 0.00358 | 0.51885 | 0.74428 | 1.41265 | 39 |
| GO_RESPONSE_TO_INTERLEUKIN_4 | 0.00143 | 0.02360 | 0.45506 | 0.77956 | 1.43428 | 21 |
| GO_RESPONSE_TO_METAL_ION | 0.00152 | 0.02461 | 0.45506 | 0.57822 | 1.15615 | 219 |
| GO_RESPONSE_TO_OXYGEN_LEVELS | 0.00000 | 0.00000 | | 0.66501 | 1.33608 | 280 |
| GO_RESPONSE_TO_TEMPERATURE_STIMULUS | 0.00038 | 0.00836 | 0.49849 | 0.61130 | 1.21525 | 163 |
| GO_RESPONSE_TO_TOPOLOGICALLY_INCORRECT_PROTEIN | 0.00023 | 0.00550 | 0.51885 | 0.60750 | 1.20843 | 171 |
| GO_RESPONSE_TO_TOXIC_SUBSTANCE | 0.00074 | 0.01408 | 0.47727 | 0.60863 | 1.20653 | 145 |
| GO_RESPONSE_TO_TUMOR_NECROSIS_FACTOR | 0.00000 | 0.00001 | 0.67496 | 0.64189 | 1.28069 | 194 |
| GO_RETINA_HOMEOSTASIS | 0.00001 | 0.00034 | 0.59333 | 0.79358 | 1.50023 | 36 |
| GO_RIBONUCLEOPROTEIN_COMPLEX_BINDING | 0.00000 | 0.00002 | 0.67496 | 0.68249 | 1.34499 | 119 |
| GO_RIBONUCLEOPROTEIN_COMPLEX_BIOGENESIS | 0.00000 | 0.00000 | | 0.67778 | 1.36859 | 397 |
| GO_RIBONUCLEOPROTEIN_COMPLEX_SUBUNIT_ORGANIZATION | 0.00000 | 0.00000 | | 0.71767 | 1.42908 | 177 |
| GO_RIBONUCLEOPROTEIN_GRANULE | 0.00003 | 0.00087 | 0.57561 | 0.61690 | 1.23097 | 193 |
| GO_RIBONUCLEOSIDE_TRIPHOSPHATE_METABOLIC_PROCESS | 0.00042 | 0.00905 | 0.49849 | 0.70431 | 1.34492 | 43 |
| GO_RIBOSE_PHOSPHATE_BIOSYNTHETIC_PROCESS | 0.00006 | 0.00175 | 0.55733 | 0.64009 | 1.26167 | 123 |
| GO_RIBOSE_PHOSPHATE_METABOLIC_PROCESS | 0.00001 | 0.00046 | 0.59333 | 0.59416 | 1.19443 | 287 |
| GO_RIBOSOMAL_LARGE_SUBUNIT_ASSEMBLY | 0.00097 | 0.01752 | 0.47727 | 0.76143 | 1.41741 | 26 |
| GO_RIBOSOMAL_LARGE_SUBUNIT_BIOGENESIS | 0.00000 | 0.00016 | 0.61053 | 0.72388 | 1.40707 | 67 |
| GO_RIBOSOMAL_SMALL_SUBUNIT_ASSEMBLY | 0.00044 | 0.00947 | 0.49849 | 0.83669 | 1.51415 | 17 |
| GO_RIBOSOMAL_SMALL_SUBUNIT_BIOGENESIS | 0.00008 | 0.00221 | 0.53843 | 0.69777 | 1.35502 | 66 |
| GO_RIBOSOMAL_SUBUNIT | 0.00000 | 0.00000 | | 0.78481 | 1.56114 | 171 |
| GO_RIBOSOME | 0.00000 | 0.00000 | | 0.77032 | 1.53895 | 209 |
| GO_RIBOSOME_ASSEMBLY | 0.00000 | 0.00001 | 0.70498 | 0.77933 | 1.50760 | 56 |
| GO_RIBOSOME_BIOGENESIS | 0.00000 | 0.00000 | | 0.66625 | 1.33849 | 278 |
| GO_RNA_3_END_PROCESSING | 0.00018 | 0.00441 | 0.51885 | 0.63085 | 1.24542 | 131 |
| GO_RNA_CATABOLIC_PROCESS | 0.00000 | 0.00000 | | 0.69999 | 1.41093 | 353 |
| GO_RNA_DEPENDENT_DNA_BIOSYNTHETIC_PROCESS | 0.00009 | 0.00239 | 0.53843 | 0.69646 | 1.35112 | 65 |
| GO_RNA_EXPORT_FROM_NUCLEUS | 0.00021 | 0.00521 | 0.51885 | 0.63237 | 1.24823 | 130 |
| GO_RNA_LOCALIZATION | 0.00001 | 0.00037 | 0.59333 | 0.61377 | 1.22678 | 213 |
| GO_RNA_SPLICING | 0.00000 | 0.00000 | | 0.64149 | 1.29546 | 398 |
| GO_RNA_SPLICING_VIA_TRANSESTERIFICATION_REACTIONS | 0.00000 | 0.00000 | | 0.65353 | 1.31529 | 320 |
| GO_ROUGH_ENDOPLASMIC_RETICULUM | 0.00001 | 0.00046 | 0.59333 | 0.71839 | 1.39228 | 61 |
| GO_RRNA_BINDING | 0.00000 | 0.00002 | 0.67496 | 0.76624 | 1.48228 | 56 |
| GO_RRNA_METABOLIC_PROCESS | 0.00000 | 0.00000 | 0.78819 | 0.66202 | 1.32275 | 211 |
| GO_SARCOLEMMA | 0.00095 | 0.01733 | 0.47727 | 0.66141 | 1.28622 | 69 |
| GO_SARCOMERE_ORGANIZATION | 0.00086 | 0.01593 | 0.47727 | 0.74755 | 1.39466 | 27 |
| GO_SCAFFOLD_PROTEIN_BINDING | 0.00239 | 0.03545 | 0.43171 | 0.69459 | 1.32411 | 42 |
| GO_SCF_DEPENDENT_PROTEASOMAL_UBIQUITIN_DEPENDENT_ | 0.00000 | 0.00000 | 0.74774 | 0.74977 | 1.46428 | 78 |
| GO_SECRETORY_GRANULE | 0.00000 | 0.00000 | | 0.63119 | 1.27645 | 491 |
| GO_SERINE_TYPE_ENDOPEPTIDASE_INHIBITOR_ACTIVITY | 0.00002 | 0.00077 | 0.57561 | 0.78427 | 1.48180 | 34 |
| GO_SINGLE_FERTILIZATION | 0.00038 | 0.00836 | 0.49849 | 0.68922 | 1.33344 | 58 |
| GO_SINGLE_STRANDED_RNA_BINDING | 0.00023 | 0.00564 | 0.51885 | 0.67937 | 1.32114 | 69 |
| GO_SKIN_DEVELOPMENT | 0.00145 | 0.02384 | 0.45506 | 0.59343 | 1.17991 | 166 |
| GO_SMALL_MOLECULE_BIOSYNTHETIC_PROCESS | 0.00000 | 0.00006 | 0.64355 | 0.58227 | 1.17685 | 455 |
| GO_SMALL_NUCLEAR_RIBONUCLEOPROTEIN_COMPLEX | 0.00001 | 0.00025 | 0.61053 | 0.73178 | 1.41694 | 60 |
| GO_SMALL_RIBOSOMAL_SUBUNIT | 0.00000 | 0.00000 | 0.78819 | 0.78637 | 1.52854 | 67 |

| | | | | | | |
|---|---|---|---|---|---|---|
| GO_SM_LIKE_PROTEIN_FAMILY_COMPLEX | 0.00003 | 0.00090 | 0.57561 | 0.70110 | 1.36388 | 70 |
| GO_SNORNA_BINDING | 0.00102 | 0.01837 | 0.45506 | 0.74516 | 1.39741 | 30 |
| GO_SNRNA_BINDING | 0.00272 | 0.03904 | 0.43171 | 0.68968 | 1.31200 | 41 |
| GO_SPERM_EGG_RECOGNITION | 0.00000 | 0.00013 | 0.62726 | 0.86639 | 1.60487 | 23 |
| GO_SPLICEOSOMAL_COMPLEX | 0.00000 | 0.00000 | 0.76146 | 0.67024 | 1.33446 | 176 |
| GO_SPLICEOSOMAL_TRI_SNRNP_COMPLEX | 0.00106 | 0.01892 | 0.45506 | 0.74408 | 1.39538 | 30 |
| GO_STEM_CELL_DIFFERENTIATION | 0.00000 | 0.00000 | 0.78819 | 0.67877 | 1.35087 | 174 |
| GO_STEROID_METABOLIC_PROCESS | 0.00143 | 0.02360 | 0.45506 | 0.58649 | 1.16981 | 191 |
| GO_STRUCTURAL_CONSTITUENT_OF_CYTOSKELETON | 0.00002 | 0.00074 | 0.57561 | 0.70996 | 1.37969 | 68 |
| GO_STRUCTURAL_CONSTITUENT_OF_MUSCLE | 0.00152 | 0.02461 | 0.45506 | 0.74625 | 1.39384 | 28 |
| GO_STRUCTURAL_CONSTITUENT_OF_RIBOSOME | 0.00000 | 0.00000 | | 0.79877 | 1.58574 | 152 |
| GO_STRUCTURAL_MOLECULE_ACTIVITY | 0.00000 | 0.00000 | | 0.72969 | 1.47335 | 421 |
| GO_SUGAR_TRANSMEMBRANE_TRANSPORTER_ACTIVITY | 0.00122 | 0.02106 | 0.45506 | 0.80911 | 1.45635 | 16 |
| GO_SYNAPSE_ORGANIZATION | 0.00383 | 0.04936 | 0.43171 | 0.57085 | 1.14142 | 219 |
| GO_TELOMERASE_HOLOENZYME_COMPLEX | 0.00183 | 0.02839 | 0.45506 | 0.78688 | 1.44595 | 20 |
| GO_TELOMERASE_RNA_LOCALIZATION | 0.00017 | 0.00433 | 0.51885 | 0.83430 | 1.52572 | 19 |
| GO_TELOMERE_MAINTENANCE_VIA_TELOMERE_LENGTHENING | 0.00009 | 0.00242 | 0.53843 | 0.68341 | 1.33084 | 72 |
| GO_THREONINE_TYPE_ENDOPEPTIDASE_ACTIVITY | 0.00000 | 0.00000 | 0.77494 | 0.95063 | 1.72034 | 17 |
| GO_THREONINE_TYPE_PEPTIDASE_ACTIVITY | 0.00000 | 0.00000 | 0.78819 | 0.93518 | 1.72719 | 22 |
| GO_TISSUE_MORPHOGENESIS | 0.00001 | 0.00025 | 0.61053 | 0.58283 | 1.17704 | 415 |
| GO_TOXIN_TRANSPORT | 0.00316 | 0.04320 | 0.43171 | 0.71373 | 1.34284 | 32 |
| GO_TRANSCRIPTION_COUPLED_NUCLEOTIDE_EXCISION_REPAIR | 0.00209 | 0.03187 | 0.43171 | 0.64495 | 1.25596 | 72 |
| GO_TRANSCRIPTION_ELONGATION_FROM_RNA_POLYMERASE_I | 0.00305 | 0.04213 | 0.43171 | 0.63599 | 1.23920 | 74 |
| GO_TRANSLATIONAL_ELONGATION | 0.00000 | 0.00000 | | 0.72824 | 1.43578 | 124 |
| GO_TRANSLATIONAL_INITIATION | 0.00000 | 0.00000 | | 0.77403 | 1.53970 | 171 |
| GO_TRANSLATIONAL_TERMINATION | 0.00000 | 0.00000 | 0.77494 | 0.72616 | 1.42508 | 101 |
| GO_TRANSLATION_FACTOR_ACTIVITY_RNA_BINDING | 0.00000 | 0.00000 | 0.71951 | 0.75726 | 1.47311 | 70 |
| GO_TRANSLATION_INITIATION_FACTOR_ACTIVITY | 0.00001 | 0.00022 | 0.61053 | 0.77199 | 1.47778 | 44 |
| GO_TRANSLATION_INITIATION_FACTOR_BINDING | 0.00216 | 0.03261 | 0.43171 | 0.73979 | 1.38177 | 28 |
| GO_TRANSLATION_PREINITIATION_COMPLEX | 0.00017 | 0.00429 | 0.51885 | 0.85099 | 1.53173 | 16 |
| GO_TRANSLATION_REGULATOR_ACTIVITY | 0.00000 | 0.00000 | 0.74774 | 0.71173 | 1.40216 | 114 |
| GO_TRANSLATION_REGULATOR_ACTIVITY_NUCLEIC_ACID_BIND | 0.00000 | 0.00000 | 0.77494 | 0.75289 | 1.47246 | 88 |
| GO_TUMOR_NECROSIS_FACTOR_MEDIATED_SIGNALING_PATHW | 0.00000 | 0.00000 | 0.76146 | 0.71080 | 1.40031 | 116 |
| GO_T_CELL_RECEPTOR_SIGNALING_PATHWAY | 0.00000 | 0.00000 | 0.73376 | 0.68978 | 1.36359 | 136 |
| GO_U12_TYPE_SPLICEOSOMAL_COMPLEX | 0.00005 | 0.00141 | 0.55733 | 0.80249 | 1.49716 | 27 |
| GO_U1_SNRNP | 0.00065 | 0.01293 | 0.47727 | 0.83035 | 1.50268 | 17 |
| GO_U2_SNRNP | 0.00079 | 0.01491 | 0.47727 | 0.81041 | 1.47426 | 18 |
| GO_U2_TYPE_SPLICEOSOMAL_COMPLEX | 0.00001 | 0.00029 | 0.59333 | 0.68735 | 1.34476 | 87 |
| GO_UBIQUITIN_LIKE_PROTEIN_LIGASE_BINDING | 0.00000 | 0.00000 | 0.71951 | 0.63008 | 1.26436 | 260 |
| GO_UNFOLDED_PROTEIN_BINDING | 0.00000 | 0.00001 | 0.69013 | 0.70218 | 1.37585 | 95 |
| GO_VACUOLAR_LUMEN | 0.00000 | 0.00001 | 0.69013 | 0.70002 | 1.37842 | 113 |
| GO_VESICLE_LUMEN | 0.00000 | 0.00000 | | 0.68680 | 1.37254 | 214 |
| GO_VESICLE_MEMBRANE | 0.00023 | 0.00546 | 0.51885 | 0.55801 | 1.12866 | 496 |
| GO_VIRAL_GENE_EXPRESSION | 0.00000 | 0.00000 | | 0.75651 | 1.50621 | 176 |
| GO_VIRAL_GENOME_REPLICATION | 0.00095 | 0.01733 | 0.47727 | 0.64761 | 1.26533 | 80 |
| GO_VIRAL_LIFE_CYCLE | 0.00000 | 0.00018 | 0.61053 | 0.61316 | 1.22797 | 237 |
| HP_ABNORMALITY_OF_ACID_BASE_HOMEOSTASIS | 0.00060 | 0.01211 | 0.47727 | 0.58131 | 1.16598 | 257 |
| HP_ABNORMAL_ABDOMEN_MORPHOLOGY | 0.00228 | 0.03396 | 0.43171 | 0.54705 | 1.10575 | 469 |
| HP_ABNORMAL_ACTIVITY_OF_MITOCHONDRIAL_RESPIRATORY_ | 0.00194 | 0.02997 | 0.45506 | 0.66077 | 1.28145 | 62 |
| HP_ABNORMAL_CNS_MYELINATION | 0.00294 | 0.04113 | 0.43171 | 0.58444 | 1.16257 | 171 |
| HP_ABNORMAL_CSF_METABOLITE_LEVEL | 0.00011 | 0.00305 | 0.53843 | 0.66094 | 1.29212 | 85 |
| HP_ABNORMAL_ERYTHROCYTE_MORPHOLOGY | 0.00009 | 0.00242 | 0.53843 | 0.57189 | 1.15486 | 392 |
| HP_ACUTE_LEUKEMIA | 0.00018 | 0.00443 | 0.51885 | 0.66998 | 1.30542 | 74 |
| HP_ANEMIA_OF_INADEQUATE_PRODUCTION | 0.00042 | 0.00898 | 0.49849 | 0.63442 | 1.24703 | 105 |
| HP_DELAYED_PUBERTY | 0.00250 | 0.03685 | 0.43171 | 0.60612 | 1.19449 | 121 |
| HP_DEPRESSED_NASAL_RIDGE | 0.00294 | 0.04113 | 0.43171 | 0.62809 | 1.22779 | 84 |
| HP_ECTOPIC_CALCIFICATION | 0.00383 | 0.04936 | 0.43171 | 0.59450 | 1.17413 | 132 |
| HP_EMOTIONAL_LABILITY | 0.00111 | 0.01975 | 0.45506 | 0.62769 | 1.22989 | 95 |
| HP_FATIGUE | 0.00124 | 0.02132 | 0.45506 | 0.58241 | 1.16439 | 221 |
| HP_HEADACHE | 0.00113 | 0.01991 | 0.45506 | 0.59380 | 1.18137 | 172 |
| HP_HYPERTROPHIC_CARDIOMYOPATHY | 0.00005 | 0.00140 | 0.55733 | 0.61019 | 1.21744 | 194 |
| HP_INCREASED_CSF_LACTATE | 0.00029 | 0.00671 | 0.49849 | 0.66992 | 1.30382 | 71 |
| HP_INCREASED_SERUM_LACTATE | 0.00045 | 0.00951 | 0.49849 | 0.61723 | 1.22000 | 135 |
| HP_LACTIC_ACIDOSIS | 0.00061 | 0.01218 | 0.47727 | 0.61959 | 1.22430 | 134 |
| HP_LEUKEMIA | 0.00279 | 0.03966 | 0.43171 | 0.61321 | 1.20680 | 108 |
| HP_LEUKODYSTROPHY | 0.00272 | 0.03904 | 0.43171 | 0.63197 | 1.23520 | 81 |
| HP_MACROCYTIC_ANEMIA | 0.00006 | 0.00179 | 0.53843 | 0.75058 | 1.42786 | 41 |
| HP_MIGRAINE | 0.00015 | 0.00390 | 0.51885 | 0.68238 | 1.33022 | 75 |
| HP_MUSCLE_SPASM | 0.00183 | 0.02839 | 0.45506 | 0.61820 | 1.21322 | 101 |
| HP_MUTISM | 0.00350 | 0.04617 | 0.43171 | 0.69676 | 1.32018 | 37 |

| | | | | | | |
|---|---|---|---|---|---|---|
| HP_OPTIC_NERVE_COLOBOMA | 0.00122 | 0.02106 | 0.45506 | 0.80841 | 1.45508 | 16 |
| HP_PALLOR | 0.00305 | 0.04213 | 0.43171 | 0.61012 | 1.19849 | 103 |
| HP_PAROXYSMAL_INVOLUNTARY_EYE_MOVEMENTS | 0.00335 | 0.04515 | 0.43171 | 0.71211 | 1.33980 | 32 |
| HP_POOR_EYE_CONTACT | 0.00339 | 0.04555 | 0.43171 | 0.65698 | 1.26722 | 53 |
| HP_POOR_HEAD_CONTROL | 0.00061 | 0.01218 | 0.47727 | 0.62565 | 1.23255 | 116 |
| HP_RENAL_TUBULAR_DYSFUNCTION | 0.00361 | 0.04735 | 0.43171 | 0.63376 | 1.23540 | 73 |
| HP_RESPIRATORY_INSUFFICIENCY | 0.00061 | 0.01218 | 0.47727 | 0.56663 | 1.14186 | 346 |
| HP_TETRAPARESIS | 0.00298 | 0.04146 | 0.43171 | 0.74965 | 1.39103 | 24 |
| **Cluster 5** | | | | | | |
| pathway | pval | padj | log2err | ES | NES | size |
| GO_ACTIVATION_OF_IMMUNE_RESPONSE | 0.00085 | 0.02168 | 0.47727 | 0.57934 | 1.17714 | 244 |
| GO_ACTIVATION_OF_INNATE_IMMUNE_RESPONSE | 0.00008 | 0.00380 | 0.53843 | 0.66093 | 1.31772 | 97 |
| GO_ALTERNATIVE_MRNA_SPLICING_VIA_SPLICEOSOME | 0.00002 | 0.00116 | 0.57561 | 0.70414 | 1.39090 | 72 |
| GO_AMINE_METABOLIC_PROCESS | 0.00010 | 0.00457 | 0.53843 | 0.65936 | 1.31459 | 97 |
| GO_ANAPHASE_PROMOTING_COMPLEX_DEPENDENT_CATABOL | 0.00000 | 0.00046 | 0.61053 | 0.71347 | 1.41102 | 78 |
| GO_ANTIGEN_PROCESSING_AND_PRESENTATION | 0.00084 | 0.02153 | 0.47727 | 0.60274 | 1.21680 | 161 |
| GO_ANTIGEN_PROCESSING_AND_PRESENTATION_OF_EXOGENC | 0.00076 | 0.02039 | 0.47727 | 0.67075 | 1.31713 | 64 |
| GO_ANTIGEN_PROCESSING_AND_PRESENTATION_OF_PEPTIDE_ | 0.00061 | 0.01696 | 0.47727 | 0.60979 | 1.22657 | 142 |
| GO_ANTIGEN_PROCESSING_AND_PRESENTATION_OF_PEPTIDE_ | 0.00191 | 0.03864 | 0.45506 | 0.64593 | 1.27693 | 77 |
| GO_APOPTOTIC_SIGNALING_PATHWAY | 0.00010 | 0.00457 | 0.53843 | 0.57134 | 1.17272 | 401 |
| GO_ATPASE_ACTIVITY | 0.00228 | 0.04417 | 0.43171 | 0.55761 | 1.14103 | 339 |
| GO_ATP_METABOLIC_PROCESS | 0.00002 | 0.00126 | 0.57561 | 0.60907 | 1.23591 | 219 |
| GO_ATP_SYNTHESIS_COUPLED_ELECTRON_TRANSPORT | 0.00216 | 0.04293 | 0.43171 | 0.64497 | 1.27554 | 78 |
| GO_ATTACHMENT_OF_SPINDLE_MICROTUBULES_TO_KINETOCH | 0.00224 | 0.04371 | 0.43171 | 0.72290 | 1.37098 | 30 |
| GO_AUTOPHAGY_OF_MITOCHONDRION | 0.00228 | 0.04417 | 0.43171 | 0.64129 | 1.26552 | 71 |
| GO_BINDING_OF_SPERM_TO_ZONA_PELLUCIDA | 0.00046 | 0.01385 | 0.49849 | 0.78250 | 1.47661 | 25 |
| GO_CADHERIN_BINDING | 0.00000 | 0.00011 | 0.65944 | 0.62087 | 1.26299 | 260 |
| GO_CAJAL_BODY | 0.00076 | 0.02032 | 0.47727 | 0.66769 | 1.30584 | 58 |
| GO_CALYX_OF_HELD | 0.00157 | 0.03409 | 0.45506 | 0.79312 | 1.46299 | 16 |
| GO_CATALYTIC_STEP_2_SPLICEOSOME | 0.00000 | 0.00000 | 0.73376 | 0.74224 | 1.47028 | 81 |
| GO_CELLULAR_COMPONENT_DISASSEMBLY | 0.00035 | 0.01129 | 0.49849 | 0.55718 | 1.14469 | 428 |
| GO_CELLULAR_KETONE_METABOLIC_PROCESS | 0.00014 | 0.00571 | 0.51885 | 0.61546 | 1.24165 | 158 |
| GO_CELLULAR_NITROGEN_COMPOUND_CATABOLIC_PROCESS | 0.00000 | 0.00000 | | 0.62598 | 1.28720 | 463 |
| GO_CELLULAR_RESPONSE_TO_HEAT | 0.00041 | 0.01283 | 0.49849 | 0.64306 | 1.28209 | 97 |
| GO_CELLULAR_RESPONSE_TO_OXYGEN_LEVELS | 0.00012 | 0.00507 | 0.53843 | 0.62048 | 1.25261 | 161 |
| GO_CELL_ADHESION_MOLECULE_BINDING | 0.00000 | 0.00032 | 0.62726 | 0.58625 | 1.20117 | 371 |
| GO_CELL_CELL_RECOGNITION | 0.00093 | 0.02341 | 0.47727 | 0.71298 | 1.37690 | 41 |
| GO_CELL_CYCLE_CHECKPOINT | 0.00170 | 0.03590 | 0.45506 | 0.58592 | 1.18498 | 184 |
| GO_CELL_CYCLE_G1_S_PHASE_TRANSITION | 0.00031 | 0.01033 | 0.49849 | 0.59089 | 1.19936 | 222 |
| GO_CELL_CYCLE_G2_M_PHASE_TRANSITION | 0.00000 | 0.00046 | 0.61053 | 0.61564 | 1.25057 | 229 |
| GO_CELL_DIVISION | 0.00000 | 0.00005 | 0.67496 | 0.58911 | 1.21132 | 465 |
| GO_CELL_SUBSTRATE_JUNCTION | 0.00000 | 0.00012 | 0.64355 | 0.60413 | 1.23505 | 322 |
| GO_CHAPERONE_BINDING | 0.00046 | 0.01385 | 0.49849 | 0.65718 | 1.30034 | 79 |
| GO_CHAPERONE_COFACTOR_DEPENDENT_PROTEIN_REFOLDIN( | 0.00152 | 0.03335 | 0.45506 | 0.75664 | 1.42782 | 25 |
| GO_CHAPERONE_COMPLEX | 0.00016 | 0.00652 | 0.51885 | 0.82163 | 1.52833 | 20 |
| GO_CHROMATIN_ASSEMBLY_OR_DISASSEMBLY | 0.00246 | 0.04690 | 0.43171 | 0.59976 | 1.20201 | 118 |
| GO_CHROMATIN_BINDING | 0.00058 | 0.01633 | 0.47727 | 0.55379 | 1.13766 | 435 |
| GO_CHROMATIN_REMODELING | 0.00042 | 0.01292 | 0.49849 | 0.60910 | 1.22748 | 152 |
| GO_CHROMOSOMAL_REGION | 0.00000 | 0.00008 | 0.65944 | 0.61418 | 1.25267 | 287 |
| GO_CHROMOSOME_CENTROMERIC_REGION | 0.00004 | 0.00216 | 0.55733 | 0.62298 | 1.25797 | 167 |
| GO_CHROMOSOME_SEGREGATION | 0.00000 | 0.00001 | 0.71951 | 0.62443 | 1.27183 | 274 |
| GO_CHROMOSOME_SEPARATION | 0.00126 | 0.02910 | 0.45506 | 0.64161 | 1.27324 | 85 |
| GO_CHROMOSOME_TELOMERIC_REGION | 0.00122 | 0.02846 | 0.45506 | 0.60670 | 1.21772 | 125 |
| GO_CONDENSED_CHROMOSOME | 0.00027 | 0.00930 | 0.49849 | 0.60791 | 1.22935 | 179 |
| GO_CONDENSED_CHROMOSOME_CENTROMERIC_REGION | 0.00154 | 0.03364 | 0.45506 | 0.62812 | 1.25374 | 100 |
| GO_COP9_SIGNALOSOME | 0.00213 | 0.04246 | 0.43171 | 0.73243 | 1.38635 | 29 |
| GO_COTRANSLATIONAL_PROTEIN_TARGETING_TO_MEMBRANE | 0.00000 | 0.00000 | 0.82666 | 0.75672 | 1.50710 | 93 |
| GO_COVALENT_CHROMATIN_MODIFICATION | 0.00079 | 0.02069 | 0.47727 | 0.56313 | 1.15398 | 378 |
| GO_CYTOKINESIS | 0.00145 | 0.03229 | 0.45506 | 0.60896 | 1.22168 | 123 |
| GO_CYTOKINE_MEDIATED_SIGNALING_PATHWAY | 0.00031 | 0.01033 | 0.49849 | 0.56308 | 1.15562 | 399 |
| GO_CYTOPLASMIC_TRANSLATION | 0.00000 | 0.00008 | 0.65944 | 0.70865 | 1.40941 | 91 |
| GO_CYTOPLASMIC_TRANSLATIONAL_INITIATION | 0.00090 | 0.02290 | 0.47727 | 0.74527 | 1.41342 | 30 |
| GO_CYTOSOLIC_LARGE_RIBOSOMAL_SUBUNIT | 0.00000 | 0.00010 | 0.65944 | 0.77113 | 1.50488 | 53 |
| GO_CYTOSOLIC_RIBOSOME | 0.00000 | 0.00000 | | 0.76249 | 1.52063 | 98 |
| GO_CYTOSOLIC_SMALL_RIBOSOMAL_SUBUNIT | 0.00003 | 0.00168 | 0.57561 | 0.76773 | 1.47996 | 40 |
| GO_DEVELOPMENT_OF_PRIMARY_SEXUAL_CHARACTERISTICS | 0.00161 | 0.03442 | 0.45506 | 0.60847 | 1.22378 | 132 |
| GO_DE_NOVO_PROTEIN_FOLDING | 0.00205 | 0.04111 | 0.43171 | 0.71878 | 1.37048 | 32 |
| GO_DNA_BINDING_TRANSCRIPTION_FACTOR_BINDING | 0.00058 | 0.01633 | 0.47727 | 0.57881 | 1.17821 | 267 |
| GO_DNA_BIOSYNTHETIC_PROCESS | 0.00148 | 0.03266 | 0.45506 | 0.59913 | 1.20602 | 148 |
| GO_DNA_CONFORMATION_CHANGE | 0.00000 | 0.00015 | 0.64355 | 0.62387 | 1.26729 | 229 |

| | | | | | | |
|---|---|---|---|---|---|---|
| GO_DNA_DAMAGE_RESPONSE_SIGNAL_TRANSDUCTION_BY_P5 | 0.00183 | 0.03774 | 0.45506 | 0.63756 | 1.26359 | 82 |
| GO_DNA_PACKAGING | 0.00013 | 0.00541 | 0.51885 | 0.63292 | 1.27265 | 131 |
| GO_DNA_REPAIR | 0.00002 | 0.00130 | 0.57561 | 0.57261 | 1.17649 | 439 |
| GO_DNA_REPLICATION | 0.00146 | 0.03236 | 0.45506 | 0.57730 | 1.17270 | 229 |
| GO_DNA_TEMPLATED_TRANSCRIPTION_ELONGATION | 0.00014 | 0.00586 | 0.51885 | 0.64574 | 1.29083 | 104 |
| GO_DNA_TEMPLATED_TRANSCRIPTION_INITIATION | 0.00019 | 0.00711 | 0.51885 | 0.60204 | 1.21899 | 196 |
| GO_DNA_TEMPLATED_TRANSCRIPTION_TERMINATION | 0.00104 | 0.02549 | 0.45506 | 0.66031 | 1.30065 | 68 |
| GO_ENDOPEPTIDASE_COMPLEX | 0.00031 | 0.01043 | 0.49849 | 0.68166 | 1.33646 | 62 |
| GO_ESTABLISHMENT_OF_PROTEIN_LOCALIZATION_TO_ENDOPL | 0.00000 | 0.00000 | 0.81404 | 0.73455 | 1.46838 | 104 |
| GO_ESTABLISHMENT_OF_PROTEIN_LOCALIZATION_TO_MEMBR | 0.00000 | 0.00000 | 0.82666 | 0.65282 | 1.32724 | 253 |
| GO_ESTABLISHMENT_OF_PROTEIN_LOCALIZATION_TO_ORGANE | 0.00000 | 0.00000 | | 0.64352 | 1.32332 | 460 |
| GO_ESTABLISHMENT_OF_RNA_LOCALIZATION | 0.00004 | 0.00197 | 0.55733 | 0.62128 | 1.25639 | 179 |
| GO_ESTABLISHMENT_OF_TISSUE_POLARITY | 0.00183 | 0.03774 | 0.45506 | 0.62744 | 1.24865 | 92 |
| GO_EXON_EXON_JUNCTION_COMPLEX | 0.00183 | 0.03774 | 0.45506 | 0.78980 | 1.46091 | 17 |
| GO_FC_EPSILON_RECEPTOR_SIGNALING_PATHWAY | 0.00198 | 0.03976 | 0.43171 | 0.62667 | 1.24809 | 93 |
| GO_FC_RECEPTOR_SIGNALING_PATHWAY | 0.00167 | 0.03536 | 0.45506 | 0.60388 | 1.21471 | 139 |
| GO_FEMALE_GAMETE_GENERATION | 0.00009 | 0.00415 | 0.53843 | 0.66650 | 1.32557 | 91 |
| GO_FIBRILLAR_CENTER | 0.00115 | 0.02725 | 0.45506 | 0.63149 | 1.26048 | 100 |
| GO_FIBROBLAST_GROWTH_FACTOR_RECEPTOR_SIGNALING_PAT | 0.00002 | 0.00116 | 0.57561 | 0.70777 | 1.39555 | 69 |
| GO_FICOLIN_1_RICH_GRANULE | 0.00024 | 0.00864 | 0.51885 | 0.64658 | 1.28946 | 98 |
| GO_GAMETE_GENERATION | 0.00016 | 0.00624 | 0.51885 | 0.56561 | 1.16173 | 422 |
| GO_GENE_SILENCING | 0.00024 | 0.00864 | 0.51885 | 0.60334 | 1.22005 | 182 |
| GO_HEAT_SHOCK_PROTEIN_BINDING | 0.00017 | 0.00671 | 0.51885 | 0.65643 | 1.30785 | 95 |
| GO_HELICASE_ACTIVITY | 0.00178 | 0.03721 | 0.45506 | 0.60017 | 1.20745 | 141 |
| GO_HEMATOPOIETIC_PROGENITOR_CELL_DIFFERENTIATION | 0.00006 | 0.00321 | 0.53843 | 0.64267 | 1.29092 | 127 |
| GO_HEMATOPOIETIC_STEM_CELL_DIFFERENTIATION | 0.00111 | 0.02667 | 0.45506 | 0.65316 | 1.29086 | 74 |
| GO_HISTONE_BINDING | 0.00122 | 0.02846 | 0.45506 | 0.58282 | 1.17971 | 190 |
| GO_IMPORT_INTO_NUCLEUS | 0.00014 | 0.00571 | 0.51885 | 0.63193 | 1.27064 | 131 |
| GO_INNATE_IMMUNE_RESPONSE | 0.00250 | 0.04718 | 0.43171 | 0.55016 | 1.12967 | 415 |
| GO_INNER_MITOCHONDRIAL_MEMBRANE_PROTEIN_COMPLEX | 0.00194 | 0.03914 | 0.45506 | 0.62153 | 1.23563 | 90 |
| GO_INTERLEUKIN_1_MEDIATED_SIGNALING_PATHWAY | 0.00010 | 0.00440 | 0.53843 | 0.67332 | 1.33286 | 80 |
| GO_INTRINSIC_APOPTOTIC_SIGNALING_PATHWAY | 0.00037 | 0.01176 | 0.49849 | 0.60048 | 1.21758 | 207 |
| GO_KINASE_REGULATOR_ACTIVITY | 0.00000 | 0.00025 | 0.62726 | 0.64434 | 1.29991 | 158 |
| GO_KINETOCHORE | 0.00079 | 0.02069 | 0.47727 | 0.61870 | 1.24034 | 119 |
| GO_LARGE_RIBOSOMAL_SUBUNIT | 0.00001 | 0.00056 | 0.61053 | 0.67092 | 1.34182 | 106 |
| GO_LEUKOCYTE_MEDIATED_IMMUNITY | 0.00111 | 0.02667 | 0.45506 | 0.55315 | 1.13634 | 427 |
| GO_MACROAUTOPHAGY | 0.00159 | 0.03426 | 0.45506 | 0.57206 | 1.16282 | 247 |
| GO_MEIOSIS_I_CELL_CYCLE_PROCESS | 0.00002 | 0.00152 | 0.57561 | 0.67248 | 1.33933 | 93 |
| GO_MEIOTIC_CELL_CYCLE | 0.00021 | 0.00777 | 0.51885 | 0.60028 | 1.21601 | 198 |
| GO_MEIOTIC_CELL_CYCLE_PROCESS | 0.00006 | 0.00322 | 0.53843 | 0.62728 | 1.26196 | 146 |
| GO_MICROTUBULE_CYTOSKELETON_ORGANIZATION | 0.00076 | 0.02039 | 0.47727 | 0.55125 | 1.13337 | 453 |
| GO_MICROTUBULE_CYTOSKELETON_ORGANIZATION_INVOLVED | 0.00047 | 0.01399 | 0.49849 | 0.62253 | 1.24891 | 123 |
| GO_MITOCHONDRIAL_TRANSPORT | 0.00051 | 0.01453 | 0.47727 | 0.59283 | 1.19974 | 189 |
| GO_MITOCHONDRION_ORGANIZATION | 0.00004 | 0.00223 | 0.55733 | 0.57647 | 1.18323 | 401 |
| GO_MITOTIC_CELL_CYCLE_CHECKPOINT | 0.00039 | 0.01216 | 0.49849 | 0.61856 | 1.24423 | 139 |
| GO_MITOTIC_NUCLEAR_DIVISION | 0.00002 | 0.00130 | 0.57561 | 0.60637 | 1.23160 | 228 |
| GO_MITOTIC_SISTER_CHROMATID_SEGREGATION | 0.00002 | 0.00157 | 0.57561 | 0.63555 | 1.27799 | 144 |
| GO_MITOTIC_SPINDLE_ASSEMBLY | 0.00121 | 0.02834 | 0.45506 | 0.66904 | 1.30588 | 54 |
| GO_MITOTIC_SPINDLE_ORGANIZATION | 0.00006 | 0.00321 | 0.53843 | 0.65933 | 1.31603 | 102 |
| GO_MORPHOGENESIS_OF_A_POLARIZED_EPITHELIUM | 0.00111 | 0.02667 | 0.45506 | 0.62548 | 1.25057 | 105 |
| GO_MRNA_3_END_PROCESSING | 0.00003 | 0.00180 | 0.57561 | 0.66356 | 1.32181 | 94 |
| GO_MRNA_BINDING | 0.00000 | 0.00000 | | 0.66560 | 1.35242 | 244 |
| GO_MRNA_EXPORT_FROM_NUCLEUS | 0.00001 | 0.00076 | 0.59333 | 0.66430 | 1.32795 | 104 |
| GO_MRNA_PROCESSING | 0.00000 | 0.00000 | | 0.66942 | 1.37599 | 445 |
| GO_MRNA_TRANSPORT | 0.00032 | 0.01057 | 0.49849 | 0.62370 | 1.25412 | 137 |
| GO_NCRNA_METABOLIC_PROCESS | 0.00008 | 0.00361 | 0.53843 | 0.57257 | 1.17396 | 390 |
| GO_NCRNA_PROCESSING | 0.00000 | 0.00022 | 0.62726 | 0.59922 | 1.22575 | 329 |
| GO_NEGATIVE_REGULATION_OF_CALCIUM_ION_TRANSMEMBR | 0.00051 | 0.01453 | 0.47727 | 0.82264 | 1.51745 | 16 |
| GO_NEGATIVE_REGULATION_OF_CALCIUM_ION_TRANSPORT | 0.00231 | 0.04475 | 0.43171 | 0.74779 | 1.40340 | 23 |
| GO_NEGATIVE_REGULATION_OF_CATALYTIC_ACTIVITY | 0.00079 | 0.02069 | 0.47727 | 0.55014 | 1.13120 | 468 |
| GO_NEGATIVE_REGULATION_OF_CELLULAR_COMPONENT_ORG | 0.00038 | 0.01193 | 0.49849 | 0.55283 | 1.13814 | 497 |
| GO_NEGATIVE_REGULATION_OF_CELL_CYCLE | 0.00001 | 0.00049 | 0.61053 | 0.57886 | 1.19095 | 481 |
| GO_NEGATIVE_REGULATION_OF_CELL_CYCLE_G2_M_PHASE_TR | 0.00012 | 0.00507 | 0.53843 | 0.65680 | 1.30444 | 88 |
| GO_NEGATIVE_REGULATION_OF_CELL_CYCLE_PHASE_TRANSITI | 0.00000 | 0.00030 | 0.62726 | 0.62773 | 1.27202 | 200 |
| GO_NEGATIVE_REGULATION_OF_CELL_CYCLE_PROCESS | 0.00000 | 0.00001 | 0.73376 | 0.63019 | 1.28279 | 267 |
| GO_NEGATIVE_REGULATION_OF_CHROMOSOME_ORGANIZATIO | 0.00002 | 0.00121 | 0.57561 | 0.65382 | 1.30928 | 115 |
| GO_NEGATIVE_REGULATION_OF_CHROMOSOME_SEGREGATION | 0.00157 | 0.03409 | 0.45506 | 0.69917 | 1.35392 | 44 |
| GO_NEGATIVE_REGULATION_OF_MITOTIC_CELL_CYCLE | 0.00000 | 0.00005 | 0.67496 | 0.62378 | 1.26923 | 262 |
| GO_NEGATIVE_REGULATION_OF_MRNA_SPLICING_VIA_SPLICEO | 0.00133 | 0.03025 | 0.45506 | 0.78250 | 1.45555 | 20 |
| GO_NEGATIVE_REGULATION_OF_MULTI_ORGANISM_PROCESS | 0.00001 | 0.00076 | 0.59333 | 0.88326 | 1.63379 | 17 |

| | | | | | | |
|---|---|---|---|---|---|---|
| GO_NEGATIVE_REGULATION_OF_NUCLEAR_DIVISION | 0.00030 | 0.01029 | 0.49849 | 0.69164 | 1.35223 | 56 |
| GO_NEGATIVE_REGULATION_OF_ORGANELLE_ORGANIZATION | 0.00003 | 0.00168 | 0.57561 | 0.59116 | 1.20702 | 297 |
| GO_NEGATIVE_REGULATION_OF_PROTEIN_MODIFICATION_PRO | 0.00135 | 0.03056 | 0.45506 | 0.55292 | 1.13566 | 422 |
| GO_NEGATIVE_REGULATION_OF_REPRODUCTIVE_PROCESS | 0.00100 | 0.02478 | 0.45506 | 0.71551 | 1.37334 | 37 |
| GO_NEGATIVE_REGULATION_OF_RNA_SPLICING | 0.00033 | 0.01075 | 0.49849 | 0.77989 | 1.46705 | 24 |
| GO_NEGATIVE_REGULATION_OF_TRANSFERASE_ACTIVITY | 0.00075 | 0.02025 | 0.47727 | 0.59277 | 1.20055 | 197 |
| GO_NIK_NF_KAPPAB_SIGNALING | 0.00004 | 0.00197 | 0.55733 | 0.64659 | 1.29589 | 118 |
| GO_NUCLEAR_CHROMOSOME_SEGREGATION | 0.00001 | 0.00060 | 0.61053 | 0.61426 | 1.24696 | 221 |
| GO_NUCLEAR_CHROMOSOME_TELOMERIC_REGION | 0.00242 | 0.04633 | 0.43171 | 0.62211 | 1.23901 | 93 |
| GO_NUCLEAR_EXPORT | 0.00000 | 0.00030 | 0.62726 | 0.63757 | 1.28934 | 181 |
| GO_NUCLEAR_MATRIX | 0.00250 | 0.04718 | 0.43171 | 0.62272 | 1.24125 | 96 |
| GO_NUCLEAR_PERIPHERY | 0.00030 | 0.01029 | 0.49849 | 0.62518 | 1.25296 | 118 |
| GO_NUCLEAR_SPECK | 0.00000 | 0.00000 | 0.82666 | 0.63944 | 1.30679 | 310 |
| GO_NUCLEAR_TRANSCRIBED_MRNA_CATABOLIC_PROCESS | 0.00000 | 0.00000 | | 0.69053 | 1.39720 | 187 |
| GO_NUCLEAR_TRANSCRIBED_MRNA_CATABOLIC_PROCESS_NON | 0.00000 | 0.00000 | | 0.75664 | 1.51504 | 112 |
| GO_NUCLEAR_TRANSPORT | 0.00000 | 0.00000 | 0.76146 | 0.63218 | 1.29061 | 298 |
| GO_NUCLEOBASE_CONTAINING_COMPOUND_TRANSPORT | 0.00003 | 0.00176 | 0.57561 | 0.60601 | 1.22999 | 220 |
| GO_NUCLEOTIDE_EXCISION_REPAIR | 0.00235 | 0.04519 | 0.43171 | 0.62334 | 1.24357 | 99 |
| GO_NUCLEUS_ORGANIZATION | 0.00001 | 0.00096 | 0.59333 | 0.66685 | 1.33381 | 107 |
| GO_OOCYTE_DIFFERENTIATION | 0.00006 | 0.00322 | 0.53843 | 0.76246 | 1.46322 | 36 |
| GO_OOCYTE_MATURATION | 0.00159 | 0.03426 | 0.45506 | 0.78827 | 1.46196 | 18 |
| GO_OOGENESIS | 0.00095 | 0.02378 | 0.47727 | 0.67396 | 1.31898 | 59 |
| GO_ORGANELLE_FISSION | 0.00000 | 0.00006 | 0.67496 | 0.60063 | 1.23005 | 362 |
| GO_ORGANIC_CYCLIC_COMPOUND_CATABOLIC_PROCESS | 0.00000 | 0.00000 | | 0.62226 | 1.28017 | 480 |
| GO_OXIDATIVE_PHOSPHORYLATION | 0.00025 | 0.00887 | 0.49849 | 0.64992 | 1.29725 | 100 |
| GO_PEPTIDASE_COMPLEX | 0.00110 | 0.02664 | 0.45506 | 0.65351 | 1.29244 | 78 |
| GO_POLYSOMAL_RIBOSOME | 0.00032 | 0.01057 | 0.49849 | 0.76699 | 1.45461 | 30 |
| GO_POLYSOME | 0.00018 | 0.00707 | 0.51885 | 0.68813 | 1.35125 | 64 |
| GO_POSITIVE_REGULATION_OF_ATPASE_ACTIVITY | 0.00079 | 0.02069 | 0.47727 | 0.71056 | 1.37619 | 43 |
| GO_POSITIVE_REGULATION_OF_CALCIUM_ION_DEPENDENT_EX | 0.00041 | 0.01275 | 0.49849 | 0.84421 | 1.55536 | 15 |
| GO_POSITIVE_REGULATION_OF_CANONICAL_WNT_SIGNALING_ | 0.00250 | 0.04718 | 0.43171 | 0.60562 | 1.21292 | 114 |
| GO_POSITIVE_REGULATION_OF_CELLULAR_AMIDE_METABOLIC_ | 0.00036 | 0.01167 | 0.49849 | 0.62398 | 1.25499 | 132 |
| GO_POSITIVE_REGULATION_OF_CELL_CYCLE | 0.00038 | 0.01193 | 0.49849 | 0.57505 | 1.17262 | 282 |
| GO_POSITIVE_REGULATION_OF_CELL_CYCLE_PROCESS | 0.00076 | 0.02032 | 0.47727 | 0.58570 | 1.18818 | 215 |
| GO_POSITIVE_REGULATION_OF_DEFENSE_RESPONSE | 0.00006 | 0.00295 | 0.55733 | 0.61440 | 1.24317 | 188 |
| GO_POSITIVE_REGULATION_OF_FIBROBLAST_PROLIFERATION | 0.00010 | 0.00451 | 0.53843 | 0.78953 | 1.49207 | 27 |
| GO_POSITIVE_REGULATION_OF_IMMUNE_RESPONSE | 0.00030 | 0.01029 | 0.49849 | 0.57552 | 1.17685 | 327 |
| GO_POSITIVE_REGULATION_OF_INNATE_IMMUNE_RESPONSE | 0.00000 | 0.00042 | 0.61053 | 0.66104 | 1.32677 | 125 |
| GO_POSITIVE_REGULATION_OF_MITOTIC_CELL_CYCLE | 0.00115 | 0.02725 | 0.45506 | 0.61353 | 1.23045 | 122 |
| GO_POSITIVE_REGULATION_OF_MRNA_PROCESSING | 0.00133 | 0.03025 | 0.45506 | 0.74425 | 1.40526 | 26 |
| GO_POSITIVE_REGULATION_OF_MRNA_SPLICING_VIA_SPLICEOS | 0.00170 | 0.03590 | 0.45506 | 0.79063 | 1.45839 | 16 |
| GO_POSITIVE_REGULATION_OF_PROTEIN_MODIFICATION_BY_S | 0.00265 | 0.04968 | 0.43171 | 0.61239 | 1.22495 | 108 |
| GO_POSITIVE_REGULATION_OF_RESPONSE_TO_BIOTIC_STIMULI | 0.00003 | 0.00176 | 0.57561 | 0.63483 | 1.27654 | 144 |
| GO_POSITIVE_REGULATION_OF_RESPONSE_TO_EXTERNAL_STIM | 0.00006 | 0.00311 | 0.55733 | 0.59661 | 1.21312 | 256 |
| GO_POSITIVE_REGULATION_OF_RNA_SPLICING | 0.00139 | 0.03128 | 0.45506 | 0.74302 | 1.40640 | 29 |
| GO_POSITIVE_REGULATION_OF_SIGNAL_TRANSDUCTION_BY_P5 | 0.00009 | 0.00418 | 0.53843 | 0.85683 | 1.58051 | 16 |
| GO_POSITIVE_REGULATION_OF_TRANSFERASE_ACTIVITY | 0.00074 | 0.02008 | 0.47727 | 0.55140 | 1.13369 | 453 |
| GO_POSITIVE_REGULATION_OF_TRANSLATION | 0.00003 | 0.00193 | 0.55733 | 0.65738 | 1.31552 | 110 |
| GO_POSITIVE_REGULATION_OF_VIRAL_PROCESS | 0.00002 | 0.00130 | 0.57561 | 0.68164 | 1.35570 | 91 |
| GO_POSTTRANSCRIPTIONAL_REGULATION_OF_GENE_EXPRESSIO | 0.00000 | 0.00000 | | 0.63749 | 1.31258 | 499 |
| GO_POST_TRANSLATIONAL_PROTEIN_MODIFICATION | 0.00009 | 0.00403 | 0.53843 | 0.59363 | 1.20678 | 251 |
| GO_PRECATALYTIC_SPLICEOSOME | 0.00032 | 0.01057 | 0.49849 | 0.70624 | 1.37133 | 48 |
| GO_PRERIBOSOME | 0.00017 | 0.00657 | 0.51885 | 0.67210 | 1.32623 | 70 |
| GO_PRE_MRNA_BINDING | 0.00020 | 0.00740 | 0.51885 | 0.75582 | 1.44588 | 34 |
| GO_PROTEASOMAL_PROTEIN_CATABOLIC_PROCESS | 0.00002 | 0.00132 | 0.57561 | 0.57648 | 1.18324 | 402 |
| GO_PROTEIN_CONTAINING_COMPLEX_DISASSEMBLY | 0.00124 | 0.02878 | 0.45506 | 0.57343 | 1.16756 | 268 |
| GO_PROTEIN_CONTAINING_COMPLEX_LOCALIZATION | 0.00021 | 0.00777 | 0.51885 | 0.59273 | 1.20434 | 241 |
| GO_PROTEIN_C_TERMINUS_BINDING | 0.00051 | 0.01453 | 0.47727 | 0.60637 | 1.22220 | 154 |
| GO_PROTEIN_DNA_COMPLEX_SUBUNIT_ORGANIZATION | 0.00001 | 0.00068 | 0.59333 | 0.62491 | 1.26386 | 180 |
| GO_PROTEIN_DOMAIN_SPECIFIC_BINDING | 0.00010 | 0.00444 | 0.53843 | 0.55999 | 1.15281 | 494 |
| GO_PROTEIN_FOLDING | 0.00003 | 0.00191 | 0.55733 | 0.61753 | 1.24809 | 176 |
| GO_PROTEIN_FOLDING_CHAPERONE | 0.00003 | 0.00190 | 0.55733 | 0.84659 | 1.56973 | 19 |
| GO_PROTEIN_IMPORT | 0.00004 | 0.00206 | 0.55733 | 0.62868 | 1.26719 | 153 |
| GO_PROTEIN_LOCALIZATION_TO_ENDOPLASMIC_RETICULUM | 0.00000 | 0.00000 | 0.82666 | 0.71898 | 1.44371 | 126 |
| GO_PROTEIN_LOCALIZATION_TO_MEMBRANE | 0.00000 | 0.00001 | 0.70498 | 0.59325 | 1.21983 | 468 |
| GO_PROTEIN_LOCALIZATION_TO_MITOCHONDRION | 0.00002 | 0.00137 | 0.57561 | 0.65077 | 1.30426 | 118 |
| GO_PROTEIN_LOCALIZATION_TO_NUCLEUS | 0.00000 | 0.00002 | 0.69013 | 0.64320 | 1.30476 | 212 |
| GO_PROTEIN_MODIFICATION_BY_SMALL_PROTEIN_REMOVAL | 0.00000 | 0.00012 | 0.65944 | 0.62408 | 1.26776 | 235 |
| GO_PROTEIN_POLYUBIQUITINATION | 0.00141 | 0.03158 | 0.45506 | 0.56871 | 1.15877 | 275 |
| GO_PROTEIN_STABILIZATION | 0.00156 | 0.03393 | 0.45506 | 0.59954 | 1.20558 | 144 |

| | | | | | | |
|---|---|---|---|---|---|---|
| GO_PROTEIN_TARGETING | 0.00000 | 0.00000 | | 0.63964 | 1.30910 | 351 |
| GO_PROTEIN_TARGETING_TO_MEMBRANE | 0.00000 | 0.00000 | 0.81404 | 0.69555 | 1.40377 | 159 |
| GO_PROTEIN_TARGETING_TO_MITOCHONDRION | 0.00056 | 0.01596 | 0.47727 | 0.65184 | 1.29281 | 83 |
| GO_PROTEIN_TRANSMEMBRANE_TRANSPORT | 0.00224 | 0.04371 | 0.43171 | 0.68053 | 1.32393 | 50 |
| GO_REGULATION_OF_ANIMAL_ORGAN_MORPHOGENESIS | 0.00051 | 0.01453 | 0.47727 | 0.60640 | 1.22228 | 154 |
| GO_REGULATION_OF_APOPTOTIC_SIGNALING_PATHWAY | 0.00028 | 0.00988 | 0.49849 | 0.58315 | 1.18704 | 267 |
| GO_REGULATION_OF_ATPASE_ACTIVITY | 0.00061 | 0.01696 | 0.47727 | 0.67333 | 1.31700 | 60 |
| GO_REGULATION_OF_ATP_METABOLIC_PROCESS | 0.00122 | 0.02846 | 0.45506 | 0.63237 | 1.25846 | 92 |
| GO_REGULATION_OF_CELLULAR_AMIDE_METABOLIC_PROCESS | 0.00000 | 0.00011 | 0.65944 | 0.59637 | 1.22029 | 348 |
| GO_REGULATION_OF_CELLULAR_AMINE_METABOLIC_PROCESS | 0.00012 | 0.00507 | 0.53843 | 0.69256 | 1.35783 | 62 |
| GO_REGULATION_OF_CELLULAR_AMINO_ACID_METABOLIC_PR( | 0.00007 | 0.00344 | 0.53843 | 0.72269 | 1.41036 | 53 |
| GO_REGULATION_OF_CELLULAR_KETONE_METABOLIC_PROCESS | 0.00025 | 0.00887 | 0.49849 | 0.63063 | 1.26432 | 120 |
| GO_REGULATION_OF_CELLULAR_RESPONSE_TO_HEAT | 0.00194 | 0.03914 | 0.45506 | 0.64504 | 1.27292 | 71 |
| GO_REGULATION_OF_CELL_CYCLE_G2_M_PHASE_TRANSITION | 0.00007 | 0.00358 | 0.53843 | 0.61032 | 1.23492 | 187 |
| GO_REGULATION_OF_CELL_CYCLE_PHASE_TRANSITION | 0.00000 | 0.00014 | 0.64355 | 0.59230 | 1.21355 | 376 |
| GO_REGULATION_OF_CHROMATIN_ORGANIZATION | 0.00080 | 0.02085 | 0.47727 | 0.60675 | 1.22008 | 144 |
| GO_REGULATION_OF_CHROMOSOME_ORGANIZATION | 0.00000 | 0.00011 | 0.65944 | 0.60942 | 1.24287 | 285 |
| GO_REGULATION_OF_CHROMOSOME_SEGREGATION | 0.00033 | 0.01066 | 0.49849 | 0.64680 | 1.28925 | 96 |
| GO_REGULATION_OF_CYCLIN_DEPENDENT_PROTEIN_KINASE_A | 0.00020 | 0.00744 | 0.51885 | 0.66290 | 1.31475 | 83 |
| GO_REGULATION_OF_CYSTEINE_TYPE_ENDOPEPTIDASE_ACTIVIT | 0.00185 | 0.03800 | 0.45506 | 0.59291 | 1.19509 | 154 |
| GO_REGULATION_OF_CYTOKINESIS | 0.00066 | 0.01832 | 0.47727 | 0.67291 | 1.32136 | 64 |
| GO_REGULATION_OF_DNA_TEMPLATED_TRANSCRIPTION_IN_RE | 0.00000 | 0.00042 | 0.61053 | 0.68595 | 1.36727 | 96 |
| GO_REGULATION_OF_FERTILIZATION | 0.00008 | 0.00395 | 0.53843 | 0.87081 | 1.60435 | 15 |
| GO_REGULATION_OF_GENE_EXPRESSION_EPIGENETIC | 0.00045 | 0.01357 | 0.49849 | 0.62323 | 1.24990 | 122 |
| GO_REGULATION_OF_HEMATOPOIETIC_PROGENITOR_CELL_DIF | 0.00242 | 0.04633 | 0.43171 | 0.64075 | 1.26673 | 75 |
| GO_REGULATION_OF_HEMATOPOIETIC_STEM_CELL_DIFFERENT | 0.00133 | 0.03025 | 0.45506 | 0.66330 | 1.30250 | 64 |
| GO_REGULATION_OF_HISTONE_MODIFICATION | 0.00187 | 0.03825 | 0.45506 | 0.61037 | 1.22227 | 115 |
| GO_REGULATION_OF_IMMUNE_RESPONSE | 0.00133 | 0.03025 | 0.45506 | 0.54866 | 1.12718 | 438 |
| GO_REGULATION_OF_INNATE_IMMUNE_RESPONSE | 0.00000 | 0.00017 | 0.64355 | 0.64123 | 1.29474 | 168 |
| GO_REGULATION_OF_INTRINSIC_APOPTOTIC_SIGNALING_PATH\ | 0.00019 | 0.00707 | 0.51885 | 0.63640 | 1.27427 | 112 |
| GO_REGULATION_OF_MORPHOGENESIS_OF_AN_EPITHELIUM | 0.00115 | 0.02725 | 0.45506 | 0.61521 | 1.23341 | 120 |
| GO_REGULATION_OF_MRNA_CATABOLIC_PROCESS | 0.00007 | 0.00344 | 0.53843 | 0.61708 | 1.24576 | 172 |
| GO_REGULATION_OF_MRNA_METABOLIC_PROCESS | 0.00000 | 0.00000 | 0.81404 | 0.64147 | 1.30824 | 285 |
| GO_REGULATION_OF_MRNA_PROCESSING | 0.00001 | 0.00063 | 0.61053 | 0.65661 | 1.31959 | 129 |
| GO_REGULATION_OF_MRNA_SPLICING_VIA_SPLICEOSOME | 0.00001 | 0.00049 | 0.61053 | 0.68510 | 1.36447 | 93 |
| GO_REGULATION_OF_MULTI_ORGANISM_PROCESS | 0.00030 | 0.01029 | 0.49849 | 0.70789 | 1.37157 | 46 |
| GO_REGULATION_OF_NUCLEAR_DIVISION | 0.00004 | 0.00216 | 0.55733 | 0.63416 | 1.27653 | 148 |
| GO_REGULATION_OF_PROTEIN_MODIFICATION_BY_SMALL_PRC | 0.00019 | 0.00707 | 0.51885 | 0.59973 | 1.21394 | 190 |
| GO_REGULATION_OF_PROTEIN_SERINE_THREONINE_KINASE_A( | 0.00004 | 0.00198 | 0.55733 | 0.57734 | 1.18198 | 358 |
| GO_REGULATION_OF_REPRODUCTIVE_PROCESS | 0.00018 | 0.00676 | 0.51885 | 0.64717 | 1.29177 | 102 |
| GO_REGULATION_OF_RESPONSE_TO_BIOTIC_STIMULUS | 0.00015 | 0.00610 | 0.51885 | 0.59663 | 1.21123 | 224 |
| GO_REGULATION_OF_RNA_SPLICING | 0.00000 | 0.00012 | 0.65944 | 0.67522 | 1.35758 | 130 |
| GO_REGULATION_OF_SIGNAL_TRANSDUCTION_BY_P53_CLASS_ | 0.00079 | 0.02069 | 0.47727 | 0.60660 | 1.22035 | 146 |
| GO_REGULATION_OF_SISTER_CHROMATID_SEGREGATION | 0.00081 | 0.02100 | 0.47727 | 0.65825 | 1.30128 | 77 |
| GO_REGULATION_OF_SMALL_MOLECULE_METABOLIC_PROCESS | 0.00017 | 0.00671 | 0.51885 | 0.57963 | 1.18274 | 291 |
| GO_REGULATION_OF_STEM_CELL_DIFFERENTIATION | 0.00146 | 0.03236 | 0.45506 | 0.62807 | 1.25301 | 99 |
| GO_REGULATION_OF_SYMBIOTIC_PROCESS | 0.00000 | 0.00013 | 0.64355 | 0.65701 | 1.32316 | 149 |
| GO_REGULATION_OF_TRANSCRIPTION_FROM_RNA_POLYMERA | 0.00012 | 0.00507 | 0.53843 | 0.68943 | 1.35624 | 66 |
| GO_REGULATION_OF_TRANSLATIONAL_INITIATION | 0.00042 | 0.01292 | 0.49849 | 0.67981 | 1.33848 | 67 |
| GO_REGULATION_OF_UBIQUITIN_PROTEIN_LIGASE_ACTIVITY | 0.00011 | 0.00475 | 0.53843 | 0.79714 | 1.49949 | 24 |
| GO_REGULATION_OF_VIRAL_GENOME_REPLICATION | 0.00052 | 0.01473 | 0.47727 | 0.68295 | 1.33525 | 56 |
| GO_REGULATION_OF_VIRAL_LIFE_CYCLE | 0.00097 | 0.02415 | 0.47727 | 0.63826 | 1.26695 | 87 |
| GO_REGULATION_OF_VIRAL_TRANSCRIPTION | 0.00086 | 0.02183 | 0.47727 | 0.68175 | 1.33167 | 55 |
| GO_REPRODUCTIVE_SYSTEM_DEVELOPMENT | 0.00016 | 0.00629 | 0.51885 | 0.58663 | 1.19379 | 265 |
| GO_RESPONSE_TO_DSRNA | 0.00220 | 0.04339 | 0.43171 | 0.77185 | 1.44450 | 22 |
| GO_RESPONSE_TO_FIBROBLAST_GROWTH_FACTOR | 0.00001 | 0.00067 | 0.59333 | 0.69552 | 1.37944 | 83 |
| GO_RESPONSE_TO_HEAT | 0.00051 | 0.01453 | 0.47727 | 0.62345 | 1.24993 | 120 |
| GO_RESPONSE_TO_INTERLEUKIN_1 | 0.00010 | 0.00457 | 0.53843 | 0.63671 | 1.27904 | 128 |
| GO_RESPONSE_TO_LEUKEMIA_INHIBITORY_FACTOR | 0.00023 | 0.00814 | 0.51885 | 0.67968 | 1.34017 | 69 |
| GO_RESPONSE_TO_OXYGEN_LEVELS | 0.00039 | 0.01224 | 0.49849 | 0.58486 | 1.18973 | 260 |
| GO_RESPONSE_TO_TEMPERATURE_STIMULUS | 0.00019 | 0.00707 | 0.51885 | 0.61199 | 1.23547 | 163 |
| GO_RESPONSE_TO_TUMOR_NECROSIS_FACTOR | 0.00110 | 0.02664 | 0.45506 | 0.59597 | 1.20523 | 181 |
| GO_RHYTHMIC_PROCESS | 0.00191 | 0.03864 | 0.45506 | 0.58163 | 1.17766 | 196 |
| GO_RIBONUCLEOPROTEIN_COMPLEX_BINDING | 0.00001 | 0.00049 | 0.61053 | 0.66220 | 1.32606 | 115 |
| GO_RIBONUCLEOPROTEIN_COMPLEX_BIOGENESIS | 0.00000 | 0.00000 | | 0.66856 | 1.37082 | 391 |
| GO_RIBONUCLEOPROTEIN_COMPLEX_SUBUNIT_ORGANIZATION | 0.00000 | 0.00000 | 0.78819 | 0.67654 | 1.36828 | 180 |
| GO_RIBONUCLEOPROTEIN_GRANULE | 0.00003 | 0.00175 | 0.57561 | 0.61996 | 1.25502 | 191 |
| GO_RIBOSOMAL_LARGE_SUBUNIT_BIOGENESIS | 0.00001 | 0.00111 | 0.59333 | 0.71632 | 1.41036 | 67 |
| GO_RIBOSOMAL_SMALL_SUBUNIT_BIOGENESIS | 0.00183 | 0.03774 | 0.45506 | 0.65521 | 1.28893 | 66 |
| GO_RIBOSOMAL_SUBUNIT | 0.00000 | 0.00001 | 0.71951 | 0.66852 | 1.35005 | 170 |

| | | | | | | |
|---|---|---|---|---|---|---|
| GO_RIBOSOME | 0.00000 | 0.00000 | 0.80122 | 0.67000 | 1.35875 | 208 |
| GO_RIBOSOME_ASSEMBLY | 0.00002 | 0.00146 | 0.57561 | 0.71525 | 1.39886 | 58 |
| GO_RIBOSOME_BINDING | 0.00224 | 0.04371 | 0.43171 | 0.67392 | 1.30573 | 46 |
| GO_RIBOSOME_BIOGENESIS | 0.00000 | 0.00000 | | 0.66460 | 1.35350 | 272 |
| GO_RNA_3_END_PROCESSING | 0.00006 | 0.00321 | 0.53843 | 0.64290 | 1.29147 | 128 |
| GO_RNA_CATABOLIC_PROCESS | 0.00000 | 0.00000 | | 0.65497 | 1.34033 | 353 |
| GO_RNA_DEPENDENT_DNA_BIOSYNTHETIC_PROCESS | 0.00183 | 0.03774 | 0.45506 | 0.65590 | 1.28595 | 62 |
| GO_RNA_EXPORT_FROM_NUCLEUS | 0.00001 | 0.00076 | 0.59333 | 0.65570 | 1.31845 | 131 |
| GO_RNA_HELICASE_ACTIVITY | 0.00050 | 0.01453 | 0.47727 | 0.67754 | 1.33400 | 67 |
| GO_RNA_LOCALIZATION | 0.00000 | 0.00002 | 0.70498 | 0.63954 | 1.29724 | 210 |
| GO_RNA_POLYMERASE_BINDING | 0.00132 | 0.03025 | 0.45506 | 0.65971 | 1.29023 | 58 |
| GO_RNA_POLYMERASE_COMPLEX | 0.00104 | 0.02549 | 0.45506 | 0.63305 | 1.26183 | 96 |
| GO_RNA_POLYMERASE_II_HOLOENZYME | 0.00178 | 0.03721 | 0.45506 | 0.64042 | 1.26372 | 70 |
| GO_RNA_SPLICING | 0.00000 | 0.00000 | | 0.67255 | 1.37919 | 393 |
| GO_RNA_SPLICING_VIA_TRANSESTERIFICATION_REACTIONS | 0.00000 | 0.00000 | | 0.69147 | 1.41368 | 316 |
| GO_RRNA_BINDING | 0.00000 | 0.00000 | 0.76146 | 0.81282 | 1.58185 | 51 |
| GO_RRNA_METABOLIC_PROCESS | 0.00000 | 0.00000 | 0.73376 | 0.65098 | 1.31990 | 206 |
| GO_SCF_DEPENDENT_PROTEASOMAL_UBIQUITIN_DEPENDENT_ | 0.00008 | 0.00382 | 0.53843 | 0.68556 | 1.35582 | 78 |
| GO_SEX_DIFFERENTIATION | 0.00146 | 0.03236 | 0.45506 | 0.59304 | 1.19643 | 158 |
| GO_SIGNAL_SEQUENCE_BINDING | 0.00111 | 0.02667 | 0.45506 | 0.71419 | 1.37082 | 37 |
| GO_SIGNAL_TRANSDUCTION_BY_P53_CLASS_MEDIATOR | 0.00003 | 0.00188 | 0.55733 | 0.60780 | 1.23302 | 215 |
| GO_SIGNAL_TRANSDUCTION_IN_RESPONSE_TO_DNA_DAMAGE | 0.00044 | 0.01328 | 0.49849 | 0.63898 | 1.27543 | 102 |
| GO_SINGLE_FERTILIZATION | 0.00082 | 0.02115 | 0.47727 | 0.65736 | 1.29847 | 76 |
| GO_SISTER_CHROMATID_SEGREGATION | 0.00001 | 0.00047 | 0.61053 | 0.63649 | 1.28521 | 169 |
| GO_SMALL_NUCLEAR_RIBONUCLEOPROTEIN_COMPLEX | 0.00004 | 0.00210 | 0.55733 | 0.70782 | 1.38431 | 58 |
| GO_SMALL_RIBOSOMAL_SUBUNIT | 0.00047 | 0.01399 | 0.49849 | 0.67878 | 1.33645 | 67 |
| GO_SM_LIKE_PROTEIN_FAMILY_COMPLEX | 0.00004 | 0.00197 | 0.55733 | 0.69870 | 1.37626 | 68 |
| GO_SNORNA_BINDING | 0.00023 | 0.00842 | 0.51885 | 0.77280 | 1.46563 | 30 |
| GO_SPERM_EGG_RECOGNITION | 0.00019 | 0.00707 | 0.51885 | 0.75782 | 1.44596 | 33 |
| GO_SPINDLE | 0.00006 | 0.00308 | 0.55733 | 0.58467 | 1.19419 | 306 |
| GO_SPINDLE_MIDZONE | 0.00064 | 0.01764 | 0.47727 | 0.74669 | 1.41884 | 31 |
| GO_SPINDLE_ORGANIZATION | 0.00012 | 0.00507 | 0.53843 | 0.61659 | 1.24394 | 158 |
| GO_SPINDLE_POLE | 0.00050 | 0.01453 | 0.47727 | 0.61192 | 1.23109 | 141 |
| GO_SPLICEOSOMAL_COMPLEX | 0.00000 | 0.00000 | | 0.70540 | 1.42406 | 172 |
| GO_SPLICEOSOMAL_COMPLEX_ASSEMBLY | 0.00048 | 0.01419 | 0.49849 | 0.70038 | 1.36085 | 49 |
| GO_STEM_CELL_DIFFERENTIATION | 0.00098 | 0.02451 | 0.45506 | 0.59330 | 1.19980 | 183 |
| GO_STRUCTURAL_CONSTITUENT_OF_RIBOSOME | 0.00000 | 0.00000 | 0.74774 | 0.68390 | 1.37770 | 151 |
| GO_STRUCTURAL_MOLECULE_ACTIVITY | 0.00000 | 0.00046 | 0.61053 | 0.58139 | 1.19396 | 420 |
| GO_TELOMERASE_RNA_LOCALIZATION | 0.00191 | 0.03864 | 0.45506 | 0.77098 | 1.42953 | 19 |
| GO_TELOMERE_MAINTENANCE_VIA_TELOMERE_LENGTHENING | 0.00045 | 0.01357 | 0.49849 | 0.67074 | 1.32254 | 69 |
| GO_TELOMERE_ORGANIZATION | 0.00010 | 0.00444 | 0.53843 | 0.63723 | 1.28008 | 128 |
| GO_TRANSCRIPTION_COREGULATOR_ACTIVITY | 0.00011 | 0.00473 | 0.53843 | 0.57190 | 1.17200 | 381 |
| GO_TRANSCRIPTION_COUPLED_NUCLEOTIDE_EXCISION_REPAIR | 0.00042 | 0.01292 | 0.49849 | 0.67128 | 1.32361 | 69 |
| GO_TRANSCRIPTION_ELONGATION_FROM_RNA_POLYMERASE_I | 0.00067 | 0.01850 | 0.47727 | 0.65794 | 1.30070 | 75 |
| GO_TRANSCRIPTION_FACTOR_BINDING | 0.00006 | 0.00308 | 0.55733 | 0.56487 | 1.16249 | 486 |
| GO_TRANSCRIPTION_INITIATION_FROM_RNA_POLYMERASE_II_ | 0.00161 | 0.03442 | 0.45506 | 0.59434 | 1.19796 | 154 |
| GO_TRANSFERASE_COMPLEX_TRANSFERRING_PHOSPHORUS_C | 0.00002 | 0.00157 | 0.57561 | 0.60808 | 1.23335 | 214 |
| GO_TRANSLATIONAL_INITIATION | 0.00000 | 0.00000 | | 0.71723 | 1.44824 | 169 |
| GO_TRANSLATION_FACTOR_ACTIVITY_RNA_BINDING | 0.00001 | 0.00071 | 0.59333 | 0.71155 | 1.40553 | 72 |
| GO_TRANSLATION_INITIATION_FACTOR_ACTIVITY | 0.00003 | 0.00180 | 0.57561 | 0.73928 | 1.43237 | 46 |
| GO_TRANSLATION_PREINITIATION_COMPLEX | 0.00121 | 0.02834 | 0.45506 | 0.79962 | 1.47499 | 16 |
| GO_TRANSLATION_REGULATOR_ACTIVITY | 0.00000 | 0.00016 | 0.64355 | 0.67945 | 1.35981 | 111 |
| GO_TRANSLATION_REGULATOR_ACTIVITY_NUCLEIC_ACID_BIND | 0.00000 | 0.00017 | 0.64355 | 0.70126 | 1.39200 | 87 |
| GO_TUMOR_NECROSIS_FACTOR_MEDIATED_SIGNALING_PATHW | 0.00017 | 0.00662 | 0.51885 | 0.63375 | 1.26958 | 113 |
| GO_T_CELL_RECEPTOR_SIGNALING_PATHWAY | 0.00092 | 0.02328 | 0.47727 | 0.60912 | 1.22258 | 125 |
| GO_U12_TYPE_SPLICEOSOMAL_COMPLEX | 0.00070 | 0.01917 | 0.47727 | 0.75854 | 1.43350 | 27 |
| GO_U2_SNRNP | 0.00216 | 0.04293 | 0.43171 | 0.78679 | 1.45534 | 17 |
| GO_U2_TYPE_CATALYTIC_STEP_2_SPLICEOSOME | 0.00261 | 0.04913 | 0.43171 | 0.71983 | 1.36517 | 30 |
| GO_U2_TYPE_SPLICEOSOMAL_COMPLEX | 0.00000 | 0.00013 | 0.64355 | 0.71358 | 1.41582 | 84 |
| GO_UBIQUITIN_LIKE_PROTEIN_LIGASE_BINDING | 0.00002 | 0.00157 | 0.57561 | 0.59780 | 1.21608 | 258 |
| GO_UNFOLDED_PROTEIN_BINDING | 0.00001 | 0.00064 | 0.59333 | 0.68201 | 1.35830 | 93 |
| GO_VESICLE_LUMEN | 0.00100 | 0.02478 | 0.45506 | 0.59330 | 1.19975 | 182 |
| GO_VIRAL_GENE_EXPRESSION | 0.00000 | 0.00000 | | 0.70527 | 1.42603 | 178 |
| GO_VIRAL_GENOME_REPLICATION | 0.00002 | 0.00128 | 0.57561 | 0.70166 | 1.38598 | 76 |
| GO_VIRAL_LIFE_CYCLE | 0.00001 | 0.00056 | 0.61053 | 0.61545 | 1.24919 | 223 |
| HP_ABNORMALITY_OF_MOUTH_SIZE | 0.00235 | 0.04519 | 0.43171 | 0.56907 | 1.15614 | 239 |
| HP_HEMATOLOGICAL_NEOPLASM | 0.00163 | 0.03470 | 0.45506 | 0.59219 | 1.19697 | 177 |
| HP_SPARSE_OR_ABSENT_EYELASHES | 0.00220 | 0.04339 | 0.43171 | 0.68365 | 1.32650 | 47 |
| **Cluster 6** | | | | | | |
| pathway | pval | padj | log2err | ES | NES | size |

| | | | | | | |
|---|---|---|---|---|---|---|
| GO_ACTIN_CYTOSKELETON | 0.00063 | 0.01100 | 0.47727 | 0.54808 | 1.15260 | 337 |
| GO_ACTIN_FILAMENT_ORGANIZATION | 0.00022 | 0.00469 | 0.51885 | 0.56103 | 1.17578 | 290 |
| GO_ACTIN_POLYMERIZATION_OR_DEPOLYMERIZATION | 0.00081 | 0.01322 | 0.47727 | 0.58702 | 1.21362 | 149 |
| GO_ACTIVATION_OF_IMMUNE_RESPONSE | 0.00000 | 0.00001 | 0.69013 | 0.60123 | 1.25894 | 281 |
| GO_ACTIVATION_OF_INNATE_IMMUNE_RESPONSE | 0.00000 | 0.00002 | 0.65944 | 0.67686 | 1.38561 | 108 |
| GO_ADP_BINDING | 0.00390 | 0.04277 | 0.43171 | 0.72644 | 1.39333 | 24 |
| GO_AEROBIC_RESPIRATION | 0.00000 | 0.00000 | 0.71951 | 0.74175 | 1.50018 | 73 |
| GO_ALTERNATIVE_MRNA_SPLICING_VIA_SPLICEOSOME | 0.00002 | 0.00071 | 0.57561 | 0.69004 | 1.39385 | 70 |
| GO_AMIDE_BINDING | 0.00003 | 0.00083 | 0.57561 | 0.59145 | 1.23104 | 217 |
| GO_AMIDE_TRANSMEMBRANE_TRANSPORTER_ACTIVITY | 0.00085 | 0.01352 | 0.47727 | 0.72504 | 1.41253 | 33 |
| GO_AMINE_METABOLIC_PROCESS | 0.00001 | 0.00028 | 0.61053 | 0.66516 | 1.35951 | 100 |
| GO_ANAPHASE_PROMOTING_COMPLEX_DEPENDENT_CATABOL | 0.00000 | 0.00000 | 0.70498 | 0.72589 | 1.47195 | 77 |
| GO_ANTIGEN_BINDING | 0.00390 | 0.04277 | 0.43171 | 0.71079 | 1.36724 | 25 |
| GO_ANTIGEN_PROCESSING_AND_PRESENTATION | 0.00000 | 0.00000 | 0.77494 | 0.66424 | 1.37711 | 169 |
| GO_ANTIGEN_PROCESSING_AND_PRESENTATION_OF_EXOGENO | 0.00000 | 0.00001 | 0.70498 | 0.74378 | 1.49735 | 65 |
| GO_ANTIGEN_PROCESSING_AND_PRESENTATION_OF_PEPTIDE_ | 0.00000 | 0.00000 | 0.74774 | 0.66345 | 1.37165 | 148 |
| GO_ANTIGEN_PROCESSING_AND_PRESENTATION_OF_PEPTIDE_ | 0.00000 | 0.00002 | 0.67496 | 0.71109 | 1.44394 | 79 |
| GO_ANTIGEN_RECEPTOR_MEDIATED_SIGNALING_PATHWAY | 0.00044 | 0.00839 | 0.49849 | 0.59156 | 1.22583 | 165 |
| GO_ANTIMICROBIAL_HUMORAL_IMMUNE_RESPONSE_MEDIATE | 0.00079 | 0.01295 | 0.47727 | 0.81759 | 1.51835 | 16 |
| GO_ANTIOXIDANT_ACTIVITY | 0.00000 | 0.00004 | 0.65944 | 0.77624 | 1.53701 | 44 |
| GO_APOPTOTIC_SIGNALING_PATHWAY | 0.00003 | 0.00087 | 0.57561 | 0.55446 | 1.17021 | 450 |
| GO_ATPASE_ACTIVITY | 0.00095 | 0.01470 | 0.47727 | 0.54634 | 1.14788 | 327 |
| GO_ATP_BIOSYNTHETIC_PROCESS | 0.00000 | 0.00002 | 0.67496 | 0.88734 | 1.69045 | 22 |
| GO_ATP_METABOLIC_PROCESS | 0.00000 | 0.00000 | | 0.71080 | 1.47976 | 218 |
| GO_ATP_SYNTHESIS_COUPLED_ELECTRON_TRANSPORT | 0.00000 | 0.00000 | | 0.82532 | 1.67455 | 78 |
| GO_AZUROPHIL_GRANULE | 0.00034 | 0.00664 | 0.49849 | 0.61384 | 1.25890 | 113 |
| GO_AZUROPHIL_GRANULE_LUMEN | 0.00059 | 0.01058 | 0.47727 | 0.65902 | 1.32621 | 63 |
| GO_BINDING_OF_SPERM_TO_ZONA_PELLUCIDA | 0.00018 | 0.00393 | 0.51885 | 0.83061 | 1.55895 | 18 |
| GO_BLOOD_MICROPARTICLE | 0.00005 | 0.00133 | 0.55733 | 0.71168 | 1.42392 | 55 |
| GO_CADHERIN_BINDING | 0.00000 | 0.00000 | | 0.65598 | 1.37213 | 264 |
| GO_CANONICAL_WNT_SIGNALING_PATHWAY | 0.00028 | 0.00572 | 0.49849 | 0.57103 | 1.19209 | 243 |
| GO_CARBOHYDRATE_BIOSYNTHETIC_PROCESS | 0.00009 | 0.00232 | 0.53843 | 0.61413 | 1.26817 | 138 |
| GO_CARBOHYDRATE_CATABOLIC_PROCESS | 0.00001 | 0.00029 | 0.59333 | 0.62140 | 1.28473 | 148 |
| GO_CATALYTIC_STEP_2_SPLICEOSOME | 0.00001 | 0.00022 | 0.61053 | 0.68046 | 1.38321 | 83 |
| GO_CELLULAR_ALDEHYDE_METABOLIC_PROCESS | 0.00420 | 0.04542 | 0.40702 | 0.66739 | 1.31871 | 42 |
| GO_CELLULAR_AMINO_ACID_METABOLIC_PROCESS | 0.00011 | 0.00277 | 0.53843 | 0.57486 | 1.19882 | 235 |
| GO_CELLULAR_COMPONENT_DISASSEMBLY | 0.00000 | 0.00000 | 0.82666 | 0.59817 | 1.26222 | 444 |
| GO_CELLULAR_KETONE_METABOLIC_PROCESS | 0.00018 | 0.00403 | 0.51885 | 0.59406 | 1.23222 | 170 |
| GO_CELLULAR_NITROGEN_COMPOUND_CATABOLIC_PROCESS | 0.00000 | 0.00000 | | 0.65936 | 1.39201 | 476 |
| GO_CELLULAR_OXIDANT_DETOXIFICATION | 0.00001 | 0.00021 | 0.61053 | 0.73447 | 1.46951 | 55 |
| GO_CELLULAR_PROTEIN_COMPLEX_DISASSEMBLY | 0.00000 | 0.00000 | 0.73376 | 0.64594 | 1.34203 | 184 |
| GO_CELLULAR_RESPIRATION | 0.00000 | 0.00000 | | 0.73417 | 1.51741 | 150 |
| GO_CELLULAR_RESPONSE_TO_CHEMICAL_STRESS | 0.00364 | 0.04067 | 0.43171 | 0.54805 | 1.14636 | 263 |
| GO_CELLULAR_RESPONSE_TO_HEAT | 0.00069 | 0.01174 | 0.47727 | 0.62374 | 1.27485 | 100 |
| GO_CELLULAR_RESPONSE_TO_ORGANIC_CYCLIC_COMPOUND | 0.00065 | 0.01123 | 0.47727 | 0.54783 | 1.15286 | 359 |
| GO_CELLULAR_RESPONSE_TO_OXYGEN_LEVELS | 0.00000 | 0.00000 | 0.76146 | 0.66017 | 1.36931 | 171 |
| GO_CELLULAR_RESPONSE_TO_REACTIVE_OXYGEN_SPECIES | 0.00213 | 0.02705 | 0.43171 | 0.59365 | 1.22045 | 122 |
| GO_CELLULAR_RESPONSE_TO_TOXIC_SUBSTANCE | 0.00017 | 0.00390 | 0.51885 | 0.67739 | 1.36393 | 64 |
| GO_CELL_ACTIVATION_INVOLVED_IN_IMMUNE_RESPONSE | 0.00000 | 0.00000 | | 0.60057 | 1.26775 | 460 |
| GO_CELL_ADHESION_MOLECULE_BINDING | 0.00000 | 0.00000 | | 0.63246 | 1.33177 | 383 |
| GO_CELL_BODY | 0.00283 | 0.03346 | 0.43171 | 0.53635 | 1.12851 | 356 |
| GO_CELL_CELL_RECOGNITION | 0.00001 | 0.00034 | 0.59333 | 0.78110 | 1.52272 | 34 |
| GO_CELL_CELL_SIGNALING_BY_WNT | 0.00084 | 0.01340 | 0.47727 | 0.54398 | 1.14541 | 379 |
| GO_CELL_COMMUNICATION_BY_ELECTRICAL_COUPLING_INVOL | 0.00178 | 0.02398 | 0.45506 | 0.78272 | 1.46907 | 18 |
| GO_CELL_CORTEX | 0.00058 | 0.01053 | 0.47727 | 0.57140 | 1.18954 | 218 |
| GO_CELL_CYCLE_G2_M_PHASE_TRANSITION | 0.00004 | 0.00116 | 0.55733 | 0.57954 | 1.20912 | 240 |
| GO_CELL_DIVISION | 0.00016 | 0.00357 | 0.51885 | 0.53967 | 1.13925 | 483 |
| GO_CELL_KILLING | 0.00046 | 0.00858 | 0.49849 | 0.65166 | 1.31424 | 68 |
| GO_CELL_RECOGNITION | 0.00005 | 0.00142 | 0.55733 | 0.68010 | 1.37549 | 72 |
| GO_CELL_REDOX_HOMEOSTASIS | 0.00002 | 0.00056 | 0.57561 | 0.74920 | 1.47550 | 40 |
| GO_CELL_SUBSTRATE_JUNCTION | 0.00000 | 0.00000 | | 0.65914 | 1.38601 | 334 |
| GO_CELL_SURFACE_RECEPTOR_SIGNALING_PATHWAY_INVOLVE | 0.00290 | 0.03403 | 0.43171 | 0.53185 | 1.12238 | 430 |
| GO_CHAPERONE_BINDING | 0.00324 | 0.03688 | 0.43171 | 0.60564 | 1.23268 | 86 |
| GO_CHAPERONE_COMPLEX | 0.00023 | 0.00477 | 0.51885 | 0.80247 | 1.52227 | 21 |
| GO_CHAPERONE_MEDIATED_PROTEIN_COMPLEX_ASSEMBLY | 0.00174 | 0.02358 | 0.45506 | 0.77526 | 1.45944 | 19 |
| GO_CHAPERONE_MEDIATED_PROTEIN_FOLDING | 0.00413 | 0.04491 | 0.40702 | 0.65068 | 1.29674 | 49 |
| GO_CHROMATIN_BINDING | 0.00309 | 0.03563 | 0.43171 | 0.52921 | 1.11704 | 437 |
| GO_CHROMATIN_REMODELING | 0.00063 | 0.01100 | 0.47727 | 0.59116 | 1.22271 | 156 |
| GO_CHROMOSOMAL_REGION | 0.00185 | 0.02452 | 0.45506 | 0.55045 | 1.15379 | 292 |
| GO_CHROMOSOME_SEGREGATION | 0.00287 | 0.03366 | 0.43171 | 0.54888 | 1.14834 | 275 |

| | | | | | | |
|---|---|---|---|---|---|---|
| GO_CLATHRIN_COATED_ENDOCYTIC_VESICLE_MEMBRANE | 0.00008 | 0.00199 | 0.53843 | 0.79948 | 1.53344 | 24 |
| GO_COLLAGEN_CONTAINING_EXTRACELLULAR_MATRIX | 0.00213 | 0.02705 | 0.43171 | 0.56325 | 1.17106 | 201 |
| GO_CONTRACTILE_FIBER | 0.00003 | 0.00082 | 0.57561 | 0.62440 | 1.28919 | 137 |
| GO_COTRANSLATIONAL_PROTEIN_TARGETING_TO_MEMBRANE | 0.00000 | 0.00000 | | 0.81897 | 1.67102 | 95 |
| GO_CYSTEINE_TYPE_ENDOPEPTIDASE_INHIBITOR_ACTIVITY | 0.00390 | 0.04277 | 0.43171 | 0.71564 | 1.38604 | 28 |
| GO_CYTOCHROME_COMPLEX | 0.00000 | 0.00000 | 0.73376 | 0.90517 | 1.71710 | 21 |
| GO_CYTOKINE_MEDIATED_SIGNALING_PATHWAY | 0.00000 | 0.00001 | 0.69013 | 0.57496 | 1.21378 | 459 |
| GO_CYTOKINE_PRODUCTION | 0.00239 | 0.02937 | 0.43171 | 0.53122 | 1.12116 | 450 |
| GO_CYTOPLASMIC_STRESS_GRANULE | 0.00183 | 0.02432 | 0.45506 | 0.64101 | 1.29047 | 65 |
| GO_CYTOPLASMIC_TRANSLATION | 0.00000 | 0.00000 | | 0.75525 | 1.53885 | 92 |
| GO_CYTOPLASMIC_TRANSLATIONAL_INITIATION | 0.00005 | 0.00145 | 0.55733 | 0.77074 | 1.49735 | 31 |
| GO_CYTOSOLIC_LARGE_RIBOSOMAL_SUBUNIT | 0.00000 | 0.00000 | | 0.83277 | 1.66542 | 54 |
| GO_CYTOSOLIC_RIBOSOME | 0.00000 | 0.00000 | | 0.82187 | 1.67916 | 98 |
| GO_CYTOSOLIC_SMALL_RIBOSOMAL_SUBUNIT | 0.00000 | 0.00000 | 0.76146 | 0.83029 | 1.63520 | 40 |
| GO_DEFENSE_RESPONSE_TO_GRAM_NEGATIVE_BACTERIUM | 0.00253 | 0.03062 | 0.43171 | 0.73936 | 1.41812 | 24 |
| GO_DENDRITE_MEMBRANE | 0.00309 | 0.03563 | 0.43171 | 0.75980 | 1.43631 | 20 |
| GO_DETOXIFICATION | 0.00005 | 0.00131 | 0.55733 | 0.67064 | 1.35857 | 75 |
| GO_DICARBOXYLIC_ACID_METABOLIC_PROCESS | 0.00472 | 0.04982 | 0.40702 | 0.62814 | 1.26528 | 66 |
| GO_DISORDERED_DOMAIN_SPECIFIC_BINDING | 0.00183 | 0.02432 | 0.45506 | 0.71730 | 1.39221 | 30 |
| GO_DNA_CONFORMATION_CHANGE | 0.00004 | 0.00111 | 0.55733 | 0.58520 | 1.22052 | 238 |
| GO_DNA_DAMAGE_RESPONSE_DETECTION_OF_DNA_DAMAGE | 0.00457 | 0.04872 | 0.40702 | 0.67946 | 1.33043 | 36 |
| GO_DNA_PACKAGING | 0.00055 | 0.01006 | 0.47727 | 0.59953 | 1.23695 | 134 |
| GO_DNA_REPAIR | 0.00298 | 0.03471 | 0.43171 | 0.53055 | 1.11995 | 468 |
| GO_ELECTRON_TRANSFER_ACTIVITY | 0.00000 | 0.00000 | 0.71951 | 0.72232 | 1.46909 | 84 |
| GO_ELECTRON_TRANSPORT_CHAIN | 0.00000 | 0.00000 | | 0.72052 | 1.48838 | 140 |
| GO_EMBRYONIC_PLACENTA_MORPHOGENESIS | 0.00154 | 0.02127 | 0.45506 | 0.78579 | 1.47483 | 18 |
| GO_ENDOCYTIC_VESICLE_MEMBRANE | 0.00228 | 0.02833 | 0.43171 | 0.60072 | 1.22975 | 106 |
| GO_ENDOLYSOSOME | 0.00128 | 0.01848 | 0.45506 | 0.81802 | 1.51468 | 15 |
| GO_ENDOPEPTIDASE_COMPLEX | 0.00000 | 0.00016 | 0.61053 | 0.71848 | 1.44668 | 64 |
| GO_ENDOPEPTIDASE_REGULATOR_ACTIVITY | 0.00012 | 0.00283 | 0.53843 | 0.65045 | 1.32221 | 83 |
| GO_ENDOPLASMIC_RETICULUM_GOLGI_INTERMEDIATE_COMPA | 0.00205 | 0.02626 | 0.43171 | 0.60469 | 1.23676 | 102 |
| GO_ENDOPLASMIC_RETICULUM_LUMEN | 0.00047 | 0.00878 | 0.49849 | 0.57773 | 1.19951 | 180 |
| GO_ENERGY_DERIVATION_BY_OXIDATION_OF_ORGANIC_COMP | 0.00000 | 0.00000 | | 0.70506 | 1.46726 | 208 |
| GO_EPHRIN_RECEPTOR_SIGNALING_PATHWAY | 0.00457 | 0.04872 | 0.40702 | 0.61559 | 1.24626 | 74 |
| GO_ESTABLISHMENT_OF_PROTEIN_LOCALIZATION_TO_ENDOPLA | 0.00000 | 0.00000 | | 0.80235 | 1.64251 | 108 |
| GO_ESTABLISHMENT_OF_PROTEIN_LOCALIZATION_TO_MEMBRA | 0.00000 | 0.00000 | | 0.68236 | 1.42780 | 277 |
| GO_ESTABLISHMENT_OF_PROTEIN_LOCALIZATION_TO_MITOCH | 0.00191 | 0.02490 | 0.45506 | 0.67355 | 1.33368 | 44 |
| GO_ESTABLISHMENT_OF_PROTEIN_LOCALIZATION_TO_ORGANE | 0.00000 | 0.00000 | | 0.65099 | 1.37415 | 480 |
| GO_ESTABLISHMENT_OF_PROTEIN_LOCALIZATION_TO_TELOME | 0.00007 | 0.00176 | 0.53843 | 0.84993 | 1.58652 | 17 |
| GO_ESTABLISHMENT_OF_RNA_LOCALIZATION | 0.00000 | 0.00002 | 0.67496 | 0.62526 | 1.29889 | 183 |
| GO_ESTABLISHMENT_OF_TISSUE_POLARITY | 0.00000 | 0.00001 | 0.70498 | 0.69004 | 1.41277 | 104 |
| GO_EUKARYOTIC_TRANSLATION_INITIATION_FACTOR_3_COMPL | 0.00165 | 0.02247 | 0.45506 | 0.81262 | 1.50468 | 15 |
| GO_EXTRACELLULAR_MATRIX_BINDING | 0.00076 | 0.01266 | 0.47727 | 0.74893 | 1.45052 | 28 |
| GO_FC_EPSILON_RECEPTOR_SIGNALING_PATHWAY | 0.00002 | 0.00072 | 0.57561 | 0.65744 | 1.34292 | 97 |
| GO_FC_RECEPTOR_SIGNALING_PATHWAY | 0.00000 | 0.00010 | 0.62726 | 0.63444 | 1.31129 | 150 |
| GO_FIBROBLAST_GROWTH_FACTOR_RECEPTOR_SIGNALING_PAT | 0.00009 | 0.00232 | 0.53843 | 0.65695 | 1.33474 | 81 |
| GO_FICOLIN_1_RICH_GRANULE | 0.00000 | 0.00000 | 0.76146 | 0.70221 | 1.43755 | 107 |
| GO_GENERATION_OF_PRECURSOR_METABOLITES_AND_ENERGY | 0.00000 | 0.00000 | | 0.66831 | 1.40728 | 378 |
| GO_GENE_SILENCING | 0.00012 | 0.00288 | 0.53843 | 0.58994 | 1.22630 | 186 |
| GO_GLUCOSE_CATABOLIC_PROCESS | 0.00064 | 0.01113 | 0.47727 | 0.75191 | 1.45380 | 27 |
| GO_GLUCOSE_METABOLIC_PROCESS | 0.00078 | 0.01287 | 0.47727 | 0.58119 | 1.20159 | 148 |
| GO_GLUTAMATERGIC_SYNAPSE | 0.00242 | 0.02977 | 0.43171 | 0.55456 | 1.15651 | 235 |
| GO_GLYCOLYTIC_PROCESS_THROUGH_FRUCTOSE_6_PHOSPHATI | 0.00059 | 0.01058 | 0.47727 | 0.78382 | 1.49325 | 22 |
| GO_GTPASE_ACTIVITY | 0.00002 | 0.00064 | 0.57561 | 0.59320 | 1.23621 | 225 |
| GO_GUANYL_NUCLEOTIDE_BINDING | 0.00003 | 0.00083 | 0.57561 | 0.57799 | 1.20890 | 273 |
| GO_HEMATOPOIETIC_PROGENITOR_CELL_DIFFERENTIATION | 0.00003 | 0.00083 | 0.57561 | 0.63424 | 1.30580 | 128 |
| GO_HEMATOPOIETIC_STEM_CELL_DIFFERENTIATION | 0.00000 | 0.00008 | 0.64355 | 0.71389 | 1.44385 | 73 |
| GO_HEPARIN_BINDING | 0.00231 | 0.02862 | 0.43171 | 0.62521 | 1.26573 | 74 |
| GO_HEXOSE_CATABOLIC_PROCESS | 0.00183 | 0.02432 | 0.45506 | 0.68357 | 1.35069 | 42 |
| GO_HOMOTYPIC_CELL_CELL_ADHESION | 0.00013 | 0.00316 | 0.51885 | 0.67923 | 1.36473 | 62 |
| GO_HOST_CELLULAR_COMPONENT | 0.00117 | 0.01719 | 0.45506 | 0.66661 | 1.33374 | 55 |
| GO_HYDROGEN_PEROXIDE_METABOLIC_PROCESS | 0.00194 | 0.02523 | 0.45506 | 0.73471 | 1.41678 | 26 |
| GO_IMMUNE_RESPONSE_REGULATING_SIGNALING_PATHWAY | 0.00009 | 0.00236 | 0.53843 | 0.57508 | 1.20273 | 262 |
| GO_IMPORT_INTO_NUCLEUS | 0.00059 | 0.01058 | 0.47727 | 0.59681 | 1.23224 | 137 |
| GO_INFLAMMATORY_RESPONSE_TO_ANTIGENIC_STIMULUS | 0.00065 | 0.01123 | 0.47727 | 0.75020 | 1.44306 | 25 |
| GO_INNATE_IMMUNE_RESPONSE | 0.00000 | 0.00000 | 0.71951 | 0.57763 | 1.21933 | 470 |
| GO_INNATE_IMMUNE_RESPONSE_ACTIVATING_SIGNAL_TRANSI | 0.00001 | 0.00043 | 0.59333 | 0.67531 | 1.37458 | 88 |
| GO_INNER_MITOCHONDRIAL_MEMBRANE_PROTEIN_COMPLEX | 0.00000 | 0.00000 | | 0.79529 | 1.62162 | 93 |
| GO_INTERACTION_WITH_SYMBIONT | 0.00042 | 0.00797 | 0.49849 | 0.67411 | 1.35300 | 59 |
| GO_INTERLEUKIN_1_BETA_PRODUCTION | 0.00383 | 0.04231 | 0.43171 | 0.65164 | 1.29866 | 49 |

| | | | | | | |
|---|---|---|---|---|---|---|
| GO_INTERLEUKIN_1_MEDIATED_SIGNALING_PATHWAY | 0.00000 | 0.00000 | 0.71951 | 0.71933 | 1.46301 | 84 |
| GO_INTERLEUKIN_1_PRODUCTION | 0.00257 | 0.03096 | 0.43171 | 0.65480 | 1.31012 | 55 |
| GO_INTRINSIC_APOPTOTIC_SIGNALING_PATHWAY | 0.00076 | 0.01266 | 0.47727 | 0.56296 | 1.17403 | 235 |
| GO_KILLING_OF_CELLS_OF_OTHER_ORGANISM | 0.00090 | 0.01415 | 0.47727 | 0.78687 | 1.48747 | 20 |
| GO_LARGE_RIBOSOMAL_SUBUNIT | 0.00000 | 0.00000 | | 0.76468 | 1.56596 | 109 |
| GO_LEADING_EDGE_MEMBRANE | 0.00253 | 0.03062 | 0.43171 | 0.59515 | 1.21879 | 109 |
| GO_LEUKOCYTE_MEDIATED_IMMUNITY | 0.00000 | 0.00000 | | 0.60252 | 1.27197 | 486 |
| GO_LOW_DENSITY_LIPOPROTEIN_RECEPTOR_PARTICLE_METABO | 0.00113 | 0.01687 | 0.45506 | 0.78221 | 1.47866 | 20 |
| GO_LUMENAL_SIDE_OF_ENDOPLASMIC_RETICULUM_MEMBRAI | 0.00368 | 0.04101 | 0.43171 | 0.75741 | 1.42584 | 19 |
| GO_LUMENAL_SIDE_OF_MEMBRANE | 0.00021 | 0.00457 | 0.51885 | 0.78419 | 1.50411 | 24 |
| GO_MACROMOLECULE_TRANSMEMBRANE_TRANSPORTER_ACT | 0.00031 | 0.00629 | 0.49849 | 0.81767 | 1.53928 | 19 |
| GO_MAINTENANCE_OF_CELL_NUMBER | 0.00012 | 0.00285 | 0.53843 | 0.60943 | 1.25868 | 139 |
| GO_MAINTENANCE_OF_LOCATION_IN_CELL | 0.00205 | 0.02626 | 0.43171 | 0.58052 | 1.19919 | 140 |
| GO_MEMBRANE_REGION | 0.00179 | 0.02408 | 0.45506 | 0.56245 | 1.17185 | 223 |
| GO_MITOCHONDRIAL_ELECTRON_TRANSPORT_NADH_TO_UBIQ | 0.00000 | 0.00000 | 0.81404 | 0.83478 | 1.65506 | 45 |
| GO_MITOCHONDRIAL_GENE_EXPRESSION | 0.00000 | 0.00000 | 0.70498 | 0.65236 | 1.34949 | 154 |
| GO_MITOCHONDRIAL_LARGE_RIBOSOMAL_SUBUNIT | 0.00000 | 0.00008 | 0.64355 | 0.74742 | 1.49474 | 54 |
| GO_MITOCHONDRIAL_MATRIX | 0.00000 | 0.00000 | 0.77494 | 0.59703 | 1.25800 | 394 |
| GO_MITOCHONDRIAL_MEMBRANE_ORGANIZATION | 0.00026 | 0.00542 | 0.49849 | 0.62341 | 1.27587 | 103 |
| GO_MITOCHONDRIAL_PROTEIN_COMPLEX | 0.00000 | 0.00000 | | 0.72735 | 1.51370 | 214 |
| GO_MITOCHONDRIAL_RESPIRATORY_CHAIN_COMPLEX_ASSEMB | 0.00001 | 0.00029 | 0.61053 | 0.67822 | 1.38064 | 87 |
| GO_MITOCHONDRIAL_SMALL_RIBOSOMAL_SUBUNIT | 0.00320 | 0.03658 | 0.43171 | 0.71973 | 1.39396 | 28 |
| GO_MITOCHONDRIAL_TRANSLATION | 0.00000 | 0.00000 | 0.71951 | 0.67234 | 1.38530 | 130 |
| GO_MITOCHONDRIAL_TRANSLATIONAL_TERMINATION | 0.00000 | 0.00000 | 0.71951 | 0.72087 | 1.46731 | 88 |
| GO_MITOCHONDRIAL_TRANSPORT | 0.00008 | 0.00208 | 0.53843 | 0.58684 | 1.22175 | 212 |
| GO_MITOCHONDRION_ORGANIZATION | 0.00000 | 0.00000 | 0.78819 | 0.59105 | 1.24745 | 440 |
| GO_MODULATION_BY_HOST_OF_SYMBIONT_PROCESS | 0.00472 | 0.04982 | 0.40702 | 0.65469 | 1.30103 | 46 |
| GO_MODULATION_BY_HOST_OF_VIRAL_PROCESS | 0.00246 | 0.03000 | 0.43171 | 0.76558 | 1.44723 | 20 |
| GO_MODULATION_OF_PROCESS_OF_OTHER_ORGANISM | 0.00141 | 0.01982 | 0.45506 | 0.62441 | 1.26616 | 77 |
| GO_MONOCARBOXYLIC_ACID_METABOLIC_PROCESS | 0.00098 | 0.01512 | 0.45506 | 0.54033 | 1.13913 | 406 |
| GO_MONOSACCHARIDE_BINDING | 0.00115 | 0.01699 | 0.45506 | 0.68281 | 1.35143 | 43 |
| GO_MONOSACCHARIDE_BIOSYNTHETIC_PROCESS | 0.00002 | 0.00081 | 0.57561 | 0.69168 | 1.39246 | 65 |
| GO_MONOSACCHARIDE_CATABOLIC_PROCESS | 0.00040 | 0.00769 | 0.49849 | 0.68342 | 1.36276 | 50 |
| GO_MONOSACCHARIDE_METABOLIC_PROCESS | 0.00302 | 0.03496 | 0.43171 | 0.56002 | 1.16434 | 201 |
| GO_MONOVALENT_INORGANIC_CATION_HOMEOSTASIS | 0.00435 | 0.04665 | 0.40702 | 0.60231 | 1.22590 | 86 |
| GO_MONOVALENT_INORGANIC_CATION_TRANSMEMBRANE_TF | 0.00361 | 0.04046 | 0.43171 | 0.57484 | 1.19158 | 167 |
| GO_MONOVALENT_INORGANIC_CATION_TRANSPORT | 0.00398 | 0.04337 | 0.40702 | 0.54655 | 1.14313 | 273 |
| GO_MORPHOGENESIS_OF_AN_EPITHELIUM | 0.00013 | 0.00316 | 0.51885 | 0.55159 | 1.16149 | 378 |
| GO_MORPHOGENESIS_OF_A_POLARIZED_EPITHELIUM | 0.00000 | 0.00001 | 0.70498 | 0.67567 | 1.38908 | 122 |
| GO_MRNA_3_END_PROCESSING | 0.00139 | 0.01969 | 0.45506 | 0.61840 | 1.26001 | 92 |
| GO_MRNA_3_UTR_BINDING | 0.00141 | 0.01982 | 0.45506 | 0.63215 | 1.27980 | 74 |
| GO_MRNA_5_UTR_BINDING | 0.00052 | 0.00950 | 0.47727 | 0.75570 | 1.45364 | 25 |
| GO_MRNA_BINDING | 0.00000 | 0.00000 | | 0.66439 | 1.38842 | 249 |
| GO_MRNA_EXPORT_FROM_NUCLEUS | 0.00004 | 0.00104 | 0.55733 | 0.64753 | 1.32573 | 104 |
| GO_MRNA_PROCESSING | 0.00000 | 0.00000 | | 0.63463 | 1.33942 | 450 |
| GO_MRNA_TRANSPORT | 0.00002 | 0.00080 | 0.57561 | 0.62118 | 1.28296 | 139 |
| GO_MULTI_ORGANISM_LOCALIZATION | 0.00077 | 0.01278 | 0.47727 | 0.66827 | 1.33968 | 57 |
| GO_MUSCLE_CELL_PROLIFERATION | 0.00028 | 0.00580 | 0.49849 | 0.61574 | 1.26279 | 113 |
| GO_MYELOID_LEUKOCYTE_ACTIVATION | 0.00000 | 0.00000 | | 0.60529 | 1.27687 | 425 |
| GO_MYELOID_LEUKOCYTE_MEDIATED_IMMUNITY | 0.00000 | 0.00000 | | 0.62135 | 1.30847 | 385 |
| GO_NADH_DEHYDROGENASE_ACTIVITY | 0.00000 | 0.00000 | 0.74774 | 0.83855 | 1.64555 | 37 |
| GO_NADH_DEHYDROGENASE_COMPLEX | 0.00000 | 0.00000 | 0.77494 | 0.83971 | 1.65375 | 40 |
| GO_NADH_DEHYDROGENASE_COMPLEX_ASSEMBLY | 0.00000 | 0.00001 | 0.69013 | 0.75772 | 1.51780 | 56 |
| GO_NADH_METABOLIC_PROCESS | 0.00004 | 0.00115 | 0.55733 | 0.77502 | 1.50817 | 32 |
| GO_NAD_BINDING | 0.00302 | 0.03496 | 0.43171 | 0.66327 | 1.31808 | 46 |
| GO_NAD_METABOLIC_PROCESS | 0.00011 | 0.00274 | 0.53843 | 0.72573 | 1.42929 | 40 |
| GO_NCRNA_EXPORT_FROM_NUCLEUS | 0.00302 | 0.03496 | 0.43171 | 0.69618 | 1.36084 | 35 |
| GO_NCRNA_METABOLIC_PROCESS | 0.00002 | 0.00066 | 0.57561 | 0.56359 | 1.18821 | 407 |
| GO_NCRNA_PROCESSING | 0.00001 | 0.00027 | 0.61053 | 0.57055 | 1.20031 | 350 |
| GO_NEGATIVE_REGULATION_OF_APOPTOTIC_SIGNALING_PATHV | 0.00048 | 0.00890 | 0.49849 | 0.58706 | 1.21767 | 171 |
| GO_NEGATIVE_REGULATION_OF_CANONICAL_WNT_SIGNALING | 0.00001 | 0.00041 | 0.59333 | 0.63146 | 1.30377 | 137 |
| GO_NEGATIVE_REGULATION_OF_CATABOLIC_PROCESS | 0.00100 | 0.01530 | 0.45506 | 0.56602 | 1.17957 | 225 |
| GO_NEGATIVE_REGULATION_OF_CELLULAR_AMIDE_METABOLIC | 0.00073 | 0.01224 | 0.47727 | 0.60158 | 1.24105 | 133 |
| GO_NEGATIVE_REGULATION_OF_CELLULAR_CATABOLIC_PROCES | 0.00435 | 0.04665 | 0.40702 | 0.56241 | 1.16943 | 189 |
| GO_NEGATIVE_REGULATION_OF_CELL_CYCLE | 0.00146 | 0.02034 | 0.45506 | 0.53214 | 1.12371 | 498 |
| GO_NEGATIVE_REGULATION_OF_CELL_CYCLE_G2_M_PHASE_TR | 0.00004 | 0.00115 | 0.55733 | 0.65987 | 1.34421 | 90 |
| GO_NEGATIVE_REGULATION_OF_CELL_CYCLE_PHASE_TRANSITIC | 0.00007 | 0.00186 | 0.53843 | 0.58784 | 1.22341 | 210 |
| GO_NEGATIVE_REGULATION_OF_CELL_CYCLE_PROCESS | 0.00007 | 0.00188 | 0.53843 | 0.57278 | 1.19972 | 283 |
| GO_NEGATIVE_REGULATION_OF_CYSTEINE_TYPE_ENDOPEPTIDA | 0.00097 | 0.01488 | 0.47727 | 0.66806 | 1.34012 | 58 |
| GO_NEGATIVE_REGULATION_OF_HYDROLASE_ACTIVITY | 0.00009 | 0.00228 | 0.53843 | 0.57544 | 1.20366 | 264 |

| | | | | | | |
|---|---|---|---|---|---|---|
| GO_NEGATIVE_REGULATION_OF_INNATE_IMMUNE_RESPONSE | 0.00191 | 0.02490 | 0.45506 | 0.68165 | 1.34247 | 40 |
| GO_NEGATIVE_REGULATION_OF_MITOTIC_CELL_CYCLE | 0.00082 | 0.01329 | 0.47727 | 0.56069 | 1.17284 | 272 |
| GO_NEGATIVE_REGULATION_OF_MRNA_METABOLIC_PROCESS | 0.00034 | 0.00664 | 0.49849 | 0.66946 | 1.34796 | 64 |
| GO_NEGATIVE_REGULATION_OF_NUCLEOCYTOPLASMIC_TRANS | 0.00283 | 0.03346 | 0.43171 | 0.76733 | 1.44019 | 18 |
| GO_NEGATIVE_REGULATION_OF_ORGANELLE_ORGANIZATION | 0.00420 | 0.04542 | 0.40702 | 0.54468 | 1.14338 | 309 |
| GO_NEGATIVE_REGULATION_OF_PEPTIDASE_ACTIVITY | 0.00002 | 0.00072 | 0.57561 | 0.63223 | 1.30165 | 126 |
| GO_NEGATIVE_REGULATION_OF_PROTEIN_CATABOLIC_PROCESS | 0.00205 | 0.02626 | 0.43171 | 0.59936 | 1.22665 | 103 |
| GO_NEGATIVE_REGULATION_OF_PROTEIN_CONTAINING_COMP | 0.00035 | 0.00689 | 0.49849 | 0.63266 | 1.29309 | 100 |
| GO_NEGATIVE_REGULATION_OF_PROTEIN_POLYMERIZATION | 0.00044 | 0.00833 | 0.49849 | 0.67390 | 1.34804 | 53 |
| GO_NEGATIVE_REGULATION_OF_PROTEOLYSIS | 0.00003 | 0.00096 | 0.55733 | 0.60060 | 1.24839 | 198 |
| GO_NEGATIVE_REGULATION_OF_RNA_CATABOLIC_PROCESS | 0.00228 | 0.02833 | 0.43171 | 0.67084 | 1.32832 | 44 |
| GO_NEGATIVE_REGULATION_OF_RNA_SPLICING | 0.00298 | 0.03471 | 0.43171 | 0.71781 | 1.38074 | 25 |
| GO_NEGATIVE_REGULATION_OF_TRANSPORT | 0.00472 | 0.04982 | 0.40702 | 0.54451 | 1.13963 | 278 |
| GO_NEGATIVE_REGULATION_OF_VIRAL_GENOME_REPLICATION | 0.00086 | 0.01360 | 0.47727 | 0.70130 | 1.37759 | 38 |
| GO_NEGATIVE_REGULATION_OF_WNT_SIGNALING_PATHWAY | 0.00012 | 0.00285 | 0.53843 | 0.60100 | 1.24403 | 158 |
| GO_NEURAL_NUCLEUS_DEVELOPMENT | 0.00012 | 0.00285 | 0.53843 | 0.73581 | 1.44739 | 39 |
| GO_NEURON_DEATH_IN_RESPONSE_TO_OXIDATIVE_STRESS | 0.00383 | 0.04231 | 0.43171 | 0.73917 | 1.40220 | 21 |
| GO_NIK_NF_KAPPAB_SIGNALING | 0.00000 | 0.00000 | 0.76146 | 0.69366 | 1.42548 | 119 |
| GO_NON_CANONICAL_WNT_SIGNALING_PATHWAY | 0.00000 | 0.00007 | 0.64355 | 0.65420 | 1.34618 | 124 |
| GO_NUCLEAR_CHROMOSOME_TELOMERIC_REGION | 0.00457 | 0.04872 | 0.40702 | 0.60069 | 1.22482 | 93 |
| GO_NUCLEAR_ENVELOPE | 0.00016 | 0.00374 | 0.51885 | 0.55358 | 1.16462 | 350 |
| GO_NUCLEAR_EXPORT | 0.00000 | 0.00002 | 0.67496 | 0.62478 | 1.29873 | 186 |
| GO_NUCLEAR_PERIPHERY | 0.00042 | 0.00801 | 0.49849 | 0.61194 | 1.25687 | 118 |
| GO_NUCLEAR_PORE | 0.00198 | 0.02561 | 0.43171 | 0.64339 | 1.29871 | 69 |
| GO_NUCLEAR_RECEPTOR_BINDING | 0.00039 | 0.00753 | 0.49849 | 0.64195 | 1.30175 | 77 |
| GO_NUCLEAR_SPECK | 0.00003 | 0.00083 | 0.57561 | 0.56710 | 1.19212 | 332 |
| GO_NUCLEAR_TRANSCRIBED_MRNA_CATABOLIC_PROCESS | 0.00000 | 0.00000 | | 0.68848 | 1.43174 | 191 |
| GO_NUCLEAR_TRANSCRIBED_MRNA_CATABOLIC_PROCESS_NON | 0.00000 | 0.00000 | | 0.79685 | 1.63434 | 115 |
| GO_NUCLEAR_TRANSPORT | 0.00000 | 0.00000 | 0.73376 | 0.60664 | 1.27351 | 308 |
| GO_NUCLEOBASE_CONTAINING_COMPOUND_TRANSPORT | 0.00000 | 0.00016 | 0.61053 | 0.60197 | 1.25450 | 225 |
| GO_NUCLEOBASE_CONTAINING_SMALL_MOLECULE_METABOLIC | 0.00014 | 0.00320 | 0.51885 | 0.54855 | 1.15678 | 419 |
| GO_NUCLEOSIDE_DIPHOSPHATE_METABOLIC_PROCESS | 0.00003 | 0.00082 | 0.57561 | 0.63672 | 1.30581 | 113 |
| GO_NUCLEOSIDE_PHOSPHATE_BIOSYNTHETIC_PROCESS | 0.00346 | 0.03914 | 0.43171 | 0.56352 | 1.16817 | 168 |
| GO_NUCLEOSIDE_TRIPHOSPHATE_BIOSYNTHETIC_PROCESS | 0.00007 | 0.00176 | 0.53843 | 0.72587 | 1.43666 | 43 |
| GO_NUCLEOSIDE_TRIPHOSPHATE_METABOLIC_PROCESS | 0.00050 | 0.00919 | 0.47727 | 0.66037 | 1.33020 | 66 |
| GO_NUCLEOSOME_ORGANIZATION | 0.00046 | 0.00864 | 0.49849 | 0.63039 | 1.28538 | 93 |
| GO_NUCLEOTIDE_PHOSPHORYLATION | 0.00002 | 0.00072 | 0.57561 | 0.65186 | 1.33176 | 99 |
| GO_ORGANELLAR_RIBOSOME | 0.00000 | 0.00000 | 0.74774 | 0.72937 | 1.48343 | 84 |
| GO_ORGANELLE_INNER_MEMBRANE | 0.00000 | 0.00000 | | 0.65111 | 1.37305 | 419 |
| GO_ORGANIC_CYCLIC_COMPOUND_CATABOLIC_PROCESS | 0.00000 | 0.00000 | | 0.65670 | 1.38705 | 500 |
| GO_OSTEOBLAST_DIFFERENTIATION | 0.00015 | 0.00348 | 0.51885 | 0.60874 | 1.25818 | 144 |
| GO_OUTER_MEMBRANE | 0.00126 | 0.01834 | 0.45506 | 0.57897 | 1.19880 | 161 |
| GO_OXIDATIVE_PHOSPHORYLATION | 0.00000 | 0.00000 | | 0.80964 | 1.65541 | 101 |
| GO_OXIDOREDUCTASE_ACTIVITY | 0.00000 | 0.00000 | | 0.61483 | 1.29782 | 480 |
| GO_OXIDOREDUCTASE_ACTIVITY_ACTING_ON_A_HEME_GROUP | 0.00001 | 0.00034 | 0.59333 | 0.86577 | 1.62983 | 19 |
| GO_OXIDOREDUCTASE_ACTIVITY_ACTING_ON_CH_OH_GROUP_ | 0.00390 | 0.04277 | 0.43171 | 0.61195 | 1.24162 | 78 |
| GO_OXIDOREDUCTASE_ACTIVITY_ACTING_ON_NAD_P_H | 0.00000 | 0.00000 | 0.76146 | 0.74655 | 1.51351 | 76 |
| GO_OXIDOREDUCTASE_ACTIVITY_ACTING_ON_NAD_P_H_QUINO | 0.00000 | 0.00000 | 0.73376 | 0.79942 | 1.58866 | 46 |
| GO_OXIDOREDUCTASE_ACTIVITY_ACTING_ON_PEROXIDE_AS_AC | 0.00008 | 0.00199 | 0.53843 | 0.78646 | 1.52060 | 27 |
| GO_OXIDOREDUCTASE_COMPLEX | 0.00000 | 0.00000 | | 0.78168 | 1.58983 | 84 |
| GO_PEPTIDASE_COMPLEX | 0.00002 | 0.00056 | 0.57561 | 0.67643 | 1.37434 | 81 |
| GO_PEPTIDASE_REGULATOR_ACTIVITY | 0.00034 | 0.00664 | 0.49849 | 0.61381 | 1.25882 | 113 |
| GO_PEPTIDE_BINDING | 0.00005 | 0.00133 | 0.55733 | 0.61092 | 1.26595 | 165 |
| GO_PEPTIDE_TRANSMEMBRANE_TRANSPORTER_ACTIVITY | 0.00003 | 0.00082 | 0.57561 | 0.81215 | 1.55773 | 24 |
| GO_PIGMENT_GRANULE | 0.00000 | 0.00000 | 0.73376 | 0.72897 | 1.48379 | 88 |
| GO_PLASMA_LIPOPROTEIN_PARTICLE_CLEARANCE | 0.00272 | 0.03256 | 0.43171 | 0.67312 | 1.32835 | 41 |
| GO_PLATELET_ACTIVATION | 0.00165 | 0.02247 | 0.45506 | 0.60323 | 1.23458 | 103 |
| GO_PLATELET_AGGREGATION | 0.00038 | 0.00738 | 0.49849 | 0.70566 | 1.39906 | 45 |
| GO_POLYSOMAL_RIBOSOME | 0.00000 | 0.00003 | 0.65944 | 0.82821 | 1.60747 | 30 |
| GO_POLYSOME | 0.00000 | 0.00000 | 0.70498 | 0.75334 | 1.51687 | 64 |
| GO_POSITIVE_REGULATION_OF_ATPASE_ACTIVITY | 0.00121 | 0.01765 | 0.45506 | 0.68374 | 1.36037 | 47 |
| GO_POSITIVE_REGULATION_OF_BINDING | 0.00102 | 0.01554 | 0.45506 | 0.59762 | 1.23288 | 133 |
| GO_POSITIVE_REGULATION_OF_CANONICAL_WNT_SIGNALING_ | 0.00005 | 0.00145 | 0.55733 | 0.63012 | 1.29510 | 121 |
| GO_POSITIVE_REGULATION_OF_CELLULAR_AMIDE_METABOLIC_ | 0.00000 | 0.00002 | 0.67496 | 0.66444 | 1.36799 | 128 |
| GO_POSITIVE_REGULATION_OF_CELLULAR_COMPONENT_BIOGE | 0.00465 | 0.04935 | 0.40702 | 0.53144 | 1.11908 | 380 |
| GO_POSITIVE_REGULATION_OF_CELLULAR_PROTEIN_LOCALIZAT | 0.00067 | 0.01155 | 0.47727 | 0.55916 | 1.16950 | 268 |
| GO_POSITIVE_REGULATION_OF_CYSTEINE_TYPE_ENDOPEPTIDAS | 0.00016 | 0.00365 | 0.51885 | 0.63642 | 1.30348 | 105 |
| GO_POSITIVE_REGULATION_OF_CYTOKINE_PRODUCTION | 0.00073 | 0.01224 | 0.47727 | 0.56256 | 1.17606 | 255 |
| GO_POSITIVE_REGULATION_OF_CYTOKINE_SECRETION | 0.00372 | 0.04128 | 0.43171 | 0.69592 | 1.35579 | 33 |
| GO_POSITIVE_REGULATION_OF_CYTOSKELETON_ORGANIZATION | 0.00069 | 0.01174 | 0.47727 | 0.58389 | 1.20894 | 160 |

| | | | | | | |
|---|---|---|---|---|---|---|
| GO_POSITIVE_REGULATION_OF_DEFENSE_RESPONSE | 0.00000 | 0.00000 | 0.74774 | 0.63773 | 1.32813 | 221 |
| GO_POSITIVE_REGULATION_OF_DNA_BIOSYNTHETIC_PROCESS | 0.00100 | 0.01530 | 0.45506 | 0.66744 | 1.33888 | 58 |
| GO_POSITIVE_REGULATION_OF_DNA_METABOLIC_PROCESS | 0.00390 | 0.04277 | 0.43171 | 0.57564 | 1.19039 | 152 |
| GO_POSITIVE_REGULATION_OF_ESTABLISHMENT_OF_PROTEIN_ | 0.00000 | 0.00008 | 0.64355 | 0.58849 | 1.23227 | 281 |
| GO_POSITIVE_REGULATION_OF_IMMUNE_RESPONSE | 0.00000 | 0.00001 | 0.70498 | 0.58709 | 1.23570 | 370 |
| GO_POSITIVE_REGULATION_OF_INFLAMMATORY_RESPONSE | 0.00027 | 0.00548 | 0.49849 | 0.66815 | 1.34658 | 67 |
| GO_POSITIVE_REGULATION_OF_INNATE_IMMUNE_RESPONSE | 0.00000 | 0.00001 | 0.69013 | 0.65610 | 1.35650 | 146 |
| GO_POSITIVE_REGULATION_OF_INTRACELLULAR_TRANSPORT | 0.00086 | 0.01360 | 0.47727 | 0.57270 | 1.18905 | 180 |
| GO_POSITIVE_REGULATION_OF_MRNA_METABOLIC_PROCESS | 0.00029 | 0.00594 | 0.49849 | 0.66033 | 1.33384 | 70 |
| GO_POSITIVE_REGULATION_OF_NITRIC_OXIDE_METABOLIC_PRO | 0.00253 | 0.03062 | 0.43171 | 0.73909 | 1.41761 | 24 |
| GO_POSITIVE_REGULATION_OF_ORGANELLE_ORGANIZATION | 0.00000 | 0.00001 | 0.70498 | 0.57548 | 1.21493 | 476 |
| GO_POSITIVE_REGULATION_OF_OSTEOBLAST_DIFFERENTIATION | 0.00316 | 0.03622 | 0.43171 | 0.68099 | 1.33638 | 37 |
| GO_POSITIVE_REGULATION_OF_PEPTIDASE_ACTIVITY | 0.00021 | 0.00454 | 0.51885 | 0.60624 | 1.25189 | 138 |
| GO_POSITIVE_REGULATION_OF_PEPTIDE_SECRETION | 0.00040 | 0.00773 | 0.49849 | 0.60921 | 1.25244 | 122 |
| GO_POSITIVE_REGULATION_OF_PROTEIN_CONTAINING_COMPL | 0.00076 | 0.01261 | 0.47727 | 0.57915 | 1.20185 | 178 |
| GO_POSITIVE_REGULATION_OF_PROTEIN_LOCALIZATION_TO_N | 0.00054 | 0.00978 | 0.47727 | 0.65376 | 1.32056 | 70 |
| GO_POSITIVE_REGULATION_OF_PROTEIN_POLYMERIZATION | 0.00220 | 0.02777 | 0.43171 | 0.61029 | 1.24440 | 93 |
| GO_POSITIVE_REGULATION_OF_REACTIVE_OXYGEN_SPECIES_BI | 0.00090 | 0.01415 | 0.47727 | 0.72961 | 1.41610 | 30 |
| GO_POSITIVE_REGULATION_OF_REACTIVE_OXYGEN_SPECIES_M | 0.00287 | 0.03366 | 0.43171 | 0.66136 | 1.32263 | 54 |
| GO_POSITIVE_REGULATION_OF_RESPONSE_TO_BIOTIC_STIMUL | 0.00000 | 0.00001 | 0.70498 | 0.64571 | 1.33934 | 170 |
| GO_POSITIVE_REGULATION_OF_RESPONSE_TO_EXTERNAL_STIM | 0.00000 | 0.00000 | 0.70498 | 0.60313 | 1.26465 | 297 |
| GO_POSITIVE_REGULATION_OF_SECRETION | 0.00239 | 0.02937 | 0.43171 | 0.56494 | 1.17449 | 199 |
| GO_POSITIVE_REGULATION_OF_SIGNAL_TRANSDUCTION_BY_P5 | 0.00194 | 0.02523 | 0.45506 | 0.77950 | 1.46304 | 18 |
| GO_POSITIVE_REGULATION_OF_SUPRAMOLECULAR_FIBER_ORG | 0.00034 | 0.00664 | 0.49849 | 0.59526 | 1.22997 | 143 |
| GO_POSITIVE_REGULATION_OF_TELOMERASE_ACTIVITY | 0.00122 | 0.01788 | 0.45506 | 0.72410 | 1.40907 | 32 |
| GO_POSITIVE_REGULATION_OF_TELOMERASE_RNA_LOCALIZATI | 0.00002 | 0.00069 | 0.57561 | 0.88966 | 1.64732 | 15 |
| GO_POSITIVE_REGULATION_OF_TELOMERE_MAINTENANCE | 0.00110 | 0.01640 | 0.45506 | 0.68712 | 1.36231 | 45 |
| GO_POSITIVE_REGULATION_OF_TELOMERE_MAINTENANCE_VIA | 0.00213 | 0.02705 | 0.43171 | 0.69414 | 1.35318 | 34 |
| GO_POSITIVE_REGULATION_OF_TRANSLATION | 0.00000 | 0.00000 | 0.71951 | 0.68748 | 1.40738 | 107 |
| GO_POSITIVE_REGULATION_OF_VIRAL_LIFE_CYCLE | 0.00224 | 0.02808 | 0.43171 | 0.65362 | 1.30520 | 52 |
| GO_POSITIVE_REGULATION_OF_VIRAL_PROCESS | 0.00002 | 0.00065 | 0.57561 | 0.67034 | 1.36700 | 94 |
| GO_POSITIVE_REGULATION_OF_VIRAL_TRANSCRIPTION | 0.00133 | 0.01915 | 0.45506 | 0.69603 | 1.36914 | 39 |
| GO_POSITIVE_REGULATION_OF_WNT_SIGNALING_PATHWAY | 0.00002 | 0.00076 | 0.57561 | 0.62062 | 1.28236 | 143 |
| GO_POSTSYNAPSE | 0.00231 | 0.02862 | 0.43171 | 0.53396 | 1.12503 | 396 |
| GO_POST_TRANSLATIONAL_PROTEIN_MODIFICATION | 0.00002 | 0.00072 | 0.57561 | 0.57986 | 1.21446 | 282 |
| GO_PRECATALYTIC_SPLICEOSOME | 0.00083 | 0.01329 | 0.47727 | 0.67485 | 1.34647 | 51 |
| GO_PROTEASOMAL_PROTEIN_CATABOLIC_PROCESS | 0.00097 | 0.01488 | 0.47727 | 0.54229 | 1.14332 | 413 |
| GO_PROTEASOMAL_UBIQUITIN_INDEPENDENT_PROTEIN_CATAE | 0.00000 | 0.00002 | 0.67496 | 0.91042 | 1.70876 | 18 |
| GO_PROTEASOME_ACCESSORY_COMPLEX | 0.00048 | 0.00890 | 0.49849 | 0.77209 | 1.48090 | 24 |
| GO_PROTEASOME_CORE_COMPLEX | 0.00000 | 0.00012 | 0.62726 | 0.89946 | 1.67039 | 16 |
| GO_PROTEIN_CONTAINING_COMPLEX_DISASSEMBLY | 0.00000 | 0.00000 | 0.80122 | 0.62773 | 1.31300 | 274 |
| GO_PROTEIN_CONTAINING_COMPLEX_LOCALIZATION | 0.00000 | 0.00018 | 0.61053 | 0.59432 | 1.23950 | 236 |
| GO_PROTEIN_DNA_COMPLEX_SUBUNIT_ORGANIZATION | 0.00005 | 0.00130 | 0.55733 | 0.59939 | 1.24627 | 187 |
| GO_PROTEIN_FOLDING | 0.00000 | 0.00000 | 0.76146 | 0.64797 | 1.34534 | 180 |
| GO_PROTEIN_FOLDING_CHAPERONE | 0.00313 | 0.03593 | 0.43171 | 0.74376 | 1.41090 | 21 |
| GO_PROTEIN_IMPORT | 0.00014 | 0.00324 | 0.51885 | 0.60190 | 1.24726 | 165 |
| GO_PROTEIN_IMPORT_INTO_MITOCHONDRIAL_MATRIX | 0.00376 | 0.04163 | 0.43171 | 0.75702 | 1.42511 | 19 |
| GO_PROTEIN_LIPID_COMPLEX_SUBUNIT_ORGANIZATION | 0.00420 | 0.04542 | 0.40702 | 0.68128 | 1.32813 | 34 |
| GO_PROTEIN_LOCALIZATION_TO_CHROMOSOME_TELOMERIC_F | 0.00090 | 0.01415 | 0.47727 | 0.74615 | 1.44513 | 28 |
| GO_PROTEIN_LOCALIZATION_TO_ENDOPLASMIC_RETICULUM | 0.00000 | 0.00000 | | 0.78432 | 1.61693 | 131 |
| GO_PROTEIN_LOCALIZATION_TO_MEMBRANE | 0.00000 | 0.00000 | | 0.61550 | 1.29961 | 491 |
| GO_PROTEIN_LOCALIZATION_TO_MITOCHONDRION | 0.00017 | 0.00382 | 0.51885 | 0.61955 | 1.27489 | 124 |
| GO_PROTEIN_LOCALIZATION_TO_NUCLEUS | 0.00004 | 0.00115 | 0.55733 | 0.59261 | 1.23474 | 226 |
| GO_PROTEIN_MODIFICATION_BY_SMALL_PROTEIN_REMOVAL | 0.00018 | 0.00395 | 0.51885 | 0.57677 | 1.20296 | 237 |
| GO_PROTEIN_N_TERMINUS_BINDING | 0.00159 | 0.02194 | 0.45506 | 0.61618 | 1.25549 | 92 |
| GO_PROTEIN_POLYMERIZATION | 0.00143 | 0.01995 | 0.45506 | 0.56820 | 1.18290 | 216 |
| GO_PROTEIN_POLYUBIQUITINATION | 0.00097 | 0.01488 | 0.47727 | 0.55401 | 1.16065 | 285 |
| GO_PROTEIN_REFOLDING | 0.00279 | 0.03332 | 0.43171 | 0.75171 | 1.43207 | 22 |
| GO_PROTEIN_STABILIZATION | 0.00194 | 0.02523 | 0.45506 | 0.58342 | 1.20648 | 152 |
| GO_PROTEIN_TARGETING | 0.00000 | 0.00000 | | 0.66973 | 1.40965 | 369 |
| GO_PROTEIN_TARGETING_TO_MEMBRANE | 0.00000 | 0.00000 | | 0.73177 | 1.51711 | 169 |
| GO_PROTEIN_TARGETING_TO_MITOCHONDRION | 0.00013 | 0.00314 | 0.51885 | 0.64596 | 1.31475 | 86 |
| GO_PROTEIN_TRANSMEMBRANE_TRANSPORT | 0.00032 | 0.00640 | 0.49849 | 0.68590 | 1.37393 | 56 |
| GO_PROTON_TRANSMEMBRANE_TRANSPORT | 0.00000 | 0.00011 | 0.62726 | 0.68544 | 1.39511 | 86 |
| GO_PROTON_TRANSMEMBRANE_TRANSPORTER_ACTIVITY | 0.00143 | 0.01995 | 0.45506 | 0.64648 | 1.30223 | 66 |
| GO_PURINE_CONTAINING_COMPOUND_BIOSYNTHETIC_PROCES | 0.00398 | 0.04337 | 0.40702 | 0.58160 | 1.19812 | 129 |
| GO_PURINE_CONTAINING_COMPOUND_METABOLIC_PROCESS | 0.00003 | 0.00087 | 0.57561 | 0.57231 | 1.20080 | 304 |
| GO_PURINE_NUCLEOSIDE_TRIPHOSPHATE_METABOLIC_PROCES | 0.00002 | 0.00081 | 0.57561 | 0.72658 | 1.44800 | 49 |
| GO_PYRUVATE_METABOLIC_PROCESS | 0.00001 | 0.00021 | 0.61053 | 0.65817 | 1.34785 | 109 |
| GO_REACTIVE_OXYGEN_SPECIES_METABOLIC_PROCESS | 0.00017 | 0.00378 | 0.51885 | 0.59818 | 1.23868 | 159 |

| | | | | | | |
|---|---|---|---|---|---|---|
| GO_RECEPTOR_MEDIATED_ENDOCYTOSIS | 0.00036 | 0.00696 | 0.49849 | 0.59032 | 1.22470 | 174 |
| GO_REGULATION_OF_ACTIN_FILAMENT_BASED_PROCESS | 0.00049 | 0.00905 | 0.47727 | 0.56208 | 1.17595 | 275 |
| GO_REGULATION_OF_ACTIN_FILAMENT_LENGTH | 0.00191 | 0.02490 | 0.45506 | 0.58922 | 1.21381 | 129 |
| GO_REGULATION_OF_ACTIN_FILAMENT_ORGANIZATION | 0.00076 | 0.01266 | 0.47727 | 0.57669 | 1.19907 | 187 |
| GO_REGULATION_OF_ACUTE_INFLAMMATORY_RESPONSE | 0.00346 | 0.03914 | 0.43171 | 0.75935 | 1.42949 | 19 |
| GO_REGULATION_OF_ANATOMICAL_STRUCTURE_SIZE | 0.00067 | 0.01155 | 0.47727 | 0.54778 | 1.15066 | 323 |
| GO_REGULATION_OF_ANIMAL_ORGAN_MORPHOGENESIS | 0.00000 | 0.00005 | 0.64355 | 0.63740 | 1.32062 | 164 |
| GO_REGULATION_OF_APOPTOTIC_SIGNALING_PATHWAY | 0.00004 | 0.00108 | 0.55733 | 0.57390 | 1.20479 | 308 |
| GO_REGULATION_OF_ATPASE_ACTIVITY | 0.00146 | 0.02034 | 0.45506 | 0.64672 | 1.30145 | 63 |
| GO_REGULATION_OF_ATP_METABOLIC_PROCESS | 0.00007 | 0.00181 | 0.53843 | 0.64898 | 1.32139 | 89 |
| GO_REGULATION_OF_BEHAVIOR | 0.00108 | 0.01616 | 0.45506 | 0.75459 | 1.44733 | 24 |
| GO_REGULATION_OF_BINDING | 0.00135 | 0.01933 | 0.45506 | 0.55637 | 1.16365 | 270 |
| GO_REGULATION_OF_CELLULAR_AMIDE_METABOLIC_PROCESS | 0.00000 | 0.00000 | 0.81404 | 0.61011 | 1.28384 | 352 |
| GO_REGULATION_OF_CELLULAR_AMINE_METABOLIC_PROCESS | 0.00000 | 0.00001 | 0.69013 | 0.74070 | 1.48822 | 62 |
| GO_REGULATION_OF_CELLULAR_AMINO_ACID_METABOLIC_PR( | 0.00000 | 0.00002 | 0.67496 | 0.76008 | 1.51779 | 52 |
| GO_REGULATION_OF_CELLULAR_COMPONENT_SIZE | 0.00179 | 0.02408 | 0.45506 | 0.55366 | 1.15794 | 262 |
| GO_REGULATION_OF_CELLULAR_KETONE_METABOLIC_PROCESS | 0.00004 | 0.00115 | 0.55733 | 0.63118 | 1.29951 | 128 |
| GO_REGULATION_OF_CELLULAR_PH | 0.00435 | 0.04665 | 0.40702 | 0.63301 | 1.27185 | 62 |
| GO_REGULATION_OF_CELLULAR_PROTEIN_LOCALIZATION | 0.00005 | 0.00129 | 0.55733 | 0.55265 | 1.16638 | 434 |
| GO_REGULATION_OF_CELLULAR_RESPONSE_TO_HEAT | 0.00106 | 0.01603 | 0.45506 | 0.64279 | 1.29840 | 70 |
| GO_REGULATION_OF_CELL_CYCLE_G2_M_PHASE_TRANSITION | 0.00003 | 0.00083 | 0.57561 | 0.59927 | 1.24621 | 193 |
| GO_REGULATION_OF_CELL_CYCLE_PHASE_TRANSITION | 0.00006 | 0.00158 | 0.55733 | 0.55860 | 1.17687 | 390 |
| GO_REGULATION_OF_CHROMOSOME_ORGANIZATION | 0.00161 | 0.02210 | 0.45506 | 0.54876 | 1.14979 | 289 |
| GO_REGULATION_OF_CYSTEINE_TYPE_ENDOPEPTIDASE_ACTIVIT | 0.00000 | 0.00013 | 0.62726 | 0.62483 | 1.29387 | 159 |
| GO_REGULATION_OF_CYTOSKELETON_ORGANIZATION | 0.00040 | 0.00773 | 0.49849 | 0.54127 | 1.14090 | 402 |
| GO_REGULATION_OF_DEFENSE_RESPONSE | 0.00000 | 0.00002 | 0.67496 | 0.57774 | 1.21799 | 406 |
| GO_REGULATION_OF_DNA_TEMPLATED_TRANSCRIPTION_IN_RI | 0.00000 | 0.00015 | 0.62726 | 0.66902 | 1.36740 | 100 |
| GO_REGULATION_OF_GENERATION_OF_PRECURSOR_METABOLI | 0.00139 | 0.01969 | 0.45506 | 0.59523 | 1.22074 | 116 |
| GO_REGULATION_OF_GENE_SILENCING | 0.00042 | 0.00801 | 0.49849 | 0.63216 | 1.28821 | 91 |
| GO_REGULATION_OF_HEMATOPOIETIC_PROGENITOR_CELL_DIF | 0.00000 | 0.00002 | 0.67496 | 0.72571 | 1.46772 | 72 |
| GO_REGULATION_OF_HEMATOPOIETIC_STEM_CELL_DIFFERENT | 0.00000 | 0.00000 | 0.73376 | 0.75777 | 1.52254 | 62 |
| GO_REGULATION_OF_HEMOPOIESIS | 0.00083 | 0.01329 | 0.47727 | 0.55217 | 1.15894 | 306 |
| GO_REGULATION_OF_IMMUNE_RESPONSE | 0.00000 | 0.00015 | 0.62726 | 0.55900 | 1.18031 | 496 |
| GO_REGULATION_OF_INNATE_IMMUNE_RESPONSE | 0.00000 | 0.00000 | 0.73376 | 0.63411 | 1.31960 | 209 |
| GO_REGULATION_OF_INTRACELLULAR_PROTEIN_TRANSPORT | 0.00033 | 0.00645 | 0.49849 | 0.57477 | 1.19616 | 214 |
| GO_REGULATION_OF_INTRACELLULAR_TRANSPORT | 0.00023 | 0.00491 | 0.51885 | 0.56524 | 1.18384 | 282 |
| GO_REGULATION_OF_INTRINSIC_APOPTOTIC_SIGNALING_PATH\ | 0.00028 | 0.00565 | 0.49849 | 0.60871 | 1.25517 | 132 |
| GO_REGULATION_OF_MORPHOGENESIS_OF_AN_EPITHELIUM | 0.00000 | 0.00000 | 0.71951 | 0.67344 | 1.38931 | 133 |
| GO_REGULATION_OF_MRNA_CATABOLIC_PROCESS | 0.00000 | 0.00000 | 0.81404 | 0.67301 | 1.39597 | 170 |
| GO_REGULATION_OF_MRNA_METABOLIC_PROCESS | 0.00000 | 0.00000 | | 0.67878 | 1.42175 | 283 |
| GO_REGULATION_OF_MRNA_PROCESSING | 0.00000 | 0.00000 | 0.74774 | 0.68424 | 1.40876 | 128 |
| GO_REGULATION_OF_MRNA_SPLICING_VIA_SPLICEOSOME | 0.00000 | 0.00002 | 0.67496 | 0.69861 | 1.42345 | 92 |
| GO_REGULATION_OF_MULTI_ORGANISM_PROCESS | 0.00179 | 0.02408 | 0.45506 | 0.68288 | 1.34488 | 40 |
| GO_REGULATION_OF_NUCLEOCYTOPLASMIC_TRANSPORT | 0.00095 | 0.01470 | 0.47727 | 0.62048 | 1.26676 | 96 |
| GO_REGULATION_OF_NUCLEOTIDE_BIOSYNTHETIC_PROCESS | 0.00126 | 0.01834 | 0.45506 | 0.75076 | 1.43999 | 24 |
| GO_REGULATION_OF_OXIDATIVE_PHOSPHORYLATION | 0.00032 | 0.00637 | 0.49849 | 0.79189 | 1.51448 | 23 |
| GO_REGULATION_OF_OXIDATIVE_STRESS_INDUCED_CELL_DEAT | 0.00053 | 0.00964 | 0.47727 | 0.69191 | 1.37501 | 46 |
| GO_REGULATION_OF_PEPTIDASE_ACTIVITY | 0.00000 | 0.00001 | 0.70498 | 0.61102 | 1.27774 | 260 |
| GO_REGULATION_OF_PEPTIDE_TRANSPORT | 0.00002 | 0.00072 | 0.57561 | 0.55369 | 1.16879 | 464 |
| GO_REGULATION_OF_PH | 0.00442 | 0.04737 | 0.40702 | 0.62443 | 1.25709 | 65 |
| GO_REGULATION_OF_PLASMA_LIPOPROTEIN_PARTICLE_LEVELS | 0.00005 | 0.00143 | 0.55733 | 0.68877 | 1.38389 | 62 |
| GO_REGULATION_OF_POSTTRANSCRIPTIONAL_GENE_SILENCINC | 0.00008 | 0.00208 | 0.53843 | 0.65824 | 1.33737 | 81 |
| GO_REGULATION_OF_PROTEIN_CONTAINING_COMPLEX_ASSEM | 0.00001 | 0.00033 | 0.59333 | 0.57619 | 1.21020 | 318 |
| GO_REGULATION_OF_PROTEIN_LOCALIZATION_TO_NUCLEUS | 0.00073 | 0.01224 | 0.47727 | 0.61816 | 1.26561 | 104 |
| GO_REGULATION_OF_PROTEIN_POLYMERIZATION | 0.00009 | 0.00222 | 0.53843 | 0.60082 | 1.24399 | 160 |
| GO_REGULATION_OF_PROTEIN_STABILITY | 0.00061 | 0.01079 | 0.47727 | 0.56719 | 1.18296 | 238 |
| GO_REGULATION_OF_PROTEOLYSIS | 0.00004 | 0.00125 | 0.55733 | 0.55200 | 1.16535 | 476 |
| GO_REGULATION_OF_REACTIVE_OXYGEN_SPECIES_METABOLIC_ | 0.00083 | 0.01329 | 0.47727 | 0.60864 | 1.24762 | 112 |
| GO_REGULATION_OF_RESPONSE_TO_BIOTIC_STIMULUS | 0.00000 | 0.00002 | 0.67496 | 0.60329 | 1.26188 | 274 |
| GO_REGULATION_OF_RNA_SPLICING | 0.00000 | 0.00000 | 0.74774 | 0.67985 | 1.40050 | 129 |
| GO_REGULATION_OF_RYANODINE_SENSITIVE_CALCIUM_RELEAS | 0.00202 | 0.02594 | 0.43171 | 0.80750 | 1.49519 | 15 |
| GO_REGULATION_OF_SMALL_MOLECULE_METABOLIC_PROCESS | 0.00001 | 0.00029 | 0.59333 | 0.57733 | 1.21187 | 312 |
| GO_REGULATION_OF_STEM_CELL_DIFFERENTIATION | 0.00000 | 0.00007 | 0.64355 | 0.68815 | 1.40213 | 92 |
| GO_REGULATION_OF_SUPRAMOLECULAR_FIBER_ORGANIZATIOI | 0.00011 | 0.00270 | 0.53843 | 0.57577 | 1.20367 | 255 |
| GO_REGULATION_OF_SYMBIOTIC_PROCESS | 0.00000 | 0.00000 | 0.71951 | 0.64728 | 1.34258 | 171 |
| GO_REGULATION_OF_TELOMERASE_ACTIVITY | 0.00183 | 0.02432 | 0.45506 | 0.68108 | 1.34406 | 41 |
| GO_REGULATION_OF_TRANSCRIPTION_FROM_RNA_POLYMERA: | 0.00000 | 0.00000 | 0.71951 | 0.75140 | 1.51269 | 65 |
| GO_REGULATION_OF_TRANSLATIONAL_INITIATION | 0.00001 | 0.00024 | 0.61053 | 0.69767 | 1.40701 | 68 |
| GO_REGULATION_OF_VIRAL_GENOME_REPLICATION | 0.00027 | 0.00548 | 0.49849 | 0.65664 | 1.32701 | 71 |
| GO_REGULATION_OF_VIRAL_LIFE_CYCLE | 0.00016 | 0.00361 | 0.51885 | 0.62793 | 1.28548 | 107 |

| | | | | | | |
|---|---|---|---|---|---|---|
| GO_REGULATION_OF_VIRAL_TRANSCRIPTION | 0.00062 | 0.01089 | 0.47727 | 0.67338 | 1.35078 | 58 |
| GO_REGULATION_OF_WNT_SIGNALING_PATHWAY | 0.00257 | 0.03096 | 0.43171 | 0.55122 | 1.15298 | 274 |
| GO_RESPIRASOME | 0.00000 | 0.00000 | | 0.82314 | 1.66645 | 74 |
| GO_RESPIRATORY_CHAIN_COMPLEX | 0.00000 | 0.00000 | | 0.83838 | 1.68449 | 62 |
| GO_RESPIRATORY_ELECTRON_TRANSPORT_CHAIN | 0.00000 | 0.00000 | | 0.78870 | 1.60702 | 92 |
| GO_RESPONSE_TO_ALCOHOL | 0.00165 | 0.02247 | 0.45506 | 0.58656 | 1.21108 | 137 |
| GO_RESPONSE_TO_CORTICOSTEROID | 0.00062 | 0.01089 | 0.47727 | 0.62780 | 1.27932 | 91 |
| GO_RESPONSE_TO_DRUG | 0.00011 | 0.00270 | 0.53843 | 0.57887 | 1.20970 | 249 |
| GO_RESPONSE_TO_ETHANOL | 0.00089 | 0.01411 | 0.47727 | 0.64686 | 1.30662 | 70 |
| GO_RESPONSE_TO_FIBROBLAST_GROWTH_FACTOR | 0.00020 | 0.00427 | 0.51885 | 0.63860 | 1.30521 | 100 |
| GO_RESPONSE_TO_GROWTH_FACTOR | 0.00032 | 0.00640 | 0.49849 | 0.53803 | 1.13576 | 471 |
| GO_RESPONSE_TO_HEAT | 0.00071 | 0.01202 | 0.47727 | 0.60008 | 1.23643 | 130 |
| GO_RESPONSE_TO_INTERFERON_BETA | 0.00309 | 0.03563 | 0.43171 | 0.76371 | 1.43769 | 19 |
| GO_RESPONSE_TO_INTERLEUKIN_1 | 0.00000 | 0.00000 | 0.74774 | 0.67276 | 1.38948 | 139 |
| GO_RESPONSE_TO_INTERLEUKIN_12 | 0.00011 | 0.00272 | 0.53843 | 0.73693 | 1.44958 | 39 |
| GO_RESPONSE_TO_INTERLEUKIN_4 | 0.00146 | 0.02034 | 0.45506 | 0.74763 | 1.43398 | 24 |
| GO_RESPONSE_TO_LEUKEMIA_INHIBITORY_FACTOR | 0.00067 | 0.01155 | 0.47727 | 0.63273 | 1.28555 | 81 |
| GO_RESPONSE_TO_NUTRIENT | 0.00350 | 0.03942 | 0.43171 | 0.59675 | 1.22162 | 108 |
| GO_RESPONSE_TO_OXIDATIVE_STRESS | 0.00191 | 0.02490 | 0.45506 | 0.54642 | 1.14776 | 319 |
| GO_RESPONSE_TO_OXYGEN_LEVELS | 0.00000 | 0.00000 | 0.78819 | 0.62495 | 1.30720 | 274 |
| GO_RESPONSE_TO_REACTIVE_OXYGEN_SPECIES | 0.00357 | 0.04012 | 0.43171 | 0.56590 | 1.17297 | 166 |
| GO_RESPONSE_TO_STEROID_HORMONE | 0.00276 | 0.03294 | 0.43171 | 0.55889 | 1.16312 | 214 |
| GO_RESPONSE_TO_TEMPERATURE_STIMULUS | 0.00117 | 0.01719 | 0.45506 | 0.57798 | 1.19801 | 166 |
| GO_RESPONSE_TO_TOXIC_SUBSTANCE | 0.00172 | 0.02337 | 0.45506 | 0.58765 | 1.21459 | 144 |
| GO_RESPONSE_TO_TUMOR_NECROSIS_FACTOR | 0.00002 | 0.00056 | 0.57561 | 0.60072 | 1.24918 | 195 |
| GO_RESPONSE_TO_VIRUS | 0.00061 | 0.01079 | 0.47727 | 0.57732 | 1.20080 | 203 |
| GO_RETINA_HOMEOSTASIS | 0.00159 | 0.02194 | 0.45506 | 0.69239 | 1.36199 | 39 |
| GO_RIBONUCLEOPROTEIN_COMPLEX_BINDING | 0.00000 | 0.00000 | 0.71951 | 0.68060 | 1.39788 | 118 |
| GO_RIBONUCLEOPROTEIN_COMPLEX_BIOGENESIS | 0.00000 | 0.00000 | | 0.63866 | 1.34633 | 403 |
| GO_RIBONUCLEOPROTEIN_COMPLEX_SUBUNIT_ORGANIZATION | 0.00000 | 0.00000 | | 0.68995 | 1.43329 | 183 |
| GO_RIBONUCLEOPROTEIN_GRANULE | 0.00000 | 0.00002 | 0.65944 | 0.62223 | 1.29358 | 199 |
| GO_RIBONUCLEOSIDE_TRIPHOSPHATE_BIOSYNTHETIC_PROCESS | 0.00002 | 0.00056 | 0.57561 | 0.77832 | 1.52141 | 35 |
| GO_RIBONUCLEOSIDE_TRIPHOSPHATE_METABOLIC_PROCESS | 0.00002 | 0.00080 | 0.57561 | 0.73492 | 1.46219 | 47 |
| GO_RIBOSE_PHOSPHATE_BIOSYNTHETIC_PROCESS | 0.00246 | 0.03000 | 0.43171 | 0.58809 | 1.21015 | 124 |
| GO_RIBOSE_PHOSPHATE_METABOLIC_PROCESS | 0.00012 | 0.00285 | 0.53843 | 0.56497 | 1.18374 | 288 |
| GO_RIBOSOMAL_LARGE_SUBUNIT_ASSEMBLY | 0.00287 | 0.03366 | 0.43171 | 0.72205 | 1.39845 | 28 |
| GO_RIBOSOMAL_LARGE_SUBUNIT_BIOGENESIS | 0.00001 | 0.00037 | 0.59333 | 0.69573 | 1.40534 | 70 |
| GO_RIBOSOMAL_SMALL_SUBUNIT_ASSEMBLY | 0.00093 | 0.01451 | 0.47727 | 0.79682 | 1.49553 | 18 |
| GO_RIBOSOMAL_SMALL_SUBUNIT_BIOGENESIS | 0.00231 | 0.02862 | 0.43171 | 0.62950 | 1.26953 | 68 |
| GO_RIBOSOMAL_SUBUNIT | 0.00000 | 0.00000 | | 0.75652 | 1.56988 | 173 |
| GO_RIBOSOME | 0.00000 | 0.00000 | | 0.75342 | 1.56826 | 211 |
| GO_RIBOSOME_ASSEMBLY | 0.00000 | 0.00007 | 0.64355 | 0.73527 | 1.47574 | 59 |
| GO_RIBOSOME_BINDING | 0.00235 | 0.02903 | 0.43171 | 0.66868 | 1.33040 | 47 |
| GO_RIBOSOME_BIOGENESIS | 0.00000 | 0.00000 | 0.77494 | 0.62197 | 1.30265 | 282 |
| GO_RNA_3_END_PROCESSING | 0.00006 | 0.00166 | 0.53843 | 0.62223 | 1.28181 | 129 |
| GO_RNA_CATABOLIC_PROCESS | 0.00000 | 0.00000 | | 0.68391 | 1.43880 | 351 |
| GO_RNA_DEPENDENT_DNA_BIOSYNTHETIC_PROCESS | 0.00253 | 0.03062 | 0.43171 | 0.63895 | 1.28378 | 62 |
| GO_RNA_EXPORT_FROM_NUCLEUS | 0.00000 | 0.00003 | 0.65944 | 0.65223 | 1.34461 | 131 |
| GO_RNA_LOCALIZATION | 0.00000 | 0.00000 | 0.76146 | 0.63998 | 1.33230 | 215 |
| GO_RNA_SPLICING | 0.00000 | 0.00000 | | 0.63865 | 1.34575 | 398 |
| GO_RNA_SPLICING_VIA_TRANSESTERIFICATION_REACTIONS | 0.00000 | 0.00000 | | 0.65628 | 1.37855 | 320 |
| GO_RNA_STABILIZATION | 0.00283 | 0.03346 | 0.43171 | 0.69055 | 1.35214 | 36 |
| GO_ROUGH_ENDOPLASMIC_RETICULUM | 0.00000 | 0.00001 | 0.69013 | 0.75367 | 1.51366 | 60 |
| GO_ROUGH_ENDOPLASMIC_RETICULUM_MEMBRANE | 0.00350 | 0.03942 | 0.43171 | 0.72106 | 1.39046 | 26 |
| GO_RRNA_BINDING | 0.00000 | 0.00017 | 0.61053 | 0.73677 | 1.47411 | 55 |
| GO_RRNA_METABOLIC_PROCESS | 0.00000 | 0.00009 | 0.62726 | 0.60757 | 1.26492 | 212 |
| GO_SCF_DEPENDENT_PROTEASOMAL_UBIQUITIN_DEPENDENT_ | 0.00000 | 0.00018 | 0.61053 | 0.69122 | 1.40360 | 79 |
| GO_SECRETORY_GRANULE_MEMBRANE | 0.00076 | 0.01261 | 0.47727 | 0.57839 | 1.20283 | 196 |
| GO_SERINE_TYPE_ENDOPEPTIDASE_INHIBITOR_ACTIVITY | 0.00045 | 0.00852 | 0.49849 | 0.72007 | 1.40374 | 34 |
| GO_SIGNAL_SEQUENCE_BINDING | 0.00069 | 0.01174 | 0.47727 | 0.70514 | 1.38514 | 38 |
| GO_SINGLE_FERTILIZATION | 0.00167 | 0.02267 | 0.45506 | 0.64169 | 1.29326 | 67 |
| GO_SINGLE_STRANDED_RNA_BINDING | 0.00074 | 0.01236 | 0.47727 | 0.65154 | 1.31312 | 67 |
| GO_SMALL_MOLECULE_BIOSYNTHETIC_PROCESS | 0.00141 | 0.01982 | 0.45506 | 0.53045 | 1.11975 | 465 |
| GO_SMALL_NUCLEAR_RIBONUCLEOPROTEIN_COMPLEX | 0.00002 | 0.00064 | 0.57561 | 0.70190 | 1.40986 | 61 |
| GO_SMALL_RIBOSOMAL_SUBUNIT | 0.00000 | 0.00000 | 0.73376 | 0.75685 | 1.52535 | 67 |
| GO_SMOOTH_MUSCLE_CELL_PROLIFERATION | 0.00287 | 0.03366 | 0.43171 | 0.61786 | 1.25463 | 79 |
| GO_SM_LIKE_PROTEIN_FAMILY_COMPLEX | 0.00009 | 0.00234 | 0.53843 | 0.66985 | 1.35370 | 71 |
| GO_SOMATIC_STEM_CELL_POPULATION_MAINTENANCE | 0.00133 | 0.01915 | 0.45506 | 0.65132 | 1.30811 | 60 |
| GO_SPERM_EGG_RECOGNITION | 0.00001 | 0.00027 | 0.61053 | 0.83003 | 1.59203 | 24 |
| GO_SPLICEOSOMAL_COMPLEX | 0.00000 | 0.00001 | 0.67496 | 0.63864 | 1.32495 | 174 |

| | | | | | | |
|---|---|---|---|---|---|---|
| GO_SPLICEOSOMAL_COMPLEX_ASSEMBLY | 0.00066 | 0.01149 | 0.47727 | 0.67341 | 1.34472 | 52 |
| GO_SPLICEOSOMAL_TRI_SNRNP_COMPLEX | 0.00128 | 0.01848 | 0.45506 | 0.72282 | 1.40293 | 30 |
| GO_STEM_CELL_DIFFERENTIATION | 0.00000 | 0.00010 | 0.62726 | 0.61427 | 1.27536 | 180 |
| GO_STEROID_HORMONE_RECEPTOR_BINDING | 0.00069 | 0.01174 | 0.47727 | 0.65761 | 1.32129 | 62 |
| GO_STRUCTURAL_CONSTITUENT_OF_CYTOSKELETON | 0.00003 | 0.00083 | 0.57561 | 0.69314 | 1.39696 | 67 |
| GO_STRUCTURAL_CONSTITUENT_OF_RIBOSOME | 0.00000 | 0.00000 | | 0.78053 | 1.61461 | 154 |
| GO_STRUCTURAL_MOLECULE_ACTIVITY | 0.00000 | 0.00000 | | 0.69109 | 1.45857 | 442 |
| GO_SUBSTANTIA_NIGRA_DEVELOPMENT | 0.00021 | 0.00445 | 0.51885 | 0.75388 | 1.46320 | 30 |
| GO_SUPRAMOLECULAR_FIBER_ORGANIZATION | 0.00000 | 0.00016 | 0.62726 | 0.56241 | 1.18733 | 478 |
| GO_TELOMERASE_RNA_LOCALIZATION | 0.00023 | 0.00483 | 0.51885 | 0.82266 | 1.54867 | 19 |
| GO_TELOMERE_MAINTENANCE_VIA_TELOMERE_LENGTHENING | 0.00083 | 0.01329 | 0.47727 | 0.65570 | 1.32357 | 69 |
| GO_TELOMERE_ORGANIZATION | 0.00420 | 0.04542 | 0.40702 | 0.57401 | 1.18363 | 132 |
| GO_THREONINE_TYPE_ENDOPEPTIDASE_ACTIVITY | 0.00000 | 0.00003 | 0.65944 | 0.91654 | 1.70211 | 16 |
| GO_THREONINE_TYPE_PEPTIDASE_ACTIVITY | 0.00000 | 0.00000 | 0.71951 | 0.89879 | 1.70500 | 21 |
| GO_TISSUE_MORPHOGENESIS | 0.00100 | 0.01530 | 0.45506 | 0.53868 | 1.13684 | 432 |
| GO_TRANSCRIPTION_COUPLED_NUCLEOTIDE_EXCISION_REPAIR | 0.00220 | 0.02777 | 0.43171 | 0.63044 | 1.27504 | 72 |
| GO_TRANSLATIONAL_ELONGATION | 0.00000 | 0.00000 | 0.82666 | 0.70531 | 1.45209 | 126 |
| GO_TRANSLATIONAL_INITIATION | 0.00000 | 0.00000 | | 0.77762 | 1.61295 | 170 |
| GO_TRANSLATIONAL_TERMINATION | 0.00000 | 0.00000 | 0.74774 | 0.70841 | 1.44888 | 102 |
| GO_TRANSLATION_FACTOR_ACTIVITY_RNA_BINDING | 0.00000 | 0.00014 | 0.62726 | 0.70317 | 1.42104 | 71 |
| GO_TRANSLATION_INITIATION_FACTOR_ACTIVITY | 0.00006 | 0.00171 | 0.53843 | 0.72658 | 1.44055 | 45 |
| GO_TRANSLATION_INITIATION_FACTOR_BINDING | 0.00398 | 0.04337 | 0.40702 | 0.71493 | 1.38229 | 27 |
| GO_TRANSLATION_PREINITIATION_COMPLEX | 0.00095 | 0.01470 | 0.47727 | 0.81423 | 1.51210 | 16 |
| GO_TRANSLATION_REGULATOR_ACTIVITY | 0.00000 | 0.00000 | 0.73376 | 0.69131 | 1.41813 | 114 |
| GO_TRANSLATION_REGULATOR_ACTIVITY_NUCLEIC_ACID_BIND | 0.00000 | 0.00002 | 0.67496 | 0.70179 | 1.42848 | 88 |
| GO_TRICARBOXYLIC_ACID_CYCLE | 0.00082 | 0.01329 | 0.47727 | 0.72594 | 1.41427 | 33 |
| GO_TRNA_TRANSPORT | 0.00143 | 0.01995 | 0.45506 | 0.70129 | 1.36712 | 34 |
| GO_TUMOR_NECROSIS_FACTOR_MEDIATED_SIGNALING_PATHW | 0.00000 | 0.00008 | 0.62726 | 0.65694 | 1.34930 | 118 |
| GO_T_CELL_RECEPTOR_SIGNALING_PATHWAY | 0.00011 | 0.00277 | 0.53843 | 0.61407 | 1.26862 | 142 |
| GO_T_CELL_SELECTION | 0.00283 | 0.03346 | 0.43171 | 0.72213 | 1.39861 | 28 |
| GO_U12_TYPE_SPLICEOSOMAL_COMPLEX | 0.00021 | 0.00450 | 0.51885 | 0.77175 | 1.48821 | 26 |
| GO_U1_SNRNP | 0.00139 | 0.01969 | 0.45506 | 0.78952 | 1.48183 | 18 |
| GO_U2_SNRNP | 0.00313 | 0.03593 | 0.43171 | 0.76468 | 1.43521 | 18 |
| GO_U2_TYPE_CATALYTIC_STEP_2_SPLICEOSOME | 0.00191 | 0.02490 | 0.45506 | 0.71672 | 1.39108 | 30 |
| GO_U2_TYPE_SPLICEOSOMAL_COMPLEX | 0.00026 | 0.00534 | 0.49849 | 0.64138 | 1.30564 | 87 |
| GO_U5_SNRNP | 0.00428 | 0.04607 | 0.40702 | 0.78973 | 1.46229 | 15 |
| GO_UBIQUITIN_LIKE_PROTEIN_LIGASE_BINDING | 0.00000 | 0.00005 | 0.64355 | 0.59865 | 1.25222 | 271 |
| GO_UNFOLDED_PROTEIN_BINDING | 0.00000 | 0.00001 | 0.67496 | 0.69648 | 1.42015 | 93 |
| GO_VACUOLAR_LUMEN | 0.00012 | 0.00288 | 0.53843 | 0.62512 | 1.28320 | 117 |
| GO_VACUOLAR_MEMBRANE | 0.00265 | 0.03173 | 0.43171 | 0.53762 | 1.12932 | 323 |
| GO_VESICLE_LUMEN | 0.00000 | 0.00000 | | 0.66338 | 1.38103 | 218 |
| GO_VIRAL_GENE_EXPRESSION | 0.00000 | 0.00000 | | 0.72567 | 1.50665 | 180 |
| GO_VIRAL_GENOME_REPLICATION | 0.00003 | 0.00081 | 0.57561 | 0.65712 | 1.33988 | 93 |
| GO_VIRAL_LIFE_CYCLE | 0.00000 | 0.00000 | 0.77494 | 0.62394 | 1.30444 | 256 |
| GO_WIDE_PORE_CHANNEL_ACTIVITY | 0.00202 | 0.02594 | 0.43171 | 0.80741 | 1.49504 | 15 |
| HP_ABNORMALITY_OF_ACID_BASE_HOMEOSTASIS | 0.00002 | 0.00068 | 0.57561 | 0.58230 | 1.21747 | 259 |
| HP_ABNORMALITY_OF_BLOOD_CIRCULATION | 0.00294 | 0.03440 | 0.43171 | 0.56716 | 1.17664 | 176 |
| HP_ABNORMALITY_OF_CARDIOVASCULAR_SYSTEM_ELECTROPH | 0.00246 | 0.03000 | 0.43171 | 0.54664 | 1.14523 | 285 |
| HP_ABNORMALITY_OF_ESOPHAGUS_PHYSIOLOGY | 0.00198 | 0.02561 | 0.43171 | 0.53842 | 1.13502 | 403 |
| HP_ABNORMALITY_OF_THE_MITOCHONDRION | 0.00021 | 0.00447 | 0.51885 | 0.61447 | 1.26495 | 125 |
| HP_ABNORMAL_ABDOMEN_MORPHOLOGY | 0.00135 | 0.01933 | 0.45506 | 0.53312 | 1.12532 | 467 |
| HP_ABNORMAL_ACTIVITY_OF_MITOCHONDRIAL_RESPIRATORY_ | 0.00013 | 0.00316 | 0.51885 | 0.67704 | 1.36247 | 63 |
| HP_ABNORMAL_BASAL_GANGLIA_MRI_SIGNAL_INTENSITY | 0.00115 | 0.01699 | 0.45506 | 0.70547 | 1.38135 | 36 |
| HP_ABNORMAL_BRAINSTEM_MRI_SIGNAL_INTENSITY | 0.00022 | 0.00469 | 0.51885 | 0.74875 | 1.45871 | 33 |
| HP_ABNORMAL_CARTILAGE_MORPHOLOGY | 0.00353 | 0.03977 | 0.43171 | 0.72038 | 1.39283 | 27 |
| HP_ABNORMAL_CIRCULATING_CARBOHYDRATE_CONCENTRATIO | 0.00115 | 0.01699 | 0.45506 | 0.68442 | 1.36173 | 47 |
| HP_ABNORMAL_CNS_MYELINATION | 0.00018 | 0.00403 | 0.51885 | 0.59357 | 1.23128 | 175 |
| HP_ABNORMAL_CSF_METABOLITE_LEVEL | 0.00000 | 0.00005 | 0.64355 | 0.70095 | 1.42414 | 81 |
| HP_ABNORMAL_ERYTHROCYTE_MORPHOLOGY | 0.00003 | 0.00082 | 0.57561 | 0.56077 | 1.18222 | 412 |
| HP_ABNORMAL_FOVEAL_MORPHOLOGY | 0.00316 | 0.03622 | 0.43171 | 0.76095 | 1.43251 | 19 |
| HP_ABNORMAL_MITOCHONDRIA_IN_MUSCLE_TISSUE | 0.00018 | 0.00406 | 0.51885 | 0.76272 | 1.47957 | 29 |
| HP_ABNORMAL_MORPHOLOGY_OF_THE_GREAT_VESSELS | 0.00104 | 0.01579 | 0.45506 | 0.54456 | 1.14622 | 363 |
| HP_ABNORMAL_MYELINATION | 0.00025 | 0.00526 | 0.49849 | 0.55290 | 1.16337 | 355 |
| HP_ABNORMAL_MYOCARDIUM_MORPHOLOGY | 0.00055 | 0.01006 | 0.47727 | 0.54419 | 1.14551 | 366 |
| HP_ABNORMAL_NUMBER_OF_INCISORS | 0.00119 | 0.01742 | 0.45506 | 0.72508 | 1.41099 | 32 |
| HP_ABNORMAL_SOCIAL_BEHAVIOR | 0.00092 | 0.01441 | 0.47727 | 0.60162 | 1.23644 | 120 |
| HP_ABNORMAL_TRICUSPID_VALVE_PHYSIOLOGY | 0.00283 | 0.03346 | 0.43171 | 0.69049 | 1.35202 | 36 |
| HP_ACUTE_LEUKEMIA | 0.00024 | 0.00499 | 0.51885 | 0.64964 | 1.31733 | 77 |
| HP_ANEMIA_OF_INADEQUATE_PRODUCTION | 0.00000 | 0.00018 | 0.61053 | 0.66256 | 1.35635 | 106 |
| HP_APHASIA | 0.00059 | 0.01058 | 0.47727 | 0.72316 | 1.41358 | 35 |

| | | | | | | |
|---|---|---|---|---|---|---|
| HP_CONGENITAL_MALFORMATION_OF_THE_GREAT_ARTERIES | 0.00130 | 0.01871 | 0.45506 | 0.55877 | 1.16848 | 260 |
| HP_DECREASED_ACTIVITY_OF_MITOCHONDRIAL_COMPLEX_I | 0.00078 | 0.01287 | 0.47727 | 0.71655 | 1.40066 | 35 |
| HP_DECREASED_ACTIVITY_OF_THE_PYRUVATE_DEHYDROGENAS | 0.00253 | 0.03062 | 0.43171 | 0.70935 | 1.38038 | 32 |
| HP_DELAYED_CRANIAL_SUTURE_CLOSURE | 0.00228 | 0.02833 | 0.43171 | 0.65806 | 1.31220 | 50 |
| HP_DEPRESSED_NASAL_RIDGE | 0.00364 | 0.04067 | 0.43171 | 0.59936 | 1.22428 | 97 |
| HP_DYSPHASIA | 0.00021 | 0.00447 | 0.51885 | 0.70964 | 1.40453 | 43 |
| HP_DYSPNEA | 0.00137 | 0.01955 | 0.45506 | 0.55796 | 1.16677 | 260 |
| HP_ECHOLALIA | 0.00004 | 0.00108 | 0.55733 | 0.77089 | 1.50184 | 33 |
| HP_EMOTIONAL_LABILITY | 0.00032 | 0.00640 | 0.49849 | 0.62711 | 1.27954 | 95 |
| HP_FATIGUE | 0.00045 | 0.00852 | 0.49849 | 0.56839 | 1.18491 | 232 |
| HP_FETAL_DISTRESS | 0.00008 | 0.00208 | 0.53843 | 0.76443 | 1.48509 | 31 |
| HP_FOCAL_T2_HYPERINTENSE_BRAINSTEM_LESION | 0.00083 | 0.01329 | 0.47727 | 0.74731 | 1.44738 | 28 |
| HP_FREQUENT_FALLS | 0.00465 | 0.04935 | 0.40702 | 0.61980 | 1.25197 | 70 |
| HP_HEADACHE | 0.00066 | 0.01149 | 0.47727 | 0.57561 | 1.19531 | 181 |
| HP_HEMATOLOGICAL_NEOPLASM | 0.00019 | 0.00420 | 0.51885 | 0.58866 | 1.22397 | 187 |
| HP_HEPATOMEGALY | 0.00324 | 0.03688 | 0.43171 | 0.53260 | 1.12056 | 348 |
| HP_HYPERTROPHIC_CARDIOMYOPATHY | 0.00004 | 0.00111 | 0.55733 | 0.59622 | 1.23952 | 199 |
| HP_IMPAIRED_SOCIAL_INTERACTIONS | 0.00023 | 0.00483 | 0.51885 | 0.65263 | 1.32208 | 75 |
| HP_IMPAIRMENT_IN_PERSONALITY_FUNCTIONING | 0.00220 | 0.02777 | 0.43171 | 0.53401 | 1.12583 | 408 |
| HP_INCOORDINATION | 0.00115 | 0.01699 | 0.45506 | 0.58552 | 1.20984 | 143 |
| HP_INCREASED_CSF_LACTATE | 0.00001 | 0.00030 | 0.59333 | 0.70715 | 1.42743 | 69 |
| HP_INCREASED_SERUM_LACTATE | 0.00012 | 0.00283 | 0.53843 | 0.61243 | 1.26355 | 134 |
| HP_INCREASED_SERUM_PYRUVATE | 0.00018 | 0.00393 | 0.51885 | 0.72500 | 1.42415 | 38 |
| HP_LACTIC_ACIDOSIS | 0.00019 | 0.00423 | 0.51885 | 0.61419 | 1.26758 | 135 |
| HP_LETHARGY | 0.00187 | 0.02466 | 0.45506 | 0.60407 | 1.23661 | 108 |
| HP_LEUKEMIA | 0.00018 | 0.00395 | 0.51885 | 0.63065 | 1.29192 | 110 |
| HP_LEUKODYSTROPHY | 0.00006 | 0.00160 | 0.55733 | 0.66291 | 1.34825 | 84 |
| HP_LEUKOENCEPHALOPATHY | 0.00161 | 0.02210 | 0.45506 | 0.58861 | 1.21548 | 138 |
| HP_LOWER_LIMB_SPASTICITY | 0.00108 | 0.01616 | 0.45506 | 0.58876 | 1.21620 | 140 |
| HP_MACROCYTIC_ANEMIA | 0.00001 | 0.00043 | 0.59333 | 0.76221 | 1.49932 | 39 |
| HP_MIGRAINE | 0.00004 | 0.00125 | 0.55733 | 0.66143 | 1.34451 | 83 |
| HP_MITOCHONDRIAL_MYOPATHY | 0.00228 | 0.02833 | 0.43171 | 0.68078 | 1.33729 | 38 |
| HP_MUSCLE_ABNORMALITY_RELATED_TO_MITOCHONDRIAL_DY | 0.00079 | 0.01295 | 0.47727 | 0.67564 | 1.34804 | 51 |
| HP_MUTISM | 0.00003 | 0.00087 | 0.57561 | 0.76972 | 1.50716 | 36 |
| HP_MYOPATHY | 0.00202 | 0.02594 | 0.43171 | 0.56699 | 1.17903 | 202 |
| HP_MYOSITIS | 0.00187 | 0.02466 | 0.45506 | 0.79960 | 1.48493 | 16 |
| HP_NAUSEA_AND_VOMITING | 0.00061 | 0.01079 | 0.47727 | 0.55780 | 1.16781 | 279 |
| HP_OPHTHALMOPARESIS | 0.00472 | 0.04982 | 0.40702 | 0.55993 | 1.16274 | 181 |
| HP_OPTIC_NERVE_COLOBOMA | 0.00224 | 0.02808 | 0.43171 | 0.77367 | 1.45209 | 18 |
| HP_OPTIC_NEUROPATHY | 0.00003 | 0.00092 | 0.55733 | 0.75496 | 1.48152 | 37 |
| HP_PALLOR | 0.00339 | 0.03843 | 0.43171 | 0.59313 | 1.21423 | 107 |
| HP_PANHYPOPITUITARISM | 0.00033 | 0.00645 | 0.49849 | 0.84657 | 1.56753 | 15 |
| HP_PARALYSIS | 0.00220 | 0.02777 | 0.43171 | 0.61041 | 1.24464 | 93 |
| HP_PARAPLEGIA_PARAPARESIS | 0.00372 | 0.04128 | 0.43171 | 0.57084 | 1.18160 | 158 |
| HP_PAROXYSMAL_INVOLUNTARY_EYE_MOVEMENTS | 0.00010 | 0.00258 | 0.53843 | 0.75986 | 1.47621 | 31 |
| HP_PATENT_DUCTUS_ARTERIOSUS | 0.00327 | 0.03724 | 0.43171 | 0.55923 | 1.16267 | 197 |
| HP_PERIPHERAL_NEUROPATHY | 0.00261 | 0.03134 | 0.43171 | 0.53075 | 1.11902 | 414 |
| HP_POOR_EYE_CONTACT | 0.00029 | 0.00594 | 0.49849 | 0.68766 | 1.37584 | 55 |
| HP_POOR_HEAD_CONTROL | 0.00128 | 0.01848 | 0.45506 | 0.59429 | 1.22353 | 126 |
| HP_PROGRESSIVE_SPASTIC_PARAPLEGIA | 0.00176 | 0.02378 | 0.45506 | 0.66396 | 1.32322 | 49 |
| HP_PROXIMAL_TUBULOPATHY | 0.00108 | 0.01616 | 0.45506 | 0.72238 | 1.40735 | 33 |
| HP_PTOSIS | 0.00060 | 0.01072 | 0.47727 | 0.53789 | 1.13525 | 454 |
| HP_REDUCED_CONSCIOUSNESS_CONFUSION | 0.00014 | 0.00333 | 0.51885 | 0.57992 | 1.20684 | 208 |
| HP_RENAL_TUBULAR_DYSFUNCTION | 0.00183 | 0.02432 | 0.45506 | 0.63346 | 1.27956 | 70 |
| HP_RESPIRATORY_INSUFFICIENCY | 0.00088 | 0.01400 | 0.47727 | 0.54579 | 1.14856 | 359 |
| HP_SINGLE_MEDIAN_MAXILLARY_INCISOR | 0.00111 | 0.01664 | 0.45506 | 0.80723 | 1.50681 | 17 |
| HP_TETRAPARESIS | 0.00108 | 0.01616 | 0.45506 | 0.74097 | 1.42530 | 25 |
| HP_TETRAPLEGIA_TETRAPARESIS | 0.00009 | 0.00224 | 0.53843 | 0.71728 | 1.42542 | 46 |
| HP_THICK_LOWER_LIP_VERMILION | 0.00148 | 0.02055 | 0.45506 | 0.59444 | 1.21913 | 116 |
| HP_TRIANGULAR_SHAPED_PHALANGES_OF_THE_HAND | 0.00364 | 0.04067 | 0.43171 | 0.58488 | 1.20212 | 121 |
| HP_TRIGONOCEPHALY | 0.00224 | 0.02808 | 0.43171 | 0.71406 | 1.38592 | 30 |
| HP_VESTIBULAR_DYSFUNCTION | 0.00428 | 0.04607 | 0.40702 | 0.61906 | 1.25408 | 75 |
| HP_WEAKNESS_DUE_TO_UPPER_MOTOR_NEURON_DYSFUNCTI | 0.00246 | 0.03000 | 0.43171 | 0.54702 | 1.14663 | 293 |
| **Cluster 7** | | | | | | |
| pathway | pval | padj | log2err | ES | NES | size |
| GO_ACTIN_BASED_CELL_PROJECTION | 0.00194 | 0.02819 | 0.45506 | 0.61302 | 1.19331 | 137 |
| GO_ACTIN_BINDING | 0.00044 | 0.01057 | 0.49849 | 0.58242 | 1.15163 | 296 |
| GO_ACTIN_CYTOSKELETON | 0.00011 | 0.00408 | 0.53843 | 0.58264 | 1.15498 | 350 |
| GO_ACTIN_FILAMENT_BASED_MOVEMENT | 0.00007 | 0.00288 | 0.53843 | 0.67079 | 1.29210 | 90 |
| GO_ACTIN_FILAMENT_BUNDLE_ORGANIZATION | 0.00007 | 0.00286 | 0.53843 | 0.65780 | 1.27333 | 112 |

| | | | | | | |
|---|---|---|---|---|---|---|
| GO_ACTIN_FILAMENT_ORGANIZATION | 0.00003 | 0.00171 | 0.55733 | 0.59945 | 1.18501 | 294 |
| GO_ACTIN_MEDIATED_CELL_CONTRACTION | 0.00119 | 0.02087 | 0.45506 | 0.67542 | 1.28377 | 63 |
| GO_ACTIVATION_OF_IMMUNE_RESPONSE | 0.00002 | 0.00125 | 0.57561 | 0.61381 | 1.20794 | 244 |
| GO_ACTOMYOSIN_STRUCTURE_ORGANIZATION | 0.00357 | 0.04144 | 0.43171 | 0.60660 | 1.18082 | 137 |
| GO_ALTERNATIVE_MRNA_SPLICING_VIA_SPLICEOSOME | 0.00007 | 0.00293 | 0.53843 | 0.69783 | 1.33279 | 70 |
| GO_AMIDE_BINDING | 0.00001 | 0.00091 | 0.59333 | 0.62480 | 1.22268 | 187 |
| GO_ANATOMICAL_STRUCTURE_HOMEOSTASIS | 0.00005 | 0.00214 | 0.55733 | 0.60709 | 1.19779 | 272 |
| GO_ANDROGEN_RECEPTOR_SIGNALING_PATHWAY | 0.00327 | 0.03902 | 0.43171 | 0.72541 | 1.33862 | 31 |
| GO_ANGIOGENESIS_INVOLVED_IN_WOUND_HEALING | 0.00137 | 0.02252 | 0.45506 | 0.82853 | 1.46954 | 15 |
| GO_ANTIMICROBIAL_HUMORAL_RESPONSE | 0.00005 | 0.00238 | 0.55733 | 0.82057 | 1.50035 | 25 |
| GO_APOPTOTIC_SIGNALING_PATHWAY | 0.00056 | 0.01245 | 0.47727 | 0.56948 | 1.13183 | 397 |
| GO_ATP_METABOLIC_PROCESS | 0.00000 | 0.00005 | 0.67496 | 0.64715 | 1.26870 | 199 |
| GO_ATP_SYNTHESIS_COUPLED_ELECTRON_TRANSPORT | 0.00007 | 0.00296 | 0.53843 | 0.69894 | 1.33557 | 71 |
| GO_AXONAL_GROWTH_CONE | 0.00405 | 0.04495 | 0.40702 | 0.75400 | 1.37168 | 23 |
| GO_AXON_DEVELOPMENT | 0.00235 | 0.03181 | 0.43171 | 0.56732 | 1.12387 | 327 |
| GO_BASEMENT_MEMBRANE | 0.00405 | 0.04495 | 0.40702 | 0.66605 | 1.26024 | 56 |
| GO_BASOLATERAL_PLASMA_MEMBRANE | 0.00077 | 0.01582 | 0.47727 | 0.61645 | 1.20221 | 150 |
| GO_BINDING_OF_SPERM_TO_ZONA_PELLUCIDA | 0.00250 | 0.03324 | 0.43171 | 0.76675 | 1.38855 | 22 |
| GO_BRUSH_BORDER | 0.00041 | 0.01020 | 0.49849 | 0.67997 | 1.29717 | 68 |
| GO_CADHERIN_BINDING | 0.00000 | 0.00000 | | 0.68447 | 1.34876 | 264 |
| GO_CADHERIN_BINDING_INVOLVED_IN_CELL_CELL_ADHESION | 0.00209 | 0.02959 | 0.43171 | 0.82163 | 1.45730 | 15 |
| GO_CARDIAC_CONDUCTION | 0.00020 | 0.00641 | 0.51885 | 0.67135 | 1.28675 | 76 |
| GO_CARDIAC_MUSCLE_CELL_ACTION_POTENTIAL | 0.00257 | 0.03380 | 0.43171 | 0.69924 | 1.30581 | 41 |
| GO_CARDIAC_MUSCLE_CELL_CONTRACTION | 0.00035 | 0.00905 | 0.49849 | 0.73639 | 1.37243 | 38 |
| GO_CARDIAC_MUSCLE_CONTRACTION | 0.00115 | 0.02051 | 0.45506 | 0.66475 | 1.27024 | 71 |
| GO_CATALYTIC_STEP_2_SPLICEOSOME | 0.00102 | 0.01880 | 0.45506 | 0.66169 | 1.26880 | 78 |
| GO_CELLULAR_COMPONENT_DISASSEMBLY | 0.00016 | 0.00511 | 0.51885 | 0.57856 | 1.15017 | 402 |
| GO_CELLULAR_NITROGEN_COMPOUND_CATABOLIC_PROCESS | 0.00000 | 0.00000 | | 0.69889 | 1.38996 | 418 |
| GO_CELLULAR_RESPIRATION | 0.00004 | 0.00197 | 0.55733 | 0.64833 | 1.26054 | 129 |
| GO_CELLULAR_RESPONSE_TO_CHEMICAL_STRESS | 0.00106 | 0.01925 | 0.45506 | 0.59050 | 1.16150 | 236 |
| GO_CELLULAR_RESPONSE_TO_HORMONE_STIMULUS | 0.00003 | 0.00150 | 0.57561 | 0.58722 | 1.16737 | 403 |
| GO_CELLULAR_RESPONSE_TO_LIPID | 0.00078 | 0.01596 | 0.47727 | 0.57876 | 1.14601 | 321 |
| GO_CELLULAR_RESPONSE_TO_NITROGEN_COMPOUND | 0.00006 | 0.00273 | 0.53843 | 0.57707 | 1.14758 | 420 |
| GO_CELLULAR_RESPONSE_TO_ORGANIC_CYCLIC_COMPOUND | 0.00000 | 0.00002 | 0.69013 | 0.61481 | 1.21843 | 335 |
| GO_CELLULAR_RESPONSE_TO_OXYGEN_LEVELS | 0.00111 | 0.01996 | 0.45506 | 0.60701 | 1.18522 | 167 |
| GO_CELLULAR_RESPONSE_TO_STEROID_HORMONE_STIMULUS | 0.00038 | 0.00968 | 0.49849 | 0.62375 | 1.21458 | 140 |
| GO_CELL_ACTIVATION_INVOLVED_IN_IMMUNE_RESPONSE | 0.00000 | 0.00000 | 0.76146 | 0.62139 | 1.23383 | 382 |
| GO_CELL_ADHESION_MEDIATOR_ACTIVITY | 0.00015 | 0.00487 | 0.51885 | 0.75780 | 1.41044 | 37 |
| GO_CELL_ADHESION_MOLECULE_BINDING | 0.00000 | 0.00000 | | 0.67293 | 1.33497 | 369 |
| GO_CELL_BODY | 0.00405 | 0.04495 | 0.40702 | 0.56271 | 1.11521 | 336 |
| GO_CELL_CELL_ADHESION | 0.00052 | 0.01189 | 0.47727 | 0.57152 | 1.13667 | 410 |
| GO_CELL_CELL_CONTACT_ZONE | 0.00228 | 0.03122 | 0.43171 | 0.68827 | 1.29447 | 47 |
| GO_CELL_CELL_JUNCTION | 0.00000 | 0.00029 | 0.62726 | 0.61134 | 1.21025 | 313 |
| GO_CELL_CELL_RECOGNITION | 0.00187 | 0.02770 | 0.45506 | 0.72130 | 1.34252 | 37 |
| GO_CELL_CELL_SIGNALING_BY_WNT | 0.00006 | 0.00259 | 0.55733 | 0.58503 | 1.16037 | 366 |
| GO_CELL_COMMUNICATION_INVOLVED_IN_CARDIAC_CONDUCT | 0.00130 | 0.02200 | 0.45506 | 0.74062 | 1.36582 | 30 |
| GO_CELL_CORTEX | 0.00009 | 0.00346 | 0.53843 | 0.61766 | 1.21104 | 202 |
| GO_CELL_CYCLE_PHASE_TRANSITION | 0.00004 | 0.00204 | 0.55733 | 0.57509 | 1.14447 | 468 |
| GO_CELL_JUNCTION_ASSEMBLY | 0.00141 | 0.02294 | 0.45506 | 0.58311 | 1.14873 | 261 |
| GO_CELL_JUNCTION_ORGANIZATION | 0.00002 | 0.00140 | 0.57561 | 0.58212 | 1.15824 | 431 |
| GO_CELL_KILLING | 0.00003 | 0.00152 | 0.57561 | 0.73469 | 1.39021 | 55 |
| GO_CELL_LEADING_EDGE | 0.00000 | 0.00010 | 0.64355 | 0.61701 | 1.22014 | 300 |
| GO_CELL_MATRIX_ADHESION | 0.00168 | 0.02608 | 0.45506 | 0.61686 | 1.19995 | 133 |
| GO_CELL_MORPHOGENESIS_INVOLVED_IN_DIFFERENTIATION | 0.00004 | 0.00184 | 0.55733 | 0.57864 | 1.15166 | 478 |
| GO_CELL_PART_MORPHOGENESIS | 0.00294 | 0.03666 | 0.43171 | 0.55666 | 1.10727 | 448 |
| GO_CELL_PROJECTION_MEMBRANE | 0.00002 | 0.00100 | 0.59333 | 0.62939 | 1.23405 | 202 |
| GO_CELL_RECOGNITION | 0.00265 | 0.03441 | 0.43171 | 0.66411 | 1.26355 | 64 |
| GO_CELL_SUBSTRATE_ADHESION | 0.00030 | 0.00835 | 0.49849 | 0.60603 | 1.18981 | 219 |
| GO_CELL_SUBSTRATE_JUNCTION | 0.00000 | 0.00000 | | 0.70232 | 1.39043 | 315 |
| GO_CELL_SUBSTRATE_JUNCTION_ORGANIZATION | 0.00033 | 0.00876 | 0.49849 | 0.66567 | 1.27585 | 76 |
| GO_CELL_SURFACE | 0.00413 | 0.04562 | 0.40702 | 0.56649 | 1.12242 | 332 |
| GO_CELL_SURFACE_RECEPTOR_SIGNALING_PATHWAY_INVOLVEI | 0.00030 | 0.00835 | 0.49849 | 0.57319 | 1.13988 | 413 |
| GO_CHAPERONE_COMPLEX | 0.00228 | 0.03122 | 0.43171 | 0.76923 | 1.39305 | 22 |
| GO_CHROMATIN_ASSEMBLY_OR_DISASSEMBLY | 0.00342 | 0.04017 | 0.43171 | 0.62174 | 1.20225 | 109 |
| GO_CHROMATIN_BINDING | 0.00000 | 0.00001 | 0.70498 | 0.60529 | 1.20379 | 423 |
| GO_CHROMATIN_REMODELING | 0.00179 | 0.02699 | 0.45506 | 0.60773 | 1.18494 | 149 |
| GO_CIRCADIAN_RHYTHM | 0.00053 | 0.01189 | 0.47727 | 0.63516 | 1.23406 | 126 |
| GO_CIRCULATORY_SYSTEM_PROCESS | 0.00183 | 0.02741 | 0.45506 | 0.58013 | 1.14347 | 266 |
| GO_CLATHRIN_COATED_PIT | 0.00253 | 0.03352 | 0.43171 | 0.67009 | 1.26948 | 58 |
| GO_CLATHRIN_COATED_VESICLE | 0.00305 | 0.03715 | 0.43171 | 0.62672 | 1.21442 | 118 |

| | | | | | | |
|---|---|---|---|---|---|---|
| GO_CLATHRIN_DEPENDENT_ENDOCYTOSIS | 0.00435 | 0.04725 | 0.40702 | 0.69736 | 1.29968 | 38 |
| GO_CLUSTER_OF_ACTIN_BASED_CELL_PROJECTIONS | 0.00035 | 0.00905 | 0.49849 | 0.65970 | 1.27232 | 94 |
| GO_COAGULATION | 0.00327 | 0.03902 | 0.43171 | 0.59911 | 1.16976 | 166 |
| GO_COLLAGEN_CONTAINING_EXTRACELLULAR_MATRIX | 0.00041 | 0.01020 | 0.49849 | 0.61509 | 1.20283 | 181 |
| GO_CONTRACTILE_FIBER | 0.00044 | 0.01057 | 0.49849 | 0.62463 | 1.21690 | 144 |
| GO_CORNIFICATION | 0.00013 | 0.00445 | 0.51885 | 0.77190 | 1.42954 | 33 |
| GO_CORTICAL_CYTOSKELETON | 0.00064 | 0.01350 | 0.47727 | 0.66445 | 1.27141 | 73 |
| GO_COTRANSLATIONAL_PROTEIN_TARGETING_TO_MEMBRANE | 0.00000 | 0.00000 | | 0.85827 | 1.65452 | 92 |
| GO_COVALENT_CHROMATIN_MODIFICATION | 0.00009 | 0.00362 | 0.53843 | 0.58693 | 1.16371 | 357 |
| GO_CYTOCHROME_COMPLEX | 0.00042 | 0.01025 | 0.49849 | 0.81293 | 1.47072 | 21 |
| GO_CYTOKINE_MEDIATED_SIGNALING_PATHWAY | 0.00014 | 0.00482 | 0.51885 | 0.57631 | 1.14517 | 395 |
| GO_CYTOKINE_PRODUCTION | 0.00016 | 0.00521 | 0.51885 | 0.58135 | 1.15452 | 385 |
| GO_CYTOKINE_PRODUCTION_INVOLVED_IN_IMMUNE_RESPONS | 0.00067 | 0.01418 | 0.47727 | 0.70741 | 1.33048 | 47 |
| GO_CYTOPLASMIC_SIDE_OF_MEMBRANE | 0.00239 | 0.03218 | 0.43171 | 0.61801 | 1.19769 | 120 |
| GO_CYTOPLASMIC_STRESS_GRANULE | 0.00001 | 0.00089 | 0.59333 | 0.72688 | 1.37751 | 59 |
| GO_CYTOPLASMIC_TRANSLATION | 0.00000 | 0.00000 | | 0.78524 | 1.51061 | 87 |
| GO_CYTOPLASMIC_TRANSLATIONAL_INITIATION | 0.00224 | 0.03105 | 0.43171 | 0.74694 | 1.37338 | 28 |
| GO_CYTOSOLIC_LARGE_RIBOSOMAL_SUBUNIT | 0.00000 | 0.00000 | | 0.87502 | 1.65295 | 52 |
| GO_CYTOSOLIC_RIBOSOME | 0.00000 | 0.00000 | | 0.86593 | 1.67022 | 93 |
| GO_CYTOSOLIC_SMALL_RIBOSOMAL_SUBUNIT | 0.00000 | 0.00000 | 0.81404 | 0.87015 | 1.62171 | 38 |
| GO_DEFENSE_RESPONSE_TO_GRAM_NEGATIVE_BACTERIUM | 0.00198 | 0.02853 | 0.43171 | 0.81838 | 1.45732 | 16 |
| GO_DENDRITIC_TREE | 0.00122 | 0.02127 | 0.45506 | 0.56868 | 1.12813 | 368 |
| GO_DESMOSOME | 0.00143 | 0.02312 | 0.45506 | 0.79385 | 1.43229 | 20 |
| GO_DETECTION_OF_STIMULUS | 0.00085 | 0.01667 | 0.47727 | 0.63910 | 1.23840 | 118 |
| GO_DETOXIFICATION | 0.00357 | 0.04144 | 0.43171 | 0.64633 | 1.23767 | 74 |
| GO_DEVELOPMENTAL_GROWTH | 0.00137 | 0.02252 | 0.45506 | 0.56530 | 1.12409 | 406 |
| GO_DISORDERED_DOMAIN_SPECIFIC_BINDING | 0.00268 | 0.03460 | 0.43171 | 0.74382 | 1.36764 | 28 |
| GO_DISTAL_AXON | 0.00450 | 0.04803 | 0.40702 | 0.58376 | 1.14237 | 187 |
| GO_DNA_BINDING_TRANSCRIPTION_FACTOR_BINDING | 0.00202 | 0.02893 | 0.43171 | 0.57994 | 1.14159 | 251 |
| GO_DNA_CONFORMATION_CHANGE | 0.00176 | 0.02669 | 0.45506 | 0.59017 | 1.15815 | 213 |
| GO_DNA_METHYLATION | 0.00465 | 0.04887 | 0.40702 | 0.67330 | 1.27095 | 51 |
| GO_DNA_METHYLATION_OR_DEMETHYLATION | 0.00052 | 0.01189 | 0.47727 | 0.68162 | 1.29531 | 62 |
| GO_DNA_MODIFICATION | 0.00121 | 0.02113 | 0.45506 | 0.65118 | 1.24960 | 79 |
| GO_DOUBLE_STRANDED_RNA_BINDING | 0.00011 | 0.00397 | 0.53843 | 0.71385 | 1.35069 | 56 |
| GO_ELECTRON_TRANSFER_ACTIVITY | 0.00335 | 0.03967 | 0.43171 | 0.64883 | 1.23747 | 67 |
| GO_ELECTRON_TRANSPORT_CHAIN | 0.00011 | 0.00408 | 0.53843 | 0.64976 | 1.25920 | 120 |
| GO_EMBRYONIC_ORGAN_DEVELOPMENT | 0.00209 | 0.02959 | 0.43171 | 0.57957 | 1.14350 | 272 |
| GO_EMBRYO_DEVELOPMENT_ENDING_IN_BIRTH_OR_EGG_HAT | 0.00213 | 0.02984 | 0.43171 | 0.55829 | 1.11047 | 451 |
| GO_EMBRYO_IMPLANTATION | 0.00287 | 0.03609 | 0.43171 | 0.71939 | 1.33228 | 33 |
| GO_ENDOCYTIC_VESICLE | 0.00290 | 0.03627 | 0.43171 | 0.59043 | 1.15861 | 209 |
| GO_ENERGY_DERIVATION_BY_OXIDATION_OF_ORGANIC_COMP | 0.00004 | 0.00196 | 0.55733 | 0.62856 | 1.23033 | 189 |
| GO_EPIDERMAL_CELL_DIFFERENTIATION | 0.00003 | 0.00147 | 0.57561 | 0.65528 | 1.27397 | 128 |
| GO_EPIDERMIS_DEVELOPMENT | 0.00239 | 0.03218 | 0.43171 | 0.59420 | 1.16466 | 197 |
| GO_EPITHELIAL_CELL_DEVELOPMENT | 0.00191 | 0.02818 | 0.45506 | 0.60753 | 1.18362 | 146 |
| GO_EPITHELIAL_CELL_DIFFERENTIATION | 0.00000 | 0.00002 | 0.69013 | 0.61246 | 1.21428 | 364 |
| GO_EPITHELIAL_TO_MESENCHYMAL_TRANSITION | 0.00376 | 0.04279 | 0.43171 | 0.63598 | 1.22669 | 93 |
| GO_EPITHELIAL_TUBE_MORPHOGENESIS | 0.00339 | 0.03988 | 0.43171 | 0.58629 | 1.15004 | 205 |
| GO_ERYTHROCYTE_HOMEOSTASIS | 0.00053 | 0.01189 | 0.47727 | 0.66862 | 1.28426 | 81 |
| GO_ESTABLISHMENT_OF_PROTEIN_LOCALIZATION_TO_ENDOPL/ | 0.00000 | 0.00000 | | 0.84349 | 1.63010 | 102 |
| GO_ESTABLISHMENT_OF_PROTEIN_LOCALIZATION_TO_MEMBR/ | 0.00000 | 0.00000 | | 0.74060 | 1.45783 | 251 |
| GO_ESTABLISHMENT_OF_PROTEIN_LOCALIZATION_TO_ORGANE | 0.00000 | 0.00000 | | 0.68345 | 1.35976 | 429 |
| GO_ESTABLISHMENT_OF_RNA_LOCALIZATION | 0.00021 | 0.00645 | 0.51885 | 0.62072 | 1.21198 | 166 |
| GO_ESTABLISHMENT_OF_TISSUE_POLARITY | 0.00079 | 0.01604 | 0.47727 | 0.64524 | 1.24579 | 100 |
| GO_ESTABLISHMENT_OR_MAINTENANCE_OF_CELL_POLARITY | 0.00098 | 0.01839 | 0.45506 | 0.60377 | 1.18022 | 172 |
| GO_EXPORT_ACROSS_PLASMA_MEMBRANE | 0.00213 | 0.02984 | 0.43171 | 0.73082 | 1.34774 | 30 |
| GO_EXTRINSIC_COMPONENT_OF_CYTOPLASMIC_SIDE_OF_PLAS | 0.00092 | 0.01761 | 0.47727 | 0.68413 | 1.29607 | 58 |
| GO_EXTRINSIC_COMPONENT_OF_PLASMA_MEMBRANE | 0.00013 | 0.00459 | 0.51885 | 0.67199 | 1.29614 | 93 |
| GO_FC_RECEPTOR_SIGNALING_PATHWAY | 0.00082 | 0.01644 | 0.47727 | 0.61972 | 1.20603 | 136 |
| GO_FIBRILLAR_CENTER | 0.00276 | 0.03526 | 0.43171 | 0.62609 | 1.21027 | 106 |
| GO_FIBROBLAST_GROWTH_FACTOR_RECEPTOR_SIGNALING_PAT | 0.00058 | 0.01272 | 0.47727 | 0.65971 | 1.26444 | 76 |
| GO_FICOLIN_1_RICH_GRANULE | 0.00024 | 0.00733 | 0.51885 | 0.66673 | 1.28381 | 89 |
| GO_FILOPODIUM | 0.00220 | 0.03074 | 0.43171 | 0.65560 | 1.25154 | 69 |
| GO_FOREBRAIN_GENERATION_OF_NEURONS | 0.00092 | 0.01761 | 0.47727 | 0.74323 | 1.37148 | 31 |
| GO_FOREBRAIN_NEURON_DIFFERENTIATION | 0.00364 | 0.04199 | 0.43171 | 0.74533 | 1.36486 | 26 |
| GO_GENERATION_OF_PRECURSOR_METABOLITES_AND_ENERGY | 0.00000 | 0.00006 | 0.65944 | 0.61642 | 1.22121 | 326 |
| GO_GENE_SILENCING | 0.00089 | 0.01730 | 0.47727 | 0.60680 | 1.18675 | 183 |
| GO_GLAND_DEVELOPMENT | 0.00001 | 0.00089 | 0.59333 | 0.61449 | 1.21240 | 272 |
| GO_GLAND_MORPHOGENESIS | 0.00163 | 0.02561 | 0.45506 | 0.65055 | 1.24896 | 80 |
| GO_GLIAL_CELL_DEVELOPMENT | 0.00290 | 0.03627 | 0.43171 | 0.65458 | 1.24872 | 68 |
| GO_GLIAL_CELL_DIFFERENTIATION | 0.00038 | 0.00968 | 0.49849 | 0.63501 | 1.23197 | 122 |

| | | | | | | |
|---|---|---|---|---|---|---|
| GO_GLIOGENESIS | 0.00003 | 0.00171 | 0.55733 | 0.63336 | 1.23665 | 166 |
| GO_GLUCOSE_CATABOLIC_PROCESS | 0.00117 | 0.02072 | 0.45506 | 0.76802 | 1.40428 | 25 |
| GO_GLUTAMATERGIC_SYNAPSE | 0.00265 | 0.03441 | 0.43171 | 0.58411 | 1.14796 | 226 |
| GO_GLYCOLYTIC_PROCESS_THROUGH_FRUCTOSE_6_PHOSPHATE | 0.00154 | 0.02447 | 0.45506 | 0.78665 | 1.42318 | 21 |
| GO_G_PROTEIN_COUPLED_RECEPTOR_SIGNALING_PATHWAY_C( | 0.00250 | 0.03324 | 0.43171 | 0.66155 | 1.26019 | 65 |
| GO_HEAD_DEVELOPMENT | 0.00011 | 0.00397 | 0.53843 | 0.57304 | 1.14050 | 478 |
| GO_HEART_PROCESS | 0.00287 | 0.03609 | 0.43171 | 0.60741 | 1.18399 | 148 |
| GO_HEMATOPOIETIC_PROGENITOR_CELL_DIFFERENTIATION | 0.00014 | 0.00472 | 0.51885 | 0.64781 | 1.25543 | 120 |
| GO_HEMATOPOIETIC_STEM_CELL_DIFFERENTIATION | 0.00106 | 0.01925 | 0.45506 | 0.66764 | 1.27285 | 66 |
| GO_HEPATICOBILIARY_SYSTEM_DEVELOPMENT | 0.00228 | 0.03122 | 0.43171 | 0.63182 | 1.21855 | 96 |
| GO_HISTONE_BINDING | 0.00137 | 0.02252 | 0.45506 | 0.60004 | 1.17384 | 184 |
| GO_HISTONE_LYSINE_N_METHYLTRANSFERASE_ACTIVITY | 0.00179 | 0.02699 | 0.45506 | 0.72958 | 1.35116 | 33 |
| GO_HISTONE_METHYLATION | 0.00005 | 0.00226 | 0.55733 | 0.66931 | 1.29380 | 104 |
| GO_HISTONE_METHYLTRANSFERASE_ACTIVITY | 0.00039 | 0.00975 | 0.49849 | 0.72752 | 1.35861 | 41 |
| GO_HORMONE_MEDIATED_SIGNALING_PATHWAY | 0.00309 | 0.03746 | 0.43171 | 0.61505 | 1.19408 | 124 |
| GO_HUMORAL_IMMUNE_RESPONSE | 0.00013 | 0.00459 | 0.51885 | 0.69341 | 1.32281 | 68 |
| GO_IMMUNE_RESPONSE_REGULATING_SIGNALING_PATHWAY | 0.00025 | 0.00740 | 0.49849 | 0.60418 | 1.18811 | 233 |
| GO_IMPORT_INTO_NUCLEUS | 0.00034 | 0.00892 | 0.49849 | 0.64066 | 1.24157 | 120 |
| GO_INFLAMMATORY_RESPONSE_TO_ANTIGENIC_STIMULUS | 0.00028 | 0.00798 | 0.49849 | 0.81055 | 1.47456 | 23 |
| GO_INNATE_IMMUNE_RESPONSE | 0.00042 | 0.01025 | 0.49849 | 0.57408 | 1.14008 | 384 |
| GO_INNER_MITOCHONDRIAL_MEMBRANE_PROTEIN_COMPLEX | 0.00027 | 0.00793 | 0.49849 | 0.66499 | 1.27718 | 83 |
| GO_INTEGRIN_BINDING | 0.00061 | 0.01314 | 0.47727 | 0.66973 | 1.28286 | 75 |
| GO_INTERACTION_WITH_HOST | 0.00148 | 0.02395 | 0.45506 | 0.61473 | 1.19713 | 143 |
| GO_INTERACTION_WITH_SYMBIONT | 0.00350 | 0.04088 | 0.43171 | 0.67262 | 1.27105 | 54 |
| GO_INTERCALATED_DISC | 0.00272 | 0.03500 | 0.43171 | 0.72088 | 1.33505 | 33 |
| GO_INTERMEDIATE_FILAMENT_BASED_PROCESS | 0.00250 | 0.03324 | 0.43171 | 0.74489 | 1.36960 | 28 |
| GO_INTERMEDIATE_FILAMENT_CYTOSKELETON | 0.00102 | 0.01880 | 0.45506 | 0.66794 | 1.27635 | 71 |
| GO_INTRINSIC_APOPTOTIC_SIGNALING_PATHWAY | 0.00357 | 0.04144 | 0.43171 | 0.58352 | 1.14509 | 213 |
| GO_ION_CHANNEL_BINDING | 0.00268 | 0.03460 | 0.43171 | 0.64049 | 1.23011 | 83 |
| GO_KERATINIZATION | 0.00097 | 0.01821 | 0.47727 | 0.73458 | 1.36923 | 39 |
| GO_KERATINOCYTE_DIFFERENTIATION | 0.00008 | 0.00323 | 0.53843 | 0.66967 | 1.29095 | 92 |
| GO_LAMELLIPODIUM | 0.00428 | 0.04685 | 0.40702 | 0.59893 | 1.16842 | 152 |
| GO_LARGE_RIBOSOMAL_SUBUNIT | 0.00000 | 0.00000 | 0.81404 | 0.74602 | 1.44173 | 102 |
| GO_LEADING_EDGE_MEMBRANE | 0.00420 | 0.04628 | 0.40702 | 0.61957 | 1.19767 | 108 |
| GO_LEUKOCYTE_MEDIATED_IMMUNITY | 0.00000 | 0.00000 | 0.81404 | 0.62623 | 1.24504 | 404 |
| GO_LEUKOCYTE_MIGRATION | 0.00006 | 0.00256 | 0.55733 | 0.62199 | 1.21913 | 197 |
| GO_LIPOPROTEIN_PARTICLE_RECEPTOR_BINDING | 0.00130 | 0.02200 | 0.45506 | 0.82946 | 1.47118 | 15 |
| GO_LYSINE_N_METHYLTRANSFERASE_ACTIVITY | 0.00457 | 0.04849 | 0.40702 | 0.70595 | 1.31587 | 39 |
| GO_LYSOSOME_LOCALIZATION | 0.00428 | 0.04685 | 0.40702 | 0.67173 | 1.26558 | 48 |
| GO_MAINTENANCE_OF_CELL_NUMBER | 0.00004 | 0.00206 | 0.55733 | 0.66395 | 1.28472 | 110 |
| GO_MAST_CELL_ACTIVATION | 0.00039 | 0.00975 | 0.49849 | 0.76874 | 1.41525 | 29 |
| GO_MAST_CELL_MEDIATED_IMMUNITY | 0.00023 | 0.00702 | 0.51885 | 0.77709 | 1.43062 | 29 |
| GO_MEGAKARYOCYTE_DIFFERENTIATION | 0.00320 | 0.03850 | 0.43171 | 0.68790 | 1.29772 | 50 |
| GO_MEMBRANE_REGION | 0.00000 | 0.00023 | 0.62726 | 0.64235 | 1.25810 | 192 |
| GO_MESENCHYMAL_CELL_DIFFERENTIATION | 0.00309 | 0.03746 | 0.43171 | 0.60866 | 1.18431 | 134 |
| GO_METHYLATION | 0.00054 | 0.01218 | 0.47727 | 0.59391 | 1.16816 | 235 |
| GO_MIDBRAIN_DEVELOPMENT | 0.00152 | 0.02442 | 0.45506 | 0.67809 | 1.28463 | 58 |
| GO_MITOCHONDRIAL_PROTEIN_COMPLEX | 0.00003 | 0.00150 | 0.57561 | 0.62454 | 1.22311 | 191 |
| GO_MITOCHONDRION_ORGANIZATION | 0.00008 | 0.00326 | 0.53843 | 0.58106 | 1.15438 | 388 |
| GO_MODULATION_OF_PROCESS_OF_OTHER_ORGANISM | 0.00194 | 0.02819 | 0.45506 | 0.65098 | 1.24801 | 77 |
| GO_MONOSACCHARIDE_BIOSYNTHETIC_PROCESS | 0.00090 | 0.01742 | 0.47727 | 0.67190 | 1.28177 | 68 |
| GO_MONOVALENT_INORGANIC_CATION_TRANSPORT | 0.00102 | 0.01880 | 0.45506 | 0.58327 | 1.14921 | 262 |
| GO_MORPHOGENESIS_OF_AN_EPITHELIUM | 0.00006 | 0.00259 | 0.55733 | 0.58898 | 1.16778 | 358 |
| GO_MORPHOGENESIS_OF_A_POLARIZED_EPITHELIUM | 0.00011 | 0.00409 | 0.53843 | 0.65571 | 1.27059 | 118 |
| GO_MRNA_3_UTR_BINDING | 0.00075 | 0.01546 | 0.47727 | 0.67139 | 1.28370 | 72 |
| GO_MRNA_5_UTR_BINDING | 0.00224 | 0.03105 | 0.43171 | 0.75455 | 1.37965 | 25 |
| GO_MRNA_BINDING | 0.00000 | 0.00000 | | 0.69815 | 1.37394 | 247 |
| GO_MRNA_PROCESSING | 0.00000 | 0.00000 | | 0.65354 | 1.29982 | 409 |
| GO_MRNA_TRANSPORT | 0.00174 | 0.02654 | 0.45506 | 0.62071 | 1.20568 | 125 |
| GO_MULTICELLULAR_ORGANISMAL_SIGNALING | 0.00043 | 0.01032 | 0.49849 | 0.64874 | 1.25172 | 98 |
| GO_MULTI_MULTICELLULAR_ORGANISM_PROCESS | 0.00290 | 0.03627 | 0.43171 | 0.62139 | 1.20235 | 110 |
| GO_MUSCLE_CELL_DIFFERENTIATION | 0.00327 | 0.03902 | 0.43171 | 0.58585 | 1.14878 | 203 |
| GO_MUSCLE_CONTRACTION | 0.00002 | 0.00112 | 0.57561 | 0.62261 | 1.21840 | 187 |
| GO_MUSCLE_STRUCTURE_DEVELOPMENT | 0.00021 | 0.00670 | 0.51885 | 0.57934 | 1.15000 | 377 |
| GO_MUSCLE_SYSTEM_PROCESS | 0.00001 | 0.00098 | 0.59333 | 0.61386 | 1.20766 | 240 |
| GO_MUSCLE_TISSUE_DEVELOPMENT | 0.00390 | 0.04377 | 0.43171 | 0.57823 | 1.13604 | 223 |
| GO_MYELOID_CELL_DIFFERENTIATION | 0.00007 | 0.00290 | 0.53843 | 0.60366 | 1.18760 | 240 |
| GO_MYELOID_CELL_HOMEOSTASIS | 0.00205 | 0.02926 | 0.43171 | 0.63836 | 1.23088 | 95 |
| GO_MYELOID_LEUKOCYTE_ACTIVATION | 0.00000 | 0.00000 | 0.77494 | 0.63009 | 1.24901 | 352 |
| GO_MYELOID_LEUKOCYTE_MEDIATED_IMMUNITY | 0.00000 | 0.00000 | 0.76146 | 0.63488 | 1.25692 | 315 |

| | | | | | | |
|---|---|---|---|---|---|---|
| GO_M_BAND | 0.00061 | 0.01314 | 0.47727 | 0.83678 | 1.49678 | 17 |
| GO_NATURAL_KILLER_CELL_MEDIATED_IMMUNITY | 0.00085 | 0.01667 | 0.47727 | 0.77170 | 1.41101 | 25 |
| GO_NCRNA_METABOLIC_PROCESS | 0.00342 | 0.04017 | 0.43171 | 0.56529 | 1.12062 | 351 |
| GO_NCRNA_PROCESSING | 0.00165 | 0.02583 | 0.45506 | 0.57783 | 1.14305 | 305 |
| GO_NEGATIVE_REGULATION_OF_CATABOLIC_PROCESS | 0.00062 | 0.01329 | 0.47727 | 0.60316 | 1.18359 | 209 |
| GO_NEGATIVE_REGULATION_OF_CELLULAR_AMIDE_METABOLIC | 0.00002 | 0.00125 | 0.57561 | 0.66272 | 1.28431 | 120 |
| GO_NEGATIVE_REGULATION_OF_CELLULAR_CATABOLIC_PROCES | 0.00124 | 0.02141 | 0.45506 | 0.60649 | 1.18544 | 174 |
| GO_NEGATIVE_REGULATION_OF_CELLULAR_COMPONENT_ORG | 0.00167 | 0.02599 | 0.45506 | 0.55650 | 1.10781 | 482 |
| GO_NEGATIVE_REGULATION_OF_CELL_CYCLE | 0.00209 | 0.02959 | 0.43171 | 0.55854 | 1.11134 | 437 |
| GO_NEGATIVE_REGULATION_OF_CELL_DIFFERENTIATION | 0.00361 | 0.04164 | 0.43171 | 0.55992 | 1.11295 | 401 |
| GO_NEGATIVE_REGULATION_OF_DENDRITE_DEVELOPMENT | 0.00405 | 0.04495 | 0.40702 | 0.80075 | 1.42592 | 16 |
| GO_NEGATIVE_REGULATION_OF_HYDROLASE_ACTIVITY | 0.00156 | 0.02470 | 0.45506 | 0.58443 | 1.14902 | 232 |
| GO_NEGATIVE_REGULATION_OF_MITOTIC_CELL_CYCLE | 0.00361 | 0.04164 | 0.43171 | 0.58395 | 1.14834 | 233 |
| GO_NEGATIVE_REGULATION_OF_MRNA_METABOLIC_PROCESS | 0.00053 | 0.01189 | 0.47727 | 0.68984 | 1.31251 | 64 |
| GO_NEGATIVE_REGULATION_OF_PROTEIN_LOCALIZATION_TO_C | 0.00361 | 0.04164 | 0.43171 | 0.80481 | 1.43315 | 16 |
| GO_NEGATIVE_REGULATION_OF_RNA_CATABOLIC_PROCESS | 0.00012 | 0.00431 | 0.53843 | 0.73078 | 1.37297 | 46 |
| GO_NEGATIVE_REGULATION_OF_RNA_SPLICING | 0.00302 | 0.03706 | 0.43171 | 0.76254 | 1.38723 | 23 |
| GO_NEGATIVE_REGULATION_OF_TRANSPORT | 0.00435 | 0.04725 | 0.40702 | 0.57243 | 1.12637 | 243 |
| GO_NEURAL_NUCLEUS_DEVELOPMENT | 0.00205 | 0.02926 | 0.43171 | 0.71959 | 1.33679 | 36 |
| GO_NEURON_DEATH | 0.00372 | 0.04245 | 0.43171 | 0.57863 | 1.13551 | 214 |
| GO_NON_CANONICAL_WNT_SIGNALING_PATHWAY | 0.00047 | 0.01114 | 0.49849 | 0.63449 | 1.22836 | 115 |
| GO_NUCLEAR_ENVELOPE | 0.00213 | 0.02984 | 0.43171 | 0.57171 | 1.13277 | 332 |
| GO_NUCLEAR_EXPORT | 0.00003 | 0.00152 | 0.57561 | 0.62821 | 1.22776 | 173 |
| GO_NUCLEAR_PERIPHERY | 0.00405 | 0.04495 | 0.40702 | 0.62028 | 1.20081 | 113 |
| GO_NUCLEAR_SPECK | 0.00000 | 0.00001 | 0.71951 | 0.63013 | 1.24622 | 298 |
| GO_NUCLEAR_TRANSCRIBED_MRNA_CATABOLIC_PROCESS | 0.00000 | 0.00000 | | 0.75296 | 1.47172 | 174 |
| GO_NUCLEAR_TRANSCRIBED_MRNA_CATABOLIC_PROCESS_NON | 0.00000 | 0.00000 | | 0.84082 | 1.62761 | 112 |
| GO_NUCLEAR_TRANSPORT | 0.00000 | 0.00000 | 0.73376 | 0.63693 | 1.25794 | 279 |
| GO_NUCLEOBASE_CONTAINING_COMPOUND_TRANSPORT | 0.00049 | 0.01152 | 0.47727 | 0.60238 | 1.18112 | 201 |
| GO_NUCLEOSOME_ORGANIZATION | 0.00246 | 0.03303 | 0.43171 | 0.64704 | 1.24352 | 84 |
| GO_N_METHYLTRANSFERASE_ACTIVITY | 0.00110 | 0.01969 | 0.45506 | 0.68101 | 1.29015 | 58 |
| GO_ORGANELLE_INNER_MEMBRANE | 0.00014 | 0.00477 | 0.51885 | 0.58067 | 1.15122 | 362 |
| GO_ORGANIC_CYCLIC_COMPOUND_CATABOLIC_PROCESS | 0.00000 | 0.00000 | | 0.69921 | 1.39129 | 439 |
| GO_OXIDATIVE_PHOSPHORYLATION | 0.00002 | 0.00136 | 0.57561 | 0.68239 | 1.31449 | 91 |
| GO_OXIDOREDUCTASE_ACTIVITY | 0.00024 | 0.00733 | 0.51885 | 0.57693 | 1.14655 | 396 |
| GO_OXIDOREDUCTASE_ACTIVITY_ACTING_ON_A_HEME_GROUP | 0.00119 | 0.02087 | 0.45506 | 0.81103 | 1.45527 | 18 |
| GO_OXIDOREDUCTASE_ACTIVITY_ACTING_ON_NAD_P_H | 0.00253 | 0.03352 | 0.43171 | 0.65282 | 1.24506 | 67 |
| GO_OXIDOREDUCTASE_COMPLEX | 0.00067 | 0.01418 | 0.47727 | 0.66802 | 1.27958 | 75 |
| GO_PEPTIDE_BINDING | 0.00028 | 0.00810 | 0.49849 | 0.63494 | 1.23639 | 141 |
| GO_PEPTIDE_BIOSYNTHETIC_PROCESS | 0.00000 | 0.00000 | | 0.67981 | 1.35341 | 500 |
| GO_PEPTIDYL_LYSINE_METHYLATION | 0.00019 | 0.00622 | 0.51885 | 0.66886 | 1.28791 | 89 |
| GO_PEPTIDYL_LYSINE_MODIFICATION | 0.00050 | 0.01170 | 0.47727 | 0.58632 | 1.15904 | 292 |
| GO_PERINUCLEAR_REGION_OF_CYTOPLASM | 0.00002 | 0.00113 | 0.57561 | 0.58073 | 1.15597 | 483 |
| GO_PHAGOCYTOSIS | 0.00168 | 0.02608 | 0.45506 | 0.60724 | 1.18454 | 155 |
| GO_PHOSPHATIDYLINOSITOL_3_KINASE_BINDING | 0.00298 | 0.03675 | 0.43171 | 0.77075 | 1.39441 | 21 |
| GO_PIGMENT_GRANULE | 0.00000 | 0.00009 | 0.65944 | 0.72664 | 1.39440 | 79 |
| GO_PLASMA_LIPOPROTEIN_PARTICLE_CLEARANCE | 0.00194 | 0.02819 | 0.45506 | 0.72092 | 1.33927 | 36 |
| GO_PLASMA_MEMBRANE_ORGANIZATION | 0.00231 | 0.03145 | 0.43171 | 0.65454 | 1.25073 | 71 |
| GO_POLYMERIC_CYTOSKELETAL_FIBER | 0.00007 | 0.00290 | 0.53843 | 0.57932 | 1.15257 | 432 |
| GO_POLYSOMAL_RIBOSOME | 0.00000 | 0.00002 | 0.69013 | 0.86609 | 1.59244 | 28 |
| GO_POLYSOME | 0.00000 | 0.00000 | 0.76146 | 0.79782 | 1.51323 | 60 |
| GO_POLY_A_BINDING | 0.00305 | 0.03715 | 0.43171 | 0.79749 | 1.42649 | 17 |
| GO_POLY_PURINE_TRACT_BINDING | 0.00265 | 0.03441 | 0.43171 | 0.76575 | 1.38674 | 22 |
| GO_POLY_PYRIMIDINE_TRACT_BINDING | 0.00154 | 0.02447 | 0.45506 | 0.77865 | 1.41654 | 23 |
| GO_POSITIVE_REGULATION_OF_BINDING | 0.00013 | 0.00459 | 0.51885 | 0.64804 | 1.25550 | 117 |
| GO_POSITIVE_REGULATION_OF_CELLULAR_AMIDE_METABOLIC_ | 0.00002 | 0.00105 | 0.57561 | 0.66580 | 1.28897 | 115 |
| GO_POSITIVE_REGULATION_OF_CELLULAR_COMPONENT_MOVI | 0.00042 | 0.01025 | 0.49849 | 0.58302 | 1.15445 | 320 |
| GO_POSITIVE_REGULATION_OF_CELLULAR_PROTEIN_LOCALIZAT | 0.00001 | 0.00089 | 0.59333 | 0.60999 | 1.20188 | 263 |
| GO_POSITIVE_REGULATION_OF_CELL_ACTIVATION | 0.00033 | 0.00876 | 0.49849 | 0.62499 | 1.21825 | 148 |
| GO_POSITIVE_REGULATION_OF_CELL_ADHESION | 0.00006 | 0.00270 | 0.53843 | 0.60969 | 1.19870 | 231 |
| GO_POSITIVE_REGULATION_OF_CELL_CYCLE | 0.00058 | 0.01272 | 0.47727 | 0.58592 | 1.15566 | 269 |
| GO_POSITIVE_REGULATION_OF_CELL_CYCLE_PROCESS | 0.00457 | 0.04849 | 0.40702 | 0.58616 | 1.14921 | 200 |
| GO_POSITIVE_REGULATION_OF_CELL_DEATH | 0.00052 | 0.01189 | 0.47727 | 0.56503 | 1.12391 | 453 |
| GO_POSITIVE_REGULATION_OF_CELL_DEVELOPMENT | 0.00006 | 0.00258 | 0.55733 | 0.58993 | 1.16917 | 336 |
| GO_POSITIVE_REGULATION_OF_CELL_MATRIX_ADHESION | 0.00420 | 0.04628 | 0.40702 | 0.70597 | 1.31397 | 37 |
| GO_POSITIVE_REGULATION_OF_CELL_POPULATION_PROLIFERAT | 0.00000 | 0.00013 | 0.64355 | 0.59094 | 1.17647 | 486 |
| GO_POSITIVE_REGULATION_OF_CELL_SUBSTRATE_ADHESION | 0.00070 | 0.01461 | 0.47727 | 0.66955 | 1.28214 | 74 |
| GO_POSITIVE_REGULATION_OF_CYTOKINE_PRODUCTION | 0.00011 | 0.00408 | 0.53843 | 0.60737 | 1.19192 | 214 |
| GO_POSITIVE_REGULATION_OF_CYTOKINE_PRODUCTION_INVO | 0.00253 | 0.03352 | 0.43171 | 0.75542 | 1.38333 | 26 |
| GO_POSITIVE_REGULATION_OF_DEFENSE_RESPONSE | 0.00028 | 0.00810 | 0.49849 | 0.60410 | 1.18337 | 193 |

| | | | | | | |
|---|---|---|---|---|---|---|
| GO_POSITIVE_REGULATION_OF_DNA_BINDING | 0.00011 | 0.00408 | 0.53843 | 0.76359 | 1.42329 | 39 |
| GO_POSITIVE_REGULATION_OF_DNA_METABOLIC_PROCESS | 0.00228 | 0.03122 | 0.43171 | 0.61192 | 1.19117 | 137 |
| GO_POSITIVE_REGULATION_OF_ESTABLISHMENT_OF_PROTEIN_ | 0.00000 | 0.00032 | 0.61053 | 0.61617 | 1.21450 | 266 |
| GO_POSITIVE_REGULATION_OF_IMMUNE_EFFECTOR_PROCESS | 0.00002 | 0.00126 | 0.57561 | 0.69152 | 1.33381 | 93 |
| GO_POSITIVE_REGULATION_OF_IMMUNE_RESPONSE | 0.00000 | 0.00008 | 0.65944 | 0.61456 | 1.21754 | 326 |
| GO_POSITIVE_REGULATION_OF_IMMUNE_SYSTEM_PROCESS | 0.00000 | 0.00000 | 0.74774 | 0.60540 | 1.20505 | 477 |
| GO_POSITIVE_REGULATION_OF_INNATE_IMMUNE_RESPONSE | 0.00168 | 0.02608 | 0.45506 | 0.62130 | 1.20682 | 125 |
| GO_POSITIVE_REGULATION_OF_INTRACELLULAR_PROTEIN_TRA | 0.00083 | 0.01646 | 0.47727 | 0.61531 | 1.19971 | 149 |
| GO_POSITIVE_REGULATION_OF_INTRACELLULAR_TRANSPORT | 0.00009 | 0.00362 | 0.53843 | 0.62384 | 1.21976 | 177 |
| GO_POSITIVE_REGULATION_OF_ION_TRANSMEMBRANE_TRANS | 0.00043 | 0.01025 | 0.49849 | 0.66250 | 1.26978 | 76 |
| GO_POSITIVE_REGULATION_OF_LEUKOCYTE_CELL_CELL_ADHES | 0.00302 | 0.03706 | 0.43171 | 0.62596 | 1.21004 | 107 |
| GO_POSITIVE_REGULATION_OF_LEUKOCYTE_MEDIATED_IMMU | 0.00159 | 0.02516 | 0.45506 | 0.68223 | 1.29094 | 55 |
| GO_POSITIVE_REGULATION_OF_LYMPHOCYTE_ACTIVATION | 0.00132 | 0.02208 | 0.45506 | 0.61946 | 1.20500 | 133 |
| GO_POSITIVE_REGULATION_OF_LYMPHOCYTE_MEDIATED_IMM | 0.00331 | 0.03931 | 0.43171 | 0.69392 | 1.29587 | 41 |
| GO_POSITIVE_REGULATION_OF_MRNA_METABOLIC_PROCESS | 0.00085 | 0.01667 | 0.47727 | 0.67534 | 1.28337 | 62 |
| GO_POSITIVE_REGULATION_OF_NERVOUS_SYSTEM_DEVELOPM | 0.00261 | 0.03414 | 0.43171 | 0.57206 | 1.13309 | 323 |
| GO_POSITIVE_REGULATION_OF_NEURAL_PRECURSOR_CELL_PR | 0.00465 | 0.04887 | 0.40702 | 0.73193 | 1.34577 | 28 |
| GO_POSITIVE_REGULATION_OF_NITRIC_OXIDE_METABOLIC_PR | 0.00450 | 0.04803 | 0.40702 | 0.75363 | 1.36480 | 22 |
| GO_POSITIVE_REGULATION_OF_NUCLEOCYTOPLASMIC_TRANSP | 0.00450 | 0.04803 | 0.40702 | 0.66373 | 1.25586 | 56 |
| GO_POSITIVE_REGULATION_OF_ORGANELLE_ORGANIZATION | 0.00003 | 0.00153 | 0.57561 | 0.58005 | 1.15416 | 463 |
| GO_POSITIVE_REGULATION_OF_PHAGOCYTOSIS | 0.00228 | 0.03122 | 0.43171 | 0.74097 | 1.36412 | 29 |
| GO_POSITIVE_REGULATION_OF_PRODUCTION_OF_MOLECULAR | 0.00013 | 0.00459 | 0.51885 | 0.73672 | 1.38186 | 45 |
| GO_POSITIVE_REGULATION_OF_PROTEIN_KINASE_B_SIGNALING | 0.00053 | 0.01189 | 0.47727 | 0.65039 | 1.25281 | 90 |
| GO_POSITIVE_REGULATION_OF_PROTEIN_LOCALIZATION_TO_N | 0.00135 | 0.02251 | 0.45506 | 0.67265 | 1.27749 | 61 |
| GO_POSITIVE_REGULATION_OF_RESPONSE_TO_BIOTIC_STIMUL | 0.00187 | 0.02770 | 0.45506 | 0.61281 | 1.19339 | 143 |
| GO_POSITIVE_REGULATION_OF_RESPONSE_TO_EXTERNAL_STIM | 0.00079 | 0.01604 | 0.47727 | 0.58882 | 1.15962 | 256 |
| GO_POSITIVE_REGULATION_OF_SODIUM_ION_TRANSPORT | 0.00287 | 0.03609 | 0.43171 | 0.76377 | 1.38315 | 22 |
| GO_POSITIVE_REGULATION_OF_TRANSFERASE_ACTIVITY | 0.00383 | 0.04325 | 0.43171 | 0.55627 | 1.10688 | 439 |
| GO_POSITIVE_REGULATION_OF_TRANSLATION | 0.00002 | 0.00110 | 0.57561 | 0.68205 | 1.31543 | 96 |
| GO_POSITIVE_REGULATION_OF_TRANSMEMBRANE_TRANSPORT | 0.00083 | 0.01646 | 0.47727 | 0.63709 | 1.23154 | 108 |
| GO_POSITIVE_REGULATION_OF_T_CELL_MEDIATED_IMMUNITY | 0.00442 | 0.04764 | 0.40702 | 0.75444 | 1.36626 | 22 |
| GO_POSITIVE_REGULATION_OF_VIRAL_PROCESS | 0.00039 | 0.00975 | 0.49849 | 0.66267 | 1.27503 | 88 |
| GO_POSITIVE_REGULATION_OF_WNT_SIGNALING_PATHWAY | 0.00141 | 0.02294 | 0.45506 | 0.62135 | 1.20829 | 130 |
| GO_POSTSYNAPSE | 0.00086 | 0.01679 | 0.47727 | 0.57243 | 1.13680 | 385 |
| GO_POSTTRANSCRIPTIONAL_REGULATION_OF_GENE_EXPRESSI | 0.00000 | 0.00000 | | 0.64046 | 1.27432 | 464 |
| GO_PRECATALYTIC_SPLICEOSOME | 0.00390 | 0.04377 | 0.43171 | 0.68308 | 1.28862 | 50 |
| GO_PRE_MRNA_BINDING | 0.00016 | 0.00511 | 0.51885 | 0.76396 | 1.41552 | 34 |
| GO_PRODUCTION_OF_MOLECULAR_MEDIATOR_OF_IMMUNE_ | 0.00276 | 0.03526 | 0.43171 | 0.62615 | 1.21035 | 103 |
| GO_PROMOTER_SPECIFIC_CHROMATIN_BINDING | 0.00213 | 0.02984 | 0.43171 | 0.71452 | 1.33671 | 42 |
| GO_PROSTATE_GLAND_DEVELOPMENT | 0.00357 | 0.04144 | 0.43171 | 0.70880 | 1.31332 | 34 |
| GO_PROTEASOMAL_PROTEIN_CATABOLIC_PROCESS | 0.00305 | 0.03715 | 0.43171 | 0.56243 | 1.11506 | 362 |
| GO_PROTEIN_CONTAINING_COMPLEX_LOCALIZATION | 0.00095 | 0.01797 | 0.47727 | 0.59194 | 1.16182 | 215 |
| GO_PROTEIN_C_TERMINUS_BINDING | 0.00026 | 0.00756 | 0.49849 | 0.62463 | 1.21844 | 151 |
| GO_PROTEIN_DNA_COMPLEX_SUBUNIT_ORGANIZATION | 0.00076 | 0.01555 | 0.47727 | 0.61151 | 1.19365 | 162 |
| GO_PROTEIN_DOMAIN_SPECIFIC_BINDING | 0.00000 | 0.00002 | 0.69013 | 0.59870 | 1.19189 | 487 |
| GO_PROTEIN_FOLDING | 0.00063 | 0.01344 | 0.47727 | 0.61759 | 1.20462 | 153 |
| GO_PROTEIN_HOMODIMERIZATION_ACTIVITY | 0.00185 | 0.02756 | 0.45506 | 0.56463 | 1.12145 | 386 |
| GO_PROTEIN_IMPORT | 0.00043 | 0.01025 | 0.49849 | 0.62508 | 1.21777 | 144 |
| GO_PROTEIN_LIPID_COMPLEX | 0.00174 | 0.02654 | 0.45506 | 0.81420 | 1.45638 | 17 |
| GO_PROTEIN_LOCALIZATION_TO_CELL_PERIPHERY | 0.00011 | 0.00408 | 0.53843 | 0.60117 | 1.18270 | 240 |
| GO_PROTEIN_LOCALIZATION_TO_ENDOPLASMIC_RETICULUM | 0.00000 | 0.00000 | | 0.81551 | 1.58327 | 124 |
| GO_PROTEIN_LOCALIZATION_TO_MEMBRANE | 0.00000 | 0.00000 | | 0.69064 | 1.37439 | 467 |
| GO_PROTEIN_LOCALIZATION_TO_MITOCHONDRION | 0.00324 | 0.03880 | 0.43171 | 0.61447 | 1.19075 | 119 |
| GO_PROTEIN_LOCALIZATION_TO_NUCLEUS | 0.00003 | 0.00171 | 0.55733 | 0.62078 | 1.21649 | 196 |
| GO_PROTEIN_LOCALIZATION_TO_PLASMA_MEMBRANE | 0.00011 | 0.00405 | 0.53843 | 0.60732 | 1.19154 | 208 |
| GO_PROTEIN_METHYLATION | 0.00004 | 0.00183 | 0.55733 | 0.65633 | 1.27333 | 122 |
| GO_PROTEIN_METHYLTRANSFERASE_ACTIVITY | 0.00450 | 0.04803 | 0.40702 | 0.66349 | 1.25540 | 56 |
| GO_PROTEIN_POLYUBIQUITINATION | 0.00457 | 0.04849 | 0.40702 | 0.57438 | 1.13118 | 256 |
| GO_PROTEIN_STABILIZATION | 0.00025 | 0.00750 | 0.49849 | 0.63507 | 1.23537 | 133 |
| GO_PROTEIN_TARGETING | 0.00000 | 0.00000 | | 0.70312 | 1.39267 | 323 |
| GO_PROTEIN_TARGETING_TO_MEMBRANE | 0.00000 | 0.00000 | | 0.78619 | 1.53246 | 148 |
| GO_PROTEIN_TARGETING_TO_MITOCHONDRION | 0.00442 | 0.04764 | 0.40702 | 0.63529 | 1.21967 | 80 |
| GO_PROTON_TRANSMEMBRANE_TRANSPORT | 0.00043 | 0.01025 | 0.49849 | 0.66675 | 1.28186 | 85 |
| GO_PROTON_TRANSMEMBRANE_TRANSPORTER_ACTIVITY | 0.00313 | 0.03776 | 0.43171 | 0.65067 | 1.24212 | 69 |
| GO_PYRUVATE_METABOLIC_PROCESS | 0.00383 | 0.04325 | 0.43171 | 0.62606 | 1.20795 | 98 |
| GO_RECEPTOR_MEDIATED_ENDOCYTOSIS | 0.00022 | 0.00685 | 0.51885 | 0.62425 | 1.21772 | 155 |
| GO_REGENERATION | 0.00198 | 0.02853 | 0.43171 | 0.61918 | 1.19871 | 115 |
| GO_REGULATION_OF_ACTIN_FILAMENT_BASED_MOVEMENT | 0.00124 | 0.02141 | 0.45506 | 0.77038 | 1.41073 | 26 |
| GO_REGULATION_OF_ACTIN_FILAMENT_BASED_PROCESS | 0.00003 | 0.00164 | 0.55733 | 0.60633 | 1.19463 | 262 |
| GO_REGULATION_OF_ACTIN_FILAMENT_BUNDLE_ASSEMBLY | 0.00383 | 0.04325 | 0.43171 | 0.64250 | 1.22942 | 73 |

| | | | | | | |
|---|---|---|---|---|---|---|
| GO_REGULATION_OF_ACTIN_FILAMENT_ORGANIZATION | 0.00137 | 0.02252 | 0.45506 | 0.59196 | 1.15842 | 187 |
| GO_REGULATION_OF_ANIMAL_ORGAN_MORPHOGENESIS | 0.00257 | 0.03380 | 0.43171 | 0.59966 | 1.17067 | 164 |
| GO_REGULATION_OF_APOPTOTIC_SIGNALING_PATHWAY | 0.00130 | 0.02200 | 0.45506 | 0.58047 | 1.14491 | 269 |
| GO_REGULATION_OF_BINDING | 0.00007 | 0.00290 | 0.53843 | 0.60729 | 1.19528 | 246 |
| GO_REGULATION_OF_CARDIAC_MUSCLE_CONTRACTION | 0.00087 | 0.01692 | 0.47727 | 0.71980 | 1.34848 | 44 |
| GO_REGULATION_OF_CELLULAR_AMIDE_METABOLIC_PROCESS | 0.00000 | 0.00000 | 0.76146 | 0.63794 | 1.26278 | 312 |
| GO_REGULATION_OF_CELLULAR_PROTEIN_CATABOLIC_PROCESS | 0.00428 | 0.04685 | 0.40702 | 0.58160 | 1.13901 | 191 |
| GO_REGULATION_OF_CELLULAR_PROTEIN_LOCALIZATION | 0.00000 | 0.00001 | 0.70498 | 0.60730 | 1.20772 | 416 |
| GO_REGULATION_OF_CELL_ACTIVATION | 0.00450 | 0.04803 | 0.40702 | 0.57183 | 1.12591 | 254 |
| GO_REGULATION_OF_CELL_ADHESION | 0.00030 | 0.00835 | 0.49849 | 0.57493 | 1.14267 | 398 |
| GO_REGULATION_OF_CELL_CYCLE_G2_M_PHASE_TRANSITION | 0.00372 | 0.04245 | 0.43171 | 0.59215 | 1.15679 | 169 |
| GO_REGULATION_OF_CELL_CYCLE_PHASE_TRANSITION | 0.00040 | 0.00994 | 0.49849 | 0.58048 | 1.15082 | 345 |
| GO_REGULATION_OF_CELL_MATRIX_ADHESION | 0.00383 | 0.04325 | 0.43171 | 0.63854 | 1.22620 | 82 |
| GO_REGULATION_OF_CELL_MORPHOGENESIS | 0.00095 | 0.01797 | 0.47727 | 0.57797 | 1.14444 | 320 |
| GO_REGULATION_OF_CELL_SHAPE | 0.00185 | 0.02756 | 0.45506 | 0.62921 | 1.21668 | 105 |
| GO_REGULATION_OF_CELL_SUBSTRATE_ADHESION | 0.00013 | 0.00459 | 0.51885 | 0.63777 | 1.24115 | 136 |
| GO_REGULATION_OF_CHROMOSOME_ORGANIZATION | 0.00178 | 0.02684 | 0.45506 | 0.57747 | 1.13899 | 269 |
| GO_REGULATION_OF_CYTOKINE_PRODUCTION_INVOLVED_IN_I | 0.00074 | 0.01532 | 0.47727 | 0.73599 | 1.36370 | 34 |
| GO_REGULATION_OF_CYTOSKELETON_ORGANIZATION | 0.00037 | 0.00956 | 0.49849 | 0.57470 | 1.14132 | 384 |
| GO_REGULATION_OF_DEFENSE_RESPONSE | 0.00324 | 0.03880 | 0.43171 | 0.56546 | 1.12096 | 351 |
| GO_REGULATION_OF_DNA_BINDING | 0.00068 | 0.01428 | 0.47727 | 0.65548 | 1.25891 | 83 |
| GO_REGULATION_OF_DNA_METABOLIC_PROCESS | 0.00331 | 0.03931 | 0.43171 | 0.57910 | 1.13964 | 244 |
| GO_REGULATION_OF_DNA_TEMPLATED_TRANSCRIPTION_IN_RE | 0.00279 | 0.03559 | 0.43171 | 0.63047 | 1.21443 | 90 |
| GO_REGULATION_OF_GENE_EXPRESSION_EPIGENETIC | 0.00030 | 0.00835 | 0.49849 | 0.63534 | 1.23326 | 123 |
| GO_REGULATION_OF_GLIOGENESIS | 0.00198 | 0.02853 | 0.43171 | 0.66717 | 1.26808 | 63 |
| GO_REGULATION_OF_HEART_RATE | 0.00020 | 0.00630 | 0.51885 | 0.74432 | 1.39442 | 44 |
| GO_REGULATION_OF_HEART_RATE_BY_CARDIAC_CONDUCTION | 0.00298 | 0.03675 | 0.43171 | 0.79301 | 1.42293 | 18 |
| GO_REGULATION_OF_HEMATOPOIETIC_PROGENITOR_CELL_DIF | 0.00031 | 0.00840 | 0.49849 | 0.68122 | 1.29874 | 66 |
| GO_REGULATION_OF_HEMATOPOIETIC_STEM_CELL_DIFFERENT | 0.00122 | 0.02127 | 0.45506 | 0.68311 | 1.29253 | 56 |
| GO_REGULATION_OF_HEMOPOIESIS | 0.00019 | 0.00621 | 0.51885 | 0.59601 | 1.17638 | 274 |
| GO_REGULATION_OF_IMMUNE_EFFECTOR_PROCESS | 0.00032 | 0.00854 | 0.49849 | 0.61247 | 1.19680 | 170 |
| GO_REGULATION_OF_IMMUNE_RESPONSE | 0.00000 | 0.00026 | 0.62726 | 0.59310 | 1.17979 | 427 |
| GO_REGULATION_OF_INNATE_IMMUNE_RESPONSE | 0.00194 | 0.02819 | 0.45506 | 0.59635 | 1.16550 | 173 |
| GO_REGULATION_OF_INTRACELLULAR_PROTEIN_TRANSPORT | 0.00002 | 0.00105 | 0.57561 | 0.62040 | 1.21721 | 208 |
| GO_REGULATION_OF_INTRACELLULAR_TRANSPORT | 0.00000 | 0.00005 | 0.67496 | 0.62489 | 1.23253 | 270 |
| GO_REGULATION_OF_LEUKOCYTE_DEGRANULATION | 0.00465 | 0.04887 | 0.40702 | 0.75282 | 1.36334 | 22 |
| GO_REGULATION_OF_LEUKOCYTE_MEDIATED_IMMUNITY | 0.00076 | 0.01555 | 0.47727 | 0.66199 | 1.27272 | 85 |
| GO_REGULATION_OF_METAL_ION_TRANSPORT | 0.00104 | 0.01908 | 0.45506 | 0.59805 | 1.17089 | 190 |
| GO_REGULATION_OF_MITOTIC_CELL_CYCLE | 0.00001 | 0.00089 | 0.59333 | 0.57902 | 1.15233 | 472 |
| GO_REGULATION_OF_MORPHOGENESIS_OF_AN_EPITHELIUM | 0.00068 | 0.01428 | 0.47727 | 0.63124 | 1.22614 | 125 |
| GO_REGULATION_OF_MRNA_CATABOLIC_PROCESS | 0.00000 | 0.00011 | 0.64355 | 0.65970 | 1.28665 | 154 |
| GO_REGULATION_OF_MRNA_METABOLIC_PROCESS | 0.00000 | 0.00000 | | 0.67186 | 1.32392 | 264 |
| GO_REGULATION_OF_MRNA_PROCESSING | 0.00000 | 0.00005 | 0.65944 | 0.68655 | 1.33355 | 125 |
| GO_REGULATION_OF_MRNA_SPLICING_VIA_SPLICEOSOME | 0.00001 | 0.00074 | 0.59333 | 0.69034 | 1.32980 | 91 |
| GO_REGULATION_OF_MUSCLE_CONTRACTION | 0.00016 | 0.00516 | 0.51885 | 0.67586 | 1.29695 | 79 |
| GO_REGULATION_OF_MUSCLE_SYSTEM_PROCESS | 0.00117 | 0.02072 | 0.45506 | 0.62401 | 1.21127 | 123 |
| GO_REGULATION_OF_NEURAL_PRECURSOR_CELL_PROLIFERATIO | 0.00030 | 0.00835 | 0.49849 | 0.72046 | 1.35731 | 49 |
| GO_REGULATION_OF_NUCLEOCYTOPLASMIC_TRANSPORT | 0.00009 | 0.00359 | 0.53843 | 0.66807 | 1.28689 | 91 |
| GO_REGULATION_OF_PEPTIDASE_ACTIVITY | 0.00113 | 0.02024 | 0.45506 | 0.58668 | 1.15388 | 234 |
| GO_REGULATION_OF_PEPTIDE_TRANSPORT | 0.00000 | 0.00020 | 0.62726 | 0.59588 | 1.18505 | 422 |
| GO_REGULATION_OF_PHAGOCYTOSIS | 0.00052 | 0.01189 | 0.47727 | 0.72987 | 1.36568 | 43 |
| GO_REGULATION_OF_PLASMA_LIPOPROTEIN_PARTICLE_LEVELS | 0.00132 | 0.02208 | 0.45506 | 0.69072 | 1.30536 | 53 |
| GO_REGULATION_OF_PRODUCTION_OF_MOLECULAR_MEDIATO | 0.00009 | 0.00353 | 0.53843 | 0.70847 | 1.34551 | 61 |
| GO_REGULATION_OF_PROTEIN_CATABOLIC_PROCESS | 0.00119 | 0.02087 | 0.45506 | 0.58167 | 1.14961 | 290 |
| GO_REGULATION_OF_PROTEIN_CONTAINING_COMPLEX_ASSEM | 0.00174 | 0.02654 | 0.45506 | 0.57730 | 1.14200 | 305 |
| GO_REGULATION_OF_PROTEIN_EXPORT_FROM_NUCLEUS | 0.00313 | 0.03776 | 0.43171 | 0.70361 | 1.31134 | 38 |
| GO_REGULATION_OF_PROTEIN_LOCALIZATION_TO_CELL_PERIPH | 0.00250 | 0.03324 | 0.43171 | 0.63221 | 1.21778 | 90 |
| GO_REGULATION_OF_PROTEIN_LOCALIZATION_TO_MEMBRANE | 0.00435 | 0.04725 | 0.40702 | 0.60498 | 1.17767 | 137 |
| GO_REGULATION_OF_PROTEIN_LOCALIZATION_TO_NUCLEUS | 0.00465 | 0.04887 | 0.40702 | 0.62620 | 1.20624 | 91 |
| GO_REGULATION_OF_PROTEIN_LOCALIZATION_TO_PLASMA_MI | 0.00108 | 0.01947 | 0.45506 | 0.65014 | 1.24610 | 76 |
| GO_REGULATION_OF_PROTEIN_STABILITY | 0.00023 | 0.00720 | 0.51885 | 0.60362 | 1.18476 | 216 |
| GO_REGULATION_OF_PROTEOLYSIS | 0.00390 | 0.04377 | 0.43171 | 0.55462 | 1.10349 | 435 |
| GO_REGULATION_OF_RNA_SPLICING | 0.00000 | 0.00011 | 0.64355 | 0.67788 | 1.31753 | 127 |
| GO_REGULATION_OF_SMALL_MOLECULE_METABOLIC_PROCESS | 0.00167 | 0.02599 | 0.45506 | 0.57985 | 1.14455 | 275 |
| GO_REGULATION_OF_STEM_CELL_DIFFERENTIATION | 0.00004 | 0.00178 | 0.55733 | 0.68412 | 1.31391 | 83 |
| GO_REGULATION_OF_STEM_CELL_PROLIFERATION | 0.00154 | 0.02447 | 0.45506 | 0.70963 | 1.32943 | 44 |
| GO_REGULATION_OF_STEROID_BIOSYNTHETIC_PROCESS | 0.00100 | 0.01868 | 0.45506 | 0.67580 | 1.28070 | 59 |
| GO_REGULATION_OF_STEROID_METABOLIC_PROCESS | 0.00126 | 0.02167 | 0.45506 | 0.66108 | 1.26591 | 74 |
| GO_REGULATION_OF_STRIATED_MUSCLE_CONTRACTION | 0.00061 | 0.01314 | 0.47727 | 0.70860 | 1.33271 | 47 |
| GO_REGULATION_OF_SUBSTRATE_ADHESION_DEPENDENT_CEL | 0.00194 | 0.02819 | 0.45506 | 0.71998 | 1.33403 | 34 |

| | | | | | | |
|---|---|---|---|---|---|---|
| GO_REGULATION_OF_SUPRAMOLECULAR_FIBER_ORGANIZATIOI | 0.00064 | 0.01350 | 0.47727 | 0.58748 | 1.15690 | 255 |
| GO_REGULATION_OF_SYMBIOTIC_PROCESS | 0.00028 | 0.00810 | 0.49849 | 0.62900 | 1.22541 | 144 |
| GO_REGULATION_OF_TRANSCRIPTION_FROM_RNA_POLYMERA! | 0.00268 | 0.03460 | 0.43171 | 0.66871 | 1.26686 | 58 |
| GO_REGULATION_OF_TRANSLATIONAL_INITIATION | 0.00056 | 0.01245 | 0.47727 | 0.68378 | 1.29582 | 59 |
| GO_REGULATION_OF_TRANSMEMBRANE_TRANSPORT | 0.00383 | 0.04325 | 0.43171 | 0.57120 | 1.12875 | 289 |
| GO_REGULATION_OF_WNT_SIGNALING_PATHWAY | 0.00045 | 0.01074 | 0.49849 | 0.58980 | 1.16145 | 255 |
| GO_REPRODUCTIVE_SYSTEM_DEVELOPMENT | 0.00013 | 0.00459 | 0.51885 | 0.59586 | 1.17527 | 269 |
| GO_RESPIRASOME | 0.00005 | 0.00237 | 0.55733 | 0.69879 | 1.33399 | 69 |
| GO_RESPIRATORY_CHAIN_COMPLEX | 0.00020 | 0.00630 | 0.51885 | 0.69730 | 1.32145 | 59 |
| GO_RESPIRATORY_ELECTRON_TRANSPORT_CHAIN | 0.00011 | 0.00408 | 0.53843 | 0.67951 | 1.30487 | 82 |
| GO_RESPONSE_TO_ACID_CHEMICAL | 0.00346 | 0.04053 | 0.43171 | 0.64281 | 1.23540 | 84 |
| GO_RESPONSE_TO_ALCOHOL | 0.00102 | 0.01880 | 0.45506 | 0.62110 | 1.20820 | 133 |
| GO_RESPONSE_TO_CATECHOLAMINE | 0.00298 | 0.03675 | 0.43171 | 0.70071 | 1.30801 | 40 |
| GO_RESPONSE_TO_DOPAMINE | 0.00178 | 0.02684 | 0.45506 | 0.73415 | 1.35473 | 31 |
| GO_RESPONSE_TO_ELECTRICAL_STIMULUS | 0.00077 | 0.01582 | 0.47727 | 0.79418 | 1.44477 | 23 |
| GO_RESPONSE_TO_FIBROBLAST_GROWTH_FACTOR | 0.00082 | 0.01644 | 0.47727 | 0.64702 | 1.24728 | 92 |
| GO_RESPONSE_TO_GROWTH_FACTOR | 0.00011 | 0.00404 | 0.53843 | 0.57773 | 1.14936 | 430 |
| GO_RESPONSE_TO_INTERLEUKIN_12 | 0.00170 | 0.02630 | 0.45506 | 0.72303 | 1.33968 | 34 |
| GO_RESPONSE_TO_INTERLEUKIN_4 | 0.00006 | 0.00262 | 0.55733 | 0.84471 | 1.52821 | 21 |
| GO_RESPONSE_TO_KETONE | 0.00042 | 0.01025 | 0.49849 | 0.63418 | 1.23035 | 122 |
| GO_RESPONSE_TO_LEUKEMIA_INHIBITORY_FACTOR | 0.00003 | 0.00150 | 0.57561 | 0.70611 | 1.35215 | 74 |
| GO_RESPONSE_TO_LIPID | 0.00008 | 0.00316 | 0.53843 | 0.57367 | 1.14192 | 481 |
| GO_RESPONSE_TO_NUTRIENT | 0.00276 | 0.03526 | 0.43171 | 0.63029 | 1.21693 | 100 |
| GO_RESPONSE_TO_OXIDATIVE_STRESS | 0.00110 | 0.01969 | 0.45506 | 0.58163 | 1.14871 | 282 |
| GO_RESPONSE_TO_OXYGEN_LEVELS | 0.00035 | 0.00905 | 0.49849 | 0.59150 | 1.16481 | 255 |
| GO_RESPONSE_TO_PEPTIDE | 0.00095 | 0.01797 | 0.47727 | 0.57429 | 1.13772 | 328 |
| GO_RESPONSE_TO_PEPTIDE_HORMONE | 0.00202 | 0.02893 | 0.43171 | 0.57775 | 1.14150 | 287 |
| GO_RESPONSE_TO_STEROID_HORMONE | 0.00010 | 0.00385 | 0.53843 | 0.60749 | 1.19188 | 208 |
| GO_RESPONSE_TO_TOPOLOGICALLY_INCORRECT_PROTEIN | 0.00472 | 0.04948 | 0.40702 | 0.60038 | 1.16967 | 144 |
| GO_RESPONSE_TO_VIRUS | 0.00083 | 0.01646 | 0.47727 | 0.60824 | 1.18822 | 169 |
| GO_RESPONSE_TO_WOUNDING | 0.00034 | 0.00892 | 0.49849 | 0.58233 | 1.15450 | 344 |
| GO_RETINOIC_ACID_RECEPTOR_BINDING | 0.00413 | 0.04562 | 0.40702 | 0.76187 | 1.37835 | 21 |
| GO_RHYTHMIC_PROCESS | 0.00002 | 0.00117 | 0.57561 | 0.63094 | 1.23309 | 173 |
| GO_RIBONUCLEOPROTEIN_COMPLEX_BINDING | 0.00033 | 0.00876 | 0.49849 | 0.64248 | 1.24235 | 109 |
| GO_RIBONUCLEOPROTEIN_COMPLEX_BIOGENESIS | 0.00000 | 0.00000 | | 0.64917 | 1.28758 | 366 |
| GO_RIBONUCLEOPROTEIN_COMPLEX_SUBUNIT_ORGANIZATION | 0.00000 | 0.00000 | 0.77494 | 0.68888 | 1.34631 | 171 |
| GO_RIBONUCLEOPROTEIN_GRANULE | 0.00000 | 0.00000 | 0.73376 | 0.67051 | 1.31043 | 173 |
| GO_RIBOSOMAL_LARGE_SUBUNIT_BIOGENESIS | 0.00028 | 0.00808 | 0.49849 | 0.69170 | 1.31446 | 62 |
| GO_RIBOSOMAL_SMALL_SUBUNIT_ASSEMBLY | 0.00235 | 0.03181 | 0.43171 | 0.81445 | 1.45033 | 16 |
| GO_RIBOSOMAL_SUBUNIT | 0.00000 | 0.00000 | | 0.75465 | 1.47307 | 162 |
| GO_RIBOSOME | 0.00000 | 0.00000 | | 0.75284 | 1.47528 | 196 |
| GO_RIBOSOME_ASSEMBLY | 0.00003 | 0.00168 | 0.55733 | 0.74715 | 1.40949 | 50 |
| GO_RIBOSOME_BIOGENESIS | 0.00000 | 0.00002 | 0.69013 | 0.63893 | 1.25739 | 247 |
| GO_RNA_CATABOLIC_PROCESS | 0.00000 | 0.00000 | | 0.71970 | 1.42510 | 321 |
| GO_RNA_EXPORT_FROM_NUCLEUS | 0.00089 | 0.01730 | 0.47727 | 0.62951 | 1.21991 | 119 |
| GO_RNA_LOCALIZATION | 0.00004 | 0.00206 | 0.55733 | 0.62409 | 1.22324 | 197 |
| GO_RNA_POLYMERASE_II_SPECIFIC_DNA_BINDING_TRANSCRIPT | 0.00159 | 0.02516 | 0.45506 | 0.59407 | 1.16415 | 196 |
| GO_RNA_SPLICING | 0.00000 | 0.00000 | | 0.65759 | 1.30481 | 371 |
| GO_RNA_SPLICING_VIA_TRANSESTERIFICATION_REACTIONS | 0.00000 | 0.00000 | | 0.67324 | 1.33141 | 299 |
| GO_RNA_STABILIZATION | 0.00009 | 0.00362 | 0.53843 | 0.76331 | 1.42071 | 37 |
| GO_ROUGH_ENDOPLASMIC_RETICULUM | 0.00001 | 0.00089 | 0.59333 | 0.73605 | 1.39315 | 57 |
| GO_RRNA_BINDING | 0.00000 | 0.00004 | 0.67496 | 0.78576 | 1.48324 | 51 |
| GO_RRNA_METABOLIC_PROCESS | 0.00001 | 0.00086 | 0.59333 | 0.62931 | 1.23245 | 191 |
| GO_RUFFLE | 0.00024 | 0.00727 | 0.51885 | 0.63531 | 1.23558 | 132 |
| GO_SARCOMERE_ORGANIZATION | 0.00163 | 0.02561 | 0.45506 | 0.77720 | 1.41389 | 23 |
| GO_SECRETORY_GRANULE | 0.00000 | 0.00036 | 0.61053 | 0.58855 | 1.17105 | 437 |
| GO_SENSORY_ORGAN_MORPHOGENESIS | 0.00435 | 0.04725 | 0.40702 | 0.60041 | 1.17013 | 147 |
| GO_SERINE_TYPE_ENDOPEPTIDASE_INHIBITOR_ACTIVITY | 0.00287 | 0.03609 | 0.43171 | 0.75289 | 1.37870 | 26 |
| GO_SIDE_OF_MEMBRANE | 0.00025 | 0.00740 | 0.49849 | 0.60077 | 1.18172 | 241 |
| GO_SIGNAL_SEQUENCE_BINDING | 0.00246 | 0.03303 | 0.43171 | 0.71655 | 1.33022 | 35 |
| GO_SINGLE_STRANDED_DNA_BINDING | 0.00405 | 0.04495 | 0.40702 | 0.64372 | 1.23642 | 81 |
| GO_SINGLE_STRANDED_RNA_BINDING | 0.00000 | 0.00029 | 0.62726 | 0.73363 | 1.39750 | 65 |
| GO_SKIN_DEVELOPMENT | 0.00026 | 0.00756 | 0.49849 | 0.62096 | 1.21249 | 165 |
| GO_SMALL_NUCLEAR_RIBONUCLEOPROTEIN_COMPLEX | 0.00028 | 0.00798 | 0.49849 | 0.69350 | 1.31424 | 59 |
| GO_SMALL_RIBOSOMAL_SUBUNIT | 0.00000 | 0.00001 | 0.70498 | 0.77350 | 1.47020 | 63 |
| GO_SM_LIKE_PROTEIN_FAMILY_COMPLEX | 0.00176 | 0.02669 | 0.45506 | 0.66142 | 1.26177 | 68 |
| GO_SOMATODENDRITIC_COMPARTMENT | 0.00194 | 0.02819 | 0.45506 | 0.55744 | 1.10963 | 491 |
| GO_SPERM_EGG_RECOGNITION | 0.00106 | 0.01925 | 0.45506 | 0.75982 | 1.39705 | 28 |
| GO_SPLICEOSOMAL_COMPLEX | 0.00001 | 0.00074 | 0.59333 | 0.64287 | 1.25500 | 164 |
| GO_STEM_CELL_DIFFERENTIATION | 0.00000 | 0.00017 | 0.64355 | 0.64942 | 1.26867 | 169 |

| | | | | | | |
|---|---|---|---|---|---|---|
| GO_STEROID_HORMONE_MEDIATED_SIGNALING_PATHWAY | 0.00435 | 0.04725 | 0.40702 | 0.61875 | 1.19515 | 101 |
| GO_STEROID_METABOLIC_PROCESS | 0.00287 | 0.03609 | 0.43171 | 0.59553 | 1.16410 | 172 |
| GO_STEROL_HOMEOSTASIS | 0.00231 | 0.03145 | 0.43171 | 0.70083 | 1.30876 | 41 |
| GO_STRESS_FIBER_ASSEMBLY | 0.00224 | 0.03105 | 0.43171 | 0.64203 | 1.23203 | 79 |
| GO_STRESS_GRANULE_ASSEMBLY | 0.00231 | 0.03145 | 0.43171 | 0.77798 | 1.40366 | 20 |
| GO_STRIATED_MUSCLE_CONTRACTION | 0.00054 | 0.01218 | 0.47727 | 0.65676 | 1.26460 | 89 |
| GO_STRUCTURAL_CONSTITUENT_OF_CYTOSKELETON | 0.00130 | 0.02200 | 0.45506 | 0.67387 | 1.27981 | 61 |
| GO_STRUCTURAL_CONSTITUENT_OF_RIBOSOME | 0.00000 | 0.00000 | | 0.77014 | 1.50160 | 149 |
| GO_STRUCTURAL_MOLECULE_ACTIVITY | 0.00000 | 0.00000 | | 0.70646 | 1.40508 | 409 |
| GO_SUBSTANTIA_NIGRA_DEVELOPMENT | 0.00287 | 0.03609 | 0.43171 | 0.73527 | 1.35362 | 29 |
| GO_SUPRAMOLECULAR_FIBER_ORGANIZATION | 0.00001 | 0.00050 | 0.61053 | 0.58498 | 1.16396 | 465 |
| GO_TAU_PROTEIN_BINDING | 0.00022 | 0.00681 | 0.51885 | 0.74424 | 1.38706 | 38 |
| GO_TISSUE_HOMEOSTASIS | 0.00442 | 0.04764 | 0.40702 | 0.60262 | 1.17201 | 132 |
| GO_TISSUE_MORPHOGENESIS | 0.00007 | 0.00290 | 0.53843 | 0.58241 | 1.15835 | 410 |
| GO_TRANSCRIPTION_COACTIVATOR_ACTIVITY | 0.00002 | 0.00142 | 0.57561 | 0.62304 | 1.22163 | 201 |
| GO_TRANSCRIPTION_COREGULATOR_ACTIVITY | 0.00000 | 0.00004 | 0.67496 | 0.60846 | 1.20731 | 371 |
| GO_TRANSCRIPTION_COREPRESSOR_ACTIVITY | 0.00150 | 0.02419 | 0.45506 | 0.60662 | 1.18180 | 145 |
| GO_TRANSCRIPTION_FACTOR_BINDING | 0.00002 | 0.00142 | 0.57561 | 0.58008 | 1.15436 | 470 |
| GO_TRANSLATIONAL_INITIATION | 0.00000 | 0.00000 | | 0.81566 | 1.59167 | 158 |
| GO_TRANSLATION_FACTOR_ACTIVITY_RNA_BINDING | 0.00003 | 0.00171 | 0.55733 | 0.71649 | 1.36183 | 63 |
| GO_TRANSLATION_INITIATION_FACTOR_ACTIVITY | 0.00132 | 0.02208 | 0.45506 | 0.71640 | 1.33729 | 40 |
| GO_TRANSLATION_INITIATION_FACTOR_BINDING | 0.00368 | 0.04218 | 0.43171 | 0.75839 | 1.37342 | 22 |
| GO_TRANSLATION_REGULATOR_ACTIVITY | 0.00000 | 0.00002 | 0.69013 | 0.70625 | 1.36523 | 106 |
| GO_TRANSLATION_REGULATOR_ACTIVITY_NUCLEIC_ACID_BIND | 0.00001 | 0.00050 | 0.61053 | 0.71293 | 1.36706 | 78 |
| GO_TRANSMEMBRANE_RECEPTOR_PROTEIN_TYROSINE_KINASE | 0.00133 | 0.02226 | 0.45506 | 0.55933 | 1.11255 | 455 |
| GO_TYPE_2_IMMUNE_RESPONSE | 0.00132 | 0.02208 | 0.45506 | 0.82906 | 1.47047 | 15 |
| GO_U1_SNRNP | 0.00290 | 0.03627 | 0.43171 | 0.80933 | 1.44121 | 16 |
| GO_U2_SNRNP | 0.00172 | 0.02646 | 0.45506 | 0.80193 | 1.43894 | 18 |
| GO_UBIQUITIN_LIKE_PROTEIN_LIGASE_BINDING | 0.00031 | 0.00840 | 0.49849 | 0.60117 | 1.18227 | 237 |
| GO_UNFOLDED_PROTEIN_BINDING | 0.00059 | 0.01288 | 0.47727 | 0.66200 | 1.27125 | 82 |
| GO_VENTRAL_SPINAL_CORD_DEVELOPMENT | 0.00257 | 0.03380 | 0.43171 | 0.76650 | 1.38811 | 22 |
| GO_VESICLE_LUMEN | 0.00000 | 0.00035 | 0.61053 | 0.64333 | 1.25812 | 179 |
| GO_VIRAL_GENE_EXPRESSION | 0.00000 | 0.00000 | | 0.76469 | 1.49387 | 169 |
| GO_VIRAL_GENOME_REPLICATION | 0.00051 | 0.01187 | 0.47727 | 0.66679 | 1.27832 | 77 |
| GO_VIRAL_LIFE_CYCLE | 0.00002 | 0.00113 | 0.57561 | 0.62076 | 1.21946 | 221 |
| GO_WOUND_HEALING | 0.00038 | 0.00968 | 0.49849 | 0.58844 | 1.16164 | 276 |
| GO_ZYMOGEN_ACTIVATION | 0.00039 | 0.00975 | 0.49849 | 0.78771 | 1.44246 | 26 |
| HP_ABNORMALITY_OF_CARDIOVASCULAR_SYSTEM_ELECTROPH | 0.00005 | 0.00215 | 0.55733 | 0.60728 | 1.19474 | 242 |
| HP_ABNORMALITY_OF_LOWER_LIP | 0.00279 | 0.03559 | 0.43171 | 0.59962 | 1.17186 | 171 |
| HP_ABNORMALITY_OF_THE_CHEEK | 0.00261 | 0.03414 | 0.43171 | 0.60739 | 1.18129 | 132 |
| HP_ABNORMALITY_OF_THE_COLUMELLA | 0.00368 | 0.04218 | 0.43171 | 0.71342 | 1.32123 | 33 |
| HP_ABNORMALITY_OF_THE_MIDFACE | 0.00091 | 0.01754 | 0.47727 | 0.57331 | 1.13642 | 353 |
| HP_ABNORMALITY_OF_THE_NASAL_DORSUM | 0.00030 | 0.00835 | 0.49849 | 0.62342 | 1.21619 | 152 |
| HP_ABNORMALITY_OF_THE_PALPEBRAL_FISSURES | 0.00465 | 0.04887 | 0.40702 | 0.55419 | 1.10268 | 464 |
| HP_ABNORMALITY_OF_THE_PHILTRUM | 0.00268 | 0.03460 | 0.43171 | 0.56300 | 1.11688 | 369 |
| HP_ABNORMALITY_OF_THE_PINNA | 0.00058 | 0.01272 | 0.47727 | 0.56966 | 1.13354 | 438 |
| HP_ABNORMALITY_OF_UPPER_LIP | 0.00031 | 0.00845 | 0.49849 | 0.56834 | 1.13135 | 488 |
| HP_ABNORMAL_CARDIAC_ATRIUM_MORPHOLOGY | 0.00080 | 0.01617 | 0.47727 | 0.58773 | 1.15692 | 251 |
| HP_ABNORMAL_CARDIAC_SEPTUM_MORPHOLOGY | 0.00064 | 0.01350 | 0.47727 | 0.57310 | 1.13859 | 391 |
| HP_ABNORMAL_CARDIAC_VENTRICLE_MORPHOLOGY | 0.00098 | 0.01839 | 0.45506 | 0.57387 | 1.13702 | 331 |
| HP_ABNORMAL_EYELASH_MORPHOLOGY | 0.00097 | 0.01821 | 0.47727 | 0.61221 | 1.19467 | 158 |
| HP_ABNORMAL_FEAR_ANXIETY_RELATED_BEHAVIOR | 0.00033 | 0.00875 | 0.49849 | 0.60880 | 1.19306 | 195 |
| HP_ABNORMAL_FINGERNAIL_MORPHOLOGY | 0.00029 | 0.00832 | 0.49849 | 0.63244 | 1.22965 | 129 |
| HP_ABNORMAL_IMMUNE_SYSTEM_MORPHOLOGY | 0.00128 | 0.02193 | 0.45506 | 0.57450 | 1.13817 | 326 |
| HP_ABNORMAL_MORPHOLOGY_OF_THE_GREAT_VESSELS | 0.00194 | 0.02819 | 0.45506 | 0.57154 | 1.13215 | 324 |
| HP_ABNORMAL_SCALP_MORPHOLOGY | 0.00220 | 0.03074 | 0.43171 | 0.58849 | 1.15485 | 210 |
| HP_ABNORMAL_T_WAVE | 0.00298 | 0.03675 | 0.43171 | 0.79198 | 1.42109 | 18 |
| HP_ABNORMAL_VENTRICULAR_SEPTUM_MORPHOLOGY | 0.00024 | 0.00727 | 0.51885 | 0.59129 | 1.16423 | 254 |
| HP_ACUTE_LEUKEMIA | 0.00000 | 0.00018 | 0.62726 | 0.72709 | 1.39391 | 77 |
| HP_ANEMIA_OF_INADEQUATE_PRODUCTION | 0.00003 | 0.00144 | 0.57561 | 0.67639 | 1.30506 | 98 |
| HP_ATRESIA_OF_THE_EXTERNAL_AUDITORY_CANAL | 0.00305 | 0.03715 | 0.43171 | 0.71828 | 1.33022 | 33 |
| HP_BRACHYDACTYLY | 0.00390 | 0.04377 | 0.43171 | 0.57823 | 1.13623 | 224 |
| HP_CARDIAC_ARREST | 0.00013 | 0.00459 | 0.51885 | 0.69808 | 1.32980 | 65 |
| HP_CLEFT_LIP | 0.00181 | 0.02720 | 0.45506 | 0.62846 | 1.21663 | 113 |
| HP_CONGENITAL_ABNORMAL_HAIR_PATTERN | 0.00205 | 0.02926 | 0.43171 | 0.61174 | 1.19131 | 143 |
| HP_CONGENITAL_DIAPHRAGMATIC_HERNIA | 0.00316 | 0.03813 | 0.43171 | 0.67334 | 1.27241 | 54 |
| HP_CONGENITAL_MALFORMATION_OF_THE_GREAT_ARTERIES | 0.00172 | 0.02646 | 0.45506 | 0.58764 | 1.15489 | 226 |
| HP_CRYPTORCHIDISM | 0.00108 | 0.01947 | 0.45506 | 0.56356 | 1.12076 | 415 |
| HP_DELAYED_PUBERTY | 0.00019 | 0.00622 | 0.51885 | 0.64100 | 1.24424 | 123 |
| HP_DELAYED_SPEECH_AND_LANGUAGE_DEVELOPMENT | 0.00133 | 0.02226 | 0.45506 | 0.55864 | 1.11199 | 490 |

| | | | | | | |
|---|---|---|---|---|---|---|
| HP_DEPRESSED_NASAL_RIDGE | 0.00004 | 0.00210 | 0.55733 | 0.68918 | 1.32374 | 81 |
| HP_DOWNSLANTED_PALPEBRAL_FISSURES | 0.00143 | 0.02312 | 0.45506 | 0.57929 | 1.14372 | 277 |
| HP_EPICANTHUS | 0.00231 | 0.03145 | 0.43171 | 0.57359 | 1.13538 | 312 |
| HP_EPISPADIAS | 0.00472 | 0.04948 | 0.40702 | 0.75445 | 1.36493 | 21 |
| HP_FATIGUE | 0.00001 | 0.00052 | 0.61053 | 0.63272 | 1.24166 | 210 |
| HP_FULL_CHEEKS | 0.00194 | 0.02819 | 0.45506 | 0.64381 | 1.23545 | 79 |
| HP_HEADACHE | 0.00054 | 0.01206 | 0.47727 | 0.61350 | 1.19790 | 167 |
| HP_HEMATOLOGICAL_NEOPLASM | 0.00001 | 0.00052 | 0.61053 | 0.64267 | 1.25549 | 169 |
| HP_IMPAIRMENT_IN_PERSONALITY_FUNCTIONING | 0.00154 | 0.02447 | 0.45506 | 0.56704 | 1.12490 | 369 |
| HP_JUVENILE_ASEPTIC_NECROSIS | 0.00098 | 0.01839 | 0.45506 | 0.79894 | 1.44541 | 21 |
| HP_LEUKEMIA | 0.00000 | 0.00029 | 0.62726 | 0.68459 | 1.32336 | 106 |
| HP_LOW_POSTERIOR_HAIRLINE | 0.00339 | 0.03988 | 0.43171 | 0.65267 | 1.24028 | 62 |
| HP_MACROCYTIC_ANEMIA | 0.00000 | 0.00026 | 0.62726 | 0.80685 | 1.50394 | 39 |
| HP_MIGRAINE | 0.00000 | 0.00004 | 0.67496 | 0.74103 | 1.41944 | 75 |
| HP_MUTISM | 0.00442 | 0.04764 | 0.40702 | 0.71921 | 1.32717 | 31 |
| HP_ORAL_CLEFT | 0.00006 | 0.00258 | 0.55733 | 0.58365 | 1.15842 | 376 |
| HP_PALLOR | 0.00055 | 0.01234 | 0.47727 | 0.65396 | 1.26126 | 94 |
| HP_PATENT_DUCTUS_ARTERIOSUS | 0.00339 | 0.03988 | 0.43171 | 0.59325 | 1.15994 | 177 |
| HP_PTOSIS | 0.00139 | 0.02276 | 0.45506 | 0.56518 | 1.12385 | 406 |
| HP_RETICULOCYTOPENIA | 0.00298 | 0.03675 | 0.43171 | 0.81225 | 1.44065 | 15 |
| HP_SHORT_DIGIT | 0.00305 | 0.03715 | 0.43171 | 0.56853 | 1.12664 | 334 |
| HP_SHORT_NOSE | 0.00003 | 0.00144 | 0.57561 | 0.62581 | 1.22524 | 190 |
| HP_SMOOTH_PHILTRUM | 0.00089 | 0.01730 | 0.47727 | 0.65303 | 1.25627 | 87 |
| HP_SPORADIC | 0.00457 | 0.04849 | 0.40702 | 0.64490 | 1.23112 | 69 |
| HP_SUDDEN_CARDIAC_DEATH | 0.00053 | 0.01189 | 0.47727 | 0.71136 | 1.34018 | 49 |
| HP_THICK_LOWER_LIP_VERMILION | 0.00036 | 0.00937 | 0.49849 | 0.64284 | 1.24387 | 110 |
| HP_THICK_VERMILION_BORDER | 0.00031 | 0.00845 | 0.49849 | 0.61330 | 1.19915 | 177 |
| HP_THIN_VERMILION_BORDER | 0.00124 | 0.02141 | 0.45506 | 0.59441 | 1.16648 | 210 |
| HP_TRIANGULAR_SHAPED_PHALANGES_OF_THE_HAND | 0.00122 | 0.02127 | 0.45506 | 0.63045 | 1.22104 | 116 |
| HP_VENTRICULAR_ARRHYTHMIA | 0.00442 | 0.04764 | 0.40702 | 0.66762 | 1.26161 | 54 |
| HP_VENTRICULAR_TACHYCARDIA | 0.00137 | 0.02252 | 0.45506 | 0.75482 | 1.38786 | 28 |
| HP_X_LINKED_RECESSIVE_INHERITANCE | 0.00368 | 0.04218 | 0.43171 | 0.62605 | 1.20873 | 100 |

### Supplementary Table S8

#### Mouse male NMF coefficient matrix

| Gene | Cluster 1 | Cluster 2 | Cluster 3 | Cluster 4 | Cluster 5 | Cluster 6 | Cluster 7 | Cluster 8 | Cluster 9 | Cluster 10 | Cluster 11 | Cluster 12 | Cluster 13 | Cluster 14 | Cluster 15 | Cluster 16 |
|---|---|---|---|---|---|---|---|---|---|---|---|---|---|---|---|---|
| A4galt | 0.05379 | 0.01966 | 0.05667 | 0.21749 | 0.01045 | 0.00610 | 0.02970 | 0.06273 | 0.19832 | 0.04475 | 0.24274 | 0.03775 | 0.03734 | 0.05182 | 0.40484 | 0.02199 |
| A630020A06 | 0.09850 | 0.01843 | 0.06304 | 2.74110 | 0.00763 | 0.07640 | 0.01772 | 0.08494 | 2.75599 | 0.26921 | 0.21385 | 0.21966 | 0.05599 | 0.43859 | 0.65147 | 0.00644 |
| AA388235 | 0.16935 | 0.00778 | 0.14572 | 0.63476 | 0.00940 | 0.04807 | 0.12110 | 0.07019 | 2.05481 | 0.44345 | 0.66872 | 0.12241 | 0.03931 | 1.90506 | 0.40670 | 0.02843 |
| AA414768 | 0.00000 | 3.00398 | 2.02521 | 2.02529 | 2.52654 | 0.00640 | 1.39723 | 0.77336 | 7.87031 | 1.68702 | 0.00000 | 0.00000 | 1.20441 | 1.48466 | 1.06174 | 0.10957 |
| AA415398 | 0.01279 | 0.00702 | 0.08796 | 8.97458 | 0.00866 | 0.00363 | 1.43255 | 0.01035 | 12.44484 | 0.00416 | 0.00000 | 0.00000 | 0.00971 | 0.00006 | 0.55378 | 0.00472 |
| AA465934 | 0.35790 | 0.00739 | 0.00596 | 0.17364 | 0.00724 | 0.00703 | 0.00732 | 0.01209 | 0.38163 | 0.01138 | 0.01032 | 0.00728 | 0.01233 | 0.00965 | 0.04522 | 0.00694 |
| AA467197 | 24.27979 | 6.89574 | 17.26449 | 0.62084 | 22.23862 | 12.00025 | 5.53252 | 5.51120 | 0.31648 | 1.97989 | 5.75353 | 0.08175 | 11.93972 | 1.40480 | 0.00000 | 12.93772 |
| AA536875 | 0.03902 | 0.00749 | 0.03265 | 0.18494 | 0.01842 | 0.01263 | 0.01746 | 0.02341 | 0.16376 | 0.05633 | 0.08698 | 0.02556 | 0.05706 | 0.12085 | 0.48176 | 0.00722 |
| AA543186 | 0.04349 | 0.04297 | 0.03398 | 0.37634 | 0.01177 | 0.01245 | 0.02897 | 0.02096 | 0.29558 | 0.03661 | 0.05397 | 0.00720 | 0.02313 | 0.00595 | 0.41162 | 0.01498 |
| AA543401 | 0.01762 | 0.00777 | 0.01151 | 2.27027 | 0.00903 | 0.00626 | 0.00738 | 0.01297 | 1.63480 | 0.01274 | 0.00000 | 0.00417 | 0.01314 | 0.00184 | 0.20515 | 0.00691 |
| AA545190 | 0.01377 | 0.00802 | 0.01225 | 5.62067 | 0.00931 | 0.00612 | 0.00732 | 0.01312 | 0.00000 | 0.01100 | 0.04191 | 0.00750 | 0.01195 | 0.00110 | 0.04445 | 0.00643 |
| AA619741 | 0.01287 | 0.00793 | 0.00826 | 4.95695 | 0.00892 | 0.00565 | 0.00721 | 0.00480 | 4.66247 | 0.49858 | 0.15285 | 0.00510 | 0.00968 | 0.00000 | 0.84013 | 0.00595 |
| AA792892 | 0.10272 | 0.14143 | 0.08863 | 33.70022 | 0.07213 | 0.04674 | 0.13755 | 0.72089 | 32.05632 | 5.19036 | 2.11182 | 0.85954 | 0.62888 | 5.26636 | 10.54997 | 0.09265 |
| AA986860 | 0.02363 | 0.00870 | 0.01226 | 0.01056 | 0.01052 | 0.00722 | 0.00784 | 0.05205 | 0.01234 | 0.01217 | 0.01417 | 0.00800 | 0.71073 | 0.01312 | 0.00965 | 0.00732 |
| Aaas | 11.76583 | 12.55330 | 11.36294 | 3.05769 | 1.25785 | 2.47491 | 9.99631 | 17.23082 | 2.37060 | 9.10665 | 10.55734 | 0.84675 | 8.14643 | 0.52815 | 3.88029 | 7.64411 |
| Aacs | 6.78200 | 2.48043 | 4.10657 | 0.08208 | 7.54879 | 5.69845 | 6.15111 | 20.38400 | 1.14080 | 19.52037 | 7.58921 | 1.18823 | 1.28134 | 1.45080 | 3.97512 | 1.43665 |
| Aaed1 | 0.08127 | 0.00562 | 0.00742 | 0.49467 | 5.43853 | 0.00688 | 0.00765 | 0.00977 | 0.66148 | 0.34423 | 0.31265 | 0.00000 | 0.00880 | 0.00767 | 0.00000 | 0.00757 |
| Aagab | 0.79780 | 2.97048 | 4.47571 | 0.48469 | 0.06134 | 0.00567 | 3.14100 | 2.20499 | 0.00576 | 1.78431 | 0.71943 | 0.02373 | 3.25270 | 0.37879 | 2.79471 | 0.43115 |
| Aak1 | 2.35932 | 2.80278 | 3.03106 | 3.39595 | 3.05530 | 0.47794 | 2.04898 | 4.68108 | 3.58525 | 2.10228 | 1.63367 | 3.29214 | 2.55738 | 4.17277 | 2.36056 | 1.76208 |
| Aamdc | 2.65090 | 0.32913 | 2.52480 | 2.28818 | 0.00697 | 0.01048 | 0.00367 | 0.83080 | 4.57962 | 0.49129 | 0.80285 | 0.26925 | 2.52857 | 0.04970 | 1.42038 | 0.00156 |
| Aamp | 17.78961 | 15.10377 | 19.39963 | 4.05193 | 14.59720 | 6.72508 | 12.20625 | 24.65539 | 6.37309 | 16.31169 | 9.86326 | 2.32375 | 17.29956 | 0.09991 | 5.31644 | 9.17935 |
| Aar2 | 7.54515 | 3.27510 | 3.46123 | 0.51357 | 0.00000 | 5.67356 | 6.70124 | 8.42407 | 2.19597 | 6.47176 | 3.88263 | 5.37948 | 4.99933 | 1.89875 | 4.11474 | 1.81145 |
| Aard | 4.75909 | 0.01552 | 0.00294 | 0.68796 | 0.01209 | 0.00402 | 0.00400 | 0.00475 | 0.00000 | 0.07097 | 0.01144 | 0.00395 | 0.06756 | 0.00720 | 0.31329 | 0.09917 |
| Aars | 16.85861 | 12.01500 | 10.51237 | 9.61801 | 6.26564 | 6.74257 | 9.08956 | 21.83323 | 15.92448 | 13.11352 | 9.57695 | 1.31746 | 14.75469 | 0.20276 | 6.03418 | 10.42344 |
| Aars2 | 1.69849 | 0.02671 | 0.00000 | 0.72110 | 0.27235 | 1.32455 | 0.00344 | 3.00012 | 0.53252 | 2.55572 | 1.96070 | 0.01043 | 3.24990 | 0.00172 | 0.00000 | 2.11660 |
| Aarsd1 | 13.83233 | 6.67650 | 10.53162 | 0.58731 | 1.00303 | 5.08862 | 4.84262 | 15.50291 | 0.46286 | 6.06169 | 3.12563 | 0.86286 | 7.67817 | 0.00918 | 0.00000 | 5.16685 |
| Aasdh | 1.82797 | 4.77932 | 2.09110 | 3.53311 | 0.00800 | 0.00232 | 4.20168 | 2.40773 | 2.82273 | 0.41280 | 1.36369 | 4.73253 | 1.12935 | 0.06387 | 2.95462 | 3.60403 |
| Aasdhppt | 8.52806 | 6.26742 | 6.09947 | 3.71320 | 5.43677 | 0.01994 | 5.60822 | 2.88501 | 5.47491 | 4.83916 | 1.97978 | 1.73996 | 4.41444 | 1.32782 | 2.29850 | 0.57347 |
| Aass | 0.01370 | 0.00766 | 0.01152 | 0.45620 | 0.00920 | 0.00662 | 0.00739 | 0.01117 | 0.00461 | 0.19992 | 0.02438 | 0.00690 | 0.01225 | 0.01327 | 0.31816 | 0.00737 |
| Aatf | 12.51278 | 8.78759 | 6.01413 | 0.68977 | 0.28300 | 0.00000 | 5.22246 | 7.50887 | 0.48425 | 11.90783 | 2.10263 | 2.79250 | 6.70833 | 3.74575 | 10.29375 | 4.33178 |
| AB124611 | 0.01367 | 0.00786 | 0.01236 | 0.09180 | 0.00933 | 0.00658 | 0.00750 | 0.01343 | 0.09793 | 0.01041 | 0.01216 | 0.00736 | 0.01260 | 0.00680 | 0.11504 | 0.01465 |
| Abat | 0.00203 | 0.00774 | 0.00000 | 0.20365 | 0.01880 | 3.71898 | 0.00710 | 1.74303 | 0.99180 | 2.23765 | 0.33123 | 0.10587 | 0.00000 | 1.47995 | 3.21259 | 0.00592 |
| Abca1 | 0.02935 | 0.00720 | 0.52370 | 0.53377 | 2.17429 | 0.00552 | 0.86679 | 0.05114 | 0.00000 | 0.00000 | 0.00000 | 0.00000 | 0.00698 | 1.86891 | 0.95838 | 0.00667 |
| Abca13 | 0.62158 | 0.00764 | 0.19793 | 2.62396 | 0.00620 | 0.00618 | 0.00731 | 0.05361 | 0.42529 | 0.04139 | 0.00581 | 0.00522 | 0.00920 | 0.31260 | 0.45304 | 0.00639 |
| Abca16 | 0.02904 | 0.00792 | 0.58317 | 0.01088 | 0.00991 | 0.00564 | 0.00729 | 0.01206 | 0.01225 | 0.01469 | 0.01220 | 0.00758 | 0.00895 | 0.01476 | 0.00189 | 0.00717 |
| Abca17 | 0.00386 | 0.06354 | 0.41441 | 0.93422 | 0.00797 | 0.36153 | 0.09312 | 0.99829 | 1.76316 | 1.09344 | 0.67599 | 0.03149 | 0.06908 | 0.15143 | 2.70846 | 0.23578 |
| Abca2 | 0.71283 | 0.00620 | 0.08618 | 0.41316 | 0.00673 | 1.22457 | 0.69354 | 0.00000 | 0.16756 | 0.00000 | 0.70765 | 0.00143 | 1.22543 | 0.00385 | 0.08994 | 0.00534 |
| Abca3 | 0.00000 | 1.70167 | 3.74872 | 0.00000 | 0.00000 | 7.01104 | 5.21856 | 5.13043 | 0.05828 | 0.99563 | 0.03972 | 0.00516 | 5.63500 | 0.02622 | 0.17155 | 6.15010 |
| Abca4 | 0.00947 | 0.00781 | 0.01074 | 0.24013 | 0.00906 | 0.00622 | 0.00676 | 0.01013 | 1.09597 | 0.00000 | 0.00024 | 0.03347 | 0.95911 | 0.03848 | 0.21304 | 0.00659 |
| Abca5 | 0.01662 | 0.00768 | 0.41177 | 0.00636 | 0.00851 | 0.00560 | 0.00721 | 0.01061 | 0.29281 | 0.00797 | 0.00974 | 0.03722 | 0.00939 | 0.03089 | 0.18058 | 0.00708 |
| Abca7 | 2.34306 | 0.00469 | 0.24847 | 0.01052 | 0.00307 | 0.00647 | 0.45180 | 0.25002 | 0.01218 | 0.00753 | 0.01103 | 0.00492 | 0.11739 | 0.01127 | 0.01529 | 0.00463 |
| Abca8b | 0.01293 | 0.00812 | 0.02402 | 0.00470 | 0.00928 | 0.00767 | 0.00763 | 0.01727 | 0.32745 | 0.01109 | 0.00972 | 0.00767 | 0.01231 | 0.03743 | 0.02549 | 0.00741 |
| Abcb10 | 3.61893 | 2.69055 | 2.22684 | 5.32652 | 0.00000 | 0.00265 | 2.42845 | 3.89925 | 1.11853 | 1.63052 | 2.94043 | 0.20253 | 0.10224 | 10.03091 | 2.88713 | 2.06213 |
| Abcb11 | 0.01371 | 0.00782 | 0.01167 | 0.00765 | 0.00933 | 0.00666 | 0.00750 | 0.01262 | 0.01140 | 0.01819 | 0.01082 | 0.01757 | 0.01318 | 0.00571 | 0.52934 | 0.00740 |
| Abcb1a | 0.00637 | 0.00700 | 0.90984 | 0.11068 | 0.02052 | 0.00875 | 0.13797 | 3.63829 | 0.00718 | 0.89608 | 1.49844 | 0.95075 | 0.10361 | 0.00508 | 0.47729 | 0.02096 |
| Abcb1b | 0.00000 | 0.03069 | 0.52915 | 1.75300 | 0.00928 | 0.00581 | 1.34435 | 1.24392 | 0.00853 | 0.63106 | 1.31171 | 0.26485 | 0.62646 | 0.10313 | 0.42464 | 0.00642 |
| Abcb4 | 0.00945 | 0.00768 | 0.03810 | 0.39621 | 0.00964 | 0.00782 | 0.06898 | 0.14920 | 0.00445 | 0.03542 | 0.01166 | 0.04854 | 0.02487 | 0.01743 | 0.15516 | 0.00723 |
| Abcb5 | 0.00000 | 0.00780 | 0.43314 | 1.24771 | 0.01033 | 0.00580 | 0.00764 | 2.85791 | 0.36551 | 0.77154 | 0.07233 | 1.10666 | 0.52438 | 0.67386 | 3.45180 | 0.00708 |
| Abcb6 | 3.51618 | 0.00676 | 12.54355 | 3.86648 | 1.12141 | 7.08725 | 7.65621 | 9.77976 | 0.59032 | 3.80803 | 6.23420 | 0.46704 | 7.52883 | 0.39110 | 1.88005 | 2.11744 |
| Abcb7 | 2.80324 | 0.12922 | 2.87794 | 4.22401 | 3.37377 | 4.47316 | 1.95645 | 2.63551 | 2.74610 | 1.75661 | 3.72795 | 9.30909 | 3.50418 | 12.15987 | 4.27443 | 3.52941 |
| Abcb8 | 6.20223 | 0.36403 | 3.68949 | 0.61015 | 0.04478 | 1.68214 | 1.81934 | 3.97901 | 0.00529 | 0.86397 | 0.28952 | 0.03147 | 6.88312 | 0.04845 | 0.00000 | 0.30864 |
| Abcb9 | 0.01087 | 0.00778 | 0.00996 | 0.22250 | 0.00957 | 0.00663 | 0.00720 | 0.12894 | 0.64156 | 0.01259 | 0.00906 | 0.01636 | 0.00817 | 0.01232 | 0.01096 | 0.00734 |
| Abcc1 | 3.65445 | 0.01285 | 1.56079 | 0.64115 | 0.00000 | 0.00239 | 2.66529 | 1.47876 | 0.08681 | 1.92949 | 0.46545 | 0.51784 | 1.32952 | 2.71836 | 0.51898 | 1.29174 |
| Abcc10 | 1.34197 | 0.00309 | 0.20173 | 1.30268 | 3.76922 | 0.00459 | 0.00415 | 5.03366 | 0.76991 | 2.87048 | 2.25927 | 0.00000 | 1.99279 | 0.01572 | 0.00000 | 3.50741 |
| Abcc2 | 0.01305 | 0.00791 | 0.01021 | 0.00000 | 0.00915 | 0.00626 | 0.00728 | 0.00777 | 0.00903 | 0.74406 | 0.08569 | 0.00227 | 0.01037 | 0.00536 | 1.60417 | 0.00722 |
| Abcc3 | 2.78799 | 0.01619 | 0.00000 | 0.00720 | 0.00488 | 0.00460 | 0.00476 | 2.96982 | 0.11567 | 1.90807 | 1.23358 | 0.02800 | 0.00000 | 0.02009 | 1.41648 | 0.00442 |
| Abcc4 | 0.04975 | 0.00803 | 0.37540 | 0.38068 | 0.00630 | 0.01839 | 0.22617 | 1.04343 | 0.00518 | 0.00000 | 0.00000 | 0.00000 | 4.74710 | 0.35052 | 0.86567 | 3.33150 |
| Abcc5 | 4.77536 | 2.32178 | 4.04151 | 3.96447 | 3.20764 | 1.04418 | 2.78748 | 9.33453 | 5.98771 | 3.36969 | 2.95366 | 1.39315 | 2.36901 | 2.39210 | 4.98224 | 3.75669 |
| Abcc6 | 0.01375 | 0.00799 | 0.01215 | 0.08936 | 0.00931 | 0.00664 | 0.00753 | 0.01377 | 0.09132 | 0.01458 | 0.01314 | 0.00794 | 0.01271 | 0.01371 | 0.01224 | 0.00740 |
| Abcc8 | 0.95252 | 0.00661 | 1.41406 | 0.01099 | 0.00368 | 0.00461 | 0.00689 | 0.01000 | 0.01218 | 0.01359 | 0.01086 | 0.00728 | 0.00732 | 0.01597 | 0.00000 | 0.00579 |

| | | | | | | | | | | | | | | | | |
|---|---|---|---|---|---|---|---|---|---|---|---|---|---|---|---|---|
| **Abcc9** | 0.14395 | 0.06049 | 0.21484 | 0.28044 | 0.12783 | 0.07237 | 0.04707 | 0.05237 | 0.26297 | 0.15427 | 1.15962 | 0.19970 | 0.56184 | 1.38977 | 0.29441 | 0.08577 |
| **Abcd1** | 2.73913 | 0.10766 | 2.73242 | 0.98420 | 3.69084 | 5.26637 | 0.01468 | 2.52182 | 3.13601 | 4.67325 | 0.83998 | 0.13054 | 0.00000 | 0.00000 | 1.06758 | 0.54477 |
| **Abcd2** | 0.11060 | 0.00815 | 0.10373 | 0.13679 | 0.02490 | 0.00658 | 0.01781 | 0.00487 | 0.15947 | 0.03274 | 0.77675 | 0.02395 | 0.07005 | 0.24224 | 0.08509 | 0.02176 |
| **Abcd3** | 10.29224 | 6.93527 | 3.48906 | 0.10844 | 0.13639 | 0.22679 | 7.00518 | 5.28403 | 0.87765 | 1.97464 | 4.09938 | 0.52882 | 3.38746 | 0.99773 | 1.18195 | 1.60686 |
| **Abcd4** | 3.68131 | 1.77862 | 10.38975 | 1.15358 | 0.18748 | 3.68381 | 4.78824 | 0.06568 | 1.40133 | 0.33111 | 2.25432 | 0.11570 | 3.24520 | 0.64524 | 0.15932 | 0.94718 |
| **Abce1** | 14.19414 | 14.24075 | 8.48476 | 4.37079 | 9.14636 | 6.46513 | 9.77576 | 14.40251 | 9.98061 | 12.39502 | 19.06948 | 14.03494 | 10.09232 | 28.44896 | 17.97990 | 2.49388 |
| **Abcf1** | 18.98990 | 9.59222 | 12.73694 | 17.96873 | 6.40865 | 6.60795 | 7.02967 | 18.14976 | 16.18189 | 25.30111 | 15.67232 | 2.89579 | 13.86934 | 1.84617 | 14.08086 | 8.86204 |
| **Abcf2** | 14.05251 | 12.86943 | 16.38724 | 19.96035 | 11.48750 | 12.26575 | 9.29846 | 26.98740 | 18.20800 | 22.99329 | 13.54681 | 7.73914 | 20.81167 | 0.47874 | 10.28797 | 8.83152 |
| **Abcf3** | 4.58444 | 10.07599 | 10.54552 | 4.12976 | 5.19891 | 0.94890 | 5.93834 | 13.05497 | 8.52380 | 8.79637 | 9.18933 | 1.10572 | 7.47535 | 2.24686 | 9.71152 | 5.19647 |
| **Abcg1** | 1.72390 | 0.00654 | 0.00000 | 0.08315 | 0.01265 | 0.01540 | 0.00686 | 0.23738 | 0.00887 | 0.00554 | 0.07127 | 0.02054 | 0.02087 | 0.00913 | 0.08815 | 0.00546 |
| **Abcg2** | 0.36471 | 0.00507 | 1.02916 | 0.20147 | 1.08212 | 0.00562 | 2.42354 | 0.59620 | 0.00421 | 0.00000 | 0.00767 | 0.00310 | 0.00000 | 0.01472 | 0.00000 | 3.63988 |
| **Abcg4** | 3.08049 | 0.00239 | 1.20219 | 0.01021 | 1.06602 | 0.00408 | 0.00306 | 2.43530 | 0.82905 | 1.16647 | 2.94343 | 2.27296 | 0.30702 | 0.00332 | 0.39465 | 0.00265 |
| **Abhd1** | 0.01335 | 0.00805 | 0.08266 | 0.01062 | 0.00918 | 0.00581 | 0.08033 | 0.01300 | 0.01235 | 0.01464 | 0.01284 | 0.00825 | 0.01774 | 0.01270 | 0.00920 | 0.00733 |
| **Abhd10** | 0.31194 | 0.15284 | 0.00000 | 0.78655 | 3.26423 | 1.87838 | 1.95472 | 0.22358 | 0.34066 | 1.62132 | 1.23823 | 0.32885 | 2.75099 | 0.88769 | 1.85441 | 2.64770 |
| **Abhd11** | 10.51729 | 5.74286 | 8.58478 | 1.59822 | 7.29705 | 6.25384 | 8.42481 | 13.98126 | 0.26799 | 6.15674 | 3.47870 | 0.58597 | 6.58230 | 0.15780 | 0.00000 | 6.01974 |
| **Abhd11os** | 0.64979 | 0.02829 | 0.66408 | 0.01575 | 0.04222 | 0.00513 | 0.01530 | 0.42577 | 0.01029 | 0.86844 | 5.13220 | 0.05838 | 0.00004 | 0.00417 | 0.00381 | 0.39953 |
| **Abhd12** | 3.02487 | 5.18357 | 5.63876 | 3.68759 | 0.04248 | 0.36716 | 5.84995 | 0.06699 | 3.30966 | 3.93455 | 0.03174 | 0.11791 | 3.35749 | 0.00000 | 0.11626 | 3.83052 |
| **Abhd12b** | 0.01215 | 0.00770 | 0.12212 | 0.01327 | 0.00849 | 0.00578 | 0.01736 | 0.00000 | 0.01042 | 2.31500 | 0.77347 | 0.05804 | 0.01063 | 0.00985 | 0.00000 | 0.00711 |
| **Abhd13** | 0.87909 | 0.00391 | 1.32793 | 6.69412 | 0.00553 | 4.37517 | 0.00244 | 2.00132 | 3.15128 | 1.56432 | 1.11440 | 9.28052 | 4.18910 | 4.90098 | 2.58512 | 0.44850 |
| **Abhd14a** | 2.37790 | 0.00475 | 0.04491 | 0.14403 | 0.02977 | 0.00657 | 0.00416 | 0.46432 | 0.00860 | 0.57966 | 0.00685 | 0.02112 | 2.28870 | 0.00478 | 0.81323 | 0.01861 |
| **Abhd14b** | 0.00806 | 0.00762 | 0.30471 | 1.05610 | 0.00679 | 0.00921 | 0.00719 | 0.00993 | 0.94241 | 2.19744 | 0.00408 | 0.01771 | 0.01832 | 0.03842 | 0.17481 | 3.05997 |
| **Abhd15** | 0.03463 | 0.12057 | 0.07508 | 0.28700 | 0.01942 | 0.01222 | 0.01723 | 0.02093 | 0.18987 | 0.05703 | 0.41997 | 0.00502 | 0.04175 | 0.03290 | 0.07825 | 0.01432 |
| **Abhd16a** | 0.00000 | 0.00564 | 3.43846 | 1.99056 | 0.07393 | 1.45317 | 2.76047 | 3.45885 | 0.32417 | 1.23896 | 1.08628 | 0.35564 | 1.41069 | 0.03051 | 0.01013 | 2.52902 |
| **Abhd17a** | 7.39541 | 0.25852 | 0.33491 | 0.01775 | 0.26568 | 0.15403 | 1.23602 | 2.52228 | 0.42728 | 15.65815 | 0.42593 | 1.02925 | 4.14842 | 0.02852 | 0.00000 | 2.36613 |
| **Abhd17b** | 0.02917 | 4.10771 | 0.60070 | 8.22382 | 0.55235 | 0.26805 | 0.00471 | 1.17283 | 5.22151 | 0.44559 | 0.78233 | 1.31096 | 0.76584 | 1.36334 | 1.45683 | 0.35751 |
| **Abhd17c** | 1.29629 | 1.45844 | 0.79270 | 4.61791 | 0.03926 | 4.08555 | 4.66146 | 3.52421 | 6.94023 | 1.78335 | 0.84772 | 0.06568 | 6.27380 | 0.68884 | 6.92689 | 0.21507 |
| **Abhd18** | 0.00000 | 0.94538 | 0.04892 | 1.45709 | 0.02558 | 0.15214 | 0.03751 | 0.02189 | 0.50227 | 0.09228 | 0.00000 | 1.53782 | 2.79302 | 1.03071 | 1.29416 | 0.19898 |
| **Abhd2** | 0.00821 | 0.00829 | 1.03561 | 1.57327 | 0.02673 | 0.00551 | 0.00719 | 0.31116 | 0.64389 | 0.00000 | 0.20881 | 0.02587 | 1.24515 | 0.00342 | 1.07754 | 0.00596 |
| **Abhd3** | 0.00176 | 0.00790 | 0.16163 | 0.18871 | 0.01139 | 0.00906 | 0.00690 | 1.64378 | 0.09277 | 0.25748 | 0.43728 | 0.00000 | 0.00292 | 2.71842 | 1.66807 | 0.00759 |
| **Abhd4** | 3.20950 | 3.99980 | 2.08437 | 4.27565 | 5.10279 | 6.74597 | 0.72511 | 0.66384 | 1.76665 | 2.00207 | 0.85596 | 0.08392 | 1.47325 | 1.44105 | 0.29629 | 3.16187 |
| **Abhd5** | 0.64757 | 0.00264 | 0.37340 | 2.56080 | 0.02119 | 1.71143 | 4.59892 | 1.66134 | 0.62218 | 1.21852 | 1.21609 | 6.92595 | 8.27162 | 0.01508 | 0.31347 | 1.99153 |
| **Abhd6** | 0.00000 | 0.02421 | 3.58160 | 15.85188 | 0.05310 | 0.00430 | 0.00000 | 16.09474 | 12.95614 | 12.51691 | 14.20173 | 2.06288 | 8.55208 | 14.21399 | 16.23278 | 0.28396 |
| **Abhd8** | 0.00875 | 0.00715 | 0.01734 | 0.07112 | 1.10596 | 0.00660 | 0.00705 | 0.02671 | 0.01030 | 0.67401 | 0.05074 | 0.02768 | 0.66858 | 0.01110 | 0.00704 | 0.98859 |
| **Abi1** | 6.36742 | 4.05519 | 2.41462 | 7.48935 | 3.20304 | 0.60812 | 8.62025 | 6.60977 | 8.11455 | 8.03490 | 4.21393 | 17.55593 | 4.27343 | 11.16391 | 5.48160 | 5.16880 |
| **Abi2** | 2.56009 | 0.00000 | 0.72775 | 2.14390 | 0.02869 | 0.01392 | 4.33107 | 0.09825 | 0.16522 | 1.11336 | 0.56004 | 8.57753 | 1.41254 | 1.68092 | 1.21931 | 0.03858 |
| **Abi3bp** | 0.01376 | 0.00806 | 0.01200 | 1.27519 | 0.00933 | 0.00652 | 0.00744 | 0.01255 | 0.00000 | 0.01112 | 0.02484 | 0.00789 | 0.01257 | 0.00439 | 0.27074 | 0.00717 |
| **Abl1** | 5.17716 | 1.35106 | 3.73165 | 6.91470 | 0.01028 | 0.31015 | 0.62255 | 5.89797 | 5.03746 | 6.23585 | 3.48110 | 0.85477 | 4.87671 | 0.51354 | 0.40089 | 6.52259 |
| **Abl2** | 5.47135 | 0.01215 | 1.09543 | 19.03354 | 0.28312 | 0.34186 | 0.08743 | 1.80097 | 10.00410 | 4.66220 | 1.56506 | 2.07717 | 1.63077 | 4.81713 | 14.83999 | 0.00376 |
| **Ablim1** | 0.00000 | 4.39643 | 1.33761 | 0.86000 | 0.01386 | 5.38401 | 2.34375 | 0.33189 | 0.79067 | 1.51827 | 0.04792 | 0.26176 | 1.67108 | 0.29364 | 1.73509 | 0.00210 |
| **Ablim2** | 0.01686 | 0.81590 | 0.00000 | 3.56438 | 0.00865 | 3.83870 | 0.00666 | 0.00508 | 0.00000 | 0.00000 | 0.19766 | 0.00680 | 0.00611 | 0.00027 | 0.05887 | 0.01221 |
| **Ablim3** | 0.01374 | 0.00816 | 0.01203 | 0.35105 | 0.00932 | 0.00663 | 0.00751 | 0.01368 | 0.17833 | 0.01408 | 0.01198 | 0.01856 | 0.01276 | 0.01149 | 0.00452 | 0.00735 |
| **Abo** | 0.01360 | 0.00808 | 0.01075 | 0.01555 | 0.00934 | 0.00652 | 0.00736 | 0.01139 | 0.01183 | 0.00772 | 0.01089 | 0.00713 | 0.01908 | 0.14463 | 0.36438 | 0.00731 |
| **Abr** | 10.41752 | 3.50748 | 1.90544 | 1.39925 | 0.73883 | 0.69748 | 4.84848 | 3.98849 | 0.41576 | 6.00049 | 1.73368 | 1.82144 | 1.81788 | 0.02504 | 1.50354 | 4.85986 |
| **Abracl** | 2.82712 | 0.00284 | 1.27401 | 0.00000 | 3.42311 | 0.00466 | 0.40411 | 0.00000 | 0.30471 | 2.11015 | 0.00000 | 0.13376 | 0.92008 | 1.95557 | 0.78143 | 0.40171 |
| **Abt1** | 3.27956 | 3.41761 | 1.17345 | 5.13274 | 3.86219 | 2.35604 | 3.01488 | 9.62470 | 8.96023 | 4.03020 | 1.07100 | 0.35154 | 0.05987 | 11.27288 | 21.36803 | 1.42260 |
| **Abtb1** | 5.32437 | 1.18399 | 0.72920 | 1.23498 | 1.88097 | 4.33749 | 3.83650 | 0.21348 | 1.73134 | 1.47118 | 0.00000 | 0.00000 | 1.51075 | 0.00265 | 0.00000 | 1.32943 |
| **Abtb2** | 0.41429 | 0.00909 | 0.00000 | 6.71543 | 0.01551 | 0.00585 | 0.00670 | 0.97131 | 5.64956 | 0.70568 | 0.43096 | 0.00000 | 1.23315 | 0.01927 | 0.01136 | 5.13529 |
| **Acaa1a** | 3.91819 | 0.70277 | 2.71420 | 0.56959 | 1.75979 | 3.80721 | 1.34985 | 0.00000 | 0.00000 | 1.05571 | 1.35901 | 0.00790 | 2.23226 | 0.00232 | 0.00000 | 3.40363 |
| **Acaa1b** | 1.70438 | 0.13879 | 0.96567 | 0.28639 | 0.72349 | 1.46615 | 0.73562 | 0.00667 | 0.00453 | 0.44650 | 0.21617 | 0.00174 | 0.73557 | 0.00631 | 0.01706 | 1.37864 |
| **Acaa2** | 13.05254 | 12.04217 | 14.35765 | 0.16507 | 9.34989 | 10.48408 | 4.63847 | 1.85655 | 0.00000 | 0.00000 | 2.43911 | 0.02704 | 4.97099 | 0.00000 | 0.01725 | 11.16812 |
| **Acaca** | 1.43167 | 4.49965 | 6.55104 | 2.08759 | 4.98835 | 0.01133 | 2.15375 | 6.65069 | 1.70152 | 3.28601 | 1.35973 | 0.09127 | 2.00621 | 0.00000 | 1.97532 | 1.65318 |
| **Acacb** | 4.13072 | 3.52472 | 0.66466 | 0.88618 | 0.00000 | 0.00162 | 0.81689 | 8.02166 | 0.93244 | 7.32850 | 3.43054 | 0.61507 | 1.80046 | 0.05636 | 0.27475 | 2.46023 |
| **Acad10** | 0.94883 | 1.14025 | 0.16153 | 0.02276 | 0.53321 | 0.80571 | 0.00551 | 1.01729 | 1.25747 | 0.04461 | 1.31965 | 0.02463 | 0.03645 | 0.00275 | 0.02657 | 2.20971 |
| **Acad11** | 6.41141 | 2.60610 | 3.91101 | 1.75384 | 9.76204 | 0.00362 | 0.10926 | 2.37226 | 1.15747 | 2.05994 | 0.54234 | 1.88753 | 1.48614 | 0.38503 | 4.31156 | 3.20971 |
| **Acad12** | 0.98790 | 3.37679 | 0.49066 | 0.00000 | 3.64326 | 0.61493 | 0.00477 | 0.81518 | 0.00933 | 0.38936 | 0.12436 | 0.00778 | 0.71767 | 0.00685 | 0.00381 | 1.49565 |
| **Acad8** | 8.19882 | 3.22073 | 12.20491 | 1.56417 | 3.23040 | 3.38022 | 5.63383 | 11.53581 | 7.10714 | 5.10392 | 6.46021 | 0.55666 | 8.84182 | 9.07870 | 8.57274 | 2.25271 |
| **Acad9** | 2.82177 | 4.97320 | 7.91444 | 0.32687 | 1.37630 | 0.05781 | 1.32021 | 7.70508 | 2.62461 | 3.45173 | 1.60872 | 1.12853 | 2.99964 | 3.98328 | 4.65168 | 0.28119 |
| **Acadl** | 18.12070 | 10.66908 | 19.74851 | 0.00000 | 12.82105 | 9.51675 | 11.06862 | 18.55855 | 0.84205 | 12.92547 | 21.27062 | 14.08147 | 16.02776 | 9.97651 | 6.20992 | 9.06393 |
| **Acadm** | 12.73706 | 13.61120 | 16.67383 | 3.46093 | 3.94790 | 16.10776 | 11.85104 | 23.58359 | 1.16021 | 23.29116 | 18.11234 | 13.95442 | 13.92948 | 25.55708 | 17.32557 | 5.37389 |
| **Acads** | 2.94318 | 0.16654 | 1.54399 | 0.12451 | 0.03086 | 1.65008 | 1.19446 | 4.38592 | 0.31932 | 8.48464 | 0.31127 | 0.23813 | 1.61519 | 0.00527 | 0.29034 | 0.88010 |
| **Acadsb** | 1.16034 | 0.00641 | 1.06766 | 2.90966 | 0.18254 | 0.05272 | 0.00659 | 0.53638 | 0.61976 | 1.30949 | 0.01400 | 0.02036 | 0.00000 | 0.10908 | 1.19328 | 0.50975 |
| **Acadvl** | 17.23505 | 7.01328 | 8.74358 | 0.66343 | 7.96942 | 10.09002 | 5.63640 | 8.44453 | 0.00000 | 0.79612 | 3.28983 | 0.26967 | 9.29839 | 0.00000 | 0.19348 | 9.57185 |

| | | | | | | | | | | | | | | | | |
|---|---|---|---|---|---|---|---|---|---|---|---|---|---|---|---|---|
| **Acan** | 0.00790 | 0.00759 | 0.03334 | 0.01083 | 0.01906 | 0.00816 | 0.00714 | 0.35613 | 0.01194 | 0.00890 | 0.01062 | 0.02618 | 0.00603 | 0.01313 | 0.00658 | 0.00739 |
| **Acap1** | 0.01360 | 0.00804 | 0.01042 | 0.35609 | 0.00921 | 0.00662 | 0.00763 | 0.01340 | 0.00623 | 0.00000 | 0.21246 | 0.00674 | 0.00956 | 0.00572 | 0.16067 | 0.00721 |
| **Acap2** | 1.34247 | 0.86024 | 1.72025 | 1.83325 | 0.95713 | 0.00378 | 0.97751 | 0.06782 | 0.02264 | 2.07383 | 0.89834 | 5.20597 | 1.80761 | 0.04738 | 0.60636 | 0.00324 |
| **Acap3** | 1.38897 | 0.00508 | 0.19261 | 2.32442 | 0.01328 | 4.01345 | 2.79737 | 0.15549 | 3.52958 | 1.14937 | 0.99467 | 0.23923 | 4.31807 | 0.00000 | 0.00000 | 1.91338 |
| **Acat1** | 2.02588 | 0.32520 | 3.69943 | 3.13811 | 3.88983 | 0.20547 | 4.95201 | 4.25638 | 0.92313 | 3.13231 | 2.10979 | 0.35501 | 3.24914 | 1.89264 | 5.31609 | 0.55618 |
| **Acat2** | 7.66166 | 3.12442 | 12.28131 | 4.38357 | 6.51775 | 2.79742 | 2.01783 | 5.22762 | 4.66964 | 6.38368 | 3.80938 | 1.25622 | 2.17362 | 1.36134 | 4.52640 | 3.42968 |
| **Acat3** | 0.74503 | 0.63548 | 2.42552 | 0.42786 | 0.85330 | 0.01248 | 0.00643 | 0.82307 | 0.39327 | 0.42510 | 0.01848 | 0.02043 | 0.31299 | 0.03179 | 1.24667 | 0.00554 |
| **Acbd3** | 2.56543 | 1.80879 | 1.01643 | 6.33289 | 0.02834 | 2.87734 | 2.11284 | 5.16400 | 3.78241 | 8.02808 | 3.89562 | 1.73110 | 0.58840 | 1.03736 | 10.28725 | 0.05941 |
| **Acbd4** | 0.53906 | 0.01745 | 0.14732 | 0.00000 | 0.00632 | 5.20691 | 0.01516 | 1.90267 | 0.44472 | 3.06847 | 2.98871 | 0.11459 | 2.42916 | 0.00080 | 0.48756 | 0.01775 |
| **Acbd5** | 1.72229 | 0.17770 | 2.24122 | 8.77269 | 3.61549 | 1.49928 | 2.98794 | 4.43928 | 9.80182 | 8.01829 | 3.10146 | 3.00783 | 2.02594 | 15.22088 | 14.32168 | 0.02906 |
| **Acbd6** | 6.25578 | 0.08351 | 0.06753 | 3.43962 | 2.07499 | 0.81720 | 0.22381 | 2.80417 | 1.44052 | 9.24962 | 5.48694 | 0.12599 | 5.27422 | 0.48727 | 0.67348 | 2.25078 |
| **Acbd7** | 0.01316 | 0.00811 | 0.01563 | 1.01567 | 0.00937 | 0.00624 | 0.12981 | 0.01176 | 0.41900 | 0.05002 | 0.01008 | 0.00713 | 0.01207 | 0.00532 | 0.10487 | 0.00719 |
| **Accs** | 0.00000 | 0.07827 | 0.91666 | 0.00459 | 0.00850 | 0.00948 | 2.88417 | 0.01120 | 0.02248 | 0.00000 | 0.13023 | 0.01248 | 1.29367 | 0.00624 | 0.13613 | 0.00649 |
| **Accsl** | 0.01219 | 0.00774 | 0.00000 | 25.41079 | 0.00961 | 0.00295 | 0.00595 | 0.25497 | 7.68069 | 2.44960 | 0.69540 | 0.14271 | 0.00367 | 0.24825 | 12.10058 | 0.00262 |
| **Acd** | 13.68880 | 1.96958 | 4.77459 | 5.48663 | 0.16526 | 6.20317 | 2.53966 | 10.97383 | 3.22484 | 4.89918 | 6.51929 | 1.10221 | 3.98193 | 0.00000 | 0.70649 | 1.82688 |
| **Ace** | 0.17686 | 0.00827 | 0.00770 | 0.11684 | 0.01503 | 0.00698 | 0.00718 | 0.01045 | 0.01025 | 0.50728 | 0.01416 | 0.00610 | 0.06546 | 0.01078 | 0.00779 | 0.00782 |
| **Ace2** | 0.01375 | 0.00811 | 0.01229 | 0.28201 | 0.00931 | 0.00662 | 0.00752 | 0.01393 | 0.00745 | 0.01543 | 0.01405 | 0.00820 | 0.01266 | 0.01130 | 0.00676 | 0.00737 |
| **Acer1** | 0.05275 | 3.00959 | 0.08301 | 0.25224 | 0.04745 | 6.23823 | 0.01453 | 0.00000 | 2.10745 | 0.00000 | 1.04748 | 0.03281 | 1.38227 | 0.19498 | 1.05760 | 4.11863 |
| **Acer2** | 9.39850 | 3.78400 | 5.57902 | 3.12869 | 4.13858 | 2.68483 | 4.75762 | 10.20299 | 2.75592 | 12.02462 | 9.78349 | 2.89082 | 4.52151 | 9.75979 | 6.75990 | 0.53283 |
| **Acer3** | 0.21902 | 0.00752 | 0.89080 | 0.35256 | 0.00594 | 1.05668 | 0.70566 | 0.27983 | 0.69561 | 0.82649 | 0.17375 | 0.29429 | 2.08947 | 4.33803 | 0.00000 | 0.00549 |
| **Acin1** | 8.41513 | 2.31468 | 2.11447 | 0.00000 | 3.01614 | 2.47253 | 1.34700 | 4.39955 | 0.15377 | 9.56375 | 0.07091 | 0.24044 | 2.73358 | 0.83186 | 2.69526 | 2.87193 |
| **Ackr4** | 0.01064 | 0.02827 | 0.69253 | 0.00652 | 0.00890 | 0.00944 | 0.00736 | 0.01181 | 0.01143 | 0.32853 | 0.01199 | 0.00444 | 0.16005 | 0.02003 | 0.51143 | 0.00754 |
| **Acly** | 12.85243 | 12.77281 | 8.06673 | 7.02804 | 10.11504 | 5.33233 | 14.64602 | 18.55121 | 9.21698 | 7.03627 | 7.32478 | 8.10509 | 6.61147 | 6.49863 | 9.42878 | 6.78071 |
| **Acmsd** | 0.04614 | 0.01881 | 0.10454 | 0.49506 | 0.04455 | 0.00758 | 0.00707 | 0.03473 | 0.55045 | 0.01688 | 0.38635 | 0.12578 | 0.03983 | 0.44106 | 0.26397 | 0.02159 |
| **Acnat1** | 0.01623 | 0.03959 | 0.07851 | 0.36881 | 0.00850 | 0.01383 | 0.11923 | 0.07626 | 0.38343 | 0.18235 | 0.02670 | 0.00000 | 0.02911 | 1.14817 | 2.20604 | 0.04974 |
| **Aco1** | 13.00357 | 12.58974 | 3.17746 | 0.68180 | 10.73485 | 4.32400 | 10.36703 | 4.79942 | 1.24431 | 2.35950 | 0.55488 | 0.16121 | 2.20031 | 0.27602 | 1.36398 | 3.83790 |
| **Aco2** | 19.55584 | 9.03164 | 18.33211 | 2.66854 | 10.23969 | 10.37295 | 12.78846 | 22.13698 | 2.14329 | 9.56300 | 9.06777 | 1.85714 | 16.09629 | 0.63342 | 2.89459 | 9.42121 |
| **Acot1** | 0.00726 | 0.00734 | 0.00304 | 0.12242 | 1.82984 | 0.00849 | 0.00817 | 0.22174 | 0.95398 | 0.86381 | 0.13075 | 0.00217 | 0.00684 | 0.01005 | 0.02903 | 0.00751 |
| **Acot10** | 0.82336 | 1.13490 | 1.27662 | 0.03274 | 0.84966 | 0.76800 | 1.21174 | 1.05032 | 0.54336 | 0.37593 | 1.55388 | 1.96028 | 0.68217 | 1.17914 | 0.60060 | 0.19980 |
| **Acot11** | 0.68045 | 0.00654 | 0.39282 | 0.71830 | 0.36187 | 0.79551 | 1.97040 | 1.80506 | 0.52742 | 0.49867 | 0.74047 | 0.25489 | 0.31298 | 0.94508 | 1.01351 | 0.00594 |
| **Acot13** | 3.22943 | 2.36146 | 3.16952 | 5.67286 | 0.00000 | 0.11277 | 1.94955 | 1.43127 | 1.17227 | 3.38185 | 6.91681 | 0.49125 | 1.94705 | 1.28499 | 0.75468 | 0.03436 |
| **Acot2** | 0.04119 | 0.04170 | 0.10511 | 0.20480 | 0.15802 | 0.15558 | 0.00756 | 0.06074 | 0.09741 | 0.11362 | 0.01174 | 0.00751 | 0.54770 | 0.04646 | 0.02069 | 0.00719 |
| **Acot3** | 0.01428 | 0.00822 | 0.00773 | 1.65374 | 0.00938 | 0.16008 | 0.00741 | 0.01310 | 0.00000 | 0.04299 | 0.01297 | 0.00813 | 0.01051 | 0.00969 | 0.00296 | 0.00709 |
| **Acot5** | 0.01444 | 0.00797 | 0.00693 | 0.43422 | 0.00931 | 1.66550 | 0.00719 | 0.01088 | 0.00330 | 0.20417 | 0.00978 | 0.00732 | 0.00465 | 0.01015 | 0.00624 | 0.00674 |
| **Acot7** | 3.93266 | 4.35985 | 1.31841 | 5.08870 | 0.01341 | 3.06350 | 0.91574 | 7.51199 | 5.08447 | 9.52414 | 7.44016 | 0.38290 | 8.72182 | 0.15415 | 1.12515 | 3.67637 |
| **Acot8** | 6.64406 | 2.58172 | 6.27422 | 1.41020 | 2.38969 | 1.80528 | 4.38281 | 2.95285 | 3.52133 | 3.45701 | 2.12901 | 0.66744 | 4.55632 | 0.00000 | 0.00000 | 5.09406 |
| **Acot9** | 6.72412 | 7.90791 | 8.50102 | 0.48531 | 6.83382 | 5.00788 | 7.42853 | 9.05838 | 3.55045 | 3.96687 | 5.94936 | 10.58016 | 6.34147 | 7.22778 | 4.52813 | 1.79413 |
| **Acox1** | 5.34435 | 4.60158 | 5.90728 | 8.66759 | 5.02147 | 0.33769 | 12.06354 | 5.07102 | 9.34466 | 4.04632 | 6.36989 | 0.15494 | 2.18109 | 0.03442 | 3.70253 | 2.53618 |
| **Acox3** | 0.01928 | 0.00817 | 0.05869 | 1.78200 | 0.01033 | 0.01171 | 0.00719 | 0.00948 | 0.41455 | 0.35417 | 0.00000 | 0.03099 | 0.01183 | 1.38845 | 1.71813 | 0.00696 |
| **Acoxl** | 0.05211 | 0.00863 | 0.00911 | 0.47664 | 0.01405 | 0.00670 | 0.00659 | 0.01392 | 0.00519 | 1.95942 | 0.05004 | 0.02361 | 1.37064 | 0.01048 | 0.00016 | 0.00679 |
| **Acp1** | 20.43530 | 15.69776 | 15.59543 | 0.00000 | 10.72578 | 9.31945 | 12.23898 | 11.72499 | 0.64893 | 11.62783 | 5.87790 | 2.04280 | 15.19464 | 5.67862 | 6.32253 | 9.15730 |
| **Acp2** | 0.15028 | 0.07607 | 1.52826 | 0.57770 | 0.00500 | 0.06193 | 2.89587 | 0.83772 | 1.21632 | 0.07682 | 0.79652 | 0.02088 | 0.48481 | 0.07805 | 0.50372 | 1.54289 |
| **Acp5** | 9.50448 | 1.81392 | 9.46454 | 0.13130 | 1.86934 | 3.30782 | 2.85878 | 7.47135 | 0.01240 | 3.72549 | 2.87371 | 0.07041 | 11.24637 | 0.05411 | 0.08088 | 3.83691 |
| **Acp6** | 9.97060 | 0.00000 | 0.00000 | 1.14160 | 0.19227 | 4.97394 | 0.95903 | 3.34667 | 0.01401 | 0.79817 | 0.07742 | 0.07454 | 0.00000 | 0.01085 | 0.11429 | 3.40926 |
| **Acp7** | 0.82963 | 0.01700 | 0.00436 | 0.44348 | 0.00615 | 0.00684 | 0.00643 | 0.19018 | 0.03450 | 0.20813 | 0.14894 | 0.00615 | 0.00615 | 0.01030 | 0.04358 | 0.00610 |
| **Acpp** | 0.11868 | 0.00481 | 1.90490 | 1.30241 | 1.17226 | 0.00541 | 2.56102 | 4.85356 | 2.85978 | 5.14601 | 1.91208 | 0.72955 | 0.86073 | 6.36860 | 7.35437 | 1.61984 |
| **Acrbp** | 0.06759 | 0.00697 | 2.95955 | 1.10335 | 0.07346 | 0.00683 | 1.52532 | 0.00000 | 0.00000 | 2.07704 | 0.73159 | 0.11313 | 0.05774 | 0.23940 | 0.55333 | 0.00602 |
| **Acsbg1** | 9.15865 | 2.88082 | 3.13899 | 0.24529 | 2.19770 | 1.72361 | 3.17818 | 3.30657 | 0.45276 | 0.00037 | 0.22310 | 0.18156 | 0.64687 | 0.01885 | 0.22963 | 0.52002 |
| **Acsbg2** | 0.00000 | 0.00717 | 0.00000 | 0.38251 | 0.01841 | 0.00611 | 0.00630 | 2.97681 | 0.76598 | 2.23076 | 3.19452 | 0.92207 | 0.00000 | 2.41561 | 1.93158 | 0.00646 |
| **Acsf2** | 0.91655 | 0.22171 | 2.05804 | 0.00230 | 0.00272 | 1.45630 | 1.03696 | 0.83656 | 0.04596 | 1.86392 | 1.81840 | 0.08777 | 1.14150 | 0.75125 | 0.15604 | 2.44486 |
| **Acsf3** | 0.87881 | 0.23566 | 0.13805 | 0.00427 | 0.00720 | 0.00494 | 0.00533 | 1.81787 | 0.00929 | 0.00000 | 0.85762 | 0.01022 | 1.97142 | 0.00837 | 0.00432 | 0.00528 |
| **Acsl1** | 1.79289 | 0.01553 | 0.15005 | 4.17363 | 0.00482 | 1.51318 | 3.20790 | 0.37890 | 3.83836 | 2.79062 | 0.04220 | 0.03074 | 0.00006 | 0.00000 | 2.52607 | 0.00416 |
| **Acsl3** | 2.75512 | 7.67519 | 4.79860 | 5.66482 | 6.42566 | 2.28616 | 8.61721 | 6.06456 | 14.72616 | 7.75768 | 5.83036 | 0.21383 | 0.60770 | 17.63918 | 11.45287 | 2.82777 |
| **Acsl4** | 2.55454 | 1.16497 | 6.00740 | 10.01118 | 6.90066 | 0.00259 | 5.70368 | 12.67482 | 20.52038 | 17.30293 | 4.43378 | 19.04475 | 5.20533 | 35.16666 | 20.35349 | 0.02969 |
| **Acsl5** | 0.21302 | 1.79428 | 0.00000 | 0.00792 | 5.11363 | 0.00685 | 0.00631 | 0.00347 | 0.01021 | 0.00216 | 0.01245 | 0.00221 | 0.00782 | 0.01317 | 0.02742 | 0.95044 |
| **Acsl6** | 0.04623 | 0.00796 | 0.00000 | 0.00178 | 0.00845 | 0.16333 | 0.00624 | 0.00000 | 0.00911 | 0.20950 | 1.51031 | 0.05181 | 0.88530 | 0.00381 | 0.96119 | 0.01388 |
| **Acsm2** | 0.02049 | 0.00809 | 0.02781 | 0.11933 | 0.00861 | 0.03667 | 0.00763 | 0.01796 | 0.01383 | 0.04373 | 0.04702 | 0.02976 | 0.02507 | 0.06593 | 0.06997 | 0.00728 |
| **Acsm3** | 0.00000 | 0.01709 | 0.21091 | 0.00957 | 0.95540 | 2.16956 | 0.00563 | 0.72468 | 0.01113 | 0.00000 | 0.00360 | 0.03321 | 3.03317 | 0.04531 | 0.00000 | 0.07455 |
| **Acss1** | 3.07121 | 2.08450 | 1.58973 | 0.23982 | 0.00987 | 0.03219 | 2.07626 | 2.02364 | 0.00426 | 0.00000 | 0.59422 | 0.03749 | 2.17519 | 0.46850 | 0.99862 | 0.34674 |
| **Acss2** | 3.40523 | 1.82755 | 0.16288 | 0.30904 | 1.76162 | 2.80315 | 0.00325 | 2.94210 | 1.92032 | 0.80597 | 0.78343 | 0.02422 | 0.75270 | 0.00000 | 2.06873 | 0.00915 |
| **Acta1** | 1.10638 | 0.34434 | 1.18109 | 0.00195 | 0.99041 | 1.25664 | 0.72354 | 1.43707 | 0.07946 | 0.22506 | 0.01093 | 0.06020 | 1.60998 | 0.01171 | 0.01504 | 0.81948 |
| **Acta2** | 0.91338 | 0.57514 | 1.40292 | 0.00000 | 1.10456 | 1.12198 | 0.70766 | 1.09247 | 0.00850 | 4.18179 | 0.00000 | 0.86807 | 1.53157 | 0.05863 | 0.00000 | 0.67151 |

| | | | | | | | | | | | | | | | | |
|---|---|---|---|---|---|---|---|---|---|---|---|---|---|---|---|---|
| **Actb** | 29.40271 | 7.66673 | 36.29374 | 1.10668 | 19.03271 | 13.84283 | 10.57997 | 13.93549 | 4.53701 | 14.21363 | 30.95542 | 2.93441 | 26.20889 | 21.09907 | 6.83150 | 20.77288 |
| **Actg1** | 55.14451 | 31.13930 | 54.74761 | 6.43094 | 42.60748 | 32.05160 | 32.58648 | 55.13984 | 3.19502 | 33.16637 | 41.11180 | 22.16416 | 59.27234 | 14.80574 | 9.65471 | 31.98801 |
| **Actg2** | 0.01289 | 0.00770 | 0.00824 | 0.00501 | 0.00878 | 0.00663 | 0.00711 | 0.00951 | 0.01049 | 1.81441 | 0.00763 | 0.33212 | 0.01051 | 0.00777 | 0.26495 | 0.00727 |
| **Actl6a** | 10.42297 | 7.05199 | 8.73270 | 5.51080 | 12.55033 | 5.18371 | 6.36289 | 14.06001 | 8.51772 | 9.28190 | 7.47482 | 2.57165 | 11.52346 | 14.91024 | 11.36707 | 8.07743 |
| **Actn1** | 3.77084 | 0.22784 | 0.51497 | 4.11852 | 3.91360 | 0.01606 | 0.98016 | 0.71492 | 1.25501 | 0.04802 | 6.02694 | 0.45182 | 0.94261 | 0.29918 | 0.00000 | 0.53756 |
| **Actn2** | 6.41131 | 4.97357 | 11.50719 | 2.32738 | 0.02924 | 4.62344 | 0.29137 | 22.67674 | 4.86927 | 16.72450 | 24.47068 | 2.59780 | 11.89708 | 1.56761 | 4.57195 | 0.66463 |
| **Actn4** | 11.40651 | 11.14681 | 15.63472 | 1.29215 | 1.82438 | 16.63877 | 14.16468 | 34.00691 | 1.11931 | 17.44595 | 4.07854 | 3.10455 | 23.84683 | 0.00000 | 0.57147 | 5.95391 |
| **Actr10** | 5.19431 | 10.81763 | 8.12804 | 5.28910 | 9.20403 | 0.73719 | 3.74245 | 11.66160 | 9.65011 | 9.17008 | 12.78999 | 1.32607 | 11.24258 | 5.20839 | 6.75801 | 4.91742 |
| **Actr1a** | 9.06135 | 0.26645 | 5.65015 | 10.96668 | 3.15325 | 7.36652 | 4.03006 | 12.34063 | 5.27443 | 12.25194 | 6.41793 | 2.65094 | 5.49826 | 1.03933 | 4.00005 | 5.07036 |
| **Actr1b** | 5.50093 | 4.49868 | 3.86001 | 2.23390 | 10.51397 | 4.14858 | 5.13555 | 5.69990 | 1.65375 | 2.29991 | 3.61186 | 0.59181 | 7.19009 | 0.34973 | 2.93647 | 3.94864 |
| **Actr2** | 8.62336 | 4.26273 | 7.43394 | 4.85487 | 6.15951 | 2.49240 | 6.20278 | 5.63684 | 5.04890 | 4.65894 | 7.52782 | 9.35408 | 6.37538 | 11.24485 | 8.52608 | 1.98426 |
| **Actr3** | 17.25851 | 16.05499 | 18.93555 | 6.90331 | 16.95021 | 13.33937 | 13.59513 | 11.10887 | 6.88391 | 11.05317 | 6.57631 | 11.61955 | 19.99518 | 15.89234 | 11.82042 | 9.51403 |
| **Actr3b** | 0.58250 | 0.00686 | 0.44226 | 0.56009 | 0.00614 | 0.00738 | 0.00662 | 0.17746 | 0.00290 | 1.39673 | 1.15690 | 0.00688 | 0.25715 | 0.00294 | 0.08562 | 1.27373 |
| **Actr5** | 0.78136 | 0.01613 | 0.00625 | 3.94776 | 0.97832 | 0.00604 | 0.01472 | 7.23277 | 9.83515 | 13.04764 | 3.65836 | 0.35710 | 2.08038 | 0.12857 | 0.14997 | 0.71270 |
| **Actr6** | 3.50352 | 1.44060 | 2.31143 | 2.03340 | 0.00998 | 1.12354 | 2.38420 | 3.37341 | 8.01935 | 0.00000 | 0.55224 | 0.95525 | 1.17470 | 15.75952 | 8.78104 | 0.01414 |
| **Actr8** | 6.66089 | 4.07716 | 5.27574 | 0.36266 | 0.03214 | 0.00000 | 1.52713 | 10.07059 | 0.98309 | 7.60582 | 4.35914 | 2.30680 | 2.59420 | 2.00773 | 5.69731 | 4.38041 |
| **Acvr1** | 0.25838 | 1.95153 | 1.11048 | 0.16560 | 0.01551 | 0.00562 | 0.02711 | 0.90400 | 0.42047 | 0.61751 | 1.00105 | 1.09738 | 0.00000 | 0.47035 | 0.29331 | 1.26264 |
| **Acvr1b** | 0.00723 | 0.10224 | 0.01056 | 0.14373 | 0.00830 | 0.01292 | 0.02772 | 0.01100 | 0.00692 | 0.00539 | 0.01236 | 0.01816 | 0.37207 | 0.01153 | 0.00591 | 0.50156 |
| **Acvr2a** | 0.17943 | 0.01649 | 0.45454 | 1.75799 | 0.01054 | 1.34999 | 2.33593 | 1.65530 | 1.19516 | 2.88119 | 1.17135 | 3.65561 | 2.05793 | 5.78340 | 1.02627 | 0.00488 |
| **Acvr2b** | 1.58794 | 0.00548 | 1.02359 | 0.29601 | 0.01066 | 3.99727 | 0.05414 | 0.07072 | 0.89766 | 0.66633 | 0.81132 | 0.01524 | 4.28512 | 0.18868 | 0.60192 | 1.77307 |
| **Acvrl1** | 0.01360 | 0.00793 | 0.01203 | 0.70271 | 0.00922 | 0.00676 | 0.00754 | 0.01322 | 0.00000 | 0.18662 | 0.01135 | 0.00785 | 0.01295 | 0.00450 | 0.26968 | 0.00730 |
| **Acy1** | 0.71335 | 3.35182 | 0.48891 | 0.20262 | 0.00619 | 0.11750 | 0.00532 | 2.18072 | 0.00730 | 0.07532 | 1.75227 | 0.03347 | 0.00000 | 0.03672 | 1.48532 | 0.34235 |
| **Acy3** | 0.97450 | 0.00634 | 0.69864 | 5.69407 | 0.00462 | 0.00709 | 0.00386 | 0.74755 | 6.70445 | 0.00000 | 1.13278 | 0.00000 | 3.23885 | 0.00000 | 0.06416 | 0.00299 |
| **Acyp1** | 1.10169 | 0.04701 | 0.36816 | 2.26453 | 3.22491 | 0.00423 | 0.01599 | 0.27053 | 0.47485 | 0.91896 | 2.14857 | 0.29345 | 0.00000 | 0.03487 | 1.01607 | 1.11705 |
| **Acyp2** | 0.08716 | 0.00642 | 0.18961 | 0.00000 | 2.10084 | 0.00785 | 0.25362 | 0.04684 | 0.28782 | 0.02365 | 0.33621 | 0.04117 | 0.04493 | 0.13038 | 0.45549 | 0.01443 |
| **Ada** | 0.02903 | 0.00748 | 0.96355 | 1.32061 | 0.01096 | 0.00640 | 0.00678 | 1.86791 | 0.26749 | 0.00000 | 0.54190 | 0.00855 | 0.00000 | 0.01726 | 0.00000 | 0.01480 |
| **Adad2** | 0.00000 | 0.00800 | 0.39501 | 0.00725 | 0.07935 | 0.00972 | 0.05112 | 0.77618 | 0.02097 | 0.37443 | 0.09080 | 0.00552 | 2.14228 | 0.01544 | 0.00000 | 0.54037 |
| **Adal** | 0.00000 | 1.43093 | 1.12990 | 0.19223 | 0.03289 | 0.00675 | 3.01167 | 0.82762 | 0.08359 | 1.87507 | 1.58861 | 0.19118 | 0.79915 | 2.17398 | 3.63177 | 0.02704 |
| **Adam10** | 2.51346 | 1.42486 | 3.13518 | 6.41708 | 0.73049 | 0.02343 | 4.58893 | 1.78701 | 4.60000 | 5.01741 | 1.83382 | 1.29254 | 3.82303 | 17.33883 | 6.83237 | 1.35715 |
| **Adam12** | 0.01353 | 0.00824 | 0.01165 | 0.58989 | 0.00927 | 0.00653 | 0.00751 | 0.01365 | 1.54355 | 0.00855 | 0.00000 | 0.01698 | 0.01316 | 0.01190 | 0.00268 | 0.00722 |
| **Adam15** | 0.44830 | 0.00572 | 0.27495 | 0.02096 | 0.05593 | 0.07757 | 1.29321 | 1.13413 | 0.37377 | 1.03627 | 1.83212 | 0.01728 | 3.32469 | 0.01296 | 0.00000 | 3.70616 |
| **Adam17** | 2.20678 | 3.35148 | 5.91309 | 8.71884 | 0.10464 | 2.81164 | 1.20600 | 8.03448 | 13.83238 | 11.46796 | 10.19640 | 8.13353 | 0.18428 | 3.38788 | 11.97989 | 2.84385 |
| **Adam19** | 6.84421 | 0.00000 | 2.39632 | 0.52511 | 6.38379 | 4.04164 | 4.73161 | 0.00000 | 0.94780 | 0.19059 | 0.03129 | 0.13383 | 2.53567 | 0.00954 | 1.38349 | 0.17716 |
| **Adam1a** | 0.00980 | 0.00779 | 0.01069 | 0.79350 | 0.00964 | 0.00625 | 0.00749 | 0.22414 | 1.08919 | 0.00000 | 0.01873 | 0.00390 | 0.00662 | 0.03118 | 0.64808 | 0.00703 |
| **Adam2** | 0.10127 | 0.00718 | 0.00538 | 0.29397 | 0.00764 | 0.00720 | 0.35539 | 0.00987 | 0.11280 | 0.00816 | 0.01121 | 0.00733 | 0.01042 | 0.01694 | 0.02052 | 0.26803 |
| **Adam22** | 0.01382 | 0.00780 | 0.01159 | 0.23391 | 0.00935 | 0.00666 | 0.00748 | 0.01295 | 0.03450 | 0.01218 | 0.01182 | 0.02798 | 0.01281 | 0.01838 | 0.04517 | 0.00738 |
| **Adam28** | 0.16099 | 0.01863 | 0.27483 | 0.30963 | 0.12105 | 0.06436 | 0.14322 | 0.02610 | 0.46126 | 0.15023 | 1.09208 | 0.03168 | 0.16724 | 0.44805 | 0.16632 | 0.07859 |
| **Adam33** | 0.01346 | 0.00823 | 0.01201 | 0.38626 | 0.00907 | 0.00642 | 0.00742 | 0.01119 | 0.87853 | 0.41874 | 0.00653 | 0.00550 | 0.01119 | 0.01029 | 0.00393 | 0.00701 |
| **Adam4** | 0.01369 | 0.00787 | 0.00985 | 0.07196 | 0.00914 | 0.00667 | 0.00742 | 0.00826 | 0.61827 | 0.00368 | 0.16442 | 0.00606 | 0.01683 | 0.09282 | 0.00000 | 0.00727 |
| **Adam8** | 1.58376 | 0.00596 | 0.20924 | 1.02454 | 0.00447 | 0.00719 | 0.00606 | 0.00968 | 0.63505 | 0.00591 | 0.02400 | 0.00501 | 0.00998 | 0.00475 | 0.13046 | 0.01945 |
| **Adam9** | 0.67147 | 0.01663 | 0.28873 | 2.40319 | 1.00065 | 1.25791 | 0.21611 | 0.48872 | 0.20960 | 0.00000 | 1.07242 | 1.47876 | 0.27584 | 2.94188 | 0.68098 | 0.99406 |
| **Adamts10** | 0.01477 | 0.00830 | 0.01717 | 0.00583 | 0.01981 | 0.00723 | 0.00752 | 0.01439 | 0.01131 | 1.97214 | 0.00971 | 0.04611 | 0.01039 | 0.01005 | 0.02595 | 0.00736 |
| **Adamts12** | 0.01371 | 0.00766 | 0.01053 | 0.00929 | 0.00913 | 0.00675 | 0.00727 | 0.01069 | 0.01203 | 0.33040 | 0.01212 | 0.09047 | 0.01253 | 0.03214 | 0.01050 | 0.00780 |
| **Adamts13** | 1.52108 | 0.23724 | 3.73532 | 3.65527 | 0.53416 | 0.50062 | 2.35937 | 2.29269 | 4.06539 | 6.06664 | 1.59572 | 7.67429 | 0.86354 | 2.15054 | 2.26354 | 0.26839 |
| **Adamts14** | 0.01385 | 0.00817 | 0.00940 | 0.01472 | 0.00918 | 0.00688 | 0.00757 | 0.00679 | 0.01083 | 0.00637 | 0.36838 | 0.01743 | 0.00924 | 0.01055 | 0.00471 | 0.00723 |
| **Adamts15** | 0.01375 | 0.00811 | 0.01225 | 0.09136 | 0.00930 | 0.00664 | 0.00753 | 0.01389 | 0.02116 | 0.01510 | 0.01373 | 0.00823 | 0.01268 | 0.01155 | 0.01292 | 0.00740 |
| **Adamts17** | 0.02352 | 0.00775 | 0.02041 | 0.31564 | 0.02376 | 0.02162 | 0.00715 | 0.01373 | 0.01219 | 0.91478 | 0.02847 | 0.00504 | 0.01663 | 0.00856 | 0.17009 | 0.00712 |
| **Adamts2** | 0.14807 | 0.18735 | 0.14134 | 22.79298 | 0.05054 | 0.15543 | 0.07603 | 0.18656 | 17.51317 | 3.61782 | 0.53852 | 0.98757 | 0.34233 | 4.15279 | 9.31668 | 0.08026 |
| **Adamts4** | 0.16081 | 0.01792 | 0.23799 | 2.12313 | 0.04395 | 0.05051 | 0.04674 | 0.00000 | 1.51017 | 0.34331 | 2.17430 | 0.10692 | 0.05075 | 0.78161 | 0.15293 | 0.05982 |
| **Adamts5** | 0.01377 | 0.00806 | 0.01152 | 0.01057 | 0.00928 | 0.00684 | 0.00756 | 0.01376 | 0.01224 | 0.03339 | 0.01234 | 0.04176 | 0.01310 | 0.01191 | 0.01085 | 0.00742 |
| **Adamts7** | 0.00815 | 0.00841 | 0.00433 | 0.01056 | 0.00952 | 0.70606 | 0.00759 | 0.55142 | 0.01181 | 0.00000 | 0.00891 | 0.00758 | 1.09352 | 0.02006 | 0.00069 | 0.02084 |
| **Adamtsl1** | 0.01524 | 0.11264 | 0.00995 | 2.59379 | 0.01000 | 0.00626 | 0.00720 | 0.01021 | 0.63191 | 0.00217 | 0.02544 | 0.01610 | 0.01210 | 0.00380 | 0.54493 | 0.00681 |
| **Adamtsl4** | 0.64857 | 0.00703 | 0.78486 | 0.92455 | 0.01092 | 0.00844 | 0.00646 | 1.01734 | 0.12230 | 2.83015 | 0.09533 | 0.06101 | 0.00190 | 0.00327 | 1.12106 | 0.00642 |
| **Adamtsl5** | 0.00000 | 0.00694 | 0.15570 | 0.00450 | 0.00734 | 0.00953 | 0.00520 | 0.72891 | 0.27485 | 0.20511 | 3.39908 | 0.10602 | 1.15849 | 0.00662 | 0.07361 | 2.87885 |
| **Adap1** | 1.28808 | 0.01786 | 0.71248 | 0.74750 | 0.00400 | 0.00811 | 0.00672 | 0.62157 | 0.20087 | 0.71280 | 0.00539 | 0.05500 | 0.19217 | 0.00988 | 0.00000 | 0.01334 |
| **Adap2** | 0.00829 | 0.00803 | 0.14430 | 0.00458 | 0.00889 | 0.46553 | 0.00689 | 0.37911 | 0.01023 | 0.00000 | 0.20153 | 0.00454 | 1.04947 | 0.09625 | 0.45602 | 0.02143 |
| **Adar** | 5.85039 | 6.80454 | 3.50501 | 2.38475 | 0.00000 | 0.00000 | 0.01955 | 5.40856 | 1.16917 | 0.98883 | 1.67418 | 1.51722 | 0.84135 | 0.37761 | 2.68151 | 5.96548 |
| **Adarb1** | 0.31233 | 0.10548 | 0.71155 | 2.73231 | 0.04787 | 0.05995 | 0.07909 | 0.04957 | 3.14749 | 0.22620 | 2.52544 | 0.21354 | 0.98864 | 0.79347 | 0.58998 | 2.21202 |
| **Adarb2** | 5.66069 | 1.33356 | 7.52648 | 9.30109 | 1.66798 | 0.91592 | 5.54742 | 1.61778 | 7.20450 | 2.96539 | 3.55325 | 1.48185 | 3.02178 | 1.70088 | 0.95811 | 4.85169 |
| **Adat1** | 0.20277 | 4.13337 | 1.03409 | 0.19387 | 0.00649 | 0.00592 | 0.00599 | 0.01338 | 0.01274 | 1.35573 | 0.31049 | 0.11130 | 0.22284 | 0.00422 | 1.19137 | 0.14893 |
| **Adat2** | 2.75505 | 0.00455 | 0.84719 | 0.00390 | 0.02040 | 0.01969 | 0.09489 | 0.83091 | 0.02309 | 0.97979 | 2.32017 | 0.00000 | 0.57825 | 0.82505 | 0.00000 | 0.03972 |
| **Adat3** | 0.17465 | 0.01869 | 0.00000 | 0.61174 | 0.12767 | 0.23161 | 0.00502 | 0.02349 | 2.71819 | 2.19972 | 1.29035 | 0.10093 | 3.20557 | 0.02182 | 0.00000 | 1.62831 |

| | | | | | | | | | | | | | | | | |
|---|---|---|---|---|---|---|---|---|---|---|---|---|---|---|---|---|
| **Adck1** | 1.21201 | 0.03105 | 3.49709 | 0.54394 | 6.25396 | 2.59623 | 6.97347 | 4.56953 | 1.23280 | 3.68900 | 1.88958 | 1.06061 | 0.17196 | 0.03122 | 1.47588 | 0.03286 |
| **Adck2** | 0.11146 | 0.00912 | 0.00000 | 0.35158 | 0.01278 | 2.51600 | 0.00783 | 0.24918 | 0.26724 | 0.00000 | 0.00433 | 0.04639 | 0.00000 | 0.00850 | 0.97812 | 2.06831 |
| **Adck5** | 7.05549 | 0.38741 | 5.13410 | 1.25747 | 2.50219 | 2.85718 | 0.87811 | 1.23134 | 1.16202 | 0.19677 | 0.98870 | 0.02977 | 2.86578 | 0.00074 | 0.22267 | 4.21343 |
| **Adcy1** | 0.01294 | 0.00803 | 0.01216 | 0.01667 | 0.01878 | 0.00666 | 0.00754 | 0.01394 | 0.07458 | 0.01491 | 0.01217 | 0.00804 | 0.01265 | 0.01250 | 0.00976 | 0.00740 |
| **Adcy10** | 0.01713 | 0.00862 | 0.00784 | 0.07814 | 0.01077 | 0.00690 | 0.00791 | 0.03459 | 0.00780 | 0.15403 | 0.00753 | 0.02791 | 0.01163 | 0.55217 | 0.21244 | 0.00747 |
| **Adcy3** | 0.01270 | 0.00796 | 0.01343 | 0.13116 | 0.01005 | 1.30457 | 0.00742 | 0.57273 | 0.00942 | 0.18508 | 0.00821 | 0.02551 | 0.00000 | 0.01014 | 0.00893 | 0.00697 |
| **Adcy5** | 0.01257 | 0.00783 | 0.13185 | 0.04864 | 0.00901 | 0.00569 | 0.00753 | 0.01178 | 1.15449 | 0.12597 | 0.00000 | 0.02536 | 0.01151 | 0.01403 | 0.14521 | 0.00763 |
| **Adcy6** | 0.00813 | 0.00753 | 0.00000 | 0.63313 | 0.00946 | 0.01348 | 0.00671 | 1.31424 | 0.02096 | 0.53165 | 1.26521 | 0.04373 | 0.00000 | 1.56176 | 0.81339 | 0.00711 |
| **Adcy7** | 0.01088 | 0.00805 | 0.35733 | 0.83757 | 0.00846 | 0.00886 | 0.00718 | 0.04051 | 0.27330 | 0.47925 | 0.00892 | 0.00592 | 0.00782 | 0.00576 | 0.07621 | 0.00683 |
| **Adcy9** | 0.43479 | 0.00422 | 8.22132 | 0.70972 | 2.58858 | 0.01277 | 0.36097 | 1.61462 | 0.08809 | 4.86558 | 0.76628 | 0.17633 | 0.00000 | 2.06185 | 1.88648 | 2.61160 |
| **Adcyap1r1** | 0.01977 | 0.00791 | 0.01963 | 0.19590 | 0.00984 | 0.01994 | 0.00725 | 0.01227 | 0.43784 | 0.00918 | 0.01865 | 0.00713 | 0.38124 | 0.01662 | 0.16324 | 0.01424 |
| **Add1** | 8.50754 | 6.00369 | 5.18976 | 7.84985 | 4.83782 | 0.02010 | 9.20430 | 2.44737 | 2.78909 | 9.68568 | 1.14644 | 5.07736 | 7.47499 | 0.01133 | 0.00000 | 1.52983 |
| **Add2** | 0.01763 | 0.00792 | 0.01207 | 0.16528 | 0.01166 | 0.00704 | 0.00758 | 0.01339 | 0.33609 | 0.00000 | 0.01122 | 0.05587 | 0.15068 | 0.01325 | 0.40598 | 0.00762 |
| **Add3** | 3.57461 | 4.98975 | 2.83494 | 0.98448 | 4.08380 | 0.00188 | 2.01937 | 2.82898 | 0.62431 | 1.05637 | 0.00000 | 0.02123 | 0.00000 | 0.00234 | 0.29359 | 1.45891 |
| **Adgra2** | 0.03748 | 0.00753 | 0.00891 | 1.75872 | 0.00846 | 0.71196 | 0.00728 | 0.05644 | 5.37344 | 0.18900 | 0.16984 | 0.09256 | 0.15087 | 0.05600 | 0.26212 | 0.01421 |
| **Adgra3** | 0.07769 | 0.00776 | 0.36144 | 4.23082 | 0.02644 | 0.35317 | 0.00679 | 0.00745 | 0.07845 | 1.99540 | 0.03739 | 0.07231 | 0.00626 | 0.00140 | 1.63476 | 1.18672 |
| **Adgre5** | 6.46742 | 0.52929 | 8.13933 | 0.00000 | 5.58724 | 6.19307 | 5.11654 | 9.66040 | 0.00060 | 0.00000 | 3.47378 | 0.00615 | 3.28642 | 0.00010 | 0.00979 | 1.24036 |
| **Adgrf3** | 0.00724 | 0.00803 | 0.00124 | 0.07920 | 0.00910 | 0.00826 | 0.00739 | 0.67314 | 0.01056 | 1.44200 | 1.62330 | 0.01705 | 0.00000 | 0.00789 | 0.48244 | 0.00709 |
| **Adgrg1** | 0.00514 | 0.00837 | 0.00285 | 0.56212 | 0.00904 | 0.00832 | 0.04120 | 0.28902 | 0.18734 | 0.42180 | 1.31260 | 0.02808 | 1.56851 | 0.00598 | 0.00000 | 0.01371 |
| **Adgrg2** | 0.01366 | 0.00818 | 0.01206 | 0.12242 | 0.00922 | 0.00686 | 0.00760 | 0.01412 | 0.07916 | 0.11391 | 0.01253 | 0.00886 | 0.01304 | 0.01122 | 0.00920 | 0.00740 |
| **Adgrg3** | 0.11215 | 0.00797 | 0.00668 | 0.00459 | 0.00774 | 0.00717 | 0.00753 | 0.01564 | 0.31283 | 0.00507 | 0.17056 | 0.06014 | 0.01152 | 0.01117 | 0.00775 | 0.00715 |
| **Adgrg6** | 4.18686 | 0.00231 | 0.00339 | 0.00915 | 1.87808 | 0.00498 | 0.00436 | 0.00553 | 0.01131 | 0.07128 | 0.01122 | 0.00436 | 0.01731 | 0.01197 | 0.01272 | 0.00292 |
| **Adgrl1** | 2.48590 | 0.00000 | 1.81852 | 1.66278 | 3.17694 | 0.01529 | 3.22604 | 0.10470 | 0.58194 | 0.10108 | 0.00532 | 0.02971 | 1.54904 | 0.00871 | 0.00000 | 0.12444 |
| **Adgrl2** | 0.00000 | 1.88953 | 0.00684 | 0.07436 | 0.00754 | 0.00637 | 1.72172 | 0.00908 | 0.00817 | 0.03134 | 0.01125 | 0.01645 | 0.46323 | 0.00714 | 0.00000 | 0.85270 |
| **Adh1** | 0.00501 | 0.01848 | 0.03024 | 0.00666 | 0.06423 | 5.34487 | 0.00661 | 0.00971 | 0.01300 | 0.19678 | 0.00774 | 0.00589 | 0.02061 | 0.01136 | 0.00983 | 14.55291 |
| **Adh5** | 15.97264 | 13.65097 | 14.66812 | 11.39314 | 13.96681 | 2.56032 | 11.94716 | 11.89144 | 1.05724 | 7.07170 | 4.75525 | 5.54461 | 11.41109 | 1.80174 | 8.37908 | 8.93751 |
| **Adhfe1** | 0.00731 | 0.20927 | 0.00869 | 0.77066 | 0.00849 | 0.00683 | 0.00813 | 0.25543 | 1.09220 | 0.46705 | 0.00000 | 0.01710 | 0.00704 | 0.01152 | 0.00000 | 0.00736 |
| **Adi1** | 16.82428 | 1.36027 | 6.65999 | 8.84244 | 3.16081 | 0.13317 | 3.69523 | 7.44051 | 5.90980 | 7.36984 | 5.10489 | 1.29098 | 10.37103 | 4.89600 | 2.99290 | 5.46980 |
| **Adipor1** | 9.58693 | 8.13582 | 9.60544 | 6.87897 | 8.61462 | 8.86649 | 14.56342 | 9.55316 | 6.44648 | 16.37717 | 8.21123 | 6.87214 | 6.35794 | 6.43455 | 9.50503 | 6.74537 |
| **Adipor2** | 10.25136 | 3.97568 | 6.38237 | 2.59432 | 7.25138 | 8.24816 | 3.93228 | 2.01153 | 4.18107 | 6.02122 | 0.77957 | 1.07912 | 8.09853 | 1.86803 | 4.42785 | 3.55922 |
| **Adk** | 2.42507 | 9.37565 | 3.23076 | 2.36219 | 3.16046 | 1.12342 | 2.43090 | 0.00000 | 4.28307 | 1.83015 | 0.00000 | 0.09049 | 1.15680 | 0.00000 | 3.83257 | 5.05199 |
| **Adm** | 0.01560 | 0.02837 | 0.17160 | 0.16118 | 0.00882 | 0.00581 | 0.00725 | 0.01119 | 0.35152 | 0.10424 | 0.00985 | 0.02508 | 0.01108 | 0.01150 | 0.00985 | 0.00759 |
| **Adm2** | 0.01377 | 0.00795 | 0.01216 | 0.73820 | 0.00925 | 0.00662 | 0.00742 | 0.01274 | 0.00666 | 0.08551 | 0.01311 | 0.00747 | 0.01299 | 0.01186 | 0.00935 | 0.00764 |
| **Adnp2** | 7.76115 | 2.96805 | 0.07253 | 2.06702 | 0.02439 | 0.00000 | 0.01738 | 0.06608 | 2.87045 | 0.97301 | 0.76852 | 5.33669 | 2.92696 | 0.01274 | 5.10273 | 1.28846 |
| **Ado** | 3.19999 | 3.02609 | 1.32423 | 0.46547 | 0.01212 | 0.01379 | 0.00255 | 1.39399 | 2.40248 | 1.04656 | 0.62763 | 0.05047 | 1.65479 | 2.30814 | 0.81161 | 0.00810 |
| **Adora1** | 1.43483 | 0.04357 | 0.89812 | 0.00925 | 2.09921 | 0.00752 | 1.48646 | 0.00000 | 0.01116 | 0.00000 | 0.01182 | 0.00173 | 1.12253 | 0.01781 | 0.00038 | 6.99420 |
| **Adpgk** | 1.69063 | 1.88348 | 4.55756 | 2.56685 | 2.19334 | 2.81681 | 3.23480 | 4.00754 | 4.87193 | 0.73612 | 1.97840 | 5.39115 | 5.51719 | 0.37897 | 4.63001 | 3.25359 |
| **Adprh** | 13.01595 | 3.90044 | 4.02139 | 2.62514 | 9.18799 | 6.29701 | 7.99111 | 16.64145 | 1.04366 | 7.29500 | 8.32006 | 1.05207 | 9.81608 | 0.00000 | 0.00000 | 6.93222 |
| **Adprhl1** | 0.00895 | 0.00741 | 0.01518 | 0.03763 | 0.00928 | 0.00624 | 0.01640 | 0.71023 | 0.17387 | 0.00000 | 0.09002 | 0.00488 | 0.83445 | 0.01354 | 0.00000 | 0.00678 |
| **Adprhl2** | 6.16370 | 3.76275 | 5.89579 | 0.06392 | 0.03535 | 0.04723 | 0.57846 | 2.28932 | 0.30256 | 6.62397 | 0.00000 | 0.44255 | 3.28342 | 0.00256 | 0.58625 | 3.81689 |
| **Adprm** | 4.85683 | 0.00016 | 3.31919 | 1.87259 | 1.17884 | 0.00008 | 0.01292 | 0.00000 | 0.44190 | 1.72135 | 0.29775 | 0.01206 | 0.66796 | 0.00523 | 0.17438 | 1.79794 |
| **Adrb3** | 0.19514 | 0.00603 | 0.08254 | 0.00457 | 0.00655 | 0.13217 | 2.14134 | 0.78260 | 0.01091 | 0.00000 | 0.19113 | 0.00267 | 0.45187 | 0.00576 | 0.09838 | 3.07653 |
| **Adrbk1** | 4.64348 | 0.52508 | 2.32703 | 0.90372 | 5.05356 | 0.19485 | 1.25978 | 6.55343 | 1.35328 | 14.63334 | 2.80642 | 0.84472 | 3.24973 | 0.00000 | 0.35170 | 0.40778 |
| **Adrm1** | 23.61496 | 1.50133 | 2.55990 | 21.61121 | 16.01623 | 2.91676 | 3.24716 | 13.55168 | 22.73638 | 37.97671 | 29.35349 | 2.76199 | 34.78437 | 0.21067 | 0.00000 | 12.92497 |
| **Adsl** | 7.04947 | 7.34639 | 9.31100 | 11.97093 | 9.23486 | 2.55037 | 2.80554 | 13.96897 | 15.26257 | 13.36837 | 9.58622 | 2.76792 | 11.06278 | 1.59960 | 7.57842 | 6.28271 |
| **Adss** | 12.23394 | 3.87291 | 11.13967 | 6.93226 | 10.06829 | 5.19096 | 8.18784 | 7.82439 | 5.10241 | 9.33280 | 8.49641 | 7.56091 | 10.76344 | 4.33698 | 10.71739 | 6.03583 |
| **Adssl1** | 0.01338 | 0.00711 | 0.00437 | 0.00469 | 0.00813 | 0.00769 | 1.04349 | 0.00000 | 0.01112 | 0.13015 | 0.41126 | 0.01683 | 0.00806 | 0.01011 | 0.00490 | 0.58600 |
| **Adtrp** | 0.80253 | 0.03903 | 8.08654 | 4.55389 | 0.03325 | 3.60307 | 1.24145 | 15.19331 | 3.48315 | 13.18242 | 10.35220 | 14.30324 | 6.08370 | 18.15070 | 7.24795 | 0.01752 |
| **Aebp1** | 0.12995 | 0.00784 | 0.40937 | 0.00796 | 0.01489 | 0.00925 | 0.00801 | 0.24956 | 0.01137 | 0.74711 | 0.00727 | 0.25420 | 0.00495 | 0.01536 | 0.00000 | 0.00730 |
| **Aebp2** | 12.95750 | 6.45855 | 13.17467 | 10.33235 | 0.38042 | 5.00351 | 5.41599 | 12.45856 | 10.09630 | 13.88441 | 19.02565 | 8.96152 | 4.87799 | 20.60444 | 16.70619 | 1.61626 |
| **Aen** | 7.83241 | 5.25434 | 7.65370 | 8.39700 | 9.94862 | 0.12305 | 7.02711 | 9.56577 | 6.38471 | 6.28700 | 6.70629 | 1.12701 | 8.46784 | 2.40732 | 2.96246 | 2.93871 |
| **Aes** | 0.88319 | 0.00583 | 0.06270 | 6.08496 | 0.00543 | 0.06564 | 0.16148 | 2.32750 | 7.11113 | 5.91167 | 4.62726 | 0.00000 | 0.98507 | 0.00000 | 0.00000 | 2.29874 |
| **AF067061** | 0.01105 | 0.00860 | 0.01815 | 7.07057 | 0.00953 | 0.04984 | 0.00788 | 0.07965 | 12.34182 | 0.25134 | 0.14975 | 0.02432 | 0.08537 | 0.01234 | 0.51311 | 0.00585 |
| **AF067063** | 0.00000 | 0.01607 | 0.00000 | 20.90326 | 0.02450 | 1.84138 | 0.00007 | 0.87349 | 29.54844 | 10.88681 | 5.32011 | 4.37930 | 4.69302 | 8.98281 | 10.03821 | 0.00516 |
| **AF357399** | 0.01360 | 0.62170 | 0.10535 | 0.00545 | 0.00777 | 0.00578 | 0.00775 | 0.03589 | 0.07965 | 0.00636 | 0.01114 | 0.00953 | 0.21807 | 0.01211 | 0.06926 | 0.02815 |
| **AF366264** | 0.08944 | 0.01726 | 0.00598 | 0.00576 | 0.00843 | 0.00787 | 0.00741 | 0.23029 | 0.10290 | 0.25100 | 0.04514 | 0.05672 | 0.08485 | 0.03644 | 0.04534 | 0.00734 |
| **Afap1** | 0.02371 | 0.00779 | 0.03475 | 0.22706 | 0.03835 | 0.05319 | 0.00728 | 0.02610 | 0.76737 | 0.23252 | 0.26918 | 0.03821 | 0.03293 | 0.13739 | 0.09423 | 0.01450 |
| **Afap1l1** | 0.03501 | 0.00681 | 0.02510 | 0.00226 | 0.00893 | 0.00662 | 2.04023 | 0.65222 | 0.44409 | 0.00000 | 0.49587 | 0.00115 | 0.18836 | 0.00886 | 0.39909 | 0.01463 |
| **Afap1l2** | 0.01319 | 0.00785 | 0.00972 | 2.59533 | 0.00918 | 0.00609 | 0.00727 | 0.01345 | 0.53350 | 0.21823 | 0.05979 | 0.02298 | 0.01062 | 0.00177 | 1.34030 | 0.01370 |
| **Aff1** | 1.35961 | 0.06737 | 0.54646 | 13.63140 | 1.44074 | 0.03274 | 2.35453 | 3.77916 | 16.92354 | 7.64912 | 7.09286 | 1.20303 | 0.00000 | 3.07648 | 6.63874 | 0.00837 |
| **Aff3** | 0.00859 | 0.00836 | 0.02374 | 2.17392 | 0.01009 | 0.00809 | 0.00811 | 0.21060 | 0.02367 | 0.26973 | 0.01087 | 0.04623 | 0.27219 | 0.01926 | 0.43624 | 0.00765 |
| **Aff4** | 6.47693 | 2.60035 | 4.14776 | 10.41900 | 0.41102 | 8.11711 | 1.87371 | 7.55238 | 9.55360 | 7.46647 | 2.19360 | 0.83182 | 4.76502 | 11.42104 | 13.62450 | 4.01780 |

| | | | | | | | | | | | | | | | | |
|---|---|---|---|---|---|---|---|---|---|---|---|---|---|---|---|---|
| **Afg3l1** | 4.71699 | 4.16826 | 7.70936 | 1.58562 | 4.04929 | 0.00674 | 6.97616 | 6.73191 | 0.82071 | 8.35315 | 3.91491 | 0.58037 | 8.34101 | 0.04158 | 1.30907 | 4.37394 |
| **Afg3l2** | 5.86464 | 1.28243 | 2.30491 | 7.56192 | 7.96792 | 3.71867 | 3.33012 | 1.49848 | 5.05598 | 2.44607 | 1.57453 | 1.79359 | 3.81914 | 0.00000 | 0.00000 | 4.97685 |
| **Afmid** | 0.88971 | 4.18257 | 10.01134 | 2.75270 | 3.85329 | 2.57462 | 7.55296 | 8.97856 | 3.16738 | 10.02823 | 8.40181 | 1.69776 | 5.66408 | 5.34227 | 7.89548 | 2.90436 |
| **Aftph** | 1.48398 | 3.68889 | 1.18723 | 8.05581 | 0.30347 | 1.70792 | 1.79186 | 3.32481 | 4.83409 | 4.92992 | 2.05623 | 6.19968 | 1.54289 | 6.62287 | 9.24296 | 0.04458 |
| **Aga** | 0.01349 | 0.00841 | 0.01179 | 0.61783 | 0.00928 | 0.00683 | 0.00774 | 0.01882 | 0.00000 | 0.05799 | 0.01252 | 0.02146 | 0.01281 | 0.01077 | 0.00510 | 0.00732 |
| **Agap1** | 0.95588 | 0.11066 | 1.54919 | 0.44075 | 2.84222 | 0.27947 | 0.00648 | 1.78202 | 1.21514 | 0.81241 | 0.37792 | 0.00000 | 0.01249 | 0.24888 | 0.24923 | 0.02812 |
| **Agap3** | 2.84885 | 0.06158 | 2.00351 | 3.89390 | 4.55120 | 4.79809 | 2.43948 | 2.56259 | 0.49325 | 2.14404 | 1.13965 | 0.00000 | 8.43674 | 0.02879 | 0.00000 | 1.51854 |
| **Agbl2** | 0.02042 | 0.00803 | 0.00219 | 0.03840 | 0.00967 | 0.00609 | 0.00715 | 0.01197 | 1.57438 | 1.38710 | 0.03295 | 0.09998 | 0.01044 | 1.32573 | 2.94407 | 0.00708 |
| **Agbl3** | 0.01374 | 0.00817 | 0.01235 | 0.92137 | 0.00934 | 0.00657 | 0.00751 | 0.01399 | 0.01466 | 0.01502 | 0.01295 | 0.00819 | 0.01267 | 0.01024 | 0.00621 | 0.00726 |
| **Agbl5** | 0.00000 | 0.00781 | 1.00425 | 0.34758 | 0.00541 | 0.00592 | 0.00752 | 0.08446 | 1.24208 | 0.14261 | 0.11073 | 0.02927 | 0.00709 | 0.01424 | 0.02837 | 3.08129 |
| **Ager** | 0.01463 | 0.00786 | 0.00673 | 0.00911 | 0.00917 | 2.79421 | 0.00716 | 0.01069 | 0.01279 | 0.00610 | 0.00963 | 0.00806 | 0.00968 | 0.01303 | 0.01019 | 0.00623 |
| **Agfg1** | 5.64015 | 8.25754 | 5.53340 | 14.53775 | 7.89595 | 8.19791 | 5.70988 | 5.30379 | 15.42388 | 3.56570 | 5.46342 | 7.69525 | 3.52333 | 6.53862 | 4.81936 | 0.32994 |
| **Agfg2** | 0.73673 | 0.01850 | 0.57423 | 1.96467 | 0.01439 | 0.00608 | 0.87536 | 3.36856 | 5.50804 | 0.15411 | 0.74726 | 0.08311 | 0.85505 | 2.84441 | 1.82705 | 0.23639 |
| **Aggf1** | 6.25615 | 4.45723 | 6.73813 | 10.02242 | 4.17338 | 2.01952 | 6.06882 | 11.20437 | 12.52281 | 15.53982 | 6.83146 | 6.35138 | 7.84200 | 15.04357 | 19.12822 | 3.46899 |
| **Agk** | 0.97990 | 0.00606 | 0.06295 | 0.25443 | 0.00585 | 0.00667 | 0.06552 | 2.40931 | 0.00523 | 0.26461 | 0.78841 | 0.18738 | 0.00000 | 0.26660 | 0.57884 | 0.07231 |
| **Agl** | 0.88303 | 0.08808 | 2.07935 | 5.61307 | 0.57457 | 0.12133 | 0.12246 | 2.06401 | 8.70650 | 0.59857 | 9.53702 | 0.39428 | 0.20281 | 3.33672 | 0.29276 | 0.63766 |
| **Agmat** | 0.11391 | 0.11772 | 1.92994 | 1.81242 | 0.00454 | 0.00540 | 0.17841 | 1.30043 | 1.47848 | 2.94188 | 2.40837 | 0.11839 | 0.90512 | 0.90380 | 2.07415 | 0.00528 |
| **Ago1** | 3.77638 | 0.00314 | 0.06069 | 0.97229 | 0.01937 | 0.00217 | 0.00262 | 2.24291 | 0.01652 | 4.30543 | 4.12379 | 0.28727 | 0.56684 | 3.58704 | 3.03927 | 0.04608 |
| **Ago2** | 2.49928 | 5.78003 | 10.45851 | 8.43244 | 1.57820 | 0.01235 | 7.10208 | 17.25121 | 13.44344 | 12.89306 | 6.88366 | 9.24890 | 2.83316 | 3.05359 | 8.61821 | 1.02418 |
| **Ago3** | 0.00000 | 0.00787 | 0.01746 | 4.19538 | 0.00795 | 0.00620 | 0.00749 | 0.72358 | 1.08215 | 1.68273 | 0.33916 | 0.68534 | 0.01847 | 3.84038 | 3.51724 | 1.74807 |
| **Ago4** | 0.00000 | 0.00603 | 0.00000 | 0.09837 | 0.01585 | 0.00465 | 0.00534 | 0.00000 | 1.21601 | 5.08371 | 1.75174 | 6.30905 | 0.55914 | 12.24866 | 5.42379 | 1.99434 |
| **Agpat1** | 1.48334 | 0.00639 | 0.04934 | 3.41712 | 0.00741 | 0.00643 | 0.00609 | 0.00521 | 0.62801 | 0.00000 | 0.17322 | 0.01374 | 0.09621 | 0.00001 | 0.20062 | 0.00512 |
| **Agpat2** | 20.19735 | 11.17312 | 10.40094 | 3.85649 | 5.54308 | 12.58886 | 11.21462 | 17.30147 | 5.46037 | 18.15447 | 13.71959 | 10.58763 | 20.95628 | 0.12828 | 4.11597 | 11.41305 |
| **Agpat3** | 1.40941 | 0.00546 | 3.22340 | 0.89279 | 0.00838 | 0.00551 | 0.00483 | 0.05149 | 0.02135 | 0.06361 | 2.62200 | 0.26895 | 0.21340 | 0.00064 | 0.39444 | 0.14533 |
| **Agpat4** | 0.05106 | 0.04596 | 2.89469 | 0.00000 | 0.01048 | 3.85766 | 4.23911 | 0.00000 | 0.10283 | 0.69126 | 0.63267 | 0.00000 | 0.25354 | 0.05448 | 0.00000 | 2.24040 |
| **Agpat5** | 4.30899 | 3.37203 | 1.65690 | 12.42507 | 2.42251 | 2.09026 | 1.26639 | 11.11745 | 19.98847 | 13.46046 | 15.71295 | 9.51087 | 5.68070 | 12.08539 | 14.23665 | 2.99301 |
| **Agpat9** | 1.64451 | 2.15072 | 3.50574 | 2.43332 | 1.10783 | 4.22079 | 1.98109 | 12.35815 | 4.25557 | 2.81191 | 6.17804 | 2.62045 | 2.41537 | 24.85866 | 12.42832 | 1.94208 |
| **Agps** | 6.03501 | 6.59141 | 6.12578 | 10.25807 | 10.71051 | 7.12513 | 5.37700 | 1.31977 | 7.62567 | 8.36505 | 7.77483 | 7.83701 | 4.16154 | 10.84796 | 10.91927 | 2.96029 |
| **Agrn** | 0.89455 | 0.00436 | 3.98070 | 0.67220 | 4.45083 | 2.04339 | 0.00457 | 1.13709 | 0.98525 | 0.00000 | 1.49352 | 0.07929 | 2.06661 | 0.45036 | 0.00000 | 4.52281 |
| **Agrp** | 0.01259 | 0.00851 | 1.16249 | 0.36918 | 0.02094 | 3.03692 | 0.00606 | 0.00000 | 0.33915 | 0.86516 | 2.84452 | 0.23160 | 1.26622 | 0.01181 | 0.08081 | 0.00468 |
| **Agtpbp1** | 0.12185 | 0.00833 | 0.57372 | 7.25150 | 0.00682 | 0.00476 | 0.00737 | 0.09285 | 1.81596 | 1.04077 | 0.69233 | 0.08786 | 0.26801 | 1.40189 | 6.76636 | 0.00541 |
| **Agtr1a** | 0.01221 | 0.00807 | 0.20696 | 0.33876 | 0.01826 | 0.00562 | 0.00742 | 0.07964 | 0.57774 | 0.00899 | 0.03614 | 0.00706 | 0.00780 | 0.01152 | 0.00000 | 0.00718 |
| **Agtrap** | 13.32140 | 1.89441 | 10.19213 | 1.90076 | 6.69688 | 5.05949 | 8.17717 | 9.25234 | 5.16958 | 3.69510 | 5.99253 | 0.93321 | 2.74798 | 0.24133 | 0.30491 | 7.14179 |
| **Ahctf1** | 3.62860 | 11.37889 | 5.28674 | 22.37861 | 1.77156 | 5.64036 | 1.45651 | 12.66446 | 24.70692 | 18.19189 | 8.59903 | 11.85161 | 2.80041 | 7.01512 | 10.54389 | 2.44041 |
| **Ahcy** | 21.78438 | 9.67101 | 16.64257 | 0.00000 | 14.65852 | 5.68474 | 11.95384 | 14.30357 | 0.57943 | 5.42600 | 5.14912 | 0.61905 | 13.85117 | 0.01797 | 1.77461 | 7.61544 |
| **Ahcyl1** | 5.58360 | 11.53538 | 5.62549 | 5.52967 | 8.59114 | 3.94699 | 5.49428 | 5.56182 | 3.58790 | 7.39934 | 4.30023 | 17.61945 | 2.35586 | 7.21710 | 5.62512 | 3.73958 |
| **Ahcyl2** | 3.95712 | 0.01469 | 1.09230 | 3.07619 | 0.01102 | 4.33301 | 0.01332 | 0.00000 | 3.14969 | 2.38880 | 1.24457 | 0.11365 | 1.68112 | 6.40753 | 1.81995 | 0.00691 |
| **Ahdc1** | 0.00000 | 0.00748 | 0.01382 | 0.49667 | 0.00700 | 0.00710 | 0.00713 | 0.35799 | 1.54305 | 0.00460 | 0.00000 | 0.01468 | 0.00000 | 0.02485 | 0.12251 | 2.55973 |
| **Ahi1** | 0.01375 | 0.00814 | 0.01230 | 0.37643 | 0.00932 | 0.00664 | 0.00751 | 0.01377 | 0.00425 | 0.02255 | 0.01293 | 0.00806 | 0.01285 | 0.01093 | 0.00615 | 0.00747 |
| **Ahnak** | 0.00000 | 0.00718 | 1.62332 | 1.14955 | 0.16006 | 0.44402 | 0.02604 | 0.00000 | 1.89152 | 0.12738 | 0.41174 | 0.29393 | 0.73494 | 3.55710 | 0.00000 | 2.42760 |
| **Ahr** | 0.01357 | 0.00753 | 0.00645 | 1.00238 | 0.00924 | 0.00682 | 0.00712 | 0.00207 | 0.00000 | 0.00000 | 0.26624 | 0.00449 | 0.00941 | 0.50279 | 0.11597 | 0.00705 |
| **Ahrr** | 0.03101 | 0.00809 | 0.04342 | 0.01035 | 0.01326 | 0.00587 | 0.00729 | 0.74798 | 0.02229 | 0.00531 | 0.00874 | 0.00660 | 0.01283 | 0.01652 | 0.00000 | 0.00747 |
| **Ahsa1** | 29.45009 | 15.31452 | 26.12790 | 13.93448 | 13.45689 | 14.07434 | 17.05972 | 23.99904 | 12.45772 | 11.61274 | 8.15263 | 2.66678 | 23.34442 | 0.40876 | 7.18103 | 15.56806 |
| **Ahsa2** | 2.97658 | 0.00422 | 1.83479 | 0.59261 | 0.00000 | 2.47622 | 0.00408 | 0.30856 | 0.47160 | 1.83226 | 0.26711 | 0.01264 | 1.05617 | 0.59756 | 0.16907 | 0.57957 |
| **Ahsg** | 0.28589 | 0.00724 | 0.00691 | 0.00572 | 0.00661 | 0.00821 | 0.00710 | 0.56764 | 0.01077 | 0.00000 | 0.36356 | 0.01426 | 0.07292 | 0.01155 | 0.00210 | 2.37465 |
| **AI314180** | 3.55091 | 5.02873 | 8.71790 | 6.88524 | 10.61664 | 0.47134 | 8.06518 | 8.67224 | 2.83598 | 8.31428 | 4.73520 | 1.52682 | 3.94706 | 13.46380 | 7.69431 | 2.27704 |
| **AI314278** | 0.00836 | 0.00768 | 0.02641 | 0.30059 | 0.00942 | 0.00781 | 0.01743 | 0.11773 | 0.36098 | 0.61406 | 0.34310 | 0.31180 | 0.04287 | 0.26968 | 0.61698 | 0.00723 |
| **AI413582** | 2.45277 | 0.20253 | 1.62630 | 3.43845 | 0.01079 | 0.05939 | 0.17367 | 1.16885 | 0.28476 | 3.53260 | 0.95473 | 0.10330 | 1.33792 | 0.20154 | 0.05444 | 0.01094 |
| **AI427809** | 0.01316 | 0.00808 | 0.01184 | 0.43030 | 0.00915 | 0.00644 | 0.00738 | 0.01361 | 0.03472 | 0.01155 | 0.01181 | 0.00817 | 0.12673 | 0.01226 | 0.00675 | 0.00715 |
| **AI450353** | 0.67789 | 0.00679 | 0.00484 | 0.01583 | 0.00657 | 0.00691 | 0.00688 | 0.02473 | 0.02797 | 0.01047 | 0.01271 | 0.00736 | 0.01103 | 0.01179 | 0.09799 | 0.02825 |
| **AI464131** | 0.49245 | 0.04433 | 4.13164 | 8.40581 | 0.06736 | 5.37412 | 3.01013 | 2.01310 | 8.92389 | 5.95394 | 7.75312 | 2.54827 | 2.82111 | 8.37657 | 13.93676 | 4.44979 |
| **AI467606** | 0.00992 | 0.00740 | 0.00760 | 0.01101 | 0.00957 | 0.00690 | 0.00707 | 0.97859 | 0.01137 | 0.00000 | 0.00761 | 0.00559 | 0.03215 | 0.03363 | 0.00000 | 0.01472 |
| **AI506816** | 0.09292 | 0.08635 | 0.25474 | 0.74983 | 0.07643 | 0.10796 | 0.06596 | 0.01606 | 0.68610 | 1.99766 | 0.30292 | 0.36489 | 0.47290 | 0.72990 | 3.19699 | 0.01339 |
| **AI507597** | 0.00441 | 0.22834 | 0.02431 | 0.01072 | 0.01762 | 0.00607 | 0.28497 | 0.37137 | 0.01198 | 0.00828 | 0.01091 | 0.00701 | 0.00601 | 0.01356 | 0.00129 | 0.00726 |
| **AI597479** | 3.55741 | 1.99814 | 4.55797 | 3.10626 | 0.00000 | 0.66700 | 2.07958 | 2.58716 | 0.24027 | 6.39618 | 2.32597 | 3.64194 | 1.47246 | 2.47563 | 8.02538 | 0.02826 |
| **AI661453** | 0.02251 | 0.00707 | 1.08685 | 6.63691 | 0.00644 | 0.29448 | 0.03654 | 0.22965 | 10.54074 | 2.05566 | 6.44122 | 0.21436 | 0.18592 | 1.31703 | 2.37555 | 0.04076 |
| **AI662270** | 8.61171 | 8.34267 | 2.97648 | 0.01824 | 1.07330 | 0.00000 | 2.43961 | 0.00000 | 0.00835 | 0.00000 | 0.00000 | 0.04101 | 1.12546 | 2.57948 | 1.89872 | 3.21091 |
| **AI837181** | 0.86563 | 0.02933 | 0.18951 | 0.68026 | 0.01010 | 7.95704 | 0.01432 | 3.76332 | 0.07414 | 1.71068 | 0.90985 | 0.01916 | 3.86521 | 0.00118 | 0.00000 | 3.56197 |
| **AI846148** | 1.78096 | 0.24317 | 0.84216 | 4.83711 | 0.55610 | 4.31281 | 2.44949 | 2.74475 | 3.83393 | 2.01095 | 0.83040 | 0.12602 | 0.90471 | 0.69565 | 5.48513 | 0.16033 |
| **AI854703** | 0.00297 | 0.00812 | 0.00371 | 2.66700 | 0.01688 | 0.00681 | 0.00785 | 0.19126 | 0.74616 | 0.00000 | 0.56446 | 0.02473 | 0.00000 | 0.00049 | 0.62688 | 0.81506 |
| **AI987944** | 0.00399 | 0.03874 | 1.47299 | 3.53950 | 0.01733 | 0.01627 | 0.02715 | 0.49612 | 0.07632 | 0.37075 | 0.06042 | 0.06455 | 0.23505 | 0.08316 | 2.99035 | 0.01393 |
| **Aicda** | 0.01374 | 0.00786 | 0.01187 | 0.00460 | 0.00932 | 0.00654 | 0.00750 | 0.01245 | 0.18187 | 0.03060 | 0.00953 | 0.01661 | 0.01236 | 0.00678 | 0.28890 | 0.00723 |

| | | | | | | | | | | | | | | | | |
|---|---|---|---|---|---|---|---|---|---|---|---|---|---|---|---|---|
| **Aida** | 4.59588 | 0.03328 | 0.25485 | 4.18768 | 1.86975 | 0.05054 | 0.02251 | 0.44693 | 3.08492 | 0.00000 | 1.06488 | 0.05877 | 0.14093 | 3.14396 | 5.06254 | 1.61381 |
| **Aif1** | 0.01272 | 0.00839 | 0.01122 | 0.00692 | 0.00932 | 0.00674 | 0.00799 | 0.03421 | 1.08410 | 0.01709 | 0.00443 | 0.00718 | 0.01186 | 0.01676 | 0.01193 | 0.00735 |
| **Aif1l** | 0.50518 | 0.00738 | 0.00079 | 0.53036 | 0.83018 | 0.00714 | 0.01572 | 0.00000 | 1.35501 | 0.73371 | 1.59148 | 0.09369 | 1.05965 | 0.00140 | 0.86910 | 0.00635 |
| **Aifm1** | 10.05675 | 7.72034 | 4.19322 | 4.95484 | 5.11602 | 7.64523 | 2.23057 | 3.21606 | 7.46399 | 11.18367 | 4.82631 | 3.83554 | 4.88935 | 11.51014 | 14.09467 | 7.49236 |
| **Aifm2** | 0.29663 | 0.00614 | 2.25002 | 0.02813 | 0.00269 | 0.00816 | 1.62335 | 0.00000 | 0.17046 | 0.38156 | 1.53799 | 0.00371 | 0.00000 | 0.39511 | 0.00000 | 1.61126 |
| **Aifm3** | 0.00671 | 0.00807 | 0.44193 | 0.00301 | 0.00821 | 0.00580 | 0.00699 | 0.37215 | 0.00923 | 3.07418 | 0.94881 | 0.05929 | 0.39901 | 0.01681 | 0.90320 | 0.00712 |
| **Aig1** | 0.01331 | 0.00770 | 0.01226 | 0.60132 | 0.00898 | 0.00648 | 0.00730 | 0.01057 | 0.00386 | 0.82367 | 0.01207 | 0.00497 | 0.01130 | 0.00470 | 0.15983 | 0.00738 |
| **Aim1** | 1.60697 | 0.00423 | 1.23162 | 1.90800 | 0.00216 | 4.61620 | 2.93596 | 2.63624 | 0.43525 | 0.00000 | 0.00000 | 0.21380 | 1.72147 | 5.00760 | 0.96414 | 1.20586 |
| **Aim1l** | 0.62680 | 1.97801 | 4.46073 | 1.40256 | 1.28640 | 5.13086 | 3.98666 | 3.44774 | 0.58239 | 0.00000 | 1.26020 | 0.00000 | 2.36836 | 0.00000 | 0.02335 | 0.05075 |
| **Aim2** | 0.01512 | 0.00773 | 0.02420 | 0.03225 | 0.00902 | 0.05859 | 0.02751 | 0.03557 | 0.01155 | 0.40600 | 0.01109 | 0.03438 | 0.03105 | 0.14356 | 0.03447 | 0.00738 |
| **Aimp1** | 23.79788 | 14.33499 | 19.78887 | 0.63288 | 9.93513 | 12.43812 | 8.79406 | 20.04987 | 0.34616 | 12.39675 | 14.44688 | 5.45499 | 14.47275 | 9.53209 | 3.82364 | 9.85943 |
| **Aimp2** | 11.79572 | 8.38891 | 14.29079 | 9.64582 | 10.86500 | 12.86277 | 6.19842 | 13.23192 | 13.32678 | 17.43189 | 8.92418 | 9.14097 | 11.48221 | 0.26700 | 7.87234 | 5.91078 |
| **Aip** | 6.48313 | 4.92391 | 5.16153 | 1.25716 | 2.24171 | 4.71153 | 5.01160 | 7.66123 | 2.81300 | 9.00044 | 1.40083 | 0.63011 | 7.79981 | 0.01602 | 0.00000 | 2.07809 |
| **Aire** | 0.01044 | 0.00776 | 0.00885 | 1.03236 | 0.00838 | 0.00655 | 0.00722 | 0.00777 | 0.58496 | 0.12249 | 0.38715 | 0.01415 | 0.00708 | 0.00551 | 0.00000 | 0.51328 |
| **Airn** | 0.01135 | 0.00756 | 0.01542 | 0.14495 | 0.00945 | 0.00734 | 0.00723 | 0.20147 | 0.02116 | 0.00887 | 0.01071 | 0.00706 | 0.00985 | 0.01926 | 0.05636 | 0.00734 |
| **Ajuba** | 1.56900 | 0.03263 | 1.86051 | 8.53559 | 4.01224 | 5.99976 | 0.00978 | 7.02561 | 2.64030 | 10.91174 | 9.95300 | 4.28309 | 2.90760 | 1.20860 | 10.94561 | 0.13513 |
| **AK010878** | 2.79217 | 1.57233 | 3.47234 | 7.51158 | 4.65200 | 0.91661 | 1.46638 | 2.40278 | 11.71877 | 4.40585 | 4.30270 | 4.69138 | 2.26305 | 1.92598 | 1.85586 | 0.17938 |
| **Ak1** | 0.93024 | 0.00647 | 0.03013 | 0.00991 | 0.00597 | 0.00772 | 0.03707 | 0.01224 | 0.01226 | 0.15992 | 0.01229 | 0.03482 | 0.01129 | 0.01137 | 0.00974 | 0.00658 |
| **Ak2** | 17.30649 | 10.18920 | 11.41613 | 2.17526 | 7.55410 | 11.13474 | 6.07011 | 7.47308 | 1.32272 | 8.21833 | 10.19078 | 2.32705 | 11.83593 | 0.97105 | 1.08513 | 6.20748 |
| **Ak3** | 0.01315 | 0.00794 | 0.01214 | 0.00704 | 0.00891 | 0.00660 | 0.00735 | 0.01053 | 0.01172 | 1.14241 | 0.01100 | 0.00518 | 0.01098 | 0.01183 | 0.00853 | 0.00733 |
| **Ak4** | 34.36381 | 19.13015 | 27.10886 | 1.10362 | 17.72652 | 12.69632 | 19.39597 | 21.07427 | 0.76166 | 19.80500 | 5.40630 | 11.33985 | 20.77982 | 16.01426 | 4.80863 | 17.41858 |
| **Ak6** | 11.04315 | 2.15514 | 7.31598 | 10.58198 | 9.75263 | 4.55432 | 1.61452 | 8.14727 | 17.65060 | 11.92907 | 6.45876 | 7.97167 | 7.89918 | 11.09271 | 8.53715 | 7.04332 |
| **Ak7** | 2.94228 | 0.01131 | 2.58257 | 1.07852 | 0.06640 | 2.21697 | 0.27809 | 13.24035 | 1.59216 | 4.78118 | 4.68823 | 1.20421 | 1.80986 | 4.30731 | 3.28536 | 0.03643 |
| **Ak8** | 0.00974 | 0.00803 | 0.31429 | 0.28997 | 0.00844 | 0.00905 | 0.00784 | 0.76288 | 0.03099 | 0.84932 | 0.01795 | 0.01322 | 0.00385 | 0.88717 | 0.15647 | 0.00695 |
| **Akap1** | 11.69212 | 2.40031 | 11.51638 | 5.38307 | 9.26207 | 9.92324 | 7.41484 | 20.87342 | 3.94077 | 19.25076 | 5.92566 | 1.89352 | 6.02657 | 1.40993 | 15.61940 | 5.22637 |
| **Akap10** | 0.01381 | 2.77737 | 0.00321 | 2.69020 | 0.00793 | 0.00601 | 0.00610 | 0.04719 | 1.37956 | 0.70848 | 0.03057 | 0.04715 | 0.00939 | 0.03173 | 1.05923 | 0.00610 |
| **Akap11** | 0.00632 | 0.00689 | 0.00000 | 3.61631 | 1.25371 | 0.01829 | 0.01692 | 0.00729 | 0.58196 | 0.82358 | 0.00063 | 0.07139 | 0.01053 | 0.01021 | 2.92758 | 0.00677 |
| **Akap12** | 13.31215 | 9.10803 | 5.96687 | 3.36749 | 13.84873 | 7.51116 | 7.90513 | 5.74060 | 3.90982 | 4.71176 | 5.13783 | 0.57451 | 7.67635 | 7.20352 | 7.11271 | 4.59654 |
| **Akap13** | 1.40195 | 0.01500 | 1.60521 | 2.66715 | 3.24627 | 0.00718 | 0.00475 | 0.08309 | 3.20324 | 2.59266 | 0.32701 | 0.14281 | 1.52634 | 1.40315 | 1.12479 | 0.00411 |
| **Akap14** | 0.27523 | 0.00787 | 0.00536 | 0.06271 | 0.00720 | 0.00713 | 0.00741 | 0.03395 | 0.00857 | 0.76230 | 0.01225 | 0.02998 | 0.00970 | 0.05552 | 0.00000 | 0.00673 |
| **Akap17b** | 0.00849 | 0.12708 | 0.00000 | 11.00908 | 0.91942 | 0.00511 | 0.00691 | 0.00798 | 2.97982 | 2.93993 | 0.09928 | 0.04921 | 0.08813 | 2.79827 | 9.49329 | 0.02802 |
| **Akap5** | 0.02128 | 0.02885 | 0.05378 | 0.27014 | 0.00854 | 0.00680 | 0.01734 | 0.00790 | 1.90053 | 0.00013 | 0.30854 | 0.00194 | 0.03691 | 0.07958 | 0.32266 | 0.00701 |
| **Akap7** | 1.03707 | 2.03770 | 0.25246 | 2.40779 | 3.56615 | 0.00523 | 0.10416 | 0.00000 | 1.86145 | 1.25958 | 0.00000 | 0.04855 | 0.16587 | 1.22809 | 4.66446 | 0.00445 |
| **Akap8** | 11.34763 | 10.98585 | 9.93421 | 22.70785 | 5.89386 | 5.46969 | 13.02338 | 18.01646 | 32.17397 | 30.73040 | 16.73928 | 21.30204 | 9.77885 | 30.84965 | 24.08147 | 5.43926 |
| **Akap8l** | 6.15963 | 0.01196 | 0.00000 | 0.05128 | 0.08463 | 0.54255 | 0.09341 | 2.00225 | 0.10685 | 5.13128 | 0.05005 | 0.23423 | 3.73869 | 0.00808 | 0.13279 | 2.35730 |
| **Akap9** | 0.52073 | 1.25985 | 0.00000 | 3.48307 | 0.00742 | 2.41360 | 0.00663 | 0.04388 | 0.19839 | 0.39020 | 0.44978 | 0.00880 | 0.27926 | 2.52191 | 3.51481 | 0.00508 |
| **Akip1** | 4.22307 | 0.45248 | 3.46720 | 2.66008 | 6.85693 | 3.57026 | 3.40148 | 3.13383 | 2.51852 | 2.11088 | 2.21767 | 0.14478 | 7.38448 | 0.97169 | 0.00000 | 0.26393 |
| **Akirin1** | 2.36062 | 6.78240 | 4.72353 | 7.79702 | 2.54630 | 3.33157 | 4.47962 | 9.67559 | 10.74792 | 14.43989 | 11.63554 | 11.27256 | 6.85619 | 5.66631 | 9.63033 | 2.22566 |
| **Akirin2** | 5.80698 | 5.23162 | 2.66439 | 19.51855 | 10.09617 | 0.51350 | 0.90998 | 5.55923 | 19.54512 | 6.99778 | 9.33406 | 0.99471 | 3.61094 | 6.76998 | 6.54370 | 1.52430 |
| **Akna** | 0.72829 | 0.00798 | 0.01128 | 0.14923 | 0.01715 | 0.00750 | 0.00687 | 0.00907 | 0.09735 | 0.00607 | 0.31084 | 0.01639 | 0.00936 | 0.00876 | 0.00416 | 0.02898 |
| **Akr1a1** | 24.83529 | 14.69078 | 21.39972 | 12.46128 | 12.90531 | 13.99443 | 12.64822 | 22.29602 | 12.24932 | 25.93541 | 18.48381 | 16.17792 | 20.97973 | 6.07106 | 10.33011 | 17.78109 |
| **Akr1b10** | 0.03626 | 0.38633 | 0.14478 | 0.00637 | 0.03523 | 0.95531 | 0.37973 | 0.06145 | 0.01150 | 0.04675 | 0.00886 | 0.00751 | 0.91008 | 0.00593 | 0.45087 | 0.16488 |
| **Akr1b7** | 0.01290 | 0.03024 | 0.00776 | 0.50748 | 0.00908 | 0.02515 | 0.00746 | 0.01659 | 1.33161 | 0.00000 | 0.14518 | 0.00495 | 0.01672 | 0.00488 | 0.24115 | 0.00703 |
| **Akr1b8** | 10.50533 | 18.18115 | 21.81494 | 1.38891 | 11.06022 | 22.59958 | 14.08789 | 13.88330 | 0.28417 | 0.00000 | 6.74652 | 0.03319 | 30.81651 | 0.35032 | 0.00000 | 9.99279 |
| **Akr1c13** | 0.09447 | 0.00742 | 0.04517 | 0.01051 | 0.00785 | 0.01200 | 0.00724 | 0.19490 | 0.01198 | 0.06995 | 0.01131 | 0.00796 | 0.00730 | 0.01199 | 0.00865 | 0.00722 |
| **Akr1c14** | 0.01091 | 0.00800 | 0.02615 | 0.00871 | 0.00957 | 0.00796 | 0.00737 | 0.01396 | 0.01203 | 0.44281 | 0.01404 | 0.00605 | 0.40479 | 0.01150 | 0.00969 | 0.00790 |
| **Akr1c21** | 4.69028 | 2.48390 | 2.18980 | 0.01210 | 0.03640 | 0.00000 | 0.00000 | 2.62978 | 0.00784 | 0.45390 | 0.00000 | 14.71638 | 0.62287 | 0.00673 | 0.00000 | 0.02240 |
| **Akr1cl** | 0.16837 | 0.00770 | 0.00554 | 0.57818 | 0.01126 | 0.00661 | 0.00721 | 0.00920 | 0.62286 | 5.09318 | 0.00426 | 0.00000 | 0.01064 | 0.00643 | 0.90932 | 0.01482 |
| **Akr1d1** | 0.01331 | 0.00795 | 0.02302 | 0.16462 | 0.00911 | 0.00764 | 0.00744 | 0.00888 | 0.43899 | 0.00700 | 0.17844 | 0.00651 | 0.01007 | 0.03517 | 0.01400 | 0.00724 |
| **Akr1e1** | 2.61516 | 1.15524 | 0.84310 | 0.20689 | 0.86033 | 0.24998 | 0.00450 | 0.00000 | 0.00460 | 0.27321 | 0.29658 | 0.01194 | 0.22360 | 0.00536 | 0.00000 | 0.29346 |
| **Akr7a5** | 3.54064 | 0.00376 | 0.00023 | 1.40682 | 0.02269 | 0.01332 | 0.03439 | 1.73204 | 0.08490 | 0.71891 | 0.21669 | 0.00000 | 0.00740 | 0.00386 | 0.02212 | 0.57529 |
| **Akt1** | 7.98510 | 2.88324 | 1.92204 | 6.83981 | 6.16964 | 7.94013 | 3.63402 | 8.85838 | 6.11756 | 9.31186 | 4.98066 | 0.60570 | 5.95976 | 0.00000 | 0.00000 | 1.55312 |
| **Akt1s1** | 9.61346 | 0.05467 | 0.00000 | 1.42619 | 8.32229 | 2.98461 | 0.49621 | 13.63903 | 1.27153 | 22.28649 | 5.05634 | 0.49934 | 9.25339 | 0.00000 | 0.00000 | 6.47185 |
| **Akt2** | 15.54251 | 2.94647 | 8.14256 | 9.61771 | 6.24668 | 7.09231 | 7.79832 | 6.81729 | 9.06565 | 8.55669 | 6.03449 | 0.89068 | 7.69874 | 0.04325 | 2.46272 | 3.30262 |
| **Akt3** | 0.00000 | 4.24940 | 4.14503 | 6.97738 | 0.87023 | 0.46481 | 0.02601 | 1.00762 | 1.96332 | 0.31012 | 0.03148 | 0.29431 | 2.63681 | 0.36440 | 4.23498 | 0.04486 |
| **Aktip** | 0.00000 | 1.43801 | 1.43281 | 0.92427 | 0.00610 | 5.44208 | 0.37475 | 0.90539 | 1.06011 | 1.57914 | 0.30821 | 0.03511 | 0.22159 | 0.77050 | 3.00818 | 0.00428 |
| **Alad** | 11.25845 | 6.73563 | 13.76315 | 1.60581 | 2.53441 | 8.82148 | 7.00631 | 17.49282 | 2.94560 | 7.93125 | 7.46685 | 0.22909 | 9.82324 | 0.04487 | 0.00000 | 8.95826 |
| **Alas1** | 2.08743 | 1.48535 | 2.12520 | 6.72581 | 0.00000 | 0.38706 | 1.08066 | 4.68308 | 6.09732 | 0.24953 | 2.64973 | 5.68986 | 0.50325 | 0.18912 | 0.80835 | 1.27472 |
| **Alas2** | 0.00000 | 2.49402 | 4.91813 | 0.00000 | 0.00000 | 0.00533 | 2.66047 | 0.00550 | 0.67017 | 17.22812 | 0.00000 | 0.00000 | 0.00410 | 0.01351 | 0.00000 | 0.00448 |
| **Aldh16a1** | 8.02510 | 0.11221 | 2.00844 | 1.30023 | 3.28712 | 6.41241 | 4.19334 | 7.67040 | 0.98820 | 4.57509 | 2.43063 | 0.15514 | 9.15940 | 0.00000 | 0.00000 | 1.11856 |
| **Aldh18a1** | 6.59511 | 3.94837 | 0.00000 | 2.99953 | 10.91305 | 4.84095 | 1.28814 | 0.00000 | 1.99081 | 1.74589 | 0.21443 | 0.06029 | 1.22226 | 0.00000 | 0.57101 | 13.70880 |
| **Aldh1a1** | 0.01254 | 0.00778 | 0.01206 | 0.00549 | 0.00874 | 0.00657 | 0.00726 | 0.00962 | 0.02456 | 2.32100 | 0.02708 | 0.00000 | 0.01085 | 0.01225 | 0.00669 | 0.00760 |

| | | | | | | | | | | | | | | | | |
|---|---|---|---|---|---|---|---|---|---|---|---|---|---|---|---|---|
| **Aldh1b1** | 0.96813 | 1.88265 | 0.70036 | 0.00000 | 0.01374 | 0.02035 | 3.05453 | 0.00356 | 0.08387 | 1.67101 | 0.02251 | 0.00000 | 0.59890 | 0.01605 | 0.00000 | 0.00487 |
| **Aldh1l1** | 0.35566 | 0.58765 | 3.17936 | 0.00526 | 0.08451 | 0.58776 | 0.00690 | 0.21303 | 0.01161 | 0.69717 | 0.02306 | 0.04922 | 0.00000 | 0.00701 | 0.06592 | 2.94466 |
| **Aldh1l2** | 0.01128 | 0.00728 | 0.00629 | 1.01544 | 0.00988 | 0.00654 | 0.00716 | 0.00875 | 2.04097 | 0.00000 | 0.00360 | 0.73269 | 0.02924 | 1.88330 | 0.00000 | 0.00713 |
| **Aldh2** | 3.94491 | 0.76512 | 3.54676 | 1.98971 | 0.00834 | 0.77541 | 0.00267 | 0.39684 | 1.67459 | 2.28255 | 0.30726 | 0.08395 | 1.05746 | 0.00000 | 0.94883 | 0.79792 |
| **Aldh3a2** | 1.11292 | 0.04994 | 0.80131 | 1.26562 | 0.01417 | 0.00442 | 0.02849 | 0.67449 | 0.01117 | 1.14527 | 0.39271 | 0.03216 | 0.00543 | 0.00299 | 0.11086 | 0.00571 |
| **Aldh3b1** | 0.01374 | 0.00783 | 0.00746 | 0.00870 | 0.00919 | 0.27635 | 0.00732 | 0.01107 | 0.01228 | 0.39421 | 0.01213 | 0.00660 | 0.00914 | 0.01166 | 0.00855 | 0.05839 |
| **Aldh3b2** | 0.00836 | 0.00770 | 0.00959 | 0.00867 | 0.00775 | 4.08129 | 0.00694 | 0.01047 | 0.01287 | 0.00000 | 0.00853 | 0.03862 | 0.00000 | 0.01431 | 0.00922 | 1.84448 |
| **Aldh3b3** | 0.01724 | 0.00811 | 0.00627 | 0.20405 | 0.01003 | 0.74888 | 0.00735 | 0.01199 | 0.00572 | 0.02094 | 0.01087 | 0.01821 | 0.00644 | 0.01121 | 0.00682 | 0.18668 |
| **Aldh4a1** | 1.02484 | 0.29747 | 7.01761 | 0.43224 | 0.67595 | 3.88321 | 0.85115 | 1.95216 | 1.20323 | 0.33649 | 3.69019 | 0.07507 | 2.39551 | 0.02576 | 0.71571 | 3.97581 |
| **Aldh5a1** | 0.01179 | 0.00790 | 0.98800 | 1.40605 | 0.01508 | 0.00552 | 0.00728 | 0.01078 | 0.00000 | 0.00000 | 0.01070 | 0.02991 | 0.00848 | 0.00619 | 1.02943 | 1.00481 |
| **Aldh6a1** | 2.15666 | 2.47638 | 1.82102 | 0.95136 | 0.05039 | 2.59484 | 3.64348 | 2.96600 | 0.37246 | 4.99862 | 1.52544 | 0.07863 | 1.69920 | 0.00205 | 1.40163 | 0.63289 |
| **Aldh7a1** | 0.34004 | 2.94519 | 0.14989 | 1.71924 | 1.43075 | 2.29791 | 5.28565 | 1.76609 | 0.92988 | 3.26502 | 0.23275 | 0.14329 | 0.42392 | 0.00000 | 1.18263 | 0.01250 |
| **Aldh9a1** | 17.33031 | 10.71590 | 5.03311 | 9.68447 | 17.98915 | 3.54634 | 7.27925 | 6.16645 | 5.60845 | 2.11288 | 1.19622 | 0.49142 | 5.79638 | 0.41827 | 5.80033 | 8.99584 |
| **Aldoa** | 36.55638 | 22.76950 | 32.27388 | 4.44893 | 26.75400 | 16.58035 | 20.08537 | 22.40173 | 13.64268 | 13.18828 | 15.11441 | 4.28042 | 29.29968 | 0.21555 | 1.56213 | 21.51718 |
| **Aldoart1** | 2.09335 | 1.58223 | 1.93075 | 0.10914 | 1.81051 | 0.84808 | 1.44490 | 0.68075 | 0.04810 | 0.25218 | 0.08607 | 0.03852 | 1.52369 | 0.00606 | 0.00000 | 1.61827 |
| **Aldoart2** | 0.08733 | 0.06274 | 0.04469 | 0.26872 | 0.01852 | 0.02699 | 0.03883 | 0.02042 | 0.73098 | 0.01744 | 0.00842 | 0.01842 | 0.07852 | 0.01130 | 0.00302 | 0.02904 |
| **Aldoc** | 0.67084 | 0.00593 | 0.00535 | 1.28111 | 0.42915 | 0.00672 | 1.59146 | 0.01261 | 0.00000 | 5.63448 | 0.39449 | 0.01541 | 0.15403 | 0.01828 | 0.75409 | 0.00657 |
| **Alg1** | 0.06134 | 0.00595 | 1.28666 | 0.00176 | 0.00534 | 0.00742 | 2.01676 | 2.69455 | 0.33125 | 0.44345 | 0.19392 | 0.82786 | 3.00202 | 0.00380 | 0.37071 | 1.16945 |
| **Alg10b** | 0.55885 | 0.00639 | 0.05649 | 1.78856 | 0.07606 | 0.07930 | 2.40198 | 1.02531 | 0.85338 | 1.27955 | 0.32709 | 0.15417 | 0.00638 | 0.00747 | 0.99316 | 0.33465 |
| **Alg11** | 0.18426 | 0.00731 | 2.47019 | 0.20825 | 0.00472 | 0.07874 | 0.00596 | 1.90477 | 0.23513 | 0.00001 | 1.99517 | 0.05359 | 0.46656 | 1.74031 | 0.53740 | 0.02004 |
| **Alg12** | 0.00000 | 2.21735 | 2.45283 | 0.62844 | 0.02327 | 0.45513 | 1.93470 | 2.00195 | 1.35149 | 1.18986 | 0.92977 | 0.94768 | 2.02075 | 0.05567 | 0.06464 | 0.50759 |
| **Alg13** | 12.96999 | 5.49728 | 9.64333 | 18.97415 | 8.29566 | 1.34209 | 0.44765 | 7.28840 | 20.36549 | 6.17654 | 15.54323 | 12.27369 | 2.55790 | 14.65831 | 5.58618 | 0.20538 |
| **Alg14** | 7.00137 | 3.95655 | 8.53449 | 0.03120 | 4.67849 | 0.17199 | 4.57212 | 4.60840 | 0.32800 | 3.44746 | 0.04389 | 1.20462 | 4.16852 | 0.56315 | 1.37875 | 2.40547 |
| **Alg2** | 0.45767 | 0.00416 | 0.70336 | 0.51579 | 0.01418 | 0.00378 | 0.09747 | 2.25368 | 0.88144 | 2.00576 | 0.40875 | 8.89812 | 0.72485 | 0.02472 | 3.30835 | 0.09875 |
| **Alg3** | 9.49585 | 0.02332 | 6.20837 | 2.57100 | 0.17399 | 5.45814 | 6.97172 | 7.69009 | 0.71033 | 1.63112 | 2.94205 | 4.24856 | 2.07454 | 0.00000 | 0.24743 | 5.43532 |
| **Alg5** | 6.38485 | 7.14014 | 4.23004 | 1.27412 | 2.11290 | 4.96418 | 2.30621 | 4.65956 | 0.18074 | 0.54788 | 2.44119 | 0.16557 | 7.25856 | 1.71066 | 0.00004 | 6.98950 |
| **Alg6** | 0.07899 | 0.03278 | 0.92952 | 0.05721 | 0.02980 | 0.04322 | 0.00723 | 0.12416 | 0.09028 | 0.48312 | 0.78441 | 0.34304 | 0.29703 | 1.63524 | 0.00105 | 0.05179 |
| **Alg8** | 0.94576 | 0.11488 | 1.58658 | 1.06587 | 0.03666 | 0.00635 | 3.72995 | 2.41626 | 0.01661 | 1.97423 | 1.99658 | 0.01043 | 3.20442 | 0.44526 | 0.09736 | 0.00383 |
| **Alg9** | 5.00379 | 0.00000 | 4.16447 | 2.08590 | 0.01293 | 0.03443 | 3.76667 | 0.82505 | 0.57024 | 1.25083 | 0.33891 | 0.00000 | 2.41643 | 0.26579 | 0.00000 | 3.94875 |
| **Alkbh1** | 5.67767 | 2.09534 | 2.98039 | 0.10801 | 6.44815 | 4.01517 | 0.68952 | 3.14097 | 3.16295 | 5.05037 | 1.47291 | 0.20935 | 1.95434 | 2.47946 | 5.55197 | 1.41654 |
| **Alkbh2** | 3.27790 | 1.30137 | 0.52090 | 3.99573 | 2.56486 | 0.01162 | 0.01360 | 1.33474 | 2.98592 | 5.12669 | 2.12700 | 0.15243 | 0.82836 | 0.00000 | 0.36667 | 0.61461 |
| **Alkbh3** | 8.32238 | 7.88645 | 4.85860 | 2.09040 | 8.38612 | 4.51297 | 6.17053 | 5.99099 | 1.25337 | 5.00384 | 2.90445 | 4.75961 | 8.58854 | 0.02057 | 2.99826 | 1.76115 |
| **Alkbh3os1** | 0.01110 | 0.00811 | 0.00370 | 0.00930 | 0.01008 | 0.69681 | 0.43644 | 0.05475 | 0.01233 | 0.00407 | 0.03099 | 0.00793 | 0.24122 | 0.01379 | 0.00889 | 0.00713 |
| **Alkbh4** | 1.94950 | 2.40156 | 1.09476 | 4.01229 | 0.09034 | 0.00461 | 2.44727 | 2.35752 | 3.99533 | 2.83765 | 3.06572 | 0.34390 | 3.78378 | 0.02952 | 0.41699 | 2.18580 |
| **Alkbh5** | 4.10745 | 0.26315 | 4.05689 | 10.97269 | 0.86007 | 1.15586 | 1.85664 | 4.23989 | 11.91177 | 5.19486 | 2.50248 | 0.37016 | 3.03313 | 2.37438 | 5.72068 | 1.13991 |
| **Alkbh6** | 4.79699 | 0.00201 | 0.02796 | 0.00330 | 1.51822 | 0.01899 | 0.00022 | 0.30990 | 0.01011 | 1.93485 | 0.06844 | 0.01805 | 4.15951 | 0.01560 | 0.00348 | 3.83836 |
| **Alkbh7** | 4.94093 | 0.01132 | 1.61305 | 1.46303 | 1.57875 | 0.57127 | 0.04181 | 1.19709 | 2.03120 | 2.05570 | 1.53058 | 0.09045 | 4.14002 | 0.00000 | 0.07921 | 3.86478 |
| **Alkbh8** | 0.01341 | 0.25507 | 0.38309 | 0.48957 | 0.09024 | 0.00757 | 0.00734 | 1.47190 | 0.93841 | 0.00000 | 0.00000 | 0.01455 | 0.32256 | 1.56064 | 1.18356 | 0.01385 |
| **Allc** | 1.73315 | 0.00591 | 1.10662 | 0.12932 | 0.00282 | 0.00913 | 0.02753 | 0.33434 | 0.02433 | 1.39916 | 0.36178 | 0.60635 | 0.44914 | 7.34289 | 0.52080 | 0.00437 |
| **Alms1** | 0.78683 | 0.05666 | 1.43259 | 2.52724 | 0.00407 | 0.00675 | 0.93385 | 1.18967 | 1.39365 | 0.70170 | 0.00000 | 0.01873 | 0.04176 | 2.88042 | 3.81611 | 0.00567 |
| **Alms1.ps2** | 0.57863 | 0.00740 | 0.10641 | 0.17099 | 0.00655 | 0.00648 | 0.00770 | 0.24318 | 0.17153 | 0.49841 | 0.00000 | 0.04346 | 0.00647 | 0.87608 | 1.71448 | 0.00682 |
| **Alox12** | 3.12654 | 0.85386 | 3.24390 | 8.81283 | 0.02731 | 0.05231 | 0.13338 | 3.28487 | 11.65848 | 3.36265 | 3.85807 | 0.30866 | 2.30124 | 4.30412 | 2.49891 | 0.03507 |
| **Alox12e** | 0.01286 | 0.00794 | 0.01194 | 0.02044 | 0.00880 | 0.00656 | 0.00733 | 0.01021 | 0.03278 | 1.67490 | 0.01021 | 0.01411 | 0.01057 | 0.01196 | 0.00689 | 0.00724 |
| **Alox15** | 0.01238 | 0.00799 | 0.01202 | 0.06840 | 0.00882 | 0.00666 | 0.00732 | 0.01556 | 0.00818 | 2.01868 | 0.01028 | 0.00339 | 0.01091 | 0.00821 | 0.05710 | 0.00745 |
| **Alox5ap** | 0.01328 | 0.00781 | 0.01199 | 0.00874 | 0.00903 | 0.00661 | 0.00725 | 0.01027 | 0.01206 | 0.44696 | 0.01265 | 0.00635 | 0.07377 | 0.01141 | 0.00992 | 0.00770 |
| **Alox8** | 0.08548 | 0.05021 | 0.07748 | 0.50120 | 0.02896 | 0.03157 | 0.06874 | 0.00734 | 0.69422 | 0.02686 | 0.55113 | 0.09010 | 0.04239 | 0.62117 | 0.18362 | 0.02148 |
| **Aloxe3** | 0.14745 | 0.00745 | 0.00000 | 1.22351 | 1.22165 | 0.00601 | 0.00711 | 0.96917 | 7.54320 | 2.99605 | 0.10951 | 0.19052 | 0.00501 | 0.01681 | 1.81074 | 0.00664 |
| **Alpi** | 0.00739 | 0.00780 | 0.07507 | 0.00906 | 0.00988 | 0.02504 | 0.16248 | 0.50158 | 0.01159 | 0.20573 | 0.03571 | 0.00612 | 0.26997 | 0.01302 | 0.01789 | 0.00739 |
| **Alpk3** | 1.64485 | 0.00681 | 0.12576 | 0.27973 | 0.00957 | 0.00733 | 0.00619 | 0.00897 | 0.90045 | 0.01137 | 0.00581 | 0.00570 | 0.01066 | 0.01251 | 0.00208 | 0.00565 |
| **Alpl** | 18.68821 | 0.00000 | 2.67381 | 2.65820 | 2.06482 | 0.29850 | 6.65910 | 0.33290 | 0.15453 | 4.44198 | 0.95995 | 0.38026 | 2.42355 | 0.01984 | 0.00000 | 9.10475 |
| **Alppl2** | 2.63109 | 1.89129 | 28.38337 | 19.01476 | 0.90368 | 10.02491 | 24.66177 | 63.99246 | 13.78556 | 49.02122 | 42.58926 | 9.18784 | 47.20259 | 10.99196 | 10.22072 | 4.36612 |
| **Als2** | 6.85279 | 9.32501 | 3.68603 | 9.33389 | 2.52245 | 0.12584 | 2.84893 | 6.44695 | 15.04033 | 12.36563 | 10.40427 | 9.26626 | 4.91446 | 14.37320 | 18.21664 | 0.98269 |
| **Als2cl** | 0.00877 | 0.00745 | 0.01106 | 0.35127 | 0.00946 | 0.01524 | 0.00704 | 0.99036 | 0.12802 | 0.23616 | 0.13051 | 0.01321 | 0.00516 | 0.02876 | 0.02677 | 0.01533 |
| **Als2cr11** | 0.01299 | 0.00808 | 0.00793 | 0.00785 | 0.00943 | 0.00689 | 0.00737 | 0.01096 | 0.01098 | 0.04209 | 0.00686 | 0.00606 | 0.01374 | 0.69653 | 0.44361 | 0.00744 |
| **Alx4** | 0.00117 | 0.00713 | 0.04408 | 0.90158 | 0.01861 | 0.00639 | 0.04678 | 0.48866 | 0.78706 | 2.14768 | 1.46584 | 1.07912 | 0.34962 | 1.23125 | 1.74220 | 0.01388 |
| **Alyref** | 2.63917 | 3.05285 | 2.12524 | 0.53886 | 0.59457 | 0.04666 | 2.26825 | 1.55002 | 3.37446 | 9.38551 | 2.11589 | 0.03403 | 3.52850 | 3.95798 | 4.54049 | 1.65625 |
| **Alyref2** | 0.12971 | 0.04113 | 0.07614 | 1.46704 | 0.02206 | 0.01205 | 0.04982 | 0.02781 | 1.95929 | 0.19459 | 0.27270 | 0.00199 | 0.12152 | 0.16333 | 0.15643 | 0.06570 |
| **Amacr** | 1.30689 | 0.00694 | 0.01338 | 0.52898 | 1.50048 | 0.01150 | 0.00710 | 0.18936 | 0.38451 | 0.01653 | 0.00682 | 0.03496 | 0.35596 | 0.00860 | 0.25722 | 0.01317 |
| **Ambra1** | 2.62461 | 0.17162 | 1.27706 | 6.08600 | 0.00287 | 0.01706 | 0.00197 | 2.53331 | 9.66765 | 2.28157 | 4.17233 | 0.08370 | 1.95567 | 1.29869 | 3.12280 | 1.72855 |
| **Amdhd2** | 4.09779 | 6.11089 | 9.46122 | 1.22374 | 2.60868 | 4.72971 | 1.09250 | 9.63281 | 0.87846 | 2.70275 | 5.92251 | 0.05800 | 11.99862 | 0.00000 | 0.00000 | 6.65719 |
| **Amer1** | 0.01380 | 0.00761 | 0.01118 | 1.40626 | 0.00932 | 0.00640 | 0.00733 | 0.01140 | 0.00000 | 0.00613 | 0.02795 | 0.01688 | 0.01887 | 0.01648 | 0.32630 | 0.00715 |
| **Amfr** | 3.73523 | 0.61025 | 3.46440 | 2.39287 | 0.11031 | 2.58750 | 3.47352 | 0.06585 | 0.02925 | 0.83961 | 0.60704 | 0.15726 | 0.14617 | 0.00208 | 3.87798 | 0.02924 |

| | | | | | | | | | | | | | | | | |
|---|---|---|---|---|---|---|---|---|---|---|---|---|---|---|---|---|
| **Amhr2** | 4.41805 | 0.85392 | 0.83685 | 9.41755 | 1.32030 | 1.90990 | 5.79914 | 5.81089 | 12.28857 | 18.36495 | 8.51297 | 0.51299 | 2.13730 | 0.00000 | 0.00000 | 0.62191 |
| **Amigo2** | 0.01073 | 0.00705 | 0.00750 | 0.00858 | 0.00937 | 0.00688 | 1.53272 | 0.00969 | 0.01216 | 0.31524 | 0.01178 | 0.00610 | 0.02366 | 0.01136 | 0.00971 | 0.01432 |
| **Amigo3** | 0.00774 | 0.00721 | 0.00429 | 2.72146 | 0.00923 | 0.00736 | 0.91387 | 0.46064 | 0.16068 | 1.24434 | 0.51027 | 0.08709 | 0.00399 | 0.00003 | 0.15620 | 0.00681 |
| **Ammecr1l** | 2.78871 | 1.54526 | 3.76386 | 19.15192 | 0.05240 | 3.05972 | 4.56847 | 6.55338 | 18.89190 | 19.79140 | 9.27743 | 4.57335 | 5.36783 | 30.94389 | 27.50192 | 1.08098 |
| **Amn** | 0.00000 | 0.00701 | 0.11276 | 0.37395 | 0.00642 | 0.00892 | 0.00510 | 0.00000 | 0.41840 | 2.38422 | 3.05367 | 0.00003 | 1.69032 | 0.00142 | 0.00000 | 3.59342 |
| **Amn1** | 1.67973 | 3.21225 | 8.07447 | 1.18885 | 0.00000 | 0.17828 | 1.22930 | 0.75721 | 1.16388 | 1.75108 | 3.82081 | 0.24626 | 0.10528 | 1.28112 | 1.51390 | 0.08398 |
| **Amot** | 7.17296 | 0.07864 | 6.12403 | 1.95164 | 9.72849 | 3.97729 | 7.43324 | 5.05067 | 0.95335 | 2.96999 | 5.84099 | 0.04722 | 5.17006 | 14.40427 | 0.00000 | 6.89630 |
| **Amotl1** | 0.01324 | 0.00812 | 0.01064 | 0.43499 | 0.00909 | 0.00681 | 0.00751 | 0.01244 | 0.04025 | 0.32750 | 0.03614 | 0.05210 | 0.06258 | 0.05093 | 0.02970 | 0.00720 |
| **Amotl2** | 0.00000 | 0.00510 | 3.74759 | 0.00000 | 0.04628 | 0.00913 | 0.00000 | 10.31040 | 0.04171 | 8.90146 | 6.39717 | 1.20092 | 8.65162 | 3.01486 | 3.54394 | 6.33380 |
| **Ampd1** | 0.01336 | 0.00800 | 0.00847 | 0.72506 | 0.00910 | 0.00633 | 0.00720 | 0.00015 | 3.89023 | 0.29291 | 0.31678 | 0.00000 | 0.00929 | 0.02020 | 0.12521 | 0.00702 |
| **Ampd2** | 1.41205 | 0.66569 | 0.00000 | 0.26687 | 1.01954 | 2.09211 | 0.02834 | 1.00393 | 0.34821 | 0.78467 | 0.03656 | 0.08841 | 2.55037 | 0.00638 | 0.00000 | 0.33465 |
| **Ampd3** | 0.01289 | 0.01841 | 0.00468 | 9.19530 | 0.00904 | 0.00545 | 0.00684 | 0.00021 | 3.69523 | 0.00000 | 0.25925 | 0.04411 | 0.00965 | 0.12814 | 1.73790 | 0.00539 |
| **Amph** | 3.04173 | 0.00506 | 0.00272 | 0.00349 | 0.17770 | 0.00549 | 0.00504 | 0.00000 | 0.29212 | 0.00000 | 0.78389 | 0.00203 | 0.00533 | 0.00556 | 0.20105 | 0.00392 |
| **Amt** | 0.04789 | 0.00762 | 0.04384 | 0.05572 | 0.01676 | 0.00834 | 0.00713 | 0.01027 | 0.00866 | 0.00462 | 0.00726 | 0.00389 | 0.01241 | 1.03756 | 0.11096 | 0.00731 |
| **Amy1** | 0.01255 | 0.00773 | 0.00782 | 0.03787 | 0.00928 | 0.00667 | 0.00726 | 0.00972 | 0.01233 | 0.32965 | 0.00786 | 0.00348 | 0.01159 | 1.02316 | 0.00000 | 0.00717 |
| **Amz1** | 0.01406 | 0.00773 | 0.00900 | 0.38860 | 0.00913 | 0.00659 | 0.00715 | 0.17032 | 1.64244 | 0.01721 | 0.00000 | 0.00405 | 0.00772 | 0.00654 | 0.15028 | 0.00739 |
| **Amz2** | 6.08141 | 6.00749 | 6.64276 | 0.67327 | 0.00000 | 3.51568 | 4.49395 | 3.97677 | 0.58254 | 6.47447 | 4.50715 | 0.21200 | 2.13102 | 0.59281 | 3.22599 | 0.00759 |
| **Anapc1** | 5.80996 | 9.46247 | 8.86872 | 8.05357 | 3.80937 | 2.60546 | 3.51767 | 4.56469 | 2.27401 | 8.46253 | 6.43413 | 6.11736 | 4.84946 | 4.60360 | 9.68955 | 0.40894 |
| **Anapc10** | 8.66645 | 7.95891 | 9.91790 | 5.19271 | 4.91474 | 5.19358 | 5.55709 | 10.47804 | 4.94280 | 16.03225 | 15.61184 | 6.32920 | 7.21851 | 18.94716 | 10.64623 | 4.42442 |
| **Anapc11** | 4.22129 | 0.48440 | 2.38441 | 6.44668 | 3.46165 | 2.16394 | 2.74074 | 6.22718 | 5.96399 | 9.71492 | 6.29312 | 1.22417 | 4.44484 | 0.56952 | 1.21851 | 2.46934 |
| **Anapc13** | 2.81029 | 0.08867 | 1.62950 | 1.84485 | 1.53381 | 0.80395 | 0.03309 | 0.05567 | 1.00334 | 0.45789 | 6.49716 | 0.12263 | 1.47195 | 0.02993 | 0.21230 | 0.99912 |
| **Anapc15** | 11.03105 | 3.59956 | 16.07862 | 13.12464 | 12.30551 | 5.18221 | 5.10868 | 11.28872 | 15.55266 | 15.59390 | 10.01686 | 2.02947 | 10.73822 | 3.54870 | 11.75568 | 1.70355 |
| **Anapc16** | 0.38108 | 0.93508 | 0.33760 | 12.35435 | 1.49970 | 0.00409 | 0.20530 | 0.59234 | 4.56506 | 1.01963 | 6.20847 | 0.45529 | 1.48298 | 0.47803 | 2.48609 | 0.00235 |
| **Anapc2** | 8.60036 | 8.81546 | 10.25646 | 2.17188 | 7.43070 | 1.44415 | 2.41650 | 12.33344 | 3.53878 | 9.85278 | 6.66450 | 0.48586 | 7.35244 | 3.82232 | 4.84878 | 8.39382 |
| **Anapc4** | 6.12509 | 3.02683 | 6.07123 | 1.37214 | 4.19456 | 4.48011 | 4.89110 | 8.79861 | 4.94358 | 8.29297 | 3.67520 | 7.32492 | 3.22677 | 12.88825 | 9.71912 | 1.25651 |
| **Anapc5** | 19.10361 | 9.75580 | 11.53681 | 12.43914 | 9.94559 | 11.56431 | 10.78173 | 22.45003 | 12.51562 | 26.42438 | 21.63343 | 10.03178 | 20.91777 | 11.56101 | 16.71008 | 11.84155 |
| **Anapc7** | 7.95882 | 6.47811 | 7.05723 | 7.70585 | 0.74964 | 3.53803 | 3.16906 | 11.54093 | 14.67052 | 19.08355 | 13.91794 | 1.63287 | 4.78979 | 0.42886 | 10.56976 | 1.81708 |
| **Angel1** | 0.03411 | 0.00752 | 1.25814 | 3.33185 | 0.01474 | 0.00888 | 0.02666 | 1.18538 | 0.98703 | 1.00467 | 0.27496 | 0.02544 | 1.47369 | 0.00000 | 0.41109 | 0.00544 |
| **Angel2** | 2.77703 | 3.27433 | 0.06779 | 19.50700 | 5.82730 | 3.20757 | 0.47303 | 0.20701 | 3.44586 | 4.18563 | 2.91310 | 0.28210 | 2.36461 | 7.38466 | 11.33995 | 1.16070 |
| **Angpt2** | 0.04377 | 0.03671 | 0.02758 | 0.00467 | 0.00808 | 0.00594 | 0.00689 | 0.07039 | 0.62882 | 0.22085 | 0.35593 | 0.00212 | 0.23750 | 0.51084 | 0.00000 | 0.00720 |
| **Angptl1** | 0.00888 | 0.00770 | 0.00860 | 0.00970 | 0.00946 | 0.00702 | 0.02744 | 0.16361 | 0.01211 | 0.19956 | 0.00903 | 0.00674 | 0.26279 | 0.32496 | 0.00000 | 0.00728 |
| **Angptl2** | 0.01626 | 0.00789 | 0.12045 | 7.35186 | 0.00901 | 0.00501 | 0.00701 | 0.01029 | 0.72531 | 0.01584 | 0.02217 | 0.11896 | 0.01068 | 0.00000 | 0.19512 | 0.00587 |
| **Angptl4** | 3.82644 | 0.06242 | 1.37711 | 0.67934 | 5.92904 | 0.00193 | 0.32859 | 3.12950 | 0.00000 | 3.28508 | 2.64397 | 0.26354 | 2.59536 | 3.08000 | 1.90164 | 0.02230 |
| **Angptl6** | 0.01610 | 0.00783 | 0.07007 | 0.16987 | 0.01126 | 0.00628 | 0.00745 | 1.11029 | 0.00837 | 0.63035 | 0.19239 | 0.00278 | 0.04439 | 0.00879 | 0.00000 | 0.00778 |
| **Ank** | 0.00000 | 3.61806 | 1.00400 | 1.45576 | 0.02429 | 11.92674 | 0.03161 | 2.11722 | 1.70217 | 0.48259 | 2.29317 | 0.05542 | 7.32188 | 0.00000 | 1.07863 | 3.93855 |
| **Ank2** | 0.00321 | 0.00478 | 0.63068 | 0.03937 | 3.08713 | 0.00554 | 1.84625 | 0.02575 | 0.02424 | 0.00588 | 0.02477 | 0.01288 | 0.00703 | 0.04416 | 0.01791 | 0.00689 |
| **Ank3** | 0.01717 | 0.00810 | 0.01720 | 0.01587 | 0.00907 | 0.00635 | 0.00769 | 0.01855 | 0.07878 | 0.02993 | 0.01183 | 0.00768 | 0.01235 | 0.00758 | 0.06933 | 0.00741 |
| **Ankar** | 0.01322 | 0.01761 | 0.01141 | 0.64820 | 0.00907 | 0.00635 | 0.00726 | 0.01034 | 0.26129 | 0.41001 | 0.01040 | 0.00559 | 0.01114 | 0.01599 | 0.47504 | 0.01403 |
| **Ankdd1b** | 0.02162 | 0.00767 | 0.91014 | 1.89172 | 0.31449 | 0.00899 | 0.53427 | 0.08630 | 0.25189 | 0.07570 | 0.66670 | 0.07498 | 0.10148 | 0.00874 | 0.54175 | 0.00719 |
| **Ankfy1** | 2.85652 | 7.98178 | 5.45657 | 5.12668 | 0.61939 | 4.46693 | 3.92344 | 4.75346 | 5.84599 | 7.85976 | 7.01378 | 7.46376 | 2.65154 | 4.72286 | 3.92855 | 4.49299 |
| **Ankhd1** | 2.00919 | 0.02290 | 2.76024 | 1.30333 | 0.01569 | 0.04486 | 1.86092 | 1.01449 | 1.43497 | 3.58391 | 0.33561 | 7.69689 | 2.94974 | 9.28361 | 5.25637 | 1.63754 |
| **Ankib1** | 2.20967 | 2.33071 | 5.94591 | 0.62758 | 6.27324 | 0.01614 | 1.18736 | 2.88822 | 1.93551 | 3.68724 | 1.72234 | 13.90517 | 1.70422 | 2.55038 | 2.69062 | 0.18870 |
| **Ankle1** | 1.07115 | 0.24382 | 1.25089 | 0.92975 | 0.02096 | 0.01752 | 0.00576 | 1.08010 | 0.98721 | 0.16614 | 0.96616 | 0.06166 | 0.50911 | 0.02975 | 0.35753 | 0.07081 |
| **Ankle2** | 2.90155 | 0.04948 | 5.64939 | 10.50101 | 2.72943 | 0.01922 | 1.31116 | 2.28366 | 12.03543 | 3.33500 | 3.63841 | 19.61701 | 2.10411 | 8.66114 | 11.80177 | 2.72895 |
| **Ankmy2** | 3.76367 | 5.81318 | 3.62902 | 0.81322 | 5.06062 | 0.00291 | 8.21732 | 8.50921 | 2.11312 | 6.79854 | 10.65798 | 1.28607 | 2.73642 | 3.67541 | 7.76796 | 1.70024 |
| **Ankra2** | 0.76776 | 0.04916 | 0.32679 | 4.30750 | 0.00753 | 0.00752 | 0.01644 | 0.00792 | 3.02716 | 0.40801 | 0.05397 | 0.00000 | 0.01061 | 0.68326 | 0.22818 | 0.00560 |
| **Ankrd1** | 0.00916 | 0.12256 | 0.01237 | 0.00953 | 0.00920 | 0.00700 | 0.00719 | 0.01149 | 0.01222 | 0.19894 | 0.01451 | 0.00653 | 0.56003 | 0.01040 | 0.00901 | 0.00777 |
| **Ankrd10** | 7.22560 | 2.50112 | 0.80771 | 21.28967 | 10.23219 | 1.79038 | 10.65385 | 15.63137 | 26.95759 | 42.27846 | 19.98957 | 22.32606 | 8.58094 | 13.69210 | 21.60613 | 0.07106 |
| **Ankrd11** | 5.96493 | 2.54970 | 2.40149 | 0.37549 | 0.02997 | 1.83107 | 6.26849 | 3.82604 | 0.01090 | 1.54313 | 0.07170 | 0.17296 | 0.32876 | 0.26745 | 1.57649 | 1.29154 |
| **Ankrd12** | 0.22286 | 0.00550 | 0.86570 | 1.00101 | 0.01413 | 0.00588 | 2.06531 | 0.71410 | 1.30169 | 2.51859 | 2.63555 | 0.83855 | 0.48008 | 10.72964 | 3.82198 | 1.03590 |
| **Ankrd13a** | 6.52414 | 0.99302 | 6.54573 | 7.20510 | 5.40565 | 5.61985 | 6.79477 | 14.38457 | 2.07755 | 9.17412 | 5.59880 | 1.00984 | 9.40550 | 0.08424 | 4.42075 | 4.50874 |
| **Ankrd13b** | 0.35781 | 0.00735 | 0.39163 | 0.05233 | 0.01615 | 0.00587 | 1.99945 | 0.04596 | 0.12133 | 0.00000 | 0.24253 | 0.00493 | 0.24201 | 0.01203 | 0.00000 | 0.00712 |
| **Ankrd13c** | 2.19258 | 1.72381 | 0.68982 | 16.20115 | 7.06651 | 2.76172 | 1.77772 | 3.76776 | 14.82621 | 12.70314 | 7.90087 | 14.15418 | 2.43026 | 14.48472 | 11.43630 | 0.08973 |
| **Ankrd13d** | 0.01028 | 0.00804 | 0.01232 | 0.08651 | 0.02712 | 0.00672 | 0.00732 | 0.00002 | 0.01095 | 0.06307 | 0.59551 | 0.03065 | 0.44480 | 0.00755 | 0.03102 | 0.00704 |
| **Ankrd16** | 1.94003 | 0.17452 | 0.00000 | 2.33610 | 0.00421 | 2.42170 | 3.09294 | 2.96283 | 0.62135 | 1.06404 | 1.03853 | 3.23875 | 0.22393 | 0.00000 | 0.92013 | 0.09741 |
| **Ankrd17** | 11.45629 | 9.16526 | 11.51920 | 16.09548 | 3.94510 | 5.42653 | 5.92776 | 8.81078 | 17.32295 | 18.61772 | 12.10707 | 16.81943 | 4.56121 | 23.49497 | 18.57937 | 5.08432 |
| **Ankrd22** | 0.01371 | 0.00830 | 0.01222 | 2.84663 | 0.00930 | 0.00637 | 0.00742 | 0.01387 | 0.32195 | 0.01240 | 0.01185 | 0.00788 | 0.01270 | 0.00959 | 0.00267 | 0.00692 |
| **Ankrd23** | 0.74077 | 0.00638 | 0.18948 | 2.17043 | 0.00599 | 0.00719 | 0.00629 | 1.16506 | 3.09542 | 0.81032 | 0.40655 | 0.01281 | 0.00379 | 0.00000 | 0.53095 | 0.00603 |
| **Ankrd24** | 1.11907 | 0.00561 | 0.29631 | 0.00489 | 0.00522 | 0.00639 | 1.01734 | 0.10862 | 0.28622 | 0.00803 | 0.02204 | 0.00541 | 0.00650 | 0.01486 | 0.00246 | 0.00600 |
| **Ankrd26** | 0.01291 | 0.00765 | 0.00511 | 0.51472 | 0.00911 | 0.00687 | 0.00716 | 0.00177 | 0.01770 | 0.69131 | 0.28062 | 0.02219 | 0.00969 | 0.44939 | 0.77882 | 0.00725 |
| **Ankrd27** | 5.76836 | 2.62860 | 1.67866 | 1.58805 | 0.00651 | 0.28510 | 0.42822 | 1.06429 | 0.53449 | 2.45508 | 0.91476 | 0.43600 | 2.73374 | 0.00170 | 1.36132 | 0.02642 |
| **Ankrd28** | 1.92465 | 1.17981 | 0.53794 | 3.68383 | 0.02977 | 1.30096 | 2.10394 | 0.10452 | 10.88980 | 1.85616 | 4.58800 | 0.20971 | 1.84232 | 5.50618 | 4.51675 | 0.04565 |

| | | | | | | | | | | | | | | | | |
|---|---|---|---|---|---|---|---|---|---|---|---|---|---|---|---|---|
| **Ankrd33b** | 0.00477 | 0.00675 | 1.43787 | 0.56171 | 1.37759 | 0.00897 | 0.00706 | 0.01386 | 0.00000 | 0.40914 | 0.01063 | 0.03164 | 0.00582 | 0.00931 | 0.00000 | 1.56981 |
| **Ankrd35** | 0.01044 | 0.00770 | 0.00353 | 5.36643 | 0.00707 | 2.54932 | 0.00662 | 0.00316 | 8.03333 | 1.15212 | 0.07976 | 0.03048 | 0.21442 | 0.00001 | 0.28494 | 3.04868 |
| **Ankrd36** | 0.01373 | 0.00819 | 1.09656 | 0.75329 | 0.00742 | 0.00553 | 0.00738 | 0.01181 | 0.01834 | 0.01111 | 0.02880 | 0.03075 | 0.00764 | 0.01041 | 0.00850 | 0.00681 |
| **Ankrd37** | 0.00954 | 0.16724 | 0.11756 | 3.50124 | 0.00905 | 0.00585 | 0.00678 | 0.00000 | 2.26011 | 0.30471 | 1.07391 | 0.06056 | 0.34922 | 0.62966 | 0.00000 | 0.00648 |
| **Ankrd39** | 1.40236 | 0.01538 | 1.04640 | 0.50664 | 0.00349 | 0.01219 | 0.38046 | 1.98918 | 1.60884 | 0.29702 | 0.11656 | 0.04032 | 3.78303 | 0.00685 | 0.00000 | 0.00395 |
| **Ankrd40** | 4.44851 | 3.16959 | 3.01027 | 10.92263 | 7.05911 | 4.10703 | 1.90494 | 3.27752 | 12.91080 | 5.62713 | 5.00018 | 3.27197 | 2.60492 | 3.22666 | 5.72216 | 0.87406 |
| **Ankrd42** | 2.00809 | 0.00552 | 0.12557 | 2.64171 | 0.00408 | 0.00306 | 0.00488 | 0.00000 | 3.45699 | 4.83574 | 2.22352 | 0.26214 | 0.00000 | 6.30298 | 3.94834 | 0.00391 |
| **Ankrd44** | 0.00753 | 0.00785 | 3.33942 | 14.10311 | 0.00431 | 0.00994 | 0.15475 | 0.03867 | 6.74730 | 0.97382 | 2.65331 | 0.34247 | 0.03331 | 0.27194 | 12.51210 | 0.00322 |
| **Ankrd45** | 2.10390 | 0.35083 | 0.00151 | 0.11585 | 1.13698 | 0.00672 | 0.00607 | 0.28193 | 1.05786 | 0.00000 | 0.41483 | 0.01241 | 0.08807 | 0.05011 | 0.02293 | 0.01157 |
| **Ankrd46** | 3.34216 | 7.99616 | 5.68950 | 6.63938 | 1.32219 | 1.28936 | 2.69567 | 4.21530 | 1.36957 | 1.01136 | 3.82757 | 0.19647 | 2.15168 | 1.19382 | 1.99052 | 1.44535 |
| **Ankrd49** | 2.60927 | 2.10562 | 2.77473 | 0.33420 | 0.00000 | 0.00285 | 0.00449 | 2.06753 | 0.00000 | 1.69230 | 0.00000 | 0.08065 | 0.29146 | 1.19424 | 7.14941 | 1.94479 |
| **Ankrd50** | 0.01033 | 0.00713 | 0.67340 | 5.27323 | 0.23314 | 0.01489 | 0.00636 | 0.07139 | 2.85366 | 0.18250 | 1.05474 | 0.58818 | 0.58257 | 0.03138 | 0.77666 | 0.00588 |
| **Ankrd52** | 0.02058 | 0.00837 | 0.05547 | 0.56238 | 0.01496 | 0.00624 | 0.00762 | 1.45281 | 0.72912 | 0.67648 | 0.01391 | 0.18551 | 0.12552 | 0.00371 | 0.59559 | 0.00757 |
| **Ankrd54** | 7.39214 | 0.08413 | 3.83671 | 2.83439 | 2.03542 | 3.72364 | 0.06013 | 1.46352 | 6.38784 | 2.56833 | 1.38732 | 0.25684 | 1.29556 | 0.00637 | 0.99001 | 3.06291 |
| **Ankrd6** | 0.30260 | 0.00720 | 2.01269 | 12.80324 | 0.01435 | 0.00702 | 0.00555 | 0.74005 | 7.40745 | 2.42150 | 0.99899 | 0.73291 | 0.47445 | 0.10403 | 7.09842 | 0.00357 |
| **Ankrd9** | 0.01435 | 0.00760 | 0.03924 | 0.42587 | 0.01076 | 0.84376 | 0.00679 | 0.20723 | 0.65601 | 1.94917 | 0.64213 | 0.02548 | 0.14013 | 0.02727 | 0.03711 | 0.16705 |
| **Anks1** | 4.98271 | 1.40564 | 3.29680 | 0.74904 | 2.16804 | 1.03688 | 2.55273 | 3.17574 | 5.63400 | 0.52375 | 0.86446 | 4.12948 | 2.06989 | 0.00000 | 0.00000 | 0.00580 |
| **Anks1b** | 0.00845 | 0.00737 | 0.01082 | 0.00740 | 0.68053 | 0.00666 | 0.00766 | 0.01934 | 0.22111 | 0.78484 | 0.00998 | 0.01467 | 0.01052 | 0.01510 | 0.05235 | 0.00762 |
| **Anks3** | 0.00000 | 0.14706 | 1.94895 | 5.32305 | 5.10637 | 4.91989 | 0.00675 | 1.34763 | 3.09191 | 2.43540 | 0.40247 | 0.10817 | 0.18630 | 0.01040 | 0.20489 | 1.61440 |
| **Anks4b** | 0.00713 | 0.00756 | 0.01139 | 0.01287 | 0.00750 | 0.00634 | 0.00734 | 0.01227 | 0.05593 | 0.00000 | 0.01148 | 0.00737 | 0.00731 | 0.02438 | 0.20123 | 1.61445 |
| **Ankzf1** | 0.78075 | 0.00596 | 0.76887 | 0.68525 | 0.06589 | 2.60868 | 1.30905 | 1.15841 | 1.42635 | 1.23567 | 0.16502 | 0.05093 | 1.10742 | 0.09377 | 0.83890 | 0.00492 |
| **Anln** | 1.76751 | 6.11316 | 0.83283 | 10.09683 | 1.86319 | 2.46555 | 0.70285 | 8.48500 | 14.53313 | 2.16645 | 7.10778 | 14.38283 | 3.49106 | 20.53701 | 14.45188 | 0.44289 |
| **Ano1** | 0.01367 | 0.00786 | 0.01217 | 0.00928 | 0.00915 | 0.00665 | 0.00735 | 0.01139 | 0.01215 | 0.29872 | 0.01317 | 0.00672 | 0.01252 | 0.01131 | 0.01042 | 0.00781 |
| **Ano10** | 0.00913 | 0.37212 | 0.03253 | 0.15579 | 0.00975 | 0.00836 | 0.02842 | 0.01196 | 0.26318 | 0.00139 | 0.03120 | 0.03685 | 0.19339 | 0.00713 | 0.14825 | 0.00743 |
| **Ano2** | 0.01178 | 0.00796 | 0.01422 | 0.00475 | 0.00991 | 0.00715 | 0.00747 | 0.01247 | 0.22887 | 0.70254 | 0.01267 | 0.00485 | 0.28945 | 0.00956 | 0.01915 | 0.00810 |
| **Ano6** | 0.00000 | 0.00539 | 2.82066 | 1.58901 | 0.00622 | 0.27280 | 3.22293 | 1.51189 | 1.12728 | 2.22240 | 5.02977 | 0.17858 | 2.18939 | 0.05227 | 0.38284 | 0.00438 |
| **Ano7** | 0.00783 | 0.00836 | 0.03965 | 0.00962 | 0.00944 | 0.00849 | 0.00822 | 0.75421 | 0.01116 | 0.36545 | 0.00691 | 0.00613 | 0.00565 | 0.01838 | 0.03105 | 0.00776 |
| **Ano9** | 0.00000 | 0.00631 | 0.00000 | 0.76843 | 0.00935 | 3.93451 | 0.00278 | 7.31406 | 1.93306 | 1.85885 | 5.45199 | 0.00000 | 3.71966 | 1.78654 | 2.05505 | 0.00324 |
| **Anp32a** | 3.84483 | 0.00190 | 0.78635 | 0.00000 | 2.91168 | 2.64319 | 0.00426 | 0.95399 | 0.00806 | 1.87802 | 0.61525 | 0.24368 | 0.34279 | 0.01864 | 0.04306 | 0.13962 |
| **Anp32b** | 25.71570 | 12.64028 | 18.09366 | 18.95230 | 17.82715 | 2.34827 | 10.75050 | 18.49849 | 18.47327 | 26.21090 | 21.70939 | 9.45821 | 21.45191 | 18.62889 | 15.50097 | 10.78548 |
| **Anp32e** | 12.17589 | 10.38225 | 15.89116 | 12.28633 | 15.97310 | 5.56663 | 6.25403 | 9.55308 | 17.72067 | 11.41229 | 14.57804 | 5.08502 | 7.35913 | 14.89926 | 15.63083 | 4.72130 |
| **Anpep** | 0.22108 | 0.00724 | 0.09029 | 0.01229 | 0.00719 | 0.00546 | 0.39752 | 1.04130 | 0.12345 | 0.50946 | 0.44244 | 0.36981 | 0.00319 | 0.39023 | 0.57526 | 0.00689 |
| **Antxr1** | 0.00836 | 0.00766 | 0.00916 | 6.14389 | 0.00894 | 0.00563 | 0.00627 | 0.00760 | 1.92326 | 0.27482 | 0.01808 | 0.10150 | 1.25139 | 0.00022 | 0.47858 | 0.00540 |
| **Antxr2** | 0.01335 | 0.00791 | 0.01095 | 0.73275 | 0.00918 | 0.00632 | 0.00731 | 0.00999 | 0.00001 | 0.55417 | 0.02411 | 0.01494 | 0.01103 | 0.00610 | 0.88693 | 0.00709 |
| **Anxa1** | 0.01246 | 0.00783 | 0.01092 | 0.00570 | 0.00868 | 0.00672 | 0.00732 | 0.01051 | 0.01113 | 2.54516 | 0.00917 | 0.06502 | 0.01085 | 0.01288 | 0.00470 | 0.00741 |
| **Anxa11** | 0.18773 | 0.37786 | 0.47999 | 1.68534 | 0.14678 | 0.01612 | 1.01527 | 1.52549 | 2.57725 | 3.17416 | 7.47250 | 0.31982 | 1.17574 | 1.20340 | 0.00000 | 0.67329 |
| **Anxa2** | 0.00000 | 6.60215 | 3.80862 | 0.30578 | 0.11267 | 16.22802 | 2.53336 | 1.38334 | 0.23869 | 4.48896 | 0.21958 | 0.08368 | 8.21899 | 1.52861 | 0.89782 | 4.84536 |
| **Anxa3** | 0.00429 | 0.00625 | 0.41958 | 0.15138 | 0.04171 | 0.02642 | 0.00466 | 0.33784 | 0.00600 | 0.63503 | 4.16800 | 1.55894 | 1.23379 | 0.00088 | 0.00000 | 1.12423 |
| **Anxa4** | 0.21006 | 5.42279 | 3.85628 | 0.76397 | 1.52074 | 0.00583 | 0.12157 | 0.00000 | 0.00000 | 3.12884 | 1.00550 | 5.27856 | 0.74758 | 0.01994 | 0.39798 | 0.01713 |
| **Anxa5** | 1.86695 | 1.62117 | 0.00000 | 0.46730 | 0.00364 | 0.01148 | 0.00508 | 0.01411 | 0.00384 | 8.27452 | 0.00736 | 0.01340 | 0.00694 | 0.00391 | 0.23585 | 1.91327 |
| **Anxa6** | 9.40871 | 6.52767 | 0.00000 | 0.50318 | 0.44081 | 11.75653 | 9.35126 | 1.77268 | 1.31823 | 0.57328 | 2.36746 | 0.01927 | 4.73595 | 0.00000 | 0.33339 | 6.38132 |
| **Anxa7** | 18.68194 | 14.60303 | 17.72155 | 19.63886 | 11.69290 | 9.99448 | 11.20535 | 9.07560 | 5.12399 | 19.20186 | 7.21969 | 2.37007 | 15.92976 | 9.52075 | 16.11277 | 7.41813 |
| **Anxa8** | 0.01338 | 0.00773 | 0.01016 | 0.00399 | 0.01031 | 0.00633 | 0.00675 | 0.00976 | 0.01011 | 0.63870 | 0.00841 | 0.03283 | 0.97853 | 0.00513 | 1.05066 | 0.00699 |
| **Anxa9** | 3.58898 | 0.00338 | 2.82197 | 0.00000 | 0.00686 | 4.25297 | 0.00416 | 2.72011 | 0.97987 | 2.58457 | 0.54709 | 0.00000 | 0.07870 | 0.90347 | 0.63490 | 0.01589 |
| **Aoc1** | 0.01332 | 0.00778 | 0.01206 | 0.00781 | 0.00899 | 0.00657 | 0.00725 | 0.01049 | 0.01188 | 0.75932 | 0.01247 | 0.00549 | 0.02509 | 0.01155 | 0.00912 | 0.00774 |
| **Aoc2** | 0.21761 | 0.00756 | 2.99218 | 0.64234 | 0.17852 | 0.16654 | 0.07549 | 0.31585 | 0.55463 | 0.63024 | 1.53322 | 0.10470 | 0.82909 | 0.26151 | 0.39556 | 0.01252 |
| **Aoc3** | 0.01383 | 0.00772 | 0.00591 | 1.15758 | 0.00897 | 0.00702 | 0.00709 | 0.00000 | 0.97138 | 0.00000 | 0.72499 | 0.14467 | 0.00588 | 0.00271 | 0.02067 | 0.00679 |
| **Aox2** | 0.01263 | 0.00794 | 0.32025 | 0.00521 | 0.00876 | 0.01209 | 0.00757 | 0.01306 | 0.17988 | 0.00971 | 0.01018 | 0.00769 | 0.00973 | 0.01563 | 0.07970 | 0.00721 |
| **Aox4** | 0.00833 | 0.00788 | 0.30254 | 0.00000 | 0.00931 | 0.00558 | 0.00738 | 0.75452 | 0.86314 | 0.00145 | 0.00000 | 0.05915 | 0.00418 | 0.00619 | 2.16806 | 0.00708 |
| **Ap1ar** | 3.40136 | 2.88697 | 3.13980 | 5.29639 | 2.17350 | 4.70567 | 0.69627 | 2.37729 | 2.47232 | 3.34197 | 1.95214 | 3.99549 | 0.59952 | 7.58150 | 13.53295 | 3.54689 |
| **Ap1b1** | 4.30298 | 6.68344 | 8.75555 | 7.73297 | 2.29038 | 0.01553 | 0.65610 | 9.77631 | 5.26503 | 8.61762 | 4.63977 | 0.32983 | 7.43567 | 0.55059 | 4.37320 | 5.95826 |
| **Ap1g1** | 5.70331 | 9.62268 | 7.57847 | 7.63656 | 6.46047 | 9.87362 | 7.58582 | 13.22792 | 12.65988 | 10.53261 | 7.28681 | 4.40737 | 7.67798 | 13.66938 | 12.66601 | 1.69019 |
| **Ap1g2** | 0.01052 | 0.00789 | 0.12614 | 0.00557 | 0.00904 | 0.00595 | 0.00669 | 0.00193 | 0.01139 | 0.00289 | 0.41778 | 0.00521 | 1.28794 | 0.01128 | 0.00246 | 0.00651 |
| **Ap1m1** | 7.76045 | 3.90571 | 1.97684 | 7.42906 | 3.33292 | 9.28314 | 2.67489 | 2.33581 | 5.25813 | 12.44285 | 4.66756 | 4.50930 | 1.57881 | 0.00000 | 0.87693 | 2.16917 |
| **Ap1m2** | 4.33892 | 8.16497 | 10.36071 | 5.72559 | 6.96227 | 2.20053 | 5.15376 | 12.84831 | 3.93618 | 9.32026 | 6.26151 | 1.53299 | 9.10368 | 2.70960 | 5.49705 | 6.56448 |
| **Ap1s1** | 12.42116 | 3.08740 | 4.81328 | 4.27597 | 0.14766 | 14.67197 | 6.30391 | 19.02017 | 3.17180 | 15.35311 | 11.31600 | 1.40436 | 17.93236 | 0.02587 | 2.03699 | 8.48371 |
| **Ap1s2** | 1.33894 | 0.13199 | 0.00000 | 2.43562 | 0.00548 | 0.00513 | 0.00678 | 0.11223 | 0.93682 | 0.45361 | 0.00000 | 0.00000 | 0.01617 | 3.37610 | 0.00000 | 0.01258 |
| **Ap1s3** | 0.11844 | 0.00684 | 2.21625 | 0.38873 | 0.01276 | 0.00721 | 0.66762 | 1.92309 | 0.19632 | 4.31846 | 0.71247 | 0.25927 | 0.05365 | 11.89373 | 3.61362 | 0.01969 |
| **Ap2a1** | 2.56527 | 0.04632 | 0.00000 | 0.03895 | 0.81716 | 0.85681 | 0.59325 | 3.01461 | 0.00891 | 3.41284 | 0.00000 | 0.08200 | 3.69027 | 0.01205 | 0.00000 | 3.09372 |
| **Ap2a2** | 6.12033 | 4.21554 | 3.50244 | 8.09739 | 0.26122 | 15.90476 | 1.72484 | 19.28838 | 4.81711 | 10.63713 | 14.19048 | 2.70588 | 19.96672 | 1.61147 | 0.76468 | 6.28256 |
| **Ap2b1** | 4.45866 | 2.76461 | 2.23873 | 3.12058 | 6.40503 | 8.64382 | 0.03775 | 5.85507 | 0.35182 | 2.41320 | 6.83522 | 5.89410 | 6.62867 | 6.49121 | 4.23846 | 3.43822 |
| **Ap2m1** | 18.43964 | 18.58833 | 22.32865 | 8.12098 | 14.42270 | 17.91425 | 15.14460 | 24.21870 | 6.93658 | 18.03523 | 19.50348 | 7.78880 | 26.42286 | 12.01491 | 12.06899 | 11.89769 |

| | | | | | | | | | | | | | | | | |
|---|---|---|---|---|---|---|---|---|---|---|---|---|---|---|---|---|
| **Ap2s1** | 11.37575 | 0.61700 | 0.00000 | 0.04004 | 9.42754 | 7.18757 | 0.62591 | 13.83972 | 0.00895 | 14.65529 | 0.50023 | 0.14068 | 13.33036 | 0.00245 | 0.00000 | 10.42847 |
| **Ap3b1** | 13.52232 | 7.77865 | 8.56169 | 3.30234 | 0.96564 | 7.32678 | 1.43794 | 8.25691 | 1.15319 | 7.80716 | 5.91828 | 7.51508 | 8.66592 | 7.78837 | 2.47139 | 5.11482 |
| **Ap3b2** | 0.50498 | 0.00797 | 0.49900 | 0.01690 | 0.00663 | 0.27576 | 0.00660 | 0.10751 | 0.01134 | 2.66949 | 3.09249 | 0.00000 | 0.28357 | 0.00193 | 0.69357 | 0.00623 |
| **Ap3d1** | 14.58658 | 9.65088 | 9.40026 | 6.63922 | 2.83544 | 10.86467 | 7.36523 | 12.89171 | 5.16817 | 14.25984 | 9.26746 | 10.39395 | 9.58576 | 9.98186 | 13.41721 | 4.63123 |
| **Ap3m1** | 5.76035 | 7.74948 | 4.56992 | 7.22580 | 0.27104 | 0.07298 | 6.67381 | 6.69829 | 1.72391 | 7.43266 | 3.00546 | 8.15466 | 2.88535 | 10.56751 | 4.77810 | 1.78960 |
| **Ap3m2** | 0.01583 | 0.00806 | 0.91616 | 0.83663 | 0.00877 | 0.20703 | 0.00739 | 0.00915 | 2.02778 | 0.59787 | 0.00000 | 0.00000 | 0.00688 | 0.10117 | 0.27939 | 0.00645 |
| **Ap3s1** | 0.90987 | 4.27279 | 0.44432 | 2.08207 | 3.36610 | 0.03293 | 0.75920 | 1.55528 | 2.13770 | 4.10347 | 1.34489 | 9.08018 | 0.79170 | 2.05677 | 0.52529 | 0.25546 |
| **Ap3s2** | 8.47314 | 6.96046 | 6.14664 | 2.92442 | 3.14181 | 2.41505 | 2.48992 | 10.08167 | 3.45697 | 10.49179 | 4.35887 | 0.26928 | 7.75221 | 1.79162 | 2.81044 | 3.96760 |
| **Ap4b1** | 4.97710 | 10.64718 | 3.13713 | 10.37033 | 2.28703 | 7.16015 | 3.07927 | 4.44603 | 9.80247 | 13.41851 | 5.97454 | 8.43016 | 2.86262 | 8.93116 | 15.22705 | 2.53068 |
| **Ap4e1** | 3.03283 | 1.76833 | 1.40101 | 3.20119 | 0.00000 | 0.02516 | 0.12670 | 0.62040 | 6.97903 | 1.48865 | 1.06293 | 1.76718 | 0.32832 | 2.28862 | 2.67239 | 0.86348 |
| **Ap4m1** | 0.25236 | 0.00736 | 0.54707 | 1.05765 | 0.01313 | 0.97274 | 1.69371 | 1.34534 | 0.69836 | 0.37170 | 1.04553 | 0.04809 | 1.86746 | 0.25013 | 3.02086 | 0.12989 |
| **Ap4s1** | 4.37540 | 0.01670 | 4.54062 | 5.45047 | 1.06991 | 1.38801 | 0.87367 | 8.17376 | 9.60298 | 13.32489 | 16.84482 | 10.82145 | 3.25363 | 8.98289 | 8.38123 | 0.00836 |
| **Ap5b1** | 0.01533 | 0.00765 | 0.01689 | 1.19494 | 0.01066 | 0.01212 | 0.00715 | 1.30306 | 0.38937 | 0.00000 | 0.01712 | 0.02459 | 0.00504 | 0.01256 | 0.00000 | 0.01442 |
| **Ap5m1** | 0.85929 | 0.41934 | 0.74688 | 1.60258 | 0.01498 | 2.30974 | 0.28969 | 1.25131 | 1.15875 | 1.75581 | 0.64289 | 3.93784 | 0.36087 | 3.83686 | 1.97108 | 1.40154 |
| **Ap5s1** | 4.00589 | 0.00000 | 6.78964 | 1.83702 | 0.00000 | 0.49793 | 1.59943 | 2.07554 | 1.60642 | 1.01504 | 0.45385 | 0.03448 | 3.16301 | 0.04688 | 0.97418 | 0.00624 |
| **Ap5z1** | 0.98086 | 0.00455 | 3.54570 | 0.67987 | 0.03404 | 3.70644 | 1.76971 | 1.90099 | 1.31674 | 2.91067 | 3.11292 | 0.31354 | 7.64393 | 0.62151 | 0.35959 | 2.89489 |
| **Apaf1** | 2.37028 | 0.00000 | 1.80501 | 8.53120 | 2.03822 | 2.55468 | 6.88944 | 1.86282 | 10.84114 | 13.29057 | 4.08963 | 8.52508 | 1.24435 | 3.00017 | 14.03470 | 3.14857 |
| **Apba1** | 0.01374 | 0.00791 | 0.01219 | 0.89241 | 0.00949 | 0.00669 | 0.00753 | 0.01380 | 0.01850 | 0.01336 | 0.01290 | 0.01806 | 0.01588 | 0.01656 | 0.02326 | 0.00734 |
| **Apba2** | 0.02089 | 0.00779 | 0.03204 | 0.41638 | 0.01776 | 0.00588 | 0.00734 | 0.01562 | 0.00416 | 0.36972 | 0.01062 | 0.00644 | 0.02868 | 0.11129 | 0.02827 | 0.01395 |
| **Apba3** | 2.29982 | 0.01474 | 0.00000 | 0.00536 | 0.94795 | 0.18867 | 0.02514 | 1.98145 | 0.24036 | 0.54020 | 0.71402 | 0.03723 | 1.90305 | 0.01631 | 0.01386 | 0.70495 |
| **Apbb1** | 1.65744 | 4.16174 | 0.00000 | 4.06123 | 4.53539 | 4.87495 | 2.51617 | 2.78904 | 0.00000 | 0.55879 | 2.16308 | 0.04143 | 1.61616 | 0.00742 | 0.14280 | 0.01520 |
| **Apbb1ip** | 0.04344 | 0.01885 | 0.11076 | 0.01064 | 0.00853 | 0.00616 | 0.00779 | 0.01470 | 0.01233 | 0.00880 | 0.01398 | 0.05095 | 0.18875 | 0.01305 | 0.00732 | 0.00744 |
| **Apbb2** | 0.09205 | 0.82594 | 1.57948 | 2.23186 | 0.06478 | 0.00513 | 3.72874 | 0.34780 | 2.35860 | 1.26301 | 2.41828 | 0.25003 | 0.00000 | 0.85725 | 2.86828 | 1.28595 |
| **Apbb3** | 0.85444 | 0.00675 | 0.07180 | 0.22075 | 0.00683 | 0.56831 | 0.32097 | 0.01920 | 0.51721 | 0.00083 | 0.01104 | 0.00569 | 1.39428 | 0.01460 | 0.00257 | 0.00580 |
| **Apc** | 2.31986 | 2.11932 | 1.53818 | 16.26738 | 5.10123 | 5.68456 | 2.83094 | 0.03199 | 8.93122 | 10.11359 | 2.89649 | 5.43831 | 0.36464 | 9.26069 | 16.25883 | 0.18104 |
| **Apc2** | 0.01376 | 0.00802 | 0.01228 | 3.59995 | 0.00934 | 0.00624 | 0.00734 | 0.01298 | 0.09948 | 0.00892 | 0.01305 | 0.00765 | 0.01262 | 0.00087 | 0.18135 | 0.00675 |
| **Apcdd1** | 0.54967 | 0.17571 | 0.86777 | 0.00916 | 0.09712 | 0.00498 | 0.20967 | 0.00727 | 0.01218 | 0.32895 | 0.01023 | 0.01627 | 0.00785 | 0.01299 | 0.00000 | 0.01314 |
| **Apeh** | 8.01205 | 0.80631 | 3.07610 | 0.48496 | 0.83431 | 0.01679 | 2.35357 | 2.98240 | 0.00000 | 2.93149 | 0.64707 | 0.02047 | 4.05142 | 0.00456 | 0.08384 | 4.00166 |
| **Apela** | 3.08018 | 1.54598 | 2.32141 | 0.00955 | 0.00000 | 0.00226 | 0.00378 | 0.00000 | 0.01240 | 0.00000 | 0.01105 | 0.00434 | 0.54471 | 0.01380 | 0.00000 | 0.00223 |
| **Apex1** | 26.69441 | 17.18034 | 21.33549 | 6.22799 | 15.14792 | 8.89455 | 14.09616 | 20.79215 | 20.26601 | 26.16683 | 20.70048 | 16.07404 | 21.51346 | 12.32858 | 21.15336 | 12.37516 |
| **Apex2** | 1.85380 | 0.02562 | 1.01169 | 2.66012 | 3.62037 | 0.23783 | 1.26063 | 2.07735 | 3.88393 | 2.78822 | 0.75632 | 0.35690 | 3.54545 | 2.81499 | 3.79583 | 2.76312 |
| **Aph1a** | 2.26086 | 0.19164 | 1.87293 | 1.69015 | 0.01381 | 0.01773 | 0.09870 | 3.42598 | 0.53416 | 2.23063 | 0.80509 | 0.06185 | 0.93564 | 1.80850 | 0.06078 | 0.57789 |
| **Aph1b** | 0.01220 | 0.00825 | 0.01331 | 0.38514 | 0.01025 | 0.00700 | 0.00756 | 0.01414 | 0.63117 | 0.03517 | 0.01049 | 0.00592 | 0.24260 | 0.09186 | 0.00000 | 0.00764 |
| **Aph1c** | 0.01704 | 0.04063 | 0.02080 | 0.22984 | 0.00951 | 0.00664 | 0.02871 | 0.01773 | 0.78386 | 0.00938 | 0.11258 | 0.00563 | 0.05994 | 0.04718 | 0.00159 | 0.03038 |
| **Api5** | 13.56267 | 9.20416 | 10.41388 | 3.83589 | 13.72083 | 8.26035 | 8.66310 | 14.70812 | 5.45768 | 18.24245 | 12.73855 | 11.46962 | 10.33833 | 16.35082 | 9.40992 | 2.44498 |
| **Apip** | 4.88976 | 4.40287 | 4.55343 | 2.62431 | 0.01025 | 0.00004 | 2.06357 | 3.22517 | 2.34601 | 0.89462 | 1.91217 | 0.27006 | 0.00000 | 0.78526 | 0.78398 | 2.79311 |
| **Apitd1** | 8.59855 | 0.05916 | 3.07771 | 12.87312 | 3.93148 | 0.00000 | 3.16730 | 5.55436 | 17.39534 | 7.65415 | 2.02299 | 0.61795 | 3.75689 | 0.09831 | 1.75998 | 0.11581 |
| **Aplf** | 5.17272 | 0.00000 | 0.33080 | 2.89414 | 3.66703 | 0.22064 | 0.08481 | 0.36374 | 1.77332 | 0.63873 | 2.38514 | 0.74887 | 2.46214 | 2.63063 | 1.45633 | 0.41384 |
| **Aplp1** | 0.01622 | 0.00778 | 0.01094 | 0.00459 | 0.00901 | 0.00657 | 0.00739 | 0.03814 | 0.43059 | 0.17911 | 0.00768 | 0.00625 | 0.01031 | 0.01494 | 0.08487 | 0.00711 |
| **Aplp2** | 9.87165 | 8.89898 | 12.21656 | 11.92264 | 8.58539 | 5.82650 | 7.74801 | 3.06729 | 14.89964 | 5.07388 | 4.31680 | 8.18573 | 6.72890 | 2.74419 | 7.58576 | 4.88320 |
| **Apmap** | 1.51820 | 3.03016 | 5.02813 | 4.54605 | 0.89239 | 0.10487 | 0.10809 | 1.83986 | 3.30683 | 2.39322 | 4.27041 | 0.73119 | 5.33823 | 1.63520 | 2.16390 | 2.04105 |
| **Apoa1** | 0.00000 | 0.00463 | 0.00000 | 0.00190 | 1.60705 | 4.25965 | 2.01499 | 7.04461 | 0.00668 | 0.00000 | 0.12779 | 0.00000 | 2.40418 | 0.01371 | 0.00000 | 0.00441 |
| **Apoa1bp** | 4.13395 | 0.06856 | 2.67820 | 4.99790 | 0.03360 | 5.12602 | 0.04078 | 1.85635 | 2.40155 | 2.70137 | 4.00843 | 0.02474 | 3.70438 | 0.18772 | 0.40160 | 1.49752 |
| **Apoa4** | 0.00545 | 0.00765 | 0.42780 | 0.00848 | 0.01809 | 0.00560 | 0.00677 | 0.69260 | 0.04042 | 0.00000 | 0.00800 | 0.00619 | 1.04844 | 0.01853 | 0.00000 | 0.00656 |
| **Apoa5** | 0.01375 | 0.00796 | 0.00799 | 1.67072 | 0.00897 | 0.00686 | 0.00732 | 0.00046 | 1.08447 | 0.26841 | 0.40371 | 0.03729 | 0.00906 | 0.00398 | 0.25920 | 0.00695 |
| **Apobec1** | 0.01347 | 0.00802 | 0.00686 | 0.63284 | 0.00893 | 0.00705 | 0.00690 | 0.00000 | 0.00000 | 0.63671 | 0.93729 | 0.00178 | 0.00518 | 0.02707 | 0.04813 | 0.00733 |
| **Apobec2** | 0.00687 | 0.00860 | 0.00816 | 0.02554 | 0.04499 | 0.00694 | 0.00833 | 0.46967 | 0.01006 | 0.68642 | 0.00735 | 0.00661 | 0.00558 | 0.01458 | 0.00206 | 0.00708 |
| **Apobec3** | 3.98014 | 0.00284 | 1.08991 | 0.56527 | 0.00000 | 0.00115 | 0.00287 | 6.85961 | 1.70858 | 3.94863 | 3.15203 | 0.05907 | 0.14938 | 6.70135 | 5.23654 | 0.03042 |
| **Apobr** | 0.76762 | 0.00658 | 0.82246 | 0.00649 | 0.02454 | 0.00562 | 0.00648 | 0.17481 | 0.10778 | 0.06182 | 0.01190 | 0.01437 | 0.75398 | 0.01439 | 0.00000 | 0.17409 |
| **Apoc1** | 5.24165 | 0.19292 | 3.67476 | 0.00000 | 7.25852 | 0.82008 | 0.80748 | 0.06672 | 1.04868 | 0.45658 | 8.38979 | 0.05230 | 2.19520 | 0.32379 | 0.00000 | 4.85904 |
| **Apoc2** | 0.01264 | 0.00784 | 0.02676 | 0.04998 | 0.00945 | 0.00604 | 0.00732 | 0.11552 | 0.00902 | 0.04816 | 0.01146 | 0.00743 | 0.00840 | 0.01086 | 0.07866 | 0.00742 |
| **Apod** | 2.80159 | 0.01568 | 0.01885 | 0.55330 | 0.00294 | 0.00482 | 0.00512 | 0.01916 | 0.02319 | 0.00257 | 0.22100 | 0.02421 | 0.01838 | 0.10605 | 0.06601 | 1.65845 |
| **Apoe** | 4.04749 | 0.02276 | 0.34932 | 3.27791 | 0.00553 | 0.00823 | 0.04191 | 0.00000 | 2.40086 | 5.45871 | 5.40221 | 0.17130 | 1.14742 | 0.01100 | 0.00000 | 13.39332 |
| **Apof** | 0.02601 | 0.00800 | 0.02723 | 0.53192 | 0.01768 | 0.01314 | 0.01804 | 0.04845 | 0.98239 | 0.05566 | 0.08191 | 0.04929 | 0.05140 | 0.07167 | 0.04458 | 0.01437 |
| **Apoh** | 0.01376 | 0.00800 | 0.01115 | 0.00444 | 0.00933 | 0.00669 | 0.00747 | 0.01276 | 0.52102 | 0.01086 | 0.00854 | 0.05885 | 0.01300 | 0.01376 | 0.00519 | 0.00741 |
| **Apol10a** | 0.01282 | 0.00779 | 0.01213 | 0.00737 | 0.00906 | 0.00654 | 0.00722 | 0.00984 | 0.01184 | 0.96978 | 0.01163 | 0.00524 | 0.09169 | 0.01200 | 0.00764 | 0.00739 |
| **Apol6** | 0.02600 | 0.01857 | 0.07353 | 0.56311 | 0.00845 | 0.02904 | 0.02931 | 0.01083 | 0.24234 | 0.06900 | 0.20876 | 0.01698 | 0.01072 | 0.16204 | 0.07819 | 0.03589 |
| **Apol7a** | 0.01254 | 0.00844 | 0.02453 | 0.01014 | 0.00996 | 0.00666 | 0.00807 | 0.02745 | 0.01231 | 0.03221 | 0.01471 | 0.00868 | 0.04344 | 0.01075 | 0.04036 | 0.00758 |
| **Apol7b** | 0.00000 | 0.00837 | 1.27240 | 3.94417 | 0.00752 | 0.00519 | 0.04332 | 0.38652 | 4.77120 | 2.10224 | 0.44243 | 0.11339 | 0.90495 | 0.31730 | 2.12352 | 0.01297 |
| **Apol7c** | 0.01247 | 0.00841 | 0.01133 | 0.01039 | 0.00941 | 0.00677 | 0.00802 | 0.04468 | 0.01227 | 0.01736 | 0.01220 | 0.02014 | 0.02626 | 0.01025 | 0.04727 | 0.00740 |
| **Apol7e** | 0.00431 | 0.00826 | 0.86159 | 3.60521 | 0.00794 | 0.00515 | 0.12264 | 0.17081 | 5.23317 | 1.55238 | 0.38058 | 0.12627 | 0.66369 | 0.25283 | 1.36329 | 0.02045 |
| **Apom** | 0.46912 | 0.00777 | 0.04409 | 5.88526 | 0.00668 | 0.00352 | 0.00695 | 0.21097 | 12.70721 | 0.66393 | 2.09430 | 0.37915 | 0.00000 | 1.95420 | 0.93397 | 0.39738 |

| | | | | | | | | | | | | | | | | |
|---|---|---|---|---|---|---|---|---|---|---|---|---|---|---|---|---|
| **Apon** | 0.01371 | 0.00741 | 0.00638 | 0.05366 | 0.01028 | 0.00662 | 0.00714 | 0.87730 | 1.66936 | 0.00000 | 0.00000 | 0.00189 | 0.01131 | 0.02227 | 0.44874 | 0.14811 |
| **Apoo** | 1.63037 | 2.05315 | 2.10504 | 1.02844 | 0.00587 | 0.84735 | 1.17825 | 1.05723 | 0.96581 | 1.97813 | 1.84611 | 2.95546 | 1.14808 | 0.63021 | 0.77000 | 0.68383 |
| **Apool** | 5.56133 | 3.12536 | 4.07151 | 0.01057 | 0.00000 | 2.31039 | 2.46393 | 0.73886 | 0.07833 | 1.63153 | 2.17148 | 0.07186 | 4.43505 | 2.62350 | 0.20942 | 4.15156 |
| **Apopt1** | 0.80845 | 0.10485 | 1.41777 | 1.75080 | 0.44217 | 1.93286 | 0.04681 | 0.61091 | 3.29216 | 0.79316 | 1.33237 | 0.24686 | 1.25910 | 1.69121 | 0.90210 | 0.05458 |
| **App** | 0.02297 | 1.58319 | 1.06551 | 0.58348 | 0.01416 | 0.00956 | 0.00737 | 0.63316 | 0.00361 | 1.54521 | 0.02039 | 0.10536 | 0.00865 | 0.00461 | 0.00000 | 0.00626 |
| **Appbp2** | 9.07431 | 5.28638 | 7.15163 | 2.03878 | 3.67749 | 0.56889 | 8.22792 | 7.14289 | 2.98084 | 6.57799 | 2.43701 | 3.88192 | 0.71233 | 12.87248 | 11.04250 | 2.08402 |
| **Appbp2os** | 0.00666 | 0.00706 | 1.09991 | 0.04186 | 0.55415 | 0.21574 | 0.54063 | 0.00984 | 0.01132 | 0.00904 | 0.01258 | 0.00648 | 0.01374 | 0.01438 | 0.00000 | 0.00693 |
| **Appl1** | 6.14017 | 2.71874 | 6.22180 | 7.22980 | 1.82665 | 0.42990 | 5.31722 | 3.30590 | 1.28904 | 6.69363 | 5.12191 | 3.40534 | 2.79877 | 8.61084 | 6.75755 | 2.15120 |
| **Appl2** | 2.01045 | 1.14559 | 1.25238 | 0.00000 | 0.43409 | 2.84125 | 1.76717 | 0.07329 | 0.50481 | 0.21381 | 0.00000 | 0.09816 | 0.00088 | 0.01431 | 0.01274 | 2.38694 |
| **Aprt** | 21.04087 | 12.03691 | 27.37216 | 15.67373 | 5.76422 | 12.08313 | 13.20986 | 36.19870 | 19.05272 | 36.55109 | 32.23144 | 11.36597 | 30.25587 | 0.55748 | 7.18593 | 10.13140 |
| **Aptx** | 5.25470 | 2.01365 | 7.00342 | 2.15052 | 0.22801 | 0.04739 | 2.49303 | 9.88327 | 1.58812 | 6.22201 | 4.77998 | 7.11630 | 6.16544 | 2.62430 | 5.05229 | 0.01614 |
| **Aqp11** | 0.13888 | 0.00754 | 0.00000 | 3.91777 | 0.00720 | 0.00635 | 0.01756 | 2.51659 | 1.03266 | 5.65978 | 0.68536 | 0.02133 | 0.00249 | 0.00000 | 2.17226 | 0.00642 |
| **Aqp3** | 2.42309 | 2.88486 | 0.98803 | 0.03435 | 1.58909 | 0.10026 | 1.47717 | 2.27321 | 0.00703 | 0.00000 | 7.81278 | 0.03905 | 1.08778 | 2.04855 | 0.00000 | 0.16098 |
| **Aqp7** | 0.05320 | 0.01885 | 0.08177 | 0.54153 | 0.02254 | 0.01332 | 0.01737 | 0.00000 | 0.79766 | 0.06877 | 0.47026 | 0.03231 | 0.11531 | 0.45788 | 0.00990 | 0.02200 |
| **Aqp8** | 12.39872 | 4.92242 | 0.00000 | 0.00665 | 14.39985 | 0.00000 | 0.00530 | 0.00000 | 0.00854 | 0.00000 | 0.00922 | 0.06905 | 0.02646 | 0.00635 | 0.01397 | 8.05361 |
| **Aqp9** | 2.31151 | 2.89540 | 7.10037 | 9.17260 | 0.02480 | 5.64611 | 5.66089 | 4.87717 | 9.31960 | 13.76619 | 11.78427 | 14.26270 | 5.99828 | 18.18264 | 16.75592 | 0.02775 |
| **Aqr** | 7.65237 | 3.78402 | 5.42281 | 3.50048 | 3.84153 | 5.02318 | 1.56295 | 6.74788 | 6.42693 | 6.55953 | 4.52369 | 5.39143 | 4.42313 | 0.41601 | 6.30586 | 0.01189 |
| **Araf** | 6.97372 | 6.82536 | 10.05239 | 3.76069 | 4.23329 | 7.77346 | 3.92566 | 6.94049 | 5.08958 | 6.86195 | 6.25067 | 1.01495 | 7.22962 | 5.65347 | 7.20989 | 3.16968 |
| **Arap1** | 0.31307 | 0.30344 | 1.25826 | 0.05572 | 0.00757 | 0.00893 | 0.00507 | 1.85909 | 0.47318 | 1.17711 | 3.70207 | 0.09267 | 1.16332 | 0.32539 | 2.19161 | 0.01307 |
| **Arap2** | 0.00760 | 2.94228 | 0.17326 | 11.06089 | 0.43919 | 0.00494 | 0.00547 | 0.37526 | 3.44041 | 1.06791 | 0.72524 | 1.67136 | 0.03291 | 0.20264 | 13.79198 | 0.00401 |
| **Arcn1** | 5.59546 | 3.41975 | 4.88570 | 8.16651 | 12.96804 | 6.60258 | 6.26623 | 6.00450 | 14.97395 | 6.83188 | 5.85369 | 6.49180 | 3.50846 | 10.06287 | 9.34161 | 4.02037 |
| **Arel1** | 2.48874 | 0.20240 | 1.98259 | 6.75704 | 0.02784 | 3.15677 | 1.90825 | 1.91687 | 3.88462 | 2.75130 | 0.37941 | 10.80652 | 2.61158 | 4.54489 | 5.42829 | 0.02904 |
| **Arf1** | 29.91912 | 17.80612 | 23.53075 | 13.54422 | 22.83018 | 18.47370 | 15.98339 | 30.38102 | 11.23569 | 29.03773 | 13.08122 | 8.00017 | 30.50475 | 0.57879 | 3.48949 | 16.97730 |
| **Arf2** | 1.72719 | 3.30351 | 7.17511 | 6.58479 | 0.02260 | 7.49148 | 8.48741 | 6.54043 | 12.86798 | 7.03029 | 4.04643 | 1.10361 | 1.14059 | 0.44449 | 4.78673 | 3.94099 |
| **Arf3** | 16.08508 | 10.22551 | 11.38994 | 6.19854 | 3.61893 | 4.08550 | 6.20649 | 10.41542 | 5.42347 | 11.29717 | 3.39542 | 1.10453 | 5.56353 | 6.42241 | 5.75217 | 7.12223 |
| **Arf4** | 12.57783 | 9.25494 | 16.43065 | 6.28566 | 14.83063 | 8.39534 | 7.50556 | 12.23198 | 5.38330 | 11.35636 | 9.18803 | 5.05162 | 15.25056 | 7.55340 | 4.61033 | 11.68959 |
| **Arf5** | 3.95688 | 0.04435 | 0.00000 | 0.10553 | 0.02274 | 1.21233 | 0.80920 | 2.61128 | 0.00732 | 0.88263 | 0.02313 | 0.00000 | 1.42286 | 0.01694 | 0.00942 | 5.76465 |
| **Arf6** | 7.40795 | 4.52214 | 4.35406 | 13.75039 | 2.93196 | 1.99890 | 6.08961 | 4.19908 | 17.63738 | 9.45521 | 10.42892 | 0.44811 | 4.83773 | 1.40758 | 4.62074 | 3.19152 |
| **Arfgap1** | 6.76629 | 2.64168 | 7.52541 | 2.78338 | 3.42075 | 4.27492 | 8.76229 | 10.45566 | 3.03194 | 6.27395 | 3.87107 | 0.45742 | 8.91302 | 0.44898 | 3.30560 | 8.14177 |
| **Arfgap2** | 0.14551 | 0.75624 | 0.55699 | 0.00168 | 0.63025 | 0.68770 | 0.03886 | 1.71017 | 0.01922 | 0.04532 | 0.30252 | 0.00236 | 1.33220 | 0.01168 | 0.05535 | 0.61768 |
| **Arfgap3** | 0.88936 | 2.69338 | 0.00000 | 1.56582 | 0.00640 | 4.85328 | 0.00518 | 0.00496 | 0.96722 | 0.90634 | 0.00000 | 0.02856 | 0.01513 | 0.07099 | 0.00000 | 1.11198 |
| **Arfgef1** | 6.76443 | 0.06449 | 1.31056 | 16.30171 | 0.49557 | 1.77552 | 2.55062 | 0.00000 | 10.20681 | 5.21117 | 0.71391 | 0.78494 | 1.69124 | 9.65867 | 17.86545 | 1.36384 |
| **Arfgef2** | 6.03952 | 2.25831 | 4.49918 | 7.20263 | 2.14899 | 3.88211 | 5.21415 | 4.55529 | 7.66589 | 7.61900 | 2.80633 | 0.77597 | 2.88999 | 0.38019 | 2.75200 | 0.00000 |
| **Arfgef3** | 0.01399 | 0.00771 | 0.00953 | 3.13301 | 0.00915 | 0.00633 | 0.00722 | 0.00343 | 0.00000 | 0.00000 | 0.31378 | 0.00509 | 0.00901 | 0.00052 | 0.45288 | 0.00676 |
| **Arfip1** | 3.62524 | 0.06859 | 4.51478 | 13.52542 | 1.64143 | 1.30346 | 2.20487 | 0.62526 | 1.92613 | 1.49423 | 1.91673 | 2.51628 | 1.98498 | 2.11976 | 15.09612 | 1.48764 |
| **Arfip2** | 2.53659 | 2.91392 | 2.07912 | 1.15679 | 5.04289 | 6.03263 | 0.64624 | 3.90298 | 0.00000 | 1.29970 | 1.78543 | 0.11059 | 3.24954 | 0.00000 | 0.53265 | 5.55615 |
| **Arfrp1** | 4.02152 | 3.37554 | 7.82450 | 10.35429 | 0.03923 | 3.65520 | 0.51254 | 6.77820 | 13.26731 | 8.91912 | 9.33513 | 4.43677 | 8.28609 | 1.42937 | 7.12316 | 1.03825 |
| **Arg1** | 0.00000 | 0.00745 | 0.64376 | 3.22117 | 0.00768 | 0.00476 | 0.00568 | 0.00000 | 2.41717 | 2.87078 | 4.03732 | 2.77081 | 0.69942 | 1.79476 | 7.37241 | 0.00534 |
| **Arg2** | 0.23311 | 1.31768 | 1.46033 | 2.85394 | 0.03158 | 0.00853 | 0.00691 | 0.14230 | 2.52211 | 0.32057 | 0.32576 | 0.06433 | 0.00000 | 2.51146 | 0.92393 | 0.00571 |
| **Arglu1** | 8.10296 | 0.51875 | 7.30807 | 12.71636 | 5.97251 | 2.17775 | 2.74395 | 4.53272 | 20.37296 | 7.74370 | 7.14820 | 0.87833 | 4.38885 | 9.42696 | 6.36560 | 7.00670 |
| **Arhgap1** | 10.81961 | 7.32369 | 12.29497 | 8.37609 | 1.83464 | 12.19384 | 6.63543 | 12.51960 | 6.31519 | 14.01153 | 14.85994 | 3.49659 | 12.44499 | 6.26666 | 15.13807 | 1.03211 |
| **Arhgap10** | 0.52622 | 0.01775 | 0.48952 | 0.57948 | 0.02060 | 0.00514 | 0.31848 | 1.08270 | 0.72121 | 1.41683 | 1.34023 | 0.06428 | 0.06069 | 1.36914 | 0.79145 | 0.55537 |
| **Arhgap11a** | 3.83291 | 2.88735 | 1.10290 | 18.59968 | 2.82455 | 4.58175 | 1.32503 | 5.62692 | 24.62558 | 9.96253 | 4.69862 | 3.59029 | 5.95011 | 7.69938 | 19.14130 | 2.51376 |
| **Arhgap12** | 0.17320 | 0.21215 | 2.42921 | 11.54981 | 3.57694 | 0.23868 | 4.79500 | 1.69157 | 7.70896 | 2.93051 | 1.73499 | 1.35910 | 0.06564 | 3.02315 | 10.71984 | 0.01024 |
| **Arhgap15** | 0.05774 | 0.00736 | 0.00802 | 0.06668 | 0.00810 | 0.00633 | 0.00732 | 0.00965 | 0.59383 | 0.68739 | 0.00425 | 0.00156 | 0.01132 | 0.00684 | 1.32547 | 0.00718 |
| **Arhgap17** | 4.65487 | 0.43796 | 0.00000 | 3.19103 | 4.37303 | 2.66605 | 2.69300 | 3.52761 | 0.83579 | 0.81503 | 1.38216 | 0.15560 | 2.35210 | 0.01393 | 0.00000 | 5.08917 |
| **Arhgap18** | 0.00000 | 0.00428 | 0.41030 | 0.68870 | 0.00768 | 3.09104 | 0.00000 | 6.94552 | 0.66429 | 7.58534 | 12.57469 | 4.60890 | 3.99166 | 13.90983 | 3.32106 | 0.00148 |
| **Arhgap19** | 3.77036 | 4.81640 | 1.44166 | 1.58873 | 2.11410 | 3.99028 | 1.67373 | 3.70752 | 1.38858 | 0.38348 | 3.02882 | 3.07177 | 0.34440 | 7.35285 | 8.05134 | 0.00542 |
| **Arhgap20** | 0.01311 | 0.00860 | 0.01157 | 0.32889 | 0.00929 | 0.00685 | 0.00792 | 0.02694 | 0.00540 | 0.07199 | 0.01166 | 0.00988 | 0.01234 | 0.00695 | 0.17237 | 0.00736 |
| **Arhgap20os** | 0.06109 | 0.01903 | 0.01957 | 8.48662 | 0.04490 | 0.11412 | 0.00687 | 0.04077 | 1.58694 | 0.70344 | 0.27972 | 0.02002 | 0.04977 | 0.00750 | 0.71010 | 0.00551 |
| **Arhgap21** | 2.85009 | 0.15458 | 2.49592 | 18.10151 | 0.56626 | 4.13509 | 6.86081 | 12.94140 | 8.36769 | 19.22172 | 9.53539 | 19.68359 | 2.67535 | 7.87120 | 3.98134 | 0.25603 |
| **Arhgap22** | 0.73827 | 0.00767 | 0.08777 | 0.43210 | 0.03597 | 0.00786 | 0.00675 | 0.01013 | 0.33424 | 0.01726 | 0.01005 | 0.00611 | 0.01596 | 0.00659 | 0.06733 | 0.74023 |
| **Arhgap23** | 0.26492 | 0.00809 | 0.00327 | 0.27038 | 0.00724 | 0.00783 | 0.00787 | 0.19891 | 2.65313 | 0.26789 | 0.00000 | 0.12924 | 0.00745 | 0.01131 | 0.00000 | 0.00686 |
| **Arhgap26** | 1.00084 | 0.00792 | 0.54149 | 0.20822 | 0.01385 | 0.00871 | 0.00704 | 0.03153 | 0.16227 | 0.43886 | 0.18277 | 0.07981 | 0.01386 | 0.00512 | 0.75160 | 0.00625 |
| **Arhgap27** | 0.00814 | 0.00840 | 0.09202 | 0.09875 | 0.00964 | 0.00887 | 0.00817 | 0.50869 | 0.04529 | 0.08576 | 0.00901 | 0.00730 | 0.00554 | 0.01807 | 0.00873 | 0.00741 |
| **Arhgap28** | 0.01265 | 0.00784 | 0.00342 | 3.56182 | 0.00883 | 0.00641 | 0.00690 | 0.00000 | 0.80773 | 0.94294 | 0.80583 | 0.02970 | 0.01268 | 0.00000 | 1.61527 | 0.01364 |
| **Arhgap29** | 0.70773 | 0.00248 | 1.93012 | 10.00344 | 0.48769 | 0.44247 | 4.85466 | 1.64735 | 13.93968 | 5.58298 | 4.81345 | 3.32290 | 0.09568 | 7.48225 | 13.93394 | 1.03948 |
| **Arhgap30** | 0.01511 | 0.01848 | 0.00697 | 0.02353 | 0.00890 | 0.00637 | 0.02696 | 0.01946 | 0.01039 | 0.46821 | 0.00000 | 0.01869 | 0.03728 | 1.24505 | 1.06840 | 0.01500 |
| **Arhgap31** | 0.08243 | 0.00864 | 0.00017 | 0.16345 | 0.01513 | 3.30812 | 0.00725 | 0.01094 | 0.00882 | 0.02242 | 0.00871 | 0.03267 | 0.02303 | 0.01134 | 0.00829 | 0.00620 |
| **Arhgap32** | 0.00000 | 0.87214 | 1.88799 | 0.94112 | 0.00609 | 0.00934 | 1.68804 | 3.09315 | 0.40047 | 2.88885 | 1.48208 | 0.07873 | 0.00000 | 1.94352 | 0.41498 | 0.00603 |
| **Arhgap33** | 0.01187 | 0.02792 | 0.01626 | 0.89539 | 0.00867 | 0.00716 | 0.07698 | 0.00688 | 0.94593 | 1.62339 | 0.09327 | 0.04763 | 0.01043 | 0.00340 | 0.12107 | 0.00725 |
| **Arhgap35** | 3.52747 | 5.14774 | 4.26336 | 8.07476 | 5.25834 | 3.05227 | 6.13546 | 5.98162 | 6.58861 | 5.31602 | 4.97597 | 0.08885 | 4.87558 | 6.14294 | 4.56274 | 0.97025 |

| | | | | | | | | | | | | | | | | |
|---|---|---|---|---|---|---|---|---|---|---|---|---|---|---|---|---|
| **Arhgap39** | 0.18784 | 0.00794 | 0.32355 | 4.24951 | 0.11325 | 0.02360 | 0.00710 | 0.00857 | 1.50349 | 0.31946 | 0.01866 | 0.01317 | 0.01712 | 0.00000 | 0.38139 | 0.00639 |
| **Arhgap4** | 0.35907 | 0.00787 | 0.00271 | 0.35492 | 0.01273 | 0.00831 | 0.00700 | 0.31730 | 0.54415 | 0.42010 | 0.00633 | 0.00592 | 0.00682 | 0.00902 | 0.08791 | 0.00707 |
| **Arhgap42** | 0.23687 | 0.01757 | 0.88860 | 4.12839 | 1.20441 | 0.00472 | 0.00713 | 2.75395 | 3.22634 | 4.87746 | 1.02919 | 1.36757 | 0.16167 | 1.28311 | 4.09108 | 0.01295 |
| **Arhgap44** | 0.07104 | 0.00751 | 0.14128 | 0.67462 | 0.00654 | 0.00655 | 0.00698 | 0.01013 | 0.77725 | 1.20915 | 0.00613 | 0.01094 | 0.01459 | 0.00523 | 0.07890 | 4.36772 |
| **Arhgap5** | 0.87613 | 4.10219 | 1.89132 | 1.31825 | 0.09399 | 2.42596 | 0.47201 | 1.23973 | 1.23057 | 0.00000 | 0.74745 | 0.00000 | 0.36282 | 14.33224 | 0.65684 | 0.08781 |
| **Arhgap8** | 5.51267 | 0.00954 | 0.97328 | 1.64067 | 0.86336 | 6.83800 | 1.25677 | 5.29593 | 2.74827 | 1.59188 | 2.26892 | 0.00893 | 5.64885 | 0.00000 | 0.00000 | 0.00960 |
| **Arhgap9** | 1.51311 | 0.01583 | 0.24242 | 0.05812 | 0.01621 | 0.14206 | 1.65718 | 1.29868 | 0.00989 | 0.57145 | 0.22524 | 0.02918 | 1.24225 | 0.01396 | 0.00208 | 0.01178 |
| **Arhgdia** | 21.96092 | 3.75664 | 8.55276 | 9.98699 | 16.63810 | 10.04189 | 6.77298 | 23.31804 | 3.70997 | 17.31270 | 9.16806 | 1.12631 | 22.52911 | 0.01847 | 0.18195 | 13.39042 |
| **Arhgdib** | 0.03661 | 0.00836 | 0.00869 | 0.00625 | 0.01806 | 1.37724 | 0.00772 | 0.02881 | 0.01205 | 0.77939 | 0.00986 | 0.01709 | 0.01843 | 0.01250 | 0.00855 | 0.03750 |
| **Arhgdig** | 0.29351 | 0.01619 | 2.17723 | 4.91922 | 0.02055 | 2.80363 | 0.10504 | 5.12658 | 2.11394 | 5.01347 | 6.84542 | 0.28990 | 0.33083 | 0.01738 | 0.56202 | 0.00303 |
| **Arhgef1** | 3.98840 | 0.80196 | 0.29102 | 1.09469 | 0.56047 | 4.24948 | 1.66708 | 6.29892 | 0.04304 | 1.42077 | 2.78104 | 0.38693 | 0.48181 | 0.01861 | 0.00000 | 0.51146 |
| **Arhgef10** | 0.02430 | 0.00775 | 0.16010 | 0.01760 | 0.00825 | 0.00885 | 0.00726 | 0.01253 | 0.01258 | 0.08618 | 0.01257 | 0.02627 | 0.01086 | 0.01205 | 0.00644 | 0.00733 |
| **Arhgef10l** | 0.01829 | 0.00771 | 0.00899 | 0.34052 | 0.00883 | 0.00677 | 0.00721 | 0.06251 | 0.46195 | 0.00022 | 0.19120 | 0.00585 | 0.00823 | 0.00946 | 0.02558 | 0.00721 |
| **Arhgef11** | 0.21255 | 0.00827 | 0.00531 | 0.58847 | 0.01698 | 0.00679 | 0.00707 | 0.00025 | 0.02657 | 0.20836 | 0.33556 | 0.00404 | 0.00886 | 0.00678 | 0.54918 | 0.00727 |
| **Arhgef12** | 5.11861 | 1.74631 | 1.10240 | 13.45385 | 6.92816 | 0.31098 | 3.66830 | 4.81465 | 13.58721 | 10.52763 | 2.54889 | 2.21138 | 1.72791 | 7.33333 | 12.97341 | 2.98383 |
| **Arhgef15** | 0.18485 | 0.00794 | 0.00690 | 0.00457 | 0.00746 | 0.00719 | 0.00743 | 0.01520 | 0.37812 | 0.05109 | 0.03819 | 0.00821 | 0.01911 | 0.01208 | 0.01018 | 0.00708 |
| **Arhgef16** | 0.02169 | 2.28532 | 0.74084 | 1.74848 | 0.03296 | 2.70959 | 1.28367 | 20.47257 | 1.56078 | 13.35839 | 5.76471 | 2.63102 | 3.67013 | 0.07100 | 0.00000 | 0.00969 |
| **Arhgef17** | 1.08164 | 0.00595 | 0.00288 | 0.00852 | 0.00587 | 0.00718 | 0.48821 | 0.00594 | 0.01215 | 0.45440 | 0.01189 | 0.05124 | 0.01104 | 0.01174 | 0.00703 | 0.00624 |
| **Arhgef18** | 4.13172 | 0.60133 | 2.25637 | 3.31961 | 0.00000 | 0.16662 | 0.00231 | 1.61957 | 1.29842 | 1.01117 | 0.95066 | 0.01883 | 2.42555 | 0.00436 | 1.98351 | 0.02920 |
| **Arhgef19** | 2.08686 | 0.00430 | 0.20522 | 0.96211 | 0.13878 | 0.00697 | 0.08462 | 4.65665 | 0.00000 | 2.39431 | 3.14276 | 0.05870 | 0.02021 | 0.00000 | 0.00000 | 0.00446 |
| **Arhgef2** | 0.25952 | 0.00676 | 2.54138 | 1.51109 | 0.00298 | 0.00714 | 1.81068 | 0.26084 | 1.88568 | 0.86699 | 0.00000 | 0.05332 | 0.07715 | 0.00812 | 0.00000 | 0.02007 |
| **Arhgef26** | 0.00439 | 0.00717 | 0.00000 | 3.24058 | 0.00910 | 5.54154 | 0.00567 | 1.76236 | 1.34788 | 1.36266 | 1.88952 | 0.46338 | 1.19518 | 0.00000 | 1.65801 | 0.00425 |
| **Arhgef28** | 0.01362 | 0.00826 | 0.01201 | 0.61518 | 0.00930 | 0.00656 | 0.00752 | 0.01391 | 0.82822 | 0.01104 | 0.00760 | 0.00705 | 0.01289 | 0.01154 | 0.00332 | 0.00727 |
| **Arhgef3** | 2.77289 | 2.58498 | 2.42371 | 2.32125 | 0.05251 | 0.41782 | 3.85980 | 6.45786 | 1.43753 | 14.92582 | 10.01905 | 17.00257 | 1.95291 | 12.43970 | 12.21543 | 0.01258 |
| **Arhgef39** | 2.20184 | 0.00000 | 4.87256 | 11.60709 | 3.29801 | 0.01886 | 2.12483 | 10.53037 | 8.17343 | 2.72031 | 4.79935 | 0.00000 | 5.15978 | 0.02475 | 0.45767 | 0.01150 |
| **Arhgef4** | 0.01459 | 0.00749 | 0.40586 | 0.67466 | 0.00826 | 0.00556 | 0.00720 | 0.81488 | 1.79979 | 0.67872 | 0.45826 | 0.00000 | 0.00479 | 0.00187 | 0.14192 | 0.02297 |
| **Arhgef40** | 0.01373 | 0.00802 | 0.01190 | 0.07675 | 0.00922 | 0.00659 | 0.00750 | 0.01264 | 0.27414 | 0.10034 | 0.01096 | 0.01726 | 0.01218 | 0.01281 | 0.01154 | 0.00730 |
| **Arhgef5** | 0.08590 | 0.00548 | 0.69328 | 0.00049 | 0.02478 | 1.21702 | 3.96744 | 0.00000 | 0.01097 | 0.00000 | 0.50686 | 0.02170 | 0.00000 | 0.01217 | 0.00000 | 3.09052 |
| **Arhgef7** | 0.48489 | 0.80181 | 1.77560 | 2.06929 | 0.14537 | 5.75537 | 0.01443 | 1.99713 | 4.76436 | 0.93475 | 1.03490 | 5.80446 | 1.70690 | 0.00000 | 1.66582 | 0.89214 |
| **Arid1a** | 12.15821 | 5.43943 | 13.00482 | 15.34420 | 5.79557 | 12.13387 | 7.12446 | 19.23040 | 12.95213 | 28.33534 | 15.89032 | 14.93082 | 7.84205 | 16.38919 | 19.83707 | 9.77308 |
| **Arid1b** | 1.00105 | 4.08139 | 1.19748 | 3.20738 | 2.35673 | 0.00589 | 0.01369 | 6.59464 | 2.94510 | 2.55606 | 2.80906 | 0.60141 | 0.99206 | 0.31807 | 1.79844 | 0.17325 |
| **Arid2** | 4.78211 | 0.42852 | 4.10121 | 11.00958 | 4.33302 | 0.31469 | 2.91362 | 5.20002 | 4.71157 | 6.19226 | 5.57230 | 1.21205 | 2.35369 | 2.59740 | 7.25711 | 5.34811 |
| **Arid3a** | 1.39662 | 0.00690 | 0.11003 | 1.01442 | 0.00664 | 0.00421 | 0.00674 | 0.00651 | 0.00000 | 0.00600 | 0.30862 | 0.00664 | 0.00761 | 0.00881 | 0.00000 | 0.23095 |
| **Arid3b** | 8.52441 | 3.01118 | 3.40502 | 1.04154 | 1.47086 | 0.83879 | 2.92998 | 3.16883 | 0.36689 | 5.08094 | 0.61933 | 0.61011 | 5.09676 | 0.00278 | 1.59775 | 0.01494 |
| **Arid3c** | 0.01097 | 0.00821 | 0.19777 | 1.97138 | 0.00886 | 0.00543 | 0.00785 | 0.07008 | 0.29407 | 0.43450 | 0.00807 | 0.17208 | 0.24043 | 0.00873 | 0.17906 | 0.00657 |
| **Arid4a** | 0.37762 | 0.00574 | 0.04922 | 1.90099 | 2.49146 | 0.02767 | 0.37599 | 0.00000 | 3.63727 | 2.11512 | 0.23657 | 0.13343 | 0.00645 | 0.00000 | 1.09491 | 1.32739 |
| **Arid4b** | 1.62919 | 2.33772 | 0.14577 | 2.59810 | 0.66015 | 0.18285 | 0.01473 | 1.80840 | 7.68757 | 0.39387 | 0.11075 | 0.27576 | 1.05304 | 6.01576 | 5.64114 | 0.66561 |
| **Arid5a** | 0.02479 | 0.00778 | 0.01743 | 6.31246 | 0.00915 | 0.00504 | 0.05877 | 1.34296 | 12.79023 | 0.00000 | 0.00000 | 0.02886 | 0.36408 | 0.05333 | 0.01153 | 0.00553 |
| **Arid5b** | 3.13770 | 5.70556 | 1.84754 | 0.86336 | 1.18465 | 0.00338 | 1.41085 | 0.51134 | 0.00000 | 0.05215 | 0.00000 | 0.02839 | 0.00000 | 1.29990 | 0.58736 | 0.00000 |
| **Arih1** | 5.86296 | 10.38118 | 6.03336 | 14.15967 | 5.98061 | 0.00000 | 4.86590 | 6.33563 | 20.37144 | 8.01515 | 4.76642 | 12.12944 | 2.99101 | 4.28500 | 14.62941 | 2.36466 |
| **Arih2** | 15.19605 | 11.16569 | 19.80546 | 8.48890 | 11.72425 | 10.50467 | 5.19117 | 34.37384 | 13.98759 | 39.25248 | 26.47051 | 20.62329 | 18.72700 | 9.03588 | 19.80880 | 6.46623 |
| **Arl1** | 10.20531 | 2.71427 | 4.73906 | 2.43935 | 6.69607 | 6.90859 | 1.07832 | 7.48796 | 1.58383 | 4.44462 | 1.57973 | 0.96493 | 9.34296 | 3.88710 | 0.00000 | 4.75439 |
| **Arl10** | 0.02580 | 0.00786 | 0.01070 | 0.03304 | 0.00846 | 0.00702 | 0.00756 | 0.02318 | 0.00997 | 0.18615 | 0.07002 | 0.00805 | 0.01259 | 0.00828 | 0.03602 | 0.11645 |
| **Arl13b** | 0.53238 | 0.00654 | 1.04098 | 0.28038 | 0.01570 | 2.27356 | 0.00621 | 0.51697 | 0.59649 | 0.17367 | 0.17501 | 2.08199 | 0.91846 | 0.00457 | 0.39204 | 0.01251 |
| **Arl14ep** | 1.17870 | 3.01897 | 3.91461 | 10.37579 | 0.01588 | 0.00203 | 0.01246 | 2.36818 | 15.65335 | 4.53271 | 5.04659 | 0.49997 | 0.60455 | 0.14949 | 3.82923 | 0.11333 |
| **Arl14epl** | 6.72387 | 4.80485 | 5.22929 | 0.57878 | 0.20650 | 0.46341 | 3.27087 | 0.00000 | 0.03785 | 3.87433 | 4.48939 | 0.09769 | 4.01859 | 0.79979 | 0.85619 | 0.01569 |
| **Arl15** | 0.25051 | 0.02821 | 1.06057 | 3.71562 | 0.03208 | 0.15056 | 0.00575 | 2.02311 | 6.85465 | 0.00000 | 2.31722 | 0.07268 | 0.08358 | 0.15094 | 1.29437 | 0.26879 |
| **Arl16** | 0.01536 | 0.36857 | 0.84776 | 1.72887 | 2.43894 | 0.00554 | 0.15027 | 1.94485 | 3.49535 | 0.04063 | 0.40867 | 0.00604 | 0.06618 | 1.90992 | 0.09252 | 0.00609 |
| **Arl2** | 10.60564 | 3.77287 | 5.80376 | 0.55403 | 4.83402 | 3.77809 | 2.62946 | 2.94225 | 0.02391 | 1.13995 | 0.15484 | 0.00000 | 3.21100 | 0.00108 | 0.00000 | 2.89900 |
| **Arl2bp** | 0.04014 | 0.00729 | 0.05671 | 7.45423 | 0.00813 | 0.01624 | 0.01637 | 0.00000 | 4.58512 | 0.51833 | 0.79942 | 0.00000 | 0.00580 | 0.11882 | 0.65295 | 0.01344 |
| **Arl3** | 1.90001 | 0.01655 | 2.93628 | 8.97727 | 0.01199 | 1.09213 | 0.00467 | 1.14165 | 2.80733 | 1.91516 | 1.25415 | 0.09738 | 0.18578 | 0.00000 | 0.85123 | 0.06689 |
| **Arl4a** | 5.95336 | 0.00000 | 1.01959 | 12.83035 | 1.45499 | 2.06730 | 0.00000 | 0.91206 | 6.35491 | 0.65697 | 3.05869 | 0.12625 | 0.41306 | 1.22503 | 8.32641 | 0.01827 |
| **Arl4c** | 0.03628 | 0.00844 | 0.01070 | 0.17268 | 0.01291 | 0.00666 | 0.00735 | 0.01086 | 0.00863 | 1.13055 | 0.01209 | 0.00521 | 0.01100 | 0.01096 | 0.00580 | 0.00750 |
| **Arl4d** | 2.66645 | 0.03711 | 0.10327 | 0.16422 | 0.02865 | 0.00291 | 0.72366 | 0.00585 | 0.19860 | 1.90552 | 0.00971 | 0.00000 | 0.01713 | 0.00549 | 0.20730 | 0.00469 |
| **Arl5a** | 8.90701 | 3.86868 | 2.41799 | 6.80293 | 7.95634 | 1.83851 | 0.63967 | 2.53063 | 3.55879 | 4.59817 | 1.62000 | 7.10761 | 4.75716 | 0.62605 | 1.26440 | 0.00532 |
| **Arl5b** | 0.89629 | 0.00729 | 0.10174 | 1.54098 | 0.00706 | 0.00534 | 0.01855 | 0.35280 | 0.56587 | 0.11836 | 0.00697 | 0.03504 | 0.13664 | 0.00243 | 0.44943 | 0.01406 |
| **Arl5c** | 0.01322 | 0.00800 | 0.01267 | 0.11115 | 0.00932 | 0.00669 | 0.00749 | 0.01090 | 0.06354 | 0.82733 | 0.02898 | 0.00553 | 0.01679 | 0.03110 | 0.00728 | 0.00725 |
| **Arl6** | 4.12330 | 0.00286 | 0.00000 | 1.63425 | 1.81047 | 0.00921 | 0.00342 | 0.23116 | 0.33964 | 0.81330 | 0.00756 | 0.09486 | 0.83865 | 1.39137 | 0.78012 | 0.00922 |
| **Arl6ip1** | 19.20366 | 8.66462 | 19.93326 | 5.77703 | 10.23441 | 4.18262 | 11.90050 | 20.84228 | 2.20116 | 15.64605 | 14.99114 | 6.99004 | 12.43591 | 18.49188 | 19.58523 | 7.88827 |
| **Arl6ip4** | 1.79196 | 3.31794 | 2.76318 | 2.56269 | 15.08971 | 1.08818 | 0.93515 | 6.06349 | 5.11903 | 10.77338 | 5.00570 | 0.56592 | 9.03962 | 0.00588 | 1.46672 | 4.60930 |
| **Arl6ip5** | 0.00000 | 1.51731 | 0.01876 | 0.00153 | 0.00756 | 0.45384 | 0.00668 | 0.00914 | 0.13193 | 0.00763 | 0.01095 | 0.02627 | 0.00659 | 0.01153 | 0.00000 | 1.09602 |
| **Arl6ip6** | 1.73820 | 0.25936 | 0.39422 | 3.48140 | 0.04487 | 0.70951 | 0.00420 | 1.47920 | 0.00000 | 2.19713 | 0.66764 | 6.52968 | 0.86586 | 1.07924 | 4.75270 | 1.24680 |

| | | | | | | | | | | | | | | | | |
|---|---|---|---|---|---|---|---|---|---|---|---|---|---|---|---|---|
| **Arl8a** | 0.97580 | 3.32496 | 0.03286 | 1.41676 | 2.06623 | 0.43589 | 1.66220 | 0.00496 | 0.73908 | 0.36155 | 0.00548 | 0.03560 | 0.93675 | 0.00147 | 0.78355 | 1.57649 |
| **Arl8b** | 9.15318 | 11.42095 | 12.93118 | 8.81792 | 10.94013 | 7.26485 | 9.21930 | 14.39923 | 9.68201 | 18.76874 | 15.93012 | 18.93512 | 10.67080 | 20.35150 | 15.10454 | 5.08910 |
| **Armc1** | 7.88952 | 3.74505 | 7.94392 | 7.78782 | 7.31675 | 2.83483 | 1.39380 | 6.22440 | 5.17887 | 8.63539 | 3.04338 | 8.71412 | 3.49578 | 2.93048 | 5.92733 | 1.77133 |
| **Armc10** | 4.43351 | 0.09578 | 1.15157 | 2.06220 | 0.10953 | 2.53995 | 0.15444 | 4.43321 | 1.02682 | 2.53931 | 2.65583 | 0.12056 | 3.23226 | 0.05358 | 1.35215 | 2.61920 |
| **Armc12** | 0.01379 | 0.00802 | 0.00865 | 0.89380 | 0.00907 | 0.00671 | 0.00750 | 0.00461 | 0.59550 | 0.00000 | 0.48666 | 0.00455 | 0.00756 | 0.00402 | 0.10744 | 0.00696 |
| **Armc2** | 0.01213 | 0.02946 | 0.01044 | 0.98288 | 0.00917 | 0.00643 | 0.14069 | 0.01341 | 1.69770 | 0.66122 | 0.00000 | 0.00237 | 0.01304 | 0.01525 | 0.19198 | 0.00733 |
| **Armc5** | 1.97954 | 0.00505 | 0.00000 | 0.55472 | 0.00417 | 0.83972 | 0.01565 | 3.49448 | 1.42094 | 1.91548 | 0.77924 | 0.05397 | 1.56585 | 0.02153 | 0.00000 | 0.00418 |
| **Armc6** | 2.56903 | 0.39893 | 3.13166 | 2.22548 | 0.84939 | 3.45654 | 2.16259 | 13.33920 | 0.22589 | 6.24896 | 1.57272 | 0.25592 | 3.51191 | 0.01613 | 0.11290 | 0.02190 |
| **Armc7** | 0.00000 | 0.00660 | 0.21215 | 6.01356 | 0.02464 | 0.00617 | 0.91364 | 1.60086 | 13.34659 | 5.05305 | 1.88576 | 0.66125 | 1.48298 | 2.65345 | 3.64745 | 1.16059 |
| **Armc8** | 8.34482 | 3.92273 | 2.07875 | 0.95900 | 6.71732 | 6.43574 | 2.87398 | 4.00934 | 1.00254 | 2.39917 | 3.73670 | 4.09928 | 2.52102 | 2.41293 | 6.03809 | 0.03605 |
| **Armc9** | 1.88814 | 0.23684 | 0.97819 | 0.38587 | 0.00758 | 0.00788 | 0.04676 | 0.32299 | 0.95844 | 0.33839 | 0.00000 | 0.03034 | 0.63096 | 0.00422 | 1.01659 | 0.00540 |
| **Armcx4** | 0.15858 | 0.08198 | 0.22025 | 0.48134 | 0.05688 | 0.07231 | 0.06786 | 0.08661 | 0.48576 | 0.14583 | 1.51539 | 0.01478 | 0.15688 | 0.71273 | 0.67537 | 0.07092 |
| **Armcx5** | 0.00000 | 3.14586 | 0.00000 | 12.19018 | 3.49759 | 0.76410 | 0.00486 | 0.91151 | 6.42683 | 1.32564 | 0.90337 | 0.42331 | 0.56164 | 4.73444 | 7.43616 | 0.44439 |
| **Armt1** | 0.81365 | 0.00209 | 6.32642 | 0.52455 | 5.47262 | 0.01335 | 2.53917 | 0.00000 | 0.86056 | 0.19077 | 0.22323 | 0.06436 | 2.75324 | 0.00432 | 0.65044 | 0.02556 |
| **Arnt** | 0.45799 | 0.12183 | 0.72009 | 3.62873 | 0.57007 | 0.06419 | 0.42936 | 1.64620 | 2.42745 | 6.92005 | 3.53439 | 6.99370 | 2.58665 | 6.28991 | 1.80142 | 2.43122 |
| **Arntl** | 4.18778 | 0.00000 | 0.17622 | 8.09259 | 7.27630 | 0.00000 | 0.00327 | 1.07399 | 6.18125 | 0.79414 | 0.82261 | 0.89080 | 0.06724 | 0.47914 | 7.00002 | 5.12378 |
| **Arntl2** | 0.04378 | 0.00766 | 0.05725 | 0.73755 | 0.00829 | 0.00581 | 0.00728 | 0.01901 | 0.00000 | 0.01354 | 0.01328 | 0.01722 | 0.01147 | 0.00933 | 0.02957 | 0.00714 |
| **Arpc1a** | 16.07424 | 6.04123 | 8.84228 | 11.44561 | 10.20199 | 9.03682 | 8.95978 | 13.02334 | 11.65156 | 17.11764 | 8.52995 | 2.40883 | 11.36490 | 1.23489 | 3.42918 | 12.29208 |
| **Arpc1b** | 21.71109 | 7.96067 | 9.36658 | 4.47398 | 16.47361 | 10.88274 | 15.88181 | 14.06904 | 2.37242 | 17.79881 | 12.81087 | 0.89477 | 18.27930 | 0.13696 | 0.00000 | 17.96748 |
| **Arpc2** | 18.70668 | 11.15630 | 22.46402 | 7.29124 | 16.46279 | 12.85269 | 7.59028 | 17.14630 | 12.18501 | 16.59533 | 7.92638 | 9.87234 | 19.27531 | 9.73729 | 7.99760 | 9.46104 |
| **Arpc3** | 2.93997 | 12.99784 | 3.01691 | 1.68703 | 5.27021 | 3.08810 | 2.76562 | 1.84564 | 2.31605 | 3.43102 | 6.53443 | 0.18638 | 3.31012 | 4.14913 | 4.93314 | 1.08363 |
| **Arpc4** | 17.87280 | 3.00647 | 5.90193 | 0.00000 | 4.86391 | 9.90798 | 3.26602 | 20.51409 | 0.12963 | 13.43186 | 1.31101 | 0.51330 | 19.02602 | 0.16895 | 0.01442 | 8.73388 |
| **Arpc5** | 14.16004 | 13.59276 | 12.72254 | 3.98406 | 15.46520 | 18.35739 | 13.09207 | 14.16985 | 6.50360 | 11.12484 | 8.78646 | 11.78859 | 19.03328 | 5.66298 | 0.75274 | 12.51391 |
| **Arpc5l** | 6.95554 | 0.31125 | 5.11738 | 13.21188 | 7.62495 | 7.87581 | 1.44114 | 4.51440 | 13.94243 | 15.42277 | 5.53519 | 3.41264 | 8.96678 | 1.12240 | 2.53719 | 6.72565 |
| **Arpin** | 1.75161 | 0.00564 | 0.00307 | 0.52544 | 0.00463 | 0.00636 | 0.04525 | 0.00000 | 1.43521 | 0.59439 | 0.49563 | 0.00896 | 0.00782 | 0.00584 | 0.10429 | 0.01246 |
| **Arpp19** | 9.45805 | 1.53474 | 6.60412 | 4.21707 | 5.56489 | 2.13904 | 2.41858 | 0.20746 | 5.97866 | 4.82784 | 11.50958 | 2.90089 | 1.64631 | 7.26160 | 1.93729 | 1.02306 |
| **Arrb1** | 1.18680 | 0.00408 | 4.92297 | 0.36627 | 5.00628 | 3.92828 | 0.00471 | 0.44260 | 0.23242 | 1.29242 | 1.61161 | 0.01836 | 1.42754 | 0.33555 | 0.36541 | 0.38291 |
| **Arrb2** | 0.58349 | 0.02700 | 0.62322 | 5.82000 | 0.00709 | 0.00431 | 0.27652 | 3.23865 | 2.93006 | 3.13492 | 2.13412 | 0.05537 | 0.09590 | 0.00000 | 0.14260 | 0.28348 |
| **Arrdc1** | 0.97435 | 0.22511 | 0.00000 | 8.32008 | 0.00546 | 0.00963 | 2.82760 | 6.11484 | 7.69032 | 12.87468 | 3.60410 | 0.47064 | 1.72842 | 0.03880 | 4.18770 | 0.01002 |
| **Arrdc2** | 0.40428 | 0.00597 | 0.45950 | 8.13305 | 0.00613 | 0.00671 | 0.00499 | 1.57333 | 11.03674 | 2.88142 | 3.13782 | 2.16031 | 0.00000 | 0.00000 | 1.00010 | 0.12224 |
| **Arrdc3** | 0.06608 | 0.00791 | 0.00840 | 0.66792 | 0.00897 | 0.00663 | 0.00751 | 0.01430 | 0.12931 | 0.00362 | 0.00889 | 0.00334 | 0.03927 | 0.00655 | 2.14968 | 0.02209 |
| **Arrdc4** | 0.01409 | 0.00797 | 0.09882 | 0.27385 | 0.00898 | 0.00567 | 0.00741 | 0.01156 | 0.54999 | 0.20046 | 0.01062 | 0.00691 | 0.01062 | 0.01631 | 0.00341 | 0.02133 |
| **Arrdc5** | 0.29934 | 0.00783 | 0.00507 | 0.79917 | 0.00719 | 0.00722 | 0.00716 | 0.02718 | 0.65550 | 0.28723 | 0.36373 | 0.00415 | 0.00832 | 0.00551 | 0.03431 | 0.00712 |
| **Arsa** | 2.50647 | 0.00562 | 0.00000 | 0.21678 | 0.00318 | 2.16977 | 0.00607 | 0.45582 | 0.00840 | 0.44520 | 0.11446 | 0.01601 | 0.00000 | 0.01072 | 0.00000 | 0.00361 |
| **Arsb** | 0.00472 | 0.02758 | 0.04921 | 1.14101 | 0.01944 | 0.00767 | 0.00702 | 0.37340 | 0.37092 | 0.00770 | 0.01415 | 0.46954 | 0.01596 | 1.70888 | 0.00000 | 0.00712 |
| **Arsg** | 0.01302 | 0.00796 | 0.01213 | 1.01393 | 0.00885 | 0.00647 | 0.00726 | 0.01027 | 0.04971 | 1.26485 | 0.01315 | 0.00571 | 0.01068 | 0.01374 | 0.00258 | 0.01409 |
| **Arsk** | 0.02439 | 0.00821 | 1.07658 | 8.59469 | 0.00759 | 0.00447 | 0.00716 | 0.00266 | 5.48326 | 0.16551 | 0.21853 | 0.00787 | 0.00730 | 0.00000 | 0.84681 | 0.00513 |
| **Art2a.ps** | 0.01371 | 0.00786 | 0.01183 | 0.33179 | 0.00927 | 0.00675 | 0.00755 | 0.01331 | 0.00625 | 0.04906 | 0.01165 | 0.00796 | 0.01324 | 0.00745 | 0.30777 | 0.00736 |
| **Art3** | 0.00000 | 0.00759 | 0.29338 | 0.31800 | 0.00861 | 0.00571 | 0.58503 | 0.00000 | 0.00506 | 0.91769 | 0.46958 | 0.07689 | 2.71365 | 2.66905 | 1.23754 | 0.01289 |
| **Art5** | 0.01284 | 0.00787 | 0.01348 | 0.79438 | 0.00994 | 0.00694 | 0.00752 | 0.01152 | 0.00000 | 0.49630 | 0.05428 | 0.00471 | 0.09778 | 0.00506 | 0.47187 | 0.00785 |
| **Artn** | 0.01383 | 0.00825 | 0.01085 | 0.01679 | 0.00920 | 0.00666 | 0.00749 | 0.00969 | 0.01232 | 0.00467 | 0.16523 | 0.00774 | 0.01027 | 0.00767 | 0.08781 | 0.00730 |
| **Arv1** | 0.04743 | 0.91795 | 0.95820 | 1.87343 | 0.00608 | 0.00544 | 0.68101 | 2.94876 | 2.49369 | 0.47614 | 2.69321 | 0.35799 | 2.35867 | 0.41965 | 0.24192 | 0.03327 |
| **Arvcf** | 5.49610 | 0.67062 | 0.54695 | 0.82509 | 5.05243 | 6.58524 | 3.03306 | 14.63341 | 1.55003 | 4.01447 | 5.14820 | 0.14000 | 8.51004 | 0.29544 | 0.23724 | 4.02633 |
| **Arx** | 0.01346 | 0.00786 | 0.01141 | 0.42898 | 0.00918 | 0.00645 | 0.00738 | 0.01100 | 1.94049 | 0.12670 | 0.01731 | 0.00287 | 0.01292 | 0.00921 | 0.17269 | 0.00759 |
| **Arxes1** | 0.01214 | 0.00810 | 0.02576 | 0.32464 | 0.01029 | 0.00662 | 0.00767 | 0.02050 | 0.00545 | 0.01504 | 0.03396 | 0.00697 | 0.16887 | 0.00550 | 0.30748 | 0.00766 |
| **Arxes2** | 0.01089 | 0.00791 | 0.02458 | 0.09047 | 0.02068 | 0.00650 | 0.00758 | 0.02449 | 0.00778 | 0.30576 | 0.03146 | 0.00279 | 0.10339 | 0.00511 | 1.77774 | 0.00776 |
| **As3mt** | 12.54003 | 4.14837 | 9.03842 | 0.10692 | 7.29485 | 3.87311 | 3.17076 | 7.05318 | 0.67911 | 0.25458 | 3.75118 | 0.03683 | 7.04040 | 0.00000 | 0.15448 | 1.35514 |
| **Asah1** | 6.89913 | 2.56157 | 6.83167 | 1.00109 | 3.98995 | 11.89921 | 8.07365 | 7.80003 | 0.89679 | 4.68139 | 11.52944 | 4.03600 | 6.39577 | 16.59432 | 8.70753 | 5.03144 |
| **Asah2** | 0.01370 | 0.00783 | 0.01117 | 0.06384 | 0.00931 | 0.00679 | 0.00754 | 0.01265 | 0.03491 | 0.05936 | 0.00930 | 0.01881 | 0.01340 | 0.01036 | 0.53566 | 0.00741 |
| **Asap1** | 2.32929 | 0.00284 | 0.14793 | 7.75750 | 3.07935 | 0.00215 | 0.28346 | 0.22456 | 2.22063 | 1.97555 | 0.28517 | 0.67022 | 0.22568 | 1.57873 | 1.27014 | 3.44008 |
| **Asap2** | 1.08263 | 0.03557 | 0.75503 | 0.00000 | 2.10420 | 0.23350 | 0.00509 | 2.27755 | 0.00584 | 1.69114 | 2.16898 | 2.64168 | 0.26812 | 0.05504 | 0.21436 | 0.04797 |
| **Asap3** | 0.03316 | 0.01753 | 0.83583 | 0.00664 | 0.02939 | 0.00869 | 0.05687 | 3.58715 | 0.10272 | 0.00000 | 1.71172 | 0.02863 | 2.31118 | 0.34649 | 0.00000 | 0.02023 |
| **Asb1** | 1.82831 | 0.02621 | 0.00000 | 0.16245 | 0.05316 | 3.31369 | 0.00548 | 0.00806 | 0.04023 | 0.00000 | 0.00946 | 0.02748 | 0.00016 | 0.01266 | 0.07805 | 2.53904 |
| **Asb10** | 0.01373 | 0.00820 | 0.01154 | 0.42464 | 0.00924 | 0.00655 | 0.00748 | 0.01205 | 0.42288 | 0.00619 | 0.06371 | 0.00755 | 0.01165 | 0.00748 | 0.07992 | 0.00727 |
| **Asb13** | 4.77513 | 0.36663 | 0.78856 | 6.40202 | 3.95875 | 8.38817 | 0.03296 | 3.16230 | 7.15864 | 3.16737 | 2.30305 | 0.40919 | 4.22091 | 1.68261 | 4.76754 | 1.67916 |
| **Asb14** | 0.01783 | 0.00786 | 0.01195 | 0.01042 | 0.00900 | 0.00664 | 0.00751 | 0.01389 | 0.01230 | 0.01260 | 0.01281 | 0.00772 | 0.01268 | 0.00999 | 0.06580 | 0.00737 |
| **Asb16** | 0.00689 | 0.00778 | 0.00897 | 0.01053 | 0.01022 | 0.00691 | 0.61543 | 0.18017 | 0.01212 | 0.00428 | 0.01262 | 0.00734 | 0.36237 | 0.01451 | 0.00706 | 0.00722 |
| **Asb17** | 0.01056 | 0.00763 | 0.38421 | 0.01030 | 0.00917 | 0.00933 | 0.27757 | 0.03563 | 0.01245 | 0.01188 | 0.01319 | 0.00690 | 0.01721 | 0.00934 | 0.02942 | 0.00742 |
| **Asb17os** | 0.05949 | 0.00735 | 0.33140 | 0.01007 | 0.00805 | 0.00835 | 0.43675 | 0.21957 | 0.01221 | 0.02649 | 0.01317 | 0.00601 | 0.08020 | 0.00811 | 0.05444 | 0.01510 |
| **Asb3** | 1.73611 | 0.00471 | 0.19861 | 0.00000 | 3.60538 | 0.00449 | 0.01856 | 0.17988 | 0.32270 | 0.27692 | 0.00637 | 0.04139 | 0.06592 | 0.00620 | 0.66374 | 0.02052 |
| **Asb4** | 2.05060 | 0.00555 | 0.25275 | 0.52871 | 0.00512 | 0.00344 | 0.00537 | 0.00000 | 0.00000 | 0.50369 | 0.01251 | 0.03078 | 0.01030 | 3.07714 | 0.82129 | 0.00506 |
| **Asb6** | 0.97844 | 2.18459 | 0.34293 | 1.18766 | 1.65616 | 0.01077 | 0.53733 | 6.95963 | 3.88735 | 0.96558 | 0.90971 | 0.23843 | 0.01834 | 0.01537 | 0.02690 | 0.24607 |

| | | | | | | | | | | | | | | | | |
|---|---|---|---|---|---|---|---|---|---|---|---|---|---|---|---|---|
| **Asb7** | 0.69149 | 0.00582 | 0.22890 | 1.27551 | 1.89442 | 0.00481 | 0.00638 | 0.32843 | 0.11030 | 0.03364 | 0.74754 | 0.04242 | 0.46197 | 0.01078 | 0.03368 | 0.52891 |
| **Asb8** | 5.65029 | 0.00000 | 1.30188 | 4.16131 | 1.35903 | 0.02131 | 1.54033 | 2.74421 | 1.00510 | 1.97561 | 2.01630 | 0.10736 | 3.09527 | 0.54295 | 3.95632 | 1.26905 |
| **Ascc1** | 4.43470 | 0.01185 | 1.72728 | 1.07554 | 5.31413 | 4.54963 | 1.14264 | 0.38811 | 0.92818 | 2.86843 | 1.00517 | 0.22185 | 4.19810 | 0.47657 | 1.42967 | 1.17498 |
| **Ascc2** | 3.10219 | 4.39336 | 4.24978 | 5.55265 | 2.42888 | 4.40343 | 4.18706 | 8.67068 | 8.27415 | 7.91368 | 7.17639 | 1.51515 | 6.19954 | 0.04519 | 3.02691 | 1.42409 |
| **Ascc3** | 2.88590 | 8.04582 | 0.41518 | 6.34432 | 0.71228 | 0.00169 | 0.00000 | 2.64542 | 4.23358 | 3.49186 | 1.52353 | 2.54333 | 1.70276 | 9.27434 | 9.33780 | 4.39485 |
| **Asf1a** | 4.87587 | 2.48038 | 3.10448 | 4.10690 | 6.68813 | 0.02339 | 1.33349 | 0.10781 | 4.59020 | 3.08435 | 5.11937 | 6.69153 | 0.05009 | 1.73142 | 0.27862 | 0.01269 |
| **Asf1b** | 7.70696 | 8.30221 | 14.23428 | 10.77459 | 5.01753 | 5.20342 | 8.09421 | 8.28670 | 10.27217 | 7.50688 | 1.63598 | 1.37822 | 4.09973 | 3.14323 | 7.30140 | 3.59558 |
| **Asgr2** | 0.01327 | 0.00780 | 0.01010 | 0.09133 | 0.00914 | 0.00671 | 0.10792 | 0.00824 | 0.58523 | 0.41481 | 0.08599 | 0.00532 | 0.01120 | 0.04864 | 0.03643 | 0.00745 |
| **Ash1l** | 0.08486 | 0.00580 | 0.37674 | 2.54479 | 0.00725 | 0.00558 | 0.01531 | 0.27936 | 5.08943 | 0.74923 | 2.18952 | 4.91420 | 0.00000 | 0.62983 | 3.94023 | 0.02005 |
| **Ash2l** | 20.61074 | 15.08734 | 16.02320 | 10.54862 | 10.76727 | 15.79701 | 15.32180 | 21.70393 | 11.22056 | 11.78504 | 15.49039 | 1.41683 | 19.14505 | 12.11920 | 9.27580 | 8.56389 |
| **Asic1** | 0.14860 | 0.00779 | 0.00282 | 0.13382 | 0.01603 | 0.01505 | 0.00716 | 0.00000 | 0.04462 | 0.25260 | 0.47920 | 0.08550 | 0.01173 | 0.08500 | 0.08599 | 0.00700 |
| **Asic3** | 0.00241 | 0.56208 | 2.13753 | 0.00820 | 0.00493 | 5.97436 | 0.00662 | 0.53851 | 0.01082 | 0.00000 | 0.18493 | 0.00397 | 0.00000 | 0.01088 | 0.00000 | 0.00436 |
| **Asl** | 1.37731 | 3.67038 | 1.34915 | 2.40947 | 2.65830 | 1.56918 | 0.00352 | 0.00000 | 1.56710 | 1.93289 | 2.95713 | 0.09905 | 0.00000 | 0.01769 | 1.08537 | 0.12769 |
| **Asna1** | 11.15425 | 10.07769 | 11.33122 | 3.18054 | 11.38777 | 8.58934 | 11.05529 | 12.72597 | 2.27417 | 15.25418 | 8.00679 | 1.88130 | 13.03687 | 0.38084 | 4.42136 | 9.92257 |
| **Asns** | 21.86554 | 12.22561 | 0.00000 | 0.57938 | 26.22617 | 0.02046 | 9.28253 | 9.03296 | 0.46945 | 0.15097 | 0.51245 | 0.94590 | 0.51863 | 0.00828 | 1.68020 | 13.03135 |
| **Asnsd1** | 8.76693 | 8.43110 | 7.66362 | 3.87119 | 13.75853 | 5.31363 | 4.09756 | 7.60010 | 3.19907 | 7.71709 | 7.08241 | 6.26621 | 5.12967 | 5.24304 | 7.63059 | 5.20350 |
| **Aspa** | 0.03875 | 0.00387 | 1.10311 | 0.00817 | 7.17939 | 0.00543 | 0.00724 | 0.01123 | 0.00951 | 0.01133 | 0.01080 | 0.00035 | 0.00695 | 0.01612 | 0.00000 | 0.00681 |
| **Aspg** | 0.00384 | 0.00475 | 0.00000 | 7.62941 | 0.02453 | 0.01195 | 0.00388 | 4.60827 | 5.15976 | 7.75326 | 7.32196 | 5.04196 | 0.00000 | 0.24684 | 2.26960 | 0.00411 |
| **Asph** | 0.00000 | 0.04490 | 1.73946 | 0.56067 | 1.26514 | 0.00503 | 3.37611 | 1.16184 | 3.12405 | 0.16552 | 0.73315 | 3.18221 | 0.35289 | 0.72943 | 3.41754 | 0.32354 |
| **Asphd1** | 0.75014 | 0.15905 | 0.00352 | 0.00555 | 0.01740 | 0.00740 | 0.00677 | 0.00172 | 0.01154 | 0.00541 | 0.30715 | 0.00619 | 0.00975 | 0.01008 | 0.00520 | 0.00667 |
| **Asphd2** | 0.01373 | 0.00812 | 0.01104 | 0.09383 | 0.00925 | 0.00665 | 0.00751 | 0.01095 | 0.41449 | 0.01543 | 0.12724 | 0.00687 | 0.01084 | 0.01142 | 0.00252 | 0.00734 |
| **Aspm** | 1.36396 | 0.02600 | 2.69620 | 6.21808 | 0.04251 | 0.97740 | 0.20950 | 0.45141 | 10.79519 | 0.82675 | 0.38004 | 2.54016 | 0.50198 | 5.33971 | 8.87169 | 0.00272 |
| **Aspn** | 0.01371 | 0.00791 | 0.00564 | 0.00617 | 0.00930 | 0.11065 | 0.00736 | 0.02247 | 0.02213 | 0.00000 | 0.16819 | 0.00699 | 0.00860 | 0.00789 | 0.15289 | 0.00726 |
| **Aspscr1** | 5.29595 | 3.62521 | 3.46424 | 2.36875 | 8.34507 | 4.86697 | 0.70532 | 9.19235 | 3.05463 | 7.11754 | 2.74564 | 0.36536 | 5.98751 | 0.00000 | 0.00000 | 4.49842 |
| **Asrgl1** | 0.73333 | 0.00796 | 0.85431 | 0.38800 | 0.02333 | 0.00477 | 0.00623 | 0.00620 | 0.00594 | 0.81755 | 0.00991 | 0.07163 | 1.07621 | 0.00886 | 0.02209 | 0.00562 |
| **Ass1** | 0.01268 | 0.00834 | 0.01175 | 0.00781 | 0.00967 | 0.00717 | 0.04068 | 0.03002 | 0.02613 | 0.04504 | 0.06347 | 0.00866 | 0.04961 | 0.01181 | 0.01031 | 0.00755 |
| **Aste1** | 1.41840 | 0.00480 | 0.68022 | 5.46056 | 0.00404 | 0.29342 | 1.04310 | 3.81354 | 6.45023 | 5.63833 | 3.20124 | 0.07136 | 0.98683 | 7.72691 | 3.00261 | 0.02440 |
| **Astl** | 0.00012 | 0.00806 | 0.75045 | 3.08051 | 0.00855 | 0.00502 | 0.00721 | 0.93480 | 3.11953 | 2.10135 | 0.00000 | 0.08518 | 0.00301 | 1.92274 | 0.00000 | 0.00626 |
| **Asun** | 7.54930 | 1.64090 | 4.80693 | 9.69045 | 6.62912 | 0.91177 | 2.30558 | 6.65023 | 3.31542 | 10.09165 | 2.74978 | 9.23828 | 2.64384 | 10.27379 | 14.52025 | 0.02091 |
| **Asxl1** | 3.64637 | 2.90257 | 1.88169 | 11.79410 | 2.77614 | 0.00000 | 0.00000 | 4.07586 | 13.82301 | 6.53792 | 3.79647 | 3.22685 | 1.71901 | 2.65438 | 7.70557 | 4.29632 |
| **Asxl2** | 0.86399 | 0.06873 | 0.50961 | 6.29723 | 0.45306 | 0.66958 | 0.41268 | 3.85275 | 11.30089 | 3.69635 | 1.22288 | 4.42374 | 1.09933 | 6.48103 | 6.08944 | 0.07569 |
| **Asz1** | 1.75987 | 0.00411 | 4.96642 | 1.74473 | 0.90364 | 0.00328 | 1.77188 | 0.48450 | 0.62464 | 1.04672 | 1.47091 | 1.10502 | 3.19573 | 4.77441 | 2.79577 | 0.00148 |
| **Atad1** | 1.57283 | 3.95735 | 5.40633 | 9.50884 | 0.00000 | 0.00799 | 5.51104 | 4.51371 | 7.67421 | 2.69944 | 2.58049 | 6.17803 | 0.15785 | 11.00613 | 9.73989 | 0.97386 |
| **Atad2** | 4.95484 | 1.91866 | 4.00007 | 11.35930 | 5.35542 | 0.09678 | 1.59376 | 2.24612 | 15.52806 | 5.50521 | 6.53080 | 2.84562 | 3.04555 | 12.25544 | 9.10813 | 1.68434 |
| **Atad2b** | 3.71134 | 2.56003 | 2.66045 | 8.32947 | 2.55820 | 4.17769 | 7.29540 | 7.35304 | 8.21757 | 10.67440 | 4.39114 | 10.14360 | 2.99844 | 9.87767 | 14.80110 | 0.83809 |
| **Atad3a** | 19.70720 | 8.40862 | 12.97615 | 6.63837 | 9.29042 | 6.92311 | 11.06517 | 15.11971 | 3.25156 | 10.33939 | 3.65222 | 0.85640 | 11.52607 | 2.17633 | 4.29194 | 4.37850 |
| **Atad5** | 2.45843 | 0.00000 | 0.00000 | 4.03703 | 0.00329 | 0.01124 | 2.01901 | 1.74586 | 4.20065 | 2.13514 | 0.36767 | 7.74379 | 0.13808 | 7.59861 | 5.07686 | 0.19682 |
| **Atat1** | 2.75097 | 0.00434 | 0.00000 | 4.69652 | 0.00208 | 0.00562 | 0.00383 | 0.53517 | 3.27856 | 1.27440 | 1.08903 | 0.01970 | 1.11301 | 0.18249 | 0.00000 | 2.08448 |
| **Atcay** | 0.01376 | 0.00765 | 0.00877 | 0.76227 | 0.00924 | 0.00671 | 0.00722 | 0.01065 | 0.87361 | 0.00348 | 0.00149 | 0.37355 | 0.01353 | 0.00985 | 0.00000 | 0.01478 |
| **Ate1** | 1.49348 | 1.76235 | 5.16756 | 2.35301 | 1.38019 | 0.00400 | 3.44097 | 2.29912 | 3.70344 | 2.13902 | 1.72967 | 1.64963 | 2.86247 | 0.70906 | 0.71446 | 1.47590 |
| **Atf1** | 6.59834 | 3.91193 | 4.71564 | 16.93756 | 2.50723 | 6.87626 | 4.52319 | 11.09608 | 18.44383 | 14.68202 | 8.40645 | 13.42093 | 7.70457 | 22.66789 | 20.18505 | 1.79106 |
| **Atf2** | 1.88224 | 1.35949 | 1.03419 | 7.11732 | 5.14300 | 0.31388 | 4.99168 | 1.26693 | 10.40928 | 6.94471 | 6.30418 | 1.68025 | 0.53860 | 15.51695 | 13.67719 | 0.04362 |
| **Atf4** | 19.93654 | 10.75299 | 18.41742 | 19.74084 | 13.96331 | 12.00241 | 10.38008 | 26.68518 | 25.82456 | 30.05025 | 16.31268 | 7.35041 | 17.25655 | 3.91694 | 8.86634 | 23.13815 |
| **Atf5** | 0.00530 | 0.03723 | 0.50079 | 0.16061 | 0.24520 | 0.00879 | 0.00698 | 0.55772 | 0.44395 | 0.97745 | 0.24732 | 0.06995 | 0.01353 | 0.00516 | 0.45218 | 0.01438 |
| **Atf6** | 0.00000 | 0.16262 | 2.69962 | 0.53382 | 1.22275 | 0.97191 | 3.01530 | 5.64476 | 0.01642 | 3.76982 | 1.21962 | 0.25776 | 1.34081 | 2.05926 | 1.94346 | 2.33100 |
| **Atf6b** | 2.44947 | 0.89191 | 2.12486 | 2.53742 | 0.42485 | 0.01130 | 0.60761 | 2.20295 | 2.50954 | 2.44574 | 3.63533 | 0.25632 | 6.51706 | 0.01679 | 0.06801 | 3.33034 |
| **Atf7** | 3.19917 | 0.00265 | 0.54660 | 1.13877 | 0.00886 | 0.12544 | 2.33567 | 0.31623 | 0.76060 | 1.42953 | 0.21496 | 0.13823 | 2.11947 | 0.65181 | 0.81361 | 1.58145 |
| **Atf7ip** | 0.81798 | 0.21649 | 3.87284 | 8.24073 | 6.62827 | 0.01378 | 1.43804 | 4.69270 | 15.63533 | 4.02634 | 1.61842 | 0.09203 | 4.45908 | 0.02751 | 0.76996 | 0.93897 |
| **Atf7ip2** | 1.33550 | 0.00179 | 0.13818 | 10.81320 | 3.36216 | 0.00124 | 0.00316 | 0.00000 | 14.48328 | 4.16239 | 4.18341 | 9.05623 | 0.47744 | 6.03534 | 6.41625 | 0.19157 |
| **Atg10** | 1.92507 | 0.00602 | 0.00000 | 0.46955 | 0.00780 | 0.00738 | 0.00594 | 0.00000 | 0.46695 | 1.64115 | 0.24286 | 0.00567 | 0.18893 | 1.13015 | 0.16311 | 0.01399 |
| **Atg101** | 9.54647 | 1.54504 | 6.50377 | 10.12366 | 8.72620 | 0.88562 | 2.41050 | 11.54991 | 12.01226 | 8.09901 | 8.40046 | 0.80653 | 7.32174 | 0.33484 | 0.47327 | 1.00127 |
| **Atg12** | 6.06869 | 7.09677 | 9.24561 | 5.26158 | 5.19432 | 5.29987 | 1.11718 | 6.02016 | 8.59035 | 16.72714 | 8.39632 | 13.15071 | 7.34698 | 6.68196 | 10.48030 | 0.00769 |
| **Atg13** | 18.20444 | 8.52425 | 8.01545 | 6.70776 | 6.77379 | 4.27881 | 11.29135 | 11.64704 | 9.57409 | 16.27245 | 8.55676 | 1.54206 | 11.96329 | 3.28296 | 13.04109 | 7.72245 |
| **Atg14** | 5.16267 | 5.43287 | 13.91625 | 12.71919 | 3.42154 | 2.70412 | 8.08948 | 18.49224 | 14.05183 | 19.29549 | 18.40795 | 4.31542 | 14.55142 | 14.78568 | 18.51132 | 0.61296 |
| **Atg16l1** | 4.72570 | 0.53168 | 5.97208 | 3.98416 | 0.09319 | 1.18980 | 0.00000 | 3.77818 | 9.31723 | 11.21693 | 1.81117 | 6.36823 | 3.29693 | 2.09482 | 14.62345 | 1.31096 |
| **Atg16l2** | 5.93166 | 1.48567 | 5.98171 | 0.06424 | 0.04878 | 5.93427 | 5.72738 | 12.10129 | 0.03773 | 5.33210 | 5.33454 | 0.04819 | 16.96866 | 0.52676 | 3.08945 | 2.46774 |
| **Atg2a** | 9.47683 | 6.60406 | 14.60088 | 6.40286 | 14.65569 | 10.65537 | 10.60865 | 21.26897 | 9.40536 | 20.27295 | 7.20182 | 3.47317 | 20.23429 | 0.00000 | 1.76918 | 10.16290 |
| **Atg2b** | 5.66443 | 5.84362 | 5.08886 | 2.13335 | 4.09347 | 4.71274 | 6.62117 | 9.44393 | 6.22726 | 10.53560 | 4.17048 | 2.95365 | 2.91615 | 7.33286 | 9.57746 | 0.04362 |
| **Atg3** | 1.70757 | 2.33854 | 1.47640 | 11.22955 | 2.31959 | 2.79751 | 1.68653 | 0.58972 | 11.63868 | 21.20199 | 9.92487 | 2.26864 | 2.67408 | 12.56303 | 11.24393 | 0.07730 |
| **Atg4a** | 0.49717 | 2.02471 | 2.77186 | 0.34993 | 3.52641 | 1.93042 | 0.00586 | 0.61670 | 0.00000 | 0.69399 | 0.00000 | 0.29102 | 0.38056 | 0.99382 | 0.78598 | 0.02588 |
| **Atg4b** | 13.77077 | 15.23212 | 32.05116 | 6.47359 | 11.25912 | 1.06665 | 20.14624 | 29.32952 | 14.43793 | 22.08555 | 25.34513 | 2.60867 | 20.31932 | 2.64919 | 14.23693 | 5.44248 |
| **Atg4c** | 0.36795 | 0.54152 | 0.03925 | 0.03256 | 0.00740 | 0.00691 | 0.00676 | 0.57112 | 0.02057 | 0.12082 | 0.03498 | 0.16073 | 0.17227 | 0.19795 | 0.12133 | 0.04346 |

| | | | | | | | | | | | | | | | | |
|---|---|---|---|---|---|---|---|---|---|---|---|---|---|---|---|---|
| **Atg4d** | 0.05577 | 0.14735 | 0.44803 | 0.74337 | 0.04456 | 4.29789 | 0.51966 | 7.53073 | 0.02064 | 5.31411 | 1.83492 | 0.08878 | 5.67746 | 0.12802 | 0.38405 | 0.01020 |
| **Atg5** | 2.94402 | 0.89930 | 3.93264 | 6.34012 | 7.40623 | 4.67515 | 4.02714 | 10.45979 | 3.29832 | 4.52393 | 2.65820 | 0.46327 | 5.57609 | 3.32505 | 2.66559 | 4.19708 |
| **Atg7** | 7.62355 | 4.56177 | 7.48482 | 1.10200 | 1.23784 | 1.77003 | 3.14104 | 16.50660 | 2.18202 | 12.89858 | 10.55595 | 1.04712 | 14.03361 | 0.45997 | 1.78862 | 0.02172 |
| **Atg9a** | 1.50047 | 2.57056 | 9.43277 | 11.45139 | 1.81859 | 0.13614 | 7.31089 | 24.21921 | 13.88586 | 25.70113 | 19.79649 | 6.75220 | 13.66341 | 4.85393 | 9.04407 | 1.38542 |
| **Atg9b** | 0.00804 | 0.00728 | 0.92553 | 0.01125 | 0.00809 | 0.00916 | 0.00685 | 1.00369 | 0.01120 | 0.00001 | 0.00646 | 0.00502 | 0.21031 | 0.01785 | 0.00000 | 0.00687 |
| **Atic** | 2.62293 | 2.83308 | 3.63356 | 0.42044 | 1.59395 | 0.70031 | 2.07411 | 5.88761 | 0.67655 | 0.42074 | 1.76210 | 0.17340 | 3.43105 | 0.60820 | 3.66101 | 2.03531 |
| **Atl1** | 0.00909 | 0.00793 | 0.41574 | 0.68805 | 0.00772 | 0.00926 | 0.00695 | 0.00000 | 0.00000 | 0.75401 | 0.45080 | 0.00980 | 0.01927 | 0.69867 | 0.95772 | 0.31015 |
| **Atl2** | 3.62981 | 3.91143 | 2.50991 | 4.86941 | 1.27627 | 4.10171 | 0.15078 | 5.42363 | 6.85021 | 0.66278 | 4.50706 | 7.16575 | 1.30301 | 7.29617 | 5.75819 | 0.97627 |
| **Atl3** | 4.12615 | 0.63991 | 0.46053 | 13.13794 | 3.06576 | 0.00656 | 1.98105 | 2.24321 | 11.65689 | 1.61965 | 1.66660 | 0.09197 | 0.13195 | 11.20418 | 6.53038 | 0.02935 |
| **Atm** | 6.14715 | 0.00000 | 5.80330 | 7.55959 | 3.47645 | 0.72758 | 4.32402 | 6.56625 | 5.07990 | 9.33517 | 6.24267 | 6.72591 | 3.23240 | 12.99556 | 10.31391 | 0.63926 |
| **Atmin** | 9.27831 | 7.49739 | 7.98420 | 10.99935 | 1.87463 | 5.88793 | 8.25816 | 7.20609 | 11.25264 | 13.10603 | 7.34981 | 0.57110 | 4.63729 | 8.37995 | 12.16178 | 0.16449 |
| **Atn1** | 3.59016 | 0.55173 | 1.34403 | 3.04376 | 2.31289 | 1.52469 | 2.18096 | 8.32770 | 4.09421 | 3.95564 | 11.53184 | 0.40752 | 2.50996 | 1.38063 | 0.95371 | 2.11225 |
| **Atox1** | 9.68385 | 0.67840 | 6.79771 | 5.35718 | 10.14370 | 2.75794 | 0.74534 | 1.17120 | 2.78194 | 6.23263 | 16.05613 | 0.57087 | 3.75474 | 0.05443 | 2.18952 | 5.99392 |
| **Atp10a** | 0.01810 | 0.00786 | 0.00963 | 0.51329 | 0.00898 | 0.00676 | 0.00732 | 0.01188 | 0.00474 | 0.01184 | 0.01266 | 0.17305 | 0.01299 | 0.01078 | 0.00000 | 0.00734 |
| **Atp10b** | 1.11181 | 0.01563 | 0.01393 | 1.23891 | 0.00575 | 0.00697 | 0.00635 | 0.01015 | 1.04421 | 0.00000 | 0.02911 | 0.00270 | 0.02992 | 0.04946 | 0.85747 | 0.00580 |
| **Atp10d** | 0.00558 | 0.00755 | 0.24108 | 0.00367 | 0.01915 | 0.00605 | 0.00692 | 0.76754 | 0.00908 | 0.15688 | 1.40289 | 0.75948 | 0.42622 | 0.06152 | 0.24182 | 0.00670 |
| **Atp11a** | 0.00343 | 0.00544 | 0.00000 | 11.11133 | 0.01863 | 0.00366 | 0.00493 | 0.58590 | 3.04319 | 2.11914 | 1.01179 | 6.66190 | 1.00951 | 0.01398 | 6.15577 | 0.01149 |
| **Atp11b** | 0.01361 | 3.86580 | 0.09808 | 3.07353 | 0.01098 | 1.25843 | 0.00334 | 1.56328 | 1.49711 | 0.46287 | 3.81572 | 5.54925 | 0.25260 | 9.03778 | 4.01089 | 0.00355 |
| **Atp11c** | 0.14555 | 3.21706 | 1.09093 | 5.99719 | 0.00531 | 0.50787 | 1.27204 | 0.34233 | 1.07920 | 0.65240 | 0.20565 | 9.57284 | 2.25093 | 9.29655 | 4.30079 | 0.63800 |
| **Atp12a** | 0.00000 | 0.00793 | 0.01105 | 0.01060 | 0.00672 | 0.01807 | 0.00686 | 0.01233 | 0.01241 | 0.00469 | 0.01250 | 0.00725 | 0.84220 | 0.01344 | 0.01020 | 3.85302 |
| **Atp13a1** | 3.47253 | 3.31549 | 7.07396 | 1.24480 | 0.08204 | 3.31695 | 6.49876 | 10.54434 | 4.77546 | 7.60146 | 2.33281 | 0.23048 | 9.68053 | 0.27642 | 0.61447 | 2.73397 |
| **Atp13a2** | 0.10056 | 0.01817 | 0.70853 | 2.82516 | 0.00794 | 0.25886 | 0.18424 | 1.32217 | 5.21230 | 1.87375 | 1.83817 | 0.71777 | 2.45989 | 0.81895 | 0.35838 | 0.24742 |
| **Atp13a3** | 13.32450 | 9.91282 | 8.67014 | 20.69028 | 4.42026 | 6.78736 | 10.40536 | 13.89150 | 19.43662 | 15.35230 | 18.52390 | 23.02707 | 7.61001 | 34.63383 | 23.43644 | 1.96599 |
| **Atp13a5** | 0.01328 | 0.00768 | 0.01237 | 0.13737 | 0.00907 | 0.00642 | 0.00730 | 0.01052 | 0.00974 | 0.86040 | 0.01182 | 0.00480 | 0.01100 | 0.00559 | 0.14362 | 0.00727 |
| **Atp1a1** | 11.69490 | 4.64581 | 11.82143 | 3.64852 | 9.79827 | 8.65794 | 14.16813 | 16.69183 | 3.46989 | 18.85826 | 8.71238 | 0.35289 | 17.85894 | 3.86826 | 4.84031 | 9.08412 |
| **Atp1a2** | 0.71469 | 0.01799 | 0.02017 | 0.05877 | 0.04752 | 0.00729 | 0.01726 | 0.00630 | 0.23875 | 0.02196 | 0.25159 | 0.00535 | 0.03440 | 0.22532 | 0.03563 | 0.00641 |
| **Atp1a3** | 0.01451 | 0.00707 | 0.25772 | 0.42337 | 0.00887 | 0.00561 | 0.74339 | 0.00952 | 0.22801 | 0.00945 | 0.02279 | 0.00595 | 0.04759 | 0.00574 | 0.27337 | 0.00704 |
| **Atp1b1** | 4.74023 | 3.89765 | 3.53245 | 4.41140 | 0.41281 | 7.14493 | 6.50973 | 2.58148 | 8.97761 | 0.42438 | 1.74827 | 0.14897 | 7.91461 | 2.74349 | 1.18550 | 5.93320 |
| **Atp1b2** | 0.89689 | 0.01767 | 0.03519 | 0.48269 | 0.00617 | 0.00483 | 0.00601 | 1.03771 | 1.35447 | 0.75522 | 2.98825 | 0.09479 | 0.00000 | 0.33728 | 1.90660 | 0.01317 |
| **Atp1b3** | 5.43537 | 7.25219 | 7.16186 | 14.48853 | 5.82241 | 4.49418 | 3.72908 | 8.62147 | 12.23507 | 19.41653 | 2.57695 | 13.40360 | 3.87078 | 13.71030 | 18.38016 | 3.72885 |
| **Atp2a1** | 0.01130 | 0.00765 | 0.01168 | 0.00475 | 0.05418 | 0.00652 | 0.00745 | 0.01284 | 0.14565 | 0.00755 | 0.01105 | 0.00698 | 0.01865 | 0.00698 | 0.22328 | 0.00728 |
| **Atp2a2** | 9.64467 | 10.03604 | 15.98573 | 18.56873 | 2.97616 | 7.65988 | 8.12517 | 11.72240 | 8.75001 | 12.08832 | 6.95468 | 4.48516 | 10.93499 | 13.58717 | 17.27290 | 5.85840 |
| **Atp2a3** | 0.05127 | 0.00749 | 0.51224 | 0.01053 | 0.00700 | 0.01121 | 0.00725 | 1.23686 | 0.02141 | 0.00271 | 0.00485 | 0.00549 | 0.00335 | 0.01694 | 0.00000 | 0.00701 |
| **Atp2b1** | 1.98870 | 0.08854 | 2.48732 | 0.23252 | 0.15504 | 0.81411 | 0.00406 | 1.54310 | 1.94396 | 1.11182 | 0.06844 | 6.17384 | 1.45983 | 8.50329 | 1.09305 | 2.93607 |
| **Atp2b2** | 0.00894 | 0.00750 | 0.00795 | 0.04944 | 0.00955 | 0.00699 | 0.00709 | 0.72576 | 0.00870 | 0.34450 | 0.00782 | 0.00491 | 0.00598 | 0.01282 | 0.00000 | 0.00782 |
| **Atp2b3** | 0.01015 | 0.00833 | 0.00830 | 0.07079 | 0.00971 | 0.00679 | 0.00810 | 0.08889 | 0.50739 | 0.00345 | 0.00451 | 0.00541 | 0.00937 | 0.64443 | 0.00000 | 0.00743 |
| **Atp2b4** | 0.00130 | 2.28459 | 0.00000 | 0.16607 | 0.13640 | 0.00608 | 0.28618 | 0.00000 | 0.65529 | 0.00000 | 2.75655 | 0.02499 | 1.31837 | 0.00000 | 2.05880 | 0.01202 |
| **Atp2c1** | 3.31163 | 1.67117 | 1.14073 | 6.08337 | 1.72714 | 0.78515 | 3.14428 | 1.92856 | 2.73455 | 5.04899 | 0.36094 | 0.28435 | 0.00000 | 0.05190 | 12.25704 | 3.32846 |
| **Atp2c2** | 0.35914 | 0.00679 | 0.14929 | 0.00764 | 1.15793 | 0.00589 | 0.00711 | 0.01066 | 0.01151 | 0.43769 | 0.01468 | 0.02289 | 0.21640 | 0.01188 | 0.00653 | 0.00744 |
| **Atp4b** | 0.00002 | 0.00734 | 0.00351 | 4.20286 | 0.00881 | 2.81754 | 0.00497 | 2.52104 | 3.08695 | 0.31491 | 0.00000 | 0.00000 | 3.73194 | 0.00000 | 0.15509 | 0.00366 |
| **Atp5a1** | 39.94049 | 22.64359 | 35.43673 | 13.27873 | 29.09977 | 21.25819 | 20.84879 | 35.77459 | 18.96589 | 32.44879 | 29.30140 | 13.77420 | 39.21827 | 16.88149 | 22.91967 | 24.53801 |
| **Atp5b** | 43.59215 | 19.94262 | 31.56107 | 24.46706 | 30.99949 | 19.73754 | 17.86213 | 29.83226 | 34.83509 | 40.15873 | 32.68041 | 18.21364 | 46.18478 | 23.84076 | 20.90557 | 29.18489 |
| **Atp5c1** | 32.51642 | 18.04962 | 26.53968 | 11.52830 | 22.98187 | 16.19415 | 14.84691 | 23.76596 | 10.35451 | 30.76655 | 17.91705 | 11.75727 | 27.99901 | 9.03843 | 7.90770 | 19.03515 |
| **Atp5d** | 15.71790 | 0.78954 | 8.28609 | 1.24068 | 12.99633 | 5.39536 | 1.43596 | 9.29160 | 0.86807 | 20.49096 | 11.77234 | 0.37216 | 27.82791 | 0.06325 | 0.00000 | 12.00390 |
| **Atp5e** | 7.69065 | 0.30949 | 4.85731 | 2.48078 | 3.42009 | 1.46436 | 1.03864 | 0.24928 | 1.47418 | 2.68094 | 14.22795 | 0.98298 | 3.01377 | 0.04900 | 0.00000 | 1.09191 |
| **Atp5f1** | 35.86409 | 15.93082 | 27.54766 | 15.76130 | 21.69596 | 18.64827 | 18.82506 | 33.73679 | 19.68186 | 36.70857 | 24.99538 | 20.91166 | 32.66624 | 17.02123 | 8.28422 | 20.13998 |
| **Atp5g1** | 26.15814 | 5.07589 | 26.95509 | 18.50578 | 19.66463 | 11.60384 | 10.69986 | 21.19457 | 27.46868 | 24.78314 | 32.75256 | 5.07741 | 24.33955 | 1.83169 | 13.05134 | 13.29943 |
| **Atp5g2** | 22.60682 | 10.68824 | 14.87677 | 1.74290 | 16.30200 | 5.05706 | 9.82367 | 13.05548 | 2.55766 | 12.01620 | 13.72811 | 1.36285 | 17.31443 | 0.62389 | 4.46641 | 13.40339 |
| **Atp5g3** | 34.83053 | 3.86051 | 16.14442 | 6.05647 | 21.50519 | 13.08515 | 7.40624 | 12.47189 | 9.84299 | 18.66615 | 18.20177 | 1.25472 | 22.33303 | 3.50598 | 1.32385 | 15.29324 |
| **Atp5h** | 21.91345 | 6.19177 | 17.69689 | 12.01050 | 14.68757 | 8.58044 | 5.25554 | 6.63995 | 10.74301 | 13.90725 | 24.93097 | 2.08699 | 19.49931 | 10.10090 | 8.32642 | 9.39431 |
| **Atp5j** | 15.58724 | 1.68270 | 16.08665 | 7.96234 | 16.30704 | 4.26433 | 2.33896 | 2.88178 | 15.90304 | 13.48382 | 25.29838 | 2.31650 | 8.01678 | 7.02266 | 4.05865 | 5.43224 |
| **Atp5j2** | 7.72653 | 0.32340 | 6.36446 | 8.46205 | 4.57516 | 1.33613 | 0.72010 | 0.47779 | 11.87727 | 5.19644 | 17.66066 | 0.60899 | 4.05367 | 3.69934 | 2.73521 | 4.37258 |
| **Atp5k** | 6.51687 | 0.35928 | 5.02650 | 4.37438 | 7.15748 | 1.59974 | 0.49989 | 1.29909 | 4.68492 | 1.43125 | 22.11217 | 0.97247 | 2.46348 | 1.44793 | 1.57897 | 3.93900 |
| **Atp5l** | 13.75705 | 1.33550 | 12.19778 | 5.82070 | 12.27814 | 4.58184 | 1.99149 | 1.09907 | 1.17673 | 7.49680 | 22.71401 | 1.70032 | 9.43716 | 5.54797 | 3.33603 | 6.14531 |
| **Atp5o** | 28.77810 | 16.71654 | 20.45717 | 6.43809 | 15.09946 | 15.49354 | 14.47947 | 16.29555 | 6.85378 | 18.97518 | 17.91145 | 6.42758 | 23.54011 | 4.47326 | 6.26578 | 17.62344 |
| **Atp5s** | 1.66487 | 0.78279 | 1.00930 | 2.69316 | 0.00549 | 1.81949 | 0.29219 | 2.13351 | 3.47941 | 1.39164 | 2.65072 | 0.28777 | 1.83159 | 1.80731 | 0.67619 | 1.85237 |
| **Atp5sl** | 6.67103 | 0.43761 | 3.68471 | 1.77368 | 3.44235 | 0.00000 | 4.31877 | 7.72957 | 3.14712 | 6.41675 | 3.32800 | 0.78353 | 1.42365 | 0.04134 | 0.00000 | 2.00868 |
| **Atp6ap1** | 1.70789 | 4.71410 | 8.29241 | 5.24974 | 5.69645 | 4.48631 | 1.20736 | 8.01846 | 3.66081 | 6.98631 | 8.38252 | 0.30558 | 10.71437 | 13.42256 | 15.18411 | 2.65017 |
| **Atp6ap1l** | 0.00000 | 0.00557 | 2.02696 | 0.21015 | 0.00694 | 1.31873 | 3.77716 | 1.65297 | 1.36497 | 0.73438 | 1.65028 | 0.06700 | 1.32599 | 2.11111 | 4.96819 | 0.00502 |
| **Atp6ap2** | 3.81491 | 3.67147 | 18.33336 | 8.06803 | 8.12384 | 5.44447 | 10.31340 | 16.09456 | 5.48722 | 9.96921 | 9.84118 | 11.82993 | 11.49781 | 22.90784 | 11.99345 | 6.47108 |
| **Atp6v0a1** | 5.79659 | 0.00000 | 2.50271 | 0.17056 | 0.47694 | 0.05413 | 6.61146 | 4.71635 | 0.02658 | 6.30096 | 4.33688 | 0.13526 | 2.16379 | 0.00295 | 0.44359 | 0.22556 |
| **Atp6v0a2** | 0.78430 | 6.97098 | 2.46071 | 6.30132 | 0.00174 | 0.43639 | 0.00221 | 1.87654 | 1.98418 | 3.10386 | 0.83479 | 0.36789 | 2.71463 | 0.23547 | 2.75646 | 1.46710 |

| | | | | | | | | | | | | | | | | |
|---|---|---|---|---|---|---|---|---|---|---|---|---|---|---|---|---|
| **Atp6v0a4** | 0.00000 | 1.81007 | 0.46986 | 0.48139 | 0.02431 | 2.15412 | 0.00468 | 0.72423 | 1.18664 | 0.81081 | 4.39603 | 0.02204 | 0.18141 | 0.61206 | 1.40580 | 0.25527 |
| **Atp6v0b** | 18.66576 | 7.11039 | 15.84351 | 7.89261 | 8.83478 | 13.73025 | 8.69080 | 15.43260 | 0.85633 | 14.12254 | 10.53211 | 11.93065 | 23.92628 | 0.05980 | 0.64360 | 8.01021 |
| **Atp6v0d1** | 9.54693 | 10.44846 | 12.60649 | 8.12330 | 12.49215 | 7.82615 | 7.21806 | 9.86831 | 7.94087 | 16.33814 | 14.67751 | 8.17090 | 13.09268 | 6.58077 | 5.09555 | 6.21065 |
| **Atp6v0e** | 15.54191 | 11.41860 | 25.61817 | 6.00647 | 4.61438 | 8.95022 | 7.33451 | 19.11347 | 12.56681 | 25.28835 | 16.10266 | 1.97994 | 19.15718 | 9.70256 | 11.59571 | 8.62904 |
| **Atp6v0e2** | 0.01227 | 0.00786 | 0.01214 | 0.33261 | 0.00946 | 0.00700 | 0.00750 | 0.01269 | 0.00612 | 0.25687 | 0.01155 | 0.00758 | 0.26264 | 0.01216 | 0.31563 | 0.00739 |
| **Atp6v1a** | 6.96691 | 9.14334 | 9.47505 | 4.25879 | 6.12041 | 4.82757 | 7.41393 | 5.33014 | 6.90901 | 5.84100 | 3.98125 | 15.63210 | 3.82031 | 11.66361 | 14.59901 | 3.27798 |
| **Atp6v1b2** | 13.33772 | 5.57585 | 6.95981 | 6.67731 | 6.50635 | 11.12685 | 9.00757 | 12.23448 | 1.14182 | 12.43455 | 8.40622 | 1.42672 | 8.67860 | 7.15223 | 6.77823 | 6.07534 |
| **Atp6v1c1** | 14.78226 | 13.17623 | 13.00047 | 3.44233 | 11.23725 | 11.55024 | 10.48498 | 15.52505 | 6.05208 | 8.74418 | 8.78838 | 11.16263 | 12.10569 | 0.20985 | 6.83112 | 6.79946 |
| **Atp6v1c2** | 0.41749 | 0.00764 | 0.00000 | 1.61959 | 0.00698 | 0.00628 | 0.00704 | 0.09950 | 0.00000 | 2.50241 | 1.36456 | 0.52951 | 0.00000 | 0.90264 | 1.87892 | 0.00623 |
| **Atp6v1d** | 12.85560 | 10.72154 | 11.20265 | 3.74541 | 3.86670 | 8.46219 | 3.29594 | 14.38186 | 8.15711 | 24.55921 | 9.39435 | 7.18390 | 11.27676 | 5.67200 | 13.05174 | 5.70845 |
| **Atp6v1e1** | 21.94367 | 16.97032 | 22.97370 | 7.47588 | 15.56366 | 11.53600 | 15.40734 | 22.14649 | 10.04857 | 23.31329 | 11.76104 | 13.81438 | 21.83168 | 2.15159 | 12.62763 | 13.46255 |
| **Atp6v1f** | 7.72579 | 1.89552 | 5.53055 | 1.18394 | 8.88496 | 2.51914 | 1.50715 | 3.69388 | 0.14217 | 4.51035 | 1.51627 | 0.37403 | 4.76288 | 0.00141 | 0.36610 | 6.60690 |
| **Atp6v1g1** | 20.24880 | 10.08092 | 14.78547 | 3.96843 | 11.90477 | 11.43285 | 9.65402 | 10.89818 | 1.17411 | 12.60077 | 9.09285 | 0.32541 | 13.13420 | 0.28614 | 0.59908 | 10.51702 |
| **Atp6v1g2** | 0.01377 | 0.00776 | 0.01045 | 0.21872 | 0.00907 | 0.00670 | 0.00725 | 0.00670 | 0.00828 | 0.36262 | 0.14997 | 0.03426 | 0.01077 | 0.01067 | 0.00512 | 0.00765 |
| **Atp6v1h** | 8.15747 | 9.33236 | 3.00644 | 7.02411 | 10.35666 | 2.81273 | 0.38251 | 2.38497 | 0.00000 | 8.24048 | 4.99788 | 4.11469 | 3.83328 | 0.94325 | 3.01130 | 0.00000 |
| **Atp7a** | 1.04310 | 0.00373 | 0.05909 | 4.75209 | 0.00619 | 0.00474 | 0.37321 | 0.00000 | 3.48760 | 0.51369 | 1.48380 | 8.94665 | 0.21864 | 4.85351 | 8.38960 | 0.00414 |
| **Atp8a1** | 1.35163 | 0.00521 | 0.76525 | 0.72134 | 0.01266 | 0.00413 | 1.46719 | 0.71323 | 1.20312 | 0.42885 | 1.77702 | 0.00000 | 0.68823 | 0.88945 | 0.00000 | 0.01277 |
| **Atp8b1** | 1.71051 | 0.00527 | 0.00000 | 0.51056 | 0.03923 | 1.17882 | 0.46925 | 0.00000 | 0.18401 | 0.26161 | 0.19082 | 0.00000 | 0.01603 | 1.54511 | 0.54166 | 0.63843 |
| **Atp8b2** | 0.44118 | 0.00767 | 0.00000 | 0.00004 | 0.03588 | 3.62388 | 0.00683 | 0.40407 | 0.14253 | 0.53827 | 0.00493 | 0.05016 | 0.00000 | 0.07733 | 0.03408 | 0.00655 |
| **Atp8b4** | 0.00000 | 0.00819 | 0.22100 | 0.01126 | 0.01119 | 0.00609 | 0.09157 | 0.01613 | 0.01085 | 1.14866 | 0.00000 | 0.20711 | 0.48430 | 6.31216 | 0.00000 | 0.00730 |
| **Atp8b5** | 0.01376 | 0.00775 | 0.01220 | 0.00899 | 0.00936 | 0.00655 | 0.00745 | 0.01258 | 0.01197 | 0.00932 | 0.01238 | 0.00705 | 0.01275 | 0.00670 | 0.28176 | 0.00741 |
| **Atp9a** | 2.21390 | 0.00281 | 1.59449 | 1.09842 | 0.82829 | 0.01369 | 1.12286 | 7.54975 | 1.62508 | 5.97271 | 2.24566 | 0.16820 | 3.61352 | 0.00000 | 0.42428 | 1.82215 |
| **Atp9b** | 4.30116 | 6.31941 | 1.69795 | 0.35980 | 0.01359 | 0.00759 | 0.04691 | 2.17922 | 0.71022 | 1.01897 | 1.46130 | 0.02024 | 0.39708 | 0.00000 | 0.19477 | 5.40885 |
| **Atpaf1** | 4.24183 | 2.80174 | 2.65593 | 0.91318 | 8.26165 | 0.06306 | 1.85882 | 0.31761 | 1.47602 | 7.30746 | 5.94041 | 5.10307 | 3.93798 | 2.48043 | 1.81400 | 1.73665 |
| **Atpaf2** | 4.62978 | 0.51973 | 1.31078 | 1.83209 | 3.74230 | 6.37154 | 4.29086 | 8.26175 | 4.73321 | 3.77121 | 7.68537 | 0.92916 | 4.76378 | 0.76696 | 3.80498 | 0.03215 |
| **Atpif1** | 11.36912 | 0.71818 | 6.93109 | 8.74808 | 7.82612 | 3.37056 | 0.88267 | 0.38883 | 14.04916 | 5.29100 | 13.40380 | 0.56241 | 5.04263 | 0.36007 | 1.29396 | 7.05870 |
| **Atr** | 1.63015 | 4.56343 | 2.12553 | 1.24648 | 2.64239 | 3.01330 | 5.79379 | 1.00288 | 0.70875 | 0.00000 | 0.47380 | 0.23346 | 0.44720 | 6.63991 | 1.39698 | 0.70275 |
| **Atraid** | 4.10696 | 2.24550 | 3.53261 | 7.38030 | 1.08903 | 0.25615 | 0.87166 | 5.06343 | 11.83879 | 9.91477 | 5.53523 | 9.11282 | 2.19452 | 1.14428 | 5.36788 | 5.31647 |
| **Atrip** | 1.86053 | 0.10215 | 1.48712 | 1.60092 | 0.02324 | 2.86856 | 0.07884 | 3.54774 | 0.73608 | 0.44499 | 1.06450 | 0.10824 | 2.34107 | 0.41560 | 0.83547 | 1.32276 |
| **Atrn** | 1.34085 | 0.01315 | 4.35295 | 1.92638 | 0.01122 | 0.00610 | 5.17457 | 1.38995 | 0.80821 | 3.91278 | 2.05311 | 0.16280 | 0.03181 | 1.18787 | 2.07140 | 0.01123 |
| **Atrnl1** | 0.01328 | 0.00774 | 0.00891 | 6.59205 | 0.00945 | 0.00587 | 0.00703 | 0.01016 | 0.68403 | 0.00000 | 0.00270 | 0.01277 | 0.01318 | 0.31890 | 0.99620 | 0.00616 |
| **Atrx** | 2.35375 | 0.89480 | 4.45040 | 10.43347 | 2.44612 | 3.74441 | 4.62335 | 4.12922 | 5.98375 | 7.60832 | 5.28318 | 9.56530 | 2.13094 | 23.15149 | 11.76296 | 0.60398 |
| **Atxn1** | 0.00000 | 2.46634 | 0.53952 | 1.83181 | 0.01605 | 0.00576 | 0.00663 | 0.35346 | 0.79548 | 0.41893 | 0.53264 | 0.01860 | 0.01425 | 1.21469 | 0.00000 | 0.00572 |
| **Atxn10** | 7.60053 | 1.07335 | 2.95783 | 3.80246 | 10.52487 | 6.60188 | 0.06527 | 5.39653 | 1.38176 | 10.56170 | 2.27918 | 1.93111 | 11.00599 | 1.28386 | 3.16522 | 11.49140 |
| **Atxn1l** | 1.59818 | 0.02805 | 0.00000 | 10.03381 | 0.33116 | 2.24849 | 0.13201 | 0.00000 | 7.33473 | 1.61596 | 3.39848 | 0.35091 | 0.54356 | 8.29486 | 2.61740 | 1.16647 |
| **Atxn2** | 2.42022 | 1.76828 | 1.88174 | 5.80441 | 2.43895 | 0.36187 | 3.50228 | 5.60811 | 1.92889 | 5.63982 | 4.31315 | 3.62159 | 0.77415 | 2.11963 | 7.66941 | 0.94231 |
| **Atxn2l** | 1.00766 | 0.88782 | 1.04681 | 9.38693 | 0.04714 | 0.49383 | 0.46462 | 1.21766 | 6.72362 | 3.00926 | 2.39590 | 0.22750 | 0.26301 | 0.15462 | 1.31182 | 0.14125 |
| **Atxn3** | 4.82355 | 0.03019 | 2.25783 | 8.64076 | 2.56397 | 1.05622 | 1.25416 | 6.62889 | 7.37953 | 6.35945 | 7.29876 | 0.93260 | 1.04660 | 9.41283 | 7.74907 | 0.11777 |
| **Atxn7** | 1.12605 | 0.65708 | 0.96898 | 5.05262 | 0.01320 | 0.62809 | 1.96895 | 1.63048 | 8.06028 | 1.05189 | 2.61911 | 5.92493 | 0.21880 | 10.87680 | 7.94042 | 0.00267 |
| **Atxn7l1** | 1.17553 | 5.67990 | 6.01726 | 0.35049 | 0.22770 | 0.31993 | 3.59805 | 0.77850 | 0.00594 | 0.15888 | 0.00113 | 0.00726 | 1.64987 | 0.00294 | 0.00000 | 4.34397 |
| **Atxn7l2** | 0.35799 | 0.00854 | 0.17037 | 0.00437 | 0.01806 | 0.00533 | 0.00735 | 0.01164 | 0.66858 | 0.07438 | 0.00882 | 0.00756 | 0.01932 | 0.02160 | 0.00181 | 0.00696 |
| **Atxn7l3** | 1.64167 | 0.66081 | 2.42069 | 1.82176 | 1.08012 | 0.14103 | 0.06635 | 5.83709 | 2.92161 | 12.37236 | 0.90728 | 0.31690 | 0.45337 | 0.01315 | 3.10385 | 0.54856 |
| **Atxn7l3b** | 2.88354 | 1.24160 | 5.63975 | 1.04369 | 0.40915 | 3.02816 | 3.35307 | 0.90087 | 4.08651 | 2.76164 | 1.40383 | 0.12274 | 3.19248 | 8.07452 | 3.27288 | 9.29939 |
| **AU015228** | 0.00000 | 0.15404 | 0.00739 | 15.57765 | 0.01739 | 0.02130 | 0.23951 | 0.10722 | 16.62205 | 0.00000 | 0.00000 | 6.91986 | 0.00854 | 0.05256 | 2.72303 | 0.00293 |
| **AU015836** | 0.00384 | 0.00830 | 1.95363 | 5.30101 | 0.00679 | 1.18253 | 0.00788 | 0.37924 | 5.13881 | 1.84386 | 0.00000 | 0.15921 | 0.09565 | 0.00000 | 0.23678 | 0.00524 |
| **AU016765** | 0.01403 | 0.00771 | 0.00846 | 7.24102 | 0.00913 | 0.00597 | 0.00697 | 0.00031 | 0.00000 | 0.00000 | 0.38240 | 0.00434 | 0.00829 | 0.03782 | 0.61177 | 0.00589 |
| **AU018091** | 6.87187 | 5.66347 | 8.28574 | 2.99115 | 2.01346 | 11.72429 | 3.19469 | 14.54682 | 11.33508 | 16.22434 | 21.52907 | 14.81392 | 16.35705 | 36.72990 | 15.03595 | 6.37021 |
| **AU019823** | 5.83292 | 0.11431 | 0.63214 | 5.54156 | 0.00000 | 0.00322 | 2.26245 | 0.00000 | 8.12154 | 2.44322 | 2.72477 | 11.66414 | 0.82582 | 8.72643 | 2.89327 | 1.32037 |
| **AU019990** | 0.01129 | 0.00793 | 0.05885 | 3.91591 | 0.00900 | 0.00798 | 0.00742 | 0.02027 | 0.39361 | 0.35104 | 0.00867 | 0.03783 | 0.17418 | 0.00758 | 0.17763 | 0.00635 |
| **AU021092** | 0.00000 | 0.00680 | 2.12888 | 0.00000 | 0.02368 | 7.36302 | 0.02295 | 2.36600 | 0.00898 | 2.40426 | 0.69483 | 0.01556 | 6.14751 | 0.01030 | 0.00000 | 0.00000 |
| **AU022252** | 9.26196 | 0.22694 | 1.99355 | 4.11156 | 4.74814 | 1.30864 | 2.39448 | 1.99481 | 3.27092 | 2.67290 | 2.34725 | 0.08472 | 6.39502 | 0.01172 | 0.80628 | 1.45020 |
| **AU022751** | 1.17559 | 0.45343 | 0.91900 | 3.56185 | 0.01548 | 0.01062 | 0.02627 | 1.05916 | 1.74692 | 0.58054 | 1.01521 | 0.00000 | 0.14957 | 0.00000 | 1.00699 | 0.00486 |
| **AU022754** | 0.07063 | 0.13234 | 0.03665 | 0.74624 | 0.00808 | 0.03062 | 0.00722 | 0.03291 | 0.73137 | 0.03631 | 0.02102 | 0.01568 | 0.08725 | 0.17149 | 0.18109 | 0.07245 |
| **AU023762** | 0.17577 | 0.00741 | 0.10229 | 1.43101 | 0.02592 | 0.02154 | 0.04749 | 0.03363 | 0.84057 | 0.05560 | 0.52251 | 0.00162 | 0.01321 | 0.52204 | 0.10773 | 0.02835 |
| **AU040320** | 3.18405 | 0.00000 | 3.47790 | 5.05287 | 0.05508 | 0.09223 | 3.84681 | 2.51553 | 0.82735 | 6.25766 | 1.39247 | 0.02853 | 0.07889 | 0.49373 | 2.31288 | 0.00001 |
| **AU041133** | 0.00000 | 0.00718 | 0.13722 | 0.39426 | 0.02092 | 0.38668 | 0.76753 | 0.92899 | 0.03144 | 0.63446 | 0.40025 | 0.02788 | 0.00000 | 4.87731 | 0.15207 | 0.00673 |
| **Auh** | 0.74090 | 0.00487 | 0.05422 | 4.94018 | 4.46654 | 0.00745 | 0.02774 | 0.01502 | 4.20880 | 1.64668 | 0.32452 | 0.04078 | 0.04732 | 0.05032 | 0.65318 | 0.02043 |
| **Aunip** | 0.39208 | 0.01615 | 2.04799 | 4.99695 | 0.00364 | 3.28272 | 0.00340 | 0.00000 | 8.87655 | 1.26501 | 5.11455 | 0.09548 | 1.42091 | 0.21392 | 2.24936 | 0.03776 |
| **Aup1** | 12.08507 | 3.41049 | 8.18947 | 0.60580 | 6.75787 | 1.73901 | 4.64565 | 6.91622 | 0.00000 | 9.30705 | 3.36798 | 0.09991 | 12.73951 | 0.01952 | 0.00000 | 6.05912 |
| **Aurka** | 10.33939 | 17.28262 | 20.70674 | 6.34807 | 9.29755 | 8.77867 | 5.64325 | 32.59828 | 17.25405 | 27.32942 | 15.76646 | 5.80462 | 16.73528 | 17.75369 | 26.42748 | 4.86095 |
| **Aurkaip1** | 14.72537 | 1.55508 | 8.58697 | 6.83177 | 8.16671 | 7.15133 | 6.12767 | 14.60913 | 6.87050 | 11.91536 | 10.94857 | 1.63799 | 12.48324 | 1.55430 | 1.11670 | 11.23599 |
| **Aurkb** | 7.08755 | 0.45310 | 8.27023 | 3.56979 | 4.55625 | 0.93470 | 1.14873 | 6.29139 | 6.04756 | 13.31073 | 3.57877 | 1.50743 | 3.72891 | 0.16482 | 6.70290 | 1.08881 |

| | | | | | | | | | | | | | | | | |
|---|---|---|---|---|---|---|---|---|---|---|---|---|---|---|---|---|
| **Aurkc** | 7.38096 | 2.96949 | 0.76950 | 5.97197 | 0.03462 | 0.00000 | 3.40327 | 6.00831 | 6.84806 | 9.74470 | 11.80388 | 1.71707 | 3.92570 | 4.67388 | 11.85741 | 0.05630 |
| **Auts2** | 1.31811 | 1.73357 | 2.04869 | 1.44437 | 2.00569 | 0.00598 | 0.05485 | 0.67126 | 1.22859 | 2.48842 | 1.04005 | 0.38203 | 0.39008 | 2.62919 | 3.64877 | 1.17153 |
| **AV051173** | 0.04143 | 0.00799 | 0.00794 | 0.32936 | 0.00800 | 0.00672 | 0.04774 | 0.01089 | 0.54816 | 0.04071 | 0.00775 | 0.01563 | 0.08143 | 0.21517 | 0.01765 | 0.11173 |
| **AV320801** | 0.01342 | 0.00761 | 0.01078 | 0.08247 | 0.00936 | 0.00631 | 0.00729 | 0.01535 | 0.13857 | 0.00393 | 0.00962 | 0.00504 | 0.01871 | 0.01037 | 1.22460 | 0.00727 |
| **Aven** | 4.05415 | 4.11946 | 7.54798 | 1.10355 | 0.41709 | 0.01004 | 0.09267 | 3.97448 | 3.16315 | 6.75790 | 5.58211 | 0.26293 | 4.49972 | 6.95457 | 1.93121 | 0.23644 |
| **Avl9** | 0.00000 | 1.91396 | 1.27368 | 8.83395 | 4.24643 | 0.41356 | 0.11929 | 2.68268 | 12.09864 | 14.12192 | 7.32257 | 17.81432 | 0.00000 | 21.12914 | 10.63674 | 0.00811 |
| **Avpi1** | 13.29513 | 0.10677 | 0.71360 | 4.54132 | 2.43123 | 6.92808 | 7.06439 | 4.75481 | 2.38987 | 4.92572 | 3.48633 | 0.15664 | 6.75194 | 0.11606 | 0.00000 | 6.65567 |
| **AW011738** | 0.01338 | 0.00818 | 0.01144 | 1.70020 | 0.00915 | 0.00662 | 0.00754 | 0.01330 | 1.96889 | 0.16883 | 0.00000 | 0.00530 | 0.01328 | 0.01125 | 0.00832 | 0.00702 |
| **AW146154** | 0.10371 | 0.27533 | 0.43126 | 0.50992 | 0.03603 | 0.16116 | 0.16331 | 0.11708 | 0.06119 | 0.75347 | 0.12936 | 0.43614 | 0.67335 | 1.50671 | 1.05658 | 0.04854 |
| **AW209491** | 0.44769 | 0.00606 | 2.45016 | 4.34077 | 0.01084 | 0.00474 | 0.00454 | 3.03229 | 1.60561 | 4.26866 | 2.53831 | 3.59071 | 1.37541 | 1.65898 | 9.56802 | 0.15786 |
| **AW495222** | 0.01190 | 0.00746 | 0.00819 | 0.01130 | 0.00943 | 0.00682 | 0.00716 | 0.30057 | 0.01183 | 0.00866 | 0.00978 | 0.00636 | 0.00642 | 0.15393 | 0.02389 | 0.00737 |
| **AW549542** | 0.01338 | 0.00806 | 0.01170 | 0.26882 | 0.00919 | 0.00683 | 0.00750 | 0.01769 | 0.00559 | 0.23147 | 0.02731 | 0.00859 | 0.01207 | 0.02964 | 0.00456 | 0.00733 |
| **AW549877** | 2.18329 | 0.00000 | 2.70932 | 1.62628 | 1.68431 | 1.37714 | 2.56410 | 0.29360 | 0.00000 | 1.28472 | 0.52520 | 11.18514 | 2.18602 | 1.53757 | 2.51430 | 2.34396 |
| **AW554918** | 0.62746 | 0.00755 | 0.05856 | 6.27018 | 0.00719 | 0.00572 | 0.01902 | 0.12973 | 0.96052 | 0.46234 | 0.00779 | 0.03814 | 0.00749 | 0.01511 | 1.20436 | 0.00540 |
| **AW822252** | 0.01223 | 0.00801 | 0.20302 | 0.31976 | 0.00877 | 0.00700 | 0.00708 | 0.01098 | 0.00654 | 0.00633 | 0.01326 | 0.00796 | 0.41762 | 0.01016 | 0.03565 | 0.00692 |
| **Awat2** | 0.15410 | 0.00785 | 0.00612 | 0.01854 | 0.31038 | 0.00716 | 0.00728 | 0.02270 | 0.01258 | 0.01854 | 0.01275 | 0.00758 | 0.01064 | 0.01195 | 0.01055 | 0.00719 |
| **Axin1** | 3.12733 | 4.21884 | 6.84362 | 9.97834 | 0.00000 | 2.58096 | 6.38543 | 6.94766 | 10.75396 | 8.49666 | 3.76533 | 0.76953 | 2.33988 | 3.51120 | 10.28106 | 5.98145 |
| **Axin2** | 1.41131 | 0.02694 | 0.03377 | 4.14635 | 0.05310 | 1.60207 | 0.00591 | 0.00408 | 2.86331 | 0.11964 | 0.01707 | 0.00000 | 0.07978 | 0.38156 | 0.33275 | 2.11504 |
| **Axl** | 0.66462 | 0.00806 | 0.00199 | 0.00855 | 0.01729 | 0.00799 | 0.00683 | 0.01374 | 0.01187 | 0.35691 | 0.00867 | 0.13723 | 0.01877 | 0.33737 | 0.07363 | 0.00692 |
| **AY074887** | 0.19774 | 0.14487 | 0.27242 | 0.03598 | 0.20723 | 0.08765 | 0.13971 | 0.10349 | 0.10649 | 0.09678 | 0.81875 | 0.02185 | 0.21023 | 0.21095 | 0.19621 | 0.17960 |
| **AY358078** | 0.33195 | 0.00748 | 0.11024 | 0.13019 | 0.01012 | 0.00595 | 0.03745 | 0.15559 | 0.75617 | 0.21522 | 0.09372 | 0.03355 | 0.01438 | 0.06054 | 0.14583 | 0.09511 |
| **AY512915** | 0.01296 | 0.00816 | 0.00872 | 5.48525 | 0.00951 | 0.00587 | 0.00730 | 0.01246 | 0.00000 | 0.00530 | 0.01061 | 0.00000 | 0.01347 | 0.11189 | 3.55912 | 0.00649 |
| **AY512931** | 0.16285 | 0.17615 | 0.28707 | 4.57715 | 0.06190 | 0.79688 | 0.13198 | 0.24468 | 2.85509 | 0.35877 | 0.58925 | 0.14995 | 0.13832 | 0.73692 | 0.92225 | 0.26420 |
| **AY761184** | 0.02009 | 0.00788 | 0.00915 | 1.68596 | 0.00879 | 0.00647 | 0.00736 | 0.00874 | 2.41500 | 0.03063 | 0.12060 | 0.05533 | 0.05612 | 0.00526 | 0.06457 | 0.00691 |
| **AY761185** | 0.01376 | 0.00815 | 0.01226 | 0.50695 | 0.00932 | 0.00659 | 0.00750 | 0.01372 | 0.00427 | 0.01364 | 0.01361 | 0.00830 | 0.01266 | 0.00940 | 0.03879 | 0.00732 |
| **Azi2** | 5.63311 | 3.89420 | 1.00261 | 1.94028 | 0.00000 | 0.14557 | 3.43609 | 3.08472 | 0.16968 | 1.68039 | 2.52353 | 7.68349 | 1.80511 | 4.09452 | 2.34444 | 1.19149 |
| **Azin1** | 6.69806 | 13.51238 | 16.21469 | 18.27377 | 9.33602 | 5.55012 | 14.53476 | 17.19907 | 20.67123 | 21.89511 | 25.37335 | 22.39122 | 7.43621 | 37.85627 | 23.12286 | 4.65436 |
| **Azin2** | 0.05037 | 0.00735 | 0.37838 | 0.00946 | 0.26111 | 1.58540 | 0.00702 | 0.01031 | 0.01257 | 0.00560 | 0.00994 | 0.04518 | 0.00433 | 0.01052 | 0.02059 | 0.00643 |
| **B2m** | 0.00000 | 6.41106 | 5.30251 | 8.54169 | 3.75534 | 4.27109 | 2.23496 | 1.76362 | 9.59414 | 4.95166 | 2.14372 | 2.12971 | 3.59991 | 4.11483 | 3.78018 | 1.97936 |
| **B3galnt1** | 0.54163 | 0.00809 | 0.05474 | 0.24378 | 0.05582 | 0.01135 | 0.02663 | 0.00576 | 0.00420 | 0.00000 | 0.00398 | 0.00000 | 0.63781 | 2.17603 | 0.13238 | 0.00645 |
| **B3galnt2** | 0.37091 | 0.00334 | 1.18571 | 4.14724 | 4.56842 | 0.31855 | 3.43595 | 0.01226 | 1.62418 | 0.60486 | 0.00000 | 0.30671 | 0.37930 | 2.48005 | 2.70844 | 0.01938 |
| **B3galt4** | 0.41307 | 0.00688 | 0.01189 | 0.05919 | 0.00683 | 0.00593 | 0.00600 | 0.80801 | 0.99096 | 0.07160 | 0.91776 | 0.00000 | 1.10514 | 0.00361 | 0.18209 | 0.00652 |
| **B3galt5** | 0.01139 | 0.00769 | 0.00478 | 0.35030 | 0.00976 | 0.00644 | 0.00706 | 0.00812 | 0.06322 | 0.00805 | 0.00484 | 0.02181 | 0.01394 | 3.80550 | 0.00000 | 0.00748 |
| **B3galt6** | 0.01838 | 0.00792 | 1.69763 | 1.78337 | 0.00593 | 0.01073 | 0.00761 | 0.54880 | 3.17536 | 0.59088 | 0.38816 | 0.02632 | 0.22044 | 0.00000 | 1.77184 | 0.00598 |
| **B3gat2** | 0.01548 | 0.00868 | 0.30591 | 0.00802 | 0.01042 | 0.00564 | 0.00776 | 0.01898 | 0.04104 | 0.00476 | 0.00735 | 0.00521 | 0.05344 | 0.98882 | 0.00000 | 0.00726 |
| **B3gat3** | 4.18504 | 0.01134 | 3.39029 | 1.40600 | 0.51862 | 0.00140 | 1.86555 | 1.67376 | 1.54005 | 6.21009 | 1.28936 | 0.10752 | 2.62150 | 0.15299 | 0.57545 | 2.11741 |
| **B3glct** | 4.71868 | 2.04934 | 2.26973 | 0.76356 | 5.94666 | 0.01170 | 5.11207 | 0.17335 | 0.32640 | 0.53594 | 0.74361 | 1.88988 | 0.07614 | 0.01546 | 0.08934 | 0.47649 |
| **B3gnt2** | 2.15062 | 0.00335 | 0.59733 | 7.67219 | 1.78284 | 0.02308 | 1.37410 | 0.71478 | 0.71973 | 0.79573 | 0.31451 | 0.00871 | 0.24627 | 0.38836 | 3.84084 | 0.01052 |
| **B3gnt3** | 0.30859 | 0.00780 | 0.00414 | 0.66698 | 0.01649 | 0.01511 | 0.00722 | 0.07074 | 0.48279 | 0.00725 | 0.01256 | 0.00690 | 0.14552 | 0.01061 | 0.01231 | 0.01448 |
| **B3gnt4** | 0.05929 | 0.00856 | 0.00985 | 0.01059 | 0.01493 | 0.00680 | 0.00751 | 0.01324 | 0.01238 | 0.01934 | 0.01285 | 0.00812 | 0.01250 | 0.01214 | 0.01072 | 0.00748 |
| **B3gnt5** | 0.09451 | 0.00584 | 0.84355 | 0.00805 | 0.10150 | 0.00905 | 3.26782 | 0.00553 | 0.01107 | 0.00000 | 0.02614 | 0.00339 | 0.62743 | 0.02276 | 0.28466 | 2.07581 |
| **B3gnt6** | 0.01202 | 0.00848 | 0.00701 | 0.01129 | 0.02198 | 0.00708 | 0.00819 | 1.19300 | 0.01110 | 0.00337 | 0.00535 | 0.00684 | 0.00567 | 0.01883 | 0.00000 | 0.00739 |
| **B3gnt7** | 4.07725 | 1.48401 | 0.19601 | 0.11711 | 2.44584 | 0.00112 | 1.36437 | 0.00000 | 1.27672 | 1.82383 | 0.06251 | 0.04648 | 0.03150 | 0.99833 | 0.00000 | 4.43199 |
| **B3gnt8** | 0.04703 | 0.00834 | 0.00400 | 0.00880 | 0.01431 | 0.00750 | 0.00673 | 0.56981 | 0.01139 | 0.15808 | 0.05808 | 0.09048 | 1.37016 | 0.01420 | 0.00000 | 0.00689 |
| **B3gntl1** | 0.01306 | 0.00768 | 0.00388 | 1.77664 | 0.00915 | 0.00648 | 0.03692 | 0.00000 | 0.00000 | 0.06758 | 0.89803 | 0.00985 | 0.00509 | 0.00116 | 1.49719 | 0.04340 |
| **B4galnt1** | 0.03207 | 0.01771 | 0.30741 | 0.09303 | 0.00799 | 0.15329 | 0.00539 | 4.70801 | 0.11864 | 1.79909 | 2.34876 | 0.28478 | 2.33459 | 0.03038 | 0.68374 | 0.00559 |
| **B4galnt2** | 1.11510 | 0.01798 | 0.44556 | 5.86489 | 0.45467 | 0.09071 | 0.09714 | 0.29046 | 7.84318 | 0.83251 | 1.48646 | 0.57834 | 0.25882 | 2.02346 | 1.03528 | 0.10037 |
| **B4galnt3** | 0.00685 | 0.23211 | 0.00803 | 0.15623 | 0.00857 | 0.00673 | 0.00703 | 0.68127 | 0.13681 | 0.21667 | 0.00685 | 0.00505 | 0.00558 | 0.00855 | 0.00000 | 0.03649 |
| **B4galnt4** | 0.23804 | 0.00762 | 0.00665 | 0.06583 | 0.00735 | 0.00706 | 0.00717 | 0.01180 | 0.01134 | 0.82197 | 0.01208 | 0.01652 | 0.01165 | 0.00968 | 0.02006 | 0.00707 |
| **B4galt1** | 9.62892 | 2.05583 | 4.18045 | 0.00000 | 1.89779 | 3.20900 | 2.42342 | 3.32763 | 0.88343 | 1.34638 | 0.39136 | 0.17336 | 1.67219 | 0.00667 | 0.00000 | 4.69906 |
| **B4galt2** | 0.00525 | 0.00761 | 0.01080 | 2.36438 | 0.00691 | 0.00601 | 0.00711 | 0.01150 | 0.32560 | 0.00309 | 0.00940 | 0.03566 | 0.02456 | 0.00575 | 0.44556 | 2.93125 |
| **B4galt3** | 5.52590 | 2.54540 | 5.70699 | 10.48515 | 0.04538 | 6.55719 | 3.28923 | 11.36495 | 13.10988 | 14.89435 | 6.70317 | 1.08860 | 9.78920 | 0.16513 | 1.72437 | 7.24453 |
| **B4galt4** | 0.01263 | 0.00763 | 0.01232 | 2.23848 | 0.00878 | 0.00624 | 0.00718 | 0.00981 | 0.22852 | 2.02006 | 0.00865 | 0.00000 | 0.01063 | 0.00152 | 0.16189 | 0.00702 |
| **B4galt5** | 4.07352 | 1.09751 | 4.74955 | 3.13114 | 0.05817 | 3.38466 | 0.05121 | 0.02295 | 3.57311 | 0.22572 | 1.03603 | 0.10511 | 3.21267 | 0.00000 | 0.23749 | 0.44167 |
| **B4galt6** | 5.09259 | 1.83486 | 0.57319 | 14.13274 | 8.37927 | 2.20725 | 2.74423 | 0.25606 | 3.34797 | 2.31078 | 1.35339 | 0.16868 | 1.41959 | 0.25274 | 3.54834 | 2.55119 |
| **B4galt7** | 0.76080 | 0.02664 | 1.30790 | 3.02692 | 0.61771 | 0.04424 | 0.28113 | 0.00000 | 4.86234 | 1.57238 | 5.01072 | 0.33479 | 1.06432 | 2.19651 | 0.65317 | 0.32832 |
| **B4gat1** | 0.12891 | 0.00712 | 0.46978 | 0.44813 | 0.03389 | 0.23041 | 0.00638 | 0.92725 | 0.16639 | 0.75832 | 0.68259 | 1.59822 | 0.60796 | 0.00860 | 0.00000 | 0.01370 |
| **B9d1** | 2.82076 | 0.24711 | 1.21858 | 4.29831 | 3.54851 | 0.00223 | 0.05016 | 0.14441 | 2.32605 | 0.00000 | 0.08906 | 0.04194 | 0.51866 | 1.17106 | 1.87362 | 1.85548 |
| **B9d2** | 3.17239 | 0.00433 | 0.00443 | 0.00436 | 0.13038 | 0.00578 | 0.00506 | 0.00066 | 0.35031 | 0.72811 | 0.02388 | 0.01171 | 0.15818 | 0.01037 | 0.01333 | 0.02021 |
| **Baat** | 0.01872 | 0.00787 | 0.00639 | 0.35603 | 0.01027 | 0.00708 | 0.00717 | 0.00084 | 0.00546 | 0.00000 | 0.34781 | 0.24531 | 0.00876 | 0.00739 | 0.00000 | 0.00719 |
| **Babam1** | 8.94342 | 0.46203 | 3.14579 | 1.68628 | 5.51744 | 3.99619 | 0.04414 | 3.79467 | 0.28240 | 3.50085 | 3.83851 | 5.51157 | 9.38126 | 0.00022 | 0.26339 | 3.80287 |
| **Bace1** | 1.21648 | 0.00000 | 4.89630 | 4.44613 | 4.51308 | 2.65717 | 2.94607 | 5.42525 | 7.81029 | 6.57423 | 4.93063 | 0.20883 | 0.45646 | 4.00048 | 9.68849 | 1.19147 |

| | | | | | | | | | | | | | | | | |
|---|---|---|---|---|---|---|---|---|---|---|---|---|---|---|---|---|
| **Bace2** | 0.01308 | 0.00755 | 0.01180 | 0.00701 | 0.00890 | 0.00659 | 0.00729 | 0.01054 | 0.01129 | 1.23533 | 0.01116 | 0.00394 | 0.01143 | 0.01425 | 0.10234 | 0.00778 |
| **Bach1** | 6.56684 | 5.48163 | 3.08864 | 6.89381 | 2.15099 | 4.52430 | 0.92860 | 8.45888 | 7.25756 | 8.13059 | 14.22640 | 4.10014 | 3.44945 | 27.97057 | 8.77469 | 1.23952 |
| **Bach2** | 0.99736 | 0.01940 | 0.07910 | 0.10531 | 0.07391 | 0.02166 | 0.02713 | 0.00675 | 0.08694 | 0.22886 | 0.28142 | 0.02376 | 0.13822 | 0.11851 | 0.45634 | 0.00644 |
| **Bad** | 10.18517 | 5.05304 | 10.51828 | 0.91114 | 4.59783 | 5.71706 | 5.69049 | 7.26775 | 2.09683 | 6.94837 | 4.11424 | 0.43771 | 7.11311 | 0.00000 | 0.19699 | 3.12409 |
| **Bag1** | 10.92273 | 3.07916 | 12.13745 | 10.98866 | 7.55166 | 7.61290 | 7.51104 | 4.89145 | 25.23020 | 17.14654 | 18.44011 | 6.16578 | 9.10953 | 8.09738 | 4.44625 | 1.64822 |
| **Bag2** | 0.08696 | 0.09688 | 0.14866 | 0.25869 | 0.00761 | 0.03550 | 0.00662 | 0.00788 | 0.00491 | 0.84310 | 0.01062 | 0.02419 | 0.89359 | 0.01098 | 0.01271 | 0.01453 |
| **Bag3** | 4.77907 | 4.30025 | 9.34533 | 3.93725 | 5.55711 | 7.03460 | 3.72283 | 10.22096 | 22.29699 | 4.32112 | 15.04101 | 0.25741 | 12.27973 | 5.81699 | 7.19836 | 3.24542 |
| **Bag4** | 8.75950 | 8.59732 | 2.46404 | 4.88161 | 4.31686 | 4.87231 | 6.17185 | 4.23207 | 4.05960 | 4.16817 | 1.86531 | 7.07088 | 6.84218 | 8.18101 | 2.86075 | 1.45576 |
| **Bag5** | 4.22049 | 3.26431 | 6.51702 | 19.16861 | 0.22680 | 4.43364 | 6.77529 | 6.86556 | 29.25787 | 9.87517 | 5.62562 | 0.94020 | 4.19888 | 2.38063 | 11.53686 | 5.86482 |
| **Bag6** | 12.28398 | 0.95648 | 3.08829 | 6.47654 | 4.73258 | 5.52069 | 3.18692 | 19.54768 | 9.16286 | 19.25586 | 13.45222 | 1.78284 | 14.67198 | 0.63144 | 3.07169 | 12.07146 |
| **Bahcc1** | 0.01367 | 0.00799 | 0.01237 | 0.41363 | 0.00929 | 0.00659 | 0.00752 | 0.01393 | 0.05322 | 0.01399 | 0.01312 | 0.00778 | 0.01260 | 0.00882 | 0.02659 | 0.01464 |
| **Bahd1** | 2.68170 | 0.00479 | 0.31100 | 5.23939 | 0.00417 | 0.05288 | 0.01370 | 0.03684 | 2.74911 | 0.53133 | 2.17929 | 0.06634 | 1.78705 | 0.35374 | 0.76785 | 0.72530 |
| **Baiap2** | 10.27282 | 0.00000 | 6.46352 | 2.07725 | 8.22923 | 0.00699 | 7.89868 | 15.53575 | 3.20579 | 4.07974 | 5.65806 | 0.17177 | 0.71172 | 0.00172 | 0.00000 | 3.61060 |
| **Baiap2l1** | 1.23473 | 0.55324 | 1.00606 | 2.47930 | 0.00345 | 0.00763 | 0.49355 | 4.33677 | 0.80550 | 1.54388 | 2.22263 | 0.25755 | 1.47446 | 0.04304 | 1.31668 | 0.01829 |
| **Baiap2l2** | 0.10686 | 0.00881 | 0.00782 | 1.04117 | 0.01534 | 0.00691 | 0.00745 | 0.01257 | 1.00648 | 0.02271 | 0.02202 | 0.01853 | 0.01269 | 0.01041 | 0.00215 | 0.00728 |
| **Baiap3** | 0.54228 | 0.00659 | 0.00972 | 0.00878 | 0.00670 | 0.00784 | 0.00662 | 0.31547 | 0.22164 | 0.02797 | 0.30713 | 0.00450 | 0.00653 | 0.01246 | 0.00017 | 0.00673 |
| **Bak1** | 0.06321 | 0.00772 | 0.01424 | 0.02023 | 0.00796 | 0.00638 | 0.00749 | 0.01712 | 0.04500 | 0.01300 | 0.03898 | 0.01837 | 0.02426 | 0.01215 | 0.00910 | 0.00725 |
| **Bambi** | 1.00805 | 0.00343 | 0.00115 | 7.00541 | 6.73992 | 0.00645 | 0.02563 | 0.98183 | 5.84576 | 0.16136 | 0.00000 | 0.11059 | 0.01663 | 0.00697 | 0.12499 | 0.98303 |
| **Bambi.ps1** | 0.01275 | 0.00812 | 0.02877 | 0.71389 | 0.00911 | 0.00709 | 0.01799 | 0.01396 | 0.04603 | 0.24665 | 0.01045 | 0.00724 | 0.01722 | 0.08108 | 0.00629 | 0.00700 |
| **Banf1** | 23.39191 | 6.90456 | 16.60330 | 7.16399 | 10.64153 | 7.72434 | 10.56059 | 16.87151 | 6.50040 | 22.24187 | 19.13185 | 3.25238 | 14.73089 | 2.23250 | 5.29947 | 9.92697 |
| **Banp** | 0.26013 | 0.03752 | 3.50558 | 2.97472 | 5.44926 | 0.07648 | 2.04748 | 2.36085 | 7.07019 | 7.21742 | 1.91487 | 0.13615 | 6.90604 | 0.20041 | 0.49716 | 1.45648 |
| **Bap1** | 7.74874 | 1.00454 | 4.88183 | 9.90165 | 6.22140 | 1.75785 | 2.64318 | 8.76292 | 5.76231 | 9.16459 | 4.63786 | 4.24655 | 5.49586 | 0.13258 | 3.11618 | 2.91259 |
| **Bard1** | 2.16659 | 1.41606 | 1.19655 | 4.38850 | 0.00326 | 0.07369 | 1.23790 | 0.82705 | 12.01448 | 1.90011 | 3.65093 | 5.80658 | 1.84278 | 0.12552 | 1.72325 | 1.19309 |
| **Barx2** | 0.01387 | 0.01878 | 0.04061 | 0.00508 | 0.00918 | 0.00616 | 0.04978 | 0.00920 | 0.01106 | 0.01993 | 0.66676 | 0.01662 | 0.00861 | 0.01053 | 0.00204 | 0.00730 |
| **Basp1** | 1.98070 | 0.11049 | 0.08527 | 0.23375 | 0.92519 | 1.44595 | 0.00500 | 0.01891 | 0.43836 | 0.12659 | 0.29598 | 0.05671 | 1.27646 | 0.00598 | 0.02589 | 1.75657 |
| **Batf** | 0.02617 | 0.00780 | 0.12663 | 0.00918 | 0.00852 | 0.00576 | 0.00735 | 0.01293 | 0.02694 | 0.01842 | 0.01239 | 0.02776 | 0.01090 | 0.01250 | 0.00796 | 0.00722 |
| **Batf3** | 0.01372 | 0.00814 | 0.01203 | 0.15097 | 0.00927 | 0.00663 | 0.00753 | 0.01354 | 0.24438 | 0.01327 | 0.02577 | 0.00781 | 0.01242 | 0.01158 | 0.00894 | 0.00737 |
| **Bax** | 1.42752 | 0.01162 | 3.69041 | 3.01347 | 8.68689 | 2.39623 | 0.01972 | 1.50662 | 1.57547 | 4.49645 | 5.63283 | 0.06419 | 7.00353 | 0.34800 | 0.36373 | 3.00261 |
| **Baz1a** | 4.02957 | 9.55149 | 8.44603 | 13.57274 | 4.55346 | 7.37058 | 7.19316 | 11.46991 | 14.24445 | 30.26625 | 15.65079 | 23.96122 | 11.20823 | 35.68636 | 26.69645 | 2.85868 |
| **Baz1b** | 8.11302 | 2.02601 | 7.48146 | 12.82597 | 0.12922 | 1.38335 | 5.34499 | 11.18417 | 9.85115 | 10.43546 | 3.93183 | 0.76406 | 4.74666 | 1.37097 | 8.85034 | 3.07361 |
| **Baz2a** | 11.82556 | 8.01211 | 6.73575 | 13.05664 | 7.33386 | 2.81203 | 12.48799 | 16.25188 | 14.42292 | 21.14999 | 11.65124 | 8.14718 | 6.65849 | 10.66875 | 20.18759 | 3.59719 |
| **Baz2b** | 6.88227 | 5.77538 | 0.08924 | 3.41191 | 2.20924 | 0.00000 | 1.70904 | 0.50092 | 3.62374 | 1.99773 | 0.32709 | 0.19717 | 0.46307 | 0.25659 | 5.75350 | 0.00000 |
| **BB283400** | 0.04979 | 0.00757 | 0.05987 | 0.23272 | 0.00806 | 0.00577 | 0.01787 | 0.00404 | 0.09305 | 0.04714 | 0.48616 | 0.03651 | 0.11055 | 0.03824 | 0.07515 | 0.02149 |
| **BB287469** | 0.01829 | 0.01877 | 0.16248 | 10.58503 | 0.03841 | 0.00403 | 0.02736 | 0.05776 | 9.04846 | 0.00000 | 0.00000 | 0.00000 | 0.03887 | 2.20913 | 1.09070 | 0.01987 |
| **BB557941** | 0.10064 | 0.00789 | 0.02783 | 3.78995 | 0.01477 | 0.00549 | 0.00701 | 0.00897 | 1.45333 | 0.09887 | 0.00000 | 0.07458 | 0.01296 | 0.01942 | 1.85040 | 0.00711 |
| **Bbc3** | 1.05826 | 0.00624 | 0.34757 | 0.01254 | 0.82455 | 0.00515 | 0.31085 | 0.03634 | 0.23214 | 0.88096 | 0.00793 | 0.02965 | 0.39237 | 0.01496 | 0.03211 | 0.00599 |
| **Bbip1** | 5.70203 | 1.15334 | 4.60569 | 3.60024 | 3.21167 | 1.25809 | 0.82107 | 2.55110 | 3.57877 | 1.16450 | 2.04033 | 3.30503 | 4.07904 | 1.17128 | 1.63813 | 0.07728 |
| **Bbof1** | 0.00388 | 0.00550 | 0.00000 | 0.26834 | 1.62194 | 0.00670 | 2.64155 | 0.00501 | 2.93774 | 0.44747 | 0.00000 | 0.01174 | 0.01002 | 1.57172 | 0.09885 | 0.03576 |
| **Bbs1** | 0.03761 | 1.73088 | 0.08100 | 3.28880 | 0.08275 | 0.02209 | 0.01657 | 0.00402 | 5.23934 | 0.95900 | 0.08228 | 0.00000 | 0.02684 | 0.02557 | 1.61527 | 0.13573 |
| **Bbs10** | 2.87629 | 0.01427 | 0.00411 | 0.50925 | 0.00271 | 0.00438 | 0.00506 | 0.00287 | 0.00482 | 0.00658 | 0.01169 | 0.00456 | 0.01109 | 0.00734 | 0.54342 | 0.00414 |
| **Bbs12** | 0.05768 | 0.00837 | 0.05418 | 0.10699 | 0.01309 | 0.01168 | 0.00735 | 0.01229 | 0.31264 | 0.01151 | 0.01182 | 0.01826 | 0.01171 | 0.00976 | 0.03785 | 0.01480 |
| **Bbs2** | 0.54982 | 0.00749 | 1.28767 | 10.92669 | 0.01004 | 0.01150 | 0.00525 | 0.00000 | 9.70623 | 6.26066 | 0.76021 | 1.41785 | 1.71170 | 1.13894 | 4.42972 | 0.00310 |
| **Bbs4** | 1.15925 | 5.18986 | 1.00448 | 1.96532 | 0.01258 | 0.00715 | 1.58301 | 0.37714 | 1.58488 | 0.52809 | 0.46509 | 0.00000 | 0.14052 | 6.48743 | 5.28675 | 1.75305 |
| **Bbs5** | 1.46089 | 4.19656 | 2.36746 | 1.84585 | 0.01073 | 2.88588 | 2.35876 | 1.98675 | 2.03969 | 1.49574 | 2.92586 | 1.22813 | 3.56655 | 5.20553 | 2.70656 | 0.00695 |
| **Bbs7** | 0.01374 | 0.00772 | 0.01183 | 0.23256 | 0.00945 | 0.00648 | 0.00739 | 0.01152 | 0.08462 | 0.01465 | 0.02134 | 0.00628 | 0.01266 | 0.00648 | 0.49201 | 0.00738 |
| **Bbs9** | 0.03102 | 0.00890 | 0.00578 | 3.28081 | 0.01270 | 0.00773 | 0.00785 | 0.11945 | 0.12053 | 0.09446 | 1.07385 | 0.00234 | 0.06290 | 0.00810 | 0.17206 | 0.00683 |
| **Bbx** | 0.00000 | 2.64813 | 0.00000 | 0.00000 | 0.03404 | 0.09613 | 0.01537 | 0.20132 | 1.02377 | 1.47617 | 0.58366 | 0.45810 | 3.10882 | 18.53137 | 5.54102 | 0.03353 |
| **BC002163** | 4.43766 | 0.24079 | 3.57172 | 0.80034 | 3.36278 | 1.15876 | 0.37511 | 0.58424 | 0.23911 | 2.32908 | 8.04445 | 0.20516 | 0.92004 | 0.00950 | 0.74303 | 0.93573 |
| **BC003331** | 2.65896 | 0.00003 | 1.18073 | 1.90209 | 0.81870 | 3.52631 | 2.69661 | 4.37426 | 1.46438 | 1.97355 | 7.89251 | 0.45464 | 3.35914 | 8.26375 | 4.55963 | 2.23997 |
| **BC003965** | 6.53396 | 0.06715 | 2.59692 | 6.71459 | 4.92306 | 5.10331 | 2.27579 | 8.35688 | 9.12475 | 10.68347 | 5.81546 | 0.66701 | 6.11165 | 3.62590 | 2.56850 | 5.01027 |
| **BC004004** | 6.18567 | 3.69486 | 2.23838 | 2.10851 | 4.53728 | 3.80916 | 2.46720 | 3.07112 | 0.49491 | 1.57129 | 1.58950 | 4.39524 | 4.13026 | 0.89966 | 1.38060 | 0.07378 |
| **BC005537** | 9.32221 | 9.82798 | 7.69174 | 6.02355 | 6.90777 | 3.35518 | 5.59089 | 10.11501 | 6.75892 | 6.13736 | 1.90632 | 2.59393 | 1.27792 | 8.82255 | 6.26511 | 0.97636 |
| **BC005561** | 0.41754 | 0.00686 | 0.50331 | 1.35972 | 0.00612 | 0.00518 | 0.00665 | 0.23213 | 0.00000 | 1.13840 | 0.96879 | 0.00000 | 0.00000 | 1.79033 | 2.39183 | 0.01375 |
| **BC005624** | 9.21560 | 4.32207 | 6.44884 | 1.25282 | 6.27213 | 0.01511 | 6.69547 | 5.31543 | 3.29861 | 5.96483 | 2.66808 | 1.34391 | 5.46503 | 3.27170 | 5.51247 | 0.48166 |
| **BC006965** | 0.01896 | 0.00765 | 0.01474 | 0.02538 | 0.00874 | 0.00719 | 0.04743 | 0.00981 | 0.02949 | 0.01658 | 0.02442 | 0.02525 | 0.02465 | 0.28355 | 0.38260 | 0.05050 |
| **BC017158** | 3.96022 | 0.00320 | 0.24808 | 0.49721 | 0.26889 | 0.52092 | 0.19648 | 0.43183 | 0.36853 | 0.00000 | 0.00758 | 0.00301 | 0.99335 | 0.01431 | 0.00000 | 2.32371 |
| **BC017643** | 1.51737 | 0.00513 | 2.16229 | 0.58781 | 0.01683 | 5.71920 | 0.00484 | 0.62309 | 10.22527 | 0.08143 | 0.64480 | 2.44236 | 0.00000 | 1.78253 | 6.23856 | 0.02411 |
| **BC018473** | 0.04167 | 0.00680 | 0.03957 | 0.29656 | 0.00890 | 0.00614 | 1.18039 | 0.02333 | 0.42105 | 0.02719 | 0.12643 | 0.06986 | 0.02223 | 0.27710 | 0.06509 | 0.00723 |
| **BC020402** | 0.01078 | 0.07754 | 0.00599 | 0.00638 | 0.00904 | 0.00668 | 0.29758 | 0.05967 | 0.14176 | 0.01223 | 0.03374 | 0.00012 | 0.00963 | 1.57376 | 0.00656 | 0.00751 |
| **BC021614** | 1.94569 | 0.02544 | 0.05749 | 1.68670 | 0.00423 | 0.00656 | 0.00473 | 0.00000 | 1.44496 | 1.25648 | 0.86812 | 0.02124 | 0.87376 | 0.08779 | 0.76015 | 0.00439 |
| **BC021767** | 1.92992 | 0.00454 | 0.61990 | 2.36694 | 0.09816 | 2.75154 | 1.50264 | 2.21494 | 1.62657 | 1.35966 | 1.15605 | 0.10792 | 0.32508 | 0.85728 | 0.69064 | 0.18615 |
| **BC021891** | 0.01368 | 0.00778 | 0.01197 | 0.58196 | 0.00938 | 0.00650 | 0.00736 | 0.01144 | 0.06193 | 0.18795 | 0.02650 | 0.00686 | 0.01211 | 0.03290 | 0.01124 | 0.00734 |

| | | | | | | | | | | | | | | | | |
|---|---|---|---|---|---|---|---|---|---|---|---|---|---|---|---|---|
| **BC022687** | 0.02040 | 0.00712 | 0.04333 | 0.37778 | 0.01708 | 0.01149 | 0.17189 | 1.06086 | 1.24636 | 5.16138 | 0.33607 | 0.08115 | 0.10738 | 0.00486 | 0.88193 | 0.00700 |
| **BC024386** | 0.29968 | 0.00726 | 0.23924 | 1.29098 | 0.05484 | 0.01791 | 0.02686 | 0.00000 | 1.75325 | 0.02785 | 1.74548 | 0.05128 | 0.07686 | 0.63494 | 0.04022 | 0.03573 |
| **BC024978** | 0.19862 | 0.07250 | 0.12944 | 0.48669 | 0.02861 | 0.02285 | 0.00695 | 0.00422 | 1.08252 | 0.31442 | 1.02415 | 0.20979 | 0.03417 | 0.22285 | 0.75864 | 0.63952 |
| **BC025920** | 0.01437 | 0.00782 | 0.00665 | 1.28318 | 0.00891 | 0.00621 | 0.00693 | 0.00783 | 2.82249 | 0.19113 | 0.00000 | 0.00000 | 0.38902 | 0.73344 | 0.76124 | 0.00681 |
| **BC026585** | 1.92558 | 1.40253 | 1.59712 | 0.69081 | 0.11295 | 0.12518 | 0.00512 | 2.07537 | 0.46670 | 2.03511 | 0.00000 | 0.02280 | 0.24592 | 0.00687 | 0.00000 | 0.55754 |
| **BC028528** | 0.32658 | 0.00693 | 0.26861 | 0.25533 | 0.37952 | 0.00684 | 0.20184 | 0.26430 | 0.04075 | 0.01510 | 0.21000 | 0.03063 | 0.28645 | 1.15025 | 0.23693 | 0.00678 |
| **BC029214** | 0.00807 | 0.00746 | 0.22950 | 0.01097 | 0.01870 | 0.00802 | 0.00707 | 0.49376 | 0.01175 | 0.02937 | 0.00977 | 0.01540 | 0.00486 | 0.01398 | 0.00132 | 0.02241 |
| **BC029722** | 0.01318 | 0.00802 | 0.01138 | 0.00913 | 0.00903 | 0.00680 | 0.00742 | 0.01182 | 0.01202 | 0.35996 | 0.01136 | 0.03102 | 0.09284 | 0.01167 | 0.01060 | 0.00731 |
| **BC030336** | 0.95412 | 0.00709 | 0.00000 | 1.00343 | 0.01599 | 0.00502 | 0.00579 | 0.21785 | 0.61342 | 1.01937 | 0.59249 | 0.27833 | 0.37769 | 4.32765 | 2.14250 | 0.00591 |
| **BC030867** | 4.25677 | 0.03932 | 10.20074 | 11.11127 | 3.62968 | 8.10702 | 2.56449 | 16.90345 | 14.60150 | 17.92691 | 8.83046 | 1.65299 | 3.34571 | 0.89116 | 11.16934 | 6.73563 |
| **BC031181** | 13.62988 | 6.72391 | 19.82377 | 9.03860 | 9.92003 | 6.25822 | 10.70337 | 17.46169 | 5.96151 | 19.11907 | 8.16949 | 10.93922 | 13.00649 | 0.74743 | 5.41375 | 8.20157 |
| **BC031361** | 0.04903 | 0.00827 | 0.00419 | 0.83502 | 0.00794 | 0.00703 | 0.00818 | 0.46292 | 2.12082 | 0.00266 | 0.00000 | 0.00429 | 0.00577 | 0.00908 | 0.43735 | 0.00683 |
| **BC037034** | 1.75793 | 0.00547 | 0.05913 | 0.93690 | 0.01115 | 0.00426 | 0.24790 | 2.55866 | 1.02535 | 0.86053 | 0.72459 | 0.00001 | 0.74665 | 0.00156 | 0.10397 | 0.00501 |
| **BC037704** | 0.01253 | 0.00809 | 0.01459 | 0.80733 | 0.00869 | 4.67704 | 0.00690 | 0.00270 | 5.60529 | 0.92719 | 0.13319 | 0.00000 | 0.00000 | 0.02399 | 0.01516 | 0.00492 |
| **BC039771** | 0.21990 | 0.03845 | 0.37014 | 0.18633 | 0.05910 | 0.01761 | 0.08794 | 0.04818 | 0.08621 | 0.10468 | 0.87293 | 0.07078 | 0.41298 | 0.49827 | 0.21029 | 0.05005 |
| **BC046251** | 0.01633 | 0.00776 | 0.03078 | 0.03144 | 0.01840 | 0.00671 | 0.00738 | 0.02782 | 0.11042 | 0.03367 | 0.03838 | 0.00715 | 0.01339 | 0.07228 | 0.02792 | 0.01509 |
| **BC048403** | 0.09356 | 0.19607 | 0.02240 | 0.02972 | 0.01224 | 0.00629 | 0.00731 | 0.07342 | 0.19940 | 0.54084 | 0.03938 | 0.00551 | 0.01689 | 0.01686 | 0.04737 | 0.00747 |
| **BC048507** | 0.48790 | 0.03825 | 0.32399 | 0.23990 | 0.42721 | 0.09663 | 0.17755 | 0.13741 | 0.63721 | 0.36801 | 0.17594 | 0.04362 | 0.13587 | 0.04597 | 0.00000 | 0.32182 |
| **BC048546** | 0.01201 | 0.00726 | 0.68245 | 0.00443 | 0.00735 | 0.01128 | 0.44442 | 0.00812 | 0.01062 | 6.22243 | 0.00826 | 0.00000 | 0.00891 | 0.03353 | 0.00000 | 0.01478 |
| **BC048679** | 0.91134 | 3.14537 | 1.88789 | 0.00149 | 0.38812 | 0.65998 | 3.45383 | 0.99801 | 0.00949 | 1.14660 | 1.90395 | 0.00761 | 6.63847 | 0.00506 | 0.00491 | 1.20228 |
| **BC049715** | 0.02621 | 0.97270 | 0.00526 | 0.03457 | 0.00854 | 0.00704 | 0.01722 | 0.06237 | 0.11274 | 0.07797 | 0.03758 | 0.01639 | 0.06895 | 0.07907 | 0.05252 | 0.00700 |
| **BC051019** | 0.00000 | 0.00569 | 0.98845 | 0.87647 | 3.96734 | 0.01721 | 0.24126 | 0.35604 | 0.35887 | 3.14548 | 0.00516 | 0.01752 | 0.87207 | 0.01100 | 0.05708 | 2.95363 |
| **BC051142** | 0.00227 | 0.03802 | 0.10937 | 0.00557 | 0.01047 | 0.00850 | 1.32971 | 1.75738 | 0.02930 | 0.00000 | 0.20509 | 0.02090 | 0.00372 | 0.01313 | 0.00000 | 0.00714 |
| **BC051226** | 0.02036 | 0.00812 | 0.01024 | 0.24295 | 0.00836 | 0.00652 | 0.00724 | 0.03056 | 0.42009 | 1.54492 | 0.00822 | 0.02565 | 0.40547 | 0.01222 | 0.00396 | 0.04267 |
| **BC051537** | 0.01154 | 0.00805 | 0.08786 | 0.02516 | 0.00903 | 0.00884 | 0.00725 | 0.02801 | 0.05097 | 0.01941 | 0.01138 | 0.00834 | 0.12006 | 0.00835 | 0.07657 | 0.00726 |
| **BC051665** | 0.01103 | 0.00767 | 0.01064 | 0.00797 | 0.02460 | 0.00638 | 0.00716 | 0.00981 | 0.01209 | 0.65904 | 0.01193 | 0.03368 | 0.01776 | 0.03251 | 0.00445 | 2.08510 |
| **BC052040** | 4.87324 | 6.33335 | 5.43854 | 0.00000 | 0.03009 | 0.25327 | 0.03966 | 6.47557 | 0.13945 | 5.61642 | 7.53387 | 1.09938 | 1.67927 | 0.66641 | 0.42412 | 0.03999 |
| **BC052688** | 8.82523 | 14.01165 | 19.58819 | 10.94618 | 9.07587 | 9.14270 | 13.41704 | 12.75338 | 5.46016 | 11.13814 | 11.03207 | 13.09076 | 16.01783 | 22.19627 | 10.97181 | 6.81691 |
| **BC053393** | 0.00000 | 10.86533 | 6.86946 | 0.21422 | 0.13257 | 11.85931 | 8.67123 | 0.00788 | 0.42839 | 0.00000 | 0.00000 | 0.05463 | 4.93951 | 0.01786 | 0.05613 | 4.57321 |
| **BC055324** | 3.45632 | 8.28994 | 3.60294 | 1.06170 | 4.65726 | 0.00226 | 0.95548 | 9.17037 | 0.49705 | 7.53881 | 2.73583 | 5.54285 | 3.21842 | 5.44633 | 10.50222 | 3.03202 |
| **BC064078** | 0.01247 | 0.00807 | 0.08807 | 0.22890 | 0.00895 | 0.00906 | 0.00739 | 0.01220 | 0.19022 | 0.06224 | 0.01025 | 0.00808 | 0.04681 | 0.00602 | 0.22280 | 0.00730 |
| **BC065397** | 0.00000 | 3.02998 | 0.87417 | 0.00483 | 0.04872 | 0.00789 | 2.66495 | 0.35375 | 0.01238 | 0.00000 | 0.21766 | 0.00399 | 0.00333 | 0.00566 | 0.01492 | 0.00605 |
| **BC068281** | 0.01500 | 0.01783 | 0.53886 | 0.54201 | 0.02636 | 0.01140 | 0.18085 | 0.10115 | 0.63900 | 0.18823 | 0.00000 | 0.41578 | 0.03033 | 0.03172 | 0.29355 | 0.00703 |
| **BC080695** | 0.01260 | 0.01828 | 0.00912 | 2.86663 | 0.00910 | 0.00604 | 0.00722 | 0.01332 | 5.66646 | 0.00000 | 0.00000 | 0.15397 | 0.01302 | 0.05755 | 0.08836 | 0.00647 |
| **BC100530** | 0.07783 | 0.00797 | 0.03232 | 0.10952 | 0.00852 | 0.04030 | 0.00759 | 0.01609 | 0.41124 | 0.02270 | 0.03880 | 0.00741 | 0.03388 | 0.01311 | 0.00405 | 0.00723 |
| **BC147527** | 0.00697 | 0.00823 | 0.00812 | 8.72124 | 0.00916 | 0.00495 | 0.02926 | 0.51913 | 11.31265 | 0.39888 | 0.00008 | 0.00000 | 0.00619 | 0.00000 | 0.45923 | 0.00504 |
| **Bcam** | 0.02122 | 0.00791 | 0.01295 | 0.06343 | 0.00932 | 0.00715 | 0.00766 | 0.01571 | 0.01136 | 0.05586 | 0.01531 | 0.00735 | 0.07642 | 0.01475 | 0.03173 | 0.01584 |
| **Bcan** | 0.01325 | 0.00731 | 0.00437 | 0.16127 | 0.00899 | 0.00701 | 0.00696 | 0.00000 | 0.60344 | 0.21554 | 0.30573 | 0.40986 | 0.01616 | 0.42564 | 0.18834 | 0.00760 |
| **Bcap29** | 5.24945 | 4.19483 | 3.50008 | 1.63407 | 0.00000 | 0.05703 | 4.47944 | 1.03377 | 3.44615 | 5.02085 | 4.40196 | 0.64080 | 2.29788 | 8.49867 | 2.87439 | 0.01057 |
| **Bcap31** | 17.69664 | 12.20695 | 26.32940 | 8.53183 | 9.19631 | 3.53018 | 11.96515 | 19.95572 | 9.13326 | 14.28190 | 15.08752 | 14.45006 | 17.34470 | 23.13972 | 19.61936 | 15.42335 |
| **Bcar1** | 1.12373 | 0.00370 | 0.09998 | 5.50635 | 5.22497 | 0.00512 | 0.00582 | 2.64278 | 1.29221 | 0.72374 | 0.48583 | 0.07284 | 0.00000 | 0.00000 | 0.00000 | 0.01920 |
| **Bcar3** | 0.01095 | 0.00772 | 0.00645 | 1.78585 | 0.00926 | 0.00629 | 0.00706 | 0.00940 | 0.00236 | 0.35146 | 0.00000 | 0.03566 | 0.01178 | 1.32298 | 0.78345 | 0.13859 |
| **Bcas1** | 0.01757 | 0.00791 | 0.01130 | 0.53322 | 0.00881 | 0.00648 | 0.00729 | 0.01094 | 1.18268 | 0.40881 | 0.00000 | 0.00385 | 0.01253 | 0.00827 | 0.24749 | 0.00760 |
| **Bcas2** | 17.85537 | 16.05001 | 16.68721 | 10.40821 | 13.03536 | 5.23912 | 16.12178 | 21.16512 | 7.00569 | 17.80592 | 17.75798 | 13.73976 | 21.25929 | 8.43122 | 1.67330 | 14.16605 |
| **Bcas3** | 10.04854 | 3.73146 | 9.21791 | 3.93431 | 3.78194 | 4.87142 | 1.27185 | 17.00423 | 6.36116 | 12.53705 | 15.68404 | 2.10287 | 8.53728 | 7.96867 | 9.33192 | 6.29722 |
| **Bcas3os1** | 0.04781 | 0.00807 | 0.03535 | 0.16658 | 0.00975 | 0.00818 | 0.01821 | 0.11915 | 0.23866 | 0.09643 | 0.27345 | 0.00605 | 0.02197 | 0.15974 | 0.11977 | 0.00734 |
| **Bcat1** | 17.31323 | 6.58585 | 20.65379 | 3.91909 | 19.20495 | 9.41488 | 11.23623 | 30.28744 | 4.10218 | 28.83802 | 29.26491 | 6.50635 | 20.86586 | 22.23090 | 18.29923 | 3.81744 |
| **Bcat2** | 5.67593 | 0.19115 | 9.36878 | 0.99814 | 2.49885 | 13.92884 | 2.51065 | 6.76811 | 1.80357 | 5.19401 | 4.72236 | 0.46911 | 9.83411 | 0.43264 | 1.56431 | 2.78484 |
| **Bccip** | 5.55445 | 8.77170 | 4.49986 | 0.00000 | 9.29807 | 5.74492 | 0.33435 | 5.82954 | 0.08021 | 1.56578 | 0.00000 | 0.09626 | 4.10860 | 8.78253 | 5.02906 | 2.81624 |
| **Bcdin3d** | 2.37437 | 2.22014 | 4.65519 | 1.12021 | 0.00059 | 0.00282 | 1.30053 | 2.53988 | 1.52809 | 1.23593 | 0.00000 | 0.02448 | 0.70643 | 0.00445 | 0.20735 | 1.11103 |
| **Bckdha** | 6.01729 | 4.01750 | 2.79597 | 0.91081 | 0.09762 | 0.08233 | 4.21870 | 9.94952 | 0.03253 | 11.35314 | 2.89826 | 0.47145 | 5.06942 | 0.08881 | 0.00000 | 1.25384 |
| **Bckdhb** | 1.17359 | 2.20770 | 3.53843 | 0.77677 | 0.37692 | 1.49326 | 3.07142 | 7.01934 | 1.69284 | 7.54492 | 0.44650 | 4.65976 | 2.39680 | 6.02964 | 3.56610 | 0.47071 |
| **Bckdk** | 12.32543 | 0.50629 | 5.37820 | 1.24515 | 0.38918 | 2.23674 | 5.85450 | 10.76024 | 1.80500 | 8.60679 | 3.59723 | 0.47554 | 7.04483 | 0.39009 | 2.18396 | 9.77835 |
| **Bcl10** | 6.04049 | 0.40040 | 1.20338 | 1.65264 | 2.11519 | 5.04028 | 3.33895 | 4.49435 | 0.49127 | 2.03590 | 0.38767 | 0.94912 | 1.45337 | 2.17724 | 2.14049 | 1.78776 |
| **Bcl11b** | 0.44806 | 0.23356 | 0.00356 | 0.02090 | 0.01670 | 0.00723 | 0.00676 | 0.00000 | 0.05940 | 0.20121 | 0.31883 | 0.06326 | 0.00845 | 0.10653 | 0.00805 | 0.00671 |
| **Bcl2** | 0.01344 | 0.00824 | 0.01156 | 0.04660 | 0.00916 | 0.00687 | 0.00770 | 0.01703 | 0.01170 | 0.17228 | 0.02457 | 0.02085 | 0.01912 | 0.01039 | 0.01881 | 0.00737 |
| **Bcl2l1** | 2.35289 | 2.08898 | 4.04548 | 2.50169 | 3.02715 | 6.96130 | 1.60096 | 6.06395 | 1.77458 | 6.87674 | 2.00660 | 0.24140 | 2.80799 | 0.00482 | 0.00000 | 6.35178 |
| **Bcl2l10** | 0.01937 | 0.00811 | 0.00326 | 24.71215 | 0.00808 | 0.00537 | 0.01726 | 0.04421 | 6.02371 | 0.37878 | 1.70741 | 0.01816 | 0.00000 | 0.08790 | 0.24785 | 0.24016 |
| **Bcl2l11** | 1.64801 | 0.00489 | 0.80452 | 7.03593 | 0.04028 | 0.08521 | 0.02591 | 1.31108 | 9.63805 | 3.58679 | 4.08471 | 3.16811 | 1.71298 | 1.94785 | 0.80170 | 2.10185 |
| **Bcl2l12** | 0.01658 | 0.00725 | 0.28895 | 0.00338 | 0.00736 | 1.90428 | 0.06526 | 0.10717 | 0.11795 | 1.73258 | 0.17001 | 0.03106 | 1.00908 | 0.01686 | 0.00020 | 0.97912 |
| **Bcl2l13** | 6.58805 | 0.16670 | 1.78106 | 13.43023 | 0.44777 | 1.87420 | 1.48967 | 5.49400 | 15.84763 | 13.94567 | 8.78790 | 9.24330 | 4.86119 | 10.64746 | 14.84287 | 3.14816 |
| **Bcl2l14** | 6.66739 | 5.84145 | 1.06624 | 1.81810 | 0.10663 | 0.00887 | 1.33888 | 9.01283 | 6.51093 | 17.98461 | 10.02535 | 3.95309 | 3.41443 | 5.87110 | 7.61808 | 0.01518 |

| | | | | | | | | | | | | | | | | |
|---|---|---|---|---|---|---|---|---|---|---|---|---|---|---|---|---|
| **Bcl2l15** | 0.00968 | 0.00792 | 0.01029 | 0.04425 | 0.00929 | 0.00681 | 0.00747 | 0.35380 | 0.45080 | 0.75172 | 0.00600 | 0.00501 | 0.00657 | 0.03055 | 0.00000 | 0.00757 |
| **Bcl2l2** | 0.67226 | 0.00699 | 0.00000 | 0.95156 | 0.22835 | 0.77237 | 0.00638 | 0.42145 | 0.02273 | 1.08054 | 0.07159 | 0.10046 | 0.75462 | 1.96922 | 4.32558 | 0.04110 |
| **Bcl3** | 5.67138 | 0.13569 | 0.00000 | 0.27049 | 5.39444 | 2.39343 | 1.90613 | 1.33435 | 1.31914 | 1.05101 | 0.90158 | 0.03487 | 0.78426 | 0.00335 | 0.00000 | 0.24380 |
| **Bcl7a** | 2.10726 | 0.00531 | 1.56841 | 0.08143 | 0.00415 | 2.24309 | 0.00458 | 1.21241 | 0.78795 | 6.30086 | 2.58495 | 0.22323 | 0.00000 | 0.35990 | 0.79850 | 1.25806 |
| **Bcl7b** | 4.79461 | 2.42293 | 2.83162 | 1.33260 | 0.02439 | 0.00182 | 0.02445 | 2.78618 | 1.04500 | 4.90200 | 1.58634 | 0.28798 | 1.43861 | 0.00810 | 0.22196 | 1.28492 |
| **Bcl7c** | 0.97796 | 0.06590 | 0.63763 | 13.76559 | 0.04841 | 0.02208 | 0.00597 | 0.22404 | 3.80405 | 2.00705 | 1.03387 | 0.10314 | 0.35445 | 0.01709 | 0.61953 | 0.00352 |
| **Bcl9** | 0.01270 | 0.00783 | 0.04230 | 2.08346 | 0.00917 | 0.01157 | 0.00744 | 0.01175 | 0.83526 | 0.28921 | 0.00684 | 0.00527 | 0.01157 | 0.00220 | 0.30532 | 0.01414 |
| **Bcl9l** | 1.44086 | 0.02948 | 0.07142 | 0.00971 | 0.01711 | 0.00425 | 0.00652 | 0.00759 | 0.01241 | 0.18111 | 0.01150 | 0.00698 | 0.01568 | 0.03233 | 0.00721 | 0.48740 |
| **Bclaf1** | 13.08742 | 13.09920 | 11.38077 | 19.14134 | 9.39946 | 7.48764 | 10.45169 | 17.81240 | 25.39272 | 19.44078 | 17.89277 | 17.04972 | 9.08616 | 32.32967 | 26.41405 | 3.90181 |
| **Bco1** | 0.00335 | 0.01731 | 0.06098 | 0.11623 | 0.02884 | 0.01318 | 0.03642 | 0.34019 | 0.02570 | 0.81031 | 0.00004 | 0.00000 | 0.02361 | 2.29147 | 0.00000 | 0.01464 |
| **Bco2** | 0.01247 | 0.00791 | 0.01164 | 0.00876 | 0.00922 | 0.00647 | 0.00713 | 0.00990 | 0.01233 | 0.39999 | 0.01230 | 0.00606 | 0.21092 | 0.07109 | 0.00383 | 0.00734 |
| **Bcor** | 4.42271 | 0.00000 | 0.28691 | 4.75723 | 4.07971 | 0.10105 | 0.86288 | 3.92035 | 3.34140 | 2.11160 | 0.38987 | 0.71828 | 1.61667 | 0.00000 | 3.91706 | 2.41451 |
| **Bcorl1** | 0.77481 | 0.00654 | 0.88000 | 7.95822 | 0.27435 | 0.00377 | 0.00664 | 1.34493 | 4.39831 | 0.03237 | 0.00000 | 0.01483 | 0.00229 | 0.00498 | 0.76945 | 0.00482 |
| **Bcr** | 4.48431 | 2.18439 | 0.00000 | 3.43794 | 4.06409 | 2.09819 | 0.01303 | 0.40350 | 2.44212 | 1.04526 | 0.00000 | 0.76540 | 0.77273 | 0.00000 | 0.45476 | 0.44962 |
| **Bcs1l** | 7.84920 | 6.36659 | 3.68581 | 1.02311 | 3.54962 | 2.07239 | 3.89483 | 1.59375 | 0.01605 | 2.84643 | 2.20895 | 1.11930 | 4.38000 | 0.08733 | 2.25275 | 1.53732 |
| **Bdh1** | 1.97738 | 1.05790 | 2.73608 | 0.82565 | 2.44362 | 1.61025 | 3.38852 | 6.14229 | 3.47927 | 2.80045 | 1.80192 | 6.27622 | 7.06484 | 0.24683 | 0.24104 | 3.10525 |
| **Bdh2** | 0.01362 | 0.00779 | 0.01216 | 0.10629 | 0.00938 | 0.00647 | 0.00740 | 0.01175 | 0.56878 | 0.15818 | 0.02482 | 0.00573 | 0.01228 | 0.00692 | 0.04951 | 0.00742 |
| **Bdp1** | 1.64459 | 0.00247 | 2.21760 | 4.59522 | 0.00000 | 1.69878 | 0.47281 | 4.06086 | 6.88596 | 6.28947 | 3.75681 | 7.03041 | 0.54163 | 12.21865 | 10.08853 | 0.56190 |
| **Becn1** | 6.22462 | 4.08813 | 1.98129 | 4.66857 | 5.83303 | 0.98553 | 2.69818 | 6.71314 | 1.94062 | 9.28779 | 0.57601 | 6.40818 | 7.14515 | 7.43241 | 5.74088 | 7.01847 |
| **Becn2** | 0.32532 | 5.53145 | 17.72167 | 8.75904 | 2.33104 | 3.15003 | 4.14739 | 17.55006 | 14.85996 | 13.94275 | 12.70382 | 2.09819 | 11.15769 | 14.98585 | 15.14524 | 0.02394 |
| **Bend3** | 1.44489 | 0.65923 | 0.22634 | 9.07877 | 0.95776 | 1.89587 | 1.00621 | 3.10674 | 13.28079 | 2.68982 | 6.57110 | 7.21931 | 1.15004 | 14.40405 | 9.29916 | 0.02998 |
| **Bend4** | 3.43949 | 0.01458 | 0.00000 | 0.61273 | 0.00000 | 7.33976 | 2.07335 | 0.32612 | 0.26001 | 9.07143 | 1.52632 | 0.31486 | 2.49856 | 0.04598 | 2.36965 | 0.05737 |
| **Bend5** | 0.04326 | 0.00849 | 0.00946 | 0.23066 | 0.01352 | 0.00658 | 0.00737 | 0.02212 | 0.14875 | 0.00982 | 0.01265 | 0.00762 | 0.00847 | 0.01107 | 0.00978 | 1.57261 |
| **Bend6** | 0.00959 | 0.00786 | 0.00563 | 0.50966 | 0.00883 | 0.00637 | 0.00705 | 0.00847 | 0.00347 | 0.00000 | 0.01053 | 0.00868 | 0.00968 | 2.35871 | 0.19333 | 0.93848 |
| **Bend7** | 0.70456 | 0.00722 | 0.03598 | 0.00487 | 0.01395 | 0.00505 | 0.03519 | 0.36671 | 0.25167 | 0.00000 | 0.02153 | 0.00534 | 1.13046 | 0.01675 | 0.00185 | 0.00602 |
| **Best2** | 0.01240 | 0.00744 | 0.00611 | 4.62635 | 0.00856 | 0.00675 | 0.00655 | 0.00000 | 0.78715 | 2.25869 | 1.25402 | 0.00000 | 0.01322 | 0.00000 | 0.04568 | 0.00633 |
| **Bet1** | 2.90338 | 0.06169 | 1.80252 | 3.36069 | 7.18800 | 0.92787 | 0.17146 | 0.86758 | 0.52395 | 1.68473 | 4.34684 | 3.92247 | 1.22646 | 0.07954 | 1.93226 | 0.36308 |
| **Bet1l** | 0.45236 | 3.85485 | 3.41502 | 3.31582 | 6.11410 | 2.31821 | 0.00472 | 4.53384 | 0.38701 | 4.00981 | 0.83795 | 0.27627 | 0.93816 | 0.11287 | 6.33140 | 2.75447 |
| **Bex1** | 0.02991 | 0.00788 | 0.00627 | 2.58970 | 0.01390 | 0.00728 | 0.00703 | 1.34397 | 0.69174 | 0.29426 | 0.00000 | 0.04738 | 0.26045 | 0.00853 | 0.00000 | 0.00736 |
| **Bex2** | 0.23034 | 0.00775 | 0.00618 | 0.38956 | 0.01652 | 0.00773 | 0.00729 | 0.10158 | 0.00615 | 0.08059 | 0.01502 | 0.00709 | 0.04565 | 0.01021 | 0.00473 | 0.00748 |
| **Bex4** | 0.24283 | 0.00808 | 0.00426 | 0.00727 | 0.07016 | 0.00747 | 0.00652 | 0.31349 | 0.01139 | 0.98749 | 0.00932 | 0.02338 | 0.85749 | 0.01184 | 0.00739 | 0.00722 |
| **Bex6** | 0.00000 | 0.00551 | 0.24860 | 8.39958 | 0.01578 | 0.00396 | 0.00180 | 6.04053 | 7.88226 | 15.33588 | 11.73588 | 1.09470 | 0.87464 | 11.44466 | 15.83707 | 0.00982 |
| **Bfar** | 2.57743 | 4.30771 | 6.08747 | 8.28568 | 1.06914 | 0.28945 | 4.65472 | 4.58801 | 8.27512 | 5.31387 | 1.91451 | 0.56871 | 0.34690 | 12.84844 | 7.33282 | 2.44209 |
| **Bfsp1** | 0.01357 | 0.00773 | 0.79358 | 0.01195 | 0.00778 | 0.00668 | 0.05829 | 0.01066 | 3.29284 | 0.00666 | 0.00000 | 0.00000 | 0.00896 | 0.01625 | 0.00000 | 0.00682 |
| **Bfsp2** | 2.39249 | 0.00597 | 0.05778 | 0.23386 | 0.00958 | 0.00325 | 0.00575 | 0.00678 | 0.00811 | 0.05127 | 0.01287 | 0.02776 | 0.02382 | 0.01357 | 0.01039 | 0.00499 |
| **Bglap** | 0.04911 | 0.00824 | 0.11246 | 0.00645 | 0.01292 | 0.00644 | 0.00737 | 0.01261 | 0.10086 | 0.01474 | 0.01187 | 0.00773 | 0.01112 | 0.01293 | 0.00807 | 0.00742 |
| **Bglap2** | 0.08315 | 0.00812 | 0.15035 | 0.00540 | 0.01195 | 0.00597 | 0.00733 | 0.01243 | 0.21040 | 0.01411 | 0.01091 | 0.00752 | 0.01080 | 0.01340 | 0.00668 | 0.00733 |
| **Bglap3** | 0.53521 | 0.00786 | 0.23239 | 0.05323 | 0.04735 | 0.22377 | 0.00695 | 0.00934 | 0.23063 | 0.01861 | 0.04045 | 0.00690 | 0.02268 | 0.01264 | 0.00335 | 0.00672 |
| **Bgn** | 0.01222 | 0.00743 | 0.01398 | 0.00593 | 0.00872 | 0.00754 | 0.00713 | 0.00982 | 0.01090 | 1.89412 | 0.00545 | 0.57669 | 0.01120 | 0.01212 | 0.00000 | 0.00752 |
| **Bhlha15** | 0.01922 | 0.00761 | 0.02051 | 0.00635 | 0.00850 | 0.02200 | 0.00723 | 0.01437 | 0.02499 | 1.26694 | 0.01216 | 0.00519 | 0.01690 | 0.02045 | 0.01592 | 0.00713 |
| **Bhlhb9** | 0.32368 | 0.00398 | 1.09915 | 1.49414 | 6.13303 | 0.00507 | 0.00650 | 0.85868 | 0.00000 | 0.83129 | 0.99300 | 0.06680 | 0.00000 | 1.03156 | 8.33767 | 0.00603 |
| **Bhlhe40** | 0.01729 | 2.56593 | 0.23478 | 0.80518 | 0.00714 | 0.04402 | 0.03722 | 0.00000 | 0.83413 | 0.00000 | 4.52578 | 0.01599 | 0.00000 | 0.00000 | 0.20287 | 0.00504 |
| **Bhlhe41** | 0.01323 | 0.00764 | 0.01095 | 0.68331 | 0.00914 | 0.00626 | 0.22737 | 0.00024 | 1.17358 | 0.00000 | 0.34007 | 0.01237 | 0.00847 | 0.00303 | 0.43950 | 0.00691 |
| **Bhmt** | 54.48173 | 33.08203 | 50.26210 | 0.58862 | 34.20086 | 23.63597 | 33.85645 | 55.41694 | 6.99901 | 55.29181 | 52.29159 | 22.13828 | 51.58990 | 24.95384 | 22.64116 | 10.89185 |
| **Bhmt2** | 4.67360 | 1.90388 | 4.28007 | 0.60438 | 0.83844 | 4.99996 | 1.74552 | 19.35663 | 0.30658 | 9.83826 | 8.34052 | 5.67548 | 8.18777 | 3.51396 | 2.60692 | 0.02944 |
| **Bicc1** | 0.00914 | 0.29227 | 0.01628 | 0.29262 | 0.00841 | 0.00728 | 0.00725 | 0.01064 | 0.52809 | 0.01014 | 0.02079 | 0.01681 | 0.01193 | 0.00866 | 0.00000 | 0.00718 |
| **Bicd2** | 1.67196 | 0.00508 | 0.00000 | 6.06614 | 0.00736 | 0.94583 | 1.35639 | 0.71610 | 3.51825 | 0.40847 | 0.00000 | 1.47203 | 0.41878 | 0.00000 | 4.60359 | 0.00410 |
| **Bid** | 2.56463 | 0.79804 | 3.82749 | 4.04647 | 0.00411 | 3.12386 | 4.08872 | 13.30588 | 1.71219 | 7.94650 | 6.14746 | 0.71463 | 5.22345 | 2.50347 | 7.07980 | 0.00000 |
| **Bik** | 0.02524 | 0.00798 | 0.22503 | 1.50722 | 0.00738 | 0.00891 | 0.00719 | 0.01215 | 1.70830 | 0.16240 | 0.00000 | 0.00379 | 0.01115 | 0.00836 | 0.00847 | 0.77801 |
| **Bin1** | 1.61553 | 0.00585 | 0.00000 | 0.01037 | 0.00398 | 0.00640 | 0.01580 | 0.00569 | 0.01243 | 0.05130 | 0.01252 | 0.03460 | 1.08728 | 0.01171 | 0.00949 | 3.02104 |
| **Bin2** | 6.87165 | 0.03624 | 0.23501 | 2.12805 | 7.30789 | 9.92323 | 2.34394 | 7.14980 | 0.02357 | 7.87021 | 2.63146 | 0.65699 | 1.39008 | 0.03139 | 0.22503 | 4.45041 |
| **Bin3** | 3.29638 | 2.24406 | 4.17870 | 2.50866 | 0.08645 | 0.00249 | 0.00219 | 0.94575 | 2.13986 | 2.50612 | 0.00000 | 0.01454 | 1.95577 | 0.00000 | 0.68744 | 0.00000 |
| **Birc2** | 2.80874 | 3.03127 | 3.20982 | 1.52559 | 4.18056 | 0.02184 | 1.09823 | 1.42428 | 5.01931 | 0.47943 | 0.61630 | 0.21460 | 0.00000 | 0.26891 | 4.23780 | 1.07551 |
| **Birc3** | 0.00000 | 0.02589 | 1.93300 | 12.09147 | 3.21354 | 0.00578 | 0.00533 | 1.77249 | 8.26714 | 4.07365 | 2.37613 | 1.92432 | 0.00000 | 10.34604 | 7.85499 | 0.45585 |
| **Birc5** | 25.97633 | 13.85529 | 22.86673 | 15.24819 | 15.77858 | 14.58937 | 12.58517 | 22.47294 | 19.14921 | 22.59412 | 13.43334 | 6.92227 | 20.36219 | 4.48041 | 11.00891 | 10.63143 |
| **Birc6** | 0.84358 | 0.67459 | 1.25712 | 0.76196 | 0.01255 | 0.00592 | 0.60258 | 1.43213 | 3.17383 | 1.54205 | 5.40072 | 0.18374 | 1.53814 | 5.27145 | 5.18316 | 0.82130 |
| **Bivm** | 0.01636 | 0.07186 | 0.01125 | 0.29619 | 0.00876 | 0.00664 | 0.00743 | 0.01284 | 0.08257 | 0.01356 | 0.03072 | 0.00782 | 0.01219 | 0.00988 | 0.00839 | 0.00729 |
| **Blcap** | 2.01769 | 1.53690 | 1.79422 | 5.58922 | 2.20016 | 3.56442 | 2.30851 | 2.12550 | 1.25784 | 2.60902 | 0.71310 | 0.15615 | 0.52672 | 5.06221 | 3.66020 | 3.34375 |
| **Blm** | 2.59459 | 1.43321 | 4.36005 | 6.46406 | 3.94032 | 1.63397 | 4.64091 | 6.62085 | 5.93061 | 4.13319 | 5.82685 | 10.85846 | 2.89526 | 11.17421 | 8.65619 | 0.01134 |
| **Blmh** | 12.96800 | 9.09875 | 4.64207 | 7.89829 | 10.13963 | 4.43121 | 5.19688 | 3.14943 | 2.53944 | 8.25397 | 3.15848 | 0.22681 | 3.51638 | 0.26088 | 4.62590 | 5.31097 |
| **Blnk** | 0.07146 | 0.00713 | 0.29535 | 0.03548 | 0.03009 | 0.31823 | 2.33316 | 0.00901 | 0.62175 | 0.00452 | 0.00755 | 0.00567 | 0.03285 | 0.01577 | 0.00105 | 0.01441 |
| **Bloc1s1** | 4.02194 | 0.46462 | 3.88978 | 0.00000 | 2.94941 | 0.86230 | 1.05417 | 0.65134 | 0.46272 | 3.02605 | 9.18836 | 0.31797 | 2.34857 | 0.00000 | 0.69757 | 1.20909 |

| | | | | | | | | | | | | | | | | |
|---|---|---|---|---|---|---|---|---|---|---|---|---|---|---|---|---|
| **Bloc1s2** | 10.39701 | 3.64818 | 4.83507 | 4.52443 | 7.88106 | 5.26239 | 5.11277 | 5.42965 | 6.82740 | 7.66222 | 5.80831 | 3.04498 | 7.31340 | 2.25206 | 0.24048 | 7.10269 |
| **Bloc1s3** | 0.27612 | 0.01804 | 0.27285 | 1.23137 | 0.26683 | 0.08498 | 0.04855 | 0.07578 | 1.80885 | 0.29989 | 1.64995 | 0.05989 | 0.45009 | 0.91479 | 0.73626 | 0.07245 |
| **Bloc1s4** | 0.17130 | 0.00766 | 0.33667 | 2.90093 | 0.01290 | 0.00521 | 0.01537 | 0.09416 | 6.81345 | 2.66210 | 0.92789 | 0.09983 | 2.21200 | 0.00000 | 0.68726 | 1.41166 |
| **Bloc1s5** | 3.15896 | 8.50839 | 3.11100 | 0.00000 | 7.98129 | 2.48131 | 4.49860 | 2.75538 | 0.07562 | 3.95377 | 2.54204 | 0.71765 | 2.78915 | 0.05017 | 0.00000 | 2.24778 |
| **Bloc1s6** | 5.52520 | 5.34320 | 5.45022 | 0.67512 | 2.60306 | 4.62606 | 6.85175 | 2.45366 | 1.11271 | 2.35736 | 4.52815 | 7.45686 | 3.45398 | 0.34441 | 2.46162 | 0.05213 |
| **Blvra** | 0.64540 | 0.03629 | 1.81157 | 0.21482 | 0.01073 | 0.43956 | 0.00546 | 1.67170 | 1.52102 | 0.69761 | 1.33879 | 0.10202 | 0.96909 | 0.00273 | 0.45582 | 2.64361 |
| **Blvrb** | 3.15669 | 0.61088 | 1.83863 | 0.09835 | 2.88238 | 2.48701 | 0.31105 | 1.25735 | 1.27331 | 1.86746 | 1.33340 | 0.02637 | 0.18846 | 0.00374 | 0.00000 | 1.24660 |
| **Blzf1** | 3.69114 | 2.12290 | 4.95013 | 0.92931 | 0.00000 | 2.63435 | 2.10529 | 3.55933 | 2.31492 | 0.61405 | 2.42516 | 4.84306 | 0.70646 | 8.47019 | 4.78703 | 0.22085 |
| **Bmf** | 1.96398 | 0.00563 | 0.52601 | 1.45512 | 0.00402 | 0.00404 | 0.00584 | 0.55953 | 0.00000 | 0.00000 | 0.05883 | 0.07807 | 0.14931 | 0.57378 | 0.58121 | 0.00481 |
| **Bmi1** | 0.00000 | 7.05777 | 2.32994 | 31.35280 | 6.32212 | 0.02336 | 2.52730 | 0.07000 | 17.80280 | 4.72525 | 1.92752 | 5.97592 | 0.83640 | 1.70760 | 31.99216 | 0.58975 |
| **Bmp1** | 0.62191 | 0.00779 | 0.00216 | 0.07166 | 0.00649 | 0.00756 | 0.01837 | 0.01199 | 0.01081 | 0.00382 | 0.78326 | 0.03896 | 0.01008 | 0.32159 | 0.10691 | 0.00631 |
| **Bmp15** | 0.01735 | 0.00732 | 0.00146 | 10.87646 | 0.00871 | 0.00512 | 0.00659 | 0.00000 | 6.00159 | 0.00000 | 0.25178 | 0.50199 | 0.00942 | 0.00000 | 1.90046 | 0.00485 |
| **Bmp2k** | 0.01402 | 0.00749 | 0.26871 | 1.48790 | 0.00853 | 0.00556 | 0.00710 | 0.00939 | 0.63168 | 0.01221 | 0.00001 | 0.03885 | 0.00946 | 0.00374 | 2.87419 | 0.14601 |
| **Bmp4** | 11.90797 | 0.00000 | 0.00475 | 0.09075 | 2.31625 | 0.00000 | 0.00000 | 0.00000 | 0.00840 | 0.44628 | 0.01200 | 0.07769 | 0.00950 | 0.01578 | 0.01333 | 0.07753 |
| **Bmp5** | 0.01214 | 0.00842 | 0.00961 | 8.31586 | 0.00951 | 0.00580 | 0.00766 | 0.04288 | 0.66813 | 0.00696 | 0.06138 | 0.00517 | 0.02625 | 0.04765 | 0.82358 | 0.01331 |
| **Bmp6** | 2.74579 | 0.00442 | 0.00000 | 10.14169 | 0.00818 | 0.00175 | 0.00408 | 0.43092 | 5.36386 | 1.12519 | 0.54232 | 1.79604 | 0.00000 | 0.00000 | 3.14477 | 2.49338 |
| **Bmp7** | 0.01340 | 0.00781 | 0.01749 | 0.00497 | 0.00926 | 0.00729 | 0.00746 | 0.01221 | 0.40153 | 0.00907 | 0.03266 | 0.02716 | 0.01223 | 0.01811 | 0.03733 | 0.00738 |
| **Bmp8a** | 0.01104 | 0.00791 | 0.01802 | 0.00981 | 0.00868 | 0.00718 | 0.00741 | 0.01844 | 0.02362 | 0.01108 | 0.01268 | 0.00798 | 0.01746 | 0.01259 | 0.01023 | 0.26929 |
| **Bmp8b** | 0.02550 | 0.00769 | 0.01009 | 0.00990 | 0.00861 | 0.00670 | 0.00738 | 0.01797 | 0.02283 | 0.00980 | 0.01120 | 0.00712 | 0.01226 | 0.11158 | 0.12324 | 0.02902 |
| **Bmper** | 0.01327 | 0.00806 | 0.01039 | 0.00665 | 0.00903 | 0.00673 | 0.00752 | 0.00933 | 0.01173 | 0.00805 | 0.19921 | 0.01780 | 0.00967 | 0.01121 | 0.00686 | 0.07187 |
| **Bmpr1a** | 4.03517 | 0.18030 | 1.41346 | 4.31595 | 0.31539 | 6.27619 | 3.06446 | 1.97437 | 5.25276 | 2.50792 | 3.15509 | 10.84069 | 1.78842 | 10.64033 | 3.24933 | 0.20881 |
| **Bmpr1b** | 0.01326 | 0.00796 | 0.00984 | 5.16683 | 0.00932 | 0.00595 | 0.00726 | 0.01058 | 1.20500 | 0.01917 | 0.01636 | 0.00198 | 0.01338 | 0.00000 | 2.19243 | 0.00644 |
| **Bmpr2** | 0.03757 | 0.00609 | 0.83730 | 6.05419 | 0.12413 | 0.92685 | 0.00567 | 0.13175 | 3.05638 | 0.98727 | 1.48293 | 6.21366 | 0.19272 | 4.79105 | 2.60724 | 1.53165 |
| **Bms1** | 2.82721 | 6.12815 | 5.36159 | 5.88890 | 11.11312 | 4.99691 | 6.45519 | 11.32270 | 12.62619 | 8.79527 | 7.96569 | 9.37631 | 6.13253 | 13.84206 | 12.28520 | 3.24152 |
| **Bmt2** | 0.00000 | 2.19159 | 2.99499 | 2.11079 | 1.86365 | 0.00779 | 0.00605 | 0.00205 | 0.83958 | 0.70549 | 0.12017 | 0.14120 | 0.13918 | 0.01043 | 0.56195 | 1.11856 |
| **Bmyc** | 0.00000 | 0.01726 | 0.70396 | 0.00342 | 0.03530 | 0.70621 | 0.13871 | 0.38280 | 0.01040 | 1.74485 | 1.24356 | 0.00000 | 16.59585 | 0.01008 | 0.00000 | 4.13825 |
| **Bnc1** | 0.01282 | 0.00763 | 0.00673 | 3.88300 | 0.00911 | 0.00587 | 0.00723 | 0.00869 | 3.09439 | 0.39782 | 0.02744 | 0.08196 | 0.01249 | 0.00000 | 2.85415 | 0.00660 |
| **Bnc2** | 0.07890 | 0.00867 | 0.00721 | 2.53557 | 0.01469 | 0.00672 | 0.00747 | 0.01308 | 0.50601 | 0.13729 | 0.02175 | 0.01925 | 0.01281 | 0.00167 | 0.76605 | 0.00706 |
| **Bnip1** | 5.18267 | 6.99249 | 4.89196 | 0.62633 | 3.00662 | 0.60669 | 2.01268 | 2.02402 | 0.10655 | 3.39525 | 0.04475 | 0.06364 | 3.07224 | 0.02180 | 0.77875 | 0.57573 |
| **Bnip2** | 0.12449 | 0.02589 | 0.23690 | 1.39002 | 0.00752 | 1.74887 | 0.21386 | 0.94830 | 3.64798 | 3.25806 | 1.16739 | 3.97309 | 0.18838 | 2.01348 | 3.06572 | 0.00548 |
| **Bnip3** | 1.53753 | 1.41151 | 1.38458 | 13.50469 | 2.93075 | 1.83176 | 2.08875 | 8.74120 | 12.57412 | 2.84301 | 23.19908 | 18.24889 | 2.18663 | 19.50365 | 14.17837 | 0.04534 |
| **Bnip3l** | 11.60232 | 12.60018 | 18.82767 | 16.30105 | 15.18248 | 7.09971 | 13.64970 | 30.02320 | 13.04792 | 32.92982 | 22.78898 | 22.65281 | 21.91633 | 26.89250 | 15.20699 | 6.05649 |
| **Bnipl** | 0.62802 | 0.00700 | 0.35243 | 0.02474 | 0.00594 | 0.00645 | 0.00673 | 0.00651 | 0.66168 | 0.57721 | 0.03240 | 0.00252 | 0.01005 | 0.28951 | 0.21420 | 0.00684 |
| **Boc** | 1.20434 | 0.00813 | 0.03901 | 0.00906 | 0.01601 | 0.00788 | 0.05156 | 0.38782 | 0.02662 | 0.06545 | 0.00853 | 0.07362 | 1.41607 | 0.01422 | 0.00433 | 0.00587 |
| **Bod1** | 8.19692 | 3.04142 | 5.04729 | 21.55492 | 1.16905 | 3.48052 | 5.28186 | 7.73020 | 9.62582 | 15.75980 | 8.30188 | 1.38225 | 3.88181 | 11.40048 | 13.69122 | 4.05080 |
| **Bod1l** | 3.44204 | 0.01336 | 3.88526 | 5.89190 | 0.00000 | 0.00000 | 0.69257 | 1.63023 | 9.58113 | 4.56183 | 2.79257 | 2.58551 | 0.38253 | 27.05918 | 19.32144 | 0.32670 |
| **Bok** | 9.33958 | 0.00000 | 2.62281 | 0.95809 | 0.05726 | 4.42756 | 1.76260 | 7.87220 | 0.00000 | 9.07970 | 11.04811 | 0.11509 | 6.00682 | 0.29416 | 0.00000 | 4.62979 |
| **Bola1** | 1.79765 | 0.01322 | 0.04336 | 3.51179 | 4.37609 | 0.90008 | 0.38362 | 2.88245 | 7.33163 | 1.72343 | 1.30767 | 0.04094 | 6.34468 | 0.00610 | 0.00000 | 0.74179 |
| **Bola2** | 4.19342 | 0.32436 | 2.52586 | 7.48497 | 3.29952 | 1.11382 | 0.38625 | 0.53517 | 13.92173 | 4.13901 | 8.58669 | 0.70553 | 2.37555 | 0.00000 | 0.00000 | 4.27814 |
| **Bola3** | 1.65457 | 0.00427 | 0.08815 | 0.44608 | 1.41324 | 1.17183 | 0.80213 | 0.02811 | 0.65323 | 0.32498 | 2.20573 | 0.02999 | 0.18904 | 0.00144 | 0.00000 | 0.01231 |
| **Bop1** | 6.97988 | 0.09039 | 0.89714 | 7.67314 | 6.00390 | 4.72947 | 1.17525 | 3.13021 | 10.49804 | 13.96513 | 10.03647 | 0.72223 | 7.58488 | 0.25455 | 2.49296 | 3.01519 |
| **Bora** | 3.08113 | 4.40502 | 4.89551 | 3.06450 | 0.04189 | 0.96786 | 5.18607 | 6.64202 | 1.82773 | 5.85779 | 4.44828 | 0.58105 | 4.03575 | 3.24989 | 8.43640 | 1.82852 |
| **Borcs5** | 1.08362 | 0.00743 | 0.76226 | 3.01884 | 0.01154 | 0.03928 | 0.36637 | 1.01140 | 0.78559 | 2.75300 | 0.01478 | 0.49818 | 1.51397 | 0.00008 | 0.33620 | 2.67839 |
| **Borcs6** | 1.36250 | 0.00612 | 0.00000 | 4.12619 | 0.00835 | 0.68201 | 0.00525 | 0.69227 | 5.18537 | 5.00466 | 1.46220 | 0.17546 | 0.00000 | 0.45175 | 0.56527 | 0.00452 |
| **Borcs7** | 2.86006 | 0.01447 | 1.13052 | 0.28669 | 0.00000 | 1.44280 | 0.12862 | 1.32399 | 1.06205 | 1.99229 | 4.20736 | 0.07800 | 0.00000 | 0.17566 | 0.05886 | 1.30812 |
| **Borcs8** | 0.00000 | 2.39315 | 2.70591 | 1.15565 | 4.23468 | 0.01717 | 0.35683 | 2.27330 | 1.55667 | 3.46249 | 1.23474 | 0.22303 | 2.51033 | 0.00506 | 1.78991 | 3.50150 |
| **Bpgm** | 0.00000 | 0.00768 | 0.86652 | 18.35104 | 0.00588 | 0.00530 | 0.00601 | 0.00000 | 1.43137 | 7.15115 | 1.07570 | 0.10398 | 0.00000 | 0.61865 | 4.07650 | 1.66764 |
| **Bphl** | 0.00690 | 0.00669 | 1.93347 | 0.74278 | 0.88892 | 0.00884 | 0.01670 | 0.00673 | 0.50278 | 1.15681 | 0.00628 | 0.00000 | 0.00691 | 0.00896 | 0.00000 | 0.00648 |
| **Bpifb9a** | 0.01345 | 0.00774 | 0.01014 | 0.01050 | 0.00940 | 0.00673 | 0.00734 | 0.01099 | 0.01265 | 0.01041 | 0.01116 | 0.00637 | 0.01310 | 0.36251 | 0.00360 | 0.00746 |
| **Bpifb9b** | 0.01355 | 0.00781 | 0.01069 | 0.01050 | 0.00938 | 0.00672 | 0.00738 | 0.01140 | 0.01263 | 0.01118 | 0.01168 | 0.00678 | 0.01305 | 0.25821 | 0.00397 | 0.00746 |
| **Bpifc** | 0.01362 | 0.00760 | 0.01119 | 0.02055 | 0.00934 | 0.00646 | 0.00742 | 0.01167 | 1.28380 | 0.00000 | 0.00000 | 0.00484 | 0.01306 | 0.02706 | 0.34707 | 0.00725 |
| **Bpnt1** | 4.24046 | 3.10181 | 3.02228 | 0.74713 | 0.29115 | 3.92005 | 0.08561 | 2.51824 | 1.69824 | 2.12034 | 3.06548 | 0.31923 | 3.86710 | 1.18200 | 3.78918 | 2.30614 |
| **Bptf** | 3.44806 | 4.64741 | 4.61832 | 15.06092 | 4.41049 | 1.55328 | 4.95661 | 3.61274 | 7.50510 | 19.77730 | 6.48727 | 8.41969 | 4.00348 | 16.84570 | 14.96029 | 0.54513 |
| **Braf** | 2.12733 | 2.84787 | 2.80596 | 1.00042 | 3.37428 | 0.43115 | 3.61799 | 0.46237 | 1.92974 | 3.66571 | 2.75570 | 9.31866 | 1.18212 | 10.84276 | 2.03476 | 0.00000 |
| **Brap** | 1.53006 | 0.10340 | 0.41720 | 11.10597 | 3.33662 | 1.99694 | 0.34179 | 4.13222 | 8.18589 | 11.75962 | 2.72431 | 0.76599 | 1.91050 | 0.00000 | 0.47774 | 2.53796 |
| **Brat1** | 0.97140 | 0.04330 | 1.15570 | 4.21671 | 0.00732 | 0.00421 | 0.54175 | 0.07886 | 1.79076 | 1.74644 | 0.37824 | 0.00000 | 4.23338 | 0.01996 | 0.90650 | 2.57008 |
| **Brca1** | 3.47084 | 7.77915 | 9.86316 | 3.93742 | 0.01313 | 6.32979 | 6.52096 | 2.20838 | 2.09770 | 7.93867 | 1.40606 | 6.81294 | 2.67374 | 1.64681 | 3.39573 | 4.39406 |
| **Brca2** | 2.81639 | 0.61137 | 2.33209 | 22.12604 | 6.45722 | 4.13889 | 2.67565 | 2.37530 | 15.20894 | 19.72616 | 7.30919 | 1.31265 | 3.55792 | 25.23167 | 17.99595 | 0.98534 |
| **Brcc3** | 1.43459 | 2.95331 | 4.59442 | 0.04593 | 2.59674 | 0.00392 | 3.31738 | 0.68042 | 0.28182 | 1.74287 | 3.26985 | 0.02410 | 2.96924 | 1.86127 | 0.21939 | 0.07821 |
| **Brd1** | 4.78407 | 3.11657 | 6.32340 | 19.26375 | 3.49956 | 0.57323 | 3.30718 | 12.78446 | 19.08558 | 8.80959 | 10.10688 | 10.04194 | 8.11191 | 0.62487 | 10.88762 | 5.72693 |
| **Brd2** | 8.30026 | 7.31967 | 9.62329 | 7.01755 | 0.86938 | 6.06512 | 5.08479 | 19.39575 | 10.53486 | 17.95072 | 9.76944 | 7.73865 | 12.07349 | 9.74521 | 16.39506 | 7.47568 |
| **Brd3** | 9.56160 | 0.33008 | 6.34933 | 2.55414 | 5.64212 | 3.58025 | 4.03278 | 17.32034 | 2.41867 | 18.54871 | 9.64150 | 1.22843 | 4.85510 | 1.28944 | 10.50179 | 1.29867 |

| | | | | | | | | | | | | | | | | |
|---|---|---|---|---|---|---|---|---|---|---|---|---|---|---|---|---|
| **Brd4** | 11.04092 | 1.08789 | 3.77536 | 4.52654 | 0.24289 | 2.10486 | 4.18980 | 12.54327 | 6.64303 | 16.94447 | 5.38471 | 0.85017 | 6.60330 | 0.48993 | 8.89147 | 4.73048 |
| **Brd7** | 12.74599 | 7.09599 | 6.56236 | 12.56677 | 6.72314 | 5.26141 | 4.85794 | 6.10816 | 9.44914 | 13.59332 | 8.25189 | 3.48111 | 7.59855 | 4.26504 | 13.51985 | 4.47358 |
| **Brd8** | 6.59122 | 0.11106 | 4.87439 | 13.10642 | 5.41952 | 3.29064 | 7.92689 | 9.24985 | 20.25100 | 8.45430 | 13.21219 | 4.56346 | 4.74354 | 9.27665 | 14.89591 | 3.02563 |
| **Brd9** | 6.66523 | 0.65036 | 8.56667 | 5.11122 | 0.02976 | 10.69707 | 0.01866 | 16.19938 | 14.74496 | 19.41172 | 15.32708 | 17.22431 | 8.40625 | 15.20310 | 16.81098 | 4.23229 |
| **Brdt** | 0.09846 | 0.00625 | 0.00000 | 16.15831 | 0.01334 | 0.00205 | 0.00544 | 0.00000 | 14.15744 | 1.31069 | 1.02444 | 5.64896 | 0.00875 | 0.11827 | 7.12813 | 0.00296 |
| **Bre** | 8.36557 | 0.00000 | 5.63671 | 0.00000 | 6.70093 | 5.28024 | 4.01330 | 4.93559 | 0.00097 | 2.64601 | 1.24468 | 0.29851 | 2.96860 | 0.02252 | 0.03681 | 3.64540 |
| **Brf1** | 2.32617 | 0.00306 | 3.10767 | 7.80113 | 3.27653 | 0.00514 | 0.00405 | 2.14330 | 6.18883 | 3.45328 | 1.96849 | 0.09833 | 1.25925 | 0.00000 | 0.14046 | 3.70587 |
| **Brf2** | 1.28287 | 0.20904 | 0.52458 | 1.68916 | 0.46873 | 0.01699 | 1.59910 | 1.49619 | 9.74278 | 4.37509 | 0.56355 | 0.13370 | 1.62785 | 0.00000 | 1.02475 | 0.03323 |
| **Bri3** | 1.94517 | 0.15229 | 0.47733 | 33.19449 | 2.30284 | 1.05589 | 0.43212 | 0.12641 | 21.16356 | 2.25490 | 9.88907 | 0.16212 | 0.76151 | 0.04505 | 0.07197 | 1.06264 |
| **Bri3bp** | 8.04175 | 0.02001 | 2.39377 | 7.12393 | 4.08526 | 4.62633 | 4.68325 | 15.97681 | 13.55550 | 24.45548 | 15.86825 | 7.70683 | 2.88856 | 3.43947 | 5.46479 | 4.42274 |
| **Brip1** | 1.43502 | 4.19376 | 4.54018 | 6.73437 | 0.18730 | 0.40888 | 4.55232 | 0.07685 | 7.37514 | 1.96792 | 3.73492 | 0.42898 | 0.10007 | 6.68758 | 3.06114 | 0.46174 |
| **Brip1os** | 3.55390 | 1.82717 | 4.62431 | 6.79599 | 1.17183 | 3.81674 | 2.22960 | 0.81330 | 10.06745 | 1.07328 | 7.33974 | 0.88765 | 2.08778 | 5.94882 | 2.00666 | 0.41659 |
| **Brix1** | 13.87780 | 13.21137 | 17.96657 | 18.91112 | 12.47811 | 10.79384 | 10.18766 | 20.99542 | 29.38979 | 29.91285 | 23.98736 | 15.07200 | 14.13648 | 19.28464 | 14.92067 | 10.11241 |
| **Brk1** | 17.25352 | 9.29254 | 19.67874 | 11.76962 | 13.27225 | 12.44270 | 11.92563 | 16.93823 | 5.24798 | 16.62137 | 8.75291 | 10.85035 | 17.49827 | 10.82409 | 10.44195 | 12.37569 |
| **Brms1** | 10.06998 | 3.78003 | 9.72230 | 10.22915 | 3.85908 | 1.64584 | 3.15477 | 12.45662 | 8.04007 | 8.89614 | 5.63253 | 1.44219 | 9.88345 | 0.08168 | 3.79299 | 5.73288 |
| **Brms1l** | 0.18293 | 1.67303 | 0.27849 | 6.13294 | 0.02176 | 2.34349 | 0.84954 | 1.57896 | 12.22430 | 1.18122 | 3.43053 | 6.38718 | 0.34402 | 3.32690 | 8.00340 | 0.10328 |
| **Brox** | 1.33724 | 4.64126 | 1.23880 | 5.46188 | 0.00208 | 3.09225 | 3.26450 | 2.47482 | 2.32328 | 3.51494 | 2.01720 | 3.55085 | 1.71011 | 1.37803 | 4.46257 | 4.33218 |
| **Brpf1** | 3.78962 | 0.09702 | 1.45338 | 2.72437 | 0.72892 | 3.84751 | 1.57716 | 4.85584 | 11.34110 | 6.47722 | 3.96608 | 0.28981 | 3.45093 | 1.35033 | 3.89731 | 0.52114 |
| **Brpf3** | 5.58704 | 1.08908 | 0.55907 | 9.96111 | 0.02479 | 0.07293 | 0.00000 | 1.49705 | 12.92318 | 2.16651 | 5.82640 | 2.78124 | 1.29864 | 0.01213 | 2.45338 | 3.58583 |
| **Brsk1** | 0.01181 | 0.00797 | 0.00963 | 0.09490 | 0.00943 | 0.00658 | 0.00721 | 0.06681 | 0.68254 | 0.00000 | 0.09083 | 0.01612 | 0.00878 | 0.01243 | 0.22031 | 0.00724 |
| **Brsk2** | 0.00777 | 0.01806 | 0.41750 | 3.83673 | 0.00931 | 0.00546 | 1.09349 | 0.11192 | 0.70357 | 0.00000 | 0.55832 | 0.00203 | 0.57177 | 0.00000 | 0.19279 | 0.00613 |
| **Brwd1** | 2.11847 | 7.02621 | 7.82492 | 5.88378 | 3.09560 | 3.25402 | 6.04181 | 9.30794 | 4.86179 | 9.87077 | 6.74450 | 6.69789 | 0.68089 | 9.21806 | 13.30950 | 1.14202 |
| **Brwd3** | 2.70617 | 0.01281 | 0.29144 | 7.54095 | 0.00584 | 1.81593 | 0.65563 | 0.32200 | 13.77016 | 5.71804 | 0.31766 | 8.88977 | 0.02116 | 14.73432 | 8.50907 | 0.74527 |
| **Bscl2** | 3.04822 | 4.41909 | 0.00000 | 9.41370 | 0.01057 | 0.00428 | 1.97542 | 0.00000 | 2.14306 | 1.26016 | 0.97974 | 0.03932 | 0.74641 | 0.00000 | 1.12945 | 0.00703 |
| **Bsdc1** | 4.21113 | 0.53996 | 1.49532 | 0.23378 | 1.36934 | 2.74204 | 2.51220 | 7.53109 | 3.61507 | 7.87830 | 2.41480 | 0.73469 | 2.27449 | 1.78218 | 6.64477 | 0.39732 |
| **Bsg** | 11.91202 | 5.48971 | 14.35138 | 4.73047 | 3.29334 | 6.35482 | 6.35127 | 11.68563 | 0.26548 | 9.81025 | 10.73399 | 1.15145 | 21.96559 | 0.19963 | 0.00000 | 15.16916 |
| **Bsn** | 0.01359 | 0.00824 | 0.01231 | 0.00580 | 0.00938 | 0.00695 | 0.00765 | 0.01426 | 0.07907 | 0.12235 | 0.01130 | 0.00879 | 0.01609 | 0.00690 | 0.14103 | 0.00748 |
| **Bspry** | 0.07218 | 0.00611 | 0.84369 | 16.33726 | 0.00598 | 3.56092 | 0.90274 | 6.18585 | 22.68398 | 8.65708 | 4.97960 | 0.39947 | 2.42238 | 0.08397 | 5.49919 | 0.00000 |
| **Bst2** | 0.87212 | 0.00608 | 0.60937 | 0.86327 | 0.00526 | 0.00824 | 0.01605 | 0.90368 | 0.00002 | 1.02731 | 1.87424 | 0.00000 | 0.00968 | 0.01882 | 0.00225 | 0.00595 |
| **Btaf1** | 3.79341 | 7.36569 | 1.98372 | 12.81973 | 0.09991 | 2.24417 | 3.55061 | 4.54072 | 13.32127 | 6.92408 | 4.03897 | 7.87145 | 1.79078 | 14.84724 | 11.92020 | 0.09718 |
| **Btbd1** | 2.20282 | 6.00439 | 7.01778 | 0.96803 | 5.32761 | 1.00428 | 4.99207 | 1.93908 | 7.10127 | 4.69038 | 7.11270 | 6.89653 | 1.82911 | 3.35640 | 3.24644 | 3.37198 |
| **Btbd10** | 8.01377 | 3.75832 | 2.71457 | 9.80502 | 4.31402 | 0.07958 | 3.66880 | 4.14151 | 4.61873 | 5.60226 | 4.19303 | 6.52795 | 2.45195 | 7.56703 | 10.53592 | 2.06977 |
| **Btbd16** | 0.06869 | 0.00778 | 0.11714 | 1.52226 | 0.00838 | 0.05150 | 0.03807 | 0.02935 | 2.42930 | 0.20798 | 0.06006 | 0.02008 | 0.01933 | 0.35874 | 0.45899 | 0.01429 |
| **Btbd18** | 0.00000 | 0.00531 | 0.51450 | 22.09754 | 0.00754 | 4.21295 | 0.00879 | 5.63391 | 22.57879 | 19.36291 | 10.11797 | 2.44581 | 6.27121 | 22.75683 | 26.43760 | 0.00254 |
| **Btbd19** | 0.97253 | 0.08537 | 1.98014 | 0.93420 | 0.02731 | 0.03532 | 0.09031 | 0.00000 | 1.43505 | 2.86140 | 3.33167 | 0.05402 | 0.15117 | 0.44184 | 0.87871 | 0.03384 |
| **Btbd2** | 1.01608 | 0.00678 | 0.00000 | 7.20015 | 0.00471 | 0.01223 | 0.00488 | 0.00213 | 1.58677 | 1.44642 | 0.13298 | 0.04056 | 2.25068 | 0.00000 | 0.19317 | 3.18730 |
| **Btbd3** | 0.01120 | 0.00507 | 0.00000 | 0.00000 | 4.94811 | 0.01414 | 0.00724 | 0.00437 | 0.49185 | 0.03722 | 0.00471 | 0.00000 | 0.02025 | 0.04819 | 1.20956 | 0.00738 |
| **Btbd6** | 0.36298 | 0.00459 | 0.05093 | 0.11083 | 3.27826 | 0.02772 | 2.56647 | 1.57857 | 0.53947 | 1.46625 | 0.28869 | 0.01971 | 0.14068 | 0.02402 | 1.18939 | 0.03115 |
| **Btbd7** | 0.16505 | 0.00648 | 0.05677 | 3.17362 | 0.00773 | 0.00883 | 1.96032 | 0.61159 | 3.00205 | 0.61126 | 0.40125 | 0.14017 | 0.44250 | 0.05584 | 2.11577 | 0.06800 |
| **Btbd8** | 0.01254 | 0.00781 | 0.01201 | 0.01000 | 0.00896 | 0.00643 | 0.00740 | 0.01342 | 0.01246 | 0.00873 | 0.01261 | 0.00761 | 0.08474 | 0.00752 | 0.08988 | 0.09963 |
| **Btbd9** | 0.92092 | 0.01745 | 1.04611 | 0.66472 | 0.01259 | 0.00618 | 0.00652 | 0.56197 | 1.32968 | 3.27870 | 0.42644 | 0.00000 | 0.00588 | 0.68624 | 0.49905 | 0.01254 |
| **Btd** | 0.00323 | 0.00667 | 2.22061 | 0.00265 | 0.00569 | 0.00520 | 0.00637 | 0.95889 | 0.02283 | 0.88631 | 0.17858 | 3.12391 | 0.47537 | 0.00518 | 0.44601 | 0.00609 |
| **Btf3** | 42.97931 | 23.39455 | 33.66318 | 15.20402 | 25.36398 | 18.04243 | 21.16883 | 35.35632 | 16.77413 | 37.08790 | 30.86620 | 23.99731 | 37.46710 | 17.71876 | 19.17507 | 22.09827 |
| **Btf3l4** | 15.49440 | 7.87385 | 10.32957 | 7.27243 | 8.52470 | 8.88988 | 5.26030 | 9.69414 | 8.11473 | 13.20814 | 13.29077 | 12.95869 | 11.87723 | 17.77704 | 9.60610 | 4.34204 |
| **Btg1** | 8.23177 | 7.48954 | 10.75209 | 5.36504 | 6.75409 | 2.68003 | 9.41641 | 14.38586 | 17.12751 | 21.09363 | 14.74302 | 14.74474 | 14.65871 | 25.41034 | 6.40831 | 1.56671 |
| **Btg2** | 6.31017 | 3.13146 | 1.24162 | 1.67739 | 0.01788 | 3.74113 | 0.04186 | 0.00000 | 1.27221 | 1.42851 | 0.41097 | 0.13416 | 0.02172 | 0.00000 | 0.32293 | 0.00000 |
| **Btg3** | 2.09509 | 0.00164 | 0.08535 | 4.74371 | 2.58047 | 0.00166 | 0.07414 | 0.32707 | 3.94451 | 0.43291 | 0.42356 | 6.87731 | 0.31211 | 0.02762 | 0.10273 | 0.04849 |
| **Btg4** | 0.00424 | 0.00849 | 0.00000 | 16.20489 | 0.00887 | 0.00446 | 0.00735 | 0.35892 | 4.54495 | 2.59952 | 0.74489 | 0.22824 | 0.00299 | 0.55909 | 4.79592 | 0.00421 |
| **Btk** | 0.00577 | 1.28884 | 0.00841 | 2.09513 | 0.00873 | 0.00673 | 0.00699 | 0.01109 | 0.78135 | 0.07910 | 0.00693 | 0.00399 | 0.10887 | 0.17052 | 0.62700 | 0.00683 |
| **Btn2a2** | 0.01660 | 0.00775 | 0.66487 | 0.42533 | 0.00805 | 0.00558 | 0.00729 | 0.01222 | 0.31469 | 0.01286 | 0.01031 | 0.00696 | 0.00874 | 0.01240 | 0.00000 | 0.00699 |
| **Btnl1** | 0.01656 | 0.00790 | 0.23424 | 0.01033 | 0.00875 | 0.00567 | 0.00737 | 0.01204 | 0.01239 | 0.09228 | 0.01175 | 0.00746 | 0.00974 | 0.01211 | 0.00683 | 0.00704 |
| **Btrc** | 16.49387 | 14.57751 | 16.39989 | 1.84493 | 14.57984 | 5.83036 | 15.13602 | 25.92243 | 6.32967 | 19.94304 | 14.20116 | 7.47132 | 14.98851 | 2.17960 | 5.51869 | 7.58866 |
| **Bub1** | 4.19371 | 9.47444 | 4.20832 | 9.19331 | 6.51803 | 4.21086 | 4.10099 | 2.93074 | 15.80371 | 13.61649 | 7.29821 | 11.06577 | 3.17811 | 12.09147 | 12.32335 | 0.84304 |
| **Bub1b** | 10.06179 | 4.35761 | 6.55608 | 0.66914 | 5.75418 | 2.22321 | 2.81116 | 7.19085 | 2.42459 | 4.65414 | 3.51522 | 7.83029 | 3.55006 | 7.05343 | 7.04274 | 1.19472 |
| **Bub3** | 18.18566 | 15.08087 | 17.27397 | 2.03322 | 9.96404 | 8.51760 | 12.05188 | 17.67320 | 11.46053 | 20.13060 | 11.53859 | 8.09396 | 13.64488 | 0.51060 | 9.43941 | 13.04350 |
| **Bud13** | 2.28908 | 0.44588 | 2.60202 | 0.04703 | 0.02314 | 0.04325 | 2.58677 | 5.21435 | 0.04416 | 1.50322 | 0.30583 | 0.20809 | 1.04524 | 2.10896 | 9.69832 | 0.73092 |
| **Bud31** | 17.82020 | 7.21916 | 10.74397 | 4.63633 | 3.56504 | 6.60614 | 6.08175 | 10.62317 | 10.07482 | 14.49093 | 5.85193 | 7.45893 | 12.01050 | 9.94654 | 7.83990 | 3.94824 |
| **Bves** | 1.34086 | 0.00637 | 0.00362 | 0.00927 | 0.00537 | 0.00657 | 0.16373 | 0.01179 | 0.01184 | 0.00920 | 0.01236 | 0.00745 | 0.01101 | 0.00792 | 0.18096 | 0.00589 |
| **Bvht** | 0.05083 | 0.04113 | 0.33808 | 0.07729 | 0.00744 | 0.00564 | 0.00754 | 0.01461 | 0.03541 | 0.01409 | 0.01144 | 0.00847 | 0.02282 | 0.01314 | 0.00203 | 0.00707 |
| **Bysl** | 10.99302 | 6.98380 | 11.14864 | 9.63459 | 8.61413 | 5.87885 | 8.23531 | 24.28898 | 10.13177 | 22.09954 | 7.61854 | 5.78271 | 15.54920 | 2.87142 | 4.58371 | 8.81958 |
| **Bzw1** | 17.14466 | 23.04586 | 20.75548 | 21.69991 | 17.91339 | 12.92034 | 19.83424 | 26.16985 | 27.27718 | 26.01238 | 13.53031 | 14.24540 | 20.04943 | 39.58483 | 32.26844 | 6.85732 |
| **Bzw2** | 12.16814 | 9.46840 | 12.55789 | 2.75547 | 13.45170 | 7.22385 | 12.11987 | 11.91565 | 5.65476 | 18.78831 | 6.37906 | 7.12486 | 9.64210 | 8.94176 | 8.72179 | 8.32967 |

| | | | | | | | | | | | | | | | | |
|---|---|---|---|---|---|---|---|---|---|---|---|---|---|---|---|---|
| **C1d** | 8.72647 | 12.17262 | 12.57959 | 6.18570 | 9.75364 | 9.94860 | 10.36676 | 8.43265 | 6.72247 | 18.32403 | 7.53628 | 11.06266 | 16.33139 | 16.63159 | 17.10928 | 3.32833 |
| **C1galt1** | 0.02092 | 0.00796 | 0.01974 | 0.56218 | 0.00864 | 0.00612 | 0.00748 | 0.01394 | 0.90131 | 0.00000 | 0.23463 | 0.00529 | 0.01775 | 0.00662 | 0.04985 | 0.03579 |
| **C1galt1c1** | 0.41242 | 1.88377 | 1.78692 | 0.00000 | 0.00384 | 6.14411 | 0.00462 | 0.85984 | 0.20163 | 1.48624 | 1.38110 | 0.00000 | 1.60382 | 4.96943 | 1.48342 | 0.00217 |
| **C1qa** | 0.16294 | 0.00782 | 0.00693 | 0.00815 | 0.00809 | 0.00781 | 0.00742 | 0.09464 | 0.01192 | 0.01990 | 0.09701 | 0.00706 | 0.10733 | 0.01129 | 0.00804 | 0.01498 |
| **C1qb** | 0.01104 | 0.00786 | 0.01199 | 0.48233 | 0.00904 | 0.00640 | 0.00695 | 0.00939 | 0.00387 | 0.35840 | 0.01415 | 0.00740 | 0.54881 | 0.01641 | 0.00332 | 0.00680 |
| **C1qbp** | 23.63984 | 12.17105 | 21.85277 | 19.73873 | 13.98744 | 3.63880 | 10.10024 | 19.47592 | 27.22680 | 30.39267 | 36.51250 | 14.97663 | 18.82659 | 14.44258 | 13.09281 | 4.45830 |
| **C1ql1** | 0.00777 | 0.00744 | 0.00433 | 6.88477 | 0.00916 | 0.00688 | 0.00679 | 0.80994 | 0.41641 | 1.51005 | 0.41042 | 0.12244 | 0.00392 | 0.00000 | 0.25985 | 0.00612 |
| **C1ql4** | 1.74643 | 0.44524 | 0.00000 | 0.01028 | 0.03031 | 0.05411 | 0.00634 | 0.05956 | 0.01241 | 0.00145 | 0.01213 | 0.00791 | 0.78859 | 0.01465 | 0.00417 | 0.96251 |
| **C1qtnf2** | 0.11721 | 0.00768 | 0.44990 | 0.00803 | 0.01662 | 0.01100 | 1.11125 | 0.00887 | 0.01237 | 0.43621 | 0.01191 | 0.00541 | 0.00921 | 0.00879 | 0.02634 | 0.00730 |
| **C1qtnf4** | 0.01000 | 0.00792 | 0.01385 | 0.01006 | 0.00887 | 0.00692 | 0.00752 | 0.01211 | 0.02678 | 2.31791 | 0.01250 | 0.00000 | 0.03765 | 0.01247 | 0.00640 | 0.43301 |
| **C1ra** | 0.01424 | 0.00780 | 0.01961 | 0.02378 | 0.00912 | 0.00613 | 0.00735 | 0.01231 | 0.01301 | 0.00801 | 0.05195 | 0.00734 | 0.10834 | 0.00767 | 0.06403 | 0.00720 |
| **C1rb** | 0.01375 | 0.00782 | 0.01236 | 0.00952 | 0.00935 | 0.00657 | 0.00748 | 0.01317 | 0.01231 | 0.00971 | 0.01276 | 0.00737 | 0.01266 | 0.00658 | 0.16724 | 0.00738 |
| **C1rl** | 0.01765 | 0.00760 | 0.02195 | 0.05562 | 0.01724 | 0.00688 | 0.00726 | 0.01788 | 0.09438 | 0.52056 | 0.06021 | 0.00570 | 0.01035 | 0.03136 | 0.01189 | 0.18897 |
| **C1s1** | 0.01200 | 0.00765 | 0.09123 | 0.28385 | 0.00892 | 0.00567 | 0.00736 | 0.01059 | 0.56400 | 0.92690 | 0.00011 | 0.30777 | 0.01113 | 0.00455 | 0.29852 | 0.00747 |
| **C1s2** | 0.01202 | 0.00798 | 0.53929 | 0.81737 | 0.00828 | 0.00548 | 0.00752 | 0.01141 | 1.75158 | 0.17993 | 0.00000 | 0.11949 | 0.01649 | 0.04826 | 0.00000 | 0.00697 |
| **C2** | 1.03920 | 0.00390 | 4.75300 | 0.02729 | 0.00000 | 0.01440 | 1.90339 | 6.80137 | 1.84686 | 7.40779 | 4.60347 | 0.33259 | 6.51869 | 9.43448 | 2.48863 | 0.00000 |
| **C2cd2** | 1.26308 | 0.00561 | 0.30952 | 1.68468 | 0.02142 | 3.23954 | 0.74763 | 0.00368 | 1.01088 | 1.02425 | 0.03238 | 0.12029 | 0.00166 | 0.55459 | 0.03845 | 0.00425 |
| **C2cd2l** | 0.00000 | 0.01585 | 0.34557 | 3.71494 | 0.01471 | 2.56383 | 0.00265 | 11.45370 | 13.46588 | 10.80064 | 9.40748 | 0.68362 | 1.10014 | 4.30418 | 6.30096 | 2.28362 |
| **C2cd3** | 0.51569 | 0.15259 | 1.49085 | 9.44106 | 2.44317 | 3.94806 | 4.75642 | 0.00000 | 7.74217 | 1.23379 | 1.72948 | 4.74896 | 0.00000 | 0.00000 | 4.75653 | 0.00908 |
| **C2cd4a** | 0.01257 | 0.00810 | 0.40834 | 0.00814 | 0.00855 | 0.00653 | 0.00754 | 0.01344 | 0.04856 | 0.01462 | 0.01169 | 0.00818 | 0.00946 | 0.01441 | 0.00381 | 0.00721 |
| **C2cd4b** | 0.01358 | 0.00825 | 0.01192 | 0.47071 | 0.00929 | 0.00656 | 0.00753 | 0.01388 | 1.08987 | 0.01019 | 0.00608 | 0.00660 | 0.01295 | 0.01184 | 0.00343 | 0.00727 |
| **C2cd5** | 1.24326 | 0.00533 | 0.77688 | 9.83701 | 1.31099 | 0.00587 | 0.01542 | 3.20945 | 3.02892 | 0.33277 | 0.32198 | 0.14413 | 1.34724 | 0.77866 | 8.57928 | 0.01818 |
| **C3** | 0.01152 | 0.00756 | 0.01183 | 0.00505 | 0.00840 | 0.00651 | 0.00720 | 0.00794 | 0.01069 | 5.58291 | 0.04865 | 0.00000 | 0.01003 | 0.01583 | 0.01628 | 0.00733 |
| **C4a** | 0.01385 | 0.00759 | 0.09066 | 0.24429 | 0.00911 | 0.00619 | 0.00706 | 0.34325 | 0.07948 | 0.01092 | 0.01047 | 0.02574 | 0.05375 | 0.01173 | 0.00000 | 0.02177 |
| **C4b** | 0.01593 | 0.00769 | 0.03386 | 0.04945 | 0.00912 | 0.00642 | 0.00717 | 0.06939 | 0.00968 | 0.02113 | 0.01225 | 0.02655 | 0.03117 | 0.01173 | 0.00748 | 0.00735 |
| **C5ar2** | 0.02667 | 0.00794 | 0.03096 | 0.12675 | 0.00941 | 0.02027 | 0.01730 | 0.01282 | 0.19576 | 0.04788 | 0.30846 | 0.00546 | 0.03534 | 0.13920 | 0.02627 | 0.02173 |
| **C6** | 0.01231 | 0.03980 | 0.08894 | 0.05297 | 0.00920 | 0.03593 | 0.12989 | 0.01112 | 0.00936 | 0.03866 | 0.17548 | 0.00592 | 0.01266 | 0.20448 | 0.04820 | 0.00757 |
| **C7** | 0.01375 | 0.00818 | 0.01225 | 0.13911 | 0.00931 | 0.00663 | 0.00752 | 0.01377 | 0.00926 | 0.01424 | 0.01384 | 0.00840 | 0.01266 | 0.01023 | 0.02994 | 0.00738 |
| **C77080** | 8.35845 | 2.71330 | 5.68377 | 0.67197 | 0.16626 | 8.52274 | 1.41852 | 9.24399 | 5.77674 | 9.00712 | 7.05240 | 0.37025 | 5.01451 | 0.75050 | 6.19648 | 2.06692 |
| **C77370** | 6.06225 | 0.00717 | 1.34611 | 0.30085 | 9.74171 | 0.00000 | 0.94036 | 0.28404 | 0.00000 | 0.78379 | 0.84627 | 0.04751 | 0.10868 | 1.23412 | 1.18423 | 1.89164 |
| **C86187** | 0.00843 | 0.00755 | 0.00000 | 20.35356 | 0.00916 | 0.00382 | 0.00586 | 0.00000 | 7.56018 | 0.00000 | 1.10861 | 0.03040 | 0.00001 | 2.66995 | 2.01611 | 0.00303 |
| **C87198** | 0.02214 | 0.00780 | 0.00694 | 2.06332 | 0.00881 | 0.04284 | 0.03806 | 0.01168 | 1.60056 | 0.04675 | 0.01285 | 0.01371 | 0.02780 | 0.03553 | 0.28491 | 0.00683 |
| **C87414** | 0.05443 | 0.03917 | 0.03694 | 10.28380 | 0.03845 | 0.01183 | 0.06882 | 0.12121 | 4.28863 | 0.42338 | 0.35566 | 0.43111 | 0.11986 | 0.77167 | 3.96364 | 0.00508 |
| **C87436** | 0.54060 | 0.15896 | 1.71459 | 8.42478 | 0.00411 | 0.00515 | 0.62175 | 1.01205 | 4.96842 | 2.00075 | 0.22676 | 0.49895 | 0.12653 | 1.45753 | 4.28913 | 1.02310 |
| **C87499** | 0.00508 | 0.00828 | 0.00000 | 8.74751 | 0.00955 | 0.00469 | 0.00751 | 0.09356 | 7.49227 | 0.32927 | 0.51386 | 0.00000 | 0.00587 | 1.44078 | 5.14497 | 0.00570 |
| **C87977** | 0.00000 | 0.00731 | 1.15214 | 5.78034 | 0.01061 | 0.00715 | 0.00472 | 0.78618 | 4.65217 | 0.00000 | 4.98380 | 0.03002 | 0.00000 | 9.63041 | 4.11506 | 0.00488 |
| **C8b** | 0.02233 | 0.00802 | 0.47052 | 0.61749 | 0.00962 | 0.00566 | 0.00738 | 0.01287 | 0.74579 | 0.01110 | 0.00727 | 0.00638 | 0.01201 | 0.01103 | 0.00000 | 0.00713 |
| **Caap1** | 0.14435 | 0.01871 | 0.06435 | 3.29246 | 0.01084 | 0.04538 | 0.00608 | 0.00000 | 1.57585 | 0.00000 | 0.86809 | 1.17345 | 0.70846 | 3.79477 | 4.45034 | 0.02017 |
| **Cab39** | 8.20821 | 6.85942 | 11.57151 | 15.56590 | 7.70076 | 3.07314 | 6.82038 | 17.06206 | 28.03282 | 21.96163 | 16.36835 | 13.87728 | 12.41010 | 31.25944 | 21.98484 | 3.98274 |
| **Cab39l** | 9.66100 | 10.70059 | 6.52658 | 3.91342 | 4.61474 | 3.05096 | 4.26279 | 3.03357 | 3.47326 | 8.42395 | 4.21303 | 8.96791 | 3.22527 | 12.75118 | 6.04160 | 3.32849 |
| **Cabin1** | 6.34670 | 0.06468 | 3.63430 | 12.00917 | 1.64087 | 0.02500 | 1.13141 | 4.93175 | 7.32496 | 11.22047 | 2.57219 | 0.42783 | 2.72989 | 0.24382 | 3.41724 | 0.48187 |
| **Cables1** | 0.24055 | 0.00739 | 0.00327 | 3.38669 | 0.00766 | 0.00627 | 0.00705 | 0.01148 | 0.25288 | 0.25357 | 0.00677 | 0.02234 | 0.01254 | 0.00000 | 1.73793 | 0.00654 |
| **Cables2** | 1.81120 | 0.06549 | 5.29620 | 5.35760 | 5.47623 | 0.01104 | 0.50741 | 3.97781 | 1.93017 | 1.90129 | 0.37266 | 4.12229 | 0.00000 | 1.03777 | 3.21688 | 1.85172 |
| **Cabp4** | 0.31316 | 0.00840 | 0.00593 | 0.22981 | 0.01741 | 0.00713 | 0.00731 | 0.00928 | 0.00809 | 0.01677 | 0.15676 | 0.00753 | 0.01052 | 0.00720 | 0.04746 | 0.00721 |
| **Cabp5** | 0.01833 | 0.00818 | 0.01157 | 0.01055 | 0.01038 | 0.00665 | 0.00745 | 0.01266 | 0.01231 | 0.02339 | 0.01277 | 0.00812 | 0.05967 | 0.01188 | 0.01232 | 0.00748 |
| **Cabp7** | 0.01362 | 0.00768 | 0.00912 | 0.01820 | 0.00923 | 0.00678 | 0.00718 | 0.20250 | 0.01234 | 0.01157 | 0.01233 | 0.00724 | 0.00710 | 0.01272 | 0.00713 | 0.00737 |
| **Cabyr** | 0.23031 | 0.00722 | 0.00583 | 0.21146 | 0.00738 | 0.00678 | 0.00725 | 0.00502 | 0.02030 | 0.00000 | 0.22482 | 0.00553 | 0.01556 | 0.00525 | 0.39173 | 0.01398 |
| **Cacfd1** | 2.79994 | 0.92441 | 0.15949 | 2.36179 | 0.00145 | 0.00409 | 0.00413 | 1.24480 | 1.24369 | 1.46455 | 0.97879 | 0.01046 | 0.21951 | 0.00000 | 1.56304 | 1.76188 |
| **Cachd1** | 1.60421 | 0.00549 | 0.19310 | 0.62590 | 0.00433 | 0.00369 | 1.50859 | 0.30229 | 3.15031 | 0.14807 | 0.14972 | 0.06991 | 0.00000 | 0.00188 | 0.55409 | 3.18965 |
| **Cacna1a** | 1.06064 | 0.00610 | 0.06560 | 0.00313 | 0.78044 | 0.00767 | 0.08896 | 0.00000 | 0.15415 | 1.22399 | 1.25894 | 0.02847 | 0.00000 | 0.00660 | 0.04152 | 0.00568 |
| **Cacna1b** | 0.01143 | 0.00787 | 0.11770 | 1.96027 | 0.00852 | 0.00806 | 0.00705 | 0.00781 | 4.03188 | 1.10019 | 0.00000 | 0.00842 | 0.02247 | 0.04641 | 0.01543 | 0.00647 |
| **Cacna1d** | 0.01144 | 0.00748 | 0.00981 | 0.00865 | 0.00952 | 0.00656 | 0.00721 | 0.09630 | 0.01137 | 0.00547 | 0.01088 | 0.01597 | 0.01638 | 0.00884 | 0.42489 | 0.00733 |
| **Cacna1h** | 0.01782 | 0.00771 | 0.01139 | 1.38104 | 0.00906 | 0.00639 | 0.00734 | 0.01205 | 0.13241 | 0.00576 | 0.01080 | 0.01636 | 0.01276 | 0.00473 | 0.49819 | 0.00708 |
| **Cacna2d1** | 0.01311 | 0.00777 | 0.01152 | 0.37653 | 0.00897 | 0.00631 | 0.00723 | 0.00988 | 0.37820 | 0.94658 | 0.00894 | 0.00407 | 0.01071 | 0.00726 | 0.60582 | 0.00711 |
| **Cacna2d3** | 0.01380 | 0.00766 | 0.01173 | 0.70385 | 0.00940 | 0.00651 | 0.00741 | 0.01214 | 0.00607 | 0.00672 | 0.01131 | 0.00716 | 0.01281 | 0.01773 | 0.29750 | 0.00727 |
| **Cacnb1** | 1.46827 | 0.00425 | 0.04364 | 0.05730 | 3.75810 | 0.01043 | 0.01604 | 1.53007 | 0.25992 | 2.59191 | 1.00224 | 0.07938 | 0.00000 | 0.00672 | 0.00000 | 5.04053 |
| **Cacnb2** | 0.01369 | 0.00798 | 0.01189 | 0.11563 | 0.00922 | 0.00688 | 0.00760 | 0.01384 | 0.00986 | 0.11732 | 0.01246 | 0.00877 | 0.01307 | 0.01432 | 0.03203 | 0.00740 |
| **Cacnb3** | 2.18588 | 0.00351 | 2.11495 | 0.73000 | 0.04572 | 8.97449 | 1.59404 | 2.90878 | 0.35999 | 3.60691 | 4.74900 | 0.04478 | 7.49641 | 0.00000 | 0.40733 | 3.54497 |
| **Cactin** | 7.10438 | 0.11445 | 3.12787 | 6.70390 | 3.14125 | 0.66333 | 3.03430 | 6.98489 | 12.84162 | 7.98780 | 2.33944 | 0.54444 | 5.70277 | 0.14814 | 0.00000 | 0.25940 |
| **Cacul1** | 9.26316 | 6.77506 | 10.53993 | 3.67666 | 7.57182 | 5.27544 | 11.10853 | 10.49980 | 0.17407 | 4.66110 | 8.15141 | 6.57608 | 7.03436 | 10.83763 | 2.20377 | 0.02753 |
| **Cacybp** | 22.50195 | 8.98698 | 17.21562 | 6.67991 | 19.27834 | 7.96790 | 9.35211 | 6.03255 | 11.46417 | 15.90842 | 7.02983 | 18.37814 | 15.36770 | 7.82564 | 4.66387 | 9.23895 |
| **Cad** | 13.78664 | 6.89499 | 11.08281 | 11.77610 | 12.05681 | 4.62441 | 11.10283 | 25.12422 | 9.91917 | 22.95142 | 15.77443 | 2.44466 | 16.06044 | 0.20121 | 2.40224 | 8.92579 |

| | | | | | | | | | | | | | | | | |
|---|---|---|---|---|---|---|---|---|---|---|---|---|---|---|---|---|
| **Cadm1** | 1.14368 | 0.16975 | 1.51710 | 3.06260 | 0.01096 | 0.05372 | 0.13152 | 3.35868 | 4.98623 | 0.00000 | 3.78625 | 0.01842 | 5.31348 | 1.91220 | 0.13018 | 0.03154 |
| **Cadm4** | 0.12740 | 0.00788 | 0.16288 | 0.63605 | 0.01560 | 0.32852 | 0.05690 | 0.19730 | 2.35724 | 0.03122 | 0.37156 | 0.01981 | 0.39167 | 0.22869 | 0.32093 | 0.00632 |
| **Cadps2** | 0.01360 | 0.00785 | 0.01133 | 0.96990 | 0.00942 | 0.00627 | 0.00730 | 0.01096 | 0.38377 | 0.00761 | 0.00866 | 0.00495 | 0.01294 | 0.00495 | 1.09576 | 0.00718 |
| **Cage1** | 0.00686 | 0.00711 | 0.20424 | 0.56025 | 1.95435 | 2.46282 | 0.00736 | 0.00829 | 0.01785 | 0.09407 | 0.00845 | 0.00638 | 0.00980 | 0.01115 | 0.00257 | 0.00596 |
| **Calb1** | 0.01130 | 0.00737 | 0.01132 | 0.32868 | 0.00840 | 0.00638 | 0.00719 | 0.00902 | 0.74648 | 5.71315 | 0.00006 | 0.00000 | 0.01068 | 0.01149 | 0.43043 | 0.00747 |
| **Calb2** | 0.15423 | 0.00747 | 0.00773 | 0.24055 | 0.00768 | 0.00674 | 0.00720 | 0.01874 | 0.00797 | 0.22232 | 0.01301 | 0.00629 | 0.01193 | 0.00559 | 0.12932 | 0.00719 |
| **Calcoco1** | 0.04754 | 0.00782 | 1.71259 | 0.18645 | 0.06824 | 0.01222 | 0.00706 | 0.24648 | 0.05677 | 1.43267 | 1.61114 | 0.19841 | 0.33039 | 0.00625 | 0.20182 | 0.01366 |
| **Calcoco2** | 57.07913 | 39.35981 | 62.37520 | 10.68133 | 43.30833 | 26.66967 | 33.36790 | 60.57298 | 26.93752 | 38.73126 | 48.51803 | 10.67922 | 58.85709 | 45.62771 | 24.64245 | 23.84294 |
| **Calcrl** | 0.02122 | 0.00773 | 0.02557 | 0.01101 | 0.00881 | 0.00678 | 0.00744 | 0.01243 | 0.01208 | 0.01232 | 0.01245 | 0.00760 | 0.01249 | 0.04871 | 0.03510 | 0.00739 |
| **Cald1** | 0.01145 | 0.00744 | 1.86446 | 0.00858 | 0.00586 | 0.00912 | 0.00700 | 0.00886 | 0.01175 | 0.54405 | 0.00658 | 0.28036 | 0.00665 | 0.01225 | 0.00000 | 0.00666 |
| **Calm1** | 29.35702 | 10.34392 | 24.08384 | 7.87335 | 25.58543 | 17.40173 | 11.27644 | 22.33201 | 13.90166 | 21.23027 | 13.10025 | 5.73731 | 24.37905 | 5.90002 | 2.80270 | 19.24451 |
| **Calm2** | 32.00314 | 17.74302 | 22.79167 | 11.75423 | 21.55219 | 14.13869 | 17.81969 | 20.13795 | 14.77334 | 16.73081 | 9.54707 | 11.83034 | 20.15033 | 20.76052 | 15.64512 | 13.97548 |
| **Calm3** | 5.78395 | 1.13279 | 2.99779 | 2.68273 | 9.85086 | 2.35764 | 0.34460 | 1.67867 | 5.82381 | 5.95483 | 6.23295 | 0.25023 | 6.36494 | 0.02225 | 0.85382 | 5.54560 |
| **Calm4** | 0.01325 | 0.00822 | 0.00972 | 0.00734 | 0.00884 | 0.00666 | 0.01888 | 0.05335 | 0.01163 | 0.46635 | 0.07830 | 0.01971 | 0.35956 | 0.02428 | 0.04299 | 0.00692 |
| **Calm5** | 0.00703 | 0.00819 | 0.00843 | 0.00721 | 0.00901 | 0.00673 | 0.04160 | 0.22252 | 0.01119 | 1.00866 | 0.00959 | 0.07934 | 0.92989 | 0.03465 | 0.08607 | 0.00670 |
| **Calml4** | 0.01078 | 0.00745 | 0.25811 | 4.07250 | 0.00856 | 0.00531 | 0.00644 | 0.00000 | 0.99838 | 1.67668 | 0.26726 | 0.09803 | 0.92149 | 0.00000 | 1.56915 | 0.00612 |
| **Caln1** | 0.01372 | 0.00792 | 0.01182 | 0.25423 | 0.00935 | 0.00666 | 0.00749 | 0.01304 | 0.15685 | 0.01261 | 0.01125 | 0.00762 | 0.01289 | 0.05667 | 0.00658 | 0.00739 |
| **Calr** | 13.63763 | 5.66839 | 22.87743 | 11.37691 | 3.74509 | 9.74497 | 6.45267 | 12.52718 | 14.50446 | 9.42497 | 10.31568 | 2.24347 | 24.79909 | 2.86165 | 6.38163 | 10.87955 |
| **Calr3** | 0.01108 | 0.00762 | 0.00633 | 0.01534 | 0.00934 | 0.00668 | 0.00745 | 0.74283 | 0.02318 | 0.21582 | 0.00536 | 0.02697 | 0.00532 | 0.04254 | 0.75738 | 0.00710 |
| **Calr4** | 1.53745 | 0.00416 | 2.10011 | 1.81762 | 0.01034 | 0.00524 | 2.18894 | 1.03377 | 0.12474 | 0.49437 | 0.45778 | 0.13822 | 0.00000 | 1.42548 | 0.83375 | 0.00461 |
| **Calu** | 6.31496 | 7.59350 | 8.48898 | 5.87199 | 7.25927 | 9.61943 | 9.95407 | 8.54635 | 4.72264 | 7.46328 | 6.82302 | 9.87179 | 8.91663 | 18.42585 | 14.22939 | 7.64650 |
| **Camk1** | 1.15506 | 0.05623 | 0.14983 | 0.00000 | 0.00551 | 5.77756 | 0.00022 | 1.00430 | 0.06878 | 0.48616 | 1.97539 | 0.03187 | 7.37561 | 0.00844 | 0.03973 | 0.00000 |
| **Camk1d** | 0.00941 | 0.00753 | 0.00654 | 1.29177 | 0.00863 | 0.00632 | 0.00706 | 0.00931 | 0.00000 | 0.00000 | 0.00005 | 0.00199 | 0.01042 | 1.32051 | 0.71091 | 0.44247 |
| **Camk2a** | 0.01552 | 0.01840 | 0.05521 | 0.03985 | 0.00911 | 0.02243 | 0.00743 | 0.02342 | 0.02730 | 0.03182 | 0.01207 | 0.00697 | 0.03739 | 0.13700 | 0.08129 | 0.00752 |
| **Camk2b** | 0.00942 | 0.00725 | 0.00335 | 0.00992 | 0.00928 | 0.00826 | 0.00666 | 0.50058 | 0.58875 | 0.95543 | 1.49594 | 0.00000 | 0.00000 | 0.00885 | 0.00000 | 0.01509 |
| **Camk2d** | 1.21996 | 1.68472 | 0.49133 | 0.00000 | 4.40993 | 3.04343 | 2.21562 | 0.81507 | 0.00706 | 0.11313 | 0.65998 | 0.03486 | 1.00974 | 0.00356 | 0.21025 | 2.47821 |
| **Camk2g** | 4.58632 | 0.63679 | 4.17724 | 8.33751 | 0.00405 | 2.07915 | 3.61626 | 3.34107 | 6.92426 | 6.33852 | 7.31253 | 0.45217 | 3.63810 | 1.94093 | 4.04747 | 1.56976 |
| **Camk2n2** | 0.01289 | 0.00794 | 0.00817 | 0.92708 | 0.00943 | 0.00693 | 0.04824 | 0.02187 | 0.40879 | 0.00000 | 0.37753 | 0.06347 | 0.01128 | 0.01760 | 0.05752 | 0.00715 |
| **Camk4** | 0.55393 | 0.01926 | 0.15220 | 0.29660 | 0.04968 | 0.02029 | 0.06906 | 0.06578 | 0.41098 | 0.19389 | 0.37786 | 0.11464 | 0.09074 | 0.39973 | 0.24362 | 0.05850 |
| **Camkk2** | 0.01366 | 0.00795 | 0.01212 | 2.13119 | 0.00922 | 0.00633 | 0.00745 | 0.01258 | 0.55222 | 0.11566 | 0.00812 | 0.00626 | 0.01196 | 0.00398 | 0.11074 | 0.00680 |
| **Camkmt** | 1.65782 | 0.00686 | 0.76404 | 0.10467 | 0.00634 | 0.00441 | 0.00690 | 0.40150 | 0.06691 | 0.29940 | 0.44202 | 0.02543 | 0.02437 | 0.00922 | 0.00000 | 1.99170 |
| **Caml** | 11.10667 | 0.04366 | 6.28044 | 6.91113 | 0.03410 | 2.01049 | 2.51333 | 5.64886 | 13.60302 | 8.98065 | 5.34481 | 7.16207 | 6.56327 | 0.63884 | 5.87508 | 1.34732 |
| **Camp** | 0.01364 | 0.00770 | 0.01070 | 0.53259 | 0.00930 | 0.00657 | 0.00734 | 0.01135 | 0.00470 | 0.25334 | 0.01108 | 0.06587 | 0.01255 | 0.00526 | 0.44887 | 0.00736 |
| **Camsap1** | 2.87628 | 3.93452 | 6.14499 | 13.97953 | 3.41720 | 4.45209 | 3.79359 | 7.80048 | 16.82628 | 7.76122 | 7.34214 | 1.74975 | 5.86483 | 4.88451 | 10.55867 | 3.71478 |
| **Camsap2** | 0.93635 | 4.98351 | 1.71826 | 12.54343 | 0.30824 | 0.56403 | 0.39617 | 0.23625 | 4.94318 | 0.20358 | 0.40658 | 12.24954 | 1.06086 | 7.22628 | 11.79609 | 1.55408 |
| **Camsap3** | 1.08081 | 0.00613 | 0.87930 | 0.12930 | 0.04798 | 2.16131 | 1.27691 | 0.38646 | 0.00680 | 0.57603 | 0.33452 | 0.03907 | 2.15068 | 0.00809 | 0.00000 | 0.27275 |
| **Camta1** | 0.09730 | 0.00862 | 0.75897 | 0.76964 | 0.01582 | 0.00907 | 0.02806 | 0.00888 | 0.26161 | 0.23875 | 0.01666 | 0.00930 | 0.03866 | 2.07453 | 0.26070 | 0.38092 |
| **Camta2** | 0.72853 | 0.00671 | 0.00000 | 0.00214 | 0.01468 | 4.61458 | 0.01715 | 0.85045 | 0.01156 | 1.18366 | 0.00011 | 0.02166 | 0.00000 | 0.01453 | 0.00012 | 2.57351 |
| **Cand1** | 23.13256 | 18.86165 | 23.01524 | 21.97063 | 13.15598 | 9.76494 | 13.00009 | 29.05168 | 23.81968 | 36.17407 | 26.20615 | 26.45270 | 16.12621 | 37.46319 | 32.94314 | 1.75266 |
| **Cand2** | 2.00336 | 0.00435 | 0.04506 | 0.08294 | 2.25241 | 0.02504 | 0.06558 | 0.01785 | 2.16931 | 0.11603 | 0.45018 | 0.03315 | 0.08314 | 0.16515 | 1.17289 | 0.01185 |
| **Cant1** | 0.75255 | 0.70270 | 3.45630 | 3.31519 | 0.58088 | 0.24673 | 0.65208 | 2.56084 | 1.64481 | 2.77959 | 0.05661 | 0.06553 | 1.64061 | 0.02170 | 0.60006 | 3.91614 |
| **Canx** | 11.82744 | 7.51816 | 24.76500 | 9.81317 | 7.95669 | 9.57581 | 12.32581 | 10.34517 | 8.09819 | 12.75186 | 11.69705 | 17.60188 | 16.35641 | 31.45549 | 15.80637 | 8.18259 |
| **Cap1** | 2.71184 | 1.94896 | 0.00000 | 0.56862 | 0.00330 | 0.00992 | 2.45252 | 0.52504 | 0.00073 | 0.73467 | 0.01767 | 0.08747 | 0.00648 | 0.32720 | 1.66491 | 0.01092 |
| **Capg** | 4.85416 | 2.51211 | 0.00000 | 0.00000 | 2.48909 | 8.68045 | 0.02130 | 0.00000 | 0.02187 | 0.40329 | 0.24867 | 0.03725 | 1.26669 | 0.00747 | 0.00232 | 7.37922 |
| **Capn1** | 1.88696 | 0.00533 | 0.11516 | 1.95303 | 0.10570 | 0.04109 | 0.02361 | 0.10585 | 1.63776 | 4.13183 | 3.93312 | 0.11463 | 1.36027 | 0.10357 | 0.48736 | 0.73431 |
| **Capn10** | 4.77227 | 0.34442 | 1.09041 | 1.68209 | 0.04944 | 0.00773 | 0.85503 | 3.73615 | 4.87288 | 2.51594 | 1.94174 | 0.07662 | 4.13078 | 0.49984 | 0.04988 | 2.95137 |
| **Capn11** | 0.32458 | 0.13648 | 0.00139 | 0.40073 | 0.00725 | 0.00828 | 0.00666 | 0.76156 | 0.91066 | 2.12000 | 1.14321 | 0.04941 | 0.05990 | 0.00465 | 0.24344 | 0.00669 |
| **Capn15** | 0.00000 | 0.63734 | 0.24654 | 0.00143 | 0.33452 | 1.97665 | 1.12873 | 0.90374 | 0.01090 | 1.47420 | 0.01936 | 0.02993 | 1.10655 | 0.00882 | 0.00426 | 1.52638 |
| **Capn2** | 0.01311 | 0.00787 | 0.14488 | 0.00743 | 0.01095 | 8.26427 | 0.00623 | 0.00742 | 0.01323 | 0.00000 | 0.00809 | 0.03627 | 0.99143 | 0.01344 | 0.01143 | 0.29025 |
| **Capn5** | 4.27690 | 0.00309 | 1.60583 | 0.23735 | 0.01732 | 3.17965 | 0.00231 | 1.04372 | 0.00741 | 0.33647 | 0.00000 | 0.00000 | 2.20875 | 0.01132 | 0.00000 | 0.00765 |
| **Capn6** | 0.00938 | 0.00767 | 2.08673 | 0.00887 | 0.00656 | 4.00384 | 0.00685 | 0.00873 | 0.01261 | 0.00000 | 0.00962 | 0.00655 | 0.47889 | 0.01777 | 0.00000 | 0.01250 |
| **Capn7** | 1.20882 | 6.04909 | 2.53555 | 5.98090 | 0.00470 | 0.38594 | 1.31012 | 1.94996 | 0.00000 | 0.00000 | 0.19128 | 0.85928 | 0.46980 | 1.96253 | 4.52211 | 0.04771 |
| **Capns1** | 23.59603 | 7.61958 | 17.49780 | 1.80543 | 15.73635 | 13.56255 | 8.98031 | 5.37546 | 0.60539 | 7.36765 | 1.73940 | 0.12481 | 16.01496 | 0.00000 | 0.33377 | 12.62119 |
| **Caprin1** | 14.51413 | 15.66901 | 16.79183 | 12.19619 | 21.23633 | 9.26703 | 10.59598 | 18.15402 | 12.85488 | 22.41276 | 12.41415 | 23.27942 | 13.64198 | 27.22315 | 19.12004 | 7.72910 |
| **Caprin2** | 7.09916 | 0.01010 | 5.14402 | 10.00082 | 0.48982 | 2.41045 | 0.00979 | 0.70166 | 7.32354 | 3.43684 | 1.90823 | 0.39101 | 0.78650 | 4.59000 | 6.39524 | 0.01708 |
| **Capsl** | 0.01235 | 0.00804 | 0.03731 | 0.25187 | 0.00882 | 0.00601 | 0.00739 | 0.01052 | 0.00765 | 1.42966 | 0.01264 | 0.00460 | 0.01122 | 0.01088 | 0.00477 | 0.00761 |
| **Capza1** | 9.72461 | 12.51825 | 10.51343 | 4.77952 | 8.47935 | 11.81028 | 6.89758 | 8.59029 | 6.90177 | 9.02460 | 8.25489 | 11.24749 | 8.93215 | 23.03218 | 9.33763 | 4.80604 |
| **Capza2** | 10.20133 | 10.01726 | 9.67428 | 4.00585 | 2.25582 | 10.15957 | 11.12281 | 14.85410 | 1.61187 | 16.05820 | 8.01419 | 11.95344 | 14.99958 | 26.56922 | 13.07362 | 7.05775 |
| **Capzb** | 19.71400 | 8.06721 | 16.07160 | 1.11870 | 16.73158 | 14.96671 | 9.60806 | 16.68772 | 0.00000 | 9.80740 | 10.30286 | 2.63971 | 17.62654 | 0.15763 | 1.01786 | 11.07505 |
| **Car2** | 0.00748 | 0.00768 | 0.01208 | 0.00503 | 0.00753 | 0.00648 | 0.00717 | 0.00952 | 0.01079 | 4.69402 | 0.00916 | 0.00000 | 0.00834 | 0.01246 | 0.00641 | 1.33810 |
| **Car4** | 0.00000 | 0.01914 | 1.90618 | 0.00095 | 0.03544 | 0.00685 | 0.00608 | 0.07957 | 0.29246 | 1.75799 | 1.06339 | 0.08291 | 2.07176 | 0.09604 | 1.07280 | 5.91498 |
| **Car5a** | 0.00838 | 0.44576 | 0.59970 | 0.09533 | 0.00725 | 0.01277 | 0.01731 | 0.00988 | 0.11968 | 0.08116 | 0.25024 | 0.00616 | 0.01470 | 0.03312 | 0.04998 | 0.02835 |

| | | | | | | | | | | | | | | | | |
|---|---|---|---|---|---|---|---|---|---|---|---|---|---|---|---|---|
| **Car5b** | 1.93316 | 0.61939 | 0.96140 | 0.51095 | 0.09916 | 0.05461 | 0.79520 | 0.47676 | 0.59852 | 0.43330 | 1.36575 | 0.74868 | 0.81588 | 1.61629 | 0.46052 | 0.05561 |
| **Car6** | 0.01266 | 0.00798 | 0.07150 | 0.00437 | 0.00920 | 0.00841 | 0.00726 | 0.01155 | 0.17995 | 0.09730 | 0.01028 | 0.00673 | 0.01832 | 0.00623 | 0.28179 | 0.00742 |
| **Car8** | 0.09949 | 0.00786 | 0.12106 | 0.39877 | 0.03003 | 0.01236 | 0.00714 | 0.00994 | 0.14874 | 0.54480 | 0.35304 | 0.03161 | 0.11828 | 0.15201 | 0.07416 | 0.03016 |
| **Car9** | 1.68251 | 0.00616 | 0.40697 | 0.60649 | 0.06143 | 0.00523 | 0.00643 | 0.21524 | 0.89391 | 0.75545 | 0.01618 | 0.01317 | 0.54572 | 0.08814 | 0.37958 | 0.00502 |
| **Card10** | 3.51393 | 1.47872 | 0.15484 | 0.26572 | 0.01812 | 0.16842 | 0.00352 | 0.81856 | 0.40116 | 2.12352 | 0.44489 | 0.00519 | 0.47733 | 0.00742 | 0.00000 | 0.00241 |
| **Card11** | 0.01356 | 0.00823 | 0.01943 | 0.01118 | 0.00917 | 0.00622 | 0.00761 | 0.04969 | 0.09787 | 0.00000 | 1.19169 | 0.00209 | 0.00000 | 0.00441 | 0.16998 | 0.00694 |
| **Card14** | 0.00842 | 0.00786 | 0.00775 | 0.00744 | 0.00932 | 0.00700 | 0.00740 | 1.06149 | 0.01068 | 0.95504 | 0.00454 | 0.03714 | 0.00582 | 0.01254 | 0.01099 | 0.00726 |
| **Card19** | 0.01176 | 0.00778 | 0.00324 | 0.00932 | 0.00952 | 1.94394 | 0.00718 | 0.20068 | 0.01226 | 0.06950 | 0.00840 | 0.01886 | 0.00000 | 0.01185 | 0.00894 | 0.00671 |
| **Card6** | 0.06622 | 0.00784 | 0.03090 | 1.60618 | 0.01635 | 0.00495 | 0.00707 | 0.15124 | 6.21733 | 3.81471 | 0.00000 | 0.07680 | 0.01533 | 7.26968 | 4.90824 | 0.01337 |
| **Card9** | 0.01374 | 0.00788 | 0.01220 | 0.05501 | 0.00920 | 0.00653 | 0.00739 | 0.01195 | 0.01144 | 0.16220 | 0.01523 | 0.00812 | 0.01163 | 0.01798 | 0.00834 | 0.00716 |
| **Carf** | 0.49466 | 0.01701 | 0.17597 | 1.23994 | 0.06198 | 0.01262 | 0.05718 | 1.05058 | 0.40144 | 0.12608 | 0.28585 | 0.05751 | 0.15159 | 0.44734 | 1.58684 | 0.04213 |
| **Carhsp1** | 6.24992 | 4.61533 | 5.29818 | 12.23152 | 2.20551 | 6.36464 | 4.18051 | 4.37432 | 10.60357 | 6.66213 | 3.53471 | 0.85034 | 5.10888 | 0.40721 | 3.44450 | 4.05377 |
| **Carkd** | 6.73822 | 1.48155 | 7.73347 | 0.55109 | 0.04856 | 4.08831 | 6.14425 | 3.08515 | 0.81594 | 3.42692 | 2.30968 | 2.35439 | 7.25357 | 0.02276 | 0.27850 | 3.29048 |
| **Carm1** | 2.80000 | 0.16207 | 0.75459 | 0.00000 | 0.89382 | 2.72749 | 1.85368 | 2.67834 | 0.05806 | 2.82051 | 0.00000 | 0.04925 | 2.04001 | 0.01042 | 0.00463 | 1.88571 |
| **Carnmt1** | 4.55798 | 4.40908 | 5.24940 | 7.39202 | 2.67828 | 2.10055 | 3.26057 | 4.26838 | 12.88207 | 2.83489 | 6.43908 | 8.70822 | 5.20540 | 1.82621 | 0.73718 | 2.87791 |
| **Carns1** | 0.06962 | 0.00745 | 0.49052 | 1.35465 | 0.01530 | 0.00856 | 0.00526 | 0.00000 | 4.22059 | 0.00000 | 2.66986 | 0.00000 | 1.48848 | 0.00000 | 0.90433 | 0.00539 |
| **Cars** | 8.19142 | 8.00398 | 4.43397 | 1.90281 | 6.08625 | 0.00000 | 4.07622 | 11.92071 | 5.46368 | 5.42066 | 3.28904 | 0.44340 | 7.08840 | 0.03077 | 0.59626 | 5.82822 |
| **Cars2** | 7.01268 | 1.33259 | 3.03274 | 2.65806 | 4.65298 | 0.00551 | 0.09163 | 2.93604 | 4.37737 | 8.53516 | 5.48202 | 1.30167 | 1.59556 | 0.04358 | 1.72856 | 0.00145 |
| **Casc1** | 0.01636 | 0.00760 | 0.13050 | 0.69717 | 0.00904 | 0.00568 | 0.00730 | 0.01221 | 0.00000 | 0.00933 | 0.01355 | 0.00706 | 0.01086 | 0.00524 | 0.17372 | 0.00715 |
| **Casc3** | 4.65710 | 5.71692 | 3.20928 | 7.88362 | 3.49925 | 0.63314 | 4.75123 | 5.51573 | 9.07195 | 13.68746 | 7.21904 | 9.63191 | 4.54128 | 7.90738 | 6.37466 | 1.89661 |
| **Casc4** | 3.84740 | 4.11968 | 5.74218 | 16.63204 | 2.66585 | 7.68892 | 4.46160 | 4.95035 | 21.70115 | 0.88279 | 22.32688 | 4.88143 | 4.48090 | 14.77814 | 5.97393 | 6.81053 |
| **Casc5** | 0.30464 | 0.00655 | 0.10334 | 2.14439 | 0.01695 | 0.00765 | 0.38414 | 0.36633 | 2.39064 | 0.00000 | 0.62839 | 3.84329 | 0.73309 | 9.67913 | 4.69014 | 0.00556 |
| **Casd1** | 1.13154 | 0.00622 | 0.00292 | 0.01095 | 0.00607 | 0.00791 | 0.00655 | 0.58257 | 0.01169 | 0.00320 | 0.01180 | 0.00576 | 0.02555 | 0.01560 | 0.00270 | 0.26309 |
| **Cask** | 0.31495 | 0.00558 | 0.00000 | 1.63179 | 2.37100 | 0.00560 | 0.16268 | 1.03969 | 2.54142 | 0.57513 | 0.32877 | 0.08031 | 0.00410 | 1.85794 | 2.92952 | 0.00685 |
| **Caskin1** | 3.44351 | 0.78702 | 2.19065 | 3.94233 | 0.06252 | 2.31467 | 0.68365 | 6.80334 | 1.85064 | 7.52440 | 5.28653 | 0.63698 | 3.32463 | 0.02705 | 0.18443 | 0.92059 |
| **Caskin2** | 1.47598 | 0.00591 | 0.23997 | 1.00059 | 0.03637 | 4.14189 | 0.49048 | 0.96493 | 1.72843 | 3.50106 | 0.31630 | 0.10927 | 1.95293 | 0.01782 | 0.23932 | 2.60023 |
| **Casp2** | 2.39999 | 0.01481 | 0.66375 | 0.37138 | 0.02288 | 0.31826 | 0.00440 | 0.59526 | 0.28461 | 2.04786 | 0.89148 | 0.03937 | 0.82210 | 0.00647 | 2.47857 | 0.53232 |
| **Casp3** | 1.81371 | 5.95164 | 6.16691 | 7.05742 | 5.48197 | 0.00262 | 4.58816 | 6.44573 | 8.94459 | 10.11388 | 5.00838 | 0.75230 | 1.71534 | 12.40362 | 7.52448 | 1.75676 |
| **Casp6** | 12.49265 | 13.52787 | 15.79194 | 1.52419 | 17.47981 | 6.74589 | 8.98792 | 12.65498 | 3.86949 | 15.25074 | 15.22877 | 1.65134 | 6.53617 | 1.87985 | 7.05211 | 2.39381 |
| **Casp7** | 3.35359 | 0.30406 | 0.77215 | 0.00000 | 3.14017 | 3.11940 | 1.38443 | 2.32843 | 0.01885 | 1.29608 | 1.57632 | 0.00000 | 2.54345 | 0.00205 | 0.47784 | 1.28329 |
| **Casp8ap2** | 1.86866 | 3.99511 | 1.41684 | 1.83224 | 1.58849 | 0.02033 | 4.53967 | 5.80629 | 4.81835 | 5.62110 | 1.21677 | 6.17867 | 0.94064 | 15.99186 | 18.53642 | 0.46808 |
| **Casp9** | 2.73206 | 0.01187 | 4.89626 | 0.88217 | 4.44263 | 0.00310 | 2.68845 | 2.56143 | 0.00000 | 1.87459 | 0.00000 | 0.05234 | 1.61184 | 0.13514 | 1.13713 | 1.30193 |
| **Casq1** | 0.01366 | 0.00800 | 0.01216 | 0.00882 | 0.00915 | 0.00664 | 0.00735 | 0.01137 | 0.01197 | 0.29748 | 0.01317 | 0.00698 | 0.01253 | 0.00832 | 0.05404 | 0.00780 |
| **Casq2** | 1.48516 | 0.00602 | 0.00354 | 0.10019 | 0.01332 | 0.00682 | 0.00633 | 0.00768 | 0.00761 | 0.01836 | 0.01156 | 0.05720 | 0.01145 | 0.01167 | 0.00778 | 0.02029 |
| **Cass4** | 0.01223 | 0.00802 | 1.20766 | 0.45471 | 0.00725 | 0.00547 | 0.00733 | 0.01015 | 0.00510 | 0.08888 | 0.01215 | 0.01952 | 0.00719 | 0.00838 | 0.53585 | 0.00659 |
| **Cast** | 0.32441 | 0.01848 | 3.48022 | 0.79140 | 0.36822 | 2.04051 | 0.21009 | 3.06154 | 1.03343 | 1.27168 | 1.83899 | 0.11081 | 3.28143 | 4.82276 | 0.59238 | 0.21899 |
| **Casz1** | 0.20091 | 0.00848 | 0.21875 | 0.00844 | 0.01667 | 0.04731 | 0.00690 | 0.00894 | 0.02370 | 0.24232 | 0.01119 | 0.02709 | 0.67583 | 0.00785 | 0.05035 | 0.02169 |
| **Cat** | 7.42567 | 5.94180 | 2.85455 | 0.00000 | 9.66949 | 7.10760 | 6.83774 | 6.07438 | 0.07020 | 2.23357 | 0.84451 | 0.02829 | 10.27236 | 0.00498 | 0.00194 | 1.88247 |
| **Catip** | 0.23843 | 0.00748 | 0.00675 | 0.00655 | 0.00732 | 0.00694 | 0.00703 | 0.00851 | 0.01159 | 0.99572 | 0.07571 | 0.00450 | 0.01100 | 0.01154 | 0.00680 | 0.00733 |
| **Catsper2** | 0.07292 | 0.35713 | 0.69719 | 0.00175 | 0.00700 | 3.17102 | 0.00796 | 0.15408 | 0.79815 | 0.21440 | 0.00000 | 0.01848 | 0.00000 | 0.07334 | 0.08275 | 0.00595 |
| **Catsperd** | 0.00824 | 0.00868 | 0.00000 | 2.18647 | 0.00971 | 0.00663 | 0.00801 | 0.25925 | 0.50782 | 3.15579 | 0.42575 | 0.09105 | 0.08303 | 0.87397 | 4.24972 | 0.00726 |
| **Catsperg1** | 0.00974 | 0.00769 | 0.08874 | 1.62676 | 0.00943 | 0.00529 | 0.00724 | 0.57860 | 2.37349 | 0.00000 | 0.67380 | 0.12081 | 0.00612 | 1.10245 | 0.29526 | 0.00653 |
| **Catsperg2** | 0.01152 | 0.00771 | 0.01631 | 0.45172 | 0.00947 | 0.00613 | 0.00739 | 0.21676 | 0.62452 | 0.00000 | 0.08207 | 0.04246 | 0.00642 | 0.38820 | 0.12581 | 0.00716 |
| **Cav1** | 0.00970 | 0.00873 | 0.00802 | 0.00660 | 0.00938 | 0.00691 | 0.00818 | 0.24708 | 0.01069 | 0.43375 | 0.00798 | 0.10487 | 0.00709 | 0.00663 | 0.42590 | 0.00768 |
| **Cav2** | 0.01210 | 0.00780 | 0.00248 | 0.00541 | 0.00949 | 0.00677 | 0.00700 | 0.00232 | 0.01128 | 0.00205 | 0.21664 | 0.09646 | 0.01021 | 2.38749 | 0.00000 | 0.00733 |
| **Cbarp** | 1.27114 | 1.13912 | 0.11220 | 0.00174 | 0.00559 | 0.00784 | 0.00562 | 1.81548 | 0.00940 | 0.00000 | 1.21984 | 0.00639 | 0.04304 | 0.00495 | 0.00578 | 0.02020 |
| **Cbfa2t2** | 6.97785 | 5.30926 | 2.96432 | 8.26398 | 5.19459 | 0.30225 | 1.66030 | 2.43562 | 15.31575 | 5.42317 | 2.19708 | 0.47381 | 1.29537 | 7.00250 | 4.47696 | 3.51441 |
| **Cbfa2t3** | 0.01747 | 0.00789 | 1.58481 | 0.47582 | 0.00937 | 0.11384 | 0.00687 | 0.31258 | 0.33472 | 0.17779 | 0.03035 | 0.00551 | 0.33932 | 0.00921 | 0.00000 | 0.00662 |
| **Cbfb** | 6.99494 | 6.77761 | 3.16228 | 4.08629 | 3.75502 | 1.37966 | 6.10865 | 3.18851 | 7.41725 | 6.97748 | 2.59459 | 6.23963 | 1.36952 | 11.91662 | 5.37052 | 0.63254 |
| **Cbl** | 0.45921 | 0.00376 | 2.87499 | 7.68396 | 0.02426 | 0.00653 | 3.03276 | 3.36533 | 5.10342 | 3.73819 | 0.77487 | 6.35541 | 0.79101 | 9.14315 | 11.49541 | 1.60835 |
| **Cblb** | 0.00000 | 1.49184 | 0.00000 | 0.00000 | 1.61343 | 0.00728 | 1.26838 | 0.00000 | 0.01027 | 0.00000 | 0.53869 | 0.01060 | 0.56262 | 0.01010 | 0.00000 | 2.18904 |
| **Cblc** | 3.82548 | 2.28620 | 0.01109 | 0.00000 | 5.49518 | 0.00594 | 0.01383 | 0.00000 | 0.00862 | 0.77557 | 0.65892 | 0.00000 | 0.21490 | 0.00789 | 0.00243 | 0.00289 |
| **Cbll1** | 4.77868 | 5.92210 | 2.53485 | 14.07435 | 0.02258 | 0.02078 | 1.33564 | 3.02713 | 18.50685 | 6.14439 | 5.67585 | 7.61707 | 3.33441 | 9.73326 | 11.68621 | 3.37233 |
| **Cbln1** | 0.03932 | 0.00683 | 0.00002 | 2.45452 | 3.18763 | 0.00781 | 0.04235 | 0.25238 | 1.68348 | 0.57656 | 0.00000 | 0.00000 | 0.00708 | 0.00300 | 0.83095 | 0.00649 |
| **Cbln3** | 0.27057 | 0.04897 | 1.06889 | 0.26114 | 0.32075 | 0.06106 | 0.09524 | 0.00000 | 0.52486 | 0.32534 | 5.15567 | 0.38837 | 1.05348 | 1.23985 | 0.27670 | 0.20535 |
| **Cbr1** | 18.44685 | 0.09536 | 3.47985 | 2.25478 | 7.43099 | 0.01633 | 5.34951 | 18.39430 | 3.58433 | 12.19395 | 11.55360 | 0.43802 | 8.83149 | 1.20150 | 0.15763 | 10.34129 |
| **Cbr2** | 0.07624 | 0.00000 | 7.76563 | 0.00000 | 4.60482 | 0.99182 | 4.66470 | 5.93398 | 0.46321 | 9.74724 | 7.37600 | 0.05195 | 9.74544 | 0.06195 | 0.99903 | 0.00000 |
| **Cbr3** | 0.06124 | 0.01817 | 0.23907 | 0.00810 | 0.00815 | 0.00569 | 0.02773 | 0.05042 | 0.03400 | 0.33159 | 0.01226 | 0.00632 | 0.02930 | 0.01234 | 0.00595 | 0.01515 |
| **Cbr4** | 2.31676 | 3.80802 | 3.25821 | 0.00345 | 1.80264 | 0.00319 | 2.26992 | 1.05266 | 0.01045 | 0.45533 | 0.00000 | 0.09892 | 1.03548 | 0.00439 | 0.05139 | 0.01629 |
| **Cbs** | 0.00000 | 0.02737 | 2.29284 | 0.71347 | 0.01257 | 7.30245 | 0.63474 | 2.98591 | 0.00000 | 2.52641 | 0.56425 | 0.03854 | 0.22105 | 0.03730 | 0.00000 | 3.31454 |
| **Cbwd1** | 0.94019 | 0.22363 | 1.21066 | 1.21231 | 2.07493 | 1.35115 | 0.82902 | 2.25076 | 5.65479 | 0.79386 | 2.71613 | 5.00063 | 0.32937 | 8.20668 | 6.81322 | 0.01133 |
| **Cbx1** | 4.89382 | 0.03382 | 0.41732 | 0.93922 | 0.02988 | 0.27811 | 0.17242 | 2.89820 | 0.98120 | 0.73469 | 0.98738 | 0.19781 | 3.43030 | 0.53797 | 0.51094 | 2.59680 |

| | | | | | | | | | | | | | | | | |
|---|---|---|---|---|---|---|---|---|---|---|---|---|---|---|---|---|
| **Cbx2** | 0.62454 | 0.00581 | 0.00000 | 7.85594 | 1.59481 | 0.00488 | 0.00669 | 1.04877 | 4.82262 | 1.12525 | 0.14366 | 0.02701 | 0.03827 | 0.00000 | 6.23354 | 0.35209 |
| **Cbx3** | 8.69809 | 6.35853 | 6.70681 | 11.52667 | 11.43758 | 4.68193 | 3.81373 | 5.75294 | 17.18585 | 10.08600 | 15.21041 | 4.25380 | 2.58334 | 19.37240 | 13.30845 | 2.12207 |
| **Cbx4** | 0.03677 | 0.00783 | 0.01030 | 0.33641 | 0.00833 | 0.00671 | 0.00739 | 0.02823 | 0.00589 | 0.01409 | 0.03044 | 0.00801 | 0.01137 | 0.01089 | 0.00700 | 0.00728 |
| **Cbx5** | 15.01701 | 4.43582 | 9.56552 | 6.13942 | 10.44253 | 10.03151 | 2.57053 | 6.92875 | 13.53616 | 16.74515 | 6.32090 | 14.66156 | 5.86127 | 25.15774 | 14.48633 | 3.00279 |
| **Cbx6** | 0.01693 | 0.00834 | 0.02938 | 0.00646 | 0.09877 | 0.00618 | 0.00754 | 0.01298 | 0.01130 | 1.03486 | 0.01055 | 0.04350 | 0.01791 | 0.00739 | 0.11263 | 0.95679 |
| **Cbx7** | 7.41423 | 4.69183 | 4.97660 | 2.33637 | 6.07096 | 0.00490 | 7.45889 | 5.20175 | 3.65078 | 4.91430 | 2.34938 | 0.88252 | 3.98979 | 2.74424 | 5.66957 | 2.13593 |
| **Cbx8** | 0.00583 | 0.00612 | 0.17324 | 0.44195 | 2.20786 | 0.00872 | 0.00699 | 0.86975 | 0.16521 | 0.02963 | 0.00493 | 0.00247 | 0.13620 | 0.03290 | 0.20691 | 0.00719 |
| **Cby1** | 1.63453 | 0.02566 | 2.31605 | 3.79314 | 0.00976 | 0.01283 | 0.00548 | 3.79325 | 0.88372 | 1.73826 | 0.66224 | 0.00000 | 0.06259 | 0.70024 | 1.31636 | 0.00429 |
| **Cc2d1a** | 3.66019 | 0.00187 | 0.05741 | 0.37162 | 1.79416 | 0.01853 | 0.48264 | 3.00237 | 0.57519 | 2.35834 | 0.03926 | 0.05713 | 8.51173 | 0.01186 | 0.01516 | 0.01469 |
| **Cc2d1b** | 6.33547 | 2.39178 | 3.23561 | 1.17792 | 0.51279 | 0.02431 | 5.98344 | 7.44964 | 1.32766 | 0.00000 | 0.63718 | 1.13355 | 6.66470 | 0.00000 | 0.46859 | 2.18682 |
| **Ccar1** | 7.45435 | 4.00880 | 6.74977 | 16.32083 | 1.36611 | 4.81899 | 5.47839 | 9.38121 | 17.13433 | 17.07157 | 10.08069 | 12.07648 | 8.50955 | 29.26608 | 23.56868 | 5.12804 |
| **Ccar2** | 1.14777 | 0.12250 | 0.00000 | 0.08085 | 0.00642 | 0.00709 | 1.41367 | 0.77390 | 0.02706 | 3.14765 | 0.00400 | 0.01657 | 1.10508 | 0.00820 | 0.59813 | 0.00565 |
| **Ccbe1** | 0.01355 | 0.00781 | 0.01132 | 0.02099 | 0.00909 | 0.00667 | 0.00728 | 0.01053 | 0.24922 | 0.51633 | 0.01032 | 0.04365 | 0.01219 | 0.01180 | 0.00662 | 0.00778 |
| **Ccdc106** | 0.68433 | 0.00621 | 0.00184 | 0.00520 | 0.00639 | 0.00799 | 1.24896 | 0.00821 | 0.07944 | 0.34431 | 0.01190 | 0.00501 | 0.01103 | 0.01131 | 0.00981 | 0.00688 |
| **Ccdc107** | 0.87105 | 0.00568 | 0.00000 | 3.52958 | 2.23510 | 1.58530 | 0.00252 | 0.20099 | 3.54230 | 1.37124 | 2.76764 | 0.03572 | 5.25731 | 0.06335 | 0.00000 | 0.01785 |
| **Ccdc109b** | 0.09284 | 0.00915 | 0.00402 | 1.51084 | 0.01586 | 0.00768 | 0.00805 | 0.54517 | 0.44370 | 0.00551 | 0.01836 | 0.00693 | 0.00572 | 0.00927 | 0.01756 | 0.00715 |
| **Ccdc112** | 1.66029 | 0.00404 | 0.23774 | 0.02329 | 0.85596 | 0.00587 | 1.45368 | 0.00150 | 0.02701 | 0.00939 | 0.01115 | 0.00379 | 0.02190 | 0.01768 | 0.24896 | 0.01246 |
| **Ccdc114** | 0.01360 | 0.00807 | 0.01198 | 0.77405 | 0.00916 | 0.00682 | 0.00753 | 0.01305 | 0.00000 | 0.28679 | 0.01224 | 0.00856 | 0.01268 | 0.01228 | 0.01284 | 0.00728 |
| **Ccdc115** | 4.20595 | 2.80443 | 8.19887 | 12.37020 | 6.35394 | 5.88514 | 5.10391 | 11.97653 | 16.42558 | 14.40905 | 7.67660 | 4.31264 | 11.76622 | 4.45133 | 6.19007 | 0.09605 |
| **Ccdc116** | 0.01561 | 0.00788 | 0.55292 | 1.16220 | 0.00808 | 0.00554 | 0.00732 | 0.08393 | 0.34554 | 0.00850 | 0.02605 | 0.03907 | 0.00627 | 0.01518 | 0.00000 | 0.00693 |
| **Ccdc117** | 1.02069 | 5.16507 | 12.31049 | 17.10938 | 0.43604 | 7.17388 | 2.15096 | 14.07063 | 18.71203 | 27.60522 | 13.75365 | 19.33006 | 3.21355 | 28.53786 | 27.17032 | 0.69033 |
| **Ccdc12** | 6.97416 | 3.95312 | 5.59019 | 9.34781 | 3.58681 | 6.03564 | 3.16289 | 7.83414 | 7.77699 | 5.99692 | 10.54937 | 1.74941 | 9.49187 | 0.22773 | 0.94525 | 9.42843 |
| **Ccdc120** | 0.19874 | 0.00822 | 0.00000 | 3.71323 | 0.01545 | 0.00678 | 0.00696 | 0.94626 | 0.00000 | 0.15030 | 0.01881 | 0.01399 | 0.00579 | 0.00583 | 1.90118 | 0.00672 |
| **Ccdc124** | 12.66939 | 0.07352 | 0.00000 | 4.32336 | 10.42172 | 11.44109 | 6.06661 | 30.21755 | 9.19448 | 25.70998 | 6.76999 | 2.43162 | 28.10654 | 0.00000 | 0.00000 | 7.25640 |
| **Ccdc125** | 0.05233 | 0.00747 | 0.07102 | 0.57590 | 0.04613 | 0.00619 | 0.02724 | 0.01342 | 3.96085 | 1.08019 | 0.10191 | 0.00000 | 0.03605 | 0.96971 | 2.46341 | 0.00692 |
| **Ccdc126** | 2.82616 | 2.85592 | 2.09554 | 7.01893 | 1.58596 | 3.43766 | 0.71454 | 0.90735 | 9.69855 | 1.04574 | 0.43411 | 0.40148 | 1.01596 | 1.11320 | 2.18599 | 0.32628 |
| **Ccdc127** | 5.44665 | 7.90206 | 5.81500 | 1.66583 | 0.67716 | 5.12457 | 4.39916 | 1.52131 | 0.07958 | 4.71701 | 2.78137 | 1.00592 | 4.76564 | 4.46365 | 3.82957 | 0.35214 |
| **Ccdc13** | 0.00619 | 0.00644 | 1.12911 | 0.06260 | 0.01192 | 0.00905 | 2.07831 | 1.39174 | 0.00775 | 0.00000 | 0.00001 | 0.00231 | 1.28067 | 0.01696 | 0.00000 | 0.12115 |
| **Ccdc130** | 1.45258 | 4.85491 | 6.55847 | 1.43474 | 0.03403 | 0.01891 | 7.91934 | 3.37590 | 2.71599 | 0.98262 | 0.43770 | 0.23859 | 2.17901 | 0.21736 | 0.00000 | 0.90941 |
| **Ccdc134** | 0.36343 | 0.48686 | 0.35360 | 4.44884 | 0.06749 | 0.12579 | 0.02542 | 2.25499 | 4.70339 | 0.00000 | 2.20788 | 0.08707 | 1.75368 | 0.00000 | 1.84572 | 0.15362 |
| **Ccdc136** | 0.01294 | 0.00774 | 0.01136 | 0.08754 | 0.00893 | 0.00662 | 0.00740 | 0.01048 | 1.19078 | 1.09819 | 0.01138 | 0.00254 | 0.01167 | 0.00515 | 0.32940 | 0.00737 |
| **Ccdc137** | 1.66272 | 4.79159 | 8.44994 | 12.04374 | 0.02647 | 5.96871 | 5.28280 | 2.90067 | 19.96711 | 17.89703 | 8.33349 | 11.46025 | 6.24503 | 14.43695 | 16.03301 | 2.63702 |
| **Ccdc138** | 1.02789 | 0.00670 | 0.19703 | 0.31608 | 0.02567 | 0.00788 | 0.78685 | 0.00209 | 0.00418 | 1.67704 | 0.28382 | 0.32685 | 0.36588 | 1.54593 | 0.74290 | 0.00596 |
| **Ccdc14** | 1.09693 | 1.12814 | 2.77640 | 1.05077 | 0.01043 | 0.00737 | 3.80587 | 0.11420 | 0.60665 | 0.93031 | 0.09091 | 0.03647 | 0.48126 | 5.60714 | 0.42468 | 0.42077 |
| **Ccdc141** | 0.01785 | 0.00791 | 0.01179 | 0.18481 | 0.00902 | 0.00667 | 0.00751 | 0.01395 | 0.00589 | 0.01426 | 0.01202 | 0.00800 | 0.01275 | 0.01622 | 0.02233 | 0.00736 |
| **Ccdc142** | 0.71905 | 0.00637 | 0.01018 | 0.14659 | 1.44768 | 0.00580 | 0.00706 | 0.00836 | 0.00819 | 0.00710 | 0.01268 | 0.03952 | 0.00868 | 0.01233 | 0.00747 | 0.59735 |
| **Ccdc146** | 0.01299 | 0.00793 | 0.01213 | 0.00662 | 0.00885 | 0.00660 | 0.00734 | 0.01037 | 0.01159 | 1.41980 | 0.01074 | 0.00449 | 0.01087 | 0.01194 | 0.00818 | 0.00734 |
| **Ccdc148** | 0.01353 | 0.00813 | 0.01110 | 0.16060 | 0.00931 | 0.00660 | 0.00756 | 0.01420 | 0.21568 | 0.02212 | 0.05279 | 0.00740 | 0.01915 | 0.04913 | 0.17612 | 0.00737 |
| **Ccdc15** | 0.01210 | 0.00810 | 0.15138 | 0.42372 | 0.00888 | 0.00565 | 0.08177 | 0.40516 | 0.09876 | 1.03356 | 0.01029 | 0.00575 | 0.00503 | 0.01610 | 0.05707 | 0.00694 |
| **Ccdc150** | 0.00962 | 0.00796 | 0.00744 | 0.00949 | 0.00961 | 0.00679 | 0.00785 | 0.65723 | 0.01131 | 0.00000 | 0.00547 | 0.23111 | 0.00630 | 0.00682 | 0.24412 | 0.00730 |
| **Ccdc152** | 0.30496 | 0.00626 | 0.00277 | 0.00274 | 2.40444 | 0.00787 | 0.00701 | 0.00300 | 0.01080 | 0.35090 | 0.16641 | 0.02210 | 0.01445 | 0.05580 | 0.06154 | 0.00667 |
| **Ccdc153** | 0.01667 | 0.00765 | 0.01507 | 0.32052 | 0.00837 | 0.00738 | 0.00708 | 0.00000 | 0.02717 | 2.74662 | 0.46323 | 0.00000 | 0.00793 | 0.02756 | 0.19795 | 0.00722 |
| **Ccdc154** | 0.00855 | 0.00778 | 0.00372 | 0.28459 | 0.00957 | 0.00775 | 0.00705 | 0.37633 | 0.75309 | 0.49305 | 0.73156 | 0.01210 | 0.00426 | 0.00469 | 0.31306 | 0.00714 |
| **Ccdc155** | 0.99036 | 0.00672 | 2.29173 | 1.27438 | 0.01093 | 0.00590 | 0.00594 | 0.00325 | 0.13673 | 1.13747 | 0.20218 | 0.06340 | 0.00522 | 0.00584 | 0.00000 | 0.00565 |
| **Ccdc157** | 0.01360 | 0.01835 | 0.65485 | 2.77132 | 0.00981 | 0.00489 | 0.01745 | 0.32240 | 4.76332 | 0.05549 | 0.00000 | 0.00611 | 0.00607 | 0.00000 | 0.49145 | 0.00647 |
| **Ccdc158** | 0.01378 | 0.00766 | 0.01161 | 0.28506 | 0.00941 | 0.00655 | 0.00743 | 0.01203 | 0.10996 | 0.00624 | 0.01034 | 0.00694 | 0.01289 | 0.01940 | 0.33870 | 0.00733 |
| **Ccdc160** | 0.15481 | 0.00766 | 0.01290 | 0.46993 | 0.00759 | 0.00619 | 0.01730 | 0.01053 | 0.70974 | 0.03715 | 0.05794 | 0.00538 | 0.01139 | 0.00558 | 0.47225 | 0.13137 |
| **Ccdc163** | 1.95643 | 0.01555 | 4.97818 | 1.23529 | 0.00000 | 0.01084 | 0.88178 | 4.96753 | 1.07942 | 6.91893 | 6.81046 | 1.21885 | 2.19177 | 6.36087 | 6.96793 | 0.00853 |
| **Ccdc166** | 0.00903 | 0.05801 | 0.02733 | 3.63875 | 0.00971 | 0.00715 | 0.00657 | 0.07522 | 2.44954 | 1.51775 | 0.00000 | 0.00532 | 0.81327 | 0.00000 | 0.61827 | 0.04323 |
| **Ccdc167** | 0.63993 | 0.59420 | 0.65335 | 0.00219 | 0.03888 | 0.00847 | 0.77166 | 0.07526 | 0.07769 | 0.93545 | 0.12759 | 0.00000 | 0.10129 | 0.47926 | 0.25272 | 0.09035 |
| **Ccdc169** | 0.02363 | 0.00745 | 0.00737 | 0.00631 | 0.00881 | 0.00667 | 0.00725 | 0.01394 | 0.20344 | 0.00467 | 0.00610 | 0.00262 | 0.01308 | 1.41623 | 0.00000 | 0.00740 |
| **Ccdc17** | 0.01550 | 0.00765 | 1.19913 | 0.00453 | 0.01621 | 0.00564 | 0.00656 | 0.00597 | 0.05148 | 0.02260 | 0.08358 | 0.01475 | 0.81622 | 0.00752 | 0.52826 | 0.00681 |
| **Ccdc170** | 0.01388 | 0.00770 | 0.53381 | 1.67298 | 0.01029 | 0.00537 | 0.46415 | 0.07683 | 0.57261 | 0.00347 | 0.02483 | 0.00587 | 0.01133 | 0.00404 | 0.19610 | 0.01401 |
| **Ccdc171** | 0.02011 | 0.00731 | 0.26604 | 0.15311 | 0.00822 | 0.00553 | 0.00688 | 0.00000 | 0.46343 | 2.27529 | 0.40697 | 0.00452 | 0.00709 | 0.20459 | 1.30185 | 0.00695 |
| **Ccdc172** | 0.01200 | 0.00772 | 0.00724 | 0.01220 | 0.00937 | 0.00658 | 0.00711 | 0.00919 | 0.01233 | 0.75135 | 0.00682 | 0.01874 | 0.01167 | 1.20555 | 0.10593 | 0.00743 |
| **Ccdc173** | 0.00974 | 0.00779 | 0.00355 | 0.84610 | 0.01992 | 1.94192 | 0.00719 | 0.03817 | 0.29660 | 0.72206 | 0.00000 | 0.01458 | 0.03110 | 1.12871 | 2.80352 | 0.00688 |
| **Ccdc174** | 1.82807 | 6.55421 | 7.44146 | 2.05287 | 5.34762 | 1.20915 | 4.34005 | 3.22126 | 1.36198 | 6.51537 | 4.20526 | 0.30693 | 2.39702 | 1.15602 | 6.03830 | 2.85504 |
| **Ccdc177** | 0.01231 | 0.00775 | 0.00648 | 0.34040 | 0.00966 | 0.00652 | 0.00716 | 0.00942 | 0.40926 | 0.00000 | 0.00000 | 0.00000 | 0.01391 | 1.40458 | 0.66504 | 0.00744 |
| **Ccdc18** | 0.00000 | 0.00761 | 0.45233 | 0.27831 | 0.07133 | 0.00555 | 0.00724 | 0.29329 | 0.50597 | 0.00000 | 0.00000 | 0.01414 | 0.00436 | 3.06048 | 1.09607 | 0.00687 |
| **Ccdc181** | 0.02010 | 0.00758 | 0.01040 | 0.78045 | 0.00856 | 0.00669 | 0.00726 | 0.00997 | 0.27969 | 1.14468 | 0.00944 | 0.05395 | 0.01158 | 0.01362 | 0.01798 | 0.00747 |
| **Ccdc183** | 0.01332 | 0.00817 | 0.02225 | 0.00564 | 0.00931 | 0.00625 | 0.00762 | 0.01389 | 0.16846 | 0.01377 | 0.02362 | 0.00805 | 0.01223 | 0.01285 | 0.00871 | 0.00739 |
| **Ccdc184** | 0.43308 | 0.00804 | 0.29761 | 0.01095 | 0.01692 | 0.00875 | 0.00702 | 0.01093 | 0.01221 | 0.01350 | 0.01203 | 0.00746 | 0.00973 | 0.01550 | 0.01193 | 0.06658 |

| | | | | | | | | | | | | | | | | |
|---|---|---|---|---|---|---|---|---|---|---|---|---|---|---|---|---|
| **Ccdc186** | 0.00620 | 0.00772 | 0.00467 | 1.59245 | 0.00948 | 0.00616 | 0.00663 | 0.64256 | 1.13598 | 0.14851 | 0.01504 | 0.02757 | 1.06133 | 0.46565 | 3.01096 | 0.00658 |
| **Ccdc189** | 0.05898 | 0.00762 | 1.34738 | 0.96964 | 0.02279 | 0.00834 | 0.00602 | 0.01545 | 0.00000 | 1.74580 | 0.01243 | 0.06736 | 1.88500 | 0.24202 | 2.14298 | 1.14874 |
| **Ccdc191** | 0.37572 | 1.99969 | 1.15627 | 0.00000 | 0.00528 | 0.00930 | 3.99167 | 0.00002 | 0.01094 | 0.10551 | 0.24556 | 0.00000 | 0.58422 | 1.65347 | 0.00000 | 0.00537 |
| **Ccdc22** | 2.69723 | 0.03763 | 5.29630 | 0.00464 | 0.02926 | 0.01085 | 1.06235 | 2.50309 | 0.00926 | 1.69163 | 0.00000 | 0.03496 | 5.23950 | 0.00536 | 0.10561 | 1.49725 |
| **Ccdc24** | 0.01374 | 0.00814 | 0.01227 | 1.09684 | 0.00941 | 0.00648 | 0.00743 | 0.01313 | 0.09724 | 0.02378 | 0.01215 | 0.00781 | 0.01281 | 0.00542 | 0.12089 | 0.00726 |
| **Ccdc25** | 4.23139 | 9.59382 | 5.89238 | 10.91681 | 8.53215 | 6.26615 | 3.56461 | 4.77596 | 14.79646 | 12.17226 | 6.11308 | 6.34982 | 1.97346 | 25.35501 | 12.68638 | 0.15080 |
| **Ccdc28a** | 6.70575 | 0.00000 | 3.96093 | 1.90363 | 0.01677 | 2.81562 | 2.47604 | 2.49764 | 4.78523 | 4.93937 | 1.03763 | 2.09131 | 0.79551 | 0.01790 | 1.69876 | 2.60892 |
| **Ccdc28b** | 3.75552 | 0.00380 | 1.50856 | 2.19431 | 0.00000 | 2.40010 | 0.00390 | 0.54698 | 1.94444 | 0.00000 | 0.28975 | 0.01947 | 0.21004 | 0.00355 | 0.01084 | 0.01530 |
| **Ccdc30** | 0.01734 | 0.00665 | 0.19715 | 0.00455 | 0.01779 | 0.00568 | 1.58503 | 0.00939 | 0.71945 | 1.16913 | 0.00585 | 0.02213 | 0.01013 | 0.01240 | 0.02511 | 0.00720 |
| **Ccdc32** | 1.95890 | 3.31786 | 4.87430 | 1.93888 | 5.45630 | 2.58528 | 5.73689 | 0.32096 | 3.90551 | 3.35976 | 1.20761 | 0.16354 | 1.44497 | 0.74507 | 2.51921 | 4.43774 |
| **Ccdc34** | 3.68401 | 3.81748 | 0.00000 | 2.67027 | 7.38018 | 2.67118 | 5.39671 | 0.28264 | 0.00000 | 3.62753 | 0.01156 | 0.02820 | 0.29938 | 2.42821 | 1.05790 | 2.82126 |
| **Ccdc36** | 0.07100 | 0.05114 | 0.11516 | 5.35929 | 0.01156 | 0.01230 | 0.05709 | 0.01389 | 3.78819 | 2.92918 | 1.08671 | 0.04938 | 0.05050 | 0.58299 | 2.22718 | 0.04271 |
| **Ccdc38** | 0.00982 | 0.00745 | 0.00976 | 0.00447 | 0.17935 | 0.00658 | 0.00739 | 0.01139 | 0.11659 | 0.00459 | 0.01036 | 0.00662 | 0.01167 | 0.01268 | 0.45857 | 0.00725 |
| **Ccdc39** | 0.00602 | 0.40791 | 0.00663 | 0.00423 | 0.00832 | 0.00627 | 0.00709 | 0.09943 | 0.11649 | 0.01145 | 0.00000 | 2.39922 | 0.00858 | 0.00747 | 0.00000 | 0.00701 |
| **Ccdc40** | 0.01369 | 0.00783 | 2.61277 | 2.01829 | 0.00403 | 0.00699 | 0.20719 | 1.01652 | 0.62068 | 0.89462 | 0.36296 | 0.12209 | 1.16163 | 0.00134 | 0.46971 | 0.00532 |
| **Ccdc42** | 0.01009 | 0.00795 | 0.11897 | 0.20877 | 0.01747 | 0.02646 | 0.00736 | 0.01260 | 0.00646 | 0.01042 | 0.01233 | 0.00795 | 0.07094 | 0.01201 | 0.00515 | 0.23377 |
| **Ccdc43** | 2.01255 | 2.71938 | 5.13090 | 0.00000 | 3.11002 | 7.58106 | 1.69746 | 12.77359 | 0.03541 | 7.32214 | 5.14236 | 1.72078 | 6.82694 | 13.01703 | 0.00000 | 0.38346 |
| **Ccdc47** | 3.78685 | 1.80476 | 9.95067 | 6.39298 | 3.31783 | 1.32431 | 6.12884 | 8.37417 | 4.21666 | 0.85873 | 4.80247 | 3.13508 | 4.57906 | 1.72097 | 8.23922 | 2.35471 |
| **Ccdc50** | 2.36474 | 2.91300 | 3.29341 | 5.43972 | 0.03032 | 2.60072 | 2.14335 | 0.27348 | 3.18503 | 3.39934 | 0.22072 | 0.10180 | 0.16452 | 2.02231 | 2.41181 | 3.02238 |
| **Ccdc51** | 4.40754 | 3.84131 | 1.91788 | 0.37710 | 0.00000 | 2.26558 | 3.59793 | 2.37409 | 0.00427 | 2.28670 | 2.03423 | 0.22667 | 0.00000 | 0.00000 | 0.97585 | 0.24952 |
| **Ccdc53** | 12.63946 | 4.19235 | 6.08283 | 0.00000 | 3.93767 | 5.46408 | 3.41200 | 5.29517 | 0.15394 | 4.60364 | 2.54932 | 0.89371 | 3.78933 | 1.02828 | 0.97823 | 0.69446 |
| **Ccdc57** | 0.89592 | 0.00647 | 0.00841 | 0.74916 | 0.00601 | 0.01128 | 0.00556 | 2.57995 | 0.34218 | 5.09611 | 0.52451 | 0.16279 | 1.67654 | 0.00498 | 2.23240 | 0.00548 |
| **Ccdc58** | 11.15769 | 6.16192 | 9.87707 | 4.22996 | 11.07530 | 3.73602 | 3.14090 | 5.72493 | 2.06151 | 7.32723 | 9.68013 | 1.75588 | 10.69003 | 10.16008 | 4.54720 | 1.71712 |
| **Ccdc59** | 10.46329 | 7.84660 | 5.20709 | 10.98061 | 0.10175 | 0.02448 | 6.91022 | 10.45770 | 13.39346 | 15.09951 | 13.15549 | 2.55315 | 9.20575 | 2.20723 | 2.11610 | 6.24708 |
| **Ccdc6** | 6.58630 | 2.48847 | 1.66556 | 24.53555 | 0.43702 | 1.01964 | 0.02981 | 1.54565 | 18.51281 | 10.21179 | 4.97680 | 2.49516 | 0.66495 | 15.30755 | 31.25139 | 3.65334 |
| **Ccdc60** | 0.00614 | 0.00861 | 0.00832 | 0.97480 | 0.00930 | 0.00664 | 0.00724 | 0.35057 | 0.01232 | 0.02411 | 0.00938 | 0.00701 | 1.61515 | 0.03732 | 0.00000 | 0.00651 |
| **Ccdc61** | 2.99490 | 0.42409 | 0.00000 | 0.86480 | 0.06986 | 1.45778 | 0.02374 | 0.04538 | 0.00000 | 2.79502 | 4.55381 | 0.02260 | 0.15463 | 0.00000 | 0.00001 | 0.19732 |
| **Ccdc62** | 0.08676 | 0.00846 | 0.00939 | 0.49287 | 0.01346 | 0.00685 | 0.00745 | 0.01343 | 0.00689 | 0.04781 | 0.01316 | 0.00811 | 0.01526 | 0.01314 | 0.00562 | 0.00739 |
| **Ccdc63** | 0.03141 | 0.00712 | 0.00195 | 0.92664 | 0.18089 | 0.00672 | 0.00687 | 0.00000 | 5.06838 | 0.61845 | 0.21644 | 1.59633 | 0.00915 | 0.00562 | 0.15728 | 0.00639 |
| **Ccdc64** | 0.00439 | 0.72487 | 0.19441 | 0.00750 | 0.00775 | 0.00602 | 0.10752 | 0.30722 | 0.11200 | 0.10656 | 0.00936 | 0.00665 | 0.40100 | 0.01021 | 0.00184 | 0.00684 |
| **Ccdc64b** | 0.01214 | 0.00803 | 0.01817 | 1.05251 | 0.00904 | 0.00726 | 0.00730 | 0.01580 | 0.87558 | 1.50615 | 0.00612 | 0.04358 | 0.01549 | 0.00847 | 0.01516 | 0.00740 |
| **Ccdc65** | 0.00442 | 0.00777 | 0.00712 | 0.10769 | 0.02023 | 0.00660 | 0.00690 | 0.36251 | 0.00958 | 0.26006 | 0.00000 | 0.01967 | 0.52592 | 0.94924 | 0.72636 | 0.00729 |
| **Ccdc66** | 0.39031 | 2.43439 | 0.82712 | 1.53945 | 5.00497 | 0.89205 | 0.00458 | 2.10008 | 0.78172 | 0.13675 | 1.18773 | 3.81474 | 0.70016 | 2.61828 | 5.73629 | 1.07369 |
| **Ccdc68** | 1.30983 | 0.00709 | 0.00275 | 0.00890 | 0.01646 | 0.00669 | 0.00638 | 0.00778 | 0.01207 | 0.01262 | 0.01139 | 0.00624 | 0.01037 | 0.00616 | 0.28089 | 0.24523 |
| **Ccdc69** | 0.01322 | 0.00741 | 0.00670 | 14.60153 | 0.00891 | 0.00514 | 0.00655 | 0.00000 | 1.20874 | 0.90561 | 0.31104 | 0.06026 | 0.00855 | 0.00000 | 1.12594 | 0.00454 |
| **Ccdc71** | 1.73555 | 0.00332 | 3.17120 | 1.68745 | 0.00786 | 0.00483 | 3.54484 | 1.33557 | 0.00000 | 3.00913 | 0.54765 | 0.00000 | 3.58369 | 0.90193 | 1.73393 | 0.00940 |
| **Ccdc71l** | 0.00743 | 0.78339 | 0.04357 | 3.00464 | 0.00811 | 0.00467 | 0.01669 | 0.00550 | 8.63610 | 0.00000 | 0.00000 | 0.19277 | 0.01095 | 0.00000 | 0.37846 | 0.00576 |
| **Ccdc73** | 1.29420 | 0.00644 | 0.00979 | 1.12605 | 0.00543 | 0.00530 | 0.00679 | 0.01343 | 0.20025 | 0.84397 | 0.01022 | 0.02849 | 0.01059 | 0.01316 | 0.25639 | 0.00562 |
| **Ccdc74a** | 0.00991 | 0.00810 | 0.08153 | 0.94828 | 0.00907 | 0.00555 | 0.00699 | 0.75186 | 0.58963 | 0.17361 | 0.37668 | 0.00314 | 0.67494 | 0.00477 | 0.04066 | 0.00655 |
| **Ccdc77** | 2.18570 | 5.10471 | 6.51059 | 5.52330 | 7.32288 | 3.47508 | 1.58946 | 5.42215 | 6.08926 | 2.03669 | 1.25187 | 0.16482 | 4.49845 | 10.36229 | 6.69650 | 2.01099 |
| **Ccdc78** | 0.04118 | 0.00810 | 0.19072 | 0.14691 | 0.03280 | 0.02119 | 0.04899 | 0.05498 | 0.21675 | 0.83945 | 0.57931 | 0.01332 | 0.11726 | 0.11124 | 0.21986 | 0.01457 |
| **Ccdc7b** | 0.04986 | 0.00786 | 0.09310 | 0.13716 | 0.00796 | 0.00577 | 0.00765 | 0.00748 | 0.09073 | 0.00916 | 0.68573 | 0.00521 | 0.00592 | 0.04635 | 0.03380 | 0.00701 |
| **Ccdc80** | 0.01361 | 0.00787 | 0.00936 | 0.00928 | 0.00917 | 0.00672 | 0.00744 | 0.01335 | 0.01211 | 0.32246 | 0.01044 | 0.23436 | 0.01145 | 0.01314 | 0.00474 | 0.00720 |
| **Ccdc81** | 0.27998 | 0.00719 | 1.35919 | 0.01079 | 0.00500 | 0.00832 | 0.00689 | 0.31095 | 0.01195 | 0.07958 | 0.00885 | 0.00565 | 0.00271 | 0.01387 | 0.00000 | 0.00638 |
| **Ccdc82** | 1.72030 | 0.62307 | 2.59968 | 44.47688 | 0.28563 | 2.92036 | 2.59037 | 2.45927 | 39.14170 | 37.01031 | 7.16566 | 7.98649 | 4.87316 | 43.73344 | 52.40767 | 0.74025 |
| **Ccdc84** | 1.39481 | 0.00604 | 0.00000 | 0.63937 | 0.00542 | 2.47975 | 0.00520 | 0.63590 | 0.22771 | 2.49777 | 1.56973 | 1.40901 | 1.61988 | 0.60735 | 0.41375 | 0.01121 |
| **Ccdc85a** | 0.01325 | 0.00756 | 0.01746 | 0.45222 | 0.00926 | 0.00714 | 0.00721 | 0.01058 | 0.02297 | 0.00865 | 0.01020 | 0.45342 | 0.01304 | 0.02593 | 0.00000 | 0.00742 |
| **Ccdc85b** | 0.02204 | 0.00827 | 0.08626 | 2.07253 | 0.03564 | 0.03056 | 0.01736 | 0.01703 | 1.37871 | 0.09949 | 0.86808 | 0.08007 | 0.04454 | 0.23952 | 0.34855 | 0.05754 |
| **Ccdc86** | 9.46090 | 0.81236 | 0.00000 | 14.25994 | 8.23570 | 5.80321 | 1.25180 | 20.15412 | 18.77238 | 18.18560 | 6.25844 | 3.16689 | 14.61927 | 2.79111 | 5.16640 | 2.10969 |
| **Ccdc87** | 0.00952 | 0.00791 | 0.02011 | 0.20328 | 0.00905 | 0.00601 | 0.00667 | 0.01084 | 0.00867 | 0.00547 | 0.01351 | 0.00736 | 1.25725 | 0.01223 | 0.00712 | 0.00653 |
| **Ccdc88a** | 0.01182 | 0.00768 | 0.00619 | 0.51632 | 0.00957 | 0.00572 | 0.05746 | 0.01392 | 1.67894 | 0.07540 | 0.01195 | 0.03349 | 0.01223 | 0.03926 | 8.33733 | 0.00715 |
| **Ccdc88b** | 0.01340 | 0.00810 | 0.01166 | 0.01342 | 0.00906 | 0.00672 | 0.00741 | 0.01137 | 0.70040 | 0.48320 | 0.00778 | 0.00586 | 0.01236 | 0.03411 | 0.00454 | 0.00754 |
| **Ccdc88c** | 1.28563 | 0.66571 | 0.98753 | 0.00000 | 0.00527 | 0.00559 | 0.32113 | 4.09716 | 0.21461 | 0.59082 | 0.00000 | 0.05550 | 1.01825 | 0.00302 | 0.62414 | 0.01307 |
| **Ccdc89** | 0.01265 | 0.00820 | 1.10782 | 0.07575 | 0.00740 | 0.00564 | 0.00764 | 0.00825 | 0.01081 | 0.00435 | 0.32673 | 0.00680 | 0.00552 | 0.01049 | 0.00000 | 0.00681 |
| **Ccdc9** | 7.35004 | 2.73879 | 1.60609 | 5.70565 | 4.44809 | 0.53214 | 0.33517 | 5.13990 | 8.12827 | 10.43172 | 6.94300 | 0.61849 | 2.78711 | 0.05148 | 2.47387 | 5.73301 |
| **Ccdc90b** | 6.38570 | 6.67161 | 3.64946 | 2.64150 | 9.53067 | 3.43740 | 3.47479 | 2.15872 | 1.14740 | 4.02519 | 5.99661 | 1.26343 | 3.01499 | 0.26107 | 1.98958 | 1.03265 |
| **Ccdc91** | 3.43160 | 2.84108 | 0.00000 | 2.04103 | 3.93757 | 6.77876 | 2.39990 | 2.07225 | 0.70925 | 0.37500 | 0.78012 | 0.37045 | 1.06659 | 0.00000 | 1.91371 | 1.53836 |
| **Ccdc92** | 0.63993 | 0.04592 | 1.01494 | 4.22889 | 2.70908 | 0.00479 | 3.99731 | 1.05635 | 0.90630 | 0.00000 | 1.66896 | 0.05955 | 2.53929 | 0.00000 | 0.24192 | 2.88792 |
| **Ccdc92b** | 3.26931 | 0.01231 | 0.11795 | 0.00000 | 3.31835 | 0.00258 | 0.00252 | 5.41318 | 4.89765 | 5.13849 | 5.92271 | 0.67638 | 0.00000 | 1.84447 | 1.30284 | 0.02435 |
| **Ccdc93** | 6.19161 | 7.00786 | 10.00801 | 0.29470 | 1.14201 | 0.00000 | 3.46551 | 8.62418 | 1.50640 | 7.25798 | 5.04184 | 9.23223 | 2.04488 | 10.36705 | 5.83637 | 6.23431 |
| **Ccdc94** | 2.43101 | 1.70507 | 1.98924 | 0.44553 | 0.01149 | 0.01814 | 5.47144 | 7.51086 | 1.27297 | 7.69690 | 1.02417 | 0.18687 | 2.60816 | 0.00827 | 0.13185 | 0.00909 |
| **Ccdc96** | 0.01627 | 0.00736 | 0.33711 | 0.75718 | 0.00831 | 0.00547 | 0.00694 | 0.00025 | 0.61704 | 1.09552 | 0.28993 | 0.00237 | 0.00746 | 0.01111 | 0.00000 | 0.01370 |

| | | | | | | | | | | | | | | | | |
|---|---|---|---|---|---|---|---|---|---|---|---|---|---|---|---|---|
| **Ccdc97** | 2.38656 | 0.01448 | 1.43458 | 2.28839 | 0.11516 | 0.37620 | 0.64277 | 6.60378 | 1.20045 | 2.68607 | 1.78757 | 1.44058 | 4.13947 | 0.59249 | 0.38746 | 1.55284 |
| **Cchcr1** | 0.00000 | 0.00687 | 0.00643 | 1.49933 | 0.02458 | 0.00865 | 0.18040 | 1.14088 | 1.66771 | 2.96678 | 3.72815 | 0.10224 | 1.00234 | 0.00000 | 0.00000 | 1.70526 |
| **Cck** | 0.01020 | 0.12165 | 0.20887 | 0.01063 | 0.00869 | 0.00647 | 0.00711 | 0.01593 | 0.01235 | 0.00580 | 0.01336 | 0.00743 | 0.46615 | 0.01313 | 0.00505 | 0.01427 |
| **Ccl22** | 0.01289 | 0.00764 | 0.01053 | 0.00512 | 0.00889 | 0.00659 | 0.00724 | 0.01135 | 0.01102 | 1.25996 | 0.12750 | 0.02551 | 0.00964 | 0.00754 | 0.23012 | 0.00714 |
| **Ccl25** | 0.00481 | 2.94972 | 1.01830 | 0.16503 | 0.00595 | 0.00564 | 0.00631 | 0.91132 | 0.00738 | 1.44376 | 0.90007 | 0.00021 | 0.00337 | 1.13817 | 1.07979 | 0.00566 |
| **Ccl27a** | 0.01056 | 0.00779 | 0.01119 | 0.01051 | 0.00909 | 0.00661 | 0.00729 | 0.06183 | 0.01233 | 0.04118 | 0.01265 | 0.00741 | 0.00922 | 0.01157 | 0.00982 | 0.15939 |
| **Ccl28** | 0.01225 | 0.00809 | 0.02383 | 0.05337 | 0.00924 | 0.00761 | 0.00776 | 0.03391 | 0.06035 | 0.09589 | 0.01273 | 0.00837 | 0.01060 | 0.01968 | 0.02660 | 0.00720 |
| **Ccl9** | 0.01345 | 0.02014 | 0.56616 | 0.11754 | 0.00870 | 0.48900 | 0.04162 | 0.85687 | 0.09303 | 1.00953 | 0.26070 | 0.05432 | 1.09828 | 0.27586 | 0.09308 | 0.02130 |
| **Ccm2** | 2.96579 | 0.02386 | 5.13620 | 0.33350 | 0.03619 | 3.49186 | 3.67156 | 2.51578 | 0.07181 | 3.04740 | 0.29246 | 0.11589 | 2.68244 | 0.00319 | 0.21152 | 0.77410 |
| **Ccm2l** | 0.01429 | 0.00863 | 0.00865 | 0.00612 | 0.00896 | 0.00688 | 0.00807 | 0.07185 | 0.01155 | 0.26854 | 0.23462 | 0.01831 | 0.00708 | 0.01186 | 0.00453 | 0.00706 |
| **Ccna2** | 9.81100 | 8.87894 | 12.12091 | 28.09333 | 7.46092 | 3.65006 | 3.51594 | 14.17998 | 38.64917 | 33.82016 | 23.49623 | 23.76735 | 9.01051 | 23.87061 | 27.00080 | 2.62167 |
| **Ccnb1** | 25.69767 | 15.32735 | 21.53973 | 22.40585 | 7.92291 | 13.00819 | 14.16666 | 35.65476 | 33.66982 | 46.33487 | 33.78595 | 31.49652 | 23.11402 | 45.29196 | 37.96821 | 8.23698 |
| **Ccnb1ip1** | 2.39755 | 3.85959 | 1.43522 | 2.94143 | 3.88723 | 2.29450 | 3.79970 | 3.91227 | 0.66915 | 6.81167 | 2.43376 | 6.46387 | 3.43996 | 2.19079 | 7.87880 | 1.02774 |
| **Ccnb2** | 12.81744 | 9.80727 | 12.81888 | 4.02228 | 13.93042 | 10.08116 | 6.47348 | 7.30080 | 7.02379 | 16.51465 | 14.46824 | 7.38934 | 12.65592 | 3.90246 | 5.24451 | 8.86988 |
| **Ccnb3** | 0.01083 | 0.00769 | 0.00435 | 0.02651 | 0.00973 | 0.00634 | 0.00707 | 0.00799 | 0.01142 | 0.25481 | 0.00123 | 0.00000 | 0.01280 | 5.04877 | 0.00461 | 0.00729 |
| **Ccnc** | 9.29991 | 4.77106 | 4.54464 | 4.74929 | 2.79719 | 0.33638 | 0.00000 | 3.55127 | 4.52878 | 6.45863 | 3.69469 | 0.89936 | 2.68748 | 6.29579 | 6.46734 | 2.20035 |
| **Ccnd1** | 0.00000 | 0.00762 | 0.00000 | 0.04512 | 0.01801 | 0.00654 | 0.00296 | 0.36520 | 0.66728 | 2.90607 | 2.26072 | 0.05209 | 7.26222 | 0.01819 | 2.33158 | 0.02615 |
| **Ccnd2** | 0.00898 | 0.00709 | 0.11246 | 0.00760 | 0.00908 | 0.00942 | 1.14926 | 0.01062 | 0.01171 | 0.55375 | 0.00686 | 0.15520 | 0.01074 | 0.63869 | 0.00000 | 0.00727 |
| **Ccnd3** | 15.79686 | 6.58334 | 13.30623 | 0.96157 | 10.55890 | 4.93593 | 11.22422 | 20.95168 | 1.31970 | 9.53143 | 4.50474 | 0.68050 | 13.91820 | 0.35591 | 0.69955 | 6.34018 |
| **Ccndbp1** | 6.32094 | 0.38580 | 6.01223 | 19.92686 | 3.55811 | 6.61509 | 2.79948 | 7.39433 | 12.52010 | 7.49555 | 2.76403 | 4.94595 | 5.07941 | 0.78313 | 10.48424 | 7.61436 |
| **Ccne1** | 2.60726 | 6.26818 | 13.42710 | 19.98333 | 4.54006 | 12.06227 | 10.75695 | 22.23347 | 32.29112 | 27.38221 | 23.50871 | 14.39734 | 13.78640 | 10.03816 | 22.04140 | 0.04171 |
| **Ccne2** | 0.00871 | 0.00692 | 0.02518 | 0.60939 | 0.00933 | 2.40074 | 0.84820 | 0.74620 | 0.13111 | 0.21230 | 0.00000 | 2.05823 | 0.00000 | 2.77084 | 1.59317 | 0.00615 |
| **Ccnf** | 4.98280 | 4.98111 | 7.28437 | 8.32379 | 1.90462 | 2.77656 | 3.27561 | 10.81364 | 12.04024 | 11.20199 | 8.39015 | 5.01413 | 3.26575 | 0.00000 | 7.94871 | 0.51170 |
| **Ccng1** | 8.24789 | 8.48334 | 4.11141 | 1.34493 | 11.89121 | 3.71742 | 10.30970 | 3.52662 | 0.00000 | 0.00000 | 4.83825 | 0.66306 | 4.24196 | 0.69523 | 0.00000 | 4.04408 |
| **Ccng2** | 0.44153 | 0.00780 | 0.52502 | 3.66547 | 0.01630 | 0.00789 | 0.00669 | 0.01028 | 3.47112 | 0.35596 | 0.00000 | 0.02224 | 0.00863 | 0.00000 | 2.89878 | 0.00582 |
| **Ccnh** | 12.67184 | 6.76443 | 2.55599 | 0.46995 | 8.01146 | 4.50514 | 1.76168 | 4.97194 | 5.94425 | 8.18939 | 7.30012 | 7.89907 | 7.04585 | 4.57831 | 2.79681 | 4.34654 |
| **Ccni** | 4.62130 | 4.72123 | 6.75793 | 1.85430 | 4.34632 | 5.25679 | 1.88740 | 1.78683 | 3.45184 | 2.90642 | 1.24488 | 7.88971 | 1.03972 | 1.30643 | 3.06881 | 1.32447 |
| **Ccnj** | 0.01050 | 0.00829 | 0.00785 | 3.17490 | 0.00941 | 0.00625 | 0.00802 | 0.14023 | 1.44372 | 0.11955 | 0.00024 | 0.04511 | 0.00838 | 0.02959 | 1.15412 | 0.00713 |
| **Ccnjl** | 0.01346 | 0.00822 | 0.01165 | 0.49958 | 0.00929 | 0.00654 | 0.00753 | 0.01346 | 1.78544 | 0.00592 | 0.00000 | 0.00491 | 0.01326 | 0.03362 | 0.00642 | 0.00725 |
| **Ccnk** | 3.23798 | 0.66795 | 1.70786 | 9.16505 | 3.97818 | 0.00927 | 1.58009 | 2.40599 | 13.21983 | 6.29698 | 7.94981 | 0.72355 | 2.43887 | 1.83630 | 0.84132 | 1.98237 |
| **Ccnl1** | 3.46940 | 8.14263 | 1.06894 | 4.41572 | 5.13487 | 0.82215 | 6.87359 | 4.79484 | 6.62891 | 10.50833 | 3.69527 | 10.13676 | 6.52447 | 9.76614 | 12.60024 | 0.23755 |
| **Ccnl2** | 8.27088 | 4.66446 | 8.72332 | 17.03305 | 2.96132 | 5.47037 | 4.42692 | 7.55679 | 20.55049 | 13.23488 | 10.25599 | 10.79605 | 11.36369 | 2.39935 | 11.37752 | 6.06991 |
| **Ccno** | 1.05182 | 0.00631 | 0.00376 | 6.41711 | 0.00581 | 0.00589 | 0.00614 | 0.00000 | 6.66440 | 2.00635 | 0.00000 | 0.00935 | 0.01065 | 0.04603 | 0.01154 | 0.00460 |
| **Ccnt1** | 0.67630 | 1.20305 | 3.32604 | 2.49925 | 0.00000 | 0.00424 | 0.31160 | 1.59993 | 3.40332 | 4.89710 | 0.83525 | 3.30627 | 1.63113 | 5.08834 | 3.67639 | 0.01766 |
| **Ccnt2** | 1.20362 | 5.41045 | 0.35452 | 3.11069 | 0.02156 | 0.10069 | 0.22507 | 2.77473 | 12.88825 | 1.07257 | 4.82206 | 0.38339 | 1.07314 | 0.64628 | 5.20546 | 0.05946 |
| **Ccny** | 1.54033 | 0.84325 | 1.55879 | 3.79349 | 0.97729 | 0.00707 | 1.25037 | 2.23880 | 1.94714 | 2.48226 | 0.00000 | 0.10273 | 1.38961 | 0.00197 | 1.09720 | 2.47110 |
| **Ccnyl1** | 5.17410 | 0.88056 | 2.65476 | 5.92846 | 1.23453 | 0.13386 | 2.32406 | 0.17169 | 9.70003 | 1.23983 | 0.16828 | 10.19575 | 2.16496 | 3.45876 | 2.27012 | 0.04670 |
| **Ccp110** | 4.81376 | 0.03972 | 3.59580 | 7.72774 | 4.58127 | 0.00783 | 2.20776 | 0.47843 | 4.85592 | 3.15446 | 3.27000 | 4.49095 | 0.07461 | 5.49256 | 7.51559 | 1.21566 |
| **Ccpg1** | 2.07485 | 2.86518 | 7.79438 | 5.03850 | 0.00603 | 6.07976 | 5.74272 | 17.81428 | 13.44755 | 12.66975 | 10.99489 | 10.47031 | 7.11970 | 14.49144 | 19.19374 | 1.29675 |
| **Ccpg1os** | 3.27320 | 0.81230 | 1.21036 | 1.44878 | 0.00000 | 0.00257 | 1.96705 | 0.26693 | 2.13743 | 0.95753 | 1.26454 | 0.63674 | 1.90403 | 2.31341 | 0.65437 | 0.02393 |
| **Ccr4** | 0.00047 | 0.01893 | 0.80234 | 0.40607 | 0.00985 | 0.00549 | 0.01728 | 0.29911 | 0.25707 | 1.02148 | 0.04647 | 0.13883 | 0.00793 | 3.29665 | 0.11914 | 0.00704 |
| **Ccr8** | 0.01337 | 0.00804 | 0.01192 | 0.00608 | 0.00907 | 0.00642 | 0.00739 | 0.01082 | 0.01117 | 0.60815 | 0.00978 | 0.00556 | 0.01085 | 0.00563 | 0.45672 | 0.00711 |
| **Ccr9** | 0.01247 | 0.00806 | 0.01195 | 0.15396 | 0.00895 | 0.00659 | 0.00750 | 0.01359 | 0.17695 | 0.01163 | 0.01212 | 0.01849 | 0.01159 | 0.01212 | 0.00600 | 0.15819 |
| **Ccs** | 7.46731 | 0.96252 | 6.31355 | 3.02272 | 0.47677 | 2.61956 | 3.81366 | 9.69933 | 3.01676 | 4.06396 | 3.95092 | 0.07342 | 4.46413 | 1.04772 | 0.54752 | 3.62099 |
| **Ccsap** | 1.36060 | 0.00586 | 0.00000 | 0.43772 | 0.00529 | 4.41856 | 0.00594 | 0.00424 | 0.04421 | 0.00000 | 0.02376 | 0.04467 | 0.01466 | 0.04061 | 0.16924 | 0.00413 |
| **Ccser1** | 0.23763 | 0.00816 | 0.01366 | 0.89250 | 0.01816 | 0.00735 | 0.00726 | 0.01142 | 1.04851 | 0.01120 | 0.00629 | 0.00521 | 0.01489 | 0.00477 | 0.24356 | 0.00712 |
| **Ccser2** | 3.23527 | 3.27612 | 0.05634 | 8.40930 | 0.47363 | 0.00372 | 0.00276 | 0.25755 | 4.15863 | 1.40087 | 1.20538 | 0.25834 | 0.00000 | 8.25365 | 8.01428 | 0.00000 |
| **Cct2** | 36.54917 | 21.33922 | 29.89120 | 9.92236 | 28.25722 | 15.45122 | 20.77832 | 30.68576 | 15.50143 | 34.26656 | 28.51208 | 25.70817 | 28.57787 | 36.93065 | 21.44836 | 15.96385 |
| **Cct3** | 37.06612 | 23.55858 | 32.84795 | 15.77797 | 23.89065 | 16.18280 | 20.48986 | 35.63079 | 27.08150 | 31.56616 | 25.34089 | 17.87351 | 32.73718 | 30.93398 | 25.13037 | 19.48556 |
| **Cct4** | 32.85969 | 17.87184 | 22.11387 | 10.43455 | 21.86891 | 11.93803 | 12.00001 | 24.98758 | 23.30603 | 37.36288 | 27.28110 | 20.34394 | 31.81080 | 30.43067 | 21.05643 | 18.52951 |
| **Cct5** | 40.09100 | 24.84740 | 29.81953 | 6.00707 | 26.84368 | 18.43765 | 19.89086 | 37.13548 | 6.59624 | 31.24437 | 26.65991 | 17.50709 | 36.43051 | 33.24418 | 21.31400 | 21.68394 |
| **Cct6a** | 32.60440 | 21.55863 | 26.68263 | 9.05466 | 25.11043 | 16.28701 | 18.31558 | 32.52247 | 14.16776 | 29.58034 | 22.52056 | 12.63511 | 26.56841 | 23.22102 | 23.58866 | 16.90434 |
| **Cct6b** | 1.40609 | 0.00602 | 0.00000 | 1.15180 | 0.00514 | 1.60598 | 0.00519 | 0.00938 | 1.12501 | 0.00000 | 0.13076 | 0.07281 | 1.49124 | 0.74956 | 2.11174 | 0.01861 |
| **Cct7** | 34.49362 | 22.08667 | 36.43739 | 4.87783 | 24.38279 | 13.42112 | 22.97650 | 41.21971 | 12.58890 | 37.37511 | 24.81902 | 14.92011 | 31.59159 | 5.92445 | 21.61436 | 18.53667 |
| **Cct8** | 35.27338 | 18.16446 | 25.42367 | 15.21417 | 24.65114 | 13.79082 | 15.87358 | 30.98852 | 25.09666 | 31.11061 | 26.35285 | 17.98046 | 31.73452 | 28.06545 | 15.75326 | 17.55912 |
| **Cct8l1** | 0.01189 | 0.00767 | 0.00630 | 2.34811 | 0.00966 | 0.00620 | 0.00706 | 0.00828 | 1.93913 | 0.01805 | 0.00000 | 0.06820 | 0.01395 | 1.68844 | 0.28834 | 0.00697 |
| **Ccz1** | 9.16057 | 5.34631 | 7.54991 | 5.67210 | 10.24391 | 7.44112 | 4.38728 | 7.72668 | 6.63166 | 16.05805 | 8.79830 | 4.29990 | 5.08179 | 5.70719 | 14.15384 | 3.48082 |
| **Cd109** | 1.53999 | 0.00461 | 0.00000 | 0.00728 | 0.21938 | 0.00805 | 2.12154 | 0.00442 | 0.01195 | 0.51314 | 0.01182 | 0.03074 | 0.01044 | 0.01178 | 0.00700 | 0.00564 |
| **Cd14** | 0.01356 | 0.00795 | 0.01212 | 0.00646 | 0.00913 | 0.00676 | 0.00744 | 0.01187 | 0.06374 | 0.39867 | 0.01173 | 0.00709 | 0.01241 | 0.01063 | 0.01510 | 0.00759 |
| **Cd151** | 12.90223 | 4.88883 | 10.59794 | 2.36906 | 11.10772 | 7.66517 | 3.88383 | 9.89339 | 0.09197 | 5.09891 | 7.86904 | 0.27157 | 7.59554 | 0.38802 | 3.65965 | 3.31200 |
| **Cd160** | 0.02953 | 0.00778 | 0.00215 | 1.26181 | 0.01210 | 2.73549 | 0.00701 | 0.07166 | 0.00000 | 0.00352 | 0.00852 | 0.00748 | 0.00000 | 0.02174 | 0.09097 | 0.00615 |
| **Cd163** | 0.01374 | 0.00823 | 0.02847 | 0.29941 | 0.00943 | 0.03893 | 0.00758 | 0.01484 | 0.15282 | 0.01509 | 0.01209 | 0.01798 | 0.02530 | 0.14744 | 0.02079 | 0.00749 |

| | | | | | | | | | | | | | | | | |
|---|---|---|---|---|---|---|---|---|---|---|---|---|---|---|---|---|
| **Cd164** | 8.74877 | 10.24419 | 9.71768 | 10.89358 | 7.29234 | 7.35569 | 10.18112 | 14.77670 | 11.96855 | 9.15901 | 12.85869 | 20.74823 | 9.98816 | 16.14374 | 13.72011 | 3.60114 |
| **Cd164l2** | 0.00469 | 0.00710 | 0.07289 | 6.30143 | 0.00861 | 0.00495 | 1.19696 | 0.28635 | 3.28930 | 4.54159 | 0.27166 | 0.13441 | 0.59509 | 0.06612 | 0.03140 | 0.00559 |
| **Cd177** | 0.01127 | 0.00783 | 0.01338 | 0.37837 | 0.00985 | 0.00712 | 0.00736 | 0.01207 | 0.13390 | 0.77783 | 0.01362 | 0.05022 | 0.39384 | 0.00541 | 0.12048 | 0.00800 |
| **Cd19** | 0.01351 | 0.00777 | 0.01158 | 0.12185 | 0.00910 | 0.00653 | 0.00739 | 0.01107 | 0.00986 | 0.46868 | 0.01095 | 0.02544 | 0.01142 | 0.00598 | 0.32074 | 0.00725 |
| **Cd1d1** | 0.01363 | 0.00785 | 0.00572 | 0.00965 | 0.00929 | 0.81233 | 0.00727 | 0.03823 | 0.01245 | 0.00985 | 0.02330 | 0.00779 | 0.00435 | 0.01232 | 0.00979 | 0.00701 |
| **Cd1d2** | 0.01436 | 0.00802 | 0.00720 | 0.01045 | 0.00924 | 0.26823 | 0.00748 | 0.01325 | 0.01241 | 0.01325 | 0.01247 | 0.00818 | 0.01612 | 0.01245 | 0.01058 | 0.00723 |
| **Cd2** | 1.76614 | 0.03663 | 0.00346 | 0.01060 | 0.00459 | 0.00664 | 0.00617 | 0.00788 | 0.01237 | 0.01532 | 0.01255 | 0.00691 | 0.01127 | 0.01220 | 0.01053 | 0.01270 |
| **Cd200** | 0.07383 | 0.00772 | 0.00897 | 0.00946 | 0.00797 | 0.00673 | 0.00727 | 0.01249 | 0.01230 | 0.26747 | 0.01258 | 0.02593 | 0.01184 | 0.01160 | 0.00840 | 0.07990 |
| **Cd209c** | 0.59849 | 0.36530 | 2.96603 | 9.26179 | 0.03644 | 0.00806 | 0.44049 | 15.24744 | 17.64895 | 12.86803 | 8.06773 | 6.02050 | 4.27372 | 12.56125 | 6.86154 | 0.74419 |
| **Cd209e** | 0.01599 | 0.00771 | 0.23373 | 0.01073 | 0.00884 | 0.00571 | 0.00725 | 0.02557 | 0.01230 | 0.01471 | 0.01268 | 0.00758 | 0.00965 | 0.01298 | 0.00652 | 0.00726 |
| **Cd209g** | 0.01372 | 0.00796 | 0.00973 | 0.82508 | 0.00919 | 0.00659 | 0.00748 | 0.00727 | 0.80701 | 0.00477 | 0.30874 | 0.00472 | 0.00908 | 0.00449 | 0.06688 | 0.00713 |
| **Cd22** | 0.17477 | 1.99198 | 1.67696 | 0.61664 | 2.13406 | 0.02197 | 0.01590 | 0.20451 | 0.89170 | 0.55528 | 0.24627 | 0.22880 | 0.05985 | 1.22299 | 0.17827 | 0.01292 |
| **Cd24a** | 4.39740 | 0.00000 | 1.66102 | 4.48767 | 5.51552 | 2.75770 | 1.22694 | 0.88683 | 2.03280 | 11.47587 | 0.76625 | 0.29091 | 0.51265 | 0.05501 | 0.67527 | 3.11581 |
| **Cd274** | 0.02153 | 0.01873 | 0.02428 | 0.07038 | 0.07878 | 0.01482 | 0.04885 | 0.03187 | 0.02481 | 0.03020 | 0.01069 | 0.01728 | 0.02053 | 0.23437 | 0.09432 | 0.01474 |
| **Cd276** | 0.01384 | 0.00794 | 0.01120 | 0.07880 | 0.00930 | 0.00670 | 0.00745 | 0.01270 | 0.01133 | 0.01371 | 0.01338 | 0.06839 | 0.01281 | 0.01173 | 0.00787 | 0.00741 |
| **Cd28** | 0.01104 | 0.00834 | 0.50149 | 0.01109 | 0.00884 | 0.00571 | 0.00812 | 0.48151 | 0.01175 | 0.00788 | 0.00908 | 0.00777 | 0.00361 | 0.01769 | 0.00000 | 0.00719 |
| **Cd2ap** | 16.21057 | 10.94668 | 10.13805 | 6.65283 | 10.79632 | 8.96949 | 3.84508 | 8.02670 | 5.16473 | 9.00627 | 8.56961 | 17.53231 | 5.68674 | 21.23022 | 12.20604 | 6.81426 |
| **Cd2bp2** | 5.53204 | 4.07342 | 8.68604 | 10.66996 | 3.44709 | 0.01070 | 4.00423 | 15.20853 | 14.62294 | 12.51385 | 11.10157 | 0.80989 | 11.23826 | 2.22009 | 6.87343 | 5.12248 |
| **Cd300a** | 0.27571 | 0.05534 | 0.24352 | 2.33691 | 0.15001 | 0.02640 | 0.18045 | 0.07117 | 4.41157 | 0.08575 | 3.49682 | 0.15506 | 0.29457 | 2.85305 | 0.81710 | 0.13179 |
| **Cd300c2** | 0.01185 | 0.00870 | 0.01162 | 0.01026 | 0.00976 | 0.00692 | 0.00817 | 0.06809 | 0.01207 | 0.01226 | 0.01302 | 0.00903 | 0.01651 | 0.01123 | 0.05205 | 0.00753 |
| **Cd300ld** | 0.90293 | 0.00792 | 0.40438 | 0.23607 | 0.01171 | 0.00467 | 0.00684 | 0.00994 | 0.00594 | 0.01298 | 0.01206 | 0.00831 | 0.00931 | 0.00815 | 0.19643 | 0.00627 |
| **Cd300lf** | 0.09339 | 0.00855 | 0.05889 | 0.38489 | 0.01230 | 0.01811 | 0.00747 | 0.01189 | 0.00967 | 0.03009 | 0.02369 | 0.00810 | 0.01119 | 0.10256 | 0.01476 | 0.00721 |
| **Cd300lg** | 0.06019 | 0.00797 | 0.00000 | 0.01046 | 0.01215 | 0.00594 | 0.00664 | 0.00000 | 0.01900 | 0.00000 | 0.24727 | 0.00000 | 0.07297 | 9.75375 | 0.37250 | 0.00695 |
| **Cd320** | 6.87667 | 3.86349 | 7.26316 | 15.79064 | 4.57781 | 1.45281 | 0.72856 | 16.07095 | 21.41955 | 26.98730 | 19.42403 | 16.48816 | 14.60354 | 7.02524 | 8.49251 | 2.85946 |
| **Cd33** | 0.03832 | 0.01826 | 0.01859 | 0.01097 | 0.03626 | 0.02276 | 0.01751 | 0.02531 | 0.05019 | 0.03522 | 0.07819 | 0.00726 | 0.06842 | 0.06866 | 0.04992 | 0.01455 |
| **Cd36** | 0.01370 | 0.00770 | 0.01216 | 0.83135 | 0.00923 | 0.00653 | 0.00734 | 0.01162 | 0.00000 | 0.15968 | 0.01249 | 0.00625 | 0.01287 | 0.00398 | 0.29388 | 0.00765 |
| **Cd38** | 2.18727 | 2.61780 | 0.24951 | 7.38286 | 0.10733 | 0.17496 | 0.33968 | 0.35499 | 11.88715 | 1.21631 | 0.64693 | 0.27173 | 0.44739 | 2.17266 | 3.81842 | 0.07579 |
| **Cd3d** | 0.02284 | 0.01815 | 0.06715 | 0.09231 | 0.00961 | 0.03337 | 0.01728 | 0.14153 | 0.16602 | 0.11819 | 0.17460 | 0.16887 | 0.01944 | 0.18625 | 0.15994 | 0.01476 |
| **Cd3eap** | 3.57534 | 0.34171 | 1.88331 | 3.73659 | 0.00733 | 0.01880 | 4.76960 | 5.58802 | 18.56431 | 5.06351 | 1.93323 | 0.14240 | 3.76787 | 4.73371 | 2.35800 | 5.41410 |
| **Cd40** | 0.01336 | 0.00792 | 0.02212 | 0.01106 | 0.00934 | 0.00625 | 0.00759 | 0.01367 | 0.01204 | 0.01302 | 0.01249 | 0.00809 | 0.01248 | 0.01864 | 0.04432 | 0.00740 |
| **Cd44** | 0.01323 | 0.00794 | 0.01081 | 0.00717 | 0.00892 | 0.00668 | 0.00736 | 0.01063 | 0.01173 | 1.08160 | 0.00999 | 0.07866 | 0.01085 | 0.01217 | 0.00608 | 0.00718 |
| **Cd47** | 0.01298 | 0.00758 | 0.00873 | 2.61852 | 0.00894 | 0.00597 | 0.00719 | 0.00865 | 3.85387 | 0.63832 | 0.00000 | 0.14456 | 0.01138 | 0.00000 | 0.62313 | 0.00660 |
| **Cd52** | 0.01171 | 0.00800 | 0.20761 | 0.00324 | 0.00874 | 0.00893 | 0.00687 | 0.00000 | 0.97302 | 0.47105 | 0.62762 | 0.07757 | 0.00670 | 0.56453 | 0.29156 | 0.00721 |
| **Cd55** | 0.01484 | 0.00772 | 0.34886 | 5.85454 | 0.02007 | 0.00839 | 0.00671 | 0.00000 | 0.73972 | 0.28304 | 0.62771 | 0.00000 | 0.00587 | 0.10960 | 2.13109 | 0.00596 |
| **Cd55b** | 0.01694 | 0.00777 | 0.03588 | 0.10666 | 0.00888 | 0.00795 | 0.00738 | 0.01240 | 0.00933 | 0.06827 | 0.01170 | 0.00696 | 0.01187 | 0.00785 | 0.21021 | 0.00727 |
| **Cd59a** | 2.17221 | 3.05254 | 6.03324 | 0.40510 | 3.89073 | 0.01022 | 3.39335 | 0.00000 | 0.62604 | 0.96554 | 0.79289 | 0.09916 | 0.63747 | 5.54373 | 0.33005 | 0.15704 |
| **Cd59b** | 0.09256 | 0.09524 | 0.56266 | 0.54571 | 0.10220 | 0.00567 | 0.20406 | 0.00484 | 1.97522 | 0.13586 | 0.30106 | 0.00000 | 0.03005 | 0.97971 | 0.00000 | 0.00711 |
| **Cd6** | 0.01405 | 0.00780 | 0.00848 | 0.23363 | 0.00908 | 0.00670 | 0.00735 | 0.00113 | 0.00786 | 0.00000 | 0.45189 | 0.02439 | 0.00782 | 0.00458 | 0.30131 | 0.00709 |
| **Cd63** | 26.83300 | 21.79151 | 45.89889 | 23.11357 | 13.37103 | 19.65402 | 21.71616 | 41.47992 | 19.87798 | 35.73023 | 41.03056 | 16.13047 | 42.50235 | 25.89546 | 24.17384 | 21.54912 |
| **Cd68** | 0.01525 | 0.00800 | 2.81922 | 0.24200 | 0.00585 | 0.00674 | 0.16318 | 0.37317 | 0.16112 | 1.40088 | 0.48908 | 0.02204 | 0.09025 | 0.06754 | 0.87793 | 0.01324 |
| **Cd7** | 0.01719 | 0.93509 | 0.00000 | 0.46088 | 0.00754 | 0.00751 | 0.19474 | 2.04519 | 0.43399 | 0.87108 | 1.28244 | 0.00389 | 0.90727 | 0.00284 | 0.33251 | 0.03546 |
| **Cd74** | 0.00779 | 0.00741 | 0.00645 | 0.11517 | 0.00839 | 0.00713 | 0.00707 | 0.13178 | 0.02145 | 5.72367 | 0.26073 | 0.24784 | 0.03347 | 0.00908 | 0.00000 | 0.00731 |
| **Cd79a** | 0.02199 | 0.00773 | 0.01256 | 0.51143 | 0.00860 | 0.00747 | 0.00702 | 0.00000 | 0.17149 | 0.00000 | 0.79904 | 0.06183 | 0.00665 | 0.00556 | 0.47549 | 0.00696 |
| **Cd79b** | 1.10619 | 0.00480 | 0.04740 | 0.00642 | 3.03007 | 0.00769 | 0.00635 | 0.00329 | 0.01083 | 0.39338 | 0.01209 | 0.00262 | 0.01030 | 0.01144 | 0.01007 | 0.00645 |
| **Cd81** | 0.01258 | 0.00784 | 0.01171 | 0.00542 | 0.00874 | 0.00662 | 0.00729 | 0.01004 | 0.03775 | 2.25858 | 0.00976 | 0.02304 | 0.01075 | 0.01267 | 0.00622 | 0.00737 |
| **Cd82** | 0.00251 | 0.00828 | 0.02478 | 0.00500 | 0.01939 | 0.01215 | 0.01826 | 2.14168 | 0.00917 | 2.56270 | 0.32053 | 0.00710 | 2.78558 | 0.01097 | 0.00431 | 0.01390 |
| **Cd84** | 30.19659 | 17.37856 | 32.42982 | 1.57827 | 21.64834 | 8.24899 | 23.10147 | 34.74057 | 1.81853 | 33.37169 | 3.81178 | 25.69132 | 19.61644 | 39.69761 | 21.65593 | 10.54927 |
| **Cd8a** | 0.80350 | 0.00753 | 0.06161 | 0.01448 | 0.00669 | 0.00541 | 0.09947 | 0.19521 | 0.14750 | 0.05482 | 0.70924 | 0.10165 | 1.73081 | 0.00841 | 0.14071 | 0.00540 |
| **Cd8b1** | 0.05244 | 0.01890 | 0.11920 | 0.08621 | 0.05821 | 0.03611 | 0.07139 | 0.09356 | 0.17640 | 0.10543 | 0.21760 | 0.03733 | 0.07840 | 0.07712 | 0.12349 | 0.01449 |
| **Cd9** | 23.95910 | 15.20882 | 36.61933 | 1.63012 | 16.04508 | 16.19507 | 19.67927 | 36.59286 | 2.21246 | 27.40437 | 18.00699 | 8.87426 | 31.70558 | 28.10040 | 15.65474 | 15.02015 |
| **Cd93** | 0.05869 | 0.00748 | 0.15391 | 0.14856 | 0.01655 | 0.01942 | 0.03741 | 0.01111 | 0.12983 | 0.06744 | 0.37426 | 0.03355 | 0.08561 | 0.31304 | 0.12688 | 0.02139 |
| **Cd99l2** | 0.23130 | 0.00847 | 0.41129 | 0.40310 | 0.00646 | 0.00681 | 0.00627 | 0.17463 | 0.40532 | 1.90155 | 0.36056 | 0.09958 | 2.50731 | 0.07424 | 0.05963 | 0.99582 |
| **Cda** | 0.01366 | 0.00807 | 0.01242 | 1.45220 | 0.00936 | 0.00640 | 0.00731 | 0.01182 | 0.00000 | 0.24435 | 0.01388 | 0.00721 | 0.01227 | 0.00943 | 0.00282 | 0.00723 |
| **Cdadc1** | 0.03924 | 3.47534 | 2.53390 | 0.43472 | 0.00856 | 0.00697 | 0.00457 | 2.48361 | 2.50375 | 3.17810 | 2.33674 | 5.99951 | 0.07179 | 0.15801 | 1.60726 | 0.03527 |
| **Cdan1** | 2.62611 | 1.78672 | 8.82236 | 8.99539 | 0.00000 | 1.63014 | 2.21564 | 12.39907 | 17.34110 | 12.73245 | 6.02720 | 9.39747 | 5.60701 | 3.59570 | 9.17385 | 0.01541 |
| **Cdc123** | 9.08224 | 8.08368 | 5.94525 | 4.33103 | 3.97907 | 6.09859 | 6.75558 | 1.89275 | 7.14247 | 0.85887 | 1.94473 | 0.18733 | 2.96822 | 1.40329 | 1.78444 | 5.92758 |
| **Cdc14a** | 0.01154 | 0.00840 | 0.00737 | 0.51938 | 0.00928 | 0.00670 | 0.00812 | 0.06520 | 0.13353 | 0.26655 | 0.17426 | 0.00502 | 0.00837 | 0.00826 | 1.32915 | 0.00716 |
| **Cdc14b** | 5.06786 | 6.82935 | 5.81604 | 18.89721 | 0.03917 | 2.11262 | 9.32946 | 7.11061 | 19.01121 | 13.62070 | 15.88652 | 5.22375 | 4.07129 | 22.83510 | 18.66019 | 0.31906 |
| **Cdc16** | 5.69744 | 8.54817 | 8.94692 | 4.37105 | 0.27074 | 4.61419 | 4.69105 | 8.56339 | 2.75939 | 8.84997 | 9.69528 | 6.92202 | 6.19555 | 7.45359 | 8.74909 | 2.99639 |
| **Cdc20** | 12.19751 | 3.79239 | 11.01409 | 1.33932 | 16.46502 | 7.98639 | 3.53883 | 25.78361 | 11.87468 | 32.85257 | 11.27138 | 10.01734 | 24.26486 | 14.87877 | 23.25335 | 8.60039 |
| **Cdc23** | 3.42380 | 5.58563 | 2.48753 | 0.15115 | 3.93675 | 0.10947 | 5.24507 | 6.19372 | 0.99707 | 3.74959 | 0.50434 | 0.43621 | 1.21070 | 0.47109 | 5.22644 | 1.71025 |
| **Cdc25a** | 2.67760 | 5.70212 | 12.10099 | 2.09375 | 6.75196 | 1.87139 | 5.43190 | 11.43399 | 13.55341 | 12.53369 | 13.35602 | 2.64443 | 5.98535 | 13.65569 | 11.61838 | 1.75966 |

| | | | | | | | | | | | | | | | | |
|---|---|---|---|---|---|---|---|---|---|---|---|---|---|---|---|---|
| **Cdc25b** | 0.00000 | 0.00594 | 2.88623 | 2.20791 | 0.01163 | 0.01822 | 0.00406 | 4.38384 | 2.11698 | 12.51699 | 6.77894 | 1.11323 | 0.45318 | 0.65851 | 9.62997 | 0.00443 |
| **Cdc25c** | 9.83107 | 2.41985 | 4.02493 | 12.06590 | 0.00000 | 1.93584 | 2.30303 | 5.04826 | 17.46913 | 10.90336 | 3.12106 | 10.66777 | 3.80590 | 0.10522 | 5.28203 | 2.70420 |
| **Cdc26** | 6.75506 | 0.13076 | 5.11169 | 0.65686 | 3.36807 | 4.07451 | 1.80759 | 2.20694 | 0.58070 | 2.53463 | 0.35187 | 0.08827 | 0.48573 | 1.57370 | 0.24271 | 2.15914 |
| **Cdc27** | 1.74037 | 7.84550 | 5.83724 | 12.58024 | 5.56968 | 4.90072 | 6.86087 | 4.78038 | 18.10272 | 7.97115 | 11.47816 | 4.60205 | 3.46820 | 32.94487 | 17.56207 | 0.04157 |
| **Cdc34** | 18.42564 | 4.03184 | 13.39574 | 17.55529 | 14.02018 | 14.70179 | 12.28904 | 27.17311 | 17.76418 | 25.12742 | 12.86042 | 1.52910 | 22.95095 | 0.00000 | 0.00000 | 13.82416 |
| **Cdc37** | 21.40245 | 9.49730 | 15.14932 | 1.30575 | 13.00163 | 8.09290 | 11.44708 | 29.57224 | 2.75187 | 22.93435 | 8.78957 | 1.22023 | 23.36280 | 0.54235 | 1.74829 | 12.73086 |
| **Cdc37l1** | 2.35480 | 2.60105 | 0.13841 | 6.89137 | 0.19327 | 0.00788 | 1.87880 | 0.00000 | 5.02716 | 0.64517 | 1.38100 | 3.59984 | 0.83467 | 4.06095 | 8.14379 | 0.33803 |
| **Cdc40** | 5.11605 | 6.39232 | 1.60353 | 7.60365 | 0.03363 | 0.00000 | 1.71411 | 5.37836 | 9.53244 | 10.05880 | 0.95758 | 7.24966 | 1.89547 | 13.45970 | 12.75265 | 1.16855 |
| **Cdc42** | 24.42664 | 13.85167 | 22.81235 | 15.38400 | 20.33387 | 15.47137 | 15.69320 | 21.31092 | 20.20415 | 25.85483 | 19.00799 | 11.57754 | 27.23420 | 25.10149 | 16.71274 | 15.85278 |
| **Cdc42bpa** | 0.02972 | 0.84650 | 0.00284 | 0.46378 | 0.01274 | 0.00678 | 1.09211 | 0.03111 | 0.44942 | 1.18048 | 0.00752 | 0.06374 | 0.00575 | 0.00554 | 0.00000 | 2.01942 |
| **Cdc42bpb** | 6.17422 | 0.01880 | 0.00000 | 1.85926 | 2.03865 | 2.45284 | 1.06309 | 9.20993 | 1.35987 | 7.89957 | 0.23982 | 0.22491 | 5.42726 | 0.00000 | 0.26445 | 2.07473 |
| **Cdc42bpg** | 0.12884 | 0.00668 | 0.00000 | 2.02467 | 0.01728 | 0.00667 | 0.09416 | 0.02088 | 2.88089 | 0.00000 | 4.55712 | 0.01160 | 0.15820 | 1.19108 | 0.71985 | 0.01282 |
| **Cdc42ep1** | 1.40971 | 0.01477 | 1.87030 | 0.28659 | 0.02019 | 4.29826 | 1.48633 | 4.37232 | 0.00902 | 0.56411 | 1.86825 | 0.05257 | 1.49264 | 0.01687 | 0.39433 | 0.02504 |
| **Cdc42ep2** | 0.00000 | 0.00672 | 2.31866 | 2.71211 | 0.00472 | 3.35247 | 0.00456 | 0.96669 | 5.54244 | 2.90933 | 2.42129 | 0.30192 | 1.96824 | 0.29508 | 1.33436 | 0.00330 |
| **Cdc42ep3** | 0.29261 | 4.35270 | 0.97341 | 0.46526 | 0.98330 | 0.09553 | 0.07744 | 9.76862 | 0.00315 | 0.24074 | 0.00000 | 0.03761 | 1.51713 | 0.00000 | 0.00000 | 2.21718 |
| **Cdc42ep4** | 2.72848 | 0.01318 | 2.92913 | 3.73154 | 0.89057 | 8.66513 | 0.00175 | 8.67352 | 1.51333 | 9.61982 | 5.23986 | 0.30825 | 0.27400 | 0.69153 | 2.16638 | 0.00000 |
| **Cdc42ep5** | 2.40485 | 0.00471 | 0.00000 | 0.07900 | 0.04423 | 5.90929 | 0.00956 | 1.75110 | 0.27762 | 4.46663 | 1.09129 | 0.03823 | 10.56435 | 0.01018 | 0.00000 | 0.03452 |
| **Cdc42se1** | 3.15212 | 1.01643 | 2.07966 | 7.30287 | 2.55921 | 4.06950 | 1.65277 | 5.83802 | 1.64199 | 4.92742 | 2.75326 | 7.05609 | 4.57413 | 0.33337 | 3.78842 | 2.45186 |
| **Cdc42se2** | 5.19349 | 2.93171 | 3.64498 | 17.60386 | 6.61358 | 0.46614 | 3.72030 | 3.09687 | 11.61123 | 6.58186 | 1.06138 | 7.42662 | 1.64153 | 4.76842 | 28.90717 | 1.88791 |
| **Cdc45** | 10.49352 | 7.53145 | 16.95164 | 10.93687 | 5.37825 | 5.84854 | 5.50520 | 12.85320 | 20.16674 | 20.38458 | 9.88426 | 11.15151 | 7.46138 | 2.19572 | 18.92865 | 1.98309 |
| **Cdc5l** | 20.20943 | 21.43187 | 18.78039 | 12.72399 | 15.81482 | 10.03331 | 14.26039 | 28.29529 | 20.65380 | 20.83467 | 18.64697 | 17.59139 | 17.61790 | 28.15331 | 13.81771 | 7.55688 |
| **Cdc6** | 6.04092 | 3.54030 | 15.30984 | 3.59861 | 4.81895 | 6.58479 | 3.37722 | 15.87012 | 14.45425 | 18.85792 | 10.13596 | 3.53622 | 5.64189 | 21.68725 | 17.73259 | 5.13399 |
| **Cdc7** | 7.55861 | 0.00601 | 0.54159 | 2.77996 | 0.77844 | 5.09860 | 5.69743 | 2.94864 | 2.59027 | 2.94194 | 3.16611 | 4.14058 | 3.83145 | 8.18136 | 8.31449 | 2.93692 |
| **Cdc73** | 9.03913 | 8.61762 | 4.32686 | 4.97622 | 5.27314 | 3.97290 | 2.06055 | 3.95251 | 1.66922 | 6.53434 | 3.22607 | 7.61661 | 2.11306 | 18.05888 | 10.93305 | 1.80892 |
| **Cdca2** | 4.11434 | 1.68881 | 4.73009 | 2.08194 | 0.16684 | 0.01035 | 0.82605 | 7.26564 | 9.48687 | 2.35059 | 1.49725 | 0.16769 | 3.92697 | 0.79853 | 2.92359 | 2.31396 |
| **Cdca3** | 21.22799 | 9.31859 | 18.47051 | 10.54266 | 12.65300 | 14.29718 | 9.65078 | 24.85248 | 10.88357 | 18.25003 | 13.05944 | 9.65485 | 16.81711 | 5.02249 | 7.17428 | 13.83009 |
| **Cdca4** | 2.13892 | 1.54801 | 5.34924 | 16.20839 | 5.36113 | 4.09803 | 1.99632 | 7.68894 | 10.05166 | 11.46272 | 2.41895 | 1.61824 | 4.34518 | 2.71031 | 7.38774 | 1.16875 |
| **Cdca5** | 9.65414 | 4.61593 | 9.13174 | 23.47157 | 10.16554 | 2.20071 | 7.81173 | 16.19266 | 28.81732 | 17.55480 | 8.52313 | 3.12563 | 10.92802 | 13.17060 | 14.05404 | 3.25991 |
| **Cdca7** | 4.37106 | 5.74242 | 3.79219 | 0.45213 | 0.56965 | 5.07008 | 4.65415 | 5.95229 | 3.21706 | 8.50001 | 6.84778 | 7.78968 | 2.89807 | 5.13894 | 5.98483 | 1.77865 |
| **Cdca7l** | 0.01328 | 0.00790 | 0.01183 | 0.42124 | 0.00905 | 0.00646 | 0.00723 | 0.01033 | 0.21504 | 0.74369 | 0.01020 | 0.00495 | 0.01848 | 0.00542 | 0.27467 | 0.00764 |
| **Cdca8** | 19.11486 | 6.37036 | 9.51730 | 18.61163 | 9.79775 | 12.59377 | 3.27555 | 20.82792 | 28.67508 | 26.78589 | 13.84032 | 11.55885 | 16.19395 | 0.73301 | 15.37790 | 9.34754 |
| **Cdcp1** | 0.01052 | 0.00836 | 0.10787 | 0.58337 | 0.00953 | 0.00576 | 0.00757 | 0.01322 | 0.00018 | 0.00450 | 0.00490 | 0.00286 | 0.01158 | 1.42093 | 0.44931 | 0.00731 |
| **Cdh1** | 6.71611 | 4.91843 | 15.77737 | 3.54378 | 0.47009 | 12.05094 | 9.52373 | 17.62968 | 7.45072 | 7.51263 | 14.14476 | 6.75649 | 15.46704 | 18.42371 | 3.41687 | 12.17780 |
| **Cdh11** | 0.59963 | 0.17380 | 1.34458 | 0.20826 | 0.04960 | 0.33436 | 0.22287 | 0.71846 | 0.18104 | 0.16467 | 0.16096 | 0.09573 | 0.42871 | 0.32773 | 0.21956 | 0.22199 |
| **Cdh15** | 0.01787 | 0.00808 | 0.01132 | 0.17921 | 0.00900 | 0.00669 | 0.00748 | 0.01357 | 0.20279 | 0.01331 | 0.01085 | 0.03911 | 0.01284 | 0.01192 | 0.00478 | 0.00736 |
| **Cdh16** | 0.09721 | 0.00856 | 0.03287 | 0.33841 | 0.08020 | 0.00626 | 0.02933 | 0.02023 | 0.26364 | 0.02443 | 0.10182 | 0.02946 | 0.05586 | 0.08148 | 0.02465 | 0.05075 |
| **Cdh17** | 0.01383 | 0.00836 | 0.00932 | 0.19732 | 0.00912 | 0.00681 | 0.00758 | 0.00670 | 0.00578 | 0.01162 | 0.41184 | 0.00674 | 0.00798 | 0.00848 | 0.00395 | 0.00712 |
| **Cdh2** | 0.62376 | 2.03923 | 0.00000 | 0.03663 | 0.01327 | 0.00630 | 0.00592 | 0.00707 | 0.04190 | 0.00022 | 0.01217 | 0.03549 | 0.01488 | 0.00653 | 0.16073 | 0.96047 |
| **Cdh23** | 0.01197 | 0.00817 | 0.01390 | 0.01793 | 0.01018 | 0.00720 | 0.00765 | 0.02769 | 0.01249 | 0.01317 | 0.01516 | 0.00794 | 0.24758 | 0.01224 | 0.01027 | 0.00764 |
| **Cdh26** | 0.01368 | 0.00809 | 0.01106 | 0.23950 | 0.00926 | 0.00662 | 0.00746 | 0.01269 | 1.03748 | 0.00859 | 0.00664 | 0.05977 | 0.01307 | 0.01246 | 0.00086 | 0.00733 |
| **Cdh3** | 0.47267 | 0.00770 | 0.00665 | 0.21882 | 0.00750 | 0.00753 | 0.00736 | 0.01754 | 0.00548 | 0.01209 | 0.01388 | 0.00772 | 0.12404 | 0.01153 | 0.00723 | 0.00706 |
| **Cdh4** | 0.01022 | 0.00786 | 0.01073 | 2.10276 | 0.01000 | 0.00681 | 0.00739 | 0.02924 | 0.00000 | 0.94588 | 0.04619 | 0.04035 | 0.42810 | 0.02265 | 2.06714 | 0.00724 |
| **Cdip1** | 4.04917 | 1.38958 | 3.98017 | 13.56033 | 4.41925 | 0.02785 | 2.35173 | 6.20531 | 14.10018 | 4.73520 | 4.87967 | 0.42849 | 3.46745 | 4.14117 | 4.61340 | 5.09740 |
| **Cdipt** | 13.13204 | 10.87937 | 21.37330 | 7.92030 | 11.90418 | 12.66094 | 12.52684 | 28.28540 | 15.88632 | 24.99960 | 20.77998 | 3.89781 | 27.44595 | 2.33755 | 11.90993 | 12.29953 |
| **Cdk1** | 22.12456 | 10.29319 | 14.24963 | 20.55986 | 8.16873 | 5.59275 | 9.63758 | 27.42857 | 28.01535 | 18.77231 | 24.71903 | 15.18093 | 13.65881 | 15.69632 | 21.16749 | 6.07757 |
| **Cdk10** | 7.56156 | 2.15800 | 0.71411 | 1.18035 | 3.17237 | 3.08218 | 3.04127 | 6.33375 | 0.95464 | 3.64504 | 1.24032 | 0.08191 | 8.34918 | 0.00466 | 0.32192 | 4.46712 |
| **Cdk11b** | 10.58673 | 7.35650 | 15.34091 | 5.21805 | 6.29825 | 10.69205 | 8.51247 | 15.46337 | 7.98784 | 15.47227 | 6.11979 | 2.86831 | 14.45425 | 0.01644 | 1.79739 | 7.45760 |
| **Cdk12** | 5.21645 | 0.41595 | 0.19480 | 12.71828 | 0.30502 | 0.00000 | 3.06040 | 1.07522 | 12.88470 | 1.66667 | 0.90770 | 1.11111 | 0.41618 | 11.56374 | 9.54763 | 0.08443 |
| **Cdk13** | 0.21580 | 0.73523 | 1.60990 | 3.05762 | 1.66554 | 1.40258 | 0.53756 | 0.79140 | 6.57629 | 3.41174 | 0.16062 | 3.23968 | 0.23982 | 3.26801 | 2.65201 | 1.21697 |
| **Cdk14** | 0.62280 | 1.81995 | 2.18752 | 0.26279 | 0.00314 | 0.65788 | 0.00589 | 0.17570 | 0.14125 | 0.00000 | 0.00675 | 0.00475 | 0.01464 | 0.04374 | 0.39542 | 0.06204 |
| **Cdk16** | 9.29301 | 3.06949 | 13.69748 | 8.76431 | 7.05319 | 6.41174 | 4.50873 | 15.08884 | 5.65502 | 14.87996 | 4.92020 | 2.00167 | 6.78247 | 1.66583 | 9.68609 | 4.27875 |
| **Cdk17** | 2.47095 | 0.02479 | 0.37328 | 1.01934 | 0.02714 | 0.01255 | 2.92072 | 0.75796 | 0.49251 | 3.22119 | 0.94144 | 0.07369 | 0.30321 | 0.57686 | 0.86320 | 0.87683 |
| **Cdk18** | 0.22767 | 4.98642 | 3.55901 | 0.02390 | 5.14278 | 0.01328 | 3.73248 | 4.82181 | 0.03976 | 0.54563 | 2.73289 | 3.14074 | 4.57259 | 0.00000 | 0.06615 | 0.00915 |
| **Cdk19** | 1.93459 | 0.63399 | 0.30283 | 1.35147 | 1.31396 | 0.00880 | 2.04257 | 2.51678 | 8.02739 | 2.99475 | 2.04135 | 1.84162 | 1.17307 | 6.38132 | 2.35208 | 0.43797 |
| **Cdk2** | 6.28027 | 1.98323 | 3.18362 | 0.76913 | 1.63587 | 2.39547 | 2.72987 | 3.09407 | 2.50441 | 9.08768 | 0.07749 | 0.32684 | 2.40667 | 13.39686 | 8.62234 | 5.59423 |
| **Cdk20** | 2.05208 | 0.02589 | 0.14078 | 2.46316 | 0.63303 | 0.01600 | 2.39299 | 0.00000 | 4.85098 | 0.94829 | 0.20923 | 0.03641 | 0.01457 | 0.00000 | 0.81616 | 0.00486 |
| **Cdk2ap1** | 10.94663 | 2.63838 | 12.24267 | 21.90024 | 5.15287 | 6.80980 | 9.79539 | 37.80992 | 24.87624 | 42.61369 | 36.75864 | 12.43130 | 21.92262 | 29.87227 | 33.82279 | 1.53358 |
| **Cdk2ap2** | 4.74134 | 0.00000 | 3.70204 | 17.76429 | 2.52068 | 2.49888 | 0.81668 | 2.30409 | 16.45262 | 11.29008 | 3.83869 | 0.81913 | 7.98467 | 0.24689 | 0.00423 | 0.02108 |
| **Cdk3.ps** | 0.01029 | 0.00807 | 0.84637 | 0.01084 | 0.00813 | 0.00577 | 0.00751 | 0.06720 | 0.01205 | 0.05564 | 0.01069 | 0.02082 | 0.00627 | 0.01507 | 0.00035 | 0.01435 |
| **Cdk4** | 26.36711 | 15.29430 | 22.24255 | 4.05271 | 19.24078 | 13.22720 | 13.19966 | 23.47251 | 4.03710 | 25.41578 | 18.44351 | 4.56287 | 26.37965 | 1.19223 | 8.93492 | 17.64301 |
| **Cdk5** | 7.62300 | 0.10101 | 3.54278 | 0.59353 | 4.33912 | 7.28021 | 1.64424 | 4.31193 | 0.00000 | 1.08871 | 0.00000 | 0.00101 | 4.87983 | 0.00687 | 0.00000 | 7.05147 |
| **Cdk5r1** | 0.00740 | 0.00764 | 0.26696 | 0.73089 | 0.00879 | 0.00679 | 0.00714 | 0.42777 | 0.36281 | 0.92957 | 0.00000 | 0.07588 | 0.28039 | 0.00647 | 0.72915 | 0.00709 |

| | | | | | | | | | | | | | | | | |
|---|---|---|---|---|---|---|---|---|---|---|---|---|---|---|---|---|
| **Cdk5r2** | 0.01082 | 0.00833 | 0.01061 | 0.35436 | 0.00957 | 0.00676 | 0.00767 | 0.10600 | 0.02195 | 0.01002 | 0.01258 | 0.00831 | 0.13538 | 0.01214 | 0.00411 | 0.00728 |
| **Cdk5rap1** | 3.51840 | 2.70697 | 2.96368 | 2.24429 | 3.42033 | 0.06290 | 0.00144 | 1.69353 | 6.74412 | 2.13986 | 4.94738 | 0.45182 | 0.11228 | 1.21734 | 2.63662 | 0.04456 |
| **Cdk5rap2** | 2.70352 | 0.04036 | 0.09212 | 0.53828 | 0.03092 | 1.98985 | 0.01312 | 3.30478 | 0.23225 | 1.14856 | 0.43865 | 0.09426 | 3.60912 | 1.20900 | 1.01600 | 0.19573 |
| **Cdk5rap3** | 8.91078 | 3.79505 | 12.39804 | 4.11213 | 4.50049 | 4.30473 | 6.69261 | 13.54214 | 7.45672 | 1.66850 | 5.37854 | 0.18297 | 9.38021 | 0.03490 | 1.57172 | 4.26320 |
| **Cdk6** | 3.66579 | 0.12227 | 1.70960 | 0.41483 | 0.01461 | 0.01400 | 0.04502 | 0.92985 | 0.52917 | 0.01498 | 0.35765 | 0.05743 | 0.75755 | 0.38518 | 0.07038 | 0.02460 |
| **Cdk7** | 9.70910 | 7.29611 | 8.60338 | 16.24304 | 7.09577 | 3.67314 | 10.03888 | 14.13507 | 21.20216 | 14.69314 | 7.68374 | 10.62922 | 3.24757 | 7.90981 | 18.39484 | 6.96219 |
| **Cdk8** | 2.32398 | 6.04018 | 2.54569 | 12.16926 | 4.73653 | 2.90065 | 4.72383 | 7.40519 | 10.81884 | 19.23537 | 7.16069 | 12.82017 | 5.44348 | 0.80583 | 12.19353 | 2.21081 |
| **Cdk9** | 10.20823 | 9.10144 | 11.79238 | 14.98019 | 7.29319 | 2.11423 | 6.86523 | 11.87374 | 16.81828 | 15.79057 | 5.38718 | 1.04384 | 9.30632 | 2.50963 | 11.53705 | 12.07274 |
| **Cdkal1** | 0.71477 | 0.05050 | 2.13030 | 2.32968 | 0.37559 | 0.12844 | 1.11385 | 0.15276 | 4.71290 | 0.46265 | 2.99514 | 0.13828 | 0.04837 | 5.29717 | 0.00000 | 1.31181 |
| **Cdkl2** | 2.60334 | 2.62807 | 0.67405 | 5.20596 | 0.00497 | 0.00258 | 0.00262 | 0.41207 | 2.32025 | 2.41786 | 1.97033 | 5.05136 | 0.00000 | 10.66227 | 7.27161 | 0.80699 |
| **Cdkl3** | 0.01331 | 0.00783 | 0.01009 | 0.02472 | 0.00921 | 0.00667 | 0.00749 | 0.01157 | 0.71524 | 0.24845 | 0.00388 | 0.00482 | 0.01310 | 0.06611 | 1.06603 | 0.00737 |
| **Cdkl4** | 0.00539 | 0.00771 | 0.02945 | 2.01644 | 0.00982 | 0.00554 | 0.00714 | 0.20297 | 1.51260 | 0.61350 | 0.11424 | 0.08515 | 0.00588 | 1.08551 | 1.20430 | 0.00665 |
| **Cdkl5** | 0.01382 | 0.00816 | 0.01135 | 1.06611 | 0.00930 | 0.00660 | 0.00742 | 0.01283 | 0.10037 | 0.01326 | 0.01321 | 0.06077 | 0.01280 | 0.01093 | 0.00000 | 0.00724 |
| **Cdkn1a** | 2.63061 | 0.03651 | 3.89089 | 2.43388 | 1.29297 | 0.01353 | 2.84057 | 8.76148 | 2.94745 | 8.41727 | 10.92903 | 0.55251 | 0.00000 | 3.21273 | 1.81470 | 0.00798 |
| **Cdkn1b** | 0.07790 | 0.00832 | 0.02435 | 0.07977 | 0.01066 | 0.00802 | 0.00758 | 0.01573 | 0.00983 | 0.01699 | 0.01394 | 0.00789 | 0.02812 | 0.01173 | 0.00886 | 0.17053 |
| **Cdkn1c** | 0.00597 | 0.00767 | 0.00973 | 8.08089 | 0.00885 | 0.00547 | 0.00655 | 0.00901 | 6.92668 | 3.11901 | 0.00000 | 0.09467 | 1.38289 | 0.03122 | 0.22181 | 0.01987 |
| **Cdkn2aip** | 0.00000 | 1.09139 | 1.90651 | 2.40553 | 0.18111 | 0.03405 | 6.01194 | 1.70946 | 4.81261 | 6.04233 | 1.68880 | 0.13709 | 0.00000 | 4.68351 | 4.15115 | 2.20989 |
| **Cdkn2aipnl** | 5.43873 | 2.63471 | 5.70044 | 4.08270 | 3.72358 | 4.26213 | 3.06509 | 6.77464 | 1.91872 | 2.76255 | 5.84804 | 0.18090 | 8.69093 | 0.00000 | 2.41539 | 2.75309 |
| **Cdkn2b** | 0.02292 | 0.00835 | 0.01103 | 0.00469 | 0.01134 | 0.00694 | 0.00754 | 0.01252 | 0.28675 | 0.33185 | 0.00933 | 0.01965 | 0.01266 | 0.01112 | 0.01361 | 0.00747 |
| **Cdkn2c** | 0.16794 | 0.00453 | 0.05692 | 1.11319 | 0.58566 | 0.00512 | 2.28286 | 0.65254 | 0.12491 | 0.06034 | 0.00000 | 4.87865 | 0.29569 | 0.02883 | 0.00000 | 0.00622 |
| **Cdkn2d** | 2.62799 | 0.00000 | 0.00000 | 6.80020 | 2.68195 | 1.58920 | 3.20781 | 1.06766 | 9.88632 | 2.71498 | 0.29971 | 0.64092 | 1.39553 | 0.00000 | 1.82408 | 4.06712 |
| **Cdkn3** | 1.33880 | 0.33667 | 1.47124 | 1.85166 | 0.00292 | 2.92585 | 0.07621 | 0.00000 | 5.46515 | 1.53455 | 0.23038 | 0.09917 | 0.00000 | 1.07603 | 0.13760 | 0.62807 |
| **Cdnf** | 0.04945 | 0.12050 | 0.04688 | 0.38152 | 0.03601 | 0.03151 | 0.03688 | 0.03629 | 0.12068 | 0.03058 | 0.08657 | 0.08314 | 0.03824 | 0.24685 | 0.15683 | 0.02908 |
| **Cdo1** | 0.01247 | 0.00809 | 0.00299 | 5.47222 | 0.00788 | 0.17623 | 0.00759 | 0.03332 | 1.38088 | 0.50460 | 0.00765 | 0.46105 | 0.00707 | 0.01283 | 1.13572 | 1.03717 |
| **Cdon** | 1.23955 | 0.00480 | 0.00000 | 0.02813 | 0.00565 | 1.18467 | 2.42852 | 0.00000 | 0.01170 | 0.39538 | 0.18949 | 0.01562 | 0.09111 | 0.01313 | 0.03605 | 0.01285 |
| **Cdpf1** | 4.54731 | 5.63433 | 6.71900 | 0.95904 | 4.83514 | 6.51799 | 6.86464 | 6.39516 | 0.19913 | 3.32024 | 1.52588 | 0.50595 | 9.51311 | 0.22052 | 0.35853 | 3.08629 |
| **Cdr2** | 0.35642 | 0.00712 | 1.51489 | 18.40916 | 0.00479 | 0.24643 | 0.00589 | 0.00000 | 15.72274 | 4.11765 | 0.92682 | 0.91220 | 0.39190 | 1.26132 | 5.69906 | 0.00934 |
| **Cdr2l** | 1.28070 | 0.37062 | 0.53683 | 0.00622 | 2.65578 | 0.00404 | 1.50483 | 0.02381 | 2.18249 | 0.00008 | 0.00000 | 0.02430 | 0.00754 | 0.03319 | 0.00443 | 0.00544 |
| **Cds1** | 1.12934 | 0.14419 | 5.83004 | 1.04753 | 0.02704 | 0.00419 | 4.91231 | 6.04758 | 0.75001 | 2.31822 | 5.68027 | 4.05001 | 0.55727 | 0.69841 | 0.89184 | 0.02436 |
| **Cds2** | 13.71315 | 5.95813 | 13.10353 | 1.91127 | 4.76483 | 8.16790 | 12.75332 | 14.88548 | 1.63070 | 17.60264 | 17.23174 | 8.84128 | 18.16707 | 1.81599 | 12.85358 | 3.46529 |
| **Cdsn** | 0.01398 | 0.00821 | 0.05079 | 2.37579 | 0.00895 | 0.00558 | 0.00776 | 0.43783 | 1.15842 | 0.98573 | 0.01665 | 0.06020 | 0.00881 | 0.00120 | 0.56242 | 0.00687 |
| **Cdt1** | 12.47385 | 3.84033 | 4.14503 | 14.11369 | 7.98640 | 3.59798 | 3.43098 | 21.76382 | 14.77890 | 32.09927 | 20.20895 | 3.75662 | 16.66556 | 0.87665 | 2.30682 | 11.46607 |
| **Cdv3** | 28.65357 | 16.31331 | 29.32372 | 25.42277 | 11.37320 | 14.63532 | 17.39382 | 19.78602 | 30.66797 | 28.55036 | 22.20995 | 18.71583 | 23.81923 | 28.02720 | 24.63893 | 11.18776 |
| **Cdx1** | 0.00914 | 0.00784 | 0.01688 | 0.01054 | 0.00781 | 0.18505 | 0.00739 | 0.01316 | 0.01241 | 0.00632 | 0.01246 | 0.00802 | 0.00701 | 0.01314 | 0.01020 | 1.22298 |
| **Cdx2** | 0.00000 | 0.00663 | 1.59737 | 0.00000 | 0.00594 | 4.37155 | 0.27554 | 3.53976 | 0.00815 | 0.00000 | 2.35485 | 0.00238 | 6.21914 | 0.01263 | 0.00000 | 1.70210 |
| **Cdyl** | 2.84800 | 0.00508 | 1.49905 | 4.04427 | 0.00934 | 0.15790 | 0.01611 | 0.31141 | 5.09071 | 1.60403 | 1.19250 | 0.05868 | 0.78258 | 0.72712 | 4.95063 | 0.74307 |
| **Cdyl2** | 5.91765 | 2.94929 | 7.90660 | 0.24475 | 5.27620 | 0.78318 | 5.05602 | 4.11512 | 0.37756 | 3.38845 | 1.13962 | 4.29050 | 2.90022 | 3.80662 | 0.00000 | 0.00000 |
| **Ceacam1** | 0.85356 | 0.00583 | 1.42724 | 0.61102 | 0.00401 | 0.00473 | 1.54085 | 0.00601 | 0.53458 | 0.00778 | 0.00750 | 0.00466 | 0.02638 | 0.07149 | 0.02321 | 0.00563 |
| **Ceacam10** | 15.00143 | 0.00000 | 10.91189 | 0.33270 | 8.02521 | 3.17444 | 5.80146 | 0.00000 | 0.00000 | 2.63109 | 3.63291 | 0.00000 | 4.08705 | 0.91621 | 0.00000 | 1.86178 |
| **Ceacam18** | 0.11656 | 0.00795 | 0.00835 | 0.23369 | 0.00768 | 0.00691 | 0.00735 | 0.01426 | 0.00761 | 0.06237 | 0.01395 | 0.00783 | 0.01285 | 0.00896 | 0.03471 | 0.01496 |
| **Ceacam19** | 0.22301 | 0.07040 | 0.10868 | 0.00656 | 0.01773 | 0.18100 | 0.06689 | 0.11302 | 0.01183 | 0.13843 | 0.11971 | 0.01556 | 0.16269 | 0.01687 | 0.07091 | 0.04992 |
| **Ceacam2** | 0.32977 | 0.01818 | 0.25075 | 0.42988 | 0.00749 | 0.00525 | 0.44434 | 0.00918 | 0.30719 | 0.02801 | 0.01043 | 0.00724 | 0.00965 | 0.00989 | 0.00984 | 0.00667 |
| **Ceacam20** | 0.30846 | 0.00759 | 0.02214 | 2.30913 | 0.00727 | 0.00727 | 0.00579 | 0.00000 | 0.08143 | 0.00000 | 0.95725 | 0.02228 | 1.88167 | 0.04518 | 0.04924 | 0.00555 |
| **Ceacam5** | 0.01363 | 0.00811 | 0.01193 | 0.00432 | 0.00932 | 0.00664 | 0.00755 | 0.01383 | 0.81716 | 0.01134 | 0.00697 | 0.00688 | 0.01295 | 0.01399 | 0.00648 | 0.00737 |
| **Cebpa** | 2.57312 | 0.00423 | 0.00000 | 0.70245 | 0.01037 | 2.02398 | 0.01394 | 0.73753 | 0.34560 | 7.19688 | 0.98157 | 0.14563 | 0.79206 | 0.76826 | 1.83835 | 3.54849 |
| **Cebpb** | 0.07972 | 0.00837 | 0.00940 | 0.00998 | 0.01607 | 0.00674 | 0.00740 | 0.01216 | 0.01255 | 0.09817 | 0.01288 | 0.00721 | 0.01247 | 0.00894 | 0.02813 | 0.00757 |
| **Cebpd** | 0.01382 | 0.00806 | 0.00816 | 0.60337 | 0.00898 | 0.00700 | 0.00745 | 0.00225 | 0.93125 | 0.15602 | 0.41305 | 0.04052 | 0.00879 | 0.00632 | 0.00000 | 0.00711 |
| **Cebpg** | 2.82849 | 0.08056 | 3.68830 | 10.26928 | 0.04715 | 0.30010 | 1.30541 | 2.91535 | 1.35264 | 7.35463 | 6.17976 | 6.26473 | 0.46657 | 11.48667 | 6.86406 | 0.01418 |
| **Cebpz** | 6.85791 | 10.36603 | 5.31828 | 4.74022 | 7.81300 | 4.59060 | 3.27140 | 8.98005 | 6.54767 | 13.55372 | 5.73132 | 10.72455 | 5.73768 | 20.28626 | 19.81234 | 3.11578 |
| **Cebpzos** | 1.97291 | 0.13526 | 1.67780 | 0.09220 | 1.51191 | 0.56172 | 0.85454 | 0.55733 | 0.00585 | 1.66626 | 4.36609 | 0.02542 | 2.29834 | 0.28844 | 1.07116 | 1.54560 |
| **Cecr2** | 9.90816 | 6.96621 | 2.01096 | 20.49587 | 7.61255 | 5.34571 | 5.39458 | 9.72957 | 20.86104 | 20.05044 | 13.23906 | 15.10207 | 1.90950 | 28.16105 | 20.64276 | 5.66477 |
| **Cecr5** | 0.01210 | 0.00826 | 0.01211 | 0.23928 | 0.00895 | 0.00661 | 0.00769 | 0.01870 | 0.00523 | 0.01275 | 0.01215 | 0.00852 | 0.01119 | 0.01187 | 0.00695 | 0.17214 |
| **Cela1** | 1.37639 | 0.00611 | 1.03443 | 1.47809 | 0.01441 | 0.00465 | 0.00606 | 0.81391 | 0.72936 | 0.75617 | 0.95717 | 0.03404 | 0.06034 | 0.04704 | 0.00000 | 0.00584 |
| **Cela2a** | 3.36052 | 0.00255 | 0.96021 | 0.01034 | 0.33425 | 0.00285 | 1.44437 | 0.40061 | 0.01153 | 0.00000 | 0.01014 | 0.00248 | 0.63655 | 0.01793 | 0.00000 | 0.00302 |
| **Cela3b** | 0.01305 | 0.00761 | 0.01198 | 0.00770 | 0.00903 | 0.00647 | 0.00724 | 0.01034 | 0.01161 | 0.91634 | 0.01339 | 0.00587 | 0.05171 | 0.03213 | 0.01181 | 0.00709 |
| **Celf1** | 19.75939 | 8.13636 | 10.98498 | 7.21022 | 3.35841 | 10.05171 | 7.49811 | 12.75189 | 9.51535 | 13.67320 | 8.85212 | 8.84198 | 14.33169 | 9.29663 | 17.00047 | 5.27233 |
| **Celf2** | 0.00554 | 3.57313 | 2.35665 | 7.85634 | 0.02826 | 0.07166 | 0.12280 | 0.02196 | 0.90746 | 2.12065 | 0.88861 | 0.01084 | 0.99616 | 0.00000 | 1.16913 | 0.00320 |
| **Celf3** | 0.01816 | 0.00746 | 0.01489 | 0.00636 | 0.00868 | 0.00740 | 0.00708 | 0.41501 | 0.01112 | 0.00000 | 0.25397 | 0.00561 | 0.00475 | 0.01250 | 0.00131 | 0.00722 |
| **Celf5** | 0.01155 | 0.00795 | 0.02976 | 0.45485 | 0.02735 | 0.00799 | 0.01777 | 0.01312 | 0.01372 | 0.03586 | 0.01261 | 0.00840 | 0.01895 | 0.06476 | 0.00908 | 0.01452 |
| **Celsr1** | 0.05353 | 0.45089 | 1.49466 | 0.26038 | 0.00535 | 0.00684 | 0.01589 | 0.56765 | 0.41843 | 0.44102 | 0.04139 | 0.06710 | 1.03750 | 0.01163 | 0.00097 | 0.00605 |
| **Celsr3** | 0.00952 | 0.00882 | 0.00834 | 1.27350 | 0.00963 | 0.00682 | 0.00815 | 0.44935 | 0.00000 | 0.00817 | 0.01065 | 0.02108 | 0.00594 | 0.01033 | 0.02565 | 0.00716 |
| **Cend1** | 0.01268 | 0.00844 | 0.70232 | 0.00445 | 0.00809 | 0.00568 | 0.00763 | 0.00907 | 0.21440 | 0.00000 | 0.22725 | 0.00749 | 0.00712 | 0.00777 | 0.06485 | 0.00699 |

| | | | | | | | | | | | | | | | | |
|---|---|---|---|---|---|---|---|---|---|---|---|---|---|---|---|---|
| **Cenpa** | 18.99564 | 0.24565 | 11.16035 | 13.67440 | 17.34506 | 8.25255 | 5.50816 | 17.97429 | 20.89101 | 19.62478 | 18.63857 | 2.79350 | 17.51848 | 3.28930 | 2.58038 | 8.44692 |
| **Cenpb** | 0.19245 | 0.01698 | 0.00000 | 0.29784 | 3.29468 | 1.13732 | 0.02609 | 0.02430 | 0.02005 | 1.45398 | 0.00870 | 0.02845 | 1.45433 | 0.00517 | 0.16649 | 1.06131 |
| **Cenpc1** | 0.54674 | 0.00682 | 1.12316 | 0.38991 | 0.00654 | 1.04954 | 0.04642 | 1.05837 | 0.95806 | 1.14877 | 0.58382 | 0.06908 | 0.55183 | 0.62283 | 3.20429 | 0.03417 |
| **Cenpe** | 5.52524 | 5.37727 | 4.97214 | 11.62425 | 4.31818 | 8.49377 | 0.55413 | 12.67347 | 14.90696 | 21.39110 | 10.72374 | 19.65464 | 4.13847 | 29.02793 | 20.36912 | 1.52379 |
| **Cenpf** | 4.05702 | 1.26938 | 6.21787 | 6.82131 | 0.29747 | 3.15454 | 1.36206 | 2.20634 | 13.69367 | 0.00000 | 5.21019 | 1.18941 | 0.00004 | 2.06293 | 3.26513 | 2.19446 |
| **Cenph** | 5.95386 | 4.65386 | 4.69716 | 4.62747 | 5.79087 | 3.05199 | 2.37858 | 7.78247 | 6.18020 | 8.61435 | 4.32361 | 8.74308 | 6.95020 | 0.16153 | 4.64165 | 2.53446 |
| **Cenpi** | 3.41041 | 0.60442 | 2.18759 | 3.56524 | 1.66283 | 0.00246 | 0.00203 | 3.37939 | 0.63946 | 5.41942 | 1.76529 | 4.49054 | 1.17206 | 12.03442 | 7.29665 | 0.00780 |
| **Cenpj** | 0.00000 | 0.13878 | 0.02690 | 5.87422 | 4.92741 | 1.67885 | 2.79285 | 0.62950 | 1.60116 | 0.00001 | 0.48206 | 6.98893 | 0.05315 | 1.01132 | 3.54486 | 0.00504 |
| **Cenpk** | 1.94867 | 2.78332 | 0.75422 | 1.76752 | 0.01158 | 1.84363 | 4.87569 | 0.99109 | 2.98927 | 2.80017 | 4.79048 | 14.18467 | 0.00000 | 4.75150 | 0.00000 | 0.01541 |
| **Cenpl** | 0.01825 | 0.09276 | 5.35489 | 2.79923 | 4.24226 | 0.01442 | 0.16644 | 3.09429 | 10.75262 | 4.06517 | 1.87979 | 1.57772 | 3.57556 | 0.01546 | 6.34690 | 0.01008 |
| **Cenpm** | 5.32765 | 1.52847 | 8.22854 | 0.30682 | 2.34320 | 0.03659 | 5.01269 | 7.41773 | 2.44628 | 3.51753 | 3.40382 | 0.33383 | 5.64505 | 0.00000 | 0.00000 | 0.06877 |
| **Cenpn** | 7.25012 | 3.38010 | 5.79422 | 18.64288 | 3.10386 | 5.28943 | 8.46452 | 6.49431 | 6.92621 | 8.31656 | 4.57792 | 1.59008 | 6.54392 | 5.95461 | 11.41857 | 4.45704 |
| **Cenpo** | 1.00345 | 8.32494 | 4.52531 | 2.29102 | 0.00741 | 0.01267 | 2.03370 | 2.16755 | 0.13023 | 0.83576 | 0.78634 | 0.06524 | 0.75089 | 0.00000 | 1.83246 | 0.00000 |
| **Cenpp** | 3.10523 | 1.90960 | 0.00000 | 3.20058 | 0.66287 | 3.61861 | 0.45386 | 3.26087 | 4.27381 | 3.73075 | 1.15138 | 0.47406 | 0.83221 | 1.40609 | 1.39936 | 2.13056 |
| **Cenpq** | 0.00000 | 2.86427 | 0.44940 | 3.81849 | 2.68897 | 0.00941 | 0.00407 | 2.76368 | 3.16353 | 4.91718 | 0.93734 | 8.59803 | 0.96424 | 4.72889 | 4.85884 | 1.50852 |
| **Cenpt** | 10.71518 | 3.08119 | 1.60363 | 9.05300 | 0.08005 | 5.82224 | 7.40029 | 10.06936 | 11.58687 | 7.63823 | 7.81639 | 3.99279 | 4.91334 | 0.23523 | 4.97271 | 6.80452 |
| **Cenpu** | 3.25731 | 1.92938 | 3.12208 | 0.00000 | 0.00000 | 0.76001 | 0.00354 | 2.22560 | 0.45397 | 2.39827 | 0.09170 | 2.30207 | 0.56090 | 0.23792 | 0.52504 | 1.75646 |
| **Cenpv** | 1.17426 | 0.27768 | 0.17166 | 4.57587 | 0.00516 | 0.00740 | 0.00460 | 0.00175 | 2.31329 | 0.22162 | 0.66621 | 0.00557 | 1.93525 | 0.00839 | 0.00000 | 0.18272 |
| **Cenpw** | 4.37367 | 0.00883 | 0.90924 | 7.04013 | 6.35649 | 1.03791 | 1.51954 | 1.89119 | 13.33632 | 4.57072 | 3.90212 | 0.49556 | 0.35821 | 0.10952 | 0.14188 | 1.59355 |
| **Cep104** | 0.94096 | 0.51575 | 2.46317 | 2.02787 | 0.03540 | 0.01113 | 0.70809 | 2.96448 | 1.31129 | 3.66782 | 0.89536 | 0.32074 | 0.57543 | 0.19693 | 1.57858 | 2.49638 |
| **Cep120** | 3.04255 | 4.44261 | 3.81406 | 11.09655 | 0.05255 | 2.07875 | 0.12567 | 0.58947 | 2.12210 | 7.41381 | 2.59620 | 0.57782 | 1.97844 | 5.47497 | 12.46668 | 3.08188 |
| **Cep126** | 0.92102 | 0.63455 | 0.26809 | 0.15474 | 0.89993 | 0.00975 | 0.00586 | 0.00000 | 0.00469 | 0.11073 | 0.16749 | 3.49378 | 0.05602 | 0.03220 | 0.03843 | 0.00601 |
| **Cep128** | 0.08632 | 0.00843 | 0.00276 | 0.38508 | 0.00894 | 0.01652 | 0.00785 | 0.97515 | 1.41803 | 0.13103 | 0.00000 | 0.00474 | 0.06368 | 0.00726 | 0.04509 | 0.01504 |
| **Cep131** | 3.12699 | 0.00423 | 0.03850 | 0.01215 | 0.00035 | 0.01506 | 0.00410 | 2.00148 | 0.03108 | 0.93437 | 0.17132 | 0.01843 | 1.90769 | 0.02244 | 0.00329 | 3.88675 |
| **Cep135** | 1.24001 | 0.14629 | 0.00000 | 3.30732 | 0.05097 | 0.00516 | 0.66709 | 0.00000 | 0.78818 | 0.30433 | 0.59321 | 0.08252 | 0.06149 | 0.00000 | 3.35961 | 0.00560 |
| **Cep152** | 1.77734 | 0.00402 | 1.09610 | 0.41689 | 0.02613 | 0.00693 | 2.68654 | 0.87816 | 3.08615 | 1.75995 | 0.00000 | 0.38626 | 1.06840 | 0.04160 | 0.04985 | 2.34768 |
| **Cep162** | 0.00000 | 0.00678 | 0.41324 | 1.31094 | 0.46924 | 0.00915 | 2.03375 | 0.58476 | 1.14114 | 1.05360 | 0.50026 | 0.12209 | 1.53509 | 0.00786 | 1.48359 | 0.02101 |
| **Cep164** | 8.57235 | 0.03207 | 3.87317 | 1.20624 | 7.93481 | 3.89699 | 6.29813 | 0.12484 | 0.00000 | 1.82656 | 0.00259 | 0.21338 | 3.77681 | 0.01011 | 1.76947 | 0.00000 |
| **Cep170** | 2.23605 | 2.71847 | 1.00428 | 2.50905 | 0.07804 | 0.02684 | 1.81338 | 0.74957 | 5.97628 | 0.00000 | 0.64219 | 5.28379 | 0.00000 | 7.49380 | 3.51964 | 0.00190 |
| **Cep170b** | 0.00000 | 0.00699 | 0.87156 | 5.49514 | 0.00644 | 0.15247 | 0.19354 | 1.25503 | 5.89931 | 0.99540 | 0.55847 | 0.08252 | 2.64307 | 0.00000 | 0.14878 | 1.85543 |
| **Cep19** | 4.08583 | 3.21638 | 0.67015 | 1.85972 | 0.57811 | 2.13367 | 0.19924 | 0.94930 | 1.14593 | 7.43579 | 0.87688 | 0.20869 | 3.24678 | 3.50126 | 0.34422 | 0.00000 |
| **Cep192** | 2.64874 | 2.06672 | 2.01595 | 7.10693 | 3.74378 | 3.26537 | 3.90088 | 3.47353 | 12.23024 | 8.05048 | 0.83155 | 8.53317 | 0.00000 | 3.51096 | 11.72071 | 1.38493 |
| **Cep250** | 3.02214 | 1.11519 | 1.32209 | 2.08546 | 0.02388 | 0.40220 | 2.23499 | 2.48353 | 3.23770 | 3.49146 | 0.75336 | 2.13344 | 0.00000 | 0.45503 | 0.14062 | 2.33982 |
| **Cep290** | 0.62718 | 0.28521 | 1.76587 | 0.13614 | 0.01194 | 0.00656 | 0.00568 | 0.95112 | 0.00367 | 0.00000 | 0.65222 | 0.00240 | 1.29983 | 2.86548 | 3.19363 | 0.28094 |
| **Cep295** | 2.88172 | 5.08158 | 1.01542 | 4.79941 | 0.02997 | 0.00410 | 3.08810 | 3.71795 | 1.70186 | 2.42333 | 1.48478 | 0.19827 | 1.42593 | 3.01920 | 2.50828 | 1.88515 |
| **Cep350** | 0.12158 | 0.00759 | 1.84215 | 1.64743 | 0.01252 | 0.94656 | 0.23891 | 0.01516 | 0.00000 | 0.00000 | 0.09472 | 0.51244 | 0.50607 | 2.75683 | 0.57119 | 0.65639 |
| **Cep41** | 1.68799 | 1.79357 | 0.00000 | 1.34914 | 0.00900 | 0.11810 | 3.18877 | 0.03584 | 0.70764 | 1.08740 | 0.00000 | 0.03480 | 0.03567 | 2.06039 | 0.00000 | 0.01223 |
| **Cep44** | 0.07942 | 0.18746 | 0.13590 | 0.00281 | 0.00759 | 0.00601 | 0.00683 | 0.77245 | 0.03854 | 0.03271 | 1.00014 | 0.00001 | 0.07651 | 0.00849 | 0.80329 | 0.06553 |
| **Cep55** | 3.83882 | 0.57306 | 0.32156 | 0.35240 | 0.00000 | 0.02295 | 0.71064 | 0.16116 | 0.22213 | 0.05712 | 0.17167 | 0.31509 | 0.72465 | 2.97862 | 2.13642 | 0.19998 |
| **Cep57** | 5.07065 | 3.74710 | 3.48011 | 3.54754 | 3.88796 | 0.80858 | 7.82595 | 4.13164 | 5.10778 | 0.97885 | 0.80961 | 0.49854 | 4.06901 | 1.61408 | 3.73856 | 4.21120 |
| **Cep57l1** | 5.32676 | 2.48667 | 3.73443 | 1.53857 | 1.65463 | 4.49093 | 8.32620 | 4.19101 | 0.00000 | 2.50924 | 3.25245 | 6.74900 | 4.53108 | 8.84721 | 3.10347 | 0.08789 |
| **Cep63** | 2.79063 | 1.10778 | 3.19197 | 5.11940 | 3.15814 | 0.01060 | 1.53767 | 3.55578 | 2.38622 | 11.16623 | 1.71023 | 5.67744 | 6.11883 | 12.67294 | 8.82635 | 0.02012 |
| **Cep68** | 1.36075 | 0.02685 | 2.55735 | 0.66441 | 0.00607 | 2.46468 | 1.00083 | 0.00000 | 2.27016 | 0.84168 | 0.64257 | 0.04927 | 0.53388 | 5.87649 | 0.31316 | 0.00378 |
| **Cep70** | 1.50560 | 2.43362 | 1.01949 | 7.53774 | 3.49547 | 5.05927 | 0.64376 | 0.39361 | 1.64762 | 0.00000 | 4.72037 | 1.06930 | 0.42989 | 2.58913 | 3.52107 | 0.00000 |
| **Cep72** | 0.01501 | 0.02881 | 1.29042 | 0.00295 | 0.01026 | 0.00890 | 0.00605 | 0.11962 | 0.00929 | 0.00000 | 1.56257 | 0.05888 | 0.51754 | 0.77829 | 0.65905 | 0.00628 |
| **Cep76** | 1.55176 | 4.23071 | 2.98167 | 2.88077 | 6.97992 | 0.93518 | 4.77068 | 3.27176 | 3.80717 | 2.42682 | 1.83318 | 0.69957 | 2.32456 | 10.47405 | 5.01196 | 0.83715 |
| **Cep78** | 1.35071 | 0.00394 | 1.24313 | 0.16043 | 2.73314 | 0.01435 | 1.63055 | 0.01062 | 0.84121 | 1.39738 | 1.01776 | 0.20195 | 0.95636 | 3.55941 | 2.19059 | 0.02618 |
| **Cep83** | 4.39745 | 3.13314 | 4.16817 | 2.00082 | 3.34796 | 1.37034 | 5.36835 | 1.61015 | 1.14733 | 2.08514 | 0.98924 | 3.77555 | 0.04537 | 1.81597 | 1.41243 | 0.86608 |
| **Cep83os** | 0.00980 | 0.11000 | 0.14678 | 0.30212 | 0.00863 | 0.00555 | 0.00732 | 0.11279 | 0.70676 | 0.00000 | 0.00322 | 0.18218 | 0.00740 | 0.00477 | 0.42273 | 0.00706 |
| **Cep85** | 2.10420 | 3.77280 | 0.00000 | 4.74202 | 0.04209 | 4.87829 | 0.69518 | 3.32163 | 3.95971 | 4.48225 | 4.18948 | 0.15348 | 0.00000 | 0.32898 | 6.34530 | 0.45112 |
| **Cep85l** | 0.01236 | 0.00787 | 0.00634 | 0.07345 | 0.00970 | 0.00648 | 0.00717 | 0.00945 | 0.36866 | 0.00000 | 0.00000 | 0.01100 | 0.01366 | 1.28802 | 0.92829 | 0.00736 |
| **Cep89** | 2.09287 | 0.09270 | 1.59544 | 1.81996 | 1.33808 | 0.00963 | 1.46104 | 2.49457 | 5.18487 | 1.69299 | 2.90559 | 0.22405 | 0.02198 | 0.09332 | 4.91580 | 1.03693 |
| **Cep95** | 3.19844 | 2.03956 | 0.53798 | 1.83104 | 0.09410 | 0.00145 | 3.24609 | 2.07104 | 0.10230 | 0.15442 | 0.49534 | 5.32805 | 0.66641 | 0.06535 | 2.89557 | 0.33006 |
| **Cep97** | 4.11322 | 0.00272 | 1.64703 | 4.34194 | 0.52411 | 0.00926 | 0.04284 | 0.08533 | 5.87415 | 0.00000 | 2.23076 | 0.08609 | 0.03978 | 1.45230 | 0.91382 | 0.02933 |
| **Cept1** | 1.67779 | 0.73998 | 5.75791 | 6.05788 | 0.67725 | 3.88411 | 2.43502 | 6.57902 | 2.19345 | 0.00000 | 2.71939 | 0.53413 | 1.52696 | 2.74231 | 1.25602 | 0.02788 |
| **Cercam** | 0.06347 | 0.01824 | 0.11975 | 1.21101 | 0.02668 | 0.01193 | 0.07842 | 0.01619 | 0.59913 | 0.09906 | 0.43733 | 0.04374 | 0.05138 | 0.15670 | 0.00000 | 0.01439 |
| **Cerk** | 0.92110 | 0.00560 | 0.13276 | 0.28325 | 0.56655 | 0.00682 | 0.82469 | 0.30367 | 0.00695 | 0.68253 | 0.11531 | 0.01061 | 0.32036 | 0.00543 | 0.08355 | 2.38731 |
| **Cers2** | 4.06687 | 10.48140 | 5.08576 | 4.31870 | 5.42397 | 7.91438 | 7.23529 | 18.92379 | 7.24997 | 10.43011 | 11.78445 | 9.13974 | 4.86837 | 17.42869 | 19.26283 | 4.48488 |
| **Cers4** | 0.01505 | 0.00785 | 0.00731 | 0.00542 | 0.00933 | 0.00679 | 0.00706 | 0.00000 | 0.01939 | 0.00000 | 0.49745 | 0.03229 | 0.45481 | 0.00694 | 1.15855 | 0.00705 |
| **Cers5** | 0.92404 | 0.09731 | 7.51670 | 7.67421 | 4.93286 | 1.24882 | 4.95789 | 2.07330 | 9.52087 | 13.49032 | 6.25720 | 7.49868 | 11.10206 | 11.40733 | 8.55516 | 3.81010 |
| **Cers6** | 2.39498 | 0.00472 | 1.46573 | 0.96521 | 0.01026 | 0.00735 | 0.24312 | 0.00000 | 0.45230 | 0.16386 | 1.26542 | 0.00757 | 0.28408 | 0.13754 | 0.87307 | 0.01181 |
| **Ces1a** | 0.01372 | 0.00819 | 0.01080 | 0.00537 | 0.00926 | 0.00670 | 0.00747 | 0.01018 | 0.01037 | 0.02304 | 0.08076 | 0.00760 | 0.01196 | 0.00623 | 0.69374 | 0.00737 |

| | | | | | | | | | | | | | | | | |
|---|---|---|---|---|---|---|---|---|---|---|---|---|---|---|---|---|
| **Ces1d** | 0.01277 | 0.00789 | 0.01231 | 0.11323 | 0.00890 | 0.00646 | 0.00725 | 0.00990 | 0.00720 | 1.84839 | 0.01044 | 0.00302 | 0.01063 | 0.01138 | 0.00477 | 0.00729 |
| **Ces1f** | 0.01239 | 0.00770 | 0.01223 | 0.00559 | 0.00868 | 0.00652 | 0.00719 | 0.00943 | 0.01117 | 2.73248 | 0.01060 | 0.00000 | 0.01089 | 0.01224 | 0.00672 | 0.00767 |
| **Cetn2** | 7.87502 | 3.94246 | 3.70633 | 0.55486 | 1.09470 | 3.99351 | 3.55626 | 0.13017 | 0.86566 | 1.67370 | 1.65464 | 0.03155 | 3.62666 | 0.00000 | 2.19068 | 0.00000 |
| **Cetn3** | 13.52497 | 3.85162 | 8.65222 | 1.58666 | 9.46722 | 5.04130 | 0.90789 | 3.69735 | 0.22421 | 0.87998 | 6.06122 | 1.04676 | 4.96176 | 7.05016 | 1.05032 | 1.16682 |
| **Cetn4** | 0.00862 | 0.00820 | 0.48812 | 0.30788 | 0.00737 | 0.00577 | 0.00761 | 0.01322 | 0.00623 | 0.26333 | 0.01171 | 0.00867 | 0.00850 | 0.01138 | 0.00000 | 0.63051 |
| **Cfap100** | 0.01374 | 0.00796 | 0.01233 | 0.01045 | 0.00941 | 0.00659 | 0.00748 | 0.01336 | 0.01240 | 0.04804 | 0.01316 | 0.00765 | 0.01275 | 0.01175 | 0.00995 | 0.00747 |
| **Cfap126** | 0.25789 | 0.00799 | 0.00647 | 1.50645 | 0.01724 | 0.00661 | 0.00695 | 0.00823 | 1.12140 | 1.74197 | 0.00020 | 0.00000 | 0.01042 | 0.00759 | 0.05284 | 0.00683 |
| **Cfap157** | 0.01011 | 0.00762 | 0.01311 | 0.09774 | 0.00917 | 0.00796 | 0.00674 | 0.00000 | 1.68526 | 1.20987 | 1.31532 | 0.03866 | 0.22432 | 0.25284 | 2.11089 | 0.00707 |
| **Cfap161** | 0.01015 | 0.00767 | 0.00742 | 0.00639 | 0.00946 | 0.00676 | 0.00709 | 0.93707 | 0.53499 | 0.00371 | 0.00000 | 0.00521 | 0.00532 | 0.00766 | 0.21748 | 0.00721 |
| **Cfap20** | 5.85928 | 2.33594 | 1.42379 | 7.83648 | 0.60300 | 2.67735 | 0.71899 | 3.44485 | 13.44313 | 3.17474 | 4.97323 | 5.15137 | 2.17315 | 3.99569 | 0.69542 | 2.95555 |
| **Cfap206** | 0.00643 | 0.00729 | 0.00504 | 0.28722 | 0.00915 | 0.00723 | 0.00637 | 0.81100 | 0.00758 | 0.70714 | 0.32459 | 0.00142 | 1.06133 | 0.00608 | 0.01494 | 0.00704 |
| **Cfap221** | 0.01372 | 0.00788 | 0.01163 | 1.29582 | 0.00933 | 0.00654 | 0.00744 | 0.01277 | 0.33365 | 0.00905 | 0.00996 | 0.00708 | 0.01293 | 0.05317 | 0.03538 | 0.00720 |
| **Cfap36** | 2.89773 | 0.00227 | 0.74365 | 1.53473 | 3.31599 | 0.00502 | 1.01426 | 0.41748 | 0.00000 | 0.20872 | 0.04533 | 0.66025 | 0.15513 | 0.92978 | 0.68492 | 0.81324 |
| **Cfap43** | 0.16301 | 4.39689 | 0.96311 | 0.43858 | 0.01317 | 0.01067 | 0.00497 | 1.23864 | 0.81029 | 0.00000 | 1.42388 | 0.00000 | 0.31657 | 1.10262 | 0.61583 | 0.01162 |
| **Cfap45** | 0.28456 | 0.00764 | 0.00415 | 0.74557 | 0.00734 | 0.01358 | 0.00714 | 0.00945 | 0.20706 | 0.00000 | 0.14139 | 0.00653 | 0.01037 | 0.00535 | 0.31287 | 0.00676 |
| **Cfap46** | 0.01340 | 0.00752 | 0.02045 | 0.01066 | 0.00926 | 0.00617 | 0.00716 | 0.24898 | 0.01210 | 0.02877 | 0.01134 | 0.00664 | 0.01672 | 0.06770 | 0.01924 | 0.00755 |
| **Cfap53** | 0.01292 | 0.00784 | 0.01164 | 1.05425 | 0.01871 | 0.00665 | 0.00751 | 0.01262 | 0.00000 | 0.04038 | 0.02707 | 0.00764 | 0.01274 | 0.00506 | 0.37878 | 0.00720 |
| **Cfap57** | 0.17466 | 0.00690 | 2.87995 | 0.23136 | 0.00274 | 0.00724 | 0.00657 | 0.14262 | 0.69524 | 0.00000 | 0.51110 | 0.00907 | 0.00000 | 0.45838 | 0.39374 | 0.00584 |
| **Cfap61** | 0.00682 | 0.33866 | 0.01421 | 0.21829 | 0.00825 | 0.00713 | 0.02700 | 0.04561 | 0.00804 | 0.31156 | 0.02370 | 0.00575 | 0.20645 | 0.15598 | 0.35735 | 0.00690 |
| **Cfap70** | 0.01142 | 0.00763 | 0.00534 | 0.00000 | 0.00936 | 0.00637 | 0.00710 | 0.00838 | 1.04514 | 0.50537 | 0.00000 | 0.02344 | 0.01259 | 1.72529 | 1.29272 | 0.00772 |
| **Cfap74** | 1.33576 | 0.26280 | 1.98451 | 6.35881 | 0.02152 | 0.27469 | 1.04859 | 2.86181 | 7.80411 | 4.78501 | 3.68042 | 2.33759 | 1.19322 | 1.27050 | 2.16495 | 0.21048 |
| **Cfap77** | 0.08199 | 0.89885 | 0.05645 | 0.00903 | 0.00752 | 0.00574 | 0.19996 | 0.01201 | 0.01262 | 0.02147 | 0.01243 | 0.00801 | 0.41560 | 0.00671 | 0.06121 | 0.00653 |
| **Cfap97** | 1.76701 | 0.00357 | 1.34721 | 8.51224 | 0.00188 | 0.00244 | 1.79696 | 0.40887 | 3.23667 | 1.69782 | 1.11931 | 2.01884 | 2.39968 | 0.71779 | 4.76085 | 1.85458 |
| **Cfb** | 0.82049 | 0.00660 | 0.31454 | 0.41407 | 0.00579 | 0.00490 | 0.00665 | 1.31213 | 0.24449 | 1.63760 | 0.20377 | 0.02845 | 0.04828 | 0.00825 | 0.22062 | 0.01351 |
| **Cfc1** | 0.01403 | 0.00798 | 0.01004 | 0.01038 | 0.00920 | 0.37968 | 0.00743 | 0.01275 | 0.01243 | 0.01284 | 0.01223 | 0.00807 | 0.00841 | 0.01243 | 0.01052 | 0.00720 |
| **Cfd** | 0.01337 | 0.00765 | 0.01230 | 0.17062 | 0.00921 | 0.00641 | 0.00727 | 0.01058 | 0.49551 | 0.63057 | 0.01000 | 0.00454 | 0.01167 | 0.00681 | 0.04698 | 0.00740 |
| **Cfdp1** | 24.34300 | 14.68915 | 20.63719 | 10.93216 | 15.85179 | 6.82020 | 11.86225 | 24.19528 | 20.53845 | 32.71622 | 21.00828 | 16.47039 | 19.74097 | 25.78138 | 22.81769 | 9.46668 |
| **Cfh** | 0.01346 | 0.00829 | 0.01194 | 0.02040 | 0.00925 | 0.00688 | 0.00777 | 0.01889 | 0.04357 | 0.07855 | 0.01224 | 0.00916 | 0.01280 | 0.01198 | 0.01130 | 0.00742 |
| **Cfi** | 0.01288 | 0.00770 | 0.01204 | 1.02667 | 0.00885 | 0.00657 | 0.00733 | 0.01078 | 0.00000 | 1.45351 | 0.01074 | 0.00379 | 0.01142 | 0.00366 | 0.28033 | 0.00733 |
| **Cfl1** | 38.74078 | 18.30677 | 29.34116 | 14.94786 | 24.98006 | 17.82518 | 19.64535 | 32.91853 | 19.47783 | 34.51141 | 26.24258 | 3.94269 | 34.66928 | 7.16159 | 4.44718 | 19.98775 |
| **Cfl2** | 8.04688 | 1.10036 | 7.62254 | 11.35741 | 0.37664 | 6.93593 | 6.84905 | 8.47154 | 11.09224 | 15.72357 | 11.62619 | 4.49173 | 5.29550 | 11.04004 | 12.92327 | 0.96867 |
| **Cflar** | 4.44369 | 0.35495 | 4.47189 | 24.18466 | 3.26488 | 0.57630 | 0.67165 | 0.00000 | 28.84864 | 2.34823 | 21.44060 | 2.97906 | 2.46364 | 12.52879 | 9.06579 | 1.37102 |
| **Cfp** | 0.93886 | 0.00602 | 0.00216 | 0.18514 | 0.00591 | 0.00720 | 0.00613 | 0.34981 | 0.50414 | 0.00000 | 0.79686 | 0.36741 | 0.00019 | 0.00764 | 0.24566 | 0.00603 |
| **Cftr** | 0.01358 | 0.00782 | 0.01098 | 1.21186 | 0.00939 | 0.00639 | 0.00737 | 0.01136 | 0.00000 | 0.01180 | 0.00964 | 0.00551 | 0.01325 | 0.00579 | 1.42014 | 0.00717 |
| **Cga** | 0.06623 | 0.00784 | 0.00864 | 0.78389 | 0.00800 | 0.00669 | 0.00726 | 0.00862 | 0.00000 | 0.12812 | 0.18944 | 0.01652 | 0.01025 | 0.00809 | 0.00000 | 0.00714 |
| **Cggbp1** | 8.10558 | 4.06617 | 7.48716 | 21.09936 | 4.04316 | 2.66775 | 5.96449 | 8.96859 | 34.43994 | 7.43282 | 7.49200 | 5.02065 | 3.71450 | 21.19763 | 3.81948 | 5.38289 |
| **Cgn** | 1.03548 | 0.00390 | 0.78623 | 4.21985 | 1.92073 | 6.37916 | 1.16507 | 7.45422 | 3.89919 | 0.46797 | 4.76945 | 0.65614 | 2.34091 | 2.81104 | 1.23874 | 1.32667 |
| **Cgnl1** | 1.00001 | 0.00579 | 0.65540 | 1.86273 | 0.07457 | 10.40752 | 3.01160 | 2.33666 | 1.17789 | 0.51867 | 0.00000 | 0.02679 | 3.20349 | 0.02557 | 0.12150 | 5.01241 |
| **Cgref1** | 0.36572 | 1.03436 | 3.31950 | 0.00864 | 0.00145 | 0.00670 | 0.00601 | 0.46401 | 0.40107 | 1.44703 | 0.83861 | 0.06734 | 0.64176 | 2.23841 | 1.00232 | 0.00478 |
| **Cgrrf1** | 0.44778 | 0.01819 | 0.73239 | 0.07044 | 0.00578 | 1.02549 | 0.00711 | 0.01185 | 0.00840 | 0.01000 | 0.00959 | 0.02983 | 0.01481 | 0.01376 | 0.00000 | 0.01331 |
| **Chac1** | 0.01686 | 0.00756 | 0.01803 | 0.54527 | 0.00859 | 0.00697 | 0.00718 | 0.01004 | 0.22573 | 1.21301 | 0.01257 | 0.00427 | 0.01039 | 0.00942 | 0.06868 | 0.01399 |
| **Chac2** | 0.43151 | 0.00818 | 0.18728 | 0.00992 | 0.02029 | 0.00594 | 0.00728 | 0.01064 | 0.01225 | 0.10861 | 0.01081 | 0.02541 | 0.25863 | 0.70919 | 0.00000 | 0.00767 |
| **Chadl** | 0.02600 | 0.00782 | 0.00982 | 0.31365 | 0.00840 | 0.00658 | 0.00715 | 0.00420 | 0.26776 | 0.57847 | 0.24688 | 0.00443 | 0.04128 | 0.00831 | 0.00000 | 0.00732 |
| **Chaf1a** | 7.60015 | 5.24704 | 5.56959 | 9.27825 | 5.55145 | 10.87203 | 2.57385 | 8.16348 | 8.50581 | 9.72537 | 2.74815 | 1.07828 | 6.38054 | 1.99710 | 10.49936 | 6.88568 |
| **Chaf1b** | 5.72707 | 0.31939 | 4.67623 | 4.62563 | 1.65645 | 5.69648 | 5.84001 | 9.70993 | 5.61337 | 11.42566 | 9.73811 | 1.11701 | 4.47458 | 1.22790 | 1.49231 | 2.83076 |
| **Champ1** | 3.27056 | 0.04435 | 3.95035 | 7.09393 | 0.08458 | 0.27156 | 0.25168 | 11.01433 | 10.61936 | 14.73003 | 7.99977 | 8.18892 | 5.05422 | 13.31560 | 14.22894 | 1.23183 |
| **Chchd1** | 8.84263 | 1.11311 | 2.99523 | 2.82461 | 7.74109 | 1.32799 | 2.15107 | 3.11473 | 3.63154 | 4.61450 | 8.66636 | 0.74466 | 5.63199 | 1.44819 | 2.40436 | 5.39674 |
| **Chchd10** | 1.92651 | 0.09430 | 0.03136 | 1.70635 | 0.05475 | 0.57667 | 0.16237 | 0.07521 | 6.27982 | 3.94471 | 5.75980 | 0.11886 | 0.85951 | 0.00000 | 0.00000 | 0.39973 |
| **Chchd2** | 22.47995 | 0.91416 | 9.03793 | 21.11546 | 20.30303 | 3.76473 | 1.59194 | 3.95607 | 38.92982 | 41.52628 | 32.37118 | 3.02783 | 29.07430 | 3.69584 | 0.56624 | 14.20186 |
| **Chchd3** | 9.93267 | 1.53495 | 6.49415 | 0.98654 | 6.21286 | 4.00191 | 5.43360 | 6.05973 | 2.16152 | 5.49299 | 0.49698 | 0.34990 | 7.71843 | 0.30349 | 0.83887 | 9.73724 |
| **Chchd4** | 15.63456 | 8.37323 | 14.17327 | 0.94180 | 16.85249 | 6.55320 | 7.52017 | 8.23904 | 1.54076 | 4.51410 | 1.75453 | 7.45013 | 16.12554 | 1.66146 | 1.93503 | 6.61231 |
| **Chchd5** | 12.05364 | 5.80494 | 13.94172 | 0.37711 | 3.32196 | 2.39195 | 7.63239 | 5.31569 | 3.22441 | 7.26170 | 7.74439 | 1.27167 | 9.45101 | 0.70257 | 2.27351 | 3.83458 |
| **Chchd6** | 4.77712 | 0.01188 | 2.14212 | 0.64962 | 0.88772 | 0.00643 | 2.02161 | 2.26329 | 0.34692 | 0.87213 | 0.52578 | 0.10552 | 4.07133 | 0.05797 | 0.00000 | 0.00698 |
| **Chchd7** | 1.47683 | 0.00625 | 0.37547 | 0.04939 | 0.03707 | 0.45620 | 0.22873 | 0.00579 | 1.08061 | 0.56034 | 0.00000 | 0.05924 | 2.85929 | 0.01698 | 0.06850 | 3.87603 |
| **Chd1** | 3.16417 | 8.17072 | 4.94854 | 11.08981 | 8.02377 | 4.32624 | 5.26889 | 8.67852 | 13.41893 | 8.97099 | 2.00360 | 7.79384 | 0.60441 | 16.19579 | 18.79179 | 2.27691 |
| **Chd1l** | 0.00000 | 0.00454 | 2.43751 | 0.00000 | 3.74425 | 0.27528 | 1.65798 | 1.85360 | 0.31088 | 2.33190 | 1.28248 | 0.10799 | 1.31033 | 0.05710 | 1.08435 | 0.01237 |
| **Chd2** | 0.06204 | 3.01644 | 0.05641 | 0.20646 | 0.01262 | 0.00562 | 1.34515 | 1.15905 | 0.27324 | 0.46546 | 0.01199 | 0.13998 | 1.88565 | 0.01323 | 2.57770 | 1.10727 |
| **Chd3** | 0.65598 | 0.00607 | 0.86918 | 0.69441 | 0.67283 | 0.10897 | 0.01647 | 0.01715 | 1.17085 | 1.02873 | 0.04952 | 0.04675 | 0.01260 | 0.00342 | 0.20173 | 2.11885 |
| **Chd3os** | 0.92227 | 0.00600 | 0.00000 | 0.66037 | 0.00592 | 0.04835 | 0.00570 | 0.78797 | 0.56682 | 1.90334 | 0.95458 | 0.03982 | 0.77717 | 0.00293 | 0.54164 | 0.00571 |
| **Chd4** | 15.00128 | 0.26597 | 1.84412 | 2.16272 | 7.71417 | 3.22366 | 7.60230 | 17.60518 | 5.46000 | 21.87101 | 5.01587 | 5.64646 | 13.07335 | 1.72672 | 0.00000 | 11.17090 |
| **Chd5** | 0.01282 | 0.03784 | 0.00690 | 0.00488 | 0.00894 | 0.00699 | 0.00689 | 0.00000 | 0.01103 | 0.68477 | 0.90881 | 0.00126 | 0.00514 | 0.01072 | 0.00000 | 0.00725 |
| **Chd6** | 2.24774 | 0.23378 | 1.37409 | 2.21111 | 3.08680 | 3.96612 | 1.93943 | 3.91240 | 3.12033 | 3.22135 | 1.74321 | 2.78911 | 0.52967 | 0.00614 | 1.34631 | 2.14521 |

| | | | | | | | | | | | | | | | | |
|---|---|---|---|---|---|---|---|---|---|---|---|---|---|---|---|---|
| **Chd7** | 2.43335 | 0.01325 | 0.18855 | 6.23302 | 0.09248 | 0.01168 | 4.56765 | 0.44553 | 3.50689 | 14.34718 | 2.70773 | 0.46017 | 0.45481 | 4.10257 | 9.27226 | 1.35406 |
| **Chd8** | 1.05338 | 0.58235 | 1.43438 | 9.32582 | 0.12968 | 1.14829 | 1.37055 | 4.43574 | 12.58411 | 4.95017 | 2.22814 | 0.24179 | 1.96332 | 6.92897 | 5.86657 | 0.80119 |
| **Chd9** | 1.59454 | 0.00526 | 0.05301 | 3.34029 | 0.01461 | 0.00428 | 1.20968 | 0.00378 | 0.81437 | 0.00000 | 0.00000 | 0.09278 | 0.23628 | 4.93391 | 1.78521 | 0.01244 |
| **Chdh** | 0.00000 | 0.00744 | 0.58223 | 0.37812 | 0.01619 | 0.00563 | 0.00723 | 0.39475 | 0.17405 | 0.10178 | 0.00728 | 0.00523 | 0.00000 | 0.20284 | 0.00000 | 1.65564 |
| **Chek1** | 7.09093 | 5.79590 | 4.04142 | 6.48808 | 9.86708 | 1.30021 | 3.26975 | 4.81981 | 4.55571 | 2.30369 | 1.39801 | 3.20828 | 1.95338 | 8.33016 | 8.35424 | 0.19866 |
| **Chek2** | 4.19200 | 1.71462 | 0.32109 | 9.66217 | 4.59389 | 1.65991 | 5.58687 | 12.73654 | 10.88883 | 10.21257 | 11.71123 | 7.58026 | 1.27465 | 18.55379 | 18.86839 | 0.02463 |
| **Cherp** | 11.47321 | 3.12265 | 5.51589 | 12.95362 | 4.85088 | 1.13639 | 4.66960 | 19.20427 | 10.37878 | 23.23528 | 9.34656 | 8.09223 | 11.38392 | 6.27560 | 9.48074 | 7.16050 |
| **Chfr** | 2.43939 | 3.94609 | 4.93892 | 2.16399 | 4.82868 | 1.83083 | 5.91336 | 3.70182 | 3.86824 | 3.70427 | 1.01500 | 4.34184 | 1.62937 | 0.79851 | 4.20001 | 1.03882 |
| **Chga** | 2.07211 | 0.23510 | 0.00803 | 0.29586 | 4.53173 | 0.01634 | 1.69335 | 0.49897 | 0.01325 | 0.03546 | 0.00653 | 0.06028 | 3.91591 | 0.00993 | 0.00000 | 0.01041 |
| **Chgb** | 0.01610 | 0.00785 | 0.07083 | 0.29337 | 0.00910 | 0.00579 | 0.00734 | 0.01251 | 0.00687 | 0.04768 | 0.01511 | 0.00817 | 0.01112 | 0.01476 | 0.00420 | 0.00713 |
| **Chic1** | 0.25898 | 0.01732 | 0.00000 | 3.11764 | 0.00763 | 0.01964 | 0.12011 | 0.00000 | 1.57452 | 4.13227 | 3.64616 | 0.90482 | 1.48966 | 13.33127 | 5.53110 | 0.22284 |
| **Chic2** | 3.76303 | 0.00000 | 0.12197 | 3.61318 | 3.09452 | 0.00125 | 0.49820 | 0.00000 | 0.00000 | 5.85454 | 4.30294 | 1.93367 | 1.43009 | 3.15344 | 0.12863 | 0.00000 |
| **Chid1** | 5.68290 | 4.17247 | 7.50703 | 0.26136 | 0.45014 | 8.22795 | 5.66797 | 2.93410 | 0.04023 | 0.46719 | 1.74677 | 0.15250 | 6.47218 | 2.53744 | 0.24768 | 0.15688 |
| **Chil1** | 0.00757 | 0.00511 | 1.17744 | 0.00877 | 4.67832 | 0.00552 | 0.00731 | 0.00874 | 0.01019 | 0.01171 | 0.01093 | 0.00280 | 0.00728 | 0.01606 | 0.00000 | 0.00675 |
| **Chil5** | 0.01348 | 0.00808 | 0.01035 | 1.44156 | 0.00937 | 0.00642 | 0.00734 | 0.01102 | 0.32739 | 0.01362 | 0.00776 | 0.00523 | 0.01270 | 0.21678 | 0.44639 | 0.00701 |
| **Chit1** | 0.00000 | 1.98419 | 0.91913 | 1.61734 | 0.02473 | 2.17913 | 1.12623 | 0.77513 | 2.27102 | 0.00406 | 0.00000 | 0.00000 | 0.00000 | 0.00165 | 0.25475 | 0.04709 |
| **Chka** | 0.00000 | 0.23397 | 9.29511 | 14.97095 | 4.55734 | 5.19103 | 8.41087 | 27.99796 | 19.67518 | 44.94642 | 21.10205 | 21.48357 | 9.14961 | 6.40681 | 19.51492 | 8.22437 |
| **Chkb** | 0.31067 | 0.07072 | 0.12642 | 0.22314 | 0.00720 | 0.03695 | 0.17494 | 0.04765 | 0.49554 | 0.09666 | 0.55056 | 0.00423 | 0.08114 | 0.01095 | 0.00438 | 0.00667 |
| **Chm** | 0.19112 | 0.01915 | 3.03934 | 0.50168 | 0.00732 | 0.00516 | 0.00637 | 1.32322 | 0.00002 | 0.03480 | 0.72918 | 0.67245 | 0.06511 | 4.17998 | 0.24041 | 0.30711 |
| **Chml** | 0.00603 | 0.00564 | 0.00198 | 1.11950 | 3.56765 | 0.00769 | 0.00722 | 0.00481 | 1.11233 | 0.01847 | 0.05756 | 0.00136 | 0.00907 | 0.01740 | 0.43505 | 0.00675 |
| **Chmp1a** | 8.00235 | 2.23199 | 6.47343 | 2.68852 | 9.42457 | 3.62837 | 0.15337 | 7.28576 | 1.94272 | 11.54380 | 2.84729 | 0.45391 | 9.10404 | 0.02071 | 3.79446 | 6.74748 |
| **Chmp1b** | 0.39184 | 0.06039 | 2.09998 | 4.29745 | 1.82430 | 0.10754 | 1.31399 | 7.73138 | 4.21969 | 1.58252 | 3.04511 | 0.48868 | 2.04625 | 3.30800 | 3.72508 | 0.56744 |
| **Chmp2a** | 12.83939 | 9.67516 | 17.27627 | 7.05803 | 4.58915 | 0.04342 | 12.31122 | 13.62786 | 9.49890 | 13.58780 | 8.31421 | 2.62658 | 8.47456 | 1.19980 | 10.02023 | 6.75000 |
| **Chmp2b** | 2.56363 | 1.65525 | 1.19816 | 1.22124 | 10.48596 | 0.01586 | 2.05572 | 0.32903 | 0.72015 | 1.26468 | 0.00000 | 0.00516 | 4.95100 | 0.00295 | 0.00000 | 3.54814 |
| **Chmp3** | 3.59080 | 6.91893 | 7.34574 | 1.50442 | 4.76558 | 7.73827 | 1.59753 | 7.06099 | 0.21076 | 5.13222 | 2.22981 | 7.08863 | 7.68758 | 6.57748 | 3.57442 | 4.43284 |
| **Chmp4b** | 3.31219 | 0.21646 | 1.84994 | 0.55720 | 1.19471 | 0.87734 | 0.11526 | 0.00000 | 4.36245 | 1.20925 | 6.30179 | 0.20882 | 0.11604 | 2.75218 | 0.25243 | 0.12721 |
| **Chmp4c** | 7.87570 | 0.50827 | 2.00240 | 0.00886 | 0.03999 | 0.00000 | 1.99013 | 5.62911 | 0.18519 | 1.57895 | 4.72699 | 0.05715 | 0.00000 | 0.58914 | 0.00000 | 0.08405 |
| **Chmp5** | 6.15172 | 6.02562 | 3.87156 | 5.93538 | 10.84472 | 2.26172 | 2.37529 | 3.65017 | 4.64942 | 2.33686 | 4.72968 | 0.05917 | 0.18238 | 9.72265 | 6.72113 | 4.07466 |
| **Chmp6** | 8.20621 | 0.00276 | 3.93397 | 0.09136 | 5.49896 | 4.55833 | 3.91758 | 2.85182 | 1.03185 | 1.62211 | 0.28797 | 0.08938 | 6.63755 | 0.17577 | 0.00000 | 5.12027 |
| **Chmp7** | 9.10654 | 2.28686 | 4.47180 | 4.93211 | 9.67268 | 4.28176 | 5.55874 | 11.51699 | 5.07075 | 11.43018 | 4.29269 | 2.10808 | 5.57663 | 0.09249 | 2.25373 | 4.17966 |
| **Chn1** | 1.88606 | 0.72239 | 0.13294 | 2.72057 | 0.79108 | 0.00916 | 0.00548 | 0.00000 | 2.26013 | 0.02649 | 0.32431 | 0.12974 | 0.08071 | 0.02250 | 1.55029 | 1.75134 |
| **Chn2** | 0.01561 | 0.00822 | 0.00778 | 0.42733 | 0.01192 | 0.00737 | 0.00739 | 0.01443 | 0.00489 | 0.33395 | 0.13503 | 0.00278 | 0.24872 | 0.91796 | 0.00000 | 0.00783 |
| **Chordc1** | 5.03912 | 8.00469 | 9.06894 | 1.26755 | 4.46752 | 3.00635 | 3.55397 | 5.33683 | 2.91782 | 4.36745 | 3.36908 | 8.81059 | 4.54658 | 11.89874 | 6.62626 | 0.92183 |
| **Chp1** | 1.73683 | 4.27201 | 12.51233 | 9.87020 | 8.65580 | 14.12560 | 6.25223 | 11.93085 | 11.35922 | 16.37140 | 13.14960 | 5.38327 | 9.97657 | 20.61259 | 16.54279 | 0.45032 |
| **Chpf** | 6.79710 | 0.00000 | 0.00000 | 0.28145 | 3.53041 | 0.06423 | 1.40612 | 3.44676 | 0.05278 | 0.00000 | 0.00000 | 0.02127 | 2.02251 | 0.01629 | 0.00000 | 4.02272 |
| **Chpf2** | 2.02173 | 0.39515 | 0.68642 | 0.28708 | 2.13217 | 0.00396 | 1.94552 | 2.29468 | 1.77195 | 0.09027 | 0.00067 | 0.02226 | 0.00159 | 0.00480 | 0.00000 | 0.00470 |
| **Chpt1** | 4.16843 | 4.70851 | 2.71178 | 1.67503 | 0.04262 | 1.61923 | 3.51096 | 1.96460 | 5.12434 | 8.42627 | 6.61988 | 5.38007 | 1.58014 | 15.98437 | 6.78476 | 0.00000 |
| **Chrac1** | 11.56685 | 4.91216 | 4.67893 | 2.70085 | 6.92482 | 0.02635 | 3.15302 | 6.85296 | 6.30913 | 3.73900 | 3.22590 | 0.23434 | 3.72927 | 0.31685 | 1.73331 | 2.11139 |
| **Chrna1** | 1.35573 | 0.05754 | 1.22018 | 7.76896 | 0.73351 | 0.18150 | 0.15329 | 0.00000 | 11.60309 | 0.00000 | 9.75886 | 0.74239 | 0.39966 | 3.66831 | 0.38677 | 0.28572 |
| **Chrna10** | 0.01108 | 0.00708 | 0.00183 | 0.00411 | 0.00909 | 0.00765 | 0.87799 | 0.00000 | 0.32353 | 0.77629 | 0.72342 | 0.03029 | 0.00647 | 0.00507 | 0.70333 | 0.00720 |
| **Chrna5** | 0.01206 | 0.00763 | 0.61402 | 0.79509 | 0.00819 | 0.00894 | 0.00709 | 0.00968 | 0.00000 | 0.51678 | 0.01113 | 0.03279 | 0.00993 | 0.00931 | 0.00000 | 0.01448 |
| **Chrna9** | 0.09990 | 0.00753 | 0.00000 | 0.18888 | 0.00759 | 0.00658 | 0.00686 | 0.00000 | 1.14327 | 0.69963 | 0.64237 | 0.00000 | 0.01139 | 2.32464 | 0.42662 | 0.00669 |
| **Chrnb1** | 0.01962 | 0.03874 | 1.25422 | 0.26464 | 0.00710 | 0.00573 | 0.00680 | 0.00000 | 0.45691 | 0.00000 | 0.68139 | 0.05549 | 0.01169 | 1.16489 | 0.53308 | 0.00661 |
| **Chrnb4** | 0.16519 | 0.03848 | 0.56881 | 0.03171 | 0.00694 | 0.00849 | 0.00722 | 0.01188 | 0.12886 | 0.05874 | 0.20970 | 0.03655 | 0.03436 | 0.05991 | 0.05548 | 0.00682 |
| **Chrnd** | 0.01393 | 0.00800 | 0.00850 | 0.48103 | 0.00904 | 0.00677 | 0.01739 | 0.00000 | 0.00274 | 0.18619 | 0.47980 | 0.00474 | 0.00830 | 0.00502 | 0.28022 | 0.00758 |
| **Chrne** | 0.01136 | 0.00834 | 0.97412 | 1.72811 | 0.00824 | 0.00572 | 0.01869 | 0.01016 | 2.24789 | 0.66288 | 0.06710 | 0.01205 | 0.05512 | 0.00705 | 0.00000 | 0.00669 |
| **Chst10** | 0.01321 | 0.00792 | 0.01226 | 0.47457 | 0.00902 | 0.00652 | 0.00723 | 0.01014 | 0.00286 | 1.06172 | 0.01172 | 0.00492 | 0.01143 | 0.01024 | 0.00416 | 0.00757 |
| **Chst11** | 0.01352 | 0.00788 | 0.01153 | 0.00484 | 0.00932 | 0.00657 | 0.00754 | 0.01337 | 1.69771 | 0.00990 | 0.00000 | 0.00492 | 0.01323 | 0.01931 | 0.02221 | 0.00733 |
| **Chst12** | 3.61174 | 0.02431 | 1.29046 | 2.50733 | 0.00669 | 2.16300 | 0.57180 | 1.47120 | 1.06889 | 2.20543 | 0.74129 | 0.10841 | 2.39909 | 0.02553 | 0.00000 | 3.13663 |
| **Chst14** | 0.07640 | 0.00851 | 0.00818 | 0.03438 | 0.01423 | 0.00716 | 0.00751 | 0.01290 | 0.00994 | 0.04280 | 0.01160 | 0.05387 | 0.01312 | 0.01170 | 0.01022 | 0.00752 |
| **Chst2** | 0.00811 | 0.00756 | 0.00611 | 0.19398 | 0.47640 | 0.01397 | 0.00748 | 0.02407 | 0.02588 | 0.04494 | 0.01113 | 0.01909 | 0.01704 | 0.01015 | 0.02697 | 0.00723 |
| **Chst3** | 0.03685 | 0.00631 | 1.20465 | 0.00994 | 0.01375 | 0.01856 | 1.91251 | 0.00000 | 0.31944 | 0.65306 | 1.68878 | 0.02546 | 0.34479 | 0.14614 | 0.39522 | 1.15050 |
| **Chst5** | 0.01321 | 0.00817 | 0.01113 | 0.39224 | 0.00917 | 0.00630 | 0.00756 | 0.01249 | 3.76019 | 0.04081 | 0.00000 | 0.01107 | 0.01259 | 0.00945 | 0.01032 | 0.00694 |
| **Chst7** | 0.01345 | 0.00810 | 0.01207 | 1.51692 | 0.00932 | 0.00630 | 0.00729 | 0.01137 | 1.20664 | 0.32185 | 0.00088 | 0.00484 | 0.01236 | 0.00925 | 0.00301 | 0.00715 |
| **Chst8** | 0.01362 | 0.00821 | 0.01197 | 0.36882 | 0.00927 | 0.00657 | 0.00753 | 0.01387 | 0.88089 | 0.01111 | 0.00823 | 0.00691 | 0.01289 | 0.01193 | 0.00362 | 0.00730 |
| **Chsy1** | 0.80397 | 0.00660 | 0.69162 | 7.01171 | 0.00506 | 0.00402 | 0.00649 | 0.00519 | 1.23278 | 1.87494 | 0.03568 | 0.03794 | 0.00822 | 0.00000 | 1.03882 | 0.33602 |
| **Chtf18** | 1.28318 | 3.14273 | 4.10770 | 3.81912 | 0.24627 | 1.38003 | 2.32043 | 5.17577 | 5.43363 | 5.29012 | 4.20659 | 0.20331 | 5.76192 | 0.01302 | 0.00000 | 3.95137 |
| **Chtf8** | 4.47807 | 1.71698 | 1.46042 | 1.42098 | 1.45778 | 0.79710 | 0.23837 | 2.79089 | 2.15077 | 4.73838 | 0.35594 | 0.63523 | 3.34300 | 0.38776 | 1.44875 | 1.32664 |
| **Chtop** | 11.22726 | 8.60583 | 10.51565 | 11.26657 | 7.79266 | 4.72033 | 2.47689 | 8.14849 | 15.68314 | 16.07351 | 4.60498 | 7.14677 | 9.99041 | 5.59820 | 10.80520 | 5.72661 |
| **Chuk** | 2.91238 | 1.64324 | 0.00000 | 4.02465 | 0.01337 | 1.72314 | 0.01351 | 0.78830 | 2.11330 | 0.41052 | 1.45854 | 1.52786 | 1.65124 | 8.07757 | 6.81625 | 0.36797 |
| **Churc1** | 3.76178 | 0.68238 | 1.25426 | 8.56777 | 1.25644 | 0.68206 | 1.13801 | 1.29237 | 13.25872 | 2.14907 | 15.47537 | 1.63474 | 2.21449 | 8.34711 | 3.41120 | 1.20923 |
| **Ciao1** | 3.48983 | 9.95622 | 9.13603 | 4.15358 | 6.07338 | 1.72818 | 3.29674 | 9.34562 | 4.95848 | 4.77707 | 7.07917 | 5.63431 | 7.60093 | 7.29603 | 3.44374 | 1.05565 |

| | | | | | | | | | | | | | | | | |
|---|---|---|---|---|---|---|---|---|---|---|---|---|---|---|---|---|
| **Ciapin1** | 8.15528 | 2.15763 | 11.09682 | 3.73044 | 0.44730 | 0.39588 | 0.25623 | 7.46701 | 0.58558 | 3.88917 | 3.17652 | 3.34080 | 11.21739 | 1.36254 | 0.66197 | 8.79683 |
| **Ciart** | 2.41008 | 1.23710 | 0.71292 | 0.77482 | 0.00216 | 0.00287 | 2.32108 | 1.55384 | 1.92056 | 2.87313 | 0.00000 | 0.00166 | 0.46990 | 0.00271 | 2.90344 | 0.00325 |
| **Cib1** | 7.10434 | 1.35006 | 5.99460 | 0.24746 | 12.07041 | 0.04214 | 2.11151 | 10.10077 | 0.18205 | 11.15004 | 1.24588 | 0.61101 | 6.38834 | 0.00000 | 0.00000 | 0.44276 |
| **Cib2** | 0.01130 | 0.06983 | 0.03596 | 0.00372 | 0.00884 | 1.16464 | 0.00716 | 0.01112 | 0.13200 | 0.06021 | 0.00966 | 0.00714 | 0.02812 | 0.01145 | 0.00766 | 0.04345 |
| **Cic** | 4.03500 | 0.28897 | 1.49119 | 7.52490 | 0.36646 | 0.00904 | 0.45436 | 1.98823 | 5.32062 | 3.86126 | 2.19369 | 0.14034 | 4.73394 | 2.06285 | 2.33722 | 5.47081 |
| **Cidea** | 0.00000 | 1.72255 | 2.33812 | 0.18626 | 0.00399 | 3.29390 | 0.98477 | 0.37165 | 0.11197 | 0.30100 | 0.83775 | 0.02310 | 0.00000 | 1.15422 | 0.46011 | 0.02025 |
| **Cideb** | 0.00839 | 0.00757 | 0.27312 | 0.00963 | 0.08499 | 0.00825 | 0.00729 | 0.00876 | 0.05590 | 0.09751 | 0.08149 | 0.00608 | 0.07500 | 0.50550 | 0.00000 | 0.00713 |
| **Cidec** | 0.01789 | 0.00771 | 0.00801 | 0.01768 | 0.00880 | 0.00716 | 0.00732 | 0.25763 | 0.01076 | 0.15695 | 0.02130 | 0.00733 | 0.00709 | 0.00640 | 0.12244 | 0.00734 |
| **Ciita** | 0.01096 | 0.00753 | 0.00602 | 1.11149 | 0.00944 | 0.00678 | 0.00723 | 1.01154 | 0.38969 | 0.06741 | 0.01002 | 0.00312 | 0.00602 | 0.00230 | 1.08173 | 0.00718 |
| **Cilp** | 0.00696 | 0.00769 | 0.99917 | 0.00000 | 0.01996 | 0.01923 | 0.00722 | 0.08747 | 0.42974 | 3.10341 | 0.47931 | 1.21220 | 0.40347 | 0.00552 | 1.20956 | 0.01470 |
| **Cilp2** | 0.19211 | 0.00766 | 0.00511 | 0.00717 | 0.00738 | 0.00724 | 0.00714 | 0.01608 | 0.01176 | 0.86540 | 0.03832 | 0.12602 | 0.01630 | 0.01199 | 0.00446 | 0.00685 |
| **Cinp** | 3.72430 | 0.31361 | 4.91624 | 0.81039 | 0.12785 | 0.78147 | 1.53297 | 4.12636 | 3.59685 | 3.80606 | 0.97950 | 1.63193 | 3.25219 | 1.02686 | 3.04644 | 2.11266 |
| **Cipc** | 1.24511 | 0.00571 | 0.65430 | 0.56878 | 0.02613 | 0.00471 | 0.00619 | 0.98843 | 0.50521 | 0.56953 | 0.00000 | 0.04406 | 0.24690 | 0.01143 | 0.63457 | 1.76872 |
| **Cir1** | 6.25669 | 7.35061 | 3.05988 | 6.11716 | 5.15269 | 1.87415 | 3.60006 | 6.41468 | 12.24973 | 5.13921 | 4.58379 | 1.39913 | 2.79878 | 8.77765 | 4.79026 | 0.48839 |
| **Cirbp** | 2.19442 | 0.01540 | 1.73284 | 0.72204 | 0.00000 | 0.03830 | 0.00374 | 5.21037 | 1.10912 | 2.31185 | 1.38449 | 0.16910 | 2.38567 | 0.00233 | 0.24396 | 2.89471 |
| **Cirh1a** | 15.90808 | 11.69034 | 10.45515 | 7.48107 | 9.88505 | 5.65007 | 5.35873 | 21.56047 | 16.10462 | 26.21582 | 16.27102 | 18.80105 | 7.02049 | 12.81164 | 20.82486 | 5.90318 |
| **Cisd1** | 8.29985 | 0.15495 | 4.38381 | 3.38962 | 8.90622 | 3.40156 | 1.77729 | 1.15053 | 0.00000 | 8.28573 | 6.46657 | 0.15575 | 10.18429 | 1.27846 | 1.22738 | 3.18523 |
| **Cisd2** | 1.72769 | 0.27681 | 0.08876 | 3.49059 | 4.16582 | 2.63231 | 0.53431 | 1.86483 | 1.61224 | 6.22587 | 1.95740 | 1.81577 | 1.73323 | 2.81139 | 1.60464 | 1.18220 |
| **Cisd3** | 0.32444 | 0.00739 | 1.21045 | 0.14872 | 0.00754 | 0.03556 | 0.00678 | 0.12408 | 0.00661 | 0.65798 | 0.19443 | 0.05464 | 0.25476 | 0.01241 | 0.00562 | 0.00663 |
| **Cish** | 0.41920 | 0.00471 | 0.38024 | 0.25861 | 1.30553 | 5.64922 | 0.00347 | 7.13458 | 0.00371 | 15.15217 | 5.43955 | 0.62956 | 2.02204 | 0.53573 | 3.15118 | 1.57754 |
| **Cit** | 0.51892 | 0.00664 | 0.22578 | 0.11440 | 0.02249 | 0.00824 | 0.00647 | 0.16926 | 0.00859 | 1.19091 | 0.62389 | 1.03568 | 0.01436 | 0.05573 | 0.94772 | 0.01396 |
| **Cited1** | 14.79199 | 5.84559 | 14.84098 | 3.38543 | 6.15013 | 0.98133 | 10.55531 | 36.86908 | 9.20176 | 47.27966 | 27.34264 | 12.62055 | 22.46079 | 1.28478 | 7.26232 | 2.73378 |
| **Cited2** | 1.41833 | 0.00714 | 0.18946 | 0.45504 | 0.01034 | 0.00746 | 0.00691 | 0.40673 | 2.05179 | 0.79233 | 0.00000 | 0.13009 | 0.00882 | 0.41354 | 0.00000 | 1.30895 |
| **Cited4** | 0.01920 | 0.00840 | 0.00580 | 1.03190 | 0.01049 | 0.95908 | 0.00738 | 0.01127 | 0.57519 | 0.18735 | 0.00743 | 0.00671 | 0.01335 | 0.04729 | 0.00000 | 0.00683 |
| **Ciz1** | 1.16169 | 1.68987 | 0.72376 | 2.47569 | 0.02159 | 0.02203 | 0.04913 | 0.82379 | 4.19006 | 1.10429 | 0.37698 | 0.12609 | 2.25389 | 0.00000 | 0.03585 | 3.85407 |
| **Ckap2** | 4.94037 | 3.02349 | 4.36549 | 7.54262 | 3.26831 | 3.17508 | 3.92597 | 6.05343 | 13.45497 | 5.89335 | 5.46967 | 3.38240 | 3.96723 | 1.26547 | 10.04402 | 2.05951 |
| **Ckap2l** | 3.90918 | 8.85163 | 3.58279 | 2.25696 | 5.42644 | 3.29697 | 2.32754 | 1.57020 | 4.00181 | 3.28879 | 4.64555 | 0.61880 | 1.84003 | 2.83728 | 5.14852 | 0.58719 |
| **Ckap4** | 2.00991 | 0.00259 | 0.42389 | 0.00000 | 6.54876 | 0.00350 | 0.57205 | 1.13429 | 0.00596 | 1.77316 | 0.01612 | 0.12606 | 0.71145 | 0.00660 | 0.22891 | 9.09644 |
| **Ckap5** | 4.29005 | 10.00011 | 5.11618 | 5.63193 | 3.31870 | 3.55874 | 2.29802 | 6.48278 | 11.70115 | 7.24249 | 2.68196 | 2.71437 | 1.85734 | 5.15206 | 12.88190 | 2.41131 |
| **Ckb** | 41.84355 | 6.81109 | 11.84601 | 7.57630 | 27.80023 | 12.00151 | 8.81979 | 28.91076 | 6.62231 | 33.46555 | 23.34044 | 12.53443 | 37.60882 | 0.00000 | 0.00000 | 16.77175 |
| **Cklf** | 0.07510 | 0.01749 | 0.27907 | 0.95712 | 0.00675 | 0.01265 | 0.00597 | 1.04476 | 3.16056 | 0.00000 | 0.88844 | 0.02107 | 1.58483 | 0.47783 | 0.00000 | 1.10505 |
| **Ckm** | 0.63632 | 0.00752 | 0.00000 | 0.01030 | 0.01771 | 0.33451 | 0.11440 | 0.29170 | 0.01208 | 0.05474 | 0.01076 | 0.00583 | 0.47792 | 0.01206 | 0.00636 | 0.00689 |
| **Ckmt1** | 16.44452 | 7.14395 | 21.13911 | 0.00000 | 7.36495 | 1.12807 | 10.66214 | 14.94947 | 0.00873 | 4.92023 | 12.49304 | 1.04287 | 19.90845 | 5.77347 | 0.00000 | 12.63969 |
| **Cks1b** | 15.83905 | 5.42702 | 15.37518 | 6.67419 | 8.58640 | 10.54457 | 8.14243 | 11.86930 | 7.02723 | 13.06430 | 8.80102 | 2.84120 | 10.30104 | 3.10048 | 6.83454 | 3.91455 |
| **Cks1brt** | 1.10072 | 0.36928 | 1.20033 | 0.84679 | 0.61139 | 0.87354 | 0.63324 | 0.56057 | 0.73497 | 0.68740 | 1.00795 | 0.11043 | 0.59784 | 0.16827 | 0.36030 | 0.26629 |
| **Cks2** | 18.28303 | 6.11436 | 15.95053 | 24.43607 | 18.14089 | 5.53856 | 2.73569 | 19.66073 | 34.12815 | 40.05129 | 39.92893 | 12.02893 | 13.79710 | 15.65604 | 21.80093 | 3.99250 |
| **Clasp1** | 2.45802 | 2.96625 | 0.99580 | 1.87072 | 2.31931 | 0.03253 | 0.00402 | 1.78116 | 0.86215 | 2.25066 | 0.05700 | 0.06862 | 0.52246 | 0.00849 | 0.35766 | 3.72015 |
| **Clasp2** | 0.00292 | 1.73601 | 0.35903 | 3.98666 | 0.07612 | 0.00619 | 0.10843 | 0.75531 | 2.56414 | 2.55724 | 1.28049 | 0.03153 | 0.21794 | 0.39221 | 0.77676 | 0.02754 |
| **Clasrp** | 2.87373 | 1.08986 | 0.75881 | 1.77262 | 0.00915 | 0.00349 | 0.01292 | 1.21310 | 3.24427 | 0.26692 | 2.95819 | 0.05147 | 1.35895 | 0.00000 | 0.00000 | 1.58913 |
| **Clca3a1** | 0.09447 | 0.05064 | 0.29233 | 0.52829 | 0.07943 | 0.10062 | 0.06935 | 0.06708 | 0.56933 | 0.02577 | 0.53671 | 0.03242 | 0.08113 | 0.52158 | 0.12116 | 0.10970 |
| **Clca3a2** | 0.02141 | 0.01829 | 0.04938 | 0.22974 | 0.01993 | 0.00656 | 0.01741 | 0.02302 | 0.20653 | 0.00585 | 0.29783 | 0.00567 | 0.01781 | 0.05805 | 0.13326 | 0.02909 |
| **Clca4c.ps** | 0.05053 | 0.00765 | 0.02044 | 0.12119 | 0.00854 | 0.01531 | 0.00729 | 0.03510 | 0.09213 | 0.02413 | 0.02521 | 0.00732 | 0.01079 | 0.05914 | 0.01517 | 0.00731 |
| **Clcc1** | 2.20771 | 0.79855 | 4.40265 | 2.63659 | 4.31072 | 4.03298 | 0.00434 | 3.50319 | 0.81431 | 3.41408 | 0.54247 | 0.14373 | 0.00000 | 0.03104 | 1.88292 | 2.05809 |
| **Clcn2** | 1.71583 | 0.00451 | 0.47442 | 1.90306 | 0.08675 | 0.01827 | 1.14302 | 4.16174 | 2.27266 | 0.21699 | 2.16988 | 0.07515 | 1.44957 | 0.04811 | 1.30014 | 0.85805 |
| **Clcn3** | 3.02882 | 1.37032 | 4.52569 | 1.19695 | 1.55021 | 0.03016 | 1.10319 | 2.92928 | 0.47686 | 1.66824 | 3.17422 | 3.57495 | 0.43739 | 2.37894 | 2.21065 | 0.45616 |
| **Clcn4** | 0.33020 | 0.00240 | 2.90024 | 0.01751 | 1.65264 | 0.00483 | 0.53789 | 0.65322 | 0.04356 | 5.74203 | 2.65418 | 11.25497 | 4.53972 | 2.63933 | 2.01573 | 0.01732 |
| **Clcn5** | 0.00982 | 0.00785 | 0.01728 | 1.72789 | 0.00974 | 0.00655 | 0.00707 | 0.05433 | 0.15530 | 1.24675 | 0.02260 | 0.00000 | 0.91386 | 0.04781 | 1.35361 | 0.00686 |
| **Clcn6** | 0.45688 | 1.43068 | 0.49337 | 3.01205 | 0.08797 | 0.48785 | 1.93356 | 1.13501 | 2.82188 | 1.63922 | 3.42511 | 0.48076 | 1.79670 | 0.21117 | 2.12096 | 0.01119 |
| **Clcn7** | 5.15696 | 0.70308 | 3.17493 | 3.88580 | 0.56833 | 0.66577 | 4.42711 | 6.11556 | 4.98672 | 1.89947 | 2.68400 | 0.40482 | 4.97415 | 0.26922 | 2.17472 | 3.85722 |
| **Clcnka** | 0.00000 | 0.00665 | 0.70320 | 0.38729 | 0.02605 | 0.00573 | 0.00529 | 6.56976 | 0.00428 | 3.40004 | 3.12699 | 0.00000 | 1.84338 | 0.00000 | 0.03126 | 0.00590 |
| **Clcnkb** | 0.82016 | 0.00638 | 0.20394 | 0.00611 | 0.00582 | 0.00511 | 0.00631 | 0.66409 | 0.01064 | 0.00000 | 0.32084 | 0.00412 | 0.51176 | 0.01565 | 0.00536 | 0.01292 |
| **Cldn10** | 0.01310 | 0.00755 | 0.01121 | 0.12553 | 0.00894 | 0.00657 | 0.00730 | 0.01026 | 0.27772 | 0.96076 | 0.00896 | 0.01362 | 0.01167 | 0.00890 | 0.44219 | 0.01512 |
| **Cldn12** | 1.76225 | 2.26845 | 1.14995 | 0.52448 | 0.03280 | 1.01265 | 1.26043 | 1.02323 | 0.43040 | 0.11723 | 2.55502 | 0.19661 | 1.85710 | 1.41476 | 2.30087 | 0.22972 |
| **Cldn15** | 0.01224 | 0.00762 | 0.00543 | 0.92969 | 0.00937 | 0.00678 | 0.00701 | 0.27784 | 0.00000 | 0.00000 | 0.23755 | 0.00440 | 0.00520 | 0.00360 | 0.69937 | 0.00700 |
| **Cldn19** | 0.01104 | 0.07918 | 0.02132 | 0.00908 | 0.00897 | 0.00682 | 0.12827 | 0.01624 | 0.01221 | 0.00822 | 0.01243 | 0.00702 | 0.01783 | 0.00629 | 0.22016 | 0.00728 |
| **Cldn22** | 0.01542 | 0.00764 | 0.54334 | 0.00975 | 0.00836 | 0.00559 | 0.00722 | 0.01054 | 0.01222 | 0.25352 | 0.01207 | 0.00649 | 0.00923 | 0.01378 | 0.00147 | 0.01452 |
| **Cldn3** | 0.73422 | 0.00772 | 0.01107 | 0.27423 | 0.05648 | 0.00723 | 0.00647 | 0.00522 | 0.24768 | 5.29722 | 0.04132 | 0.00000 | 0.08567 | 0.01045 | 0.00212 | 0.25476 |
| **Cldn34b1** | 0.01351 | 0.00825 | 0.01106 | 0.11716 | 0.00924 | 0.00694 | 0.00753 | 0.01265 | 0.00768 | 0.16072 | 0.01123 | 0.00816 | 0.01309 | 0.15189 | 0.00435 | 0.00740 |
| **Cldn34b2** | 0.01271 | 0.00810 | 0.00805 | 0.81627 | 0.00924 | 0.00697 | 0.00736 | 0.01055 | 0.00000 | 0.36263 | 0.02122 | 0.00542 | 0.01258 | 0.77224 | 0.01517 | 0.00726 |
| **Cldn34c1** | 0.77522 | 0.00762 | 0.00000 | 0.78814 | 0.03766 | 0.09557 | 0.01730 | 0.01159 | 0.00595 | 0.12554 | 0.01020 | 0.00577 | 0.01206 | 0.08789 | 0.48152 | 0.04461 |
| **Cldn34d** | 0.01004 | 0.00841 | 0.00847 | 0.47732 | 0.00958 | 0.00665 | 0.00816 | 0.25945 | 0.28938 | 0.00423 | 0.00864 | 0.01870 | 0.00654 | 0.00611 | 0.42517 | 0.00718 |
| **Cldn4** | 0.00050 | 0.10381 | 6.97553 | 0.00000 | 0.53731 | 17.80297 | 2.56043 | 23.45781 | 0.53138 | 9.97935 | 12.89922 | 0.01702 | 25.73508 | 4.13357 | 0.68542 | 12.76557 |

| | | | | | | | | | | | | | | | | |
|---|---|---|---|---|---|---|---|---|---|---|---|---|---|---|---|---|
| **Cldn5** | 0.01318 | 0.00772 | 0.00902 | 0.00467 | 0.00875 | 0.00684 | 0.00716 | 0.00000 | 0.01099 | 1.48451 | 0.49232 | 0.00000 | 0.00766 | 0.00600 | 0.11533 | 0.00723 |
| **Cldn6** | 0.00000 | 7.10709 | 6.01159 | 0.63681 | 0.10489 | 12.86469 | 0.00000 | 5.52615 | 0.00000 | 5.50385 | 1.01719 | 0.14726 | 11.00933 | 1.45041 | 2.39804 | 1.68983 |
| **Cldn7** | 3.85230 | 5.08215 | 13.01790 | 0.88356 | 0.08659 | 12.23946 | 4.51560 | 19.61171 | 1.87190 | 8.33644 | 10.95029 | 0.70621 | 19.90824 | 0.07848 | 0.00838 | 9.63030 |
| **Cldn9** | 0.00000 | 0.00730 | 0.44454 | 1.10415 | 0.00841 | 0.00890 | 0.00554 | 0.93122 | 1.82269 | 0.00000 | 0.01811 | 0.25162 | 2.84351 | 0.00887 | 0.00000 | 0.00560 |
| **Cldnd1** | 2.17048 | 0.12315 | 1.40516 | 2.67152 | 7.75273 | 3.72819 | 4.50166 | 1.75082 | 6.58025 | 2.28128 | 2.18459 | 6.03258 | 0.22024 | 4.63922 | 2.20249 | 0.74060 |
| **Clec10a** | 0.00305 | 0.00770 | 0.05894 | 0.25686 | 0.00848 | 0.00596 | 0.00358 | 1.76937 | 1.44113 | 0.91027 | 0.00000 | 0.02106 | 7.26441 | 0.56700 | 0.42327 | 0.00433 |
| **Clec11a** | 0.01571 | 0.00830 | 0.00868 | 0.21097 | 0.00991 | 0.28969 | 0.00761 | 0.01604 | 0.00648 | 0.00724 | 0.01461 | 0.00795 | 0.40897 | 0.01195 | 0.00703 | 0.00734 |
| **Clec16a** | 3.26205 | 0.00549 | 2.17408 | 0.44093 | 0.00431 | 2.21975 | 0.06103 | 0.02606 | 0.46896 | 1.00993 | 0.16408 | 0.03636 | 0.93034 | 0.00471 | 0.90695 | 0.00190 |
| **Clec2e** | 0.01365 | 0.00818 | 0.01199 | 0.36454 | 0.00924 | 0.00655 | 0.00750 | 0.01393 | 0.46263 | 0.01255 | 0.01065 | 0.00760 | 0.01914 | 0.01158 | 0.00572 | 0.00729 |
| **Clec2f** | 2.29163 | 0.00578 | 0.04072 | 0.21849 | 0.00547 | 0.00392 | 0.00578 | 0.01297 | 0.00545 | 0.33948 | 0.01307 | 0.01528 | 0.02839 | 0.01068 | 0.00601 | 0.00558 |
| **Clec2g** | 0.07582 | 0.00766 | 0.03109 | 0.83766 | 0.00809 | 0.08116 | 0.00704 | 0.02263 | 0.73943 | 0.03509 | 0.05938 | 0.04066 | 0.01179 | 3.43303 | 0.00000 | 0.01454 |
| **Clec2h** | 0.07304 | 0.00805 | 0.01682 | 0.67241 | 0.00795 | 0.02159 | 0.02832 | 0.03906 | 0.14908 | 0.29718 | 0.09417 | 0.00719 | 0.02236 | 0.04879 | 0.25179 | 0.00720 |
| **Clec2i** | 0.00935 | 0.00789 | 0.03364 | 0.03865 | 0.00981 | 0.00599 | 0.00735 | 0.88617 | 0.12477 | 0.01109 | 0.00000 | 0.00409 | 0.00610 | 0.66295 | 0.20276 | 0.00742 |
| **Clec2l** | 0.01377 | 0.00781 | 0.01124 | 0.17642 | 0.00934 | 0.00662 | 0.00744 | 0.01253 | 0.43100 | 0.00753 | 0.00892 | 0.06537 | 0.01290 | 0.00664 | 0.09326 | 0.00735 |
| **Clec3b** | 0.01330 | 0.00780 | 0.01138 | 0.00785 | 0.00934 | 0.00668 | 0.00744 | 0.01711 | 0.03391 | 0.01343 | 0.07457 | 0.01713 | 0.02001 | 0.01228 | 0.04743 | 0.02936 |
| **Clec4a1** | 0.16042 | 0.00805 | 0.08347 | 0.32602 | 0.04040 | 0.00573 | 0.00770 | 0.04090 | 0.79691 | 0.49377 | 0.00599 | 0.00717 | 0.01665 | 0.01501 | 0.02206 | 0.00697 |
| **Clgn** | 1.19938 | 0.00698 | 0.00254 | 0.00392 | 0.03355 | 0.00675 | 0.00641 | 0.00747 | 0.34689 | 0.00529 | 0.00880 | 0.00538 | 0.01148 | 0.00719 | 0.42845 | 0.00625 |
| **Clhc1** | 0.02885 | 0.00766 | 0.00098 | 0.01259 | 0.00836 | 0.02485 | 0.00705 | 0.01323 | 2.26419 | 0.60116 | 0.00000 | 0.02316 | 0.14491 | 2.95457 | 0.49381 | 0.00709 |
| **Clic1** | 3.97352 | 2.12545 | 0.00000 | 0.07903 | 0.03272 | 0.01038 | 2.12589 | 0.00000 | 0.12373 | 1.66024 | 0.75844 | 0.06018 | 0.92273 | 0.00405 | 0.46339 | 7.18305 |
| **Clic3** | 0.00000 | 0.26215 | 2.56483 | 0.00170 | 0.02064 | 0.07640 | 2.20201 | 0.79226 | 0.00990 | 0.00000 | 2.57102 | 0.05758 | 3.51105 | 0.00506 | 0.00961 | 0.45693 |
| **Clic4** | 7.20165 | 13.42566 | 14.90958 | 4.99758 | 8.06232 | 16.90092 | 4.73224 | 8.51512 | 14.36160 | 3.29149 | 4.29321 | 4.64865 | 11.28962 | 4.90122 | 1.28868 | 5.10217 |
| **Clint1** | 6.23081 | 8.03057 | 1.56170 | 1.79431 | 4.40769 | 3.29985 | 5.48257 | 6.10441 | 2.46742 | 4.00121 | 1.29660 | 1.96546 | 1.56015 | 7.70649 | 6.28286 | 5.69638 |
| **Clip1** | 0.83718 | 5.13145 | 1.04855 | 5.38811 | 0.00436 | 0.68779 | 0.37736 | 0.48225 | 4.87366 | 2.42131 | 4.75582 | 0.78390 | 0.31148 | 2.92239 | 4.51825 | 1.13319 |
| **Clip2** | 0.00878 | 0.00803 | 0.00997 | 0.45538 | 0.00890 | 0.00640 | 0.00669 | 0.00865 | 0.00532 | 0.54103 | 0.03859 | 0.10539 | 1.05455 | 0.03238 | 0.00000 | 0.02738 |
| **Clip3** | 0.01378 | 0.00789 | 0.01108 | 0.26558 | 0.00932 | 0.00655 | 0.00733 | 0.00882 | 0.04906 | 0.18414 | 0.11822 | 0.02596 | 0.01094 | 0.00923 | 0.00529 | 0.00739 |
| **Clip4** | 0.01379 | 0.00766 | 0.00967 | 1.32189 | 0.00924 | 0.00630 | 0.00724 | 0.00478 | 0.00000 | 0.08861 | 0.20889 | 0.00446 | 0.00979 | 0.00437 | 0.91616 | 0.00701 |
| **Clk1** | 7.54861 | 10.39900 | 1.24553 | 9.80388 | 10.37963 | 10.20844 | 10.43465 | 6.76855 | 18.00432 | 11.85835 | 6.89271 | 18.77604 | 5.30575 | 20.47603 | 11.85069 | 6.77242 |
| **Clk2** | 2.24157 | 1.81900 | 0.36388 | 5.76910 | 2.53677 | 3.53796 | 2.52789 | 2.14398 | 2.23212 | 3.00012 | 5.41014 | 0.32264 | 2.38398 | 0.71709 | 2.01231 | 2.15127 |
| **Clk3** | 3.85231 | 7.65322 | 5.69913 | 6.37521 | 7.17394 | 6.37189 | 5.43945 | 9.85492 | 10.35608 | 5.23090 | 2.41101 | 0.60199 | 5.59241 | 0.01847 | 1.04510 | 0.92224 |
| **Clk4** | 2.00029 | 2.32352 | 0.52821 | 2.29419 | 1.81690 | 5.04198 | 1.63962 | 0.95454 | 7.37465 | 2.62460 | 2.74383 | 0.83112 | 1.03191 | 2.85239 | 5.25858 | 0.73504 |
| **Clmn** | 1.64421 | 0.00508 | 1.33311 | 1.32484 | 0.01874 | 0.04623 | 0.04204 | 0.69957 | 1.87889 | 2.36182 | 1.56816 | 0.14655 | 3.25196 | 0.99857 | 0.10802 | 1.01580 |
| **Clmp** | 0.00443 | 0.00780 | 0.32402 | 2.49398 | 0.00829 | 0.00815 | 0.00603 | 0.00000 | 2.87681 | 1.47037 | 1.05002 | 0.07503 | 1.48068 | 0.34578 | 0.98294 | 0.00608 |
| **Cln3** | 1.97832 | 1.89684 | 11.04574 | 1.92497 | 0.02036 | 7.25828 | 0.13325 | 13.38749 | 1.67334 | 11.75076 | 5.98418 | 0.59344 | 4.28281 | 3.86159 | 3.74900 | 1.89349 |
| **Cln5** | 6.17054 | 6.46679 | 0.59275 | 0.00000 | 0.22444 | 3.51018 | 5.11640 | 0.55605 | 0.00689 | 1.71181 | 1.83195 | 0.03833 | 0.00000 | 0.00270 | 3.16466 | 1.58411 |
| **Cln6** | 0.01943 | 0.02863 | 0.00000 | 0.72592 | 0.02221 | 1.26478 | 0.00683 | 0.00000 | 0.28361 | 0.63593 | 0.78478 | 0.07496 | 0.03954 | 0.17299 | 0.00000 | 0.00690 |
| **Cln8** | 2.34086 | 2.80742 | 5.84197 | 1.05212 | 0.06758 | 6.13578 | 5.26611 | 15.27977 | 1.28296 | 5.55635 | 6.09915 | 0.60817 | 10.89074 | 7.82099 | 5.66288 | 4.07320 |
| **Clns1a** | 17.60093 | 1.75040 | 8.31253 | 10.39131 | 15.34496 | 9.85526 | 6.81172 | 21.30102 | 15.61269 | 32.37813 | 15.20313 | 15.62479 | 16.35961 | 15.58792 | 7.19777 | 8.13719 |
| **Clock** | 1.18899 | 0.00592 | 1.52246 | 5.76250 | 0.00730 | 0.04206 | 1.29914 | 0.30688 | 1.80545 | 0.98957 | 0.00000 | 0.07876 | 0.02395 | 6.74986 | 6.63551 | 0.00458 |
| **Clp1** | 3.70190 | 6.87486 | 8.09113 | 9.59797 | 4.42721 | 4.69026 | 5.51148 | 12.44655 | 8.46770 | 6.86608 | 3.52570 | 7.64984 | 1.82862 | 9.33588 | 12.48899 | 1.66765 |
| **Clpb** | 3.72289 | 0.04303 | 5.50171 | 0.55065 | 2.39174 | 4.33777 | 0.01739 | 9.38441 | 0.39804 | 3.63600 | 3.03132 | 1.03617 | 11.22954 | 0.36552 | 0.88402 | 1.29147 |
| **Clpp** | 19.77703 | 6.39841 | 19.14880 | 7.16401 | 14.35152 | 7.01419 | 10.93589 | 20.91507 | 13.79246 | 18.33329 | 14.54317 | 5.61648 | 16.16996 | 0.00000 | 0.35572 | 8.89095 |
| **Clps** | 0.01264 | 0.00829 | 0.26177 | 0.00568 | 0.00874 | 0.00569 | 0.00767 | 0.00733 | 0.63641 | 0.00000 | 0.34861 | 0.00636 | 0.05229 | 0.00609 | 0.14671 | 0.00711 |
| **Clptm1** | 11.15292 | 0.06215 | 0.00000 | 4.94473 | 4.98922 | 6.48941 | 0.76348 | 17.70557 | 4.82169 | 11.64578 | 4.40530 | 0.51618 | 15.49561 | 0.00000 | 0.00000 | 4.87074 |
| **Clptm1l** | 12.18365 | 6.51323 | 8.01540 | 6.44578 | 5.12826 | 1.23697 | 7.02645 | 11.47202 | 2.12974 | 11.88292 | 7.04531 | 9.23141 | 8.29486 | 6.33836 | 6.58847 | 0.45033 |
| **Clpx** | 8.59667 | 5.90836 | 8.27195 | 16.03461 | 13.22398 | 7.55461 | 6.31628 | 22.15835 | 26.62006 | 38.42058 | 26.92534 | 19.78279 | 5.82143 | 30.46639 | 28.95019 | 2.56204 |
| **Clrn1** | 0.04931 | 0.00737 | 0.04560 | 0.02286 | 0.00792 | 0.00857 | 0.00727 | 0.01453 | 0.04755 | 0.07713 | 0.02408 | 0.27640 | 0.01183 | 0.04916 | 0.01340 | 0.01476 |
| **Clspn** | 4.08759 | 4.99707 | 5.75854 | 7.69838 | 0.00000 | 0.03124 | 1.09537 | 3.68566 | 15.08094 | 4.49604 | 2.34171 | 1.30205 | 5.54129 | 4.70934 | 11.41621 | 0.37955 |
| **Clstn1** | 7.25124 | 0.02807 | 7.88795 | 2.38119 | 0.02195 | 6.05630 | 2.43640 | 0.00000 | 3.02860 | 2.44315 | 1.88054 | 0.02048 | 4.46686 | 0.66176 | 0.50285 | 7.33856 |
| **Clstn3** | 0.03976 | 0.00725 | 0.76746 | 0.51023 | 0.02227 | 0.00871 | 0.00648 | 2.35545 | 0.77028 | 0.00000 | 0.32059 | 0.01680 | 0.58412 | 0.17242 | 0.26678 | 1.73283 |
| **Clta** | 19.46823 | 5.77952 | 10.19150 | 13.75802 | 13.60397 | 7.67500 | 4.35936 | 5.02857 | 12.74053 | 28.78490 | 23.43958 | 3.04054 | 23.14157 | 5.47049 | 5.48707 | 14.40958 |
| **Cltb** | 0.56861 | 0.00645 | 0.21171 | 8.00270 | 0.00652 | 1.63416 | 0.01649 | 2.44888 | 5.74374 | 4.91346 | 2.49706 | 0.43538 | 0.18850 | 1.54375 | 1.03161 | 0.19346 |
| **Cltc** | 16.40578 | 15.33054 | 9.23245 | 11.57511 | 16.40579 | 10.18408 | 9.98271 | 9.89903 | 12.73748 | 11.97648 | 7.83548 | 9.94619 | 12.40505 | 23.55912 | 9.37773 | 7.69169 |
| **Clu** | 0.21601 | 0.00788 | 0.00736 | 0.33388 | 0.03427 | 0.00673 | 0.00705 | 0.02118 | 0.27142 | 6.07131 | 0.00611 | 0.03094 | 0.01003 | 0.01317 | 0.12601 | 0.00736 |
| **Cluap1** | 1.00198 | 4.83675 | 2.07698 | 0.29468 | 2.70367 | 0.00583 | 2.21926 | 3.63214 | 0.06544 | 2.86003 | 3.61055 | 0.55044 | 2.41116 | 0.04203 | 2.20040 | 0.11329 |
| **Cluh** | 16.69352 | 10.05549 | 14.87855 | 4.67261 | 7.49430 | 10.19158 | 7.42202 | 17.63468 | 4.18338 | 13.69187 | 12.32647 | 1.37368 | 13.18426 | 3.36313 | 3.17622 | 5.47001 |
| **Clvs2** | 0.01337 | 0.00769 | 0.00631 | 1.82950 | 0.00905 | 0.00638 | 0.00703 | 0.00000 | 0.12971 | 0.27760 | 0.48611 | 0.01039 | 0.00828 | 0.00170 | 1.77778 | 0.01488 |
| **Clybl** | 4.66594 | 0.00000 | 0.00000 | 4.27434 | 0.37510 | 4.50592 | 0.00000 | 0.54930 | 2.98358 | 3.83840 | 2.71632 | 5.01680 | 1.00118 | 4.15990 | 3.00187 | 0.00000 |
| **Cmah** | 7.40838 | 6.38292 | 8.91693 | 17.61257 | 1.77835 | 1.51409 | 8.09528 | 3.42327 | 16.13330 | 5.89698 | 2.96929 | 12.01178 | 2.96814 | 16.28606 | 8.37550 | 5.29299 |
| **Cmas** | 10.18409 | 2.55850 | 2.98546 | 6.64468 | 1.80933 | 0.00000 | 5.07189 | 1.12245 | 4.47603 | 2.42430 | 1.26575 | 0.99831 | 2.98485 | 0.00000 | 0.21018 | 1.24475 |
| **Cmbl** | 1.96843 | 3.02921 | 6.70078 | 0.00000 | 0.01038 | 0.16240 | 0.17472 | 3.73616 | 0.00771 | 0.78759 | 0.78983 | 0.00000 | 2.55144 | 0.01741 | 0.69855 | 4.16851 |
| **Cmc1** | 2.96059 | 0.13282 | 0.30418 | 0.15503 | 0.61165 | 0.02032 | 0.10298 | 0.34656 | 0.83225 | 0.64724 | 0.33909 | 0.00595 | 0.27443 | 0.25193 | 0.83337 | 0.49785 |
| **Cmc2** | 8.28928 | 1.77280 | 5.89829 | 1.29377 | 3.22026 | 2.06533 | 1.89506 | 5.00165 | 4.03023 | 3.53262 | 5.43354 | 0.65433 | 7.50013 | 0.33965 | 1.31564 | 2.18792 |

| | | | | | | | | | | | | | | | | |
|---|---|---|---|---|---|---|---|---|---|---|---|---|---|---|---|---|
| **Cmc4** | 0.00652 | 0.00701 | 0.24643 | 0.97338 | 1.58457 | 0.05111 | 0.00716 | 0.14780 | 0.00000 | 0.23952 | 0.34034 | 0.01294 | 0.00000 | 0.00437 | 0.04233 | 1.01261 |
| **Cmip** | 3.61411 | 1.22302 | 2.02189 | 0.12592 | 6.11995 | 6.08258 | 0.00000 | 9.79775 | 0.00000 | 2.31302 | 0.80123 | 0.71357 | 6.50578 | 0.43587 | 0.45519 | 0.94325 |
| **Cmklr1** | 0.05993 | 0.00798 | 0.05736 | 1.54978 | 0.00829 | 0.00794 | 0.13940 | 0.02066 | 0.00000 | 0.01012 | 0.01238 | 0.00757 | 0.02593 | 0.01763 | 0.17568 | 0.00694 |
| **Cmpk1** | 4.26963 | 4.14860 | 2.26186 | 5.63762 | 5.70224 | 4.09303 | 1.23602 | 2.56841 | 4.14317 | 5.63076 | 3.35286 | 3.77615 | 3.82667 | 5.94705 | 5.88978 | 3.33239 |
| **Cmpk2** | 0.01361 | 0.00777 | 0.01118 | 3.33101 | 0.00944 | 0.00606 | 0.00722 | 0.01080 | 0.00000 | 0.00644 | 0.01009 | 0.00500 | 0.01280 | 0.00175 | 1.43071 | 0.00675 |
| **Cmss1** | 5.59463 | 2.35305 | 4.76203 | 4.22678 | 0.01811 | 4.14474 | 0.00000 | 3.12479 | 4.70229 | 7.11287 | 2.70447 | 7.86336 | 6.53089 | 2.99227 | 0.44904 | 2.36507 |
| **Cmtm3** | 0.01244 | 0.00785 | 0.01156 | 0.00633 | 0.00877 | 0.00651 | 0.06750 | 0.01014 | 0.01156 | 1.62261 | 0.01056 | 0.00355 | 0.02278 | 0.01194 | 0.00760 | 0.03600 |
| **Cmtm4** | 3.50449 | 0.01978 | 6.71869 | 14.72895 | 2.29574 | 3.69473 | 2.64100 | 8.16335 | 16.25745 | 11.45962 | 7.40589 | 1.73529 | 3.66993 | 2.99313 | 5.04733 | 0.02971 |
| **Cmtm6** | 3.62330 | 5.18530 | 7.81293 | 3.40951 | 1.03300 | 5.62665 | 10.48861 | 10.43020 | 9.43919 | 6.61132 | 6.18637 | 4.58544 | 7.30259 | 1.42736 | 3.72182 | 4.78816 |
| **Cmtm7** | 11.02215 | 1.86285 | 6.47797 | 0.00000 | 9.74632 | 8.95646 | 4.92089 | 6.99440 | 0.75239 | 11.73151 | 4.10148 | 3.58782 | 12.87620 | 0.48347 | 0.00000 | 8.33829 |
| **Cmtm8** | 0.00696 | 0.00841 | 0.01124 | 0.74904 | 0.00808 | 0.00653 | 0.00801 | 0.05423 | 0.02782 | 1.35059 | 0.01067 | 0.00512 | 0.00719 | 0.01247 | 0.00993 | 0.66406 |
| **Cmtr1** | 0.01125 | 0.00775 | 0.03278 | 0.01065 | 0.00969 | 0.00715 | 0.04751 | 0.09676 | 0.01225 | 0.01316 | 0.01314 | 0.00740 | 0.01135 | 0.01240 | 0.00872 | 0.00747 |
| **Cmtr2** | 0.00634 | 0.00709 | 0.70441 | 2.22402 | 0.00936 | 0.00541 | 2.35916 | 0.43275 | 0.41844 | 0.00000 | 0.03096 | 0.02758 | 0.07587 | 0.00355 | 0.31174 | 0.00644 |
| **Cmya5** | 0.01377 | 0.00790 | 0.01238 | 5.80639 | 0.00936 | 0.00604 | 0.00725 | 0.01278 | 0.00000 | 0.00845 | 0.01264 | 0.00709 | 0.01264 | 0.00188 | 0.24266 | 0.00638 |
| **Cnbd2** | 1.16220 | 1.75223 | 4.47323 | 9.03729 | 1.41251 | 2.67126 | 4.38608 | 2.73731 | 6.09977 | 13.71490 | 5.24977 | 5.52673 | 3.04475 | 3.97774 | 6.43663 | 0.43454 |
| **Cnbp** | 33.24468 | 25.35375 | 34.61772 | 22.69148 | 26.07511 | 15.67076 | 24.92979 | 42.46917 | 28.97080 | 46.53930 | 39.76008 | 19.17530 | 35.77401 | 31.85373 | 29.31387 | 20.74959 |
| **Cndp1** | 0.01179 | 0.00778 | 1.69832 | 1.35424 | 0.00631 | 0.00800 | 0.00720 | 0.01033 | 0.49312 | 0.29353 | 0.00691 | 0.00580 | 0.00734 | 0.00913 | 0.00000 | 0.00651 |
| **Cndp2** | 20.01179 | 20.16913 | 23.65534 | 3.29803 | 12.28457 | 19.02927 | 12.04791 | 25.14806 | 8.34641 | 17.36784 | 18.85104 | 12.49051 | 25.81700 | 11.32827 | 8.92969 | 13.23375 |
| **Cnep1r1** | 0.55094 | 2.17386 | 1.14620 | 7.73677 | 0.91022 | 0.90204 | 0.20712 | 1.65896 | 8.96434 | 0.88954 | 1.14362 | 4.66289 | 0.41938 | 7.15748 | 4.74040 | 0.22423 |
| **Cnga1** | 0.01361 | 0.00780 | 0.01147 | 0.88629 | 0.00923 | 0.00668 | 0.00747 | 0.01220 | 0.00000 | 0.22526 | 0.01070 | 0.00736 | 0.01286 | 0.00967 | 0.44910 | 0.00728 |
| **Cnga3** | 0.65573 | 0.01698 | 0.23731 | 0.03152 | 0.07937 | 0.01538 | 0.00668 | 0.00957 | 0.12272 | 0.02311 | 0.14591 | 0.00578 | 0.02228 | 0.18287 | 0.02847 | 0.00645 |
| **Cnga4** | 0.02808 | 0.00786 | 0.00865 | 0.51777 | 0.20918 | 0.00671 | 0.00738 | 0.01100 | 0.00348 | 0.00559 | 0.02422 | 0.00709 | 0.01140 | 0.01756 | 0.24071 | 0.00735 |
| **Cngb1** | 0.01305 | 0.00786 | 0.01032 | 2.89602 | 0.00910 | 0.00626 | 0.00743 | 0.01110 | 4.11206 | 0.20797 | 0.00000 | 0.01068 | 0.01333 | 0.00000 | 0.46460 | 0.00668 |
| **Cnih1** | 12.22330 | 8.52422 | 12.46237 | 11.09746 | 11.15299 | 9.71964 | 5.26686 | 7.50882 | 11.02187 | 11.07568 | 3.93332 | 7.42399 | 9.35466 | 5.31778 | 2.88627 | 2.61102 |
| **Cnih2** | 0.01375 | 0.00805 | 0.01225 | 0.01061 | 0.00932 | 0.00666 | 0.00754 | 0.01390 | 0.01235 | 0.01531 | 0.01310 | 0.00811 | 0.01270 | 0.01234 | 0.01062 | 0.00741 |
| **Cnih4** | 7.29439 | 1.98873 | 7.69119 | 3.23632 | 2.12679 | 4.81527 | 2.63518 | 2.57848 | 4.28752 | 5.63401 | 9.87283 | 0.59044 | 4.77499 | 2.98077 | 3.99321 | 2.27906 |
| **Cnksr1** | 0.01745 | 0.00752 | 0.47546 | 0.01011 | 0.01969 | 0.00636 | 0.00622 | 0.00000 | 0.36535 | 0.29937 | 2.75677 | 0.02119 | 0.13628 | 0.00224 | 0.18155 | 0.00645 |
| **Cnksr3** | 0.36272 | 1.70899 | 0.13428 | 6.37344 | 3.90218 | 0.00762 | 0.70724 | 0.73062 | 3.87354 | 0.42400 | 0.59535 | 1.40933 | 1.27089 | 0.06022 | 0.48435 | 2.03823 |
| **Cnn1** | 0.01393 | 0.00807 | 0.00958 | 4.04971 | 0.00912 | 0.00653 | 0.00730 | 0.00686 | 0.00000 | 0.03681 | 0.26007 | 0.03740 | 0.00982 | 0.00378 | 0.00000 | 0.00684 |
| **Cnn2** | 0.29547 | 0.01942 | 1.15661 | 0.48447 | 0.08657 | 13.97512 | 0.04395 | 4.37667 | 0.33773 | 5.45284 | 1.27476 | 0.04931 | 5.77655 | 0.04135 | 0.20059 | 9.19361 |
| **Cnn3** | 12.27156 | 10.23804 | 14.81739 | 10.40674 | 9.93605 | 13.17043 | 10.57731 | 15.99314 | 17.60297 | 4.54640 | 6.89034 | 7.52372 | 22.12210 | 5.74875 | 2.10180 | 7.45228 |
| **Cnnm1** | 0.01347 | 0.00772 | 0.00661 | 0.00456 | 0.00911 | 0.00699 | 0.00698 | 0.00000 | 0.34941 | 0.51380 | 0.64279 | 0.04717 | 0.00686 | 0.11029 | 0.00000 | 0.00729 |
| **Cnnm2** | 5.40366 | 1.52416 | 4.00099 | 5.38356 | 0.00813 | 0.01492 | 2.27426 | 5.86327 | 11.18160 | 5.42590 | 2.17464 | 0.14956 | 6.06561 | 0.00000 | 5.24342 | 0.91398 |
| **Cnnm3** | 0.01224 | 0.00629 | 3.47656 | 7.15799 | 1.03926 | 0.02006 | 0.01613 | 2.26822 | 3.82876 | 1.67064 | 0.75130 | 0.30163 | 1.66896 | 0.06680 | 2.94636 | 0.01111 |
| **Cnnm4** | 0.02505 | 1.85106 | 4.46974 | 3.31077 | 0.01620 | 1.68157 | 0.68389 | 10.59950 | 2.61452 | 5.65860 | 10.43897 | 5.99109 | 5.57226 | 3.48949 | 5.97210 | 0.49378 |
| **Cnot1** | 16.51356 | 15.74564 | 11.77436 | 12.46431 | 12.52160 | 7.87461 | 11.72378 | 26.51271 | 10.50446 | 26.93715 | 15.12870 | 18.55935 | 11.13656 | 29.33825 | 25.42861 | 8.09071 |
| **Cnot10** | 7.04741 | 8.26511 | 6.98225 | 3.37857 | 10.51250 | 9.33264 | 6.74403 | 7.45069 | 7.18038 | 10.43185 | 7.21311 | 1.40268 | 3.11419 | 0.68404 | 5.61608 | 3.29151 |
| **Cnot11** | 1.25684 | 2.32053 | 0.71567 | 4.96761 | 2.08544 | 2.69232 | 0.12824 | 3.05057 | 1.77490 | 1.20505 | 7.51294 | 0.90743 | 0.00000 | 4.23786 | 1.75879 | 0.02398 |
| **Cnot2** | 9.52792 | 11.57932 | 12.85779 | 9.01444 | 9.17298 | 2.98646 | 2.58440 | 14.25072 | 5.10251 | 15.02440 | 7.69266 | 6.05872 | 1.27734 | 21.94619 | 13.53437 | 7.14068 |
| **Cnot3** | 1.98602 | 1.33504 | 0.43889 | 0.16484 | 2.15861 | 0.26085 | 1.66130 | 4.45799 | 0.23307 | 1.93451 | 0.00000 | 0.07913 | 1.51530 | 0.00451 | 0.00944 | 1.70802 |
| **Cnot4** | 1.43753 | 2.11068 | 3.67178 | 2.79113 | 0.21517 | 1.67430 | 0.32187 | 3.95767 | 1.96872 | 4.52846 | 2.30951 | 1.90267 | 1.07402 | 4.48317 | 4.42805 | 0.79202 |
| **Cnot6** | 2.05173 | 2.24580 | 3.76963 | 5.59130 | 2.71442 | 0.00855 | 6.72373 | 1.81738 | 2.71527 | 5.74439 | 4.24627 | 8.06227 | 0.82289 | 9.59987 | 8.46598 | 0.36171 |
| **Cnot6l** | 1.96887 | 2.88288 | 1.41512 | 9.00221 | 0.01004 | 0.58051 | 1.11303 | 0.00000 | 2.29157 | 1.95407 | 0.44656 | 0.54833 | 2.92051 | 5.22655 | 3.40564 | 0.41244 |
| **Cnot7** | 1.67425 | 3.32836 | 1.60849 | 18.18728 | 1.32279 | 0.17381 | 4.05578 | 2.15012 | 9.58676 | 7.67345 | 5.83670 | 13.88191 | 1.34231 | 16.75395 | 8.10309 | 0.01140 |
| **Cnot8** | 12.64838 | 2.67410 | 7.33075 | 17.87529 | 7.45666 | 10.84906 | 3.65282 | 12.78811 | 18.59807 | 19.86117 | 16.17631 | 9.55398 | 10.21893 | 6.26976 | 20.18820 | 1.60932 |
| **Cnot9** | 9.21379 | 2.49023 | 12.45283 | 3.05854 | 5.52984 | 6.77653 | 9.28456 | 6.98877 | 2.34776 | 13.69096 | 1.10579 | 3.20253 | 6.99112 | 11.01606 | 6.26261 | 3.26184 |
| **Cnp** | 1.16873 | 0.00619 | 0.00000 | 0.15264 | 0.00539 | 0.16477 | 1.12686 | 1.99678 | 0.71619 | 4.83672 | 1.60545 | 0.07267 | 0.28043 | 0.01674 | 0.02394 | 0.01258 |
| **Cnppd1** | 5.71653 | 0.82930 | 7.89214 | 21.87473 | 0.01461 | 3.07415 | 0.45287 | 12.20131 | 25.28324 | 18.44940 | 6.92612 | 1.35762 | 5.83566 | 1.92885 | 5.05136 | 0.00983 |
| **Cnpy1** | 0.03461 | 0.00726 | 0.00000 | 1.12445 | 0.00730 | 0.00661 | 0.11484 | 0.00000 | 0.28560 | 0.00000 | 0.56447 | 0.48868 | 0.00375 | 0.00352 | 1.50511 | 1.47043 |
| **Cnpy2** | 14.95463 | 8.37061 | 11.06678 | 0.26929 | 11.31152 | 3.01386 | 3.02241 | 3.54932 | 0.00000 | 1.49584 | 4.16755 | 0.10300 | 3.70643 | 0.04286 | 0.63812 | 7.29649 |
| **Cnpy3** | 2.38597 | 3.78488 | 6.25313 | 1.51122 | 0.00000 | 0.00991 | 1.63580 | 2.87748 | 4.02269 | 3.72850 | 2.23135 | 0.08899 | 1.92847 | 0.01239 | 0.77025 | 2.73661 |
| **Cnpy4** | 0.70534 | 2.89543 | 0.00000 | 6.78869 | 0.37845 | 2.12634 | 0.06268 | 0.63455 | 3.97173 | 0.40793 | 1.08022 | 0.24466 | 0.49769 | 4.26631 | 4.88939 | 0.19932 |
| **Cnr2** | 0.09003 | 0.01817 | 0.14807 | 0.30043 | 0.02564 | 0.09561 | 0.03916 | 0.02866 | 0.32505 | 0.11653 | 0.40403 | 0.20412 | 0.04623 | 0.67667 | 0.47167 | 0.04358 |
| **Cnrip1** | 0.01374 | 0.00763 | 0.00940 | 0.29656 | 0.00924 | 0.00656 | 0.00730 | 0.01088 | 0.00712 | 0.22460 | 0.00946 | 0.18059 | 0.01198 | 0.00469 | 0.43961 | 0.00721 |
| **Cnst** | 3.75826 | 2.98808 | 2.25215 | 10.10213 | 0.07801 | 4.49091 | 0.60977 | 1.06212 | 3.05624 | 4.64113 | 0.35671 | 0.24614 | 1.72355 | 3.18302 | 8.59189 | 0.67287 |
| **Cntd1** | 0.44814 | 0.00739 | 0.00663 | 0.01056 | 0.00747 | 0.00747 | 0.00750 | 0.01418 | 0.01234 | 0.01408 | 0.01453 | 0.00775 | 0.05306 | 0.01241 | 0.01065 | 0.00717 |
| **Cntf** | 0.01197 | 0.00777 | 0.01180 | 0.01102 | 0.03681 | 0.00667 | 0.00752 | 0.01350 | 0.01207 | 0.01368 | 0.01261 | 0.00790 | 0.01251 | 0.01742 | 0.03252 | 0.00738 |
| **Cntln** | 0.01401 | 0.00623 | 0.00276 | 4.38357 | 0.00878 | 0.00490 | 0.00640 | 0.00745 | 1.97496 | 1.02555 | 0.00000 | 4.24332 | 0.01178 | 2.05295 | 2.70167 | 0.00617 |
| **Cntn1** | 0.01374 | 0.00782 | 0.01163 | 1.13120 | 0.00942 | 0.00636 | 0.00733 | 0.01137 | 0.00000 | 0.00583 | 0.01142 | 0.00663 | 0.01279 | 0.00683 | 0.71118 | 0.00716 |
| **Cntn2** | 0.04979 | 0.01852 | 0.14167 | 0.50716 | 0.05263 | 0.01928 | 0.04773 | 0.00680 | 0.13918 | 0.08786 | 0.31422 | 0.02354 | 0.10624 | 0.14984 | 0.14418 | 0.04363 |
| **Cntn4** | 0.01372 | 0.00766 | 0.01168 | 0.61664 | 0.00936 | 0.00644 | 0.00740 | 0.01189 | 0.48363 | 0.00435 | 0.00838 | 0.00609 | 0.01279 | 0.01007 | 0.40266 | 0.00720 |
| **Cntn6** | 0.01375 | 0.00779 | 0.01185 | 1.01196 | 0.00937 | 0.00655 | 0.00747 | 0.01310 | 0.28101 | 0.00935 | 0.00988 | 0.00725 | 0.01283 | 0.01922 | 0.07502 | 0.00722 |

| | | | | | | | | | | | | | | | | |
|---|---|---|---|---|---|---|---|---|---|---|---|---|---|---|---|---|
| **Cntnap1** | 0.01375 | 0.00785 | 0.01123 | 5.48844 | 0.00925 | 0.00604 | 0.00720 | 0.01128 | 0.70120 | 0.03278 | 0.02671 | 0.02842 | 0.01199 | 0.00337 | 0.22532 | 0.00625 |
| **Cntrl** | 0.02400 | 0.00844 | 0.00000 | 2.68560 | 0.00839 | 0.04452 | 0.00767 | 0.22679 | 0.27797 | 0.58164 | 0.29103 | 0.02644 | 0.35868 | 0.02764 | 0.23568 | 0.00700 |
| **Cntrob** | 0.51753 | 2.00320 | 2.39022 | 1.90205 | 1.55940 | 1.31488 | 2.83530 | 1.45546 | 0.36087 | 0.00893 | 0.00236 | 0.03729 | 2.70722 | 0.00616 | 0.04179 | 0.01030 |
| **Coa3** | 10.49524 | 7.94164 | 9.00219 | 0.30225 | 5.17067 | 1.69628 | 4.65446 | 5.52023 | 0.19262 | 4.40550 | 3.98835 | 0.29086 | 9.20290 | 0.00767 | 0.33661 | 3.01743 |
| **Coa4** | 6.83360 | 5.31381 | 7.81417 | 0.00000 | 0.90293 | 5.70597 | 4.36557 | 3.30721 | 0.73421 | 2.55630 | 1.64271 | 2.00042 | 1.78555 | 0.15426 | 0.00000 | 3.05144 |
| **Coa5** | 7.08898 | 4.60458 | 13.13353 | 0.81909 | 4.20919 | 6.92944 | 9.42099 | 8.04711 | 0.51968 | 7.50843 | 3.99793 | 1.52435 | 7.38863 | 5.53011 | 5.15704 | 2.26749 |
| **Coa6** | 3.26242 | 0.00057 | 0.00000 | 3.60690 | 3.99683 | 0.80837 | 0.14718 | 3.49855 | 3.61501 | 3.66720 | 1.65710 | 0.16742 | 1.50857 | 0.00827 | 0.06353 | 1.41550 |
| **Coa7** | 8.49834 | 4.64793 | 6.85156 | 3.03249 | 4.93330 | 0.00896 | 3.73214 | 5.84369 | 1.06598 | 10.53337 | 5.16767 | 0.33751 | 6.73673 | 0.81789 | 8.11176 | 2.24877 |
| **Coasy** | 11.59501 | 8.61166 | 6.06892 | 0.50214 | 6.15241 | 4.51246 | 1.45096 | 5.60122 | 0.02507 | 3.49573 | 2.52208 | 0.08624 | 6.49371 | 0.20465 | 0.00000 | 3.06917 |
| **Cobl** | 0.01737 | 0.00776 | 0.01015 | 2.56659 | 0.00898 | 0.00638 | 0.00740 | 0.01195 | 0.28312 | 0.12691 | 0.00633 | 0.00459 | 0.01337 | 0.00446 | 1.62363 | 0.00694 |
| **Cobll1** | 0.01041 | 0.00842 | 0.00233 | 1.37027 | 0.00956 | 0.00698 | 0.00788 | 0.12778 | 0.07698 | 0.00000 | 0.60801 | 0.09248 | 0.00560 | 0.00339 | 1.29590 | 0.00687 |
| **Coch** | 0.01849 | 0.00837 | 0.01186 | 0.56030 | 0.01054 | 0.00671 | 0.00755 | 0.01413 | 0.04085 | 0.02454 | 0.01360 | 0.00864 | 0.01289 | 0.01082 | 0.00532 | 0.00737 |
| **Cog1** | 0.66483 | 3.11752 | 0.62725 | 0.10955 | 0.91371 | 0.02444 | 0.31280 | 4.67255 | 0.01857 | 0.72729 | 2.13443 | 0.06193 | 2.74430 | 0.03472 | 1.93813 | 2.15507 |
| **Cog2** | 4.00810 | 5.92188 | 4.14059 | 1.26473 | 9.52212 | 3.39777 | 1.77660 | 3.78875 | 2.72784 | 5.12342 | 3.61535 | 0.28128 | 1.24372 | 0.50196 | 2.10837 | 1.47493 |
| **Cog3** | 0.00000 | 0.00701 | 0.29345 | 2.78183 | 0.69358 | 0.00562 | 0.20395 | 1.58738 | 0.63722 | 0.78803 | 0.51129 | 0.21766 | 0.20521 | 0.30727 | 4.63353 | 2.96400 |
| **Cog4** | 9.70462 | 9.14774 | 8.57893 | 20.34117 | 0.61599 | 4.79658 | 7.25069 | 7.31467 | 20.13174 | 11.52001 | 9.20927 | 1.79710 | 7.12154 | 6.58599 | 15.61993 | 4.93534 |
| **Cog5** | 3.32462 | 2.64565 | 0.80688 | 0.40735 | 0.01141 | 4.08938 | 0.05223 | 0.77471 | 0.29756 | 0.96111 | 2.87919 | 0.04581 | 0.00000 | 0.92687 | 1.83840 | 0.00000 |
| **Cog6** | 1.72805 | 0.64315 | 0.00000 | 0.50146 | 0.00780 | 0.00367 | 3.69036 | 0.00000 | 0.00581 | 2.39535 | 1.64294 | 5.95606 | 2.72182 | 0.03534 | 0.30268 | 0.89082 |
| **Cog7** | 6.42390 | 6.02572 | 2.87050 | 2.34459 | 0.75211 | 2.87516 | 3.73671 | 3.81417 | 1.75696 | 3.22214 | 3.55657 | 0.01792 | 1.55434 | 0.05540 | 0.09395 | 2.02736 |
| **Cog8** | 0.00000 | 0.00688 | 0.04180 | 0.39301 | 1.39470 | 1.26325 | 0.26984 | 1.39295 | 0.27620 | 2.36043 | 0.01666 | 0.04295 | 1.31281 | 0.00965 | 0.00000 | 5.67205 |
| **Coil** | 5.15725 | 7.80943 | 13.40967 | 4.24961 | 5.79028 | 6.66224 | 7.62173 | 3.29219 | 7.69595 | 4.64241 | 3.64564 | 3.16431 | 13.68597 | 0.00000 | 0.00000 | 5.27627 |
| **Col10a1** | 0.05574 | 0.00824 | 0.20016 | 0.00350 | 0.01337 | 0.44371 | 0.00732 | 0.16948 | 0.51883 | 0.00000 | 0.14957 | 0.23064 | 0.00626 | 0.63012 | 0.00000 | 0.00722 |
| **Col11a2** | 0.02000 | 0.03976 | 0.01143 | 0.00611 | 0.00877 | 0.00670 | 0.00752 | 0.01317 | 0.09328 | 0.01463 | 0.01218 | 0.00795 | 0.01267 | 0.01239 | 0.00930 | 0.00737 |
| **Col12a1** | 0.01377 | 0.00761 | 0.00775 | 0.01060 | 0.00921 | 0.00682 | 0.00726 | 0.01115 | 0.01201 | 0.03313 | 0.00754 | 0.73668 | 0.01265 | 0.01078 | 0.00544 | 0.00722 |
| **Col13a1** | 0.15182 | 0.00877 | 0.00786 | 0.76831 | 0.01709 | 0.00689 | 0.00740 | 0.01298 | 0.03745 | 0.01479 | 0.01289 | 0.00832 | 0.01237 | 0.01091 | 0.00426 | 0.02237 |
| **Col16a1** | 0.01038 | 0.00764 | 0.00955 | 0.00895 | 0.00940 | 0.00681 | 0.00715 | 0.16231 | 0.01193 | 0.31076 | 0.02485 | 0.00605 | 0.00775 | 0.01140 | 0.00794 | 0.00781 |
| **Col18a1** | 0.01123 | 0.00759 | 0.00824 | 0.00721 | 0.00943 | 0.00690 | 0.00712 | 0.42016 | 0.07648 | 0.03408 | 0.00985 | 0.01572 | 0.00616 | 0.01287 | 0.00444 | 0.00758 |
| **Col19a1** | 0.01372 | 0.00824 | 0.01236 | 0.29755 | 0.00924 | 0.00655 | 0.00752 | 0.01293 | 0.00564 | 0.12559 | 0.01204 | 0.00784 | 0.01177 | 0.01043 | 0.00643 | 0.00713 |
| **Col1a1** | 0.01248 | 0.00696 | 0.00640 | 0.01744 | 0.00872 | 0.00660 | 0.00689 | 0.00962 | 0.01157 | 1.54639 | 0.00000 | 1.86197 | 0.01142 | 0.01059 | 0.00000 | 0.00755 |
| **Col1a2** | 0.01303 | 0.00730 | 0.00717 | 0.00650 | 0.00893 | 0.00669 | 0.00702 | 0.00947 | 0.01100 | 1.22595 | 0.00535 | 0.90225 | 0.01101 | 0.03422 | 0.00000 | 0.00730 |
| **Col20a1** | 0.00776 | 0.00782 | 0.01154 | 0.84550 | 0.00910 | 0.00696 | 0.00717 | 0.03494 | 0.00000 | 0.17020 | 0.01418 | 0.09382 | 0.50438 | 0.01161 | 0.00433 | 0.35252 |
| **Col22a1** | 0.01360 | 0.00829 | 0.01199 | 1.67520 | 0.00929 | 0.00645 | 0.00748 | 0.01385 | 0.98475 | 0.00965 | 0.00712 | 0.00677 | 0.01288 | 0.01062 | 0.00288 | 0.00707 |
| **Col23a1** | 0.01346 | 0.00827 | 0.01195 | 0.03485 | 0.00922 | 0.00689 | 0.00775 | 0.01789 | 0.01237 | 0.14692 | 0.01255 | 0.00910 | 0.01271 | 0.01171 | 0.01133 | 0.00741 |
| **Col24a1** | 0.01376 | 0.00798 | 0.01213 | 0.45052 | 0.00932 | 0.00662 | 0.00752 | 0.01375 | 0.05395 | 0.01444 | 0.01263 | 0.00801 | 0.01272 | 0.01442 | 0.01293 | 0.00734 |
| **Col26a1** | 0.74452 | 0.23446 | 0.00000 | 0.18278 | 0.54866 | 1.84379 | 0.00641 | 0.00537 | 0.00803 | 0.04724 | 0.02341 | 0.04021 | 0.03830 | 0.54325 | 0.04952 | 1.71933 |
| **Col2a1** | 0.01383 | 0.00797 | 0.01117 | 0.11621 | 0.00930 | 0.00669 | 0.00745 | 0.01268 | 0.12190 | 0.01304 | 0.01274 | 0.06866 | 0.01284 | 0.01187 | 0.00561 | 0.00740 |
| **Col3a1** | 0.01357 | 0.00775 | 0.00998 | 0.00879 | 0.00925 | 0.00670 | 0.00734 | 0.01191 | 0.01207 | 0.45622 | 0.01110 | 0.15876 | 0.01197 | 0.01202 | 0.00504 | 0.00747 |
| **Col4a1** | 9.96599 | 0.01158 | 4.67291 | 0.21457 | 7.96170 | 4.04079 | 1.94310 | 0.36988 | 0.27862 | 0.00000 | 1.53356 | 0.45037 | 3.52000 | 0.00353 | 0.00000 | 10.85144 |
| **Col4a2** | 12.22565 | 1.72598 | 5.54294 | 0.04029 | 5.48241 | 5.57596 | 4.58600 | 2.55419 | 0.15412 | 0.00000 | 1.35348 | 0.34566 | 10.12848 | 0.57402 | 0.00000 | 11.88478 |
| **Col4a3bp** | 1.74572 | 0.53094 | 2.75177 | 13.68021 | 3.79480 | 2.18832 | 1.85701 | 2.45966 | 10.71010 | 0.30752 | 3.39366 | 12.46011 | 1.27590 | 6.21643 | 7.54314 | 0.54832 |
| **Col5a1** | 0.44707 | 0.00767 | 0.00262 | 0.01072 | 0.01624 | 0.00787 | 0.00682 | 0.00855 | 0.01212 | 0.00859 | 0.00953 | 0.48319 | 0.01265 | 0.01113 | 0.00631 | 0.00725 |
| **Col5a2** | 0.00888 | 0.00824 | 0.00747 | 0.01103 | 0.00960 | 0.00689 | 0.00813 | 0.53217 | 0.01144 | 0.00315 | 0.00521 | 0.57335 | 0.00595 | 0.01370 | 0.00000 | 0.00749 |
| **Col5a3** | 0.01414 | 0.00727 | 0.54156 | 1.54836 | 0.02178 | 0.02314 | 0.00368 | 0.01043 | 0.72596 | 0.54757 | 7.48398 | 0.09788 | 1.68872 | 0.01257 | 0.25782 | 0.00512 |
| **Col6a1** | 0.01382 | 0.00781 | 0.01086 | 0.01000 | 0.00921 | 0.00675 | 0.00734 | 0.01151 | 0.01220 | 0.14088 | 0.01265 | 0.08296 | 0.01293 | 0.01134 | 0.00923 | 0.00776 |
| **Col6a2** | 0.01370 | 0.00803 | 0.01170 | 0.00950 | 0.00920 | 0.00658 | 0.00744 | 0.01175 | 0.01232 | 0.25800 | 0.01161 | 0.03726 | 0.01157 | 0.01143 | 0.00909 | 0.00720 |
| **Col6a3** | 0.01366 | 0.00762 | 0.00858 | 1.41557 | 0.00926 | 0.00649 | 0.00719 | 0.01003 | 1.90712 | 0.00000 | 0.00000 | 0.36397 | 0.01329 | 0.01650 | 0.12872 | 0.00697 |
| **Col6a4** | 0.00786 | 0.00785 | 0.00000 | 0.01144 | 0.01117 | 0.00766 | 0.00745 | 2.27866 | 0.32778 | 5.26550 | 0.85177 | 0.00000 | 0.00502 | 0.81439 | 0.70445 | 0.00727 |
| **Col6a5** | 0.00748 | 0.00852 | 1.21649 | 0.37500 | 0.00860 | 0.00581 | 0.00843 | 0.41860 | 0.06038 | 0.00000 | 0.00713 | 0.00736 | 0.00452 | 0.03350 | 0.31857 | 0.00696 |
| **Col6a6** | 0.01379 | 0.00827 | 0.01004 | 0.03526 | 0.00911 | 0.00690 | 0.00750 | 0.00730 | 0.09566 | 0.09531 | 0.21985 | 0.00793 | 0.01041 | 0.00616 | 0.26737 | 0.00733 |
| **Col7a1** | 2.18020 | 0.00472 | 0.22670 | 0.00190 | 1.46537 | 0.00641 | 0.00528 | 0.00429 | 0.85571 | 0.00000 | 0.00621 | 0.03477 | 0.95060 | 0.01796 | 0.00127 | 0.01157 |
| **Col8a2** | 0.01359 | 0.00775 | 0.00833 | 0.18787 | 0.00915 | 0.00660 | 0.00726 | 0.00636 | 0.59054 | 0.12413 | 0.13934 | 0.04329 | 0.01125 | 0.06464 | 0.95822 | 0.00755 |
| **Col9a2** | 0.27912 | 0.00722 | 0.11772 | 0.00316 | 0.04763 | 0.00559 | 0.48363 | 2.08341 | 0.00856 | 0.10695 | 5.79438 | 0.00000 | 2.07855 | 0.46357 | 0.18744 | 0.00521 |
| **Col9a3** | 0.01376 | 0.00820 | 0.01212 | 0.79972 | 0.00930 | 0.00657 | 0.00750 | 0.01350 | 0.00000 | 0.01448 | 0.03101 | 0.00832 | 0.01226 | 0.01064 | 0.00361 | 0.00727 |
| **Colec12** | 0.62271 | 0.21288 | 1.33667 | 24.91075 | 0.06710 | 0.18635 | 0.66787 | 0.30730 | 16.55414 | 4.86957 | 1.00010 | 1.25404 | 0.53806 | 10.24194 | 15.12690 | 0.21860 |
| **Colgalt1** | 5.10321 | 4.48566 | 3.43048 | 5.51970 | 0.40090 | 0.01518 | 1.22678 | 1.77637 | 3.00958 | 6.15910 | 3.41840 | 0.24044 | 2.99870 | 3.09672 | 6.28338 | 6.71445 |
| **Colgalt2** | 0.01317 | 0.00797 | 0.00985 | 2.97698 | 0.00928 | 0.00621 | 0.00722 | 0.00945 | 2.62133 | 0.00000 | 0.01713 | 0.00000 | 0.01272 | 0.28483 | 0.10532 | 0.00664 |
| **Colq** | 0.01322 | 0.00816 | 0.01039 | 0.23314 | 0.00904 | 0.01321 | 0.00756 | 0.01277 | 1.03050 | 0.53604 | 0.02125 | 0.00549 | 0.01203 | 0.00480 | 0.47022 | 0.00729 |
| **Commd1** | 9.72703 | 2.68974 | 2.77725 | 0.56393 | 8.03143 | 3.25858 | 2.19020 | 2.79102 | 0.40289 | 2.86134 | 1.98330 | 0.16377 | 2.21351 | 1.07582 | 0.14712 | 6.25798 |
| **Commd10** | 11.97475 | 2.25057 | 7.42835 | 0.00000 | 7.64223 | 3.24785 | 0.96628 | 3.20484 | 0.58052 | 4.97277 | 2.25102 | 0.75076 | 6.90504 | 3.39738 | 2.88386 | 2.77404 |
| **Commd2** | 11.96751 | 6.42275 | 7.17010 | 8.76967 | 8.27815 | 7.13892 | 4.94830 | 8.97678 | 9.58385 | 13.59431 | 6.91108 | 1.54185 | 11.54824 | 2.31657 | 3.22802 | 7.93682 |
| **Commd3** | 11.76436 | 5.69222 | 6.76107 | 3.65554 | 4.96265 | 7.59496 | 3.01947 | 1.55397 | 0.00000 | 3.32055 | 2.03966 | 0.31454 | 8.91951 | 0.00000 | 0.00000 | 3.66160 |
| **Commd4** | 16.72419 | 7.04833 | 5.29125 | 2.22833 | 2.75422 | 3.31761 | 7.59433 | 3.85885 | 0.34607 | 4.97348 | 4.90491 | 0.31090 | 4.53194 | 0.02843 | 0.36843 | 7.02084 |

| | | | | | | | | | | | | | | | | |
|---|---|---|---|---|---|---|---|---|---|---|---|---|---|---|---|---|
| **Commd5** | 10.99913 | 0.42196 | 1.79556 | 1.35772 | 4.51082 | 1.17956 | 5.07481 | 19.64299 | 11.53198 | 17.78997 | 8.58891 | 0.77312 | 14.46063 | 0.47164 | 1.79636 | 6.77553 |
| **Commd6** | 5.32081 | 0.57793 | 3.16561 | 3.67215 | 2.69703 | 2.05138 | 0.18278 | 2.49573 | 0.61612 | 3.92599 | 1.89913 | 0.33166 | 2.10055 | 1.23664 | 2.66398 | 3.55384 |
| **Commd7** | 2.81884 | 1.03186 | 0.91531 | 9.77155 | 0.42249 | 2.49512 | 0.30615 | 2.73418 | 10.94159 | 9.79453 | 8.85189 | 5.60591 | 3.24246 | 1.66911 | 4.43682 | 2.57836 |
| **Commd8** | 6.80824 | 10.87614 | 8.43549 | 0.00000 | 6.67098 | 5.92316 | 7.04838 | 6.19540 | 1.13102 | 4.55258 | 2.01810 | 0.31886 | 5.66609 | 2.65685 | 1.48164 | 5.07277 |
| **Commd9** | 5.05444 | 3.78122 | 10.20090 | 0.28589 | 3.86512 | 0.35106 | 7.35354 | 4.37188 | 0.70503 | 6.07875 | 2.36107 | 0.31935 | 4.72492 | 0.07104 | 0.34840 | 0.00000 |
| **Comp** | 0.01140 | 0.01974 | 0.46374 | 0.02554 | 0.01167 | 0.00602 | 0.00760 | 1.66850 | 0.38704 | 0.03281 | 0.00000 | 0.20626 | 0.01498 | 0.00790 | 0.06109 | 0.00757 |
| **Comt** | 5.03179 | 0.62379 | 1.92503 | 7.76087 | 2.63828 | 2.67722 | 0.47479 | 2.89245 | 12.23368 | 1.87511 | 10.63665 | 1.22574 | 2.72343 | 4.94091 | 3.60293 | 0.66462 |
| **Comtd1** | 5.62805 | 0.01112 | 0.02302 | 0.01229 | 0.00785 | 0.01038 | 0.00046 | 0.21207 | 0.84701 | 0.03366 | 1.27288 | 0.00000 | 1.22537 | 0.00906 | 0.00000 | 1.55410 |
| **Copa** | 7.97783 | 3.06908 | 9.75525 | 7.51228 | 7.74769 | 10.22134 | 2.51819 | 5.08488 | 6.01023 | 12.33765 | 8.76985 | 13.54706 | 8.54002 | 11.02188 | 7.90516 | 9.48208 |
| **Copb1** | 2.23522 | 8.27560 | 3.44481 | 1.55965 | 4.72802 | 1.56539 | 0.06474 | 2.39634 | 1.94013 | 2.00879 | 0.52946 | 3.04117 | 1.10902 | 1.61695 | 0.91570 | 0.21633 |
| **Copb2** | 12.76603 | 12.58202 | 13.31452 | 6.84797 | 17.68732 | 5.55028 | 10.74035 | 12.63356 | 5.47346 | 14.70021 | 6.78657 | 4.80369 | 8.79543 | 12.27209 | 10.88810 | 6.99523 |
| **Cope** | 16.63204 | 10.16972 | 16.11112 | 2.47601 | 12.96441 | 3.97669 | 9.04344 | 15.92356 | 3.14677 | 18.51455 | 9.10587 | 1.04071 | 18.98867 | 0.08889 | 0.00000 | 7.62390 |
| **Copg1** | 13.54399 | 12.43043 | 11.07654 | 2.62839 | 11.57531 | 8.87039 | 7.67719 | 10.01902 | 0.81178 | 8.27843 | 5.04472 | 3.06521 | 9.80546 | 4.98474 | 4.36346 | 6.21262 |
| **Copg2** | 3.56388 | 0.19217 | 1.61139 | 2.70492 | 0.00000 | 0.26462 | 0.00418 | 0.00000 | 0.26039 | 0.00000 | 0.67578 | 0.01042 | 0.03380 | 0.00656 | 0.06918 | 0.00957 |
| **Coprs** | 3.01585 | 0.65750 | 0.90595 | 1.72957 | 0.01498 | 0.00296 | 0.93917 | 1.86885 | 1.94515 | 6.73071 | 6.53409 | 0.60114 | 5.32603 | 0.34105 | 0.00000 | 4.68445 |
| **Cops2** | 12.54927 | 9.72510 | 13.82709 | 5.51502 | 12.96621 | 0.35634 | 6.08406 | 10.12780 | 9.53725 | 14.65601 | 10.25869 | 11.90905 | 11.00294 | 23.34994 | 13.80964 | 4.47744 |
| **Cops3** | 12.69641 | 11.41997 | 13.36310 | 1.07803 | 5.12059 | 10.49761 | 7.23497 | 13.65686 | 1.67205 | 10.14295 | 2.36370 | 10.50368 | 9.47201 | 4.09472 | 9.54496 | 8.32503 |
| **Cops4** | 22.98032 | 16.08729 | 15.61882 | 7.31890 | 17.68646 | 6.22682 | 9.12139 | 16.54825 | 11.20120 | 25.19038 | 16.72286 | 18.01830 | 22.11781 | 5.52389 | 16.90904 | 10.20455 |
| **Cops5** | 24.83277 | 15.58041 | 22.21875 | 0.93892 | 17.00248 | 13.34884 | 14.64121 | 19.47619 | 4.17835 | 14.67865 | 13.93791 | 10.60372 | 17.76105 | 15.76980 | 6.78065 | 14.63444 |
| **Cops6** | 17.63522 | 8.27594 | 12.07431 | 6.32957 | 11.96746 | 4.07516 | 6.07749 | 9.41109 | 19.53711 | 14.49221 | 16.68382 | 9.78208 | 12.05947 | 6.47285 | 7.97927 | 6.13788 |
| **Cops7a** | 18.88998 | 8.77985 | 11.23845 | 12.78502 | 19.19116 | 12.73443 | 8.13256 | 26.81982 | 16.76358 | 12.66867 | 15.78605 | 2.94448 | 13.71729 | 6.87765 | 12.40598 | 9.33094 |
| **Cops7b** | 0.69324 | 0.02658 | 0.11707 | 0.56344 | 3.05697 | 0.02277 | 2.60025 | 6.51109 | 0.46954 | 8.09838 | 1.56957 | 0.17400 | 4.76967 | 0.55226 | 1.26475 | 0.64220 |
| **Cops8** | 13.92547 | 7.63348 | 11.82866 | 7.85647 | 2.29695 | 6.94021 | 9.18723 | 13.12489 | 6.16511 | 12.10107 | 8.72388 | 11.23543 | 12.00908 | 2.90595 | 11.02691 | 6.03725 |
| **Copz1** | 20.16456 | 15.47970 | 25.03487 | 3.89080 | 9.10442 | 8.36157 | 12.31750 | 19.82528 | 4.94999 | 17.23888 | 12.70079 | 8.02956 | 18.82694 | 17.97955 | 9.57118 | 10.76073 |
| **Copz2** | 0.01132 | 0.00779 | 0.09540 | 0.62810 | 0.00928 | 0.00667 | 0.00728 | 0.04613 | 0.11863 | 0.13364 | 0.01111 | 0.04582 | 0.01730 | 0.00877 | 0.06200 | 0.00735 |
| **Coq10a** | 0.12935 | 0.00853 | 0.00788 | 1.64886 | 0.01586 | 0.00693 | 0.00725 | 0.00631 | 0.05678 | 0.07113 | 0.19671 | 0.03973 | 0.39535 | 0.24413 | 1.02556 | 0.00720 |
| **Coq10b** | 1.82274 | 0.00000 | 0.91262 | 3.54649 | 3.13490 | 0.01560 | 3.62387 | 5.62385 | 2.17213 | 4.25570 | 1.79147 | 2.46568 | 2.71101 | 0.00000 | 2.00339 | 2.07689 |
| **Coq2** | 2.60455 | 2.10632 | 1.83843 | 2.74171 | 0.01096 | 4.55976 | 0.43296 | 1.76714 | 0.64533 | 0.53279 | 0.64144 | 1.61148 | 2.81319 | 0.00000 | 1.20918 | 1.95251 |
| **Coq3** | 3.91803 | 5.00984 | 7.10201 | 4.57060 | 7.36061 | 1.27712 | 8.68645 | 3.71578 | 2.63214 | 7.62945 | 7.38876 | 0.18961 | 4.57763 | 4.94807 | 5.41375 | 4.14638 |
| **Coq4** | 5.21450 | 0.06090 | 2.17012 | 8.73058 | 4.95024 | 2.70085 | 0.17219 | 3.86054 | 15.02867 | 12.27922 | 4.28821 | 0.07662 | 4.54212 | 0.00000 | 0.54041 | 3.95257 |
| **Coq5** | 3.19521 | 1.56250 | 3.54742 | 1.11447 | 5.05438 | 6.66447 | 5.06862 | 2.21045 | 1.24222 | 5.74992 | 4.96348 | 0.49510 | 5.53967 | 0.70864 | 1.35563 | 2.18891 |
| **Coq6** | 9.51334 | 7.60679 | 6.70860 | 2.67583 | 3.41346 | 7.94936 | 3.62738 | 13.17692 | 6.57753 | 7.52842 | 4.83873 | 0.65943 | 9.40941 | 0.07881 | 1.49397 | 2.89313 |
| **Coq7** | 11.56072 | 7.35499 | 9.85718 | 1.94105 | 1.70804 | 7.85199 | 2.45775 | 6.62984 | 1.19704 | 8.15145 | 7.83183 | 0.43290 | 10.03307 | 0.76553 | 0.46017 | 8.19705 |
| **Coq8a** | 0.02337 | 0.00736 | 0.00318 | 0.70470 | 0.00918 | 0.00679 | 0.74447 | 0.02597 | 0.00000 | 0.00000 | 0.19600 | 0.00394 | 1.65403 | 0.00568 | 0.78879 | 0.01403 |
| **Coq8b** | 2.07602 | 0.00450 | 0.10675 | 1.44614 | 0.00303 | 2.91007 | 0.07677 | 4.09014 | 0.15749 | 2.45106 | 1.50612 | 0.07166 | 1.25903 | 0.00000 | 0.52349 | 2.40645 |
| **Coq9** | 5.80866 | 6.80962 | 4.57416 | 1.95338 | 4.46428 | 1.61454 | 0.68555 | 5.75488 | 0.30462 | 3.86344 | 1.99367 | 0.20791 | 6.24996 | 3.03338 | 4.01586 | 5.14402 |
| **Coro1a** | 0.01368 | 0.00746 | 0.02393 | 0.22704 | 0.00860 | 0.09055 | 0.00705 | 0.00309 | 0.00800 | 3.30551 | 0.22554 | 0.18846 | 0.02852 | 0.00791 | 0.02138 | 0.00735 |
| **Coro1b** | 11.91347 | 5.11613 | 8.69580 | 2.29810 | 4.48925 | 6.03620 | 8.79053 | 4.93721 | 2.97936 | 5.46136 | 2.72424 | 0.13475 | 7.00539 | 0.01267 | 0.00000 | 7.29927 |
| **Coro1c** | 16.34384 | 7.96528 | 19.94434 | 3.29651 | 10.34304 | 4.41856 | 8.11714 | 21.18275 | 4.83582 | 12.22491 | 8.78686 | 1.23991 | 15.44932 | 0.18420 | 0.22117 | 11.94528 |
| **Coro2a** | 1.28108 | 0.00657 | 0.67005 | 0.00275 | 0.03416 | 0.60906 | 1.06025 | 1.60375 | 0.04375 | 0.20546 | 1.07337 | 0.30275 | 0.87896 | 0.10232 | 0.57206 | 0.00550 |
| **Coro2b** | 1.42093 | 0.00448 | 0.65118 | 0.96071 | 3.39836 | 0.00749 | 0.00598 | 0.02975 | 0.27141 | 0.02030 | 0.00884 | 0.02207 | 0.00699 | 0.00976 | 0.00000 | 0.00525 |
| **Coro7** | 2.78275 | 0.44136 | 0.65044 | 0.33981 | 0.00000 | 0.91374 | 0.00341 | 2.38001 | 0.00524 | 0.00000 | 0.26760 | 0.19402 | 2.47586 | 0.00460 | 0.07558 | 6.11011 |
| **Cort** | 0.01320 | 0.00787 | 0.00901 | 0.20057 | 0.01831 | 0.00680 | 0.00736 | 0.00288 | 0.37802 | 0.12965 | 0.39195 | 0.01483 | 0.00884 | 0.00795 | 0.00000 | 0.00752 |
| **Cotl1** | 4.90087 | 0.03575 | 0.54316 | 3.13486 | 0.02106 | 0.00000 | 0.00213 | 1.75614 | 1.82957 | 1.29763 | 2.79186 | 0.14660 | 1.68987 | 0.00000 | 0.00000 | 0.63507 |
| **Cox10** | 1.14945 | 0.09641 | 5.86354 | 6.96750 | 0.14172 | 2.68792 | 3.56867 | 6.04035 | 2.00286 | 5.26349 | 2.84964 | 0.22019 | 3.43011 | 0.51149 | 1.34658 | 3.03040 |
| **Cox11** | 1.49080 | 0.03931 | 0.00000 | 2.40092 | 0.31960 | 0.00457 | 0.56446 | 0.19515 | 2.84744 | 2.65544 | 2.12422 | 1.15864 | 1.85744 | 1.46046 | 1.11835 | 0.01906 |
| **Cox14** | 10.21008 | 0.08917 | 1.45749 | 6.74409 | 7.43822 | 3.09000 | 0.24492 | 2.70398 | 7.72962 | 5.76319 | 5.56947 | 0.24693 | 6.99622 | 0.02520 | 0.15099 | 2.05888 |
| **Cox15** | 0.85260 | 2.47413 | 1.83487 | 0.90212 | 0.02863 | 0.37171 | 0.02514 | 2.56853 | 0.37934 | 1.01354 | 1.54503 | 2.94099 | 1.75292 | 3.58755 | 3.36429 | 2.49379 |
| **Cox16** | 1.14888 | 0.01602 | 1.03433 | 0.08690 | 0.65588 | 0.31806 | 0.20774 | 0.38928 | 0.02726 | 1.13385 | 0.63679 | 0.07705 | 0.26500 | 0.00800 | 0.00000 | 1.19749 |
| **Cox17** | 8.02627 | 0.18859 | 2.66996 | 0.30582 | 4.92831 | 2.22708 | 0.42799 | 0.32050 | 1.89887 | 1.53498 | 10.22684 | 0.37013 | 2.90046 | 0.00000 | 0.10446 | 2.75845 |
| **Cox18** | 2.75894 | 4.29821 | 6.46857 | 4.52204 | 0.93549 | 1.25323 | 1.70715 | 2.32915 | 1.22047 | 6.45044 | 5.17885 | 0.81946 | 3.30014 | 0.04271 | 0.78989 | 3.34159 |
| **Cox19** | 5.71297 | 3.95061 | 2.52522 | 0.00000 | 0.22659 | 0.85878 | 1.57738 | 2.30613 | 0.33794 | 5.15317 | 1.62787 | 0.83620 | 1.99866 | 0.04466 | 0.43801 | 0.01209 |
| **Cox20** | 2.98629 | 0.08522 | 2.17572 | 1.14383 | 2.61750 | 0.31569 | 0.04341 | 0.40009 | 0.26516 | 0.72407 | 2.20733 | 3.46262 | 1.80412 | 0.87741 | 0.22136 | 1.47136 |
| **Cox4i1** | 31.59342 | 11.40002 | 24.66997 | 14.79043 | 20.82856 | 12.85572 | 13.99030 | 21.50438 | 18.84510 | 23.55651 | 23.96660 | 4.48825 | 30.72279 | 2.02257 | 3.08023 | 20.52233 |
| **Cox4i2** | 0.00754 | 0.00766 | 0.00164 | 2.29373 | 0.00931 | 0.00738 | 0.00693 | 0.19324 | 2.35786 | 2.24498 | 0.91710 | 0.22941 | 0.00934 | 0.03826 | 0.88803 | 0.00698 |
| **Cox5a** | 12.14909 | 3.44435 | 11.77826 | 17.38536 | 11.93703 | 4.68203 | 4.40928 | 0.20376 | 27.87095 | 23.25399 | 23.87168 | 1.12063 | 12.64024 | 6.33096 | 1.64330 | 6.73288 |
| **Cox5b** | 6.44982 | 0.09941 | 3.53836 | 7.91980 | 7.61471 | 2.18758 | 0.10355 | 0.00000 | 11.60768 | 7.42763 | 10.93243 | 0.07321 | 10.27912 | 0.00000 | 0.00000 | 6.86516 |
| **Cox6a1** | 14.63895 | 0.27823 | 6.97045 | 13.78761 | 13.44454 | 6.10528 | 0.53177 | 2.63852 | 18.76172 | 16.53885 | 23.05064 | 0.82491 | 13.53466 | 0.93405 | 0.00000 | 11.45074 |
| **Cox6b1** | 15.28463 | 1.43437 | 11.13156 | 6.79580 | 5.96561 | 4.77225 | 1.85431 | 0.84726 | 10.99133 | 10.27655 | 16.64879 | 0.86515 | 10.68495 | 5.44067 | 4.85100 | 9.56581 |
| **Cox6b2** | 1.68789 | 0.00563 | 0.13980 | 16.33374 | 0.00523 | 0.00226 | 0.00498 | 0.00000 | 3.25152 | 0.50776 | 1.51525 | 0.11547 | 0.00000 | 0.31480 | 2.76842 | 0.00219 |
| **Cox6c** | 12.91490 | 0.66912 | 13.28857 | 13.48316 | 13.10580 | 4.21906 | 1.42344 | 1.37875 | 13.18566 | 6.28338 | 27.79697 | 1.97644 | 6.12935 | 3.89299 | 1.89511 | 6.68972 |
| **Cox7a1** | 1.01555 | 0.00669 | 0.05909 | 0.76456 | 0.00648 | 0.22040 | 0.00620 | 0.00000 | 0.67635 | 0.00000 | 1.35572 | 0.00964 | 0.23460 | 0.02134 | 0.07357 | 0.54006 |

| | | | | | | | | | | | | | | | | |
|---|---|---|---|---|---|---|---|---|---|---|---|---|---|---|---|---|
| **Cox7a2** | 10.27101 | 0.65700 | 9.74543 | 13.06445 | 7.26521 | 1.51687 | 1.27140 | 0.24754 | 14.33622 | 5.95595 | 18.31562 | 1.20070 | 4.56832 | 5.06970 | 2.91785 | 4.24068 |
| **Cox7a2l** | 24.83908 | 13.48869 | 23.53745 | 3.84680 | 7.85859 | 9.20758 | 14.37920 | 24.11580 | 3.09742 | 24.67049 | 12.23197 | 4.68269 | 23.42887 | 0.96807 | 7.93274 | 10.74579 |
| **Cox7b** | 11.57372 | 0.94247 | 10.75756 | 10.95299 | 8.09967 | 1.87824 | 0.77542 | 0.72568 | 6.92595 | 3.41070 | 7.81517 | 0.35725 | 5.03368 | 4.13697 | 2.25537 | 3.24940 |
| **Cox7b2** | 0.01216 | 0.00755 | 0.01074 | 0.01079 | 0.00951 | 0.00664 | 0.00728 | 0.06099 | 0.01174 | 0.00898 | 0.01205 | 0.00726 | 0.01006 | 0.01905 | 0.12379 | 0.00739 |
| **Cox7c** | 4.73596 | 0.21581 | 3.57142 | 0.46160 | 2.97479 | 0.87945 | 0.78197 | 0.50764 | 5.42497 | 1.27948 | 11.95976 | 0.48706 | 2.56863 | 0.47693 | 0.65359 | 3.60649 |
| **Cox8a** | 11.99679 | 0.03695 | 3.71897 | 13.62593 | 12.16462 | 2.10991 | 0.16307 | 0.33055 | 17.02065 | 18.03093 | 19.17695 | 0.37803 | 11.03536 | 0.46464 | 0.00000 | 7.55891 |
| **Cox8c** | 0.01348 | 0.00773 | 0.01199 | 0.00913 | 0.00924 | 0.00657 | 0.00742 | 0.01272 | 0.01193 | 0.45198 | 0.01217 | 0.00629 | 0.01193 | 0.01573 | 0.03150 | 0.00757 |
| **Cp** | 0.01213 | 0.00797 | 0.08201 | 0.00973 | 0.00901 | 0.00919 | 0.00752 | 0.01998 | 0.01183 | 0.18486 | 0.01077 | 0.00797 | 0.01149 | 0.08299 | 0.08302 | 0.00739 |
| **Cpa1** | 0.00333 | 0.00793 | 0.00000 | 15.72877 | 0.00905 | 0.00965 | 0.23313 | 1.65527 | 7.18243 | 1.31351 | 2.01884 | 0.24859 | 0.00000 | 0.00000 | 11.41724 | 0.00371 |
| **Cpa2** | 0.05070 | 0.00784 | 0.00626 | 0.35531 | 0.00791 | 0.00692 | 0.00723 | 0.00000 | 0.34567 | 0.00000 | 0.61692 | 0.04425 | 0.00663 | 0.00423 | 0.22708 | 0.00687 |
| **Cpa4** | 0.01358 | 0.00806 | 0.01080 | 0.05708 | 0.00935 | 0.00670 | 0.00739 | 0.01148 | 0.29478 | 0.00953 | 0.01095 | 0.00652 | 0.01311 | 0.20560 | 0.00000 | 0.00744 |
| **Cpb2** | 0.01383 | 0.00823 | 0.00918 | 1.54772 | 0.00911 | 0.00671 | 0.00754 | 0.00693 | 0.00000 | 0.00609 | 0.42343 | 0.00641 | 0.00791 | 0.00548 | 0.00000 | 0.00691 |
| **Cpd** | 3.24492 | 0.00414 | 0.23689 | 8.25702 | 0.01153 | 0.19499 | 0.04638 | 0.00000 | 6.06904 | 1.54560 | 0.22569 | 0.02858 | 1.25408 | 0.00000 | 4.85632 | 0.09142 |
| **Cpe** | 3.02959 | 0.00455 | 1.15205 | 0.32767 | 0.23674 | 0.00243 | 0.00464 | 0.00000 | 0.01068 | 1.79737 | 0.89799 | 0.00746 | 0.00868 | 0.00521 | 0.00000 | 0.01880 |
| **Cpeb1** | 0.00534 | 0.00808 | 0.37939 | 4.86571 | 0.00927 | 0.00535 | 0.00708 | 0.11506 | 1.56275 | 1.00774 | 0.64232 | 0.49018 | 0.01055 | 0.85191 | 2.34510 | 0.00659 |
| **Cpeb2** | 2.11266 | 0.00000 | 1.33325 | 2.47443 | 0.33334 | 3.31516 | 1.55471 | 1.60970 | 1.57412 | 0.05074 | 1.56757 | 13.44886 | 0.44618 | 6.10219 | 3.98299 | 0.37617 |
| **Cpeb3** | 1.29146 | 0.00559 | 0.22586 | 2.33377 | 0.01346 | 0.19641 | 0.00451 | 1.36887 | 2.63712 | 3.69181 | 2.62183 | 1.45287 | 1.02512 | 6.33783 | 3.94164 | 0.01114 |
| **Cpeb4** | 0.11343 | 0.10153 | 0.69388 | 8.62833 | 0.00696 | 0.00418 | 0.35383 | 0.12946 | 7.93208 | 0.00000 | 1.80068 | 1.71104 | 0.12133 | 4.27197 | 1.62372 | 0.00456 |
| **Cplx1** | 0.00704 | 2.15672 | 0.00326 | 0.39943 | 0.00809 | 0.89438 | 0.00637 | 0.00912 | 0.00767 | 0.32309 | 0.01081 | 0.00573 | 0.00622 | 0.00380 | 0.17721 | 0.00614 |
| **Cplx2** | 0.01806 | 0.02827 | 0.05253 | 0.18615 | 0.00879 | 0.02322 | 0.07820 | 0.02470 | 0.23375 | 0.08116 | 0.02151 | 0.02443 | 0.03016 | 0.31029 | 0.10005 | 0.00726 |
| **Cpm** | 0.00557 | 0.00785 | 0.00000 | 0.91015 | 0.02998 | 0.30785 | 0.00579 | 0.37436 | 3.15663 | 0.59406 | 1.34684 | 0.13672 | 1.52628 | 0.50652 | 0.54873 | 0.00640 |
| **Cpn1** | 7.73299 | 8.15698 | 6.60933 | 1.66455 | 8.35402 | 0.03441 | 5.45226 | 7.06083 | 5.64881 | 2.41702 | 0.37974 | 0.37034 | 10.80739 | 0.00000 | 0.84525 | 10.34929 |
| **Cpne1** | 1.30384 | 1.19422 | 1.06221 | 7.31088 | 1.08571 | 0.02393 | 1.98133 | 1.84352 | 8.33061 | 0.77325 | 2.35834 | 12.47587 | 0.26794 | 9.96618 | 7.53117 | 0.69831 |
| **Cpne2** | 0.01382 | 0.00813 | 0.01029 | 0.08054 | 0.00914 | 0.00671 | 0.00754 | 0.00901 | 0.01082 | 0.00852 | 0.24651 | 0.00735 | 0.00945 | 0.01046 | 0.00353 | 0.00725 |
| **Cpne3** | 2.39169 | 0.00257 | 4.07499 | 0.17020 | 5.38698 | 1.89464 | 0.08536 | 1.03968 | 0.45556 | 1.14424 | 0.02087 | 0.06691 | 0.27564 | 0.01322 | 0.00000 | 0.00256 |
| **Cpne5** | 1.19194 | 0.00716 | 0.74805 | 2.35288 | 0.00473 | 0.00471 | 0.00718 | 0.31223 | 0.33071 | 0.57960 | 0.00628 | 0.03245 | 0.01450 | 0.00155 | 0.31055 | 0.00557 |
| **Cpne7** | 1.35426 | 0.00686 | 0.00225 | 2.17956 | 0.01600 | 0.00714 | 0.00592 | 0.88811 | 1.46221 | 2.76016 | 0.02284 | 0.11900 | 0.00544 | 0.00575 | 0.01921 | 0.01334 |
| **Cpne8** | 3.43156 | 5.43260 | 3.20702 | 5.48773 | 2.67865 | 0.34455 | 5.89103 | 2.61316 | 10.57944 | 6.89224 | 1.98392 | 5.82026 | 1.65644 | 4.30927 | 2.95674 | 3.02845 |
| **Cpne9** | 1.87116 | 0.04365 | 0.01319 | 2.60250 | 0.03704 | 0.00586 | 0.01418 | 0.38567 | 1.74440 | 0.00000 | 1.53283 | 4.25568 | 0.00000 | 0.49134 | 0.29585 | 0.03201 |
| **Cpox** | 1.96065 | 2.53072 | 1.00018 | 0.46624 | 0.05415 | 0.00386 | 0.67904 | 0.05061 | 0.37165 | 0.79756 | 1.38298 | 0.09397 | 1.84891 | 0.40727 | 0.49839 | 0.01056 |
| **Cpped1** | 0.29090 | 0.00652 | 0.00000 | 0.02515 | 2.08650 | 2.47134 | 0.00648 | 1.63607 | 0.00968 | 0.00770 | 0.00000 | 0.02452 | 1.41486 | 0.00623 | 0.16861 | 0.00554 |
| **Cpsf1** | 9.61995 | 8.54481 | 9.35109 | 3.32960 | 5.44923 | 4.32384 | 4.96514 | 11.99400 | 1.37440 | 6.27011 | 8.07940 | 0.22568 | 5.57599 | 0.01694 | 2.01994 | 2.37077 |
| **Cpsf2** | 6.53171 | 2.88133 | 3.35670 | 4.55698 | 2.34823 | 2.41156 | 3.82354 | 8.87357 | 2.13014 | 5.46740 | 4.32162 | 0.72929 | 2.04645 | 7.34675 | 10.43276 | 2.22842 |
| **Cpsf3** | 13.73107 | 15.36305 | 16.51571 | 1.70280 | 8.67616 | 5.88464 | 12.52072 | 17.14824 | 5.80237 | 13.42418 | 16.53075 | 4.02894 | 14.39446 | 14.65299 | 12.27548 | 8.65566 |
| **Cpsf3l** | 8.36972 | 6.85157 | 15.49474 | 5.48496 | 7.25069 | 7.39073 | 8.13978 | 13.77241 | 7.92285 | 11.23176 | 8.34703 | 4.11612 | 12.29073 | 1.40856 | 9.85179 | 5.04071 |
| **Cpsf4** | 6.41066 | 0.79962 | 4.91267 | 2.62832 | 4.73355 | 0.62525 | 4.86398 | 6.25637 | 4.98421 | 4.73017 | 0.29741 | 0.28962 | 2.72253 | 0.00000 | 0.12237 | 2.86636 |
| **Cpsf4l** | 0.00819 | 0.00892 | 0.16434 | 2.39821 | 0.00915 | 0.00544 | 0.00825 | 0.18915 | 0.56387 | 1.00330 | 0.01507 | 0.03054 | 0.00931 | 0.00223 | 0.51323 | 0.01418 |
| **Cpsf6** | 5.35808 | 7.35918 | 3.05209 | 14.34815 | 5.96417 | 2.21779 | 9.01323 | 13.27860 | 21.86871 | 9.10086 | 7.85777 | 10.20940 | 10.07714 | 12.95506 | 4.02494 | 2.63567 |
| **Cpsf7** | 4.16036 | 0.03127 | 6.71852 | 1.15741 | 2.68666 | 3.68846 | 0.27783 | 2.40234 | 0.63046 | 1.22548 | 1.94938 | 7.36042 | 1.79889 | 3.89495 | 1.97221 | 2.81319 |
| **Cpt1a** | 9.50871 | 1.48514 | 2.96301 | 1.09198 | 4.19424 | 2.70740 | 1.56900 | 9.84597 | 6.31377 | 7.82739 | 7.55540 | 4.26882 | 2.48211 | 7.90201 | 8.34301 | 2.34707 |
| **Cpt1c** | 0.00554 | 0.00746 | 0.00734 | 1.15285 | 0.00947 | 0.00649 | 0.00631 | 0.71745 | 0.88027 | 0.73752 | 0.01843 | 0.03644 | 1.51905 | 0.10203 | 0.49515 | 0.00665 |
| **Cpt2** | 4.06529 | 0.21751 | 2.20148 | 5.78240 | 0.02909 | 2.63851 | 0.01398 | 0.01030 | 4.22250 | 0.13317 | 0.00000 | 0.01863 | 0.03279 | 0.00000 | 0.62290 | 1.18470 |
| **Cptp** | 2.64266 | 1.75245 | 3.40802 | 9.95597 | 0.00000 | 7.37661 | 2.98329 | 2.48757 | 11.67890 | 2.78016 | 0.53065 | 0.13911 | 2.95876 | 0.00000 | 2.47734 | 0.00000 |
| **Cpxm1** | 4.56095 | 0.96720 | 0.00000 | 0.01242 | 0.00550 | 0.00265 | 5.27499 | 1.33015 | 0.00936 | 0.00434 | 0.00000 | 0.00750 | 2.95507 | 0.00856 | 0.42439 | 7.04213 |
| **Cpxm2** | 0.13670 | 0.00841 | 0.00765 | 0.00944 | 0.01697 | 0.00699 | 0.00741 | 0.01246 | 0.02667 | 0.01401 | 0.01246 | 0.01829 | 0.01253 | 0.01573 | 0.02079 | 0.01501 |
| **Cr1l** | 3.62210 | 2.42774 | 4.51325 | 0.00000 | 2.99390 | 0.29238 | 0.68244 | 1.04159 | 0.01936 | 2.42828 | 1.48049 | 2.05591 | 1.05995 | 0.41284 | 0.34122 | 1.69893 |
| **Crabp2** | 0.01769 | 0.00800 | 0.00936 | 1.10696 | 0.00883 | 0.00658 | 0.00742 | 0.00822 | 1.63326 | 0.06897 | 0.13853 | 0.00439 | 0.01129 | 0.00446 | 0.51252 | 0.00706 |
| **Cracr2a** | 0.00844 | 0.00800 | 0.01196 | 0.25296 | 0.00905 | 0.00640 | 0.00668 | 0.00986 | 0.04735 | 0.00514 | 0.01313 | 0.00743 | 1.33127 | 0.01230 | 0.00643 | 0.00654 |
| **Cracr2b** | 0.12066 | 0.01738 | 0.10470 | 0.20029 | 0.73085 | 0.02784 | 0.08723 | 0.24983 | 0.71129 | 2.37650 | 2.90055 | 0.05417 | 0.16655 | 0.52808 | 0.06010 | 0.02763 |
| **Cradd** | 1.56548 | 0.00724 | 0.05454 | 0.00553 | 0.01209 | 0.00439 | 0.00679 | 0.34560 | 0.01090 | 0.20047 | 0.35213 | 0.00544 | 0.01129 | 0.03171 | 0.00000 | 0.00595 |
| **Cramp1l** | 3.00020 | 0.00713 | 5.27143 | 3.50966 | 6.00299 | 1.21023 | 4.60116 | 2.32238 | 4.22224 | 4.94079 | 1.54318 | 5.16355 | 0.26011 | 0.11135 | 7.33524 | 0.03632 |
| **Crat** | 3.77339 | 0.00000 | 0.37075 | 1.56436 | 5.83133 | 4.43407 | 1.97889 | 2.00494 | 0.00000 | 0.65695 | 0.91682 | 0.06357 | 0.00000 | 0.00008 | 0.12869 | 0.00000 |
| **Crb3** | 0.00000 | 0.11694 | 0.00000 | 17.55227 | 0.04107 | 7.70376 | 0.01544 | 20.06048 | 15.84611 | 25.85000 | 12.82208 | 1.50809 | 11.16083 | 0.02067 | 0.00000 | 0.00000 |
| **Crbn** | 2.35686 | 6.60314 | 3.38880 | 2.31027 | 6.16140 | 4.15455 | 0.81784 | 4.92090 | 1.71596 | 4.91624 | 3.75164 | 4.25614 | 1.36675 | 9.52996 | 3.98215 | 2.15520 |
| **Crcp** | 3.56348 | 4.91924 | 6.01545 | 4.43525 | 5.67590 | 0.22586 | 2.89142 | 6.86415 | 7.04837 | 6.20288 | 4.84613 | 2.28510 | 4.44994 | 2.19602 | 7.63354 | 0.00996 |
| **Creb1** | 3.99380 | 0.08613 | 2.16175 | 0.87229 | 3.10499 | 1.61635 | 3.91422 | 0.24870 | 7.31557 | 2.27474 | 0.77289 | 1.44142 | 0.95318 | 0.17373 | 2.45181 | 0.44551 |
| **Creb3** | 0.14689 | 3.42942 | 7.69407 | 0.61120 | 0.00000 | 3.56166 | 3.23694 | 5.64689 | 10.36343 | 3.16383 | 1.33639 | 0.46328 | 3.34626 | 4.33217 | 6.00157 | 3.75623 |
| **Creb3l1** | 0.01320 | 0.00794 | 0.00840 | 0.95873 | 0.00984 | 0.00747 | 0.00743 | 0.00000 | 0.78513 | 0.11538 | 0.88106 | 0.01972 | 0.10240 | 0.00214 | 0.12950 | 0.00778 |
| **Creb3l2** | 1.42990 | 0.00584 | 0.00000 | 0.18460 | 0.00316 | 0.00563 | 0.38146 | 0.00407 | 0.03713 | 0.36994 | 0.01000 | 0.05354 | 0.00714 | 0.00994 | 0.00957 | 5.67502 |
| **Creb3l3** | 0.01162 | 0.00772 | 0.00409 | 1.59065 | 0.01809 | 0.00676 | 0.01730 | 0.00000 | 2.99509 | 0.00000 | 0.94906 | 0.00689 | 0.14982 | 0.03106 | 0.66569 | 0.00656 |
| **Creb3l4** | 0.00805 | 0.00769 | 0.41488 | 6.15406 | 0.00680 | 0.01032 | 0.00677 | 0.00000 | 3.15894 | 0.92253 | 0.64579 | 0.00403 | 0.00000 | 0.00000 | 0.79984 | 1.61581 |
| **Crebbp** | 0.41935 | 0.01823 | 1.21446 | 3.34275 | 1.35965 | 0.01415 | 0.00592 | 1.29820 | 3.29611 | 1.93031 | 0.00000 | 0.01994 | 1.41629 | 7.32359 | 4.49113 | 0.00517 |

| | | | | | | | | | | | | | | | | |
|---|---|---|---|---|---|---|---|---|---|---|---|---|---|---|---|---|
| **Crebl2** | 0.79235 | 0.00895 | 0.00297 | 0.93369 | 0.01694 | 0.00575 | 0.00760 | 0.28504 | 3.35397 | 0.00000 | 0.00000 | 0.02002 | 0.00807 | 0.71297 | 0.78308 | 0.00617 |
| **Crebrf** | 0.00000 | 0.00000 | 2.52171 | 6.96248 | 0.00346 | 0.00229 | 0.74464 | 1.16420 | 9.78368 | 10.91546 | 8.47733 | 20.06248 | 0.00000 | 12.53186 | 16.07585 | 0.00130 |
| **Crebzf** | 0.75677 | 0.61678 | 0.85727 | 6.02494 | 1.64037 | 0.37022 | 0.02619 | 1.85225 | 1.65425 | 1.52092 | 0.18672 | 2.39115 | 0.25800 | 2.11989 | 2.34568 | 0.27804 |
| **Creg1** | 8.63119 | 0.59921 | 6.75660 | 1.48833 | 2.88902 | 0.17661 | 2.19652 | 4.92187 | 0.06509 | 9.45879 | 6.26755 | 1.23488 | 3.59527 | 3.22103 | 3.05341 | 0.00000 |
| **Creld1** | 0.01164 | 0.00798 | 0.18680 | 1.64510 | 0.01000 | 0.00759 | 0.00630 | 0.00116 | 1.39353 | 0.06503 | 0.25673 | 0.00284 | 1.42093 | 0.00319 | 0.21070 | 0.00634 |
| **Creld2** | 3.14420 | 3.00970 | 5.90503 | 12.32324 | 0.03471 | 0.04763 | 4.81851 | 5.90735 | 4.49914 | 1.20460 | 6.28670 | 3.13961 | 11.24334 | 1.35473 | 3.04908 | 3.40458 |
| **Crem** | 1.65638 | 0.00533 | 3.43101 | 2.25548 | 0.00000 | 0.00486 | 0.01633 | 0.33195 | 1.47714 | 0.00000 | 0.41350 | 0.03469 | 0.00000 | 0.00285 | 0.89721 | 0.17492 |
| **Crim1** | 0.02649 | 0.00828 | 0.17706 | 0.19943 | 0.01199 | 0.00897 | 0.00733 | 0.01224 | 0.02755 | 0.01460 | 0.01311 | 0.02810 | 0.01058 | 0.01181 | 0.00337 | 0.00739 |
| **Crip1** | 1.63123 | 0.03447 | 0.10731 | 0.41338 | 0.01342 | 3.58911 | 0.02300 | 0.01071 | 0.00468 | 1.58020 | 3.79615 | 0.00000 | 1.87166 | 0.00000 | 0.00000 | 3.01860 |
| **Crip2** | 16.45450 | 4.86120 | 12.47531 | 0.69371 | 15.17254 | 14.70591 | 6.48500 | 38.36844 | 0.38504 | 18.09841 | 17.85370 | 0.64010 | 33.72514 | 0.07697 | 0.00000 | 9.77308 |
| **Cript** | 10.41101 | 10.46111 | 10.19523 | 17.89536 | 10.57021 | 8.40886 | 3.50689 | 7.23804 | 18.99114 | 14.57354 | 10.56779 | 12.75101 | 10.79465 | 11.21833 | 10.90392 | 3.52638 |
| **Crispld2** | 0.49790 | 0.00768 | 0.10263 | 1.18948 | 0.00672 | 0.00513 | 0.09056 | 0.01151 | 0.80732 | 0.00727 | 0.03667 | 0.03173 | 0.01085 | 0.00436 | 0.31543 | 0.00642 |
| **Crk** | 5.91386 | 6.30485 | 6.53411 | 12.68184 | 8.41209 | 5.21225 | 6.21740 | 1.69781 | 4.81641 | 11.84043 | 0.38182 | 9.24489 | 1.61145 | 17.10084 | 14.94355 | 0.90312 |
| **Crkl** | 6.56886 | 1.36878 | 2.34025 | 10.95475 | 4.08456 | 5.66714 | 2.12219 | 2.15952 | 3.42594 | 2.52531 | 2.43136 | 3.50284 | 2.75046 | 3.58202 | 4.14781 | 2.64689 |
| **Crlf1** | 0.01328 | 0.00800 | 0.00859 | 3.26886 | 0.00904 | 0.00649 | 0.00722 | 0.00453 | 0.53365 | 0.00000 | 0.28932 | 0.06751 | 0.12892 | 0.00408 | 0.00000 | 0.00655 |
| **Crlf2** | 0.05562 | 0.01793 | 0.00704 | 0.88624 | 0.00799 | 0.00711 | 0.00669 | 0.00000 | 0.41651 | 0.36010 | 1.58268 | 0.00744 | 0.01306 | 0.00802 | 0.00000 | 0.01479 |
| **Crlf3** | 5.47533 | 0.00682 | 3.86407 | 2.70706 | 2.40386 | 4.96049 | 3.66694 | 7.50598 | 5.13768 | 10.84426 | 4.84095 | 3.06558 | 4.24140 | 4.61509 | 9.20763 | 2.03879 |
| **Crls1** | 2.06107 | 1.95183 | 0.30492 | 1.13751 | 0.06035 | 0.07440 | 1.34787 | 1.97141 | 1.63442 | 3.51448 | 9.51100 | 1.51124 | 1.27705 | 11.07469 | 1.46035 | 0.21960 |
| **Crmp1** | 0.00620 | 0.00788 | 0.01120 | 0.23913 | 0.01813 | 0.00635 | 0.00633 | 0.00946 | 0.00769 | 0.02836 | 0.01206 | 0.03721 | 1.93015 | 0.01188 | 0.00869 | 0.00623 |
| **Crnkl1** | 2.23686 | 8.14650 | 7.64421 | 6.49211 | 5.09746 | 0.00000 | 2.60124 | 22.96210 | 18.86095 | 33.08924 | 12.16841 | 14.15018 | 4.09074 | 25.82274 | 31.30187 | 1.86368 |
| **Crocc** | 1.13103 | 0.02970 | 0.47301 | 0.92302 | 0.03290 | 0.00579 | 0.38398 | 0.00000 | 0.00000 | 3.36745 | 1.12544 | 0.03253 | 0.03216 | 0.00379 | 0.00000 | 3.48043 |
| **Crot** | 9.30880 | 5.17582 | 5.60109 | 1.57825 | 2.58218 | 5.97456 | 6.63687 | 2.21135 | 0.00000 | 4.66957 | 0.84633 | 0.08067 | 1.31486 | 1.48561 | 2.78301 | 0.00000 |
| **Crtac1** | 0.01368 | 0.00850 | 0.01096 | 0.00509 | 0.00975 | 0.00712 | 0.00768 | 0.00436 | 0.01119 | 0.18549 | 0.41325 | 0.00610 | 0.03225 | 0.00695 | 0.05776 | 0.00735 |
| **Crtam** | 0.01017 | 0.00828 | 0.02386 | 0.00934 | 0.01056 | 0.00660 | 0.00755 | 0.01160 | 0.02420 | 0.00687 | 0.00876 | 0.00180 | 0.11684 | 1.76043 | 0.00297 | 0.00773 |
| **Crtap** | 5.43309 | 5.11515 | 5.82884 | 2.64090 | 5.61911 | 0.10117 | 3.00726 | 8.66988 | 2.65766 | 4.75479 | 5.63393 | 8.33395 | 6.13648 | 1.77927 | 2.58958 | 2.64212 |
| **Crtc1** | 0.00462 | 0.00795 | 0.15556 | 0.12315 | 0.30631 | 0.01251 | 0.00764 | 0.23117 | 0.00654 | 0.44019 | 0.13127 | 0.01637 | 0.83678 | 0.01453 | 0.00352 | 0.00716 |
| **Crtc2** | 0.08678 | 0.11535 | 0.00000 | 0.34554 | 2.22669 | 0.07995 | 0.00701 | 0.34269 | 0.43320 | 0.03220 | 0.48192 | 0.02023 | 0.02556 | 0.01356 | 0.65794 | 0.02111 |
| **Crtc3** | 1.86326 | 0.01661 | 0.36731 | 1.06369 | 0.03075 | 0.25859 | 0.00585 | 0.00000 | 2.38300 | 0.63524 | 0.04744 | 0.01928 | 0.01611 | 0.07762 | 0.01548 | 0.83131 |
| **Crx** | 0.03685 | 0.01776 | 0.23872 | 6.59849 | 0.01653 | 0.02372 | 0.02666 | 0.02818 | 3.41457 | 3.13426 | 0.61656 | 0.19385 | 0.03538 | 1.88962 | 3.63247 | 0.07085 |
| **Crxos** | 27.54326 | 14.54120 | 31.40792 | 14.36798 | 22.14591 | 13.08025 | 15.14107 | 22.89954 | 33.45477 | 28.92922 | 41.93466 | 16.11519 | 28.64284 | 23.43484 | 20.85308 | 11.16334 |
| **Cry1** | 2.79588 | 1.93291 | 3.45442 | 16.43797 | 0.00566 | 0.00000 | 2.70580 | 6.71882 | 12.67027 | 11.44352 | 2.74367 | 9.99122 | 3.20275 | 1.68955 | 14.97965 | 1.70514 |
| **Cry2** | 1.69694 | 0.00531 | 0.68905 | 3.44518 | 0.01937 | 0.09429 | 0.01416 | 3.04652 | 3.41562 | 0.58329 | 1.38824 | 0.02865 | 2.43989 | 0.02809 | 2.09146 | 0.01005 |
| **Cryab** | 0.00000 | 0.03157 | 4.96970 | 0.00802 | 0.03671 | 0.15856 | 0.00744 | 0.15349 | 0.02194 | 1.86649 | 0.00323 | 0.29092 | 0.22899 | 0.03193 | 0.00000 | 2.84661 |
| **Cryba1** | 0.01292 | 0.00812 | 0.00631 | 0.00470 | 0.00924 | 0.00711 | 0.00772 | 0.02059 | 0.01053 | 0.00248 | 0.71254 | 0.02667 | 0.00584 | 0.01554 | 0.23075 | 0.00723 |
| **Crybg3** | 0.11247 | 0.00605 | 0.30314 | 1.25801 | 2.28116 | 0.00477 | 0.69502 | 0.30927 | 5.10711 | 0.90968 | 0.39324 | 0.07245 | 0.92672 | 0.00465 | 2.58310 | 0.00571 |
| **Cryga** | 0.05382 | 0.00796 | 0.05522 | 0.24393 | 0.00994 | 0.00589 | 0.00737 | 0.01248 | 0.00757 | 0.01753 | 0.01349 | 0.03893 | 0.01192 | 0.01129 | 0.00528 | 0.00732 |
| **Crygc** | 1.07692 | 0.00770 | 0.00379 | 0.01047 | 0.01624 | 0.00732 | 0.00673 | 0.01055 | 0.01222 | 0.05216 | 0.01205 | 0.00817 | 0.01171 | 0.01180 | 0.01219 | 0.08882 |
| **Crygd** | 2.09001 | 0.00595 | 0.02399 | 0.01025 | 0.00926 | 0.00452 | 0.00574 | 0.01223 | 0.01249 | 0.00980 | 0.01248 | 0.00628 | 0.01547 | 0.00858 | 0.05226 | 1.23025 |
| **Cryge** | 1.20733 | 0.00666 | 0.06271 | 0.00904 | 0.12325 | 0.00469 | 0.00645 | 0.00848 | 0.01206 | 0.08286 | 0.01145 | 0.01699 | 0.01072 | 0.00600 | 0.18461 | 0.21674 |
| **Crygf** | 0.31303 | 0.00782 | 0.01623 | 0.01029 | 0.02330 | 0.01227 | 0.00722 | 0.01260 | 0.01249 | 0.01935 | 0.01283 | 0.00755 | 0.01145 | 0.00880 | 0.03901 | 0.11677 |
| **Crygs** | 1.47640 | 0.33114 | 0.00000 | 0.07308 | 0.00495 | 0.25298 | 0.00422 | 0.26632 | 0.76420 | 1.07731 | 4.44234 | 0.14570 | 0.47243 | 0.48911 | 1.32583 | 0.00464 |
| **Cryl1** | 0.00227 | 0.00373 | 0.00000 | 1.63099 | 4.49838 | 0.00591 | 0.00655 | 0.02328 | 0.64654 | 0.94381 | 0.00000 | 4.34502 | 0.00933 | 0.00610 | 0.78982 | 0.00663 |
| **Crym** | 1.88278 | 0.02618 | 0.23850 | 0.01072 | 0.00278 | 0.00663 | 0.00561 | 0.00482 | 0.01238 | 0.00401 | 0.01196 | 0.01652 | 0.56512 | 0.01454 | 0.00562 | 3.40893 |
| **Cryz** | 7.11817 | 2.98013 | 3.86521 | 0.40485 | 5.68439 | 0.01869 | 0.00000 | 0.83874 | 0.11542 | 0.15480 | 1.36131 | 0.00830 | 1.38194 | 0.00199 | 0.00000 | 1.35145 |
| **Cryzl1** | 3.33675 | 2.84001 | 6.94933 | 0.00000 | 3.42678 | 5.18756 | 4.62631 | 0.15778 | 0.56575 | 1.87976 | 0.77518 | 0.02214 | 0.13065 | 0.00369 | 0.74529 | 1.93617 |
| **Cs** | 16.74591 | 12.94762 | 16.05676 | 13.61127 | 16.43898 | 9.18026 | 16.77362 | 21.56382 | 17.43514 | 10.56671 | 12.11671 | 7.12835 | 15.98202 | 11.64155 | 10.11410 | 10.98065 |
| **Csad** | 0.13837 | 0.08656 | 0.79971 | 1.98120 | 3.29641 | 0.00928 | 0.01667 | 1.74056 | 0.15708 | 0.09765 | 2.11163 | 0.40827 | 0.13205 | 0.20008 | 1.27591 | 0.00622 |
| **Csdc2** | 1.13615 | 0.00612 | 0.32470 | 0.93471 | 0.01302 | 0.00433 | 0.19414 | 3.42998 | 0.56643 | 4.23863 | 0.30278 | 0.13515 | 0.02756 | 0.39648 | 0.41497 | 0.00559 |
| **Csde1** | 14.59809 | 15.19969 | 12.41322 | 16.20095 | 12.59236 | 9.23465 | 10.19570 | 20.10001 | 21.96779 | 30.28366 | 20.36888 | 14.82686 | 11.15809 | 44.90874 | 32.15552 | 6.27281 |
| **Cse1l** | 10.00882 | 17.04987 | 6.92278 | 12.43854 | 11.17633 | 4.05373 | 6.28091 | 12.34530 | 8.99718 | 21.61723 | 14.77392 | 19.15170 | 7.16457 | 26.75321 | 21.27407 | 6.30929 |
| **Csf1** | 0.72534 | 0.00772 | 0.00340 | 2.49705 | 0.01778 | 0.00722 | 0.00659 | 0.00762 | 0.14423 | 0.01355 | 0.02554 | 0.23467 | 0.01194 | 0.00907 | 0.00001 | 0.00650 |
| **Csf2ra** | 0.00867 | 0.02977 | 0.19106 | 1.18421 | 1.81507 | 0.00590 | 0.50325 | 3.44984 | 0.63564 | 2.14100 | 2.10774 | 0.06671 | 0.36511 | 0.00455 | 0.03138 | 3.20199 |
| **Csf2rb** | 0.01761 | 0.00806 | 0.03419 | 0.13997 | 0.00996 | 0.01312 | 0.01784 | 0.01829 | 0.13478 | 1.23222 | 0.23067 | 0.01607 | 0.00930 | 0.13132 | 0.01455 | 0.00731 |
| **Csf2rb2** | 0.01655 | 0.00738 | 0.13120 | 0.47122 | 0.00954 | 0.01719 | 0.04703 | 0.21153 | 0.90768 | 1.04828 | 0.80829 | 0.85520 | 0.10500 | 0.49492 | 0.91958 | 0.00693 |
| **Csf3r** | 1.45087 | 1.77802 | 7.07055 | 0.00000 | 9.56935 | 10.00940 | 9.59647 | 5.54076 | 0.48554 | 1.57980 | 1.61577 | 0.16555 | 12.37945 | 0.57800 | 1.42210 | 0.50567 |
| **Csgalnact2** | 0.00000 | 0.00800 | 0.16694 | 0.34747 | 0.00999 | 1.68028 | 0.02888 | 2.41244 | 0.32810 | 0.29189 | 0.62254 | 0.02015 | 0.00000 | 3.61177 | 1.06879 | 0.22944 |
| **Csk** | 18.00624 | 2.51906 | 5.75227 | 3.34258 | 6.26348 | 6.42079 | 3.83763 | 14.10243 | 3.85752 | 10.54908 | 3.99865 | 0.91338 | 7.78242 | 0.00000 | 0.00000 | 3.55276 |
| **Csl** | 0.77934 | 0.76210 | 1.17272 | 0.65183 | 1.43825 | 0.65679 | 1.44691 | 1.33403 | 0.80627 | 0.56488 | 0.66303 | 0.08800 | 1.17163 | 0.35044 | 0.35532 | 0.58873 |
| **Csnk1a1** | 4.80065 | 9.81500 | 5.84464 | 14.74727 | 6.01679 | 0.14400 | 6.20431 | 6.82462 | 18.66655 | 8.70143 | 6.50718 | 0.67796 | 1.80678 | 6.14128 | 13.69416 | 2.62186 |
| **Csnk1d** | 7.73651 | 4.48442 | 1.44714 | 5.21043 | 0.04954 | 0.76575 | 4.46615 | 7.03072 | 3.37002 | 6.37839 | 2.97118 | 0.51143 | 4.46969 | 1.33380 | 4.32085 | 3.10708 |
| **Csnk1e** | 7.89482 | 3.43331 | 1.51945 | 13.39135 | 6.51584 | 5.45454 | 7.31325 | 22.67129 | 21.56211 | 26.44657 | 10.97685 | 9.47218 | 7.97019 | 8.71767 | 14.53004 | 2.14747 |
| **Csnk1g1** | 3.15702 | 0.00000 | 6.42081 | 2.71914 | 1.99230 | 0.00319 | 4.28256 | 1.94171 | 3.81292 | 7.09665 | 2.16155 | 1.19274 | 0.00000 | 15.08009 | 7.08351 | 2.20382 |

| | | | | | | | | | | | | | | | | |
|---|---|---|---|---|---|---|---|---|---|---|---|---|---|---|---|---|
| **Csnk1g2** | 10.19098 | 0.03535 | 0.00000 | 1.96946 | 4.14093 | 7.26383 | 0.38510 | 6.79622 | 0.40574 | 7.43062 | 0.87682 | 0.19051 | 6.60933 | 0.00000 | 0.00000 | 8.89146 |
| **Csnk1g3** | 1.52924 | 0.20279 | 0.00000 | 8.13379 | 3.48820 | 1.79008 | 4.45120 | 3.99895 | 12.52876 | 10.36758 | 2.26052 | 10.16205 | 1.43067 | 16.88481 | 11.51753 | 0.02194 |
| **Csnk2a1** | 6.59881 | 6.05613 | 8.10949 | 4.79435 | 3.35931 | 3.97130 | 1.77179 | 6.69070 | 11.42874 | 9.60931 | 8.30992 | 2.02990 | 4.80081 | 15.53482 | 10.59543 | 4.03076 |
| **Csnk2a2** | 1.63503 | 8.09933 | 1.23298 | 6.54415 | 1.45987 | 1.03559 | 2.33647 | 5.17264 | 10.01421 | 3.67545 | 6.54645 | 5.09581 | 0.00000 | 4.05423 | 6.28265 | 0.31594 |
| **Csnk2b** | 18.66107 | 13.18982 | 19.87279 | 23.16663 | 17.75373 | 12.80448 | 13.01373 | 21.79026 | 21.03401 | 21.18367 | 22.25903 | 6.09309 | 20.86623 | 4.79254 | 8.50398 | 14.80820 |
| **Cspg5** | 0.00574 | 0.00726 | 1.66809 | 0.00413 | 0.01687 | 0.00915 | 0.00693 | 0.74098 | 0.63002 | 0.00000 | 0.00000 | 0.01381 | 0.09196 | 0.02005 | 0.17962 | 0.00660 |
| **Cspp1** | 0.81856 | 2.07121 | 1.47518 | 0.78019 | 0.01224 | 3.68524 | 2.71765 | 1.94191 | 1.40462 | 0.27616 | 1.50648 | 7.72102 | 1.73137 | 2.18569 | 0.70886 | 0.00922 |
| **Csrnp1** | 6.39361 | 0.16583 | 7.67860 | 3.92535 | 0.02111 | 5.16937 | 0.00013 | 6.96407 | 6.98925 | 13.00357 | 10.54803 | 4.62630 | 5.89222 | 0.14767 | 9.97928 | 3.13067 |
| **Csrnp2** | 0.83052 | 0.01439 | 0.87495 | 6.61676 | 0.04710 | 1.08652 | 0.00257 | 1.18796 | 10.69900 | 1.02526 | 5.09494 | 7.93644 | 1.18821 | 11.51973 | 6.62389 | 0.00905 |
| **Csrnp3** | 0.04268 | 0.01922 | 0.04694 | 2.28006 | 0.00937 | 0.01207 | 0.01698 | 0.00000 | 0.88167 | 0.70500 | 0.49703 | 0.00000 | 0.07279 | 2.75892 | 1.86927 | 0.02047 |
| **Csrp1** | 0.53869 | 0.98819 | 2.45031 | 0.07869 | 0.22625 | 8.50500 | 3.22124 | 5.70803 | 0.05714 | 7.76881 | 1.68434 | 0.32958 | 6.29949 | 0.00619 | 0.00000 | 1.80113 |
| **Csrp2** | 1.53635 | 1.23954 | 2.38215 | 0.03103 | 0.16545 | 4.45784 | 5.80302 | 2.23061 | 0.18154 | 2.00042 | 1.48354 | 0.19725 | 6.24602 | 0.05487 | 0.00000 | 1.80047 |
| **Csrp2bp** | 0.23337 | 3.44878 | 4.78449 | 3.67585 | 2.03495 | 1.94382 | 2.80899 | 10.71018 | 7.39017 | 9.21636 | 3.08027 | 1.64459 | 6.98889 | 0.97552 | 9.38827 | 3.79061 |
| **Cst13** | 0.01383 | 0.00820 | 0.00872 | 0.00537 | 0.00909 | 0.00687 | 0.00760 | 0.00592 | 0.01128 | 0.00593 | 0.49707 | 0.00637 | 0.00743 | 0.01119 | 0.00285 | 0.00715 |
| **Cst3** | 10.20738 | 0.01974 | 2.15867 | 0.00000 | 8.83476 | 5.15877 | 2.92660 | 6.80350 | 0.12708 | 8.91049 | 4.18317 | 0.22495 | 9.57796 | 0.00083 | 0.20905 | 8.51396 |
| **Cst6** | 0.01023 | 0.00885 | 0.04895 | 0.48106 | 0.01905 | 0.00613 | 0.00831 | 0.06443 | 0.00413 | 0.05632 | 0.01196 | 0.02384 | 0.01160 | 0.01221 | 0.00906 | 0.00735 |
| **Cst7** | 1.33528 | 0.00331 | 1.39554 | 8.30758 | 0.00291 | 2.05411 | 0.00342 | 2.66218 | 8.83326 | 4.02344 | 2.71835 | 6.75613 | 1.02425 | 3.94976 | 2.32205 | 0.00856 |
| **Cst8** | 0.01333 | 0.00793 | 0.01222 | 0.00768 | 0.00906 | 0.00653 | 0.00735 | 0.01069 | 0.01195 | 0.82330 | 0.01139 | 0.00579 | 0.01105 | 0.01176 | 0.00832 | 0.00726 |
| **Csta1** | 0.78330 | 0.00771 | 1.55719 | 4.60899 | 0.05193 | 1.69013 | 0.01561 | 0.01560 | 6.00169 | 0.15070 | 0.71971 | 0.00000 | 1.52021 | 0.25130 | 0.07277 | 0.52861 |
| **Cstb** | 7.58200 | 1.54041 | 10.07369 | 6.77707 | 9.14761 | 9.96434 | 2.85665 | 3.33656 | 9.99955 | 3.99505 | 25.91259 | 1.82744 | 11.07664 | 0.61270 | 0.88448 | 5.34821 |
| **Cstf1** | 6.08372 | 1.82732 | 10.90680 | 11.16009 | 5.42370 | 2.67639 | 6.45717 | 9.28059 | 16.97939 | 7.71049 | 6.34802 | 0.59542 | 7.71533 | 0.00730 | 2.58955 | 3.88450 |
| **Cstf2** | 7.02765 | 1.81588 | 3.50357 | 3.99868 | 8.18504 | 4.42554 | 5.17331 | 9.83596 | 3.08408 | 7.24055 | 3.45946 | 1.31661 | 3.07287 | 6.38904 | 3.46211 | 0.28985 |
| **Cstf2t** | 3.31396 | 2.22285 | 3.25203 | 7.95813 | 3.49675 | 0.11902 | 0.13257 | 1.50682 | 12.72427 | 3.66888 | 1.85677 | 12.81126 | 1.02270 | 16.50987 | 14.08598 | 1.40670 |
| **Cstf3** | 5.08829 | 2.24083 | 2.09794 | 4.94909 | 9.06199 | 1.17780 | 3.55761 | 4.12500 | 8.80657 | 6.68303 | 1.65580 | 6.81380 | 6.28518 | 7.03882 | 0.88744 | 1.32524 |
| **Ctbp1** | 5.21213 | 1.02940 | 2.19124 | 0.06341 | 4.56631 | 4.99135 | 1.71451 | 2.40370 | 0.00995 | 5.64298 | 0.12286 | 0.95867 | 7.21013 | 0.00626 | 0.00000 | 1.96642 |
| **Ctbp2** | 9.02826 | 5.37465 | 8.51768 | 2.43827 | 3.72376 | 6.20671 | 7.98289 | 3.22208 | 4.00420 | 4.23045 | 4.12242 | 1.36531 | 4.87289 | 0.01835 | 0.99673 | 8.92544 |
| **Ctbs** | 0.00448 | 0.21071 | 0.13160 | 0.28248 | 2.71348 | 2.63110 | 0.35732 | 1.05454 | 0.11237 | 1.49387 | 0.00446 | 0.00711 | 0.00000 | 0.00652 | 0.58604 | 0.01337 |
| **Ctc1** | 6.23126 | 3.55649 | 4.65785 | 10.02059 | 2.64631 | 5.10438 | 5.76079 | 4.24301 | 6.37537 | 8.09824 | 4.09739 | 0.41622 | 2.74674 | 2.15494 | 5.25250 | 3.10791 |
| **Ctcf** | 8.02087 | 5.21618 | 6.65136 | 19.65717 | 7.77438 | 0.41937 | 2.34598 | 11.17336 | 13.55962 | 19.91990 | 5.44968 | 9.26927 | 9.25309 | 8.14275 | 12.14338 | 3.02728 |
| **Ctcfl** | 0.10799 | 0.00758 | 0.27624 | 0.04758 | 0.04900 | 0.04137 | 0.06842 | 0.01553 | 0.19027 | 0.05264 | 0.58925 | 0.04354 | 0.27503 | 0.16180 | 0.15987 | 0.02113 |
| **Ctdnep1** | 7.75915 | 4.77340 | 3.77785 | 4.51176 | 0.07361 | 5.97279 | 5.01970 | 4.00240 | 8.59555 | 6.53270 | 2.02699 | 0.21313 | 4.26852 | 0.37455 | 3.98070 | 5.71380 |
| **Ctdp1** | 5.45574 | 0.00000 | 0.78595 | 1.40234 | 1.83295 | 1.41025 | 3.09047 | 8.40283 | 2.51402 | 9.79646 | 0.39816 | 5.86761 | 8.03273 | 0.02760 | 0.00000 | 1.98626 |
| **Ctdsp1** | 1.44984 | 0.90341 | 0.27423 | 3.88670 | 2.19155 | 0.76165 | 0.00539 | 0.06276 | 1.09523 | 0.75712 | 0.00000 | 0.02551 | 0.34538 | 0.00001 | 1.12438 | 3.09159 |
| **Ctdsp2** | 7.94023 | 0.19075 | 1.42342 | 2.06118 | 2.06245 | 7.63224 | 1.10222 | 2.24668 | 5.90138 | 8.26758 | 2.90292 | 0.59342 | 5.90687 | 2.11410 | 3.81973 | 6.57011 |
| **Ctdspl** | 0.00000 | 0.00731 | 0.07598 | 2.55141 | 0.01563 | 0.88678 | 0.00569 | 0.00315 | 0.17854 | 0.75745 | 0.34215 | 0.02092 | 1.97958 | 0.00302 | 0.47079 | 1.85702 |
| **Ctdspl2** | 2.28965 | 1.66079 | 2.71159 | 7.86595 | 1.60739 | 0.08294 | 2.73286 | 5.52284 | 8.02901 | 4.96064 | 3.07703 | 2.68964 | 0.41359 | 18.32799 | 8.22487 | 0.53112 |
| **Ctf1** | 0.01287 | 0.00775 | 0.01180 | 0.24638 | 0.00938 | 0.00658 | 0.00736 | 0.03559 | 0.04932 | 0.01120 | 0.01239 | 0.00730 | 0.01124 | 0.00675 | 0.08855 | 0.00734 |
| **Ctf2** | 0.01100 | 0.00756 | 0.00799 | 0.67340 | 0.00933 | 0.00672 | 0.00708 | 0.58641 | 1.07584 | 0.00483 | 0.00000 | 0.00455 | 0.00567 | 0.01155 | 0.00000 | 0.00715 |
| **Ctgf** | 0.01829 | 0.00743 | 0.00701 | 0.00485 | 0.01045 | 0.00695 | 0.00713 | 0.01845 | 0.17904 | 0.04147 | 0.00632 | 0.57443 | 0.01298 | 0.01199 | 0.13849 | 0.00776 |
| **Cth** | 5.80252 | 9.76747 | 11.80778 | 6.95979 | 0.15383 | 7.88937 | 6.95653 | 25.82305 | 13.56359 | 27.85546 | 14.74876 | 11.71481 | 15.08398 | 29.13679 | 20.45892 | 3.48510 |
| **Cthrc1** | 0.00000 | 0.00468 | 0.87824 | 0.00146 | 0.02280 | 0.00854 | 0.09619 | 7.49508 | 0.00523 | 6.60728 | 8.44032 | 3.05256 | 4.43953 | 2.57016 | 0.00000 | 0.00343 |
| **Ctif** | 0.01826 | 0.00803 | 0.15917 | 0.49227 | 0.07940 | 0.00604 | 0.28120 | 0.01130 | 0.20294 | 0.02089 | 0.01120 | 0.06291 | 0.01056 | 0.01129 | 0.00043 | 0.00722 |
| **Ctla2a** | 0.04815 | 0.03855 | 0.04823 | 0.10292 | 0.01645 | 0.00805 | 0.05806 | 0.07346 | 0.24036 | 0.07006 | 0.02299 | 0.02486 | 0.05850 | 0.47563 | 0.08249 | 0.02907 |
| **Ctnna1** | 16.54503 | 19.23922 | 15.40836 | 11.57622 | 10.71458 | 13.28192 | 13.62406 | 23.78814 | 17.64146 | 13.50467 | 15.26606 | 9.01159 | 16.33533 | 8.01776 | 5.53796 | 9.10307 |
| **Ctnnal1** | 0.83821 | 3.14354 | 3.84650 | 7.15305 | 0.00744 | 4.32098 | 2.25274 | 6.23214 | 12.17616 | 8.18777 | 3.19666 | 12.08628 | 2.11734 | 15.31318 | 6.21467 | 0.52244 |
| **Ctnnb1** | 16.06403 | 14.24440 | 14.04912 | 11.96139 | 15.58458 | 8.45851 | 12.78929 | 12.09867 | 18.53514 | 13.32874 | 9.14789 | 15.54413 | 10.55351 | 6.28024 | 16.00299 | 10.14475 |
| **Ctnnbip1** | 2.61870 | 0.00321 | 1.06471 | 5.80984 | 0.60497 | 3.65078 | 0.00267 | 4.51663 | 6.74720 | 2.04275 | 4.18493 | 0.24676 | 0.39922 | 0.17801 | 0.99768 | 1.75945 |
| **Ctnnbl1** | 4.16637 | 3.86220 | 6.69594 | 10.55040 | 11.48299 | 4.45787 | 7.13566 | 10.49862 | 13.44671 | 11.42839 | 6.78716 | 5.09830 | 8.94971 | 4.82817 | 9.91902 | 5.40373 |
| **Ctnnd1** | 13.59918 | 5.17290 | 9.00985 | 2.99112 | 7.30719 | 5.95706 | 2.44046 | 9.88413 | 4.59032 | 8.52108 | 7.76608 | 2.59049 | 7.34308 | 13.26366 | 13.61261 | 5.90872 |
| **Ctns** | 0.61009 | 3.64707 | 1.30474 | 0.19750 | 0.00670 | 0.00707 | 0.60232 | 1.69080 | 0.00597 | 0.00000 | 0.00000 | 0.05023 | 0.69368 | 1.86949 | 1.69474 | 1.09503 |
| **Ctps** | 8.78975 | 8.63851 | 9.32784 | 15.29804 | 8.37267 | 6.79288 | 9.75496 | 14.15775 | 10.91776 | 18.18069 | 11.14385 | 4.08716 | 8.11908 | 8.64394 | 18.96049 | 1.21391 |
| **Ctps2** | 4.21318 | 0.00051 | 1.05582 | 5.06220 | 2.33648 | 0.00176 | 0.00333 | 1.51635 | 4.57340 | 2.36085 | 0.69944 | 0.02148 | 0.00000 | 8.57216 | 2.12320 | 0.02311 |
| **Ctr9** | 8.26560 | 6.90225 | 6.42412 | 4.60866 | 6.89998 | 1.90502 | 3.12745 | 5.70909 | 6.73288 | 6.80238 | 1.64609 | 0.27263 | 0.89051 | 2.04668 | 6.15891 | 0.00559 |
| **Ctrb1** | 0.01211 | 0.00825 | 0.01371 | 0.26775 | 0.01000 | 0.00729 | 0.00779 | 0.01677 | 1.24004 | 0.14886 | 0.00003 | 0.00604 | 0.23463 | 0.00678 | 0.04875 | 0.00758 |
| **Ctrc** | 0.00659 | 1.56082 | 0.00702 | 0.10019 | 0.00796 | 0.00655 | 0.00669 | 0.00934 | 0.00871 | 0.00906 | 0.01303 | 0.00722 | 0.01084 | 0.00637 | 0.04476 | 0.00670 |
| **Ctrcos** | 0.02346 | 0.00807 | 0.04454 | 0.71470 | 0.00858 | 0.00702 | 0.01807 | 0.00801 | 0.65342 | 0.00717 | 0.23079 | 0.04689 | 0.05242 | 0.28927 | 0.06307 | 0.01428 |
| **Ctrl** | 0.00739 | 0.00784 | 0.00167 | 0.00529 | 0.00860 | 0.00839 | 0.00571 | 0.00000 | 0.69682 | 2.91960 | 4.59179 | 0.04058 | 0.94274 | 0.01089 | 0.23842 | 0.00625 |
| **Cts8** | 0.05226 | 0.00795 | 0.07838 | 0.03043 | 0.02661 | 0.01585 | 0.04978 | 0.02438 | 0.11391 | 0.05598 | 0.04848 | 0.01689 | 0.07367 | 0.17075 | 0.08646 | 0.02874 |
| **Ctsa** | 16.53916 | 11.35369 | 26.99828 | 12.24106 | 7.70233 | 6.41862 | 14.34682 | 34.73893 | 10.86347 | 27.71796 | 30.17443 | 6.83734 | 31.34335 | 4.05063 | 23.33190 | 15.53239 |
| **Ctsb** | 22.37089 | 7.38908 | 22.84284 | 19.59105 | 12.50455 | 9.30843 | 14.90363 | 14.93867 | 29.33806 | 25.54874 | 21.12230 | 18.46898 | 9.63368 | 20.73225 | 26.38693 | 11.11636 |
| **Ctsc** | 5.79595 | 4.60091 | 15.14216 | 9.99141 | 7.18335 | 7.67192 | 12.65820 | 25.95496 | 21.07014 | 30.43570 | 21.83597 | 17.79046 | 10.80122 | 36.83653 | 27.02730 | 2.39884 |
| **Ctsd** | 19.87404 | 10.27040 | 33.66962 | 1.23986 | 9.05422 | 9.33363 | 15.90117 | 31.15209 | 0.56973 | 21.49859 | 15.90744 | 1.32504 | 28.02635 | 10.87752 | 6.99194 | 8.11341 |

| | | | | | | | | | | | | | | | | |
|---|---|---|---|---|---|---|---|---|---|---|---|---|---|---|---|---|
| **Ctsf** | 0.00855 | 0.43942 | 0.00939 | 0.00864 | 0.00821 | 0.00692 | 0.00731 | 0.01168 | 0.01231 | 0.25005 | 0.01179 | 0.00801 | 0.01885 | 0.00680 | 0.04380 | 0.00717 |
| **Ctsh** | 0.00000 | 0.07676 | 0.73923 | 9.97068 | 0.02428 | 0.03421 | 3.10494 | 4.67351 | 6.37720 | 9.02471 | 9.57301 | 8.43268 | 2.13550 | 3.79222 | 4.47688 | 0.00160 |
| **Ctsk** | 0.01314 | 0.00778 | 0.01060 | 0.14813 | 0.00909 | 0.00687 | 0.13032 | 0.01155 | 0.49570 | 0.48866 | 0.00771 | 0.00656 | 0.01215 | 0.01337 | 0.06105 | 0.00737 |
| **Ctsl** | 24.25766 | 14.03602 | 34.23804 | 27.17517 | 13.81881 | 12.26668 | 21.33944 | 37.69822 | 38.81977 | 42.82503 | 38.04343 | 30.23881 | 27.35634 | 53.31677 | 45.86616 | 19.72853 |
| **Ctss** | 0.01342 | 0.00806 | 0.01204 | 0.38771 | 0.00907 | 0.00668 | 0.00737 | 0.01114 | 0.31028 | 0.60412 | 0.00971 | 0.00621 | 0.01212 | 0.01050 | 0.00486 | 0.00755 |
| **Ctsz** | 2.85627 | 0.01830 | 4.59582 | 4.01062 | 0.09948 | 0.01172 | 1.72903 | 12.16379 | 5.40561 | 18.75349 | 17.53562 | 8.72261 | 15.75618 | 1.50686 | 6.34559 | 8.75885 |
| **Cttn** | 13.05502 | 7.25229 | 13.51576 | 7.11660 | 7.76043 | 13.73651 | 9.63138 | 21.41295 | 8.49876 | 20.76672 | 15.71656 | 2.38677 | 19.04760 | 1.19658 | 11.06700 | 10.96671 |
| **Cttnbp2nl** | 1.64307 | 0.00428 | 0.40288 | 13.89688 | 0.00688 | 0.16541 | 3.12527 | 0.23686 | 8.35880 | 2.45048 | 0.00000 | 0.11427 | 0.54577 | 0.17087 | 5.89443 | 0.18205 |
| **Ctu1** | 1.22334 | 0.44692 | 1.23869 | 1.68382 | 0.01274 | 0.02072 | 0.27539 | 0.67189 | 4.07782 | 6.17922 | 2.50029 | 0.20946 | 2.46381 | 2.68601 | 2.10531 | 0.03314 |
| **Ctu2** | 5.95199 | 0.94908 | 0.26633 | 1.07704 | 3.04986 | 2.35131 | 2.13017 | 6.81292 | 2.65695 | 6.19014 | 4.21556 | 0.14485 | 5.29669 | 0.00000 | 0.00000 | 3.22206 |
| **Ctxn1** | 0.09281 | 0.04867 | 0.67176 | 0.77305 | 0.01411 | 0.00623 | 0.00649 | 0.01890 | 1.22700 | 1.83364 | 0.00000 | 0.50068 | 1.21535 | 0.02946 | 0.14730 | 2.18226 |
| **Cubn** | 2.20869 | 0.00286 | 0.00217 | 0.13807 | 4.36992 | 0.00692 | 0.01583 | 0.00000 | 0.73684 | 0.52247 | 0.39776 | 0.05160 | 0.04354 | 0.00738 | 0.15164 | 0.09288 |
| **Cuedc1** | 0.01376 | 0.00794 | 0.01189 | 0.01039 | 0.00937 | 0.00666 | 0.00746 | 0.01297 | 0.01235 | 0.06154 | 0.01290 | 0.02766 | 0.01286 | 0.01166 | 0.00981 | 0.00750 |
| **Cuedc2** | 7.55359 | 4.73578 | 3.50994 | 1.70042 | 4.70628 | 6.13385 | 1.44213 | 3.90080 | 1.64371 | 0.85239 | 3.93616 | 0.25639 | 4.44931 | 0.00000 | 0.75281 | 4.22601 |
| **Cul1** | 6.17479 | 9.99482 | 5.52005 | 31.07637 | 8.28141 | 2.36264 | 3.78054 | 14.17083 | 16.93205 | 17.72787 | 9.22618 | 23.88669 | 5.59839 | 19.38545 | 23.07395 | 2.28710 |
| **Cul2** | 7.68793 | 10.73480 | 9.53151 | 0.72292 | 9.02482 | 2.70682 | 6.24743 | 12.07403 | 3.24689 | 9.95047 | 3.43111 | 7.39221 | 4.80842 | 10.86861 | 9.61838 | 0.62433 |
| **Cul3** | 9.93971 | 11.20227 | 9.38115 | 25.25543 | 13.27163 | 6.65095 | 7.81429 | 8.45716 | 31.04835 | 14.68836 | 9.12745 | 12.21258 | 5.74466 | 21.80185 | 21.65154 | 2.20293 |
| **Cul4a** | 2.58914 | 2.95307 | 5.39335 | 1.50965 | 5.11197 | 5.45680 | 0.00000 | 1.21389 | 1.00294 | 6.14566 | 3.66476 | 4.08065 | 2.68230 | 0.23942 | 3.22654 | 0.25288 |
| **Cul4b** | 7.34129 | 18.12663 | 11.49788 | 21.85994 | 10.33319 | 3.62675 | 11.13854 | 7.90578 | 19.98572 | 6.37720 | 4.30173 | 8.66620 | 5.98858 | 18.29670 | 16.30723 | 5.19482 |
| **Cul5** | 6.56448 | 5.52809 | 7.80611 | 2.36720 | 6.99645 | 1.44588 | 7.94431 | 19.12945 | 7.04995 | 20.60520 | 14.55160 | 16.21587 | 8.48724 | 34.07608 | 12.88213 | 0.87254 |
| **Cul7** | 7.03129 | 0.00000 | 0.00000 | 0.34608 | 3.63499 | 2.03946 | 1.57966 | 1.60270 | 0.00420 | 1.23734 | 0.00000 | 0.01999 | 0.00000 | 0.00562 | 0.05820 | 2.50872 |
| **Cul9** | 0.01213 | 0.00813 | 0.01107 | 0.76109 | 0.00916 | 0.00652 | 0.00730 | 0.04573 | 0.00000 | 0.52257 | 0.02330 | 0.02687 | 0.00929 | 0.00975 | 0.00195 | 0.00699 |
| **Cuta** | 11.26940 | 2.36200 | 8.52996 | 2.66333 | 6.23191 | 5.38591 | 2.89758 | 7.29257 | 0.83992 | 11.24362 | 5.28985 | 0.33654 | 13.36689 | 0.00000 | 0.53946 | 3.48240 |
| **Cutc** | 2.70707 | 2.99110 | 1.00369 | 0.40295 | 4.85128 | 0.01363 | 0.01554 | 1.44091 | 0.81274 | 1.21850 | 0.43960 | 2.67741 | 0.02204 | 1.30588 | 1.87424 | 2.60308 |
| **Cux1** | 0.58537 | 0.52616 | 2.41974 | 0.27701 | 3.51150 | 0.01315 | 3.15053 | 1.32200 | 0.00000 | 2.97183 | 1.62109 | 0.24553 | 1.35459 | 1.22595 | 0.38313 | 0.07084 |
| **Cux2** | 0.00000 | 0.26875 | 1.10656 | 2.73447 | 0.00578 | 0.00615 | 0.00456 | 7.84514 | 1.79088 | 4.33626 | 5.37993 | 0.26292 | 1.17603 | 0.55869 | 1.30875 | 0.00482 |
| **Cuzd1** | 0.01370 | 0.00814 | 0.01214 | 0.08056 | 0.00934 | 0.00665 | 0.00754 | 0.01391 | 0.33001 | 0.01358 | 0.01026 | 0.00770 | 0.01281 | 0.01263 | 0.00649 | 0.00739 |
| **Cwc15** | 17.08240 | 11.30289 | 11.45166 | 13.49402 | 5.56166 | 10.76444 | 7.35586 | 14.22390 | 22.39698 | 21.62742 | 20.42583 | 12.70871 | 13.57726 | 15.80191 | 17.42529 | 8.13249 |
| **Cwc22** | 8.83962 | 7.23721 | 9.87182 | 20.79755 | 1.48016 | 4.19330 | 3.42576 | 8.80547 | 15.96720 | 12.83938 | 8.07167 | 5.58399 | 8.06694 | 14.33931 | 14.84748 | 3.78547 |
| **Cwc25** | 0.48457 | 0.42900 | 1.48327 | 4.03839 | 1.40318 | 1.20075 | 0.07676 | 3.69931 | 8.22936 | 9.81698 | 2.60981 | 0.92380 | 4.20681 | 0.62179 | 5.48628 | 1.48360 |
| **Cwc27** | 7.21902 | 5.57868 | 1.89885 | 0.30383 | 10.68064 | 7.76135 | 4.12984 | 2.49984 | 0.00000 | 2.52950 | 1.51837 | 9.63263 | 7.43809 | 6.51838 | 2.48021 | 1.04703 |
| **Cwf19l1** | 6.99788 | 3.05123 | 5.62859 | 0.00807 | 0.00000 | 1.14075 | 6.31480 | 6.23376 | 3.21977 | 4.84368 | 1.23733 | 5.07297 | 4.04971 | 3.58257 | 2.75424 | 0.00863 |
| **Cwf19l2** | 0.00000 | 4.23098 | 0.58638 | 0.07861 | 1.49973 | 0.00758 | 2.05793 | 1.62989 | 1.08029 | 1.53281 | 0.62761 | 0.00000 | 1.59170 | 5.55925 | 1.03367 | 0.01146 |
| **Cwh43** | 0.01456 | 0.00670 | 0.00302 | 0.00375 | 0.00865 | 0.00799 | 1.45598 | 0.00000 | 0.01071 | 1.31511 | 0.78144 | 0.00000 | 0.00560 | 0.01394 | 0.00000 | 0.00702 |
| **Cx3cl1** | 0.01295 | 0.00778 | 0.01196 | 1.13859 | 0.00891 | 0.00634 | 0.00717 | 0.01441 | 0.00000 | 1.09512 | 0.01457 | 0.00577 | 0.01618 | 0.01878 | 0.00260 | 0.00691 |
| **Cx3cr1** | 0.01491 | 0.04995 | 0.04911 | 0.10610 | 0.02716 | 0.00750 | 0.00736 | 0.01154 | 0.17184 | 0.03657 | 0.01078 | 0.00648 | 0.04285 | 0.28963 | 0.05358 | 0.00736 |
| **Cxadr** | 1.76932 | 0.20682 | 1.31548 | 5.32248 | 0.67845 | 7.05513 | 1.72699 | 3.06306 | 15.08282 | 4.17343 | 12.19441 | 4.87578 | 3.18570 | 22.24447 | 9.06526 | 1.00125 |
| **Cxcl10** | 0.01299 | 0.00773 | 0.01186 | 0.52046 | 0.01881 | 0.00659 | 0.00748 | 0.01311 | 0.00444 | 0.01140 | 0.01304 | 0.00772 | 0.01265 | 0.01874 | 0.07606 | 0.00731 |
| **Cxcl11** | 0.01352 | 0.00807 | 0.01113 | 1.41804 | 0.00906 | 0.00632 | 0.00739 | 0.00977 | 1.26835 | 0.24496 | 0.06863 | 0.00473 | 0.01676 | 0.01061 | 0.01507 | 0.00679 |
| **Cxcl12** | 0.01338 | 0.00801 | 0.01211 | 0.00767 | 0.00904 | 0.00670 | 0.00736 | 0.01104 | 0.01177 | 0.72276 | 0.01204 | 0.00650 | 0.01190 | 0.00953 | 0.03069 | 0.00758 |
| **Cxcl16** | 0.00610 | 0.00782 | 1.54237 | 8.90660 | 0.00714 | 0.00842 | 0.00700 | 0.30156 | 5.60192 | 1.15683 | 0.00000 | 0.00000 | 0.01471 | 0.00839 | 0.10252 | 0.00531 |
| **Cxcl9** | 0.01737 | 0.00787 | 0.01576 | 0.19525 | 0.00906 | 0.00627 | 0.00738 | 0.01181 | 0.00822 | 0.00928 | 0.01180 | 0.03743 | 0.01271 | 0.02673 | 0.54981 | 0.02197 |
| **Cxcr2** | 1.11119 | 0.12238 | 1.22364 | 5.82578 | 0.67735 | 0.22588 | 0.18499 | 0.00000 | 10.67512 | 0.00000 | 9.35408 | 0.62105 | 0.51696 | 3.63373 | 1.04792 | 0.14034 |
| **Cxx1a** | 0.04561 | 0.00769 | 0.00990 | 0.06867 | 0.00823 | 0.00675 | 0.00742 | 0.01118 | 0.09659 | 0.14026 | 0.02740 | 0.02685 | 0.01239 | 0.01144 | 0.00691 | 0.00751 |
| **Cxx1b** | 0.02660 | 0.00769 | 0.01028 | 0.11412 | 0.00860 | 0.00673 | 0.00730 | 0.03164 | 0.11999 | 0.13761 | 0.01155 | 0.03550 | 0.01165 | 0.01105 | 0.00595 | 0.00762 |
| **Cxx1c** | 0.04567 | 0.00769 | 0.00948 | 0.03731 | 0.00824 | 0.00680 | 0.00744 | 0.01178 | 0.29752 | 0.07771 | 0.01040 | 0.04741 | 0.01281 | 0.01297 | 0.00564 | 0.00744 |
| **Cxxc1** | 6.87682 | 1.53290 | 7.02816 | 1.74820 | 4.57259 | 8.92525 | 3.50819 | 9.02553 | 4.20423 | 4.85821 | 2.82807 | 0.80514 | 10.90505 | 0.02165 | 1.07958 | 2.89560 |
| **Cxxc5** | 0.27251 | 2.31286 | 0.45166 | 0.13178 | 0.02532 | 2.82436 | 0.00576 | 0.78991 | 0.87889 | 0.77561 | 0.34263 | 0.00915 | 0.10795 | 0.00368 | 0.00000 | 0.01996 |
| **Cyb561** | 0.14079 | 0.00745 | 0.70658 | 0.07499 | 0.01542 | 0.00719 | 0.16598 | 0.00876 | 0.23740 | 1.47211 | 0.00978 | 0.02122 | 1.88234 | 0.03791 | 0.00000 | 0.00647 |
| **Cyb561a3** | 2.75240 | 1.26188 | 3.83860 | 4.84793 | 0.37792 | 1.64215 | 2.68592 | 7.04962 | 2.85888 | 2.40808 | 1.75289 | 0.90477 | 2.35086 | 1.36077 | 3.75816 | 0.04383 |
| **Cyb561d1** | 0.01594 | 0.00808 | 0.01621 | 0.00817 | 0.01029 | 0.00644 | 0.00730 | 0.01214 | 0.01182 | 0.26689 | 0.01061 | 0.01916 | 0.40504 | 0.00591 | 0.22963 | 0.00721 |
| **Cyb561d2** | 5.54430 | 2.12886 | 1.97351 | 0.34724 | 0.03151 | 0.16241 | 4.96371 | 2.09288 | 1.17532 | 0.59216 | 1.51450 | 3.41346 | 2.56384 | 0.04212 | 0.00000 | 0.05451 |
| **Cyb5a** | 19.76599 | 4.15569 | 8.35099 | 0.00000 | 14.42997 | 3.10551 | 0.00000 | 2.13796 | 0.79539 | 3.47935 | 2.06694 | 1.68497 | 4.99869 | 0.97138 | 0.80094 | 4.30594 |
| **Cyb5b** | 3.43164 | 4.31691 | 7.29002 | 1.14098 | 3.02206 | 0.01651 | 5.66790 | 0.00000 | 3.03708 | 14.05273 | 0.84893 | 0.43750 | 1.23694 | 7.59905 | 7.36564 | 6.00999 |
| **Cyb5d1** | 0.01749 | 0.00788 | 0.00574 | 0.11488 | 0.00901 | 0.00668 | 0.02701 | 0.00711 | 0.22625 | 0.00000 | 0.08673 | 0.00303 | 0.01145 | 0.93532 | 0.39018 | 0.00724 |
| **Cyb5d2** | 1.70434 | 0.16784 | 0.71443 | 1.37173 | 0.02944 | 0.58176 | 0.05400 | 0.14217 | 0.65279 | 0.44381 | 0.94571 | 0.05839 | 1.50150 | 1.40352 | 0.79971 | 2.24159 |
| **Cyb5r1** | 11.95660 | 5.16337 | 4.68469 | 0.00000 | 5.64379 | 7.89297 | 8.07311 | 3.73815 | 0.01056 | 0.00000 | 1.67521 | 0.14590 | 5.97353 | 0.52846 | 0.12753 | 8.85214 |
| **Cyb5r2** | 0.63307 | 0.05008 | 0.23862 | 0.52850 | 0.01047 | 0.00487 | 4.35161 | 0.01104 | 0.18599 | 0.27650 | 0.01053 | 0.02269 | 0.01531 | 0.00874 | 0.21178 | 0.00642 |
| **Cyb5r3** | 4.79528 | 4.73030 | 5.63567 | 1.33792 | 4.88099 | 14.23466 | 5.35395 | 1.64431 | 1.35575 | 6.39938 | 1.42878 | 0.17149 | 4.05758 | 0.00000 | 0.21442 | 3.90626 |
| **Cyb5r4** | 3.67466 | 0.25792 | 1.27449 | 6.53466 | 3.68454 | 4.02564 | 2.19451 | 5.59167 | 3.29415 | 10.35396 | 6.33001 | 6.67043 | 6.14865 | 12.09872 | 7.22444 | 1.55055 |
| **Cyb5rl** | 0.12738 | 0.00790 | 2.77534 | 0.09179 | 0.00986 | 0.00553 | 0.00712 | 0.00985 | 0.01015 | 0.04539 | 0.00934 | 0.02943 | 0.00605 | 0.00995 | 0.12951 | 0.01373 |
| **Cyba** | 0.00538 | 0.00709 | 0.12689 | 0.06028 | 0.00942 | 0.01870 | 0.00446 | 1.97264 | 0.01934 | 2.06875 | 3.97073 | 0.00000 | 2.41374 | 0.00146 | 0.09557 | 0.00583 |

| | | | | | | | | | | | | | | | | |
|---|---|---|---|---|---|---|---|---|---|---|---|---|---|---|---|---|
| **Cybb** | 0.35348 | 0.00743 | 0.00081 | 0.00545 | 0.00718 | 0.00752 | 0.00692 | 0.01046 | 0.01010 | 0.14000 | 0.00796 | 0.21145 | 0.01292 | 0.00582 | 0.89914 | 0.00733 |
| **Cybrd1** | 0.02307 | 0.01886 | 0.14322 | 0.25424 | 0.03640 | 0.00680 | 0.06923 | 0.00752 | 0.14230 | 0.04543 | 0.36878 | 0.09181 | 0.10197 | 0.09493 | 0.05430 | 0.01477 |
| **Cyc1** | 19.12319 | 6.12586 | 9.48627 | 8.87916 | 16.71515 | 12.63904 | 6.88359 | 6.85958 | 0.15434 | 14.32385 | 9.45382 | 2.64599 | 16.29643 | 0.10062 | 2.66964 | 12.67696 |
| **Cycs** | 24.12400 | 10.03802 | 20.29714 | 9.13388 | 12.65995 | 10.05806 | 11.71109 | 15.27953 | 17.57166 | 18.58573 | 26.20862 | 17.06183 | 19.86145 | 14.13300 | 12.29358 | 8.77848 |
| **Cyct** | 2.00640 | 0.55007 | 0.44174 | 0.00246 | 0.00658 | 0.00954 | 0.00476 | 0.00000 | 0.01059 | 0.99326 | 1.84203 | 0.00000 | 0.00000 | 0.02217 | 0.18015 | 0.00481 |
| **Cyfip1** | 0.40386 | 2.70808 | 4.92368 | 5.05917 | 10.22950 | 5.70834 | 6.72406 | 4.41625 | 1.53311 | 6.15039 | 4.95607 | 11.32744 | 1.82153 | 3.87425 | 5.63274 | 3.59684 |
| **Cyfip2** | 1.91998 | 0.00569 | 0.00000 | 0.64003 | 0.00427 | 0.00576 | 0.00565 | 0.00001 | 2.27090 | 0.24623 | 0.00000 | 0.01531 | 0.01116 | 1.60252 | 0.00000 | 0.01171 |
| **Cyhr1** | 4.26358 | 2.44057 | 3.40852 | 3.23566 | 1.49255 | 3.67362 | 3.61088 | 0.61444 | 3.29406 | 2.14062 | 2.67785 | 0.02965 | 3.47524 | 0.06721 | 0.00000 | 1.84432 |
| **Cyld** | 0.28961 | 0.20342 | 0.69606 | 3.87959 | 0.23640 | 0.05531 | 0.22239 | 0.17297 | 4.06805 | 0.54590 | 3.84302 | 0.52796 | 0.23047 | 2.30792 | 1.99793 | 0.09978 |
| **Cyp11a1** | 0.01298 | 0.00757 | 0.25459 | 0.00472 | 0.00790 | 0.00570 | 0.00715 | 0.00913 | 0.01032 | 9.48099 | 0.00749 | 0.00000 | 0.00966 | 0.01610 | 0.00029 | 0.00724 |
| **Cyp17a1** | 0.00422 | 0.00820 | 1.95752 | 0.00512 | 0.00629 | 0.00586 | 0.00666 | 0.08855 | 0.18891 | 0.27852 | 0.00709 | 0.00645 | 1.47460 | 0.00619 | 0.46975 | 0.00613 |
| **Cyp1a2** | 0.01073 | 0.00792 | 0.01420 | 0.10450 | 0.00990 | 0.00715 | 0.00732 | 0.01200 | 0.00994 | 0.70947 | 0.01581 | 0.00516 | 0.53114 | 0.01212 | 0.01143 | 0.00803 |
| **Cyp1b1** | 0.01368 | 0.00766 | 0.01198 | 0.35775 | 0.00918 | 0.00662 | 0.00734 | 0.01125 | 0.00658 | 0.27893 | 0.01174 | 0.00636 | 0.01261 | 0.00918 | 0.14061 | 0.00774 |
| **Cyp20a1** | 3.63065 | 0.07659 | 2.59311 | 1.89562 | 0.23630 | 1.14790 | 0.65288 | 5.76104 | 0.75248 | 3.13551 | 5.55361 | 0.41111 | 2.17288 | 1.90710 | 2.30197 | 0.30576 |
| **Cyp26a1** | 4.86620 | 0.00263 | 0.00367 | 0.00838 | 0.00000 | 0.00433 | 0.00373 | 0.00351 | 0.02631 | 0.23778 | 0.01098 | 0.00532 | 0.01139 | 0.01176 | 0.01227 | 0.00231 |
| **Cyp26b1** | 0.01364 | 0.00777 | 0.01007 | 0.11062 | 0.00921 | 0.00671 | 0.00723 | 0.10817 | 0.00998 | 0.01310 | 0.01300 | 0.01739 | 0.00856 | 0.01173 | 0.00652 | 0.00736 |
| **Cyp2ab1** | 0.01402 | 0.00791 | 0.00799 | 0.35738 | 0.00905 | 0.00676 | 0.00741 | 0.00031 | 0.13020 | 0.00000 | 0.60057 | 0.01409 | 0.00670 | 0.00418 | 0.22877 | 0.00700 |
| **Cyp2b10** | 0.01848 | 0.00749 | 0.01013 | 0.09974 | 0.00861 | 0.00643 | 0.00705 | 0.00985 | 1.21727 | 0.00000 | 0.00000 | 0.02557 | 0.54416 | 0.02717 | 0.10761 | 0.00684 |
| **Cyp2b13** | 0.05155 | 0.00767 | 0.00989 | 0.01074 | 0.00803 | 0.00672 | 0.00748 | 0.01228 | 0.01232 | 0.01314 | 0.01258 | 0.00801 | 0.05845 | 0.01298 | 0.02111 | 0.00724 |
| **Cyp2b23** | 1.54224 | 0.00596 | 0.00361 | 0.02666 | 0.00499 | 0.00675 | 0.00625 | 0.01301 | 0.01061 | 0.01872 | 0.01202 | 0.01674 | 0.01821 | 0.01106 | 0.01524 | 0.02018 |
| **Cyp2c44** | 0.57462 | 2.25323 | 0.00000 | 0.94432 | 0.03616 | 0.01154 | 0.00560 | 0.05947 | 0.79622 | 0.50842 | 0.25925 | 0.03833 | 0.19954 | 0.86655 | 2.79799 | 0.00616 |
| **Cyp2c55** | 0.01391 | 0.00775 | 0.00952 | 0.00903 | 0.00931 | 0.00679 | 0.00732 | 0.01123 | 0.26826 | 0.00913 | 0.00969 | 0.21789 | 0.01316 | 0.01288 | 0.00324 | 0.00745 |
| **Cyp2d10** | 0.01951 | 0.00822 | 0.01065 | 0.01894 | 0.00881 | 0.00687 | 0.00784 | 0.05333 | 0.01109 | 0.03081 | 0.01196 | 0.00882 | 0.01074 | 0.01247 | 0.01051 | 0.00742 |
| **Cyp2d12** | 0.01047 | 0.00818 | 0.00948 | 0.03914 | 0.00957 | 0.00707 | 0.00767 | 0.17095 | 0.03577 | 0.06100 | 0.02292 | 0.02022 | 0.01719 | 0.01214 | 0.00784 | 0.00742 |
| **Cyp2d22** | 0.01329 | 0.00778 | 0.01119 | 0.00631 | 0.00891 | 0.00672 | 0.00725 | 0.00757 | 0.01146 | 1.05681 | 0.09717 | 0.01445 | 0.01091 | 0.01134 | 0.00679 | 0.00766 |
| **Cyp2d40** | 0.01305 | 0.00826 | 0.01128 | 0.09752 | 0.00930 | 0.00670 | 0.00770 | 0.02787 | 0.49456 | 0.02240 | 0.02679 | 0.01853 | 0.01173 | 0.01245 | 0.00480 | 0.00746 |
| **Cyp2d9** | 0.00803 | 0.00782 | 0.00735 | 0.07200 | 0.00941 | 0.00722 | 0.00772 | 2.21698 | 0.06423 | 0.73472 | 0.00000 | 0.01427 | 0.00571 | 0.01274 | 0.00000 | 0.00740 |
| **Cyp2e1** | 0.39454 | 0.02848 | 0.47283 | 3.25907 | 0.29624 | 0.06508 | 0.07731 | 0.00000 | 4.90283 | 0.00000 | 3.92583 | 0.28541 | 0.11049 | 1.29955 | 0.21132 | 0.16236 |
| **Cyp2f2** | 0.00789 | 0.00769 | 0.04543 | 0.17769 | 0.00896 | 0.00588 | 0.00746 | 0.39315 | 0.15174 | 2.09760 | 0.00436 | 0.04896 | 0.00607 | 0.00491 | 0.81943 | 0.00753 |
| **Cyp2j13** | 0.01909 | 0.00794 | 0.09400 | 0.12040 | 0.00858 | 0.00618 | 0.06031 | 0.04484 | 0.12504 | 0.08507 | 0.03913 | 0.29381 | 0.00937 | 0.13309 | 0.18694 | 0.05096 |
| **Cyp2j6** | 0.48153 | 0.00746 | 0.00542 | 0.01062 | 0.00688 | 0.00711 | 0.00712 | 0.01438 | 0.01235 | 0.01534 | 0.01280 | 0.00778 | 0.01215 | 0.01229 | 0.01080 | 0.00680 |
| **Cyp2s1** | 0.30099 | 9.62306 | 17.52741 | 2.13716 | 2.59319 | 12.86903 | 8.54753 | 25.74402 | 4.97392 | 7.58433 | 19.66168 | 5.21529 | 27.71331 | 2.57340 | 1.47803 | 8.00525 |
| **Cyp2t4** | 0.15213 | 0.00689 | 1.13106 | 0.00892 | 0.00593 | 0.01873 | 0.00665 | 2.32389 | 0.09928 | 0.00000 | 0.84019 | 0.00533 | 0.20578 | 0.00260 | 0.28932 | 0.00691 |
| **Cyp39a1** | 0.01317 | 0.00765 | 0.01007 | 4.47119 | 0.00912 | 0.00585 | 0.00709 | 0.00926 | 1.74977 | 0.39540 | 0.01053 | 0.00169 | 0.01129 | 0.06861 | 0.95613 | 0.00626 |
| **Cyp46a1** | 0.01126 | 0.00838 | 0.01778 | 0.93828 | 0.00966 | 0.00641 | 0.01808 | 0.01350 | 0.78776 | 0.00129 | 0.04085 | 0.00689 | 0.42359 | 0.01167 | 0.00154 | 0.00705 |
| **Cyp4f13** | 0.00662 | 0.00800 | 0.01257 | 0.01018 | 0.00906 | 0.10882 | 0.00638 | 0.00897 | 0.01247 | 0.07433 | 0.01159 | 0.00659 | 1.95809 | 0.01133 | 0.01076 | 0.00630 |
| **Cyp4f16** | 2.99189 | 0.00395 | 0.00301 | 0.02818 | 0.00363 | 0.00510 | 1.37352 | 0.00000 | 0.12346 | 0.46228 | 0.01050 | 0.00342 | 0.01644 | 0.01262 | 0.01519 | 0.00389 |
| **Cyp4f17** | 0.01678 | 0.01795 | 0.00836 | 0.01036 | 0.00889 | 0.00683 | 0.60325 | 0.01027 | 0.01234 | 0.01571 | 0.01281 | 0.01719 | 0.01797 | 0.01173 | 0.01125 | 0.01449 |
| **Cyp4f37** | 0.07395 | 0.00789 | 0.00830 | 0.01040 | 0.00788 | 0.00688 | 0.10987 | 0.01329 | 0.01243 | 0.05209 | 0.01218 | 0.00775 | 0.01171 | 0.01145 | 0.01077 | 0.00707 |
| **Cyp4f39** | 2.32940 | 0.00532 | 1.46984 | 1.20710 | 0.03325 | 2.93229 | 0.00303 | 0.00000 | 0.31178 | 0.00000 | 0.07921 | 0.01278 | 3.55774 | 0.02525 | 0.04755 | 0.01520 |
| **Cyp4f41.ps** | 0.01297 | 0.00810 | 0.08584 | 0.01058 | 0.00989 | 0.01629 | 0.01834 | 0.01563 | 0.01232 | 0.01291 | 0.01472 | 0.00773 | 0.08108 | 0.01257 | 0.00910 | 0.00767 |
| **Cyp4v3** | 0.01268 | 0.00765 | 0.00728 | 0.41108 | 0.00946 | 0.00672 | 0.00730 | 0.01045 | 0.00510 | 0.22410 | 0.01706 | 0.00354 | 0.01272 | 0.82994 | 0.55758 | 0.00737 |
| **Cyp51** | 3.48902 | 4.62058 | 5.36164 | 2.59686 | 0.07513 | 0.00900 | 6.11258 | 2.54603 | 1.44664 | 0.38092 | 1.20389 | 0.11565 | 3.08019 | 2.78458 | 0.00000 | 2.45049 |
| **Cypt3** | 0.01369 | 0.00805 | 0.01132 | 0.50437 | 0.00933 | 0.00652 | 0.00740 | 0.01215 | 0.45840 | 0.00500 | 0.00958 | 0.03964 | 0.01275 | 0.00694 | 0.22513 | 0.01452 |
| **Cyr61** | 0.99420 | 0.00511 | 0.10970 | 0.18186 | 0.00564 | 0.00807 | 1.58550 | 2.12954 | 0.31557 | 0.03529 | 0.52955 | 0.12040 | 0.98985 | 0.39906 | 1.14076 | 0.11105 |
| **Cys1** | 0.01373 | 0.00782 | 0.01196 | 0.00963 | 0.00919 | 0.00654 | 0.00737 | 0.01156 | 0.01213 | 0.21270 | 0.01435 | 0.01866 | 0.01151 | 0.01884 | 0.00740 | 0.00715 |
| **Cysrt1** | 0.00961 | 0.00804 | 0.00729 | 0.01030 | 0.00912 | 0.37278 | 0.00689 | 0.01178 | 0.01230 | 0.02866 | 0.01148 | 0.00836 | 1.12905 | 0.01198 | 0.01197 | 0.00668 |
| **Cystm1** | 7.23400 | 2.89962 | 6.05793 | 1.00584 | 1.74983 | 7.02509 | 4.42643 | 7.08802 | 0.00030 | 7.72017 | 3.27713 | 1.13078 | 7.42721 | 0.01793 | 0.08304 | 3.66842 |
| **Cyth1** | 0.00935 | 0.00794 | 0.05646 | 0.00554 | 0.00871 | 0.00749 | 0.00730 | 0.01199 | 0.35775 | 0.06278 | 0.00929 | 0.00735 | 0.35385 | 0.01227 | 0.02114 | 0.20891 |
| **Cyth2** | 7.94995 | 1.08754 | 9.42143 | 5.43711 | 5.83780 | 4.62306 | 4.02075 | 14.13113 | 4.43791 | 13.99324 | 12.76005 | 1.72237 | 5.75554 | 4.54433 | 9.21006 | 2.68515 |
| **Cyth3** | 2.17129 | 0.00420 | 0.43106 | 3.79291 | 0.00741 | 0.00419 | 0.00439 | 2.13218 | 0.00000 | 1.39675 | 0.62377 | 3.77048 | 0.52389 | 1.78906 | 2.56044 | 0.00416 |
| **D10Jhu81e** | 4.00526 | 0.00277 | 1.61658 | 0.28878 | 0.20954 | 4.51718 | 0.17054 | 0.85513 | 0.30701 | 1.01136 | 0.56515 | 0.00000 | 0.00000 | 0.00492 | 0.00000 | 2.12096 |
| **D10Wsu102e** | 6.88131 | 7.71714 | 3.02926 | 14.59020 | 8.71904 | 4.35617 | 4.74777 | 10.40243 | 13.28943 | 11.01004 | 4.45757 | 4.57092 | 6.08945 | 5.23422 | 14.65361 | 6.53787 |
| **D11Wsu47e** | 2.89621 | 0.05037 | 5.02759 | 3.73615 | 4.39488 | 0.06585 | 0.00175 | 0.00000 | 4.56698 | 3.13344 | 2.56786 | 0.36896 | 2.56164 | 2.50249 | 2.85947 | 0.43889 |
| **D16Ertd472e** | 6.83052 | 0.00000 | 3.97094 | 13.08563 | 0.03320 | 0.00315 | 0.84914 | 2.12757 | 7.67345 | 5.29105 | 0.00000 | 0.30320 | 1.70226 | 0.24322 | 8.66604 | 0.96784 |
| **D16Ertd519e** | 0.01375 | 0.00831 | 0.01234 | 2.84991 | 0.00932 | 0.00640 | 0.00743 | 0.01392 | 0.00000 | 0.01445 | 0.01353 | 0.00845 | 0.01263 | 0.00947 | 0.00269 | 0.00693 |
| **D17Ertd648e** | 0.14352 | 0.16195 | 0.44872 | 43.74434 | 0.03635 | 0.16401 | 0.32430 | 2.98255 | 44.52124 | 12.14586 | 4.98029 | 3.23232 | 1.81080 | 11.12885 | 19.09024 | 0.13194 |
| **D17H6S53E** | 0.29330 | 0.04825 | 0.30617 | 1.10106 | 0.07082 | 0.03713 | 0.10794 | 0.00000 | 1.19814 | 1.13195 | 3.72308 | 0.07985 | 0.44699 | 0.70976 | 0.33173 | 0.11363 |
| **D17H6S56E.5** | 1.92132 | 0.87733 | 5.53989 | 6.21742 | 1.35921 | 1.98551 | 4.28205 | 2.94172 | 4.98239 | 6.06878 | 4.32942 | 1.77397 | 1.57512 | 2.57902 | 7.07143 | 0.70754 |
| **D17Wsu92e** | 5.78831 | 3.15117 | 6.33534 | 14.25972 | 2.40075 | 4.82375 | 4.76889 | 9.34657 | 14.93627 | 9.14917 | 5.25171 | 0.44508 | 10.45192 | 0.56016 | 5.79319 | 3.87348 |
| **D1Ertd622e** | 1.44552 | 0.00358 | 0.86270 | 17.20015 | 0.86121 | 0.01228 | 0.00485 | 0.00000 | 5.21400 | 1.34001 | 0.00000 | 6.07853 | 0.01556 | 0.01591 | 6.09825 | 0.00185 |
| **D1Pas1** | 0.00745 | 0.00577 | 0.53472 | 7.45242 | 0.00784 | 0.00394 | 0.00616 | 0.00817 | 1.29545 | 0.00000 | 0.00000 | 7.06371 | 0.02331 | 1.06020 | 1.85749 | 0.01255 |

| | | | | | | | | | | | | | | | | |
|---|---|---|---|---|---|---|---|---|---|---|---|---|---|---|---|---|
| **D2hgdh** | 3.01636 | 2.67091 | 1.93248 | 0.53329 | 0.66126 | 0.01627 | 2.87575 | 1.49474 | 2.30258 | 0.00000 | 1.70879 | 0.05484 | 1.77918 | 0.00000 | 2.00782 | 1.40736 |
| **D2Wsu81e** | 6.73154 | 6.02661 | 8.53523 | 3.00036 | 4.75427 | 4.39555 | 1.54490 | 6.05905 | 3.45696 | 5.31866 | 0.87137 | 0.39779 | 1.89236 | 0.00000 | 0.69835 | 6.22938 |
| **D3Ertd254e** | 0.41687 | 0.98122 | 1.02546 | 2.49339 | 0.19090 | 1.29138 | 0.05727 | 1.96981 | 4.64629 | 0.40771 | 4.16708 | 0.35558 | 0.07372 | 5.05752 | 5.94876 | 0.03370 |
| **D3Ertd751e** | 0.01253 | 0.00786 | 0.01250 | 0.28160 | 0.01023 | 0.00710 | 0.05028 | 0.01374 | 0.00749 | 0.00528 | 0.01203 | 0.04676 | 0.20527 | 0.01562 | 0.31253 | 0.00753 |
| **D4Ertd617e** | 0.01376 | 0.00812 | 0.01205 | 0.34268 | 0.00931 | 0.00658 | 0.00745 | 0.01294 | 0.00559 | 0.02296 | 0.01400 | 0.00855 | 0.01243 | 0.03312 | 0.04962 | 0.00726 |
| **D5Ertd577e** | 0.42563 | 0.33202 | 0.57261 | 4.72961 | 0.15176 | 0.10578 | 0.35729 | 1.43231 | 11.36017 | 4.19790 | 4.86464 | 7.32662 | 1.84238 | 8.01913 | 4.67476 | 0.00289 |
| **D5Ertd579e** | 1.48002 | 1.41246 | 0.29245 | 0.41828 | 0.05351 | 0.13061 | 0.09280 | 1.20315 | 0.69993 | 1.40225 | 0.15703 | 0.07854 | 2.44440 | 0.00438 | 0.28055 | 3.44443 |
| **D5Ertd605e** | 0.02060 | 0.03790 | 1.56276 | 0.75475 | 0.00577 | 3.14678 | 0.22834 | 0.93767 | 1.81228 | 2.54278 | 1.66442 | 0.10103 | 0.00000 | 0.08682 | 1.72777 | 0.71929 |
| **D5Ertd615e** | 0.08825 | 0.06901 | 0.16821 | 0.42297 | 0.01595 | 0.03055 | 0.11896 | 0.24991 | 0.61050 | 0.35185 | 0.12971 | 0.10112 | 0.08102 | 0.73886 | 0.38898 | 0.15070 |
| **D6Ertd474e** | 0.01053 | 0.00751 | 0.00312 | 11.47199 | 0.00902 | 0.00483 | 0.00665 | 0.00563 | 1.81219 | 1.65990 | 0.07257 | 0.09845 | 0.00997 | 0.88932 | 4.90679 | 0.00499 |
| **D6Ertd527e** | 0.01257 | 0.00802 | 0.01090 | 2.22501 | 0.00977 | 0.00700 | 0.00758 | 0.00910 | 3.02156 | 0.49524 | 0.12909 | 0.06529 | 0.09563 | 0.00388 | 0.06211 | 0.00714 |
| **D6Wsu163e** | 2.20954 | 3.10680 | 2.81327 | 0.00000 | 0.02095 | 0.00358 | 0.02186 | 4.96967 | 0.51946 | 6.97058 | 5.82705 | 8.81389 | 0.23393 | 3.79597 | 8.75587 | 0.84353 |
| **D7Ertd143e** | 1.85888 | 0.20271 | 2.46366 | 1.07770 | 0.01244 | 0.00376 | 0.18585 | 2.48473 | 3.47269 | 0.55499 | 1.89647 | 0.03430 | 0.16740 | 0.01122 | 0.00000 | 2.15755 |
| **D7Ertd443e** | 0.00358 | 0.00763 | 0.00429 | 0.17286 | 0.00920 | 0.00564 | 0.00690 | 0.01417 | 0.81440 | 3.45364 | 0.00000 | 0.06007 | 0.01022 | 7.98934 | 14.18706 | 0.00692 |
| **D8Ertd738e** | 16.65863 | 0.83977 | 5.89743 | 13.68134 | 4.23883 | 5.06671 | 3.39836 | 11.98944 | 27.21024 | 23.37469 | 17.20754 | 2.56554 | 14.11756 | 0.24577 | 0.00000 | 6.19487 |
| **Daam1** | 2.81849 | 0.03294 | 2.34052 | 5.19894 | 3.59581 | 0.74691 | 0.92334 | 1.60415 | 3.03246 | 1.37112 | 0.95643 | 0.12231 | 0.33515 | 0.01240 | 0.49357 | 3.67257 |
| **Dab1** | 0.01246 | 0.00801 | 0.31073 | 0.28737 | 0.00873 | 0.00553 | 0.00760 | 0.01330 | 1.18194 | 0.00632 | 0.00010 | 0.00589 | 0.00975 | 0.00618 | 0.13470 | 0.00705 |
| **Dab2** | 0.76076 | 6.21336 | 11.20358 | 5.46799 | 2.73615 | 20.93419 | 7.97230 | 12.92800 | 2.14252 | 8.84365 | 4.37752 | 4.74108 | 23.01347 | 4.46185 | 0.00000 | 14.09656 |
| **Dab2ip** | 1.02454 | 0.20553 | 0.84800 | 0.00000 | 0.82991 | 4.65665 | 3.17677 | 2.46870 | 0.00868 | 0.00000 | 0.26027 | 0.00746 | 1.86732 | 0.01860 | 0.03723 | 1.49539 |
| **Dact1** | 0.01357 | 0.00764 | 0.01181 | 0.95381 | 0.00920 | 0.00645 | 0.00729 | 0.01090 | 0.00000 | 0.27145 | 0.01093 | 0.00515 | 0.01266 | 0.00342 | 0.64263 | 0.00764 |
| **Dact3** | 0.00368 | 0.00646 | 0.00732 | 0.15351 | 0.00948 | 0.00649 | 2.01382 | 0.00731 | 1.44086 | 1.03560 | 0.01163 | 0.00860 | 0.75447 | 0.00594 | 0.26641 | 0.00685 |
| **Dad1** | 7.72040 | 3.66655 | 11.26886 | 3.33680 | 2.89289 | 4.97324 | 4.43925 | 5.44899 | 4.76559 | 9.28476 | 9.80676 | 2.29109 | 10.67419 | 0.64318 | 2.37023 | 7.93270 |
| **Dag1** | 3.36320 | 0.00000 | 7.04317 | 4.79283 | 0.07134 | 8.49163 | 7.02098 | 4.60090 | 3.60300 | 5.98544 | 2.16097 | 0.46827 | 9.72176 | 0.49199 | 1.26990 | 4.74521 |
| **Dagla** | 0.00836 | 0.00814 | 0.01788 | 2.11897 | 0.00790 | 0.00686 | 0.00735 | 0.01376 | 0.00000 | 0.00933 | 0.01293 | 0.00838 | 0.00854 | 0.01034 | 0.00261 | 0.94348 |
| **Daglb** | 0.39240 | 0.00699 | 0.00253 | 0.57965 | 0.00771 | 0.00781 | 0.42468 | 0.00000 | 2.00807 | 0.00000 | 0.70335 | 0.01642 | 0.12479 | 0.00196 | 0.53395 | 0.00671 |
| **Dalrd3** | 7.04980 | 1.01514 | 3.90197 | 0.19331 | 2.98325 | 4.74036 | 1.56260 | 8.49728 | 0.10613 | 7.54285 | 1.19241 | 0.55671 | 9.63167 | 0.18968 | 0.69063 | 2.75051 |
| **Dancr** | 1.39070 | 2.19425 | 1.10493 | 6.39092 | 1.20525 | 0.04723 | 0.00196 | 0.00000 | 11.50286 | 4.27336 | 4.37532 | 0.29526 | 2.73053 | 0.12211 | 0.27120 | 0.01560 |
| **Dand5** | 0.06765 | 0.00789 | 0.00833 | 1.71484 | 0.02603 | 0.00664 | 0.00744 | 0.01245 | 1.88640 | 0.01249 | 0.00000 | 0.01604 | 0.02639 | 0.01169 | 0.00406 | 0.00695 |
| **Dap** | 10.72314 | 0.67819 | 6.42222 | 0.15676 | 13.02075 | 15.84468 | 6.45023 | 13.60426 | 0.00000 | 1.03428 | 8.69624 | 0.20049 | 14.50099 | 0.08611 | 0.00000 | 4.57863 |
| **Dap3** | 16.87388 | 6.98262 | 15.35341 | 3.41243 | 7.70803 | 8.65787 | 5.94878 | 9.93261 | 6.26829 | 9.49033 | 5.72961 | 7.20419 | 12.59502 | 0.34350 | 4.01028 | 8.37549 |
| **Dapk1** | 0.00871 | 0.00726 | 0.13349 | 0.00264 | 0.00970 | 1.27675 | 1.52999 | 0.01579 | 0.01129 | 2.30431 | 0.00601 | 0.04200 | 1.09937 | 0.17587 | 0.00000 | 0.00639 |
| **Dapk2** | 0.49920 | 0.00749 | 0.00537 | 0.10014 | 0.00687 | 0.00712 | 0.00710 | 0.01439 | 0.00760 | 0.01538 | 0.01213 | 0.00783 | 0.01214 | 0.01181 | 0.00902 | 0.00677 |
| **Dapk3** | 0.03104 | 0.01945 | 0.01136 | 0.00621 | 0.01467 | 0.00724 | 0.00754 | 0.05571 | 0.42426 | 0.01545 | 0.01135 | 0.00684 | 0.16888 | 0.01772 | 0.02765 | 0.00769 |
| **Dapp1** | 0.02765 | 0.00846 | 0.00791 | 0.31959 | 0.01240 | 0.00721 | 0.00747 | 0.00599 | 0.00439 | 0.01145 | 0.60706 | 0.00531 | 0.01047 | 0.00740 | 0.21639 | 0.02226 |
| **Dars** | 17.35898 | 11.32103 | 12.28751 | 4.57070 | 9.88929 | 7.98795 | 11.08525 | 15.82302 | 4.23720 | 10.58650 | 11.03977 | 12.83871 | 11.60180 | 8.95634 | 15.10284 | 4.68761 |
| **Dars2** | 3.44299 | 0.07376 | 5.87264 | 1.34208 | 5.68849 | 0.38423 | 3.93307 | 5.97760 | 2.53693 | 3.85444 | 4.19764 | 6.51857 | 0.94172 | 3.26415 | 5.43865 | 4.23389 |
| **Daw1** | 0.01352 | 0.00767 | 0.01142 | 1.15230 | 0.00928 | 0.00627 | 0.00738 | 0.01164 | 1.85507 | 0.00400 | 0.00000 | 0.00358 | 0.01289 | 0.00566 | 0.40957 | 0.00697 |
| **Daxx** | 1.55132 | 0.19446 | 6.15207 | 1.21449 | 4.68666 | 0.38881 | 3.40973 | 3.95441 | 2.83853 | 7.99721 | 0.29789 | 3.02017 | 3.34245 | 0.08121 | 0.89433 | 3.71247 |
| **Dazap1** | 5.58183 | 0.00000 | 0.00000 | 1.92968 | 0.57108 | 0.40103 | 0.03055 | 1.37619 | 2.88997 | 2.65348 | 2.55701 | 0.06161 | 2.68677 | 0.00000 | 0.00000 | 3.45374 |
| **Dazap2** | 10.46800 | 6.74107 | 13.73750 | 13.14970 | 7.39977 | 5.21511 | 7.20003 | 15.42236 | 11.27516 | 18.79587 | 8.71039 | 8.61788 | 12.24933 | 13.34683 | 11.68230 | 5.50391 |
| **Dazl** | 0.00000 | 0.01401 | 0.00000 | 3.27234 | 0.00800 | 0.00325 | 0.00406 | 0.00000 | 0.65793 | 2.69960 | 3.18678 | 15.02005 | 0.26693 | 2.06899 | 1.19538 | 0.06143 |
| **Dbf4** | 4.10957 | 5.71788 | 1.61941 | 20.07050 | 6.69343 | 2.73182 | 3.17019 | 7.97282 | 9.44300 | 12.08690 | 7.84321 | 5.26080 | 3.22051 | 16.99764 | 26.27574 | 0.57486 |
| **Dbhos** | 0.22487 | 0.09191 | 1.00745 | 6.69646 | 0.00677 | 0.28643 | 0.11808 | 3.55080 | 11.12779 | 3.77978 | 2.93956 | 1.75075 | 0.75821 | 2.63351 | 1.58569 | 0.13964 |
| **Dbi** | 6.76364 | 0.23002 | 2.86855 | 9.46911 | 7.60828 | 1.10039 | 1.13975 | 0.66671 | 4.60589 | 1.20207 | 2.61373 | 0.38466 | 1.05663 | 0.13626 | 2.78301 | 3.59275 |
| **Dbil5** | 0.05002 | 0.01920 | 0.00832 | 0.77633 | 0.00799 | 0.00674 | 0.00739 | 0.00748 | 0.00000 | 0.00000 | 0.31435 | 0.00715 | 0.00885 | 0.00617 | 0.10112 | 0.00697 |
| **Dbn1** | 0.00838 | 0.00747 | 0.00854 | 0.00629 | 0.00934 | 0.00678 | 0.00711 | 0.46822 | 0.02488 | 1.43997 | 0.00923 | 0.00308 | 0.00593 | 0.01799 | 0.00000 | 0.00729 |
| **Dbndd1** | 0.01323 | 0.00798 | 0.01169 | 2.25995 | 0.00945 | 0.00645 | 0.00748 | 0.01112 | 1.76630 | 0.43540 | 0.01626 | 0.03331 | 0.02042 | 0.00499 | 0.17582 | 0.00681 |
| **Dbndd2** | 0.00000 | 0.00726 | 0.14101 | 3.20091 | 0.21549 | 0.02817 | 0.00646 | 0.03153 | 4.04664 | 1.79118 | 0.45582 | 0.00000 | 1.16762 | 0.00000 | 0.66362 | 0.52520 |
| **Dbnl** | 9.77777 | 2.82533 | 7.25154 | 2.91096 | 7.66356 | 6.88386 | 7.16037 | 15.99727 | 7.62116 | 13.04522 | 9.48299 | 3.72457 | 10.35150 | 3.47408 | 3.23285 | 6.53106 |
| **Dbp** | 0.61946 | 0.00732 | 0.00497 | 0.00961 | 0.00780 | 0.00704 | 0.00693 | 0.01025 | 0.01226 | 0.22458 | 0.01254 | 0.00688 | 0.01153 | 0.01228 | 0.01012 | 0.00675 |
| **Dbr1** | 4.81447 | 2.02760 | 0.60512 | 4.09788 | 2.24514 | 4.02648 | 1.14514 | 0.25515 | 2.40833 | 1.95037 | 2.76080 | 0.04926 | 3.04639 | 1.55974 | 4.77339 | 1.78060 |
| **Dbt** | 1.37029 | 4.94282 | 0.67403 | 2.89637 | 1.58953 | 1.64512 | 0.93400 | 2.40092 | 5.42397 | 1.02252 | 4.94984 | 3.50064 | 1.86978 | 14.11866 | 9.60821 | 0.13194 |
| **Dcaf10** | 6.80564 | 0.07008 | 3.84500 | 7.56848 | 0.07133 | 2.25343 | 4.36448 | 1.00014 | 13.82421 | 4.63064 | 5.22235 | 8.05234 | 1.65301 | 14.33913 | 8.03916 | 0.78112 |
| **Dcaf11** | 1.61418 | 9.20552 | 6.38645 | 4.47366 | 6.03182 | 6.92018 | 8.93821 | 6.40599 | 7.97359 | 10.22086 | 3.76146 | 2.32869 | 4.07698 | 5.03271 | 7.15269 | 1.17307 |
| **Dcaf12** | 10.40354 | 5.45269 | 7.23862 | 11.04336 | 5.57235 | 7.60674 | 4.97519 | 8.80430 | 17.56789 | 8.58953 | 8.02198 | 7.13162 | 4.41450 | 4.60577 | 11.96911 | 7.27932 |
| **Dcaf12l1** | 14.32307 | 0.00001 | 1.68908 | 0.00000 | 12.31357 | 0.00000 | 0.00510 | 1.79070 | 1.85125 | 0.84472 | 0.00000 | 0.00000 | 1.21805 | 0.51317 | 0.25742 | 0.00000 |
| **Dcaf13** | 8.31604 | 12.02302 | 16.31693 | 13.95350 | 12.59954 | 3.01628 | 10.25063 | 19.98132 | 16.34550 | 24.92348 | 17.98626 | 17.27528 | 10.77086 | 27.55864 | 19.95457 | 9.15690 |
| **Dcaf15** | 4.87243 | 1.28925 | 1.14748 | 0.09182 | 1.59059 | 1.58066 | 1.31732 | 8.25949 | 0.24256 | 7.09794 | 1.04593 | 0.76557 | 7.24267 | 0.00446 | 0.00000 | 3.06930 |
| **Dcaf17** | 2.09965 | 0.03837 | 2.61586 | 4.69778 | 0.95166 | 0.06639 | 0.11072 | 1.58311 | 6.98757 | 2.51869 | 6.28013 | 0.39329 | 1.76927 | 2.77106 | 4.88938 | 2.89332 |
| **Dcaf4** | 9.40015 | 5.47953 | 12.55688 | 3.36240 | 7.23822 | 5.52537 | 7.86172 | 21.94536 | 2.90311 | 27.27243 | 14.49316 | 1.62496 | 7.33634 | 0.02115 | 2.55260 | 5.59317 |
| **Dcaf5** | 1.19530 | 0.01275 | 0.58337 | 7.14824 | 4.39432 | 0.07143 | 2.67161 | 3.36231 | 16.40348 | 6.29291 | 2.67333 | 0.54280 | 0.22621 | 2.40607 | 5.70865 | 1.77956 |
| **Dcaf6** | 0.01209 | 0.00832 | 0.00733 | 0.67680 | 0.00964 | 0.00629 | 0.00777 | 0.03045 | 1.31512 | 0.00000 | 0.00000 | 0.02037 | 0.01534 | 0.22172 | 3.23400 | 0.00720 |

| | | | | | | | | | | | | | | | | |
|---|---|---|---|---|---|---|---|---|---|---|---|---|---|---|---|---|
| **Dcaf7** | 10.32057 | 6.77109 | 6.44486 | 7.14595 | 4.10115 | 2.48231 | 5.44116 | 4.10588 | 12.58577 | 9.35129 | 6.70552 | 1.44888 | 5.38452 | 5.28655 | 7.83644 | 0.23561 |
| **Dcaf8** | 8.63099 | 6.29939 | 3.15580 | 11.00764 | 3.61715 | 1.27747 | 3.90559 | 4.45397 | 11.71869 | 4.94246 | 4.34236 | 0.52173 | 3.40336 | 3.23762 | 5.13383 | 3.54303 |
| **Dcakd** | 3.50646 | 0.48629 | 2.15752 | 5.45393 | 1.97644 | 0.01312 | 4.52283 | 2.35619 | 6.77941 | 1.70976 | 1.88168 | 0.20528 | 3.11688 | 0.01209 | 0.00000 | 3.34998 |
| **Dcbld1** | 5.48117 | 4.91018 | 2.94099 | 4.13521 | 6.02220 | 3.66280 | 5.74230 | 10.95788 | 4.30229 | 4.87993 | 5.99467 | 1.85788 | 4.36910 | 2.64591 | 2.88845 | 5.37206 |
| **Dcbld2** | 0.58198 | 0.07307 | 0.00000 | 4.27971 | 0.00660 | 0.00597 | 4.56607 | 2.02724 | 1.98428 | 0.00000 | 3.43875 | 0.05578 | 0.83791 | 2.96565 | 1.71759 | 0.00494 |
| **Dcc** | 0.01587 | 0.00792 | 0.01174 | 3.68089 | 0.00891 | 0.01321 | 0.00699 | 0.00932 | 2.15035 | 0.00000 | 0.00000 | 0.00000 | 0.12838 | 0.38511 | 1.26212 | 0.00636 |
| **Dcdc2a** | 0.02574 | 0.00770 | 0.01845 | 2.09175 | 0.01937 | 0.00743 | 0.01711 | 0.01900 | 0.27447 | 0.00000 | 0.03329 | 0.02148 | 0.01780 | 0.71550 | 0.51344 | 0.02156 |
| **Dcdc2b** | 0.62304 | 0.19942 | 0.66422 | 1.09228 | 0.01594 | 0.55644 | 0.18755 | 0.38212 | 1.36172 | 0.49997 | 0.72296 | 0.04624 | 0.38577 | 0.08524 | 0.16894 | 0.17300 |
| **Dcdc2c** | 0.68322 | 1.42497 | 2.18007 | 67.96443 | 1.03069 | 0.98758 | 1.17714 | 6.13381 | 65.23173 | 45.50991 | 15.23888 | 18.82159 | 2.45764 | 40.49353 | 49.25253 | 0.29428 |
| **Dck** | 0.80825 | 2.27413 | 2.11906 | 4.16221 | 1.05057 | 3.14865 | 0.56560 | 3.86926 | 7.20904 | 3.10319 | 0.90692 | 5.57463 | 2.24012 | 7.95697 | 6.59299 | 0.02281 |
| **Dclk2** | 0.65970 | 0.00660 | 0.43454 | 10.92122 | 0.00567 | 0.00343 | 0.00553 | 0.00000 | 5.38601 | 1.15437 | 1.16554 | 0.09038 | 0.20466 | 0.00000 | 0.10801 | 0.84427 |
| **Dclk3** | 0.01778 | 0.00761 | 0.01136 | 1.64300 | 0.00908 | 0.00632 | 0.00730 | 0.01164 | 0.01049 | 0.00592 | 0.01122 | 0.00611 | 0.01278 | 0.00579 | 0.72497 | 0.00705 |
| **Dclre1a** | 0.96415 | 0.00653 | 0.71415 | 1.41618 | 0.40415 | 0.00706 | 0.00643 | 0.30456 | 0.03221 | 0.64199 | 0.42902 | 0.02002 | 0.67337 | 1.25195 | 0.76414 | 0.00554 |
| **Dclre1b** | 0.26099 | 3.22941 | 0.61541 | 8.08987 | 2.87296 | 0.73302 | 0.35416 | 1.70331 | 7.93873 | 1.69217 | 5.57089 | 0.38114 | 1.16917 | 4.77849 | 7.68241 | 1.85664 |
| **Dclre1c** | 3.46123 | 0.10065 | 3.75611 | 9.85596 | 3.95761 | 0.17865 | 1.48167 | 1.39502 | 16.43417 | 2.89523 | 16.29635 | 6.73677 | 0.89985 | 10.56202 | 1.28498 | 0.52891 |
| **Dcn** | 0.01168 | 0.00775 | 0.01212 | 0.00503 | 0.00844 | 0.00650 | 0.00723 | 0.00940 | 0.01080 | 4.82481 | 0.00896 | 0.00565 | 0.01023 | 0.01421 | 0.00502 | 0.00729 |
| **Dcp1a** | 0.00000 | 5.75385 | 4.41710 | 9.79235 | 0.58935 | 0.30412 | 5.28720 | 2.21559 | 3.81013 | 2.86063 | 2.45482 | 0.25523 | 3.49088 | 0.40139 | 3.55954 | 3.58855 |
| **Dcp1b** | 0.01366 | 0.00823 | 0.01174 | 0.15721 | 0.00928 | 0.00671 | 0.00757 | 0.01353 | 0.52644 | 0.01881 | 0.03578 | 0.00768 | 0.01243 | 0.01188 | 0.00455 | 0.00735 |
| **Dcp2** | 2.40916 | 2.57224 | 6.65938 | 8.37524 | 1.83934 | 2.87438 | 2.89892 | 1.37243 | 3.62511 | 3.70492 | 2.51474 | 5.62695 | 0.68842 | 1.61751 | 9.05369 | 4.77683 |
| **Dcps** | 12.11859 | 3.68036 | 2.74701 | 1.48997 | 0.00000 | 0.00000 | 0.00000 | 0.28336 | 2.58204 | 1.08972 | 1.47600 | 0.09893 | 3.67420 | 0.10663 | 0.91245 | 2.50284 |
| **Dcst1** | 0.01373 | 0.00790 | 0.01239 | 0.03507 | 0.00941 | 0.00656 | 0.00745 | 0.01305 | 0.01276 | 0.07588 | 0.01372 | 0.00738 | 0.01268 | 0.01048 | 0.01404 | 0.00747 |
| **Dctd** | 9.71688 | 0.02437 | 12.17745 | 5.39802 | 11.27555 | 14.13009 | 6.57956 | 8.10886 | 3.36757 | 23.84396 | 6.25333 | 12.51220 | 12.80600 | 13.61043 | 16.45874 | 3.02272 |
| **Dctn1** | 6.23670 | 2.62294 | 2.73509 | 0.62827 | 5.17880 | 4.21872 | 0.33490 | 4.83558 | 0.33018 | 3.74133 | 2.14973 | 0.11427 | 7.37235 | 0.00015 | 0.00000 | 2.57809 |
| **Dctn2** | 16.75911 | 4.47808 | 6.30702 | 1.22265 | 6.50571 | 9.91433 | 3.49719 | 9.18257 | 0.19902 | 6.55392 | 8.20719 | 4.88213 | 15.35743 | 0.82319 | 1.07557 | 10.16159 |
| **Dctn3** | 8.55301 | 1.91272 | 9.86405 | 2.64430 | 9.57550 | 3.69785 | 2.78588 | 3.12704 | 1.52041 | 9.39709 | 4.58706 | 0.21295 | 5.67495 | 0.01254 | 0.09257 | 7.64553 |
| **Dctn4** | 1.46902 | 3.48362 | 3.25133 | 3.81050 | 1.61388 | 0.00422 | 0.16358 | 3.47258 | 1.81904 | 3.50077 | 4.46698 | 1.43700 | 1.35949 | 10.81241 | 2.42272 | 0.03900 |
| **Dctn5** | 8.84487 | 2.19793 | 7.07733 | 0.71928 | 3.85352 | 3.01444 | 0.00464 | 7.18574 | 1.49058 | 8.30418 | 4.35154 | 1.06320 | 4.56354 | 8.01378 | 7.13635 | 2.71382 |
| **Dctn6** | 9.23940 | 8.09134 | 5.27959 | 0.80576 | 3.19906 | 2.99503 | 2.64427 | 5.62262 | 0.59887 | 8.82243 | 4.11620 | 4.29358 | 1.88663 | 0.54606 | 1.10584 | 2.84291 |
| **Dctpp1** | 12.38967 | 0.61666 | 4.24477 | 3.03694 | 7.11495 | 2.96843 | 1.22095 | 5.96859 | 2.21773 | 5.87118 | 9.51850 | 0.66201 | 9.15232 | 0.00451 | 0.38014 | 8.07895 |
| **Dcun1d1** | 2.01724 | 7.60581 | 4.31353 | 11.05096 | 2.65105 | 0.06168 | 0.42407 | 2.37986 | 1.66056 | 0.00000 | 0.73897 | 0.03761 | 3.59315 | 4.44359 | 10.51574 | 1.43540 |
| **Dcun1d2** | 0.14444 | 0.00720 | 2.13475 | 8.56976 | 0.01268 | 1.25657 | 0.00645 | 2.69618 | 3.30506 | 3.05132 | 0.17458 | 0.43013 | 0.13639 | 0.07270 | 4.24622 | 0.81942 |
| **Dcun1d3** | 1.49777 | 3.08497 | 1.82850 | 5.29228 | 0.02666 | 0.02394 | 0.00183 | 0.12584 | 6.56158 | 5.77280 | 3.40495 | 5.51523 | 0.85777 | 13.50085 | 12.00226 | 0.01598 |
| **Dcun1d4** | 1.82674 | 0.00567 | 0.00000 | 0.72039 | 0.00627 | 0.00537 | 0.00575 | 0.15914 | 0.00000 | 0.41411 | 0.00000 | 0.21402 | 0.01959 | 2.42681 | 0.21686 | 1.32581 |
| **Dcun1d5** | 6.81900 | 4.57623 | 5.62642 | 21.49592 | 6.76805 | 5.25710 | 5.79556 | 4.24515 | 18.38724 | 9.58655 | 10.24301 | 6.87242 | 5.49770 | 7.48557 | 2.33978 | 2.23533 |
| **Dcx** | 0.02998 | 0.01800 | 0.02063 | 0.03633 | 0.02613 | 0.02625 | 0.01770 | 0.04234 | 0.05832 | 0.08118 | 0.06801 | 0.04792 | 0.03040 | 0.00955 | 0.06139 | 0.01433 |
| **Dcxr** | 0.26539 | 1.04926 | 0.30659 | 0.22918 | 2.37908 | 0.00733 | 1.16042 | 0.93392 | 0.00206 | 4.13721 | 1.86085 | 0.00000 | 1.05887 | 0.00980 | 0.00000 | 3.07364 |
| **Dda1** | 3.95130 | 0.84290 | 0.95935 | 11.53958 | 8.16040 | 1.39941 | 3.93458 | 5.73486 | 9.07930 | 8.62614 | 1.02505 | 4.81583 | 8.55121 | 0.46768 | 1.60507 | 6.30013 |
| **Ddah1** | 9.54872 | 6.37094 | 5.81273 | 0.00000 | 3.38862 | 8.29773 | 5.78926 | 5.14593 | 0.01157 | 5.17927 | 5.10486 | 0.61565 | 9.86025 | 13.91019 | 1.67762 | 8.20931 |
| **Ddah2** | 2.79139 | 0.00101 | 1.86024 | 0.00000 | 3.09563 | 0.00914 | 1.79462 | 2.32564 | 0.00587 | 4.87818 | 3.63504 | 0.08172 | 2.91711 | 0.00384 | 0.00265 | 3.39266 |
| **Ddb1** | 22.85531 | 15.07117 | 14.85991 | 15.49082 | 19.26063 | 8.35645 | 16.17068 | 15.27051 | 11.19777 | 19.01745 | 21.03875 | 4.31457 | 22.67748 | 25.36959 | 17.89828 | 7.63779 |
| **Ddb2** | 0.00000 | 0.00744 | 4.04961 | 0.00000 | 0.00469 | 0.33427 | 0.26727 | 2.05921 | 0.00558 | 2.57966 | 0.00000 | 0.41841 | 2.72059 | 2.68270 | 3.19725 | 2.27627 |
| **Ddc** | 0.65981 | 0.00801 | 0.19833 | 0.01273 | 0.01357 | 0.00499 | 0.02988 | 0.08474 | 0.81230 | 0.00499 | 0.00000 | 0.15380 | 0.00785 | 0.02273 | 0.10412 | 0.00667 |
| **Ddhd1** | 4.86337 | 0.30712 | 4.34103 | 5.88645 | 1.05805 | 0.00000 | 3.00002 | 1.25030 | 3.73864 | 1.32982 | 0.28216 | 0.22970 | 0.16290 | 0.00000 | 1.65444 | 0.04153 |
| **Ddhd2** | 4.45927 | 3.84256 | 3.44495 | 7.53040 | 0.02846 | 0.05030 | 4.50832 | 4.93760 | 3.78156 | 7.38519 | 3.83117 | 0.58298 | 3.69815 | 4.74163 | 7.19824 | 2.22245 |
| **Ddi2** | 0.02308 | 0.43516 | 0.01462 | 0.00106 | 0.02580 | 0.67330 | 0.71995 | 1.40944 | 0.74915 | 0.28142 | 0.00000 | 0.02239 | 0.04673 | 0.00535 | 0.33702 | 0.00689 |
| **Ddias** | 3.77945 | 4.74648 | 1.84716 | 11.83893 | 0.16698 | 0.45737 | 7.17639 | 7.28475 | 21.98094 | 15.33995 | 10.04222 | 13.50275 | 0.65335 | 24.78846 | 24.75799 | 2.91187 |
| **Ddit3** | 0.02819 | 0.02625 | 0.35019 | 0.30108 | 0.00770 | 1.80337 | 2.87221 | 1.44668 | 3.00163 | 2.41196 | 0.87765 | 0.87267 | 1.71139 | 0.00204 | 0.00000 | 1.36715 |
| **Ddit4** | 0.30171 | 0.00603 | 0.21963 | 2.18581 | 0.00698 | 0.00471 | 0.10475 | 2.07358 | 4.79505 | 4.22655 | 5.86979 | 0.17356 | 0.01782 | 1.12945 | 1.43911 | 0.01997 |
| **Ddit4l** | 1.01005 | 0.00711 | 0.48225 | 10.98536 | 0.01669 | 0.00224 | 0.00492 | 1.48480 | 13.72009 | 0.41740 | 1.74348 | 0.48729 | 1.22252 | 0.11850 | 0.00000 | 0.00266 |
| **Ddo** | 0.11265 | 0.00835 | 0.03443 | 0.36874 | 0.01224 | 0.01774 | 0.00748 | 0.02510 | 0.08910 | 0.08591 | 0.08730 | 0.00757 | 0.04209 | 0.00870 | 0.01589 | 0.00728 |
| **Ddost** | 9.67740 | 0.62307 | 15.83349 | 1.10880 | 5.74789 | 7.79508 | 4.88152 | 9.89764 | 0.25199 | 6.43002 | 3.66337 | 0.54325 | 7.76484 | 3.96388 | 1.68540 | 8.86185 |
| **Ddr1** | 0.58896 | 0.00511 | 0.02789 | 2.36046 | 0.01642 | 0.00503 | 3.35418 | 5.04614 | 1.20559 | 4.10923 | 5.43676 | 0.12911 | 0.49161 | 0.51886 | 1.13755 | 0.01268 |
| **Ddr2** | 2.64870 | 0.00381 | 0.00000 | 8.82837 | 2.53326 | 0.00780 | 0.00382 | 0.30099 | 3.74229 | 4.87814 | 2.76258 | 0.06178 | 0.19016 | 12.83294 | 2.19431 | 3.92305 |
| **Ddrgk1** | 8.46105 | 2.51948 | 9.01917 | 0.53740 | 10.25886 | 0.25913 | 4.01404 | 4.55122 | 0.35297 | 4.77475 | 3.76677 | 0.15713 | 10.50822 | 0.05424 | 0.00000 | 4.68430 |
| **Ddt** | 9.42108 | 2.77308 | 2.66267 | 2.88345 | 3.06333 | 0.87433 | 1.43979 | 1.80052 | 6.07432 | 3.42488 | 1.42072 | 0.10709 | 2.36279 | 0.00000 | 0.36961 | 4.90592 |
| **Ddx1** | 13.39767 | 13.29372 | 7.48901 | 2.33968 | 8.79024 | 8.48823 | 7.02793 | 12.46275 | 1.43068 | 6.06036 | 7.73721 | 5.56817 | 6.65296 | 7.42108 | 10.22108 | 10.40551 |
| **Ddx10** | 5.52396 | 8.07757 | 5.08731 | 3.57338 | 9.54087 | 0.00000 | 4.49514 | 6.18811 | 6.22712 | 6.78369 | 3.72086 | 4.94344 | 2.77253 | 19.14330 | 13.73408 | 0.60149 |
| **Ddx11** | 0.55230 | 0.00708 | 0.14029 | 0.77611 | 0.00755 | 0.02140 | 0.18158 | 0.02114 | 0.19256 | 0.00000 | 0.25766 | 0.01911 | 3.00583 | 0.27917 | 1.00653 | 1.76966 |
| **Ddx17** | 23.90172 | 15.02901 | 17.80968 | 9.83853 | 21.05149 | 7.71412 | 8.65937 | 14.97691 | 8.48182 | 24.44621 | 17.08235 | 15.35671 | 14.68065 | 26.26050 | 15.43551 | 5.61680 |
| **Ddx18** | 20.91376 | 9.47097 | 14.02594 | 11.60244 | 12.92150 | 10.88750 | 10.51324 | 24.98444 | 16.93287 | 20.02144 | 12.11314 | 14.61966 | 14.51343 | 24.79810 | 22.67027 | 9.61580 |
| **Ddx19a** | 0.32696 | 0.01945 | 0.41003 | 4.90874 | 0.38028 | 0.01438 | 0.64859 | 1.14036 | 6.91953 | 2.09166 | 1.93861 | 0.29126 | 0.52774 | 0.53350 | 0.57873 | 0.37247 |
| **Ddx19b** | 2.85863 | 3.08794 | 2.25882 | 3.52259 | 0.02247 | 3.39046 | 5.33511 | 7.99553 | 5.29506 | 7.49045 | 5.59016 | 0.98746 | 3.36524 | 11.27939 | 4.18677 | 2.01963 |

| | | | | | | | | | | | | | | | | |
|---|---|---|---|---|---|---|---|---|---|---|---|---|---|---|---|---|
| **Ddx20** | 3.37436 | 6.49825 | 4.27969 | 9.64328 | 9.89743 | 5.90163 | 3.79611 | 9.49918 | 17.52693 | 7.88103 | 7.54335 | 2.15870 | 5.75876 | 16.44994 | 14.81102 | 2.64082 |
| **Ddx21** | 14.50108 | 14.64603 | 10.91887 | 12.37821 | 10.01910 | 7.73689 | 8.48534 | 26.50668 | 20.68102 | 28.43753 | 18.50040 | 11.32919 | 11.69569 | 25.07572 | 21.02314 | 4.46267 |
| **Ddx23** | 2.25823 | 0.00602 | 0.00000 | 0.00712 | 0.00867 | 0.01387 | 0.00556 | 0.02707 | 0.01139 | 1.22347 | 0.00997 | 0.02397 | 0.98988 | 0.01793 | 0.06421 | 0.62459 |
| **Ddx24** | 8.04464 | 6.08314 | 7.22902 | 7.57127 | 7.84435 | 4.18781 | 10.51017 | 6.33002 | 19.50064 | 16.08549 | 4.60734 | 9.71595 | 8.03741 | 2.45732 | 15.40338 | 2.67915 |
| **Ddx25** | 0.01319 | 0.00813 | 0.01047 | 0.00443 | 0.00900 | 0.00674 | 0.00743 | 0.01140 | 0.07177 | 0.84911 | 0.00838 | 0.05523 | 0.01175 | 0.00605 | 0.55486 | 0.00733 |
| **Ddx26b** | 2.84864 | 3.15150 | 0.00000 | 4.76873 | 4.15204 | 0.00181 | 0.69637 | 0.00000 | 6.45283 | 2.18605 | 0.00000 | 0.00000 | 0.00966 | 3.26858 | 2.34574 | 0.00173 |
| **Ddx27** | 16.55596 | 9.82471 | 8.04844 | 13.03462 | 6.65273 | 5.49398 | 10.43580 | 14.73346 | 13.82354 | 13.67734 | 6.91834 | 10.39805 | 10.72573 | 5.12253 | 8.72892 | 4.35338 |
| **Ddx28** | 1.59331 | 0.00193 | 4.89910 | 1.93885 | 2.68523 | 0.02828 | 1.39490 | 5.82888 | 3.99846 | 6.30274 | 0.61157 | 2.97490 | 5.96872 | 4.00235 | 4.97562 | 1.70647 |
| **Ddx31** | 1.28907 | 6.06158 | 2.37069 | 4.18130 | 3.04924 | 0.00452 | 5.29310 | 6.69562 | 3.64322 | 10.87644 | 1.70958 | 0.40598 | 1.53896 | 3.71574 | 6.57642 | 2.80650 |
| **Ddx39** | 24.64444 | 19.79745 | 24.58308 | 18.74776 | 14.09966 | 16.08380 | 17.66378 | 34.27838 | 29.67936 | 23.43879 | 23.58067 | 8.94280 | 25.63440 | 9.32001 | 14.65007 | 9.61416 |
| **Ddx39b** | 24.11633 | 18.22448 | 15.52924 | 4.91298 | 16.76090 | 6.58356 | 7.41579 | 21.29435 | 8.39960 | 17.14617 | 16.14466 | 8.11790 | 16.53633 | 2.47134 | 13.82264 | 11.63393 |
| **Ddx3x** | 12.10246 | 13.10925 | 25.01921 | 27.82790 | 14.24190 | 8.34138 | 14.46320 | 26.08647 | 25.10806 | 24.23187 | 24.56497 | 22.96769 | 16.91578 | 46.64340 | 26.70864 | 5.69707 |
| **Ddx3y** | 0.00000 | 5.97461 | 0.00000 | 0.00000 | 0.00767 | 1.72471 | 2.28822 | 0.00000 | 0.00712 | 0.18951 | 2.51013 | 7.80195 | 0.73672 | 0.01735 | 0.00000 | 0.00320 |
| **Ddx4** | 0.00831 | 0.00710 | 3.34554 | 4.33204 | 0.00358 | 0.00628 | 0.00611 | 0.00000 | 0.50696 | 1.14841 | 0.48561 | 0.01314 | 0.61497 | 0.00009 | 2.52672 | 0.00480 |
| **Ddx41** | 9.00333 | 3.50081 | 8.55588 | 5.26969 | 10.11778 | 4.90821 | 6.98137 | 15.71408 | 9.26991 | 13.07840 | 7.99283 | 3.03987 | 11.60834 | 0.15802 | 5.80302 | 4.86281 |
| **Ddx42** | 6.73495 | 8.61723 | 7.43807 | 4.25039 | 7.24039 | 2.31323 | 2.29903 | 7.14881 | 6.24272 | 3.61534 | 2.69820 | 0.52048 | 1.42669 | 10.30474 | 8.16913 | 1.94783 |
| **Ddx43** | 0.21236 | 0.00594 | 0.00000 | 0.82850 | 0.01554 | 0.00557 | 1.12460 | 0.80639 | 1.63043 | 0.61531 | 1.72097 | 2.30316 | 0.00078 | 4.99532 | 0.00000 | 0.00622 |
| **Ddx46** | 11.06864 | 9.54813 | 2.84376 | 17.65908 | 7.16269 | 8.07127 | 5.64627 | 12.55755 | 19.39516 | 16.57024 | 10.31644 | 13.01216 | 4.75508 | 15.93891 | 19.08854 | 5.52398 |
| **Ddx47** | 9.07932 | 14.34467 | 13.01990 | 7.54705 | 8.66178 | 6.02033 | 10.32823 | 17.86720 | 10.25931 | 8.93245 | 5.65190 | 1.18404 | 10.40797 | 13.70816 | 19.62800 | 4.80463 |
| **Ddx49** | 4.31929 | 3.76987 | 3.58132 | 0.16128 | 1.56036 | 0.04266 | 3.88644 | 13.31313 | 0.32658 | 12.79381 | 0.59214 | 0.42413 | 6.56767 | 0.01604 | 1.75799 | 3.76369 |
| **Ddx5** | 28.78631 | 23.09906 | 21.12622 | 22.77645 | 21.45022 | 15.25643 | 18.44768 | 35.29257 | 31.22029 | 41.78961 | 29.94317 | 27.64491 | 25.84413 | 39.37996 | 30.30426 | 16.29427 |
| **Ddx50** | 6.08475 | 5.77519 | 5.14741 | 3.04610 | 5.92391 | 0.00602 | 4.26211 | 3.07431 | 2.01290 | 4.89219 | 1.59768 | 10.68808 | 2.52014 | 0.55616 | 4.97567 | 2.50012 |
| **Ddx51** | 6.03119 | 2.04793 | 3.02397 | 7.11864 | 3.29064 | 0.00000 | 4.24361 | 6.54858 | 16.48267 | 5.98305 | 2.59778 | 1.16213 | 6.94039 | 3.27211 | 4.91212 | 0.00000 |
| **Ddx52** | 1.93105 | 8.24002 | 11.21529 | 11.07512 | 6.85236 | 1.33601 | 9.10092 | 7.64308 | 21.39152 | 13.41386 | 9.43995 | 8.84890 | 5.63494 | 19.41719 | 18.76339 | 2.57899 |
| **Ddx54** | 13.21919 | 0.11382 | 0.00000 | 17.38428 | 7.16863 | 5.65180 | 2.04268 | 20.70454 | 20.40066 | 21.58799 | 9.23072 | 0.88719 | 11.48664 | 0.02191 | 0.48909 | 6.67639 |
| **Ddx55** | 8.37013 | 0.01777 | 0.54169 | 5.07363 | 1.07518 | 0.67196 | 0.73693 | 8.23503 | 8.69955 | 3.63691 | 3.65997 | 1.69112 | 4.07449 | 3.46418 | 5.65492 | 3.35074 |
| **Ddx56** | 15.66965 | 9.31785 | 15.11162 | 7.87803 | 10.33347 | 7.68843 | 9.26653 | 20.82071 | 10.50564 | 11.29175 | 12.44768 | 6.70362 | 16.64625 | 5.41103 | 11.75086 | 15.63530 |
| **Ddx58** | 1.23384 | 0.00649 | 0.03315 | 0.87711 | 0.01344 | 0.00756 | 0.01612 | 0.00608 | 0.02974 | 1.13321 | 0.40104 | 0.05560 | 0.04895 | 0.30422 | 0.37377 | 0.00563 |
| **Ddx59** | 1.99910 | 0.01578 | 0.98255 | 4.05147 | 0.00570 | 0.43044 | 0.12725 | 0.71420 | 0.60703 | 1.22972 | 1.31842 | 0.03990 | 0.57365 | 0.00000 | 0.40714 | 0.00373 |
| **Ddx6** | 4.92519 | 0.32837 | 7.64725 | 3.41836 | 0.23969 | 3.23023 | 0.14561 | 1.42052 | 3.08015 | 5.08922 | 1.82063 | 0.07235 | 4.56914 | 2.38521 | 3.22800 | 0.03845 |
| **Ddx60** | 0.01475 | 0.00639 | 0.00315 | 1.43163 | 0.00899 | 0.00542 | 0.00656 | 0.00774 | 1.13005 | 0.55065 | 0.00000 | 3.79475 | 0.01428 | 2.91010 | 1.03065 | 0.00680 |
| **Deaf1** | 8.26331 | 0.35226 | 2.46092 | 3.61650 | 0.02096 | 3.59631 | 0.03827 | 6.89460 | 4.43885 | 5.16968 | 5.53153 | 6.06140 | 1.11462 | 0.00695 | 1.11361 | 0.05292 |
| **Deb1** | 4.50086 | 0.04402 | 1.66517 | 7.28965 | 0.03041 | 0.41354 | 0.69438 | 0.55820 | 6.87065 | 1.87569 | 5.86298 | 0.22597 | 2.30761 | 3.25114 | 2.20460 | 1.54369 |
| **Decr1** | 4.80287 | 5.06675 | 0.00000 | 0.57042 | 4.76343 | 0.00893 | 5.23995 | 0.00000 | 0.00000 | 1.05653 | 2.13426 | 2.85268 | 1.03628 | 0.75724 | 0.21513 | 0.00658 |
| **Decr2** | 0.01752 | 0.01732 | 0.53698 | 4.56535 | 1.28921 | 0.03045 | 1.80626 | 2.56034 | 5.62435 | 1.07987 | 4.89409 | 0.07187 | 0.86445 | 3.03513 | 2.40878 | 0.09258 |
| **Dedd** | 2.26828 | 1.80910 | 2.06001 | 2.99668 | 0.00000 | 4.18532 | 0.54405 | 1.08057 | 6.41609 | 4.81866 | 3.99520 | 0.41741 | 0.90803 | 9.38039 | 3.17001 | 2.19066 |
| **Dedd2** | 4.17629 | 1.92255 | 0.88926 | 8.61151 | 0.01957 | 0.21684 | 0.29552 | 2.40132 | 8.77701 | 8.14764 | 6.82338 | 0.17695 | 1.72771 | 0.06344 | 1.05306 | 0.03530 |
| **Def6** | 2.78179 | 0.00497 | 0.65283 | 1.01572 | 0.00007 | 0.00276 | 0.00565 | 0.10737 | 0.65754 | 0.42989 | 0.74511 | 0.00236 | 0.00600 | 0.00426 | 0.00000 | 0.00383 |
| **Def8** | 5.71548 | 0.16436 | 8.48892 | 6.16413 | 0.08882 | 0.01355 | 7.96817 | 15.30285 | 8.01188 | 12.67379 | 14.32551 | 1.74724 | 10.91234 | 12.61589 | 13.03231 | 3.10574 |
| **Defa29** | 0.01311 | 0.00801 | 0.00967 | 1.83786 | 0.00911 | 0.00628 | 0.00731 | 0.00908 | 2.59083 | 0.02109 | 0.12986 | 0.02372 | 0.07401 | 0.00373 | 0.08132 | 0.00684 |
| **Defb23** | 0.01227 | 0.00831 | 0.39090 | 1.31713 | 0.00854 | 0.00536 | 0.00761 | 0.01366 | 2.02197 | 0.00731 | 0.00000 | 0.00511 | 0.00956 | 0.01100 | 0.00000 | 0.00684 |
| **Defb30** | 0.01255 | 0.00759 | 0.00671 | 0.65115 | 0.00958 | 0.00659 | 0.00714 | 0.00949 | 0.00000 | 0.00823 | 0.00595 | 0.12664 | 0.01321 | 1.24444 | 0.31223 | 0.00722 |
| **Defb42** | 0.01406 | 0.00758 | 0.01730 | 2.30190 | 0.00888 | 0.00574 | 0.00677 | 0.00000 | 3.27614 | 0.63003 | 1.05277 | 0.09041 | 0.00419 | 0.29975 | 0.97206 | 0.00653 |
| **Degs1** | 7.78520 | 2.44080 | 0.00000 | 17.52134 | 2.22701 | 8.14710 | 2.55354 | 8.26630 | 8.61654 | 1.04625 | 4.85827 | 1.02575 | 6.36132 | 1.17123 | 2.67060 | 5.95383 |
| **Degs2** | 0.01241 | 0.00830 | 0.01621 | 0.01793 | 0.00929 | 0.00646 | 0.00798 | 0.03714 | 0.01118 | 0.14427 | 0.01204 | 0.01781 | 0.01138 | 0.00716 | 0.08518 | 0.00777 |
| **Dek** | 4.85033 | 3.80495 | 3.37777 | 2.29879 | 4.72005 | 2.72809 | 2.51921 | 5.13843 | 1.01862 | 12.43744 | 1.34405 | 1.13290 | 1.71196 | 12.17027 | 17.43439 | 1.73420 |
| **Dennd1a** | 0.18976 | 0.07264 | 0.68538 | 3.27311 | 0.01568 | 0.91735 | 0.01768 | 1.07144 | 0.96444 | 0.47373 | 0.75715 | 0.00000 | 0.00000 | 0.01498 | 0.33820 | 2.74510 |
| **Dennd1b** | 0.00810 | 0.00620 | 0.00000 | 0.20805 | 0.00841 | 3.66369 | 0.00523 | 0.53495 | 0.75667 | 1.29757 | 0.39930 | 4.60571 | 2.33354 | 4.17342 | 0.61418 | 0.01211 |
| **Dennd1c** | 0.01311 | 0.00741 | 0.06955 | 1.08611 | 0.00834 | 0.00865 | 0.00637 | 0.76474 | 0.27656 | 1.84224 | 1.84464 | 0.57227 | 0.32951 | 0.09861 | 0.20715 | 0.00660 |
| **Dennd2a** | 0.60636 | 0.00645 | 0.85284 | 5.55577 | 0.01300 | 0.00431 | 0.00561 | 0.66103 | 9.11177 | 4.48492 | 1.88442 | 0.03423 | 0.35055 | 0.16364 | 1.68326 | 1.49710 |
| **Dennd2c** | 1.54723 | 0.03422 | 1.79587 | 1.75569 | 0.20534 | 0.00365 | 3.36007 | 0.12911 | 1.07802 | 0.26928 | 0.24158 | 0.04764 | 0.02580 | 0.00127 | 0.15508 | 0.02485 |
| **Dennd3** | 0.00675 | 0.00804 | 0.00000 | 2.30878 | 0.01053 | 5.19282 | 0.00649 | 0.57392 | 1.59635 | 2.71351 | 0.60825 | 0.00735 | 0.00000 | 1.34047 | 0.73771 | 0.00476 |
| **Dennd4a** | 0.92148 | 0.00616 | 0.00000 | 3.40500 | 0.00597 | 0.00476 | 1.12587 | 0.00000 | 1.12257 | 0.00000 | 0.63824 | 0.03060 | 0.00615 | 0.60696 | 4.78986 | 0.00528 |
| **Dennd4b** | 1.61426 | 0.00569 | 0.00234 | 1.25713 | 0.00496 | 0.00555 | 0.00603 | 0.00457 | 1.02070 | 0.54528 | 0.00000 | 0.06360 | 0.01172 | 0.00280 | 1.08869 | 0.00541 |
| **Dennd4c** | 1.07295 | 1.61602 | 1.04386 | 8.88380 | 0.98900 | 0.00554 | 8.17069 | 2.45202 | 4.22732 | 2.07469 | 3.56921 | 0.44627 | 0.11686 | 6.71593 | 8.15151 | 0.04007 |
| **Dennd5a** | 5.72922 | 3.43279 | 2.35749 | 6.03308 | 9.52313 | 0.63271 | 6.28058 | 5.23714 | 6.22275 | 14.01844 | 1.27261 | 3.99196 | 0.24576 | 1.93944 | 0.52377 | 4.25836 |
| **Dennd5b** | 4.40343 | 4.70793 | 1.31627 | 8.31406 | 0.98404 | 0.45388 | 0.05691 | 2.57463 | 12.38907 | 1.25403 | 11.24789 | 1.58709 | 2.94662 | 16.65241 | 1.27030 | 1.75738 |
| **Dennd6a** | 0.44339 | 0.10648 | 1.51994 | 3.39297 | 2.11525 | 1.05065 | 1.25618 | 0.05685 | 4.01454 | 0.68543 | 2.44211 | 0.24149 | 0.78641 | 1.03907 | 3.83597 | 0.16488 |
| **Dennd6b** | 2.43259 | 0.00227 | 3.03234 | 0.16844 | 1.21961 | 0.00300 | 2.63782 | 1.26298 | 0.00716 | 0.99822 | 0.45165 | 0.01997 | 0.98690 | 0.00482 | 0.21071 | 3.74224 |
| **Denr** | 19.02335 | 3.94642 | 5.71432 | 13.59474 | 12.07060 | 5.45227 | 4.79111 | 12.38449 | 14.02576 | 18.10297 | 10.75000 | 2.34098 | 13.78080 | 9.18485 | 2.80919 | 8.16639 |
| **Depdc1a** | 0.06782 | 0.00760 | 0.00181 | 0.84062 | 1.08068 | 0.00753 | 0.33655 | 0.00893 | 0.98722 | 0.00000 | 0.09162 | 0.00339 | 0.01580 | 0.00428 | 0.55721 | 0.00705 |
| **Depdc1b** | 1.12044 | 0.11413 | 0.54496 | 0.00422 | 0.00488 | 0.00740 | 0.00600 | 0.40884 | 0.18920 | 0.73251 | 0.19832 | 0.00000 | 0.15389 | 0.51838 | 0.00000 | 0.00592 |

| | | | | | | | | | | | | | | | | |
|---|---|---|---|---|---|---|---|---|---|---|---|---|---|---|---|---|
| **Depdc5** | 3.29819 | 0.21010 | 2.26888 | 3.56644 | 0.07580 | 1.07156 | 2.59956 | 0.69447 | 4.60420 | 0.61778 | 0.25370 | 0.40274 | 0.07604 | 0.18284 | 2.93615 | 0.94743 |
| **Depdc7** | 6.91351 | 0.62154 | 2.43270 | 11.40209 | 1.27250 | 5.29810 | 2.16699 | 8.27931 | 19.48298 | 20.69208 | 9.25741 | 14.12608 | 7.07105 | 15.16425 | 13.80325 | 1.80434 |
| **Deptor** | 4.00846 | 1.65245 | 3.43260 | 4.64056 | 7.57038 | 0.02894 | 2.40559 | 4.14265 | 5.27500 | 5.33200 | 2.37018 | 6.18817 | 1.21274 | 15.99487 | 4.02511 | 0.08547 |
| **Dera** | 4.90988 | 0.14428 | 1.26709 | 0.07896 | 8.66443 | 4.39136 | 0.29602 | 1.54175 | 0.49003 | 2.34617 | 3.02188 | 0.36165 | 0.41160 | 0.00000 | 0.80235 | 4.40696 |
| **Derl1** | 7.29748 | 5.85777 | 8.04254 | 1.95112 | 5.33657 | 7.69978 | 0.67000 | 4.21126 | 4.22316 | 6.25207 | 5.48699 | 1.10357 | 7.50052 | 3.74308 | 0.99437 | 4.88967 |
| **Derl2** | 10.47666 | 6.75076 | 12.55903 | 1.56868 | 3.68536 | 2.41255 | 7.35012 | 5.10285 | 0.03929 | 4.39055 | 3.04984 | 0.28212 | 6.55111 | 0.13938 | 2.30957 | 1.78029 |
| **Derl3** | 0.03291 | 0.00759 | 3.09692 | 3.54632 | 0.01671 | 0.00556 | 0.00686 | 0.00726 | 2.81941 | 1.03483 | 0.00000 | 0.00000 | 0.00842 | 0.02199 | 0.00000 | 0.00523 |
| **Des** | 0.01304 | 0.00789 | 0.01042 | 0.00870 | 0.00913 | 0.00678 | 0.21944 | 0.01113 | 0.01209 | 0.43942 | 0.01161 | 0.00714 | 0.01128 | 0.01159 | 0.01015 | 0.00724 |
| **Desi1** | 5.57308 | 3.62989 | 2.29082 | 4.98229 | 3.07130 | 1.21660 | 3.36623 | 2.88535 | 2.17537 | 7.30790 | 2.55454 | 0.23607 | 2.77982 | 4.46818 | 0.47249 | 3.95221 |
| **Desi2** | 2.74187 | 1.68637 | 2.17207 | 6.03146 | 0.45707 | 0.00947 | 0.01468 | 0.35442 | 5.68075 | 0.85172 | 0.14537 | 0.09929 | 1.52256 | 1.86938 | 4.43147 | 0.23523 |
| **Det1** | 0.71882 | 0.09590 | 4.41539 | 0.03294 | 0.67262 | 0.01221 | 2.26001 | 2.82400 | 2.06030 | 3.86003 | 0.00000 | 0.15997 | 1.77109 | 0.00475 | 2.03598 | 0.01877 |
| **Dexi** | 0.00460 | 0.00748 | 0.32344 | 1.76548 | 0.01549 | 0.00553 | 0.00707 | 0.04283 | 0.75445 | 0.96822 | 0.01853 | 0.00291 | 0.00484 | 0.00361 | 0.05665 | 2.17244 |
| **Dffa** | 1.64944 | 1.08581 | 3.45020 | 5.53633 | 0.52632 | 2.78390 | 3.36941 | 2.77167 | 8.44967 | 3.73760 | 2.70496 | 0.81648 | 3.43802 | 1.11448 | 2.29366 | 0.76798 |
| **Dffb** | 3.95464 | 2.43385 | 2.15106 | 0.23638 | 0.00823 | 0.00902 | 4.55772 | 8.00673 | 0.00459 | 6.74448 | 1.37784 | 1.17804 | 1.58909 | 0.24114 | 1.70613 | 2.01069 |
| **Dfna5** | 0.01835 | 0.00831 | 0.01173 | 0.01827 | 0.01064 | 0.00665 | 0.00745 | 0.01858 | 0.01099 | 0.01237 | 0.01289 | 0.00835 | 0.01231 | 0.00895 | 0.06682 | 0.00744 |
| **Dgat1** | 3.43020 | 0.00000 | 4.94528 | 2.88176 | 0.18826 | 0.03119 | 5.58957 | 1.97754 | 2.53996 | 0.39580 | 0.46807 | 0.14568 | 2.60020 | 0.08231 | 1.26386 | 2.56728 |
| **Dgat2** | 4.60516 | 2.61158 | 4.17296 | 3.12711 | 1.64845 | 0.00566 | 3.23221 | 6.58830 | 2.09220 | 16.47361 | 15.02569 | 2.02114 | 0.14727 | 0.34449 | 6.56200 | 2.80980 |
| **Dgcr14** | 3.69104 | 0.07600 | 2.88317 | 8.58110 | 1.86396 | 0.01089 | 2.77297 | 9.00986 | 10.01949 | 7.25915 | 4.94332 | 2.08538 | 3.28064 | 0.07015 | 3.36603 | 5.29408 |
| **Dgcr2** | 6.87056 | 3.14019 | 2.53215 | 6.81405 | 3.76969 | 3.33011 | 6.12881 | 4.28319 | 4.40709 | 1.67968 | 2.23060 | 0.27634 | 3.51140 | 1.01965 | 5.37876 | 5.81140 |
| **Dgcr6** | 5.36689 | 0.52326 | 4.20794 | 0.22262 | 3.20303 | 0.00941 | 0.44076 | 0.79919 | 0.29060 | 2.00387 | 3.08978 | 0.16405 | 4.98078 | 0.00281 | 0.01169 | 0.01202 |
| **Dgcr8** | 3.53604 | 0.00000 | 1.85543 | 3.17412 | 2.62050 | 1.91311 | 3.04124 | 4.55233 | 3.68919 | 3.11865 | 0.00000 | 4.21547 | 1.25864 | 4.18484 | 9.19061 | 3.49634 |
| **Dgka** | 3.52112 | 0.18771 | 7.76299 | 0.69307 | 1.20505 | 2.51999 | 7.56877 | 5.86284 | 0.76215 | 5.06543 | 4.64189 | 0.47509 | 1.52856 | 3.72957 | 2.14027 | 0.57554 |
| **Dgkd** | 0.98046 | 0.91030 | 2.38539 | 7.28395 | 5.47695 | 0.03208 | 2.55934 | 3.54581 | 2.99487 | 2.76024 | 1.41962 | 2.10917 | 2.71892 | 1.09950 | 5.01300 | 1.12396 |
| **Dgke** | 0.20780 | 0.10282 | 0.22011 | 15.13429 | 0.05541 | 0.01895 | 0.52470 | 2.18549 | 9.59304 | 4.51915 | 1.93091 | 8.30440 | 1.12604 | 10.63360 | 15.78096 | 0.05350 |
| **Dgkeos** | 0.07773 | 0.00757 | 0.00131 | 0.29829 | 0.00782 | 0.00706 | 0.00711 | 0.01447 | 1.13230 | 0.62284 | 0.00000 | 0.00000 | 0.01178 | 0.82463 | 0.80842 | 0.00733 |
| **Dgkh** | 0.01758 | 0.00811 | 0.00605 | 0.03541 | 0.01091 | 0.00658 | 0.00742 | 0.01271 | 0.94198 | 0.00000 | 0.00000 | 0.00000 | 0.01341 | 1.18292 | 0.57180 | 0.00740 |
| **Dgkk** | 0.42147 | 0.00739 | 0.00361 | 0.18996 | 0.00951 | 0.00754 | 0.00698 | 0.00929 | 0.00865 | 0.00976 | 0.01189 | 0.21337 | 0.01889 | 0.01209 | 0.02184 | 0.01442 |
| **Dgkq** | 0.42616 | 0.00673 | 0.06658 | 0.00703 | 0.00705 | 0.00537 | 0.69644 | 0.71532 | 0.01101 | 0.64255 | 0.00702 | 0.01217 | 1.59690 | 0.00713 | 0.19874 | 0.00623 |
| **Dgkz** | 3.31555 | 1.68992 | 4.30108 | 0.05747 | 5.68695 | 4.06381 | 1.63297 | 6.05991 | 0.71246 | 6.75952 | 0.69304 | 0.20938 | 2.05639 | 0.17352 | 1.27373 | 1.28507 |
| **Dguok** | 2.10670 | 0.02198 | 2.86287 | 0.57177 | 2.93197 | 2.88918 | 0.07483 | 1.72476 | 0.00000 | 0.88216 | 2.32371 | 3.39964 | 1.83491 | 0.00379 | 0.14695 | 1.03084 |
| **Dhcr24** | 1.44423 | 0.06603 | 5.10213 | 0.45416 | 1.18957 | 0.01006 | 6.52762 | 0.85263 | 0.06912 | 3.14572 | 1.11828 | 0.13335 | 6.07335 | 1.85467 | 4.31078 | 1.90511 |
| **Dhcr7** | 1.85690 | 0.83135 | 1.79883 | 2.96063 | 0.97002 | 1.26027 | 3.03838 | 0.73887 | 1.63734 | 2.66032 | 2.00975 | 1.38211 | 2.16718 | 0.11745 | 0.00000 | 1.98534 |
| **Dhdds** | 1.68740 | 2.66917 | 7.26198 | 3.11300 | 1.48186 | 2.82848 | 5.04916 | 6.61998 | 8.10944 | 6.14326 | 5.86760 | 1.62042 | 1.65417 | 3.22919 | 3.73798 | 0.79525 |
| **Dhdh** | 0.01355 | 0.00783 | 0.00986 | 0.67767 | 0.00916 | 0.01272 | 0.00737 | 0.01084 | 0.12477 | 0.34534 | 0.00827 | 0.03524 | 0.01131 | 0.08224 | 0.48748 | 0.00702 |
| **Dhfr** | 8.20934 | 6.59313 | 5.80273 | 2.30690 | 4.45410 | 6.09710 | 3.68722 | 3.61200 | 8.76471 | 5.46522 | 6.93268 | 4.25578 | 4.58094 | 3.58605 | 4.99435 | 0.18009 |
| **Dhh** | 0.27232 | 0.00781 | 0.00475 | 0.01266 | 0.01721 | 0.00723 | 0.00667 | 0.21860 | 0.01263 | 0.00000 | 0.77272 | 0.03045 | 0.00267 | 0.00479 | 0.06904 | 0.00698 |
| **Dhodh** | 7.79029 | 5.16758 | 6.22554 | 2.36354 | 5.72015 | 7.01854 | 5.29117 | 7.35616 | 1.47356 | 5.63629 | 1.90742 | 0.77091 | 3.33162 | 0.99864 | 0.00000 | 3.36039 |
| **Dhps** | 3.45995 | 0.69883 | 4.03193 | 2.51803 | 3.27807 | 5.13343 | 1.79398 | 0.72594 | 5.32772 | 5.88381 | 6.82608 | 0.14128 | 8.09259 | 3.91510 | 1.10478 | 4.39041 |
| **Dhrs1** | 4.11667 | 3.12643 | 1.06572 | 4.11006 | 0.00000 | 3.12578 | 0.32740 | 3.46250 | 1.01657 | 2.05820 | 2.42848 | 0.11966 | 1.24119 | 1.24793 | 0.00000 | 0.02138 |
| **Dhrs11** | 0.99215 | 0.01642 | 0.93187 | 0.57906 | 0.55442 | 5.73790 | 2.30672 | 0.06233 | 0.56485 | 0.19677 | 0.00000 | 0.00818 | 0.86547 | 0.71160 | 0.52406 | 0.82280 |
| **Dhrs13** | 1.92151 | 1.34871 | 0.10414 | 1.55251 | 0.01288 | 2.99927 | 0.22235 | 0.54392 | 1.65849 | 3.72309 | 1.22906 | 0.37845 | 2.53733 | 0.00000 | 0.91736 | 0.04488 |
| **Dhrs2** | 0.01364 | 0.00776 | 0.01150 | 0.28026 | 0.00942 | 0.00637 | 0.00735 | 0.01118 | 0.40087 | 0.00415 | 0.00791 | 0.00536 | 0.01279 | 0.00620 | 0.87821 | 0.00724 |
| **Dhrs3** | 4.00908 | 0.42097 | 0.00001 | 0.08666 | 1.00234 | 0.00594 | 0.00354 | 0.00000 | 0.00987 | 1.37403 | 0.57762 | 0.01588 | 0.00496 | 0.00693 | 0.00000 | 1.03069 |
| **Dhrs4** | 0.04381 | 0.00842 | 0.00999 | 0.01061 | 0.01394 | 0.00680 | 0.02819 | 0.01274 | 0.01235 | 0.01493 | 0.01300 | 0.01841 | 0.01260 | 0.01228 | 0.01041 | 0.00753 |
| **Dhrs7** | 2.63595 | 4.89359 | 7.18751 | 0.00000 | 0.05830 | 11.51170 | 2.39776 | 1.19263 | 0.58004 | 1.42441 | 3.05008 | 0.11335 | 3.77374 | 0.42609 | 0.54563 | 0.00000 |
| **Dhrs7b** | 11.22135 | 5.31506 | 12.81320 | 1.21839 | 5.16613 | 3.00042 | 1.20746 | 6.97975 | 0.76368 | 10.19246 | 4.95658 | 1.86622 | 8.82787 | 4.47575 | 9.27975 | 0.45003 |
| **Dhtkd1** | 0.00504 | 0.00817 | 2.27011 | 2.21023 | 0.01068 | 0.00586 | 0.00721 | 1.06790 | 0.00000 | 1.37809 | 0.71183 | 0.07454 | 0.09338 | 2.01089 | 2.14578 | 0.00655 |
| **Dhx15** | 11.40828 | 12.64640 | 11.33750 | 10.02082 | 16.51946 | 9.00833 | 10.64319 | 15.81513 | 10.56712 | 21.53958 | 9.90329 | 26.49174 | 9.85346 | 35.14568 | 19.03337 | 7.34624 |
| **Dhx16** | 7.27502 | 11.46074 | 10.93019 | 5.58245 | 16.59696 | 5.57561 | 16.24104 | 14.46373 | 4.33774 | 13.49093 | 12.25727 | 9.29485 | 6.53949 | 2.04926 | 5.15598 | 6.62114 |
| **Dhx29** | 2.63549 | 6.68288 | 4.10726 | 4.30348 | 5.31236 | 5.66627 | 0.12163 | 4.44248 | 4.34211 | 7.24045 | 2.63368 | 0.87965 | 3.41240 | 2.07996 | 7.55161 | 2.05596 |
| **Dhx30** | 6.77004 | 8.05403 | 7.01423 | 10.76639 | 0.65888 | 2.05184 | 2.89606 | 9.60698 | 10.11227 | 17.42591 | 9.65050 | 3.53644 | 5.76714 | 0.39130 | 11.31900 | 3.49564 |
| **Dhx32** | 12.12467 | 5.92370 | 6.80329 | 6.02150 | 3.84604 | 5.81326 | 3.49682 | 12.83179 | 2.33348 | 6.05970 | 2.38194 | 0.92202 | 2.81080 | 1.13822 | 9.62578 | 6.75198 |
| **Dhx33** | 4.40422 | 2.86755 | 7.69106 | 7.06558 | 5.28909 | 4.19810 | 5.84420 | 8.48762 | 12.04783 | 11.99963 | 5.16103 | 9.31248 | 5.30088 | 18.65005 | 11.87976 | 1.46726 |
| **Dhx34** | 2.35278 | 0.78272 | 0.42442 | 0.33110 | 1.70402 | 1.18598 | 0.72207 | 1.75923 | 0.03779 | 0.06249 | 1.15146 | 0.04395 | 2.54897 | 0.00338 | 0.00000 | 0.91603 |
| **Dhx35** | 3.16798 | 0.00391 | 0.67673 | 0.90257 | 0.00000 | 0.01145 | 0.07745 | 1.84567 | 0.06533 | 4.42428 | 0.63870 | 0.07072 | 1.75255 | 0.00412 | 0.09247 | 1.90955 |
| **Dhx36** | 3.86539 | 2.14579 | 1.34930 | 3.90729 | 2.49913 | 0.00000 | 4.92753 | 2.58982 | 10.01675 | 3.86807 | 3.49135 | 4.30740 | 1.02096 | 12.85829 | 8.76841 | 0.74540 |
| **Dhx37** | 5.33197 | 1.67074 | 0.00000 | 0.00000 | 0.00638 | 0.50941 | 0.09079 | 6.16538 | 2.99744 | 4.02793 | 0.72479 | 0.32700 | 1.59152 | 0.92065 | 2.63293 | 0.01467 |
| **Dhx38** | 6.32451 | 0.35796 | 3.41664 | 2.70466 | 6.41120 | 1.21634 | 0.09258 | 7.21011 | 7.42373 | 2.75131 | 1.81103 | 0.08680 | 3.98907 | 0.00650 | 0.28410 | 3.09459 |
| **Dhx40** | 5.88470 | 5.08321 | 3.83766 | 4.26949 | 9.02778 | 0.00288 | 7.03010 | 7.48858 | 11.29683 | 12.67050 | 5.72434 | 11.58954 | 3.33540 | 12.63151 | 12.08791 | 0.69528 |
| **Dhx57** | 0.06773 | 2.23189 | 0.41253 | 2.70418 | 0.02529 | 0.56729 | 1.75168 | 3.92429 | 2.73830 | 8.70718 | 1.38843 | 0.12091 | 2.26167 | 0.28790 | 0.66739 | 2.84128 |
| **Dhx58** | 0.01179 | 0.00774 | 0.03935 | 0.14699 | 0.00908 | 0.00561 | 0.00629 | 0.00000 | 0.98950 | 0.00000 | 2.51438 | 0.32657 | 0.00000 | 0.84179 | 0.55648 | 0.00653 |
| **Dhx8** | 0.79710 | 8.05335 | 5.25198 | 6.27156 | 4.21045 | 2.39935 | 4.03793 | 11.93097 | 12.04934 | 13.29444 | 3.49568 | 1.15770 | 3.29281 | 3.33116 | 12.35054 | 1.26405 |

| | | | | | | | | | | | | | | | | |
|---|---|---|---|---|---|---|---|---|---|---|---|---|---|---|---|---|
| **Dhx9** | 3.04647 | 0.64323 | 2.68114 | 12.49633 | 2.00995 | 1.19400 | 0.47343 | 0.44017 | 21.03511 | 2.24477 | 13.99977 | 0.39658 | 1.89762 | 6.92881 | 4.47902 | 2.18091 |
| **Diablo** | 11.47648 | 9.03275 | 11.79981 | 3.42384 | 12.95909 | 6.25196 | 7.96758 | 16.18432 | 10.35735 | 13.78589 | 5.69674 | 1.45922 | 13.07241 | 2.34916 | 8.57836 | 6.00668 |
| **Diaph1** | 9.45298 | 1.51902 | 4.31844 | 5.74440 | 0.76658 | 10.15856 | 4.84634 | 3.04602 | 1.99068 | 3.16097 | 3.50796 | 0.43090 | 0.93912 | 0.06305 | 1.77355 | 2.94312 |
| **Diaph2** | 0.17647 | 4.77532 | 1.39107 | 0.19166 | 0.08239 | 0.04283 | 0.04770 | 0.03699 | 0.33304 | 0.40637 | 0.59718 | 0.09248 | 0.19244 | 2.89089 | 0.00000 | 1.34957 |
| **Diaph3** | 0.00762 | 0.04031 | 1.04186 | 5.50409 | 0.27102 | 3.23585 | 0.00585 | 0.00000 | 6.68027 | 3.70278 | 2.58222 | 0.28254 | 0.00000 | 7.58362 | 1.75604 | 0.38410 |
| **Dicer1** | 2.87336 | 2.25369 | 4.48814 | 11.45340 | 0.04712 | 4.43982 | 3.16594 | 2.92522 | 11.96988 | 8.25721 | 6.29027 | 7.07835 | 6.46962 | 6.03466 | 8.44218 | 4.26091 |
| **Dido1** | 9.76206 | 3.59013 | 3.27979 | 9.13989 | 3.22660 | 5.36869 | 3.93131 | 3.36218 | 10.57594 | 8.54932 | 2.74222 | 2.98743 | 1.62088 | 7.07406 | 10.67072 | 3.20187 |
| **Diexf** | 2.93492 | 2.51377 | 4.78259 | 2.34086 | 0.00001 | 2.85158 | 3.61478 | 4.21968 | 0.71086 | 5.77942 | 0.88446 | 0.23407 | 0.89646 | 1.55779 | 5.97623 | 0.15614 |
| **Dimt1** | 2.58154 | 0.97970 | 0.12413 | 8.05990 | 2.91920 | 0.00796 | 0.98463 | 8.76468 | 14.25093 | 2.82336 | 8.00525 | 0.87018 | 2.19926 | 8.69155 | 4.89020 | 0.43588 |
| **Dio1** | 0.00000 | 0.00721 | 0.00000 | 0.58209 | 0.00929 | 0.00661 | 0.00348 | 1.54734 | 2.78637 | 2.25477 | 3.96123 | 0.06925 | 4.00150 | 1.50774 | 0.16827 | 0.00472 |
| **Dio3** | 1.90452 | 0.00542 | 0.00288 | 3.08580 | 0.00425 | 0.00634 | 0.61876 | 0.00000 | 0.35835 | 0.00285 | 0.31329 | 0.00378 | 0.02197 | 0.00360 | 0.00002 | 0.01160 |
| **Dio3os** | 0.07393 | 0.00771 | 0.00924 | 0.34325 | 0.00788 | 0.00683 | 0.00733 | 0.01320 | 0.00555 | 0.14371 | 0.01330 | 0.00710 | 0.01285 | 0.01315 | 0.01771 | 0.00758 |
| **Dip2a** | 0.01265 | 0.12596 | 0.12190 | 1.15014 | 0.01029 | 2.31150 | 0.00721 | 0.00000 | 0.68697 | 0.88300 | 0.37099 | 0.00990 | 0.11040 | 0.00314 | 0.74171 | 0.00632 |
| **Dip2b** | 4.48201 | 2.94775 | 0.75738 | 3.22869 | 6.28057 | 0.56421 | 4.10241 | 2.66487 | 7.47045 | 4.43978 | 4.34225 | 0.09305 | 1.06764 | 1.75903 | 3.48380 | 2.54533 |
| **Dip2c** | 0.21756 | 4.71611 | 2.06714 | 2.50926 | 0.01680 | 2.26449 | 0.25013 | 0.00599 | 0.00000 | 0.00505 | 0.00969 | 0.01443 | 0.00595 | 0.00229 | 0.21093 | 1.63551 |
| **Diras2** | 0.01376 | 0.00805 | 0.00737 | 1.49032 | 0.00897 | 0.00698 | 0.00742 | 0.00006 | 0.82024 | 0.09801 | 0.71830 | 0.00303 | 0.00664 | 0.02084 | 0.00000 | 0.01464 |
| **Dirc2** | 9.02031 | 1.17222 | 12.95770 | 8.78334 | 0.00000 | 4.95968 | 6.18649 | 15.61708 | 5.29995 | 20.78791 | 15.79701 | 16.08977 | 6.67026 | 11.53675 | 19.23539 | 1.89067 |
| **Dis3** | 3.50071 | 4.69493 | 0.42693 | 5.95674 | 0.09912 | 3.19457 | 4.66906 | 1.26609 | 7.95507 | 7.56766 | 3.30757 | 4.99251 | 1.98891 | 13.60829 | 13.56253 | 0.01915 |
| **Dis3l** | 0.00000 | 0.35506 | 1.76735 | 9.14964 | 0.00474 | 0.00266 | 0.00622 | 0.07507 | 20.78098 | 3.80483 | 0.87564 | 0.03886 | 0.04853 | 5.04064 | 4.27048 | 2.09766 |
| **Dis3l2** | 0.38033 | 0.00780 | 0.00000 | 0.03926 | 0.00757 | 1.66824 | 0.00768 | 0.01842 | 0.00957 | 0.36586 | 0.01001 | 0.00861 | 0.04182 | 0.01174 | 0.01072 | 0.00667 |
| **Disp1** | 0.01066 | 0.04623 | 0.00571 | 1.30065 | 0.00877 | 0.00612 | 0.00655 | 0.00864 | 0.14578 | 0.00000 | 0.00000 | 2.73912 | 0.26480 | 0.00960 | 0.00000 | 0.00675 |
| **Dixdc1** | 0.01543 | 0.00737 | 0.04406 | 0.46917 | 0.00902 | 0.00592 | 0.00716 | 0.00980 | 0.00379 | 0.62273 | 0.01097 | 0.01373 | 0.01174 | 0.01129 | 0.42535 | 0.00768 |
| **Dkc1** | 19.79672 | 19.07185 | 16.28253 | 3.28404 | 14.60202 | 5.74763 | 8.82062 | 17.78796 | 4.63920 | 26.05577 | 6.20469 | 14.64275 | 17.17549 | 34.10845 | 25.73857 | 7.83403 |
| **Dkk1** | 0.68317 | 0.00682 | 0.01360 | 0.03186 | 0.04401 | 0.01530 | 0.01695 | 0.01358 | 0.02395 | 0.01242 | 0.02515 | 0.00636 | 0.01270 | 0.01408 | 0.02034 | 8.80517 |
| **Dkk3** | 0.01324 | 0.00778 | 0.01118 | 0.00620 | 0.00897 | 0.00671 | 0.00729 | 0.01044 | 0.03902 | 1.04691 | 0.01091 | 0.05474 | 0.01158 | 0.01105 | 0.01094 | 0.00759 |
| **Dkkl1** | 1.59608 | 1.87377 | 1.46977 | 2.91097 | 0.01853 | 8.29367 | 2.08300 | 1.70993 | 0.21943 | 1.18413 | 0.38598 | 0.24888 | 4.31193 | 0.00000 | 0.10776 | 2.09257 |
| **Dlat** | 8.26973 | 5.76794 | 5.86178 | 1.39164 | 11.49340 | 3.90764 | 4.12308 | 5.20235 | 0.33094 | 3.22059 | 3.88579 | 0.05305 | 2.87389 | 0.29069 | 3.29525 | 3.98959 |
| **Dlc1** | 5.55300 | 3.68939 | 1.17891 | 2.22109 | 1.15607 | 0.07346 | 2.71927 | 3.72789 | 0.39149 | 0.00000 | 4.07594 | 0.01713 | 1.99509 | 0.00000 | 0.00000 | 3.03246 |
| **Dld** | 7.97285 | 3.38900 | 10.76054 | 4.04422 | 11.24935 | 9.14811 | 7.24689 | 8.52628 | 4.49807 | 6.88903 | 2.84161 | 12.74047 | 6.77902 | 9.80090 | 9.50762 | 4.22285 |
| **Dlec1** | 0.01377 | 0.00792 | 0.01223 | 0.08669 | 0.00924 | 0.00655 | 0.00744 | 0.01264 | 0.01053 | 0.08464 | 0.01516 | 0.00828 | 0.01190 | 0.01682 | 0.00718 | 0.00719 |
| **Dleu2** | 0.63180 | 0.00735 | 0.00000 | 0.47798 | 0.00683 | 0.53408 | 0.00696 | 0.01615 | 0.40733 | 0.52468 | 0.00891 | 0.01580 | 0.00742 | 0.01139 | 0.00385 | 0.00638 |
| **Dlg1** | 0.99473 | 0.07264 | 1.27092 | 4.75519 | 3.68581 | 0.01598 | 0.33612 | 0.22245 | 0.89818 | 0.98179 | 2.91852 | 0.59027 | 0.27538 | 0.59735 | 2.23191 | 1.33102 |
| **Dlg3** | 0.13852 | 0.00693 | 0.27253 | 1.28905 | 0.01745 | 5.86523 | 0.38370 | 0.67521 | 1.48794 | 5.76832 | 4.12520 | 0.59954 | 2.30086 | 0.51033 | 2.51068 | 0.01031 |
| **Dlg5** | 2.15807 | 4.21526 | 0.00000 | 5.30684 | 0.16450 | 0.00197 | 0.12634 | 0.00000 | 1.99621 | 8.27524 | 1.89220 | 5.75354 | 0.18350 | 5.55714 | 10.23312 | 2.36038 |
| **Dlgap1** | 0.63016 | 0.01738 | 0.00506 | 1.38292 | 0.00657 | 0.00597 | 0.00679 | 0.00811 | 2.21211 | 1.22577 | 0.00990 | 0.09731 | 0.02596 | 0.16584 | 0.81082 | 0.00628 |
| **Dlgap4** | 8.39921 | 0.25999 | 2.54538 | 2.21824 | 0.03230 | 3.19077 | 2.06415 | 9.09326 | 2.27513 | 10.75411 | 1.58524 | 0.35381 | 7.48943 | 0.00000 | 0.00000 | 2.44551 |
| **Dlgap5** | 8.27868 | 6.77213 | 8.41974 | 8.22360 | 6.97101 | 5.72302 | 2.25349 | 7.29136 | 7.43463 | 4.46565 | 8.20665 | 5.28191 | 4.86666 | 6.79876 | 7.52452 | 4.44320 |
| **Dlk2** | 0.01313 | 0.00822 | 0.01024 | 1.61154 | 0.00907 | 0.00678 | 0.00740 | 0.00702 | 0.00000 | 0.09325 | 0.25667 | 0.00769 | 0.13815 | 0.00696 | 0.00000 | 0.00700 |
| **Dll3** | 0.55997 | 0.01715 | 0.30982 | 1.27845 | 0.00665 | 0.00522 | 0.00562 | 0.00000 | 2.30942 | 0.74410 | 4.32439 | 0.12651 | 0.10093 | 0.00000 | 0.10200 | 0.00575 |
| **Dlst** | 7.73155 | 5.50142 | 6.68881 | 1.26570 | 7.60630 | 10.68910 | 4.40500 | 2.56171 | 4.31595 | 2.83950 | 2.90497 | 0.09970 | 4.32275 | 0.30474 | 2.02178 | 8.66580 |
| **Dlx2** | 0.01377 | 0.00813 | 0.01207 | 0.73096 | 0.00940 | 0.00648 | 0.00741 | 0.01240 | 0.00000 | 0.00757 | 0.01241 | 0.00794 | 0.01270 | 0.00599 | 0.31138 | 0.00725 |
| **Dlx3** | 0.01361 | 0.00853 | 0.00365 | 0.01837 | 0.00893 | 0.00804 | 0.00699 | 0.00000 | 0.02917 | 0.02358 | 2.90922 | 0.00736 | 0.00000 | 0.00286 | 0.06339 | 0.00665 |
| **Dmap1** | 7.62737 | 0.06557 | 0.00000 | 0.05855 | 8.49779 | 2.70204 | 0.78733 | 11.18034 | 0.91930 | 16.08063 | 2.86059 | 0.13265 | 10.20796 | 0.00387 | 0.03152 | 6.09129 |
| **Dmbx1** | 0.00000 | 0.61597 | 0.00000 | 0.24113 | 0.00792 | 0.00638 | 0.00630 | 0.69234 | 0.00312 | 0.34847 | 1.58610 | 0.21256 | 0.16001 | 2.11708 | 2.41295 | 0.01378 |
| **Dmc1** | 3.26587 | 3.22107 | 1.97318 | 0.28117 | 0.01221 | 4.15114 | 6.21647 | 3.23482 | 0.39455 | 0.27568 | 1.26966 | 1.45688 | 3.74275 | 1.46367 | 0.32821 | 3.93180 |
| **Dmd** | 0.30356 | 6.32392 | 0.00000 | 0.51280 | 0.51669 | 0.00582 | 2.24843 | 0.69774 | 0.58172 | 1.01271 | 0.00000 | 0.27719 | 0.01706 | 3.15565 | 0.99151 | 1.32720 |
| **Dmgdh** | 3.13387 | 7.88385 | 6.32441 | 2.48351 | 2.34346 | 8.17102 | 5.66359 | 10.80257 | 0.44707 | 6.41619 | 16.86183 | 3.31699 | 9.66714 | 4.09722 | 5.61834 | 3.57072 |
| **Dmkn** | 1.20831 | 0.00586 | 0.80084 | 0.00813 | 0.00423 | 2.44519 | 0.00530 | 0.00278 | 0.01257 | 0.23859 | 0.00882 | 0.00455 | 1.26214 | 0.01439 | 0.00001 | 0.00447 |
| **Dmpk** | 0.51936 | 0.00695 | 0.00370 | 0.41121 | 0.00666 | 0.00739 | 0.02663 | 0.00000 | 1.82817 | 0.31845 | 0.62410 | 0.05702 | 0.03509 | 0.00602 | 0.00000 | 0.00672 |
| **Dmrt1** | 1.74052 | 0.00542 | 0.08458 | 0.29388 | 0.01469 | 0.00379 | 0.00549 | 0.58590 | 0.00695 | 0.62881 | 0.24255 | 0.00194 | 0.20599 | 0.00783 | 0.00000 | 0.00526 |
| **Dmrta2** | 0.26944 | 0.00809 | 0.00000 | 0.05832 | 0.00777 | 0.05390 | 0.00773 | 0.02466 | 0.06280 | 0.01796 | 0.01425 | 0.00818 | 0.03737 | 0.01216 | 0.00903 | 0.00720 |
| **Dmrtb1** | 0.00570 | 0.00706 | 0.00000 | 8.07245 | 0.00810 | 0.00458 | 0.00532 | 0.00000 | 13.22123 | 5.54379 | 4.09895 | 0.95075 | 0.39645 | 0.16558 | 2.14532 | 0.01157 |
| **Dmtf1** | 5.16833 | 0.41839 | 2.54247 | 4.91883 | 0.03789 | 3.22740 | 0.39027 | 7.57588 | 7.27758 | 9.76064 | 4.36874 | 9.98657 | 3.42602 | 19.03650 | 10.49667 | 1.21408 |
| **Dmtn** | 3.91093 | 0.09926 | 1.89148 | 3.06220 | 0.00990 | 0.01630 | 0.77196 | 2.29265 | 1.05300 | 1.77828 | 2.95737 | 0.58555 | 2.12190 | 0.02859 | 0.97341 | 0.05716 |
| **Dmwd** | 6.58684 | 0.01915 | 5.08152 | 11.59239 | 0.19936 | 5.22989 | 2.55850 | 5.69894 | 8.06597 | 7.61043 | 3.18605 | 0.27223 | 3.90212 | 0.00000 | 0.11801 | 1.14042 |
| **Dmxl1** | 1.68148 | 0.00536 | 0.00000 | 4.38534 | 0.00545 | 0.00340 | 0.27429 | 0.43703 | 3.38787 | 0.00000 | 0.46764 | 1.12776 | 0.01265 | 4.76502 | 2.77089 | 0.00428 |
| **Dmxl2** | 1.23750 | 0.00227 | 2.00139 | 5.49033 | 0.00929 | 0.02469 | 0.00337 | 4.04174 | 1.37684 | 1.42445 | 1.24421 | 11.09415 | 1.24894 | 5.72227 | 7.46356 | 3.75550 |
| **Dna2** | 2.82701 | 3.29867 | 1.91732 | 7.90281 | 0.08187 | 0.00000 | 2.29740 | 2.27551 | 9.20465 | 3.92962 | 4.49096 | 2.23545 | 0.59505 | 3.86177 | 2.96411 | 2.66800 |
| **Dnaaf1** | 0.00000 | 0.00513 | 2.94701 | 1.61195 | 0.00549 | 0.00604 | 2.56127 | 5.31947 | 0.38220 | 2.78043 | 0.78485 | 1.86134 | 2.38806 | 0.00612 | 0.16916 | 0.45830 |
| **Dnaaf2** | 4.80800 | 1.04432 | 1.05646 | 4.24720 | 8.86059 | 3.03566 | 2.52060 | 9.68658 | 9.73324 | 10.60264 | 5.55792 | 2.48163 | 2.51163 | 0.84363 | 6.10410 | 2.75195 |
| **Dnaaf3** | 0.01317 | 0.00809 | 0.02500 | 0.17303 | 0.00939 | 0.00771 | 0.00752 | 0.01395 | 0.08527 | 0.01461 | 0.01299 | 0.01819 | 0.01529 | 0.01170 | 0.00637 | 0.00745 |
| **Dnaaf5** | 0.39884 | 1.33794 | 1.56771 | 4.04946 | 5.21374 | 0.31931 | 0.09923 | 5.38663 | 4.75668 | 1.80469 | 1.90387 | 0.30714 | 0.81091 | 0.07531 | 1.95800 | 0.00405 |

| | | | | | | | | | | | | | | | | |
|---|---|---|---|---|---|---|---|---|---|---|---|---|---|---|---|---|
| **Dnah1** | 0.02970 | 0.00809 | 0.01089 | 0.01028 | 0.00836 | 0.00664 | 0.00741 | 0.01358 | 0.01222 | 0.01584 | 0.01290 | 0.00829 | 0.05237 | 0.01025 | 0.03405 | 0.00732 |
| **Dnah10** | 0.01325 | 0.00797 | 0.00968 | 0.63784 | 0.00904 | 0.00659 | 0.03737 | 0.00383 | 0.08084 | 0.33800 | 0.21221 | 0.00503 | 0.08392 | 0.00447 | 0.43786 | 0.00745 |
| **Dnah11** | 0.01057 | 0.00753 | 0.00690 | 0.10190 | 0.00961 | 0.00651 | 0.00712 | 0.17902 | 0.00858 | 0.00000 | 0.09242 | 0.00392 | 0.00666 | 0.01728 | 1.18940 | 0.00724 |
| **Dnah17** | 0.00737 | 0.00793 | 0.00197 | 0.27778 | 0.00929 | 0.00808 | 0.00711 | 0.66294 | 0.54070 | 0.94403 | 1.41160 | 0.00000 | 0.08180 | 0.00647 | 0.30977 | 0.00720 |
| **Dnah2** | 1.39477 | 0.65261 | 0.22614 | 1.57620 | 0.40096 | 0.00566 | 0.00619 | 0.45187 | 1.14533 | 0.60519 | 0.00000 | 0.05447 | 1.35665 | 0.00415 | 0.82602 | 0.04193 |
| **Dnah5** | 0.01366 | 0.00798 | 0.01217 | 0.00903 | 0.00916 | 0.00666 | 0.00736 | 0.01146 | 0.01205 | 0.29853 | 0.01317 | 0.00699 | 0.01252 | 0.00963 | 0.02789 | 0.00778 |
| **Dnah6** | 0.24478 | 0.00767 | 0.00764 | 0.16589 | 0.00795 | 0.00747 | 0.00747 | 0.01758 | 0.51750 | 0.00495 | 0.01165 | 0.00664 | 0.22060 | 0.02042 | 0.03602 | 0.00721 |
| **Dnah7a** | 0.09895 | 0.00733 | 0.00746 | 0.24375 | 0.00769 | 0.00657 | 0.00716 | 0.00832 | 0.14652 | 0.95617 | 0.02158 | 0.01522 | 0.07277 | 0.01068 | 0.51544 | 0.00690 |
| **Dnah7b** | 0.11428 | 0.00726 | 0.00653 | 0.80384 | 0.00764 | 0.00651 | 0.00710 | 0.00612 | 0.21968 | 0.16093 | 0.10311 | 0.00491 | 0.16061 | 0.00902 | 0.80325 | 0.00691 |
| **Dnah7c** | 0.16858 | 0.00733 | 0.00563 | 0.57773 | 0.00754 | 0.00645 | 0.00714 | 0.00885 | 0.14655 | 0.35981 | 0.00896 | 0.00426 | 0.06446 | 0.00598 | 1.16869 | 0.00708 |
| **Dnah8** | 0.01887 | 0.00832 | 0.06102 | 0.53596 | 0.00855 | 0.00573 | 0.00770 | 0.00081 | 0.00138 | 0.40315 | 0.87353 | 0.00163 | 0.00457 | 0.26335 | 0.66880 | 0.00684 |
| **Dnah9** | 0.01346 | 0.00758 | 0.01019 | 1.37847 | 0.00916 | 0.00631 | 0.00723 | 0.00845 | 1.20534 | 0.22601 | 0.05726 | 0.01136 | 0.01210 | 0.00275 | 0.89876 | 0.00748 |
| **Dnaic1** | 0.00981 | 0.00746 | 0.00362 | 0.00458 | 0.01037 | 0.00873 | 0.02729 | 0.59471 | 0.01016 | 0.00000 | 1.94399 | 0.00000 | 0.02579 | 0.01411 | 0.00000 | 0.00733 |
| **Dnaic2** | 0.01345 | 0.00800 | 0.01156 | 2.18829 | 0.00921 | 0.00650 | 0.00750 | 0.01342 | 1.36668 | 0.08286 | 0.00000 | 0.02708 | 0.01305 | 0.00222 | 0.08637 | 0.01417 |
| **Dnaja1** | 39.21310 | 23.24173 | 43.52372 | 13.76758 | 29.45377 | 16.73918 | 24.07083 | 51.07403 | 19.76967 | 44.62897 | 45.16602 | 32.97282 | 46.98273 | 41.80200 | 23.49896 | 12.79608 |
| **Dnaja2** | 17.76526 | 12.90803 | 17.54845 | 20.43238 | 14.60748 | 6.65804 | 10.04016 | 19.71606 | 26.63928 | 28.25130 | 17.31499 | 19.01503 | 16.98114 | 27.96413 | 22.66865 | 8.38124 |
| **Dnaja3** | 9.20797 | 2.38389 | 6.30533 | 5.54157 | 4.69061 | 1.40307 | 7.70522 | 7.62298 | 5.22481 | 4.83627 | 6.23173 | 0.30931 | 7.47030 | 7.17888 | 6.13092 | 5.00771 |
| **Dnaja4** | 0.00305 | 0.00865 | 0.05636 | 5.79861 | 0.00877 | 0.00530 | 0.00700 | 0.51075 | 1.44184 | 0.48616 | 0.01881 | 0.08112 | 2.44574 | 0.00000 | 1.18382 | 0.01233 |
| **Dnajb1** | 10.76570 | 10.61269 | 11.43359 | 9.96687 | 2.84243 | 7.39772 | 7.44791 | 7.54836 | 21.79561 | 15.49721 | 8.48630 | 3.36865 | 12.43578 | 0.09521 | 6.03937 | 3.02626 |
| **Dnajb11** | 5.89775 | 5.44173 | 10.66450 | 3.47515 | 6.26749 | 7.59212 | 4.15366 | 7.63146 | 5.65070 | 8.90222 | 4.87220 | 6.61346 | 13.86920 | 12.60040 | 10.51473 | 10.89491 |
| **Dnajb12** | 0.05049 | 0.00756 | 0.00000 | 0.10492 | 0.00807 | 0.03246 | 0.00731 | 0.21984 | 0.90999 | 0.00000 | 0.15419 | 0.16455 | 0.03031 | 0.01430 | 0.69319 | 0.02125 |
| **Dnajb13** | 0.01343 | 0.00803 | 0.01186 | 0.40238 | 0.00908 | 0.00677 | 0.00752 | 0.01253 | 0.58851 | 0.39932 | 0.00821 | 0.00694 | 0.01255 | 0.00602 | 0.10309 | 0.00730 |
| **Dnajb14** | 0.00000 | 0.06705 | 0.32065 | 9.45547 | 0.00763 | 0.00285 | 0.00572 | 1.17450 | 15.41572 | 9.24167 | 1.77154 | 2.30938 | 0.00000 | 2.81164 | 11.54481 | 0.00402 |
| **Dnajb2** | 2.90172 | 3.72523 | 6.51042 | 1.10378 | 0.02826 | 3.60346 | 4.21427 | 2.95229 | 2.11432 | 5.67293 | 3.47692 | 0.13071 | 4.02930 | 0.49547 | 1.31098 | 0.00416 |
| **Dnajb3** | 0.00971 | 0.00754 | 0.01923 | 0.24457 | 0.00917 | 0.00607 | 0.00666 | 0.01877 | 0.49136 | 0.00000 | 0.00878 | 0.05445 | 1.11914 | 0.05023 | 0.02228 | 0.00662 |
| **Dnajb4** | 0.01897 | 0.00777 | 0.00435 | 0.65536 | 0.01027 | 2.35680 | 0.00710 | 0.00881 | 0.61296 | 0.59346 | 0.03294 | 0.05141 | 0.00513 | 0.00583 | 0.51893 | 0.00687 |
| **Dnajb6** | 21.47813 | 10.79921 | 20.49983 | 23.17589 | 16.83788 | 9.74669 | 10.48804 | 26.05094 | 25.67941 | 43.40434 | 31.71771 | 22.07013 | 28.54371 | 31.78883 | 23.70541 | 13.81058 |
| **Dnajb9** | 1.64304 | 2.91259 | 5.74282 | 22.37411 | 0.43633 | 4.42611 | 0.00000 | 7.51210 | 23.56476 | 19.39536 | 11.56704 | 30.10492 | 6.81380 | 27.40490 | 17.31943 | 0.29523 |
| **Dnajc1** | 0.77771 | 4.54130 | 4.01710 | 1.89774 | 0.11546 | 0.00525 | 0.18061 | 1.42056 | 5.23042 | 2.80430 | 2.72892 | 0.62251 | 0.00000 | 0.55862 | 1.51187 | 2.43964 |
| **Dnajc10** | 2.96294 | 0.00224 | 4.26452 | 1.10019 | 0.07646 | 3.51895 | 2.43374 | 4.10782 | 1.77115 | 1.06671 | 1.63490 | 0.08809 | 5.18987 | 0.34379 | 1.25954 | 7.70304 |
| **Dnajc11** | 12.19672 | 12.43040 | 11.56322 | 3.31680 | 4.45521 | 10.81842 | 9.65331 | 18.22879 | 3.03240 | 9.48303 | 6.76103 | 0.52180 | 18.12180 | 1.62167 | 5.59041 | 7.01958 |
| **Dnajc12** | 1.33162 | 0.00569 | 0.85142 | 5.27731 | 0.01320 | 0.00311 | 2.98816 | 0.00000 | 5.78981 | 1.84161 | 0.84094 | 0.35181 | 1.18977 | 2.09243 | 6.77149 | 0.01079 |
| **Dnajc13** | 10.06880 | 8.42340 | 6.09682 | 6.94058 | 6.74875 | 3.87750 | 7.29788 | 7.38085 | 3.95743 | 11.90347 | 3.16811 | 8.07943 | 5.57208 | 11.23601 | 8.05736 | 1.80793 |
| **Dnajc14** | 2.52435 | 1.76532 | 0.42449 | 1.09828 | 0.01040 | 7.50504 | 1.74591 | 5.26440 | 1.18462 | 4.09759 | 0.66438 | 2.90650 | 4.07643 | 0.06731 | 4.70022 | 1.49463 |
| **Dnajc15** | 11.67974 | 1.73205 | 7.11557 | 3.03509 | 2.41188 | 0.54672 | 5.77299 | 7.25733 | 12.80080 | 15.16268 | 16.94176 | 5.05302 | 10.00328 | 1.27747 | 0.31554 | 4.19226 |
| **Dnajc16** | 2.37818 | 0.00484 | 2.16860 | 0.90074 | 0.00000 | 0.00294 | 0.00544 | 2.15655 | 0.01732 | 1.52604 | 0.00000 | 0.09450 | 0.41972 | 0.00333 | 0.26692 | 0.00345 |
| **Dnajc17** | 3.58861 | 3.05979 | 4.33474 | 0.94394 | 0.85379 | 4.01238 | 0.16812 | 6.25538 | 0.20400 | 6.69923 | 1.42848 | 0.11903 | 0.85708 | 0.03763 | 0.05476 | 2.13048 |
| **Dnajc18** | 2.25629 | 0.01549 | 0.58979 | 1.36219 | 0.00291 | 0.00303 | 0.04396 | 0.00000 | 0.38250 | 0.18628 | 0.00000 | 0.00000 | 0.00908 | 2.84949 | 0.73584 | 0.00412 |
| **Dnajc19** | 11.62507 | 0.00000 | 0.83277 | 0.40392 | 7.51181 | 2.04612 | 0.84464 | 2.28147 | 0.00000 | 2.58476 | 9.60635 | 0.30291 | 6.67241 | 0.08435 | 0.34107 | 5.30567 |
| **Dnajc2** | 11.84923 | 3.72664 | 10.17205 | 4.91013 | 12.22066 | 7.59939 | 4.94139 | 11.98350 | 3.20449 | 25.14888 | 22.33372 | 19.05998 | 13.17383 | 25.34408 | 15.13990 | 3.83708 |
| **Dnajc21** | 10.25154 | 4.96198 | 7.23107 | 0.90537 | 6.12878 | 9.17804 | 6.74955 | 21.61875 | 1.15347 | 30.21220 | 14.69193 | 6.18614 | 8.46974 | 33.13436 | 31.37337 | 5.98235 |
| **Dnajc22** | 4.47396 | 0.16950 | 0.00000 | 0.00236 | 0.71027 | 0.00342 | 0.36580 | 0.00000 | 0.68523 | 0.63476 | 0.00515 | 0.01010 | 0.00635 | 0.04431 | 0.00000 | 3.46602 |
| **Dnajc24** | 3.21038 | 0.00000 | 0.71651 | 0.56693 | 8.70951 | 2.49359 | 0.06833 | 0.31528 | 0.03693 | 7.15291 | 6.80695 | 0.32265 | 1.51278 | 0.40560 | 0.00000 | 0.80204 |
| **Dnajc25** | 0.05249 | 0.02862 | 0.46164 | 2.99221 | 0.01659 | 0.32494 | 0.00572 | 2.12850 | 0.38252 | 1.42253 | 1.80308 | 0.55185 | 2.30683 | 0.00535 | 0.86661 | 0.01199 |
| **Dnajc27** | 0.60181 | 0.00616 | 0.00000 | 1.86376 | 0.01028 | 0.00547 | 2.15361 | 3.31599 | 1.96380 | 1.07531 | 0.33020 | 0.10045 | 3.19642 | 0.00038 | 2.15325 | 0.00485 |
| **Dnajc28** | 0.10066 | 0.00882 | 0.00954 | 0.44027 | 0.01761 | 0.00714 | 0.00761 | 0.01466 | 0.00575 | 0.01798 | 0.01511 | 0.00834 | 0.02639 | 0.01097 | 0.00582 | 0.00764 |
| **Dnajc3** | 1.03221 | 4.36980 | 6.43961 | 4.84231 | 0.00000 | 0.00396 | 0.72759 | 1.18376 | 4.44865 | 12.24435 | 2.49414 | 2.31632 | 0.29795 | 16.94540 | 15.10575 | 0.08685 |
| **Dnajc30** | 0.71779 | 1.22733 | 0.00000 | 2.92775 | 3.50857 | 2.14217 | 0.03497 | 1.95865 | 5.63561 | 0.51335 | 0.45481 | 0.23138 | 1.89152 | 0.00000 | 0.13882 | 0.03347 |
| **Dnajc4** | 3.23981 | 0.41149 | 1.06810 | 9.82433 | 0.00000 | 2.50222 | 0.76019 | 1.00275 | 9.74383 | 1.00949 | 1.70390 | 0.09325 | 1.05889 | 0.00000 | 0.24724 | 2.14453 |
| **Dnajc5** | 7.48084 | 0.20911 | 8.22283 | 7.37393 | 2.71991 | 6.50719 | 0.59578 | 4.99375 | 7.49300 | 9.54736 | 4.67361 | 9.80843 | 2.94339 | 6.47238 | 5.29464 | 4.38062 |
| **Dnajc5g** | 1.31860 | 0.00478 | 0.00000 | 1.54848 | 0.00545 | 0.00408 | 0.00492 | 0.21980 | 2.89361 | 2.91945 | 1.74622 | 3.91722 | 0.00000 | 4.08705 | 0.87700 | 0.00518 |
| **Dnajc6** | 0.56429 | 0.00830 | 0.00280 | 0.00422 | 0.03089 | 0.00821 | 0.00755 | 0.30792 | 0.99298 | 0.00000 | 0.28140 | 0.15165 | 0.00959 | 0.01267 | 0.00000 | 0.00688 |
| **Dnajc7** | 2.81204 | 5.57381 | 5.40025 | 1.92806 | 4.42017 | 3.90803 | 3.61013 | 11.30433 | 8.38921 | 5.56518 | 13.00182 | 6.86040 | 2.56738 | 12.54681 | 9.80683 | 2.09243 |
| **Dnajc8** | 17.18863 | 18.85386 | 18.37146 | 4.32687 | 9.66695 | 3.76391 | 11.71198 | 17.44054 | 9.65814 | 14.66502 | 10.52388 | 6.43611 | 16.66737 | 10.36068 | 14.03995 | 6.27077 |
| **Dnajc9** | 0.48074 | 2.84010 | 3.57318 | 4.50929 | 0.00008 | 0.35111 | 0.04560 | 0.12121 | 4.56668 | 3.02886 | 5.55400 | 0.18472 | 0.00000 | 2.26618 | 2.14616 | 0.00948 |
| **Dnal1** | 0.09943 | 0.00761 | 0.11936 | 0.49105 | 0.00777 | 0.00826 | 0.71341 | 0.03067 | 0.03092 | 0.04668 | 1.13944 | 0.00631 | 0.23166 | 1.84570 | 0.51650 | 0.02176 |
| **Dnal4** | 1.48689 | 1.75857 | 1.85807 | 11.39681 | 0.29495 | 0.65602 | 0.35523 | 3.95505 | 2.45179 | 5.88970 | 0.31176 | 0.08916 | 2.61755 | 0.00000 | 0.00000 | 0.08796 |
| **Dnase1** | 0.04848 | 0.00765 | 0.00596 | 0.13074 | 0.00796 | 0.00743 | 0.00725 | 0.27005 | 0.21055 | 0.23188 | 0.00938 | 0.00725 | 0.01652 | 0.01428 | 0.01416 | 0.00718 |
| **Dnase1l1** | 0.01330 | 0.00823 | 0.00563 | 0.03189 | 0.00907 | 0.00693 | 0.00757 | 0.00933 | 0.25123 | 0.00000 | 0.87896 | 0.01406 | 0.00464 | 0.00641 | 0.48999 | 0.00699 |
| **Dnase1l2** | 0.09351 | 0.00714 | 1.24824 | 0.00539 | 0.00564 | 0.03893 | 0.43044 | 0.00000 | 0.01078 | 0.60134 | 0.55747 | 0.00244 | 0.00583 | 0.01016 | 0.23998 | 0.00657 |
| **Dnase1l3** | 0.01343 | 0.00752 | 0.02595 | 1.11638 | 0.00924 | 0.00620 | 0.00724 | 0.01448 | 3.44193 | 0.20692 | 0.00000 | 0.01847 | 0.01217 | 0.00540 | 0.14178 | 0.00706 |
| **Dnase2a** | 1.51809 | 0.02557 | 0.99478 | 0.07287 | 0.02887 | 0.01531 | 2.21106 | 0.15425 | 0.27495 | 0.51434 | 1.24213 | 0.03369 | 0.18803 | 0.33895 | 0.55121 | 1.11849 |

| | | | | | | | | | | | | | | | | |
|---|---|---|---|---|---|---|---|---|---|---|---|---|---|---|---|---|
| **Dnd1** | 0.04620 | 0.00884 | 0.39585 | 0.87698 | 0.01187 | 2.98185 | 0.00673 | 0.52156 | 1.11776 | 0.32928 | 0.14440 | 0.00279 | 1.82503 | 0.03678 | 0.00924 | 0.18741 |
| **Dner** | 0.01362 | 0.00816 | 0.01245 | 1.42694 | 0.00917 | 0.00642 | 0.00745 | 0.01238 | 0.00000 | 0.20993 | 0.01226 | 0.00736 | 0.01147 | 0.00784 | 0.01207 | 0.01398 |
| **Dnlz** | 0.76510 | 0.00691 | 0.51060 | 1.02289 | 0.28373 | 0.01069 | 0.05766 | 0.48282 | 0.47082 | 1.01428 | 0.97662 | 0.03908 | 1.97885 | 0.00533 | 0.13866 | 0.00541 |
| **Dnm1** | 0.01294 | 0.00809 | 0.03255 | 0.65525 | 0.00908 | 0.00601 | 0.00738 | 0.01104 | 0.00000 | 0.53038 | 0.01351 | 0.00653 | 0.01186 | 0.00990 | 0.00578 | 0.00769 |
| **Dnm1l** | 3.04724 | 8.98818 | 5.05944 | 8.32686 | 4.75470 | 2.51100 | 3.99882 | 4.91607 | 2.87425 | 5.05458 | 3.81201 | 1.36680 | 4.97605 | 0.04917 | 8.35477 | 5.63717 |
| **Dnm2** | 13.27191 | 0.12946 | 0.79896 | 2.02468 | 2.63486 | 7.29741 | 2.92475 | 11.38831 | 3.62757 | 8.41236 | 2.53749 | 0.35393 | 2.40444 | 0.00000 | 0.34637 | 3.78117 |
| **Dnm3** | 0.01263 | 0.00785 | 0.00843 | 0.03156 | 0.00917 | 0.00664 | 0.00728 | 0.00985 | 0.01286 | 0.52553 | 0.00832 | 0.00403 | 0.01108 | 0.77506 | 0.00000 | 0.00710 |
| **Dnmbp** | 1.64555 | 1.14997 | 1.48453 | 1.63063 | 0.01097 | 3.34177 | 0.01340 | 0.00000 | 1.81897 | 0.00000 | 1.37732 | 0.02443 | 2.06350 | 0.00262 | 0.03036 | 0.00181 |
| **Dnmt1** | 9.23716 | 1.19778 | 1.03515 | 6.74210 | 8.06483 | 2.09916 | 3.38232 | 7.96916 | 6.43919 | 10.41833 | 5.61169 | 0.58107 | 5.14933 | 0.03759 | 0.39066 | 11.76896 |
| **Dnmt3a** | 2.70510 | 4.61068 | 9.73648 | 6.79238 | 2.14314 | 0.00278 | 7.45781 | 7.07847 | 7.58413 | 5.64991 | 6.02491 | 1.34802 | 3.32639 | 3.88405 | 3.98353 | 4.51419 |
| **Dnmt3b** | 8.01421 | 3.84482 | 5.72926 | 1.92090 | 0.15658 | 11.54509 | 6.41934 | 13.59505 | 0.25811 | 9.04526 | 5.51497 | 1.67224 | 7.44427 | 3.73239 | 6.03627 | 20.52531 |
| **Dnmt3l** | 9.89699 | 3.01136 | 6.95181 | 3.62080 | 9.48183 | 16.92009 | 1.95018 | 4.90299 | 1.21829 | 3.08336 | 5.99656 | 5.20407 | 14.40535 | 1.35165 | 1.06776 | 22.57146 |
| **Dnpep** | 13.55052 | 3.10534 | 13.44132 | 7.47155 | 11.02133 | 12.34647 | 11.18210 | 16.47539 | 9.21415 | 8.86305 | 6.86470 | 4.60019 | 13.44120 | 0.26435 | 1.76028 | 8.33134 |
| **Dnph1** | 3.35729 | 0.01226 | 1.49487 | 0.91451 | 1.71841 | 0.00842 | 0.26757 | 5.15069 | 0.35500 | 3.09213 | 5.23998 | 0.73897 | 3.57979 | 0.07028 | 0.41591 | 0.50323 |
| **Dnttip1** | 4.23813 | 2.93603 | 4.80389 | 2.13969 | 1.96997 | 2.91932 | 0.02169 | 3.45208 | 3.93800 | 5.05654 | 2.06773 | 4.43532 | 0.00000 | 0.00000 | 0.46743 | 0.02000 |
| **Dnttip2** | 16.20605 | 13.60431 | 12.50150 | 11.76276 | 12.67207 | 4.32853 | 8.41680 | 17.14617 | 20.08184 | 23.17511 | 17.16191 | 9.36324 | 12.71057 | 28.58979 | 21.63548 | 6.39719 |
| **Doc2a** | 3.92014 | 2.93582 | 3.49813 | 2.60157 | 4.32427 | 3.13738 | 3.91603 | 5.65995 | 5.89112 | 7.29212 | 1.99517 | 0.13228 | 3.99090 | 0.00000 | 0.00000 | 2.25113 |
| **Doc2b** | 0.01080 | 0.00752 | 0.01021 | 0.15077 | 0.00944 | 0.00658 | 0.00717 | 0.12643 | 0.00964 | 0.25459 | 0.01479 | 0.00687 | 0.00753 | 0.01124 | 0.05356 | 0.00711 |
| **Doc2g** | 0.01016 | 0.00734 | 0.00388 | 0.01128 | 0.00963 | 0.05452 | 0.00707 | 1.11499 | 0.01119 | 0.00335 | 0.00648 | 0.00538 | 0.00468 | 0.01651 | 0.00000 | 0.00734 |
| **Dock1** | 1.43124 | 0.01536 | 2.61211 | 0.02159 | 0.37970 | 0.00772 | 0.00532 | 1.04617 | 0.04989 | 0.86097 | 0.04202 | 0.04343 | 0.73646 | 0.76555 | 0.02816 | 0.14600 |
| **Dock10** | 0.00726 | 0.00766 | 0.00827 | 0.01716 | 0.38690 | 0.00706 | 0.00736 | 0.01168 | 0.01220 | 0.03274 | 0.02431 | 0.00821 | 0.35136 | 0.01182 | 0.01147 | 0.00712 |
| **Dock11** | 0.00000 | 0.00777 | 0.29851 | 1.06827 | 0.00770 | 0.00880 | 1.02842 | 0.17911 | 1.18907 | 0.41746 | 0.50750 | 0.00000 | 0.00002 | 0.00229 | 0.23867 | 0.64868 |
| **Dock3** | 0.01064 | 0.00839 | 0.00837 | 1.35549 | 0.00923 | 0.00663 | 0.00788 | 0.08736 | 1.45424 | 0.13688 | 0.16881 | 0.02434 | 0.08696 | 0.03563 | 0.09816 | 0.00732 |
| **Dock4** | 0.15214 | 0.01812 | 1.33646 | 0.01752 | 0.00867 | 0.00576 | 0.00701 | 0.33434 | 0.61814 | 0.00000 | 0.06666 | 0.04543 | 0.00000 | 0.01628 | 0.40127 | 0.00660 |
| **Dock5** | 4.92113 | 1.09372 | 2.52539 | 5.04238 | 0.00747 | 0.00157 | 2.21403 | 3.23240 | 0.63988 | 3.27976 | 4.30740 | 0.03378 | 2.66868 | 2.12320 | 2.03026 | 0.29819 |
| **Dock6** | 3.34208 | 0.20996 | 1.30341 | 0.80164 | 0.00905 | 0.00549 | 0.00363 | 3.75678 | 0.90969 | 2.48261 | 1.76014 | 0.01790 | 0.16429 | 0.00000 | 0.49793 | 1.17094 |
| **Dock7** | 0.10250 | 3.79591 | 0.10663 | 10.54006 | 0.00747 | 0.32765 | 0.00622 | 0.13394 | 0.00000 | 0.29066 | 0.12863 | 0.04761 | 0.00222 | 0.71269 | 2.31526 | 0.00386 |
| **Dock8** | 0.29529 | 0.85220 | 0.98928 | 0.02605 | 0.00557 | 0.00550 | 0.00664 | 0.01867 | 0.01129 | 0.00313 | 0.01157 | 0.01793 | 0.24175 | 0.01195 | 0.00000 | 0.01316 |
| **Dock9** | 3.34993 | 0.01300 | 0.68921 | 6.89013 | 0.22956 | 0.50599 | 0.26621 | 5.65565 | 1.93102 | 2.08437 | 5.15545 | 0.19888 | 0.89477 | 2.60277 | 4.55718 | 2.55459 |
| **Dohh** | 8.84918 | 0.35648 | 0.06236 | 0.42857 | 4.77356 | 0.60833 | 0.04595 | 2.11992 | 2.05742 | 12.08860 | 2.23922 | 0.06036 | 10.66883 | 0.00000 | 0.00000 | 4.81161 |
| **Dok1** | 1.13373 | 0.00591 | 1.14718 | 0.18066 | 0.14480 | 0.90422 | 0.00594 | 2.94564 | 0.00770 | 0.96130 | 0.35291 | 0.00504 | 0.22239 | 0.00603 | 0.00000 | 0.01275 |
| **Dok2** | 0.00000 | 1.18725 | 2.95026 | 0.00000 | 0.00000 | 4.47977 | 2.43288 | 3.30445 | 0.00866 | 0.00000 | 0.40568 | 0.02150 | 5.96405 | 0.01600 | 0.00000 | 8.39075 |
| **Dok3** | 0.01364 | 0.00818 | 0.01195 | 0.24648 | 0.00924 | 0.00683 | 0.00761 | 0.01433 | 0.34783 | 0.07356 | 0.00998 | 0.00844 | 0.01318 | 0.01223 | 0.00891 | 0.00737 |
| **Dok4** | 0.97049 | 0.00666 | 0.00000 | 1.33283 | 0.03206 | 2.62264 | 1.37655 | 0.05206 | 1.51726 | 0.12321 | 0.00000 | 0.04135 | 0.89705 | 0.00244 | 0.32823 | 0.24959 |
| **Dok5** | 0.01181 | 0.00809 | 0.04278 | 0.01365 | 0.00986 | 0.01565 | 0.00757 | 0.02502 | 0.01253 | 0.57288 | 0.01390 | 0.00599 | 0.12002 | 0.04323 | 0.03710 | 0.03853 |
| **Dolk** | 2.31339 | 0.43550 | 2.49770 | 4.02719 | 2.73077 | 0.00351 | 3.69741 | 3.13848 | 8.38161 | 3.56949 | 2.38463 | 0.39331 | 1.51783 | 2.04201 | 3.95411 | 1.41846 |
| **Dolpp1** | 4.09461 | 1.80770 | 0.99865 | 0.17323 | 0.23996 | 2.61275 | 1.11047 | 5.69837 | 0.38157 | 0.50205 | 2.34181 | 0.21514 | 6.36526 | 0.00028 | 0.55261 | 4.31436 |
| **Donson** | 3.36889 | 2.44944 | 0.00000 | 11.90523 | 0.00588 | 0.02274 | 0.80695 | 1.84987 | 8.90003 | 3.32013 | 2.77020 | 0.77299 | 0.51232 | 2.86993 | 7.35468 | 1.17618 |
| **Dopey1** | 0.00000 | 0.00708 | 0.11526 | 1.06248 | 0.01121 | 0.10115 | 2.12360 | 1.38709 | 0.73940 | 0.79991 | 0.05776 | 0.08037 | 0.11153 | 1.25324 | 1.88170 | 0.00711 |
| **Dopey2** | 2.72728 | 0.00430 | 0.01133 | 1.56381 | 0.00491 | 0.00438 | 0.00410 | 0.60358 | 0.67901 | 0.60615 | 0.65624 | 0.03921 | 0.95543 | 0.00054 | 0.85318 | 0.00405 |
| **Dot1l** | 0.60751 | 0.02527 | 1.19406 | 0.54849 | 2.92895 | 0.01753 | 0.54568 | 3.17314 | 1.13996 | 2.31812 | 0.22945 | 0.10377 | 2.13562 | 0.12112 | 0.91082 | 1.10705 |
| **Dpagt1** | 1.51281 | 5.44093 | 4.08582 | 2.83813 | 2.45388 | 7.49155 | 0.79833 | 2.63161 | 3.94598 | 2.88313 | 1.51219 | 0.67549 | 3.26437 | 2.22273 | 2.98595 | 6.61602 |
| **Dpcd** | 5.19823 | 8.07482 | 3.04073 | 0.36452 | 5.80006 | 0.00094 | 1.37156 | 0.62732 | 0.35756 | 4.40493 | 2.38398 | 0.29187 | 3.40178 | 0.10809 | 1.55205 | 0.37381 |
| **Dpcr1** | 0.01510 | 0.00822 | 0.01741 | 0.03355 | 0.03093 | 0.00725 | 0.00764 | 0.01663 | 0.60702 | 0.02909 | 0.04501 | 0.00699 | 0.01139 | 0.01534 | 0.01457 | 0.00733 |
| **Dpep1** | 0.37307 | 0.00748 | 0.83667 | 0.00993 | 0.00558 | 0.00508 | 0.00698 | 0.45801 | 2.16098 | 0.66404 | 0.00000 | 0.06267 | 0.00382 | 0.00882 | 0.02183 | 0.00683 |
| **Dpep2** | 0.00000 | 0.00768 | 0.26145 | 3.93505 | 0.00860 | 0.00792 | 0.00694 | 0.24942 | 8.31119 | 2.13351 | 0.00000 | 0.04739 | 0.33100 | 1.90532 | 5.70445 | 0.00583 |
| **Dpep3** | 0.00000 | 0.01799 | 2.48412 | 1.14879 | 0.01968 | 1.25969 | 0.84644 | 0.03591 | 1.33702 | 2.32340 | 4.19372 | 0.14269 | 3.45515 | 0.15905 | 0.89613 | 0.01146 |
| **Dpf1** | 0.05755 | 0.00838 | 0.02590 | 0.56924 | 0.01411 | 0.00604 | 0.00752 | 0.01288 | 0.00000 | 0.01236 | 0.01176 | 0.01780 | 0.01216 | 0.00548 | 0.16232 | 0.00734 |
| **Dpf2** | 8.11637 | 10.71617 | 13.25513 | 10.43923 | 8.03927 | 9.24409 | 6.36928 | 17.85200 | 6.31057 | 22.83337 | 14.76872 | 2.83822 | 13.36899 | 3.97686 | 14.47712 | 3.16738 |
| **Dpf3** | 2.62707 | 0.06816 | 2.03705 | 1.06459 | 0.00000 | 0.00358 | 2.40893 | 6.75603 | 4.16152 | 3.29956 | 5.78424 | 0.03824 | 1.07305 | 0.39330 | 0.38102 | 0.00885 |
| **Dph1** | 1.66050 | 0.01605 | 0.00000 | 0.00188 | 0.12079 | 1.13197 | 1.06233 | 7.62324 | 0.00687 | 6.64922 | 0.03276 | 0.14217 | 4.81579 | 0.01218 | 0.00000 | 0.21958 |
| **Dph2** | 2.18653 | 2.48015 | 5.46853 | 5.54408 | 3.90352 | 0.00839 | 0.00000 | 4.49125 | 11.75185 | 6.32792 | 3.92351 | 3.89448 | 2.64761 | 0.02831 | 4.34062 | 1.04783 |
| **Dph3** | 7.11869 | 3.53352 | 8.47504 | 12.32710 | 8.78390 | 0.26303 | 6.93493 | 7.19509 | 11.84290 | 18.52784 | 15.36148 | 1.90275 | 13.06143 | 3.98890 | 2.91002 | 0.03555 |
| **Dph5** | 11.19041 | 5.20390 | 5.70556 | 0.52669 | 1.20629 | 0.09063 | 6.76259 | 5.00999 | 0.46489 | 3.64786 | 1.84510 | 3.56349 | 5.08457 | 8.09837 | 0.47072 | 0.24644 |
| **Dph6** | 3.04284 | 2.76425 | 3.46141 | 0.65315 | 2.05108 | 0.03829 | 2.60939 | 3.26727 | 0.30827 | 0.27569 | 1.81340 | 0.28294 | 1.76025 | 2.93093 | 2.84127 | 0.37719 |
| **Dph7** | 3.13993 | 0.60259 | 0.25031 | 1.42479 | 0.01244 | 0.00234 | 0.52527 | 0.32289 | 2.74164 | 0.77569 | 0.72766 | 0.10962 | 0.00000 | 0.28255 | 7.40964 | 1.48510 |
| **Dpm1** | 0.53376 | 0.40871 | 0.49676 | 2.17243 | 0.33275 | 0.02726 | 0.15206 | 0.26692 | 2.84387 | 0.40846 | 0.87120 | 0.24417 | 0.35439 | 0.11967 | 0.14247 | 0.44743 |
| **Dpm2** | 5.85471 | 5.53884 | 12.76369 | 3.68259 | 9.02955 | 0.89321 | 3.66437 | 7.28732 | 1.29748 | 9.24824 | 3.94368 | 1.85375 | 7.60967 | 2.48124 | 2.82278 | 5.53324 |
| **Dpm3** | 3.70619 | 0.06326 | 1.61718 | 0.13264 | 1.92108 | 0.37639 | 0.17436 | 0.12695 | 0.20839 | 0.60750 | 0.69847 | 0.04526 | 1.44727 | 0.00937 | 0.00000 | 0.46868 |
| **Dpp10** | 0.01370 | 0.00777 | 0.01136 | 2.16276 | 0.00943 | 0.00623 | 0.00727 | 0.01074 | 0.00000 | 0.00484 | 0.02035 | 0.00583 | 0.01258 | 0.00453 | 0.97799 | 0.00698 |
| **Dpp3** | 11.67048 | 1.07056 | 1.47629 | 0.74242 | 1.94396 | 2.23919 | 1.70048 | 7.38057 | 0.31070 | 4.81402 | 3.79766 | 0.06495 | 11.29474 | 0.00000 | 0.03956 | 6.79340 |
| **Dpp4** | 2.13742 | 0.00233 | 0.00363 | 0.00442 | 2.34314 | 0.00492 | 4.39035 | 0.00000 | 0.00954 | 0.12740 | 0.00865 | 0.20140 | 0.01207 | 0.00616 | 0.41155 | 0.00509 |

| | | | | | | | | | | | | | | | | |
|---|---|---|---|---|---|---|---|---|---|---|---|---|---|---|---|---|
| **Dpp7** | 6.70282 | 3.33518 | 13.07246 | 0.00000 | 0.11730 | 6.13378 | 9.55003 | 7.94690 | 0.81258 | 0.37483 | 8.62411 | 0.09589 | 16.67109 | 0.10500 | 1.59221 | 2.19005 |
| **Dpp8** | 2.27741 | 0.00553 | 0.00000 | 0.66187 | 0.00595 | 0.00467 | 0.00534 | 1.56765 | 0.20307 | 0.23391 | 1.07696 | 0.02206 | 0.11325 | 2.00154 | 1.25139 | 0.00486 |
| **Dpp9** | 1.75061 | 2.46408 | 0.87746 | 5.28406 | 4.76473 | 0.45723 | 0.01348 | 2.69000 | 6.47441 | 3.20467 | 0.41371 | 0.63098 | 2.12302 | 0.00000 | 0.64719 | 1.97497 |
| **Dppa1** | 0.00000 | 0.01781 | 0.01970 | 0.00000 | 0.28252 | 12.28919 | 0.00556 | 0.00447 | 1.03395 | 0.00000 | 0.00000 | 0.00486 | 1.51524 | 0.01827 | 0.00441 | 2.19480 |
| **Dppa2** | 24.77857 | 20.06955 | 16.44406 | 14.28150 | 22.59207 | 7.83483 | 14.64659 | 29.50402 | 23.85140 | 17.88023 | 27.84472 | 14.90004 | 14.66049 | 43.82085 | 31.46315 | 4.56420 |
| **Dppa3** | 43.43137 | 30.19215 | 49.17332 | 31.96539 | 32.10870 | 20.67482 | 27.56540 | 45.66849 | 42.10666 | 55.38266 | 50.55017 | 22.55218 | 46.02930 | 43.41861 | 37.84792 | 16.25833 |
| **Dppa4** | 19.29090 | 9.62832 | 19.57113 | 17.38752 | 16.77631 | 8.48787 | 8.56788 | 20.21633 | 26.72493 | 28.41034 | 21.70930 | 13.57555 | 18.83032 | 22.63153 | 21.75564 | 8.09088 |
| **Dppa5a** | 44.26855 | 15.27019 | 34.41819 | 16.46749 | 27.14855 | 15.43145 | 13.43581 | 21.89725 | 29.06733 | 22.87828 | 38.36327 | 8.75707 | 28.28143 | 17.81818 | 17.01832 | 22.58817 |
| **Dpy19l1** | 2.48217 | 3.76332 | 0.15814 | 0.46996 | 0.43634 | 0.00598 | 1.36324 | 0.02823 | 0.00381 | 0.00000 | 0.02088 | 0.01061 | 1.14843 | 0.76865 | 0.27347 | 2.46690 |
| **Dpy19l2** | 0.00953 | 0.00856 | 0.00836 | 0.01095 | 0.00965 | 0.00695 | 0.00822 | 0.44588 | 0.01187 | 0.00897 | 0.00980 | 0.00829 | 0.00596 | 0.01625 | 0.00344 | 0.00739 |
| **Dpy19l3** | 0.01702 | 0.01768 | 0.00669 | 0.90953 | 0.00886 | 0.00691 | 0.05815 | 0.01125 | 0.49033 | 0.46081 | 0.00082 | 1.66978 | 0.01718 | 0.00949 | 0.00000 | 0.00721 |
| **Dpy19l4** | 0.10670 | 0.00756 | 0.36980 | 3.83076 | 0.00790 | 0.03339 | 0.00738 | 0.25690 | 0.82810 | 2.89490 | 0.02276 | 0.01755 | 0.05096 | 0.41065 | 0.51042 | 0.00675 |
| **Dpy30** | 18.42754 | 4.28077 | 9.51449 | 11.99667 | 9.04378 | 4.09577 | 5.45495 | 8.62784 | 11.18906 | 11.12084 | 11.95588 | 2.42264 | 12.15249 | 2.77396 | 5.44281 | 6.09159 |
| **Dpys** | 0.05036 | 0.00800 | 0.00751 | 0.45426 | 0.00808 | 0.00728 | 0.00743 | 0.01376 | 0.00384 | 0.11632 | 0.01089 | 0.15740 | 0.01323 | 0.01064 | 0.00299 | 0.00723 |
| **Dpysl2** | 0.00120 | 0.05067 | 0.01808 | 0.01632 | 2.25587 | 0.00861 | 0.00812 | 0.08822 | 0.01101 | 0.13578 | 0.01385 | 0.01846 | 0.02453 | 0.02051 | 0.00545 | 0.02142 |
| **Dpysl3** | 0.01349 | 0.00763 | 0.00845 | 2.66991 | 0.00926 | 0.00608 | 0.00725 | 0.00937 | 1.75373 | 0.00000 | 0.05192 | 0.06940 | 0.01180 | 0.00490 | 1.31081 | 0.00669 |
| **Dpysl4** | 1.55114 | 0.90201 | 0.01313 | 0.00291 | 1.02399 | 0.00443 | 0.12449 | 3.52893 | 0.00816 | 0.35946 | 0.45426 | 0.00306 | 2.43948 | 0.01617 | 0.00000 | 1.95576 |
| **Dpysl5** | 0.02258 | 0.00776 | 0.00736 | 0.38506 | 0.00966 | 0.00671 | 0.71501 | 0.00982 | 0.09428 | 0.40071 | 0.01005 | 0.00628 | 0.15544 | 0.00678 | 0.07577 | 0.00691 |
| **Dqx1** | 0.12423 | 0.02927 | 0.36175 | 0.35003 | 0.05215 | 0.04741 | 0.04838 | 0.01804 | 0.77004 | 0.06905 | 0.77598 | 0.10875 | 0.22814 | 0.54810 | 0.07304 | 0.05066 |
| **Dr1** | 1.99444 | 0.06343 | 0.82808 | 18.91862 | 2.61884 | 1.52234 | 3.04747 | 4.96811 | 16.71251 | 2.34536 | 0.36242 | 10.32400 | 2.35826 | 9.77435 | 10.58260 | 0.43680 |
| **Dram1** | 3.96494 | 0.01350 | 1.28385 | 0.00000 | 1.52395 | 0.00140 | 1.60436 | 0.03475 | 0.03943 | 0.52173 | 0.00000 | 1.54069 | 0.38938 | 0.19589 | 1.46502 | 0.00284 |
| **Dram2** | 0.11538 | 0.09984 | 1.29004 | 1.09104 | 4.92548 | 7.15156 | 5.88180 | 5.80361 | 0.11016 | 1.29175 | 2.39541 | 1.78348 | 1.91647 | 5.47535 | 0.00000 | 0.71049 |
| **Drap1** | 4.13588 | 0.04170 | 2.13672 | 0.89747 | 0.77953 | 1.05388 | 1.40783 | 1.26573 | 1.00629 | 1.43040 | 4.12216 | 0.35207 | 1.28022 | 0.03450 | 0.00000 | 1.35041 |
| **Draxin** | 0.01310 | 0.00788 | 0.01167 | 0.57126 | 0.00912 | 0.00643 | 0.03714 | 0.00956 | 0.10043 | 0.99740 | 0.02910 | 0.01474 | 0.01108 | 0.00899 | 0.01523 | 0.01468 |
| **Drc1** | 0.00000 | 0.00601 | 0.29619 | 0.44461 | 0.00911 | 0.00555 | 2.92154 | 1.76648 | 0.03429 | 2.74447 | 1.11569 | 0.25683 | 0.00000 | 2.35254 | 6.88076 | 0.00682 |
| **Drc7** | 0.01330 | 0.00767 | 0.00883 | 0.24677 | 0.00913 | 0.00658 | 0.00726 | 0.01472 | 1.37544 | 0.00000 | 0.21820 | 0.02279 | 0.00978 | 0.01286 | 0.31028 | 0.00738 |
| **Drd3** | 0.00604 | 0.00779 | 0.47966 | 0.00594 | 0.00863 | 0.00520 | 0.00775 | 0.16576 | 0.00937 | 0.00000 | 0.00000 | 2.59433 | 0.10674 | 0.00587 | 0.80293 | 0.00682 |
| **Drg1** | 19.77251 | 10.78171 | 18.53039 | 8.31851 | 16.28106 | 6.15408 | 9.07605 | 22.39937 | 11.94168 | 22.80105 | 16.98329 | 9.42860 | 17.79623 | 14.38207 | 16.51703 | 10.58626 |
| **Drg2** | 11.73747 | 7.80593 | 15.49188 | 4.48548 | 6.84252 | 5.16866 | 10.97315 | 31.38267 | 6.12864 | 26.56343 | 15.81105 | 2.31404 | 14.76976 | 6.90590 | 18.44395 | 5.06442 |
| **Drosha** | 2.60088 | 4.27682 | 3.86204 | 7.19088 | 0.06515 | 5.46796 | 5.37202 | 5.54165 | 5.83146 | 7.25141 | 9.44577 | 3.08283 | 6.43577 | 5.80347 | 7.65782 | 6.60084 |
| **Dsc2** | 0.00000 | 0.00856 | 0.03345 | 1.85171 | 0.00752 | 0.00802 | 0.00773 | 0.20509 | 0.12617 | 0.00212 | 0.01032 | 0.00854 | 0.73397 | 0.01382 | 0.00000 | 1.46700 |
| **Dscam** | 0.01360 | 0.00820 | 0.01192 | 0.30857 | 0.00927 | 0.00657 | 0.00753 | 0.01385 | 1.01154 | 0.01062 | 0.00758 | 0.00668 | 0.01292 | 0.01209 | 0.00379 | 0.00731 |
| **Dscc1** | 5.44696 | 0.43886 | 1.99828 | 2.93551 | 2.09377 | 0.70565 | 1.19299 | 2.30496 | 2.99498 | 12.24511 | 9.03048 | 11.74397 | 9.35604 | 5.49773 | 4.53759 | 6.17164 |
| **Dscr3** | 6.50666 | 1.99555 | 9.15701 | 0.13663 | 0.01428 | 5.42341 | 6.09328 | 3.42416 | 1.44091 | 4.30344 | 0.81845 | 5.39564 | 0.83153 | 2.16769 | 0.17393 | 1.55444 |
| **Dse** | 1.60044 | 0.00592 | 0.00345 | 0.78679 | 0.00508 | 0.00630 | 0.00617 | 0.00727 | 0.81039 | 0.01149 | 0.00723 | 0.02613 | 0.01779 | 0.00830 | 0.15535 | 0.00543 |
| **Dsel** | 0.01364 | 0.00779 | 0.01138 | 0.36650 | 0.00924 | 0.00653 | 0.00747 | 0.01308 | 0.90244 | 0.00781 | 0.00717 | 0.01659 | 0.01931 | 0.01928 | 0.03445 | 0.00727 |
| **Dsg1b** | 0.03180 | 0.00788 | 0.10649 | 0.37070 | 0.03343 | 0.01866 | 0.00712 | 0.01676 | 0.10991 | 0.06819 | 0.36488 | 0.03351 | 0.12212 | 0.33151 | 0.08677 | 0.01467 |
| **Dsg2** | 2.40950 | 3.32666 | 1.70287 | 0.00540 | 0.19879 | 5.63165 | 0.01194 | 0.00831 | 0.02524 | 0.00000 | 0.00475 | 0.02036 | 2.86666 | 0.17012 | 0.01773 | 2.82128 |
| **Dsn1** | 3.28027 | 9.14756 | 3.83750 | 11.89445 | 0.00821 | 0.47870 | 2.37758 | 6.58283 | 14.53472 | 10.13982 | 3.72055 | 0.74376 | 1.93174 | 9.86463 | 18.45838 | 0.00000 |
| **Dsp** | 0.42840 | 0.01992 | 4.49760 | 1.42037 | 0.04853 | 11.60056 | 1.00977 | 1.16305 | 0.00000 | 0.64195 | 1.15206 | 0.01563 | 4.63796 | 0.05030 | 0.00000 | 0.02165 |
| **Dst** | 2.09616 | 0.00324 | 0.00000 | 11.59794 | 0.01409 | 3.86856 | 4.68426 | 0.09889 | 2.18335 | 1.06252 | 0.97167 | 0.10404 | 0.89196 | 0.00000 | 3.70227 | 0.00803 |
| **Dstn** | 16.17043 | 16.20321 | 19.09147 | 8.21437 | 15.96411 | 16.62857 | 14.35256 | 12.58765 | 8.28284 | 14.78442 | 9.52808 | 15.78464 | 11.74270 | 14.22208 | 10.33597 | 16.02828 |
| **Dstyk** | 0.21151 | 0.00756 | 0.02661 | 3.21063 | 0.00731 | 3.12408 | 0.00681 | 0.00000 | 1.53563 | 0.95237 | 0.40106 | 0.01805 | 0.00645 | 0.01051 | 2.21987 | 0.00498 |
| **Dtd1** | 1.78837 | 1.30250 | 0.75960 | 0.64406 | 0.01403 | 3.51531 | 0.00422 | 2.07172 | 0.02376 | 1.81149 | 2.82222 | 0.18515 | 0.00000 | 0.66884 | 1.06215 | 1.72627 |
| **Dtd2** | 0.64732 | 0.12223 | 1.55973 | 0.20294 | 0.61508 | 0.59045 | 0.85687 | 0.83788 | 0.02595 | 0.00000 | 0.17794 | 0.03874 | 0.40749 | 0.00783 | 0.00000 | 3.16768 |
| **Dtl** | 4.45820 | 7.41921 | 5.98145 | 10.70995 | 2.44112 | 7.01040 | 9.61449 | 8.09304 | 18.42258 | 19.72061 | 15.27808 | 9.20297 | 4.53865 | 26.79511 | 18.72017 | 1.53935 |
| **Dtnb** | 3.32632 | 3.42783 | 2.43957 | 1.02734 | 0.00963 | 0.00808 | 1.92114 | 1.60776 | 2.99806 | 6.02568 | 4.18694 | 1.48776 | 1.65783 | 2.36321 | 1.77034 | 1.49470 |
| **Dtnbp1** | 1.96880 | 3.00887 | 5.19059 | 2.54845 | 0.11029 | 0.01136 | 4.59450 | 1.10609 | 1.44563 | 2.42449 | 3.88562 | 0.08452 | 0.47574 | 0.80599 | 2.81698 | 0.01554 |
| **Dtwd1** | 2.64598 | 0.48146 | 1.60419 | 0.00227 | 0.00000 | 0.00305 | 0.00474 | 0.73883 | 0.02317 | 0.81779 | 0.76732 | 0.00817 | 1.43881 | 0.01396 | 0.81457 | 0.64068 |
| **Dtwd2** | 0.43151 | 1.63030 | 1.01431 | 3.79016 | 0.00604 | 0.01076 | 0.47696 | 0.08849 | 3.26004 | 2.21430 | 0.30625 | 0.57866 | 0.07284 | 2.77102 | 0.75308 | 0.01198 |
| **Dtx2** | 1.41221 | 0.00247 | 0.17587 | 25.22986 | 4.71180 | 0.01908 | 1.07723 | 6.59347 | 13.51581 | 14.62224 | 2.72837 | 0.59672 | 4.92857 | 0.07795 | 0.00000 | 5.30493 |
| **Dtx3** | 1.54263 | 0.00628 | 0.00363 | 0.59060 | 0.00498 | 0.00679 | 0.00609 | 0.02614 | 0.01433 | 0.07007 | 0.02734 | 0.00693 | 0.01085 | 0.01016 | 0.00442 | 0.00561 |
| **Dtx3l** | 0.04974 | 0.00767 | 0.07946 | 0.37734 | 0.04227 | 0.00611 | 0.03737 | 0.01701 | 1.06534 | 0.01352 | 0.49427 | 0.05926 | 0.10900 | 0.19999 | 0.46009 | 0.02147 |
| **Dtx4** | 0.01319 | 0.00776 | 0.01047 | 0.91839 | 0.00927 | 0.00638 | 0.00752 | 0.01510 | 0.96583 | 0.38791 | 0.03702 | 0.04283 | 0.01141 | 0.00628 | 0.30645 | 0.00722 |
| **Dtymk** | 13.86918 | 2.78503 | 7.03280 | 2.45012 | 9.72894 | 8.88717 | 0.44652 | 15.62565 | 3.60998 | 17.16217 | 8.49478 | 1.42373 | 14.82093 | 3.77350 | 0.52929 | 11.18673 |
| **Dubr** | 0.85103 | 0.00793 | 0.00425 | 0.00476 | 0.01709 | 0.00694 | 0.00673 | 0.00816 | 0.50892 | 0.32140 | 0.00737 | 0.01597 | 0.01704 | 0.01202 | 0.00724 | 0.00643 |
| **Duox1** | 0.49547 | 0.00828 | 0.22290 | 0.01070 | 0.01540 | 0.00510 | 0.12170 | 0.00994 | 0.01238 | 0.00436 | 0.01234 | 0.00799 | 0.00604 | 0.01471 | 0.00613 | 2.64939 |
| **Duoxa2** | 0.06619 | 0.02772 | 0.00473 | 0.00416 | 0.00874 | 0.00737 | 0.85522 | 0.00943 | 0.24111 | 0.01063 | 0.01077 | 0.02625 | 0.01088 | 0.01280 | 0.00735 | 0.00727 |
| **Dus1l** | 9.49165 | 2.02951 | 8.23210 | 1.39959 | 4.25251 | 1.75841 | 7.95435 | 9.67743 | 3.89152 | 13.45341 | 9.49696 | 0.24262 | 7.32068 | 0.15686 | 2.41084 | 4.63880 |
| **Dus2** | 6.20979 | 2.93115 | 2.73879 | 1.70532 | 1.69992 | 5.03949 | 5.20204 | 4.56425 | 2.97125 | 4.54373 | 1.37353 | 3.15281 | 3.06231 | 0.00000 | 2.34957 | 1.67232 |
| **Dus3l** | 9.62869 | 0.18777 | 0.00000 | 3.97441 | 0.66299 | 5.02831 | 2.15461 | 10.88778 | 3.55885 | 12.48567 | 7.38433 | 0.53929 | 9.99234 | 0.01068 | 0.67118 | 3.46727 |
| **Dus4l** | 0.01158 | 0.00775 | 1.43731 | 0.34480 | 0.00743 | 0.00582 | 0.00741 | 0.00954 | 0.00517 | 0.02958 | 0.02425 | 0.00517 | 0.01508 | 0.58652 | 0.00000 | 0.00722 |

| | | | | | | | | | | | | | | | | |
|---|---|---|---|---|---|---|---|---|---|---|---|---|---|---|---|---|
| **Dusp1** | 0.37177 | 1.68442 | 0.00000 | 3.67081 | 0.01511 | 0.07639 | 0.00607 | 0.30792 | 8.63645 | 0.19870 | 0.00000 | 0.00927 | 0.81597 | 0.00000 | 2.08305 | 0.00472 |
| **Dusp10** | 0.00000 | 0.00091 | 0.54783 | 1.89835 | 8.80258 | 0.01118 | 1.53056 | 1.28849 | 1.56585 | 4.59553 | 1.16449 | 2.47798 | 1.68920 | 1.61809 | 6.65559 | 0.93017 |
| **Dusp11** | 7.41378 | 1.32031 | 6.97277 | 2.21361 | 4.40798 | 1.25743 | 6.08312 | 12.51431 | 4.07708 | 5.11713 | 5.73086 | 5.49046 | 7.18408 | 2.64578 | 2.38048 | 8.47960 |
| **Dusp12** | 8.70782 | 0.02831 | 2.01527 | 3.28855 | 0.04581 | 0.02399 | 1.33626 | 8.40988 | 3.54049 | 18.61352 | 9.19956 | 1.72731 | 7.70877 | 10.34609 | 1.96389 | 7.52316 |
| **Dusp14** | 0.00000 | 2.48005 | 8.04441 | 19.20721 | 0.17458 | 15.53303 | 11.59362 | 18.13127 | 27.44581 | 7.32270 | 9.46211 | 1.26502 | 14.65764 | 3.41753 | 7.49300 | 1.72668 |
| **Dusp16** | 5.13087 | 1.92944 | 0.00000 | 8.12852 | 1.22209 | 0.86680 | 0.00000 | 2.43871 | 0.95804 | 2.14165 | 2.14054 | 0.19385 | 1.09759 | 0.90660 | 8.90916 | 0.06207 |
| **Dusp18** | 0.02422 | 0.00791 | 0.53495 | 2.05612 | 0.00833 | 0.26623 | 0.00743 | 0.05877 | 0.29063 | 1.63758 | 0.00659 | 0.00000 | 0.09339 | 0.07473 | 1.47590 | 0.00693 |
| **Dusp19** | 1.17942 | 0.19552 | 3.11362 | 0.00993 | 0.05221 | 0.01061 | 0.21989 | 0.30965 | 0.00904 | 1.51909 | 1.95536 | 0.19836 | 0.00000 | 0.98374 | 0.34205 | 0.59739 |
| **Dusp2** | 0.00872 | 0.00827 | 0.00301 | 1.01811 | 0.00930 | 0.00717 | 0.00773 | 0.28276 | 0.28203 | 0.80620 | 0.49110 | 0.11949 | 0.00426 | 0.00335 | 0.80910 | 0.00722 |
| **Dusp22** | 0.09571 | 0.02836 | 0.15350 | 1.78726 | 0.02860 | 0.04524 | 0.05742 | 0.01144 | 2.55571 | 1.38450 | 0.38316 | 0.11834 | 0.17484 | 0.17207 | 2.96664 | 0.03751 |
| **Dusp23** | 0.01297 | 0.00796 | 0.01227 | 0.10179 | 0.00889 | 0.00644 | 0.00730 | 0.01008 | 0.01808 | 1.47160 | 0.01097 | 0.00406 | 0.01042 | 0.01125 | 0.00548 | 0.00716 |
| **Dusp26** | 0.00000 | 0.00662 | 0.00169 | 0.21966 | 0.02020 | 2.37574 | 2.17687 | 2.03172 | 0.00695 | 6.35163 | 0.90186 | 0.25234 | 0.00000 | 0.02514 | 1.10275 | 0.00629 |
| **Dusp28** | 0.00974 | 0.00785 | 0.11456 | 0.09337 | 0.00910 | 0.00901 | 0.00747 | 0.01244 | 1.14879 | 0.42255 | 0.00816 | 0.01280 | 0.14133 | 0.00574 | 0.08709 | 0.25546 |
| **Dusp3** | 0.05623 | 0.00829 | 0.00000 | 0.10772 | 0.03232 | 4.10816 | 0.01814 | 1.72899 | 0.08978 | 1.46836 | 0.98539 | 0.08738 | 0.80413 | 0.17183 | 0.17413 | 0.01288 |
| **Dusp4** | 4.29342 | 2.11627 | 0.79722 | 0.34042 | 0.09067 | 0.01341 | 0.00000 | 0.77511 | 1.22549 | 0.00000 | 0.00000 | 0.18554 | 6.07841 | 0.00675 | 0.11804 | 12.28998 |
| **Dusp5** | 0.52560 | 0.00697 | 0.11000 | 0.17174 | 0.00896 | 0.00528 | 0.88733 | 0.15704 | 0.05318 | 0.02204 | 0.01072 | 0.00692 | 0.00580 | 0.00791 | 0.04503 | 0.33354 |
| **Dusp6** | 1.07251 | 0.02708 | 0.00000 | 1.63421 | 0.00556 | 0.05898 | 0.00631 | 0.00000 | 0.17098 | 0.00000 | 0.27641 | 0.01233 | 0.00728 | 0.82317 | 0.00000 | 0.54030 |
| **Dusp7** | 0.51262 | 0.00672 | 0.42229 | 5.89349 | 0.35694 | 0.00508 | 0.00588 | 0.00301 | 1.45323 | 0.68964 | 0.05678 | 0.01552 | 1.40280 | 0.01864 | 2.23994 | 0.30179 |
| **Dusp8** | 0.01330 | 0.00798 | 0.01173 | 0.00978 | 0.00934 | 0.00674 | 0.00746 | 0.02452 | 0.02350 | 0.02076 | 0.01264 | 0.01877 | 0.01218 | 0.01207 | 0.01055 | 0.00742 |
| **Dusp9** | 2.19866 | 0.05214 | 2.22306 | 0.97077 | 0.53341 | 0.00369 | 0.70156 | 0.13490 | 1.10211 | 0.05588 | 1.44687 | 0.00638 | 0.82613 | 0.00431 | 0.11660 | 0.13357 |
| **Dut** | 15.87532 | 5.26299 | 5.25640 | 18.14672 | 12.30521 | 2.50513 | 1.31549 | 0.70160 | 12.85372 | 12.85606 | 15.05803 | 0.46656 | 17.57077 | 1.23330 | 1.76286 | 17.57280 |
| **Dvl1** | 3.66581 | 0.12491 | 0.80474 | 0.94566 | 0.04185 | 0.22370 | 0.00000 | 5.93991 | 2.46086 | 3.39167 | 4.65901 | 0.06993 | 2.79803 | 0.00000 | 0.78774 | 2.20878 |
| **Dvl2** | 0.67193 | 0.41264 | 0.87034 | 1.76424 | 0.78056 | 0.11675 | 0.00510 | 0.74077 | 1.07944 | 2.14780 | 3.08288 | 0.02190 | 0.89491 | 0.25543 | 0.41863 | 3.01518 |
| **Dvl3** | 0.00274 | 0.00730 | 0.08566 | 0.00561 | 0.74299 | 0.17621 | 0.13116 | 0.94802 | 0.11970 | 0.52147 | 0.06565 | 0.01251 | 0.14470 | 0.21160 | 0.15607 | 0.40362 |
| **Dxo** | 6.34244 | 0.01025 | 0.00000 | 6.10626 | 0.09581 | 3.81229 | 0.01792 | 2.75994 | 9.55346 | 4.12847 | 3.74657 | 0.52197 | 4.09420 | 0.00000 | 0.94621 | 7.97146 |
| **Dydc1** | 0.01053 | 0.00790 | 0.00890 | 0.54579 | 0.00942 | 0.00660 | 0.00749 | 0.12991 | 0.00535 | 0.11261 | 0.05355 | 0.00658 | 0.12153 | 0.00986 | 0.33658 | 0.00702 |
| **Dydc2** | 0.00772 | 0.00768 | 0.00000 | 1.41587 | 0.00919 | 0.00815 | 0.00679 | 0.69281 | 0.00000 | 0.57962 | 1.09245 | 0.05763 | 0.48854 | 0.00147 | 0.67407 | 0.00658 |
| **Dym** | 6.29859 | 1.83697 | 4.62164 | 6.08942 | 3.54549 | 0.03398 | 3.21703 | 3.78658 | 3.64249 | 8.15406 | 2.69595 | 0.19698 | 4.01883 | 0.30985 | 2.97355 | 5.78055 |
| **Dynap** | 0.01024 | 0.00742 | 0.00000 | 0.00000 | 0.00916 | 0.00620 | 0.00694 | 0.00318 | 0.00709 | 0.20838 | 0.28786 | 1.81848 | 0.00982 | 2.31339 | 1.44997 | 0.00753 |
| **Dync1h1** | 9.44449 | 12.80693 | 9.84085 | 6.08608 | 9.87850 | 4.58656 | 7.94237 | 10.53222 | 5.85411 | 5.61299 | 2.23016 | 0.45368 | 10.50051 | 0.00000 | 2.38545 | 5.79939 |
| **Dync1i1** | 2.81138 | 0.00573 | 0.00574 | 0.23828 | 0.01275 | 0.00458 | 0.00504 | 0.00000 | 0.00521 | 1.49519 | 0.00981 | 0.00239 | 0.02256 | 0.01070 | 0.00529 | 0.00435 |
| **Dync1i2** | 4.25438 | 6.38897 | 5.96976 | 4.14176 | 8.90845 | 1.40258 | 5.16890 | 6.77637 | 2.75804 | 3.34393 | 2.68013 | 6.82019 | 2.28012 | 5.76864 | 5.08411 | 1.34449 |
| **Dync1li1** | 3.34492 | 3.84280 | 2.73100 | 0.22522 | 7.40522 | 2.06314 | 0.89456 | 5.59190 | 0.15586 | 6.67301 | 0.06962 | 0.28002 | 1.97949 | 3.00471 | 1.03162 | 1.64429 |
| **Dync1li2** | 2.27521 | 3.36959 | 1.42136 | 0.63748 | 6.25804 | 2.94601 | 5.55855 | 4.00915 | 0.34212 | 6.53805 | 3.80633 | 0.95714 | 0.00000 | 4.72879 | 6.23869 | 4.97153 |
| **Dync2h1** | 2.24038 | 3.47343 | 0.11076 | 0.40066 | 3.00414 | 0.59567 | 1.56717 | 1.43974 | 0.05296 | 0.09089 | 0.20423 | 0.03522 | 0.65015 | 0.01520 | 0.26839 | 0.48443 |
| **Dync2li1** | 0.31597 | 0.00822 | 0.00629 | 0.00769 | 0.01775 | 0.00696 | 0.00709 | 0.00990 | 0.01191 | 0.80628 | 0.01223 | 0.00535 | 0.01145 | 0.01208 | 0.00820 | 0.00753 |
| **Dynll1** | 29.20151 | 5.15229 | 20.67266 | 24.52400 | 22.36891 | 8.43297 | 12.34432 | 15.86712 | 38.02388 | 29.90517 | 23.73964 | 6.59645 | 15.24057 | 4.59049 | 2.32403 | 17.74399 |
| **Dynll2** | 5.17901 | 0.01027 | 0.57501 | 13.56801 | 2.94304 | 0.08098 | 1.81019 | 0.19645 | 4.63508 | 7.51210 | 4.85776 | 0.32191 | 1.16012 | 1.47649 | 3.40921 | 0.64207 |
| **Dynlrb1** | 9.31594 | 3.99683 | 9.49586 | 7.02339 | 9.51371 | 6.87633 | 3.61229 | 3.63963 | 9.20793 | 10.17702 | 13.02809 | 1.61654 | 5.91081 | 2.50472 | 2.42560 | 4.32694 |
| **Dynlrb2** | 0.14732 | 0.00784 | 0.00796 | 0.01022 | 0.00762 | 0.00688 | 0.00736 | 0.01418 | 0.01239 | 0.09450 | 0.01245 | 0.00754 | 0.01219 | 0.01144 | 0.01045 | 0.00714 |
| **Dynlt1a** | 6.68203 | 2.14663 | 1.39350 | 2.26722 | 4.96164 | 3.59122 | 1.76737 | 1.47486 | 3.47371 | 0.57543 | 3.71825 | 0.35593 | 1.77749 | 2.48465 | 1.74685 | 4.49170 |
| **Dynlt1b** | 6.96939 | 2.36917 | 1.40076 | 0.14130 | 5.28220 | 3.38945 | 1.65312 | 1.47925 | 0.42763 | 0.59185 | 4.29586 | 0.16136 | 1.72219 | 1.64460 | 1.25542 | 4.13530 |
| **Dynlt1f** | 0.00935 | 0.02915 | 0.01996 | 0.06360 | 0.00870 | 0.31495 | 0.36745 | 0.00983 | 0.01720 | 0.00285 | 0.10214 | 0.00735 | 0.10027 | 0.01103 | 0.00994 | 0.25352 |
| **Dynlt3** | 5.62171 | 2.12116 | 4.30441 | 7.42386 | 3.77714 | 6.01100 | 1.48192 | 1.21482 | 1.20313 | 1.77490 | 0.07655 | 6.63682 | 1.38667 | 0.99797 | 4.57994 | 3.19233 |
| **Dyrk1a** | 4.51954 | 4.75338 | 6.36087 | 15.70754 | 2.46065 | 0.83252 | 7.01430 | 6.37883 | 16.19839 | 18.00321 | 6.91437 | 12.79607 | 3.50275 | 4.00661 | 7.55876 | 4.15795 |
| **Dyrk1b** | 0.01267 | 0.00804 | 0.01990 | 0.10364 | 0.00903 | 0.00642 | 0.00725 | 0.01121 | 0.16024 | 0.29832 | 0.01066 | 0.00801 | 0.38926 | 0.01132 | 0.00843 | 0.00717 |
| **Dyrk3** | 7.16474 | 2.96058 | 6.35521 | 8.48109 | 0.18979 | 0.75193 | 6.83574 | 3.06578 | 7.30452 | 1.92674 | 2.46380 | 2.49703 | 3.55296 | 0.00000 | 7.10504 | 0.26894 |
| **Dyrk4** | 0.77392 | 0.00714 | 0.00269 | 0.00324 | 0.00637 | 0.00691 | 0.00672 | 0.01037 | 0.75126 | 0.00286 | 0.00608 | 0.02772 | 0.01191 | 0.00709 | 0.51574 | 0.00635 |
| **Dysf** | 1.22620 | 0.00425 | 0.69318 | 0.84985 | 2.88314 | 0.00625 | 0.00356 | 0.00000 | 1.49220 | 9.51966 | 2.93318 | 2.42925 | 3.45288 | 4.88316 | 5.85675 | 0.05426 |
| **Dyx1c1** | 0.00855 | 0.00800 | 0.00332 | 0.18648 | 0.01000 | 0.00681 | 0.00691 | 0.06131 | 0.00506 | 0.11646 | 0.25404 | 0.00000 | 0.08818 | 1.73939 | 0.09752 | 0.00732 |
| **Dzank1** | 0.00000 | 0.02921 | 0.00000 | 0.41070 | 0.06607 | 0.01529 | 0.01742 | 0.48168 | 1.28842 | 0.34090 | 1.13571 | 0.03063 | 0.03516 | 7.62188 | 1.14482 | 0.00685 |
| **Dzip1** | 0.01718 | 0.00806 | 0.01035 | 0.01408 | 0.00899 | 0.00683 | 0.00747 | 0.01411 | 0.01272 | 0.71909 | 0.01215 | 0.03311 | 0.01525 | 0.20292 | 0.00000 | 0.00775 |
| **Dzip3** | 1.14918 | 2.47968 | 0.00000 | 2.44335 | 0.01405 | 0.46724 | 0.00505 | 0.00391 | 0.31934 | 0.48482 | 0.00701 | 0.01051 | 0.48431 | 0.00242 | 1.41245 | 0.00468 |
| **E2f1** | 4.26268 | 0.38317 | 2.62624 | 13.83246 | 4.44076 | 0.02222 | 1.07245 | 3.26254 | 9.27554 | 13.43814 | 2.46894 | 0.67400 | 3.27710 | 0.31922 | 9.67179 | 4.34994 |
| **E2f2** | 0.07276 | 0.00615 | 1.82723 | 1.22503 | 2.75567 | 0.01443 | 0.00633 | 0.00000 | 3.70723 | 0.46437 | 0.59633 | 0.04667 | 0.57057 | 0.02384 | 1.17558 | 0.16651 |
| **E2f3** | 0.02185 | 0.00600 | 0.00044 | 0.00000 | 0.00875 | 2.10967 | 2.81513 | 0.01032 | 0.53331 | 0.00332 | 0.00626 | 0.00443 | 0.00313 | 0.00663 | 0.26242 | 0.06230 |
| **E2f4** | 4.98623 | 5.31960 | 5.40601 | 17.08484 | 4.81603 | 2.19962 | 3.59841 | 9.02447 | 13.06081 | 8.41216 | 6.74385 | 2.01017 | 8.98989 | 1.17143 | 7.49363 | 4.94801 |
| **E2f5** | 0.21774 | 1.93852 | 0.79376 | 21.11555 | 0.00615 | 0.00191 | 0.00467 | 1.05209 | 9.34912 | 1.57473 | 0.41005 | 4.76529 | 0.00674 | 1.94107 | 5.56120 | 0.00142 |
| **E2f6** | 0.78369 | 6.78118 | 2.67789 | 0.57502 | 5.05543 | 0.89944 | 3.86451 | 1.11010 | 0.00000 | 0.89832 | 0.31518 | 7.71053 | 1.46302 | 0.00242 | 2.29385 | 0.00000 |
| **E2f7** | 0.01222 | 0.00703 | 0.00301 | 1.14481 | 0.62921 | 0.00820 | 0.00645 | 0.74227 | 0.40498 | 0.29400 | 0.46807 | 0.02666 | 1.05800 | 0.00289 | 0.03947 | 0.00709 |
| **E2f8** | 0.00333 | 0.00725 | 1.09496 | 1.32516 | 0.00948 | 2.04979 | 0.00622 | 0.71935 | 0.44694 | 0.84427 | 0.92835 | 4.99828 | 1.72265 | 0.00244 | 0.39369 | 0.00495 |
| **E4f1** | 1.59406 | 0.02010 | 3.02279 | 7.76242 | 5.71557 | 5.16668 | 3.82131 | 5.94860 | 10.11835 | 4.76879 | 4.99046 | 0.37864 | 5.08019 | 0.00000 | 0.54714 | 2.29113 |

| | | | | | | | | | | | | | | | | |
|---|---|---|---|---|---|---|---|---|---|---|---|---|---|---|---|---|
| **Eaf1** | 0.64183 | 1.40636 | 4.91976 | 8.79209 | 1.54895 | 1.06135 | 5.11745 | 5.46825 | 6.28157 | 3.41327 | 3.87468 | 6.98526 | 3.51298 | 10.06795 | 6.11796 | 2.53502 |
| **Eaf2** | 0.14303 | 0.01846 | 0.48593 | 3.38011 | 0.45310 | 0.43558 | 0.01684 | 0.44201 | 5.63065 | 1.82623 | 0.37896 | 0.59365 | 0.59404 | 0.94316 | 0.55678 | 0.03513 |
| **Eapp** | 10.94961 | 8.33611 | 9.97313 | 5.02933 | 4.41053 | 5.87697 | 5.64360 | 8.26280 | 6.90734 | 7.21705 | 12.07579 | 3.09046 | 4.92722 | 6.85583 | 4.76219 | 3.47036 |
| **Ears2** | 3.44832 | 6.88887 | 4.52176 | 3.64204 | 0.28084 | 0.01424 | 0.77132 | 3.36207 | 7.26653 | 3.23365 | 4.15474 | 0.45067 | 3.82780 | 0.74474 | 3.65166 | 3.99185 |
| **Ebag9** | 2.89614 | 4.03183 | 0.24612 | 0.30481 | 2.67371 | 1.62400 | 1.38838 | 0.65859 | 0.00671 | 1.12727 | 1.03222 | 0.00000 | 0.07468 | 0.03316 | 2.02165 | 2.11077 |
| **Ebf3** | 0.01338 | 0.00778 | 0.01121 | 0.69041 | 0.00936 | 0.00644 | 0.00736 | 0.01723 | 1.17942 | 0.00285 | 0.00002 | 0.00509 | 0.01282 | 0.02759 | 0.30808 | 0.00725 |
| **Ebf4** | 0.01313 | 0.00810 | 0.01213 | 0.01057 | 0.00912 | 0.00677 | 0.00759 | 0.01437 | 0.01226 | 0.03081 | 0.01259 | 0.00868 | 0.01249 | 0.01199 | 0.01153 | 0.07293 |
| **Ebna1bp2** | 20.52178 | 11.86518 | 16.36241 | 17.06951 | 14.02536 | 10.35123 | 13.76243 | 23.63263 | 28.25185 | 20.53975 | 17.06300 | 9.86070 | 15.69508 | 11.43374 | 11.72978 | 7.45844 |
| **Ebp** | 10.90007 | 5.78974 | 25.72837 | 3.31599 | 7.39235 | 6.16357 | 8.41645 | 10.37605 | 6.49687 | 5.23201 | 4.92197 | 2.06893 | 3.25986 | 2.73445 | 10.22012 | 0.07299 |
| **Ebpl** | 0.90833 | 0.00668 | 0.08916 | 8.25934 | 0.06262 | 0.00602 | 0.00565 | 0.12821 | 0.77139 | 1.04958 | 1.12235 | 0.13484 | 0.33944 | 0.18449 | 0.23294 | 2.06355 |
| **Ecd** | 6.60865 | 8.84692 | 4.38894 | 3.52906 | 5.08618 | 7.39730 | 4.43095 | 12.47527 | 4.58260 | 11.97048 | 9.65367 | 2.34964 | 3.08230 | 2.69034 | 9.09299 | 4.89682 |
| **Ece1** | 0.02182 | 0.00782 | 0.00740 | 1.03768 | 0.00843 | 0.00699 | 0.00703 | 0.00226 | 0.32214 | 0.85917 | 0.55327 | 0.03290 | 0.01498 | 0.02148 | 0.04614 | 0.00723 |
| **Ece2** | 1.04909 | 2.91438 | 1.76398 | 3.89689 | 0.02125 | 1.11726 | 4.83827 | 3.12741 | 8.08097 | 8.54041 | 3.63872 | 4.76011 | 1.37372 | 0.57465 | 4.07598 | 0.31282 |
| **Ecel1** | 0.00735 | 0.00797 | 0.01255 | 0.07864 | 0.01881 | 0.00660 | 0.00673 | 0.01050 | 0.01137 | 0.00518 | 0.01397 | 0.00725 | 1.46374 | 0.01274 | 0.00893 | 0.00663 |
| **Ech1** | 11.06737 | 1.99209 | 11.53504 | 0.45947 | 3.51171 | 6.15302 | 5.56959 | 4.51299 | 0.00000 | 3.90377 | 0.70433 | 0.00956 | 6.18822 | 0.00590 | 0.40124 | 7.18470 |
| **Echdc1** | 0.03518 | 0.00827 | 0.79303 | 0.01050 | 0.00693 | 0.00727 | 0.01916 | 0.03614 | 0.01228 | 0.00884 | 0.00997 | 0.00726 | 0.00927 | 0.25616 | 0.00000 | 0.00712 |
| **Echdc2** | 30.42771 | 4.35068 | 19.62947 | 5.73533 | 21.81564 | 12.22523 | 11.62422 | 28.08668 | 13.80028 | 26.92008 | 24.46780 | 6.01703 | 25.91479 | 2.92835 | 0.27409 | 17.98749 |
| **Echdc3** | 0.01316 | 0.00833 | 0.01313 | 1.37430 | 0.00985 | 0.00675 | 0.00769 | 0.01485 | 1.31031 | 0.03168 | 0.00000 | 0.00557 | 0.04337 | 0.02518 | 0.12924 | 0.01461 |
| **Echs1** | 16.05249 | 5.86370 | 12.82102 | 3.81318 | 9.11816 | 0.66661 | 6.73467 | 3.67111 | 1.48456 | 1.72517 | 1.20929 | 0.04531 | 10.30329 | 1.50341 | 1.23247 | 9.65169 |
| **Eci1** | 2.63454 | 0.74884 | 3.63834 | 6.55034 | 0.03507 | 1.94469 | 0.05989 | 1.82659 | 15.30862 | 8.85774 | 11.47958 | 4.03514 | 2.23009 | 2.34565 | 3.35281 | 5.20561 |
| **Eci2** | 8.48990 | 5.31195 | 2.20723 | 0.61390 | 7.90292 | 4.77966 | 6.88197 | 5.67200 | 3.70207 | 3.51236 | 7.55308 | 0.41830 | 1.43611 | 0.00000 | 1.34641 | 7.13802 |
| **Eci3** | 0.56155 | 0.10096 | 0.06911 | 0.00665 | 0.20811 | 0.09155 | 0.13319 | 0.12172 | 0.06737 | 0.04742 | 0.01101 | 0.00713 | 0.03122 | 0.00992 | 0.03174 | 0.19393 |
| **Ecm1** | 0.00640 | 0.00766 | 0.04156 | 0.83385 | 0.00814 | 0.03835 | 0.00718 | 1.39221 | 0.12280 | 2.78716 | 0.15395 | 0.11834 | 0.70179 | 0.00941 | 0.07882 | 0.43760 |
| **Ecm2** | 0.06974 | 0.01775 | 0.13767 | 0.47633 | 0.01028 | 0.04038 | 0.04681 | 0.63745 | 0.77583 | 0.28893 | 0.62090 | 0.84556 | 0.13326 | 0.35574 | 0.35735 | 0.04259 |
| **Ecscr** | 0.01349 | 0.00816 | 0.01167 | 0.00427 | 0.00929 | 0.00658 | 0.00756 | 0.01378 | 1.76037 | 0.00863 | 0.00000 | 0.00535 | 0.01318 | 0.01378 | 0.00891 | 0.00731 |
| **Ecsit** | 6.05551 | 4.97166 | 9.05256 | 4.92166 | 3.57436 | 3.03653 | 5.22783 | 7.11952 | 7.08917 | 12.12295 | 7.35653 | 2.10387 | 6.29360 | 1.15913 | 8.00127 | 1.96827 |
| **Ect2** | 0.50950 | 7.99698 | 1.44646 | 4.06803 | 6.90562 | 0.00437 | 3.41445 | 5.89742 | 15.25489 | 10.72548 | 6.20858 | 4.88769 | 0.79950 | 25.38127 | 14.47273 | 2.14426 |
| **Eda** | 2.02103 | 0.00382 | 0.07585 | 3.93704 | 0.00534 | 4.53627 | 0.06317 | 5.15662 | 12.31264 | 9.31472 | 0.94722 | 4.41362 | 4.62648 | 13.23259 | 6.22891 | 0.00682 |
| **Eda2r** | 0.54400 | 0.00797 | 0.48411 | 0.01056 | 0.01654 | 0.00863 | 0.00678 | 0.00841 | 0.01233 | 0.01118 | 0.01111 | 0.07425 | 0.00941 | 0.07389 | 0.00000 | 0.00690 |
| **Edar** | 0.01373 | 0.00821 | 0.01217 | 0.00524 | 0.00933 | 0.00661 | 0.00751 | 0.01348 | 0.17660 | 0.01073 | 0.01148 | 0.00815 | 0.01272 | 0.00857 | 0.08594 | 0.00739 |
| **Edaradd** | 1.26710 | 0.00619 | 0.00000 | 0.00145 | 0.01627 | 0.00512 | 1.44053 | 0.00000 | 0.03660 | 0.00000 | 0.34479 | 0.00000 | 0.02669 | 3.33393 | 0.48101 | 0.00587 |
| **Edc3** | 5.90465 | 0.00000 | 0.46360 | 1.23033 | 3.08046 | 4.59211 | 0.98685 | 2.15935 | 0.43231 | 3.85150 | 0.95738 | 0.28058 | 2.57031 | 1.08693 | 3.86393 | 0.01986 |
| **Edc4** | 10.02371 | 3.98984 | 3.23980 | 4.39520 | 4.47539 | 5.61554 | 2.89625 | 10.94504 | 2.88870 | 9.07868 | 5.70044 | 1.07850 | 2.63525 | 0.11013 | 2.87685 | 5.77432 |
| **Edem1** | 3.00932 | 0.61893 | 3.15506 | 4.44139 | 2.04481 | 0.76828 | 0.47216 | 1.51005 | 4.33896 | 1.90374 | 0.98903 | 0.02089 | 3.00391 | 0.02231 | 4.10453 | 4.81375 |
| **Edem2** | 5.44039 | 0.00000 | 0.41660 | 1.04039 | 3.78394 | 0.83592 | 6.12849 | 3.13994 | 0.30112 | 3.48131 | 0.81874 | 0.17359 | 7.32154 | 0.00696 | 0.00000 | 4.50033 |
| **Edem3** | 0.36614 | 2.25283 | 0.00000 | 0.83918 | 0.00699 | 1.75654 | 1.26739 | 1.71564 | 2.42012 | 0.00000 | 0.91534 | 5.72942 | 1.88842 | 7.34738 | 3.87215 | 0.00350 |
| **Edf1** | 12.60414 | 0.51881 | 4.79579 | 9.26380 | 7.44517 | 4.34431 | 2.42688 | 9.66389 | 5.29769 | 23.22136 | 15.15750 | 1.18013 | 20.13444 | 0.02920 | 0.00000 | 6.16579 |
| **Edn2** | 1.01038 | 0.00603 | 0.00223 | 0.01126 | 0.00576 | 0.00781 | 0.00634 | 0.93183 | 0.01136 | 0.00451 | 0.00725 | 0.00506 | 0.00540 | 0.01640 | 0.00000 | 0.00626 |
| **Edn3** | 0.01335 | 0.00776 | 0.01156 | 0.00297 | 0.00920 | 0.00645 | 0.00734 | 0.01074 | 0.40885 | 0.49535 | 0.00774 | 0.00484 | 0.01169 | 0.00647 | 0.62019 | 0.00728 |
| **Ednra** | 0.32783 | 0.00847 | 0.00434 | 0.23935 | 0.01639 | 0.00670 | 0.00684 | 0.00981 | 0.00528 | 0.00532 | 0.01177 | 0.00753 | 0.56752 | 0.01163 | 0.01158 | 2.25261 |
| **Ednrb** | 0.01687 | 0.00788 | 0.03022 | 0.39911 | 0.00899 | 0.00799 | 0.02794 | 0.01193 | 0.11721 | 0.03100 | 0.02665 | 0.00721 | 0.03255 | 0.07794 | 0.02698 | 0.00740 |
| **Edrf1** | 1.76176 | 3.85102 | 0.52007 | 2.32514 | 0.00401 | 0.00386 | 4.03487 | 0.91023 | 0.30079 | 1.09282 | 1.27225 | 2.06915 | 1.66045 | 2.04949 | 3.80588 | 0.33583 |
| **Eea1** | 3.65636 | 0.00000 | 1.30509 | 8.02829 | 0.00000 | 5.31605 | 0.20704 | 0.47171 | 3.10115 | 0.00000 | 0.66470 | 6.33138 | 1.53612 | 6.42352 | 6.97580 | 1.57206 |
| **Eed** | 16.31256 | 14.62607 | 15.44079 | 6.95726 | 12.65641 | 7.20952 | 11.15713 | 14.32302 | 5.82267 | 16.92989 | 8.26146 | 7.57310 | 12.71284 | 19.05807 | 12.09880 | 9.69664 |
| **Eef1a1** | 69.78303 | 43.64737 | 65.23476 | 20.36106 | 49.82072 | 32.30229 | 37.57081 | 67.94116 | 28.28754 | 62.50518 | 51.54959 | 32.15995 | 64.71997 | 58.40761 | 40.56137 | 40.06266 |
| **Eef1akmt1** | 11.79667 | 5.10159 | 9.40855 | 4.90914 | 0.31845 | 2.59886 | 4.72727 | 9.96479 | 7.05133 | 15.64947 | 13.78636 | 2.56030 | 7.28153 | 6.76825 | 9.53816 | 0.99604 |
| **Eef1b2** | 49.56292 | 27.29162 | 40.00142 | 13.70375 | 30.95040 | 17.71359 | 24.92137 | 44.77663 | 20.87978 | 46.01499 | 44.54728 | 20.99167 | 39.84460 | 13.01457 | 9.80809 | 24.34535 |
| **Eef1d** | 20.72687 | 17.65129 | 23.78617 | 10.37205 | 14.68614 | 7.96275 | 13.79385 | 15.27823 | 10.50457 | 15.97440 | 24.73174 | 11.61532 | 20.19897 | 5.13776 | 10.43813 | 15.80399 |
| **Eef1e1** | 17.20347 | 10.31405 | 18.14682 | 3.25692 | 13.52788 | 7.16889 | 7.00325 | 17.84262 | 4.71072 | 23.65616 | 14.71817 | 12.67334 | 16.70101 | 22.44259 | 12.07593 | 6.90375 |
| **Eef1g** | 49.87631 | 29.12437 | 43.58570 | 3.80897 | 31.53903 | 24.34191 | 26.61684 | 50.63878 | 8.53335 | 42.33679 | 35.31920 | 14.30629 | 46.33719 | 24.77649 | 23.18612 | 30.21435 |
| **Eef2** | 38.74268 | 23.87693 | 34.94108 | 14.98821 | 24.73652 | 17.69242 | 23.18076 | 51.71832 | 21.47609 | 40.49158 | 35.63027 | 9.80102 | 39.40588 | 3.24284 | 13.27953 | 24.19761 |
| **Eef2k** | 1.97239 | 0.33391 | 0.02861 | 0.88718 | 1.72397 | 0.00644 | 0.01500 | 0.75532 | 0.48773 | 0.08226 | 0.77465 | 0.01570 | 0.00000 | 0.04855 | 0.62442 | 3.14779 |
| **Eef2kmt** | 5.32680 | 4.79893 | 2.82173 | 0.07593 | 1.73998 | 0.03045 | 2.97392 | 3.90364 | 0.20715 | 1.11158 | 3.65183 | 0.01474 | 3.10391 | 0.21961 | 0.41033 | 0.26453 |
| **Eefsec** | 8.74701 | 0.14697 | 1.13478 | 2.58327 | 3.72502 | 0.00571 | 0.44908 | 1.69467 | 2.83163 | 2.74059 | 0.63027 | 0.12951 | 2.31252 | 0.00201 | 0.00000 | 1.97885 |
| **Eepd1** | 0.01368 | 0.00798 | 0.04801 | 1.02915 | 0.00842 | 0.00851 | 0.00717 | 0.22638 | 1.22666 | 0.16901 | 1.40846 | 0.00000 | 0.77038 | 0.00944 | 0.00000 | 0.04820 |
| **Efcab10** | 1.13880 | 0.01673 | 0.00175 | 0.00971 | 0.95359 | 0.00700 | 0.00642 | 0.00774 | 0.01179 | 0.05108 | 0.01334 | 0.00638 | 0.10393 | 0.01662 | 0.00935 | 1.41771 |
| **Efcab11** | 0.08334 | 0.00723 | 0.52480 | 0.30688 | 0.00741 | 0.00555 | 0.00707 | 0.00977 | 0.00642 | 0.03147 | 0.01117 | 0.00547 | 0.00949 | 0.00696 | 0.65056 | 0.00723 |
| **Efcab12** | 0.25305 | 0.00758 | 0.03766 | 2.17338 | 0.00747 | 0.00530 | 0.01696 | 0.00983 | 1.86954 | 0.00000 | 0.00000 | 0.00000 | 0.01194 | 0.20806 | 0.90343 | 0.00638 |
| **Efcab14** | 3.02406 | 0.02428 | 3.02258 | 5.43266 | 4.36248 | 0.04001 | 0.54334 | 0.01087 | 2.72435 | 3.34897 | 0.07600 | 0.02941 | 1.31016 | 0.84215 | 2.78662 | 3.51464 |
| **Efcab2** | 0.49465 | 1.55820 | 0.02744 | 0.08094 | 0.00693 | 0.00566 | 0.02602 | 0.00836 | 0.07310 | 0.02737 | 0.01087 | 0.02848 | 0.69016 | 0.00740 | 0.00000 | 0.04155 |
| **Efcab3** | 0.01016 | 0.00801 | 0.00000 | 0.72471 | 0.00926 | 0.00639 | 0.00655 | 0.00000 | 0.00000 | 0.29737 | 1.42901 | 0.00950 | 0.00000 | 2.20365 | 1.57665 | 0.00669 |
| **Efcab5** | 0.00631 | 0.00839 | 0.23546 | 1.18413 | 0.00895 | 0.00576 | 0.00732 | 0.07870 | 1.40514 | 1.06305 | 0.00000 | 0.01528 | 1.13256 | 0.00760 | 0.34227 | 0.00643 |

| | | | | | | | | | | | | | | | | |
|---|---|---|---|---|---|---|---|---|---|---|---|---|---|---|---|---|
| **Efcab6** | 0.01369 | 0.00820 | 0.01182 | 1.67446 | 0.00939 | 0.00643 | 0.00740 | 0.01299 | 0.54550 | 0.03838 | 0.00948 | 0.02788 | 0.01292 | 0.00861 | 0.01208 | 0.00716 |
| **Efcab7** | 0.43836 | 0.00748 | 0.09279 | 0.49997 | 0.00703 | 0.00528 | 0.00702 | 0.01286 | 0.04748 | 0.04985 | 0.01253 | 0.00741 | 0.01086 | 0.01050 | 0.00347 | 0.00661 |
| **Efcc1** | 0.11951 | 0.00773 | 0.06210 | 2.26279 | 0.02652 | 0.02240 | 0.00605 | 0.00000 | 3.49106 | 0.00000 | 2.97282 | 0.00000 | 0.02178 | 0.75081 | 0.00000 | 0.01324 |
| **Efemp1** | 0.01945 | 0.00799 | 0.00628 | 0.00920 | 0.01046 | 1.97577 | 0.00721 | 0.01095 | 0.01270 | 0.05562 | 0.01012 | 0.00768 | 0.01125 | 0.01152 | 0.01020 | 0.00673 |
| **Efemp2** | 0.01134 | 0.00803 | 0.66959 | 1.32463 | 0.01712 | 0.00555 | 0.00746 | 0.01091 | 0.00000 | 0.71153 | 0.00993 | 0.09710 | 0.01516 | 0.01297 | 0.00000 | 0.00670 |
| **Efhb** | 0.31740 | 0.00728 | 0.00000 | 0.59028 | 0.00705 | 0.00547 | 0.00605 | 0.00000 | 1.23422 | 1.90722 | 2.46001 | 0.17308 | 0.00000 | 6.23121 | 2.78236 | 0.00625 |
| **Efhc2** | 0.12430 | 0.00732 | 0.36936 | 0.33190 | 0.00727 | 0.00551 | 0.00713 | 0.01191 | 0.00601 | 0.00658 | 0.01238 | 0.00645 | 0.01022 | 0.00560 | 0.33010 | 0.00690 |
| **Efhd1** | 1.78667 | 2.26705 | 0.97432 | 0.95733 | 0.07051 | 0.01023 | 0.02415 | 5.44415 | 1.14207 | 5.72557 | 2.31592 | 0.71828 | 1.39081 | 1.84223 | 1.50403 | 0.00324 |
| **Efhd2** | 1.35766 | 0.05814 | 3.89181 | 0.00000 | 7.48998 | 7.68803 | 3.68518 | 3.95705 | 0.01042 | 3.98290 | 2.00325 | 0.18408 | 9.27707 | 0.04982 | 0.00000 | 2.29037 |
| **Efl1** | 3.17530 | 0.87528 | 2.35212 | 0.06682 | 2.28857 | 1.89987 | 0.92012 | 1.07617 | 0.80964 | 0.27129 | 0.56326 | 0.10355 | 2.85123 | 0.40609 | 1.23351 | 1.88179 |
| **Efna1** | 0.18424 | 0.00634 | 3.85236 | 1.41155 | 0.00000 | 0.02256 | 0.00325 | 1.40556 | 1.17187 | 4.98466 | 5.05504 | 0.26288 | 2.77902 | 0.68393 | 1.94612 | 1.51685 |
| **Efna2** | 2.17428 | 0.00532 | 0.00000 | 0.05256 | 0.03440 | 1.86310 | 0.88021 | 2.47614 | 0.00744 | 2.02929 | 1.04752 | 0.13043 | 0.16067 | 0.01370 | 0.00280 | 0.04133 |
| **Efna3** | 0.36341 | 0.00700 | 0.00347 | 0.00889 | 0.05658 | 0.00720 | 0.16589 | 0.00765 | 0.01209 | 0.35289 | 0.01264 | 0.00569 | 0.55058 | 0.01111 | 0.01178 | 1.02746 |
| **Efna4** | 1.12665 | 0.00756 | 0.00000 | 0.10912 | 0.01410 | 0.00612 | 0.00567 | 0.00597 | 0.47114 | 0.00076 | 0.00945 | 0.01731 | 1.16080 | 0.00996 | 0.11121 | 4.02818 |
| **Efna5** | 0.01054 | 0.01809 | 0.02258 | 0.38812 | 0.02858 | 0.02348 | 0.02683 | 0.01108 | 0.03436 | 0.20991 | 0.10392 | 0.00000 | 0.01060 | 2.20289 | 0.00000 | 0.00728 |
| **Efnb1** | 0.00687 | 0.00862 | 0.00925 | 0.29340 | 0.01851 | 0.00698 | 0.00781 | 0.26310 | 0.00716 | 0.24519 | 0.19353 | 0.00803 | 0.83277 | 0.01123 | 0.00000 | 0.00690 |
| **Efnb2** | 1.50163 | 2.58965 | 0.25260 | 6.11004 | 1.53602 | 6.53994 | 0.00360 | 1.34079 | 5.87525 | 0.26749 | 1.45258 | 0.53543 | 0.39340 | 4.54731 | 4.56703 | 0.61280 |
| **Efnb3** | 3.15498 | 0.00401 | 0.00000 | 0.90031 | 0.01905 | 0.01355 | 0.00466 | 1.05844 | 0.13971 | 0.58803 | 0.00000 | 0.00152 | 0.00559 | 0.00771 | 0.12843 | 0.07274 |
| **Efr3a** | 4.51257 | 0.00000 | 0.65198 | 4.68881 | 0.08232 | 0.08019 | 2.98282 | 1.08446 | 0.45608 | 1.15696 | 0.03441 | 0.41156 | 1.19522 | 4.23361 | 3.98411 | 0.42363 |
| **Efr3b** | 0.05116 | 0.00804 | 0.10465 | 3.91614 | 0.01246 | 0.00578 | 0.05737 | 0.00000 | 2.62204 | 1.28940 | 0.76022 | 0.33181 | 0.06676 | 0.27802 | 0.73124 | 0.07219 |
| **Efs** | 0.01375 | 0.00813 | 0.01230 | 0.36292 | 0.00931 | 0.00662 | 0.00752 | 0.01393 | 0.00657 | 0.01541 | 0.01413 | 0.00822 | 0.01266 | 0.01118 | 0.00614 | 0.00735 |
| **Eftud2** | 15.27751 | 11.53767 | 11.11303 | 1.00928 | 14.57485 | 1.41995 | 11.51143 | 19.38463 | 0.35661 | 17.04526 | 8.24060 | 4.79694 | 13.31932 | 2.91942 | 9.27004 | 6.31283 |
| **Egf** | 0.31227 | 0.01841 | 0.00000 | 1.36891 | 0.00783 | 0.00821 | 0.00720 | 0.07068 | 0.01159 | 0.01758 | 0.11471 | 0.03410 | 0.18221 | 0.96657 | 0.69308 | 0.00702 |
| **Egfl7** | 2.72028 | 1.95270 | 1.97493 | 0.23220 | 0.04010 | 0.00433 | 3.25755 | 0.37102 | 0.00740 | 0.08045 | 1.09288 | 0.02946 | 2.84704 | 0.00548 | 0.08137 | 2.22399 |
| **Egfl8** | 0.01280 | 0.00742 | 0.00000 | 0.15253 | 0.01089 | 2.99764 | 0.00677 | 0.57115 | 0.11956 | 1.04829 | 0.32733 | 0.07900 | 0.00000 | 0.01126 | 0.00000 | 0.00612 |
| **Egflam** | 0.01261 | 0.00765 | 0.01212 | 0.14952 | 0.01865 | 0.00640 | 0.00728 | 0.01036 | 0.85994 | 0.76673 | 0.00583 | 0.00352 | 0.01156 | 0.00652 | 0.04973 | 0.00737 |
| **Egfr** | 0.01703 | 0.00781 | 0.01081 | 2.99849 | 0.00900 | 0.00624 | 0.00725 | 0.02289 | 1.41552 | 0.00167 | 0.01608 | 0.02424 | 0.01182 | 0.00928 | 0.19616 | 0.00670 |
| **Egln1** | 6.34449 | 11.57576 | 10.63597 | 6.87596 | 4.32232 | 12.13974 | 12.00017 | 18.37455 | 7.96268 | 10.83563 | 12.45085 | 4.87606 | 11.99324 | 0.66801 | 4.82286 | 3.25170 |
| **Egln2** | 9.23125 | 2.28479 | 3.06747 | 5.28893 | 5.44593 | 2.56377 | 0.43795 | 7.24686 | 5.95173 | 7.71552 | 2.02985 | 2.35443 | 6.64302 | 0.00000 | 0.15809 | 2.85283 |
| **Egln3** | 0.01651 | 0.01986 | 0.01012 | 0.09180 | 0.00959 | 0.00743 | 0.00739 | 0.00499 | 0.20032 | 0.00000 | 0.48252 | 0.02550 | 0.74593 | 0.02497 | 0.00000 | 0.00716 |
| **Egr1** | 5.05247 | 0.00225 | 0.00293 | 1.55563 | 0.00000 | 0.00356 | 0.00314 | 0.00000 | 0.00000 | 0.55023 | 0.01175 | 0.02156 | 0.00967 | 0.00789 | 0.01123 | 0.91639 |
| **Egr3** | 0.01306 | 0.00777 | 0.01187 | 0.00469 | 0.00888 | 0.00642 | 0.00722 | 0.00963 | 0.59368 | 1.23625 | 0.00881 | 0.00430 | 0.01039 | 0.02132 | 0.00502 | 0.00707 |
| **Ehbp1** | 0.01571 | 0.00786 | 0.00000 | 5.61022 | 0.00945 | 0.16371 | 0.00706 | 0.00000 | 6.92491 | 0.04738 | 0.71907 | 0.00000 | 0.20089 | 0.00000 | 0.90252 | 0.02254 |
| **Ehbp1l1** | 1.88983 | 0.06940 | 0.45761 | 0.64422 | 0.01450 | 0.06455 | 0.14123 | 0.09402 | 0.70542 | 0.79068 | 0.90699 | 0.05701 | 0.23811 | 0.46275 | 0.16847 | 0.06645 |
| **Ehd1** | 8.44508 | 6.28472 | 8.81445 | 1.17496 | 0.57623 | 5.48121 | 4.66663 | 15.31399 | 1.89696 | 10.54635 | 9.09363 | 0.27327 | 14.20461 | 0.00000 | 1.77300 | 11.19048 |
| **Ehd2** | 0.01357 | 0.00781 | 0.00394 | 0.12757 | 0.00909 | 0.00648 | 0.00710 | 0.00000 | 0.00862 | 0.02657 | 0.58942 | 0.06253 | 0.01209 | 0.00585 | 1.80134 | 0.00697 |
| **Ehd4** | 3.77793 | 8.37379 | 9.63398 | 4.97467 | 9.70842 | 3.30555 | 5.17589 | 5.09143 | 5.14667 | 1.13898 | 2.49010 | 4.15482 | 5.35013 | 2.34003 | 0.00000 | 3.05216 |
| **Ehf** | 0.01353 | 0.00768 | 0.01021 | 2.38263 | 0.00909 | 0.00652 | 0.00729 | 0.00689 | 0.34301 | 0.47574 | 0.14633 | 0.00497 | 0.01082 | 0.00688 | 0.43004 | 0.00695 |
| **Ehmt1** | 3.51092 | 0.43667 | 4.79010 | 4.37588 | 3.86914 | 2.98150 | 2.05630 | 2.76758 | 4.75619 | 9.52799 | 2.34022 | 2.32254 | 4.11886 | 0.65095 | 6.95345 | 3.10079 |
| **Ehmt2** | 3.83307 | 0.27603 | 0.00000 | 1.72595 | 4.46692 | 6.22714 | 0.28629 | 14.34448 | 1.98718 | 18.98841 | 6.17177 | 0.86380 | 10.62630 | 0.04584 | 0.16464 | 9.37488 |
| **Ei24** | 12.65220 | 6.05914 | 9.49914 | 21.80748 | 5.99540 | 0.11392 | 9.08272 | 14.61313 | 20.96674 | 25.28278 | 14.85829 | 7.73410 | 11.41420 | 22.83053 | 22.26230 | 0.47337 |
| **Eid1** | 0.01580 | 0.00761 | 0.09575 | 0.00714 | 0.00888 | 0.00581 | 0.00726 | 0.01038 | 0.01181 | 0.71509 | 0.01074 | 0.02446 | 0.01074 | 0.00608 | 0.16966 | 0.00729 |
| **Eid2** | 1.74398 | 0.26590 | 1.04184 | 12.55048 | 3.52648 | 2.43597 | 0.02477 | 0.05498 | 21.63492 | 1.58223 | 1.02400 | 0.00000 | 0.21572 | 0.25986 | 0.50654 | 0.85912 |
| **Eid2b** | 0.00000 | 0.00606 | 0.88062 | 1.31060 | 0.00849 | 0.00826 | 1.57356 | 0.36535 | 3.03001 | 0.00000 | 0.40190 | 2.83920 | 0.08352 | 0.00240 | 0.02819 | 0.00618 |
| **Eid3** | 0.01347 | 0.00787 | 0.01748 | 0.35495 | 0.00938 | 0.00624 | 0.00744 | 0.01225 | 0.02547 | 0.00677 | 0.01294 | 0.00706 | 0.01253 | 0.00576 | 0.42668 | 0.00729 |
| **Eif1** | 24.54950 | 7.59836 | 13.61329 | 37.55791 | 17.70441 | 7.94040 | 6.52911 | 17.81391 | 44.55763 | 39.73742 | 38.53285 | 8.46804 | 20.36753 | 17.77321 | 9.86203 | 11.32611 |
| **Eif1a** | 13.82694 | 17.05912 | 18.47677 | 9.65701 | 11.47493 | 10.54284 | 11.16429 | 18.80156 | 16.85141 | 25.84414 | 13.94800 | 15.07812 | 10.53513 | 29.17889 | 15.89177 | 8.89039 |
| **Eif1ad** | 14.62695 | 3.03241 | 11.32698 | 8.56970 | 9.09957 | 10.38663 | 5.36454 | 17.33078 | 19.55469 | 16.21668 | 12.50379 | 5.36432 | 14.54017 | 1.54977 | 4.99595 | 6.54190 |
| **Eif1ax** | 23.19782 | 3.69070 | 6.06279 | 8.39472 | 8.01014 | 6.95258 | 1.64358 | 14.65445 | 13.61661 | 18.85285 | 9.46521 | 1.45367 | 15.71806 | 5.48338 | 0.00000 | 5.28477 |
| **Eif1b** | 5.77757 | 0.01735 | 2.57249 | 8.38863 | 8.53736 | 0.14609 | 0.00000 | 1.28612 | 4.89553 | 8.62632 | 5.98436 | 2.03343 | 0.63026 | 2.77912 | 2.06020 | 5.14054 |
| **Eif2a** | 9.27829 | 9.32000 | 6.93034 | 6.12877 | 15.09660 | 4.48257 | 5.31116 | 11.46419 | 7.43606 | 16.56633 | 9.15875 | 8.81020 | 5.85407 | 23.00319 | 15.07627 | 2.57849 |
| **Eif2ak1** | 2.01491 | 0.01484 | 1.10695 | 3.39280 | 0.21068 | 1.09963 | 0.02401 | 1.22570 | 5.49771 | 3.46929 | 2.60291 | 5.50059 | 1.66013 | 0.68354 | 3.31177 | 0.51967 |
| **Eif2ak2** | 0.22620 | 0.00731 | 0.33909 | 0.06468 | 0.00706 | 0.01145 | 0.04911 | 0.05891 | 0.35616 | 0.05633 | 0.11602 | 0.07269 | 0.00935 | 0.20463 | 0.05350 | 0.00698 |
| **Eif2ak3** | 0.01454 | 0.02656 | 0.75376 | 0.82627 | 1.10208 | 0.00749 | 1.29525 | 0.64314 | 1.12675 | 0.07037 | 0.00000 | 0.88834 | 0.47953 | 0.75673 | 3.29428 | 0.01343 |
| **Eif2ak4** | 0.92102 | 2.32383 | 0.14925 | 6.13701 | 1.23920 | 4.34494 | 0.00371 | 0.00000 | 0.93632 | 0.12817 | 1.88208 | 3.69301 | 0.63171 | 0.00000 | 1.84569 | 2.24123 |
| **Eif2b1** | 18.17664 | 14.16980 | 13.25842 | 7.34668 | 5.42558 | 4.34798 | 12.25289 | 24.89500 | 9.82775 | 24.55394 | 9.95114 | 12.72005 | 13.52391 | 3.68895 | 18.09363 | 11.25108 |
| **Eif2b2** | 10.12457 | 0.45670 | 11.00459 | 5.97510 | 0.45634 | 4.36867 | 5.51220 | 12.27161 | 14.58429 | 13.24314 | 9.27163 | 4.05610 | 7.15016 | 0.00000 | 4.15379 | 9.93204 |
| **Eif2b3** | 6.08219 | 12.19652 | 11.39314 | 3.25920 | 10.61656 | 8.86264 | 8.92249 | 11.07402 | 8.69011 | 13.39218 | 17.55542 | 5.75016 | 10.18963 | 13.55104 | 13.63272 | 4.23637 |
| **Eif2b4** | 10.93180 | 7.23086 | 9.43511 | 14.06328 | 7.47139 | 4.85921 | 5.85705 | 14.53269 | 21.23071 | 18.23916 | 12.63950 | 7.84060 | 13.88299 | 0.00000 | 1.14090 | 8.34555 |
| **Eif2b5** | 12.25447 | 9.88647 | 14.25256 | 8.08048 | 8.98760 | 9.86940 | 6.72209 | 20.07701 | 6.09161 | 18.89739 | 13.28967 | 4.04282 | 18.45813 | 13.03819 | 16.68311 | 11.16175 |
| **Eif2d** | 16.02352 | 8.77954 | 13.39821 | 5.69860 | 10.75699 | 3.56885 | 13.26688 | 17.87622 | 1.74937 | 17.56060 | 7.97287 | 2.87021 | 13.83724 | 1.36445 | 10.58890 | 3.70707 |
| **Eif2s1** | 27.54814 | 19.10566 | 22.77688 | 11.31767 | 21.56123 | 13.25630 | 12.79381 | 22.24749 | 20.41568 | 28.00790 | 17.46522 | 11.69110 | 19.56242 | 28.65406 | 19.54991 | 11.69566 |

| | | | | | | | | | | | | | | | | |
|---|---|---|---|---|---|---|---|---|---|---|---|---|---|---|---|---|
| **Eif2s2** | 46.68192 | 31.49928 | 38.57171 | 16.35664 | 34.38997 | 19.18241 | 24.55576 | 44.53640 | 30.71569 | 43.96765 | 39.31482 | 26.55256 | 35.45776 | 44.22434 | 29.98625 | 23.36122 |
| **Eif2s3x** | 19.14385 | 9.84722 | 16.40378 | 11.93706 | 15.28690 | 9.66585 | 10.58695 | 25.41552 | 10.11540 | 29.18551 | 19.03302 | 6.64166 | 17.66323 | 30.94417 | 29.33158 | 12.81585 |
| **Eif2s3y** | 0.00509 | 8.18490 | 0.00000 | 0.97519 | 0.00780 | 2.29461 | 1.70967 | 0.00000 | 2.00450 | 2.71252 | 5.13813 | 5.73444 | 1.40930 | 0.00000 | 0.00000 | 0.00919 |
| **Eif3a** | 19.80134 | 19.95734 | 16.82930 | 17.38766 | 13.67815 | 9.77948 | 12.79112 | 24.56321 | 28.44235 | 32.05918 | 23.42355 | 27.62029 | 16.73758 | 41.45475 | 34.86402 | 10.53568 |
| **Eif3b** | 22.88505 | 19.32347 | 21.09961 | 11.50669 | 14.53333 | 12.77647 | 15.48237 | 36.02174 | 19.83298 | 29.18064 | 18.39059 | 5.11454 | 22.53837 | 0.43649 | 7.96850 | 13.06757 |
| **Eif3c** | 25.41574 | 19.09624 | 23.66282 | 14.80617 | 18.26539 | 14.86759 | 20.04202 | 36.96488 | 29.05044 | 42.11566 | 29.40128 | 18.85484 | 27.88815 | 6.51775 | 13.89025 | 16.26887 |
| **Eif3d** | 23.98625 | 19.18822 | 23.86245 | 4.00423 | 18.07223 | 12.19081 | 16.36598 | 33.31907 | 6.62954 | 28.20345 | 24.22350 | 5.00863 | 22.42979 | 6.55014 | 20.55339 | 15.92413 |
| **Eif3e** | 27.28805 | 17.45547 | 19.97198 | 3.52203 | 22.04485 | 15.77863 | 14.01091 | 25.62616 | 9.65191 | 26.76203 | 21.33347 | 16.30201 | 21.07881 | 29.89960 | 22.83074 | 15.80110 |
| **Eif3f** | 16.60740 | 10.86606 | 10.55331 | 11.07901 | 14.47054 | 8.63250 | 8.90997 | 15.36067 | 23.70145 | 18.40280 | 25.59528 | 11.89743 | 11.05300 | 8.59917 | 3.69741 | 6.00718 |
| **Eif3g** | 32.72628 | 9.08771 | 19.80817 | 13.02862 | 15.43891 | 11.42313 | 16.53553 | 32.74664 | 16.13961 | 42.16239 | 19.52035 | 14.82928 | 33.45418 | 0.00000 | 0.00000 | 19.79536 |
| **Eif3h** | 33.07467 | 14.60200 | 27.62771 | 20.02241 | 19.97517 | 14.63389 | 16.65881 | 35.45166 | 32.17632 | 37.47156 | 30.24189 | 15.11862 | 33.16752 | 15.56048 | 11.42681 | 15.16587 |
| **Eif3i** | 30.22368 | 18.07279 | 25.92633 | 8.26964 | 18.80042 | 10.91678 | 16.76552 | 35.43436 | 8.70784 | 32.01858 | 18.34526 | 12.64167 | 27.41627 | 13.05108 | 17.38193 | 12.05183 |
| **Eif3k** | 17.60087 | 8.92936 | 17.86964 | 9.47675 | 5.64332 | 9.72555 | 10.42116 | 19.52360 | 8.04491 | 22.78090 | 21.63586 | 8.33817 | 27.83756 | 3.67454 | 7.21262 | 12.56677 |
| **Eif3l** | 24.35482 | 17.61992 | 19.82759 | 11.69103 | 19.56692 | 12.68931 | 15.40361 | 31.86987 | 11.10765 | 31.65587 | 24.47726 | 4.15454 | 23.64019 | 1.44890 | 14.76287 | 13.06421 |
| **Eif3m** | 24.96482 | 16.18361 | 26.97935 | 8.42239 | 18.99474 | 9.83577 | 12.20806 | 27.83995 | 13.53852 | 31.39223 | 21.41358 | 18.44860 | 24.00808 | 24.92641 | 19.98452 | 12.44821 |
| **Eif4a1** | 35.77397 | 28.56613 | 36.27921 | 16.73247 | 24.32095 | 18.13816 | 21.45622 | 43.17769 | 22.63623 | 42.30119 | 34.61161 | 11.99395 | 32.77261 | 23.39418 | 25.08586 | 20.90570 |
| **Eif4a2** | 13.97183 | 21.88639 | 31.09626 | 9.97246 | 7.17382 | 9.95707 | 17.98318 | 25.19510 | 10.33273 | 22.94375 | 22.64585 | 16.11194 | 23.40648 | 36.56475 | 17.11448 | 10.83556 |
| **Eif4a3** | 25.08237 | 14.94161 | 23.04259 | 17.92933 | 11.48126 | 11.21692 | 16.11019 | 37.08677 | 25.99772 | 35.11052 | 26.37073 | 13.71628 | 23.86790 | 6.84546 | 12.19693 | 10.92786 |
| **Eif4b** | 28.26226 | 16.82528 | 19.37781 | 13.14432 | 13.62239 | 11.66572 | 10.57685 | 26.34978 | 15.76686 | 27.24427 | 20.07540 | 10.51723 | 21.09055 | 24.68329 | 21.28212 | 15.19331 |
| **Eif4e** | 22.86821 | 16.23938 | 21.29121 | 27.46300 | 14.20950 | 11.05462 | 11.53171 | 31.58069 | 31.91825 | 45.12897 | 32.42149 | 30.68719 | 16.30425 | 46.31638 | 43.26046 | 6.40416 |
| **Eif4e1b** | 0.08172 | 1.27934 | 0.07820 | 3.17509 | 0.01636 | 0.00549 | 0.01686 | 0.00665 | 3.04671 | 0.23613 | 0.31609 | 0.05494 | 0.42716 | 0.00000 | 0.00000 | 1.15470 |
| **Eif4e2** | 18.78043 | 11.04969 | 11.95590 | 12.91624 | 9.18124 | 7.62833 | 7.62798 | 12.60492 | 10.72298 | 19.64585 | 16.22738 | 3.46272 | 13.44962 | 13.75893 | 14.79118 | 11.18687 |
| **Eif4e3** | 0.18613 | 0.00818 | 0.02125 | 10.35116 | 0.01451 | 0.00565 | 0.00670 | 0.00361 | 3.83892 | 1.48666 | 0.11992 | 0.05321 | 0.02269 | 0.00029 | 2.26578 | 0.00515 |
| **Eif4ebp1** | 22.06311 | 4.51224 | 14.13253 | 3.65866 | 15.71999 | 0.88833 | 7.68497 | 9.91894 | 0.30537 | 9.20106 | 15.66435 | 2.06248 | 26.60883 | 0.09305 | 0.00000 | 12.41020 |
| **Eif4ebp2** | 0.60686 | 0.03762 | 0.92294 | 0.71584 | 0.03172 | 0.00499 | 1.14781 | 0.19199 | 0.00000 | 6.01847 | 0.00485 | 0.03862 | 0.07851 | 0.01682 | 2.15106 | 0.20356 |
| **Eif4ebp3** | 1.73210 | 0.07074 | 0.78385 | 1.38034 | 0.00402 | 0.02103 | 0.00385 | 0.34101 | 2.57047 | 1.34255 | 1.53140 | 1.41335 | 2.31489 | 1.26556 | 0.65825 | 1.13768 |
| **Eif4enif1** | 6.50286 | 6.78383 | 5.88537 | 19.36305 | 1.69990 | 6.61913 | 3.13714 | 5.86721 | 16.84607 | 10.43819 | 3.10900 | 4.05776 | 2.34033 | 0.09493 | 13.42681 | 4.94897 |
| **Eif4g1** | 15.84960 | 0.20273 | 0.00000 | 4.50142 | 12.07012 | 9.22905 | 3.28210 | 22.86514 | 2.25672 | 23.33835 | 4.86465 | 0.89698 | 22.58964 | 2.50933 | 0.87556 | 11.29065 |
| **Eif4g2** | 21.29456 | 26.60150 | 27.09143 | 31.72455 | 23.17436 | 16.21036 | 20.41950 | 34.48577 | 37.06119 | 36.05911 | 28.15682 | 35.96657 | 25.94043 | 48.03146 | 36.34753 | 15.66978 |
| **Eif4g3** | 10.08709 | 7.43315 | 1.15109 | 4.69026 | 6.09227 | 7.28705 | 0.94244 | 11.43655 | 5.01087 | 8.50652 | 9.87449 | 4.26531 | 3.68528 | 5.10763 | 11.63802 | 2.15728 |
| **Eif4h** | 20.02557 | 12.08723 | 18.45381 | 13.00641 | 18.52957 | 16.30817 | 15.30095 | 36.20959 | 15.84569 | 45.50677 | 24.58546 | 11.90600 | 26.26414 | 8.47313 | 9.39296 | 14.58407 |
| **Eif5** | 15.19682 | 20.38734 | 27.00507 | 24.83876 | 16.49650 | 14.34660 | 16.89038 | 32.42591 | 33.18854 | 41.52279 | 36.28336 | 34.55694 | 19.83839 | 38.62902 | 31.39010 | 5.40640 |
| **Eif5a** | 46.88536 | 21.70137 | 40.56566 | 19.11828 | 30.59526 | 22.35643 | 22.61157 | 44.18563 | 16.18361 | 47.20896 | 31.37361 | 7.44778 | 47.46401 | 8.20090 | 21.35678 | 27.27967 |
| **Eif5a2** | 0.01323 | 0.00690 | 0.24612 | 1.33405 | 0.00904 | 0.00533 | 0.00670 | 0.00126 | 0.01179 | 0.00639 | 0.23289 | 1.27690 | 0.00965 | 2.61684 | 0.20915 | 0.00694 |
| **Eif5b** | 15.64280 | 14.36477 | 13.33717 | 13.21734 | 13.94325 | 3.79670 | 9.63570 | 15.95243 | 16.06669 | 24.47003 | 17.72511 | 22.76255 | 13.96495 | 35.55705 | 27.28031 | 7.08267 |
| **Eif6** | 21.09302 | 3.80007 | 13.37459 | 21.63877 | 13.00982 | 7.90749 | 6.33684 | 28.30643 | 24.61987 | 35.55646 | 17.87536 | 4.50459 | 19.74626 | 2.98662 | 4.24434 | 12.37639 |
| **Elac1** | 1.74941 | 4.68360 | 1.59215 | 0.58978 | 0.01442 | 0.00431 | 1.65610 | 0.53920 | 0.03999 | 0.85356 | 0.58167 | 0.04931 | 0.37276 | 0.07466 | 0.32200 | 0.01064 |
| **Elac2** | 4.38191 | 3.39771 | 5.69657 | 0.36559 | 1.63830 | 1.89589 | 5.94247 | 8.83044 | 0.37685 | 5.20385 | 0.27557 | 1.11509 | 9.42959 | 0.03653 | 2.33167 | 5.41919 |
| **Elavl1** | 10.23243 | 6.76424 | 14.89544 | 1.00554 | 3.38927 | 1.18593 | 3.94639 | 8.22794 | 2.50305 | 10.98246 | 5.29057 | 4.12633 | 6.52109 | 3.80009 | 4.83600 | 5.84416 |
| **Elavl2** | 0.24671 | 0.00805 | 0.00000 | 12.90499 | 0.01830 | 0.00471 | 0.00655 | 0.00608 | 2.20463 | 0.21116 | 0.00000 | 0.02774 | 0.01901 | 4.32581 | 5.46195 | 0.00516 |
| **Elavl3** | 0.34483 | 0.00640 | 0.54640 | 3.58423 | 0.00631 | 0.10207 | 0.00514 | 2.62577 | 1.08363 | 1.12320 | 2.98874 | 0.05361 | 0.66057 | 0.00083 | 1.44917 | 0.00534 |
| **Eldr** | 0.01283 | 0.00797 | 0.82176 | 0.00698 | 0.00790 | 0.00563 | 0.00752 | 0.00887 | 0.02268 | 0.00702 | 0.13595 | 0.01817 | 0.00758 | 0.07835 | 0.00000 | 0.00701 |
| **Elf1** | 0.48702 | 0.00578 | 1.02596 | 8.55856 | 0.04977 | 0.00542 | 1.86567 | 0.00000 | 14.20725 | 0.14692 | 0.79075 | 2.65056 | 0.25280 | 4.31687 | 3.05304 | 0.02638 |
| **Elf2** | 2.01903 | 2.46777 | 1.50596 | 1.06185 | 0.15142 | 0.02347 | 3.09995 | 3.64972 | 0.10169 | 2.54465 | 0.87556 | 0.07248 | 1.32503 | 1.52194 | 3.83822 | 0.02438 |
| **Elf3** | 8.64130 | 14.38786 | 15.52360 | 1.13699 | 19.01711 | 19.55228 | 18.66411 | 27.39984 | 0.57254 | 5.01188 | 5.81106 | 0.17300 | 29.72227 | 0.52353 | 2.75742 | 16.22996 |
| **Elf4** | 2.38714 | 0.00472 | 0.55122 | 0.40149 | 0.02306 | 0.03507 | 0.00515 | 0.87881 | 0.01688 | 2.27766 | 1.01833 | 0.65442 | 0.00000 | 0.05682 | 2.13509 | 0.02665 |
| **Elf5** | 0.84285 | 0.00705 | 0.00436 | 0.01062 | 0.00620 | 0.00709 | 0.00681 | 0.01334 | 0.01235 | 0.01501 | 0.01267 | 0.00751 | 0.01169 | 0.01231 | 0.01086 | 0.02793 |
| **Elk1** | 0.96273 | 1.30717 | 0.35229 | 2.49099 | 0.00714 | 0.04725 | 0.03648 | 2.24139 | 1.35554 | 1.96119 | 2.35856 | 0.12136 | 0.12618 | 4.18131 | 2.08245 | 0.12787 |
| **Elk3** | 0.44706 | 0.00757 | 0.00388 | 0.00533 | 0.00686 | 0.00772 | 0.00768 | 0.07620 | 0.01112 | 0.01936 | 0.51262 | 0.01719 | 0.01427 | 0.01219 | 0.00194 | 0.00698 |
| **Elk4** | 0.31841 | 0.00815 | 0.01267 | 1.45719 | 0.01544 | 0.00584 | 0.00696 | 0.00842 | 1.00220 | 0.24874 | 0.02037 | 0.00259 | 0.04544 | 0.12509 | 1.15072 | 0.00706 |
| **Ell** | 3.61103 | 0.36542 | 0.17161 | 5.50583 | 0.00000 | 6.09505 | 0.25460 | 7.25724 | 2.63895 | 1.92725 | 3.20878 | 0.22159 | 0.21452 | 0.85300 | 1.99189 | 1.79684 |
| **Ell2** | 5.74108 | 0.00718 | 0.17053 | 10.58348 | 8.85192 | 0.27157 | 6.81431 | 7.96650 | 19.43756 | 18.70124 | 10.91648 | 19.35587 | 0.00000 | 23.79378 | 19.19460 | 0.00000 |
| **Ell3** | 0.00000 | 1.12391 | 0.00000 | 0.66649 | 0.01605 | 5.64527 | 0.03030 | 5.10319 | 1.05543 | 10.12921 | 10.68309 | 0.45248 | 6.54894 | 2.73568 | 7.59579 | 0.00000 |
| **Elmo2** | 2.15742 | 3.20223 | 0.74755 | 1.19850 | 2.95218 | 0.00383 | 2.62937 | 0.45155 | 1.78586 | 2.48004 | 0.20573 | 0.15633 | 2.09657 | 0.01995 | 1.25241 | 1.20424 |
| **Elmo3** | 1.73043 | 0.00489 | 0.02269 | 0.00576 | 0.00550 | 0.23236 | 1.67644 | 0.08135 | 0.01177 | 0.13120 | 0.35438 | 0.01215 | 1.58462 | 0.00577 | 0.10405 | 0.01174 |
| **Elmod1** | 0.01383 | 0.00820 | 0.00872 | 0.00536 | 0.00909 | 0.00690 | 0.00760 | 0.00589 | 0.01129 | 0.00591 | 0.50249 | 0.00616 | 0.00739 | 0.01095 | 0.00298 | 0.00715 |
| **Elmod2** | 0.67618 | 1.44470 | 1.90037 | 10.88028 | 0.01127 | 0.01138 | 0.68095 | 0.00000 | 3.81817 | 2.66404 | 2.89608 | 2.00985 | 0.41204 | 7.29441 | 8.36744 | 1.19451 |
| **Elmod3** | 1.13809 | 0.00507 | 0.25103 | 0.66668 | 1.66942 | 0.00414 | 0.01654 | 2.60719 | 2.64226 | 0.27707 | 0.00000 | 0.00299 | 0.00339 | 0.12354 | 0.00000 | 0.22321 |
| **Elmsan1** | 0.00669 | 0.00732 | 0.41572 | 1.04494 | 0.00862 | 0.02830 | 0.00656 | 1.67961 | 0.80514 | 0.81953 | 1.36437 | 0.12682 | 0.63573 | 0.25969 | 0.13948 | 0.05000 |
| **Eln** | 0.01333 | 0.00828 | 0.02186 | 1.28167 | 0.00928 | 0.02001 | 0.00750 | 0.01394 | 0.00000 | 0.01498 | 0.01359 | 0.00847 | 0.01217 | 0.01055 | 0.00314 | 0.00718 |
| **Elof1** | 13.60122 | 5.12468 | 13.83972 | 12.48032 | 6.04979 | 8.28500 | 4.01859 | 14.80211 | 18.79525 | 24.34013 | 11.02538 | 8.49267 | 10.67581 | 0.98770 | 15.85950 | 5.68827 |
| **Elovl1** | 4.59310 | 2.58214 | 4.48629 | 1.72979 | 2.55373 | 7.69468 | 2.16551 | 1.46712 | 0.68547 | 0.00000 | 0.68411 | 0.17200 | 5.72839 | 0.00034 | 0.10478 | 4.39935 |

| | | | | | | | | | | | | | | | | |
|---|---|---|---|---|---|---|---|---|---|---|---|---|---|---|---|---|
| **Elovl2** | 0.01376 | 0.00818 | 0.01214 | 0.06045 | 0.00938 | 0.00657 | 0.00747 | 0.01289 | 0.03539 | 0.00876 | 0.01204 | 0.00817 | 0.01270 | 0.00752 | 0.19438 | 0.00737 |
| **Elovl3** | 0.01362 | 0.00724 | 0.00731 | 0.40167 | 0.00928 | 0.00671 | 0.00705 | 0.00979 | 0.00547 | 0.28243 | 0.00752 | 0.76382 | 0.01251 | 0.04923 | 0.04043 | 0.00736 |
| **Elovl5** | 5.90522 | 4.11634 | 7.64989 | 0.11128 | 0.02429 | 1.71214 | 7.38798 | 5.99979 | 0.00331 | 1.18131 | 7.11216 | 0.00000 | 1.24304 | 7.77566 | 0.00000 | 1.84161 |
| **Elovl6** | 13.25463 | 12.74455 | 15.39927 | 7.46646 | 10.53238 | 14.33882 | 12.57105 | 19.03607 | 16.09876 | 15.60958 | 16.64536 | 7.92083 | 13.61736 | 34.23491 | 18.93537 | 9.10128 |
| **Elovl7** | 3.42150 | 6.28667 | 6.96011 | 1.44738 | 5.90103 | 0.13608 | 3.94097 | 0.54575 | 0.48829 | 0.00000 | 0.59954 | 0.04134 | 2.86650 | 2.37770 | 0.88638 | 1.51202 |
| **Elp2** | 10.88802 | 8.99823 | 10.67721 | 1.13576 | 1.45753 | 0.00000 | 7.35815 | 9.88938 | 4.39351 | 5.46183 | 9.31663 | 7.29782 | 6.61997 | 8.81672 | 7.88275 | 5.39030 |
| **Elp3** | 9.33581 | 12.02303 | 12.87955 | 0.00000 | 6.52641 | 0.21891 | 10.57644 | 12.62685 | 0.01674 | 4.52809 | 6.72227 | 0.22151 | 8.66848 | 2.22734 | 5.43907 | 2.64215 |
| **Elp4** | 0.19815 | 1.58567 | 3.89772 | 0.22369 | 0.01114 | 0.01207 | 1.08405 | 1.64559 | 0.10891 | 2.38002 | 3.50108 | 7.75469 | 2.23781 | 0.75141 | 0.00000 | 0.15248 |
| **Elp5** | 18.79399 | 10.08750 | 14.22193 | 13.89681 | 10.30660 | 1.26410 | 8.88268 | 9.88922 | 23.18363 | 17.27390 | 7.17662 | 4.45292 | 10.76564 | 9.36571 | 12.16927 | 9.54017 |
| **Elp6** | 4.48090 | 1.50899 | 2.94432 | 8.13225 | 0.04903 | 4.74294 | 2.69279 | 8.17588 | 8.77493 | 10.48435 | 3.95330 | 4.22127 | 6.00094 | 0.11595 | 0.00000 | 0.02302 |
| **Emb** | 0.54199 | 0.00841 | 0.01111 | 1.70631 | 0.03333 | 0.00790 | 0.00731 | 0.15617 | 1.08197 | 6.36537 | 0.00000 | 0.00000 | 0.00000 | 0.94575 | 0.35216 | 6.93388 |
| **Emc1** | 3.79529 | 3.54516 | 6.17294 | 4.36142 | 0.03432 | 0.01418 | 0.12984 | 5.52168 | 5.05168 | 6.13710 | 12.49488 | 1.96887 | 2.12017 | 14.23248 | 6.47643 | 3.83409 |
| **Emc10** | 9.75253 | 2.96002 | 4.26736 | 0.05168 | 5.61146 | 1.97319 | 1.74688 | 2.65536 | 0.01848 | 3.56311 | 8.51068 | 3.37984 | 13.05807 | 0.00000 | 0.00000 | 5.80904 |
| **Emc2** | 8.66916 | 8.08146 | 6.37332 | 1.50258 | 5.00353 | 8.22105 | 7.00881 | 2.00597 | 1.39802 | 5.36279 | 7.45858 | 6.92521 | 7.90888 | 9.62009 | 6.52581 | 7.27432 |
| **Emc3** | 5.17946 | 4.72808 | 12.31739 | 15.22533 | 3.85473 | 10.94124 | 7.88015 | 20.72003 | 20.60414 | 29.46971 | 19.41276 | 11.76030 | 15.06273 | 16.99696 | 22.26326 | 1.91519 |
| **Emc4** | 15.57844 | 6.30474 | 11.68086 | 6.74735 | 7.40288 | 9.43378 | 8.44948 | 20.51404 | 9.49637 | 20.48078 | 9.09384 | 5.57420 | 15.60423 | 4.49350 | 2.38960 | 9.11370 |
| **Emc6** | 2.70268 | 2.58310 | 5.69647 | 6.82336 | 8.47370 | 0.49690 | 1.75515 | 1.36782 | 2.40656 | 12.34008 | 7.59681 | 0.83715 | 7.53817 | 0.00000 | 3.25636 | 2.15653 |
| **Emc7** | 6.24905 | 6.73566 | 10.92804 | 7.11263 | 6.22834 | 6.79883 | 7.06399 | 9.11930 | 11.26975 | 7.69350 | 10.12325 | 10.92642 | 6.84910 | 4.52237 | 4.67765 | 3.39567 |
| **Emc8** | 8.19566 | 0.00565 | 3.87408 | 6.77448 | 2.71749 | 4.54925 | 1.53097 | 3.44705 | 13.00670 | 6.82529 | 6.45675 | 0.48863 | 10.87569 | 0.05461 | 1.50731 | 2.48586 |
| **Emc9** | 3.46865 | 0.00458 | 0.00000 | 0.01808 | 0.01560 | 3.59631 | 0.00420 | 0.66569 | 0.84306 | 0.68650 | 1.15489 | 0.00636 | 1.00094 | 0.00952 | 0.00000 | 0.36379 |
| **Emd** | 0.35262 | 3.34998 | 3.17594 | 1.25701 | 0.04516 | 0.00706 | 0.06593 | 0.01217 | 0.68088 | 1.03108 | 2.72171 | 0.08399 | 1.03901 | 0.50965 | 1.08796 | 0.30706 |
| **Eme1** | 7.17137 | 4.38143 | 3.47820 | 4.74279 | 1.89101 | 4.16768 | 5.18746 | 3.20179 | 2.96220 | 10.48724 | 0.25163 | 1.83870 | 3.63977 | 0.53012 | 8.52193 | 5.52447 |
| **Eme2** | 0.75878 | 0.00462 | 0.64009 | 0.06712 | 4.04018 | 0.01564 | 0.00548 | 0.95808 | 0.05950 | 0.00186 | 0.00652 | 0.00025 | 2.04447 | 0.00963 | 0.06194 | 1.15615 |
| **Emg1** | 11.64739 | 14.64635 | 12.20490 | 0.40534 | 9.70903 | 3.69407 | 10.30601 | 19.24749 | 1.52463 | 23.21788 | 4.22238 | 9.00139 | 13.85347 | 5.76275 | 16.22202 | 9.48164 |
| **Emid1** | 0.01367 | 0.00799 | 0.01165 | 0.37729 | 0.00919 | 0.00686 | 0.00756 | 0.01350 | 0.04664 | 0.15802 | 0.01200 | 0.03115 | 0.01299 | 0.00817 | 0.05041 | 0.00735 |
| **Emilin1** | 0.01802 | 0.00808 | 0.01958 | 0.01058 | 0.01826 | 0.00638 | 0.00765 | 0.01847 | 0.01228 | 0.02503 | 0.01272 | 0.01915 | 0.01239 | 0.01226 | 0.01067 | 0.00742 |
| **Emilin2** | 0.00946 | 0.00824 | 0.01151 | 4.25137 | 0.00893 | 0.00605 | 0.00711 | 0.01493 | 0.94665 | 1.35075 | 0.02663 | 0.01270 | 0.70926 | 0.00000 | 0.22969 | 0.00618 |
| **Eml1** | 0.11206 | 0.00713 | 0.18789 | 0.00581 | 0.00759 | 0.00563 | 0.26155 | 0.00907 | 0.09232 | 0.23821 | 0.01212 | 0.00615 | 0.01070 | 0.01195 | 0.00701 | 0.00746 |
| **Eml2** | 0.66507 | 0.02845 | 1.99276 | 0.09498 | 0.01319 | 0.02931 | 0.80663 | 0.91941 | 0.00757 | 0.00000 | 0.80287 | 0.00871 | 0.00000 | 0.00623 | 0.00000 | 0.00594 |
| **Eml3** | 1.04677 | 0.01697 | 0.00000 | 1.50188 | 0.00657 | 4.36371 | 0.00500 | 1.42641 | 6.25334 | 2.76466 | 0.64948 | 0.01733 | 1.88984 | 0.00000 | 1.78969 | 0.01818 |
| **Eml4** | 3.10866 | 0.02252 | 0.31161 | 1.91437 | 2.25752 | 4.50944 | 2.07922 | 1.60456 | 1.97930 | 0.85275 | 0.00000 | 0.56925 | 0.75656 | 2.81141 | 1.69114 | 1.84157 |
| **Eml5** | 0.24842 | 0.00726 | 0.03529 | 3.05703 | 0.00724 | 0.00546 | 0.00710 | 0.36974 | 0.39881 | 0.10950 | 0.29253 | 0.05557 | 0.00479 | 0.82535 | 0.95670 | 0.00633 |
| **Eml6** | 0.01484 | 0.00748 | 0.18808 | 0.05229 | 0.00943 | 0.01749 | 0.31076 | 0.01158 | 0.04998 | 0.03493 | 0.06076 | 0.00544 | 0.07686 | 0.00552 | 0.51541 | 0.00741 |
| **Emp1** | 0.01370 | 0.00774 | 0.01069 | 0.00905 | 0.00912 | 0.00674 | 0.00727 | 0.01070 | 0.01206 | 0.36660 | 0.01233 | 0.09032 | 0.01244 | 0.01145 | 0.00808 | 0.00780 |
| **Emp2** | 0.09451 | 0.00614 | 5.01526 | 1.10498 | 0.09370 | 3.78798 | 2.58198 | 1.74414 | 2.05001 | 0.69540 | 0.76645 | 0.09941 | 1.64229 | 0.51519 | 0.09842 | 4.15397 |
| **Emp3** | 0.01295 | 0.00797 | 0.01140 | 0.47100 | 0.00888 | 0.00665 | 0.00729 | 0.01035 | 0.05475 | 1.40116 | 0.00991 | 0.02605 | 0.01127 | 0.04306 | 0.04373 | 0.00744 |
| **Emsy** | 1.77413 | 0.31303 | 0.72048 | 0.59421 | 0.19819 | 1.91867 | 2.77431 | 1.09473 | 1.60628 | 4.08523 | 0.00000 | 4.50712 | 1.43862 | 3.20851 | 3.55930 | 0.00343 |
| **Emx2** | 0.01349 | 0.00812 | 0.01165 | 0.00464 | 0.00922 | 0.00660 | 0.00753 | 0.01301 | 1.71960 | 0.08160 | 0.00000 | 0.00488 | 0.01311 | 0.01246 | 0.00681 | 0.00743 |
| **Emx2os** | 0.03525 | 0.00777 | 0.05247 | 0.05580 | 0.00871 | 0.02663 | 0.00731 | 0.01044 | 0.10463 | 0.57224 | 0.03260 | 0.06271 | 0.03951 | 0.09466 | 0.02075 | 0.00729 |
| **En2** | 0.00838 | 0.00762 | 0.00583 | 0.45072 | 0.77776 | 0.00727 | 0.00748 | 0.00876 | 0.09293 | 0.00395 | 0.17798 | 0.00682 | 0.01607 | 0.00772 | 0.05644 | 0.00710 |
| **Enah** | 22.41586 | 17.85592 | 17.80800 | 6.12131 | 18.96433 | 13.03535 | 14.81634 | 20.20897 | 3.69346 | 21.42863 | 21.29973 | 20.27785 | 19.50439 | 27.09138 | 10.39312 | 8.04835 |
| **Enc1** | 1.10590 | 2.34972 | 2.51649 | 1.97989 | 0.06011 | 0.01060 | 3.29556 | 0.66996 | 0.20447 | 0.77529 | 0.65695 | 0.14899 | 0.00000 | 0.35545 | 1.19421 | 5.78233 |
| **Endod1** | 0.28847 | 0.00631 | 1.18406 | 1.68167 | 0.03379 | 1.08053 | 2.13559 | 0.00000 | 0.00000 | 0.77570 | 0.72158 | 0.13500 | 0.98985 | 0.00223 | 1.71979 | 0.00588 |
| **Endog** | 0.41152 | 0.05875 | 0.05591 | 0.22721 | 0.00918 | 0.00531 | 0.00627 | 0.25784 | 0.57776 | 2.52580 | 1.12316 | 0.09191 | 0.30968 | 0.00391 | 0.18206 | 0.69487 |
| **Endov** | 1.11823 | 0.04945 | 2.03693 | 0.68012 | 0.24154 | 0.07562 | 2.32636 | 2.36251 | 1.01895 | 0.10862 | 1.07021 | 0.22521 | 0.10809 | 0.13340 | 0.19087 | 0.08201 |
| **Eng** | 2.63666 | 0.00499 | 0.07083 | 0.00793 | 0.00532 | 0.00623 | 0.00507 | 0.00000 | 0.02625 | 5.22895 | 0.50159 | 0.07624 | 0.02031 | 0.00764 | 0.10521 | 0.00447 |
| **Engase** | 0.20084 | 0.54375 | 1.19148 | 0.00689 | 0.01450 | 0.00790 | 0.00666 | 0.00779 | 0.02302 | 0.70983 | 0.01063 | 0.00418 | 0.01510 | 0.01138 | 0.00000 | 2.43620 |
| **Enho** | 0.00902 | 0.00739 | 0.00654 | 0.83555 | 0.00944 | 0.01334 | 0.00703 | 1.29576 | 0.00000 | 0.44212 | 0.00487 | 0.03012 | 0.05865 | 0.00908 | 0.00000 | 0.00752 |
| **Enkd1** | 2.56810 | 0.08482 | 0.00000 | 1.15814 | 0.42697 | 3.67978 | 0.01538 | 0.31924 | 1.39294 | 1.39591 | 0.05823 | 2.82205 | 1.44242 | 0.00763 | 0.00000 | 0.01850 |
| **Eno1** | 0.14323 | 0.37197 | 0.33370 | 0.01009 | 0.07696 | 0.07777 | 0.15818 | 0.09006 | 0.01231 | 0.02435 | 0.01183 | 0.01698 | 0.34815 | 0.00905 | 0.01019 | 0.21229 |
| **Eno1b** | 8.70227 | 7.38973 | 10.80995 | 2.34428 | 7.95152 | 3.82013 | 4.57661 | 2.71286 | 0.19059 | 2.28439 | 1.08774 | 0.15872 | 8.49218 | 0.00000 | 0.00000 | 4.55686 |
| **Eno2** | 0.04515 | 0.00826 | 0.00398 | 0.05643 | 0.02328 | 0.00775 | 0.00700 | 0.00257 | 0.07310 | 0.00336 | 2.33825 | 0.00828 | 0.06152 | 0.00958 | 0.00315 | 0.01388 |
| **Eno3** | 0.00069 | 0.00781 | 0.81811 | 0.40023 | 0.00794 | 3.51452 | 0.01745 | 0.76866 | 0.00479 | 1.73472 | 1.28784 | 0.19673 | 2.01222 | 0.00955 | 0.09110 | 1.52338 |
| **Eno4** | 0.75654 | 0.00791 | 0.59267 | 0.00815 | 0.00600 | 0.00487 | 0.00797 | 0.30707 | 0.01169 | 0.74063 | 0.00775 | 0.00675 | 0.00331 | 0.01628 | 0.00000 | 0.00608 |
| **Enoph1** | 3.72169 | 1.71726 | 6.41046 | 3.57940 | 5.51715 | 3.85437 | 6.71303 | 18.56722 | 8.14309 | 12.95601 | 3.88180 | 2.36930 | 10.49008 | 0.64563 | 0.02691 | 8.19506 |
| **Enox1** | 0.00000 | 0.00515 | 0.00000 | 2.48670 | 0.00974 | 0.00472 | 0.00560 | 1.20032 | 0.00000 | 1.64466 | 1.69262 | 8.60032 | 0.00000 | 1.58803 | 1.79776 | 0.01362 |
| **Enox2** | 0.01374 | 0.00784 | 0.01231 | 0.00539 | 0.00929 | 0.00665 | 0.00747 | 0.01305 | 0.12027 | 0.04407 | 0.01190 | 0.00699 | 0.01305 | 0.00680 | 0.08667 | 0.00768 |
| **Enpep** | 0.00000 | 6.41287 | 3.66107 | 0.00000 | 0.07423 | 7.41976 | 1.53366 | 1.46379 | 0.07295 | 0.00000 | 0.00000 | 0.00000 | 3.62334 | 0.02380 | 0.39633 | 1.79400 |
| **Enpp1** | 0.00000 | 0.00776 | 0.47120 | 0.00948 | 0.01837 | 0.00952 | 0.00647 | 0.00851 | 0.01207 | 0.28327 | 0.01197 | 0.00564 | 1.34227 | 0.01280 | 0.00419 | 1.02509 |
| **Enpp2** | 0.01317 | 0.00761 | 0.01172 | 0.19492 | 0.00899 | 0.00646 | 0.00728 | 0.01019 | 0.68324 | 0.83104 | 0.00626 | 0.00298 | 0.01173 | 0.00436 | 0.40743 | 0.00763 |
| **Enpp3** | 0.00000 | 0.00469 | 0.00000 | 1.14187 | 0.00806 | 0.01009 | 0.00560 | 1.75255 | 1.53307 | 4.34903 | 1.13730 | 9.02410 | 0.00000 | 5.65167 | 5.91812 | 0.00612 |
| **Enpp4** | 0.29375 | 0.01727 | 0.16062 | 0.54448 | 0.00742 | 0.49943 | 0.00664 | 0.27999 | 0.00033 | 1.60050 | 0.31630 | 0.84213 | 0.49529 | 0.00482 | 1.27421 | 0.12696 |

| | | | | | | | | | | | | | | | | |
|---|---|---|---|---|---|---|---|---|---|---|---|---|---|---|---|---|
| **Enpp5** | 0.01160 | 0.00804 | 0.00974 | 1.24420 | 0.01022 | 0.12705 | 0.00754 | 0.01407 | 0.35905 | 0.05022 | 0.00985 | 0.01537 | 0.48298 | 0.00369 | 0.47128 | 0.00711 |
| **Ensa** | 0.75649 | 0.04941 | 0.09268 | 2.49461 | 0.02401 | 0.01999 | 0.00693 | 0.62217 | 4.34258 | 1.16616 | 0.10608 | 0.45327 | 0.07890 | 2.49074 | 1.48367 | 0.01340 |
| **Enthd2** | 0.38115 | 0.41057 | 3.41538 | 1.78710 | 0.00859 | 0.00509 | 2.34113 | 4.62240 | 1.70028 | 3.65740 | 3.13844 | 0.12947 | 0.51346 | 0.00038 | 0.09274 | 5.27632 |
| **Entpd1** | 0.01192 | 0.00793 | 0.01608 | 0.22134 | 0.01928 | 0.01403 | 0.00723 | 0.01058 | 0.00534 | 1.16056 | 0.01113 | 0.00458 | 0.01437 | 0.01190 | 0.00782 | 1.26110 |
| **Entpd2** | 0.00000 | 0.00583 | 0.00000 | 0.83563 | 0.01085 | 1.56439 | 3.76080 | 3.27975 | 0.33489 | 1.46741 | 1.84847 | 0.13178 | 2.72794 | 0.00229 | 0.28950 | 0.01306 |
| **Entpd3** | 0.43053 | 0.00649 | 0.13514 | 0.00000 | 0.00629 | 1.61649 | 0.00639 | 6.60378 | 1.67321 | 8.86599 | 2.24688 | 0.18211 | 0.00000 | 6.45211 | 3.35263 | 0.00562 |
| **Entpd4** | 0.01332 | 0.00769 | 0.00902 | 1.16892 | 0.00884 | 0.00676 | 0.00715 | 0.00000 | 0.65460 | 1.10937 | 0.45190 | 0.00000 | 0.00844 | 0.00256 | 0.12773 | 0.00715 |
| **Entpd5** | 1.58323 | 0.00659 | 0.41245 | 4.94801 | 0.03455 | 0.70746 | 0.02643 | 0.39766 | 1.53544 | 2.34617 | 0.31102 | 1.26730 | 0.84835 | 0.02816 | 3.67354 | 0.01149 |
| **Entpd6** | 2.83559 | 3.08658 | 4.64124 | 3.49606 | 0.43052 | 1.52264 | 4.15125 | 0.00000 | 1.20468 | 0.26537 | 1.25621 | 0.28042 | 3.86945 | 0.10024 | 2.60379 | 3.41701 |
| **Entpd7** | 0.76604 | 0.00802 | 0.68346 | 0.60972 | 0.01379 | 0.00753 | 0.01797 | 1.12886 | 0.41462 | 0.97309 | 0.00000 | 0.04021 | 0.05718 | 0.69577 | 0.31523 | 0.28510 |
| **Eny2** | 5.06596 | 0.24095 | 2.86932 | 17.17980 | 8.23839 | 4.58905 | 1.27192 | 0.22282 | 21.63548 | 3.31640 | 9.37880 | 0.91743 | 2.60741 | 5.37198 | 2.70561 | 2.12657 |
| **Eogt** | 0.61304 | 0.00488 | 0.06922 | 0.44571 | 0.00616 | 0.00850 | 3.22347 | 2.89016 | 0.45200 | 1.10168 | 0.00000 | 0.01059 | 0.85023 | 4.56189 | 0.21592 | 0.02866 |
| **Eomes** | 8.79155 | 2.74133 | 3.16413 | 0.77274 | 6.83242 | 10.08196 | 0.78081 | 4.98572 | 0.00000 | 2.62364 | 1.06499 | 0.08647 | 1.01293 | 2.19076 | 0.31458 | 6.30457 |
| **Ep300** | 0.00019 | 0.00548 | 0.07175 | 17.74314 | 3.25816 | 0.00242 | 0.00369 | 0.00000 | 17.23658 | 0.26854 | 6.41800 | 0.19511 | 0.88311 | 1.26638 | 2.95865 | 0.00174 |
| **Ep400** | 5.63502 | 4.51979 | 8.70624 | 11.91341 | 9.21299 | 0.73277 | 7.07205 | 9.20706 | 10.50333 | 10.05816 | 5.21446 | 12.70369 | 4.84022 | 7.26676 | 6.07460 | 3.54300 |
| **Epas1** | 1.34309 | 2.16950 | 2.80597 | 0.40866 | 0.08266 | 12.63949 | 7.24555 | 18.99565 | 0.14405 | 10.39530 | 14.15342 | 2.83689 | 10.89886 | 0.70516 | 0.00000 | 0.40740 |
| **Epb41** | 3.13457 | 3.79687 | 4.33435 | 7.86138 | 3.69857 | 7.87928 | 3.61476 | 4.21431 | 7.18266 | 6.96896 | 0.00000 | 0.79997 | 1.91104 | 0.00000 | 6.20874 | 5.01107 |
| **Epb41l1** | 5.28923 | 2.69849 | 5.09749 | 0.29162 | 0.01968 | 2.96788 | 3.51659 | 0.37058 | 0.04983 | 0.00000 | 0.68371 | 0.03388 | 2.33995 | 0.00329 | 0.28056 | 0.30994 |
| **Epb41l2** | 5.67035 | 7.49085 | 6.74793 | 4.13539 | 8.07930 | 5.72552 | 3.46175 | 6.20713 | 3.17717 | 0.62071 | 3.33180 | 0.38574 | 3.70006 | 7.99875 | 3.30033 | 0.50769 |
| **Epb41l3** | 0.00000 | 0.00767 | 0.01314 | 0.51969 | 0.00820 | 0.00689 | 0.00742 | 0.01382 | 0.09661 | 0.24147 | 0.01454 | 0.05769 | 0.27873 | 0.03704 | 0.39229 | 1.86389 |
| **Epb41l4a** | 0.00240 | 1.72770 | 0.10923 | 0.76174 | 0.00807 | 0.00628 | 0.06247 | 0.01643 | 0.00000 | 0.19867 | 0.00947 | 0.01735 | 0.14470 | 0.55446 | 0.00000 | 0.01433 |
| **Epb41l4aos** | 0.76394 | 0.00757 | 0.21575 | 1.04646 | 0.02836 | 0.01180 | 0.22430 | 0.01386 | 1.82735 | 1.07909 | 4.33948 | 0.07197 | 0.87017 | 0.15385 | 0.05426 | 0.27158 |
| **Epb41l4b** | 10.88851 | 5.92267 | 4.61904 | 6.28578 | 6.09585 | 3.39099 | 10.50061 | 10.51855 | 3.25479 | 3.43362 | 3.62470 | 8.40698 | 5.73326 | 6.60732 | 5.46120 | 5.15400 |
| **Epb41l5** | 0.25851 | 3.91629 | 0.19381 | 5.62850 | 0.02324 | 3.01031 | 1.54545 | 5.15739 | 1.71760 | 0.00000 | 0.11934 | 0.93939 | 2.26821 | 0.83018 | 1.03783 | 0.25808 |
| **Epb42** | 0.04020 | 0.00788 | 0.08788 | 0.08922 | 0.03858 | 0.00695 | 0.01702 | 0.01512 | 0.07220 | 0.07983 | 0.46218 | 0.00000 | 0.05609 | 1.16287 | 0.05366 | 0.03576 |
| **Epc1** | 0.75147 | 0.21316 | 1.03720 | 6.25703 | 0.01661 | 2.71040 | 0.38927 | 0.91857 | 5.64079 | 1.90446 | 0.75814 | 0.36878 | 0.06847 | 1.36726 | 0.77021 | 2.21033 |
| **Epc2** | 0.92921 | 1.56656 | 0.65048 | 12.24789 | 5.19525 | 3.27526 | 1.94513 | 3.89065 | 3.77149 | 5.42371 | 5.96490 | 6.59372 | 1.74837 | 6.15888 | 15.20854 | 1.80065 |
| **Epcam** | 10.49859 | 8.80256 | 11.77248 | 6.19242 | 4.76827 | 18.32685 | 13.57079 | 7.37950 | 5.08700 | 2.29364 | 2.19026 | 0.18159 | 21.11244 | 0.86141 | 3.98022 | 4.33820 |
| **Epg5** | 2.68310 | 0.02520 | 7.70691 | 7.90854 | 0.03024 | 1.40827 | 0.79661 | 6.38540 | 3.19498 | 3.45055 | 3.08269 | 2.11546 | 3.62048 | 0.50143 | 3.82835 | 0.01956 |
| **Epha1** | 0.01327 | 0.00817 | 0.02666 | 0.00848 | 0.00915 | 2.32770 | 0.02780 | 0.01112 | 0.01244 | 0.00586 | 0.01010 | 0.00873 | 0.00347 | 0.00910 | 0.08637 | 0.02064 |
| **Epha10** | 0.61792 | 0.00613 | 0.21755 | 2.19261 | 0.00722 | 0.00806 | 1.36273 | 0.00000 | 1.00165 | 1.56435 | 1.01811 | 0.10412 | 0.89681 | 0.05970 | 1.13153 | 0.00568 |
| **Epha2** | 0.00000 | 0.00735 | 0.55293 | 0.01110 | 0.00474 | 0.00941 | 0.00709 | 1.03293 | 0.01119 | 0.04143 | 0.00545 | 0.03326 | 0.17456 | 0.01460 | 0.00000 | 5.48749 |
| **Epha3** | 0.01353 | 0.00811 | 0.01187 | 0.20351 | 0.00921 | 0.00676 | 0.00762 | 0.01434 | 1.03917 | 0.06461 | 0.00615 | 0.00681 | 0.01330 | 0.00891 | 0.02189 | 0.00733 |
| **Epha4** | 6.40013 | 6.27237 | 6.80830 | 1.05216 | 6.52962 | 0.00000 | 0.75416 | 3.10114 | 1.43887 | 0.28966 | 1.03778 | 2.16653 | 0.19283 | 5.43802 | 1.22875 | 5.75830 |
| **Epha7** | 0.01350 | 0.01834 | 0.01197 | 0.01794 | 0.00923 | 0.00668 | 0.00745 | 0.01287 | 0.01254 | 0.03551 | 0.01330 | 0.00750 | 0.01298 | 0.03144 | 0.01743 | 0.00764 |
| **Ephb2** | 0.00866 | 0.00523 | 0.00000 | 0.56788 | 4.46269 | 0.99226 | 0.02692 | 0.16039 | 1.52131 | 0.45289 | 0.49208 | 0.29457 | 0.77995 | 0.53301 | 0.56561 | 0.15000 |
| **Ephb3** | 4.39385 | 4.88998 | 8.05969 | 1.97559 | 2.76662 | 0.50935 | 5.88344 | 3.54239 | 2.52465 | 0.00000 | 1.12457 | 0.00000 | 6.11291 | 0.00000 | 0.00000 | 8.63297 |
| **Ephb4** | 3.10910 | 0.23142 | 0.66191 | 1.37272 | 0.02144 | 0.00309 | 0.01452 | 1.96270 | 0.99227 | 0.15100 | 0.12681 | 0.03181 | 0.00447 | 0.03328 | 1.88991 | 1.46098 |
| **Ephb6** | 4.62066 | 0.00000 | 1.69460 | 0.23833 | 0.00487 | 3.37385 | 2.83587 | 0.76376 | 0.00692 | 0.11591 | 0.00001 | 0.02663 | 0.24447 | 0.00531 | 0.36061 | 0.00000 |
| **Ephx1** | 0.56563 | 3.62232 | 2.91871 | 0.00000 | 0.01181 | 0.01421 | 5.89113 | 0.40825 | 0.01179 | 0.75123 | 0.02842 | 0.00939 | 0.08441 | 0.00517 | 0.02628 | 0.00465 |
| **Ephx2** | 7.58040 | 0.00000 | 0.00000 | 0.22443 | 3.54256 | 0.00000 | 0.15303 | 0.01920 | 1.00887 | 0.00222 | 0.00000 | 0.04212 | 0.00428 | 0.00683 | 0.55091 | 6.54012 |
| **Ephx3** | 0.06022 | 0.00791 | 0.00815 | 0.00464 | 0.00798 | 0.00687 | 0.01787 | 0.00857 | 0.66661 | 0.01036 | 0.24284 | 0.00594 | 0.02480 | 0.03165 | 0.00208 | 0.00716 |
| **Ephx4** | 0.02087 | 0.00816 | 0.48221 | 1.04832 | 0.00794 | 0.00563 | 0.05055 | 0.00862 | 0.00000 | 0.00383 | 0.26025 | 0.00688 | 0.00871 | 0.00587 | 0.02980 | 0.00688 |
| **Epm2a** | 0.01350 | 0.00807 | 0.01194 | 0.51620 | 0.00926 | 0.00677 | 0.00772 | 0.01756 | 0.01672 | 0.06494 | 0.01169 | 0.00835 | 0.01293 | 0.00545 | 0.22783 | 0.00733 |
| **Epm2aip1** | 0.00372 | 0.00852 | 1.28768 | 3.55021 | 0.02167 | 0.14562 | 0.00748 | 1.21019 | 6.62009 | 3.63433 | 0.50877 | 0.04522 | 0.00000 | 3.00564 | 3.25468 | 0.02037 |
| **Epn1** | 0.28326 | 0.04000 | 0.00000 | 6.02671 | 0.70989 | 1.81657 | 0.12638 | 0.15005 | 0.26028 | 0.44984 | 2.85884 | 0.11070 | 0.12107 | 0.04760 | 0.04284 | 0.14283 |
| **Epn2** | 2.34662 | 3.75342 | 1.70585 | 14.04613 | 0.01218 | 3.74799 | 1.64015 | 6.44673 | 8.37811 | 7.69445 | 8.78737 | 4.30672 | 1.02434 | 1.90746 | 11.32796 | 0.00348 |
| **Epn3** | 0.04149 | 0.00837 | 0.06475 | 0.01065 | 0.01375 | 0.00850 | 0.00734 | 0.01274 | 0.01235 | 0.01415 | 0.01261 | 0.00788 | 0.04349 | 0.01252 | 0.00967 | 0.00743 |
| **Eprs** | 19.54487 | 16.49795 | 15.95379 | 4.66405 | 13.72060 | 13.73636 | 12.79397 | 20.25204 | 6.51413 | 13.18811 | 9.81504 | 14.00260 | 14.32230 | 19.27184 | 17.15020 | 10.31709 |
| **Eps15** | 7.26770 | 2.46538 | 6.47191 | 7.44829 | 7.51424 | 5.69010 | 4.13956 | 6.69002 | 0.76697 | 3.95733 | 2.96832 | 0.49249 | 9.18132 | 11.85592 | 2.91155 | 1.30479 |
| **Eps15l1** | 3.44821 | 4.02381 | 2.89816 | 2.54685 | 5.31946 | 0.39409 | 0.24507 | 7.25034 | 3.52926 | 2.18285 | 3.53090 | 0.31792 | 0.67750 | 1.30992 | 3.85655 | 0.31961 |
| **Eps8** | 1.03931 | 1.13824 | 0.00000 | 7.24724 | 0.00606 | 0.00408 | 0.00569 | 0.00186 | 2.10758 | 0.00000 | 0.05572 | 0.02750 | 0.01144 | 3.18700 | 7.61450 | 0.00451 |
| **Eps8l1** | 0.01412 | 0.00806 | 0.03554 | 0.00499 | 0.01013 | 0.00832 | 0.00737 | 0.00318 | 0.01092 | 2.05815 | 0.51779 | 0.00000 | 0.28840 | 0.01487 | 0.00000 | 0.00721 |
| **Eps8l2** | 1.11388 | 0.00527 | 0.23959 | 0.44516 | 0.00456 | 3.91676 | 0.26975 | 5.31167 | 0.00365 | 0.69192 | 3.85479 | 0.00000 | 3.09759 | 0.00000 | 0.00000 | 1.17209 |
| **Ept1** | 2.98450 | 2.97349 | 2.37278 | 0.74989 | 2.32956 | 0.04768 | 3.05880 | 3.71100 | 0.16496 | 1.83959 | 2.84545 | 0.41283 | 2.50324 | 11.12593 | 4.87085 | 0.25684 |
| **Eqtn** | 0.01213 | 0.00780 | 0.01099 | 0.04332 | 0.00947 | 0.00664 | 0.00731 | 0.05750 | 0.62906 | 0.01050 | 0.00795 | 0.00664 | 0.01034 | 0.01529 | 0.01015 | 0.00737 |
| **Eral1** | 2.21602 | 2.17679 | 0.00000 | 0.87701 | 0.02024 | 3.45127 | 1.57113 | 2.68097 | 0.95938 | 0.96944 | 0.51660 | 0.11768 | 2.27989 | 1.83891 | 4.14744 | 0.02308 |
| **Erap1** | 0.01475 | 0.00702 | 0.00000 | 0.61022 | 0.00880 | 0.00625 | 0.00615 | 0.00000 | 0.77600 | 0.04683 | 1.62924 | 1.45354 | 0.00000 | 2.55220 | 0.23968 | 0.00682 |
| **Eras** | 0.75673 | 0.00688 | 0.00196 | 1.54090 | 0.00641 | 0.00757 | 0.00622 | 0.81273 | 0.35375 | 0.00301 | 0.00695 | 0.00487 | 0.56564 | 0.00529 | 0.21563 | 0.00591 |
| **Erbb2** | 0.00000 | 0.00660 | 1.01445 | 0.00000 | 0.02091 | 7.13510 | 1.12496 | 1.12968 | 0.01035 | 0.58684 | 0.52292 | 0.00000 | 2.72849 | 0.00379 | 0.09500 | 0.00341 |
| **Erbb3** | 0.01940 | 0.00856 | 0.13644 | 0.00706 | 0.00983 | 0.00918 | 0.00791 | 0.22250 | 0.01141 | 0.60109 | 0.00907 | 0.00577 | 0.09381 | 0.01051 | 0.26296 | 0.02897 |
| **Erbin** | 0.00000 | 4.40912 | 0.78323 | 3.93377 | 0.00799 | 3.02781 | 1.87537 | 2.80253 | 2.78532 | 2.09028 | 0.56579 | 10.31404 | 0.00000 | 1.15445 | 2.71850 | 0.00244 |

| | | | | | | | | | | | | | | | | |
|---|---|---|---|---|---|---|---|---|---|---|---|---|---|---|---|---|
| **Erc1** | 1.50088 | 0.01736 | 2.12446 | 0.55889 | 0.20786 | 0.01425 | 0.00533 | 0.40257 | 1.79704 | 1.51003 | 1.39749 | 0.01820 | 1.10552 | 1.07272 | 0.00000 | 0.04185 |
| **Erc2** | 0.01316 | 0.00826 | 0.00967 | 1.47897 | 0.00922 | 0.00673 | 0.00769 | 0.02064 | 0.78864 | 0.27719 | 0.00491 | 0.16318 | 0.01206 | 0.01064 | 0.00000 | 0.00708 |
| **Ercc1** | 4.48995 | 0.00747 | 4.57879 | 13.76907 | 7.35974 | 2.89615 | 2.10580 | 4.41246 | 5.12549 | 4.25293 | 4.65863 | 1.76398 | 5.98144 | 0.03005 | 3.01312 | 1.22575 |
| **Ercc2** | 1.13364 | 0.46074 | 1.00003 | 1.69534 | 0.00942 | 0.32320 | 0.00468 | 4.04658 | 2.24390 | 3.07401 | 2.94488 | 0.14393 | 0.91627 | 0.00000 | 0.00000 | 2.51372 |
| **Ercc3** | 3.76213 | 8.07829 | 4.09158 | 3.66193 | 9.71658 | 3.13087 | 2.17834 | 8.01513 | 7.63550 | 8.05168 | 5.36246 | 1.76110 | 2.32517 | 0.30775 | 5.53280 | 0.14712 |
| **Ercc4** | 1.26154 | 0.01689 | 1.83393 | 3.40585 | 0.01080 | 2.00308 | 0.01453 | 1.80241 | 4.48332 | 1.10500 | 2.19599 | 0.03149 | 2.05825 | 0.19912 | 0.01887 | 1.79693 |
| **Ercc5** | 0.79440 | 1.44083 | 2.06620 | 0.62879 | 0.00262 | 0.00585 | 2.17723 | 0.35004 | 0.42308 | 0.00000 | 1.53578 | 0.14754 | 0.95308 | 0.00245 | 1.31857 | 0.53810 |
| **Ercc6** | 1.12329 | 0.00604 | 0.29117 | 5.72871 | 0.02742 | 0.03145 | 0.07526 | 1.32280 | 7.30332 | 0.18358 | 4.35876 | 0.09727 | 4.04445 | 5.31361 | 9.51428 | 0.07515 |
| **Ercc6l** | 2.90891 | 3.28425 | 0.83699 | 10.75110 | 0.01127 | 0.00000 | 2.87560 | 1.45144 | 11.49752 | 5.76235 | 6.29861 | 14.45795 | 0.71448 | 6.70669 | 14.02571 | 1.58914 |
| **Ercc6l2** | 0.93462 | 0.54016 | 0.02613 | 1.93839 | 0.00613 | 0.00717 | 3.59598 | 0.59380 | 0.86131 | 1.06173 | 0.79261 | 0.12148 | 1.01429 | 8.99068 | 3.84579 | 0.00505 |
| **Ercc8** | 5.23264 | 1.23871 | 3.34223 | 1.33325 | 0.00000 | 3.58464 | 0.00000 | 4.94996 | 0.00000 | 3.46202 | 2.09641 | 0.23490 | 3.18409 | 2.45351 | 2.46801 | 1.81877 |
| **Erdr1** | 1.20046 | 0.04667 | 1.01548 | 2.38059 | 0.10397 | 2.21713 | 0.36070 | 0.25419 | 4.51402 | 11.12799 | 9.45092 | 1.32675 | 3.48231 | 0.00000 | 0.00000 | 1.86469 |
| **Erf** | 1.43770 | 0.03940 | 0.57786 | 0.15751 | 0.86671 | 0.51033 | 0.09424 | 0.33355 | 0.91148 | 0.25907 | 2.88631 | 0.88749 | 0.07386 | 0.00550 | 0.16313 | 3.38982 |
| **Erg** | 0.01344 | 0.00793 | 0.01168 | 1.37697 | 0.00915 | 0.00664 | 0.00750 | 0.01237 | 1.10169 | 0.26297 | 0.01457 | 0.00554 | 0.01274 | 0.00258 | 0.17990 | 0.00714 |
| **Ergic1** | 1.63943 | 0.00382 | 1.87458 | 0.63072 | 1.31988 | 0.08583 | 1.82165 | 1.61178 | 1.21648 | 2.26177 | 0.56664 | 0.03173 | 2.57774 | 0.16011 | 0.20471 | 0.14643 |
| **Ergic2** | 6.39296 | 4.30356 | 4.72842 | 2.58285 | 2.70025 | 0.15669 | 5.39634 | 5.82951 | 0.70317 | 8.95856 | 0.96980 | 2.30032 | 2.15299 | 8.76757 | 9.89539 | 1.27374 |
| **Ergic3** | 14.50260 | 4.52220 | 15.33425 | 7.29981 | 2.32719 | 11.13128 | 4.80045 | 10.71695 | 6.39166 | 10.27690 | 8.69374 | 7.94530 | 18.69724 | 0.71675 | 6.41375 | 9.63194 |
| **Erh** | 24.13325 | 11.21110 | 24.72927 | 15.12687 | 13.87665 | 11.18092 | 8.32592 | 18.53548 | 19.38907 | 24.30768 | 22.01977 | 6.33703 | 19.98474 | 14.20225 | 12.48021 | 11.90794 |
| **Eri1** | 2.89170 | 4.66168 | 3.07268 | 7.12868 | 3.91388 | 0.00000 | 0.89904 | 4.48707 | 6.18851 | 9.12071 | 6.16480 | 13.29479 | 3.50774 | 6.66412 | 8.54285 | 2.28425 |
| **Eri2** | 1.43082 | 3.40143 | 1.33661 | 1.60303 | 4.00172 | 0.65329 | 0.05830 | 1.51244 | 1.06921 | 0.81691 | 0.88512 | 0.08724 | 0.12909 | 4.00341 | 6.06537 | 0.16618 |
| **Eri3** | 5.79585 | 4.64260 | 4.72081 | 0.14922 | 0.16611 | 4.00501 | 0.47907 | 3.79389 | 0.78527 | 3.47829 | 1.05582 | 0.02735 | 3.40733 | 0.00226 | 0.02859 | 0.05282 |
| **Erich1** | 2.35807 | 2.25047 | 5.74947 | 0.18594 | 1.14460 | 0.00461 | 3.16381 | 2.78842 | 0.71627 | 2.42965 | 1.11069 | 0.13294 | 0.53668 | 0.17379 | 1.65538 | 0.02421 |
| **Erich5** | 0.36686 | 0.15534 | 0.84875 | 1.26542 | 0.13788 | 0.09049 | 0.04630 | 0.00000 | 0.47783 | 0.08665 | 3.33708 | 0.12205 | 0.75146 | 0.37197 | 1.07447 | 0.11337 |
| **Erlec1** | 2.50897 | 0.00401 | 0.00000 | 0.99217 | 0.00394 | 0.00462 | 2.12205 | 0.00000 | 0.31832 | 1.42154 | 1.05172 | 0.09487 | 0.02121 | 1.24852 | 0.68428 | 0.00434 |
| **Erlin1** | 0.38385 | 1.06616 | 1.43990 | 3.19162 | 0.66380 | 0.00666 | 2.47997 | 5.05037 | 1.34511 | 4.04409 | 3.31770 | 7.34306 | 1.93961 | 11.66869 | 4.43584 | 2.79921 |
| **Erlin2** | 3.64123 | 2.55278 | 5.62005 | 0.00000 | 3.91883 | 3.73706 | 6.59691 | 3.92286 | 1.13322 | 3.50186 | 3.11285 | 1.07794 | 3.84632 | 10.14449 | 8.26892 | 0.48814 |
| **Ermap** | 0.02057 | 0.00791 | 0.00786 | 0.16537 | 0.00911 | 0.03849 | 0.01818 | 0.02175 | 0.06018 | 0.02933 | 0.01281 | 0.03898 | 0.01719 | 0.05509 | 0.10170 | 0.00743 |
| **Ermard** | 0.59667 | 0.47578 | 3.65105 | 0.35057 | 0.57755 | 0.01406 | 0.62243 | 1.55837 | 0.14457 | 0.26534 | 0.40423 | 1.27345 | 0.95279 | 0.00618 | 1.40826 | 1.18546 |
| **Ermn** | 0.02943 | 0.00787 | 0.03959 | 0.00574 | 0.00918 | 0.05755 | 0.04905 | 0.01889 | 0.04621 | 0.01194 | 0.23788 | 0.00652 | 0.03288 | 0.06287 | 0.03150 | 0.05759 |
| **Ermp1** | 1.18102 | 6.75916 | 11.52786 | 32.13809 | 0.00000 | 10.82692 | 8.32283 | 7.46195 | 2.58255 | 9.73622 | 7.33844 | 10.68560 | 11.04586 | 21.71610 | 18.94165 | 3.20675 |
| **Ern1** | 0.59811 | 0.00743 | 0.00000 | 2.64819 | 0.00722 | 0.00637 | 0.00774 | 0.21895 | 5.12165 | 0.22642 | 0.29630 | 0.03428 | 0.06024 | 0.01446 | 2.17225 | 0.15053 |
| **Ero1l** | 0.04247 | 0.19455 | 1.77036 | 3.55587 | 0.00972 | 0.00547 | 0.49799 | 0.36084 | 1.37447 | 2.55846 | 0.91709 | 0.11464 | 0.09694 | 4.05132 | 1.53361 | 1.55532 |
| **Ero1lb** | 0.00214 | 0.01909 | 0.16785 | 4.34325 | 0.01014 | 0.00518 | 0.00698 | 0.42814 | 1.34975 | 0.65083 | 0.56524 | 0.08389 | 0.00258 | 1.65927 | 3.13114 | 0.00620 |
| **Erp29** | 10.00464 | 7.39963 | 12.38871 | 4.15482 | 6.56934 | 7.74151 | 4.89188 | 10.00815 | 6.55304 | 8.59032 | 8.96988 | 0.47687 | 11.38264 | 0.00000 | 1.14157 | 10.93134 |
| **Erp44** | 5.10765 | 0.20510 | 3.54193 | 1.66002 | 6.44521 | 5.26119 | 6.89287 | 4.51648 | 0.53603 | 5.74289 | 4.60442 | 10.86005 | 4.95243 | 1.52465 | 2.37963 | 0.87278 |
| **Errfi1** | 0.08451 | 0.98771 | 0.15693 | 15.32765 | 0.04839 | 1.16915 | 0.00625 | 0.28361 | 12.63652 | 2.53011 | 0.86309 | 0.00000 | 0.44550 | 3.10504 | 5.14787 | 1.66886 |
| **Esam** | 2.01469 | 0.00590 | 0.00205 | 0.00319 | 0.73531 | 0.00698 | 0.00584 | 0.10212 | 0.48568 | 0.74194 | 1.82785 | 0.00000 | 0.08920 | 0.00291 | 0.26347 | 0.00522 |
| **Esco1** | 1.29088 | 2.27607 | 1.10302 | 24.17012 | 1.63962 | 0.32477 | 0.00000 | 1.76111 | 17.88291 | 1.39094 | 1.73271 | 12.95025 | 3.33155 | 19.17315 | 12.03281 | 0.00347 |
| **Esco2** | 3.45576 | 6.18240 | 6.48797 | 6.92768 | 1.46035 | 1.47887 | 8.95569 | 8.37863 | 21.71792 | 10.97593 | 9.59541 | 19.96956 | 3.08261 | 31.73943 | 17.28088 | 1.01669 |
| **Esd** | 10.13283 | 1.76634 | 3.37136 | 0.15766 | 0.64742 | 4.02194 | 4.58723 | 4.95129 | 1.49455 | 2.42222 | 2.87355 | 0.62856 | 4.13259 | 0.19429 | 2.70580 | 3.34114 |
| **Esf1** | 6.38275 | 5.01700 | 2.01234 | 6.04486 | 3.67916 | 3.61903 | 0.83078 | 4.69439 | 8.60455 | 3.63034 | 2.42157 | 12.47086 | 0.35951 | 19.19000 | 16.39078 | 0.50070 |
| **Esp24** | 0.01576 | 0.01753 | 0.00590 | 2.76609 | 0.00878 | 0.01825 | 0.01733 | 0.02391 | 8.00593 | 0.19126 | 0.00058 | 0.07432 | 0.01103 | 0.20580 | 0.58112 | 0.00619 |
| **Espl1** | 10.25146 | 0.01407 | 0.00000 | 18.74856 | 1.49790 | 3.83889 | 0.15222 | 13.14967 | 21.58030 | 27.44786 | 11.47618 | 0.67149 | 11.49930 | 0.19044 | 0.00000 | 2.11841 |
| **Esr1** | 0.01292 | 0.00804 | 0.01123 | 2.74890 | 0.00901 | 0.00621 | 0.00741 | 0.01316 | 0.49307 | 0.88158 | 0.00651 | 0.00342 | 0.01128 | 0.00035 | 0.82167 | 0.00694 |
| **Esr2** | 0.05656 | 0.00744 | 0.02067 | 1.00683 | 0.00822 | 0.00678 | 0.01749 | 0.01171 | 1.12152 | 0.01348 | 0.10910 | 0.06038 | 0.02445 | 0.15269 | 0.35693 | 0.06557 |
| **Esrp1** | 1.92104 | 0.00232 | 2.12608 | 6.55079 | 0.00000 | 1.31933 | 4.17928 | 1.86698 | 6.63240 | 6.99711 | 2.61194 | 1.96418 | 0.08667 | 7.96475 | 8.83293 | 0.19403 |
| **Esrp2** | 0.32103 | 0.87176 | 4.04711 | 0.29114 | 0.00883 | 6.17903 | 0.00368 | 7.48498 | 0.00857 | 7.95826 | 0.33327 | 1.35032 | 3.81148 | 2.61614 | 4.79529 | 1.84415 |
| **Esrra** | 5.83363 | 0.00000 | 2.29135 | 0.74819 | 0.04212 | 1.43274 | 1.42348 | 9.37503 | 3.03059 | 6.12513 | 3.30155 | 0.24233 | 2.24903 | 0.01965 | 0.36315 | 1.95468 |
| **Esrrb** | 19.19795 | 8.54005 | 13.67928 | 5.73643 | 6.32198 | 0.00000 | 6.17047 | 13.54081 | 10.77573 | 17.67601 | 19.31350 | 5.10845 | 4.09050 | 6.23331 | 5.41357 | 2.22626 |
| **Esrrg** | 0.01378 | 0.00763 | 0.01186 | 2.30927 | 0.00940 | 0.00632 | 0.00732 | 0.01185 | 0.00000 | 0.01914 | 0.01194 | 0.00659 | 0.01295 | 0.00599 | 0.42232 | 0.00709 |
| **Esyt1** | 0.13909 | 0.00605 | 2.67985 | 5.25906 | 0.87968 | 5.92778 | 0.38450 | 2.70822 | 7.84171 | 2.11933 | 1.21082 | 0.21944 | 1.20030 | 0.00432 | 1.08807 | 3.31652 |
| **Esyt2** | 2.09501 | 2.64328 | 2.79437 | 7.02639 | 1.10028 | 0.27518 | 5.56655 | 2.36499 | 6.53413 | 3.85064 | 4.22840 | 11.49679 | 1.49642 | 7.91878 | 4.47622 | 0.63849 |
| **Esyt3** | 1.80050 | 0.17029 | 1.09601 | 3.26485 | 1.31602 | 0.00639 | 0.13428 | 0.00977 | 0.47409 | 0.58482 | 0.15603 | 0.08761 | 0.01253 | 0.25440 | 2.17959 | 0.00414 |
| **Etaa1** | 1.64570 | 7.61918 | 2.55535 | 14.18598 | 3.03538 | 0.00710 | 2.98368 | 1.96161 | 17.45809 | 11.87328 | 7.62531 | 9.08740 | 2.36325 | 8.88479 | 9.84757 | 1.69901 |
| **Etaa1os** | 0.01597 | 0.00809 | 0.07080 | 0.52523 | 0.00913 | 0.00582 | 0.00743 | 0.01344 | 0.54927 | 0.01575 | 0.00948 | 0.00742 | 0.01179 | 0.01122 | 0.00433 | 0.00723 |
| **Etd** | 0.01375 | 0.00814 | 0.01230 | 0.40144 | 0.00931 | 0.00661 | 0.00752 | 0.01394 | 0.00587 | 0.01545 | 0.01399 | 0.00824 | 0.01266 | 0.01115 | 0.00578 | 0.00735 |
| **Etf1** | 20.05352 | 16.30691 | 21.37642 | 21.46591 | 19.14004 | 15.17057 | 15.42634 | 22.64421 | 25.78478 | 30.62665 | 14.23136 | 21.98052 | 17.19485 | 21.10368 | 26.66666 | 11.83751 |
| **Etfa** | 18.97341 | 8.33049 | 14.81997 | 5.92597 | 12.82781 | 11.52366 | 6.47561 | 13.72805 | 6.99986 | 12.69755 | 10.61089 | 7.79558 | 13.68226 | 4.56874 | 4.97487 | 7.74626 |
| **Etfb** | 25.02012 | 12.64355 | 22.53099 | 0.70519 | 9.55650 | 12.23899 | 8.35460 | 15.82816 | 0.38942 | 9.55298 | 9.08382 | 0.13204 | 14.81878 | 0.04177 | 0.34405 | 9.83277 |
| **Etfdh** | 1.08849 | 0.12472 | 0.26407 | 0.27021 | 3.23412 | 0.00453 | 2.78989 | 0.03099 | 0.00717 | 1.08674 | 0.05309 | 0.01406 | 0.88778 | 2.21499 | 0.83586 | 1.62509 |
| **Ethe1** | 1.68760 | 0.00298 | 7.03340 | 1.72766 | 1.20094 | 1.80810 | 1.73398 | 2.71026 | 0.64506 | 2.27396 | 7.36556 | 0.11738 | 5.83382 | 0.00000 | 0.00016 | 6.66831 |
| **Etl4** | 3.15189 | 0.14105 | 1.41344 | 0.38399 | 5.54587 | 2.65779 | 0.78592 | 2.60119 | 0.31596 | 2.16484 | 3.53772 | 0.02920 | 2.76442 | 5.19303 | 1.23457 | 0.32056 |

| | | | | | | | | | | | | | | | | |
|---|---|---|---|---|---|---|---|---|---|---|---|---|---|---|---|---|
| **Etnk1** | 8.32607 | 0.06638 | 3.52321 | 2.80137 | 0.08195 | 5.28588 | 2.74459 | 1.37749 | 2.61717 | 17.67910 | 4.02445 | 9.63463 | 1.52184 | 10.50134 | 22.44218 | 0.37801 |
| **Etnk2** | 1.52397 | 0.00567 | 0.07151 | 0.00511 | 0.00714 | 0.00416 | 1.18806 | 0.00000 | 0.01145 | 0.00000 | 0.29859 | 0.00514 | 0.00837 | 0.01146 | 0.00385 | 0.01286 |
| **Etohd2** | 0.00595 | 3.40060 | 0.24318 | 0.00978 | 0.00746 | 0.54945 | 0.00685 | 0.06964 | 0.26502 | 0.30574 | 0.00300 | 0.49846 | 0.00342 | 0.00905 | 0.04843 | 0.00580 |
| **Ets2** | 1.06378 | 0.00648 | 0.02215 | 2.62583 | 0.00514 | 0.00693 | 0.00645 | 0.00698 | 0.00000 | 0.25213 | 0.00780 | 0.09417 | 0.01062 | 0.00131 | 1.35464 | 2.10440 |
| **Etv1** | 1.27686 | 0.00629 | 0.00000 | 0.63298 | 0.00737 | 0.01494 | 0.00637 | 0.00740 | 0.00000 | 0.00707 | 0.01133 | 0.03607 | 0.01153 | 0.01269 | 0.27060 | 0.00596 |
| **Etv3** | 0.32960 | 0.00632 | 0.00000 | 0.61211 | 0.00705 | 0.00602 | 0.04570 | 0.16677 | 0.22305 | 0.00000 | 0.29264 | 2.07687 | 0.00571 | 0.00584 | 1.63762 | 0.01346 |
| **Etv4** | 1.21387 | 0.00756 | 0.00231 | 0.00873 | 0.01583 | 0.00681 | 0.00652 | 0.00856 | 0.01220 | 0.00518 | 0.07121 | 0.00698 | 0.01931 | 0.01225 | 0.00958 | 1.46879 |
| **Etv5** | 20.59300 | 9.05285 | 3.10467 | 3.05642 | 10.79629 | 0.00000 | 7.13856 | 4.85685 | 5.58896 | 2.22362 | 0.66403 | 0.08504 | 0.00000 | 2.56065 | 1.41704 | 7.78680 |
| **Etv6** | 4.07372 | 1.33390 | 0.00000 | 0.51875 | 0.02994 | 0.21105 | 0.00280 | 1.56799 | 3.93072 | 2.76850 | 0.43422 | 2.31010 | 0.11532 | 8.77499 | 6.55211 | 0.06005 |
| **EU599041** | 0.93974 | 0.17859 | 0.56646 | 3.48379 | 0.05135 | 0.00447 | 2.17327 | 0.68104 | 0.47881 | 0.67237 | 1.66901 | 0.66908 | 0.45546 | 1.05560 | 2.19901 | 0.01929 |
| **Eva1a** | 3.40550 | 3.29654 | 3.64600 | 3.69205 | 9.26696 | 5.11928 | 6.66806 | 7.57989 | 2.72578 | 6.79737 | 6.77794 | 0.69013 | 3.26702 | 2.24016 | 0.53301 | 4.64658 |
| **Eva1b** | 0.01308 | 0.00799 | 0.02953 | 0.13437 | 0.00975 | 0.00721 | 0.00748 | 0.01272 | 0.00982 | 0.43154 | 0.01528 | 0.02614 | 0.15430 | 0.01082 | 0.00659 | 0.00789 |
| **Evi5** | 4.54340 | 7.71772 | 7.15410 | 15.72406 | 5.14060 | 6.14396 | 5.96063 | 6.47014 | 10.02815 | 10.71852 | 6.72157 | 8.72044 | 4.18769 | 23.63759 | 23.91803 | 2.45134 |
| **Evi5l** | 0.00446 | 2.01672 | 0.00725 | 0.51641 | 0.00846 | 0.00688 | 0.00612 | 0.01421 | 0.11031 | 0.00000 | 0.01216 | 0.04426 | 1.24582 | 0.01865 | 0.06214 | 0.00605 |
| **Evl** | 0.03911 | 0.00755 | 0.29156 | 0.41749 | 0.00785 | 0.00915 | 0.35572 | 0.01025 | 0.99325 | 0.47873 | 0.02179 | 0.04860 | 0.32929 | 0.01248 | 0.86351 | 0.12608 |
| **Evpl** | 0.01937 | 0.00737 | 0.43921 | 0.13669 | 0.00756 | 0.00896 | 0.00633 | 0.70519 | 0.00682 | 0.00116 | 0.00817 | 0.00517 | 1.23677 | 0.01090 | 0.02972 | 0.00642 |
| **Evx1** | 0.01324 | 0.00770 | 0.03107 | 0.69435 | 0.00893 | 0.00588 | 0.00729 | 0.00985 | 0.00000 | 1.45562 | 0.02180 | 0.00323 | 0.01033 | 0.00435 | 0.17091 | 0.00708 |
| **Evx1os** | 0.12854 | 0.00833 | 0.00780 | 0.09528 | 0.01253 | 0.00678 | 0.00741 | 0.01292 | 0.01043 | 0.01013 | 0.01330 | 0.00821 | 0.01672 | 0.01223 | 0.00886 | 0.41947 |
| **Ewsr1** | 15.18350 | 5.62322 | 9.22038 | 7.09521 | 8.23063 | 11.67635 | 9.56708 | 23.44245 | 8.72196 | 10.00088 | 14.42374 | 4.96092 | 13.39236 | 8.13133 | 2.95076 | 11.39390 |
| **Exd1** | 0.00384 | 0.00698 | 0.00000 | 0.61647 | 0.00893 | 0.00583 | 0.00485 | 1.65143 | 4.41555 | 2.27887 | 6.22876 | 0.20957 | 0.55156 | 0.08808 | 4.57514 | 0.00583 |
| **Exd2** | 5.35811 | 2.92055 | 0.16829 | 9.12262 | 1.31258 | 0.00000 | 3.95351 | 5.45510 | 7.27874 | 6.54124 | 3.73556 | 9.43777 | 0.74487 | 0.14217 | 4.32929 | 2.69387 |
| **Exo1** | 7.46244 | 6.48260 | 2.50569 | 5.15443 | 4.69294 | 6.45314 | 5.36807 | 4.74332 | 7.48469 | 15.52898 | 7.03901 | 10.18851 | 3.62796 | 1.41042 | 3.92981 | 0.84988 |
| **Exo5** | 4.25708 | 0.67284 | 1.42654 | 0.85900 | 0.01556 | 1.12585 | 0.00279 | 4.37653 | 0.43068 | 5.49961 | 1.88222 | 0.11291 | 0.80324 | 1.07778 | 1.77119 | 2.13459 |
| **Exoc1** | 3.61836 | 6.72510 | 3.12939 | 3.70402 | 0.49458 | 3.21398 | 2.11931 | 1.08832 | 3.07099 | 5.59774 | 2.53379 | 0.67984 | 2.77613 | 6.00711 | 3.98507 | 0.16356 |
| **Exoc2** | 2.45382 | 1.46582 | 2.31290 | 2.11101 | 5.42989 | 0.01103 | 4.09151 | 0.62705 | 0.63129 | 2.92003 | 0.84897 | 6.25802 | 1.61217 | 2.15312 | 2.83046 | 1.59933 |
| **Exoc3** | 2.44596 | 4.21269 | 5.60239 | 1.36414 | 7.06310 | 3.86835 | 2.75710 | 5.32091 | 3.77226 | 5.81830 | 9.82215 | 8.07235 | 3.59374 | 5.66317 | 6.41210 | 0.00000 |
| **Exoc3l** | 0.01357 | 0.00804 | 0.01185 | 0.00871 | 0.00909 | 0.00654 | 0.00742 | 0.01118 | 0.01221 | 0.46594 | 0.01122 | 0.02686 | 0.01103 | 0.01149 | 0.00879 | 0.00712 |
| **Exoc3l4** | 0.00849 | 0.00851 | 0.00003 | 0.00488 | 0.00949 | 0.01763 | 0.00799 | 0.51651 | 0.01031 | 0.33842 | 0.92032 | 0.00442 | 0.00000 | 0.01119 | 0.00186 | 0.00727 |
| **Exoc4** | 1.40689 | 0.24754 | 0.11271 | 0.00551 | 0.00548 | 0.00747 | 0.05859 | 0.68677 | 0.01011 | 0.19293 | 0.04884 | 0.00000 | 0.01662 | 1.79280 | 0.42022 | 0.33092 |
| **Exoc5** | 4.44171 | 3.85114 | 4.68720 | 4.02438 | 2.70829 | 5.84650 | 4.50787 | 5.35921 | 6.02513 | 6.57947 | 2.00177 | 12.77158 | 3.35978 | 13.74778 | 4.45136 | 0.80262 |
| **Exoc6** | 1.31383 | 0.00526 | 0.61516 | 0.14214 | 0.84847 | 1.54105 | 0.00448 | 1.89036 | 0.79118 | 2.53989 | 4.09892 | 0.19810 | 0.34092 | 0.23354 | 1.39557 | 0.23457 |
| **Exoc6b** | 0.44745 | 2.92771 | 0.12950 | 0.00000 | 0.01740 | 0.00884 | 0.00637 | 0.52811 | 0.01104 | 0.61817 | 0.02125 | 0.02539 | 0.00527 | 0.01333 | 0.47355 | 0.01356 |
| **Exoc7** | 3.85039 | 2.26345 | 1.45973 | 2.26346 | 0.21938 | 3.93015 | 0.00218 | 0.13059 | 0.59996 | 2.16774 | 2.41877 | 0.06873 | 0.19945 | 0.00000 | 0.57566 | 2.12522 |
| **Exoc8** | 2.03035 | 1.87268 | 4.24462 | 3.15810 | 0.02398 | 0.02258 | 2.26461 | 4.04241 | 11.96080 | 2.60336 | 1.80515 | 0.12259 | 1.22299 | 1.32929 | 3.82426 | 3.26978 |
| **Exog** | 1.27513 | 4.04764 | 0.68648 | 2.94413 | 0.03841 | 2.01449 | 0.01500 | 0.00570 | 0.82517 | 0.34876 | 0.60051 | 0.11562 | 0.03694 | 0.04786 | 2.05688 | 0.01011 |
| **Exosc1** | 12.91623 | 10.99906 | 15.13226 | 5.22571 | 0.48268 | 4.19493 | 2.40560 | 7.37757 | 4.03345 | 12.91655 | 6.89397 | 7.27792 | 8.77150 | 1.65072 | 9.03104 | 3.96134 |
| **Exosc10** | 11.06000 | 0.25675 | 8.58828 | 10.89903 | 1.78497 | 1.96101 | 4.03381 | 9.43257 | 10.30405 | 16.07495 | 10.42794 | 7.21512 | 0.15515 | 0.68142 | 7.97451 | 3.31906 |
| **Exosc2** | 11.72146 | 5.79508 | 5.56453 | 8.86030 | 7.59372 | 3.84532 | 5.07698 | 7.69891 | 15.23709 | 1.63958 | 12.24990 | 0.85609 | 4.40680 | 7.80394 | 6.51484 | 1.36373 |
| **Exosc3** | 2.35228 | 1.71755 | 1.20353 | 6.33536 | 8.39383 | 0.03510 | 0.35897 | 0.19290 | 11.47149 | 14.93787 | 7.96958 | 0.25452 | 3.77032 | 0.67237 | 2.44040 | 4.73540 |
| **Exosc4** | 9.22865 | 0.03696 | 0.00000 | 0.69424 | 7.91320 | 2.32272 | 0.08716 | 6.45789 | 2.90620 | 8.44087 | 0.36377 | 0.36127 | 13.31812 | 0.09346 | 0.33716 | 2.83919 |
| **Exosc5** | 8.51185 | 0.18117 | 0.56451 | 0.52331 | 4.98786 | 1.07636 | 6.04421 | 17.29642 | 0.59252 | 17.31151 | 0.83414 | 5.18031 | 11.81471 | 0.03808 | 0.00000 | 1.91349 |
| **Exosc6** | 0.75443 | 0.06174 | 0.20458 | 13.97255 | 0.42405 | 1.20492 | 0.07697 | 0.18051 | 21.21515 | 4.08240 | 6.42279 | 0.17042 | 0.27352 | 0.36018 | 0.32017 | 0.47584 |
| **Exosc7** | 14.78484 | 8.71902 | 16.23466 | 1.16608 | 6.55703 | 4.62600 | 5.49911 | 18.32903 | 2.35421 | 15.49706 | 9.71168 | 3.26651 | 10.73916 | 5.10528 | 7.17784 | 4.87900 |
| **Exosc8** | 19.45314 | 12.48062 | 13.59650 | 10.83432 | 10.02226 | 4.21181 | 8.96443 | 11.15491 | 6.54205 | 14.83170 | 12.53035 | 11.15125 | 16.65099 | 12.19379 | 6.73259 | 7.49911 |
| **Exosc9** | 5.83786 | 3.35372 | 11.30246 | 0.04006 | 9.06445 | 3.00186 | 8.99402 | 10.72783 | 1.64085 | 7.10121 | 4.91546 | 2.02412 | 5.63365 | 6.36904 | 6.78204 | 3.41083 |
| **Exph5** | 0.00568 | 0.00835 | 0.63129 | 0.00525 | 0.00915 | 0.00955 | 0.00727 | 1.25717 | 0.01008 | 0.30200 | 1.45753 | 0.00000 | 0.47811 | 0.00531 | 0.01466 | 0.01407 |
| **Ext1** | 2.11647 | 0.55715 | 0.28249 | 5.36484 | 0.00414 | 0.00310 | 1.09081 | 1.73381 | 0.88993 | 4.05910 | 0.49250 | 2.60756 | 0.00251 | 0.00000 | 4.51066 | 0.00361 |
| **Ext2** | 7.98966 | 0.27457 | 0.00000 | 0.45772 | 0.42561 | 3.20797 | 2.36325 | 4.66527 | 0.05597 | 0.48117 | 0.89128 | 0.01424 | 1.09292 | 0.01109 | 0.06376 | 0.00481 |
| **Extl1** | 0.08417 | 0.00774 | 2.29665 | 1.95474 | 0.02561 | 0.23961 | 0.02717 | 0.07704 | 1.79302 | 0.17601 | 0.00000 | 0.03826 | 0.00608 | 0.00067 | 0.15307 | 0.00602 |
| **Extl2** | 0.01793 | 0.00800 | 0.01095 | 0.01059 | 0.00898 | 0.01312 | 0.00750 | 0.01384 | 0.01234 | 0.01853 | 0.01307 | 0.01821 | 0.01270 | 0.01212 | 0.01052 | 0.00745 |
| **Extl3** | 4.00152 | 0.00386 | 0.94688 | 0.36886 | 0.02612 | 3.01363 | 0.02591 | 0.67537 | 0.77698 | 1.15512 | 0.12057 | 0.01083 | 1.29490 | 0.00424 | 0.63322 | 2.78688 |
| **Eya1** | 0.15873 | 0.00679 | 0.00333 | 1.46849 | 0.64927 | 0.00711 | 0.04778 | 0.00751 | 0.88170 | 0.00000 | 0.03946 | 0.02425 | 0.01021 | 0.00843 | 0.44197 | 0.00670 |
| **Eya2** | 0.01278 | 0.00825 | 0.01419 | 0.15427 | 0.01012 | 0.00710 | 0.00773 | 0.01643 | 0.00953 | 0.01117 | 0.01624 | 0.00795 | 0.16358 | 0.01171 | 0.00785 | 0.00763 |
| **Eya3** | 4.32979 | 4.98803 | 6.85359 | 1.82283 | 0.07853 | 0.10717 | 5.83441 | 7.72697 | 4.80010 | 1.95052 | 4.15184 | 6.16119 | 1.54222 | 10.06842 | 6.22718 | 0.00299 |
| **Ezh1** | 2.72977 | 1.67347 | 0.92572 | 9.70560 | 1.14955 | 0.41120 | 8.44219 | 1.18987 | 6.84496 | 8.86747 | 2.01147 | 0.34319 | 0.10908 | 2.77384 | 6.04499 | 3.70488 |
| **Ezh2** | 13.84431 | 18.96398 | 17.60416 | 15.69917 | 10.28348 | 7.01450 | 12.97703 | 18.19321 | 18.21996 | 21.44413 | 10.76382 | 19.66948 | 9.49353 | 24.91474 | 17.93561 | 4.94543 |
| **Ezr** | 18.52806 | 7.93000 | 5.65329 | 0.35012 | 12.59826 | 10.65716 | 10.48215 | 1.61768 | 0.18980 | 6.71583 | 0.00000 | 0.20388 | 8.23113 | 0.00000 | 0.00000 | 13.06797 |
| **F10** | 0.03611 | 0.01883 | 0.03342 | 0.38335 | 0.03513 | 0.00642 | 0.04874 | 0.02764 | 0.06007 | 0.02150 | 0.13365 | 0.00582 | 0.15037 | 0.35719 | 0.02739 | 0.00722 |
| **F11r** | 9.20203 | 7.80055 | 15.04752 | 3.64546 | 3.11241 | 13.20086 | 9.95680 | 19.88525 | 5.92177 | 15.56728 | 13.31893 | 2.56949 | 22.57490 | 27.35800 | 14.33907 | 9.26734 |
| **F12** | 0.01201 | 0.00839 | 0.01461 | 0.21146 | 0.01004 | 0.00708 | 0.00766 | 0.01369 | 0.00561 | 0.58240 | 0.01261 | 0.00632 | 0.25300 | 0.01056 | 0.00583 | 0.00740 |
| **F13a1** | 0.01376 | 0.00772 | 0.01132 | 0.47894 | 0.00939 | 0.00644 | 0.00736 | 0.01073 | 0.16115 | 0.00303 | 0.03225 | 0.00623 | 0.01232 | 0.01021 | 0.60692 | 0.00726 |
| **F2** | 0.00705 | 0.00731 | 0.17442 | 0.91389 | 0.00928 | 0.00879 | 0.00683 | 1.47039 | 2.08210 | 0.45084 | 0.73161 | 0.01534 | 0.00000 | 0.00436 | 0.13049 | 0.00705 |

| | | | | | | | | | | | | | | | | |
|---|---|---|---|---|---|---|---|---|---|---|---|---|---|---|---|---|
| **F2r** | 1.38190 | 0.00532 | 0.00000 | 1.26723 | 0.00506 | 0.00745 | 1.22330 | 0.00671 | 1.44157 | 0.00920 | 0.00000 | 0.09580 | 0.00873 | 0.05306 | 0.05708 | 0.96407 |
| **F2rl1** | 1.48624 | 1.03502 | 0.02431 | 0.00970 | 0.45903 | 0.04088 | 0.02694 | 0.01277 | 0.01227 | 0.00148 | 0.01311 | 0.00644 | 0.38826 | 0.01390 | 0.00000 | 0.82986 |
| **F3** | 0.09370 | 0.00802 | 0.00578 | 0.01022 | 0.01461 | 0.00725 | 0.00729 | 0.03210 | 0.01144 | 0.01858 | 0.01040 | 0.10029 | 0.01170 | 0.01536 | 0.21669 | 0.00745 |
| **F8** | 0.01180 | 0.00828 | 0.11899 | 0.01020 | 0.00911 | 0.00887 | 0.00761 | 0.01894 | 0.01247 | 0.00801 | 0.01064 | 0.00665 | 0.01085 | 0.30746 | 0.05169 | 0.00738 |
| **F8a** | 0.70043 | 0.00747 | 0.42160 | 0.29948 | 0.01299 | 0.00493 | 0.00667 | 0.00318 | 0.38839 | 1.15483 | 0.17633 | 0.01334 | 0.02774 | 0.00999 | 0.00000 | 0.01355 |
| **Fa2h** | 0.01118 | 0.00833 | 0.00952 | 8.69436 | 0.00914 | 0.00540 | 0.00768 | 0.04928 | 6.68881 | 0.00000 | 0.03925 | 0.00590 | 0.00984 | 0.02879 | 0.08893 | 0.00534 |
| **Faah** | 1.06208 | 0.54599 | 2.11245 | 0.02593 | 0.01040 | 6.69476 | 0.00477 | 1.87938 | 0.81032 | 0.00000 | 0.00000 | 0.00000 | 1.71991 | 0.04171 | 0.01046 | 0.43078 |
| **Faap100** | 1.77828 | 0.00547 | 0.02816 | 0.07959 | 0.01371 | 0.00617 | 0.43521 | 0.41070 | 0.04772 | 2.55810 | 0.09935 | 0.01853 | 0.96153 | 0.01401 | 0.00000 | 1.65843 |
| **Faap20** | 1.70285 | 0.04309 | 0.64282 | 1.72559 | 4.07196 | 5.69117 | 0.85146 | 3.78089 | 0.84009 | 1.53815 | 3.04629 | 0.04991 | 3.92694 | 0.10453 | 0.00000 | 0.00847 |
| **Faap24** | 4.15762 | 0.90800 | 0.90743 | 6.25294 | 2.08126 | 0.00800 | 3.01918 | 3.14019 | 4.80389 | 10.60482 | 1.36604 | 0.23274 | 0.32918 | 0.00926 | 0.68330 | 4.78391 |
| **Fabp3** | 8.79590 | 7.22354 | 19.88338 | 0.06132 | 11.33711 | 18.23637 | 11.31470 | 4.92598 | 2.80739 | 1.88544 | 17.66010 | 1.49494 | 27.40737 | 0.00000 | 0.00000 | 14.76446 |
| **Fabp4** | 0.15585 | 0.00679 | 0.18359 | 2.95253 | 0.00713 | 0.00857 | 0.60142 | 0.00000 | 0.94331 | 2.80580 | 2.29483 | 0.37321 | 0.00000 | 4.79072 | 2.52552 | 0.00594 |
| **Fabp5** | 29.22517 | 14.91194 | 15.51012 | 3.39941 | 31.69057 | 2.82273 | 10.47511 | 4.25932 | 0.02216 | 0.58317 | 5.71782 | 0.02094 | 13.45074 | 0.00001 | 1.29349 | 14.41137 |
| **Fabp6** | 0.01191 | 0.00825 | 0.01174 | 1.09365 | 0.00912 | 0.00642 | 0.00719 | 0.01140 | 0.05313 | 0.00642 | 0.01266 | 0.00800 | 0.39096 | 0.03384 | 0.00161 | 0.00688 |
| **Fabp9** | 0.00465 | 1.01856 | 0.00739 | 0.00242 | 0.00810 | 0.00656 | 0.00686 | 0.00909 | 0.20537 | 0.00391 | 0.00846 | 0.00504 | 0.01144 | 0.58385 | 0.00000 | 0.01435 |
| **Fadd** | 0.96104 | 1.29387 | 0.93454 | 2.28070 | 0.00680 | 3.37170 | 0.06489 | 0.85240 | 2.20569 | 4.13275 | 0.57168 | 0.31362 | 2.45559 | 0.39071 | 1.38232 | 1.88859 |
| **Fads1** | 2.57337 | 2.36432 | 0.58914 | 0.37568 | 0.01969 | 0.00434 | 1.62299 | 0.00000 | 0.39313 | 3.28683 | 1.97241 | 0.04759 | 0.02291 | 0.00390 | 2.89274 | 0.00334 |
| **Fads2** | 0.01352 | 0.00827 | 0.01145 | 0.01046 | 0.00935 | 0.00670 | 0.00760 | 0.01565 | 0.01249 | 0.01272 | 0.01253 | 0.00766 | 0.01269 | 0.11374 | 0.00631 | 0.00744 |
| **Fads3** | 0.51599 | 0.00761 | 0.35931 | 0.00359 | 0.00677 | 0.00574 | 0.00667 | 0.00000 | 0.14773 | 0.01778 | 1.14729 | 0.38227 | 0.77133 | 0.09795 | 0.32455 | 0.00647 |
| **Faf1** | 8.76296 | 7.84562 | 12.10200 | 5.96296 | 8.14002 | 10.65787 | 6.52935 | 12.19442 | 13.71103 | 14.08590 | 9.68143 | 12.49454 | 9.40764 | 14.72148 | 12.42018 | 5.28914 |
| **Faf2** | 4.37279 | 4.89620 | 7.14544 | 6.92244 | 3.09922 | 1.34614 | 0.17996 | 3.42625 | 7.46799 | 4.39600 | 3.04045 | 7.26695 | 2.01101 | 6.30429 | 10.16661 | 1.47747 |
| **Fah** | 0.00674 | 0.00849 | 0.00616 | 0.00417 | 0.00901 | 0.00714 | 0.00827 | 0.64839 | 0.03391 | 2.26898 | 0.12001 | 0.00000 | 0.00526 | 0.23911 | 0.35314 | 0.00731 |
| **Fahd1** | 0.14949 | 0.01707 | 0.00000 | 0.36390 | 4.40880 | 2.98049 | 0.00650 | 0.00708 | 1.22261 | 0.00000 | 0.00000 | 0.04281 | 0.67454 | 0.01372 | 0.00167 | 0.01293 |
| **Fahd2a** | 0.26012 | 0.16063 | 0.03431 | 0.37872 | 0.00674 | 0.00571 | 0.10030 | 0.65402 | 0.05722 | 0.42184 | 0.77964 | 0.02043 | 1.24182 | 0.00451 | 0.18513 | 2.30204 |
| **Faim** | 0.92408 | 0.11075 | 0.58297 | 0.66406 | 0.11491 | 0.00464 | 0.07398 | 0.40576 | 1.33658 | 1.49797 | 1.34432 | 1.71396 | 1.13655 | 0.09991 | 0.38165 | 0.53611 |
| **Faiml** | 0.95114 | 1.03293 | 1.09686 | 0.14725 | 0.16031 | 0.00590 | 0.36030 | 0.93861 | 1.59407 | 0.52198 | 2.11065 | 0.12346 | 0.03358 | 0.88293 | 1.66235 | 0.04151 |
| **Fam101a** | 0.01298 | 0.00786 | 0.01147 | 1.08063 | 0.00944 | 0.00646 | 0.00729 | 0.03272 | 0.00000 | 0.00774 | 0.01333 | 0.00758 | 0.01136 | 0.00726 | 0.25867 | 0.00718 |
| **Fam101b** | 3.43234 | 0.93455 | 10.02936 | 0.25933 | 0.03259 | 7.05047 | 3.74624 | 13.19664 | 0.38896 | 14.36249 | 14.63236 | 12.83038 | 8.98238 | 3.89493 | 5.95895 | 6.37274 |
| **Fam102a** | 4.81658 | 4.36878 | 4.91584 | 4.32614 | 5.82155 | 3.68157 | 2.93821 | 9.11057 | 4.81803 | 8.56525 | 3.62120 | 0.42403 | 4.80627 | 4.36350 | 3.05192 | 7.09991 |
| **Fam102b** | 0.01360 | 0.00773 | 0.01059 | 1.40448 | 0.00944 | 0.00624 | 0.00727 | 0.01073 | 0.59397 | 0.00984 | 0.00643 | 0.02431 | 0.01314 | 0.00498 | 1.16413 | 0.00713 |
| **Fam103a1** | 6.30206 | 10.80433 | 11.85857 | 7.12954 | 2.39260 | 6.14037 | 6.40669 | 12.82975 | 16.37189 | 20.56132 | 11.77074 | 17.04543 | 4.37923 | 21.06841 | 16.32893 | 2.48142 |
| **Fam104a** | 2.33647 | 0.03419 | 2.62176 | 11.05192 | 4.10647 | 2.77474 | 1.37755 | 1.93621 | 11.81460 | 10.36096 | 7.42840 | 1.38139 | 1.75507 | 1.18546 | 0.65434 | 1.93095 |
| **Fam105a** | 0.01270 | 0.00752 | 0.00535 | 0.01088 | 0.00919 | 0.00682 | 0.00713 | 0.00424 | 2.33046 | 0.00000 | 0.31134 | 0.00000 | 0.00905 | 0.72772 | 0.00000 | 0.00713 |
| **Fam107a** | 0.01158 | 0.00809 | 0.46861 | 0.15947 | 0.00818 | 0.00564 | 0.00746 | 0.01537 | 0.50873 | 1.15721 | 0.00798 | 0.00450 | 0.00925 | 0.01161 | 0.00000 | 0.00719 |
| **Fam107b** | 0.48092 | 3.29077 | 0.00000 | 4.99650 | 0.07347 | 7.26824 | 0.03465 | 0.77135 | 2.55423 | 0.39100 | 0.41933 | 0.16110 | 4.69361 | 0.00000 | 1.25581 | 2.79570 |
| **Fam109a** | 1.41340 | 0.01516 | 0.13790 | 11.38575 | 2.57142 | 4.86741 | 1.40107 | 2.59115 | 18.20096 | 6.13704 | 1.58733 | 0.13388 | 2.95085 | 0.01810 | 0.33342 | 0.19325 |
| **Fam110a** | 0.01027 | 0.00800 | 0.01188 | 0.20064 | 0.00918 | 0.00648 | 0.00695 | 0.01054 | 0.25591 | 0.02960 | 0.01051 | 0.00692 | 0.77969 | 0.01135 | 0.00562 | 0.00713 |
| **Fam110b** | 0.84545 | 0.00769 | 0.00335 | 0.00642 | 0.01663 | 0.00682 | 0.00618 | 0.01318 | 0.09413 | 0.00438 | 0.01147 | 0.01702 | 0.89259 | 0.01243 | 0.01411 | 0.36566 |
| **Fam110c** | 0.01377 | 0.00823 | 0.01190 | 0.99144 | 0.00930 | 0.00658 | 0.00750 | 0.01293 | 0.00000 | 0.01330 | 0.05290 | 0.00820 | 0.01183 | 0.01063 | 0.00293 | 0.00722 |
| **Fam111a** | 0.01213 | 0.00794 | 0.73910 | 0.22655 | 0.00778 | 0.00971 | 0.00732 | 0.01129 | 0.00525 | 0.36896 | 0.01009 | 0.00762 | 0.00917 | 0.01131 | 0.00000 | 0.00707 |
| **Fam114a1** | 0.01306 | 0.00786 | 0.01158 | 0.00671 | 0.00890 | 0.00655 | 0.00729 | 0.01010 | 0.01166 | 1.35689 | 0.01058 | 0.03394 | 0.01073 | 0.01211 | 0.00650 | 0.00731 |
| **Fam114a2** | 7.49271 | 4.04451 | 3.84186 | 1.55722 | 2.70565 | 2.64972 | 6.88427 | 5.27311 | 0.64501 | 4.39073 | 3.80325 | 5.21546 | 3.02494 | 1.84286 | 3.95221 | 1.37488 |
| **Fam117a** | 0.00000 | 0.01966 | 3.43975 | 8.94018 | 0.01120 | 1.96700 | 0.00368 | 0.82711 | 4.91603 | 1.91870 | 0.79724 | 0.06291 | 6.43538 | 0.07765 | 1.20736 | 1.30145 |
| **Fam117b** | 2.77706 | 1.83646 | 3.63182 | 9.51408 | 5.57402 | 0.00754 | 2.70910 | 3.92356 | 14.75431 | 1.80398 | 2.90624 | 1.85794 | 2.13621 | 5.10488 | 5.79403 | 0.92523 |
| **Fam118a** | 1.40214 | 4.37512 | 4.60353 | 11.02418 | 4.09053 | 2.09279 | 1.27398 | 8.74511 | 17.16640 | 13.83302 | 9.57190 | 1.49185 | 4.02155 | 18.77778 | 12.39508 | 2.37014 |
| **Fam118b** | 7.36092 | 1.30269 | 5.27341 | 1.73549 | 5.53716 | 9.63204 | 0.47627 | 4.85003 | 7.85058 | 13.04434 | 4.27411 | 9.84656 | 2.87193 | 5.94441 | 9.26992 | 2.87152 |
| **Fam120a** | 5.59569 | 1.95181 | 6.51445 | 4.16530 | 8.86907 | 5.98581 | 8.47300 | 11.07236 | 4.15764 | 10.69754 | 8.82313 | 18.61720 | 6.61361 | 21.12798 | 12.11748 | 2.67484 |
| **Fam120aos** | 0.01338 | 0.00793 | 0.41762 | 0.53336 | 0.00840 | 0.00560 | 0.17606 | 0.00973 | 0.13271 | 0.60392 | 0.01077 | 0.01507 | 0.02318 | 0.05549 | 0.09866 | 0.00746 |
| **Fam120b** | 0.79681 | 0.00624 | 0.00000 | 0.49604 | 0.01805 | 0.00591 | 2.71547 | 0.00000 | 1.32498 | 0.79428 | 1.11670 | 0.03865 | 1.73857 | 0.18977 | 1.68265 | 0.00573 |
| **Fam120c** | 0.01783 | 0.00786 | 0.01081 | 0.60146 | 0.01023 | 0.00650 | 0.00738 | 0.01062 | 0.15056 | 0.65599 | 0.00829 | 0.01550 | 0.01119 | 0.01128 | 0.46686 | 0.00712 |
| **Fam122a** | 2.65547 | 0.00480 | 0.23921 | 2.83850 | 0.00409 | 0.00341 | 0.03424 | 0.38748 | 4.92621 | 1.57093 | 0.23398 | 0.03525 | 0.65033 | 0.15345 | 0.94010 | 0.01016 |
| **Fam122b** | 0.00446 | 0.00603 | 1.02954 | 0.70303 | 1.22528 | 0.37226 | 0.57728 | 0.00000 | 0.43772 | 1.10855 | 2.09951 | 4.15704 | 0.68582 | 3.88069 | 3.73263 | 0.01266 |
| **Fam124a** | 0.03153 | 0.01861 | 0.02736 | 0.06316 | 0.01030 | 0.00595 | 0.00722 | 0.01563 | 0.05964 | 0.71027 | 0.01192 | 0.02436 | 0.01758 | 0.06469 | 0.02044 | 0.00742 |
| **Fam124b** | 0.01345 | 0.00661 | 0.76702 | 0.18585 | 0.00801 | 0.00562 | 1.52131 | 0.00818 | 0.00821 | 0.34994 | 0.01287 | 0.00477 | 0.00874 | 0.01302 | 0.00000 | 0.00735 |
| **Fam126a** | 0.59424 | 0.00745 | 0.12296 | 1.50954 | 0.01648 | 0.00808 | 1.77139 | 0.41378 | 1.97388 | 0.00000 | 0.79119 | 0.59515 | 0.00638 | 0.02205 | 1.31147 | 0.00606 |
| **Fam126b** | 0.62341 | 0.00186 | 2.17880 | 10.36501 | 0.00258 | 0.00342 | 2.62869 | 1.03929 | 6.96064 | 1.73905 | 1.61865 | 9.86015 | 0.15018 | 11.59303 | 5.59413 | 1.02134 |
| **Fam129a** | 4.33995 | 8.17203 | 4.89903 | 0.00000 | 0.04679 | 10.53376 | 4.43724 | 4.64038 | 0.23371 | 0.95719 | 1.47378 | 1.69126 | 5.70938 | 1.72454 | 0.00000 | 0.45114 |
| **Fam129b** | 0.00000 | 0.00633 | 2.53860 | 0.17378 | 0.01429 | 0.01921 | 0.00467 | 3.67521 | 0.00548 | 0.34328 | 3.76869 | 0.06085 | 1.02820 | 0.00057 | 0.00000 | 4.68004 |
| **Fam129c** | 0.01332 | 0.00757 | 0.01102 | 0.74701 | 0.00929 | 0.00633 | 0.00723 | 0.01027 | 0.90707 | 0.58079 | 0.00146 | 0.03150 | 0.01190 | 0.00632 | 0.40062 | 0.00728 |
| **Fam131a** | 0.17794 | 0.00767 | 0.00515 | 0.99012 | 0.00753 | 0.00670 | 0.00714 | 0.00590 | 2.34010 | 0.00807 | 0.21588 | 0.00000 | 0.01012 | 0.03538 | 0.33244 | 0.00680 |
| **Fam131c** | 0.00855 | 0.00777 | 0.00454 | 0.00948 | 0.00945 | 0.00759 | 0.00733 | 0.63606 | 0.88043 | 0.53395 | 0.57223 | 0.00000 | 0.00322 | 0.01066 | 0.00066 | 0.00714 |
| **Fam132a** | 4.63656 | 0.04954 | 1.59924 | 4.43340 | 0.00000 | 5.22373 | 2.52918 | 5.39670 | 3.12741 | 1.52758 | 4.31761 | 1.55593 | 2.33187 | 0.19295 | 0.00000 | 3.25125 |

| | | | | | | | | | | | | | | | | |
|---|---|---|---|---|---|---|---|---|---|---|---|---|---|---|---|---|
| **Fam132b** | 0.09588 | 0.00678 | 0.92515 | 0.28371 | 0.00528 | 0.30529 | 0.00543 | 2.66721 | 0.01127 | 0.00000 | 0.96645 | 0.00000 | 1.86045 | 0.40518 | 0.00000 | 2.15830 |
| **Fam133b** | 5.86353 | 7.91152 | 5.06059 | 4.51233 | 5.18457 | 0.00570 | 0.12354 | 5.27980 | 5.31983 | 1.49127 | 1.03440 | 0.80051 | 0.96503 | 0.09515 | 3.76917 | 1.23546 |
| **Fam134a** | 6.84801 | 2.97043 | 5.35236 | 6.92545 | 0.04528 | 4.46227 | 0.30085 | 11.04492 | 6.74809 | 12.97189 | 9.91491 | 1.63449 | 5.50530 | 2.06927 | 13.80464 | 1.82680 |
| **Fam134b** | 0.01471 | 0.00812 | 0.02051 | 0.01022 | 0.00930 | 0.00629 | 0.00748 | 0.01350 | 0.01219 | 0.01368 | 0.01328 | 0.00831 | 0.01249 | 0.00996 | 0.04065 | 0.00739 |
| **Fam134c** | 0.81547 | 2.74194 | 0.98769 | 2.54104 | 0.02870 | 0.15774 | 0.28123 | 3.99006 | 4.61632 | 3.22639 | 4.61596 | 0.11030 | 1.68876 | 4.80230 | 7.95725 | 1.13542 |
| **Fam135a** | 0.00000 | 0.24549 | 0.35475 | 1.94941 | 0.12081 | 0.01043 | 0.00615 | 0.10304 | 1.36439 | 1.16987 | 0.36523 | 6.90489 | 0.36766 | 1.75106 | 4.84854 | 0.00611 |
| **Fam136a** | 11.83453 | 8.44826 | 12.01766 | 0.00000 | 8.99567 | 9.08332 | 5.49978 | 8.01608 | 0.00000 | 2.22030 | 1.92747 | 0.14873 | 10.50827 | 3.36689 | 0.00000 | 6.06305 |
| **Fam13b** | 1.09077 | 4.08003 | 0.45044 | 1.13015 | 1.96959 | 0.00750 | 0.02426 | 1.94723 | 1.42468 | 1.81352 | 0.00000 | 0.09375 | 1.39180 | 5.91737 | 1.95147 | 0.01075 |
| **Fam13c** | 0.01379 | 0.00765 | 0.01086 | 0.05464 | 0.00940 | 0.00649 | 0.00734 | 0.01119 | 0.02105 | 0.00483 | 0.01129 | 0.03625 | 0.01289 | 0.01113 | 0.66800 | 0.00736 |
| **Fam149b** | 3.65658 | 0.16780 | 1.42493 | 3.15725 | 0.25665 | 0.02378 | 1.55093 | 1.32186 | 1.04573 | 0.00000 | 0.00000 | 0.12778 | 0.28767 | 0.77711 | 1.24346 | 0.04408 |
| **Fam151a** | 1.44470 | 0.00380 | 1.90797 | 0.00250 | 1.00255 | 0.49969 | 1.65071 | 4.00227 | 0.00765 | 1.38971 | 0.00000 | 0.00000 | 3.58019 | 0.14981 | 0.20370 | 1.99438 |
| **Fam159a** | 0.07473 | 0.00805 | 0.05288 | 0.00940 | 0.00782 | 0.00580 | 0.00732 | 0.02092 | 0.98630 | 0.20529 | 0.21916 | 0.00515 | 0.62227 | 0.01096 | 0.08212 | 0.00677 |
| **Fam159b** | 2.54151 | 0.00545 | 0.28792 | 0.20501 | 0.02718 | 0.00296 | 0.00461 | 0.00000 | 0.09854 | 0.72069 | 2.25186 | 0.20129 | 0.15717 | 0.45683 | 0.18441 | 0.45861 |
| **Fam160a1** | 0.00046 | 0.23861 | 1.26839 | 5.45167 | 3.60811 | 0.00875 | 0.00702 | 0.10560 | 0.95211 | 0.16330 | 0.29080 | 0.02210 | 1.04747 | 0.00000 | 1.08409 | 0.00521 |
| **Fam160a2** | 2.65581 | 2.37921 | 2.05820 | 1.18461 | 0.28977 | 0.00395 | 0.67467 | 0.25735 | 0.65973 | 1.54472 | 0.59132 | 0.08649 | 2.78966 | 0.01996 | 0.06131 | 3.24679 |
| **Fam160b1** | 0.28874 | 1.16342 | 1.60527 | 1.08757 | 0.03795 | 0.00668 | 0.33802 | 0.41589 | 0.31524 | 0.12552 | 0.30967 | 6.27851 | 0.61444 | 1.56304 | 0.27731 | 0.01243 |
| **Fam160b2** | 0.01689 | 2.97215 | 0.82595 | 2.14498 | 5.67705 | 3.97741 | 0.94659 | 9.25988 | 1.34715 | 4.84363 | 7.03727 | 0.12258 | 9.54681 | 0.31964 | 2.38121 | 2.63110 |
| **Fam161a** | 0.01901 | 2.85553 | 2.05994 | 0.39935 | 0.01562 | 0.24883 | 0.01589 | 0.00000 | 5.73565 | 1.79352 | 0.49332 | 0.22274 | 0.03325 | 0.00001 | 0.06612 | 0.01866 |
| **Fam161b** | 0.11347 | 0.01737 | 0.19390 | 1.53185 | 0.00735 | 0.01675 | 0.01627 | 0.52846 | 4.58932 | 4.05786 | 3.59154 | 2.57106 | 1.14611 | 0.50550 | 1.79357 | 0.00528 |
| **Fam162a** | 2.37818 | 0.00333 | 0.14105 | 0.22505 | 4.40477 | 0.15517 | 0.04012 | 0.02214 | 0.00201 | 0.55851 | 0.01308 | 0.04672 | 0.06234 | 0.18618 | 0.00000 | 1.77747 |
| **Fam163a** | 0.01366 | 0.00807 | 0.01211 | 0.20502 | 0.00928 | 0.00660 | 0.00753 | 0.01391 | 0.58364 | 0.01219 | 0.01021 | 0.00710 | 0.01281 | 0.01108 | 0.00976 | 0.00735 |
| **Fam166a** | 0.01115 | 0.00848 | 0.01039 | 0.05536 | 0.00949 | 0.00674 | 0.00816 | 0.10850 | 0.19130 | 0.01077 | 0.00997 | 0.00814 | 0.00857 | 0.00876 | 0.05862 | 0.00735 |
| **Fam167a** | 0.01260 | 0.00754 | 0.00760 | 2.46832 | 0.00941 | 0.00636 | 0.00711 | 0.00876 | 1.23070 | 0.01499 | 0.01606 | 0.00153 | 0.01268 | 1.04110 | 0.11014 | 0.00676 |
| **Fam167b** | 0.01252 | 0.00774 | 0.00972 | 0.29839 | 0.00922 | 0.00675 | 0.00714 | 0.14565 | 0.11217 | 0.26337 | 0.01297 | 0.00621 | 0.00813 | 0.01042 | 0.00363 | 0.00773 |
| **Fam168a** | 9.39740 | 1.24005 | 5.57577 | 4.79725 | 8.83922 | 0.00000 | 1.35889 | 3.15434 | 4.62238 | 4.40093 | 5.05761 | 3.92362 | 1.35097 | 9.63171 | 4.34296 | 4.70015 |
| **Fam168b** | 6.38183 | 1.72686 | 7.64026 | 9.81599 | 7.12448 | 4.48428 | 3.15259 | 2.76614 | 6.04701 | 6.75860 | 5.41305 | 10.44690 | 2.17240 | 0.44950 | 6.38391 | 1.95960 |
| **Fam169a** | 0.03842 | 0.01758 | 0.00000 | 1.37736 | 0.01888 | 3.55813 | 1.96656 | 0.11167 | 3.51071 | 1.13540 | 2.51580 | 0.74188 | 0.00000 | 3.38861 | 3.11403 | 0.00504 |
| **Fam169b** | 1.99846 | 0.22909 | 0.23179 | 8.33984 | 0.10696 | 0.18253 | 0.52166 | 1.12519 | 12.18795 | 7.22525 | 1.53828 | 0.62585 | 0.34859 | 2.65309 | 5.93888 | 1.10804 |
| **Fam171a1** | 0.01013 | 0.00748 | 0.01654 | 0.00474 | 0.00937 | 0.00625 | 1.60105 | 0.02362 | 0.18711 | 0.68971 | 0.00936 | 0.00463 | 0.00958 | 0.11357 | 0.00000 | 0.00718 |
| **Fam171a2** | 0.01388 | 0.00812 | 0.00865 | 0.00920 | 0.00908 | 0.00691 | 0.00755 | 0.00524 | 0.11378 | 0.00634 | 0.43662 | 0.02841 | 0.00809 | 0.00874 | 0.01093 | 0.00720 |
| **Fam172a** | 1.06815 | 0.93548 | 0.22092 | 5.53475 | 3.75577 | 0.00713 | 0.00566 | 1.88125 | 2.22467 | 0.17137 | 0.00038 | 0.00000 | 0.00380 | 1.62132 | 0.40221 | 0.00509 |
| **Fam173a** | 1.92161 | 0.01519 | 0.71748 | 0.17459 | 0.29749 | 2.53122 | 0.11624 | 4.34513 | 0.00590 | 2.40103 | 1.49532 | 0.01419 | 2.30254 | 0.00561 | 0.03035 | 2.20237 |
| **Fam173b** | 0.09296 | 0.97724 | 3.88346 | 0.18717 | 0.11160 | 2.25588 | 0.22207 | 3.67184 | 0.00753 | 0.72445 | 1.15344 | 0.25609 | 3.05556 | 0.01452 | 0.19554 | 0.09645 |
| **Fam174a** | 0.05838 | 0.01950 | 0.19638 | 0.33488 | 0.01429 | 0.00618 | 0.00732 | 0.02048 | 0.00436 | 0.00001 | 0.00745 | 0.06348 | 0.07581 | 2.12133 | 0.00676 | 0.00770 |
| **Fam174b** | 0.02154 | 0.00724 | 0.00618 | 0.47179 | 0.00851 | 0.00710 | 0.00704 | 0.85533 | 0.01363 | 0.02685 | 0.00684 | 0.00525 | 0.00567 | 0.01465 | 0.10179 | 0.00749 |
| **Fam175a** | 5.43913 | 2.96672 | 0.97347 | 1.91139 | 3.86214 | 0.00000 | 3.86995 | 2.58558 | 4.30034 | 5.42205 | 2.34229 | 0.16771 | 0.46750 | 1.98120 | 3.79265 | 0.00585 |
| **Fam175b** | 1.95513 | 8.59358 | 3.62404 | 5.84047 | 5.73695 | 1.81016 | 2.08638 | 4.40830 | 3.25047 | 6.97496 | 3.33530 | 4.68514 | 0.76196 | 9.51092 | 7.14691 | 1.99435 |
| **Fam178a** | 1.42094 | 2.14603 | 3.25523 | 8.06108 | 0.04923 | 0.81336 | 6.80664 | 0.05874 | 8.12366 | 4.46976 | 2.13793 | 1.69400 | 1.75640 | 5.25770 | 10.57514 | 2.86138 |
| **Fam178b** | 0.44297 | 0.00603 | 0.85508 | 3.28584 | 0.01386 | 0.00489 | 1.79979 | 0.00000 | 2.80350 | 0.75458 | 0.69881 | 0.14529 | 2.05056 | 0.00000 | 0.27937 | 0.00481 |
| **Fam179b** | 0.07062 | 2.61977 | 0.10788 | 8.41563 | 0.07402 | 0.00373 | 2.25371 | 1.70609 | 6.14828 | 2.11654 | 0.13759 | 8.78665 | 0.78416 | 8.29061 | 11.97908 | 0.13129 |
| **Fam183b** | 0.01298 | 0.00810 | 0.01388 | 0.00882 | 0.00990 | 0.00718 | 0.00765 | 0.01392 | 0.01204 | 0.41785 | 0.01418 | 0.00687 | 0.08761 | 0.01152 | 0.00994 | 0.00791 |
| **Fam184a** | 0.01086 | 0.00846 | 0.00769 | 2.94928 | 0.00945 | 0.00624 | 0.00806 | 0.13477 | 0.27811 | 0.11032 | 0.01239 | 0.00373 | 0.00830 | 0.00163 | 1.83769 | 0.00717 |
| **Fam184b** | 0.01782 | 0.00790 | 0.01205 | 0.01025 | 0.00897 | 0.00667 | 0.00749 | 0.01392 | 0.01247 | 0.02408 | 0.01318 | 0.00757 | 0.01287 | 0.00863 | 0.03968 | 0.00753 |
| **Fam185a** | 0.60444 | 2.49938 | 0.35103 | 1.97286 | 2.29816 | 0.39170 | 4.07096 | 0.00278 | 0.90364 | 0.00703 | 0.25980 | 0.01632 | 0.96278 | 0.00000 | 0.00000 | 0.00443 |
| **Fam187b** | 0.00776 | 0.00788 | 0.01188 | 0.27276 | 0.00766 | 0.00648 | 0.00733 | 0.01219 | 0.00482 | 0.11494 | 0.02469 | 0.00715 | 0.00840 | 0.01036 | 0.00672 | 1.34260 |
| **Fam188a** | 0.47613 | 1.33577 | 0.00000 | 8.02085 | 6.67177 | 4.85141 | 0.97548 | 0.85504 | 0.71204 | 3.59480 | 1.65673 | 0.07209 | 1.08246 | 6.91254 | 1.30627 | 0.86466 |
| **Fam188b** | 0.01375 | 0.00816 | 0.01232 | 0.11771 | 0.00927 | 0.00660 | 0.00755 | 0.01370 | 0.00993 | 0.03860 | 0.01301 | 0.00804 | 0.01222 | 0.01115 | 0.00860 | 0.00725 |
| **Fam189a2** | 8.63930 | 0.00000 | 0.00459 | 0.45883 | 2.06885 | 0.00974 | 0.00000 | 0.04555 | 0.00000 | 1.46806 | 0.53344 | 0.04965 | 0.02659 | 0.16537 | 0.09737 | 3.13441 |
| **Fam189b** | 0.73972 | 0.00686 | 0.87196 | 3.01290 | 0.01275 | 0.00447 | 0.00629 | 0.03374 | 2.30163 | 0.99724 | 0.49274 | 0.00713 | 0.22232 | 0.00000 | 0.07057 | 0.98980 |
| **Fam192a** | 8.85975 | 5.03721 | 7.80650 | 9.66907 | 11.20685 | 7.76618 | 6.06817 | 9.64214 | 10.14118 | 12.58336 | 2.52019 | 1.99257 | 7.28786 | 2.37620 | 6.68785 | 5.13804 |
| **Fam193a** | 2.05574 | 4.39228 | 0.71988 | 3.99238 | 0.00764 | 1.63854 | 3.00087 | 0.59370 | 1.71768 | 1.70821 | 0.01382 | 0.03439 | 1.14191 | 0.01564 | 2.36075 | 0.31702 |
| **Fam193b** | 3.10819 | 0.00269 | 0.51835 | 0.00507 | 2.52688 | 3.09728 | 1.41379 | 0.40618 | 0.02261 | 4.60080 | 0.12501 | 0.25130 | 2.33926 | 0.01160 | 0.01724 | 0.68984 |
| **Fam195a** | 2.36186 | 0.01214 | 1.31773 | 5.19511 | 1.24538 | 0.03618 | 0.48380 | 1.68275 | 2.09216 | 17.04657 | 15.50368 | 0.22414 | 12.33376 | 0.01278 | 0.19362 | 0.02503 |
| **Fam195b** | 1.36160 | 0.55846 | 1.71627 | 0.72344 | 0.00331 | 0.36156 | 0.56445 | 0.77981 | 0.86862 | 1.79824 | 0.47094 | 0.00005 | 0.94483 | 0.00190 | 0.26188 | 0.03358 |
| **Fam196a** | 0.01323 | 0.00752 | 0.01010 | 0.05499 | 0.00939 | 0.00661 | 0.26573 | 0.01095 | 0.00887 | 0.00779 | 0.01177 | 0.00674 | 0.01170 | 0.00609 | 0.28463 | 0.00732 |
| **Fam198b** | 0.04131 | 0.02903 | 1.49503 | 0.08636 | 0.14247 | 1.37021 | 0.03941 | 0.02762 | 0.11603 | 0.54129 | 0.27243 | 0.05337 | 0.79845 | 0.17324 | 0.00319 | 1.73404 |
| **Fam199x** | 0.00904 | 0.00735 | 0.00280 | 16.79644 | 0.00948 | 0.00444 | 0.66080 | 0.00415 | 2.48801 | 0.00000 | 0.12906 | 0.01176 | 0.45740 | 0.06960 | 4.23263 | 0.00395 |
| **Fam19a4** | 0.01365 | 0.00784 | 0.01177 | 0.00313 | 0.00937 | 0.00645 | 0.00741 | 0.01182 | 0.74249 | 0.00417 | 0.00676 | 0.00571 | 0.01276 | 0.00633 | 0.47099 | 0.00726 |
| **Fam19a5** | 0.00000 | 0.00648 | 0.00436 | 0.28600 | 3.47372 | 0.00824 | 0.01995 | 0.64249 | 0.03349 | 0.00438 | 0.00963 | 0.00437 | 0.03027 | 0.01412 | 0.00000 | 0.00741 |
| **Fam204a** | 4.03866 | 1.39188 | 0.78735 | 1.53064 | 2.06586 | 0.05968 | 0.02359 | 1.22700 | 3.03720 | 1.38588 | 1.49896 | 6.50572 | 0.10581 | 3.68097 | 1.62247 | 0.10665 |
| **Fam206a** | 7.28636 | 0.48857 | 4.83349 | 0.50129 | 0.07546 | 0.07976 | 0.85598 | 2.57657 | 0.08274 | 4.17323 | 1.45631 | 2.81491 | 1.40450 | 0.07804 | 2.41819 | 3.42878 |
| **Fam207a** | 3.29549 | 0.25521 | 0.79770 | 4.88594 | 3.00103 | 1.84128 | 0.90985 | 3.44153 | 9.69795 | 10.19534 | 4.13759 | 0.53667 | 3.52776 | 1.24979 | 2.15732 | 3.07328 |

| | | | | | | | | | | | | | | | | |
|---|---|---|---|---|---|---|---|---|---|---|---|---|---|---|---|---|
| **Fam208a** | 1.43418 | 4.34118 | 2.61477 | 6.91393 | 3.49928 | 0.59867 | 1.80525 | 0.02451 | 2.52049 | 6.53769 | 3.58064 | 10.02845 | 2.18807 | 10.11789 | 6.13974 | 2.00263 |
| **Fam208b** | 2.54147 | 0.02351 | 1.28390 | 2.11168 | 1.32314 | 2.65503 | 1.18505 | 3.36573 | 7.99731 | 5.58822 | 0.66929 | 3.19139 | 2.99570 | 13.15688 | 9.95749 | 0.77116 |
| **Fam209** | 0.01339 | 0.00787 | 0.01073 | 1.42208 | 0.00927 | 0.00648 | 0.00744 | 0.01197 | 2.36001 | 0.00000 | 0.00000 | 0.01419 | 0.01363 | 0.09247 | 0.02844 | 0.00709 |
| **Fam20b** | 1.54211 | 2.32060 | 1.41364 | 2.53034 | 0.15012 | 1.61086 | 0.08099 | 2.49073 | 1.39523 | 1.20280 | 0.58194 | 0.20173 | 0.67807 | 4.01459 | 5.67200 | 1.13038 |
| **Fam21** | 8.09221 | 5.02330 | 3.67694 | 9.11926 | 6.60890 | 8.25796 | 7.91445 | 9.71691 | 13.39236 | 15.42389 | 14.55230 | 13.10193 | 2.52520 | 3.61476 | 14.73232 | 3.66909 |
| **Fam210a** | 7.67495 | 1.90003 | 0.64306 | 4.69910 | 2.17382 | 1.41261 | 0.00593 | 0.98772 | 9.60474 | 4.45058 | 3.30230 | 12.15503 | 3.40652 | 7.29089 | 8.21959 | 1.72441 |
| **Fam210b** | 1.04057 | 0.00761 | 0.00643 | 1.14117 | 0.02038 | 0.00643 | 0.00682 | 0.00892 | 0.76676 | 0.40432 | 0.00608 | 0.01352 | 0.11868 | 0.00639 | 0.43402 | 0.00640 |
| **Fam212a** | 0.00925 | 0.00735 | 0.00000 | 0.97948 | 0.00806 | 0.00666 | 0.00630 | 0.00000 | 7.65258 | 7.71955 | 2.00093 | 0.29456 | 0.00000 | 0.09200 | 1.15243 | 0.00621 |
| **Fam212b** | 0.00799 | 0.00727 | 0.08151 | 0.00897 | 0.00914 | 0.01861 | 0.98829 | 0.00000 | 0.01184 | 0.31810 | 0.41217 | 0.01622 | 0.40704 | 0.03255 | 0.00894 | 0.00708 |
| **Fam213a** | 0.00000 | 1.17937 | 7.71975 | 1.58566 | 1.07465 | 13.64644 | 4.75209 | 0.39406 | 0.07722 | 0.00680 | 0.00234 | 0.02397 | 3.20762 | 0.01380 | 0.18454 | 11.54813 |
| **Fam213b** | 0.01207 | 0.00806 | 0.38775 | 0.01079 | 0.02129 | 0.00934 | 0.00711 | 0.01170 | 0.01228 | 0.00543 | 0.01288 | 0.02763 | 0.68939 | 0.01467 | 0.00341 | 0.00704 |
| **Fam214a** | 0.00000 | 0.00486 | 0.00000 | 7.04249 | 0.00803 | 0.00315 | 0.20118 | 3.35744 | 11.53333 | 6.21513 | 5.49964 | 5.08854 | 0.00000 | 12.80572 | 8.82302 | 0.00375 |
| **Fam214b** | 1.60839 | 1.90754 | 2.12490 | 4.36527 | 1.07510 | 2.59263 | 4.96159 | 7.26120 | 11.36102 | 4.32395 | 9.29265 | 3.28446 | 0.12540 | 8.77026 | 6.35825 | 0.04976 |
| **Fam216a** | 5.02501 | 0.00000 | 1.17953 | 6.78554 | 2.65176 | 0.01135 | 0.00011 | 0.59725 | 9.86358 | 2.71074 | 1.75344 | 0.04931 | 2.34438 | 0.02546 | 0.92608 | 3.21663 |
| **Fam217a** | 0.06669 | 0.00730 | 0.91393 | 0.01020 | 0.00642 | 0.00557 | 0.40201 | 0.00939 | 0.01214 | 0.01563 | 0.02815 | 0.00701 | 0.00818 | 0.01521 | 0.00000 | 0.00694 |
| **Fam219a** | 0.01980 | 0.00744 | 1.30651 | 0.06241 | 0.00657 | 0.00639 | 0.11565 | 0.00000 | 0.01061 | 0.00000 | 0.44679 | 0.00340 | 0.00428 | 0.00500 | 0.54296 | 0.00654 |
| **Fam219b** | 5.15249 | 1.53282 | 1.33300 | 0.97564 | 0.02002 | 0.03045 | 0.14835 | 1.82550 | 0.33645 | 1.68657 | 1.81845 | 0.00000 | 1.95340 | 0.30322 | 0.47043 | 0.05140 |
| **Fam220a** | 2.27654 | 3.08855 | 1.95345 | 13.60500 | 0.02054 | 0.24052 | 2.85384 | 1.74037 | 4.21875 | 2.56515 | 1.37259 | 0.64676 | 2.12727 | 0.85661 | 2.73765 | 0.26337 |
| **Fam221a** | 2.22497 | 0.04757 | 0.95814 | 0.42589 | 0.00342 | 0.06045 | 1.58066 | 1.79847 | 0.13014 | 0.57660 | 0.64250 | 0.13748 | 1.86244 | 0.00350 | 0.51962 | 0.01829 |
| **Fam222a** | 0.38186 | 0.00857 | 0.00490 | 1.17968 | 0.01556 | 0.00707 | 0.00728 | 0.01106 | 4.56829 | 0.07560 | 0.00000 | 0.00000 | 0.01273 | 0.00788 | 0.00835 | 0.00670 |
| **Fam222b** | 2.23520 | 3.68820 | 2.72497 | 8.72440 | 0.01756 | 3.24971 | 4.55801 | 0.09797 | 5.54187 | 4.70343 | 1.97545 | 0.31169 | 1.72280 | 0.00000 | 5.65019 | 0.67995 |
| **Fam227a** | 0.12500 | 0.03958 | 0.13784 | 1.79419 | 0.01089 | 0.07188 | 0.03668 | 0.03890 | 4.07424 | 0.56098 | 0.50886 | 0.02487 | 0.17888 | 2.57426 | 0.34888 | 0.06796 |
| **Fam228b** | 0.00921 | 0.00772 | 0.00000 | 1.40714 | 0.00900 | 0.01426 | 0.00637 | 0.26487 | 0.56458 | 1.65395 | 1.48422 | 0.01213 | 0.55786 | 0.57368 | 0.82700 | 0.00661 |
| **Fam234a** | 10.44205 | 3.98725 | 14.47014 | 1.59922 | 8.20320 | 1.65218 | 8.11806 | 21.58951 | 4.67452 | 12.91600 | 7.80813 | 1.00603 | 16.64607 | 0.07180 | 6.37804 | 4.41745 |
| **Fam234b** | 0.01270 | 0.00761 | 0.01163 | 0.00438 | 0.00887 | 0.00642 | 0.00729 | 0.01045 | 0.01061 | 1.65805 | 0.00938 | 0.01100 | 0.01088 | 0.00513 | 0.58668 | 0.00751 |
| **Fam25c** | 4.60469 | 0.38367 | 0.03461 | 0.00807 | 5.46461 | 0.00731 | 0.03391 | 0.00000 | 0.01033 | 0.00451 | 0.01084 | 0.05017 | 0.07279 | 0.00984 | 0.01070 | 2.37416 |
| **Fam26f** | 0.01323 | 0.00803 | 0.00671 | 1.57934 | 0.00887 | 0.00680 | 0.00719 | 0.00000 | 3.67771 | 0.21573 | 0.68719 | 0.00000 | 0.00688 | 0.03386 | 0.02214 | 0.00691 |
| **Fam32a** | 15.90144 | 6.55237 | 16.62937 | 8.36536 | 8.74701 | 8.20552 | 9.47854 | 16.21159 | 12.47535 | 14.98019 | 7.72049 | 8.06138 | 13.15920 | 4.44896 | 12.58184 | 4.51505 |
| **Fam35a** | 2.32388 | 0.77599 | 1.90861 | 0.61976 | 0.59518 | 0.00459 | 0.00500 | 0.68541 | 1.34885 | 0.07016 | 0.00000 | 1.62176 | 0.00000 | 1.20866 | 0.68066 | 0.00329 |
| **Fam3a** | 0.01711 | 0.00693 | 2.25763 | 0.96064 | 0.01488 | 0.73998 | 0.74425 | 2.49845 | 1.53430 | 0.92813 | 0.86763 | 0.00071 | 3.28480 | 0.53518 | 2.08918 | 0.00453 |
| **Fam3b** | 0.36323 | 0.00678 | 1.81219 | 0.08378 | 0.00397 | 0.00742 | 0.31176 | 0.00000 | 0.01083 | 0.02524 | 0.60587 | 0.00303 | 0.00897 | 0.01326 | 0.02082 | 0.00590 |
| **Fam3c** | 6.35203 | 3.65788 | 4.29456 | 3.81656 | 10.00567 | 2.30510 | 0.17794 | 1.34165 | 1.19078 | 2.88469 | 2.90842 | 0.49926 | 3.82009 | 2.81166 | 1.25769 | 1.32873 |
| **Fam43a** | 0.01319 | 0.00781 | 0.01210 | 0.19059 | 0.00889 | 0.00642 | 0.00723 | 0.00986 | 0.12250 | 1.09821 | 0.01360 | 0.00554 | 0.01039 | 0.01790 | 0.00302 | 0.00705 |
| **Fam43b** | 0.29638 | 0.00746 | 0.00416 | 6.01013 | 0.00723 | 0.00603 | 0.00697 | 0.00970 | 10.04636 | 0.00000 | 0.00000 | 0.00000 | 0.01260 | 0.02534 | 0.20100 | 0.00521 |
| **Fam45a** | 0.26833 | 6.83799 | 6.63924 | 5.94305 | 0.21196 | 5.12083 | 3.67420 | 6.19096 | 3.71641 | 9.22291 | 10.13634 | 9.34025 | 8.47880 | 17.98444 | 16.34190 | 0.02999 |
| **Fam46a** | 1.85770 | 0.00534 | 0.00000 | 0.12004 | 0.00550 | 0.00571 | 0.00577 | 0.00358 | 0.12986 | 0.05963 | 0.00878 | 0.02081 | 0.01861 | 0.00646 | 1.40234 | 0.00576 |
| **Fam46b** | 0.00747 | 0.00755 | 0.00505 | 4.54069 | 0.00883 | 0.00576 | 0.00639 | 0.00000 | 8.67925 | 0.57359 | 0.43945 | 0.00000 | 1.06725 | 0.00000 | 0.73356 | 0.00560 |
| **Fam46c** | 5.64450 | 5.73743 | 0.00000 | 3.51570 | 5.53371 | 0.01775 | 1.69362 | 10.54423 | 0.13556 | 2.10963 | 8.75073 | 0.41202 | 1.37235 | 0.00000 | 1.13544 | 0.00000 |
| **Fam47e** | 0.14532 | 0.00756 | 0.00773 | 0.21282 | 0.00764 | 0.00684 | 0.00726 | 0.00920 | 0.57822 | 0.86385 | 0.01962 | 0.00501 | 0.01119 | 0.01251 | 0.00354 | 0.00708 |
| **Fam49a** | 0.00957 | 0.00681 | 0.22447 | 2.84325 | 0.00843 | 0.00620 | 0.00675 | 0.01024 | 0.53557 | 0.00782 | 0.00000 | 4.22304 | 0.01111 | 0.00028 | 0.72110 | 0.00651 |
| **Fam49b** | 5.20410 | 3.06144 | 4.27138 | 0.00000 | 2.53760 | 2.33882 | 7.94594 | 0.00000 | 0.69982 | 3.49130 | 2.04394 | 7.91778 | 0.52207 | 1.76042 | 1.19110 | 2.53075 |
| **Fam50a** | 11.95034 | 4.96026 | 11.79997 | 7.42591 | 3.26176 | 7.92334 | 2.43203 | 11.36071 | 4.98930 | 11.70976 | 8.17024 | 1.73797 | 12.02262 | 10.41414 | 6.58363 | 4.74317 |
| **Fam53a** | 1.78949 | 0.18439 | 0.06466 | 3.73733 | 3.27959 | 0.01366 | 1.90626 | 1.03902 | 6.47640 | 6.76188 | 0.55575 | 0.05299 | 2.07760 | 0.00000 | 0.00000 | 0.73446 |
| **Fam53b** | 1.56596 | 0.00624 | 0.20143 | 0.76756 | 0.02493 | 0.94598 | 0.00583 | 0.00418 | 0.31257 | 1.10719 | 0.00813 | 0.02239 | 0.01587 | 0.03386 | 0.00104 | 0.02033 |
| **Fam53c** | 2.28278 | 2.51148 | 0.00000 | 11.37255 | 0.03817 | 3.04131 | 5.14534 | 2.37039 | 4.24181 | 3.87831 | 2.99400 | 0.32059 | 0.36139 | 0.00285 | 2.51846 | 0.02279 |
| **Fam57a** | 2.33352 | 0.00396 | 0.13177 | 0.16453 | 1.77289 | 0.00641 | 0.00522 | 0.53595 | 0.00384 | 0.00295 | 0.00843 | 0.00354 | 0.08767 | 0.00813 | 0.19007 | 0.00463 |
| **Fam57b** | 0.06934 | 0.01650 | 0.24733 | 0.85267 | 0.01635 | 0.07484 | 0.63652 | 8.43158 | 0.63689 | 12.39465 | 1.39734 | 0.30341 | 5.25215 | 0.00000 | 0.00000 | 0.00452 |
| **Fam58b** | 2.11559 | 0.15572 | 5.34839 | 7.03271 | 3.10173 | 0.71733 | 0.50989 | 5.44647 | 15.89752 | 7.49511 | 0.84817 | 0.30588 | 2.52636 | 0.00000 | 0.76804 | 4.45724 |
| **Fam60a** | 8.48381 | 4.78080 | 4.97743 | 1.65134 | 4.63821 | 2.27357 | 4.71386 | 5.34721 | 1.76903 | 5.72971 | 4.13852 | 0.12158 | 0.73207 | 11.52494 | 8.20520 | 1.71870 |
| **Fam63a** | 7.63690 | 7.09790 | 10.27246 | 14.09944 | 8.58291 | 10.04893 | 3.72553 | 15.63291 | 13.16713 | 19.74768 | 12.70135 | 5.98357 | 9.42277 | 3.41241 | 10.08738 | 2.85006 |
| **Fam63b** | 0.00871 | 0.00748 | 0.20734 | 1.81065 | 0.00842 | 0.00837 | 0.04623 | 0.00901 | 0.11457 | 1.63207 | 0.00774 | 0.02034 | 0.46131 | 0.10724 | 0.89299 | 0.00678 |
| **Fam65a** | 1.36333 | 0.05352 | 1.42300 | 1.58300 | 1.45525 | 8.15702 | 4.19007 | 8.23098 | 1.36911 | 7.88949 | 6.26204 | 0.41881 | 7.17027 | 0.00000 | 1.18101 | 2.36518 |
| **Fam65b** | 0.06729 | 0.01810 | 0.24229 | 0.64526 | 0.01875 | 0.01293 | 0.04775 | 0.01008 | 0.28489 | 0.58228 | 0.59987 | 0.09695 | 0.08780 | 0.53646 | 0.12036 | 0.01410 |
| **Fam65c** | 0.01266 | 0.00811 | 0.01178 | 0.00651 | 0.00893 | 0.00665 | 0.00737 | 0.01673 | 0.01149 | 1.34535 | 0.01132 | 0.00511 | 0.01079 | 0.03295 | 0.02180 | 0.00734 |
| **Fam69a** | 1.20241 | 2.96454 | 0.00000 | 1.07118 | 0.02260 | 6.13973 | 4.73240 | 4.14996 | 2.37146 | 1.91339 | 0.61368 | 6.39861 | 2.84074 | 2.08552 | 4.09642 | 0.08023 |
| **Fam69b** | 0.05090 | 0.00752 | 0.41373 | 0.42936 | 0.00716 | 0.00901 | 0.00705 | 0.01001 | 0.51384 | 1.56729 | 0.00720 | 0.03075 | 0.00979 | 0.00532 | 0.09733 | 0.00711 |
| **Fam72a** | 0.75916 | 0.32030 | 0.26887 | 5.47500 | 1.10390 | 6.19863 | 0.01600 | 0.53529 | 15.76586 | 0.36145 | 0.81780 | 0.78088 | 0.50261 | 0.00000 | 0.39792 | 0.00201 |
| **Fam73a** | 0.02608 | 0.00631 | 1.18741 | 2.40337 | 2.48408 | 3.48060 | 0.00522 | 1.52546 | 2.42596 | 4.20985 | 2.88566 | 1.13625 | 0.87130 | 2.28198 | 4.26208 | 1.39256 |
| **Fam73b** | 0.19767 | 0.00895 | 1.78888 | 1.88154 | 0.01373 | 1.63659 | 0.00740 | 0.72239 | 0.53741 | 0.56169 | 0.88810 | 0.04374 | 0.00000 | 0.00642 | 0.15978 | 0.15105 |
| **Fam76a** | 5.05546 | 0.10030 | 2.96289 | 5.34698 | 0.02826 | 6.20164 | 4.41865 | 7.32766 | 5.02348 | 15.47614 | 8.98924 | 6.66987 | 4.40414 | 7.38018 | 15.26006 | 1.53018 |
| **Fam76b** | 0.04749 | 2.25609 | 1.82641 | 3.65404 | 0.00583 | 1.50177 | 0.69252 | 0.46526 | 6.62817 | 2.67994 | 1.00342 | 2.15363 | 1.80528 | 2.51386 | 5.24311 | 0.02473 |
| **Fam78a** | 0.11871 | 0.00802 | 0.04484 | 0.00959 | 0.00758 | 0.00576 | 0.00732 | 0.15830 | 0.01197 | 0.03755 | 0.01096 | 0.00792 | 0.00715 | 0.00703 | 0.12554 | 0.00697 |

| | | | | | | | | | | | | | | | | |
|---|---|---|---|---|---|---|---|---|---|---|---|---|---|---|---|---|
| **Fam83a** | 0.31086 | 0.04064 | 0.36638 | 0.52044 | 0.05841 | 0.60083 | 0.10594 | 0.05394 | 0.69741 | 0.01611 | 0.66262 | 0.05774 | 0.29669 | 0.40805 | 0.03617 | 0.13097 |
| **Fam83b** | 0.00923 | 0.00851 | 0.00809 | 0.34139 | 0.00962 | 0.00689 | 0.00820 | 0.58056 | 0.25053 | 0.07108 | 0.00725 | 0.00803 | 0.00551 | 0.01786 | 0.00000 | 0.00710 |
| **Fam83d** | 1.19719 | 0.65249 | 0.99746 | 2.75578 | 0.27319 | 0.00376 | 0.01640 | 1.46727 | 7.63572 | 0.15730 | 0.75635 | 1.02059 | 0.25099 | 0.00000 | 1.31677 | 0.02018 |
| **Fam83e** | 0.00921 | 0.00753 | 0.31352 | 0.06583 | 0.00901 | 0.00915 | 0.00705 | 0.48724 | 0.02578 | 0.00460 | 0.04422 | 0.02485 | 0.00424 | 0.01300 | 0.00000 | 0.00730 |
| **Fam83g** | 0.99981 | 0.00720 | 0.05255 | 0.96516 | 0.01656 | 0.36241 | 0.00527 | 2.11382 | 0.07712 | 1.80969 | 2.24771 | 0.05367 | 0.59369 | 0.00000 | 0.12909 | 0.00558 |
| **Fam83h** | 1.09255 | 0.00526 | 0.00000 | 0.02777 | 0.03586 | 1.06561 | 2.39259 | 0.39786 | 0.01172 | 0.04634 | 1.33191 | 0.02303 | 0.82389 | 0.12553 | 0.00000 | 0.00577 |
| **Fam84a** | 0.01282 | 0.00794 | 0.01107 | 0.51419 | 0.00881 | 0.00643 | 0.00724 | 0.00933 | 0.38446 | 1.57545 | 0.02423 | 0.02331 | 0.01079 | 0.00474 | 0.40533 | 0.00736 |
| **Fam84b** | 0.00000 | 3.13261 | 0.83281 | 5.08688 | 0.01378 | 0.04696 | 0.02873 | 0.15117 | 1.59098 | 0.47829 | 0.03855 | 0.04595 | 1.69291 | 0.00000 | 3.24225 | 1.29915 |
| **Fam89a** | 0.00827 | 0.00902 | 0.01516 | 1.14533 | 0.00922 | 0.00716 | 0.00826 | 0.29791 | 2.20304 | 0.67109 | 0.00000 | 0.00261 | 0.03935 | 0.03425 | 0.00000 | 0.00678 |
| **Fam89b** | 1.70427 | 1.75481 | 0.00000 | 2.07212 | 0.12032 | 2.06596 | 0.00529 | 1.76121 | 4.94832 | 2.51414 | 0.74126 | 0.29098 | 0.00000 | 0.23179 | 0.27378 | 0.00400 |
| **Fam8a1** | 0.01314 | 0.00847 | 0.00526 | 1.37103 | 0.00987 | 0.01529 | 0.00744 | 0.00483 | 0.27729 | 0.01523 | 1.41148 | 0.00126 | 0.08201 | 0.00219 | 0.05550 | 0.00749 |
| **Fam90a1a** | 0.01358 | 0.00829 | 0.01179 | 1.11447 | 0.00927 | 0.00650 | 0.00750 | 0.01355 | 1.17869 | 0.00853 | 0.01784 | 0.00638 | 0.01264 | 0.01092 | 0.00260 | 0.00715 |
| **Fam91a1** | 4.68084 | 6.73643 | 5.06300 | 7.55979 | 6.95297 | 8.07855 | 6.36763 | 5.82327 | 7.19071 | 10.77427 | 8.20073 | 19.67966 | 2.83027 | 21.83782 | 10.68623 | 0.30406 |
| **Fam92a** | 0.96194 | 0.02889 | 1.00301 | 0.92622 | 0.20670 | 0.00806 | 1.04503 | 0.03337 | 0.44505 | 0.20255 | 0.02054 | 0.02104 | 1.09501 | 0.00885 | 0.00000 | 1.30150 |
| **Fam92b** | 0.01347 | 0.00779 | 0.01215 | 0.00811 | 0.00905 | 0.00661 | 0.00728 | 0.01049 | 0.01195 | 0.64346 | 0.01270 | 0.00578 | 0.01194 | 0.01149 | 0.00933 | 0.00778 |
| **Fam96a** | 28.14843 | 17.73567 | 27.45271 | 1.55749 | 10.28230 | 15.54554 | 15.31806 | 22.36568 | 2.21466 | 18.11556 | 14.99579 | 5.70737 | 23.40341 | 3.88257 | 8.06751 | 12.35802 |
| **Fam96b** | 10.42372 | 2.00600 | 6.07861 | 6.11306 | 4.82590 | 2.56469 | 4.33765 | 2.66651 | 2.34809 | 5.62976 | 6.69084 | 1.03613 | 8.05251 | 0.13019 | 0.00000 | 2.48153 |
| **Fam98a** | 8.80815 | 4.16373 | 3.07031 | 7.12922 | 5.07030 | 2.49990 | 2.98680 | 9.26442 | 4.92942 | 8.00704 | 9.50661 | 2.87743 | 6.76192 | 7.75316 | 6.21289 | 3.15252 |
| **Fam98b** | 4.01579 | 0.29487 | 1.64763 | 9.63457 | 0.16739 | 0.68709 | 1.09212 | 0.29461 | 11.59056 | 2.92077 | 10.80388 | 0.25834 | 3.40089 | 5.56287 | 6.03424 | 1.26962 |
| **Fam98c** | 0.05699 | 0.00727 | 0.38247 | 0.00559 | 1.26662 | 0.00573 | 0.00697 | 0.64171 | 0.01061 | 0.39836 | 0.32166 | 0.01184 | 0.66189 | 0.00720 | 0.02585 | 1.96441 |
| **Fan1** | 0.01355 | 0.00791 | 0.01232 | 0.08913 | 0.00946 | 0.00661 | 0.00751 | 0.01243 | 0.01098 | 0.06327 | 0.01199 | 0.01647 | 0.02563 | 0.00556 | 0.38976 | 0.01458 |
| **Fanca** | 0.57452 | 1.50909 | 0.88185 | 0.36000 | 0.00435 | 0.95383 | 2.07059 | 2.24014 | 0.16447 | 1.74089 | 0.00000 | 0.07473 | 1.68080 | 0.17335 | 0.92964 | 1.88481 |
| **Fancb** | 0.00000 | 3.36118 | 2.27966 | 1.03164 | 6.74810 | 0.00640 | 0.05671 | 1.16821 | 0.00000 | 1.95776 | 1.27428 | 0.75915 | 0.33608 | 2.24441 | 0.81127 | 1.44736 |
| **Fancc** | 3.61552 | 0.00321 | 1.05673 | 3.80879 | 0.30565 | 1.06427 | 1.48800 | 2.56570 | 0.36951 | 1.54079 | 0.59387 | 0.10987 | 0.00000 | 1.13581 | 4.62270 | 0.00208 |
| **Fancd2** | 4.03494 | 3.09639 | 2.87561 | 1.89983 | 0.01000 | 0.08461 | 0.96873 | 0.05199 | 0.70337 | 0.68217 | 1.67924 | 0.85651 | 4.59354 | 1.19477 | 6.00221 | 0.03500 |
| **Fancd2os** | 0.30643 | 0.00679 | 0.37421 | 3.69776 | 0.00655 | 0.00728 | 0.40076 | 0.12281 | 2.55205 | 0.80047 | 0.68406 | 0.04866 | 0.54623 | 1.19204 | 3.53163 | 0.00587 |
| **Fance** | 6.35151 | 0.06167 | 1.59680 | 1.90578 | 0.00000 | 6.16264 | 2.27761 | 1.89352 | 3.27729 | 3.51176 | 4.05815 | 0.11472 | 1.19464 | 0.04423 | 0.57923 | 3.00701 |
| **Fancf** | 0.02019 | 0.00811 | 0.00000 | 1.35930 | 0.00856 | 0.29881 | 0.00760 | 0.00303 | 3.67890 | 0.02220 | 0.66990 | 0.00000 | 0.00328 | 0.00377 | 0.00000 | 0.00666 |
| **Fancg** | 4.68482 | 0.00000 | 2.97549 | 1.92762 | 0.00901 | 4.86211 | 1.26231 | 1.71828 | 2.39374 | 0.54449 | 0.88091 | 0.06348 | 3.30133 | 0.01429 | 1.59887 | 0.01332 |
| **Fanci** | 2.63010 | 0.18655 | 2.68431 | 1.62626 | 4.08483 | 0.01916 | 2.18107 | 3.50516 | 1.09476 | 2.78451 | 1.09659 | 0.14638 | 1.56159 | 5.59504 | 2.83266 | 2.05646 |
| **Fancl** | 1.96124 | 4.27214 | 2.12618 | 0.00000 | 0.00994 | 0.01581 | 0.01382 | 4.33416 | 0.08023 | 3.86360 | 1.67372 | 0.92798 | 0.18029 | 0.67514 | 2.85339 | 3.41098 |
| **Fancm** | 6.98398 | 4.68283 | 0.31696 | 9.66752 | 0.02424 | 0.00000 | 2.15136 | 1.03735 | 13.44359 | 7.71749 | 7.12921 | 8.56725 | 0.76457 | 5.96719 | 8.58566 | 0.29910 |
| **Fank1** | 1.39319 | 2.52299 | 0.00000 | 4.81532 | 0.00499 | 0.26944 | 0.23887 | 0.00000 | 0.00000 | 0.26583 | 0.07087 | 0.13691 | 1.62045 | 0.00000 | 1.51445 | 0.91318 |
| **Far1** | 0.68242 | 2.42386 | 1.34405 | 4.47160 | 2.40300 | 2.85576 | 0.38641 | 0.98801 | 10.39774 | 0.00000 | 1.91109 | 0.14286 | 1.06217 | 3.61578 | 0.24509 | 0.39692 |
| **Far2** | 0.12121 | 0.00883 | 0.00379 | 0.01088 | 0.01536 | 0.00775 | 0.00809 | 0.27150 | 0.01153 | 0.00634 | 0.00979 | 0.00815 | 0.00659 | 0.02623 | 0.12238 | 0.00741 |
| **Farp1** | 0.04413 | 0.00818 | 1.88117 | 3.45995 | 0.01423 | 0.04377 | 0.00641 | 0.00657 | 2.71023 | 0.35278 | 0.18102 | 0.16328 | 1.51553 | 0.47985 | 2.30822 | 0.01227 |
| **Farp2** | 1.24696 | 0.00574 | 0.74403 | 0.24462 | 0.00409 | 0.00424 | 1.13601 | 1.51381 | 0.29646 | 0.98226 | 0.00000 | 0.07745 | 0.00449 | 0.75534 | 1.67106 | 0.00543 |
| **Fars2** | 0.45610 | 2.20015 | 3.39340 | 12.70944 | 1.98723 | 0.18958 | 0.00244 | 1.89533 | 0.00000 | 2.46946 | 1.03997 | 5.73106 | 3.93382 | 0.00000 | 1.24482 | 0.00829 |
| **Farsa** | 11.53106 | 5.10300 | 13.18465 | 2.55957 | 3.56571 | 0.43366 | 4.66347 | 12.63872 | 1.07091 | 11.13827 | 8.58084 | 1.87002 | 9.85730 | 4.23115 | 4.23299 | 2.09513 |
| **Farsb** | 20.33021 | 13.68696 | 17.11194 | 7.08924 | 16.77800 | 8.92927 | 13.65701 | 21.48948 | 6.81116 | 13.52227 | 14.50279 | 10.22837 | 19.41947 | 19.17117 | 11.60573 | 11.81955 |
| **Fasn** | 13.14685 | 6.66744 | 10.25928 | 3.08282 | 10.02227 | 2.10535 | 13.10758 | 22.69856 | 2.68220 | 21.85426 | 4.88851 | 2.45793 | 7.97263 | 1.50413 | 0.00000 | 10.77701 |
| **Fastk** | 6.76924 | 2.37041 | 5.37599 | 0.13367 | 2.44109 | 6.94531 | 5.88132 | 7.04560 | 0.40903 | 2.60628 | 1.22023 | 0.18221 | 8.70692 | 0.01336 | 0.00000 | 4.46875 |
| **Fastkd1** | 0.26152 | 0.10870 | 0.01151 | 1.45128 | 0.61784 | 0.00696 | 0.00633 | 0.72972 | 2.24692 | 0.50474 | 0.65757 | 0.17889 | 0.01970 | 10.73263 | 4.72031 | 0.00643 |
| **Fastkd2** | 5.97928 | 0.00000 | 6.52243 | 0.92323 | 3.11859 | 2.32414 | 0.67970 | 1.92932 | 0.14500 | 5.83983 | 2.76083 | 7.61967 | 4.90739 | 8.59473 | 3.77879 | 0.01618 |
| **Fastkd3** | 6.06736 | 2.47063 | 5.44824 | 12.00606 | 0.06220 | 1.14295 | 4.48618 | 2.18924 | 9.95491 | 5.65495 | 3.18200 | 12.43413 | 5.26107 | 0.00000 | 9.00153 | 1.43580 |
| **Fastkd5** | 1.69998 | 0.18708 | 2.15597 | 5.77644 | 0.01150 | 3.06768 | 4.15179 | 1.37199 | 5.93706 | 4.02864 | 0.61734 | 0.04345 | 1.05760 | 2.68032 | 4.52060 | 0.00028 |
| **Fat1** | 2.20755 | 1.79074 | 1.14013 | 2.37286 | 2.91651 | 0.25545 | 3.41167 | 1.87396 | 0.00000 | 0.04470 | 0.56695 | 0.20272 | 0.77504 | 0.72141 | 1.95622 | 0.04776 |
| **Fau** | 32.72855 | 4.67374 | 25.17728 | 18.30706 | 19.17026 | 9.64039 | 8.05121 | 18.43110 | 25.32398 | 34.23591 | 34.30803 | 4.58846 | 29.56085 | 0.17508 | 1.12184 | 18.16736 |
| **Faxc** | 0.00859 | 0.00785 | 0.00000 | 2.66319 | 0.00947 | 0.00622 | 0.00685 | 0.03365 | 1.95318 | 0.98272 | 0.58289 | 0.01804 | 0.00763 | 1.24371 | 1.91768 | 0.00665 |
| **Fbf1** | 0.06324 | 0.03702 | 1.35040 | 0.98464 | 0.00589 | 2.06881 | 0.01522 | 0.00000 | 0.70474 | 0.02258 | 0.76476 | 0.01784 | 2.48915 | 0.00433 | 0.57032 | 0.01180 |
| **Fbl** | 26.45791 | 14.09028 | 24.17513 | 19.17038 | 15.14633 | 9.03690 | 15.24944 | 27.17254 | 24.92524 | 27.66393 | 28.09752 | 5.00858 | 25.45051 | 7.49388 | 16.42940 | 14.11588 |
| **Fblim1** | 6.78176 | 10.70998 | 9.73944 | 3.98432 | 8.96947 | 2.45457 | 7.73238 | 14.44480 | 4.80342 | 10.62153 | 12.40953 | 2.41898 | 6.09064 | 1.92936 | 4.61723 | 4.99810 |
| **Fbln1** | 1.15869 | 2.29622 | 0.25287 | 0.05704 | 2.31708 | 2.88818 | 0.75545 | 0.84724 | 0.07492 | 0.56712 | 0.00000 | 0.01050 | 1.62331 | 0.06469 | 0.00860 | 5.55502 |
| **Fbln2** | 0.01385 | 0.00794 | 0.00969 | 0.24319 | 0.00935 | 0.00660 | 0.00730 | 0.01117 | 0.22616 | 0.00332 | 0.01037 | 0.14364 | 0.01306 | 0.00617 | 0.39626 | 0.00734 |
| **Fbln5** | 0.00459 | 0.00786 | 0.01097 | 0.01065 | 0.00667 | 0.00628 | 0.01752 | 0.01821 | 0.01242 | 0.00879 | 0.01208 | 0.02828 | 0.00522 | 0.01256 | 0.00992 | 3.41678 |
| **Fbln7** | 0.01001 | 0.00830 | 0.81432 | 0.00000 | 0.00741 | 0.00555 | 0.00748 | 0.01098 | 0.00857 | 2.92698 | 0.00615 | 0.02042 | 0.00803 | 0.06449 | 1.53805 | 0.00687 |
| **Fbn1** | 0.01366 | 0.00789 | 0.00969 | 0.00995 | 0.00919 | 0.00685 | 0.00749 | 0.01483 | 0.01213 | 0.15872 | 0.01169 | 0.17936 | 0.01275 | 0.01163 | 0.00762 | 0.00777 |
| **Fbp1** | 2.43238 | 0.00277 | 0.81491 | 0.31450 | 1.52560 | 0.00909 | 2.82525 | 0.14633 | 0.11792 | 0.00000 | 0.00950 | 0.01068 | 0.02994 | 0.06040 | 0.02603 | 3.06680 |
| **Fbp2** | 32.53176 | 20.72002 | 29.24078 | 0.26492 | 23.56631 | 15.81056 | 15.52719 | 24.04644 | 0.00000 | 11.66082 | 13.23383 | 0.96949 | 27.26825 | 7.43834 | 0.00000 | 18.93261 |
| **Fbrs** | 3.41081 | 0.14328 | 0.75274 | 6.76619 | 1.28264 | 0.23297 | 0.60660 | 4.28959 | 5.38378 | 1.92225 | 4.11649 | 0.07792 | 2.82748 | 0.42049 | 1.85753 | 4.71049 |
| **Fbrsl1** | 7.35149 | 3.49211 | 5.19838 | 13.28991 | 0.67911 | 2.20879 | 2.46969 | 9.91359 | 17.67880 | 18.86335 | 6.32549 | 1.62021 | 4.31080 | 1.23736 | 10.85226 | 2.53177 |
| **Fbxl12** | 1.65560 | 0.02606 | 0.14837 | 4.66702 | 0.01200 | 0.01074 | 0.00538 | 0.27853 | 7.74390 | 0.54777 | 0.53231 | 0.01591 | 0.39465 | 0.00629 | 0.66883 | 1.85661 |

| | | | | | | | | | | | | | | | | |
|---|---|---|---|---|---|---|---|---|---|---|---|---|---|---|---|---|
| **Fbxl12os** | 0.00915 | 0.00776 | 0.01286 | 1.77049 | 0.00943 | 0.08531 | 0.05793 | 0.10128 | 2.25788 | 0.03906 | 0.00000 | 0.04270 | 0.03573 | 0.30366 | 0.02933 | 0.03605 |
| **Fbxl13** | 0.01059 | 0.00771 | 0.00457 | 1.34203 | 0.00916 | 0.00663 | 0.00702 | 0.00973 | 0.00000 | 1.63707 | 0.10425 | 0.05200 | 0.01037 | 1.94579 | 0.00000 | 0.00712 |
| **Fbxl14** | 0.16410 | 0.01731 | 0.90074 | 3.05541 | 0.03062 | 2.07349 | 0.92325 | 1.45981 | 0.40517 | 1.71432 | 0.96400 | 0.06955 | 1.35118 | 0.03833 | 2.09276 | 0.83361 |
| **Fbxl15** | 0.01233 | 0.00721 | 0.01882 | 0.00461 | 0.98898 | 0.00755 | 0.00734 | 0.01398 | 0.01089 | 1.91021 | 0.01232 | 0.00000 | 0.27854 | 0.01304 | 0.00647 | 0.15009 |
| **Fbxl16** | 0.32305 | 0.00696 | 0.00000 | 0.00253 | 0.00816 | 0.05435 | 0.01613 | 0.65491 | 0.57557 | 0.42746 | 1.59095 | 0.02329 | 0.52022 | 0.00586 | 0.85178 | 0.01400 |
| **Fbxl17** | 1.75758 | 0.03486 | 1.57642 | 5.63951 | 0.09489 | 1.41976 | 0.32014 | 1.49763 | 3.97665 | 0.29234 | 0.30185 | 0.07467 | 0.06671 | 0.60224 | 1.15983 | 0.00324 |
| **Fbxl18** | 0.42768 | 0.00611 | 3.77009 | 0.67484 | 0.02125 | 0.00591 | 0.00515 | 4.95580 | 0.09737 | 3.77743 | 1.48609 | 0.04944 | 1.35064 | 0.00043 | 0.00000 | 2.33536 |
| **Fbxl19** | 1.56517 | 0.65101 | 0.72759 | 2.38195 | 2.41060 | 0.00469 | 0.00439 | 1.29750 | 1.73862 | 2.01148 | 1.35647 | 0.00711 | 1.83842 | 0.00000 | 0.13845 | 0.83876 |
| **Fbxl2** | 0.01653 | 0.00755 | 0.23946 | 0.78449 | 0.00883 | 0.00556 | 0.00723 | 0.01098 | 0.58335 | 0.00238 | 0.00790 | 0.02529 | 0.01029 | 0.04250 | 0.14231 | 0.00708 |
| **Fbxl20** | 6.57267 | 5.52788 | 7.81179 | 1.55116 | 5.72604 | 0.00866 | 7.33379 | 19.36763 | 6.05824 | 16.14780 | 14.37473 | 2.86054 | 4.23109 | 10.88964 | 17.49733 | 0.00071 |
| **Fbxl21** | 0.01347 | 0.00800 | 0.01219 | 0.17838 | 0.00934 | 0.00660 | 0.00745 | 0.01829 | 0.01713 | 0.02351 | 0.01389 | 0.00787 | 0.01234 | 0.01119 | 0.00737 | 0.01475 |
| **Fbxl22** | 0.00739 | 0.00798 | 0.01184 | 0.07677 | 0.00760 | 0.00638 | 0.00744 | 0.01387 | 0.24900 | 0.00480 | 0.01058 | 0.00786 | 0.00754 | 0.01366 | 0.00661 | 1.43833 |
| **Fbxl3** | 0.33374 | 3.02979 | 1.67469 | 2.10510 | 1.22603 | 0.03732 | 3.73052 | 0.93617 | 4.19416 | 0.15135 | 3.47035 | 5.89805 | 0.77964 | 9.89147 | 11.34090 | 1.05181 |
| **Fbxl4** | 0.96417 | 0.00612 | 1.99343 | 1.30927 | 0.39705 | 0.00432 | 0.00631 | 0.00313 | 1.16193 | 0.56828 | 0.02168 | 0.00577 | 0.00726 | 0.44645 | 2.10654 | 0.00551 |
| **Fbxl5** | 3.90540 | 0.00084 | 1.27178 | 12.25956 | 0.17662 | 0.00528 | 1.53786 | 0.00000 | 3.67115 | 7.25703 | 4.68046 | 2.07782 | 1.40824 | 1.19198 | 9.18775 | 2.64938 |
| **Fbxl6** | 3.54991 | 1.50441 | 2.52170 | 0.21198 | 1.09783 | 3.01437 | 7.57503 | 6.78147 | 0.26690 | 6.79577 | 0.07190 | 0.00000 | 3.46522 | 0.10198 | 1.35320 | 0.08539 |
| **Fbxl7** | 0.01381 | 0.00806 | 0.01008 | 0.28131 | 0.00914 | 0.00667 | 0.00754 | 0.00805 | 0.15911 | 0.00237 | 0.28611 | 0.00650 | 0.00908 | 0.00620 | 0.05533 | 0.00718 |
| **Fbxl8** | 0.55616 | 0.00798 | 0.01223 | 0.58391 | 0.01710 | 0.01418 | 0.00658 | 0.00702 | 0.19095 | 0.85353 | 0.01129 | 0.00482 | 0.28294 | 0.01055 | 0.00345 | 0.00680 |
| **Fbxo10** | 0.01133 | 0.00774 | 0.01033 | 0.02444 | 0.01878 | 0.00662 | 0.00731 | 0.04942 | 0.99422 | 0.16557 | 0.00441 | 0.03397 | 0.01093 | 0.00776 | 0.19075 | 0.00773 |
| **Fbxo11** | 3.36466 | 6.31004 | 4.93764 | 20.24733 | 3.33880 | 0.95313 | 8.00659 | 3.66945 | 17.40668 | 7.34665 | 3.71525 | 0.46424 | 2.84141 | 21.50520 | 15.42330 | 0.17574 |
| **Fbxo15** | 32.38671 | 25.23479 | 42.12146 | 26.64958 | 23.67793 | 17.39557 | 30.30802 | 53.53515 | 37.83159 | 61.09934 | 52.34212 | 33.96273 | 40.61745 | 62.46051 | 46.41962 | 17.51902 |
| **Fbxo16** | 0.00000 | 1.09671 | 0.42539 | 4.55975 | 0.00787 | 0.00844 | 0.00569 | 0.00000 | 5.60793 | 0.54895 | 1.78394 | 0.00000 | 1.58922 | 0.39626 | 0.00000 | 0.00520 |
| **Fbxo17** | 1.97108 | 1.79542 | 1.15415 | 2.28634 | 1.89681 | 0.03506 | 0.04494 | 1.03031 | 3.21161 | 0.67995 | 1.93906 | 0.01399 | 0.55028 | 1.55945 | 1.35467 | 0.02482 |
| **Fbxo18** | 4.44096 | 3.38008 | 4.10179 | 7.44756 | 0.79407 | 4.18471 | 3.56157 | 0.14571 | 3.51241 | 3.22572 | 2.10018 | 0.22491 | 1.96883 | 0.04801 | 6.75236 | 0.18508 |
| **Fbxo2** | 4.43044 | 0.00282 | 0.16961 | 1.27915 | 0.15304 | 0.01227 | 0.00339 | 0.50079 | 0.37746 | 0.48934 | 0.01931 | 0.11714 | 0.44080 | 0.00891 | 0.00000 | 0.00201 |
| **Fbxo21** | 8.41770 | 4.85329 | 6.21398 | 0.53403 | 2.15940 | 1.53878 | 2.48448 | 4.22584 | 0.56755 | 0.48655 | 0.00000 | 2.92749 | 0.96495 | 0.17733 | 0.27213 | 3.62094 |
| **Fbxo22** | 10.20105 | 2.07185 | 4.96822 | 3.25580 | 5.25644 | 0.31811 | 7.22755 | 5.93826 | 1.90436 | 12.27202 | 4.43176 | 1.31594 | 7.30750 | 1.69313 | 7.13536 | 5.37028 |
| **Fbxo25** | 1.62635 | 0.00614 | 0.87491 | 1.08137 | 0.43812 | 0.00426 | 0.31533 | 0.27670 | 0.32860 | 1.19716 | 0.04273 | 0.02200 | 0.07045 | 0.01933 | 0.08162 | 0.01280 |
| **Fbxo27** | 1.92882 | 0.00515 | 0.52094 | 0.19687 | 0.00871 | 1.67873 | 0.94580 | 0.00186 | 0.18376 | 0.49217 | 0.01787 | 0.01352 | 0.03530 | 0.01061 | 0.16652 | 0.51109 |
| **Fbxo28** | 2.70613 | 5.39900 | 2.56396 | 5.44471 | 0.61923 | 0.00257 | 1.03202 | 1.17579 | 7.20667 | 1.91225 | 0.70456 | 2.50247 | 2.71290 | 2.59142 | 3.99406 | 1.36580 |
| **Fbxo3** | 6.73238 | 8.28095 | 4.38224 | 0.09532 | 8.91237 | 0.00000 | 2.68229 | 0.46031 | 0.19367 | 0.00000 | 0.00002 | 0.06301 | 2.72212 | 0.02011 | 0.03400 | 2.14694 |
| **Fbxo30** | 1.60453 | 6.94486 | 2.33223 | 2.28348 | 0.72899 | 1.54127 | 0.00344 | 1.78365 | 2.08106 | 2.40985 | 0.00000 | 0.16609 | 0.21214 | 7.41125 | 2.10378 | 0.33485 |
| **Fbxo31** | 5.94498 | 0.08133 | 2.14489 | 0.08314 | 4.89949 | 2.14847 | 0.64541 | 1.01538 | 1.19537 | 19.42211 | 6.16020 | 0.69785 | 6.06687 | 0.25768 | 0.00000 | 4.03727 |
| **Fbxo32** | 0.11236 | 0.00774 | 0.15087 | 3.57687 | 0.02975 | 0.00592 | 0.00676 | 0.00000 | 2.35575 | 0.00000 | 1.32975 | 0.06732 | 0.03808 | 0.47009 | 0.05103 | 0.02800 |
| **Fbxo33** | 3.29299 | 1.36341 | 1.44825 | 20.77480 | 0.19045 | 0.75698 | 4.72003 | 5.35155 | 27.00068 | 9.11763 | 10.58999 | 10.53878 | 2.03685 | 14.43143 | 18.60615 | 0.71017 |
| **Fbxo34** | 2.89084 | 0.00190 | 0.18269 | 12.15629 | 0.60695 | 0.01307 | 0.00190 | 11.92344 | 10.02124 | 13.09863 | 4.84540 | 1.71104 | 1.20166 | 3.81178 | 12.39803 | 0.00000 |
| **Fbxo36** | 0.01595 | 0.00758 | 0.70449 | 5.06969 | 0.00735 | 0.00508 | 0.00664 | 0.00000 | 7.28293 | 2.03564 | 0.90295 | 0.06939 | 0.00000 | 1.67211 | 1.78770 | 0.00547 |
| **Fbxo38** | 5.30566 | 7.60602 | 1.89302 | 4.08085 | 3.03652 | 3.29622 | 4.30454 | 4.50619 | 8.22186 | 7.41807 | 3.87647 | 10.07307 | 2.83363 | 5.76619 | 2.50546 | 3.96670 |
| **Fbxo4** | 0.01303 | 0.00825 | 0.01175 | 0.16418 | 0.00915 | 0.00650 | 0.00739 | 0.02025 | 0.44843 | 0.45802 | 0.00796 | 0.00593 | 0.01068 | 0.03211 | 0.00284 | 0.00710 |
| **Fbxo41** | 0.01276 | 0.00770 | 0.29373 | 0.71972 | 0.00866 | 0.63154 | 0.00710 | 0.00865 | 0.24454 | 0.13191 | 0.07744 | 0.01568 | 0.00631 | 0.01107 | 0.00000 | 0.01426 |
| **Fbxo42** | 2.45083 | 0.14773 | 3.63385 | 9.88093 | 11.48841 | 5.56698 | 1.09486 | 5.55124 | 7.21878 | 5.42301 | 2.63256 | 1.13011 | 3.88836 | 0.80475 | 2.54735 | 5.18052 |
| **Fbxo43** | 0.02637 | 0.00781 | 0.19756 | 6.05369 | 0.02807 | 0.00624 | 0.00684 | 0.01268 | 2.65642 | 0.52621 | 0.42409 | 0.01548 | 0.11190 | 2.66103 | 3.34067 | 0.02027 |
| **Fbxo44** | 1.80758 | 0.00669 | 0.19766 | 0.63152 | 0.00814 | 1.65435 | 0.00658 | 0.26874 | 1.12226 | 0.59426 | 0.00000 | 0.01427 | 0.00000 | 0.00507 | 0.14803 | 0.00501 |
| **Fbxo45** | 0.18138 | 0.00435 | 1.39600 | 11.42505 | 0.30646 | 2.85687 | 1.65146 | 2.12786 | 11.92709 | 6.66831 | 2.67316 | 5.13382 | 0.00000 | 4.41938 | 13.08348 | 3.57759 |
| **Fbxo46** | 0.31839 | 0.00805 | 0.01274 | 0.50422 | 0.03512 | 0.01574 | 0.00669 | 0.00000 | 2.01841 | 0.36411 | 0.37056 | 0.07402 | 0.00863 | 0.53245 | 0.16567 | 0.01460 |
| **Fbxo47** | 0.04374 | 0.00717 | 0.00150 | 2.53327 | 0.00826 | 0.00631 | 0.00701 | 0.32630 | 1.88695 | 0.13231 | 0.02868 | 0.01669 | 0.00638 | 0.02128 | 1.87905 | 0.00680 |
| **Fbxo5** | 3.15523 | 6.42392 | 7.89740 | 2.77716 | 7.81604 | 4.09165 | 5.61084 | 10.08240 | 12.09784 | 7.86608 | 8.90776 | 6.28825 | 7.00391 | 8.33861 | 8.96925 | 3.43389 |
| **Fbxo6** | 19.78399 | 10.93930 | 15.80641 | 5.19835 | 8.38596 | 5.76804 | 6.49543 | 15.86795 | 14.41743 | 15.53300 | 6.44621 | 2.66729 | 14.85247 | 1.32113 | 3.87385 | 6.87309 |
| **Fbxo7** | 0.00000 | 2.91658 | 0.00000 | 0.41874 | 2.83671 | 0.00699 | 3.85560 | 0.03809 | 0.01667 | 1.93848 | 0.00721 | 0.15676 | 0.65019 | 0.00411 | 1.07448 | 0.02825 |
| **Fbxo8** | 4.12689 | 0.00000 | 2.57040 | 7.36851 | 5.84592 | 0.00000 | 0.02265 | 2.84503 | 4.87810 | 8.01222 | 0.21371 | 8.06631 | 1.75008 | 3.05411 | 1.65474 | 3.88622 |
| **Fbxo9** | 0.19462 | 3.00003 | 2.67126 | 2.43202 | 1.96223 | 7.23559 | 0.41456 | 2.98456 | 5.50975 | 2.14575 | 3.61506 | 0.84476 | 4.10918 | 2.75042 | 5.89330 | 2.02010 |
| **Fbxw11** | 6.03909 | 0.09139 | 8.56814 | 11.63362 | 6.15285 | 4.72330 | 6.84497 | 7.27759 | 14.72075 | 7.31532 | 5.52451 | 0.63450 | 6.38356 | 0.84018 | 6.90782 | 4.17954 |
| **Fbxw13** | 0.01373 | 0.00789 | 0.00908 | 0.57786 | 0.00932 | 0.00654 | 0.00732 | 0.01096 | 0.86847 | 0.01883 | 0.00000 | 0.20336 | 0.01301 | 0.02865 | 0.41817 | 0.00714 |
| **Fbxw14** | 0.01058 | 0.00750 | 0.00018 | 12.04177 | 0.00900 | 0.00451 | 0.00661 | 0.15586 | 6.50872 | 1.01722 | 0.28220 | 0.02729 | 0.00593 | 0.00000 | 4.56466 | 0.00491 |
| **Fbxw15** | 0.00125 | 0.00748 | 0.00002 | 9.21779 | 0.01095 | 0.00485 | 0.00657 | 0.64835 | 2.59539 | 0.88512 | 0.23309 | 0.21543 | 0.00482 | 3.45195 | 5.83918 | 0.00560 |
| **Fbxw16** | 0.01133 | 0.00761 | 0.00433 | 8.10030 | 0.00920 | 0.00508 | 0.00694 | 0.02183 | 1.76602 | 0.81523 | 0.06448 | 0.10185 | 0.01036 | 0.04635 | 8.17595 | 0.00590 |
| **Fbxw17** | 0.01372 | 0.00771 | 0.01225 | 0.24507 | 0.00937 | 0.00648 | 0.00740 | 0.01221 | 0.00818 | 0.03570 | 0.01298 | 0.00678 | 0.01989 | 0.00529 | 0.25051 | 0.00740 |
| **Fbxw18** | 0.01346 | 0.00768 | 0.00793 | 4.38190 | 0.00925 | 0.00591 | 0.00712 | 0.00540 | 0.83792 | 0.00000 | 0.15898 | 0.02131 | 0.01066 | 0.00000 | 2.23368 | 0.00646 |
| **Fbxw19** | 0.00813 | 0.00763 | 0.00000 | 8.84096 | 0.00868 | 0.00459 | 0.00693 | 0.02490 | 6.04000 | 1.43796 | 1.33819 | 1.29278 | 0.00000 | 0.04181 | 5.34828 | 0.00504 |
| **Fbxw2** | 3.23352 | 0.29785 | 3.61216 | 4.40166 | 5.09028 | 2.84390 | 0.74331 | 5.26330 | 9.36453 | 5.71603 | 8.67255 | 2.19404 | 0.00000 | 3.48697 | 8.19705 | 0.64881 |
| **Fbxw20** | 0.00850 | 0.01898 | 0.00711 | 1.67756 | 0.01039 | 0.00627 | 0.00729 | 0.01460 | 4.77066 | 0.00000 | 0.00000 | 0.02753 | 0.22618 | 1.69452 | 1.45728 | 0.01452 |
| **Fbxw21** | 0.01145 | 0.00799 | 0.00468 | 3.30923 | 0.00945 | 0.00593 | 0.00709 | 0.01306 | 1.07594 | 0.24504 | 0.02318 | 0.15769 | 0.01137 | 1.39259 | 1.97623 | 0.00651 |

| | | | | | | | | | | | | | | | | |
|---|---|---|---|---|---|---|---|---|---|---|---|---|---|---|---|---|
| **Fbxw22** | 0.01329 | 0.00792 | 0.00746 | 3.90196 | 0.00915 | 0.00612 | 0.00741 | 0.01032 | 1.87279 | 0.01283 | 0.22777 | 0.05408 | 0.00954 | 0.00062 | 0.86270 | 0.00652 |
| **Fbxw24** | 0.00002 | 0.00754 | 0.00088 | 18.78660 | 0.01926 | 0.00326 | 0.00603 | 0.00000 | 6.97822 | 1.27696 | 0.18836 | 1.01987 | 0.41850 | 3.79919 | 10.28679 | 0.00336 |
| **Fbxw26** | 0.01154 | 0.00741 | 0.00463 | 0.62895 | 0.00947 | 0.00680 | 0.00702 | 0.45896 | 1.00290 | 0.00000 | 0.20598 | 0.02055 | 0.00544 | 0.00921 | 0.81583 | 0.00720 |
| **Fbxw27** | 0.01057 | 0.00789 | 0.00363 | 4.13222 | 0.00932 | 0.00593 | 0.00722 | 0.32157 | 1.77629 | 0.25168 | 0.12472 | 0.00000 | 0.00580 | 0.00000 | 2.63203 | 0.00663 |
| **Fbxw28** | 0.00913 | 0.00748 | 0.00000 | 9.63979 | 0.00945 | 0.00491 | 0.00671 | 0.22705 | 4.22260 | 0.45251 | 0.30754 | 0.04042 | 0.00593 | 0.00000 | 5.36846 | 0.00545 |
| **Fbxw4** | 0.01748 | 0.00769 | 0.01762 | 6.32631 | 0.00889 | 0.00655 | 0.00704 | 0.00811 | 0.43646 | 0.30826 | 0.12065 | 0.00496 | 0.01081 | 0.01372 | 0.03207 | 0.00636 |
| **Fbxw5** | 0.14834 | 0.02858 | 0.02733 | 3.01781 | 1.17022 | 0.03986 | 1.72799 | 0.26228 | 0.99583 | 0.00000 | 0.63378 | 0.03866 | 1.66360 | 0.00034 | 0.03391 | 2.52307 |
| **Fbxw7** | 1.76620 | 0.07585 | 1.74985 | 0.93944 | 0.03434 | 0.00461 | 0.00635 | 0.01985 | 0.69506 | 0.26343 | 0.00437 | 0.02566 | 0.02705 | 0.00674 | 0.61093 | 0.24551 |
| **Fbxw8** | 5.30462 | 2.57054 | 5.20940 | 0.30053 | 5.43999 | 0.01580 | 2.21866 | 4.35111 | 0.39213 | 4.08392 | 1.68919 | 0.21372 | 2.82823 | 0.06363 | 0.21250 | 4.38844 |
| **Fbxw9** | 0.91478 | 0.10420 | 4.68064 | 6.27802 | 0.96734 | 4.31824 | 5.56450 | 7.70334 | 8.26631 | 6.12145 | 3.76029 | 1.67611 | 5.96028 | 0.24442 | 0.50570 | 0.10220 |
| **Fcer1g** | 0.01287 | 0.00784 | 0.01216 | 0.18342 | 0.00882 | 0.00651 | 0.00723 | 0.00981 | 0.00581 | 1.68793 | 0.01175 | 0.00402 | 0.01059 | 0.01475 | 0.00384 | 0.00728 |
| **Fcer2a** | 0.17372 | 0.00773 | 0.16265 | 0.09199 | 0.00954 | 0.00624 | 0.05870 | 0.04603 | 0.44773 | 0.09138 | 1.22131 | 0.08915 | 0.23121 | 0.37216 | 0.11025 | 0.05011 |
| **Fcf1** | 11.26535 | 8.98618 | 10.46095 | 6.68881 | 3.98194 | 2.14482 | 3.35264 | 5.38255 | 9.54455 | 21.02621 | 15.78093 | 3.12276 | 10.13290 | 16.00773 | 12.90401 | 3.61156 |
| **Fcgbp** | 0.01290 | 0.00813 | 0.01022 | 0.88504 | 0.00906 | 0.00669 | 0.00748 | 0.00707 | 0.09520 | 1.24187 | 0.22572 | 0.00313 | 0.01271 | 0.00569 | 0.55990 | 0.00728 |
| **Fcgr1** | 0.01337 | 0.00831 | 0.01103 | 0.22346 | 0.00921 | 0.00671 | 0.00760 | 0.01356 | 2.45989 | 0.05184 | 0.00000 | 0.02868 | 0.01361 | 0.01127 | 0.00332 | 0.00725 |
| **Fcgr2b** | 0.63919 | 0.27651 | 0.00000 | 0.00001 | 0.01447 | 0.00519 | 0.14489 | 1.78967 | 0.00548 | 5.66088 | 0.00000 | 0.72134 | 0.80577 | 6.58646 | 4.46011 | 0.74765 |
| **Fcgr3** | 1.53227 | 3.07864 | 0.00000 | 21.07981 | 0.13988 | 0.00000 | 1.52104 | 15.56529 | 23.12919 | 30.96545 | 12.22982 | 10.51533 | 0.67173 | 29.15465 | 21.25071 | 6.29878 |
| **Fcgrt** | 0.00042 | 0.00672 | 0.65880 | 1.65693 | 0.00907 | 0.00861 | 0.87216 | 1.63211 | 1.59601 | 2.53624 | 1.84898 | 0.20451 | 0.03541 | 0.00104 | 0.11679 | 0.00698 |
| **Fcho1** | 0.45301 | 0.00760 | 0.00549 | 0.28247 | 0.00693 | 0.00708 | 0.00713 | 0.01450 | 0.33037 | 0.01361 | 0.01139 | 0.00746 | 0.01222 | 0.01172 | 0.00504 | 0.00677 |
| **Fcho2** | 0.03195 | 1.06954 | 0.07958 | 4.23192 | 0.29361 | 3.64826 | 3.50136 | 0.65453 | 1.06277 | 3.17733 | 2.56365 | 8.52311 | 0.00000 | 5.04998 | 2.99909 | 1.00380 |
| **Fchsd1** | 0.37627 | 0.00729 | 0.00365 | 0.12129 | 0.00699 | 0.00751 | 0.00689 | 0.00000 | 0.44845 | 0.06126 | 0.92635 | 0.08283 | 0.02392 | 0.03401 | 0.00000 | 0.06913 |
| **Fchsd2** | 0.04399 | 0.00762 | 0.00000 | 11.80081 | 0.01535 | 0.87384 | 0.68266 | 0.13977 | 2.06147 | 0.00000 | 0.18332 | 0.03820 | 0.00000 | 2.05074 | 7.08353 | 0.07972 |
| **Fcrl5** | 0.08928 | 0.09960 | 0.01783 | 0.20409 | 0.00781 | 0.17588 | 0.16035 | 0.03014 | 0.19582 | 0.02448 | 0.37577 | 0.39061 | 0.10965 | 0.06490 | 0.06800 | 0.00709 |
| **Fcrl6** | 0.03141 | 0.00781 | 0.02333 | 0.01523 | 0.00822 | 0.06910 | 0.03957 | 0.02492 | 0.03659 | 0.02680 | 0.02493 | 0.00783 | 0.01632 | 0.01180 | 0.04022 | 0.07904 |
| **Fdft1** | 4.56301 | 9.73143 | 14.10743 | 19.14474 | 9.06872 | 5.40877 | 11.67622 | 26.82060 | 24.85839 | 25.47581 | 19.98618 | 12.59752 | 5.70548 | 12.67501 | 17.75891 | 2.61835 |
| **Fdps** | 20.74781 | 16.22580 | 32.15902 | 3.35839 | 19.98935 | 6.72648 | 16.53013 | 24.78334 | 5.40104 | 22.87446 | 17.35099 | 10.64873 | 11.53252 | 18.50068 | 8.65179 | 2.93253 |
| **Fdx1** | 0.03896 | 1.35372 | 0.73392 | 0.36216 | 2.25139 | 0.63974 | 0.04582 | 0.71106 | 1.00564 | 1.98909 | 6.62073 | 0.12970 | 1.40803 | 0.41598 | 0.06993 | 0.01221 |
| **Fdx1l** | 0.11194 | 0.00799 | 0.01438 | 0.00917 | 0.01061 | 0.00796 | 0.01753 | 0.09792 | 0.01200 | 0.33052 | 0.01341 | 0.00643 | 0.69598 | 0.01281 | 0.00828 | 0.48923 |
| **Fdxacb1** | 0.37380 | 2.67819 | 3.26340 | 0.28192 | 0.00305 | 0.00782 | 4.22134 | 0.35305 | 0.00319 | 0.60658 | 0.33343 | 0.01110 | 0.07826 | 1.71403 | 1.79210 | 0.01251 |
| **Fdxr** | 4.61234 | 3.86426 | 1.70168 | 0.83428 | 7.92589 | 0.00184 | 4.70996 | 1.56297 | 0.10792 | 1.58819 | 1.51834 | 0.09297 | 0.01202 | 0.00000 | 0.00000 | 0.00780 |
| **Fech** | 1.95476 | 2.56410 | 2.95110 | 2.28381 | 2.95065 | 0.01038 | 1.19733 | 4.09637 | 0.78855 | 6.47471 | 0.37350 | 0.50682 | 0.93772 | 0.17269 | 2.24812 | 0.03240 |
| **Fem1a** | 0.02128 | 1.52221 | 1.33673 | 6.10607 | 0.02198 | 0.00669 | 0.00585 | 0.00000 | 15.08789 | 0.23680 | 0.97634 | 0.00000 | 0.08433 | 0.10129 | 2.08493 | 0.01969 |
| **Fem1b** | 6.00816 | 6.14593 | 5.90448 | 14.45327 | 9.15956 | 6.32557 | 3.68006 | 4.65108 | 14.17459 | 7.60349 | 2.70141 | 5.15034 | 2.83234 | 12.98340 | 16.64848 | 5.18873 |
| **Fem1c** | 1.10974 | 3.25470 | 1.34405 | 16.20256 | 1.00796 | 0.00096 | 0.96609 | 2.13908 | 20.52806 | 4.46158 | 2.45691 | 3.84085 | 0.89782 | 6.17636 | 7.33584 | 2.53973 |
| **Fen1** | 5.19901 | 11.70098 | 12.02665 | 4.28268 | 5.31321 | 7.90446 | 4.19172 | 12.01615 | 4.98304 | 4.66742 | 4.29056 | 0.75788 | 7.09372 | 0.29178 | 7.41332 | 6.14237 |
| **Fendrr** | 0.01939 | 0.00780 | 0.00829 | 0.12052 | 0.01775 | 0.00684 | 0.00730 | 0.01188 | 0.03498 | 0.00549 | 0.35322 | 0.00595 | 0.00829 | 0.13101 | 0.00000 | 0.00716 |
| **Fer** | 2.82540 | 0.00551 | 1.34792 | 4.75709 | 0.00260 | 0.00823 | 1.29598 | 0.21656 | 0.00000 | 2.27360 | 0.02516 | 0.03651 | 0.90540 | 0.02727 | 0.04327 | 0.01007 |
| **Fer1l4** | 0.01169 | 0.00788 | 0.00723 | 0.50350 | 0.00947 | 0.03722 | 0.00723 | 0.10292 | 0.00768 | 0.01238 | 0.03034 | 0.00768 | 0.00811 | 0.01079 | 0.00299 | 0.00729 |
| **Fer1l5** | 0.01243 | 0.00786 | 0.00930 | 0.58024 | 0.00931 | 0.00672 | 0.00722 | 0.04460 | 0.04355 | 0.00650 | 0.22257 | 0.02710 | 0.00836 | 0.00824 | 0.00000 | 0.00718 |
| **Fermt2** | 2.96365 | 0.01042 | 2.64140 | 4.14853 | 0.40325 | 2.27154 | 2.49704 | 8.01308 | 5.52304 | 9.09638 | 8.83869 | 7.63381 | 2.94968 | 17.61379 | 4.60447 | 0.39961 |
| **Fermt3** | 0.00442 | 0.00747 | 1.46600 | 0.06022 | 0.00567 | 0.01121 | 0.21971 | 0.55020 | 0.37000 | 1.09898 | 0.41365 | 0.14210 | 0.99312 | 0.02720 | 0.04159 | 0.64542 |
| **Fes** | 0.00799 | 0.00724 | 0.00254 | 0.00527 | 0.00919 | 0.12436 | 0.00620 | 1.00898 | 0.12962 | 0.42907 | 0.02879 | 0.05658 | 1.53469 | 0.01393 | 0.00000 | 0.00668 |
| **Fetub** | 1.15768 | 0.00627 | 0.00000 | 0.10242 | 0.00559 | 1.49732 | 0.00572 | 0.00000 | 1.30916 | 0.00000 | 0.69037 | 0.05779 | 0.43091 | 0.00687 | 0.19312 | 0.00501 |
| **Fez1** | 0.01031 | 0.00776 | 0.05058 | 0.00000 | 0.00822 | 0.00548 | 0.00698 | 0.00000 | 0.23056 | 5.20457 | 0.85521 | 1.83368 | 0.00507 | 2.25453 | 3.24772 | 0.00695 |
| **Fez2** | 2.30922 | 3.22111 | 4.11433 | 0.28955 | 5.38181 | 4.01477 | 0.49797 | 5.54311 | 1.30874 | 0.57191 | 2.71804 | 0.10013 | 4.32152 | 0.51881 | 1.26613 | 7.50251 |
| **Ffar2** | 0.01216 | 0.00715 | 0.01857 | 0.04666 | 1.19908 | 0.00713 | 0.00732 | 0.00889 | 0.02658 | 0.01427 | 0.02501 | 0.04562 | 0.01894 | 0.19724 | 0.01843 | 0.00733 |
| **Ffar3** | 0.00894 | 0.00747 | 3.60539 | 0.01056 | 0.00354 | 0.00911 | 0.00674 | 0.00911 | 0.01244 | 0.00364 | 0.00997 | 0.00529 | 0.45575 | 0.01066 | 0.02336 | 0.00580 |
| **Fgd1** | 2.39667 | 0.00381 | 0.01184 | 0.35426 | 0.01192 | 0.00380 | 1.62145 | 0.14406 | 0.17539 | 0.66609 | 0.00903 | 0.01934 | 0.00723 | 0.02085 | 0.46205 | 0.06371 |
| **Fgd2** | 0.01364 | 0.00818 | 0.01112 | 0.33056 | 0.00918 | 0.00687 | 0.00749 | 0.01256 | 0.00681 | 0.24918 | 0.01262 | 0.05372 | 0.01275 | 0.03161 | 0.00449 | 0.00736 |
| **Fgd3** | 0.01295 | 0.00808 | 0.02322 | 0.00692 | 0.01020 | 0.00673 | 0.00754 | 0.01272 | 0.01133 | 0.00000 | 0.07278 | 0.00684 | 0.29980 | 0.00661 | 0.35038 | 0.00751 |
| **Fgd4** | 0.02741 | 0.00784 | 0.10449 | 0.05064 | 0.01205 | 0.00728 | 0.00715 | 2.60062 | 0.04576 | 0.00000 | 1.11261 | 0.00000 | 0.13462 | 1.75860 | 0.14315 | 0.02172 |
| **Fgd6** | 2.28956 | 1.44132 | 0.19778 | 6.59847 | 0.18866 | 0.00567 | 1.80236 | 0.00433 | 2.57362 | 1.11768 | 0.06795 | 0.00882 | 1.20158 | 0.00000 | 3.78612 | 0.44093 |
| **Fgf1** | 0.00506 | 0.00743 | 0.01126 | 4.75471 | 0.00845 | 0.00637 | 0.00671 | 0.95984 | 4.07694 | 3.59577 | 0.00000 | 0.08304 | 0.92143 | 0.00000 | 1.13976 | 0.00580 |
| **Fgf10** | 8.10863 | 2.52230 | 1.08850 | 0.20524 | 5.00441 | 0.00780 | 6.03443 | 0.00000 | 0.19085 | 0.04024 | 0.02263 | 0.03666 | 0.01297 | 0.05476 | 0.51165 | 0.73381 |
| **Fgf11** | 0.59099 | 0.00825 | 0.00529 | 0.01747 | 0.03664 | 0.02027 | 0.00714 | 0.01088 | 0.01259 | 0.19310 | 0.01193 | 0.00870 | 0.02618 | 0.02992 | 0.01036 | 0.00700 |
| **Fgf13** | 0.06675 | 0.01776 | 0.53647 | 0.01556 | 0.00810 | 0.06024 | 0.04642 | 0.02576 | 0.02467 | 0.48211 | 0.06274 | 0.01486 | 0.06768 | 0.32508 | 0.04365 | 0.02087 |
| **Fgf17** | 0.01366 | 0.00823 | 0.01209 | 0.37229 | 0.00932 | 0.00661 | 0.00753 | 0.01395 | 0.55662 | 0.01237 | 0.00904 | 0.00747 | 0.01284 | 0.01180 | 0.00395 | 0.00733 |
| **Fgf18** | 0.01327 | 0.00791 | 0.01037 | 0.25627 | 0.00932 | 0.00677 | 0.23929 | 0.01167 | 0.17288 | 0.02696 | 0.01157 | 0.00785 | 0.01212 | 0.01113 | 0.00605 | 0.00735 |
| **Fgf3** | 0.00567 | 0.00801 | 0.00232 | 0.17670 | 0.00938 | 0.00766 | 0.00625 | 1.03643 | 3.25939 | 0.47916 | 3.84566 | 0.04403 | 0.43795 | 0.14122 | 0.13399 | 0.00621 |
| **Fgf4** | 7.05660 | 0.00000 | 0.00000 | 6.19990 | 0.00000 | 0.00286 | 0.00000 | 1.78964 | 2.99549 | 1.94550 | 7.00558 | 2.22769 | 0.00000 | 8.59771 | 4.20683 | 0.07470 |
| **Fgf5** | 0.01375 | 0.00791 | 0.01163 | 0.03723 | 0.00937 | 0.00652 | 0.00741 | 0.01175 | 0.01129 | 0.00862 | 0.01185 | 0.00715 | 0.01284 | 0.00875 | 0.59624 | 0.00735 |
| **Fgf7** | 0.01364 | 0.00816 | 0.01171 | 1.29775 | 0.00929 | 0.00651 | 0.00747 | 0.01307 | 0.78685 | 0.00848 | 0.02352 | 0.00697 | 0.01257 | 0.02986 | 0.00068 | 0.00716 |

| | | | | | | | | | | | | | | | | |
|---|---|---|---|---|---|---|---|---|---|---|---|---|---|---|---|---|
| **Fgf9** | 0.01359 | 0.00791 | 0.01157 | 1.29697 | 0.00931 | 0.00648 | 0.00741 | 0.01236 | 0.42976 | 0.00989 | 0.02577 | 0.00718 | 0.03520 | 0.04122 | 0.05627 | 0.00710 |
| **Fgfbp1** | 0.01564 | 0.00854 | 0.16709 | 0.00986 | 0.02942 | 0.00880 | 0.00756 | 0.18096 | 0.01208 | 0.00019 | 0.03694 | 0.00785 | 1.18514 | 0.01659 | 0.00398 | 0.00674 |
| **Fgfbp3** | 0.01280 | 0.00812 | 0.30878 | 0.57265 | 0.00875 | 0.00616 | 0.00753 | 0.01332 | 0.05093 | 0.01388 | 0.01211 | 0.00820 | 0.00986 | 0.02845 | 0.00000 | 0.00716 |
| **Fgfr1** | 10.76459 | 0.61870 | 2.58819 | 3.39855 | 5.25728 | 6.71288 | 4.63834 | 6.51383 | 0.54362 | 9.30123 | 4.35509 | 0.69379 | 2.56352 | 0.77619 | 0.24341 | 6.53182 |
| **Fgfr1op** | 1.04199 | 4.78799 | 1.34562 | 5.63979 | 4.15012 | 2.50294 | 0.01184 | 0.41679 | 4.67264 | 6.05782 | 4.83709 | 11.18146 | 1.28369 | 5.68573 | 7.13635 | 2.59474 |
| **Fgfr1op2** | 4.39429 | 2.19020 | 2.76929 | 4.33688 | 8.72356 | 0.58181 | 0.63208 | 4.30298 | 1.90245 | 4.79872 | 4.47623 | 9.69460 | 4.22829 | 1.05872 | 4.19660 | 4.65565 |
| **Fgfr2** | 2.67500 | 4.60184 | 1.18229 | 0.69330 | 5.97121 | 6.45933 | 0.62133 | 2.19524 | 0.15450 | 6.25481 | 6.27634 | 1.90416 | 0.23069 | 8.10240 | 0.13347 | 4.28433 |
| **Fgfr3** | 0.41863 | 0.17467 | 0.20112 | 0.24414 | 0.83700 | 0.01186 | 1.04744 | 1.56436 | 1.12822 | 1.56566 | 0.28978 | 0.25038 | 0.38826 | 0.84135 | 1.21373 | 2.19534 |
| **Fgfr4** | 0.68831 | 1.69081 | 0.30634 | 0.05896 | 0.01435 | 1.44462 | 1.98191 | 0.83623 | 2.47254 | 3.05123 | 0.70329 | 0.19300 | 0.55858 | 0.00759 | 1.32367 | 0.00515 |
| **Fgfrl1** | 0.31802 | 0.00725 | 0.00000 | 5.61110 | 0.00702 | 0.63801 | 0.00680 | 0.01550 | 1.73546 | 0.00000 | 0.00000 | 0.03185 | 0.00678 | 0.25581 | 0.72520 | 0.22132 |
| **Fgl1** | 0.77523 | 0.08070 | 0.00416 | 0.33180 | 0.00630 | 0.00706 | 0.00673 | 0.01159 | 0.00687 | 0.03930 | 0.01477 | 0.00775 | 0.01139 | 0.01325 | 0.00393 | 0.00626 |
| **Fgl2** | 0.01341 | 0.00790 | 0.01191 | 0.00437 | 0.00912 | 0.00661 | 0.00734 | 0.01080 | 0.65359 | 0.56990 | 0.00749 | 0.00521 | 0.01208 | 0.01216 | 0.00724 | 0.00758 |
| **Fgr** | 0.00364 | 0.00717 | 0.00000 | 0.46003 | 0.00883 | 0.00600 | 0.00655 | 0.74873 | 1.88371 | 5.65889 | 3.10080 | 2.17045 | 0.00000 | 0.15265 | 6.03700 | 0.00675 |
| **Fh1** | 7.91409 | 4.82697 | 8.62839 | 0.45725 | 7.40487 | 7.66109 | 5.04137 | 4.11239 | 1.36522 | 0.73070 | 3.21910 | 0.09746 | 6.17837 | 0.03501 | 0.65430 | 5.63716 |
| **Fhad1** | 0.63793 | 0.00725 | 0.00378 | 0.09275 | 0.00652 | 0.00703 | 0.00673 | 0.01538 | 0.59779 | 0.42432 | 0.00860 | 0.00451 | 0.01183 | 0.00854 | 0.30149 | 0.00699 |
| **Fhl1** | 0.01220 | 0.00791 | 0.01123 | 0.00546 | 0.00867 | 0.00663 | 0.00724 | 0.01001 | 0.01109 | 2.88779 | 0.00953 | 0.04172 | 0.01730 | 0.03551 | 0.00000 | 0.01485 |
| **Fhl2** | 0.80312 | 0.99338 | 4.95077 | 0.40997 | 4.44347 | 3.54104 | 3.74936 | 0.04074 | 0.00000 | 0.00000 | 0.00633 | 0.03517 | 2.24125 | 0.02348 | 0.23286 | 5.30158 |
| **Fhl3** | 0.06839 | 0.00790 | 0.00329 | 2.54219 | 0.00770 | 0.00734 | 0.00643 | 0.00000 | 2.68122 | 0.00000 | 2.14702 | 0.06578 | 0.05681 | 0.05559 | 0.09343 | 0.00607 |
| **Fhl4** | 0.07494 | 0.00790 | 0.00636 | 0.35164 | 0.01533 | 0.00682 | 0.24486 | 0.01473 | 0.41210 | 0.00568 | 0.01055 | 0.00599 | 0.01768 | 0.00540 | 0.30350 | 0.00731 |
| **Fhod1** | 2.82170 | 0.55739 | 1.50099 | 0.64859 | 1.33401 | 4.28716 | 0.00337 | 0.00000 | 0.00000 | 0.22671 | 1.07339 | 0.04604 | 1.26351 | 0.00869 | 0.00099 | 8.74608 |
| **Fhod3** | 1.33954 | 0.00507 | 1.48091 | 22.03935 | 0.00281 | 0.00727 | 3.02036 | 0.23787 | 4.78427 | 5.53697 | 0.10090 | 0.97819 | 0.00606 | 0.14969 | 3.53619 | 0.00000 |
| **Fibcd1** | 0.01376 | 0.00782 | 0.01236 | 0.40998 | 0.00933 | 0.00654 | 0.00748 | 0.01331 | 0.00629 | 0.01055 | 0.01375 | 0.00746 | 0.01263 | 0.00627 | 0.13062 | 0.00732 |
| **Fibp** | 10.05940 | 6.98904 | 0.00000 | 0.19286 | 1.53636 | 4.14774 | 4.77409 | 5.32865 | 0.00548 | 1.46649 | 2.43200 | 0.12513 | 6.43599 | 0.00000 | 0.56961 | 0.01859 |
| **Ficd** | 1.42912 | 0.01421 | 1.81687 | 9.36507 | 0.00367 | 0.00347 | 0.02148 | 6.50669 | 5.55041 | 12.71316 | 7.30429 | 4.23845 | 1.84698 | 7.04189 | 17.28710 | 0.01508 |
| **Fig4** | 3.29713 | 3.13762 | 4.78109 | 0.19069 | 0.00000 | 3.41022 | 0.55374 | 11.67548 | 0.58358 | 13.42126 | 7.34028 | 0.73333 | 2.34221 | 3.62182 | 1.01764 | 0.03480 |
| **Fignl1** | 4.05010 | 6.10356 | 3.23015 | 4.19880 | 1.38671 | 0.44845 | 1.91980 | 3.36327 | 2.92160 | 5.26737 | 1.42642 | 11.04715 | 1.90656 | 11.23342 | 9.18488 | 0.33284 |
| **Filip1l** | 0.00178 | 0.00670 | 0.13716 | 0.71439 | 0.01131 | 0.00948 | 2.04860 | 0.38780 | 0.00000 | 0.09377 | 0.00000 | 0.47450 | 0.00626 | 0.03111 | 1.03191 | 0.01462 |
| **Fip1l1** | 7.93522 | 5.12663 | 5.53184 | 4.67332 | 0.47053 | 2.36414 | 3.43428 | 8.80469 | 7.06325 | 7.05841 | 5.32918 | 4.29300 | 4.99714 | 0.12158 | 1.46392 | 4.38556 |
| **Firre** | 0.12987 | 0.00695 | 0.70769 | 0.00000 | 0.01505 | 2.94962 | 1.74953 | 1.32094 | 0.00936 | 0.00000 | 1.06948 | 0.00000 | 1.03102 | 0.71622 | 0.10286 | 0.00496 |
| **Fis1** | 12.58317 | 6.66609 | 9.44338 | 2.35620 | 9.51885 | 1.01381 | 7.21764 | 9.94308 | 1.69761 | 8.36139 | 4.56546 | 1.03208 | 7.57286 | 0.00585 | 0.51279 | 6.60786 |
| **Fitm2** | 0.00603 | 0.00857 | 0.16246 | 0.00483 | 0.00923 | 0.00831 | 0.00765 | 0.37731 | 0.62168 | 0.13253 | 0.05020 | 0.00575 | 0.68862 | 0.30044 | 0.00000 | 0.00701 |
| **Fiz1** | 8.15266 | 0.00000 | 0.84131 | 15.23852 | 4.05936 | 7.26502 | 2.13142 | 3.91179 | 21.82480 | 20.72884 | 8.88570 | 4.86958 | 8.23911 | 4.96032 | 11.35964 | 3.88202 |
| **Fjx1** | 1.93837 | 0.54635 | 0.00170 | 0.22342 | 0.43333 | 0.00662 | 0.00555 | 0.00256 | 0.00744 | 0.01207 | 0.01328 | 0.00591 | 0.01074 | 0.00797 | 0.00000 | 0.00513 |
| **Fkbp10** | 0.00454 | 0.00705 | 0.00000 | 0.02969 | 0.00800 | 0.00735 | 0.01622 | 1.62887 | 0.00964 | 8.60283 | 3.79961 | 0.16662 | 0.00000 | 0.03347 | 1.24171 | 0.00665 |
| **Fkbp11** | 1.25841 | 0.01203 | 7.44369 | 0.25020 | 0.00000 | 0.01985 | 3.69063 | 10.04373 | 0.02138 | 1.65328 | 7.48837 | 0.09728 | 7.10623 | 0.43728 | 0.61168 | 2.85181 |
| **Fkbp14** | 0.01357 | 0.00783 | 0.01136 | 1.98141 | 0.00909 | 0.00671 | 0.00744 | 0.01177 | 0.00000 | 0.41060 | 0.02717 | 0.01944 | 0.01228 | 0.01121 | 0.10409 | 0.00706 |
| **Fkbp15** | 1.48982 | 5.10311 | 5.88128 | 1.86599 | 4.76616 | 0.00502 | 2.59525 | 1.47600 | 1.92207 | 0.58552 | 1.25568 | 0.10583 | 3.80217 | 0.57389 | 0.99041 | 1.36448 |
| **Fkbp1a** | 10.34071 | 4.47761 | 4.21581 | 11.60384 | 4.90696 | 2.32036 | 5.46150 | 5.16667 | 10.19602 | 6.53698 | 11.49984 | 1.42756 | 4.69467 | 5.75326 | 3.45220 | 4.29804 |
| **Fkbp1b** | 1.93940 | 0.00423 | 0.03322 | 2.74139 | 0.00648 | 0.00297 | 0.00436 | 0.00000 | 3.91836 | 2.04786 | 2.26280 | 4.25942 | 0.35289 | 0.00000 | 0.21420 | 0.00427 |
| **Fkbp2** | 4.63321 | 0.99015 | 4.46915 | 1.75487 | 0.20082 | 3.23417 | 2.28831 | 1.33633 | 2.20492 | 4.28739 | 6.33920 | 0.11986 | 5.93085 | 0.07231 | 1.31042 | 1.53027 |
| **Fkbp3** | 20.25417 | 12.52986 | 13.00010 | 1.06845 | 9.59126 | 10.16670 | 8.96671 | 10.50699 | 1.91908 | 12.99910 | 9.63383 | 7.99602 | 12.24217 | 1.68157 | 2.32758 | 9.97399 |
| **Fkbp4** | 28.75872 | 13.55923 | 29.02292 | 11.94944 | 14.74054 | 13.10911 | 17.03696 | 20.12772 | 15.40473 | 23.57301 | 18.00070 | 7.45533 | 23.87930 | 3.39373 | 8.95716 | 14.18304 |
| **Fkbp5** | 13.31390 | 4.59025 | 5.93081 | 5.87628 | 2.18983 | 1.17095 | 1.65735 | 2.04373 | 4.68905 | 3.58595 | 0.54309 | 7.13126 | 1.52561 | 3.59755 | 8.73459 | 0.01620 |
| **Fkbp6** | 1.34724 | 10.12323 | 15.97149 | 1.60009 | 0.01808 | 4.20662 | 7.00180 | 11.33524 | 6.76503 | 12.13050 | 9.42861 | 4.02242 | 12.41277 | 3.18462 | 8.43297 | 3.75199 |
| **Fkbp8** | 13.79818 | 0.64568 | 6.49655 | 2.75274 | 4.77273 | 8.28392 | 4.97409 | 12.36164 | 1.65967 | 7.11386 | 6.95196 | 0.48871 | 13.72630 | 0.11766 | 0.00000 | 13.75733 |
| **Fkbp9** | 5.24731 | 1.59382 | 1.82902 | 0.00000 | 1.68825 | 2.94695 | 2.11527 | 2.25141 | 0.00517 | 2.22061 | 1.24307 | 0.04705 | 1.90089 | 2.80083 | 0.71374 | 3.03768 |
| **Fkbpl** | 2.31554 | 0.00564 | 0.45734 | 1.07214 | 0.00733 | 0.00316 | 1.38787 | 1.25164 | 2.97761 | 1.41240 | 0.44214 | 0.07250 | 1.28300 | 0.00366 | 0.04478 | 2.90378 |
| **Fkrp** | 0.29611 | 0.07466 | 0.58962 | 0.26576 | 3.26428 | 0.00524 | 0.00628 | 1.02238 | 0.00342 | 0.11722 | 0.02367 | 0.00068 | 0.86561 | 0.00818 | 0.09585 | 0.01327 |
| **Fktn** | 0.36500 | 0.15101 | 0.18981 | 0.67166 | 0.04327 | 0.04944 | 0.16324 | 0.38765 | 0.38264 | 0.21081 | 0.28632 | 0.15476 | 0.30928 | 1.14587 | 0.30692 | 0.02845 |
| **Flad1** | 5.09300 | 5.15317 | 3.70628 | 2.67143 | 0.39335 | 0.00000 | 4.10004 | 2.85754 | 1.68700 | 0.95933 | 2.42168 | 0.40372 | 4.51409 | 0.88026 | 1.48412 | 3.54738 |
| **Flcn** | 1.16132 | 4.40218 | 4.78820 | 11.19819 | 4.62715 | 4.26488 | 1.72749 | 5.98894 | 9.12934 | 1.22757 | 3.29039 | 0.22051 | 4.69151 | 0.21683 | 5.56653 | 0.58651 |
| **Flii** | 6.28576 | 4.17255 | 2.44791 | 0.28634 | 0.46769 | 9.87248 | 3.74895 | 10.35062 | 0.21192 | 2.82861 | 1.87041 | 0.68407 | 4.61590 | 0.00704 | 0.00000 | 0.15847 |
| **Flna** | 0.00000 | 0.00684 | 0.00000 | 0.58110 | 2.06869 | 0.17644 | 0.00394 | 1.35350 | 7.31050 | 3.55481 | 2.30962 | 0.94752 | 5.63485 | 1.83972 | 0.81188 | 4.28808 |
| **Flnb** | 8.07301 | 9.22697 | 9.20921 | 4.47335 | 2.73110 | 11.40459 | 9.39400 | 13.01404 | 5.18106 | 5.90392 | 4.08534 | 9.69552 | 8.54104 | 5.02593 | 3.33104 | 9.60686 |
| **Flot1** | 2.23957 | 0.02449 | 0.00000 | 3.23539 | 7.41819 | 2.53475 | 0.00384 | 2.22212 | 1.96891 | 1.65937 | 0.97956 | 0.01318 | 1.92816 | 0.00000 | 0.14246 | 0.01024 |
| **Flot2** | 1.99792 | 0.01817 | 3.25828 | 0.31814 | 0.02380 | 5.00828 | 0.01279 | 3.18851 | 1.66468 | 2.02816 | 0.64562 | 0.06194 | 6.98866 | 0.04992 | 1.58344 | 3.44298 |
| **Flrt1** | 0.01467 | 0.03999 | 2.22107 | 0.21715 | 0.00499 | 0.03707 | 0.00729 | 0.01482 | 0.00959 | 0.02764 | 0.03979 | 0.00638 | 0.00592 | 0.01528 | 0.00000 | 0.01360 |
| **Flrt2** | 0.68964 | 0.00709 | 0.00428 | 0.22716 | 0.00655 | 0.00723 | 0.00698 | 0.00991 | 0.35507 | 0.01136 | 0.00944 | 0.04844 | 0.01214 | 0.03450 | 0.00156 | 0.00661 |
| **Flrt3** | 0.24332 | 0.00871 | 0.00593 | 0.31781 | 0.01691 | 0.00684 | 0.00730 | 0.01206 | 0.36554 | 0.00501 | 0.02164 | 0.00771 | 0.00868 | 0.01244 | 0.00401 | 0.86455 |
| **Flt1** | 0.01377 | 0.00803 | 0.01205 | 0.01062 | 0.00932 | 0.00667 | 0.00752 | 0.01365 | 0.01235 | 0.01500 | 0.01302 | 0.01842 | 0.01272 | 0.01231 | 0.01038 | 0.00742 |
| **Flt3** | 0.01358 | 0.00783 | 0.01156 | 0.12011 | 0.00915 | 0.00683 | 0.00754 | 0.01265 | 0.24103 | 0.25881 | 0.00972 | 0.00764 | 0.01284 | 0.01316 | 0.18775 | 0.00738 |
| **Flt3l** | 0.00352 | 0.00654 | 0.08883 | 2.46343 | 0.00871 | 0.00520 | 0.33630 | 1.98131 | 3.05971 | 2.33260 | 5.19549 | 0.12766 | 1.11653 | 0.12714 | 0.00000 | 0.00533 |

| | | | | | | | | | | | | | | | | |
|---|---|---|---|---|---|---|---|---|---|---|---|---|---|---|---|---|
| Flywch1 | 0.11750 | 0.01849 | 0.02525 | 0.10430 | 0.00841 | 0.01232 | 0.00746 | 0.03778 | 0.74889 | 0.26450 | 0.00684 | 0.04214 | 0.01105 | 0.01122 | 0.00607 | 0.00721 |
| Flywch2 | 0.01255 | 0.00786 | 0.00880 | 1.16145 | 0.00899 | 0.00659 | 0.00704 | 0.01629 | 1.02576 | 0.89972 | 0.40695 | 0.00003 | 0.00796 | 0.00432 | 0.04129 | 0.00719 |
| Fmc1 | 2.19251 | 0.00496 | 0.39691 | 0.74923 | 0.02177 | 1.03435 | 0.00473 | 0.07817 | 0.00000 | 1.53600 | 0.78638 | 0.13250 | 0.80159 | 0.28744 | 0.00000 | 0.83603 |
| Fmn1 | 0.82267 | 0.00705 | 0.00292 | 0.41179 | 0.00619 | 0.00727 | 0.23477 | 0.00870 | 0.10956 | 0.45392 | 0.00887 | 0.03431 | 0.01058 | 0.18282 | 0.00444 | 0.00609 |
| Fmn2 | 0.10314 | 0.00774 | 0.00000 | 4.02473 | 0.00910 | 0.00586 | 0.00682 | 0.00711 | 2.00014 | 0.33016 | 0.00000 | 0.00000 | 0.01313 | 3.18739 | 1.62266 | 0.00686 |
| Fmnl1 | 0.01846 | 0.00828 | 0.01171 | 0.01607 | 0.01052 | 0.00679 | 0.00758 | 0.01414 | 0.21417 | 0.02819 | 0.01148 | 0.00832 | 0.01302 | 0.01240 | 0.00936 | 0.00746 |
| Fmnl2 | 1.77375 | 2.42844 | 0.00000 | 3.01972 | 0.00591 | 6.83839 | 0.00468 | 1.20998 | 0.18544 | 0.00000 | 0.00000 | 1.09164 | 0.23639 | 3.42402 | 0.56183 | 0.01650 |
| Fmnl3 | 0.60088 | 0.00638 | 1.64668 | 18.15119 | 0.00396 | 0.01432 | 0.00529 | 0.60403 | 12.79494 | 0.08054 | 0.46276 | 0.16586 | 1.07152 | 0.00000 | 2.70372 | 0.00204 |
| Fmo1 | 1.22622 | 1.77523 | 3.50100 | 2.88773 | 0.35275 | 1.59304 | 2.71915 | 4.43120 | 4.46622 | 9.27721 | 0.78129 | 5.21321 | 2.18266 | 11.50553 | 8.26648 | 0.87034 |
| Fmo5 | 0.01746 | 0.00809 | 0.01540 | 0.01117 | 0.01020 | 0.01237 | 0.00736 | 0.01009 | 0.03539 | 0.65115 | 0.04645 | 0.00695 | 0.02343 | 0.07127 | 0.02258 | 0.00713 |
| Fmr1 | 0.65431 | 5.42883 | 5.84029 | 6.04489 | 3.60888 | 3.79030 | 1.22929 | 3.74100 | 14.26424 | 0.30904 | 2.50970 | 9.88470 | 1.72759 | 20.17668 | 5.72249 | 2.31289 |
| Fmr1nb | 4.20322 | 1.93551 | 8.01492 | 11.34332 | 1.52256 | 4.52780 | 0.00000 | 8.80377 | 0.00000 | 10.13697 | 11.12177 | 0.92723 | 11.55681 | 8.66451 | 6.97595 | 13.27375 |
| Fn1 | 19.39125 | 1.58446 | 2.37418 | 1.01610 | 4.06681 | 0.00000 | 0.00000 | 4.16740 | 1.31560 | 4.11899 | 0.20086 | 1.93349 | 2.58802 | 0.01563 | 0.41669 | 11.06675 |
| Fnbp1 | 3.23801 | 0.01053 | 2.93195 | 1.13226 | 7.10260 | 0.11357 | 1.45689 | 2.95160 | 3.00337 | 0.11516 | 1.81191 | 0.14590 | 1.75186 | 2.03312 | 1.60482 | 0.44232 |
| Fnbp1l | 0.00000 | 2.61494 | 0.03432 | 10.11841 | 0.05721 | 0.11402 | 2.57437 | 0.54904 | 2.86443 | 0.00000 | 2.45523 | 1.55704 | 2.06460 | 3.38545 | 0.93182 | 0.48230 |
| Fnbp4 | 7.79218 | 4.86201 | 6.01175 | 10.04581 | 4.19195 | 4.77853 | 2.57159 | 8.58326 | 11.13208 | 11.59393 | 9.33585 | 15.07749 | 6.96284 | 5.89181 | 17.19143 | 3.38427 |
| Fndc10 | 0.05084 | 0.00820 | 0.00741 | 1.61821 | 0.00814 | 0.03508 | 0.00760 | 0.01643 | 0.00000 | 0.57636 | 0.01279 | 0.03185 | 0.16671 | 0.00799 | 0.04941 | 0.88384 |
| Fndc3a | 3.95146 | 2.98256 | 0.65313 | 4.22608 | 7.24285 | 1.56197 | 3.36087 | 0.00000 | 2.99319 | 0.00000 | 0.00000 | 0.00000 | 0.31748 | 1.76737 | 1.72530 | 2.81645 |
| Fndc3b | 0.32564 | 2.92479 | 0.10868 | 2.82869 | 4.19379 | 0.00501 | 1.07263 | 0.03663 | 3.96376 | 0.25464 | 0.00875 | 0.30652 | 0.03113 | 2.31613 | 2.12824 | 0.01243 |
| Fndc4 | 0.00940 | 0.00749 | 0.61249 | 0.16726 | 0.01886 | 0.00598 | 0.00718 | 0.77191 | 0.31394 | 0.10991 | 0.26568 | 0.01534 | 0.07204 | 0.00840 | 0.00000 | 0.00719 |
| Fndc5 | 0.01181 | 0.00767 | 0.57814 | 0.00761 | 0.00794 | 0.00931 | 0.00718 | 0.01035 | 0.01174 | 0.92977 | 0.01088 | 0.01452 | 0.00936 | 0.01387 | 0.00152 | 0.00739 |
| Fndc7 | 0.01355 | 0.00798 | 0.01212 | 0.00890 | 0.00916 | 0.00675 | 0.00745 | 0.01201 | 0.01204 | 0.41432 | 0.01213 | 0.00746 | 0.01231 | 0.01169 | 0.01033 | 0.00748 |
| Fndc8 | 0.01236 | 0.00787 | 0.00872 | 0.45579 | 0.00905 | 0.00645 | 0.00722 | 0.01419 | 0.06620 | 0.89326 | 0.00762 | 0.00274 | 0.01045 | 0.47788 | 0.47851 | 0.00702 |
| Fnip1 | 4.27514 | 0.22196 | 0.12843 | 6.97971 | 1.45729 | 0.30074 | 0.45879 | 0.37639 | 6.09725 | 0.49659 | 1.35506 | 6.61170 | 0.37751 | 6.74194 | 3.01190 | 0.00758 |
| Fnip2 | 0.59372 | 2.27791 | 0.37008 | 1.57383 | 2.43487 | 0.07142 | 0.82059 | 0.45729 | 1.60835 | 1.94624 | 2.17281 | 0.06350 | 0.08253 | 0.84296 | 4.44570 | 0.01953 |
| Fnta | 8.42133 | 5.69073 | 5.05470 | 7.55130 | 6.53820 | 2.37471 | 3.33966 | 3.93373 | 3.17568 | 9.45848 | 7.92403 | 5.21006 | 6.42047 | 11.01397 | 2.78164 | 0.46392 |
| Fntb | 8.20498 | 0.00000 | 3.04048 | 2.21775 | 3.77099 | 3.74104 | 0.00000 | 2.83100 | 4.01380 | 6.45898 | 1.74321 | 0.11580 | 2.87780 | 0.10854 | 0.69705 | 1.75753 |
| Focad | 5.64313 | 0.10925 | 2.52390 | 0.75952 | 2.43402 | 0.00000 | 0.00000 | 2.55839 | 0.00000 | 1.94446 | 0.45651 | 0.01787 | 2.38584 | 0.00301 | 0.26605 | 4.38622 |
| Folr1 | 14.34889 | 12.62329 | 28.61415 | 12.58249 | 4.62514 | 10.93068 | 16.39186 | 27.39269 | 18.95451 | 35.53499 | 24.84791 | 12.82216 | 28.62525 | 2.11517 | 18.08742 | 9.37540 |
| Folr2 | 0.00000 | 0.00687 | 0.89087 | 0.83880 | 0.87398 | 0.08478 | 0.02612 | 1.03886 | 0.94651 | 0.88617 | 0.07599 | 0.00749 | 1.80425 | 3.54938 | 0.16268 | 0.01345 |
| Fopnl | 4.82570 | 6.98755 | 6.52295 | 3.11999 | 6.12057 | 0.26163 | 3.07694 | 4.14356 | 3.78918 | 9.39160 | 6.03685 | 1.00880 | 3.56212 | 4.39657 | 6.45108 | 3.28452 |
| Fos | 0.00000 | 3.14466 | 0.51439 | 0.12573 | 0.03023 | 0.00577 | 0.21229 | 0.01009 | 0.00935 | 0.04243 | 0.01068 | 0.03234 | 0.00724 | 0.01072 | 0.16625 | 5.76960 |
| Fosb | 0.01350 | 0.00811 | 0.01207 | 0.67042 | 0.00927 | 0.00658 | 0.00751 | 0.01381 | 0.39073 | 0.01231 | 0.01007 | 0.00759 | 0.01261 | 0.01321 | 0.00629 | 0.02920 |
| Fosl2 | 0.01731 | 0.00787 | 0.03541 | 0.08969 | 0.01033 | 0.00824 | 0.00736 | 0.01160 | 0.01094 | 0.24153 | 0.01175 | 0.03687 | 0.01231 | 0.00549 | 0.26073 | 0.00757 |
| Foxa1 | 1.24942 | 0.00000 | 0.13807 | 0.55584 | 0.01138 | 0.00237 | 0.00000 | 5.82828 | 0.04903 | 17.98126 | 12.63345 | 11.66245 | 0.00000 | 6.75598 | 8.87764 | 0.00285 |
| Foxa3 | 0.01361 | 0.00823 | 0.01221 | 0.36650 | 0.00932 | 0.00658 | 0.00749 | 0.01344 | 0.06860 | 0.01087 | 0.01191 | 0.00832 | 0.01257 | 0.00787 | 0.09087 | 0.02191 |
| Foxc1 | 0.78119 | 0.06968 | 0.53737 | 1.32711 | 0.00535 | 0.03182 | 0.03919 | 0.43946 | 0.73849 | 2.22115 | 0.32910 | 0.45631 | 0.51213 | 0.01114 | 0.00000 | 0.01226 |
| Foxc2 | 0.01296 | 0.00751 | 0.00547 | 0.01914 | 0.00895 | 0.00693 | 0.00701 | 0.00000 | 0.01203 | 0.99061 | 0.36537 | 0.12832 | 0.00845 | 0.36042 | 0.22897 | 0.00718 |
| Foxe1 | 0.00917 | 0.00775 | 0.02790 | 0.00687 | 0.00925 | 0.00614 | 0.00706 | 0.19613 | 0.01141 | 1.28622 | 0.01046 | 0.00394 | 0.51108 | 0.01330 | 0.00515 | 0.00721 |
| Foxf1 | 0.01353 | 0.00795 | 0.00379 | 0.05459 | 0.00897 | 0.00737 | 0.00706 | 0.00000 | 0.01063 | 0.19217 | 1.23188 | 0.04705 | 0.00000 | 0.00437 | 0.66160 | 0.00708 |
| Foxf2 | 0.75130 | 0.01993 | 0.00497 | 0.01888 | 0.01952 | 0.00743 | 0.03877 | 0.01183 | 0.01119 | 0.01405 | 0.01365 | 0.00748 | 0.04415 | 0.01213 | 0.01035 | 0.01482 |
| Foxg1 | 0.01679 | 0.00809 | 0.01407 | 0.09874 | 0.00894 | 0.00648 | 0.00763 | 0.02226 | 0.07537 | 0.01013 | 0.06975 | 0.01834 | 0.01845 | 0.13923 | 0.02501 | 0.00734 |
| Foxh1 | 6.02274 | 0.01916 | 0.88639 | 0.07953 | 3.98426 | 0.41607 | 0.33379 | 1.03885 | 0.00408 | 0.12157 | 0.03853 | 0.07119 | 0.25682 | 0.00600 | 0.07222 | 2.84136 |
| Foxi3 | 0.01374 | 0.00821 | 0.01202 | 0.83467 | 0.00931 | 0.00658 | 0.00749 | 0.01366 | 0.20265 | 0.01371 | 0.01234 | 0.01872 | 0.01274 | 0.01097 | 0.00301 | 0.00727 |
| Foxj1 | 0.01336 | 0.00789 | 0.01223 | 0.00775 | 0.00906 | 0.00651 | 0.00732 | 0.01051 | 0.01198 | 0.79288 | 0.01165 | 0.00563 | 0.01111 | 0.01169 | 0.00829 | 0.00734 |
| Foxj2 | 0.01008 | 0.00765 | 0.01253 | 0.47214 | 0.00800 | 3.44671 | 0.00700 | 0.00000 | 1.19605 | 0.00000 | 0.29589 | 0.00299 | 0.00000 | 0.00317 | 0.55229 | 1.14174 |
| Foxj3 | 1.98344 | 0.40383 | 3.53536 | 23.37398 | 0.95520 | 0.28504 | 2.34261 | 2.56988 | 26.32975 | 3.75777 | 3.02210 | 3.85136 | 1.38633 | 4.16157 | 14.53096 | 1.19301 |
| Foxk1 | 0.60902 | 0.00726 | 0.40946 | 2.77430 | 0.01483 | 0.22441 | 0.55125 | 1.82980 | 0.85103 | 0.40845 | 0.00000 | 0.03725 | 0.11610 | 0.23496 | 0.64363 | 0.00671 |
| Foxk2 | 1.62037 | 0.00405 | 3.73340 | 8.62313 | 3.04798 | 0.01217 | 0.04784 | 0.00000 | 2.81043 | 2.80747 | 0.32195 | 0.20600 | 0.07702 | 4.13294 | 15.02612 | 2.52012 |
| Foxl1 | 0.03866 | 0.00790 | 0.04778 | 0.06012 | 0.02755 | 0.00616 | 0.00731 | 0.01113 | 0.20998 | 0.00736 | 0.26770 | 0.04792 | 0.04389 | 0.10205 | 0.05550 | 0.00711 |
| Foxm1 | 3.60684 | 0.03201 | 1.01876 | 17.08004 | 2.27597 | 7.75811 | 1.59364 | 4.14536 | 16.39202 | 4.99648 | 0.99988 | 0.38282 | 0.70076 | 0.60864 | 6.25739 | 1.51752 |
| Foxn1 | 0.01385 | 0.00798 | 0.01050 | 0.01025 | 0.00914 | 0.00675 | 0.00746 | 0.00851 | 0.01221 | 0.03880 | 0.21335 | 0.00676 | 0.01017 | 0.01076 | 0.00691 | 0.00760 |
| Foxn2 | 1.71229 | 1.82004 | 0.59600 | 10.49043 | 3.11548 | 1.34240 | 2.10176 | 1.78126 | 8.23920 | 2.76699 | 5.04120 | 4.77784 | 2.53382 | 14.93488 | 6.61508 | 1.04033 |
| Foxn3 | 0.00780 | 0.00782 | 0.54130 | 0.72979 | 0.01054 | 0.00715 | 0.00714 | 0.42147 | 1.39776 | 1.25071 | 0.12790 | 0.25465 | 0.00337 | 1.27057 | 1.67112 | 0.01423 |
| Foxo1 | 1.63239 | 0.03027 | 4.78088 | 16.94613 | 7.49782 | 2.52731 | 5.65275 | 2.47141 | 8.78379 | 5.43064 | 4.56897 | 0.55980 | 2.79943 | 3.31325 | 7.49730 | 0.01937 |
| Foxo3 | 0.52453 | 3.27283 | 0.00841 | 0.10500 | 0.00681 | 0.00666 | 0.00506 | 1.96670 | 0.01761 | 4.03490 | 1.57698 | 0.04776 | 0.97200 | 0.00413 | 0.16482 | 0.01890 |
| Foxo4 | 0.03393 | 0.01795 | 0.01805 | 0.22133 | 0.01133 | 0.00607 | 1.27419 | 0.00954 | 0.38332 | 0.27781 | 0.01030 | 0.00493 | 0.01060 | 0.02848 | 0.02117 | 0.00736 |
| Foxo6 | 0.01810 | 0.00798 | 0.01173 | 0.01779 | 0.01030 | 0.00660 | 0.00729 | 0.01103 | 0.01106 | 0.39563 | 0.01264 | 0.00647 | 0.02561 | 0.01131 | 0.00978 | 0.00779 |
| Foxp1 | 0.01334 | 0.00791 | 0.01126 | 0.01692 | 0.00898 | 0.00671 | 0.00736 | 0.01081 | 0.08923 | 0.84092 | 0.01093 | 0.04775 | 0.01143 | 0.01191 | 0.00714 | 0.00738 |
| Foxp4 | 0.65619 | 0.00627 | 0.14151 | 0.67250 | 0.02354 | 0.00663 | 0.29088 | 3.11048 | 0.00000 | 1.85741 | 0.30822 | 0.04257 | 0.34779 | 0.00599 | 0.00000 | 0.28882 |
| Foxq1 | 0.02425 | 0.00836 | 0.01079 | 0.01791 | 0.01143 | 0.00650 | 0.00744 | 0.01214 | 0.01296 | 0.11208 | 0.01194 | 0.01772 | 0.00870 | 0.01034 | 0.01531 | 0.78957 |
| Foxr1 | 4.31387 | 1.16764 | 1.02126 | 14.52155 | 4.43222 | 0.00257 | 2.60185 | 3.70656 | 31.18933 | 6.79857 | 4.11092 | 1.24155 | 1.30990 | 0.93159 | 3.08800 | 1.27128 |

| | | | | | | | | | | | | | | | | |
|---|---|---|---|---|---|---|---|---|---|---|---|---|---|---|---|---|
| **Foxred1** | 5.53759 | 0.80386 | 1.08984 | 0.84766 | 2.17591 | 3.68593 | 1.77113 | 2.11049 | 0.94845 | 0.00000 | 2.01355 | 0.16583 | 2.84593 | 0.15373 | 3.14895 | 0.64207 |
| **Foxred2** | 0.94612 | 0.00556 | 0.00000 | 0.29421 | 0.01480 | 3.19746 | 1.71872 | 0.14325 | 0.02514 | 0.00000 | 0.68750 | 0.00881 | 0.03645 | 1.56040 | 0.04328 | 0.08632 |
| **Fpgs** | 8.98651 | 2.02702 | 2.09903 | 4.18408 | 0.01974 | 0.00000 | 0.14062 | 5.15472 | 1.39410 | 12.78479 | 1.80239 | 0.40297 | 3.18382 | 0.00000 | 3.78202 | 3.65235 |
| **Fpgt** | 1.42743 | 0.00631 | 0.43690 | 3.11646 | 0.01052 | 0.00592 | 0.00569 | 0.00000 | 0.21802 | 0.19931 | 0.70286 | 0.02412 | 0.00876 | 0.02172 | 2.86256 | 0.00550 |
| **Fra10ac1** | 5.97959 | 4.33364 | 1.10645 | 1.17108 | 4.44410 | 3.79541 | 0.00000 | 1.11445 | 0.00012 | 0.20749 | 2.67157 | 0.10148 | 0.47303 | 2.20265 | 1.29768 | 2.01166 |
| **Frat1** | 0.19750 | 0.00680 | 2.08097 | 0.33213 | 0.00401 | 0.28222 | 0.22676 | 0.41727 | 2.41053 | 0.91389 | 0.00000 | 0.01574 | 0.00000 | 0.00430 | 0.55088 | 0.02082 |
| **Frat2** | 1.69052 | 0.00442 | 0.00000 | 1.74840 | 0.05118 | 1.68214 | 0.00447 | 1.36455 | 0.48626 | 2.60326 | 0.37594 | 4.67997 | 1.49181 | 0.00374 | 1.98587 | 0.00386 |
| **Frem2** | 0.00617 | 2.43073 | 0.00563 | 0.34716 | 0.00784 | 0.00625 | 0.00636 | 0.01289 | 1.16294 | 0.53998 | 0.00000 | 0.12940 | 0.01089 | 0.00352 | 0.75935 | 0.00655 |
| **Frg1** | 6.51738 | 4.29338 | 8.98305 | 7.37930 | 3.70116 | 0.09808 | 5.94754 | 12.40691 | 14.15959 | 8.06007 | 6.87552 | 0.96756 | 7.35501 | 1.35486 | 8.71703 | 5.52619 |
| **Frk** | 0.00000 | 0.03310 | 0.00000 | 0.11043 | 0.01149 | 3.22480 | 0.00716 | 0.40184 | 0.00843 | 1.68716 | 0.25550 | 0.03777 | 1.20573 | 3.30540 | 0.00000 | 0.61341 |
| **Frmd3** | 0.02309 | 0.01740 | 0.02170 | 0.17439 | 0.00863 | 0.03730 | 0.00728 | 0.01528 | 0.02525 | 0.00364 | 0.00979 | 0.00610 | 0.02834 | 0.01968 | 0.68816 | 0.02904 |
| **Frmd4a** | 0.03978 | 0.01974 | 0.02185 | 0.22948 | 0.01146 | 0.00610 | 0.45598 | 0.02709 | 0.91122 | 0.54541 | 0.00519 | 0.03540 | 0.01070 | 0.03088 | 0.08581 | 0.00770 |
| **Frmd4b** | 0.58247 | 0.12886 | 10.01329 | 0.21954 | 0.03035 | 0.00701 | 2.41343 | 4.18178 | 0.55675 | 5.30514 | 0.33751 | 4.35614 | 6.37520 | 0.24072 | 5.05356 | 1.85064 |
| **Frmd6** | 2.79112 | 2.51971 | 3.29792 | 1.06246 | 0.36871 | 2.75225 | 3.06882 | 5.16489 | 0.56043 | 0.82726 | 1.08172 | 0.22951 | 0.00000 | 0.33859 | 0.07418 | 0.03069 |
| **Frmd8** | 10.06697 | 2.53980 | 9.78509 | 8.35020 | 0.49940 | 10.33244 | 2.18086 | 15.74880 | 4.86694 | 4.65623 | 9.53936 | 5.40168 | 11.05000 | 1.10882 | 2.76922 | 4.80366 |
| **Frmpd1** | 0.00000 | 0.00698 | 2.35204 | 1.38821 | 0.26397 | 0.82305 | 0.00377 | 3.96648 | 1.65047 | 4.97789 | 0.48909 | 0.27284 | 5.62322 | 0.11506 | 0.25486 | 0.82406 |
| **Frmpd3** | 0.01365 | 0.00799 | 0.01185 | 0.00663 | 0.00930 | 0.00663 | 0.00754 | 0.01370 | 0.79108 | 0.01092 | 0.00737 | 0.00677 | 0.01294 | 0.01609 | 0.01197 | 0.00738 |
| **Frrs1** | 6.35105 | 6.26598 | 10.38239 | 7.70195 | 1.05064 | 6.56956 | 9.18520 | 10.96063 | 9.93352 | 8.00264 | 8.66622 | 0.97770 | 10.51994 | 17.02518 | 11.19123 | 2.16199 |
| **Frrs1l** | 0.01689 | 0.00768 | 0.69122 | 0.08107 | 0.00780 | 0.00709 | 0.00730 | 0.01190 | 0.02208 | 0.07914 | 0.01082 | 0.00695 | 0.00881 | 0.01725 | 0.01478 | 0.00712 |
| **Frs2** | 1.04404 | 3.02071 | 0.63451 | 4.87687 | 1.77487 | 1.28247 | 0.84771 | 0.93375 | 2.69601 | 0.45720 | 1.73393 | 8.97761 | 1.27167 | 0.00000 | 1.90837 | 0.00202 |
| **Fry** | 1.14634 | 0.00802 | 0.11335 | 0.90335 | 0.01081 | 0.00812 | 0.00627 | 0.86482 | 1.05850 | 0.20979 | 0.00000 | 0.01356 | 1.90809 | 0.57543 | 0.58090 | 0.00543 |
| **Fryl** | 2.46873 | 0.00427 | 0.44299 | 3.48105 | 0.00284 | 0.00342 | 0.02473 | 1.05370 | 2.85590 | 0.20750 | 0.50268 | 1.81622 | 0.00000 | 4.41739 | 3.53322 | 1.32202 |
| **Fsbp** | 0.00600 | 0.00758 | 0.95781 | 0.00619 | 0.00722 | 0.00733 | 0.12174 | 0.42236 | 0.01035 | 1.11015 | 0.00605 | 0.00101 | 1.70744 | 0.01362 | 0.37502 | 0.00641 |
| **Fscn1** | 5.42086 | 0.02154 | 0.00000 | 5.36821 | 2.84945 | 0.00000 | 0.19024 | 0.09031 | 14.51398 | 7.06094 | 7.91686 | 0.76874 | 0.09194 | 0.00000 | 0.00000 | 4.13318 |
| **Fscn2** | 0.01375 | 0.00805 | 0.01226 | 0.06152 | 0.00931 | 0.00665 | 0.00753 | 0.01390 | 0.01182 | 0.01536 | 0.01380 | 0.00812 | 0.01268 | 0.01197 | 0.00960 | 0.00740 |
| **Fsd1** | 0.01074 | 0.00837 | 0.01004 | 5.38573 | 0.00948 | 0.00618 | 0.00747 | 0.12851 | 1.24980 | 0.00652 | 0.00000 | 0.01763 | 0.00816 | 0.00954 | 0.00780 | 0.00639 |
| **Fsd1l** | 0.06582 | 0.00768 | 0.41645 | 0.43065 | 0.04646 | 0.14206 | 0.03636 | 0.00000 | 0.01679 | 0.48341 | 1.47491 | 0.07418 | 0.52950 | 1.55806 | 0.30822 | 0.01359 |
| **Fsd2** | 0.01997 | 0.00756 | 0.61794 | 0.68617 | 0.00793 | 0.00545 | 0.00696 | 0.01192 | 0.78601 | 0.00000 | 0.00000 | 0.00935 | 0.00886 | 1.57376 | 0.00000 | 0.00687 |
| **Fsip1** | 0.40825 | 0.00894 | 0.00088 | 0.00702 | 0.01715 | 0.00797 | 0.00781 | 0.22979 | 0.01067 | 0.13368 | 0.00795 | 0.13849 | 0.00686 | 0.00809 | 0.56757 | 0.00724 |
| **Fstl1** | 0.01354 | 0.00791 | 0.00883 | 0.25736 | 0.00908 | 0.00708 | 0.00745 | 0.01263 | 0.00490 | 0.48606 | 0.00871 | 0.28603 | 0.01241 | 0.01225 | 0.00298 | 0.00737 |
| **Fstl3** | 0.00000 | 0.00844 | 0.58257 | 0.00449 | 0.00875 | 0.86525 | 0.08191 | 0.22464 | 0.02150 | 2.20970 | 0.00747 | 0.00000 | 3.01354 | 0.01253 | 0.00878 | 0.59770 |
| **Fth1** | 30.19265 | 8.34431 | 12.82661 | 22.73191 | 26.76484 | 8.98063 | 8.21312 | 5.26437 | 37.99140 | 34.27989 | 28.07510 | 3.19621 | 28.79996 | 5.04288 | 0.00000 | 16.06732 |
| **Fthl17.ps1** | 2.44463 | 0.00585 | 1.83643 | 1.64329 | 0.00945 | 0.18446 | 0.00549 | 1.42960 | 8.14095 | 1.70406 | 0.00000 | 0.04538 | 0.00000 | 0.31858 | 0.37717 | 0.27337 |
| **Fthl17a** | 0.01332 | 0.00802 | 0.22218 | 0.00492 | 0.00862 | 0.29324 | 0.00737 | 0.00189 | 0.01132 | 0.04685 | 0.51673 | 0.00535 | 0.00432 | 0.01144 | 0.00129 | 0.00688 |
| **Fthl17b** | 1.82023 | 0.00632 | 1.76902 | 0.96500 | 0.00969 | 0.20289 | 0.00591 | 1.38404 | 8.11842 | 1.35451 | 0.00000 | 0.04573 | 0.00000 | 0.23476 | 0.33645 | 0.19684 |
| **Fthl17d** | 1.16764 | 0.00695 | 1.76923 | 0.52413 | 0.00921 | 0.16610 | 0.00648 | 1.44248 | 7.12878 | 1.37896 | 0.00000 | 0.04794 | 0.00000 | 0.28790 | 0.39352 | 0.16245 |
| **Fthl17e** | 1.44967 | 0.00669 | 1.63998 | 3.14933 | 0.00956 | 0.18040 | 0.00620 | 1.81169 | 8.29729 | 2.01918 | 0.00000 | 0.03139 | 0.00000 | 0.58333 | 0.63922 | 0.31621 |
| **Fthl17f** | 2.10289 | 0.00574 | 3.12839 | 1.88610 | 0.04309 | 1.74158 | 0.00540 | 2.89385 | 11.80584 | 2.64914 | 0.00000 | 0.05970 | 0.00000 | 0.33720 | 0.77556 | 1.05831 |
| **Ftl1** | 58.32151 | 32.92937 | 53.85800 | 13.85057 | 34.28309 | 29.43612 | 33.10425 | 49.88310 | 14.71463 | 51.61982 | 38.78860 | 18.45228 | 55.25262 | 0.24809 | 3.23031 | 30.41021 |
| **Fto** | 1.96572 | 0.01907 | 2.85790 | 0.65719 | 0.02153 | 0.00959 | 1.37627 | 1.24604 | 0.94146 | 1.22428 | 0.00000 | 0.02315 | 0.86090 | 0.52270 | 0.41353 | 4.16470 |
| **Ftsj1** | 9.34526 | 1.71546 | 12.93739 | 3.14864 | 7.05611 | 3.34265 | 7.54826 | 10.93128 | 3.17026 | 9.60053 | 5.86266 | 0.97557 | 2.84617 | 4.18010 | 7.91425 | 2.11815 |
| **Ftsj3** | 13.21861 | 13.61589 | 9.49723 | 8.49807 | 8.84499 | 7.20963 | 6.25734 | 20.40687 | 5.61197 | 11.56871 | 11.93135 | 4.43684 | 9.09339 | 21.00131 | 14.28366 | 8.85324 |
| **Ftx** | 0.43524 | 0.01693 | 0.04618 | 0.01377 | 0.00711 | 0.00594 | 0.02684 | 0.00485 | 0.46905 | 0.00000 | 0.10230 | 0.01187 | 0.04713 | 0.18220 | 1.46476 | 0.53957 |
| **Fubp1** | 8.79095 | 13.42417 | 8.27053 | 7.68278 | 8.90012 | 0.85989 | 5.08567 | 12.37914 | 6.65145 | 8.18881 | 0.00001 | 7.76883 | 2.73888 | 4.87821 | 5.20333 | 5.10053 |
| **Fubp3** | 5.40256 | 5.79669 | 1.58204 | 2.39489 | 0.01788 | 2.83094 | 1.67653 | 4.63123 | 4.86101 | 0.67367 | 2.43838 | 4.98890 | 1.83558 | 4.33367 | 2.49922 | 2.75374 |
| **Fuca1** | 6.26763 | 0.00000 | 4.29624 | 2.00121 | 4.92971 | 1.88909 | 2.03269 | 0.23565 | 0.39190 | 2.62698 | 0.54485 | 0.09436 | 3.06828 | 0.00380 | 1.24509 | 10.15787 |
| **Fuca2** | 2.00646 | 0.00388 | 4.25153 | 0.29782 | 1.47530 | 0.10151 | 0.07306 | 1.25518 | 0.56440 | 0.39125 | 0.94143 | 0.02341 | 5.10583 | 1.43443 | 0.00000 | 6.08460 |
| **Fuk** | 1.63381 | 0.05468 | 0.65539 | 0.31948 | 0.01174 | 0.15319 | 0.00345 | 3.92885 | 1.84581 | 2.19916 | 1.65294 | 0.18850 | 3.24657 | 0.04678 | 0.08947 | 0.10710 |
| **Fundc1** | 6.68215 | 5.41387 | 7.88713 | 12.74907 | 8.41627 | 4.95040 | 2.32514 | 0.60539 | 3.86942 | 3.79547 | 0.00000 | 5.36383 | 3.48722 | 4.13273 | 1.29990 | 2.30213 |
| **Fundc2** | 1.06338 | 0.00698 | 0.00000 | 0.02867 | 0.00570 | 0.44655 | 0.00621 | 0.09003 | 1.24985 | 2.78697 | 1.13265 | 0.00000 | 0.68451 | 0.00415 | 1.02685 | 0.02766 |
| **Fuom** | 3.33585 | 0.08606 | 0.40728 | 0.13047 | 2.60634 | 0.00828 | 0.49229 | 0.00000 | 0.22242 | 0.54872 | 0.01058 | 0.03187 | 1.45409 | 0.01085 | 0.00000 | 0.17481 |
| **Furin** | 1.41960 | 0.47388 | 2.46020 | 1.55087 | 0.01750 | 1.46522 | 0.08655 | 2.59908 | 0.58440 | 1.96197 | 2.28494 | 0.02876 | 2.32072 | 0.40222 | 0.45105 | 1.11803 |
| **Fus** | 2.18176 | 0.02412 | 2.45204 | 5.39553 | 1.87078 | 0.89776 | 0.18287 | 0.00000 | 16.20415 | 0.06347 | 20.61338 | 0.48349 | 1.27731 | 2.69509 | 1.00624 | 1.43361 |
| **Fut10** | 0.20445 | 0.00776 | 0.35670 | 2.76775 | 0.01336 | 0.00843 | 0.00687 | 0.01346 | 0.11335 | 0.00437 | 0.00850 | 0.01366 | 0.00934 | 0.01793 | 1.17622 | 0.00659 |
| **Fut11** | 0.02710 | 0.00810 | 0.00501 | 0.96141 | 0.01047 | 0.07390 | 0.00744 | 0.01103 | 2.08902 | 0.35113 | 0.00000 | 0.02664 | 0.01178 | 0.00488 | 0.93963 | 0.05842 |
| **Fut2** | 1.27418 | 0.28061 | 0.00656 | 0.45953 | 0.00556 | 0.00545 | 0.00600 | 0.06637 | 0.31299 | 0.54026 | 0.00904 | 0.00339 | 0.17182 | 0.00424 | 0.36629 | 0.00574 |
| **Fut4** | 1.26524 | 0.00724 | 0.00000 | 0.10762 | 0.01656 | 0.89593 | 0.00579 | 0.00493 | 0.00980 | 0.01989 | 0.01130 | 0.00638 | 0.71726 | 0.00693 | 0.21498 | 0.00565 |
| **Fut8** | 3.00005 | 5.84785 | 1.53894 | 1.35908 | 0.01721 | 1.42654 | 0.14393 | 0.65201 | 0.07441 | 3.25247 | 0.81662 | 1.04968 | 2.05483 | 2.96000 | 2.09421 | 1.79886 |
| **Fut9** | 0.01354 | 0.00803 | 0.01123 | 0.00710 | 0.00917 | 0.00682 | 0.00752 | 0.01249 | 0.01116 | 0.29520 | 0.01015 | 0.03134 | 0.01276 | 0.00588 | 0.42730 | 0.00741 |
| **Fuz** | 0.01113 | 0.00842 | 0.35098 | 0.01059 | 0.00912 | 0.00563 | 0.00795 | 0.38768 | 0.01191 | 0.03439 | 0.00993 | 0.00777 | 0.07252 | 0.01307 | 0.01103 | 0.00721 |
| **Fv1** | 0.01352 | 0.00803 | 0.01167 | 0.43346 | 0.00927 | 0.00647 | 0.00750 | 0.01320 | 1.70733 | 0.00581 | 0.00000 | 0.00518 | 0.01302 | 0.00980 | 0.08415 | 0.00719 |
| **Fxn** | 0.96946 | 0.02839 | 1.09168 | 0.11691 | 0.00457 | 0.00539 | 0.01642 | 0.00000 | 2.36842 | 1.88491 | 1.40769 | 0.07195 | 0.18803 | 0.03188 | 0.67161 | 0.28700 |

| | | | | | | | | | | | | | | | | |
|---|---|---|---|---|---|---|---|---|---|---|---|---|---|---|---|---|
| **Fxr1** | 17.32667 | 15.78905 | 8.45241 | 12.43360 | 7.62881 | 1.32561 | 10.15535 | 9.82779 | 19.52417 | 5.36389 | 7.60726 | 5.06610 | 8.56282 | 6.50146 | 8.20762 | 4.23238 |
| **Fxr2** | 6.58811 | 5.74038 | 9.58237 | 10.24266 | 5.02918 | 4.78321 | 3.50335 | 10.04032 | 8.26881 | 5.38869 | 5.02138 | 0.58351 | 9.64755 | 1.27445 | 6.78280 | 3.67368 |
| **Fxyd3** | 0.01355 | 0.00791 | 0.01216 | 0.00873 | 0.00914 | 0.00667 | 0.00738 | 0.01137 | 0.01205 | 0.45445 | 0.01241 | 0.00678 | 0.01214 | 0.01153 | 0.00990 | 0.00758 |
| **Fxyd4** | 0.51632 | 0.00546 | 0.70591 | 0.00368 | 0.00564 | 0.21002 | 0.76373 | 0.93074 | 0.01861 | 4.31026 | 4.58705 | 0.00906 | 1.14638 | 0.00880 | 0.00000 | 0.01983 |
| **Fxyd5** | 0.01614 | 0.00763 | 0.25713 | 0.00796 | 0.00871 | 0.00572 | 0.00719 | 0.01006 | 0.01188 | 0.74911 | 0.01157 | 0.01567 | 0.01034 | 0.01267 | 0.00466 | 0.00729 |
| **Fxyd6** | 0.00000 | 0.00762 | 0.03281 | 0.10588 | 0.03185 | 9.26987 | 0.00506 | 0.00468 | 0.00710 | 0.75322 | 0.00000 | 0.00927 | 2.58192 | 0.90073 | 0.04670 | 4.33471 |
| **Fyb** | 0.01605 | 0.00779 | 0.16532 | 0.24156 | 0.00903 | 0.00576 | 0.00725 | 0.01136 | 0.00756 | 0.19776 | 0.01353 | 0.00701 | 0.01080 | 0.01074 | 0.00436 | 0.00732 |
| **Fyco1** | 1.43129 | 2.09554 | 3.22403 | 10.81539 | 0.84559 | 0.02189 | 1.40245 | 0.70674 | 2.25000 | 2.19614 | 1.86426 | 0.33204 | 0.02816 | 3.06039 | 3.24858 | 2.45381 |
| **Fyn** | 0.00897 | 0.00766 | 0.00662 | 4.04214 | 0.01059 | 0.00678 | 0.00733 | 0.75920 | 1.98943 | 0.00000 | 0.09287 | 0.00000 | 0.07710 | 0.01875 | 1.26190 | 0.00682 |
| **Fyttd1** | 3.03614 | 0.00000 | 2.67635 | 8.95948 | 5.51218 | 6.05576 | 1.12300 | 1.53514 | 6.02157 | 7.69472 | 2.08678 | 8.61673 | 3.34253 | 5.18034 | 5.80244 | 0.05627 |
| **Fzd2** | 0.01326 | 0.00813 | 0.01192 | 0.00989 | 0.00933 | 0.00654 | 0.00761 | 0.01715 | 0.59450 | 0.30704 | 0.00868 | 0.00618 | 0.01196 | 0.01278 | 0.00609 | 0.00743 |
| **Fzd3** | 0.67091 | 0.00628 | 0.50786 | 5.52689 | 0.01439 | 0.00409 | 0.00464 | 1.97891 | 5.56043 | 2.40719 | 4.00144 | 0.13258 | 1.03372 | 0.30274 | 2.24365 | 0.02659 |
| **Fzd4** | 0.01284 | 0.00787 | 0.00343 | 3.13196 | 0.00889 | 0.00628 | 0.00687 | 0.00000 | 3.00768 | 0.34707 | 0.94260 | 0.00000 | 0.00427 | 0.00000 | 1.18812 | 0.00621 |
| **Fzd5** | 1.54196 | 1.24466 | 0.33519 | 3.73414 | 1.06816 | 0.77472 | 0.00566 | 0.63756 | 0.49816 | 0.07914 | 1.16062 | 0.03506 | 0.15101 | 0.27114 | 0.07718 | 0.01851 |
| **Fzd6** | 0.24167 | 0.00798 | 0.00134 | 0.00943 | 0.01596 | 0.00831 | 0.00691 | 0.51094 | 0.01153 | 0.22059 | 0.02487 | 0.01590 | 0.00000 | 0.01260 | 0.00413 | 1.42881 |
| **Fzd7** | 0.84853 | 0.00666 | 0.01260 | 2.96096 | 0.01446 | 0.00468 | 0.00675 | 0.00810 | 4.10343 | 0.00000 | 0.00000 | 0.03091 | 0.01778 | 0.00967 | 1.20741 | 0.00546 |
| **Fzd8** | 0.02203 | 0.00773 | 0.00670 | 0.01630 | 0.00849 | 0.00693 | 0.00717 | 0.47213 | 0.01171 | 0.15797 | 0.00803 | 0.01617 | 0.00563 | 0.00778 | 0.18264 | 0.00706 |
| **Fzd9** | 0.01163 | 0.00778 | 0.00761 | 0.98492 | 0.00937 | 0.00674 | 0.00690 | 0.01308 | 5.70204 | 0.00000 | 0.46700 | 0.04140 | 0.83035 | 0.00218 | 0.00000 | 0.01390 |
| **Fzr1** | 7.70671 | 2.28320 | 5.48911 | 1.40037 | 5.48329 | 5.81185 | 2.81869 | 4.38925 | 2.08252 | 4.91655 | 2.01123 | 0.15345 | 7.15802 | 0.03010 | 0.00000 | 4.39787 |
| **G2e3** | 0.00532 | 0.00658 | 0.43375 | 3.24809 | 0.00889 | 1.29048 | 0.00666 | 0.53472 | 11.76468 | 1.46928 | 1.21126 | 5.37039 | 0.00000 | 0.00000 | 5.42277 | 0.00482 |
| **G3bp1** | 25.22897 | 11.13574 | 17.92519 | 14.14944 | 19.66481 | 10.20762 | 14.13278 | 29.69035 | 17.96885 | 23.24044 | 18.29709 | 4.38627 | 24.94682 | 11.12119 | 15.06273 | 13.42471 |
| **G3bp2** | 20.97919 | 15.44451 | 21.88771 | 20.01393 | 17.90044 | 19.10960 | 16.40513 | 29.81194 | 20.12859 | 33.69302 | 26.83494 | 17.00896 | 32.78623 | 35.34228 | 28.80268 | 13.23992 |
| **G6pc** | 0.01376 | 0.00782 | 0.01176 | 0.20618 | 0.00933 | 0.00664 | 0.00751 | 0.01292 | 0.24066 | 0.01001 | 0.02188 | 0.00734 | 0.01254 | 0.01903 | 0.05897 | 0.00736 |
| **G6pc3** | 0.01561 | 0.00556 | 2.73933 | 0.21706 | 3.54582 | 0.00686 | 0.01768 | 4.56433 | 1.20882 | 5.88275 | 1.37219 | 0.22681 | 0.00000 | 0.08923 | 0.95295 | 6.10738 |
| **G6pd2** | 0.00663 | 0.23673 | 0.15499 | 0.00953 | 0.23032 | 0.23142 | 0.05041 | 0.09771 | 0.01191 | 0.00507 | 0.01193 | 0.00744 | 0.04375 | 0.05989 | 0.10740 | 0.07253 |
| **G6pdx** | 0.14774 | 5.60922 | 3.52316 | 11.84388 | 9.31038 | 6.28020 | 4.42808 | 3.25382 | 2.34954 | 0.36286 | 1.39985 | 0.12716 | 3.01132 | 2.35304 | 3.26795 | 3.32736 |
| **Gaa** | 12.44352 | 5.33056 | 20.36670 | 4.61077 | 7.48181 | 7.30644 | 11.08297 | 27.38754 | 1.34017 | 16.84722 | 17.44035 | 5.21412 | 19.52416 | 0.01152 | 1.10016 | 6.59495 |
| **Gab1** | 6.12918 | 5.38573 | 5.24328 | 0.00000 | 4.67965 | 1.03082 | 3.69942 | 5.67350 | 1.55453 | 4.09089 | 3.42294 | 7.22918 | 3.23017 | 5.34359 | 4.12229 | 2.72445 |
| **Gab2** | 0.55093 | 3.22634 | 2.17356 | 1.64860 | 0.00290 | 0.00822 | 4.05933 | 0.45837 | 1.50591 | 0.00000 | 0.01292 | 0.00860 | 0.57044 | 0.02253 | 0.16203 | 0.00373 |
| **Gab3** | 0.02042 | 0.00798 | 0.01070 | 0.32889 | 0.00886 | 0.01372 | 0.00761 | 0.01313 | 0.08823 | 0.27225 | 0.01109 | 0.00831 | 0.01522 | 0.04690 | 0.04072 | 0.00739 |
| **Gabarap** | 7.31469 | 5.57962 | 3.12495 | 10.08660 | 2.81481 | 5.77949 | 1.90550 | 5.91222 | 9.39358 | 12.79361 | 7.23369 | 4.20705 | 7.15003 | 0.84492 | 5.58100 | 3.26152 |
| **Gabarapl1** | 8.51497 | 0.15215 | 7.23393 | 0.24601 | 10.96927 | 5.08489 | 2.08665 | 0.27485 | 0.00000 | 2.99906 | 0.37702 | 0.00000 | 2.65600 | 0.42921 | 0.00000 | 0.00000 |
| **Gabarapl2** | 45.70508 | 23.56372 | 40.73470 | 17.33871 | 32.21975 | 22.95576 | 23.24157 | 40.18810 | 25.09320 | 41.57961 | 44.24605 | 22.09509 | 43.54083 | 25.82968 | 20.41980 | 25.33494 |
| **Gabbr1** | 1.10644 | 0.00676 | 0.28250 | 1.04760 | 0.09435 | 0.01032 | 0.07650 | 0.01673 | 0.81322 | 0.11945 | 0.00000 | 0.02501 | 0.01689 | 0.69048 | 0.00000 | 0.01316 |
| **Gabpa** | 9.37939 | 10.79143 | 9.69277 | 18.46685 | 14.30121 | 4.94845 | 5.25145 | 14.70169 | 20.72292 | 11.51904 | 12.52732 | 12.10294 | 7.73819 | 18.31541 | 22.46354 | 3.54052 |
| **Gabpb1** | 1.83091 | 0.41749 | 2.84439 | 10.23905 | 3.14018 | 0.95552 | 5.01958 | 3.18643 | 10.43386 | 4.32631 | 5.41089 | 3.55422 | 0.43360 | 17.67074 | 6.05842 | 1.38006 |
| **Gabpb2** | 2.73108 | 0.10449 | 2.57864 | 4.04987 | 0.27079 | 1.00750 | 1.01008 | 2.23530 | 7.31135 | 0.55750 | 2.83534 | 8.03338 | 1.09464 | 11.11990 | 11.21700 | 0.06403 |
| **Gabra3** | 0.02164 | 0.00787 | 0.01618 | 0.96220 | 0.00876 | 0.00634 | 0.00746 | 0.01327 | 0.00000 | 0.01796 | 0.01270 | 0.00761 | 0.01283 | 0.05046 | 0.04937 | 0.00733 |
| **Gabrb3** | 0.29829 | 0.00784 | 0.00086 | 0.01911 | 0.01778 | 0.00779 | 0.00695 | 0.00821 | 0.01272 | 0.00719 | 0.00854 | 0.00387 | 0.01931 | 1.06734 | 0.00521 | 0.00733 |
| **Gabrd** | 0.00014 | 0.00726 | 1.76334 | 0.06513 | 0.00775 | 0.01305 | 0.01614 | 2.13216 | 0.75565 | 0.00000 | 1.25240 | 0.14793 | 1.45463 | 0.14331 | 0.01192 | 0.00577 |
| **Gabre** | 0.01663 | 0.00817 | 0.07340 | 0.11026 | 0.00878 | 0.00721 | 0.00752 | 0.01407 | 0.55755 | 0.01234 | 0.00924 | 0.00745 | 0.01163 | 0.01243 | 0.00750 | 0.00730 |
| **Gabrp** | 0.00976 | 0.00815 | 0.01132 | 0.89948 | 0.00916 | 0.00658 | 0.00696 | 0.01207 | 0.00464 | 0.03836 | 0.01142 | 0.03261 | 0.92347 | 0.00678 | 0.09111 | 0.00679 |
| **Gad2** | 0.10742 | 0.01835 | 0.15318 | 0.44920 | 0.07413 | 0.02940 | 0.03868 | 0.08188 | 0.37063 | 0.08461 | 0.44800 | 0.07376 | 0.18104 | 0.39787 | 0.19354 | 0.01407 |
| **Gadd45a** | 0.09358 | 0.00742 | 0.01774 | 0.00504 | 0.00769 | 0.00604 | 0.00718 | 0.20870 | 0.31525 | 1.72227 | 0.00707 | 0.01254 | 0.00912 | 0.02616 | 0.07016 | 0.00702 |
| **Gadd45b** | 0.00023 | 0.02981 | 0.21681 | 0.94010 | 0.03066 | 2.63000 | 0.00520 | 4.18755 | 6.17611 | 1.07579 | 0.38531 | 0.02146 | 3.86675 | 0.01942 | 0.00000 | 0.70663 |
| **Gadd45g** | 0.00000 | 0.00479 | 0.00000 | 0.00000 | 4.18292 | 1.46036 | 0.02702 | 2.79294 | 0.27589 | 2.76317 | 0.53594 | 0.00000 | 0.00000 | 0.01141 | 0.00000 | 0.01447 |
| **Gadd45gip1** | 2.98156 | 0.05404 | 0.96094 | 9.75128 | 0.91609 | 0.66324 | 0.32861 | 0.71265 | 13.08493 | 17.86281 | 20.26769 | 0.43745 | 6.45459 | 0.29080 | 0.33810 | 6.00648 |
| **Gak** | 4.21117 | 4.97375 | 6.31400 | 6.79308 | 7.72624 | 8.07522 | 5.09039 | 6.82479 | 9.90578 | 6.57509 | 6.84946 | 1.18307 | 9.61037 | 1.57116 | 3.15134 | 7.16503 |
| **Gal** | 0.01365 | 0.00827 | 0.01212 | 0.76540 | 0.00921 | 0.00681 | 0.00757 | 0.01396 | 0.00000 | 0.14740 | 0.01299 | 0.00914 | 0.01296 | 0.01026 | 0.00530 | 0.00729 |
| **Gal3st1** | 0.01263 | 0.00819 | 0.01115 | 0.21870 | 0.01902 | 0.00665 | 0.00767 | 0.01967 | 0.00546 | 0.05086 | 0.01163 | 0.06029 | 0.01232 | 0.00921 | 0.04455 | 0.00742 |
| **Gal3st2** | 0.02098 | 0.00800 | 1.37813 | 7.69676 | 0.00687 | 0.00890 | 0.01917 | 1.00608 | 2.06637 | 0.00168 | 0.01308 | 0.02275 | 0.35388 | 0.00052 | 0.53880 | 0.00547 |
| **Gal3st3** | 0.17988 | 0.03902 | 0.48573 | 0.54452 | 0.54095 | 0.04587 | 0.02788 | 0.03658 | 0.47780 | 0.30145 | 0.46830 | 0.07732 | 0.13712 | 0.46283 | 0.23602 | 0.08501 |
| **Galc** | 0.01264 | 0.00768 | 0.00607 | 0.83860 | 0.00879 | 0.00693 | 0.01742 | 0.00000 | 2.28020 | 1.00912 | 0.60572 | 0.00000 | 0.00726 | 0.24005 | 0.03994 | 0.00721 |
| **Gale** | 7.53649 | 1.77332 | 3.45994 | 2.02049 | 3.74038 | 2.48426 | 3.75523 | 2.42072 | 0.41608 | 1.54992 | 1.80178 | 0.11865 | 3.40475 | 0.00000 | 0.00000 | 0.01634 |
| **Galk1** | 10.92700 | 0.54630 | 4.88484 | 0.42991 | 12.22524 | 7.06323 | 5.55100 | 19.32460 | 0.00000 | 22.47617 | 19.23863 | 2.81980 | 26.43894 | 0.00000 | 0.00000 | 13.41054 |
| **Galk2** | 3.06742 | 1.07572 | 0.89997 | 0.30933 | 1.51766 | 5.11309 | 1.97083 | 2.11132 | 0.00187 | 2.53951 | 1.76662 | 0.10876 | 2.72107 | 0.02561 | 2.48766 | 3.28645 |
| **Galm** | 2.86490 | 0.17523 | 0.02136 | 0.43133 | 0.39506 | 0.00298 | 5.51936 | 0.28734 | 2.07507 | 3.87735 | 0.38189 | 0.00000 | 0.02303 | 0.47490 | 1.03189 | 0.00404 |
| **Galns** | 0.01214 | 0.00760 | 0.01004 | 2.39334 | 0.01869 | 3.46103 | 0.00701 | 0.03723 | 0.16403 | 0.00367 | 0.00755 | 0.02683 | 0.00000 | 0.00206 | 0.31394 | 0.00555 |
| **Galnt1** | 5.85938 | 4.56373 | 5.37007 | 3.70983 | 5.17761 | 4.29009 | 5.45994 | 2.04965 | 0.20968 | 4.36711 | 1.04957 | 7.34878 | 2.00029 | 3.92217 | 4.93538 | 4.54711 |
| **Galnt10** | 0.01324 | 0.00804 | 0.01211 | 0.27177 | 0.00896 | 0.00647 | 0.00737 | 0.01078 | 0.00738 | 0.71425 | 0.01178 | 0.01607 | 0.01062 | 0.00989 | 0.00816 | 0.02808 |
| **Galnt11** | 2.83357 | 0.00000 | 2.96569 | 3.32875 | 3.75692 | 1.42407 | 2.41128 | 6.34021 | 4.51534 | 2.88091 | 3.50648 | 0.70325 | 0.03924 | 4.47832 | 6.05720 | 2.60961 |
| **Galnt12** | 0.01371 | 0.00761 | 0.01097 | 1.13428 | 0.00925 | 0.00648 | 0.00731 | 0.01111 | 0.47130 | 0.08523 | 0.00762 | 0.03323 | 0.01321 | 0.00559 | 0.49623 | 0.00752 |

| | | | | | | | | | | | | | | | | |
|---|---|---|---|---|---|---|---|---|---|---|---|---|---|---|---|---|
| **Galnt16** | 0.00000 | 0.00824 | 1.20878 | 0.28396 | 0.00738 | 0.00557 | 0.00712 | 0.62982 | 0.27716 | 2.13335 | 0.00000 | 0.03849 | 0.91363 | 1.43887 | 1.06985 | 0.00627 |
| **Galnt2** | 1.15406 | 2.15157 | 3.13329 | 0.52016 | 0.00002 | 3.39305 | 4.81740 | 5.46217 | 0.92152 | 4.94787 | 2.17973 | 0.11236 | 0.55163 | 0.37265 | 0.97544 | 3.09182 |
| **Galnt3** | 0.01026 | 0.00737 | 0.22107 | 0.41014 | 0.03026 | 0.00738 | 0.54858 | 0.01184 | 0.00348 | 0.02217 | 0.01119 | 0.04234 | 0.07847 | 0.00605 | 0.60798 | 0.00749 |
| **Galnt4** | 0.04802 | 0.00791 | 0.07383 | 1.12623 | 0.01260 | 0.08867 | 0.00710 | 0.00929 | 0.49968 | 0.00362 | 0.00693 | 0.00412 | 0.02350 | 0.00587 | 1.31364 | 0.01434 |
| **Galnt6** | 4.39704 | 0.00201 | 2.33473 | 0.57945 | 0.26251 | 4.30418 | 0.00000 | 2.13167 | 2.44028 | 4.00374 | 2.24248 | 0.01988 | 5.08842 | 0.02435 | 0.85995 | 0.84611 |
| **Galnt7** | 1.38590 | 0.00591 | 0.00288 | 0.00872 | 0.00529 | 0.00669 | 0.00638 | 0.00743 | 0.01198 | 0.01059 | 0.01190 | 0.00598 | 0.01154 | 0.00608 | 0.31253 | 0.00593 |
| **Galnt9** | 0.00644 | 0.00854 | 0.00693 | 0.00434 | 0.01855 | 0.00695 | 0.00824 | 0.40620 | 0.33877 | 1.65852 | 0.10672 | 0.02411 | 0.00534 | 0.31609 | 0.00000 | 0.00714 |
| **Galntl5** | 0.00838 | 0.00787 | 0.52305 | 1.47754 | 0.00898 | 0.00542 | 0.00708 | 0.15618 | 0.68562 | 0.00000 | 0.51638 | 0.00245 | 0.02259 | 0.26557 | 0.18723 | 0.00663 |
| **Galntl6** | 0.03078 | 0.01852 | 0.11369 | 0.18799 | 0.01051 | 0.01236 | 0.03799 | 0.01269 | 0.09773 | 0.04958 | 0.12312 | 0.02538 | 0.04005 | 0.21246 | 0.00735 | 0.01486 |
| **Galt** | 5.08248 | 0.86731 | 4.42969 | 1.06734 | 3.39516 | 3.57695 | 1.30769 | 1.35123 | 4.01885 | 3.55370 | 1.70479 | 1.69553 | 2.47919 | 0.03151 | 0.19041 | 2.15879 |
| **Gamt** | 0.80467 | 0.01650 | 0.80981 | 0.00486 | 0.00621 | 0.00801 | 0.00562 | 0.26111 | 0.25275 | 1.56667 | 2.43865 | 0.01625 | 0.18438 | 0.00362 | 0.23336 | 0.00594 |
| **Gan** | 1.30772 | 0.00712 | 0.25758 | 1.68630 | 0.01528 | 0.00396 | 0.00612 | 0.00000 | 2.43081 | 0.74959 | 0.52230 | 0.16093 | 0.06273 | 1.75276 | 0.00000 | 0.00544 |
| **Ganab** | 10.34328 | 0.01812 | 8.65299 | 2.50286 | 5.06955 | 7.31463 | 8.12878 | 12.09552 | 3.01953 | 6.22348 | 3.46890 | 0.60335 | 4.80932 | 5.39504 | 9.59146 | 4.23644 |
| **Ganc** | 0.00947 | 0.00727 | 0.97340 | 0.13665 | 0.00742 | 0.00734 | 0.00695 | 0.19502 | 0.39718 | 1.30580 | 0.00705 | 0.01239 | 0.00986 | 0.02851 | 0.02841 | 0.02837 |
| **Gapdh** | 40.96248 | 21.49199 | 28.30940 | 11.72569 | 28.67127 | 19.55495 | 17.02734 | 4.77568 | 15.77732 | 8.01435 | 2.16543 | 5.14605 | 31.19249 | 0.07348 | 4.30225 | 27.60821 |
| **Gapdhs** | 0.00783 | 0.00772 | 0.27724 | 0.00913 | 0.53068 | 0.00611 | 0.01863 | 0.34854 | 0.01161 | 0.25222 | 0.01058 | 0.00643 | 0.02572 | 0.01188 | 0.00817 | 0.41981 |
| **Gapt** | 0.00584 | 0.00584 | 0.00513 | 1.29220 | 0.00886 | 0.00495 | 0.00680 | 0.17796 | 0.68227 | 0.00872 | 0.00000 | 7.23518 | 0.00722 | 0.23103 | 0.69533 | 0.00641 |
| **Gapvd1** | 1.47434 | 2.72486 | 2.97602 | 5.54533 | 2.44936 | 3.34494 | 1.85749 | 7.19759 | 6.86432 | 7.02509 | 2.42284 | 0.69084 | 2.09740 | 7.55684 | 10.01910 | 1.61440 |
| **Gar1** | 4.50684 | 1.80017 | 4.35181 | 1.60142 | 7.42890 | 2.91128 | 3.49468 | 3.93235 | 4.24290 | 2.13258 | 6.42172 | 0.62418 | 3.93530 | 1.66654 | 4.50501 | 1.47069 |
| **Garem** | 0.00700 | 0.00803 | 0.09515 | 0.01131 | 0.01053 | 0.00605 | 0.00788 | 1.66072 | 2.93419 | 0.00000 | 0.26589 | 0.00737 | 0.29386 | 0.00331 | 0.35091 | 0.00724 |
| **Gareml** | 0.01103 | 0.00872 | 0.01794 | 0.00980 | 0.00946 | 0.00627 | 0.00821 | 0.19243 | 0.01179 | 0.10037 | 0.00945 | 0.09246 | 0.00685 | 0.01063 | 0.09283 | 0.00717 |
| **Garnl3** | 0.06176 | 0.00890 | 0.89118 | 1.45793 | 0.01579 | 0.00947 | 0.00775 | 0.04507 | 0.20072 | 1.06774 | 0.00076 | 0.34237 | 0.08844 | 0.00349 | 1.30527 | 0.00721 |
| **Gars** | 19.43104 | 14.44560 | 17.82430 | 4.86069 | 18.39616 | 8.93123 | 11.74680 | 16.29258 | 5.63364 | 13.71557 | 10.54701 | 4.09723 | 13.47779 | 14.69442 | 8.54337 | 9.87255 |
| **Gart** | 11.11282 | 12.15411 | 14.96371 | 5.93087 | 9.69819 | 4.22416 | 12.72264 | 19.68906 | 4.56912 | 13.95142 | 10.72175 | 1.31742 | 11.20306 | 0.44915 | 10.70533 | 10.51645 |
| **Gas1** | 0.01336 | 0.00784 | 0.01156 | 0.00672 | 0.00895 | 0.00669 | 0.00730 | 0.00861 | 0.01158 | 0.87490 | 0.07487 | 0.00548 | 0.01112 | 0.01164 | 0.00774 | 0.00760 |
| **Gas2** | 0.12326 | 0.00703 | 0.30901 | 0.29347 | 0.01549 | 0.01786 | 0.01692 | 0.00000 | 0.82226 | 2.26453 | 0.35209 | 0.01776 | 0.00989 | 0.01068 | 0.00000 | 0.01393 |
| **Gas2l1** | 5.29528 | 0.00118 | 0.18431 | 4.60332 | 0.00000 | 2.27865 | 0.00309 | 2.38937 | 6.92713 | 2.83693 | 0.00000 | 0.05886 | 0.00000 | 0.00974 | 0.26525 | 0.07680 |
| **Gas2l3** | 1.77989 | 1.05776 | 2.58839 | 10.26422 | 0.99491 | 0.55192 | 0.42608 | 0.00000 | 12.94771 | 2.23978 | 13.31570 | 1.06125 | 2.36462 | 4.98361 | 2.69077 | 0.42075 |
| **Gas5** | 13.54321 | 1.70742 | 15.27607 | 10.86344 | 13.21060 | 7.11134 | 9.49267 | 11.90780 | 22.22504 | 17.43886 | 30.52905 | 14.30255 | 11.42451 | 11.07703 | 10.39365 | 6.13315 |
| **Gas6** | 0.00984 | 0.00756 | 0.01353 | 0.07670 | 0.00914 | 0.01433 | 0.00698 | 0.08851 | 0.07696 | 0.90701 | 0.27054 | 0.01295 | 0.14620 | 0.01000 | 0.00000 | 0.00740 |
| **Gas8** | 0.57916 | 0.00773 | 1.33375 | 0.06458 | 0.01888 | 0.00501 | 0.00573 | 0.87042 | 0.52364 | 0.86476 | 1.76678 | 0.01432 | 1.17897 | 0.48252 | 0.57781 | 2.94440 |
| **Gata1** | 0.00000 | 0.00444 | 3.91009 | 12.28659 | 0.00950 | 4.52613 | 0.00232 | 13.94069 | 16.34015 | 16.16546 | 10.18901 | 0.58593 | 0.00000 | 1.02607 | 7.79336 | 0.00000 |
| **Gata2** | 0.01331 | 0.00799 | 0.01585 | 2.27700 | 0.21208 | 4.52640 | 0.00693 | 0.00906 | 0.52299 | 1.51288 | 0.00429 | 0.01411 | 0.01467 | 0.00157 | 0.18352 | 0.53688 |
| **Gata3** | 0.00000 | 4.40789 | 2.87154 | 0.00000 | 0.10077 | 6.80107 | 0.40428 | 2.18096 | 0.04808 | 1.44327 | 0.34463 | 0.00000 | 10.90905 | 1.08873 | 0.00000 | 1.86139 |
| **Gata4** | 4.90511 | 3.43219 | 0.00000 | 16.21447 | 8.44153 | 0.00000 | 0.00000 | 0.25767 | 12.22178 | 12.67000 | 3.46490 | 0.87378 | 0.00000 | 4.87330 | 19.08872 | 9.81466 |
| **Gata6** | 14.50544 | 17.49312 | 12.12391 | 0.20352 | 14.18921 | 8.56907 | 15.43088 | 22.77773 | 0.04134 | 28.83693 | 16.72868 | 9.23258 | 11.32362 | 7.93634 | 3.63183 | 13.88427 |
| **Gatad1** | 4.40240 | 4.67197 | 1.19471 | 0.36991 | 0.00713 | 0.78098 | 1.01247 | 0.78091 | 0.00339 | 0.53924 | 0.11392 | 0.02379 | 2.26518 | 0.73135 | 0.31027 | 2.14472 |
| **Gatad2a** | 11.09083 | 3.76503 | 6.50413 | 21.58237 | 8.32603 | 10.39536 | 7.06735 | 16.73889 | 19.28870 | 20.16802 | 8.77676 | 6.82123 | 14.72017 | 2.23060 | 5.60244 | 9.30251 |
| **Gatad2b** | 0.00704 | 0.00616 | 1.02284 | 0.64408 | 0.00954 | 0.00553 | 2.76901 | 0.86827 | 0.65999 | 2.23699 | 0.19084 | 0.10801 | 0.00000 | 3.89848 | 3.76993 | 0.00658 |
| **Gatb** | 9.37950 | 1.06136 | 2.84590 | 0.52198 | 1.20766 | 3.74558 | 2.62012 | 0.00000 | 0.01975 | 0.72594 | 0.02448 | 0.07469 | 1.94918 | 0.00436 | 1.96659 | 2.59053 |
| **Gatc** | 3.68658 | 0.83458 | 0.00000 | 0.27288 | 0.02868 | 2.38698 | 0.00289 | 1.17966 | 1.70910 | 2.70071 | 0.59277 | 1.35002 | 1.48693 | 0.67545 | 0.84071 | 0.13741 |
| **Gatm** | 0.01280 | 0.00778 | 0.00988 | 0.00473 | 0.00877 | 0.00676 | 0.00740 | 0.00604 | 0.01023 | 1.71665 | 0.19209 | 0.00252 | 0.00991 | 0.02067 | 0.38984 | 0.00735 |
| **Gatsl2** | 0.01350 | 0.00798 | 0.01220 | 0.46010 | 0.00907 | 0.00646 | 0.00740 | 0.01108 | 0.15636 | 0.52880 | 0.01041 | 0.00616 | 0.01094 | 0.01058 | 0.01794 | 0.00703 |
| **Gatsl3** | 4.87683 | 3.17425 | 14.22812 | 3.14918 | 8.29617 | 6.54725 | 12.58746 | 11.93277 | 6.22790 | 10.91321 | 7.22894 | 2.63229 | 10.66528 | 0.01024 | 3.37995 | 4.43390 |
| **Gba** | 11.41842 | 7.26616 | 18.97178 | 0.00000 | 0.95679 | 2.05384 | 8.69855 | 12.37506 | 2.19652 | 8.86438 | 10.84798 | 7.94209 | 12.31486 | 5.09539 | 6.48837 | 1.16966 |
| **Gba2** | 0.01739 | 0.00791 | 1.00632 | 0.92975 | 0.02093 | 4.56681 | 0.00681 | 1.07702 | 2.09730 | 0.07224 | 0.34508 | 0.81547 | 0.72785 | 0.00182 | 0.06878 | 0.01958 |
| **Gbas** | 1.12241 | 0.03602 | 0.00000 | 9.45906 | 1.87016 | 2.89330 | 0.00517 | 0.00000 | 4.56610 | 0.65027 | 1.15280 | 0.03404 | 0.63970 | 0.75592 | 1.71628 | 0.01668 |
| **Gbe1** | 0.00826 | 0.00752 | 2.60557 | 0.00492 | 0.02759 | 0.19121 | 0.00671 | 0.25005 | 0.01080 | 0.78894 | 0.35185 | 0.02934 | 0.14059 | 0.05206 | 0.00000 | 0.01437 |
| **Gbf1** | 2.71172 | 0.04042 | 1.47888 | 7.94096 | 0.02599 | 0.01519 | 0.01389 | 4.70116 | 3.40190 | 2.61213 | 0.50004 | 0.22207 | 1.58799 | 0.00000 | 0.55754 | 4.28042 |
| **Gbp11** | 0.01685 | 0.00818 | 0.01120 | 0.00849 | 0.01817 | 0.00667 | 0.00767 | 0.01663 | 0.01213 | 0.07854 | 0.07235 | 0.00794 | 0.01101 | 0.01161 | 0.00909 | 0.00725 |
| **Gbp4** | 0.58757 | 0.00750 | 0.00000 | 0.00702 | 0.00654 | 0.00812 | 0.00756 | 0.46299 | 0.01112 | 0.87231 | 0.01345 | 0.00012 | 0.01374 | 1.06144 | 0.00000 | 0.00666 |
| **Gbp7** | 0.01372 | 0.00760 | 0.00976 | 0.00331 | 0.00948 | 0.01266 | 0.00730 | 0.01070 | 0.18227 | 0.09004 | 0.00774 | 0.04316 | 0.01289 | 0.01036 | 0.97658 | 0.00747 |
| **Gbp9** | 0.00929 | 0.00846 | 0.00749 | 0.02106 | 0.00950 | 0.00689 | 0.00821 | 0.39807 | 0.01273 | 0.30645 | 0.06721 | 0.00620 | 0.00558 | 0.28304 | 0.03616 | 0.00716 |
| **Gbx1** | 0.01382 | 0.00822 | 0.00884 | 0.06223 | 0.00910 | 0.00686 | 0.00760 | 0.00619 | 0.00868 | 0.00624 | 0.48134 | 0.00646 | 0.00754 | 0.01026 | 0.00000 | 0.00715 |
| **Gbx2** | 0.03661 | 0.00831 | 0.00706 | 4.77495 | 0.01470 | 0.00656 | 0.00721 | 0.00887 | 4.40506 | 0.00000 | 0.00000 | 0.02377 | 0.13936 | 0.69141 | 0.00000 | 0.00653 |
| **Gc** | 0.01490 | 0.00710 | 0.01278 | 0.45039 | 0.00909 | 0.00637 | 0.06586 | 0.00000 | 0.00819 | 0.00455 | 4.07789 | 0.00581 | 0.00000 | 0.45110 | 0.00000 | 0.00655 |
| **Gca** | 3.10243 | 5.72075 | 2.94827 | 0.93703 | 0.11102 | 4.53454 | 4.21600 | 4.67219 | 2.25158 | 0.66860 | 0.15740 | 8.12648 | 3.58049 | 8.33856 | 6.32536 | 1.54798 |
| **Gcat** | 15.55010 | 2.74371 | 2.36363 | 0.26199 | 5.21726 | 0.23944 | 4.22283 | 4.70325 | 0.00019 | 2.04086 | 4.23004 | 0.04786 | 8.33222 | 0.00000 | 0.00000 | 2.18525 |
| **Gcc1** | 0.69394 | 0.00475 | 0.00000 | 0.65793 | 2.09672 | 0.00567 | 0.00587 | 0.84285 | 0.18178 | 0.23896 | 1.22966 | 3.21881 | 0.00582 | 0.00282 | 0.63645 | 0.35786 |
| **Gcc2** | 0.90273 | 5.01783 | 2.23743 | 3.08571 | 0.04648 | 0.55878 | 0.00239 | 1.22271 | 0.93897 | 4.32018 | 3.24116 | 5.38519 | 1.08994 | 0.00000 | 3.46899 | 0.08011 |
| **Gcdh** | 2.37092 | 4.33580 | 3.30523 | 1.68834 | 4.44917 | 4.14398 | 3.71798 | 7.61752 | 3.88937 | 6.25590 | 7.11395 | 2.37792 | 8.39877 | 1.28492 | 7.19368 | 0.02484 |
| **Gcfc2** | 0.40217 | 0.00783 | 0.01191 | 0.00736 | 0.00705 | 0.00598 | 0.69967 | 0.14615 | 0.01155 | 0.41244 | 0.03068 | 0.00000 | 0.00812 | 1.98454 | 0.00000 | 0.00683 |

| | | | | | | | | | | | | | | | | |
|---|---|---|---|---|---|---|---|---|---|---|---|---|---|---|---|---|
| **Gch1** | 2.77666 | 0.07057 | 3.86443 | 8.12636 | 1.64238 | 4.81209 | 3.51494 | 4.99823 | 8.22973 | 11.23303 | 13.13348 | 2.33736 | 3.02770 | 4.53080 | 1.10879 | 0.95581 |
| **Gchfr** | 0.27553 | 0.00713 | 0.00000 | 0.07373 | 0.01500 | 1.07491 | 0.00707 | 0.02957 | 0.47714 | 0.00751 | 0.00939 | 0.00655 | 0.00979 | 0.00835 | 0.10181 | 1.81472 |
| **Gck** | 0.01383 | 0.00792 | 0.01147 | 0.40017 | 0.00933 | 0.00660 | 0.00743 | 0.01258 | 0.00574 | 0.00984 | 0.01323 | 0.03912 | 0.01276 | 0.01035 | 0.11450 | 0.00733 |
| **Gckr** | 0.01373 | 0.00814 | 0.01197 | 1.47531 | 0.00931 | 0.00651 | 0.00748 | 0.01338 | 0.24970 | 0.01229 | 0.02768 | 0.00781 | 0.01236 | 0.01218 | 0.00473 | 0.00714 |
| **Gclc** | 1.13174 | 0.14248 | 1.72608 | 5.13011 | 0.00475 | 0.00525 | 6.43018 | 3.79332 | 4.39315 | 1.88568 | 0.70505 | 9.32645 | 0.00000 | 0.32508 | 2.89761 | 1.70337 |
| **Gclm** | 1.09970 | 4.01089 | 2.27471 | 2.44292 | 7.28002 | 4.57172 | 3.17269 | 0.57950 | 3.87139 | 4.46097 | 0.61143 | 2.25859 | 3.60256 | 1.41477 | 2.70778 | 0.01428 |
| **Gcm1** | 0.01279 | 0.00753 | 0.00633 | 0.00000 | 0.00940 | 0.02990 | 0.00715 | 0.01018 | 0.00907 | 0.00000 | 0.00422 | 0.00188 | 0.11943 | 0.39416 | 2.01590 | 0.00719 |
| **Gcm2** | 0.01346 | 0.00819 | 0.01147 | 2.44757 | 0.00926 | 0.00627 | 0.00742 | 0.01301 | 2.19558 | 0.00512 | 0.00000 | 0.01570 | 0.01300 | 0.00639 | 0.05346 | 0.00681 |
| **Gcn1l1** | 8.20322 | 0.23074 | 0.91668 | 0.41444 | 0.16956 | 0.20626 | 3.85966 | 15.03052 | 1.10822 | 7.93665 | 0.56989 | 0.33544 | 9.56144 | 0.00000 | 0.00000 | 3.11800 |
| **Gcnt1** | 0.00000 | 0.09859 | 7.91364 | 1.19547 | 2.27575 | 6.86894 | 0.02284 | 4.50105 | 0.00001 | 0.66346 | 4.85705 | 0.67918 | 4.22057 | 5.57677 | 0.54308 | 3.98992 |
| **Gcnt2** | 0.31373 | 3.87592 | 0.00000 | 5.02114 | 0.01518 | 0.00525 | 0.01509 | 0.42321 | 3.65662 | 2.04021 | 0.03549 | 0.00434 | 0.00588 | 5.02387 | 2.72763 | 0.00420 |
| **Gcnt3** | 0.01329 | 0.00782 | 0.01182 | 0.00723 | 0.00897 | 0.00670 | 0.00732 | 0.01014 | 0.01175 | 0.88487 | 0.02794 | 0.00565 | 0.01161 | 0.02968 | 0.00523 | 0.00764 |
| **Gcnt4** | 0.01381 | 0.00758 | 0.00831 | 0.70472 | 0.00926 | 0.00677 | 0.00719 | 0.01034 | 0.60058 | 0.00438 | 0.00543 | 0.45127 | 0.01345 | 0.01041 | 0.00000 | 0.01483 |
| **Gcnt7** | 0.01373 | 0.00817 | 0.01196 | 0.10689 | 0.00933 | 0.00666 | 0.00749 | 0.01315 | 0.00938 | 0.01390 | 0.01323 | 0.00783 | 0.01280 | 0.05359 | 0.00682 | 0.00742 |
| **Gcsh** | 18.37219 | 4.74923 | 15.72874 | 20.67094 | 14.30275 | 9.31566 | 6.21174 | 34.83236 | 30.52631 | 53.55512 | 38.24339 | 26.20241 | 27.60095 | 28.49156 | 19.59671 | 12.34094 |
| **Gda** | 0.01708 | 0.00750 | 0.01549 | 7.58264 | 0.01721 | 0.00617 | 0.00666 | 0.00000 | 1.10781 | 2.93391 | 0.85120 | 0.00000 | 0.02046 | 0.00987 | 4.86318 | 0.00594 |
| **Gdap1** | 0.00000 | 0.00779 | 0.00101 | 12.72100 | 0.00919 | 0.00366 | 0.00640 | 0.00000 | 9.13161 | 3.34593 | 0.18775 | 0.00000 | 0.00932 | 11.22199 | 13.97716 | 0.00409 |
| **Gdap10** | 0.00000 | 0.00682 | 0.01296 | 0.00876 | 0.01040 | 0.00727 | 2.01908 | 0.00749 | 0.01176 | 0.00000 | 0.01351 | 0.00609 | 1.33379 | 0.01074 | 0.15418 | 0.02075 |
| **Gdap2** | 2.16025 | 10.62354 | 4.58876 | 14.25156 | 0.08468 | 12.23427 | 3.46829 | 19.85329 | 17.30398 | 21.80274 | 15.66561 | 15.50179 | 4.69654 | 22.85289 | 20.21738 | 0.10916 |
| **Gde1** | 14.00326 | 6.13143 | 8.10999 | 17.55566 | 2.40131 | 3.43066 | 6.10761 | 15.17178 | 14.89174 | 17.74517 | 12.94880 | 5.81917 | 7.24724 | 5.19560 | 5.80771 | 3.28398 |
| **Gdf11** | 0.01554 | 0.00811 | 0.10566 | 0.04042 | 0.00937 | 0.00638 | 0.01842 | 0.01307 | 0.01157 | 0.94906 | 0.01492 | 0.00589 | 0.11446 | 0.01789 | 0.06993 | 0.00737 |
| **Gdf15** | 0.05366 | 0.00811 | 0.00997 | 0.74044 | 0.00801 | 0.00669 | 0.00744 | 0.01503 | 0.08121 | 0.01308 | 0.01297 | 0.00823 | 0.01186 | 0.01103 | 0.00405 | 0.10732 |
| **Gdf3** | 15.19883 | 0.97025 | 6.85161 | 0.99463 | 6.52686 | 0.04947 | 4.89190 | 7.22856 | 0.70136 | 2.54644 | 1.74385 | 0.18883 | 6.41351 | 0.82598 | 0.54831 | 5.13938 |
| **Gdf5** | 0.01372 | 0.00806 | 0.01229 | 0.29909 | 0.00927 | 0.00656 | 0.00752 | 0.01294 | 0.00896 | 0.07924 | 0.01128 | 0.00756 | 0.01216 | 0.00562 | 0.17788 | 0.00718 |
| **Gdf9** | 0.77431 | 0.13792 | 0.49354 | 19.04854 | 0.85457 | 0.00242 | 0.04775 | 0.05536 | 12.06746 | 3.64316 | 2.30348 | 0.36487 | 0.39343 | 0.05570 | 6.16140 | 0.00210 |
| **Gdi1** | 1.43083 | 0.00659 | 0.00003 | 1.45202 | 0.00372 | 1.37628 | 0.01598 | 0.16783 | 1.81564 | 2.30801 | 0.01448 | 0.06736 | 1.92177 | 0.24021 | 3.83025 | 4.40953 |
| **Gdi2** | 20.48264 | 19.18741 | 16.77204 | 18.79670 | 19.41438 | 17.09641 | 14.19514 | 27.78715 | 21.17761 | 33.02070 | 20.17967 | 26.85192 | 24.31424 | 29.14062 | 20.49689 | 12.15482 |
| **Gdpd1** | 1.09329 | 0.00437 | 1.30202 | 1.65331 | 0.00346 | 0.25471 | 2.67957 | 0.00000 | 1.65821 | 1.91332 | 0.22570 | 0.15342 | 0.00467 | 0.00588 | 1.92173 | 0.00500 |
| **Gdpd2** | 0.01334 | 0.00813 | 0.01192 | 0.00672 | 0.00903 | 0.00660 | 0.00734 | 0.01071 | 0.01133 | 0.72862 | 0.01136 | 0.00598 | 0.01160 | 0.01113 | 0.21893 | 0.00752 |
| **Gdpd3** | 0.02052 | 0.00771 | 0.01127 | 0.00767 | 0.00886 | 0.00665 | 0.00753 | 0.01294 | 0.55853 | 0.00874 | 0.00874 | 0.00685 | 0.01289 | 0.01977 | 0.06335 | 0.00736 |
| **Gdpd5** | 2.47529 | 0.00552 | 0.00000 | 0.00834 | 0.02355 | 3.63991 | 0.00505 | 0.00780 | 0.01299 | 0.04925 | 0.00883 | 0.00528 | 0.00336 | 0.01042 | 0.01561 | 4.34965 |
| **Gdpgp1** | 1.76449 | 0.00554 | 0.31681 | 0.36252 | 0.03156 | 0.02990 | 0.03735 | 1.45581 | 1.20948 | 0.70015 | 0.83885 | 0.14079 | 0.08165 | 0.24732 | 1.61795 | 1.11977 |
| **Gemin2** | 4.50537 | 3.46081 | 5.50610 | 0.96246 | 4.14062 | 4.59532 | 2.56394 | 9.05473 | 4.46661 | 14.89248 | 11.81445 | 1.96266 | 9.29319 | 3.27112 | 8.73103 | 5.26307 |
| **Gemin4** | 8.91698 | 8.89267 | 6.64430 | 2.73487 | 0.00000 | 0.21547 | 6.82532 | 4.82130 | 2.92763 | 10.30452 | 3.74760 | 0.90207 | 3.95034 | 3.09676 | 10.93212 | 0.89368 |
| **Gemin5** | 10.68331 | 11.88416 | 14.98607 | 7.12722 | 10.31123 | 2.88324 | 7.25680 | 21.87873 | 13.14567 | 24.20672 | 15.83669 | 7.75802 | 6.30497 | 15.57752 | 22.63698 | 4.08647 |
| **Gemin6** | 5.55626 | 5.11195 | 4.91708 | 1.52698 | 3.92742 | 4.04981 | 0.69924 | 5.82383 | 1.52296 | 1.40905 | 0.00000 | 1.93787 | 1.48546 | 3.74583 | 2.51364 | 0.03154 |
| **Gemin7** | 3.92480 | 0.00153 | 3.79002 | 9.21783 | 0.36531 | 0.02069 | 1.92813 | 0.58071 | 4.99466 | 5.14208 | 3.58490 | 0.27696 | 2.13190 | 0.00000 | 0.55226 | 3.81822 |
| **Gemin8** | 2.33574 | 0.02397 | 4.08596 | 1.09426 | 4.96116 | 0.00944 | 0.07043 | 2.10441 | 1.86194 | 6.24044 | 1.68575 | 0.34397 | 0.78468 | 1.18742 | 3.67062 | 0.05337 |
| **Gen1** | 2.24117 | 0.72462 | 0.48751 | 0.62399 | 0.01341 | 1.08464 | 0.62006 | 0.06485 | 0.31666 | 0.64830 | 0.00000 | 0.14312 | 0.00287 | 3.25003 | 1.17552 | 0.00399 |
| **Get4** | 5.78760 | 0.57620 | 4.70856 | 2.10467 | 5.42008 | 7.71622 | 5.62723 | 9.09970 | 0.67377 | 12.67619 | 2.75726 | 0.57197 | 11.10572 | 0.00000 | 0.00000 | 3.33726 |
| **Gfer** | 3.23349 | 1.80456 | 3.58747 | 12.72328 | 5.55703 | 0.00154 | 2.74987 | 1.95785 | 10.31831 | 5.83247 | 8.43536 | 0.11218 | 2.12919 | 0.20356 | 2.27859 | 4.24798 |
| **Gfi1** | 0.01362 | 0.00802 | 0.01132 | 0.02651 | 0.00929 | 0.00655 | 0.00740 | 0.01200 | 1.17384 | 0.02208 | 0.00006 | 0.02618 | 0.01350 | 0.00578 | 0.27593 | 0.00758 |
| **Gfi1b** | 0.01877 | 0.00752 | 0.00959 | 0.01074 | 0.00880 | 0.00671 | 0.00722 | 0.10028 | 0.01206 | 0.04063 | 0.01308 | 0.01791 | 0.00837 | 0.01971 | 0.01169 | 0.00720 |
| **Gfm1** | 7.01215 | 4.91605 | 3.52818 | 1.34458 | 5.09971 | 0.15281 | 4.18846 | 4.61793 | 3.13159 | 0.00000 | 3.84093 | 3.34104 | 3.81094 | 4.05489 | 8.20827 | 1.86932 |
| **Gfm2** | 3.43002 | 2.25031 | 3.98248 | 1.55158 | 0.34486 | 0.97097 | 2.53775 | 3.24709 | 1.01307 | 3.15693 | 1.18250 | 0.30062 | 1.44586 | 2.48303 | 4.39613 | 1.40520 |
| **Gfod1** | 0.01319 | 0.00781 | 0.00922 | 0.97650 | 0.00932 | 0.00627 | 0.00726 | 0.00943 | 2.17209 | 0.00000 | 0.01077 | 0.00000 | 0.01280 | 0.25750 | 0.67444 | 0.00695 |
| **Gfod2** | 0.01127 | 0.00773 | 1.08929 | 1.16311 | 0.00865 | 0.00615 | 0.03708 | 0.88244 | 3.13718 | 1.62749 | 0.00000 | 0.06622 | 0.01089 | 0.03107 | 1.14743 | 0.00674 |
| **Gfpt1** | 4.40329 | 4.62619 | 1.54973 | 9.34887 | 2.57379 | 0.00000 | 2.37844 | 4.89105 | 11.94219 | 10.20299 | 9.96511 | 8.77004 | 3.37943 | 12.70263 | 13.42814 | 1.25296 |
| **Gfpt2** | 0.01019 | 0.00757 | 0.02859 | 0.01719 | 0.42804 | 0.00803 | 0.00738 | 0.01134 | 0.01248 | 0.06271 | 0.02996 | 0.04569 | 0.01341 | 0.01126 | 0.01051 | 0.00748 |
| **Gfra1** | 0.01246 | 0.00819 | 0.00624 | 1.20595 | 0.00953 | 0.00640 | 0.00724 | 0.00982 | 0.00000 | 0.07797 | 0.00000 | 0.01171 | 0.01261 | 1.15488 | 1.14664 | 0.00696 |
| **Gfra2** | 0.03147 | 0.02891 | 0.01267 | 0.01324 | 0.01897 | 0.01886 | 0.02798 | 0.01665 | 0.03948 | 0.03126 | 0.04215 | 0.02812 | 0.02711 | 0.07235 | 0.02854 | 0.00734 |
| **Gfra3** | 0.44327 | 0.00627 | 0.96850 | 2.14651 | 0.01483 | 0.00423 | 0.00562 | 1.47114 | 1.48530 | 4.94389 | 2.14040 | 4.42079 | 0.00000 | 0.03072 | 0.14209 | 0.01312 |
| **Gfra4** | 0.00641 | 1.93161 | 0.00666 | 1.30872 | 0.00811 | 0.00641 | 0.00651 | 0.00909 | 0.25501 | 0.47585 | 0.01076 | 0.01423 | 0.01236 | 0.00562 | 0.00000 | 0.00646 |
| **Gga1** | 6.12387 | 0.06795 | 2.73962 | 4.00253 | 4.09458 | 3.90698 | 2.73469 | 8.18410 | 3.61161 | 8.14086 | 2.97663 | 0.29623 | 6.60311 | 0.00000 | 0.00000 | 5.72961 |
| **Gga2** | 19.95757 | 10.23279 | 15.03377 | 1.56222 | 10.85524 | 6.92021 | 8.22069 | 16.96326 | 1.09665 | 11.62023 | 15.33605 | 1.16501 | 13.21149 | 5.73980 | 2.02001 | 8.58738 |
| **Gga3** | 10.87791 | 5.67539 | 9.12139 | 10.99694 | 5.67925 | 0.05267 | 5.28491 | 1.81210 | 10.58657 | 5.90501 | 2.24194 | 0.43366 | 5.80494 | 2.70638 | 6.87493 | 0.94546 |
| **Ggact** | 0.28536 | 0.00783 | 0.03554 | 0.39964 | 0.01691 | 0.00827 | 0.00687 | 0.01332 | 0.39968 | 0.48420 | 0.31520 | 0.02284 | 0.00873 | 0.07163 | 0.39126 | 0.00719 |
| **Ggct** | 0.21931 | 2.15023 | 3.48033 | 11.89407 | 3.68127 | 0.00384 | 1.57089 | 2.29943 | 3.88589 | 3.27098 | 0.00000 | 0.04454 | 0.00000 | 0.00000 | 1.48786 | 0.00984 |
| **Ggcx** | 1.67192 | 0.00593 | 0.40097 | 0.68652 | 0.07017 | 0.07907 | 0.01577 | 0.55743 | 0.06352 | 0.13106 | 0.72226 | 0.02528 | 1.45417 | 1.56166 | 0.04587 | 2.60135 |
| **Ggh** | 0.01639 | 0.00809 | 0.53131 | 0.00858 | 0.00934 | 0.00832 | 0.00734 | 0.01065 | 0.01193 | 0.56243 | 0.01054 | 0.00692 | 0.00929 | 0.01336 | 0.00338 | 0.00713 |
| **Ggnbp1** | 1.14143 | 0.00640 | 0.01842 | 0.12808 | 0.00596 | 0.00747 | 0.00595 | 0.00000 | 0.02840 | 1.30413 | 0.60745 | 0.04027 | 0.47254 | 0.01074 | 0.00000 | 0.00588 |
| **Ggnbp2** | 3.17234 | 3.92662 | 3.44847 | 6.18909 | 2.28405 | 2.63827 | 5.75081 | 0.89590 | 3.84181 | 4.84579 | 1.34515 | 1.39703 | 0.07876 | 9.35676 | 2.30501 | 3.60051 |

| | | | | | | | | | | | | | | | | |
|---|---|---|---|---|---|---|---|---|---|---|---|---|---|---|---|---|
| **Ggnbp2os** | 0.01631 | 0.00785 | 0.21662 | 1.35868 | 0.00871 | 0.00569 | 0.00748 | 0.01171 | 0.00000 | 0.76878 | 0.01157 | 0.00599 | 0.01312 | 0.01540 | 0.04527 | 0.00717 |
| **Ggps1** | 3.26882 | 2.19821 | 3.23469 | 2.28377 | 1.06774 | 0.05604 | 1.51881 | 0.34244 | 11.97461 | 1.87377 | 2.72867 | 0.11965 | 0.17530 | 2.21578 | 1.80100 | 3.16940 |
| **Ggt1** | 10.05387 | 5.36822 | 13.73196 | 0.11699 | 2.76785 | 1.03372 | 5.23370 | 4.23777 | 0.00232 | 0.33036 | 1.53245 | 0.23165 | 2.55918 | 0.00000 | 0.60418 | 2.24651 |
| **Ggt5** | 0.01098 | 0.00780 | 0.00924 | 0.12691 | 0.00950 | 0.00680 | 0.00720 | 0.11786 | 0.23750 | 0.01304 | 0.11346 | 0.00681 | 0.00768 | 0.01099 | 0.00214 | 0.00740 |
| **Ggt6** | 2.41711 | 0.00459 | 0.06455 | 2.49783 | 0.01152 | 0.00567 | 0.06577 | 2.55756 | 6.54497 | 0.00000 | 2.96930 | 0.00000 | 0.00000 | 0.00000 | 0.53895 | 0.00351 |
| **Ggt7** | 0.01347 | 0.00807 | 0.01148 | 0.00782 | 0.00930 | 0.00676 | 0.00752 | 0.01798 | 0.01102 | 0.04510 | 0.01060 | 0.00814 | 0.01296 | 0.00706 | 0.49798 | 0.00742 |
| **Ggta1** | 0.00000 | 0.00799 | 0.00728 | 4.10475 | 0.01729 | 3.42893 | 0.00693 | 0.09001 | 0.75577 | 0.00000 | 1.30009 | 0.01895 | 0.29598 | 0.00000 | 1.22018 | 1.02951 |
| **Ghdc** | 0.83143 | 2.56854 | 3.16161 | 1.48363 | 0.37557 | 3.56155 | 2.07832 | 8.24956 | 1.64142 | 0.12054 | 1.44020 | 0.14048 | 2.90880 | 1.65677 | 0.49433 | 3.92468 |
| **Ghitm** | 21.35021 | 17.59802 | 23.71740 | 9.79572 | 18.42859 | 13.35260 | 16.43131 | 20.71959 | 17.69365 | 13.72643 | 15.27339 | 13.64325 | 22.73698 | 12.45861 | 10.89068 | 8.73739 |
| **Ghrh** | 0.01374 | 0.00813 | 0.01230 | 0.18447 | 0.00934 | 0.00665 | 0.00754 | 0.01398 | 0.00597 | 0.01538 | 0.01241 | 0.00820 | 0.01271 | 0.01157 | 0.00767 | 0.00739 |
| **Ghrl** | 0.01348 | 0.00827 | 0.01722 | 0.01030 | 0.00932 | 0.00637 | 0.00757 | 0.01353 | 0.14254 | 0.01209 | 0.02530 | 0.00825 | 0.01228 | 0.00983 | 0.04226 | 0.00738 |
| **Gid4** | 7.38169 | 3.85697 | 6.58539 | 6.10150 | 3.75731 | 0.02984 | 2.12295 | 3.13197 | 3.36325 | 6.47423 | 3.22304 | 9.48920 | 0.12827 | 7.42091 | 7.70011 | 0.26853 |
| **Gid8** | 8.80545 | 2.02970 | 4.92842 | 2.85592 | 2.53534 | 3.53795 | 3.53215 | 2.42257 | 4.80199 | 2.87379 | 1.24935 | 0.15250 | 2.34758 | 3.64652 | 1.66153 | 3.97357 |
| **Gigyf1** | 1.29646 | 0.01584 | 0.26006 | 0.16870 | 0.10012 | 0.00465 | 0.00600 | 0.64077 | 0.16603 | 0.09997 | 0.68022 | 0.02951 | 0.03104 | 0.00610 | 0.12489 | 0.13518 |
| **Gigyf2** | 3.42123 | 5.64945 | 2.82497 | 2.20169 | 1.05280 | 0.01806 | 2.83289 | 5.96234 | 2.69672 | 3.67756 | 3.94364 | 0.10657 | 2.30091 | 4.00487 | 5.50153 | 0.26734 |
| **Gimap1** | 0.06019 | 0.00833 | 0.25099 | 1.64405 | 0.00940 | 0.13302 | 0.05111 | 0.21867 | 2.01937 | 0.14626 | 0.24476 | 0.05711 | 0.26706 | 0.48171 | 0.32135 | 0.07835 |
| **Gimap3** | 0.01316 | 0.00785 | 0.00994 | 0.07359 | 0.00897 | 0.00645 | 0.00734 | 0.01015 | 1.54556 | 0.76179 | 0.00000 | 0.10939 | 0.01098 | 0.02852 | 0.23220 | 0.00703 |
| **Gimap8** | 0.01092 | 0.03959 | 0.03127 | 0.14347 | 0.00881 | 0.00709 | 0.01743 | 0.03399 | 0.11229 | 0.02207 | 0.01152 | 0.00781 | 0.04252 | 0.05657 | 0.08472 | 0.12932 |
| **Gimap9** | 1.69608 | 0.00604 | 0.45879 | 0.00389 | 0.00429 | 0.00379 | 0.00584 | 0.00983 | 0.18472 | 0.54588 | 0.00779 | 0.01435 | 0.00879 | 0.00572 | 0.50945 | 0.00505 |
| **Gin1** | 1.98910 | 4.63418 | 1.00004 | 0.00000 | 0.02996 | 0.03331 | 2.10058 | 4.00873 | 2.12737 | 3.70234 | 0.36382 | 1.43763 | 0.30138 | 6.92116 | 4.91377 | 0.39500 |
| **Ginm1** | 1.28219 | 1.34191 | 0.00000 | 0.05385 | 0.01328 | 0.46638 | 1.70593 | 0.60929 | 0.00901 | 1.45539 | 0.07164 | 0.06138 | 1.91203 | 0.00760 | 0.53077 | 2.17108 |
| **Gins1** | 15.00202 | 6.05389 | 9.38359 | 2.64949 | 10.78620 | 0.91488 | 9.89886 | 12.25167 | 7.67461 | 18.57394 | 9.56763 | 3.56290 | 7.48776 | 8.74291 | 2.93942 | 2.25103 |
| **Gins2** | 12.50045 | 2.89482 | 10.01141 | 5.91768 | 4.46627 | 3.11172 | 2.30481 | 4.54201 | 9.56664 | 9.83153 | 3.72515 | 0.21529 | 1.85052 | 0.08591 | 4.07272 | 5.58402 |
| **Gins3** | 2.03691 | 0.04876 | 1.56012 | 7.09810 | 0.00272 | 0.53917 | 1.29335 | 2.36680 | 7.41157 | 7.32325 | 3.12277 | 0.97270 | 4.49235 | 0.29227 | 10.82634 | 0.00736 |
| **Gins4** | 14.07667 | 1.49838 | 9.93109 | 5.48292 | 10.01963 | 1.57435 | 3.32538 | 5.57682 | 12.57819 | 14.27730 | 14.60390 | 9.22310 | 16.04033 | 5.36752 | 4.31804 | 7.38952 |
| **Gip** | 0.01306 | 0.00785 | 0.00604 | 0.23786 | 0.00955 | 0.00769 | 0.00746 | 0.00000 | 2.33145 | 0.36758 | 0.61832 | 0.11295 | 0.04433 | 0.03709 | 0.38522 | 0.00739 |
| **Gipc1** | 8.31814 | 0.01627 | 0.00000 | 0.00000 | 3.70102 | 8.66364 | 0.04586 | 5.71301 | 0.16923 | 1.58503 | 2.18013 | 0.06781 | 7.69304 | 0.00435 | 0.00228 | 0.76895 |
| **Gipc2** | 4.05919 | 0.01246 | 1.69464 | 1.51832 | 0.02913 | 0.02012 | 1.37954 | 0.00000 | 0.22887 | 0.00000 | 0.19563 | 0.01655 | 1.54884 | 0.03770 | 0.73382 | 0.02280 |
| **Gipr** | 0.00844 | 0.00774 | 0.05317 | 0.01084 | 0.00970 | 0.00690 | 0.00735 | 0.80248 | 0.01136 | 0.07620 | 0.00808 | 0.00690 | 0.00615 | 0.01755 | 0.00000 | 0.00733 |
| **Git1** | 7.63714 | 1.01902 | 0.22896 | 1.62087 | 5.54319 | 4.69276 | 0.24302 | 6.76703 | 0.93034 | 2.08355 | 0.47440 | 1.33063 | 6.03651 | 0.00000 | 0.00000 | 2.48238 |
| **Git2** | 4.23436 | 1.49412 | 3.39935 | 3.03755 | 1.90512 | 0.13985 | 3.69699 | 0.84240 | 2.62865 | 2.81386 | 4.21111 | 6.35472 | 1.54047 | 0.54883 | 1.84749 | 0.00533 |
| **Gja1** | 7.62973 | 6.41313 | 10.21861 | 0.74876 | 4.99794 | 4.70029 | 5.20537 | 11.53109 | 2.90939 | 0.00000 | 12.55813 | 6.08912 | 4.18637 | 33.60755 | 12.28981 | 0.44849 |
| **Gja10** | 0.01373 | 0.00807 | 0.01218 | 0.06027 | 0.00929 | 0.00664 | 0.00753 | 0.01390 | 0.19646 | 0.01438 | 0.01235 | 0.00788 | 0.01273 | 0.01245 | 0.00770 | 0.00740 |
| **Gja3** | 0.01371 | 0.00800 | 0.01186 | 0.43442 | 0.00943 | 0.00644 | 0.00738 | 0.01183 | 0.18632 | 0.01456 | 0.01068 | 0.00681 | 0.01292 | 0.00574 | 0.49072 | 0.00734 |
| **Gja4** | 0.01153 | 0.00816 | 0.00999 | 0.72123 | 0.01013 | 0.00723 | 0.00723 | 0.19653 | 0.20297 | 0.00000 | 0.08783 | 0.01669 | 0.55488 | 0.01056 | 0.34391 | 0.00724 |
| **Gjb3** | 17.99900 | 6.03223 | 14.95927 | 0.00000 | 10.59303 | 8.94982 | 15.64911 | 19.44452 | 0.01045 | 2.38037 | 9.61287 | 0.40181 | 11.11739 | 0.25633 | 0.00000 | 8.46321 |
| **Gjb4** | 7.33561 | 0.00803 | 0.01355 | 0.09112 | 1.19763 | 0.00586 | 1.82184 | 0.00000 | 0.00697 | 0.00000 | 0.01171 | 0.01969 | 0.00791 | 0.01338 | 0.01158 | 4.42341 |
| **Gjb5** | 17.69795 | 7.90644 | 9.74623 | 0.00823 | 8.65957 | 3.87854 | 12.27187 | 1.05525 | 0.00464 | 0.00000 | 0.00000 | 0.09066 | 1.22676 | 0.00856 | 0.00000 | 11.01038 |
| **Gjc1** | 0.00946 | 0.01683 | 0.00872 | 0.08647 | 0.00945 | 0.00646 | 2.31778 | 0.00927 | 1.31113 | 0.00636 | 0.00004 | 0.00387 | 0.01008 | 0.01501 | 0.00924 | 0.02155 |
| **Gjc3** | 0.09477 | 0.02858 | 0.26200 | 0.44996 | 0.01871 | 0.01474 | 0.11833 | 0.04770 | 0.34087 | 0.08178 | 0.47755 | 0.02175 | 0.20651 | 0.72439 | 0.15219 | 0.03621 |
| **Gje1** | 0.23302 | 0.04080 | 0.26398 | 0.15327 | 0.14126 | 0.03733 | 0.01679 | 0.00559 | 0.24106 | 0.09707 | 1.04153 | 0.09933 | 0.22454 | 0.37007 | 0.05410 | 0.05854 |
| **Gk** | 0.03111 | 0.92474 | 0.93271 | 2.68762 | 0.01002 | 4.16797 | 0.00581 | 0.00000 | 2.00576 | 2.32283 | 0.45593 | 0.09945 | 0.62083 | 2.51279 | 1.02755 | 0.02675 |
| **Gk5** | 0.18103 | 0.00826 | 0.21387 | 0.04717 | 0.01460 | 0.00677 | 0.01726 | 2.52586 | 0.01705 | 0.93948 | 0.00000 | 0.02610 | 0.14980 | 9.67049 | 0.22305 | 0.00691 |
| **Gkap1** | 4.49282 | 3.96418 | 3.11006 | 6.86208 | 4.39135 | 0.42218 | 0.83174 | 2.84433 | 4.27599 | 3.40141 | 0.23127 | 0.80691 | 3.13275 | 2.01337 | 1.63101 | 1.69255 |
| **Gla** | 4.89817 | 0.12955 | 18.87476 | 7.32417 | 6.02987 | 7.59705 | 9.15641 | 8.86245 | 14.47318 | 7.84297 | 5.44580 | 3.77942 | 13.02285 | 13.34121 | 13.01839 | 3.81440 |
| **Glb1** | 9.51369 | 3.69108 | 5.80014 | 2.49160 | 3.40745 | 8.22236 | 3.75579 | 15.85262 | 2.69928 | 13.70044 | 14.14675 | 2.23404 | 5.85880 | 6.84658 | 5.70378 | 4.73750 |
| **Glb1l** | 0.01779 | 0.00748 | 0.42953 | 0.29124 | 0.00858 | 0.00600 | 0.00706 | 0.01119 | 0.42678 | 0.04196 | 0.00969 | 0.00470 | 0.36494 | 0.00447 | 0.47776 | 0.00710 |
| **Glcci1** | 0.58733 | 0.95257 | 1.54537 | 14.34961 | 1.92755 | 0.61239 | 0.80679 | 1.41479 | 5.88259 | 3.15762 | 1.46675 | 0.89356 | 1.19809 | 2.59894 | 5.70085 | 0.53761 |
| **Glce** | 3.44994 | 0.00000 | 3.58682 | 6.24030 | 3.78914 | 0.29300 | 1.47654 | 2.28501 | 3.18832 | 1.48770 | 0.16492 | 0.13179 | 0.02297 | 0.03487 | 3.05992 | 0.86812 |
| **Gldc** | 19.43881 | 7.83282 | 17.96098 | 7.33146 | 9.31381 | 10.01666 | 9.42679 | 35.79494 | 16.09199 | 38.94771 | 25.34855 | 21.03517 | 23.29136 | 20.11985 | 27.68972 | 11.89222 |
| **Gle1** | 7.81714 | 1.38431 | 10.84470 | 17.84710 | 6.77569 | 1.50946 | 5.65031 | 15.23618 | 12.68108 | 19.10710 | 9.16249 | 4.71600 | 8.61977 | 12.16721 | 15.81339 | 5.72835 |
| **Glg1** | 4.90284 | 0.00000 | 6.08986 | 1.15107 | 1.93211 | 0.00186 | 1.46035 | 3.49649 | 0.28163 | 3.41574 | 0.00000 | 0.07931 | 1.97710 | 0.18156 | 2.11861 | 4.94989 |
| **Gli1** | 0.01625 | 0.00756 | 0.55128 | 0.51974 | 0.00824 | 0.00553 | 0.00716 | 0.01571 | 0.85224 | 0.05443 | 0.00635 | 0.00524 | 0.00921 | 0.00996 | 0.08097 | 0.00728 |
| **Gli2** | 0.01312 | 0.00802 | 0.01124 | 1.19056 | 0.00922 | 0.00622 | 0.00739 | 0.01073 | 0.71335 | 0.53639 | 0.00432 | 0.00267 | 0.01284 | 0.00263 | 1.24918 | 0.00693 |
| **Gli3** | 0.01313 | 0.00755 | 0.00893 | 0.00286 | 0.00923 | 0.00610 | 0.00731 | 0.00795 | 5.17027 | 0.00005 | 0.03227 | 0.06353 | 0.01249 | 0.00395 | 0.65738 | 0.00699 |
| **Glipr1** | 0.63297 | 0.00710 | 0.00000 | 0.12432 | 0.03505 | 2.27582 | 1.16286 | 0.00596 | 0.00693 | 0.00000 | 0.00875 | 0.03661 | 0.81216 | 0.01381 | 0.00655 | 2.12490 |
| **Glipr2** | 6.15696 | 1.54190 | 1.40404 | 0.64883 | 12.20557 | 5.71313 | 0.73180 | 1.85796 | 0.25967 | 1.81215 | 1.51459 | 0.05333 | 2.10544 | 0.03845 | 0.23652 | 6.23572 |
| **Glis1** | 0.01069 | 0.00823 | 0.01175 | 3.68046 | 0.00951 | 0.00661 | 0.00747 | 0.07959 | 0.77496 | 1.03591 | 0.00645 | 0.00414 | 0.01772 | 0.00432 | 0.02983 | 0.00675 |
| **Glis2** | 1.65986 | 0.00303 | 0.76330 | 7.76792 | 3.91411 | 0.00576 | 0.01470 | 0.98544 | 5.86077 | 4.11286 | 0.62060 | 0.66091 | 0.00000 | 5.20092 | 1.74315 | 5.62759 |
| **Glis3** | 0.01290 | 0.00792 | 0.00744 | 2.93643 | 0.00931 | 0.00590 | 0.00726 | 0.00963 | 2.06147 | 0.14611 | 0.00000 | 0.03780 | 0.01269 | 0.13604 | 2.81994 | 0.00664 |
| **Glmn** | 2.70965 | 1.75847 | 0.46365 | 0.25134 | 1.50936 | 0.00486 | 0.00386 | 2.50447 | 0.11660 | 1.50490 | 0.43400 | 5.26007 | 0.00326 | 0.43349 | 2.73970 | 0.18962 |
| **Glmp** | 8.43963 | 3.82905 | 13.87315 | 4.77922 | 0.95003 | 13.20068 | 6.62166 | 19.46362 | 6.81409 | 11.32506 | 8.17750 | 1.66204 | 14.46365 | 6.11166 | 8.35556 | 8.78274 |

| | | | | | | | | | | | | | | | | |
|---|---|---|---|---|---|---|---|---|---|---|---|---|---|---|---|---|
| **Glo1** | 10.50411 | 10.72417 | 6.87985 | 0.11908 | 6.10290 | 3.93361 | 6.32726 | 0.90018 | 0.05340 | 2.10213 | 1.39589 | 0.16703 | 3.21452 | 2.16452 | 1.26848 | 8.68913 |
| **Glod4** | 8.34690 | 14.09119 | 14.00395 | 3.97925 | 17.43271 | 10.89540 | 9.88385 | 9.57197 | 6.15159 | 1.78743 | 7.29258 | 2.47799 | 9.56037 | 1.97677 | 3.82725 | 5.77033 |
| **Glod5** | 0.87043 | 0.00672 | 0.00000 | 3.55856 | 0.00608 | 0.00612 | 0.00678 | 0.00823 | 1.88605 | 0.40742 | 0.19704 | 0.00796 | 0.00933 | 1.19888 | 0.61867 | 0.00557 |
| **Glrb** | 0.01375 | 0.00807 | 0.01225 | 0.01058 | 0.00932 | 0.00666 | 0.00754 | 0.01389 | 0.01234 | 0.01523 | 0.01311 | 0.00815 | 0.01270 | 0.01220 | 0.01187 | 0.00741 |
| **Glrx** | 31.75589 | 9.95980 | 10.01822 | 1.05430 | 19.14032 | 6.89979 | 7.79641 | 0.00000 | 0.34467 | 0.00000 | 0.00717 | 0.07226 | 1.75469 | 0.11582 | 0.07550 | 13.14147 |
| **Glrx2** | 0.96054 | 0.01564 | 1.45219 | 8.73509 | 0.03752 | 0.16204 | 0.69354 | 0.17717 | 19.79811 | 3.87508 | 7.93027 | 0.41232 | 2.13449 | 4.09438 | 2.87034 | 0.86364 |
| **Glrx3** | 5.75980 | 2.35409 | 4.79648 | 3.91086 | 6.37150 | 3.47916 | 2.58982 | 5.43001 | 7.84035 | 6.82777 | 12.03752 | 7.07098 | 8.63737 | 3.79642 | 1.13048 | 4.13315 |
| **Glrx5** | 4.03297 | 0.20358 | 0.87943 | 5.30173 | 6.95532 | 4.17917 | 1.10339 | 2.64113 | 3.67062 | 3.55048 | 0.88061 | 0.56441 | 0.32983 | 0.03337 | 0.00000 | 0.00684 |
| **Gls** | 0.32927 | 0.08080 | 2.09391 | 7.73553 | 3.26044 | 0.32878 | 3.13305 | 4.85003 | 12.80482 | 7.00675 | 4.36637 | 13.62052 | 4.77439 | 7.80236 | 1.76598 | 4.00502 |
| **Gls2** | 4.67028 | 0.25235 | 4.18894 | 0.64561 | 0.00000 | 0.73328 | 1.46771 | 0.39811 | 1.23270 | 8.87472 | 6.35210 | 0.74982 | 0.96254 | 1.85152 | 3.28534 | 0.02101 |
| **Glt1d1** | 0.05925 | 0.00801 | 0.04428 | 0.12388 | 0.00907 | 0.00767 | 0.00742 | 0.00406 | 0.01750 | 0.00219 | 0.49350 | 0.01331 | 0.19657 | 0.52048 | 0.00000 | 0.00734 |
| **Glt28d2** | 0.05062 | 0.03993 | 0.10172 | 0.30369 | 0.03145 | 0.01275 | 0.00725 | 0.02910 | 0.06380 | 0.47022 | 0.21635 | 0.00651 | 0.02482 | 0.00962 | 0.02876 | 0.00724 |
| **Glt8d1** | 1.09787 | 3.29317 | 2.09937 | 5.54452 | 0.00270 | 3.08583 | 2.74146 | 0.87289 | 2.93098 | 1.55059 | 1.66389 | 0.53958 | 0.95250 | 2.00487 | 3.43615 | 0.01634 |
| **Glt8d2** | 0.01178 | 0.00676 | 0.00000 | 0.00358 | 0.00939 | 0.02146 | 1.59700 | 0.00000 | 0.13885 | 0.00000 | 0.39881 | 0.00441 | 0.00769 | 0.01113 | 0.00243 | 0.00709 |
| **Gltp** | 3.19310 | 2.37795 | 5.69444 | 3.37157 | 1.50645 | 10.00562 | 2.94213 | 2.15734 | 1.18436 | 1.07287 | 0.79027 | 0.08321 | 5.21900 | 0.02183 | 0.14906 | 0.26941 |
| **Gltscr1** | 0.18550 | 0.00715 | 0.00560 | 0.36237 | 0.00747 | 0.00815 | 1.23172 | 0.12155 | 0.05425 | 1.39084 | 0.04113 | 0.01128 | 0.01143 | 0.07911 | 0.97953 | 0.00714 |
| **Gltscr1l** | 3.17023 | 1.63801 | 0.81133 | 7.91388 | 0.00000 | 0.06267 | 0.00236 | 3.58398 | 8.70603 | 7.45878 | 3.11569 | 0.63385 | 0.00000 | 12.21549 | 10.44135 | 0.00760 |
| **Gltscr2** | 14.69528 | 0.12382 | 0.00000 | 7.62434 | 9.47767 | 2.27776 | 0.11570 | 11.54344 | 13.33607 | 30.60319 | 13.20678 | 0.76954 | 20.60023 | 0.00000 | 0.00000 | 11.69792 |
| **Glud1** | 23.27444 | 19.78983 | 14.30612 | 12.84527 | 18.97615 | 2.77426 | 16.13863 | 8.49710 | 10.21183 | 14.53682 | 7.37895 | 16.16517 | 9.82600 | 3.84999 | 13.77804 | 14.14409 |
| **Glul** | 10.83897 | 11.55083 | 16.15989 | 19.90070 | 6.47316 | 5.49941 | 12.47971 | 21.31414 | 30.20472 | 16.56533 | 21.18956 | 2.08002 | 18.96766 | 9.51416 | 13.95148 | 5.97395 |
| **Glyctk** | 0.63456 | 0.00623 | 0.32807 | 0.08465 | 0.01456 | 0.00498 | 0.00573 | 1.13521 | 0.07733 | 0.08514 | 0.06035 | 0.01271 | 1.38863 | 0.05030 | 0.33997 | 0.00563 |
| **Glyr1** | 7.12113 | 9.62145 | 8.28983 | 3.43208 | 11.53578 | 4.79855 | 7.87661 | 7.18995 | 8.18403 | 9.57337 | 8.64012 | 1.98762 | 4.79984 | 8.89590 | 11.20314 | 7.24142 |
| **Gm10012** | 5.31180 | 0.20715 | 4.41049 | 2.21272 | 3.23098 | 0.95879 | 0.67591 | 0.03043 | 8.56265 | 1.21725 | 17.58407 | 0.48385 | 2.21543 | 1.27951 | 0.72242 | 2.29906 |
| **Gm10033** | 0.22695 | 0.11579 | 0.45564 | 1.81978 | 0.08229 | 0.10215 | 1.23602 | 0.15319 | 2.24560 | 0.59511 | 0.87731 | 0.21926 | 0.17494 | 0.17938 | 0.48380 | 0.05244 |
| **Gm10046** | 0.03068 | 0.00777 | 0.01609 | 0.01035 | 0.00814 | 0.00731 | 0.00732 | 0.01087 | 0.01247 | 0.02903 | 0.01243 | 0.00676 | 0.01837 | 0.18457 | 0.00547 | 0.22170 |
| **Gm10069** | 3.91917 | 2.83368 | 3.10304 | 1.50591 | 1.52177 | 1.57190 | 1.00070 | 2.16124 | 0.29225 | 2.46055 | 1.38906 | 0.58316 | 2.55045 | 2.28545 | 2.66660 | 1.81038 |
| **Gm10190** | 0.16621 | 0.00774 | 0.07289 | 0.94484 | 0.04278 | 0.00635 | 0.02811 | 0.00000 | 2.08168 | 0.00000 | 0.80359 | 0.05230 | 0.04054 | 0.33482 | 0.02720 | 0.02871 |
| **Gm10319** | 0.01376 | 0.00812 | 0.01224 | 1.40691 | 0.00931 | 0.00653 | 0.00748 | 0.01384 | 0.00000 | 0.01481 | 0.01378 | 0.00826 | 0.01266 | 0.01192 | 0.00537 | 0.00718 |
| **Gm10324** | 2.57904 | 3.54272 | 3.06864 | 0.10390 | 1.94602 | 1.52740 | 2.28065 | 5.76011 | 0.23649 | 4.34974 | 5.51109 | 0.29215 | 2.46453 | 14.73753 | 9.39816 | 0.51500 |
| **Gm10334** | 0.01368 | 0.00832 | 0.01179 | 0.24159 | 0.00927 | 0.00687 | 0.00759 | 0.01388 | 0.06960 | 0.09796 | 0.01090 | 0.02163 | 0.01316 | 0.00773 | 0.07506 | 0.00738 |
| **Gm10336** | 0.25504 | 0.00736 | 0.26704 | 2.65842 | 0.01168 | 0.19179 | 0.48850 | 2.00319 | 0.33921 | 0.00000 | 0.63934 | 0.27476 | 1.18393 | 2.55441 | 0.78524 | 1.53730 |
| **Gm10354** | 0.01172 | 0.00805 | 0.07035 | 0.04896 | 0.00931 | 0.00600 | 0.00760 | 0.04203 | 0.00984 | 0.08440 | 0.01131 | 0.00840 | 0.01070 | 0.00777 | 0.16281 | 0.00745 |
| **Gm10364** | 0.14474 | 0.11615 | 0.01996 | 0.27349 | 0.12431 | 0.11337 | 0.04854 | 0.02421 | 1.12943 | 0.02332 | 0.00000 | 0.01489 | 0.07870 | 0.00491 | 0.31806 | 0.10204 |
| **Gm10389** | 0.04828 | 0.04858 | 0.08714 | 0.21600 | 0.00857 | 0.01515 | 0.03736 | 0.04327 | 0.70330 | 0.07191 | 0.17844 | 0.05136 | 0.06437 | 0.34711 | 0.09353 | 0.02956 |
| **Gm10406** | 0.04395 | 0.00787 | 0.03847 | 1.46822 | 0.00822 | 0.00587 | 0.00719 | 0.00968 | 0.12243 | 0.01563 | 0.06326 | 0.00669 | 0.02584 | 0.21421 | 0.00000 | 0.00707 |
| **Gm10408** | 0.01431 | 0.00802 | 0.02460 | 0.23620 | 0.00923 | 0.01981 | 0.01762 | 0.01184 | 0.03709 | 0.03009 | 0.02559 | 0.00730 | 0.01209 | 0.05450 | 0.02280 | 0.00748 |
| **Gm10409** | 0.09814 | 0.00790 | 0.01274 | 0.48136 | 0.00779 | 0.00652 | 0.00736 | 0.01391 | 0.01115 | 0.02701 | 0.02621 | 0.00812 | 0.01982 | 0.02800 | 0.00771 | 0.00722 |
| **Gm10416** | 1.14246 | 1.26821 | 0.76699 | 0.03135 | 0.27723 | 0.01216 | 0.01618 | 0.47017 | 0.00993 | 0.17210 | 0.23272 | 0.42277 | 0.79249 | 0.27703 | 0.02681 | 2.34625 |
| **Gm10432** | 0.03877 | 0.00789 | 0.00834 | 0.04057 | 0.00821 | 0.00713 | 0.03970 | 0.01546 | 0.02344 | 0.09449 | 0.01023 | 0.02000 | 0.02732 | 0.19528 | 0.02047 | 0.00732 |
| **Gm10433** | 0.00892 | 0.00776 | 0.01968 | 0.42228 | 0.00910 | 0.00816 | 0.00721 | 0.90029 | 0.57644 | 0.69445 | 0.00000 | 0.00396 | 0.01292 | 0.00832 | 0.03213 | 0.01482 |
| **Gm10436** | 0.00000 | 0.00753 | 0.00000 | 23.08931 | 0.00858 | 0.00349 | 0.00580 | 0.38361 | 12.16573 | 4.72848 | 0.66509 | 0.13785 | 0.00434 | 13.89717 | 17.80091 | 0.00219 |
| **Gm10439** | 0.01369 | 0.00780 | 0.01211 | 0.09203 | 0.00956 | 0.00655 | 0.00743 | 0.01173 | 0.03186 | 0.01914 | 0.02511 | 0.00632 | 0.01560 | 0.00619 | 0.55600 | 0.00748 |
| **Gm10509** | 0.53044 | 0.12413 | 0.51461 | 0.35089 | 0.00569 | 0.23278 | 0.00692 | 0.05190 | 1.87531 | 0.07094 | 0.12679 | 0.29990 | 0.01860 | 0.59699 | 1.05035 | 0.87037 |
| **Gm10512** | 0.01521 | 0.00783 | 0.06423 | 7.49929 | 0.00912 | 0.00495 | 0.00673 | 0.00096 | 1.17282 | 0.00000 | 0.25233 | 0.00000 | 0.00897 | 0.16222 | 3.36537 | 0.00570 |
| **Gm10548** | 0.10400 | 0.06203 | 0.26019 | 0.02987 | 0.03474 | 0.05033 | 0.11315 | 0.13279 | 0.15738 | 0.11668 | 0.22962 | 0.01771 | 0.04662 | 0.04348 | 0.01672 | 0.04279 |
| **Gm10636** | 0.13251 | 0.00809 | 0.08007 | 0.09267 | 0.09897 | 0.00625 | 0.06980 | 0.01777 | 0.10445 | 0.02671 | 0.30949 | 0.11152 | 0.03420 | 0.22818 | 0.06994 | 0.00739 |
| **Gm10638** | 0.01288 | 0.00818 | 0.01072 | 0.20298 | 0.00922 | 0.00615 | 0.00751 | 0.01214 | 5.97905 | 0.02119 | 0.00000 | 0.00000 | 0.01338 | 0.01045 | 0.00397 | 0.00706 |
| **Gm10639** | 0.18533 | 0.00741 | 0.08211 | 0.02897 | 0.00725 | 0.39659 | 0.09903 | 0.42519 | 0.00929 | 0.40401 | 0.35319 | 0.03380 | 0.67051 | 0.29402 | 0.17404 | 0.00653 |
| **Gm10653** | 12.49611 | 1.82009 | 7.46141 | 39.90341 | 9.51170 | 3.18747 | 2.04891 | 2.39834 | 38.77015 | 18.99856 | 19.00193 | 3.80707 | 8.37822 | 10.21077 | 15.75432 | 5.46635 |
| **Gm10658** | 0.01155 | 0.00828 | 0.83639 | 0.26690 | 0.00777 | 0.00911 | 0.00785 | 0.09947 | 0.45911 | 0.00000 | 0.01642 | 0.03941 | 0.00462 | 0.03221 | 0.61887 | 0.00686 |
| **Gm10696** | 0.02296 | 0.00868 | 0.67094 | 15.84472 | 0.01082 | 0.00298 | 0.00709 | 0.09976 | 11.88547 | 3.09931 | 0.48043 | 0.38130 | 0.03891 | 3.47003 | 7.55364 | 0.00349 |
| **Gm10767** | 1.02024 | 3.57163 | 2.99398 | 1.51632 | 0.47059 | 1.53087 | 1.39100 | 3.19422 | 5.05776 | 5.49101 | 2.03773 | 5.26288 | 1.05985 | 6.52850 | 5.03567 | 0.41315 |
| **Gm10768** | 0.14121 | 0.00811 | 0.00712 | 0.00395 | 0.01662 | 0.00679 | 0.00732 | 0.01108 | 0.34791 | 0.00469 | 0.00893 | 0.00639 | 0.01263 | 0.01311 | 0.36708 | 0.00739 |
| **Gm10778** | 0.15243 | 0.41717 | 2.08186 | 3.66405 | 0.74295 | 0.65974 | 1.20323 | 3.94025 | 4.75009 | 2.11975 | 5.82064 | 0.38782 | 2.46825 | 6.27142 | 3.32820 | 0.08342 |
| **Gm10785** | 0.00934 | 0.00851 | 0.00751 | 0.00540 | 0.00958 | 0.00706 | 0.03086 | 0.26169 | 0.03127 | 0.01107 | 0.48591 | 0.00671 | 0.02171 | 0.01505 | 0.03230 | 0.00720 |
| **Gm10789** | 0.03549 | 0.00796 | 0.10961 | 0.89404 | 0.01024 | 0.00741 | 0.00757 | 0.06852 | 1.73783 | 0.09339 | 0.03808 | 0.76947 | 0.02487 | 0.43131 | 0.26366 | 0.00721 |
| **Gm10791** | 0.02397 | 0.03846 | 0.09461 | 0.34556 | 0.02598 | 0.01451 | 0.01740 | 0.03296 | 0.16105 | 0.06298 | 0.16186 | 0.16146 | 0.11696 | 0.16372 | 0.09008 | 0.01437 |
| **Gm10814** | 0.01370 | 0.00761 | 0.01137 | 0.85514 | 0.00933 | 0.00642 | 0.00739 | 0.01175 | 0.75823 | 0.00186 | 0.00745 | 0.00575 | 0.01288 | 0.02216 | 0.33255 | 0.00716 |
| **Gm10845** | 1.22722 | 2.51962 | 7.96906 | 8.38153 | 0.84741 | 4.06719 | 3.93781 | 12.04996 | 5.09370 | 14.28411 | 2.04098 | 14.98243 | 2.90537 | 15.55234 | 13.29481 | 1.27187 |
| **Gm10857** | 0.84507 | 0.00710 | 0.48295 | 14.36288 | 0.00571 | 0.07919 | 0.02772 | 0.10053 | 2.39893 | 0.00000 | 0.05509 | 0.02549 | 0.07093 | 0.28374 | 4.23743 | 0.00360 |
| **Gm10865** | 0.05886 | 0.03885 | 0.43500 | 10.78869 | 0.00731 | 0.02229 | 0.14335 | 0.79356 | 8.57297 | 0.50190 | 0.28375 | 0.23855 | 0.46400 | 0.78791 | 1.22790 | 0.03381 |
| **Gm10941** | 0.02931 | 0.00777 | 0.02164 | 0.07034 | 0.00858 | 0.00706 | 0.02821 | 0.02645 | 0.19687 | 0.01431 | 0.05241 | 0.00685 | 0.01503 | 0.12160 | 0.05113 | 0.00734 |

| | | | | | | | | | | | | | | | | |
|---|---|---|---|---|---|---|---|---|---|---|---|---|---|---|---|---|
| **Gm1110** | 0.80358 | 0.00740 | 0.00000 | 0.00352 | 0.01620 | 0.00670 | 0.00595 | 0.67068 | 0.35278 | 0.85428 | 1.28631 | 0.00978 | 0.00000 | 2.11474 | 0.00000 | 0.00674 |
| **Gm11127** | 0.75969 | 1.11325 | 4.06483 | 3.88935 | 0.53158 | 1.91965 | 1.70620 | 8.64468 | 7.74366 | 9.00334 | 6.85748 | 9.50822 | 1.35244 | 5.25714 | 4.19790 | 0.73562 |
| **Gm11166** | 0.01368 | 0.00813 | 0.01208 | 0.12904 | 0.00928 | 0.00662 | 0.00753 | 0.01389 | 0.48635 | 0.01292 | 0.01061 | 0.00748 | 0.01281 | 0.01247 | 0.00533 | 0.00737 |
| **Gm1123** | 0.01229 | 0.00792 | 0.00668 | 0.11728 | 0.00962 | 0.00668 | 0.00718 | 0.00927 | 0.00954 | 0.00614 | 0.00815 | 0.00213 | 0.01343 | 1.68241 | 0.00305 | 0.00742 |
| **Gm11238** | 0.01112 | 0.00813 | 0.01031 | 4.27702 | 0.00941 | 0.00617 | 0.00717 | 0.07833 | 3.13146 | 0.03174 | 0.00000 | 0.00213 | 0.00996 | 0.03147 | 0.00597 | 0.00652 |
| **Gm1140** | 0.20155 | 0.00845 | 0.01525 | 0.00975 | 0.20842 | 0.00622 | 0.00785 | 0.03442 | 0.01193 | 0.01065 | 0.01232 | 0.02112 | 0.01031 | 0.00925 | 0.08828 | 0.00720 |
| **Gm11413** | 0.00906 | 0.00815 | 0.22626 | 2.06478 | 0.00906 | 0.00533 | 0.00735 | 0.00897 | 1.15728 | 0.47377 | 0.00000 | 0.02743 | 0.00976 | 1.89963 | 1.39488 | 0.00656 |
| **Gm11423** | 0.00000 | 0.00648 | 0.00268 | 1.08720 | 0.46574 | 0.00746 | 1.74641 | 0.35900 | 1.91221 | 0.30300 | 0.00000 | 0.02442 | 0.00779 | 0.00620 | 0.28074 | 0.00690 |
| **Gm11487** | 0.01072 | 0.00773 | 0.00873 | 2.48450 | 0.00938 | 0.00602 | 0.00716 | 0.08585 | 4.09912 | 0.16630 | 0.00000 | 0.03594 | 0.00938 | 0.02503 | 0.70850 | 0.00678 |
| **Gm11517** | 4.39457 | 0.68650 | 7.55408 | 17.73581 | 1.34060 | 0.31760 | 5.88986 | 28.86866 | 30.11750 | 24.58784 | 38.37708 | 6.43287 | 7.70923 | 29.83406 | 25.48425 | 0.79908 |
| **Gm11532** | 0.01590 | 0.00799 | 0.05395 | 0.22623 | 0.00915 | 0.00588 | 0.00737 | 0.01264 | 0.00763 | 0.03755 | 0.01475 | 0.00849 | 0.01152 | 0.01126 | 0.04948 | 0.00719 |
| **Gm11544** | 0.00000 | 0.00720 | 0.21265 | 23.04041 | 0.00848 | 0.07178 | 0.19493 | 1.16235 | 22.42695 | 5.12022 | 1.34276 | 1.21484 | 1.70682 | 1.70721 | 10.02910 | 0.00000 |
| **Gm11545** | 5.09630 | 3.37677 | 17.92498 | 13.38501 | 0.44368 | 8.51737 | 11.84898 | 26.59109 | 14.43026 | 20.31376 | 17.88220 | 11.22349 | 15.72379 | 33.95933 | 19.24505 | 6.32885 |
| **Gm11549** | 0.10497 | 0.01864 | 0.08946 | 0.17194 | 0.00987 | 0.00582 | 0.01738 | 0.00003 | 0.23255 | 0.01610 | 0.77936 | 0.04438 | 0.07034 | 0.17699 | 0.00426 | 0.01431 |
| **Gm11665** | 0.01382 | 0.00789 | 0.01083 | 0.85016 | 0.00927 | 0.00649 | 0.00734 | 0.00737 | 0.07393 | 0.10983 | 0.18039 | 0.00585 | 0.01019 | 0.01072 | 0.15327 | 0.00720 |
| **Gm11696** | 1.36585 | 0.00611 | 0.05699 | 0.10300 | 0.01331 | 0.03774 | 0.00638 | 0.02726 | 0.20533 | 0.03712 | 0.42822 | 0.01412 | 0.03087 | 0.16080 | 0.03796 | 0.01995 |
| **Gm11744** | 0.01203 | 0.00822 | 0.01430 | 0.01055 | 0.01019 | 0.00713 | 0.00767 | 0.01614 | 0.01235 | 0.00859 | 0.01518 | 0.00783 | 0.31969 | 0.01272 | 0.01022 | 0.00756 |
| **Gm11944** | 0.01282 | 0.00756 | 0.00919 | 0.02547 | 0.00905 | 0.00664 | 0.00723 | 0.00894 | 0.01264 | 0.84917 | 0.04433 | 0.00409 | 0.01077 | 0.39511 | 0.15456 | 0.00721 |
| **Gm11961** | 0.01626 | 0.00763 | 0.17117 | 0.00281 | 0.00905 | 0.00572 | 0.00722 | 0.01002 | 0.52974 | 0.00525 | 0.01735 | 0.00496 | 0.01199 | 0.00885 | 0.83708 | 0.00731 |
| **Gm11992** | 0.01364 | 0.00822 | 0.01199 | 0.74586 | 0.00931 | 0.00650 | 0.00748 | 0.01343 | 0.80227 | 0.00813 | 0.00812 | 0.00706 | 0.01280 | 0.00750 | 0.07295 | 0.00721 |
| **Gm12018** | 0.00649 | 0.00764 | 0.01291 | 0.20599 | 0.06269 | 0.01268 | 0.07945 | 0.06933 | 0.15660 | 0.04746 | 0.02321 | 0.00625 | 0.06095 | 0.49346 | 0.00935 | 0.02158 |
| **Gm12060** | 0.80125 | 0.33553 | 0.40067 | 0.82798 | 0.45699 | 0.67713 | 0.21010 | 0.43216 | 1.33576 | 0.51801 | 0.36661 | 0.35030 | 0.56882 | 0.49916 | 0.62309 | 0.14973 |
| **Gm12070** | 32.50846 | 17.43914 | 21.25269 | 9.14701 | 22.96614 | 15.94727 | 13.32680 | 2.67507 | 13.51944 | 3.68784 | 2.19985 | 3.12464 | 23.14425 | 0.12121 | 2.98567 | 22.82521 |
| **Gm12169** | 0.00546 | 0.09412 | 0.47861 | 0.00913 | 0.00861 | 3.11194 | 0.13084 | 0.00902 | 0.01269 | 0.00000 | 0.00909 | 0.00761 | 1.44745 | 0.01824 | 0.00032 | 0.26816 |
| **Gm12171** | 0.01306 | 0.00803 | 0.02481 | 0.08572 | 0.00917 | 0.06039 | 0.00745 | 0.01318 | 0.00799 | 0.01443 | 0.01208 | 0.00804 | 0.01838 | 0.01196 | 0.00882 | 0.02191 |
| **Gm12185** | 0.18062 | 0.02776 | 0.30373 | 7.61388 | 0.00827 | 0.01031 | 0.27241 | 0.00941 | 5.19161 | 0.54931 | 0.52156 | 0.18439 | 0.16246 | 1.52069 | 2.63525 | 0.04824 |
| **Gm12238** | 0.01353 | 0.00789 | 0.01213 | 0.01057 | 0.00935 | 0.00664 | 0.00747 | 0.01899 | 0.01237 | 0.01409 | 0.01300 | 0.00778 | 0.01232 | 0.01098 | 0.03092 | 0.00741 |
| **Gm12250** | 0.01272 | 0.00760 | 0.23604 | 0.05850 | 0.00931 | 0.00564 | 0.00696 | 0.00722 | 0.00683 | 0.00000 | 0.01911 | 0.00411 | 0.01722 | 3.22969 | 1.01692 | 0.00722 |
| **Gm12253** | 0.04105 | 0.00805 | 0.01579 | 0.27736 | 0.01990 | 0.00637 | 0.02858 | 0.00423 | 0.38824 | 0.00978 | 0.39712 | 0.00447 | 0.01086 | 0.37442 | 0.02860 | 0.01445 |
| **Gm12338** | 3.73517 | 0.07440 | 1.61805 | 0.97570 | 1.19478 | 0.60650 | 0.38958 | 0.15051 | 2.52569 | 1.09796 | 8.18966 | 0.21608 | 1.99447 | 0.38049 | 0.87662 | 2.57588 |
| **Gm12504** | 21.23292 | 8.89322 | 19.76616 | 0.75713 | 13.66863 | 10.29075 | 10.66534 | 24.56782 | 1.83027 | 20.59881 | 10.25843 | 6.10413 | 22.74332 | 9.61023 | 7.34121 | 11.02203 |
| **Gm12505** | 0.08677 | 0.00746 | 0.19652 | 0.44830 | 0.17153 | 0.03017 | 0.28537 | 0.10226 | 0.23110 | 0.19313 | 0.00775 | 0.02519 | 0.06438 | 0.23331 | 0.25884 | 0.00713 |
| **Gm12530** | 0.01375 | 0.00805 | 0.01225 | 0.01061 | 0.00932 | 0.00666 | 0.00754 | 0.01390 | 0.01235 | 0.01531 | 0.01310 | 0.00811 | 0.01270 | 0.01234 | 0.01062 | 0.00741 |
| **Gm12596** | 0.01053 | 0.00792 | 0.01301 | 0.59879 | 0.00963 | 0.00734 | 0.00755 | 0.05094 | 0.54527 | 0.15753 | 0.00000 | 0.02399 | 0.02280 | 0.50276 | 1.34033 | 0.00743 |
| **Gm12657** | 15.65689 | 5.60200 | 15.07437 | 8.60983 | 9.47375 | 7.50404 | 8.01784 | 21.38758 | 12.96852 | 24.04604 | 12.02172 | 6.65857 | 15.51107 | 8.31027 | 3.01206 | 6.84616 |
| **Gm12789** | 0.01651 | 0.05003 | 0.16098 | 11.75887 | 0.00845 | 0.01127 | 0.16066 | 0.13482 | 7.55002 | 0.14596 | 0.05563 | 0.00000 | 0.13867 | 0.16898 | 0.68466 | 0.02693 |
| **Gm12794** | 0.01500 | 0.01915 | 0.02990 | 2.62877 | 0.00902 | 0.00613 | 0.04968 | 0.05117 | 2.62971 | 0.01922 | 0.00000 | 0.00307 | 0.02738 | 0.00556 | 0.05738 | 0.00676 |
| **Gm128** | 0.00991 | 0.00735 | 0.06195 | 3.06417 | 0.00976 | 0.00570 | 0.00689 | 1.35967 | 1.29470 | 0.12416 | 0.85032 | 0.00000 | 0.00416 | 0.00000 | 0.27642 | 0.00675 |
| **Gm12992** | 0.01140 | 0.00811 | 0.03750 | 2.10688 | 0.00937 | 0.00577 | 0.00739 | 0.04663 | 0.70503 | 0.08577 | 0.00766 | 0.02711 | 0.01080 | 0.09153 | 0.09830 | 0.00736 |
| **Gm13023** | 0.00000 | 0.00742 | 0.15984 | 18.64440 | 0.00925 | 0.00529 | 0.00586 | 0.12380 | 12.90572 | 0.72496 | 0.93659 | 0.56592 | 0.00000 | 8.08204 | 6.39991 | 0.00317 |
| **Gm13034** | 0.00253 | 0.22281 | 0.01665 | 1.00325 | 0.02015 | 0.01199 | 0.14183 | 0.10460 | 0.27256 | 0.31758 | 0.01326 | 0.00000 | 0.02471 | 1.91058 | 0.49883 | 0.01416 |
| **Gm13078** | 0.05651 | 0.24276 | 0.58330 | 7.14732 | 0.09753 | 0.12935 | 0.39192 | 1.53260 | 10.36395 | 0.18997 | 0.00000 | 0.00000 | 0.22574 | 0.87601 | 0.08390 | 0.01245 |
| **Gm13083** | 0.16345 | 0.41618 | 0.87223 | 2.97446 | 0.08279 | 0.23661 | 0.59341 | 2.01804 | 5.64323 | 0.36012 | 0.00000 | 0.61076 | 0.30542 | 1.47877 | 0.25564 | 0.03525 |
| **Gm13084** | 0.01422 | 0.00751 | 0.00000 | 7.80919 | 0.00882 | 0.00513 | 0.00637 | 0.00000 | 1.78396 | 0.24252 | 1.09975 | 0.00000 | 0.00004 | 4.38444 | 5.67057 | 0.00544 |
| **Gm13088** | 0.01304 | 0.00769 | 0.00648 | 1.04679 | 0.00944 | 0.00665 | 0.00718 | 0.00706 | 0.00000 | 0.04153 | 0.09967 | 0.00369 | 0.01145 | 0.75143 | 0.54394 | 0.00717 |
| **Gm13102** | 5.41859 | 13.84198 | 22.76193 | 6.09640 | 9.97307 | 5.56540 | 8.25749 | 26.43495 | 16.19560 | 22.72993 | 13.43230 | 18.47579 | 17.39487 | 30.29791 | 14.64443 | 1.44859 |
| **Gm13103** | 0.03586 | 0.00751 | 0.00097 | 2.21833 | 0.00838 | 0.00675 | 0.00714 | 0.00605 | 0.01712 | 0.05062 | 0.10393 | 0.00000 | 0.01135 | 1.58392 | 0.94550 | 0.00682 |
| **Gm13119** | 0.00263 | 0.38424 | 0.00516 | 0.72073 | 0.00837 | 0.00460 | 0.00630 | 0.03029 | 4.92616 | 0.00000 | 0.00000 | 5.58017 | 0.01079 | 0.49431 | 4.00882 | 0.00616 |
| **Gm13125** | 0.01315 | 0.00755 | 0.01201 | 1.16209 | 0.00896 | 0.00627 | 0.00720 | 0.00999 | 0.00000 | 1.02783 | 0.01230 | 0.00451 | 0.01045 | 0.00597 | 0.40815 | 0.00690 |
| **Gm13128** | 0.41078 | 1.49875 | 3.48556 | 0.68008 | 0.00513 | 0.02202 | 1.33346 | 0.88224 | 2.22476 | 0.00000 | 0.00000 | 0.00000 | 1.56592 | 3.96577 | 3.29308 | 0.51336 |
| **Gm13212** | 2.16041 | 5.79676 | 4.36607 | 5.92330 | 2.75570 | 1.63023 | 4.03595 | 7.56030 | 12.34690 | 4.20329 | 6.11148 | 23.19021 | 5.54815 | 28.95720 | 2.77878 | 2.53158 |
| **Gm13238** | 0.01284 | 0.00760 | 0.00997 | 0.16909 | 0.00981 | 0.00707 | 0.39144 | 0.01038 | 0.14601 | 0.00873 | 0.05962 | 0.00691 | 0.02420 | 0.01637 | 0.02685 | 0.00754 |
| **Gm13315** | 0.02497 | 0.05956 | 0.01625 | 0.08867 | 0.03726 | 0.00737 | 0.00735 | 0.01647 | 0.10696 | 0.01452 | 0.01222 | 0.02814 | 0.02599 | 0.01501 | 0.02322 | 0.02186 |
| **Gm13375** | 0.62266 | 0.00688 | 0.00944 | 0.01244 | 0.01454 | 0.00793 | 0.00686 | 0.25778 | 0.01229 | 0.57666 | 0.05168 | 0.00472 | 0.19841 | 0.00669 | 0.09228 | 0.07448 |
| **Gm13498** | 7.73595 | 5.95601 | 10.97833 | 0.00000 | 5.85800 | 2.41800 | 5.27757 | 12.02816 | 0.00921 | 7.15796 | 0.16004 | 3.50978 | 7.81050 | 7.46539 | 4.94114 | 3.24028 |
| **Gm13648** | 0.01193 | 0.00774 | 0.00297 | 1.32961 | 0.00987 | 0.00726 | 0.04885 | 0.00000 | 2.21193 | 0.83277 | 1.34491 | 0.01150 | 0.08355 | 0.00000 | 1.39250 | 0.00721 |
| **Gm13752** | 0.00886 | 0.00788 | 0.00692 | 1.85771 | 0.01010 | 0.00728 | 0.00731 | 0.51874 | 0.00000 | 0.30472 | 0.27131 | 0.01423 | 0.01048 | 0.00521 | 0.00000 | 0.00699 |
| **Gm13830** | 0.02027 | 0.01918 | 0.04236 | 0.06530 | 0.00994 | 0.01750 | 0.00741 | 0.02592 | 0.67770 | 0.06369 | 0.02122 | 0.05003 | 0.04461 | 0.02944 | 0.02162 | 0.00726 |
| **Gm13884** | 0.00895 | 0.00751 | 0.06656 | 0.00950 | 0.14396 | 0.00877 | 0.00729 | 0.01115 | 0.01216 | 0.19077 | 0.01302 | 0.00645 | 0.10888 | 0.00931 | 0.03714 | 0.00772 |
| **Gm13985** | 0.08476 | 0.03915 | 0.14212 | 0.30609 | 0.09129 | 0.03308 | 0.02753 | 0.04644 | 0.24867 | 0.13655 | 0.45985 | 0.09926 | 0.15768 | 0.21212 | 0.19836 | 0.05894 |
| **Gm13986** | 0.04065 | 0.00785 | 0.01484 | 0.00975 | 0.01053 | 0.00737 | 0.03750 | 0.01088 | 0.01187 | 0.02130 | 0.02444 | 0.11612 | 0.01843 | 0.03796 | 0.11116 | 0.01459 |
| **Gm14005** | 1.53575 | 0.14099 | 0.80234 | 4.90230 | 1.83512 | 0.08490 | 0.18776 | 0.00448 | 7.16623 | 0.00000 | 4.71818 | 0.29620 | 0.22294 | 2.48090 | 0.40088 | 0.14524 |

| | | | | | | | | | | | | | | | | |
|---|---|---|---|---|---|---|---|---|---|---|---|---|---|---|---|---|
| **Gm14023** | 0.00764 | 1.56826 | 0.74618 | 0.00433 | 0.00677 | 0.00559 | 0.00653 | 0.00797 | 0.06006 | 0.00512 | 0.02392 | 0.00660 | 0.01511 | 0.02784 | 0.00000 | 0.01365 |
| **Gm14057** | 0.00067 | 0.00778 | 0.13995 | 1.68178 | 0.02045 | 0.00892 | 0.27395 | 0.85395 | 1.57704 | 0.82075 | 0.67762 | 0.10274 | 0.01164 | 1.37175 | 0.00000 | 0.00675 |
| **Gm14085** | 0.01364 | 0.00790 | 0.01222 | 0.32348 | 0.00914 | 0.00648 | 0.00732 | 0.01108 | 0.00503 | 0.35433 | 0.01439 | 0.00773 | 0.01119 | 0.01733 | 0.00379 | 0.00708 |
| **Gm14124** | 0.21832 | 0.04863 | 0.36223 | 0.17908 | 0.02371 | 0.01744 | 1.05637 | 0.43159 | 0.13244 | 0.06644 | 0.31236 | 0.05160 | 0.11354 | 1.75244 | 0.57442 | 0.01384 |
| **Gm14137** | 0.06259 | 0.00747 | 0.00372 | 0.68865 | 0.00789 | 0.00735 | 1.06827 | 0.06807 | 0.00000 | 0.00530 | 0.04143 | 0.00685 | 0.00854 | 0.00540 | 0.24371 | 0.00701 |
| **Gm14139** | 0.06233 | 0.01763 | 1.85173 | 0.02019 | 0.00602 | 0.00595 | 1.54641 | 0.29231 | 0.49127 | 0.00000 | 0.81977 | 0.04663 | 1.18544 | 1.78794 | 0.82907 | 0.01283 |
| **Gm14164** | 0.00562 | 0.00851 | 0.00627 | 0.64302 | 0.01080 | 0.00652 | 0.00807 | 0.20579 | 0.00000 | 0.00431 | 0.00340 | 0.01123 | 0.00736 | 2.61897 | 0.07971 | 0.00731 |
| **Gm14204** | 0.05192 | 0.01781 | 0.07731 | 0.96574 | 0.03540 | 0.01965 | 0.11017 | 0.05004 | 1.20474 | 0.09183 | 0.11718 | 0.05197 | 0.07389 | 0.81866 | 0.10898 | 0.01424 |
| **Gm14288** | 0.07955 | 0.09440 | 0.10026 | 0.06001 | 0.01025 | 0.11050 | 0.03866 | 0.02628 | 0.06252 | 0.01207 | 0.04112 | 0.00740 | 0.07558 | 0.04540 | 0.04783 | 0.05085 |
| **Gm14295** | 2.87968 | 2.15712 | 9.51084 | 0.72908 | 0.18822 | 3.15323 | 1.35606 | 0.55316 | 0.57083 | 1.14877 | 0.67044 | 0.03562 | 3.44934 | 4.30337 | 0.00000 | 1.36964 |
| **Gm14305** | 0.13250 | 0.03953 | 0.24109 | 0.03914 | 0.02568 | 0.00694 | 0.01736 | 0.02967 | 0.03715 | 0.04709 | 0.27729 | 0.03452 | 0.06044 | 0.48700 | 0.03410 | 0.14391 |
| **Gm14322** | 1.57915 | 2.43660 | 6.80010 | 2.69490 | 0.42071 | 0.46209 | 1.56796 | 0.89177 | 1.63248 | 1.51611 | 1.64758 | 0.46980 | 5.77299 | 2.56405 | 3.22158 | 2.08009 |
| **Gm14325** | 3.98540 | 4.87114 | 8.95820 | 1.50116 | 0.34856 | 1.17872 | 2.02662 | 2.21191 | 0.81295 | 4.26999 | 1.63560 | 0.59534 | 3.64852 | 9.49719 | 1.98536 | 1.92261 |
| **Gm14326** | 0.66159 | 0.71760 | 1.29566 | 0.60834 | 0.05763 | 0.29311 | 0.56447 | 0.15725 | 0.06549 | 0.85506 | 0.83220 | 0.33876 | 0.54340 | 2.06523 | 0.42062 | 0.48409 |
| **Gm14327** | 0.01268 | 0.01835 | 0.02949 | 0.00701 | 0.00910 | 0.00790 | 0.02794 | 0.01196 | 0.51073 | 0.01036 | 0.02088 | 0.00687 | 0.02596 | 0.01671 | 0.01828 | 0.00733 |
| **Gm14391** | 0.01379 | 0.00783 | 0.01135 | 0.18788 | 0.00923 | 0.00661 | 0.00740 | 0.00967 | 0.09670 | 0.10172 | 0.12004 | 0.00647 | 0.01107 | 0.00797 | 0.03558 | 0.00750 |
| **Gm14393** | 0.17015 | 0.23381 | 0.38926 | 0.29667 | 0.11461 | 0.05311 | 0.27044 | 0.17019 | 0.05813 | 0.08521 | 0.33637 | 0.01599 | 0.22075 | 0.33539 | 0.04290 | 0.47613 |
| **Gm14403** | 2.08121 | 3.56602 | 5.00866 | 2.07151 | 0.14209 | 0.13662 | 0.60113 | 0.14501 | 4.97732 | 1.92417 | 0.40609 | 3.49846 | 2.61617 | 6.60579 | 9.05762 | 2.01089 |
| **Gm14405** | 0.21718 | 0.21566 | 0.42497 | 0.05469 | 0.01151 | 0.33842 | 0.22925 | 0.05081 | 0.11112 | 0.11132 | 0.02636 | 0.01089 | 0.12979 | 1.80764 | 0.18721 | 0.03582 |
| **Gm14440** | 0.34131 | 2.04055 | 2.08371 | 0.44051 | 0.00489 | 1.92447 | 0.87785 | 0.13296 | 0.70701 | 0.19389 | 0.10295 | 0.13638 | 0.83153 | 3.65716 | 0.35934 | 0.56170 |
| **Gm14461** | 0.13092 | 0.17698 | 0.16748 | 0.38552 | 0.01795 | 0.06574 | 0.07022 | 0.10790 | 0.40655 | 0.29115 | 0.53910 | 0.33165 | 0.14053 | 0.20181 | 0.27097 | 0.07313 |
| **Gm14634** | 0.01360 | 0.00827 | 0.01198 | 0.30068 | 0.00922 | 0.00684 | 0.00760 | 0.01396 | 0.36304 | 0.13788 | 0.00980 | 0.00849 | 0.01307 | 0.01101 | 0.00578 | 0.00735 |
| **Gm14635** | 0.01375 | 0.00819 | 0.01231 | 0.56226 | 0.00934 | 0.00661 | 0.00752 | 0.01397 | 0.00058 | 0.01535 | 0.01319 | 0.00831 | 0.01268 | 0.01102 | 0.00498 | 0.00732 |
| **Gm14827** | 0.01803 | 0.00808 | 0.06308 | 0.11279 | 0.01038 | 0.00611 | 0.00700 | 0.02768 | 0.28813 | 0.02291 | 0.02165 | 0.01431 | 0.24054 | 0.59196 | 0.10031 | 0.00712 |
| **Gm14851** | 0.02031 | 0.00800 | 0.01021 | 0.96197 | 0.00872 | 0.00644 | 0.00745 | 0.01105 | 1.64384 | 0.00556 | 0.07648 | 0.01557 | 0.02409 | 0.00797 | 0.05150 | 0.00702 |
| **Gm15056** | 0.01369 | 0.00786 | 0.01139 | 0.00421 | 0.00931 | 0.00657 | 0.00740 | 0.01171 | 0.70403 | 0.02501 | 0.00684 | 0.01577 | 0.01333 | 0.00808 | 0.34922 | 0.00763 |
| **Gm15107** | 0.01390 | 0.00788 | 0.00991 | 0.00852 | 0.00921 | 0.00668 | 0.00745 | 0.00667 | 0.03378 | 0.00683 | 0.27024 | 0.00640 | 0.00941 | 0.01068 | 0.13446 | 0.00731 |
| **Gm1527** | 0.01323 | 0.00782 | 0.00913 | 0.80599 | 0.00938 | 0.00633 | 0.00727 | 0.00993 | 1.73793 | 0.00000 | 0.00000 | 0.01089 | 0.01316 | 0.26093 | 0.79447 | 0.00703 |
| **Gm15319** | 0.01371 | 0.00792 | 0.01234 | 0.35884 | 0.00946 | 0.00668 | 0.00755 | 0.01360 | 0.14908 | 0.01071 | 0.02595 | 0.00746 | 0.01530 | 0.01037 | 0.06514 | 0.00739 |
| **Gm15401** | 0.07252 | 0.07303 | 0.15298 | 0.07959 | 0.07513 | 0.01236 | 0.01741 | 0.01443 | 0.61813 | 0.06061 | 0.53079 | 0.04419 | 0.19766 | 0.10172 | 0.08298 | 0.05070 |
| **Gm15417** | 0.02674 | 0.00813 | 0.01791 | 0.08318 | 0.00974 | 0.00635 | 0.00757 | 0.02029 | 0.20667 | 0.02765 | 0.06744 | 0.00754 | 0.01963 | 0.05282 | 0.02662 | 0.00734 |
| **Gm15421** | 4.27010 | 0.36064 | 2.65379 | 1.24029 | 4.39692 | 0.25669 | 0.21235 | 0.26132 | 2.61832 | 0.85754 | 15.11639 | 0.50785 | 2.57162 | 2.13140 | 0.96270 | 2.01080 |
| **Gm15446** | 0.33806 | 0.61432 | 0.37820 | 0.25957 | 0.02475 | 0.22069 | 0.07787 | 0.34452 | 0.07682 | 0.25778 | 4.04067 | 0.79190 | 0.42463 | 5.48455 | 0.15178 | 1.17959 |
| **Gm15455** | 3.31594 | 0.90915 | 0.36738 | 11.73733 | 0.16377 | 1.35239 | 0.80894 | 1.43952 | 7.66324 | 1.60583 | 0.40030 | 0.19820 | 1.22158 | 0.54639 | 4.07841 | 0.71928 |
| **Gm15545** | 0.01207 | 0.00798 | 0.01113 | 0.63453 | 0.00942 | 0.00661 | 0.00729 | 0.06204 | 0.19991 | 0.01994 | 0.01214 | 0.00763 | 0.01014 | 0.01071 | 0.00325 | 0.00730 |
| **Gm15645** | 3.30320 | 1.56322 | 6.02774 | 7.70559 | 1.42803 | 2.54346 | 4.05533 | 5.94635 | 12.46445 | 10.12642 | 13.56894 | 8.34104 | 3.97746 | 4.70662 | 5.56634 | 1.46597 |
| **Gm15694** | 0.29214 | 0.00730 | 0.15606 | 1.93631 | 0.04426 | 0.03503 | 0.16985 | 0.64378 | 1.10196 | 0.05767 | 0.50868 | 0.08845 | 0.09286 | 0.03682 | 0.32589 | 0.07139 |
| **Gm15698** | 17.62846 | 11.41590 | 24.04569 | 17.60641 | 13.76557 | 11.36718 | 11.02310 | 16.47380 | 15.31873 | 9.61210 | 9.87783 | 6.65125 | 12.25022 | 12.53437 | 9.49543 | 8.06743 |
| **Gm15706** | 0.01374 | 0.00816 | 0.01228 | 0.30555 | 0.00932 | 0.00667 | 0.00755 | 0.01414 | 0.00544 | 0.01852 | 0.01297 | 0.00840 | 0.01279 | 0.01123 | 0.00668 | 0.00737 |
| **Gm15708** | 0.01337 | 0.00797 | 0.01202 | 0.07638 | 0.00922 | 0.00685 | 0.00755 | 0.01955 | 0.01140 | 0.17260 | 0.01254 | 0.00828 | 0.01253 | 0.00726 | 0.06693 | 0.00740 |
| **Gm15760** | 0.01123 | 0.02789 | 0.11881 | 1.60503 | 0.00895 | 0.45117 | 0.00699 | 0.01769 | 0.30591 | 1.26742 | 0.01394 | 0.01735 | 0.01458 | 1.80412 | 0.06636 | 0.00695 |
| **Gm15787** | 0.01314 | 0.00799 | 0.00894 | 0.00530 | 0.00946 | 0.00676 | 0.00729 | 0.01028 | 0.17222 | 0.00751 | 0.00867 | 0.00510 | 0.01329 | 0.64309 | 0.00000 | 0.00746 |
| **Gm15800** | 3.62492 | 5.31278 | 2.53832 | 5.45505 | 0.93862 | 0.00231 | 1.05012 | 3.67076 | 6.19053 | 6.51322 | 0.85823 | 0.30959 | 0.16085 | 0.40596 | 1.55134 | 1.68359 |
| **Gm15910** | 0.05549 | 0.00772 | 0.09687 | 0.04881 | 0.00788 | 0.00648 | 0.00744 | 0.04349 | 0.26419 | 0.02652 | 0.44688 | 0.00503 | 0.09179 | 0.11185 | 0.05626 | 0.01417 |
| **Gm15915** | 0.01120 | 0.00776 | 0.01034 | 0.01069 | 0.00956 | 0.00670 | 0.00724 | 0.11208 | 0.01224 | 0.01309 | 0.01266 | 0.00750 | 0.00847 | 0.01241 | 0.00888 | 0.00740 |
| **Gm15941** | 0.01363 | 0.00817 | 0.01167 | 2.57802 | 0.00926 | 0.00632 | 0.00742 | 0.01264 | 1.00052 | 0.00478 | 0.03973 | 0.00639 | 0.01205 | 0.00425 | 0.06626 | 0.00687 |
| **Gm1600** | 0.01622 | 0.00810 | 0.01716 | 0.05612 | 0.01805 | 0.00728 | 0.01838 | 0.01920 | 0.12160 | 0.08021 | 0.02313 | 0.03897 | 0.01127 | 0.03064 | 0.05357 | 0.00716 |
| **Gm16023** | 0.01345 | 0.00828 | 0.01185 | 0.32803 | 0.00925 | 0.00652 | 0.00762 | 0.01719 | 0.74257 | 0.04657 | 0.01021 | 0.00765 | 0.01196 | 0.01417 | 0.02270 | 0.00714 |
| **Gm16062** | 0.01346 | 0.01880 | 0.01200 | 0.00900 | 0.00925 | 0.00666 | 0.01792 | 0.01352 | 0.12484 | 0.01456 | 0.01218 | 0.00792 | 0.01267 | 0.01258 | 0.00927 | 0.00740 |
| **Gm16070** | 0.05961 | 0.00759 | 0.06394 | 8.59351 | 0.00903 | 0.06453 | 0.04662 | 0.09771 | 5.38331 | 3.48413 | 0.00000 | 0.16953 | 0.19648 | 6.84702 | 13.48244 | 0.00507 |
| **Gm16157** | 0.02672 | 0.00746 | 0.01051 | 0.27937 | 0.00849 | 0.00664 | 0.00729 | 0.00848 | 0.08863 | 0.56791 | 0.06803 | 0.00601 | 0.01199 | 0.05970 | 0.03675 | 0.00722 |
| **Gm16223** | 0.09499 | 0.28137 | 0.24670 | 0.03753 | 0.05770 | 0.22743 | 0.24239 | 0.19597 | 0.01768 | 0.40589 | 0.06774 | 0.23015 | 0.03521 | 0.78478 | 0.47771 | 0.01424 |
| **Gm16287** | 0.11361 | 0.01890 | 0.26209 | 0.04030 | 0.00921 | 0.01258 | 0.03879 | 0.01778 | 0.73334 | 0.06680 | 0.95378 | 0.04389 | 0.19991 | 0.29781 | 0.03737 | 0.00688 |
| **Gm16381** | 0.00991 | 0.00755 | 0.00421 | 25.75119 | 0.00901 | 0.00230 | 0.00622 | 0.00000 | 19.36304 | 0.00000 | 0.11015 | 0.00000 | 0.01316 | 0.64956 | 2.34730 | 0.00871 |
| **Gm16386** | 0.00963 | 0.06162 | 0.05809 | 0.69388 | 0.17814 | 0.02226 | 0.04851 | 0.03541 | 0.13129 | 0.03198 | 0.02263 | 0.01598 | 0.00984 | 0.52423 | 0.01406 | 0.00720 |
| **Gm16513** | 3.59773 | 3.06552 | 5.31308 | 8.41271 | 2.31089 | 0.20910 | 4.06896 | 4.11767 | 10.55515 | 9.94350 | 12.84178 | 6.98218 | 3.80043 | 20.83497 | 8.08123 | 0.23241 |
| **Gm16523** | 0.04162 | 0.02908 | 0.12115 | 0.18729 | 0.00957 | 0.06646 | 0.01710 | 0.00974 | 0.10964 | 0.08618 | 0.40971 | 0.05440 | 0.14727 | 2.12717 | 0.06744 | 0.03705 |
| **Gm1653** | 0.01347 | 0.00770 | 0.01189 | 0.86535 | 0.00907 | 0.00665 | 0.00733 | 0.01091 | 0.00000 | 0.62957 | 0.01141 | 0.00618 | 0.01207 | 0.01370 | 0.05059 | 0.00751 |
| **Gm16551** | 0.01352 | 0.00777 | 0.01007 | 2.26716 | 0.00934 | 0.00610 | 0.00724 | 0.01041 | 1.41864 | 0.00177 | 0.00000 | 0.03543 | 0.01300 | 0.00518 | 1.17944 | 0.00678 |
| **Gm16702** | 2.97168 | 0.43823 | 2.38600 | 18.71433 | 1.68772 | 0.33651 | 0.52783 | 0.00000 | 21.32065 | 0.89064 | 16.36592 | 1.85119 | 1.16961 | 8.94339 | 6.19440 | 0.61164 |
| **Gm16740** | 0.19514 | 0.00859 | 0.00000 | 0.06431 | 0.01745 | 0.37275 | 0.00739 | 0.01194 | 0.01139 | 0.03455 | 0.01273 | 0.00807 | 0.00861 | 0.01160 | 0.00908 | 0.00728 |
| **Gm16796** | 0.08004 | 0.00862 | 0.00922 | 0.01057 | 0.01575 | 0.00683 | 0.00750 | 0.01311 | 0.01239 | 0.02324 | 0.01269 | 0.00810 | 0.01238 | 0.01200 | 0.01077 | 0.00745 |

| | | | | | | | | | | | | | | | | |
|---|---|---|---|---|---|---|---|---|---|---|---|---|---|---|---|---|
| **Gm16845** | 0.01167 | 0.01963 | 0.01266 | 1.51295 | 0.00996 | 0.00729 | 0.00765 | 0.01521 | 0.63099 | 0.11394 | 0.02653 | 0.04418 | 0.29715 | 0.04381 | 0.01014 | 0.00737 |
| **Gm16853** | 0.01325 | 0.00828 | 0.01140 | 3.42336 | 0.00922 | 0.00624 | 0.00743 | 0.01297 | 3.07150 | 0.09791 | 0.00000 | 0.00263 | 0.01333 | 0.00765 | 0.00520 | 0.00669 |
| **Gm16907** | 0.01370 | 0.00820 | 0.01216 | 0.40984 | 0.00929 | 0.00660 | 0.00752 | 0.01395 | 0.35105 | 0.01344 | 0.01110 | 0.00779 | 0.01277 | 0.01156 | 0.00421 | 0.00733 |
| **Gm16938** | 0.34929 | 0.01923 | 0.14504 | 8.06829 | 0.03922 | 0.12493 | 0.20701 | 0.60597 | 8.39016 | 1.43012 | 0.21016 | 0.42244 | 0.84692 | 1.46512 | 1.94573 | 0.07209 |
| **Gm16973** | 1.76361 | 1.89810 | 0.09137 | 6.02257 | 0.00454 | 0.00389 | 0.00600 | 0.13687 | 7.11878 | 0.79261 | 0.00000 | 0.03445 | 0.04021 | 0.00000 | 0.00000 | 0.01844 |
| **Gm17019** | 0.01098 | 0.00799 | 0.13687 | 0.06744 | 0.00927 | 0.00585 | 0.00755 | 0.06260 | 0.13622 | 0.06366 | 0.01023 | 0.01905 | 0.00933 | 0.00747 | 0.13504 | 0.00741 |
| **Gm17026** | 0.02892 | 0.00780 | 0.03229 | 0.07937 | 0.00866 | 0.00614 | 0.01806 | 0.01256 | 0.12873 | 0.01197 | 0.04882 | 0.00748 | 0.01381 | 0.01138 | 0.02120 | 0.00736 |
| **Gm17066** | 1.07016 | 0.69489 | 0.00000 | 3.59701 | 0.00670 | 1.49551 | 0.04540 | 0.48664 | 1.74270 | 0.00000 | 1.49211 | 5.78934 | 0.62364 | 5.43913 | 0.00000 | 0.00358 |
| **Gm17644** | 2.52132 | 0.33520 | 0.76750 | 0.00000 | 0.00966 | 2.08783 | 0.31638 | 1.06354 | 0.12058 | 7.54545 | 0.03408 | 0.09181 | 1.91348 | 4.69984 | 1.49757 | 2.25430 |
| **Gm17751** | 0.01352 | 0.00753 | 0.01066 | 3.56338 | 0.00924 | 0.00613 | 0.00715 | 0.00973 | 0.00000 | 0.19714 | 0.02674 | 0.00369 | 0.01264 | 0.00307 | 1.42304 | 0.00720 |
| **Gm17762** | 0.06727 | 0.00773 | 0.11629 | 0.00755 | 0.00777 | 0.00565 | 0.00711 | 0.02019 | 0.01194 | 0.00537 | 0.13139 | 0.00704 | 0.25303 | 0.01182 | 0.00628 | 0.06294 |
| **Gm17801** | 0.03921 | 0.02993 | 0.10232 | 0.05724 | 0.04846 | 0.01186 | 0.01844 | 0.00462 | 0.09563 | 0.02337 | 0.48400 | 0.03650 | 0.03939 | 0.15380 | 0.12738 | 0.02183 |
| **Gm1821** | 34.48891 | 23.04935 | 34.23132 | 15.95158 | 23.73089 | 10.88701 | 19.99742 | 22.05799 | 27.98440 | 20.56178 | 18.62087 | 10.16974 | 29.90762 | 4.51031 | 16.30327 | 18.82883 |
| **Gm1943** | 1.55131 | 2.89510 | 6.33049 | 10.59831 | 7.04879 | 5.45467 | 2.55760 | 6.65308 | 11.76463 | 11.76929 | 8.21172 | 4.06955 | 4.15280 | 5.89493 | 8.17651 | 1.54350 |
| **Gm19434** | 0.16474 | 0.03820 | 0.45116 | 3.02987 | 0.01702 | 0.01190 | 0.33291 | 0.05896 | 2.59524 | 0.44572 | 0.33516 | 0.15340 | 0.04529 | 0.56329 | 0.82565 | 0.05643 |
| **Gm19510** | 1.62506 | 0.00609 | 0.01103 | 0.06032 | 0.00394 | 0.01278 | 0.00630 | 0.00899 | 0.05290 | 0.00535 | 0.01259 | 0.00722 | 0.00651 | 0.01306 | 0.01284 | 2.40461 |
| **Gm19557** | 0.01375 | 0.00807 | 0.01225 | 0.01058 | 0.00932 | 0.00666 | 0.00754 | 0.01389 | 0.01234 | 0.01523 | 0.01311 | 0.00815 | 0.01270 | 0.01220 | 0.01187 | 0.00741 |
| **Gm19583** | 0.01589 | 0.00844 | 0.90592 | 0.34446 | 0.00739 | 0.00574 | 0.00751 | 0.01151 | 0.00588 | 0.06687 | 0.00798 | 0.00708 | 0.00875 | 0.22166 | 0.72678 | 0.00694 |
| **Gm19589** | 0.09727 | 0.09033 | 0.07564 | 0.98009 | 0.00779 | 0.01224 | 0.00727 | 0.06410 | 0.88403 | 0.11692 | 0.09778 | 0.02468 | 0.01884 | 0.02501 | 0.08100 | 0.06637 |
| **Gm19619** | 0.01758 | 0.00800 | 0.05165 | 0.02657 | 0.01116 | 0.00765 | 0.18254 | 0.01153 | 0.04105 | 0.01487 | 0.02456 | 0.00678 | 0.02642 | 0.14296 | 0.06073 | 0.00757 |
| **Gm1965** | 0.01021 | 0.00736 | 0.00000 | 22.38147 | 0.00866 | 0.00365 | 0.00592 | 0.00000 | 11.62217 | 0.80684 | 1.01768 | 0.01395 | 0.00873 | 0.27952 | 2.19428 | 0.00241 |
| **Gm19705** | 0.45410 | 0.00789 | 0.03866 | 0.57914 | 0.00905 | 0.00823 | 0.00725 | 0.03012 | 0.61993 | 0.01328 | 0.51862 | 0.00421 | 0.01228 | 0.00874 | 0.03416 | 0.00684 |
| **Gm19710** | 0.01100 | 0.00784 | 0.01089 | 0.04126 | 0.00982 | 0.00738 | 0.24868 | 0.01359 | 0.04948 | 0.14543 | 0.01121 | 0.00672 | 0.09580 | 0.26606 | 0.07895 | 0.12953 |
| **Gm19757** | 0.71461 | 1.72716 | 2.63034 | 3.68673 | 0.02528 | 1.44900 | 1.87768 | 2.97167 | 2.47140 | 6.87279 | 0.48555 | 0.41725 | 1.50816 | 11.68869 | 6.74451 | 0.52676 |
| **Gm1976** | 0.08047 | 0.01916 | 0.09118 | 0.37545 | 0.02057 | 0.00692 | 0.00721 | 0.02841 | 0.49126 | 0.08312 | 0.65582 | 0.01165 | 0.05258 | 0.43935 | 0.10984 | 0.01434 |
| **Gm19784** | 0.01380 | 0.00800 | 0.01166 | 0.62585 | 0.00934 | 0.00653 | 0.00739 | 0.01174 | 0.00000 | 0.01263 | 0.02884 | 0.01783 | 0.01256 | 0.00570 | 0.29745 | 0.00741 |
| **Gm1995** | 0.03653 | 0.00778 | 0.59976 | 19.33900 | 0.00703 | 0.00111 | 0.01785 | 0.46462 | 27.73276 | 1.76127 | 0.83216 | 0.06566 | 0.03233 | 0.00000 | 4.02845 | 0.02393 |
| **Gm2011** | 0.00957 | 0.06446 | 0.01685 | 0.05238 | 0.00896 | 0.00724 | 0.05206 | 0.05108 | 0.09135 | 0.00778 | 0.01124 | 0.00835 | 0.22343 | 0.01104 | 0.02226 | 0.02146 |
| **Gm20110** | 0.01736 | 0.00798 | 0.02924 | 0.00955 | 0.01060 | 0.00717 | 0.00736 | 0.02295 | 0.02360 | 0.05155 | 0.01370 | 0.00785 | 0.01126 | 0.01386 | 0.01348 | 0.00732 |
| **Gm20125** | 0.02638 | 0.04117 | 0.03256 | 0.01595 | 0.01077 | 0.00610 | 0.01796 | 0.01964 | 0.02392 | 0.09006 | 0.01191 | 0.01769 | 0.02099 | 0.11386 | 0.09632 | 0.00751 |
| **Gm20139** | 0.01379 | 0.00800 | 0.02920 | 0.01667 | 0.00929 | 0.00608 | 0.01789 | 0.01070 | 0.02467 | 0.03709 | 0.09403 | 0.00745 | 0.05552 | 0.03939 | 0.05266 | 0.01454 |
| **Gm2016** | 0.04402 | 0.05074 | 1.13735 | 8.07685 | 0.01648 | 0.00374 | 0.04839 | 0.02445 | 10.72129 | 0.32216 | 0.00000 | 0.49703 | 0.29558 | 1.74310 | 0.93477 | 0.02731 |
| **Gm20172** | 0.00364 | 0.00728 | 0.07805 | 1.05678 | 0.00899 | 0.00784 | 0.00666 | 0.42682 | 0.71573 | 1.90200 | 1.06798 | 1.08933 | 0.15042 | 0.87697 | 1.66118 | 0.02132 |
| **Gm20187** | 0.22869 | 0.00749 | 0.03214 | 0.21203 | 0.00845 | 0.03113 | 0.00720 | 0.06084 | 0.13277 | 0.00961 | 0.02520 | 0.00684 | 0.07771 | 0.05274 | 0.05887 | 0.04364 |
| **Gm20199** | 0.08888 | 0.00784 | 0.02479 | 0.00655 | 0.00834 | 0.00832 | 0.01806 | 0.01418 | 0.02197 | 0.55141 | 0.02836 | 0.00615 | 0.18566 | 0.05022 | 0.24684 | 0.00737 |
| **Gm20219** | 0.01371 | 0.00781 | 0.00647 | 0.14064 | 0.00896 | 1.14313 | 0.00726 | 0.01479 | 0.49580 | 0.99488 | 0.00613 | 0.00445 | 0.00542 | 0.01244 | 0.01675 | 0.00702 |
| **Gm2022** | 0.17537 | 0.02938 | 0.13930 | 11.86403 | 0.02595 | 0.02083 | 0.01751 | 0.03758 | 11.84379 | 0.14533 | 0.00000 | 0.00000 | 0.04697 | 0.45547 | 1.74302 | 0.01905 |
| **Gm2027** | 0.03568 | 0.04983 | 0.08284 | 4.85934 | 0.11954 | 0.09925 | 0.03776 | 0.14681 | 6.58032 | 0.26225 | 0.30294 | 1.47960 | 0.23015 | 0.86403 | 0.72623 | 0.02060 |
| **Gm20300** | 0.47983 | 2.72421 | 0.85958 | 0.05363 | 0.03860 | 2.83537 | 0.04571 | 0.40778 | 0.13652 | 0.53514 | 0.03367 | 0.17777 | 0.00000 | 1.17380 | 0.00000 | 1.63912 |
| **Gm20324** | 0.01362 | 0.00813 | 0.01282 | 0.01059 | 0.00967 | 0.00693 | 0.00767 | 0.01506 | 0.01231 | 0.01773 | 0.01371 | 0.01868 | 0.02318 | 0.01219 | 0.01056 | 0.00758 |
| **Gm20337** | 0.19337 | 0.02792 | 0.86097 | 3.35327 | 0.09076 | 0.02324 | 0.03640 | 0.10648 | 2.50889 | 0.45405 | 2.46414 | 0.09363 | 0.21899 | 0.61580 | 0.57246 | 0.09819 |
| **Gm20356** | 0.01751 | 0.00842 | 0.01348 | 3.54168 | 0.00906 | 0.01256 | 0.00817 | 0.34801 | 0.00000 | 0.00000 | 0.00950 | 0.00745 | 0.01748 | 0.10709 | 0.22703 | 0.00684 |
| **Gm20362** | 22.90131 | 7.88086 | 5.87447 | 0.54917 | 9.98188 | 6.03175 | 2.95395 | 5.68934 | 0.34929 | 0.39304 | 0.46003 | 6.98785 | 5.22895 | 1.97297 | 0.08221 | 17.70128 |
| **Gm2042** | 0.00000 | 0.00762 | 0.03392 | 17.27647 | 0.00915 | 0.00457 | 0.00635 | 0.18313 | 6.59393 | 3.12116 | 0.19675 | 0.15477 | 0.00599 | 7.04153 | 14.61657 | 0.00380 |
| **Gm20594** | 9.68874 | 11.12449 | 7.87143 | 15.03884 | 3.96011 | 3.09272 | 8.01756 | 18.56727 | 11.46087 | 19.69037 | 12.03269 | 12.91331 | 3.20458 | 38.98987 | 29.74824 | 2.26549 |
| **Gm20597** | 0.03297 | 0.04072 | 0.02693 | 0.18470 | 0.02059 | 0.00607 | 0.00739 | 0.01137 | 0.22302 | 0.00596 | 0.01032 | 0.00700 | 0.01246 | 0.15847 | 0.07841 | 0.00739 |
| **Gm20605** | 0.03433 | 0.00819 | 0.01116 | 0.00994 | 0.00839 | 0.24724 | 0.07277 | 0.04444 | 0.01232 | 0.11404 | 0.01198 | 0.00785 | 0.06100 | 0.01234 | 0.00966 | 0.00734 |
| **Gm20611** | 0.01290 | 0.00766 | 0.00822 | 0.00755 | 0.00930 | 0.00690 | 0.00715 | 0.23693 | 0.02607 | 0.00665 | 0.08686 | 0.00670 | 0.00639 | 0.01233 | 0.00549 | 0.00735 |
| **Gm20750** | 0.01275 | 0.00780 | 0.00742 | 0.81113 | 0.01823 | 0.00644 | 0.00717 | 0.00462 | 0.00000 | 0.39328 | 0.37100 | 0.03416 | 0.00794 | 0.00412 | 0.99591 | 0.00688 |
| **Gm20752** | 0.03696 | 0.00794 | 0.04933 | 0.34657 | 0.00819 | 0.38133 | 0.00732 | 0.01594 | 0.41617 | 0.09489 | 0.07041 | 0.04880 | 0.01512 | 0.01057 | 0.01484 | 0.08567 |
| **Gm20759** | 0.01373 | 0.00831 | 0.01211 | 1.14774 | 0.00932 | 0.00656 | 0.00747 | 0.01350 | 0.06636 | 0.01373 | 0.01279 | 0.00812 | 0.01275 | 0.03306 | 0.00159 | 0.00723 |
| **Gm20767** | 0.07014 | 0.04874 | 0.26756 | 40.23747 | 0.02803 | 0.08835 | 0.22601 | 1.57714 | 40.81483 | 9.01722 | 3.89969 | 1.97196 | 0.82831 | 9.25575 | 16.01234 | 0.03288 |
| **Gm20939** | 3.65985 | 5.18758 | 2.37758 | 1.28014 | 1.48500 | 1.81878 | 3.28242 | 5.80663 | 2.42843 | 6.20031 | 5.74126 | 2.56888 | 1.78047 | 21.36329 | 7.12944 | 0.88131 |
| **Gm21119** | 0.01374 | 0.00800 | 0.01229 | 0.26654 | 0.00933 | 0.00661 | 0.00752 | 0.01378 | 0.10078 | 0.01316 | 0.01241 | 0.00778 | 0.01269 | 0.00926 | 0.04006 | 0.00736 |
| **Gm21190** | 0.01528 | 0.00795 | 0.09629 | 0.10561 | 0.00890 | 0.00599 | 0.00751 | 0.03570 | 0.25229 | 0.08797 | 0.01017 | 0.00735 | 0.01065 | 0.00814 | 0.11391 | 0.00749 |
| **Gm21283** | 2.17247 | 0.00557 | 0.01150 | 0.01031 | 0.19481 | 0.00687 | 0.12217 | 0.00852 | 0.01230 | 0.01240 | 0.01441 | 0.00600 | 0.06286 | 0.01128 | 0.01579 | 0.03565 |
| **Gm21284** | 0.22544 | 0.00758 | 0.24869 | 0.09871 | 0.00912 | 0.06299 | 0.02732 | 0.00075 | 0.48770 | 0.12350 | 0.94117 | 0.06817 | 0.20139 | 0.47636 | 0.11155 | 0.00678 |
| **Gm21319** | 0.03522 | 0.02921 | 0.08437 | 6.91073 | 0.00881 | 0.01192 | 0.04791 | 0.03805 | 5.38534 | 0.03617 | 0.21703 | 0.00000 | 0.04952 | 1.15900 | 1.54528 | 0.03519 |
| **Gm21637** | 0.22586 | 0.00764 | 0.00968 | 0.53979 | 0.00750 | 0.00632 | 0.00744 | 0.01959 | 1.15309 | 0.14080 | 0.00000 | 0.03533 | 0.01217 | 0.00795 | 0.20447 | 0.00697 |
| **Gm2381** | 0.75958 | 1.65833 | 0.72289 | 1.33075 | 0.16032 | 0.01080 | 0.40481 | 0.37907 | 5.52971 | 2.45781 | 1.36931 | 0.41206 | 0.88320 | 4.05131 | 2.44130 | 0.03313 |
| **Gm2518** | 0.06635 | 0.00768 | 0.71289 | 0.01942 | 0.00683 | 0.00563 | 0.00730 | 0.01310 | 0.78674 | 0.03394 | 0.00663 | 0.00657 | 0.01536 | 0.01303 | 0.00400 | 0.01407 |
| **Gm2560** | 0.01344 | 0.00829 | 0.08210 | 0.24372 | 0.00928 | 1.98094 | 0.00792 | 0.11268 | 0.00533 | 0.00595 | 0.00831 | 0.01921 | 0.00000 | 0.00801 | 0.08497 | 0.00647 |

| | | | | | | | | | | | | | | | | |
|---|---|---|---|---|---|---|---|---|---|---|---|---|---|---|---|---|
| **Gm26705** | 0.38771 | 0.00757 | 0.00981 | 0.17845 | 0.00720 | 0.00642 | 0.03821 | 0.03198 | 0.26295 | 0.03973 | 0.00894 | 0.05968 | 0.02254 | 0.17312 | 0.09381 | 0.00682 |
| **Gm2694** | 0.16646 | 0.00849 | 0.00617 | 5.01813 | 0.01131 | 0.00660 | 0.00736 | 0.01523 | 4.22389 | 0.71648 | 0.00000 | 0.03249 | 0.02125 | 0.00000 | 0.13540 | 0.00627 |
| **Gm28042** | 0.01373 | 0.00813 | 0.01219 | 0.01041 | 0.00927 | 0.00660 | 0.01799 | 0.01332 | 0.01243 | 0.05663 | 0.01225 | 0.00792 | 0.01204 | 0.01153 | 0.01071 | 0.00723 |
| **Gm2837** | 0.41446 | 0.00745 | 0.01468 | 1.99211 | 0.00712 | 0.02339 | 0.00726 | 0.01736 | 3.37402 | 0.20018 | 0.01709 | 0.08154 | 0.01142 | 0.02738 | 0.41488 | 0.00637 |
| **Gm2897** | 0.10619 | 0.00789 | 0.01775 | 0.39169 | 0.00784 | 0.00625 | 0.00732 | 0.01387 | 0.00516 | 0.01758 | 0.01335 | 0.00776 | 0.01259 | 0.04913 | 0.00590 | 0.00723 |
| **Gm29443** | 0.03713 | 0.01815 | 0.03749 | 0.38743 | 0.00889 | 0.00669 | 0.01734 | 0.01269 | 0.66999 | 0.02245 | 0.30027 | 0.02337 | 0.02527 | 0.27883 | 0.07273 | 0.01469 |
| **Gm29683** | 0.05751 | 0.01866 | 0.01920 | 0.04761 | 0.00784 | 0.00801 | 0.16485 | 0.03866 | 0.13638 | 0.01429 | 0.04353 | 0.00672 | 0.04879 | 0.36523 | 0.03405 | 0.00718 |
| **Gm29811** | 0.01288 | 0.00784 | 0.00994 | 5.60137 | 0.00899 | 0.00567 | 0.00705 | 0.00839 | 4.57095 | 0.51462 | 0.00000 | 0.00000 | 0.01250 | 0.00000 | 0.72422 | 0.00659 |
| **Gm29968** | 0.16421 | 0.15119 | 0.04827 | 0.21112 | 0.02396 | 0.02186 | 0.02719 | 0.14132 | 0.07586 | 0.12465 | 0.00699 | 0.00491 | 0.02046 | 0.58397 | 0.18833 | 0.02828 |
| **Gm2a** | 2.68469 | 4.26961 | 15.94398 | 4.72191 | 2.06555 | 16.55269 | 5.76842 | 14.54239 | 5.84442 | 13.87511 | 22.10944 | 3.37899 | 19.82008 | 14.57867 | 10.53411 | 5.92115 |
| **Gm3002** | 0.05378 | 0.00817 | 0.00977 | 0.24197 | 0.00810 | 0.00690 | 0.00747 | 0.01465 | 0.09401 | 0.02091 | 0.01293 | 0.00782 | 0.01491 | 0.05323 | 0.01008 | 0.10296 |
| **Gm30505** | 0.01352 | 0.00807 | 0.01217 | 0.25455 | 0.00935 | 0.00663 | 0.00746 | 0.01913 | 0.00639 | 0.01525 | 0.01332 | 0.00810 | 0.01232 | 0.01130 | 0.00688 | 0.00737 |
| **Gm30551** | 0.01367 | 0.01839 | 0.07380 | 0.42257 | 0.00932 | 0.00607 | 0.01769 | 0.02173 | 0.19762 | 0.06236 | 0.06178 | 0.00726 | 0.02310 | 0.01158 | 0.03826 | 0.00717 |
| **Gm3086** | 0.02039 | 0.00781 | 0.01275 | 0.01862 | 0.00941 | 0.00703 | 0.00772 | 0.01479 | 0.02476 | 0.01029 | 0.01324 | 0.00764 | 0.04397 | 0.01958 | 0.08027 | 0.00763 |
| **Gm3139** | 17.03241 | 11.39869 | 24.53123 | 21.11049 | 12.32468 | 7.56475 | 13.22957 | 25.66089 | 21.26907 | 32.89178 | 22.80720 | 16.35447 | 21.60369 | 37.86133 | 27.21709 | 2.58295 |
| **Gm3143** | 0.42866 | 0.00733 | 0.42245 | 6.46713 | 0.00652 | 0.00453 | 0.03802 | 0.04956 | 0.79395 | 0.07541 | 0.04624 | 0.04580 | 0.02520 | 0.01580 | 0.30605 | 0.02602 |
| **Gm3219** | 0.74158 | 0.74883 | 0.42573 | 13.44827 | 0.55161 | 0.01857 | 0.00584 | 0.16991 | 3.88202 | 0.99262 | 0.83799 | 0.03555 | 0.12504 | 0.00000 | 0.40869 | 0.03376 |
| **Gm32511** | 0.53431 | 0.00735 | 0.00000 | 0.00709 | 0.00674 | 0.02707 | 0.00615 | 0.00861 | 0.02243 | 0.54887 | 0.00957 | 0.00439 | 1.39562 | 0.16607 | 0.20564 | 0.00619 |
| **Gm3259** | 0.01317 | 0.00782 | 0.01071 | 3.56421 | 0.00937 | 0.00623 | 0.00733 | 0.01134 | 0.81045 | 0.18245 | 0.02487 | 0.00319 | 0.04336 | 0.02284 | 1.17500 | 0.00695 |
| **Gm3264** | 0.04614 | 0.00806 | 0.00986 | 0.01027 | 0.00811 | 0.00682 | 0.00738 | 0.01329 | 0.01231 | 0.04538 | 0.01309 | 0.00759 | 0.02035 | 0.05243 | 0.01215 | 0.00753 |
| **Gm3286** | 0.98422 | 1.06531 | 3.32707 | 9.50469 | 1.08215 | 0.71837 | 1.78197 | 5.10870 | 8.96429 | 8.54295 | 7.78006 | 4.51958 | 3.91209 | 14.58471 | 9.76655 | 0.18080 |
| **Gm3336** | 0.01345 | 0.00811 | 0.00621 | 0.00427 | 0.00952 | 0.00747 | 0.00748 | 0.00600 | 0.01107 | 0.06133 | 1.42519 | 0.00080 | 0.00282 | 0.00408 | 0.10368 | 0.00733 |
| **Gm3383** | 0.24920 | 0.00775 | 0.00569 | 0.98590 | 0.00732 | 0.00712 | 0.00714 | 0.01204 | 0.00000 | 0.01857 | 0.02592 | 0.01758 | 0.01968 | 0.11782 | 0.00000 | 0.01435 |
| **Gm3414** | 0.02108 | 0.01818 | 0.30010 | 0.08487 | 0.01105 | 0.00830 | 0.00707 | 0.00047 | 0.03858 | 0.00984 | 0.34721 | 0.04205 | 0.11889 | 0.05086 | 0.57468 | 0.01463 |
| **Gm3435** | 0.08350 | 0.41342 | 1.06171 | 0.21023 | 0.00624 | 0.00704 | 0.32141 | 0.16688 | 0.00778 | 0.13166 | 0.15616 | 0.00591 | 0.29370 | 0.00838 | 0.41761 | 0.79217 |
| **Gm3500** | 0.16499 | 0.00783 | 0.00980 | 0.50828 | 0.00755 | 0.00664 | 0.00727 | 0.01338 | 0.01036 | 0.01948 | 0.01283 | 0.00790 | 0.02686 | 0.06573 | 0.00389 | 0.00711 |
| **Gm3558** | 0.12290 | 0.00786 | 0.01560 | 0.62670 | 0.00775 | 0.00631 | 0.00724 | 0.01210 | 0.00418 | 0.03678 | 0.03987 | 0.01707 | 0.01235 | 0.08756 | 0.00000 | 0.00730 |
| **Gm3604** | 0.68962 | 7.81834 | 2.05395 | 2.52207 | 0.74129 | 1.43497 | 0.98077 | 0.44122 | 2.96918 | 0.72634 | 0.34426 | 6.36051 | 0.92987 | 4.59224 | 0.52508 | 0.07369 |
| **Gm3636** | 0.04308 | 0.00791 | 0.03233 | 1.00557 | 0.00828 | 0.00597 | 0.00729 | 0.01248 | 0.02725 | 0.01975 | 0.02665 | 0.00753 | 0.01223 | 0.08114 | 0.00000 | 0.00725 |
| **Gm364** | 0.00814 | 0.00748 | 2.07109 | 0.59059 | 0.00630 | 0.00767 | 0.00648 | 0.53008 | 0.00000 | 0.24977 | 1.08947 | 0.01168 | 0.00000 | 2.11697 | 0.69953 | 0.00674 |
| **Gm3696** | 0.08561 | 0.00802 | 0.00866 | 0.26289 | 0.00784 | 0.00690 | 0.00737 | 0.01399 | 0.05795 | 0.03194 | 0.01280 | 0.00747 | 0.01299 | 0.05249 | 0.00458 | 0.00742 |
| **Gm3776** | 0.50526 | 0.00699 | 0.00000 | 0.01940 | 0.00676 | 0.40591 | 0.07801 | 0.76137 | 0.01836 | 0.83900 | 1.16862 | 0.06961 | 0.85847 | 1.19590 | 0.00000 | 0.00595 |
| **Gm38407** | 0.01368 | 0.00825 | 0.01214 | 1.03712 | 0.00933 | 0.00655 | 0.00750 | 0.01391 | 0.43336 | 0.01248 | 0.00984 | 0.00767 | 0.01280 | 0.01135 | 0.00360 | 0.00722 |
| **Gm38415** | 0.02147 | 0.00741 | 0.02624 | 0.80280 | 0.00885 | 0.00596 | 0.03747 | 0.02560 | 1.12211 | 0.30402 | 0.00000 | 0.08269 | 0.01366 | 0.06582 | 1.42641 | 0.00755 |
| **Gm38416** | 0.02000 | 0.00753 | 0.01717 | 0.95733 | 0.00881 | 0.00710 | 0.00733 | 0.01611 | 0.00000 | 0.00720 | 0.01168 | 0.00714 | 0.01930 | 0.01338 | 0.25986 | 0.00718 |
| **Gm38426** | 0.33308 | 0.00766 | 0.60204 | 0.01088 | 0.01314 | 0.00526 | 0.00689 | 0.00842 | 0.01214 | 0.02190 | 0.00977 | 0.22813 | 0.00891 | 0.01337 | 0.00000 | 0.00685 |
| **Gm3893** | 0.00923 | 0.00739 | 0.01097 | 0.76980 | 0.00892 | 0.00637 | 0.00669 | 0.00806 | 0.06688 | 0.95417 | 0.01117 | 0.00463 | 0.86837 | 0.10239 | 0.10520 | 0.00659 |
| **Gm40893** | 0.12127 | 0.05347 | 0.06391 | 0.07638 | 0.78991 | 0.07777 | 0.01747 | 0.02801 | 0.17467 | 0.01654 | 0.16151 | 0.01523 | 0.11824 | 0.30938 | 0.03902 | 0.11622 |
| **Gm4224** | 0.01363 | 0.00815 | 0.01184 | 0.10181 | 0.00925 | 0.00664 | 0.00747 | 0.01285 | 0.20982 | 0.01557 | 0.01147 | 0.00764 | 0.01993 | 0.03516 | 0.01806 | 0.01482 |
| **Gm4262** | 0.94711 | 1.56454 | 7.00913 | 3.69007 | 0.51436 | 2.10652 | 2.63797 | 9.44309 | 9.04340 | 10.36186 | 6.38364 | 10.88127 | 1.39302 | 7.45071 | 4.44472 | 0.76469 |
| **Gm4285** | 0.08927 | 0.00763 | 0.10191 | 0.16031 | 0.01765 | 0.00615 | 0.01739 | 0.00788 | 0.36829 | 0.01478 | 0.57231 | 0.04810 | 0.06122 | 1.63948 | 0.07667 | 0.02169 |
| **Gm4349** | 0.81326 | 0.00631 | 1.09343 | 3.39785 | 0.00453 | 0.00594 | 0.00621 | 0.19166 | 3.12280 | 1.17476 | 0.03262 | 0.08533 | 0.00921 | 1.60089 | 0.91651 | 0.00518 |
| **Gm4371** | 0.72320 | 0.00738 | 0.00280 | 0.01939 | 0.01695 | 0.00777 | 0.00653 | 0.36865 | 0.01221 | 0.00893 | 0.01104 | 0.00631 | 0.00614 | 0.01544 | 0.01073 | 0.00686 |
| **Gm4425** | 0.21377 | 0.08125 | 0.60197 | 0.01883 | 0.02798 | 0.16283 | 0.12813 | 0.42460 | 0.25738 | 0.16320 | 0.70215 | 0.02305 | 0.43767 | 0.18488 | 0.28065 | 0.11255 |
| **Gm4432** | 0.01835 | 0.00773 | 0.00885 | 0.00827 | 0.01079 | 0.00667 | 0.00727 | 0.01000 | 0.01192 | 0.00269 | 0.02266 | 0.00520 | 0.01295 | 0.41326 | 0.32287 | 0.00751 |
| **Gm4532** | 0.01817 | 0.00836 | 0.01159 | 0.76182 | 0.01065 | 0.00645 | 0.00745 | 0.01324 | 1.47161 | 0.03977 | 0.00000 | 0.00539 | 0.01316 | 0.01039 | 0.00319 | 0.00731 |
| **Gm4541** | 0.01521 | 0.10499 | 0.00968 | 0.08627 | 0.01006 | 0.00674 | 0.02758 | 0.01077 | 0.02923 | 0.01036 | 0.01248 | 0.00668 | 0.01265 | 0.25042 | 0.00252 | 0.00743 |
| **Gm4566** | 0.01378 | 0.00786 | 0.01203 | 0.15284 | 0.00932 | 0.00664 | 0.00751 | 0.01350 | 0.00921 | 0.01387 | 0.01338 | 0.00793 | 0.01272 | 0.01694 | 0.02967 | 0.00739 |
| **Gm4567** | 0.01920 | 0.00855 | 0.02875 | 0.06572 | 0.00875 | 0.00613 | 0.00805 | 0.12275 | 0.15397 | 0.03642 | 0.05033 | 0.00733 | 0.10170 | 0.13155 | 0.09210 | 0.00740 |
| **Gm4598** | 0.01565 | 0.00759 | 0.13544 | 0.01065 | 0.00906 | 0.00575 | 0.00738 | 0.01206 | 0.01172 | 0.00869 | 0.01195 | 0.00738 | 0.01090 | 0.01964 | 0.14837 | 0.00729 |
| **Gm4737** | 17.15883 | 7.52888 | 14.10380 | 0.00000 | 12.34763 | 4.26249 | 10.53438 | 12.00802 | 0.00000 | 3.47339 | 1.91495 | 0.12638 | 11.05587 | 0.12120 | 0.78131 | 6.15565 |
| **Gm4791** | 0.01356 | 0.00808 | 0.01236 | 0.06459 | 0.00979 | 0.00692 | 0.00758 | 0.01290 | 0.04265 | 0.02204 | 0.03958 | 0.00705 | 0.02550 | 0.07916 | 0.03839 | 0.00766 |
| **Gm4827** | 0.01722 | 0.00836 | 0.01594 | 0.74521 | 0.00889 | 0.00642 | 0.00763 | 0.01436 | 1.72004 | 0.07801 | 0.01322 | 0.00619 | 0.01304 | 0.01067 | 0.00431 | 0.00717 |
| **Gm4832** | 0.09058 | 0.00865 | 0.12102 | 0.08821 | 0.09652 | 0.02353 | 0.03028 | 0.09616 | 0.96743 | 0.01722 | 0.87118 | 0.00298 | 0.05604 | 0.08644 | 0.12238 | 0.05148 |
| **Gm4841** | 0.05194 | 0.02863 | 0.04931 | 0.04146 | 0.03390 | 0.05618 | 0.00743 | 0.11618 | 0.08712 | 0.08271 | 0.12905 | 0.13069 | 0.05386 | 0.05304 | 0.05829 | 0.00719 |
| **Gm4850** | 0.01037 | 0.00760 | 0.06773 | 12.41640 | 0.00880 | 0.00404 | 0.00682 | 0.00004 | 5.80358 | 0.97634 | 0.19394 | 0.01452 | 0.00932 | 0.00891 | 4.78509 | 0.01191 |
| **Gm4858** | 0.01233 | 0.00792 | 0.00937 | 1.69477 | 0.00940 | 0.00626 | 0.00723 | 0.00924 | 0.82313 | 0.14145 | 0.02953 | 0.02153 | 0.01212 | 0.88211 | 0.45219 | 0.00702 |
| **Gm4871** | 0.01365 | 0.00829 | 0.01164 | 0.47830 | 0.00928 | 0.00676 | 0.00752 | 0.01332 | 0.20683 | 0.02870 | 0.01199 | 0.00810 | 0.01316 | 0.07543 | 0.00000 | 0.00734 |
| **Gm4922** | 0.16883 | 0.05220 | 0.10389 | 0.00880 | 0.01617 | 0.01265 | 0.00757 | 0.06928 | 0.01215 | 0.05594 | 0.04707 | 0.05010 | 0.04079 | 0.18779 | 0.04167 | 0.19568 |
| **Gm4925** | 0.03221 | 0.01840 | 0.21961 | 0.03074 | 0.03507 | 0.03088 | 0.02822 | 0.03693 | 0.60731 | 0.05600 | 0.68968 | 0.06275 | 0.08781 | 0.16656 | 0.08292 | 0.02864 |
| **Gm4926** | 0.00000 | 0.62659 | 2.03124 | 0.65440 | 1.78411 | 2.00495 | 0.16248 | 0.62187 | 0.09497 | 0.40960 | 0.36011 | 0.23327 | 0.72361 | 0.61845 | 0.30498 | 1.17655 |
| **Gm4951** | 0.04321 | 0.01794 | 0.16421 | 0.08251 | 0.01764 | 0.01306 | 0.01692 | 0.02260 | 0.14055 | 0.03788 | 0.64246 | 0.04009 | 0.18861 | 0.50230 | 0.04616 | 0.04355 |

| | | | | | | | | | | | | | | | | |
|---|---|---|---|---|---|---|---|---|---|---|---|---|---|---|---|---|
| **Gm4956** | 0.01313 | 0.00803 | 0.01055 | 0.00356 | 0.00916 | 0.00637 | 0.00725 | 0.01152 | 0.67203 | 0.00154 | 0.00696 | 0.05113 | 0.13108 | 0.00758 | 0.27717 | 0.00711 |
| **Gm4961** | 1.10958 | 0.10750 | 0.00000 | 5.44137 | 0.42814 | 0.41421 | 0.01628 | 3.87109 | 1.22638 | 4.80116 | 1.78169 | 0.87961 | 1.14355 | 2.21785 | 4.41469 | 0.00441 |
| **Gm4971** | 0.01236 | 0.02908 | 0.01658 | 1.40912 | 0.00939 | 0.00624 | 0.00758 | 0.02221 | 0.85629 | 0.00437 | 0.00659 | 0.00605 | 0.01466 | 0.00879 | 0.25900 | 0.01437 |
| **Gm4984** | 0.38299 | 0.00740 | 0.01130 | 0.22917 | 0.00687 | 0.00731 | 0.00647 | 0.01213 | 0.16349 | 0.00000 | 0.06072 | 0.02618 | 1.22536 | 0.00622 | 0.17201 | 0.01299 |
| **Gm5** | 0.01064 | 0.00778 | 0.00492 | 9.69642 | 0.00871 | 0.00427 | 0.00672 | 0.00000 | 18.03860 | 0.61231 | 0.25515 | 0.00000 | 0.00959 | 0.24754 | 0.77619 | 0.00467 |
| **Gm5039** | 0.11052 | 0.39781 | 0.32318 | 11.60741 | 0.15661 | 0.11667 | 0.16201 | 0.40330 | 11.02799 | 0.40671 | 0.00000 | 0.00000 | 0.10931 | 1.64044 | 0.83105 | 0.20289 |
| **Gm5065** | 0.04693 | 0.00794 | 0.00571 | 0.56805 | 0.00805 | 0.00727 | 0.00743 | 0.04239 | 0.15092 | 0.28972 | 0.04804 | 0.26149 | 0.01049 | 0.00699 | 0.14812 | 0.00721 |
| **Gm5069** | 5.72260 | 2.64225 | 2.93836 | 0.29186 | 4.25927 | 2.87021 | 1.68389 | 0.14582 | 0.51062 | 0.13209 | 0.00703 | 0.12976 | 2.87531 | 0.00811 | 0.15142 | 5.55592 |
| **Gm5071** | 0.01259 | 0.00826 | 0.01074 | 0.00983 | 0.00939 | 0.00670 | 0.00798 | 0.04318 | 0.01233 | 0.00951 | 0.01173 | 0.06128 | 0.01092 | 0.00729 | 0.11951 | 0.00738 |
| **Gm5072** | 0.01074 | 0.00845 | 0.00957 | 0.00960 | 0.00952 | 0.00675 | 0.00819 | 0.14607 | 0.01218 | 0.01329 | 0.01052 | 0.04121 | 0.00768 | 0.00693 | 0.16182 | 0.00729 |
| **Gm5088** | 10.81589 | 0.64443 | 2.20219 | 11.16710 | 8.79722 | 5.42338 | 4.29405 | 20.25354 | 9.62789 | 29.62865 | 21.38486 | 3.16592 | 25.94093 | 3.33048 | 2.94979 | 4.31739 |
| **Gm5113** | 0.15934 | 0.05105 | 0.60790 | 0.74085 | 0.06992 | 0.05942 | 0.11118 | 0.15222 | 0.94163 | 0.00003 | 0.82132 | 0.02774 | 1.14704 | 0.94110 | 1.20575 | 0.05604 |
| **Gm5114** | 0.01375 | 0.00789 | 0.01239 | 0.01005 | 0.00933 | 0.00661 | 0.00752 | 0.01366 | 0.01247 | 0.01231 | 0.01298 | 0.00764 | 0.01266 | 0.00756 | 0.07766 | 0.00739 |
| **Gm5127** | 0.01226 | 0.00739 | 0.54131 | 0.90631 | 0.00823 | 0.00880 | 0.00703 | 0.00955 | 0.00000 | 0.00326 | 0.00837 | 0.41285 | 0.00926 | 0.00477 | 0.12152 | 0.00704 |
| **Gm5129** | 0.01595 | 0.00829 | 0.01227 | 0.39289 | 0.01021 | 0.00672 | 0.00755 | 0.01459 | 0.13714 | 0.02454 | 0.01375 | 0.00824 | 0.04435 | 0.01177 | 0.00516 | 0.19038 |
| **Gm5134** | 0.24620 | 0.02794 | 0.25164 | 3.04685 | 0.05942 | 0.95410 | 0.00616 | 1.56618 | 2.73442 | 1.04321 | 0.80275 | 6.41180 | 0.00000 | 3.18137 | 2.02055 | 0.00550 |
| **Gm5141** | 0.30506 | 0.76666 | 0.75118 | 1.76060 | 0.12270 | 0.15332 | 0.45692 | 0.47062 | 0.99802 | 0.22163 | 0.82841 | 1.05682 | 0.11294 | 2.14794 | 0.95136 | 0.10720 |
| **Gm5148** | 24.87101 | 2.86325 | 17.40314 | 8.65704 | 15.75061 | 6.39201 | 3.88464 | 7.12456 | 18.19470 | 24.06428 | 32.98443 | 2.68374 | 16.70704 | 4.49144 | 7.90348 | 10.62623 |
| **Gm5176** | 1.33786 | 0.72263 | 1.51784 | 0.65800 | 1.35790 | 0.41497 | 0.85279 | 1.23807 | 2.86801 | 1.69146 | 0.58761 | 0.15450 | 0.87957 | 2.62850 | 0.92565 | 0.62545 |
| **Gm5177** | 2.74776 | 1.27211 | 1.29802 | 4.00280 | 2.20761 | 1.34547 | 0.73424 | 0.05690 | 1.19857 | 0.07688 | 0.09223 | 0.03437 | 1.34808 | 0.00000 | 0.34260 | 2.90459 |
| **Gm525** | 0.01366 | 0.00783 | 0.01128 | 0.26662 | 0.00914 | 0.00664 | 0.00730 | 0.01102 | 0.04261 | 0.31280 | 0.01261 | 0.04549 | 0.01254 | 0.00565 | 0.19627 | 0.00772 |
| **Gm527** | 2.42715 | 0.00556 | 0.00016 | 6.44729 | 0.01130 | 0.00560 | 0.00565 | 0.11792 | 1.03941 | 1.31873 | 0.38136 | 0.00970 | 0.00570 | 0.05559 | 0.81924 | 0.00351 |
| **Gm53** | 0.65934 | 0.00702 | 0.00477 | 0.01061 | 0.00658 | 0.00712 | 0.00708 | 0.01107 | 0.01238 | 0.01911 | 0.01253 | 0.01804 | 0.01185 | 0.01220 | 0.01066 | 0.00663 |
| **Gm5415** | 0.02481 | 0.00750 | 0.00922 | 0.00911 | 0.00897 | 0.00712 | 0.00738 | 0.18381 | 0.02670 | 0.02769 | 0.01360 | 0.01701 | 0.01706 | 0.01443 | 0.03598 | 0.00742 |
| **Gm5431** | 0.05995 | 0.00841 | 0.06985 | 0.06465 | 0.00905 | 0.00758 | 0.00788 | 0.03094 | 0.04631 | 0.09558 | 0.01199 | 0.02975 | 0.01716 | 0.00923 | 0.06226 | 0.01523 |
| **Gm5434** | 0.22248 | 0.10985 | 0.63358 | 2.13061 | 0.30882 | 0.21735 | 0.02877 | 0.27882 | 0.00000 | 0.29801 | 0.58902 | 0.01453 | 0.44644 | 0.13768 | 0.00000 | 0.18819 |
| **Gm5458** | 0.01506 | 0.00798 | 0.03597 | 0.63066 | 0.00916 | 0.00599 | 0.00719 | 0.00997 | 0.19760 | 0.12710 | 0.01040 | 0.01573 | 0.01996 | 0.33467 | 0.00721 | 0.00758 |
| **Gm5475** | 0.01303 | 0.00803 | 0.01781 | 0.19471 | 0.00925 | 0.00745 | 0.00745 | 0.01185 | 0.31669 | 0.36705 | 0.00869 | 0.00592 | 0.01481 | 0.05573 | 0.22984 | 0.00757 |
| **Gm5485** | 0.01270 | 0.00877 | 0.01086 | 0.00511 | 0.00922 | 0.00668 | 0.00802 | 0.11746 | 0.01094 | 0.08126 | 0.35061 | 0.00698 | 0.00632 | 0.00704 | 0.12142 | 0.00752 |
| **Gm5512** | 0.51836 | 1.19686 | 0.45900 | 1.55188 | 0.07137 | 1.12115 | 1.23661 | 0.01148 | 0.32494 | 0.58492 | 0.46135 | 0.01192 | 0.30752 | 0.00337 | 0.40947 | 1.01144 |
| **Gm5523** | 7.61272 | 3.00540 | 4.05085 | 0.93688 | 6.29213 | 3.72760 | 2.13292 | 0.14582 | 0.82819 | 0.30501 | 0.00000 | 0.35155 | 3.67797 | 0.00401 | 0.16996 | 6.75061 |
| **Gm5532** | 0.01164 | 0.00785 | 0.00750 | 0.12245 | 0.00892 | 0.00649 | 0.00741 | 0.01276 | 0.00599 | 1.87749 | 0.00307 | 0.03128 | 0.01082 | 0.48226 | 1.26980 | 0.00734 |
| **Gm5547** | 0.09692 | 0.00782 | 0.00477 | 0.00480 | 0.00762 | 0.00747 | 0.00784 | 0.06556 | 0.01087 | 0.29916 | 0.69373 | 0.01750 | 0.00444 | 0.07155 | 0.00000 | 0.00686 |
| **Gm5577** | 0.02903 | 0.00815 | 0.03964 | 2.98933 | 0.00841 | 0.00738 | 0.01793 | 0.02285 | 0.38254 | 0.14223 | 0.04810 | 0.00765 | 0.02383 | 0.05291 | 0.01884 | 0.00687 |
| **Gm5607** | 0.15020 | 0.00750 | 0.34919 | 0.30802 | 0.15185 | 0.03861 | 0.13750 | 0.08262 | 0.62715 | 0.19688 | 0.54754 | 0.04864 | 0.21255 | 0.62825 | 0.15779 | 0.05071 |
| **Gm561** | 1.36698 | 0.02598 | 0.32707 | 3.69782 | 0.91489 | 0.24286 | 0.00474 | 0.05241 | 4.63462 | 0.49619 | 2.86602 | 0.17770 | 0.18791 | 1.20794 | 1.56828 | 0.07180 |
| **Gm5617** | 0.08792 | 0.00767 | 0.01496 | 0.15215 | 0.00819 | 0.00637 | 0.00690 | 0.13691 | 0.00949 | 0.36178 | 0.11032 | 0.00548 | 0.41941 | 0.00996 | 0.00337 | 0.02213 |
| **Gm5622** | 0.02371 | 0.28213 | 0.04665 | 1.09656 | 0.01355 | 0.23563 | 0.35541 | 0.43795 | 0.14602 | 0.06806 | 0.01906 | 0.26984 | 0.00000 | 0.13592 | 0.41659 | 1.53606 |
| **Gm5640** | 0.67989 | 0.00691 | 0.00314 | 0.46831 | 0.01062 | 0.00767 | 0.37052 | 0.18277 | 0.00544 | 0.05616 | 0.01324 | 0.00578 | 0.10114 | 0.00681 | 0.04170 | 0.01443 |
| **Gm5643** | 9.35542 | 6.48127 | 7.41474 | 1.44955 | 5.14794 | 3.86488 | 5.76365 | 5.74120 | 5.57768 | 8.62594 | 11.14838 | 0.77143 | 6.24027 | 22.75800 | 11.10803 | 4.74243 |
| **Gm5662** | 0.05796 | 0.19916 | 1.44466 | 11.88683 | 0.09995 | 0.01476 | 0.07876 | 0.12197 | 13.18712 | 0.42275 | 0.07749 | 0.62779 | 0.28161 | 2.30129 | 1.49127 | 0.01155 |
| **Gm5712** | 0.05405 | 0.01822 | 0.05938 | 0.00989 | 0.04977 | 0.02416 | 0.05817 | 0.05820 | 0.01235 | 0.01943 | 0.02591 | 0.00729 | 0.04102 | 0.01007 | 0.02004 | 0.02930 |
| **Gm572** | 0.01066 | 0.00773 | 0.00977 | 0.00694 | 0.00962 | 0.00673 | 0.00721 | 0.16146 | 0.06743 | 0.01186 | 0.01165 | 0.00726 | 0.00757 | 0.01285 | 0.00758 | 0.00740 |
| **Gm5766** | 0.40774 | 0.24735 | 0.17447 | 0.02640 | 0.20225 | 0.08033 | 0.13225 | 0.30678 | 0.17762 | 0.05827 | 0.56338 | 0.05588 | 0.18129 | 0.19828 | 0.11255 | 0.26681 |
| **Gm5779** | 1.64211 | 0.62562 | 1.28944 | 0.00179 | 0.63763 | 0.67845 | 1.27068 | 2.03583 | 0.02924 | 1.50818 | 0.00000 | 0.40515 | 1.70402 | 0.00605 | 0.02411 | 1.05463 |
| **Gm5795** | 0.01791 | 0.00809 | 0.01067 | 0.36510 | 0.00890 | 0.00672 | 0.00734 | 0.01108 | 0.10525 | 0.05708 | 0.03960 | 0.01675 | 0.01259 | 0.10795 | 0.00742 | 0.00763 |
| **Gm5796** | 0.08939 | 0.00789 | 0.00883 | 0.22412 | 0.00783 | 0.00687 | 0.00741 | 0.01479 | 0.02261 | 0.01819 | 0.01328 | 0.00797 | 0.01276 | 0.02901 | 0.00676 | 0.00728 |
| **Gm5797** | 0.01428 | 0.00801 | 0.01882 | 1.44010 | 0.00918 | 0.00612 | 0.00717 | 0.00987 | 0.74702 | 0.12281 | 0.00716 | 0.00489 | 0.02035 | 0.30856 | 0.00000 | 0.00747 |
| **Gm5801** | 0.76983 | 0.00593 | 0.12290 | 3.27266 | 1.21133 | 0.10146 | 0.49147 | 2.04065 | 3.58594 | 3.80610 | 1.51650 | 0.14894 | 0.70660 | 0.00000 | 0.00000 | 1.17053 |
| **Gm5803** | 1.51095 | 2.53480 | 2.17876 | 0.04914 | 1.10201 | 1.50205 | 2.54765 | 2.91867 | 0.31813 | 1.33015 | 0.32637 | 0.00627 | 1.28760 | 16.31594 | 3.81199 | 1.64406 |
| **Gm5820** | 0.01348 | 0.00778 | 0.01082 | 0.39928 | 0.00938 | 0.00654 | 0.00741 | 0.01167 | 0.66016 | 0.00555 | 0.00619 | 0.00563 | 0.02905 | 0.14112 | 0.21468 | 0.00723 |
| **Gm5860** | 2.10840 | 0.66286 | 3.13520 | 3.05091 | 0.49809 | 0.42566 | 1.72756 | 0.60247 | 4.03498 | 1.20754 | 1.55603 | 0.92977 | 0.56235 | 0.74804 | 0.45561 | 2.21160 |
| **Gm5861** | 0.02255 | 0.00787 | 0.05106 | 0.05800 | 0.00852 | 0.00718 | 0.00740 | 0.01028 | 0.17200 | 0.00733 | 0.15761 | 0.01701 | 0.01007 | 0.00869 | 0.02200 | 0.00723 |
| **Gm5868** | 0.01349 | 0.00784 | 0.01107 | 0.21568 | 0.00944 | 0.00664 | 0.00751 | 0.01398 | 0.00588 | 0.00616 | 0.01082 | 0.00678 | 0.01288 | 0.17411 | 0.14349 | 0.00741 |
| **Gm5907** | 0.01093 | 0.00722 | 1.04588 | 0.00563 | 0.00704 | 0.00921 | 0.00643 | 0.12276 | 0.01117 | 0.28550 | 0.37384 | 0.00312 | 0.76878 | 0.01192 | 0.00000 | 0.68376 |
| **Gm5936** | 0.65853 | 0.00677 | 0.00264 | 0.38453 | 0.00986 | 0.00762 | 0.43089 | 0.24267 | 0.01707 | 0.04888 | 0.01215 | 0.00564 | 0.13287 | 0.00716 | 0.03744 | 0.03642 |
| **Gm5941** | 0.01308 | 0.00800 | 0.01717 | 1.91990 | 0.00906 | 0.00683 | 0.00738 | 0.00990 | 2.60807 | 0.04187 | 0.10658 | 0.00282 | 0.01051 | 0.01037 | 0.00795 | 0.00669 |
| **Gm595** | 0.00925 | 0.00840 | 0.00455 | 0.00507 | 0.00957 | 0.00770 | 0.00797 | 0.29754 | 0.01070 | 0.00000 | 0.71053 | 0.06114 | 0.00292 | 0.01211 | 0.00000 | 0.00710 |
| **Gm6083** | 20.37422 | 4.71217 | 14.82164 | 6.49520 | 12.14688 | 5.66979 | 3.99058 | 6.33775 | 13.34707 | 6.51405 | 19.62416 | 1.95058 | 9.37399 | 4.76299 | 4.87178 | 9.53133 |
| **Gm6086** | 0.03923 | 0.00836 | 0.09666 | 0.12090 | 0.00916 | 0.00914 | 0.00799 | 0.10791 | 0.09904 | 0.01652 | 0.05482 | 0.02712 | 0.04596 | 0.09005 | 0.13751 | 0.03045 |
| **Gm6093** | 0.12756 | 0.00775 | 0.00676 | 0.38311 | 0.00765 | 0.00701 | 0.01815 | 0.01239 | 0.05140 | 0.02068 | 0.07400 | 0.01790 | 0.01122 | 0.10234 | 0.02623 | 0.00712 |
| **Gm6194** | 0.03302 | 0.00773 | 0.04639 | 0.24499 | 0.05150 | 0.00753 | 0.05864 | 0.12457 | 0.23988 | 0.19774 | 0.20641 | 0.00509 | 0.11000 | 0.07685 | 0.09652 | 0.01481 |

| | | | | | | | | | | | | | | | | |
|---|---|---|---|---|---|---|---|---|---|---|---|---|---|---|---|---|
| **Gm6225** | 0.01407 | 0.00811 | 0.02500 | 0.86864 | 0.00935 | 0.00608 | 0.00760 | 0.01640 | 0.34826 | 0.01398 | 0.00997 | 0.00711 | 0.01239 | 0.06528 | 0.02559 | 0.00727 |
| **Gm6297** | 0.01253 | 0.00843 | 1.76546 | 0.02366 | 0.00647 | 0.00557 | 0.00778 | 0.04011 | 0.01178 | 0.00317 | 0.01066 | 0.02068 | 0.02267 | 0.00864 | 0.29498 | 0.00659 |
| **Gm6313** | 0.25982 | 0.03874 | 0.25499 | 6.90751 | 0.12858 | 0.18483 | 0.22739 | 0.08332 | 4.41582 | 0.59455 | 0.41333 | 0.16445 | 0.14022 | 2.17845 | 2.90771 | 0.21670 |
| **Gm6367** | 0.02564 | 0.01787 | 0.01163 | 8.12657 | 0.00862 | 0.01343 | 0.00711 | 0.01562 | 2.16264 | 0.15112 | 0.02265 | 0.07980 | 0.05179 | 0.09498 | 1.48017 | 0.00594 |
| **Gm6402** | 4.18398 | 0.24891 | 3.38300 | 3.11144 | 3.21483 | 0.83937 | 0.87797 | 0.13843 | 6.42555 | 1.85079 | 10.12269 | 0.30857 | 2.89165 | 3.15724 | 0.91034 | 2.15812 |
| **Gm6455** | 0.03174 | 0.00790 | 0.13114 | 0.03736 | 0.00807 | 0.00710 | 0.00740 | 0.00864 | 0.30828 | 0.00448 | 0.28828 | 0.00622 | 0.00855 | 0.00951 | 0.00814 | 0.00711 |
| **Gm6460** | 0.01352 | 0.00789 | 0.07775 | 0.01848 | 0.00910 | 0.00771 | 0.00743 | 0.01335 | 0.02190 | 0.01451 | 0.01240 | 0.00767 | 0.01150 | 0.01012 | 0.02356 | 0.00736 |
| **Gm6498** | 4.21980 | 1.69843 | 2.49709 | 1.05666 | 3.27218 | 2.55385 | 1.25883 | 0.23131 | 0.69595 | 0.15756 | 0.00005 | 0.16958 | 2.37414 | 0.00299 | 0.19052 | 4.73446 |
| **Gm6524** | 0.13365 | 0.28083 | 0.17240 | 0.15500 | 0.05437 | 0.12203 | 0.16086 | 0.13424 | 0.31686 | 0.24206 | 0.07632 | 0.19714 | 0.19125 | 0.37289 | 0.49060 | 0.10168 |
| **Gm6525** | 3.12719 | 0.16043 | 2.07699 | 1.08481 | 1.21137 | 0.29453 | 0.19233 | 0.08511 | 2.85765 | 0.76141 | 8.76644 | 0.07695 | 1.70997 | 1.85342 | 0.88650 | 0.96356 |
| **Gm6548** | 38.04270 | 24.86627 | 37.10702 | 3.60047 | 28.15698 | 17.11310 | 20.25091 | 36.13867 | 6.07803 | 27.23757 | 18.33284 | 13.80799 | 35.18348 | 30.52357 | 17.80960 | 22.88146 |
| **Gm6566** | 0.01049 | 0.00789 | 0.04573 | 0.34227 | 0.00920 | 0.01163 | 0.06866 | 0.08526 | 0.04013 | 0.49858 | 0.06398 | 0.17207 | 0.31568 | 0.00675 | 0.11836 | 0.01431 |
| **Gm6568** | 0.06296 | 0.36081 | 0.42189 | 9.79355 | 0.00957 | 0.51973 | 0.00603 | 0.62415 | 8.78443 | 2.03659 | 0.81190 | 2.27159 | 0.58100 | 1.59275 | 2.14804 | 0.01158 |
| **Gm6578** | 0.51000 | 0.08102 | 0.26885 | 0.91592 | 0.10591 | 0.11517 | 0.07860 | 0.21324 | 0.23042 | 0.67252 | 0.35891 | 0.07118 | 0.17110 | 0.06623 | 0.42513 | 0.03617 |
| **Gm6602** | 0.01258 | 0.03001 | 0.02061 | 0.01896 | 0.00926 | 0.01244 | 0.00769 | 0.01634 | 0.01115 | 0.01918 | 0.01191 | 0.00811 | 0.01233 | 0.07700 | 0.03153 | 0.00742 |
| **Gm6607** | 0.04920 | 0.00825 | 0.02028 | 0.04085 | 0.02337 | 0.00617 | 0.00747 | 0.00974 | 0.11008 | 0.02524 | 0.16508 | 0.00750 | 0.01647 | 0.01213 | 0.00609 | 0.01467 |
| **Gm6623** | 0.44045 | 1.57671 | 1.19298 | 1.41400 | 0.36915 | 0.68721 | 1.07034 | 2.38710 | 2.42588 | 3.62033 | 2.48948 | 3.68623 | 1.65075 | 1.09813 | 2.65250 | 0.36144 |
| **Gm6639** | 0.56750 | 0.00677 | 0.10092 | 2.24428 | 0.00731 | 0.01441 | 0.00657 | 0.23051 | 1.70089 | 0.26431 | 0.00000 | 1.30197 | 0.00611 | 1.62512 | 0.75619 | 0.00604 |
| **Gm6654** | 18.70583 | 1.85303 | 14.19236 | 11.06040 | 11.70098 | 4.06761 | 3.59983 | 4.44380 | 19.99132 | 11.21937 | 29.99706 | 2.14252 | 11.94152 | 0.97207 | 3.06451 | 7.08225 |
| **Gm6710** | 0.48247 | 0.46542 | 2.45865 | 0.14255 | 0.11803 | 0.33795 | 1.55577 | 0.52227 | 0.00554 | 0.98237 | 0.00000 | 0.06713 | 0.97297 | 2.73157 | 1.82211 | 0.60643 |
| **Gm6712** | 0.63546 | 5.20255 | 1.66101 | 0.92764 | 0.64631 | 3.14181 | 2.05751 | 3.67450 | 0.63146 | 6.66318 | 3.62870 | 16.71060 | 1.85963 | 17.39324 | 2.87720 | 1.68176 |
| **Gm6756** | 1.78717 | 2.04761 | 5.49388 | 0.12224 | 1.20057 | 1.68490 | 3.01251 | 9.50478 | 0.18324 | 4.46093 | 2.38110 | 0.74261 | 6.07989 | 0.03876 | 0.00000 | 1.46311 |
| **Gm6787** | 0.02135 | 0.02857 | 0.00934 | 0.45462 | 0.00871 | 0.01926 | 0.00755 | 0.01573 | 0.13068 | 0.01481 | 0.01138 | 0.00740 | 0.01231 | 0.02117 | 0.27616 | 0.00723 |
| **Gm6792** | 1.87617 | 0.01861 | 0.00064 | 1.34991 | 0.01216 | 0.01466 | 0.00594 | 0.00753 | 0.00000 | 0.01280 | 0.01235 | 0.01811 | 0.01757 | 0.00756 | 0.04419 | 0.00536 |
| **Gm6815** | 1.24863 | 0.44713 | 0.69898 | 0.58062 | 0.00590 | 0.67531 | 0.48608 | 0.86185 | 0.18418 | 1.26102 | 0.00000 | 0.11808 | 0.29169 | 6.16585 | 4.21455 | 0.23085 |
| **Gm6880** | 0.50792 | 0.00866 | 0.50401 | 0.50096 | 0.01192 | 0.00725 | 0.00703 | 0.15796 | 0.05760 | 2.33074 | 0.10987 | 0.02301 | 1.14780 | 0.00812 | 0.82588 | 0.02090 |
| **Gm6890** | 0.35019 | 0.00893 | 0.87720 | 0.77451 | 0.01271 | 0.00720 | 0.00712 | 0.03934 | 0.26597 | 1.43247 | 0.17593 | 0.00000 | 0.83990 | 0.02088 | 0.59383 | 0.00630 |
| **Gm6902** | 0.00000 | 0.00870 | 0.34491 | 9.92149 | 0.00892 | 0.00341 | 0.00761 | 0.70992 | 13.96386 | 0.00000 | 0.08745 | 0.00000 | 0.74950 | 0.65700 | 0.00000 | 0.00419 |
| **Gm6904** | 0.02351 | 0.12859 | 0.02529 | 0.01075 | 0.01883 | 0.00686 | 0.00782 | 0.04298 | 0.01223 | 0.01273 | 0.01250 | 0.00845 | 0.03753 | 0.01468 | 0.01357 | 0.00735 |
| **Gm694** | 0.12627 | 0.02957 | 0.08228 | 0.01388 | 0.00881 | 0.09213 | 0.00744 | 0.05954 | 0.04312 | 0.01691 | 0.01157 | 0.00822 | 0.11237 | 0.01166 | 0.01342 | 0.20014 |
| **Gm6981** | 4.42032 | 1.48272 | 2.18412 | 0.84614 | 2.96521 | 1.62848 | 0.76044 | 0.18532 | 1.04525 | 0.13784 | 0.05006 | 0.11936 | 1.77564 | 0.02245 | 0.11297 | 3.09704 |
| **Gm7008** | 0.01100 | 0.16179 | 0.01012 | 0.01053 | 0.00874 | 0.00673 | 0.00731 | 0.01141 | 0.01236 | 0.01242 | 0.01251 | 0.08603 | 0.01248 | 0.01093 | 0.00738 | 0.00735 |
| **Gm7030** | 0.08544 | 0.00761 | 0.02634 | 0.01016 | 0.04185 | 0.14095 | 0.01733 | 0.03635 | 0.01226 | 0.06016 | 0.01260 | 0.00751 | 0.12200 | 0.01123 | 0.01520 | 0.13715 |
| **Gm7056** | 0.00988 | 0.00762 | 0.00554 | 19.15024 | 0.00905 | 0.00389 | 0.00672 | 0.00291 | 10.41818 | 4.36418 | 0.08927 | 0.06973 | 0.03295 | 0.00000 | 4.32214 | 0.00380 |
| **Gm7102** | 0.75651 | 1.27411 | 1.75127 | 62.15898 | 0.43587 | 0.76637 | 0.94742 | 4.97058 | 58.38779 | 40.75876 | 11.49194 | 16.21901 | 2.01220 | 37.86173 | 46.14028 | 0.28444 |
| **Gm7104** | 0.00409 | 0.00793 | 0.41165 | 15.60988 | 0.00820 | 0.00202 | 0.00691 | 0.29214 | 22.33026 | 0.88636 | 0.66065 | 0.00000 | 0.00100 | 0.00000 | 2.74646 | 0.00316 |
| **Gm715** | 0.09113 | 0.00672 | 0.00407 | 0.71903 | 1.35801 | 0.00773 | 0.00745 | 0.01010 | 0.00000 | 0.01030 | 0.01271 | 0.00653 | 0.01115 | 0.01859 | 0.04657 | 0.00709 |
| **Gm7244** | 0.01373 | 0.00814 | 0.01219 | 2.58254 | 0.00932 | 0.00638 | 0.00742 | 0.01352 | 0.26543 | 0.01037 | 0.01153 | 0.00778 | 0.01268 | 0.00576 | 0.06700 | 0.00694 |
| **Gm7325** | 1.77064 | 0.00641 | 0.02769 | 0.25210 | 0.01176 | 0.25789 | 0.00576 | 0.04412 | 0.00791 | 0.00785 | 0.01292 | 0.00653 | 0.03095 | 0.01170 | 0.00604 | 0.47891 |
| **Gm7361** | 0.01585 | 0.00811 | 0.01290 | 0.01833 | 0.00905 | 0.05960 | 0.00744 | 0.04651 | 0.01120 | 0.04846 | 0.01156 | 0.00801 | 0.00953 | 0.00845 | 0.04625 | 0.00718 |
| **Gm7534** | 0.00815 | 0.00802 | 0.00497 | 0.00369 | 0.01030 | 0.00770 | 0.00738 | 1.19015 | 1.86923 | 0.12054 | 0.42589 | 0.00952 | 0.01864 | 0.00325 | 0.28003 | 0.00745 |
| **Gm7609** | 0.00960 | 0.00786 | 0.07068 | 1.24777 | 0.01174 | 0.01261 | 0.02732 | 0.38099 | 0.80269 | 1.16034 | 0.27145 | 0.02443 | 0.28904 | 1.95994 | 1.36826 | 0.01430 |
| **Gm7616** | 0.04325 | 0.00779 | 0.02699 | 0.32333 | 0.02961 | 0.00667 | 0.03842 | 0.00044 | 0.46810 | 0.00852 | 0.39800 | 0.01472 | 0.01912 | 0.15208 | 0.01710 | 0.00718 |
| **Gm7694** | 0.01356 | 0.00775 | 0.01172 | 1.02850 | 0.00933 | 0.00652 | 0.00740 | 0.01776 | 0.00000 | 0.01119 | 0.01163 | 0.00770 | 0.01870 | 0.01842 | 0.06920 | 0.00720 |
| **Gm7788** | 0.02269 | 0.00782 | 0.03533 | 0.23876 | 0.00878 | 0.01177 | 0.00730 | 0.02306 | 0.05057 | 0.03015 | 0.01307 | 0.00780 | 0.02373 | 0.01107 | 0.01264 | 0.01442 |
| **Gm7827** | 0.01349 | 0.00771 | 0.01026 | 1.16219 | 0.00941 | 0.00638 | 0.00739 | 0.01196 | 0.49033 | 0.00000 | 0.02266 | 0.00491 | 0.01251 | 0.13585 | 0.62018 | 0.00712 |
| **Gm7849** | 0.01370 | 0.00821 | 0.01212 | 0.16177 | 0.00927 | 0.00674 | 0.00758 | 0.01432 | 0.16836 | 0.02755 | 0.01234 | 0.00853 | 0.01299 | 0.01103 | 0.01353 | 0.00739 |
| **Gm7904** | 0.03921 | 0.00810 | 0.01055 | 0.00541 | 0.00898 | 0.00676 | 0.00749 | 0.01442 | 0.20699 | 0.02573 | 0.01144 | 0.00811 | 0.01260 | 0.01413 | 0.00984 | 0.00728 |
| **Gm7977** | 0.97747 | 0.64212 | 0.70642 | 0.00000 | 1.52430 | 2.00137 | 0.79516 | 4.00236 | 0.00596 | 2.94543 | 0.00000 | 2.87444 | 2.73587 | 0.90835 | 0.42555 | 0.62536 |
| **Gm805** | 0.07966 | 0.00791 | 0.40788 | 0.01074 | 0.00780 | 0.01662 | 0.00743 | 0.01691 | 0.01227 | 0.00763 | 0.01448 | 0.00734 | 0.32233 | 0.01446 | 0.00358 | 0.01473 |
| **Gm8096** | 1.38523 | 1.25429 | 3.73559 | 0.39041 | 0.91904 | 1.60939 | 2.10505 | 6.39407 | 0.19065 | 3.77620 | 3.49983 | 0.37936 | 4.44645 | 0.00016 | 0.00000 | 0.78480 |
| **Gm813** | 0.01363 | 0.00817 | 0.01182 | 0.25006 | 0.00932 | 0.00657 | 0.00751 | 0.01314 | 0.90866 | 0.00714 | 0.01852 | 0.00666 | 0.01257 | 0.00868 | 0.06756 | 0.00729 |
| **Gm8234** | 2.17722 | 0.69211 | 1.13240 | 18.65729 | 2.29484 | 0.24019 | 0.38678 | 3.43531 | 17.60799 | 4.18222 | 1.07392 | 1.35028 | 2.67603 | 6.62303 | 9.47885 | 1.51168 |
| **Gm8267** | 0.28217 | 0.44023 | 0.07162 | 1.31327 | 0.01794 | 0.01107 | 0.12966 | 0.90909 | 0.55085 | 0.17626 | 0.00000 | 1.13796 | 0.00694 | 0.26983 | 0.55746 | 0.02170 |
| **Gm8300** | 0.17810 | 0.33238 | 0.51793 | 15.88096 | 0.22111 | 0.07287 | 0.11094 | 0.24576 | 15.15140 | 0.84789 | 0.00000 | 0.00000 | 0.06510 | 1.85928 | 3.13037 | 0.12096 |
| **Gm8363** | 0.05060 | 0.00824 | 0.04463 | 0.03269 | 0.01905 | 0.03837 | 0.01799 | 0.10472 | 0.00992 | 0.05411 | 0.07575 | 0.03816 | 0.07337 | 0.02617 | 0.03907 | 0.03628 |
| **Gm839** | 0.01374 | 0.00826 | 0.01228 | 0.43406 | 0.00927 | 0.00658 | 0.00753 | 0.01378 | 0.05343 | 0.02601 | 0.01303 | 0.00829 | 0.01237 | 0.00982 | 0.01944 | 0.00723 |
| **Gm8579** | 0.35136 | 0.00819 | 0.08601 | 0.10174 | 0.03359 | 1.09416 | 0.00695 | 0.39225 | 0.01054 | 0.39154 | 0.00714 | 0.07677 | 0.00000 | 0.01153 | 0.00213 | 0.00650 |
| **Gm8580** | 13.46111 | 3.22694 | 9.19858 | 3.39665 | 6.57673 | 2.98227 | 4.59158 | 10.29015 | 5.43479 | 11.58862 | 14.18006 | 1.60780 | 10.23806 | 0.00000 | 0.31749 | 6.23878 |
| **Gm8615** | 5.23808 | 4.81542 | 3.13921 | 0.11086 | 3.22869 | 4.94605 | 4.20785 | 4.14863 | 1.35482 | 2.38605 | 4.80003 | 6.26628 | 2.15597 | 3.59198 | 7.89549 | 1.35808 |
| **Gm8624** | 11.22145 | 0.89148 | 8.62815 | 5.07782 | 7.32466 | 2.11882 | 1.51973 | 1.23301 | 11.78405 | 9.10217 | 21.49770 | 0.76372 | 6.76887 | 2.15419 | 2.25433 | 4.41167 |
| **Gm867** | 0.01656 | 0.00798 | 0.30599 | 0.00536 | 0.00867 | 0.00572 | 0.00733 | 0.01216 | 0.14630 | 0.01501 | 0.01079 | 0.01911 | 0.01004 | 0.00770 | 0.12300 | 0.00720 |

| | | | | | | | | | | | | | | | | |
|---|---|---|---|---|---|---|---|---|---|---|---|---|---|---|---|---|
| **Gm8709** | 0.04043 | 0.01949 | 0.01931 | 0.22241 | 0.06919 | 0.03272 | 0.02753 | 0.02188 | 0.05589 | 0.02448 | 0.01213 | 0.00763 | 0.02438 | 0.01082 | 0.00788 | 0.02237 |
| **Gm8801** | 1.79001 | 0.07638 | 1.39989 | 6.86304 | 1.08873 | 2.79918 | 0.00399 | 1.70568 | 1.68492 | 4.33068 | 0.40539 | 2.29211 | 1.86953 | 0.17595 | 3.42689 | 0.78889 |
| **Gm884** | 0.01361 | 0.00779 | 0.01215 | 0.00885 | 0.00912 | 0.00652 | 0.00732 | 0.01089 | 0.01202 | 0.40783 | 0.01409 | 0.00725 | 0.01125 | 0.01805 | 0.00721 | 0.00721 |
| **Gm8909** | 0.16155 | 0.00708 | 0.07943 | 0.01238 | 0.33054 | 0.12990 | 0.44236 | 0.41571 | 0.00997 | 0.13859 | 0.78272 | 0.00238 | 0.40818 | 0.01135 | 0.00338 | 0.39363 |
| **Gm8994** | 5.53857 | 2.92813 | 5.09952 | 7.65476 | 2.09678 | 2.43342 | 3.82945 | 10.45608 | 10.35377 | 9.57145 | 10.11192 | 3.87015 | 4.75930 | 1.31093 | 2.89299 | 1.75027 |
| **Gm9** | 0.47030 | 0.00836 | 0.00001 | 4.38487 | 0.01374 | 0.00513 | 0.00654 | 0.06390 | 0.51040 | 0.00000 | 0.76030 | 0.00107 | 1.06706 | 4.29339 | 3.69294 | 0.00531 |
| **Gm9079** | 0.10077 | 0.06032 | 0.10727 | 0.13964 | 0.05363 | 0.10514 | 0.16330 | 0.10987 | 0.02898 | 0.16905 | 0.04125 | 0.01684 | 0.16769 | 0.14946 | 0.07284 | 0.07317 |
| **Gm9125** | 0.00678 | 0.00738 | 0.00000 | 14.43844 | 0.00887 | 0.00324 | 0.00648 | 0.00000 | 14.38035 | 1.09840 | 0.41790 | 1.08335 | 0.19034 | 0.68704 | 4.92977 | 0.00375 |
| **Gm9159** | 0.05122 | 0.01753 | 0.09644 | 0.10372 | 0.18286 | 0.04403 | 0.07805 | 0.06231 | 0.19973 | 0.12330 | 0.18067 | 0.06161 | 0.15280 | 0.72322 | 0.10403 | 0.02160 |
| **Gm9199** | 2.78871 | 0.49808 | 2.85518 | 2.52155 | 4.03881 | 1.67997 | 1.04851 | 5.80456 | 6.88127 | 18.17757 | 10.02593 | 9.40938 | 5.52565 | 7.26335 | 4.07254 | 2.28200 |
| **Gm9268** | 0.00823 | 0.00765 | 0.00704 | 0.01266 | 0.00936 | 0.00650 | 0.00673 | 0.00747 | 0.04994 | 0.00000 | 0.04577 | 0.00336 | 0.81112 | 1.10607 | 0.00135 | 0.00677 |
| **Gm9376** | 8.70883 | 2.73356 | 12.16091 | 4.09043 | 8.20467 | 4.06988 | 3.99904 | 5.80275 | 18.88685 | 8.49070 | 23.27932 | 4.52122 | 13.16191 | 9.05859 | 6.18951 | 2.63486 |
| **Gm960** | 0.02098 | 0.00822 | 0.02123 | 0.04833 | 0.00928 | 0.00698 | 0.01916 | 0.02370 | 0.00967 | 0.47122 | 0.01217 | 0.00822 | 0.05230 | 0.01151 | 0.08746 | 0.00765 |
| **Gm973** | 0.00294 | 3.09758 | 1.22263 | 0.02814 | 0.04200 | 0.08375 | 0.00654 | 0.00810 | 0.26451 | 0.00000 | 0.08577 | 0.01494 | 0.11920 | 0.08551 | 0.27716 | 0.00576 |
| **Gm9731** | 0.01146 | 0.00803 | 0.10182 | 0.34429 | 0.07631 | 0.01780 | 0.00755 | 0.04412 | 1.19063 | 0.08758 | 0.02107 | 0.00541 | 0.03645 | 0.02913 | 0.00956 | 0.06513 |
| **Gm9758** | 0.01141 | 0.00807 | 0.18652 | 0.07205 | 0.00912 | 0.00583 | 0.00765 | 0.04596 | 0.05155 | 0.04840 | 0.01093 | 0.01950 | 0.00956 | 0.00756 | 0.15533 | 0.00739 |
| **Gm9776** | 0.86813 | 0.02679 | 0.19991 | 3.01446 | 0.11638 | 0.04216 | 0.04553 | 0.00000 | 4.47802 | 0.22875 | 3.09374 | 0.09107 | 0.11285 | 1.30036 | 0.11663 | 0.08496 |
| **Gm9833** | 0.22239 | 0.91322 | 0.53382 | 0.16571 | 0.91763 | 0.72176 | 0.10223 | 0.03905 | 0.19174 | 0.21994 | 0.25364 | 0.02285 | 0.49698 | 0.40036 | 0.84908 | 0.24571 |
| **Gm9895** | 0.01365 | 0.00761 | 0.51414 | 0.62812 | 0.00832 | 0.00577 | 0.00707 | 0.30805 | 0.35252 | 0.18607 | 0.00460 | 0.02538 | 0.16802 | 0.01568 | 0.80339 | 0.00702 |
| **Gm9926** | 0.08370 | 0.00761 | 0.06880 | 0.08604 | 0.00824 | 0.00609 | 0.00725 | 0.01807 | 0.17487 | 0.05491 | 0.12853 | 0.02636 | 0.04525 | 0.26517 | 0.00874 | 0.00722 |
| **Gm9958** | 0.01234 | 0.00806 | 0.29254 | 0.27651 | 0.00875 | 0.00559 | 0.00757 | 0.01337 | 0.45663 | 0.00907 | 0.00642 | 0.28635 | 0.02316 | 0.01123 | 0.00000 | 0.00720 |
| **Gm996** | 0.01375 | 0.00805 | 0.01225 | 0.01061 | 0.00932 | 0.00666 | 0.00754 | 0.01390 | 0.01235 | 0.01531 | 0.01310 | 0.00811 | 0.01270 | 0.01234 | 0.01062 | 0.00741 |
| **Gm9992** | 0.42321 | 1.06849 | 0.00000 | 0.00719 | 0.00771 | 3.18454 | 0.00622 | 0.01002 | 0.01289 | 0.04070 | 0.00820 | 0.00745 | 0.55171 | 0.00611 | 0.09237 | 0.00500 |
| **Gm9994** | 0.01127 | 0.00848 | 0.06980 | 0.01055 | 0.00960 | 0.00889 | 0.00807 | 0.05219 | 0.01234 | 0.01421 | 0.01303 | 0.00840 | 0.01796 | 0.01215 | 0.01516 | 0.00754 |
| **Gmcl1** | 4.17922 | 4.27713 | 2.41539 | 4.98783 | 4.32811 | 3.62543 | 0.90922 | 1.03594 | 3.45739 | 1.74871 | 1.30512 | 0.01420 | 0.30815 | 4.13782 | 0.20184 | 0.06437 |
| **Gmds** | 1.87212 | 0.00531 | 0.54131 | 0.12010 | 0.62019 | 0.01599 | 0.00589 | 0.01025 | 0.63078 | 0.05285 | 0.02372 | 0.00433 | 0.01525 | 0.01548 | 0.00890 | 0.00517 |
| **Gmeb1** | 2.42325 | 0.19030 | 4.07457 | 13.18663 | 5.90871 | 0.41467 | 0.30775 | 0.31328 | 17.32468 | 2.41818 | 14.07576 | 1.25513 | 3.14177 | 7.78030 | 5.09752 | 0.50128 |
| **Gmeb2** | 1.66206 | 0.03423 | 0.45825 | 4.89673 | 1.35062 | 0.00629 | 2.03506 | 1.16859 | 3.59511 | 2.86349 | 0.87810 | 0.58922 | 0.69501 | 0.04896 | 3.72007 | 0.01790 |
| **Gmfb** | 4.84520 | 4.15286 | 2.31502 | 3.40123 | 0.31701 | 2.46815 | 1.61076 | 2.40369 | 1.70402 | 4.17659 | 3.92862 | 8.50169 | 2.23965 | 9.95481 | 2.33279 | 1.50877 |
| **Gmfg** | 1.03655 | 0.87219 | 1.30980 | 0.00000 | 1.72634 | 0.00545 | 0.00512 | 0.00000 | 0.00984 | 0.03902 | 2.59915 | 0.00000 | 0.00000 | 0.00783 | 0.01680 | 0.01224 |
| **Gmip** | 0.01505 | 0.00774 | 0.74884 | 0.55037 | 0.02535 | 0.00957 | 1.26555 | 0.00714 | 0.45283 | 0.00000 | 0.11451 | 0.07408 | 0.27549 | 0.02179 | 0.05925 | 0.01491 |
| **Gml2** | 0.86638 | 0.00711 | 0.00368 | 0.00428 | 0.41215 | 0.00733 | 0.00681 | 0.00438 | 0.01085 | 0.00480 | 0.94868 | 0.00306 | 0.00432 | 0.01171 | 0.00000 | 0.00608 |
| **Gmnn** | 21.99162 | 7.37899 | 22.31717 | 24.54752 | 8.78492 | 8.93210 | 9.38217 | 15.76108 | 32.56965 | 28.24652 | 20.62537 | 10.17184 | 15.51194 | 15.55919 | 16.78468 | 8.89105 |
| **Gmppa** | 5.24969 | 0.58988 | 7.12036 | 0.00000 | 2.29075 | 0.26616 | 0.00000 | 1.50326 | 0.01980 | 2.05545 | 2.11223 | 0.09573 | 3.27360 | 0.00200 | 0.49105 | 3.60332 |
| **Gmppb** | 4.69109 | 6.13728 | 7.93691 | 0.05283 | 4.87332 | 4.12723 | 0.18543 | 6.81700 | 0.20680 | 3.85642 | 2.85171 | 0.02582 | 6.17386 | 0.00007 | 1.43035 | 5.16733 |
| **Gmpr** | 0.01107 | 0.00708 | 0.21059 | 0.06590 | 0.00878 | 0.00576 | 1.59816 | 2.39223 | 0.00803 | 0.84037 | 0.24782 | 0.00000 | 0.00396 | 0.00835 | 0.00600 | 0.00726 |
| **Gmpr2** | 6.64149 | 5.90200 | 4.50954 | 0.63675 | 4.98308 | 0.06777 | 3.66636 | 0.00000 | 0.37434 | 0.83940 | 0.71231 | 0.02497 | 0.09496 | 0.00271 | 0.78257 | 1.10677 |
| **Gmps** | 12.95833 | 14.37559 | 8.36698 | 7.68497 | 13.94076 | 5.93103 | 9.73579 | 13.32006 | 14.11183 | 19.84939 | 10.46733 | 11.42721 | 11.33789 | 14.63245 | 13.30096 | 4.73984 |
| **Gna11** | 6.62020 | 0.00000 | 6.58690 | 3.29459 | 2.78232 | 6.54667 | 0.48390 | 6.33823 | 0.75596 | 10.40809 | 1.20645 | 0.33844 | 5.55646 | 0.00561 | 0.00000 | 2.20007 |
| **Gna12** | 0.11170 | 0.27555 | 0.01109 | 0.38485 | 0.04169 | 0.00652 | 0.53021 | 0.37676 | 0.35010 | 0.34606 | 0.00876 | 0.01371 | 0.15617 | 0.00519 | 0.06127 | 1.12523 |
| **Gna13** | 9.46364 | 7.50905 | 8.46889 | 2.50029 | 7.90727 | 3.71444 | 9.62144 | 4.29503 | 2.03120 | 6.11153 | 9.14920 | 0.39769 | 3.60036 | 6.19354 | 0.39408 | 4.45281 |
| **Gna14** | 0.01470 | 0.00789 | 0.01252 | 1.26261 | 0.00983 | 0.00715 | 0.00749 | 0.01354 | 0.13967 | 0.00000 | 0.05874 | 0.00678 | 0.47915 | 0.05443 | 0.28015 | 0.00730 |
| **Gna15** | 0.58896 | 0.05041 | 0.04277 | 0.33245 | 0.00762 | 0.00849 | 0.00666 | 0.00775 | 0.56956 | 0.00000 | 0.56521 | 0.00297 | 0.82516 | 0.00588 | 0.00000 | 0.00637 |
| **Gnai1** | 0.88633 | 0.01667 | 1.41912 | 2.63849 | 0.00382 | 0.00460 | 0.00654 | 0.00537 | 0.19736 | 0.14629 | 0.00400 | 0.05490 | 0.03574 | 0.05298 | 3.98032 | 0.78849 |
| **Gnai2** | 3.70046 | 0.10407 | 5.03948 | 0.39061 | 1.34324 | 3.56004 | 1.34748 | 2.34281 | 0.72297 | 6.23789 | 0.80117 | 0.37413 | 2.05020 | 0.01019 | 0.17729 | 3.07856 |
| **Gnai3** | 5.25772 | 11.07582 | 7.45788 | 7.15601 | 13.86732 | 5.74394 | 8.66038 | 6.65118 | 8.92343 | 9.92794 | 4.49509 | 18.70504 | 2.96492 | 11.98434 | 7.31899 | 1.64621 |
| **Gnal** | 1.21534 | 0.18322 | 1.16043 | 0.44907 | 1.11178 | 2.42496 | 1.45853 | 0.79896 | 0.77729 | 3.04254 | 2.01160 | 0.11487 | 0.37896 | 2.14603 | 0.79983 | 0.09203 |
| **Gnaq** | 2.80686 | 0.04610 | 1.43472 | 0.15134 | 1.53733 | 0.13839 | 2.60996 | 0.01009 | 1.06616 | 2.83111 | 0.00000 | 0.09350 | 0.01227 | 5.27470 | 2.17587 | 2.73280 |
| **Gnas** | 7.78156 | 7.86742 | 7.51184 | 6.00520 | 8.39454 | 2.58292 | 10.12426 | 10.37341 | 1.44924 | 16.48105 | 15.54961 | 5.73979 | 5.78588 | 4.11253 | 0.85086 | 7.02786 |
| **Gnb1** | 14.91139 | 9.09292 | 19.02551 | 10.11952 | 7.24062 | 8.11231 | 12.75729 | 13.33252 | 2.62035 | 10.60200 | 13.68726 | 3.44505 | 14.29254 | 9.09107 | 4.93901 | 9.97689 |
| **Gnb1l** | 1.35416 | 4.54120 | 5.30847 | 2.12928 | 0.00604 | 0.00418 | 0.00238 | 3.64562 | 0.59755 | 7.29410 | 0.97537 | 0.12800 | 2.98311 | 0.10019 | 0.53812 | 0.48520 |
| **Gnb2** | 5.18067 | 0.03231 | 3.31006 | 4.31098 | 1.19844 | 5.73524 | 0.38719 | 3.76119 | 0.77226 | 5.02058 | 1.28013 | 1.63366 | 7.24504 | 0.00000 | 0.00000 | 4.76637 |
| **Gnb3** | 0.00996 | 0.00803 | 0.01027 | 0.01019 | 0.01061 | 0.00735 | 0.00745 | 0.41016 | 0.01173 | 0.00502 | 0.01394 | 0.00708 | 0.12814 | 0.00998 | 0.04770 | 0.00769 |
| **Gnb4** | 0.01325 | 0.00748 | 0.00751 | 3.79154 | 0.00907 | 0.00605 | 0.00696 | 0.00000 | 1.76655 | 0.71158 | 0.33104 | 0.01687 | 0.00904 | 0.00086 | 0.90146 | 0.00663 |
| **Gnb5** | 0.01248 | 0.00788 | 0.42135 | 0.01082 | 0.00842 | 0.00920 | 0.00735 | 0.01268 | 0.01228 | 0.01485 | 0.01206 | 0.00769 | 0.00943 | 0.01402 | 0.00393 | 0.00721 |
| **Gne** | 2.61775 | 0.00508 | 0.00000 | 1.11799 | 0.02758 | 2.34781 | 0.00514 | 0.00392 | 0.42507 | 0.44025 | 0.00674 | 0.02177 | 0.02043 | 0.00361 | 0.13261 | 0.00380 |
| **Gng10** | 0.94797 | 2.76050 | 6.71934 | 0.00000 | 3.73699 | 0.00661 | 3.02455 | 0.01812 | 0.08363 | 1.00830 | 0.32510 | 0.01937 | 1.27030 | 0.33571 | 0.00000 | 0.17359 |
| **Gng12** | 2.73046 | 0.00446 | 0.00848 | 7.58547 | 0.00423 | 0.00378 | 0.00421 | 0.00000 | 2.36514 | 2.01560 | 0.64856 | 1.45652 | 0.88738 | 0.00000 | 0.77326 | 0.41459 |
| **Gng13** | 0.00848 | 0.03071 | 0.01144 | 0.27585 | 0.00826 | 0.00653 | 0.00787 | 0.03281 | 0.31004 | 0.00682 | 0.01096 | 0.00833 | 0.00866 | 0.01217 | 0.00843 | 0.57553 |
| **Gng2** | 0.00000 | 1.01502 | 1.03151 | 0.06184 | 0.01880 | 3.73342 | 0.00528 | 0.00000 | 0.07369 | 0.32440 | 0.20119 | 0.01229 | 1.84327 | 0.02167 | 0.00000 | 4.50286 |
| **Gng3** | 0.01359 | 0.00795 | 0.00967 | 6.03382 | 0.00905 | 0.00600 | 0.00716 | 0.00625 | 2.06322 | 0.08917 | 0.19499 | 0.00323 | 0.00957 | 0.04009 | 0.00000 | 0.00597 |
| **Gng5** | 10.47129 | 0.07470 | 2.30488 | 8.41200 | 8.89622 | 1.76802 | 0.47293 | 0.83691 | 5.54836 | 6.36855 | 7.68655 | 0.42335 | 5.01992 | 0.01711 | 0.37580 | 5.88369 |

| | | | | | | | | | | | | | | | | |
|---|---|---|---|---|---|---|---|---|---|---|---|---|---|---|---|---|
| **Gng7** | 0.01322 | 0.00784 | 0.01141 | 0.00420 | 0.00908 | 0.00651 | 0.00726 | 0.01032 | 0.18481 | 0.86985 | 0.00927 | 0.02454 | 0.01163 | 0.00572 | 0.38280 | 0.00757 |
| **Gngt1** | 0.03122 | 0.05892 | 0.12850 | 0.05827 | 0.00823 | 0.03734 | 0.01725 | 0.00000 | 0.07254 | 0.16462 | 0.62990 | 0.04095 | 0.14993 | 0.22337 | 0.60000 | 0.02152 |
| **Gnl1** | 9.51614 | 5.51872 | 3.37379 | 6.52899 | 1.47402 | 6.28162 | 8.81191 | 12.95706 | 6.47719 | 13.75366 | 6.97191 | 0.73653 | 12.37832 | 0.21456 | 1.12512 | 7.28690 |
| **Gnl2** | 9.58679 | 13.73547 | 12.65918 | 3.67619 | 5.15313 | 0.31676 | 9.89280 | 18.55046 | 10.34507 | 13.58703 | 9.99614 | 8.60727 | 5.53508 | 12.36900 | 11.49312 | 4.95123 |
| **Gnl3** | 11.21732 | 18.17308 | 14.54776 | 18.64042 | 12.71801 | 9.03108 | 15.35708 | 22.36068 | 22.08984 | 28.18550 | 22.01337 | 20.51189 | 16.23734 | 31.57960 | 26.33059 | 14.37161 |
| **Gnl3l** | 6.93970 | 7.03883 | 6.27938 | 9.36395 | 3.66511 | 0.11483 | 4.00599 | 4.24997 | 8.87205 | 12.36596 | 12.01872 | 11.09157 | 6.68704 | 15.16006 | 12.67177 | 2.20567 |
| **Gnmt** | 0.01289 | 0.00813 | 0.03640 | 0.28298 | 0.00915 | 0.00802 | 0.00746 | 0.01356 | 0.72569 | 0.01177 | 0.00907 | 0.00698 | 0.01228 | 0.01203 | 0.00365 | 0.00732 |
| **Gnpat** | 7.41801 | 10.44768 | 5.89266 | 3.02947 | 9.32735 | 2.54500 | 3.11455 | 1.31569 | 2.30828 | 3.91386 | 4.22865 | 5.14775 | 3.72684 | 2.55451 | 3.71831 | 2.34314 |
| **Gnpda1** | 5.25404 | 5.26352 | 3.36745 | 0.29793 | 2.26118 | 5.88136 | 5.62658 | 4.95183 | 1.49886 | 3.16621 | 4.87581 | 5.46231 | 2.27079 | 4.19177 | 9.00067 | 1.66055 |
| **Gnpda2** | 1.78030 | 0.63019 | 0.00000 | 0.10549 | 0.00720 | 2.95201 | 0.95405 | 1.01466 | 0.79401 | 2.68902 | 0.58719 | 1.20961 | 0.00000 | 0.49996 | 0.91303 | 0.00405 |
| **Gnpnat1** | 9.10841 | 8.16591 | 12.02019 | 10.10887 | 1.50029 | 5.85005 | 12.12157 | 15.88359 | 14.77609 | 12.59746 | 17.31196 | 19.35734 | 11.05315 | 27.00708 | 14.33588 | 0.16879 |
| **Gnptab** | 4.03921 | 4.99214 | 7.26998 | 1.63178 | 4.79580 | 4.20813 | 5.04702 | 12.43450 | 8.71933 | 12.68033 | 7.48132 | 6.54476 | 5.75624 | 21.47492 | 11.13350 | 4.03873 |
| **Gnptg** | 0.23256 | 0.24053 | 6.20256 | 4.28088 | 2.62989 | 3.87059 | 3.37951 | 6.33839 | 6.99894 | 9.23435 | 4.67072 | 1.43812 | 7.07361 | 3.61153 | 6.66920 | 3.60260 |
| **Gnrh1** | 0.01312 | 0.00836 | 0.07031 | 5.89442 | 0.00889 | 0.00803 | 0.00726 | 0.00490 | 0.63687 | 0.01306 | 0.44257 | 0.01598 | 0.00740 | 0.00000 | 0.04289 | 0.00605 |
| **Gns** | 1.67883 | 5.61626 | 4.82657 | 1.17279 | 5.17626 | 0.90282 | 0.45320 | 3.09642 | 0.11657 | 1.14749 | 1.80487 | 2.35595 | 4.13560 | 0.00354 | 1.53864 | 0.45394 |
| **Golga1** | 1.23307 | 1.75157 | 1.32109 | 2.07168 | 0.00482 | 0.00605 | 0.85867 | 1.36131 | 6.06051 | 3.37821 | 1.36016 | 0.27287 | 0.46474 | 7.74679 | 3.18062 | 0.00385 |
| **Golga2** | 3.59577 | 0.12829 | 5.28788 | 7.56712 | 0.01842 | 0.00159 | 2.63902 | 6.39233 | 5.43831 | 2.16291 | 4.96418 | 0.24482 | 3.52899 | 2.74005 | 3.96438 | 3.89594 |
| **Golga3** | 2.55422 | 4.44829 | 2.30589 | 1.42571 | 2.78532 | 0.01015 | 2.13544 | 2.63352 | 2.30186 | 5.00297 | 0.98204 | 0.21090 | 0.37965 | 1.79519 | 2.88887 | 0.00000 |
| **Golga4** | 6.53348 | 2.41270 | 2.92804 | 3.28193 | 1.19153 | 7.38418 | 4.16234 | 2.65245 | 1.05826 | 3.72364 | 0.38068 | 0.20777 | 3.69491 | 4.57541 | 2.26494 | 2.12626 |
| **Golga5** | 3.76527 | 1.34016 | 4.91114 | 3.71019 | 4.62811 | 1.75592 | 0.97481 | 4.34120 | 3.52681 | 2.21532 | 3.10198 | 0.05245 | 0.00000 | 2.73984 | 6.13117 | 0.02944 |
| **Golga7** | 12.52783 | 4.52667 | 10.58724 | 1.62200 | 10.59348 | 7.39667 | 5.80249 | 13.49602 | 2.97189 | 14.53077 | 9.34258 | 6.39523 | 9.43730 | 0.78145 | 4.52186 | 4.31896 |
| **Golga7b** | 0.01215 | 0.00805 | 0.01377 | 0.25170 | 0.01011 | 0.00703 | 0.00765 | 0.01335 | 0.00772 | 0.19964 | 0.04043 | 0.02494 | 0.26914 | 0.00481 | 0.36929 | 0.00739 |
| **Golgb1** | 5.18581 | 2.19890 | 0.00000 | 3.05403 | 2.25898 | 2.26219 | 1.64885 | 2.04858 | 1.93669 | 1.61280 | 0.36958 | 0.15870 | 0.88867 | 1.26957 | 3.02461 | 1.04873 |
| **Golm1** | 7.85672 | 2.15352 | 3.54958 | 1.71040 | 6.78038 | 2.13734 | 1.92356 | 0.80312 | 0.50744 | 1.46021 | 0.00000 | 0.69300 | 0.91755 | 0.00438 | 0.61399 | 2.58274 |
| **Golph3** | 4.34255 | 3.46836 | 3.17935 | 12.07120 | 13.60542 | 7.36478 | 1.98482 | 3.25162 | 9.31202 | 7.31944 | 0.26724 | 10.55699 | 2.57528 | 10.05715 | 9.67009 | 3.19839 |
| **Golph3l** | 3.47059 | 0.01548 | 0.00000 | 0.58907 | 0.01256 | 1.42146 | 0.96300 | 0.18446 | 2.58693 | 1.07017 | 0.64407 | 0.18508 | 1.47590 | 0.00335 | 2.24987 | 2.00136 |
| **Golt1a** | 0.00414 | 0.00624 | 0.00000 | 0.00289 | 0.01059 | 0.00896 | 2.05731 | 0.42224 | 0.01004 | 0.38107 | 1.43318 | 0.00666 | 0.00000 | 0.01222 | 0.10319 | 0.00720 |
| **Golt1b** | 5.56493 | 3.10266 | 6.29844 | 4.63846 | 4.02134 | 0.16791 | 7.68970 | 4.70014 | 7.79975 | 7.12986 | 3.12006 | 6.14266 | 5.22716 | 12.15573 | 5.53310 | 0.99936 |
| **Gon4l** | 5.61131 | 3.11740 | 5.51000 | 3.95366 | 0.74698 | 4.86840 | 1.76954 | 6.25821 | 2.92592 | 4.02933 | 2.12121 | 0.24997 | 0.95916 | 0.61230 | 3.44749 | 3.56606 |
| **Gopc** | 0.47679 | 0.19024 | 0.00000 | 0.66308 | 0.50590 | 0.00712 | 1.10857 | 0.00000 | 0.00000 | 0.00000 | 0.69960 | 0.00923 | 0.10366 | 0.00501 | 1.45844 | 0.94586 |
| **Gorab** | 0.00000 | 2.85440 | 0.95393 | 0.15730 | 0.00618 | 0.00610 | 0.00626 | 1.41345 | 0.37476 | 0.47937 | 0.54162 | 0.08100 | 0.44216 | 0.00672 | 1.12362 | 0.24482 |
| **Gorasp1** | 3.64056 | 1.67002 | 1.05165 | 3.96230 | 6.30185 | 0.01334 | 3.92625 | 8.17883 | 4.50556 | 25.03275 | 9.16756 | 1.41188 | 5.34816 | 0.00000 | 6.90114 | 2.41584 |
| **Gorasp2** | 12.45710 | 7.22517 | 10.68906 | 7.53184 | 5.31685 | 4.84019 | 0.45267 | 8.50033 | 3.91899 | 9.04585 | 4.43207 | 2.64978 | 9.90814 | 0.16232 | 2.81560 | 8.55016 |
| **Gosr1** | 2.84112 | 5.55978 | 2.43005 | 1.09023 | 3.54380 | 0.00800 | 1.92328 | 4.45196 | 1.67823 | 5.63810 | 1.94792 | 0.19736 | 4.92600 | 0.00237 | 4.90120 | 0.00000 |
| **Gosr2** | 4.85347 | 3.63636 | 6.06188 | 4.67441 | 3.37675 | 1.12230 | 4.73836 | 7.05755 | 2.45214 | 10.60066 | 5.32937 | 0.48375 | 5.67491 | 7.66708 | 9.35472 | 1.37336 |
| **Got1** | 14.88726 | 16.69316 | 25.49907 | 12.04395 | 13.82781 | 15.03483 | 15.38985 | 20.73180 | 10.81757 | 8.50883 | 12.37810 | 3.34556 | 23.76490 | 1.94265 | 7.26552 | 11.28383 |
| **Got1l1** | 1.22287 | 4.06138 | 5.77022 | 0.00000 | 0.00000 | 0.58238 | 2.41287 | 2.65293 | 0.00796 | 0.08482 | 3.66013 | 0.01796 | 0.62121 | 1.33669 | 0.00000 | 0.00825 |
| **Got2** | 10.51113 | 2.50491 | 8.53353 | 4.80205 | 5.19843 | 2.95128 | 3.62318 | 10.89123 | 1.58081 | 11.25275 | 3.77212 | 1.38367 | 2.48483 | 1.40648 | 4.56583 | 3.84772 |
| **Gp1ba** | 0.32385 | 0.00731 | 0.15762 | 0.00434 | 0.00848 | 0.00539 | 0.00698 | 0.05437 | 0.60497 | 0.13776 | 0.00661 | 0.00514 | 0.02370 | 0.01254 | 0.22643 | 0.12051 |
| **Gpa33** | 0.99834 | 0.00642 | 0.21162 | 2.17548 | 0.16705 | 1.40562 | 0.00529 | 0.01077 | 0.41146 | 0.00155 | 0.01000 | 0.00439 | 1.59937 | 0.01025 | 0.00812 | 5.94513 |
| **Gpaa1** | 3.59062 | 0.08627 | 5.09428 | 2.95316 | 0.09516 | 2.10647 | 2.33467 | 3.99080 | 1.62884 | 2.60608 | 1.33017 | 0.05809 | 9.59797 | 0.00000 | 0.00000 | 6.72336 |
| **Gpalpp1** | 3.33770 | 6.95841 | 4.14757 | 3.50916 | 1.61341 | 0.09648 | 2.47056 | 6.35182 | 10.13438 | 4.56025 | 4.66181 | 1.91264 | 4.20382 | 7.81191 | 3.85508 | 1.74079 |
| **Gpam** | 0.01107 | 0.00792 | 0.04044 | 1.31329 | 0.00875 | 0.00571 | 0.07791 | 0.01007 | 0.25885 | 1.71028 | 0.01038 | 0.00304 | 0.01042 | 0.04158 | 0.01611 | 0.12339 |
| **Gpank1** | 7.83131 | 1.68534 | 4.52614 | 5.79313 | 6.86115 | 0.15959 | 6.18334 | 7.48668 | 11.21236 | 13.78848 | 6.16886 | 0.88121 | 8.07065 | 0.00000 | 1.86631 | 3.59578 |
| **Gpat2** | 0.41831 | 0.00755 | 0.00000 | 1.99350 | 0.01427 | 0.24076 | 0.54294 | 0.29802 | 1.65237 | 1.73709 | 2.56940 | 0.08620 | 3.44895 | 0.08178 | 1.08538 | 1.97207 |
| **Gpat4** | 7.96250 | 8.11502 | 0.55739 | 7.78349 | 2.17982 | 6.24546 | 2.58399 | 1.65232 | 3.17928 | 3.70301 | 0.80519 | 0.12056 | 5.97472 | 0.04542 | 1.07506 | 1.83222 |
| **Gpatch1** | 2.04462 | 0.33541 | 3.50880 | 0.15106 | 0.01344 | 0.00565 | 3.22609 | 1.67025 | 3.12452 | 4.42869 | 2.75136 | 0.18775 | 0.00000 | 0.21006 | 0.59359 | 0.01082 |
| **Gpatch11** | 1.41871 | 0.03815 | 6.62295 | 0.26655 | 0.01909 | 0.39395 | 1.60540 | 4.98349 | 0.97897 | 6.09549 | 3.62100 | 0.85858 | 1.07356 | 0.22593 | 2.53690 | 2.59343 |
| **Gpatch2** | 1.99083 | 4.71083 | 2.99894 | 2.07517 | 0.81310 | 0.00346 | 3.14225 | 1.21081 | 3.33265 | 0.12546 | 1.10980 | 0.06106 | 2.21887 | 0.23525 | 1.13431 | 0.03594 |
| **Gpatch2l** | 0.24805 | 1.34200 | 0.00000 | 3.03276 | 0.03258 | 0.49465 | 0.39396 | 1.29085 | 0.66115 | 0.42660 | 0.30711 | 0.02787 | 2.31213 | 0.83161 | 1.81485 | 0.41987 |
| **Gpatch3** | 0.79431 | 0.46640 | 0.00000 | 8.24550 | 0.00622 | 0.14417 | 0.00450 | 2.30541 | 15.95552 | 2.21891 | 1.06492 | 0.03861 | 2.23069 | 0.04030 | 0.10881 | 0.02482 |
| **Gpatch4** | 9.72883 | 1.67837 | 4.55378 | 2.71267 | 0.14601 | 4.85393 | 0.12675 | 10.44200 | 3.44184 | 3.81390 | 2.95667 | 0.25986 | 4.42552 | 2.07809 | 7.58883 | 5.27700 |
| **Gpatch8** | 2.68873 | 7.32477 | 2.52527 | 4.17095 | 0.02711 | 4.96405 | 3.04170 | 1.32975 | 4.50537 | 4.87015 | 1.78325 | 2.92464 | 0.00000 | 4.90924 | 5.17181 | 0.81630 |
| **Gpbp1** | 8.62141 | 7.54943 | 5.93668 | 33.55019 | 9.89435 | 3.66973 | 7.66040 | 13.93309 | 35.08698 | 20.93000 | 14.09140 | 18.68788 | 6.47064 | 30.87579 | 31.94881 | 5.10199 |
| **Gpbp1l1** | 1.53211 | 0.31195 | 0.24792 | 3.56824 | 3.85724 | 0.20869 | 2.11616 | 0.58811 | 9.44512 | 2.32051 | 2.89817 | 0.12296 | 0.71640 | 4.95059 | 7.76192 | 1.66673 |
| **Gpc2** | 0.01374 | 0.00808 | 0.01202 | 0.09285 | 0.00932 | 0.00661 | 0.00749 | 0.01316 | 0.00781 | 0.02701 | 0.01213 | 0.01824 | 0.01970 | 0.01139 | 0.00973 | 0.00737 |
| **Gpc3** | 0.01175 | 0.00711 | 0.00784 | 0.01010 | 0.00941 | 0.00686 | 1.13982 | 0.00966 | 0.01227 | 0.03069 | 0.01331 | 0.00659 | 0.01089 | 0.01149 | 0.01058 | 0.00754 |
| **Gpc4** | 1.51344 | 0.00577 | 0.00000 | 0.08823 | 0.78794 | 0.00463 | 0.95323 | 1.33939 | 2.90567 | 1.84085 | 1.86382 | 1.73028 | 0.16898 | 5.25148 | 0.79547 | 3.89215 |
| **Gpc6** | 0.03556 | 0.00740 | 0.01484 | 0.08995 | 0.00824 | 0.00742 | 0.07736 | 0.01544 | 0.12863 | 0.18472 | 0.01030 | 0.03413 | 0.04142 | 0.17981 | 0.08036 | 0.01516 |
| **Gpcpd1** | 4.40716 | 8.05527 | 8.92175 | 12.26707 | 6.83373 | 1.68494 | 12.85856 | 13.43574 | 17.84920 | 14.45483 | 17.98402 | 25.94085 | 2.83266 | 21.62958 | 21.66007 | 2.97815 |
| **Gpd1** | 0.01053 | 0.00779 | 0.01032 | 3.16673 | 0.00975 | 0.00641 | 0.00725 | 0.01071 | 1.96233 | 1.56660 | 0.00000 | 0.06939 | 0.33005 | 0.03138 | 2.13455 | 0.00729 |
| **Gpd1l** | 14.71683 | 12.10146 | 27.63109 | 21.16304 | 12.07225 | 16.27910 | 17.13303 | 49.79588 | 27.18047 | 44.82636 | 43.51721 | 25.38044 | 34.49545 | 42.62153 | 31.18325 | 4.67195 |

| | | | | | | | | | | | | | | | | |
|---|---|---|---|---|---|---|---|---|---|---|---|---|---|---|---|---|
| **Gpd2** | 9.52259 | 11.22344 | 14.02869 | 3.46517 | 14.35449 | 6.87571 | 11.53423 | 15.87275 | 4.91955 | 23.89027 | 18.99239 | 15.99205 | 16.01332 | 29.75338 | 14.69999 | 9.07794 |
| **Gpha2** | 0.00832 | 0.04109 | 0.91961 | 0.55254 | 0.00994 | 0.00665 | 0.00559 | 0.17433 | 0.12577 | 0.05255 | 0.84459 | 0.02513 | 2.68504 | 0.00466 | 0.00000 | 0.01317 |
| **Gphn** | 0.62873 | 0.12323 | 0.00000 | 1.26860 | 3.86724 | 4.85806 | 1.25568 | 0.05479 | 0.62209 | 0.61644 | 0.68130 | 0.04956 | 0.01291 | 0.01933 | 1.00393 | 0.31729 |
| **Gpi1** | 23.00942 | 13.02844 | 13.44027 | 5.16739 | 10.26410 | 5.76118 | 7.48104 | 10.85566 | 4.59484 | 7.55909 | 9.48648 | 0.97795 | 17.67966 | 0.94201 | 3.97790 | 9.60999 |
| **Gpkow** | 0.58860 | 1.21820 | 2.70727 | 5.40866 | 1.22924 | 0.45298 | 0.49472 | 5.93377 | 11.25757 | 0.56218 | 0.12205 | 2.24521 | 1.84701 | 5.90981 | 12.86009 | 1.06599 |
| **Gpld1** | 0.01404 | 0.00786 | 0.00495 | 0.04971 | 0.00853 | 0.01291 | 0.00679 | 0.38303 | 0.11887 | 0.00987 | 0.00642 | 0.06940 | 1.41289 | 0.00805 | 0.99987 | 0.07963 |
| **Gpm6a** | 0.01858 | 0.00773 | 0.01085 | 1.57710 | 0.01066 | 0.00633 | 0.00729 | 0.01006 | 0.00000 | 0.00310 | 0.04585 | 0.00606 | 0.01206 | 0.00769 | 0.69460 | 0.00713 |
| **Gpn1** | 13.09341 | 9.48220 | 10.55151 | 6.13318 | 0.04970 | 4.07380 | 3.99003 | 6.52761 | 6.63156 | 12.39985 | 5.33892 | 1.43996 | 4.18995 | 5.01219 | 12.99811 | 3.32714 |
| **Gpn2** | 0.33883 | 1.31416 | 0.77847 | 1.23776 | 1.76222 | 0.40396 | 0.24861 | 1.84463 | 0.49633 | 0.06706 | 0.95211 | 0.07285 | 0.54184 | 0.20998 | 1.86547 | 0.14873 |
| **Gpn3** | 5.74278 | 2.71323 | 7.25888 | 12.33365 | 3.66528 | 0.57516 | 2.06439 | 14.15506 | 17.89070 | 10.47544 | 10.99015 | 10.01754 | 4.97010 | 0.55223 | 3.94969 | 0.09112 |
| **Gpnmb** | 0.01379 | 0.00801 | 0.01187 | 0.76022 | 0.00933 | 0.00661 | 0.00748 | 0.01339 | 0.01693 | 0.01423 | 0.01281 | 0.02848 | 0.01274 | 0.01166 | 0.01019 | 0.00729 |
| **Gpr1** | 0.01377 | 0.00778 | 0.01197 | 0.65518 | 0.00935 | 0.00645 | 0.00740 | 0.01180 | 0.04871 | 0.00578 | 0.02955 | 0.00690 | 0.01230 | 0.00516 | 0.35897 | 0.00724 |
| **Gpr107** | 5.75581 | 0.00000 | 6.67264 | 1.63006 | 0.00000 | 1.74833 | 2.98313 | 6.84572 | 1.77103 | 4.60072 | 0.90238 | 0.15841 | 3.45397 | 0.00000 | 3.73171 | 1.58696 |
| **Gpr108** | 5.95408 | 1.95441 | 6.67014 | 0.00000 | 6.36258 | 6.36334 | 3.89481 | 0.62993 | 0.00546 | 2.40137 | 0.71442 | 0.01521 | 6.41023 | 0.00570 | 0.21839 | 6.07752 |
| **Gpr137** | 0.19240 | 0.00789 | 3.77332 | 0.68306 | 0.00000 | 0.28636 | 0.00730 | 1.06621 | 0.25285 | 0.59934 | 0.41091 | 0.10776 | 0.40983 | 0.00594 | 0.00000 | 3.03463 |
| **Gpr137b** | 2.36051 | 2.83356 | 12.88404 | 4.42330 | 2.04457 | 9.29404 | 5.97591 | 18.52658 | 7.69588 | 20.27109 | 14.70407 | 13.32042 | 16.90993 | 14.74287 | 7.21200 | 2.81145 |
| **Gpr137b.ps** | 1.63828 | 2.03615 | 8.87138 | 5.88422 | 1.25618 | 8.92825 | 4.46151 | 15.07233 | 5.18163 | 15.31520 | 12.87914 | 9.37632 | 14.06428 | 8.30017 | 3.86707 | 0.71118 |
| **Gpr143** | 0.01393 | 0.00795 | 0.00921 | 1.26073 | 0.00909 | 0.00662 | 0.00744 | 0.00444 | 0.00000 | 0.00000 | 0.43222 | 0.00555 | 0.00785 | 0.00356 | 0.14834 | 0.00693 |
| **Gpr146** | 1.81585 | 0.00570 | 0.00000 | 1.13887 | 0.00443 | 0.01407 | 0.00586 | 0.00200 | 1.11137 | 0.00000 | 0.25961 | 0.00219 | 0.00896 | 0.00334 | 0.36677 | 0.01888 |
| **Gpr155** | 0.01500 | 0.00761 | 0.88277 | 0.09623 | 0.00768 | 0.00546 | 0.00716 | 0.00938 | 1.24738 | 0.00000 | 0.00000 | 0.23229 | 0.00867 | 0.01233 | 0.00000 | 0.00709 |
| **Gpr156** | 0.00956 | 0.00709 | 1.88191 | 0.28494 | 0.00661 | 0.00799 | 1.41713 | 0.00804 | 1.09961 | 0.21786 | 0.00000 | 0.00016 | 0.00944 | 0.00428 | 0.91623 | 0.00646 |
| **Gpr157** | 0.18886 | 0.07238 | 0.34483 | 1.19127 | 0.12224 | 0.02716 | 0.09896 | 0.29524 | 1.94774 | 0.87651 | 2.09577 | 0.64396 | 0.22340 | 1.40653 | 0.72613 | 0.08576 |
| **Gpr160** | 0.00000 | 2.20709 | 2.31929 | 0.21649 | 0.00402 | 0.01785 | 0.01550 | 0.64864 | 0.35037 | 3.60659 | 1.17447 | 0.17119 | 1.03128 | 0.73325 | 3.35260 | 0.01210 |
| **Gpr161** | 2.41641 | 2.22249 | 8.57389 | 9.66456 | 3.57008 | 1.69419 | 9.93237 | 17.58475 | 13.37363 | 9.74490 | 17.15351 | 10.68072 | 7.90487 | 16.60134 | 13.78874 | 2.19038 |
| **Gpr171** | 0.00021 | 0.00739 | 0.00843 | 0.01735 | 0.02300 | 2.90313 | 0.00687 | 1.36149 | 0.01050 | 0.00000 | 0.00000 | 0.01999 | 0.11531 | 3.39418 | 0.13880 | 0.00623 |
| **Gpr173** | 0.01102 | 0.01883 | 0.01516 | 0.02174 | 0.03658 | 0.01254 | 0.00739 | 0.02117 | 0.04084 | 0.04754 | 0.05442 | 0.00729 | 0.02469 | 0.04683 | 0.02870 | 0.00743 |
| **Gpr174** | 0.07340 | 0.00787 | 0.02731 | 0.03250 | 0.01780 | 0.03989 | 0.02800 | 0.02412 | 0.01135 | 0.05099 | 0.04098 | 0.00733 | 0.03004 | 0.05915 | 0.03953 | 0.02153 |
| **Gpr176** | 0.01380 | 0.00775 | 0.01176 | 0.06302 | 0.00934 | 0.00658 | 0.00744 | 0.01262 | 0.01146 | 0.00874 | 0.01300 | 0.03639 | 0.01272 | 0.00632 | 0.18168 | 0.00737 |
| **Gpr180** | 0.33146 | 0.00726 | 0.47664 | 2.38834 | 0.01593 | 0.00503 | 0.04838 | 3.84201 | 1.19529 | 1.13376 | 0.04925 | 0.00000 | 0.04247 | 0.01405 | 1.78434 | 0.00632 |
| **Gpr19** | 2.40930 | 1.59122 | 2.01076 | 1.87944 | 0.00362 | 0.95048 | 0.01460 | 0.59137 | 0.00000 | 0.24184 | 1.31302 | 0.00000 | 0.00000 | 0.20346 | 0.45211 | 0.12693 |
| **Gpr20** | 0.01551 | 0.00834 | 0.03973 | 0.01056 | 0.01167 | 0.01697 | 0.00755 | 0.01538 | 0.01235 | 0.00774 | 0.01501 | 0.00764 | 0.40470 | 0.01285 | 0.00973 | 0.00755 |
| **Gpr3** | 0.01088 | 0.00765 | 0.00586 | 3.10659 | 0.00913 | 0.00625 | 0.00693 | 0.05145 | 4.96972 | 0.23081 | 0.40230 | 0.05483 | 0.00678 | 0.02845 | 0.38352 | 0.00643 |
| **Gpr35** | 0.01540 | 0.00781 | 0.05999 | 1.14319 | 0.00931 | 0.00610 | 0.00735 | 0.01119 | 0.02755 | 0.01709 | 0.05276 | 0.00729 | 0.01311 | 0.07119 | 0.01166 | 0.01453 |
| **Gpr4** | 0.00000 | 0.00717 | 0.18706 | 0.04624 | 0.02839 | 0.04658 | 1.67548 | 1.28793 | 0.01048 | 0.60184 | 0.08027 | 0.02690 | 1.68967 | 0.02131 | 0.00000 | 2.82562 |
| **Gpr50** | 0.01161 | 0.00755 | 0.00219 | 6.82424 | 0.00877 | 0.00534 | 0.00692 | 0.00000 | 12.48282 | 0.32513 | 0.28137 | 0.72082 | 0.01112 | 0.00000 | 1.37552 | 0.01267 |
| **Gpr68** | 0.03893 | 0.00784 | 0.04739 | 1.41420 | 0.00897 | 0.00632 | 0.01786 | 0.01485 | 0.68081 | 0.00532 | 0.14717 | 0.00469 | 0.05531 | 0.03576 | 0.53786 | 0.00688 |
| **Gpr75** | 0.01369 | 0.00792 | 0.01145 | 0.54931 | 0.00939 | 0.00638 | 0.00734 | 0.01122 | 0.35634 | 0.00410 | 0.00962 | 0.00623 | 0.01277 | 0.00752 | 0.73314 | 0.00721 |
| **Gpr85** | 0.01215 | 0.00686 | 0.01634 | 0.00869 | 0.00933 | 0.00629 | 1.35515 | 0.00932 | 0.01179 | 0.25572 | 0.01272 | 0.00560 | 0.01089 | 0.01017 | 0.08191 | 0.00769 |
| **Gpr89** | 5.51910 | 3.87935 | 4.10424 | 0.41672 | 2.50682 | 4.03040 | 5.42578 | 3.39121 | 0.31344 | 0.65611 | 1.64973 | 0.04648 | 5.57323 | 2.18187 | 0.66767 | 1.26005 |
| **Gprasp1** | 1.97300 | 0.00629 | 0.91202 | 0.56871 | 0.00709 | 0.00394 | 0.00630 | 0.67510 | 0.95927 | 1.12364 | 0.35586 | 0.88333 | 0.00000 | 0.00898 | 1.49185 | 0.00474 |
| **Gprasp2** | 0.01775 | 0.00817 | 0.01178 | 0.35736 | 0.00895 | 0.00661 | 0.00751 | 0.01421 | 0.51469 | 0.01244 | 0.01069 | 0.00752 | 0.01279 | 0.01144 | 0.00724 | 0.00730 |
| **Gprc5a** | 4.16309 | 1.71503 | 15.04283 | 0.18029 | 2.71446 | 9.42106 | 8.98785 | 17.27628 | 0.01213 | 7.61485 | 5.06835 | 0.06901 | 13.32946 | 2.62594 | 1.62543 | 5.00660 |
| **Gprc5b** | 0.02016 | 0.00782 | 0.05885 | 0.01037 | 0.00888 | 0.00591 | 0.00744 | 0.01365 | 0.01249 | 0.01415 | 0.01306 | 0.00767 | 0.01176 | 0.00929 | 0.03336 | 0.00733 |
| **Gprc5c** | 0.00000 | 0.05785 | 4.10398 | 2.44468 | 0.12468 | 0.00601 | 1.54962 | 8.50380 | 1.31556 | 8.03276 | 6.21901 | 0.38550 | 3.39309 | 7.72070 | 8.97689 | 0.00994 |
| **Gprin1** | 0.01356 | 0.00798 | 0.01211 | 0.01059 | 0.00932 | 0.00666 | 0.00747 | 0.01832 | 0.01234 | 0.01873 | 0.01314 | 0.00792 | 0.01246 | 0.01208 | 0.01067 | 0.00748 |
| **Gps1** | 15.31262 | 11.23053 | 12.00101 | 16.37326 | 11.29485 | 11.87555 | 9.71445 | 10.52415 | 16.50015 | 15.15693 | 10.65035 | 10.30441 | 9.80123 | 0.16714 | 12.41390 | 10.44545 |
| **Gps2** | 7.58188 | 0.75018 | 3.86423 | 7.08430 | 3.73181 | 0.50069 | 1.14417 | 9.60144 | 4.66443 | 9.16141 | 4.52461 | 6.37299 | 9.99831 | 0.02821 | 1.10359 | 1.74397 |
| **Gpsm1** | 0.00000 | 0.00673 | 0.40234 | 0.00352 | 0.00751 | 0.00909 | 0.00587 | 6.05798 | 0.00699 | 3.88744 | 2.62804 | 0.06496 | 0.95899 | 0.43179 | 0.36003 | 0.02064 |
| **Gpsm2** | 2.53091 | 0.00000 | 0.28707 | 2.59823 | 3.44965 | 4.94598 | 0.92814 | 5.51557 | 1.45362 | 3.38694 | 1.58875 | 3.08076 | 3.39878 | 2.42010 | 1.43140 | 0.01441 |
| **Gpsm3** | 0.00829 | 0.00769 | 0.04747 | 0.01562 | 0.02068 | 0.00793 | 0.05923 | 0.53289 | 0.01196 | 0.31281 | 0.02591 | 0.00525 | 0.07583 | 0.01351 | 0.00091 | 0.00776 |
| **Gpt** | 0.00649 | 0.00788 | 1.41726 | 0.61685 | 0.00715 | 0.00529 | 0.03736 | 0.71058 | 1.24386 | 0.38773 | 0.00000 | 0.02782 | 0.54890 | 0.01485 | 0.02386 | 0.00617 |
| **Gpt2** | 0.16589 | 2.73102 | 0.91679 | 4.98191 | 6.66739 | 0.00703 | 1.27348 | 0.87786 | 6.69402 | 3.33112 | 0.92532 | 1.32439 | 0.13541 | 1.18241 | 10.69019 | 0.00441 |
| **Gpx1** | 32.76585 | 4.83781 | 17.39982 | 15.35907 | 19.58406 | 15.44386 | 7.76758 | 28.24476 | 4.84302 | 30.92400 | 25.88794 | 3.51474 | 29.61838 | 0.13963 | 0.98530 | 16.06435 |
| **Gpx2** | 6.06170 | 0.01758 | 0.64827 | 0.20888 | 0.00000 | 0.00265 | 8.56555 | 0.00000 | 0.00959 | 0.00000 | 1.33269 | 0.02260 | 0.08266 | 0.01489 | 0.05949 | 1.25601 |
| **Gpx2.ps1** | 1.36030 | 0.00484 | 0.13952 | 0.03065 | 0.00613 | 0.00764 | 2.74469 | 0.01880 | 0.28987 | 0.00000 | 0.23102 | 0.00344 | 0.03440 | 0.00996 | 0.04503 | 0.36337 |
| **Gpx3** | 1.72164 | 0.03154 | 0.07833 | 6.76060 | 0.21376 | 0.00440 | 0.01743 | 0.98833 | 0.19308 | 5.40890 | 0.18695 | 0.31775 | 0.55910 | 0.00000 | 0.62176 | 2.45865 |
| **Gpx4** | 28.07105 | 9.73527 | 19.27339 | 0.27005 | 16.67145 | 13.68015 | 13.52144 | 28.53013 | 1.28736 | 22.98575 | 7.35125 | 2.77320 | 29.70650 | 0.00000 | 0.00000 | 13.44559 |
| **Gpx5** | 0.03343 | 0.00840 | 0.39750 | 0.01077 | 0.01391 | 0.00941 | 0.00718 | 0.01208 | 0.01230 | 0.00596 | 0.01326 | 0.00734 | 0.57050 | 0.01479 | 0.00365 | 0.00719 |
| **Gpx6** | 0.01021 | 0.00725 | 0.00686 | 0.05024 | 0.00959 | 0.00674 | 0.00709 | 0.77627 | 0.02204 | 0.00000 | 0.00556 | 0.00506 | 0.00542 | 0.02428 | 0.46694 | 0.00725 |
| **Gpx7** | 1.30261 | 0.00470 | 1.77292 | 3.44619 | 0.01101 | 0.02757 | 1.59319 | 0.00000 | 11.60108 | 0.21570 | 1.60193 | 0.00000 | 1.99077 | 0.79568 | 0.00000 | 0.01731 |
| **Gramd1a** | 1.74541 | 0.02520 | 0.72599 | 0.00799 | 0.00333 | 0.00630 | 1.52734 | 0.00452 | 0.01189 | 0.47810 | 0.01072 | 0.00431 | 0.00914 | 0.01134 | 0.02150 | 0.01963 |
| **Gramd1b** | 0.12901 | 0.00738 | 0.00301 | 0.08757 | 0.00754 | 0.00751 | 0.00704 | 0.00000 | 0.88383 | 0.00000 | 0.45233 | 0.00186 | 0.00790 | 0.56744 | 0.00000 | 0.00685 |

| | | | | | | | | | | | | | | | | |
|---|---|---|---|---|---|---|---|---|---|---|---|---|---|---|---|---|
| **Gramd1c** | 0.01166 | 0.00746 | 0.00324 | 0.08525 | 0.00957 | 0.00645 | 0.00703 | 0.00483 | 0.85969 | 0.00000 | 0.09988 | 0.00440 | 0.01133 | 3.23538 | 0.00000 | 0.00715 |
| **Gramd2** | 0.01202 | 0.00775 | 1.49757 | 1.19145 | 0.00682 | 0.00537 | 0.00736 | 0.01369 | 0.66037 | 0.00000 | 0.09635 | 0.01639 | 0.00612 | 0.01627 | 0.13870 | 0.00646 |
| **Gramd3** | 0.01236 | 0.00790 | 0.01221 | 0.00561 | 0.00864 | 0.00659 | 0.00732 | 0.01005 | 0.01117 | 2.71529 | 0.00965 | 0.00000 | 0.01056 | 0.01230 | 0.00730 | 0.00731 |
| **Gramd4** | 1.78661 | 2.66344 | 0.50757 | 0.93208 | 0.00440 | 0.00720 | 2.10716 | 0.00000 | 0.00000 | 0.06134 | 0.51088 | 0.01920 | 0.38629 | 0.12754 | 0.12266 | 0.00399 |
| **Grap** | 0.01439 | 0.00791 | 0.02019 | 0.45980 | 0.00925 | 0.00620 | 0.00744 | 0.01781 | 0.74942 | 0.01520 | 0.02039 | 0.00646 | 0.01205 | 0.00863 | 0.03978 | 0.00723 |
| **Grasp** | 0.40737 | 0.00694 | 0.30649 | 0.29415 | 0.00727 | 0.02644 | 0.00678 | 0.31311 | 0.35336 | 0.00755 | 0.00818 | 0.00562 | 0.00460 | 0.00898 | 0.02490 | 0.00674 |
| **Grb10** | 1.43296 | 0.00345 | 0.26489 | 2.08118 | 4.06163 | 0.00716 | 2.24555 | 0.00000 | 0.45540 | 1.11854 | 0.30542 | 0.04730 | 0.01877 | 3.23862 | 2.30801 | 0.00512 |
| **Grb14** | 0.01437 | 0.00775 | 0.00383 | 0.00906 | 0.00912 | 2.81950 | 0.00707 | 0.01031 | 0.01261 | 0.02756 | 0.00813 | 0.13458 | 0.17230 | 0.01443 | 0.00801 | 0.00616 |
| **Grb2** | 7.56841 | 2.73672 | 5.02088 | 8.45524 | 1.18791 | 6.68246 | 3.59026 | 15.18000 | 12.73375 | 17.03584 | 8.40351 | 2.29024 | 4.89242 | 7.36703 | 15.10569 | 2.68863 |
| **Grb7** | 1.61239 | 0.36026 | 0.00000 | 0.00000 | 0.00566 | 2.69949 | 0.00488 | 0.00000 | 0.01120 | 0.00000 | 1.18296 | 0.03973 | 1.21579 | 0.01000 | 0.00000 | 0.18472 |
| **Grcc10** | 4.58624 | 0.01248 | 0.36228 | 1.88308 | 0.23217 | 3.17524 | 0.55603 | 0.00000 | 0.18483 | 2.65811 | 0.27331 | 0.05786 | 1.19911 | 0.02775 | 0.08446 | 1.28218 |
| **Greb1** | 0.01208 | 0.00761 | 2.71201 | 0.43987 | 0.00494 | 0.00543 | 0.02841 | 0.01372 | 0.01962 | 0.49214 | 0.00870 | 0.01445 | 0.00563 | 0.03278 | 0.20167 | 0.00635 |
| **Greb1l** | 0.15002 | 0.00628 | 3.75576 | 0.33536 | 0.00280 | 0.00627 | 0.84354 | 4.75258 | 0.01941 | 4.22240 | 1.60884 | 0.67148 | 3.40781 | 15.38548 | 5.72329 | 0.01845 |
| **Grhl1** | 0.01536 | 0.00768 | 0.01031 | 1.93297 | 0.00905 | 4.88204 | 0.00684 | 0.00000 | 0.05191 | 0.14153 | 0.28301 | 0.01520 | 0.02100 | 0.00598 | 1.17020 | 0.01219 |
| **Grhl2** | 3.96991 | 1.14988 | 5.93148 | 10.00790 | 0.16484 | 4.61167 | 0.02107 | 3.00369 | 7.45203 | 2.01442 | 1.92759 | 0.28125 | 3.23630 | 4.95612 | 6.31849 | 2.59480 |
| **Grhl3** | 0.02148 | 0.00827 | 0.06456 | 0.00264 | 0.00880 | 0.00575 | 0.22500 | 0.11571 | 0.39678 | 0.10818 | 0.59157 | 0.01412 | 0.00466 | 0.06473 | 0.88055 | 0.00720 |
| **Grhpr** | 9.72631 | 0.02224 | 6.50371 | 0.07302 | 3.87106 | 5.04448 | 5.43298 | 7.02591 | 0.50230 | 5.15822 | 4.18880 | 0.98660 | 4.46480 | 0.00000 | 0.09966 | 3.53185 |
| **Gria4** | 0.01372 | 0.00763 | 0.01136 | 0.99737 | 0.00940 | 0.00638 | 0.00731 | 0.01104 | 0.00000 | 0.01056 | 0.01040 | 0.00565 | 0.01310 | 0.00775 | 0.92528 | 0.00735 |
| **Grid1** | 0.01349 | 0.01911 | 0.01167 | 0.12095 | 0.00927 | 0.00667 | 0.00749 | 0.01275 | 0.00711 | 0.01347 | 0.02754 | 0.01824 | 0.01242 | 0.03258 | 0.00674 | 0.00739 |
| **Grid2** | 0.01154 | 0.00784 | 0.00820 | 0.00905 | 0.00921 | 0.00667 | 0.00700 | 0.00878 | 0.01200 | 0.08959 | 0.00899 | 0.00442 | 0.28846 | 0.65768 | 0.17205 | 0.00751 |
| **Grid2ip** | 0.00846 | 0.00770 | 0.00813 | 0.00749 | 0.00930 | 0.00683 | 0.00720 | 0.71160 | 0.01129 | 0.93748 | 0.00632 | 0.00380 | 0.00530 | 0.01073 | 0.01032 | 0.00715 |
| **Grik1** | 0.01384 | 0.00795 | 0.01001 | 0.00572 | 0.00900 | 0.00668 | 0.00733 | 0.00309 | 0.01155 | 0.38734 | 0.28795 | 0.01530 | 0.00878 | 0.00924 | 0.00872 | 0.00715 |
| **Grik5** | 0.01356 | 0.00792 | 0.01223 | 0.27287 | 0.00928 | 0.00644 | 0.00732 | 0.01175 | 0.05639 | 0.28390 | 0.01328 | 0.00686 | 0.02518 | 0.01056 | 0.00509 | 0.00738 |
| **Grin1** | 0.00729 | 0.00808 | 0.01693 | 0.00576 | 0.00828 | 0.00621 | 0.00738 | 0.01754 | 0.01118 | 2.38388 | 0.01078 | 0.00002 | 0.51488 | 0.01777 | 0.00498 | 0.35316 |
| **Grin2d** | 0.00963 | 0.00781 | 0.00741 | 0.01696 | 0.00921 | 0.00744 | 0.00772 | 1.24560 | 0.11956 | 2.55940 | 0.07319 | 0.01916 | 0.01158 | 0.02900 | 0.10652 | 0.00752 |
| **Grin3a** | 0.01378 | 0.00792 | 0.01153 | 0.33735 | 0.00931 | 0.01289 | 0.00750 | 0.01377 | 0.00424 | 0.01281 | 0.01208 | 0.00777 | 0.01892 | 0.00862 | 0.05465 | 0.00731 |
| **Grin3b** | 0.00702 | 0.00787 | 0.06391 | 0.01017 | 0.00889 | 0.00831 | 0.01667 | 0.00965 | 0.01238 | 0.00905 | 0.02400 | 0.00685 | 1.48280 | 0.01251 | 0.00934 | 0.00647 |
| **Grina** | 0.51040 | 2.84801 | 0.00000 | 4.38263 | 0.03580 | 0.01619 | 0.02634 | 0.84051 | 1.06271 | 2.68183 | 0.18761 | 0.05570 | 0.80296 | 0.00000 | 0.70397 | 0.01362 |
| **Grip1** | 0.17937 | 0.00772 | 0.00398 | 0.34795 | 0.50677 | 0.00699 | 0.00713 | 0.01364 | 0.00411 | 0.07795 | 0.01030 | 0.00527 | 0.01086 | 0.00628 | 0.59179 | 0.00741 |
| **Gripap1** | 0.00000 | 0.00701 | 0.21031 | 1.54973 | 0.01173 | 0.81346 | 2.39458 | 2.40499 | 3.21287 | 0.16848 | 2.12125 | 0.17381 | 0.22480 | 0.00000 | 2.93378 | 0.00612 |
| **Grk1** | 0.12956 | 0.01814 | 0.04504 | 0.63259 | 0.03320 | 0.04229 | 0.06833 | 0.13834 | 0.52076 | 0.01751 | 0.15065 | 0.00406 | 0.01848 | 0.32837 | 0.40977 | 0.02127 |
| **Grk4** | 0.32070 | 0.62095 | 0.64112 | 2.25948 | 0.36734 | 0.04540 | 0.36884 | 0.18360 | 2.88866 | 1.20539 | 0.18649 | 1.02854 | 0.47321 | 3.87928 | 0.73449 | 0.15949 |
| **Grk5** | 2.20171 | 0.00389 | 1.44318 | 3.28619 | 0.38059 | 0.00617 | 1.71245 | 3.74201 | 6.53258 | 8.80976 | 1.39544 | 0.42645 | 0.05649 | 4.03011 | 7.75549 | 0.01086 |
| **Grk6** | 6.30455 | 5.52420 | 5.71082 | 0.32793 | 10.63646 | 5.20676 | 4.09513 | 7.43999 | 0.49269 | 4.69913 | 7.24822 | 5.48101 | 3.66109 | 0.55841 | 0.95702 | 3.44579 |
| **Grm2** | 0.01374 | 0.00788 | 0.01183 | 1.19508 | 0.00936 | 0.00645 | 0.00741 | 0.01252 | 0.33733 | 0.00652 | 0.01020 | 0.01727 | 0.01274 | 0.00598 | 0.22390 | 0.00714 |
| **Grm4** | 0.01347 | 0.00811 | 0.01118 | 0.07771 | 0.00916 | 0.00655 | 0.00747 | 0.01271 | 0.78142 | 0.00805 | 0.04265 | 0.01750 | 0.03725 | 0.01283 | 0.00374 | 0.01451 |
| **Grm8** | 0.01378 | 0.00781 | 0.01195 | 0.15255 | 0.00939 | 0.00654 | 0.00743 | 0.01231 | 0.00710 | 0.00734 | 0.01196 | 0.00734 | 0.01274 | 0.00974 | 0.30573 | 0.00735 |
| **Grn** | 8.70077 | 5.96232 | 12.62140 | 1.63666 | 5.78672 | 11.55198 | 16.90099 | 33.59950 | 2.64778 | 24.57674 | 29.81184 | 11.79529 | 26.92291 | 6.00703 | 4.62397 | 2.44347 |
| **Grpel1** | 24.05106 | 12.65919 | 19.06692 | 6.16436 | 9.03270 | 10.27577 | 10.10276 | 19.35129 | 7.98505 | 21.81081 | 13.63845 | 7.59107 | 21.86931 | 3.05773 | 5.47850 | 8.65303 |
| **Grpel2** | 2.98155 | 4.36800 | 6.76358 | 2.62860 | 0.17273 | 4.18289 | 4.47726 | 7.81756 | 1.72596 | 4.71681 | 3.92711 | 1.50963 | 0.81363 | 0.73538 | 1.79766 | 0.78901 |
| **Grrp1** | 0.01364 | 0.00818 | 0.01199 | 0.33095 | 0.00927 | 0.00659 | 0.00751 | 0.01376 | 0.78175 | 0.01520 | 0.00888 | 0.00692 | 0.01295 | 0.01166 | 0.00395 | 0.00739 |
| **Grsf1** | 14.79728 | 9.41043 | 7.72450 | 20.08601 | 8.77474 | 8.76477 | 1.96124 | 5.67072 | 18.25954 | 13.13380 | 7.24392 | 14.97797 | 8.25877 | 17.69764 | 21.23311 | 0.40489 |
| **Grtp1** | 8.28145 | 0.30916 | 3.78163 | 0.41150 | 7.35721 | 6.30457 | 3.21123 | 6.06865 | 0.00000 | 6.59832 | 3.76189 | 6.60547 | 5.84120 | 0.00000 | 0.00000 | 2.33889 |
| **Grwd1** | 15.25731 | 10.91156 | 15.36977 | 9.73380 | 13.73632 | 6.76439 | 9.98794 | 18.52446 | 12.16801 | 21.42976 | 8.43453 | 3.49768 | 14.73778 | 1.35030 | 8.61478 | 6.43235 |
| **Grxcr2** | 0.17767 | 0.00829 | 0.00561 | 0.00491 | 0.01718 | 0.00727 | 0.20841 | 0.01035 | 0.30461 | 0.01283 | 0.01041 | 0.00728 | 0.01191 | 0.01299 | 0.00829 | 0.00742 |
| **Gsap** | 0.00908 | 0.35812 | 0.00972 | 0.00497 | 0.00839 | 0.00670 | 0.00724 | 0.01088 | 0.15937 | 0.01511 | 0.01124 | 0.02725 | 0.01195 | 0.01078 | 0.00469 | 0.00719 |
| **Gsc** | 2.46449 | 0.00487 | 0.00333 | 0.00969 | 0.00332 | 0.00586 | 0.08744 | 0.00304 | 0.02381 | 0.00719 | 0.01216 | 0.00601 | 0.42303 | 0.01201 | 0.01571 | 0.01856 |
| **Gsdma** | 0.83863 | 0.00789 | 0.01152 | 0.52544 | 0.01993 | 0.00803 | 0.00667 | 0.42248 | 0.10836 | 0.93742 | 0.08819 | 0.01352 | 0.72515 | 0.00425 | 0.36278 | 0.03788 |
| **Gsdma2** | 0.01709 | 0.00888 | 0.01111 | 0.03181 | 0.01092 | 0.00684 | 0.00806 | 0.04551 | 1.10836 | 0.00737 | 0.00636 | 0.00689 | 0.02659 | 0.03988 | 0.00160 | 0.00751 |
| **Gsdma3** | 0.04369 | 0.00943 | 0.00666 | 0.01866 | 0.01579 | 0.00745 | 0.00807 | 0.58377 | 0.01078 | 0.00000 | 0.00794 | 0.00690 | 0.43422 | 0.22919 | 0.02436 | 0.00743 |
| **Gsdmc** | 0.30904 | 0.00784 | 0.14217 | 0.41278 | 0.00906 | 0.00621 | 0.08782 | 0.08984 | 0.08187 | 0.00015 | 0.13330 | 0.00656 | 0.17835 | 0.05282 | 0.01290 | 0.00678 |
| **Gsdmc2** | 1.90444 | 0.00545 | 1.45793 | 2.48974 | 0.00260 | 0.00387 | 0.41937 | 0.86694 | 1.08587 | 0.26991 | 0.25717 | 0.03527 | 1.97144 | 0.80866 | 0.65690 | 0.01032 |
| **Gsdmc3** | 0.30884 | 0.00769 | 0.40650 | 1.37224 | 0.00775 | 0.00581 | 0.13809 | 0.31666 | 0.36063 | 0.74021 | 0.07209 | 0.02959 | 0.51477 | 0.64815 | 1.23432 | 0.00634 |
| **Gsdmc4** | 0.46897 | 0.00760 | 0.42143 | 1.57711 | 0.00773 | 0.00559 | 0.14796 | 0.18961 | 0.31224 | 0.05093 | 0.41188 | 0.02421 | 0.48643 | 0.10264 | 0.14069 | 0.00596 |
| **Gsdmd** | 0.50326 | 0.64229 | 0.53510 | 0.46854 | 0.20876 | 0.00852 | 0.01637 | 0.00000 | 0.84803 | 0.00000 | 0.33988 | 0.00368 | 0.01427 | 0.00567 | 0.00000 | 0.04314 |
| **Gse1** | 0.71804 | 0.00682 | 0.80572 | 4.80962 | 0.01150 | 0.00639 | 0.00533 | 3.43497 | 2.08629 | 4.53930 | 1.62376 | 0.27850 | 1.06226 | 0.00000 | 0.57701 | 0.01921 |
| **Gsg1** | 0.01693 | 0.00796 | 0.01093 | 0.00451 | 0.00916 | 0.00679 | 0.00744 | 0.02778 | 0.44177 | 0.01435 | 0.00905 | 0.05770 | 0.01464 | 0.01325 | 0.00550 | 0.00740 |
| **Gsg1l** | 0.03341 | 0.01804 | 0.10925 | 0.00895 | 0.02591 | 0.02429 | 0.00724 | 0.02314 | 0.19646 | 0.02787 | 0.00830 | 0.30317 | 0.03118 | 0.13835 | 0.10344 | 0.00744 |
| **Gsg2** | 0.00000 | 0.39585 | 2.58538 | 15.70550 | 3.67206 | 0.06419 | 4.16147 | 3.96786 | 10.14299 | 2.85814 | 2.16296 | 3.88755 | 3.08245 | 0.00000 | 8.59374 | 0.01549 |
| **Gsk3a** | 1.11027 | 0.05750 | 0.08690 | 7.65556 | 3.76015 | 0.05928 | 0.00483 | 0.00000 | 11.47047 | 0.76342 | 3.70866 | 0.33969 | 0.15964 | 0.26237 | 0.00000 | 0.01070 |
| **Gsk3b** | 0.78639 | 1.02789 | 2.18786 | 9.37710 | 0.24279 | 0.08549 | 1.99005 | 3.68755 | 8.20705 | 6.60978 | 3.30011 | 3.51811 | 2.64762 | 9.15696 | 7.08468 | 0.08824 |
| **Gskip** | 1.59789 | 0.15802 | 0.00000 | 4.02870 | 0.01463 | 1.59352 | 0.00452 | 0.41516 | 3.89932 | 1.73959 | 1.90309 | 0.19461 | 1.15592 | 3.96602 | 1.89883 | 0.01079 |

| | | | | | | | | | | | | | | | | |
|---|---|---|---|---|---|---|---|---|---|---|---|---|---|---|---|---|
| **Gsn** | 0.02694 | 3.34823 | 2.79528 | 0.00000 | 0.03918 | 5.79790 | 0.00000 | 0.01197 | 0.01065 | 4.62413 | 0.52284 | 0.05071 | 9.16080 | 0.00860 | 0.00000 | 0.02890 |
| **Gspt1** | 18.79631 | 16.78715 | 11.50834 | 7.13173 | 13.32252 | 9.70003 | 9.40987 | 15.13250 | 11.35096 | 19.98223 | 11.07634 | 19.38460 | 12.11954 | 28.81804 | 19.50039 | 6.24578 |
| **Gspt2** | 3.44353 | 2.41823 | 0.00000 | 2.25401 | 0.00001 | 0.00317 | 0.00373 | 0.27182 | 0.00000 | 2.26357 | 0.12030 | 0.13077 | 1.30845 | 0.01936 | 3.67418 | 0.00171 |
| **Gsr** | 6.49992 | 4.30022 | 2.51824 | 1.56124 | 4.60703 | 1.27529 | 2.69541 | 2.92146 | 0.80078 | 9.58019 | 6.27086 | 0.58436 | 2.81497 | 2.64177 | 4.56853 | 0.94808 |
| **Gss** | 12.32376 | 0.87122 | 8.77481 | 7.42926 | 13.02771 | 11.09366 | 7.58826 | 7.05787 | 3.26343 | 2.46580 | 0.62938 | 1.72338 | 9.21957 | 0.00674 | 0.06061 | 3.97673 |
| **Gsta1** | 0.59478 | 0.00680 | 0.00000 | 0.01860 | 0.00657 | 0.45664 | 0.10827 | 0.69928 | 0.00906 | 0.97848 | 1.41500 | 0.08648 | 1.25386 | 1.45510 | 0.00000 | 0.01276 |
| **Gsta2** | 0.62163 | 0.00673 | 0.00000 | 0.02604 | 0.01519 | 0.42845 | 0.41603 | 0.40703 | 0.00882 | 0.48831 | 1.10148 | 0.04456 | 0.91434 | 1.41266 | 0.22721 | 0.00595 |
| **Gsta3** | 2.22253 | 0.00559 | 0.00328 | 0.00999 | 0.00372 | 0.00660 | 0.00569 | 0.00773 | 0.01221 | 0.15229 | 0.01141 | 0.01787 | 0.06907 | 0.01154 | 0.01156 | 0.00490 |
| **Gsta4** | 10.29024 | 4.17826 | 6.46341 | 0.50731 | 6.35453 | 13.54514 | 3.22400 | 0.10937 | 0.36845 | 1.13511 | 2.10866 | 0.08769 | 10.94424 | 1.15551 | 2.36925 | 11.01591 |
| **Gstcd** | 0.03512 | 0.00817 | 0.29023 | 0.58109 | 0.00853 | 2.03384 | 0.00788 | 0.04683 | 1.68066 | 0.32139 | 0.17444 | 0.01521 | 0.03769 | 0.00409 | 0.19593 | 0.02149 |
| **Gstk1** | 0.00721 | 1.99996 | 0.00307 | 0.19198 | 0.00796 | 1.94759 | 0.00653 | 0.00980 | 0.00574 | 0.09400 | 0.00876 | 0.00808 | 0.00447 | 0.00554 | 0.00000 | 0.00585 |
| **Gstm1** | 7.55462 | 0.00000 | 0.48546 | 0.01759 | 4.45126 | 0.01980 | 2.49196 | 0.00000 | 0.23729 | 16.36803 | 0.00408 | 0.00000 | 0.01521 | 0.01488 | 0.00000 | 0.01362 |
| **Gstm2** | 2.75086 | 0.00486 | 0.00000 | 0.00000 | 0.00531 | 3.31735 | 0.00450 | 0.00299 | 0.18309 | 13.64261 | 0.00500 | 0.00000 | 0.60921 | 0.01401 | 0.00467 | 0.01040 |
| **Gstm3** | 1.38245 | 0.00471 | 0.05422 | 0.00362 | 0.44016 | 0.00438 | 2.52568 | 0.00429 | 0.01104 | 1.91252 | 0.01044 | 0.00000 | 0.01006 | 0.01303 | 0.00637 | 0.00579 |
| **Gstm4** | 3.71475 | 0.49839 | 2.12295 | 0.00342 | 0.00000 | 0.00723 | 0.00403 | 0.00000 | 0.78125 | 0.61247 | 0.00477 | 0.00000 | 0.00982 | 0.01152 | 0.00000 | 0.17822 |
| **Gstm5** | 0.01317 | 0.00764 | 0.01011 | 7.30781 | 0.00911 | 0.00578 | 0.00703 | 0.01022 | 0.00000 | 0.64118 | 0.00811 | 0.03132 | 0.01185 | 0.01176 | 1.54353 | 0.00636 |
| **Gstm6** | 2.65577 | 0.00464 | 0.19751 | 0.14150 | 0.82971 | 0.00936 | 0.15982 | 0.00252 | 0.00576 | 0.01752 | 0.01133 | 0.00557 | 0.00987 | 0.01215 | 0.00481 | 0.00439 |
| **Gstm7** | 1.60589 | 1.31127 | 0.77884 | 0.00000 | 2.99815 | 0.00385 | 0.00563 | 0.00000 | 0.00952 | 4.28674 | 0.00836 | 0.00000 | 0.00878 | 0.01108 | 0.00000 | 0.02000 |
| **Gsto1** | 15.86243 | 11.21661 | 19.05876 | 0.23920 | 23.34211 | 23.50333 | 16.97780 | 36.89981 | 1.75575 | 16.98852 | 11.71822 | 0.65946 | 40.29738 | 1.98509 | 0.00000 | 17.07542 |
| **Gstp1** | 26.37735 | 10.26765 | 20.95686 | 1.13851 | 16.93479 | 13.24004 | 7.78344 | 12.60348 | 0.00741 | 7.38724 | 11.86414 | 0.33676 | 19.43473 | 0.00000 | 0.83392 | 13.97593 |
| **Gstp2** | 22.50629 | 12.23862 | 23.92448 | 1.12989 | 16.88821 | 13.55358 | 8.94330 | 12.37483 | 0.00000 | 5.19268 | 11.12774 | 0.22118 | 17.35369 | 0.00000 | 0.27622 | 14.93033 |
| **Gstt2** | 0.01141 | 0.00803 | 0.06597 | 0.00581 | 0.00953 | 0.00642 | 0.01850 | 0.01288 | 0.01144 | 1.84895 | 0.01204 | 0.00274 | 0.08661 | 0.00703 | 0.09780 | 0.00783 |
| **Gstz1** | 5.67529 | 4.29059 | 5.78822 | 0.42416 | 0.04025 | 3.63462 | 0.91931 | 3.55570 | 0.16594 | 1.20023 | 1.52638 | 0.40059 | 6.50111 | 2.90076 | 0.25069 | 0.11009 |
| **Gt.ROSA.26S** | 0.21437 | 0.00781 | 0.78342 | 5.78349 | 0.00628 | 0.03857 | 0.00621 | 0.56582 | 2.12044 | 0.06419 | 0.70346 | 0.56071 | 2.41066 | 0.00000 | 1.56052 | 0.00460 |
| **Gtdc1** | 0.62912 | 2.46638 | 1.03786 | 14.49361 | 0.01138 | 0.03763 | 0.18999 | 0.96193 | 12.81086 | 2.76191 | 0.71052 | 0.70104 | 0.07016 | 1.97100 | 7.63146 | 0.05357 |
| **Gtf2a1** | 3.94567 | 0.21432 | 2.87869 | 15.56995 | 0.10702 | 2.94836 | 6.01874 | 4.30798 | 13.29298 | 0.00000 | 2.62544 | 5.40853 | 2.38424 | 13.02508 | 13.56620 | 0.15585 |
| **Gtf2a1l** | 0.01375 | 0.00812 | 0.01199 | 3.20223 | 0.00930 | 0.00636 | 0.00740 | 0.01329 | 0.22808 | 0.01176 | 0.02542 | 0.01777 | 0.01238 | 0.00802 | 0.00728 | 0.00685 |
| **Gtf2a2** | 12.76885 | 4.08893 | 3.77394 | 3.35290 | 5.68867 | 3.97403 | 4.31732 | 4.35107 | 0.66044 | 1.59085 | 2.21375 | 1.71078 | 6.22484 | 1.81736 | 0.59202 | 7.18729 |
| **Gtf2b** | 17.67007 | 8.83110 | 10.34055 | 6.53452 | 6.76632 | 2.24788 | 3.46831 | 5.33987 | 11.76493 | 4.07888 | 1.77456 | 1.37733 | 6.21273 | 2.56076 | 1.04757 | 8.99576 |
| **Gtf2e1** | 5.65817 | 8.91505 | 6.08905 | 5.75561 | 6.97077 | 0.64722 | 7.27939 | 5.80284 | 2.81694 | 3.91036 | 3.23547 | 1.29371 | 3.46253 | 1.96440 | 5.92389 | 3.34893 |
| **Gtf2e2** | 11.21792 | 7.79013 | 7.51844 | 3.50020 | 11.47587 | 3.45300 | 6.25381 | 7.59224 | 6.34910 | 5.80834 | 3.28441 | 8.09831 | 7.72456 | 6.91226 | 7.02227 | 7.57184 |
| **Gtf2f1** | 12.74452 | 9.44574 | 9.91555 | 12.22338 | 8.49822 | 8.08388 | 5.94468 | 21.88017 | 16.99848 | 20.68741 | 7.76343 | 5.48787 | 18.60047 | 0.00000 | 0.00000 | 12.15860 |
| **Gtf2f2** | 12.17457 | 6.94625 | 7.56859 | 10.59639 | 9.29414 | 2.54266 | 3.62041 | 16.90051 | 20.60663 | 13.60708 | 11.50708 | 8.09075 | 8.50099 | 4.11766 | 7.49499 | 4.94823 |
| **Gtf2h1** | 12.63561 | 8.08594 | 8.87822 | 10.88546 | 3.19832 | 2.98710 | 3.39382 | 4.43578 | 14.05258 | 12.44260 | 10.62489 | 1.97768 | 8.67427 | 18.56477 | 17.94427 | 4.83858 |
| **Gtf2h2** | 9.47509 | 8.33037 | 6.66990 | 8.03457 | 5.74505 | 9.16059 | 7.03897 | 13.49317 | 5.98526 | 15.08291 | 12.55962 | 19.04157 | 7.99822 | 10.33487 | 13.61538 | 6.68244 |
| **Gtf2h3** | 2.54048 | 2.07670 | 3.43665 | 0.88218 | 5.07707 | 0.01039 | 1.22858 | 4.67684 | 0.45051 | 4.13849 | 1.13794 | 0.17411 | 0.57909 | 3.30265 | 8.74684 | 2.00250 |
| **Gtf2h4** | 5.99584 | 0.00000 | 0.00000 | 0.01518 | 8.98878 | 3.57004 | 0.33295 | 6.52521 | 0.05009 | 16.15738 | 0.00000 | 0.12414 | 10.96849 | 0.00549 | 0.05883 | 4.18360 |
| **Gtf2h5** | 11.92652 | 8.17849 | 8.79905 | 9.95710 | 5.20628 | 4.48407 | 5.47923 | 8.00251 | 9.13185 | 6.64922 | 14.20784 | 3.31485 | 7.98639 | 6.87416 | 8.45908 | 1.05683 |
| **Gtf2i** | 5.76053 | 3.72670 | 2.00183 | 6.97734 | 0.01924 | 0.00000 | 0.61206 | 1.92000 | 4.63080 | 11.61060 | 3.15038 | 0.21362 | 3.54467 | 0.01758 | 4.06540 | 4.69544 |
| **Gtf2ird1** | 6.11587 | 6.25988 | 5.53875 | 3.05399 | 5.71658 | 3.49296 | 6.40283 | 3.45566 | 3.38588 | 2.67232 | 10.31061 | 3.65701 | 4.33971 | 1.32148 | 1.05731 | 2.86607 |
| **Gtf2ird2** | 0.01126 | 0.00751 | 0.00245 | 0.41260 | 0.00902 | 0.00741 | 0.00698 | 1.02041 | 1.54394 | 0.90973 | 0.61516 | 0.00545 | 0.00286 | 0.01789 | 0.51203 | 0.00714 |
| **Gtf3a** | 0.46937 | 0.11735 | 0.25559 | 0.00714 | 0.00674 | 0.00639 | 0.18708 | 0.11125 | 0.01167 | 0.96206 | 0.01084 | 0.00531 | 0.66891 | 0.01327 | 0.00369 | 0.01411 |
| **Gtf3c1** | 3.55049 | 1.02128 | 1.41757 | 14.04378 | 2.24463 | 5.61802 | 1.35807 | 6.26646 | 14.31924 | 15.27443 | 11.49631 | 1.28841 | 0.29518 | 0.00000 | 3.78552 | 2.03143 |
| **Gtf3c2** | 6.31207 | 7.85605 | 9.66785 | 16.66985 | 1.59730 | 0.82634 | 7.40545 | 10.18118 | 17.08733 | 13.20531 | 7.13087 | 1.15193 | 9.15872 | 17.76901 | 19.46958 | 5.93232 |
| **Gtf3c3** | 2.82680 | 3.67388 | 3.02010 | 6.60807 | 2.98685 | 0.00208 | 3.69404 | 3.12679 | 10.05999 | 2.92374 | 3.24331 | 1.05918 | 0.85708 | 12.79883 | 9.45903 | 0.02812 |
| **Gtf3c4** | 2.46056 | 3.85985 | 2.99150 | 12.66380 | 3.45692 | 2.85190 | 4.81665 | 6.35629 | 12.36619 | 6.79578 | 5.53662 | 8.97576 | 1.58109 | 11.41717 | 8.93242 | 0.01030 |
| **Gtf3c5** | 3.52872 | 2.23909 | 2.66291 | 14.94178 | 6.57074 | 0.00000 | 0.10227 | 1.77330 | 16.85360 | 8.10166 | 3.73326 | 2.29229 | 4.72065 | 0.56397 | 10.62373 | 1.24083 |
| **Gtf3c6** | 8.49022 | 3.31485 | 4.16317 | 9.22569 | 1.53570 | 0.52893 | 1.70879 | 2.35439 | 19.84924 | 5.59387 | 6.35588 | 0.93175 | 5.37730 | 4.67822 | 3.51600 | 1.59610 |
| **Gtpbp1** | 5.71140 | 4.83181 | 1.61601 | 1.26607 | 0.03809 | 2.19482 | 2.58856 | 2.53052 | 0.82454 | 3.01965 | 0.23554 | 0.08210 | 5.27248 | 0.01612 | 0.00000 | 0.16377 |
| **Gtpbp10** | 1.20949 | 1.71891 | 1.69357 | 2.10129 | 0.88417 | 0.16194 | 0.64540 | 2.34441 | 0.65822 | 1.03229 | 2.66224 | 4.56440 | 0.00000 | 0.84214 | 12.23124 | 0.11173 |
| **Gtpbp2** | 0.11107 | 0.00741 | 0.03127 | 0.78221 | 0.00793 | 0.00717 | 0.02755 | 0.10719 | 0.00000 | 0.05857 | 0.19206 | 0.28310 | 0.01604 | 0.15099 | 0.07304 | 0.06142 |
| **Gtpbp3** | 2.98346 | 0.00326 | 1.04100 | 1.02900 | 2.03072 | 0.00910 | 0.18018 | 1.43526 | 4.22435 | 1.74970 | 0.22446 | 0.07126 | 1.38793 | 0.26601 | 0.27205 | 0.34513 |
| **Gtpbp4** | 11.32072 | 18.46756 | 14.03327 | 14.90939 | 10.59029 | 6.60990 | 12.40337 | 15.55862 | 17.85837 | 18.05375 | 11.54000 | 8.27981 | 9.18334 | 25.88862 | 19.51335 | 7.58687 |
| **Gtpbp6** | 0.69696 | 0.00713 | 0.00000 | 0.00571 | 0.13578 | 0.50030 | 0.01699 | 0.99466 | 0.01045 | 1.63505 | 0.00320 | 0.04186 | 0.71875 | 0.01492 | 0.00000 | 0.18213 |
| **Gtpbp8** | 3.30400 | 4.99042 | 0.00000 | 0.65884 | 1.86264 | 4.01569 | 0.00000 | 2.31422 | 0.44313 | 0.13446 | 2.71740 | 6.63319 | 1.04359 | 3.39583 | 1.34080 | 0.08043 |
| **Gtse1** | 8.65521 | 7.29301 | 9.13782 | 5.38326 | 6.16215 | 1.15857 | 2.34111 | 2.60888 | 8.07555 | 2.55214 | 2.98248 | 0.52792 | 4.83612 | 0.00000 | 2.54880 | 1.40853 |
| **Gtsf1** | 1.62812 | 0.14810 | 1.32082 | 1.30290 | 0.00709 | 0.00513 | 0.00276 | 0.21993 | 1.87218 | 1.97182 | 4.32542 | 0.00000 | 2.51684 | 0.00000 | 1.43581 | 0.00310 |
| **Gtsf1l** | 14.90029 | 1.71386 | 3.89593 | 0.84908 | 5.65324 | 2.62758 | 0.00605 | 0.93423 | 0.14787 | 2.45616 | 0.38641 | 0.96861 | 0.61279 | 0.13829 | 0.68723 | 0.02530 |
| **Guca1a** | 16.81445 | 0.17659 | 0.10528 | 0.76695 | 6.10056 | 10.71268 | 6.29687 | 27.51000 | 3.26430 | 33.89529 | 1.78614 | 2.26454 | 24.36497 | 0.00000 | 0.41215 | 3.70131 |
| **Guca1b** | 0.00819 | 0.00784 | 0.01361 | 1.10831 | 0.00910 | 0.00668 | 0.00750 | 0.01342 | 1.68697 | 0.05926 | 0.00000 | 0.00326 | 0.26083 | 0.01133 | 0.55045 | 0.55805 |
| **Gucd1** | 2.12849 | 4.61176 | 3.98404 | 1.36555 | 0.96771 | 0.01128 | 0.00179 | 1.49616 | 1.84934 | 2.39746 | 2.19923 | 5.46835 | 0.82552 | 1.64171 | 4.17232 | 1.33307 |
| **Gucy1a2** | 0.05294 | 0.01828 | 0.06110 | 0.02686 | 0.00797 | 0.01307 | 0.00741 | 0.01807 | 0.02028 | 0.04139 | 0.01052 | 0.02718 | 0.01831 | 0.18231 | 0.07783 | 0.01446 |

| | | | | | | | | | | | | | | | | |
|---|---|---|---|---|---|---|---|---|---|---|---|---|---|---|---|---|
| **Gucy1b2** | 0.00997 | 0.00756 | 0.02071 | 0.01090 | 0.00952 | 0.00780 | 0.00712 | 0.51435 | 0.01187 | 0.00769 | 0.01024 | 0.00647 | 0.00573 | 0.01303 | 0.00680 | 0.00739 |
| **Gucy2c** | 0.01377 | 0.00780 | 0.01194 | 0.80171 | 0.00935 | 0.00657 | 0.00748 | 0.01321 | 0.14847 | 0.01077 | 0.01138 | 0.00751 | 0.01277 | 0.01722 | 0.06707 | 0.00726 |
| **Gucy2e** | 0.00000 | 0.01817 | 0.00000 | 2.29463 | 0.00791 | 2.76989 | 0.00437 | 2.88030 | 2.43288 | 5.72933 | 4.16050 | 0.46852 | 2.29695 | 1.16001 | 3.91590 | 0.01122 |
| **Guf1** | 0.20483 | 0.00536 | 0.00000 | 0.90794 | 0.05841 | 0.00648 | 3.52811 | 0.00661 | 0.06217 | 1.29155 | 0.99166 | 0.09277 | 1.63254 | 0.39947 | 0.94444 | 0.01311 |
| **Guk1** | 10.94284 | 3.59928 | 8.13953 | 0.56480 | 6.11665 | 5.70549 | 1.88775 | 4.10697 | 0.33163 | 7.92401 | 9.38959 | 5.26193 | 10.63383 | 0.01427 | 0.53192 | 8.69741 |
| **Gulo** | 17.02431 | 5.71508 | 19.95769 | 2.32932 | 7.31648 | 7.52440 | 20.07667 | 33.69207 | 3.09322 | 29.89379 | 25.81134 | 11.98621 | 19.20430 | 6.65662 | 13.84579 | 4.21753 |
| **Gusb** | 1.17506 | 3.02961 | 0.00000 | 2.28810 | 0.07717 | 0.73030 | 0.00264 | 4.18616 | 3.93101 | 1.36948 | 1.87972 | 0.11716 | 3.74180 | 1.31565 | 3.69072 | 0.00218 |
| **Gxylt1** | 1.66626 | 0.01089 | 0.53158 | 9.41408 | 0.05330 | 1.63591 | 6.58066 | 2.60340 | 8.69338 | 3.95844 | 2.22535 | 6.42518 | 1.42681 | 7.51004 | 5.76191 | 1.32186 |
| **Gyg** | 0.41168 | 2.34455 | 0.51229 | 11.77740 | 2.44642 | 0.00554 | 0.02849 | 0.34340 | 8.51348 | 1.01189 | 1.01824 | 0.19661 | 1.88230 | 3.24352 | 9.15449 | 3.89127 |
| **Gyltl1b** | 0.01104 | 0.00769 | 0.12561 | 0.00592 | 0.01199 | 1.42726 | 0.00633 | 3.74437 | 0.00913 | 0.63135 | 0.05567 | 0.00000 | 1.81216 | 0.01390 | 0.00000 | 0.38903 |
| **Gys1** | 3.40756 | 1.05405 | 3.99124 | 2.02993 | 3.93171 | 0.28891 | 0.87871 | 4.43253 | 4.40332 | 1.92500 | 1.65570 | 0.42701 | 1.67192 | 0.64122 | 0.76603 | 3.74014 |
| **Gzf1** | 4.15161 | 3.51969 | 5.79325 | 6.15512 | 0.29060 | 0.65811 | 0.31411 | 0.48379 | 2.19512 | 3.19751 | 0.70141 | 0.00949 | 0.00000 | 0.55175 | 4.14396 | 0.53156 |
| **Gzmm** | 1.27242 | 0.00576 | 1.55472 | 1.03445 | 0.00282 | 0.00470 | 0.00617 | 0.21541 | 0.00010 | 0.91259 | 0.23949 | 0.37760 | 0.00696 | 0.08252 | 0.00000 | 0.00547 |
| **H13** | 7.43151 | 0.36929 | 6.79054 | 3.66987 | 5.20189 | 2.67538 | 1.69151 | 6.30593 | 4.67633 | 7.31457 | 5.96033 | 5.65153 | 6.09850 | 0.39011 | 0.43202 | 4.90627 |
| **H19** | 0.00447 | 0.00796 | 0.01101 | 0.01063 | 0.00582 | 0.00615 | 0.00727 | 0.01405 | 0.01241 | 0.00484 | 0.01252 | 0.00757 | 0.00463 | 0.01347 | 0.01019 | 6.02312 |
| **H1f0** | 1.45848 | 0.03034 | 0.55402 | 5.13519 | 0.03704 | 0.01183 | 0.15216 | 0.05346 | 0.80630 | 4.35406 | 0.69740 | 0.17463 | 0.71384 | 2.04343 | 1.67351 | 1.88876 |
| **H1foo** | 0.00768 | 0.06817 | 0.10300 | 8.85431 | 0.00924 | 3.74448 | 0.27700 | 0.90451 | 4.51933 | 2.73311 | 2.11182 | 0.03922 | 0.00000 | 0.62699 | 1.12706 | 0.00409 |
| **H1fx** | 0.01348 | 0.00798 | 0.01268 | 0.06394 | 0.00949 | 0.00665 | 0.00750 | 0.01358 | 0.01140 | 0.06767 | 0.01406 | 0.00752 | 0.01538 | 0.01130 | 0.00856 | 0.02983 |
| **H2.Aa** | 0.01223 | 0.00779 | 0.01224 | 0.02176 | 0.00860 | 0.00665 | 0.00731 | 0.01013 | 0.01114 | 3.02163 | 0.01005 | 0.00000 | 0.01092 | 0.01059 | 0.01372 | 0.00747 |
| **H2.Ab1** | 0.00879 | 0.00793 | 0.02160 | 0.00508 | 0.00767 | 0.00633 | 0.00716 | 0.01341 | 0.01080 | 4.23239 | 0.04962 | 0.00000 | 0.41518 | 0.01255 | 0.00597 | 0.99413 |
| **H2.Bl** | 0.80029 | 0.01778 | 0.42438 | 2.61200 | 0.21488 | 4.86046 | 0.08728 | 0.10881 | 0.88278 | 0.34729 | 0.39273 | 0.01140 | 2.30068 | 0.00349 | 0.00097 | 0.11535 |
| **H2.D1** | 0.46467 | 0.08238 | 0.86646 | 2.74198 | 0.51992 | 0.73901 | 0.16227 | 0.35969 | 2.39516 | 1.52842 | 1.64929 | 0.02605 | 0.70379 | 0.69146 | 0.50256 | 3.11181 |
| **H2.DMa** | 0.00134 | 0.94495 | 0.09863 | 1.71212 | 0.00780 | 0.00563 | 0.10552 | 0.65879 | 0.80061 | 1.01783 | 1.58493 | 0.01243 | 3.05037 | 0.03783 | 0.17607 | 0.00522 |
| **H2.DMb1** | 0.09924 | 0.00731 | 0.00809 | 0.23918 | 0.00789 | 0.00653 | 0.00730 | 0.00980 | 0.00812 | 0.32722 | 0.02295 | 0.00498 | 0.01183 | 0.00494 | 0.65864 | 0.00725 |
| **H2.DMb2** | 0.02053 | 0.00773 | 0.01193 | 0.00912 | 0.00891 | 0.00655 | 0.00746 | 0.01232 | 0.01230 | 0.09005 | 0.01223 | 0.00695 | 0.01249 | 0.00600 | 0.16867 | 0.00738 |
| **H2.Eb1** | 1.20094 | 0.01882 | 0.00881 | 0.89864 | 0.00540 | 0.00755 | 0.01822 | 0.50123 | 3.60815 | 5.03458 | 0.07716 | 0.01206 | 0.00574 | 0.00629 | 0.03029 | 0.01305 |
| **H2.K1** | 7.60147 | 0.04401 | 9.85157 | 10.06679 | 4.09080 | 11.35612 | 2.50842 | 6.31658 | 6.00114 | 5.79311 | 7.24991 | 0.12719 | 15.88256 | 0.04164 | 1.49056 | 4.87129 |
| **H2.K2** | 6.81665 | 0.00408 | 1.14637 | 0.21962 | 10.58757 | 0.01820 | 1.91968 | 0.26103 | 1.88087 | 0.84069 | 0.79976 | 0.03667 | 0.96615 | 0.12057 | 0.30150 | 0.04270 |
| **H2.Ke6** | 1.39304 | 0.00556 | 1.15716 | 0.51436 | 0.00301 | 0.00823 | 0.08480 | 1.05532 | 0.00351 | 0.06477 | 1.52466 | 0.03910 | 0.00000 | 0.00260 | 0.00000 | 0.02656 |
| **H2.M5** | 0.85342 | 0.00683 | 0.00401 | 0.01842 | 0.00613 | 0.00710 | 0.00662 | 0.02270 | 0.01241 | 0.01273 | 0.01248 | 0.01729 | 0.05731 | 0.01409 | 0.01508 | 0.01347 |
| **H2.Oa** | 1.26024 | 0.00568 | 2.04881 | 0.08240 | 0.01004 | 0.00791 | 0.00477 | 1.55122 | 0.00771 | 0.24160 | 0.34927 | 0.00000 | 1.97212 | 0.00815 | 0.00000 | 0.01189 |
| **H2.Q1** | 0.39678 | 0.00710 | 0.19399 | 0.51768 | 0.48299 | 0.34134 | 0.16044 | 0.28249 | 0.31783 | 0.30540 | 0.03484 | 0.01419 | 0.71753 | 0.01573 | 0.12978 | 0.15051 |
| **H2.Q10** | 0.17091 | 0.00744 | 0.12421 | 5.27012 | 0.31104 | 0.17825 | 0.22330 | 0.57064 | 10.27623 | 2.32877 | 0.21872 | 0.08988 | 0.87687 | 1.50453 | 1.26486 | 0.10846 |
| **H2.Q2** | 1.09397 | 0.00607 | 0.06891 | 0.00814 | 0.99473 | 0.09420 | 0.16970 | 0.11421 | 0.02360 | 0.10841 | 0.01100 | 0.00534 | 0.20943 | 0.00933 | 0.02636 | 0.09182 |
| **H2.Q4** | 0.31577 | 0.00728 | 0.09100 | 0.16234 | 0.11112 | 0.19641 | 0.13827 | 0.30359 | 0.21923 | 0.56215 | 0.00000 | 0.00786 | 1.10316 | 1.04487 | 2.08002 | 0.21892 |
| **H2.Q5** | 0.02110 | 0.00766 | 0.10864 | 2.41447 | 0.04225 | 0.04233 | 0.03946 | 2.84278 | 8.39645 | 1.98264 | 0.34172 | 0.03485 | 1.24589 | 0.09039 | 0.07618 | 0.01309 |
| **H2.T.ps** | 0.46414 | 0.00719 | 0.28298 | 0.48376 | 0.40663 | 0.26974 | 0.13977 | 0.39280 | 0.03016 | 0.14276 | 0.04809 | 0.00496 | 1.35505 | 0.04028 | 0.06918 | 0.27371 |
| **H2.T10** | 0.06865 | 0.17250 | 0.06030 | 2.04405 | 0.09845 | 1.07168 | 0.05652 | 0.13447 | 2.85440 | 0.00000 | 0.24228 | 0.01575 | 0.32122 | 0.66103 | 1.01227 | 0.02061 |
| **H2.T23** | 2.01654 | 0.16567 | 1.24988 | 7.43622 | 1.13737 | 0.43296 | 0.26839 | 0.00000 | 12.75852 | 0.22409 | 12.22807 | 1.01291 | 1.03941 | 4.95435 | 0.69886 | 0.49149 |
| **H2.T24** | 0.18499 | 0.01762 | 0.29640 | 2.38169 | 0.17936 | 0.03769 | 0.03739 | 0.00000 | 3.20046 | 0.00000 | 2.07747 | 0.10586 | 0.08640 | 1.27029 | 0.46646 | 0.06330 |
| **H2.T3** | 0.05248 | 0.03961 | 0.06777 | 0.09825 | 0.00864 | 0.01965 | 0.00734 | 0.00760 | 0.08325 | 0.05116 | 0.17288 | 0.02691 | 0.05797 | 0.05626 | 0.04751 | 0.00727 |
| **H2afj** | 0.21042 | 0.07562 | 2.88662 | 0.41537 | 0.63633 | 0.04132 | 0.11542 | 0.04399 | 0.69211 | 4.23982 | 0.49321 | 0.05327 | 0.34816 | 0.04932 | 0.25979 | 0.06398 |
| **H2afv** | 1.35387 | 0.06383 | 2.85890 | 8.49826 | 2.14674 | 2.55362 | 2.88152 | 2.40058 | 15.63922 | 14.85018 | 16.23864 | 0.62177 | 5.54181 | 1.59930 | 2.06599 | 1.09830 |
| **H2afx** | 5.21179 | 0.55124 | 1.91490 | 0.00000 | 2.48341 | 0.02555 | 0.91054 | 0.72370 | 7.70796 | 22.60505 | 6.97132 | 0.68530 | 11.41553 | 0.96183 | 3.17414 | 2.39845 |
| **H2afy** | 22.94358 | 8.58354 | 12.32594 | 0.00000 | 12.27400 | 12.37823 | 2.12277 | 8.67746 | 0.14334 | 1.66181 | 4.11255 | 0.03863 | 4.49385 | 2.09834 | 0.00000 | 1.10998 |
| **H2afy2** | 0.01605 | 0.00787 | 0.19780 | 0.02352 | 0.00846 | 0.00689 | 0.00750 | 0.01471 | 0.34907 | 0.41993 | 0.01098 | 0.00613 | 0.00971 | 0.00814 | 0.22640 | 0.00704 |
| **H2afy3** | 0.01296 | 0.00788 | 0.04281 | 0.07768 | 0.00923 | 0.00813 | 0.00741 | 0.01323 | 0.01157 | 0.02704 | 0.01361 | 0.00747 | 0.01209 | 0.00865 | 0.03643 | 0.00742 |
| **H2afz** | 55.69449 | 23.64024 | 38.01529 | 27.18616 | 33.56518 | 20.14050 | 23.18473 | 34.93476 | 42.57274 | 43.80811 | 26.62843 | 24.77626 | 37.99328 | 9.34287 | 1.57278 | 27.92578 |
| **H3f3a** | 32.24050 | 13.61988 | 30.45568 | 22.21944 | 20.17427 | 15.82698 | 17.73667 | 40.09516 | 31.56202 | 45.40683 | 29.85004 | 15.66410 | 31.91092 | 21.05936 | 9.64938 | 15.89499 |
| **H3f3b** | 30.38786 | 18.35493 | 25.98202 | 26.16459 | 21.46646 | 19.67266 | 22.96001 | 32.56240 | 40.78679 | 52.43823 | 33.76977 | 24.50596 | 35.23264 | 27.61345 | 22.66683 | 21.30798 |
| **H60b** | 0.02324 | 0.00666 | 0.41207 | 0.00703 | 0.06970 | 0.00535 | 0.00677 | 0.45839 | 0.01024 | 0.00770 | 0.10912 | 1.83107 | 0.00373 | 0.04686 | 0.00000 | 0.00716 |
| **H6pd** | 0.02069 | 0.00799 | 1.69432 | 0.00888 | 0.00652 | 0.00630 | 0.02950 | 0.01401 | 0.01175 | 0.49428 | 0.01082 | 0.05387 | 0.07767 | 0.01439 | 0.00000 | 0.00708 |
| **Habp4** | 0.29790 | 0.01791 | 0.10729 | 0.72860 | 0.00726 | 1.90516 | 0.03817 | 0.00795 | 0.41383 | 0.09406 | 0.02787 | 0.02808 | 0.45084 | 0.03222 | 0.21939 | 0.00609 |
| **Hacd1** | 0.63404 | 0.11315 | 0.12795 | 0.67585 | 0.00669 | 1.94935 | 0.03634 | 0.00677 | 0.00000 | 1.48643 | 0.00875 | 0.11605 | 0.00536 | 0.01592 | 0.00000 | 0.01257 |
| **Hacd2** | 3.64517 | 0.61708 | 3.66293 | 2.26767 | 5.07470 | 0.62554 | 3.35476 | 0.00000 | 6.02540 | 1.54970 | 2.35465 | 3.37157 | 1.10475 | 0.62833 | 1.05363 | 0.87118 |
| **Hacd3** | 2.89875 | 0.02216 | 5.99870 | 11.18273 | 2.51281 | 0.00000 | 2.24263 | 2.09960 | 19.04816 | 6.74261 | 13.03015 | 0.19506 | 2.67806 | 2.71652 | 10.42783 | 0.04519 |
| **Hace1** | 0.20895 | 2.18209 | 0.00000 | 0.00000 | 0.14047 | 2.09107 | 0.05594 | 0.24813 | 0.03623 | 0.00000 | 0.28972 | 1.32958 | 0.00000 | 0.00609 | 0.23745 | 0.00527 |
| **Hacl1** | 0.28681 | 0.00594 | 1.76290 | 0.78724 | 0.00432 | 0.00523 | 2.56859 | 0.10529 | 0.00000 | 0.00000 | 0.00760 | 0.00000 | 1.49791 | 0.03954 | 2.13520 | 0.00535 |
| **Hadh** | 12.61371 | 5.32666 | 13.35220 | 0.00000 | 14.87438 | 13.72235 | 6.13231 | 18.55950 | 0.00000 | 5.00532 | 8.15143 | 0.56946 | 16.48899 | 0.30636 | 0.65797 | 7.44769 |
| **Hadha** | 10.50563 | 11.52118 | 10.48799 | 2.09016 | 7.05846 | 9.96219 | 11.05370 | 6.56269 | 0.55738 | 3.48431 | 5.33444 | 0.27060 | 7.90456 | 3.14525 | 4.50740 | 7.44432 |
| **Hadhb** | 6.75467 | 10.03967 | 8.60766 | 7.15263 | 11.53840 | 5.79095 | 6.81926 | 12.08290 | 4.42929 | 9.73745 | 8.11651 | 3.86870 | 6.27378 | 7.32885 | 13.57149 | 5.38838 |
| **Hagh** | 3.58543 | 1.06116 | 4.67319 | 1.68731 | 4.72208 | 0.84727 | 1.07789 | 0.05950 | 0.86415 | 0.00780 | 0.00000 | 0.01544 | 2.34218 | 0.03065 | 0.21851 | 4.42443 |

| | | | | | | | | | | | | | | | | |
|---|---|---|---|---|---|---|---|---|---|---|---|---|---|---|---|---|
| **Haghl** | 0.01367 | 0.00859 | 0.01157 | 0.01029 | 0.01062 | 0.00663 | 0.00738 | 0.02826 | 0.01236 | 0.07326 | 0.01223 | 0.00764 | 0.98488 | 0.01205 | 0.01056 | 0.00701 |
| **Hal** | 2.80781 | 0.00132 | 0.98686 | 0.00431 | 0.05025 | 0.01385 | 5.59206 | 0.01342 | 0.01157 | 0.96477 | 0.01048 | 0.00000 | 0.03947 | 0.01434 | 0.00000 | 0.00411 |
| **Hamp** | 0.01378 | 0.00795 | 0.01201 | 0.46975 | 0.00931 | 0.00661 | 0.00748 | 0.01345 | 0.00527 | 0.01390 | 0.01366 | 0.02778 | 0.01269 | 0.00866 | 0.02626 | 0.00733 |
| **Hand1** | 1.01254 | 0.00412 | 0.28132 | 0.81353 | 0.04669 | 0.00432 | 0.00000 | 4.78458 | 0.15524 | 24.66952 | 16.51394 | 1.63927 | 0.00000 | 1.31068 | 2.98972 | 0.00328 |
| **Hap1** | 0.19782 | 0.00834 | 0.28444 | 0.00518 | 0.01371 | 0.00838 | 0.09181 | 0.12665 | 0.01102 | 0.23909 | 0.58430 | 0.00431 | 0.00303 | 0.01321 | 0.00772 | 0.00687 |
| **Hapln4** | 0.00000 | 0.00738 | 0.00491 | 0.33683 | 0.00853 | 0.00677 | 0.00642 | 1.00763 | 0.03161 | 1.01320 | 0.17352 | 0.01495 | 1.75483 | 0.95519 | 0.37613 | 0.70244 |
| **Harbi1** | 1.63521 | 3.30865 | 4.61882 | 0.51570 | 2.76089 | 0.88343 | 2.31316 | 0.98525 | 0.07055 | 1.02659 | 0.65101 | 0.05790 | 1.70491 | 0.01665 | 3.09423 | 0.00853 |
| **Hars** | 13.30705 | 9.63023 | 10.94262 | 3.62858 | 13.52063 | 7.12316 | 9.64328 | 12.74380 | 10.16613 | 15.25583 | 8.39318 | 3.83357 | 10.18362 | 0.21814 | 8.36653 | 11.83184 |
| **Hars2** | 5.49356 | 5.40406 | 4.98600 | 3.10144 | 1.47817 | 1.14837 | 3.38191 | 2.62567 | 4.39808 | 5.86432 | 1.54712 | 4.28790 | 5.42197 | 6.40813 | 6.00011 | 2.32165 |
| **Has3** | 0.05497 | 0.00832 | 0.00000 | 2.45431 | 0.01463 | 0.00586 | 0.00709 | 0.00822 | 1.05838 | 0.00000 | 0.00000 | 0.01831 | 0.01335 | 1.83052 | 4.21052 | 0.00682 |
| **Hat1** | 25.16818 | 17.32705 | 24.85140 | 2.89467 | 21.59092 | 7.33397 | 10.70169 | 15.42069 | 4.56861 | 17.93412 | 14.03826 | 10.94876 | 14.34423 | 13.11622 | 14.01368 | 8.27574 |
| **Haus1** | 17.57268 | 7.02157 | 7.45836 | 0.68246 | 9.89369 | 3.13771 | 6.93547 | 4.89660 | 1.36266 | 3.43688 | 3.54423 | 0.71836 | 4.66015 | 3.44531 | 2.09856 | 2.91576 |
| **Haus2** | 3.99413 | 4.21302 | 6.21586 | 4.62533 | 5.20063 | 0.67821 | 2.17785 | 6.24947 | 6.16285 | 7.94122 | 9.17769 | 5.58769 | 5.98873 | 9.21135 | 12.90229 | 0.35745 |
| **Haus3** | 5.23797 | 10.22921 | 6.04381 | 10.15635 | 2.70894 | 1.04890 | 5.48745 | 4.99213 | 18.27491 | 10.31365 | 3.08512 | 13.99106 | 5.09343 | 11.27817 | 11.57542 | 1.03331 |
| **Haus4** | 5.90400 | 4.20336 | 5.80824 | 13.35642 | 0.04939 | 3.46544 | 3.48131 | 4.92063 | 10.32346 | 13.39763 | 12.29012 | 0.85974 | 9.82398 | 0.42830 | 2.64365 | 0.25533 |
| **Haus5** | 9.31804 | 9.79353 | 7.82724 | 1.79645 | 7.05304 | 7.69288 | 4.86862 | 11.41780 | 1.03543 | 7.44108 | 3.55061 | 2.43929 | 6.98150 | 0.05772 | 0.80669 | 1.67087 |
| **Haus6** | 1.79205 | 3.67439 | 0.80526 | 6.91927 | 3.71807 | 2.20544 | 3.92850 | 2.49212 | 9.25162 | 5.12392 | 1.79985 | 1.22714 | 2.46502 | 20.54865 | 16.37363 | 1.35399 |
| **Haus7** | 6.06215 | 6.69296 | 7.40064 | 1.82743 | 5.80832 | 0.01323 | 3.72209 | 0.27097 | 3.54290 | 2.28706 | 2.30805 | 0.02002 | 1.99128 | 1.26343 | 1.07370 | 2.72097 |
| **Haus8** | 3.67359 | 7.32065 | 7.62484 | 1.21531 | 7.46253 | 9.70332 | 9.96620 | 6.60999 | 1.62324 | 5.70661 | 2.64254 | 1.43718 | 9.69408 | 0.00000 | 0.37074 | 4.37577 |
| **Havcr1** | 2.79214 | 4.24703 | 10.14440 | 0.14976 | 5.92580 | 8.66956 | 8.57520 | 6.53628 | 1.88086 | 6.62775 | 7.06617 | 9.57760 | 7.27219 | 14.76534 | 6.38959 | 5.51007 |
| **Havcr2** | 0.02010 | 0.04012 | 0.01021 | 0.00840 | 0.02014 | 0.00662 | 0.00745 | 0.01950 | 0.01175 | 0.00691 | 0.01170 | 0.03610 | 0.01216 | 0.01322 | 0.37056 | 0.00740 |
| **Hax1** | 13.33290 | 14.48435 | 18.26707 | 18.80783 | 7.72867 | 8.73481 | 10.21654 | 27.50791 | 21.04372 | 28.58432 | 27.25076 | 8.47879 | 22.46336 | 24.58060 | 24.58255 | 8.87995 |
| **Hbegf** | 0.45967 | 0.00704 | 0.01117 | 2.06867 | 0.00684 | 0.00708 | 0.00679 | 0.00712 | 1.66613 | 0.63655 | 0.03262 | 0.03483 | 0.01658 | 0.06325 | 0.14060 | 0.00611 |
| **Hbp1** | 1.11566 | 9.57754 | 3.09966 | 14.90363 | 0.02773 | 1.26086 | 5.40851 | 9.36083 | 18.82358 | 16.91794 | 10.69505 | 9.80661 | 2.38072 | 14.52758 | 16.06916 | 1.46251 |
| **Hbs1l** | 9.09658 | 1.91912 | 7.26039 | 2.66674 | 2.83947 | 2.91853 | 8.28995 | 4.03101 | 0.49984 | 3.13754 | 4.49914 | 4.75445 | 9.60902 | 5.59587 | 3.07212 | 4.58140 |
| **Hccs** | 0.35788 | 2.77366 | 2.23660 | 8.87612 | 3.69043 | 0.79175 | 2.41824 | 3.94906 | 9.30472 | 6.14350 | 2.33360 | 8.89514 | 1.64538 | 13.42986 | 10.46536 | 1.27616 |
| **Hcfc1** | 11.93375 | 10.02126 | 6.07659 | 6.65813 | 9.00886 | 1.65896 | 7.61686 | 13.07168 | 12.12168 | 16.02284 | 9.75436 | 1.97192 | 7.05104 | 9.50024 | 19.11643 | 4.12486 |
| **Hcfc1r1** | 8.42001 | 0.60816 | 4.31303 | 0.99330 | 5.66863 | 1.92667 | 3.32171 | 6.27787 | 0.28951 | 7.44523 | 1.57750 | 0.10584 | 3.70318 | 0.00000 | 0.18830 | 1.84524 |
| **Hcfc2** | 0.94715 | 0.00769 | 0.00000 | 1.20029 | 0.01600 | 3.35923 | 0.00628 | 0.00618 | 0.00000 | 0.70923 | 0.02440 | 0.00547 | 0.00077 | 0.00539 | 0.11736 | 0.00492 |
| **Hck** | 0.47262 | 0.00503 | 0.00000 | 0.00855 | 4.14060 | 0.01407 | 0.00680 | 0.00263 | 0.01082 | 0.00635 | 0.00706 | 0.00887 | 0.02780 | 1.09318 | 0.00497 | 0.00665 |
| **Hcn3** | 0.00137 | 0.00630 | 0.00000 | 0.00186 | 0.00955 | 0.22187 | 0.59632 | 1.37998 | 0.23359 | 2.57660 | 0.40627 | 3.06512 | 0.24283 | 0.01532 | 0.25416 | 0.00703 |
| **Hcrt** | 0.01199 | 0.00676 | 0.01006 | 0.02011 | 0.00800 | 0.00657 | 0.00642 | 4.63229 | 0.01676 | 5.88352 | 0.98071 | 0.00001 | 1.07847 | 0.00807 | 0.00000 | 0.00678 |
| **Hdac1** | 22.95355 | 15.46995 | 18.06926 | 13.70053 | 14.49764 | 8.74096 | 14.61592 | 32.60723 | 17.57886 | 26.93883 | 21.45768 | 15.47413 | 15.92637 | 30.30307 | 26.00452 | 4.58550 |
| **Hdac10** | 1.69837 | 0.00599 | 0.00148 | 0.84031 | 0.01387 | 0.00661 | 0.00595 | 0.00231 | 1.19471 | 0.20564 | 0.08440 | 0.03145 | 0.01116 | 0.18370 | 0.38285 | 0.01341 |
| **Hdac11** | 0.01276 | 0.00799 | 0.01412 | 0.00736 | 0.00996 | 0.00705 | 0.00755 | 0.01275 | 0.03383 | 0.63433 | 0.01423 | 0.00567 | 0.10829 | 0.01165 | 0.00829 | 0.00796 |
| **Hdac2** | 10.80820 | 13.67962 | 10.32471 | 6.20746 | 9.61727 | 10.36674 | 7.94794 | 8.25258 | 2.65829 | 8.96013 | 5.72744 | 5.01947 | 7.13326 | 4.63515 | 3.85003 | 4.59737 |
| **Hdac3** | 3.86647 | 4.21836 | 1.74472 | 0.00000 | 0.00000 | 0.13144 | 1.28088 | 0.88244 | 0.01043 | 1.28987 | 0.04104 | 0.12841 | 2.42215 | 0.11711 | 0.37347 | 1.67879 |
| **Hdac4** | 1.11386 | 0.00631 | 2.19993 | 0.26055 | 0.01079 | 0.00601 | 0.00520 | 1.00178 | 1.90665 | 0.00000 | 0.00000 | 0.05518 | 1.26184 | 0.01336 | 0.00000 | 0.00464 |
| **Hdac5** | 0.74907 | 0.00579 | 0.35336 | 0.03990 | 0.00586 | 0.93892 | 2.10709 | 1.70250 | 0.59147 | 5.66357 | 0.87129 | 0.10301 | 3.24689 | 0.02961 | 0.00000 | 0.00445 |
| **Hdac6** | 4.56468 | 2.64201 | 0.98959 | 8.44328 | 0.12722 | 0.79906 | 3.04163 | 6.66851 | 8.91977 | 7.88928 | 4.40775 | 1.23201 | 5.92869 | 2.64202 | 10.80948 | 2.63140 |
| **Hdac7** | 0.02067 | 0.00804 | 0.05173 | 0.00817 | 0.00852 | 0.00596 | 0.00747 | 0.01209 | 0.01191 | 0.63494 | 0.01206 | 0.00673 | 0.01836 | 0.01187 | 0.00914 | 0.00757 |
| **Hdac8** | 0.00484 | 0.00725 | 0.00454 | 0.19112 | 0.00973 | 0.00645 | 0.42759 | 0.00186 | 0.68862 | 0.00000 | 0.18865 | 0.00745 | 1.02287 | 1.74320 | 0.21419 | 0.00657 |
| **Hdac9** | 0.01224 | 0.00834 | 0.01308 | 1.24606 | 0.01023 | 0.01376 | 0.00755 | 0.01415 | 0.04859 | 0.00760 | 0.01311 | 0.00719 | 0.27735 | 0.00636 | 0.59971 | 0.00741 |
| **Hdc** | 0.20645 | 0.00925 | 0.06363 | 1.42209 | 0.01677 | 0.00541 | 0.00778 | 1.11882 | 1.47383 | 2.38689 | 0.93871 | 0.03033 | 0.00000 | 0.50310 | 0.00000 | 1.68621 |
| **Hddc2** | 0.55838 | 0.00683 | 3.15287 | 0.23248 | 0.21794 | 0.04755 | 0.00655 | 0.98570 | 1.05961 | 2.98231 | 0.06351 | 1.74729 | 0.01255 | 0.00456 | 0.42210 | 0.05891 |
| **Hddc3** | 1.17633 | 0.00655 | 1.11120 | 0.32954 | 0.01422 | 3.25251 | 0.00396 | 5.08199 | 0.00856 | 1.62028 | 1.36101 | 0.00702 | 3.16872 | 0.02793 | 0.00000 | 0.45675 |
| **Hdgf** | 30.44539 | 18.86174 | 31.28059 | 14.88024 | 19.25780 | 17.03599 | 20.78900 | 41.98847 | 22.43287 | 43.62192 | 38.24719 | 13.50433 | 34.27636 | 33.77428 | 29.95831 | 13.94821 |
| **Hdgfrp2** | 11.30326 | 6.84255 | 7.28685 | 5.92106 | 6.42840 | 1.01096 | 7.71245 | 16.90325 | 8.68413 | 14.78253 | 5.57376 | 0.90119 | 9.02641 | 0.02849 | 2.66207 | 8.09888 |
| **Hdgfrp3** | 1.37446 | 0.07501 | 0.09883 | 0.01838 | 0.00504 | 0.00747 | 0.34090 | 0.00524 | 0.01263 | 0.06156 | 0.01254 | 0.04492 | 0.01758 | 0.01108 | 0.00772 | 0.00603 |
| **Hdhd2** | 4.07555 | 0.29725 | 4.19348 | 0.39037 | 0.00932 | 0.05978 | 0.00364 | 0.71631 | 0.19277 | 1.22492 | 1.30544 | 0.03952 | 0.72747 | 0.01120 | 0.35513 | 1.91906 |
| **Hdhd3** | 2.10846 | 0.00598 | 0.00459 | 0.00483 | 0.01525 | 0.00581 | 0.00587 | 0.00978 | 0.44550 | 0.01329 | 0.00884 | 0.00617 | 0.01124 | 0.01353 | 0.00785 | 0.00515 |
| **Hdlbp** | 21.13835 | 16.83209 | 15.97203 | 11.29650 | 16.18290 | 11.88647 | 15.24648 | 29.22033 | 9.80384 | 29.39877 | 16.92012 | 11.97919 | 20.63840 | 8.03912 | 8.69580 | 9.44340 |
| **Hdx** | 0.01453 | 0.00772 | 0.02007 | 0.12335 | 0.00917 | 0.00612 | 0.00739 | 0.01101 | 0.26513 | 0.20614 | 0.00853 | 0.00528 | 0.01153 | 0.00672 | 0.62130 | 0.00710 |
| **Heatr1** | 6.95396 | 8.97216 | 9.97115 | 4.55809 | 4.10904 | 0.13515 | 3.36863 | 15.68947 | 12.35123 | 25.19909 | 6.75772 | 11.07088 | 4.67526 | 7.41372 | 11.20796 | 0.63062 |
| **Heatr3** | 1.71204 | 1.25793 | 4.07200 | 5.08194 | 0.00854 | 0.00346 | 0.02312 | 7.01933 | 8.75678 | 7.27030 | 4.33013 | 1.45228 | 0.86972 | 16.64014 | 11.71211 | 2.90373 |
| **Heatr5a** | 1.98650 | 8.50111 | 2.29656 | 0.81368 | 2.79827 | 0.01264 | 4.70095 | 6.84913 | 2.38296 | 0.46235 | 0.50250 | 0.16720 | 2.08939 | 0.09982 | 2.81918 | 0.21987 |
| **Heatr5b** | 1.81604 | 2.15424 | 2.01549 | 1.84135 | 0.00392 | 0.02086 | 2.97559 | 1.50460 | 0.21596 | 3.35075 | 1.03920 | 0.35351 | 0.00000 | 0.01879 | 2.68036 | 0.85009 |
| **Heatr6** | 2.13252 | 0.45477 | 0.81842 | 0.56098 | 0.01103 | 2.74022 | 0.08405 | 0.99834 | 0.48486 | 1.99282 | 2.21458 | 0.57553 | 1.44665 | 0.50862 | 1.21057 | 0.74439 |
| **Hebp1** | 0.98796 | 0.00617 | 0.00000 | 0.00216 | 0.01398 | 2.02969 | 0.00630 | 0.00000 | 0.01101 | 0.00000 | 0.43737 | 0.01419 | 0.00000 | 0.01266 | 0.19287 | 0.01990 |
| **Hebp2** | 0.01308 | 0.00802 | 0.01790 | 0.00782 | 0.00920 | 0.00643 | 0.00745 | 0.01121 | 0.01195 | 0.75600 | 0.01215 | 0.00584 | 0.01440 | 0.01163 | 0.00882 | 0.00753 |
| **Heca** | 0.01720 | 0.00898 | 0.07021 | 1.20485 | 0.01250 | 0.53522 | 0.04323 | 0.63297 | 0.00000 | 0.08464 | 0.00484 | 0.00568 | 0.00000 | 0.00412 | 0.96223 | 0.00740 |
| **Hectd1** | 5.74479 | 10.40962 | 7.92188 | 9.80581 | 2.76799 | 11.12684 | 5.76313 | 4.84680 | 11.91527 | 14.24525 | 5.61028 | 9.52864 | 4.70706 | 10.13273 | 16.09237 | 2.23377 |

| | | | | | | | | | | | | | | | | |
|---|---|---|---|---|---|---|---|---|---|---|---|---|---|---|---|---|
| **Hectd2** | 0.02187 | 0.00788 | 0.01140 | 0.22801 | 0.00872 | 0.00662 | 0.00747 | 0.01415 | 0.00801 | 0.01396 | 0.01344 | 0.00801 | 0.01905 | 0.01504 | 0.02449 | 0.00730 |
| **Hectd3** | 0.93144 | 0.25851 | 4.09654 | 1.68942 | 4.40517 | 1.17987 | 0.25091 | 2.81218 | 1.05061 | 0.32814 | 0.80347 | 0.01285 | 3.50750 | 0.28784 | 1.92334 | 0.01733 |
| **Hecw2** | 0.00151 | 0.00788 | 0.00550 | 1.20383 | 0.01156 | 0.00593 | 1.13656 | 0.19877 | 0.18163 | 0.66780 | 0.00000 | 0.00000 | 0.04454 | 4.47364 | 3.62042 | 0.00691 |
| **Heg1** | 1.64431 | 0.09550 | 0.00206 | 0.00832 | 0.17903 | 0.00673 | 0.00603 | 0.00495 | 0.02578 | 0.00561 | 0.01102 | 0.01559 | 0.01105 | 0.01989 | 0.27745 | 0.00562 |
| **Helb** | 0.34608 | 1.65476 | 0.90383 | 2.31546 | 0.02821 | 0.03990 | 0.87311 | 1.21072 | 1.57309 | 1.39742 | 0.80513 | 0.12077 | 0.04154 | 0.21139 | 1.45115 | 0.87790 |
| **Hells** | 2.82845 | 5.45601 | 1.13071 | 6.50916 | 3.14223 | 2.25730 | 3.69442 | 2.44945 | 4.72043 | 0.82004 | 2.34615 | 11.99428 | 1.02603 | 14.30516 | 9.64796 | 0.59607 |
| **Helq** | 0.25372 | 0.27936 | 0.43784 | 5.54222 | 0.00721 | 0.00472 | 0.10924 | 1.33051 | 3.66614 | 0.00000 | 1.18730 | 0.13209 | 0.64869 | 3.12779 | 6.89465 | 0.00492 |
| **Helz** | 1.77051 | 1.96943 | 0.00370 | 0.37077 | 0.06778 | 0.01607 | 0.71291 | 0.01596 | 0.00571 | 5.36810 | 0.06375 | 0.03719 | 1.87498 | 0.00413 | 1.46370 | 0.80165 |
| **Hemk1** | 1.87507 | 0.00375 | 1.43162 | 0.01676 | 0.01858 | 0.01614 | 3.33106 | 0.00000 | 0.04513 | 0.58971 | 1.31477 | 0.06963 | 3.93931 | 0.00452 | 0.11452 | 0.00998 |
| **Henmt1** | 0.01381 | 0.00768 | 0.01173 | 0.47330 | 0.00939 | 0.00653 | 0.00742 | 0.01226 | 0.00383 | 0.00740 | 0.01197 | 0.01705 | 0.01277 | 0.01071 | 0.27189 | 0.00730 |
| **Hepacam** | 0.00367 | 0.00806 | 0.00641 | 0.00767 | 0.00797 | 0.06654 | 0.00799 | 0.10243 | 0.01185 | 0.71050 | 0.00945 | 0.05750 | 0.00469 | 0.22028 | 0.00000 | 1.01007 |
| **Herc1** | 2.72057 | 0.22469 | 1.66683 | 1.19319 | 1.01331 | 3.35770 | 3.91034 | 3.95046 | 0.90095 | 1.91557 | 0.72979 | 2.21418 | 2.32314 | 0.61421 | 1.25753 | 0.12171 |
| **Herc2** | 1.72419 | 1.49975 | 0.57486 | 12.89183 | 1.03690 | 0.01219 | 1.13930 | 2.77235 | 4.68521 | 2.96445 | 0.21629 | 0.30123 | 3.53401 | 1.63880 | 7.76583 | 0.11043 |
| **Herc3** | 2.56564 | 0.00519 | 0.00000 | 0.04076 | 0.00791 | 0.16567 | 0.00519 | 0.00365 | 0.01871 | 0.00393 | 0.01101 | 0.00532 | 0.24953 | 0.00683 | 0.28788 | 0.00448 |
| **Herc4** | 6.49454 | 4.27980 | 6.23466 | 6.12513 | 7.79814 | 5.88714 | 8.20228 | 5.39474 | 13.82003 | 8.70720 | 6.55694 | 8.29818 | 4.18993 | 17.16773 | 7.47406 | 3.46644 |
| **Herpud1** | 0.00000 | 2.96587 | 1.01025 | 0.00489 | 0.00483 | 0.01869 | 0.01587 | 1.84185 | 0.01074 | 0.68146 | 0.00000 | 0.00000 | 0.00000 | 0.00907 | 0.00000 | 3.03446 |
| **Herpud2** | 0.70821 | 1.03493 | 0.35297 | 15.03009 | 0.29179 | 2.27836 | 0.30442 | 1.52890 | 3.23566 | 3.09851 | 0.41133 | 0.63409 | 0.77974 | 2.16632 | 7.86519 | 0.00953 |
| **Hes1** | 0.91120 | 0.53943 | 0.39775 | 0.87953 | 0.00522 | 0.00783 | 0.02152 | 0.07490 | 0.01514 | 4.51968 | 11.20284 | 0.40203 | 0.00000 | 0.04351 | 0.46097 | 0.01102 |
| **Hes6** | 1.98890 | 0.00579 | 0.08739 | 1.42209 | 0.15204 | 0.00674 | 0.00541 | 0.13426 | 0.00000 | 1.66187 | 0.01172 | 0.01124 | 0.00817 | 0.01108 | 0.00000 | 1.00680 |
| **Hesx1** | 7.07610 | 1.81128 | 0.95785 | 1.03537 | 0.00000 | 0.00000 | 2.51474 | 0.00000 | 4.49951 | 0.00000 | 0.09142 | 0.75855 | 0.32202 | 0.77661 | 0.14261 | 0.00000 |
| **Hexa** | 15.49576 | 7.75657 | 9.63923 | 0.33156 | 13.08986 | 13.25942 | 10.33572 | 10.56112 | 0.50970 | 5.04475 | 4.11072 | 0.13482 | 16.19805 | 0.00000 | 1.56105 | 6.45345 |
| **Hexb** | 9.71150 | 0.92599 | 9.37321 | 5.95294 | 6.44543 | 4.68700 | 8.70968 | 10.20794 | 3.72428 | 15.56036 | 12.74511 | 6.41738 | 13.08306 | 11.83462 | 9.41750 | 5.02613 |
| **Hexdc** | 1.83021 | 0.00504 | 0.77833 | 0.13616 | 0.14565 | 0.00495 | 0.42696 | 1.59644 | 0.49480 | 0.40679 | 0.34676 | 0.00000 | 2.17639 | 0.04037 | 0.06780 | 0.03194 |
| **Hexim1** | 4.50032 | 0.03960 | 0.00000 | 6.56785 | 4.20613 | 1.78830 | 0.15877 | 4.52937 | 13.41124 | 5.79442 | 2.97256 | 0.31251 | 2.67754 | 0.16966 | 0.00000 | 2.74681 |
| **Hexim2** | 1.10657 | 0.00693 | 0.00000 | 1.65538 | 0.01666 | 1.29539 | 0.00525 | 0.55713 | 1.35273 | 1.51623 | 1.10244 | 0.03135 | 1.06698 | 0.00000 | 0.26874 | 0.00489 |
| **Hey1** | 0.00974 | 0.00834 | 0.00819 | 0.59136 | 0.01893 | 0.00679 | 0.00809 | 0.12485 | 0.72315 | 0.00550 | 0.01800 | 0.32344 | 0.00819 | 0.01289 | 0.00000 | 0.00725 |
| **Heyl** | 1.78244 | 0.00567 | 0.00355 | 0.24182 | 0.01276 | 0.00659 | 0.00615 | 0.00792 | 0.00750 | 0.04261 | 0.01301 | 0.00661 | 0.01147 | 0.01316 | 0.01255 | 0.00553 |
| **Hfm1** | 0.04729 | 0.01849 | 0.20092 | 0.49205 | 0.01733 | 0.00693 | 0.01808 | 0.08110 | 1.32182 | 0.19395 | 0.09686 | 0.02631 | 0.01362 | 0.24035 | 0.07994 | 0.00689 |
| **Hgh1** | 5.29552 | 4.68975 | 5.07240 | 1.71075 | 0.19225 | 0.00138 | 0.00766 | 3.66623 | 0.51694 | 2.10300 | 0.29364 | 1.71567 | 5.39241 | 0.08432 | 3.28849 | 4.03097 |
| **Hgs** | 4.63523 | 0.22944 | 0.58960 | 7.17175 | 9.30231 | 3.44800 | 0.45378 | 7.73682 | 8.04032 | 7.88685 | 6.70528 | 0.92156 | 10.99616 | 0.00000 | 0.00000 | 4.77100 |
| **Hgsnat** | 0.25299 | 0.00601 | 1.81482 | 1.21834 | 1.53077 | 0.00675 | 0.00543 | 3.58172 | 1.10189 | 0.33981 | 0.37762 | 0.11714 | 2.27444 | 0.01927 | 1.52771 | 0.01237 |
| **Hhex** | 3.63260 | 0.02665 | 0.67529 | 10.12257 | 0.01253 | 0.05575 | 0.00215 | 2.14241 | 1.77670 | 0.47211 | 1.69253 | 0.07176 | 3.29964 | 0.00000 | 0.88059 | 0.00000 |
| **Hhipl2** | 0.02102 | 0.00781 | 0.01036 | 0.09194 | 0.01080 | 0.00657 | 0.00713 | 0.00574 | 0.01003 | 1.72086 | 0.12472 | 0.01302 | 0.00970 | 0.03647 | 0.00000 | 0.00729 |
| **Hibadh** | 4.07296 | 0.00918 | 2.45652 | 2.50472 | 4.70703 | 1.59846 | 3.68188 | 2.02827 | 0.14384 | 0.54597 | 0.00000 | 0.29119 | 1.87498 | 1.14642 | 0.36524 | 2.99399 |
| **Hibch** | 7.91990 | 10.33682 | 6.04652 | 1.62354 | 7.71690 | 7.43181 | 3.58762 | 1.57017 | 1.67342 | 6.43707 | 0.38623 | 0.88404 | 1.66386 | 5.14164 | 4.98906 | 0.53893 |
| **Hic1** | 0.02444 | 0.00791 | 0.00912 | 0.56935 | 0.01199 | 0.00676 | 0.01692 | 0.00000 | 0.58592 | 0.68066 | 1.22986 | 0.00997 | 0.35294 | 0.00531 | 0.00000 | 0.00702 |
| **Hic2** | 5.54554 | 0.02911 | 7.79927 | 16.39969 | 3.04915 | 10.49017 | 3.51385 | 5.86959 | 20.15290 | 7.93842 | 4.46244 | 0.77821 | 7.04699 | 6.25683 | 10.07207 | 4.64922 |
| **Hif1a** | 2.40777 | 3.45427 | 0.92221 | 2.24522 | 2.99570 | 0.50452 | 0.56302 | 0.72864 | 0.00000 | 0.67054 | 0.02280 | 2.51871 | 0.68073 | 0.00352 | 0.84315 | 0.13788 |
| **Hif1an** | 4.20722 | 4.78600 | 1.49215 | 3.02222 | 4.92844 | 3.67620 | 1.73811 | 2.88121 | 3.08490 | 6.86312 | 5.10869 | 0.16460 | 0.00000 | 4.25869 | 4.30289 | 1.36668 |
| **Hif3a** | 0.00774 | 0.00687 | 0.00000 | 1.15019 | 1.15639 | 0.07726 | 0.00687 | 0.00600 | 1.80900 | 1.08204 | 0.01990 | 0.05484 | 0.56518 | 0.00264 | 0.11105 | 0.01377 |
| **Higd1a** | 4.57645 | 0.32828 | 4.75375 | 15.50675 | 5.97888 | 0.79161 | 0.43129 | 0.87408 | 6.03001 | 2.88473 | 4.87554 | 0.41269 | 5.12128 | 1.01746 | 1.52769 | 0.85799 |
| **Higd2a** | 6.85259 | 1.31971 | 4.41870 | 5.01528 | 2.34620 | 0.04999 | 1.93606 | 1.68811 | 0.95089 | 4.86254 | 6.09839 | 0.34389 | 7.46993 | 0.03020 | 3.92223 | 5.46779 |
| **Hilpda** | 1.62712 | 4.24262 | 3.09821 | 4.23852 | 0.00234 | 0.00458 | 0.10213 | 4.66438 | 3.04198 | 1.39994 | 1.94023 | 0.06216 | 7.62709 | 0.02131 | 1.72799 | 0.00000 |
| **Hinfp** | 0.62770 | 0.01308 | 2.72226 | 6.33547 | 0.00962 | 0.00762 | 5.85509 | 5.48028 | 15.02862 | 4.75641 | 4.84367 | 5.08121 | 3.91549 | 0.04916 | 8.48461 | 0.00000 |
| **Hint1** | 15.04328 | 0.52666 | 7.46663 | 5.38189 | 11.53100 | 7.56467 | 1.36973 | 3.76059 | 12.45280 | 9.81157 | 13.13847 | 1.11238 | 7.32907 | 0.23429 | 0.00000 | 7.16545 |
| **Hint2** | 0.54367 | 0.00666 | 0.62415 | 0.12401 | 1.22913 | 1.40832 | 0.02730 | 0.00000 | 0.00556 | 0.02755 | 0.46644 | 0.01424 | 2.10021 | 0.00877 | 0.00000 | 2.17878 |
| **Hint3** | 1.69334 | 0.36709 | 1.40859 | 0.08249 | 0.47340 | 2.81779 | 1.16090 | 0.23061 | 1.63755 | 2.50117 | 8.73945 | 0.05699 | 0.97949 | 1.61723 | 0.00000 | 3.08967 |
| **Hip1** | 11.11934 | 7.23024 | 12.39148 | 1.08298 | 0.55475 | 10.02457 | 6.63909 | 10.72243 | 0.52621 | 10.34936 | 10.76571 | 3.87400 | 8.94437 | 9.61639 | 3.45075 | 6.54913 |
| **Hip1r** | 0.00508 | 0.00745 | 0.00270 | 2.28012 | 0.01967 | 0.88800 | 0.00532 | 1.21020 | 1.61781 | 1.43464 | 0.00000 | 0.05042 | 3.07505 | 0.00760 | 0.00000 | 0.78219 |
| **Hipk1** | 7.48046 | 3.21344 | 2.55975 | 19.60393 | 3.71098 | 5.38059 | 8.35792 | 8.14210 | 26.69770 | 14.71240 | 5.76530 | 12.47878 | 2.31677 | 16.63907 | 10.18774 | 2.26638 |
| **Hipk2** | 3.03309 | 0.02064 | 0.67819 | 0.69927 | 2.97554 | 0.00590 | 2.75767 | 0.00000 | 0.38366 | 1.13115 | 0.02256 | 0.03185 | 1.53444 | 0.01004 | 0.36609 | 0.01807 |
| **Hipk3** | 4.27902 | 2.81861 | 7.19675 | 11.33005 | 0.07769 | 4.28122 | 7.82665 | 11.27185 | 26.47542 | 13.11747 | 10.80737 | 19.38835 | 2.29886 | 25.36131 | 8.73264 | 2.48678 |
| **Hipk4** | 0.01250 | 0.00760 | 0.00933 | 0.00546 | 0.00896 | 0.00666 | 0.00710 | 0.02290 | 0.01120 | 0.83307 | 0.33072 | 0.00464 | 0.00747 | 0.02065 | 0.00000 | 0.00703 |
| **Hira** | 0.84168 | 2.25925 | 3.44880 | 1.33438 | 3.20405 | 0.02071 | 3.06593 | 5.41852 | 1.22851 | 9.48659 | 4.90462 | 1.55429 | 0.00000 | 0.10435 | 9.23132 | 1.63324 |
| **Hirip3** | 12.47889 | 6.63304 | 12.97389 | 16.95762 | 5.89227 | 1.08045 | 4.35256 | 10.79170 | 4.12313 | 5.34000 | 5.99862 | 1.46379 | 9.15546 | 0.00000 | 3.18364 | 0.25782 |
| **Hist1h1a** | 0.02542 | 0.00815 | 0.01129 | 0.01417 | 0.01756 | 0.00663 | 0.00753 | 0.01393 | 0.23642 | 0.07929 | 0.01000 | 0.00754 | 0.01194 | 0.01161 | 0.01013 | 0.00713 |
| **Hist1h1b** | 0.77744 | 0.00683 | 0.00000 | 0.00979 | 0.00617 | 1.22995 | 0.00666 | 0.00753 | 0.01280 | 0.00286 | 0.00896 | 0.00645 | 0.00528 | 0.22136 | 0.00574 | 0.33873 |
| **Hist1h1c** | 0.68312 | 0.00726 | 0.00000 | 0.00865 | 0.01850 | 0.05019 | 0.00660 | 0.01214 | 0.01208 | 0.46205 | 0.01136 | 0.00628 | 0.46828 | 0.01148 | 0.01036 | 0.00651 |
| **Hist1h1d** | 0.02072 | 0.00702 | 0.00370 | 0.00941 | 1.98495 | 0.00801 | 0.00744 | 0.01065 | 0.01140 | 0.01265 | 0.01228 | 0.00622 | 0.02304 | 0.01252 | 0.01114 | 0.00727 |
| **Hist1h2ab** | 0.02720 | 0.00764 | 0.01894 | 0.00886 | 0.02681 | 0.00631 | 0.00735 | 0.01102 | 0.01211 | 0.41046 | 0.01324 | 0.01658 | 0.03936 | 0.01323 | 0.00804 | 0.06551 |
| **Hist1h2ac** | 0.01311 | 0.00801 | 0.01758 | 0.00998 | 0.00934 | 0.00635 | 0.00749 | 0.01314 | 0.01231 | 0.14375 | 0.01312 | 0.00755 | 0.04493 | 0.01204 | 0.00975 | 0.02223 |
| **Hist1h2ad** | 0.03814 | 0.00760 | 0.00930 | 0.00836 | 0.03480 | 0.01321 | 0.00736 | 0.01516 | 0.01201 | 0.42694 | 0.03008 | 0.01673 | 0.04531 | 0.01337 | 0.00763 | 0.04378 |

| | | | | | | | | | | | | | | | | |
|---|---|---|---|---|---|---|---|---|---|---|---|---|---|---|---|---|
| **Hist1h2ae** | 0.01703 | 0.00829 | 0.05361 | 0.00993 | 0.01062 | 0.00596 | 0.00760 | 0.01305 | 0.01228 | 0.15575 | 0.01328 | 0.00801 | 0.01332 | 0.01345 | 0.00858 | 0.03680 |
| **Hist1h2af** | 0.01643 | 0.00794 | 0.01177 | 0.00887 | 0.02067 | 0.00655 | 0.00731 | 0.01090 | 0.01211 | 0.41035 | 0.01288 | 0.00687 | 0.02486 | 0.01336 | 0.00852 | 0.02926 |
| **Hist1h2ag** | 0.02326 | 0.00791 | 0.01140 | 0.01019 | 0.02958 | 0.00663 | 0.00746 | 0.01247 | 0.01230 | 0.09456 | 0.01343 | 0.00778 | 0.01219 | 0.01338 | 0.00955 | 0.00736 |
| **Hist1h2ah** | 0.01786 | 0.00785 | 0.01187 | 0.01035 | 0.00897 | 0.00661 | 0.00743 | 0.01282 | 0.01221 | 0.10760 | 0.01312 | 0.00755 | 0.01237 | 0.01492 | 0.01464 | 0.00740 |
| **Hist1h2ai** | 0.03197 | 0.00769 | 0.01667 | 0.00852 | 0.01750 | 0.00643 | 0.00743 | 0.01571 | 0.01212 | 0.49520 | 0.01322 | 0.00659 | 0.04981 | 0.01248 | 0.01227 | 0.03662 |
| **Hist1h2al** | 0.18708 | 0.00790 | 0.00000 | 0.21755 | 0.02191 | 0.10801 | 0.01765 | 0.09579 | 0.36341 | 0.15962 | 0.07574 | 0.00651 | 0.11120 | 0.04592 | 0.04596 | 0.08041 |
| **Hist1h2an** | 0.01458 | 0.00787 | 0.02059 | 0.01002 | 0.00929 | 0.00628 | 0.00741 | 0.01242 | 0.01225 | 0.13194 | 0.01360 | 0.00767 | 0.02858 | 0.01358 | 0.00937 | 0.00737 |
| **Hist1h2bb** | 0.01846 | 0.00824 | 0.01174 | 0.00482 | 0.01059 | 0.00668 | 0.00754 | 0.01371 | 0.33896 | 0.01349 | 0.01022 | 0.00766 | 0.01281 | 0.01327 | 0.00800 | 0.00745 |
| **Hist1h2bc** | 1.76221 | 0.00584 | 0.01470 | 0.00205 | 1.30179 | 0.01033 | 0.05659 | 0.00000 | 3.70581 | 1.34352 | 0.00000 | 0.00000 | 0.17720 | 0.01151 | 0.00650 | 0.08058 |
| **Hist1h2be** | 0.14466 | 0.00820 | 0.01112 | 0.00872 | 0.12551 | 0.01303 | 0.00747 | 0.01187 | 0.12533 | 0.10371 | 0.01159 | 0.00790 | 0.01918 | 0.01186 | 0.01027 | 0.02253 |
| **Hist1h4c** | 0.02574 | 0.00818 | 0.02336 | 0.01062 | 0.01154 | 0.01218 | 0.00737 | 0.02198 | 0.01235 | 0.01396 | 0.01292 | 0.00793 | 0.02432 | 0.01233 | 0.01027 | 0.21830 |
| **Hist1h4d** | 0.05773 | 0.01939 | 0.00936 | 0.01058 | 0.01109 | 0.00697 | 0.01795 | 0.03339 | 0.01231 | 0.01763 | 0.01358 | 0.00808 | 0.03470 | 0.01333 | 0.00969 | 0.03697 |
| **Hist1h4f** | 0.01719 | 0.00808 | 0.01004 | 0.01065 | 0.01055 | 0.01308 | 0.00739 | 0.04840 | 0.01232 | 0.01352 | 0.01268 | 0.00791 | 0.02477 | 0.01242 | 0.01000 | 0.02936 |
| **Hist1h4h** | 0.04307 | 0.00862 | 0.01332 | 0.01056 | 0.01393 | 0.02485 | 0.00758 | 0.03145 | 0.01239 | 0.02637 | 0.01231 | 0.00823 | 0.01080 | 0.01206 | 0.01035 | 0.01473 |
| **Hist1h4i** | 0.60540 | 0.00752 | 0.24248 | 0.11222 | 0.01043 | 0.03918 | 0.05799 | 0.35023 | 0.90908 | 0.00000 | 0.25417 | 0.01389 | 0.46945 | 0.01018 | 0.00000 | 0.44241 |
| **Hist2h2bb** | 0.41257 | 0.00827 | 0.00264 | 0.01034 | 0.01791 | 0.02084 | 0.00723 | 0.01210 | 0.01250 | 0.01418 | 0.01285 | 0.00770 | 0.01199 | 0.00932 | 0.03472 | 0.00720 |
| **Hist2h2be** | 0.00632 | 2.40563 | 0.00597 | 0.20610 | 0.00790 | 0.00679 | 0.00659 | 0.01011 | 0.03950 | 0.23786 | 0.01090 | 0.00798 | 0.01191 | 0.00694 | 0.00000 | 0.00638 |
| **Hist3h2a** | 0.18829 | 0.00794 | 0.00000 | 0.00948 | 0.03080 | 0.50162 | 0.00702 | 0.48320 | 0.02285 | 0.06689 | 0.00907 | 0.00666 | 0.00648 | 0.01516 | 0.00268 | 0.01459 |
| **Hivep1** | 2.06285 | 1.21662 | 0.76811 | 2.52936 | 0.61491 | 0.00492 | 0.03531 | 0.53772 | 2.64280 | 2.51094 | 2.64912 | 0.07349 | 1.34877 | 0.00000 | 0.06307 | 2.03774 |
| **Hivep2** | 0.37962 | 0.00676 | 0.00514 | 0.01038 | 0.00712 | 0.00654 | 0.00696 | 0.00868 | 0.01159 | 0.06577 | 0.00432 | 1.42766 | 0.01196 | 0.01036 | 0.00451 | 0.00668 |
| **Hivep3** | 0.01248 | 0.01799 | 0.03655 | 0.01047 | 0.00940 | 0.00604 | 0.00724 | 0.10947 | 0.01231 | 0.01985 | 0.01243 | 0.00753 | 0.00850 | 0.00968 | 0.02366 | 0.00736 |
| **Hjurp** | 1.05102 | 0.06553 | 0.04539 | 0.74142 | 2.09332 | 0.02964 | 0.01658 | 0.25711 | 2.92608 | 0.05947 | 0.00000 | 0.12322 | 0.01202 | 0.02841 | 0.71925 | 0.00564 |
| **Hk1** | 0.64918 | 0.01488 | 0.01409 | 0.75019 | 0.01428 | 0.01278 | 0.00000 | 5.37906 | 0.84444 | 5.54518 | 11.12189 | 0.35025 | 2.38614 | 0.06565 | 0.07285 | 0.00354 |
| **Hk1os** | 0.01381 | 0.00783 | 0.01115 | 0.28533 | 0.00940 | 0.00660 | 0.00735 | 0.01193 | 0.00711 | 0.09126 | 0.01325 | 0.07438 | 0.01275 | 0.01158 | 0.00555 | 0.00743 |
| **Hk2** | 2.83061 | 1.12652 | 7.71380 | 0.23050 | 1.28763 | 0.06370 | 2.46202 | 1.63976 | 0.17277 | 0.00000 | 0.00000 | 0.02237 | 0.97637 | 0.03547 | 0.15161 | 2.84438 |
| **Hkdc1** | 2.51673 | 0.02703 | 11.05545 | 0.23031 | 0.30687 | 0.00422 | 10.59754 | 5.96571 | 0.82066 | 2.77223 | 4.06573 | 1.39658 | 1.80381 | 0.08789 | 0.51283 | 0.01234 |
| **Hlcs** | 0.54054 | 3.75125 | 3.41269 | 0.81608 | 2.20974 | 0.00709 | 1.21859 | 1.14523 | 0.54564 | 2.57782 | 2.42403 | 0.05091 | 0.51716 | 0.75482 | 1.89861 | 0.00308 |
| **Hlf** | 0.40654 | 0.00621 | 0.01165 | 0.24183 | 0.00696 | 0.00601 | 1.29372 | 0.03562 | 0.00565 | 0.68285 | 0.20374 | 0.01416 | 0.00813 | 0.01554 | 0.09639 | 0.00689 |
| **Hltf** | 1.77674 | 3.58513 | 1.60057 | 0.35795 | 0.16410 | 0.54774 | 3.99824 | 1.91838 | 0.16928 | 0.00000 | 1.10657 | 1.25002 | 0.08963 | 0.05663 | 2.34071 | 0.00279 |
| **Hmbox1** | 0.00000 | 2.46260 | 0.06848 | 2.12170 | 0.00750 | 0.00459 | 0.00589 | 1.60787 | 3.54403 | 4.25795 | 0.46446 | 3.53719 | 0.33494 | 0.75789 | 3.72324 | 0.00510 |
| **Hmbs** | 9.25850 | 3.70695 | 7.26097 | 11.21328 | 8.28038 | 6.89612 | 3.32556 | 4.52483 | 14.52157 | 14.15364 | 7.92282 | 4.09662 | 13.33668 | 1.62716 | 6.26493 | 5.44128 |
| **Hmces** | 5.57837 | 0.00534 | 0.77079 | 11.56234 | 10.46279 | 3.81821 | 2.69534 | 1.10077 | 7.50188 | 3.91740 | 0.73190 | 0.17878 | 0.11592 | 0.00000 | 1.08248 | 1.99548 |
| **Hmg20a** | 1.92529 | 1.62687 | 2.54474 | 1.87125 | 5.28206 | 4.30798 | 0.04455 | 3.36713 | 0.00000 | 4.67794 | 0.85522 | 0.08931 | 2.72917 | 2.89192 | 2.19664 | 1.54177 |
| **Hmg20b** | 1.39338 | 0.04770 | 0.01233 | 0.07250 | 2.34506 | 0.40068 | 0.03429 | 1.52603 | 0.02926 | 7.13631 | 0.19825 | 0.06294 | 5.87078 | 0.01063 | 0.00000 | 5.66255 |
| **Hmga2** | 17.27296 | 16.21270 | 16.37351 | 0.34546 | 12.49503 | 5.29449 | 13.51384 | 23.44830 | 0.15207 | 9.16585 | 13.00374 | 18.70371 | 12.65915 | 31.09772 | 2.72103 | 9.36100 |
| **Hmga2.ps1** | 0.01365 | 0.00757 | 0.00818 | 0.35426 | 0.04110 | 0.01278 | 0.00718 | 0.26144 | 0.07554 | 0.08373 | 0.00813 | 0.10855 | 0.00670 | 0.02077 | 0.56891 | 0.00735 |
| **Hmgb1** | 21.97254 | 13.53041 | 24.10313 | 6.27584 | 18.70268 | 9.13661 | 15.61065 | 22.99650 | 21.54049 | 28.14530 | 14.17101 | 15.15578 | 17.27441 | 37.66324 | 19.78082 | 12.45721 |
| **Hmgb1.rs17** | 2.28881 | 1.14932 | 2.34011 | 0.45527 | 2.26929 | 0.70713 | 1.86929 | 2.26744 | 2.90860 | 2.84107 | 1.15093 | 0.15611 | 1.23429 | 4.43889 | 1.90302 | 1.02938 |
| **Hmgb2** | 13.61732 | 14.03265 | 7.99578 | 1.45771 | 8.91883 | 7.17737 | 6.09782 | 20.47599 | 3.76512 | 5.45956 | 6.68214 | 6.87805 | 9.81218 | 17.92972 | 16.66474 | 6.82254 |
| **Hmgb3** | 10.71749 | 8.91611 | 12.39247 | 12.59989 | 9.76079 | 0.71092 | 1.84178 | 15.36137 | 21.33593 | 15.10571 | 6.71057 | 2.89454 | 7.09354 | 19.98363 | 21.53091 | 2.09410 |
| **Hmgcl** | 8.47888 | 5.24359 | 12.12015 | 3.23653 | 4.40699 | 6.56903 | 10.50896 | 5.65289 | 5.42842 | 6.42355 | 3.32849 | 0.23618 | 4.90019 | 0.08381 | 3.93416 | 6.96643 |
| **Hmgcll1** | 0.01372 | 0.00779 | 0.01119 | 5.47581 | 0.00927 | 0.00601 | 0.00724 | 0.01074 | 0.80019 | 0.00701 | 0.04660 | 0.01525 | 0.01177 | 0.00000 | 0.36513 | 0.00628 |
| **Hmgcr** | 2.76717 | 5.23129 | 6.80951 | 0.82903 | 0.84661 | 0.79602 | 6.08787 | 1.08089 | 1.60407 | 0.55081 | 1.20660 | 0.08600 | 0.00000 | 3.00184 | 0.87908 | 0.54559 |
| **Hmgcs1** | 6.31423 | 5.31935 | 8.45417 | 9.60240 | 0.46401 | 5.17995 | 0.69587 | 3.27200 | 9.31813 | 9.73084 | 2.97204 | 5.53650 | 3.07430 | 6.18526 | 8.89751 | 2.73276 |
| **Hmgcs2** | 0.38091 | 0.00744 | 0.00523 | 0.00807 | 0.00700 | 0.00740 | 0.00725 | 0.01407 | 0.01140 | 0.71160 | 0.01035 | 0.00690 | 0.01190 | 0.01418 | 0.11232 | 0.00692 |
| **Hmgn1** | 23.50380 | 8.65301 | 13.80074 | 13.79960 | 11.34589 | 8.43078 | 8.29418 | 4.52675 | 8.38504 | 33.32367 | 24.63822 | 9.79334 | 25.55026 | 12.19381 | 6.37830 | 15.73475 |
| **Hmgn2** | 18.35379 | 6.64394 | 16.94190 | 17.85249 | 14.41175 | 7.94062 | 11.32939 | 23.19439 | 25.66858 | 30.05963 | 23.68728 | 7.07158 | 16.02628 | 14.14007 | 8.99758 | 11.24495 |
| **Hmgn3** | 0.01353 | 0.00807 | 0.01222 | 1.92426 | 0.00925 | 0.00636 | 0.00725 | 0.01100 | 0.38688 | 0.41444 | 0.01048 | 0.00599 | 0.01212 | 0.00895 | 0.00288 | 0.00724 |
| **Hmgn5** | 11.06274 | 11.94085 | 12.26545 | 5.92743 | 18.50853 | 7.26718 | 7.07196 | 1.89076 | 21.01743 | 0.96395 | 0.00314 | 5.01922 | 2.36886 | 5.38159 | 7.03427 | 1.62992 |
| **Hmgxb3** | 3.45910 | 2.31091 | 4.63036 | 8.96901 | 2.91364 | 0.01440 | 5.44776 | 5.90291 | 11.00583 | 7.03924 | 2.39730 | 0.23072 | 5.69820 | 0.17323 | 2.24526 | 3.92701 |
| **Hmgxb4** | 3.33605 | 5.25978 | 5.26857 | 4.70390 | 6.78225 | 0.03828 | 3.37631 | 4.54872 | 5.37119 | 3.86149 | 4.47700 | 0.83926 | 1.08871 | 13.02800 | 9.04511 | 1.40292 |
| **Hmha1** | 3.06184 | 0.01353 | 0.96457 | 0.05085 | 0.01683 | 0.01411 | 0.00296 | 0.00000 | 0.01167 | 0.57244 | 2.86792 | 0.00000 | 0.96167 | 0.00788 | 0.06080 | 0.27247 |
| **Hmmr** | 2.80723 | 3.70768 | 3.46332 | 10.26362 | 4.08906 | 2.54174 | 0.92826 | 4.89701 | 14.64799 | 4.03729 | 9.89294 | 10.31370 | 1.46636 | 23.65482 | 18.92274 | 1.61548 |
| **Hmox1** | 0.00794 | 0.00734 | 0.00622 | 0.34091 | 0.00919 | 0.00691 | 0.00652 | 1.13587 | 0.09013 | 0.00000 | 0.09793 | 0.11392 | 0.89930 | 0.00953 | 0.00000 | 0.00670 |
| **Hmox2** | 9.40436 | 4.23657 | 3.24441 | 0.63852 | 5.67095 | 4.54982 | 6.30992 | 4.53388 | 0.74784 | 6.99134 | 0.12161 | 7.06282 | 2.59121 | 0.62459 | 5.91142 | 4.58203 |
| **Hn1** | 15.87414 | 10.58357 | 11.92039 | 15.35085 | 1.64976 | 9.68117 | 6.84741 | 18.53585 | 22.27785 | 21.28669 | 13.18737 | 3.18182 | 14.96005 | 0.15245 | 2.08263 | 6.76417 |
| **Hn1l** | 12.09459 | 6.13765 | 6.45870 | 6.23530 | 9.13839 | 5.51524 | 2.49340 | 2.98179 | 5.57758 | 2.99051 | 1.06617 | 1.54333 | 1.50482 | 3.13245 | 6.54793 | 4.22112 |
| **Hnf1a** | 0.19442 | 0.00756 | 0.02042 | 1.08531 | 0.01631 | 0.00602 | 0.03696 | 0.01380 | 0.28826 | 0.17418 | 0.34357 | 0.04791 | 1.75829 | 0.00541 | 0.02642 | 0.00610 |
| **Hnf1b** | 0.00634 | 0.97852 | 0.01194 | 0.64954 | 0.00843 | 0.00748 | 0.00683 | 0.98788 | 0.00000 | 0.01274 | 0.00598 | 0.00577 | 0.00537 | 0.00485 | 0.00000 | 0.01402 |
| **Hnf4a** | 6.68794 | 3.46321 | 3.70935 | 0.00682 | 10.50255 | 0.00000 | 0.74223 | 0.00000 | 0.00854 | 0.00000 | 0.01816 | 0.01103 | 0.01486 | 0.01222 | 0.00000 | 0.01180 |
| **Hnf4aos** | 0.01237 | 0.00799 | 0.01981 | 0.70486 | 0.00910 | 0.00599 | 0.00710 | 0.01215 | 0.29537 | 0.00368 | 0.01033 | 0.00747 | 0.48615 | 0.03140 | 0.00165 | 0.00682 |
| **Hnmt** | 0.01363 | 0.00766 | 0.01115 | 2.38864 | 0.00931 | 0.00626 | 0.00733 | 0.01131 | 1.38023 | 0.00279 | 0.01417 | 0.01445 | 0.01266 | 0.00715 | 0.35002 | 0.00685 |

| | | | | | | | | | | | | | | | | |
|---|---|---|---|---|---|---|---|---|---|---|---|---|---|---|---|---|
| **Hnrnpa0** | 15.12692 | 12.15198 | 7.65664 | 8.22095 | 7.82737 | 8.21751 | 10.16372 | 2.71016 | 9.66955 | 13.91194 | 9.96986 | 6.63980 | 12.04205 | 25.94400 | 11.23129 | 9.75773 |
| **Hnrnpa1** | 0.53454 | 0.73816 | 1.48809 | 0.38001 | 0.86970 | 0.01361 | 0.61099 | 1.25088 | 0.22403 | 0.93176 | 2.10420 | 0.00000 | 0.74275 | 6.30947 | 1.20733 | 0.66433 |
| **Hnrnpa2b1** | 16.85378 | 14.14499 | 17.28732 | 10.33642 | 11.75883 | 11.98700 | 5.82919 | 24.64239 | 12.66962 | 10.54407 | 13.70760 | 13.99845 | 13.34777 | 24.51457 | 13.18039 | 12.53303 |
| **Hnrnpa3** | 18.07159 | 14.76658 | 19.64484 | 13.58500 | 13.76690 | 11.67811 | 9.16380 | 11.36887 | 10.02870 | 20.71751 | 19.03821 | 21.14965 | 13.91291 | 31.34764 | 16.14382 | 10.29074 |
| **Hnrnpab** | 31.76484 | 19.67667 | 25.15915 | 27.19944 | 23.29654 | 12.50219 | 17.52642 | 29.81239 | 34.50634 | 45.46617 | 36.55526 | 23.71565 | 24.62454 | 30.91638 | 28.22571 | 12.23153 |
| **Hnrnpc** | 34.22168 | 23.20034 | 32.41307 | 15.51894 | 23.35530 | 12.37867 | 19.51418 | 38.09457 | 23.38967 | 42.96462 | 30.51351 | 17.35849 | 31.04891 | 44.05131 | 35.38794 | 19.72748 |
| **Hnrnpd** | 23.67529 | 14.59755 | 12.30803 | 7.74864 | 17.36312 | 12.76045 | 13.00593 | 15.36542 | 11.98930 | 19.75660 | 17.28739 | 11.89524 | 16.57044 | 21.17137 | 14.67708 | 10.53464 |
| **Hnrnpdl** | 18.23278 | 12.36662 | 8.17077 | 18.24873 | 13.01659 | 6.02105 | 7.71951 | 11.29410 | 22.51671 | 26.44289 | 17.18669 | 16.96563 | 18.76050 | 17.86464 | 18.07363 | 15.01943 |
| **Hnrnpf** | 38.06716 | 23.74673 | 35.41632 | 22.11878 | 26.01931 | 16.57218 | 20.55126 | 40.13729 | 27.36288 | 41.08345 | 26.86345 | 21.53256 | 32.81596 | 15.64413 | 31.49733 | 16.13325 |
| **Hnrnph1** | 12.84109 | 12.97310 | 15.32748 | 6.06056 | 11.58056 | 9.78701 | 10.40880 | 21.83695 | 5.49195 | 30.48582 | 13.27927 | 17.04535 | 18.70663 | 29.81771 | 24.37454 | 9.75790 |
| **Hnrnph2** | 7.95127 | 11.02708 | 7.51113 | 4.53620 | 2.46623 | 3.12510 | 7.15758 | 10.15927 | 2.23332 | 7.65710 | 4.05794 | 0.74084 | 6.01984 | 26.29819 | 14.69806 | 2.00144 |
| **Hnrnph3** | 0.07737 | 0.00773 | 0.14966 | 0.12608 | 0.01626 | 0.00641 | 0.00704 | 0.09705 | 0.39491 | 0.18250 | 0.02121 | 0.00513 | 0.58267 | 0.10500 | 0.30425 | 0.85917 |
| **Hnrnpk** | 28.36681 | 24.74617 | 25.96798 | 17.58046 | 20.51426 | 13.52352 | 18.17608 | 32.75590 | 18.37923 | 28.67791 | 22.10088 | 7.51013 | 26.19020 | 24.59128 | 24.69597 | 13.22232 |
| **Hnrnpl** | 0.83545 | 1.08893 | 1.12974 | 2.03644 | 0.38130 | 0.04546 | 0.19993 | 0.00000 | 5.04845 | 0.00000 | 7.01991 | 0.09509 | 1.76845 | 11.20466 | 3.78621 | 2.08861 |
| **Hnrnpll** | 3.51800 | 9.88656 | 4.65670 | 4.86908 | 8.65378 | 3.71159 | 6.78163 | 3.30153 | 14.35204 | 12.34547 | 6.79687 | 18.75210 | 5.20807 | 16.81637 | 15.59175 | 1.86599 |
| **Hnrnpm** | 23.85765 | 4.22209 | 5.28571 | 12.89781 | 12.84075 | 10.19340 | 5.12717 | 23.97647 | 12.65088 | 26.98135 | 20.24650 | 4.52798 | 19.88436 | 1.09851 | 5.98901 | 14.61812 |
| **Hnrnpr** | 13.69808 | 11.33533 | 5.25021 | 11.21616 | 5.64499 | 7.56156 | 7.93681 | 10.42797 | 12.20431 | 15.83793 | 4.12586 | 14.77811 | 5.34119 | 4.31217 | 11.71168 | 4.70384 |
| **Hnrnpu** | 23.88685 | 23.70133 | 23.40038 | 16.03215 | 18.91093 | 7.41557 | 13.20277 | 25.86394 | 25.76246 | 28.37593 | 24.20604 | 8.22090 | 23.82907 | 33.73645 | 27.29163 | 12.44144 |
| **Hnrnpul1** | 1.30494 | 0.11104 | 0.91518 | 1.58350 | 0.11875 | 0.87746 | 0.90402 | 1.75024 | 1.20718 | 0.28777 | 0.00000 | 0.12786 | 2.79464 | 0.20511 | 0.94436 | 0.80379 |
| **Hnrnpul2** | 4.52713 | 3.36679 | 3.42660 | 14.01786 | 0.10632 | 3.16932 | 5.21502 | 2.75883 | 13.89575 | 5.35397 | 6.12730 | 6.09628 | 3.74381 | 15.25820 | 10.87762 | 1.18253 |
| **Hoga1** | 0.40460 | 0.00772 | 0.00324 | 0.08938 | 0.01738 | 0.00766 | 0.00666 | 0.22321 | 0.01064 | 0.00765 | 0.01205 | 0.00665 | 0.26341 | 0.01476 | 0.02644 | 0.00698 |
| **Homer1** | 0.98111 | 0.00652 | 0.00000 | 4.09509 | 0.03044 | 1.14201 | 0.00589 | 0.05885 | 0.05334 | 0.78466 | 0.67043 | 0.02125 | 0.00000 | 0.00000 | 3.34755 | 0.00537 |
| **Homer2** | 4.11663 | 2.57877 | 0.00000 | 4.63263 | 2.66492 | 3.94955 | 2.36085 | 1.90138 | 4.19878 | 1.97014 | 0.90131 | 0.06936 | 0.46182 | 1.53281 | 0.46213 | 2.01136 |
| **Homer3** | 0.01264 | 0.00813 | 0.18397 | 0.00566 | 0.00907 | 0.00600 | 0.01835 | 0.01179 | 0.01117 | 0.10995 | 0.93666 | 0.01743 | 0.03149 | 0.01146 | 0.00000 | 0.00722 |
| **Homez** | 0.35063 | 0.00759 | 0.42330 | 0.51344 | 0.01344 | 0.00538 | 0.03663 | 0.00769 | 0.70022 | 0.69154 | 0.00568 | 0.02120 | 0.00917 | 0.00403 | 0.45723 | 0.00673 |
| **Hook1** | 0.01182 | 0.00775 | 0.00694 | 0.78794 | 0.00998 | 0.00647 | 0.00745 | 0.00954 | 0.64789 | 0.41108 | 0.05695 | 0.00000 | 0.04393 | 0.29638 | 4.99861 | 0.00765 |
| **Hook2** | 9.12381 | 0.05860 | 1.51147 | 1.84680 | 4.04663 | 0.00000 | 4.05847 | 0.87409 | 1.02873 | 3.18312 | 4.46274 | 4.42050 | 8.31371 | 0.00000 | 0.00000 | 3.05180 |
| **Hook3** | 0.43191 | 0.00460 | 0.07400 | 0.61422 | 0.02175 | 0.00607 | 2.21299 | 0.00000 | 0.05160 | 0.46732 | 0.39474 | 4.68349 | 0.00979 | 2.03057 | 0.99856 | 0.00647 |
| **Hopx** | 0.01364 | 0.00798 | 0.01166 | 0.14046 | 0.00928 | 0.00659 | 0.00751 | 0.01336 | 0.98709 | 0.00955 | 0.00710 | 0.02674 | 0.01296 | 0.01044 | 0.01531 | 0.00733 |
| **Hormad1** | 0.00000 | 3.97535 | 5.64711 | 5.49246 | 0.00655 | 0.00246 | 5.44645 | 0.96490 | 7.85923 | 5.44090 | 1.69785 | 17.09658 | 0.43219 | 8.81335 | 6.89970 | 0.00000 |
| **Hormad2** | 1.15868 | 0.00626 | 0.72842 | 0.00845 | 0.00525 | 0.00567 | 0.00640 | 0.00653 | 0.01103 | 0.00000 | 0.00778 | 0.01306 | 0.27225 | 0.63244 | 0.41125 | 0.01344 |
| **Hotairm1** | 0.01370 | 0.00808 | 0.01262 | 0.16392 | 0.00943 | 0.00666 | 0.00754 | 0.01361 | 0.03826 | 0.05935 | 0.01391 | 0.00820 | 0.01486 | 0.01419 | 0.00650 | 0.00726 |
| **Hoxa1** | 0.00969 | 0.00766 | 0.00770 | 0.00444 | 0.00939 | 0.00693 | 0.00708 | 0.78247 | 0.26194 | 0.35922 | 0.00548 | 0.01399 | 0.00580 | 0.00665 | 0.08853 | 0.00779 |
| **Hoxa10** | 0.00896 | 0.00779 | 0.00759 | 0.32102 | 0.00968 | 0.00702 | 0.00740 | 1.15542 | 0.00529 | 0.01087 | 0.00560 | 0.00610 | 0.00544 | 0.01190 | 0.00000 | 0.00733 |
| **Hoxa7** | 0.01368 | 0.00798 | 0.01196 | 0.75429 | 0.00918 | 0.00665 | 0.00737 | 0.01177 | 0.00000 | 0.26210 | 0.01294 | 0.01729 | 0.01263 | 0.01080 | 0.00610 | 0.00760 |
| **Hoxb13** | 1.77231 | 0.00612 | 0.71667 | 0.14744 | 0.00468 | 0.00753 | 0.00603 | 0.00377 | 0.00939 | 0.00510 | 0.15062 | 0.00630 | 0.02683 | 0.00688 | 0.16568 | 0.01245 |
| **Hoxb7** | 1.21533 | 0.00719 | 0.00371 | 0.00643 | 0.03271 | 0.00717 | 0.00630 | 0.00573 | 0.01145 | 1.53904 | 0.01079 | 0.00343 | 0.01136 | 0.01192 | 0.00884 | 0.00654 |
| **Hoxb9** | 0.01360 | 0.00791 | 0.01215 | 0.02839 | 0.00913 | 0.00651 | 0.00746 | 0.01117 | 0.02526 | 0.25400 | 0.03535 | 0.00655 | 0.01102 | 0.00588 | 0.13668 | 0.01418 |
| **Hoxc13** | 0.06394 | 0.00794 | 0.00614 | 0.01044 | 0.01392 | 0.00718 | 0.47423 | 0.00999 | 0.01234 | 0.01351 | 0.01312 | 0.00742 | 0.01134 | 0.01212 | 0.01048 | 0.00746 |
| **Hoxd11** | 0.01341 | 0.00793 | 0.01671 | 0.02644 | 0.00939 | 0.01253 | 0.00756 | 0.01294 | 0.06882 | 0.03151 | 0.01083 | 0.01694 | 0.02144 | 0.02421 | 0.28065 | 0.00725 |
| **Hoxd8** | 0.01373 | 0.00785 | 0.01208 | 0.14927 | 0.00929 | 0.00649 | 0.00748 | 0.01222 | 0.00965 | 0.04669 | 0.01145 | 0.00672 | 0.01218 | 0.00578 | 0.40605 | 0.00719 |
| **Hoxd9** | 0.00985 | 0.00858 | 0.00886 | 0.01053 | 0.00959 | 0.00689 | 0.00823 | 0.30839 | 0.01220 | 0.02083 | 0.01031 | 0.00819 | 0.00636 | 0.01076 | 0.02469 | 0.00737 |
| **Hp** | 0.00688 | 0.00743 | 0.00788 | 1.34492 | 0.00898 | 0.00639 | 0.00642 | 0.67881 | 2.59249 | 0.58832 | 0.00000 | 0.00000 | 1.14268 | 0.01311 | 0.00000 | 0.00631 |
| **Hp1bp3** | 1.97829 | 0.10472 | 1.24932 | 1.79665 | 3.69355 | 3.27295 | 2.42234 | 1.60958 | 2.93612 | 4.88463 | 2.17303 | 3.71877 | 3.16588 | 3.61388 | 1.44412 | 3.37289 |
| **Hpcal1** | 0.11842 | 0.19621 | 1.34451 | 5.42767 | 0.01623 | 0.19337 | 0.16736 | 0.04265 | 2.97622 | 0.91986 | 0.05897 | 0.02815 | 0.65008 | 0.02005 | 0.04374 | 4.52334 |
| **Hpd** | 0.01331 | 0.00789 | 0.01171 | 0.71227 | 0.00906 | 0.00632 | 0.00738 | 0.01074 | 0.97905 | 0.52115 | 0.00397 | 0.00374 | 0.01107 | 0.02555 | 0.27500 | 0.00694 |
| **Hpdl** | 0.07260 | 2.52530 | 1.82634 | 2.79967 | 2.44205 | 7.41255 | 5.03422 | 11.89876 | 7.28045 | 12.94371 | 6.55976 | 0.70897 | 8.16072 | 0.03278 | 2.54309 | 1.27239 |
| **Hpf1** | 8.89665 | 5.64147 | 8.52925 | 19.15222 | 6.81186 | 1.22155 | 4.27829 | 8.07006 | 10.50689 | 10.32481 | 12.82295 | 7.13013 | 8.65892 | 6.92405 | 2.15916 | 7.71939 |
| **Hpgds** | 0.71112 | 0.00706 | 0.14806 | 0.26306 | 0.02856 | 0.00609 | 0.05655 | 0.01172 | 0.36642 | 0.67192 | 1.49837 | 0.03263 | 0.11784 | 0.74001 | 0.35806 | 0.04458 |
| **Hprt** | 22.09427 | 20.01838 | 35.86885 | 16.66580 | 17.33062 | 20.31810 | 18.24539 | 41.61418 | 20.06227 | 40.95460 | 32.64043 | 19.62429 | 32.13485 | 33.88437 | 21.25586 | 16.60624 |
| **Hps1** | 1.41560 | 0.00707 | 0.00325 | 0.31846 | 0.01449 | 0.00784 | 0.00595 | 0.28700 | 0.28416 | 0.60849 | 0.01176 | 0.00366 | 0.80183 | 0.01075 | 0.00052 | 0.00647 |
| **Hps3** | 1.72578 | 0.00488 | 1.43506 | 0.47396 | 0.02375 | 0.00502 | 2.10001 | 0.93843 | 0.17399 | 0.84848 | 1.31162 | 0.52172 | 1.34544 | 0.38689 | 1.61551 | 0.74032 |
| **Hps4** | 2.43169 | 0.22998 | 1.46039 | 0.61390 | 0.00000 | 1.90090 | 0.63711 | 3.61683 | 0.14219 | 2.53772 | 0.62216 | 0.30806 | 2.14705 | 0.00298 | 0.00000 | 5.25191 |
| **Hps5** | 3.94480 | 5.40308 | 1.87550 | 2.03129 | 0.15356 | 0.00150 | 4.21434 | 0.82231 | 5.99425 | 4.46377 | 2.54623 | 0.47168 | 0.00000 | 3.76884 | 3.38424 | 0.00000 |
| **Hps6** | 1.15828 | 0.10761 | 1.04966 | 3.00107 | 0.01090 | 0.00392 | 0.80445 | 1.06429 | 2.65827 | 3.93043 | 0.49409 | 0.08487 | 0.24982 | 0.30051 | 0.36487 | 0.01259 |
| **Hpse** | 0.00000 | 0.00499 | 0.00000 | 1.91303 | 1.55128 | 0.00747 | 3.48337 | 0.11623 | 0.79486 | 0.00000 | 0.47569 | 0.02390 | 1.12232 | 0.00802 | 0.44072 | 0.00606 |
| **Hr** | 0.01211 | 0.00767 | 0.05939 | 0.18530 | 0.00917 | 0.00845 | 0.30293 | 0.01079 | 0.00855 | 0.01403 | 0.01355 | 0.00741 | 0.01115 | 0.01066 | 0.01120 | 0.00733 |
| **Hras** | 6.54069 | 3.40711 | 1.64397 | 8.51806 | 5.74378 | 4.15596 | 4.75100 | 6.02849 | 12.83588 | 6.07902 | 9.03862 | 2.08142 | 5.70095 | 0.16371 | 2.14642 | 0.45563 |
| **Hs1bp3** | 5.41298 | 1.53350 | 4.55229 | 0.00000 | 0.90293 | 0.12125 | 2.37899 | 5.32519 | 0.00572 | 2.10627 | 1.82934 | 0.10793 | 7.01707 | 0.00000 | 1.87328 | 0.71054 |
| **Hs2st1** | 4.61948 | 3.12453 | 2.63543 | 7.06126 | 1.78919 | 0.90933 | 5.36473 | 4.18163 | 2.98238 | 3.26151 | 2.16870 | 1.49783 | 2.77654 | 4.05261 | 4.80593 | 4.16215 |
| **Hs3st1** | 4.78980 | 0.04509 | 0.00359 | 0.27596 | 0.01504 | 0.00392 | 0.00327 | 0.25606 | 0.00705 | 0.96091 | 0.01079 | 0.01044 | 0.00673 | 0.01029 | 0.00120 | 0.00275 |
| **Hs3st3b1** | 0.17617 | 0.42838 | 0.67340 | 0.43992 | 0.04814 | 2.23896 | 0.00630 | 0.08798 | 0.00554 | 0.00000 | 0.37924 | 0.00000 | 1.34413 | 1.13628 | 0.00000 | 1.98649 |

| | | | | | | | | | | | | | | | | |
|---|---|---|---|---|---|---|---|---|---|---|---|---|---|---|---|---|
| **Hs3st5** | 0.01847 | 0.00833 | 0.01184 | 0.25215 | 0.01057 | 0.00665 | 0.00753 | 0.01380 | 0.23866 | 0.01407 | 0.01121 | 0.00793 | 0.01276 | 0.01177 | 0.00535 | 0.00741 |
| **Hs3st6** | 0.01276 | 0.00789 | 0.00847 | 0.61613 | 0.00923 | 0.00663 | 0.00705 | 0.01934 | 0.84470 | 0.35664 | 0.47692 | 0.00309 | 0.00741 | 0.00524 | 0.02946 | 0.00720 |
| **Hs6st1** | 3.14276 | 0.58646 | 4.23051 | 6.03756 | 0.01045 | 0.00283 | 1.48596 | 6.01785 | 7.40144 | 11.22408 | 6.20983 | 0.59677 | 2.34290 | 4.30675 | 2.77379 | 2.75676 |
| **Hs6st2** | 0.03200 | 0.00778 | 0.01568 | 2.54644 | 0.01776 | 0.00601 | 0.02737 | 0.01209 | 0.42117 | 0.00172 | 0.12350 | 0.02515 | 0.01239 | 0.08002 | 0.62714 | 0.01426 |
| **Hsbp1** | 11.38374 | 7.66584 | 10.91772 | 7.91776 | 6.49178 | 5.35050 | 6.16015 | 5.87884 | 0.00005 | 4.85443 | 11.50345 | 0.99455 | 8.19402 | 4.40567 | 2.25062 | 6.97125 |
| **Hsbp1l1** | 0.40497 | 0.33601 | 0.11986 | 0.27595 | 0.01650 | 0.04536 | 0.90863 | 0.20637 | 0.39448 | 1.49186 | 0.07720 | 0.05857 | 0.04216 | 0.01317 | 0.57596 | 0.01385 |
| **Hscb** | 5.03446 | 0.78825 | 3.24766 | 1.80619 | 5.40340 | 2.39762 | 1.39398 | 0.74041 | 2.06739 | 0.23763 | 1.25003 | 0.01788 | 0.63649 | 0.81908 | 0.04080 | 0.00624 |
| **Hsd11b2** | 0.19657 | 0.00735 | 0.03795 | 0.01057 | 0.00709 | 0.00830 | 0.00708 | 0.01518 | 0.01231 | 0.02420 | 0.01261 | 0.00742 | 0.26391 | 0.01199 | 0.01063 | 0.49032 |
| **Hsd17b1** | 0.07021 | 0.06522 | 0.00267 | 0.00595 | 0.00770 | 0.00801 | 0.00682 | 1.80008 | 0.02410 | 0.00000 | 0.24692 | 0.02069 | 0.22167 | 0.01221 | 0.00000 | 0.00704 |
| **Hsd17b10** | 5.11340 | 4.52768 | 9.42140 | 0.35912 | 2.28057 | 2.65855 | 4.58402 | 2.42200 | 1.09623 | 3.33662 | 1.34321 | 0.25338 | 2.62449 | 1.15761 | 0.60205 | 4.86532 |
| **Hsd17b11** | 0.33249 | 0.00716 | 0.60305 | 0.00470 | 0.00631 | 0.00542 | 0.00617 | 0.00000 | 0.35926 | 0.35094 | 1.41781 | 0.00722 | 0.98470 | 0.28113 | 0.00000 | 0.00647 |
| **Hsd17b12** | 7.09061 | 5.07484 | 11.41476 | 2.68332 | 6.43456 | 6.00980 | 3.08436 | 7.03988 | 6.41246 | 3.06114 | 3.58993 | 0.49841 | 6.46154 | 3.99253 | 1.87482 | 2.60346 |
| **Hsd17b13** | 0.02094 | 0.00770 | 0.01398 | 10.21572 | 0.00869 | 0.01693 | 0.00686 | 0.01324 | 1.50969 | 0.69241 | 0.04761 | 0.07996 | 0.02415 | 1.90077 | 1.20870 | 0.01221 |
| **Hsd17b14** | 0.00000 | 0.01295 | 3.61558 | 6.10466 | 0.01979 | 0.09849 | 1.14071 | 9.74647 | 13.49559 | 21.03357 | 15.36105 | 5.95542 | 3.35478 | 0.41196 | 1.14256 | 0.01424 |
| **Hsd17b2** | 0.03880 | 0.00761 | 0.00773 | 0.00452 | 0.00824 | 0.00703 | 0.00755 | 0.01747 | 0.97748 | 0.23887 | 0.00000 | 0.01669 | 0.01244 | 0.23888 | 0.11841 | 0.00724 |
| **Hsd17b4** | 9.42740 | 6.78527 | 6.51908 | 0.28785 | 4.01217 | 0.00001 | 7.73915 | 8.69549 | 0.82276 | 2.38943 | 6.48125 | 0.13945 | 4.51697 | 3.26873 | 0.52672 | 1.00495 |
| **Hsd17b6** | 1.10803 | 0.00648 | 0.00000 | 1.13231 | 0.01395 | 12.80281 | 0.00581 | 0.00000 | 1.60639 | 0.85658 | 0.66307 | 0.20991 | 0.00000 | 0.47055 | 0.86767 | 0.00000 |
| **Hsd17b7** | 2.64596 | 0.01574 | 2.06676 | 0.54558 | 0.00000 | 0.01475 | 0.00497 | 1.20981 | 0.85082 | 1.10419 | 0.48444 | 0.07195 | 0.63398 | 4.14568 | 0.00000 | 0.01874 |
| **Hsd3b1** | 0.01157 | 0.00778 | 0.01216 | 0.00463 | 0.00843 | 0.00657 | 0.00728 | 0.01006 | 0.11187 | 5.00515 | 0.00820 | 0.00000 | 0.01068 | 0.01269 | 0.00608 | 0.00754 |
| **Hsd3b7** | 0.30334 | 0.03924 | 0.07197 | 1.63842 | 0.00708 | 2.92296 | 0.00677 | 0.00000 | 0.02382 | 1.72249 | 0.86210 | 0.00000 | 0.07273 | 0.00509 | 0.00000 | 1.58057 |
| **Hsdl1** | 0.59809 | 5.19146 | 0.46994 | 1.89907 | 2.80591 | 2.75620 | 0.00464 | 2.08698 | 0.00000 | 0.00000 | 0.30920 | 0.06234 | 0.00000 | 0.95493 | 2.60491 | 0.45723 |
| **Hsdl2** | 3.06702 | 1.30054 | 1.59192 | 2.24696 | 0.06763 | 0.93472 | 2.04226 | 1.54743 | 3.28596 | 3.14212 | 1.96045 | 6.27125 | 1.49393 | 0.00000 | 0.00000 | 1.50336 |
| **Hsf1** | 4.50031 | 0.06283 | 1.16262 | 16.09980 | 0.18684 | 0.00000 | 2.00324 | 2.26096 | 13.69355 | 2.62427 | 2.60246 | 0.16002 | 2.26493 | 0.51899 | 2.40300 | 2.03810 |
| **Hsf2** | 3.32421 | 3.21826 | 0.44217 | 0.71411 | 0.03079 | 3.28273 | 0.47363 | 0.26348 | 0.02442 | 0.00000 | 3.80495 | 0.15259 | 0.23714 | 0.14403 | 0.00000 | 0.00783 |
| **Hsf2bp** | 5.75247 | 1.01743 | 3.43209 | 5.44348 | 4.19896 | 0.01049 | 4.44708 | 2.25662 | 1.89851 | 3.90681 | 0.00000 | 0.09371 | 1.39596 | 0.33670 | 0.60358 | 2.35367 |
| **Hsf5** | 0.01717 | 0.00800 | 0.03984 | 0.47763 | 0.01155 | 0.00855 | 0.00763 | 0.01424 | 0.00275 | 0.00687 | 0.02572 | 0.00726 | 0.07401 | 0.01893 | 0.13079 | 0.00765 |
| **Hsh2d** | 0.01528 | 0.00816 | 0.00434 | 0.00434 | 0.01028 | 0.00747 | 0.00700 | 0.03388 | 0.01051 | 1.19984 | 0.72607 | 0.18863 | 0.00554 | 0.15244 | 0.08491 | 0.00732 |
| **Hsp90aa1** | 53.12343 | 40.47319 | 57.51495 | 17.30512 | 42.73428 | 26.82772 | 29.13402 | 61.60745 | 31.13670 | 49.45409 | 50.46280 | 40.39495 | 52.03360 | 54.45097 | 31.79987 | 24.21778 |
| **Hsp90ab1** | 56.45178 | 37.24382 | 60.12696 | 17.77811 | 41.64058 | 24.87951 | 34.67133 | 67.94667 | 30.20247 | 59.72398 | 56.03662 | 27.97231 | 55.28660 | 52.94024 | 39.66574 | 29.58575 |
| **Hsp90b1** | 16.21606 | 12.92870 | 29.44046 | 8.95140 | 13.57285 | 18.67018 | 13.52846 | 7.08350 | 3.23752 | 5.92144 | 4.17614 | 12.04818 | 19.31055 | 3.07796 | 12.17332 | 12.97207 |
| **Hspa12a** | 0.19169 | 0.00712 | 0.00224 | 0.90132 | 0.00749 | 0.00731 | 0.00698 | 0.51605 | 0.17810 | 0.09444 | 0.03427 | 0.01382 | 0.00583 | 0.00338 | 0.60732 | 0.00702 |
| **Hspa12b** | 0.01372 | 0.00812 | 0.01219 | 0.07947 | 0.00932 | 0.00665 | 0.00754 | 0.01396 | 0.20479 | 0.01424 | 0.01109 | 0.00791 | 0.01277 | 0.01239 | 0.00732 | 0.00740 |
| **Hspa13** | 0.00241 | 0.00840 | 0.64076 | 0.34072 | 0.00728 | 0.00624 | 0.00823 | 0.33480 | 0.00409 | 0.01731 | 0.00908 | 0.01956 | 0.00000 | 0.01226 | 0.00000 | 0.74779 |
| **Hspa14** | 12.59827 | 9.91561 | 8.54076 | 7.86791 | 7.88585 | 6.83070 | 6.36196 | 11.52305 | 17.54911 | 9.37446 | 10.36668 | 3.63837 | 9.18550 | 6.47978 | 9.10295 | 6.35429 |
| **Hspa1a** | 0.11839 | 0.00801 | 0.07920 | 4.66758 | 0.00739 | 0.00481 | 0.00740 | 0.71588 | 9.61889 | 0.18523 | 0.24852 | 0.01827 | 0.00913 | 0.46463 | 0.19458 | 0.00555 |
| **Hspa1b** | 0.02716 | 0.04025 | 0.25396 | 1.81393 | 0.00769 | 0.00564 | 0.00635 | 0.20851 | 17.06998 | 0.00000 | 3.46043 | 0.00000 | 0.00000 | 0.03532 | 0.00000 | 0.00515 |
| **Hspa1l** | 0.01388 | 0.00815 | 0.01037 | 0.21497 | 0.00930 | 0.02540 | 0.00753 | 0.01390 | 0.25234 | 0.01390 | 0.01175 | 0.00791 | 0.01241 | 0.01188 | 0.00552 | 0.00735 |
| **Hspa2** | 2.10836 | 0.00445 | 0.01906 | 2.99505 | 0.00459 | 0.00637 | 1.81890 | 1.17014 | 0.08551 | 2.17416 | 0.95377 | 0.04208 | 0.55520 | 0.21225 | 1.30315 | 0.01142 |
| **Hspa4** | 18.85741 | 14.10759 | 12.42832 | 8.98625 | 14.13827 | 7.98797 | 9.73698 | 16.53514 | 12.12426 | 18.05372 | 6.97627 | 17.02328 | 8.77075 | 14.01618 | 15.06763 | 7.51590 |
| **Hspa4l** | 0.02315 | 0.00812 | 0.00456 | 3.67240 | 0.01113 | 2.68182 | 0.00707 | 0.00996 | 0.39329 | 0.56372 | 0.00498 | 0.04089 | 0.00336 | 0.00045 | 4.16541 | 0.00563 |
| **Hspa5** | 15.67733 | 12.37543 | 33.57085 | 19.05873 | 8.37425 | 12.77392 | 13.64232 | 15.87177 | 20.25418 | 7.45541 | 7.71685 | 4.05362 | 30.33644 | 17.63346 | 22.47952 | 13.65133 |
| **Hspa8** | 56.46366 | 33.02301 | 65.28548 | 16.36821 | 43.45648 | 27.73293 | 33.96195 | 71.96007 | 32.57189 | 67.82710 | 63.83474 | 34.31835 | 67.29732 | 33.68902 | 28.79418 | 32.96211 |
| **Hspa9** | 24.48572 | 18.47567 | 15.96361 | 17.06487 | 18.51408 | 13.08208 | 13.12449 | 21.20535 | 23.29314 | 19.81562 | 22.00688 | 16.48925 | 21.61524 | 29.05950 | 21.03982 | 12.02609 |
| **Hspb1** | 0.88341 | 0.00675 | 0.29224 | 22.00428 | 0.01781 | 0.02491 | 0.00303 | 0.00000 | 15.08547 | 5.01183 | 7.40432 | 0.57673 | 0.00000 | 0.00000 | 0.24353 | 0.39699 |
| **Hspb11** | 0.57128 | 0.31556 | 0.07003 | 0.23247 | 0.01512 | 0.55167 | 0.33706 | 1.02366 | 0.27315 | 0.45205 | 3.07987 | 0.09467 | 0.41636 | 0.70189 | 0.35604 | 0.09925 |
| **Hspb2** | 0.01239 | 0.00812 | 0.09611 | 0.01063 | 0.01043 | 0.00877 | 0.00681 | 0.01040 | 0.01240 | 0.00995 | 0.01244 | 0.00719 | 1.08112 | 0.01280 | 0.00893 | 0.00681 |
| **Hspb6** | 0.54010 | 1.96430 | 1.23426 | 0.39388 | 0.01537 | 0.00488 | 0.00620 | 0.01135 | 0.66207 | 0.00498 | 0.00781 | 0.01639 | 0.00735 | 0.00765 | 0.00000 | 0.00568 |
| **Hspb7** | 0.01901 | 0.00793 | 0.03311 | 0.02634 | 0.00872 | 0.00717 | 0.01784 | 0.03316 | 0.03056 | 0.04879 | 0.02323 | 0.00847 | 0.01757 | 0.01063 | 0.03243 | 0.00736 |
| **Hspb8** | 1.35760 | 5.75803 | 4.72949 | 0.00000 | 0.00551 | 0.00399 | 0.00380 | 0.94455 | 0.01166 | 1.12617 | 0.00000 | 0.03932 | 0.00000 | 0.00870 | 0.00000 | 0.02389 |
| **Hspb9** | 0.01270 | 0.00826 | 0.07256 | 0.00863 | 0.00902 | 0.00578 | 0.00758 | 0.01166 | 0.01221 | 0.49159 | 0.01119 | 0.00710 | 0.01041 | 0.01185 | 0.00825 | 0.00709 |
| **Hspbap1** | 1.89201 | 0.98531 | 0.82293 | 2.11835 | 2.10031 | 2.72186 | 2.18500 | 1.56795 | 2.70717 | 0.38808 | 1.46040 | 4.84253 | 0.59202 | 0.04359 | 1.23818 | 0.61112 |
| **Hspbp1** | 9.28170 | 0.08627 | 2.13257 | 1.79038 | 5.43069 | 4.13105 | 4.95538 | 19.56144 | 3.49594 | 15.09510 | 8.47464 | 0.98230 | 15.41124 | 0.00000 | 0.09475 | 6.06719 |
| **Hspd1** | 27.48433 | 18.87697 | 30.09299 | 7.06426 | 20.48956 | 19.89142 | 19.28290 | 27.51602 | 5.28338 | 20.45869 | 15.52383 | 6.41314 | 21.59637 | 15.89280 | 18.42721 | 17.10554 |
| **Hspe1** | 22.82783 | 2.50281 | 18.11948 | 9.89642 | 19.33180 | 6.62170 | 3.81820 | 4.71623 | 5.70977 | 10.27012 | 17.11633 | 1.55264 | 12.20264 | 1.38933 | 2.03822 | 8.66587 |
| **Hspg2** | 0.01585 | 0.00749 | 0.00197 | 0.41948 | 0.00776 | 0.99702 | 0.00709 | 0.00699 | 0.02418 | 0.06016 | 0.11533 | 0.23854 | 0.00481 | 0.00952 | 0.00000 | 1.13027 |
| **Hsph1** | 28.49946 | 21.44082 | 29.77312 | 11.90986 | 19.46378 | 9.44332 | 14.20921 | 13.25883 | 22.58925 | 8.72910 | 6.16756 | 8.23638 | 22.27970 | 17.49114 | 11.26501 | 14.78339 |
| **Htatip2** | 5.01268 | 3.25422 | 7.72831 | 0.33089 | 4.31486 | 1.28695 | 2.89812 | 2.39670 | 0.00000 | 0.00000 | 2.83188 | 0.08170 | 8.91340 | 0.32049 | 0.12712 | 4.75494 |
| **Htatsf1** | 3.65466 | 3.02042 | 3.09883 | 6.49892 | 2.95243 | 0.00154 | 2.74162 | 4.63728 | 3.75410 | 11.06586 | 6.32000 | 2.93201 | 2.36642 | 15.33657 | 13.79239 | 2.80836 |
| **Htr5b** | 0.00000 | 0.03996 | 0.00457 | 0.00330 | 0.00799 | 0.00698 | 1.34515 | 0.01661 | 0.00966 | 0.60986 | 0.24845 | 0.09464 | 0.07004 | 0.02611 | 1.15121 | 2.00858 |
| **Htr7** | 0.01375 | 0.00806 | 0.01226 | 0.07866 | 0.00931 | 0.00664 | 0.00753 | 0.01390 | 0.01133 | 0.01537 | 0.01385 | 0.00813 | 0.01267 | 0.01188 | 0.00929 | 0.00740 |
| **Htra1** | 0.56644 | 0.00615 | 3.46707 | 1.60183 | 0.17697 | 0.16711 | 2.06417 | 0.23208 | 2.42373 | 2.33775 | 0.51660 | 0.13935 | 1.51490 | 0.39730 | 1.49280 | 3.26757 |
| **Htra2** | 2.04871 | 2.71972 | 2.14910 | 0.00000 | 2.06708 | 5.23301 | 3.42621 | 4.69868 | 0.00533 | 1.90751 | 0.00000 | 0.23949 | 2.85676 | 0.41460 | 0.56191 | 4.03448 |

| | | | | | | | | | | | | | | | | |
|---|---|---|---|---|---|---|---|---|---|---|---|---|---|---|---|---|
| **Htra3** | 0.01362 | 0.00784 | 0.01232 | 0.00919 | 0.00932 | 0.00653 | 0.00734 | 0.01140 | 0.01225 | 0.33062 | 0.01283 | 0.00665 | 0.01213 | 0.01159 | 0.00880 | 0.00750 |
| **Htt** | 2.51729 | 3.96182 | 0.92670 | 1.24564 | 1.37488 | 4.46808 | 0.00254 | 2.65732 | 1.46500 | 0.90431 | 0.74440 | 0.08790 | 1.89675 | 0.09890 | 0.11317 | 2.54514 |
| **Hunk** | 0.57282 | 0.00749 | 0.00512 | 0.29289 | 0.00666 | 0.00707 | 0.00703 | 0.01326 | 0.00698 | 0.10886 | 0.01277 | 0.00777 | 0.01154 | 0.01106 | 0.00675 | 0.01352 |
| **Hus1** | 1.34289 | 0.00569 | 0.81703 | 3.01215 | 0.11788 | 2.16705 | 1.04736 | 2.96429 | 1.26676 | 1.14237 | 1.16823 | 0.26132 | 1.69492 | 5.88372 | 3.57103 | 0.02571 |
| **Hus1b** | 0.13207 | 0.00852 | 0.00745 | 0.01062 | 0.01619 | 0.00673 | 0.00673 | 0.01059 | 0.01235 | 0.00823 | 0.01236 | 0.00775 | 1.12434 | 0.01287 | 0.01084 | 0.00677 |
| **Huwe1** | 14.13622 | 11.75275 | 11.43454 | 13.13934 | 7.41270 | 8.02297 | 8.01787 | 23.32590 | 14.52727 | 17.24349 | 15.07132 | 18.12854 | 14.69994 | 30.25027 | 11.48504 | 2.76959 |
| **Hvcn1** | 1.60422 | 0.00625 | 2.01655 | 6.00882 | 0.01826 | 0.34251 | 0.00521 | 1.38842 | 3.58115 | 2.79210 | 1.61563 | 0.07099 | 0.44934 | 0.00000 | 0.50898 | 0.64395 |
| **Hyal1** | 0.73408 | 0.00571 | 0.17585 | 0.09826 | 0.08056 | 0.00519 | 1.93385 | 0.08184 | 0.32549 | 0.04344 | 0.21036 | 0.01270 | 0.03257 | 0.22190 | 0.08284 | 0.02721 |
| **Hyal2** | 2.69137 | 0.00363 | 1.48314 | 0.52648 | 2.53456 | 0.13133 | 0.35007 | 0.80993 | 0.00000 | 0.31532 | 0.25549 | 0.00126 | 0.10585 | 0.00812 | 0.00000 | 1.10357 |
| **Hyal3** | 0.24864 | 0.00714 | 0.61634 | 0.59010 | 0.14642 | 0.04387 | 0.12976 | 0.06975 | 0.94252 | 0.31802 | 2.32552 | 0.11753 | 0.42514 | 0.66407 | 0.34068 | 0.08600 |
| **Hydin** | 0.14234 | 0.00641 | 0.00000 | 0.13557 | 0.00760 | 0.00817 | 1.89480 | 0.00925 | 0.00639 | 0.00333 | 0.01057 | 0.00537 | 0.01648 | 0.06835 | 0.35241 | 0.01425 |
| **Hyi** | 0.08438 | 0.00780 | 0.62479 | 0.00942 | 0.00693 | 0.74343 | 0.00732 | 0.01258 | 0.01235 | 0.19794 | 0.01096 | 0.00711 | 0.00783 | 0.01389 | 0.00183 | 0.00683 |
| **Hykk** | 1.51246 | 0.66161 | 1.18514 | 0.39481 | 0.02814 | 0.00943 | 0.02552 | 0.69570 | 1.07300 | 0.77808 | 0.44012 | 0.04917 | 0.07862 | 0.28203 | 0.07086 | 0.01851 |
| **Hyls1** | 0.76309 | 0.33094 | 0.00769 | 1.10203 | 0.01560 | 0.00741 | 0.00701 | 0.06932 | 2.16053 | 0.12854 | 0.00000 | 0.02081 | 0.01292 | 0.03767 | 0.61915 | 0.00621 |
| **Hyou1** | 1.76026 | 4.30434 | 3.30865 | 0.90581 | 0.00595 | 0.09318 | 0.70581 | 1.28334 | 1.80871 | 0.85120 | 1.06101 | 0.07815 | 4.07948 | 4.30506 | 1.95073 | 4.60854 |
| **Hypk** | 11.04384 | 1.59357 | 10.35191 | 17.21107 | 7.22665 | 2.84412 | 3.01148 | 2.03850 | 16.74200 | 7.98480 | 18.65887 | 1.28436 | 5.87401 | 4.64310 | 8.74204 | 9.78133 |
| **Iah1** | 2.77661 | 2.63295 | 2.71713 | 1.96848 | 0.02523 | 0.30658 | 1.14856 | 2.28166 | 1.19370 | 0.32994 | 1.00407 | 0.13928 | 1.58632 | 0.06370 | 1.05626 | 0.05484 |
| **Iars** | 13.87844 | 11.93073 | 10.83692 | 13.75292 | 18.69875 | 6.43021 | 7.15044 | 14.38793 | 15.99064 | 6.54387 | 6.94659 | 15.65195 | 6.61445 | 7.14529 | 8.86713 | 7.34177 |
| **Iars2** | 4.84007 | 4.40734 | 0.84147 | 0.58678 | 5.50590 | 3.71625 | 2.56099 | 4.33562 | 0.26883 | 0.00000 | 1.84324 | 0.04894 | 0.18075 | 0.01409 | 1.55018 | 0.09637 |
| **Iba57** | 0.00000 | 0.00714 | 0.00032 | 0.06521 | 0.01637 | 0.03166 | 0.01633 | 0.96543 | 0.04285 | 0.31865 | 1.30937 | 0.00606 | 0.00000 | 0.01087 | 0.28579 | 2.06163 |
| **Ibtk** | 6.76166 | 0.77414 | 1.80378 | 9.68845 | 2.79323 | 4.44800 | 0.73686 | 8.55970 | 4.35300 | 7.15760 | 4.66670 | 12.82032 | 2.96880 | 15.05033 | 13.93391 | 4.75383 |
| **Ica1** | 0.01674 | 0.00764 | 0.00568 | 5.20001 | 0.00900 | 0.00568 | 0.00699 | 0.01270 | 2.44230 | 0.18160 | 0.01706 | 0.07710 | 0.01272 | 0.23234 | 2.63850 | 0.00665 |
| **Ica1l** | 0.74208 | 0.00664 | 0.00000 | 1.32197 | 0.00560 | 0.00559 | 0.00630 | 0.45950 | 1.01539 | 2.06283 | 0.12663 | 0.01057 | 0.03517 | 0.45144 | 2.15813 | 2.11062 |
| **Icam1** | 0.00000 | 0.00505 | 0.81613 | 2.79577 | 0.00494 | 0.91209 | 1.31490 | 7.19684 | 13.64567 | 1.75963 | 3.91513 | 3.07742 | 0.00000 | 1.25817 | 3.39533 | 3.22128 |
| **Icam2** | 0.01191 | 0.00823 | 0.05347 | 0.06941 | 0.00997 | 0.00642 | 0.00733 | 0.01251 | 0.02856 | 0.53220 | 0.01415 | 0.01612 | 0.63620 | 0.00920 | 0.02774 | 0.00726 |
| **Icam4** | 1.13707 | 0.00560 | 3.33056 | 0.22131 | 0.00000 | 0.84455 | 0.04647 | 3.58768 | 0.58865 | 0.76684 | 0.75927 | 0.09580 | 3.01434 | 0.00333 | 0.00000 | 0.00329 |
| **Ice1** | 3.79471 | 7.33240 | 3.73253 | 20.71979 | 0.14890 | 2.08855 | 4.91071 | 5.67876 | 12.68337 | 6.07849 | 6.71930 | 11.37576 | 3.45717 | 3.13702 | 12.28039 | 2.44264 |
| **Ice2** | 3.42925 | 1.08095 | 1.63555 | 2.81000 | 3.95791 | 0.00185 | 0.00436 | 0.42842 | 0.96977 | 2.50554 | 1.05825 | 0.22882 | 0.11823 | 1.12816 | 1.03560 | 2.65854 |
| **Ick** | 2.11634 | 0.15995 | 1.29459 | 1.10878 | 0.00426 | 1.37152 | 0.00591 | 0.45529 | 0.00000 | 0.10744 | 0.00000 | 0.07415 | 0.53036 | 0.61259 | 0.30930 | 1.07094 |
| **Icmt** | 1.68835 | 0.42509 | 4.01224 | 4.15708 | 5.79357 | 5.47574 | 2.61011 | 6.07287 | 2.97478 | 2.72159 | 1.37071 | 0.52674 | 1.08396 | 0.05751 | 1.52158 | 2.01963 |
| **Icosl** | 0.00000 | 1.02285 | 0.16066 | 0.59397 | 0.44818 | 0.01818 | 0.00663 | 0.00766 | 0.24235 | 0.21397 | 0.04420 | 0.10867 | 0.13074 | 0.01149 | 0.23532 | 7.78226 |
| **Ict1** | 14.00888 | 0.72833 | 3.34160 | 2.88575 | 3.57997 | 4.95434 | 1.11026 | 9.71599 | 2.35322 | 12.99541 | 11.30037 | 0.74606 | 9.74360 | 0.00000 | 0.49021 | 5.84420 |
| **Id1** | 8.58850 | 0.00000 | 0.13282 | 1.98027 | 6.21078 | 0.00000 | 0.02950 | 5.55710 | 11.99331 | 4.69365 | 4.55589 | 0.04770 | 0.63949 | 0.23756 | 0.00000 | 0.00448 |
| **Id2** | 0.00000 | 1.28468 | 3.22914 | 1.32234 | 0.11075 | 0.06200 | 0.00600 | 0.03991 | 0.36377 | 0.00000 | 0.65283 | 0.00096 | 0.15290 | 0.00271 | 0.05356 | 0.01903 |
| **Id3** | 26.20049 | 1.76293 | 1.66990 | 0.01550 | 0.00000 | 0.02984 | 0.00000 | 23.41880 | 2.06982 | 14.48484 | 20.82396 | 0.36854 | 0.00000 | 0.01991 | 0.00458 | 1.29876 |
| **Id4** | 0.01377 | 0.00767 | 0.01166 | 0.20757 | 0.00936 | 0.00659 | 0.00742 | 0.01208 | 0.25327 | 0.02425 | 0.00955 | 0.00656 | 0.01314 | 0.01550 | 0.25190 | 0.00755 |
| **Ide** | 18.72121 | 13.95891 | 10.11882 | 2.78721 | 19.03138 | 8.84519 | 10.74385 | 9.47829 | 3.28352 | 4.97515 | 6.72444 | 7.77023 | 4.96858 | 7.65235 | 0.59719 | 3.11286 |
| **Idh1** | 18.79287 | 13.33546 | 8.46859 | 0.00000 | 13.19618 | 0.81138 | 7.14249 | 7.04905 | 0.37323 | 6.07379 | 7.22584 | 0.96377 | 8.97605 | 18.47092 | 2.87498 | 8.84485 |
| **Idh2** | 7.89024 | 5.78347 | 13.49668 | 2.87130 | 0.82166 | 0.91300 | 0.67264 | 10.08930 | 1.28520 | 10.31288 | 11.81833 | 0.47582 | 15.77587 | 0.84078 | 3.10602 | 4.48880 |
| **Idh3a** | 9.62596 | 18.37287 | 18.76192 | 5.24382 | 11.09870 | 17.93481 | 12.22337 | 11.34563 | 3.55535 | 5.63331 | 5.27257 | 6.05840 | 17.37907 | 1.22382 | 2.79955 | 9.41386 |
| **Idh3b** | 20.97440 | 11.73428 | 17.18725 | 4.86391 | 11.91765 | 11.05509 | 11.67121 | 19.55679 | 7.35571 | 17.82011 | 12.91502 | 2.08036 | 11.53381 | 4.19809 | 6.76882 | 11.96997 |
| **Idh3g** | 27.03527 | 11.00609 | 24.26184 | 8.47624 | 16.25111 | 12.62167 | 10.54415 | 25.80361 | 8.01911 | 24.37773 | 13.77879 | 13.62144 | 22.00694 | 14.73871 | 12.84347 | 17.94842 |
| **Idi1** | 2.05440 | 0.33080 | 0.86248 | 0.94094 | 0.01138 | 1.89841 | 1.81538 | 1.06008 | 0.27308 | 0.37978 | 1.51208 | 0.05713 | 0.16829 | 0.78865 | 2.32945 | 0.66957 |
| **Idi2** | 0.64769 | 0.08651 | 1.59495 | 0.05185 | 0.01175 | 0.01437 | 1.19170 | 3.49391 | 0.02509 | 1.10339 | 0.01727 | 0.02508 | 0.00000 | 2.69280 | 1.76640 | 0.00587 |
| **Idnk** | 0.20971 | 0.02634 | 0.50506 | 0.17449 | 3.47857 | 0.48112 | 1.06418 | 0.01716 | 0.40719 | 2.05645 | 0.37949 | 0.20076 | 0.06523 | 0.08924 | 1.44886 | 0.00661 |
| **Ids** | 3.67072 | 4.60482 | 3.09137 | 0.40496 | 0.01272 | 0.00257 | 0.01148 | 3.88739 | 3.26615 | 3.41774 | 5.71557 | 1.19603 | 0.00017 | 1.66515 | 4.11876 | 1.09993 |
| **Idua** | 1.12658 | 0.00344 | 0.20565 | 3.25461 | 0.00562 | 0.00648 | 0.00434 | 1.34411 | 0.94579 | 4.65837 | 2.64799 | 7.97147 | 0.03883 | 0.00000 | 2.19884 | 0.00460 |
| **Ier2** | 2.71400 | 0.02526 | 0.17039 | 0.29858 | 0.00959 | 0.02017 | 0.02359 | 0.02246 | 0.44871 | 3.97426 | 1.12389 | 0.10276 | 2.64855 | 0.00755 | 0.00000 | 4.42457 |
| **Ier3** | 3.29944 | 0.00262 | 0.63310 | 1.62384 | 2.65148 | 0.00581 | 0.00429 | 0.00000 | 0.14708 | 0.86768 | 1.11163 | 0.03367 | 0.01110 | 0.00136 | 0.05716 | 0.00315 |
| **Ier3ip1** | 10.31305 | 2.69667 | 4.68047 | 5.64151 | 3.31009 | 3.33904 | 2.83321 | 1.71688 | 10.00795 | 4.21133 | 5.59619 | 0.58158 | 2.43890 | 0.02056 | 0.00000 | 3.66917 |
| **Ier5** | 0.16971 | 0.00871 | 0.00381 | 4.68741 | 0.01718 | 0.01350 | 0.00730 | 0.01087 | 6.81083 | 0.01498 | 0.00000 | 0.00483 | 0.01244 | 0.00214 | 0.03141 | 0.00608 |
| **Ier5l** | 0.03340 | 0.00775 | 0.01696 | 0.07008 | 0.00824 | 0.00722 | 0.00699 | 0.00886 | 0.00862 | 0.34356 | 0.07487 | 0.00681 | 0.47515 | 0.01702 | 0.00446 | 0.00691 |
| **Iffo1** | 1.95598 | 0.01485 | 0.00000 | 0.82341 | 0.34441 | 0.15893 | 2.52575 | 0.00602 | 1.07316 | 0.01269 | 0.00523 | 0.02297 | 0.00916 | 0.00965 | 0.01033 | 0.00503 |
| **Iffo2** | 0.00000 | 0.00739 | 2.07599 | 20.78306 | 0.01320 | 0.34794 | 0.05468 | 0.00000 | 16.04096 | 1.91434 | 1.49513 | 0.09934 | 2.17597 | 4.98205 | 8.88961 | 3.30021 |
| **Ifi203** | 0.01463 | 0.00770 | 0.04974 | 0.68539 | 0.00863 | 0.00659 | 0.01785 | 0.30661 | 0.47795 | 1.13058 | 0.57266 | 0.37024 | 0.03605 | 0.46789 | 0.71322 | 0.02137 |
| **Ifi27** | 0.00585 | 0.00759 | 0.00000 | 0.04229 | 0.00852 | 0.00676 | 0.00708 | 0.34685 | 3.89863 | 3.78428 | 1.13750 | 0.44974 | 0.00000 | 0.04498 | 1.52621 | 0.00689 |
| **Ifi30** | 6.98640 | 0.06978 | 12.08299 | 2.21052 | 2.77096 | 9.08500 | 1.91157 | 6.73246 | 1.64379 | 1.31254 | 3.59718 | 0.05772 | 7.14280 | 0.64553 | 1.26505 | 2.82104 |
| **Ifi35** | 1.33324 | 1.69312 | 0.34372 | 6.42635 | 0.00440 | 0.00307 | 1.56473 | 4.74117 | 11.26028 | 11.00855 | 5.63029 | 0.50711 | 1.48850 | 1.13799 | 8.56695 | 0.04461 |
| **Ifit1bl2** | 0.02836 | 0.00800 | 0.11680 | 1.41977 | 0.00825 | 0.03049 | 0.02870 | 0.08006 | 0.95709 | 0.49725 | 0.00000 | 0.36620 | 0.02605 | 0.07822 | 0.01224 | 0.02223 |
| **Ifitm1** | 0.01209 | 0.00764 | 0.01126 | 0.56495 | 0.00860 | 0.00625 | 0.00721 | 0.00885 | 1.90949 | 2.96590 | 0.00000 | 0.00000 | 0.01048 | 0.00640 | 0.21361 | 0.00714 |
| **Ifitm10** | 0.01086 | 0.00765 | 0.00874 | 0.00828 | 0.00939 | 0.00669 | 0.00709 | 0.27365 | 0.04072 | 0.00700 | 0.01105 | 0.00702 | 0.12674 | 0.01372 | 0.00585 | 0.00723 |
| **Ifitm2** | 3.10174 | 0.27349 | 0.96151 | 0.38747 | 0.00913 | 0.11224 | 0.00401 | 0.07278 | 0.08088 | 6.13477 | 0.00345 | 0.11989 | 1.95538 | 0.15941 | 0.55788 | 0.59481 |

| | | | | | | | | | | | | | | | | |
|---|---|---|---|---|---|---|---|---|---|---|---|---|---|---|---|---|
| **Ifitm3** | 0.03246 | 0.00791 | 0.03134 | 2.64777 | 0.01081 | 0.00620 | 0.00701 | 0.00891 | 0.09081 | 5.18407 | 0.00791 | 0.03282 | 0.01055 | 0.01899 | 0.14437 | 0.00708 |
| **Ifitm6** | 0.01361 | 0.00795 | 0.01170 | 4.53927 | 0.00918 | 0.00652 | 0.00739 | 0.01285 | 0.00000 | 0.25036 | 0.01137 | 0.00838 | 0.01283 | 0.03390 | 0.07208 | 0.00665 |
| **Ifitm7** | 0.01741 | 0.00802 | 0.01151 | 0.23362 | 0.00896 | 0.00658 | 0.01771 | 0.01897 | 0.20984 | 0.02078 | 0.02431 | 0.00750 | 0.01824 | 0.01030 | 0.01189 | 0.00727 |
| **Ifnar1** | 2.03245 | 0.61691 | 5.96137 | 2.28788 | 2.12176 | 0.58516 | 1.47340 | 2.57181 | 7.36271 | 3.60066 | 14.03179 | 1.54164 | 6.08682 | 3.72821 | 4.72626 | 1.03687 |
| **Ifnar2** | 0.14966 | 0.00843 | 0.00729 | 0.00617 | 0.01640 | 0.00697 | 0.00724 | 0.01017 | 0.06575 | 0.53753 | 0.01149 | 0.00659 | 0.01018 | 0.01166 | 0.00999 | 0.58827 |
| **Ifngr1** | 0.30278 | 0.00720 | 0.75776 | 0.00000 | 0.03782 | 0.00865 | 0.00676 | 0.00469 | 1.26966 | 2.14642 | 0.09261 | 0.03230 | 0.00711 | 0.03215 | 2.05484 | 1.59915 |
| **Ifngr2** | 0.18399 | 0.00747 | 0.32032 | 0.18254 | 0.01510 | 0.03063 | 0.02757 | 0.27137 | 0.07137 | 0.44984 | 0.03766 | 0.15541 | 0.00776 | 1.46234 | 0.14084 | 0.01422 |
| **Ifnlr1** | 0.01307 | 0.00756 | 0.01078 | 0.01361 | 0.00915 | 0.00642 | 0.00725 | 0.01036 | 0.62380 | 0.00000 | 0.00741 | 0.01606 | 0.14954 | 0.02482 | 0.33095 | 0.00714 |
| **Ifrd1** | 6.85333 | 10.69200 | 7.86275 | 14.29541 | 3.22719 | 2.23285 | 6.91302 | 9.12023 | 22.76055 | 12.00911 | 14.25072 | 18.98739 | 1.60796 | 9.20386 | 5.47579 | 4.29580 |
| **Ifrd2** | 2.03677 | 1.38152 | 0.53792 | 0.27126 | 0.24302 | 0.34120 | 0.57497 | 4.19941 | 1.08589 | 1.65557 | 0.32924 | 0.14829 | 1.28346 | 0.03883 | 0.14050 | 0.52397 |
| **Ift122** | 0.58329 | 0.00698 | 1.97582 | 0.17883 | 0.96095 | 0.00485 | 0.13424 | 0.50676 | 0.75859 | 0.96389 | 0.41180 | 0.15157 | 0.00000 | 0.00952 | 0.11819 | 1.19667 |
| **Ift140** | 1.29174 | 1.60998 | 1.89981 | 1.15782 | 0.00617 | 2.57900 | 4.62136 | 0.87550 | 0.32185 | 1.21895 | 0.00000 | 0.03228 | 0.09030 | 0.00331 | 0.43802 | 0.00367 |
| **Ift172** | 1.41519 | 0.01637 | 0.94410 | 1.10368 | 0.00370 | 0.00434 | 1.50256 | 1.73602 | 1.04203 | 0.93594 | 1.08610 | 0.01979 | 2.20326 | 0.03108 | 0.73855 | 0.01139 |
| **Ift20** | 13.79436 | 3.59891 | 7.86496 | 9.71278 | 1.41868 | 2.52151 | 6.61714 | 6.71411 | 18.55209 | 9.10567 | 4.28423 | 1.78156 | 4.09705 | 0.25293 | 1.10230 | 2.83262 |
| **Ift22** | 2.80214 | 0.00417 | 3.95138 | 0.01774 | 0.03879 | 0.24613 | 0.00415 | 1.65685 | 0.01102 | 0.62870 | 1.46222 | 0.00000 | 0.00000 | 0.03198 | 0.00987 | 0.00305 |
| **Ift27** | 0.32189 | 0.00696 | 0.00000 | 0.65383 | 0.06054 | 0.00928 | 1.29959 | 0.96045 | 0.00000 | 0.79087 | 0.04231 | 0.01948 | 0.49529 | 0.00879 | 0.12404 | 0.03135 |
| **Ift43** | 4.86002 | 1.04027 | 1.46080 | 9.27889 | 0.00000 | 0.00722 | 0.03147 | 0.99793 | 2.94088 | 3.35556 | 2.42370 | 0.08559 | 2.28294 | 1.53886 | 0.85668 | 0.02707 |
| **Ift46** | 11.09190 | 1.13777 | 11.02805 | 6.36664 | 6.35824 | 6.07709 | 4.22969 | 5.73849 | 4.14458 | 5.22950 | 2.88398 | 0.77643 | 7.27330 | 5.64001 | 5.14316 | 4.53394 |
| **Ift52** | 6.00100 | 8.76807 | 4.89947 | 3.67936 | 3.85631 | 2.66688 | 4.05658 | 0.36652 | 4.22972 | 0.66314 | 0.51876 | 0.07084 | 2.44666 | 2.44039 | 2.09987 | 1.80439 |
| **Ift74** | 0.71489 | 4.03704 | 2.04386 | 1.77072 | 0.00870 | 0.00442 | 0.00479 | 0.00405 | 1.54594 | 1.49779 | 0.00000 | 0.03921 | 0.60520 | 2.30794 | 1.89854 | 0.51350 |
| **Ift80** | 1.63017 | 0.00599 | 0.77997 | 0.29835 | 1.17951 | 0.00706 | 0.00568 | 0.06953 | 0.19907 | 0.54571 | 0.00741 | 0.04678 | 0.18540 | 1.39286 | 0.97330 | 0.00505 |
| **Ift81** | 0.18923 | 0.00610 | 0.00000 | 1.52668 | 2.89017 | 0.00620 | 0.00694 | 0.00549 | 0.05522 | 0.49038 | 0.00000 | 0.04527 | 0.00951 | 1.58118 | 1.49541 | 0.00646 |
| **Ift88** | 2.35220 | 3.57602 | 0.27724 | 2.76474 | 0.02046 | 1.72065 | 0.00388 | 0.77298 | 2.27175 | 0.97934 | 0.00000 | 0.08568 | 0.88809 | 0.05117 | 2.41744 | 0.42890 |
| **Igbp1** | 14.64181 | 10.18405 | 15.01376 | 0.00000 | 2.73018 | 6.00710 | 7.87800 | 13.72038 | 0.01080 | 8.40832 | 5.89547 | 9.96336 | 8.94308 | 9.99890 | 3.27091 | 13.47623 |
| **Igdcc3** | 0.01372 | 0.00764 | 0.01026 | 1.33992 | 0.00936 | 0.01255 | 0.00729 | 0.01144 | 0.72800 | 0.02262 | 0.00701 | 0.04285 | 0.01942 | 0.00317 | 0.48278 | 0.00714 |
| **Igf1** | 0.01332 | 0.00805 | 0.01211 | 0.12133 | 0.00907 | 0.00676 | 0.00744 | 0.01158 | 0.00697 | 0.76910 | 0.01103 | 0.00693 | 0.01184 | 0.01126 | 0.00738 | 0.00738 |
| **Igf1r** | 0.68641 | 0.00683 | 0.00655 | 0.85919 | 0.00661 | 0.00650 | 0.00677 | 0.01061 | 0.05642 | 0.35187 | 0.01286 | 0.06082 | 0.01720 | 0.00432 | 0.27673 | 0.26554 |
| **Igf2** | 0.71725 | 0.00802 | 0.00478 | 0.05299 | 0.02099 | 0.00771 | 0.00704 | 0.01142 | 0.01206 | 0.00858 | 0.01473 | 0.10618 | 0.14501 | 0.01184 | 0.00780 | 0.01505 |
| **Igf2bp1** | 17.89128 | 11.18443 | 7.89151 | 2.03288 | 0.09253 | 5.50609 | 3.07314 | 13.90469 | 5.15517 | 4.68415 | 5.31133 | 0.52276 | 14.35988 | 0.51895 | 1.68362 | 4.15426 |
| **Igf2bp2** | 15.41751 | 15.01582 | 14.32157 | 7.25323 | 11.39026 | 11.29549 | 9.28187 | 13.96146 | 2.36759 | 18.79526 | 9.29386 | 5.62914 | 8.84480 | 7.09185 | 10.94183 | 8.31500 |
| **Igf2bp3** | 14.64387 | 8.13663 | 9.96629 | 12.77269 | 3.34572 | 5.78614 | 7.07965 | 0.36535 | 9.59787 | 4.84525 | 2.02296 | 0.43125 | 8.21206 | 5.15710 | 11.86597 | 5.15749 |
| **Igf2r** | 2.92376 | 0.12670 | 3.91436 | 4.33215 | 0.17677 | 0.55074 | 0.92587 | 4.16070 | 10.82812 | 3.84747 | 13.60119 | 1.18491 | 1.91094 | 4.90785 | 10.64325 | 0.65402 |
| **Igfbp2** | 0.78216 | 0.00658 | 0.15342 | 1.34980 | 0.00591 | 0.00809 | 0.00627 | 0.00000 | 1.14659 | 1.02376 | 0.63462 | 0.34848 | 0.00706 | 0.00181 | 0.07107 | 0.00610 |
| **Igfbp3** | 0.01160 | 0.00808 | 0.00904 | 0.00287 | 0.00951 | 0.00685 | 0.00741 | 0.05133 | 0.00923 | 0.64836 | 0.14294 | 0.01614 | 0.01506 | 0.00610 | 1.09496 | 0.00754 |
| **Igfbp4** | 0.01210 | 0.00779 | 0.01220 | 0.00539 | 0.00856 | 0.00657 | 0.00725 | 0.00967 | 0.02128 | 3.42964 | 0.00953 | 0.00000 | 0.01067 | 0.01238 | 0.00675 | 0.00749 |
| **Igfbp5** | 0.01228 | 0.00771 | 0.01088 | 0.00547 | 0.00866 | 0.00666 | 0.00723 | 0.00979 | 0.01102 | 3.06450 | 0.00902 | 0.07063 | 0.01085 | 0.01312 | 0.00355 | 0.00747 |
| **Igfbp6** | 0.01624 | 0.00795 | 0.01107 | 0.31220 | 0.00875 | 0.00655 | 0.00723 | 0.02541 | 0.00438 | 1.40020 | 0.01043 | 0.03370 | 0.01014 | 0.01070 | 0.00247 | 0.00729 |
| **Igfbp7** | 0.01354 | 0.00778 | 0.01064 | 0.00837 | 0.00915 | 0.00672 | 0.00730 | 0.01083 | 0.01196 | 0.58014 | 0.01138 | 0.09531 | 0.01195 | 0.01193 | 0.00647 | 0.00749 |
| **Igflr1** | 0.42945 | 0.00586 | 1.09740 | 0.07498 | 0.00551 | 3.20443 | 1.85589 | 0.55385 | 0.01069 | 0.38929 | 0.72052 | 0.05635 | 0.17106 | 0.00721 | 0.00000 | 0.00587 |
| **Ighmbp2** | 2.85976 | 0.01330 | 0.57067 | 0.80754 | 0.12189 | 0.01336 | 0.76710 | 6.16427 | 1.90177 | 1.71865 | 1.51118 | 0.04474 | 3.35456 | 0.35650 | 0.03244 | 1.76418 |
| **Igip** | 1.23515 | 0.00621 | 0.00000 | 0.24694 | 0.00619 | 0.00628 | 0.00644 | 0.01075 | 0.00637 | 0.67671 | 0.00398 | 0.01787 | 0.15569 | 2.29088 | 0.65687 | 0.00613 |
| **Igsf1** | 0.01137 | 0.00762 | 0.00687 | 0.01053 | 0.00939 | 0.00652 | 0.00706 | 0.00936 | 0.01259 | 0.00014 | 0.00741 | 0.00302 | 0.19135 | 1.39032 | 0.00322 | 0.00718 |
| **Igsf23** | 0.01375 | 0.00793 | 0.01218 | 1.40552 | 0.00928 | 0.00641 | 0.00747 | 0.01242 | 0.04250 | 0.03836 | 0.02635 | 0.00704 | 0.01188 | 0.00313 | 0.23394 | 0.00698 |
| **Igsf3** | 0.03068 | 0.00658 | 1.78907 | 2.98812 | 0.01356 | 4.43058 | 0.75746 | 3.58843 | 2.04596 | 0.00000 | 1.07049 | 0.06279 | 1.22717 | 0.38017 | 0.71809 | 0.00359 |
| **Igsf5** | 0.01375 | 0.00805 | 0.01225 | 0.01061 | 0.00932 | 0.00666 | 0.00754 | 0.01390 | 0.01235 | 0.01531 | 0.01310 | 0.00811 | 0.01270 | 0.01234 | 0.01062 | 0.00741 |
| **Igsf8** | 0.01375 | 0.00815 | 0.01125 | 0.00646 | 0.00919 | 0.00684 | 0.00758 | 0.01175 | 0.04985 | 0.05547 | 0.10507 | 0.00821 | 0.01132 | 0.01133 | 0.00939 | 0.00734 |
| **Igsf9** | 0.01130 | 0.01769 | 1.17955 | 0.05905 | 0.02649 | 0.02760 | 0.00624 | 0.00384 | 0.01072 | 0.96427 | 0.34976 | 0.28788 | 1.58190 | 0.02070 | 0.14854 | 0.34826 |
| **Igsf9b** | 0.01627 | 0.00866 | 0.85243 | 0.39413 | 0.00868 | 0.00568 | 0.00732 | 0.07779 | 0.05917 | 0.00290 | 0.01217 | 0.00771 | 1.45458 | 0.01252 | 0.01907 | 0.00631 |
| **Ihh** | 0.01334 | 0.00760 | 0.01206 | 0.08200 | 0.00907 | 0.00653 | 0.00725 | 0.01016 | 0.01049 | 0.86588 | 0.01262 | 0.00507 | 0.01155 | 0.01441 | 0.02107 | 0.00759 |
| **Ik** | 10.16266 | 10.93507 | 9.42117 | 9.82412 | 2.15190 | 9.53651 | 6.10805 | 10.88057 | 19.78940 | 6.59605 | 9.15385 | 1.40872 | 8.30483 | 13.16134 | 7.98548 | 3.94185 |
| **Ikbip** | 1.10254 | 0.03966 | 0.94827 | 1.73708 | 0.06371 | 0.06355 | 0.16439 | 0.37585 | 2.27359 | 1.58788 | 0.89057 | 0.17431 | 1.95467 | 0.41674 | 2.10445 | 0.07738 |
| **Ikbkap** | 5.64401 | 12.05971 | 7.96143 | 2.47328 | 2.75018 | 0.00703 | 8.85524 | 8.13414 | 1.31392 | 8.38971 | 3.99454 | 4.05852 | 2.96151 | 2.96813 | 5.64474 | 4.38770 |
| **Ikbkb** | 0.00769 | 0.00805 | 0.00941 | 0.00553 | 0.00749 | 0.00682 | 0.00747 | 0.00695 | 0.01130 | 0.11322 | 0.30262 | 0.00738 | 0.00618 | 0.00715 | 0.05815 | 1.55101 |
| **Ikbkg** | 1.63334 | 0.33791 | 6.95406 | 6.78585 | 2.62210 | 0.15265 | 0.20423 | 4.04987 | 12.98134 | 5.08297 | 11.90078 | 5.13047 | 1.70793 | 14.26873 | 5.27320 | 0.58840 |
| **Ikzf2** | 0.01960 | 0.00765 | 0.00600 | 0.00711 | 0.01035 | 3.48111 | 0.00707 | 0.01046 | 0.01163 | 0.00766 | 0.00842 | 0.00694 | 0.00006 | 0.01798 | 0.37688 | 0.00622 |
| **Ikzf4** | 0.01315 | 0.00731 | 0.00736 | 0.00727 | 0.00925 | 0.00690 | 0.00709 | 0.63099 | 0.06629 | 0.00301 | 0.00821 | 0.00594 | 0.00563 | 0.02234 | 0.08885 | 0.00732 |
| **Ikzf5** | 0.87806 | 2.51628 | 0.44439 | 3.03095 | 3.81316 | 2.93582 | 0.00518 | 0.80883 | 0.85232 | 1.06663 | 0.13634 | 2.24107 | 0.00000 | 1.56523 | 0.88945 | 0.52260 |
| **Il10rb** | 1.60257 | 0.00591 | 0.62416 | 0.15442 | 0.00358 | 0.00785 | 0.01504 | 0.00383 | 0.05875 | 0.88498 | 0.00985 | 0.00315 | 1.33935 | 0.01145 | 0.00000 | 0.37290 |
| **Il11** | 0.14196 | 0.06088 | 0.46569 | 0.92848 | 0.25442 | 0.01796 | 0.09915 | 0.00746 | 2.08687 | 0.25790 | 1.67640 | 0.10853 | 0.48679 | 0.68867 | 0.54194 | 0.03546 |
| **Il11ra1** | 0.00572 | 0.00815 | 0.00822 | 0.41518 | 0.00773 | 2.39146 | 0.01789 | 0.01225 | 1.29435 | 0.42471 | 0.00000 | 0.00477 | 0.08598 | 0.01056 | 0.00431 | 2.34082 |
| **Il12rb1** | 0.01373 | 0.00794 | 0.01173 | 0.01074 | 0.00932 | 0.00660 | 0.00745 | 0.01290 | 0.01207 | 0.01633 | 0.01251 | 0.00772 | 0.01923 | 0.04182 | 0.05020 | 0.00741 |
| **Il12rb2** | 0.03757 | 0.00783 | 0.00251 | 0.01816 | 0.01182 | 0.00677 | 0.00691 | 0.00839 | 0.01206 | 0.01458 | 0.00753 | 0.01188 | 0.22101 | 1.88824 | 0.00779 | 0.00709 |

| | | | | | | | | | | | | | | | | |
|---|---|---|---|---|---|---|---|---|---|---|---|---|---|---|---|---|
| **Il13ra1** | 0.01370 | 0.00762 | 0.01133 | 1.56956 | 0.00932 | 0.00633 | 0.00734 | 0.01184 | 0.60545 | 0.00254 | 0.00765 | 0.00603 | 0.01913 | 0.02079 | 0.32078 | 0.00702 |
| **Il17b** | 0.00972 | 0.00775 | 0.02101 | 0.00916 | 0.00948 | 0.00801 | 0.00724 | 0.34942 | 0.01171 | 0.21612 | 0.03414 | 0.00684 | 0.00793 | 0.01219 | 0.00558 | 0.00738 |
| **Il17d** | 0.01774 | 0.00800 | 0.01173 | 0.00922 | 0.00891 | 0.00685 | 0.00759 | 0.01414 | 0.01219 | 0.07420 | 0.02751 | 0.00842 | 0.01279 | 0.00696 | 0.09091 | 0.00739 |
| **Il17ra** | 1.00037 | 0.00564 | 1.95017 | 6.92736 | 0.04423 | 0.00531 | 0.57254 | 1.25229 | 12.43797 | 1.45625 | 0.25275 | 0.18786 | 0.00686 | 0.59364 | 1.71511 | 0.07487 |
| **Il17rb** | 1.86129 | 0.00576 | 0.52381 | 0.97476 | 0.00363 | 0.00349 | 0.00573 | 0.41956 | 0.47159 | 0.81516 | 0.06634 | 0.00007 | 0.24176 | 0.20429 | 0.11533 | 0.00482 |
| **Il17rc** | 0.02206 | 0.00797 | 0.01729 | 0.10249 | 0.00948 | 0.00707 | 0.00669 | 0.01154 | 0.01019 | 0.03107 | 0.01401 | 0.00731 | 1.20849 | 0.01334 | 0.00790 | 0.00674 |
| **Il17rd** | 2.94495 | 0.01354 | 1.14881 | 5.49608 | 0.01717 | 0.00167 | 0.04406 | 1.26285 | 5.95414 | 0.13169 | 0.77416 | 3.14804 | 0.00037 | 3.79497 | 3.04485 | 0.02286 |
| **Il17re** | 0.00876 | 0.00772 | 0.11543 | 0.00509 | 0.00887 | 0.00898 | 0.00672 | 0.01487 | 0.14051 | 0.04872 | 0.16912 | 0.00584 | 0.86348 | 0.01144 | 0.00532 | 0.00722 |
| **Il18** | 0.01124 | 0.00865 | 0.68443 | 0.00555 | 0.00839 | 0.00566 | 0.00800 | 0.24106 | 0.01106 | 0.51184 | 0.20509 | 0.00593 | 0.00254 | 0.00745 | 0.09127 | 0.00698 |
| **Il18bp** | 0.44011 | 0.79903 | 0.00000 | 0.00722 | 0.00768 | 0.00853 | 0.00670 | 0.01104 | 0.01185 | 0.63682 | 0.01132 | 0.01597 | 0.04195 | 0.08451 | 0.02708 | 0.01402 |
| **Il1bos** | 0.01115 | 0.00803 | 0.03502 | 0.02457 | 0.00942 | 0.00596 | 0.10104 | 0.04131 | 0.01198 | 0.03499 | 0.01095 | 0.00719 | 0.01602 | 0.21023 | 0.21572 | 0.00728 |
| **Il1f9** | 0.02152 | 0.04117 | 0.48637 | 7.22092 | 0.04240 | 2.11241 | 0.00710 | 0.02738 | 6.68450 | 0.03843 | 0.43275 | 0.38242 | 0.25106 | 0.58277 | 1.13133 | 0.21778 |
| **Il1r1** | 0.02179 | 0.00790 | 0.01091 | 0.47352 | 0.00858 | 0.00650 | 0.00722 | 0.01057 | 0.26841 | 0.67533 | 0.01121 | 0.02805 | 0.01109 | 0.00988 | 0.11005 | 0.00714 |
| **Il1rap** | 0.01543 | 0.00712 | 0.00450 | 0.41049 | 0.01807 | 0.00727 | 1.34981 | 0.00000 | 0.30048 | 0.00000 | 0.37605 | 0.01509 | 0.00799 | 0.00708 | 0.01725 | 0.00701 |
| **Il1rapl2** | 0.02165 | 0.00764 | 0.03892 | 0.03194 | 0.03548 | 0.03261 | 0.00729 | 0.01951 | 0.10566 | 0.04641 | 0.02261 | 0.01561 | 0.04133 | 0.45031 | 0.07951 | 0.00734 |
| **Il1rl1** | 0.17315 | 0.02997 | 0.18835 | 0.04320 | 0.23299 | 0.03508 | 0.00752 | 0.02120 | 0.14123 | 0.00696 | 0.60498 | 0.09421 | 0.39596 | 0.06132 | 0.03404 | 0.04945 |
| **Il20rb** | 0.01170 | 0.00803 | 0.00881 | 0.00394 | 0.00919 | 0.00664 | 0.00721 | 0.00939 | 0.23634 | 0.79373 | 0.00427 | 0.00000 | 0.01204 | 1.20832 | 0.12409 | 0.00765 |
| **Il23a** | 0.17261 | 0.00821 | 0.00479 | 0.01968 | 0.00738 | 0.00743 | 0.00772 | 0.09544 | 0.01106 | 0.02148 | 0.26095 | 0.03242 | 0.01090 | 0.01474 | 0.00995 | 0.00689 |
| **Il2ra** | 0.02787 | 0.00779 | 0.03012 | 0.00816 | 0.00841 | 0.00714 | 0.01742 | 0.02558 | 0.11251 | 0.02225 | 0.05408 | 0.01715 | 0.01725 | 0.15194 | 0.04080 | 0.00731 |
| **Il2rg** | 0.01008 | 0.01846 | 2.55776 | 0.42313 | 0.00464 | 0.00948 | 0.00654 | 0.00000 | 0.12576 | 1.85526 | 0.81838 | 0.71263 | 0.76101 | 1.83969 | 1.17400 | 0.00598 |
| **Il31** | 0.01374 | 0.00781 | 0.01185 | 0.00514 | 0.00931 | 0.00666 | 0.00750 | 0.01316 | 0.32609 | 0.01749 | 0.00990 | 0.00714 | 0.01300 | 0.01792 | 0.05160 | 0.00751 |
| **Il33** | 0.00957 | 0.00852 | 0.35035 | 0.00815 | 0.00960 | 0.00649 | 0.00790 | 0.02113 | 0.02661 | 0.22177 | 0.01276 | 0.02072 | 0.55978 | 0.00868 | 0.03932 | 0.00745 |
| **Il34** | 0.71136 | 0.00703 | 2.78239 | 0.69412 | 0.00151 | 0.00789 | 0.00540 | 0.00000 | 0.18748 | 0.00886 | 0.45888 | 0.08603 | 1.50432 | 0.02046 | 0.16257 | 0.01829 |
| **Il3ra** | 0.01171 | 0.00763 | 0.01333 | 0.01125 | 0.02159 | 0.00747 | 0.00703 | 0.89381 | 0.01143 | 0.00933 | 0.00704 | 0.00581 | 0.17534 | 0.01559 | 0.00000 | 0.01435 |
| **Il4** | 0.01362 | 0.00815 | 0.01207 | 0.10849 | 0.00918 | 0.00652 | 0.01789 | 0.01241 | 0.49461 | 0.15897 | 0.00929 | 0.00672 | 0.01169 | 0.01062 | 0.00854 | 0.00712 |
| **Il4ra** | 0.01133 | 0.00715 | 0.05499 | 0.00546 | 0.00937 | 0.00590 | 0.00666 | 0.78687 | 0.01035 | 0.34955 | 1.06195 | 0.00860 | 0.00000 | 0.00904 | 0.00105 | 0.00757 |
| **Il5ra** | 0.02558 | 0.04111 | 0.12176 | 0.03878 | 0.00927 | 0.00618 | 0.00748 | 0.02121 | 0.02225 | 0.03156 | 0.22592 | 0.01682 | 0.08792 | 0.02183 | 0.10097 | 0.01440 |
| **Il6** | 0.01374 | 0.00806 | 0.01221 | 0.00657 | 0.00932 | 0.00666 | 0.00754 | 0.01392 | 0.09824 | 0.01479 | 0.01210 | 0.00801 | 0.01273 | 0.01264 | 0.00978 | 0.00741 |
| **Il6ra** | 1.85991 | 0.00463 | 0.97838 | 1.27067 | 0.01082 | 0.00714 | 3.10758 | 0.15150 | 0.43681 | 0.71255 | 0.00000 | 0.05649 | 1.84950 | 0.11755 | 4.13315 | 0.01119 |
| **Il6st** | 5.44385 | 0.00000 | 0.00042 | 2.45447 | 1.24657 | 0.00195 | 0.00286 | 0.00492 | 2.01319 | 0.21818 | 0.00000 | 0.03876 | 0.01013 | 0.00136 | 1.33896 | 0.23291 |
| **Il7** | 0.01361 | 0.00758 | 0.01106 | 5.49999 | 0.00943 | 0.00586 | 0.00712 | 0.01036 | 0.00000 | 0.00499 | 0.01912 | 0.00430 | 0.01260 | 0.00000 | 1.52905 | 0.00639 |
| **Ildr1** | 3.56929 | 0.00023 | 0.00000 | 0.52686 | 4.24094 | 0.00981 | 0.12176 | 0.00000 | 0.41797 | 0.00000 | 0.00859 | 0.05007 | 3.31872 | 0.01109 | 0.17494 | 3.57573 |
| **Ilf2** | 13.90875 | 11.96592 | 12.34535 | 0.24286 | 7.43677 | 3.49792 | 5.35779 | 12.78979 | 1.52978 | 12.78000 | 3.03097 | 6.29147 | 5.70800 | 26.38885 | 18.27439 | 5.25481 |
| **Ilf3** | 8.20641 | 8.70891 | 11.14505 | 2.32002 | 1.36330 | 6.96156 | 6.87195 | 11.62962 | 2.43878 | 11.09378 | 4.05677 | 1.99294 | 7.22289 | 1.61780 | 13.98157 | 3.67804 |
| **Ilk** | 1.78597 | 0.56120 | 3.34111 | 1.53650 | 3.88628 | 8.35649 | 0.71200 | 3.18510 | 1.81209 | 4.10193 | 0.43783 | 0.02375 | 1.32599 | 2.30045 | 1.69770 | 0.03727 |
| **Ilkap** | 4.41538 | 7.04250 | 4.61280 | 0.99942 | 2.61841 | 0.00175 | 2.45813 | 0.64016 | 0.27542 | 2.48738 | 0.88074 | 0.16901 | 2.56111 | 0.00000 | 1.18255 | 2.43890 |
| **Ilvbl** | 0.00000 | 0.00786 | 0.02278 | 0.00769 | 0.00735 | 0.01527 | 0.00690 | 0.01038 | 0.01167 | 0.67356 | 0.01108 | 0.06505 | 1.05804 | 0.00957 | 0.10130 | 2.80733 |
| **Immp1l** | 2.13776 | 3.48100 | 1.86237 | 9.05140 | 0.00515 | 2.38916 | 0.01253 | 0.30828 | 1.21494 | 1.93391 | 2.08547 | 1.30809 | 1.70762 | 0.00000 | 1.70489 | 2.98629 |
| **Immp2l** | 0.00824 | 0.00819 | 0.01438 | 0.21991 | 0.00894 | 0.00704 | 0.00773 | 0.01682 | 0.00840 | 0.00740 | 0.01627 | 0.00798 | 0.15994 | 0.01188 | 0.00696 | 0.70830 |
| **Immt** | 14.22969 | 11.26948 | 7.22546 | 4.73265 | 14.61291 | 6.70392 | 5.38129 | 8.18444 | 4.31076 | 4.86621 | 5.04319 | 3.53445 | 8.21323 | 5.29528 | 12.28294 | 8.60356 |
| **Imp3** | 4.40155 | 0.75900 | 0.27533 | 22.50170 | 4.49211 | 3.29052 | 0.00000 | 1.99162 | 31.68638 | 15.21136 | 12.59082 | 0.51661 | 1.60397 | 1.72938 | 1.70818 | 0.23417 |
| **Imp4** | 19.10162 | 10.48356 | 17.94522 | 13.79245 | 7.77738 | 8.35952 | 9.74979 | 22.39734 | 20.71894 | 25.02550 | 12.63572 | 13.13459 | 19.00243 | 2.73364 | 11.13701 | 11.19080 |
| **Impa1** | 4.97346 | 10.38358 | 5.99627 | 0.00000 | 3.99889 | 0.00000 | 2.87933 | 0.00000 | 1.67703 | 3.86090 | 1.21134 | 1.21691 | 0.35614 | 0.49567 | 4.45939 | 1.46694 |
| **Impa2** | 4.39616 | 2.81292 | 5.19035 | 3.64850 | 3.22980 | 3.02269 | 3.14021 | 0.14344 | 5.94614 | 4.17944 | 3.59271 | 0.15738 | 8.14959 | 0.03695 | 1.02499 | 7.34241 |
| **Impact** | 8.57038 | 4.17503 | 3.34178 | 0.62550 | 3.46980 | 0.00000 | 8.39346 | 1.70707 | 0.03905 | 8.76592 | 2.30407 | 14.65128 | 7.57239 | 17.69941 | 11.59181 | 0.02680 |
| **Impad1** | 1.36421 | 0.19495 | 0.11188 | 0.00003 | 3.22209 | 0.61661 | 0.03558 | 0.18179 | 1.14900 | 0.23292 | 0.12358 | 1.18314 | 0.26005 | 0.55039 | 0.31077 | 0.01303 |
| **Impdh1** | 14.96840 | 1.95120 | 7.36576 | 9.77856 | 11.22029 | 6.94934 | 8.96629 | 18.15166 | 10.69071 | 16.68770 | 16.36270 | 1.74457 | 16.06998 | 0.18791 | 0.34881 | 6.45058 |
| **Impdh2** | 30.35209 | 23.24129 | 30.54070 | 20.26822 | 19.87651 | 15.79988 | 23.25704 | 57.85254 | 40.59877 | 61.34086 | 55.96853 | 29.02838 | 37.92826 | 34.90808 | 34.11602 | 22.59084 |
| **Impg2** | 0.00979 | 0.00793 | 0.00872 | 0.16571 | 0.17561 | 0.00677 | 0.00760 | 0.03025 | 0.19755 | 0.28578 | 0.00922 | 0.09922 | 0.00997 | 0.00926 | 0.03364 | 0.00705 |
| **Ina** | 1.37365 | 0.00631 | 0.00000 | 1.32560 | 0.00784 | 0.90304 | 0.00608 | 0.00489 | 0.00000 | 2.19587 | 0.02332 | 0.22040 | 0.30480 | 0.01695 | 1.90411 | 0.00527 |
| **Inadl** | 2.35651 | 3.23675 | 0.19885 | 1.59899 | 0.00450 | 3.83064 | 0.00317 | 0.25837 | 0.12602 | 0.29266 | 0.07361 | 0.03728 | 1.36348 | 0.02365 | 2.26968 | 2.18819 |
| **Inafm1** | 0.00747 | 0.00806 | 1.02788 | 0.46494 | 0.00979 | 0.00581 | 0.60444 | 0.19321 | 0.00544 | 0.00000 | 2.77244 | 0.04898 | 1.30173 | 0.00122 | 0.30049 | 0.00585 |
| **Incenp** | 12.09499 | 3.09840 | 7.34992 | 17.27053 | 3.12727 | 0.49835 | 0.69834 | 15.84231 | 23.22070 | 9.87154 | 7.61280 | 1.23964 | 7.46619 | 6.15735 | 7.31806 | 4.19820 |
| **Inf2** | 0.01793 | 0.03830 | 0.00000 | 0.05150 | 0.00946 | 5.58485 | 0.00581 | 0.42141 | 0.01144 | 1.25141 | 2.14458 | 0.17716 | 0.00000 | 0.00865 | 0.00736 | 4.23210 |
| **Ing1** | 10.52969 | 1.56527 | 10.46372 | 18.31027 | 0.35154 | 4.68825 | 2.98403 | 8.50837 | 21.02732 | 16.36527 | 7.50349 | 3.12648 | 6.44584 | 8.43601 | 11.94367 | 3.94251 |
| **Ing2** | 8.75847 | 3.14861 | 1.71819 | 5.58215 | 4.55444 | 2.93339 | 3.87033 | 4.05079 | 4.85392 | 5.03332 | 1.92097 | 0.48746 | 5.22106 | 2.01706 | 3.10426 | 5.20122 |
| **Ing3** | 0.11848 | 0.47921 | 1.06784 | 0.53855 | 0.61814 | 0.22764 | 0.00631 | 0.55484 | 0.07007 | 0.63639 | 0.62586 | 0.05658 | 0.50946 | 0.60171 | 2.06270 | 1.44503 |
| **Ing4** | 8.81652 | 3.69317 | 10.60955 | 4.72486 | 1.73389 | 0.02288 | 5.95392 | 4.39022 | 6.47946 | 13.57053 | 4.89610 | 1.23675 | 6.68822 | 1.03401 | 6.56928 | 4.58401 |
| **Ing5** | 4.11498 | 3.20472 | 6.23055 | 2.29436 | 3.16390 | 1.06323 | 2.12813 | 8.71640 | 1.03600 | 4.59425 | 3.38122 | 0.70553 | 6.14924 | 1.04408 | 4.99963 | 2.97581 |
| **Inha** | 0.01221 | 0.00799 | 0.01217 | 0.00494 | 0.00869 | 0.00655 | 0.00742 | 0.01165 | 0.08877 | 2.82334 | 0.00894 | 0.00000 | 0.01041 | 0.01268 | 0.00654 | 0.00725 |
| **Inhba** | 0.04787 | 0.04007 | 0.02561 | 0.02777 | 0.03741 | 0.01426 | 0.00731 | 0.01230 | 0.04463 | 0.02451 | 0.01064 | 0.26617 | 0.04791 | 0.05007 | 0.04165 | 0.02956 |
| **Inhbb** | 0.67681 | 0.00706 | 0.00903 | 0.00594 | 0.36488 | 0.00613 | 0.00674 | 0.00754 | 0.01150 | 0.93421 | 0.31920 | 0.03305 | 0.00876 | 0.00567 | 0.11515 | 0.01403 |

| | | | | | | | | | | | | | | | | |
|---|---|---|---|---|---|---|---|---|---|---|---|---|---|---|---|---|
| **Inip** | 0.50981 | 0.02561 | 2.52265 | 4.23419 | 0.04559 | 2.60653 | 3.36988 | 3.84837 | 5.72492 | 4.23795 | 1.99573 | 0.40865 | 5.27342 | 0.91365 | 0.44158 | 1.83197 |
| **Ino80** | 5.24087 | 0.03657 | 3.38903 | 4.38259 | 2.74810 | 0.00000 | 6.29543 | 6.51309 | 6.50337 | 7.03614 | 3.42125 | 0.23523 | 2.63248 | 0.59863 | 3.33276 | 2.62512 |
| **Ino80b** | 0.01508 | 0.00812 | 0.02547 | 0.09810 | 0.00913 | 0.00615 | 0.00761 | 0.01416 | 0.01030 | 0.04917 | 0.20179 | 0.00740 | 0.00979 | 0.01047 | 0.00381 | 0.00736 |
| **Ino80c** | 12.06062 | 9.94640 | 6.80965 | 8.11586 | 0.40119 | 9.33714 | 9.19285 | 9.46194 | 6.71487 | 8.51178 | 9.43344 | 6.80303 | 9.03658 | 4.66201 | 13.48213 | 4.88198 |
| **Ino80d** | 4.43844 | 1.62078 | 1.17419 | 7.14338 | 2.70529 | 5.17572 | 3.14874 | 2.09104 | 3.69048 | 6.78426 | 6.67223 | 2.37029 | 2.86774 | 9.20629 | 10.64086 | 1.62280 |
| **Ino80dos** | 0.01508 | 0.00768 | 0.04974 | 0.03898 | 0.00992 | 0.00628 | 0.00750 | 0.01279 | 0.11303 | 0.10738 | 0.01098 | 0.01569 | 0.06031 | 0.00992 | 0.37513 | 0.00765 |
| **Ino80e** | 6.64585 | 0.62991 | 1.37900 | 0.57325 | 3.56614 | 1.33794 | 1.71441 | 6.43416 | 1.01021 | 5.33286 | 0.77181 | 0.18497 | 9.72226 | 0.00216 | 0.00000 | 5.13104 |
| **Inpp1** | 0.01836 | 0.00814 | 0.00811 | 7.28077 | 0.00953 | 0.02020 | 0.07408 | 0.02412 | 4.25336 | 0.00000 | 0.00000 | 0.03781 | 0.18639 | 0.27529 | 0.00000 | 0.00602 |
| **Inpp4a** | 0.02117 | 0.00797 | 0.99933 | 1.97864 | 0.01295 | 0.00878 | 0.00746 | 1.62555 | 0.58110 | 0.00110 | 0.01812 | 0.01012 | 0.01323 | 0.00621 | 1.42163 | 0.01439 |
| **Inpp5a** | 0.04140 | 0.01701 | 0.46261 | 1.09542 | 0.02450 | 0.00890 | 0.06635 | 0.59776 | 0.25469 | 0.51333 | 0.66230 | 1.57201 | 0.02409 | 0.00221 | 0.46487 | 0.30182 |
| **Inpp5b** | 3.32699 | 3.49306 | 0.00000 | 1.48654 | 3.47552 | 0.00348 | 0.00296 | 2.78427 | 1.49909 | 0.36036 | 0.91379 | 0.13416 | 1.00344 | 0.00000 | 2.15897 | 0.02356 |
| **Inpp5d** | 3.61427 | 5.96718 | 1.17050 | 0.37405 | 0.63494 | 1.50371 | 1.01330 | 2.49798 | 0.98313 | 0.00000 | 4.88621 | 0.12949 | 0.19697 | 0.46976 | 0.11051 | 0.37019 |
| **Inpp5e** | 0.92075 | 0.00553 | 0.00000 | 1.26281 | 0.01397 | 0.00575 | 1.21144 | 0.24276 | 0.08572 | 0.00000 | 0.21028 | 0.02577 | 0.16071 | 1.35489 | 0.63987 | 0.89543 |
| **Inpp5f** | 0.00000 | 1.06968 | 4.32575 | 2.58943 | 1.60275 | 1.34221 | 0.00582 | 1.00240 | 1.09855 | 1.05160 | 0.30616 | 1.23510 | 0.54361 | 3.92855 | 0.95019 | 2.87793 |
| **Inpp5k** | 3.56969 | 0.00000 | 2.24086 | 0.28900 | 4.60042 | 4.90166 | 2.27452 | 7.03513 | 1.78790 | 9.01476 | 4.25224 | 0.46882 | 5.13267 | 0.02273 | 0.45109 | 0.75422 |
| **Inppl1** | 0.28560 | 0.00797 | 0.00237 | 0.50995 | 0.12619 | 0.00816 | 0.00665 | 0.36264 | 0.08311 | 0.10457 | 0.77939 | 0.04299 | 0.01080 | 0.02447 | 0.13106 | 0.00701 |
| **Insig1** | 1.62960 | 3.69039 | 1.50609 | 4.17961 | 0.01099 | 6.54255 | 5.88637 | 3.01594 | 0.68935 | 2.92427 | 3.91911 | 0.06755 | 1.85944 | 5.50009 | 4.85442 | 0.73592 |
| **Insig2** | 2.34650 | 1.16478 | 0.87265 | 0.00000 | 0.01303 | 0.00294 | 1.35417 | 0.76916 | 0.07580 | 0.36836 | 0.39493 | 5.08988 | 1.21534 | 0.00463 | 0.95970 | 0.01003 |
| **Insm1** | 0.05528 | 0.00824 | 0.03122 | 0.41036 | 0.02717 | 0.04578 | 0.00744 | 0.00919 | 0.38638 | 0.00755 | 0.11321 | 0.00679 | 0.03034 | 0.04607 | 0.02370 | 0.02225 |
| **Insr** | 0.27731 | 0.00758 | 0.20680 | 0.07977 | 0.01317 | 0.00577 | 0.74522 | 0.50425 | 0.01623 | 0.00000 | 0.01078 | 0.05182 | 0.02739 | 1.92525 | 1.15849 | 1.54623 |
| **Ints1** | 9.74707 | 0.58461 | 3.81701 | 9.44573 | 0.30459 | 3.46699 | 2.61240 | 12.07021 | 7.65241 | 18.56287 | 4.22940 | 0.84617 | 13.35366 | 0.00000 | 0.75177 | 6.58602 |
| **Ints10** | 2.65768 | 4.84133 | 6.57095 | 1.85142 | 1.90153 | 1.01802 | 0.03501 | 1.81777 | 3.68989 | 5.54230 | 1.10190 | 0.54049 | 1.34108 | 0.81575 | 7.83940 | 0.26417 |
| **Ints12** | 9.23105 | 7.02254 | 3.01299 | 3.92105 | 1.50298 | 0.89599 | 3.41392 | 7.97489 | 8.98761 | 5.53534 | 7.14519 | 7.12984 | 4.30410 | 0.57684 | 6.15386 | 3.25034 |
| **Ints2** | 0.96732 | 1.38357 | 2.65542 | 3.70361 | 1.54373 | 0.80042 | 0.00272 | 1.11043 | 5.90683 | 5.42230 | 0.64615 | 8.25707 | 2.78092 | 12.55839 | 13.71511 | 0.03786 |
| **Ints3** | 14.23720 | 6.26203 | 14.63067 | 9.78064 | 11.35039 | 7.44309 | 5.02876 | 14.04932 | 9.92860 | 21.63495 | 3.12672 | 2.03355 | 8.36920 | 1.80224 | 15.46424 | 4.64202 |
| **Ints4** | 7.25884 | 10.64327 | 4.09765 | 5.31642 | 1.54614 | 4.76376 | 2.24084 | 9.20185 | 11.44514 | 13.23030 | 7.67403 | 5.86771 | 6.73561 | 21.20449 | 11.72994 | 0.02949 |
| **Ints5** | 8.47160 | 2.05015 | 2.89990 | 10.23653 | 3.91342 | 7.57219 | 4.04250 | 5.13916 | 9.16666 | 6.35688 | 2.33744 | 0.55095 | 2.53030 | 0.00000 | 0.60612 | 4.62869 |
| **Ints6** | 0.36911 | 4.14075 | 2.51230 | 3.53110 | 0.01733 | 2.99091 | 0.00388 | 0.00000 | 5.63315 | 8.39496 | 1.75772 | 3.03230 | 1.42043 | 1.69608 | 6.64957 | 0.01628 |
| **Ints7** | 9.65799 | 4.55137 | 5.71325 | 3.01142 | 0.61394 | 0.84760 | 5.86096 | 7.24717 | 2.60726 | 12.69747 | 5.59922 | 1.19810 | 2.87693 | 0.78823 | 5.52454 | 6.36311 |
| **Ints8** | 4.20359 | 1.33485 | 0.24304 | 1.00155 | 0.82137 | 0.77110 | 4.22115 | 2.75197 | 1.58628 | 1.99076 | 0.82992 | 0.42532 | 0.00000 | 15.47938 | 5.71147 | 1.88029 |
| **Ints9** | 3.99577 | 3.47401 | 2.72817 | 1.37898 | 3.42135 | 3.12288 | 5.48704 | 5.50823 | 2.43550 | 6.72580 | 1.70387 | 0.79731 | 1.11100 | 4.85213 | 4.84052 | 3.03939 |
| **Intu** | 0.29598 | 0.14882 | 0.32807 | 36.39591 | 0.03452 | 0.11422 | 0.42117 | 0.13402 | 32.45018 | 6.30898 | 2.69157 | 1.15109 | 0.63970 | 13.13038 | 20.04110 | 0.11058 |
| **Invs** | 2.32215 | 1.69927 | 2.01777 | 0.97503 | 1.88053 | 0.67335 | 1.45516 | 3.55406 | 0.91436 | 2.04711 | 1.23527 | 1.87478 | 1.68120 | 0.45126 | 1.98689 | 0.68336 |
| **Ip6k1** | 1.06075 | 0.09158 | 1.56460 | 2.90048 | 0.01690 | 0.01005 | 2.92133 | 2.95387 | 6.56485 | 1.94463 | 2.49323 | 0.03182 | 0.26911 | 1.47120 | 2.01071 | 0.00451 |
| **Ip6k2** | 0.88714 | 0.00757 | 0.97301 | 0.01039 | 0.01688 | 0.00537 | 0.00661 | 0.05416 | 0.01241 | 0.03944 | 0.01072 | 0.00678 | 1.06637 | 0.01069 | 0.01964 | 0.51717 |
| **Ipcef1** | 0.01319 | 0.01813 | 0.01696 | 0.27050 | 0.00932 | 0.00625 | 0.00746 | 0.01216 | 0.35031 | 0.00519 | 0.00932 | 0.00647 | 0.01258 | 0.00843 | 0.38267 | 0.00727 |
| **Ipmk** | 2.50695 | 0.02367 | 1.37483 | 14.07394 | 4.54867 | 0.00285 | 0.00336 | 3.07442 | 3.29854 | 1.52856 | 2.24519 | 1.02977 | 0.76623 | 3.38485 | 14.37991 | 3.52958 |
| **Ipo11** | 2.21711 | 8.42803 | 6.10608 | 3.35479 | 12.31610 | 0.61799 | 0.49652 | 10.92429 | 9.12371 | 12.83186 | 4.62156 | 2.85584 | 2.62533 | 14.63115 | 9.01987 | 0.01036 |
| **Ipo13** | 2.50305 | 0.02487 | 2.83312 | 2.32011 | 3.77320 | 4.62783 | 0.00260 | 0.31800 | 1.20455 | 3.50984 | 0.00000 | 2.44976 | 4.14410 | 0.00219 | 0.89977 | 4.10658 |
| **Ipo4** | 13.17577 | 3.02397 | 4.33925 | 12.25209 | 10.79163 | 7.50694 | 10.13506 | 28.87888 | 11.93182 | 28.27792 | 21.21962 | 5.08262 | 15.95431 | 0.43415 | 1.96822 | 5.30722 |
| **Ipo5** | 18.07591 | 14.49229 | 19.58868 | 16.47843 | 9.55087 | 9.59792 | 10.13214 | 36.32201 | 25.03919 | 42.06680 | 29.21778 | 25.16502 | 21.59900 | 39.94276 | 31.29528 | 13.05761 |
| **Ipo7** | 2.00990 | 4.89690 | 2.22617 | 0.92801 | 0.97493 | 0.15694 | 0.45559 | 3.89142 | 1.53291 | 0.65832 | 0.96574 | 5.83014 | 0.21811 | 10.30936 | 4.68535 | 0.01559 |
| **Ipo8** | 2.08795 | 6.15735 | 3.59160 | 15.78376 | 6.27637 | 5.38765 | 2.43717 | 3.43404 | 6.59991 | 3.65700 | 4.47274 | 0.06690 | 3.42652 | 7.00655 | 7.55096 | 3.65449 |
| **Ipo9** | 7.83854 | 6.24636 | 8.21783 | 7.52498 | 1.53381 | 2.60420 | 2.91103 | 4.78107 | 4.64262 | 4.32562 | 0.89895 | 0.17605 | 4.01519 | 6.63444 | 4.74818 | 4.70743 |
| **Ipp** | 1.36682 | 0.00597 | 0.58564 | 1.35475 | 0.00568 | 0.00773 | 0.00522 | 0.00000 | 0.36788 | 0.28476 | 0.88886 | 0.14304 | 0.79248 | 2.04370 | 3.25766 | 0.00515 |
| **Ippk** | 3.23565 | 3.33496 | 3.03095 | 5.65166 | 0.07662 | 0.00929 | 5.02425 | 0.35466 | 6.11277 | 7.81743 | 5.38553 | 0.10897 | 1.19138 | 5.69219 | 4.88772 | 0.01305 |
| **Iqca** | 0.01015 | 0.00603 | 0.19667 | 13.87824 | 0.00933 | 0.00427 | 2.75667 | 0.00000 | 1.27333 | 0.56971 | 0.32842 | 0.07882 | 0.02134 | 0.00000 | 2.98256 | 0.00472 |
| **Iqcb1** | 0.11635 | 0.05440 | 0.00543 | 0.22622 | 0.03152 | 0.01316 | 0.00731 | 0.01190 | 0.00798 | 0.00573 | 0.01322 | 0.00799 | 0.00732 | 0.01222 | 0.01397 | 1.58351 |
| **Iqcc** | 0.87513 | 0.00500 | 0.00000 | 0.56779 | 0.00633 | 4.01085 | 2.62616 | 0.00490 | 0.02140 | 0.91884 | 0.02931 | 0.06287 | 0.02302 | 0.01518 | 0.00000 | 0.00467 |
| **Iqcd** | 0.35198 | 0.00788 | 0.00365 | 0.01264 | 0.02106 | 0.00812 | 1.25235 | 0.01057 | 0.01052 | 0.01990 | 0.84188 | 0.00299 | 0.04314 | 0.01112 | 0.00000 | 0.02321 |
| **Iqce** | 1.61255 | 0.00463 | 0.57408 | 0.00322 | 0.01249 | 0.01546 | 3.01286 | 0.00000 | 0.01069 | 0.89975 | 0.22051 | 0.00000 | 1.18504 | 0.01126 | 0.15858 | 0.02078 |
| **Iqcf1** | 0.00909 | 0.00765 | 0.00879 | 2.32631 | 0.00911 | 0.00618 | 0.00704 | 0.18084 | 2.11854 | 1.33334 | 0.00000 | 0.06945 | 0.01424 | 0.00584 | 0.35266 | 0.00672 |
| **Iqcf5** | 0.01240 | 0.00838 | 0.00492 | 1.76860 | 0.00898 | 0.00742 | 0.00732 | 0.00758 | 0.12116 | 0.12400 | 1.71297 | 0.01127 | 0.15320 | 0.00081 | 0.00000 | 0.00645 |
| **Iqcg** | 2.98276 | 3.02621 | 5.88068 | 6.70450 | 0.00000 | 0.15974 | 0.26329 | 2.44940 | 8.36370 | 11.09223 | 3.79853 | 5.42248 | 0.78057 | 10.16116 | 8.91838 | 0.00468 |
| **Iqgap1** | 6.22121 | 4.26279 | 7.74393 | 3.21272 | 4.20922 | 6.98490 | 7.44737 | 8.75881 | 4.02570 | 3.20536 | 2.85365 | 0.95391 | 2.60243 | 7.87680 | 3.59498 | 8.13689 |
| **Iqgap2** | 0.88788 | 0.00784 | 0.00000 | 2.19774 | 0.01682 | 0.00575 | 0.44065 | 0.35153 | 0.74377 | 0.25497 | 0.00041 | 0.00000 | 0.28092 | 1.26720 | 1.02966 | 0.00603 |
| **Iqgap3** | 4.63146 | 1.34479 | 3.61056 | 0.76655 | 2.23803 | 3.46677 | 3.79817 | 16.37193 | 0.34162 | 5.73572 | 7.04659 | 1.66312 | 4.55782 | 0.00000 | 3.07502 | 6.06591 |
| **Iqsec1** | 1.42446 | 0.05095 | 0.13558 | 2.36835 | 0.21125 | 1.55944 | 0.00418 | 0.50270 | 0.75403 | 2.93739 | 1.84915 | 0.14244 | 3.41227 | 0.02117 | 0.03698 | 0.48567 |
| **Iqsec2** | 0.01219 | 0.00805 | 0.02809 | 0.35670 | 0.01009 | 0.00805 | 0.00766 | 0.01567 | 0.20950 | 0.00746 | 0.01345 | 0.02662 | 0.17928 | 0.00779 | 0.04863 | 0.00757 |
| **Iqub** | 0.00000 | 0.89238 | 1.90585 | 4.10716 | 0.00442 | 0.00370 | 0.00543 | 3.77787 | 10.74897 | 4.08042 | 1.76260 | 3.01464 | 0.53429 | 9.28945 | 3.50332 | 0.00394 |
| **Irak1** | 6.54497 | 1.74734 | 4.93898 | 18.08205 | 5.93535 | 0.66614 | 1.00350 | 0.00000 | 22.48907 | 0.99477 | 22.12789 | 2.47864 | 5.25770 | 11.48892 | 2.46049 | 1.24715 |
| **Irak1bp1** | 0.01541 | 0.00751 | 0.04912 | 1.07861 | 0.00930 | 0.00575 | 0.00723 | 0.00927 | 0.18582 | 0.03220 | 0.05888 | 0.00538 | 0.01140 | 0.00922 | 0.67888 | 0.00717 |

| | | | | | | | | | | | | | | | | |
|---|---|---|---|---|---|---|---|---|---|---|---|---|---|---|---|---|
| **Irak2** | 1.28414 | 0.10510 | 2.83190 | 1.83172 | 0.21559 | 0.11557 | 0.39860 | 4.79911 | 3.67828 | 4.79672 | 6.23948 | 0.51155 | 4.28756 | 2.72832 | 3.18115 | 0.22771 |
| **Irak3** | 0.01105 | 0.00737 | 0.01424 | 0.51821 | 0.55812 | 0.01237 | 0.00728 | 0.69186 | 0.79030 | 1.02637 | 0.19780 | 0.12079 | 0.01385 | 0.04284 | 0.88737 | 0.00721 |
| **Irak4** | 0.29584 | 0.00827 | 0.00451 | 0.00734 | 0.01778 | 0.00725 | 0.23320 | 0.00857 | 0.01183 | 0.77082 | 0.01156 | 0.01519 | 0.01088 | 0.02770 | 0.04418 | 0.00724 |
| **Ireb2** | 5.66433 | 3.87562 | 3.20728 | 18.68543 | 8.55639 | 0.23296 | 2.72932 | 2.74244 | 9.45617 | 6.93011 | 5.24559 | 17.27314 | 0.73737 | 18.29532 | 16.36322 | 2.85882 |
| **Irf1** | 0.56336 | 0.00743 | 0.00000 | 0.75752 | 0.01368 | 0.00656 | 0.00659 | 0.96145 | 5.12853 | 0.03933 | 0.00000 | 0.03152 | 0.00569 | 0.00000 | 1.22554 | 0.00684 |
| **Irf2bp1** | 2.68682 | 0.09891 | 0.88286 | 0.00047 | 1.50458 | 0.03075 | 0.40991 | 1.52111 | 5.42289 | 1.90425 | 0.52782 | 0.01778 | 5.16484 | 0.00265 | 0.05993 | 0.60279 |
| **Irf2bp2** | 1.49939 | 0.00550 | 0.19803 | 0.14565 | 0.00476 | 0.00744 | 0.00599 | 0.00449 | 0.04708 | 0.75258 | 0.01025 | 0.00347 | 0.01047 | 0.01372 | 0.28534 | 0.00572 |
| **Irf2bpl** | 2.46641 | 0.03506 | 1.15208 | 5.48146 | 0.00453 | 6.17254 | 1.64196 | 8.99685 | 4.37971 | 10.30865 | 3.74571 | 3.02224 | 4.36917 | 5.79980 | 9.17774 | 0.24656 |
| **Irf3** | 5.93445 | 0.02273 | 0.00000 | 3.48985 | 0.43563 | 5.75264 | 1.42239 | 3.54461 | 4.80958 | 4.18987 | 0.43381 | 0.13306 | 6.11957 | 0.03186 | 0.00000 | 7.44301 |
| **Irf5** | 0.12838 | 0.00920 | 0.00248 | 0.00498 | 0.01621 | 0.01512 | 0.00793 | 0.12228 | 0.24275 | 0.54932 | 0.44609 | 0.00512 | 0.00859 | 0.01265 | 0.00111 | 0.00718 |
| **Irf6** | 0.72265 | 0.00697 | 0.42191 | 0.28946 | 0.00813 | 0.00905 | 0.00592 | 0.51360 | 0.00670 | 1.98462 | 1.95432 | 0.00000 | 0.07765 | 0.06311 | 0.00000 | 0.00641 |
| **Irf7** | 0.01112 | 0.00793 | 0.56026 | 0.70948 | 0.00807 | 0.00554 | 0.00739 | 0.00945 | 0.79286 | 0.94497 | 0.01436 | 0.04594 | 0.13477 | 0.00405 | 0.20463 | 0.00680 |
| **Irf8** | 0.01309 | 0.00812 | 0.01133 | 0.93789 | 0.00898 | 0.00654 | 0.00739 | 0.01062 | 0.00000 | 0.95901 | 0.02597 | 0.00555 | 0.01126 | 0.00444 | 0.64752 | 0.00713 |
| **Irf9** | 1.01081 | 0.00000 | 0.87225 | 2.76846 | 0.18370 | 0.00475 | 6.38476 | 2.78328 | 4.33042 | 3.39448 | 3.13623 | 7.31095 | 0.45387 | 1.13515 | 4.53291 | 1.52949 |
| **Irgm1** | 0.05345 | 0.00803 | 0.02175 | 0.46835 | 0.03796 | 0.01200 | 0.01767 | 0.00044 | 0.80831 | 0.00000 | 0.40240 | 0.00421 | 0.00813 | 0.12098 | 0.02555 | 0.01433 |
| **Irgm2** | 0.05057 | 0.01868 | 0.04828 | 0.07533 | 0.01903 | 0.02033 | 0.02799 | 0.03612 | 0.08720 | 0.03765 | 0.10558 | 0.00602 | 0.03579 | 0.28637 | 0.09942 | 0.00735 |
| **Irgq** | 0.31751 | 0.02891 | 5.24227 | 10.62336 | 0.01462 | 3.12211 | 0.00205 | 2.92278 | 19.32340 | 6.59764 | 3.17956 | 0.84669 | 5.21374 | 0.00000 | 3.68453 | 1.56881 |
| **Irs3** | 0.92035 | 0.00774 | 0.00420 | 0.57350 | 0.01712 | 0.00702 | 0.00660 | 0.02448 | 0.00425 | 0.01518 | 0.01375 | 0.00744 | 0.01099 | 0.01083 | 0.00492 | 0.00658 |
| **Irx2** | 0.92482 | 0.00668 | 0.00390 | 0.97130 | 0.37954 | 0.00710 | 0.00682 | 0.01002 | 0.01186 | 0.01454 | 0.01262 | 0.00746 | 0.01081 | 0.01089 | 0.00419 | 0.00619 |
| **Isca1** | 8.50093 | 8.05162 | 5.38478 | 4.52938 | 12.35762 | 9.81128 | 3.96654 | 9.72669 | 6.19604 | 10.59400 | 3.16220 | 7.19869 | 5.01693 | 0.29230 | 3.79140 | 3.42884 |
| **Isca2** | 3.84673 | 2.65374 | 1.87515 | 0.93718 | 5.25551 | 2.36822 | 0.00000 | 2.18090 | 2.30415 | 2.36272 | 2.89280 | 3.19616 | 1.82279 | 0.00000 | 2.15660 | 0.02000 |
| **Iscu** | 4.64188 | 2.23581 | 1.18329 | 6.16904 | 2.92739 | 2.54398 | 1.94072 | 0.73421 | 6.38670 | 4.81507 | 8.90658 | 6.38105 | 1.86357 | 0.41026 | 0.95142 | 1.40693 |
| **Isg15** | 0.01306 | 0.00777 | 0.01087 | 0.88634 | 0.00899 | 0.00631 | 0.01735 | 0.00887 | 3.58377 | 0.50349 | 0.01826 | 0.00000 | 0.01217 | 0.02218 | 0.00835 | 0.00742 |
| **Isg20** | 0.01146 | 0.00755 | 0.00960 | 0.99937 | 0.02625 | 1.18646 | 0.00713 | 0.00952 | 0.20196 | 4.39994 | 0.00756 | 0.00000 | 0.01253 | 0.00412 | 0.09651 | 0.00676 |
| **Isg20l2** | 1.83094 | 1.23347 | 1.21781 | 4.40707 | 0.72970 | 0.01276 | 0.20717 | 1.69886 | 0.40241 | 3.58080 | 1.02543 | 0.25651 | 2.35146 | 12.63387 | 10.45802 | 0.99383 |
| **Isl2** | 0.40937 | 0.00709 | 0.00350 | 2.89437 | 0.00731 | 0.00779 | 0.00671 | 0.23806 | 0.04917 | 3.83885 | 0.55170 | 0.00006 | 0.72650 | 0.00439 | 0.00000 | 0.00605 |
| **Islr2** | 0.00777 | 0.00801 | 0.01109 | 0.12901 | 0.00763 | 0.00644 | 0.00747 | 0.01189 | 0.00972 | 0.00500 | 0.10767 | 0.00777 | 0.00700 | 0.01111 | 0.00574 | 1.32847 |
| **Ism2** | 0.03409 | 0.00787 | 0.01030 | 0.34787 | 0.00842 | 0.00653 | 0.00727 | 0.01089 | 0.16534 | 0.32447 | 0.03976 | 0.01624 | 0.01164 | 0.00619 | 0.19895 | 0.01455 |
| **Isoc1** | 4.04649 | 4.67049 | 2.63523 | 0.25729 | 0.09208 | 0.97959 | 1.62757 | 0.60660 | 0.25030 | 1.51053 | 1.75850 | 3.19616 | 2.66687 | 3.06136 | 0.05531 | 0.02856 |
| **Isoc2b** | 0.11071 | 0.89736 | 0.86638 | 0.42345 | 0.02399 | 0.00579 | 0.72838 | 0.00000 | 0.58006 | 0.94575 | 2.09955 | 0.01394 | 1.04707 | 0.00263 | 0.10994 | 0.00567 |
| **Israa** | 0.01802 | 0.00744 | 0.00722 | 0.00461 | 0.00886 | 0.00670 | 0.13736 | 0.00246 | 0.01007 | 0.00000 | 0.28321 | 0.01436 | 0.00905 | 0.01351 | 0.60846 | 0.00713 |
| **Ist1** | 9.69976 | 10.18347 | 13.03622 | 14.96703 | 8.38788 | 7.75459 | 6.67896 | 15.10486 | 21.48199 | 17.76861 | 10.04505 | 8.38620 | 10.01816 | 15.50496 | 16.95474 | 0.04830 |
| **Isy1** | 11.09948 | 5.90703 | 6.40461 | 2.97657 | 6.36541 | 5.45133 | 3.19086 | 17.40972 | 2.42463 | 6.19966 | 5.81624 | 5.09836 | 10.82213 | 0.61217 | 2.85603 | 5.27936 |
| **Isyna1** | 36.34319 | 13.26649 | 32.47032 | 6.52954 | 24.28348 | 17.17990 | 22.33391 | 49.72426 | 4.52707 | 50.61146 | 38.16139 | 17.50849 | 53.00295 | 0.19095 | 0.00000 | 17.08185 |
| **Itch** | 2.74318 | 5.24627 | 6.34217 | 3.01298 | 1.53608 | 6.59231 | 7.88684 | 8.96056 | 1.78295 | 12.52823 | 3.99141 | 8.59527 | 2.11388 | 17.52864 | 14.11813 | 1.82681 |
| **Itfg1** | 2.63776 | 3.13141 | 3.54207 | 0.83656 | 2.18457 | 1.02381 | 0.37888 | 1.13400 | 0.20620 | 1.68359 | 1.02762 | 0.01473 | 1.17361 | 0.73839 | 1.29373 | 0.01523 |
| **Itfg2** | 1.68307 | 3.43560 | 3.88126 | 1.85937 | 4.66340 | 0.00302 | 5.17651 | 6.43500 | 3.80455 | 4.19165 | 0.25996 | 5.08754 | 2.56233 | 3.51906 | 1.96649 | 4.08024 |
| **Itga1** | 0.00934 | 0.00734 | 0.00597 | 0.01199 | 0.00956 | 0.00695 | 0.00681 | 0.50382 | 0.49774 | 0.02071 | 0.22534 | 0.03121 | 0.37513 | 0.05799 | 0.00000 | 0.00716 |
| **Itga10** | 0.01376 | 0.00780 | 0.01187 | 0.59884 | 0.00934 | 0.00658 | 0.00749 | 0.01322 | 0.25315 | 0.01050 | 0.01072 | 0.00737 | 0.01281 | 0.01911 | 0.05752 | 0.00730 |
| **Itga11** | 0.01396 | 0.00796 | 0.00488 | 0.00236 | 0.00837 | 0.02276 | 0.00750 | 0.02701 | 0.35986 | 3.04089 | 1.08367 | 0.25713 | 0.00705 | 0.00574 | 0.72262 | 0.00689 |
| **Itga2b** | 0.44911 | 0.00741 | 0.00475 | 0.09166 | 0.00680 | 0.00723 | 0.00674 | 0.02218 | 0.02013 | 0.23195 | 0.10564 | 0.00709 | 0.45552 | 0.01054 | 0.00688 | 0.00662 |
| **Itga3** | 6.98567 | 4.00154 | 5.01557 | 0.00000 | 0.04360 | 3.96188 | 8.85978 | 3.01336 | 0.02210 | 0.89840 | 2.94624 | 0.04988 | 3.19901 | 0.00657 | 0.00104 | 4.18783 |
| **Itga4** | 0.12792 | 0.01899 | 0.05870 | 0.48968 | 0.06311 | 0.00602 | 0.07108 | 0.03662 | 0.71557 | 0.74154 | 0.02616 | 0.03816 | 0.02716 | 0.53474 | 0.48160 | 0.01467 |
| **Itga5** | 4.84962 | 0.01044 | 3.87425 | 0.00007 | 0.51686 | 0.02952 | 5.29445 | 0.98450 | 0.14106 | 0.31451 | 1.14611 | 0.25988 | 5.89362 | 0.00880 | 0.00000 | 6.08923 |
| **Itga6** | 7.27787 | 3.55686 | 9.02764 | 5.89611 | 9.93319 | 2.08581 | 7.47585 | 1.75652 | 8.71319 | 3.13323 | 2.53817 | 3.05776 | 1.21064 | 4.11216 | 8.29670 | 2.12004 |
| **Itga7** | 0.00765 | 0.00793 | 0.74060 | 0.29965 | 0.00786 | 0.13261 | 0.00671 | 0.00403 | 0.00441 | 0.15304 | 0.21396 | 0.00657 | 1.03172 | 0.03512 | 0.00000 | 0.00657 |
| **Itga9** | 0.01367 | 0.00815 | 0.01141 | 4.00898 | 0.00922 | 0.00643 | 0.00739 | 0.01179 | 0.20964 | 0.08626 | 0.04882 | 0.00774 | 0.01249 | 0.00000 | 0.40030 | 0.00674 |
| **Itgae** | 0.11609 | 1.22847 | 1.35691 | 1.26784 | 0.00521 | 0.01171 | 1.25352 | 1.76017 | 0.47364 | 0.46637 | 2.38915 | 0.03049 | 1.67150 | 1.72652 | 2.25794 | 0.01168 |
| **Itgal** | 0.00636 | 1.76634 | 0.00594 | 6.87607 | 0.00788 | 0.00542 | 0.00634 | 0.00801 | 2.57541 | 1.12022 | 0.00000 | 0.00769 | 0.01018 | 0.00000 | 1.28571 | 0.00507 |
| **Itgam** | 0.01334 | 0.01858 | 0.01166 | 0.00467 | 0.00916 | 0.00686 | 0.00757 | 0.01300 | 0.34376 | 0.21806 | 0.00875 | 0.00776 | 0.01286 | 0.01365 | 0.05739 | 0.00738 |
| **Itgav** | 2.72068 | 0.00367 | 0.42658 | 5.90568 | 0.00214 | 2.32852 | 0.05562 | 1.19608 | 3.76057 | 5.10025 | 1.07825 | 2.41181 | 0.78230 | 11.01359 | 6.88874 | 0.00005 |
| **Itgb1** | 6.83284 | 1.66865 | 4.26908 | 3.57207 | 3.01104 | 0.04219 | 8.11391 | 7.32819 | 0.31963 | 2.53075 | 3.28188 | 0.78148 | 6.58420 | 5.74248 | 3.31264 | 3.92734 |
| **Itgb1bp1** | 6.42085 | 5.31734 | 11.10069 | 7.82502 | 7.05895 | 0.00000 | 5.48453 | 12.84091 | 9.90239 | 6.31602 | 9.48394 | 0.88242 | 9.23455 | 0.79454 | 0.00000 | 6.57869 |
| **Itgb3** | 0.17856 | 0.01836 | 0.00747 | 2.39569 | 0.00761 | 0.00635 | 0.00706 | 0.02251 | 0.11450 | 0.34659 | 0.02756 | 0.02547 | 0.01806 | 0.00131 | 0.75354 | 0.01466 |
| **Itgb3bp** | 0.59093 | 0.52182 | 0.01420 | 0.12003 | 0.00947 | 0.00578 | 0.00744 | 0.24816 | 0.00937 | 0.00825 | 0.01100 | 0.01886 | 0.00666 | 0.03588 | 0.00000 | 0.01393 |
| **Itgb4** | 0.00544 | 0.00705 | 0.16426 | 1.56230 | 0.00779 | 0.00900 | 0.65199 | 2.01551 | 1.01445 | 6.80303 | 0.58976 | 0.10229 | 0.00002 | 0.00346 | 0.00000 | 0.00678 |
| **Itgb5** | 6.43684 | 0.00001 | 5.41467 | 7.09148 | 0.01169 | 6.18883 | 6.38401 | 14.96187 | 5.14543 | 16.36811 | 6.30975 | 2.96636 | 7.64509 | 0.83179 | 10.26172 | 2.43129 |
| **Itgb7** | 1.94171 | 0.00447 | 1.78671 | 0.46210 | 0.00965 | 0.00310 | 1.06984 | 2.71660 | 4.96179 | 2.92449 | 2.27156 | 0.07066 | 0.36054 | 0.05150 | 0.81219 | 0.00366 |
| **Itih5** | 0.01320 | 0.00816 | 0.01217 | 0.00671 | 0.00895 | 0.00652 | 0.00738 | 0.01066 | 0.01161 | 1.01021 | 0.01064 | 0.00577 | 0.01065 | 0.00735 | 0.09066 | 0.00714 |
| **Itln1** | 0.01086 | 0.00783 | 0.00779 | 0.13542 | 0.00951 | 0.00687 | 0.00711 | 0.31013 | 0.00701 | 0.00000 | 0.08592 | 0.01710 | 0.00599 | 0.00733 | 0.11293 | 0.00735 |
| **Itm2a** | 0.01371 | 0.00756 | 0.00571 | 0.00624 | 0.00926 | 0.00672 | 0.00702 | 0.00781 | 0.01039 | 0.00000 | 0.07187 | 1.03288 | 0.01200 | 0.00790 | 0.41704 | 0.00735 |
| **Itm2b** | 3.42941 | 2.62481 | 2.07088 | 1.53102 | 2.97214 | 8.62480 | 0.16131 | 0.00000 | 3.27212 | 9.89659 | 4.99466 | 0.07531 | 2.69666 | 1.96922 | 0.00000 | 2.74075 |

| | | | | | | | | | | | | | | | | |
|---|---|---|---|---|---|---|---|---|---|---|---|---|---|---|---|---|
| **Itm2c** | 1.11464 | 0.00731 | 0.17422 | 0.28311 | 0.01436 | 0.00483 | 0.00638 | 0.01355 | 0.14079 | 4.31242 | 0.00781 | 0.06396 | 0.96639 | 0.02572 | 0.45817 | 0.02097 |
| **Itpa** | 12.96160 | 10.48044 | 13.73493 | 5.46386 | 16.20409 | 9.47634 | 7.05336 | 17.80005 | 9.10949 | 17.56325 | 16.42249 | 1.65704 | 18.61809 | 6.07025 | 11.77846 | 6.14308 |
| **Itpk1** | 10.25397 | 0.02924 | 1.83750 | 6.56479 | 0.11653 | 0.00000 | 2.34613 | 0.08011 | 4.60885 | 1.90067 | 2.13772 | 0.68050 | 0.88676 | 0.00000 | 0.21029 | 4.27261 |
| **Itpka** | 0.43403 | 0.00696 | 0.00000 | 0.62650 | 0.00697 | 0.01594 | 0.00700 | 2.22664 | 0.00143 | 1.19699 | 0.71588 | 0.41695 | 0.00545 | 0.00350 | 0.00000 | 0.01381 |
| **Itpkb** | 0.01375 | 0.00812 | 0.01230 | 0.30812 | 0.00931 | 0.00662 | 0.00752 | 0.01394 | 0.00678 | 0.01549 | 0.01388 | 0.00821 | 0.01266 | 0.01128 | 0.00642 | 0.00736 |
| **Itpkc** | 1.58643 | 0.00293 | 0.33871 | 8.30667 | 1.60527 | 0.02728 | 2.02438 | 2.33397 | 14.46985 | 5.09081 | 6.12795 | 0.43150 | 0.58439 | 0.40774 | 8.11254 | 0.52875 |
| **Itpr1** | 0.01143 | 0.00773 | 0.00576 | 9.86184 | 0.00915 | 0.00477 | 0.00700 | 0.01233 | 4.68671 | 0.66811 | 0.05623 | 0.00000 | 0.01108 | 0.00544 | 6.46562 | 0.00532 |
| **Itpr2** | 0.01086 | 0.00776 | 0.00377 | 0.84889 | 0.00902 | 0.00711 | 0.01667 | 0.12751 | 2.01983 | 0.80535 | 0.67243 | 0.02632 | 0.00427 | 0.19103 | 0.35626 | 0.00716 |
| **Itpr3** | 4.23473 | 0.03417 | 2.09523 | 1.44410 | 0.00000 | 0.00000 | 1.45886 | 3.60437 | 2.79677 | 3.81325 | 2.10845 | 0.20692 | 2.51377 | 0.00000 | 0.00000 | 0.00742 |
| **Itpripl1** | 0.45770 | 0.00792 | 0.83801 | 0.00882 | 0.01168 | 0.00924 | 0.00713 | 0.01321 | 0.02323 | 0.17551 | 0.01017 | 0.01894 | 0.00877 | 0.00784 | 0.07294 | 0.00676 |
| **Itpripl2** | 0.01884 | 0.00783 | 0.25689 | 0.00693 | 0.02170 | 1.42870 | 0.00700 | 0.00924 | 0.01216 | 0.59703 | 0.00973 | 0.03319 | 0.00511 | 0.01220 | 0.00523 | 0.01483 |
| **Itsn1** | 6.02384 | 2.75667 | 1.93059 | 8.43871 | 4.63071 | 0.06748 | 5.82728 | 3.56839 | 9.02980 | 8.15581 | 8.48587 | 3.46433 | 0.13448 | 3.05688 | 8.14218 | 1.71736 |
| **Itsn2** | 1.66584 | 0.95175 | 1.85904 | 2.57050 | 0.00944 | 1.57705 | 0.02367 | 1.23394 | 0.00000 | 0.97558 | 0.81033 | 0.04434 | 1.64275 | 3.28804 | 0.69559 | 0.32515 |
| **Ivd** | 1.61732 | 3.57679 | 1.75270 | 6.23621 | 4.30090 | 2.84911 | 0.93558 | 2.94607 | 7.18682 | 0.77967 | 2.48792 | 0.11455 | 2.63835 | 0.00000 | 2.27691 | 5.99522 |
| **Ivns1abp** | 6.50063 | 12.71570 | 17.33124 | 12.61512 | 10.57590 | 5.22907 | 12.07525 | 16.64378 | 14.59473 | 21.60389 | 21.57782 | 16.25072 | 14.72609 | 30.84441 | 20.46238 | 4.74949 |
| **Iws1** | 6.16171 | 3.91458 | 2.15567 | 11.52747 | 8.69901 | 4.58748 | 3.23945 | 5.95160 | 14.70382 | 11.67155 | 8.68712 | 3.97960 | 4.69124 | 15.92915 | 14.96305 | 1.79674 |
| **Izumo1** | 0.04418 | 0.00803 | 0.00986 | 0.01056 | 0.00803 | 0.00666 | 0.00728 | 0.01276 | 0.01243 | 0.00866 | 0.01247 | 0.00787 | 0.21856 | 0.03410 | 0.00913 | 0.00708 |
| **Izumo1r** | 0.01369 | 0.00816 | 0.01057 | 1.60623 | 0.00930 | 0.00661 | 0.00750 | 0.01472 | 0.00000 | 0.01068 | 0.03815 | 0.08739 | 0.01208 | 0.02873 | 0.01353 | 0.00714 |
| **Izumo4** | 0.01921 | 0.00807 | 0.07271 | 0.57636 | 0.00857 | 0.02619 | 0.00746 | 0.01347 | 0.27885 | 0.02318 | 0.01095 | 0.00808 | 0.01158 | 0.01090 | 0.00502 | 0.00724 |
| **Jade1** | 4.73109 | 5.97500 | 10.41071 | 3.77778 | 5.05400 | 2.05196 | 1.50610 | 2.94572 | 9.49868 | 8.56806 | 1.36732 | 0.25434 | 1.14480 | 1.52766 | 7.59526 | 6.10347 |
| **Jade2** | 0.01070 | 0.00807 | 0.00569 | 3.68304 | 0.00984 | 0.00708 | 0.00721 | 0.00000 | 2.52612 | 0.24965 | 0.49732 | 0.10147 | 0.47651 | 0.00000 | 1.30995 | 0.01440 |
| **Jade3** | 0.37662 | 4.69714 | 2.29842 | 1.93608 | 2.02998 | 0.75213 | 0.00333 | 0.19892 | 2.27108 | 0.47381 | 0.06813 | 9.98517 | 1.29035 | 3.03795 | 0.00000 | 2.26781 |
| **Jag1** | 0.01394 | 0.00763 | 0.00923 | 0.86006 | 0.00929 | 0.00661 | 0.00728 | 0.00868 | 0.00000 | 0.00369 | 0.08985 | 0.09275 | 0.01141 | 0.03157 | 0.29822 | 0.00723 |
| **Jag2** | 0.52431 | 0.05289 | 0.03215 | 0.33694 | 0.00687 | 0.19037 | 0.02966 | 0.10921 | 0.09410 | 2.42364 | 0.00941 | 0.00000 | 1.89464 | 0.01410 | 0.00000 | 0.00579 |
| **Jagn1** | 10.43786 | 2.47387 | 8.56753 | 3.89866 | 7.13781 | 10.65048 | 2.02060 | 12.85721 | 2.88530 | 10.60715 | 4.29866 | 0.23613 | 10.25713 | 0.05817 | 3.59091 | 5.45909 |
| **Jak1** | 5.85029 | 2.13195 | 5.84161 | 5.56474 | 1.63733 | 4.84817 | 4.25277 | 4.22424 | 2.33190 | 3.43110 | 2.83420 | 7.41304 | 0.37214 | 8.15961 | 5.04871 | 1.45962 |
| **Jak2** | 0.83239 | 0.00564 | 1.66450 | 19.49471 | 1.31138 | 0.00101 | 0.00542 | 0.00000 | 17.17910 | 9.02192 | 0.83953 | 1.63601 | 0.04581 | 7.76877 | 26.31581 | 0.02245 |
| **Jak3** | 0.62903 | 0.01773 | 0.01140 | 0.00465 | 0.04684 | 0.00636 | 0.00684 | 0.04424 | 0.76618 | 1.39979 | 0.00618 | 0.00000 | 0.36784 | 0.01516 | 0.04398 | 0.01438 |
| **Jakmip1** | 0.61874 | 0.00643 | 1.37129 | 0.02917 | 0.00424 | 0.00472 | 0.01656 | 2.18038 | 0.10000 | 4.32301 | 1.16406 | 1.27228 | 0.00000 | 0.02991 | 0.62362 | 0.00564 |
| **Jakmip3** | 0.01359 | 0.00819 | 0.01164 | 0.17513 | 0.00924 | 0.00681 | 0.00762 | 0.01391 | 0.76062 | 0.05834 | 0.02136 | 0.00769 | 0.01296 | 0.01289 | 0.00833 | 0.00735 |
| **Jam2** | 7.90364 | 2.10228 | 1.55966 | 2.28221 | 11.29169 | 0.00223 | 0.02847 | 0.75476 | 0.81481 | 2.39452 | 2.65153 | 0.79296 | 1.61095 | 0.00000 | 2.63550 | 0.05585 |
| **Jam3** | 0.01512 | 0.00761 | 0.00468 | 4.84052 | 0.01003 | 0.00468 | 0.33246 | 0.00849 | 11.92722 | 0.33185 | 0.00000 | 0.07519 | 0.01751 | 0.00651 | 3.02963 | 0.00562 |
| **Jarid2** | 16.98446 | 12.73639 | 21.53959 | 6.72540 | 9.69858 | 8.28996 | 12.01782 | 25.67937 | 3.94646 | 15.04030 | 24.63800 | 8.05381 | 12.12135 | 17.95077 | 11.56590 | 9.36030 |
| **Jazf1** | 0.01068 | 0.00829 | 0.00677 | 9.24716 | 0.00938 | 0.00514 | 0.00775 | 0.09660 | 3.61328 | 0.00000 | 0.00000 | 0.01779 | 0.01314 | 0.07515 | 2.59714 | 0.00535 |
| **Jchain** | 0.01187 | 0.00786 | 0.01125 | 0.05746 | 0.00911 | 0.00643 | 0.00714 | 0.01189 | 0.02763 | 0.01876 | 0.01320 | 0.02904 | 0.40808 | 0.01539 | 0.03042 | 0.00705 |
| **Jdp2** | 0.00971 | 0.00774 | 0.08937 | 0.01063 | 0.00922 | 0.02578 | 0.00719 | 0.10549 | 0.01230 | 0.03190 | 0.01170 | 0.00721 | 0.00751 | 0.01203 | 0.00761 | 0.04340 |
| **Jkamp** | 8.34279 | 1.13622 | 4.45061 | 1.03730 | 2.39722 | 0.03069 | 1.41364 | 4.12070 | 1.64101 | 4.88955 | 2.15649 | 2.44165 | 5.19267 | 2.26552 | 1.12122 | 4.46069 |
| **Jmjd1c** | 3.91006 | 0.82246 | 0.36298 | 2.84739 | 0.00000 | 0.00785 | 0.99441 | 2.46179 | 4.80930 | 1.37649 | 1.96154 | 5.92238 | 0.17720 | 7.12037 | 4.64685 | 2.21776 |
| **Jmjd4** | 1.51748 | 0.00557 | 1.77945 | 4.00254 | 0.01289 | 2.61812 | 0.05818 | 2.69245 | 3.72392 | 0.96474 | 0.54041 | 0.03424 | 0.84189 | 0.13177 | 0.56106 | 1.99607 |
| **Jmjd6** | 7.74046 | 3.53918 | 8.55718 | 10.49401 | 5.58476 | 4.01587 | 4.82010 | 14.99823 | 21.64169 | 20.80504 | 8.12425 | 2.14736 | 4.16895 | 2.36951 | 10.52154 | 4.04091 |
| **Jmjd7** | 0.14841 | 0.05907 | 0.15952 | 0.11844 | 0.00784 | 0.06002 | 0.02887 | 0.04406 | 0.32012 | 0.22332 | 0.11715 | 0.00576 | 0.02382 | 0.08112 | 0.02017 | 0.14886 |
| **Jmjd8** | 0.08529 | 0.26933 | 0.36936 | 0.28064 | 0.01597 | 0.17820 | 0.62322 | 0.52819 | 0.22567 | 0.65974 | 0.67306 | 0.02036 | 0.80035 | 0.00443 | 0.19790 | 0.13460 |
| **Jmy** | 1.38569 | 0.00000 | 2.48731 | 16.06219 | 0.00000 | 0.37378 | 2.20679 | 6.80325 | 17.40849 | 9.69696 | 6.47958 | 6.97294 | 0.00000 | 22.14442 | 16.70696 | 0.04207 |
| **Josd1** | 7.30354 | 2.50308 | 4.37089 | 8.36962 | 4.19098 | 0.83818 | 6.42521 | 3.56706 | 5.08045 | 5.14946 | 0.32210 | 0.80551 | 0.39813 | 0.54348 | 3.33716 | 3.24812 |
| **Josd2** | 8.97608 | 0.31228 | 2.09533 | 0.20904 | 6.49935 | 4.79032 | 2.95528 | 7.04934 | 0.00001 | 0.26825 | 1.10711 | 0.06951 | 6.06031 | 0.00023 | 0.00000 | 4.16038 |
| **Jph1** | 0.94689 | 0.00718 | 0.00000 | 0.00552 | 0.01075 | 0.00631 | 0.00627 | 0.00000 | 0.01140 | 0.00000 | 0.22235 | 0.01837 | 0.01460 | 1.91202 | 0.01252 | 0.53840 |
| **Jpx** | 0.10380 | 0.01808 | 0.23677 | 0.24788 | 0.02704 | 0.02012 | 0.03783 | 0.00000 | 0.44415 | 0.05867 | 1.24361 | 0.08928 | 0.23710 | 0.38691 | 0.19391 | 0.41683 |
| **Jrk** | 0.00181 | 0.01897 | 2.86937 | 0.89661 | 0.00371 | 0.43430 | 0.00578 | 0.01270 | 0.00000 | 0.00000 | 0.02158 | 0.05018 | 2.30554 | 0.00622 | 0.35021 | 0.46512 |
| **Jsrp1** | 0.01102 | 0.00770 | 0.01014 | 0.02539 | 0.00951 | 0.00675 | 0.00719 | 0.12327 | 0.02451 | 0.06310 | 0.01321 | 0.00695 | 0.00854 | 0.01137 | 0.00885 | 0.00772 |
| **Jtb** | 7.68643 | 0.01337 | 1.13395 | 6.13991 | 5.32814 | 0.07207 | 2.59293 | 11.92009 | 9.48870 | 17.20174 | 14.91844 | 0.84584 | 11.64492 | 2.28441 | 0.26444 | 4.19035 |
| **Jun** | 0.00759 | 0.00778 | 0.04080 | 0.67064 | 0.01826 | 0.05259 | 0.00698 | 0.52140 | 0.00000 | 0.08164 | 0.90028 | 0.03509 | 0.55448 | 0.00542 | 0.00000 | 0.02811 |
| **Junb** | 0.90012 | 0.00678 | 0.00613 | 0.08074 | 0.00690 | 0.00635 | 0.02694 | 0.06494 | 0.08405 | 0.34206 | 0.35970 | 0.04810 | 0.74295 | 0.00969 | 0.00000 | 0.86600 |
| **Jund** | 0.20518 | 0.00678 | 0.00000 | 1.75090 | 0.92833 | 0.29356 | 0.00621 | 0.02461 | 2.34009 | 4.09877 | 2.02905 | 0.06505 | 0.07966 | 0.13827 | 0.33114 | 1.02589 |
| **Jup** | 0.25948 | 3.01266 | 0.61943 | 0.84954 | 4.46175 | 6.24543 | 3.66579 | 4.95161 | 0.33136 | 3.32159 | 3.86811 | 0.10497 | 2.75278 | 0.84045 | 0.10925 | 6.13304 |
| **Kalrn** | 0.29736 | 0.00500 | 0.99048 | 4.34459 | 4.21551 | 1.02306 | 0.00664 | 0.01318 | 1.34381 | 0.17955 | 0.14671 | 0.01181 | 0.38552 | 1.64964 | 0.65135 | 0.16693 |
| **Kank1** | 1.21194 | 0.00575 | 1.41097 | 1.35237 | 0.03416 | 2.67888 | 2.27981 | 0.14857 | 0.83435 | 2.69626 | 0.55649 | 0.01280 | 0.83887 | 0.01319 | 0.16371 | 0.00423 |
| **Kank2** | 1.27967 | 0.00601 | 2.01739 | 2.50118 | 0.01165 | 0.43238 | 0.00456 | 0.60952 | 8.93083 | 1.61404 | 5.46917 | 0.14432 | 0.43150 | 0.58278 | 1.01484 | 0.78664 |
| **Kank3** | 0.00000 | 0.00726 | 2.13968 | 1.12559 | 0.21923 | 0.00641 | 1.09261 | 0.09829 | 1.39944 | 0.61288 | 0.02356 | 0.01115 | 0.25443 | 0.00489 | 0.17324 | 0.00670 |
| **Kank4** | 0.00910 | 0.00762 | 0.00000 | 3.65868 | 0.00900 | 0.00628 | 0.01645 | 0.00000 | 3.70423 | 1.45655 | 2.76715 | 0.58068 | 0.00000 | 1.53420 | 0.83278 | 0.00612 |
| **Kank4os** | 0.00000 | 5.11433 | 0.19086 | 0.63937 | 0.00725 | 0.00838 | 1.36232 | 1.03774 | 1.04618 | 0.00000 | 0.25478 | 0.09558 | 0.00334 | 0.04385 | 1.01342 | 0.00488 |
| **Kansl1** | 3.05539 | 5.51193 | 1.76632 | 12.51825 | 4.00808 | 0.00000 | 4.87192 | 5.06996 | 12.88024 | 11.16358 | 4.20450 | 6.86047 | 1.98447 | 5.16808 | 6.05125 | 4.31585 |
| **Kansl1l** | 0.22282 | 0.00763 | 0.00000 | 1.12412 | 0.00872 | 0.57261 | 0.00707 | 0.01924 | 0.18097 | 0.00000 | 0.10224 | 0.00570 | 0.00593 | 0.00621 | 0.68434 | 0.00658 |

| | | | | | | | | | | | | | | | | |
|---|---|---|---|---|---|---|---|---|---|---|---|---|---|---|---|---|
| **Kansl2** | 11.33537 | 8.97527 | 11.95999 | 15.03999 | 10.61949 | 0.05879 | 8.27336 | 12.30133 | 19.36174 | 17.38796 | 12.67807 | 8.33922 | 8.64905 | 11.80188 | 19.84861 | 9.47255 |
| **Kansl3** | 5.41795 | 0.85867 | 6.03832 | 8.52941 | 0.01186 | 0.05818 | 2.77618 | 6.97442 | 14.90052 | 10.13835 | 8.57720 | 4.65563 | 5.58382 | 9.35960 | 9.86131 | 3.36869 |
| **Kantr** | 0.73342 | 0.00616 | 0.00000 | 0.01477 | 0.00656 | 0.00625 | 0.67077 | 0.00482 | 0.40007 | 0.00000 | 0.02764 | 0.04390 | 0.01117 | 2.38703 | 0.08970 | 0.00661 |
| **Kars** | 21.24774 | 16.40869 | 21.50451 | 9.73279 | 15.20007 | 13.61695 | 10.09308 | 29.58119 | 13.96735 | 33.54849 | 19.51207 | 17.82112 | 18.76393 | 7.21295 | 20.02613 | 15.36363 |
| **Kat2a** | 14.86085 | 7.24182 | 6.70750 | 4.77900 | 8.65899 | 3.79185 | 6.46329 | 12.24321 | 5.37760 | 7.75437 | 4.79989 | 2.27719 | 5.43781 | 0.72590 | 7.28994 | 4.19822 |
| **Kat2b** | 5.13605 | 0.00691 | 0.00000 | 3.90465 | 2.82344 | 0.61981 | 0.00000 | 0.44087 | 4.25415 | 0.35979 | 2.86209 | 11.90849 | 0.00000 | 4.33403 | 3.55011 | 0.28376 |
| **Kat5** | 1.89504 | 2.54524 | 5.97974 | 2.46032 | 6.23134 | 0.01202 | 0.00940 | 8.49115 | 2.11785 | 7.82239 | 3.52726 | 1.50369 | 6.87734 | 2.32222 | 4.08282 | 4.17646 |
| **Kat6a** | 3.77391 | 1.35977 | 0.65937 | 15.13278 | 4.67337 | 0.00057 | 0.83264 | 0.00000 | 11.38810 | 5.50184 | 4.67015 | 0.80356 | 0.55427 | 2.87745 | 3.84065 | 1.59222 |
| **Kat6b** | 5.52363 | 2.88941 | 5.48998 | 6.44038 | 0.57074 | 2.59629 | 0.57108 | 8.54218 | 5.05384 | 7.91935 | 3.15238 | 7.29603 | 3.04551 | 9.71848 | 9.76865 | 4.40564 |
| **Kat7** | 9.99605 | 8.77916 | 9.84305 | 13.85833 | 7.19141 | 5.96722 | 7.60485 | 12.20546 | 13.68631 | 16.71520 | 9.61010 | 5.69560 | 11.51168 | 14.63021 | 14.06776 | 6.52819 |
| **Kat8** | 9.12531 | 0.44961 | 3.77968 | 4.38624 | 0.63823 | 0.00670 | 2.26840 | 9.56372 | 9.00008 | 11.39770 | 7.72976 | 2.06299 | 6.88558 | 0.95010 | 7.81231 | 1.09134 |
| **Katna1** | 9.33598 | 11.76576 | 10.87171 | 17.60831 | 4.40675 | 5.45029 | 9.86861 | 8.91577 | 30.08853 | 14.61539 | 8.89223 | 10.06289 | 7.53961 | 14.99473 | 17.39883 | 6.96961 |
| **Katnal1** | 0.02536 | 0.01708 | 0.29773 | 0.33376 | 0.00829 | 0.00540 | 0.00706 | 0.00910 | 1.60977 | 0.21443 | 0.00000 | 0.03612 | 0.01024 | 0.14105 | 0.99024 | 0.02164 |
| **Katnal2** | 0.05811 | 0.00763 | 0.00911 | 0.01767 | 0.00809 | 0.00672 | 0.00738 | 0.01323 | 0.01172 | 0.00715 | 0.01204 | 0.01741 | 0.01277 | 0.01482 | 0.26320 | 0.00726 |
| **Katnb1** | 3.76866 | 0.03558 | 2.20583 | 0.78631 | 0.47386 | 2.90523 | 0.00298 | 4.68847 | 3.76023 | 3.57737 | 0.31881 | 0.30331 | 1.87341 | 0.00000 | 0.06939 | 1.71381 |
| **Katnbl1** | 1.16576 | 3.14648 | 0.28101 | 9.47840 | 2.63727 | 3.92329 | 0.00272 | 1.65526 | 3.45609 | 4.48523 | 4.07359 | 1.16018 | 0.92193 | 2.65621 | 6.11921 | 0.02256 |
| **Kazald1** | 0.01172 | 0.00787 | 0.01163 | 0.26241 | 0.00861 | 0.00647 | 0.00737 | 0.01068 | 0.00764 | 0.11029 | 0.07287 | 0.00809 | 0.00966 | 0.01611 | 0.00336 | 0.28298 |
| **Kazn** | 2.69693 | 1.63139 | 3.12555 | 0.80192 | 0.17870 | 0.05954 | 1.50054 | 4.43906 | 1.37387 | 3.61904 | 2.28862 | 1.30605 | 0.03779 | 0.00000 | 1.07462 | 2.10956 |
| **Kbtbd11** | 0.01191 | 0.01882 | 0.01100 | 0.03353 | 0.00924 | 0.00651 | 0.00736 | 0.04723 | 0.99004 | 0.19697 | 0.00468 | 0.00523 | 0.00987 | 0.03122 | 0.03184 | 0.00708 |
| **Kbtbd13** | 0.01956 | 0.00867 | 0.00000 | 0.13003 | 0.01187 | 3.77858 | 0.00784 | 0.45896 | 1.67287 | 0.00000 | 0.17428 | 0.02526 | 0.18266 | 0.00380 | 0.40409 | 0.00574 |
| **Kbtbd2** | 1.10604 | 3.41470 | 0.55109 | 2.23894 | 0.00676 | 0.03986 | 0.38463 | 0.56527 | 4.97579 | 0.44218 | 2.56038 | 8.68804 | 0.14870 | 0.36650 | 5.23388 | 0.56331 |
| **Kbtbd3** | 0.58637 | 0.00720 | 0.07140 | 1.03527 | 0.00672 | 0.00488 | 0.00714 | 0.00665 | 2.69200 | 0.00507 | 0.31444 | 0.00000 | 0.00829 | 0.00593 | 0.00000 | 0.03435 |
| **Kbtbd4** | 6.10901 | 4.55532 | 0.80683 | 4.21078 | 0.00510 | 3.45053 | 0.09365 | 2.60825 | 6.51834 | 6.80386 | 3.13595 | 0.27494 | 1.82787 | 0.02311 | 2.85228 | 1.50048 |
| **Kbtbd6** | 0.02723 | 0.00739 | 0.00370 | 0.17643 | 0.00842 | 0.01980 | 0.00721 | 0.36233 | 2.50143 | 0.00000 | 0.00000 | 0.02100 | 0.00606 | 0.10117 | 0.35984 | 0.00703 |
| **Kbtbd7** | 0.06943 | 3.12899 | 0.72168 | 8.39606 | 0.02390 | 0.38158 | 0.00550 | 0.08591 | 10.62025 | 0.45700 | 0.63430 | 0.04179 | 0.50918 | 0.02011 | 3.69699 | 0.00325 |
| **Kbtbd8** | 0.00000 | 0.62882 | 0.00000 | 23.07362 | 0.88999 | 0.00841 | 0.93217 | 0.54775 | 9.56594 | 3.05139 | 3.00274 | 2.32878 | 0.00000 | 12.31035 | 8.45797 | 0.00156 |
| **Kcmf1** | 15.87034 | 13.29345 | 12.14805 | 21.58303 | 12.26228 | 8.27477 | 12.66494 | 15.31313 | 23.21920 | 18.07058 | 18.06304 | 19.06999 | 11.63498 | 4.23480 | 6.93932 | 7.27205 |
| **Kcna2** | 0.02014 | 0.00800 | 0.00947 | 0.03518 | 0.00891 | 0.01341 | 0.00759 | 0.01423 | 0.10144 | 0.03833 | 0.01115 | 0.00773 | 0.03940 | 0.20026 | 0.00825 | 0.00731 |
| **Kcna7** | 0.01259 | 0.00811 | 0.16561 | 0.40038 | 0.00887 | 0.00890 | 0.00736 | 0.01277 | 0.31272 | 0.01056 | 0.01068 | 0.00762 | 0.01062 | 0.00835 | 0.06147 | 0.00722 |
| **Kcnab1** | 0.01215 | 0.08277 | 0.01156 | 0.49318 | 0.00896 | 0.00662 | 0.00742 | 0.01253 | 0.00338 | 0.01459 | 0.01282 | 0.00812 | 0.01958 | 0.01103 | 0.00468 | 0.00727 |
| **Kcnab2** | 0.07445 | 0.00755 | 0.00611 | 0.20432 | 0.00773 | 0.00747 | 0.00729 | 0.00000 | 0.03603 | 0.31045 | 0.63738 | 0.05088 | 0.00731 | 0.00652 | 0.02933 | 0.01434 |
| **Kcnb1** | 0.01304 | 0.00691 | 0.00393 | 0.18829 | 2.47293 | 0.00909 | 0.00749 | 0.01273 | 0.00291 | 0.24770 | 0.03774 | 0.00552 | 0.60293 | 0.01060 | 0.00812 | 0.00738 |
| **Kcnc3** | 0.07442 | 0.00835 | 0.00903 | 0.06821 | 0.01595 | 0.00674 | 0.00729 | 0.02150 | 0.05714 | 0.09764 | 0.02981 | 0.00722 | 0.01164 | 0.01136 | 0.00727 | 0.00756 |
| **Kcnd1** | 0.01814 | 0.00810 | 0.00659 | 0.33004 | 0.00866 | 2.13881 | 0.00720 | 0.01039 | 0.01065 | 1.24177 | 0.00899 | 0.00503 | 0.00362 | 0.00987 | 0.01066 | 0.00627 |
| **Kcnd3** | 0.01363 | 0.00816 | 0.01173 | 0.10692 | 0.00930 | 0.00663 | 0.00753 | 0.01362 | 0.95875 | 0.01043 | 0.00601 | 0.01758 | 0.01300 | 0.01306 | 0.00415 | 0.00735 |
| **Kcne1** | 0.01235 | 0.00837 | 0.29462 | 0.40506 | 0.00875 | 0.00551 | 0.00768 | 0.01356 | 1.85712 | 0.00725 | 0.01516 | 0.00565 | 0.00992 | 0.01370 | 0.00000 | 0.00706 |
| **Kcne3** | 0.01229 | 0.00751 | 0.00915 | 0.00483 | 0.00853 | 0.00675 | 0.00701 | 0.00000 | 0.02374 | 3.56363 | 0.46607 | 0.00000 | 0.01379 | 0.01203 | 0.00000 | 0.00744 |
| **Kcnf1** | 0.00000 | 0.00616 | 1.96933 | 0.00000 | 0.01564 | 0.00916 | 0.00534 | 6.53294 | 0.00559 | 1.61287 | 3.00788 | 0.03396 | 0.72818 | 1.14627 | 2.79497 | 0.00570 |
| **Kcng3** | 0.01378 | 0.00769 | 0.01193 | 0.14560 | 0.00935 | 0.00654 | 0.00743 | 0.01225 | 0.00932 | 0.00906 | 0.01269 | 0.00701 | 0.01282 | 0.00881 | 0.34436 | 0.00739 |
| **Kcng4** | 0.01209 | 0.00787 | 0.41467 | 1.19173 | 0.00854 | 0.00894 | 0.00750 | 0.01903 | 0.07293 | 0.01177 | 0.01075 | 0.00690 | 0.00910 | 0.00506 | 0.24322 | 0.00699 |
| **Kcnh1** | 0.00662 | 0.00830 | 0.00255 | 5.02838 | 0.00982 | 0.00632 | 0.00785 | 0.24558 | 0.00000 | 0.64222 | 0.26489 | 0.01226 | 0.00507 | 1.04838 | 1.11825 | 0.00621 |
| **Kcnh2** | 0.01197 | 0.00863 | 0.01101 | 0.10040 | 0.00924 | 0.00676 | 0.00801 | 0.05396 | 0.04766 | 0.30614 | 0.01193 | 0.00746 | 0.01047 | 0.03391 | 0.00747 | 0.00776 |
| **Kcnh3** | 0.00841 | 0.00791 | 0.00862 | 0.06514 | 0.00943 | 0.00702 | 0.00719 | 0.67556 | 0.01727 | 0.56333 | 0.91594 | 0.02449 | 0.00000 | 0.00596 | 0.04444 | 0.00733 |
| **Kcnh4** | 0.01383 | 0.00815 | 0.00958 | 0.00576 | 0.00913 | 0.00679 | 0.00757 | 0.00749 | 0.01144 | 0.00699 | 0.34640 | 0.00696 | 0.00859 | 0.01114 | 0.00463 | 0.00721 |
| **Kcnip2** | 0.01197 | 0.00778 | 0.00773 | 1.28786 | 0.00906 | 0.00631 | 0.00722 | 0.01459 | 0.00000 | 1.13425 | 0.00360 | 0.00000 | 0.01029 | 0.58267 | 0.96461 | 0.00686 |
| **Kcnip3** | 0.10668 | 0.00773 | 0.01500 | 1.74057 | 0.00823 | 0.00728 | 0.00713 | 0.01048 | 0.00000 | 1.29050 | 0.02214 | 0.00406 | 0.01194 | 0.03692 | 0.04016 | 0.00693 |
| **Kcnip4** | 0.01338 | 0.00798 | 0.01240 | 0.49528 | 0.00940 | 0.00662 | 0.00744 | 0.01111 | 0.00368 | 0.50517 | 0.02527 | 0.00624 | 0.01683 | 0.00725 | 0.43509 | 0.00719 |
| **Kcnj10** | 0.01566 | 0.01775 | 0.01958 | 0.50411 | 0.00850 | 0.01936 | 0.00698 | 0.03721 | 0.02397 | 1.15679 | 0.01370 | 0.00461 | 0.41929 | 0.06191 | 0.05451 | 0.01368 |
| **Kcnj13** | 0.00954 | 0.00751 | 0.20815 | 0.37307 | 0.00910 | 0.00887 | 0.00689 | 0.00835 | 0.04950 | 0.07500 | 0.00737 | 0.00404 | 0.30218 | 0.84812 | 0.00903 | 0.00718 |
| **Kcnj14** | 0.01007 | 0.00711 | 0.94869 | 0.00671 | 0.00752 | 0.00786 | 0.00699 | 0.14835 | 0.04198 | 0.15409 | 0.00000 | 1.39200 | 0.00387 | 0.00634 | 0.28617 | 0.00691 |
| **Kcnj15** | 0.05244 | 0.03943 | 0.05359 | 0.01192 | 0.02658 | 0.07504 | 0.01800 | 0.08795 | 0.12033 | 0.03542 | 0.03645 | 0.01682 | 0.05996 | 0.08896 | 0.07448 | 0.02202 |
| **Kcnj16** | 0.01055 | 0.01870 | 0.02797 | 0.00823 | 0.01869 | 0.00730 | 0.01819 | 0.04493 | 0.03752 | 0.02914 | 0.02633 | 0.01808 | 0.02556 | 0.03701 | 0.04077 | 0.01454 |
| **Kcnj2** | 0.01780 | 0.01775 | 0.00600 | 0.19054 | 0.00882 | 0.00690 | 0.00711 | 0.00000 | 0.20811 | 0.00000 | 0.33769 | 0.01335 | 0.02319 | 0.37162 | 0.24477 | 0.00713 |
| **Kcnk4** | 0.01462 | 0.00751 | 0.02545 | 0.01085 | 0.00920 | 0.00605 | 0.00710 | 0.44931 | 0.01194 | 0.00832 | 0.01081 | 0.00657 | 0.00586 | 0.01269 | 0.00769 | 0.00734 |
| **Kcnk5** | 2.66772 | 0.19616 | 2.66017 | 4.12681 | 0.03852 | 0.52114 | 0.00954 | 9.35762 | 6.13040 | 18.28962 | 8.89949 | 2.45707 | 3.03265 | 1.82349 | 12.51442 | 0.35837 |
| **Kcnk6** | 1.16821 | 0.02402 | 1.96023 | 13.70729 | 0.04638 | 3.95130 | 0.00000 | 7.32670 | 16.42597 | 15.08454 | 19.59640 | 2.28914 | 6.54328 | 18.33159 | 11.80797 | 0.02368 |
| **Kcnmb3** | 0.01353 | 0.00820 | 0.01210 | 0.01062 | 0.00932 | 0.00668 | 0.00771 | 0.01864 | 0.01234 | 0.01531 | 0.01293 | 0.00836 | 0.01238 | 0.01264 | 0.01053 | 0.00741 |
| **Kcnn1** | 0.09796 | 0.02905 | 0.35780 | 0.16588 | 0.04167 | 0.04492 | 0.08643 | 0.00000 | 0.40188 | 0.74412 | 1.29520 | 0.02343 | 1.81284 | 0.55793 | 0.26008 | 0.08940 |
| **Kcnn2** | 0.32345 | 0.68375 | 0.00274 | 0.00343 | 0.00771 | 0.00822 | 0.00712 | 0.01513 | 0.35233 | 0.00770 | 0.01163 | 0.00686 | 0.08659 | 0.01305 | 0.00000 | 0.00700 |
| **Kcnn4** | 0.60406 | 0.00944 | 0.14572 | 0.59513 | 0.03625 | 3.12898 | 0.00775 | 0.20644 | 0.08346 | 0.05236 | 0.00647 | 0.01078 | 0.10500 | 0.01231 | 0.00000 | 0.01311 |
| **Kcnq1** | 0.01322 | 0.00756 | 0.01054 | 1.17834 | 0.00906 | 0.00658 | 0.00718 | 0.01007 | 0.01283 | 0.64701 | 0.01120 | 0.00506 | 0.01873 | 0.20097 | 0.00000 | 0.00758 |
| **Kcnq1ot1** | 0.88801 | 0.52648 | 4.94341 | 4.53770 | 0.55769 | 0.39756 | 1.02393 | 2.06985 | 8.33518 | 4.26347 | 1.92194 | 2.45390 | 1.99720 | 11.27291 | 1.74812 | 0.43805 |

| | | | | | | | | | | | | | | | | |
|---|---|---|---|---|---|---|---|---|---|---|---|---|---|---|---|---|
| **Kcnq5** | 0.01353 | 0.00788 | 0.01084 | 0.00217 | 0.00938 | 0.00633 | 0.00734 | 0.01089 | 1.01584 | 0.00296 | 0.00000 | 0.01508 | 0.01296 | 0.00665 | 1.09685 | 0.00721 |
| **Kcnrg** | 0.01521 | 0.00822 | 0.01383 | 0.00968 | 0.02300 | 0.00713 | 0.00768 | 0.01428 | 0.01212 | 0.26754 | 0.01348 | 0.00694 | 0.22307 | 0.01387 | 0.02109 | 0.02270 |
| **Kcns3** | 4.39394 | 4.08036 | 6.93628 | 0.70374 | 5.26538 | 6.27923 | 0.43470 | 1.96891 | 0.74878 | 3.76557 | 3.05772 | 0.42200 | 0.30580 | 0.12730 | 3.96797 | 0.46629 |
| **Kcnu1** | 0.00569 | 0.00761 | 2.05034 | 0.58512 | 0.00701 | 0.00921 | 0.00688 | 0.35705 | 1.52073 | 0.00470 | 0.00000 | 0.00000 | 0.37112 | 0.00689 | 1.34000 | 0.00637 |
| **Kcnv2** | 0.01690 | 0.00775 | 0.01131 | 0.00493 | 0.02117 | 0.00639 | 0.00734 | 0.01153 | 0.05425 | 0.00557 | 0.01083 | 0.00615 | 0.02508 | 0.00604 | 0.64869 | 0.00731 |
| **Kcp** | 0.00577 | 0.00773 | 0.01196 | 0.28903 | 0.00909 | 0.00753 | 0.00604 | 0.25769 | 0.00452 | 0.00000 | 0.53233 | 0.01347 | 1.63728 | 0.00563 | 0.06666 | 0.00632 |
| **Kctd1** | 0.17331 | 0.00729 | 0.01700 | 0.27694 | 0.00781 | 0.00706 | 0.02678 | 0.00309 | 0.00796 | 0.66236 | 0.17208 | 0.05103 | 0.75277 | 0.01016 | 0.01289 | 0.00719 |
| **Kctd10** | 4.40709 | 1.40488 | 1.30737 | 0.12338 | 0.26666 | 1.66970 | 2.23672 | 6.56163 | 0.00533 | 6.72199 | 0.35430 | 0.21398 | 2.59965 | 0.04254 | 2.05619 | 2.78297 |
| **Kctd12** | 0.01359 | 0.00780 | 0.01057 | 2.03953 | 0.00915 | 0.01261 | 0.00735 | 0.01062 | 0.69762 | 0.29870 | 0.01666 | 0.00485 | 0.01144 | 0.01891 | 0.38445 | 0.00680 |
| **Kctd12b** | 0.00898 | 0.00871 | 0.00922 | 0.39894 | 0.01064 | 0.00757 | 0.00830 | 0.37045 | 0.00612 | 0.00000 | 0.14654 | 0.00736 | 0.19136 | 0.01066 | 0.00000 | 0.00757 |
| **Kctd13** | 1.84280 | 0.00451 | 0.00000 | 8.20669 | 1.05008 | 0.20078 | 0.21274 | 3.36339 | 6.72929 | 0.56700 | 2.80156 | 0.32123 | 0.00000 | 0.03261 | 1.51927 | 1.08071 |
| **Kctd14** | 0.02117 | 0.01812 | 0.12431 | 0.64066 | 0.02736 | 0.00696 | 0.01707 | 0.02245 | 1.09223 | 3.42125 | 0.21155 | 0.04414 | 0.01512 | 0.18471 | 0.03667 | 0.01480 |
| **Kctd15** | 0.74560 | 0.01530 | 0.00000 | 1.84685 | 0.01451 | 2.12237 | 2.28674 | 1.03062 | 1.26302 | 2.68949 | 0.56940 | 0.03163 | 0.46843 | 1.52689 | 0.63655 | 0.16748 |
| **Kctd17** | 0.01345 | 0.00805 | 0.01189 | 0.30883 | 0.00931 | 0.00659 | 0.00766 | 0.01763 | 0.34207 | 0.00927 | 0.01056 | 0.00751 | 0.01239 | 0.01095 | 0.09257 | 0.01461 |
| **Kctd18** | 0.00000 | 3.30013 | 1.01955 | 5.82260 | 1.22669 | 2.10158 | 0.01212 | 4.34152 | 5.82995 | 5.20436 | 0.00000 | 0.61624 | 7.00581 | 1.97718 | 1.37173 | 3.47080 |
| **Kctd2** | 0.53870 | 0.82709 | 1.62293 | 0.72879 | 0.37984 | 0.01424 | 0.00611 | 0.22403 | 0.44744 | 1.63250 | 0.00000 | 0.13796 | 0.24031 | 0.00372 | 0.52597 | 2.47036 |
| **Kctd20** | 2.81968 | 3.63926 | 6.09208 | 9.42441 | 5.14652 | 0.27678 | 4.40265 | 8.01523 | 8.11268 | 9.18497 | 2.22686 | 1.45922 | 3.32900 | 1.42126 | 8.67765 | 1.56330 |
| **Kctd21** | 0.24944 | 0.00789 | 0.00765 | 0.80999 | 0.00796 | 0.00735 | 0.00759 | 0.01413 | 0.76179 | 0.00387 | 0.01122 | 0.00669 | 0.11341 | 0.02669 | 0.06354 | 0.00720 |
| **Kctd3** | 2.25526 | 0.42771 | 3.00585 | 12.52958 | 3.78766 | 1.26822 | 3.83329 | 7.22832 | 8.87828 | 4.94006 | 3.71436 | 1.68479 | 0.55412 | 0.98956 | 9.31850 | 2.58611 |
| **Kctd4** | 0.01278 | 0.00764 | 0.00777 | 0.01067 | 0.00951 | 0.00675 | 0.00725 | 0.00983 | 0.01245 | 0.01117 | 0.00844 | 0.00409 | 0.01314 | 1.06483 | 0.00420 | 0.00738 |
| **Kctd5** | 8.19742 | 0.00000 | 3.35010 | 2.24846 | 4.58111 | 0.01105 | 4.78227 | 3.44805 | 3.54756 | 5.19729 | 6.55790 | 8.13671 | 7.46693 | 2.36888 | 2.32934 | 5.24844 |
| **Kctd6** | 2.41285 | 2.32300 | 0.59322 | 2.84192 | 0.00239 | 0.01477 | 0.02355 | 0.28072 | 5.78029 | 0.70977 | 1.46997 | 0.01915 | 0.42146 | 2.67543 | 1.36578 | 0.01762 |
| **Kctd7** | 0.01351 | 0.00768 | 0.00497 | 0.86218 | 0.00903 | 0.00651 | 0.00700 | 0.00000 | 1.16734 | 0.00000 | 0.65177 | 0.03919 | 0.02312 | 0.00346 | 0.99376 | 0.00677 |
| **Kctd9** | 1.24648 | 3.66978 | 3.00453 | 4.81664 | 2.50814 | 4.43051 | 5.43120 | 7.09855 | 6.25226 | 3.96728 | 4.24089 | 0.81470 | 1.97472 | 11.25498 | 1.38314 | 2.77791 |
| **Kdelc1** | 3.55387 | 2.78377 | 0.00000 | 0.00555 | 1.23738 | 0.00296 | 0.00333 | 0.00000 | 0.01150 | 0.00060 | 0.01257 | 0.01427 | 0.00847 | 0.00754 | 0.27710 | 0.86797 |
| **Kdelc2** | 0.46693 | 0.08853 | 0.46350 | 0.41467 | 2.40072 | 0.09500 | 0.11763 | 0.01763 | 2.07627 | 0.45206 | 2.41464 | 0.14901 | 0.43699 | 0.47365 | 0.35406 | 0.09240 |
| **Kdelr1** | 3.05816 | 0.04408 | 2.46279 | 1.67438 | 2.05577 | 5.38972 | 2.60277 | 2.46768 | 0.43293 | 3.01562 | 1.32751 | 0.01529 | 9.82151 | 0.00277 | 0.14372 | 7.63753 |
| **Kdelr2** | 3.81599 | 2.42557 | 6.79852 | 0.44982 | 2.13585 | 0.28098 | 0.09044 | 1.10870 | 0.00000 | 4.57427 | 3.18138 | 0.23370 | 4.62029 | 0.36966 | 0.69103 | 4.00579 |
| **Kdelr3** | 1.80279 | 3.20561 | 0.30695 | 0.28170 | 0.02109 | 0.00393 | 0.62119 | 0.00000 | 0.00410 | 3.40433 | 2.90301 | 0.04767 | 0.04417 | 0.55943 | 0.40179 | 3.04241 |
| **Kdf1** | 3.03016 | 0.00202 | 0.08183 | 0.78147 | 4.31158 | 0.03998 | 0.01322 | 0.04814 | 2.01469 | 5.39004 | 0.62441 | 0.00000 | 1.78379 | 0.02602 | 0.00000 | 6.28790 |
| **Kdm1a** | 7.29246 | 5.15747 | 1.47318 | 0.60748 | 3.50352 | 3.88873 | 2.80754 | 2.00740 | 0.25289 | 1.10233 | 0.47173 | 3.54850 | 1.63237 | 12.74827 | 14.57658 | 0.29596 |
| **Kdm1b** | 1.05321 | 0.01995 | 0.54319 | 8.39137 | 0.01480 | 0.13844 | 0.01673 | 0.17142 | 4.04627 | 0.19296 | 0.33535 | 0.11520 | 0.00580 | 2.66505 | 2.65887 | 0.00483 |
| **Kdm2a** | 1.67813 | 2.69186 | 2.19849 | 2.75231 | 0.00000 | 0.00628 | 0.00471 | 1.04408 | 2.20480 | 1.52956 | 1.15081 | 0.03282 | 0.00000 | 0.56333 | 3.84887 | 0.00298 |
| **Kdm2b** | 9.05182 | 5.22558 | 10.23752 | 15.52603 | 6.12699 | 6.17936 | 9.75955 | 17.82168 | 18.69623 | 24.76665 | 17.45454 | 17.73328 | 12.47264 | 17.60799 | 17.67784 | 0.53631 |
| **Kdm3a** | 3.60730 | 5.76216 | 3.88522 | 7.63063 | 0.20751 | 1.82204 | 6.35770 | 7.41004 | 11.03395 | 6.68109 | 3.45887 | 9.84881 | 0.72302 | 14.50487 | 8.47412 | 3.63577 |
| **Kdm3b** | 5.88635 | 5.26024 | 3.37753 | 1.59871 | 7.09097 | 5.27331 | 10.77744 | 5.89985 | 1.39493 | 12.07737 | 1.51739 | 0.83361 | 5.63105 | 6.80862 | 9.79247 | 0.45645 |
| **Kdm4a** | 0.29793 | 0.01849 | 0.04938 | 3.71460 | 0.01321 | 0.02472 | 0.00657 | 0.00000 | 6.19717 | 0.00000 | 1.37576 | 0.38344 | 0.05664 | 0.13070 | 0.22627 | 0.05040 |
| **Kdm4b** | 0.41978 | 0.95836 | 0.00000 | 1.50971 | 7.98484 | 2.68688 | 2.12011 | 2.08233 | 3.64599 | 1.75223 | 0.68041 | 0.01773 | 0.00000 | 0.03469 | 0.00000 | 3.15969 |
| **Kdm4c** | 0.14006 | 5.40385 | 0.07961 | 2.83795 | 0.01235 | 1.00210 | 0.00380 | 1.84006 | 6.00693 | 4.18020 | 1.58469 | 3.51167 | 0.52125 | 15.92559 | 13.71537 | 0.65029 |
| **Kdm4d** | 0.01190 | 0.00778 | 0.01078 | 0.94794 | 0.00932 | 0.00663 | 0.00719 | 0.06593 | 0.13362 | 0.14140 | 0.01261 | 0.01628 | 0.01016 | 0.00901 | 0.02641 | 0.00761 |
| **Kdm5a** | 1.41670 | 3.28035 | 5.47833 | 4.22712 | 0.08790 | 1.97141 | 0.09661 | 5.88546 | 5.00608 | 2.49219 | 1.10828 | 3.43332 | 1.75005 | 10.03948 | 13.32879 | 0.00705 |
| **Kdm5b** | 9.00977 | 12.83064 | 3.38952 | 19.74124 | 5.70873 | 3.62316 | 0.11151 | 13.46238 | 22.68060 | 13.12016 | 12.91153 | 10.85467 | 2.29400 | 27.38308 | 20.18742 | 1.85126 |
| **Kdm5c** | 12.98436 | 2.20509 | 4.30944 | 3.25429 | 0.89178 | 0.87800 | 4.20696 | 5.50286 | 2.90037 | 6.03068 | 2.03402 | 0.15217 | 6.71294 | 7.89665 | 5.03364 | 1.35860 |
| **Kdm5d** | 0.01455 | 0.00780 | 0.00000 | 0.00074 | 0.00914 | 2.47302 | 0.12949 | 0.00000 | 0.22510 | 0.00000 | 0.43282 | 0.00585 | 0.00000 | 0.01127 | 0.00258 | 0.00625 |
| **Kdm6a** | 1.86072 | 0.85561 | 2.30578 | 1.55052 | 3.90863 | 0.00424 | 1.64701 | 5.84532 | 0.23867 | 2.58139 | 0.00001 | 0.14326 | 0.67302 | 16.72195 | 11.73717 | 0.16786 |
| **Kdm6b** | 3.43809 | 0.06674 | 0.35302 | 1.05755 | 0.07772 | 1.45868 | 0.71236 | 1.13633 | 0.64667 | 1.40737 | 0.54181 | 0.03769 | 1.58740 | 0.02363 | 0.00000 | 2.62311 |
| **Kdm7a** | 0.51797 | 0.71489 | 0.12258 | 11.80143 | 0.06523 | 4.15367 | 0.32317 | 2.07973 | 10.22427 | 6.63190 | 2.64283 | 10.82914 | 1.01781 | 12.15074 | 18.98075 | 0.03649 |
| **Kdm8** | 0.18015 | 0.00724 | 0.00204 | 0.89974 | 0.00746 | 0.00726 | 0.02689 | 0.11343 | 0.55668 | 0.06262 | 0.49052 | 0.00000 | 0.00528 | 0.91010 | 0.09059 | 0.00688 |
| **Kdsr** | 1.12563 | 0.19675 | 1.89348 | 2.62379 | 0.00277 | 6.60857 | 3.45880 | 0.90170 | 1.20657 | 0.30759 | 2.97794 | 4.79979 | 2.09364 | 2.43497 | 3.42920 | 0.30202 |
| **Keap1** | 6.86154 | 0.04206 | 3.60620 | 2.07015 | 0.05753 | 4.15434 | 1.15305 | 4.39745 | 1.16596 | 3.40468 | 4.73013 | 0.13125 | 3.33282 | 0.11066 | 1.50960 | 2.56044 |
| **Kel** | 0.14549 | 0.00762 | 0.00801 | 0.01061 | 0.00766 | 0.00694 | 0.00751 | 0.01298 | 0.01235 | 0.01522 | 0.01301 | 0.00811 | 0.01250 | 0.01243 | 0.01072 | 0.01449 |
| **Khdc1a** | 2.95691 | 2.49334 | 5.97800 | 19.90923 | 3.13781 | 4.10640 | 3.90807 | 6.23440 | 19.18302 | 32.60888 | 24.20273 | 18.09002 | 5.44657 | 32.80042 | 22.84919 | 2.89997 |
| **Khdc1b** | 0.03323 | 0.11726 | 0.64622 | 20.50489 | 0.00707 | 0.01457 | 0.12648 | 1.12394 | 9.22988 | 6.56195 | 3.73361 | 0.85057 | 0.03507 | 11.29795 | 29.67687 | 0.00190 |
| **Khdc1c** | 0.84487 | 1.11971 | 0.63403 | 5.94818 | 0.00509 | 0.16714 | 0.71867 | 0.59009 | 10.69233 | 2.96623 | 11.86758 | 0.78663 | 0.41849 | 5.00253 | 5.73860 | 0.00169 |
| **Khdc3** | 18.42051 | 3.27512 | 7.50777 | 0.00000 | 21.22458 | 1.23398 | 2.99214 | 4.77013 | 19.56049 | 2.92262 | 26.10422 | 0.37264 | 12.12151 | 0.16653 | 0.00000 | 7.51415 |
| **Khdrbs1** | 7.44376 | 7.74770 | 6.35921 | 17.44002 | 3.57813 | 1.66092 | 6.77439 | 6.54315 | 19.86251 | 9.60593 | 13.49255 | 2.20639 | 6.78344 | 11.87766 | 13.24340 | 0.80598 |
| **Khdrbs3** | 0.01344 | 0.00776 | 0.01135 | 0.88581 | 0.00903 | 0.00647 | 0.00722 | 0.00974 | 0.00000 | 0.76293 | 0.02510 | 0.03778 | 0.01067 | 0.01932 | 0.00318 | 0.00702 |
| **Khk** | 0.01200 | 0.00788 | 0.01060 | 1.20817 | 0.00988 | 0.00631 | 0.00611 | 0.00521 | 0.17997 | 0.27574 | 0.13333 | 0.00453 | 2.10921 | 0.00378 | 0.10950 | 0.00613 |
| **Khnyn** | 2.20912 | 0.01375 | 1.14138 | 0.12888 | 0.01115 | 1.32298 | 0.86089 | 4.40200 | 0.00782 | 0.83469 | 3.72675 | 0.20548 | 7.13336 | 0.03718 | 1.40669 | 0.29892 |
| **Khsrp** | 2.82870 | 0.25418 | 2.09249 | 8.16462 | 0.81555 | 0.11585 | 0.06582 | 1.06435 | 9.50652 | 1.00490 | 11.21883 | 0.50685 | 1.79086 | 2.54566 | 2.29377 | 1.33711 |
| **Kidins220** | 0.00000 | 0.00387 | 8.07394 | 2.26067 | 0.02293 | 0.08791 | 2.48700 | 3.59586 | 3.02762 | 5.58292 | 6.37776 | 2.14873 | 3.66386 | 13.61746 | 7.65683 | 0.02971 |
| **Kif11** | 0.03870 | 6.54641 | 4.31588 | 7.17391 | 5.73431 | 0.61291 | 0.50983 | 6.54691 | 14.05813 | 3.79641 | 4.97117 | 6.56597 | 1.69902 | 16.11413 | 12.18420 | 0.67383 |

| | | | | | | | | | | | | | | | | |
|---|---|---|---|---|---|---|---|---|---|---|---|---|---|---|---|---|
| **Kif13a** | 0.58069 | 4.75719 | 0.19100 | 0.77030 | 0.05573 | 0.00657 | 4.22095 | 0.00555 | 2.25427 | 1.45440 | 0.00000 | 0.00603 | 0.59859 | 0.00804 | 0.34086 | 0.00417 |
| **Kif13b** | 0.00089 | 0.00825 | 0.01217 | 0.29991 | 0.01246 | 0.20671 | 0.01821 | 2.73858 | 0.84926 | 0.31511 | 0.16199 | 0.02002 | 0.01582 | 3.47869 | 0.00000 | 0.00727 |
| **Kif14** | 0.21212 | 0.00859 | 1.23875 | 3.14442 | 0.03427 | 0.19108 | 0.14205 | 0.65228 | 1.44560 | 0.29478 | 0.42313 | 0.06126 | 0.08736 | 3.48189 | 1.39389 | 0.01489 |
| **Kif15** | 1.01596 | 2.76331 | 0.00000 | 0.12281 | 1.06722 | 1.55682 | 0.84235 | 0.65647 | 0.66244 | 0.51297 | 1.92712 | 0.07708 | 0.88358 | 4.15660 | 1.41853 | 0.41341 |
| **Kif16b** | 4.74046 | 5.58408 | 3.79711 | 0.56693 | 3.95899 | 1.99392 | 0.02937 | 5.53736 | 0.01523 | 10.13750 | 7.58732 | 2.13089 | 1.64051 | 8.09836 | 9.35752 | 0.00000 |
| **Kif17** | 0.01332 | 0.00782 | 0.00704 | 4.07641 | 0.00894 | 0.00604 | 0.00710 | 0.00000 | 2.69797 | 0.28936 | 0.43463 | 0.00000 | 0.00779 | 0.00000 | 0.82390 | 0.00620 |
| **Kif18a** | 0.17619 | 5.63082 | 1.55108 | 1.16287 | 2.71918 | 0.00497 | 1.65113 | 1.62131 | 15.25239 | 4.70471 | 2.56691 | 12.42807 | 0.01425 | 25.72735 | 11.71887 | 0.00884 |
| **Kif18b** | 1.46516 | 1.57326 | 2.86509 | 4.29769 | 0.00877 | 0.00379 | 0.13351 | 2.30839 | 11.31814 | 4.12163 | 2.45378 | 0.32735 | 0.69297 | 0.00000 | 3.19617 | 0.25016 |
| **Kif1a** | 0.02173 | 0.00779 | 0.01460 | 0.09461 | 0.00857 | 0.00631 | 0.00726 | 0.00000 | 0.01049 | 0.39373 | 0.43552 | 0.01470 | 0.00758 | 0.00868 | 0.00000 | 0.00697 |
| **Kif1b** | 4.31574 | 0.00000 | 4.51178 | 7.80606 | 4.81364 | 0.00637 | 4.49417 | 3.83661 | 15.08136 | 10.51034 | 6.64119 | 7.83783 | 2.21873 | 14.82338 | 14.16519 | 0.01732 |
| **Kif1bp** | 3.36934 | 1.66996 | 3.03263 | 4.63394 | 5.17104 | 0.04279 | 1.41209 | 6.17912 | 5.31548 | 6.56082 | 2.24887 | 9.02353 | 0.77677 | 2.34860 | 8.06877 | 0.02844 |
| **Kif1c** | 0.39753 | 1.29256 | 1.51901 | 0.84875 | 0.19245 | 2.28167 | 0.00579 | 0.27656 | 1.86697 | 0.99060 | 0.66681 | 0.16606 | 0.23266 | 0.01745 | 0.00000 | 0.25353 |
| **Kif20a** | 5.62002 | 8.16013 | 5.24061 | 0.21615 | 7.84191 | 6.48870 | 6.40098 | 19.93088 | 0.19580 | 15.94774 | 0.00000 | 4.77643 | 2.74249 | 0.06094 | 6.46153 | 1.90540 |
| **Kif20b** | 0.00000 | 1.80216 | 1.59169 | 5.28836 | 0.03158 | 0.00369 | 0.01500 | 0.17500 | 11.58646 | 1.46212 | 0.40545 | 6.77724 | 0.00000 | 14.06524 | 7.02402 | 0.00347 |
| **Kif21a** | 0.01170 | 0.00809 | 0.01081 | 0.21756 | 0.01006 | 0.00755 | 0.46511 | 0.01406 | 0.00768 | 0.24318 | 0.01432 | 0.02019 | 0.11473 | 0.00805 | 0.05613 | 0.00767 |
| **Kif21b** | 0.68251 | 0.00440 | 1.66556 | 0.74188 | 0.01590 | 0.00601 | 3.85335 | 0.00850 | 0.64067 | 1.60630 | 0.02118 | 0.00572 | 0.01038 | 0.00417 | 0.58407 | 0.47856 |
| **Kif22** | 7.77686 | 7.75982 | 9.93049 | 11.41464 | 12.19435 | 9.80225 | 7.75167 | 21.61069 | 23.20406 | 25.59704 | 11.48149 | 7.91578 | 6.00013 | 0.45483 | 16.93651 | 3.23238 |
| **Kif23** | 6.25780 | 11.97715 | 2.76732 | 12.51285 | 9.79906 | 6.30557 | 3.80421 | 20.49040 | 15.99567 | 18.14756 | 17.23088 | 13.18185 | 4.24344 | 23.70363 | 20.37161 | 4.63793 |
| **Kif24** | 0.01927 | 0.00697 | 0.63206 | 1.72719 | 0.03641 | 0.02380 | 1.65928 | 0.44194 | 1.01325 | 0.62916 | 0.44749 | 0.03358 | 0.23577 | 2.76780 | 2.38087 | 0.02902 |
| **Kif26a** | 0.75368 | 0.05804 | 0.00401 | 0.37988 | 0.00647 | 0.00699 | 0.04735 | 0.00576 | 0.12627 | 0.18918 | 0.09294 | 0.00613 | 0.01089 | 0.00678 | 0.06400 | 0.00652 |
| **Kif27** | 0.00984 | 0.00830 | 0.02497 | 0.88949 | 0.00946 | 0.00562 | 0.00814 | 0.49563 | 2.82207 | 0.00000 | 0.00000 | 0.05878 | 0.00560 | 0.00597 | 1.97510 | 0.00683 |
| **Kif2a** | 2.52014 | 1.95494 | 3.45397 | 13.26357 | 2.22351 | 4.80606 | 6.23527 | 5.62457 | 12.25254 | 7.16624 | 1.36057 | 6.09661 | 2.86154 | 15.57018 | 12.47710 | 1.89033 |
| **Kif2c** | 1.71817 | 3.85206 | 2.86571 | 12.44365 | 4.70611 | 5.46466 | 2.49614 | 8.23113 | 15.96830 | 7.24794 | 1.69132 | 0.47228 | 0.84149 | 0.53124 | 8.28593 | 0.07107 |
| **Kif3a** | 4.81484 | 0.06830 | 1.66527 | 2.59898 | 0.03117 | 1.98276 | 1.14868 | 2.74966 | 3.60370 | 0.67819 | 3.17287 | 0.58983 | 1.02493 | 1.93177 | 1.78427 | 4.13923 |
| **Kif3b** | 1.24119 | 0.00248 | 0.43738 | 1.88623 | 6.15933 | 0.00346 | 0.09127 | 1.11674 | 4.93736 | 0.00000 | 0.00000 | 1.60612 | 1.45074 | 0.00000 | 2.93493 | 1.00915 |
| **Kif3c** | 0.00175 | 0.00664 | 0.66020 | 2.39736 | 0.00693 | 0.00464 | 0.00641 | 6.46398 | 6.67829 | 5.90239 | 2.45438 | 0.31808 | 0.00000 | 0.37619 | 9.89577 | 0.00567 |
| **Kif4** | 5.27367 | 2.87187 | 3.39197 | 11.63655 | 4.13372 | 0.34692 | 2.91573 | 11.19208 | 16.64761 | 14.79649 | 5.55617 | 5.91098 | 0.23666 | 14.22191 | 19.49964 | 3.83928 |
| **Kif5a** | 0.01598 | 0.00787 | 0.01020 | 0.44517 | 0.00910 | 2.62615 | 0.00702 | 0.00393 | 0.07983 | 0.00000 | 0.22523 | 0.04571 | 0.00000 | 0.02724 | 0.00000 | 0.00613 |
| **Kif5b** | 8.28656 | 8.38997 | 8.33929 | 2.33382 | 4.75332 | 2.70519 | 4.23968 | 7.31530 | 5.69787 | 18.25829 | 6.19472 | 13.96961 | 8.68789 | 17.42830 | 19.52769 | 5.24385 |
| **Kif5c** | 0.01006 | 0.00765 | 0.00346 | 6.58343 | 0.00991 | 0.00526 | 0.00687 | 0.00633 | 2.34924 | 0.02456 | 0.03125 | 0.00000 | 0.01291 | 2.65212 | 4.50897 | 0.00607 |
| **Kif7** | 0.04859 | 0.00842 | 0.00915 | 0.56451 | 0.01404 | 0.00692 | 0.00750 | 0.00955 | 0.28551 | 0.06185 | 0.14325 | 0.00717 | 0.01109 | 0.00513 | 0.14506 | 0.00737 |
| **Kif9** | 0.17448 | 2.21721 | 0.00000 | 3.48345 | 0.11504 | 0.02294 | 2.74755 | 1.25233 | 5.10811 | 2.09130 | 0.28204 | 4.16414 | 1.97179 | 0.61777 | 2.57202 | 0.00470 |
| **Kifap3** | 0.01124 | 0.60145 | 1.93903 | 0.00193 | 0.01466 | 0.00744 | 0.39780 | 0.04108 | 0.00921 | 3.09554 | 1.71917 | 1.79694 | 0.00000 | 0.72443 | 0.00000 | 0.01297 |
| **Kifc1** | 1.57609 | 2.22863 | 2.08508 | 3.65351 | 3.86716 | 6.41401 | 1.95222 | 5.75108 | 9.06730 | 3.67712 | 5.04435 | 0.34875 | 3.38038 | 0.04770 | 1.74761 | 0.09213 |
| **Kifc2** | 4.62471 | 0.05176 | 0.54146 | 5.90257 | 2.65431 | 0.09845 | 0.52738 | 2.51122 | 4.26746 | 2.60001 | 0.88802 | 0.16449 | 0.09613 | 0.00000 | 0.46060 | 1.03315 |
| **Kifc3** | 4.91254 | 0.00582 | 1.02191 | 2.48436 | 0.38880 | 0.02705 | 3.77530 | 19.19290 | 4.37791 | 13.09241 | 9.02445 | 4.53921 | 5.78161 | 0.35501 | 1.04522 | 2.45178 |
| **Kifc5b** | 1.03856 | 0.70514 | 1.44255 | 1.86576 | 1.37725 | 3.63975 | 1.60640 | 3.23165 | 6.04091 | 1.98865 | 2.30350 | 0.12445 | 1.67468 | 0.08823 | 1.32151 | 0.02423 |
| **Kin** | 0.00000 | 4.62038 | 5.49923 | 2.04886 | 4.41293 | 0.25559 | 5.20947 | 7.11735 | 5.21193 | 5.91768 | 0.45015 | 0.67723 | 0.48820 | 5.26086 | 6.94060 | 0.01711 |
| **Kirrel** | 0.00000 | 0.00731 | 0.00748 | 0.01110 | 0.00580 | 0.00625 | 0.00689 | 0.58509 | 0.01167 | 0.00000 | 0.00930 | 0.02547 | 0.00000 | 0.01726 | 0.00028 | 6.05103 |
| **Kirrel2** | 0.01311 | 0.00768 | 0.01966 | 0.00833 | 0.00981 | 0.00735 | 0.00748 | 0.01251 | 0.01183 | 0.27795 | 0.01273 | 0.00592 | 0.02725 | 0.00732 | 0.24351 | 0.00769 |
| **Kiss1r** | 0.01327 | 0.00811 | 0.01196 | 0.00960 | 0.00910 | 0.00652 | 0.00744 | 0.01217 | 0.02378 | 0.05903 | 0.01170 | 0.00789 | 0.11132 | 0.01158 | 0.01065 | 0.00711 |
| **Kit** | 1.26019 | 0.00522 | 0.00000 | 4.14896 | 0.01604 | 0.00390 | 2.73390 | 0.00134 | 2.79115 | 0.00000 | 0.04969 | 0.03662 | 0.00973 | 0.44721 | 8.16112 | 0.00498 |
| **Kitl** | 0.01375 | 0.00814 | 0.01231 | 1.25509 | 0.00931 | 0.00650 | 0.00747 | 0.01368 | 0.04064 | 0.01260 | 0.01374 | 0.00815 | 0.01263 | 0.00689 | 0.05348 | 0.00719 |
| **Kiz** | 0.97932 | 0.00246 | 0.97487 | 1.67274 | 5.23274 | 0.03133 | 1.20601 | 1.14231 | 3.95186 | 5.50955 | 1.84849 | 1.32914 | 1.44319 | 1.65095 | 3.28268 | 0.15554 |
| **Kl** | 0.02897 | 0.00774 | 0.05554 | 0.20293 | 0.07844 | 0.01405 | 0.01803 | 0.03686 | 1.12082 | 0.11484 | 0.24254 | 0.09188 | 0.08156 | 0.47190 | 0.41952 | 0.20162 |
| **Klb** | 0.57479 | 6.87926 | 4.61056 | 4.37361 | 0.03470 | 0.51241 | 3.48653 | 2.41257 | 7.42388 | 3.40599 | 3.06131 | 3.38943 | 0.11185 | 0.00000 | 3.65303 | 0.00750 |
| **Klc1** | 3.88493 | 0.00314 | 0.00983 | 1.57083 | 0.00000 | 0.01070 | 0.43527 | 0.84864 | 1.50066 | 1.09363 | 0.25696 | 0.06548 | 1.91636 | 0.00727 | 0.04290 | 2.27895 |
| **Klc2** | 0.01324 | 0.00796 | 0.01177 | 0.07079 | 0.00923 | 0.00654 | 0.00745 | 0.01244 | 0.43610 | 0.02446 | 0.02716 | 0.01616 | 0.01214 | 0.02140 | 0.06145 | 0.05862 |
| **Klc3** | 0.01206 | 0.00791 | 0.01216 | 0.00805 | 0.00889 | 0.00669 | 0.00740 | 0.01128 | 0.01197 | 0.67498 | 0.01193 | 0.02608 | 0.01360 | 0.01170 | 0.00832 | 0.18946 |
| **Klc4** | 4.21801 | 0.36375 | 1.24795 | 2.95186 | 0.04813 | 3.39019 | 0.01001 | 5.82998 | 0.23491 | 8.28902 | 3.74021 | 0.84932 | 3.71803 | 0.01547 | 0.47590 | 0.01280 |
| **Klf1** | 0.01375 | 0.00795 | 0.01237 | 0.12039 | 0.00931 | 0.00662 | 0.00752 | 0.01387 | 0.01046 | 0.01424 | 0.01387 | 0.00782 | 0.01265 | 0.00906 | 0.03253 | 0.00739 |
| **Klf10** | 1.67164 | 0.01550 | 1.10144 | 1.10945 | 0.08061 | 0.28745 | 2.62151 | 7.18706 | 0.09650 | 2.22945 | 5.63299 | 0.16280 | 2.24674 | 3.13042 | 1.26239 | 0.02515 |
| **Klf11** | 2.19525 | 0.00435 | 0.21823 | 7.44529 | 0.01078 | 0.04044 | 0.00324 | 4.06890 | 13.90020 | 9.57604 | 4.88797 | 0.59254 | 0.44030 | 0.00000 | 6.34701 | 1.51444 |
| **Klf13** | 5.98990 | 2.29419 | 4.47748 | 0.55955 | 2.15069 | 5.31276 | 1.34030 | 4.19923 | 4.00318 | 5.81312 | 5.28946 | 0.62845 | 2.31613 | 5.37684 | 3.82885 | 1.81673 |
| **Klf15** | 0.52614 | 0.00773 | 0.23775 | 0.00506 | 0.03583 | 4.33588 | 0.00643 | 0.00729 | 0.26432 | 0.00279 | 0.00697 | 0.00689 | 0.00000 | 0.01562 | 0.00408 | 0.00527 |
| **Klf16** | 2.61665 | 0.01572 | 2.61604 | 7.50956 | 0.03223 | 0.02727 | 0.00299 | 2.07623 | 11.23955 | 3.06063 | 3.92689 | 0.22369 | 0.92374 | 3.04628 | 6.75188 | 2.10314 |
| **Klf17** | 0.00000 | 0.00258 | 0.00000 | 25.41376 | 0.00735 | 0.00000 | 0.00000 | 3.90117 | 21.66711 | 31.73864 | 9.33571 | 11.35540 | 0.00000 | 24.89650 | 40.82393 | 0.00000 |
| **Klf2** | 5.50860 | 0.00000 | 0.32914 | 2.08005 | 0.00000 | 0.00000 | 0.95509 | 5.01547 | 13.42005 | 17.81908 | 14.50790 | 4.79971 | 1.10455 | 0.07703 | 1.37404 | 1.50004 |
| **Klf3** | 2.56479 | 2.78394 | 0.17392 | 1.70969 | 0.05294 | 0.00580 | 0.00387 | 0.75083 | 3.69122 | 0.93566 | 1.17319 | 0.03047 | 1.23146 | 0.00000 | 0.00000 | 2.92291 |
| **Klf4** | 18.21371 | 1.74529 | 3.95950 | 4.22969 | 7.75119 | 2.00739 | 3.06696 | 15.39786 | 7.93424 | 13.06190 | 8.73108 | 3.36040 | 7.47582 | 10.14990 | 6.32545 | 2.16763 |
| **Klf5** | 10.59863 | 0.00000 | 0.00000 | 0.09194 | 4.56910 | 6.34933 | 0.93177 | 10.12040 | 0.78334 | 29.45608 | 0.00000 | 0.42131 | 12.53285 | 0.23928 | 0.13197 | 11.04562 |
| **Klf6** | 5.57983 | 0.27947 | 1.12827 | 0.00014 | 3.26538 | 0.01038 | 0.09191 | 0.47951 | 0.00982 | 1.27950 | 0.00622 | 0.06870 | 0.01106 | 0.03340 | 0.00000 | 5.50046 |

| | | | | | | | | | | | | | | | | |
|---|---|---|---|---|---|---|---|---|---|---|---|---|---|---|---|---|
| **Klf7** | 0.01184 | 0.00810 | 0.01328 | 0.00664 | 0.00996 | 0.00710 | 0.12454 | 0.01300 | 0.01168 | 0.86662 | 0.01214 | 0.01491 | 0.17314 | 0.00629 | 0.17242 | 0.00744 |
| **Klf8** | 1.93308 | 1.41214 | 3.01535 | 1.52004 | 4.47187 | 0.00394 | 0.00414 | 2.53082 | 0.53827 | 4.25133 | 1.74613 | 0.58204 | 0.32673 | 9.93213 | 0.06419 | 0.15330 |
| **Klf9** | 5.47203 | 4.47618 | 3.47524 | 6.21126 | 0.80032 | 4.98023 | 5.19290 | 5.81338 | 5.20145 | 4.87257 | 3.58403 | 0.15890 | 2.90295 | 5.27116 | 6.69819 | 3.72379 |
| **Klhdc1** | 0.01005 | 0.00745 | 0.00597 | 0.19500 | 0.00951 | 0.00634 | 0.00715 | 0.26258 | 0.26014 | 0.05195 | 0.00000 | 0.27260 | 0.00660 | 0.00595 | 1.88166 | 0.01459 |
| **Klhdc10** | 4.34349 | 1.94312 | 3.60399 | 6.10741 | 1.22019 | 0.00753 | 2.76835 | 3.54684 | 3.39958 | 7.00806 | 5.02246 | 6.30228 | 1.95441 | 3.69639 | 3.42243 | 1.10085 |
| **Klhdc2** | 4.02835 | 0.53288 | 0.87068 | 25.33538 | 10.68870 | 0.61351 | 0.01045 | 2.69213 | 11.36025 | 4.26614 | 3.86165 | 1.97226 | 2.04153 | 0.00000 | 8.02933 | 2.16368 |
| **Klhdc3** | 8.71099 | 3.28754 | 3.55375 | 2.74114 | 5.61103 | 6.27346 | 3.81513 | 2.75842 | 2.01993 | 3.63434 | 0.03192 | 0.30910 | 6.77258 | 0.00000 | 0.00000 | 9.03078 |
| **Klhdc4** | 8.64707 | 6.96233 | 13.20515 | 7.55486 | 10.35358 | 7.73266 | 4.49889 | 19.59310 | 8.72170 | 5.90365 | 7.78250 | 0.72631 | 12.73908 | 0.25355 | 5.20914 | 9.12804 |
| **Klhdc9** | 2.14225 | 0.52065 | 1.08880 | 0.07124 | 1.20425 | 1.55046 | 1.98259 | 0.52181 | 0.23091 | 2.95903 | 0.45531 | 0.13075 | 0.40998 | 4.30618 | 0.12932 | 1.10161 |
| **Klhl10** | 0.00000 | 0.09945 | 0.00000 | 0.90802 | 0.00799 | 0.00577 | 0.63850 | 2.04833 | 0.01516 | 7.11178 | 6.13419 | 0.42919 | 1.40869 | 2.66256 | 8.46657 | 0.00538 |
| **Klhl11** | 0.17995 | 0.75143 | 0.00000 | 18.78203 | 2.16752 | 0.00277 | 0.00485 | 3.16617 | 26.00001 | 1.84230 | 3.61958 | 0.21281 | 0.02546 | 0.85982 | 4.15736 | 0.00885 |
| **Klhl12** | 5.59179 | 4.37522 | 3.03745 | 7.65901 | 4.13273 | 0.01367 | 7.77215 | 4.21080 | 12.08246 | 11.78995 | 4.64387 | 3.08883 | 3.56989 | 11.72686 | 12.08863 | 1.58220 |
| **Klhl13** | 0.04321 | 0.00802 | 0.00903 | 0.68318 | 0.00891 | 0.00643 | 0.00729 | 0.01167 | 0.59077 | 0.00321 | 0.00756 | 0.01688 | 0.01280 | 0.00609 | 0.82053 | 0.00709 |
| **Klhl15** | 2.01246 | 4.21504 | 0.57929 | 6.65879 | 0.32918 | 2.43039 | 0.49868 | 1.68810 | 1.01885 | 3.13592 | 1.54361 | 1.87244 | 0.12188 | 0.62600 | 3.09795 | 1.50478 |
| **Klhl17** | 1.40250 | 0.00601 | 0.52507 | 0.22108 | 0.00678 | 0.00427 | 0.00618 | 1.27280 | 0.45875 | 0.77389 | 0.01282 | 0.00000 | 0.00313 | 0.03015 | 0.15571 | 0.11308 |
| **Klhl18** | 0.99761 | 0.92892 | 0.26350 | 4.13129 | 0.00567 | 0.00499 | 0.00581 | 1.79030 | 4.88818 | 2.04366 | 0.56960 | 0.09126 | 0.57329 | 0.50743 | 0.36521 | 0.00467 |
| **Klhl2** | 1.12299 | 0.00509 | 1.29882 | 2.27460 | 0.36528 | 0.00446 | 1.34342 | 4.96885 | 0.48007 | 1.58765 | 2.35732 | 0.06170 | 1.65892 | 1.84146 | 4.52362 | 0.21539 |
| **Klhl20** | 3.86504 | 1.02139 | 0.95728 | 0.92729 | 0.58553 | 2.22696 | 0.49401 | 0.41503 | 0.24827 | 1.02858 | 0.12728 | 0.02267 | 1.05711 | 0.84387 | 1.28798 | 1.85039 |
| **Klhl21** | 6.27902 | 1.24843 | 5.49106 | 9.02035 | 4.88941 | 3.51791 | 1.71151 | 8.55344 | 10.18575 | 8.25834 | 4.92088 | 2.03773 | 1.06700 | 7.47205 | 13.91974 | 0.03667 |
| **Klhl22** | 1.12735 | 0.01558 | 6.50479 | 4.56673 | 0.03466 | 0.00458 | 3.55601 | 1.45252 | 1.35538 | 3.41236 | 1.34933 | 0.05908 | 9.49318 | 0.00000 | 1.99801 | 0.04296 |
| **Klhl23** | 0.01230 | 0.00769 | 0.89034 | 0.58511 | 0.00769 | 0.96651 | 0.03719 | 0.00976 | 0.34765 | 0.00548 | 0.01782 | 0.01614 | 0.00530 | 0.01074 | 0.00000 | 0.00652 |
| **Klhl24** | 0.42021 | 0.00318 | 0.71744 | 0.88712 | 2.79261 | 0.00703 | 0.00542 | 0.02341 | 3.74252 | 1.40286 | 1.04392 | 7.75493 | 0.23998 | 0.55307 | 3.63355 | 2.00589 |
| **Klhl25** | 0.00000 | 0.05709 | 4.11169 | 1.07268 | 0.00230 | 3.61398 | 0.22958 | 0.00439 | 0.76995 | 2.17250 | 0.00000 | 0.03515 | 1.32148 | 0.00798 | 0.00000 | 0.01097 |
| **Klhl26** | 0.48792 | 0.01975 | 2.09625 | 0.45636 | 0.01417 | 0.00646 | 0.17285 | 0.01217 | 0.00361 | 0.32546 | 0.00933 | 0.03033 | 0.02203 | 0.00967 | 0.00000 | 0.08905 |
| **Klhl28** | 0.06891 | 0.00800 | 0.18287 | 0.22820 | 0.00812 | 0.00623 | 0.00779 | 0.37197 | 1.30493 | 0.26555 | 0.00000 | 0.00315 | 0.08934 | 0.00664 | 1.18123 | 0.00739 |
| **Klhl3** | 0.01344 | 0.00824 | 0.01357 | 0.00946 | 0.00972 | 0.00689 | 0.00769 | 0.01382 | 0.01233 | 0.25694 | 0.01284 | 0.00725 | 0.02934 | 0.01133 | 0.00996 | 0.00738 |
| **Klhl30** | 0.00850 | 0.00756 | 0.28950 | 0.44915 | 0.00920 | 0.00901 | 0.00706 | 0.53662 | 0.00308 | 0.00685 | 0.00830 | 0.03443 | 0.00443 | 0.01104 | 0.00000 | 0.00725 |
| **Klhl32** | 0.01375 | 0.00805 | 0.01225 | 0.01061 | 0.00932 | 0.00666 | 0.00754 | 0.01390 | 0.01235 | 0.01531 | 0.01310 | 0.00811 | 0.01270 | 0.01234 | 0.01062 | 0.00741 |
| **Klhl35** | 0.01351 | 0.00797 | 0.01228 | 0.14188 | 0.00922 | 0.00655 | 0.00734 | 0.01120 | 0.00926 | 0.50136 | 0.01316 | 0.00673 | 0.01189 | 0.01011 | 0.01614 | 0.00742 |
| **Klhl36** | 0.82199 | 0.00747 | 0.04053 | 0.27594 | 0.00767 | 0.00507 | 0.00675 | 0.00963 | 0.04287 | 0.27914 | 0.10045 | 0.00630 | 0.33926 | 0.01113 | 0.00558 | 0.02863 |
| **Klhl41** | 1.23394 | 0.20407 | 3.50824 | 0.20727 | 0.67174 | 0.25995 | 0.40964 | 0.00000 | 2.65355 | 0.81399 | 9.58582 | 0.72777 | 3.65271 | 1.53605 | 0.85489 | 0.73573 |
| **Klhl42** | 0.15226 | 1.79748 | 1.26370 | 1.00702 | 1.87215 | 0.00519 | 1.07067 | 0.00000 | 1.21310 | 0.70419 | 0.57063 | 0.02953 | 0.58181 | 0.01239 | 0.31822 | 0.00543 |
| **Klhl5** | 0.00830 | 0.00742 | 0.00676 | 0.00610 | 0.00903 | 0.00698 | 0.00716 | 0.97258 | 0.01033 | 1.60738 | 0.08806 | 0.02911 | 0.00477 | 0.01415 | 0.05665 | 0.00708 |
| **Klhl6** | 0.01375 | 0.00791 | 0.01194 | 0.29820 | 0.00933 | 0.00670 | 0.00754 | 0.01372 | 0.10410 | 0.01820 | 0.01156 | 0.00803 | 0.01293 | 0.01668 | 0.04024 | 0.00736 |
| **Klhl7** | 0.89072 | 1.15799 | 0.80665 | 4.96454 | 0.27075 | 1.20755 | 3.53918 | 0.29604 | 5.74658 | 1.02888 | 5.64427 | 3.71638 | 0.31758 | 3.25798 | 2.06677 | 0.14306 |
| **Klhl8** | 0.95578 | 0.00702 | 0.67985 | 19.02848 | 0.01274 | 0.00201 | 0.00546 | 0.00596 | 8.46697 | 1.73887 | 0.59616 | 0.29306 | 0.00449 | 0.00610 | 9.42758 | 0.00224 |
| **Klhl9** | 0.85076 | 0.05722 | 1.13318 | 0.61782 | 0.00519 | 2.18338 | 0.03564 | 0.00000 | 0.51721 | 0.63596 | 1.65102 | 0.04111 | 0.74994 | 0.41911 | 2.63965 | 0.41236 |
| **Klk1b24** | 0.01038 | 0.00853 | 0.00910 | 0.00766 | 0.00966 | 0.00685 | 0.00814 | 0.17117 | 0.02629 | 0.06378 | 0.07118 | 0.00786 | 0.00720 | 0.01320 | 0.00587 | 0.00743 |
| **Klrb1a** | 0.01291 | 0.00778 | 0.02396 | 0.01006 | 0.00914 | 0.00764 | 0.01744 | 0.01672 | 0.01231 | 0.11290 | 0.01300 | 0.00710 | 0.01897 | 0.02881 | 0.00798 | 0.00774 |
| **Klrg2** | 1.19354 | 1.91184 | 0.06185 | 9.84977 | 2.98420 | 0.00271 | 5.35357 | 1.82187 | 7.39879 | 7.44685 | 3.86183 | 0.47026 | 0.12023 | 0.52007 | 7.35987 | 0.20952 |
| **Klri2** | 0.01331 | 0.00782 | 0.01784 | 0.10286 | 0.00928 | 0.00728 | 0.04868 | 0.01279 | 0.15806 | 0.01972 | 0.01118 | 0.00768 | 0.01240 | 0.01512 | 0.03674 | 0.00734 |
| **Kmt2a** | 1.92372 | 0.01601 | 0.00000 | 0.04091 | 0.01367 | 0.82462 | 0.00467 | 0.00257 | 0.58110 | 0.20432 | 0.00752 | 0.02280 | 1.96212 | 0.00754 | 0.05084 | 1.87329 |
| **Kmt2b** | 6.92413 | 0.12870 | 5.21416 | 12.08647 | 0.00000 | 3.98217 | 3.73259 | 5.78309 | 2.54420 | 3.50453 | 1.99001 | 5.23087 | 4.92081 | 0.00000 | 0.70314 | 3.26297 |
| **Kmt2c** | 1.10718 | 2.84672 | 5.59971 | 10.75836 | 0.27545 | 4.02162 | 3.67176 | 8.50327 | 12.12664 | 12.50792 | 9.60645 | 18.16377 | 1.30712 | 21.32999 | 18.52869 | 2.79833 |
| **Kmt2d** | 3.83538 | 0.18987 | 1.92260 | 5.35428 | 0.24066 | 1.03742 | 0.00203 | 0.59298 | 5.75770 | 0.65916 | 0.63916 | 0.02932 | 3.04472 | 1.51929 | 0.22563 | 2.59145 |
| **Kmt2e** | 0.82871 | 2.88833 | 0.00985 | 0.00000 | 0.01303 | 0.00717 | 0.00567 | 0.59336 | 0.17594 | 0.78817 | 0.10999 | 0.02283 | 0.33499 | 4.15744 | 0.23120 | 0.00539 |
| **Kmt5a** | 11.11852 | 6.41097 | 9.94519 | 9.87380 | 10.43570 | 4.54955 | 9.69828 | 19.32396 | 11.60434 | 17.40635 | 12.04732 | 2.01729 | 8.76128 | 4.42392 | 6.89418 | 3.34679 |
| **Kncn** | 0.01331 | 0.00839 | 0.01175 | 0.00933 | 0.00933 | 0.00653 | 0.00770 | 0.02124 | 0.01188 | 0.01880 | 0.01242 | 0.00846 | 0.02476 | 0.00753 | 0.17365 | 0.00737 |
| **Kndc1** | 1.76337 | 0.00643 | 0.02395 | 0.78171 | 0.00701 | 0.00411 | 0.00619 | 0.00835 | 0.57660 | 0.04547 | 0.00956 | 0.00664 | 0.01721 | 0.01325 | 0.00262 | 0.01252 |
| **Kng1** | 0.00000 | 5.20091 | 5.04023 | 0.00000 | 0.00704 | 0.01970 | 9.92907 | 7.33188 | 0.64646 | 1.30193 | 3.46525 | 6.61247 | 4.86991 | 0.00000 | 1.10235 | 0.00675 |
| **Kng2** | 0.00000 | 0.61256 | 1.13388 | 0.05039 | 0.00594 | 0.00703 | 2.25342 | 1.80077 | 0.35425 | 0.22774 | 0.95331 | 0.92392 | 1.45209 | 0.06922 | 0.00023 | 0.00552 |
| **Knop1** | 14.44511 | 7.06361 | 14.71457 | 8.81896 | 9.26179 | 7.71278 | 10.43124 | 11.42815 | 15.86987 | 19.85881 | 12.32435 | 8.06958 | 10.47230 | 21.88315 | 16.71190 | 3.20964 |
| **Knstrn** | 7.83950 | 0.00669 | 1.90530 | 15.84865 | 5.39540 | 5.08317 | 0.00000 | 3.95726 | 4.45478 | 5.89833 | 2.31766 | 0.13636 | 1.81945 | 2.59119 | 1.62885 | 4.03823 |
| **Kntc1** | 2.35828 | 0.11632 | 4.83816 | 5.76377 | 5.14978 | 2.45936 | 4.89076 | 1.72209 | 8.69267 | 3.02354 | 2.60577 | 4.14194 | 0.97097 | 4.75021 | 6.18475 | 1.65101 |
| **Kpna1** | 3.70969 | 10.36553 | 9.43168 | 13.27572 | 7.76123 | 1.16650 | 0.55060 | 8.25018 | 12.98857 | 16.60760 | 8.83149 | 6.74227 | 6.44164 | 32.54362 | 20.76141 | 3.13398 |
| **Kpna2** | 24.54105 | 22.27845 | 24.28693 | 20.83171 | 14.82530 | 10.16826 | 13.11871 | 38.48489 | 27.18217 | 38.19802 | 25.16210 | 25.33314 | 16.31312 | 43.89174 | 39.22468 | 6.72913 |
| **Kpna3** | 5.73935 | 5.27543 | 4.70639 | 9.57117 | 7.86772 | 1.32710 | 2.95397 | 2.94060 | 3.43825 | 2.13101 | 1.29579 | 8.05325 | 6.89506 | 7.10590 | 13.43633 | 2.06293 |
| **Kpna4** | 1.49330 | 4.10128 | 0.44901 | 6.53278 | 4.12999 | 2.26532 | 1.83240 | 1.66921 | 10.61661 | 4.69767 | 4.75673 | 5.80815 | 1.96990 | 10.42392 | 2.66696 | 0.63281 |
| **Kpna6** | 9.88190 | 5.97057 | 6.97945 | 7.55865 | 7.72143 | 7.66413 | 7.62882 | 11.85445 | 13.02911 | 7.81360 | 3.20617 | 0.67990 | 5.67093 | 5.69251 | 8.60673 | 7.63684 |
| **Kpna7** | 0.01169 | 0.11247 | 0.01104 | 2.65565 | 0.00887 | 0.00640 | 0.00732 | 0.01167 | 0.00000 | 0.02842 | 0.02349 | 0.02695 | 0.01185 | 0.00197 | 0.13237 | 0.00675 |
| **Kpnb1** | 25.75640 | 18.66526 | 20.54634 | 17.44485 | 21.02657 | 11.73475 | 17.56101 | 25.53772 | 18.78628 | 23.99662 | 16.09363 | 8.30476 | 22.08069 | 28.10607 | 22.93652 | 13.81247 |
| **Kptn** | 3.55387 | 0.09338 | 0.10832 | 0.12290 | 0.07205 | 0.00326 | 3.20291 | 6.32865 | 0.02628 | 4.45444 | 2.03213 | 0.05529 | 2.29918 | 0.13964 | 0.00000 | 3.29815 |

| | | | | | | | | | | | | | | | | |
|---|---|---|---|---|---|---|---|---|---|---|---|---|---|---|---|---|
| **Kras** | 3.35892 | 2.39909 | 0.49008 | 2.72960 | 3.33902 | 2.37570 | 2.20964 | 0.63344 | 1.51665 | 3.48585 | 2.40167 | 12.61840 | 0.59970 | 3.76780 | 4.40742 | 0.09976 |
| **Krba1** | 0.07312 | 0.00878 | 0.91870 | 0.57399 | 0.01366 | 0.00744 | 0.00610 | 0.41256 | 0.00000 | 0.00000 | 1.21503 | 0.00879 | 2.58431 | 0.00381 | 0.83433 | 0.04915 |
| **Krcc1** | 1.94822 | 0.11753 | 0.70553 | 3.29022 | 3.69967 | 7.71956 | 1.70200 | 1.29082 | 0.00000 | 2.83696 | 0.00000 | 1.05872 | 0.69367 | 5.47779 | 1.47283 | 0.00823 |
| **Kremen1** | 0.29485 | 0.00830 | 0.52323 | 2.53958 | 0.00899 | 0.00922 | 0.00687 | 0.00000 | 0.60945 | 1.40591 | 0.62064 | 0.08385 | 0.31037 | 0.10748 | 0.18016 | 3.22462 |
| **Kremen2** | 0.11205 | 0.08861 | 0.33224 | 0.11146 | 0.01962 | 5.25603 | 0.16018 | 0.75683 | 0.36337 | 4.50018 | 2.01745 | 0.16376 | 5.79671 | 1.08155 | 0.17645 | 0.07517 |
| **Kri1** | 3.02387 | 3.88094 | 2.59662 | 0.00000 | 2.33336 | 2.48849 | 2.56991 | 1.23145 | 0.14359 | 2.49005 | 0.02875 | 0.05203 | 2.04357 | 2.22762 | 3.56780 | 3.15789 |
| **Krit1** | 0.45137 | 3.67560 | 1.28874 | 0.90688 | 0.00429 | 0.15955 | 3.03253 | 1.50775 | 0.74136 | 3.15160 | 1.34293 | 15.18303 | 1.77801 | 7.94721 | 3.31293 | 1.74222 |
| **Krr1** | 7.08061 | 4.78988 | 3.96662 | 8.12345 | 3.88915 | 3.03025 | 5.19826 | 10.22142 | 12.28310 | 9.38052 | 6.98606 | 13.12421 | 3.05439 | 24.10702 | 13.82308 | 3.41732 |
| **Krt10** | 0.12631 | 0.00794 | 0.30181 | 0.74174 | 0.03324 | 0.00786 | 0.00640 | 0.02387 | 0.43509 | 0.48545 | 0.54939 | 0.02078 | 0.78407 | 0.00410 | 0.02723 | 0.00684 |
| **Krt12** | 0.01377 | 0.00810 | 0.01219 | 1.35856 | 0.00936 | 0.00650 | 0.00745 | 0.01331 | 0.00000 | 0.01096 | 0.01254 | 0.00820 | 0.01268 | 0.00771 | 0.10073 | 0.00717 |
| **Krt18** | 0.52551 | 0.11232 | 6.39765 | 0.00000 | 0.85786 | 30.35846 | 3.59175 | 6.88247 | 0.59928 | 11.07762 | 6.30096 | 0.00000 | 30.57578 | 0.08994 | 0.00000 | 21.03458 |
| **Krt19** | 0.01087 | 0.00741 | 0.16621 | 0.00494 | 0.00794 | 0.00887 | 0.00695 | 0.00000 | 0.06104 | 6.04983 | 0.41508 | 0.00000 | 0.00761 | 0.01208 | 0.00000 | 0.00730 |
| **Krt222** | 0.00101 | 0.01980 | 0.04190 | 0.12557 | 0.04534 | 0.00847 | 0.00811 | 0.53863 | 0.30113 | 0.76529 | 0.07088 | 0.01067 | 0.00595 | 1.26083 | 0.52701 | 0.00768 |
| **Krt25** | 0.00982 | 0.00806 | 0.06487 | 0.00809 | 0.00927 | 0.00860 | 0.00754 | 0.10687 | 0.01180 | 0.33914 | 0.05617 | 0.00734 | 0.02448 | 0.01472 | 0.00683 | 0.00725 |
| **Krt27** | 0.01163 | 0.00794 | 1.12553 | 0.00858 | 0.00795 | 0.00953 | 0.00746 | 0.01164 | 0.01225 | 0.45380 | 0.01091 | 0.08133 | 0.04442 | 0.00920 | 0.01906 | 0.00707 |
| **Krt28** | 0.01114 | 0.00721 | 0.00000 | 0.79734 | 0.00844 | 0.00825 | 0.00644 | 2.75455 | 5.56771 | 7.12621 | 1.90399 | 0.25195 | 0.64610 | 0.21312 | 0.03599 | 0.00616 |
| **Krt7** | 0.03441 | 0.00771 | 0.00268 | 0.00479 | 0.01021 | 0.00806 | 0.00616 | 0.65383 | 0.19008 | 0.16159 | 0.71525 | 0.01232 | 1.37046 | 0.00670 | 0.01460 | 0.00676 |
| **Krt8** | 5.02824 | 8.20894 | 11.62984 | 1.41454 | 8.19874 | 28.17821 | 5.58374 | 5.46887 | 0.74121 | 0.63948 | 0.04655 | 0.02402 | 27.54653 | 0.00000 | 0.00000 | 23.78959 |
| **Krt84** | 0.01360 | 0.00787 | 0.01167 | 1.23490 | 0.00925 | 0.00640 | 0.00746 | 0.01246 | 1.23373 | 0.00360 | 0.03858 | 0.00528 | 0.01206 | 0.00441 | 0.10124 | 0.00706 |
| **Krtcap2** | 5.37073 | 0.38789 | 6.13815 | 0.53850 | 5.36819 | 3.16130 | 2.03884 | 2.80457 | 0.62756 | 5.99856 | 9.04470 | 0.39330 | 8.98722 | 0.06121 | 0.00000 | 0.70627 |
| **Krtcap3** | 1.03891 | 0.01639 | 0.07677 | 2.41300 | 0.00561 | 0.00785 | 0.00509 | 0.01779 | 0.00000 | 1.18767 | 1.37446 | 0.01446 | 1.10636 | 0.00000 | 0.29685 | 0.00538 |
| **Krtdap** | 0.01001 | 0.00774 | 0.04096 | 0.01058 | 0.00921 | 0.20459 | 0.00702 | 0.07012 | 0.01234 | 0.00694 | 0.01192 | 0.00742 | 0.29629 | 0.01299 | 0.00879 | 0.00704 |
| **Ksr1** | 0.59409 | 0.00617 | 0.00172 | 0.21092 | 0.00709 | 0.00846 | 1.52267 | 0.00839 | 0.00545 | 0.00892 | 0.01348 | 0.01640 | 0.02848 | 0.00977 | 0.06100 | 0.00688 |
| **Kti12** | 2.75221 | 1.76638 | 0.46998 | 11.13117 | 3.79339 | 2.24116 | 1.68222 | 7.16214 | 16.98836 | 14.74433 | 5.76688 | 2.06931 | 3.17305 | 2.32413 | 5.68310 | 1.25852 |
| **Ktn1** | 1.15408 | 1.85821 | 4.76916 | 8.67093 | 6.22846 | 4.80942 | 7.39359 | 3.69966 | 2.17678 | 1.06067 | 2.20387 | 0.74538 | 2.59116 | 8.27475 | 12.69621 | 4.32784 |
| **Kxd1** | 16.26531 | 8.74301 | 12.64825 | 1.91159 | 1.42878 | 9.18778 | 9.16897 | 15.90158 | 2.53896 | 14.51262 | 13.15011 | 6.66894 | 13.89996 | 2.47538 | 5.16393 | 7.16977 |
| **Kyat1** | 3.70975 | 0.00301 | 0.52670 | 1.72961 | 0.53098 | 0.00256 | 0.01341 | 1.52058 | 0.80096 | 5.37229 | 1.15219 | 1.13191 | 0.51895 | 0.01857 | 1.84118 | 1.06327 |
| **Kyat3** | 11.55664 | 7.31677 | 7.88077 | 4.99189 | 13.85245 | 0.00000 | 6.94201 | 0.88413 | 11.02157 | 1.91261 | 0.53053 | 0.05349 | 7.11716 | 0.03573 | 4.26829 | 2.18579 |
| **L1cam** | 0.02013 | 0.00828 | 0.13798 | 3.09397 | 0.00941 | 0.05936 | 0.00719 | 0.08082 | 2.39985 | 0.75920 | 0.76226 | 0.04781 | 0.89833 | 0.00000 | 1.64685 | 0.00622 |
| **L1td1** | 20.68785 | 14.49730 | 15.43223 | 13.75060 | 19.91244 | 9.89403 | 13.68757 | 2.14139 | 19.90382 | 0.00128 | 1.57314 | 0.21998 | 4.04686 | 5.76219 | 0.51330 | 12.95121 |
| **L2hgdh** | 5.34648 | 2.42986 | 1.60327 | 11.93557 | 0.62641 | 1.79100 | 1.65097 | 5.87472 | 8.30735 | 13.17954 | 12.99973 | 18.85973 | 7.53156 | 22.37797 | 18.34733 | 0.51513 |
| **L3hypdh** | 1.04889 | 0.00694 | 0.00379 | 0.07198 | 0.01144 | 0.00684 | 0.00656 | 0.00829 | 0.01130 | 0.05543 | 0.01284 | 0.01539 | 0.01207 | 0.00570 | 0.21047 | 0.00677 |
| **L3mbtl2** | 5.53298 | 4.62986 | 6.10062 | 13.05524 | 5.25841 | 0.87102 | 5.71898 | 10.24340 | 17.36848 | 10.53637 | 5.50231 | 0.70951 | 5.15169 | 0.18021 | 1.65364 | 10.63476 |
| **L3mbtl3** | 0.00000 | 0.14450 | 0.90514 | 0.29645 | 0.00675 | 0.00565 | 1.01347 | 0.15078 | 0.00408 | 0.16984 | 0.10832 | 0.05729 | 0.11323 | 0.00587 | 1.21945 | 0.74646 |
| **l7Rn6** | 19.43614 | 5.37831 | 12.82715 | 5.62647 | 15.39658 | 8.54999 | 4.72544 | 12.23352 | 3.67716 | 15.23252 | 8.81577 | 3.57814 | 13.98080 | 2.06962 | 0.00000 | 4.04714 |
| **Lacc1** | 0.01033 | 0.00739 | 0.00750 | 0.08431 | 0.00974 | 0.00673 | 1.00468 | 0.02904 | 0.00807 | 0.00684 | 0.01003 | 0.01399 | 0.01014 | 0.52494 | 0.00002 | 0.00740 |
| **Lace1** | 0.00536 | 2.26218 | 0.57018 | 2.43685 | 0.00754 | 0.00709 | 0.00645 | 0.00000 | 0.00000 | 0.21760 | 0.75061 | 0.01403 | 0.17385 | 0.00000 | 0.16579 | 0.00606 |
| **Lactb** | 3.06428 | 0.00213 | 3.53039 | 2.85191 | 0.00000 | 6.47252 | 2.08174 | 1.30249 | 4.55050 | 3.61640 | 2.69509 | 0.76561 | 3.06564 | 5.29357 | 3.62461 | 0.02660 |
| **Lactb2** | 7.02911 | 2.24108 | 3.06353 | 2.53290 | 9.55584 | 3.98875 | 5.01013 | 5.54359 | 3.76920 | 4.15243 | 12.45273 | 6.13842 | 1.51510 | 3.58750 | 2.76979 | 0.34560 |
| **Lad1** | 0.54254 | 0.00677 | 0.00000 | 0.00000 | 0.00664 | 2.90326 | 0.00530 | 3.18866 | 0.00846 | 0.53983 | 3.39422 | 0.00419 | 1.38183 | 0.00865 | 0.00000 | 0.00447 |
| **Lag3** | 0.00710 | 0.00698 | 0.00539 | 0.01477 | 1.31489 | 0.01373 | 0.00729 | 0.01351 | 0.03513 | 0.25470 | 0.02734 | 0.01493 | 0.01705 | 0.06315 | 0.01229 | 0.00746 |
| **Lage3** | 10.50733 | 2.82676 | 4.58813 | 8.22459 | 1.48892 | 0.01846 | 4.47148 | 5.43682 | 1.17245 | 5.98857 | 3.80452 | 0.37915 | 3.91288 | 3.19759 | 3.55627 | 5.14182 |
| **Lama1** | 7.28020 | 2.56380 | 0.55706 | 0.00674 | 0.02171 | 0.01014 | 0.00000 | 0.01664 | 0.01285 | 0.00113 | 0.01085 | 0.00000 | 0.00460 | 0.02409 | 0.18760 | 23.16235 |
| **Lama2** | 0.01274 | 0.00770 | 0.04263 | 1.18266 | 0.00942 | 0.00578 | 0.00738 | 0.01086 | 0.00000 | 0.00463 | 0.00953 | 0.00445 | 0.01210 | 0.00821 | 1.67441 | 0.00711 |
| **Lama3** | 2.26882 | 2.04440 | 0.09086 | 2.74394 | 2.84722 | 0.01552 | 0.92525 | 0.00000 | 0.16069 | 0.00000 | 0.23672 | 0.00925 | 0.90634 | 0.00000 | 0.00000 | 0.00322 |
| **Lama4** | 0.01623 | 0.00766 | 0.96638 | 0.93811 | 0.00736 | 0.00552 | 0.00729 | 0.01024 | 0.00000 | 0.42052 | 0.00827 | 0.03435 | 0.00841 | 0.00413 | 1.11322 | 0.00686 |
| **Lama5** | 5.50232 | 1.67893 | 1.83085 | 0.48815 | 6.44929 | 8.91466 | 2.19857 | 7.29322 | 0.00000 | 1.15469 | 4.49478 | 0.10765 | 5.06486 | 0.00000 | 0.00000 | 11.74659 |
| **Lamb1** | 14.35090 | 2.15577 | 10.49768 | 3.03721 | 4.21937 | 0.89284 | 14.56411 | 2.35915 | 0.07189 | 1.61703 | 1.85374 | 0.65706 | 5.02721 | 4.84813 | 3.18666 | 18.12575 |
| **Lamb2** | 0.00000 | 0.00727 | 1.56804 | 0.65669 | 0.00604 | 0.36612 | 0.00429 | 0.82062 | 0.98454 | 1.09035 | 0.84959 | 0.05090 | 5.11210 | 0.00308 | 0.00000 | 0.01186 |
| **Lamc1** | 5.05592 | 0.01561 | 1.84425 | 2.74298 | 4.15109 | 9.23071 | 6.12284 | 1.29236 | 0.46580 | 0.00000 | 0.00000 | 4.72076 | 3.11496 | 0.00000 | 0.29016 | 8.39794 |
| **Lamc2** | 0.74994 | 0.00759 | 0.00000 | 0.53508 | 0.00687 | 0.00660 | 0.03782 | 0.00000 | 0.00000 | 0.12844 | 0.59453 | 0.04886 | 0.45930 | 0.47580 | 2.01761 | 0.01344 |
| **Lamc3** | 0.01307 | 0.00774 | 0.00931 | 1.41811 | 0.00899 | 0.00639 | 0.00717 | 0.00919 | 0.75969 | 1.05067 | 0.03951 | 0.10840 | 0.01122 | 0.01896 | 0.46741 | 0.00735 |
| **Lamp1** | 28.19441 | 16.33826 | 28.54420 | 8.11297 | 12.20633 | 15.30827 | 19.20490 | 27.85608 | 0.93880 | 24.85612 | 25.51809 | 9.91344 | 28.41137 | 2.29464 | 13.14903 | 9.59593 |
| **Lamp2** | 4.32860 | 4.22875 | 8.33862 | 16.80795 | 1.20729 | 2.48440 | 8.07296 | 9.39103 | 20.03905 | 19.94973 | 6.34629 | 16.66820 | 8.59857 | 30.81769 | 17.31355 | 0.00783 |
| **Lamp3** | 1.51989 | 1.35668 | 1.69361 | 0.92129 | 2.27551 | 0.63693 | 1.00489 | 1.16650 | 1.21798 | 0.74805 | 0.44090 | 0.97794 | 0.66065 | 0.75603 | 0.69879 | 0.76969 |
| **Lamtor1** | 16.26145 | 4.49562 | 19.84285 | 9.29718 | 12.87992 | 9.78273 | 2.22499 | 16.35883 | 6.73306 | 17.16541 | 10.72564 | 0.83275 | 14.58216 | 2.99455 | 2.80611 | 5.74512 |
| **Lamtor2** | 4.00008 | 0.27396 | 1.52389 | 2.92018 | 1.17258 | 0.33033 | 0.76688 | 0.66425 | 0.62303 | 1.28937 | 0.88769 | 0.07805 | 2.28832 | 0.20088 | 0.87552 | 0.02947 |
| **Lamtor3** | 1.72032 | 6.33346 | 9.52815 | 3.65891 | 7.72086 | 6.69428 | 0.66534 | 6.15268 | 3.53515 | 6.80645 | 3.37185 | 6.23494 | 5.92382 | 0.64187 | 5.74188 | 4.13870 |
| **Lamtor4** | 3.13240 | 0.84038 | 1.80387 | 0.50707 | 2.97398 | 2.65113 | 1.28193 | 2.56970 | 0.68353 | 2.27113 | 2.56842 | 0.06018 | 4.32284 | 0.09059 | 0.43554 | 2.70887 |
| **Lamtor5** | 17.16808 | 10.57014 | 6.47436 | 2.60352 | 12.54023 | 3.18478 | 5.24709 | 2.42720 | 0.00000 | 6.98909 | 3.41460 | 1.32937 | 3.68452 | 1.80294 | 0.74945 | 7.76914 |
| **Lancl1** | 5.68699 | 1.97839 | 4.44388 | 1.12716 | 6.56728 | 0.62923 | 7.72896 | 1.93178 | 1.47625 | 2.85816 | 1.85509 | 0.09362 | 3.17030 | 1.50937 | 0.96499 | 3.62549 |
| **Lancl2** | 3.07416 | 1.69285 | 0.00000 | 0.70952 | 0.00364 | 2.47259 | 0.00406 | 0.00785 | 0.00000 | 1.25299 | 0.11753 | 0.00000 | 0.01532 | 0.00297 | 0.36096 | 1.60425 |

| | | | | | | | | | | | | | | | | |
|---|---|---|---|---|---|---|---|---|---|---|---|---|---|---|---|---|
| **Lap3** | 24.41827 | 15.40590 | 16.59601 | 5.56695 | 20.25417 | 14.22556 | 15.15048 | 16.02182 | 6.58010 | 8.51565 | 6.03659 | 3.58990 | 15.32630 | 0.78012 | 8.68750 | 15.11745 |
| **Laptm4a** | 0.76420 | 3.29463 | 4.95269 | 8.02883 | 6.61589 | 4.90293 | 4.64301 | 9.68715 | 5.81215 | 2.28869 | 13.06205 | 0.51087 | 5.74617 | 1.93393 | 2.10307 | 4.53736 |
| **Laptm4b** | 5.06160 | 3.66036 | 2.99034 | 22.10353 | 0.01060 | 3.37644 | 2.61095 | 0.27534 | 14.81584 | 7.32128 | 0.49604 | 0.35823 | 7.96815 | 3.93999 | 7.55442 | 1.53617 |
| **Laptm5** | 0.01724 | 0.00828 | 0.01143 | 0.42241 | 0.01004 | 0.00667 | 0.00734 | 0.01083 | 0.37667 | 1.07889 | 0.00976 | 0.00523 | 0.01829 | 0.03202 | 0.00274 | 0.00740 |
| **Large** | 0.82043 | 0.00630 | 1.04248 | 0.54278 | 0.00468 | 0.08827 | 0.00634 | 0.91712 | 0.00014 | 0.00199 | 0.00552 | 0.02526 | 0.00000 | 0.00656 | 0.46953 | 0.00586 |
| **Larp1** | 18.01568 | 8.36699 | 9.79652 | 14.69918 | 10.84213 | 7.42619 | 11.01420 | 19.14982 | 17.83724 | 23.47680 | 21.93771 | 3.42767 | 14.11687 | 8.82292 | 16.84848 | 9.23349 |
| **Larp1b** | 0.00000 | 2.33581 | 1.04872 | 29.50454 | 2.61435 | 3.97207 | 0.57398 | 9.30122 | 31.64384 | 35.08275 | 16.34631 | 20.10551 | 8.76972 | 19.25768 | 32.59999 | 0.03937 |
| **Larp4** | 4.04907 | 6.70085 | 4.04331 | 18.37414 | 1.72167 | 3.07442 | 2.50002 | 19.56735 | 22.66685 | 17.16120 | 34.48672 | 27.72624 | 6.57027 | 48.67540 | 26.46372 | 1.66519 |
| **Larp4b** | 11.78702 | 7.22391 | 9.60228 | 11.14842 | 7.56961 | 9.65978 | 5.89724 | 10.71973 | 12.99789 | 12.25912 | 7.37560 | 5.89445 | 6.82781 | 17.16991 | 15.03570 | 6.03837 |
| **Larp7** | 16.45040 | 13.47188 | 12.09730 | 8.48875 | 12.59848 | 6.77646 | 10.19080 | 14.08145 | 9.49713 | 15.74835 | 8.69161 | 2.97818 | 6.06283 | 12.52590 | 16.17397 | 6.35733 |
| **Lars** | 7.92128 | 9.15065 | 5.44405 | 0.14173 | 16.83751 | 3.52537 | 9.02440 | 8.64151 | 0.00000 | 10.08979 | 1.88164 | 1.22894 | 6.63840 | 6.86177 | 5.03460 | 1.78917 |
| **Lars2** | 3.47572 | 2.20545 | 0.23412 | 1.86660 | 4.64614 | 3.49535 | 0.00409 | 0.10819 | 0.82727 | 0.39819 | 0.45805 | 0.10208 | 0.39766 | 1.37619 | 1.07622 | 1.07697 |
| **Las1l** | 11.74562 | 4.41342 | 10.29902 | 7.94287 | 3.54410 | 6.36486 | 6.07678 | 3.55703 | 19.49242 | 8.97464 | 8.07175 | 1.26630 | 7.72260 | 6.76369 | 8.79801 | 3.80092 |
| **Lasp1** | 11.21760 | 9.04719 | 10.40690 | 0.04563 | 15.03291 | 7.83993 | 7.44867 | 17.85141 | 0.74411 | 11.40531 | 16.28352 | 0.91182 | 12.83324 | 1.11645 | 0.00000 | 8.47373 |
| **Lat** | 0.01418 | 0.07147 | 0.48224 | 0.82654 | 0.00821 | 0.00704 | 1.13023 | 0.00000 | 0.69862 | 2.76674 | 1.54373 | 0.20927 | 0.05494 | 0.01052 | 0.68207 | 0.02199 |
| **Lat2** | 0.00000 | 0.00676 | 1.35487 | 0.92814 | 0.00681 | 0.01227 | 0.00462 | 5.15764 | 1.82265 | 0.74874 | 3.62699 | 0.00000 | 2.04655 | 0.19766 | 0.76078 | 0.01267 |
| **Lats1** | 0.00000 | 0.00313 | 0.51061 | 2.39156 | 0.81255 | 0.14201 | 1.42474 | 1.22798 | 3.58056 | 3.10264 | 1.33027 | 10.60658 | 0.00000 | 7.60122 | 6.67801 | 2.71149 |
| **Lats2** | 0.00000 | 1.35844 | 1.20312 | 1.06199 | 0.00643 | 2.30121 | 0.00182 | 3.92855 | 3.59274 | 3.72686 | 6.78419 | 10.03138 | 1.52213 | 1.11379 | 1.86707 | 0.01032 |
| **Lbh** | 0.12099 | 0.00489 | 0.01498 | 0.00000 | 5.27829 | 0.00852 | 0.00707 | 0.00000 | 3.01933 | 0.74480 | 0.07792 | 0.03643 | 0.01649 | 0.01071 | 0.00713 | 0.00706 |
| **Lbhd1** | 0.00952 | 0.00866 | 0.00507 | 0.00000 | 0.00952 | 0.00628 | 0.00813 | 0.41071 | 0.00692 | 0.51591 | 0.00038 | 0.10382 | 0.00618 | 0.01616 | 3.53961 | 0.00739 |
| **Lbp** | 0.01212 | 0.00812 | 0.01654 | 0.96255 | 0.00871 | 0.01228 | 0.00732 | 0.00982 | 0.84701 | 2.54045 | 0.01554 | 0.00000 | 0.01093 | 0.00901 | 0.00723 | 0.00717 |
| **Lbr** | 5.83232 | 2.08141 | 2.89025 | 11.41643 | 3.05859 | 8.32314 | 5.08262 | 12.54060 | 11.91316 | 12.19702 | 13.54954 | 14.59082 | 3.50749 | 21.10878 | 14.41954 | 0.06881 |
| **Lbx2** | 0.01289 | 0.00819 | 0.01036 | 1.05360 | 0.00891 | 0.00632 | 0.00738 | 0.00822 | 3.48613 | 0.82793 | 0.08368 | 0.00000 | 0.01069 | 0.00842 | 0.00119 | 0.00693 |
| **Lca5** | 0.01309 | 0.00776 | 0.01176 | 0.19032 | 0.00915 | 0.00671 | 0.00738 | 0.02090 | 0.00586 | 0.41953 | 0.02202 | 0.00628 | 0.01173 | 0.00538 | 0.20955 | 0.00757 |
| **Lca5l** | 0.01371 | 0.00820 | 0.01195 | 0.76050 | 0.00930 | 0.00657 | 0.00749 | 0.01362 | 0.39800 | 0.01264 | 0.01155 | 0.01844 | 0.01278 | 0.01114 | 0.00285 | 0.00727 |
| **Lcat** | 0.00941 | 0.00756 | 0.00592 | 0.00682 | 0.00959 | 0.00701 | 0.00702 | 1.63935 | 0.00998 | 0.00000 | 0.12258 | 0.03407 | 0.00460 | 0.00609 | 0.15100 | 0.00721 |
| **Lck** | 0.67506 | 0.00307 | 1.17393 | 1.55828 | 3.43687 | 0.00549 | 3.09062 | 3.21683 | 5.43799 | 2.90216 | 0.40885 | 0.12157 | 0.00000 | 0.87136 | 2.70066 | 0.01253 |
| **Lclat1** | 0.00000 | 0.42308 | 4.99658 | 14.87039 | 0.01676 | 0.10376 | 3.88569 | 5.23267 | 11.59134 | 14.08024 | 11.66477 | 12.93954 | 3.75530 | 13.62944 | 16.75963 | 1.45913 |
| **Lcmt1** | 6.98106 | 3.05830 | 7.26554 | 1.49478 | 7.30295 | 4.89389 | 1.86394 | 2.44205 | 0.00000 | 2.35584 | 1.43594 | 0.18945 | 4.22390 | 0.15630 | 0.49355 | 0.03619 |
| **Lcmt2** | 2.21799 | 1.98028 | 1.08594 | 9.20409 | 0.79359 | 0.07551 | 1.17288 | 1.22738 | 11.94502 | 4.13859 | 9.87232 | 0.82443 | 0.46517 | 4.22356 | 2.58104 | 0.33985 |
| **Lcn10** | 0.02390 | 0.01924 | 0.00641 | 0.00885 | 0.01199 | 3.26775 | 0.00731 | 0.01144 | 0.01283 | 0.00516 | 0.01055 | 0.00801 | 0.02384 | 0.01304 | 0.00993 | 0.00637 |
| **Lcn2** | 0.01043 | 0.00829 | 0.00987 | 0.08926 | 0.00846 | 0.00669 | 0.00771 | 0.02019 | 0.00759 | 4.60321 | 0.00565 | 0.00000 | 0.00991 | 0.36228 | 0.00000 | 0.00726 |
| **Lcn9** | 0.00762 | 0.00846 | 0.74280 | 0.52986 | 0.00854 | 0.00928 | 0.00810 | 0.45107 | 0.49346 | 0.00668 | 0.00601 | 0.00677 | 0.00250 | 0.01241 | 0.00000 | 0.00702 |
| **Lcor** | 0.35736 | 0.00688 | 0.00142 | 1.06765 | 0.65612 | 0.00825 | 0.00714 | 0.00853 | 0.00000 | 0.45864 | 0.00891 | 0.00305 | 0.28730 | 0.77336 | 0.17257 | 0.00697 |
| **Lcorl** | 0.01109 | 0.00727 | 0.56484 | 0.03144 | 0.12189 | 0.00550 | 0.00704 | 0.50820 | 0.01540 | 0.43836 | 0.00000 | 1.18176 | 0.20996 | 0.00703 | 0.69967 | 0.00734 |
| **Lcp1** | 5.74896 | 5.61515 | 1.23529 | 0.50300 | 1.33824 | 14.49863 | 5.47007 | 0.09867 | 0.96616 | 0.68981 | 0.09459 | 0.02551 | 0.08457 | 0.00248 | 0.00000 | 8.19690 |
| **Lct** | 0.00510 | 0.00820 | 0.44640 | 0.68174 | 0.00805 | 0.80314 | 0.00670 | 0.01722 | 1.50212 | 0.49941 | 0.06284 | 0.03345 | 1.75016 | 0.00314 | 0.41584 | 0.00573 |
| **Lctl** | 0.00000 | 1.95053 | 2.15766 | 1.26362 | 0.00606 | 0.01486 | 0.03510 | 4.97799 | 0.76063 | 1.65713 | 1.54113 | 4.40604 | 0.32742 | 2.02205 | 2.38171 | 0.00489 |
| **Ldah** | 5.98599 | 6.23125 | 9.92082 | 1.02556 | 6.72528 | 0.64884 | 2.24234 | 10.75553 | 1.63030 | 11.09916 | 3.99247 | 2.07236 | 5.23429 | 14.33530 | 4.97261 | 1.87574 |
| **Ldb1** | 1.35950 | 1.97005 | 0.91688 | 1.03738 | 2.31656 | 1.43564 | 1.94183 | 4.03284 | 1.65408 | 0.84175 | 5.03684 | 0.63239 | 0.36571 | 0.20167 | 0.58373 | 1.13812 |
| **Ldha** | 16.89623 | 13.81224 | 10.60901 | 6.94016 | 12.82024 | 4.08920 | 0.66207 | 3.13324 | 5.28874 | 6.96748 | 1.32795 | 8.78435 | 4.99453 | 0.00000 | 1.43530 | 6.20953 |
| **Ldhb** | 0.00000 | 0.00665 | 0.86370 | 22.53793 | 0.01684 | 0.00233 | 0.00240 | 0.77634 | 13.34882 | 11.28856 | 5.01758 | 1.98615 | 3.43953 | 7.52104 | 17.57153 | 0.00000 |
| **Ldhc** | 0.02312 | 0.00802 | 0.00763 | 3.68359 | 0.00930 | 0.00622 | 0.00745 | 0.01178 | 3.96260 | 0.25287 | 0.00000 | 0.07528 | 0.19956 | 0.01562 | 2.77028 | 0.00657 |
| **Ldhd** | 0.19687 | 0.00771 | 0.00645 | 0.15077 | 0.00742 | 0.00711 | 0.00714 | 0.01140 | 0.46598 | 0.52437 | 0.00863 | 0.03518 | 0.01220 | 0.01094 | 0.00588 | 0.00741 |
| **Ldlr** | 0.80509 | 0.00317 | 3.51263 | 0.21382 | 0.01027 | 2.06248 | 4.33096 | 2.70080 | 1.89524 | 0.52764 | 4.55498 | 0.06025 | 0.00000 | 1.50914 | 0.34909 | 0.04064 |
| **Ldlrad2** | 0.22788 | 0.07252 | 0.63372 | 0.14743 | 0.19998 | 0.05521 | 0.12036 | 0.01199 | 0.12848 | 0.18936 | 1.42674 | 0.10745 | 0.71377 | 0.46137 | 0.15893 | 0.12928 |
| **Ldlrad3** | 0.01346 | 0.00782 | 0.01178 | 0.18241 | 0.00907 | 0.00664 | 0.01765 | 0.01203 | 0.03680 | 0.47929 | 0.02692 | 0.00613 | 0.01190 | 0.01260 | 0.01401 | 0.00776 |
| **Ldlrad4** | 0.01379 | 0.00786 | 0.01176 | 0.67044 | 0.00939 | 0.00650 | 0.00740 | 0.01216 | 0.00000 | 0.00659 | 0.01300 | 0.00736 | 0.01279 | 0.02391 | 0.27132 | 0.00728 |
| **Ldlrap1** | 3.87032 | 0.00000 | 3.80493 | 3.11703 | 3.15422 | 3.85017 | 4.94102 | 6.67028 | 4.02661 | 5.58476 | 4.34262 | 4.41793 | 4.30278 | 0.02310 | 2.36577 | 0.01862 |
| **Ldoc1** | 0.02061 | 0.00777 | 0.01431 | 0.00562 | 0.01982 | 5.85958 | 0.00660 | 0.00813 | 0.01314 | 0.06735 | 0.00721 | 0.00657 | 0.58917 | 0.00522 | 0.21660 | 0.00482 |
| **Ldoc1l** | 0.14030 | 0.00748 | 0.01925 | 1.58581 | 0.00787 | 0.00846 | 0.02750 | 1.68800 | 0.36825 | 0.21502 | 0.37264 | 0.07221 | 0.22799 | 0.00141 | 1.86743 | 0.00693 |
| **Lect1** | 0.00864 | 0.00838 | 0.00728 | 0.24784 | 0.01023 | 0.00757 | 0.00802 | 0.21532 | 0.25494 | 0.51866 | 0.46163 | 0.01385 | 0.22900 | 0.00747 | 0.00000 | 0.01520 |
| **Lef1** | 0.01376 | 0.00795 | 0.01172 | 0.28907 | 0.00940 | 0.00657 | 0.00737 | 0.01186 | 0.00593 | 0.11779 | 0.02618 | 0.03639 | 0.01236 | 0.01043 | 0.00421 | 0.00742 |
| **Lekr1** | 0.01332 | 0.00762 | 1.51063 | 0.00996 | 0.00646 | 0.00658 | 0.00711 | 0.00806 | 0.57990 | 0.69586 | 0.02111 | 0.00380 | 0.00670 | 0.01368 | 0.00000 | 0.00648 |
| **Lemd1** | 0.01337 | 0.00760 | 0.00964 | 0.00855 | 0.00952 | 0.00661 | 0.00729 | 0.01066 | 0.01188 | 0.00324 | 0.00943 | 0.00531 | 0.01330 | 0.38171 | 0.35711 | 0.00748 |
| **Lemd2** | 0.42532 | 0.33936 | 0.39965 | 0.00000 | 2.32709 | 1.70236 | 1.32470 | 2.54191 | 0.72854 | 2.19690 | 0.46838 | 0.00000 | 2.37060 | 0.00716 | 0.00478 | 5.10643 |
| **Lemd3** | 0.00907 | 0.05552 | 0.07692 | 7.67815 | 0.00891 | 0.00576 | 0.00503 | 0.92900 | 6.35783 | 0.00000 | 1.20131 | 8.78256 | 0.00000 | 8.52874 | 1.49162 | 0.00471 |
| **Lenep** | 0.18344 | 1.19135 | 0.30487 | 0.46733 | 0.01490 | 0.00822 | 1.40590 | 0.03581 | 0.23047 | 0.31826 | 0.73260 | 0.02234 | 0.09732 | 0.00375 | 0.06482 | 0.00623 |
| **Leng1** | 1.38124 | 0.00346 | 1.48197 | 7.74386 | 2.55096 | 2.60453 | 0.24013 | 6.22154 | 24.08429 | 4.36856 | 2.36671 | 0.32198 | 0.00000 | 0.07513 | 0.56145 | 4.06394 |
| **Leng8** | 1.34825 | 0.00572 | 0.73895 | 5.86481 | 0.10478 | 1.00741 | 0.01436 | 1.55196 | 4.70021 | 3.52017 | 1.23484 | 0.15861 | 2.18385 | 0.37915 | 0.00000 | 0.23297 |
| **Leng9** | 0.01326 | 0.00828 | 0.01671 | 0.26161 | 0.00934 | 0.00631 | 0.00752 | 0.01288 | 0.44021 | 0.01754 | 0.04583 | 0.00724 | 0.02796 | 0.03362 | 0.00114 | 0.01469 |
| **Leo1** | 10.59517 | 8.33213 | 6.10644 | 10.59480 | 10.03968 | 3.36123 | 7.85591 | 15.40992 | 20.57503 | 17.57088 | 11.97444 | 11.42091 | 10.10654 | 3.86324 | 3.58653 | 5.57590 |

| | | | | | | | | | | | | | | | | |
|---|---|---|---|---|---|---|---|---|---|---|---|---|---|---|---|---|
| **Lepr** | 0.45362 | 0.18521 | 0.15447 | 5.39135 | 0.22773 | 0.11018 | 0.11742 | 0.27062 | 10.48576 | 0.00000 | 4.27878 | 0.32201 | 0.20729 | 2.46140 | 1.50475 | 0.09672 |
| **Leprot** | 6.01475 | 0.01888 | 3.87859 | 0.00000 | 0.10370 | 4.20639 | 7.10200 | 0.67608 | 0.53738 | 3.14148 | 0.50976 | 0.05982 | 4.24241 | 0.95617 | 1.54474 | 6.10194 |
| **Leprotl1** | 1.95826 | 0.09159 | 2.67663 | 0.16758 | 0.34406 | 0.00457 | 0.18936 | 1.21261 | 1.74660 | 2.10354 | 1.56580 | 0.11388 | 1.05059 | 1.34092 | 1.28588 | 1.85845 |
| **Letm1** | 9.00370 | 0.22500 | 7.10071 | 5.54587 | 7.15938 | 5.19084 | 8.68358 | 10.76625 | 4.40938 | 3.59086 | 4.29100 | 0.74533 | 10.03018 | 1.11629 | 4.37723 | 0.09443 |
| **Letm2** | 0.00577 | 0.00755 | 0.00934 | 2.21219 | 0.00893 | 0.00580 | 0.00653 | 0.34522 | 5.18233 | 0.84088 | 0.00000 | 0.03277 | 1.19993 | 0.00000 | 0.48723 | 0.00605 |
| **Letmd1** | 0.64156 | 2.84303 | 0.59301 | 0.16223 | 0.01606 | 0.00535 | 3.16342 | 0.00000 | 0.02591 | 6.21255 | 2.13060 | 0.18008 | 2.03983 | 0.00734 | 1.49731 | 0.00404 |
| **Lexm** | 0.01064 | 0.00775 | 0.01355 | 0.28468 | 0.00937 | 0.00737 | 0.00715 | 0.56100 | 0.00679 | 0.11215 | 0.00807 | 0.00573 | 0.00558 | 0.00577 | 0.35021 | 0.00722 |
| **Lgals1** | 0.00000 | 0.37839 | 0.49137 | 3.89261 | 0.65973 | 3.39607 | 0.00500 | 0.00000 | 0.61809 | 2.92031 | 2.18316 | 0.10509 | 1.40742 | 0.00000 | 0.67531 | 3.76455 |
| **Lgals12** | 0.00943 | 0.00753 | 0.01066 | 1.02115 | 0.00980 | 2.44724 | 0.00685 | 0.60083 | 0.00000 | 0.00001 | 0.17124 | 0.00546 | 0.00000 | 0.00760 | 0.00000 | 0.00639 |
| **Lgals2** | 0.00790 | 0.00763 | 0.42082 | 0.97068 | 0.00889 | 0.00526 | 0.00725 | 0.44339 | 2.02170 | 0.00000 | 0.00000 | 0.01376 | 0.10919 | 0.02464 | 0.09177 | 0.02094 |
| **Lgals3** | 2.88275 | 0.12694 | 1.76479 | 1.29613 | 0.02154 | 6.13431 | 0.01389 | 0.04287 | 3.52575 | 4.96148 | 2.65882 | 0.15860 | 0.75955 | 0.06700 | 0.41139 | 0.03003 |
| **Lgals3bp** | 0.01327 | 0.00792 | 0.01216 | 0.00739 | 0.00896 | 0.00657 | 0.00734 | 0.01048 | 0.01184 | 0.95010 | 0.01137 | 0.00543 | 0.01106 | 0.01169 | 0.00868 | 0.00738 |
| **Lgals4** | 0.00901 | 0.00768 | 0.24011 | 0.00942 | 0.00821 | 2.37807 | 0.00689 | 0.01414 | 0.01266 | 0.00266 | 0.00977 | 0.00754 | 0.31186 | 0.01491 | 0.00571 | 0.27679 |
| **Lgals7** | 0.00689 | 0.06897 | 0.01148 | 0.00973 | 0.00914 | 0.00646 | 0.33847 | 0.09943 | 0.01219 | 0.16634 | 0.01246 | 0.00666 | 0.43463 | 0.01335 | 0.00647 | 0.00709 |
| **Lgals8** | 6.45831 | 8.12229 | 6.95419 | 0.54432 | 5.62754 | 8.17690 | 6.82700 | 11.21823 | 0.92672 | 8.59690 | 19.97254 | 6.41095 | 5.28588 | 12.76350 | 4.17526 | 1.82026 |
| **Lgals9** | 5.69046 | 11.84790 | 10.37342 | 0.34011 | 0.03237 | 0.47224 | 3.45552 | 0.00000 | 0.21868 | 2.48451 | 2.29370 | 0.18037 | 4.05472 | 1.52079 | 0.97543 | 0.02801 |
| **Lgalsl** | 3.14392 | 0.05315 | 0.00000 | 4.93038 | 4.35895 | 1.94659 | 0.04294 | 4.46621 | 3.92004 | 3.31969 | 3.67289 | 4.95357 | 1.27753 | 8.26210 | 1.10227 | 0.44832 |
| **Lgi2** | 0.01366 | 0.00785 | 0.01169 | 0.47238 | 0.00938 | 0.00642 | 0.00738 | 0.01175 | 0.51982 | 0.00183 | 0.00855 | 0.00601 | 0.01286 | 0.02435 | 0.46402 | 0.00725 |
| **Lgi3** | 0.03421 | 0.00801 | 0.30346 | 0.12182 | 0.00978 | 0.00582 | 0.00739 | 0.00966 | 0.00990 | 0.05718 | 0.14987 | 0.00780 | 0.00938 | 0.01210 | 0.00866 | 0.00721 |
| **Lgmn** | 4.92564 | 3.87288 | 8.07261 | 3.80953 | 1.57753 | 4.09640 | 8.54569 | 4.35281 | 7.52098 | 20.65441 | 4.98412 | 6.46790 | 11.13859 | 0.07004 | 1.50245 | 4.56891 |
| **Lgr4** | 0.43898 | 2.44975 | 0.47800 | 1.06909 | 2.96183 | 0.96796 | 0.00567 | 0.49870 | 0.44832 | 1.10784 | 0.01292 | 0.61815 | 0.00000 | 5.48918 | 0.00000 | 0.01287 |
| **Lgr5** | 0.06129 | 0.01904 | 0.06884 | 0.01947 | 0.02058 | 0.01263 | 0.03713 | 0.00223 | 0.64067 | 0.03640 | 0.41679 | 0.02974 | 0.11089 | 1.33079 | 0.32131 | 0.02899 |
| **Lhb** | 0.01323 | 0.00803 | 0.01133 | 0.14557 | 0.00930 | 0.00687 | 0.00751 | 0.02515 | 0.13272 | 0.06367 | 0.01108 | 0.03125 | 0.01246 | 0.01251 | 0.01134 | 0.00739 |
| **Lhfp** | 0.01404 | 0.00697 | 0.00000 | 1.44801 | 0.00889 | 3.09417 | 0.00653 | 0.00000 | 0.00000 | 0.00000 | 0.35701 | 1.46138 | 0.00000 | 0.00460 | 0.00000 | 0.00565 |
| **Lhfpl2** | 5.53765 | 7.34617 | 2.91105 | 2.67993 | 5.88475 | 0.01651 | 7.14677 | 0.00000 | 1.93188 | 0.88956 | 0.00000 | 0.03783 | 1.24899 | 0.00000 | 0.67774 | 2.79692 |
| **Lhfpl5** | 0.01713 | 0.00781 | 1.21390 | 0.00432 | 0.00860 | 0.01171 | 0.00719 | 1.76730 | 0.02287 | 0.15766 | 0.88442 | 0.00056 | 0.00000 | 0.00557 | 0.13647 | 0.00683 |
| **Lhpp** | 0.01335 | 0.00819 | 0.01219 | 2.99683 | 0.00915 | 0.00665 | 0.00743 | 0.01243 | 0.00000 | 0.61767 | 0.01148 | 0.00785 | 0.01405 | 0.00193 | 0.16330 | 0.00694 |
| **Lhx2** | 0.00948 | 0.00847 | 0.08570 | 0.66141 | 0.00919 | 0.00795 | 0.00800 | 0.12927 | 0.01938 | 0.06611 | 0.00881 | 0.06582 | 0.26789 | 0.01515 | 0.06141 | 0.00695 |
| **Lhx8** | 0.01307 | 0.00818 | 0.01597 | 2.20423 | 0.00926 | 0.00602 | 0.00750 | 0.01337 | 2.70951 | 0.00258 | 0.00000 | 0.00350 | 0.01303 | 0.02783 | 0.00271 | 0.00686 |
| **Lias** | 9.13181 | 5.26426 | 9.33702 | 6.17755 | 0.01491 | 10.74140 | 6.66599 | 10.60095 | 5.73458 | 16.60044 | 13.47656 | 9.48552 | 5.45951 | 23.38441 | 17.62006 | 4.80446 |
| **Lif** | 0.01427 | 0.00768 | 0.00598 | 0.01477 | 0.03499 | 0.00694 | 0.00709 | 1.69225 | 0.01047 | 0.30092 | 0.00000 | 0.06665 | 0.00914 | 0.02778 | 0.26574 | 0.00735 |
| **Lifr** | 1.83443 | 0.55678 | 0.45583 | 1.26870 | 4.24572 | 0.33869 | 0.12837 | 0.26450 | 0.79214 | 0.11215 | 0.68775 | 0.08569 | 0.21597 | 0.97137 | 0.22334 | 0.25043 |
| **Lig1** | 13.43036 | 1.74054 | 7.38168 | 0.81615 | 8.29104 | 0.01044 | 8.72636 | 12.37278 | 0.00000 | 3.51124 | 7.38270 | 0.86727 | 8.64830 | 0.12047 | 1.90741 | 11.36991 |
| **Lig3** | 7.89742 | 7.65169 | 6.44066 | 1.69704 | 4.84316 | 1.99921 | 7.43362 | 13.74126 | 1.75154 | 12.43855 | 3.09090 | 2.06851 | 4.84185 | 10.86208 | 13.24568 | 3.97249 |
| **Lig4** | 0.01783 | 0.00800 | 0.00520 | 1.12188 | 0.00937 | 0.01284 | 0.00735 | 0.07223 | 1.24707 | 0.00000 | 0.00000 | 0.35920 | 0.02538 | 2.77487 | 3.18815 | 0.00727 |
| **Lilra6** | 0.34485 | 0.00728 | 0.02591 | 0.30912 | 0.00733 | 0.00580 | 0.00708 | 0.00911 | 0.00677 | 0.01435 | 0.01183 | 0.00625 | 0.01237 | 0.38173 | 0.00000 | 0.00697 |
| **Lim2** | 0.01366 | 0.00813 | 0.01213 | 0.05596 | 0.00915 | 0.00654 | 0.00749 | 0.01246 | 0.01169 | 0.09321 | 0.01233 | 0.00783 | 0.03153 | 0.01116 | 0.00955 | 0.00715 |
| **Lima1** | 3.98876 | 0.00000 | 6.21669 | 1.77210 | 4.60690 | 2.99032 | 2.44773 | 4.75366 | 4.39269 | 2.13150 | 2.52354 | 0.17748 | 1.35001 | 0.00000 | 4.31243 | 0.00559 |
| **Limch1** | 0.01315 | 0.00795 | 0.00333 | 3.57057 | 0.00979 | 0.08923 | 0.00755 | 0.00750 | 6.17312 | 0.00000 | 0.18080 | 0.06487 | 0.09501 | 0.00000 | 0.13637 | 0.00650 |
| **Limd1** | 2.86157 | 0.01402 | 2.85126 | 8.33921 | 3.22046 | 2.48968 | 0.51240 | 0.98274 | 3.28465 | 1.93556 | 0.76701 | 0.13545 | 0.32984 | 0.00000 | 5.09376 | 0.00000 |
| **Limd2** | 1.56367 | 0.01652 | 1.71045 | 5.33444 | 0.01911 | 0.13997 | 0.14584 | 3.10717 | 10.40149 | 3.00799 | 3.58607 | 0.05394 | 0.90587 | 2.88295 | 1.66901 | 1.17941 |
| **Lime1** | 1.38971 | 0.00586 | 2.92740 | 3.86953 | 0.00844 | 1.35140 | 0.02861 | 1.78155 | 5.03448 | 0.86145 | 1.40623 | 0.45860 | 0.56001 | 0.02368 | 0.67561 | 1.61712 |
| **Limk1** | 0.05828 | 0.04180 | 0.06236 | 2.48068 | 0.01351 | 2.18480 | 0.00698 | 0.00000 | 1.94544 | 0.02228 | 0.48870 | 0.04409 | 0.01786 | 0.00415 | 0.07617 | 0.00612 |
| **Limk2** | 7.99737 | 0.17647 | 6.53372 | 6.28681 | 2.16164 | 10.90914 | 4.21131 | 11.09513 | 7.74808 | 5.09011 | 8.00496 | 0.42827 | 6.88184 | 1.40947 | 2.69124 | 7.29839 |
| **Lims1** | 1.75042 | 2.17820 | 2.49090 | 6.74495 | 1.07473 | 2.72338 | 3.53021 | 0.35707 | 3.34343 | 3.44749 | 2.16814 | 6.21668 | 4.19669 | 8.44091 | 4.56150 | 1.11726 |
| **Lims2** | 1.53111 | 0.00383 | 3.27777 | 0.16019 | 0.00000 | 3.46393 | 1.97782 | 2.86767 | 0.75202 | 0.94863 | 1.66539 | 0.05169 | 1.00448 | 0.00731 | 0.00000 | 0.00989 |
| **Lin28a** | 0.00000 | 0.00727 | 0.25367 | 1.41296 | 0.00682 | 0.33300 | 0.02583 | 1.97224 | 0.31008 | 1.78091 | 2.29418 | 0.08790 | 2.23256 | 2.47773 | 2.06787 | 1.59753 |
| **Lin28b** | 1.64622 | 5.24448 | 2.64778 | 1.48130 | 7.30694 | 0.00592 | 4.81577 | 0.52333 | 0.67422 | 0.00000 | 1.04382 | 0.01552 | 0.00000 | 6.19983 | 1.05842 | 0.00196 |
| **Lin37** | 8.23326 | 0.54253 | 3.53472 | 2.45732 | 5.82748 | 0.59248 | 1.85671 | 6.65717 | 0.00000 | 4.60515 | 3.57238 | 0.09346 | 4.89699 | 0.06016 | 0.00000 | 5.68388 |
| **Lin52** | 0.26788 | 1.26120 | 0.96324 | 1.39422 | 0.03412 | 1.07312 | 0.04674 | 0.00000 | 5.71945 | 0.11027 | 1.17089 | 6.29260 | 0.00000 | 5.57814 | 6.99155 | 0.33626 |
| **Lin54** | 0.10468 | 2.18468 | 2.26735 | 8.00012 | 3.61469 | 0.00554 | 0.58008 | 2.11864 | 10.05029 | 1.26727 | 1.41156 | 1.06091 | 0.00000 | 0.18838 | 9.51672 | 0.00331 |
| **Lin7b** | 0.01372 | 0.00817 | 0.00969 | 1.51084 | 0.00910 | 0.00661 | 0.00750 | 0.00852 | 0.66873 | 0.00299 | 0.28785 | 0.00583 | 0.00919 | 0.00621 | 0.00000 | 0.00703 |
| **Lin7c** | 1.35345 | 6.22590 | 7.35976 | 9.43232 | 7.76208 | 5.00797 | 6.42098 | 8.24043 | 5.24945 | 14.26250 | 8.22301 | 14.20018 | 3.83131 | 27.87840 | 14.83463 | 3.36113 |
| **Lin9** | 0.07944 | 0.02953 | 0.00000 | 1.45790 | 0.02939 | 0.31934 | 0.04631 | 1.37967 | 1.07931 | 0.00000 | 0.28966 | 0.03932 | 0.00000 | 7.24923 | 3.38766 | 0.00658 |
| **Lincred1** | 0.01379 | 0.00802 | 0.01189 | 0.01049 | 0.00932 | 0.00667 | 0.00750 | 0.01342 | 0.01236 | 0.01426 | 0.01297 | 0.02861 | 0.01274 | 0.01098 | 0.01889 | 0.00741 |
| **Lingo1** | 0.55767 | 0.00822 | 0.00515 | 0.14788 | 0.01718 | 0.00710 | 0.00708 | 0.01170 | 0.00917 | 0.01539 | 0.01349 | 0.00789 | 0.01197 | 0.01157 | 0.00970 | 0.01454 |
| **Lingo2** | 0.01346 | 0.00797 | 0.00971 | 1.23899 | 0.00942 | 0.00666 | 0.00728 | 0.01062 | 0.00000 | 0.01009 | 0.01103 | 0.00624 | 0.01314 | 0.39410 | 0.00000 | 0.00725 |
| **Lins1** | 0.00667 | 0.00748 | 1.61272 | 0.65464 | 1.93674 | 0.00559 | 0.00726 | 0.37490 | 0.00000 | 0.07081 | 0.00716 | 0.00396 | 1.02187 | 0.09027 | 0.09630 | 0.02208 |
| **Lipe** | 0.27858 | 0.03883 | 0.00590 | 0.03198 | 0.00719 | 0.00695 | 0.00708 | 0.00795 | 0.09992 | 1.00439 | 0.09222 | 0.00446 | 0.00991 | 0.01394 | 0.02375 | 0.00674 |
| **Lipg** | 0.01796 | 0.00822 | 0.01036 | 0.01048 | 0.01054 | 0.00674 | 0.00750 | 0.01356 | 0.01258 | 0.01085 | 0.01184 | 0.00709 | 0.01259 | 0.22486 | 0.00468 | 0.03703 |
| **Liph** | 3.13757 | 0.06098 | 1.23913 | 0.44671 | 0.00000 | 1.59961 | 0.04974 | 0.34781 | 0.25947 | 0.33446 | 0.11785 | 0.03568 | 0.00000 | 0.07831 | 0.00332 | 0.02590 |
| **Lipo2** | 0.01275 | 0.00801 | 0.01216 | 0.08401 | 0.01002 | 0.00719 | 0.16447 | 0.01227 | 0.14043 | 0.07023 | 0.05246 | 0.00679 | 0.09936 | 0.00716 | 0.09400 | 0.00803 |

| | | | | | | | | | | | | | | | | |
|---|---|---|---|---|---|---|---|---|---|---|---|---|---|---|---|---|
| **Lipo3** | 0.48612 | 0.01593 | 0.19696 | 2.22123 | 0.12129 | 0.00454 | 2.31225 | 0.00000 | 4.99739 | 0.72451 | 0.61585 | 0.02190 | 0.02089 | 0.00000 | 0.28556 | 0.00621 |
| **Lipt1** | 1.54376 | 0.68929 | 2.27635 | 1.09610 | 0.00179 | 1.85595 | 0.00342 | 1.33255 | 0.00000 | 2.44903 | 1.12700 | 0.07740 | 3.60999 | 1.10259 | 2.45691 | 0.01598 |
| **Lipt2** | 0.76512 | 0.00782 | 0.00497 | 0.00481 | 0.01702 | 0.00685 | 0.05784 | 0.00940 | 0.41783 | 0.01165 | 0.00914 | 0.03810 | 0.01834 | 0.01308 | 0.00815 | 0.00679 |
| **Litaf** | 5.49180 | 4.80279 | 14.09004 | 8.31242 | 8.49628 | 12.18366 | 7.11668 | 10.75182 | 3.13889 | 19.23715 | 10.58005 | 4.15417 | 9.52114 | 10.67288 | 14.20354 | 7.24917 |
| **Lix1** | 0.24200 | 0.05881 | 0.63500 | 0.42535 | 0.08652 | 0.05551 | 0.23217 | 0.04925 | 0.36012 | 0.17289 | 1.04476 | 0.13780 | 0.08248 | 0.61478 | 0.33474 | 0.16614 |
| **Lix1l** | 0.01377 | 0.00809 | 0.01183 | 0.20981 | 0.00929 | 0.00672 | 0.00753 | 0.01373 | 0.00810 | 0.02093 | 0.01341 | 0.02986 | 0.01289 | 0.01129 | 0.00707 | 0.00739 |
| **Llgl1** | 4.96918 | 1.22690 | 6.24539 | 1.13021 | 6.22181 | 4.28767 | 3.64266 | 12.52092 | 3.00174 | 2.33140 | 3.55193 | 0.50049 | 1.34561 | 0.11136 | 4.83691 | 1.35403 |
| **Llgl2** | 0.83371 | 0.00445 | 0.00000 | 1.27411 | 2.01408 | 6.92638 | 0.86776 | 5.67162 | 3.33558 | 8.83696 | 0.31683 | 0.08106 | 3.79529 | 0.00253 | 0.00000 | 8.02233 |
| **Llph** | 22.58106 | 11.11157 | 18.17945 | 14.47539 | 14.84713 | 6.25220 | 7.87234 | 19.82263 | 20.68465 | 26.41947 | 21.91059 | 9.68497 | 23.68320 | 20.03742 | 13.49046 | 5.19146 |
| **Lman1** | 4.72034 | 0.60042 | 3.98709 | 0.13488 | 4.21627 | 3.10741 | 1.34319 | 2.29379 | 0.47788 | 0.76150 | 1.57063 | 1.81374 | 2.34499 | 2.75187 | 0.85557 | 1.57155 |
| **Lman1l** | 0.01273 | 0.02980 | 0.04365 | 0.02782 | 0.00882 | 0.00575 | 0.00740 | 0.00060 | 0.72423 | 0.52691 | 0.35458 | 0.01436 | 0.00801 | 0.00634 | 0.12689 | 0.00695 |
| **Lman2** | 13.84097 | 8.09978 | 17.79003 | 7.49391 | 7.72188 | 7.45045 | 8.94462 | 16.30774 | 14.96165 | 12.04201 | 10.81697 | 3.03292 | 16.39250 | 3.96140 | 6.83337 | 9.67067 |
| **Lman2l** | 3.13746 | 3.36583 | 0.39949 | 2.98945 | 0.01263 | 0.00130 | 0.00219 | 1.94995 | 0.08732 | 0.26385 | 0.11681 | 5.21621 | 1.06086 | 1.36784 | 1.02469 | 0.00000 |
| **Lmbr1** | 1.14649 | 0.00720 | 1.03933 | 0.85201 | 0.00934 | 0.01064 | 0.00595 | 0.00980 | 1.61154 | 3.55441 | 0.09191 | 0.01172 | 1.07680 | 0.43670 | 1.69353 | 0.87090 |
| **Lmbr1l** | 0.05276 | 0.00823 | 0.05762 | 3.23956 | 0.02824 | 0.00669 | 0.04776 | 0.00000 | 3.85676 | 0.64831 | 1.78354 | 0.05231 | 0.17575 | 0.10430 | 0.46567 | 0.30197 |
| **Lmbrd1** | 2.06061 | 0.01966 | 4.57513 | 0.96390 | 0.08156 | 0.03317 | 1.68912 | 3.27152 | 0.32482 | 4.42179 | 5.41319 | 14.08531 | 3.88230 | 10.24630 | 10.26968 | 0.53245 |
| **Lmbrd2** | 0.00129 | 0.00819 | 0.13746 | 1.31423 | 0.00734 | 0.00559 | 0.00793 | 0.19523 | 0.11882 | 0.00000 | 0.01055 | 0.03233 | 0.00336 | 0.00993 | 0.16274 | 1.80797 |
| **Lmf1** | 0.14075 | 1.76527 | 0.58006 | 0.00000 | 0.22680 | 0.00559 | 1.44418 | 0.00000 | 0.01091 | 0.00000 | 0.74016 | 0.00167 | 0.52816 | 0.00987 | 0.00000 | 0.06885 |
| **Lmf2** | 6.30020 | 1.68023 | 0.80643 | 6.25812 | 0.00000 | 0.52758 | 0.00000 | 3.80467 | 1.34575 | 2.83489 | 1.32150 | 0.11885 | 3.23446 | 0.00000 | 0.00000 | 2.44887 |
| **Lmln** | 0.01376 | 0.00787 | 0.01116 | 0.33035 | 0.00942 | 0.01266 | 0.00736 | 0.01144 | 0.06233 | 0.00557 | 0.01085 | 0.00652 | 0.01265 | 0.00653 | 0.71477 | 0.00728 |
| **Lmna** | 15.24860 | 0.10090 | 1.09471 | 3.88526 | 3.40274 | 9.44924 | 0.13432 | 8.45679 | 12.98581 | 7.85660 | 4.06664 | 0.43329 | 10.18150 | 0.00000 | 0.00000 | 12.87875 |
| **Lmnb1** | 3.63285 | 3.70200 | 3.24831 | 2.70738 | 2.67408 | 2.87000 | 0.04325 | 2.03204 | 5.45639 | 0.00000 | 2.95922 | 0.15202 | 1.58080 | 3.89274 | 4.03765 | 3.64884 |
| **Lmnb2** | 2.95826 | 1.65440 | 3.65572 | 0.43092 | 5.93355 | 0.91677 | 1.63881 | 3.61642 | 0.06890 | 1.80773 | 2.79029 | 0.28440 | 3.32209 | 0.26016 | 2.75996 | 6.94779 |
| **Lmntd2** | 0.42132 | 0.00709 | 0.00312 | 0.31679 | 0.01852 | 0.00813 | 1.89431 | 0.00687 | 0.00664 | 0.00000 | 0.09680 | 0.00565 | 0.19030 | 0.01056 | 0.00306 | 0.00731 |
| **Lmo1** | 0.01178 | 0.00788 | 0.01140 | 1.06405 | 0.00987 | 0.00720 | 0.07011 | 0.00387 | 0.41411 | 0.90124 | 0.27446 | 0.01196 | 0.25752 | 0.00617 | 0.00000 | 0.00780 |
| **Lmo2** | 1.72071 | 2.92068 | 0.78204 | 0.00000 | 8.64857 | 0.00365 | 0.26828 | 0.56994 | 0.00718 | 0.85343 | 0.19793 | 0.00373 | 0.00000 | 0.00997 | 0.00000 | 0.04890 |
| **Lmo4** | 0.44637 | 0.00574 | 0.00000 | 1.18277 | 1.83040 | 0.98837 | 0.10644 | 0.00396 | 0.67045 | 1.07510 | 0.01620 | 0.00000 | 0.00594 | 0.52062 | 0.00000 | 0.00623 |
| **Lmo7** | 0.56520 | 0.00261 | 3.04838 | 5.24929 | 0.18059 | 0.00727 | 2.75096 | 12.60252 | 3.45005 | 8.63103 | 6.23774 | 0.56422 | 2.37668 | 5.76756 | 3.51703 | 0.00957 |
| **Lmod1** | 0.01322 | 0.00792 | 0.00796 | 1.22059 | 0.00878 | 0.00679 | 0.00706 | 0.00000 | 0.41867 | 1.32338 | 0.53090 | 0.04330 | 0.00723 | 0.00557 | 0.00462 | 0.00689 |
| **Lmod3** | 0.01009 | 0.00776 | 0.00352 | 0.00001 | 0.00953 | 0.00694 | 0.00722 | 0.80541 | 0.02305 | 0.73978 | 0.20884 | 0.02963 | 0.00592 | 0.02114 | 1.29814 | 0.00755 |
| **Lmtk2** | 1.61023 | 0.01519 | 0.00000 | 4.09353 | 0.01312 | 4.84980 | 1.46054 | 2.00802 | 1.68964 | 2.83863 | 2.08026 | 1.25883 | 1.28550 | 0.52240 | 1.89501 | 0.00205 |
| **Lmx1a** | 0.01275 | 0.00747 | 0.01033 | 4.99956 | 0.00903 | 0.00576 | 0.00706 | 0.01410 | 3.55683 | 0.56119 | 0.00001 | 0.00901 | 0.01091 | 0.04299 | 0.33574 | 0.00606 |
| **Lmx1b** | 0.01179 | 0.00807 | 0.01447 | 0.06369 | 0.01028 | 0.00704 | 0.00750 | 0.01360 | 0.00874 | 0.33511 | 0.01420 | 0.00640 | 0.33172 | 0.01119 | 0.00731 | 0.00774 |
| **Lncenc1** | 1.28056 | 3.26036 | 8.17702 | 0.00000 | 0.09084 | 0.00519 | 1.89578 | 0.49162 | 0.42504 | 0.00177 | 0.00000 | 0.00861 | 0.00000 | 0.01266 | 0.00000 | 0.00911 |
| **Lnpep** | 0.01501 | 0.00811 | 0.04031 | 3.87439 | 0.00891 | 0.00787 | 0.00715 | 0.00298 | 0.36919 | 0.01304 | 0.37680 | 0.00000 | 0.00834 | 0.86000 | 2.88105 | 0.00649 |
| **Lnpk1** | 0.57507 | 0.04271 | 1.50029 | 2.67721 | 0.02298 | 0.01189 | 0.01551 | 2.41033 | 2.36242 | 0.00000 | 1.54666 | 2.84441 | 1.12858 | 5.44646 | 0.57115 | 0.12729 |
| **Lnx1** | 0.02913 | 1.52472 | 0.77471 | 0.08371 | 0.01004 | 0.45726 | 0.38575 | 0.00733 | 0.12003 | 0.00000 | 0.01721 | 0.05312 | 0.37636 | 1.06626 | 0.13693 | 0.15770 |
| **Lnx2** | 0.52688 | 0.05758 | 1.00210 | 2.45836 | 0.12355 | 3.57093 | 0.00575 | 1.38144 | 2.88019 | 0.04106 | 0.57535 | 0.15296 | 1.06761 | 0.00000 | 0.10048 | 0.00406 |
| **LOC10050470** | 0.01378 | 0.00789 | 0.01206 | 0.02729 | 0.00933 | 0.00666 | 0.00753 | 0.01359 | 0.01118 | 0.01419 | 0.01264 | 0.00797 | 0.01274 | 0.01656 | 0.02563 | 0.00741 |
| **LOC10050502** | 0.01619 | 0.01888 | 0.08098 | 0.54773 | 0.02123 | 0.00617 | 0.01766 | 0.01612 | 0.68849 | 0.01154 | 0.17822 | 0.04586 | 0.01278 | 0.21757 | 0.00640 | 0.00726 |
| **LOC10086161** | 0.07763 | 0.00803 | 0.00888 | 0.31811 | 0.00787 | 0.00689 | 0.00744 | 0.01449 | 0.00956 | 0.01669 | 0.03853 | 0.01893 | 0.01215 | 0.01078 | 0.00701 | 0.01441 |
| **LOC10105607** | 0.62326 | 2.14289 | 0.51738 | 0.31369 | 1.06910 | 0.22349 | 0.69184 | 0.76838 | 0.12976 | 0.53930 | 0.24672 | 2.24768 | 0.40271 | 5.50255 | 0.77382 | 0.29266 |
| **LOC10263481** | 1.30536 | 0.07713 | 1.14442 | 3.06404 | 0.41897 | 0.12765 | 0.07298 | 0.00000 | 5.28177 | 0.17048 | 8.40633 | 0.10352 | 0.42704 | 2.51502 | 0.16544 | 0.46466 |
| **LOC10524239** | 0.01345 | 0.00814 | 0.26459 | 2.08269 | 0.01067 | 0.00560 | 0.00751 | 0.16225 | 0.84112 | 0.31598 | 0.00426 | 0.00494 | 0.00708 | 0.00270 | 0.36555 | 0.00719 |
| **LOC10524257** | 0.23791 | 0.13591 | 0.36876 | 0.22692 | 0.14967 | 0.07858 | 0.09701 | 0.05418 | 0.44789 | 0.25492 | 1.47726 | 0.17203 | 0.43557 | 0.70867 | 0.29539 | 0.07838 |
| **LOC10524586** | 0.09152 | 0.17244 | 0.00375 | 0.00257 | 0.01495 | 0.00696 | 0.00727 | 0.54554 | 0.79256 | 0.22787 | 0.58974 | 0.31519 | 0.00344 | 0.18799 | 0.46790 | 0.00757 |
| **LOC10524605** | 0.01856 | 0.00801 | 0.01167 | 0.01103 | 0.01059 | 0.00668 | 0.00751 | 0.01333 | 0.01209 | 0.01376 | 0.01265 | 0.00794 | 0.01273 | 0.01740 | 0.03251 | 0.00746 |
| **LOC10524650** | 3.93553 | 0.27897 | 0.33798 | 0.03790 | 4.00253 | 0.01882 | 0.00501 | 0.21896 | 0.04852 | 0.36222 | 0.05589 | 0.01235 | 0.00972 | 0.03007 | 0.00000 | 1.10911 |
| **LOC10602923** | 0.00877 | 0.00819 | 0.01149 | 9.32401 | 0.00914 | 0.00409 | 0.00751 | 0.04006 | 19.48173 | 0.01217 | 0.26298 | 0.00000 | 0.04876 | 0.02554 | 0.26128 | 0.00458 |
| **LOC106740** | 0.56081 | 0.01993 | 0.09088 | 0.38559 | 0.01002 | 0.00548 | 0.00773 | 0.37095 | 0.24454 | 0.88919 | 0.00829 | 0.00487 | 0.01213 | 0.01077 | 0.54378 | 0.00680 |
| **LOC215458** | 0.26713 | 0.00772 | 0.02928 | 0.01421 | 0.00815 | 0.00569 | 0.00723 | 0.09630 | 0.03744 | 0.36000 | 0.02001 | 0.03840 | 0.07853 | 0.18640 | 0.54506 | 0.00696 |
| **LOC666331** | 0.01373 | 0.00791 | 0.01236 | 0.00576 | 0.00933 | 0.00661 | 0.00752 | 0.01370 | 0.13078 | 0.01167 | 0.01171 | 0.00746 | 0.01269 | 0.00760 | 0.07498 | 0.00739 |
| **Lockd** | 0.86960 | 0.00759 | 0.00415 | 0.00495 | 0.01781 | 0.00709 | 0.00678 | 0.00983 | 0.51048 | 0.05432 | 0.00828 | 0.00620 | 0.01204 | 0.01632 | 0.02460 | 0.00676 |
| **Lonp1** | 11.24581 | 4.12825 | 10.83755 | 8.28661 | 10.93357 | 7.19571 | 8.34036 | 14.30725 | 9.42181 | 18.85757 | 10.62671 | 6.94524 | 9.24548 | 0.19230 | 9.55048 | 9.45450 |
| **Lonp2** | 4.65699 | 0.37313 | 4.71929 | 9.20173 | 1.60801 | 0.00331 | 6.12616 | 10.89626 | 8.07181 | 8.47270 | 7.63376 | 8.30590 | 3.98381 | 5.24462 | 6.06918 | 5.63611 |
| **Lonrf1** | 1.91951 | 0.92048 | 1.14334 | 4.54061 | 0.00687 | 1.22642 | 2.47995 | 0.13328 | 0.00000 | 0.73095 | 0.01642 | 0.01784 | 0.00000 | 0.00000 | 0.80389 | 0.01898 |
| **Lonrf2** | 0.00921 | 0.00742 | 0.01128 | 0.02737 | 0.00966 | 0.02171 | 0.00715 | 0.50288 | 0.01151 | 0.00838 | 0.00945 | 0.00654 | 0.00565 | 0.01898 | 0.05388 | 0.00738 |
| **Lonrf3** | 0.86300 | 0.00560 | 0.00000 | 10.56488 | 0.00570 | 0.00318 | 1.09370 | 5.39754 | 11.92260 | 4.80296 | 3.50488 | 0.07247 | 0.00000 | 2.34273 | 3.15502 | 0.00318 |
| **Loxhd1** | 0.01290 | 0.00800 | 0.01382 | 0.12586 | 0.00990 | 0.00706 | 0.00764 | 0.01359 | 0.00950 | 0.56412 | 0.01386 | 0.00612 | 0.07386 | 0.00597 | 0.26284 | 0.00771 |
| **Loxl1** | 0.01025 | 0.00799 | 0.01086 | 0.63619 | 0.00948 | 0.00728 | 0.00727 | 0.00946 | 0.01778 | 3.66153 | 0.26804 | 0.09854 | 0.31312 | 0.00848 | 0.01295 | 0.00751 |
| **Loxl2** | 1.20273 | 0.08000 | 1.02577 | 5.96154 | 0.37584 | 0.12180 | 0.16474 | 0.00000 | 10.66053 | 0.00000 | 9.75425 | 1.35307 | 0.50332 | 4.12647 | 0.40289 | 0.15300 |
| **Lpar1** | 3.11005 | 3.23238 | 2.18527 | 0.00000 | 0.00000 | 0.72052 | 0.00000 | 2.07364 | 0.56733 | 2.50581 | 3.69375 | 2.46180 | 1.44546 | 4.99226 | 2.56377 | 0.22517 |

| | | | | | | | | | | | | | | | | |
|---|---|---|---|---|---|---|---|---|---|---|---|---|---|---|---|---|
| **Lpar2** | 0.21417 | 0.00753 | 0.19250 | 0.01927 | 0.00710 | 0.00549 | 0.00739 | 0.43153 | 0.01173 | 0.00000 | 0.00946 | 0.00714 | 0.57823 | 0.02837 | 0.02895 | 0.50010 |
| **Lpar5** | 0.02200 | 0.00813 | 0.10574 | 0.00847 | 0.01263 | 0.20120 | 0.00756 | 0.46941 | 0.01162 | 0.33694 | 0.01091 | 0.00614 | 0.35464 | 0.00743 | 0.09811 | 0.00753 |
| **Lpar6** | 0.00758 | 0.00751 | 0.01054 | 1.64143 | 0.00770 | 0.00592 | 0.00722 | 0.01047 | 2.89298 | 0.00272 | 0.00000 | 0.01143 | 0.00795 | 0.00421 | 0.53306 | 1.29829 |
| **Lpcat1** | 2.51410 | 1.76483 | 0.00000 | 0.00000 | 0.00323 | 2.90435 | 0.03617 | 1.06705 | 0.01053 | 0.64381 | 0.00000 | 0.02105 | 1.15201 | 0.00678 | 0.58435 | 1.05333 |
| **Lpcat2** | 0.01052 | 0.00881 | 0.00672 | 0.84841 | 0.01003 | 0.00816 | 0.00784 | 0.04567 | 0.56153 | 0.26791 | 1.05770 | 0.03132 | 0.26184 | 0.00375 | 0.09049 | 0.00735 |
| **Lpcat3** | 16.88330 | 6.51817 | 15.43220 | 0.54704 | 6.39619 | 6.90283 | 11.05521 | 15.39013 | 1.14546 | 9.19998 | 11.58445 | 2.90272 | 16.19145 | 4.02426 | 4.40015 | 6.00759 |
| **Lpcat4** | 0.95453 | 0.00743 | 0.73235 | 0.00480 | 0.17187 | 6.21671 | 0.00528 | 0.27506 | 0.12028 | 1.87291 | 0.03041 | 0.01200 | 1.56806 | 0.05134 | 0.01477 | 0.00346 |
| **Lpgat1** | 2.60505 | 0.00000 | 3.38865 | 6.29666 | 0.22548 | 9.15266 | 6.69633 | 3.62616 | 1.21108 | 9.14818 | 5.43520 | 4.42584 | 6.64975 | 9.93538 | 10.93296 | 3.02555 |
| **Lpin1** | 1.79463 | 0.00306 | 0.21103 | 0.43278 | 0.01200 | 2.23441 | 5.46926 | 0.48756 | 0.00304 | 3.46728 | 0.70440 | 0.09616 | 0.00000 | 0.02079 | 0.00000 | 0.91925 |
| **Lpin2** | 0.69990 | 0.00742 | 0.00000 | 2.20057 | 0.01653 | 0.01148 | 0.00610 | 0.00986 | 2.21033 | 1.68611 | 0.00000 | 0.04693 | 0.83934 | 0.17780 | 2.93815 | 0.00626 |
| **Lpin3** | 1.65000 | 0.00252 | 1.89433 | 0.99056 | 0.00508 | 3.21502 | 4.93483 | 3.65452 | 0.77578 | 0.00000 | 1.06039 | 0.07961 | 0.00000 | 0.00517 | 0.62131 | 0.00311 |
| **Lpl** | 0.07459 | 0.00809 | 0.04043 | 1.90784 | 0.00798 | 0.01409 | 0.00733 | 0.03712 | 0.99654 | 0.06677 | 0.09054 | 0.02694 | 0.05212 | 0.12241 | 0.37384 | 0.03593 |
| **Lpo** | 0.01335 | 0.00806 | 0.01228 | 0.00735 | 0.00907 | 0.00648 | 0.00734 | 0.01062 | 0.01189 | 0.79145 | 0.01137 | 0.00591 | 0.01089 | 0.00827 | 0.04948 | 0.00721 |
| **Lpp** | 5.16969 | 0.03162 | 3.02075 | 0.86929 | 2.87663 | 1.75265 | 0.15551 | 0.94904 | 0.80199 | 2.26924 | 0.70091 | 0.18918 | 2.64103 | 0.38407 | 0.36547 | 3.52967 |
| **Lppos** | 0.97361 | 0.00725 | 0.14144 | 0.91304 | 0.01694 | 0.00483 | 0.00649 | 0.18203 | 1.34178 | 0.43543 | 0.05106 | 0.08753 | 0.00663 | 0.00259 | 0.23728 | 0.00633 |
| **Lrba** | 1.69729 | 1.71213 | 1.99723 | 2.17603 | 0.31320 | 5.07125 | 1.64113 | 4.01905 | 2.15610 | 4.13826 | 2.77613 | 1.18501 | 6.73287 | 0.62475 | 1.17960 | 0.42021 |
| **Lrch1** | 0.82565 | 0.00670 | 0.64371 | 0.99867 | 0.00517 | 0.00463 | 0.00685 | 0.00899 | 0.73291 | 0.19895 | 0.01277 | 0.04405 | 0.00868 | 0.00535 | 0.57288 | 0.09259 |
| **Lrch3** | 0.44459 | 1.04485 | 0.00000 | 0.12353 | 0.01690 | 0.00806 | 0.00652 | 0.01313 | 0.00938 | 0.02348 | 0.01254 | 0.01728 | 0.01756 | 0.01063 | 0.04538 | 0.00673 |
| **Lrcol1** | 0.01276 | 0.00817 | 0.01166 | 0.00577 | 0.00888 | 0.00645 | 0.00747 | 0.01384 | 0.01095 | 1.35402 | 0.01237 | 0.00544 | 0.01010 | 0.02967 | 0.20245 | 0.00708 |
| **Lrfn1** | 0.22378 | 0.03993 | 0.00000 | 0.04726 | 0.08093 | 0.04478 | 0.00717 | 0.01452 | 0.01211 | 0.27441 | 0.06580 | 0.00653 | 0.01749 | 0.01058 | 0.00788 | 0.00752 |
| **Lrfn4** | 0.03680 | 0.00829 | 0.00870 | 0.10313 | 0.01460 | 0.06732 | 0.00727 | 0.01611 | 0.01076 | 0.16559 | 0.25241 | 0.00588 | 0.22212 | 0.00998 | 0.00000 | 0.00761 |
| **Lrguk** | 0.01628 | 0.00792 | 0.12022 | 0.08328 | 0.00897 | 0.00596 | 0.00747 | 0.01322 | 0.00961 | 0.12328 | 0.01220 | 0.00840 | 0.01156 | 0.00989 | 0.01908 | 0.00731 |
| **Lrif1** | 0.00292 | 0.00341 | 1.58526 | 1.90108 | 0.00760 | 0.00475 | 0.00421 | 0.33640 | 2.71824 | 0.00000 | 1.58882 | 13.22588 | 0.55825 | 9.05774 | 3.51625 | 0.00485 |
| **Lrig1** | 0.26044 | 0.00683 | 0.98343 | 0.95149 | 0.01650 | 0.62891 | 0.33468 | 0.95638 | 0.18435 | 0.13782 | 0.03284 | 0.02821 | 0.00000 | 0.00403 | 1.34379 | 0.01378 |
| **Lrig2** | 0.04351 | 0.07706 | 0.44451 | 2.10275 | 0.00787 | 0.00876 | 0.00722 | 0.00000 | 7.38228 | 0.04902 | 0.48134 | 0.07043 | 0.41514 | 3.43569 | 1.83311 | 0.00617 |
| **Lrig3** | 0.01039 | 0.00648 | 0.00659 | 0.00219 | 0.00955 | 0.00674 | 2.05077 | 0.01001 | 0.51935 | 0.00928 | 0.00536 | 0.20010 | 0.01052 | 0.00532 | 0.32995 | 0.00722 |
| **Lrmp** | 0.16491 | 0.00793 | 0.00782 | 0.01023 | 0.00953 | 0.00688 | 0.00736 | 0.01391 | 0.01256 | 0.03923 | 0.01284 | 0.00742 | 0.01252 | 0.00878 | 0.03335 | 0.00727 |
| **Lrp1** | 0.00834 | 0.00584 | 0.00696 | 4.22457 | 3.14064 | 1.96103 | 0.00688 | 0.00349 | 0.50842 | 0.46515 | 0.11735 | 0.25988 | 0.30303 | 0.00000 | 0.37526 | 0.00555 |
| **Lrp10** | 2.77351 | 0.03432 | 1.98297 | 0.45265 | 0.06592 | 0.05632 | 2.36264 | 2.62645 | 1.21515 | 3.56089 | 0.74649 | 0.09985 | 4.14894 | 0.11682 | 0.04874 | 6.95347 |
| **Lrp11** | 1.61965 | 2.76176 | 0.99774 | 0.22099 | 4.95157 | 0.08257 | 1.93533 | 2.76787 | 0.01613 | 1.13406 | 1.12256 | 7.47900 | 1.29607 | 0.90622 | 0.70462 | 0.90685 |
| **Lrp12** | 0.22397 | 0.00783 | 0.30643 | 0.55912 | 0.00669 | 0.40189 | 0.00712 | 0.00955 | 0.04242 | 0.50584 | 0.00003 | 0.01823 | 0.00696 | 2.09708 | 1.23820 | 0.35662 |
| **Lrp2** | 0.00000 | 5.59585 | 9.21626 | 0.44649 | 0.04388 | 15.54434 | 10.41246 | 2.46805 | 0.30068 | 0.00224 | 0.00000 | 0.01890 | 19.90247 | 0.00527 | 0.62053 | 4.54809 |
| **Lrp3** | 0.01203 | 0.00771 | 0.34910 | 0.00864 | 0.00840 | 0.00918 | 0.00716 | 0.01479 | 0.01201 | 0.51422 | 0.01174 | 0.00582 | 0.00967 | 0.01246 | 0.00482 | 0.00747 |
| **Lrp4** | 0.01215 | 0.01871 | 0.08626 | 0.13708 | 0.00893 | 0.00842 | 0.01731 | 0.00761 | 0.32749 | 0.04384 | 0.28843 | 0.03714 | 0.03615 | 0.15225 | 0.19238 | 0.00715 |
| **Lrp5** | 0.01335 | 0.00872 | 0.01205 | 0.57052 | 0.00857 | 0.23349 | 0.00793 | 0.05802 | 1.13206 | 0.63378 | 0.00000 | 0.02850 | 0.00642 | 0.01106 | 0.00152 | 0.15411 |
| **Lrp6** | 0.99584 | 0.00628 | 1.56594 | 9.32549 | 0.01103 | 0.66177 | 0.17841 | 0.85553 | 2.23056 | 2.94777 | 2.40823 | 1.02440 | 0.87136 | 8.95062 | 6.57354 | 1.63265 |
| **Lrp8** | 3.35263 | 0.01380 | 0.51301 | 0.52881 | 3.81891 | 2.99596 | 0.00440 | 0.65415 | 0.17473 | 0.00000 | 0.40684 | 0.04138 | 0.88585 | 0.00568 | 0.87492 | 0.00881 |
| **Lrpap1** | 13.13297 | 10.93424 | 34.96778 | 4.69095 | 10.76116 | 10.62368 | 21.04426 | 40.94514 | 6.12764 | 24.17563 | 27.82008 | 13.73731 | 42.67399 | 12.05368 | 4.43220 | 14.94268 |
| **Lrpprc** | 9.06137 | 12.76069 | 13.72600 | 0.46750 | 19.34580 | 6.56382 | 7.27156 | 11.65893 | 1.02782 | 10.00276 | 7.80029 | 8.61262 | 9.62824 | 7.38594 | 7.98038 | 4.19628 |
| **Lrr1** | 6.41725 | 6.61640 | 10.32590 | 16.80245 | 7.55423 | 7.62623 | 3.44819 | 19.09360 | 13.11042 | 39.02812 | 21.70382 | 21.91976 | 16.92145 | 15.61938 | 14.06492 | 0.26341 |
| **Lrrc1** | 1.30981 | 0.00605 | 0.00361 | 1.03183 | 0.00540 | 0.00692 | 0.00641 | 0.01762 | 0.00000 | 0.01585 | 0.01174 | 0.00702 | 0.01116 | 0.01667 | 0.02978 | 0.00581 |
| **Lrrc10b** | 0.02251 | 0.00859 | 0.01811 | 0.00502 | 0.01168 | 0.00731 | 0.01838 | 0.01675 | 0.32597 | 0.10467 | 0.00945 | 0.00739 | 0.01400 | 0.01225 | 0.00811 | 0.00732 |
| **Lrrc14** | 3.96342 | 1.44955 | 3.10624 | 2.31047 | 0.15502 | 0.00208 | 2.89116 | 2.85516 | 2.45108 | 0.26653 | 0.37023 | 0.04864 | 2.87051 | 0.00000 | 1.68776 | 1.27709 |
| **Lrrc14b** | 0.51287 | 0.07873 | 0.26667 | 0.39860 | 0.04121 | 0.01373 | 0.06093 | 0.05151 | 0.24861 | 0.35522 | 0.79251 | 0.02080 | 0.20233 | 0.95474 | 0.14511 | 0.03554 |
| **Lrrc15** | 1.23535 | 0.00697 | 1.32485 | 3.54124 | 0.01341 | 0.00394 | 0.00596 | 0.00282 | 0.00000 | 1.23131 | 0.13912 | 0.16902 | 0.01394 | 2.23401 | 4.66235 | 0.01307 |
| **Lrrc16a** | 0.69280 | 3.87589 | 0.76865 | 7.37649 | 3.87337 | 0.03207 | 2.70408 | 0.62585 | 1.28232 | 0.25463 | 0.90728 | 0.03489 | 0.33473 | 0.00000 | 0.00000 | 0.10835 |
| **Lrrc17** | 0.03482 | 0.06808 | 0.04739 | 2.48167 | 0.00867 | 0.00597 | 0.00670 | 0.00000 | 1.16712 | 1.09877 | 0.94870 | 0.11055 | 0.00524 | 0.16379 | 2.73264 | 0.05087 |
| **Lrrc2** | 2.43587 | 3.37743 | 1.33225 | 3.34867 | 2.53994 | 0.95870 | 4.03674 | 0.00000 | 6.18177 | 0.08740 | 3.70041 | 0.20516 | 0.44977 | 2.08298 | 1.25929 | 0.12902 |
| **Lrrc20** | 1.90184 | 0.00516 | 1.25145 | 0.54490 | 0.05132 | 0.01244 | 0.02539 | 0.04944 | 0.22458 | 0.40534 | 0.26698 | 0.00086 | 0.67284 | 0.18968 | 0.27543 | 0.02746 |
| **Lrrc23** | 0.01002 | 0.00644 | 0.00254 | 0.00417 | 1.94873 | 0.00850 | 0.00691 | 0.65181 | 0.00986 | 1.39243 | 0.00767 | 0.01023 | 0.51287 | 0.01726 | 0.00000 | 0.00704 |
| **Lrrc25** | 0.00890 | 0.00749 | 0.00852 | 0.09511 | 0.00938 | 0.00673 | 0.00711 | 0.38770 | 0.32398 | 0.83590 | 0.00823 | 0.03566 | 0.00574 | 0.01883 | 0.00000 | 0.00709 |
| **Lrrc27** | 0.05546 | 0.00702 | 0.00465 | 0.23888 | 0.00848 | 0.00731 | 1.19777 | 0.01012 | 0.00692 | 0.01649 | 0.01330 | 0.00672 | 0.01067 | 0.01214 | 0.00879 | 0.00720 |
| **Lrrc28** | 0.07271 | 0.68345 | 2.08158 | 3.92503 | 1.17425 | 0.55931 | 0.74829 | 2.77531 | 1.74708 | 1.20581 | 0.41457 | 0.06504 | 0.00000 | 1.14480 | 0.28288 | 0.00521 |
| **Lrrc3** | 0.05458 | 0.02925 | 0.23450 | 0.17822 | 0.03290 | 0.02632 | 0.03766 | 0.01293 | 0.29601 | 0.13093 | 0.48602 | 0.03242 | 0.13652 | 0.26400 | 0.17550 | 0.01447 |
| **Lrrc31** | 0.08834 | 0.01842 | 0.09272 | 0.05467 | 0.02055 | 0.03210 | 0.06826 | 0.00255 | 0.17165 | 0.03952 | 0.61174 | 0.07134 | 0.16221 | 0.26863 | 0.10863 | 0.02889 |
| **Lrrc32** | 0.01375 | 0.00796 | 0.01208 | 0.53333 | 0.00932 | 0.00660 | 0.00751 | 0.01368 | 0.14875 | 0.01377 | 0.01246 | 0.00781 | 0.01274 | 0.01487 | 0.01306 | 0.00732 |
| **Lrrc34** | 0.75883 | 0.00768 | 0.00451 | 0.09285 | 0.01177 | 0.00709 | 0.01715 | 0.01602 | 0.00786 | 0.02286 | 0.01176 | 0.00740 | 0.01140 | 0.01061 | 0.01425 | 0.00664 |
| **Lrrc39** | 0.50674 | 0.69306 | 0.63989 | 0.70094 | 0.55204 | 0.30037 | 0.16747 | 0.27975 | 1.07368 | 0.38088 | 0.55965 | 1.01325 | 1.05755 | 0.80565 | 0.72246 | 0.28811 |
| **Lrrc4** | 0.01612 | 0.00748 | 0.32541 | 1.77710 | 0.00871 | 0.00538 | 0.00706 | 0.01048 | 1.22466 | 0.00000 | 0.13324 | 0.02132 | 0.00979 | 0.00174 | 0.95951 | 0.00668 |
| **Lrrc40** | 5.09882 | 5.54215 | 5.88899 | 10.17530 | 5.21531 | 0.01878 | 5.96438 | 8.21745 | 2.75146 | 5.15864 | 5.56707 | 1.32214 | 6.05869 | 1.01801 | 10.88303 | 3.12016 |
| **Lrrc41** | 1.43532 | 1.66183 | 2.96464 | 2.83121 | 0.03038 | 0.40757 | 3.35357 | 2.33949 | 0.93493 | 3.89986 | 0.59940 | 2.61439 | 1.41052 | 1.91270 | 0.45852 | 2.09254 |
| **Lrrc42** | 4.05899 | 2.09645 | 5.92594 | 11.28643 | 0.26803 | 3.72042 | 1.80435 | 22.38334 | 23.08863 | 20.83035 | 15.29884 | 3.63761 | 6.09575 | 8.34013 | 11.13847 | 0.17940 |

| | | | | | | | | | | | | | | | | |
|---|---|---|---|---|---|---|---|---|---|---|---|---|---|---|---|---|
| **Lrrc43** | 0.01361 | 0.00795 | 0.01214 | 0.23698 | 0.00910 | 0.00660 | 0.00731 | 0.01109 | 0.10685 | 0.36382 | 0.01318 | 0.00653 | 0.01239 | 0.00774 | 0.05137 | 0.00776 |
| **Lrrc45** | 0.00000 | 1.38754 | 3.34713 | 1.10183 | 3.01503 | 2.21286 | 2.43899 | 2.09932 | 0.47922 | 0.70989 | 1.27089 | 0.00620 | 2.58314 | 0.01634 | 0.00000 | 2.00973 |
| **Lrrc46** | 0.01909 | 0.00779 | 0.24018 | 0.59786 | 0.00914 | 0.00581 | 0.00731 | 0.03407 | 0.93140 | 0.00000 | 0.00678 | 0.00386 | 0.02320 | 0.02589 | 0.78510 | 0.00718 |
| **Lrrc47** | 1.39723 | 0.38056 | 1.73809 | 0.20043 | 0.02431 | 0.00463 | 0.51249 | 2.02070 | 0.35725 | 0.49647 | 1.32921 | 0.05740 | 0.69501 | 0.04237 | 2.45052 | 0.67525 |
| **Lrrc48** | 0.01267 | 0.00807 | 0.01367 | 0.40212 | 0.01015 | 0.00713 | 0.00763 | 0.01431 | 0.00828 | 0.18943 | 0.02932 | 0.01745 | 0.17326 | 0.01179 | 0.03144 | 0.00770 |
| **Lrrc49** | 0.01245 | 0.00792 | 0.09308 | 0.64511 | 0.00901 | 0.00570 | 0.00739 | 0.01032 | 0.00000 | 0.86573 | 0.02882 | 0.00532 | 0.01083 | 0.00759 | 0.02327 | 0.00741 |
| **Lrrc51** | 0.25017 | 0.00763 | 1.19431 | 0.07174 | 0.00535 | 3.00509 | 0.00700 | 0.11451 | 0.24513 | 0.10866 | 0.00000 | 0.50649 | 0.00013 | 0.83181 | 1.39228 | 0.00553 |
| **Lrrc56** | 0.04800 | 0.00788 | 0.00000 | 1.24980 | 0.01085 | 3.11937 | 0.01725 | 0.01260 | 3.43101 | 0.35705 | 0.02033 | 0.00000 | 0.06160 | 0.00151 | 0.09264 | 0.00588 |
| **Lrrc57** | 2.00333 | 1.86401 | 1.95574 | 2.31720 | 1.75852 | 3.30761 | 1.21553 | 5.29016 | 3.43663 | 2.16879 | 1.36589 | 0.18427 | 0.13662 | 4.49308 | 7.94657 | 0.00124 |
| **Lrrc58** | 7.11636 | 4.96761 | 11.02240 | 4.42092 | 3.87177 | 2.94089 | 3.84636 | 2.54912 | 3.66085 | 12.41174 | 11.02795 | 14.55988 | 7.35445 | 20.91902 | 17.03577 | 5.90007 |
| **Lrrc59** | 28.04224 | 16.48544 | 22.28567 | 6.17859 | 19.74906 | 14.71820 | 13.66339 | 24.34190 | 11.00547 | 13.72989 | 9.54837 | 6.26545 | 21.71308 | 10.19159 | 10.33481 | 15.39372 |
| **Lrrc61** | 0.06731 | 0.04973 | 0.18060 | 0.16460 | 0.02690 | 0.13007 | 0.06773 | 0.04004 | 0.02798 | 0.43407 | 0.13582 | 0.02340 | 0.77307 | 0.06232 | 0.13133 | 0.03625 |
| **Lrrc73** | 0.01116 | 0.00832 | 0.01317 | 0.60534 | 0.01016 | 0.00686 | 0.00748 | 0.01408 | 2.07697 | 0.01737 | 0.01726 | 0.02443 | 0.44662 | 0.00996 | 0.00669 | 0.00739 |
| **Lrrc74a** | 0.01375 | 0.00794 | 0.01194 | 0.00658 | 0.00929 | 0.00662 | 0.00749 | 0.01305 | 0.07958 | 0.01200 | 0.01186 | 0.00792 | 0.01975 | 0.01442 | 0.05594 | 0.00738 |
| **Lrrc74b** | 0.01941 | 0.04113 | 0.03459 | 0.27411 | 0.00973 | 0.00824 | 0.03753 | 0.01795 | 0.18391 | 0.58034 | 0.09956 | 0.00548 | 0.01928 | 0.04472 | 0.18187 | 0.00736 |
| **Lrrc75a** | 0.01551 | 0.00769 | 0.04098 | 0.15213 | 0.01031 | 4.17992 | 0.00657 | 0.00706 | 0.00636 | 0.00000 | 0.03254 | 0.00704 | 0.65671 | 0.02144 | 0.04146 | 0.00543 |
| **Lrrc75b** | 2.01388 | 0.00413 | 1.27103 | 0.00267 | 0.15833 | 0.01513 | 0.03423 | 0.33890 | 0.01627 | 1.73692 | 0.74754 | 3.28954 | 0.00000 | 0.02810 | 0.05209 | 2.51388 |
| **Lrrc8a** | 1.00868 | 0.00515 | 0.16363 | 3.28919 | 0.01374 | 2.29520 | 0.65789 | 2.34063 | 6.10829 | 1.22324 | 1.28635 | 1.78159 | 0.00000 | 0.82062 | 1.77187 | 0.00391 |
| **Lrrc8b** | 0.06727 | 0.00769 | 0.28895 | 0.23324 | 0.00732 | 0.00790 | 0.00745 | 0.01293 | 0.00585 | 0.03304 | 0.00692 | 0.07320 | 0.01045 | 0.00762 | 1.80069 | 0.00709 |
| **Lrrc8d** | 1.19068 | 0.20186 | 1.92356 | 3.47146 | 2.00222 | 2.02992 | 0.36310 | 0.74505 | 0.45827 | 0.37897 | 1.26814 | 0.26603 | 0.45732 | 1.25103 | 1.03888 | 0.00360 |
| **Lrrc8e** | 0.07975 | 0.00761 | 0.02969 | 4.52862 | 0.00876 | 0.02247 | 0.00641 | 0.18625 | 2.76113 | 1.50628 | 1.31539 | 0.20555 | 0.05749 | 1.11978 | 1.73082 | 0.04098 |
| **Lrrcc1** | 1.25855 | 0.00553 | 1.41066 | 1.29868 | 0.00368 | 0.00476 | 0.04716 | 1.30091 | 0.55422 | 0.43114 | 0.71497 | 0.00000 | 0.00848 | 0.00345 | 1.43822 | 0.01294 |
| **Lrrfip1** | 0.46583 | 3.93598 | 3.97897 | 7.26403 | 0.02661 | 8.04421 | 0.03506 | 0.03992 | 0.04360 | 0.00000 | 0.44224 | 0.08083 | 3.14981 | 2.62086 | 7.27852 | 2.04548 |
| **Lrrfip2** | 4.73649 | 5.04026 | 5.94853 | 2.53838 | 7.54825 | 7.38438 | 4.12875 | 7.07243 | 2.11139 | 6.68842 | 4.54180 | 1.24762 | 1.51323 | 5.00932 | 6.58289 | 0.02358 |
| **Lrriq3** | 0.01379 | 0.00774 | 0.01188 | 1.00735 | 0.00941 | 0.00647 | 0.00740 | 0.01227 | 0.01272 | 0.00719 | 0.01147 | 0.00725 | 0.01277 | 0.01130 | 0.29527 | 0.00721 |
| **Lrrn4** | 0.00206 | 0.00818 | 0.93619 | 0.50091 | 0.01682 | 0.00933 | 0.01876 | 0.29356 | 0.39943 | 2.26256 | 1.30744 | 0.00546 | 0.00000 | 0.09803 | 0.00000 | 0.66127 |
| **Lrrn4cl** | 0.11465 | 0.02910 | 0.32318 | 0.59525 | 0.06078 | 0.01825 | 0.06805 | 0.02089 | 0.34449 | 0.11893 | 0.88580 | 0.08040 | 0.26112 | 0.47706 | 0.21070 | 0.05003 |
| **Lrrtm2** | 0.03801 | 0.00795 | 0.01079 | 0.01054 | 0.00830 | 0.00674 | 0.00749 | 0.01491 | 0.01241 | 0.01505 | 0.01304 | 0.00799 | 0.01268 | 0.01126 | 0.01579 | 0.00732 |
| **Lrsam1** | 0.58590 | 0.09062 | 0.12875 | 1.55706 | 0.40099 | 0.00504 | 3.25041 | 1.94019 | 0.00000 | 0.00041 | 1.65151 | 1.28623 | 2.57447 | 0.00000 | 0.00000 | 0.01251 |
| **Lrwd1** | 10.19742 | 0.66147 | 3.10556 | 4.25975 | 2.00885 | 0.00239 | 0.40333 | 9.15336 | 5.88318 | 4.85308 | 5.70001 | 0.40990 | 8.82861 | 0.00719 | 0.27261 | 4.38466 |
| **Lsg1** | 11.14725 | 8.11346 | 10.65312 | 2.75542 | 8.79596 | 5.93498 | 7.93449 | 13.42377 | 5.12973 | 11.19824 | 4.11890 | 6.91919 | 8.24048 | 0.60825 | 0.37105 | 4.76342 |
| **Lsm1** | 7.88064 | 0.14798 | 2.39962 | 5.24849 | 4.50354 | 3.73901 | 1.32241 | 2.77569 | 6.64839 | 8.30441 | 2.64652 | 0.94622 | 4.74959 | 0.75660 | 0.56612 | 4.90363 |
| **Lsm10** | 4.04313 | 0.00000 | 1.68802 | 5.90434 | 4.81685 | 5.02214 | 0.52255 | 0.78957 | 3.58140 | 3.52382 | 1.21226 | 0.03725 | 0.00000 | 0.00000 | 2.90615 | 2.27988 |
| **Lsm11** | 0.99814 | 0.01662 | 0.05376 | 2.07751 | 0.01422 | 2.37952 | 3.02818 | 0.09478 | 0.44005 | 0.77407 | 2.26823 | 0.05778 | 0.10698 | 1.49330 | 0.36546 | 1.85503 |
| **Lsm12** | 1.02812 | 0.00000 | 5.35440 | 3.70883 | 5.21532 | 3.15362 | 3.88627 | 8.02562 | 7.37301 | 14.12452 | 15.04740 | 1.75687 | 9.95059 | 0.00000 | 3.80567 | 5.53762 |
| **Lsm14a** | 10.49669 | 7.97514 | 3.20456 | 22.40358 | 10.42913 | 4.60667 | 1.24905 | 5.67926 | 12.42597 | 4.72932 | 4.88789 | 12.86016 | 8.34576 | 7.27210 | 10.64737 | 6.24517 |
| **Lsm14b** | 1.07510 | 0.01952 | 0.00000 | 10.03258 | 0.01241 | 0.58729 | 0.03911 | 0.16555 | 7.05435 | 1.61972 | 1.54587 | 0.10665 | 0.14839 | 0.33326 | 1.26949 | 0.35400 |
| **Lsm2** | 3.51049 | 0.72977 | 5.58543 | 7.34906 | 3.21227 | 0.00738 | 0.48640 | 1.37916 | 9.44871 | 15.77962 | 6.61061 | 0.43406 | 3.98034 | 3.38408 | 3.11327 | 0.48541 |
| **Lsm3** | 11.31615 | 5.40929 | 6.98528 | 0.21148 | 10.98453 | 5.23826 | 2.69255 | 3.31177 | 0.14370 | 5.20371 | 5.34236 | 0.43809 | 8.65817 | 7.58194 | 2.11578 | 2.97750 |
| **Lsm4** | 3.50082 | 0.38682 | 0.83780 | 1.05349 | 1.41151 | 0.02695 | 0.08229 | 0.03697 | 2.96886 | 13.80042 | 6.90197 | 0.09809 | 19.15460 | 0.00000 | 0.00000 | 10.94333 |
| **Lsm5** | 6.78175 | 0.81666 | 4.20528 | 0.00000 | 6.28350 | 1.59869 | 0.96007 | 1.01651 | 1.13165 | 6.02999 | 12.37053 | 1.27465 | 5.30125 | 9.42378 | 4.54155 | 3.54735 |
| **Lsm6** | 8.83921 | 0.57091 | 7.57983 | 5.05709 | 3.96233 | 2.21517 | 1.71127 | 1.68522 | 5.09333 | 2.58996 | 5.07767 | 0.45702 | 6.21742 | 2.08902 | 1.11697 | 3.57608 |
| **Lsm7** | 6.00296 | 0.49525 | 5.31327 | 5.55629 | 5.26769 | 1.18685 | 0.24653 | 0.42742 | 2.51518 | 4.82229 | 10.02069 | 0.46164 | 2.47576 | 0.05463 | 0.31664 | 0.92075 |
| **Lsm8** | 8.08647 | 3.96808 | 5.80558 | 8.01611 | 8.05437 | 1.72718 | 3.06140 | 6.15044 | 1.88958 | 5.44622 | 9.85563 | 3.74041 | 9.56292 | 4.06596 | 6.14075 | 7.31370 |
| **Lsp1** | 0.01192 | 0.00769 | 0.01064 | 5.74977 | 0.00858 | 0.00587 | 0.00700 | 0.00885 | 1.96915 | 3.39669 | 0.00000 | 0.03461 | 0.01079 | 0.00000 | 0.36238 | 0.00640 |
| **Lsr** | 8.37062 | 4.37849 | 7.35294 | 1.14642 | 1.89721 | 4.22186 | 3.02078 | 3.57050 | 0.29027 | 0.00000 | 1.32528 | 0.09474 | 6.47726 | 0.00000 | 1.09239 | 7.97114 |
| **Lss** | 1.45163 | 6.25048 | 4.02636 | 4.66750 | 0.29311 | 0.41849 | 2.05308 | 7.28000 | 1.39105 | 5.62278 | 4.09185 | 0.09241 | 1.75653 | 4.73394 | 6.02855 | 0.30602 |
| **Lst1** | 0.01049 | 0.00727 | 0.00001 | 2.66968 | 0.00906 | 0.00715 | 0.00510 | 0.00000 | 4.12820 | 0.30023 | 5.07179 | 0.07662 | 0.10822 | 0.63625 | 0.70397 | 0.00590 |
| **Lta** | 0.01303 | 0.00769 | 0.01111 | 0.81461 | 0.00889 | 0.00633 | 0.00731 | 0.00999 | 0.91097 | 1.17928 | 0.00001 | 0.02235 | 0.01047 | 0.01089 | 0.23486 | 0.00690 |
| **Lta4h** | 9.26263 | 4.98897 | 5.00186 | 0.18915 | 5.49971 | 5.55252 | 3.70748 | 3.11140 | 0.00105 | 0.10426 | 0.00000 | 0.06426 | 6.36606 | 0.08285 | 0.72782 | 6.41948 |
| **Ltb** | 0.01530 | 0.00748 | 0.03897 | 0.04177 | 0.00878 | 0.00595 | 0.00695 | 0.00000 | 0.15479 | 1.78251 | 0.78560 | 0.00000 | 0.01152 | 0.00895 | 0.00000 | 0.00719 |
| **Ltbp1** | 0.01380 | 0.00790 | 0.01164 | 0.01031 | 0.00925 | 0.00671 | 0.00742 | 0.01243 | 0.01227 | 0.07154 | 0.01317 | 0.03645 | 0.01305 | 0.01143 | 0.01038 | 0.00774 |
| **Ltbp2** | 0.00000 | 0.00562 | 0.64904 | 0.41725 | 3.56036 | 0.00878 | 0.00652 | 0.00449 | 0.00000 | 0.93086 | 0.02147 | 0.16155 | 0.86690 | 0.00986 | 0.00598 | 0.00661 |
| **Ltbp3** | 0.01227 | 0.00782 | 0.00778 | 0.85033 | 0.00862 | 0.00678 | 0.00692 | 0.00000 | 2.92878 | 1.70960 | 1.25591 | 0.05082 | 0.00911 | 0.00266 | 0.03013 | 0.01413 |
| **Ltbp4** | 2.62144 | 0.00534 | 0.00000 | 4.26926 | 0.01904 | 1.38803 | 0.01373 | 1.49250 | 1.46623 | 4.31295 | 0.57694 | 0.22023 | 2.53961 | 0.66282 | 0.07145 | 2.43683 |
| **Ltf** | 0.01039 | 0.00767 | 0.01233 | 0.00456 | 0.00808 | 0.00648 | 0.00721 | 0.00880 | 0.01013 | 12.57147 | 0.00686 | 0.00000 | 0.01022 | 0.01361 | 0.00433 | 0.00735 |
| **Ltk** | 0.01186 | 0.00693 | 0.00343 | 0.57091 | 0.00930 | 0.00715 | 1.05331 | 0.00000 | 0.06749 | 0.00000 | 0.47722 | 0.01224 | 0.00723 | 0.00469 | 0.47068 | 0.00694 |
| **Ltn1** | 4.65674 | 4.65977 | 4.79241 | 4.07487 | 3.81948 | 0.03157 | 0.77609 | 7.76938 | 5.32943 | 0.83733 | 4.63251 | 15.81282 | 0.73690 | 5.70023 | 9.56439 | 0.01762 |
| **Ltv1** | 12.85364 | 10.68319 | 15.21429 | 15.32265 | 11.03526 | 8.92110 | 9.39456 | 22.15230 | 20.41046 | 26.05977 | 21.21644 | 15.41479 | 16.29610 | 21.47067 | 19.75970 | 9.57515 |
| **Luc7l** | 9.47368 | 6.03939 | 11.48892 | 8.05962 | 4.00094 | 1.67334 | 8.95290 | 17.27069 | 7.89978 | 18.39975 | 11.61877 | 5.38310 | 11.15388 | 17.42615 | 9.18419 | 2.79632 |
| **Luc7l2** | 7.19890 | 1.52811 | 2.22668 | 12.08956 | 6.71641 | 3.25116 | 1.18238 | 1.74808 | 17.46869 | 5.66880 | 13.25341 | 4.77536 | 3.10035 | 11.81462 | 10.41225 | 1.43946 |
| **Luc7l3** | 9.01977 | 9.82666 | 9.82587 | 7.81870 | 8.46206 | 5.25601 | 5.22192 | 12.56639 | 11.80936 | 11.85077 | 4.91473 | 14.22293 | 5.68824 | 16.02767 | 12.32659 | 4.14917 |

| | | | | | | | | | | | | | | | | |
|---|---|---|---|---|---|---|---|---|---|---|---|---|---|---|---|---|
| **Lurap1** | 0.16680 | 0.00755 | 0.00576 | 0.00468 | 0.00744 | 0.00704 | 0.00715 | 0.00122 | 0.30542 | 0.13517 | 0.49730 | 0.01386 | 0.00717 | 0.02821 | 0.00000 | 0.29749 |
| **Lurap1l** | 0.00778 | 0.00764 | 0.11903 | 0.26344 | 0.02064 | 0.02652 | 0.00603 | 0.57778 | 0.41176 | 0.00066 | 0.00589 | 0.00485 | 1.92664 | 0.01651 | 0.00000 | 0.32693 |
| **Luzp1** | 2.25058 | 0.87541 | 6.41713 | 10.23874 | 7.15712 | 2.14085 | 2.04107 | 6.62015 | 9.49484 | 8.00679 | 1.55235 | 2.71245 | 2.51655 | 1.52559 | 7.04015 | 0.02649 |
| **Luzp2** | 0.00751 | 0.00707 | 0.00661 | 0.52064 | 0.00942 | 0.00642 | 0.00694 | 0.73848 | 0.00484 | 0.09919 | 0.01307 | 1.70689 | 0.00529 | 0.00927 | 0.00000 | 0.00694 |
| **Luzp4** | 0.01372 | 0.00799 | 0.01251 | 0.08335 | 0.00949 | 0.00667 | 0.00754 | 0.01359 | 0.02136 | 0.00985 | 0.01380 | 0.00782 | 0.01577 | 0.00853 | 0.14342 | 0.00744 |
| **Lxn** | 1.44037 | 0.00469 | 1.21187 | 0.06463 | 0.74395 | 0.00796 | 0.94275 | 0.00178 | 0.06628 | 0.45536 | 0.01830 | 0.01240 | 0.00677 | 0.02015 | 0.29049 | 1.65692 |
| **Ly6a** | 9.27321 | 2.07287 | 12.41221 | 4.36362 | 2.59895 | 0.53312 | 2.83797 | 0.27170 | 1.69443 | 3.23685 | 0.04019 | 0.16169 | 7.06396 | 0.01522 | 0.00000 | 4.51299 |
| **Ly6c1** | 2.11276 | 0.00582 | 0.35738 | 0.00400 | 0.02044 | 0.00386 | 0.15230 | 0.03857 | 0.73394 | 0.37371 | 0.26150 | 0.03299 | 1.67693 | 0.09041 | 0.00000 | 0.18772 |
| **Ly6c2** | 0.31223 | 0.00795 | 0.00682 | 0.02697 | 0.00706 | 0.00671 | 0.00740 | 0.03083 | 0.01158 | 0.02904 | 0.01167 | 0.00822 | 0.22229 | 0.01223 | 0.01030 | 0.10360 |
| **Ly6e** | 4.85327 | 2.83708 | 7.36363 | 1.47618 | 5.75546 | 0.00000 | 4.61201 | 4.14188 | 0.51867 | 3.75452 | 0.00000 | 0.20608 | 1.76448 | 0.00000 | 0.03306 | 0.72394 |
| **Ly6f** | 0.20665 | 0.00690 | 1.13319 | 0.01726 | 1.64992 | 0.34731 | 0.00222 | 0.52117 | 0.01070 | 0.00000 | 0.00782 | 0.00000 | 9.90710 | 0.01948 | 0.00000 | 2.06863 |
| **Ly6g** | 0.04668 | 0.00830 | 0.02909 | 0.07146 | 0.01665 | 0.00800 | 0.01862 | 0.04258 | 0.00990 | 0.00846 | 0.02865 | 0.00839 | 0.18151 | 0.01280 | 0.00916 | 0.11399 |
| **Ly6g5b** | 1.80972 | 0.00571 | 0.60565 | 0.34479 | 0.00368 | 0.00767 | 0.00569 | 0.00539 | 2.13275 | 0.79183 | 0.00000 | 0.45970 | 0.30777 | 0.03615 | 0.00000 | 0.01223 |
| **Ly6g5c** | 0.01209 | 0.00836 | 0.01002 | 0.34227 | 0.00923 | 0.00688 | 0.00812 | 0.05100 | 0.04099 | 0.31678 | 0.00895 | 0.04603 | 0.01083 | 0.00636 | 0.48221 | 0.00733 |
| **Ly6g6c** | 0.27045 | 0.00714 | 1.25735 | 0.01101 | 0.00500 | 0.00764 | 0.00613 | 0.08158 | 0.01219 | 0.00055 | 0.01115 | 0.00597 | 1.30254 | 0.01797 | 0.01056 | 0.01276 |
| **Ly6g6e** | 2.89105 | 0.32720 | 0.33405 | 0.14409 | 1.67638 | 0.00295 | 0.00505 | 0.61809 | 0.24623 | 0.43178 | 0.31101 | 0.10213 | 0.28638 | 0.01026 | 0.05186 | 0.24471 |
| **Ly6i** | 0.27413 | 0.00831 | 0.16101 | 0.69569 | 0.00715 | 0.00657 | 0.07424 | 0.29560 | 0.22388 | 0.03686 | 0.00927 | 0.00850 | 0.06245 | 0.00952 | 0.11710 | 0.01361 |
| **Ly75** | 0.17812 | 0.00827 | 0.00641 | 0.27014 | 0.01754 | 0.00674 | 0.00724 | 0.01062 | 0.00743 | 0.00610 | 0.01257 | 0.01711 | 0.01252 | 0.00684 | 0.52163 | 0.00735 |
| **Ly96** | 0.01132 | 0.89991 | 0.06694 | 0.11450 | 0.00804 | 0.05658 | 1.13315 | 0.00000 | 0.35554 | 0.09979 | 3.35637 | 0.19017 | 0.02966 | 3.11827 | 0.00000 | 0.00615 |
| **Lyar** | 18.58910 | 6.49810 | 12.34033 | 20.40349 | 11.01479 | 10.03032 | 9.53108 | 22.00323 | 24.06699 | 31.40369 | 21.25899 | 10.85804 | 18.86561 | 3.11412 | 6.70713 | 9.64039 |
| **Lyn** | 1.23067 | 0.00652 | 0.56321 | 0.48442 | 0.00421 | 0.00816 | 0.00682 | 0.55505 | 0.05583 | 1.04381 | 0.86103 | 0.01026 | 0.01633 | 0.00489 | 0.03581 | 0.57741 |
| **Lypd1** | 0.33615 | 0.03888 | 0.23824 | 2.44004 | 0.22269 | 0.03011 | 0.07742 | 0.00000 | 4.03531 | 0.02557 | 2.75381 | 0.15292 | 0.05730 | 1.04588 | 0.07095 | 0.06987 |
| **Lypd3** | 0.00946 | 0.00828 | 0.95380 | 1.96538 | 0.00707 | 0.06573 | 0.00741 | 0.01348 | 0.00000 | 0.34870 | 0.01125 | 0.02910 | 0.02087 | 0.00885 | 0.00424 | 0.35492 |
| **Lypd5** | 0.01367 | 0.00781 | 0.01147 | 0.01045 | 0.00935 | 0.00668 | 0.00744 | 0.01222 | 0.01261 | 0.01187 | 0.01223 | 0.00726 | 0.01294 | 0.13505 | 0.01480 | 0.00745 |
| **Lypd6** | 0.01358 | 0.00753 | 0.01100 | 0.95895 | 0.00955 | 0.00627 | 0.00728 | 0.01068 | 0.01215 | 0.05460 | 0.00846 | 0.00426 | 0.01309 | 0.00866 | 1.38872 | 0.00732 |
| **Lypd8** | 0.01478 | 0.00854 | 0.01063 | 0.01066 | 0.00911 | 0.00681 | 0.00809 | 0.07132 | 0.01226 | 0.01717 | 0.01211 | 0.00901 | 0.00982 | 0.01344 | 0.00985 | 0.00737 |
| **Lypla1** | 5.08615 | 5.64836 | 2.92139 | 4.03111 | 2.70428 | 1.57898 | 7.00950 | 11.68265 | 3.19965 | 13.93474 | 6.75231 | 18.31726 | 3.28814 | 12.22637 | 10.90380 | 1.86692 |
| **Lypla2** | 5.09292 | 0.01241 | 0.00000 | 0.18785 | 0.05625 | 0.72460 | 0.00000 | 4.80525 | 0.05367 | 6.27722 | 1.92446 | 0.12775 | 6.06934 | 0.01011 | 0.01228 | 4.91260 |
| **Lyplal1** | 0.05005 | 0.00769 | 0.00932 | 1.58626 | 0.00808 | 0.00676 | 0.00739 | 0.01284 | 0.00000 | 0.16539 | 0.02088 | 0.00717 | 0.01281 | 0.01015 | 0.37563 | 0.00714 |
| **Lyrm1** | 1.35495 | 2.09989 | 4.26143 | 0.47433 | 2.70173 | 0.01998 | 2.38070 | 1.05893 | 2.01481 | 1.26456 | 0.46867 | 0.04419 | 3.59191 | 1.06617 | 0.07847 | 0.00890 |
| **Lyrm2** | 0.38453 | 0.00190 | 3.01757 | 1.65050 | 4.00409 | 4.72133 | 2.75260 | 0.07682 | 0.03637 | 0.28755 | 9.37283 | 0.02908 | 0.84971 | 1.25495 | 3.68786 | 0.19967 |
| **Lyrm4** | 1.25725 | 1.63656 | 2.65429 | 2.94992 | 3.01722 | 0.10048 | 2.43943 | 1.27696 | 0.06349 | 0.75891 | 1.97860 | 0.43326 | 3.05750 | 1.01796 | 0.00000 | 0.04485 |
| **Lyrm5** | 3.08406 | 0.46868 | 3.83881 | 0.88543 | 3.43218 | 3.18506 | 7.06044 | 1.78438 | 0.92494 | 0.20963 | 2.23088 | 3.38462 | 2.24483 | 0.24555 | 2.34095 | 2.38313 |
| **Lyrm7** | 0.61042 | 0.00786 | 0.94985 | 0.00672 | 0.03403 | 0.13005 | 0.00633 | 0.46817 | 0.20595 | 0.81898 | 1.08107 | 0.00000 | 0.00389 | 1.09676 | 0.04078 | 0.00640 |
| **Lyrm9** | 1.19114 | 0.07951 | 1.47248 | 1.01789 | 0.02533 | 0.26108 | 0.08608 | 0.05185 | 0.27420 | 0.12349 | 0.73166 | 0.00978 | 1.15692 | 0.66744 | 0.00000 | 0.00450 |
| **Lysmd1** | 0.00000 | 0.55120 | 0.07924 | 7.28711 | 0.00813 | 4.14531 | 0.00577 | 0.00549 | 12.10917 | 0.47805 | 0.00000 | 0.27824 | 1.38782 | 0.71936 | 0.52054 | 1.48261 |
| **Lysmd2** | 0.01200 | 0.02764 | 0.32700 | 0.46515 | 0.07002 | 0.00947 | 1.36078 | 0.00000 | 0.00466 | 0.47715 | 0.88732 | 0.15345 | 0.01007 | 0.01429 | 0.00000 | 0.00707 |
| **Lysmd3** | 0.01601 | 0.00801 | 0.55773 | 0.53977 | 0.01041 | 3.96706 | 0.00749 | 0.86735 | 0.06939 | 0.07355 | 0.26140 | 0.01686 | 0.00000 | 0.01199 | 0.44766 | 0.01329 |
| **Lysmd4** | 1.24908 | 0.00465 | 0.01816 | 3.26293 | 0.00550 | 0.00569 | 2.55048 | 0.70509 | 0.63943 | 0.82727 | 0.10663 | 0.27540 | 2.30362 | 0.00719 | 0.47852 | 0.35855 |
| **Lyst** | 0.01061 | 0.00754 | 0.01613 | 0.36908 | 0.00962 | 0.00624 | 0.00724 | 0.17678 | 0.00607 | 0.04010 | 0.01014 | 0.00596 | 0.00747 | 0.37209 | 0.05126 | 0.02208 |
| **Lyz1** | 0.01311 | 0.00796 | 0.01179 | 0.84163 | 0.00907 | 0.00642 | 0.00724 | 0.01023 | 0.62860 | 1.05490 | 0.00772 | 0.01452 | 0.01142 | 0.00533 | 0.09457 | 0.00727 |
| **Lyz2** | 0.01296 | 0.00803 | 0.01162 | 0.05924 | 0.00888 | 0.00671 | 0.00737 | 0.01082 | 0.14491 | 1.33940 | 0.01003 | 0.01667 | 0.01150 | 0.00643 | 0.18849 | 0.00745 |
| **Lzic** | 2.49456 | 2.87492 | 1.73263 | 4.53578 | 1.67658 | 0.00208 | 1.50399 | 2.83679 | 6.96635 | 0.00000 | 1.51213 | 0.24802 | 0.49501 | 0.20397 | 1.94677 | 0.46278 |
| **Lztfl1** | 0.96826 | 1.43254 | 2.23958 | 1.58728 | 0.00376 | 0.00387 | 0.18900 | 0.90997 | 7.63750 | 1.56336 | 0.94719 | 11.97119 | 0.00000 | 6.73317 | 7.29979 | 0.01790 |
| **Lztr1** | 1.84450 | 1.50644 | 1.82683 | 3.84976 | 4.76764 | 0.02365 | 1.60150 | 7.76627 | 4.13803 | 3.76022 | 7.56104 | 0.04858 | 5.22607 | 0.04823 | 2.91977 | 2.47761 |
| **Lzts2** | 0.20826 | 0.00840 | 0.00000 | 0.10220 | 0.01677 | 0.55096 | 0.03955 | 1.54453 | 0.70010 | 7.52840 | 0.93859 | 0.00000 | 0.01035 | 0.00380 | 0.02895 | 0.01459 |
| **Lzts3** | 0.05621 | 0.02904 | 0.01869 | 0.25020 | 0.00812 | 0.02675 | 0.02759 | 0.01055 | 0.58613 | 0.03184 | 0.04880 | 0.00527 | 0.02117 | 0.14505 | 0.14557 | 0.01450 |
| **M6pr** | 11.63514 | 8.47065 | 12.35403 | 6.33532 | 11.06229 | 4.68768 | 10.88321 | 16.02788 | 9.24984 | 10.27898 | 9.14447 | 13.43528 | 9.44205 | 16.48919 | 13.02579 | 2.51130 |
| **Maats1** | 1.70705 | 0.00603 | 0.00349 | 0.01055 | 0.00466 | 0.00667 | 0.00608 | 0.01010 | 0.01239 | 0.01436 | 0.01225 | 0.03703 | 0.01139 | 0.01110 | 0.01677 | 0.00550 |
| **Mab21l3** | 0.00979 | 0.00780 | 0.04555 | 0.01046 | 0.00955 | 0.01666 | 0.00725 | 0.14715 | 0.01222 | 0.06335 | 0.01175 | 0.00702 | 0.00906 | 0.01164 | 0.00816 | 0.00747 |
| **Macf1** | 10.37543 | 3.66875 | 6.77463 | 4.60011 | 4.13386 | 3.86318 | 3.39933 | 13.61127 | 10.11535 | 14.13704 | 8.98729 | 0.95702 | 5.13520 | 11.91251 | 4.45848 | 2.71407 |
| **Macrod1** | 0.01323 | 0.00810 | 0.01756 | 0.00789 | 0.00909 | 0.00751 | 0.00752 | 0.01213 | 0.03389 | 0.06759 | 0.05849 | 0.00857 | 0.04726 | 0.01149 | 0.01080 | 0.00736 |
| **Macrod2** | 2.10208 | 2.07124 | 0.48861 | 2.56954 | 0.99098 | 0.00303 | 0.09567 | 0.01001 | 1.19825 | 0.00389 | 0.00000 | 0.03397 | 0.08898 | 0.00009 | 0.21685 | 0.00359 |
| **Mad1l1** | 7.18521 | 1.22721 | 4.51947 | 3.54960 | 10.53037 | 3.91905 | 2.32296 | 8.55762 | 1.61604 | 1.90654 | 7.56702 | 0.60331 | 2.86400 | 0.00190 | 0.10489 | 2.98694 |
| **Mad2l1** | 10.26076 | 11.34290 | 8.78504 | 12.38838 | 12.68324 | 12.65020 | 4.46657 | 14.03239 | 10.65331 | 13.57899 | 9.85455 | 10.14448 | 4.86637 | 16.67967 | 11.78075 | 4.76656 |
| **Mad2l1bp** | 4.17804 | 6.24559 | 10.19763 | 21.24817 | 3.10031 | 5.49610 | 6.73564 | 20.54944 | 30.43064 | 31.16987 | 12.09300 | 15.56040 | 15.73507 | 14.50003 | 30.25491 | 7.58894 |
| **Mad2l2** | 12.41523 | 7.31025 | 6.96355 | 6.54899 | 6.48450 | 9.05664 | 8.72662 | 7.82544 | 11.46386 | 27.85759 | 10.05284 | 4.91779 | 5.77499 | 0.00000 | 1.26140 | 1.37494 |
| **Madd** | 0.10600 | 0.00700 | 0.23792 | 1.90518 | 0.01545 | 0.22871 | 0.01719 | 1.87286 | 0.26615 | 1.82362 | 1.09985 | 0.11567 | 0.08070 | 0.04144 | 0.24796 | 1.16816 |
| **Maea** | 12.95055 | 6.89936 | 6.52875 | 6.44172 | 6.54725 | 7.45338 | 9.58385 | 14.21327 | 6.30490 | 9.78739 | 5.52203 | 4.14161 | 12.34254 | 0.00000 | 0.00000 | 8.60930 |
| **Mael** | 0.00768 | 0.00627 | 0.06979 | 3.83366 | 0.00979 | 0.00597 | 2.43292 | 0.00000 | 0.84133 | 2.36418 | 0.97523 | 0.17043 | 0.17377 | 0.00000 | 1.37936 | 0.00686 |
| **Maf1** | 10.39327 | 2.88467 | 3.00609 | 3.11441 | 11.93819 | 3.59990 | 3.53714 | 2.35124 | 2.91019 | 6.68624 | 0.78951 | 0.28178 | 7.13582 | 0.38454 | 3.23622 | 1.38803 |
| **Mafb** | 0.01139 | 0.00774 | 0.00966 | 0.13024 | 0.00942 | 0.01332 | 0.00733 | 0.06895 | 0.01283 | 0.60283 | 0.01019 | 0.00599 | 0.01076 | 0.07564 | 0.06408 | 0.00745 |

| | | | | | | | | | | | | | | | | |
|---|---|---|---|---|---|---|---|---|---|---|---|---|---|---|---|---|
| **Maff** | 0.74234 | 0.00682 | 0.01176 | 0.31521 | 0.01538 | 0.02880 | 0.00586 | 0.00007 | 0.44641 | 0.31010 | 1.40455 | 0.00312 | 0.31357 | 0.53932 | 1.36962 | 0.00642 |
| **Mafg** | 0.82705 | 0.23389 | 1.82019 | 7.61630 | 0.00345 | 0.00371 | 2.69825 | 3.16754 | 7.05462 | 1.64459 | 3.13173 | 0.25980 | 0.74009 | 0.29196 | 4.25182 | 2.10202 |
| **Mafk** | 0.40745 | 0.33137 | 0.12736 | 2.86801 | 0.00858 | 0.01587 | 0.79215 | 0.58282 | 7.17377 | 1.41076 | 0.93244 | 0.08259 | 0.88861 | 0.01513 | 0.30398 | 0.00518 |
| **Magea1** | 0.32195 | 0.00728 | 0.06001 | 0.43270 | 0.00716 | 0.07583 | 0.00692 | 0.01220 | 0.00521 | 0.06523 | 0.01200 | 0.00631 | 0.15147 | 0.00793 | 0.34342 | 0.00686 |
| **Magea10** | 0.01367 | 0.00769 | 0.01138 | 1.09790 | 0.00940 | 0.00629 | 0.00730 | 0.01098 | 0.06813 | 0.00511 | 0.01096 | 0.00551 | 0.01279 | 0.00590 | 1.06790 | 0.00715 |
| **Magea2** | 0.36563 | 0.00715 | 0.02282 | 0.17306 | 0.00717 | 0.11990 | 0.00679 | 0.01104 | 0.00800 | 0.07972 | 0.01015 | 0.04290 | 0.26930 | 0.00845 | 0.78547 | 0.00680 |
| **Magea3** | 0.19476 | 0.00766 | 0.00667 | 0.01675 | 0.00761 | 0.72696 | 0.00713 | 0.01873 | 0.01200 | 0.00675 | 0.01168 | 0.00722 | 0.14485 | 0.01954 | 0.23767 | 0.00681 |
| **Magea4** | 0.03781 | 0.00785 | 0.01033 | 0.01067 | 0.00822 | 0.00662 | 0.00741 | 0.01443 | 0.01211 | 0.01207 | 0.01250 | 0.00800 | 0.03763 | 0.01324 | 0.05249 | 0.00723 |
| **Magea5** | 1.41461 | 0.00590 | 0.04397 | 0.00000 | 0.00525 | 0.04182 | 0.01515 | 0.01109 | 0.00758 | 0.00000 | 0.00718 | 0.03790 | 1.56486 | 0.01283 | 3.36979 | 0.01208 |
| **Magea6** | 0.17451 | 0.00760 | 0.03152 | 0.00815 | 0.00740 | 0.00791 | 0.00708 | 0.01242 | 0.01115 | 0.00424 | 0.01147 | 0.01754 | 0.08314 | 0.00876 | 0.45791 | 0.00696 |
| **Magea8** | 3.78776 | 0.00341 | 0.31298 | 0.52414 | 0.00000 | 0.00527 | 0.00415 | 0.00600 | 0.00372 | 0.00000 | 0.00930 | 0.06712 | 0.29969 | 0.00991 | 2.64769 | 0.00323 |
| **Mageb16** | 8.26007 | 6.41751 | 8.39602 | 0.00000 | 8.45977 | 3.38184 | 5.11738 | 3.30911 | 1.64179 | 4.77456 | 0.45964 | 0.18920 | 3.08960 | 8.74185 | 0.00000 | 7.42736 |
| **Mageb16.ps1** | 1.10108 | 0.66930 | 0.96163 | 0.06346 | 0.91688 | 0.41778 | 0.52423 | 0.32857 | 0.08825 | 0.46372 | 0.01132 | 0.09536 | 0.30627 | 1.21921 | 0.05756 | 0.92070 |
| **Mageb3** | 0.01458 | 0.00813 | 0.05134 | 0.00444 | 0.00896 | 0.00579 | 0.00776 | 0.02134 | 0.65396 | 0.86800 | 0.10411 | 0.01352 | 0.00915 | 0.01041 | 0.45128 | 0.00705 |
| **Mageb4** | 0.01244 | 0.00751 | 0.01239 | 1.40193 | 0.00891 | 0.00683 | 0.00715 | 0.00950 | 1.51863 | 0.71644 | 0.00000 | 0.55548 | 0.01089 | 0.00191 | 1.01320 | 0.00674 |
| **Maged1** | 1.93104 | 1.03190 | 0.00000 | 2.43483 | 6.63419 | 3.76814 | 0.00577 | 1.13903 | 1.08839 | 4.04998 | 0.00000 | 0.10763 | 0.00000 | 0.32590 | 2.32992 | 12.61584 |
| **Maged2** | 0.13369 | 0.00795 | 0.00000 | 0.00618 | 0.03350 | 0.00865 | 0.00706 | 1.24118 | 0.01008 | 1.94205 | 0.00000 | 0.09950 | 0.00000 | 0.01385 | 0.00000 | 2.27383 |
| **Magee2** | 0.00959 | 0.00885 | 0.00935 | 0.00702 | 0.01049 | 0.00717 | 0.00835 | 0.32777 | 0.01051 | 0.00453 | 0.01081 | 0.00771 | 0.06748 | 0.00786 | 0.64426 | 0.00761 |
| **Magef1** | 0.01372 | 0.00825 | 0.01110 | 0.02406 | 0.00926 | 0.01347 | 0.01811 | 0.01390 | 0.02611 | 0.07231 | 0.01214 | 0.00875 | 0.01299 | 0.03049 | 0.01942 | 0.00742 |
| **Mageh1** | 0.01335 | 0.00805 | 0.01279 | 0.29448 | 0.00949 | 0.00704 | 0.01820 | 0.01344 | 0.00526 | 0.44330 | 0.01265 | 0.00778 | 0.02254 | 0.01248 | 0.01724 | 0.00748 |
| **Magi1** | 2.77743 | 0.00575 | 0.00000 | 1.69330 | 0.01873 | 0.01354 | 0.00567 | 0.35178 | 1.05188 | 0.30143 | 0.00000 | 0.06852 | 0.64812 | 0.00722 | 0.55153 | 0.54336 |
| **Magi2** | 0.21280 | 0.00682 | 0.00968 | 1.12997 | 0.00734 | 0.04719 | 0.00698 | 0.78636 | 0.73063 | 0.11437 | 0.18288 | 2.30272 | 0.01315 | 0.00694 | 0.00000 | 0.02116 |
| **Magi3** | 1.18468 | 0.00564 | 0.00000 | 0.59873 | 0.00542 | 0.00762 | 0.01608 | 0.00506 | 0.91185 | 0.00000 | 0.00000 | 0.85151 | 0.00994 | 0.04675 | 0.11584 | 0.47867 |
| **Magoh** | 14.37306 | 9.53347 | 11.48673 | 20.75281 | 13.68111 | 4.91207 | 6.33285 | 9.90670 | 33.81367 | 21.02706 | 24.20900 | 4.29480 | 17.62737 | 25.25955 | 18.88411 | 5.16267 |
| **Magohb** | 2.61687 | 0.01331 | 0.36942 | 1.54696 | 2.45276 | 0.02017 | 1.26545 | 0.83445 | 0.49974 | 4.51465 | 3.39575 | 0.19845 | 1.57611 | 3.36621 | 2.99145 | 1.12658 |
| **Magt1** | 4.02047 | 4.38952 | 9.62380 | 7.26306 | 7.46535 | 0.18050 | 7.20531 | 2.68282 | 5.14644 | 6.32401 | 7.11080 | 9.12562 | 5.19964 | 12.85335 | 7.61763 | 2.35141 |
| **Majin** | 0.76778 | 0.00770 | 0.50951 | 0.00000 | 0.04134 | 0.00484 | 0.00428 | 1.35687 | 0.00741 | 2.45136 | 3.71608 | 1.09468 | 3.18910 | 0.44340 | 2.12793 | 0.01174 |
| **Mak** | 0.01379 | 0.00765 | 0.01176 | 0.60597 | 0.00938 | 0.00646 | 0.00738 | 0.01194 | 0.00363 | 0.00665 | 0.01270 | 0.01655 | 0.01274 | 0.00719 | 0.40569 | 0.00726 |
| **Mak16** | 13.52442 | 6.04538 | 10.28856 | 15.82540 | 4.25935 | 1.62333 | 5.26586 | 9.04795 | 13.19016 | 8.54731 | 7.34286 | 1.26468 | 5.03652 | 10.99942 | 17.49397 | 4.81482 |
| **Mal** | 0.03046 | 0.00763 | 0.00825 | 0.00941 | 0.00838 | 0.00696 | 0.00722 | 0.01114 | 0.01205 | 0.28308 | 0.01152 | 0.19026 | 0.01269 | 0.01139 | 0.00683 | 0.00770 |
| **Malat1** | 2.15620 | 4.54094 | 3.35323 | 4.89101 | 0.13763 | 0.28337 | 0.14017 | 2.55886 | 4.29935 | 0.14361 | 2.45915 | 3.53865 | 3.26478 | 10.21994 | 9.69434 | 2.32960 |
| **Malsu1** | 2.51338 | 0.12395 | 0.67954 | 1.15100 | 4.56097 | 0.00362 | 0.02084 | 0.00000 | 1.94369 | 3.79778 | 11.71268 | 0.05426 | 0.24343 | 0.52996 | 0.00000 | 0.51078 |
| **Malt1** | 0.25337 | 2.57645 | 1.00047 | 0.91858 | 0.05461 | 0.19479 | 0.15527 | 1.19356 | 3.21323 | 1.46568 | 2.26348 | 0.15220 | 0.48937 | 1.20618 | 1.51870 | 0.01912 |
| **Mamdc2** | 0.02286 | 0.00752 | 2.68366 | 0.02544 | 0.00570 | 0.00553 | 0.00709 | 0.01057 | 0.01214 | 0.01100 | 0.02757 | 0.00621 | 0.01266 | 0.01423 | 0.00000 | 0.00638 |
| **Mamdc4** | 0.01074 | 0.01795 | 1.03159 | 0.30489 | 0.00783 | 0.00566 | 0.00709 | 0.13408 | 0.30679 | 0.00624 | 0.00829 | 0.00619 | 0.00806 | 0.07713 | 0.00000 | 0.00697 |
| **Maml1** | 1.87453 | 2.66113 | 0.47318 | 4.05617 | 0.05275 | 0.00288 | 0.01396 | 1.21544 | 9.84581 | 1.42872 | 0.00000 | 3.79511 | 0.99214 | 0.00000 | 0.04792 | 0.16145 |
| **Maml2** | 0.01656 | 0.00676 | 0.00293 | 0.00350 | 0.00892 | 0.04911 | 1.79096 | 0.01008 | 0.13270 | 0.49376 | 0.00973 | 0.00481 | 0.00966 | 0.00623 | 0.12358 | 0.00702 |
| **Man1a2** | 2.26111 | 1.44821 | 3.76681 | 4.56592 | 3.40872 | 0.44631 | 3.27637 | 5.51725 | 3.47993 | 2.96109 | 3.03153 | 1.29452 | 1.84589 | 11.83012 | 4.92341 | 0.70095 |
| **Man1b1** | 3.29280 | 0.91681 | 1.53547 | 7.86870 | 0.00794 | 2.23893 | 2.76628 | 3.48714 | 8.23192 | 3.96370 | 5.54378 | 0.99273 | 2.65923 | 2.15704 | 1.19233 | 1.65290 |
| **Man1c1** | 0.30370 | 0.00745 | 0.01011 | 1.67152 | 0.00722 | 0.00615 | 0.00756 | 0.01883 | 0.34275 | 0.00822 | 0.05808 | 0.00693 | 0.01692 | 0.00397 | 0.23866 | 0.01371 |
| **Man2a1** | 3.73891 | 1.51225 | 2.40328 | 2.56262 | 6.76010 | 0.00177 | 6.10662 | 0.01111 | 0.00000 | 0.00000 | 0.01704 | 0.00180 | 1.49350 | 0.02044 | 0.36620 | 2.29291 |
| **Man2a2** | 1.17855 | 0.08617 | 0.00000 | 13.77424 | 0.00573 | 0.10184 | 1.75351 | 0.61077 | 11.93978 | 0.14415 | 0.16397 | 0.30482 | 0.01796 | 0.16895 | 0.08373 | 0.02467 |
| **Man2b1** | 6.25208 | 2.48341 | 5.82211 | 3.19265 | 2.47830 | 7.54388 | 6.56994 | 6.28899 | 2.09050 | 1.94353 | 9.37885 | 0.51916 | 5.33074 | 0.58799 | 1.39385 | 0.03421 |
| **Man2b2** | 0.44387 | 0.00694 | 0.00171 | 0.00478 | 0.00686 | 0.00847 | 0.08595 | 1.24354 | 0.01004 | 0.41225 | 0.56049 | 0.00125 | 0.14824 | 0.00488 | 0.05498 | 0.00655 |
| **Man2c1** | 4.95360 | 0.01024 | 3.15515 | 0.10632 | 0.09634 | 0.53827 | 2.43558 | 1.43903 | 3.11989 | 1.35403 | 0.67159 | 0.17205 | 5.00014 | 0.00380 | 0.11391 | 0.01331 |
| **Man2c1os** | 0.01421 | 0.00797 | 0.13299 | 2.15368 | 0.00881 | 0.00653 | 0.00714 | 0.01020 | 0.82404 | 0.44571 | 0.00777 | 0.00509 | 0.01068 | 0.00950 | 0.00092 | 0.00697 |
| **Manbal** | 4.67487 | 2.79696 | 9.58419 | 0.43347 | 0.00000 | 2.40584 | 6.19543 | 8.66619 | 2.00796 | 8.83564 | 0.96764 | 0.45130 | 2.03904 | 0.36024 | 0.86039 | 2.10720 |
| **Manea** | 0.59298 | 0.00776 | 0.01714 | 0.23508 | 0.02754 | 0.00736 | 0.03015 | 0.12855 | 0.11992 | 0.06240 | 0.83723 | 0.04492 | 0.30819 | 1.06636 | 0.03548 | 0.00630 |
| **Maneal** | 0.99054 | 0.00701 | 0.49503 | 0.85682 | 0.00513 | 0.00449 | 0.00706 | 0.01695 | 0.77834 | 0.46768 | 0.44058 | 0.06387 | 0.00622 | 3.89070 | 3.20142 | 0.00609 |
| **Manf** | 7.12411 | 0.70896 | 7.82124 | 9.00159 | 0.55375 | 8.96281 | 2.57159 | 7.64113 | 12.08849 | 15.07079 | 10.17630 | 1.08302 | 18.29620 | 1.59744 | 2.40391 | 9.10460 |
| **Mansc1** | 0.01374 | 0.00797 | 0.01208 | 0.23159 | 0.00927 | 0.00672 | 0.00756 | 0.01379 | 0.01471 | 0.02395 | 0.02620 | 0.00811 | 0.01264 | 0.00879 | 0.05189 | 0.00737 |
| **Mansc4** | 0.55379 | 0.00708 | 0.00470 | 0.01064 | 0.00664 | 0.00694 | 0.00665 | 0.00889 | 0.01240 | 0.00612 | 0.01240 | 0.00740 | 0.77836 | 0.01325 | 0.01050 | 0.00623 |
| **Maoa** | 2.31446 | 0.00530 | 0.00000 | 0.29740 | 0.02597 | 0.08235 | 0.03538 | 0.02629 | 0.67865 | 0.00000 | 0.04786 | 0.00196 | 0.75769 | 0.56530 | 0.13634 | 1.36792 |
| **Map10** | 0.13368 | 0.00727 | 0.00529 | 1.57719 | 0.00770 | 0.00647 | 0.00733 | 0.00897 | 5.60435 | 0.00281 | 0.00000 | 0.00674 | 0.01278 | 0.00580 | 0.39221 | 0.01356 |
| **Map1a** | 0.49100 | 0.00713 | 0.72999 | 0.00535 | 0.00564 | 0.00511 | 0.00680 | 0.00000 | 0.01113 | 0.86668 | 0.40928 | 0.02265 | 0.00654 | 0.01208 | 0.00000 | 0.00665 |
| **Map1b** | 0.08305 | 0.00682 | 0.00195 | 1.38161 | 1.63237 | 0.00790 | 0.00738 | 0.00948 | 0.31747 | 4.07386 | 0.00694 | 0.01162 | 0.05240 | 0.00323 | 0.92890 | 0.00724 |
| **Map1lc3a** | 0.79877 | 1.91912 | 4.34824 | 4.54213 | 4.70355 | 3.81479 | 2.87240 | 8.88381 | 10.57932 | 24.58643 | 22.50402 | 1.53094 | 8.48476 | 0.06251 | 5.05459 | 0.29073 |
| **Map1lc3b** | 46.44801 | 29.47417 | 49.21889 | 35.32413 | 33.29238 | 19.75516 | 29.43826 | 52.02937 | 45.12898 | 62.73587 | 54.46611 | 27.33657 | 46.77590 | 27.42221 | 38.06402 | 20.51227 |
| **Map1s** | 2.97661 | 0.00407 | 0.23911 | 1.82368 | 0.76787 | 0.03187 | 0.00357 | 0.79295 | 0.67744 | 8.10180 | 1.52175 | 0.09319 | 1.66110 | 0.00457 | 0.00000 | 2.70454 |
| **Map2** | 0.01261 | 0.00790 | 0.00903 | 0.76639 | 0.00894 | 0.00649 | 0.00724 | 0.00860 | 0.00000 | 1.24996 | 0.04506 | 0.00303 | 0.01009 | 0.35991 | 0.26158 | 0.00694 |
| **Map2k1** | 9.23219 | 3.82227 | 3.06680 | 7.69765 | 11.80419 | 2.39485 | 1.34078 | 6.89809 | 8.19724 | 10.73698 | 3.25404 | 1.66430 | 1.71753 | 9.20817 | 11.30896 | 2.99262 |
| **Map2k2** | 13.36569 | 0.90926 | 2.84054 | 3.54729 | 1.59754 | 9.95765 | 3.18843 | 26.06566 | 4.65796 | 22.55324 | 8.82807 | 4.29540 | 15.32946 | 0.02929 | 0.33601 | 8.96320 |

| | | | | | | | | | | | | | | | | |
|---|---|---|---|---|---|---|---|---|---|---|---|---|---|---|---|---|
| **Map2k3** | 8.52915 | 5.86369 | 4.64687 | 2.21206 | 4.48860 | 3.73711 | 8.47595 | 6.71085 | 1.32882 | 5.59618 | 3.88612 | 0.53442 | 5.95781 | 0.00000 | 2.83330 | 2.10038 |
| **Map2k3os** | 0.45903 | 0.00714 | 0.05922 | 0.36610 | 0.00710 | 0.00640 | 0.00695 | 0.27904 | 0.00390 | 0.01366 | 0.01054 | 0.00710 | 0.00617 | 0.01079 | 0.00681 | 0.00674 |
| **Map2k4** | 3.19644 | 3.50159 | 6.75059 | 5.14695 | 8.39922 | 4.73763 | 4.34010 | 3.79663 | 3.98280 | 4.37273 | 1.81226 | 12.97298 | 2.11054 | 0.16368 | 2.19161 | 2.19507 |
| **Map2k5** | 0.68809 | 2.70952 | 0.00000 | 0.09781 | 0.00801 | 0.35849 | 0.00569 | 0.00759 | 0.06346 | 0.01290 | 0.01020 | 0.01643 | 0.00626 | 0.00641 | 0.00000 | 1.54697 |
| **Map2k6** | 6.83135 | 2.85058 | 3.09181 | 0.33204 | 3.11427 | 4.51897 | 2.61091 | 1.40021 | 0.00000 | 0.98402 | 0.00000 | 0.18903 | 1.92989 | 0.00680 | 1.36827 | 0.03219 |
| **Map2k7** | 5.72235 | 0.00000 | 1.27866 | 2.76015 | 0.06200 | 4.41966 | 2.57234 | 4.40962 | 2.83869 | 6.83657 | 3.66900 | 0.35277 | 2.30679 | 0.98000 | 3.63300 | 2.75445 |
| **Map3k1** | 0.05828 | 1.95535 | 0.05539 | 0.27917 | 0.00773 | 0.00834 | 0.01605 | 0.00966 | 0.00671 | 0.44751 | 0.00541 | 0.00000 | 0.57331 | 1.63107 | 0.20540 | 0.21357 |
| **Map3k10** | 0.01678 | 0.00799 | 0.01597 | 0.34046 | 0.00878 | 0.00626 | 0.00716 | 0.06198 | 0.48949 | 0.12599 | 0.35624 | 0.03606 | 0.13356 | 0.00763 | 0.01364 | 0.02149 |
| **Map3k11** | 2.63468 | 0.03553 | 0.20453 | 2.77145 | 0.02919 | 0.00316 | 0.00431 | 0.38723 | 0.06484 | 2.19228 | 1.27402 | 0.39568 | 0.66082 | 0.00000 | 0.20892 | 0.05401 |
| **Map3k12** | 0.71201 | 0.92763 | 0.43148 | 0.48089 | 0.00650 | 0.00543 | 1.32846 | 0.01329 | 0.12882 | 0.06378 | 0.01483 | 0.03754 | 0.17372 | 0.02827 | 0.17157 | 0.11651 |
| **Map3k13** | 0.04632 | 0.00728 | 0.00592 | 0.03388 | 0.00810 | 0.00717 | 0.77287 | 0.01070 | 0.01001 | 0.01265 | 0.01256 | 0.00715 | 0.01097 | 0.01201 | 0.00973 | 0.00724 |
| **Map3k14** | 0.01321 | 0.00778 | 0.01658 | 0.59110 | 0.00933 | 0.00609 | 0.00739 | 0.01123 | 1.15754 | 0.00549 | 0.00000 | 0.00394 | 0.01293 | 0.00454 | 0.93837 | 0.00718 |
| **Map3k15** | 1.47499 | 0.00574 | 0.00179 | 0.00965 | 0.03363 | 0.00741 | 0.00635 | 0.09221 | 0.01156 | 0.02407 | 0.00980 | 0.04950 | 0.06757 | 0.40477 | 0.24619 | 0.00622 |
| **Map3k2** | 1.06688 | 0.00468 | 0.72614 | 0.00000 | 0.00436 | 0.67952 | 1.63329 | 0.66006 | 1.53039 | 0.72501 | 0.68007 | 1.72654 | 1.09655 | 0.62255 | 3.39268 | 0.57706 |
| **Map3k3** | 4.12020 | 0.04621 | 0.58064 | 0.47742 | 0.78492 | 0.09347 | 1.84018 | 1.86365 | 4.08795 | 5.19559 | 0.00000 | 0.05610 | 1.07738 | 3.38782 | 5.65710 | 0.67312 |
| **Map3k4** | 1.88032 | 2.26422 | 3.65073 | 0.98847 | 0.85433 | 0.53578 | 0.18003 | 3.00959 | 0.08071 | 0.35887 | 0.00000 | 0.10472 | 4.33034 | 1.60130 | 0.43729 | 2.85066 |
| **Map3k5** | 1.28183 | 4.86591 | 0.00000 | 2.01598 | 0.29065 | 3.02344 | 0.49690 | 1.49662 | 1.02278 | 1.67139 | 1.37319 | 0.05462 | 2.57378 | 1.91666 | 4.21690 | 0.03063 |
| **Map3k6** | 1.85185 | 0.39812 | 0.00000 | 4.16048 | 5.99426 | 0.00456 | 0.00620 | 0.35388 | 2.28091 | 0.00000 | 0.00000 | 0.01923 | 0.00942 | 0.10309 | 0.01840 | 5.19505 |
| **Map3k7** | 2.38170 | 1.24323 | 0.22256 | 3.38056 | 8.37420 | 2.36971 | 0.34292 | 3.43668 | 3.45625 | 6.43768 | 4.63383 | 7.84210 | 2.12347 | 5.31680 | 7.90064 | 3.38890 |
| **Map3k9** | 0.00428 | 0.01893 | 1.35080 | 1.01353 | 0.00603 | 0.00898 | 0.04127 | 0.94364 | 0.00000 | 1.51714 | 1.83367 | 0.03683 | 0.00000 | 0.51438 | 0.65935 | 0.94713 |
| **Map4** | 10.36605 | 5.41804 | 13.66813 | 11.16182 | 12.80301 | 11.25653 | 6.97595 | 12.17579 | 10.07413 | 7.70640 | 12.34603 | 0.70416 | 16.74442 | 7.45833 | 2.40887 | 5.46744 |
| **Map4k1** | 0.69649 | 0.02667 | 0.83195 | 1.51536 | 0.00751 | 0.00488 | 0.55698 | 0.72606 | 1.91373 | 2.28536 | 4.10574 | 0.01298 | 1.95674 | 0.00000 | 0.03879 | 0.00473 |
| **Map4k2** | 3.40252 | 0.00400 | 0.25062 | 0.81133 | 0.00646 | 0.01425 | 0.23991 | 0.94034 | 0.17255 | 0.14982 | 1.02191 | 0.00482 | 2.21133 | 0.00523 | 0.19306 | 0.73603 |
| **Map4k3** | 1.44856 | 3.76889 | 2.04791 | 1.02482 | 0.60878 | 0.52993 | 2.08453 | 1.23954 | 1.37194 | 0.59846 | 3.36762 | 1.10784 | 0.00000 | 7.86758 | 2.63854 | 0.14941 |
| **Map4k4** | 10.36924 | 1.34834 | 3.61431 | 0.72045 | 8.45626 | 5.49033 | 5.06667 | 3.08364 | 0.52680 | 1.19296 | 2.89563 | 2.70506 | 2.89863 | 0.08416 | 0.30136 | 8.75048 |
| **Map4k5** | 2.51533 | 5.11853 | 0.35891 | 17.18061 | 3.45226 | 2.52610 | 3.41201 | 4.09043 | 10.79096 | 13.52265 | 6.22950 | 7.15960 | 1.94820 | 9.22407 | 18.32318 | 2.75625 |
| **Map6** | 0.42161 | 0.00773 | 0.00486 | 1.24234 | 0.00867 | 0.00702 | 0.01725 | 0.01184 | 1.30710 | 0.04472 | 0.00000 | 0.01523 | 0.01290 | 0.00493 | 0.13093 | 0.00700 |
| **Map6d1** | 0.01220 | 0.00766 | 0.00091 | 0.10653 | 0.00897 | 0.00657 | 0.00682 | 0.00000 | 0.15610 | 1.19065 | 1.16078 | 0.00361 | 0.00000 | 0.00416 | 2.78161 | 0.00724 |
| **Map7** | 9.10225 | 7.06831 | 8.81271 | 2.45580 | 5.27066 | 7.29257 | 10.36724 | 18.68007 | 3.65444 | 16.62567 | 17.73573 | 18.58826 | 10.08044 | 12.12981 | 10.36732 | 4.85667 |
| **Map7d1** | 0.67666 | 0.00746 | 0.79227 | 1.30408 | 0.02236 | 2.96064 | 0.15275 | 0.00725 | 0.00000 | 0.01720 | 0.00842 | 0.06434 | 0.19619 | 0.00966 | 0.00000 | 0.00523 |
| **Map7d2** | 0.97997 | 0.00379 | 2.39237 | 0.71057 | 0.01160 | 0.00573 | 0.00284 | 1.78135 | 1.65064 | 4.90032 | 2.76182 | 7.28181 | 2.56960 | 12.89433 | 12.52019 | 0.00288 |
| **Map9** | 0.01621 | 0.00725 | 1.68782 | 0.30769 | 0.00632 | 0.00542 | 0.00660 | 0.00000 | 0.00384 | 0.00000 | 0.74116 | 0.41254 | 0.00000 | 0.00651 | 1.51578 | 0.00627 |
| **Mapk1** | 13.63608 | 9.58694 | 10.96660 | 5.89375 | 8.95745 | 2.00337 | 5.09062 | 7.12427 | 10.12539 | 8.35803 | 6.65675 | 6.39709 | 5.16262 | 7.96968 | 11.39651 | 5.49964 |
| **Mapk10** | 0.01304 | 0.00796 | 0.01019 | 0.00606 | 0.01848 | 0.00667 | 0.00750 | 0.00868 | 0.01175 | 0.00474 | 0.23274 | 0.01724 | 0.01566 | 0.00717 | 0.05171 | 0.00724 |
| **Mapk11** | 0.16688 | 0.00866 | 0.00651 | 3.29539 | 0.01710 | 0.00681 | 0.00717 | 0.01073 | 1.34314 | 0.09409 | 0.00000 | 0.00489 | 0.01315 | 0.08087 | 0.00958 | 0.00712 |
| **Mapk12** | 1.52434 | 0.00577 | 0.00308 | 0.38635 | 0.00502 | 0.00701 | 0.00631 | 0.00686 | 0.14759 | 0.19257 | 0.01076 | 0.00641 | 0.01185 | 0.01434 | 0.11875 | 0.00571 |
| **Mapk13** | 0.01737 | 0.00556 | 0.15162 | 2.24092 | 2.50768 | 0.00788 | 0.01477 | 2.57056 | 0.46285 | 2.15035 | 3.98358 | 0.06233 | 2.46547 | 0.12832 | 0.48940 | 0.01949 |
| **Mapk14** | 5.24302 | 2.94887 | 8.46166 | 16.23156 | 8.36944 | 0.01893 | 6.64656 | 8.59043 | 10.26985 | 7.21575 | 1.34568 | 1.40313 | 2.85124 | 8.20197 | 10.76334 | 4.36055 |
| **Mapk1ip1** | 1.88976 | 1.04948 | 2.01230 | 1.68571 | 0.00375 | 2.47700 | 3.89063 | 1.30732 | 0.51019 | 0.00000 | 1.06411 | 5.92826 | 0.03448 | 1.00763 | 0.16003 | 3.07968 |
| **Mapk1ip1l** | 15.17719 | 6.43385 | 8.55683 | 10.76870 | 5.73374 | 4.87058 | 6.75839 | 11.44168 | 13.15270 | 14.35112 | 11.46286 | 2.08654 | 8.85673 | 18.36092 | 9.29342 | 3.91306 |
| **Mapk3** | 5.39487 | 0.00000 | 1.18351 | 5.00787 | 0.50741 | 0.58896 | 0.28171 | 6.51165 | 0.10767 | 3.94257 | 0.64578 | 0.56689 | 5.35575 | 0.00043 | 0.34141 | 1.30050 |
| **Mapk6** | 3.91587 | 8.33028 | 8.22879 | 16.76508 | 0.61144 | 6.92901 | 6.93191 | 13.22674 | 16.93900 | 22.57836 | 14.93309 | 20.58860 | 8.26636 | 11.20325 | 13.45515 | 3.93623 |
| **Mapk7** | 5.48889 | 2.38284 | 0.38311 | 1.01034 | 0.07543 | 0.90355 | 0.60306 | 0.38821 | 3.18139 | 3.10125 | 1.18404 | 0.00853 | 1.84655 | 0.00000 | 0.60125 | 0.46445 |
| **Mapk8** | 2.14143 | 0.02345 | 0.54648 | 11.65711 | 1.86366 | 2.99195 | 0.01373 | 0.53465 | 5.82269 | 0.71816 | 0.92864 | 7.09268 | 0.63512 | 3.46843 | 3.99973 | 3.40547 |
| **Mapk8ip1** | 0.00966 | 0.00734 | 0.00000 | 0.25217 | 0.00964 | 0.38087 | 0.47509 | 0.06538 | 0.39290 | 0.83703 | 0.41292 | 0.02966 | 0.00743 | 0.00705 | 0.01263 | 0.00746 |
| **Mapk8ip3** | 3.56534 | 0.00173 | 2.53202 | 1.60039 | 0.13928 | 1.03426 | 3.05699 | 1.81230 | 5.61627 | 1.03221 | 0.00000 | 0.01250 | 0.36526 | 0.03192 | 0.00005 | 0.08315 |
| **Mapk9** | 1.21235 | 2.42984 | 1.50522 | 2.25215 | 0.01382 | 2.58557 | 2.38841 | 0.64448 | 4.72885 | 0.42492 | 1.77527 | 0.03812 | 1.43820 | 0.00000 | 4.28616 | 1.12104 |
| **Mapkap1** | 1.30452 | 0.00000 | 2.19041 | 11.14028 | 4.57480 | 0.89383 | 4.10582 | 5.14248 | 4.94663 | 4.14331 | 1.40081 | 0.19778 | 1.25604 | 0.09601 | 1.47697 | 0.00167 |
| **Mapkapk2** | 1.88284 | 0.05339 | 0.85054 | 4.16389 | 0.00326 | 3.57922 | 3.06206 | 1.03769 | 4.41584 | 2.51389 | 0.25031 | 0.65350 | 2.82174 | 0.00000 | 0.82172 | 0.07368 |
| **Mapkapk3** | 0.00679 | 0.00822 | 1.24887 | 0.19584 | 0.02246 | 0.00923 | 0.00595 | 2.33965 | 0.19865 | 2.16598 | 1.50109 | 0.08008 | 2.14215 | 0.10428 | 1.97150 | 0.00593 |
| **Mapkapk5** | 6.01925 | 2.95137 | 4.95980 | 14.04229 | 7.59116 | 0.02100 | 2.52239 | 4.83943 | 11.65619 | 8.93181 | 6.75350 | 1.99626 | 4.34183 | 1.12684 | 8.14932 | 3.92222 |
| **Mapkbp1** | 0.00696 | 0.00797 | 1.10205 | 5.66534 | 0.00721 | 0.01598 | 0.00603 | 0.00000 | 5.43116 | 1.03962 | 1.38577 | 0.03878 | 1.16745 | 0.06962 | 5.07640 | 0.00491 |
| **Mapre1** | 20.03581 | 15.73641 | 20.86305 | 15.48056 | 14.63607 | 13.10521 | 18.88241 | 26.21799 | 14.10827 | 25.90605 | 16.71360 | 19.37367 | 18.28625 | 23.92639 | 18.79122 | 7.68932 |
| **Mapre2** | 1.21142 | 0.00784 | 0.81809 | 2.54100 | 0.25378 | 0.16449 | 0.00655 | 0.69869 | 0.83809 | 0.44482 | 0.36903 | 0.13769 | 1.05517 | 0.27885 | 0.30365 | 0.25115 |
| **Mapre3** | 0.08310 | 0.00789 | 0.23008 | 0.20001 | 0.00739 | 0.00907 | 0.03749 | 0.01796 | 0.00711 | 0.00481 | 0.01253 | 0.02728 | 0.54484 | 0.03421 | 0.00106 | 0.12028 |
| **Mapt** | 4.48085 | 1.82772 | 2.76539 | 0.03136 | 1.52194 | 0.01550 | 2.02911 | 5.03305 | 0.00584 | 5.10585 | 4.70851 | 0.72366 | 1.95957 | 1.95099 | 1.42248 | 1.38279 |
| **X2.Mar** | 9.22504 | 0.02844 | 0.00000 | 9.42473 | 0.04068 | 2.23604 | 0.00000 | 1.84582 | 7.89367 | 6.77205 | 8.13257 | 2.15840 | 0.12216 | 2.99722 | 3.04520 | 0.00000 |
| **X2.Mar.1** | 2.42067 | 0.02718 | 1.73597 | 3.77476 | 0.00000 | 0.00546 | 0.00576 | 0.38711 | 0.75878 | 4.82430 | 0.36695 | 0.05169 | 0.02375 | 0.05512 | 1.38347 | 0.03984 |
| **X3.Mar** | 0.00868 | 0.00772 | 0.00766 | 0.46765 | 0.51386 | 0.00696 | 0.00749 | 0.01300 | 0.41745 | 0.01229 | 0.01021 | 0.00724 | 0.01085 | 0.01160 | 0.00414 | 0.00718 |
| **X5.Mar** | 7.79810 | 4.39815 | 5.17521 | 22.42224 | 5.66666 | 2.60288 | 5.77580 | 7.89141 | 13.40466 | 11.02698 | 15.03387 | 9.79140 | 5.40804 | 11.26835 | 12.93016 | 0.21289 |
| **X6.Mar** | 4.70492 | 4.28075 | 5.74849 | 6.65903 | 3.71001 | 7.33791 | 2.55620 | 6.29383 | 5.51228 | 5.29675 | 9.76271 | 9.81619 | 5.33749 | 12.12362 | 7.15771 | 1.90435 |
| **X7.Mar** | 0.52837 | 0.70095 | 2.24631 | 0.94352 | 0.06813 | 0.41230 | 0.25033 | 1.03225 | 0.61944 | 0.00000 | 0.00000 | 3.68748 | 0.65151 | 6.01020 | 3.50321 | 0.26731 |

| | | | | | | | | | | | | | | | | |
|---|---|---|---|---|---|---|---|---|---|---|---|---|---|---|---|---|
| **X8.Mar** | 0.89850 | 0.00725 | 0.53989 | 3.06661 | 0.01084 | 0.00436 | 0.00593 | 0.00000 | 0.01219 | 0.08973 | 0.26616 | 0.08093 | 0.62063 | 0.01126 | 0.50239 | 0.00531 |
| **X9.Mar** | 0.06990 | 0.00828 | 0.00396 | 0.57035 | 0.00779 | 0.00744 | 0.00755 | 0.08839 | 1.45758 | 0.59158 | 0.93771 | 0.02229 | 0.00129 | 0.00276 | 0.23745 | 0.00692 |
| **Marcks** | 8.42458 | 17.12675 | 11.78899 | 0.40477 | 14.67232 | 7.24924 | 7.28921 | 0.29554 | 0.41334 | 1.33357 | 7.57342 | 5.06658 | 4.83723 | 0.08313 | 0.00000 | 6.51492 |
| **Marcksl1** | 17.88358 | 6.19925 | 14.76564 | 2.29828 | 3.82311 | 14.13621 | 12.48384 | 21.74997 | 3.36118 | 16.86281 | 16.43216 | 1.74881 | 21.75672 | 8.05350 | 8.65004 | 12.99411 |
| **Marf1** | 2.58684 | 0.08897 | 2.99642 | 7.08225 | 2.37614 | 1.10146 | 1.10449 | 3.85391 | 4.03894 | 1.21189 | 3.96093 | 1.17573 | 0.69864 | 4.25015 | 5.09945 | 0.00835 |
| **Mark1** | 0.01150 | 0.00591 | 0.00000 | 1.70707 | 0.01276 | 1.02423 | 2.96033 | 3.51170 | 0.76312 | 5.50641 | 8.66228 | 0.34890 | 0.00000 | 0.04607 | 1.45249 | 0.09568 |
| **Mark2** | 6.84341 | 0.50171 | 2.01308 | 4.12185 | 2.17931 | 4.66103 | 0.80971 | 4.56268 | 4.28821 | 15.72665 | 1.65399 | 0.28076 | 3.37192 | 0.01570 | 3.56246 | 4.58827 |
| **Mark3** | 8.31704 | 7.47884 | 7.36862 | 7.62026 | 10.40216 | 7.64302 | 6.53290 | 15.88972 | 13.52963 | 16.30728 | 7.02382 | 6.26682 | 5.44883 | 0.92112 | 9.10861 | 1.64370 |
| **Mark4** | 0.04092 | 0.00836 | 0.01824 | 0.17606 | 0.01556 | 0.00667 | 0.00766 | 0.01448 | 0.02923 | 0.06325 | 0.01493 | 0.00695 | 0.04813 | 0.00589 | 0.12418 | 0.08549 |
| **Mars** | 18.43649 | 13.70197 | 14.51811 | 3.38741 | 11.76818 | 6.76699 | 11.09547 | 18.75214 | 3.28689 | 12.40790 | 10.76950 | 7.07430 | 14.87340 | 0.59278 | 5.99887 | 9.40144 |
| **Mars2** | 0.23563 | 0.01646 | 0.00000 | 3.45031 | 0.01488 | 0.02923 | 1.12002 | 0.88883 | 6.26247 | 3.20424 | 0.42976 | 0.20231 | 0.22975 | 4.17226 | 7.66307 | 1.79305 |
| **Marveld1** | 0.01599 | 0.00777 | 0.01064 | 0.33829 | 0.00980 | 0.00642 | 0.00730 | 0.01011 | 0.00645 | 2.13472 | 0.00859 | 0.01058 | 0.01050 | 0.05269 | 0.75331 | 0.02153 |
| **Marveld2** | 0.22601 | 0.00716 | 0.00000 | 3.49384 | 0.00713 | 0.13610 | 0.00655 | 1.61503 | 4.50737 | 2.64693 | 0.85109 | 0.05889 | 0.16544 | 5.38901 | 0.60569 | 0.20239 |
| **Marveld3** | 0.01009 | 0.00754 | 0.04286 | 0.00654 | 0.00936 | 0.00842 | 0.00701 | 0.00770 | 0.02498 | 0.54738 | 0.10389 | 0.01412 | 0.55606 | 0.01780 | 0.08831 | 0.00748 |
| **Masp2** | 0.02438 | 0.00748 | 0.25504 | 0.54504 | 0.00830 | 0.55480 | 0.00713 | 0.01005 | 1.27287 | 0.00000 | 0.00000 | 0.01420 | 0.00683 | 0.01972 | 0.33752 | 0.00671 |
| **Mast1** | 1.32737 | 0.00516 | 0.00313 | 4.99780 | 0.00505 | 0.00591 | 0.00314 | 1.92284 | 3.87021 | 7.94692 | 7.18311 | 0.08250 | 0.08075 | 0.03711 | 0.00000 | 0.03934 |
| **Mast2** | 4.07914 | 3.62115 | 0.13549 | 6.40396 | 0.04185 | 0.06022 | 6.94592 | 4.63751 | 5.76383 | 4.86423 | 2.10337 | 0.33747 | 3.83638 | 0.00000 | 3.10776 | 2.70873 |
| **Mast3** | 0.15731 | 0.00604 | 0.35648 | 1.83368 | 1.34257 | 4.26062 | 0.00000 | 2.37342 | 0.68404 | 7.58226 | 8.98925 | 0.39223 | 5.80069 | 0.10753 | 2.83483 | 0.00002 |
| **Mast4** | 0.93322 | 0.00523 | 0.44821 | 1.76605 | 0.00550 | 0.00485 | 2.25097 | 0.69776 | 0.30995 | 0.05700 | 0.65377 | 0.02304 | 0.01678 | 0.00456 | 0.80983 | 0.00575 |
| **Mastl** | 0.11577 | 0.00421 | 1.91479 | 5.46167 | 4.97202 | 0.87859 | 0.02470 | 5.08543 | 0.52101 | 3.87583 | 3.09480 | 0.10534 | 2.54421 | 11.30720 | 6.79509 | 0.03959 |
| **Mat1a** | 0.01375 | 0.00772 | 0.01200 | 0.48288 | 0.00941 | 0.00645 | 0.00739 | 0.01194 | 0.00357 | 0.00663 | 0.01168 | 0.00669 | 0.01273 | 0.00610 | 0.48892 | 0.00728 |
| **Mat2a** | 12.87707 | 17.78948 | 18.60297 | 14.52660 | 7.09612 | 15.28231 | 16.15284 | 36.86554 | 23.65105 | 40.57692 | 20.47703 | 12.59777 | 21.58494 | 39.83224 | 37.46540 | 11.37329 |
| **Mat2b** | 2.26321 | 4.09525 | 5.49619 | 24.81428 | 1.70969 | 1.07554 | 2.14550 | 5.59808 | 20.82270 | 18.17783 | 9.61148 | 9.79672 | 2.14432 | 12.37285 | 18.51536 | 2.66121 |
| **Matk** | 0.01374 | 0.00808 | 0.01225 | 0.08387 | 0.00931 | 0.00665 | 0.00753 | 0.01394 | 0.06476 | 0.01509 | 0.01285 | 0.00808 | 0.01271 | 0.01203 | 0.00847 | 0.00740 |
| **Matn2** | 0.01332 | 0.00787 | 0.00976 | 0.00423 | 0.00919 | 0.00673 | 0.00724 | 0.00948 | 0.49740 | 0.30721 | 0.02306 | 0.01370 | 0.01247 | 0.28953 | 0.00000 | 0.00778 |
| **Matr3** | 6.98509 | 10.72923 | 8.53650 | 4.29272 | 5.07821 | 3.96760 | 5.87948 | 15.03874 | 8.58062 | 9.91078 | 8.76559 | 15.69752 | 5.19512 | 28.47227 | 12.44286 | 3.23403 |
| **Mau2** | 3.75358 | 0.11428 | 4.00791 | 1.44056 | 0.07695 | 0.14698 | 3.66358 | 6.28809 | 0.89825 | 4.81641 | 2.91945 | 0.22248 | 1.77043 | 5.56886 | 6.37836 | 0.11679 |
| **Mavs** | 3.28713 | 2.45113 | 6.23756 | 1.79719 | 0.00000 | 7.77420 | 0.05266 | 1.09863 | 6.75583 | 4.80178 | 1.90429 | 0.35670 | 0.00000 | 0.00000 | 3.18513 | 1.21662 |
| **Max** | 9.75353 | 3.41215 | 3.20606 | 3.33836 | 3.56129 | 0.18810 | 6.24659 | 7.66662 | 3.98540 | 3.71829 | 3.33518 | 0.53025 | 3.21756 | 0.88726 | 3.39855 | 2.02081 |
| **Maz** | 3.06779 | 0.24728 | 0.06585 | 0.26413 | 0.01007 | 0.00323 | 0.04971 | 0.03705 | 1.23266 | 0.31152 | 0.00000 | 0.00402 | 0.26484 | 0.00987 | 0.04136 | 2.30536 |
| **Mb21d1** | 0.12988 | 0.00757 | 1.29666 | 1.46620 | 0.02924 | 0.05662 | 0.07863 | 0.02658 | 0.93856 | 0.65004 | 0.83944 | 0.04467 | 0.17267 | 0.50699 | 0.75231 | 0.02935 |
| **Mb21d2** | 0.01649 | 0.00788 | 0.00841 | 1.73752 | 0.00850 | 0.00639 | 0.00797 | 0.07942 | 0.00007 | 2.13574 | 0.03455 | 0.33779 | 0.00792 | 0.01390 | 0.50789 | 0.16596 |
| **Mbd1** | 1.10247 | 0.00574 | 0.85943 | 4.42345 | 0.69186 | 0.00585 | 2.10376 | 0.21095 | 0.08482 | 0.10751 | 1.32532 | 0.10554 | 1.66314 | 0.00434 | 1.67205 | 0.73109 |
| **Mbd2** | 0.49645 | 1.62253 | 0.29922 | 2.50903 | 0.00647 | 0.00484 | 1.08687 | 0.00000 | 1.53247 | 0.21757 | 0.57317 | 0.09689 | 1.43480 | 0.00000 | 0.00000 | 0.12408 |
| **Mbd3** | 14.84628 | 5.27235 | 11.08828 | 9.17326 | 13.90001 | 6.90091 | 4.17635 | 13.83124 | 6.77046 | 19.55132 | 9.83930 | 1.06055 | 14.31327 | 0.00000 | 0.33587 | 6.08422 |
| **Mbd3l2** | 0.00000 | 0.00729 | 0.00000 | 14.70152 | 0.00849 | 0.00314 | 0.00537 | 0.00000 | 10.22167 | 2.89616 | 2.05952 | 1.45120 | 0.73408 | 13.03835 | 10.80260 | 0.00308 |
| **Mbd4** | 1.40908 | 0.02743 | 0.50639 | 1.18292 | 0.21418 | 0.65407 | 0.07657 | 0.47133 | 0.94024 | 0.39957 | 1.14177 | 0.01295 | 0.21814 | 0.66466 | 1.78750 | 0.48095 |
| **Mbd5** | 0.76196 | 0.00785 | 0.00150 | 2.17778 | 0.01039 | 0.00689 | 0.00768 | 0.22076 | 0.83401 | 0.22872 | 0.00006 | 0.00236 | 0.05591 | 0.00179 | 1.47206 | 0.00689 |
| **Mbd6** | 3.26906 | 0.00285 | 1.75612 | 2.76822 | 2.43493 | 0.40139 | 0.02383 | 1.15891 | 0.83365 | 1.76866 | 2.55655 | 0.03102 | 2.59458 | 0.00000 | 0.07134 | 2.14462 |
| **Mbip** | 5.43054 | 4.09521 | 1.88830 | 2.56413 | 3.59111 | 2.52066 | 1.33301 | 1.64367 | 1.95177 | 8.64495 | 0.92000 | 1.29643 | 0.45840 | 0.86447 | 3.65615 | 2.55920 |
| **Mbl2** | 0.00000 | 0.00651 | 0.00000 | 2.09748 | 0.01024 | 0.00599 | 2.03431 | 1.36211 | 0.33038 | 1.95809 | 0.51032 | 0.08237 | 0.01024 | 2.52096 | 2.80971 | 0.00670 |
| **Mblac1** | 1.75300 | 0.00633 | 0.00000 | 0.00655 | 0.00506 | 0.20525 | 0.00649 | 0.05153 | 0.01158 | 1.28544 | 0.01099 | 0.04586 | 0.04310 | 0.01303 | 0.00571 | 0.00577 |
| **Mblac2** | 1.11438 | 0.00638 | 0.00000 | 0.00274 | 0.01655 | 0.95126 | 1.10144 | 0.00562 | 0.01033 | 0.96698 | 0.00748 | 0.01173 | 0.02603 | 0.02890 | 0.77921 | 0.00589 |
| **Mbnl1** | 0.01250 | 0.00796 | 0.00791 | 0.14395 | 0.00915 | 0.01238 | 0.00732 | 0.01051 | 0.00472 | 0.98539 | 0.00561 | 0.06628 | 0.01156 | 0.00569 | 3.52503 | 0.01468 |
| **Mbnl2** | 0.00450 | 0.43275 | 1.44370 | 0.00978 | 0.00561 | 0.66965 | 0.00668 | 0.00747 | 0.01252 | 0.00310 | 0.00916 | 0.01571 | 0.39255 | 0.06073 | 0.00000 | 0.00611 |
| **Mbnl3** | 0.00000 | 0.00845 | 1.14964 | 0.18739 | 0.05382 | 2.51142 | 0.00645 | 0.37366 | 0.03350 | 2.61900 | 1.26061 | 0.12395 | 1.32724 | 0.80102 | 1.36780 | 4.65660 |
| **Mboat1** | 1.28222 | 0.03061 | 0.85142 | 0.00137 | 0.08678 | 0.35865 | 0.66365 | 0.99140 | 0.00857 | 0.97637 | 1.59073 | 0.02628 | 0.34124 | 2.69228 | 0.23842 | 0.01210 |
| **Mboat2** | 3.76005 | 6.96716 | 3.47109 | 0.96795 | 3.88199 | 1.24791 | 5.47149 | 3.57787 | 3.05820 | 5.81144 | 7.69243 | 1.14562 | 6.83613 | 7.59083 | 5.19862 | 0.02301 |
| **Mboat4** | 0.03200 | 0.00820 | 2.03428 | 0.21262 | 0.05878 | 0.00600 | 0.03828 | 0.00858 | 0.01735 | 1.71376 | 0.02745 | 0.03420 | 0.01064 | 0.01037 | 0.00000 | 0.01373 |
| **Mboat7** | 5.63992 | 0.11773 | 0.81092 | 3.43867 | 0.88051 | 3.01116 | 0.02933 | 4.81635 | 3.17205 | 8.90693 | 2.83711 | 0.41406 | 4.67580 | 0.09072 | 0.71884 | 3.62115 |
| **Mbp** | 0.16653 | 0.01753 | 0.08328 | 0.14759 | 0.02286 | 4.18964 | 0.00581 | 0.00969 | 0.59577 | 0.00000 | 0.08301 | 0.02333 | 1.66930 | 0.14080 | 0.04153 | 2.98217 |
| **Mbtd1** | 10.87289 | 3.47837 | 10.32369 | 22.91087 | 0.11029 | 0.27487 | 7.81055 | 7.44573 | 15.96486 | 11.33342 | 4.30287 | 9.75072 | 2.79604 | 21.24008 | 21.65449 | 0.60179 |
| **Mbtps1** | 7.68449 | 2.47406 | 5.01398 | 7.86562 | 4.98921 | 5.52355 | 3.91232 | 7.11918 | 8.75624 | 13.69923 | 6.36710 | 5.30913 | 2.56846 | 1.37686 | 2.64001 | 3.28763 |
| **Mbtps2** | 4.61301 | 0.15871 | 1.06593 | 2.79568 | 0.02676 | 0.00646 | 1.12952 | 1.70711 | 3.04257 | 2.30878 | 0.32097 | 8.60518 | 3.15833 | 2.71020 | 1.58098 | 1.26625 |
| **Mc1r** | 0.02317 | 0.01908 | 0.11148 | 0.36768 | 0.01097 | 0.00726 | 0.03736 | 0.04904 | 0.19979 | 0.03995 | 0.33602 | 0.04085 | 0.12634 | 1.30405 | 0.10865 | 0.01452 |
| **Mcam** | 0.00962 | 0.00786 | 0.00814 | 0.00991 | 0.00965 | 0.00691 | 0.00743 | 0.39855 | 0.01181 | 0.00678 | 0.03847 | 0.02727 | 0.00593 | 0.01414 | 0.00343 | 0.00740 |
| **Mcat** | 7.62873 | 2.49288 | 3.60594 | 1.60526 | 6.91968 | 0.02055 | 0.27609 | 2.01516 | 0.82158 | 0.55312 | 0.32565 | 0.02273 | 1.97141 | 0.00223 | 0.00000 | 0.41169 |
| **Mcc** | 0.35642 | 0.05974 | 0.60186 | 4.27951 | 0.08113 | 0.12100 | 0.17747 | 0.02368 | 1.41338 | 0.36446 | 1.75372 | 0.24846 | 0.65091 | 3.65052 | 0.59880 | 1.08413 |
| **Mccc1** | 3.79237 | 3.59400 | 0.58213 | 0.00000 | 3.99963 | 3.14151 | 2.92360 | 5.39566 | 1.92285 | 0.51541 | 1.81099 | 8.00580 | 1.59613 | 0.00000 | 0.76914 | 5.30942 |
| **Mccc2** | 4.84248 | 2.14513 | 6.04210 | 0.00000 | 0.21111 | 6.37989 | 3.59125 | 4.20078 | 0.70593 | 3.22899 | 0.63656 | 0.05540 | 3.85082 | 1.53924 | 1.19468 | 8.19284 |
| **Mcee** | 3.82385 | 1.53095 | 2.58712 | 2.68637 | 4.15052 | 1.08148 | 2.84642 | 1.30266 | 0.79475 | 1.81138 | 0.97583 | 0.68789 | 2.24129 | 0.05516 | 0.69173 | 0.93052 |
| **Mcf2** | 2.54505 | 0.00359 | 0.00000 | 2.07486 | 1.97181 | 0.00375 | 0.00505 | 0.00000 | 1.09033 | 1.71166 | 0.00000 | 0.08184 | 0.00986 | 0.02590 | 2.33479 | 0.00391 |

| | | | | | | | | | | | | | | | | |
|---|---|---|---|---|---|---|---|---|---|---|---|---|---|---|---|---|
| **Mcfd2** | 7.06695 | 6.77659 | 8.85053 | 14.38744 | 2.10650 | 6.16090 | 6.40585 | 4.97518 | 6.46810 | 8.17202 | 5.50847 | 0.72131 | 6.57595 | 5.49455 | 8.19256 | 7.33391 |
| **Mcl1** | 12.49806 | 11.82700 | 18.22562 | 17.48066 | 8.40438 | 4.41723 | 9.31900 | 19.59627 | 23.12871 | 31.98765 | 25.61234 | 15.96488 | 16.12527 | 27.15468 | 27.12355 | 9.15543 |
| **Mcm10** | 13.13333 | 15.17631 | 9.86910 | 7.60841 | 11.28896 | 5.57365 | 3.91724 | 17.21433 | 11.76923 | 18.82415 | 12.77317 | 7.77453 | 12.37941 | 11.52413 | 16.43952 | 4.53363 |
| **Mcm2** | 21.37665 | 12.96063 | 16.59888 | 14.80257 | 12.88850 | 8.68814 | 13.65282 | 22.99169 | 15.47008 | 16.62050 | 14.93504 | 3.59354 | 13.64571 | 4.51288 | 14.98071 | 7.37582 |
| **Mcm3** | 17.26389 | 14.49312 | 11.74393 | 10.37237 | 6.71195 | 4.55960 | 12.08189 | 20.37285 | 10.02420 | 16.08175 | 8.41401 | 8.51014 | 8.54958 | 3.84746 | 14.97749 | 9.41618 |
| **Mcm3ap** | 3.23580 | 6.48625 | 4.43375 | 6.20813 | 9.44363 | 5.56017 | 4.18995 | 3.05589 | 7.59426 | 6.90602 | 2.33789 | 0.42885 | 0.57390 | 3.93301 | 1.64180 | 0.02481 |
| **Mcm4** | 11.12590 | 12.58709 | 9.49812 | 8.61846 | 6.66325 | 1.98076 | 9.31230 | 14.90789 | 12.17688 | 7.74036 | 7.34814 | 8.29261 | 5.74379 | 13.13537 | 18.50880 | 6.16102 |
| **Mcm5** | 17.85355 | 14.44858 | 16.87934 | 4.01248 | 14.82700 | 9.26262 | 11.31171 | 21.58874 | 6.76901 | 20.11317 | 10.23667 | 7.57176 | 10.57317 | 3.56117 | 10.28303 | 13.27660 |
| **Mcm6** | 17.00298 | 12.74304 | 17.22485 | 15.75122 | 11.96258 | 3.48517 | 10.29851 | 19.38615 | 26.05312 | 28.76009 | 17.88220 | 13.55688 | 10.90186 | 23.25649 | 25.94703 | 9.78082 |
| **Mcm7** | 18.92906 | 14.22099 | 19.49873 | 10.39455 | 10.48569 | 3.89233 | 16.08608 | 25.54494 | 9.38793 | 19.91328 | 21.07259 | 3.96517 | 19.07664 | 4.16160 | 9.69545 | 16.36857 |
| **Mcm8** | 4.98287 | 7.48709 | 7.08489 | 3.24289 | 7.90612 | 0.06339 | 3.99656 | 3.20546 | 1.06915 | 4.51425 | 2.73710 | 8.49695 | 0.48483 | 3.14108 | 4.17269 | 4.02975 |
| **Mcm9** | 0.48836 | 0.65424 | 1.34273 | 0.78364 | 0.01153 | 0.76746 | 0.00671 | 0.02050 | 0.65015 | 0.31604 | 0.00000 | 0.06713 | 0.19820 | 0.32814 | 0.12279 | 0.00598 |
| **Mcmbp** | 8.34841 | 5.50151 | 5.17537 | 13.01524 | 9.55210 | 10.06252 | 0.18789 | 9.71552 | 19.89329 | 12.85305 | 9.46448 | 7.13403 | 6.73472 | 6.97432 | 8.93383 | 2.54828 |
| **Mcmdc2** | 0.22644 | 0.01800 | 0.11903 | 0.44046 | 0.00740 | 0.03052 | 0.24566 | 0.06798 | 0.62404 | 0.26087 | 0.05759 | 0.10656 | 0.04800 | 0.70600 | 0.16974 | 0.04330 |
| **Mcoln1** | 0.00000 | 2.12652 | 1.35066 | 0.82165 | 0.01468 | 0.12658 | 0.00604 | 4.79115 | 0.00000 | 0.00000 | 1.10371 | 0.02792 | 0.01165 | 0.00106 | 0.97440 | 0.02060 |
| **Mcph1** | 2.44651 | 1.65597 | 3.27706 | 5.02695 | 3.01630 | 0.01091 | 0.09578 | 4.83158 | 3.37670 | 4.23948 | 2.79039 | 0.14860 | 1.90406 | 1.26296 | 11.99717 | 0.01537 |
| **Mcrs1** | 14.22075 | 12.56669 | 14.49878 | 13.91374 | 8.11209 | 5.67059 | 8.08706 | 28.12703 | 7.71093 | 25.25371 | 13.45569 | 2.06791 | 19.46390 | 0.02215 | 5.80826 | 10.23067 |
| **Mctp1** | 0.01365 | 0.00811 | 0.01202 | 0.66050 | 0.00929 | 0.00656 | 0.00752 | 0.01387 | 0.69281 | 0.01134 | 0.00844 | 0.00704 | 0.01286 | 0.01224 | 0.00666 | 0.00727 |
| **Mctp2** | 0.01270 | 0.00770 | 0.00672 | 2.15586 | 0.00954 | 0.00658 | 0.00711 | 0.00857 | 0.00000 | 0.00588 | 0.03202 | 0.01253 | 0.01275 | 1.27021 | 0.00000 | 0.00706 |
| **Mcts1** | 6.54183 | 4.18785 | 5.80273 | 3.42545 | 3.00557 | 0.10626 | 4.37514 | 4.24089 | 3.36805 | 5.69960 | 6.73023 | 0.00000 | 7.62369 | 8.30854 | 3.82187 | 7.20422 |
| **Mcts2** | 2.06850 | 0.44376 | 0.94174 | 4.85049 | 0.01749 | 2.66381 | 1.91411 | 3.84019 | 2.85395 | 6.94201 | 1.42981 | 0.57620 | 2.27984 | 1.31894 | 2.45927 | 0.00877 |
| **Mcu** | 0.08130 | 0.03917 | 0.19707 | 2.72679 | 1.45787 | 0.00875 | 0.01754 | 0.01070 | 0.32968 | 0.02144 | 0.06792 | 0.07285 | 0.00960 | 0.00759 | 0.47966 | 0.00657 |
| **Mcur1** | 4.82312 | 2.43465 | 0.46756 | 3.94229 | 0.90141 | 2.79011 | 2.68199 | 0.55693 | 1.90897 | 3.44611 | 6.38517 | 0.14402 | 2.64625 | 2.49378 | 0.78806 | 0.00000 |
| **Mdc1** | 7.13862 | 2.52840 | 5.77465 | 9.08002 | 1.03820 | 2.81879 | 1.82779 | 1.61470 | 13.56127 | 3.37069 | 2.33556 | 0.26282 | 4.36817 | 4.42483 | 9.67588 | 3.57970 |
| **Mdfic** | 0.01087 | 0.00783 | 0.01062 | 0.01068 | 0.00846 | 0.00661 | 0.00738 | 0.01218 | 0.01232 | 0.00654 | 0.01207 | 0.09908 | 0.01019 | 0.01234 | 0.00972 | 0.44553 |
| **Mdga1** | 0.01354 | 0.00802 | 0.01703 | 0.00698 | 0.00915 | 0.00731 | 0.00749 | 0.00977 | 0.01168 | 0.01137 | 0.17421 | 0.00742 | 0.01022 | 0.01245 | 0.01726 | 0.00737 |
| **Mdh1** | 13.73692 | 9.63461 | 25.24977 | 2.74039 | 8.48123 | 1.05992 | 6.86656 | 4.48023 | 2.42191 | 7.43506 | 0.45239 | 0.70688 | 8.49658 | 2.64258 | 2.17094 | 9.14282 |
| **Mdh2** | 36.24715 | 8.64001 | 21.64987 | 4.95123 | 21.69101 | 17.21373 | 12.96109 | 35.48682 | 8.94186 | 30.97258 | 19.03592 | 7.30956 | 31.83267 | 3.78485 | 9.66643 | 15.44672 |
| **Mdk** | 8.89933 | 0.00000 | 0.00000 | 0.38688 | 2.48507 | 0.91147 | 0.09969 | 1.14917 | 0.00000 | 0.11184 | 0.49615 | 0.06510 | 0.00000 | 0.00444 | 0.12417 | 3.08581 |
| **Mdm1** | 1.67631 | 0.01357 | 0.00333 | 0.87045 | 2.02032 | 0.00562 | 1.42595 | 0.00000 | 0.11971 | 0.65721 | 0.34701 | 0.04981 | 0.20590 | 0.00461 | 0.53086 | 0.01254 |
| **Mdm2** | 4.42745 | 1.63937 | 5.11497 | 19.62788 | 6.10992 | 0.52724 | 5.05108 | 6.69422 | 15.23785 | 10.04530 | 3.38052 | 9.57487 | 2.25461 | 7.34240 | 16.55128 | 2.52472 |
| **Mdm4** | 7.93436 | 4.00415 | 6.28488 | 28.33421 | 6.33268 | 2.22189 | 8.16790 | 3.67042 | 10.87807 | 11.00136 | 8.66011 | 5.16193 | 5.70455 | 22.14668 | 29.62864 | 2.78558 |
| **Mdn1** | 9.93421 | 15.89494 | 8.57662 | 11.01530 | 0.03205 | 4.27033 | 2.53957 | 10.73030 | 11.95786 | 23.66048 | 11.28876 | 15.93233 | 7.41740 | 26.43892 | 19.08898 | 3.83734 |
| **Mdp1** | 0.80508 | 4.46118 | 6.89517 | 2.29898 | 4.64847 | 0.05253 | 1.49874 | 1.57451 | 7.95322 | 10.45127 | 7.99499 | 0.33538 | 2.10542 | 1.67981 | 3.79813 | 1.60388 |
| **Mdrl** | 0.86271 | 3.54015 | 0.00000 | 1.13810 | 0.00632 | 0.00486 | 0.96081 | 0.02284 | 1.52865 | 1.47325 | 1.04019 | 1.86600 | 0.69073 | 0.12310 | 0.41549 | 0.00405 |
| **Me1** | 19.17904 | 17.98605 | 9.51308 | 6.46482 | 17.36872 | 6.44538 | 8.22153 | 1.19678 | 1.51871 | 10.19252 | 8.79786 | 4.59989 | 6.09830 | 3.32757 | 8.29050 | 2.68378 |
| **Me2** | 0.80062 | 3.25053 | 1.70582 | 6.45052 | 0.03313 | 0.00407 | 0.29782 | 2.07087 | 2.92738 | 3.61884 | 0.41270 | 0.43494 | 0.48011 | 0.00000 | 3.36171 | 0.17940 |
| **Me3** | 0.02232 | 0.01954 | 0.01062 | 0.01912 | 0.01120 | 0.00682 | 0.01783 | 0.01177 | 0.01167 | 0.44640 | 0.01065 | 0.00762 | 0.01240 | 0.00672 | 0.45654 | 0.00747 |
| **Mea1** | 11.27585 | 6.10171 | 8.16284 | 3.42437 | 4.83405 | 8.58997 | 3.62303 | 5.70849 | 1.46902 | 7.02870 | 2.56589 | 0.78953 | 5.73315 | 0.18635 | 2.97950 | 8.10375 |
| **Meaf6** | 10.34322 | 3.27814 | 9.55740 | 6.74780 | 0.24767 | 5.95746 | 2.61566 | 8.91819 | 7.39399 | 7.46910 | 10.50306 | 6.74207 | 6.61210 | 1.58767 | 2.78798 | 2.17834 |
| **Mecom** | 0.01340 | 0.00764 | 0.01163 | 0.03455 | 0.00916 | 0.00666 | 0.00739 | 0.01109 | 0.00930 | 0.62613 | 0.01014 | 0.00601 | 0.01197 | 0.01370 | 0.26685 | 0.00743 |
| **Mecp2** | 0.09035 | 0.00820 | 0.74497 | 0.29350 | 0.00679 | 0.00748 | 0.13043 | 0.51096 | 0.19514 | 0.37951 | 0.18416 | 0.02306 | 0.00220 | 0.76560 | 0.07640 | 0.01421 |
| **Mecr** | 5.46269 | 0.13348 | 0.00000 | 1.36951 | 1.27349 | 5.62277 | 2.27256 | 5.59630 | 1.93077 | 2.93681 | 1.59562 | 0.13691 | 2.41774 | 0.00000 | 0.61845 | 4.93980 |
| **Med1** | 6.07222 | 4.87805 | 6.35800 | 8.45596 | 3.23553 | 0.71355 | 2.27865 | 4.82110 | 10.44932 | 5.68814 | 5.79378 | 1.29649 | 1.94874 | 8.52748 | 9.30091 | 1.08288 |
| **Med10** | 15.71582 | 8.77622 | 12.50257 | 11.95251 | 7.99936 | 8.68189 | 6.52009 | 12.17617 | 17.73875 | 17.66411 | 8.66020 | 9.03712 | 14.39111 | 2.13919 | 6.95069 | 2.27261 |
| **Med11** | 9.76067 | 0.61660 | 4.30775 | 2.78450 | 6.01292 | 0.00796 | 2.96951 | 6.69788 | 7.79024 | 6.94773 | 3.29370 | 0.78535 | 2.19976 | 0.00000 | 0.80811 | 1.44292 |
| **Med12** | 3.17803 | 0.01486 | 0.00000 | 3.64993 | 1.56383 | 1.62124 | 0.11137 | 0.65947 | 5.12318 | 4.75903 | 1.48805 | 0.00000 | 2.76488 | 0.00000 | 0.00000 | 3.48171 |
| **Med12l** | 0.78391 | 0.00581 | 0.00000 | 0.10080 | 1.94716 | 4.05016 | 0.00605 | 1.19068 | 0.06383 | 0.18820 | 0.67612 | 0.00587 | 1.19716 | 1.67428 | 0.21679 | 0.00422 |
| **Med13** | 0.65515 | 3.32240 | 0.96807 | 13.98363 | 2.28604 | 0.00255 | 2.39168 | 1.06007 | 16.95935 | 0.64584 | 5.27112 | 3.40162 | 0.00000 | 11.81753 | 8.51742 | 1.59348 |
| **Med13l** | 2.59396 | 2.90259 | 3.01210 | 4.54482 | 0.01486 | 0.00200 | 0.81424 | 3.02018 | 2.72396 | 2.16598 | 1.68581 | 0.14438 | 0.58247 | 1.45901 | 3.07403 | 0.09162 |
| **Med14** | 0.51678 | 1.73244 | 2.48208 | 10.64953 | 0.14829 | 0.06993 | 0.03302 | 6.34184 | 12.41884 | 12.78888 | 2.97489 | 13.56431 | 0.47883 | 27.65881 | 21.13296 | 0.05153 |
| **Med15** | 8.72207 | 0.87978 | 2.61760 | 8.80225 | 4.43046 | 1.45636 | 1.27405 | 8.56658 | 2.97852 | 2.36494 | 1.94502 | 0.18442 | 2.82801 | 1.11373 | 2.56889 | 3.26556 |
| **Med16** | 5.64550 | 0.01081 | 0.73206 | 2.92441 | 2.98874 | 0.01141 | 0.06323 | 0.06480 | 4.08733 | 7.41602 | 3.14172 | 0.00000 | 3.00012 | 0.15187 | 0.35649 | 4.30987 |
| **Med17** | 8.51399 | 10.53099 | 8.87816 | 2.05567 | 6.54125 | 0.52412 | 3.26159 | 10.62403 | 6.14146 | 13.92943 | 7.86917 | 5.77809 | 7.31798 | 8.26839 | 12.54518 | 3.59522 |
| **Med18** | 0.29349 | 0.01887 | 0.03553 | 0.01522 | 0.01364 | 0.00542 | 0.02665 | 0.01487 | 0.94252 | 0.71397 | 0.05270 | 0.03301 | 1.91076 | 0.23714 | 0.51117 | 0.02037 |
| **Med19** | 8.69006 | 3.65130 | 3.62305 | 1.73108 | 3.91402 | 2.42150 | 1.04924 | 6.87210 | 4.71815 | 9.97118 | 4.65164 | 3.17748 | 3.29386 | 1.80147 | 4.15892 | 0.82657 |
| **Med20** | 6.32472 | 5.18360 | 6.53805 | 6.53536 | 3.61256 | 4.15712 | 6.75755 | 7.78014 | 6.77664 | 12.42553 | 7.85722 | 3.29093 | 8.03228 | 0.18865 | 5.61610 | 4.46680 |
| **Med21** | 10.22144 | 4.68896 | 7.22560 | 9.29027 | 10.26170 | 6.91230 | 2.91885 | 4.76393 | 14.26384 | 14.72652 | 11.26546 | 2.74845 | 7.32949 | 14.72374 | 9.45825 | 0.67475 |
| **Med22** | 5.14633 | 0.80936 | 6.39760 | 5.59695 | 0.09298 | 3.48646 | 3.83936 | 6.56571 | 3.16815 | 15.30215 | 2.77914 | 0.96850 | 6.12317 | 0.34711 | 5.09634 | 0.22974 |
| **Med23** | 5.32364 | 5.28648 | 2.27394 | 7.92285 | 0.00000 | 1.98003 | 5.67640 | 2.33586 | 8.18861 | 11.41709 | 3.38852 | 7.16105 | 1.90868 | 9.76707 | 12.41399 | 2.15510 |
| **Med24** | 0.99216 | 2.36488 | 4.52526 | 6.07069 | 5.80514 | 0.16329 | 4.41871 | 9.31894 | 6.24897 | 9.56855 | 7.34979 | 8.18276 | 5.30950 | 0.20528 | 0.31067 | 6.46931 |
| **Med25** | 5.08788 | 0.02122 | 0.00000 | 0.11734 | 3.88795 | 0.89714 | 0.03864 | 0.61699 | 0.21071 | 11.03583 | 0.31444 | 0.00000 | 13.44902 | 0.00906 | 0.00000 | 2.01913 |

| | | | | | | | | | | | | | | | | |
|---|---|---|---|---|---|---|---|---|---|---|---|---|---|---|---|---|
| **Med26** | 3.00921 | 2.48263 | 3.69344 | 11.94325 | 1.83759 | 3.14553 | 1.20773 | 1.39278 | 11.56917 | 9.98946 | 1.84851 | 0.52957 | 1.63873 | 3.55673 | 3.60779 | 0.07677 |
| **Med27** | 10.76066 | 8.41510 | 11.29022 | 16.44874 | 7.44330 | 0.00000 | 6.92072 | 12.73220 | 9.24715 | 26.10518 | 12.86306 | 3.22277 | 12.59001 | 2.43446 | 11.43647 | 5.65497 |
| **Med28** | 4.67566 | 1.70074 | 5.95222 | 12.94330 | 3.88719 | 1.31289 | 0.22369 | 4.38970 | 9.03582 | 20.67488 | 19.69636 | 1.10462 | 13.18344 | 2.02237 | 3.87138 | 5.90326 |
| **Med29** | 5.57062 | 0.05901 | 1.56650 | 20.50589 | 0.84728 | 0.38790 | 3.54275 | 7.03144 | 30.71079 | 18.10901 | 13.67751 | 1.55371 | 7.96484 | 1.21643 | 2.49690 | 1.39776 |
| **Med30** | 1.13311 | 0.29924 | 0.68788 | 2.17596 | 3.01444 | 0.25631 | 0.04708 | 0.00000 | 0.00000 | 0.16098 | 0.73311 | 0.48081 | 0.05410 | 0.00542 | 0.05144 | 0.59219 |
| **Med31** | 2.25099 | 1.10855 | 1.69234 | 7.47936 | 4.34092 | 0.74431 | 0.06202 | 1.35785 | 16.58496 | 6.85097 | 5.32531 | 0.54119 | 1.26659 | 3.87119 | 7.45865 | 0.97043 |
| **Med4** | 9.92721 | 11.95378 | 13.31387 | 8.87132 | 10.10018 | 1.81156 | 5.77337 | 10.28064 | 15.34003 | 16.67637 | 6.09859 | 10.90731 | 8.65301 | 7.70810 | 15.77506 | 1.24267 |
| **Med6** | 2.87946 | 5.34890 | 2.36911 | 5.80596 | 2.71364 | 1.81827 | 0.07831 | 7.60703 | 10.73423 | 5.61519 | 7.26568 | 9.65898 | 3.37538 | 13.85748 | 13.72835 | 1.19028 |
| **Med7** | 4.57867 | 6.22674 | 7.14034 | 9.88334 | 0.37810 | 1.98600 | 2.30845 | 1.72521 | 16.55734 | 9.36202 | 5.42087 | 3.02143 | 2.85077 | 6.30171 | 3.26579 | 0.55292 |
| **Med8** | 13.95023 | 13.07063 | 11.93183 | 4.03699 | 6.03421 | 0.63975 | 8.15848 | 8.79542 | 4.20072 | 10.89684 | 4.04582 | 7.07115 | 9.90266 | 5.46665 | 4.89657 | 2.85297 |
| **Med9** | 5.73575 | 1.76713 | 0.61554 | 5.89601 | 1.63573 | 0.00698 | 0.08049 | 0.63038 | 9.74261 | 6.00578 | 3.63110 | 0.83015 | 0.40733 | 0.17606 | 0.45617 | 0.75893 |
| **Medag** | 0.02148 | 0.01856 | 0.07502 | 0.11829 | 0.00867 | 0.00640 | 0.00763 | 0.01740 | 0.08812 | 0.02052 | 0.11227 | 0.00716 | 0.01013 | 0.06485 | 0.02133 | 0.00730 |
| **Mef2a** | 1.77026 | 0.00512 | 0.85575 | 4.88516 | 0.00303 | 0.00673 | 0.00578 | 0.03210 | 1.90898 | 0.33985 | 0.00000 | 2.77212 | 0.20033 | 0.13296 | 0.00003 | 0.00388 |
| **Mef2d** | 0.86965 | 0.54230 | 1.73242 | 0.68692 | 0.00247 | 4.45418 | 2.15275 | 4.84910 | 1.55818 | 5.83575 | 7.60679 | 0.04687 | 5.94574 | 4.65454 | 0.27985 | 1.43620 |
| **Meg3** | 3.26007 | 0.00444 | 0.01562 | 0.57664 | 0.01587 | 0.01334 | 0.01541 | 0.00637 | 0.00512 | 0.01225 | 0.10739 | 0.01528 | 0.02510 | 0.01272 | 0.05465 | 0.02570 |
| **Megf11** | 0.01358 | 0.00733 | 1.46413 | 0.53771 | 0.01370 | 0.21011 | 0.04672 | 0.93576 | 0.69724 | 0.32868 | 1.25001 | 0.00000 | 1.18976 | 0.00757 | 0.82786 | 0.92216 |
| **Megf6** | 0.00728 | 0.00770 | 0.01004 | 0.00422 | 0.00884 | 0.00833 | 0.00701 | 0.59666 | 0.11029 | 2.64896 | 1.37326 | 0.00000 | 0.00000 | 0.08681 | 0.00000 | 0.00692 |
| **Megf8** | 0.60899 | 0.00708 | 0.91538 | 0.49503 | 0.06385 | 0.00642 | 0.00515 | 0.28062 | 1.02807 | 1.26794 | 0.38204 | 0.00689 | 2.56113 | 0.13692 | 0.01129 | 0.01228 |
| **Megf9** | 0.01359 | 0.00761 | 0.01189 | 1.99156 | 0.00942 | 0.00625 | 0.00728 | 0.01133 | 0.55517 | 0.15627 | 0.00750 | 0.00491 | 0.01270 | 0.00246 | 0.42860 | 0.00712 |
| **Mei1** | 0.04541 | 0.00712 | 1.04805 | 3.50393 | 0.01388 | 0.00889 | 1.90774 | 0.51334 | 1.47486 | 3.56522 | 3.13381 | 0.13194 | 0.00000 | 0.06987 | 2.56405 | 0.00579 |
| **Mei4** | 0.03117 | 0.00794 | 0.06197 | 0.10769 | 0.00920 | 0.00717 | 0.00750 | 0.01272 | 0.12391 | 0.06439 | 0.10955 | 0.03608 | 0.05412 | 0.16028 | 0.02790 | 0.03623 |
| **Meikin** | 0.00461 | 0.00775 | 0.00000 | 1.03738 | 0.01033 | 0.00717 | 0.00658 | 0.37336 | 0.76129 | 0.61391 | 0.63427 | 0.00401 | 0.84502 | 0.86625 | 1.26497 | 0.00699 |
| **Meioc** | 0.01103 | 0.00755 | 0.00718 | 0.48259 | 0.01948 | 0.00660 | 0.00723 | 0.00923 | 1.38946 | 0.00000 | 0.00000 | 0.00000 | 0.01694 | 1.60153 | 0.05218 | 0.00723 |
| **Meis1** | 0.42055 | 0.00415 | 1.71974 | 0.27966 | 4.26223 | 0.00877 | 1.74712 | 0.02203 | 0.00181 | 0.54510 | 0.01000 | 0.01032 | 0.00409 | 0.01271 | 0.00000 | 2.74279 |
| **Meis2** | 0.00901 | 0.00778 | 0.00414 | 4.73479 | 0.00946 | 0.00550 | 0.00694 | 0.00766 | 1.27675 | 1.11504 | 0.00000 | 0.02302 | 0.01124 | 2.90689 | 4.01706 | 0.00633 |
| **Meis3** | 0.01362 | 0.00823 | 0.01208 | 0.43230 | 0.00917 | 0.00684 | 0.00757 | 0.01371 | 0.07671 | 0.18733 | 0.01229 | 0.00888 | 0.01289 | 0.01060 | 0.00625 | 0.00734 |
| **Melk** | 6.41256 | 9.94112 | 5.23177 | 4.64268 | 8.13610 | 5.42633 | 6.71166 | 10.13017 | 6.66973 | 9.93313 | 9.51681 | 14.34468 | 7.74689 | 13.11382 | 10.51820 | 5.04810 |
| **Memo1** | 5.54483 | 4.04784 | 1.28182 | 15.92047 | 5.56708 | 5.90943 | 2.01119 | 10.12863 | 19.38238 | 18.27028 | 14.17161 | 7.15274 | 4.08683 | 7.64335 | 10.29487 | 1.28466 |
| **Men1** | 8.13619 | 0.69321 | 3.90793 | 3.13287 | 1.98945 | 4.58371 | 5.30683 | 6.61214 | 0.00000 | 6.07389 | 1.77977 | 0.22211 | 4.48702 | 0.00000 | 2.00873 | 0.09992 |
| **Mep1b** | 6.36242 | 0.02097 | 0.00000 | 2.19745 | 0.00000 | 0.00456 | 0.00000 | 4.12432 | 0.78063 | 7.70138 | 3.30073 | 1.13099 | 0.58873 | 8.90220 | 5.12593 | 0.00000 |
| **Mepce** | 5.93305 | 3.52251 | 8.74911 | 13.42442 | 5.97403 | 1.52109 | 6.53413 | 6.42345 | 15.58428 | 10.68805 | 2.70634 | 3.32506 | 3.45268 | 0.11901 | 9.34306 | 0.02703 |
| **Mertk** | 0.07172 | 0.00626 | 0.44288 | 2.80832 | 0.00702 | 0.00700 | 0.00615 | 0.70465 | 3.91808 | 1.18653 | 0.31544 | 5.96793 | 0.67598 | 3.65606 | 3.32093 | 0.00551 |
| **Mesdc1** | 2.68025 | 0.06451 | 1.95446 | 1.13397 | 0.05129 | 0.00299 | 0.78251 | 1.24738 | 6.62491 | 1.48831 | 0.48228 | 0.21676 | 0.45776 | 0.16133 | 0.83283 | 3.55444 |
| **Mesdc2** | 15.95651 | 10.96173 | 23.74380 | 5.71693 | 14.08582 | 9.69018 | 14.01587 | 17.47918 | 5.91675 | 9.43631 | 8.79849 | 13.32187 | 16.17707 | 5.88946 | 10.02231 | 16.16544 |
| **Mesp2** | 0.04836 | 0.00795 | 0.57991 | 10.94651 | 0.01373 | 0.00415 | 0.00678 | 3.12118 | 6.89514 | 1.67887 | 0.64692 | 0.02083 | 0.00000 | 0.00000 | 0.19387 | 0.00530 |
| **Mest** | 0.52206 | 0.04643 | 0.00000 | 0.24268 | 0.01740 | 1.14680 | 0.00741 | 0.38681 | 0.00568 | 0.34191 | 0.00436 | 0.04621 | 0.70759 | 0.00786 | 1.15752 | 0.17003 |
| **Metap1** | 9.28740 | 5.59421 | 9.18259 | 3.95712 | 9.10744 | 6.20886 | 3.34860 | 14.43176 | 8.02650 | 10.40839 | 8.25827 | 6.71352 | 3.81327 | 9.79939 | 11.66860 | 3.16045 |
| **Metap1d** | 4.42662 | 5.63560 | 1.04634 | 0.17971 | 4.18104 | 1.59439 | 0.95268 | 1.60997 | 0.00138 | 3.30547 | 0.77805 | 0.18003 | 1.66456 | 1.39329 | 2.28351 | 0.03499 |
| **Metap2** | 21.32978 | 18.42017 | 26.02806 | 21.67600 | 19.15269 | 9.71341 | 16.00296 | 32.81056 | 24.45817 | 40.97060 | 34.81501 | 18.39117 | 24.97227 | 34.44104 | 31.64027 | 8.40780 |
| **Metrn** | 0.01193 | 0.00801 | 0.04682 | 0.09529 | 0.00907 | 0.00781 | 0.00732 | 0.01220 | 0.01042 | 0.01008 | 0.01313 | 0.00796 | 0.21367 | 0.01227 | 0.00806 | 0.00718 |
| **Mettl1** | 1.37408 | 0.02432 | 1.75904 | 0.11158 | 0.92691 | 0.65050 | 1.64415 | 1.85910 | 0.20868 | 3.23376 | 0.29200 | 3.21579 | 1.59100 | 0.41524 | 0.42362 | 0.46775 |
| **Mettl10** | 1.11246 | 0.78402 | 0.25781 | 0.37161 | 0.00772 | 0.00470 | 0.09937 | 0.33814 | 0.00320 | 0.44566 | 0.69215 | 0.06690 | 1.55320 | 0.97345 | 0.09122 | 2.97605 |
| **Mettl13** | 5.20201 | 3.43198 | 6.65063 | 3.31176 | 4.23561 | 0.95962 | 2.66905 | 8.49358 | 3.87534 | 4.35365 | 1.25746 | 0.45255 | 2.51187 | 1.74718 | 6.99048 | 2.07302 |
| **Mettl14** | 0.44631 | 0.01851 | 2.64799 | 0.18487 | 0.02214 | 0.00644 | 0.00466 | 3.41550 | 0.64351 | 3.67504 | 4.24268 | 0.32793 | 0.96255 | 2.14250 | 4.55002 | 0.35958 |
| **Mettl15** | 0.83191 | 0.00867 | 0.00083 | 0.01026 | 0.02172 | 0.00783 | 0.08793 | 0.25438 | 0.86821 | 0.87480 | 0.11975 | 0.09826 | 0.24594 | 0.02940 | 0.84519 | 0.02315 |
| **Mettl16** | 7.43996 | 9.55167 | 8.52094 | 3.41326 | 1.10690 | 0.00271 | 9.43742 | 6.17523 | 5.78103 | 9.42414 | 4.68451 | 3.86309 | 6.68944 | 0.64896 | 5.98175 | 3.48784 |
| **Mettl17** | 2.41509 | 0.02200 | 5.67833 | 2.71683 | 0.65691 | 4.31079 | 5.17480 | 6.33617 | 0.65938 | 4.22382 | 3.31591 | 0.13479 | 4.44112 | 0.04312 | 0.41073 | 1.95657 |
| **Mettl18** | 0.62270 | 0.01693 | 2.15908 | 0.25374 | 0.01149 | 0.01531 | 3.12017 | 0.06964 | 0.68074 | 1.22807 | 1.27076 | 0.07513 | 2.78445 | 0.00380 | 2.50693 | 0.02560 |
| **Mettl2** | 8.63960 | 2.13930 | 3.10030 | 3.32394 | 4.45307 | 3.15487 | 3.40939 | 6.86679 | 10.24208 | 9.21119 | 5.85103 | 5.31875 | 4.67096 | 3.06502 | 7.46516 | 0.74530 |
| **Mettl20** | 0.12167 | 0.00810 | 0.65118 | 0.61276 | 0.00590 | 0.00723 | 0.00786 | 0.46718 | 0.08108 | 0.21737 | 0.02021 | 0.01706 | 0.00000 | 0.09307 | 1.45387 | 1.41129 |
| **Mettl21a** | 2.24854 | 2.04418 | 0.84784 | 0.38395 | 0.01300 | 0.00694 | 0.00393 | 1.15343 | 0.18983 | 1.21925 | 0.31134 | 0.00000 | 2.30033 | 0.00377 | 1.40358 | 0.28613 |
| **Mettl21c** | 0.01359 | 0.00801 | 0.01213 | 0.32796 | 0.00912 | 0.00660 | 0.00732 | 0.01106 | 0.16691 | 0.39299 | 0.01129 | 0.00644 | 0.01237 | 0.00803 | 0.04169 | 0.00775 |
| **Mettl22** | 4.94944 | 0.17248 | 2.17799 | 0.56322 | 0.00000 | 1.85819 | 3.08839 | 2.35771 | 0.72012 | 0.39380 | 0.39929 | 0.16738 | 2.43511 | 0.02240 | 0.08614 | 0.86760 |
| **Mettl23** | 1.51140 | 0.10423 | 0.65162 | 0.47260 | 2.84905 | 0.00456 | 1.81990 | 0.45281 | 2.01335 | 2.28190 | 1.00475 | 0.05692 | 1.29211 | 0.24518 | 0.57269 | 1.89764 |
| **Mettl25** | 0.42464 | 0.11256 | 1.18427 | 0.10994 | 0.43882 | 0.00827 | 0.00603 | 0.00000 | 0.43350 | 0.59767 | 1.49601 | 0.02541 | 0.69964 | 0.01996 | 0.53137 | 0.51151 |
| **Mettl26** | 6.02487 | 0.04927 | 1.93133 | 0.63221 | 1.82347 | 0.28539 | 1.08710 | 2.87692 | 0.96746 | 0.76795 | 4.75666 | 0.01174 | 5.75447 | 0.00000 | 0.00000 | 3.62330 |
| **Mettl3** | 3.69841 | 3.64054 | 3.59720 | 2.07753 | 0.95953 | 0.66511 | 3.78676 | 2.02432 | 1.96164 | 2.84194 | 0.91152 | 0.26818 | 1.33372 | 2.86788 | 4.69838 | 2.73229 |
| **Mettl4** | 1.32965 | 4.85892 | 0.72427 | 5.58037 | 0.00406 | 0.00544 | 0.00364 | 1.75454 | 1.83608 | 1.78459 | 1.17491 | 0.55087 | 0.39720 | 1.64825 | 4.64752 | 0.00204 |
| **Mettl5** | 5.91573 | 3.91210 | 3.23721 | 0.51528 | 4.28627 | 0.86381 | 0.56978 | 0.85120 | 0.57764 | 3.37845 | 1.66352 | 0.19995 | 2.98611 | 3.32013 | 0.49507 | 4.38089 |
| **Mettl6** | 3.07867 | 2.72685 | 4.89984 | 4.23630 | 0.09258 | 0.01122 | 2.33108 | 2.47805 | 2.17109 | 2.61852 | 1.66528 | 0.57050 | 0.33324 | 0.00000 | 3.44164 | 3.60781 |
| **Mettl7a1** | 0.24776 | 0.00764 | 0.00370 | 0.78662 | 0.00693 | 0.04933 | 0.07161 | 0.01268 | 0.14425 | 0.15670 | 0.09199 | 0.00670 | 0.06361 | 0.01100 | 0.01137 | 1.51348 |
| **Mettl7a2** | 0.05081 | 0.00778 | 0.00267 | 1.26005 | 0.00815 | 0.11988 | 0.00764 | 0.00752 | 0.79032 | 0.02486 | 0.25083 | 0.00533 | 0.01241 | 0.01378 | 0.04042 | 0.23623 |

| | | | | | | | | | | | | | | | | |
|---|---|---|---|---|---|---|---|---|---|---|---|---|---|---|---|---|
| **Mettl7a3** | 0.01437 | 0.00799 | 0.01125 | 0.01057 | 0.00926 | 0.01334 | 0.08025 | 0.01282 | 0.01237 | 0.00655 | 0.01377 | 0.00769 | 0.51037 | 0.01296 | 0.01023 | 0.03623 |
| **Mettl8** | 4.41661 | 0.00348 | 0.83369 | 1.63612 | 0.00000 | 0.00172 | 0.24648 | 0.57920 | 0.01156 | 1.77261 | 0.00708 | 0.03363 | 0.72974 | 0.00435 | 1.01321 | 0.00980 |
| **Mettl9** | 6.60499 | 3.09480 | 4.31426 | 23.46282 | 5.61068 | 0.30141 | 4.08822 | 1.09127 | 11.60931 | 7.40873 | 4.17341 | 0.21675 | 0.80627 | 3.74203 | 4.85151 | 0.55080 |
| **Mex3a** | 0.09821 | 0.00786 | 0.09078 | 0.15597 | 0.01028 | 1.41108 | 0.00713 | 0.00000 | 1.16274 | 0.00000 | 0.43649 | 0.01164 | 0.01656 | 0.00507 | 0.44102 | 0.00647 |
| **Mex3b** | 0.01009 | 0.01757 | 0.52137 | 0.03842 | 2.69962 | 0.00573 | 0.00708 | 0.01897 | 0.82177 | 0.00000 | 1.54109 | 0.13113 | 0.00000 | 0.00725 | 0.65808 | 0.01417 |
| **Mex3c** | 0.49335 | 3.39177 | 0.99675 | 10.86102 | 5.43095 | 0.09234 | 0.68768 | 4.59615 | 11.17839 | 1.75464 | 4.78318 | 5.13804 | 0.76219 | 25.66683 | 7.85220 | 0.02196 |
| **Mex3d** | 0.29575 | 0.53942 | 0.00000 | 0.87897 | 0.00989 | 0.00750 | 0.01607 | 0.18041 | 0.04645 | 1.06387 | 1.34733 | 0.02874 | 0.33370 | 0.00712 | 0.69446 | 0.00635 |
| **Mfap1a** | 2.40062 | 6.78977 | 5.10165 | 4.88143 | 1.49789 | 0.33320 | 4.75643 | 2.77542 | 9.27576 | 8.39881 | 2.52046 | 0.98737 | 3.54492 | 8.81005 | 10.13989 | 1.86802 |
| **Mfap1b** | 0.72370 | 1.75254 | 1.24779 | 0.00000 | 0.48664 | 0.03974 | 1.53171 | 0.85088 | 0.17245 | 1.92125 | 0.00000 | 0.10713 | 0.86389 | 2.01194 | 3.32332 | 0.48431 |
| **Mfap2** | 0.01301 | 0.00821 | 0.00988 | 2.70449 | 0.00893 | 0.00641 | 0.00735 | 0.00893 | 3.82286 | 0.50452 | 0.05943 | 0.03724 | 0.01864 | 0.00462 | 0.02391 | 0.00667 |
| **Mfap3** | 4.12525 | 1.54218 | 3.49030 | 1.44689 | 1.20529 | 0.00000 | 4.09088 | 0.00000 | 0.59299 | 1.03776 | 0.92619 | 3.77013 | 0.10813 | 3.79000 | 2.47221 | 1.51660 |
| **Mfap3l** | 0.01387 | 0.00782 | 0.00824 | 0.76120 | 0.00932 | 0.00662 | 0.00719 | 0.01006 | 0.51465 | 0.00000 | 0.01972 | 0.33512 | 0.01286 | 0.00470 | 0.31603 | 0.00721 |
| **Mfap4** | 0.01461 | 2.73481 | 0.08238 | 1.47442 | 0.00752 | 0.00558 | 0.00642 | 2.55130 | 0.63947 | 0.34937 | 0.00000 | 0.01263 | 0.02411 | 0.00172 | 0.69352 | 0.00572 |
| **Mfap5** | 0.37872 | 0.00646 | 0.30668 | 4.82631 | 0.02764 | 0.00570 | 0.33157 | 1.86050 | 1.65980 | 2.10044 | 3.58802 | 0.68711 | 1.39088 | 1.01078 | 2.15195 | 0.00477 |
| **Mff** | 6.31794 | 2.21117 | 4.17793 | 10.66244 | 9.33087 | 8.27812 | 2.50032 | 2.73178 | 2.99410 | 3.90580 | 0.07711 | 0.33757 | 2.59959 | 1.81354 | 2.43604 | 0.11154 |
| **Mfge8** | 10.22776 | 2.15405 | 7.78022 | 0.52362 | 7.12400 | 3.20757 | 7.52424 | 11.85050 | 0.02526 | 10.67225 | 7.08547 | 0.25927 | 15.10138 | 0.13800 | 0.05369 | 12.74567 |
| **Mfhas1** | 0.01120 | 0.06157 | 0.20662 | 0.17566 | 0.00855 | 0.00567 | 0.00739 | 0.01054 | 0.19621 | 0.76634 | 0.01021 | 0.01553 | 0.01053 | 0.01045 | 0.00236 | 0.00751 |
| **Mfi2** | 0.00789 | 0.00726 | 0.88578 | 0.01157 | 0.00830 | 0.00914 | 0.00696 | 2.06962 | 0.01042 | 0.00000 | 0.00000 | 0.00337 | 0.00747 | 0.01912 | 0.00000 | 0.00700 |
| **Mfn1** | 17.24894 | 13.77738 | 17.56211 | 8.84917 | 11.44353 | 10.18929 | 9.59207 | 22.41371 | 9.36041 | 16.65738 | 11.69678 | 10.32604 | 8.45984 | 6.79465 | 10.69384 | 3.85470 |
| **Mfn2** | 2.10735 | 4.65411 | 4.26538 | 9.26705 | 3.58292 | 2.27601 | 5.59759 | 1.85867 | 1.40805 | 10.70569 | 6.75087 | 0.42735 | 4.44159 | 0.21645 | 4.87171 | 1.77761 |
| **Mfng** | 0.28006 | 0.00680 | 0.00267 | 0.01042 | 0.84948 | 0.00825 | 0.00689 | 0.47290 | 0.01133 | 0.00899 | 0.01002 | 0.00579 | 0.00567 | 0.01409 | 0.00389 | 0.00696 |
| **Mfsd1** | 19.06529 | 7.18646 | 21.29429 | 7.84700 | 16.49742 | 9.68610 | 14.18217 | 13.81430 | 1.47048 | 13.66687 | 16.23000 | 12.55272 | 17.72167 | 14.59564 | 13.62269 | 13.19706 |
| **Mfsd10** | 5.90813 | 0.00000 | 1.38289 | 0.63278 | 1.66192 | 0.35403 | 2.09947 | 10.22759 | 0.73556 | 3.30801 | 0.73365 | 0.99500 | 8.00750 | 0.00000 | 0.00000 | 4.82764 |
| **Mfsd11** | 2.79682 | 0.12446 | 6.99765 | 8.36198 | 1.22379 | 5.58343 | 4.30303 | 0.39925 | 10.39916 | 0.21311 | 10.59495 | 0.98450 | 3.09878 | 3.81471 | 3.31657 | 2.28901 |
| **Mfsd12** | 0.01275 | 0.00796 | 0.04206 | 0.06592 | 0.00917 | 0.00812 | 0.00738 | 0.01251 | 0.57941 | 0.04188 | 0.00857 | 0.00663 | 0.03300 | 0.01151 | 0.00766 | 0.00737 |
| **Mfsd13a** | 4.48030 | 0.00357 | 0.05923 | 2.36730 | 0.15818 | 0.45562 | 0.05916 | 0.34451 | 0.18979 | 1.03964 | 0.00767 | 0.01000 | 2.89858 | 0.00864 | 0.00419 | 1.36476 |
| **Mfsd14a** | 3.41918 | 3.09834 | 4.20202 | 15.31878 | 2.44820 | 1.55904 | 7.84486 | 13.00793 | 14.83611 | 6.07904 | 7.45653 | 11.67428 | 4.22308 | 25.03588 | 23.43365 | 0.01373 |
| **Mfsd14b** | 1.10915 | 0.00000 | 3.33984 | 0.86161 | 1.10284 | 0.89946 | 3.33585 | 6.20030 | 2.60387 | 2.70577 | 4.96439 | 12.20549 | 1.53588 | 28.66554 | 13.56076 | 1.19485 |
| **Mfsd2a** | 0.01272 | 0.00796 | 0.00608 | 6.52593 | 0.00891 | 0.00546 | 0.00711 | 0.00000 | 8.92207 | 0.01382 | 0.21198 | 0.03285 | 0.01042 | 0.00000 | 0.99970 | 0.00559 |
| **Mfsd2b** | 0.43029 | 0.00741 | 0.02914 | 1.34226 | 0.01520 | 0.00616 | 0.00574 | 0.02177 | 2.14604 | 0.00000 | 2.94570 | 0.00000 | 1.70729 | 0.32381 | 0.05905 | 0.01199 |
| **Mfsd3** | 0.14950 | 0.00691 | 1.02902 | 0.00933 | 0.00607 | 0.00930 | 0.54729 | 0.96575 | 0.02652 | 0.00278 | 0.00718 | 0.00409 | 0.62723 | 0.01525 | 0.00211 | 0.00668 |
| **Mfsd4a** | 0.08840 | 0.02939 | 0.14942 | 0.09862 | 0.05211 | 0.00605 | 0.01764 | 0.01338 | 0.27565 | 0.02456 | 0.32815 | 0.01499 | 0.05021 | 0.16304 | 0.07052 | 0.03646 |
| **Mfsd4b1** | 1.75932 | 0.00632 | 0.07642 | 0.00418 | 0.00506 | 0.00748 | 0.00614 | 0.00770 | 0.42849 | 0.00249 | 0.00868 | 0.00459 | 0.07968 | 0.38546 | 0.02692 | 0.00578 |
| **Mfsd4b2** | 0.07033 | 0.00801 | 0.00951 | 0.04100 | 0.00798 | 0.00680 | 0.00747 | 0.01448 | 0.00954 | 0.01286 | 0.01247 | 0.00829 | 0.01265 | 0.00897 | 0.05857 | 0.00726 |
| **Mfsd4b3** | 0.07980 | 0.00810 | 0.01005 | 0.01035 | 0.00822 | 0.00710 | 0.00761 | 0.01665 | 0.01224 | 0.01407 | 0.01413 | 0.00824 | 0.02658 | 0.01098 | 0.02410 | 0.00744 |
| **Mfsd4b5** | 0.05434 | 0.00795 | 0.03689 | 0.09795 | 0.07786 | 0.00775 | 0.01807 | 0.02801 | 0.15189 | 0.06692 | 0.01130 | 0.00738 | 0.04382 | 0.04011 | 0.16963 | 0.00743 |
| **Mfsd5** | 5.80431 | 0.07278 | 2.53732 | 10.11388 | 0.03862 | 1.61162 | 2.18845 | 19.50164 | 19.21866 | 22.49513 | 12.50900 | 2.71693 | 6.33910 | 0.08814 | 7.81143 | 0.14516 |
| **Mfsd6** | 0.01180 | 0.00747 | 0.01291 | 1.38919 | 0.00912 | 0.00698 | 0.00710 | 0.00793 | 1.79734 | 0.50627 | 0.02578 | 0.00000 | 0.01209 | 0.65356 | 0.79837 | 0.00745 |
| **Mfsd7a** | 0.00751 | 0.00776 | 1.41171 | 0.00427 | 0.00787 | 0.00562 | 0.00651 | 0.47585 | 0.21274 | 0.03653 | 2.24689 | 0.00000 | 0.00000 | 0.17663 | 0.14022 | 0.00638 |
| **Mfsd7b** | 1.71586 | 0.50528 | 0.17141 | 4.14213 | 0.00538 | 0.05944 | 0.00580 | 0.32637 | 5.49320 | 0.83954 | 0.96167 | 0.01144 | 0.43324 | 0.00378 | 1.79973 | 0.86054 |
| **Mfsd7c** | 5.65240 | 6.50114 | 12.00394 | 2.43485 | 0.14950 | 3.63370 | 9.82543 | 24.86569 | 4.25719 | 16.41592 | 18.51620 | 2.77960 | 19.39539 | 15.17261 | 16.93711 | 0.36410 |
| **Mfsd8** | 0.85101 | 0.00409 | 0.00000 | 0.74474 | 2.17633 | 2.57786 | 0.06578 | 0.81099 | 0.00000 | 1.94426 | 1.30029 | 3.33239 | 0.20387 | 0.09320 | 1.56871 | 0.00503 |
| **Mfsd9** | 0.01341 | 0.00751 | 0.01057 | 2.14413 | 0.00925 | 0.00612 | 0.00714 | 0.00995 | 0.00000 | 0.00323 | 0.00982 | 0.01512 | 0.07615 | 0.00875 | 1.17087 | 0.00686 |
| **Mga** | 2.57480 | 4.16749 | 2.26344 | 13.77754 | 2.06624 | 0.03449 | 0.38204 | 3.70574 | 13.74870 | 3.35490 | 5.41289 | 0.84979 | 1.85035 | 21.57330 | 15.57193 | 0.11597 |
| **Mgarp** | 22.51104 | 8.99422 | 13.96071 | 0.15036 | 12.02446 | 0.00000 | 9.24652 | 4.35020 | 0.69178 | 4.12688 | 4.86444 | 0.22646 | 1.61713 | 0.32347 | 0.00000 | 7.05695 |
| **Mgat1** | 0.10736 | 0.00737 | 0.89697 | 0.88693 | 0.00765 | 0.33182 | 0.00699 | 0.27598 | 0.00000 | 0.72154 | 0.56885 | 0.06000 | 0.03512 | 0.00305 | 0.44961 | 0.00701 |
| **Mgat2** | 1.46923 | 3.29134 | 4.60526 | 14.80849 | 3.77498 | 4.80595 | 2.91085 | 7.87464 | 22.28284 | 9.62106 | 4.67393 | 5.14876 | 3.81646 | 8.60417 | 13.31957 | 3.05393 |
| **Mgat3** | 0.48454 | 0.00681 | 0.00000 | 0.85416 | 0.00657 | 1.68095 | 0.00522 | 1.67508 | 1.36934 | 0.00000 | 2.40414 | 0.03370 | 0.79057 | 2.64862 | 3.17299 | 0.42954 |
| **Mgat4a** | 0.00000 | 0.00686 | 1.51844 | 3.01642 | 0.01581 | 2.10116 | 0.00585 | 3.64370 | 1.18800 | 2.69915 | 2.03956 | 0.05895 | 0.57945 | 1.50855 | 2.80997 | 0.72650 |
| **Mgat4b** | 0.07046 | 0.00770 | 0.11131 | 0.45724 | 0.02064 | 0.04055 | 0.03833 | 0.01208 | 1.03711 | 0.46177 | 0.00272 | 0.02212 | 0.08400 | 0.00384 | 0.24709 | 0.21350 |
| **Mgat4e** | 0.01381 | 0.00807 | 0.01102 | 0.26007 | 0.00932 | 0.00660 | 0.00741 | 0.00879 | 0.00505 | 0.07843 | 0.17926 | 0.00685 | 0.01032 | 0.00964 | 0.00000 | 0.00736 |
| **Mgat5** | 0.01575 | 0.00795 | 0.04547 | 0.09456 | 0.00923 | 0.01189 | 0.00744 | 0.01341 | 0.00820 | 0.01532 | 0.01269 | 0.00796 | 0.01192 | 0.01188 | 0.00827 | 0.00735 |
| **Mgea5** | 8.92168 | 1.96875 | 0.50219 | 10.91502 | 3.17734 | 3.94271 | 2.88143 | 6.60052 | 10.02074 | 21.34790 | 8.40982 | 13.60751 | 0.67569 | 16.06494 | 18.01365 | 2.21233 |
| **Mgl2** | 0.24527 | 0.19904 | 0.54092 | 1.19572 | 0.01637 | 0.05295 | 0.17682 | 4.84813 | 1.11892 | 1.36496 | 0.21048 | 0.21237 | 4.28953 | 0.62388 | 0.91471 | 0.32707 |
| **Mgll** | 1.05430 | 0.94443 | 2.43060 | 0.00835 | 4.22582 | 0.01663 | 2.23319 | 0.00140 | 0.00999 | 0.00000 | 0.01081 | 0.01934 | 0.09932 | 0.01358 | 0.00000 | 0.42505 |
| **Mgme1** | 1.50952 | 3.35618 | 7.69187 | 2.80864 | 0.00515 | 1.68781 | 0.10630 | 8.71320 | 1.23289 | 7.27539 | 2.71653 | 1.06920 | 2.26839 | 2.63283 | 12.13884 | 1.30504 |
| **Mgmt** | 0.07080 | 0.00423 | 1.00081 | 0.00691 | 5.57323 | 0.00541 | 0.00686 | 0.36851 | 0.02230 | 0.00480 | 0.00922 | 0.01952 | 0.00000 | 0.01585 | 0.00000 | 0.02108 |
| **Mgp** | 0.01357 | 0.00801 | 0.01219 | 0.00893 | 0.00916 | 0.00665 | 0.00744 | 0.01177 | 0.01213 | 0.40494 | 0.01192 | 0.00720 | 0.01184 | 0.01156 | 0.00991 | 0.00734 |
| **Mgrn1** | 3.64157 | 0.00000 | 1.64253 | 6.10400 | 0.00000 | 1.46178 | 5.43475 | 7.52073 | 2.46014 | 10.70706 | 8.94526 | 0.52028 | 0.78269 | 0.03213 | 3.56643 | 1.55777 |
| **Mgst1** | 0.01295 | 0.00790 | 0.01206 | 0.00657 | 0.00885 | 0.00672 | 0.00735 | 0.01063 | 0.01148 | 1.46672 | 0.01093 | 0.00469 | 0.01137 | 0.01202 | 0.00869 | 0.00750 |
| **Mgst2** | 0.59079 | 0.79992 | 0.00000 | 0.00950 | 0.02271 | 1.43165 | 0.00565 | 0.00708 | 0.01269 | 0.00000 | 0.01087 | 0.02618 | 1.29299 | 0.01368 | 0.00000 | 0.44496 |
| **Mgst3** | 0.11795 | 0.33938 | 0.39171 | 0.21324 | 0.02242 | 4.13668 | 0.00632 | 0.01724 | 0.40004 | 0.09354 | 0.02927 | 0.01312 | 0.55608 | 0.00483 | 0.59194 | 5.56515 |

| | | | | | | | | | | | | | | | | |
|---|---|---|---|---|---|---|---|---|---|---|---|---|---|---|---|---|
| **Mia** | 0.01349 | 0.00779 | 0.00910 | 0.00635 | 0.00915 | 0.00665 | 0.00723 | 0.00608 | 0.01186 | 0.30068 | 0.16362 | 0.04297 | 0.02853 | 0.17052 | 0.00000 | 0.00720 |
| **Mia2** | 0.01361 | 0.00814 | 0.01108 | 0.00367 | 0.00938 | 0.00663 | 0.00744 | 0.01195 | 0.32777 | 0.00571 | 0.00815 | 0.00708 | 0.01333 | 0.07921 | 0.48590 | 0.00741 |
| **Mia3** | 5.72263 | 2.64620 | 7.42414 | 2.15738 | 1.36355 | 3.95180 | 7.86228 | 7.89372 | 3.35532 | 3.25949 | 4.71134 | 1.58614 | 2.01408 | 0.83801 | 2.60210 | 1.78337 |
| **Mib1** | 0.09704 | 0.00662 | 0.08805 | 0.08701 | 0.00829 | 0.94221 | 0.16660 | 1.91340 | 2.83317 | 1.49549 | 0.68792 | 1.84256 | 0.00000 | 0.56741 | 7.27071 | 0.09597 |
| **Mib2** | 2.14843 | 0.01493 | 0.00000 | 0.01155 | 1.12620 | 0.01122 | 0.03460 | 0.26756 | 0.09707 | 0.77195 | 0.00937 | 0.00000 | 3.28923 | 0.00821 | 0.03446 | 7.65762 |
| **Mical1** | 3.14331 | 2.96148 | 0.66499 | 0.09859 | 2.27843 | 0.04370 | 1.29824 | 1.96752 | 0.00660 | 0.00000 | 0.66089 | 0.00139 | 3.83573 | 0.00438 | 0.00000 | 5.42769 |
| **Mical2** | 0.28104 | 0.00718 | 0.00519 | 0.15420 | 0.00731 | 0.00715 | 0.00715 | 0.00946 | 0.00645 | 0.30186 | 0.01106 | 0.05484 | 0.01229 | 0.00886 | 0.18729 | 0.00736 |
| **Mical3** | 2.61283 | 3.09814 | 5.34184 | 2.29400 | 0.01950 | 0.00354 | 0.00138 | 8.14875 | 1.03907 | 5.11914 | 4.24185 | 0.11154 | 0.96149 | 2.80427 | 8.00374 | 0.00696 |
| **Micall1** | 0.82605 | 0.01629 | 0.00000 | 8.72076 | 0.00771 | 0.00615 | 0.00552 | 1.59824 | 9.92346 | 3.58638 | 2.46798 | 0.15386 | 0.00000 | 0.00000 | 0.95544 | 0.00428 |
| **Micall2** | 0.04167 | 0.00776 | 0.71754 | 0.73328 | 0.00713 | 0.00553 | 0.01724 | 0.01014 | 0.00000 | 0.00958 | 0.08098 | 0.04800 | 0.02192 | 0.00728 | 0.04614 | 0.01411 |
| **Micu1** | 2.84352 | 0.08464 | 5.59237 | 4.32161 | 0.17821 | 0.01173 | 0.07153 | 0.57354 | 4.83753 | 0.60804 | 1.70034 | 0.18402 | 1.73108 | 0.99391 | 0.00001 | 2.31368 |
| **Micu2** | 3.94595 | 0.00218 | 0.00000 | 2.53280 | 1.86387 | 2.70465 | 0.00380 | 0.91793 | 1.98705 | 0.83635 | 0.68000 | 0.00000 | 0.00000 | 0.09217 | 1.44989 | 0.00947 |
| **Mid1** | 0.33507 | 0.00867 | 0.00465 | 0.38205 | 0.01954 | 1.27942 | 0.00733 | 0.01190 | 0.18195 | 0.17386 | 0.01008 | 0.00657 | 0.23508 | 0.00528 | 0.25636 | 0.11339 |
| **Mid1ip1** | 0.00000 | 0.00819 | 0.43161 | 2.14155 | 0.01292 | 0.23406 | 0.00659 | 0.45232 | 7.66261 | 3.67150 | 0.19561 | 0.00000 | 2.14363 | 0.00509 | 0.00000 | 6.35256 |
| **Midn** | 0.17708 | 0.68835 | 0.82575 | 4.45658 | 0.31480 | 0.31513 | 0.29747 | 0.00000 | 1.88968 | 0.13864 | 1.79137 | 0.00000 | 0.36912 | 0.68629 | 0.01216 | 0.20925 |
| **Mief1** | 3.41477 | 1.78679 | 3.02634 | 2.78960 | 2.21853 | 4.30464 | 0.14110 | 2.72396 | 4.83271 | 6.97978 | 2.64843 | 5.09665 | 4.04705 | 5.95241 | 9.81367 | 2.33025 |
| **Mief2** | 0.69267 | 0.02893 | 1.06523 | 0.51914 | 0.04447 | 0.00530 | 0.17870 | 2.14150 | 1.50080 | 0.67481 | 1.20221 | 0.00796 | 0.50308 | 0.45020 | 1.62021 | 0.99861 |
| **Mien1** | 15.73031 | 6.52972 | 9.08280 | 1.33148 | 6.49198 | 8.74599 | 7.53562 | 8.87998 | 0.57771 | 8.45728 | 7.18746 | 0.41012 | 19.10762 | 0.02321 | 0.25192 | 7.59059 |
| **Mier1** | 2.11585 | 2.12820 | 0.31199 | 1.04537 | 0.00394 | 3.06543 | 0.07636 | 0.15446 | 1.12600 | 1.23304 | 0.93093 | 6.84374 | 0.20155 | 6.64681 | 1.11161 | 0.32052 |
| **Mier2** | 3.75910 | 0.00395 | 0.11815 | 0.60095 | 0.02855 | 0.09534 | 0.19601 | 0.42542 | 0.56142 | 1.33831 | 0.10714 | 0.04867 | 3.20696 | 0.00846 | 0.00000 | 3.65530 |
| **Mier3** | 4.50609 | 6.01900 | 11.34619 | 13.73155 | 1.25689 | 1.18849 | 12.34509 | 17.81131 | 19.98520 | 33.56780 | 24.99625 | 30.44471 | 7.86096 | 44.35948 | 36.42490 | 1.74257 |
| **Mif** | 12.52828 | 0.17802 | 4.87356 | 4.53913 | 11.71003 | 2.37821 | 2.59116 | 5.27633 | 3.98804 | 11.49969 | 9.60715 | 0.67111 | 9.60836 | 0.00000 | 0.00000 | 4.44625 |
| **Mif4gd** | 7.41332 | 0.45301 | 0.67772 | 5.32616 | 0.04176 | 1.17745 | 3.66130 | 7.48291 | 5.09723 | 4.32530 | 6.23142 | 0.35625 | 2.19126 | 0.00000 | 0.37477 | 4.31776 |
| **Miip** | 6.90174 | 0.52985 | 7.71365 | 0.30329 | 11.20458 | 5.65346 | 3.37810 | 17.81686 | 3.05624 | 2.04921 | 4.13764 | 0.10850 | 10.99948 | 0.00693 | 0.25450 | 5.16329 |
| **Milr1** | 1.07604 | 0.46247 | 0.29915 | 0.30863 | 0.00679 | 0.03439 | 0.00622 | 0.00000 | 0.43171 | 0.67954 | 1.00277 | 0.02543 | 0.01553 | 3.65715 | 0.00000 | 0.00544 |
| **Mina** | 0.45854 | 0.00556 | 0.47417 | 0.20161 | 1.03933 | 1.66017 | 1.05732 | 3.99113 | 0.79790 | 3.84577 | 0.00000 | 0.17249 | 2.61261 | 3.37172 | 5.84618 | 0.01933 |
| **Mink1** | 0.17326 | 0.01800 | 0.21015 | 0.73941 | 0.01716 | 0.08205 | 0.06796 | 0.80480 | 0.71148 | 2.70905 | 3.88663 | 0.23386 | 0.28629 | 0.00000 | 0.00000 | 0.00633 |
| **Minos1** | 9.52063 | 1.27392 | 12.69965 | 13.92114 | 7.26011 | 1.73553 | 3.13247 | 0.69958 | 14.27723 | 7.03186 | 16.30322 | 1.14892 | 7.44553 | 1.81780 | 3.54805 | 3.62866 |
| **Minpp1** | 3.72569 | 0.14932 | 3.75080 | 0.65354 | 4.38688 | 0.08253 | 0.00189 | 0.96615 | 0.14916 | 3.91798 | 0.03467 | 0.17797 | 4.16195 | 0.00591 | 9.58835 | 2.03206 |
| **Mios** | 4.70505 | 4.53151 | 0.00000 | 8.49256 | 2.08052 | 0.46567 | 0.01009 | 2.87612 | 2.70247 | 5.39750 | 2.99924 | 2.75787 | 0.35633 | 4.18955 | 13.40768 | 0.14651 |
| **Miox** | 0.00090 | 0.00625 | 0.00000 | 6.15969 | 3.22652 | 0.00648 | 0.00775 | 0.75143 | 6.19699 | 0.45724 | 0.00000 | 0.09887 | 0.09006 | 0.25176 | 2.39038 | 0.00623 |
| **Mip** | 0.01843 | 0.00850 | 0.01015 | 0.01012 | 0.01057 | 0.00671 | 0.00740 | 0.01111 | 0.01231 | 0.03730 | 0.01137 | 0.00724 | 0.01244 | 0.24867 | 0.02527 | 0.00730 |
| **Mipep** | 1.55467 | 0.00268 | 1.71261 | 1.04719 | 5.79957 | 3.38843 | 0.01403 | 1.74184 | 1.28534 | 0.44972 | 1.90576 | 0.18102 | 1.97833 | 1.06093 | 3.20692 | 1.16034 |
| **Mipol1** | 0.05370 | 0.00814 | 0.01050 | 0.02102 | 0.00853 | 0.00721 | 0.00768 | 0.01522 | 0.35715 | 0.00800 | 0.11430 | 0.00698 | 0.03802 | 0.01222 | 0.00523 | 0.00748 |
| **Mir1291** | 0.01118 | 0.00850 | 0.07266 | 0.00965 | 0.02802 | 0.00585 | 0.00790 | 0.02645 | 0.01199 | 0.03714 | 0.01164 | 0.00807 | 0.06138 | 0.06338 | 0.14621 | 0.00734 |
| **Mir17hg** | 0.00000 | 1.29362 | 0.13994 | 0.21728 | 3.33300 | 0.00561 | 0.01632 | 0.80090 | 0.17686 | 1.21558 | 0.90497 | 0.00000 | 0.61919 | 0.00875 | 0.38446 | 0.92403 |
| **Mir1949** | 0.02853 | 0.00810 | 0.00869 | 0.01047 | 0.01242 | 0.00684 | 0.00732 | 0.01054 | 0.01258 | 0.01022 | 0.01114 | 0.00639 | 0.01310 | 0.38155 | 0.00460 | 0.00755 |
| **Mir1982** | 0.04632 | 0.06017 | 0.02239 | 0.00890 | 0.00802 | 0.00769 | 0.01735 | 0.01650 | 0.01225 | 0.11619 | 0.04897 | 0.01655 | 0.10010 | 0.01076 | 0.00817 | 0.02878 |
| **Mir1a.1hg** | 0.03119 | 0.01869 | 0.01486 | 0.10732 | 0.01740 | 0.01263 | 0.01825 | 0.01724 | 0.13279 | 0.03197 | 0.01167 | 0.00771 | 0.01217 | 0.03251 | 0.04516 | 0.00734 |
| **Mir22hg** | 0.00867 | 0.00771 | 0.01547 | 0.33897 | 0.00906 | 0.00683 | 0.00663 | 0.00844 | 0.20421 | 0.00000 | 0.00791 | 0.01416 | 1.08816 | 0.05068 | 1.00777 | 0.01365 |
| **Mir25** | 0.00937 | 0.00800 | 0.11779 | 0.01045 | 0.01010 | 0.00596 | 0.00753 | 0.17788 | 0.01219 | 0.07522 | 0.01279 | 0.00722 | 0.01458 | 0.01253 | 0.00535 | 0.00765 |
| **Mir3064** | 0.07089 | 0.00787 | 0.00359 | 0.00863 | 0.10891 | 0.36457 | 0.14450 | 0.03937 | 0.01168 | 0.07358 | 0.00844 | 0.05089 | 0.10159 | 0.36957 | 0.20425 | 0.11158 |
| **Mir3068** | 0.01637 | 0.00787 | 0.11185 | 0.01032 | 0.00907 | 0.00579 | 0.00737 | 0.01264 | 0.01205 | 0.01212 | 0.01286 | 0.00792 | 0.01110 | 0.01088 | 0.06811 | 0.00730 |
| **Mir3091** | 0.20896 | 0.00754 | 0.09677 | 0.02610 | 0.00773 | 0.00630 | 0.01830 | 0.01021 | 0.02394 | 0.18450 | 0.09608 | 0.00726 | 0.05437 | 0.01624 | 0.00447 | 0.08863 |
| **Mir3471.1** | 0.09417 | 0.03805 | 0.06133 | 0.43988 | 0.00776 | 0.01234 | 0.01732 | 0.01551 | 0.41182 | 0.10463 | 0.00840 | 0.00588 | 0.02435 | 0.24107 | 0.07480 | 0.00701 |
| **Mir3471.2** | 0.01376 | 0.00785 | 0.01203 | 0.19197 | 0.00932 | 0.00661 | 0.00750 | 0.01327 | 0.19548 | 0.01390 | 0.01157 | 0.00739 | 0.01284 | 0.01231 | 0.07422 | 0.00742 |
| **Mir3473f** | 0.01205 | 0.00763 | 0.03305 | 0.15696 | 0.03663 | 0.02027 | 0.01749 | 0.00512 | 0.16820 | 0.02387 | 0.21341 | 0.00568 | 0.01811 | 0.17381 | 0.04471 | 0.00728 |
| **Mir425** | 0.01465 | 0.00807 | 0.02040 | 0.01170 | 0.00927 | 0.00627 | 0.00749 | 0.01361 | 0.43715 | 0.01262 | 0.01052 | 0.00749 | 0.01261 | 0.01207 | 0.01473 | 0.00737 |
| **Mir6236** | 0.04023 | 0.11269 | 0.02702 | 0.01440 | 0.10557 | 0.09196 | 0.01718 | 0.03045 | 0.03980 | 0.02983 | 0.01298 | 0.00725 | 0.00976 | 0.01088 | 0.00870 | 0.00736 |
| **Mir6385** | 0.01598 | 0.00763 | 0.03358 | 0.00701 | 0.01776 | 0.00818 | 0.00701 | 0.05666 | 0.01163 | 0.14487 | 0.15128 | 0.00544 | 0.11035 | 0.16595 | 0.00757 | 0.00754 |
| **Mir6516** | 0.01314 | 0.00806 | 0.01145 | 0.00444 | 0.09060 | 0.08135 | 0.00745 | 0.03577 | 0.43924 | 0.01100 | 0.00902 | 0.01797 | 0.00992 | 0.01254 | 0.01259 | 0.00734 |
| **Mir677** | 0.01375 | 0.00818 | 0.01224 | 0.01030 | 0.00932 | 0.00664 | 0.00753 | 0.01378 | 0.01223 | 0.01415 | 0.01322 | 0.00839 | 0.01268 | 0.01053 | 0.03136 | 0.00741 |
| **Mir680.2** | 0.05471 | 0.00781 | 0.01514 | 0.13471 | 0.00820 | 0.00636 | 0.00737 | 0.01601 | 0.06020 | 0.03198 | 0.03699 | 0.03714 | 0.01178 | 0.03947 | 0.08332 | 0.00726 |
| **Mir682** | 0.05381 | 0.00809 | 0.02279 | 0.06815 | 0.04110 | 0.01968 | 0.01807 | 0.00520 | 0.30794 | 0.01605 | 0.38728 | 0.00629 | 0.03354 | 0.01056 | 0.03338 | 0.03607 |
| **Mir684.1** | 0.23964 | 0.24449 | 0.71155 | 0.60250 | 0.24752 | 0.15418 | 0.40634 | 0.53177 | 0.75924 | 1.06870 | 3.87210 | 0.97984 | 0.45455 | 1.36995 | 0.20925 | 0.29410 |
| **Mir684.2** | 0.11198 | 0.01894 | 0.17285 | 0.15738 | 0.06428 | 0.04373 | 0.03855 | 0.11111 | 0.09669 | 0.30455 | 0.47696 | 0.22338 | 0.07680 | 0.28413 | 0.06762 | 0.01444 |
| **Mir6910** | 0.01119 | 0.00802 | 0.01025 | 0.01015 | 0.00940 | 0.00661 | 0.00742 | 0.10501 | 0.01203 | 0.06167 | 0.01333 | 0.00845 | 0.02316 | 0.01527 | 0.02282 | 0.00717 |
| **Mir692.1** | 1.71281 | 0.63815 | 1.17258 | 0.16642 | 0.87103 | 1.04436 | 0.74576 | 0.68039 | 0.09348 | 0.53181 | 0.67791 | 0.10309 | 1.60421 | 0.00653 | 0.00000 | 0.54052 |
| **Mir6935** | 0.04314 | 0.00778 | 0.10822 | 0.01027 | 0.00794 | 0.08396 | 0.00735 | 0.03657 | 0.01221 | 0.01259 | 0.01267 | 0.00815 | 0.01617 | 0.01047 | 0.03149 | 0.00723 |
| **Mir6951** | 0.01318 | 0.00778 | 0.08466 | 0.01758 | 0.00931 | 0.00579 | 0.27428 | 0.03617 | 0.01240 | 0.01263 | 0.01343 | 0.00780 | 0.00987 | 0.01051 | 0.02809 | 0.00732 |
| **Mir697** | 0.01270 | 0.00807 | 0.11336 | 0.00968 | 0.00905 | 0.00878 | 0.00735 | 0.01254 | 0.01201 | 0.01537 | 0.01260 | 0.00794 | 0.01111 | 0.00781 | 0.11775 | 0.00735 |
| **Mir7013** | 0.01284 | 0.00793 | 0.04824 | 0.10124 | 0.00907 | 0.00816 | 0.00731 | 0.01151 | 0.33312 | 0.16506 | 0.01199 | 0.00753 | 0.01111 | 0.01758 | 0.01182 | 0.00711 |

| | | | | | | | | | | | | | | | | |
|---|---|---|---|---|---|---|---|---|---|---|---|---|---|---|---|---|
| **Mir7014** | 0.01315 | 0.00774 | 0.00990 | 0.01071 | 0.00925 | 0.00673 | 0.00722 | 0.12772 | 0.01223 | 0.01281 | 0.01227 | 0.00745 | 0.00812 | 0.01245 | 0.00863 | 0.00738 |
| **Mir7024** | 0.01263 | 0.00822 | 0.01718 | 0.01005 | 0.01853 | 0.00626 | 0.00755 | 0.01332 | 0.01237 | 0.10296 | 0.01192 | 0.00805 | 0.02387 | 0.01075 | 0.01693 | 0.00713 |
| **Mir703** | 15.72767 | 9.22024 | 15.04841 | 0.98959 | 12.59559 | 5.11577 | 6.57794 | 8.06564 | 4.09362 | 8.88567 | 17.00820 | 4.86606 | 11.43050 | 10.31249 | 4.91944 | 7.81702 |
| **Mir7036** | 0.12609 | 0.00753 | 0.11211 | 0.01059 | 0.00736 | 0.00882 | 0.34632 | 0.04936 | 0.01228 | 0.00889 | 0.01263 | 0.00740 | 0.08528 | 0.01327 | 0.00781 | 0.12795 |
| **Mir7036b** | 0.01369 | 0.00817 | 0.01216 | 0.02030 | 0.00924 | 0.00673 | 0.00759 | 0.01379 | 0.10926 | 0.08349 | 0.01127 | 0.00826 | 0.01254 | 0.01171 | 0.01009 | 0.00731 |
| **Mir7051** | 0.01070 | 0.06181 | 0.05331 | 0.03549 | 0.00916 | 0.00662 | 0.00769 | 0.03724 | 0.00984 | 0.02001 | 0.01219 | 0.00804 | 0.02188 | 0.01616 | 0.02463 | 0.00753 |
| **Mir7067** | 0.02805 | 0.00771 | 0.04848 | 0.11346 | 0.00818 | 0.00764 | 0.09613 | 0.10189 | 0.10563 | 0.09255 | 0.01266 | 0.01765 | 0.00780 | 0.01199 | 0.01632 | 0.12106 |
| **Mir7069** | 0.01335 | 0.00767 | 0.03637 | 0.01060 | 0.00933 | 0.00592 | 0.00719 | 0.07576 | 0.01235 | 0.01282 | 0.01273 | 0.00739 | 0.03198 | 0.01125 | 0.01400 | 0.00731 |
| **Mir7079** | 0.05036 | 0.17587 | 0.20349 | 0.19489 | 0.52655 | 0.00680 | 0.00714 | 0.23519 | 0.64494 | 0.20005 | 0.29365 | 0.09885 | 0.21349 | 0.00548 | 0.15286 | 0.26770 |
| **Mir763** | 0.01233 | 0.00764 | 0.01469 | 0.01051 | 0.00947 | 0.00735 | 0.00724 | 0.01000 | 0.01261 | 0.00803 | 0.00915 | 0.00456 | 0.01308 | 0.88457 | 0.00299 | 0.00745 |
| **Mir8091** | 0.01161 | 0.00844 | 0.01250 | 0.02548 | 0.01025 | 0.00713 | 0.00824 | 0.06485 | 0.07271 | 0.00569 | 0.01365 | 0.00801 | 0.08058 | 0.01214 | 0.22494 | 0.00766 |
| **Mir8112** | 0.06962 | 0.00838 | 0.00152 | 0.00972 | 0.14420 | 0.13334 | 0.00760 | 0.01499 | 0.01217 | 0.15084 | 0.01328 | 0.01974 | 0.01760 | 0.01678 | 0.00847 | 0.00716 |
| **Mira** | 0.01368 | 0.00790 | 0.01136 | 1.90846 | 0.00922 | 0.00669 | 0.00749 | 0.01281 | 0.18736 | 0.12298 | 0.02126 | 0.03002 | 0.01289 | 0.00373 | 0.21302 | 0.00708 |
| **Mirc35hg** | 0.30861 | 0.02782 | 0.54650 | 0.07301 | 0.15127 | 0.05924 | 0.08614 | 0.00000 | 2.17801 | 0.06951 | 2.26275 | 0.17528 | 0.68964 | 1.52615 | 0.48425 | 0.12058 |
| **Mirt1** | 0.09812 | 0.00744 | 0.21879 | 0.28679 | 0.06483 | 0.03121 | 0.07827 | 0.04534 | 0.88963 | 0.09270 | 0.34403 | 0.09133 | 0.11232 | 0.39077 | 0.11827 | 0.02158 |
| **Mis12** | 1.09538 | 4.15810 | 4.41550 | 7.65829 | 0.00000 | 0.72412 | 2.59826 | 3.00860 | 3.89058 | 2.73521 | 0.15952 | 2.07642 | 0.00000 | 5.62312 | 9.16361 | 1.09978 |
| **Mis18a** | 9.78952 | 0.57911 | 2.99225 | 13.01840 | 2.00711 | 0.00000 | 0.96574 | 7.98827 | 15.19026 | 18.84549 | 6.16126 | 3.55622 | 8.54880 | 2.51242 | 6.58965 | 3.16911 |
| **Mis18bp1** | 2.12612 | 2.62413 | 2.36907 | 7.63736 | 0.00341 | 5.12407 | 0.05135 | 1.27839 | 9.22697 | 3.45689 | 2.67588 | 14.26361 | 2.08028 | 5.55264 | 5.86978 | 0.02007 |
| **Misp** | 0.19219 | 0.00708 | 0.17803 | 0.43860 | 0.00727 | 0.00877 | 0.00696 | 0.57682 | 0.09616 | 0.00699 | 0.00894 | 0.00598 | 0.00488 | 0.01137 | 0.00000 | 0.00700 |
| **Mitd1** | 1.77417 | 4.91200 | 4.23337 | 0.00000 | 2.74101 | 2.15980 | 2.63959 | 0.92627 | 0.66217 | 2.72283 | 0.78331 | 4.57137 | 2.15958 | 4.05714 | 1.64014 | 2.66842 |
| **Mitf** | 1.78430 | 0.00708 | 0.20499 | 0.14263 | 0.01959 | 0.00436 | 0.00651 | 0.01009 | 0.03422 | 0.02073 | 0.01247 | 0.00600 | 0.12931 | 0.00558 | 0.29987 | 0.00586 |
| **Mixl1** | 0.26714 | 0.00628 | 0.00000 | 0.35172 | 0.00711 | 0.00586 | 0.00615 | 1.16971 | 0.00363 | 1.27435 | 1.33548 | 1.67692 | 0.00000 | 1.41093 | 1.66651 | 0.00663 |
| **Mki67** | 6.84866 | 5.25063 | 4.74744 | 19.57566 | 11.96662 | 0.57446 | 0.02547 | 4.76790 | 20.96991 | 6.91084 | 3.55327 | 5.12368 | 4.87171 | 12.31630 | 16.22458 | 0.04023 |
| **Mkks** | 5.60597 | 0.49679 | 4.51554 | 8.79511 | 7.70351 | 4.97268 | 3.74406 | 3.03591 | 3.60522 | 5.75881 | 6.88528 | 9.83889 | 3.90008 | 6.44367 | 9.71565 | 3.03268 |
| **Mkl1** | 1.19999 | 0.94379 | 0.29285 | 1.87527 | 0.01198 | 0.00606 | 2.35478 | 3.76703 | 1.08929 | 0.28624 | 0.40404 | 0.05389 | 1.44512 | 0.00012 | 0.05859 | 0.01213 |
| **Mkl2** | 0.06407 | 0.00740 | 0.30228 | 2.50752 | 0.04587 | 0.03063 | 0.23902 | 0.51056 | 0.13482 | 0.38170 | 0.89033 | 0.14842 | 0.10186 | 0.18421 | 0.63406 | 0.19251 |
| **Mkln1** | 3.65844 | 5.61645 | 1.99402 | 3.90408 | 1.06681 | 3.50552 | 0.55522 | 4.25623 | 1.44195 | 0.57873 | 4.44152 | 6.68132 | 1.00774 | 3.04684 | 2.28542 | 1.91650 |
| **Mknk1** | 1.44880 | 0.00552 | 0.95036 | 4.25207 | 0.00290 | 2.30873 | 0.00513 | 2.84570 | 8.36287 | 4.63609 | 1.26086 | 0.57325 | 0.00000 | 0.11807 | 4.42202 | 1.13247 |
| **Mknk2** | 4.24926 | 0.32855 | 0.48171 | 7.67016 | 0.64362 | 0.33687 | 0.98400 | 5.42185 | 11.06021 | 10.60117 | 8.65174 | 0.49562 | 0.84506 | 0.65362 | 2.32021 | 2.03199 |
| **Mkrn1** | 21.90979 | 13.33158 | 21.22295 | 16.10314 | 20.97321 | 8.62630 | 21.13364 | 21.90482 | 27.55099 | 33.11440 | 35.87794 | 17.45094 | 22.20912 | 12.73970 | 10.30246 | 5.35947 |
| **Mkrn2** | 5.10612 | 2.38814 | 7.46550 | 15.18471 | 0.01309 | 0.96717 | 1.13941 | 10.18035 | 15.57859 | 11.46942 | 9.28277 | 0.84778 | 4.53135 | 6.73110 | 17.13359 | 2.80896 |
| **Mkrn2os** | 0.01587 | 0.00732 | 0.00000 | 3.02900 | 0.01822 | 0.00695 | 0.01622 | 0.00000 | 3.93823 | 5.49374 | 5.25833 | 0.38690 | 0.00000 | 0.69637 | 3.36681 | 0.04268 |
| **Mks1** | 1.77804 | 0.02787 | 1.97721 | 0.00433 | 0.01842 | 0.00607 | 0.63767 | 0.06590 | 0.01088 | 0.00000 | 0.31047 | 0.04700 | 2.12488 | 0.02427 | 0.23454 | 0.01114 |
| **Mlec** | 8.28847 | 0.36158 | 9.28943 | 2.02263 | 11.10252 | 9.24467 | 7.16009 | 6.12054 | 1.20399 | 3.53901 | 0.98531 | 2.50042 | 3.63468 | 1.68260 | 3.29266 | 7.26074 |
| **Mlf1** | 0.01295 | 0.00801 | 0.02082 | 0.75826 | 0.00919 | 0.00595 | 0.00760 | 0.01246 | 2.12640 | 0.09304 | 0.00000 | 0.00372 | 0.01197 | 0.01415 | 0.06683 | 0.01403 |
| **Mlf2** | 24.38399 | 6.11258 | 32.07277 | 17.01498 | 19.57434 | 11.27966 | 15.19482 | 37.17365 | 14.06107 | 40.08944 | 33.41572 | 4.49999 | 30.37183 | 1.61337 | 3.97201 | 15.65774 |
| **Mlh1** | 1.44774 | 3.25831 | 1.10045 | 0.01020 | 0.20045 | 0.01629 | 4.48909 | 3.34615 | 1.22903 | 1.08541 | 0.89763 | 1.46882 | 3.89391 | 0.00909 | 0.08871 | 0.00189 |
| **Mlh3** | 1.50103 | 0.00549 | 1.62650 | 0.23783 | 0.08463 | 0.01268 | 0.04538 | 0.01116 | 0.35646 | 0.43247 | 0.15285 | 0.04785 | 0.10516 | 3.80069 | 0.89199 | 0.01936 |
| **Mlkl** | 0.01124 | 0.00709 | 0.00744 | 0.21313 | 0.00951 | 0.00680 | 1.49258 | 0.00961 | 0.07204 | 0.00933 | 0.01158 | 0.05433 | 0.01052 | 0.03249 | 0.00476 | 0.00732 |
| **Mllt1** | 0.78693 | 0.00542 | 0.32520 | 3.06701 | 0.10095 | 0.00453 | 2.54994 | 0.35526 | 4.80027 | 1.73074 | 2.68045 | 0.14388 | 1.73260 | 0.00000 | 0.42297 | 0.01208 |
| **Mllt10** | 2.07032 | 2.83639 | 4.70243 | 7.74561 | 0.01289 | 0.01207 | 1.05864 | 5.87211 | 2.63487 | 3.43313 | 2.93053 | 4.74100 | 2.28437 | 9.06139 | 11.22132 | 0.98051 |
| **Mllt11** | 1.38797 | 0.00602 | 0.45804 | 6.26788 | 0.00560 | 0.00318 | 0.04614 | 0.00000 | 4.66311 | 0.52726 | 0.46526 | 0.00000 | 0.00775 | 0.14238 | 1.29675 | 0.01147 |
| **Mllt3** | 0.87437 | 0.00653 | 0.81056 | 8.29064 | 0.67309 | 0.00378 | 0.11591 | 0.01947 | 3.11196 | 0.70850 | 0.54317 | 0.04162 | 0.01656 | 1.41960 | 2.51470 | 0.71824 |
| **Mllt4** | 6.05562 | 0.05187 | 2.51095 | 9.71311 | 1.77554 | 3.66094 | 2.66207 | 2.54383 | 2.51028 | 8.52467 | 2.08509 | 8.59705 | 3.59969 | 9.28766 | 4.60900 | 0.63382 |
| **Mllt6** | 5.38165 | 5.86556 | 3.78125 | 2.48239 | 0.01460 | 5.92753 | 6.07672 | 6.93529 | 1.46235 | 5.08709 | 6.34027 | 0.69615 | 6.66113 | 2.15804 | 2.40255 | 8.41888 |
| **Mlph** | 0.02936 | 0.01813 | 0.02840 | 0.21127 | 0.00834 | 0.00722 | 0.01760 | 0.02138 | 0.03005 | 0.03258 | 0.01255 | 0.01743 | 0.01879 | 0.00909 | 0.02840 | 0.00732 |
| **Mlst8** | 4.01955 | 6.39977 | 7.84881 | 0.98492 | 3.84769 | 0.40580 | 4.04253 | 7.34509 | 2.10248 | 3.56522 | 4.09880 | 0.32447 | 2.58903 | 5.68996 | 4.42806 | 0.03244 |
| **Mlx** | 5.57973 | 3.80636 | 0.23938 | 0.35253 | 8.31305 | 3.09483 | 0.29376 | 4.90565 | 1.20270 | 0.71291 | 1.29054 | 0.53177 | 1.65631 | 2.73713 | 4.10328 | 2.30923 |
| **Mlxip** | 2.84058 | 0.00344 | 0.22427 | 3.09974 | 2.48758 | 2.34984 | 0.00465 | 1.71829 | 1.46226 | 2.82978 | 0.00000 | 0.39180 | 0.48873 | 2.22964 | 5.51684 | 0.27943 |
| **Mlxipl** | 1.59139 | 0.00512 | 0.90471 | 7.39692 | 0.01082 | 2.04823 | 0.66960 | 0.29316 | 7.53158 | 1.37602 | 6.34936 | 0.00000 | 0.38412 | 0.00000 | 0.47934 | 0.00183 |
| **Mlycd** | 0.28932 | 1.22079 | 0.48732 | 0.19709 | 2.04687 | 0.01707 | 0.23326 | 0.80752 | 0.00312 | 0.00571 | 0.00715 | 0.00408 | 0.00538 | 0.00944 | 0.00000 | 0.00639 |
| **Mmaa** | 3.05967 | 4.00046 | 3.53770 | 0.23792 | 2.52530 | 0.70843 | 1.59377 | 2.58949 | 2.97980 | 1.18670 | 1.30133 | 0.27644 | 1.66366 | 0.82404 | 1.00843 | 0.03592 |
| **Mmab** | 1.35635 | 0.89389 | 1.68560 | 0.45269 | 0.00834 | 0.00414 | 0.00498 | 0.31680 | 0.02041 | 0.92673 | 0.00681 | 0.00000 | 1.37473 | 0.00392 | 0.36985 | 3.17615 |
| **Mmachc** | 4.97889 | 4.41475 | 6.53204 | 0.03499 | 3.73255 | 3.44084 | 4.08158 | 1.70002 | 0.82530 | 2.82534 | 1.97431 | 0.13594 | 0.11342 | 4.62797 | 4.32480 | 0.05413 |
| **Mmadhc** | 15.58909 | 8.76991 | 9.44666 | 3.52235 | 11.41933 | 8.67956 | 3.60092 | 8.19066 | 0.52858 | 4.22968 | 3.49632 | 9.70039 | 6.18386 | 1.40941 | 3.40202 | 2.25205 |
| **Mmd** | 0.01351 | 0.00782 | 0.01069 | 0.21133 | 0.00922 | 0.00674 | 0.00752 | 0.01209 | 0.58990 | 0.19317 | 0.00492 | 0.01771 | 0.01320 | 0.01069 | 0.70392 | 0.00735 |
| **Mmd2** | 1.08095 | 0.00618 | 0.00315 | 0.00476 | 0.00586 | 0.00738 | 0.00649 | 0.00000 | 0.24933 | 0.19605 | 0.50971 | 0.01284 | 0.01126 | 0.01197 | 0.16351 | 0.00652 |
| **Mme** | 0.25728 | 1.63065 | 0.82750 | 0.00857 | 0.00732 | 0.00557 | 2.29437 | 0.92331 | 0.01168 | 0.06425 | 0.00711 | 0.00270 | 0.00000 | 0.02492 | 0.00000 | 1.56412 |
| **Mmel1** | 0.01394 | 0.00785 | 0.00703 | 0.00900 | 0.00917 | 3.05771 | 0.00706 | 0.00951 | 0.01281 | 0.00406 | 0.00947 | 0.00806 | 0.13663 | 0.01350 | 0.01013 | 0.00608 |
| **Mmgt1** | 1.25416 | 0.11334 | 2.81820 | 8.68817 | 1.62546 | 1.65015 | 2.97623 | 1.80874 | 5.73586 | 5.40467 | 4.72024 | 5.76928 | 6.03077 | 10.07638 | 9.63858 | 0.82196 |
| **Mmgt2** | 0.00329 | 0.00730 | 1.47028 | 0.92915 | 0.01544 | 0.01772 | 0.01578 | 0.58061 | 0.08392 | 1.84073 | 1.63638 | 0.08035 | 1.26321 | 0.00144 | 0.71779 | 0.01270 |
| **Mmp11** | 1.78660 | 0.00631 | 0.00354 | 0.13546 | 0.00723 | 0.00659 | 0.00607 | 0.00951 | 0.00689 | 0.01466 | 0.01171 | 0.01680 | 0.01132 | 0.00982 | 0.02309 | 0.00554 |

| | | | | | | | | | | | | | | | | |
|---|---|---|---|---|---|---|---|---|---|---|---|---|---|---|---|---|
| **Mmp12** | 2.28470 | 3.66486 | 0.05837 | 0.82839 | 0.00323 | 0.00351 | 0.00409 | 0.00000 | 0.00000 | 0.00076 | 0.01561 | 0.00542 | 0.01290 | 0.00519 | 0.00000 | 5.14883 |
| **Mmp14** | 0.02359 | 0.00784 | 0.03421 | 0.01774 | 0.01915 | 0.01363 | 0.00722 | 0.01878 | 0.10183 | 0.77243 | 0.04812 | 0.07071 | 0.01072 | 0.10890 | 0.02604 | 0.00768 |
| **Mmp15** | 2.42021 | 0.00160 | 0.93142 | 0.47264 | 6.37739 | 0.70437 | 0.00537 | 0.16171 | 0.01553 | 0.05172 | 0.00909 | 0.00003 | 0.00000 | 0.00572 | 0.11813 | 0.07611 |
| **Mmp16** | 0.04600 | 0.00771 | 0.03631 | 0.29217 | 0.00811 | 0.01252 | 0.00726 | 0.01456 | 0.19756 | 0.02938 | 0.15616 | 0.00648 | 0.00973 | 0.00726 | 0.07658 | 0.00710 |
| **Mmp17** | 0.01805 | 0.00792 | 0.00688 | 0.14759 | 0.01036 | 0.00698 | 0.00731 | 0.00000 | 0.46339 | 0.03342 | 0.72147 | 0.01371 | 0.05992 | 0.01361 | 0.14223 | 0.00724 |
| **Mmp19** | 0.01465 | 0.00772 | 0.11208 | 1.38218 | 0.00861 | 0.00826 | 0.00708 | 0.00726 | 5.32922 | 0.00000 | 0.00000 | 0.01176 | 0.01204 | 0.53364 | 1.47555 | 0.00709 |
| **Mmp1b** | 2.47473 | 0.00554 | 0.49258 | 0.00767 | 0.01011 | 0.00297 | 0.00520 | 0.00728 | 0.01117 | 0.00500 | 0.01141 | 0.00520 | 0.01548 | 0.00709 | 0.50166 | 0.00464 |
| **Mmp2** | 0.01367 | 0.00812 | 0.01122 | 1.96758 | 0.00940 | 0.00638 | 0.00744 | 0.01425 | 0.00000 | 0.00614 | 0.01171 | 0.04003 | 0.01262 | 0.00335 | 0.46282 | 0.00703 |
| **Mmp21** | 0.14551 | 0.00775 | 0.00873 | 0.01100 | 0.00815 | 0.00731 | 0.00732 | 0.01643 | 0.01196 | 0.00365 | 0.01392 | 0.00738 | 0.58347 | 0.02155 | 0.06076 | 0.00710 |
| **Mmp23** | 1.50237 | 0.00572 | 0.00000 | 0.07817 | 0.00452 | 2.15923 | 0.00587 | 0.45888 | 0.01080 | 1.96294 | 0.04881 | 0.01073 | 0.00000 | 0.01196 | 0.00000 | 1.08370 |
| **Mmp24** | 0.09241 | 0.00752 | 0.00727 | 0.01356 | 0.00780 | 0.00678 | 0.00707 | 0.01027 | 0.17793 | 0.85598 | 0.01002 | 0.06941 | 0.11673 | 0.00617 | 0.11133 | 0.00718 |
| **Mmp25** | 0.03313 | 0.00750 | 0.13734 | 0.03042 | 0.00789 | 0.00887 | 0.02777 | 0.00625 | 0.28340 | 1.23590 | 0.43967 | 0.02887 | 0.06062 | 0.94665 | 0.45662 | 0.00714 |
| **Mmp9** | 0.01206 | 0.00771 | 0.01693 | 0.22887 | 0.09484 | 0.00722 | 0.00726 | 0.04460 | 0.00768 | 0.01758 | 0.01365 | 0.00763 | 0.02248 | 0.01103 | 0.00802 | 0.00732 |
| **Mmrn2** | 0.00603 | 0.00740 | 0.00534 | 0.19442 | 0.00716 | 0.00628 | 0.00701 | 0.00473 | 1.60315 | 0.95456 | 0.36465 | 0.09700 | 0.00423 | 0.00284 | 0.81267 | 2.51215 |
| **Mms19** | 7.96636 | 6.62298 | 8.69804 | 3.15894 | 1.60414 | 4.78581 | 2.89998 | 7.01032 | 7.83213 | 8.97688 | 5.33682 | 1.83601 | 1.57959 | 2.12139 | 3.69510 | 6.69285 |
| **Mms22l** | 3.81893 | 2.82258 | 2.94834 | 6.55852 | 5.40669 | 0.22076 | 0.70279 | 5.41152 | 2.93830 | 9.67222 | 4.02959 | 9.63297 | 2.05789 | 7.95317 | 4.58609 | 1.37505 |
| **Mn1** | 0.01267 | 0.00765 | 0.00670 | 1.85633 | 0.00934 | 0.00628 | 0.00726 | 0.00869 | 2.72818 | 0.00000 | 0.02410 | 0.03899 | 0.01331 | 0.55566 | 1.27150 | 0.00688 |
| **Mnat1** | 10.13975 | 8.59073 | 7.46997 | 3.39586 | 7.26295 | 3.59681 | 2.75137 | 8.02440 | 5.42174 | 3.16650 | 5.66488 | 1.29569 | 3.67376 | 6.06908 | 4.21385 | 7.22518 |
| **Mnd1** | 0.95536 | 0.03569 | 1.93747 | 0.55292 | 0.02678 | 3.48510 | 2.83975 | 0.00000 | 1.29724 | 0.72168 | 2.31158 | 0.04939 | 0.80456 | 0.00211 | 0.87069 | 6.86188 |
| **Mnd1.ps** | 0.12704 | 0.00696 | 0.56236 | 0.36285 | 0.00623 | 1.06945 | 0.82797 | 0.00553 | 0.50177 | 0.19254 | 0.81281 | 0.03309 | 0.14103 | 0.00424 | 0.24151 | 2.12366 |
| **Mns1** | 0.00071 | 0.00590 | 0.35137 | 0.15406 | 0.01027 | 0.00546 | 2.36944 | 1.26104 | 0.34034 | 1.09660 | 0.03857 | 0.84989 | 0.00396 | 0.59360 | 1.70574 | 0.00707 |
| **Mnt** | 1.64073 | 0.01614 | 0.25262 | 1.46337 | 0.01270 | 4.90313 | 0.03595 | 4.88534 | 2.22958 | 3.49903 | 1.23851 | 0.69095 | 1.30192 | 0.32679 | 3.20672 | 0.02432 |
| **Mob1a** | 0.03906 | 0.00728 | 0.01308 | 5.16768 | 0.04522 | 0.15771 | 0.01735 | 0.15585 | 1.95368 | 0.66690 | 0.90252 | 0.92469 | 0.06884 | 0.77671 | 0.39592 | 0.00638 |
| **Mob1b** | 0.74441 | 0.00655 | 0.27275 | 7.60734 | 0.00568 | 0.01540 | 0.00522 | 0.14347 | 1.88262 | 1.24564 | 2.61513 | 0.24861 | 0.50090 | 7.21568 | 1.49141 | 0.71162 |
| **Mob2** | 3.64749 | 3.21766 | 6.03781 | 2.74195 | 1.70104 | 0.69732 | 2.29425 | 4.45021 | 2.32116 | 9.20786 | 2.16126 | 0.37031 | 3.52675 | 0.00000 | 0.53593 | 1.20893 |
| **Mob3a** | 3.80488 | 0.00343 | 0.00000 | 0.36216 | 0.01214 | 3.09240 | 0.00311 | 0.00000 | 2.00077 | 0.00000 | 0.16210 | 0.04658 | 1.36901 | 0.00342 | 0.23821 | 2.94479 |
| **Mob3b** | 0.20056 | 0.01848 | 0.60343 | 0.27354 | 0.02596 | 0.03272 | 0.07792 | 0.00000 | 0.95782 | 0.35281 | 2.92204 | 0.05040 | 0.60922 | 0.40505 | 0.12218 | 2.04901 |
| **Mob3c** | 0.81645 | 0.01679 | 2.83443 | 0.30383 | 0.00167 | 0.05677 | 0.17949 | 0.06103 | 0.35125 | 0.75977 | 0.12774 | 0.02985 | 0.12918 | 0.26246 | 0.22115 | 0.00515 |
| **Mob4** | 6.72048 | 5.13392 | 6.25095 | 9.87900 | 4.34233 | 5.04759 | 4.54309 | 6.92499 | 12.99357 | 13.96992 | 8.25345 | 12.36316 | 8.91343 | 7.66406 | 8.19402 | 1.49420 |
| **Mocos** | 2.82368 | 0.00457 | 0.00000 | 0.00403 | 1.75304 | 2.86467 | 0.00483 | 0.85183 | 0.31726 | 0.48382 | 2.77111 | 0.02616 | 0.00000 | 0.00470 | 0.09489 | 0.00265 |
| **Mocs1** | 4.44674 | 0.00302 | 0.03896 | 0.04918 | 0.00963 | 0.00401 | 0.03437 | 0.00000 | 0.00956 | 0.01924 | 0.02205 | 0.19842 | 0.97145 | 0.01166 | 0.00712 | 2.20524 |
| **Mocs2** | 0.00000 | 0.00780 | 0.16846 | 1.07701 | 0.01080 | 0.00601 | 0.03795 | 1.06092 | 0.12686 | 0.29390 | 0.03904 | 0.00000 | 0.06643 | 1.84427 | 0.00000 | 1.78715 |
| **Mocs3** | 0.47539 | 0.89897 | 0.03492 | 4.37223 | 2.03902 | 2.34283 | 0.00491 | 0.00000 | 9.55394 | 2.46012 | 1.44409 | 0.04631 | 2.21035 | 0.02249 | 1.06117 | 0.17739 |
| **Mog** | 0.01616 | 0.00739 | 0.36415 | 0.14705 | 0.00854 | 0.00544 | 0.00718 | 0.00995 | 1.51256 | 0.00324 | 0.00000 | 0.00344 | 0.00948 | 0.00796 | 0.57929 | 0.00692 |
| **Mogat1** | 0.01366 | 0.00807 | 0.01227 | 0.47517 | 0.00938 | 0.00648 | 0.00738 | 0.01235 | 0.31060 | 0.14264 | 0.01196 | 0.00698 | 0.01256 | 0.00894 | 0.02218 | 0.00738 |
| **Mogat2** | 1.68219 | 0.00420 | 0.75260 | 0.00374 | 0.00226 | 0.00374 | 2.47052 | 0.00000 | 0.01106 | 0.00000 | 0.38268 | 0.03252 | 0.02494 | 0.01254 | 0.00000 | 4.02363 |
| **Mogs** | 2.97055 | 0.01163 | 4.12287 | 2.41089 | 0.01842 | 0.06007 | 2.91651 | 6.11969 | 6.49726 | 4.03455 | 3.53875 | 0.15917 | 7.24195 | 0.18004 | 1.92900 | 5.05329 |
| **Mok** | 0.80623 | 0.00681 | 1.17465 | 0.00506 | 0.00637 | 0.47623 | 0.00608 | 1.63464 | 0.03789 | 1.49023 | 0.03941 | 0.07844 | 1.01610 | 0.01305 | 0.00000 | 0.11101 |
| **Mon1a** | 3.17980 | 0.27120 | 0.81180 | 9.72213 | 0.02483 | 4.15332 | 0.09900 | 2.20443 | 12.78898 | 25.58490 | 12.24069 | 2.32442 | 13.34131 | 0.00000 | 1.08117 | 0.01423 |
| **Mon1b** | 0.12474 | 2.15514 | 7.11818 | 2.94455 | 3.29398 | 5.89969 | 0.05759 | 1.66757 | 2.68854 | 2.67913 | 0.41723 | 0.10936 | 0.50176 | 0.99847 | 1.03923 | 0.00794 |
| **Mon2** | 6.78635 | 2.16470 | 2.92797 | 10.55718 | 3.35059 | 4.45130 | 4.42841 | 6.36338 | 4.38867 | 6.71309 | 4.42049 | 2.88206 | 3.51909 | 5.34262 | 7.05463 | 2.86083 |
| **Morc1** | 4.63097 | 0.40336 | 0.00000 | 0.98415 | 1.27294 | 0.37529 | 0.01379 | 1.56125 | 0.58791 | 0.81235 | 0.59102 | 0.02624 | 1.01836 | 2.72807 | 0.00000 | 3.12506 |
| **Morc2a** | 2.48538 | 1.25551 | 4.23197 | 6.96963 | 0.80134 | 6.17135 | 2.32065 | 7.50513 | 5.01721 | 7.93413 | 3.59032 | 8.68864 | 3.10612 | 11.71878 | 15.69545 | 2.45316 |
| **Morc3** | 7.54591 | 7.47442 | 3.55534 | 2.26487 | 2.36319 | 4.60861 | 1.79831 | 11.92207 | 4.62043 | 8.67056 | 5.00715 | 8.42188 | 2.19856 | 10.65072 | 14.29108 | 0.45346 |
| **Morc4** | 1.12708 | 0.00661 | 0.00403 | 0.40801 | 0.00572 | 0.00694 | 0.00670 | 0.01020 | 0.25133 | 0.01426 | 0.01190 | 0.00719 | 0.01161 | 0.01159 | 0.00480 | 0.00609 |
| **Morf4l1** | 22.75051 | 17.26788 | 17.45941 | 5.28166 | 15.23208 | 10.79057 | 12.53109 | 12.52453 | 4.85103 | 8.36398 | 9.42554 | 1.63398 | 10.73956 | 20.58893 | 15.06690 | 9.97518 |
| **Morf4l1.ps1** | 14.38892 | 11.30858 | 10.06841 | 2.21434 | 9.13605 | 6.72102 | 7.79368 | 7.15089 | 2.00700 | 4.05631 | 4.84621 | 0.66142 | 5.48601 | 12.99860 | 8.76110 | 5.69390 |
| **Morf4l1b** | 13.71732 | 11.04173 | 9.15499 | 2.08282 | 8.72046 | 6.27099 | 7.61866 | 5.95818 | 2.33160 | 3.69439 | 4.95773 | 0.54900 | 5.19295 | 12.61978 | 8.81634 | 5.15221 |
| **Morf4l2** | 24.66010 | 16.87151 | 29.05960 | 9.71531 | 19.24418 | 8.74757 | 10.54520 | 13.68486 | 8.32115 | 9.24772 | 5.42181 | 1.02160 | 18.05601 | 9.19069 | 10.01099 | 16.95299 |
| **Morn1** | 0.00873 | 0.00787 | 0.01447 | 0.49327 | 0.00995 | 0.00789 | 0.00734 | 0.84956 | 0.27711 | 0.43131 | 0.21510 | 0.00334 | 0.01206 | 0.00615 | 0.01991 | 0.00727 |
| **Morn2** | 2.51310 | 0.33000 | 0.52619 | 0.02248 | 0.01073 | 0.00944 | 0.29035 | 0.61320 | 0.81901 | 0.32356 | 0.69821 | 0.01553 | 0.00578 | 0.33392 | 0.00000 | 1.14474 |
| **Morn4** | 1.12961 | 0.00683 | 1.22830 | 1.85768 | 0.00422 | 0.00776 | 0.00604 | 0.00000 | 0.59384 | 0.31751 | 0.43852 | 0.01200 | 0.71437 | 0.02154 | 0.54296 | 0.00514 |
| **Morn5** | 0.01382 | 0.00819 | 0.00624 | 0.00536 | 0.00903 | 0.00720 | 0.00752 | 0.00094 | 0.02450 | 0.01984 | 1.04619 | 0.02563 | 0.00880 | 0.01115 | 0.00000 | 0.00699 |
| **Mos** | 0.01356 | 0.00791 | 0.01162 | 0.14762 | 0.00928 | 0.00649 | 0.00729 | 0.00959 | 0.41029 | 0.41983 | 0.05318 | 0.01544 | 0.01144 | 0.01093 | 0.00250 | 0.00740 |
| **Mospd1** | 2.60442 | 2.86035 | 2.01404 | 15.73090 | 3.38364 | 2.54731 | 1.28192 | 8.40207 | 11.31060 | 12.13030 | 9.91735 | 7.05788 | 1.99815 | 35.47696 | 23.59053 | 0.00000 |
| **Mospd2** | 1.39967 | 0.07464 | 0.39917 | 1.23918 | 0.08823 | 2.07806 | 2.96362 | 1.54541 | 0.00000 | 0.00000 | 0.41486 | 0.00000 | 0.66441 | 0.00247 | 0.73404 | 0.45850 |
| **Mospd3** | 4.52799 | 1.13428 | 1.63919 | 2.59684 | 3.00586 | 0.04875 | 0.68381 | 1.66946 | 1.56571 | 8.50263 | 2.02550 | 1.32319 | 6.14225 | 0.00000 | 0.09128 | 5.02723 |
| **Mospd4** | 0.01238 | 0.00822 | 0.01286 | 0.71916 | 0.00910 | 0.00657 | 0.00735 | 0.01035 | 0.08109 | 1.31273 | 0.01109 | 0.00452 | 0.12653 | 0.00989 | 0.00358 | 0.00703 |
| **Mov10** | 8.29353 | 1.60410 | 1.30122 | 0.00000 | 3.80983 | 0.13389 | 0.00983 | 4.82812 | 0.13186 | 10.40664 | 1.89034 | 0.22653 | 0.00000 | 0.01994 | 2.77616 | 2.96536 |
| **Mov10l1** | 0.07504 | 2.28613 | 1.79559 | 0.68918 | 0.00369 | 0.00518 | 0.00610 | 0.00000 | 2.05188 | 1.19461 | 0.54590 | 0.01483 | 0.00000 | 0.00181 | 0.70112 | 2.31215 |
| **Mpc1** | 11.76379 | 7.73220 | 13.48115 | 14.12033 | 11.35642 | 6.05039 | 8.16981 | 13.20453 | 23.43521 | 24.74633 | 22.51609 | 17.33447 | 15.08077 | 7.43832 | 6.24850 | 2.16093 |
| **Mpc2** | 5.00101 | 0.11432 | 1.30181 | 15.47559 | 0.04418 | 0.65933 | 0.12696 | 0.64442 | 26.96434 | 29.48520 | 23.58917 | 1.37428 | 18.15402 | 3.05176 | 0.00000 | 2.28950 |

| | | | | | | | | | | | | | | | | |
|---|---|---|---|---|---|---|---|---|---|---|---|---|---|---|---|---|
| **Mpdu1** | 9.57034 | 4.26908 | 16.03733 | 2.64661 | 4.34757 | 10.02082 | 11.16512 | 19.33206 | 4.45924 | 10.79889 | 6.05416 | 0.57493 | 9.50476 | 0.05056 | 5.99368 | 6.59187 |
| **Mpdz** | 0.60155 | 0.00567 | 0.00000 | 9.50952 | 0.22788 | 0.00425 | 1.50838 | 0.00364 | 2.84090 | 0.83603 | 0.17062 | 0.00000 | 0.00569 | 2.06563 | 5.42094 | 2.06235 |
| **Mpeg1** | 0.01624 | 0.00797 | 0.01040 | 0.00848 | 0.01046 | 0.00680 | 0.00729 | 0.07047 | 0.01191 | 0.53752 | 0.01197 | 0.00629 | 0.00970 | 0.01160 | 0.00920 | 0.00775 |
| **Mpg** | 4.94289 | 0.00953 | 1.67211 | 0.58588 | 6.00217 | 0.48121 | 0.16783 | 2.19900 | 0.08761 | 0.17195 | 0.42909 | 0.01881 | 2.36773 | 0.00460 | 0.02325 | 3.89445 |
| **Mphosph10** | 7.07722 | 7.16850 | 7.21568 | 12.74094 | 9.35657 | 2.51112 | 2.38584 | 9.41085 | 15.41434 | 9.76933 | 7.09053 | 14.42520 | 3.42401 | 16.23773 | 13.89884 | 3.51081 |
| **Mphosph6** | 8.33746 | 2.89355 | 4.96238 | 15.77402 | 10.22081 | 2.60898 | 6.42574 | 9.66012 | 7.11566 | 8.52354 | 6.02313 | 1.93295 | 4.03567 | 16.56402 | 12.74431 | 7.07519 |
| **Mphosph8** | 3.28045 | 7.05223 | 3.37130 | 0.14378 | 2.50890 | 6.14607 | 1.91338 | 3.65026 | 0.50633 | 1.82592 | 0.47924 | 0.56908 | 4.55887 | 1.36810 | 6.37568 | 3.23376 |
| **Mphosph9** | 2.24820 | 2.02862 | 0.00000 | 7.36511 | 3.09538 | 2.45365 | 2.59410 | 1.35973 | 3.55523 | 2.92006 | 0.63564 | 8.28296 | 0.07306 | 0.32236 | 9.07065 | 0.10987 |
| **Mpi** | 4.69526 | 0.00000 | 2.82038 | 0.72507 | 3.91509 | 0.00093 | 0.00000 | 0.00000 | 0.80879 | 2.76931 | 1.42278 | 0.06163 | 3.92376 | 3.08694 | 1.96645 | 3.89392 |
| **Mpl** | 0.00815 | 0.00829 | 0.00908 | 1.23379 | 0.00869 | 0.00635 | 0.00786 | 0.12182 | 0.00000 | 3.20801 | 0.00822 | 0.00000 | 0.36413 | 0.00854 | 0.77077 | 0.00691 |
| **Mplkip** | 2.22083 | 1.20043 | 0.60406 | 5.51521 | 0.10739 | 0.01873 | 1.39727 | 0.07458 | 8.98706 | 3.54432 | 2.92288 | 0.33782 | 1.54948 | 1.42811 | 1.65496 | 1.00955 |
| **Mpnd** | 4.27165 | 0.01203 | 0.05165 | 0.21183 | 2.61021 | 0.01523 | 0.01272 | 0.80215 | 0.00517 | 0.00923 | 0.00630 | 0.01803 | 2.04238 | 0.00533 | 0.48859 | 0.00901 |
| **Mpp1** | 13.81952 | 6.91662 | 16.66091 | 7.59045 | 9.91028 | 6.39072 | 6.50741 | 3.66123 | 7.46912 | 14.76784 | 1.60127 | 7.93263 | 5.39332 | 11.10280 | 15.07879 | 8.72761 |
| **Mpp3** | 0.02449 | 0.00741 | 0.00425 | 0.08377 | 0.00840 | 0.00702 | 0.01675 | 0.10588 | 0.00975 | 0.40911 | 0.57160 | 0.01320 | 0.00785 | 0.08558 | 0.52462 | 0.00692 |
| **Mpp4** | 0.02670 | 0.00777 | 0.23773 | 0.17076 | 0.00794 | 0.00552 | 0.00614 | 0.12763 | 0.30253 | 0.55780 | 1.82007 | 0.10828 | 1.24577 | 0.56814 | 0.00000 | 0.00641 |
| **Mpp5** | 2.10924 | 0.00311 | 0.64600 | 4.43599 | 0.00336 | 1.05848 | 0.01484 | 0.00657 | 3.81140 | 0.53470 | 0.00000 | 6.46775 | 0.08149 | 0.02139 | 4.39291 | 0.32228 |
| **Mpp6** | 4.49701 | 5.99970 | 0.53303 | 1.94022 | 0.06633 | 2.60078 | 1.44647 | 1.37195 | 3.74695 | 6.98882 | 2.49225 | 7.97132 | 0.95094 | 4.31169 | 2.56502 | 3.17525 |
| **Mpp7** | 4.62328 | 1.52066 | 0.18188 | 0.15272 | 0.07086 | 0.60119 | 0.18038 | 1.15257 | 0.25801 | 0.01764 | 0.68008 | 0.19204 | 2.61746 | 1.33219 | 0.18413 | 1.36340 |
| **Mppe1** | 0.31660 | 0.07611 | 1.30482 | 0.01606 | 2.56679 | 0.58434 | 1.32823 | 1.65444 | 0.00584 | 1.37050 | 0.03179 | 0.12398 | 0.17712 | 0.04693 | 0.66613 | 0.17318 |
| **Mpped1** | 0.01373 | 0.00817 | 0.01176 | 0.75684 | 0.00943 | 0.00642 | 0.00735 | 0.01166 | 0.00000 | 0.00539 | 0.01204 | 0.00717 | 0.01281 | 0.01606 | 0.54477 | 0.00725 |
| **Mpped2** | 2.14880 | 3.69420 | 1.81386 | 0.18892 | 0.98972 | 0.13446 | 0.36918 | 0.35872 | 0.00459 | 0.00124 | 0.00814 | 0.00356 | 0.00000 | 0.00999 | 0.00000 | 1.30465 |
| **Mprip** | 10.07595 | 5.81981 | 9.03217 | 14.39680 | 4.73124 | 8.08925 | 11.07602 | 17.82465 | 8.45799 | 13.27799 | 7.19866 | 5.88801 | 11.35670 | 2.70591 | 8.58381 | 7.25162 |
| **Mpst** | 0.01773 | 0.00786 | 0.01197 | 0.02606 | 0.00902 | 0.00650 | 0.00733 | 0.01193 | 0.01056 | 0.29403 | 0.01240 | 0.00681 | 0.01873 | 0.01130 | 0.00968 | 0.00742 |
| **Mpv17** | 3.74796 | 0.77717 | 2.62906 | 4.34070 | 0.30527 | 2.91368 | 0.12143 | 2.02885 | 7.41349 | 0.00000 | 5.38338 | 5.57418 | 0.64090 | 1.85669 | 2.67959 | 0.34760 |
| **Mpv17l** | 2.68379 | 0.12872 | 2.57150 | 11.37481 | 1.30874 | 0.29964 | 0.31317 | 0.00000 | 16.97210 | 0.17134 | 14.03228 | 1.64670 | 1.06690 | 6.46777 | 1.47141 | 0.56289 |
| **Mpv17l2** | 1.26655 | 0.00623 | 3.31067 | 1.20020 | 0.21821 | 0.04106 | 0.22478 | 0.00000 | 0.98129 | 3.80279 | 1.94084 | 0.05115 | 1.48996 | 0.26071 | 0.06367 | 2.27496 |
| **Mpzl1** | 0.98238 | 3.50037 | 0.98470 | 2.85897 | 0.01305 | 0.00497 | 3.72017 | 0.00297 | 0.10213 | 2.10582 | 0.14290 | 0.01796 | 0.15878 | 0.00000 | 0.82668 | 0.03609 |
| **Mpzl2** | 3.23878 | 0.90759 | 5.41749 | 0.19084 | 3.25931 | 5.22315 | 4.87183 | 5.22457 | 1.82505 | 1.55923 | 2.12667 | 7.33367 | 4.44626 | 4.34185 | 0.00000 | 1.87791 |
| **Mpzl3** | 0.01808 | 0.00817 | 0.01829 | 0.09273 | 0.01058 | 0.00725 | 0.00751 | 0.01362 | 0.00789 | 0.01859 | 0.01212 | 0.00784 | 0.01240 | 0.00971 | 0.02352 | 0.00740 |
| **Mrap** | 0.38837 | 0.00718 | 0.00000 | 0.40584 | 0.01718 | 0.00840 | 1.40298 | 0.00811 | 0.03372 | 0.80285 | 0.00992 | 0.02391 | 0.01088 | 0.00485 | 0.42262 | 0.00729 |
| **Mras** | 8.48342 | 4.52296 | 5.12816 | 1.19816 | 3.45193 | 5.56982 | 5.76640 | 2.51215 | 0.64149 | 0.65204 | 0.37585 | 0.49644 | 7.99498 | 0.00175 | 0.10712 | 6.29549 |
| **Mrc1** | 0.01365 | 0.00811 | 0.00840 | 0.50760 | 0.01005 | 0.15676 | 0.00766 | 0.01351 | 0.55416 | 0.00848 | 0.00706 | 0.01709 | 0.05646 | 0.00752 | 0.90646 | 0.00746 |
| **Mre11a** | 6.50042 | 2.22053 | 2.52988 | 2.01486 | 1.39846 | 0.09472 | 7.03709 | 3.02092 | 1.56745 | 3.51573 | 4.33153 | 2.30965 | 1.20107 | 2.92767 | 1.36899 | 2.83480 |
| **Mreg** | 6.57378 | 3.80657 | 10.20701 | 0.88768 | 3.68806 | 4.46435 | 6.51714 | 8.18449 | 0.63877 | 6.70641 | 8.44919 | 0.49281 | 7.80380 | 20.26213 | 6.91668 | 4.20604 |
| **Mrfap1** | 22.86793 | 8.41185 | 18.78246 | 6.65902 | 7.72576 | 8.80965 | 11.37350 | 11.08941 | 11.51228 | 21.03283 | 20.58616 | 2.91557 | 17.90025 | 12.97556 | 9.20384 | 11.44258 |
| **Mrgbp** | 0.21101 | 1.97767 | 0.78129 | 0.00901 | 0.01091 | 0.00658 | 0.00627 | 0.21834 | 0.01221 | 0.03208 | 0.01097 | 0.00672 | 0.46501 | 0.05345 | 0.01376 | 0.00621 |
| **Mrgpra6** | 0.86754 | 0.73915 | 2.80396 | 1.16482 | 0.51568 | 0.33629 | 2.34071 | 2.71240 | 3.01360 | 2.42231 | 0.99414 | 0.64640 | 1.10283 | 16.06625 | 2.36476 | 0.29077 |
| **Mrgprb1** | 0.03674 | 0.00796 | 0.05693 | 0.07202 | 0.01833 | 0.00693 | 0.01807 | 0.01046 | 0.14066 | 0.03773 | 0.18146 | 0.04633 | 0.06761 | 0.24224 | 0.02961 | 0.00717 |
| **Mrgpre** | 0.01111 | 0.00800 | 0.03289 | 0.02454 | 0.05912 | 0.01510 | 0.01814 | 0.04056 | 0.02439 | 0.04589 | 0.01257 | 0.00749 | 0.02840 | 0.08582 | 0.03480 | 0.03017 |
| **Mri1** | 3.38176 | 0.63200 | 0.93050 | 0.85994 | 2.44917 | 0.97646 | 1.13357 | 1.02204 | 0.98433 | 2.42625 | 3.11387 | 0.48962 | 0.42663 | 0.00812 | 0.00000 | 1.96877 |
| **Mrln** | 0.01370 | 0.00811 | 0.01209 | 0.30772 | 0.00930 | 0.00664 | 0.00751 | 0.01370 | 0.00578 | 0.01496 | 0.01323 | 0.01865 | 0.01264 | 0.01127 | 0.00613 | 0.01468 |
| **Mrm1** | 0.00000 | 0.01551 | 4.28510 | 2.83078 | 0.39185 | 0.03631 | 2.58476 | 1.00454 | 3.02770 | 0.65088 | 0.00000 | 0.11392 | 0.91199 | 0.00000 | 1.59977 | 2.26651 |
| **Mrm2** | 4.43737 | 0.01010 | 4.88804 | 0.00000 | 2.35830 | 3.96419 | 1.52027 | 5.19689 | 0.22689 | 4.75837 | 1.82376 | 0.31391 | 4.70028 | 0.00002 | 0.62971 | 1.63182 |
| **Mro** | 0.02669 | 0.00803 | 0.03797 | 0.51086 | 0.00849 | 0.00768 | 0.00768 | 0.01556 | 0.50537 | 0.20373 | 0.03478 | 0.00776 | 0.01391 | 0.02953 | 0.00432 | 0.01462 |
| **Mroh1** | 2.45821 | 3.44001 | 2.84288 | 0.25010 | 1.59037 | 5.08732 | 2.99692 | 0.00000 | 0.51529 | 0.89412 | 1.23142 | 0.01106 | 1.03809 | 0.31733 | 0.52097 | 0.00769 |
| **Mroh5** | 0.01331 | 0.00803 | 0.01217 | 0.17316 | 0.00901 | 0.00652 | 0.00733 | 0.01046 | 0.18370 | 0.83967 | 0.00993 | 0.00537 | 0.01105 | 0.01086 | 0.00541 | 0.00732 |
| **Mroh8** | 0.00898 | 0.00749 | 0.00000 | 0.31998 | 0.00932 | 0.00610 | 0.00629 | 0.00000 | 1.56397 | 0.30487 | 1.69977 | 0.18023 | 0.00000 | 3.36425 | 1.37471 | 0.00700 |
| **Mrpl1** | 7.36577 | 8.15155 | 1.75437 | 0.03735 | 2.04374 | 4.15733 | 4.17036 | 2.71385 | 0.21208 | 1.47923 | 2.11386 | 8.25303 | 4.85039 | 6.58646 | 3.14644 | 4.01686 |
| **Mrpl10** | 8.05588 | 1.60212 | 8.13149 | 4.07058 | 2.84759 | 1.90342 | 3.56929 | 8.80665 | 2.86946 | 5.68675 | 3.63924 | 0.24943 | 5.88338 | 1.32093 | 3.58350 | 4.26460 |
| **Mrpl11** | 15.92629 | 4.15972 | 10.34195 | 2.81705 | 10.81958 | 8.17422 | 8.89472 | 7.69524 | 3.09268 | 5.21679 | 8.61169 | 2.93539 | 12.68752 | 0.95019 | 1.12856 | 4.23796 |
| **Mrpl12** | 20.38600 | 3.03222 | 10.12564 | 8.81565 | 9.85239 | 10.01107 | 5.49698 | 14.15686 | 14.84438 | 27.18480 | 13.27736 | 2.34143 | 17.88861 | 0.67170 | 1.01327 | 14.03745 |
| **Mrpl13** | 10.42522 | 9.99687 | 11.08881 | 1.97311 | 9.83253 | 5.39539 | 7.65985 | 8.55121 | 2.32760 | 8.63565 | 8.55604 | 3.18206 | 6.16627 | 3.59698 | 2.17692 | 6.07057 |
| **Mrpl14** | 7.58324 | 0.16412 | 2.48824 | 0.19746 | 7.71102 | 2.12708 | 2.36881 | 5.54513 | 0.20636 | 9.14928 | 5.08878 | 0.09558 | 3.28657 | 0.00000 | 0.16348 | 3.56607 |
| **Mrpl15** | 23.76298 | 4.98265 | 11.68071 | 2.47535 | 12.31235 | 8.14082 | 4.77471 | 19.74789 | 5.99344 | 24.58284 | 22.23528 | 11.30880 | 13.20133 | 5.81570 | 2.50744 | 12.21557 |
| **Mrpl16** | 9.86878 | 7.09130 | 6.56108 | 0.79841 | 9.62776 | 3.23259 | 1.38740 | 9.80593 | 2.00461 | 9.11017 | 6.73325 | 2.04998 | 4.07203 | 1.29757 | 0.94840 | 4.41995 |
| **Mrpl17** | 21.92933 | 1.46707 | 9.31869 | 0.63623 | 18.02450 | 8.15174 | 7.14900 | 22.20321 | 0.64156 | 13.87459 | 14.32384 | 1.39329 | 24.47617 | 3.90418 | 0.00000 | 15.35293 |
| **Mrpl18** | 18.44902 | 11.98190 | 20.25813 | 7.38125 | 14.47948 | 8.20968 | 9.16726 | 19.14758 | 7.51538 | 22.39091 | 11.83405 | 9.67916 | 18.14421 | 5.07663 | 7.45748 | 10.53120 |
| **Mrpl19** | 12.36150 | 4.87552 | 7.79422 | 1.14083 | 10.85217 | 5.66273 | 4.63381 | 8.83082 | 0.84722 | 10.50791 | 4.49678 | 1.17155 | 11.64203 | 7.52377 | 1.97711 | 1.13417 |
| **Mrpl2** | 10.35267 | 0.95919 | 7.68348 | 1.21963 | 6.20116 | 6.16192 | 5.01729 | 18.86898 | 2.03682 | 6.38560 | 6.33690 | 0.18840 | 14.99631 | 0.01085 | 0.00000 | 9.42860 |
| **Mrpl20** | 15.29765 | 5.30622 | 13.39461 | 9.61944 | 7.54972 | 5.94719 | 2.35353 | 13.16500 | 13.17774 | 23.79465 | 15.78819 | 2.49629 | 16.73444 | 2.09292 | 3.77762 | 7.85350 |
| **Mrpl21** | 10.33875 | 4.23612 | 5.75144 | 0.29884 | 11.44671 | 3.26446 | 1.16272 | 2.86123 | 0.71934 | 9.60636 | 6.82566 | 3.57798 | 9.00144 | 0.68942 | 5.24305 | 4.33855 |
| **Mrpl22** | 12.25545 | 7.01434 | 13.04792 | 0.74225 | 6.86656 | 5.99818 | 7.60736 | 19.29551 | 0.23273 | 23.92150 | 18.31530 | 6.51020 | 19.17632 | 12.96212 | 14.09845 | 3.82494 |

| | | | | | | | | | | | | | | | | |
|---|---|---|---|---|---|---|---|---|---|---|---|---|---|---|---|---|
| **Mrpl23** | 8.61600 | 0.67646 | 4.34198 | 2.28370 | 4.46485 | 2.21310 | 1.54851 | 1.12154 | 1.10023 | 5.20921 | 14.65979 | 0.33214 | 5.81801 | 2.59018 | 1.16381 | 2.68558 |
| **Mrpl24** | 11.24510 | 3.19795 | 14.51157 | 0.05200 | 5.59407 | 1.41224 | 3.47755 | 10.13018 | 0.09752 | 5.54379 | 2.13392 | 0.17308 | 6.72439 | 5.91698 | 4.38578 | 6.12260 |
| **Mrpl27** | 10.38579 | 5.00455 | 2.05135 | 3.12511 | 0.08619 | 3.09096 | 3.36490 | 6.94357 | 5.50099 | 5.51051 | 9.45729 | 0.80633 | 5.58642 | 0.98600 | 1.08374 | 2.93865 |
| **Mrpl28** | 18.33186 | 3.88170 | 11.19306 | 3.17682 | 7.44222 | 8.89658 | 9.92601 | 17.34913 | 0.89183 | 10.69160 | 5.01204 | 1.26580 | 16.51887 | 0.00061 | 0.00000 | 12.15575 |
| **Mrpl3** | 6.94464 | 5.96636 | 4.12397 | 3.51886 | 4.83882 | 5.13849 | 2.62718 | 2.93100 | 0.97798 | 9.37698 | 5.25468 | 7.10023 | 5.47838 | 3.25634 | 3.46666 | 4.24779 |
| **Mrpl30** | 14.94303 | 3.70764 | 12.62698 | 1.75369 | 5.93817 | 4.84618 | 6.97378 | 10.02509 | 0.36266 | 12.43909 | 12.35661 | 2.20891 | 10.04450 | 3.81552 | 3.62172 | 3.30870 |
| **Mrpl32** | 11.23687 | 2.40921 | 4.67765 | 3.97714 | 4.30579 | 2.90875 | 3.63531 | 5.49166 | 5.41060 | 9.73556 | 8.33600 | 1.68577 | 4.69862 | 1.30693 | 3.16641 | 1.44175 |
| **Mrpl33** | 6.94690 | 0.36550 | 4.94248 | 5.44828 | 2.38169 | 1.27291 | 0.45973 | 0.50387 | 3.09226 | 1.89809 | 15.36949 | 0.43913 | 1.65810 | 4.55179 | 1.50551 | 0.64470 |
| **Mrpl34** | 10.29127 | 0.00000 | 0.00000 | 9.52875 | 3.15254 | 2.87358 | 0.15656 | 3.94079 | 11.43286 | 10.65171 | 2.10125 | 0.14435 | 5.64695 | 0.01929 | 0.00000 | 4.41903 |
| **Mrpl35** | 14.34040 | 0.54203 | 8.51227 | 1.82664 | 7.44427 | 5.23020 | 4.42218 | 10.60944 | 2.98301 | 14.00762 | 12.20122 | 2.34724 | 5.70751 | 7.36251 | 4.87947 | 4.84836 |
| **Mrpl36** | 16.98384 | 1.25822 | 5.70201 | 0.00000 | 9.18655 | 6.77323 | 2.62262 | 7.84800 | 0.07118 | 6.66387 | 4.68381 | 1.54974 | 11.14979 | 0.78947 | 0.00000 | 5.00797 |
| **Mrpl37** | 7.88511 | 4.44517 | 3.80700 | 5.06403 | 6.32119 | 4.59181 | 2.33450 | 8.94896 | 0.83104 | 9.48960 | 6.11517 | 6.58488 | 10.14785 | 0.11308 | 1.41583 | 3.78333 |
| **Mrpl38** | 8.28772 | 5.02571 | 3.17583 | 3.53644 | 2.86485 | 3.84251 | 0.44555 | 3.42538 | 2.72679 | 5.67034 | 4.26774 | 0.51521 | 6.77247 | 0.11579 | 1.15228 | 6.56216 |
| **Mrpl39** | 11.88113 | 4.17508 | 2.58491 | 0.80547 | 8.20108 | 5.56936 | 5.25034 | 10.03010 | 2.01224 | 6.26832 | 10.90136 | 6.01183 | 7.17462 | 2.93198 | 0.74325 | 4.18684 |
| **Mrpl4** | 19.48572 | 1.69859 | 7.03025 | 7.55961 | 13.88862 | 11.95802 | 6.39991 | 21.27727 | 12.04364 | 21.64120 | 7.54096 | 3.31233 | 19.00093 | 0.92007 | 1.80066 | 11.24991 |
| **Mrpl40** | 19.21387 | 10.38813 | 15.54986 | 14.55394 | 9.03137 | 7.58507 | 7.75829 | 14.86421 | 24.79763 | 26.86156 | 21.16248 | 9.00607 | 13.81309 | 7.42877 | 5.45686 | 6.43143 |
| **Mrpl41** | 8.65072 | 4.66081 | 9.25014 | 7.69185 | 8.07613 | 0.58723 | 5.15159 | 5.26737 | 10.47057 | 8.44400 | 6.51409 | 1.05693 | 3.64469 | 1.05280 | 5.19121 | 6.30406 |
| **Mrpl42** | 14.41278 | 6.28460 | 11.63926 | 4.66637 | 6.02468 | 7.46102 | 3.60829 | 2.56969 | 4.67655 | 10.06124 | 13.13578 | 1.88293 | 8.85591 | 6.24765 | 4.36553 | 5.58930 |
| **Mrpl43** | 13.89859 | 0.93653 | 1.10478 | 4.61121 | 12.11794 | 6.54767 | 1.31334 | 11.95132 | 12.10958 | 16.47150 | 6.81827 | 1.97631 | 6.34593 | 1.20162 | 0.61579 | 5.00780 |
| **Mrpl44** | 14.52694 | 2.26338 | 10.46711 | 3.36942 | 16.27839 | 6.61223 | 8.06228 | 20.20857 | 1.20201 | 18.86967 | 9.27607 | 1.46866 | 12.90120 | 7.93916 | 5.39061 | 6.55092 |
| **Mrpl45** | 13.32960 | 12.79452 | 13.24195 | 6.43859 | 12.67774 | 6.78097 | 9.89916 | 12.68784 | 5.64094 | 14.22565 | 7.04082 | 1.58030 | 8.73436 | 1.51883 | 8.28397 | 6.98203 |
| **Mrpl46** | 13.93745 | 10.52927 | 10.55255 | 2.42346 | 10.20623 | 4.66585 | 7.30661 | 14.68339 | 3.71754 | 19.24333 | 8.44922 | 1.21654 | 12.14718 | 1.02682 | 0.97670 | 8.84117 |
| **Mrpl47** | 10.84159 | 8.24718 | 9.34420 | 0.93084 | 9.28267 | 8.39028 | 5.86000 | 7.54337 | 2.06272 | 8.24482 | 10.09889 | 3.33765 | 9.08976 | 5.83547 | 6.24361 | 6.06005 |
| **Mrpl48** | 11.80093 | 4.24613 | 7.03166 | 11.65262 | 8.90549 | 3.33304 | 3.36456 | 8.70659 | 11.21471 | 7.97592 | 9.44727 | 7.04281 | 8.17795 | 5.34305 | 5.87350 | 3.02844 |
| **Mrpl49** | 9.94736 | 4.81040 | 13.73627 | 4.85338 | 7.21138 | 3.36634 | 6.41969 | 10.71471 | 8.84616 | 13.81992 | 9.48627 | 4.12868 | 12.98900 | 0.57361 | 4.85958 | 5.48710 |
| **Mrpl50** | 8.43166 | 5.35651 | 5.26926 | 0.81287 | 6.84899 | 2.50956 | 0.00000 | 2.49262 | 1.55959 | 3.90599 | 2.49730 | 2.58057 | 6.41949 | 2.57198 | 5.50480 | 1.00974 |
| **Mrpl51** | 18.10760 | 6.94644 | 9.73265 | 10.31198 | 9.52411 | 6.83221 | 1.08487 | 11.31717 | 19.70456 | 14.54674 | 15.23136 | 1.06048 | 10.38088 | 1.26637 | 6.97341 | 6.26406 |
| **Mrpl52** | 3.45117 | 0.02371 | 0.20322 | 0.43673 | 3.57530 | 0.27778 | 0.11255 | 0.32633 | 0.65309 | 1.19774 | 2.78051 | 0.09734 | 3.42956 | 0.19098 | 0.00000 | 3.21401 |
| **Mrpl53** | 12.64057 | 3.11560 | 3.91508 | 1.75174 | 9.49882 | 3.27771 | 3.55285 | 2.62101 | 4.25684 | 10.97867 | 3.42571 | 0.50868 | 8.42740 | 0.03229 | 0.00000 | 5.69045 |
| **Mrpl54** | 5.09473 | 0.02999 | 1.95230 | 3.78432 | 5.89429 | 0.14395 | 0.26918 | 0.00000 | 6.91734 | 4.96659 | 8.30391 | 0.16144 | 6.84299 | 0.22638 | 0.00000 | 0.59866 |
| **Mrpl55** | 8.23557 | 1.70970 | 9.34202 | 1.47192 | 6.32238 | 3.55557 | 3.21591 | 9.53044 | 4.10784 | 5.44991 | 6.57460 | 3.59708 | 1.71693 | 0.03650 | 2.11218 | 1.38687 |
| **Mrpl57** | 12.99718 | 0.11300 | 0.57191 | 3.76798 | 5.90301 | 1.37866 | 0.50599 | 1.65586 | 6.16675 | 11.73642 | 7.14137 | 0.56126 | 8.75133 | 0.00000 | 0.35969 | 1.74940 |
| **Mrpl9** | 4.17519 | 1.85590 | 2.16720 | 5.40849 | 4.45160 | 1.98504 | 4.20132 | 7.22926 | 5.54011 | 2.09113 | 6.07436 | 1.88960 | 2.59058 | 2.52483 | 4.13188 | 1.19963 |
| **Mrps10** | 13.48626 | 0.27104 | 0.00000 | 0.06799 | 0.79166 | 5.78402 | 1.51890 | 8.67709 | 0.66785 | 14.87552 | 2.81780 | 1.84449 | 11.60795 | 0.00000 | 0.00000 | 3.11648 |
| **Mrps11** | 11.09782 | 7.11011 | 8.44042 | 2.92756 | 7.05994 | 1.16469 | 6.41622 | 7.89577 | 1.54560 | 7.29982 | 2.10311 | 0.76597 | 8.69045 | 0.00000 | 0.00000 | 4.47681 |
| **Mrps12** | 14.42610 | 0.11705 | 1.40748 | 2.65207 | 9.30276 | 6.48304 | 3.42503 | 7.80278 | 5.32343 | 8.19062 | 4.47422 | 0.38467 | 9.60708 | 0.00000 | 0.00000 | 8.09791 |
| **Mrps14** | 14.08037 | 5.04707 | 7.94819 | 5.75083 | 4.14348 | 4.36022 | 5.15096 | 4.74055 | 7.59350 | 8.41541 | 9.62585 | 1.02993 | 8.06697 | 2.75903 | 3.33259 | 1.23623 |
| **Mrps15** | 2.31765 | 1.61830 | 1.54005 | 3.79510 | 4.16650 | 0.06649 | 0.73617 | 0.00000 | 4.52360 | 8.37406 | 11.44114 | 1.20248 | 10.88752 | 0.46131 | 0.00000 | 5.71983 |
| **Mrps16** | 7.64161 | 1.11492 | 6.36273 | 3.26512 | 4.36902 | 2.16646 | 3.18468 | 1.12314 | 2.46254 | 3.02140 | 10.26241 | 0.19320 | 8.19738 | 0.47973 | 0.30633 | 3.04829 |
| **Mrps17** | 11.82689 | 5.57947 | 9.10163 | 3.47164 | 1.52663 | 6.01093 | 4.33046 | 4.10846 | 4.81110 | 8.01554 | 6.73695 | 1.00790 | 4.32723 | 0.50345 | 5.38846 | 2.44671 |
| **Mrps18a** | 10.07745 | 2.82804 | 7.14987 | 0.36338 | 6.60160 | 6.17883 | 5.01291 | 10.25948 | 1.30616 | 13.57168 | 11.51374 | 4.89052 | 11.24923 | 0.40766 | 0.54386 | 5.66429 |
| **Mrps18b** | 20.26546 | 9.74093 | 27.61645 | 7.93230 | 13.37475 | 5.12153 | 13.73467 | 28.99402 | 9.83085 | 27.66474 | 23.58326 | 13.87423 | 26.26132 | 16.16626 | 15.57593 | 13.69952 |
| **Mrps18c** | 7.22893 | 0.60192 | 2.82797 | 0.96870 | 4.94609 | 2.32803 | 1.06629 | 0.00000 | 5.38341 | 5.60846 | 17.11568 | 0.41526 | 3.47162 | 3.59032 | 2.13233 | 1.59641 |
| **Mrps2** | 14.03332 | 3.54446 | 8.55193 | 4.64488 | 9.02530 | 5.51292 | 8.70289 | 6.68527 | 5.34403 | 12.75700 | 8.23414 | 0.35041 | 12.98815 | 3.69095 | 9.96426 | 5.67891 |
| **Mrps21** | 6.56598 | 0.24602 | 4.19766 | 2.93133 | 5.60225 | 2.79381 | 0.47369 | 0.83727 | 2.73767 | 3.46613 | 10.77904 | 0.58315 | 1.83029 | 1.77983 | 1.19057 | 1.10257 |
| **Mrps22** | 6.50928 | 3.30088 | 5.09100 | 2.39286 | 1.53320 | 0.28859 | 1.72264 | 1.94690 | 7.98058 | 15.90512 | 11.53714 | 2.79516 | 8.59679 | 4.26485 | 1.30150 | 8.22157 |
| **Mrps23** | 15.93782 | 9.35266 | 16.31046 | 5.67675 | 9.48621 | 9.84464 | 6.54446 | 12.63749 | 3.82104 | 15.66352 | 6.46206 | 2.07970 | 20.01806 | 0.12011 | 0.57674 | 10.60674 |
| **Mrps24** | 5.09794 | 0.46173 | 2.36445 | 8.12486 | 5.10005 | 0.73607 | 0.38516 | 0.12382 | 9.97298 | 13.42838 | 19.16583 | 0.29403 | 3.41619 | 0.39904 | 0.08115 | 0.51388 |
| **Mrps25** | 7.85714 | 2.10144 | 6.68406 | 2.35269 | 2.48803 | 0.38943 | 3.74168 | 3.47179 | 3.59614 | 3.04461 | 4.29390 | 0.27775 | 3.60495 | 1.38738 | 2.30090 | 2.72920 |
| **Mrps26** | 7.51076 | 0.00000 | 0.00000 | 12.52044 | 4.95437 | 3.16760 | 0.44113 | 4.33526 | 10.77490 | 15.02898 | 10.50265 | 0.93725 | 6.29866 | 0.79849 | 0.22221 | 3.37982 |
| **Mrps27** | 6.58306 | 8.88613 | 7.26217 | 0.70701 | 4.25201 | 4.58037 | 0.94778 | 8.28293 | 0.92927 | 5.29913 | 6.30859 | 0.29562 | 7.68967 | 0.05058 | 3.63509 | 0.04776 |
| **Mrps28** | 5.53404 | 0.26238 | 0.00000 | 3.87734 | 2.57505 | 0.75702 | 1.33986 | 3.34425 | 4.65379 | 0.68758 | 4.86240 | 0.32316 | 1.42349 | 0.06294 | 0.00000 | 0.65177 |
| **Mrps30** | 10.10415 | 3.27749 | 2.78587 | 8.31642 | 1.01317 | 0.00000 | 2.11724 | 1.79559 | 11.90534 | 11.43913 | 14.78054 | 2.13537 | 0.95563 | 1.62262 | 2.47553 | 2.78312 |
| **Mrps31** | 9.42475 | 4.80688 | 5.78813 | 0.00000 | 4.83719 | 1.62320 | 3.58675 | 6.86916 | 0.00013 | 4.15070 | 3.73219 | 3.39973 | 5.46289 | 6.54448 | 8.43344 | 4.68958 |
| **Mrps33** | 8.06181 | 2.37858 | 6.01817 | 3.06087 | 10.44729 | 0.84618 | 1.68450 | 4.63786 | 2.45373 | 5.47958 | 6.02023 | 0.74567 | 4.13384 | 5.66918 | 2.95874 | 2.82169 |
| **Mrps34** | 1.33177 | 0.28890 | 0.74345 | 0.98115 | 4.96855 | 0.78717 | 0.27070 | 0.76348 | 3.45247 | 3.57568 | 6.14242 | 0.21057 | 3.52477 | 0.00000 | 0.34652 | 2.69718 |
| **Mrps35** | 10.69622 | 1.97684 | 3.97865 | 1.17563 | 1.91356 | 0.10640 | 0.00402 | 3.19189 | 0.39672 | 6.07271 | 3.48180 | 4.47265 | 3.98948 | 0.17999 | 0.73303 | 3.67783 |
| **Mrps36** | 8.80718 | 2.69646 | 4.23474 | 4.43929 | 5.81396 | 3.22964 | 2.16680 | 1.87652 | 2.95332 | 3.72837 | 5.61210 | 0.39172 | 5.25427 | 0.49083 | 0.82608 | 6.14446 |
| **Mrps5** | 12.01843 | 6.24429 | 7.99154 | 0.87069 | 1.10037 | 0.65402 | 3.36196 | 10.35906 | 0.00000 | 3.19987 | 3.52533 | 0.31240 | 2.62537 | 0.01622 | 1.82449 | 6.68730 |
| **Mrps6** | 5.69596 | 0.02010 | 1.98479 | 9.29536 | 1.84692 | 1.59651 | 2.59618 | 4.70871 | 17.33657 | 10.06007 | 7.77755 | 0.31920 | 9.65522 | 2.60183 | 0.62537 | 2.47978 |
| **Mrps7** | 16.60912 | 6.60966 | 10.19758 | 17.30484 | 10.12873 | 7.14056 | 6.43258 | 14.35783 | 9.36665 | 25.20713 | 12.92084 | 2.91548 | 16.21375 | 1.64435 | 7.72009 | 8.20701 |
| **Mrps9** | 9.81312 | 5.47881 | 8.59634 | 1.49808 | 7.68387 | 8.61490 | 3.23579 | 9.95273 | 1.52664 | 10.71459 | 5.29421 | 2.87717 | 9.30386 | 0.00000 | 2.65716 | 8.71145 |

| | | | | | | | | | | | | | | | | |
|---|---|---|---|---|---|---|---|---|---|---|---|---|---|---|---|---|
| **Mrrf** | 6.06954 | 1.01537 | 12.22168 | 1.09642 | 0.94973 | 0.84948 | 2.76485 | 5.38205 | 2.16253 | 5.85825 | 3.64491 | 5.82585 | 5.21812 | 0.48023 | 3.99163 | 0.00691 |
| **Mrs2** | 0.00000 | 0.12562 | 2.52984 | 1.10960 | 0.51600 | 3.46604 | 0.00411 | 1.43473 | 0.37943 | 2.91481 | 4.88333 | 0.20264 | 2.49438 | 0.00000 | 1.12569 | 0.01013 |
| **Mrto4** | 22.04693 | 9.07480 | 22.30651 | 14.25045 | 15.67714 | 4.12826 | 6.06056 | 28.08280 | 15.64840 | 25.78588 | 24.49052 | 3.88926 | 23.38793 | 6.79592 | 11.22966 | 9.89307 |
| **Ms4a1** | 0.01377 | 0.00761 | 0.01152 | 0.08112 | 0.00941 | 0.00650 | 0.00738 | 0.01143 | 0.00742 | 0.00740 | 0.01041 | 0.00638 | 0.01296 | 0.01341 | 0.60828 | 0.00743 |
| **Ms4a6c** | 0.01284 | 0.00848 | 0.03496 | 0.08331 | 0.00963 | 0.00629 | 0.03010 | 0.02183 | 0.11105 | 0.24610 | 0.05218 | 0.01789 | 0.02068 | 0.17772 | 0.04569 | 0.00733 |
| **Msantd1** | 0.01445 | 0.00778 | 0.02023 | 0.00514 | 0.00931 | 0.00611 | 0.00727 | 0.01071 | 0.71225 | 0.40849 | 0.00748 | 0.01469 | 0.01197 | 0.01248 | 0.00541 | 0.00742 |
| **Msantd2** | 4.76220 | 6.74868 | 2.56363 | 5.99853 | 0.00520 | 0.67366 | 5.82279 | 5.64974 | 3.29150 | 9.06884 | 4.10209 | 2.00936 | 0.45331 | 16.32292 | 15.15079 | 0.00190 |
| **Msantd3** | 3.76955 | 0.00235 | 0.00344 | 0.46759 | 2.29419 | 0.00473 | 0.01424 | 0.07190 | 0.36754 | 0.01346 | 0.00870 | 0.00203 | 0.00790 | 0.00991 | 0.06571 | 0.01753 |
| **Msantd4** | 3.02525 | 0.00242 | 0.85022 | 2.51717 | 0.00544 | 0.55976 | 0.01313 | 1.70108 | 0.00000 | 0.00000 | 1.95119 | 7.73484 | 0.94233 | 0.56731 | 3.06102 | 0.32368 |
| **Msc** | 4.01562 | 1.55131 | 0.76734 | 0.00000 | 0.00000 | 2.86054 | 2.08844 | 0.66615 | 0.02023 | 0.33215 | 0.65218 | 0.00725 | 0.06959 | 0.01008 | 0.00000 | 1.46930 |
| **Msh2** | 12.71971 | 7.17736 | 8.78575 | 1.65157 | 16.87528 | 2.36026 | 6.60623 | 11.35409 | 4.42710 | 6.72553 | 6.88724 | 0.93389 | 6.25950 | 7.29305 | 4.20980 | 10.90893 |
| **Msh3** | 0.96552 | 0.72984 | 1.23649 | 3.66768 | 4.10784 | 3.37236 | 2.58396 | 0.77153 | 2.47178 | 3.77391 | 1.13672 | 6.48023 | 0.24297 | 2.44770 | 7.40448 | 0.00268 |
| **Msh4** | 1.58360 | 7.21114 | 1.00179 | 1.39579 | 0.00142 | 3.26357 | 2.38618 | 7.33098 | 0.55370 | 1.13145 | 1.63816 | 0.06746 | 0.74243 | 3.33447 | 1.68973 | 4.08655 |
| **Msh5** | 0.00139 | 0.36219 | 0.09722 | 2.05241 | 0.00808 | 0.00558 | 0.81871 | 0.03869 | 1.03978 | 1.85085 | 0.00000 | 0.05373 | 0.01056 | 0.00382 | 0.00000 | 0.15629 |
| **Msh6** | 17.18532 | 16.27996 | 11.24407 | 2.74283 | 13.65445 | 7.28659 | 6.06703 | 12.07326 | 8.15832 | 11.46158 | 7.49490 | 5.76287 | 7.07257 | 25.50046 | 18.24370 | 5.91542 |
| **Msi2** | 0.01343 | 0.01878 | 0.00993 | 0.35092 | 0.00897 | 0.00693 | 0.00744 | 0.00650 | 0.29980 | 0.35584 | 0.20826 | 0.01900 | 0.01045 | 0.01118 | 0.00469 | 0.00726 |
| **Msl1** | 2.92410 | 0.00868 | 5.68320 | 4.36220 | 0.20813 | 1.92499 | 3.47066 | 2.51626 | 2.49384 | 8.79881 | 2.49251 | 9.56264 | 5.42385 | 6.23637 | 6.54745 | 2.02027 |
| **Msl2** | 2.03385 | 1.36136 | 0.51961 | 9.77997 | 2.66731 | 2.55598 | 4.09753 | 1.61180 | 10.02200 | 1.49094 | 0.97936 | 0.11664 | 1.65020 | 2.66895 | 10.21340 | 0.01447 |
| **Msl3** | 1.91212 | 0.05906 | 6.66351 | 10.21215 | 0.00707 | 0.13558 | 1.67492 | 7.39268 | 2.19157 | 6.26172 | 3.11515 | 2.74586 | 2.90036 | 0.18335 | 5.76766 | 0.79538 |
| **Msln** | 0.01364 | 0.00797 | 0.01275 | 0.01839 | 0.00946 | 0.00683 | 0.00749 | 0.01279 | 0.01111 | 0.17532 | 0.01361 | 0.00706 | 0.01874 | 0.01124 | 0.01052 | 0.00793 |
| **Mslnl** | 0.00753 | 0.00752 | 0.06267 | 2.27712 | 0.00854 | 0.00823 | 0.00656 | 0.00770 | 0.77512 | 2.02122 | 0.00425 | 0.02621 | 0.94016 | 0.00069 | 0.69879 | 0.00672 |
| **Msmo1** | 0.00000 | 3.93986 | 6.03140 | 0.32147 | 0.01298 | 0.10727 | 3.25348 | 11.65616 | 0.31635 | 3.10966 | 8.79511 | 10.74985 | 3.98630 | 18.22597 | 3.99782 | 0.02820 |
| **Msn** | 1.96145 | 0.00645 | 0.00000 | 0.12802 | 0.00375 | 10.66230 | 0.00431 | 0.38910 | 0.95967 | 1.20165 | 0.40403 | 0.15851 | 2.51546 | 1.00749 | 0.31935 | 1.70523 |
| **Msra** | 1.44723 | 0.00727 | 0.00000 | 0.42287 | 0.01447 | 0.31113 | 0.00607 | 0.00651 | 0.72016 | 0.00141 | 0.00734 | 0.00583 | 0.03707 | 0.00871 | 0.18218 | 4.16975 |
| **Msrb1** | 0.28344 | 0.31722 | 0.37028 | 6.61561 | 0.01543 | 0.02573 | 0.04900 | 0.19762 | 18.62135 | 0.09035 | 2.91711 | 0.06854 | 0.02064 | 0.06334 | 0.06942 | 0.03983 |
| **Msrb2** | 2.15395 | 0.00242 | 0.88958 | 0.00000 | 3.47813 | 0.04683 | 0.01515 | 0.00000 | 0.04131 | 0.00000 | 0.00000 | 3.23887 | 0.03318 | 0.00610 | 0.38005 | 1.84128 |
| **Msrb3** | 0.38402 | 0.00841 | 0.00745 | 0.50352 | 0.01840 | 0.00683 | 0.00700 | 0.00000 | 0.82161 | 0.13372 | 0.42343 | 0.06239 | 0.20719 | 0.00520 | 0.61508 | 0.00706 |
| **Mss51** | 0.01587 | 0.00791 | 0.12447 | 0.21467 | 0.00895 | 0.00574 | 0.00733 | 0.01224 | 0.00801 | 0.01305 | 0.01346 | 0.00792 | 0.08244 | 0.01178 | 0.00495 | 0.00720 |
| **Mst1r** | 0.05409 | 0.00800 | 0.00888 | 0.00643 | 0.00799 | 0.00686 | 0.00747 | 0.01088 | 0.13471 | 0.00812 | 0.19844 | 0.00711 | 0.00996 | 0.01154 | 0.00594 | 0.00717 |
| **Msto1** | 3.89814 | 0.01038 | 1.97558 | 1.01876 | 2.23950 | 2.84221 | 3.85545 | 0.39689 | 1.56613 | 3.49349 | 0.62885 | 0.08739 | 4.58221 | 0.02387 | 1.94332 | 4.53754 |
| **Msx2** | 2.44183 | 0.00352 | 0.38769 | 0.02130 | 4.26801 | 0.01608 | 0.02901 | 0.12070 | 0.02130 | 0.00953 | 0.03694 | 0.00261 | 0.00649 | 0.07116 | 0.00103 | 0.09106 |
| **Mt1** | 4.86703 | 0.74453 | 5.52482 | 8.58908 | 0.19299 | 10.99499 | 3.98084 | 1.54221 | 20.40807 | 28.84238 | 35.71204 | 4.47520 | 15.26843 | 0.30150 | 0.00000 | 6.88796 |
| **Mt2** | 3.03402 | 0.62749 | 6.85350 | 20.04316 | 1.32518 | 9.92383 | 4.24140 | 0.00000 | 37.76954 | 19.31845 | 52.89436 | 5.76360 | 13.92113 | 1.02781 | 9.13605 | 6.28284 |
| **Mt4** | 0.08519 | 0.00809 | 0.00612 | 3.18410 | 0.00766 | 0.00690 | 0.00729 | 0.00466 | 0.00000 | 0.00400 | 0.96085 | 0.00397 | 0.00431 | 0.00027 | 0.03965 | 0.00630 |
| **Mta1** | 2.10202 | 0.09036 | 0.00000 | 0.46606 | 0.04160 | 0.49857 | 0.00548 | 0.60487 | 0.04146 | 0.67052 | 0.00488 | 0.32657 | 0.16703 | 0.00888 | 0.00555 | 1.25844 |
| **Mta2** | 0.14787 | 4.07870 | 2.50195 | 15.62665 | 0.61697 | 0.03173 | 2.64073 | 3.62422 | 18.20934 | 7.85149 | 6.86506 | 0.75833 | 7.87869 | 4.41413 | 11.28528 | 5.00892 |
| **Mta3** | 15.37943 | 9.49058 | 18.96157 | 4.33228 | 14.39536 | 15.84330 | 11.17980 | 37.98635 | 2.88910 | 38.49686 | 20.37813 | 13.99263 | 24.94258 | 0.23631 | 0.00000 | 10.24967 |
| **Mtap** | 8.94628 | 2.74893 | 0.00000 | 0.50186 | 12.65359 | 2.10427 | 0.07646 | 0.78854 | 0.00000 | 0.57219 | 0.12859 | 1.30438 | 3.25711 | 0.00307 | 0.58506 | 4.39288 |
| **Mtap7d3** | 0.01335 | 0.00814 | 0.01086 | 0.80901 | 0.00914 | 0.00664 | 0.00745 | 0.01189 | 0.00000 | 0.42443 | 0.00949 | 0.01927 | 0.01254 | 0.00449 | 0.97321 | 0.00726 |
| **Mtbp** | 0.99009 | 3.80326 | 0.22772 | 0.49248 | 0.04797 | 1.98538 | 1.25580 | 0.62857 | 0.02233 | 0.11241 | 1.30676 | 0.95974 | 0.00000 | 0.13815 | 3.30512 | 0.62030 |
| **Mtch1** | 9.71067 | 0.27189 | 7.13283 | 1.30647 | 2.24598 | 4.25009 | 3.80066 | 3.11801 | 0.96773 | 4.41236 | 5.02296 | 0.36231 | 0.00000 | 0.08278 | 0.61743 | 3.23321 |
| **Mtch2** | 27.78808 | 17.43280 | 28.43960 | 6.61377 | 17.18957 | 13.48268 | 14.21167 | 24.65559 | 8.48549 | 23.43653 | 22.69680 | 3.84610 | 26.57359 | 6.14383 | 6.31402 | 13.59441 |
| **Mtcl1** | 4.19510 | 0.22320 | 0.95486 | 0.66784 | 3.78616 | 4.25765 | 4.31288 | 0.01795 | 0.21959 | 0.19701 | 0.01515 | 0.00899 | 0.07548 | 1.05193 | 0.93686 | 2.92351 |
| **Mtcp1** | 0.01570 | 0.00790 | 0.02257 | 0.01804 | 0.00905 | 0.00690 | 0.00736 | 0.03978 | 0.01252 | 0.01463 | 0.01321 | 0.00785 | 0.01092 | 0.01221 | 0.00974 | 0.00738 |
| **Mtdh** | 6.24582 | 9.56900 | 5.96069 | 9.81280 | 2.36450 | 2.96899 | 8.26094 | 5.40369 | 7.52540 | 4.46377 | 1.56056 | 2.54361 | 1.49472 | 7.56206 | 10.51437 | 2.53492 |
| **Mterf1a** | 0.06372 | 0.01824 | 0.00483 | 0.00977 | 0.00851 | 0.00711 | 0.00720 | 0.01076 | 0.01202 | 0.00460 | 0.00918 | 0.00507 | 0.01331 | 0.64809 | 0.15651 | 0.00735 |
| **Mterf1b** | 0.04753 | 0.01832 | 0.00678 | 0.00931 | 0.00894 | 0.00708 | 0.00731 | 0.01147 | 0.02352 | 0.00570 | 0.01025 | 0.03407 | 0.01658 | 0.45345 | 0.09513 | 0.00745 |
| **Mterf2** | 1.63421 | 0.03773 | 0.00000 | 0.00000 | 0.00493 | 3.40150 | 0.03631 | 0.00000 | 0.53694 | 0.06236 | 0.73346 | 0.00093 | 1.09037 | 0.00434 | 0.41247 | 0.72730 |
| **Mterf3** | 1.23149 | 5.30595 | 0.49736 | 1.24519 | 2.81763 | 1.68465 | 1.48024 | 0.99077 | 0.17117 | 0.06589 | 0.18245 | 0.18681 | 1.22830 | 3.16653 | 2.16853 | 0.73781 |
| **Mterf4** | 7.28188 | 3.67116 | 1.71980 | 1.64372 | 0.15568 | 0.00000 | 0.00000 | 2.21266 | 0.73338 | 6.71574 | 3.34062 | 0.59409 | 3.62129 | 5.38152 | 3.23587 | 4.95518 |
| **Mtf1** | 3.70759 | 3.22881 | 0.00000 | 9.91124 | 0.07320 | 4.72923 | 2.50962 | 6.67712 | 10.70886 | 9.58506 | 6.83807 | 1.68450 | 2.74016 | 4.84880 | 10.96161 | 0.02521 |
| **Mtf2** | 9.83285 | 13.23418 | 19.96184 | 11.88031 | 8.44002 | 8.99137 | 6.98899 | 22.37290 | 21.12488 | 20.72781 | 22.52836 | 26.31273 | 10.27105 | 40.64113 | 20.02072 | 4.07642 |
| **Mtfmt** | 2.33084 | 0.01575 | 0.32985 | 1.22970 | 1.50268 | 1.02236 | 0.48152 | 1.44997 | 1.33431 | 1.15474 | 1.13984 | 0.05831 | 0.41682 | 0.43451 | 0.70703 | 0.92794 |
| **Mtfp1** | 4.18874 | 0.13520 | 0.00000 | 0.19544 | 0.01184 | 4.41504 | 2.24338 | 0.00667 | 0.00818 | 0.00000 | 0.00797 | 0.05938 | 3.36795 | 0.01244 | 0.01029 | 0.87437 |
| **Mtfr1** | 4.38619 | 8.80377 | 13.18624 | 5.45336 | 14.45574 | 5.46407 | 7.19009 | 6.18415 | 4.67378 | 3.06360 | 2.61331 | 7.19618 | 7.99624 | 2.98574 | 4.20727 | 3.54201 |
| **Mtfr1l** | 12.78702 | 6.19255 | 9.99322 | 8.47331 | 5.98388 | 0.26783 | 8.56903 | 4.12669 | 1.84147 | 6.78153 | 0.88769 | 0.94401 | 5.74408 | 0.00001 | 2.80302 | 1.52144 |
| **Mtfr2** | 0.01350 | 0.00808 | 0.01272 | 0.26061 | 0.00989 | 0.00697 | 0.00765 | 0.01386 | 0.03867 | 0.00777 | 0.02538 | 0.00732 | 0.03456 | 0.06354 | 0.08404 | 0.00760 |
| **Mtg1** | 12.41069 | 2.43172 | 8.99537 | 2.03746 | 3.64672 | 4.18205 | 6.48456 | 5.93596 | 0.24098 | 5.06923 | 3.98902 | 0.20268 | 8.97761 | 0.00000 | 1.15291 | 2.26508 |
| **Mtg2** | 6.18504 | 0.00000 | 0.01957 | 0.02916 | 0.11395 | 0.00202 | 0.27682 | 3.78085 | 0.04471 | 1.48784 | 0.09324 | 0.02915 | 4.07888 | 0.00742 | 0.02423 | 1.58129 |
| **Mthfd1** | 20.97461 | 15.29921 | 17.22485 | 2.80900 | 14.06892 | 9.21606 | 14.10821 | 22.30037 | 4.48194 | 20.05489 | 15.44660 | 8.07253 | 16.98488 | 2.23732 | 2.89940 | 10.66845 |
| **Mthfd1l** | 5.75160 | 0.25402 | 2.15894 | 0.08684 | 0.20390 | 0.01373 | 0.47568 | 0.10749 | 0.11478 | 2.08076 | 0.02308 | 0.03053 | 1.80762 | 0.68749 | 1.54976 | 0.00799 |
| **Mthfd2** | 1.20783 | 2.90318 | 1.98973 | 11.03164 | 2.45059 | 0.01251 | 2.59208 | 10.39293 | 1.56399 | 3.11077 | 0.40716 | 4.26305 | 6.12912 | 0.25733 | 2.46089 | 6.54812 |

| | | | | | | | | | | | | | | | | |
|---|---|---|---|---|---|---|---|---|---|---|---|---|---|---|---|---|
| **Mthfr** | 3.97707 | 2.77476 | 0.58783 | 8.57843 | 4.51316 | 3.83887 | 0.05885 | 10.31507 | 10.37472 | 14.97159 | 5.84015 | 4.69901 | 3.03753 | 1.27766 | 8.75097 | 2.03540 |
| **Mthfs** | 1.43287 | 1.01478 | 0.47304 | 1.96700 | 3.02109 | 0.21765 | 0.16731 | 1.06946 | 0.69575 | 0.15911 | 1.24607 | 0.22757 | 0.51590 | 0.12363 | 0.00000 | 0.09344 |
| **Mthfsd** | 0.26617 | 0.28454 | 5.42364 | 0.00709 | 0.00000 | 0.63581 | 3.71862 | 5.16812 | 0.00720 | 2.49818 | 3.52941 | 0.19780 | 5.01596 | 0.02962 | 0.00000 | 0.02351 |
| **Mthfsl** | 1.05151 | 0.85759 | 0.48219 | 1.78427 | 3.20773 | 0.32928 | 0.20768 | 1.18724 | 0.52129 | 0.28591 | 1.18047 | 0.16210 | 0.25651 | 0.32998 | 0.00000 | 0.11502 |
| **Mtif2** | 2.58160 | 1.40328 | 3.59733 | 1.85168 | 4.09422 | 2.86043 | 0.76632 | 0.00000 | 2.98436 | 2.80235 | 3.82249 | 0.37878 | 2.88670 | 6.74921 | 2.62045 | 0.14577 |
| **Mtif3** | 0.51473 | 0.11221 | 0.21013 | 3.06893 | 0.00644 | 2.29550 | 0.01463 | 4.07300 | 2.85917 | 4.85424 | 1.66609 | 0.09033 | 4.34354 | 0.00000 | 0.00000 | 1.05190 |
| **Mtl5** | 0.01786 | 0.00790 | 0.01185 | 0.01025 | 0.00894 | 0.00669 | 0.00744 | 0.01319 | 0.01216 | 0.05815 | 0.01317 | 0.00748 | 0.01303 | 0.01028 | 0.04626 | 0.00770 |
| **Mtm1** | 0.18697 | 2.99444 | 0.44374 | 0.03724 | 0.61229 | 0.00894 | 0.79675 | 0.54931 | 0.00685 | 0.95724 | 1.81925 | 0.02318 | 0.00000 | 2.13701 | 0.97356 | 0.08023 |
| **Mtmr1** | 0.75732 | 0.00198 | 0.41290 | 1.26145 | 1.81326 | 0.00582 | 0.00476 | 0.01397 | 1.39224 | 1.75573 | 0.08580 | 13.23972 | 1.09070 | 3.15947 | 0.38194 | 0.00451 |
| **Mtmr10** | 0.00000 | 2.03347 | 0.37566 | 1.58131 | 4.26723 | 5.15851 | 1.84905 | 0.86976 | 0.52765 | 3.30938 | 3.26504 | 0.20307 | 1.22307 | 0.08148 | 3.48291 | 2.01958 |
| **Mtmr11** | 0.03977 | 0.00816 | 1.17242 | 0.81792 | 0.01173 | 0.00607 | 0.00744 | 0.01285 | 0.09512 | 0.01859 | 0.03340 | 0.00667 | 0.13405 | 0.03472 | 0.00000 | 0.00711 |
| **Mtmr12** | 4.78430 | 0.16998 | 3.84816 | 2.18352 | 0.00000 | 3.70508 | 2.88534 | 2.31586 | 3.64891 | 3.08983 | 1.78142 | 9.32864 | 0.99412 | 2.83025 | 6.22174 | 2.08816 |
| **Mtmr14** | 1.16116 | 3.29461 | 3.47139 | 16.94773 | 0.06817 | 1.82255 | 0.03269 | 6.94719 | 8.62108 | 7.60660 | 5.47012 | 0.58885 | 1.58729 | 0.02267 | 0.65647 | 0.33629 |
| **Mtmr2** | 2.79299 | 0.04932 | 0.10043 | 6.62909 | 2.17533 | 0.52243 | 0.03718 | 0.09221 | 1.42094 | 0.60064 | 2.56518 | 0.04614 | 0.93114 | 0.20265 | 5.02944 | 1.08944 |
| **Mtmr3** | 2.80437 | 1.00287 | 7.23991 | 9.33475 | 2.24037 | 0.00924 | 1.87399 | 5.95261 | 6.93148 | 7.27930 | 3.07811 | 1.88495 | 4.79500 | 5.21483 | 6.17280 | 0.28993 |
| **Mtmr4** | 3.08591 | 6.36574 | 6.97447 | 4.19580 | 4.50368 | 0.16990 | 6.37637 | 3.47898 | 2.87871 | 1.57470 | 1.39370 | 11.48048 | 5.01259 | 0.40637 | 5.39812 | 4.74328 |
| **Mtmr6** | 0.82241 | 1.94055 | 0.00000 | 2.07542 | 0.03475 | 2.46835 | 0.00545 | 0.01204 | 1.86570 | 3.24508 | 0.00000 | 0.14300 | 1.02372 | 8.93618 | 7.57771 | 1.40773 |
| **Mtmr7** | 5.74812 | 0.01103 | 3.50199 | 0.25525 | 3.14789 | 0.00000 | 0.00000 | 0.79423 | 0.01875 | 0.00000 | 1.05805 | 0.10008 | 2.31574 | 0.03500 | 2.48716 | 5.37622 |
| **Mtmr9** | 1.56572 | 0.03531 | 2.48360 | 1.74266 | 0.00000 | 0.84583 | 0.00533 | 2.21293 | 1.21998 | 0.09888 | 0.57868 | 0.07780 | 0.18951 | 0.00686 | 0.51953 | 0.01200 |
| **Mto1** | 4.05181 | 0.23533 | 0.77528 | 0.80296 | 1.43117 | 0.00000 | 0.11246 | 0.15735 | 1.49026 | 2.09812 | 1.93589 | 8.58091 | 2.24348 | 0.92960 | 1.32973 | 0.05882 |
| **Mtor** | 2.27282 | 4.35628 | 5.20707 | 6.09123 | 0.02546 | 3.35577 | 0.00000 | 4.52316 | 6.48270 | 4.60473 | 3.49223 | 4.78366 | 3.09047 | 4.97731 | 6.09252 | 2.66892 |
| **Mtpap** | 5.48977 | 2.96457 | 5.49898 | 3.63971 | 7.37198 | 0.01618 | 7.98003 | 5.61576 | 0.00000 | 9.73606 | 2.74755 | 0.53524 | 2.11964 | 2.91946 | 3.72152 | 1.58969 |
| **Mtpn** | 13.06612 | 9.02278 | 8.98511 | 11.84073 | 13.78505 | 6.46337 | 7.81685 | 7.47091 | 11.54580 | 5.30734 | 6.26097 | 11.11078 | 7.40032 | 14.52456 | 6.66586 | 3.34369 |
| **Mtr** | 2.67522 | 5.46805 | 1.61847 | 0.05223 | 0.01069 | 3.47718 | 0.83699 | 3.44880 | 2.85293 | 3.75934 | 0.37735 | 0.08240 | 0.57958 | 0.05000 | 1.23279 | 1.39593 |
| **Mtrf1** | 0.01526 | 0.49370 | 0.40511 | 1.36931 | 0.31264 | 0.02148 | 0.01814 | 0.02305 | 3.61640 | 0.02327 | 0.00000 | 0.00000 | 0.42609 | 0.00916 | 0.18238 | 0.00664 |
| **Mtrf1l** | 1.69832 | 0.66238 | 1.48397 | 1.01707 | 1.90039 | 0.00439 | 3.75984 | 0.61677 | 1.59561 | 0.90737 | 2.29119 | 0.09240 | 0.48945 | 0.04610 | 1.74599 | 0.01153 |
| **Mtrr** | 3.15612 | 6.32324 | 4.57832 | 3.23121 | 4.47831 | 3.00133 | 5.50039 | 7.76922 | 3.35965 | 5.44070 | 7.25918 | 4.89113 | 3.53014 | 4.83464 | 6.95895 | 3.96854 |
| **Mtss1** | 0.00000 | 0.00687 | 0.00000 | 2.31210 | 0.00696 | 0.45221 | 0.00610 | 4.30909 | 0.87284 | 1.69616 | 1.95822 | 0.15084 | 0.68274 | 0.20383 | 0.88030 | 1.86594 |
| **Mtss1l** | 0.01655 | 0.00810 | 0.00806 | 0.57219 | 0.02427 | 0.00654 | 0.00709 | 0.15365 | 0.08612 | 0.00407 | 0.01214 | 0.00737 | 0.00000 | 0.01287 | 0.00266 | 1.89887 |
| **Mttp** | 0.00371 | 0.00878 | 0.00000 | 1.19384 | 0.01223 | 0.00625 | 0.00779 | 0.61950 | 0.56606 | 1.88647 | 0.66343 | 0.04113 | 0.00290 | 3.31242 | 1.59636 | 0.00716 |
| **Mturn** | 0.00451 | 0.00762 | 0.31209 | 0.01085 | 0.00873 | 0.00916 | 0.00611 | 0.95774 | 0.01122 | 0.00000 | 0.00780 | 0.02511 | 2.04400 | 0.01541 | 0.00861 | 0.01350 |
| **Mtus1** | 0.00152 | 3.13461 | 4.17032 | 24.64892 | 0.00754 | 7.23159 | 3.12029 | 10.22884 | 16.54471 | 18.53823 | 13.39464 | 10.35484 | 4.36973 | 18.95347 | 24.87497 | 0.34883 |
| **Mtus2** | 2.22815 | 0.01676 | 0.00000 | 3.60085 | 0.00467 | 0.00394 | 0.00514 | 0.00000 | 8.08429 | 0.32993 | 1.60116 | 0.19751 | 0.20708 | 0.03186 | 1.83251 | 0.00349 |
| **Mtx1** | 18.94206 | 0.92883 | 2.29371 | 0.12219 | 7.51973 | 5.76049 | 6.37916 | 11.37735 | 0.01587 | 10.69398 | 1.23753 | 0.52417 | 14.12631 | 0.01213 | 0.00776 | 5.68312 |
| **Mtx2** | 3.19638 | 2.57691 | 4.81519 | 2.90835 | 8.94927 | 2.14951 | 4.68716 | 0.00000 | 1.64761 | 4.45332 | 3.31945 | 9.91781 | 1.12069 | 3.67029 | 0.00000 | 0.22718 |
| **Mtx3** | 0.00000 | 2.26247 | 0.54721 | 0.52363 | 0.00750 | 0.00500 | 0.00574 | 0.98614 | 0.77212 | 2.24872 | 0.15772 | 6.00171 | 0.01877 | 1.25734 | 3.50435 | 0.00594 |
| **Muc1** | 0.90127 | 0.02471 | 2.42351 | 0.31584 | 0.01018 | 0.00549 | 2.92948 | 1.77578 | 0.00413 | 0.68004 | 0.00000 | 0.00746 | 0.00000 | 0.00883 | 0.00000 | 1.34323 |
| **Muc13** | 1.06882 | 0.00626 | 0.00321 | 1.32937 | 1.12306 | 0.00700 | 0.00648 | 0.01675 | 1.83789 | 0.00568 | 0.00000 | 0.01430 | 0.02210 | 0.00965 | 0.02108 | 0.00584 |
| **Muc4** | 0.01783 | 0.00795 | 0.01205 | 2.20097 | 0.00903 | 0.00642 | 0.00741 | 0.01371 | 0.00000 | 0.01084 | 0.01258 | 0.00775 | 0.01264 | 0.00424 | 0.11141 | 0.00699 |
| **Mul1** | 6.55285 | 3.18187 | 5.18519 | 5.74091 | 1.09050 | 0.56672 | 2.47846 | 7.38742 | 3.29363 | 7.70144 | 1.60905 | 0.10435 | 6.05370 | 0.02697 | 0.62931 | 1.44127 |
| **Mum1** | 6.09315 | 5.00684 | 11.87570 | 2.02863 | 2.26063 | 0.38569 | 0.73219 | 14.49062 | 3.03973 | 7.75572 | 5.23093 | 0.32563 | 16.39956 | 0.86934 | 0.05458 | 9.49296 |
| **Mum1l1** | 0.01333 | 0.00785 | 0.01148 | 1.86294 | 0.00933 | 0.00617 | 0.00721 | 0.01044 | 0.37851 | 0.45536 | 0.00781 | 0.00408 | 0.01200 | 0.00187 | 0.86490 | 0.00713 |
| **Murc** | 0.01033 | 0.00646 | 0.00766 | 0.00983 | 0.00964 | 0.00670 | 2.36804 | 0.00944 | 0.01225 | 0.01028 | 0.01334 | 0.00603 | 0.01014 | 0.01249 | 0.00935 | 0.00729 |
| **Mus81** | 0.50629 | 0.00451 | 1.16966 | 1.78456 | 4.45739 | 0.64743 | 0.30147 | 2.00605 | 1.27657 | 1.71034 | 0.15936 | 0.08642 | 0.35929 | 0.00613 | 0.66757 | 1.93520 |
| **Musk** | 0.01375 | 0.00794 | 0.01245 | 1.58294 | 0.00935 | 0.00647 | 0.00746 | 0.01366 | 0.00000 | 0.01193 | 0.01272 | 0.00766 | 0.01263 | 0.00479 | 0.08076 | 0.00713 |
| **Mustn1** | 0.03226 | 0.00812 | 0.01050 | 0.00818 | 0.01256 | 0.00690 | 0.00743 | 0.01110 | 0.01148 | 0.69731 | 0.01087 | 0.00679 | 0.01205 | 0.01421 | 0.08898 | 0.00757 |
| **Mut** | 2.60400 | 3.16809 | 0.72658 | 0.14202 | 0.00000 | 0.09444 | 1.19518 | 1.08745 | 0.00486 | 0.25004 | 1.01173 | 0.03269 | 0.02626 | 2.92465 | 0.57193 | 2.64030 |
| **Mutyh** | 2.44122 | 0.27126 | 3.89087 | 0.45732 | 9.98397 | 1.38145 | 0.54843 | 7.14091 | 1.36892 | 1.42427 | 7.01342 | 0.12078 | 2.18852 | 0.28544 | 0.00000 | 2.79492 |
| **Mvb12a** | 2.86630 | 0.01467 | 1.09172 | 3.82018 | 0.00000 | 0.03550 | 0.00000 | 3.16523 | 2.63837 | 6.61020 | 2.64882 | 0.04115 | 4.62678 | 0.00931 | 0.00000 | 4.58881 |
| **Mvb12b** | 0.00000 | 0.00804 | 0.00000 | 6.76672 | 0.00811 | 0.00736 | 0.00763 | 0.74942 | 4.21851 | 0.94114 | 1.26679 | 0.00000 | 0.00000 | 0.02369 | 1.03807 | 1.32114 |
| **Mvd** | 2.19722 | 0.00000 | 4.82061 | 0.38447 | 0.05770 | 0.08694 | 6.85768 | 5.89303 | 0.00774 | 3.81675 | 1.16127 | 0.08993 | 2.97124 | 0.00151 | 0.00006 | 0.30431 |
| **Mvk** | 6.70899 | 0.00000 | 0.32059 | 3.76582 | 0.00497 | 6.26145 | 3.84109 | 2.46703 | 2.01155 | 0.00000 | 0.00000 | 0.03064 | 2.86570 | 0.01059 | 0.00000 | 0.00959 |
| **Mvp** | 5.91857 | 3.74221 | 11.25726 | 4.12731 | 0.00000 | 3.09070 | 1.90469 | 4.27750 | 5.59469 | 5.66889 | 4.94639 | 0.27208 | 1.82490 | 0.00000 | 0.00000 | 3.31838 |
| **Mxd1** | 0.01258 | 0.00728 | 0.00000 | 3.52230 | 0.00927 | 2.14431 | 0.00679 | 1.53192 | 0.93090 | 0.16063 | 0.32656 | 0.06624 | 0.00000 | 0.00014 | 0.82643 | 0.00601 |
| **Mxd3** | 3.63301 | 0.16486 | 2.44085 | 0.16283 | 0.00000 | 4.84170 | 0.00189 | 4.53708 | 0.16560 | 2.96974 | 1.30626 | 0.13160 | 2.32905 | 0.03121 | 0.05206 | 1.92762 |
| **Mxd4** | 0.01332 | 0.00832 | 0.12023 | 0.67033 | 0.00897 | 0.00563 | 0.00767 | 0.00899 | 1.15181 | 0.00000 | 0.30083 | 0.00513 | 0.00849 | 0.00684 | 0.00341 | 0.00702 |
| **Mxi1** | 2.58170 | 0.64984 | 1.67129 | 1.58771 | 0.00343 | 0.00422 | 0.00416 | 1.15316 | 1.35108 | 3.37117 | 0.04410 | 3.20187 | 1.19874 | 0.00186 | 0.64231 | 0.05479 |
| **Mxra7** | 0.01214 | 0.00774 | 0.13553 | 0.12004 | 0.00875 | 0.00675 | 0.00672 | 0.00000 | 0.41429 | 0.31511 | 0.86495 | 0.01298 | 0.45927 | 0.12511 | 0.22213 | 0.01393 |
| **Mxra8** | 0.00879 | 0.01719 | 0.00864 | 1.77399 | 0.00892 | 0.00625 | 0.44816 | 0.00769 | 1.42460 | 0.77116 | 0.00000 | 0.00963 | 1.23941 | 0.00118 | 0.40474 | 0.00651 |
| **Myadm** | 3.04127 | 0.08979 | 1.61579 | 1.57901 | 1.47152 | 1.42555 | 1.59851 | 3.33074 | 4.08426 | 1.40874 | 5.22853 | 1.35423 | 0.30682 | 9.09112 | 5.28810 | 0.13360 |
| **Myadml2** | 0.01357 | 0.00821 | 0.01207 | 3.10656 | 0.00937 | 0.00623 | 0.00736 | 0.01321 | 1.12450 | 0.04930 | 0.00684 | 0.00584 | 0.01295 | 0.00841 | 0.00312 | 0.00689 |
| **Mybbp1a** | 19.36321 | 12.72545 | 12.68023 | 17.31758 | 18.57823 | 5.30399 | 12.06176 | 31.99685 | 25.27929 | 31.39799 | 30.05568 | 11.54885 | 16.32395 | 15.09300 | 14.81140 | 9.76591 |

| | | | | | | | | | | | | | | | | |
|---|---|---|---|---|---|---|---|---|---|---|---|---|---|---|---|---|
| **Mybl1** | 0.01294 | 0.00751 | 0.00709 | 6.37231 | 0.00916 | 0.00575 | 0.04681 | 0.00588 | 2.55486 | 0.00000 | 0.08064 | 0.00885 | 0.04383 | 0.26403 | 1.00884 | 0.00592 |
| **Mybl2** | 23.10452 | 9.52798 | 15.44625 | 12.88270 | 13.08368 | 9.28969 | 12.00874 | 20.29320 | 12.07496 | 14.74670 | 10.46093 | 1.25629 | 15.59612 | 1.61913 | 6.96629 | 13.60899 |
| **Mybpc1** | 0.93777 | 0.82899 | 0.00000 | 0.36827 | 2.55041 | 0.00669 | 0.08684 | 0.07184 | 0.20364 | 0.00000 | 0.00801 | 0.04220 | 0.01720 | 0.47483 | 0.10205 | 1.03418 |
| **Mybpc2** | 0.00000 | 0.02410 | 4.57572 | 0.03508 | 4.21374 | 0.25923 | 0.97181 | 11.39085 | 0.43512 | 3.53327 | 4.96584 | 0.25897 | 4.91960 | 0.27682 | 1.48800 | 0.00241 |
| **Myc** | 9.30214 | 2.88126 | 0.00000 | 1.41751 | 3.34840 | 2.28013 | 0.61374 | 9.02062 | 0.29328 | 2.96681 | 1.45584 | 0.68906 | 0.06357 | 1.74126 | 3.13436 | 3.62809 |
| **Mycbp** | 5.52172 | 5.39690 | 10.96771 | 4.83742 | 10.01668 | 8.27562 | 7.59379 | 7.66534 | 4.87975 | 9.72591 | 8.08134 | 2.88284 | 5.58694 | 5.70773 | 4.86372 | 6.76515 |
| **Mycbp2** | 5.34976 | 7.34792 | 5.02255 | 7.18454 | 2.08723 | 0.00000 | 5.18855 | 3.48879 | 3.71052 | 5.21140 | 0.46051 | 0.37072 | 5.71401 | 6.26882 | 11.29760 | 1.10781 |
| **Mycbpap** | 0.00348 | 0.00722 | 0.07524 | 0.00178 | 0.09860 | 1.98731 | 0.44238 | 0.15131 | 0.01134 | 0.00075 | 0.45203 | 0.01314 | 0.45427 | 0.04273 | 0.00000 | 0.01417 |
| **Mycl** | 0.95513 | 0.25265 | 2.53633 | 0.00320 | 0.00208 | 0.00489 | 1.19048 | 0.00465 | 0.02414 | 0.23092 | 0.38649 | 0.05493 | 0.02380 | 0.42419 | 0.00000 | 0.01342 |
| **Mycn** | 3.53795 | 0.00344 | 0.00000 | 0.00970 | 1.23714 | 0.11620 | 0.00501 | 0.00954 | 0.01176 | 0.01014 | 0.01362 | 0.00449 | 0.05713 | 0.01276 | 0.01102 | 0.15176 |
| **Mycs** | 0.01366 | 0.00795 | 0.01188 | 0.40574 | 0.00928 | 0.00674 | 0.00758 | 0.01403 | 0.45772 | 0.03235 | 0.00867 | 0.00758 | 0.01321 | 0.01023 | 0.07973 | 0.00733 |
| **Myd88** | 1.89914 | 0.09295 | 0.88509 | 3.06559 | 0.05165 | 0.00660 | 0.01631 | 0.02572 | 0.04466 | 2.55810 | 0.06632 | 0.15904 | 1.05025 | 0.00054 | 0.49050 | 0.01957 |
| **Mydgf** | 14.26521 | 1.50413 | 4.51618 | 0.56271 | 6.12038 | 0.11395 | 1.70238 | 5.09341 | 0.33039 | 5.39939 | 2.65209 | 0.08705 | 7.65898 | 0.41142 | 0.40070 | 7.84012 |
| **Myef2** | 2.49231 | 5.03389 | 3.45534 | 1.62616 | 4.09625 | 2.25238 | 1.74419 | 1.04713 | 2.37388 | 1.57166 | 7.80556 | 4.75437 | 2.11695 | 4.03294 | 4.72521 | 1.49319 |
| **Myeov2** | 2.56061 | 0.07693 | 1.41069 | 16.73243 | 0.18016 | 0.66899 | 0.70706 | 0.00000 | 7.52860 | 0.12657 | 8.28580 | 0.08142 | 0.48434 | 1.07556 | 1.93039 | 0.41649 |
| **Myg1** | 3.44610 | 2.63799 | 6.28677 | 6.60163 | 2.43337 | 1.98982 | 1.85686 | 7.57427 | 6.29290 | 8.55155 | 3.44516 | 0.92807 | 2.47243 | 0.00000 | 2.02442 | 3.43838 |
| **Myh10** | 4.20076 | 6.94367 | 5.83486 | 2.32675 | 2.62025 | 13.90360 | 7.80639 | 16.52732 | 1.94142 | 8.45406 | 7.27064 | 2.69080 | 10.91144 | 6.97761 | 2.22716 | 1.31320 |
| **Myh11** | 0.01043 | 0.00760 | 0.00853 | 2.77773 | 0.00979 | 0.00650 | 0.00729 | 0.03251 | 1.37960 | 1.23336 | 0.01367 | 0.02495 | 0.02977 | 0.62674 | 0.92707 | 0.00720 |
| **Myh13** | 0.53187 | 0.01627 | 0.00000 | 0.01023 | 0.00586 | 0.00726 | 1.59709 | 0.18396 | 0.01205 | 0.00000 | 0.01239 | 0.00585 | 0.00000 | 0.01594 | 0.00581 | 2.49776 |
| **Myh14** | 0.98631 | 0.01990 | 0.00000 | 0.33465 | 0.01024 | 0.01635 | 0.00686 | 0.16499 | 0.00662 | 0.95960 | 0.79480 | 0.01388 | 0.39540 | 0.00622 | 0.00000 | 0.00611 |
| **Myh15** | 0.01334 | 0.00795 | 0.01200 | 0.01062 | 0.00935 | 0.00665 | 0.00744 | 0.02350 | 0.01234 | 0.01503 | 0.01308 | 0.00795 | 0.01204 | 0.01224 | 0.01045 | 0.00741 |
| **Myh3** | 0.01336 | 0.00792 | 0.00922 | 0.28925 | 0.00903 | 0.00676 | 0.05873 | 0.00731 | 0.27507 | 0.00456 | 0.17687 | 0.06781 | 0.01001 | 0.00949 | 0.00000 | 0.05770 |
| **Myh6** | 0.01083 | 0.00862 | 0.01025 | 0.04897 | 0.00984 | 0.00720 | 0.00829 | 0.23460 | 0.01050 | 0.50245 | 0.01324 | 0.01920 | 0.02193 | 0.01695 | 0.00698 | 0.00761 |
| **Myh7** | 0.00855 | 0.00870 | 0.01042 | 0.25272 | 0.01017 | 0.00734 | 0.00830 | 0.27851 | 0.91368 | 0.85261 | 0.04981 | 0.02804 | 0.14851 | 0.01408 | 0.00784 | 0.00767 |
| **Myh7b** | 1.69805 | 0.19340 | 0.64920 | 0.00606 | 0.00862 | 0.93270 | 0.00507 | 0.00000 | 0.01197 | 0.00000 | 0.14247 | 0.00518 | 1.60174 | 0.01370 | 0.00000 | 0.00418 |
| **Myh9** | 0.46611 | 0.24869 | 0.91193 | 0.89259 | 0.00692 | 4.20361 | 0.05706 | 1.35439 | 1.63067 | 0.00000 | 0.37327 | 0.44178 | 1.13217 | 0.19232 | 0.05259 | 0.07764 |
| **Myl12a** | 20.41898 | 13.60890 | 21.06066 | 4.84719 | 11.64834 | 16.73761 | 16.07254 | 18.44997 | 2.86240 | 18.08896 | 13.53693 | 8.75881 | 20.21890 | 3.43543 | 7.07585 | 12.96060 |
| **Myl12b** | 10.89856 | 10.76627 | 15.26511 | 2.25859 | 6.63790 | 13.73899 | 10.95920 | 13.92346 | 0.99719 | 9.38059 | 7.22374 | 2.51625 | 20.04641 | 2.32425 | 4.44385 | 10.47968 |
| **Myl2** | 0.01372 | 0.00785 | 0.01244 | 0.31460 | 0.00935 | 0.00651 | 0.00746 | 0.01280 | 0.00622 | 0.07672 | 0.01237 | 0.00704 | 0.01235 | 0.00551 | 0.14071 | 0.00729 |
| **Myl4** | 0.01254 | 0.00781 | 1.65068 | 0.07488 | 0.00636 | 0.00865 | 0.00730 | 0.00590 | 0.01122 | 0.00921 | 0.42745 | 0.00498 | 0.00403 | 0.00954 | 0.00000 | 0.00678 |
| **Myl6** | 23.76172 | 10.52550 | 20.02605 | 9.80836 | 13.84168 | 14.02390 | 6.84504 | 17.31536 | 5.90356 | 23.28497 | 21.48608 | 2.24818 | 30.43045 | 9.67022 | 7.12536 | 18.66266 |
| **Myl6b** | 0.12128 | 0.02821 | 0.08122 | 0.31315 | 0.02885 | 0.07912 | 0.01722 | 0.05819 | 0.06302 | 0.10599 | 0.08947 | 0.00620 | 0.20923 | 0.17083 | 0.01681 | 0.09471 |
| **Myl9** | 0.02989 | 0.00886 | 0.73747 | 0.00556 | 0.01037 | 0.00884 | 0.09675 | 0.15586 | 0.01098 | 2.50618 | 0.00678 | 0.09378 | 0.00413 | 0.01387 | 0.00478 | 0.00716 |
| **Mylip** | 0.02491 | 0.01703 | 0.54408 | 1.24632 | 0.00717 | 0.00902 | 1.26232 | 0.01748 | 1.91392 | 0.57064 | 0.01286 | 0.00000 | 1.18385 | 0.00457 | 0.55109 | 0.00635 |
| **Mylk** | 1.85201 | 0.00558 | 0.00350 | 0.36044 | 0.00434 | 0.00651 | 0.00572 | 0.00458 | 0.41745 | 1.34168 | 0.00967 | 0.00168 | 0.01139 | 0.00762 | 0.02773 | 0.00557 |
| **Mylk2** | 0.01368 | 0.00785 | 0.01101 | 0.67391 | 0.00894 | 0.00655 | 0.00715 | 0.00000 | 0.61200 | 0.22440 | 0.96077 | 0.00153 | 0.00510 | 0.00391 | 0.00000 | 0.00741 |
| **Mylk3** | 0.01366 | 0.00783 | 0.01212 | 0.04020 | 0.00923 | 0.00660 | 0.00745 | 0.01271 | 0.62615 | 0.07216 | 0.00986 | 0.00594 | 0.01310 | 0.00695 | 0.05843 | 0.00771 |
| **Mylk4** | 0.01914 | 0.00758 | 0.03499 | 0.11308 | 0.00919 | 0.01205 | 0.00719 | 0.01503 | 0.10998 | 0.03179 | 0.03972 | 0.22829 | 0.09075 | 0.21509 | 0.02917 | 0.00737 |
| **Mylpf** | 6.18642 | 4.02674 | 12.43201 | 0.73133 | 2.90788 | 2.06985 | 3.55154 | 8.03237 | 1.47484 | 6.71460 | 20.05518 | 1.42857 | 11.32452 | 2.11290 | 1.84470 | 3.80379 |
| **Mynn** | 1.04936 | 4.20089 | 1.72639 | 4.38811 | 2.60939 | 1.10398 | 2.31994 | 1.49048 | 3.68237 | 4.10150 | 2.98708 | 0.80951 | 0.98508 | 3.43908 | 8.51949 | 0.46271 |
| **Myo10** | 0.19986 | 0.00694 | 0.77744 | 5.77879 | 0.01374 | 0.00515 | 1.07284 | 0.01442 | 2.00989 | 5.82254 | 0.09701 | 0.06161 | 0.15285 | 0.00000 | 4.72395 | 2.28712 |
| **Myo15** | 0.01006 | 0.00845 | 0.00910 | 0.99411 | 0.00954 | 0.00671 | 0.00812 | 0.17327 | 0.14169 | 0.02534 | 0.00873 | 0.01882 | 0.02860 | 0.15619 | 0.02372 | 0.00707 |
| **Myo16** | 0.01358 | 0.00827 | 0.01201 | 1.05967 | 0.00911 | 0.00641 | 0.00742 | 0.01114 | 0.46899 | 0.32984 | 0.02411 | 0.00652 | 0.01109 | 0.00978 | 0.00330 | 0.00693 |
| **Myo18a** | 0.67862 | 0.00601 | 0.06983 | 2.04216 | 0.01286 | 0.00776 | 0.31159 | 3.88551 | 2.33740 | 0.92955 | 1.61945 | 0.02271 | 0.87177 | 0.00000 | 0.07843 | 3.26466 |
| **Myo19** | 2.04093 | 1.72650 | 2.92467 | 3.70240 | 0.40234 | 0.12090 | 1.81100 | 2.57563 | 3.21112 | 0.41334 | 0.67442 | 0.02396 | 0.57009 | 0.00003 | 0.76404 | 0.01658 |
| **Myo1a** | 0.02514 | 0.00821 | 0.05817 | 0.82561 | 0.02224 | 0.01152 | 0.00735 | 0.01107 | 0.88349 | 0.00603 | 0.06793 | 0.02778 | 0.01740 | 0.01097 | 0.00000 | 0.00718 |
| **Myo1b** | 7.20577 | 10.27830 | 7.52230 | 7.31365 | 2.30618 | 0.13087 | 9.89179 | 15.01845 | 10.92405 | 19.04874 | 11.49753 | 10.62758 | 8.80733 | 16.07089 | 13.63946 | 5.16513 |
| **Myo1c** | 2.80893 | 0.39696 | 6.08615 | 9.34506 | 0.05965 | 7.80822 | 0.65920 | 11.77914 | 7.85950 | 6.95063 | 5.67872 | 5.02392 | 9.27383 | 1.41743 | 2.75882 | 1.42634 |
| **Myo1d** | 2.19889 | 0.00547 | 0.68101 | 2.32804 | 0.02595 | 0.01627 | 0.00577 | 2.52261 | 2.05315 | 1.15894 | 0.00000 | 0.02240 | 0.19061 | 0.00000 | 0.42515 | 0.01232 |
| **Myo1e** | 0.11357 | 0.00732 | 0.91125 | 0.93030 | 0.01316 | 0.02601 | 0.02682 | 0.00606 | 0.04585 | 0.05812 | 0.28963 | 0.07205 | 0.08322 | 0.27580 | 0.03870 | 4.57371 |
| **Myo1g** | 0.01076 | 0.00768 | 0.99201 | 0.50945 | 0.00773 | 0.00570 | 0.00694 | 0.46317 | 0.88982 | 0.43364 | 0.49515 | 0.01397 | 0.14899 | 0.00662 | 0.00000 | 0.00673 |
| **Myo1h** | 0.01314 | 0.00760 | 0.00758 | 0.09465 | 0.02189 | 0.00686 | 0.00710 | 0.55531 | 0.01989 | 0.00578 | 0.01049 | 0.00628 | 0.00568 | 0.00786 | 0.08946 | 0.00735 |
| **Myo3b** | 0.01367 | 0.00770 | 0.00754 | 1.16685 | 0.00914 | 0.00614 | 0.00712 | 0.00041 | 0.74203 | 0.00000 | 0.31804 | 0.00120 | 0.00876 | 0.00356 | 1.74868 | 0.00685 |
| **Myo5a** | 0.24339 | 0.01757 | 1.54449 | 5.63985 | 0.04600 | 0.00732 | 0.01576 | 0.93325 | 0.62518 | 0.69897 | 2.56281 | 0.09980 | 1.18124 | 1.09149 | 4.87535 | 0.00459 |
| **Myo5b** | 0.55112 | 0.35982 | 2.81776 | 6.18135 | 0.01195 | 1.15862 | 0.00595 | 1.56540 | 1.71617 | 0.33093 | 0.20805 | 0.01116 | 0.50147 | 0.62455 | 6.46446 | 0.30703 |
| **Myo5c** | 0.28366 | 0.00709 | 0.21637 | 0.01681 | 0.42354 | 0.00532 | 0.02684 | 0.01486 | 0.03489 | 0.00000 | 0.01041 | 0.10733 | 0.00710 | 0.03349 | 0.07956 | 1.26560 |
| **Myo6** | 2.57316 | 0.00421 | 0.16518 | 2.16728 | 0.01140 | 3.97584 | 0.24868 | 1.21045 | 3.59407 | 3.15997 | 2.55116 | 0.01294 | 0.00000 | 0.00000 | 0.83033 | 0.00219 |
| **Myo7a** | 0.02539 | 0.27520 | 2.69262 | 0.57719 | 0.00372 | 2.08011 | 0.00693 | 0.00924 | 0.00000 | 1.08390 | 0.00765 | 0.00353 | 0.00701 | 0.00767 | 0.00000 | 0.98091 |
| **Myo7b** | 0.00816 | 0.01796 | 0.02433 | 0.00815 | 0.00962 | 0.00683 | 2.48455 | 0.01090 | 0.01166 | 0.04969 | 0.01151 | 0.00739 | 0.05847 | 0.00680 | 0.19386 | 0.00724 |
| **Myo9a** | 2.86344 | 1.43577 | 0.96125 | 3.69791 | 0.00330 | 0.56577 | 0.46778 | 4.94324 | 2.55846 | 9.63439 | 6.09834 | 8.71419 | 1.92038 | 26.34059 | 8.62533 | 3.65727 |
| **Myo9b** | 0.98279 | 0.02454 | 0.00725 | 5.09433 | 0.00402 | 4.11822 | 2.49626 | 1.39130 | 2.48942 | 3.33982 | 0.00000 | 0.08664 | 2.18587 | 0.00000 | 0.84644 | 5.23342 |
| **Myod1** | 2.86533 | 0.00434 | 0.00310 | 0.00912 | 1.57382 | 0.00581 | 0.00537 | 0.00498 | 0.01143 | 0.01106 | 0.01207 | 0.00518 | 0.01012 | 0.00982 | 0.05382 | 0.00436 |

| | | | | | | | | | | | | | | | | |
|---|---|---|---|---|---|---|---|---|---|---|---|---|---|---|---|---|
| **Myof** | 0.07681 | 0.00841 | 0.00546 | 0.24875 | 0.01324 | 3.32533 | 0.00709 | 0.01545 | 0.00510 | 0.65012 | 0.00720 | 0.04016 | 0.00000 | 0.01070 | 0.00502 | 0.16582 |
| **Myom1** | 0.01354 | 0.00756 | 0.00992 | 0.00651 | 0.00953 | 0.00649 | 0.00730 | 0.01077 | 0.01070 | 0.00224 | 0.00875 | 0.02336 | 0.01331 | 0.17078 | 0.85606 | 0.00747 |
| **Myom2** | 0.01009 | 0.00745 | 0.17536 | 2.17493 | 0.00913 | 0.00867 | 0.00694 | 0.00831 | 0.05964 | 0.00000 | 0.01452 | 0.01959 | 0.10590 | 1.23644 | 0.65595 | 0.00679 |
| **Myom3** | 0.01182 | 0.00810 | 0.01162 | 0.49058 | 0.00910 | 0.00645 | 0.00719 | 0.01141 | 0.26746 | 0.00572 | 0.01175 | 0.01854 | 0.41354 | 0.01209 | 0.00344 | 0.00695 |
| **Myoz1** | 0.00904 | 0.00759 | 0.00872 | 0.01117 | 0.01031 | 0.00720 | 0.00731 | 0.84581 | 0.01143 | 0.00482 | 0.00946 | 0.00588 | 0.01288 | 0.01620 | 0.00000 | 0.00764 |
| **Myoz3** | 0.06418 | 0.02759 | 0.07258 | 0.72708 | 0.02454 | 0.03125 | 0.01702 | 0.00000 | 1.10403 | 0.04664 | 0.65341 | 0.00999 | 0.03203 | 0.33542 | 0.32408 | 0.01409 |
| **Mypn** | 2.16896 | 0.01464 | 4.11595 | 0.72670 | 0.00000 | 3.37439 | 0.07402 | 3.80988 | 0.61717 | 2.90261 | 0.64417 | 0.56385 | 2.09089 | 1.64371 | 3.75096 | 0.04525 |
| **Mypop** | 0.01170 | 0.00813 | 0.20846 | 2.95743 | 0.01798 | 0.01724 | 0.00720 | 0.00000 | 0.66803 | 0.10150 | 1.08278 | 0.00196 | 0.13757 | 0.00215 | 0.34928 | 0.00647 |
| **Myrf** | 0.01103 | 0.00774 | 0.01097 | 0.07750 | 0.01862 | 0.00683 | 0.00740 | 0.05853 | 0.00808 | 0.40644 | 0.01051 | 0.00683 | 0.01030 | 0.00605 | 0.15841 | 0.00748 |
| **Myrfl** | 0.22517 | 0.00716 | 0.25353 | 0.00552 | 0.00705 | 0.00532 | 0.93478 | 0.01004 | 0.63583 | 0.05056 | 0.00455 | 0.06141 | 0.01649 | 0.57116 | 0.00000 | 0.00690 |
| **Myrip** | 0.01359 | 0.00778 | 0.01064 | 1.11019 | 0.00924 | 0.00664 | 0.00747 | 0.01211 | 0.40905 | 0.13183 | 0.00708 | 0.02882 | 0.01327 | 0.00686 | 0.73925 | 0.00720 |
| **Mysm1** | 0.54175 | 0.23483 | 0.90327 | 4.58676 | 0.81169 | 0.18425 | 2.38369 | 8.42589 | 4.26482 | 9.85505 | 7.32066 | 8.90101 | 0.92983 | 19.26860 | 17.46491 | 1.64967 |
| **Myt1** | 0.01377 | 0.00805 | 0.01216 | 0.96926 | 0.00935 | 0.00653 | 0.00746 | 0.01329 | 0.01924 | 0.01093 | 0.01255 | 0.00809 | 0.01269 | 0.00871 | 0.09937 | 0.00723 |
| **Myt1l** | 0.00769 | 2.08159 | 0.09187 | 0.09245 | 0.00784 | 0.00574 | 0.00609 | 0.51802 | 0.38508 | 0.00000 | 0.89056 | 0.00083 | 0.00000 | 0.05457 | 0.05033 | 0.00612 |
| **Myzap** | 0.01772 | 0.00778 | 0.01135 | 0.00440 | 0.00886 | 0.00651 | 0.00743 | 0.01132 | 0.32428 | 0.35676 | 0.00775 | 0.00580 | 0.01131 | 0.01355 | 0.23135 | 0.00709 |
| **Mzb1** | 0.01368 | 0.00815 | 0.01083 | 0.85101 | 0.00898 | 0.00654 | 0.00733 | 0.00558 | 0.00000 | 0.51342 | 0.19729 | 0.00622 | 0.00940 | 0.00802 | 0.00000 | 0.00696 |
| **Mzf1** | 0.11871 | 0.00715 | 0.00549 | 0.00787 | 0.01369 | 0.00708 | 1.27357 | 0.00165 | 0.17054 | 0.18610 | 0.23792 | 0.00433 | 0.00954 | 0.00743 | 0.02549 | 0.00769 |
| **Mzt1** | 5.76397 | 2.67771 | 5.79154 | 7.24032 | 3.82759 | 1.12719 | 1.67460 | 2.54052 | 0.92218 | 6.10139 | 2.48011 | 0.95583 | 2.60888 | 6.79959 | 3.19756 | 1.41609 |
| **Mzt2** | 4.61123 | 0.24356 | 0.35515 | 0.21976 | 1.08235 | 3.99712 | 1.04359 | 5.58034 | 0.40093 | 3.54642 | 3.28235 | 0.20556 | 0.50995 | 0.00193 | 0.04611 | 0.01493 |
| **N4bp1** | 6.62095 | 4.88050 | 0.69346 | 15.16560 | 10.68943 | 5.84026 | 6.97427 | 7.40814 | 3.29974 | 8.85035 | 3.52996 | 2.95284 | 4.83111 | 2.42257 | 6.54726 | 2.39429 |
| **N4bp2** | 1.27794 | 0.00581 | 0.00000 | 5.50506 | 0.00570 | 0.00396 | 0.04564 | 0.00737 | 3.04161 | 0.80006 | 0.00000 | 0.11108 | 0.01125 | 0.05378 | 5.55040 | 0.00492 |
| **N4bp2l1** | 0.18192 | 0.00729 | 0.00000 | 0.00998 | 0.00827 | 0.00781 | 0.00713 | 0.00842 | 0.01239 | 0.08986 | 0.00738 | 0.02169 | 0.01223 | 1.45620 | 0.01114 | 0.00699 |
| **N4bp2l2** | 3.39962 | 8.51215 | 7.43342 | 14.41145 | 2.25853 | 0.02124 | 4.46892 | 8.14006 | 17.07145 | 15.07428 | 9.77748 | 8.82099 | 6.77721 | 19.82670 | 22.88883 | 5.04968 |
| **N4bp3** | 0.17094 | 0.08212 | 0.04901 | 0.42742 | 0.03105 | 0.00800 | 0.00691 | 2.64444 | 0.00391 | 0.64530 | 0.39380 | 0.00000 | 0.00267 | 0.00423 | 0.01983 | 0.47326 |
| **N6amt1** | 3.19278 | 0.02590 | 0.51893 | 0.98117 | 0.01667 | 2.11143 | 1.33565 | 1.68102 | 2.47034 | 2.45513 | 4.12005 | 1.57920 | 0.38855 | 2.52297 | 0.62281 | 0.03098 |
| **Naa10** | 20.50080 | 6.10036 | 14.03636 | 5.07080 | 8.21664 | 2.82083 | 8.13009 | 13.87964 | 7.59459 | 11.89248 | 5.49977 | 1.68588 | 8.12842 | 0.21279 | 1.44058 | 11.04958 |
| **Naa11** | 3.90353 | 4.42073 | 6.31487 | 2.47684 | 7.17492 | 2.07464 | 2.88448 | 1.52009 | 0.07327 | 2.43066 | 1.13614 | 4.94557 | 0.06157 | 0.15333 | 0.80546 | 6.98509 |
| **Naa15** | 9.14831 | 11.64579 | 9.42021 | 9.33424 | 9.23488 | 9.62666 | 7.74863 | 11.11213 | 8.67482 | 9.61211 | 8.89426 | 10.04987 | 3.98950 | 28.27211 | 14.64242 | 7.15333 |
| **Naa16** | 5.10011 | 9.87391 | 2.85820 | 4.59475 | 0.76445 | 1.40252 | 2.59370 | 4.92652 | 10.44656 | 7.55240 | 4.18026 | 1.65000 | 4.23711 | 6.35948 | 6.83610 | 4.27657 |
| **Naa20** | 3.44255 | 4.58448 | 0.00000 | 6.99877 | 3.85099 | 1.65916 | 0.43243 | 2.42582 | 5.16028 | 0.46582 | 0.67457 | 3.54960 | 0.64919 | 6.49459 | 0.00000 | 0.19020 |
| **Naa25** | 3.65130 | 9.08609 | 6.78287 | 4.16448 | 0.01528 | 4.94647 | 5.83928 | 5.94936 | 3.55136 | 7.29477 | 1.46979 | 13.73472 | 3.41104 | 10.27914 | 5.28163 | 0.02129 |
| **Naa30** | 1.13368 | 4.99311 | 2.36609 | 21.19399 | 2.47728 | 2.48020 | 0.03182 | 2.59049 | 4.91786 | 1.48283 | 2.64528 | 4.20911 | 2.74309 | 10.51373 | 6.06315 | 0.07628 |
| **Naa35** | 10.53050 | 10.67518 | 5.84169 | 8.71790 | 5.99424 | 5.83847 | 6.75454 | 16.18300 | 9.73571 | 15.01581 | 10.31260 | 7.43268 | 6.42095 | 13.34293 | 11.79874 | 2.25578 |
| **Naa38** | 5.03145 | 0.83051 | 2.34903 | 4.02921 | 6.01911 | 1.14341 | 0.72019 | 1.23986 | 1.69489 | 2.66529 | 4.27506 | 0.28295 | 4.96458 | 0.00000 | 0.00000 | 3.19628 |
| **Naa40** | 7.00805 | 5.33103 | 6.56175 | 4.43299 | 7.96336 | 0.08260 | 8.07313 | 2.04550 | 2.69909 | 4.05097 | 0.97347 | 5.95249 | 7.30533 | 0.08883 | 3.66547 | 5.89627 |
| **Naa50** | 13.86543 | 12.98175 | 13.44687 | 17.82402 | 6.29641 | 7.24395 | 10.17330 | 10.26067 | 17.97133 | 17.36636 | 15.68557 | 15.55539 | 11.47720 | 27.58216 | 8.67308 | 3.25588 |
| **Naa60** | 3.18483 | 4.28756 | 1.08119 | 5.18503 | 5.85559 | 7.78751 | 6.56202 | 6.86440 | 1.34315 | 3.81695 | 2.49468 | 1.46749 | 4.08896 | 0.00000 | 0.86622 | 0.00001 |
| **Naaa** | 2.09725 | 1.47921 | 0.56225 | 0.13965 | 0.05918 | 6.42886 | 3.45683 | 3.41334 | 0.91584 | 3.55913 | 4.79216 | 0.06449 | 3.38249 | 2.36811 | 1.12119 | 3.88054 |
| **Naalad2** | 1.66603 | 3.09787 | 7.49353 | 7.37359 | 0.03170 | 2.61916 | 0.04365 | 22.88409 | 5.45302 | 15.06355 | 19.59249 | 19.49099 | 3.82786 | 27.92294 | 19.42159 | 0.49288 |
| **Naaladl1** | 0.01521 | 0.00870 | 0.00799 | 0.00938 | 0.01044 | 0.00704 | 0.00796 | 0.25163 | 0.01182 | 0.08961 | 0.00949 | 0.00888 | 0.42755 | 0.00684 | 0.14994 | 0.00719 |
| **Nab1** | 1.47922 | 3.56249 | 0.47032 | 7.88747 | 0.08182 | 0.00337 | 1.97508 | 0.00000 | 0.66869 | 0.42667 | 0.08514 | 0.00799 | 0.15212 | 0.01521 | 2.08559 | 0.00259 |
| **Nab2** | 0.37778 | 0.00740 | 0.00000 | 1.21778 | 0.01523 | 0.03334 | 0.00709 | 0.02035 | 0.06986 | 0.38689 | 0.02837 | 0.00586 | 0.01785 | 0.00422 | 0.09762 | 0.00706 |
| **Nabp1** | 0.33528 | 0.37780 | 2.95967 | 3.29165 | 0.26138 | 3.18894 | 1.68004 | 7.21570 | 17.70515 | 5.78560 | 3.69170 | 12.12572 | 5.38782 | 19.83994 | 17.17194 | 0.00588 |
| **Nabp2** | 2.84635 | 0.11738 | 0.00000 | 0.93676 | 4.98120 | 3.73853 | 0.15759 | 3.36764 | 1.11880 | 4.16758 | 1.22227 | 0.08648 | 0.73393 | 0.01654 | 0.05405 | 1.64568 |
| **Naca** | 46.17900 | 27.48332 | 40.48891 | 17.79562 | 30.35640 | 20.09737 | 22.17904 | 37.86927 | 27.03181 | 43.36190 | 40.82920 | 19.54800 | 45.12242 | 23.58332 | 21.14898 | 25.93257 |
| **Nacad** | 0.00963 | 0.00747 | 0.00690 | 0.00610 | 0.00977 | 0.00702 | 0.00702 | 0.25007 | 0.01127 | 0.19721 | 0.26512 | 0.04119 | 0.00552 | 0.01158 | 0.00119 | 0.00742 |
| **Nacc1** | 2.69695 | 2.97079 | 2.26175 | 5.14938 | 0.54686 | 2.04794 | 4.85298 | 2.75317 | 6.62017 | 3.79351 | 2.05833 | 0.09554 | 5.12029 | 0.03660 | 0.05873 | 1.57247 |
| **Nacc2** | 3.34228 | 0.01172 | 5.07706 | 7.77401 | 0.01660 | 0.08943 | 2.43633 | 7.33432 | 4.93571 | 11.07777 | 6.04622 | 1.15078 | 3.69813 | 5.02577 | 5.28440 | 0.84548 |
| **Nadk** | 10.26828 | 10.41173 | 12.78741 | 8.32983 | 8.16186 | 7.70842 | 6.78268 | 9.74095 | 4.66783 | 15.20130 | 11.11568 | 14.31327 | 11.08022 | 10.21837 | 10.35112 | 6.31583 |
| **Nadk2** | 2.33995 | 2.33329 | 0.94747 | 4.26651 | 0.22164 | 0.87277 | 0.00414 | 0.17581 | 1.01665 | 1.84409 | 0.59237 | 0.02097 | 0.58028 | 4.38429 | 2.31977 | 0.00936 |
| **Nadsyn1** | 0.84693 | 0.03746 | 0.08322 | 0.01050 | 0.00626 | 0.01075 | 0.00634 | 0.00866 | 0.01193 | 0.09236 | 0.01215 | 0.04618 | 0.99972 | 0.01557 | 0.05347 | 0.00636 |
| **Nae1** | 4.53707 | 7.36767 | 2.61292 | 3.09226 | 1.91646 | 0.01631 | 5.09053 | 5.01868 | 3.17317 | 5.55636 | 5.87862 | 2.54301 | 3.63468 | 3.58783 | 9.23847 | 4.76169 |
| **Naf1** | 0.93423 | 0.61843 | 0.13580 | 3.74789 | 2.46668 | 0.00266 | 0.22778 | 0.99507 | 5.28436 | 2.81589 | 5.96516 | 9.09465 | 0.22279 | 2.44098 | 5.10080 | 0.37916 |
| **Naga** | 1.94950 | 0.24479 | 0.65037 | 0.00000 | 0.01974 | 4.18948 | 1.87129 | 2.77099 | 0.09394 | 0.33841 | 0.00000 | 0.00681 | 2.18244 | 0.01247 | 0.00000 | 0.03061 |
| **Nagk** | 5.97157 | 5.87301 | 7.56742 | 0.00000 | 5.06888 | 2.76784 | 0.00123 | 4.04093 | 0.58746 | 2.91397 | 2.89569 | 0.38698 | 9.63928 | 0.03359 | 0.07090 | 7.83464 |
| **Naglu** | 0.06127 | 0.00792 | 0.01873 | 0.00710 | 0.00869 | 0.00787 | 0.00735 | 0.01229 | 0.01180 | 0.43003 | 0.10424 | 0.00584 | 0.41762 | 0.01132 | 0.00659 | 0.00754 |
| **Nagpa** | 2.53447 | 1.04429 | 2.44693 | 2.44647 | 0.46859 | 6.06026 | 2.15187 | 6.29459 | 0.16268 | 1.80094 | 3.89536 | 0.29074 | 11.19317 | 0.00000 | 1.35642 | 3.27257 |
| **Nags** | 0.34886 | 0.00749 | 0.00255 | 0.00295 | 0.00700 | 0.00700 | 0.00625 | 0.00758 | 1.55836 | 0.34047 | 0.04531 | 0.07047 | 1.17697 | 0.00633 | 0.48183 | 0.00666 |
| **Naif1** | 0.01042 | 0.03068 | 0.22427 | 0.00694 | 0.00935 | 0.00954 | 0.00803 | 0.03735 | 0.06690 | 0.06081 | 0.01112 | 0.00758 | 0.12402 | 0.01590 | 0.08432 | 0.00733 |
| **Naip2** | 0.05037 | 0.01842 | 0.03127 | 0.18857 | 0.00918 | 0.03837 | 0.01770 | 0.01410 | 0.30088 | 0.04575 | 0.16708 | 0.01554 | 0.06215 | 0.28161 | 0.05023 | 0.00725 |
| **Naip5** | 0.18820 | 0.02771 | 0.28097 | 1.83422 | 0.10471 | 0.06062 | 0.03687 | 0.06926 | 2.90998 | 0.53989 | 1.40510 | 0.13661 | 0.09977 | 1.00811 | 0.25822 | 0.05154 |
| **Nampt** | 5.38802 | 2.45517 | 1.64162 | 8.45834 | 2.50601 | 3.52563 | 1.02896 | 2.67378 | 15.57221 | 3.91830 | 4.44683 | 4.07706 | 2.66054 | 4.84686 | 10.36780 | 3.52219 |

| | | | | | | | | | | | | | | | | |
|---|---|---|---|---|---|---|---|---|---|---|---|---|---|---|---|---|
| **Nanog** | 19.07460 | 9.93228 | 21.02989 | 12.57850 | 1.03580 | 5.54358 | 14.54913 | 7.93302 | 19.62326 | 3.05676 | 18.29695 | 1.55574 | 8.81598 | 11.19488 | 2.44013 | 4.15383 |
| **Nanos1** | 0.01156 | 0.00866 | 1.13645 | 7.54286 | 0.00817 | 0.01199 | 0.00726 | 0.39233 | 6.84290 | 0.28631 | 0.19054 | 0.48788 | 1.23995 | 0.00000 | 4.08290 | 0.00461 |
| **Nanos2** | 0.00944 | 0.00841 | 0.00796 | 3.25156 | 0.00872 | 0.00624 | 0.00802 | 0.09058 | 0.47313 | 2.71382 | 0.15813 | 0.00000 | 0.01089 | 0.00000 | 0.98599 | 0.00646 |
| **Nanos3** | 0.01285 | 0.00772 | 0.01086 | 1.49349 | 0.00909 | 0.00624 | 0.00732 | 0.01166 | 0.61334 | 1.06080 | 0.00600 | 0.02181 | 0.01136 | 0.00213 | 0.86940 | 0.00713 |
| **Nanp** | 1.53565 | 0.00186 | 0.00629 | 2.06799 | 7.48434 | 0.00675 | 0.00520 | 0.03050 | 0.00000 | 0.00126 | 0.01146 | 0.00835 | 1.44236 | 0.00508 | 0.72260 | 0.01154 |
| **Nans** | 2.33062 | 0.06127 | 0.33371 | 3.49167 | 2.89660 | 0.01022 | 1.30939 | 0.00000 | 5.29198 | 4.35182 | 0.32090 | 0.17643 | 0.37077 | 0.00359 | 0.00542 | 5.68758 |
| **Nap1l1** | 30.63744 | 20.01769 | 26.07023 | 20.77023 | 23.87153 | 14.92538 | 14.25026 | 25.73667 | 28.53359 | 42.62229 | 34.31363 | 26.34394 | 23.87926 | 36.50610 | 29.48656 | 13.28037 |
| **Nap1l2** | 0.01255 | 0.00818 | 0.01394 | 0.13056 | 0.01017 | 0.00715 | 0.00770 | 0.01593 | 0.00687 | 0.01197 | 0.01400 | 0.01793 | 0.21417 | 0.01328 | 0.01122 | 0.00768 |
| **Nap1l3** | 0.00644 | 0.01654 | 0.00452 | 0.04047 | 2.15164 | 0.00736 | 0.00741 | 0.00915 | 0.00753 | 0.00309 | 0.01094 | 0.00551 | 0.00975 | 0.01611 | 0.20999 | 0.00713 |
| **Nap1l4** | 4.90288 | 2.30724 | 7.57934 | 16.57751 | 7.10997 | 5.94536 | 4.91040 | 10.68558 | 15.45973 | 12.71862 | 11.26329 | 0.39559 | 8.92483 | 1.15901 | 7.76049 | 5.93576 |
| **Napa** | 12.38263 | 4.72819 | 14.67412 | 2.95668 | 12.90872 | 10.68203 | 9.81664 | 16.77656 | 4.06485 | 10.93647 | 5.44307 | 0.59536 | 10.97274 | 0.01086 | 3.37878 | 6.56661 |
| **Napb** | 0.01745 | 0.00794 | 0.01051 | 0.11085 | 0.00859 | 0.00742 | 0.14657 | 0.01047 | 0.00633 | 0.37184 | 0.00107 | 0.01296 | 0.31524 | 1.30046 | 0.53003 | 0.03570 |
| **Napepld** | 0.99818 | 0.00413 | 1.80419 | 1.60537 | 3.99985 | 2.37719 | 0.00442 | 0.92247 | 1.55337 | 4.21373 | 2.65546 | 0.06894 | 1.12293 | 0.00000 | 0.99677 | 1.30137 |
| **Napg** | 6.00876 | 1.99263 | 6.82766 | 3.70773 | 4.40420 | 3.97923 | 2.37646 | 3.77723 | 3.77158 | 2.77745 | 4.82891 | 11.94417 | 2.13709 | 2.15840 | 8.96606 | 4.83990 |
| **Naprt** | 2.96660 | 0.00404 | 0.00000 | 0.00274 | 0.01916 | 0.98446 | 0.00471 | 0.56255 | 0.65356 | 0.02605 | 0.00000 | 0.00328 | 0.04250 | 0.04196 | 0.00000 | 0.00372 |
| **Napsa** | 0.00000 | 0.00683 | 1.54529 | 0.00468 | 0.01749 | 0.00571 | 2.53633 | 0.00656 | 0.01124 | 0.52357 | 0.10111 | 0.05491 | 0.99702 | 0.00759 | 0.07517 | 0.01353 |
| **Narf** | 0.10890 | 0.00750 | 0.67665 | 1.75457 | 0.01635 | 0.00552 | 1.43381 | 2.00008 | 1.94537 | 0.11279 | 0.10528 | 0.00000 | 0.04374 | 0.83742 | 0.28970 | 1.97047 |
| **Narfl** | 4.21880 | 7.60238 | 8.66920 | 4.22722 | 6.42181 | 7.41431 | 3.79746 | 9.99298 | 8.32803 | 13.83780 | 2.61222 | 3.55165 | 11.47476 | 0.00000 | 2.17593 | 5.04986 |
| **Nars** | 22.89023 | 19.33025 | 18.80494 | 3.49633 | 15.79206 | 8.31535 | 15.53380 | 22.67425 | 5.29002 | 14.29474 | 8.05903 | 8.37350 | 16.73887 | 16.91110 | 13.66399 | 11.71709 |
| **Nars2** | 0.27580 | 0.00845 | 0.00877 | 0.00000 | 0.03512 | 0.00806 | 0.00664 | 0.50356 | 0.20240 | 0.08721 | 0.17586 | 0.27228 | 0.86643 | 1.06908 | 2.30078 | 0.01442 |
| **Nasp** | 26.70205 | 22.37761 | 20.33619 | 24.13854 | 12.84075 | 14.19040 | 18.71256 | 29.30812 | 40.61431 | 40.12860 | 31.59164 | 23.25909 | 20.93769 | 35.08779 | 32.22305 | 13.67710 |
| **Nat1** | 0.01254 | 0.00826 | 0.57368 | 0.22655 | 0.00835 | 0.00565 | 0.00763 | 0.01376 | 0.00542 | 0.01466 | 0.01131 | 0.00847 | 0.00891 | 0.01232 | 0.00000 | 0.00711 |
| **Nat10** | 10.12356 | 6.55163 | 8.57804 | 4.23614 | 0.77593 | 3.87619 | 8.33612 | 15.26134 | 7.01835 | 10.84601 | 6.32907 | 4.64081 | 5.84268 | 0.78984 | 3.81819 | 2.65599 |
| **Nat14** | 0.00861 | 0.00833 | 0.03819 | 0.56949 | 0.02738 | 0.00854 | 0.00795 | 0.22852 | 0.32385 | 0.07946 | 0.42732 | 0.04903 | 0.00710 | 0.00897 | 0.00000 | 0.00742 |
| **Nat6** | 2.79891 | 0.00387 | 2.25388 | 0.20820 | 0.00000 | 3.12254 | 0.20635 | 1.91599 | 0.19483 | 1.64613 | 0.00000 | 0.00488 | 1.52571 | 0.00925 | 0.00000 | 0.01644 |
| **Nat8** | 0.01351 | 0.00782 | 0.01563 | 0.04181 | 0.00931 | 0.13583 | 0.00737 | 0.01234 | 0.06161 | 0.05071 | 0.01261 | 0.00734 | 0.01077 | 0.04530 | 0.00489 | 0.00739 |
| **Nat8f1** | 0.01578 | 0.00753 | 0.21287 | 0.13533 | 0.00999 | 3.13862 | 0.00687 | 0.00833 | 1.84419 | 0.54018 | 0.00000 | 0.00000 | 0.00312 | 0.27361 | 0.08209 | 0.00615 |
| **Nat8f2** | 0.26603 | 0.00738 | 0.03589 | 6.57403 | 0.00714 | 0.00462 | 0.00708 | 1.46852 | 4.34930 | 0.53812 | 0.10387 | 0.06369 | 0.00672 | 0.55076 | 0.98959 | 0.01276 |
| **Nat8f3** | 0.01378 | 0.00805 | 0.01203 | 0.01616 | 0.00933 | 0.00655 | 0.00744 | 0.01250 | 0.01211 | 0.01287 | 0.01338 | 0.00811 | 0.01253 | 0.00890 | 0.24049 | 0.00732 |
| **Nat8f4** | 0.02069 | 0.00788 | 0.93300 | 0.21582 | 0.00707 | 0.00546 | 0.00724 | 0.07308 | 0.48111 | 0.01996 | 0.00757 | 0.03806 | 0.31514 | 0.00932 | 0.05421 | 0.00674 |
| **Nat8f7** | 0.01378 | 0.00796 | 0.01207 | 0.01062 | 0.00933 | 0.00664 | 0.00751 | 0.01342 | 0.01204 | 0.01251 | 0.01277 | 0.00806 | 0.01272 | 0.01355 | 0.06110 | 0.00740 |
| **Nat8l** | 0.01256 | 0.00786 | 0.10072 | 0.74632 | 0.00904 | 0.00877 | 0.00748 | 0.01559 | 0.00000 | 0.01199 | 0.01263 | 0.05767 | 0.01108 | 0.01271 | 0.03791 | 0.00724 |
| **Nat9** | 6.74669 | 0.04168 | 6.02687 | 0.00268 | 2.08542 | 0.02074 | 0.00833 | 2.50753 | 0.00781 | 0.18297 | 0.21254 | 0.00013 | 6.65894 | 0.02633 | 0.00783 | 3.15639 |
| **Natd1** | 4.78558 | 2.89647 | 9.68298 | 9.94956 | 0.00887 | 5.58239 | 6.03122 | 5.62754 | 13.05420 | 9.18529 | 5.80140 | 0.65631 | 2.26995 | 5.19104 | 7.62822 | 4.38105 |
| **Nav1** | 1.33595 | 2.89042 | 0.88687 | 1.23330 | 0.29645 | 1.68819 | 0.12605 | 0.70069 | 8.22394 | 2.22654 | 2.19165 | 0.35429 | 3.34994 | 16.00279 | 0.00000 | 0.12100 |
| **Nav2** | 0.00000 | 0.00795 | 0.03618 | 2.94989 | 0.00859 | 2.95714 | 0.02602 | 1.19642 | 1.24549 | 1.07374 | 0.12872 | 0.03919 | 3.69696 | 0.00830 | 0.38774 | 0.00394 |
| **Nav3** | 0.01129 | 0.00747 | 0.47701 | 0.12401 | 0.00859 | 0.00572 | 0.00727 | 0.15444 | 0.38463 | 1.06402 | 0.00494 | 0.00000 | 0.00685 | 0.01040 | 2.49599 | 0.00698 |
| **Nbas** | 0.00000 | 0.00519 | 1.71010 | 0.78680 | 3.74029 | 0.00814 | 0.00671 | 2.03467 | 0.05266 | 0.67036 | 0.89717 | 0.29230 | 0.00000 | 0.03849 | 0.00000 | 2.12453 |
| **Nbea** | 0.01137 | 0.00801 | 0.00535 | 0.00893 | 0.00946 | 0.00685 | 0.00730 | 0.00988 | 0.01152 | 0.33966 | 0.00461 | 0.00000 | 0.01305 | 2.72772 | 0.00435 | 0.00737 |
| **Nbeal1** | 1.02293 | 0.00626 | 0.00000 | 0.89390 | 0.00586 | 0.01764 | 0.00609 | 0.00000 | 0.86857 | 0.00000 | 0.14046 | 0.66944 | 0.01031 | 2.18776 | 0.95560 | 0.00573 |
| **Nbeal2** | 1.07337 | 0.00586 | 0.82415 | 0.49971 | 0.18903 | 0.21429 | 0.16404 | 2.86429 | 0.00375 | 4.17804 | 0.56469 | 0.00000 | 0.75478 | 0.03520 | 0.00000 | 1.21621 |
| **Nbl1** | 0.00599 | 0.06885 | 0.08671 | 0.35042 | 0.01703 | 0.00871 | 0.00644 | 0.00613 | 0.00613 | 1.99797 | 0.06068 | 0.02071 | 1.21209 | 0.01067 | 0.00000 | 0.00686 |
| **Nbn** | 3.19131 | 0.00424 | 1.83534 | 1.96971 | 0.00970 | 0.00504 | 0.00232 | 0.00000 | 4.17770 | 7.15380 | 5.19823 | 0.48396 | 1.24422 | 5.29084 | 5.16718 | 1.74388 |
| **Nbr1** | 6.07114 | 0.11592 | 7.02688 | 13.95287 | 0.51000 | 7.00838 | 5.62853 | 6.71426 | 11.76330 | 7.33138 | 4.86698 | 7.24868 | 7.81973 | 5.02596 | 12.68307 | 2.60448 |
| **Ncald** | 0.01333 | 0.00781 | 0.01317 | 0.21747 | 0.01002 | 0.00688 | 0.00762 | 0.01370 | 0.39779 | 0.01163 | 0.00958 | 0.00573 | 0.05394 | 0.00741 | 0.50334 | 0.00768 |
| **Ncam1** | 0.78439 | 0.00704 | 0.00815 | 0.00636 | 0.00636 | 0.00632 | 0.00704 | 0.01340 | 0.03614 | 1.01213 | 0.00996 | 0.10134 | 0.02833 | 0.06885 | 0.02056 | 0.00676 |
| **Ncan** | 0.02380 | 0.00818 | 0.04979 | 0.02933 | 0.02640 | 0.01167 | 0.00758 | 0.01121 | 0.10198 | 0.01881 | 0.19548 | 0.02883 | 0.05016 | 0.01013 | 0.26249 | 0.02164 |
| **Ncapd2** | 17.00474 | 11.00631 | 11.81240 | 6.23634 | 9.92021 | 5.48334 | 9.03055 | 11.06925 | 8.88831 | 10.19378 | 2.04101 | 4.83950 | 8.88478 | 5.19587 | 12.64610 | 6.66409 |
| **Ncapd3** | 0.84992 | 1.92821 | 1.46788 | 2.49351 | 0.10129 | 0.00455 | 2.56063 | 2.29762 | 3.11563 | 0.29053 | 2.50006 | 7.70128 | 0.48061 | 0.76030 | 3.15718 | 0.64410 |
| **Ncapg** | 2.62685 | 5.88216 | 4.33343 | 1.25417 | 0.28494 | 0.54572 | 3.00128 | 2.71375 | 0.94571 | 1.05272 | 0.33865 | 0.36289 | 1.93693 | 0.00000 | 5.55048 | 1.43747 |
| **Ncapg2** | 1.07964 | 1.95962 | 2.97791 | 10.84491 | 0.04067 | 3.55660 | 3.39183 | 3.65974 | 9.47939 | 0.74729 | 3.26783 | 10.87686 | 0.78607 | 2.55469 | 10.84074 | 0.39928 |
| **Ncaph** | 9.20179 | 1.66013 | 2.64460 | 1.24619 | 0.73526 | 0.00649 | 3.54606 | 9.28169 | 2.05965 | 2.57153 | 2.19993 | 0.20108 | 4.87259 | 0.13057 | 5.13325 | 1.33587 |
| **Ncaph2** | 14.23428 | 2.33634 | 14.57177 | 4.01763 | 5.80775 | 10.16263 | 8.70396 | 18.54285 | 4.40261 | 12.33078 | 9.54014 | 0.83415 | 20.88467 | 0.49621 | 7.85415 | 11.39508 |
| **Ncbp1** | 5.05763 | 7.94682 | 7.86304 | 10.80654 | 0.38584 | 7.48070 | 11.99055 | 15.64429 | 15.77524 | 22.25445 | 10.38809 | 5.08508 | 13.15713 | 21.51219 | 23.20190 | 4.64808 |
| **Ncbp2** | 13.09157 | 13.04774 | 13.95160 | 15.79177 | 1.97049 | 5.90236 | 7.59064 | 12.04263 | 19.80874 | 24.35233 | 16.41775 | 11.78984 | 12.20661 | 15.27202 | 20.90560 | 3.62062 |
| **Ncbp3** | 4.60274 | 4.90681 | 4.58752 | 10.19082 | 5.69410 | 4.77483 | 4.72117 | 3.48420 | 14.07600 | 7.33898 | 9.37094 | 4.56311 | 1.54071 | 13.95236 | 12.28429 | 1.94426 |
| **Ncdn** | 5.84358 | 0.04361 | 0.46488 | 1.95501 | 0.68556 | 0.15401 | 0.00977 | 0.80857 | 1.70590 | 6.14760 | 0.80394 | 0.06363 | 4.81440 | 0.04523 | 0.17237 | 0.11429 |
| **Nceh1** | 0.00651 | 0.00823 | 1.78859 | 5.48861 | 0.00687 | 0.00717 | 0.00695 | 0.01871 | 3.03844 | 0.91262 | 0.00000 | 0.03117 | 0.40235 | 0.32442 | 3.15528 | 0.00546 |
| **Ncf1** | 0.03282 | 0.00777 | 0.02783 | 0.08820 | 0.00805 | 0.03364 | 0.00706 | 0.00000 | 0.20473 | 0.22517 | 0.84810 | 0.02350 | 0.02603 | 0.07112 | 1.24281 | 0.00705 |
| **Ncf2** | 0.66730 | 0.01951 | 0.24591 | 0.05106 | 0.01502 | 0.00507 | 0.00657 | 0.00263 | 0.01887 | 1.80487 | 0.50378 | 0.02121 | 0.01051 | 0.02905 | 0.01643 | 1.77081 |
| **Nck1** | 2.48416 | 5.37647 | 0.16305 | 0.16677 | 0.02145 | 0.01378 | 1.85670 | 1.34084 | 2.43876 | 5.55562 | 1.13588 | 10.00170 | 2.97576 | 0.79081 | 1.28480 | 0.01517 |
| **Nck2** | 0.01420 | 0.04949 | 0.00908 | 0.11465 | 0.01057 | 0.00665 | 0.00715 | 0.08616 | 0.00744 | 0.00144 | 0.16386 | 0.11896 | 0.00775 | 0.32654 | 0.00129 | 0.00735 |

| | | | | | | | | | | | | | | | | |
|---|---|---|---|---|---|---|---|---|---|---|---|---|---|---|---|---|
| **Nckap1** | 7.27133 | 6.13966 | 8.19587 | 19.64701 | 5.42906 | 8.44978 | 3.69904 | 7.02269 | 9.34955 | 12.96509 | 11.14288 | 18.86992 | 7.83032 | 20.27523 | 17.13835 | 0.82550 |
| **Nckap1l** | 0.00734 | 0.00788 | 0.01180 | 0.36680 | 0.00760 | 0.00636 | 0.00740 | 0.01207 | 0.36617 | 0.93363 | 0.00702 | 0.00443 | 0.00777 | 0.00472 | 0.31763 | 1.40087 |
| **Nckap5** | 0.01368 | 0.00789 | 0.01016 | 0.50222 | 0.00900 | 0.00686 | 0.00742 | 0.00517 | 0.00452 | 0.40667 | 0.25446 | 0.00648 | 0.01006 | 0.00476 | 0.17568 | 0.00724 |
| **Nckap5l** | 0.80406 | 0.00659 | 0.00342 | 0.09820 | 0.00628 | 0.00737 | 0.00656 | 0.15853 | 0.58729 | 0.01520 | 0.00745 | 0.00580 | 0.04936 | 0.01261 | 0.00304 | 0.00644 |
| **Nckipsd** | 3.32142 | 0.00346 | 1.16443 | 0.41282 | 0.00504 | 0.01531 | 1.49988 | 0.27703 | 0.00405 | 0.00000 | 0.42831 | 0.03768 | 3.47446 | 0.02258 | 0.01951 | 2.00275 |
| **Ncl** | 34.87694 | 28.35505 | 32.97174 | 17.87219 | 25.84932 | 14.50073 | 20.40271 | 39.87150 | 26.27602 | 30.07491 | 34.63108 | 23.19390 | 31.79202 | 31.53150 | 34.05280 | 19.59727 |
| **Ncln** | 5.62937 | 0.03669 | 0.06566 | 0.01304 | 5.09725 | 0.61468 | 3.04376 | 8.45408 | 0.53743 | 3.99516 | 0.58323 | 0.32723 | 10.52280 | 0.01567 | 0.04307 | 5.74573 |
| **Ncmap** | 0.67081 | 0.01791 | 0.00202 | 4.46819 | 0.01455 | 0.00662 | 0.08289 | 0.01473 | 5.69537 | 0.95739 | 0.00000 | 0.03680 | 0.01708 | 0.05848 | 0.41590 | 0.00531 |
| **Ncoa1** | 2.19079 | 0.00466 | 1.38605 | 6.18233 | 0.78400 | 4.79839 | 0.02583 | 0.79693 | 2.67293 | 1.48638 | 0.74532 | 0.03425 | 0.26137 | 0.00000 | 1.17282 | 2.04075 |
| **Ncoa2** | 0.00000 | 0.00354 | 0.40858 | 13.21808 | 4.07180 | 4.17925 | 3.23090 | 2.38230 | 2.83128 | 0.60787 | 0.51138 | 0.12716 | 2.38651 | 0.00000 | 1.34947 | 0.23012 |
| **Ncoa3** | 11.31546 | 10.10467 | 9.62117 | 16.41097 | 9.96565 | 8.06673 | 7.47426 | 21.25571 | 17.92156 | 22.30630 | 18.82627 | 15.78083 | 8.26544 | 13.17076 | 18.49726 | 5.86064 |
| **Ncoa4** | 6.88130 | 8.93477 | 12.45160 | 7.68607 | 10.06316 | 7.02376 | 4.13444 | 9.71912 | 4.06411 | 9.10697 | 10.48443 | 16.65068 | 9.26705 | 14.03442 | 6.18631 | 4.86677 |
| **Ncoa5** | 0.81260 | 2.85747 | 2.99790 | 1.53950 | 4.81282 | 0.01240 | 0.00426 | 0.86279 | 0.33042 | 2.15962 | 0.69003 | 0.04163 | 2.94602 | 0.32469 | 1.69032 | 0.03960 |
| **Ncoa6** | 2.28663 | 5.07579 | 3.16806 | 19.27545 | 5.06919 | 3.67727 | 4.91156 | 1.61286 | 7.70696 | 2.78318 | 4.41958 | 1.77582 | 2.43671 | 11.41124 | 16.61969 | 2.12407 |
| **Ncoa7** | 0.38145 | 0.00696 | 0.00000 | 0.00903 | 0.00714 | 0.40553 | 0.00717 | 0.00977 | 0.01125 | 0.01206 | 0.01032 | 0.00688 | 0.00843 | 0.01805 | 0.39450 | 0.00677 |
| **Ncor1** | 9.28016 | 2.52140 | 5.42376 | 4.95298 | 4.50326 | 5.87817 | 0.07964 | 4.18712 | 4.64438 | 5.67184 | 2.36036 | 0.53142 | 2.96874 | 7.36723 | 10.25365 | 7.01787 |
| **Ncor2** | 2.08687 | 0.00391 | 0.60194 | 0.07004 | 1.62758 | 0.01388 | 1.25241 | 0.64577 | 0.15772 | 0.22119 | 0.16043 | 0.11492 | 0.37303 | 0.01136 | 0.00816 | 2.67194 |
| **Ncs1** | 0.00854 | 0.00771 | 0.37376 | 0.00525 | 0.00747 | 0.00567 | 0.00735 | 0.00984 | 0.22487 | 1.07549 | 0.00878 | 0.06404 | 0.02124 | 0.01408 | 0.01988 | 0.66006 |
| **Ncstn** | 0.19647 | 1.63544 | 1.71107 | 0.13861 | 0.00453 | 0.00749 | 0.56551 | 4.23051 | 0.29347 | 1.16856 | 0.93131 | 0.00000 | 0.84468 | 6.96272 | 3.58591 | 2.41830 |
| **Ndc1** | 2.31659 | 3.02533 | 9.88620 | 0.85816 | 7.06314 | 1.54748 | 6.61683 | 7.38857 | 0.76604 | 6.27270 | 8.55141 | 3.46385 | 4.68598 | 5.61840 | 8.83641 | 0.34046 |
| **Ndc80** | 3.82525 | 5.50418 | 5.27388 | 2.68701 | 6.13138 | 6.37911 | 6.58307 | 10.93607 | 13.64598 | 15.62894 | 8.61225 | 3.58235 | 3.60463 | 17.78134 | 11.06352 | 2.65881 |
| **Nde1** | 2.98447 | 0.01119 | 5.69767 | 5.85883 | 3.67863 | 2.09993 | 2.57820 | 0.00000 | 3.00487 | 1.98269 | 1.89202 | 0.09412 | 0.00000 | 0.00000 | 1.94596 | 1.95087 |
| **Ndel1** | 6.32745 | 0.52812 | 2.31070 | 8.37928 | 7.04289 | 1.90808 | 1.53903 | 0.78755 | 7.54620 | 11.38955 | 5.20787 | 4.95201 | 5.26935 | 0.00555 | 1.96590 | 5.00530 |
| **Ndfip1** | 2.44302 | 3.53480 | 1.08305 | 1.47261 | 7.36078 | 3.39122 | 0.32358 | 0.75677 | 0.79555 | 0.45811 | 2.78523 | 1.09263 | 1.63548 | 1.09082 | 0.65593 | 0.87475 |
| **Ndfip2** | 0.92260 | 5.03114 | 0.13528 | 6.30018 | 0.54903 | 0.43703 | 3.71513 | 0.92896 | 1.12536 | 0.54290 | 3.04045 | 0.18431 | 1.18838 | 2.73912 | 0.85134 | 0.00229 |
| **Ndnl2** | 2.68435 | 5.05961 | 2.34319 | 3.46652 | 6.45214 | 0.03064 | 0.05529 | 4.12853 | 4.84463 | 5.98022 | 2.58855 | 0.99706 | 0.89270 | 2.91981 | 2.95659 | 0.06446 |
| **Ndor1** | 3.59171 | 0.01084 | 5.14396 | 0.73401 | 0.00558 | 0.78581 | 4.03036 | 9.50488 | 5.73210 | 10.26988 | 4.23143 | 1.25163 | 4.95430 | 0.26570 | 4.88082 | 3.24102 |
| **Ndrg1** | 0.03035 | 0.00396 | 3.30466 | 0.00000 | 0.06041 | 8.03575 | 4.72807 | 0.00000 | 0.05131 | 0.00000 | 0.27786 | 1.85328 | 0.02623 | 0.00693 | 0.00000 | 4.11514 |
| **Ndrg2** | 0.00421 | 0.00711 | 5.32637 | 0.00668 | 0.00000 | 0.00715 | 0.00666 | 0.75275 | 0.01065 | 1.47912 | 0.00271 | 0.00000 | 0.00000 | 0.01313 | 0.00000 | 0.00568 |
| **Ndrg3** | 6.81001 | 5.73983 | 11.38116 | 1.26675 | 10.55161 | 8.21003 | 6.84623 | 12.64346 | 0.66452 | 5.75261 | 8.72669 | 4.74109 | 5.89353 | 10.20321 | 2.01331 | 8.39260 |
| **Ndrg4** | 0.01367 | 0.00830 | 0.01094 | 1.19099 | 0.00920 | 0.00651 | 0.00748 | 0.01094 | 0.88625 | 0.00574 | 0.11756 | 0.00637 | 0.01077 | 0.00667 | 0.04320 | 0.00705 |
| **Ndst1** | 2.45007 | 0.00508 | 1.21764 | 1.78943 | 0.00000 | 0.00274 | 0.00556 | 1.29052 | 1.15362 | 2.99803 | 0.26430 | 0.42027 | 0.17083 | 1.72710 | 2.66315 | 0.02525 |
| **Ndst2** | 0.01019 | 0.00782 | 0.76482 | 0.18907 | 0.00753 | 0.00945 | 0.00712 | 0.00000 | 0.00777 | 0.00000 | 0.53627 | 0.00334 | 0.00931 | 0.00524 | 0.78906 | 0.85683 |
| **Ndufa1** | 1.39852 | 0.07719 | 1.65987 | 2.26639 | 4.92623 | 0.41655 | 0.24731 | 0.04954 | 1.64500 | 0.58064 | 1.99729 | 0.11061 | 0.35401 | 0.28710 | 0.81043 | 1.42852 |
| **Ndufa10** | 14.16378 | 2.71480 | 12.90717 | 0.67852 | 3.47315 | 7.70224 | 6.07528 | 4.27264 | 1.00083 | 9.07051 | 3.84712 | 2.69516 | 12.15836 | 0.23585 | 2.69469 | 8.44406 |
| **Ndufa11** | 7.38403 | 2.07917 | 3.33332 | 6.96884 | 9.55022 | 3.09544 | 2.61720 | 4.69431 | 1.84552 | 8.53582 | 7.86452 | 0.89604 | 9.58937 | 0.16090 | 2.93987 | 4.15321 |
| **Ndufa12** | 11.46405 | 0.32638 | 6.01756 | 2.21327 | 7.78186 | 3.00072 | 1.19568 | 1.95595 | 0.39532 | 8.38621 | 12.28177 | 0.41769 | 8.65798 | 0.23111 | 0.00000 | 6.09530 |
| **Ndufa13** | 9.84333 | 0.54253 | 5.40273 | 3.05181 | 7.87865 | 1.69441 | 0.65019 | 1.25216 | 2.69198 | 4.98633 | 7.23090 | 0.38226 | 2.99963 | 0.13089 | 0.23199 | 5.61917 |
| **Ndufa2** | 3.14538 | 0.08672 | 2.05239 | 12.03155 | 6.67917 | 1.74164 | 1.01696 | 0.09384 | 18.84652 | 2.65758 | 5.05848 | 0.31062 | 1.93776 | 0.00000 | 0.02187 | 3.34245 |
| **Ndufa3** | 3.32107 | 0.44478 | 2.47602 | 0.11543 | 1.14390 | 0.55834 | 0.12183 | 0.57387 | 0.80116 | 1.90141 | 11.01030 | 0.18422 | 1.81343 | 1.45563 | 0.38580 | 3.06969 |
| **Ndufa4** | 11.14697 | 1.81517 | 10.80181 | 2.08312 | 13.70962 | 2.58924 | 1.99045 | 1.29955 | 0.67598 | 2.41948 | 7.04352 | 0.80418 | 6.63299 | 3.97377 | 5.06243 | 3.83066 |
| **Ndufa5** | 5.54250 | 0.41586 | 5.70556 | 0.98526 | 10.16849 | 1.42648 | 0.45845 | 0.00000 | 0.84734 | 1.25496 | 12.55663 | 0.43813 | 1.87758 | 0.89793 | 0.73714 | 2.03301 |
| **Ndufa6** | 8.08954 | 1.08945 | 1.37767 | 4.66126 | 1.49240 | 4.34973 | 0.20872 | 0.95850 | 4.83198 | 4.79819 | 3.14239 | 0.01914 | 2.94971 | 0.05787 | 0.00000 | 3.45894 |
| **Ndufa7** | 8.36865 | 0.97699 | 5.80513 | 4.69865 | 3.93007 | 2.27897 | 1.05421 | 0.68459 | 9.43495 | 4.02828 | 11.37346 | 0.35244 | 3.28975 | 0.11215 | 1.37382 | 5.53979 |
| **Ndufa8** | 9.95568 | 8.03747 | 9.21457 | 0.13313 | 10.75712 | 3.14282 | 4.96917 | 7.16383 | 0.00000 | 7.99890 | 7.75709 | 0.41277 | 11.53597 | 0.21299 | 0.18652 | 7.26367 |
| **Ndufa9** | 17.16394 | 10.81027 | 10.27852 | 4.41855 | 8.78751 | 6.27193 | 11.38243 | 17.92029 | 7.37614 | 10.96036 | 16.20470 | 5.92525 | 17.51476 | 10.09811 | 4.02578 | 6.92160 |
| **Ndufab1** | 3.21044 | 1.19577 | 1.80241 | 9.59312 | 5.65315 | 0.49525 | 0.14585 | 0.00000 | 8.50796 | 4.38811 | 5.20937 | 0.18736 | 1.57524 | 0.40007 | 0.07118 | 2.64252 |
| **Ndufaf1** | 3.81440 | 1.69705 | 6.73072 | 3.77442 | 1.85436 | 0.00733 | 1.53193 | 2.79578 | 3.19774 | 7.86544 | 3.96099 | 3.18012 | 2.46433 | 4.62797 | 4.04519 | 1.37257 |
| **Ndufaf2** | 4.63516 | 2.76702 | 2.08541 | 6.22446 | 6.81533 | 1.92659 | 2.23180 | 1.73560 | 3.69874 | 4.62449 | 2.67647 | 0.47046 | 3.04079 | 0.53867 | 1.57104 | 3.34290 |
| **Ndufaf3** | 3.71161 | 0.85261 | 2.68738 | 0.43378 | 1.38569 | 0.08291 | 0.44232 | 0.00000 | 0.00010 | 0.33361 | 0.77321 | 0.00000 | 2.29503 | 0.00898 | 0.00000 | 0.37202 |
| **Ndufaf4** | 8.53988 | 6.00811 | 4.25270 | 7.47337 | 4.35596 | 6.98145 | 3.14707 | 8.23787 | 3.02324 | 4.18895 | 4.42359 | 1.32550 | 2.70803 | 5.26108 | 6.09570 | 2.91472 |
| **Ndufaf5** | 3.42314 | 0.60519 | 2.15349 | 3.09949 | 0.00761 | 0.00089 | 1.18037 | 0.10573 | 7.67320 | 5.42639 | 1.52236 | 4.81943 | 0.70986 | 2.55195 | 1.34479 | 0.91622 |
| **Ndufaf6** | 1.71116 | 0.09565 | 1.64633 | 0.01753 | 0.63583 | 0.01724 | 0.00535 | 0.00000 | 1.35003 | 2.33891 | 1.78377 | 0.02732 | 0.09926 | 0.09225 | 0.33684 | 0.00486 |
| **Ndufaf7** | 8.03610 | 3.85888 | 11.03686 | 4.45751 | 0.28343 | 8.62866 | 6.58358 | 6.14576 | 7.90652 | 11.28420 | 11.72859 | 2.04631 | 7.90330 | 15.30329 | 10.60201 | 4.33475 |
| **Ndufb10** | 18.21422 | 5.65696 | 10.12362 | 4.59307 | 6.47680 | 5.46655 | 6.07207 | 7.61801 | 11.28206 | 13.29679 | 14.42210 | 0.77265 | 10.66817 | 1.92068 | 0.36225 | 6.67854 |
| **Ndufb11** | 13.41221 | 2.94376 | 10.51965 | 11.54815 | 10.26574 | 4.72224 | 3.39394 | 10.40200 | 20.05247 | 18.65983 | 14.84769 | 2.38186 | 10.30714 | 1.46182 | 1.21194 | 8.27533 |
| **Ndufb2** | 2.92473 | 0.33122 | 0.93443 | 4.56173 | 2.88483 | 0.07434 | 0.01217 | 0.00000 | 7.01373 | 5.37070 | 5.19000 | 0.06389 | 2.26909 | 0.00000 | 0.00000 | 1.95962 |
| **Ndufb3** | 7.94933 | 0.91838 | 4.43260 | 3.03278 | 5.32498 | 1.13168 | 0.78231 | 1.49433 | 2.96341 | 1.23099 | 8.58892 | 0.61795 | 2.82132 | 1.91205 | 2.51895 | 4.43742 |
| **Ndufb4** | 7.37243 | 0.52030 | 2.71647 | 0.26000 | 3.32111 | 1.75189 | 0.98614 | 0.54160 | 0.05000 | 3.16757 | 4.34825 | 0.28025 | 3.50597 | 0.34379 | 0.10728 | 3.52470 |
| **Ndufb5** | 16.38840 | 1.75128 | 6.90790 | 0.89955 | 12.03869 | 9.12085 | 4.44039 | 8.91098 | 3.39595 | 11.74883 | 7.10668 | 0.84363 | 8.64408 | 0.53255 | 0.00000 | 10.12570 |
| **Ndufb6** | 9.95011 | 0.70839 | 8.08699 | 6.62465 | 9.67626 | 1.83885 | 2.07227 | 2.44926 | 6.61689 | 7.34017 | 11.36777 | 0.77142 | 6.87887 | 0.23152 | 0.00000 | 5.47477 |
| **Ndufb7** | 10.19391 | 1.32811 | 4.16983 | 2.38301 | 3.43617 | 0.99762 | 0.89366 | 0.48909 | 2.25933 | 4.56251 | 11.29730 | 0.26970 | 2.12706 | 0.00912 | 0.00011 | 0.57214 |

| | | | | | | | | | | | | | | | | |
|---|---|---|---|---|---|---|---|---|---|---|---|---|---|---|---|---|
| **Ndufb8** | 19.16856 | 4.73527 | 8.36971 | 6.45609 | 12.87662 | 7.06483 | 5.86131 | 3.38154 | 0.00000 | 5.67174 | 8.12730 | 0.40152 | 9.23466 | 0.00000 | 0.58113 | 13.06034 |
| **Ndufb9** | 19.65451 | 2.27992 | 7.76785 | 4.08572 | 13.51179 | 9.09139 | 5.69694 | 11.82043 | 3.28747 | 8.90259 | 11.97903 | 0.74156 | 15.72260 | 0.91480 | 0.41519 | 14.32274 |
| **Ndufc1** | 6.03861 | 0.18475 | 3.75525 | 2.90835 | 10.53309 | 1.01411 | 0.62172 | 0.03343 | 6.22450 | 3.11569 | 10.97846 | 0.24184 | 2.62039 | 0.38031 | 0.00000 | 2.40348 |
| **Ndufc2** | 11.78298 | 1.20086 | 4.86804 | 0.49414 | 5.39338 | 3.91850 | 2.42893 | 3.45502 | 0.22035 | 6.42895 | 5.90369 | 0.25995 | 4.57656 | 0.00000 | 0.00000 | 4.82246 |
| **Ndufs1** | 12.45214 | 12.12561 | 9.99010 | 4.30614 | 3.92087 | 5.95955 | 4.45029 | 9.31537 | 3.97218 | 13.55357 | 11.59114 | 14.19718 | 6.85855 | 11.44261 | 11.53347 | 6.46856 |
| **Ndufs2** | 18.57058 | 6.75787 | 10.94905 | 7.09945 | 12.58455 | 11.31017 | 8.29374 | 10.95074 | 7.75653 | 9.59387 | 7.56171 | 9.56033 | 13.03575 | 2.24651 | 1.25392 | 12.77348 |
| **Ndufs3** | 19.79834 | 5.07740 | 9.86234 | 4.35537 | 15.04747 | 9.66343 | 6.93069 | 16.23953 | 3.61532 | 17.98185 | 16.28779 | 3.51014 | 20.81154 | 0.28324 | 0.46427 | 11.57962 |
| **Ndufs4** | 7.87100 | 3.20469 | 5.05440 | 0.58359 | 0.11367 | 5.10972 | 2.99252 | 0.85332 | 1.15965 | 3.73764 | 3.22558 | 0.76468 | 4.17334 | 1.57042 | 0.00000 | 8.07142 |
| **Ndufs6** | 14.21596 | 0.76623 | 5.29376 | 1.16621 | 3.31858 | 4.06842 | 1.17978 | 1.64965 | 0.00000 | 5.79270 | 16.78492 | 0.63575 | 7.53205 | 0.18724 | 0.58909 | 7.04939 |
| **Ndufs7** | 16.46640 | 0.93784 | 9.46115 | 0.68199 | 7.77536 | 6.90958 | 4.98554 | 14.10766 | 2.22296 | 12.77922 | 10.24048 | 0.54223 | 17.43597 | 0.08896 | 0.00000 | 10.51534 |
| **Ndufs8** | 15.03273 | 10.43694 | 12.64178 | 6.34940 | 10.20104 | 4.50776 | 5.87886 | 11.41757 | 9.09210 | 11.94916 | 12.99941 | 1.34880 | 8.67458 | 0.07070 | 0.81876 | 6.27888 |
| **Ndufv1** | 15.75369 | 13.37523 | 19.70121 | 0.86467 | 9.80352 | 3.95399 | 10.48934 | 13.93895 | 0.02693 | 7.12010 | 4.48696 | 0.56465 | 15.93179 | 0.00000 | 3.08080 | 10.38750 |
| **Ndufv2** | 19.41046 | 6.18778 | 9.43928 | 6.18975 | 12.00188 | 8.93082 | 5.95865 | 3.55147 | 8.07954 | 9.81620 | 10.19293 | 3.92128 | 10.09248 | 2.65951 | 0.00000 | 14.68892 |
| **Ndufv3** | 3.29143 | 0.06427 | 2.86916 | 0.94100 | 3.06245 | 0.16058 | 0.53380 | 1.43703 | 1.78419 | 3.37251 | 8.13612 | 0.14572 | 2.81206 | 0.82831 | 1.29241 | 3.20567 |
| **Neat1** | 0.14001 | 0.01681 | 5.01426 | 0.84007 | 1.14118 | 0.85775 | 2.49477 | 0.05189 | 0.80008 | 2.02516 | 3.77188 | 0.21358 | 4.70419 | 0.46972 | 2.41575 | 0.22290 |
| **Neb** | 0.02378 | 0.00754 | 0.00671 | 1.03664 | 0.00852 | 0.00634 | 0.00712 | 0.01615 | 0.73838 | 0.28283 | 0.08706 | 0.06790 | 0.03955 | 0.00362 | 1.66188 | 0.01515 |
| **Necab2** | 0.80248 | 0.00746 | 0.00336 | 1.80745 | 0.22983 | 0.00722 | 0.00741 | 0.05279 | 1.57093 | 1.85584 | 0.04121 | 0.07270 | 0.00850 | 0.02720 | 0.00226 | 0.00588 |
| **Necab3** | 1.33191 | 0.00655 | 2.15445 | 0.43778 | 0.00486 | 0.00765 | 0.01632 | 0.00658 | 0.06958 | 0.00386 | 0.01098 | 0.00597 | 0.16002 | 0.00828 | 0.14317 | 0.00507 |
| **Necap1** | 2.29644 | 6.21348 | 7.77754 | 2.05662 | 8.08332 | 2.93802 | 2.72221 | 12.52772 | 7.83218 | 11.25018 | 10.01676 | 10.89810 | 7.72995 | 8.96602 | 11.10798 | 2.20599 |
| **Necap2** | 8.77979 | 1.02998 | 3.60154 | 8.52009 | 6.15646 | 4.93286 | 6.01228 | 6.53784 | 9.90669 | 11.67932 | 0.43645 | 1.37241 | 5.98306 | 0.00000 | 0.00000 | 6.98803 |
| **Nectin1** | 1.86547 | 1.98863 | 2.57322 | 0.04222 | 0.01673 | 0.00373 | 1.85280 | 3.31886 | 0.00763 | 0.05898 | 0.16175 | 0.02350 | 1.18744 | 0.00963 | 0.00000 | 1.26748 |
| **Nectin2** | 9.31495 | 0.12965 | 4.38578 | 3.24127 | 0.06960 | 6.88556 | 2.10345 | 5.60836 | 10.77858 | 0.02538 | 6.93897 | 0.36739 | 5.32992 | 0.00000 | 0.00000 | 4.99082 |
| **Nectin3** | 1.08349 | 0.00218 | 0.48133 | 5.07643 | 4.26942 | 0.00426 | 2.16372 | 0.02563 | 0.44317 | 0.89289 | 0.01789 | 1.51383 | 0.00803 | 0.90894 | 2.77956 | 0.03449 |
| **Nedd1** | 1.45789 | 2.28818 | 0.00000 | 6.83361 | 0.05172 | 0.13010 | 1.72302 | 4.99616 | 5.66074 | 4.85528 | 9.17257 | 15.52841 | 0.00000 | 14.52389 | 16.27827 | 0.16553 |
| **Nedd4** | 5.17503 | 7.67300 | 4.37541 | 1.12810 | 0.38827 | 0.03715 | 0.14058 | 0.00000 | 2.83251 | 5.37145 | 1.09848 | 9.60975 | 0.00000 | 0.00000 | 2.05845 | 3.94553 |
| **Nedd4l** | 2.15836 | 1.62852 | 1.04240 | 2.39327 | 0.06364 | 2.03287 | 1.54488 | 15.46124 | 0.83961 | 11.28368 | 2.33214 | 0.28979 | 4.24748 | 0.00000 | 0.40487 | 4.66061 |
| **Nedd8** | 18.47867 | 6.08726 | 11.75891 | 15.13508 | 12.21789 | 2.08470 | 5.99743 | 7.92878 | 19.20275 | 12.36778 | 14.71163 | 2.06997 | 11.96588 | 2.11764 | 2.35639 | 6.07722 |
| **Nedd9** | 0.01358 | 0.00761 | 0.01166 | 2.34790 | 0.00944 | 0.00625 | 0.00723 | 0.01096 | 0.00000 | 0.22917 | 0.01051 | 0.00547 | 0.01244 | 0.03078 | 0.51430 | 0.00708 |
| **Nefh** | 1.88953 | 3.85415 | 0.08207 | 0.83219 | 0.00544 | 0.00762 | 0.00426 | 0.28215 | 0.00000 | 0.64979 | 0.70343 | 0.05694 | 1.48461 | 0.00172 | 0.84205 | 0.00363 |
| **Nefl** | 1.36941 | 0.00710 | 0.00065 | 0.01062 | 0.01040 | 0.00659 | 0.00636 | 0.00935 | 0.01244 | 0.01014 | 0.01203 | 0.00699 | 0.01293 | 0.01273 | 0.01083 | 2.97706 |
| **Nefm** | 0.71273 | 0.00799 | 0.00507 | 0.01051 | 0.03679 | 0.00702 | 0.00696 | 0.01133 | 0.01230 | 0.03706 | 0.01246 | 0.00787 | 0.01864 | 0.01188 | 0.01125 | 0.00689 |
| **Neil1** | 0.70376 | 0.00607 | 0.58086 | 0.00686 | 0.00552 | 0.00583 | 1.40149 | 0.86707 | 0.01085 | 1.03727 | 0.00493 | 0.02033 | 0.94445 | 0.01738 | 0.01136 | 1.41829 |
| **Neil2** | 0.01050 | 0.00832 | 0.01314 | 0.79454 | 0.01006 | 0.00675 | 0.00747 | 0.01385 | 3.65464 | 0.00000 | 0.00000 | 0.00000 | 0.53576 | 0.04020 | 0.00126 | 0.00706 |
| **Neil3** | 1.63207 | 0.06773 | 0.31534 | 1.62253 | 0.01235 | 0.00359 | 0.00646 | 0.15073 | 0.00000 | 0.03471 | 0.04070 | 1.52132 | 0.00600 | 0.00355 | 1.79773 | 0.01282 |
| **Nek1** | 0.08699 | 0.00709 | 0.19530 | 3.91495 | 0.85074 | 0.01598 | 0.00709 | 1.04228 | 0.61185 | 0.24232 | 0.16692 | 0.42863 | 0.01743 | 2.93054 | 3.62279 | 0.01396 |
| **Nek11** | 0.00988 | 0.00766 | 0.00888 | 0.01080 | 0.00975 | 0.00683 | 0.00717 | 0.31985 | 0.01200 | 0.00984 | 0.01158 | 0.00700 | 0.00670 | 0.01313 | 0.00678 | 0.00742 |
| **Nek2** | 4.87058 | 2.94218 | 6.67238 | 14.79963 | 5.18987 | 5.14133 | 3.98457 | 18.43146 | 22.86376 | 11.60297 | 11.88214 | 11.63505 | 7.53118 | 12.38569 | 12.62932 | 0.78715 |
| **Nek3** | 0.79861 | 0.00467 | 0.68656 | 1.05075 | 0.01204 | 0.00912 | 0.00390 | 0.25977 | 0.26274 | 0.27421 | 2.09326 | 7.18515 | 1.87774 | 0.02163 | 0.81454 | 3.91382 |
| **Nek4** | 2.18049 | 0.05855 | 0.58802 | 2.07457 | 0.59443 | 0.03411 | 0.01521 | 0.00000 | 3.01860 | 0.10686 | 1.98717 | 0.11057 | 0.11288 | 0.78192 | 0.36705 | 0.04004 |
| **Nek5** | 0.01666 | 0.00782 | 0.01289 | 0.44367 | 0.00936 | 0.00706 | 0.00752 | 0.01241 | 0.00388 | 1.02890 | 0.01218 | 0.00443 | 0.08782 | 0.00886 | 0.34562 | 0.00788 |
| **Nek6** | 1.45755 | 3.49719 | 4.14233 | 0.00000 | 0.00000 | 8.69972 | 2.28567 | 0.00000 | 0.01059 | 0.35088 | 0.51711 | 0.07464 | 2.29757 | 0.00874 | 0.06975 | 1.79243 |
| **Nek7** | 0.28547 | 0.03669 | 0.55249 | 1.06138 | 2.19715 | 0.16091 | 0.06689 | 0.64477 | 0.60376 | 0.78474 | 1.35727 | 0.24353 | 1.16607 | 0.72223 | 0.60288 | 0.27019 |
| **Nek8** | 0.00000 | 0.00685 | 0.53728 | 1.54002 | 0.01760 | 0.00817 | 1.00095 | 2.26529 | 1.27747 | 2.36601 | 0.06735 | 0.85193 | 1.40085 | 0.46590 | 2.12526 | 0.00632 |
| **Nek9** | 3.66734 | 0.87239 | 3.02846 | 0.98286 | 5.07098 | 3.87915 | 1.78206 | 3.64810 | 0.35932 | 2.76729 | 2.53838 | 2.16439 | 3.39513 | 2.03352 | 2.93179 | 3.12714 |
| **Nelfa** | 4.86458 | 4.76313 | 2.96459 | 7.15931 | 2.42886 | 0.70898 | 0.33598 | 7.84728 | 8.00338 | 9.52992 | 7.60262 | 2.30462 | 6.40381 | 3.87567 | 10.11788 | 4.76236 |
| **Nelfb** | 9.15856 | 2.67016 | 10.19312 | 6.81398 | 2.41036 | 3.99174 | 7.17696 | 7.12014 | 6.59256 | 8.64996 | 5.31759 | 0.31064 | 8.19698 | 0.91197 | 9.13158 | 4.21429 |
| **Nelfcd** | 10.62800 | 5.16631 | 3.69217 | 8.84060 | 0.05736 | 2.51132 | 4.42589 | 13.63186 | 2.77549 | 12.97814 | 12.00474 | 1.52984 | 10.22758 | 0.46496 | 7.56667 | 3.42574 |
| **Nelfe** | 9.10155 | 4.15702 | 12.42101 | 10.40146 | 2.61826 | 5.07508 | 4.34325 | 12.77053 | 13.77113 | 17.51428 | 13.43591 | 2.87859 | 12.35930 | 0.01975 | 5.14836 | 7.78845 |
| **Nell1** | 0.01693 | 0.00760 | 0.01119 | 2.56519 | 0.01842 | 0.00617 | 0.00727 | 0.01144 | 1.97875 | 0.19267 | 0.00000 | 0.00263 | 0.01271 | 0.02176 | 0.14459 | 0.00690 |
| **Nemf** | 1.86550 | 4.69457 | 3.88081 | 6.93012 | 1.70632 | 4.54990 | 3.08195 | 2.44913 | 6.32788 | 3.67770 | 4.00775 | 9.18660 | 4.36824 | 8.88498 | 6.11260 | 0.74443 |
| **Nemp1** | 0.72588 | 0.01218 | 1.93156 | 8.05908 | 1.44461 | 0.00202 | 2.02680 | 1.05204 | 10.25327 | 10.02167 | 4.23987 | 8.56927 | 1.24714 | 16.82307 | 17.03849 | 1.10407 |
| **Nemp2** | 0.01576 | 0.00724 | 0.09639 | 0.01507 | 0.00915 | 0.00561 | 0.00707 | 0.00981 | 0.45711 | 0.11408 | 0.00487 | 0.57383 | 0.01160 | 0.00530 | 0.17167 | 0.00734 |
| **Nenf** | 0.60260 | 0.00683 | 0.00365 | 0.01042 | 0.00827 | 0.00743 | 0.00639 | 0.00000 | 2.23681 | 0.67409 | 1.13341 | 0.00000 | 0.00001 | 0.00917 | 0.00000 | 0.00664 |
| **Neo1** | 0.02834 | 1.40294 | 0.21365 | 2.74176 | 0.01119 | 4.57855 | 0.20437 | 0.11016 | 1.11053 | 0.32954 | 0.02647 | 0.03495 | 0.00002 | 0.00676 | 0.07212 | 0.02077 |
| **Nepro** | 0.32061 | 1.36803 | 0.12495 | 12.00306 | 0.03536 | 0.00274 | 0.00307 | 2.45839 | 15.35466 | 8.16920 | 5.25229 | 2.30221 | 2.11993 | 2.30605 | 10.32132 | 1.37028 |
| **Nes** | 1.22459 | 0.04163 | 0.00106 | 0.72296 | 0.01114 | 0.00680 | 0.00619 | 0.00665 | 0.94420 | 0.00000 | 0.00012 | 0.14441 | 0.00753 | 0.00719 | 0.07763 | 1.72727 |
| **Nespas** | 0.00642 | 0.67776 | 0.24096 | 0.01867 | 0.00766 | 0.00565 | 0.00693 | 1.04800 | 0.01142 | 0.00359 | 0.00697 | 0.00553 | 0.00426 | 0.01247 | 0.00000 | 0.00692 |
| **Net1** | 1.82965 | 3.04636 | 4.04281 | 2.56963 | 0.18870 | 2.74883 | 0.04618 | 1.09534 | 0.97260 | 2.69215 | 5.86524 | 0.82588 | 1.59717 | 4.64871 | 4.77472 | 4.16222 |
| **Neto2** | 1.57423 | 0.00461 | 2.00538 | 1.46656 | 0.01552 | 2.69309 | 0.38822 | 0.38954 | 3.99199 | 1.32838 | 2.10981 | 4.61566 | 0.35173 | 7.93877 | 6.42673 | 0.01743 |
| **Neu1** | 1.37658 | 1.66470 | 6.68133 | 1.86358 | 0.02682 | 8.13646 | 1.59403 | 4.83926 | 0.64746 | 7.53693 | 3.06340 | 1.21372 | 12.94176 | 0.09008 | 2.25759 | 1.35212 |
| **Neu3** | 1.03257 | 2.83533 | 0.73653 | 1.87198 | 0.09247 | 0.09312 | 0.24248 | 0.00200 | 3.58956 | 0.10115 | 2.86537 | 0.24646 | 0.18203 | 1.15450 | 0.27675 | 0.12061 |
| **Neurl1a** | 0.04340 | 0.00767 | 0.05262 | 0.03782 | 0.00862 | 0.00888 | 0.00733 | 0.00000 | 0.00935 | 0.34477 | 0.66784 | 0.01429 | 0.08865 | 0.01589 | 0.00000 | 0.00725 |

| | | | | | | | | | | | | | | | | |
|---|---|---|---|---|---|---|---|---|---|---|---|---|---|---|---|---|
| **Neurl2** | 0.59530 | 0.00802 | 0.00000 | 4.01976 | 0.01651 | 0.16571 | 0.00663 | 1.49815 | 0.86175 | 0.76392 | 0.73061 | 0.00000 | 0.00000 | 0.01234 | 0.08501 | 0.00586 |
| **Neurl3** | 0.01734 | 0.00783 | 0.01007 | 0.14151 | 0.00895 | 0.00672 | 0.00745 | 0.01970 | 0.09888 | 0.02578 | 0.01067 | 0.03721 | 0.01961 | 0.11706 | 0.09682 | 0.00741 |
| **Neurl4** | 4.10753 | 4.47952 | 7.07664 | 0.34445 | 3.79705 | 3.19351 | 7.12094 | 5.74198 | 0.69690 | 2.10571 | 1.88943 | 0.13814 | 8.57463 | 0.00000 | 1.65963 | 4.73048 |
| **Nexn** | 1.69520 | 0.00096 | 0.00000 | 16.76041 | 0.00469 | 0.00000 | 2.97568 | 0.00000 | 6.77607 | 6.94831 | 4.62555 | 7.11261 | 0.00000 | 12.44932 | 9.63493 | 0.00000 |
| **Nf1** | 0.07228 | 0.00842 | 0.00318 | 1.15744 | 0.01589 | 0.00753 | 0.90411 | 0.06814 | 4.49367 | 2.59636 | 0.21121 | 0.03867 | 0.02224 | 0.47732 | 2.44067 | 0.06045 |
| **Nf2** | 5.26089 | 4.60687 | 5.47222 | 11.93520 | 4.44813 | 3.75306 | 0.27924 | 5.10939 | 13.90268 | 17.05241 | 10.46748 | 2.35866 | 3.09913 | 7.02448 | 15.68135 | 4.71627 |
| **Nfat5** | 0.07340 | 0.00798 | 0.82985 | 3.51837 | 0.01581 | 0.00917 | 0.00781 | 0.39386 | 1.79722 | 0.83187 | 0.00000 | 0.21710 | 0.03882 | 0.00549 | 0.32096 | 0.01447 |
| **Nfatc1** | 0.01082 | 0.00777 | 0.00744 | 0.11686 | 0.00923 | 0.00691 | 0.00727 | 0.77723 | 0.01321 | 0.48271 | 0.00449 | 0.18855 | 0.00546 | 0.05589 | 0.00000 | 0.00719 |
| **Nfatc2** | 0.00885 | 0.00717 | 0.00334 | 0.07282 | 0.00922 | 0.00799 | 0.00621 | 0.17415 | 0.00787 | 0.92117 | 2.36091 | 0.00363 | 0.00000 | 0.01934 | 0.00000 | 0.00669 |
| **Nfatc2ip** | 6.16954 | 6.11264 | 6.86442 | 4.37009 | 0.42303 | 0.04324 | 0.44531 | 3.09848 | 9.61873 | 5.78213 | 17.17401 | 0.30244 | 4.56460 | 8.36325 | 2.83972 | 5.90249 |
| **Nfatc3** | 1.76374 | 2.74844 | 6.20782 | 7.62416 | 1.37583 | 0.48374 | 3.17876 | 5.14841 | 9.84171 | 11.16660 | 7.05095 | 6.36158 | 2.88153 | 7.43718 | 11.97265 | 4.54584 |
| **Nfatc4** | 1.20095 | 2.13482 | 1.14264 | 0.00666 | 0.01173 | 0.00977 | 1.12782 | 0.00416 | 0.02742 | 0.00095 | 0.01196 | 0.00601 | 0.03087 | 0.01301 | 0.00000 | 1.04618 |
| **Nfe2** | 2.84655 | 0.00391 | 1.75830 | 3.93506 | 0.00797 | 0.00301 | 0.00468 | 0.18993 | 7.83775 | 0.00000 | 0.07045 | 0.83359 | 0.00390 | 0.00000 | 0.41622 | 0.00233 |
| **Nfe2l1** | 4.65159 | 6.53214 | 6.27571 | 5.46284 | 1.02166 | 6.97842 | 9.81196 | 8.45382 | 6.54396 | 7.49034 | 4.02495 | 0.48858 | 5.88244 | 0.20774 | 5.25408 | 4.85374 |
| **Nfe2l2** | 6.34929 | 6.21901 | 4.74692 | 3.23707 | 9.20432 | 8.12872 | 3.86573 | 0.64953 | 3.44207 | 3.07489 | 0.20614 | 0.06586 | 1.68547 | 2.39280 | 0.82705 | 6.52280 |
| **Nfe2l3** | 0.01357 | 0.00769 | 0.01109 | 1.82003 | 0.00932 | 0.00625 | 0.00734 | 0.01133 | 1.64237 | 0.00000 | 0.00000 | 0.00425 | 0.01300 | 0.01934 | 0.43886 | 0.00689 |
| **Nfib** | 0.01337 | 0.00804 | 0.00875 | 0.51808 | 0.00897 | 0.00670 | 0.00725 | 0.01029 | 0.01051 | 0.82771 | 0.00792 | 0.30995 | 0.01081 | 0.04597 | 0.01318 | 0.00702 |
| **Nfic** | 1.41269 | 0.00521 | 0.00000 | 0.00000 | 1.87192 | 2.07252 | 0.00617 | 0.60498 | 0.55930 | 0.27792 | 0.15421 | 0.07833 | 0.03939 | 0.01229 | 0.00243 | 0.15333 |
| **Nfil3** | 0.00000 | 0.07049 | 1.14149 | 4.02428 | 0.00667 | 0.00524 | 0.00441 | 1.97113 | 2.37595 | 0.00000 | 3.12997 | 0.01159 | 2.84878 | 0.00790 | 0.64592 | 0.01112 |
| **Nfix** | 0.01928 | 0.00758 | 0.01450 | 0.21237 | 0.00840 | 0.02075 | 0.01735 | 0.00949 | 0.32431 | 2.09304 | 0.00802 | 0.03170 | 0.01017 | 0.01510 | 0.03882 | 0.00709 |
| **Nfkb1** | 1.46298 | 0.00603 | 0.07122 | 0.43853 | 0.00519 | 0.00420 | 0.00607 | 0.00478 | 0.00897 | 0.91295 | 0.01226 | 0.01407 | 0.01045 | 0.01027 | 0.00315 | 0.01279 |
| **Nfkb2** | 1.08960 | 0.00556 | 0.00475 | 0.26859 | 0.47232 | 0.00687 | 0.80323 | 0.71030 | 0.73382 | 0.00003 | 0.02496 | 0.02291 | 0.04683 | 0.01250 | 0.00000 | 0.00632 |
| **Nfkbia** | 0.00175 | 0.00545 | 0.00432 | 1.41680 | 1.98694 | 0.00679 | 0.00454 | 0.13414 | 3.97491 | 3.70159 | 6.47864 | 0.03132 | 0.00000 | 0.00000 | 0.27296 | 0.01287 |
| **Nfkbib** | 5.40592 | 0.01838 | 0.00000 | 0.91800 | 8.04968 | 6.34509 | 0.04924 | 4.88146 | 3.13034 | 6.76485 | 1.96503 | 0.17525 | 9.67751 | 0.03373 | 0.14784 | 4.67413 |
| **Nfkbie** | 0.65672 | 0.00828 | 0.00471 | 0.00740 | 0.01754 | 0.00718 | 0.00719 | 0.01622 | 0.05432 | 0.01521 | 0.01214 | 0.00793 | 0.01169 | 0.01284 | 0.01032 | 0.00695 |
| **Nfkbil1** | 0.07409 | 0.00734 | 0.05637 | 0.00442 | 0.00849 | 0.00705 | 0.00629 | 0.35700 | 0.15400 | 0.75568 | 0.36530 | 0.01036 | 1.41721 | 0.00532 | 0.07930 | 0.04516 |
| **Nfkbiz** | 0.02087 | 0.00814 | 0.01054 | 0.33092 | 0.02399 | 0.00679 | 0.00695 | 0.01411 | 0.01858 | 0.47780 | 0.03918 | 0.00673 | 0.87199 | 0.00545 | 0.16885 | 0.00703 |
| **Nfrkb** | 0.33417 | 0.00585 | 0.68578 | 1.69425 | 0.00628 | 0.04103 | 1.52738 | 3.97287 | 5.88261 | 2.40355 | 3.04803 | 0.58253 | 0.45648 | 0.33039 | 4.11486 | 0.19246 |
| **Nfs1** | 7.30394 | 2.27022 | 8.35201 | 4.32138 | 9.58037 | 0.86358 | 3.88463 | 9.17995 | 6.89658 | 5.98716 | 1.87601 | 1.23805 | 3.09737 | 3.86654 | 3.12878 | 4.01416 |
| **Nfu1** | 22.16190 | 6.30278 | 7.17116 | 8.16016 | 11.11262 | 3.35889 | 6.13508 | 6.91366 | 6.36262 | 9.73565 | 10.05839 | 0.75794 | 7.82447 | 0.82851 | 2.24857 | 3.81003 |
| **Nfx1** | 4.73703 | 14.02334 | 7.15297 | 2.14057 | 5.87955 | 7.09954 | 9.24064 | 11.64766 | 3.63706 | 5.48714 | 5.83106 | 6.27486 | 8.00418 | 5.58204 | 10.06136 | 6.85093 |
| **Nfxl1** | 2.19549 | 0.00302 | 1.56354 | 3.96949 | 0.21797 | 3.62641 | 1.97249 | 0.00000 | 1.72742 | 1.27034 | 0.51886 | 2.27628 | 0.00000 | 5.41395 | 9.55528 | 4.34414 |
| **Nfya** | 3.73025 | 0.14762 | 0.00000 | 6.40122 | 0.00344 | 0.19420 | 0.59827 | 4.24480 | 8.20255 | 0.09657 | 1.82404 | 0.14800 | 0.03163 | 0.00000 | 0.29558 | 0.02460 |
| **Nfyb** | 8.27475 | 4.84495 | 3.62896 | 10.75063 | 4.57166 | 0.30566 | 1.85855 | 7.50260 | 15.69496 | 10.76985 | 9.60328 | 9.12930 | 3.93153 | 17.92438 | 14.93171 | 2.26544 |
| **Nfyc** | 13.82359 | 5.76795 | 10.57805 | 12.04018 | 10.19816 | 4.21792 | 7.34192 | 18.91897 | 6.39739 | 23.89231 | 12.53340 | 1.92363 | 12.33867 | 3.55884 | 14.79635 | 8.35927 |
| **Ngdn** | 20.72354 | 17.74788 | 16.22216 | 5.87308 | 12.47282 | 12.23683 | 12.33407 | 13.81363 | 12.33852 | 16.67808 | 16.87195 | 9.02028 | 18.97156 | 2.00005 | 10.56855 | 7.68833 |
| **Ngef** | 0.00725 | 0.00753 | 0.79440 | 0.34001 | 0.00741 | 0.00815 | 0.00648 | 0.00000 | 1.59957 | 1.23757 | 0.47788 | 0.00000 | 0.90380 | 0.00295 | 0.21861 | 0.00622 |
| **Ngfr** | 0.02620 | 0.00798 | 0.02133 | 0.35804 | 0.03504 | 0.00696 | 0.04939 | 0.01359 | 0.03999 | 0.03823 | 0.05428 | 0.00754 | 0.02071 | 0.07503 | 0.09261 | 0.00728 |
| **Ngfrap1** | 0.72540 | 0.00649 | 0.19311 | 0.00431 | 0.68704 | 0.00844 | 0.00674 | 1.07610 | 0.00951 | 3.61493 | 0.00355 | 0.00002 | 0.27050 | 0.01440 | 0.00000 | 0.00683 |
| **Ngly1** | 0.05871 | 0.42427 | 0.31609 | 1.94260 | 2.91595 | 0.00576 | 0.00641 | 1.33035 | 0.11296 | 1.95331 | 0.02354 | 0.12243 | 1.30552 | 0.00396 | 3.03341 | 0.01384 |
| **Ngrn** | 10.40867 | 2.88394 | 2.78505 | 7.28387 | 3.29285 | 3.90529 | 2.36952 | 4.37244 | 8.35648 | 6.97043 | 3.99842 | 0.32332 | 3.33322 | 0.16991 | 1.69508 | 2.90211 |
| **Nhej1** | 0.07268 | 0.01850 | 1.61550 | 2.02616 | 0.00625 | 0.00602 | 0.00574 | 1.73653 | 0.27960 | 1.34685 | 0.69493 | 2.33834 | 2.49708 | 0.90511 | 2.03866 | 0.01213 |
| **Nhlrc2** | 2.43914 | 0.06808 | 0.13958 | 2.81894 | 0.07798 | 0.00921 | 0.58501 | 0.90191 | 1.81469 | 1.55935 | 1.46962 | 0.15349 | 1.15699 | 3.31177 | 0.00000 | 0.49270 |
| **Nhlrc3** | 0.01366 | 0.01868 | 0.01189 | 0.01064 | 0.00808 | 0.00659 | 0.00751 | 0.01441 | 0.01237 | 0.00780 | 0.01307 | 0.00812 | 0.01102 | 0.01285 | 0.01020 | 0.80473 |
| **Nhlrc4** | 0.01851 | 0.00815 | 0.01170 | 0.01077 | 0.01051 | 0.00662 | 0.00751 | 0.01359 | 0.01226 | 0.01861 | 0.01262 | 0.00807 | 0.01887 | 0.01372 | 0.01712 | 0.00738 |
| **Nhp2** | 29.74625 | 7.72654 | 19.27629 | 19.76832 | 16.26324 | 6.78042 | 13.85995 | 32.32070 | 31.86963 | 37.01667 | 29.44994 | 7.01413 | 27.98264 | 1.21425 | 3.86062 | 13.09391 |
| **Nhp2l1** | 27.94532 | 13.06680 | 27.38967 | 26.40160 | 15.39776 | 12.47185 | 14.68770 | 34.94875 | 33.84199 | 42.04594 | 29.82961 | 20.26675 | 26.45448 | 26.56487 | 32.97400 | 14.63115 |
| **Nhsl1** | 1.38997 | 0.00601 | 0.00000 | 5.67774 | 0.00561 | 0.01199 | 0.06756 | 0.62448 | 3.94126 | 2.06148 | 0.00000 | 0.01848 | 0.03480 | 0.11531 | 2.20663 | 0.00471 |
| **Nhsl2** | 0.02440 | 0.00794 | 0.01785 | 0.00842 | 0.00984 | 0.00639 | 0.01757 | 0.01045 | 0.02344 | 0.02933 | 0.05300 | 0.00718 | 0.01945 | 0.13390 | 0.01869 | 0.00751 |
| **Nicn1** | 4.78267 | 0.00000 | 1.83082 | 4.97490 | 5.77458 | 0.00560 | 0.75041 | 3.02832 | 1.61160 | 7.81127 | 0.00000 | 0.21249 | 1.87664 | 4.83819 | 4.45048 | 1.72249 |
| **Nid1** | 3.56500 | 0.00891 | 5.22571 | 0.00000 | 0.00663 | 3.49821 | 6.81919 | 8.31613 | 0.26096 | 2.17201 | 2.82324 | 0.14382 | 3.30248 | 5.35989 | 2.30866 | 6.24575 |
| **Nid2** | 0.00000 | 0.00641 | 1.45225 | 1.64717 | 0.00638 | 0.00911 | 2.84259 | 0.62267 | 1.11042 | 0.26676 | 0.10111 | 0.04947 | 3.07579 | 0.07391 | 1.23653 | 0.11746 |
| **Nif3l1** | 4.67358 | 7.34398 | 11.93824 | 6.39805 | 3.57654 | 6.81592 | 4.24234 | 9.65489 | 3.86000 | 7.10128 | 4.46414 | 1.11427 | 1.51156 | 4.78104 | 5.96830 | 7.01818 |
| **Nifk** | 17.27971 | 15.92592 | 20.95997 | 13.96056 | 10.57435 | 2.39151 | 11.32438 | 20.57705 | 19.64058 | 24.26404 | 11.97531 | 14.05348 | 16.04893 | 23.52672 | 22.67649 | 11.16177 |
| **Nim1k** | 0.00929 | 0.00861 | 0.00840 | 0.50343 | 0.00959 | 0.00680 | 0.00820 | 0.46107 | 1.10172 | 0.00559 | 0.00000 | 0.00648 | 0.00592 | 0.01364 | 0.00000 | 0.00720 |
| **Nin** | 1.67601 | 2.15320 | 2.92174 | 16.79391 | 0.01734 | 0.67104 | 0.25237 | 11.88168 | 15.63558 | 22.13737 | 17.29039 | 18.40524 | 1.46835 | 18.98505 | 26.03860 | 3.00465 |
| **Ninj1** | 0.83542 | 0.00570 | 0.00000 | 12.26320 | 0.01493 | 2.94807 | 1.35300 | 0.94939 | 0.54027 | 1.35913 | 0.26196 | 0.00000 | 2.65427 | 0.00000 | 0.21259 | 0.00204 |
| **Ninj2** | 0.01382 | 0.00812 | 0.01122 | 1.81028 | 0.00925 | 0.00645 | 0.00741 | 0.01033 | 0.00000 | 0.00461 | 0.12221 | 0.00724 | 0.01077 | 0.01524 | 0.10661 | 0.00702 |
| **Ninl** | 0.00000 | 0.00598 | 2.06688 | 0.89362 | 0.00751 | 0.00575 | 2.79281 | 3.29281 | 0.71270 | 1.96711 | 2.63148 | 0.02402 | 0.00000 | 0.52365 | 4.65313 | 0.00627 |
| **Nip7** | 17.11493 | 14.75901 | 22.22094 | 15.55907 | 12.96114 | 4.85118 | 12.25315 | 19.72281 | 26.87830 | 22.23291 | 21.59671 | 4.54270 | 13.61511 | 18.24932 | 23.32388 | 8.23959 |
| **Nipa2** | 5.49777 | 4.08619 | 0.77943 | 6.05244 | 1.67134 | 4.16222 | 0.00000 | 1.39001 | 6.67440 | 4.43350 | 0.89324 | 4.27926 | 4.23128 | 6.16544 | 9.68381 | 0.16030 |
| **Nipal1** | 2.63205 | 0.02252 | 4.14806 | 2.38009 | 0.03191 | 0.00113 | 0.00000 | 5.35097 | 8.25740 | 14.12740 | 10.05958 | 5.82815 | 3.69301 | 14.20509 | 16.72888 | 0.00000 |

| | | | | | | | | | | | | | | | | |
|---|---|---|---|---|---|---|---|---|---|---|---|---|---|---|---|---|
| **Nipal3** | 1.63490 | 0.00622 | 0.11330 | 2.25681 | 0.00438 | 0.01677 | 0.04009 | 0.99412 | 0.36508 | 1.25528 | 1.45615 | 0.05624 | 0.00000 | 0.00113 | 0.34516 | 0.01206 |
| **Nipbl** | 5.72734 | 7.12543 | 3.98540 | 4.20428 | 0.00000 | 4.00550 | 1.27590 | 5.48261 | 2.72071 | 12.49927 | 3.92739 | 7.59729 | 2.11400 | 11.21196 | 10.85840 | 2.07681 |
| **Nipsnap1** | 0.52626 | 0.00630 | 6.14757 | 3.23381 | 0.00000 | 1.45833 | 0.01521 | 0.82292 | 0.00000 | 2.99542 | 2.22859 | 0.09239 | 0.00000 | 0.26436 | 3.55116 | 3.11396 |
| **Nipsnap3b** | 8.97312 | 5.03568 | 0.99538 | 0.00000 | 5.84894 | 9.57326 | 1.96400 | 1.00840 | 0.00094 | 5.81518 | 0.96100 | 0.98753 | 3.66337 | 4.06018 | 1.10620 | 1.63382 |
| **Nisch** | 2.91259 | 1.13396 | 0.68038 | 3.54560 | 0.22339 | 0.00406 | 2.10325 | 0.00000 | 1.36546 | 2.48938 | 0.04161 | 0.07359 | 3.71715 | 0.02781 | 0.35834 | 5.52216 |
| **Nit1** | 5.62044 | 0.24769 | 4.54629 | 0.64147 | 1.31726 | 0.32853 | 1.12753 | 2.50993 | 0.38807 | 1.14123 | 0.85401 | 0.03153 | 1.50313 | 0.02265 | 1.70789 | 1.03750 |
| **Nit2** | 9.21794 | 2.49247 | 3.59670 | 0.38479 | 1.95006 | 5.19608 | 5.16385 | 2.75526 | 0.29354 | 1.51424 | 0.58491 | 3.84664 | 3.77968 | 0.11781 | 1.16343 | 0.05004 |
| **Nkain1** | 1.32689 | 0.19977 | 0.02386 | 2.99181 | 0.03982 | 0.14653 | 0.01655 | 0.01156 | 1.53054 | 0.24041 | 0.00000 | 0.07552 | 0.14780 | 0.00017 | 0.79544 | 1.75525 |
| **Nkain3** | 0.03161 | 0.00781 | 0.01447 | 0.13536 | 0.00829 | 0.00638 | 0.01761 | 0.03276 | 0.15887 | 0.02984 | 0.06190 | 0.00733 | 0.02269 | 0.03046 | 0.03844 | 0.00719 |
| **Nkap** | 1.85205 | 4.82715 | 3.04975 | 2.15971 | 3.03912 | 1.92665 | 0.97533 | 5.76959 | 0.86389 | 4.09527 | 2.01092 | 1.80286 | 1.71113 | 7.34122 | 9.20458 | 5.00240 |
| **Nkapl** | 0.01360 | 0.00786 | 0.01206 | 0.37187 | 0.00931 | 0.00650 | 0.00750 | 0.01335 | 1.05398 | 0.00699 | 0.00607 | 0.00570 | 0.01281 | 0.00613 | 0.10401 | 0.00724 |
| **Nkd1** | 0.01331 | 0.00751 | 0.01548 | 1.95772 | 0.00890 | 0.00713 | 0.04692 | 0.00055 | 0.11868 | 0.66201 | 0.32457 | 0.02113 | 0.00934 | 0.00120 | 0.29908 | 0.00737 |
| **Nkd2** | 0.01382 | 0.00807 | 0.01092 | 0.45061 | 0.00917 | 0.00660 | 0.00750 | 0.00940 | 0.00552 | 0.03815 | 0.18783 | 0.00694 | 0.00981 | 0.00742 | 0.01745 | 0.00715 |
| **Nkiras1** | 3.92583 | 4.05996 | 6.46328 | 9.99994 | 3.35224 | 0.00000 | 0.11248 | 6.99406 | 8.04693 | 8.14456 | 7.90153 | 2.53524 | 1.89410 | 6.19874 | 5.57154 | 2.28273 |
| **Nkiras2** | 0.28397 | 0.02623 | 1.88411 | 0.13040 | 0.66157 | 1.42125 | 3.66383 | 3.28514 | 0.52886 | 1.38369 | 0.07758 | 0.01180 | 0.85950 | 0.00505 | 0.05548 | 0.00552 |
| **Nkpd1** | 0.01184 | 0.00813 | 0.01194 | 0.66931 | 0.00912 | 0.00644 | 0.00720 | 0.01168 | 0.02709 | 0.00782 | 0.01348 | 0.00814 | 0.41331 | 0.01146 | 0.00438 | 0.00693 |
| **Nkrf** | 4.93683 | 0.00000 | 0.00000 | 6.11584 | 0.00000 | 0.00000 | 0.00179 | 1.91974 | 0.66904 | 5.66718 | 0.75193 | 8.88803 | 0.00124 | 7.83627 | 12.10762 | 0.01445 |
| **Nktr** | 4.68830 | 0.15612 | 3.38897 | 13.83307 | 2.56425 | 3.19849 | 4.75064 | 3.76628 | 13.93297 | 6.47547 | 6.61662 | 4.18572 | 4.06455 | 7.70442 | 12.33190 | 2.95288 |
| **Nkx2.3** | 0.00901 | 0.00846 | 0.00889 | 0.00658 | 0.01059 | 0.00763 | 0.00823 | 0.23742 | 0.01110 | 0.00000 | 0.23745 | 0.00675 | 0.31949 | 0.01928 | 0.04801 | 0.00751 |
| **Nkx6.2** | 1.11970 | 0.00550 | 0.01462 | 0.01317 | 1.89511 | 0.00508 | 0.00641 | 0.05135 | 0.01152 | 0.43059 | 0.05140 | 0.00387 | 0.01837 | 0.01275 | 0.00729 | 1.48510 |
| **Nle1** | 3.32167 | 0.18877 | 1.42053 | 2.94819 | 2.31489 | 0.21552 | 2.57901 | 6.08021 | 6.43921 | 5.31373 | 7.22136 | 0.48336 | 3.20457 | 0.19602 | 0.48525 | 1.67636 |
| **Nlgn1** | 0.01203 | 0.00757 | 0.00594 | 0.18494 | 0.00971 | 0.00662 | 0.00715 | 0.00880 | 0.01061 | 0.00490 | 0.01266 | 0.00979 | 0.01338 | 2.16004 | 0.00897 | 0.00740 |
| **Nlgn2** | 0.20073 | 0.00859 | 0.00607 | 0.00480 | 0.01682 | 0.00695 | 0.00709 | 0.00607 | 0.20334 | 0.00710 | 0.21753 | 0.01736 | 0.43814 | 0.01186 | 0.00461 | 0.00694 |
| **Nlk** | 4.41696 | 5.01316 | 0.40606 | 2.18881 | 5.10133 | 3.76934 | 1.77047 | 1.12732 | 4.44032 | 3.73124 | 3.05920 | 0.24126 | 0.55364 | 3.47305 | 1.28666 | 0.08574 |
| **Nln** | 5.66656 | 3.92826 | 3.51547 | 3.03923 | 2.71695 | 0.11115 | 1.45948 | 6.43367 | 0.50115 | 1.29063 | 1.41893 | 0.96090 | 1.06067 | 0.17366 | 1.28899 | 2.46244 |
| **Nlrc3** | 0.01377 | 0.00811 | 0.08198 | 0.00683 | 0.00945 | 0.86915 | 0.01775 | 0.01150 | 0.01119 | 0.00411 | 0.01145 | 0.00743 | 0.03628 | 0.01006 | 0.52833 | 0.00705 |
| **Nlrc4** | 1.13856 | 0.56484 | 0.00000 | 1.92131 | 0.00872 | 0.00661 | 0.00560 | 0.00309 | 1.76098 | 0.51443 | 0.01220 | 0.00000 | 0.70025 | 0.04722 | 0.44205 | 0.00506 |
| **Nlrp12** | 1.02253 | 0.00635 | 0.46124 | 0.17275 | 0.01397 | 0.00433 | 1.47287 | 0.00469 | 5.99099 | 0.62326 | 0.00000 | 0.05001 | 0.04039 | 0.31132 | 1.17559 | 0.00591 |
| **Nlrp14** | 0.00000 | 0.01884 | 0.00000 | 18.81753 | 0.00854 | 0.00320 | 0.00540 | 0.08559 | 8.24182 | 3.49267 | 1.05259 | 0.90570 | 1.87651 | 11.55610 | 14.70844 | 0.00236 |
| **Nlrp1a** | 1.96479 | 0.49288 | 3.16422 | 3.55868 | 0.47344 | 0.56433 | 1.73717 | 0.91839 | 4.67811 | 1.64909 | 1.97755 | 2.35316 | 1.06702 | 1.45920 | 0.65210 | 2.07055 |
| **Nlrp2** | 0.01744 | 0.00778 | 0.00997 | 1.37395 | 0.00904 | 0.00612 | 0.00726 | 0.01046 | 1.50899 | 0.00272 | 0.00000 | 0.00251 | 0.01297 | 0.00802 | 1.49207 | 0.00689 |
| **Nlrp3** | 0.03838 | 0.00851 | 0.01050 | 0.03675 | 0.01352 | 0.00676 | 0.00752 | 0.01308 | 0.23692 | 0.01334 | 0.02297 | 0.00775 | 0.01241 | 0.01262 | 0.00760 | 0.01495 |
| **Nlrp4a** | 0.04055 | 0.09077 | 0.01984 | 2.52595 | 0.07012 | 0.00677 | 0.05790 | 0.00709 | 2.03649 | 0.00000 | 0.08038 | 0.01511 | 0.02695 | 0.06625 | 3.16103 | 0.00666 |
| **Nlrp4b** | 0.01035 | 0.00822 | 0.00769 | 1.69925 | 0.00946 | 0.00648 | 0.00808 | 0.16148 | 1.35957 | 0.00000 | 0.06433 | 0.04914 | 0.00699 | 0.01336 | 0.52665 | 0.00684 |
| **Nlrp4c** | 7.42433 | 10.55705 | 0.46254 | 0.78044 | 9.67809 | 2.80804 | 5.45406 | 0.00000 | 1.22067 | 1.21999 | 4.79684 | 8.13188 | 0.98847 | 10.60508 | 4.00955 | 0.00631 |
| **Nlrp4e** | 0.19359 | 0.53321 | 0.00000 | 3.00527 | 0.44254 | 0.08989 | 0.13946 | 0.00675 | 1.98500 | 0.04086 | 0.01456 | 0.32548 | 0.05555 | 1.25834 | 0.17122 | 0.00630 |
| **Nlrp4f** | 0.07668 | 2.55010 | 0.00000 | 12.75395 | 0.00768 | 0.01030 | 0.15015 | 0.00000 | 3.24267 | 0.88747 | 0.27216 | 1.10752 | 0.00924 | 0.00000 | 4.03574 | 0.00392 |
| **Nlrp4g** | 0.01371 | 0.00785 | 0.01159 | 2.14828 | 0.00940 | 0.00626 | 0.00729 | 0.01129 | 0.00000 | 0.00681 | 0.01116 | 0.00634 | 0.01285 | 0.00367 | 0.85615 | 0.00700 |
| **Nlrp5** | 0.00000 | 0.00764 | 0.61316 | 9.64921 | 0.00752 | 0.39445 | 0.00685 | 1.38075 | 11.47240 | 4.94144 | 0.82477 | 0.21377 | 0.03353 | 3.78542 | 6.90467 | 0.00435 |
| **Nlrp5.ps** | 0.00294 | 0.00767 | 0.00866 | 1.21827 | 0.00952 | 0.46242 | 0.00692 | 0.07370 | 0.46801 | 1.89436 | 0.28688 | 0.23239 | 0.24494 | 2.37051 | 0.43065 | 0.01389 |
| **Nlrp6** | 0.01381 | 0.00777 | 0.01173 | 0.55876 | 0.00937 | 0.00661 | 0.00747 | 0.01292 | 0.00005 | 0.01106 | 0.01210 | 0.01780 | 0.01278 | 0.01875 | 0.07821 | 0.00732 |
| **Nlrp9a** | 0.01363 | 0.00744 | 0.00675 | 4.35181 | 0.00945 | 0.00633 | 0.00698 | 0.00836 | 0.87603 | 0.06015 | 0.05705 | 0.67305 | 0.01482 | 0.00000 | 0.48216 | 0.00670 |
| **Nlrp9b** | 0.00972 | 0.00813 | 0.00997 | 1.03866 | 0.01041 | 0.00684 | 0.00726 | 0.01050 | 2.39450 | 0.00000 | 0.00000 | 0.00000 | 0.47673 | 0.72923 | 0.30873 | 0.00723 |
| **Nlrp9c** | 0.01704 | 0.00780 | 0.00879 | 3.00906 | 0.00910 | 0.00590 | 0.00713 | 0.00933 | 0.57902 | 0.32100 | 0.02961 | 0.01949 | 0.01204 | 0.01596 | 2.49723 | 0.00689 |
| **Nlrx1** | 0.00721 | 0.00792 | 0.37259 | 0.21580 | 0.02059 | 0.00937 | 0.15723 | 0.07751 | 1.24001 | 0.45311 | 0.00000 | 0.01546 | 1.38415 | 0.01253 | 0.00000 | 0.00663 |
| **Nmb** | 0.15620 | 0.01985 | 1.10675 | 2.20510 | 0.02856 | 0.00961 | 0.15630 | 0.38195 | 14.78606 | 0.30980 | 0.86579 | 0.00000 | 0.00000 | 0.12298 | 0.06684 | 0.01299 |
| **Nmd3** | 8.55977 | 12.01976 | 11.64453 | 11.59330 | 8.52219 | 3.91877 | 6.96108 | 17.36251 | 16.96071 | 15.62911 | 11.99644 | 11.46645 | 8.33352 | 15.45799 | 13.42409 | 7.94865 |
| **Nme1** | 34.55186 | 16.56165 | 35.11430 | 2.92610 | 24.84287 | 14.37702 | 14.54480 | 25.81429 | 14.02603 | 32.16398 | 31.60511 | 4.99285 | 29.97034 | 7.76699 | 15.23084 | 14.90753 |
| **Nme2** | 41.79779 | 17.90810 | 39.45112 | 24.17972 | 27.43327 | 14.69412 | 20.26422 | 37.76373 | 35.60338 | 45.16602 | 49.21640 | 9.26897 | 42.85397 | 9.54018 | 22.07022 | 23.34245 |
| **Nme3** | 1.01632 | 0.00506 | 0.76361 | 2.71133 | 0.84508 | 0.02539 | 1.58041 | 1.12254 | 0.00000 | 0.34142 | 0.75086 | 0.00675 | 2.70925 | 0.00101 | 0.00000 | 1.12592 |
| **Nme4** | 5.28564 | 0.89138 | 1.95303 | 0.25249 | 2.03540 | 0.00000 | 0.40747 | 0.67181 | 1.19807 | 1.91921 | 2.75877 | 0.34688 | 3.89817 | 0.59765 | 0.00000 | 0.00622 |
| **Nme5** | 0.92836 | 0.00633 | 0.36847 | 3.76788 | 0.00532 | 0.00763 | 0.00580 | 0.18288 | 3.29910 | 1.16682 | 1.53506 | 0.05609 | 0.02326 | 0.88765 | 0.90858 | 0.00519 |
| **Nme6** | 8.15469 | 2.68087 | 8.72301 | 4.53190 | 2.58657 | 4.50201 | 6.11044 | 10.54007 | 3.79492 | 6.30957 | 4.35616 | 0.31857 | 5.09483 | 0.94976 | 3.25090 | 3.98626 |
| **Nme7** | 6.37035 | 4.04501 | 2.91769 | 0.99228 | 6.39671 | 2.92341 | 0.65183 | 2.36043 | 0.99765 | 0.10270 | 0.97990 | 3.07051 | 1.76993 | 0.22762 | 1.44238 | 2.11737 |
| **Nmi** | 0.01362 | 0.00817 | 0.01195 | 0.02555 | 0.00921 | 0.00678 | 0.00759 | 0.01379 | 0.46207 | 0.10709 | 0.00956 | 0.00792 | 0.01284 | 0.01207 | 0.00839 | 0.00734 |
| **Nmnat1** | 0.30354 | 0.03896 | 1.89657 | 4.84061 | 0.00582 | 0.13795 | 0.00606 | 0.00000 | 2.62202 | 1.71396 | 1.06156 | 0.04415 | 0.37003 | 0.00000 | 2.90976 | 0.00488 |
| **Nmnat2** | 0.01959 | 0.00784 | 0.88687 | 0.10589 | 0.01756 | 0.02397 | 0.00729 | 0.01166 | 0.26496 | 0.00341 | 0.01224 | 0.00646 | 0.03014 | 0.00739 | 0.42039 | 0.00703 |
| **Nmnat3** | 0.02770 | 0.00845 | 0.01065 | 0.23477 | 0.01197 | 0.00694 | 0.00747 | 0.01021 | 0.00790 | 0.42716 | 0.07549 | 0.00795 | 0.01163 | 0.01030 | 0.00582 | 0.00743 |
| **Nmral1** | 2.35864 | 0.01425 | 0.87991 | 0.71988 | 0.69906 | 0.00362 | 2.03193 | 1.25746 | 2.09370 | 1.65174 | 3.32748 | 0.10928 | 1.66259 | 0.53797 | 1.16327 | 3.82365 |
| **Nmrk1** | 3.88315 | 0.12217 | 4.05845 | 6.33189 | 2.41894 | 2.63395 | 3.10885 | 0.66902 | 7.49522 | 0.00000 | 6.51841 | 0.32998 | 1.78981 | 2.61697 | 0.72845 | 0.23458 |
| **Nmrk2** | 0.13411 | 0.02797 | 0.41798 | 0.15034 | 0.40147 | 0.06293 | 0.12046 | 0.00939 | 0.76251 | 0.12733 | 1.00379 | 0.04939 | 0.56857 | 0.40376 | 0.15526 | 0.07141 |
| **Nmt1** | 3.52878 | 1.24655 | 2.54069 | 1.08024 | 0.57551 | 0.06877 | 0.01306 | 2.81241 | 1.32307 | 0.20355 | 1.40559 | 0.11963 | 1.14644 | 2.83094 | 2.25238 | 0.00829 |

| | | | | | | | | | | | | | | | | |
|---|---|---|---|---|---|---|---|---|---|---|---|---|---|---|---|---|
| **Nmt2** | 4.06048 | 2.29694 | 1.82837 | 2.01458 | 4.22768 | 5.71119 | 5.36694 | 4.16239 | 10.81346 | 5.70677 | 3.46809 | 7.95011 | 3.09662 | 9.02418 | 3.76468 | 1.48916 |
| **Nmu** | 0.01369 | 0.00799 | 0.01197 | 0.17232 | 0.00914 | 0.00661 | 0.00733 | 0.01082 | 0.00899 | 0.26501 | 0.03253 | 0.00697 | 0.01227 | 0.00744 | 0.07106 | 0.00777 |
| **Nnat** | 0.01340 | 0.00785 | 0.01189 | 0.26980 | 0.00924 | 0.00647 | 0.03736 | 0.01001 | 0.00755 | 0.60479 | 0.03128 | 0.00595 | 0.01140 | 0.01031 | 0.00557 | 0.00740 |
| **Nnt** | 0.08744 | 3.16626 | 2.16723 | 0.00603 | 0.94988 | 0.80587 | 0.00680 | 0.69601 | 0.01127 | 1.36544 | 0.00418 | 0.00000 | 0.04980 | 0.01015 | 0.00000 | 2.34687 |
| **Noa1** | 0.90424 | 4.50727 | 1.42826 | 7.18308 | 1.43556 | 0.00177 | 0.10774 | 1.24118 | 12.26716 | 12.63177 | 5.69276 | 7.58616 | 0.01048 | 2.95575 | 10.78421 | 0.00000 |
| **Nob1** | 11.28721 | 9.88253 | 15.33144 | 12.63640 | 6.69101 | 10.01582 | 9.55675 | 22.23574 | 17.60606 | 19.73084 | 11.65885 | 3.10628 | 12.76305 | 3.69371 | 15.76690 | 9.50889 |
| **Nobox** | 0.01359 | 0.00790 | 0.01166 | 0.60090 | 0.00927 | 0.00649 | 0.00753 | 0.01317 | 1.29864 | 0.02705 | 0.00000 | 0.00544 | 0.01269 | 0.01673 | 0.05270 | 0.00710 |
| **Noc2l** | 8.94167 | 8.72653 | 8.23119 | 0.18410 | 10.17849 | 8.28808 | 3.08417 | 18.73083 | 0.51804 | 17.59295 | 1.57943 | 1.87528 | 12.10638 | 6.26056 | 11.23142 | 11.11211 |
| **Noc3l** | 1.97065 | 7.18188 | 1.27064 | 1.63320 | 1.54811 | 0.02326 | 0.04484 | 2.66852 | 4.13598 | 2.45654 | 1.80196 | 0.63461 | 1.60489 | 7.60122 | 9.20479 | 0.96797 |
| **Noc4l** | 15.87781 | 8.54337 | 8.69123 | 8.94572 | 10.00699 | 0.06258 | 3.04692 | 15.84301 | 11.19399 | 12.48448 | 7.21693 | 1.59587 | 16.03395 | 0.31610 | 7.10200 | 8.09903 |
| **Noct** | 2.98230 | 5.01311 | 3.68759 | 20.88732 | 8.95495 | 2.29868 | 1.89629 | 5.60817 | 18.29425 | 7.35434 | 5.01982 | 5.62055 | 7.38352 | 5.39051 | 11.05610 | 6.51539 |
| **Nod1** | 0.00981 | 1.50139 | 0.00898 | 0.00749 | 0.02055 | 0.00643 | 0.11755 | 0.00300 | 0.79688 | 0.57091 | 0.21847 | 0.02694 | 0.02919 | 0.00580 | 0.07499 | 0.00672 |
| **Nol10** | 1.27657 | 3.97101 | 2.21157 | 4.10073 | 2.04010 | 0.06495 | 0.86986 | 1.63767 | 4.64070 | 2.84242 | 3.95364 | 3.51418 | 1.22620 | 2.50090 | 3.39673 | 0.16923 |
| **Nol11** | 3.06376 | 6.35656 | 9.47621 | 6.02316 | 7.86435 | 8.04936 | 9.24925 | 14.55518 | 14.80573 | 15.60150 | 3.93686 | 7.77085 | 6.10290 | 25.65255 | 17.37577 | 1.67000 |
| **Nol12** | 13.35614 | 8.45521 | 11.67745 | 5.78765 | 2.96362 | 5.76456 | 9.69602 | 11.48671 | 16.52000 | 21.04717 | 12.15244 | 7.77447 | 13.40628 | 0.48908 | 9.22209 | 3.35149 |
| **Nol3** | 0.06445 | 0.00858 | 0.00951 | 0.00913 | 0.01505 | 0.00689 | 0.00751 | 0.01315 | 0.02684 | 0.02147 | 0.01253 | 0.00843 | 0.01996 | 0.01222 | 0.01102 | 0.00752 |
| **Nol4l** | 0.52354 | 0.00241 | 0.00005 | 7.80165 | 0.39522 | 4.38418 | 3.47133 | 8.82271 | 10.03109 | 7.08911 | 12.80250 | 0.73951 | 3.60862 | 9.84790 | 9.53721 | 2.26046 |
| **Nol6** | 7.45942 | 2.68051 | 5.67869 | 1.83796 | 4.38574 | 3.61225 | 4.83978 | 8.41530 | 1.86116 | 9.19702 | 4.61470 | 0.26604 | 6.92753 | 0.28038 | 4.18158 | 5.00092 |
| **Nol7** | 4.66108 | 4.43461 | 6.14526 | 22.69081 | 10.14833 | 2.21658 | 0.98105 | 3.17113 | 23.06423 | 26.07048 | 25.74289 | 3.04964 | 5.93718 | 10.64293 | 4.90118 | 3.19573 |
| **Nol8** | 3.70107 | 5.40026 | 6.05827 | 7.91354 | 0.00000 | 0.00559 | 3.45000 | 4.06050 | 14.76292 | 6.05614 | 6.36978 | 12.56333 | 2.12656 | 18.54043 | 15.92680 | 0.03035 |
| **Nol9** | 7.31415 | 6.72925 | 2.56479 | 14.57604 | 9.77553 | 2.24260 | 1.83330 | 11.15563 | 9.88712 | 4.11554 | 5.80929 | 2.41069 | 2.37022 | 3.61632 | 9.56895 | 0.66794 |
| **Nolc1** | 15.43954 | 13.29656 | 21.32976 | 14.89684 | 14.51591 | 10.85599 | 16.13850 | 39.63295 | 16.34428 | 38.61598 | 19.91867 | 15.82969 | 19.26557 | 24.61882 | 32.57073 | 10.37221 |
| **Nom1** | 0.70024 | 3.29080 | 3.46651 | 8.99941 | 1.89636 | 0.02193 | 0.03076 | 2.96647 | 18.89100 | 7.48334 | 8.88595 | 1.56082 | 2.49278 | 5.93459 | 10.62896 | 2.47760 |
| **Nomo1** | 8.02901 | 3.27106 | 4.88304 | 4.00720 | 0.46467 | 5.43181 | 5.03866 | 4.67909 | 3.91317 | 2.80091 | 2.82650 | 0.15159 | 9.05144 | 0.08602 | 0.75250 | 4.68673 |
| **Nono** | 21.92574 | 12.81238 | 18.51462 | 10.46060 | 11.42176 | 6.18704 | 9.96431 | 20.88215 | 20.03131 | 29.86842 | 19.11344 | 6.80957 | 12.45389 | 28.93797 | 26.22296 | 11.13636 |
| **Nop10** | 6.47964 | 1.21034 | 9.78894 | 4.29577 | 5.87461 | 1.94431 | 0.99070 | 0.93842 | 10.29761 | 4.64398 | 12.59946 | 1.13337 | 5.33131 | 4.91987 | 7.93146 | 3.68265 |
| **Nop14** | 12.12046 | 9.46104 | 9.06406 | 8.95229 | 3.76964 | 3.43716 | 7.88861 | 17.06449 | 9.29192 | 9.83635 | 8.59774 | 7.11285 | 8.98212 | 3.65107 | 8.90773 | 8.43479 |
| **Nop16** | 13.20436 | 13.13683 | 18.66590 | 10.90427 | 5.69343 | 6.72917 | 10.56300 | 16.09553 | 19.54325 | 16.70230 | 6.45051 | 2.37003 | 8.44431 | 2.59188 | 12.98621 | 7.91185 |
| **Nop2** | 13.86496 | 9.69446 | 8.64639 | 19.96003 | 13.48963 | 6.04407 | 10.46724 | 21.60753 | 23.58483 | 23.88779 | 14.98664 | 8.14060 | 11.01067 | 1.49864 | 16.10782 | 7.46148 |
| **Nop56** | 28.46720 | 21.68376 | 26.37917 | 17.25500 | 20.92166 | 14.47888 | 16.83559 | 35.03289 | 23.19077 | 27.61683 | 21.96058 | 11.63042 | 28.64424 | 22.56654 | 24.21604 | 16.03115 |
| **Nop58** | 7.90895 | 9.87373 | 8.72725 | 10.62443 | 11.97650 | 3.68925 | 2.81657 | 11.79262 | 18.31946 | 9.04473 | 17.42608 | 9.35276 | 4.26909 | 25.98961 | 12.80999 | 3.19067 |
| **Nop9** | 12.39467 | 4.33517 | 6.81639 | 8.88790 | 5.88416 | 0.00000 | 2.99965 | 12.85171 | 17.98239 | 20.16931 | 13.34996 | 2.98130 | 6.42550 | 5.73269 | 5.59951 | 4.41113 |
| **Nos1** | 1.01132 | 1.22146 | 3.22639 | 2.57032 | 0.01102 | 3.08172 | 2.26965 | 4.17759 | 4.31188 | 4.24252 | 1.86940 | 0.23664 | 2.83651 | 1.08963 | 2.30504 | 2.80700 |
| **Nos3** | 0.00884 | 0.03106 | 0.14467 | 0.01035 | 0.00924 | 0.00581 | 0.00798 | 0.24828 | 0.01200 | 0.10303 | 0.01111 | 0.00785 | 0.00634 | 0.01345 | 0.00490 | 0.00767 |
| **Nosip** | 4.48294 | 0.82821 | 3.26036 | 2.41982 | 0.05125 | 1.26647 | 4.61624 | 16.38765 | 4.94738 | 11.26626 | 4.74989 | 0.40315 | 10.29560 | 1.69369 | 0.42016 | 3.69239 |
| **Nostrin** | 2.03392 | 0.00576 | 0.01608 | 0.81295 | 0.00402 | 0.00397 | 0.00642 | 0.02843 | 1.08343 | 0.04333 | 0.00000 | 0.00438 | 0.00902 | 0.00385 | 0.36888 | 0.28653 |
| **Notch1** | 3.13157 | 0.00383 | 0.00000 | 0.41776 | 0.00229 | 0.08500 | 0.02519 | 0.64774 | 0.00490 | 1.19664 | 1.77013 | 0.49845 | 0.00000 | 0.00258 | 0.69784 | 0.00375 |
| **Notch2** | 0.00783 | 0.00633 | 0.04361 | 1.84223 | 0.00953 | 0.00823 | 2.51918 | 0.00953 | 0.64744 | 0.13054 | 0.00707 | 0.04157 | 0.07162 | 0.00211 | 0.36519 | 0.01471 |
| **Notch3** | 0.01265 | 0.00805 | 0.01096 | 0.00729 | 0.00948 | 0.00651 | 0.00738 | 0.04103 | 0.01091 | 0.00665 | 0.01159 | 0.00734 | 0.01097 | 0.00702 | 0.58155 | 0.00735 |
| **Notch4** | 0.30003 | 0.00706 | 0.00222 | 0.00504 | 0.00726 | 0.00748 | 0.00666 | 0.17367 | 0.01106 | 0.13543 | 0.28800 | 0.00354 | 0.31597 | 0.22275 | 0.22232 | 0.00686 |
| **Noto** | 0.00839 | 0.00784 | 0.00081 | 0.00326 | 0.00989 | 0.00817 | 0.00694 | 0.15386 | 0.01929 | 0.81748 | 1.02992 | 0.00000 | 0.02729 | 1.00155 | 0.39815 | 0.00778 |
| **Notum** | 0.00929 | 0.00733 | 0.00467 | 0.61195 | 0.00954 | 0.00743 | 0.00690 | 0.95431 | 0.01315 | 0.00000 | 0.42896 | 0.00335 | 0.00342 | 0.00781 | 0.00000 | 0.00705 |
| **Noxo1** | 0.01069 | 0.17130 | 0.36468 | 1.96232 | 0.00803 | 0.00531 | 0.00751 | 0.04692 | 2.46535 | 1.24138 | 0.86945 | 0.15093 | 0.04149 | 0.00767 | 0.05761 | 0.00639 |
| **Noxred1** | 0.82200 | 0.23214 | 0.74762 | 0.30780 | 0.03602 | 0.03489 | 0.86791 | 0.00000 | 0.00586 | 0.00008 | 1.69025 | 0.06123 | 0.64061 | 0.00566 | 0.34913 | 1.08719 |
| **Npas1** | 0.01364 | 0.00792 | 0.01213 | 0.36301 | 0.00914 | 0.00660 | 0.00734 | 0.01136 | 0.17833 | 0.24179 | 0.01257 | 0.00657 | 0.01994 | 0.00978 | 0.00846 | 0.00771 |
| **Npas2** | 0.06017 | 0.00795 | 0.00000 | 0.00355 | 0.01449 | 0.00750 | 0.00649 | 1.11130 | 0.25967 | 1.77594 | 0.00000 | 0.00000 | 0.95869 | 0.13729 | 0.51380 | 1.48473 |
| **Npat** | 0.16149 | 1.52061 | 1.45602 | 12.25268 | 0.11960 | 0.53034 | 5.15573 | 2.29395 | 13.67989 | 4.44877 | 2.55454 | 17.66384 | 1.48523 | 12.54301 | 10.51791 | 1.36959 |
| **Npc1** | 5.03983 | 0.11488 | 5.09633 | 1.19915 | 0.02481 | 4.07715 | 3.31209 | 6.31280 | 3.98721 | 3.80084 | 3.40618 | 0.80892 | 5.21901 | 1.06722 | 1.85364 | 0.93124 |
| **Npc1l1** | 0.00000 | 0.01774 | 0.00000 | 1.12858 | 0.00920 | 0.00529 | 0.00655 | 1.27766 | 4.05771 | 2.32402 | 0.86513 | 0.54129 | 0.00000 | 6.43756 | 7.88547 | 0.00657 |
| **Npc2** | 15.88160 | 13.26433 | 30.20378 | 12.95104 | 9.65132 | 15.19879 | 16.60991 | 37.48325 | 9.48145 | 27.50486 | 28.00922 | 15.25331 | 32.67008 | 10.33192 | 11.82516 | 11.34737 |
| **Npdc1** | 0.84518 | 0.01969 | 0.00000 | 0.83015 | 0.01620 | 4.36240 | 0.02625 | 1.89181 | 1.45824 | 8.59881 | 2.88000 | 0.00000 | 1.71103 | 0.00444 | 0.04285 | 0.58511 |
| **Npepl1** | 0.00000 | 0.00699 | 0.28414 | 1.03217 | 0.02812 | 2.79280 | 1.32693 | 0.00788 | 0.56402 | 0.43011 | 0.00000 | 0.47959 | 2.19760 | 0.00949 | 0.00000 | 0.01337 |
| **Npepps** | 6.06000 | 3.98674 | 0.44197 | 4.01365 | 6.05423 | 3.96553 | 0.36607 | 0.30232 | 5.91572 | 2.58306 | 1.07607 | 1.41315 | 4.80057 | 0.70579 | 2.51441 | 3.07455 |
| **Npff** | 0.01370 | 0.00828 | 0.01190 | 0.17032 | 0.00930 | 0.00682 | 0.00758 | 0.01375 | 0.00613 | 0.03405 | 0.02384 | 0.00857 | 0.01282 | 0.03229 | 0.00758 | 0.00739 |
| **Nphp1** | 0.12665 | 0.00858 | 0.00780 | 0.40782 | 0.01679 | 0.00670 | 0.00731 | 0.01167 | 0.00761 | 0.00770 | 0.01271 | 0.00785 | 0.01885 | 0.00644 | 0.28004 | 0.00735 |
| **Nphp3** | 1.74760 | 0.05442 | 0.03423 | 1.21169 | 0.00486 | 0.00384 | 1.31125 | 0.00278 | 1.02719 | 0.00000 | 0.00000 | 0.00999 | 0.01058 | 0.44976 | 0.29315 | 0.01231 |
| **Nphp4** | 0.01384 | 0.00795 | 0.01076 | 0.03923 | 0.00921 | 0.00666 | 0.00750 | 0.00902 | 0.00972 | 0.00263 | 0.19737 | 0.00675 | 0.00992 | 0.00626 | 0.10758 | 0.00728 |
| **Nphs1** | 0.04161 | 0.00751 | 0.07964 | 0.17919 | 0.01673 | 0.01988 | 0.08848 | 0.04789 | 0.59331 | 0.05235 | 0.58282 | 0.02138 | 0.03144 | 0.73853 | 0.17721 | 0.05720 |
| **Nphs2** | 0.00989 | 0.00806 | 0.00755 | 0.45724 | 0.00916 | 0.00641 | 0.00693 | 0.01298 | 0.00320 | 0.49175 | 0.00398 | 0.06096 | 0.52508 | 0.56509 | 0.75147 | 0.00678 |
| **Npl** | 6.71488 | 8.02596 | 1.59346 | 15.41189 | 6.58309 | 3.04055 | 3.10159 | 8.12482 | 18.17265 | 3.68142 | 9.75863 | 7.71553 | 0.93244 | 24.78458 | 22.80604 | 0.23739 |
| **Nploc4** | 10.28799 | 3.39455 | 6.12703 | 3.33801 | 2.53772 | 6.22602 | 2.15044 | 6.85136 | 8.13351 | 8.16510 | 6.94798 | 4.54572 | 5.78644 | 0.96449 | 7.32626 | 3.90753 |
| **Npm1** | 56.05784 | 38.04144 | 51.20099 | 24.68040 | 39.38403 | 22.66807 | 32.65889 | 60.52009 | 35.11850 | 60.76445 | 56.00672 | 35.79581 | 55.25421 | 62.30026 | 44.62516 | 31.40150 |

| | | | | | | | | | | | | | | | | |
|---|---|---|---|---|---|---|---|---|---|---|---|---|---|---|---|---|
| **Npm2** | 2.80501 | 0.00484 | 0.56285 | 5.30875 | 0.00209 | 0.00167 | 0.00492 | 1.21043 | 8.38535 | 5.61240 | 0.27168 | 0.03386 | 0.14542 | 2.04378 | 3.56191 | 0.00278 |
| **Npm3** | 0.48478 | 0.07118 | 0.09158 | 2.03131 | 0.00942 | 0.85017 | 0.45175 | 0.14938 | 1.21557 | 0.20569 | 3.05022 | 0.06339 | 0.40795 | 0.00231 | 0.08780 | 0.39415 |
| **Npm3.ps1** | 0.29654 | 0.01786 | 0.06674 | 0.99441 | 0.00901 | 0.30102 | 0.18338 | 0.14377 | 0.65581 | 0.07209 | 1.33888 | 0.02067 | 0.24736 | 0.02716 | 0.01022 | 0.10645 |
| **Npnt** | 0.01358 | 0.00816 | 0.01172 | 0.01063 | 0.00932 | 0.00670 | 0.00767 | 0.01776 | 0.01234 | 0.01469 | 0.01277 | 0.02945 | 0.01245 | 0.01258 | 0.01006 | 0.00741 |
| **Nppa** | 0.00939 | 0.00884 | 0.00780 | 0.25478 | 0.00940 | 0.00731 | 0.00827 | 0.29907 | 0.06883 | 0.40283 | 0.11015 | 0.00907 | 0.00637 | 0.01185 | 0.00000 | 0.00733 |
| **Nppb** | 0.00758 | 0.29676 | 0.01504 | 0.01102 | 0.00922 | 0.00632 | 0.00739 | 0.93186 | 0.01151 | 0.00896 | 0.00699 | 0.01647 | 0.00752 | 0.01368 | 0.00000 | 0.02162 |
| **Npr1** | 1.89529 | 1.23151 | 5.09840 | 0.67355 | 0.00516 | 0.26990 | 1.86191 | 3.11003 | 1.73165 | 2.49035 | 2.07345 | 0.06029 | 3.20036 | 0.37325 | 1.77732 | 0.02322 |
| **Npr2** | 0.79210 | 0.00789 | 0.25999 | 0.00461 | 0.00788 | 0.00545 | 0.00732 | 0.01343 | 0.01071 | 0.00000 | 0.45063 | 0.01727 | 0.17107 | 0.00623 | 0.20751 | 0.00654 |
| **Npr3** | 0.01227 | 0.00515 | 0.09226 | 0.35367 | 2.00308 | 0.00585 | 2.77345 | 0.00000 | 0.01609 | 0.73010 | 0.43629 | 0.00000 | 0.00747 | 0.46490 | 0.00000 | 0.00692 |
| **Nprl2** | 1.76744 | 0.01405 | 4.19140 | 1.62288 | 0.70531 | 3.17136 | 1.45190 | 3.72925 | 0.23652 | 1.90402 | 2.21283 | 0.05134 | 3.17275 | 0.34251 | 0.82676 | 1.71312 |
| **Nprl3** | 0.48790 | 0.00757 | 0.08270 | 0.35983 | 0.01727 | 0.01671 | 0.03849 | 0.23799 | 0.34937 | 0.18349 | 1.35358 | 0.01963 | 0.96232 | 0.23652 | 1.19913 | 0.12564 |
| **Npsr1** | 0.01364 | 0.00793 | 0.01083 | 0.01454 | 0.00935 | 0.00640 | 0.00735 | 0.01096 | 0.88801 | 0.00200 | 0.00638 | 0.02711 | 0.01283 | 0.00757 | 0.74246 | 0.00723 |
| **Nptn** | 0.79508 | 0.16023 | 0.23319 | 11.17824 | 0.30519 | 0.01084 | 1.20193 | 0.92132 | 9.58403 | 0.00000 | 4.79198 | 4.69733 | 0.00000 | 2.04301 | 0.73483 | 0.00256 |
| **Nptx1** | 0.53502 | 0.00721 | 0.00177 | 1.36494 | 0.01500 | 0.00685 | 0.00714 | 0.02072 | 0.00000 | 1.07628 | 0.03560 | 0.04254 | 0.01028 | 0.00509 | 0.89395 | 0.11417 |
| **Nptx2** | 0.29982 | 0.00831 | 0.00430 | 14.07385 | 0.00755 | 0.00607 | 0.00713 | 0.11431 | 9.10233 | 0.31261 | 0.08742 | 0.01052 | 0.35374 | 0.01226 | 0.08812 | 0.00415 |
| **Npw** | 0.01336 | 0.00802 | 0.01167 | 0.28972 | 0.00906 | 0.00661 | 0.00736 | 0.01092 | 0.12901 | 0.70964 | 0.01041 | 0.02584 | 0.01149 | 0.02796 | 0.01537 | 0.00733 |
| **Npy** | 0.00316 | 0.00724 | 0.06349 | 2.88710 | 0.00838 | 0.00758 | 0.00540 | 0.00000 | 3.69773 | 3.79809 | 0.91858 | 0.10481 | 2.42320 | 0.05655 | 0.00000 | 0.00548 |
| **Npy1r** | 0.09611 | 0.01833 | 0.20378 | 0.23174 | 0.09181 | 0.02251 | 0.13888 | 0.03832 | 0.39136 | 0.17668 | 0.39250 | 0.05004 | 0.30160 | 0.71703 | 0.38106 | 0.06547 |
| **Nqo1** | 2.14595 | 0.10507 | 1.24572 | 0.00288 | 0.01952 | 0.00701 | 0.03994 | 0.36373 | 0.16320 | 0.25844 | 0.00610 | 0.03258 | 0.39610 | 0.01119 | 0.67770 | 0.00436 |
| **Nqo2** | 5.65411 | 1.76338 | 4.19597 | 0.04385 | 0.00000 | 0.27860 | 3.48036 | 4.39990 | 0.23637 | 1.81023 | 1.83496 | 0.06712 | 1.44962 | 1.74555 | 0.10606 | 3.05158 |
| **Nr0b2** | 0.01384 | 0.00816 | 0.00927 | 0.00755 | 0.00911 | 0.00684 | 0.00758 | 0.00683 | 0.01187 | 0.00655 | 0.40276 | 0.00657 | 0.00814 | 0.01089 | 0.00395 | 0.00719 |
| **Nr1d1** | 0.00448 | 0.00721 | 0.07553 | 0.22384 | 0.00891 | 0.01797 | 0.23676 | 3.45870 | 0.13945 | 2.25102 | 0.29277 | 0.12931 | 0.00450 | 0.02897 | 0.00000 | 0.00723 |
| **Nr1d2** | 0.00000 | 0.10114 | 0.15129 | 5.68932 | 0.00778 | 0.00475 | 3.48361 | 3.33899 | 3.21271 | 4.69236 | 3.73166 | 0.76000 | 0.77996 | 14.77110 | 7.07193 | 0.00447 |
| **Nr1h2** | 6.33204 | 2.74302 | 1.86230 | 4.68422 | 3.91326 | 6.57204 | 3.43820 | 20.75807 | 2.73594 | 15.63742 | 7.59458 | 0.73739 | 6.92317 | 0.03212 | 0.00000 | 3.66521 |
| **Nr1h3** | 0.36335 | 0.00735 | 0.00000 | 6.49325 | 0.01764 | 0.00511 | 0.00543 | 4.55833 | 4.91125 | 11.29807 | 4.58300 | 1.35862 | 0.16866 | 0.00000 | 2.25357 | 0.00496 |
| **Nr1h4** | 2.97425 | 0.08014 | 0.05413 | 0.00808 | 0.02515 | 0.00265 | 0.00477 | 0.00000 | 0.01211 | 0.62744 | 0.01214 | 0.00362 | 0.01101 | 0.01133 | 0.00614 | 0.00444 |
| **Nr2c1** | 1.12568 | 0.00663 | 0.68824 | 0.64515 | 0.00451 | 0.00440 | 0.58041 | 0.17769 | 0.10605 | 2.60174 | 0.51532 | 0.09703 | 0.69708 | 0.00614 | 0.36114 | 0.00546 |
| **Nr2c2** | 0.31739 | 0.00664 | 0.33720 | 8.05962 | 0.00641 | 0.00304 | 0.00577 | 2.01020 | 13.43088 | 1.64227 | 2.43693 | 0.97657 | 0.00000 | 3.04473 | 6.33681 | 0.00396 |
| **Nr2c2ap** | 2.22193 | 1.32390 | 4.04573 | 2.46598 | 2.78398 | 1.70865 | 0.78924 | 1.58142 | 9.24284 | 2.77422 | 3.82806 | 0.10136 | 1.79187 | 0.09692 | 0.90158 | 0.05168 |
| **Nr2f2** | 0.01200 | 0.01744 | 0.25320 | 0.01607 | 0.01824 | 0.02361 | 0.00709 | 0.08381 | 0.01240 | 0.27789 | 0.01208 | 0.01528 | 0.00776 | 0.04657 | 0.01723 | 0.00760 |
| **Nr2f6** | 0.08972 | 0.00814 | 0.00000 | 0.00708 | 0.01531 | 0.84989 | 0.00712 | 0.06449 | 0.01188 | 0.75495 | 0.01044 | 0.01492 | 0.00029 | 0.01297 | 0.01466 | 0.01546 |
| **Nr3c1** | 0.88002 | 0.23722 | 0.11185 | 1.00529 | 0.29024 | 0.00475 | 0.00649 | 0.00777 | 0.06249 | 0.19464 | 0.02514 | 0.05969 | 0.01073 | 0.00690 | 0.94864 | 0.00665 |
| **Nr4a1** | 1.23907 | 0.92081 | 0.17207 | 0.00000 | 0.02184 | 3.95005 | 0.00456 | 0.79639 | 0.27720 | 0.00000 | 2.67528 | 0.01349 | 0.18961 | 0.00650 | 0.00178 | 0.00383 |
| **Nr4a2** | 0.01021 | 0.00799 | 0.00787 | 0.00623 | 0.00939 | 0.00719 | 0.00729 | 0.59669 | 0.07880 | 0.15576 | 0.00750 | 0.00729 | 0.00604 | 0.03065 | 0.02063 | 0.01469 |
| **Nr5a2** | 0.12989 | 2.71683 | 1.27164 | 11.34142 | 0.94952 | 1.09150 | 1.67972 | 10.75355 | 19.88821 | 10.50330 | 13.84139 | 6.01476 | 0.00000 | 15.07077 | 15.48471 | 0.02070 |
| **Nr6a1** | 2.39632 | 7.13138 | 10.31147 | 16.75688 | 0.00900 | 6.31269 | 2.40920 | 3.53066 | 5.05748 | 4.79880 | 1.37306 | 0.38811 | 5.69795 | 0.03282 | 8.41377 | 0.66584 |
| **Nradd** | 0.27723 | 0.00486 | 0.51514 | 0.01196 | 3.71764 | 0.04433 | 0.81233 | 3.04226 | 0.00541 | 0.00000 | 0.12793 | 0.13783 | 0.01301 | 0.01351 | 0.00000 | 0.00655 |
| **Nrap** | 0.88101 | 0.00558 | 0.05339 | 0.00932 | 2.00572 | 0.00792 | 0.01708 | 0.01088 | 0.01146 | 0.01494 | 0.01222 | 0.00528 | 0.00969 | 0.01147 | 0.01520 | 0.00633 |
| **Nrarp** | 0.01144 | 0.00784 | 0.01475 | 0.62122 | 0.00923 | 0.00612 | 0.00728 | 0.05238 | 0.20441 | 0.20555 | 0.00841 | 0.01649 | 0.12394 | 0.00869 | 0.31795 | 0.00691 |
| **Nras** | 7.92519 | 11.93665 | 9.93120 | 7.22550 | 5.66772 | 9.08724 | 4.84615 | 4.69769 | 6.86524 | 5.29462 | 3.85133 | 0.72327 | 4.06667 | 2.68499 | 6.68104 | 5.09434 |
| **Nrbf2** | 1.52566 | 6.42080 | 2.97366 | 10.99893 | 7.09275 | 1.40864 | 7.04879 | 2.98853 | 4.22921 | 6.50699 | 1.50442 | 9.29250 | 3.16737 | 2.23509 | 3.74420 | 2.23507 |
| **Nrbp1** | 4.66863 | 4.53340 | 7.71232 | 7.37223 | 1.39627 | 5.73089 | 9.58753 | 17.78452 | 7.76859 | 28.10865 | 5.88123 | 8.00363 | 8.73989 | 0.04477 | 4.06888 | 5.27876 |
| **Nrbp2** | 0.00660 | 0.00735 | 0.00587 | 0.09694 | 0.00902 | 0.00706 | 0.00630 | 1.13768 | 0.01009 | 0.81847 | 0.14335 | 0.02908 | 1.27606 | 0.01033 | 0.00000 | 0.00708 |
| **Nrcam** | 0.02592 | 0.00806 | 0.20971 | 0.05468 | 0.00869 | 0.00911 | 0.01814 | 0.02105 | 0.01924 | 0.10486 | 0.77421 | 0.00317 | 0.20684 | 0.10451 | 0.04793 | 0.00717 |
| **Nrd1** | 7.65932 | 11.72127 | 10.06035 | 2.25086 | 10.33420 | 9.24597 | 9.40369 | 5.96036 | 4.18415 | 6.75059 | 6.03513 | 2.29556 | 5.36040 | 0.66807 | 4.67505 | 1.45329 |
| **Nrde2** | 3.55532 | 0.00000 | 1.57663 | 0.74606 | 5.06641 | 1.78866 | 0.91454 | 4.95567 | 0.47860 | 4.58368 | 0.34989 | 0.34679 | 2.46296 | 0.08567 | 0.00000 | 1.23800 |
| **Nrep** | 0.01377 | 0.00814 | 0.01160 | 0.74849 | 0.00930 | 0.00660 | 0.00748 | 0.01298 | 0.21883 | 0.01218 | 0.02634 | 0.02925 | 0.01245 | 0.01171 | 0.00710 | 0.00727 |
| **Nrf1** | 5.37908 | 2.78416 | 2.84577 | 5.29291 | 2.59887 | 2.02047 | 6.47443 | 11.61153 | 8.03726 | 12.45509 | 3.43721 | 0.63749 | 5.40417 | 0.55343 | 6.65594 | 4.64514 |
| **Nrg1** | 0.00000 | 0.00632 | 0.47234 | 0.18713 | 2.67500 | 0.00559 | 1.04389 | 0.14686 | 0.00423 | 0.37547 | 0.00991 | 0.08893 | 0.00932 | 0.05014 | 0.04364 | 0.00689 |
| **Nrg2** | 0.03256 | 0.00766 | 0.04613 | 0.00558 | 0.00814 | 0.00586 | 0.00728 | 0.03753 | 0.01135 | 0.01084 | 0.41293 | 0.00607 | 0.00630 | 0.01123 | 0.00291 | 0.14726 |
| **Nrg4** | 0.01433 | 0.00778 | 0.06134 | 2.55207 | 0.00936 | 0.00559 | 0.00715 | 0.01001 | 0.63575 | 0.00000 | 0.01078 | 0.01088 | 0.01199 | 0.86857 | 1.03912 | 0.00682 |
| **Nrip1** | 0.00570 | 0.00737 | 0.26211 | 0.14046 | 0.06981 | 0.00891 | 0.00704 | 0.79888 | 0.01014 | 0.00569 | 0.00000 | 0.19103 | 0.00433 | 0.00582 | 0.95060 | 0.01447 |
| **Nrip2** | 0.01057 | 0.00822 | 0.01066 | 0.01050 | 0.00916 | 0.00649 | 0.00716 | 0.01333 | 0.01256 | 0.00439 | 0.01134 | 0.00719 | 0.62344 | 0.17014 | 0.00581 | 0.00692 |
| **Nrip3** | 0.74204 | 0.00509 | 0.68632 | 0.54944 | 0.48150 | 0.00472 | 2.60452 | 0.00989 | 0.17662 | 0.00279 | 0.01123 | 0.00495 | 0.00784 | 0.00912 | 0.30648 | 0.00593 |
| **Nrl** | 0.00945 | 0.00781 | 0.01311 | 0.64186 | 0.12108 | 0.00705 | 0.00770 | 0.01599 | 0.06249 | 0.00814 | 0.01429 | 0.00723 | 0.13693 | 0.00622 | 0.09378 | 0.00746 |
| **Nrm** | 0.01039 | 1.97007 | 0.40142 | 0.48966 | 0.00774 | 0.36739 | 0.02572 | 2.71578 | 0.17953 | 0.00000 | 0.22933 | 0.05394 | 2.31820 | 0.21455 | 0.01049 | 0.01212 |
| **Nrn1l** | 0.01323 | 0.00811 | 0.49584 | 0.25105 | 0.00842 | 0.00558 | 0.00747 | 0.01103 | 0.15949 | 0.25572 | 0.02604 | 0.01735 | 0.00928 | 0.01073 | 0.00000 | 0.00698 |
| **Nrp** | 0.01859 | 0.00863 | 0.00966 | 0.18687 | 0.01334 | 0.01431 | 0.00785 | 0.08871 | 0.44965 | 0.04703 | 0.00980 | 0.03751 | 0.66923 | 0.14205 | 0.05700 | 0.00755 |
| **Nrp1** | 0.01206 | 0.00808 | 0.01094 | 0.46052 | 0.02769 | 0.00656 | 0.01818 | 0.01693 | 0.92108 | 0.01136 | 0.00607 | 0.03901 | 0.01868 | 0.01287 | 0.01683 | 0.00719 |
| **Nrp2** | 1.17853 | 0.01560 | 0.01559 | 0.02448 | 0.00564 | 0.00726 | 0.60458 | 0.01099 | 0.03329 | 0.34115 | 0.01168 | 0.02334 | 0.02342 | 0.01136 | 0.04746 | 0.03508 |
| **Nrros** | 0.20767 | 0.00820 | 0.00259 | 1.49154 | 0.01548 | 0.00824 | 0.00747 | 0.52079 | 0.01921 | 0.47411 | 0.79366 | 0.05025 | 0.00008 | 0.00431 | 0.05212 | 0.00668 |
| **Nrsn2** | 0.01281 | 0.00744 | 1.37344 | 0.55038 | 0.00717 | 0.00570 | 0.00693 | 0.12131 | 0.00485 | 0.08564 | 0.47769 | 0.01481 | 0.00000 | 0.00477 | 0.05023 | 0.00670 |

| | | | | | | | | | | | | | | | | |
|---|---|---|---|---|---|---|---|---|---|---|---|---|---|---|---|---|
| **Nrxn2** | 0.02193 | 0.00784 | 0.01162 | 0.01009 | 0.00886 | 0.00660 | 0.00740 | 0.01310 | 0.01246 | 0.10695 | 0.01299 | 0.01683 | 0.01264 | 0.01055 | 0.01413 | 0.00744 |
| **Nsa2** | 23.28361 | 15.54177 | 20.65555 | 12.91970 | 19.69191 | 11.89716 | 14.65043 | 22.92598 | 21.75462 | 28.86217 | 21.73748 | 16.52392 | 21.87612 | 26.21178 | 22.12087 | 14.84143 |
| **Nsd1** | 11.11308 | 8.53655 | 4.69528 | 8.41154 | 1.05850 | 6.63081 | 1.90091 | 6.99810 | 5.77400 | 7.18964 | 5.93531 | 0.76313 | 5.85601 | 11.56609 | 10.40381 | 5.75581 |
| **Nsdhl** | 3.47686 | 7.17751 | 5.26274 | 0.39078 | 1.53656 | 1.47337 | 1.99237 | 0.00000 | 0.39573 | 1.53621 | 0.39411 | 0.04020 | 0.12906 | 0.00248 | 0.00000 | 1.74570 |
| **Nsf** | 9.57925 | 11.92665 | 8.09450 | 4.61895 | 12.34112 | 6.51642 | 9.83457 | 4.86105 | 3.51721 | 4.35089 | 4.16025 | 0.51450 | 5.38484 | 1.61581 | 7.20119 | 2.85970 |
| **Nsfl1c** | 19.95327 | 9.93122 | 20.42074 | 2.20852 | 16.38305 | 9.46608 | 12.37294 | 16.94055 | 1.01009 | 8.50848 | 14.49788 | 6.07730 | 19.68158 | 2.90068 | 6.02962 | 6.27139 |
| **Nsl1** | 1.13831 | 0.00594 | 4.68177 | 2.36896 | 0.11431 | 3.16582 | 0.96261 | 0.23538 | 1.51197 | 1.54178 | 3.19658 | 0.97001 | 0.43544 | 1.24573 | 2.15589 | 0.03239 |
| **Nsmaf** | 3.22765 | 0.08943 | 2.53037 | 0.00000 | 9.07729 | 0.00388 | 4.21395 | 1.06141 | 0.00429 | 0.08743 | 0.32863 | 0.00952 | 1.25489 | 0.00445 | 0.33526 | 0.02371 |
| **Nsmce1** | 14.02038 | 8.31793 | 18.62353 | 6.36740 | 8.74714 | 8.37432 | 9.71877 | 16.27558 | 6.49273 | 13.38512 | 14.22326 | 7.77034 | 19.14353 | 1.75116 | 3.97318 | 6.18340 |
| **Nsmce2** | 15.40652 | 6.92035 | 11.50907 | 6.23668 | 8.64985 | 5.17986 | 4.67752 | 5.25191 | 12.54441 | 12.19428 | 7.87194 | 6.66835 | 8.25159 | 2.58579 | 7.58867 | 6.01828 |
| **Nsmce4a** | 15.48529 | 5.45048 | 13.85714 | 17.72899 | 9.55065 | 2.85854 | 5.52303 | 10.27832 | 19.04130 | 27.03517 | 28.56340 | 19.35153 | 8.30234 | 7.60267 | 8.33363 | 2.48312 |
| **Nsmf** | 0.01047 | 0.00779 | 0.01082 | 0.02076 | 0.00976 | 0.00661 | 0.00721 | 0.30114 | 0.30682 | 0.05878 | 0.00942 | 0.02435 | 0.01046 | 0.07182 | 0.00000 | 0.00748 |
| **Nsrp1** | 7.17118 | 0.46511 | 4.98147 | 8.22635 | 0.01709 | 1.23417 | 4.40179 | 5.66358 | 9.56292 | 8.83155 | 4.08423 | 4.84762 | 5.22146 | 10.73677 | 7.17500 | 3.20602 |
| **Nsun2** | 18.36494 | 11.69401 | 16.65847 | 14.56462 | 12.44433 | 4.70930 | 11.36626 | 19.16877 | 14.80486 | 26.45346 | 19.87661 | 17.69128 | 13.77325 | 7.77986 | 22.04229 | 11.31747 |
| **Nsun3** | 0.25408 | 0.00802 | 0.00513 | 0.00931 | 0.01239 | 0.00708 | 0.10777 | 0.01573 | 0.01167 | 0.00787 | 0.01236 | 0.00756 | 0.01882 | 0.00906 | 0.23106 | 0.00720 |
| **Nsun4** | 3.17690 | 3.15410 | 1.49380 | 10.66485 | 0.01016 | 0.00291 | 1.14013 | 1.24575 | 4.92256 | 3.01445 | 1.40335 | 0.12984 | 2.78913 | 4.47601 | 6.53500 | 0.92609 |
| **Nsun5** | 8.70404 | 4.52900 | 5.28099 | 14.05735 | 4.13363 | 4.30323 | 1.71708 | 16.62981 | 14.44036 | 12.56350 | 8.09288 | 1.36888 | 5.10992 | 0.03341 | 1.57699 | 3.46253 |
| **Nsun6** | 0.91618 | 0.00187 | 3.82591 | 0.00000 | 0.79925 | 0.00482 | 2.99595 | 0.00000 | 0.00685 | 2.04600 | 0.51578 | 6.90696 | 2.90366 | 0.85140 | 1.41702 | 0.01020 |
| **Nsun7** | 0.00805 | 0.00800 | 0.00560 | 1.06483 | 0.00999 | 0.00565 | 0.00708 | 0.01160 | 2.22795 | 0.22736 | 0.00000 | 0.02951 | 0.01916 | 3.52274 | 4.97706 | 0.00698 |
| **Nt5c** | 2.70710 | 0.41401 | 0.88395 | 0.00002 | 0.04243 | 2.88070 | 1.57654 | 5.96977 | 0.00798 | 7.72096 | 15.50539 | 0.49584 | 9.91882 | 0.19810 | 1.11270 | 3.06290 |
| **Nt5c2** | 7.85782 | 8.46475 | 3.14846 | 3.06499 | 4.20326 | 0.74901 | 0.65391 | 0.51203 | 2.93902 | 3.40672 | 0.15148 | 0.11682 | 1.17365 | 2.64463 | 2.80817 | 3.97314 |
| **Nt5c3** | 2.83523 | 0.93786 | 0.00000 | 1.20455 | 0.01115 | 0.00339 | 0.00338 | 0.01064 | 0.00000 | 2.16253 | 1.23604 | 5.75172 | 0.03452 | 0.00295 | 1.51144 | 0.08158 |
| **Nt5c3b** | 5.32141 | 4.99555 | 5.18160 | 0.96219 | 0.09273 | 0.00000 | 6.02965 | 14.90256 | 1.07360 | 11.59830 | 14.10252 | 2.04679 | 9.87687 | 0.09747 | 1.34952 | 5.39991 |
| **Nt5dc1** | 2.82362 | 0.00226 | 0.38658 | 0.00000 | 0.01242 | 2.68232 | 0.54941 | 0.92161 | 3.73131 | 0.12560 | 0.80081 | 6.50702 | 2.24320 | 1.57689 | 1.51005 | 0.00952 |
| **Nt5dc2** | 0.00454 | 0.00573 | 0.00137 | 0.00073 | 1.44964 | 0.00837 | 2.00860 | 0.00683 | 0.30470 | 0.66423 | 0.00917 | 0.04877 | 0.01716 | 0.01092 | 0.01226 | 0.00750 |
| **Nt5dc3** | 0.66362 | 0.02635 | 0.28852 | 4.26957 | 3.57094 | 0.02494 | 0.13568 | 0.00000 | 2.16209 | 0.04532 | 1.27274 | 0.07993 | 1.90742 | 0.42950 | 0.09370 | 3.52911 |
| **Nt5e** | 1.32222 | 0.00740 | 0.00164 | 0.01064 | 0.01521 | 0.00659 | 0.00640 | 0.00898 | 0.01241 | 0.00587 | 0.01230 | 0.00719 | 0.01598 | 0.01336 | 0.01061 | 2.29298 |
| **Nt5m** | 1.63218 | 0.27800 | 2.53958 | 0.20376 | 0.00000 | 0.01165 | 1.79565 | 0.66655 | 0.00949 | 0.00080 | 0.00594 | 0.01207 | 0.00000 | 0.01004 | 0.14825 | 1.66045 |
| **Ntan1** | 9.50135 | 0.81518 | 1.46585 | 3.74807 | 1.45774 | 2.03862 | 2.36648 | 0.13413 | 2.94017 | 0.59284 | 1.69403 | 0.31075 | 5.08794 | 0.01400 | 0.60694 | 1.38000 |
| **Ntf5** | 0.08948 | 0.00759 | 0.04149 | 0.04341 | 0.00788 | 0.02020 | 0.00728 | 0.01579 | 0.53845 | 0.12440 | 0.27489 | 0.01602 | 0.01956 | 0.12416 | 0.07275 | 0.02174 |
| **Nthl1** | 3.25483 | 0.67259 | 3.17799 | 1.00338 | 2.10890 | 0.04581 | 1.93056 | 0.00000 | 0.40485 | 0.58460 | 0.63765 | 0.06600 | 0.39920 | 0.00285 | 0.20312 | 0.25403 |
| **Ntmt1** | 2.89631 | 1.67013 | 2.05770 | 9.28215 | 7.62900 | 0.35603 | 2.30700 | 2.56674 | 13.48251 | 13.83684 | 3.21543 | 0.78482 | 6.25589 | 0.00000 | 0.00000 | 0.03397 |
| **Ntn1** | 0.01358 | 0.00776 | 0.01160 | 0.15461 | 0.00942 | 0.00653 | 0.00737 | 0.01672 | 0.01649 | 0.03074 | 0.01236 | 0.01702 | 0.01256 | 0.00932 | 0.28001 | 0.00743 |
| **Ntn3** | 0.01787 | 0.00770 | 0.01141 | 0.05820 | 0.00904 | 0.00656 | 0.00741 | 0.01234 | 0.01079 | 0.00619 | 0.01228 | 0.00701 | 0.01282 | 0.02499 | 0.30873 | 0.00736 |
| **Ntpcr** | 2.27240 | 0.00552 | 0.00349 | 0.15666 | 0.00370 | 0.00623 | 0.00567 | 0.00995 | 0.00646 | 0.01343 | 0.01177 | 0.00646 | 0.06913 | 0.01097 | 0.01331 | 0.01200 |
| **Ntrk2** | 0.06244 | 0.00742 | 0.01312 | 0.61567 | 0.00796 | 0.01478 | 0.02782 | 0.01527 | 0.06990 | 0.01448 | 0.01108 | 0.01732 | 0.01946 | 0.15951 | 0.04046 | 0.00720 |
| **Nuak1** | 3.65694 | 2.16884 | 0.00000 | 0.00824 | 2.83908 | 0.06648 | 1.39776 | 0.00836 | 0.01104 | 0.00014 | 0.01199 | 0.05846 | 0.01492 | 0.01110 | 0.00000 | 0.09002 |
| **Nuak2** | 0.01365 | 0.00801 | 0.01172 | 0.37505 | 0.00918 | 0.00662 | 0.00734 | 0.01141 | 0.42091 | 0.24858 | 0.00967 | 0.02588 | 0.01270 | 0.01051 | 0.00348 | 0.00771 |
| **Nub1** | 9.39498 | 10.54610 | 23.66485 | 19.21266 | 5.98436 | 9.49582 | 7.71187 | 30.03320 | 30.23624 | 35.95025 | 34.17938 | 12.99148 | 11.96951 | 19.51556 | 25.23015 | 6.77461 |
| **Nubp1** | 4.57094 | 0.33483 | 1.89678 | 0.07483 | 1.88302 | 0.00729 | 0.11994 | 1.36302 | 1.52744 | 5.21981 | 9.96492 | 6.71131 | 1.79411 | 0.54818 | 1.92009 | 2.50603 |
| **Nubp2** | 20.18397 | 13.08155 | 19.66115 | 6.62831 | 8.83126 | 5.57174 | 11.71501 | 20.79331 | 14.00802 | 21.02036 | 7.21616 | 1.94232 | 18.55025 | 1.33378 | 12.58223 | 8.45051 |
| **Nubpl** | 0.88853 | 0.00771 | 0.76522 | 0.80589 | 0.03116 | 1.40631 | 0.01756 | 0.12809 | 0.00000 | 1.41474 | 0.33111 | 0.02114 | 1.19452 | 0.04165 | 0.54693 | 0.02110 |
| **Nucb1** | 10.00329 | 1.86494 | 6.08791 | 1.00854 | 5.44087 | 3.07641 | 6.79891 | 2.96468 | 5.46041 | 2.17738 | 0.72799 | 0.51581 | 10.66306 | 0.00000 | 0.59678 | 2.33955 |
| **Nucb2** | 4.26964 | 0.00000 | 9.81826 | 1.03486 | 2.13424 | 0.00000 | 7.07081 | 0.85805 | 1.93273 | 3.33947 | 0.60586 | 3.71684 | 2.97883 | 3.37100 | 5.06446 | 0.03914 |
| **Nucks1** | 25.19663 | 11.60064 | 14.91576 | 6.39551 | 16.57568 | 7.66160 | 12.44392 | 10.49818 | 16.84238 | 13.02711 | 11.35930 | 7.56951 | 13.47818 | 21.69550 | 12.33620 | 9.19173 |
| **Nudc** | 26.26964 | 4.64368 | 13.01025 | 11.80835 | 14.51956 | 8.42591 | 5.77552 | 17.52227 | 16.72912 | 22.04436 | 20.24776 | 7.90017 | 20.08329 | 2.01938 | 3.93656 | 12.55953 |
| **Nudcd1** | 2.93996 | 9.38098 | 3.26230 | 2.97958 | 8.67405 | 0.00000 | 5.19084 | 2.88995 | 4.84816 | 6.17028 | 4.42576 | 15.48652 | 3.87831 | 6.41009 | 3.25917 | 3.30545 |
| **Nudcd2** | 11.93284 | 4.47567 | 0.00000 | 7.80711 | 8.67094 | 5.60994 | 1.93982 | 12.36063 | 4.51419 | 16.65387 | 7.42549 | 1.69734 | 7.87801 | 7.42325 | 2.26029 | 8.45050 |
| **Nudcd3** | 5.15224 | 2.19440 | 3.87279 | 4.42319 | 3.93898 | 7.46704 | 6.30118 | 10.27042 | 1.38552 | 2.23746 | 2.49424 | 2.30146 | 5.02543 | 3.13957 | 1.89128 | 1.29767 |
| **Nudt1** | 4.40293 | 5.90447 | 2.28992 | 0.52135 | 0.00000 | 3.70383 | 3.58125 | 0.84152 | 0.08032 | 0.61071 | 3.16363 | 0.13367 | 2.50889 | 0.00000 | 0.76063 | 2.44767 |
| **Nudt10** | 0.96094 | 0.33346 | 0.00000 | 0.00771 | 2.94782 | 0.00793 | 1.46326 | 0.16108 | 0.01040 | 0.00074 | 0.01067 | 0.04876 | 0.00734 | 0.01033 | 0.14280 | 0.00607 |
| **Nudt11** | 0.24817 | 0.00609 | 0.24529 | 0.00818 | 1.91144 | 0.00721 | 0.18379 | 0.01244 | 0.01138 | 0.23056 | 0.01215 | 0.01360 | 0.01617 | 0.01241 | 0.00500 | 0.00704 |
| **Nudt12** | 2.17407 | 0.00500 | 0.22137 | 0.00777 | 0.30105 | 0.07608 | 0.15875 | 0.02729 | 0.03527 | 0.00000 | 0.00572 | 0.00001 | 0.02186 | 1.29701 | 0.11108 | 0.00491 |
| **Nudt13** | 2.15301 | 1.70228 | 1.25174 | 0.00369 | 0.00491 | 0.02174 | 1.26317 | 0.49127 | 0.02561 | 0.46688 | 1.65362 | 0.01390 | 0.50484 | 0.56141 | 0.10521 | 0.01062 |
| **Nudt14** | 1.60543 | 0.03389 | 1.26564 | 1.76268 | 1.94511 | 0.44888 | 0.06513 | 1.50932 | 0.42466 | 2.36795 | 1.05343 | 0.13128 | 0.00000 | 0.67481 | 0.05961 | 0.03374 |
| **Nudt15** | 2.70266 | 2.02805 | 0.68851 | 0.29936 | 0.47235 | 0.01150 | 0.91553 | 1.81588 | 0.00294 | 3.11962 | 0.00000 | 0.08424 | 0.24716 | 1.96160 | 1.80125 | 0.00366 |
| **Nudt16l1** | 4.06349 | 2.30213 | 4.93500 | 0.24208 | 7.49279 | 1.43288 | 4.39099 | 1.81694 | 0.01562 | 3.40543 | 4.53720 | 0.84956 | 3.89348 | 0.00232 | 0.27472 | 4.48208 |
| **Nudt17** | 1.16080 | 0.01851 | 0.07136 | 0.00400 | 0.04474 | 0.00546 | 0.87051 | 0.27849 | 0.01046 | 0.40292 | 1.19450 | 0.03223 | 0.21610 | 0.01017 | 0.00000 | 1.74105 |
| **Nudt18** | 0.00677 | 0.00574 | 0.96472 | 0.00000 | 2.93051 | 0.52760 | 0.99193 | 0.01954 | 0.24516 | 0.15406 | 0.00680 | 0.04843 | 0.01279 | 0.01723 | 0.02422 | 0.01410 |
| **Nudt19** | 6.44855 | 3.05853 | 1.43358 | 1.28665 | 5.07544 | 2.38853 | 1.56954 | 3.05771 | 0.69683 | 4.65848 | 4.62460 | 1.91263 | 9.29575 | 0.00000 | 1.57871 | 0.02238 |
| **Nudt2** | 6.83366 | 7.43110 | 9.30965 | 1.16681 | 2.64939 | 1.81464 | 1.84347 | 3.95840 | 0.40501 | 5.99503 | 3.59978 | 2.63644 | 5.21665 | 5.13878 | 1.41291 | 2.35813 |
| **Nudt21** | 16.43801 | 6.60717 | 2.51734 | 3.07390 | 10.24134 | 6.04808 | 0.60457 | 9.71578 | 9.04646 | 22.35812 | 14.57973 | 4.15116 | 14.72163 | 6.49986 | 4.04362 | 9.38317 |

| | | | | | | | | | | | | | | | | |
|---|---|---|---|---|---|---|---|---|---|---|---|---|---|---|---|---|
| **Nudt22** | 3.35988 | 0.01388 | 1.04759 | 3.84357 | 2.66092 | 1.98330 | 0.00283 | 1.66063 | 1.72440 | 1.82923 | 1.25522 | 0.36726 | 2.27812 | 0.00100 | 0.11490 | 2.15234 |
| **Nudt3** | 3.92787 | 1.36961 | 10.23391 | 2.69257 | 0.24472 | 0.44805 | 0.95426 | 9.14508 | 3.59826 | 9.58819 | 5.95501 | 0.59778 | 3.44190 | 2.97142 | 6.47587 | 0.82879 |
| **Nudt4** | 17.50583 | 14.00197 | 21.98696 | 23.76063 | 10.20121 | 11.04943 | 15.31947 | 23.44518 | 32.72657 | 40.06753 | 39.46636 | 34.07982 | 14.66174 | 38.33989 | 28.69942 | 5.37050 |
| **Nudt5** | 12.39273 | 9.57989 | 12.89464 | 6.66700 | 7.41706 | 0.00000 | 5.53899 | 1.29008 | 2.60752 | 0.00000 | 0.32827 | 0.05450 | 2.87682 | 0.52492 | 0.03655 | 4.75933 |
| **Nudt6** | 0.12369 | 0.00774 | 1.96632 | 0.17729 | 0.01187 | 0.00641 | 0.00678 | 0.02561 | 0.22376 | 1.92045 | 0.22836 | 0.01874 | 0.00340 | 0.00896 | 0.00000 | 0.00648 |
| **Nudt7** | 0.01837 | 0.00773 | 0.10637 | 1.56834 | 0.00825 | 0.01140 | 0.00714 | 0.00865 | 0.00000 | 0.48553 | 0.08201 | 0.02518 | 0.08281 | 0.00941 | 0.02462 | 0.31916 |
| **Nudt8** | 0.02268 | 0.00785 | 0.01263 | 0.00408 | 0.40286 | 0.00665 | 0.00702 | 0.01077 | 0.56016 | 0.16581 | 0.00769 | 0.00677 | 0.92539 | 0.01227 | 0.00906 | 1.41176 |
| **Nudt9** | 6.90465 | 1.25385 | 4.08723 | 0.96525 | 0.35414 | 1.54339 | 9.38163 | 14.15647 | 0.22729 | 7.22966 | 5.53424 | 0.82316 | 7.75846 | 0.08084 | 0.35414 | 2.55628 |
| **Nuf2** | 4.93408 | 3.34961 | 1.10664 | 2.98958 | 4.04985 | 3.79651 | 0.15516 | 4.23419 | 0.91620 | 4.45338 | 1.99388 | 0.65889 | 2.36681 | 6.06002 | 5.08691 | 0.01208 |
| **Nufip1** | 7.34239 | 5.69814 | 8.03573 | 12.02363 | 1.91399 | 0.02107 | 5.17195 | 9.19853 | 17.59215 | 24.94033 | 8.91114 | 12.19599 | 8.88484 | 7.76852 | 19.19818 | 5.45346 |
| **Nufip2** | 1.66479 | 1.59311 | 2.89267 | 6.91621 | 0.01796 | 1.94033 | 1.25481 | 3.00719 | 13.03202 | 1.62107 | 7.09723 | 7.50127 | 1.61758 | 15.15341 | 9.43262 | 3.00737 |
| **Numa1** | 9.10914 | 1.60602 | 4.26849 | 5.08737 | 1.34274 | 12.57156 | 4.35978 | 7.49922 | 4.09033 | 6.36557 | 2.79214 | 0.29267 | 9.25271 | 0.97766 | 3.45063 | 2.58985 |
| **Numb** | 0.11201 | 0.03573 | 0.00000 | 3.21887 | 2.91563 | 0.03159 | 0.00654 | 0.05801 | 5.38646 | 0.77243 | 0.56829 | 0.58983 | 0.16595 | 0.12256 | 0.14410 | 0.09437 |
| **Numbl** | 0.94200 | 0.00732 | 0.28460 | 0.74616 | 0.01169 | 0.00835 | 0.00642 | 0.01512 | 0.09854 | 2.09070 | 0.27638 | 0.01040 | 0.28196 | 0.00982 | 0.01759 | 0.32510 |
| **Nup107** | 2.63994 | 3.39090 | 0.76878 | 5.78566 | 3.09850 | 2.43574 | 0.12372 | 2.99270 | 6.19315 | 4.45155 | 3.14110 | 7.49780 | 0.45697 | 9.82195 | 17.05750 | 0.20759 |
| **Nup133** | 10.41727 | 7.77524 | 9.36613 | 8.68286 | 4.97879 | 4.97818 | 6.02028 | 13.39757 | 9.57744 | 14.83568 | 12.69435 | 4.98167 | 10.36727 | 9.55451 | 10.59201 | 7.74612 |
| **Nup153** | 1.65740 | 5.18118 | 5.33590 | 2.59378 | 7.95895 | 3.14347 | 8.55160 | 9.51607 | 5.68950 | 14.20026 | 3.25757 | 20.22108 | 3.62330 | 15.26682 | 14.66031 | 3.85942 |
| **Nup155** | 5.02564 | 5.77031 | 2.90071 | 8.96234 | 1.68024 | 0.02634 | 3.82853 | 9.04448 | 16.92719 | 8.82417 | 7.00464 | 8.77116 | 3.45954 | 20.33815 | 13.63334 | 0.89470 |
| **Nup160** | 1.92720 | 8.25163 | 4.04461 | 2.58957 | 3.69979 | 3.68837 | 3.42614 | 7.22173 | 5.20147 | 9.67033 | 2.58134 | 11.29642 | 1.99022 | 14.75391 | 15.05652 | 0.80308 |
| **Nup188** | 5.08434 | 0.05417 | 0.00769 | 6.35538 | 3.60840 | 4.12809 | 1.04693 | 5.70976 | 3.41927 | 7.81914 | 3.05218 | 0.21184 | 1.05343 | 0.04033 | 0.32350 | 2.12811 |
| **Nup205** | 10.03897 | 7.87701 | 8.58115 | 2.12365 | 8.39954 | 0.50212 | 5.82264 | 12.86148 | 3.98968 | 10.65508 | 10.18461 | 6.51965 | 6.91334 | 16.57427 | 12.18852 | 3.82094 |
| **Nup210** | 3.98021 | 2.22038 | 0.39920 | 0.36772 | 0.00000 | 0.00428 | 0.00291 | 1.29634 | 0.62542 | 0.00000 | 0.00000 | 0.18334 | 0.71402 | 0.09529 | 0.00000 | 0.00177 |
| **Nup210l** | 0.01344 | 0.00804 | 0.00792 | 0.45221 | 0.00905 | 0.00669 | 0.00740 | 0.00578 | 2.26668 | 0.17224 | 0.17360 | 0.06194 | 0.01098 | 0.00689 | 0.63810 | 0.00714 |
| **Nup214** | 2.75520 | 3.97678 | 3.78134 | 14.44478 | 5.64944 | 0.80200 | 4.24886 | 6.19468 | 14.04101 | 10.05906 | 4.60184 | 0.35760 | 5.63872 | 0.00000 | 3.76143 | 3.12605 |
| **Nup35** | 12.64860 | 10.66070 | 14.64073 | 18.42459 | 9.24399 | 9.33780 | 5.54086 | 17.96814 | 17.72317 | 25.72783 | 14.53904 | 17.25546 | 10.98049 | 24.44537 | 25.48387 | 2.56472 |
| **Nup37** | 11.28834 | 3.90638 | 8.84159 | 2.84224 | 5.36753 | 2.76559 | 8.25279 | 8.41445 | 1.67385 | 8.52356 | 5.51053 | 6.62042 | 7.26435 | 5.48595 | 5.88389 | 3.75110 |
| **Nup43** | 11.53763 | 12.91295 | 6.58812 | 7.21411 | 3.98304 | 4.15535 | 5.71936 | 10.48858 | 20.76493 | 20.36941 | 12.70033 | 13.77071 | 14.81607 | 7.45529 | 13.41114 | 2.32737 |
| **Nup50** | 11.27109 | 7.28325 | 9.92960 | 14.25280 | 6.63964 | 3.28772 | 5.35913 | 8.10225 | 15.68226 | 14.75019 | 10.30507 | 8.83402 | 11.78014 | 11.86487 | 9.63405 | 6.29182 |
| **Nup54** | 7.46193 | 7.05770 | 1.64511 | 2.62741 | 7.00017 | 0.09896 | 1.24319 | 7.54008 | 2.62820 | 5.26118 | 2.40156 | 5.05179 | 1.27382 | 6.79891 | 9.82603 | 0.24956 |
| **Nup62** | 14.14169 | 2.82059 | 12.84403 | 20.22420 | 8.88412 | 7.50555 | 6.60704 | 12.42630 | 16.53186 | 8.71964 | 10.03891 | 1.42333 | 12.90816 | 0.00000 | 4.24911 | 12.02716 |
| **Nup62cl** | 0.00000 | 6.68208 | 0.01424 | 1.64651 | 0.00867 | 0.01698 | 0.98073 | 0.00677 | 1.67924 | 1.05297 | 0.00000 | 0.00000 | 0.44662 | 1.28125 | 0.00000 | 2.03729 |
| **Nup85** | 14.72123 | 13.67436 | 7.99008 | 7.32933 | 6.14943 | 5.86245 | 8.28645 | 16.93210 | 1.77977 | 18.59768 | 15.86769 | 15.17831 | 9.41676 | 7.46506 | 17.84351 | 9.95345 |
| **Nup88** | 7.48603 | 1.85372 | 3.76332 | 2.22639 | 7.51297 | 3.49642 | 5.62370 | 6.48123 | 4.54722 | 2.43201 | 9.40918 | 0.96813 | 3.92771 | 11.79460 | 8.72211 | 1.44550 |
| **Nup93** | 7.86159 | 4.65060 | 6.91348 | 1.87775 | 8.12543 | 5.62949 | 6.02253 | 9.78599 | 5.32630 | 8.01856 | 2.44396 | 9.04683 | 5.63642 | 3.89588 | 7.74471 | 6.16689 |
| **Nup98** | 2.40074 | 4.55159 | 2.10626 | 13.47101 | 3.56735 | 3.42711 | 0.21312 | 1.13653 | 18.38742 | 2.40632 | 11.25421 | 2.11678 | 0.64113 | 9.55668 | 11.31568 | 0.27465 |
| **Nupl1** | 1.34491 | 2.01176 | 1.13958 | 0.92640 | 1.26003 | 0.99971 | 0.33695 | 1.38965 | 0.05637 | 0.32765 | 0.06406 | 4.63872 | 0.38101 | 0.86852 | 2.96693 | 0.44534 |
| **Nupl2** | 1.10488 | 0.01325 | 1.71439 | 2.48234 | 2.90448 | 0.04477 | 2.62775 | 1.81535 | 3.33949 | 2.86325 | 1.12705 | 0.17108 | 0.94687 | 0.16990 | 1.02831 | 1.38303 |
| **Nupr1** | 0.00537 | 0.00724 | 0.00000 | 0.59620 | 0.00718 | 0.00633 | 0.00608 | 0.00000 | 3.49499 | 2.87542 | 2.31576 | 0.53211 | 0.03868 | 0.08735 | 1.36865 | 1.85298 |
| **Nupr1l** | 0.01368 | 0.00808 | 0.01020 | 0.82244 | 0.00916 | 0.00652 | 0.00738 | 0.00996 | 1.15252 | 0.13615 | 0.05213 | 0.08562 | 0.01127 | 0.00957 | 0.00000 | 0.00698 |
| **Nus1** | 6.71225 | 6.67801 | 4.65045 | 16.52357 | 11.45706 | 0.75985 | 1.19106 | 3.03137 | 17.62272 | 3.02937 | 8.60318 | 3.68999 | 3.41121 | 9.59333 | 10.83132 | 1.35605 |
| **Nusap1** | 4.29332 | 4.29380 | 4.86130 | 5.50115 | 0.00771 | 0.01632 | 0.00000 | 4.15122 | 10.25964 | 5.35637 | 3.27697 | 5.83223 | 0.99141 | 8.41860 | 6.10846 | 2.33482 |
| **Nutf2** | 2.60200 | 0.09190 | 2.29919 | 0.00377 | 0.00764 | 0.00450 | 0.42003 | 0.56348 | 0.33794 | 0.10819 | 0.00005 | 0.03862 | 0.05395 | 0.00561 | 0.19706 | 0.01104 |
| **Nutm1** | 0.10727 | 0.00796 | 0.00429 | 0.56641 | 0.00764 | 0.00696 | 0.00774 | 0.07086 | 2.92011 | 1.39812 | 0.14140 | 0.02910 | 0.03149 | 0.00347 | 0.41303 | 0.00695 |
| **Nvl** | 5.58262 | 9.45080 | 6.32492 | 6.70682 | 9.25288 | 0.06701 | 5.32845 | 8.47913 | 8.86695 | 11.80669 | 7.81462 | 6.32116 | 5.12327 | 15.09506 | 14.25461 | 1.91038 |
| **Nwd1** | 0.01320 | 0.00794 | 0.07661 | 0.04418 | 0.00986 | 0.00794 | 0.00758 | 0.01158 | 0.04242 | 0.04664 | 0.37535 | 0.02665 | 0.07877 | 0.01531 | 0.02546 | 0.00747 |
| **Nxf1** | 13.43827 | 7.35108 | 6.86344 | 10.77773 | 9.54864 | 8.66367 | 6.69587 | 15.63968 | 24.06631 | 26.12842 | 13.31845 | 12.72495 | 9.31324 | 22.59495 | 24.09891 | 7.28161 |
| **Nxf2** | 0.01295 | 0.00782 | 0.00879 | 0.00000 | 0.00958 | 0.00609 | 0.00726 | 0.01020 | 0.00661 | 0.14648 | 0.00649 | 0.03840 | 0.01286 | 0.00612 | 4.16575 | 0.00745 |
| **Nxf7** | 0.01143 | 0.00774 | 0.90121 | 0.01099 | 0.01683 | 0.00927 | 0.00729 | 0.01222 | 0.01220 | 0.01418 | 0.01140 | 0.00739 | 0.00817 | 0.01569 | 0.00000 | 0.00702 |
| **Nxn** | 2.63690 | 0.00468 | 0.23193 | 0.87311 | 0.00001 | 0.00600 | 0.00517 | 0.00997 | 0.00000 | 3.73340 | 0.03419 | 0.14770 | 0.14616 | 0.01223 | 0.06045 | 4.09193 |
| **Nxnl1** | 0.01778 | 0.00816 | 0.01174 | 0.11246 | 0.00897 | 0.00658 | 0.00747 | 0.01354 | 0.45956 | 0.00846 | 0.01079 | 0.00759 | 0.01273 | 0.00790 | 0.11172 | 0.00731 |
| **Nxnl2** | 0.01827 | 0.00828 | 0.01169 | 0.00961 | 0.01040 | 0.00691 | 0.00757 | 0.01320 | 0.01207 | 0.23947 | 0.01196 | 0.00871 | 0.01279 | 0.01163 | 0.01182 | 0.00746 |
| **Nxpe3** | 0.01730 | 0.81023 | 0.17900 | 0.41473 | 0.00797 | 1.13423 | 0.63666 | 0.00000 | 0.01130 | 0.00000 | 0.96060 | 0.04096 | 0.01384 | 0.04297 | 0.09501 | 0.00613 |
| **Nxpe4** | 0.85309 | 1.76234 | 0.00000 | 1.56015 | 0.00607 | 0.00654 | 0.02531 | 0.00687 | 0.00000 | 0.77816 | 0.02673 | 0.01490 | 0.01592 | 0.00522 | 0.00000 | 0.00516 |
| **Nxpe5** | 0.01334 | 0.00814 | 0.01199 | 0.01896 | 0.00934 | 0.00667 | 0.00763 | 0.02325 | 0.01119 | 0.01515 | 0.01260 | 0.00824 | 0.01207 | 0.01249 | 0.01024 | 0.00741 |
| **Nxt1** | 7.32301 | 0.21673 | 1.76667 | 11.43970 | 7.59159 | 0.07711 | 2.25012 | 9.67559 | 17.28812 | 14.92599 | 4.29351 | 2.56414 | 6.00726 | 1.74587 | 0.38058 | 5.59205 |
| **Nxt2** | 0.09592 | 1.31905 | 0.01134 | 2.87889 | 0.01073 | 0.02350 | 0.07649 | 0.00016 | 4.24750 | 0.00000 | 0.34832 | 0.05860 | 0.07212 | 0.47661 | 4.54899 | 0.00568 |
| **Nyap1** | 0.00929 | 0.00787 | 0.00624 | 0.15099 | 0.00951 | 0.00683 | 0.00718 | 0.24948 | 0.20716 | 0.41897 | 0.20831 | 0.00340 | 0.03875 | 0.07413 | 0.50573 | 0.00712 |
| **Nynrin** | 6.16802 | 6.92343 | 3.06626 | 0.00648 | 6.10379 | 2.19591 | 0.05947 | 0.75828 | 0.00897 | 0.00000 | 0.00331 | 0.05537 | 4.56443 | 0.00546 | 0.04433 | 3.96141 |
| **Nyx** | 0.01378 | 0.00756 | 0.00743 | 0.07122 | 0.00915 | 0.00682 | 0.00709 | 0.18017 | 0.25977 | 0.00540 | 0.00639 | 0.54433 | 0.00760 | 0.03115 | 0.01870 | 0.00741 |
| **Oacyl** | 0.00953 | 0.00850 | 0.01131 | 1.73602 | 0.01056 | 0.00721 | 0.00823 | 0.20286 | 0.00000 | 0.00306 | 0.02805 | 0.02903 | 0.24681 | 0.00467 | 0.11540 | 0.00732 |
| **Oaf** | 4.12962 | 0.00336 | 0.00242 | 3.12323 | 0.00000 | 0.00452 | 0.00418 | 0.00000 | 0.47643 | 0.02536 | 0.00742 | 0.01511 | 0.01151 | 0.00360 | 0.33679 | 0.00248 |
| **Oard1** | 12.23473 | 7.93148 | 12.24101 | 1.82536 | 10.36217 | 3.86727 | 5.91399 | 6.24787 | 1.03238 | 4.23822 | 2.98125 | 0.30728 | 9.93116 | 0.01596 | 3.89987 | 8.41985 |

| | | | | | | | | | | | | | | | | |
|---|---|---|---|---|---|---|---|---|---|---|---|---|---|---|---|---|
| **Oas1a** | 2.70875 | 3.16849 | 2.37023 | 0.08472 | 0.88844 | 1.95879 | 2.80390 | 0.39320 | 0.00648 | 0.00000 | 0.33154 | 0.03702 | 1.38072 | 0.01120 | 0.00000 | 5.03639 |
| **Oas1b** | 0.01710 | 0.04903 | 0.04333 | 0.41682 | 0.03491 | 0.02112 | 0.07048 | 0.02614 | 0.44105 | 0.39120 | 0.01822 | 0.00634 | 0.03940 | 0.00952 | 0.32322 | 0.04331 |
| **Oas1c** | 0.02743 | 0.01752 | 0.04460 | 3.06378 | 0.00861 | 0.22601 | 0.02713 | 0.01013 | 0.63218 | 0.78066 | 0.02933 | 0.03558 | 0.01064 | 0.01849 | 2.49891 | 0.02113 |
| **Oas1d** | 0.37990 | 0.10483 | 0.97317 | 13.57691 | 0.01478 | 2.31550 | 0.99054 | 0.42127 | 5.80701 | 1.73814 | 2.21667 | 0.20115 | 0.00000 | 0.01030 | 7.20487 | 0.01759 |
| **Oas1e** | 0.02772 | 0.00750 | 0.06258 | 2.60625 | 0.00837 | 0.14836 | 0.00674 | 0.00000 | 0.24266 | 0.30762 | 0.49149 | 0.03684 | 0.33433 | 0.46121 | 1.53092 | 0.00623 |
| **Oas1f** | 0.01217 | 0.00777 | 1.14260 | 0.65250 | 0.00744 | 0.01845 | 0.00727 | 0.01227 | 0.00000 | 0.01581 | 0.01092 | 0.00702 | 0.00927 | 0.01246 | 0.01639 | 0.00691 |
| **Oas1g** | 2.02709 | 2.71756 | 2.17503 | 0.11478 | 0.63574 | 1.39861 | 2.42752 | 0.23106 | 0.00615 | 0.00000 | 0.51545 | 0.00776 | 1.96621 | 0.01692 | 0.00000 | 4.39544 |
| **Oas1h** | 0.01368 | 0.00775 | 0.01140 | 2.46695 | 0.00944 | 0.00619 | 0.00726 | 0.01101 | 0.02986 | 0.00597 | 0.01004 | 0.00569 | 0.01280 | 0.00410 | 1.05472 | 0.00691 |
| **Oas3** | 0.17923 | 0.06150 | 0.20533 | 1.36528 | 0.01897 | 0.05346 | 0.08107 | 0.09298 | 1.59719 | 0.11978 | 0.07909 | 0.45863 | 0.05479 | 0.36376 | 0.17647 | 0.04380 |
| **Oasl1** | 0.01364 | 0.00823 | 0.01207 | 1.07779 | 0.00930 | 0.00651 | 0.00750 | 0.01388 | 0.74590 | 0.01096 | 0.00870 | 0.00707 | 0.01284 | 0.01057 | 0.00549 | 0.00719 |
| **Oasl2** | 0.03845 | 0.00695 | 0.00000 | 0.29860 | 0.23222 | 0.00814 | 0.00705 | 3.06778 | 0.49865 | 2.11169 | 0.00000 | 1.49131 | 0.01572 | 0.05291 | 0.00000 | 0.01501 |
| **Oat** | 0.07074 | 0.01532 | 0.00000 | 6.02190 | 0.08823 | 2.50033 | 5.14327 | 0.00000 | 10.89506 | 5.98965 | 0.00000 | 0.03919 | 0.91662 | 0.00001 | 6.23928 | 0.01912 |
| **Oaz1** | 1.86252 | 0.28482 | 0.53055 | 4.37122 | 1.12123 | 0.36765 | 0.36654 | 0.46035 | 2.07061 | 1.10596 | 1.13342 | 0.05507 | 1.26888 | 0.05753 | 0.00000 | 1.27444 |
| **Oaz2** | 7.97822 | 2.21675 | 2.80046 | 10.07851 | 1.96795 | 5.17776 | 3.12271 | 6.17308 | 15.75778 | 7.32904 | 5.60524 | 0.39476 | 5.07438 | 0.71575 | 2.14752 | 5.72949 |
| **Oaz3** | 0.01373 | 0.00800 | 0.01184 | 0.77489 | 0.00935 | 0.00646 | 0.00740 | 0.01219 | 0.29637 | 0.00637 | 0.01102 | 0.00724 | 0.01277 | 0.00699 | 0.34754 | 0.00721 |
| **Obfc1** | 1.14198 | 1.44202 | 1.45936 | 2.09311 | 4.27972 | 1.04150 | 1.79570 | 1.81955 | 2.35953 | 3.26113 | 2.35115 | 0.35047 | 0.81147 | 0.00000 | 1.15435 | 0.01055 |
| **Obox1** | 0.00778 | 0.00831 | 0.00000 | 9.04955 | 0.00965 | 0.00546 | 0.00725 | 0.03069 | 2.73561 | 0.00000 | 0.75126 | 0.02044 | 0.00498 | 2.66247 | 1.32222 | 0.00533 |
| **Obox2** | 0.00944 | 0.02962 | 0.00000 | 8.29575 | 0.00919 | 0.00540 | 0.00671 | 0.00000 | 2.55967 | 0.00000 | 0.78111 | 0.00000 | 0.00544 | 1.99049 | 2.12839 | 0.00538 |
| **Obox3** | 0.00664 | 0.00757 | 0.00228 | 29.88221 | 0.00840 | 0.00178 | 0.01585 | 0.00000 | 33.24919 | 0.66957 | 0.65180 | 0.00000 | 0.01438 | 0.40544 | 3.25393 | 0.00000 |
| **Obox5** | 0.01116 | 0.01763 | 0.00000 | 19.59177 | 0.00880 | 0.00403 | 0.00629 | 0.00000 | 4.94202 | 1.31465 | 0.58623 | 0.00000 | 0.00672 | 0.00154 | 4.41432 | 0.00343 |
| **Obox6** | 0.00000 | 7.34359 | 8.90102 | 20.46999 | 0.02811 | 0.00000 | 2.22162 | 18.02580 | 31.82791 | 39.96998 | 29.70760 | 22.01784 | 3.45179 | 39.71804 | 31.43383 | 0.03680 |
| **Obox7** | 0.00819 | 0.46780 | 0.00522 | 5.26106 | 0.00821 | 0.00590 | 0.00692 | 0.00354 | 2.86931 | 0.05735 | 0.18931 | 0.00000 | 0.01004 | 0.00000 | 1.25878 | 0.00606 |
| **Obp2a** | 0.01244 | 0.00794 | 0.41590 | 0.01079 | 0.00869 | 0.00936 | 0.00744 | 0.01347 | 0.01227 | 0.01474 | 0.01264 | 0.00766 | 0.01341 | 0.01397 | 0.00398 | 0.00733 |
| **Obsl1** | 0.04783 | 0.01824 | 0.00975 | 0.00607 | 0.00807 | 0.00682 | 0.00694 | 0.01069 | 0.07930 | 0.34383 | 0.01219 | 0.01553 | 0.63475 | 0.01137 | 0.00937 | 0.00746 |
| **Oc90** | 0.00821 | 0.03670 | 0.00757 | 0.01005 | 0.00948 | 0.00683 | 0.00705 | 1.54995 | 0.01098 | 0.22504 | 0.00000 | 0.00341 | 0.01209 | 0.04249 | 0.00000 | 0.00744 |
| **Ocel1** | 1.49398 | 0.02729 | 0.00429 | 1.16069 | 0.00613 | 0.00644 | 0.00585 | 0.00000 | 0.00000 | 0.41673 | 0.58162 | 0.03249 | 0.02323 | 0.00529 | 0.00000 | 0.00580 |
| **Ociad1** | 17.92040 | 10.81629 | 17.15913 | 6.71648 | 9.31784 | 4.57519 | 7.54341 | 19.24390 | 7.43180 | 12.51617 | 10.98757 | 2.30556 | 11.21902 | 5.85095 | 10.40425 | 7.97664 |
| **Ociad2** | 1.89875 | 0.06852 | 0.20595 | 2.10247 | 0.17763 | 0.04900 | 0.13838 | 0.06190 | 3.32550 | 0.09451 | 2.19522 | 0.08459 | 0.12823 | 1.74769 | 0.15699 | 1.29422 |
| **Ocln** | 0.74063 | 0.00435 | 0.00000 | 0.25224 | 0.01414 | 0.79135 | 3.36958 | 2.72090 | 1.28378 | 1.99096 | 0.17426 | 0.38852 | 0.00000 | 4.47099 | 0.71561 | 0.00591 |
| **Ocm** | 1.50563 | 0.00348 | 2.82467 | 2.40605 | 0.71286 | 0.00442 | 0.20500 | 4.39053 | 0.00000 | 2.85348 | 5.35490 | 2.60783 | 0.69818 | 3.00529 | 1.76871 | 0.00290 |
| **Ocrl** | 0.81450 | 0.05736 | 1.88691 | 0.73715 | 3.59471 | 3.45387 | 2.77532 | 2.13489 | 0.00000 | 2.61588 | 3.49692 | 9.42048 | 3.99020 | 10.52953 | 0.71691 | 3.00990 |
| **Odc1** | 24.43373 | 15.34287 | 36.06267 | 30.34730 | 13.84901 | 18.55954 | 23.04900 | 41.04216 | 38.80300 | 57.41895 | 49.43665 | 34.10814 | 33.34066 | 11.92751 | 34.58604 | 14.47777 |
| **Odf2** | 2.83106 | 0.33099 | 2.79698 | 6.95618 | 7.71630 | 0.44501 | 0.58612 | 2.10122 | 8.78853 | 2.24040 | 2.27824 | 0.15412 | 2.95675 | 0.00000 | 1.09820 | 0.02930 |
| **Odf2l** | 0.16835 | 0.03612 | 0.00000 | 0.20730 | 0.66883 | 0.00562 | 0.26127 | 0.05386 | 0.00532 | 0.00000 | 0.02896 | 3.31291 | 0.01149 | 0.00729 | 3.30749 | 0.00670 |
| **Odf3b** | 0.01361 | 0.00794 | 0.01238 | 0.23968 | 0.00934 | 0.00649 | 0.00732 | 0.01139 | 0.00676 | 0.34330 | 0.01320 | 0.00677 | 0.01207 | 0.01039 | 0.00822 | 0.00743 |
| **Ofd1** | 4.61934 | 1.13856 | 0.99230 | 6.09799 | 0.73679 | 0.78418 | 3.45936 | 2.20333 | 2.61849 | 1.65022 | 2.06007 | 0.11922 | 3.67667 | 6.43978 | 7.67362 | 0.02009 |
| **Ogdh** | 13.53252 | 4.75166 | 15.76207 | 17.88086 | 10.59621 | 6.61919 | 7.55072 | 12.00028 | 27.15384 | 12.41920 | 24.42247 | 7.56542 | 12.60438 | 14.51959 | 10.37573 | 6.92770 |
| **Ogdhl** | 0.02207 | 0.01872 | 0.21747 | 0.17766 | 0.00758 | 0.00923 | 0.01777 | 0.01429 | 0.00607 | 0.00535 | 0.01292 | 0.00747 | 0.46255 | 0.01277 | 0.00290 | 1.13471 |
| **Ogfod1** | 6.12600 | 3.44987 | 1.85420 | 1.69614 | 2.97870 | 1.64738 | 2.89576 | 3.24862 | 0.22047 | 9.53075 | 1.66687 | 4.21800 | 1.62346 | 1.74880 | 5.83483 | 0.00000 |
| **Ogfod2** | 0.08468 | 0.00685 | 0.22781 | 0.09075 | 0.00688 | 0.37105 | 0.00546 | 0.60116 | 0.75144 | 1.27525 | 2.95112 | 0.07843 | 0.73934 | 0.23784 | 0.00000 | 1.93819 |
| **Ogfod3** | 3.27559 | 0.27337 | 4.85722 | 0.00298 | 0.01333 | 0.01082 | 0.61316 | 2.43602 | 0.90987 | 1.97894 | 0.82816 | 0.15876 | 2.02890 | 0.24531 | 0.04839 | 0.00840 |
| **Ogfr** | 1.69448 | 0.44028 | 0.77604 | 0.00000 | 2.65054 | 3.62462 | 4.19298 | 0.03780 | 0.33464 | 4.05002 | 0.25349 | 0.27121 | 4.56778 | 0.01185 | 0.00000 | 5.39248 |
| **Ogfrl1** | 0.86161 | 0.31667 | 1.85099 | 2.09681 | 0.00391 | 0.00585 | 4.06197 | 0.07437 | 1.01906 | 0.11160 | 0.00000 | 0.04018 | 0.04415 | 4.49746 | 0.53921 | 0.01249 |
| **Ogg1** | 0.68948 | 0.52247 | 0.80387 | 0.18711 | 1.45457 | 0.00688 | 0.00583 | 0.41583 | 0.15666 | 0.18198 | 0.11094 | 0.01408 | 1.01875 | 0.00894 | 0.00000 | 0.18961 |
| **Ogn** | 0.01340 | 0.00769 | 0.00594 | 0.00805 | 0.00892 | 0.00712 | 0.00718 | 0.00000 | 0.59270 | 0.49998 | 0.76293 | 0.00197 | 0.00654 | 0.01019 | 0.58776 | 0.00719 |
| **Ogt** | 6.29606 | 7.83187 | 9.55709 | 6.33237 | 5.02662 | 0.72992 | 8.58408 | 11.31971 | 5.68078 | 5.34065 | 2.94045 | 0.35424 | 3.28597 | 18.00942 | 10.47507 | 2.86273 |
| **Oip5** | 0.41635 | 0.01650 | 0.10564 | 3.13322 | 0.07829 | 0.03443 | 0.61315 | 0.67402 | 4.03134 | 1.02519 | 3.32990 | 0.05997 | 2.93716 | 1.33825 | 0.12365 | 1.45304 |
| **Ola1** | 7.55844 | 5.25441 | 5.40010 | 4.94409 | 8.03064 | 5.79954 | 4.25999 | 9.84894 | 8.14946 | 16.56127 | 11.30608 | 9.96725 | 13.63980 | 15.88551 | 11.33451 | 8.26960 |
| **Olah** | 0.01372 | 0.00798 | 0.01164 | 0.54187 | 0.00944 | 0.00640 | 0.00734 | 0.01137 | 0.00022 | 0.00579 | 0.01139 | 0.00687 | 0.01278 | 0.00662 | 0.74129 | 0.00726 |
| **Olfm1** | 0.01647 | 0.00786 | 0.00000 | 4.30120 | 0.00896 | 1.09074 | 0.00724 | 0.61482 | 0.35509 | 0.78568 | 0.05818 | 0.03830 | 0.00000 | 0.94531 | 1.88811 | 0.00622 |
| **Olfml1** | 0.25320 | 0.00842 | 0.00672 | 0.19132 | 0.01832 | 0.00694 | 0.00721 | 0.01120 | 0.00591 | 0.11828 | 0.01223 | 0.01716 | 0.01236 | 0.01077 | 0.00637 | 0.00742 |
| **Olfml2a** | 0.01280 | 0.00783 | 0.01108 | 0.40032 | 0.00911 | 0.00670 | 0.00745 | 0.02278 | 0.00579 | 0.57914 | 0.02742 | 0.00594 | 0.01148 | 0.00466 | 0.66567 | 0.00732 |
| **Olfr1033** | 0.01483 | 0.00796 | 0.05309 | 0.02359 | 0.00893 | 0.04632 | 0.00764 | 0.04185 | 0.01229 | 0.23952 | 0.00958 | 0.23022 | 0.00984 | 0.00731 | 0.16996 | 0.00744 |
| **Olfr1055** | 0.10894 | 0.11043 | 0.53887 | 1.35283 | 0.18159 | 0.29580 | 0.14698 | 1.55963 | 1.79666 | 1.68895 | 1.07927 | 2.89808 | 0.29001 | 1.15075 | 0.75001 | 0.06397 |
| **Olfr118** | 0.01377 | 0.00816 | 0.01210 | 0.62219 | 0.00933 | 0.00661 | 0.00750 | 0.01370 | 0.00001 | 0.01507 | 0.01355 | 0.01883 | 0.01269 | 0.01096 | 0.00423 | 0.00731 |
| **Olfr1300.ps1** | 0.01380 | 0.00784 | 0.01161 | 0.49588 | 0.00927 | 0.00654 | 0.00745 | 0.01106 | 0.02259 | 0.00503 | 0.09151 | 0.00696 | 0.01119 | 0.00538 | 0.15478 | 0.00726 |
| **Olfr1372.ps1** | 0.01160 | 0.01853 | 0.08879 | 0.01696 | 0.00895 | 0.00589 | 0.00747 | 0.46889 | 0.04680 | 0.37503 | 0.03347 | 0.17306 | 0.07069 | 0.37979 | 0.22859 | 0.00718 |
| **Olfr287** | 0.01466 | 0.00801 | 0.03509 | 0.39500 | 0.00939 | 0.01306 | 0.00749 | 0.01347 | 0.18026 | 0.01440 | 0.01140 | 0.00752 | 0.01499 | 0.03539 | 0.02917 | 0.00738 |
| **Olfr376** | 0.01367 | 0.00828 | 0.01213 | 1.08610 | 0.00931 | 0.00654 | 0.00750 | 0.01395 | 0.50992 | 0.01210 | 0.00949 | 0.00761 | 0.01281 | 0.01092 | 0.00313 | 0.00721 |
| **Olfr456** | 0.01382 | 0.00800 | 0.01112 | 0.18303 | 0.00921 | 0.00665 | 0.00752 | 0.01068 | 0.00724 | 0.01354 | 0.13253 | 0.00738 | 0.01053 | 0.01036 | 0.03782 | 0.00725 |
| **Olfr613** | 0.30899 | 0.24686 | 0.42722 | 0.93744 | 0.03943 | 0.10216 | 0.34317 | 0.29613 | 1.38988 | 0.21719 | 0.01444 | 0.44491 | 0.24310 | 1.41846 | 0.52698 | 0.05630 |
| **Olfr787** | 0.01358 | 0.00822 | 0.01189 | 0.33470 | 0.00928 | 0.00657 | 0.00753 | 0.01386 | 1.13694 | 0.01017 | 0.00613 | 0.00649 | 0.01296 | 0.01210 | 0.00366 | 0.00729 |

| | | | | | | | | | | | | | | | | |
|---|---|---|---|---|---|---|---|---|---|---|---|---|---|---|---|---|
| **Olfr815** | 0.01360 | 0.00824 | 0.01158 | 0.91676 | 0.00930 | 0.00654 | 0.00748 | 0.01341 | 1.18064 | 0.00884 | 0.00273 | 0.02840 | 0.01310 | 0.01143 | 0.00187 | 0.00720 |
| **Olfr856.ps1** | 0.04031 | 0.04943 | 0.07749 | 0.65609 | 0.00793 | 0.05788 | 0.08035 | 0.39374 | 0.41349 | 0.14575 | 0.47460 | 0.01059 | 0.08506 | 0.71877 | 0.42150 | 0.02102 |
| **Olfr976** | 0.01372 | 0.00762 | 0.01137 | 0.77494 | 0.00940 | 0.00644 | 0.00738 | 0.01162 | 0.56196 | 0.00220 | 0.00718 | 0.00593 | 0.01291 | 0.02199 | 0.40806 | 0.00719 |
| **Oma1** | 3.49492 | 3.46055 | 5.40719 | 0.62008 | 5.23096 | 0.01509 | 1.50325 | 0.17371 | 0.67802 | 0.85965 | 0.00000 | 0.38023 | 1.60241 | 1.27795 | 0.80669 | 6.61574 |
| **Omd** | 0.77885 | 0.06960 | 0.61809 | 4.23434 | 0.36118 | 0.10695 | 0.10576 | 0.00000 | 7.36720 | 0.00000 | 7.29437 | 0.37638 | 0.29793 | 2.25465 | 0.09774 | 0.13873 |
| **Omt2a** | 0.00000 | 0.00717 | 0.00000 | 49.27958 | 0.00825 | 0.00000 | 0.00435 | 0.00000 | 28.40761 | 4.48718 | 2.49264 | 0.30188 | 0.00000 | 0.68926 | 17.24816 | 0.00394 |
| **Omt2b** | 0.00000 | 0.00724 | 0.00000 | 49.04464 | 0.00853 | 0.00000 | 0.00423 | 0.00000 | 25.13610 | 2.39586 | 3.04744 | 0.27611 | 0.00000 | 0.85694 | 17.42256 | 0.00000 |
| **Onecut2** | 0.01387 | 0.00764 | 0.01066 | 0.04875 | 0.00930 | 0.00667 | 0.00740 | 0.01145 | 0.05763 | 0.01552 | 0.02445 | 0.05828 | 0.01240 | 0.02271 | 0.17413 | 0.00731 |
| **Onecut3** | 0.02622 | 0.00822 | 0.02235 | 0.06182 | 0.00969 | 0.00678 | 0.00739 | 0.01182 | 0.00913 | 0.00655 | 0.02391 | 0.00740 | 0.01215 | 0.01004 | 0.44807 | 0.00732 |
| **Ooep** | 6.89113 | 1.33113 | 2.60152 | 2.43480 | 1.82671 | 0.04634 | 0.88555 | 0.51544 | 0.70361 | 0.10178 | 1.99830 | 0.04521 | 1.71984 | 0.39569 | 0.56094 | 6.19377 |
| **Oog2** | 0.01550 | 0.00747 | 0.00000 | 7.21579 | 0.00887 | 0.00556 | 0.00674 | 0.00000 | 0.12923 | 0.71146 | 0.28705 | 0.01148 | 0.00895 | 2.26848 | 3.39705 | 0.00580 |
| **Oog3** | 0.00000 | 0.00751 | 0.00000 | 19.61923 | 0.00914 | 0.00327 | 0.00589 | 0.00000 | 6.26012 | 2.83618 | 1.20888 | 0.37434 | 0.00000 | 1.56507 | 27.20399 | 0.00314 |
| **Oog4** | 0.00792 | 0.00789 | 0.00169 | 2.56224 | 0.01906 | 0.08967 | 0.00710 | 0.04136 | 3.00750 | 0.66472 | 0.80884 | 0.34995 | 0.07046 | 1.11889 | 1.16845 | 0.07231 |
| **Oosp1** | 0.00000 | 0.07328 | 0.00000 | 27.19239 | 0.00874 | 0.00257 | 0.00449 | 0.15509 | 8.32876 | 0.00000 | 2.49904 | 0.59966 | 2.68189 | 2.78394 | 19.42095 | 0.00000 |
| **Oosp2** | 0.01341 | 0.00788 | 0.02201 | 3.85663 | 0.00937 | 0.00581 | 0.00733 | 0.01081 | 0.00000 | 0.00348 | 0.05747 | 0.00710 | 0.01160 | 0.00294 | 0.41333 | 0.00669 |
| **Oosp3** | 0.01340 | 0.00765 | 0.00604 | 10.49013 | 0.00910 | 0.00524 | 0.00685 | 0.00076 | 2.83000 | 0.00000 | 0.26158 | 0.02654 | 0.00924 | 0.00000 | 1.85914 | 0.00514 |
| **Opa1** | 8.19267 | 6.97008 | 2.30365 | 2.60738 | 6.09070 | 5.86749 | 1.17756 | 1.87518 | 0.51107 | 2.94367 | 3.19050 | 0.52471 | 8.40834 | 6.92948 | 4.45271 | 2.16626 |
| **Opa3** | 2.86067 | 0.04232 | 1.08005 | 0.18314 | 0.00995 | 0.01043 | 0.01604 | 0.57822 | 0.94720 | 3.10753 | 0.00000 | 0.08545 | 1.39210 | 4.16513 | 3.16278 | 0.00346 |
| **Ophn1** | 0.01321 | 0.00769 | 0.01075 | 1.54991 | 0.00909 | 0.00620 | 0.00726 | 0.01014 | 1.15033 | 0.54992 | 0.00000 | 0.01152 | 0.01172 | 0.00277 | 0.93168 | 0.00713 |
| **Oplah** | 2.64688 | 0.79654 | 0.18940 | 0.00701 | 0.67983 | 0.02445 | 0.00397 | 0.93412 | 0.01081 | 0.58561 | 0.00511 | 0.01893 | 1.59609 | 0.01045 | 0.00000 | 0.03253 |
| **Opn3** | 0.01345 | 0.00788 | 0.01218 | 0.48744 | 0.00903 | 0.00643 | 0.00725 | 0.01032 | 0.00568 | 0.68585 | 0.01424 | 0.00693 | 0.01072 | 0.01735 | 0.00267 | 0.00703 |
| **Oprd1** | 0.02367 | 0.00760 | 0.06066 | 0.47902 | 0.00875 | 0.00577 | 0.00737 | 0.01125 | 0.42562 | 0.00967 | 0.05544 | 0.00624 | 0.01115 | 0.00553 | 0.12790 | 0.00720 |
| **Optn** | 1.10566 | 5.10233 | 4.08641 | 0.36379 | 0.02050 | 0.06628 | 0.01470 | 2.36089 | 0.06570 | 1.34274 | 1.19729 | 2.65509 | 0.00000 | 0.28454 | 1.50742 | 0.07938 |
| **Orai1** | 0.00854 | 0.00739 | 0.39894 | 6.51148 | 0.01865 | 0.00441 | 0.00665 | 0.76590 | 6.28608 | 0.05150 | 0.00000 | 0.01109 | 0.36370 | 0.00651 | 0.00000 | 0.00556 |
| **Orai2** | 1.51902 | 0.00532 | 2.44779 | 0.32526 | 0.00000 | 0.00458 | 0.01599 | 0.00000 | 0.40460 | 0.00000 | 0.12945 | 0.02178 | 0.00538 | 0.00637 | 0.43188 | 0.00455 |
| **Orai3** | 0.01678 | 0.00760 | 0.01720 | 1.11397 | 0.00864 | 0.00695 | 0.00724 | 0.01023 | 0.06479 | 1.28908 | 0.00940 | 0.00346 | 0.01053 | 0.00814 | 0.35006 | 0.00696 |
| **Oraov1** | 1.37086 | 0.00238 | 3.57931 | 4.86245 | 1.10496 | 0.00312 | 1.95859 | 2.97708 | 9.54587 | 8.62718 | 8.40880 | 1.88778 | 1.77681 | 1.72007 | 6.25566 | 0.00798 |
| **Orc1** | 2.30868 | 4.45877 | 2.20297 | 9.42667 | 3.25848 | 0.01136 | 0.08169 | 2.17770 | 11.69751 | 5.80960 | 3.67023 | 6.51488 | 1.11478 | 0.50934 | 4.16095 | 3.91949 |
| **Orc2** | 3.06119 | 7.62081 | 5.94252 | 9.83058 | 3.34557 | 0.27968 | 3.61781 | 5.72541 | 8.22475 | 9.58501 | 3.74388 | 8.00712 | 2.13447 | 13.32038 | 10.09950 | 6.08894 |
| **Orc3** | 7.60803 | 6.42595 | 10.18664 | 1.19144 | 4.08294 | 3.39836 | 6.64254 | 6.03031 | 1.76684 | 5.16874 | 4.25207 | 2.81227 | 5.72646 | 7.34342 | 5.60908 | 4.26402 |
| **Orc4** | 4.93998 | 6.79390 | 10.30611 | 4.84002 | 6.92576 | 7.28506 | 4.01455 | 5.44046 | 11.57532 | 10.13928 | 5.17757 | 14.19750 | 2.21206 | 11.47074 | 5.45501 | 4.19500 |
| **Orc5** | 5.80823 | 9.53682 | 6.22455 | 4.98022 | 8.70105 | 3.13425 | 4.14130 | 6.09767 | 0.56300 | 3.81666 | 2.60386 | 12.07537 | 3.75272 | 6.55447 | 4.68315 | 4.43004 |
| **Orc6** | 6.22056 | 10.67395 | 8.58548 | 5.11963 | 9.96712 | 7.68088 | 5.85983 | 14.60428 | 6.16581 | 9.45036 | 3.15744 | 4.67816 | 8.80270 | 14.89770 | 20.04102 | 7.11776 |
| **Ormdl1** | 1.38285 | 5.32697 | 5.04429 | 0.00000 | 4.16109 | 3.74112 | 4.92911 | 2.33359 | 0.01354 | 0.89081 | 4.25499 | 9.18459 | 1.53168 | 2.94582 | 0.73933 | 0.00575 |
| **Ormdl2** | 3.81664 | 3.37273 | 6.89261 | 0.00000 | 1.43424 | 1.65787 | 3.07608 | 8.23959 | 0.01680 | 1.79250 | 0.00000 | 3.15043 | 3.60344 | 4.96352 | 2.89292 | 2.54097 |
| **Ormdl3** | 3.45220 | 0.04496 | 3.45165 | 7.55004 | 0.33456 | 2.51598 | 2.29782 | 6.36543 | 11.76344 | 11.84874 | 4.94628 | 0.82161 | 6.62336 | 0.01584 | 4.25575 | 0.48531 |
| **Os9** | 2.04314 | 0.00559 | 2.11337 | 3.74658 | 0.00915 | 1.28551 | 0.33782 | 0.64775 | 1.72057 | 0.93673 | 0.22415 | 0.18222 | 0.03333 | 0.00000 | 0.35166 | 0.37875 |
| **Osbp** | 3.33082 | 1.17391 | 3.96302 | 5.32462 | 0.16492 | 0.43020 | 0.71325 | 0.27506 | 3.13298 | 0.23048 | 1.80573 | 0.73774 | 0.01570 | 7.43834 | 1.79369 | 0.11907 |
| **Osbp2** | 0.03121 | 0.00882 | 0.01838 | 0.34027 | 0.01431 | 0.00678 | 0.00778 | 0.01604 | 0.17862 | 0.00909 | 0.01276 | 0.00808 | 0.06082 | 0.01190 | 0.00467 | 0.31631 |
| **Osbpl10** | 0.03647 | 0.00521 | 1.06568 | 0.18042 | 3.41806 | 0.00922 | 0.00386 | 4.15180 | 0.48742 | 0.51360 | 1.12572 | 0.01295 | 5.15114 | 0.00150 | 0.06667 | 0.00494 |
| **Osbpl11** | 2.16613 | 8.36889 | 1.18637 | 13.36840 | 0.06894 | 2.73868 | 3.00604 | 6.66335 | 13.30200 | 10.99932 | 4.90614 | 10.60448 | 1.12818 | 0.00000 | 6.26227 | 1.10895 |
| **Osbpl1a** | 7.11968 | 0.00866 | 3.71882 | 1.04712 | 2.90662 | 0.00917 | 0.00000 | 0.50738 | 6.76714 | 0.81299 | 0.04449 | 0.20077 | 0.11882 | 0.00000 | 1.20600 | 0.35696 |
| **Osbpl2** | 3.20487 | 0.03161 | 3.29694 | 5.92340 | 0.06032 | 4.39733 | 3.85666 | 11.07545 | 9.94960 | 7.77219 | 7.06141 | 1.80092 | 1.97016 | 0.21654 | 9.02294 | 3.05926 |
| **Osbpl3** | 0.62603 | 0.00639 | 0.00000 | 0.00000 | 0.00738 | 0.01642 | 1.06118 | 0.53797 | 0.83785 | 0.73118 | 0.07114 | 0.79656 | 1.46550 | 0.00644 | 0.84633 | 0.00562 |
| **Osbpl5** | 0.00761 | 0.00792 | 0.00625 | 0.79994 | 0.00746 | 3.05571 | 0.00698 | 0.01039 | 0.19694 | 0.14974 | 0.00694 | 0.05925 | 0.01674 | 0.00717 | 0.04221 | 2.48126 |
| **Osbpl6** | 0.00000 | 2.31808 | 1.68512 | 8.67891 | 0.02893 | 2.50395 | 3.60607 | 11.90976 | 6.53228 | 7.51167 | 8.72313 | 0.84880 | 1.83345 | 11.22432 | 11.25750 | 0.00756 |
| **Osbpl7** | 1.91277 | 2.28457 | 6.60228 | 0.43655 | 0.39488 | 7.09868 | 0.09277 | 13.06007 | 2.08232 | 6.32510 | 4.83753 | 0.31504 | 4.98324 | 0.54316 | 0.75720 | 0.01081 |
| **Osbpl8** | 1.61936 | 4.55194 | 2.04540 | 15.75556 | 1.54568 | 3.74592 | 1.87043 | 5.44297 | 15.00026 | 2.01664 | 6.44319 | 7.28358 | 1.08889 | 10.64225 | 8.13522 | 1.26650 |
| **Osbpl9** | 4.59797 | 7.01797 | 4.37393 | 3.20738 | 4.44957 | 2.68432 | 3.21964 | 12.30920 | 4.06149 | 7.95848 | 9.45604 | 1.11105 | 1.35659 | 13.06717 | 11.34666 | 3.59351 |
| **Oscar** | 0.00659 | 0.18931 | 0.00020 | 0.10405 | 0.00952 | 0.00834 | 0.00719 | 0.58903 | 0.04760 | 1.18245 | 0.85319 | 0.04776 | 0.24445 | 0.40286 | 0.21642 | 0.00685 |
| **Oscp1** | 0.28617 | 0.00802 | 1.37468 | 0.51706 | 0.00684 | 0.00519 | 0.01720 | 0.00000 | 0.81198 | 1.34100 | 0.88829 | 0.02155 | 0.00000 | 0.00356 | 0.18397 | 0.18661 |
| **Oser1** | 10.61600 | 6.88885 | 10.61906 | 21.37369 | 12.56253 | 8.05880 | 5.51675 | 15.79295 | 28.19244 | 19.52517 | 11.45184 | 7.72375 | 10.11363 | 11.00018 | 9.83282 | 2.05146 |
| **Osgep** | 3.76352 | 0.02174 | 7.23604 | 0.93249 | 3.87037 | 5.88284 | 0.24426 | 1.12227 | 2.21075 | 10.36640 | 8.42093 | 0.82513 | 3.80564 | 5.89435 | 0.72589 | 3.10151 |
| **Osgepl1** | 3.46222 | 3.60205 | 7.56363 | 0.99142 | 6.35820 | 0.00313 | 0.01207 | 0.28765 | 0.34093 | 3.15836 | 3.38385 | 1.27127 | 0.23273 | 8.66177 | 3.45838 | 0.74853 |
| **Osgin1** | 0.01367 | 0.00805 | 0.01185 | 0.43732 | 0.00920 | 0.00661 | 0.00738 | 0.01199 | 0.55126 | 0.15304 | 0.01019 | 0.01632 | 0.01294 | 0.01043 | 0.00376 | 0.00768 |
| **Osgin2** | 2.82755 | 4.26420 | 2.01867 | 0.97370 | 0.00000 | 0.02477 | 1.48327 | 0.91511 | 4.33203 | 1.89588 | 2.32454 | 0.02347 | 0.00000 | 0.00000 | 2.03116 | 0.87005 |
| **Osm** | 0.86576 | 0.00671 | 0.00715 | 0.62998 | 0.00620 | 0.00750 | 0.00636 | 0.19630 | 0.36336 | 0.49999 | 0.00468 | 0.01874 | 0.00724 | 0.93756 | 0.00000 | 0.00630 |
| **Osr2** | 1.55393 | 0.00424 | 0.05946 | 1.37767 | 4.98499 | 0.00479 | 0.04899 | 0.00000 | 0.90525 | 0.25688 | 0.57872 | 0.36446 | 0.16365 | 0.00332 | 0.00000 | 0.19119 |
| **Ost4** | 1.80083 | 0.13022 | 0.24916 | 3.39552 | 0.39561 | 0.38937 | 0.12474 | 0.71571 | 3.65002 | 0.27172 | 3.21166 | 0.37165 | 0.38996 | 0.72254 | 1.79088 | 0.37187 |
| **Ostc** | 12.87038 | 9.98073 | 18.19148 | 2.69967 | 13.36568 | 11.55403 | 9.56154 | 8.51906 | 1.86401 | 11.44083 | 3.91877 | 9.09759 | 12.61966 | 5.88538 | 8.04062 | 8.62557 |
| **Ostf1** | 8.14152 | 6.52424 | 10.28154 | 2.27800 | 12.95930 | 6.55753 | 5.67118 | 4.96801 | 2.01872 | 6.44242 | 5.88903 | 0.47401 | 12.63543 | 1.44684 | 0.84030 | 9.80425 |
| **Ostm1** | 2.24094 | 1.72909 | 6.66751 | 2.95676 | 5.92545 | 2.88308 | 8.13185 | 6.95139 | 3.08451 | 2.00200 | 6.77266 | 4.24376 | 0.68213 | 8.69865 | 6.63159 | 2.31618 |

| | | | | | | | | | | | | | | | | |
|---|---|---|---|---|---|---|---|---|---|---|---|---|---|---|---|---|
| **Otoa** | 0.00919 | 0.00850 | 0.00656 | 0.00701 | 0.00966 | 0.00721 | 0.00817 | 0.52531 | 0.01116 | 0.00000 | 0.18024 | 0.01881 | 0.00500 | 0.01581 | 0.00000 | 0.00732 |
| **Otog** | 0.00726 | 0.00781 | 0.00878 | 0.12549 | 0.03081 | 0.01432 | 0.00743 | 1.10891 | 0.00905 | 0.01039 | 0.00964 | 0.00545 | 0.17327 | 0.00744 | 0.10907 | 0.01546 |
| **Otop1** | 0.01021 | 0.00793 | 0.01752 | 0.01048 | 0.00927 | 0.00606 | 0.05666 | 0.00923 | 0.01259 | 0.00000 | 0.00895 | 0.01413 | 0.67140 | 0.73408 | 0.00422 | 0.00684 |
| **Otub1** | 3.36342 | 0.49820 | 2.70992 | 1.16304 | 0.03285 | 2.76573 | 2.15769 | 3.61373 | 3.04004 | 10.99037 | 3.83603 | 0.23418 | 2.60271 | 0.04995 | 3.32923 | 0.10016 |
| **Otub2** | 1.63280 | 0.00532 | 0.02711 | 1.72289 | 0.00502 | 0.00433 | 0.88378 | 0.00490 | 0.28786 | 0.21561 | 0.02829 | 0.03263 | 0.01630 | 0.00406 | 0.53085 | 0.00572 |
| **Otud1** | 7.75388 | 0.27399 | 0.79113 | 2.27369 | 5.81266 | 0.85164 | 1.32174 | 8.57305 | 0.43064 | 2.39410 | 0.77155 | 0.19727 | 0.59467 | 5.45702 | 1.58341 | 1.80638 |
| **Otud3** | 0.00000 | 0.00744 | 0.00000 | 1.82218 | 0.00830 | 0.02602 | 0.00550 | 2.20820 | 5.14726 | 3.58648 | 2.36465 | 0.10077 | 1.94806 | 0.05398 | 0.41990 | 0.00558 |
| **Otud4** | 2.21539 | 7.14079 | 3.46829 | 17.87905 | 8.25802 | 1.31571 | 3.06563 | 6.26030 | 17.89865 | 7.22478 | 2.65539 | 8.19744 | 1.68398 | 20.67621 | 12.05259 | 3.51674 |
| **Otud5** | 10.48628 | 1.04448 | 11.04084 | 13.54932 | 2.43480 | 0.00689 | 5.17203 | 14.74192 | 17.54982 | 20.50594 | 8.68867 | 2.51343 | 13.26646 | 2.09063 | 10.64478 | 8.71031 |
| **Otud6b** | 7.66238 | 11.44557 | 15.08378 | 9.54911 | 8.31523 | 8.37624 | 11.44731 | 15.87240 | 18.87567 | 16.90264 | 11.72533 | 22.12145 | 12.04131 | 21.89226 | 24.38208 | 3.11837 |
| **Otud7a** | 0.01369 | 0.00805 | 0.01217 | 0.00567 | 0.00929 | 0.00677 | 0.00759 | 0.01416 | 0.12894 | 0.03157 | 0.01103 | 0.00819 | 0.01310 | 0.00695 | 0.10849 | 0.00741 |
| **Otud7b** | 0.52723 | 3.43575 | 0.46695 | 0.87323 | 3.79569 | 0.01690 | 0.01453 | 1.31071 | 0.91585 | 0.79965 | 0.83320 | 5.71346 | 1.75831 | 0.10823 | 0.09582 | 1.26818 |
| **Otulin** | 5.67905 | 0.24124 | 3.16948 | 1.32748 | 1.20777 | 0.00000 | 0.00000 | 1.68488 | 3.44402 | 8.87689 | 5.00524 | 4.08983 | 1.58246 | 1.20044 | 4.19523 | 0.04998 |
| **Otx1** | 0.01273 | 0.00845 | 0.01122 | 0.01427 | 0.00933 | 0.00670 | 0.00797 | 0.03802 | 0.02785 | 0.01701 | 0.04520 | 0.00878 | 0.01054 | 0.01387 | 0.00995 | 0.00734 |
| **Otx2** | 0.00000 | 0.00596 | 0.00000 | 0.00000 | 2.31008 | 7.15482 | 0.00560 | 1.68154 | 0.00783 | 0.00000 | 0.72969 | 0.00000 | 2.24902 | 0.01120 | 0.00000 | 0.01095 |
| **Ovca2** | 1.44007 | 0.25590 | 1.36153 | 0.00306 | 0.14666 | 0.03801 | 0.00550 | 1.48054 | 0.07934 | 0.38076 | 0.65279 | 0.01797 | 0.51012 | 0.20449 | 0.55317 | 1.00389 |
| **Ovgp1** | 0.00451 | 0.00754 | 0.35596 | 0.26897 | 0.00707 | 0.00562 | 0.00740 | 1.09950 | 0.43491 | 35.47751 | 0.00000 | 0.00000 | 0.00376 | 0.03129 | 0.00000 | 0.02160 |
| **Ovol1** | 1.47006 | 0.00487 | 0.08341 | 4.17826 | 0.02090 | 0.00689 | 0.00482 | 5.82311 | 3.42383 | 3.48326 | 2.63078 | 0.10467 | 0.00000 | 0.02987 | 5.05049 | 0.00437 |
| **Ovol2** | 0.00000 | 0.00520 | 0.15645 | 4.70312 | 0.02625 | 3.35286 | 0.00016 | 1.53474 | 4.76604 | 13.68162 | 7.88482 | 4.27616 | 3.24685 | 7.45753 | 8.86244 | 0.56754 |
| **Ovol3** | 0.01054 | 0.00805 | 0.01164 | 0.22084 | 0.00911 | 0.00645 | 0.00702 | 0.01071 | 0.25380 | 0.00425 | 0.01108 | 0.01821 | 0.71800 | 0.01291 | 0.00458 | 0.00682 |
| **Oxa1l** | 12.34962 | 0.94650 | 8.95532 | 2.91154 | 6.95584 | 0.84137 | 2.21468 | 9.23515 | 4.50636 | 7.24357 | 3.71873 | 0.34376 | 9.43513 | 0.00000 | 0.89146 | 8.54340 |
| **Oxct1** | 0.00000 | 5.47485 | 1.75286 | 0.41031 | 0.01952 | 8.95044 | 2.65084 | 1.34735 | 2.42304 | 4.40076 | 1.93145 | 3.74032 | 4.36984 | 2.79567 | 0.22410 | 6.61251 |
| **Oxgr1** | 0.03391 | 0.01934 | 0.02242 | 0.04924 | 0.04120 | 0.01960 | 0.02797 | 0.02516 | 0.05179 | 0.07137 | 0.14011 | 0.00699 | 0.03467 | 0.07519 | 0.02496 | 0.01478 |
| **Oxld1** | 2.51673 | 0.00539 | 0.09157 | 1.01602 | 0.05803 | 0.00926 | 0.02495 | 0.03368 | 1.28709 | 0.24227 | 0.02391 | 0.00221 | 0.98070 | 0.00644 | 0.03721 | 0.00430 |
| **Oxnad1** | 1.15661 | 4.02694 | 3.56477 | 2.03802 | 0.00000 | 1.30587 | 1.79082 | 0.66647 | 0.74114 | 0.85829 | 1.29682 | 0.03657 | 0.81087 | 1.07071 | 1.59914 | 0.00926 |
| **Oxr1** | 1.38879 | 5.07221 | 0.00000 | 1.27277 | 0.01424 | 0.01305 | 0.00435 | 0.01370 | 0.78425 | 0.00000 | 0.00000 | 5.02936 | 0.03916 | 0.00288 | 1.11483 | 0.01157 |
| **Oxsm** | 0.08071 | 0.00762 | 0.48394 | 0.04474 | 0.00791 | 0.01830 | 0.03767 | 0.17884 | 0.19872 | 0.00502 | 0.11474 | 0.00306 | 0.01047 | 0.86361 | 0.00969 | 0.00721 |
| **Oxsr1** | 4.52667 | 3.58455 | 9.72881 | 4.11077 | 4.81981 | 9.28175 | 4.18372 | 5.85081 | 4.59913 | 3.40144 | 3.67253 | 0.58567 | 4.41686 | 12.97578 | 4.58404 | 4.36158 |
| **P2rx2** | 0.01359 | 0.00778 | 0.01084 | 0.32665 | 0.00921 | 0.00659 | 0.00734 | 0.01127 | 0.00471 | 0.00768 | 0.01133 | 0.03743 | 0.06507 | 0.04979 | 0.07521 | 0.00728 |
| **P2rx3** | 0.03457 | 0.00800 | 0.13588 | 0.21667 | 0.00834 | 0.03307 | 0.00655 | 0.01153 | 0.25743 | 0.01401 | 0.07704 | 0.01628 | 1.50328 | 0.00962 | 0.01589 | 0.17787 |
| **P2rx4** | 0.15388 | 0.34662 | 5.10130 | 1.12327 | 0.28635 | 6.71565 | 0.00361 | 5.68681 | 3.92229 | 0.81880 | 4.95122 | 0.16706 | 1.63671 | 0.40389 | 0.98342 | 0.10879 |
| **P2rx5** | 0.01686 | 0.00782 | 0.40130 | 0.00341 | 0.00836 | 0.00581 | 0.00694 | 0.00000 | 0.00988 | 0.41109 | 0.89958 | 0.00267 | 0.00132 | 0.00491 | 0.48761 | 0.00706 |
| **P2rx7** | 0.07954 | 0.00886 | 0.49447 | 0.00987 | 0.15877 | 0.00622 | 0.00776 | 0.07894 | 0.75469 | 0.14240 | 0.00002 | 0.04450 | 0.07827 | 0.18614 | 0.57391 | 0.00763 |
| **P2ry14** | 0.00844 | 0.00750 | 2.14096 | 0.00919 | 0.00631 | 0.00903 | 0.00680 | 0.32624 | 0.02583 | 0.00000 | 0.00000 | 0.00000 | 0.00000 | 2.81586 | 0.00000 | 0.00647 |
| **P2ry4** | 0.04244 | 0.00745 | 0.84238 | 0.07177 | 0.00722 | 0.01776 | 0.03740 | 0.00799 | 0.09747 | 0.02497 | 0.21023 | 0.02489 | 0.02927 | 0.20381 | 0.07689 | 0.04290 |
| **P3h1** | 0.48736 | 0.00501 | 0.61807 | 0.81578 | 4.04749 | 0.00517 | 0.00679 | 0.65225 | 1.12438 | 2.50027 | 0.20868 | 0.13327 | 0.00000 | 0.00441 | 0.57346 | 2.44603 |
| **P3h3** | 0.49591 | 0.21722 | 0.31860 | 3.36185 | 0.10382 | 0.01687 | 0.06723 | 0.00000 | 4.86198 | 0.09469 | 4.06587 | 0.27395 | 0.15994 | 1.37449 | 0.06458 | 0.07064 |
| **P3h4** | 0.18758 | 0.00756 | 0.01323 | 0.00299 | 0.00740 | 0.01625 | 0.00661 | 0.00000 | 0.06289 | 2.57934 | 1.25337 | 0.09980 | 0.00000 | 0.00397 | 0.75027 | 0.00681 |
| **P4ha1** | 2.22707 | 2.26779 | 2.96732 | 0.17069 | 0.26562 | 0.00370 | 8.05664 | 3.28628 | 2.59845 | 1.53351 | 1.30000 | 0.20943 | 3.12794 | 0.02172 | 0.35059 | 3.74250 |
| **P4ha2** | 2.45803 | 3.08054 | 5.23110 | 3.23905 | 0.11630 | 0.00354 | 8.94838 | 4.56310 | 2.35984 | 0.00000 | 0.32561 | 0.55201 | 6.71256 | 0.78490 | 0.60773 | 6.82686 |
| **P4ha3** | 0.00000 | 7.62876 | 4.76695 | 0.00000 | 0.00000 | 0.00527 | 0.00393 | 0.00000 | 6.70765 | 2.22654 | 1.00485 | 0.30845 | 0.52987 | 0.64993 | 0.98304 | 0.00181 |
| **P4hb** | 13.61209 | 7.08287 | 17.98714 | 15.31375 | 6.04541 | 14.25381 | 12.03347 | 18.46699 | 18.33428 | 23.88689 | 12.16484 | 8.41055 | 18.68987 | 1.07727 | 11.70563 | 14.45501 |
| **P4htm** | 0.25154 | 0.04961 | 0.93516 | 0.00509 | 0.01372 | 0.00548 | 0.01577 | 2.24806 | 1.82861 | 0.98468 | 1.36275 | 0.73069 | 1.03720 | 0.01999 | 0.00000 | 0.02149 |
| **Pa2g4** | 23.12800 | 19.45103 | 26.30036 | 23.63282 | 13.83466 | 14.67544 | 19.74834 | 34.40232 | 21.46699 | 47.43905 | 27.65946 | 21.84378 | 27.31156 | 23.16946 | 21.36651 | 12.54512 |
| **Pabpc1** | 29.41908 | 21.17068 | 27.18677 | 22.75423 | 23.36968 | 15.96153 | 23.39589 | 37.51028 | 27.01694 | 34.44611 | 25.48073 | 31.51089 | 24.54990 | 33.46297 | 32.51141 | 14.20605 |
| **Pabpc1l** | 0.01369 | 0.00815 | 0.01138 | 0.93575 | 0.00938 | 0.00659 | 0.00739 | 0.01207 | 0.00000 | 0.00969 | 0.01230 | 0.00778 | 0.01293 | 0.12267 | 0.04985 | 0.00729 |
| **Pabpc4** | 19.54945 | 1.03660 | 4.56207 | 7.32627 | 15.30400 | 10.02597 | 6.47682 | 30.32906 | 7.50570 | 45.13936 | 26.88502 | 5.55350 | 37.79868 | 3.74958 | 4.77596 | 8.28284 |
| **Pabpn1** | 18.15787 | 10.53568 | 12.21510 | 5.60917 | 10.14857 | 9.51505 | 7.71010 | 14.96501 | 9.62405 | 25.68717 | 12.28252 | 1.53831 | 15.25730 | 10.72095 | 20.06901 | 9.95368 |
| **Pabpn1l** | 0.01384 | 0.00810 | 0.01037 | 0.57320 | 0.00918 | 0.00666 | 0.00750 | 0.00919 | 0.04644 | 0.00735 | 0.20532 | 0.01792 | 0.00985 | 0.00918 | 0.01392 | 0.00718 |
| **Pacrgl** | 4.23857 | 2.52031 | 1.72475 | 5.14550 | 0.34728 | 0.03601 | 0.01182 | 0.91551 | 4.84908 | 4.30708 | 1.00562 | 0.18208 | 1.02029 | 2.64958 | 2.67688 | 2.32155 |
| **Pacs1** | 1.12308 | 0.00381 | 1.11084 | 3.49407 | 5.60478 | 0.01606 | 0.00641 | 0.00000 | 2.46764 | 1.79823 | 0.00000 | 0.03160 | 0.06752 | 0.00000 | 1.24332 | 0.01973 |
| **Pacs2** | 3.06465 | 0.01324 | 0.08058 | 12.64192 | 0.03677 | 0.00744 | 2.08462 | 0.01141 | 4.98999 | 3.04514 | 0.05048 | 0.25432 | 1.47819 | 0.02599 | 4.93405 | 1.59895 |
| **Pacsin1** | 0.33980 | 0.00692 | 0.56508 | 1.04738 | 0.00628 | 0.00508 | 0.00627 | 0.80687 | 0.73156 | 0.00000 | 0.00000 | 0.03350 | 0.94480 | 0.01213 | 0.00000 | 0.00588 |
| **Pacsin2** | 7.08266 | 1.60033 | 4.34100 | 20.12327 | 3.47894 | 7.66306 | 4.18906 | 3.26650 | 9.77320 | 2.80598 | 5.40339 | 3.05242 | 3.80702 | 4.00256 | 17.16385 | 3.10312 |
| **Pacsin3** | 0.04406 | 0.00681 | 0.19031 | 1.37057 | 4.77168 | 1.79058 | 0.00614 | 0.11239 | 0.00000 | 2.30834 | 0.29461 | 0.01927 | 3.11854 | 0.00390 | 0.24344 | 1.15108 |
| **Padi1** | 0.01011 | 0.00763 | 0.00616 | 0.00520 | 0.00957 | 0.00719 | 0.12690 | 0.11533 | 0.01115 | 0.07678 | 0.55257 | 0.00444 | 0.00470 | 0.01050 | 0.00161 | 0.00713 |
| **Padi2** | 0.35333 | 0.02729 | 0.32019 | 2.46806 | 0.04162 | 0.02438 | 0.14904 | 0.75933 | 1.35147 | 0.24581 | 1.54535 | 0.05333 | 0.05720 | 0.85376 | 0.53780 | 0.04159 |
| **Padi3** | 0.01069 | 0.00832 | 0.13695 | 0.32214 | 0.06155 | 0.00574 | 0.01864 | 0.00744 | 0.02309 | 0.00602 | 0.38622 | 0.00716 | 0.00768 | 0.00828 | 0.00000 | 0.00705 |
| **Padi4** | 0.11084 | 0.01770 | 0.01302 | 0.34629 | 0.00809 | 0.00613 | 0.01723 | 0.03363 | 0.75572 | 0.26151 | 0.72137 | 0.04716 | 0.00482 | 0.22053 | 0.01957 | 0.00675 |
| **Padi6** | 0.00000 | 4.84039 | 1.84429 | 9.85741 | 0.02185 | 0.00345 | 0.00000 | 4.72862 | 9.31188 | 10.71699 | 5.17468 | 1.98175 | 4.94366 | 6.43058 | 11.11186 | 0.00000 |
| **Paf1** | 7.55259 | 2.54612 | 2.56881 | 10.69123 | 5.80618 | 4.63829 | 3.52976 | 5.89787 | 15.56217 | 7.56835 | 4.43791 | 1.18541 | 8.42171 | 0.00000 | 2.31123 | 7.61863 |
| **Pafah1b1** | 10.37733 | 6.66455 | 12.10132 | 10.58914 | 5.93825 | 7.08717 | 8.89383 | 10.21381 | 6.15236 | 11.56493 | 16.08167 | 6.35674 | 12.42388 | 19.07937 | 13.23453 | 4.69862 |

| | | | | | | | | | | | | | | | | |
|---|---|---|---|---|---|---|---|---|---|---|---|---|---|---|---|---|
| **Pafah1b2** | 5.01808 | 0.70533 | 6.60956 | 11.00346 | 0.01779 | 5.79229 | 4.21099 | 3.77742 | 13.81084 | 13.58313 | 9.72014 | 12.44237 | 5.69410 | 15.36050 | 14.64181 | 3.71997 |
| **Pafah1b3** | 5.08787 | 0.79148 | 4.82240 | 5.20123 | 4.39413 | 0.55202 | 0.02778 | 7.37751 | 6.22756 | 10.29331 | 7.32405 | 0.40073 | 6.07937 | 0.25036 | 0.40589 | 5.13422 |
| **Pafah2** | 2.71016 | 1.62253 | 3.81636 | 0.02294 | 0.16742 | 4.75635 | 0.02634 | 1.23743 | 1.78806 | 0.00000 | 0.27719 | 0.00486 | 2.42726 | 0.00545 | 0.34582 | 2.80055 |
| **Pag1** | 0.57832 | 0.00615 | 1.13042 | 5.48107 | 0.02185 | 0.00798 | 1.41809 | 1.14617 | 2.69947 | 0.40719 | 0.00000 | 0.09405 | 0.00831 | 0.03568 | 0.46106 | 0.18932 |
| **Pagr1a** | 9.74878 | 0.06214 | 1.58095 | 6.40661 | 0.06287 | 6.79918 | 3.30290 | 7.43449 | 3.51822 | 6.57539 | 2.37033 | 0.27209 | 3.89684 | 0.40600 | 0.59533 | 6.55632 |
| **Paics** | 20.64322 | 16.34972 | 20.57942 | 11.96286 | 17.04120 | 16.06152 | 15.98468 | 26.42387 | 20.76428 | 30.60229 | 20.40158 | 17.39428 | 20.97974 | 33.17501 | 25.36152 | 14.46474 |
| **Paip1** | 0.48305 | 0.12844 | 1.47323 | 5.07338 | 0.01300 | 0.13930 | 0.59632 | 0.00000 | 2.00590 | 0.11944 | 2.07260 | 1.49677 | 0.06377 | 2.67899 | 0.69437 | 0.27634 |
| **Paip2** | 13.72195 | 5.82583 | 9.91103 | 8.34030 | 9.44265 | 9.73521 | 5.88190 | 16.70760 | 13.69655 | 17.61475 | 15.31710 | 16.09299 | 8.29104 | 22.31915 | 13.21069 | 1.73635 |
| **Paip2b** | 2.05963 | 2.00125 | 4.05842 | 19.39002 | 3.38490 | 10.47386 | 2.11946 | 5.04368 | 14.25870 | 14.94186 | 9.49874 | 7.52790 | 5.11048 | 19.51352 | 17.18104 | 0.85707 |
| **Pak1** | 0.00000 | 0.00461 | 2.95127 | 8.79788 | 0.03189 | 0.01260 | 3.75209 | 5.11068 | 6.19418 | 5.57387 | 0.78673 | 0.58594 | 1.88059 | 8.89450 | 7.72477 | 0.01032 |
| **Pak1ip1** | 14.47634 | 9.52535 | 14.40591 | 8.00819 | 17.40999 | 7.15329 | 8.09190 | 23.27736 | 20.59019 | 25.31308 | 16.53480 | 10.33542 | 12.75549 | 12.29225 | 10.58610 | 7.16951 |
| **Pak2** | 3.97113 | 0.42936 | 4.38299 | 5.97075 | 3.19205 | 4.10999 | 2.23627 | 2.64243 | 8.35190 | 8.62260 | 6.47035 | 0.69520 | 2.31830 | 15.79135 | 7.65521 | 6.28175 |
| **Pak4** | 7.51990 | 0.17474 | 0.00000 | 10.51827 | 7.97755 | 6.55858 | 0.91301 | 6.23137 | 2.50414 | 12.15438 | 8.22167 | 0.41730 | 3.63003 | 0.12698 | 1.34503 | 5.88303 |
| **Pak6** | 0.00000 | 0.01834 | 0.01889 | 0.00847 | 0.06434 | 3.00273 | 0.00566 | 0.01338 | 0.03967 | 0.55747 | 0.17304 | 0.00341 | 2.90369 | 0.02730 | 0.11956 | 1.40925 |
| **Pak7** | 0.01356 | 0.00764 | 0.00862 | 6.03760 | 0.00944 | 0.00583 | 0.00703 | 0.00843 | 0.00000 | 0.00000 | 0.06250 | 0.03088 | 0.01191 | 0.09451 | 1.73721 | 0.00632 |
| **Palb2** | 0.92680 | 3.20973 | 0.60288 | 0.51217 | 4.31164 | 0.00543 | 0.34400 | 1.65169 | 0.39586 | 1.50519 | 0.57517 | 0.08218 | 0.06102 | 0.00303 | 3.38840 | 0.00451 |
| **Pald1** | 1.98480 | 0.00641 | 0.42254 | 0.63496 | 0.01149 | 0.00347 | 0.00610 | 0.02823 | 0.09476 | 0.52115 | 0.60863 | 0.01268 | 0.01626 | 0.01603 | 0.04066 | 0.00515 |
| **Palld** | 1.85144 | 2.59504 | 2.02018 | 3.84283 | 0.13049 | 0.00587 | 1.84539 | 0.05458 | 2.73537 | 3.95954 | 0.15285 | 0.21547 | 1.84395 | 2.31508 | 2.62303 | 0.76030 |
| **Palm** | 2.82125 | 0.00422 | 0.00000 | 0.01221 | 0.00508 | 2.62365 | 0.00434 | 1.91815 | 1.38787 | 2.08110 | 1.39397 | 0.05474 | 0.00000 | 0.00651 | 0.00095 | 1.31259 |
| **Palm3** | 7.25108 | 3.00365 | 11.31938 | 1.80105 | 7.51776 | 7.90921 | 2.45445 | 7.55456 | 0.55425 | 1.96861 | 2.71001 | 0.11119 | 4.75639 | 0.00000 | 0.00018 | 0.00000 |
| **Pam16** | 7.09595 | 0.07140 | 1.89438 | 6.96943 | 5.19370 | 1.01904 | 1.09557 | 5.28347 | 3.93978 | 12.89140 | 8.79183 | 0.48203 | 9.21861 | 0.09961 | 0.00000 | 3.71927 |
| **Pan2** | 2.22935 | 7.27396 | 3.93394 | 6.85862 | 4.62683 | 5.94824 | 2.87753 | 6.90864 | 1.41600 | 3.57299 | 1.87813 | 0.26700 | 0.27534 | 2.77907 | 4.69617 | 0.57939 |
| **Pan3** | 0.69840 | 0.01680 | 1.44890 | 6.54372 | 0.08952 | 0.21394 | 0.73471 | 1.48960 | 9.27404 | 2.81291 | 5.10936 | 0.24599 | 0.44046 | 8.38640 | 3.07738 | 0.02521 |
| **Panct2** | 0.04957 | 0.01610 | 2.22152 | 1.85898 | 4.13578 | 2.00446 | 0.09082 | 0.02501 | 1.78976 | 0.00005 | 1.20881 | 0.05430 | 0.11877 | 0.16078 | 0.45046 | 0.01207 |
| **Pank1** | 0.01894 | 0.01785 | 0.07972 | 0.00798 | 0.00843 | 0.00834 | 0.00698 | 0.00253 | 1.32810 | 0.45223 | 0.19248 | 0.00000 | 0.03105 | 1.96645 | 0.00000 | 0.00693 |
| **Pank2** | 2.89554 | 5.40051 | 0.21159 | 4.76223 | 0.00997 | 0.50021 | 2.15069 | 0.00002 | 3.32552 | 3.67262 | 0.12765 | 0.00001 | 0.11115 | 4.64307 | 2.44734 | 0.00000 |
| **Pank3** | 1.16646 | 6.73423 | 9.38333 | 22.75937 | 0.02921 | 0.81686 | 2.43930 | 13.66876 | 27.62324 | 25.25395 | 23.80226 | 26.28121 | 6.73618 | 39.45265 | 30.40581 | 1.35784 |
| **Pank4** | 0.15416 | 0.00843 | 0.01834 | 0.00493 | 0.01653 | 0.00789 | 0.00721 | 0.09672 | 0.01108 | 0.25323 | 0.43119 | 0.01566 | 0.00909 | 0.07302 | 0.09138 | 0.00710 |
| **Panx1** | 1.62415 | 0.00627 | 0.00000 | 0.18811 | 0.00485 | 0.64414 | 0.28621 | 0.36489 | 0.03191 | 0.05980 | 0.60692 | 0.01588 | 0.00000 | 1.74220 | 1.81278 | 0.00526 |
| **Paox** | 2.68479 | 0.00172 | 0.76978 | 2.43508 | 2.21251 | 0.00610 | 3.09492 | 0.00000 | 1.47983 | 0.70779 | 0.00000 | 0.02470 | 0.02175 | 0.00183 | 0.51018 | 0.59713 |
| **Papd4** | 1.40890 | 3.41733 | 3.31010 | 5.48981 | 5.27311 | 2.12073 | 1.34978 | 4.63864 | 7.31747 | 7.18822 | 5.50916 | 12.39698 | 2.37057 | 14.24068 | 4.23935 | 2.72925 |
| **Papd5** | 4.23887 | 5.83016 | 4.91600 | 14.38102 | 1.81809 | 6.42834 | 5.07983 | 11.85151 | 9.70474 | 21.00871 | 11.19123 | 18.79040 | 4.75983 | 28.76814 | 20.29773 | 0.47252 |
| **Papd7** | 2.36764 | 0.00143 | 4.50461 | 25.94441 | 4.64271 | 5.01673 | 0.90754 | 1.99163 | 13.89449 | 5.90230 | 0.92657 | 0.88437 | 0.00000 | 0.00000 | 15.59942 | 0.94760 |
| **Papln** | 0.00796 | 0.00792 | 0.00868 | 0.49324 | 0.00882 | 0.00665 | 0.00645 | 0.00000 | 0.00385 | 0.99491 | 0.56003 | 0.00193 | 1.23077 | 0.00593 | 0.00000 | 0.00639 |
| **Papola** | 23.32862 | 13.11754 | 15.72903 | 17.31543 | 17.39953 | 11.73885 | 10.38917 | 18.77513 | 22.25211 | 32.92320 | 23.61114 | 19.22530 | 14.84008 | 25.59199 | 19.44366 | 7.71863 |
| **Papolb** | 0.00683 | 0.00778 | 0.01947 | 3.12030 | 0.00958 | 0.03641 | 0.00716 | 0.03741 | 4.90305 | 0.51563 | 0.00000 | 0.00000 | 0.01161 | 2.28518 | 0.45297 | 0.00648 |
| **Papolg** | 0.89530 | 0.00627 | 0.40635 | 0.05642 | 1.02873 | 0.00494 | 0.00583 | 0.42252 | 1.33364 | 0.83638 | 2.17089 | 0.65974 | 0.00000 | 4.84432 | 0.98814 | 0.94877 |
| **Pappa** | 0.01346 | 0.00784 | 0.01473 | 0.00791 | 0.00930 | 0.00642 | 0.00743 | 0.01161 | 0.04851 | 0.00969 | 0.02391 | 0.05701 | 0.01901 | 0.10468 | 0.02213 | 0.00740 |
| **Papss1** | 0.01298 | 0.75186 | 0.00000 | 6.01226 | 0.00802 | 2.60862 | 0.00579 | 0.82763 | 4.25697 | 5.67804 | 1.36840 | 0.25467 | 0.01297 | 7.30674 | 10.90312 | 0.01858 |
| **Papss2** | 0.05289 | 0.01868 | 0.04512 | 0.33888 | 0.01134 | 0.01922 | 0.00717 | 0.01371 | 0.02560 | 0.01722 | 0.03794 | 0.03430 | 0.01808 | 0.54726 | 0.04913 | 0.00744 |
| **Paqr4** | 0.69835 | 0.00682 | 1.05230 | 0.36667 | 0.01390 | 3.84445 | 0.00367 | 2.66111 | 0.00579 | 3.81770 | 2.08567 | 0.00831 | 3.37557 | 0.00470 | 0.10851 | 0.00318 |
| **Paqr6** | 0.01328 | 0.00816 | 0.01173 | 1.42436 | 0.00931 | 0.00649 | 0.00738 | 0.02230 | 0.58283 | 0.01035 | 0.02580 | 0.00716 | 0.01183 | 0.01008 | 0.00306 | 0.00713 |
| **Paqr7** | 0.01841 | 0.00774 | 0.14443 | 0.38656 | 0.00840 | 0.00806 | 0.00729 | 0.00000 | 2.52562 | 0.00000 | 0.44704 | 0.00000 | 0.03917 | 0.14483 | 0.00000 | 0.04291 |
| **Pard3** | 5.35353 | 1.67357 | 3.59834 | 12.16807 | 5.15110 | 1.65628 | 3.98049 | 2.87941 | 3.97123 | 2.89336 | 0.78971 | 8.28128 | 0.32732 | 0.24409 | 4.12490 | 5.82036 |
| **Pard3b** | 0.02714 | 0.01849 | 0.01021 | 0.01788 | 0.01107 | 0.00674 | 0.00739 | 0.01799 | 0.03485 | 0.01950 | 0.01147 | 0.03695 | 0.02227 | 0.01396 | 0.16885 | 0.00750 |
| **Pard6a** | 4.62534 | 0.05897 | 4.85396 | 0.01222 | 0.00000 | 0.01298 | 0.74560 | 0.00000 | 0.69946 | 2.50239 | 0.50066 | 0.13829 | 1.49194 | 0.00999 | 0.65672 | 0.00000 |
| **Pard6b** | 0.17895 | 0.00339 | 3.50071 | 5.38990 | 0.04734 | 2.42745 | 3.44569 | 10.28823 | 0.20417 | 4.37975 | 6.81117 | 0.06098 | 3.20764 | 0.03355 | 0.58562 | 1.61249 |
| **Pard6g** | 0.00655 | 0.00752 | 0.06588 | 0.23042 | 1.14293 | 0.00572 | 0.00806 | 0.31173 | 1.01793 | 0.00052 | 0.01949 | 0.07436 | 0.00562 | 0.01646 | 0.00000 | 0.00717 |
| **Parg** | 2.22266 | 2.51923 | 4.00551 | 1.35029 | 6.70817 | 2.67877 | 1.52224 | 7.17486 | 1.37098 | 7.75483 | 4.11764 | 2.59969 | 1.73500 | 17.11795 | 11.05551 | 3.18220 |
| **Park2** | 0.38288 | 0.03907 | 0.32201 | 2.80339 | 0.21082 | 0.08114 | 0.08484 | 0.00000 | 4.35509 | 0.50683 | 4.28889 | 0.20175 | 0.13477 | 3.32632 | 1.12692 | 0.13874 |
| **Park7** | 16.39167 | 8.27396 | 18.35772 | 5.35288 | 7.69365 | 2.42797 | 8.27588 | 8.00588 | 8.38501 | 10.88543 | 8.27109 | 0.33976 | 8.53476 | 0.00000 | 2.01094 | 9.95687 |
| **Parl** | 7.15019 | 5.70157 | 9.45687 | 9.05528 | 10.41147 | 6.95778 | 3.80842 | 7.77728 | 4.33777 | 7.56312 | 12.14366 | 7.65341 | 11.55410 | 2.85048 | 3.44369 | 10.51665 |
| **Parn** | 4.04171 | 3.48593 | 6.27537 | 2.73519 | 6.05842 | 4.00213 | 8.11580 | 7.46007 | 2.10755 | 7.23483 | 2.52359 | 0.33291 | 3.27512 | 4.14278 | 5.48453 | 0.26377 |
| **Parp1** | 23.28969 | 17.91225 | 17.57425 | 11.21073 | 15.84808 | 9.32494 | 13.77251 | 26.84637 | 11.50850 | 27.19240 | 21.17374 | 12.93365 | 18.54264 | 5.27728 | 7.06989 | 9.73696 |
| **Parp10** | 0.01645 | 0.00788 | 0.49017 | 3.74649 | 0.00830 | 0.00524 | 0.00721 | 0.01221 | 1.08955 | 0.00948 | 0.01844 | 0.00562 | 0.00918 | 0.00844 | 0.00000 | 0.00644 |
| **Parp11** | 1.11211 | 0.00671 | 0.00673 | 0.01405 | 0.00578 | 0.00582 | 0.00646 | 0.00337 | 0.03115 | 0.20399 | 0.22208 | 0.00477 | 0.00994 | 0.00729 | 0.33428 | 0.00586 |
| **Parp12** | 0.09917 | 0.00735 | 0.00000 | 25.07725 | 0.00760 | 0.00185 | 0.00580 | 0.83670 | 17.11955 | 0.28333 | 0.79473 | 0.15484 | 0.00236 | 0.98620 | 15.78202 | 0.00177 |
| **Parp14** | 0.01301 | 0.00840 | 0.01083 | 0.15722 | 0.00906 | 0.00663 | 0.00752 | 0.01136 | 3.15782 | 0.37867 | 0.00000 | 0.00000 | 0.01280 | 0.03505 | 0.11802 | 0.01451 |
| **Parp16** | 3.44870 | 0.00000 | 0.38453 | 0.00000 | 3.54144 | 0.00446 | 2.53105 | 0.64926 | 0.67708 | 0.59972 | 1.38175 | 0.00670 | 0.92573 | 0.00280 | 1.29653 | 1.10191 |
| **Parp2** | 8.09349 | 3.56403 | 7.02660 | 7.44518 | 5.70286 | 7.24850 | 7.90994 | 9.52204 | 6.49650 | 6.91233 | 8.01350 | 13.39924 | 4.87705 | 11.18900 | 10.78928 | 2.78437 |
| **Parp4** | 0.00000 | 0.01825 | 0.33461 | 0.96907 | 0.01113 | 0.02788 | 0.63077 | 0.74805 | 2.76503 | 0.91190 | 1.11297 | 1.82179 | 0.81726 | 4.53274 | 2.07651 | 0.00647 |
| **Parp6** | 1.81923 | 0.08423 | 2.00566 | 2.72459 | 1.89032 | 0.00519 | 2.94890 | 0.52465 | 1.47931 | 0.20103 | 1.84329 | 0.14228 | 1.31837 | 0.41659 | 3.94199 | 0.00328 |

| | | | | | | | | | | | | | | | | |
|---|---|---|---|---|---|---|---|---|---|---|---|---|---|---|---|---|
| **Parp8** | 2.97889 | 6.69639 | 5.67028 | 2.14206 | 0.00976 | 0.00270 | 6.79789 | 17.74719 | 3.90341 | 20.32778 | 14.20073 | 7.08897 | 9.05129 | 15.00860 | 7.77008 | 0.95400 |
| **Parp9** | 1.14415 | 0.00271 | 0.41096 | 0.53428 | 0.67031 | 0.00257 | 2.45200 | 1.30617 | 7.80188 | 3.11781 | 0.00000 | 5.96628 | 0.18585 | 13.97075 | 9.03098 | 0.00419 |
| **Parpbp** | 3.37977 | 5.53280 | 1.87706 | 0.16338 | 2.13016 | 3.87739 | 4.88257 | 0.36029 | 0.02185 | 2.53921 | 0.43691 | 0.26099 | 0.00000 | 0.10451 | 1.79613 | 0.61284 |
| **Pars2** | 1.03664 | 0.01508 | 2.39494 | 1.09717 | 0.14635 | 0.00602 | 1.63046 | 2.72454 | 0.08881 | 0.00000 | 0.49015 | 0.00641 | 0.26943 | 0.00787 | 0.27918 | 0.00489 |
| **Particl** | 0.01338 | 0.00838 | 0.01709 | 0.91417 | 0.00924 | 0.00645 | 0.00763 | 0.01473 | 0.36118 | 0.04699 | 0.01084 | 0.00875 | 0.01300 | 0.01047 | 0.00395 | 0.00725 |
| **Parva** | 0.00609 | 0.00705 | 2.10669 | 0.00291 | 0.92045 | 0.01914 | 0.00710 | 0.00830 | 0.15147 | 0.90387 | 0.00529 | 0.23333 | 0.05646 | 0.32890 | 0.06140 | 0.00710 |
| **Parvb** | 8.96828 | 0.02545 | 3.05273 | 0.03128 | 5.17050 | 5.12123 | 5.03869 | 4.99263 | 0.00342 | 6.63742 | 2.40161 | 0.77944 | 5.10921 | 0.46492 | 0.07898 | 3.21447 |
| **Pask** | 1.78228 | 0.04264 | 0.23999 | 0.87051 | 0.01195 | 3.81526 | 2.75823 | 2.00854 | 2.17960 | 4.27537 | 2.42891 | 6.68802 | 1.92704 | 0.05344 | 1.46864 | 2.89404 |
| **Pate2** | 0.01395 | 0.00839 | 0.00817 | 0.00450 | 0.00907 | 0.00683 | 0.00749 | 0.00196 | 0.01074 | 0.00000 | 0.59245 | 0.00572 | 0.00674 | 0.00608 | 0.16849 | 0.00708 |
| **Pate3** | 0.02188 | 0.00797 | 0.01168 | 0.01785 | 0.00880 | 0.00661 | 0.00746 | 0.01364 | 0.01098 | 0.00965 | 0.01252 | 0.00780 | 0.01267 | 0.00711 | 0.16404 | 0.00734 |
| **Pate4** | 0.01386 | 0.00816 | 0.01037 | 0.01494 | 0.00921 | 0.00665 | 0.00747 | 0.00805 | 0.01052 | 0.00000 | 0.20710 | 0.00746 | 0.00995 | 0.00749 | 0.18257 | 0.00730 |
| **Patl1** | 3.58560 | 1.93792 | 1.03781 | 1.90918 | 3.23363 | 0.70276 | 3.08603 | 3.36327 | 4.69034 | 3.97284 | 2.97676 | 14.26593 | 0.26770 | 3.29757 | 5.30426 | 0.00714 |
| **Patl2** | 0.01931 | 0.00724 | 2.92726 | 0.22809 | 0.00568 | 0.05441 | 0.00583 | 0.11889 | 0.87742 | 0.00074 | 0.00484 | 0.00294 | 1.67987 | 0.01318 | 0.00000 | 0.01245 |
| **Patz1** | 2.45888 | 0.01424 | 4.81272 | 3.80743 | 1.78606 | 0.50185 | 0.01273 | 6.40383 | 2.90579 | 9.35201 | 1.98794 | 0.36660 | 2.43028 | 3.78090 | 6.45162 | 0.46100 |
| **Pawr** | 3.48742 | 5.65857 | 0.00000 | 0.21444 | 5.58783 | 4.90464 | 2.40818 | 3.49384 | 0.00000 | 0.93448 | 0.13503 | 0.09070 | 1.87186 | 2.69547 | 0.00000 | 2.47077 |
| **Pax3** | 0.02212 | 0.02897 | 0.02549 | 0.56792 | 0.01735 | 0.02304 | 0.01739 | 0.02132 | 0.18158 | 0.02613 | 0.02655 | 0.02829 | 0.03139 | 0.01047 | 0.00766 | 0.01447 |
| **Pax6** | 0.00874 | 0.00778 | 0.00000 | 0.17807 | 0.00818 | 0.00583 | 0.00661 | 0.00000 | 0.09932 | 8.63932 | 0.75346 | 0.25452 | 0.00574 | 9.34274 | 2.31408 | 0.00683 |
| **Pax8** | 0.01313 | 0.00777 | 0.01216 | 0.00695 | 0.00896 | 0.00658 | 0.00725 | 0.01009 | 0.01167 | 1.19085 | 0.01183 | 0.00452 | 0.01140 | 0.01187 | 0.00801 | 0.00764 |
| **Pax9** | 0.02365 | 0.05042 | 0.08374 | 0.21368 | 0.09217 | 0.00668 | 0.00764 | 0.03074 | 0.13095 | 0.00000 | 0.27388 | 0.05791 | 0.35788 | 0.07122 | 0.16241 | 0.01471 |
| **Paxbp1** | 3.72921 | 8.58626 | 2.84511 | 5.10232 | 1.45617 | 7.41538 | 3.76105 | 1.26099 | 13.33974 | 3.64980 | 3.62108 | 8.41749 | 1.22786 | 7.46817 | 10.02459 | 2.44236 |
| **Paxip1** | 7.40849 | 2.93257 | 4.90424 | 6.12558 | 1.66088 | 3.39998 | 9.18251 | 8.85412 | 5.24050 | 6.47327 | 5.42042 | 8.31990 | 4.13809 | 1.34225 | 6.41564 | 3.65582 |
| **Pbdc1** | 18.98446 | 9.33053 | 21.72554 | 3.82695 | 19.31589 | 6.35962 | 10.81425 | 17.39546 | 5.20702 | 21.59268 | 6.32566 | 5.69208 | 11.47398 | 15.37659 | 17.94262 | 6.28397 |
| **Pbk** | 11.45164 | 9.50362 | 10.08317 | 10.12057 | 9.39610 | 8.08881 | 4.39365 | 10.96538 | 4.81633 | 20.13881 | 12.98080 | 9.80530 | 7.27360 | 16.30271 | 7.70095 | 4.70052 |
| **Pbld1** | 1.49774 | 0.10141 | 1.15973 | 8.83381 | 1.10371 | 0.18359 | 0.14327 | 0.00000 | 12.87429 | 0.69706 | 12.32624 | 1.11609 | 0.66592 | 4.16992 | 1.10365 | 0.54585 |
| **Pbld2** | 0.00326 | 0.00703 | 0.34506 | 2.01959 | 1.28276 | 0.00654 | 0.00688 | 0.06726 | 2.77047 | 0.08580 | 1.02813 | 0.07227 | 0.00000 | 0.00000 | 1.04356 | 0.00631 |
| **Pbrm1** | 2.06400 | 0.09973 | 2.34866 | 5.06397 | 2.46377 | 0.00970 | 0.01306 | 1.76254 | 7.19570 | 5.02636 | 3.14319 | 0.58300 | 1.55146 | 5.92063 | 6.51153 | 5.47006 |
| **Pbx1** | 0.01193 | 0.00813 | 0.51561 | 0.00721 | 0.00811 | 0.00563 | 0.00750 | 0.01044 | 0.01178 | 1.12738 | 0.01000 | 0.00543 | 0.00877 | 0.01394 | 0.00190 | 0.00695 |
| **Pbx2** | 0.82259 | 0.51129 | 0.58890 | 0.03725 | 0.00520 | 0.01643 | 0.04802 | 1.02548 | 0.05781 | 0.16855 | 0.40993 | 0.00295 | 0.27048 | 0.00896 | 0.00000 | 0.20276 |
| **Pbx3** | 0.01443 | 0.08138 | 0.19185 | 2.39373 | 0.00987 | 0.00532 | 1.69254 | 0.00891 | 1.00294 | 0.62254 | 0.00000 | 0.13372 | 0.01625 | 0.10808 | 1.77280 | 0.00657 |
| **Pbx4** | 0.16269 | 0.00735 | 1.02895 | 0.54122 | 0.01474 | 0.00619 | 0.00652 | 0.09521 | 0.02696 | 1.94846 | 0.09693 | 0.01084 | 0.98443 | 0.03211 | 0.02449 | 0.00643 |
| **Pbxip1** | 0.87070 | 0.00685 | 0.34354 | 1.00814 | 0.01278 | 0.00487 | 0.00540 | 0.00159 | 0.27510 | 0.18411 | 0.17656 | 0.00401 | 1.87802 | 0.09933 | 0.44203 | 3.46397 |
| **Pcbd1** | 1.25362 | 0.00587 | 1.15083 | 2.58455 | 0.12638 | 0.00547 | 0.00481 | 0.00000 | 0.22070 | 0.96253 | 0.40944 | 0.01904 | 1.88381 | 0.00554 | 0.00000 | 0.01879 |
| **Pcbd2** | 0.09213 | 0.00643 | 0.44770 | 1.66818 | 1.39796 | 0.00544 | 0.00643 | 0.03454 | 2.32124 | 0.28961 | 1.50473 | 0.08371 | 0.00000 | 0.88560 | 0.00000 | 0.00662 |
| **Pcbp1** | 35.94368 | 16.51614 | 24.40944 | 21.60534 | 27.10373 | 8.00274 | 10.00657 | 30.75962 | 21.50177 | 49.02290 | 33.17477 | 16.57101 | 38.96321 | 14.09123 | 20.97329 | 17.69900 |
| **Pcbp2** | 14.38049 | 9.75557 | 10.61160 | 10.67290 | 12.10303 | 3.04036 | 7.96702 | 8.27653 | 10.20933 | 14.46988 | 2.41006 | 6.25257 | 1.99074 | 3.75472 | 8.79062 | 7.79036 |
| **Pcbp3** | 0.02009 | 0.00700 | 0.24851 | 2.51759 | 0.00830 | 0.00530 | 0.00678 | 1.38740 | 1.80320 | 0.48873 | 0.44189 | 0.09112 | 0.00300 | 0.02728 | 0.54492 | 0.00694 |
| **Pcbp4** | 0.12522 | 0.00893 | 0.00146 | 0.23165 | 0.01589 | 3.77554 | 0.00727 | 0.01207 | 0.02712 | 0.99084 | 0.00781 | 0.01802 | 0.02474 | 0.01098 | 0.00522 | 0.37990 |
| **Pcca** | 1.48101 | 2.14757 | 0.00000 | 0.79989 | 0.00541 | 0.00529 | 0.00471 | 0.43893 | 0.07677 | 0.24787 | 0.03959 | 0.08309 | 0.76193 | 0.00894 | 0.37927 | 0.04992 |
| **Pccb** | 7.39108 | 0.17031 | 2.01669 | 4.48225 | 0.04535 | 2.07806 | 2.16627 | 5.55531 | 2.02109 | 13.95711 | 9.85450 | 8.20807 | 3.51885 | 4.54924 | 5.25242 | 0.88831 |
| **Pcdh1** | 0.00861 | 0.00750 | 0.00724 | 0.01024 | 0.55932 | 0.00708 | 0.01782 | 0.01241 | 0.01206 | 0.01393 | 0.01259 | 0.01792 | 0.01076 | 0.01237 | 0.01060 | 0.00727 |
| **Pcdh10** | 0.01366 | 0.00816 | 0.01203 | 0.20305 | 0.00928 | 0.00660 | 0.00753 | 0.01389 | 0.63647 | 0.01215 | 0.00982 | 0.00727 | 0.01284 | 0.01223 | 0.00455 | 0.00735 |
| **Pcdh15** | 0.01331 | 0.00777 | 0.00901 | 1.02549 | 0.00951 | 0.00650 | 0.00723 | 0.01025 | 0.29463 | 0.00000 | 0.00708 | 0.00456 | 0.01343 | 0.41987 | 0.46607 | 0.00728 |
| **Pcdh19** | 0.53594 | 0.00712 | 0.34444 | 0.28669 | 0.00551 | 0.00508 | 0.00698 | 0.01759 | 0.00462 | 0.00000 | 0.01036 | 0.03851 | 0.10389 | 0.02574 | 0.08838 | 2.43484 |
| **Pcdh8** | 0.00996 | 0.00845 | 0.00888 | 1.66215 | 0.00960 | 0.00671 | 0.00816 | 0.25849 | 0.41133 | 0.00713 | 0.00743 | 0.00774 | 0.00667 | 0.01824 | 0.04075 | 0.00707 |
| **Pcdh9** | 0.01347 | 0.00762 | 0.01044 | 0.06758 | 0.00941 | 0.00628 | 0.00733 | 0.01050 | 0.97981 | 0.00000 | 0.00000 | 0.00316 | 0.01308 | 0.01869 | 1.49869 | 0.00721 |
| **Pced1a** | 0.31114 | 2.59302 | 0.34029 | 1.76426 | 0.01378 | 0.01644 | 1.85084 | 1.16443 | 0.00000 | 1.08494 | 0.55827 | 0.00594 | 0.16303 | 0.00000 | 0.78391 | 2.26204 |
| **Pced1b** | 0.73081 | 0.00604 | 0.13992 | 0.43745 | 0.05785 | 0.20867 | 0.57672 | 0.00000 | 1.68967 | 1.53254 | 3.84494 | 0.06123 | 0.00000 | 1.76710 | 1.66219 | 0.26536 |
| **Pcf11** | 2.14093 | 6.63095 | 0.37368 | 1.69600 | 1.98028 | 5.80099 | 2.87560 | 2.66696 | 3.29326 | 1.01399 | 1.07950 | 10.86702 | 1.11507 | 13.35478 | 4.37464 | 2.40025 |
| **Pcgf1** | 1.42926 | 0.02428 | 1.71938 | 12.49838 | 0.02434 | 0.81438 | 0.21918 | 3.47050 | 17.69504 | 12.90171 | 6.30223 | 5.02572 | 2.24719 | 6.20239 | 8.75960 | 0.61762 |
| **Pcgf2** | 0.24442 | 0.00690 | 0.00000 | 0.01022 | 0.00740 | 0.01542 | 0.83401 | 0.00989 | 0.01237 | 0.03623 | 0.01228 | 0.05297 | 0.51165 | 0.01157 | 0.00872 | 0.00676 |
| **Pcgf3** | 0.21764 | 0.01881 | 1.05276 | 2.54640 | 0.68191 | 0.54326 | 0.17119 | 1.40399 | 1.07477 | 1.87915 | 1.19652 | 0.76460 | 0.72247 | 0.84170 | 1.05154 | 1.03243 |
| **Pcgf5** | 0.06011 | 0.13430 | 0.83238 | 10.89836 | 0.01525 | 5.48387 | 0.10879 | 2.50573 | 4.57044 | 8.53295 | 2.50537 | 10.70777 | 2.83635 | 7.48029 | 14.11582 | 0.02183 |
| **Pcgf6** | 4.96691 | 2.22547 | 5.72210 | 15.72432 | 8.43963 | 0.00643 | 5.67118 | 4.09760 | 16.72490 | 15.83663 | 2.01090 | 13.13760 | 3.66519 | 7.80080 | 16.35121 | 0.14213 |
| **Pcid2** | 0.57258 | 1.74690 | 1.75761 | 0.38110 | 0.62510 | 0.00599 | 2.22686 | 3.33141 | 0.07555 | 1.05620 | 0.10867 | 0.36008 | 0.00000 | 0.31103 | 1.67121 | 4.05289 |
| **Pcif1** | 1.84997 | 0.02546 | 1.21293 | 1.58759 | 2.02829 | 0.60344 | 1.39930 | 1.10155 | 0.10730 | 1.91680 | 0.17161 | 0.07351 | 1.57568 | 0.00694 | 0.00000 | 1.08515 |
| **Pck2** | 5.56003 | 5.32619 | 2.91337 | 0.96620 | 1.30323 | 0.00710 | 8.94410 | 6.26662 | 0.68288 | 2.70749 | 3.98656 | 0.06335 | 3.73531 | 0.00000 | 0.06106 | 2.54259 |
| **Pclo** | 0.08270 | 0.00748 | 0.13600 | 0.04327 | 0.02484 | 0.03460 | 0.02757 | 0.02715 | 0.18829 | 0.04858 | 0.54140 | 0.05237 | 0.08331 | 0.17507 | 0.47587 | 0.05733 |
| **Pcm1** | 3.67485 | 4.76731 | 1.28194 | 8.08012 | 6.60114 | 1.01898 | 1.93182 | 1.88203 | 1.59852 | 3.40337 | 1.23256 | 3.65235 | 0.83360 | 17.63389 | 9.55845 | 0.29986 |
| **Pcmt1** | 8.58237 | 3.05609 | 3.60294 | 9.34226 | 7.14574 | 3.55873 | 1.12287 | 11.26627 | 10.98475 | 14.13471 | 6.14578 | 1.88198 | 6.78707 | 5.73784 | 7.89915 | 6.84114 |
| **Pcmtd1** | 1.98187 | 5.40371 | 7.97255 | 7.97829 | 4.56480 | 0.72620 | 1.85202 | 2.20199 | 5.04553 | 0.90681 | 1.90107 | 2.45657 | 0.85228 | 4.91622 | 0.76139 | 1.28852 |
| **Pcmtd2** | 1.13382 | 1.50083 | 0.06083 | 0.04976 | 0.00640 | 0.00540 | 0.00604 | 0.00000 | 0.00667 | 0.16392 | 0.62883 | 0.01292 | 0.11488 | 0.00457 | 2.00401 | 0.00577 |
| **Pcna** | 24.61001 | 13.19160 | 20.69767 | 15.62744 | 19.80687 | 11.12571 | 9.45790 | 23.07298 | 27.53670 | 18.67511 | 16.26856 | 18.45018 | 18.01334 | 12.09451 | 19.59854 | 18.78270 |

| | | | | | | | | | | | | | | | | |
|---|---|---|---|---|---|---|---|---|---|---|---|---|---|---|---|---|
| **Pcnp** | 10.07455 | 9.06601 | 9.64449 | 2.86462 | 7.20079 | 5.50011 | 3.11941 | 4.61272 | 4.84492 | 4.95221 | 3.70637 | 9.60759 | 6.84282 | 9.73420 | 4.33152 | 5.52387 |
| **Pcnt** | 4.30152 | 0.80117 | 2.49614 | 8.28591 | 4.36336 | 0.65738 | 4.56330 | 6.68298 | 8.40766 | 21.12502 | 5.14011 | 10.00893 | 3.37615 | 0.00000 | 1.24689 | 3.52030 |
| **Pcnx** | 1.52809 | 0.00380 | 0.41625 | 1.34512 | 1.06754 | 0.00719 | 1.54481 | 3.31927 | 1.03603 | 2.15260 | 0.62475 | 0.71641 | 1.34032 | 0.00134 | 0.31344 | 0.82257 |
| **Pcnx2** | 0.00687 | 0.00747 | 0.00683 | 0.30888 | 0.00969 | 0.00653 | 0.00705 | 1.37749 | 0.09487 | 0.96556 | 0.00000 | 0.01644 | 0.00549 | 0.44881 | 0.55856 | 0.00741 |
| **Pcnx3** | 1.59967 | 0.20123 | 0.00000 | 0.23526 | 0.02447 | 1.94525 | 1.19883 | 1.63215 | 0.48414 | 2.89066 | 0.00000 | 0.15393 | 2.70009 | 0.00743 | 0.01450 | 6.26633 |
| **Pcnx4** | 0.92249 | 0.00649 | 0.88315 | 1.12957 | 0.01293 | 0.00425 | 0.00651 | 0.01005 | 2.91142 | 0.00031 | 0.00000 | 0.05244 | 0.02187 | 0.00447 | 1.80611 | 0.32067 |
| **Pcolce** | 0.01482 | 0.00751 | 0.01944 | 0.28375 | 0.00867 | 0.00799 | 0.00690 | 0.15271 | 0.01227 | 1.22766 | 0.76044 | 0.02874 | 0.00355 | 0.00536 | 0.00000 | 0.00704 |
| **Pcolce2** | 1.18260 | 0.00751 | 0.00059 | 0.00470 | 0.01509 | 0.01540 | 0.00679 | 0.01387 | 0.31064 | 0.00456 | 0.16284 | 0.01683 | 0.00969 | 0.07680 | 0.01996 | 0.01374 |
| **Pcp2** | 0.01314 | 0.00822 | 0.01379 | 0.01018 | 0.00996 | 0.00729 | 0.00784 | 0.01720 | 0.01231 | 0.05398 | 0.01388 | 0.00845 | 0.08190 | 0.00909 | 0.03267 | 0.00771 |
| **Pcp4l1** | 0.01353 | 0.00783 | 0.00965 | 1.84355 | 0.00907 | 0.00650 | 0.07732 | 0.00549 | 0.00000 | 0.32773 | 0.17706 | 0.00531 | 0.01700 | 0.00302 | 0.36779 | 0.00738 |
| **Pcsk1n** | 0.03251 | 0.00800 | 0.01038 | 0.07019 | 0.00834 | 0.00681 | 0.00738 | 0.01734 | 0.03506 | 0.12576 | 0.07698 | 0.00773 | 0.12899 | 0.01082 | 0.00766 | 0.00722 |
| **Pcsk6** | 4.83008 | 0.80504 | 8.15464 | 0.19022 | 5.59271 | 5.52380 | 5.87357 | 9.59558 | 0.14420 | 3.70571 | 1.35987 | 2.01625 | 4.08820 | 0.17703 | 0.67376 | 7.02664 |
| **Pcsk7** | 4.33473 | 0.00000 | 3.43536 | 2.46867 | 0.00000 | 3.07270 | 0.00000 | 6.17851 | 1.84418 | 5.72551 | 3.56574 | 1.77450 | 4.31579 | 0.85812 | 2.04482 | 0.00000 |
| **Pctp** | 0.03141 | 0.00773 | 0.62569 | 0.21369 | 0.00841 | 0.02332 | 0.00725 | 0.01628 | 0.04191 | 0.43055 | 0.06646 | 0.01717 | 0.05910 | 0.09183 | 0.00011 | 0.00705 |
| **Pcx** | 1.27860 | 1.42990 | 0.46439 | 0.72371 | 0.00451 | 0.00507 | 0.00602 | 1.36681 | 2.81460 | 1.70317 | 0.47206 | 0.03988 | 0.04753 | 0.00235 | 0.00000 | 0.00486 |
| **Pcyox1** | 1.83323 | 0.00529 | 0.68160 | 1.70847 | 0.01230 | 0.00374 | 0.04603 | 1.02797 | 0.44061 | 0.91625 | 0.00657 | 0.10687 | 0.00631 | 0.04110 | 1.08522 | 0.02708 |
| **Pcyox1l** | 0.09129 | 0.00771 | 0.08381 | 0.00496 | 0.00726 | 0.00591 | 0.00686 | 0.05889 | 0.22932 | 0.78426 | 0.67149 | 0.00171 | 1.42768 | 0.01362 | 0.00847 | 1.64274 |
| **Pcyt1a** | 2.12916 | 6.25694 | 3.36302 | 12.63685 | 7.01205 | 4.30669 | 4.31941 | 3.83301 | 19.48906 | 15.91800 | 4.79212 | 4.24383 | 3.89855 | 12.23646 | 16.58677 | 1.03772 |
| **Pcyt1b** | 1.60253 | 0.00190 | 3.06979 | 0.71798 | 6.84704 | 0.78403 | 0.00577 | 4.26989 | 1.02611 | 1.35345 | 0.64289 | 0.00669 | 0.27645 | 0.07247 | 2.85647 | 0.42005 |
| **Pcyt2** | 6.00682 | 7.14852 | 9.53061 | 5.84296 | 3.54800 | 8.86760 | 9.05670 | 25.53100 | 4.47603 | 14.38423 | 13.38380 | 0.50258 | 14.49794 | 0.13966 | 4.36211 | 3.99805 |
| **Pdap1** | 6.92412 | 1.96589 | 4.51311 | 7.57290 | 2.36281 | 0.67399 | 2.08149 | 0.72268 | 9.87477 | 3.79640 | 12.80488 | 0.74895 | 4.04338 | 1.62699 | 2.21873 | 2.08521 |
| **Pdcd10** | 10.93666 | 8.28125 | 8.82806 | 9.14201 | 6.94404 | 2.64146 | 8.22996 | 9.87389 | 12.98522 | 8.41326 | 5.31600 | 9.19107 | 9.01964 | 8.82932 | 5.80912 | 2.68455 |
| **Pdcd11** | 5.37270 | 0.07028 | 0.00000 | 3.28978 | 6.12598 | 6.25995 | 2.06611 | 13.76501 | 4.04579 | 15.32459 | 0.70264 | 0.45231 | 7.43709 | 0.55300 | 0.44953 | 4.11504 |
| **Pdcd2** | 1.97883 | 1.04545 | 1.37968 | 9.42434 | 4.80109 | 0.98103 | 1.91884 | 3.29218 | 10.64367 | 0.91955 | 4.35017 | 0.90957 | 1.82175 | 0.45474 | 1.71706 | 0.59584 |
| **Pdcd2l** | 5.48829 | 0.18289 | 7.23091 | 9.97019 | 0.77982 | 0.21062 | 0.02520 | 7.16217 | 7.59570 | 11.73309 | 10.26888 | 0.76646 | 6.93399 | 2.82796 | 2.41881 | 4.71099 |
| **Pdcd4** | 1.90258 | 0.12470 | 0.57316 | 0.56675 | 0.03893 | 0.00499 | 0.35843 | 0.17118 | 0.67059 | 1.01503 | 0.38091 | 0.06830 | 0.05688 | 0.73352 | 0.39561 | 1.38299 |
| **Pdcd5** | 2.85937 | 0.28951 | 3.31635 | 1.97587 | 4.68693 | 0.53921 | 1.16378 | 0.03162 | 3.71868 | 2.03702 | 9.91249 | 0.15793 | 3.28023 | 0.23261 | 0.00000 | 1.38084 |
| **Pdcd6** | 14.84110 | 2.49729 | 7.73150 | 2.27866 | 10.27120 | 9.30910 | 8.56652 | 18.62861 | 0.00000 | 15.61939 | 8.59781 | 1.18964 | 19.58091 | 5.57885 | 1.06928 | 9.96056 |
| **Pdcd6ip** | 6.48618 | 2.36108 | 0.78380 | 8.11624 | 5.56060 | 2.20260 | 3.48713 | 4.44595 | 5.34404 | 10.17585 | 7.56333 | 2.59674 | 3.84165 | 0.52662 | 2.62492 | 3.85099 |
| **Pdcd7** | 4.51911 | 2.81430 | 1.97345 | 5.29942 | 0.66038 | 0.99883 | 2.18496 | 6.35626 | 6.14469 | 12.42240 | 3.14389 | 0.89222 | 3.75021 | 12.33358 | 13.71194 | 0.03315 |
| **Pdcl** | 2.49081 | 2.71026 | 4.27780 | 1.57908 | 2.57356 | 3.59242 | 0.65184 | 2.82847 | 3.47798 | 8.61860 | 2.07101 | 0.66773 | 3.62943 | 17.96913 | 9.44769 | 1.66734 |
| **Pdcl3** | 4.73529 | 7.77171 | 4.15910 | 1.50813 | 4.33280 | 9.51924 | 8.62560 | 7.15682 | 0.32273 | 4.92833 | 5.18455 | 5.94279 | 5.10107 | 8.42405 | 3.67992 | 6.82508 |
| **Pddc1** | 0.40531 | 0.22256 | 2.68938 | 0.45879 | 0.00252 | 0.06709 | 0.00563 | 0.07037 | 0.63882 | 1.43933 | 0.14195 | 0.04149 | 1.91819 | 0.27545 | 1.71711 | 1.10149 |
| **Pde12** | 1.71327 | 2.51426 | 2.38803 | 10.39830 | 0.23438 | 4.57261 | 0.00000 | 4.95861 | 17.80948 | 7.86405 | 1.63562 | 8.00125 | 2.77062 | 1.54899 | 5.44736 | 0.46345 |
| **Pde1b** | 0.00000 | 2.26370 | 0.00669 | 0.31087 | 0.00692 | 0.00642 | 0.00656 | 0.01053 | 0.11224 | 0.04311 | 0.00928 | 0.03934 | 0.00692 | 0.00428 | 0.30531 | 2.65706 |
| **Pde2a** | 0.01346 | 0.00790 | 0.01185 | 0.02225 | 0.00908 | 0.00684 | 0.00751 | 0.01241 | 0.01266 | 0.48372 | 0.01154 | 0.01869 | 0.01233 | 0.00633 | 0.13044 | 0.00739 |
| **Pde3a** | 0.01764 | 0.00779 | 0.01134 | 0.21434 | 0.00903 | 0.00660 | 0.00759 | 0.01638 | 0.00783 | 0.00780 | 0.01250 | 0.00757 | 0.01252 | 0.02000 | 0.20916 | 0.00733 |
| **Pde3b** | 0.00794 | 0.00792 | 0.01211 | 8.26128 | 0.00999 | 0.00500 | 0.00689 | 0.01030 | 2.34153 | 0.00000 | 0.04477 | 0.02185 | 0.01166 | 3.10938 | 4.21629 | 0.00557 |
| **Pde4a** | 0.04703 | 0.00774 | 0.07213 | 1.52439 | 0.00870 | 0.00560 | 0.00710 | 0.00000 | 1.47411 | 0.49277 | 0.43543 | 0.02336 | 0.01555 | 0.00466 | 0.00000 | 0.00688 |
| **Pde4c** | 0.12167 | 0.02828 | 0.19813 | 0.56098 | 0.04917 | 0.01886 | 0.15130 | 0.01458 | 0.43553 | 0.09221 | 0.79342 | 1.21662 | 0.27607 | 0.58148 | 0.21987 | 0.05066 |
| **Pde4d** | 0.01326 | 0.00814 | 0.01130 | 0.40155 | 0.00905 | 0.00668 | 0.00748 | 0.01179 | 0.76358 | 0.54488 | 0.00580 | 0.00608 | 0.01237 | 0.00498 | 0.47459 | 0.00728 |
| **Pde4dip** | 1.26214 | 0.00430 | 0.11282 | 4.26995 | 3.17722 | 4.00722 | 1.41780 | 0.09986 | 1.21827 | 0.86637 | 0.81093 | 0.01201 | 0.65541 | 0.00000 | 1.33290 | 0.84126 |
| **Pde5a** | 0.14079 | 0.05099 | 0.22628 | 0.39077 | 0.08498 | 0.04220 | 0.12308 | 0.07612 | 0.25465 | 0.81725 | 0.35228 | 0.01259 | 0.09272 | 0.35070 | 0.11686 | 0.07227 |
| **Pde6a** | 0.40363 | 0.00700 | 0.03687 | 1.57127 | 0.00685 | 0.00780 | 0.00596 | 0.00000 | 3.17710 | 0.99636 | 2.01251 | 0.03262 | 0.00000 | 0.30067 | 1.10187 | 0.00609 |
| **Pde6b** | 0.01166 | 0.00812 | 0.01012 | 0.00959 | 0.01032 | 0.00717 | 0.05967 | 0.01193 | 0.01183 | 0.00000 | 0.01081 | 0.00551 | 0.19291 | 0.67496 | 0.13399 | 0.00764 |
| **Pde6d** | 2.95064 | 0.02488 | 0.42305 | 0.73988 | 0.40535 | 0.00845 | 0.04582 | 1.32597 | 1.60114 | 3.09942 | 0.32242 | 0.04739 | 0.77299 | 0.00184 | 1.47628 | 0.00360 |
| **Pde6g** | 0.01022 | 0.00758 | 0.00461 | 0.56877 | 0.00948 | 0.00663 | 0.00699 | 0.94089 | 0.02101 | 0.00000 | 0.12151 | 0.01320 | 0.02659 | 0.00467 | 0.99945 | 0.00706 |
| **Pde7a** | 0.23563 | 3.20829 | 0.61513 | 0.85137 | 0.03477 | 1.50340 | 2.20870 | 0.01167 | 0.59981 | 1.70409 | 0.31048 | 2.14394 | 1.85532 | 0.85549 | 1.79635 | 0.19501 |
| **Pde8a** | 0.49794 | 0.94287 | 0.33154 | 1.35022 | 0.00656 | 0.00650 | 0.00614 | 0.27244 | 0.16912 | 1.30243 | 0.00788 | 0.02182 | 0.34596 | 0.00394 | 0.46276 | 0.68509 |
| **Pdf** | 0.53625 | 0.09100 | 1.19426 | 2.35337 | 3.44640 | 0.11720 | 0.09615 | 2.26426 | 3.90375 | 7.31943 | 4.30464 | 3.75470 | 0.31747 | 0.30460 | 3.85825 | 0.85286 |
| **Pdgfa** | 1.81690 | 1.31787 | 2.77906 | 0.00000 | 0.07878 | 2.57380 | 1.83213 | 3.51474 | 0.00850 | 2.06418 | 0.39364 | 0.00000 | 3.87517 | 0.00500 | 0.01253 | 0.85488 |
| **Pdgfb** | 2.15143 | 0.00549 | 0.00000 | 0.02432 | 0.00389 | 0.00573 | 0.00541 | 0.05310 | 0.21601 | 0.15915 | 0.00765 | 0.02097 | 0.01553 | 0.09650 | 1.07595 | 0.00522 |
| **Pdgfra** | 9.14059 | 4.01808 | 0.06701 | 0.00000 | 7.83613 | 0.00000 | 7.98364 | 0.00000 | 0.04700 | 0.19629 | 0.03271 | 0.07995 | 0.02438 | 1.95497 | 1.34445 | 4.61272 |
| **Pdgfrb** | 0.01382 | 0.00787 | 0.01129 | 0.01042 | 0.00940 | 0.00664 | 0.00740 | 0.01230 | 0.01238 | 0.05899 | 0.01259 | 0.06552 | 0.01278 | 0.01163 | 0.00866 | 0.00745 |
| **Pdha1** | 14.51340 | 5.23868 | 19.95343 | 5.04939 | 10.90386 | 10.31439 | 8.92538 | 28.57279 | 9.09482 | 21.97694 | 12.62477 | 10.32586 | 20.92507 | 21.73310 | 12.31320 | 6.33226 |
| **Pdha2** | 0.35140 | 0.00739 | 0.01270 | 0.00982 | 0.00712 | 0.00757 | 0.00732 | 0.01140 | 0.95706 | 0.00944 | 0.03509 | 0.00630 | 0.01186 | 0.01383 | 0.00558 | 0.00693 |
| **Pdhb** | 21.96547 | 15.10873 | 21.58796 | 13.74065 | 17.54117 | 14.62638 | 16.36046 | 25.32709 | 18.64835 | 22.65742 | 21.47975 | 9.48010 | 23.70728 | 15.77033 | 10.01907 | 12.81795 |
| **Pdhx** | 4.04374 | 1.88093 | 6.70837 | 4.81880 | 6.11216 | 2.93967 | 1.93566 | 10.66424 | 3.30352 | 8.88871 | 6.40943 | 0.69927 | 6.28311 | 1.29469 | 5.93460 | 3.02732 |
| **Pdia2** | 0.01225 | 0.00739 | 0.83407 | 0.00692 | 0.00776 | 0.00917 | 0.00697 | 0.00000 | 0.01159 | 1.08061 | 0.34433 | 0.02002 | 0.00929 | 0.01184 | 0.00000 | 0.00716 |
| **Pdia3** | 3.05579 | 6.08700 | 5.09877 | 6.65842 | 1.90399 | 6.62541 | 2.53561 | 1.33851 | 10.92253 | 3.64629 | 10.10367 | 8.14687 | 4.07550 | 15.04602 | 8.72479 | 1.40774 |
| **Pdia4** | 0.77006 | 0.01555 | 0.45517 | 1.99810 | 0.01445 | 1.59650 | 1.95565 | 0.98696 | 3.00958 | 2.48381 | 0.00000 | 0.03499 | 1.62782 | 1.06489 | 4.10093 | 0.01193 |
| **Pdia5** | 6.98764 | 5.53877 | 13.00852 | 1.24735 | 7.58365 | 1.29282 | 10.90958 | 8.82472 | 1.12418 | 3.52488 | 15.31085 | 0.09432 | 14.58116 | 4.73514 | 1.99474 | 5.13203 |

| | | | | | | | | | | | | | | | | |
|---|---|---|---|---|---|---|---|---|---|---|---|---|---|---|---|---|
| **Pdia6** | 11.86447 | 9.26919 | 28.99791 | 7.68924 | 3.84559 | 13.31102 | 10.93591 | 9.95934 | 3.07462 | 8.06452 | 4.91710 | 2.03563 | 15.74593 | 1.00500 | 4.69035 | 15.03582 |
| **Pdik1l** | 1.18661 | 0.00532 | 0.15799 | 0.24526 | 0.00935 | 0.00839 | 1.67663 | 0.00709 | 0.43680 | 0.62969 | 0.00000 | 1.41473 | 0.29457 | 0.00774 | 0.84253 | 0.01371 |
| **Pdk1** | 1.02758 | 0.00701 | 0.00000 | 7.01326 | 0.01236 | 0.01092 | 0.00613 | 0.00876 | 1.99323 | 0.17062 | 0.04591 | 0.04425 | 0.03838 | 0.03009 | 3.40628 | 0.00544 |
| **Pdk2** | 2.75924 | 0.00445 | 0.00253 | 0.05610 | 0.01394 | 0.00567 | 0.00502 | 0.00049 | 0.01088 | 0.38535 | 0.01295 | 0.00487 | 0.17758 | 0.04132 | 0.10234 | 0.38250 |
| **Pdk3** | 0.00711 | 0.00669 | 0.06474 | 2.98956 | 0.00843 | 0.00804 | 2.70989 | 0.00905 | 2.44707 | 3.51423 | 0.02766 | 0.47075 | 0.00937 | 2.53234 | 3.32835 | 0.00625 |
| **Pdlim1** | 15.21133 | 8.48721 | 13.57838 | 5.95158 | 16.38241 | 9.24389 | 14.84730 | 5.57361 | 10.23954 | 7.93929 | 3.67651 | 6.65989 | 13.65383 | 0.58034 | 0.80215 | 9.36329 |
| **Pdlim2** | 0.01214 | 0.01827 | 0.00738 | 1.19092 | 0.00921 | 0.00682 | 0.00705 | 0.83039 | 0.00000 | 0.00685 | 0.02212 | 0.02604 | 0.00944 | 0.00877 | 0.01921 | 0.00721 |
| **Pdlim3** | 0.18962 | 0.00773 | 1.15506 | 0.00788 | 0.00555 | 0.00544 | 0.01883 | 0.04695 | 0.01114 | 2.95130 | 0.02504 | 0.12102 | 0.00568 | 0.00806 | 0.29371 | 0.00669 |
| **Pdlim4** | 1.35852 | 0.00411 | 0.00058 | 0.00000 | 5.13853 | 0.00724 | 0.00604 | 0.00000 | 0.68046 | 1.87940 | 0.00441 | 0.02706 | 0.00757 | 0.01246 | 0.00617 | 2.01307 |
| **Pdlim5** | 0.97611 | 2.85259 | 1.97811 | 3.37454 | 0.07522 | 5.58677 | 1.47939 | 2.79134 | 4.68139 | 3.47513 | 2.85822 | 11.24391 | 2.25531 | 0.01128 | 0.80698 | 1.17317 |
| **Pdlim7** | 4.17535 | 0.02393 | 0.24976 | 2.60538 | 0.03598 | 0.97021 | 0.04163 | 2.82583 | 0.00000 | 2.35582 | 1.38166 | 0.13250 | 3.46364 | 0.00000 | 0.00000 | 0.89758 |
| **Pdp1** | 0.00837 | 0.00724 | 0.00000 | 2.39918 | 0.01881 | 0.52923 | 0.01605 | 0.47844 | 0.65623 | 0.23177 | 0.13301 | 0.01077 | 1.25388 | 0.00000 | 0.44434 | 0.01351 |
| **Pdp2** | 0.13495 | 0.73621 | 4.46790 | 10.02873 | 0.20484 | 0.02069 | 1.53603 | 4.69463 | 13.96188 | 10.67228 | 5.09673 | 1.33225 | 1.67383 | 17.74419 | 8.59771 | 1.84936 |
| **Pdpk1** | 10.26509 | 1.37562 | 3.53612 | 4.88157 | 5.00985 | 3.75256 | 1.75870 | 2.75771 | 5.80762 | 11.87017 | 3.09366 | 10.35444 | 4.07781 | 20.30761 | 8.29217 | 0.52560 |
| **Pdpn** | 11.22335 | 6.05011 | 11.44493 | 1.63928 | 3.27899 | 0.00000 | 8.79500 | 0.28964 | 4.68075 | 0.73401 | 6.67013 | 0.44282 | 8.59199 | 3.17764 | 1.68352 | 1.98770 |
| **Pdpr** | 0.23007 | 0.00738 | 0.00000 | 1.12586 | 0.00741 | 0.00664 | 0.00723 | 0.13929 | 4.30561 | 0.51763 | 0.00000 | 0.00513 | 0.00976 | 0.00000 | 1.82904 | 0.00678 |
| **Pdrg1** | 6.82282 | 0.00000 | 9.36282 | 14.64189 | 9.46840 | 3.86041 | 3.05466 | 1.70519 | 24.38278 | 10.52762 | 8.04219 | 1.19106 | 5.58920 | 1.87777 | 7.18770 | 0.00000 |
| **Pds5a** | 0.57443 | 0.31227 | 1.05518 | 2.83693 | 0.58148 | 1.54442 | 1.42412 | 3.21038 | 5.06627 | 2.71265 | 0.00000 | 3.42743 | 0.55803 | 12.12647 | 7.85657 | 0.30835 |
| **Pds5b** | 0.20292 | 4.33890 | 0.49497 | 5.96905 | 1.48519 | 6.04294 | 0.54756 | 0.00000 | 10.55463 | 1.16085 | 1.31896 | 11.07286 | 0.35322 | 13.04183 | 5.49832 | 0.21922 |
| **Pdss1** | 5.28914 | 2.00830 | 3.75100 | 0.23624 | 0.01504 | 0.97518 | 1.88339 | 1.32610 | 0.63859 | 3.15484 | 1.97498 | 0.27721 | 3.99576 | 2.23941 | 1.25624 | 4.89373 |
| **Pdss2** | 0.63265 | 0.21141 | 0.80482 | 3.51552 | 0.01229 | 3.33250 | 0.00614 | 1.76110 | 1.03922 | 2.64693 | 0.53949 | 0.07701 | 0.36585 | 2.52930 | 1.00264 | 0.29785 |
| **Pdxdc1** | 3.56105 | 6.03177 | 3.10263 | 3.41936 | 0.03863 | 1.87150 | 2.15985 | 2.63419 | 1.72268 | 1.28918 | 2.97180 | 0.73171 | 1.10789 | 1.43988 | 2.91483 | 3.98729 |
| **Pdxk** | 6.17171 | 5.43896 | 19.85095 | 16.28664 | 7.36993 | 3.72356 | 12.63106 | 39.67593 | 25.31774 | 51.94072 | 34.45878 | 9.91344 | 26.42799 | 1.24338 | 9.71347 | 0.21129 |
| **Pdxk.ps** | 0.01849 | 0.00783 | 0.02027 | 0.08954 | 0.00884 | 0.00699 | 0.01817 | 0.04383 | 0.20726 | 0.03956 | 0.12651 | 0.00626 | 0.03276 | 0.14239 | 0.15732 | 0.00730 |
| **Pdxp** | 0.89779 | 0.01547 | 0.00000 | 10.07590 | 0.03873 | 4.66171 | 2.96633 | 4.89153 | 7.71259 | 4.45811 | 3.47301 | 1.04747 | 1.23408 | 0.00000 | 2.96651 | 0.03178 |
| **Pdzd11** | 10.59403 | 7.59208 | 13.97921 | 7.20826 | 0.81153 | 3.32847 | 6.08779 | 6.41329 | 11.70896 | 12.34194 | 9.99889 | 8.77427 | 10.84631 | 7.25035 | 9.72827 | 4.17386 |
| **Pdzd2** | 0.10083 | 0.00714 | 2.95576 | 3.37044 | 0.05451 | 0.03296 | 0.00641 | 0.00000 | 4.35245 | 0.48038 | 0.68706 | 0.12125 | 0.10232 | 0.01537 | 1.67313 | 0.01871 |
| **Pdzd3** | 1.64360 | 5.27950 | 10.27313 | 0.00000 | 0.10776 | 4.32905 | 2.67798 | 8.38392 | 0.01044 | 1.22205 | 8.09359 | 0.18730 | 12.13982 | 0.07911 | 0.00000 | 3.61045 |
| **Pdzd4** | 0.12928 | 0.00700 | 0.00000 | 1.69006 | 0.00742 | 0.00746 | 0.00561 | 0.00000 | 0.32514 | 1.21945 | 3.63059 | 0.00000 | 0.00000 | 0.29998 | 0.99794 | 0.01353 |
| **Pdzd8** | 1.41485 | 0.06374 | 4.06012 | 5.65485 | 4.30220 | 0.05739 | 2.73433 | 3.06923 | 2.74773 | 2.62202 | 6.78714 | 2.80802 | 3.31362 | 12.75007 | 8.77222 | 2.70216 |
| **Pdzk1** | 0.32559 | 0.08636 | 1.75481 | 1.06670 | 0.05957 | 8.70551 | 0.01569 | 0.87175 | 0.28427 | 0.72662 | 0.14023 | 0.00883 | 1.35211 | 0.00196 | 1.05492 | 0.07558 |
| **Pdzk1ip1** | 0.00805 | 0.00839 | 0.02379 | 0.00237 | 0.01056 | 3.45631 | 0.00637 | 0.01067 | 0.01216 | 1.33007 | 0.00845 | 0.00382 | 2.18450 | 0.01211 | 0.00816 | 0.01286 |
| **Pdzph1** | 0.01347 | 0.00771 | 0.01055 | 1.70070 | 0.00908 | 0.00647 | 0.00722 | 0.01024 | 1.55314 | 0.41382 | 0.00000 | 0.06958 | 0.01260 | 0.01112 | 0.02510 | 0.00743 |
| **Pdzrn3** | 0.01341 | 0.00810 | 0.01319 | 0.30397 | 0.00958 | 0.00689 | 0.00748 | 0.01222 | 0.00443 | 0.42369 | 0.01325 | 0.00653 | 0.02666 | 0.00693 | 0.08141 | 0.00797 |
| **Pea15a** | 0.18091 | 0.00752 | 2.62116 | 2.25886 | 0.00701 | 0.00565 | 0.00590 | 0.09092 | 0.00000 | 5.05680 | 0.01014 | 0.04826 | 1.81301 | 0.00133 | 1.74244 | 0.00522 |
| **Peak1** | 2.33644 | 0.00522 | 1.60335 | 10.09045 | 0.01495 | 0.00433 | 0.18580 | 0.58285 | 5.25918 | 1.40214 | 1.75682 | 0.05861 | 2.63180 | 4.17331 | 1.88912 | 0.96916 |
| **Pebp1** | 22.99961 | 5.08897 | 11.27424 | 14.01500 | 12.56680 | 6.13238 | 8.67722 | 16.39986 | 20.73371 | 26.78162 | 10.17412 | 4.24682 | 19.86938 | 2.98428 | 6.89858 | 11.42574 |
| **Pecam1** | 4.49762 | 1.32810 | 5.93993 | 1.26780 | 0.00000 | 0.00000 | 2.04725 | 3.21602 | 0.78246 | 11.16789 | 4.10650 | 0.72031 | 0.37354 | 10.59731 | 13.70115 | 0.01260 |
| **Pecr** | 0.99761 | 2.20254 | 0.00000 | 6.48410 | 0.34435 | 0.00436 | 0.56275 | 0.13097 | 3.20919 | 0.62047 | 0.33575 | 0.44825 | 1.53404 | 1.44272 | 1.31203 | 0.01865 |
| **Pef1** | 13.61463 | 7.06959 | 10.89387 | 0.23648 | 0.45884 | 2.47553 | 9.39970 | 14.02708 | 0.03625 | 11.94807 | 3.41274 | 0.41386 | 12.16494 | 2.00881 | 3.41964 | 7.53578 |
| **Peg13** | 0.58674 | 0.00704 | 0.00442 | 0.00922 | 0.00666 | 0.00754 | 0.00707 | 0.00963 | 0.01202 | 0.33389 | 0.01147 | 0.05209 | 0.01222 | 0.01169 | 0.01028 | 0.00683 |
| **Peg3** | 0.01585 | 0.00810 | 0.01668 | 0.00996 | 0.06370 | 0.00637 | 0.00754 | 0.01340 | 0.01224 | 0.01152 | 0.01288 | 0.00798 | 0.01213 | 0.00822 | 0.09048 | 0.00740 |
| **Peli1** | 0.46483 | 3.44281 | 5.44594 | 2.09228 | 3.33292 | 1.89641 | 4.08094 | 4.84190 | 5.11954 | 0.48001 | 0.36112 | 4.05020 | 0.00000 | 4.47118 | 2.12156 | 0.06850 |
| **Peli2** | 0.15090 | 0.00820 | 0.81316 | 0.03478 | 0.03390 | 0.00903 | 0.00715 | 0.01072 | 0.28571 | 0.02291 | 0.02225 | 0.00653 | 0.00839 | 0.01396 | 0.00000 | 0.00713 |
| **Pelo** | 4.51907 | 1.15639 | 4.04808 | 0.71367 | 7.92980 | 2.02223 | 2.04604 | 9.70699 | 0.55333 | 4.34523 | 3.78049 | 0.34347 | 0.77150 | 0.64658 | 0.43777 | 3.93908 |
| **Pelp1** | 8.14798 | 1.77995 | 1.31424 | 10.25588 | 4.27885 | 0.03781 | 2.17552 | 4.02233 | 12.09584 | 14.59154 | 16.77519 | 1.22755 | 2.78553 | 0.22509 | 1.91943 | 0.37909 |
| **Pemt** | 1.70841 | 8.27395 | 40.20038 | 17.72312 | 0.53147 | 6.46996 | 14.77608 | 42.95349 | 19.06032 | 37.38647 | 40.41718 | 17.69265 | 30.67427 | 15.42534 | 18.53813 | 3.89492 |
| **Peo1** | 5.79387 | 2.53639 | 3.18647 | 3.64864 | 4.48997 | 4.26219 | 1.88802 | 3.24767 | 8.20432 | 7.74691 | 2.81952 | 0.77090 | 2.79534 | 0.52154 | 9.97606 | 1.12769 |
| **Pepd** | 3.68491 | 5.77703 | 6.15120 | 0.62952 | 6.35979 | 6.01188 | 4.21078 | 6.45105 | 0.00000 | 0.91606 | 1.59194 | 0.07659 | 10.20809 | 0.00000 | 0.00487 | 5.69604 |
| **Per1** | 0.00000 | 0.00826 | 0.68242 | 0.15367 | 0.01470 | 0.00843 | 0.04161 | 0.39430 | 0.94715 | 2.40971 | 0.19017 | 0.07586 | 0.55048 | 0.52577 | 0.00000 | 1.58364 |
| **Per2** | 0.00000 | 0.00421 | 0.00000 | 0.00000 | 0.00943 | 0.00550 | 2.01075 | 0.50090 | 0.02187 | 0.21923 | 0.42932 | 6.33126 | 0.00456 | 0.01095 | 0.60967 | 0.00686 |
| **Per3** | 0.00283 | 0.47302 | 0.00549 | 1.67728 | 0.00809 | 0.00637 | 0.46714 | 0.00000 | 3.17304 | 0.00000 | 0.25069 | 0.00754 | 0.75992 | 0.00000 | 0.21276 | 0.07568 |
| **Perm1** | 0.01857 | 0.00823 | 0.01853 | 1.14192 | 0.00886 | 0.00591 | 0.00748 | 0.01242 | 3.85393 | 0.00139 | 0.04795 | 0.00000 | 0.01193 | 0.01093 | 0.00186 | 0.00689 |
| **Perp** | 0.39401 | 4.33774 | 4.51878 | 0.26033 | 0.53450 | 11.15952 | 2.22547 | 2.87292 | 0.00632 | 1.51959 | 10.65861 | 0.17314 | 7.66618 | 0.00000 | 0.00000 | 4.39794 |
| **Pes1** | 10.74393 | 2.94073 | 9.48562 | 10.36459 | 7.83730 | 1.81432 | 4.72395 | 10.99718 | 13.48063 | 12.40642 | 11.37795 | 1.19910 | 9.83184 | 13.00850 | 9.31431 | 3.48176 |
| **Pet100** | 0.38087 | 0.00563 | 1.25711 | 0.00829 | 0.48013 | 0.27834 | 2.19251 | 0.00000 | 1.38985 | 0.40371 | 2.29194 | 0.12993 | 0.09779 | 0.28572 | 0.09604 | 0.05015 |
| **Pet2** | 0.32729 | 0.00838 | 1.53401 | 0.00446 | 0.01502 | 0.00519 | 0.00730 | 0.01136 | 0.93778 | 0.01030 | 0.00207 | 0.00617 | 0.00717 | 0.01647 | 0.00000 | 0.00662 |
| **Pex1** | 1.61488 | 0.04848 | 2.43654 | 1.97788 | 1.45854 | 0.12633 | 2.14561 | 1.92810 | 0.06892 | 4.25702 | 1.70168 | 0.27117 | 0.09290 | 1.78331 | 2.96080 | 0.01152 |
| **Pex10** | 1.36374 | 0.00691 | 0.00000 | 0.30617 | 0.01333 | 0.11188 | 0.01704 | 0.00000 | 0.00669 | 0.26505 | 0.71367 | 0.01383 | 0.31345 | 0.02816 | 0.23706 | 0.36779 |
| **Pex11a** | 0.08914 | 0.00719 | 0.12302 | 0.73605 | 0.00781 | 0.00574 | 0.00713 | 0.49907 | 0.29523 | 2.15692 | 0.00456 | 0.08451 | 0.00657 | 0.00496 | 0.93977 | 0.11483 |
| **Pex11b** | 7.61420 | 0.00000 | 1.47488 | 1.53583 | 0.00000 | 0.00000 | 0.01821 | 1.01633 | 3.16287 | 2.78508 | 0.01553 | 0.07444 | 4.55459 | 0.00563 | 1.00849 | 4.67968 |
| **Pex11g** | 0.00841 | 0.62866 | 1.06619 | 0.18967 | 0.00639 | 0.00559 | 0.00671 | 0.00003 | 0.27173 | 0.36062 | 0.29865 | 0.00369 | 0.00663 | 0.00560 | 0.00566 | 0.00692 |

| | | | | | | | | | | | | | | | | |
|---|---|---|---|---|---|---|---|---|---|---|---|---|---|---|---|---|
| **Pex12** | 0.65705 | 0.00394 | 2.45809 | 4.27904 | 4.62275 | 0.68048 | 1.58984 | 1.19584 | 3.18929 | 5.57182 | 0.35376 | 0.55208 | 0.42385 | 1.43967 | 7.78373 | 0.01192 |
| **Pex13** | 2.84231 | 2.44353 | 1.49126 | 4.99086 | 0.18197 | 0.15602 | 1.88402 | 0.00000 | 4.13649 | 0.00000 | 0.82780 | 4.07863 | 1.64597 | 0.56041 | 1.61137 | 0.18591 |
| **Pex14** | 4.53121 | 0.36530 | 1.05270 | 1.10762 | 3.54415 | 0.19882 | 0.31909 | 2.49298 | 1.75445 | 5.16384 | 2.36222 | 0.10879 | 2.71173 | 0.03382 | 0.00000 | 0.20746 |
| **Pex16** | 7.27085 | 0.87452 | 6.32850 | 6.22135 | 2.14083 | 4.48682 | 7.29858 | 19.18798 | 7.19072 | 11.96474 | 10.80182 | 2.57607 | 14.32755 | 0.00000 | 2.18587 | 6.02022 |
| **Pex19** | 7.57964 | 6.18085 | 6.43607 | 8.95537 | 9.72085 | 7.82030 | 7.04268 | 10.63214 | 19.98112 | 16.94628 | 4.70069 | 2.23539 | 8.85456 | 11.76830 | 18.44622 | 4.06964 |
| **Pex2** | 4.19214 | 8.19036 | 5.78307 | 2.02604 | 1.75606 | 3.11278 | 3.73804 | 3.37663 | 1.67020 | 3.67720 | 0.44654 | 2.76230 | 1.95842 | 0.02003 | 1.57513 | 1.97800 |
| **Pex26** | 1.39361 | 0.00390 | 6.31569 | 0.15950 | 0.01121 | 0.00472 | 3.53689 | 0.79326 | 2.19407 | 0.00000 | 0.94687 | 0.11469 | 3.11817 | 0.00303 | 0.46487 | 0.03712 |
| **Pex3** | 7.63198 | 0.94102 | 5.01340 | 2.49886 | 6.33977 | 0.00000 | 2.30380 | 2.32321 | 6.41300 | 9.88910 | 7.06613 | 3.26421 | 2.83561 | 4.87867 | 8.28474 | 0.03010 |
| **Pex5** | 3.58440 | 0.09797 | 0.25390 | 0.05940 | 0.89914 | 7.68935 | 0.03892 | 5.03245 | 0.00732 | 2.19504 | 1.53550 | 0.13641 | 0.88778 | 1.48309 | 0.32775 | 4.65402 |
| **Pex6** | 4.16491 | 0.04559 | 1.52320 | 0.06250 | 0.02090 | 0.06339 | 0.39590 | 4.40009 | 0.55876 | 3.41344 | 0.23662 | 0.14850 | 5.27998 | 0.00840 | 0.00000 | 2.00317 |
| **Pex7** | 1.77374 | 1.48581 | 1.34185 | 0.91238 | 0.01012 | 0.01382 | 1.64846 | 0.24014 | 0.71126 | 1.89159 | 0.74964 | 3.78603 | 1.81005 | 0.70899 | 0.00000 | 1.36668 |
| **Pfas** | 4.15900 | 2.04143 | 1.74977 | 1.48900 | 0.08520 | 0.67667 | 0.98752 | 1.36677 | 2.68141 | 0.44376 | 3.03054 | 0.25715 | 1.71802 | 3.12661 | 0.82105 | 2.28492 |
| **Pfdn1** | 11.08122 | 6.46539 | 7.95524 | 1.00807 | 7.73634 | 6.03345 | 6.63693 | 3.95845 | 0.19039 | 6.45182 | 9.80789 | 1.00341 | 12.16984 | 0.41216 | 1.94041 | 8.60901 |
| **Pfdn2** | 7.44199 | 1.73813 | 6.11674 | 0.00919 | 8.43041 | 1.98893 | 2.55957 | 4.30838 | 0.56004 | 13.33605 | 3.46547 | 0.40093 | 6.98811 | 9.50647 | 6.59692 | 2.19869 |
| **Pfdn4** | 11.40508 | 3.65997 | 3.86935 | 2.45457 | 6.10668 | 3.83798 | 1.85805 | 3.96952 | 0.35054 | 7.03417 | 5.74062 | 0.72815 | 4.79308 | 4.64431 | 3.23968 | 2.50243 |
| **Pfdn5** | 12.86173 | 3.39814 | 7.58058 | 15.97814 | 4.58503 | 2.56858 | 5.09981 | 6.74805 | 19.19335 | 16.17472 | 26.63651 | 1.91298 | 8.21824 | 2.89415 | 9.08958 | 7.34542 |
| **Pfdn6** | 11.66762 | 5.06669 | 6.99724 | 7.82563 | 8.58405 | 0.98084 | 4.63537 | 9.44872 | 12.46551 | 12.70824 | 14.48634 | 0.91446 | 9.55851 | 0.53471 | 7.54299 | 4.93487 |
| **Pfkfb1** | 1.89815 | 0.00612 | 1.34337 | 0.32193 | 0.03168 | 0.00450 | 0.96438 | 0.40528 | 0.44982 | 0.26822 | 0.25753 | 0.00000 | 0.27610 | 0.00582 | 1.13498 | 0.00568 |
| **Pfkfb2** | 1.66875 | 1.76818 | 1.04126 | 0.96125 | 1.88116 | 3.00117 | 0.04959 | 0.65328 | 0.04686 | 2.60241 | 2.53884 | 0.15230 | 2.74001 | 3.55083 | 1.00865 | 1.40047 |
| **Pfkfb3** | 3.02516 | 0.00000 | 3.67937 | 6.64179 | 0.03344 | 0.00671 | 0.06976 | 8.64982 | 9.64392 | 14.42427 | 17.97960 | 3.35054 | 1.23515 | 19.59576 | 16.84408 | 0.04035 |
| **Pfkfb4** | 0.00000 | 0.01664 | 3.20748 | 0.10641 | 0.01323 | 1.55173 | 2.71952 | 4.58475 | 0.00675 | 0.00000 | 0.47644 | 0.04288 | 2.41015 | 0.00998 | 0.30079 | 0.00418 |
| **Pfkl** | 3.50258 | 5.11504 | 2.64803 | 2.79263 | 1.30379 | 1.71647 | 0.60343 | 2.17418 | 1.30690 | 5.01841 | 2.13892 | 0.04139 | 8.00516 | 0.01586 | 0.27036 | 7.26073 |
| **Pfkm** | 3.11837 | 0.00414 | 0.00656 | 0.27106 | 0.00208 | 0.00555 | 1.22008 | 0.11032 | 0.00745 | 0.73984 | 0.00849 | 0.00236 | 0.01097 | 0.00546 | 0.55893 | 0.01056 |
| **Pfkp** | 4.17037 | 2.14945 | 0.00000 | 1.43432 | 5.79277 | 0.01656 | 3.11315 | 0.00000 | 3.70367 | 1.32504 | 0.71645 | 0.06727 | 0.07732 | 0.00000 | 1.19610 | 0.01601 |
| **Pfn1** | 10.05332 | 4.84167 | 1.74935 | 0.00036 | 6.50619 | 2.26804 | 2.31064 | 4.85975 | 0.00000 | 25.72945 | 0.24483 | 0.09596 | 12.48389 | 3.12693 | 4.80738 | 7.91559 |
| **Pfn2** | 4.67999 | 0.00000 | 0.00253 | 1.10295 | 4.82668 | 0.00427 | 0.00447 | 0.16094 | 0.62720 | 0.62736 | 0.00532 | 0.06055 | 0.02800 | 0.00812 | 0.38859 | 0.00952 |
| **Pfn4** | 0.06419 | 2.01920 | 0.00000 | 0.54792 | 0.00825 | 0.02709 | 3.74392 | 0.01170 | 0.00421 | 0.14110 | 0.02990 | 0.02273 | 0.10190 | 0.06742 | 0.27740 | 0.00624 |
| **Pga5** | 1.17527 | 0.00654 | 0.00122 | 0.34530 | 0.00493 | 0.00690 | 0.00640 | 0.00000 | 0.00533 | 0.04187 | 0.47860 | 0.02464 | 0.00407 | 0.01329 | 0.16390 | 1.66618 |
| **Pgam1** | 30.40834 | 16.60213 | 22.17634 | 11.79114 | 19.30393 | 8.32777 | 14.36352 | 27.15265 | 15.32550 | 30.85239 | 28.60263 | 7.70504 | 27.06571 | 12.03802 | 11.53548 | 15.19158 |
| **Pgam2** | 2.63307 | 2.45387 | 4.21283 | 0.00000 | 1.90824 | 0.08803 | 1.05018 | 0.66010 | 0.01916 | 1.31989 | 1.93634 | 0.02684 | 4.16397 | 0.00326 | 0.50570 | 10.77067 |
| **Pgam5** | 9.35475 | 8.56241 | 10.77172 | 0.90008 | 5.88094 | 1.34767 | 6.99168 | 7.42063 | 1.54896 | 5.57800 | 1.44616 | 11.49257 | 10.48655 | 1.64235 | 9.09159 | 7.60659 |
| **Pgap1** | 0.00861 | 0.00715 | 0.03339 | 1.14058 | 0.53804 | 1.88998 | 0.00723 | 0.01565 | 0.18162 | 0.00000 | 0.00946 | 0.01535 | 0.02708 | 0.03912 | 0.30275 | 0.01380 |
| **Pgap2** | 4.37986 | 0.31473 | 0.00000 | 6.05387 | 1.74019 | 6.55468 | 0.00806 | 4.42106 | 3.54306 | 5.87508 | 6.99172 | 6.94653 | 3.05767 | 7.67465 | 10.64674 | 0.29313 |
| **Pgap3** | 0.07573 | 0.00796 | 0.00913 | 0.01534 | 0.00843 | 0.00713 | 0.00738 | 0.05952 | 0.01226 | 0.21136 | 0.01410 | 0.00688 | 0.05671 | 0.00693 | 0.11976 | 0.00757 |
| **Pgbd5** | 0.54144 | 4.68173 | 0.00000 | 1.85775 | 0.03483 | 0.00837 | 0.01615 | 0.09176 | 1.49309 | 0.12182 | 0.58363 | 0.00000 | 1.65909 | 0.00000 | 0.25668 | 0.00387 |
| **Pgc** | 0.02724 | 0.00813 | 0.02022 | 0.01005 | 0.01024 | 0.01801 | 0.00706 | 0.01174 | 0.01227 | 0.11948 | 0.01286 | 0.00712 | 0.60875 | 0.01142 | 0.01085 | 0.02266 |
| **Pgd** | 25.45097 | 19.52974 | 26.62444 | 10.70514 | 25.48900 | 13.09381 | 15.55199 | 29.33676 | 14.55946 | 21.08627 | 18.27529 | 1.78860 | 20.17357 | 0.71125 | 5.24804 | 18.49673 |
| **Pggt1b** | 0.56853 | 0.00679 | 0.30771 | 0.32015 | 0.01396 | 0.01495 | 0.20625 | 0.98991 | 0.00523 | 0.44994 | 0.28992 | 0.02989 | 0.85376 | 3.23951 | 0.83647 | 0.00576 |
| **Pgk1** | 7.72686 | 20.02419 | 18.67335 | 9.56563 | 13.16253 | 12.14677 | 11.59213 | 2.42109 | 0.47926 | 9.39920 | 3.35633 | 1.45301 | 16.63397 | 2.82302 | 8.83642 | 7.34144 |
| **Pgls** | 0.40507 | 0.00716 | 0.00112 | 0.10873 | 1.77991 | 0.40583 | 0.05725 | 0.02054 | 0.03244 | 0.95121 | 0.04237 | 0.04214 | 0.49055 | 0.01127 | 0.00352 | 1.66092 |
| **Pglyrp1** | 0.90682 | 0.09756 | 0.11007 | 3.32731 | 0.00594 | 0.00602 | 0.64356 | 1.48109 | 2.66004 | 2.26301 | 2.38090 | 0.13643 | 1.94816 | 0.00147 | 0.00000 | 0.00426 |
| **Pgm1** | 6.25340 | 2.49799 | 1.93298 | 0.55721 | 5.12759 | 0.00000 | 2.28071 | 2.31375 | 0.33923 | 0.62127 | 0.00000 | 0.03494 | 3.05111 | 0.27483 | 1.46645 | 1.68812 |
| **Pgm2** | 3.88852 | 10.94181 | 3.13673 | 0.24424 | 2.62260 | 10.82084 | 0.13012 | 0.00000 | 0.49363 | 1.18599 | 0.70105 | 0.10320 | 1.69871 | 0.00000 | 0.06027 | 0.00861 |
| **Pgm2l1** | 0.20293 | 0.00583 | 0.70929 | 4.17624 | 2.07364 | 0.00516 | 0.00672 | 0.00000 | 3.09205 | 1.56705 | 0.18457 | 2.46928 | 0.38543 | 9.49924 | 4.25625 | 0.00569 |
| **Pgm3** | 4.77852 | 4.54447 | 3.06771 | 1.88690 | 4.68974 | 0.00571 | 2.95877 | 5.31772 | 0.53379 | 8.01269 | 1.57212 | 4.99405 | 2.48407 | 1.40463 | 9.07344 | 5.80008 |
| **Pgm5** | 0.01226 | 0.01751 | 0.01683 | 0.01631 | 0.00910 | 0.00726 | 0.00722 | 0.02561 | 0.01195 | 0.37008 | 0.01183 | 0.01505 | 0.01803 | 0.04706 | 0.15744 | 0.00777 |
| **Pgp** | 0.95364 | 0.04899 | 0.53159 | 6.09933 | 3.04266 | 0.37856 | 0.16598 | 0.00000 | 7.93911 | 8.70522 | 6.97986 | 0.14563 | 4.43035 | 0.00000 | 0.00000 | 1.44446 |
| **Pgpep1** | 1.80837 | 0.02831 | 0.00394 | 1.54079 | 0.02224 | 0.00653 | 0.00599 | 1.70071 | 0.17399 | 1.21784 | 0.00000 | 0.11875 | 0.09654 | 0.01107 | 0.00000 | 0.30705 |
| **Pgpep1l** | 0.01353 | 0.00797 | 0.01273 | 0.27136 | 0.00979 | 0.00679 | 0.00758 | 0.01285 | 0.40976 | 0.00370 | 0.04023 | 0.00635 | 0.02894 | 0.00652 | 0.34025 | 0.00752 |
| **Pgrmc1** | 2.65567 | 7.40004 | 7.86973 | 9.90297 | 2.42639 | 1.71778 | 4.89870 | 6.52967 | 8.89389 | 8.55444 | 8.39498 | 7.32870 | 11.68908 | 11.66829 | 0.84095 | 2.22425 |
| **Pgrmc2** | 1.67936 | 3.20534 | 2.49551 | 5.56301 | 6.45505 | 1.83939 | 3.46243 | 4.46855 | 10.96074 | 8.66839 | 5.15134 | 1.38816 | 3.13075 | 9.47373 | 5.39498 | 2.49145 |
| **Pgs1** | 3.38242 | 1.43445 | 0.79747 | 13.82922 | 7.98166 | 1.64090 | 2.63186 | 9.45360 | 9.33459 | 11.65912 | 7.73441 | 3.30741 | 2.66981 | 0.73685 | 1.33349 | 0.29806 |
| **Phactr1** | 0.47997 | 0.00823 | 0.00455 | 0.14382 | 0.03602 | 0.00712 | 0.04763 | 0.00861 | 0.00664 | 0.43552 | 0.01025 | 0.00605 | 0.01068 | 0.09660 | 0.01658 | 0.03571 |
| **Phactr2** | 0.67550 | 0.00699 | 0.07336 | 0.01010 | 0.00637 | 0.00868 | 0.00704 | 0.01066 | 0.01213 | 0.13541 | 0.01176 | 0.00823 | 0.01168 | 0.01184 | 0.01110 | 0.00663 |
| **Phactr4** | 2.52448 | 0.15669 | 0.09866 | 4.01699 | 0.02156 | 0.00664 | 0.40454 | 0.14586 | 1.84611 | 0.72851 | 1.24586 | 0.08078 | 0.17091 | 0.00000 | 0.00000 | 3.44788 |
| **Phax** | 8.94655 | 7.62148 | 7.79150 | 3.38538 | 5.32610 | 0.32329 | 6.89455 | 12.84427 | 7.83273 | 19.66581 | 8.65287 | 8.91845 | 6.20786 | 1.92025 | 9.75611 | 1.06298 |
| **Phb** | 24.02117 | 1.09227 | 1.04012 | 1.58489 | 17.15525 | 5.81019 | 4.79242 | 19.53808 | 0.23069 | 19.68826 | 2.89440 | 2.38801 | 17.26206 | 0.55976 | 0.00000 | 10.90884 |
| **Phb2** | 21.03813 | 11.93013 | 15.32029 | 5.08419 | 15.71319 | 8.63992 | 9.25268 | 20.27381 | 1.48245 | 21.63601 | 7.14004 | 1.39051 | 24.42474 | 3.40888 | 11.32867 | 12.52485 |
| **Phc1** | 3.44189 | 2.34403 | 0.00000 | 3.88148 | 2.18543 | 1.96610 | 3.00439 | 2.60173 | 1.15813 | 6.39705 | 2.85445 | 0.25569 | 4.06129 | 2.85237 | 5.80920 | 5.25820 |
| **Phc2** | 2.83219 | 2.42475 | 1.75843 | 22.86257 | 5.80485 | 0.00671 | 0.00201 | 3.98011 | 12.50018 | 5.36250 | 2.03677 | 0.34284 | 1.23142 | 0.00000 | 9.60917 | 0.83451 |
| **Phc3** | 0.04494 | 0.00704 | 0.07914 | 0.17843 | 0.00780 | 0.00567 | 1.30484 | 0.63802 | 0.73582 | 0.36746 | 0.01731 | 0.01335 | 0.01244 | 0.00932 | 0.01456 | 0.00723 |
| **Phex** | 0.02399 | 0.03893 | 0.01659 | 0.70818 | 0.00873 | 0.00640 | 0.02835 | 0.01092 | 0.07384 | 0.04313 | 0.02381 | 0.05630 | 0.02604 | 0.69772 | 0.00789 | 0.00732 |

| | | | | | | | | | | | | | | | | |
|---|---|---|---|---|---|---|---|---|---|---|---|---|---|---|---|---|
| **Phf1** | 0.53140 | 0.00735 | 0.53167 | 13.37092 | 0.01443 | 0.00400 | 0.01776 | 0.37405 | 5.63200 | 1.32588 | 0.86113 | 0.13815 | 0.00000 | 0.00127 | 1.58100 | 0.01152 |
| **Phf10** | 3.45325 | 3.01067 | 1.08603 | 0.95613 | 2.86735 | 0.01156 | 1.78847 | 0.00000 | 0.31827 | 1.12618 | 0.75415 | 0.43962 | 0.10199 | 0.13825 | 3.11891 | 0.00949 |
| **Phf11a** | 0.89975 | 2.28895 | 0.88902 | 0.27672 | 0.93293 | 1.36915 | 0.21621 | 0.66664 | 0.26341 | 0.34118 | 0.43498 | 0.91185 | 0.84520 | 0.03937 | 0.36168 | 0.13723 |
| **Phf11b** | 0.35163 | 0.67139 | 0.39523 | 0.19517 | 0.25701 | 0.24348 | 0.04995 | 0.19318 | 0.03646 | 0.12671 | 0.06677 | 0.17890 | 0.16505 | 0.00973 | 0.02336 | 0.02915 |
| **Phf11c** | 0.01155 | 0.00823 | 0.12077 | 0.01069 | 0.00907 | 0.00575 | 0.00751 | 0.01271 | 0.01237 | 0.00889 | 0.01248 | 0.00850 | 0.26602 | 0.01353 | 0.00827 | 0.00712 |
| **Phf11d** | 5.60941 | 8.41315 | 5.81660 | 0.21477 | 4.36547 | 4.49239 | 0.92416 | 1.21809 | 0.11148 | 0.16037 | 0.77271 | 0.12904 | 3.24860 | 0.04252 | 0.09638 | 0.95143 |
| **Phf12** | 1.71642 | 2.79329 | 2.30204 | 6.75446 | 4.20693 | 5.25073 | 3.22667 | 4.30651 | 4.36841 | 6.85980 | 3.22173 | 1.15164 | 3.07861 | 0.33694 | 2.73415 | 3.32750 |
| **Phf13** | 9.36129 | 6.88362 | 6.38172 | 12.44653 | 5.19657 | 7.66591 | 5.38122 | 8.36759 | 9.50724 | 17.39948 | 6.41775 | 1.60746 | 8.21803 | 14.65614 | 11.44244 | 4.49699 |
| **Phf14** | 2.30923 | 3.66749 | 3.35643 | 7.44804 | 1.23111 | 2.48374 | 3.11393 | 1.37357 | 1.12938 | 3.68705 | 1.52731 | 0.14834 | 2.09625 | 1.30603 | 12.37769 | 4.46193 |
| **Phf19** | 0.02281 | 0.00813 | 0.00579 | 0.41575 | 0.00860 | 0.69216 | 0.00737 | 0.01111 | 0.00554 | 0.42924 | 0.02428 | 0.00714 | 0.00625 | 0.02686 | 0.00263 | 0.00672 |
| **Phf2** | 0.06363 | 0.00668 | 0.00938 | 0.25880 | 1.52779 | 0.00918 | 0.00740 | 0.97646 | 0.69822 | 0.21138 | 0.00000 | 0.01420 | 0.02209 | 0.01076 | 0.00000 | 0.18946 |
| **Phf20** | 4.60077 | 6.35334 | 9.26351 | 10.25374 | 5.66814 | 1.37885 | 1.78657 | 0.39288 | 12.56698 | 6.69231 | 2.41275 | 13.14666 | 3.21015 | 6.39821 | 11.56783 | 1.13787 |
| **Phf20l1** | 0.03201 | 0.02798 | 1.45188 | 2.12951 | 0.00640 | 0.00860 | 0.35770 | 0.00000 | 1.96743 | 2.50902 | 2.44082 | 0.08031 | 0.00000 | 0.19843 | 4.17899 | 0.01305 |
| **Phf21a** | 0.54636 | 0.00779 | 0.00171 | 2.15701 | 0.03707 | 0.00720 | 0.00672 | 0.72453 | 1.27373 | 2.24349 | 0.64360 | 0.06663 | 0.01145 | 0.00319 | 0.77354 | 0.00662 |
| **Phf21b** | 0.78294 | 0.00842 | 0.06691 | 0.13570 | 0.01304 | 0.52391 | 0.00752 | 0.68603 | 0.00891 | 1.09518 | 0.29436 | 0.11394 | 0.42114 | 0.01694 | 0.00000 | 0.02110 |
| **Phf23** | 13.33465 | 8.10086 | 8.29263 | 16.72413 | 11.49735 | 8.47741 | 5.44523 | 14.54782 | 20.35001 | 14.08092 | 6.01052 | 0.75644 | 15.20948 | 0.06038 | 11.34176 | 9.38241 |
| **Phf3** | 6.30830 | 6.59020 | 3.95805 | 8.57359 | 9.79719 | 10.22088 | 8.33913 | 9.19856 | 8.22407 | 13.39077 | 6.78520 | 11.70293 | 7.91982 | 19.94161 | 16.38950 | 1.75393 |
| **Phf5a** | 25.06201 | 10.47644 | 23.07191 | 13.07802 | 13.50169 | 9.14988 | 8.40312 | 18.79863 | 14.38165 | 21.95381 | 16.06799 | 8.25544 | 22.95572 | 13.74550 | 11.94712 | 9.98794 |
| **Phf6** | 7.92009 | 7.13655 | 4.54921 | 3.12522 | 0.00000 | 0.46950 | 0.00664 | 0.00000 | 1.07454 | 0.85336 | 0.36822 | 0.05454 | 3.22607 | 2.58982 | 0.00234 | 1.03808 |
| **Phf7** | 2.85602 | 0.00449 | 1.07919 | 3.57501 | 0.01454 | 0.00269 | 0.77364 | 0.24810 | 0.58951 | 0.79572 | 0.62472 | 0.00000 | 0.04914 | 0.01482 | 1.03805 | 0.00387 |
| **Phf8** | 0.62333 | 0.00599 | 3.32131 | 3.80844 | 1.41276 | 0.00535 | 0.03544 | 3.60539 | 5.27477 | 2.38684 | 2.71220 | 0.27051 | 1.94406 | 7.14488 | 3.19066 | 0.69374 |
| **Phgdh** | 20.96271 | 16.67132 | 34.00366 | 3.82924 | 14.56104 | 15.85544 | 20.16678 | 45.57754 | 3.70791 | 39.79025 | 24.66064 | 9.45037 | 36.88665 | 0.35393 | 1.68106 | 12.86685 |
| **Phip** | 1.33974 | 2.18208 | 0.27273 | 1.30137 | 0.05488 | 0.39449 | 2.98017 | 0.47825 | 0.41593 | 0.05084 | 0.25437 | 7.18782 | 1.83150 | 4.54651 | 1.33840 | 1.23093 |
| **Phka1** | 0.01375 | 0.00801 | 0.01180 | 0.10474 | 0.00942 | 0.00649 | 0.00739 | 0.01174 | 0.00688 | 0.00618 | 0.01117 | 0.00731 | 0.01278 | 0.00774 | 0.54173 | 0.00734 |
| **Phka2** | 0.01517 | 0.00726 | 0.51914 | 1.06354 | 0.62400 | 0.00926 | 0.00742 | 0.01317 | 0.00000 | 0.00439 | 0.01254 | 0.00603 | 0.04742 | 0.00610 | 0.29653 | 0.00706 |
| **Phkb** | 0.00000 | 0.44897 | 1.10225 | 1.30473 | 0.04952 | 2.75351 | 0.01552 | 0.19163 | 1.89342 | 0.09961 | 0.31932 | 0.01598 | 1.19296 | 0.00237 | 0.81928 | 1.99715 |
| **Phkg1** | 0.76691 | 0.00674 | 0.00000 | 3.43900 | 0.00656 | 0.00637 | 0.00656 | 0.88914 | 4.61336 | 0.75562 | 0.13655 | 0.05057 | 0.01262 | 0.01567 | 1.40267 | 0.00557 |
| **Phkg2** | 0.07978 | 0.35965 | 0.13651 | 0.00266 | 0.00709 | 1.91642 | 0.34106 | 1.29357 | 0.01022 | 1.04227 | 0.00000 | 0.07505 | 1.08270 | 0.00902 | 0.31685 | 1.58575 |
| **Phlda1** | 2.17617 | 1.02184 | 0.59238 | 1.77168 | 0.13614 | 0.04144 | 1.38157 | 2.51021 | 1.17278 | 0.90081 | 5.09400 | 0.08744 | 0.28755 | 0.17649 | 0.00000 | 3.30109 |
| **Phlda2** | 0.06575 | 0.00773 | 0.00454 | 0.20264 | 0.00778 | 0.00757 | 0.00712 | 0.53720 | 0.00814 | 3.89501 | 0.00654 | 0.03816 | 0.02710 | 0.00973 | 0.01622 | 0.00717 |
| **Phlda3** | 0.01248 | 0.00638 | 0.83053 | 0.00000 | 3.32254 | 0.00586 | 1.85611 | 0.96252 | 0.00797 | 0.87966 | 0.44280 | 0.00000 | 0.01478 | 0.01387 | 0.00000 | 0.00691 |
| **Phldb1** | 0.01315 | 0.00789 | 0.01122 | 0.00438 | 0.00909 | 0.00652 | 0.00728 | 0.01179 | 0.62084 | 0.12761 | 0.00796 | 0.02569 | 0.11423 | 0.01228 | 0.00694 | 0.00761 |
| **Phldb2** | 0.00860 | 0.00795 | 0.18468 | 0.00786 | 0.00772 | 0.00950 | 0.00732 | 0.01166 | 0.01134 | 0.32481 | 0.00966 | 0.10392 | 0.00952 | 0.00659 | 0.22987 | 0.63241 |
| **Phldb3** | 0.00313 | 0.00714 | 0.00187 | 0.00688 | 0.00834 | 0.00832 | 0.00579 | 0.85296 | 0.09122 | 5.85431 | 1.40202 | 0.03108 | 1.82910 | 0.01180 | 0.02805 | 0.06426 |
| **Phlpp1** | 0.07647 | 0.00548 | 0.00000 | 0.14355 | 0.00779 | 0.00586 | 1.08709 | 0.02653 | 0.00576 | 0.84131 | 0.00000 | 5.27836 | 0.24539 | 0.01256 | 0.00000 | 0.00658 |
| **Phlpp2** | 0.28123 | 0.41674 | 0.44643 | 3.53947 | 0.00640 | 0.00707 | 0.10270 | 0.91492 | 2.76814 | 0.10502 | 0.00000 | 0.98944 | 0.04895 | 0.00000 | 1.69770 | 0.00590 |
| **Phospho2** | 1.92971 | 1.21304 | 0.00000 | 19.20806 | 0.03068 | 0.00172 | 0.02447 | 2.52078 | 9.89074 | 9.42357 | 1.47949 | 0.86321 | 1.12172 | 1.37012 | 10.18581 | 0.84875 |
| **Phox2a** | 0.01375 | 0.00815 | 0.01230 | 0.36219 | 0.00932 | 0.00662 | 0.00752 | 0.01396 | 0.00536 | 0.01540 | 0.01345 | 0.00824 | 0.01267 | 0.01121 | 0.00610 | 0.00735 |
| **Phpt1** | 7.33721 | 2.44270 | 3.84904 | 0.03950 | 4.86614 | 0.77583 | 0.99521 | 1.05783 | 0.29651 | 1.98373 | 3.76040 | 0.16234 | 3.49529 | 0.07976 | 0.71423 | 2.44800 |
| **Phrf1** | 6.34769 | 4.15957 | 8.51262 | 10.39125 | 3.99657 | 0.58725 | 6.49183 | 14.75479 | 19.41374 | 15.17319 | 12.95642 | 1.70489 | 8.42150 | 1.50359 | 8.72561 | 4.26680 |
| **Phtf1** | 0.00000 | 0.00505 | 1.15348 | 4.36990 | 2.49032 | 0.00536 | 1.93526 | 0.00000 | 3.53686 | 1.48623 | 0.47804 | 0.10241 | 1.04769 | 1.75014 | 5.52755 | 0.96565 |
| **Phtf2** | 1.25054 | 3.76621 | 0.72993 | 13.10184 | 0.01650 | 2.61080 | 2.43329 | 5.22800 | 13.79134 | 1.87625 | 0.87152 | 0.55123 | 0.07691 | 3.14990 | 14.33022 | 1.12761 |
| **Phyh** | 10.61563 | 4.19840 | 7.69210 | 2.27605 | 3.44684 | 3.80938 | 4.93422 | 8.03817 | 1.94537 | 8.47951 | 10.86674 | 2.05842 | 7.28329 | 1.22101 | 1.23546 | 2.98689 |
| **Phyhd1** | 0.00001 | 0.00709 | 0.00000 | 1.36166 | 1.90272 | 0.01529 | 0.00794 | 0.12443 | 0.03065 | 0.44086 | 0.00847 | 0.01513 | 0.22366 | 0.16917 | 0.32802 | 0.36908 |
| **Phyhipl** | 0.01262 | 0.00802 | 0.00663 | 1.19768 | 0.00937 | 0.00620 | 0.00751 | 0.01468 | 1.84366 | 0.02045 | 0.02207 | 0.04051 | 0.01251 | 0.40545 | 1.80414 | 0.00716 |
| **Phykpl** | 1.75395 | 0.00586 | 0.81792 | 0.10262 | 0.00486 | 0.00591 | 0.09950 | 0.57760 | 0.40019 | 0.34898 | 0.21533 | 0.38527 | 0.17414 | 1.25996 | 0.71963 | 0.00525 |
| **Pi15** | 0.00770 | 0.01767 | 0.03101 | 0.03576 | 0.18101 | 0.01924 | 0.01751 | 0.04146 | 1.09178 | 0.01884 | 0.00000 | 0.01456 | 0.02022 | 0.19675 | 0.09526 | 0.00734 |
| **Pi16** | 0.06418 | 0.00771 | 0.46682 | 0.00465 | 0.00704 | 0.00595 | 0.00695 | 0.45146 | 0.01043 | 2.03030 | 0.87635 | 0.05880 | 0.00000 | 0.02102 | 0.00000 | 0.00677 |
| **Pi4k2a** | 5.26607 | 2.80864 | 6.84876 | 14.97464 | 0.11778 | 6.50518 | 1.80169 | 3.17934 | 17.26709 | 8.22084 | 2.95657 | 1.47280 | 4.89069 | 0.03698 | 3.78533 | 6.37531 |
| **Pi4k2b** | 3.95447 | 0.54779 | 5.69830 | 9.00400 | 2.50833 | 0.39903 | 1.47128 | 3.73958 | 9.36205 | 6.11274 | 2.58350 | 5.15296 | 1.34881 | 10.14184 | 8.24969 | 0.47479 |
| **Pi4ka** | 4.78833 | 3.04067 | 6.50667 | 6.64059 | 0.02194 | 1.00837 | 2.02462 | 6.38533 | 3.42782 | 2.81785 | 1.13409 | 0.27909 | 0.74690 | 4.79691 | 3.71545 | 4.61696 |
| **Pi4kb** | 0.07024 | 0.01816 | 0.03305 | 0.60909 | 0.01241 | 0.02182 | 0.00706 | 0.00753 | 0.63013 | 0.03359 | 0.03646 | 0.00000 | 0.05074 | 1.60843 | 1.77456 | 0.03663 |
| **Pianp** | 0.09203 | 0.00725 | 2.92076 | 0.28690 | 0.00414 | 0.00802 | 0.69312 | 0.01461 | 0.00537 | 0.00000 | 0.01013 | 0.01570 | 1.15041 | 0.02073 | 0.00000 | 0.00557 |
| **Pias1** | 0.23966 | 4.12503 | 2.22875 | 2.85163 | 0.59373 | 0.06524 | 0.04566 | 0.68190 | 4.15051 | 6.33795 | 0.97033 | 1.17690 | 1.07121 | 0.63034 | 5.16537 | 0.48294 |
| **Pias2** | 10.17539 | 9.93769 | 7.80606 | 4.36937 | 8.70272 | 4.32080 | 9.13239 | 5.39722 | 3.17071 | 5.32099 | 4.73191 | 4.44141 | 6.45532 | 7.41630 | 4.05156 | 2.43352 |
| **Pias3** | 0.40787 | 0.00451 | 1.18866 | 5.74399 | 4.47814 | 0.00533 | 0.56768 | 1.66837 | 3.24990 | 1.39500 | 0.46987 | 0.01366 | 1.40983 | 0.00000 | 1.37809 | 0.00467 |
| **Pias4** | 0.35934 | 0.02820 | 0.20212 | 0.16026 | 0.00757 | 0.02187 | 0.02533 | 0.03091 | 0.15503 | 13.57640 | 0.17758 | 0.00000 | 3.60615 | 0.00998 | 0.00000 | 0.30877 |
| **Pibf1** | 1.73343 | 5.01994 | 2.45731 | 1.82696 | 6.31500 | 0.04045 | 0.00000 | 3.63323 | 2.20297 | 1.24274 | 3.31435 | 10.88262 | 0.00000 | 3.70651 | 3.78532 | 0.79277 |
| **Picalm** | 5.42223 | 2.40002 | 3.79112 | 9.02445 | 3.14023 | 3.09403 | 3.66813 | 9.42671 | 7.54985 | 5.43345 | 8.46277 | 11.41503 | 2.82626 | 14.71293 | 5.52208 | 0.46632 |
| **Pick1** | 1.97113 | 1.88003 | 2.90657 | 0.00000 | 0.02351 | 0.77677 | 0.00298 | 1.89445 | 0.00989 | 1.50609 | 0.52187 | 0.05253 | 3.13377 | 0.00372 | 0.03224 | 2.63028 |
| **Pid1** | 0.01361 | 0.00794 | 0.01149 | 1.29840 | 0.00929 | 0.00656 | 0.00745 | 0.01230 | 0.43640 | 0.11342 | 0.00787 | 0.00679 | 0.01316 | 0.00588 | 0.45711 | 0.00720 |
| **Pidd1** | 0.56492 | 0.00650 | 2.59500 | 0.09181 | 0.10905 | 0.01171 | 1.48425 | 1.84653 | 0.05110 | 1.91261 | 0.00000 | 0.06757 | 1.55985 | 0.00512 | 0.76750 | 1.12604 |

| | | | | | | | | | | | | | | | | |
|---|---|---|---|---|---|---|---|---|---|---|---|---|---|---|---|---|
| **Piezo1** | 1.66212 | 0.01523 | 0.10156 | 0.17810 | 0.00425 | 1.80110 | 1.80698 | 1.47836 | 0.00876 | 0.08477 | 0.00000 | 0.10018 | 1.29128 | 0.00707 | 0.04623 | 1.25298 |
| **Pif1** | 2.35071 | 1.42195 | 0.00000 | 2.07690 | 0.22827 | 0.02254 | 4.06400 | 0.06657 | 5.95574 | 0.02729 | 0.48494 | 0.05851 | 0.64825 | 0.00000 | 2.24425 | 0.02547 |
| **Piga** | 0.23845 | 2.53733 | 1.70441 | 5.25828 | 3.00737 | 0.00581 | 0.00566 | 1.13199 | 5.90134 | 4.51337 | 0.43564 | 1.59804 | 1.08704 | 0.80436 | 4.95139 | 0.90690 |
| **Pigb** | 0.00000 | 0.00737 | 0.95499 | 0.02185 | 0.01826 | 0.00827 | 0.00689 | 2.05212 | 0.01023 | 1.16567 | 0.00000 | 0.00000 | 0.40893 | 0.00968 | 0.39729 | 0.86498 |
| **Pigc** | 0.01960 | 2.99213 | 1.78636 | 0.29712 | 0.00528 | 1.42651 | 0.00570 | 0.16221 | 0.03372 | 0.24647 | 1.44770 | 0.14423 | 1.38310 | 0.89725 | 0.18473 | 0.01163 |
| **Pigf** | 1.02581 | 5.39890 | 4.16980 | 2.00168 | 1.06467 | 2.42073 | 1.23818 | 2.69016 | 0.23241 | 0.12991 | 0.42273 | 8.17678 | 2.70895 | 0.15704 | 3.00501 | 2.43586 |
| **Pigg** | 0.00844 | 0.00819 | 0.10183 | 0.48417 | 0.00984 | 0.00585 | 0.00746 | 0.30241 | 0.00495 | 0.19051 | 0.14857 | 0.01444 | 0.00775 | 0.08184 | 0.59787 | 0.00745 |
| **Pigh** | 0.04976 | 0.00668 | 0.40229 | 0.74831 | 1.76106 | 0.00621 | 0.32605 | 0.68313 | 1.43875 | 1.08650 | 0.43222 | 0.00664 | 0.05318 | 0.00476 | 0.06446 | 0.00716 |
| **Pigk** | 3.23643 | 1.93644 | 4.16274 | 0.37419 | 2.21003 | 0.16497 | 5.31652 | 3.80605 | 0.01767 | 4.74852 | 5.61525 | 0.44786 | 6.54967 | 0.15339 | 1.74775 | 2.34779 |
| **Pigl** | 6.38600 | 2.25522 | 2.23892 | 1.91451 | 1.24630 | 0.02108 | 4.83414 | 2.11004 | 1.39122 | 0.00000 | 0.00000 | 6.35796 | 0.03414 | 0.07072 | 1.28885 | 0.01921 |
| **Pigm** | 1.60277 | 0.04612 | 2.49309 | 4.39185 | 0.13347 | 0.03806 | 1.89078 | 1.49865 | 6.71647 | 3.45310 | 1.52127 | 0.07424 | 0.82098 | 8.01144 | 5.44780 | 0.01014 |
| **Pign** | 1.77895 | 0.00350 | 1.57645 | 2.70746 | 0.00000 | 0.30673 | 3.72508 | 0.70108 | 0.98365 | 0.00000 | 0.00000 | 0.00000 | 0.85916 | 2.67294 | 0.88697 | 1.50004 |
| **Pigo** | 4.33460 | 0.26873 | 3.20764 | 4.19655 | 2.41360 | 0.27441 | 2.20217 | 2.53435 | 3.42381 | 3.63336 | 2.95790 | 4.21242 | 4.37838 | 0.40936 | 3.33816 | 3.70849 |
| **Pigp** | 14.29894 | 2.97097 | 11.44397 | 9.00668 | 8.35279 | 3.34970 | 2.64840 | 3.01188 | 10.01013 | 4.01494 | 15.55148 | 1.12284 | 5.29963 | 3.12423 | 2.62325 | 6.11095 |
| **Pigq** | 6.15557 | 1.37478 | 6.18244 | 11.36140 | 0.02462 | 6.18286 | 4.33087 | 7.53662 | 7.20988 | 7.21162 | 6.26598 | 0.39606 | 8.91260 | 0.26259 | 10.13280 | 2.55823 |
| **Pigs** | 6.01608 | 1.97863 | 6.96141 | 7.86993 | 0.15846 | 0.69503 | 7.21312 | 12.18313 | 6.51130 | 17.55114 | 7.98216 | 9.74798 | 7.55622 | 1.47189 | 3.25257 | 1.36660 |
| **Pigt** | 0.61509 | 0.01776 | 0.02070 | 0.00604 | 0.00733 | 0.01301 | 0.00620 | 0.17044 | 0.06426 | 0.89342 | 0.59587 | 0.05617 | 0.89516 | 0.00598 | 0.23735 | 0.07216 |
| **Pigu** | 9.22331 | 0.02615 | 8.61324 | 0.00741 | 0.01443 | 0.00397 | 4.32277 | 0.50358 | 0.01134 | 2.22771 | 0.47835 | 0.01495 | 0.29107 | 0.00632 | 0.13215 | 0.00000 |
| **Pigv** | 0.67334 | 0.22169 | 2.40225 | 0.17085 | 0.32772 | 0.21514 | 0.25708 | 0.15377 | 0.48447 | 1.23770 | 1.05668 | 0.05969 | 0.37298 | 0.78232 | 0.13404 | 0.05774 |
| **Pigw** | 0.01046 | 0.00794 | 0.06319 | 0.00509 | 0.00990 | 0.00937 | 0.00754 | 0.01329 | 0.78197 | 0.14879 | 0.00454 | 0.00358 | 0.38195 | 0.00797 | 1.38605 | 0.00755 |
| **Pigx** | 0.01300 | 0.72148 | 3.75910 | 4.76853 | 1.54674 | 0.10530 | 0.04948 | 0.76067 | 8.91026 | 11.93097 | 17.64277 | 1.99725 | 2.54057 | 3.50447 | 1.07447 | 0.00000 |
| **Pigyl** | 7.97165 | 0.88995 | 2.03875 | 7.84397 | 0.00000 | 3.67267 | 1.66705 | 5.70688 | 9.64681 | 7.79040 | 1.87564 | 0.35639 | 7.93715 | 0.00000 | 0.00000 | 5.32096 |
| **Pigz** | 0.00420 | 0.00792 | 1.17071 | 0.01085 | 0.00709 | 0.05380 | 0.00644 | 0.01811 | 0.01223 | 0.00039 | 0.01106 | 0.00662 | 2.01675 | 0.01862 | 0.00000 | 0.00583 |
| **Pih1d1** | 8.06973 | 1.44110 | 3.59286 | 4.09053 | 2.37842 | 7.79030 | 0.47246 | 5.40388 | 3.57843 | 5.82715 | 4.10873 | 0.33873 | 11.35471 | 0.00000 | 0.62214 | 3.51787 |
| **Pih1d2** | 0.00844 | 0.00713 | 0.01063 | 0.14343 | 0.60039 | 0.00641 | 0.00739 | 0.01039 | 0.06073 | 0.00278 | 0.01145 | 0.00615 | 0.01057 | 0.01346 | 0.53134 | 0.00715 |
| **Pik3ap1** | 0.01374 | 0.00805 | 0.01226 | 0.01015 | 0.00924 | 0.00662 | 0.00749 | 0.01283 | 0.01235 | 0.10611 | 0.01259 | 0.00757 | 0.01229 | 0.01138 | 0.01075 | 0.00743 |
| **Pik3c2a** | 0.34701 | 0.11589 | 0.19343 | 11.15348 | 0.18590 | 3.40558 | 1.50917 | 3.83197 | 11.27284 | 2.71751 | 7.23758 | 8.26003 | 1.92865 | 23.32808 | 12.59146 | 0.32715 |
| **Pik3c2b** | 1.65397 | 0.01651 | 0.47262 | 1.02236 | 0.10913 | 0.03756 | 0.03621 | 0.00000 | 1.46729 | 0.70732 | 0.90508 | 0.02658 | 0.20964 | 2.35027 | 2.70815 | 0.01184 |
| **Pik3c2g** | 0.00587 | 2.61619 | 0.01081 | 0.00610 | 0.00784 | 0.00621 | 0.00641 | 0.00908 | 0.03583 | 0.00818 | 0.01288 | 0.00751 | 0.01050 | 0.01076 | 0.00000 | 0.00632 |
| **Pik3c3** | 2.39470 | 4.61126 | 2.52479 | 1.45270 | 5.18150 | 5.26707 | 3.85742 | 10.35230 | 4.39540 | 7.40422 | 6.48055 | 7.45847 | 4.85083 | 10.36712 | 9.86567 | 0.58844 |
| **Pik3ca** | 0.80658 | 0.18838 | 4.09440 | 3.03061 | 0.23948 | 0.34691 | 0.91159 | 2.44803 | 4.20120 | 1.90233 | 0.74523 | 0.11337 | 0.00000 | 6.18554 | 5.21423 | 1.52408 |
| **Pik3cb** | 0.00000 | 0.15253 | 1.77770 | 8.79924 | 0.01519 | 0.03889 | 0.61391 | 1.35988 | 14.50449 | 3.10330 | 12.35455 | 3.57345 | 0.00000 | 4.86590 | 2.61664 | 0.93953 |
| **Pik3cd** | 0.00893 | 0.01792 | 1.03708 | 5.07357 | 0.00720 | 0.11799 | 0.03506 | 1.30170 | 2.91312 | 4.44040 | 1.47399 | 0.08977 | 1.60357 | 0.00588 | 1.53621 | 0.00502 |
| **Pik3ip1** | 0.01669 | 0.00838 | 0.00967 | 0.75182 | 0.00954 | 0.00751 | 0.00760 | 0.00501 | 0.66194 | 0.20241 | 0.36638 | 0.00469 | 0.15740 | 0.10537 | 0.28649 | 0.00746 |
| **Pik3r1** | 1.39933 | 0.00637 | 2.06153 | 3.56443 | 0.03457 | 1.31269 | 0.02732 | 0.01360 | 3.13993 | 0.08256 | 0.63815 | 0.00000 | 0.00000 | 2.78636 | 4.50251 | 0.01115 |
| **Pik3r2** | 5.31071 | 0.00000 | 0.00000 | 0.54626 | 1.16414 | 5.36766 | 0.22294 | 5.29182 | 0.06872 | 4.44029 | 0.61320 | 0.04344 | 6.94683 | 0.00262 | 0.00000 | 3.54373 |
| **Pik3r3** | 0.01492 | 0.00711 | 0.00433 | 5.06276 | 0.00926 | 0.11674 | 1.09503 | 0.00969 | 2.21727 | 4.29258 | 0.00000 | 0.02324 | 0.06124 | 2.42649 | 4.25067 | 0.00643 |
| **Pik3r4** | 5.78737 | 2.14247 | 3.95836 | 7.24400 | 0.05442 | 0.00000 | 0.82762 | 5.16939 | 2.36219 | 2.91317 | 4.76164 | 0.18321 | 2.72669 | 0.81023 | 0.81018 | 0.08408 |
| **Pik3r6** | 0.68535 | 0.00535 | 2.60835 | 0.01018 | 0.01261 | 0.01428 | 2.60606 | 2.92036 | 0.00935 | 0.00000 | 0.00000 | 1.28659 | 3.24098 | 0.00954 | 0.00000 | 0.01153 |
| **Pikfyve** | 1.83082 | 0.43112 | 3.73377 | 5.31590 | 2.90802 | 0.30310 | 0.72416 | 1.44787 | 2.56713 | 1.77408 | 0.66374 | 13.86150 | 2.05205 | 1.24859 | 3.31083 | 0.45429 |
| **Pilra** | 0.35775 | 0.00701 | 0.13954 | 1.53249 | 0.05025 | 0.01695 | 0.00595 | 0.00000 | 1.85970 | 0.32285 | 2.57268 | 0.06403 | 0.07150 | 0.68709 | 0.21538 | 0.02090 |
| **Pilrb2** | 0.01306 | 0.00819 | 0.04270 | 0.01061 | 0.00928 | 0.00601 | 0.00765 | 0.01423 | 0.01235 | 0.01883 | 0.01300 | 0.00836 | 0.01210 | 0.01244 | 0.01010 | 0.00746 |
| **Pim1** | 2.50176 | 1.45323 | 2.05739 | 1.37103 | 0.00389 | 2.51065 | 1.34366 | 4.57194 | 3.69605 | 0.96356 | 4.48245 | 0.12749 | 0.85932 | 2.32555 | 2.14614 | 0.16190 |
| **Pim2** | 2.34525 | 0.00457 | 3.27564 | 5.91434 | 0.04392 | 0.01033 | 0.00335 | 3.24568 | 6.33052 | 4.85613 | 0.38885 | 0.23564 | 2.26597 | 0.92293 | 5.85711 | 7.41697 |
| **Pim3** | 2.83249 | 0.06611 | 0.91688 | 9.80936 | 0.04151 | 1.95179 | 0.78014 | 12.67741 | 11.45366 | 7.69689 | 5.61041 | 0.48043 | 4.76473 | 2.52540 | 1.08800 | 2.87628 |
| **Pin1** | 25.15240 | 5.60177 | 11.56743 | 5.57118 | 5.86442 | 7.11224 | 6.52161 | 17.92958 | 9.94255 | 17.18897 | 13.01309 | 2.13946 | 9.69645 | 0.17619 | 0.43015 | 9.66285 |
| **Pin1rt1** | 0.40088 | 0.05907 | 0.15885 | 8.15991 | 0.03545 | 0.16029 | 0.09838 | 4.86405 | 8.26062 | 6.69890 | 4.00602 | 0.17657 | 1.54524 | 0.46344 | 0.53337 | 0.12124 |
| **Pin4** | 5.18338 | 0.50034 | 3.39912 | 4.98012 | 1.30232 | 0.37610 | 0.29249 | 0.65078 | 14.14292 | 2.47475 | 13.18834 | 0.53561 | 2.62287 | 4.31533 | 3.19375 | 1.18052 |
| **Pink1** | 0.02715 | 1.41815 | 2.13506 | 1.56891 | 0.06292 | 0.72714 | 0.57988 | 13.27310 | 1.80466 | 11.29973 | 14.80391 | 0.22270 | 4.87680 | 1.58224 | 2.02533 | 3.13081 |
| **Pinlyp** | 0.00000 | 1.12952 | 1.53743 | 3.62114 | 0.00556 | 2.64051 | 1.52515 | 1.08857 | 3.60960 | 2.54601 | 2.25218 | 0.19203 | 4.00895 | 0.07210 | 1.09017 | 0.00258 |
| **Pinx1** | 2.25282 | 5.72391 | 3.62747 | 0.27530 | 0.07968 | 1.81205 | 3.89423 | 5.85659 | 0.84039 | 0.69540 | 3.88391 | 0.98789 | 1.44488 | 1.69347 | 5.45528 | 1.01258 |
| **Pip4k2a** | 3.00279 | 0.00387 | 0.07276 | 1.89936 | 0.05998 | 0.01542 | 0.47457 | 0.01554 | 4.46679 | 1.65299 | 0.00000 | 2.03159 | 0.26195 | 0.01838 | 0.70948 | 0.25709 |
| **Pip4k2b** | 2.22448 | 0.07535 | 0.07465 | 0.43969 | 1.89052 | 0.00426 | 1.00676 | 0.17089 | 1.66279 | 0.29217 | 0.82618 | 0.03794 | 0.03307 | 0.00298 | 0.61412 | 0.00472 |
| **Pip4k2c** | 2.25221 | 2.44983 | 9.22578 | 2.34204 | 0.34363 | 6.05752 | 7.28164 | 15.33115 | 2.07046 | 8.85607 | 5.06268 | 1.36209 | 10.50510 | 3.27905 | 8.13414 | 3.35948 |
| **Pip5k1a** | 0.04195 | 0.04997 | 2.25013 | 1.90601 | 0.79016 | 0.01190 | 0.01712 | 1.53465 | 2.97163 | 9.18027 | 0.25588 | 0.12852 | 0.32871 | 0.31492 | 7.98757 | 0.09844 |
| **Pip5k1b** | 0.02662 | 0.00771 | 0.01012 | 0.78705 | 0.00950 | 0.00648 | 0.00733 | 0.01104 | 2.35292 | 0.19732 | 0.00000 | 0.00217 | 0.01304 | 0.01149 | 0.13055 | 0.00747 |
| **Pip5k1c** | 1.77922 | 0.02673 | 0.62685 | 2.41575 | 0.01372 | 1.67829 | 0.01518 | 3.40144 | 6.70663 | 0.96219 | 0.28875 | 0.47451 | 1.69961 | 0.00000 | 0.56713 | 0.01039 |
| **Pipox** | 0.09314 | 0.00752 | 0.00562 | 0.00278 | 0.00776 | 0.00680 | 0.00718 | 0.01089 | 0.68327 | 1.12705 | 0.00355 | 0.07578 | 0.02622 | 0.00755 | 0.66422 | 0.00736 |
| **Pir** | 5.59677 | 0.00000 | 0.10304 | 0.00948 | 0.01594 | 0.00000 | 1.99330 | 0.00000 | 0.01249 | 0.03977 | 0.01283 | 0.00912 | 0.50899 | 0.40040 | 0.00000 | 0.00959 |
| **Pira6** | 0.02868 | 0.00757 | 0.24662 | 0.23787 | 0.00826 | 0.00567 | 0.00710 | 0.00854 | 0.00750 | 0.00808 | 0.00814 | 0.01239 | 0.01050 | 1.31773 | 0.00000 | 0.00719 |
| **Pirb** | 0.03136 | 0.00758 | 0.07453 | 0.02839 | 0.00840 | 0.00599 | 0.00735 | 0.01085 | 0.01132 | 0.36145 | 0.01042 | 0.00623 | 0.01185 | 0.02730 | 0.53501 | 0.00737 |
| **Pisd** | 5.40090 | 1.16201 | 3.56741 | 2.70240 | 1.17387 | 2.24815 | 2.03771 | 8.30504 | 6.05688 | 7.18259 | 4.72592 | 1.57532 | 3.28207 | 3.90303 | 3.80006 | 6.08232 |

| | | | | | | | | | | | | | | | | |
|---|---|---|---|---|---|---|---|---|---|---|---|---|---|---|---|---|
| **Pisd.ps1** | 5.66007 | 1.13584 | 5.28709 | 5.20198 | 0.11938 | 2.85304 | 5.62843 | 4.42654 | 7.44971 | 2.25164 | 3.32658 | 2.77320 | 4.01563 | 1.18317 | 5.06103 | 4.71751 |
| **Pisd.ps2** | 0.79032 | 0.37210 | 0.37535 | 0.01020 | 0.03211 | 0.98377 | 0.43579 | 0.82107 | 0.34014 | 0.47184 | 0.01618 | 0.09004 | 0.90466 | 0.12067 | 0.45507 | 1.14303 |
| **Pithd1** | 5.34265 | 0.74261 | 4.38859 | 7.53916 | 4.00690 | 2.66855 | 0.74833 | 1.52819 | 8.59441 | 4.68850 | 3.95679 | 2.10028 | 3.92483 | 0.14259 | 1.49989 | 2.15737 |
| **Pitpna** | 8.87628 | 5.79549 | 7.10111 | 8.79476 | 11.83381 | 5.63985 | 5.87768 | 13.79813 | 13.41173 | 18.65010 | 19.14206 | 13.00480 | 5.22718 | 9.80011 | 9.00545 | 2.95330 |
| **Pitpnb** | 6.25832 | 7.27062 | 13.15417 | 10.42691 | 7.83414 | 5.18677 | 5.63430 | 15.57179 | 11.44077 | 22.71633 | 13.55421 | 6.87621 | 9.42376 | 3.93303 | 16.65086 | 3.74282 |
| **Pitpnc1** | 0.38055 | 2.56120 | 3.01500 | 3.64571 | 4.85383 | 1.19811 | 4.46991 | 4.60127 | 2.37427 | 0.00000 | 0.38323 | 0.01509 | 1.26233 | 9.27450 | 0.00000 | 0.79978 |
| **Pitpnm1** | 0.01250 | 0.00793 | 0.36757 | 0.06735 | 0.00852 | 0.00917 | 0.00735 | 0.01279 | 0.00974 | 0.01502 | 0.01211 | 0.00776 | 0.00962 | 0.01316 | 0.00288 | 0.00722 |
| **Pitpnm2** | 2.28016 | 0.00392 | 0.99865 | 0.72036 | 1.46978 | 0.00676 | 0.00506 | 0.00000 | 0.12640 | 2.67125 | 0.63000 | 0.00435 | 0.02799 | 0.02613 | 0.00000 | 0.00419 |
| **Pitrm1** | 4.13352 | 7.96706 | 2.53395 | 0.95415 | 1.42730 | 4.94073 | 5.04965 | 1.38198 | 1.87213 | 3.84295 | 0.56503 | 0.11290 | 7.15769 | 0.11174 | 3.09946 | 0.00807 |
| **Pitx1** | 0.07023 | 0.00758 | 0.00510 | 0.04127 | 1.11332 | 0.00802 | 0.00702 | 0.29266 | 0.01067 | 1.05369 | 0.10195 | 0.00215 | 0.54323 | 0.01158 | 0.00215 | 0.00796 |
| **Piwil1** | 0.02529 | 0.00784 | 0.00896 | 0.76378 | 0.00853 | 0.01294 | 0.00721 | 0.06036 | 0.00000 | 0.04951 | 0.01246 | 0.00705 | 0.10200 | 0.02442 | 0.04493 | 0.00713 |
| **Piwil2** | 0.54543 | 4.12105 | 5.93084 | 6.42212 | 5.57957 | 0.01161 | 0.45191 | 4.16049 | 4.59058 | 13.53261 | 11.55975 | 0.89608 | 3.56993 | 7.88748 | 3.54651 | 0.97109 |
| **Pja1** | 0.00000 | 2.87969 | 0.00516 | 0.30989 | 0.00719 | 0.00700 | 2.09693 | 3.09366 | 0.00631 | 0.33501 | 0.06635 | 0.40596 | 0.97075 | 0.25255 | 0.22208 | 3.29135 |
| **Pja2** | 0.07447 | 0.01825 | 3.83575 | 0.81502 | 0.24163 | 0.00607 | 0.00673 | 0.01222 | 0.67310 | 0.70528 | 0.00850 | 0.02130 | 0.10149 | 0.00429 | 1.01216 | 1.57524 |
| **Pkd1** | 0.05945 | 0.00697 | 0.08982 | 0.20093 | 0.39616 | 0.44712 | 0.00693 | 0.01907 | 0.20894 | 0.25461 | 0.00000 | 1.56331 | 0.16233 | 0.00508 | 0.76315 | 1.69470 |
| **Pkd1l2** | 0.13632 | 0.00583 | 0.00000 | 1.10801 | 1.40540 | 0.00635 | 1.59229 | 0.29724 | 0.74898 | 1.47641 | 0.89174 | 0.16366 | 0.03371 | 0.19687 | 2.15590 | 0.00679 |
| **Pkd1l3** | 0.00938 | 0.00585 | 0.00268 | 0.20339 | 0.00915 | 0.39138 | 1.58112 | 0.01595 | 0.45018 | 0.60745 | 0.00000 | 3.87981 | 0.01363 | 0.11991 | 0.35741 | 0.00685 |
| **Pkd2** | 0.04941 | 0.01878 | 0.17154 | 0.66973 | 0.00995 | 0.00636 | 0.04860 | 0.03442 | 0.00000 | 2.24166 | 0.11492 | 0.04889 | 0.06044 | 0.05493 | 2.47683 | 0.00716 |
| **Pkd2l1** | 0.21142 | 0.00653 | 0.08647 | 0.08136 | 1.38644 | 0.00593 | 0.17462 | 0.00666 | 0.00642 | 0.00232 | 0.01142 | 0.00566 | 0.00966 | 0.00780 | 0.57058 | 0.01388 |
| **Pkd2l2** | 0.16761 | 0.00774 | 0.39257 | 0.65914 | 0.00700 | 0.00552 | 0.00725 | 0.03204 | 0.43136 | 0.01050 | 0.01320 | 0.01377 | 0.00908 | 1.13462 | 0.49308 | 0.00686 |
| **Pkig** | 12.45112 | 0.88039 | 9.20087 | 5.86196 | 6.88288 | 5.86562 | 7.73085 | 8.29788 | 2.75759 | 17.44020 | 7.21592 | 2.17513 | 8.29577 | 0.22421 | 0.00000 | 3.85429 |
| **Pklr** | 0.00848 | 0.00739 | 0.10065 | 0.12370 | 0.00939 | 0.00854 | 0.00706 | 1.45237 | 0.00916 | 0.05145 | 0.00559 | 0.01531 | 0.00462 | 0.02328 | 0.02237 | 0.00713 |
| **Pkm** | 29.74638 | 12.69049 | 12.38920 | 16.87618 | 17.61319 | 12.05540 | 13.59113 | 0.68960 | 17.24483 | 7.15535 | 4.44493 | 0.84395 | 15.50374 | 0.05032 | 1.86205 | 15.63403 |
| **Pkmyt1** | 1.34122 | 0.00639 | 0.00000 | 0.61214 | 0.00683 | 1.93378 | 0.08775 | 1.98314 | 0.66201 | 5.01987 | 0.27684 | 0.20754 | 2.55702 | 0.00730 | 0.00000 | 1.50596 |
| **Pkn1** | 1.51391 | 0.86986 | 1.37187 | 0.00327 | 0.01603 | 0.00412 | 0.01367 | 0.08139 | 0.01125 | 0.14921 | 0.45873 | 0.00037 | 3.26437 | 0.01023 | 0.00000 | 0.66237 |
| **Pkn2** | 3.47431 | 6.42073 | 1.33532 | 9.70724 | 5.36675 | 1.36512 | 0.50117 | 6.65249 | 7.15578 | 5.81219 | 3.95528 | 12.08201 | 0.88450 | 9.45958 | 9.77060 | 1.59156 |
| **Pkn3** | 3.69096 | 0.00336 | 0.38977 | 0.94648 | 0.00464 | 0.00114 | 1.07624 | 0.86990 | 4.10725 | 2.44457 | 1.43318 | 0.17300 | 0.43275 | 0.03145 | 0.83018 | 0.00959 |
| **Pknox1** | 6.54870 | 0.10585 | 6.12116 | 1.09635 | 0.02548 | 0.00397 | 3.71946 | 10.22258 | 6.58569 | 16.51896 | 3.71791 | 7.00283 | 5.18430 | 5.93654 | 6.97607 | 1.91537 |
| **Pknox2** | 0.01359 | 0.00808 | 0.01205 | 0.00937 | 0.00915 | 0.00686 | 0.00754 | 0.01293 | 0.01205 | 0.29515 | 0.01200 | 0.00834 | 0.01266 | 0.01161 | 0.01152 | 0.00745 |
| **Pkp2** | 0.00000 | 3.68761 | 4.29325 | 0.40845 | 0.00000 | 0.80861 | 3.85250 | 4.77269 | 0.67696 | 1.62650 | 1.89743 | 6.56423 | 2.80856 | 0.00000 | 0.00000 | 2.30272 |
| **Pkp3** | 0.77566 | 0.00721 | 0.00228 | 0.17731 | 0.01765 | 0.00857 | 0.00591 | 0.62436 | 1.04969 | 0.48870 | 1.98795 | 0.00418 | 0.09481 | 0.00458 | 0.00000 | 0.00645 |
| **Pkp4** | 4.72617 | 3.92784 | 1.71391 | 13.63809 | 4.55501 | 0.78867 | 0.09309 | 2.35200 | 3.61771 | 9.35689 | 2.42396 | 0.39868 | 2.23462 | 1.75909 | 10.24388 | 5.11529 |
| **Pla1a** | 1.75024 | 0.29470 | 1.53703 | 0.35638 | 0.01030 | 0.02194 | 0.00295 | 0.00000 | 0.18923 | 0.94611 | 3.26951 | 2.46829 | 2.56170 | 3.06156 | 0.38939 | 0.01050 |
| **Pla2g10** | 3.07120 | 1.26105 | 7.67793 | 0.00838 | 0.02601 | 0.02940 | 2.32236 | 1.69095 | 0.13270 | 0.00000 | 3.18754 | 0.01373 | 1.77896 | 0.00487 | 0.00884 | 1.70429 |
| **Pla2g10os** | 0.02144 | 0.00824 | 0.00996 | 0.13389 | 0.00862 | 0.00672 | 0.00766 | 0.01602 | 0.31870 | 0.00947 | 0.19046 | 0.00699 | 0.00964 | 0.00897 | 0.03329 | 0.01433 |
| **Pla2g12a** | 0.50948 | 2.34363 | 2.53723 | 3.70186 | 0.00311 | 3.28085 | 5.51133 | 4.41199 | 2.58303 | 9.79440 | 9.84578 | 2.06962 | 3.43137 | 1.46198 | 1.49109 | 1.38511 |
| **Pla2g12b** | 2.16979 | 0.00550 | 0.00343 | 0.00430 | 0.01228 | 0.00632 | 0.00595 | 0.00777 | 1.69519 | 0.00868 | 0.01164 | 0.00399 | 0.01122 | 0.01458 | 0.00594 | 0.00497 |
| **Pla2g15** | 3.55842 | 1.79765 | 6.10971 | 10.40425 | 0.92603 | 7.99510 | 5.48867 | 12.21225 | 11.55587 | 12.77740 | 14.62960 | 0.59227 | 5.23217 | 5.01470 | 13.55130 | 1.82800 |
| **Pla2g16** | 1.83018 | 0.77689 | 2.10720 | 14.91573 | 0.03624 | 3.12993 | 2.01020 | 6.26021 | 16.61333 | 20.26127 | 11.61695 | 4.17392 | 4.06361 | 5.09698 | 15.32544 | 0.03941 |
| **Pla2g1b** | 0.51768 | 0.00744 | 0.00000 | 0.18041 | 0.00997 | 0.02138 | 0.00659 | 0.00000 | 0.00569 | 0.16236 | 0.53262 | 0.00254 | 0.00966 | 0.61791 | 0.00000 | 0.00717 |
| **Pla2g2d** | 0.03115 | 0.03804 | 0.10961 | 0.17893 | 0.14758 | 0.04409 | 0.01731 | 0.04909 | 0.11869 | 0.56636 | 0.12040 | 0.03156 | 0.15042 | 0.23255 | 0.13929 | 0.07649 |
| **Pla2g2f** | 0.27754 | 0.00827 | 0.00659 | 0.01061 | 0.01423 | 0.00705 | 0.00734 | 0.01251 | 0.01236 | 0.01539 | 0.01295 | 0.00802 | 0.01237 | 0.01236 | 0.01076 | 0.00730 |
| **Pla2g4a** | 0.01357 | 0.00765 | 0.01199 | 0.00777 | 0.00912 | 0.00660 | 0.00731 | 0.01092 | 0.01194 | 0.41232 | 0.01217 | 0.01489 | 0.01232 | 0.00570 | 0.19247 | 0.00780 |
| **Pla2g4c** | 0.00646 | 0.01619 | 0.76444 | 5.22049 | 0.00720 | 0.07088 | 0.00595 | 0.00000 | 2.79200 | 0.38705 | 0.16561 | 6.44169 | 0.00760 | 2.26347 | 0.94387 | 0.00538 |
| **Pla2g4d** | 0.01356 | 0.00773 | 0.00599 | 1.32800 | 0.00888 | 0.00720 | 0.00719 | 0.00199 | 1.32585 | 0.50312 | 0.28030 | 0.34235 | 0.00995 | 0.00699 | 0.06005 | 0.00699 |
| **Pla2g4e** | 0.00000 | 0.00501 | 0.00000 | 0.05516 | 0.00996 | 0.00485 | 1.01800 | 0.93672 | 0.85473 | 9.85899 | 5.80013 | 5.18804 | 0.00000 | 6.27767 | 7.93631 | 0.00574 |
| **Pla2g4f** | 0.00911 | 0.00777 | 1.12774 | 0.00321 | 0.26850 | 0.74299 | 0.00740 | 0.00000 | 0.27766 | 0.48046 | 0.41394 | 0.03215 | 0.00000 | 0.01027 | 0.04429 | 0.00653 |
| **Pla2g5** | 0.85101 | 0.05147 | 0.12379 | 0.39922 | 0.01453 | 3.48380 | 0.00586 | 1.91927 | 0.00845 | 0.44812 | 0.68374 | 0.01750 | 0.07416 | 0.00269 | 0.59007 | 0.00528 |
| **Pla2g6** | 3.44073 | 3.91499 | 4.10835 | 0.53587 | 1.48251 | 3.28700 | 3.52828 | 3.42291 | 0.58049 | 0.36654 | 0.93896 | 0.06030 | 2.54841 | 0.00000 | 0.00000 | 0.00585 |
| **Pla2g7** | 0.00630 | 0.26058 | 0.84430 | 0.00413 | 0.00708 | 0.00576 | 0.00693 | 3.20804 | 0.46534 | 0.01076 | 0.00000 | 0.00000 | 0.00219 | 0.02951 | 0.00000 | 0.00719 |
| **Plaa** | 15.00543 | 11.42202 | 14.26244 | 5.80649 | 11.99398 | 8.70394 | 7.18854 | 13.02033 | 10.36706 | 20.67964 | 10.42755 | 19.99195 | 9.00486 | 27.22749 | 18.85960 | 3.60388 |
| **Plac1** | 0.43430 | 0.48618 | 0.42531 | 0.52340 | 0.01639 | 0.00928 | 0.00676 | 0.00856 | 0.01641 | 0.28557 | 0.01051 | 0.00751 | 0.01006 | 0.00655 | 0.00000 | 0.00674 |
| **Plac8** | 0.94535 | 0.00714 | 0.00000 | 5.50611 | 0.00859 | 0.01317 | 0.00581 | 0.00245 | 2.45292 | 0.78522 | 0.07188 | 0.00000 | 0.64786 | 0.02573 | 0.01413 | 2.72181 |
| **Plac9b** | 6.59059 | 1.40701 | 6.96630 | 2.60170 | 0.36874 | 1.18035 | 4.19282 | 2.87906 | 1.81685 | 3.55741 | 5.35468 | 0.96367 | 3.77285 | 0.28972 | 1.16611 | 1.68128 |
| **Plag1** | 0.01443 | 0.00787 | 0.00442 | 2.86066 | 0.00909 | 0.83497 | 0.00731 | 0.01042 | 2.45198 | 0.04694 | 0.02542 | 0.03584 | 0.00675 | 0.02093 | 0.41718 | 0.00640 |
| **Plagl1** | 0.52030 | 0.00814 | 0.03803 | 0.46247 | 0.03780 | 0.00792 | 0.00695 | 0.01081 | 0.00559 | 0.05201 | 0.01386 | 0.00729 | 0.01162 | 0.01049 | 0.00461 | 0.00706 |
| **Plagl2** | 0.19059 | 0.02652 | 1.16199 | 0.39901 | 0.46652 | 0.00931 | 0.01651 | 0.11626 | 0.12937 | 0.40448 | 1.36840 | 0.04383 | 0.27052 | 0.18341 | 1.44123 | 0.00639 |
| **Plat** | 0.01283 | 0.00777 | 0.01203 | 2.14575 | 0.00899 | 2.00313 | 0.00692 | 0.00000 | 2.00729 | 0.65932 | 0.61076 | 0.02856 | 0.00000 | 0.06339 | 0.73224 | 0.00637 |
| **Platr10** | 1.49771 | 0.00650 | 0.00000 | 0.62094 | 0.00696 | 0.03738 | 0.00602 | 0.26032 | 1.06355 | 0.21427 | 0.26482 | 0.02385 | 0.04199 | 0.08383 | 0.22521 | 0.03459 |
| **Platr14** | 5.18850 | 4.79458 | 2.75882 | 0.37164 | 0.07136 | 5.69041 | 0.00000 | 2.27286 | 0.21535 | 0.47757 | 0.24077 | 5.37968 | 2.02219 | 1.58645 | 3.69323 | 1.28859 |
| **Platr17** | 0.19721 | 0.09731 | 3.23190 | 13.87226 | 0.03128 | 3.14319 | 2.06903 | 13.80447 | 14.65617 | 15.62192 | 16.85066 | 4.03877 | 2.64668 | 1.87989 | 5.59130 | 0.67344 |
| **Platr20** | 0.01787 | 0.00768 | 0.01165 | 0.45766 | 0.00887 | 0.00665 | 0.00734 | 0.01178 | 0.00472 | 0.22685 | 0.01322 | 0.00678 | 0.01270 | 0.01436 | 0.04118 | 0.00769 |

| | | | | | | | | | | | | | | | | |
|---|---|---|---|---|---|---|---|---|---|---|---|---|---|---|---|---|
| **Platr21** | 0.01237 | 0.00783 | 0.67304 | 0.01094 | 0.00800 | 0.00925 | 0.00731 | 0.01240 | 0.01224 | 0.01453 | 0.01170 | 0.00755 | 0.00870 | 0.01524 | 0.00100 | 0.00712 |
| **Platr22** | 0.01823 | 0.00755 | 0.00809 | 0.09675 | 0.00881 | 0.00658 | 0.00721 | 0.00200 | 0.00960 | 0.00000 | 0.32003 | 0.02396 | 0.00878 | 0.01227 | 0.65204 | 0.00710 |
| **Platr26** | 2.68956 | 0.24148 | 0.00159 | 0.00838 | 1.18611 | 0.00589 | 0.00508 | 0.00000 | 0.01093 | 0.00350 | 0.01164 | 0.00456 | 0.01035 | 0.01261 | 0.25975 | 0.00428 |
| **Platr4** | 0.09871 | 0.00524 | 0.62699 | 0.46725 | 0.00637 | 0.06844 | 3.34454 | 1.18541 | 0.22593 | 0.00000 | 0.00715 | 0.52409 | 0.00000 | 0.05149 | 0.42581 | 0.94651 |
| **Platr5** | 0.01271 | 0.00807 | 0.04047 | 0.03789 | 0.00984 | 0.00849 | 0.01823 | 0.01329 | 0.02281 | 0.01692 | 0.07157 | 0.00679 | 0.05650 | 0.01520 | 0.11262 | 0.00784 |
| **Platr9** | 0.04008 | 0.00750 | 0.03266 | 1.06165 | 0.00894 | 0.02356 | 0.01718 | 0.26350 | 0.49989 | 0.00000 | 0.21037 | 0.01482 | 0.02547 | 0.04812 | 0.15030 | 0.00709 |
| **Plaur** | 0.01399 | 0.07081 | 0.89204 | 0.01960 | 0.00727 | 0.09391 | 0.01771 | 0.85047 | 0.01169 | 0.00000 | 1.17938 | 0.02061 | 0.00000 | 0.00898 | 0.00000 | 0.01352 |
| **Plb1** | 1.72503 | 0.09908 | 0.56229 | 1.56523 | 0.00508 | 0.00376 | 1.38560 | 0.00576 | 2.35740 | 0.94830 | 0.00004 | 0.03502 | 0.29810 | 0.34636 | 0.40723 | 0.89655 |
| **Plbd1** | 1.73062 | 3.04367 | 4.39093 | 0.21084 | 5.12396 | 2.04423 | 2.18087 | 5.10314 | 1.64873 | 9.74365 | 10.83078 | 7.15754 | 8.35386 | 9.50155 | 4.80469 | 0.02482 |
| **Plbd2** | 4.25400 | 0.03071 | 5.37246 | 0.14071 | 0.33115 | 0.79565 | 4.49753 | 6.54348 | 0.02549 | 1.61346 | 2.74257 | 0.04435 | 5.45152 | 0.78739 | 0.83653 | 0.00000 |
| **Plcb1** | 0.01784 | 0.00770 | 0.01078 | 0.50545 | 0.00894 | 0.00660 | 0.00741 | 0.01138 | 0.00435 | 0.00865 | 0.04959 | 0.01698 | 0.01180 | 0.04625 | 0.15275 | 0.02169 |
| **Plcb3** | 4.46800 | 0.01381 | 0.68932 | 1.32908 | 0.02282 | 4.76347 | 0.12054 | 2.74649 | 0.36262 | 1.71921 | 2.74511 | 0.03334 | 4.33345 | 0.00001 | 0.00000 | 2.01480 |
| **Plcd1** | 4.43316 | 3.08747 | 6.83459 | 0.00000 | 2.61480 | 7.24981 | 2.81956 | 9.20440 | 0.00610 | 3.48514 | 5.36363 | 0.74043 | 4.12371 | 0.04769 | 1.83453 | 3.62400 |
| **Plcd3** | 0.02632 | 0.00562 | 0.00000 | 0.00000 | 0.01070 | 5.66747 | 0.01217 | 13.28038 | 1.04046 | 11.53400 | 7.28210 | 0.61442 | 4.28575 | 0.13717 | 2.20172 | 0.00980 |
| **Plcd4** | 0.00000 | 0.00762 | 0.00000 | 4.82601 | 0.00930 | 0.00513 | 0.00721 | 1.47570 | 9.39656 | 4.81196 | 0.65147 | 0.65724 | 0.23035 | 6.18939 | 5.05704 | 0.00606 |
| **Plce1** | 0.01128 | 0.00795 | 0.00609 | 2.59873 | 0.00913 | 0.00641 | 0.00705 | 0.00877 | 0.73624 | 1.00371 | 0.00000 | 0.02786 | 0.01193 | 1.73654 | 0.23742 | 0.00713 |
| **Plcg1** | 1.32479 | 0.01532 | 0.51605 | 0.06839 | 0.00444 | 0.75798 | 3.08231 | 1.62265 | 0.00944 | 0.58467 | 0.00000 | 0.03810 | 0.32152 | 1.65309 | 0.33712 | 0.00515 |
| **Plcg2** | 3.08889 | 1.04096 | 4.06673 | 0.90700 | 0.25340 | 0.20952 | 0.02131 | 3.50137 | 0.13477 | 3.50376 | 4.69709 | 0.59290 | 3.12034 | 4.54033 | 1.57153 | 0.00000 |
| **Plch1** | 0.01692 | 0.00806 | 0.05256 | 0.01067 | 0.00893 | 0.00593 | 0.00764 | 0.01450 | 0.01230 | 0.01377 | 0.01265 | 0.00823 | 0.01183 | 0.01219 | 0.03608 | 0.00735 |
| **Plcxd1** | 0.01095 | 0.00783 | 0.01716 | 0.19374 | 0.01024 | 0.09977 | 0.00744 | 0.17798 | 1.85980 | 0.00146 | 0.02299 | 0.00392 | 0.05630 | 0.02954 | 0.00237 | 0.00748 |
| **Pld1** | 0.76045 | 0.00667 | 0.00000 | 15.80504 | 0.00651 | 0.00298 | 0.00566 | 0.00000 | 7.73425 | 1.11253 | 0.50357 | 0.00000 | 0.00768 | 2.17289 | 5.73282 | 0.00320 |
| **Pld2** | 0.74034 | 0.00656 | 0.02151 | 5.53979 | 0.01650 | 0.00620 | 0.00609 | 1.09241 | 4.34765 | 1.56388 | 0.85507 | 0.06218 | 0.00000 | 0.18775 | 2.18252 | 0.27946 |
| **Pld3** | 5.63237 | 1.58070 | 14.93288 | 0.87617 | 0.05097 | 2.77222 | 5.20752 | 11.10090 | 0.65314 | 7.32480 | 9.31098 | 1.23013 | 9.71363 | 0.08542 | 1.98200 | 1.29502 |
| **Pld4** | 0.01323 | 0.00805 | 0.01168 | 0.52159 | 0.00901 | 0.00671 | 0.00746 | 0.01176 | 0.41019 | 0.72110 | 0.00865 | 0.00635 | 0.01207 | 0.00491 | 0.28613 | 0.00727 |
| **Pld5** | 0.01374 | 0.00800 | 0.01166 | 0.45600 | 0.00920 | 0.00661 | 0.00733 | 0.01151 | 0.00504 | 0.16407 | 0.01349 | 0.02737 | 0.01283 | 0.00585 | 0.17097 | 0.00771 |
| **Pld6** | 0.01167 | 0.00790 | 0.11023 | 3.34484 | 0.00861 | 0.00768 | 0.00715 | 0.00000 | 8.77337 | 0.00000 | 0.24304 | 0.00000 | 0.00899 | 0.15312 | 0.00000 | 0.00600 |
| **Plec** | 2.44579 | 0.00359 | 1.38130 | 2.15535 | 0.05959 | 6.41581 | 2.65982 | 0.06528 | 0.96038 | 0.88526 | 0.87761 | 0.11212 | 1.56745 | 0.00001 | 0.00000 | 3.58647 |
| **Plek** | 0.02835 | 0.00803 | 0.05080 | 1.41399 | 0.02079 | 0.00692 | 0.00724 | 0.00000 | 0.22156 | 0.00853 | 1.13316 | 0.01178 | 0.04425 | 0.13127 | 0.38613 | 0.01392 |
| **Plek2** | 3.41113 | 0.09573 | 0.65471 | 0.01380 | 0.00000 | 0.41821 | 0.03597 | 2.37466 | 0.57091 | 5.61222 | 1.57201 | 0.21809 | 0.00000 | 1.48286 | 2.23411 | 1.10465 |
| **Plekha1** | 3.93904 | 1.60677 | 5.63393 | 3.58843 | 2.43182 | 5.19105 | 1.99887 | 4.57266 | 6.03590 | 4.12398 | 4.14629 | 7.01222 | 6.14113 | 9.47364 | 6.00179 | 1.29334 |
| **Plekha2** | 0.07083 | 0.00766 | 0.00915 | 1.90204 | 0.00776 | 0.00644 | 0.02766 | 0.02736 | 1.49990 | 0.37135 | 0.42274 | 0.03089 | 0.01437 | 0.06471 | 0.01364 | 0.06005 |
| **Plekha3** | 1.23026 | 1.18544 | 0.62898 | 11.75063 | 0.53297 | 0.00276 | 1.94435 | 1.97542 | 4.52199 | 4.16167 | 1.48885 | 1.40773 | 1.02592 | 1.70086 | 9.29044 | 0.15837 |
| **Plekha4** | 4.46138 | 0.58416 | 1.86026 | 4.04753 | 0.00573 | 0.00000 | 0.74063 | 0.75887 | 3.07797 | 2.07785 | 3.31781 | 0.36302 | 1.76392 | 0.09730 | 0.29620 | 0.00748 |
| **Plekha5** | 1.39087 | 0.61462 | 0.49067 | 1.60607 | 0.67358 | 0.07806 | 0.42259 | 1.23682 | 2.29241 | 1.63841 | 1.46854 | 0.52638 | 0.40649 | 5.11218 | 7.39216 | 0.47390 |
| **Plekha6** | 2.52328 | 0.00340 | 0.00000 | 3.08453 | 0.00321 | 0.00307 | 1.81943 | 0.01088 | 1.82059 | 2.81990 | 1.79330 | 0.26395 | 0.21759 | 1.18222 | 2.96016 | 0.00340 |
| **Plekha7** | 1.38679 | 0.00517 | 3.66935 | 0.88781 | 0.38577 | 0.01527 | 0.65954 | 0.75403 | 3.26831 | 0.77644 | 1.72252 | 0.01414 | 1.63214 | 0.00000 | 0.64300 | 1.17318 |
| **Plekha8** | 0.59829 | 6.02510 | 4.41600 | 13.64643 | 0.11462 | 0.00380 | 0.33507 | 1.74499 | 4.15678 | 2.04669 | 1.36586 | 1.34667 | 0.01935 | 6.48854 | 15.10372 | 2.40605 |
| **Plekhb1** | 0.25472 | 0.00780 | 0.02645 | 1.25964 | 0.00730 | 0.00571 | 0.00717 | 0.02202 | 0.04706 | 1.64186 | 0.00730 | 0.02258 | 0.01068 | 0.16175 | 0.90640 | 0.00681 |
| **Plekhb2** | 0.06679 | 0.03692 | 2.12197 | 3.96898 | 0.11067 | 2.65255 | 4.34592 | 0.00000 | 2.10709 | 1.75453 | 1.27062 | 0.08071 | 1.97338 | 3.17774 | 1.60604 | 0.26456 |
| **Plekhd1** | 0.00250 | 0.00737 | 0.81076 | 0.27780 | 0.00980 | 0.19545 | 1.03777 | 0.76635 | 0.18373 | 0.10521 | 0.00504 | 0.00472 | 0.44698 | 0.01026 | 0.00000 | 0.00684 |
| **Plekhf1** | 8.62101 | 1.37938 | 1.67521 | 0.00000 | 2.13171 | 6.93031 | 2.56477 | 5.76815 | 0.50297 | 2.05016 | 2.20372 | 0.07238 | 8.10936 | 0.00698 | 0.00757 | 6.26754 |
| **Plekhf2** | 7.35342 | 8.15144 | 10.64720 | 10.74298 | 5.53471 | 1.32417 | 3.15369 | 8.83555 | 14.45353 | 16.13895 | 15.59739 | 2.78641 | 8.33120 | 22.39868 | 15.89039 | 2.57023 |
| **Plekhg1** | 0.01335 | 0.00816 | 0.01124 | 0.68524 | 0.00910 | 0.00671 | 0.01791 | 0.01172 | 1.05429 | 0.37286 | 0.01957 | 0.00596 | 0.01244 | 0.00969 | 0.23113 | 0.00722 |
| **Plekhg2** | 2.17176 | 0.00464 | 0.00000 | 1.02098 | 0.01202 | 1.98357 | 0.01580 | 3.61503 | 0.23117 | 3.05143 | 0.53752 | 0.08735 | 0.29820 | 0.00275 | 0.00000 | 0.48799 |
| **Plekhg3** | 3.49174 | 2.97108 | 2.11965 | 3.18531 | 0.73513 | 1.21821 | 6.84144 | 7.39315 | 6.94702 | 9.55916 | 5.70764 | 0.53172 | 2.25465 | 0.02580 | 2.32916 | 2.92001 |
| **Plekhg4** | 0.76827 | 0.00687 | 0.02400 | 0.00335 | 1.39455 | 0.00530 | 0.00702 | 0.01494 | 0.13052 | 0.01271 | 0.01110 | 0.00621 | 0.00990 | 0.01212 | 0.01422 | 0.00657 |
| **Plekhg5** | 2.92645 | 0.00320 | 0.00000 | 8.76103 | 0.01750 | 0.16487 | 0.00000 | 5.49023 | 9.23639 | 10.13455 | 8.70947 | 0.52347 | 0.15347 | 0.69513 | 1.39239 | 0.21636 |
| **Plekhg6** | 0.00868 | 0.00732 | 0.02833 | 1.93078 | 0.00888 | 0.00856 | 0.00650 | 0.89882 | 1.36582 | 1.34571 | 1.55335 | 0.02266 | 0.00000 | 0.00000 | 0.17852 | 0.00683 |
| **Plekhh1** | 0.48919 | 0.14891 | 3.74987 | 0.43695 | 0.31373 | 2.37552 | 1.55932 | 0.67691 | 2.59139 | 1.57696 | 1.91904 | 0.69970 | 0.44241 | 1.04375 | 1.10995 | 0.10670 |
| **Plekhh2** | 1.09963 | 0.04279 | 0.02622 | 0.08094 | 0.01715 | 0.00738 | 0.03792 | 0.01418 | 0.12868 | 0.10903 | 0.06663 | 0.03966 | 0.04492 | 0.09754 | 0.04676 | 0.00643 |
| **Plekhh3** | 0.01336 | 0.00787 | 0.13241 | 0.08322 | 0.01081 | 0.00586 | 0.00716 | 0.40636 | 0.06594 | 0.43165 | 0.43450 | 0.00354 | 0.00383 | 0.00868 | 0.00000 | 0.00772 |
| **Plekhj1** | 2.99775 | 0.21292 | 1.30599 | 4.08492 | 0.22007 | 2.80647 | 0.07145 | 2.44570 | 4.77296 | 6.21717 | 2.79274 | 0.13456 | 8.35774 | 0.00000 | 1.33435 | 5.48763 |
| **Plekhm1** | 1.63741 | 2.89579 | 4.48645 | 7.55946 | 0.06001 | 0.07934 | 2.55649 | 9.86061 | 10.09490 | 10.94052 | 6.44580 | 0.67061 | 2.36295 | 1.38131 | 9.65474 | 0.03665 |
| **Plekhm2** | 0.10187 | 1.28762 | 0.00014 | 0.29603 | 5.11641 | 3.08002 | 0.22849 | 3.57104 | 1.86956 | 2.64745 | 2.55228 | 0.12073 | 1.75702 | 0.91150 | 0.00000 | 0.01135 |
| **Plekhm3** | 0.73903 | 2.09338 | 1.88778 | 2.04398 | 0.00254 | 0.00593 | 3.50984 | 0.76728 | 3.47936 | 9.20006 | 4.72339 | 0.92896 | 0.58514 | 10.27266 | 10.22833 | 1.69820 |
| **Plekhn1** | 0.02765 | 0.00822 | 0.00902 | 0.01072 | 0.02967 | 0.00755 | 0.00742 | 0.28674 | 0.01200 | 0.00664 | 0.01409 | 0.00703 | 0.21061 | 0.01363 | 0.00587 | 0.01551 |
| **Plekho1** | 0.07980 | 0.43986 | 0.00350 | 0.00452 | 0.02470 | 0.01373 | 0.00631 | 0.67623 | 0.11407 | 0.46389 | 0.00719 | 0.03999 | 1.26446 | 0.00963 | 0.06550 | 0.00633 |
| **Plekho2** | 0.05174 | 2.16766 | 3.83496 | 6.25436 | 3.50123 | 0.00481 | 0.00408 | 2.84386 | 9.54514 | 1.85535 | 1.12828 | 0.00000 | 2.54015 | 0.03501 | 1.84988 | 1.98268 |
| **Plekhs1** | 0.01130 | 0.00773 | 0.01222 | 0.00484 | 0.00830 | 0.00655 | 0.00723 | 0.00933 | 0.01058 | 6.36588 | 0.00842 | 0.00000 | 0.01041 | 0.01383 | 0.00545 | 0.00741 |
| **Plet1** | 4.84331 | 6.30017 | 4.73850 | 0.00000 | 0.04045 | 5.68110 | 2.50953 | 0.00000 | 0.01112 | 4.99421 | 0.00577 | 0.00000 | 6.56706 | 0.03108 | 0.00000 | 5.25762 |
| **Plgrkt** | 0.00328 | 0.31748 | 0.54744 | 0.10970 | 0.00735 | 0.00694 | 0.00705 | 0.18478 | 0.00726 | 0.55045 | 0.95742 | 0.00150 | 1.44471 | 0.00476 | 0.07808 | 0.00644 |
| **Plin1** | 0.01304 | 0.00786 | 0.00885 | 0.18937 | 0.00882 | 0.00709 | 0.00721 | 0.00000 | 0.00848 | 0.92000 | 0.49405 | 0.00429 | 0.09086 | 0.00772 | 0.00000 | 0.00722 |

| | | | | | | | | | | | | | | | | |
|---|---|---|---|---|---|---|---|---|---|---|---|---|---|---|---|---|
| **Plin2** | 26.02195 | 14.31017 | 17.78261 | 7.72989 | 18.04738 | 11.61851 | 12.32440 | 3.72229 | 8.91535 | 1.17044 | 4.48155 | 0.21029 | 7.68191 | 3.87214 | 4.03059 | 7.71232 |
| **Plin3** | 14.75009 | 1.41193 | 2.68994 | 4.94417 | 4.28295 | 5.36405 | 7.36560 | 6.32967 | 6.25197 | 5.82225 | 6.23710 | 0.17420 | 8.03609 | 0.43913 | 0.16893 | 9.91321 |
| **Plin5** | 0.30854 | 0.00746 | 0.00550 | 0.00832 | 0.00708 | 0.00702 | 0.00688 | 0.00938 | 0.01201 | 0.49252 | 0.02398 | 0.03782 | 0.32878 | 0.01447 | 0.00712 | 0.00657 |
| **Plk1** | 9.04939 | 0.29929 | 0.01293 | 1.60982 | 5.75648 | 2.91949 | 1.16849 | 27.57376 | 8.80305 | 22.52798 | 5.89463 | 1.65893 | 6.65084 | 0.21715 | 0.00000 | 3.44615 |
| **Plk2** | 0.28745 | 0.00767 | 0.05437 | 0.11719 | 0.10725 | 0.00562 | 0.28714 | 0.54679 | 0.52590 | 0.00000 | 0.00000 | 0.07961 | 0.00699 | 0.00670 | 0.45719 | 0.00687 |
| **Plk3** | 1.32564 | 0.00579 | 0.00201 | 1.76197 | 0.31217 | 0.00672 | 0.00504 | 0.57835 | 2.46429 | 0.00000 | 0.49317 | 0.02524 | 1.30012 | 0.03494 | 0.00000 | 0.00457 |
| **Plk4** | 1.51988 | 2.48472 | 1.32001 | 5.64415 | 2.89200 | 3.02988 | 0.39402 | 3.71409 | 2.20332 | 1.18587 | 2.90518 | 0.52475 | 0.00000 | 9.25647 | 9.01594 | 2.32617 |
| **Plk5** | 0.00000 | 0.00709 | 0.00000 | 0.00422 | 0.02392 | 0.00824 | 0.00414 | 4.19913 | 0.17971 | 0.14566 | 2.73834 | 0.32310 | 4.27842 | 0.00399 | 0.00000 | 0.00575 |
| **Pllp** | 0.01314 | 0.00789 | 0.01214 | 0.79114 | 0.00891 | 0.00640 | 0.00720 | 0.00982 | 0.10213 | 1.19160 | 0.01285 | 0.00522 | 0.01060 | 0.01361 | 0.01297 | 0.00711 |
| **Plod1** | 1.80747 | 0.02936 | 4.98498 | 0.50963 | 4.06414 | 0.30703 | 7.15817 | 7.20526 | 0.37358 | 0.69474 | 2.62772 | 1.16590 | 1.79196 | 0.00000 | 0.39083 | 1.88039 |
| **Plod2** | 8.47644 | 0.00000 | 0.11983 | 0.00858 | 1.43072 | 0.00000 | 4.03293 | 0.00000 | 0.01149 | 0.00000 | 0.01267 | 0.00827 | 0.06014 | 0.01306 | 0.02220 | 3.24454 |
| **Plod3** | 3.16257 | 2.11025 | 2.95748 | 0.43863 | 0.01659 | 1.02954 | 2.34104 | 6.58451 | 4.02468 | 3.38616 | 1.64512 | 0.28817 | 6.44739 | 0.08586 | 1.24989 | 2.64575 |
| **Plp1** | 0.03433 | 0.01782 | 0.22841 | 0.02720 | 1.73586 | 0.01223 | 0.01788 | 0.01510 | 0.08227 | 0.06411 | 0.35731 | 0.02661 | 0.03993 | 0.51555 | 0.07414 | 0.01430 |
| **Plp2** | 17.83162 | 4.50488 | 13.66204 | 0.47528 | 7.12542 | 11.11180 | 7.51724 | 19.05298 | 0.72393 | 8.76610 | 5.40963 | 7.99376 | 16.73139 | 2.63034 | 0.00000 | 5.20493 |
| **Plpp1** | 13.66730 | 4.14882 | 5.04206 | 0.79996 | 11.49493 | 5.79516 | 2.00019 | 7.61554 | 1.37983 | 4.56970 | 5.05400 | 6.48647 | 3.52841 | 0.00126 | 4.90093 | 5.80050 |
| **Plpp2** | 5.27429 | 0.00000 | 4.18866 | 0.00993 | 0.00021 | 3.57617 | 4.33325 | 0.90173 | 0.33086 | 0.78224 | 1.13014 | 2.30206 | 2.41490 | 0.05014 | 0.36063 | 4.69892 |
| **Plpp3** | 0.26065 | 0.00654 | 2.44770 | 1.30425 | 0.02011 | 0.04190 | 0.02618 | 0.03901 | 3.24806 | 1.17207 | 0.04032 | 1.06872 | 1.65464 | 1.53033 | 1.08726 | 0.02035 |
| **Plpp5** | 1.71154 | 1.57532 | 2.54559 | 9.61763 | 0.01572 | 0.00222 | 0.14370 | 5.16107 | 13.70812 | 13.25494 | 7.20858 | 11.91504 | 2.94607 | 4.77699 | 13.51038 | 2.90856 |
| **Plpp6** | 0.01001 | 0.01723 | 0.00724 | 0.28491 | 0.00953 | 0.00664 | 0.00711 | 0.49263 | 0.28377 | 0.00541 | 0.00651 | 0.00494 | 0.01263 | 0.01707 | 0.52230 | 0.00716 |
| **Plppr3** | 0.64167 | 0.12753 | 0.47161 | 0.09527 | 0.00607 | 0.00882 | 0.00700 | 0.07213 | 0.08260 | 0.00362 | 0.01309 | 0.01757 | 0.53038 | 0.01322 | 0.00043 | 0.00627 |
| **Plppr4** | 0.03340 | 0.02909 | 0.02232 | 0.02679 | 0.00846 | 0.00621 | 0.00741 | 0.01265 | 0.01135 | 0.02661 | 0.01203 | 0.03751 | 0.02880 | 0.07518 | 0.07477 | 0.01459 |
| **Plrg1** | 13.59207 | 6.16329 | 9.84752 | 10.51266 | 10.71811 | 0.45740 | 5.55009 | 7.12886 | 21.02932 | 10.20051 | 8.33018 | 8.29418 | 8.98797 | 12.09469 | 11.36001 | 5.13093 |
| **Pls1** | 0.01511 | 0.00858 | 1.02366 | 1.53178 | 0.01024 | 0.00942 | 0.00700 | 0.16252 | 0.68664 | 0.00000 | 0.84536 | 0.06170 | 0.80512 | 0.05825 | 0.16081 | 0.01353 |
| **Pls3** | 1.85735 | 6.73437 | 2.48249 | 0.08294 | 3.18437 | 0.24119 | 0.20840 | 0.14439 | 1.63357 | 0.54833 | 0.00000 | 0.04135 | 3.58512 | 3.58910 | 1.82056 | 1.68050 |
| **Plscr1** | 8.93994 | 0.00000 | 0.15694 | 0.00756 | 3.10216 | 0.00940 | 1.05685 | 0.00000 | 0.01062 | 0.12595 | 0.01035 | 0.04025 | 0.42523 | 0.00950 | 0.03214 | 1.89003 |
| **Plscr3** | 1.10370 | 0.00718 | 0.00237 | 0.00966 | 0.21790 | 0.00744 | 0.00686 | 0.42639 | 0.01177 | 0.23317 | 0.00819 | 0.00618 | 0.51990 | 0.01375 | 0.00443 | 0.00560 |
| **Pltp** | 1.07492 | 0.01664 | 0.64031 | 0.00781 | 0.02100 | 0.00479 | 0.76075 | 0.32453 | 0.01163 | 0.36377 | 0.00829 | 0.02813 | 0.00648 | 0.00599 | 0.15466 | 2.32992 |
| **Plvap** | 0.01196 | 0.00767 | 0.01038 | 0.76354 | 0.00930 | 0.00659 | 0.00732 | 0.05851 | 0.53465 | 0.20052 | 0.00890 | 0.04268 | 0.01046 | 0.00453 | 0.20013 | 0.00761 |
| **Plxdc1** | 0.01377 | 0.00803 | 0.01204 | 0.00978 | 0.00931 | 0.00667 | 0.00752 | 0.01366 | 0.02361 | 0.01492 | 0.01286 | 0.01842 | 0.01273 | 0.01236 | 0.01026 | 0.00742 |
| **Plxdc2** | 0.01348 | 0.00797 | 0.01173 | 0.49181 | 0.00908 | 0.00656 | 0.00728 | 0.01060 | 0.35313 | 0.50864 | 0.00937 | 0.01568 | 0.01222 | 0.00546 | 0.16453 | 0.00770 |
| **Plxna1** | 3.76896 | 0.00175 | 2.83558 | 5.29600 | 0.50052 | 0.47929 | 2.94887 | 0.29044 | 2.05058 | 1.60034 | 0.30500 | 0.23102 | 1.42652 | 0.40050 | 0.22049 | 1.60438 |
| **Plxna2** | 0.01782 | 0.00770 | 0.01065 | 0.84032 | 0.00907 | 0.00642 | 0.00732 | 0.01155 | 0.26573 | 0.00386 | 0.00915 | 0.03610 | 0.01287 | 0.00679 | 0.63610 | 0.00716 |
| **Plxna3** | 0.26103 | 0.00802 | 0.00154 | 0.11908 | 0.01537 | 0.00810 | 0.00801 | 0.40369 | 0.00656 | 0.00000 | 0.11773 | 0.05510 | 0.00544 | 0.00789 | 0.39603 | 0.00689 |
| **Plxna4** | 0.01362 | 0.00764 | 0.01123 | 2.79233 | 0.00939 | 0.00609 | 0.00721 | 0.01093 | 0.00000 | 0.00556 | 0.00987 | 0.00520 | 0.01907 | 0.00293 | 1.21431 | 0.00682 |
| **Plxnb1** | 0.01834 | 0.00735 | 1.23351 | 0.87511 | 0.00653 | 0.00904 | 0.00678 | 0.00000 | 0.14147 | 0.00000 | 0.55416 | 0.37294 | 0.00422 | 0.01822 | 0.00000 | 0.00664 |
| **Plxnb2** | 6.15958 | 0.50903 | 7.76549 | 0.29153 | 3.45751 | 5.08655 | 7.00010 | 10.50085 | 0.25617 | 5.11467 | 4.00968 | 0.09780 | 7.23663 | 0.00000 | 0.00000 | 3.60085 |
| **Plxnb3** | 0.00936 | 0.00667 | 1.65373 | 0.88899 | 0.00706 | 0.00545 | 2.15382 | 0.00842 | 0.07291 | 0.00836 | 0.01145 | 0.00557 | 0.00665 | 0.00998 | 0.00000 | 0.00656 |
| **Plxnc1** | 0.00920 | 0.05259 | 0.00637 | 4.45089 | 0.00913 | 0.00594 | 0.00800 | 0.20363 | 1.23931 | 0.40852 | 0.01559 | 0.00000 | 0.00732 | 0.00001 | 2.40220 | 0.00682 |
| **Plxnd1** | 0.02184 | 0.00857 | 0.00998 | 0.18258 | 0.01142 | 0.00672 | 0.01870 | 0.03002 | 0.00665 | 0.59802 | 0.03446 | 0.02804 | 0.02323 | 0.03057 | 0.06239 | 0.00745 |
| **Pm20d2** | 0.01379 | 0.00789 | 0.01192 | 0.29461 | 0.00934 | 0.00665 | 0.00751 | 0.01319 | 0.00453 | 0.01308 | 0.02371 | 0.00787 | 0.01247 | 0.01721 | 0.03111 | 0.00736 |
| **Pmaip1** | 5.58550 | 0.00000 | 3.01407 | 1.92289 | 2.25851 | 0.00000 | 0.08054 | 0.05412 | 1.67313 | 0.00000 | 0.01771 | 0.37636 | 0.00000 | 0.00186 | 0.69206 | 2.74944 |
| **Pmch** | 0.02528 | 0.00738 | 0.08391 | 0.01074 | 0.00860 | 0.00567 | 0.00724 | 0.01418 | 0.01165 | 0.00792 | 0.00614 | 1.15909 | 0.01181 | 0.01099 | 0.00298 | 0.00737 |
| **Pmel** | 0.00999 | 0.04060 | 0.86994 | 1.60273 | 0.02898 | 0.24811 | 0.73108 | 4.96362 | 0.44439 | 3.00304 | 2.38193 | 0.15922 | 2.34216 | 0.14058 | 5.63522 | 0.93755 |
| **Pmepa1** | 0.69773 | 0.00515 | 1.23775 | 0.01831 | 3.33128 | 0.07507 | 0.30151 | 0.00365 | 0.01063 | 0.01120 | 0.00980 | 0.07739 | 0.00685 | 0.01450 | 0.00000 | 0.24358 |
| **Pmf1** | 15.86245 | 0.40078 | 8.36560 | 5.39692 | 6.75195 | 5.26306 | 5.41767 | 12.11560 | 2.57695 | 6.26931 | 4.28496 | 2.08044 | 7.33084 | 0.00849 | 0.00000 | 7.95258 |
| **Pmfbp1** | 0.00586 | 0.00779 | 0.00551 | 1.82222 | 0.00918 | 0.00591 | 0.00620 | 0.01358 | 3.12183 | 0.00000 | 0.00000 | 0.00000 | 1.59099 | 1.76301 | 1.29940 | 0.00618 |
| **Pmis2** | 0.10443 | 0.13892 | 0.31555 | 0.18557 | 0.01591 | 0.00556 | 0.00745 | 0.00341 | 0.00845 | 0.00447 | 0.34460 | 0.00704 | 0.00753 | 0.00818 | 0.00000 | 0.00711 |
| **Pml** | 5.74748 | 0.96935 | 2.05905 | 2.64723 | 0.01316 | 1.95180 | 1.64155 | 1.12954 | 0.19928 | 6.42203 | 3.09815 | 4.10954 | 1.73024 | 0.89896 | 4.94395 | 0.01177 |
| **Pmm1** | 32.22055 | 10.00068 | 27.50734 | 9.49124 | 23.62374 | 9.10631 | 16.17307 | 43.45465 | 20.26951 | 34.45555 | 24.04505 | 15.27482 | 30.22039 | 0.28229 | 2.72082 | 13.34945 |
| **Pmm2** | 9.00881 | 6.69032 | 9.73204 | 0.68507 | 7.42666 | 8.52010 | 5.94597 | 12.23796 | 2.17181 | 11.04671 | 6.54369 | 1.53815 | 7.08976 | 1.84281 | 8.66718 | 7.65518 |
| **Pmp22** | 0.01319 | 0.00774 | 0.01208 | 0.00714 | 0.00913 | 0.00647 | 0.00723 | 0.01010 | 0.01182 | 1.09181 | 0.01174 | 0.01415 | 0.01121 | 0.01213 | 0.00652 | 0.00744 |
| **Pmpca** | 7.94614 | 0.96970 | 5.26661 | 0.76580 | 3.20117 | 3.89406 | 2.19510 | 5.03590 | 1.80100 | 4.33886 | 3.73012 | 0.35768 | 6.30728 | 1.59127 | 3.96650 | 2.97835 |
| **Pmpcb** | 12.67309 | 7.38354 | 11.06790 | 2.89389 | 1.40664 | 6.90489 | 7.04223 | 11.44763 | 5.23893 | 7.39670 | 9.43723 | 9.08056 | 7.07294 | 6.08526 | 2.87306 | 6.04317 |
| **Pms1** | 1.38231 | 1.79638 | 0.26602 | 0.38222 | 0.40440 | 0.03867 | 0.00595 | 0.02558 | 0.00798 | 0.22850 | 0.47471 | 0.15019 | 0.04109 | 0.02892 | 0.69719 | 0.05332 |
| **Pms2** | 3.16543 | 3.40703 | 1.68105 | 5.08484 | 0.01077 | 0.00176 | 6.74279 | 1.97861 | 6.57600 | 1.73678 | 0.74341 | 2.75148 | 0.44849 | 0.66682 | 8.39770 | 0.42420 |
| **Pmvk** | 8.46202 | 3.54000 | 15.85099 | 0.24608 | 9.95949 | 6.72971 | 4.60693 | 3.11395 | 0.57784 | 0.23482 | 0.99268 | 0.03461 | 6.31995 | 0.00460 | 0.12724 | 0.06548 |
| **Pnck** | 1.07373 | 0.00682 | 0.00404 | 0.01060 | 0.00578 | 0.00700 | 0.00663 | 0.01269 | 0.01238 | 0.01942 | 0.01242 | 0.00732 | 0.01153 | 0.01209 | 0.01095 | 0.00612 |
| **Pnisr** | 3.23865 | 0.06680 | 2.29376 | 0.72337 | 0.35873 | 0.01054 | 1.60966 | 3.47960 | 3.21369 | 8.24728 | 0.11735 | 7.17579 | 0.31395 | 7.06924 | 13.37057 | 0.97430 |
| **Pnkd** | 9.60219 | 2.05246 | 7.08624 | 4.42607 | 10.23040 | 3.05541 | 3.79601 | 0.52838 | 3.31151 | 4.78011 | 2.33079 | 0.30351 | 3.86915 | 0.41440 | 0.23353 | 2.54997 |
| **Pnkp** | 4.13521 | 2.14678 | 5.73299 | 0.88039 | 2.27992 | 0.04325 | 2.50618 | 5.49009 | 0.93696 | 6.65830 | 4.58862 | 0.32298 | 5.73494 | 0.02658 | 1.12773 | 0.70882 |
| **Pnldc1** | 2.02694 | 0.00000 | 1.55821 | 4.87027 | 0.09373 | 4.13601 | 4.81129 | 1.84943 | 4.90936 | 2.67888 | 2.13823 | 5.74184 | 0.00000 | 0.65903 | 3.60629 | 0.73040 |
| **Pnlip** | 0.01371 | 0.00795 | 0.01196 | 0.12557 | 0.00935 | 0.00657 | 0.00747 | 0.01267 | 0.15360 | 0.00909 | 0.01094 | 0.00724 | 0.01273 | 0.04149 | 0.17137 | 0.00734 |

| | | | | | | | | | | | | | | | | |
|---|---|---|---|---|---|---|---|---|---|---|---|---|---|---|---|---|
| **Pnliprp2** | 2.99204 | 0.01231 | 5.44195 | 0.00244 | 0.00000 | 0.01693 | 1.96776 | 2.36772 | 0.00862 | 0.97376 | 1.71782 | 0.03350 | 6.93798 | 0.02813 | 0.00000 | 3.96645 |
| **Pnma1** | 0.01177 | 0.00797 | 0.01081 | 0.01034 | 0.00949 | 0.00673 | 0.00731 | 0.07782 | 0.01213 | 0.01778 | 0.01273 | 0.00815 | 0.00959 | 0.01022 | 0.03001 | 0.00740 |
| **Pnma2** | 0.01080 | 0.00808 | 0.01172 | 0.00951 | 0.00912 | 0.00634 | 0.00701 | 0.01156 | 0.01189 | 0.00274 | 0.01285 | 0.00817 | 0.65537 | 0.00901 | 0.14905 | 0.00686 |
| **Pnma3** | 0.41028 | 0.01929 | 0.02368 | 0.02760 | 0.01049 | 1.12305 | 0.15304 | 0.82037 | 0.94777 | 0.15793 | 0.13439 | 0.00000 | 1.47579 | 0.00765 | 0.61781 | 0.32583 |
| **Pnma5** | 3.61102 | 0.14825 | 0.82321 | 7.28140 | 0.05989 | 8.29583 | 2.18020 | 1.43542 | 15.25065 | 1.71119 | 1.26353 | 0.05989 | 2.84361 | 0.83856 | 2.10721 | 0.31134 |
| **Pnmal2** | 0.08272 | 0.00811 | 0.24897 | 0.00454 | 0.00805 | 0.00597 | 0.00765 | 0.01407 | 0.76648 | 0.75403 | 0.00671 | 0.00445 | 0.04775 | 0.01079 | 0.04635 | 0.00723 |
| **Pnn** | 3.94391 | 3.20427 | 6.42288 | 15.73671 | 3.42362 | 1.93462 | 1.25420 | 10.48992 | 26.21471 | 10.85204 | 5.75573 | 11.04173 | 4.27903 | 6.08241 | 11.99240 | 7.08793 |
| **Pno1** | 9.24940 | 7.84509 | 6.07506 | 12.37575 | 11.59002 | 8.68379 | 4.64412 | 18.66675 | 13.23443 | 20.17283 | 11.46855 | 17.77987 | 10.09830 | 13.55127 | 7.54674 | 10.73873 |
| **Pnoc** | 0.01378 | 0.00801 | 0.01143 | 0.02912 | 0.00928 | 0.00667 | 0.00749 | 0.01192 | 0.05981 | 0.01060 | 0.05323 | 0.00763 | 0.01193 | 0.04141 | 0.03224 | 0.00739 |
| **Pnp** | 15.79107 | 9.10227 | 18.25650 | 11.37372 | 3.81786 | 5.71771 | 8.65105 | 15.04852 | 22.64295 | 11.74347 | 6.74563 | 3.43686 | 15.47090 | 14.18467 | 6.39541 | 4.81245 |
| **Pnp2** | 4.57831 | 1.66124 | 4.48665 | 5.22788 | 0.35474 | 2.84266 | 3.18465 | 6.79014 | 8.18041 | 5.38500 | 3.05102 | 0.58284 | 3.59051 | 8.66581 | 3.22204 | 1.41723 |
| **Pnpla1** | 0.00507 | 0.01833 | 1.12855 | 0.52108 | 0.00798 | 0.64717 | 0.10766 | 0.16073 | 1.93877 | 0.33784 | 0.16725 | 0.27444 | 0.20174 | 0.36541 | 0.84223 | 0.00645 |
| **Pnpla2** | 1.11995 | 0.16900 | 0.00000 | 1.51955 | 5.20517 | 3.48831 | 0.32010 | 9.90268 | 0.28330 | 6.76815 | 3.59204 | 0.05929 | 3.05328 | 0.00000 | 0.00000 | 0.09236 |
| **Pnpla3** | 0.02730 | 0.27585 | 2.60445 | 2.74314 | 3.98667 | 0.01160 | 0.02348 | 2.92695 | 3.11271 | 4.45458 | 5.04098 | 0.26077 | 4.02594 | 5.99554 | 4.74443 | 0.03871 |
| **Pnpla6** | 10.09593 | 2.86121 | 12.38519 | 1.08248 | 6.82287 | 1.49008 | 6.13356 | 6.69323 | 0.24021 | 1.31289 | 5.31015 | 0.40209 | 5.82915 | 0.00000 | 1.57145 | 4.38674 |
| **Pnpla7** | 0.94902 | 0.00594 | 0.96947 | 3.12264 | 0.00330 | 0.00633 | 0.00538 | 0.16809 | 1.98654 | 2.19599 | 1.90825 | 0.04724 | 0.00000 | 0.01354 | 3.49341 | 3.07059 |
| **Pnpla8** | 2.56997 | 1.26789 | 4.96260 | 9.55823 | 4.82444 | 5.25519 | 7.97994 | 10.28620 | 8.09073 | 7.87382 | 6.34505 | 24.55582 | 3.60935 | 20.54681 | 10.90907 | 0.03025 |
| **Pnpo** | 1.29538 | 3.32266 | 6.47653 | 0.27815 | 0.00000 | 0.72104 | 0.72205 | 2.04410 | 0.34999 | 0.44452 | 1.10783 | 0.00245 | 0.13950 | 0.25664 | 0.00000 | 2.25386 |
| **Pnpt1** | 2.93757 | 1.12272 | 0.00000 | 0.00000 | 2.23551 | 2.07219 | 0.39748 | 0.50453 | 0.61953 | 0.00000 | 0.34707 | 2.68709 | 0.23388 | 2.88163 | 1.18507 | 0.00251 |
| **Pnrc1** | 1.74400 | 0.13511 | 0.00000 | 14.95008 | 2.54178 | 0.00265 | 0.03490 | 0.00000 | 20.72218 | 2.40340 | 0.00000 | 0.00000 | 0.01055 | 0.00000 | 1.42020 | 0.00897 |
| **Pnrc2** | 1.95520 | 0.87742 | 5.03493 | 0.23436 | 0.10227 | 0.01024 | 2.92488 | 1.05874 | 0.37443 | 1.17281 | 1.36410 | 0.16851 | 3.01630 | 2.37980 | 7.76229 | 0.61397 |
| **Poc1a** | 6.84049 | 0.74985 | 2.40002 | 3.19854 | 4.72209 | 3.59733 | 4.83290 | 5.43323 | 6.70429 | 6.42333 | 2.45688 | 0.22017 | 3.33172 | 0.64330 | 5.32930 | 5.07972 |
| **Poc1b** | 0.33141 | 0.00873 | 0.56952 | 0.80769 | 0.01517 | 0.00925 | 0.00787 | 0.43374 | 0.10675 | 1.10164 | 0.00792 | 0.03669 | 0.00949 | 0.01672 | 0.00000 | 0.00720 |
| **Poc5** | 3.86108 | 2.43644 | 2.24196 | 7.87644 | 1.05245 | 0.02456 | 2.24216 | 4.43419 | 6.67109 | 4.00953 | 2.24138 | 7.66514 | 1.47198 | 1.73754 | 1.78219 | 0.70971 |
| **Podnl1** | 0.01240 | 0.00797 | 0.00696 | 1.12104 | 0.00911 | 0.00709 | 0.00719 | 0.03254 | 0.81119 | 0.06389 | 0.54063 | 0.02900 | 0.00701 | 0.00449 | 0.12572 | 0.00704 |
| **Podxl** | 0.01552 | 0.00798 | 0.01724 | 0.01030 | 0.00898 | 0.00727 | 0.00742 | 0.04324 | 0.01233 | 0.08708 | 0.01182 | 0.03771 | 0.01040 | 0.01152 | 0.00944 | 0.01447 |
| **Pofut1** | 1.23784 | 0.00414 | 0.56056 | 0.80773 | 3.36951 | 0.00706 | 0.00623 | 1.57609 | 0.01557 | 0.83879 | 0.44003 | 0.11194 | 0.10741 | 0.00682 | 0.60031 | 0.02005 |
| **Pofut2** | 2.10352 | 0.02545 | 1.21577 | 4.92644 | 1.16356 | 0.66705 | 1.02791 | 2.05562 | 2.60585 | 2.29061 | 0.47138 | 0.51566 | 4.47624 | 0.00000 | 0.19319 | 3.87659 |
| **Pogk** | 0.00000 | 0.30392 | 0.05944 | 0.05706 | 0.00889 | 0.00635 | 2.37277 | 0.65325 | 1.18044 | 0.92594 | 0.01218 | 0.39556 | 0.16018 | 0.02263 | 0.24705 | 0.22731 |
| **Poglut1** | 7.72299 | 9.62733 | 11.28949 | 1.98716 | 9.91110 | 3.33463 | 7.91623 | 4.18703 | 3.90368 | 2.62886 | 1.98001 | 7.36118 | 3.17323 | 2.79965 | 3.20824 | 3.73003 |
| **Pogz** | 3.93468 | 0.14800 | 4.16065 | 4.01597 | 3.09423 | 1.61855 | 0.00000 | 3.15950 | 2.54039 | 3.84183 | 3.59804 | 0.44493 | 5.47282 | 3.32210 | 6.51476 | 1.07698 |
| **Pola1** | 1.00102 | 1.86372 | 2.10852 | 2.49708 | 0.09645 | 0.44722 | 0.01405 | 5.49878 | 1.17357 | 6.63933 | 4.13609 | 2.34239 | 0.15382 | 17.80401 | 5.26580 | 2.85823 |
| **Pola2** | 3.28618 | 0.17392 | 0.28998 | 3.14920 | 0.18028 | 0.38313 | 0.66876 | 4.95353 | 5.93430 | 2.53955 | 6.06573 | 0.40029 | 3.20327 | 1.33971 | 0.34219 | 0.07286 |
| **Polb** | 7.02604 | 3.94067 | 4.80794 | 3.36595 | 4.29546 | 4.34941 | 2.33254 | 10.79996 | 3.83878 | 8.32641 | 5.85196 | 4.82662 | 8.83237 | 1.81404 | 4.45528 | 4.57479 |
| **Pold1** | 9.25601 | 1.49759 | 7.00322 | 0.22445 | 1.68998 | 5.77389 | 0.39354 | 11.47362 | 0.00583 | 7.38864 | 6.90460 | 0.38675 | 8.40777 | 0.00000 | 0.00728 | 6.03032 |
| **Pold2** | 16.36977 | 6.10098 | 10.66505 | 0.27999 | 7.62389 | 3.93221 | 5.80298 | 8.61990 | 0.14898 | 8.14870 | 4.39797 | 0.46447 | 5.71317 | 0.02434 | 0.00000 | 11.18064 |
| **Pold3** | 7.70121 | 7.20794 | 12.34443 | 5.66348 | 7.03441 | 5.02177 | 9.29311 | 19.09899 | 4.40545 | 22.83947 | 15.22487 | 14.22303 | 14.12756 | 13.01687 | 16.20475 | 4.91158 |
| **Pold4** | 0.27897 | 0.51063 | 0.41959 | 0.00953 | 0.01530 | 0.42140 | 0.00687 | 0.01170 | 0.01235 | 0.00391 | 0.01332 | 0.00757 | 0.29144 | 0.00967 | 0.02442 | 0.00650 |
| **Poldip2** | 10.77158 | 3.21480 | 7.51133 | 0.89550 | 5.44754 | 5.32440 | 2.91006 | 7.52602 | 1.61073 | 2.02751 | 0.82924 | 0.23398 | 3.19430 | 0.10233 | 1.02534 | 1.44814 |
| **Poldip3** | 11.25634 | 9.10933 | 20.83384 | 12.88515 | 10.61278 | 8.70726 | 7.89892 | 14.64578 | 17.62311 | 12.69918 | 9.55593 | 1.65111 | 15.97874 | 12.33509 | 14.71363 | 9.38921 |
| **Pole** | 4.78630 | 0.01656 | 2.37162 | 6.02024 | 0.00569 | 2.13456 | 8.23453 | 7.45478 | 4.91632 | 5.90538 | 4.03595 | 3.87933 | 4.97091 | 7.38816 | 2.77285 | 4.24568 |
| **Pole2** | 3.76913 | 0.53424 | 1.18096 | 0.77955 | 1.80154 | 2.48454 | 1.47439 | 3.89646 | 4.60216 | 0.64364 | 4.88548 | 0.64529 | 2.95466 | 4.02264 | 4.89912 | 3.91627 |
| **Pole3** | 5.20399 | 4.44315 | 8.43727 | 5.27874 | 2.66511 | 3.61758 | 4.89485 | 9.93896 | 10.28875 | 8.88209 | 11.74360 | 0.54706 | 10.92548 | 0.96451 | 1.74356 | 7.08269 |
| **Pole4** | 12.15846 | 6.08176 | 1.73763 | 19.80205 | 11.76721 | 1.86935 | 0.92938 | 4.75399 | 18.81117 | 21.34243 | 11.31050 | 10.90516 | 7.69004 | 7.08937 | 10.52956 | 2.00892 |
| **Polg** | 1.23643 | 3.18348 | 5.71183 | 0.75167 | 0.06595 | 3.45669 | 3.67278 | 7.70100 | 0.31059 | 1.52993 | 1.56436 | 0.06923 | 4.12680 | 0.05494 | 0.62969 | 1.51609 |
| **Polg2** | 0.04487 | 0.28133 | 0.33440 | 0.83890 | 0.00726 | 0.00647 | 0.12028 | 0.02741 | 0.99691 | 0.33318 | 0.42795 | 0.19595 | 0.26350 | 0.10203 | 0.71280 | 0.50677 |
| **Polh** | 0.01258 | 0.00781 | 0.00791 | 1.88746 | 0.00939 | 0.00619 | 0.00717 | 0.00910 | 2.31659 | 0.24853 | 0.00000 | 0.00000 | 0.01290 | 0.56912 | 0.76459 | 0.00700 |
| **Poli** | 1.78903 | 0.00544 | 0.10693 | 0.71803 | 0.01337 | 0.01254 | 0.82444 | 0.00000 | 0.42414 | 0.54092 | 0.36064 | 0.08534 | 1.70942 | 0.20360 | 0.02275 | 0.04130 |
| **Polk** | 0.48673 | 0.00818 | 1.19370 | 0.34067 | 0.04778 | 0.00497 | 0.41733 | 0.97069 | 0.02019 | 0.80498 | 0.28941 | 0.89483 | 0.00000 | 1.81006 | 1.95023 | 0.02155 |
| **Poll** | 3.61659 | 4.71255 | 1.33370 | 1.72132 | 0.00847 | 0.00943 | 0.63272 | 3.65029 | 1.26938 | 5.53418 | 1.17288 | 0.09146 | 3.16308 | 0.04876 | 0.41102 | 0.20776 |
| **Polm** | 0.12241 | 0.00695 | 0.34512 | 10.90188 | 0.03885 | 4.00955 | 0.60752 | 2.78817 | 6.48654 | 0.84006 | 0.45088 | 0.07741 | 1.52550 | 0.01208 | 2.85987 | 9.64532 |
| **Poln** | 0.01165 | 0.10093 | 0.01122 | 0.01208 | 0.00897 | 0.00647 | 0.00727 | 0.01055 | 1.02853 | 0.39032 | 0.00518 | 0.00461 | 0.01202 | 0.01195 | 0.00346 | 0.00736 |
| **Polq** | 3.00862 | 3.63724 | 2.09497 | 4.11227 | 0.00967 | 0.05463 | 1.99307 | 1.24910 | 9.04848 | 13.89844 | 5.37557 | 14.98792 | 0.90402 | 17.45401 | 15.57109 | 2.21258 |
| **Polr1a** | 13.19523 | 9.46431 | 12.77009 | 7.77099 | 3.51381 | 3.86436 | 6.64774 | 19.32351 | 7.23827 | 16.08931 | 13.46581 | 1.74425 | 10.40600 | 2.80716 | 10.35149 | 2.50168 |
| **Polr1b** | 16.60889 | 13.53662 | 12.38420 | 7.51235 | 12.68320 | 1.12474 | 10.90545 | 21.77257 | 10.68996 | 19.15036 | 10.60367 | 10.47718 | 13.31098 | 17.78523 | 14.35194 | 6.61416 |
| **Polr1c** | 20.42839 | 10.42262 | 17.24288 | 19.20686 | 12.66476 | 4.85955 | 12.91511 | 26.09631 | 19.85284 | 28.00150 | 14.29006 | 10.93665 | 18.18039 | 14.33990 | 20.96268 | 10.27324 |
| **Polr1d** | 20.88144 | 10.31078 | 14.09311 | 14.24869 | 10.95760 | 9.98862 | 7.42596 | 19.44188 | 22.68774 | 26.36169 | 20.68506 | 3.97934 | 18.52884 | 19.41714 | 15.25847 | 12.09034 |
| **Polr1e** | 5.00905 | 0.11751 | 0.10869 | 10.78988 | 2.51742 | 0.04490 | 1.09080 | 8.07790 | 19.27101 | 14.49916 | 7.52316 | 0.99548 | 2.58322 | 9.60271 | 19.27926 | 2.79577 |
| **Polr2a** | 19.67910 | 7.70048 | 4.99317 | 5.71337 | 3.52568 | 3.62854 | 5.18436 | 15.28752 | 8.17473 | 14.93829 | 7.32648 | 15.26484 | 12.10138 | 2.01111 | 10.96713 | 10.39013 |
| **Polr2b** | 10.38556 | 10.89724 | 7.27358 | 9.76427 | 11.42274 | 4.10784 | 7.99457 | 16.05607 | 15.65306 | 16.78508 | 12.01910 | 13.46244 | 3.44040 | 26.49408 | 16.60403 | 4.29827 |
| **Polr2c** | 13.66160 | 9.88316 | 16.19632 | 20.22109 | 3.71769 | 7.12447 | 10.11065 | 17.61645 | 13.41116 | 30.04966 | 11.51599 | 2.99312 | 15.23773 | 1.95639 | 11.61452 | 9.80984 |
| **Polr2d** | 9.69108 | 7.34328 | 8.56615 | 0.90717 | 5.29256 | 3.34080 | 3.98625 | 14.56725 | 4.81140 | 12.48316 | 1.33001 | 0.69837 | 5.69798 | 12.12324 | 11.64671 | 2.25845 |

| | | | | | | | | | | | | | | | | |
|---|---|---|---|---|---|---|---|---|---|---|---|---|---|---|---|---|
| **Polr2e** | 28.47478 | 10.18100 | 24.08719 | 13.81820 | 18.73211 | 13.86064 | 12.87341 | 32.13015 | 16.22571 | 30.76004 | 19.44201 | 14.31645 | 25.45003 | 0.62954 | 12.25951 | 15.74046 |
| **Polr2f** | 11.16279 | 0.75431 | 7.26362 | 12.25644 | 10.40748 | 1.36120 | 3.42013 | 9.56108 | 13.42474 | 28.06195 | 13.27465 | 1.89719 | 12.24550 | 0.00000 | 0.00000 | 3.86086 |
| **Polr2g** | 13.22829 | 2.74688 | 11.69079 | 4.18840 | 4.69968 | 5.97686 | 5.36413 | 7.58237 | 12.69454 | 5.12919 | 11.85808 | 1.56895 | 5.81021 | 6.94657 | 6.88809 | 3.44075 |
| **Polr2h** | 15.56990 | 8.55424 | 10.72423 | 5.16864 | 8.09534 | 3.86411 | 7.21024 | 16.65064 | 13.90139 | 20.75344 | 8.73230 | 3.20356 | 9.92676 | 8.86077 | 9.97285 | 3.97657 |
| **Polr2i** | 5.86798 | 0.15732 | 2.10502 | 4.90585 | 5.24990 | 0.78165 | 0.16974 | 0.88310 | 9.28106 | 4.12398 | 8.44593 | 0.51591 | 2.88134 | 0.00785 | 0.06403 | 2.24751 |
| **Polr2j** | 12.38767 | 0.23846 | 7.11256 | 24.56404 | 12.37758 | 2.33129 | 0.82922 | 3.45494 | 22.04761 | 15.08829 | 16.54743 | 2.40846 | 6.09949 | 8.52082 | 11.82566 | 2.59772 |
| **Polr2k** | 3.77683 | 0.13448 | 2.33652 | 10.73130 | 1.90162 | 0.06491 | 1.04713 | 0.00000 | 13.18082 | 1.67465 | 21.53238 | 0.86692 | 3.49139 | 10.70411 | 4.60926 | 0.03861 |
| **Polr2l** | 7.85276 | 0.25154 | 5.60819 | 23.40126 | 6.16199 | 0.75879 | 1.03151 | 1.68505 | 33.78312 | 10.10183 | 30.97638 | 2.43601 | 6.23725 | 2.97785 | 3.58923 | 2.04267 |
| **Polr2m** | 14.85674 | 11.86706 | 5.99297 | 4.89300 | 13.12627 | 4.57330 | 7.85523 | 5.43053 | 3.22526 | 12.52189 | 10.49126 | 7.99153 | 10.99706 | 6.75092 | 13.77462 | 7.62650 |
| **Polr3a** | 7.79736 | 5.24163 | 4.23540 | 14.49272 | 0.00000 | 3.85574 | 1.10756 | 5.77063 | 10.32578 | 12.59166 | 3.06744 | 5.65819 | 3.14610 | 2.41737 | 17.20812 | 4.01784 |
| **Polr3b** | 4.92785 | 4.23653 | 3.88750 | 3.54021 | 2.25916 | 0.00000 | 2.20801 | 2.45390 | 8.40601 | 8.53122 | 4.40067 | 0.59299 | 3.13109 | 1.52381 | 8.08231 | 4.47260 |
| **Polr3c** | 4.70995 | 5.19430 | 3.84369 | 1.05277 | 0.87130 | 1.68839 | 2.15751 | 7.13683 | 1.15548 | 8.62577 | 1.65074 | 4.61355 | 5.49173 | 5.74534 | 8.22960 | 0.00000 |
| **Polr3d** | 2.86089 | 0.12621 | 7.03291 | 9.57913 | 0.68563 | 0.00108 | 1.47387 | 3.51108 | 15.66646 | 11.19492 | 7.34582 | 1.02684 | 8.62293 | 0.82551 | 7.90736 | 8.45627 |
| **Polr3e** | 8.24819 | 4.13927 | 4.99348 | 10.41987 | 1.02756 | 3.68856 | 6.72622 | 7.20675 | 12.71565 | 5.84937 | 1.29581 | 1.21764 | 3.71609 | 0.26280 | 2.04816 | 3.09416 |
| **Polr3f** | 2.63418 | 4.68950 | 2.21733 | 3.68495 | 1.73510 | 0.01912 | 0.99198 | 5.28534 | 4.25970 | 4.89532 | 2.44211 | 7.37028 | 1.75075 | 6.35577 | 7.38985 | 0.27068 |
| **Polr3g** | 1.93778 | 1.56272 | 6.31281 | 4.14401 | 8.09528 | 3.99010 | 4.03773 | 9.02759 | 7.23430 | 13.16189 | 8.50245 | 19.49740 | 12.42661 | 12.30597 | 16.01874 | 0.20288 |
| **Polr3gl** | 0.44219 | 0.03996 | 0.01443 | 0.50216 | 0.11119 | 2.70678 | 0.12998 | 0.05719 | 0.05586 | 0.10513 | 0.19284 | 0.07750 | 0.09263 | 0.12262 | 0.05639 | 0.09148 |
| **Polr3h** | 3.57278 | 4.10830 | 3.19909 | 15.51650 | 3.39291 | 5.11116 | 3.12728 | 6.97430 | 15.23534 | 8.83980 | 0.97396 | 0.21425 | 5.99306 | 0.48346 | 1.26056 | 1.48625 |
| **Polr3k** | 1.55462 | 0.67587 | 2.11275 | 4.60342 | 3.27531 | 0.78431 | 2.52340 | 1.61508 | 11.57981 | 4.01835 | 4.32114 | 0.41209 | 1.01408 | 5.98543 | 8.86509 | 0.07272 |
| **Polrmt** | 2.97554 | 0.00728 | 5.10491 | 9.42766 | 4.44036 | 2.44694 | 2.97208 | 12.33248 | 13.57187 | 17.42530 | 13.64745 | 1.09199 | 7.63142 | 0.14670 | 4.61130 | 2.54543 |
| **Pom121** | 0.47416 | 0.02745 | 3.05434 | 8.12982 | 0.04328 | 0.53490 | 1.48686 | 9.14554 | 9.85287 | 1.24006 | 1.26858 | 0.41200 | 2.27468 | 0.54983 | 12.14724 | 1.83416 |
| **Pomc** | 0.03815 | 0.00795 | 0.26391 | 2.38975 | 0.00857 | 0.00676 | 0.17593 | 0.01186 | 0.52178 | 0.73676 | 0.01125 | 0.00493 | 0.31199 | 0.00871 | 0.00114 | 0.01483 |
| **Pomgnt1** | 8.04761 | 4.16818 | 4.53412 | 0.34037 | 3.03957 | 6.51334 | 7.36479 | 4.52641 | 1.20727 | 3.33060 | 3.13456 | 0.34441 | 7.91907 | 2.01294 | 1.65460 | 2.77088 |
| **Pomgnt2** | 3.45831 | 0.01374 | 0.00550 | 0.00802 | 0.00256 | 0.00405 | 0.00441 | 0.95872 | 0.01099 | 0.50660 | 0.04521 | 0.00204 | 0.00569 | 0.01719 | 0.00000 | 0.01054 |
| **Pomk** | 3.18518 | 0.17228 | 2.05167 | 0.01933 | 0.00691 | 0.00483 | 2.30660 | 0.86036 | 0.89030 | 0.00000 | 0.00000 | 0.18956 | 0.12737 | 0.04866 | 0.03886 | 0.00287 |
| **Pomp** | 21.85126 | 15.59241 | 26.56376 | 5.91108 | 14.17644 | 7.01968 | 11.11577 | 18.71449 | 11.75669 | 17.63271 | 26.20199 | 5.77385 | 22.22943 | 13.69793 | 11.05504 | 11.08649 |
| **Pomt1** | 3.96779 | 0.00201 | 3.17291 | 1.49347 | 0.00508 | 0.01029 | 1.40484 | 0.51053 | 0.39398 | 0.12383 | 0.23619 | 0.00000 | 1.96910 | 0.00246 | 0.07923 | 0.13861 |
| **Pomt2** | 0.74844 | 0.00223 | 0.75360 | 0.16870 | 0.00448 | 0.00382 | 4.92589 | 1.82593 | 0.71146 | 0.33633 | 1.20206 | 6.75719 | 0.36031 | 1.20284 | 0.07115 | 1.42684 |
| **Pon2** | 2.30562 | 0.00503 | 0.06966 | 0.00000 | 0.01538 | 5.19038 | 0.01534 | 0.31035 | 0.21966 | 0.57669 | 0.23743 | 0.03883 | 0.00000 | 0.20858 | 0.00000 | 1.88918 |
| **Pop1** | 4.49817 | 1.14239 | 3.65353 | 5.74849 | 2.85018 | 0.00000 | 4.27343 | 7.34496 | 6.60064 | 2.94587 | 1.87207 | 1.03606 | 3.12175 | 0.26337 | 1.07786 | 3.40107 |
| **Pop4** | 17.45389 | 14.78794 | 20.34996 | 1.69837 | 10.48338 | 9.51851 | 11.37495 | 16.65160 | 6.80764 | 19.86172 | 8.20198 | 8.45972 | 17.58620 | 11.28143 | 14.13962 | 11.20958 |
| **Pop5** | 3.55058 | 0.42246 | 3.02447 | 6.04956 | 0.03930 | 0.01249 | 0.58648 | 3.08406 | 12.90300 | 17.68940 | 18.42421 | 1.36735 | 2.83394 | 3.64318 | 3.99890 | 1.63239 |
| **Pop7** | 9.49703 | 6.81285 | 11.10794 | 3.88530 | 4.11325 | 0.00000 | 3.74413 | 7.37742 | 5.36412 | 9.67747 | 5.01847 | 0.98697 | 12.26617 | 0.24588 | 0.00000 | 3.62198 |
| **Popdc3** | 3.71298 | 0.15508 | 0.07381 | 0.25089 | 3.43180 | 0.00190 | 0.00435 | 0.92117 | 0.97992 | 0.00000 | 0.10137 | 0.11106 | 0.08640 | 0.00581 | 0.61280 | 0.04620 |
| **Por** | 2.27491 | 0.01515 | 2.84485 | 1.46201 | 3.40376 | 5.63608 | 10.99660 | 14.45672 | 1.80135 | 1.75836 | 3.04740 | 1.90989 | 2.97647 | 0.00377 | 0.00000 | 0.80951 |
| **Porcn** | 4.60045 | 1.35225 | 7.52796 | 2.02146 | 0.00000 | 0.18455 | 0.69255 | 3.24078 | 0.05621 | 5.90458 | 5.82744 | 0.71584 | 3.72300 | 2.26638 | 1.30346 | 1.35077 |
| **Postn** | 0.00931 | 0.00857 | 0.00794 | 0.00878 | 0.00937 | 0.00697 | 0.00824 | 0.27740 | 0.01161 | 0.41078 | 0.01848 | 0.34484 | 0.00635 | 0.01406 | 0.00000 | 0.00718 |
| **Pot1a** | 2.04703 | 0.00422 | 1.44027 | 0.92579 | 0.00454 | 0.00335 | 2.41338 | 1.41828 | 0.45045 | 5.01385 | 0.98614 | 1.57165 | 1.01306 | 1.82908 | 4.34414 | 0.01113 |
| **Pot1b** | 0.59379 | 0.00716 | 0.22359 | 0.72033 | 0.19595 | 0.01948 | 0.03894 | 0.01471 | 1.74600 | 0.58058 | 0.19603 | 0.52216 | 0.24913 | 0.60303 | 0.48017 | 0.00661 |
| **Pou2f1** | 6.81522 | 0.90883 | 0.80971 | 0.84651 | 2.04338 | 0.69227 | 2.10244 | 0.16893 | 0.20257 | 3.09415 | 0.34275 | 8.44557 | 0.87042 | 1.09032 | 0.24261 | 0.01298 |
| **Pou2f2** | 0.14862 | 0.00810 | 0.00493 | 1.96213 | 0.01664 | 0.00672 | 0.00708 | 0.01171 | 4.59140 | 0.53272 | 0.00000 | 0.07066 | 0.01174 | 0.00000 | 0.18812 | 0.00705 |
| **Pou2f3** | 4.97562 | 0.00207 | 0.62530 | 0.01211 | 0.01720 | 0.00166 | 0.00277 | 0.00000 | 1.06513 | 3.65995 | 0.87196 | 0.08370 | 0.00916 | 0.71461 | 3.90574 | 0.00143 |
| **Pou3f1** | 0.01207 | 0.00831 | 0.01072 | 0.70783 | 0.00944 | 0.00661 | 0.00805 | 0.05996 | 0.33020 | 0.00750 | 0.00934 | 0.01886 | 0.01012 | 0.00991 | 0.14208 | 0.00722 |
| **Pou3f3** | 0.01364 | 0.00786 | 0.01209 | 0.10237 | 0.00933 | 0.00649 | 0.00733 | 0.01177 | 0.01080 | 0.24374 | 0.01367 | 0.01654 | 0.01881 | 0.01098 | 0.00692 | 0.00743 |
| **Pou4f1** | 0.00865 | 0.00744 | 0.00262 | 9.90142 | 0.00998 | 1.34754 | 0.00660 | 0.16507 | 1.03887 | 0.00000 | 0.02752 | 0.00000 | 0.00000 | 1.59242 | 1.80748 | 0.00510 |
| **Pou4f2** | 0.01161 | 0.00787 | 0.01200 | 0.01041 | 0.00926 | 0.00647 | 0.00713 | 0.01110 | 0.01242 | 0.05556 | 0.01269 | 0.00729 | 0.45747 | 0.01170 | 0.00985 | 0.00721 |
| **Pou5f1** | 18.60843 | 11.20198 | 23.83468 | 6.30154 | 19.53587 | 10.97595 | 12.12648 | 28.73622 | 2.67177 | 17.97269 | 21.74485 | 1.04672 | 20.89366 | 2.45248 | 5.01491 | 11.68538 |
| **Pou6f1** | 1.61184 | 0.00147 | 1.00873 | 3.00942 | 1.95999 | 0.00256 | 0.29170 | 1.65458 | 1.22420 | 5.40468 | 1.19426 | 10.49791 | 1.96318 | 2.05440 | 8.19506 | 0.00299 |
| **Pp2d1** | 0.38909 | 0.00720 | 0.00451 | 0.05700 | 0.00696 | 0.00739 | 0.00721 | 0.00118 | 0.64794 | 0.09618 | 0.35686 | 0.01629 | 0.00884 | 0.07634 | 0.00000 | 0.00672 |
| **Ppa1** | 19.32137 | 8.01959 | 13.40730 | 6.11275 | 11.88478 | 3.57994 | 6.99548 | 3.75233 | 7.24778 | 15.07929 | 4.50756 | 8.92653 | 12.10595 | 5.64622 | 3.94768 | 10.49198 |
| **Ppa2** | 0.86145 | 0.01324 | 1.95799 | 1.71661 | 3.66446 | 2.75948 | 2.41673 | 1.16334 | 0.01939 | 2.83991 | 2.38127 | 2.85191 | 2.21397 | 1.66003 | 0.89453 | 0.01018 |
| **Ppan** | 17.70058 | 1.60246 | 9.29708 | 16.88978 | 11.66557 | 4.28116 | 8.34527 | 21.51190 | 16.74706 | 27.12003 | 10.34800 | 2.06773 | 17.59921 | 0.02710 | 0.00000 | 15.12574 |
| **Ppara** | 0.01389 | 0.00763 | 0.00822 | 0.59390 | 0.00936 | 0.00678 | 0.00716 | 0.01039 | 0.00103 | 0.02261 | 0.01005 | 0.49256 | 0.01328 | 0.02951 | 0.00000 | 0.00737 |
| **Ppard** | 1.84302 | 1.65015 | 0.76823 | 6.15442 | 0.00437 | 0.02845 | 0.03428 | 1.96383 | 0.39846 | 0.58196 | 0.29557 | 0.08491 | 0.29387 | 0.00000 | 0.19111 | 0.00311 |
| **Ppargc1b** | 0.00000 | 0.50283 | 2.00230 | 0.01017 | 0.02615 | 0.01299 | 0.20988 | 1.20112 | 0.01104 | 0.00000 | 0.01755 | 0.00311 | 1.86692 | 0.01443 | 0.00000 | 0.02008 |
| **Ppat** | 9.75033 | 7.71236 | 9.57325 | 19.68502 | 12.78498 | 4.52811 | 10.34452 | 21.40082 | 32.76148 | 19.67897 | 15.29207 | 17.89504 | 10.15541 | 25.52601 | 22.18414 | 8.08543 |
| **Ppcdc** | 2.26984 | 2.23705 | 9.30485 | 0.00000 | 0.06210 | 0.01134 | 0.00216 | 3.71596 | 0.15877 | 2.89080 | 2.54212 | 0.23375 | 2.13319 | 0.62877 | 0.52095 | 2.40197 |
| **Ppcs** | 3.90682 | 4.26369 | 4.87179 | 5.89041 | 14.53776 | 0.01929 | 1.23084 | 2.63946 | 9.10157 | 8.04070 | 0.95041 | 0.18669 | 6.35013 | 0.86420 | 2.76003 | 4.98952 |
| **Ppdpf** | 5.56937 | 0.00000 | 0.49249 | 7.26163 | 0.10931 | 0.36995 | 0.00898 | 0.58242 | 3.31894 | 5.47768 | 4.55567 | 0.02362 | 3.49249 | 0.18331 | 0.00000 | 3.66431 |
| **Ppef1** | 0.01350 | 0.01761 | 0.01107 | 0.34524 | 0.00939 | 0.00643 | 0.00732 | 0.01068 | 0.00573 | 0.01372 | 0.02213 | 0.00579 | 0.01271 | 0.01575 | 0.76307 | 0.00736 |
| **Ppef2** | 0.01305 | 0.00797 | 0.01028 | 0.01053 | 0.00933 | 0.00675 | 0.24536 | 0.01606 | 0.01234 | 0.01436 | 0.01294 | 0.00803 | 0.01160 | 0.01258 | 0.01045 | 0.00738 |
| **Ppfia1** | 5.00145 | 7.53663 | 7.13364 | 23.54501 | 5.24058 | 3.18132 | 6.74457 | 13.54393 | 13.69621 | 11.96751 | 12.68334 | 5.98450 | 6.43896 | 3.21362 | 15.26936 | 0.25616 |

| | | | | | | | | | | | | | | | | |
|---|---|---|---|---|---|---|---|---|---|---|---|---|---|---|---|---|
| **Ppfia3** | 0.84535 | 0.00730 | 0.00414 | 0.00881 | 0.01056 | 0.00700 | 0.00642 | 0.00728 | 0.01218 | 0.34369 | 0.02406 | 0.00601 | 0.36120 | 0.01147 | 0.00910 | 0.00650 |
| **Ppfia4** | 0.00926 | 0.00725 | 0.05705 | 0.51227 | 0.00917 | 0.00743 | 0.00649 | 0.79480 | 0.00347 | 0.73674 | 1.73979 | 0.01646 | 0.02584 | 0.03800 | 0.00000 | 0.01430 |
| **Ppfibp1** | 1.38270 | 3.36559 | 0.00000 | 6.42899 | 3.01491 | 2.02392 | 0.09645 | 0.27836 | 4.90539 | 0.00000 | 1.50124 | 0.66812 | 0.08667 | 1.05964 | 7.74771 | 0.61220 |
| **Ppfibp2** | 2.69558 | 0.74722 | 7.05685 | 0.49502 | 0.02656 | 0.03396 | 1.56447 | 18.09138 | 0.21851 | 16.50437 | 15.01222 | 11.85275 | 4.27396 | 16.70431 | 8.59563 | 0.00329 |
| **Pphln1** | 4.45076 | 3.84087 | 9.59155 | 10.89145 | 7.35750 | 8.54886 | 4.45851 | 16.26139 | 19.57332 | 18.78312 | 14.46469 | 6.51317 | 7.87515 | 19.14349 | 13.67716 | 3.61712 |
| **Ppia** | 52.03405 | 29.29199 | 41.90479 | 21.54226 | 34.97404 | 18.80363 | 23.81622 | 42.00430 | 28.49156 | 41.19130 | 43.49912 | 20.50402 | 44.28213 | 19.89646 | 25.56316 | 27.73739 |
| **Ppib** | 5.18488 | 4.65020 | 9.01113 | 0.00000 | 0.82296 | 4.06847 | 5.23473 | 1.95598 | 0.02199 | 2.07828 | 0.00000 | 0.19668 | 6.08404 | 0.27268 | 0.65980 | 5.96465 |
| **Ppic** | 0.00000 | 5.14510 | 0.03729 | 0.26534 | 0.59195 | 0.01227 | 1.98617 | 0.72347 | 0.00663 | 1.09659 | 2.46391 | 0.11560 | 1.03016 | 0.00000 | 0.00000 | 0.02693 |
| **Ppid** | 19.01731 | 15.22770 | 19.32873 | 10.74600 | 20.90575 | 7.63602 | 11.62855 | 17.17041 | 14.40639 | 12.76720 | 10.63330 | 11.63672 | 12.45673 | 10.68608 | 11.98969 | 9.57176 |
| **Ppie** | 8.55252 | 9.59727 | 11.26109 | 0.00000 | 4.85976 | 7.02032 | 8.37569 | 9.23882 | 0.20414 | 14.70124 | 1.37956 | 0.49803 | 13.39626 | 0.15393 | 5.30455 | 7.30172 |
| **Ppif** | 2.76745 | 0.11589 | 6.96698 | 2.34210 | 9.29254 | 0.02600 | 5.36784 | 3.67224 | 3.33444 | 3.91646 | 5.14232 | 3.86155 | 5.85842 | 1.28253 | 0.68475 | 0.07880 |
| **Ppig** | 8.12055 | 10.96414 | 15.96678 | 5.30451 | 10.17531 | 4.07568 | 9.76238 | 16.78342 | 6.02261 | 23.31113 | 6.68597 | 18.74405 | 10.20397 | 36.26854 | 26.28986 | 5.08339 |
| **Ppih** | 14.71545 | 10.56141 | 12.00267 | 5.59811 | 8.14422 | 7.45940 | 6.68284 | 10.69940 | 3.34808 | 8.88205 | 6.31313 | 4.14809 | 12.11194 | 9.15474 | 10.12118 | 8.27394 |
| **Ppil1** | 15.42972 | 15.85556 | 18.28400 | 4.65677 | 14.37665 | 4.02408 | 12.50457 | 14.03185 | 4.11197 | 18.49094 | 9.92282 | 1.28401 | 11.96101 | 3.73898 | 11.25820 | 7.45107 |
| **Ppil2** | 8.40379 | 8.07008 | 7.92055 | 5.51883 | 8.79948 | 0.05544 | 3.17521 | 10.68955 | 7.37083 | 13.26708 | 4.29718 | 2.22503 | 5.99629 | 0.00000 | 1.51560 | 4.41820 |
| **Ppil3** | 3.78326 | 5.79429 | 2.21762 | 0.00000 | 7.29632 | 4.86217 | 0.45333 | 1.75655 | 0.77452 | 2.03928 | 1.55085 | 0.07803 | 1.08739 | 2.84717 | 0.45598 | 3.62717 |
| **Ppil4** | 1.88543 | 5.33496 | 5.67233 | 4.87557 | 9.01931 | 0.01538 | 3.07489 | 3.58083 | 5.46265 | 12.32528 | 3.12531 | 5.91097 | 3.04309 | 5.23598 | 10.71231 | 0.00477 |
| **Ppil6** | 0.01372 | 0.00821 | 0.00789 | 0.51956 | 0.00904 | 0.00694 | 0.00758 | 0.00560 | 0.13777 | 0.00558 | 0.65544 | 0.00550 | 0.01197 | 0.00595 | 0.00000 | 0.00703 |
| **Ppip5k1** | 0.73320 | 0.11491 | 2.40099 | 0.14413 | 0.00237 | 0.01221 | 0.01849 | 0.03873 | 0.41353 | 0.40278 | 0.02509 | 0.13489 | 0.01166 | 0.07115 | 0.48855 | 0.00534 |
| **Ppip5k2** | 1.25497 | 6.21197 | 2.20995 | 3.67474 | 1.47608 | 0.94651 | 5.13652 | 3.72733 | 1.99224 | 1.97126 | 0.85044 | 2.07699 | 0.76401 | 5.55759 | 6.21798 | 1.70809 |
| **Ppl** | 0.00413 | 0.00750 | 0.64491 | 0.00475 | 0.00811 | 0.00582 | 0.00541 | 0.00000 | 0.02071 | 0.17348 | 2.44764 | 0.00414 | 2.25028 | 0.00910 | 0.00000 | 0.04479 |
| **Ppm1a** | 17.60178 | 21.46721 | 19.24210 | 25.50560 | 15.69902 | 3.83273 | 15.67643 | 32.55164 | 19.02210 | 23.58550 | 28.46249 | 19.07298 | 14.81941 | 25.64659 | 21.63810 | 0.95311 |
| **Ppm1b** | 8.96872 | 6.39691 | 6.33628 | 16.56149 | 7.25238 | 7.47787 | 3.87569 | 6.81777 | 14.10844 | 13.64288 | 10.76161 | 11.12521 | 3.78201 | 19.25385 | 16.32051 | 4.31761 |
| **Ppm1d** | 0.72942 | 2.51870 | 1.00370 | 5.68001 | 0.02089 | 3.04594 | 2.45835 | 0.10195 | 10.71113 | 2.91791 | 0.00000 | 1.68828 | 1.38250 | 0.97315 | 2.73259 | 1.22261 |
| **Ppm1e** | 0.01172 | 0.00787 | 0.00676 | 0.90102 | 0.00917 | 0.00677 | 0.00728 | 0.00950 | 1.27310 | 0.68035 | 0.00000 | 0.00000 | 0.01976 | 1.34131 | 0.08055 | 0.00713 |
| **Ppm1f** | 2.88438 | 0.00016 | 1.96227 | 0.29510 | 4.55825 | 0.03280 | 0.97865 | 2.18870 | 1.04609 | 2.01852 | 2.85739 | 0.19468 | 1.58798 | 1.42279 | 1.41413 | 1.69533 |
| **Ppm1g** | 15.89230 | 8.44085 | 14.27587 | 12.61858 | 11.93967 | 10.08464 | 11.88506 | 23.82405 | 15.19409 | 32.80398 | 11.69493 | 12.58833 | 13.39739 | 2.68160 | 21.79259 | 7.76200 |
| **Ppm1h** | 0.01484 | 0.00723 | 0.01321 | 8.06414 | 0.00892 | 0.03352 | 0.65708 | 0.00000 | 2.95067 | 0.62727 | 0.65233 | 0.00783 | 0.01982 | 1.49707 | 1.11090 | 0.01291 |
| **Ppm1j** | 0.27235 | 0.00862 | 0.01012 | 0.00811 | 0.01720 | 0.00634 | 0.00718 | 0.02795 | 0.01206 | 0.58106 | 0.01089 | 0.01710 | 0.02155 | 0.00999 | 0.02188 | 0.00702 |
| **Ppm1k** | 1.80170 | 0.16709 | 3.36580 | 10.19074 | 4.19937 | 0.26431 | 2.38603 | 0.00000 | 16.78977 | 0.00000 | 16.03773 | 4.68356 | 1.17371 | 10.59882 | 0.52537 | 1.08095 |
| **Ppm1l** | 1.35191 | 0.01635 | 0.21609 | 1.34849 | 0.05610 | 2.83873 | 0.00461 | 1.68048 | 0.46009 | 3.31472 | 1.79225 | 0.40888 | 1.94679 | 4.82826 | 0.92936 | 0.00351 |
| **Ppm1m** | 0.35601 | 0.00647 | 0.72574 | 0.00908 | 1.74815 | 0.00848 | 0.00607 | 0.88691 | 0.00990 | 0.00000 | 0.00248 | 0.00155 | 1.05394 | 0.29347 | 0.13412 | 0.85086 |
| **Ppm1n** | 0.00410 | 0.00913 | 0.07979 | 0.06686 | 0.00941 | 1.25699 | 0.00752 | 0.40597 | 0.02260 | 0.18504 | 0.04919 | 0.00755 | 1.42102 | 0.37568 | 0.11807 | 0.00634 |
| **Ppme1** | 6.12970 | 0.00000 | 3.32325 | 0.99926 | 10.09314 | 1.75746 | 5.04766 | 9.50546 | 1.34051 | 3.85973 | 0.35327 | 0.63974 | 6.09743 | 0.00000 | 0.28814 | 5.53378 |
| **Ppox** | 0.70099 | 0.00681 | 1.37682 | 0.25166 | 0.01642 | 0.02310 | 0.00554 | 0.79555 | 0.04678 | 0.03685 | 1.28183 | 0.00000 | 0.83039 | 0.80706 | 0.00000 | 0.01277 |
| **Ppp1ca** | 27.73465 | 11.26638 | 22.57951 | 4.51748 | 17.19992 | 16.88499 | 10.24772 | 24.52529 | 4.67785 | 26.66137 | 4.81735 | 3.75060 | 25.76756 | 0.20682 | 5.71111 | 15.01175 |
| **Ppp1cb** | 6.16179 | 14.97400 | 5.00572 | 9.48448 | 6.09399 | 6.87123 | 7.51647 | 9.34289 | 9.62765 | 6.01734 | 6.76061 | 0.13700 | 5.10872 | 14.23181 | 7.67918 | 3.69889 |
| **Ppp1cc** | 24.31896 | 11.75528 | 12.09238 | 24.66633 | 18.97195 | 17.66793 | 10.58644 | 27.03170 | 31.87903 | 45.58791 | 22.45662 | 20.64898 | 20.71320 | 17.99668 | 19.92123 | 10.50888 |
| **Ppp1r10** | 4.13062 | 0.36984 | 3.89960 | 6.01440 | 2.29252 | 6.42281 | 0.00941 | 3.80727 | 4.93940 | 9.95163 | 1.34881 | 4.37277 | 5.06822 | 1.83280 | 5.57559 | 1.44140 |
| **Ppp1r11** | 13.25409 | 1.19175 | 9.84792 | 0.95148 | 9.58329 | 2.16554 | 4.01347 | 4.21932 | 2.84934 | 2.76201 | 3.96301 | 0.26625 | 8.11441 | 0.00000 | 2.28089 | 1.30382 |
| **Ppp1r12a** | 3.03939 | 4.16049 | 0.00000 | 7.62379 | 5.02103 | 2.29238 | 0.54353 | 2.88120 | 4.77258 | 5.18969 | 1.51277 | 0.27917 | 0.47333 | 4.78810 | 6.17220 | 0.11974 |
| **Ppp1r12b** | 0.23280 | 0.26977 | 0.52876 | 4.88017 | 0.00678 | 0.04277 | 0.24642 | 0.07470 | 3.94271 | 0.88894 | 0.13579 | 0.20302 | 0.20354 | 1.78879 | 1.31787 | 0.02670 |
| **Ppp1r12c** | 7.52129 | 1.20933 | 0.83101 | 8.57191 | 3.98359 | 6.23292 | 0.57500 | 4.50120 | 4.22785 | 5.52292 | 2.15440 | 0.43818 | 5.91767 | 0.61277 | 1.08244 | 6.20757 |
| **Ppp1r13b** | 2.14440 | 2.36833 | 3.04471 | 1.41155 | 2.44791 | 0.00306 | 4.57745 | 0.70405 | 5.02367 | 1.66968 | 0.19305 | 0.44355 | 0.03210 | 0.00000 | 0.82278 | 0.00252 |
| **Ppp1r13l** | 0.03837 | 0.00710 | 1.46850 | 1.04661 | 0.01104 | 0.00873 | 1.56096 | 0.00711 | 0.21546 | 0.91282 | 0.00930 | 0.00175 | 0.72221 | 0.00700 | 0.26249 | 0.00648 |
| **Ppp1r14a** | 0.64984 | 0.00815 | 0.08143 | 0.00846 | 0.01945 | 0.00567 | 0.00701 | 0.01494 | 0.01202 | 0.43377 | 0.02656 | 0.00655 | 0.22564 | 0.01251 | 0.00765 | 0.12340 |
| **Ppp1r14b** | 8.28989 | 1.00094 | 4.99245 | 7.86807 | 8.97890 | 5.48236 | 2.92897 | 4.73350 | 8.16610 | 15.54911 | 9.97254 | 0.47765 | 12.87099 | 0.00000 | 0.00000 | 2.02931 |
| **Ppp1r14c** | 0.01056 | 0.00819 | 0.01599 | 0.09278 | 0.00975 | 0.00751 | 0.02836 | 0.06399 | 0.10576 | 0.02343 | 0.02431 | 0.01606 | 0.01335 | 0.43760 | 0.00000 | 0.00739 |
| **Ppp1r14d** | 10.91275 | 0.00000 | 0.13952 | 0.00000 | 4.97394 | 0.00000 | 0.00000 | 0.00000 | 0.00881 | 0.55417 | 0.33934 | 0.02354 | 0.64031 | 0.01256 | 0.10421 | 5.76708 |
| **Ppp1r15a** | 1.15883 | 0.09989 | 1.73294 | 20.14352 | 0.91341 | 0.00280 | 0.00446 | 7.47069 | 18.00516 | 3.08849 | 2.47862 | 0.58091 | 0.13341 | 0.00000 | 5.10714 | 0.82420 |
| **Ppp1r15b** | 3.74556 | 6.70166 | 3.72208 | 14.36671 | 3.63948 | 0.80674 | 2.45989 | 4.14579 | 16.60849 | 0.00000 | 5.43966 | 5.44897 | 1.86621 | 23.69990 | 17.21372 | 1.47731 |
| **Ppp1r16a** | 2.49158 | 0.00449 | 0.92708 | 3.21394 | 0.14550 | 0.00891 | 1.15494 | 0.66800 | 6.05533 | 5.01184 | 1.47888 | 0.05346 | 1.41884 | 0.00000 | 0.00687 | 0.01733 |
| **Ppp1r16b** | 1.29462 | 0.07900 | 0.96753 | 8.51885 | 0.75501 | 0.08824 | 0.21344 | 0.00000 | 12.28181 | 5.51892 | 10.38341 | 1.06060 | 0.47809 | 6.66901 | 10.63448 | 0.26956 |
| **Ppp1r18** | 6.17989 | 0.00000 | 0.00000 | 0.00000 | 2.64427 | 5.44764 | 1.54380 | 0.36098 | 0.24673 | 0.77876 | 0.46718 | 0.06554 | 2.45279 | 0.25577 | 0.00000 | 0.78224 |
| **Ppp1r1b** | 0.04353 | 0.00854 | 0.01042 | 0.19868 | 0.01376 | 0.00684 | 0.00748 | 0.01256 | 0.00820 | 0.14133 | 0.01322 | 0.00807 | 0.01266 | 0.01073 | 0.00804 | 0.00759 |
| **Ppp1r2** | 7.64749 | 2.28183 | 3.66025 | 7.84764 | 12.71043 | 1.77540 | 2.35668 | 4.80583 | 15.23186 | 11.77618 | 8.83560 | 10.79416 | 3.22904 | 5.84309 | 5.43634 | 3.40557 |
| **Ppp1r2.ps3** | 1.42405 | 0.02669 | 0.15241 | 1.36725 | 2.27856 | 0.04061 | 0.38178 | 0.56944 | 1.39737 | 2.07920 | 0.62257 | 0.74448 | 0.28390 | 0.29735 | 0.85097 | 0.23776 |
| **Ppp1r2.ps7** | 0.01338 | 0.00812 | 0.00768 | 4.18982 | 0.00899 | 0.00643 | 0.00743 | 0.00648 | 2.40181 | 0.00000 | 0.55915 | 0.00000 | 0.00683 | 0.00000 | 0.00000 | 0.00624 |
| **Ppp1r21** | 2.60312 | 0.35121 | 2.81073 | 2.17767 | 1.07366 | 2.60814 | 0.03406 | 4.72543 | 1.03413 | 3.10759 | 1.24285 | 0.11718 | 3.73323 | 0.06627 | 0.26918 | 0.07968 |
| **Ppp1r26** | 1.06847 | 0.02312 | 1.89210 | 0.31453 | 3.52769 | 1.58638 | 2.50838 | 0.93073 | 0.83481 | 0.87620 | 1.31679 | 0.99999 | 0.45033 | 0.32613 | 1.24547 | 0.01963 |
| **Ppp1r35** | 0.35420 | 0.00800 | 0.02291 | 0.79029 | 0.01113 | 0.00674 | 0.00682 | 0.00398 | 0.25181 | 1.36653 | 0.21383 | 0.00229 | 0.59137 | 0.00922 | 0.01939 | 1.12728 |
| **Ppp1r36** | 0.01294 | 0.00757 | 0.01100 | 2.66658 | 0.00903 | 0.00614 | 0.00715 | 0.01002 | 0.00000 | 1.28342 | 0.00972 | 0.03104 | 0.01095 | 0.00159 | 0.72908 | 0.00689 |

| | | | | | | | | | | | | | | | | |
|---|---|---|---|---|---|---|---|---|---|---|---|---|---|---|---|---|
| **Ppp1r37** | 0.01523 | 0.17378 | 0.55275 | 0.44192 | 0.01080 | 0.00567 | 0.06804 | 0.67916 | 0.13144 | 0.70937 | 1.09850 | 0.01035 | 0.14219 | 0.00537 | 0.00000 | 0.00705 |
| **Ppp1r3b** | 0.56484 | 1.13405 | 5.87817 | 0.60869 | 0.02341 | 0.01354 | 3.33993 | 2.13767 | 0.18414 | 1.62495 | 0.95965 | 0.04490 | 4.53619 | 0.00322 | 0.80474 | 0.26024 |
| **Ppp1r3c** | 0.01375 | 0.00771 | 0.01217 | 0.96363 | 0.00938 | 0.00641 | 0.00738 | 0.01215 | 0.00000 | 0.00712 | 0.01265 | 0.00672 | 0.01268 | 0.00437 | 0.42637 | 0.00719 |
| **Ppp1r3d** | 0.01353 | 0.00758 | 0.00680 | 3.53983 | 0.00932 | 0.00609 | 0.00707 | 0.00000 | 0.88695 | 0.00000 | 0.35812 | 0.02739 | 0.01119 | 0.00000 | 1.92088 | 0.01365 |
| **Ppp1r3e** | 0.01369 | 0.00823 | 0.01212 | 0.29814 | 0.00930 | 0.00660 | 0.00752 | 0.01382 | 0.41644 | 0.01194 | 0.01041 | 0.00776 | 0.01278 | 0.01019 | 0.02559 | 0.00734 |
| **Ppp1r3f** | 0.01077 | 0.00809 | 0.06879 | 0.68658 | 0.00926 | 0.00815 | 0.00716 | 0.01178 | 0.00000 | 0.01270 | 0.01208 | 0.00730 | 0.43455 | 0.17336 | 0.00000 | 0.00708 |
| **Ppp1r3fos** | 0.01275 | 0.00789 | 0.08186 | 0.00807 | 0.00915 | 0.00829 | 0.00730 | 0.01136 | 0.01114 | 0.00618 | 0.01165 | 0.00717 | 0.01142 | 0.00841 | 0.46776 | 0.00731 |
| **Ppp1r3g** | 0.01366 | 0.00824 | 0.01210 | 0.71166 | 0.00930 | 0.00656 | 0.00751 | 0.01390 | 0.55958 | 0.01215 | 0.01003 | 0.00747 | 0.01281 | 0.01128 | 0.00340 | 0.00727 |
| **Ppp1r7** | 7.94319 | 9.40326 | 8.68971 | 5.44167 | 6.86262 | 4.90174 | 7.73494 | 5.14758 | 11.26305 | 4.25850 | 8.75770 | 5.69128 | 5.25452 | 4.30125 | 6.24756 | 6.65019 |
| **Ppp1r8** | 11.36803 | 8.05767 | 7.95642 | 13.61451 | 0.51739 | 7.16456 | 2.51762 | 6.14277 | 17.34090 | 10.85579 | 9.79960 | 1.50937 | 5.86594 | 5.24172 | 8.68201 | 6.10459 |
| **Ppp1r9a** | 0.02968 | 0.00746 | 0.00355 | 0.70081 | 0.00809 | 0.00732 | 0.00686 | 0.00501 | 0.03252 | 2.80935 | 0.29650 | 0.83004 | 0.04818 | 0.27152 | 2.37188 | 0.02136 |
| **Ppp1r9b** | 0.58746 | 0.00723 | 0.50900 | 1.07010 | 0.01040 | 0.00740 | 0.00653 | 0.00709 | 0.01897 | 0.64461 | 0.00822 | 0.31593 | 0.00937 | 0.00920 | 0.00000 | 0.00658 |
| **Ppp2ca** | 15.84217 | 10.99253 | 13.48001 | 12.94205 | 16.96907 | 7.96806 | 4.54000 | 9.81959 | 14.08193 | 14.44786 | 12.78562 | 17.49827 | 6.40608 | 11.00299 | 14.19834 | 4.73135 |
| **Ppp2cb** | 0.03744 | 0.12242 | 0.21747 | 13.51218 | 0.22652 | 0.55124 | 0.08469 | 1.07488 | 15.04557 | 0.00000 | 3.25302 | 1.84509 | 0.00000 | 6.51144 | 4.79859 | 0.03124 |
| **Ppp2r1a** | 21.63763 | 19.50379 | 20.08065 | 5.11502 | 17.49842 | 10.32720 | 13.61210 | 21.39663 | 2.89724 | 12.06486 | 6.77044 | 0.86696 | 17.83031 | 4.90852 | 10.17551 | 11.82794 |
| **Ppp2r1b** | 4.63815 | 5.17271 | 2.30664 | 2.06884 | 12.98319 | 3.09725 | 1.24697 | 1.63877 | 2.33345 | 1.71656 | 1.38815 | 7.31986 | 3.10945 | 0.00000 | 2.80250 | 1.43281 |
| **Ppp2r2a** | 9.22354 | 10.45721 | 7.62413 | 14.28076 | 12.35664 | 5.13130 | 6.83693 | 10.10321 | 17.78074 | 20.43238 | 9.16332 | 16.92049 | 4.60238 | 16.25513 | 17.62946 | 3.73224 |
| **Ppp2r2b** | 0.00405 | 0.00748 | 1.31632 | 2.06996 | 0.00670 | 0.00531 | 0.03775 | 0.48515 | 1.07446 | 2.28434 | 2.11309 | 0.04854 | 0.93640 | 0.00000 | 1.68763 | 0.01297 |
| **Ppp2r2c** | 1.92417 | 0.18554 | 0.03960 | 0.00748 | 0.00926 | 0.01568 | 2.27011 | 2.05090 | 0.71033 | 3.15624 | 0.31993 | 0.01907 | 0.74877 | 0.92029 | 0.42185 | 0.82191 |
| **Ppp2r2d** | 10.41100 | 0.99970 | 2.17934 | 6.84115 | 5.49320 | 2.67890 | 1.32941 | 6.35775 | 6.32096 | 11.17034 | 4.16134 | 6.11216 | 3.19040 | 1.71586 | 3.76383 | 4.29620 |
| **Ppp2r3a** | 0.01573 | 0.00772 | 0.01174 | 1.03792 | 0.00985 | 0.00678 | 0.00744 | 0.01268 | 0.58834 | 0.00000 | 0.00851 | 0.03339 | 0.27917 | 0.01401 | 1.26132 | 0.00726 |
| **Ppp2r3c** | 7.61176 | 3.26878 | 6.22951 | 1.55616 | 0.01761 | 4.14238 | 0.00000 | 2.46289 | 1.45081 | 4.43056 | 2.63625 | 10.64595 | 2.80302 | 0.32429 | 2.61976 | 0.00000 |
| **Ppp2r3d** | 3.39762 | 0.10229 | 2.75936 | 0.96085 | 2.20339 | 3.18965 | 4.42124 | 2.42332 | 0.62824 | 2.13724 | 4.05025 | 7.52122 | 3.55577 | 3.98665 | 0.09671 | 0.15159 |
| **Ppp2r5a** | 1.61575 | 0.08675 | 0.73060 | 1.14760 | 2.93284 | 1.42576 | 0.49727 | 1.44253 | 0.41846 | 0.16136 | 0.00001 | 0.05893 | 0.66677 | 0.00778 | 0.00000 | 0.00428 |
| **Ppp2r5b** | 0.71684 | 0.10439 | 0.19282 | 0.84158 | 0.09521 | 4.31723 | 3.34261 | 0.07839 | 0.21808 | 1.11867 | 0.02359 | 0.04199 | 2.09768 | 0.00791 | 0.00000 | 2.34552 |
| **Ppp2r5c** | 13.01657 | 11.49496 | 11.82974 | 14.02560 | 4.44389 | 8.64074 | 9.25017 | 10.20807 | 5.90177 | 6.10541 | 6.38377 | 0.66468 | 4.42668 | 1.50522 | 9.45441 | 6.32782 |
| **Ppp2r5d** | 2.89436 | 0.00000 | 2.71865 | 4.86288 | 6.22280 | 4.47633 | 0.74369 | 3.49566 | 4.97716 | 2.59503 | 1.40638 | 0.14640 | 3.00162 | 0.00000 | 0.64624 | 2.10889 |
| **Ppp2r5e** | 1.94064 | 6.42862 | 5.20118 | 6.88221 | 3.54728 | 0.98638 | 4.98230 | 10.29248 | 5.22879 | 3.47488 | 3.72772 | 0.31996 | 0.00000 | 3.04476 | 3.85468 | 2.06942 |
| **Ppp3ca** | 1.29681 | 1.96473 | 0.91445 | 0.47111 | 3.61903 | 1.99297 | 1.68424 | 3.08032 | 0.35870 | 0.06714 | 0.00000 | 0.02283 | 0.70664 | 0.00162 | 0.32845 | 2.31675 |
| **Ppp3cb** | 3.29544 | 2.57752 | 0.19335 | 2.51172 | 0.01047 | 4.12896 | 0.53442 | 0.11489 | 1.22143 | 0.20252 | 0.00000 | 0.00000 | 0.37278 | 3.42898 | 1.90249 | 0.01572 |
| **Ppp3cc** | 0.04195 | 0.00767 | 0.00480 | 0.06885 | 0.00841 | 0.00689 | 0.00721 | 0.00959 | 0.06289 | 0.00227 | 0.00771 | 0.01444 | 0.01354 | 0.80577 | 0.42042 | 0.00740 |
| **Ppp3r1** | 10.12963 | 7.12270 | 9.54823 | 3.11374 | 5.54097 | 1.40974 | 5.45179 | 2.78775 | 1.28817 | 0.61236 | 1.99422 | 7.79896 | 3.89245 | 7.36113 | 3.64038 | 0.26209 |
| **Ppp4c** | 10.35995 | 1.97249 | 4.05756 | 0.00000 | 5.02412 | 5.50279 | 0.90853 | 8.91429 | 0.00061 | 12.69280 | 0.17432 | 0.37393 | 8.26030 | 0.00701 | 0.32363 | 5.65115 |
| **Ppp4r1** | 7.22696 | 11.95303 | 5.52627 | 2.62007 | 14.82266 | 7.98153 | 5.81135 | 12.78862 | 2.77481 | 0.94406 | 2.95128 | 0.23210 | 8.04022 | 0.11984 | 0.65736 | 5.11107 |
| **Ppp4r1l.ps** | 0.02031 | 0.00787 | 0.06178 | 0.02716 | 0.02862 | 1.55514 | 0.02826 | 0.03234 | 0.03856 | 0.24044 | 0.05067 | 0.01585 | 0.05966 | 0.11459 | 0.02379 | 0.00702 |
| **Ppp4r2** | 8.29350 | 6.24679 | 1.71484 | 26.60734 | 6.05023 | 5.79141 | 4.41370 | 11.19717 | 27.48560 | 22.19480 | 13.56868 | 22.46594 | 4.02018 | 32.53216 | 25.94328 | 2.99571 |
| **Ppp4r4** | 0.01350 | 0.00767 | 0.01147 | 0.04268 | 0.00930 | 0.00634 | 0.00740 | 0.01147 | 1.76256 | 0.00395 | 0.00000 | 0.00344 | 0.01288 | 0.00537 | 0.52125 | 0.00715 |
| **Ppp5c** | 13.54664 | 0.85029 | 2.66860 | 2.96284 | 5.65223 | 4.06338 | 3.89474 | 9.79290 | 3.34756 | 18.23807 | 6.20130 | 0.69212 | 12.79922 | 0.00000 | 0.00000 | 7.69015 |
| **Ppp6c** | 4.31624 | 5.78321 | 3.97427 | 3.79126 | 5.97087 | 3.57771 | 2.76239 | 13.05822 | 8.08583 | 7.51949 | 10.54458 | 3.16957 | 4.78430 | 10.03998 | 7.25695 | 0.09447 |
| **Ppp6r1** | 0.72486 | 0.02718 | 0.28723 | 2.36030 | 0.01506 | 0.18183 | 0.14529 | 1.18159 | 1.83974 | 0.14218 | 1.94327 | 0.10466 | 0.86282 | 0.00000 | 0.04336 | 0.76622 |
| **Ppp6r2** | 0.01635 | 0.00804 | 0.10588 | 1.91924 | 0.00861 | 0.00846 | 0.00720 | 0.00988 | 1.90395 | 0.00641 | 0.04572 | 0.00348 | 0.01101 | 0.00492 | 0.37833 | 0.00701 |
| **Ppp6r3** | 13.74888 | 7.58548 | 3.41911 | 6.02171 | 3.71105 | 7.99901 | 5.23308 | 23.06003 | 4.67887 | 17.01906 | 12.53179 | 0.92403 | 13.80260 | 2.17648 | 3.95409 | 4.08582 |
| **Pprc1** | 4.48336 | 7.05123 | 8.09670 | 18.11917 | 3.85443 | 0.87719 | 8.83303 | 16.45163 | 24.15067 | 29.93668 | 14.60464 | 5.01204 | 11.25362 | 3.74528 | 10.05144 | 6.92273 |
| **Ppt1** | 4.18079 | 2.78329 | 1.40157 | 1.32400 | 0.00802 | 0.00179 | 2.55403 | 0.00000 | 0.51524 | 0.52032 | 0.54995 | 7.14293 | 0.08665 | 0.00203 | 1.35457 | 0.40912 |
| **Ppt2** | 0.76867 | 0.04531 | 1.15556 | 6.50994 | 0.01395 | 0.29106 | 0.71911 | 1.05267 | 5.24626 | 16.49897 | 24.87044 | 2.11076 | 0.57829 | 2.13920 | 3.59448 | 0.01436 |
| **Pptc7** | 1.40048 | 0.35805 | 5.13425 | 9.03434 | 0.00000 | 5.10332 | 0.00444 | 0.00000 | 7.08544 | 4.13462 | 0.25607 | 0.18893 | 1.24691 | 3.51305 | 5.10670 | 0.00682 |
| **Ppwd1** | 1.67335 | 0.00465 | 0.15387 | 0.07490 | 3.83242 | 1.14694 | 0.00662 | 0.08812 | 0.00797 | 0.45327 | 0.42959 | 0.05518 | 0.04312 | 0.84493 | 4.43170 | 0.02016 |
| **Pqbp1** | 6.56418 | 4.48389 | 11.39277 | 0.82471 | 0.01696 | 4.78702 | 4.39733 | 17.61544 | 2.72785 | 20.03972 | 7.71218 | 0.92062 | 10.20605 | 0.01562 | 0.56905 | 2.97958 |
| **Pqlc1** | 0.15464 | 1.73815 | 6.14546 | 2.36293 | 0.05767 | 9.96558 | 0.00602 | 17.19556 | 3.41183 | 16.39759 | 11.71868 | 3.95010 | 8.57443 | 3.27453 | 3.37950 | 0.25749 |
| **Pqlc2** | 3.32511 | 0.49420 | 0.83700 | 4.23235 | 3.20659 | 2.37395 | 1.98499 | 5.76994 | 2.23796 | 2.44763 | 1.11134 | 0.00000 | 2.26859 | 0.00000 | 0.00000 | 1.00004 |
| **Pqlc3** | 3.06978 | 0.72474 | 2.05111 | 0.00159 | 0.00715 | 0.58000 | 0.00430 | 0.00000 | 0.61174 | 0.35421 | 0.57084 | 0.00080 | 0.00004 | 0.01085 | 0.00008 | 0.00272 |
| **Pradc1** | 4.64219 | 0.00000 | 4.31413 | 6.36352 | 0.04371 | 0.58743 | 3.99943 | 3.11450 | 10.46212 | 5.50449 | 2.11147 | 0.21618 | 3.59636 | 3.02619 | 1.92165 | 0.31365 |
| **Praf2** | 10.62087 | 0.00614 | 0.16287 | 1.48718 | 0.02322 | 0.00000 | 0.00000 | 3.00092 | 1.70124 | 4.64552 | 0.99450 | 0.73125 | 0.33266 | 0.78576 | 0.90729 | 11.25343 |
| **Pram1** | 0.34881 | 0.00739 | 0.19770 | 0.22845 | 0.00676 | 0.00711 | 0.00729 | 0.01190 | 0.00824 | 0.01533 | 0.01336 | 0.00799 | 0.01033 | 0.01198 | 0.00354 | 0.00685 |
| **Pramef12** | 5.72071 | 0.26748 | 7.63627 | 16.46955 | 0.12504 | 0.11924 | 1.01913 | 0.50217 | 12.34256 | 9.78506 | 8.00626 | 2.93394 | 4.86165 | 12.81174 | 7.95813 | 0.02064 |
| **Pramef17** | 0.62332 | 2.43084 | 2.33559 | 0.58399 | 0.00708 | 0.01399 | 0.66742 | 0.91630 | 1.56822 | 0.07098 | 0.00000 | 0.01029 | 1.24215 | 2.45415 | 2.58021 | 0.57449 |
| **Pramef25** | 0.01151 | 0.04914 | 0.00665 | 4.88880 | 0.00902 | 0.00577 | 0.00688 | 0.01324 | 4.21104 | 0.00000 | 0.00000 | 1.08723 | 0.01283 | 0.20215 | 0.41111 | 0.00608 |
| **Pramef6** | 0.03287 | 0.28814 | 0.72442 | 0.51979 | 0.07227 | 0.23931 | 0.49376 | 1.68407 | 1.25244 | 0.17336 | 0.00000 | 0.11083 | 0.23616 | 1.54331 | 0.29850 | 0.04382 |
| **Pramef8** | 1.13839 | 4.34299 | 1.79163 | 3.10076 | 4.24710 | 0.50882 | 4.08382 | 0.68417 | 4.18402 | 2.25393 | 0.70723 | 2.53732 | 2.34277 | 0.00743 | 7.78750 | 2.91420 |
| **Pramel1** | 0.02855 | 0.00759 | 0.05868 | 0.42348 | 0.00821 | 0.00564 | 0.00733 | 0.01005 | 0.78110 | 0.97868 | 0.01976 | 0.00410 | 0.01017 | 0.07583 | 0.11241 | 0.00689 |
| **Pramel3** | 0.01373 | 0.00769 | 0.01163 | 0.00563 | 0.00941 | 0.00646 | 0.00738 | 0.01144 | 0.05311 | 0.00545 | 0.01067 | 0.00632 | 0.01281 | 0.00909 | 0.68475 | 0.00735 |
| **Pramel4** | 5.17919 | 14.26373 | 23.45252 | 6.84315 | 9.23067 | 5.78674 | 8.53998 | 26.81060 | 17.03250 | 22.00886 | 14.19457 | 18.44914 | 17.62786 | 30.26673 | 14.73218 | 1.49596 |

| | | | | | | | | | | | | | | | | |
|---|---|---|---|---|---|---|---|---|---|---|---|---|---|---|---|---|
| **Pramel5** | 8.09844 | 8.58512 | 26.73133 | 7.41627 | 14.03423 | 12.47454 | 14.80611 | 39.73855 | 14.65381 | 24.14979 | 25.58247 | 12.61083 | 21.75422 | 31.27152 | 10.06606 | 3.53554 |
| **Pramel6** | 14.23564 | 18.09156 | 35.42808 | 14.27501 | 20.48395 | 17.85345 | 16.01206 | 26.03710 | 19.53109 | 32.65290 | 35.19213 | 16.95931 | 27.78733 | 26.38731 | 28.49742 | 9.84647 |
| **Pramel7** | 1.36600 | 7.09150 | 0.83469 | 3.21602 | 5.22772 | 3.34099 | 4.80757 | 12.51284 | 3.78800 | 0.00000 | 3.98831 | 15.59997 | 1.76709 | 42.40682 | 26.16683 | 0.45042 |
| **Prc1** | 4.51710 | 5.68063 | 5.59797 | 12.47138 | 1.55783 | 3.53865 | 1.82714 | 17.72350 | 16.51655 | 10.59923 | 16.84904 | 3.85350 | 1.10991 | 26.61521 | 21.00638 | 0.91612 |
| **Prcc** | 0.02795 | 1.52051 | 0.00678 | 0.00000 | 0.11020 | 0.00574 | 0.00655 | 0.01792 | 0.36269 | 1.19608 | 0.00000 | 0.04473 | 0.02414 | 0.92905 | 7.99578 | 0.00661 |
| **Prcp** | 3.52705 | 4.18422 | 2.97885 | 0.12070 | 0.00546 | 0.00845 | 4.28171 | 2.94548 | 0.67878 | 5.90331 | 0.93066 | 0.93556 | 3.43133 | 4.43121 | 5.10116 | 0.02077 |
| **Prdm10** | 0.00143 | 0.33095 | 0.29816 | 0.46793 | 0.01621 | 0.00820 | 0.00745 | 1.02212 | 1.36805 | 2.39842 | 0.11281 | 0.26186 | 0.42939 | 0.54298 | 0.46510 | 0.00664 |
| **Prdm14** | 0.93485 | 0.00629 | 0.00000 | 1.08164 | 0.00637 | 0.01114 | 0.01595 | 1.27963 | 8.56428 | 4.87769 | 2.49185 | 0.21110 | 0.01888 | 3.22790 | 1.54958 | 0.00527 |
| **Prdm15** | 0.89028 | 0.13571 | 1.70998 | 6.06791 | 0.00315 | 0.00504 | 3.82356 | 1.30259 | 4.62353 | 0.88588 | 1.55810 | 0.03544 | 0.92930 | 0.00000 | 0.76729 | 0.00355 |
| **Prdm16** | 0.00629 | 0.00624 | 0.65468 | 0.07020 | 0.00939 | 0.00547 | 2.19279 | 1.68973 | 1.38999 | 0.00000 | 0.00000 | 0.00000 | 0.00212 | 0.03204 | 0.16129 | 0.00679 |
| **Prdm2** | 3.79003 | 7.15857 | 7.95585 | 5.35852 | 4.20095 | 1.52287 | 4.60762 | 9.43761 | 11.70216 | 5.62058 | 3.03567 | 7.77380 | 1.43966 | 7.67250 | 7.87428 | 1.07773 |
| **Prdm4** | 5.04718 | 2.08982 | 3.31286 | 8.51626 | 3.11149 | 0.00398 | 5.93331 | 5.82510 | 10.54843 | 9.43024 | 3.98212 | 8.42961 | 2.13419 | 11.45724 | 13.78345 | 2.15183 |
| **Prdm6** | 0.01320 | 0.00782 | 0.01374 | 0.05938 | 0.01015 | 0.00698 | 0.00761 | 0.01380 | 0.00877 | 0.22360 | 0.01333 | 0.00668 | 0.07185 | 0.01640 | 0.03803 | 0.00779 |
| **Prdm9** | 0.06261 | 0.03091 | 0.49498 | 5.46464 | 0.16891 | 0.02890 | 0.00735 | 0.20395 | 0.25556 | 0.08276 | 0.57618 | 0.04716 | 0.20355 | 0.18324 | 2.16178 | 0.02769 |
| **Prdx1** | 50.39159 | 27.21835 | 26.93733 | 8.41068 | 33.40896 | 15.93563 | 18.12984 | 7.97082 | 5.23385 | 25.59646 | 19.07104 | 17.01156 | 24.36593 | 22.48174 | 16.97078 | 20.37211 |
| **Prdx2** | 22.77206 | 7.92313 | 6.60350 | 18.26976 | 11.68516 | 6.06755 | 4.52333 | 3.77371 | 3.45933 | 9.06762 | 7.14102 | 4.76362 | 9.09032 | 0.04733 | 4.92294 | 13.02291 |
| **Prdx3** | 5.29254 | 2.46877 | 6.60404 | 0.00000 | 8.90223 | 2.81718 | 6.31165 | 11.90331 | 0.09131 | 5.15527 | 1.40635 | 0.47797 | 1.42378 | 16.06191 | 4.70516 | 1.99833 |
| **Prdx4** | 10.44178 | 0.53858 | 8.72779 | 9.67905 | 3.28181 | 6.08281 | 4.06098 | 10.53073 | 12.42904 | 11.60705 | 9.30892 | 2.45844 | 9.34300 | 5.52044 | 1.35975 | 7.21966 |
| **Prdx5** | 6.51832 | 1.80613 | 10.32016 | 3.52201 | 12.61523 | 7.04096 | 4.65250 | 12.39722 | 6.14084 | 13.66919 | 10.33459 | 2.33952 | 11.71761 | 0.96125 | 1.71908 | 8.77076 |
| **Prdx6** | 36.08766 | 20.34897 | 26.11136 | 2.87339 | 21.70264 | 12.14665 | 15.05115 | 23.95695 | 9.11727 | 21.32649 | 18.21410 | 10.72185 | 30.91062 | 10.61642 | 1.54587 | 15.67589 |
| **Prdx6b** | 0.35283 | 0.25315 | 0.20869 | 0.02526 | 0.14847 | 0.07404 | 0.16696 | 0.14967 | 0.03086 | 0.07184 | 0.01108 | 0.00714 | 0.28148 | 0.03018 | 0.01198 | 0.16109 |
| **Preb** | 12.43090 | 0.92931 | 6.35525 | 3.75126 | 8.56725 | 6.37935 | 0.00000 | 9.16944 | 1.92069 | 7.88220 | 5.28111 | 0.28212 | 14.05009 | 0.71570 | 4.21973 | 5.85711 |
| **Prelid1** | 19.53714 | 7.73164 | 14.42417 | 9.63773 | 15.21819 | 12.24204 | 8.37973 | 13.23394 | 8.29999 | 27.47092 | 12.35858 | 11.91931 | 26.91953 | 0.00000 | 0.49961 | 16.04811 |
| **Prelid2** | 6.84121 | 9.21550 | 12.52090 | 3.76971 | 0.51277 | 7.53612 | 6.82664 | 12.74045 | 4.09416 | 6.56183 | 13.53872 | 2.73744 | 17.35279 | 11.54259 | 0.65448 | 1.93732 |
| **Prelp** | 0.01034 | 0.00824 | 0.00928 | 2.59788 | 0.00955 | 0.00653 | 0.00804 | 0.14716 | 0.00000 | 0.41636 | 0.02568 | 0.02837 | 0.00773 | 0.01466 | 0.11310 | 0.00702 |
| **Prep** | 6.99478 | 6.12708 | 8.70341 | 0.71044 | 12.76835 | 7.20343 | 7.84444 | 14.84840 | 1.48071 | 1.43394 | 3.65098 | 0.68769 | 11.69233 | 0.26578 | 0.00000 | 9.60905 |
| **Prepl** | 0.20460 | 2.04011 | 1.42119 | 0.24633 | 2.79987 | 0.11181 | 1.66350 | 2.65470 | 0.03980 | 0.38849 | 0.89288 | 0.83453 | 0.74776 | 3.58977 | 0.77135 | 0.52503 |
| **Prex1** | 1.36365 | 0.00683 | 0.00000 | 0.67923 | 0.01342 | 3.56100 | 0.07557 | 1.60728 | 0.57659 | 0.00000 | 0.55851 | 0.00000 | 0.00000 | 0.33142 | 0.07962 | 1.79792 |
| **Prex2** | 1.52352 | 1.21153 | 0.53584 | 29.85935 | 1.06276 | 0.33184 | 0.87662 | 0.74186 | 26.19152 | 3.74384 | 2.45186 | 1.79117 | 1.09358 | 4.90267 | 7.90035 | 0.82645 |
| **Prg4** | 0.01419 | 0.14084 | 0.00575 | 0.50181 | 0.00842 | 0.00649 | 0.00698 | 0.00801 | 1.27162 | 0.06487 | 0.09146 | 1.58692 | 0.08158 | 0.00612 | 0.15642 | 0.00691 |
| **Prickle1** | 0.00071 | 0.00575 | 0.00485 | 23.41911 | 0.00877 | 0.00251 | 0.00579 | 1.31687 | 3.84551 | 1.88137 | 0.00000 | 5.80816 | 0.00554 | 0.39319 | 17.68769 | 0.00265 |
| **Prickle2** | 0.01354 | 0.00818 | 0.01184 | 0.07944 | 0.00931 | 0.00660 | 0.00755 | 0.01396 | 1.34371 | 0.00992 | 0.00000 | 0.00614 | 0.01315 | 0.01354 | 0.00484 | 0.00733 |
| **Prickle3** | 0.51550 | 0.00712 | 0.04378 | 0.17574 | 0.00670 | 0.00813 | 0.00703 | 0.01026 | 0.87536 | 0.06143 | 0.00792 | 0.01552 | 0.01200 | 0.01141 | 0.00384 | 0.00703 |
| **Prim1** | 14.36609 | 3.31459 | 4.18470 | 5.75052 | 4.73514 | 6.36759 | 2.53921 | 6.02025 | 2.88663 | 10.41221 | 7.92790 | 9.68649 | 6.06318 | 12.24280 | 7.12268 | 6.70936 |
| **Prim2** | 4.44025 | 4.67538 | 9.34290 | 1.24676 | 3.38736 | 0.01342 | 4.23819 | 5.46286 | 3.40585 | 10.36414 | 5.24237 | 9.20423 | 2.97878 | 2.05146 | 10.45275 | 3.05820 |
| **Primpol** | 1.94626 | 2.63754 | 1.13685 | 0.69205 | 0.41112 | 0.04233 | 0.95219 | 0.42245 | 0.21250 | 1.29081 | 0.61496 | 4.94939 | 1.29171 | 2.66994 | 2.14921 | 1.14606 |
| **Prkaa1** | 0.00022 | 7.02539 | 3.61002 | 2.50026 | 3.55953 | 1.86698 | 6.02031 | 10.15614 | 6.33436 | 8.66765 | 10.36029 | 4.29324 | 6.12702 | 11.53454 | 9.11000 | 1.22706 |
| **Prkaa2** | 2.42119 | 0.00470 | 0.00000 | 0.66465 | 0.00472 | 3.36420 | 1.40456 | 1.01324 | 0.00000 | 1.72887 | 0.08099 | 0.07904 | 2.91083 | 3.14689 | 0.51742 | 0.13776 |
| **Prkab1** | 13.04698 | 5.68842 | 14.64982 | 8.32390 | 6.50405 | 1.15236 | 12.27965 | 13.71792 | 14.01791 | 24.66134 | 14.13138 | 5.76656 | 17.51724 | 3.42176 | 6.92857 | 12.50630 |
| **Prkab2** | 0.00576 | 0.00793 | 0.01680 | 0.20487 | 0.00854 | 0.11073 | 0.00780 | 0.55519 | 0.00826 | 1.38032 | 0.00623 | 0.03320 | 0.01148 | 0.00492 | 0.31236 | 0.46417 |
| **Prkaca** | 3.08045 | 2.58711 | 0.00000 | 1.03402 | 4.52226 | 2.97602 | 0.40458 | 0.90016 | 0.89676 | 2.00863 | 0.00000 | 0.00000 | 0.60675 | 0.00233 | 1.27544 | 2.08041 |
| **Prkacb** | 0.91641 | 1.61508 | 2.26451 | 1.32101 | 1.77762 | 0.47381 | 0.37991 | 2.84643 | 3.38217 | 4.20214 | 1.66032 | 6.72677 | 1.68528 | 5.86535 | 2.45012 | 0.34340 |
| **Prkag1** | 7.45239 | 6.75954 | 14.36601 | 6.51746 | 13.69091 | 0.29571 | 4.01545 | 11.52887 | 3.37429 | 5.59311 | 4.22343 | 0.33030 | 12.20349 | 0.23079 | 0.64373 | 3.04498 |
| **Prkag2** | 0.02231 | 0.00784 | 0.01524 | 0.09650 | 0.01004 | 1.19879 | 0.00701 | 0.00983 | 0.04727 | 0.32906 | 0.01124 | 0.00606 | 0.23981 | 0.00742 | 0.04490 | 0.61383 |
| **Prkag2os1** | 0.01349 | 0.00833 | 0.01177 | 0.41017 | 0.00919 | 0.00680 | 0.00759 | 0.01366 | 1.03461 | 0.16810 | 0.00476 | 0.00733 | 0.01314 | 0.01078 | 0.00466 | 0.00730 |
| **Prkar1a** | 11.65883 | 10.58424 | 9.11286 | 1.57970 | 9.49449 | 0.79220 | 4.61578 | 6.14931 | 3.16870 | 7.35408 | 4.13951 | 0.69176 | 7.86555 | 0.85798 | 6.06881 | 5.71903 |
| **Prkar1b** | 0.01353 | 0.00782 | 0.01216 | 0.00845 | 0.00910 | 0.00662 | 0.00730 | 0.01074 | 0.01202 | 0.53180 | 0.01279 | 0.00611 | 0.01207 | 0.01145 | 0.00956 | 0.00775 |
| **Prkar2a** | 3.44990 | 0.00242 | 5.21326 | 2.18615 | 1.01742 | 1.37227 | 0.23712 | 3.16388 | 1.68461 | 5.63405 | 0.54203 | 0.70495 | 7.11063 | 0.34102 | 6.70546 | 0.45599 |
| **Prkar2b** | 0.01337 | 0.00783 | 0.01050 | 2.62855 | 0.00916 | 0.00613 | 0.00730 | 0.01010 | 0.28889 | 0.37848 | 0.02225 | 0.01401 | 0.01124 | 0.00302 | 1.25952 | 0.00669 |
| **Prkca** | 0.01377 | 0.00820 | 0.01204 | 0.07750 | 0.00934 | 0.00666 | 0.00751 | 0.01355 | 0.00812 | 0.01367 | 0.01241 | 0.01920 | 0.01273 | 0.01018 | 0.03355 | 0.00740 |
| **Prkcb** | 0.02142 | 0.00802 | 0.02658 | 0.07342 | 0.00973 | 0.01630 | 0.01766 | 0.01178 | 0.05026 | 0.26711 | 0.01098 | 0.00775 | 0.03358 | 0.12660 | 0.03141 | 0.00738 |
| **Prkcd** | 0.74006 | 0.00378 | 0.11447 | 18.07795 | 1.46606 | 0.00489 | 0.02340 | 3.38675 | 9.77370 | 3.02376 | 4.68486 | 3.83564 | 0.18255 | 1.67305 | 14.68103 | 0.45944 |
| **Prkce** | 0.00256 | 0.00805 | 0.00000 | 0.00982 | 0.01056 | 0.00756 | 0.13245 | 1.27061 | 0.02596 | 0.51140 | 0.59450 | 0.01760 | 0.25846 | 1.61697 | 0.05738 | 0.00758 |
| **Prkcg** | 0.47623 | 0.36701 | 3.05215 | 0.93189 | 0.03605 | 4.23771 | 2.13556 | 7.36475 | 0.53261 | 6.81538 | 4.31068 | 2.58094 | 3.51257 | 0.32515 | 4.22131 | 0.00900 |
| **Prkch** | 0.24829 | 0.00746 | 0.00000 | 0.09683 | 0.00723 | 1.89756 | 0.00684 | 0.00000 | 0.01028 | 1.50818 | 0.44345 | 0.00000 | 0.00000 | 0.00841 | 0.00000 | 0.00628 |
| **Prkci** | 5.36953 | 5.36527 | 1.24626 | 4.13257 | 3.53019 | 2.40317 | 1.64595 | 2.78515 | 0.25379 | 0.10585 | 0.23917 | 5.10609 | 1.29480 | 0.24014 | 1.79862 | 2.84502 |
| **Prkcq** | 0.01379 | 0.00775 | 0.01145 | 0.22394 | 0.00935 | 0.00663 | 0.00747 | 0.01269 | 0.23823 | 0.00886 | 0.01011 | 0.02748 | 0.01287 | 0.01918 | 0.08844 | 0.00736 |
| **Prkcsh** | 8.55808 | 5.27283 | 12.53301 | 1.36029 | 3.37301 | 1.69317 | 9.71233 | 12.69161 | 3.75271 | 7.34336 | 6.68355 | 4.23764 | 9.54342 | 0.03089 | 0.00000 | 10.46436 |
| **Prkcz** | 4.26196 | 0.10305 | 10.15746 | 3.04292 | 3.89657 | 6.07047 | 0.01704 | 4.56194 | 5.61421 | 3.95292 | 4.93353 | 0.51452 | 7.23941 | 0.00573 | 0.70464 | 2.30012 |
| **Prkd1** | 0.00950 | 0.00843 | 0.00566 | 3.89758 | 0.00961 | 0.00609 | 0.00805 | 0.55780 | 0.24747 | 0.00002 | 0.00270 | 0.05458 | 0.00560 | 0.00654 | 1.99829 | 0.02077 |
| **Prkd2** | 0.66690 | 0.00677 | 1.11352 | 0.86653 | 0.04744 | 0.00474 | 0.00689 | 0.00238 | 0.58678 | 0.02654 | 0.37356 | 0.04473 | 0.00574 | 0.00584 | 0.00000 | 0.00600 |
| **Prkd3** | 0.34346 | 0.00797 | 0.00000 | 1.71445 | 0.01610 | 0.00661 | 0.00691 | 0.00794 | 0.90399 | 0.67896 | 0.01284 | 0.03040 | 0.00745 | 0.00329 | 0.76184 | 3.58529 |

| | | | | | | | | | | | | | | | | |
|---|---|---|---|---|---|---|---|---|---|---|---|---|---|---|---|---|
| **Prkdc** | 3.81209 | 0.08387 | 3.72082 | 2.74337 | 0.00000 | 0.00291 | 2.53716 | 0.94650 | 2.42018 | 2.12502 | 1.85615 | 3.76306 | 0.00000 | 0.53889 | 6.13383 | 1.38829 |
| **Prkg1** | 0.01265 | 0.00794 | 0.00685 | 0.58398 | 0.00968 | 0.00654 | 0.00718 | 0.00971 | 0.00000 | 0.00289 | 0.00471 | 0.01208 | 0.01363 | 1.07232 | 0.69501 | 0.00736 |
| **Prkra** | 0.00301 | 1.05523 | 0.19872 | 0.15677 | 0.00748 | 0.00944 | 0.00638 | 2.40926 | 0.00838 | 0.81991 | 0.68029 | 0.00000 | 0.66685 | 0.00365 | 0.00000 | 0.00700 |
| **Prkrip1** | 7.71365 | 7.39819 | 9.41884 | 8.09712 | 8.08823 | 3.01883 | 5.07655 | 12.71230 | 14.96917 | 13.68144 | 10.96351 | 1.65726 | 9.33999 | 1.44383 | 7.23672 | 2.99252 |
| **Prkrir** | 2.25080 | 1.09620 | 1.42913 | 8.37619 | 5.29662 | 0.02012 | 2.65756 | 0.00000 | 7.00346 | 3.45782 | 2.47278 | 5.96445 | 1.51526 | 4.05257 | 8.10904 | 2.06945 |
| **Prkx** | 0.04925 | 0.00840 | 0.13384 | 0.15985 | 0.02343 | 1.66254 | 0.51760 | 0.61454 | 0.65867 | 0.19505 | 0.17396 | 0.08934 | 1.40116 | 2.89674 | 1.04673 | 0.11274 |
| **Prl3b1** | 0.00766 | 0.00716 | 0.43463 | 0.00000 | 0.00865 | 0.00522 | 0.00652 | 0.25277 | 0.00593 | 4.37549 | 0.75025 | 3.89391 | 1.57532 | 1.96982 | 2.57709 | 0.00605 |
| **Prlr** | 0.00599 | 0.00772 | 0.00743 | 4.26591 | 0.01056 | 0.00645 | 0.01760 | 0.02682 | 0.05655 | 1.41135 | 0.00000 | 0.00000 | 0.06913 | 2.32474 | 6.83403 | 0.00679 |
| **Prm2** | 0.01026 | 0.00827 | 0.22804 | 0.12378 | 0.00917 | 0.00576 | 0.00740 | 0.02317 | 0.56961 | 0.00333 | 0.00859 | 0.01883 | 0.41942 | 0.01471 | 0.00020 | 0.00704 |
| **Prm3** | 0.01320 | 0.00820 | 0.01394 | 0.01056 | 0.01005 | 0.00709 | 0.00776 | 0.01634 | 0.01233 | 0.01305 | 0.01487 | 0.00796 | 0.08356 | 0.01238 | 0.01041 | 0.00769 |
| **Prmt1** | 0.08164 | 0.06035 | 0.08615 | 0.28126 | 0.04372 | 0.03845 | 1.49860 | 0.08017 | 0.70462 | 0.80103 | 0.32371 | 0.05434 | 0.15722 | 0.02462 | 0.33299 | 0.02847 |
| **Prmt2** | 0.01301 | 0.00753 | 0.00862 | 3.82754 | 0.00881 | 0.00632 | 0.00700 | 0.00388 | 0.00000 | 1.59567 | 0.19772 | 0.07690 | 0.00993 | 0.00000 | 0.52484 | 0.00694 |
| **Prmt3** | 6.96161 | 6.25836 | 5.82501 | 2.96674 | 8.75520 | 0.04962 | 6.41296 | 7.09054 | 0.61106 | 7.74253 | 5.61950 | 8.33090 | 4.20560 | 9.60410 | 6.96388 | 0.25684 |
| **Prmt5** | 16.57284 | 14.01866 | 17.22648 | 6.00714 | 11.58368 | 12.50462 | 8.92690 | 24.62417 | 7.67596 | 21.37025 | 11.71471 | 5.38993 | 11.56933 | 8.70717 | 15.00734 | 10.67792 |
| **Prmt6** | 6.11155 | 3.48936 | 1.66436 | 12.89064 | 1.68557 | 0.35992 | 0.45544 | 3.12950 | 21.34412 | 4.75022 | 7.62875 | 1.12802 | 0.57144 | 13.90286 | 10.35297 | 0.07978 |
| **Prmt7** | 8.92020 | 8.37298 | 11.64292 | 0.74344 | 4.12474 | 2.86841 | 5.20794 | 7.93873 | 2.69945 | 3.18355 | 6.59841 | 0.00000 | 10.32703 | 0.75949 | 2.47037 | 5.63663 |
| **Prmt9** | 0.22611 | 0.02869 | 3.86849 | 0.00000 | 0.01921 | 0.00582 | 0.00321 | 3.64216 | 0.02880 | 6.97496 | 7.36702 | 4.26019 | 0.44140 | 6.22301 | 6.05415 | 2.75338 |
| **Prnp** | 14.18733 | 14.59250 | 34.49788 | 9.77948 | 9.89363 | 17.43605 | 19.51882 | 44.65810 | 15.93605 | 47.24031 | 14.65374 | 31.18042 | 18.57565 | 40.03347 | 33.55697 | 11.50550 |
| **Prob1** | 0.43437 | 0.00710 | 0.01012 | 1.04244 | 0.00690 | 0.00737 | 0.00685 | 0.01036 | 0.06272 | 0.33181 | 0.01001 | 0.03178 | 0.01227 | 0.00325 | 0.77356 | 0.00706 |
| **Proc** | 0.01366 | 0.00799 | 0.01123 | 0.06200 | 0.00921 | 0.00657 | 0.00729 | 0.00863 | 0.31105 | 0.33282 | 0.09055 | 0.01609 | 0.01126 | 0.00802 | 0.04478 | 0.00756 |
| **Proca1** | 0.52875 | 0.00563 | 0.00000 | 0.77357 | 0.00733 | 0.00716 | 2.30701 | 0.40996 | 1.60927 | 0.65085 | 0.00000 | 0.10724 | 0.11181 | 0.09757 | 1.32755 | 0.00717 |
| **Procr** | 0.19821 | 0.45818 | 0.95805 | 0.00360 | 0.88397 | 0.00529 | 0.00683 | 0.00000 | 0.22317 | 0.08478 | 0.42969 | 0.00376 | 0.05043 | 0.01139 | 0.00000 | 0.00677 |
| **Prodh** | 2.20551 | 0.02694 | 0.75655 | 0.47084 | 0.04456 | 0.01095 | 4.73301 | 3.38343 | 0.80682 | 0.62638 | 0.27686 | 0.10614 | 2.37441 | 0.00539 | 0.00000 | 0.02550 |
| **Prokr2** | 0.00635 | 0.00823 | 0.00000 | 1.86295 | 0.00969 | 0.01397 | 0.00659 | 0.00000 | 1.58625 | 1.54506 | 0.49872 | 0.00000 | 1.05611 | 2.47086 | 0.67625 | 0.00683 |
| **Prom1** | 0.39646 | 0.00897 | 0.00223 | 0.44794 | 0.02121 | 0.00852 | 0.00796 | 0.28100 | 0.00368 | 0.00598 | 0.01060 | 0.00758 | 0.30435 | 0.01288 | 0.49072 | 0.00735 |
| **Prom2** | 0.00583 | 0.00770 | 0.00000 | 0.00113 | 0.00910 | 2.07861 | 0.00600 | 0.73219 | 0.01025 | 0.47256 | 0.85578 | 0.05066 | 1.33175 | 0.00466 | 0.09734 | 0.00567 |
| **Prorsd1** | 6.15980 | 4.96217 | 2.17025 | 2.19829 | 4.64408 | 2.90685 | 3.35629 | 2.14475 | 3.96281 | 1.37967 | 3.03154 | 0.53749 | 0.94039 | 3.17069 | 4.70796 | 0.00000 |
| **Pros1** | 6.40919 | 0.24470 | 2.16302 | 0.00000 | 3.70818 | 0.00544 | 0.00000 | 1.53947 | 0.38401 | 0.00000 | 1.41302 | 0.02638 | 0.37578 | 0.00278 | 0.05953 | 2.23060 |
| **Prosc** | 2.00337 | 5.42394 | 4.31120 | 8.13953 | 9.75784 | 2.73993 | 6.10527 | 10.48537 | 10.37749 | 23.81244 | 9.54832 | 12.40710 | 3.33022 | 12.88457 | 19.53071 | 0.50037 |
| **Proser1** | 3.07620 | 0.00378 | 1.22485 | 0.00000 | 0.01309 | 0.01386 | 0.01460 | 3.16167 | 0.01277 | 0.88271 | 0.53021 | 0.18965 | 0.73702 | 0.00667 | 1.54147 | 0.01883 |
| **Proser2** | 0.06372 | 0.00830 | 1.37472 | 0.40906 | 0.00611 | 0.00693 | 0.02900 | 0.05673 | 1.44720 | 4.43087 | 1.99152 | 0.15310 | 1.04047 | 0.02176 | 0.01677 | 0.21661 |
| **Proser3** | 0.89926 | 0.02559 | 0.93731 | 0.39864 | 0.00460 | 0.00458 | 1.90849 | 0.01029 | 0.00475 | 0.00375 | 0.01140 | 0.02519 | 0.00779 | 0.01105 | 0.21539 | 0.00581 |
| **Prox2** | 0.02374 | 0.00802 | 0.03360 | 0.04699 | 0.00864 | 0.00599 | 0.00737 | 0.00515 | 0.02649 | 0.03179 | 0.25031 | 0.01671 | 0.01734 | 0.13346 | 0.00636 | 0.00711 |
| **Prpf18** | 4.54041 | 3.27560 | 7.37489 | 3.51707 | 3.59966 | 1.31266 | 4.39504 | 10.15262 | 4.77667 | 7.41050 | 4.96807 | 7.67591 | 4.68443 | 7.56893 | 13.10548 | 2.18987 |
| **Prpf19** | 2.33981 | 0.18261 | 1.51402 | 5.48697 | 1.11858 | 0.38043 | 0.43063 | 0.86849 | 6.70571 | 1.34229 | 7.47414 | 0.25248 | 1.15577 | 0.34106 | 0.69550 | 1.43185 |
| **Prpf3** | 3.29271 | 4.15625 | 3.97212 | 0.79941 | 0.00000 | 1.95214 | 4.11094 | 5.13437 | 1.74804 | 5.59129 | 1.82891 | 3.02922 | 1.81403 | 7.65103 | 8.52449 | 1.02526 |
| **Prpf31** | 9.11442 | 7.92491 | 8.70299 | 1.43976 | 7.45772 | 1.82578 | 6.85165 | 10.00712 | 1.39666 | 4.62345 | 5.58289 | 0.49731 | 10.74647 | 3.76701 | 4.28961 | 6.49039 |
| **Prpf38a** | 10.41339 | 6.64502 | 12.68517 | 10.79167 | 5.61600 | 7.28207 | 10.15884 | 10.46238 | 17.17018 | 11.74163 | 7.44679 | 9.45989 | 12.63818 | 14.58216 | 9.68518 | 5.89441 |
| **Prpf38b** | 5.10822 | 5.04028 | 5.19379 | 7.71746 | 8.20369 | 8.75322 | 2.79664 | 6.04032 | 11.92239 | 8.50429 | 2.26669 | 0.39781 | 4.88911 | 1.72599 | 8.02413 | 0.76810 |
| **Prpf39** | 0.02730 | 0.00609 | 1.81988 | 2.81950 | 0.00457 | 0.56324 | 1.87876 | 1.25407 | 11.53665 | 1.17023 | 1.77796 | 0.45838 | 1.19208 | 7.17191 | 2.94954 | 1.68380 |
| **Prpf4** | 8.86125 | 9.43464 | 6.07091 | 8.42151 | 7.62396 | 2.98196 | 5.11927 | 6.27600 | 12.16592 | 7.82205 | 3.52383 | 1.05726 | 4.22575 | 5.07534 | 13.21883 | 0.98515 |
| **Prpf40a** | 6.99692 | 3.77722 | 8.52547 | 8.97770 | 2.85744 | 2.81114 | 10.43896 | 5.23543 | 21.99876 | 12.09001 | 8.83370 | 17.62245 | 3.09510 | 23.34717 | 16.63928 | 2.67172 |
| **Prpf40b** | 2.55238 | 0.00454 | 1.49071 | 0.00452 | 0.05017 | 0.00276 | 0.02732 | 0.00000 | 0.00991 | 0.55443 | 0.28891 | 1.69288 | 0.01188 | 0.00851 | 0.00000 | 1.64172 |
| **Prpf4b** | 1.10518 | 0.82291 | 3.24433 | 4.35798 | 4.54714 | 5.45020 | 8.16729 | 5.33617 | 9.16591 | 5.38302 | 3.22218 | 7.26483 | 1.78507 | 12.99813 | 13.34249 | 1.76825 |
| **Prpf6** | 10.80957 | 5.19885 | 7.33887 | 2.79133 | 0.02335 | 6.12808 | 5.20069 | 7.85901 | 6.88593 | 7.20049 | 8.31430 | 0.93410 | 4.11886 | 1.86306 | 8.30510 | 3.68873 |
| **Prpf8** | 18.47843 | 17.91355 | 16.20646 | 4.72892 | 9.50027 | 6.16104 | 10.34163 | 21.36048 | 6.82711 | 19.60535 | 12.11682 | 12.54472 | 12.15599 | 9.76038 | 14.12081 | 4.05369 |
| **Prph** | 0.13979 | 0.00762 | 0.00824 | 0.13994 | 0.00765 | 0.00684 | 0.00753 | 0.01279 | 0.00962 | 0.03728 | 0.01286 | 0.00772 | 0.01210 | 0.00860 | 0.03219 | 0.00706 |
| **Prph2** | 0.01346 | 0.00814 | 0.01162 | 0.78060 | 0.00925 | 0.00648 | 0.00752 | 0.01369 | 1.96390 | 0.00751 | 0.00000 | 0.00492 | 0.01314 | 0.01367 | 0.00489 | 0.00715 |
| **Prps1** | 30.23491 | 18.30603 | 33.10683 | 29.46193 | 10.45892 | 15.73621 | 17.28638 | 30.37867 | 32.74834 | 39.80165 | 26.75321 | 28.02424 | 23.07461 | 40.43912 | 40.62453 | 10.60668 |
| **Prps1l3** | 8.92507 | 6.35462 | 10.23645 | 16.47890 | 4.20068 | 2.92384 | 5.10610 | 7.99149 | 21.18591 | 12.21587 | 6.30405 | 8.84965 | 7.02302 | 15.14873 | 19.14715 | 3.05590 |
| **Prps2** | 1.68148 | 2.93840 | 0.19989 | 0.00000 | 0.01971 | 0.01708 | 0.00526 | 0.00620 | 0.49281 | 0.54035 | 0.00000 | 1.31268 | 0.21809 | 0.00951 | 1.21202 | 0.01321 |
| **Prpsap1** | 12.52460 | 1.01245 | 9.89820 | 18.43427 | 13.21311 | 7.72527 | 8.64741 | 11.73722 | 18.24715 | 19.37539 | 9.48237 | 7.82769 | 3.76618 | 0.74015 | 10.54459 | 5.56296 |
| **Prpsap2** | 21.23805 | 12.23851 | 20.54357 | 5.39724 | 9.52968 | 11.81594 | 13.58050 | 21.21978 | 7.67234 | 26.00292 | 14.60161 | 11.93029 | 12.13268 | 9.80754 | 14.81589 | 4.06323 |
| **Prr11** | 0.53222 | 0.16246 | 1.12662 | 10.03787 | 1.75121 | 0.00314 | 0.01382 | 1.70776 | 12.80609 | 4.77235 | 8.28307 | 0.70211 | 0.00000 | 9.24104 | 10.95881 | 0.03759 |
| **Prr12** | 0.84335 | 0.00651 | 0.20685 | 3.11032 | 0.40909 | 0.00760 | 0.04588 | 0.63941 | 5.41028 | 0.81118 | 1.67236 | 0.00166 | 0.15729 | 0.00000 | 0.09283 | 0.74245 |
| **Prr13** | 18.13810 | 6.24880 | 16.66747 | 10.28029 | 5.35064 | 7.92326 | 14.28090 | 24.29623 | 13.50283 | 23.74874 | 24.52836 | 12.15894 | 14.37968 | 8.86943 | 16.20719 | 7.86016 |
| **Prr14** | 4.89573 | 4.25703 | 0.03200 | 2.02474 | 0.04442 | 4.16362 | 5.76285 | 7.74136 | 1.33391 | 2.53859 | 1.16315 | 1.10777 | 2.47495 | 0.28844 | 2.08624 | 3.05995 |
| **Prr14l** | 2.66284 | 4.56487 | 2.97474 | 9.86693 | 2.74101 | 4.93640 | 2.80263 | 1.17035 | 13.36731 | 2.86569 | 0.24708 | 9.40034 | 0.84983 | 1.29283 | 3.46684 | 2.04493 |
| **Prr15l** | 0.02066 | 0.00808 | 0.01402 | 0.01614 | 0.01228 | 0.00709 | 0.00723 | 0.00000 | 0.01104 | 0.78216 | 0.86914 | 0.01093 | 0.19468 | 0.00576 | 0.01997 | 0.01597 |
| **Prr18** | 0.00767 | 0.00780 | 0.05641 | 0.65508 | 0.00931 | 0.00588 | 0.00611 | 0.00869 | 0.02067 | 0.00000 | 0.01088 | 0.01546 | 2.13669 | 0.02723 | 0.77711 | 0.00622 |
| **Prr19** | 0.01228 | 0.00792 | 0.01101 | 0.24301 | 0.00963 | 0.00700 | 0.00749 | 0.02657 | 0.10277 | 0.77582 | 0.27398 | 0.01356 | 0.02592 | 0.02371 | 0.04103 | 0.00728 |
| **Prr22** | 0.01609 | 0.00797 | 0.01069 | 0.00897 | 0.01916 | 0.00748 | 0.30059 | 0.01427 | 0.01196 | 0.35246 | 0.01296 | 0.01939 | 0.04503 | 0.01156 | 0.01100 | 0.00763 |

| | | | | | | | | | | | | | | | | |
|---|---|---|---|---|---|---|---|---|---|---|---|---|---|---|---|---|
| **Prr23a1** | 0.00000 | 0.00728 | 1.10184 | 7.30555 | 0.00685 | 0.00390 | 0.06826 | 0.46449 | 10.17524 | 3.90718 | 0.76673 | 1.25914 | 0.01628 | 0.28148 | 0.20626 | 0.00487 |
| **Prr23a2** | 0.00969 | 0.00806 | 0.14119 | 0.37545 | 0.00920 | 0.00573 | 0.01798 | 0.09476 | 0.27193 | 0.50957 | 0.03240 | 0.10717 | 0.01503 | 0.00689 | 0.04711 | 0.00721 |
| **Prr23a3** | 2.77610 | 0.00403 | 0.00000 | 0.43418 | 0.00268 | 0.00371 | 0.01409 | 1.17123 | 1.24999 | 1.95369 | 0.30901 | 0.23476 | 1.18768 | 0.00632 | 1.88204 | 0.00343 |
| **Prr3** | 2.76680 | 0.01940 | 0.96880 | 3.43073 | 6.69154 | 0.00329 | 5.66924 | 4.28689 | 1.34945 | 4.35884 | 1.23847 | 2.22528 | 1.20667 | 0.58638 | 2.90825 | 3.34024 |
| **Prr36** | 0.01123 | 0.00746 | 0.00748 | 0.50706 | 0.00937 | 0.00683 | 0.00706 | 0.83583 | 0.62212 | 0.00294 | 0.01832 | 0.00468 | 0.00539 | 0.01134 | 0.00000 | 0.00721 |
| **Prr5** | 1.40319 | 0.21317 | 0.76070 | 0.51625 | 0.08171 | 0.90996 | 0.70048 | 1.66904 | 0.55079 | 2.75705 | 0.79238 | 0.02878 | 2.60382 | 0.00509 | 0.00000 | 1.55317 |
| **Prr5l** | 0.03372 | 0.00749 | 0.06099 | 1.86021 | 0.01703 | 0.01209 | 0.03685 | 0.01812 | 1.45253 | 1.01019 | 0.17498 | 0.02148 | 0.01624 | 0.27132 | 0.00000 | 0.01365 |
| **Prrc1** | 10.57272 | 6.16174 | 4.66891 | 0.06777 | 2.19244 | 0.00001 | 4.24752 | 0.00000 | 0.02478 | 1.70630 | 8.10675 | 0.03825 | 8.28173 | 0.00000 | 0.80269 | 6.41211 |
| **Prrc2a** | 4.09806 | 0.68815 | 0.33008 | 1.21357 | 1.23304 | 3.97083 | 2.57962 | 9.70749 | 1.73182 | 7.91040 | 1.83029 | 0.44748 | 4.78774 | 0.10741 | 0.00000 | 3.03299 |
| **Prrc2b** | 7.81057 | 0.85826 | 1.22601 | 2.33495 | 1.53781 | 1.01660 | 1.60399 | 5.56203 | 1.63480 | 4.71767 | 2.41264 | 0.97786 | 3.05154 | 1.11370 | 2.72204 | 5.68936 |
| **Prrc2c** | 3.68172 | 0.69146 | 4.52586 | 17.22826 | 0.02082 | 1.55071 | 0.11898 | 5.80337 | 19.08550 | 9.92492 | 4.60258 | 3.75257 | 8.36671 | 19.41335 | 11.71320 | 0.02843 |
| **Prrg1** | 0.01379 | 0.00794 | 0.01192 | 1.33927 | 0.00931 | 0.00652 | 0.00746 | 0.01306 | 0.00000 | 0.01238 | 0.02987 | 0.00799 | 0.01230 | 0.01372 | 0.03423 | 0.00717 |
| **Prrg2** | 2.37686 | 0.00362 | 2.12330 | 2.23987 | 0.05172 | 4.43522 | 1.76732 | 2.17561 | 0.81250 | 0.94289 | 0.16722 | 0.06726 | 2.27663 | 0.00380 | 0.00000 | 0.00000 |
| **Prrg4** | 0.01363 | 0.00820 | 0.01214 | 0.27573 | 0.00923 | 0.00681 | 0.00760 | 0.01448 | 0.00745 | 0.05635 | 0.01329 | 0.00898 | 0.01301 | 0.01088 | 0.00791 | 0.01471 |
| **Prrt3** | 0.16269 | 0.00787 | 0.00653 | 0.15876 | 0.00756 | 0.00680 | 0.00723 | 0.01016 | 0.00863 | 0.00000 | 0.11651 | 0.01823 | 0.01092 | 0.00648 | 0.33230 | 0.00706 |
| **Prrt4** | 0.01644 | 0.00629 | 0.00062 | 0.00292 | 1.41227 | 0.00804 | 0.00685 | 0.00000 | 0.00931 | 0.04715 | 0.27755 | 1.09504 | 0.02848 | 0.00845 | 0.19952 | 0.04324 |
| **Prrx1** | 0.02054 | 0.00789 | 0.01080 | 0.01727 | 0.00878 | 0.00666 | 0.00747 | 0.01172 | 0.03568 | 0.16106 | 0.01082 | 0.04830 | 0.01184 | 0.03167 | 0.03152 | 0.00717 |
| **Prrx2** | 0.01258 | 0.00812 | 0.02351 | 0.42522 | 0.00907 | 0.00748 | 0.00734 | 0.01047 | 2.80779 | 0.36531 | 0.00000 | 0.01222 | 0.01247 | 0.03443 | 0.00174 | 0.00732 |
| **Prss16** | 0.72294 | 0.00757 | 0.34559 | 0.01105 | 0.00692 | 0.00497 | 0.00744 | 0.36270 | 0.01190 | 0.00941 | 0.00994 | 0.00750 | 0.00464 | 0.01731 | 0.00029 | 0.00644 |
| **Prss23** | 0.01384 | 0.00789 | 0.01115 | 0.01047 | 0.00926 | 0.00676 | 0.00742 | 0.01236 | 0.01227 | 0.04608 | 0.01281 | 0.06639 | 0.01312 | 0.01156 | 0.00987 | 0.00766 |
| **Prss3** | 0.01360 | 0.00835 | 0.01221 | 0.13781 | 0.00936 | 0.00690 | 0.00763 | 0.01433 | 0.12066 | 0.05880 | 0.01100 | 0.00913 | 0.02401 | 0.00830 | 0.06345 | 0.00743 |
| **Prss32** | 0.01384 | 0.00791 | 0.01138 | 0.01041 | 0.00933 | 0.01303 | 0.00751 | 0.01353 | 0.01241 | 0.01328 | 0.01288 | 0.01783 | 0.01259 | 0.00962 | 0.04475 | 0.00740 |
| **Prss35** | 3.52956 | 0.15007 | 0.12151 | 0.35447 | 3.28080 | 0.01351 | 0.00447 | 0.05996 | 0.22116 | 0.00000 | 0.10660 | 0.21901 | 0.17928 | 0.07228 | 0.26488 | 0.22346 |
| **Prss36** | 4.94743 | 0.01264 | 2.62271 | 0.32373 | 0.00725 | 0.03215 | 2.33848 | 1.33112 | 0.46802 | 1.21214 | 0.45995 | 0.01268 | 0.55483 | 0.10508 | 0.39604 | 0.00000 |
| **Prss38** | 0.01128 | 0.00744 | 3.11168 | 0.00364 | 0.01348 | 0.00760 | 0.00685 | 0.00000 | 0.02374 | 0.00000 | 0.74798 | 0.01116 | 0.00000 | 0.00603 | 0.41481 | 0.01386 |
| **Prss41** | 0.43823 | 0.00711 | 0.10311 | 0.14070 | 0.00661 | 0.00839 | 0.00681 | 0.01401 | 0.00935 | 0.00727 | 0.01297 | 0.00739 | 0.40443 | 0.01249 | 0.00610 | 0.13950 |
| **Prss45** | 0.01199 | 0.00804 | 0.88625 | 2.44679 | 0.00756 | 0.00530 | 0.00723 | 0.00959 | 0.09380 | 0.72367 | 0.01215 | 0.05566 | 0.00775 | 0.00833 | 0.11312 | 0.00633 |
| **Prss48** | 0.01345 | 0.00823 | 0.01043 | 0.00676 | 0.00918 | 0.00669 | 0.00768 | 0.01600 | 0.01174 | 0.00887 | 0.18841 | 0.00777 | 0.02204 | 0.01172 | 0.00701 | 0.00726 |
| **Prss53** | 0.01718 | 0.00838 | 0.71810 | 0.14993 | 0.00945 | 0.00563 | 0.00761 | 0.01277 | 0.00648 | 0.01299 | 0.04072 | 0.00831 | 0.00835 | 0.01238 | 0.00000 | 0.00711 |
| **Prss57** | 0.01014 | 0.00772 | 0.00931 | 0.00455 | 0.00963 | 0.00670 | 0.00719 | 0.21399 | 0.87124 | 0.00717 | 0.00556 | 0.00581 | 0.00700 | 0.01489 | 0.00360 | 0.00734 |
| **Prss8** | 3.23792 | 2.49483 | 2.38416 | 4.33375 | 0.02986 | 12.88946 | 0.01138 | 0.56522 | 1.39414 | 0.48660 | 0.30309 | 0.03263 | 1.80729 | 0.00000 | 0.51758 | 0.00000 |
| **Prtg** | 1.32613 | 0.00660 | 1.11308 | 0.07993 | 0.00626 | 0.00686 | 0.00631 | 0.00887 | 0.62627 | 0.00141 | 0.00815 | 0.00535 | 0.00715 | 0.01544 | 0.00131 | 2.03184 |
| **Prtn3** | 0.00809 | 0.00853 | 0.04365 | 0.00465 | 0.00967 | 0.00898 | 0.00789 | 0.19849 | 0.21048 | 1.45646 | 0.72117 | 0.02450 | 0.05575 | 0.00802 | 0.01911 | 0.00738 |
| **Prune** | 1.87659 | 3.25879 | 5.04787 | 8.05778 | 5.03752 | 4.02813 | 4.00263 | 9.66345 | 6.73507 | 8.80870 | 4.29296 | 2.39164 | 5.80519 | 9.45329 | 8.44498 | 5.50167 |
| **Prune2** | 1.97588 | 0.00614 | 0.00351 | 0.19422 | 0.00635 | 0.00586 | 0.00589 | 0.00038 | 0.26472 | 0.10045 | 0.18600 | 0.05691 | 0.00833 | 0.00623 | 0.17114 | 0.97175 |
| **Prx** | 0.98574 | 0.00639 | 1.12840 | 0.01727 | 0.00405 | 0.00458 | 0.00648 | 0.47385 | 0.51504 | 0.21515 | 0.01455 | 0.03285 | 0.00982 | 0.00560 | 0.29935 | 0.00578 |
| **Psap** | 14.27350 | 10.32353 | 26.56439 | 9.15113 | 8.11639 | 19.30721 | 17.10715 | 23.04871 | 9.38965 | 10.36732 | 33.07497 | 1.51100 | 28.97905 | 2.19178 | 8.82644 | 13.45927 |
| **Psat1** | 23.10202 | 20.31622 | 33.61128 | 9.41282 | 24.26504 | 7.50278 | 23.19724 | 46.22133 | 20.66501 | 45.15552 | 35.25225 | 29.21415 | 33.14977 | 42.70439 | 27.83986 | 6.50096 |
| **Psd** | 0.01217 | 0.00798 | 0.01307 | 0.00975 | 0.00960 | 0.00688 | 0.00738 | 0.01293 | 0.01220 | 0.17895 | 0.01424 | 0.00689 | 0.29700 | 0.01127 | 0.01077 | 0.00786 |
| **Psd2** | 0.01302 | 0.00775 | 0.00339 | 0.78708 | 0.00891 | 0.00707 | 0.00685 | 0.00000 | 1.83306 | 0.22001 | 1.09165 | 0.00000 | 0.00011 | 0.13288 | 0.68757 | 0.00679 |
| **Psd3** | 0.01531 | 0.00775 | 0.02472 | 3.25833 | 0.02122 | 0.01503 | 0.00718 | 0.01906 | 0.76679 | 0.00299 | 0.07766 | 0.04320 | 0.03514 | 0.06881 | 1.07654 | 0.00671 |
| **Psd4** | 2.29630 | 0.56974 | 2.83265 | 0.01821 | 0.01262 | 2.65683 | 1.54982 | 2.77659 | 0.52379 | 1.25211 | 2.09907 | 0.05114 | 3.67375 | 0.37063 | 0.42556 | 7.26507 |
| **Psen1** | 0.30566 | 4.17307 | 12.89855 | 2.63026 | 9.00073 | 6.01824 | 2.32564 | 9.27915 | 1.25883 | 1.65364 | 2.15566 | 4.45474 | 9.08405 | 0.32271 | 4.72928 | 2.85926 |
| **Psen2** | 1.48999 | 0.07191 | 1.12944 | 1.01504 | 0.01205 | 0.01583 | 0.00600 | 0.00000 | 0.40262 | 0.50388 | 0.62277 | 0.00144 | 0.03829 | 0.00687 | 0.69344 | 0.00494 |
| **Psenen** | 6.70499 | 0.76504 | 5.20685 | 3.30674 | 5.46622 | 4.53779 | 2.30127 | 1.35293 | 0.25103 | 3.31110 | 4.66014 | 0.27989 | 7.22988 | 3.32693 | 0.61539 | 2.87892 |
| **Psg16** | 3.42688 | 3.68712 | 7.06228 | 25.33884 | 2.56379 | 2.38830 | 3.93251 | 14.28345 | 33.14197 | 31.28268 | 3.34447 | 7.17540 | 6.52760 | 39.72626 | 45.85714 | 1.84156 |
| **Psg17** | 0.01373 | 0.00819 | 0.01216 | 0.04790 | 0.00931 | 0.00661 | 0.00751 | 0.01361 | 0.20844 | 0.01199 | 0.01267 | 0.00807 | 0.01270 | 0.00942 | 0.05093 | 0.00738 |
| **Psg23** | 0.84147 | 0.42479 | 2.24119 | 6.71837 | 0.17104 | 0.73802 | 1.18490 | 3.44385 | 13.96049 | 4.64291 | 3.06922 | 5.87911 | 0.82461 | 2.13530 | 3.01829 | 0.53747 |
| **Psg28** | 0.01155 | 0.00788 | 0.01634 | 0.05072 | 0.00946 | 0.01847 | 0.07907 | 0.04589 | 0.01935 | 0.00889 | 0.02327 | 0.00704 | 0.02345 | 0.19293 | 0.01402 | 0.00738 |
| **Psg29** | 0.16822 | 0.12945 | 0.44554 | 1.36881 | 0.07991 | 0.14681 | 0.13778 | 1.00355 | 1.59642 | 0.82356 | 0.74253 | 1.08739 | 0.11292 | 0.49588 | 0.46265 | 0.20651 |
| **Psip1** | 4.06145 | 4.24577 | 1.35792 | 17.40204 | 0.00836 | 0.16656 | 3.77115 | 1.17553 | 8.43978 | 2.40130 | 0.00052 | 2.08292 | 1.11553 | 15.32636 | 22.09539 | 0.54075 |
| **Pskh1** | 2.40205 | 0.00320 | 1.90884 | 2.66761 | 0.00374 | 0.02439 | 3.21232 | 0.00000 | 1.62606 | 1.76932 | 0.76546 | 0.04787 | 0.15031 | 0.31199 | 4.41973 | 2.00145 |
| **Psma1** | 33.90032 | 20.83124 | 30.47741 | 8.05041 | 24.49640 | 14.52299 | 14.59730 | 19.89840 | 15.19130 | 23.40880 | 18.12025 | 13.10079 | 25.91998 | 17.40824 | 10.91025 | 13.84651 |
| **Psma2** | 35.59149 | 22.92874 | 25.63579 | 8.89877 | 24.15453 | 11.96603 | 17.26653 | 22.07636 | 14.66549 | 18.71012 | 21.82270 | 11.79998 | 26.88631 | 20.62999 | 11.62044 | 13.29868 |
| **Psma3** | 27.54550 | 17.38064 | 30.88530 | 8.72634 | 21.88890 | 12.07531 | 14.53381 | 27.03048 | 16.46672 | 23.10908 | 21.21811 | 9.13532 | 23.86683 | 12.49757 | 10.87054 | 12.38910 |
| **Psma4** | 36.00738 | 24.25116 | 31.41300 | 12.76478 | 26.33871 | 16.39249 | 20.02120 | 26.64356 | 17.65929 | 23.17357 | 21.51776 | 12.66357 | 27.09782 | 24.52037 | 16.07767 | 13.61473 |
| **Psma5** | 34.52595 | 17.03393 | 24.56487 | 2.93004 | 20.94788 | 9.37143 | 14.02833 | 20.67097 | 1.90249 | 17.31173 | 11.01895 | 5.32706 | 17.45749 | 14.23198 | 10.86631 | 8.52701 |
| **Psma6** | 35.87590 | 20.05491 | 37.76034 | 12.78549 | 24.72772 | 15.71473 | 20.38381 | 32.48875 | 19.19415 | 38.06698 | 33.84210 | 21.41678 | 31.01180 | 29.17438 | 22.28916 | 12.99183 |
| **Psma7** | 37.70862 | 18.25413 | 27.50221 | 0.25958 | 23.62073 | 16.66974 | 15.40225 | 32.92872 | 0.05028 | 24.38119 | 15.16672 | 6.68977 | 28.41061 | 6.96815 | 6.46096 | 16.52320 |
| **Psma8** | 0.58823 | 3.42557 | 0.00000 | 0.08817 | 0.01740 | 0.00615 | 0.00555 | 0.00682 | 0.01543 | 0.00999 | 0.01192 | 0.00709 | 0.01053 | 0.00661 | 0.00000 | 0.00567 |
| **Psmb1** | 34.82922 | 21.04407 | 24.25754 | 12.94464 | 21.26883 | 11.83145 | 17.28941 | 24.91936 | 15.16051 | 25.48903 | 23.57744 | 9.01288 | 27.42607 | 11.29297 | 14.15209 | 13.48387 |
| **Psmb10** | 1.41126 | 0.20022 | 0.90352 | 2.85537 | 0.03371 | 0.14759 | 0.38474 | 1.43279 | 0.08512 | 5.22754 | 10.05990 | 0.36717 | 2.45105 | 0.00000 | 0.19484 | 0.02517 |

| | | | | | | | | | | | | | | | | |
|---|---|---|---|---|---|---|---|---|---|---|---|---|---|---|---|---|
| **Psmb11** | 0.13022 | 0.02907 | 0.27653 | 0.51861 | 0.08598 | 0.12080 | 0.08913 | 0.06884 | 0.73719 | 0.58158 | 1.23928 | 0.07160 | 0.60384 | 0.53112 | 0.17847 | 0.04460 |
| **Psmb2** | 28.49553 | 17.25767 | 19.85560 | 0.73865 | 19.52760 | 12.58065 | 16.53208 | 23.10506 | 1.51256 | 24.01188 | 3.80437 | 3.37825 | 21.68160 | 5.54434 | 11.37085 | 11.97382 |
| **Psmb3** | 29.49689 | 11.30232 | 22.95426 | 0.30044 | 20.88728 | 8.20505 | 12.42681 | 13.21243 | 4.14770 | 17.82064 | 17.35094 | 2.25656 | 24.59233 | 4.03341 | 5.01479 | 10.57734 |
| **Psmb4** | 34.38205 | 8.82183 | 18.85153 | 14.89147 | 25.59839 | 12.01661 | 4.64237 | 22.66229 | 19.56374 | 32.91907 | 25.66457 | 4.13252 | 32.21703 | 3.88311 | 4.35998 | 17.45208 |
| **Psmb5** | 28.26472 | 8.12220 | 23.58912 | 14.82292 | 19.99340 | 11.66620 | 13.16741 | 26.58300 | 21.88176 | 26.27513 | 24.62649 | 4.92962 | 23.39362 | 0.92108 | 1.25123 | 13.63834 |
| **Psmb6** | 31.55496 | 18.98317 | 25.91465 | 3.94792 | 22.56725 | 12.30868 | 16.50087 | 21.70271 | 3.60025 | 20.87305 | 20.18879 | 2.10767 | 26.64571 | 1.80826 | 6.65678 | 14.77429 |
| **Psmb7** | 24.33927 | 19.59747 | 19.35243 | 7.80373 | 16.34756 | 11.22744 | 14.13503 | 24.83188 | 13.82351 | 20.66262 | 20.74621 | 16.91424 | 20.17030 | 22.01833 | 13.67592 | 10.73532 |
| **Psmb8** | 0.01351 | 0.00782 | 0.01228 | 0.00846 | 0.00923 | 0.00652 | 0.00731 | 0.01086 | 0.01211 | 0.53771 | 0.01246 | 0.00616 | 0.01176 | 0.01169 | 0.00837 | 0.00748 |
| **Psmb9** | 4.93291 | 0.19782 | 0.00422 | 0.00999 | 0.01225 | 0.00915 | 0.77700 | 0.24534 | 0.01215 | 0.08959 | 0.01094 | 0.00290 | 0.01443 | 0.01117 | 0.00367 | 0.00222 |
| **Psmc1** | 24.02439 | 18.11742 | 26.45441 | 3.16202 | 20.91382 | 14.43547 | 14.67958 | 25.15491 | 11.08759 | 25.75151 | 21.13574 | 6.68805 | 23.93903 | 18.24273 | 17.58638 | 11.06986 |
| **Psmc2** | 29.83930 | 20.70547 | 28.91326 | 6.09803 | 18.93582 | 14.93811 | 18.16342 | 29.12398 | 9.05665 | 25.28996 | 19.39616 | 11.38589 | 25.93134 | 15.06590 | 17.78293 | 12.94891 |
| **Psmc3** | 29.18654 | 17.99838 | 25.17970 | 1.27439 | 22.90720 | 16.52386 | 14.24833 | 25.93494 | 5.02648 | 13.30238 | 17.11712 | 2.34450 | 26.64176 | 4.92605 | 9.29518 | 17.14123 |
| **Psmc3ip** | 1.76444 | 3.66111 | 2.94942 | 10.22876 | 3.07347 | 0.00938 | 2.51532 | 5.95352 | 8.52332 | 4.78933 | 3.64181 | 1.46226 | 2.02507 | 1.31659 | 5.83938 | 1.64831 |
| **Psmc4** | 29.65668 | 17.64060 | 28.47685 | 5.03761 | 19.88353 | 12.36428 | 16.70921 | 29.92489 | 9.62954 | 24.60424 | 24.17862 | 11.32318 | 30.83272 | 3.72238 | 8.46537 | 15.25697 |
| **Psmc5** | 26.68362 | 15.43209 | 26.03345 | 4.16702 | 17.78945 | 17.04499 | 15.73613 | 30.36718 | 2.25154 | 16.52378 | 21.23230 | 8.21189 | 30.83768 | 8.10875 | 12.99705 | 10.72061 |
| **Psmc6** | 28.72470 | 22.93073 | 23.10744 | 3.89495 | 23.74328 | 13.37322 | 15.83169 | 23.94641 | 7.92877 | 13.69091 | 14.10731 | 14.08166 | 20.65348 | 26.74295 | 11.49539 | 12.31094 |
| **Psmd1** | 24.39467 | 20.81860 | 18.98183 | 6.45469 | 19.80194 | 9.12638 | 12.79871 | 23.10998 | 8.18661 | 16.73758 | 13.71397 | 4.69879 | 15.21725 | 23.76842 | 12.09987 | 7.23892 |
| **Psmd10** | 3.26196 | 8.37114 | 9.96672 | 0.82115 | 5.69565 | 3.63119 | 4.00099 | 0.32425 | 0.00000 | 2.61237 | 0.05495 | 0.01499 | 4.04592 | 3.15454 | 0.00000 | 4.13388 |
| **Psmd11** | 24.14794 | 15.93844 | 28.45823 | 10.44965 | 18.04080 | 11.99139 | 15.39682 | 26.50033 | 14.40167 | 22.15616 | 18.40210 | 2.81137 | 22.41261 | 19.29511 | 20.24796 | 6.92960 |
| **Psmd12** | 25.19421 | 21.85232 | 19.15790 | 10.70226 | 22.64501 | 14.75643 | 13.87919 | 27.62729 | 14.91276 | 26.20206 | 19.70130 | 11.50452 | 23.67660 | 14.14187 | 12.00144 | 9.55669 |
| **Psmd13** | 21.46396 | 17.18306 | 21.12156 | 0.86115 | 13.85231 | 8.07289 | 14.24328 | 21.08232 | 3.43739 | 19.11590 | 10.14925 | 4.93504 | 18.86300 | 5.60513 | 10.35841 | 10.69419 |
| **Psmd14** | 29.98783 | 19.85631 | 28.33706 | 6.68464 | 21.54635 | 16.60199 | 16.22699 | 24.01188 | 19.15042 | 24.78968 | 17.65892 | 19.99351 | 19.72746 | 11.64611 | 15.82010 | 14.33102 |
| **Psmd2** | 27.17287 | 21.65508 | 25.00782 | 8.54017 | 17.98610 | 11.96768 | 16.60108 | 25.75151 | 9.01640 | 13.14842 | 15.15461 | 11.42452 | 19.18788 | 10.95836 | 7.50655 | 12.62063 |
| **Psmd3** | 28.62218 | 13.62899 | 24.44093 | 10.57276 | 11.53157 | 8.93689 | 15.63659 | 33.60240 | 11.65918 | 27.88873 | 14.65189 | 2.82728 | 24.94629 | 0.06786 | 5.12555 | 12.60427 |
| **Psmd4** | 31.50469 | 18.13787 | 32.38156 | 14.22118 | 18.95799 | 16.16576 | 16.23410 | 33.50423 | 21.97543 | 31.16574 | 25.08443 | 6.87577 | 32.35710 | 2.09011 | 3.34994 | 16.59007 |
| **Psmd5** | 7.15148 | 5.73751 | 5.98704 | 8.60818 | 10.68637 | 5.30078 | 6.92321 | 3.89994 | 7.37866 | 12.66911 | 4.51028 | 1.73282 | 6.09177 | 2.91661 | 6.65281 | 2.46264 |
| **Psmd6** | 26.18213 | 9.61983 | 12.13718 | 3.09005 | 21.03998 | 13.05711 | 5.99830 | 27.26093 | 13.00614 | 16.25277 | 21.01642 | 4.94100 | 20.80261 | 11.52295 | 6.33473 | 13.58546 |
| **Psmd7** | 30.58598 | 14.95826 | 20.33486 | 8.22512 | 23.11121 | 12.69080 | 12.78642 | 29.87521 | 11.79576 | 34.22697 | 19.23300 | 13.74996 | 26.34531 | 19.13718 | 18.58151 | 10.28213 |
| **Psmd8** | 18.67553 | 7.38129 | 16.30385 | 11.50610 | 16.71408 | 10.16696 | 7.34988 | 23.25753 | 10.20351 | 32.35586 | 20.80407 | 10.66545 | 31.17836 | 6.75134 | 6.67019 | 12.72512 |
| **Psmd9** | 7.74312 | 2.08128 | 7.40151 | 3.39700 | 7.04861 | 5.50303 | 5.51088 | 9.08453 | 3.01206 | 6.02048 | 4.42447 | 0.96468 | 5.23401 | 0.42919 | 0.95620 | 7.19312 |
| **Psme1** | 10.10348 | 4.29178 | 8.99673 | 2.49638 | 5.90985 | 3.84122 | 2.84186 | 6.71512 | 2.57538 | 13.43013 | 9.51605 | 6.24519 | 7.43696 | 0.00000 | 0.45889 | 2.45639 |
| **Psme2** | 4.73017 | 3.92471 | 4.26382 | 1.97039 | 2.84864 | 5.05298 | 3.03330 | 0.52998 | 5.69722 | 2.48531 | 1.07660 | 0.26196 | 3.04497 | 0.57122 | 2.49864 | 2.06015 |
| **Psme2b** | 1.09998 | 0.85083 | 0.76287 | 0.63980 | 0.63232 | 1.11122 | 0.59378 | 0.12650 | 1.71341 | 0.43056 | 0.42388 | 0.01823 | 0.57414 | 0.04608 | 0.48793 | 0.40859 |
| **Psme3** | 15.59021 | 10.47486 | 11.49193 | 15.06504 | 6.77200 | 6.58212 | 10.48021 | 15.19952 | 18.08186 | 18.42145 | 12.86963 | 7.85578 | 11.53104 | 1.67933 | 6.93746 | 5.37256 |
| **Psme4** | 7.95765 | 6.70844 | 1.77511 | 20.01278 | 4.46903 | 2.32269 | 5.25370 | 5.76942 | 9.95482 | 5.56275 | 3.52263 | 14.93923 | 2.11860 | 15.94969 | 14.71349 | 2.24884 |
| **Psmf1** | 5.07092 | 2.97150 | 3.87268 | 2.49871 | 6.12341 | 0.00744 | 3.21334 | 5.23997 | 1.61514 | 6.21131 | 4.13545 | 2.82110 | 8.06632 | 0.01510 | 4.37059 | 6.13030 |
| **Psmg1** | 10.10830 | 0.36622 | 2.72928 | 0.81867 | 9.25574 | 0.73481 | 3.74521 | 9.43007 | 0.81895 | 4.08993 | 2.59836 | 0.25350 | 6.21274 | 3.45015 | 0.21844 | 3.19735 |
| **Psmg2** | 17.80281 | 9.91738 | 10.95220 | 2.15213 | 9.96769 | 10.60255 | 9.04111 | 16.19315 | 1.62133 | 10.74354 | 11.96189 | 4.67877 | 12.92929 | 2.90367 | 6.16736 | 6.72384 |
| **Psmg3** | 10.58340 | 11.00481 | 9.30920 | 1.62190 | 6.83789 | 0.01480 | 4.27480 | 9.97931 | 2.38258 | 3.82598 | 3.02602 | 0.82733 | 5.57549 | 0.77499 | 5.24179 | 2.04194 |
| **Psmg4** | 3.79348 | 0.03253 | 0.42925 | 0.01083 | 1.77842 | 0.00311 | 0.19827 | 0.20662 | 0.93650 | 2.38124 | 3.27998 | 0.00000 | 2.09093 | 0.01089 | 0.00000 | 2.31368 |
| **Psors1c2** | 0.01362 | 0.00788 | 0.01018 | 0.68931 | 0.00927 | 0.00656 | 0.00730 | 0.01083 | 0.22934 | 0.10309 | 0.25385 | 0.00590 | 0.00948 | 0.00746 | 0.00000 | 0.00726 |
| **Pspc1** | 6.03703 | 0.93170 | 1.53620 | 1.49044 | 3.58310 | 4.06663 | 2.68179 | 1.06291 | 3.56373 | 3.69228 | 5.02606 | 0.32156 | 2.87466 | 0.42728 | 3.71670 | 0.17351 |
| **Psph** | 2.35952 | 6.33842 | 8.77290 | 0.93540 | 3.77177 | 3.89367 | 1.26497 | 12.08733 | 0.15255 | 6.51851 | 8.54878 | 0.35218 | 5.68416 | 3.21237 | 1.45509 | 0.06483 |
| **Psrc1** | 12.38788 | 2.10667 | 9.32378 | 0.48910 | 10.17433 | 0.02028 | 6.77386 | 8.71116 | 0.79178 | 0.98251 | 4.30446 | 0.19164 | 0.00000 | 0.38654 | 0.00000 | 7.98104 |
| **Pstk** | 2.53031 | 0.02354 | 1.52242 | 0.62981 | 2.75143 | 2.17234 | 0.00001 | 0.08580 | 0.42619 | 2.01738 | 4.87891 | 0.12092 | 3.77018 | 0.00000 | 0.00000 | 3.43635 |
| **Pstpip1** | 1.26217 | 0.00616 | 0.00254 | 0.01109 | 0.00533 | 0.00720 | 0.03540 | 0.19859 | 0.01201 | 0.00579 | 0.01061 | 0.02545 | 0.66548 | 0.01662 | 0.01824 | 0.00550 |
| **Pstpip2** | 0.01317 | 0.00798 | 0.01599 | 0.43390 | 0.00925 | 0.00615 | 0.00745 | 0.01106 | 0.35992 | 0.19316 | 0.00565 | 0.02497 | 0.01171 | 0.00527 | 1.09668 | 0.00707 |
| **Ptafr** | 0.13345 | 0.02782 | 0.16685 | 1.14786 | 0.02598 | 0.01169 | 0.72595 | 0.00000 | 2.01541 | 0.09051 | 0.88208 | 0.08874 | 0.17279 | 0.22206 | 0.22056 | 0.04968 |
| **Ptar1** | 0.80219 | 0.00770 | 0.23168 | 2.64459 | 0.04020 | 0.00521 | 0.00679 | 0.00839 | 0.00000 | 0.53128 | 0.03275 | 0.05227 | 0.33719 | 0.02504 | 0.99098 | 0.00699 |
| **Ptbp1** | 29.05860 | 13.60468 | 21.85208 | 16.52524 | 16.34504 | 13.25924 | 15.54779 | 36.36095 | 17.22413 | 37.61896 | 23.02679 | 17.16861 | 22.65525 | 11.43981 | 13.77965 | 16.30229 |
| **Ptbp2** | 0.45187 | 0.00679 | 0.00000 | 4.45735 | 0.01490 | 0.00531 | 0.32347 | 0.00000 | 1.45254 | 0.34054 | 0.28108 | 0.16019 | 0.35045 | 0.85695 | 2.96775 | 0.00550 |
| **Ptbp3** | 14.20631 | 8.47423 | 7.62814 | 2.75359 | 16.99962 | 9.82060 | 8.44474 | 7.96161 | 2.74518 | 2.03063 | 2.05295 | 0.60385 | 5.58775 | 3.12499 | 5.09138 | 6.97501 |
| **Ptcd1** | 2.40333 | 0.01488 | 1.90532 | 1.06957 | 0.00000 | 3.75180 | 0.28036 | 0.32565 | 1.48380 | 2.02015 | 0.39876 | 0.18348 | 0.07696 | 0.00122 | 1.63673 | 0.00240 |
| **Ptcd2** | 2.76158 | 9.72929 | 4.69572 | 0.79551 | 7.96583 | 6.25680 | 4.25069 | 4.74712 | 0.44053 | 9.76886 | 1.58709 | 4.38616 | 6.48864 | 1.72626 | 1.38180 | 0.26279 |
| **Ptcd3** | 10.67762 | 10.65428 | 10.53377 | 2.40246 | 11.09287 | 6.54512 | 10.43641 | 13.93673 | 4.01604 | 12.12075 | 6.61531 | 2.03574 | 8.07713 | 14.92391 | 12.45464 | 3.51550 |
| **Ptch1** | 2.86597 | 1.63023 | 1.91996 | 0.31235 | 2.91076 | 0.01011 | 0.00384 | 0.00000 | 0.00310 | 0.00000 | 0.00000 | 4.00385 | 0.02239 | 0.06996 | 0.38584 | 0.03132 |
| **Ptch2** | 0.00832 | 0.00722 | 0.03681 | 0.00499 | 0.00969 | 0.00856 | 0.01664 | 1.41770 | 0.00960 | 0.00000 | 0.92486 | 0.00014 | 0.00000 | 0.01025 | 0.00000 | 0.00707 |
| **Ptcra** | 0.12587 | 0.03944 | 0.06048 | 3.05124 | 0.08214 | 0.01955 | 0.06735 | 0.00830 | 1.43331 | 0.03064 | 1.64662 | 0.07223 | 0.04735 | 0.81711 | 0.16470 | 0.05689 |
| **Ptdss1** | 14.87522 | 8.93740 | 11.74832 | 3.51255 | 4.87523 | 8.98914 | 11.09420 | 17.06104 | 1.48386 | 11.52251 | 9.27157 | 9.47039 | 11.15449 | 8.43548 | 7.52747 | 9.23944 |
| **Ptdss2** | 7.56678 | 3.58084 | 13.08372 | 17.85400 | 1.50325 | 6.20246 | 8.30157 | 12.54433 | 16.65495 | 26.30238 | 19.69649 | 8.49190 | 11.38160 | 4.84757 | 17.88104 | 5.96054 |
| **Pten** | 2.91787 | 0.48851 | 2.96040 | 9.72182 | 3.94774 | 2.58220 | 3.89880 | 5.20435 | 11.73973 | 10.06604 | 7.46117 | 5.22927 | 3.38844 | 11.86963 | 9.44290 | 0.30884 |
| **Pter** | 0.01009 | 0.00608 | 0.00139 | 1.32509 | 3.22873 | 0.00798 | 0.00725 | 0.00592 | 0.42476 | 0.12767 | 0.02352 | 0.00256 | 0.01130 | 0.00325 | 0.52588 | 0.00706 |

| | | | | | | | | | | | | | | | | |
|---|---|---|---|---|---|---|---|---|---|---|---|---|---|---|---|---|
| **Ptger1** | 0.74148 | 0.00718 | 0.01609 | 0.00607 | 0.00887 | 0.00812 | 0.00661 | 0.27869 | 0.06540 | 0.66670 | 0.02039 | 0.00520 | 0.01447 | 0.01244 | 0.00536 | 0.00667 |
| **Ptger3** | 0.00914 | 0.00762 | 0.01189 | 0.00775 | 0.00904 | 0.00651 | 0.00669 | 0.00861 | 0.02598 | 0.52416 | 0.01196 | 0.00537 | 1.02098 | 0.01352 | 0.02253 | 0.00724 |
| **Ptger4** | 0.01292 | 0.00758 | 0.01013 | 2.88500 | 0.00922 | 0.00589 | 0.00726 | 0.01539 | 4.58310 | 0.00358 | 0.00000 | 0.00811 | 0.01256 | 0.00299 | 0.55241 | 0.00647 |
| **Ptges** | 10.78333 | 16.93121 | 35.11079 | 2.22789 | 10.70596 | 19.69829 | 23.09076 | 27.38473 | 3.39412 | 0.00000 | 14.64802 | 1.40031 | 32.07763 | 0.60998 | 0.00000 | 10.08089 |
| **Ptges2** | 6.58840 | 3.43890 | 4.01008 | 2.44873 | 3.03859 | 2.86711 | 2.94085 | 3.51914 | 1.42394 | 4.86615 | 3.41741 | 0.43323 | 5.37287 | 0.21109 | 0.91214 | 1.70665 |
| **Ptges3** | 18.46802 | 14.97556 | 22.80154 | 9.53766 | 10.61866 | 5.02000 | 7.12770 | 15.91572 | 6.61823 | 18.87757 | 17.13906 | 10.10711 | 19.54936 | 36.42894 | 12.15500 | 8.89348 |
| **Ptges3l** | 0.00940 | 0.00701 | 0.79843 | 0.46311 | 0.00770 | 0.00501 | 0.00636 | 0.10544 | 1.35025 | 0.59370 | 1.65214 | 2.67642 | 0.00000 | 0.11719 | 0.91226 | 0.00623 |
| **Ptgfrn** | 6.50027 | 1.38280 | 3.58307 | 0.49520 | 2.73876 | 0.00000 | 0.33863 | 1.78831 | 0.00000 | 0.01230 | 0.00000 | 0.02194 | 0.00000 | 0.00409 | 0.00000 | 2.50024 |
| **Ptgis** | 4.32069 | 3.72530 | 7.86983 | 0.00011 | 0.02631 | 1.53998 | 6.43551 | 13.43117 | 0.22779 | 15.24986 | 11.76135 | 0.81960 | 6.22408 | 2.49403 | 4.36840 | 4.77279 |
| **Ptgr1** | 22.74850 | 20.48891 | 18.74588 | 0.05665 | 11.83235 | 15.94581 | 14.51306 | 9.79780 | 0.33303 | 0.00000 | 2.78129 | 0.18193 | 14.74210 | 0.81810 | 0.00000 | 15.38876 |
| **Ptgr2** | 2.29604 | 1.58943 | 2.87291 | 0.86083 | 4.45759 | 0.04462 | 2.28236 | 1.35468 | 0.90242 | 0.00000 | 2.61584 | 0.01770 | 2.04078 | 1.31245 | 0.78641 | 0.54771 |
| **Ptgs1** | 0.05403 | 0.00796 | 0.00952 | 0.00831 | 0.00797 | 0.00703 | 0.00744 | 0.01307 | 0.01186 | 0.59903 | 0.01137 | 0.01915 | 0.01219 | 0.01176 | 0.01038 | 0.00731 |
| **Pth1r** | 8.91248 | 0.00000 | 0.00206 | 0.00853 | 4.16334 | 0.00840 | 0.00000 | 0.00869 | 0.01068 | 0.00994 | 0.01046 | 0.00933 | 0.02106 | 0.01366 | 0.01098 | 5.58844 |
| **Ptk2** | 5.16285 | 2.85053 | 2.98935 | 7.24859 | 5.49631 | 2.89208 | 5.01028 | 0.97762 | 8.42962 | 4.37960 | 3.32059 | 0.15342 | 1.90817 | 3.61897 | 8.33478 | 3.01326 |
| **Ptk2b** | 0.01276 | 0.00790 | 0.01210 | 0.02551 | 0.00888 | 0.00670 | 0.00737 | 0.01117 | 0.01036 | 0.58551 | 0.01191 | 0.00655 | 0.01172 | 0.01146 | 0.00958 | 0.09757 |
| **Ptk6** | 0.67142 | 0.00770 | 0.04038 | 0.24625 | 0.01599 | 0.00596 | 0.02742 | 0.01251 | 0.67217 | 0.04852 | 0.09898 | 0.01581 | 0.05361 | 0.02855 | 0.07040 | 0.01424 |
| **Ptk7** | 0.06343 | 0.00845 | 0.02380 | 0.01010 | 0.01503 | 0.00601 | 0.00751 | 0.01247 | 0.02314 | 0.04102 | 0.01115 | 0.02900 | 0.03621 | 0.01674 | 0.05828 | 0.00727 |
| **Ptma** | 55.89922 | 28.31471 | 50.65593 | 11.25226 | 36.80595 | 27.25295 | 29.74387 | 61.10727 | 17.79517 | 58.95068 | 41.49341 | 23.16356 | 55.18411 | 35.70519 | 28.64102 | 29.63418 |
| **Ptms** | 3.60826 | 0.40069 | 1.39130 | 0.14071 | 0.12136 | 2.65358 | 0.09500 | 1.64681 | 0.14986 | 2.50678 | 0.46760 | 0.07770 | 0.94234 | 0.00591 | 0.00000 | 1.71930 |
| **Ptov1** | 13.81449 | 0.00000 | 0.00000 | 2.72327 | 5.81406 | 3.59758 | 1.19055 | 3.96489 | 0.86993 | 3.55156 | 0.27162 | 0.08954 | 8.08876 | 0.02257 | 0.00000 | 1.60732 |
| **Ptp4a1** | 0.30517 | 1.69894 | 1.16691 | 0.00000 | 0.01470 | 0.01203 | 1.52900 | 0.33468 | 0.51630 | 0.47646 | 1.55944 | 3.99709 | 0.40579 | 4.25570 | 0.60048 | 0.53031 |
| **Ptp4a2** | 19.24102 | 16.67159 | 24.55744 | 17.91307 | 8.72220 | 12.11552 | 14.20643 | 19.83704 | 17.36050 | 22.50128 | 21.85466 | 14.45228 | 18.83462 | 32.79238 | 19.79266 | 7.36839 |
| **Ptp4a3** | 2.93936 | 0.27710 | 0.76115 | 1.26652 | 0.05647 | 0.06361 | 0.00360 | 0.00000 | 0.43162 | 0.70881 | 1.96717 | 0.30959 | 0.97263 | 0.34719 | 0.00000 | 0.04515 |
| **Ptpa** | 6.89726 | 3.09414 | 6.77165 | 2.46716 | 2.12343 | 6.80952 | 0.40043 | 8.56849 | 2.38230 | 11.37053 | 3.79894 | 0.37054 | 3.11276 | 0.00621 | 3.45606 | 2.37785 |
| **Ptpdc1** | 0.00004 | 0.00716 | 0.00781 | 3.64921 | 0.01216 | 0.00670 | 2.97001 | 0.36277 | 0.00000 | 1.14045 | 0.01517 | 0.03488 | 0.02007 | 0.00061 | 1.02774 | 0.00676 |
| **Ptpmt1** | 3.06085 | 0.10454 | 0.57751 | 1.05696 | 0.02524 | 4.59703 | 0.42062 | 0.61222 | 2.03574 | 1.52640 | 10.51224 | 0.01810 | 6.30126 | 0.18719 | 0.29405 | 5.05387 |
| **Ptpn1** | 7.05978 | 2.71239 | 4.16708 | 8.27470 | 5.59546 | 2.64048 | 4.34384 | 4.62195 | 14.85372 | 6.23625 | 2.91879 | 0.50876 | 3.73051 | 0.08339 | 1.70657 | 2.10123 |
| **Ptpn11** | 12.38726 | 5.08121 | 8.74168 | 17.88606 | 9.09099 | 6.21628 | 8.14117 | 12.82141 | 17.73049 | 10.32283 | 6.89790 | 0.99814 | 6.96228 | 8.49844 | 11.86544 | 1.04913 |
| **Ptpn12** | 3.95106 | 10.37132 | 5.09977 | 1.04981 | 6.76642 | 4.69664 | 6.30306 | 5.37543 | 6.03428 | 8.04394 | 1.37901 | 15.09427 | 5.29497 | 5.72258 | 4.68550 | 3.14834 |
| **Ptpn13** | 4.54692 | 2.18415 | 2.52688 | 5.83716 | 0.11194 | 0.38177 | 2.93101 | 5.19285 | 1.28946 | 13.06860 | 7.30616 | 10.38185 | 1.12813 | 2.00169 | 5.99986 | 4.41901 |
| **Ptpn14** | 0.03052 | 0.00900 | 0.00822 | 0.21162 | 0.01410 | 0.00731 | 0.00826 | 0.23862 | 0.00564 | 0.02388 | 0.01104 | 0.00788 | 0.01437 | 0.00808 | 0.05387 | 0.01544 |
| **Ptpn18** | 0.25885 | 0.00667 | 0.01567 | 0.06423 | 1.03753 | 0.00651 | 0.00548 | 0.06006 | 0.25880 | 0.28149 | 0.60429 | 0.00028 | 2.21311 | 0.00690 | 0.00720 | 3.13390 |
| **Ptpn2** | 8.53541 | 9.08599 | 4.99773 | 15.20852 | 7.12896 | 7.98663 | 5.93265 | 14.78903 | 17.48976 | 16.81033 | 14.25546 | 9.14580 | 6.47770 | 22.44692 | 20.71678 | 3.11106 |
| **Ptpn21** | 1.51707 | 0.02372 | 0.58755 | 14.60596 | 0.20455 | 2.95468 | 0.02035 | 7.72260 | 3.97416 | 16.43977 | 9.06770 | 11.19842 | 0.91269 | 4.45662 | 12.55153 | 0.00660 |
| **Ptpn22** | 0.00000 | 0.00500 | 0.00000 | 0.98943 | 0.00801 | 0.00569 | 2.19004 | 5.90955 | 1.64389 | 4.83682 | 6.73496 | 0.68436 | 2.09538 | 11.10622 | 1.08630 | 0.02644 |
| **Ptpn23** | 1.15026 | 0.32629 | 2.69094 | 7.65093 | 2.82123 | 0.02168 | 2.52956 | 8.96151 | 3.14189 | 5.52932 | 4.70833 | 2.53983 | 4.05441 | 0.00895 | 0.99423 | 6.63423 |
| **Ptpn3** | 1.70389 | 0.00473 | 3.14261 | 1.27077 | 2.34044 | 0.00608 | 0.08882 | 0.54344 | 0.00000 | 1.64651 | 0.64698 | 0.04144 | 1.56590 | 5.71548 | 2.06107 | 0.02562 |
| **Ptpn4** | 1.83821 | 0.00454 | 0.00000 | 1.87466 | 0.00732 | 0.00558 | 0.00518 | 0.00839 | 0.88059 | 0.63833 | 0.61501 | 2.97540 | 0.01972 | 0.00962 | 0.23674 | 0.01290 |
| **Ptpn6** | 5.57975 | 1.10794 | 2.89230 | 0.17949 | 6.35739 | 0.00000 | 1.54349 | 7.63067 | 0.75781 | 0.16701 | 4.33286 | 0.00003 | 4.93229 | 0.00000 | 0.99638 | 0.01145 |
| **Ptpn7** | 0.09835 | 0.01838 | 0.24082 | 0.13603 | 0.01763 | 0.02630 | 0.08023 | 0.03184 | 0.53778 | 0.14799 | 0.67580 | 0.01254 | 0.27090 | 0.33191 | 0.12714 | 0.11570 |
| **Ptpn9** | 0.83725 | 0.00528 | 0.00069 | 5.68472 | 1.99400 | 0.00687 | 0.00495 | 0.00000 | 7.89771 | 4.85070 | 1.07604 | 0.11563 | 1.53409 | 6.26630 | 7.43623 | 0.03992 |
| **Ptpra** | 5.50181 | 0.05965 | 2.34633 | 3.92950 | 4.32547 | 4.29843 | 6.75087 | 1.87487 | 2.09606 | 2.05273 | 2.61319 | 0.38998 | 1.48396 | 0.00000 | 1.44313 | 1.13949 |
| **Ptprb** | 0.05996 | 0.00482 | 1.32676 | 0.25204 | 3.15971 | 0.00533 | 1.53829 | 0.00463 | 0.00069 | 0.64097 | 0.00504 | 0.26671 | 0.00692 | 0.01048 | 0.31455 | 0.01349 |
| **Ptprcap** | 0.01681 | 0.00745 | 0.02092 | 0.19220 | 0.00872 | 0.00770 | 0.00716 | 0.58232 | 0.18647 | 0.02761 | 0.04132 | 0.29269 | 0.01477 | 0.01204 | 0.00000 | 0.00714 |
| **Ptpre** | 0.01326 | 0.01836 | 0.01037 | 1.18473 | 0.00933 | 0.00629 | 0.00729 | 0.01080 | 0.81920 | 0.00000 | 0.00359 | 0.01442 | 0.01305 | 0.06138 | 1.01138 | 0.00707 |
| **Ptprf** | 1.14260 | 0.02593 | 0.28310 | 0.99971 | 0.04575 | 3.76240 | 0.13532 | 0.99292 | 0.00000 | 3.27169 | 0.39873 | 0.04253 | 2.36782 | 0.00420 | 0.05176 | 4.51967 |
| **Ptprg** | 1.09503 | 0.00457 | 0.00000 | 8.93925 | 3.50864 | 0.00444 | 0.00564 | 0.00000 | 0.24312 | 1.56837 | 0.71576 | 0.34797 | 0.02554 | 0.13515 | 3.70196 | 0.04955 |
| **Ptprh** | 0.01254 | 0.00774 | 0.04491 | 0.82113 | 0.00910 | 0.00783 | 0.00715 | 0.00439 | 2.84154 | 0.06373 | 0.16627 | 0.00000 | 0.01044 | 0.00293 | 0.90197 | 0.00711 |
| **Ptprj** | 0.01453 | 0.00762 | 0.08278 | 0.36780 | 0.00897 | 0.00576 | 0.00711 | 0.42796 | 0.91901 | 0.67995 | 0.53039 | 0.02506 | 0.00417 | 1.28032 | 1.23422 | 0.01436 |
| **Ptprk** | 0.11133 | 0.00748 | 0.09324 | 0.00253 | 1.16718 | 0.00916 | 0.00696 | 0.03276 | 0.40075 | 0.25008 | 0.32722 | 0.00295 | 1.24259 | 0.01172 | 0.00327 | 0.00706 |
| **Ptprm** | 0.01366 | 0.00786 | 0.01141 | 0.06122 | 0.00918 | 0.00661 | 0.00743 | 0.01231 | 0.11123 | 0.01223 | 0.01233 | 0.02827 | 0.04597 | 0.01546 | 0.01746 | 0.00733 |
| **Ptprn** | 0.45617 | 0.00710 | 0.02678 | 0.25590 | 0.30630 | 0.01709 | 0.00600 | 1.38891 | 0.00674 | 0.95716 | 0.11478 | 0.00000 | 2.20942 | 0.01077 | 0.00000 | 0.01303 |
| **Ptpro** | 0.01381 | 0.00771 | 0.01183 | 0.89452 | 0.00940 | 0.00653 | 0.00744 | 0.01258 | 0.00000 | 0.00830 | 0.01144 | 0.00746 | 0.01279 | 0.01873 | 0.17999 | 0.00725 |
| **Ptprr** | 0.01767 | 0.00785 | 0.01076 | 3.92680 | 0.00894 | 0.00613 | 0.00731 | 0.01170 | 1.98343 | 0.00069 | 0.02831 | 0.00408 | 0.01223 | 0.00852 | 0.19264 | 0.00648 |
| **Ptprs** | 1.15180 | 0.00590 | 0.01702 | 0.45112 | 0.03817 | 0.00710 | 0.02580 | 0.00000 | 0.00521 | 0.63213 | 0.52221 | 0.02921 | 0.67405 | 0.00479 | 0.09154 | 0.01307 |
| **Ptprt** | 0.02369 | 0.01747 | 0.00805 | 0.54744 | 0.00843 | 0.01870 | 0.00719 | 0.01384 | 0.18776 | 0.99250 | 0.00751 | 0.01998 | 0.01859 | 0.00664 | 1.54461 | 0.01529 |
| **Ptpru** | 0.00588 | 0.00837 | 0.00846 | 0.11358 | 0.01914 | 0.00718 | 0.00698 | 0.02121 | 0.01045 | 0.00240 | 0.57116 | 0.01550 | 1.86427 | 0.00850 | 0.00000 | 0.00647 |
| **Ptprz1** | 0.01808 | 0.00771 | 0.00855 | 0.54288 | 0.00889 | 0.00679 | 0.01735 | 0.00917 | 0.03412 | 0.00687 | 0.07071 | 0.17042 | 0.01889 | 0.08633 | 0.00651 | 0.00728 |
| **Ptrf** | 2.10601 | 0.00505 | 0.02176 | 0.17640 | 0.00442 | 0.02357 | 0.00468 | 1.17053 | 0.11859 | 3.32934 | 0.69203 | 0.03916 | 1.49740 | 0.02556 | 0.29598 | 0.00452 |
| **Ptrh1** | 2.95719 | 0.21335 | 0.23300 | 2.55169 | 1.76542 | 0.00318 | 0.96928 | 1.73078 | 8.50128 | 12.39672 | 6.79263 | 0.11517 | 6.12343 | 0.18818 | 0.77333 | 0.00000 |
| **Ptrh2** | 6.70271 | 4.35261 | 7.16766 | 4.02967 | 2.56259 | 2.57699 | 2.88986 | 7.93171 | 6.77957 | 11.18289 | 9.09519 | 1.93594 | 4.31648 | 7.41573 | 7.82184 | 0.04240 |
| **Ptrhd1** | 1.18467 | 0.07850 | 1.50830 | 2.18045 | 1.99960 | 1.35650 | 0.09847 | 1.46849 | 3.16324 | 1.35131 | 3.21494 | 0.32615 | 1.42781 | 0.46147 | 0.15090 | 0.12683 |

| | | | | | | | | | | | | | | | | |
|---|---|---|---|---|---|---|---|---|---|---|---|---|---|---|---|---|
| **Pts** | 0.19870 | 0.00578 | 0.00000 | 0.30275 | 3.50841 | 2.58154 | 0.04635 | 0.00000 | 0.05614 | 1.18201 | 0.24021 | 0.00000 | 0.45615 | 0.13944 | 0.02437 | 0.02119 |
| **Pttg1** | 22.41959 | 5.10839 | 12.05015 | 20.53423 | 17.94188 | 1.83386 | 6.69877 | 10.90552 | 26.25281 | 18.96481 | 25.43808 | 6.56158 | 10.49563 | 15.41143 | 14.88595 | 6.76761 |
| **Pttg1ip** | 11.66758 | 7.09870 | 18.04925 | 19.96480 | 4.82758 | 8.01197 | 13.24808 | 33.77437 | 25.58051 | 40.09285 | 29.88543 | 23.00202 | 21.84071 | 24.72120 | 29.96174 | 1.56260 |
| **Puf60** | 20.55238 | 1.53771 | 3.59666 | 13.30487 | 14.36401 | 8.29557 | 5.42839 | 19.30333 | 9.09396 | 24.99963 | 15.80310 | 4.46508 | 20.19424 | 4.73293 | 5.48193 | 8.08557 |
| **Pum1** | 8.50232 | 4.33468 | 7.02539 | 7.56265 | 3.03800 | 0.01199 | 5.06782 | 5.79001 | 11.14924 | 14.04238 | 8.28076 | 2.99102 | 0.00000 | 7.68631 | 9.45428 | 0.24221 |
| **Pum2** | 6.49849 | 10.39424 | 9.01291 | 13.76435 | 2.77261 | 4.97493 | 7.25086 | 8.79598 | 16.08948 | 22.86777 | 9.72538 | 27.81080 | 6.04957 | 29.32103 | 22.50309 | 2.61718 |
| **Pum3** | 12.62512 | 9.96071 | 11.45999 | 10.10044 | 12.61890 | 6.83146 | 9.93918 | 15.94262 | 17.94393 | 15.44678 | 10.17481 | 15.36643 | 10.07739 | 6.33303 | 17.08388 | 5.04577 |
| **Pura** | 2.19206 | 0.08123 | 0.00000 | 0.73671 | 0.00499 | 2.43263 | 0.07843 | 1.21789 | 0.08430 | 1.66117 | 0.13268 | 0.09257 | 0.00000 | 0.17510 | 0.13292 | 2.58493 |
| **Purb** | 3.75750 | 3.12462 | 8.19783 | 13.22950 | 3.42559 | 6.57388 | 3.16482 | 9.91677 | 17.94308 | 20.02947 | 5.12475 | 12.27969 | 3.01229 | 32.08788 | 15.09592 | 4.65390 |
| **Purg** | 1.57100 | 0.00518 | 0.93288 | 0.42644 | 1.91427 | 0.00398 | 0.00646 | 0.00161 | 0.09750 | 0.00000 | 0.70925 | 0.00138 | 0.00316 | 0.01557 | 0.09337 | 0.00504 |
| **Pus1** | 13.94069 | 3.81673 | 8.38792 | 8.15225 | 9.08312 | 6.33993 | 8.90287 | 6.23769 | 10.42862 | 10.32400 | 1.30039 | 1.49764 | 8.45132 | 0.08782 | 3.50433 | 6.00226 |
| **Pus10** | 3.97701 | 1.68646 | 4.71288 | 0.00000 | 6.64809 | 0.00230 | 5.18002 | 0.08957 | 0.00008 | 6.25993 | 2.22391 | 0.28127 | 1.59245 | 6.63649 | 1.71545 | 3.10948 |
| **Pus3** | 3.05916 | 3.59075 | 4.94105 | 1.46634 | 2.52802 | 0.02063 | 3.27367 | 3.10237 | 0.00000 | 5.49130 | 3.34838 | 2.09591 | 2.24197 | 11.27156 | 9.52433 | 3.59016 |
| **Pus7** | 3.06580 | 0.91281 | 3.90620 | 5.41997 | 5.09027 | 0.13040 | 3.28435 | 6.19989 | 6.16208 | 3.31457 | 4.63552 | 7.59723 | 1.59549 | 5.93202 | 7.40474 | 1.08541 |
| **Pus7l** | 1.60857 | 0.00544 | 1.70114 | 0.42847 | 0.00195 | 1.69862 | 0.22183 | 1.71862 | 0.25482 | 3.31259 | 0.20868 | 0.10682 | 0.35663 | 0.00341 | 1.00197 | 0.00395 |
| **Pusl1** | 7.24610 | 0.00000 | 0.00000 | 0.55928 | 3.74177 | 4.13766 | 0.06152 | 7.10557 | 3.43801 | 6.25375 | 0.44253 | 0.09334 | 3.96667 | 0.77941 | 0.30954 | 5.00280 |
| **Pvr** | 0.02052 | 0.00759 | 0.04797 | 0.11299 | 0.00842 | 0.00627 | 0.00704 | 0.27481 | 0.88891 | 0.92230 | 0.72173 | 0.08556 | 0.00282 | 0.02147 | 0.54647 | 0.00690 |
| **Pvt1** | 2.33371 | 0.00280 | 3.47280 | 1.37765 | 0.51306 | 0.00504 | 2.54347 | 1.60812 | 1.36737 | 1.91895 | 4.25443 | 0.01947 | 0.00000 | 0.61427 | 0.57830 | 0.00259 |
| **Pwp1** | 10.52403 | 15.06551 | 7.38270 | 8.97724 | 7.95845 | 7.53959 | 9.84320 | 17.05241 | 19.63113 | 19.18955 | 13.44952 | 8.86658 | 15.10107 | 13.31042 | 19.39619 | 3.06937 |
| **Pwp2** | 5.66956 | 6.02401 | 5.89859 | 4.98070 | 0.60021 | 1.05783 | 5.72590 | 11.24174 | 11.62692 | 10.87521 | 4.50516 | 0.66708 | 6.15092 | 0.88399 | 9.47526 | 3.15633 |
| **Pwwp2a** | 0.02543 | 1.19496 | 1.06746 | 5.31557 | 4.23575 | 0.41702 | 7.99112 | 2.57281 | 5.83988 | 5.87945 | 1.84392 | 0.50758 | 0.99333 | 4.55794 | 6.18379 | 2.45366 |
| **Pwwp2b** | 0.00000 | 0.00670 | 0.88859 | 0.00455 | 0.29439 | 0.00923 | 0.00677 | 3.14861 | 0.00814 | 0.00000 | 0.86637 | 0.00000 | 0.00000 | 0.01409 | 0.00350 | 1.82992 |
| **Pxdc1** | 0.01370 | 0.00790 | 0.01220 | 0.13569 | 0.00916 | 0.00664 | 0.00736 | 0.01163 | 0.00985 | 0.24674 | 0.01423 | 0.00691 | 0.01260 | 0.01068 | 0.00827 | 0.00778 |
| **Pxdn** | 2.90112 | 0.00356 | 0.00000 | 0.00352 | 0.01067 | 0.00597 | 0.01473 | 0.00000 | 0.97151 | 0.77862 | 0.00000 | 2.01304 | 0.01166 | 0.01049 | 0.00000 | 0.00434 |
| **Pxk** | 1.58340 | 0.01083 | 6.52166 | 5.85386 | 2.56371 | 7.80492 | 3.78766 | 5.43528 | 4.65869 | 5.38048 | 6.82940 | 2.94592 | 6.44510 | 4.58388 | 3.61852 | 1.14456 |
| **Pxmp2** | 4.69301 | 1.83573 | 0.58889 | 5.08600 | 0.81178 | 6.15754 | 0.58930 | 1.73423 | 8.53975 | 5.80220 | 7.37839 | 4.23196 | 2.51900 | 3.68376 | 2.20233 | 2.99684 |
| **Pxmp4** | 2.90390 | 2.76655 | 3.32628 | 0.62115 | 0.04784 | 6.87142 | 0.06341 | 1.31791 | 1.51946 | 1.48419 | 1.91242 | 1.40003 | 2.22098 | 0.28816 | 0.00000 | 4.06003 |
| **Pxn** | 2.46258 | 0.01638 | 0.65667 | 4.07415 | 0.07693 | 2.09281 | 1.72700 | 0.97255 | 3.34474 | 0.00000 | 2.49960 | 0.04203 | 1.83508 | 0.27612 | 0.32283 | 0.19293 |
| **Pxt1** | 0.04548 | 0.00535 | 1.19144 | 10.89803 | 4.32340 | 0.00345 | 0.00638 | 0.66495 | 10.34855 | 0.00513 | 0.33158 | 0.07941 | 0.00000 | 0.00000 | 0.75801 | 1.55266 |
| **Pxylp1** | 0.01374 | 0.00798 | 0.01169 | 0.10594 | 0.00940 | 0.00645 | 0.00736 | 0.01149 | 0.00907 | 0.00586 | 0.01226 | 0.00706 | 0.01276 | 0.00746 | 0.65096 | 0.00734 |
| **Pycard** | 1.32577 | 0.00372 | 3.51472 | 0.00283 | 0.01518 | 0.01297 | 1.77773 | 0.54856 | 0.01920 | 0.54533 | 5.16347 | 0.01158 | 3.93837 | 0.00485 | 0.07518 | 2.04351 |
| **Pycr1** | 0.36364 | 0.02724 | 0.72967 | 0.05467 | 0.00654 | 0.04306 | 1.34680 | 1.64696 | 0.11172 | 0.14883 | 0.49304 | 0.02734 | 0.94754 | 0.24215 | 0.01500 | 0.01982 |
| **Pycr2** | 9.92051 | 0.35561 | 2.79756 | 0.18911 | 3.23761 | 0.01948 | 2.02826 | 4.67476 | 5.28000 | 0.30423 | 1.23551 | 0.00757 | 8.63460 | 0.03288 | 0.00000 | 10.17492 |
| **Pycrl** | 2.10353 | 0.64032 | 9.44485 | 2.65390 | 0.91774 | 0.30667 | 5.91774 | 3.59767 | 0.15733 | 2.06285 | 0.05794 | 0.47402 | 5.72434 | 0.05155 | 0.00000 | 1.84147 |
| **Pydc3** | 0.02032 | 0.00771 | 0.01120 | 0.13742 | 0.00885 | 0.00667 | 0.01775 | 0.01804 | 0.07390 | 0.01233 | 0.01292 | 0.01732 | 0.01238 | 0.04319 | 0.02320 | 0.00738 |
| **Pygb** | 7.61194 | 1.84193 | 6.65351 | 0.89789 | 3.65485 | 0.41199 | 5.03000 | 5.52790 | 0.73600 | 1.57654 | 1.66250 | 0.10779 | 2.31518 | 0.34477 | 0.00000 | 2.32360 |
| **Pygl** | 0.00000 | 0.00685 | 0.11604 | 0.00000 | 2.12818 | 0.00898 | 0.00740 | 0.48264 | 0.58782 | 0.31695 | 1.08509 | 0.08151 | 0.43310 | 0.12503 | 2.05878 | 0.02219 |
| **Pygm** | 0.00743 | 0.00748 | 0.39605 | 0.93975 | 0.00917 | 0.00547 | 0.00724 | 0.63294 | 0.44639 | 0.58031 | 0.00000 | 0.06987 | 0.00479 | 0.15385 | 0.00000 | 0.00714 |
| **Pygo1** | 0.01032 | 0.00752 | 0.00626 | 1.72409 | 0.00956 | 0.00687 | 0.00694 | 0.29031 | 1.55765 | 0.03287 | 0.34328 | 0.00204 | 0.00492 | 0.00432 | 0.00000 | 0.00698 |
| **Pygo2** | 2.61323 | 3.29663 | 6.25864 | 8.06815 | 0.63588 | 2.16667 | 0.94312 | 6.50801 | 5.92776 | 5.42093 | 3.70017 | 0.97658 | 3.58725 | 0.57367 | 1.96298 | 7.00531 |
| **Pym1** | 0.08480 | 0.10349 | 0.07944 | 0.58349 | 0.00996 | 0.01732 | 0.05890 | 0.06792 | 0.02216 | 0.19253 | 0.01058 | 0.01708 | 0.02786 | 0.02284 | 0.21274 | 0.00706 |
| **Pyroxd1** | 1.68215 | 0.00476 | 1.14001 | 0.17388 | 0.01007 | 0.00881 | 0.00394 | 0.84136 | 0.17945 | 2.50455 | 0.87072 | 4.00671 | 2.09814 | 0.01771 | 0.07464 | 0.00386 |
| **Pyurf** | 0.00000 | 0.03884 | 0.21799 | 0.05299 | 0.01604 | 0.09006 | 0.00686 | 0.00985 | 0.01102 | 0.51714 | 0.01162 | 0.07165 | 0.61086 | 0.05759 | 0.00010 | 2.54385 |
| **Pyy** | 0.03141 | 0.16190 | 0.01253 | 0.01012 | 0.64963 | 0.00628 | 0.00727 | 0.01017 | 0.01210 | 0.00506 | 0.01260 | 0.01759 | 0.01202 | 0.01219 | 0.00796 | 1.68209 |
| **Pzp** | 1.73979 | 0.54517 | 0.00000 | 0.30314 | 1.81365 | 1.07719 | 0.00346 | 1.04833 | 3.36108 | 1.82605 | 5.52495 | 0.10046 | 0.00000 | 4.40486 | 1.36257 | 0.01108 |
| **Qars** | 13.33683 | 13.65635 | 18.55377 | 13.21968 | 15.33436 | 10.55479 | 11.14639 | 29.05409 | 18.44116 | 16.26796 | 16.72034 | 6.21186 | 21.47203 | 16.91660 | 10.48449 | 12.90319 |
| **Qdpr** | 7.76136 | 0.27789 | 6.71517 | 4.36460 | 3.13834 | 0.00000 | 4.03780 | 0.11216 | 1.03610 | 1.25439 | 1.36464 | 0.53024 | 5.23960 | 0.00000 | 0.07318 | 2.40078 |
| **Qk** | 10.50792 | 8.59632 | 7.69934 | 8.54714 | 8.12086 | 3.05038 | 6.51327 | 8.51625 | 12.20593 | 6.33936 | 3.71614 | 5.76465 | 4.95662 | 18.67254 | 5.93282 | 2.85407 |
| **Qpctl** | 0.38782 | 0.00594 | 0.35635 | 2.46187 | 0.00663 | 0.00525 | 2.31887 | 0.23076 | 4.56863 | 1.06901 | 0.00000 | 0.03584 | 0.00691 | 0.02001 | 0.00000 | 0.00602 |
| **Qrich1** | 5.62686 | 5.59646 | 3.12683 | 2.07026 | 1.57878 | 0.78559 | 3.91459 | 4.43222 | 2.45997 | 2.69482 | 0.09200 | 0.01193 | 1.06231 | 12.09666 | 4.79018 | 3.03866 |
| **Qrich2** | 0.01385 | 0.00826 | 0.01042 | 0.44682 | 0.00912 | 0.00665 | 0.00746 | 0.00805 | 0.00448 | 0.04520 | 0.23297 | 0.00729 | 0.00965 | 0.00785 | 0.02290 | 0.00732 |
| **Qrsl1** | 2.00578 | 0.00453 | 0.33310 | 1.17786 | 0.02196 | 3.33400 | 1.93980 | 3.37877 | 0.80864 | 2.33695 | 2.56547 | 1.15137 | 1.29428 | 1.40945 | 0.80339 | 1.85652 |
| **Qser1** | 1.09223 | 0.01632 | 0.39660 | 3.81074 | 0.00639 | 0.00387 | 2.38361 | 0.00969 | 2.23093 | 0.00000 | 0.00000 | 0.00000 | 0.00860 | 0.71182 | 4.22159 | 0.00489 |
| **Qsox1** | 0.04940 | 0.00670 | 1.78123 | 2.44418 | 0.55341 | 0.79005 | 0.00650 | 0.00000 | 1.22523 | 0.84139 | 0.44449 | 0.04336 | 0.16360 | 0.00465 | 0.36019 | 0.03466 |
| **Qsox2** | 0.00000 | 0.08001 | 2.45710 | 2.45472 | 0.02031 | 0.02448 | 0.05032 | 1.69889 | 4.30664 | 7.25582 | 0.49994 | 7.28796 | 0.36096 | 0.25611 | 10.05884 | 1.76651 |
| **Qtrt1** | 10.12948 | 0.02399 | 3.42929 | 1.90892 | 3.91129 | 4.16983 | 1.95780 | 8.25267 | 2.91184 | 7.29705 | 3.05829 | 0.48752 | 8.00496 | 0.11105 | 2.73628 | 3.94907 |
| **Qtrtd1** | 2.57335 | 2.12346 | 0.70691 | 0.36566 | 0.01153 | 0.47777 | 0.69865 | 1.74292 | 0.18998 | 0.69538 | 1.28791 | 0.45137 | 2.92919 | 3.15641 | 5.67047 | 0.06738 |
| **R3hcc1** | 4.36372 | 1.40394 | 9.62807 | 5.05216 | 8.65703 | 1.30685 | 4.63031 | 18.01778 | 2.78493 | 8.64774 | 11.09025 | 0.70930 | 14.39881 | 0.02486 | 0.72098 | 2.43632 |
| **R3hcc1l** | 0.89661 | 2.59469 | 0.00000 | 0.76610 | 0.02158 | 0.01316 | 0.76247 | 1.88329 | 0.40968 | 0.99261 | 1.11427 | 0.04065 | 0.01856 | 1.85237 | 1.27109 | 2.15055 |
| **R3hdm1** | 7.44649 | 0.55617 | 1.06581 | 11.46015 | 4.50112 | 3.97738 | 3.10445 | 8.40205 | 15.02943 | 12.41385 | 6.45029 | 6.36159 | 2.61823 | 13.48608 | 14.79299 | 0.24896 |
| **R3hdm2** | 7.85411 | 0.03000 | 0.34720 | 10.98631 | 2.11670 | 2.60282 | 0.86362 | 6.49018 | 7.71337 | 10.88009 | 5.70583 | 0.50777 | 4.44099 | 1.63452 | 2.05835 | 2.94634 |
| **R3hdm4** | 6.01397 | 0.00892 | 3.04913 | 1.96152 | 2.61521 | 6.70163 | 1.53696 | 4.91198 | 0.18131 | 1.14437 | 1.44738 | 0.17782 | 3.29359 | 0.39944 | 0.61554 | 2.42768 |

| | | | | | | | | | | | | | | | | |
|---|---|---|---|---|---|---|---|---|---|---|---|---|---|---|---|---|
| **Rab10** | 6.52823 | 7.59048 | 3.29120 | 8.12093 | 6.93582 | 6.19540 | 5.19498 | 11.80640 | 15.87698 | 16.83324 | 13.68795 | 10.29057 | 4.28709 | 18.19415 | 12.60130 | 0.52460 |
| **Rab10os** | 1.64227 | 0.04842 | 0.08039 | 7.34414 | 0.00614 | 0.16614 | 0.02497 | 1.45553 | 9.82787 | 1.97085 | 6.05824 | 0.19181 | 0.00000 | 0.00000 | 0.00000 | 0.00288 |
| **Rab11a** | 10.35281 | 6.18956 | 8.46837 | 11.47738 | 9.57243 | 3.80704 | 2.73143 | 3.94283 | 13.45761 | 10.02240 | 14.98721 | 12.24132 | 12.76851 | 10.74616 | 3.82845 | 9.04067 |
| **Rab11b** | 12.01925 | 1.45849 | 7.24563 | 6.91179 | 1.73197 | 6.79065 | 3.84371 | 17.54984 | 11.78238 | 21.36512 | 7.62284 | 1.92267 | 3.41480 | 0.10442 | 2.01033 | 0.06579 |
| **Rab11fip1** | 1.13878 | 0.33080 | 2.61730 | 3.50043 | 0.47562 | 2.89552 | 0.38801 | 8.10774 | 7.94176 | 10.05575 | 7.08035 | 1.29070 | 3.46632 | 15.52789 | 12.07958 | 0.08635 |
| **Rab11fip2** | 0.03758 | 0.00781 | 0.00754 | 0.81224 | 0.00813 | 0.00640 | 0.00722 | 0.00870 | 0.09710 | 0.36410 | 0.25912 | 0.00323 | 0.00869 | 0.00533 | 1.11761 | 0.00682 |
| **Rab11fip3** | 0.01698 | 0.00760 | 0.00292 | 1.19383 | 0.00990 | 0.01408 | 0.00651 | 0.29103 | 1.32713 | 0.81748 | 0.36432 | 0.03571 | 1.07486 | 0.00306 | 0.41418 | 0.01434 |
| **Rab11fip4** | 0.17292 | 0.00811 | 0.17451 | 0.29236 | 0.00965 | 1.68685 | 0.00684 | 0.00392 | 0.50924 | 0.00000 | 0.26952 | 0.04573 | 0.59477 | 0.00636 | 0.05772 | 0.00625 |
| **Rab11fip5** | 2.46067 | 1.99966 | 4.08910 | 11.87363 | 0.48051 | 1.75061 | 0.14152 | 3.94713 | 13.33302 | 7.33845 | 6.51400 | 1.21951 | 4.84632 | 5.85777 | 6.39586 | 3.59275 |
| **Rab12** | 3.29973 | 0.22659 | 1.05116 | 1.07813 | 2.48064 | 0.01975 | 0.00268 | 1.26671 | 5.31432 | 3.36922 | 1.25427 | 0.00000 | 2.45767 | 0.99950 | 3.34005 | 0.00007 |
| **Rab13** | 1.86051 | 0.01581 | 0.41771 | 1.80110 | 0.01495 | 2.34362 | 1.84320 | 0.83239 | 6.18708 | 1.08469 | 0.00000 | 0.00000 | 0.00000 | 0.25281 | 0.17726 | 0.00351 |
| **Rab14** | 10.72367 | 8.21320 | 9.92441 | 16.31208 | 5.14701 | 5.91983 | 6.25869 | 5.45757 | 9.75500 | 9.92720 | 5.34500 | 10.26975 | 7.62749 | 2.66050 | 12.32015 | 3.31073 |
| **Rab15** | 0.01646 | 0.00746 | 0.00488 | 0.72992 | 0.01946 | 0.00651 | 1.45997 | 0.00884 | 0.00000 | 0.07910 | 0.01186 | 0.02602 | 0.00866 | 0.10499 | 0.04104 | 3.52466 |
| **Rab17** | 0.25671 | 0.01799 | 7.03140 | 0.00000 | 0.21793 | 4.60058 | 0.02277 | 2.28788 | 0.22680 | 0.00000 | 0.87141 | 0.00000 | 6.50125 | 0.00791 | 0.00516 | 0.00746 |
| **Rab18** | 14.98809 | 12.57196 | 14.27869 | 4.20198 | 15.08789 | 11.07052 | 8.53300 | 13.97781 | 3.49397 | 4.14120 | 9.90286 | 9.83978 | 7.84539 | 5.09995 | 4.80288 | 2.47580 |
| **Rab19** | 0.08135 | 0.00744 | 0.00753 | 0.00799 | 0.00786 | 0.00651 | 0.00718 | 0.01049 | 0.23048 | 1.68777 | 0.00756 | 0.01992 | 0.01076 | 0.00760 | 0.66336 | 0.00716 |
| **Rab1a** | 24.00089 | 15.72921 | 18.60491 | 10.79790 | 19.38727 | 11.85996 | 14.51074 | 21.87697 | 15.69218 | 14.88680 | 13.19320 | 5.81435 | 16.82509 | 14.86832 | 9.51447 | 10.36730 |
| **Rab1b** | 8.69730 | 3.85314 | 1.15060 | 0.00000 | 6.46634 | 3.58311 | 0.18726 | 5.03243 | 0.01231 | 3.82282 | 1.81605 | 0.10403 | 10.80310 | 0.09128 | 0.08417 | 2.75662 |
| **Rab20** | 4.22992 | 0.20936 | 5.58104 | 0.98883 | 1.30134 | 4.78328 | 2.99082 | 12.91618 | 1.24986 | 17.72935 | 1.39846 | 1.24272 | 8.10281 | 0.23518 | 5.80317 | 0.81134 |
| **Rab21** | 4.18286 | 0.01304 | 0.61298 | 6.16155 | 0.01798 | 0.03499 | 0.29836 | 0.00000 | 1.72004 | 1.19662 | 0.06346 | 0.15910 | 0.46303 | 0.00000 | 0.66375 | 1.25407 |
| **Rab22a** | 2.56755 | 2.50898 | 0.57751 | 4.43516 | 1.98862 | 0.45594 | 0.01345 | 2.75002 | 1.87837 | 0.30021 | 0.35557 | 5.07332 | 0.32156 | 0.00000 | 1.29553 | 1.25501 |
| **Rab23** | 3.58682 | 1.42478 | 1.31434 | 5.13474 | 0.04019 | 0.00000 | 2.26096 | 1.88844 | 2.66957 | 3.62569 | 2.20494 | 2.67199 | 0.60325 | 0.14657 | 1.85486 | 0.00000 |
| **Rab24** | 4.67431 | 2.97927 | 5.66875 | 9.34962 | 6.21001 | 2.17455 | 1.53736 | 2.28270 | 8.83045 | 4.11319 | 6.08895 | 1.01408 | 4.02189 | 1.63033 | 1.79007 | 0.01597 |
| **Rab25** | 0.00540 | 0.00767 | 0.01153 | 0.00742 | 0.00656 | 0.00630 | 0.00710 | 0.01030 | 0.01179 | 0.90596 | 0.01237 | 0.00489 | 0.00618 | 0.01300 | 0.00814 | 3.67213 |
| **Rab26** | 0.47505 | 0.01806 | 0.26530 | 0.32514 | 0.01603 | 0.01187 | 0.00716 | 0.00354 | 0.26018 | 1.41921 | 0.21784 | 0.08463 | 0.23661 | 0.00698 | 0.03658 | 0.01519 |
| **Rab26os** | 0.01402 | 0.00797 | 0.31216 | 0.65907 | 0.00853 | 0.00538 | 0.00728 | 0.00902 | 2.20857 | 0.04269 | 0.50358 | 0.03269 | 0.06165 | 0.12928 | 0.21726 | 0.03626 |
| **Rab27a** | 0.70389 | 0.04072 | 0.35327 | 1.80979 | 0.47905 | 0.00480 | 0.03875 | 0.03197 | 1.22641 | 0.01487 | 1.02178 | 0.08506 | 0.69190 | 0.62249 | 0.50447 | 0.27375 |
| **Rab27b** | 0.02723 | 0.01817 | 0.04050 | 0.04573 | 0.01781 | 0.01363 | 0.01773 | 0.01424 | 0.02098 | 0.04542 | 0.13997 | 0.00636 | 0.05683 | 0.10378 | 0.03203 | 0.01494 |
| **Rab28** | 0.69068 | 0.03629 | 0.59731 | 0.00000 | 0.02331 | 0.00407 | 1.44563 | 0.36032 | 0.53407 | 0.25918 | 0.08406 | 7.48871 | 0.77661 | 0.01034 | 0.94757 | 0.27073 |
| **Rab29** | 1.98050 | 1.01503 | 10.76917 | 0.00000 | 7.58974 | 0.01924 | 4.01710 | 7.48360 | 1.66337 | 3.60305 | 7.20405 | 0.00898 | 4.02157 | 1.47604 | 0.40488 | 0.01233 |
| **Rab2a** | 13.27946 | 10.29350 | 10.19826 | 7.88868 | 10.06546 | 6.06940 | 5.72585 | 5.96111 | 4.32800 | 12.42374 | 11.96225 | 0.82342 | 11.30492 | 5.82398 | 6.85121 | 0.63816 |
| **Rab2b** | 0.00000 | 0.00653 | 1.99121 | 5.67476 | 0.26922 | 0.00545 | 0.14725 | 0.82484 | 4.85238 | 1.53314 | 2.73393 | 1.02447 | 0.00000 | 5.28483 | 2.36590 | 0.00474 |
| **Rab30** | 0.00000 | 4.94880 | 1.19121 | 1.41273 | 0.01352 | 0.00546 | 2.92822 | 0.31059 | 0.39303 | 0.85968 | 0.00763 | 0.07687 | 0.11977 | 1.19604 | 1.34256 | 1.91469 |
| **Rab31** | 1.36842 | 0.00499 | 0.88914 | 0.27767 | 0.02709 | 0.01741 | 3.05157 | 1.80987 | 0.61795 | 0.22916 | 0.98788 | 0.04624 | 0.94983 | 1.29566 | 0.40216 | 0.01223 |
| **Rab32** | 0.01350 | 0.00806 | 0.01230 | 1.99273 | 0.00922 | 0.00642 | 0.00728 | 0.01125 | 0.00000 | 0.51427 | 0.01294 | 0.00686 | 0.01192 | 0.00938 | 0.00332 | 0.00711 |
| **Rab33b** | 0.81236 | 1.03620 | 0.83711 | 11.13077 | 2.75596 | 0.24333 | 0.14531 | 0.61526 | 12.71507 | 0.62642 | 12.23662 | 1.01795 | 0.74313 | 6.03621 | 2.30132 | 0.19760 |
| **Rab34** | 0.31577 | 0.00622 | 0.64981 | 4.00001 | 0.01630 | 0.00496 | 2.35625 | 1.71993 | 0.87855 | 9.71222 | 2.46448 | 0.42559 | 0.13360 | 2.02661 | 3.70523 | 0.00569 |
| **Rab35** | 9.85420 | 2.48717 | 5.18482 | 12.91878 | 5.68223 | 2.04663 | 3.97606 | 6.12706 | 15.26880 | 8.40777 | 4.05320 | 4.46852 | 5.60099 | 0.03184 | 1.23044 | 3.71807 |
| **Rab38** | 8.26347 | 0.00000 | 0.00000 | 0.46618 | 0.01932 | 0.00830 | 0.00000 | 0.15423 | 0.69532 | 0.23762 | 0.00478 | 0.04006 | 1.90308 | 0.00466 | 1.02463 | 0.00000 |
| **Rab39** | 1.80605 | 0.84372 | 0.38807 | 5.89912 | 0.01311 | 0.29007 | 1.29364 | 3.07113 | 4.95602 | 4.18776 | 3.59607 | 7.72738 | 0.56248 | 6.74064 | 5.39523 | 0.00185 |
| **Rab39b** | 0.01365 | 0.00811 | 0.01197 | 0.00464 | 0.00931 | 0.00664 | 0.00755 | 0.01386 | 0.67795 | 0.01190 | 0.00790 | 0.00713 | 0.01291 | 0.01381 | 0.00679 | 0.00738 |
| **Rab3a** | 0.74671 | 0.01670 | 0.00000 | 1.30839 | 0.00705 | 3.48739 | 0.00646 | 0.00000 | 2.34903 | 1.42500 | 0.68566 | 0.04395 | 0.05145 | 0.00407 | 0.00545 | 0.00531 |
| **Rab3b** | 2.03476 | 0.00000 | 2.09808 | 1.51556 | 9.08392 | 0.10115 | 0.00436 | 3.88264 | 1.75402 | 1.19928 | 0.68675 | 0.00000 | 1.66982 | 0.02854 | 0.17347 | 0.00304 |
| **Rab3c** | 0.01158 | 0.00787 | 0.02482 | 0.06029 | 0.01925 | 0.00787 | 0.03869 | 0.01887 | 0.08895 | 0.03090 | 0.01170 | 0.00727 | 0.03092 | 0.11821 | 0.01622 | 0.02212 |
| **Rab3d** | 0.07354 | 0.02951 | 0.12479 | 13.05007 | 0.01194 | 2.98548 | 0.00496 | 1.19803 | 8.51254 | 0.00000 | 1.57214 | 0.04223 | 1.44341 | 1.11334 | 5.21156 | 0.09516 |
| **Rab3gap1** | 1.89218 | 0.00397 | 1.94954 | 1.77412 | 0.24010 | 0.00478 | 0.82501 | 3.70847 | 3.73869 | 1.49884 | 2.28777 | 1.53676 | 0.00000 | 1.29684 | 2.45114 | 1.15776 |
| **Rab3gap2** | 1.77549 | 0.06515 | 0.12188 | 2.20476 | 0.33771 | 2.52999 | 0.03871 | 0.62788 | 0.93545 | 0.90085 | 0.64514 | 0.20950 | 0.17053 | 0.28834 | 1.40418 | 0.01103 |
| **Rab3il1** | 6.86661 | 4.24180 | 11.22348 | 2.31973 | 0.09823 | 2.93818 | 7.60861 | 18.15704 | 0.28085 | 20.85436 | 9.89918 | 8.45865 | 11.51014 | 2.89549 | 11.85125 | 1.68376 |
| **Rab3ip** | 2.62171 | 4.54848 | 1.21163 | 9.11901 | 6.61395 | 4.31237 | 1.37524 | 8.59994 | 9.03996 | 4.98002 | 3.20578 | 0.64628 | 5.93240 | 0.00967 | 3.28716 | 1.49979 |
| **Rab40b** | 0.00190 | 0.00730 | 0.00371 | 0.06447 | 0.01025 | 0.00639 | 0.00698 | 1.27993 | 0.45017 | 1.39398 | 0.14354 | 0.01126 | 0.01124 | 2.49499 | 0.61249 | 0.00714 |
| **Rab40c** | 3.02817 | 6.08161 | 6.63556 | 5.45879 | 0.07189 | 3.17875 | 4.64395 | 6.87655 | 11.52269 | 10.07655 | 2.47480 | 2.14376 | 3.61039 | 0.13746 | 7.69615 | 3.01880 |
| **Rab42** | 0.08152 | 0.00771 | 0.00880 | 0.59451 | 0.00786 | 0.00669 | 0.00748 | 0.01163 | 0.11747 | 0.11231 | 0.01124 | 0.01742 | 0.01184 | 0.02298 | 0.12899 | 0.00696 |
| **Rab43** | 0.00000 | 0.02834 | 0.07080 | 10.48613 | 0.23760 | 0.71015 | 0.01666 | 0.02375 | 12.55489 | 2.50446 | 0.97862 | 0.02304 | 0.00000 | 0.00000 | 3.04153 | 1.16477 |
| **Rab4a** | 0.00000 | 0.75606 | 1.51534 | 2.14717 | 0.00546 | 0.23995 | 0.00628 | 0.43175 | 1.11571 | 1.34448 | 0.10345 | 0.00000 | 2.62420 | 0.00198 | 0.00000 | 0.00491 |
| **Rab4b** | 2.30615 | 1.00072 | 0.00000 | 0.20794 | 0.00373 | 2.36879 | 0.00384 | 0.77302 | 0.23356 | 0.88205 | 0.88972 | 0.15904 | 1.07303 | 0.01701 | 0.00000 | 2.26640 |
| **Rab5a** | 7.32228 | 10.62078 | 5.22937 | 14.43584 | 5.52858 | 2.77387 | 3.50349 | 17.19002 | 18.04815 | 19.21205 | 17.15076 | 26.84672 | 7.16643 | 22.83312 | 16.30853 | 1.05028 |
| **Rab5b** | 2.46111 | 3.45815 | 5.14480 | 2.56470 | 0.63451 | 2.91427 | 3.30658 | 4.62948 | 2.06019 | 5.84491 | 4.82948 | 0.29706 | 3.19883 | 8.11646 | 4.38342 | 0.01981 |
| **Rab5c** | 4.86293 | 1.79630 | 6.00133 | 2.62107 | 0.57570 | 4.33402 | 2.84598 | 6.43440 | 1.40863 | 9.87575 | 5.97140 | 0.58639 | 7.74091 | 0.07885 | 0.00000 | 1.57668 |
| **Rab6a** | 8.59265 | 9.23197 | 6.40115 | 16.90850 | 8.27143 | 7.25555 | 2.81722 | 6.34322 | 17.23104 | 5.35944 | 7.31684 | 8.33770 | 3.88286 | 6.88827 | 8.58143 | 2.17374 |
| **Rab6b** | 0.00929 | 0.00780 | 0.68565 | 0.00660 | 0.01751 | 0.01724 | 0.00719 | 0.01007 | 0.03882 | 0.84107 | 0.01099 | 0.00540 | 0.35807 | 0.01394 | 0.00189 | 0.01490 |
| **Rab7** | 22.52250 | 14.16003 | 17.85556 | 15.73264 | 20.15444 | 15.71886 | 16.39088 | 27.96847 | 22.75097 | 36.89125 | 23.77844 | 18.24758 | 22.68973 | 5.44150 | 21.35570 | 10.71540 |
| **Rab8a** | 8.81367 | 5.37207 | 7.44759 | 5.47464 | 12.02058 | 9.13828 | 7.97600 | 13.85863 | 5.44587 | 14.23341 | 4.93000 | 0.88029 | 9.07016 | 0.14847 | 2.31107 | 0.77774 |

| | | | | | | | | | | | | | | | | |
|---|---|---|---|---|---|---|---|---|---|---|---|---|---|---|---|---|
| **Rab8b** | 0.93985 | 0.02521 | 0.07931 | 3.13487 | 0.00762 | 0.00407 | 4.59350 | 0.01010 | 2.00565 | 1.01278 | 0.00000 | 3.46902 | 0.01580 | 0.42875 | 1.03840 | 0.02615 |
| **Rab9** | 7.91022 | 2.37958 | 10.95005 | 8.02994 | 9.55474 | 4.76973 | 3.52298 | 5.30000 | 10.62481 | 6.63153 | 4.90637 | 0.54740 | 11.59842 | 2.78499 | 10.09536 | 1.28703 |
| **Rabac1** | 14.09732 | 0.11856 | 2.43568 | 2.47505 | 6.83588 | 3.96816 | 1.54246 | 9.10989 | 1.21013 | 9.87487 | 10.56407 | 0.10774 | 12.18748 | 0.00000 | 0.00000 | 5.26219 |
| **Rabep1** | 2.16140 | 2.43105 | 1.03490 | 6.48589 | 4.51678 | 0.19848 | 1.21975 | 3.40606 | 8.44028 | 7.93538 | 8.23922 | 6.06698 | 1.20674 | 17.72171 | 11.27063 | 1.19602 |
| **Rabep2** | 0.49918 | 0.06184 | 0.22326 | 2.84238 | 7.11900 | 3.71764 | 0.46948 | 2.01891 | 6.76527 | 11.67843 | 5.40459 | 0.82574 | 2.98713 | 0.02116 | 1.07329 | 1.32255 |
| **Rabepk** | 3.05214 | 6.27927 | 4.00774 | 0.20924 | 3.81224 | 0.03615 | 5.34654 | 2.77620 | 1.83431 | 3.63578 | 1.06616 | 0.85020 | 2.74057 | 0.00000 | 1.13694 | 4.47172 |
| **Rabgap1** | 0.00000 | 0.04874 | 1.46195 | 3.05523 | 0.00642 | 0.00570 | 0.01548 | 1.38437 | 2.22112 | 1.91053 | 2.86626 | 2.52143 | 0.59982 | 4.03740 | 3.06093 | 0.01239 |
| **Rabgap1l** | 0.01050 | 0.01672 | 0.00494 | 1.50089 | 0.00941 | 0.00667 | 1.81315 | 0.00381 | 0.73636 | 0.00000 | 0.17791 | 0.01177 | 0.00902 | 0.00356 | 0.40235 | 0.00679 |
| **Rabgef1** | 0.04731 | 2.08775 | 2.31629 | 8.52206 | 1.67779 | 2.99963 | 1.20605 | 4.46676 | 9.91561 | 4.11726 | 3.76883 | 0.60921 | 3.39438 | 4.97476 | 16.55182 | 0.13768 |
| **Rabggta** | 1.23430 | 0.04416 | 6.26208 | 1.71965 | 6.68966 | 0.03316 | 1.26366 | 7.89490 | 1.36496 | 6.71511 | 8.60001 | 0.48272 | 5.40792 | 0.35510 | 0.64980 | 3.17430 |
| **Rabggtb** | 12.56577 | 6.69320 | 6.62023 | 3.40902 | 8.37395 | 9.31529 | 7.39280 | 17.57782 | 6.13426 | 2.73517 | 13.35422 | 15.03917 | 4.95371 | 18.92426 | 11.81493 | 7.07465 |
| **Rabif** | 7.91235 | 5.72452 | 7.03387 | 6.82876 | 6.26388 | 7.24249 | 3.61597 | 7.23712 | 8.27500 | 9.16156 | 5.22864 | 7.83128 | 3.56634 | 2.78276 | 7.07491 | 4.25474 |
| **Rabl2** | 2.66652 | 1.23047 | 2.98256 | 1.24666 | 2.88049 | 0.01568 | 2.36077 | 1.92763 | 0.69361 | 1.77165 | 1.06382 | 1.65205 | 0.68276 | 1.12616 | 0.65524 | 0.68202 |
| **Rabl3** | 2.00775 | 6.64022 | 3.60814 | 1.01126 | 5.03927 | 0.00488 | 6.95964 | 0.00000 | 0.03094 | 2.72473 | 2.18930 | 0.11564 | 2.62037 | 2.54871 | 1.25326 | 1.26273 |
| **Rabl6** | 3.19909 | 4.66975 | 5.42657 | 0.04570 | 3.97532 | 3.82015 | 0.63996 | 1.50442 | 0.72899 | 2.59680 | 0.52028 | 0.14433 | 2.76265 | 0.00379 | 0.06691 | 0.16204 |
| **Rac1** | 11.45108 | 8.71080 | 7.27644 | 19.99726 | 4.85914 | 7.22888 | 8.16215 | 7.21652 | 18.86513 | 10.57202 | 14.10503 | 1.34031 | 6.45693 | 7.74542 | 6.23766 | 1.82102 |
| **Rac2** | 0.02858 | 0.01860 | 0.12796 | 0.22822 | 0.00979 | 0.01325 | 0.01729 | 0.01980 | 0.14386 | 0.63230 | 0.47343 | 0.04953 | 0.03246 | 0.38394 | 0.04418 | 0.06826 |
| **Rac3** | 0.01329 | 0.00750 | 0.07783 | 0.99986 | 0.01124 | 0.02112 | 0.00618 | 4.36285 | 0.55140 | 0.44352 | 0.83994 | 0.00000 | 1.54729 | 0.01304 | 0.00000 | 0.02130 |
| **Racgap1** | 12.20529 | 3.23123 | 12.69283 | 19.64119 | 8.52861 | 7.26367 | 6.29099 | 17.89924 | 28.04387 | 29.75868 | 24.14294 | 14.71377 | 10.06933 | 14.86807 | 23.05379 | 2.91174 |
| **Rack1** | 55.89185 | 30.31810 | 49.16913 | 13.76866 | 35.68493 | 23.55302 | 30.29252 | 56.46687 | 28.39606 | 62.07029 | 47.87979 | 27.51606 | 57.48267 | 20.97111 | 29.58094 | 33.20950 |
| **Rad1** | 4.83330 | 0.23739 | 8.80598 | 6.28196 | 5.06062 | 2.74302 | 3.14225 | 2.75276 | 6.69160 | 9.02535 | 6.37549 | 2.68233 | 4.42062 | 6.37501 | 7.11861 | 2.30492 |
| **Rad17** | 3.26875 | 0.47729 | 1.89196 | 12.81321 | 1.54805 | 1.68469 | 3.32760 | 7.19938 | 10.88124 | 12.87090 | 4.04301 | 3.18128 | 0.16172 | 1.31485 | 0.95250 | 0.10057 |
| **Rad18** | 1.09238 | 3.04572 | 2.47735 | 0.42960 | 0.00450 | 0.00599 | 0.02615 | 1.19423 | 1.70934 | 0.80222 | 0.39689 | 0.47964 | 0.00000 | 0.33268 | 0.99532 | 0.03401 |
| **Rad21** | 21.95815 | 15.00762 | 11.79609 | 18.72656 | 22.49678 | 13.09508 | 11.98072 | 24.16408 | 25.39159 | 32.59473 | 16.92863 | 23.76677 | 13.90224 | 35.22683 | 33.09998 | 7.87971 |
| **Rad23a** | 0.63411 | 0.13022 | 0.06448 | 0.00788 | 0.40284 | 0.31721 | 0.02717 | 1.02532 | 0.01073 | 0.00000 | 0.03941 | 0.01643 | 0.17820 | 0.00757 | 0.16406 | 0.01341 |
| **Rad23b** | 15.34323 | 7.94951 | 9.27523 | 15.35407 | 9.24533 | 4.08958 | 3.87007 | 10.70389 | 16.20616 | 7.70845 | 17.35113 | 4.94050 | 10.48969 | 15.68233 | 5.78268 | 5.15538 |
| **Rad50** | 1.94615 | 6.41371 | 2.95710 | 6.03562 | 5.27153 | 0.00291 | 3.91123 | 0.04621 | 3.77649 | 2.59906 | 1.58688 | 0.08523 | 0.12134 | 3.32247 | 5.30447 | 1.82946 |
| **Rad51** | 10.29725 | 5.15573 | 7.16843 | 8.09838 | 13.75866 | 7.09937 | 7.95875 | 3.94388 | 4.68306 | 12.20076 | 6.49790 | 1.16818 | 6.49371 | 7.78686 | 10.66206 | 4.52714 |
| **Rad51ap1** | 6.70082 | 1.66622 | 7.49410 | 7.40444 | 7.53139 | 6.04038 | 5.12273 | 3.79176 | 18.61313 | 11.80893 | 6.66206 | 13.46208 | 4.88544 | 0.75896 | 10.78795 | 3.53241 |
| **Rad51b** | 0.08751 | 4.88098 | 1.00129 | 0.01084 | 0.38427 | 0.11909 | 0.00584 | 0.32963 | 0.26349 | 0.30215 | 0.09990 | 0.11578 | 0.00000 | 0.61572 | 0.90660 | 0.00503 |
| **Rad51c** | 0.08560 | 5.99319 | 0.98132 | 2.90301 | 0.03845 | 0.03882 | 0.00499 | 2.11595 | 3.36679 | 1.59945 | 0.67204 | 0.12781 | 0.62306 | 3.36572 | 2.81876 | 1.16587 |
| **Rad51d** | 4.30441 | 0.03505 | 5.22822 | 4.30777 | 0.20906 | 1.14464 | 0.77276 | 5.55614 | 0.59399 | 3.65489 | 3.48249 | 0.48581 | 2.97282 | 2.54745 | 1.04726 | 3.16459 |
| **Rad52** | 1.63235 | 1.96114 | 1.10043 | 2.86729 | 7.70529 | 4.01548 | 1.06378 | 4.59957 | 2.10614 | 0.00000 | 3.96880 | 0.11428 | 4.04691 | 0.00000 | 0.00000 | 2.95057 |
| **Rad54b** | 1.11901 | 1.96447 | 3.97976 | 4.98042 | 0.18558 | 0.01893 | 5.51315 | 4.75671 | 5.16742 | 3.40992 | 5.02737 | 8.81705 | 4.17096 | 7.60082 | 1.92816 | 0.81331 |
| **Rad54l** | 6.54781 | 3.98061 | 5.44668 | 1.31211 | 3.01954 | 1.86071 | 4.43095 | 3.03176 | 3.26270 | 4.54302 | 1.00548 | 0.14783 | 1.01555 | 3.67212 | 4.38405 | 2.79346 |
| **Rad54l2** | 3.40162 | 2.35868 | 4.45471 | 6.04608 | 1.90862 | 1.94473 | 1.14379 | 5.02691 | 6.58195 | 8.72188 | 7.15783 | 0.66803 | 1.98354 | 2.68648 | 7.81954 | 2.08908 |
| **Rad9a** | 3.88134 | 3.32745 | 5.80940 | 1.94379 | 0.00668 | 4.32425 | 0.16403 | 4.16695 | 2.47362 | 4.51890 | 0.94189 | 4.89570 | 6.25553 | 0.06615 | 8.17105 | 0.52647 |
| **Rad9b** | 0.35679 | 0.00821 | 0.35919 | 0.97236 | 0.00723 | 0.00557 | 0.00778 | 0.46276 | 3.08672 | 0.19011 | 0.00000 | 0.00000 | 0.21993 | 0.28717 | 0.28383 | 0.00711 |
| **Radil** | 0.00448 | 0.00830 | 0.00357 | 0.06741 | 0.01836 | 0.00791 | 0.00707 | 0.18577 | 0.45850 | 0.91782 | 0.89542 | 0.00000 | 1.55304 | 0.17408 | 0.03971 | 0.00688 |
| **Rae1** | 14.56978 | 2.45156 | 13.48253 | 8.28880 | 5.44435 | 0.74511 | 5.13497 | 11.90972 | 7.60792 | 8.85947 | 10.92432 | 8.10729 | 10.28953 | 1.31154 | 4.56134 | 4.61142 |
| **Raet1a** | 0.64108 | 0.00567 | 0.00289 | 0.00896 | 2.14271 | 0.00771 | 0.00668 | 0.08516 | 0.01127 | 0.07950 | 0.01178 | 0.00467 | 0.05379 | 0.01156 | 0.00938 | 0.00662 |
| **Raet1d** | 0.19859 | 0.00717 | 0.00844 | 0.01020 | 0.60044 | 0.00662 | 0.00726 | 0.01603 | 0.01206 | 0.01631 | 0.01265 | 0.00720 | 0.04765 | 0.01226 | 0.01058 | 0.00707 |
| **Raet1e** | 1.06627 | 0.00461 | 0.01009 | 0.00000 | 3.60714 | 0.00541 | 0.01670 | 0.00915 | 0.01032 | 0.01077 | 0.27578 | 0.01158 | 0.12196 | 0.00730 | 0.03646 | 0.00606 |
| **Raf1** | 3.29405 | 1.95172 | 3.06082 | 13.49119 | 4.23053 | 2.36322 | 2.42674 | 5.71675 | 8.05670 | 1.35392 | 1.48943 | 0.38484 | 1.95766 | 4.18241 | 15.16274 | 0.95291 |
| **Rai1** | 0.41912 | 0.00702 | 0.02752 | 0.18513 | 0.10732 | 0.00775 | 0.17945 | 0.00947 | 0.00837 | 0.01826 | 0.01387 | 0.00727 | 0.01279 | 0.01128 | 0.00745 | 0.00698 |
| **Rai14** | 0.89949 | 0.00534 | 2.97805 | 3.92417 | 0.77035 | 0.15627 | 1.22930 | 0.00000 | 0.00000 | 2.58070 | 0.82786 | 0.01027 | 1.64379 | 0.00152 | 1.21049 | 0.35419 |
| **Rala** | 3.90517 | 1.52893 | 0.55077 | 4.80156 | 1.83556 | 1.91714 | 0.59960 | 1.75072 | 1.23355 | 1.03395 | 1.55047 | 0.12951 | 1.61058 | 0.04243 | 4.29835 | 0.83022 |
| **Ralb** | 8.02478 | 2.11047 | 3.30129 | 0.00000 | 0.99730 | 2.09097 | 2.32643 | 1.36439 | 0.10117 | 0.49284 | 0.56215 | 0.00000 | 0.75321 | 0.00381 | 0.26778 | 0.84251 |
| **Ralbp1** | 4.77615 | 1.10323 | 4.09116 | 9.62153 | 8.17622 | 9.84297 | 2.42567 | 6.70950 | 3.85250 | 11.28669 | 3.97772 | 6.30805 | 2.56580 | 4.69144 | 6.35593 | 3.88747 |
| **Ralgapa1** | 1.01838 | 0.00251 | 1.98309 | 1.50334 | 0.00321 | 4.85821 | 2.61945 | 0.61762 | 1.59728 | 0.97757 | 0.28127 | 8.13136 | 0.20219 | 0.34310 | 1.17307 | 0.04057 |
| **Ralgapa2** | 1.97745 | 0.00529 | 0.35092 | 0.39718 | 0.01264 | 0.90222 | 0.22912 | 2.01959 | 0.07782 | 0.34697 | 0.46484 | 0.00762 | 1.70301 | 0.02153 | 0.24638 | 0.00436 |
| **Ralgapb** | 5.43372 | 0.66243 | 1.25655 | 4.83610 | 0.01660 | 0.00000 | 0.30965 | 2.23353 | 1.37713 | 4.19642 | 0.79392 | 4.18139 | 1.07675 | 3.75871 | 5.67988 | 0.00000 |
| **Ralgds** | 0.05100 | 0.01767 | 2.42057 | 2.26568 | 0.82986 | 7.14951 | 0.00226 | 6.34532 | 4.04397 | 6.10060 | 2.13293 | 0.17964 | 6.27383 | 0.08481 | 0.60037 | 2.47321 |
| **Ralgps1** | 1.32338 | 0.00590 | 0.00137 | 2.12767 | 0.00526 | 0.00736 | 0.00586 | 1.04643 | 1.76371 | 0.47424 | 0.45033 | 0.00000 | 0.01152 | 0.00000 | 0.26343 | 0.00532 |
| **Ralgps2** | 6.10202 | 0.00000 | 4.93466 | 12.38753 | 1.96061 | 0.00479 | 2.47891 | 3.99708 | 6.33162 | 5.67966 | 2.09242 | 7.55665 | 3.33513 | 2.43977 | 3.83254 | 3.27598 |
| **Raly** | 2.75528 | 0.11232 | 0.96016 | 2.77580 | 11.01540 | 3.31396 | 0.62811 | 5.87422 | 1.12905 | 4.36603 | 1.85092 | 0.23147 | 0.64598 | 0.00950 | 0.17897 | 3.47911 |
| **Ramp1** | 0.01193 | 0.00775 | 0.12783 | 0.00760 | 0.00998 | 0.00628 | 0.00739 | 0.10665 | 0.01170 | 0.86445 | 0.01400 | 0.00464 | 0.09558 | 0.01219 | 0.00544 | 0.00800 |
| **Ramp2** | 1.24373 | 0.00408 | 0.00266 | 1.42597 | 2.25464 | 0.00508 | 0.03418 | 0.00000 | 1.23749 | 6.30598 | 7.02537 | 0.30432 | 0.00000 | 3.05034 | 3.48898 | 0.00443 |
| **Ramp3** | 0.01375 | 0.00794 | 0.01248 | 2.84270 | 0.00934 | 0.00636 | 0.00742 | 0.01376 | 0.03192 | 0.01221 | 0.01296 | 0.00756 | 0.01261 | 0.00345 | 0.06175 | 0.00691 |
| **Ran** | 44.88417 | 18.34032 | 38.88586 | 16.73261 | 33.29278 | 20.17854 | 22.00432 | 41.41357 | 27.04201 | 41.66510 | 36.44800 | 19.99499 | 37.21246 | 22.68130 | 16.17348 | 21.21447 |
| **Ranbp1** | 18.24193 | 8.96372 | 13.27385 | 3.25461 | 12.77758 | 8.21101 | 6.25322 | 10.44632 | 5.70345 | 30.14928 | 20.47710 | 7.76257 | 21.63100 | 3.44506 | 2.95318 | 15.79414 |
| **Ranbp10** | 5.37005 | 2.00501 | 3.76582 | 14.06930 | 2.13697 | 4.35657 | 1.14975 | 10.42438 | 16.30220 | 19.90526 | 7.88082 | 1.91036 | 2.98801 | 4.61143 | 11.50235 | 1.47447 |

| | | | | | | | | | | | | | | | | |
|---|---|---|---|---|---|---|---|---|---|---|---|---|---|---|---|---|
| **Ranbp17** | 4.72326 | 2.53748 | 0.00000 | 0.93596 | 2.61684 | 0.00201 | 1.22918 | 0.89689 | 1.45545 | 3.99122 | 1.07238 | 0.27839 | 0.85984 | 3.59202 | 2.83285 | 2.05371 |
| **Ranbp2** | 2.77495 | 3.49080 | 0.00000 | 16.33241 | 9.67901 | 7.58258 | 3.76664 | 4.76700 | 19.00612 | 7.78918 | 2.85222 | 8.92034 | 4.81430 | 20.65349 | 13.30047 | 0.96857 |
| **Ranbp3** | 10.02439 | 1.44968 | 2.30943 | 11.87474 | 4.87498 | 5.89368 | 1.27411 | 13.48677 | 8.86771 | 19.44264 | 8.13412 | 1.87854 | 8.66682 | 0.18117 | 5.12374 | 7.53814 |
| **Ranbp3l** | 0.06575 | 0.01789 | 0.31465 | 0.18841 | 0.04403 | 0.03916 | 0.10920 | 0.04549 | 0.18254 | 0.18135 | 0.56685 | 0.06143 | 0.24761 | 0.44536 | 0.06840 | 0.04305 |
| **Ranbp6** | 0.01055 | 0.01872 | 0.89587 | 0.05262 | 0.00756 | 0.00562 | 0.00715 | 0.09275 | 0.02071 | 0.95629 | 0.64537 | 0.01802 | 0.00000 | 0.00514 | 2.42031 | 0.01430 |
| **Ranbp9** | 4.47722 | 5.73147 | 5.13681 | 15.74607 | 3.87350 | 7.07633 | 6.67143 | 5.62826 | 7.45108 | 6.11546 | 1.35061 | 1.37570 | 5.77323 | 9.01613 | 13.11211 | 3.54018 |
| **Rangap1** | 15.56692 | 16.04591 | 20.16689 | 6.34055 | 13.03908 | 7.74681 | 13.06138 | 23.91673 | 7.59218 | 19.94376 | 11.75611 | 1.74853 | 20.10596 | 10.25000 | 16.25478 | 9.20361 |
| **Rangrf** | 11.11018 | 3.00292 | 5.11126 | 3.06125 | 5.78351 | 0.03373 | 0.35195 | 1.25446 | 1.48313 | 3.68565 | 3.42758 | 0.34659 | 3.59108 | 0.00000 | 0.00000 | 1.82414 |
| **Rap1a** | 8.85147 | 8.88244 | 9.23570 | 1.41569 | 10.84247 | 4.56239 | 10.95466 | 4.14805 | 2.05189 | 13.32424 | 12.82298 | 18.14653 | 7.91099 | 16.62088 | 4.65944 | 6.14674 |
| **Rap1b** | 6.00507 | 7.48909 | 6.35098 | 15.09817 | 4.37390 | 6.91110 | 8.61065 | 3.54361 | 25.87864 | 8.18483 | 9.05322 | 13.33333 | 6.90409 | 15.50598 | 2.68324 | 0.52725 |
| **Rap1gap** | 0.71779 | 0.00859 | 0.01220 | 0.00491 | 0.01234 | 0.00608 | 0.00727 | 0.27023 | 0.10045 | 0.19712 | 1.04044 | 0.00352 | 0.00020 | 0.02487 | 0.00000 | 0.00676 |
| **Rap1gap2** | 0.07461 | 0.00630 | 0.84247 | 1.63210 | 2.74570 | 0.00574 | 0.00742 | 0.00681 | 1.26777 | 0.64590 | 0.07845 | 0.03852 | 0.11082 | 0.16363 | 0.40263 | 0.01482 |
| **Rap1gds1** | 1.23873 | 3.35985 | 0.09832 | 6.93704 | 3.41063 | 0.00655 | 0.46846 | 0.05271 | 3.53818 | 1.48563 | 0.00000 | 0.30327 | 0.49398 | 0.05068 | 2.56933 | 0.01707 |
| **Rap2a** | 0.10603 | 0.00760 | 0.21832 | 4.19651 | 1.04169 | 0.01236 | 0.01846 | 1.06689 | 7.72569 | 1.79079 | 0.36521 | 0.89608 | 0.05572 | 1.00867 | 2.19189 | 0.02062 |
| **Rap2b** | 1.13213 | 0.00687 | 1.18490 | 0.20686 | 0.00559 | 0.01152 | 0.03895 | 0.01993 | 0.00717 | 0.00507 | 0.01346 | 0.01851 | 0.04450 | 0.00774 | 0.17702 | 0.01342 |
| **Rap2c** | 1.12454 | 7.03653 | 0.70374 | 0.42086 | 3.39494 | 0.00996 | 0.33506 | 1.32188 | 0.76115 | 5.57387 | 1.15431 | 0.89820 | 1.33020 | 8.56298 | 1.81022 | 0.00205 |
| **Rapgef1** | 8.06843 | 4.87397 | 0.94913 | 1.04336 | 0.00000 | 3.36385 | 5.95302 | 3.05488 | 1.12479 | 0.00000 | 2.13093 | 1.63806 | 2.96451 | 0.00000 | 1.01809 | 5.78090 |
| **Rapgef2** | 0.03973 | 0.01824 | 0.98907 | 1.76289 | 0.02713 | 1.26369 | 0.84155 | 0.72383 | 4.90474 | 0.85604 | 0.00000 | 5.24354 | 0.00000 | 0.01206 | 0.44405 | 0.02740 |
| **Rapgef4** | 0.05824 | 0.01869 | 0.46062 | 0.29273 | 0.02907 | 0.06073 | 0.02692 | 0.00519 | 0.41094 | 0.44233 | 1.77465 | 0.13716 | 0.60707 | 0.41809 | 0.07736 | 0.11195 |
| **Rapgef5** | 0.30000 | 0.00729 | 0.00000 | 0.94650 | 0.00727 | 0.00619 | 0.00704 | 0.01002 | 1.75989 | 0.00000 | 0.02286 | 0.05881 | 0.01224 | 0.00564 | 2.58006 | 0.00660 |
| **Rapgef6** | 4.75999 | 0.09298 | 2.95918 | 4.37879 | 0.07105 | 4.30577 | 1.21008 | 1.33683 | 4.13018 | 1.97514 | 1.66210 | 3.55359 | 1.10304 | 2.63549 | 7.92393 | 2.96289 |
| **Rapgefl1** | 0.01547 | 0.00753 | 0.24897 | 3.14658 | 0.02507 | 0.06727 | 0.00646 | 1.25113 | 0.74373 | 1.51993 | 0.94339 | 0.01249 | 0.36229 | 0.00000 | 0.11751 | 0.00673 |
| **Raph1** | 0.14008 | 0.06364 | 2.05380 | 1.14713 | 0.04535 | 5.10885 | 0.01489 | 0.47689 | 2.10431 | 0.94138 | 1.00424 | 0.45873 | 1.87780 | 5.43092 | 0.88373 | 0.27018 |
| **Rapsn** | 0.01819 | 0.00842 | 0.01714 | 0.68788 | 0.01060 | 0.00633 | 0.00755 | 0.01393 | 0.07177 | 0.01493 | 0.01356 | 0.00837 | 0.01253 | 0.01104 | 0.00407 | 0.00734 |
| **Rarb** | 1.72008 | 0.17742 | 0.31901 | 0.00000 | 0.00459 | 2.68640 | 0.09730 | 0.66889 | 0.55400 | 0.96794 | 0.00000 | 0.00695 | 1.53946 | 1.19265 | 0.35794 | 0.00353 |
| **Rarg** | 0.00000 | 0.00557 | 0.18143 | 8.43321 | 0.00827 | 0.00802 | 0.13577 | 8.55490 | 4.20489 | 8.27987 | 8.80304 | 0.32427 | 0.00088 | 0.00000 | 0.34820 | 0.00425 |
| **Rarres2** | 0.01363 | 0.00805 | 0.01211 | 0.00945 | 0.00916 | 0.00678 | 0.00750 | 0.01260 | 0.01211 | 0.27059 | 0.01221 | 0.00791 | 0.01252 | 0.01152 | 0.01118 | 0.00748 |
| **Rars** | 21.01259 | 14.89456 | 19.11807 | 13.58830 | 16.47025 | 8.52139 | 12.13799 | 19.84043 | 22.81844 | 22.34296 | 17.56398 | 12.61083 | 16.81327 | 24.88737 | 22.62870 | 8.24016 |
| **Rars2** | 10.75998 | 3.52528 | 6.56556 | 1.04741 | 5.18532 | 1.39375 | 2.07553 | 5.42311 | 0.12225 | 2.66100 | 3.71520 | 0.14896 | 4.48915 | 6.75265 | 0.63673 | 2.44791 |
| **Rasa1** | 3.28510 | 6.48834 | 4.70849 | 3.66099 | 8.31052 | 3.17366 | 3.51188 | 1.98632 | 2.10163 | 3.67045 | 1.31283 | 2.33564 | 1.16764 | 7.34046 | 11.72693 | 1.10104 |
| **Rasa2** | 0.25169 | 0.00483 | 0.03459 | 13.49683 | 0.00749 | 0.05657 | 3.34220 | 0.00000 | 2.83630 | 1.19682 | 1.76049 | 2.18828 | 0.01318 | 5.30956 | 6.66673 | 0.01128 |
| **Rasa3** | 0.01377 | 0.00812 | 0.01193 | 0.34312 | 0.00935 | 0.00652 | 0.00742 | 0.01215 | 0.07131 | 0.00668 | 0.02627 | 0.00783 | 0.01237 | 0.00667 | 0.25479 | 0.00730 |
| **Rasa4** | 0.02107 | 0.00841 | 0.19529 | 3.26296 | 0.00856 | 0.00553 | 0.00761 | 0.00748 | 0.31167 | 0.17332 | 0.28577 | 0.00611 | 0.01303 | 0.00387 | 0.00000 | 0.00653 |
| **Rasal1** | 0.97790 | 0.00737 | 0.00303 | 0.00556 | 0.01643 | 0.00708 | 0.00595 | 0.08113 | 0.16168 | 0.01445 | 0.00985 | 0.01559 | 0.90430 | 0.00879 | 0.03456 | 0.01324 |
| **Rasal2** | 2.50445 | 0.00550 | 0.00292 | 1.40473 | 0.02409 | 0.01884 | 0.00455 | 1.22457 | 0.25168 | 2.03494 | 1.57832 | 0.16579 | 0.93290 | 2.37195 | 0.77618 | 0.13786 |
| **Rasal3** | 0.02855 | 0.00668 | 0.14102 | 0.00909 | 2.82587 | 0.00563 | 0.00742 | 0.01003 | 0.01109 | 0.00681 | 0.01197 | 0.00579 | 0.38270 | 0.01400 | 0.00807 | 0.00686 |
| **Rasd1** | 0.01443 | 0.00805 | 0.00485 | 0.04137 | 0.00913 | 0.78064 | 0.00747 | 0.00808 | 0.38855 | 0.05290 | 0.16193 | 0.00709 | 0.00556 | 0.00805 | 0.10292 | 0.00693 |
| **Rasd2** | 0.01799 | 0.01755 | 0.04759 | 1.64171 | 0.02707 | 0.00807 | 0.00725 | 0.01568 | 0.00000 | 0.00509 | 0.01151 | 0.03391 | 0.01386 | 0.00411 | 0.76467 | 0.00707 |
| **Rasef** | 0.01372 | 0.00830 | 0.01185 | 1.30151 | 0.00932 | 0.00654 | 0.00745 | 0.01319 | 0.21460 | 0.01237 | 0.01225 | 0.01852 | 0.01281 | 0.03470 | 0.00000 | 0.00720 |
| **Rasgef1a** | 0.01372 | 0.00814 | 0.01230 | 0.01003 | 0.00923 | 0.00657 | 0.00753 | 0.01279 | 0.01241 | 0.13591 | 0.01195 | 0.00768 | 0.01174 | 0.01137 | 0.01047 | 0.00717 |
| **Rasgef1b** | 0.01369 | 0.00776 | 0.01212 | 0.58455 | 0.00935 | 0.00645 | 0.00743 | 0.01253 | 0.47137 | 0.00602 | 0.00974 | 0.00627 | 0.01269 | 0.00506 | 0.28168 | 0.00721 |
| **Rasgrf1** | 0.01405 | 0.00783 | 0.00770 | 0.30403 | 0.00929 | 0.13302 | 0.00744 | 0.01782 | 0.25936 | 0.01001 | 0.01169 | 0.00716 | 0.01041 | 0.00679 | 0.08263 | 0.00723 |
| **Rasgrp2** | 0.01195 | 0.00869 | 0.01016 | 0.11686 | 0.00925 | 0.00681 | 0.00794 | 0.05593 | 0.03186 | 0.21574 | 0.11943 | 0.00817 | 0.00867 | 0.00965 | 0.02118 | 0.00717 |
| **Rasgrp3** | 0.01267 | 0.00783 | 0.01221 | 0.61664 | 0.00965 | 0.00694 | 0.00741 | 0.02681 | 0.03936 | 0.37741 | 0.04268 | 0.00595 | 0.02662 | 0.07474 | 0.00283 | 0.00792 |
| **Rasgrp4** | 3.00450 | 2.91256 | 5.72748 | 0.55238 | 0.07671 | 2.83780 | 0.21339 | 4.94543 | 1.72897 | 0.56145 | 1.41446 | 0.04451 | 2.77958 | 0.00448 | 0.83562 | 0.00602 |
| **Rasip1** | 0.00971 | 0.00884 | 0.00841 | 0.00610 | 0.02645 | 0.00755 | 0.00828 | 0.71472 | 0.09436 | 0.81155 | 0.00775 | 0.01630 | 0.17979 | 0.01594 | 0.00001 | 1.69427 |
| **Rasl10a** | 0.05777 | 0.00741 | 0.00000 | 1.50528 | 0.03961 | 0.12529 | 0.00626 | 0.59888 | 0.51894 | 4.82059 | 3.14597 | 0.00000 | 0.00341 | 0.00000 | 0.00000 | 0.16174 |
| **Rasl11a** | 0.23064 | 0.00752 | 0.00387 | 0.04687 | 0.00741 | 0.00671 | 0.00712 | 0.00683 | 0.30617 | 1.76299 | 0.06466 | 0.03148 | 0.01052 | 0.01377 | 1.00475 | 0.00710 |
| **Rasl11b** | 0.02033 | 0.00792 | 0.02683 | 1.65048 | 0.00860 | 0.00568 | 0.00700 | 0.01041 | 1.91037 | 0.00194 | 0.00000 | 0.00478 | 0.56066 | 0.01386 | 0.00270 | 0.00651 |
| **Rasl12** | 0.01051 | 0.00768 | 0.01776 | 0.05723 | 0.00952 | 0.00617 | 0.01718 | 0.21062 | 0.02336 | 0.08626 | 0.04966 | 0.01585 | 0.03354 | 0.04833 | 0.06308 | 0.02830 |
| **Rasl2.9** | 5.63365 | 1.35133 | 5.18626 | 0.81112 | 5.29635 | 2.27221 | 2.37529 | 4.15919 | 1.82726 | 4.00313 | 4.52782 | 1.76415 | 4.07982 | 1.34278 | 0.45462 | 1.87852 |
| **Rassf1** | 4.16367 | 0.00000 | 2.17518 | 1.63195 | 1.72808 | 0.08604 | 1.87172 | 1.78276 | 0.00000 | 2.99164 | 0.90322 | 0.39766 | 0.02758 | 0.94134 | 0.93253 | 1.35423 |
| **Rassf3** | 2.26703 | 3.21857 | 1.65244 | 4.79568 | 2.71613 | 0.03116 | 0.45704 | 1.83648 | 10.28496 | 3.46315 | 6.55898 | 0.21141 | 0.00000 | 3.89352 | 3.26471 | 0.00000 |
| **Rassf4** | 0.00855 | 0.00791 | 0.01197 | 0.36818 | 0.00906 | 0.00649 | 0.00662 | 0.00953 | 0.00624 | 0.04553 | 0.01384 | 0.00684 | 1.29278 | 0.01046 | 0.00652 | 0.00705 |
| **Rassf5** | 0.01099 | 0.00798 | 0.02056 | 4.05549 | 0.00874 | 0.00714 | 0.00755 | 0.02066 | 1.27009 | 0.00000 | 0.31962 | 0.03259 | 0.00753 | 0.01487 | 0.58588 | 0.16801 |
| **Rassf7** | 4.56953 | 0.10061 | 0.00000 | 1.36319 | 2.92489 | 9.57857 | 0.78361 | 12.41272 | 2.61772 | 4.56533 | 7.04766 | 0.26635 | 10.37970 | 0.00323 | 0.00000 | 3.47846 |
| **Rassf8** | 0.02650 | 0.00777 | 0.63352 | 0.05917 | 0.00807 | 0.46380 | 0.00741 | 0.74839 | 0.15147 | 0.87998 | 0.18777 | 0.01149 | 0.10214 | 0.00844 | 0.02172 | 0.00716 |
| **Raver1** | 7.39613 | 1.17113 | 2.99481 | 3.96873 | 2.10779 | 6.17441 | 0.28692 | 7.59017 | 4.70584 | 6.56453 | 5.45768 | 0.87901 | 4.25896 | 0.25200 | 2.30527 | 3.00671 |
| **Raver2** | 0.01268 | 0.00803 | 0.01335 | 0.18093 | 0.00997 | 0.00739 | 0.00778 | 0.01533 | 0.00572 | 0.28782 | 0.01155 | 0.00780 | 0.14975 | 0.01312 | 0.20217 | 0.00769 |
| **Rb1** | 0.01166 | 0.00768 | 0.00644 | 10.72563 | 0.00909 | 0.00501 | 0.00692 | 0.01319 | 0.85315 | 1.16423 | 0.04612 | 0.05981 | 0.01046 | 0.00000 | 6.33361 | 0.00534 |
| **Rb1cc1** | 4.03284 | 5.88182 | 5.87213 | 16.93489 | 4.13120 | 0.00000 | 3.79950 | 11.20298 | 19.38646 | 36.15169 | 21.69623 | 21.05069 | 5.77990 | 45.84961 | 33.51084 | 0.00773 |
| **Rbak** | 0.39511 | 0.02878 | 0.48014 | 0.83470 | 0.31733 | 0.04476 | 0.09746 | 0.00000 | 0.67590 | 0.37389 | 1.71573 | 0.09611 | 0.40907 | 0.53926 | 1.40657 | 0.08290 |

| | | | | | | | | | | | | | | | | |
|---|---|---|---|---|---|---|---|---|---|---|---|---|---|---|---|---|
| **Rbakdn** | 0.01274 | 0.00811 | 0.09571 | 0.01011 | 0.00905 | 0.00877 | 0.00740 | 0.01302 | 0.01215 | 0.01281 | 0.01280 | 0.00816 | 0.01125 | 0.00932 | 0.05572 | 0.00735 |
| **Rbbp4** | 2.60237 | 4.03962 | 6.22150 | 6.93504 | 0.22913 | 3.22264 | 0.18164 | 1.93021 | 12.27820 | 2.23877 | 13.30920 | 0.70275 | 2.01005 | 10.01243 | 5.12916 | 2.68986 |
| **Rbbp5** | 4.41803 | 5.48802 | 5.89156 | 2.46193 | 0.00000 | 4.14795 | 3.95343 | 3.81588 | 2.25711 | 7.24012 | 1.85973 | 13.72285 | 1.89645 | 10.76319 | 7.27742 | 0.03033 |
| **Rbbp6** | 3.28794 | 8.74833 | 4.57443 | 20.00965 | 6.55938 | 1.76969 | 5.35661 | 8.44568 | 22.95249 | 8.66544 | 2.52812 | 5.61147 | 2.36710 | 6.34986 | 9.97947 | 0.44573 |
| **Rbbp7** | 30.60783 | 17.19037 | 28.06407 | 10.91475 | 24.73708 | 13.82865 | 20.24671 | 35.48119 | 15.38228 | 26.79411 | 12.12326 | 17.95545 | 16.88187 | 29.01091 | 19.36950 | 13.90436 |
| **Rbbp8** | 2.63555 | 6.24482 | 0.10952 | 6.77643 | 3.04648 | 1.21962 | 2.37226 | 7.51043 | 4.71752 | 12.43695 | 7.40037 | 9.39562 | 1.79702 | 16.82114 | 13.32661 | 1.56200 |
| **Rbbp9** | 1.34066 | 0.00183 | 0.42291 | 0.00000 | 2.52753 | 2.55964 | 0.68387 | 1.83647 | 0.44611 | 0.38593 | 0.48211 | 7.31938 | 0.00000 | 0.00666 | 2.21971 | 0.00442 |
| **Rbck1** | 3.97952 | 0.10278 | 0.81983 | 2.77491 | 2.14159 | 0.78259 | 1.66231 | 6.90548 | 0.34473 | 4.06639 | 2.01338 | 0.17040 | 3.62900 | 0.00000 | 0.13321 | 0.04463 |
| **Rbfa** | 0.86615 | 0.00605 | 0.09562 | 0.00429 | 1.12901 | 0.00579 | 0.04725 | 1.26204 | 0.00929 | 2.24995 | 0.00000 | 0.00758 | 0.00835 | 0.01530 | 0.00000 | 0.04479 |
| **Rbfox2** | 2.81871 | 0.01523 | 5.24194 | 5.35286 | 0.38953 | 0.00938 | 0.56824 | 1.49566 | 6.16955 | 4.14486 | 1.04126 | 0.07876 | 1.93853 | 0.93301 | 0.72004 | 1.21496 |
| **Rbks** | 2.64113 | 0.00497 | 0.02295 | 0.00480 | 0.00268 | 0.00984 | 0.16036 | 0.77553 | 0.01017 | 0.00000 | 0.90069 | 0.00000 | 1.84126 | 0.01142 | 0.00000 | 1.83746 |
| **Rbl1** | 1.59869 | 3.18664 | 0.00000 | 1.47170 | 0.00601 | 0.76328 | 0.00461 | 0.40875 | 1.04945 | 0.00000 | 0.83396 | 1.79404 | 0.06646 | 0.33293 | 1.08010 | 0.00363 |
| **Rbl2** | 0.16681 | 3.16958 | 0.14786 | 0.01049 | 0.01421 | 0.00895 | 1.96562 | 0.06678 | 0.01099 | 0.39241 | 0.00830 | 0.00416 | 0.00847 | 0.00629 | 1.09145 | 0.01324 |
| **Rbm10** | 10.01572 | 4.12100 | 3.06590 | 11.51345 | 1.48897 | 6.37270 | 4.89566 | 9.44265 | 11.51727 | 4.99714 | 3.85847 | 0.79817 | 6.35318 | 0.59955 | 7.54605 | 4.66089 |
| **Rbm11** | 0.14943 | 0.02955 | 0.09011 | 0.19410 | 0.03485 | 0.03230 | 0.03780 | 0.02084 | 0.47802 | 0.04486 | 0.59594 | 0.06162 | 0.07232 | 0.14176 | 0.18130 | 0.02197 |
| **Rbm12** | 1.51786 | 0.00134 | 1.11007 | 4.66282 | 4.87862 | 0.06552 | 1.11272 | 3.89884 | 6.97264 | 5.00383 | 1.58942 | 2.26170 | 2.26965 | 0.01255 | 3.75830 | 2.69758 |
| **Rbm12b1** | 0.57064 | 0.02755 | 0.08440 | 2.76846 | 0.01132 | 0.00475 | 0.08428 | 0.27799 | 1.52432 | 2.00045 | 0.74415 | 0.02859 | 0.00011 | 0.08327 | 2.86189 | 0.00601 |
| **Rbm12b2** | 0.42242 | 0.99426 | 0.02684 | 3.57448 | 0.00715 | 0.00507 | 2.13947 | 0.89219 | 2.33457 | 0.20610 | 1.00025 | 0.04176 | 0.01394 | 1.65648 | 4.15171 | 0.00524 |
| **Rbm14** | 5.52253 | 2.85201 | 3.16418 | 3.89038 | 0.06328 | 5.57775 | 3.69097 | 11.84862 | 6.66701 | 13.85778 | 10.56694 | 1.72630 | 4.61711 | 1.03818 | 0.86121 | 1.14373 |
| **Rbm15** | 1.95015 | 0.13088 | 0.10365 | 9.10589 | 0.01960 | 1.73954 | 2.54302 | 0.09473 | 10.38669 | 1.08009 | 2.59435 | 6.44395 | 0.58535 | 0.16827 | 1.21894 | 0.00000 |
| **Rbm15b** | 1.46133 | 0.07488 | 0.00000 | 2.26167 | 0.00489 | 0.46553 | 0.41122 | 2.55173 | 1.46632 | 0.57290 | 0.00000 | 0.08071 | 0.22505 | 0.00077 | 0.60700 | 0.04147 |
| **Rbm17** | 12.93767 | 7.10218 | 12.70553 | 12.40574 | 3.51423 | 6.01999 | 9.88311 | 11.17391 | 14.17359 | 16.23415 | 12.35748 | 1.66890 | 7.43191 | 17.11678 | 17.72599 | 3.69932 |
| **Rbm18** | 1.74561 | 2.16468 | 2.72771 | 16.53048 | 0.18859 | 4.95361 | 0.52518 | 2.72529 | 6.41212 | 8.23482 | 3.13343 | 7.34443 | 4.07851 | 5.53084 | 14.17621 | 5.20005 |
| **Rbm19** | 9.28491 | 3.05326 | 5.21120 | 6.33435 | 6.17406 | 0.01650 | 6.65430 | 7.08840 | 6.00601 | 6.37350 | 1.59443 | 2.91770 | 8.66437 | 0.03876 | 1.37279 | 0.07633 |
| **Rbm20** | 1.30926 | 0.00585 | 0.14086 | 0.01503 | 0.00532 | 0.00492 | 0.00627 | 0.00654 | 0.01235 | 0.19306 | 0.01234 | 0.00516 | 0.03273 | 0.00566 | 0.24945 | 0.00626 |
| **Rbm22** | 1.64175 | 3.18257 | 3.09443 | 14.20355 | 3.24519 | 0.00122 | 1.41058 | 3.17581 | 19.28670 | 12.08321 | 8.00632 | 4.33589 | 3.29934 | 2.50147 | 5.41003 | 1.45225 |
| **Rbm25** | 11.04396 | 9.99779 | 13.65036 | 11.87925 | 4.88588 | 0.04319 | 4.57942 | 10.38179 | 11.58882 | 10.01021 | 6.22512 | 12.38247 | 8.01192 | 21.39311 | 21.36932 | 4.58913 |
| **Rbm26** | 4.59327 | 2.02396 | 3.81922 | 4.56602 | 0.60623 | 1.71419 | 5.48199 | 2.50645 | 2.33323 | 6.33090 | 1.39463 | 9.75084 | 2.97677 | 5.77030 | 5.47418 | 1.06006 |
| **Rbm27** | 3.16252 | 0.10668 | 1.72001 | 12.51777 | 0.87881 | 2.60548 | 4.02997 | 2.51024 | 19.60868 | 5.44960 | 1.42612 | 6.49073 | 1.54151 | 8.37541 | 9.27758 | 2.04713 |
| **Rbm28** | 6.69906 | 9.76747 | 8.74113 | 10.65072 | 1.56326 | 8.08466 | 6.32720 | 14.73787 | 17.92755 | 17.04273 | 5.97491 | 5.94465 | 5.24322 | 9.66941 | 16.77040 | 3.73850 |
| **Rbm3** | 24.54399 | 19.36035 | 17.14545 | 9.24352 | 18.31299 | 8.93196 | 14.02317 | 26.24343 | 17.84501 | 27.01200 | 18.97363 | 16.17844 | 19.92490 | 32.29820 | 24.87181 | 16.78603 |
| **Rbm33** | 7.41558 | 4.91226 | 4.11523 | 8.53372 | 4.84577 | 4.85356 | 6.11857 | 5.60173 | 12.59060 | 9.32767 | 6.21118 | 5.87397 | 1.13769 | 5.28085 | 9.96438 | 1.48873 |
| **Rbm34** | 5.50271 | 4.73879 | 4.67694 | 1.67481 | 7.40928 | 0.03220 | 3.36972 | 3.95774 | 3.29899 | 9.55081 | 5.44395 | 8.06457 | 4.23070 | 7.05879 | 6.80341 | 2.94663 |
| **Rbm38** | 0.38760 | 0.00771 | 0.12762 | 12.68973 | 0.01556 | 0.72075 | 0.67178 | 4.21550 | 9.53675 | 0.02860 | 0.59893 | 0.01263 | 0.02189 | 0.00363 | 0.00000 | 0.04245 |
| **Rbm39** | 14.79064 | 15.08833 | 11.21809 | 14.75328 | 12.40917 | 4.60323 | 7.41781 | 13.82878 | 22.98228 | 22.35441 | 14.90719 | 24.13604 | 15.16935 | 22.87070 | 21.86781 | 8.80070 |
| **Rbm3os** | 0.00929 | 0.00750 | 0.00755 | 1.80392 | 0.00970 | 0.00651 | 0.00703 | 0.43534 | 0.81153 | 0.00000 | 0.00000 | 0.02183 | 0.00592 | 0.37756 | 0.12901 | 0.00696 |
| **Rbm4** | 0.01827 | 0.17057 | 0.23535 | 0.00986 | 0.00841 | 0.00608 | 0.00740 | 0.18647 | 0.01178 | 0.20465 | 0.01348 | 0.00721 | 0.02881 | 0.02331 | 0.02293 | 0.12974 |
| **Rbm41** | 1.38841 | 1.74280 | 0.00000 | 2.99993 | 0.82587 | 0.00199 | 0.04359 | 0.76885 | 0.74969 | 9.00710 | 4.93774 | 15.51729 | 0.10156 | 12.08693 | 7.74579 | 0.02430 |
| **Rbm42** | 15.76552 | 0.28675 | 2.26476 | 21.80791 | 10.74676 | 2.58260 | 2.67189 | 10.60036 | 29.59237 | 30.98034 | 20.84309 | 2.20483 | 27.38097 | 1.68628 | 2.34024 | 12.71981 |
| **Rbm44** | 3.48756 | 0.16767 | 0.78910 | 1.40716 | 0.04558 | 0.00231 | 0.00353 | 0.33457 | 0.58669 | 2.39699 | 0.45459 | 0.09837 | 1.63271 | 0.47678 | 2.52337 | 0.00960 |
| **Rbm45** | 0.00000 | 0.00693 | 0.00000 | 0.05486 | 0.06299 | 0.05641 | 0.00624 | 1.73685 | 0.38498 | 2.16303 | 2.35529 | 0.00000 | 0.21176 | 2.80759 | 0.76592 | 0.32828 |
| **Rbm46os** | 0.01318 | 0.00793 | 0.01125 | 0.54030 | 0.00947 | 0.00666 | 0.00743 | 0.02145 | 0.60303 | 0.01650 | 0.03409 | 0.00619 | 0.01427 | 0.07527 | 0.10436 | 0.00730 |
| **Rbm47** | 1.46445 | 0.70801 | 2.00695 | 0.02292 | 0.46958 | 2.10512 | 0.96866 | 11.46219 | 1.47745 | 1.16517 | 3.69664 | 0.20005 | 1.71194 | 11.84576 | 4.62275 | 0.30752 |
| **Rbm48** | 0.00000 | 0.00296 | 0.72747 | 2.83621 | 2.98209 | 0.04550 | 4.18248 | 2.98504 | 2.44012 | 8.88912 | 3.73730 | 0.27530 | 2.88565 | 1.35210 | 8.61329 | 0.16819 |
| **Rbm4b** | 0.01354 | 0.50459 | 0.36141 | 0.84309 | 0.07782 | 0.00604 | 1.33928 | 0.36653 | 1.61442 | 0.53312 | 0.85413 | 0.15107 | 0.06603 | 0.80569 | 0.24592 | 1.32158 |
| **Rbm5** | 6.87875 | 8.28072 | 7.74828 | 14.94735 | 3.63113 | 5.30728 | 5.08880 | 8.65908 | 16.38382 | 9.89342 | 4.36440 | 5.94765 | 8.10633 | 17.63987 | 11.88618 | 4.68388 |
| **Rbm6** | 6.27624 | 4.15086 | 7.95008 | 7.64487 | 6.57225 | 0.01985 | 7.42702 | 5.67840 | 10.63419 | 18.85243 | 12.92866 | 14.96846 | 8.53921 | 14.95272 | 17.35152 | 5.22758 |
| **Rbm7** | 6.97374 | 11.54079 | 6.64745 | 3.14748 | 4.26807 | 5.61305 | 6.43653 | 11.49194 | 4.00503 | 7.76848 | 3.58593 | 7.74289 | 5.82310 | 3.72839 | 7.27296 | 2.33121 |
| **Rbm8a** | 24.44320 | 10.45842 | 16.63589 | 15.49997 | 11.07117 | 7.81271 | 8.76884 | 17.35273 | 24.54308 | 24.04532 | 20.93701 | 3.33845 | 23.00845 | 4.48631 | 13.81669 | 9.55221 |
| **Rbms1** | 0.01002 | 0.00815 | 1.03947 | 0.00498 | 0.01110 | 0.00950 | 0.98047 | 0.39245 | 0.01080 | 2.13459 | 0.02758 | 0.01195 | 0.00241 | 0.03451 | 0.00000 | 0.01458 |
| **Rbms2** | 10.85332 | 3.03946 | 5.89026 | 1.39112 | 0.57853 | 3.96458 | 1.82012 | 5.36067 | 1.36942 | 11.69350 | 6.53592 | 0.86031 | 6.09835 | 6.06818 | 5.94069 | 0.26508 |
| **Rbmx** | 3.83027 | 3.66127 | 4.30698 | 7.31547 | 0.82067 | 0.12345 | 3.06288 | 5.21989 | 8.08435 | 2.21468 | 9.16466 | 0.49402 | 5.70248 | 10.94628 | 7.97752 | 0.13174 |
| **Rbmx2** | 3.05498 | 0.00000 | 5.11335 | 8.97763 | 4.93451 | 0.47889 | 3.81466 | 3.48945 | 10.00290 | 5.72589 | 3.86035 | 0.38075 | 5.12796 | 1.16653 | 8.23449 | 2.82585 |
| **Rbmxl1** | 21.80287 | 18.38722 | 22.55058 | 6.52500 | 20.25101 | 8.47783 | 10.76735 | 22.99769 | 13.02088 | 23.10989 | 15.14724 | 2.96334 | 16.36366 | 11.50198 | 11.26672 | 9.49381 |
| **Rbmxl2** | 0.80163 | 0.00763 | 0.09578 | 3.30536 | 0.01338 | 0.00484 | 0.00672 | 0.00706 | 0.00000 | 0.00382 | 0.52582 | 0.02487 | 0.01092 | 0.00151 | 0.00000 | 0.35838 |
| **Rbp1** | 0.01245 | 0.00772 | 0.01215 | 0.01491 | 0.00870 | 0.00660 | 0.00728 | 0.01019 | 0.01059 | 2.50214 | 0.01027 | 0.00000 | 0.01097 | 0.01233 | 0.01762 | 0.00761 |
| **Rbp7** | 0.00379 | 0.00725 | 0.00755 | 0.72884 | 0.00915 | 0.00642 | 0.00670 | 0.47491 | 2.55858 | 0.48396 | 0.00000 | 0.00000 | 0.64459 | 0.72321 | 0.00000 | 0.11962 |
| **Rbpj** | 14.70255 | 13.79132 | 13.38272 | 8.96463 | 13.27438 | 3.71170 | 11.90128 | 12.91208 | 11.34998 | 14.98582 | 16.15143 | 18.44067 | 5.88460 | 13.78750 | 10.72481 | 5.34459 |
| **Rbpms** | 7.03196 | 4.65281 | 3.98496 | 11.22404 | 0.84251 | 5.88441 | 4.30429 | 8.76737 | 16.84525 | 16.46160 | 20.01036 | 12.96642 | 2.59114 | 5.74422 | 10.03653 | 3.17923 |
| **Rbpms2** | 23.66555 | 8.30728 | 16.79043 | 20.33513 | 14.70992 | 9.18402 | 13.16872 | 22.34593 | 18.63245 | 24.83798 | 18.45679 | 14.35170 | 9.88002 | 1.65188 | 13.01846 | 9.68174 |
| **Rbsn** | 2.81322 | 2.43878 | 0.27530 | 16.85201 | 0.01912 | 1.17145 | 0.05089 | 1.81396 | 13.65128 | 14.36312 | 9.97205 | 11.22736 | 1.09725 | 23.27114 | 28.87411 | 0.90670 |
| **Rbx1** | 10.85126 | 5.67720 | 10.31412 | 17.13399 | 10.01052 | 5.57998 | 2.66041 | 3.31100 | 21.04903 | 11.60733 | 24.81677 | 2.34872 | 8.96839 | 7.47044 | 9.54395 | 6.27451 |

| | | | | | | | | | | | | | | | | |
|---|---|---|---|---|---|---|---|---|---|---|---|---|---|---|---|---|
| **Rc3h1** | 2.50020 | 3.41715 | 3.46048 | 10.33740 | 1.49214 | 0.00000 | 0.12894 | 4.69209 | 11.58857 | 6.53051 | 3.35016 | 8.66634 | 0.37158 | 8.23154 | 6.47062 | 0.69866 |
| **Rc3h2** | 3.67412 | 0.00934 | 0.54675 | 2.69352 | 0.03357 | 0.03649 | 1.02364 | 2.11935 | 1.66199 | 3.16790 | 2.98699 | 14.63744 | 0.10179 | 20.56571 | 5.64886 | 0.02378 |
| **Rcan1** | 3.83625 | 2.69348 | 5.73440 | 2.97502 | 0.35056 | 6.48445 | 0.00000 | 11.52572 | 1.60257 | 8.24115 | 6.82144 | 0.74925 | 8.94993 | 3.52446 | 0.86072 | 1.23690 |
| **Rcan3** | 0.61956 | 0.00710 | 0.59360 | 13.31931 | 0.01136 | 0.00386 | 0.00592 | 1.01584 | 2.53808 | 5.25970 | 0.85668 | 0.38511 | 0.00000 | 2.69239 | 10.04190 | 0.00382 |
| **Rcbtb1** | 1.78419 | 0.00549 | 0.00000 | 0.39908 | 0.45368 | 0.00505 | 0.00563 | 0.00193 | 0.53492 | 0.00000 | 0.00000 | 1.05972 | 0.01263 | 1.99668 | 0.14320 | 0.01253 |
| **Rcbtb2** | 0.01622 | 0.00813 | 0.13560 | 1.77153 | 0.00888 | 0.00566 | 0.00731 | 0.01128 | 0.25996 | 0.16022 | 0.02650 | 0.00767 | 0.03788 | 0.00929 | 0.00077 | 0.00690 |
| **Rcc1** | 11.90934 | 3.87399 | 8.77425 | 5.00419 | 5.05855 | 0.00000 | 7.66466 | 22.01931 | 6.32182 | 21.84349 | 11.39893 | 1.54479 | 10.27897 | 7.71364 | 14.34204 | 8.19769 |
| **Rcc2** | 22.67733 | 11.90541 | 18.31466 | 2.52654 | 15.95562 | 4.92805 | 15.23343 | 27.85914 | 3.75384 | 20.00445 | 16.34554 | 1.61830 | 19.98915 | 0.26728 | 5.07099 | 6.21120 |
| **Rccd1** | 0.01139 | 0.00698 | 1.19698 | 0.24943 | 0.00746 | 0.00546 | 0.45917 | 0.60747 | 0.77963 | 0.34859 | 0.66850 | 0.00000 | 0.00035 | 0.00952 | 0.28075 | 0.01348 |
| **Rce1** | 6.08701 | 1.40120 | 3.53364 | 1.59749 | 5.83807 | 2.72197 | 0.91401 | 5.70637 | 0.65278 | 7.49379 | 3.01281 | 0.23310 | 7.82160 | 0.62972 | 2.30685 | 5.37024 |
| **Rchy1** | 8.27084 | 7.27643 | 9.91590 | 16.32318 | 8.72595 | 4.01139 | 5.23164 | 7.98090 | 8.48405 | 14.14546 | 6.34352 | 3.96195 | 6.06982 | 8.29412 | 5.98598 | 5.65676 |
| **Rcl1** | 3.07349 | 5.48438 | 5.29398 | 6.45438 | 5.33280 | 5.01852 | 0.25147 | 12.51966 | 13.32586 | 23.52885 | 8.12283 | 0.95029 | 12.51163 | 4.73092 | 12.19923 | 2.28344 |
| **Rcn1** | 8.52792 | 0.00000 | 0.02240 | 0.00784 | 5.41219 | 0.00624 | 0.11852 | 0.00000 | 0.26629 | 1.79584 | 0.00440 | 0.03799 | 0.00619 | 0.43180 | 0.00000 | 4.69147 |
| **Rcn2** | 5.03708 | 0.01046 | 4.68212 | 2.99958 | 0.00000 | 3.04578 | 2.31649 | 1.47808 | 1.50033 | 2.02270 | 0.59653 | 0.01646 | 0.00000 | 1.30636 | 3.03150 | 1.58397 |
| **Rcn3** | 0.02656 | 0.00742 | 0.01108 | 1.36928 | 0.03676 | 0.00851 | 0.00607 | 0.91126 | 0.00000 | 2.58579 | 2.98600 | 0.06467 | 0.00000 | 0.00000 | 0.00006 | 0.00719 |
| **Rcor1** | 0.81356 | 0.00699 | 0.78842 | 4.81710 | 0.00504 | 0.24221 | 0.03808 | 0.41337 | 3.20016 | 2.74783 | 0.32391 | 0.05976 | 0.00000 | 0.00000 | 4.58491 | 0.00491 |
| **Rcor2** | 1.69799 | 0.00560 | 0.14951 | 8.48218 | 0.00993 | 0.02319 | 0.00192 | 0.56255 | 6.03670 | 4.06829 | 4.97767 | 0.07610 | 3.32709 | 0.02441 | 1.49546 | 0.00177 |
| **Rcor3** | 0.27824 | 0.00803 | 0.30064 | 0.62310 | 0.01782 | 0.00885 | 0.06297 | 0.57061 | 0.03989 | 1.08188 | 0.99140 | 0.21734 | 0.92819 | 0.00407 | 0.00000 | 1.08802 |
| **Rcsd1** | 0.01044 | 0.00669 | 0.00700 | 0.84138 | 0.00955 | 0.00667 | 2.23966 | 0.00923 | 0.53597 | 0.01096 | 0.01018 | 0.00537 | 0.01025 | 0.00873 | 0.02758 | 0.00717 |
| **Rd3** | 0.01277 | 0.00805 | 0.01328 | 0.01460 | 0.00988 | 0.00680 | 0.00748 | 0.01266 | 0.61949 | 0.25562 | 0.01055 | 0.00576 | 0.14184 | 0.01133 | 0.01036 | 0.00763 |
| **Rdh1** | 0.03452 | 0.01833 | 0.15762 | 0.13975 | 0.02750 | 0.04563 | 0.06743 | 0.05139 | 0.36922 | 0.05478 | 0.14323 | 0.05364 | 0.04086 | 0.25603 | 0.10043 | 0.05031 |
| **Rdh10** | 1.41820 | 2.92127 | 7.04356 | 4.30836 | 2.41415 | 0.07850 | 4.99254 | 6.80901 | 5.92278 | 5.40319 | 4.24423 | 8.01715 | 2.46099 | 1.36770 | 11.52575 | 0.39372 |
| **Rdh11** | 0.35335 | 3.61907 | 6.54276 | 1.76465 | 0.00422 | 0.00509 | 7.93021 | 0.00486 | 0.51200 | 0.00000 | 0.00000 | 0.02876 | 0.00391 | 4.09710 | 0.44110 | 0.00906 |
| **Rdh12** | 0.00970 | 0.00781 | 0.09253 | 2.76148 | 0.00958 | 0.00841 | 0.00697 | 0.01289 | 0.24420 | 0.00000 | 0.00866 | 0.03549 | 0.01138 | 1.79066 | 0.99277 | 0.00685 |
| **Rdh13** | 2.34895 | 0.00333 | 4.63038 | 0.18286 | 0.00000 | 0.00273 | 0.03252 | 2.23004 | 1.81406 | 5.88922 | 4.01192 | 2.30224 | 5.23279 | 4.46860 | 0.83231 | 4.06563 |
| **Rdh14** | 1.08711 | 0.00325 | 0.03567 | 0.87609 | 6.02864 | 0.00723 | 0.01605 | 0.00547 | 0.00000 | 0.04894 | 0.00802 | 0.00761 | 0.01602 | 0.62861 | 0.01498 | 0.01370 |
| **Rdh16** | 0.01232 | 0.00803 | 0.01976 | 0.56497 | 0.00924 | 0.08172 | 0.00744 | 0.03113 | 1.01991 | 0.00650 | 0.07968 | 0.00611 | 0.00960 | 0.03056 | 0.01261 | 0.00719 |
| **Rdh18.ps** | 0.12205 | 0.00773 | 0.06009 | 0.13581 | 0.03379 | 0.42024 | 0.00739 | 0.02328 | 0.44538 | 0.27275 | 0.06698 | 0.00595 | 0.02232 | 0.13035 | 0.19207 | 0.00706 |
| **Rdh19** | 0.04226 | 0.00791 | 0.03595 | 0.01430 | 0.01036 | 0.27846 | 0.01754 | 0.01144 | 0.30786 | 0.01342 | 0.00923 | 0.00710 | 0.00904 | 0.05120 | 0.01116 | 0.00722 |
| **Rdh5** | 0.00624 | 0.85015 | 0.04708 | 0.01012 | 0.00788 | 0.01671 | 0.00689 | 0.00951 | 0.01235 | 0.00789 | 0.01064 | 0.28789 | 0.01110 | 0.00901 | 0.00000 | 0.00707 |
| **Rdh7** | 0.06460 | 0.00781 | 0.00410 | 0.00351 | 0.00799 | 0.56218 | 0.00744 | 0.01152 | 0.54507 | 0.00619 | 0.00750 | 0.00714 | 0.00744 | 0.03707 | 0.02019 | 0.00701 |
| **Rdh9** | 0.00809 | 0.00745 | 0.00938 | 1.55128 | 0.00953 | 0.00620 | 0.00712 | 0.84027 | 3.04266 | 0.09040 | 0.01844 | 0.00000 | 0.03095 | 0.00224 | 0.17820 | 0.02104 |
| **Rdm1** | 0.00000 | 1.39368 | 8.69102 | 6.85834 | 5.08343 | 3.11483 | 7.96380 | 5.27735 | 5.94811 | 8.36841 | 6.63873 | 0.44059 | 10.45556 | 0.02202 | 2.20394 | 6.08154 |
| **Rdx** | 7.08829 | 10.45881 | 7.16586 | 11.24475 | 10.12044 | 2.98552 | 7.83002 | 3.86891 | 5.37970 | 1.50649 | 2.30362 | 3.63429 | 2.23624 | 8.13385 | 9.42010 | 3.47010 |
| **Rec114** | 2.34672 | 0.47790 | 1.09081 | 0.28256 | 0.00203 | 0.00336 | 0.00579 | 0.39499 | 0.11301 | 0.21644 | 0.42807 | 0.00323 | 0.00348 | 0.00564 | 0.01157 | 0.00420 |
| **Rec8** | 6.34216 | 2.01367 | 6.11830 | 0.00000 | 0.33428 | 0.00600 | 3.10667 | 1.56436 | 0.01768 | 1.11747 | 2.40365 | 0.00000 | 4.18058 | 0.00485 | 0.00351 | 0.01644 |
| **Reck** | 0.35985 | 0.00829 | 0.00557 | 0.18854 | 0.01103 | 0.00714 | 0.00724 | 0.00953 | 0.08626 | 0.01105 | 0.18733 | 0.00774 | 0.01010 | 0.00920 | 0.01975 | 0.00700 |
| **Recql** | 1.26379 | 2.12741 | 0.60586 | 0.86724 | 0.00856 | 0.01286 | 0.01459 | 0.89521 | 0.75914 | 0.97554 | 0.04033 | 0.00000 | 1.99689 | 0.31177 | 1.03889 | 0.00392 |
| **Recql4** | 0.08085 | 1.14809 | 0.53350 | 0.01025 | 0.02398 | 0.01403 | 3.00718 | 0.36918 | 0.01049 | 1.55249 | 0.00452 | 0.02044 | 2.56186 | 0.33698 | 0.08162 | 0.01221 |
| **Recql5** | 5.96087 | 7.34474 | 5.90474 | 6.37039 | 2.56593 | 2.61984 | 5.19562 | 5.30966 | 2.99645 | 2.66553 | 1.78957 | 0.21764 | 2.52288 | 0.00000 | 1.09551 | 0.00000 |
| **Reep1** | 4.53879 | 7.87515 | 15.60933 | 0.05650 | 7.70553 | 12.09245 | 10.30742 | 16.02427 | 0.02296 | 7.42136 | 8.03442 | 0.25153 | 7.91955 | 7.06081 | 4.27868 | 1.59974 |
| **Reep2** | 0.01288 | 0.00769 | 0.01093 | 4.37559 | 0.00896 | 0.00592 | 0.00722 | 0.01048 | 3.05985 | 0.74941 | 0.00000 | 0.00000 | 0.01175 | 0.00000 | 0.41504 | 0.00667 |
| **Reep3** | 4.55836 | 10.00481 | 6.91173 | 0.95820 | 3.62719 | 0.25711 | 1.21540 | 1.76017 | 3.77651 | 5.75797 | 3.71769 | 0.43534 | 1.64149 | 2.23056 | 2.41984 | 0.05280 |
| **Reep4** | 0.19534 | 1.91883 | 2.20319 | 5.11886 | 6.45117 | 5.62144 | 0.00279 | 4.70592 | 0.44897 | 0.53907 | 2.15261 | 0.02668 | 3.94944 | 0.00649 | 0.23711 | 0.01704 |
| **Reep5** | 1.44655 | 0.01532 | 1.50374 | 3.58142 | 0.36170 | 2.33779 | 0.65615 | 0.57952 | 0.44479 | 1.13252 | 0.47448 | 0.04354 | 0.09371 | 0.13955 | 2.09360 | 2.10652 |
| **Reep6** | 0.02648 | 0.00835 | 0.08613 | 1.16525 | 0.01266 | 0.01447 | 0.00662 | 0.03040 | 1.45563 | 2.00784 | 0.18096 | 0.00000 | 1.59297 | 0.01043 | 0.00000 | 0.15483 |
| **Reg1** | 0.01253 | 0.00758 | 0.01300 | 0.00403 | 0.00893 | 0.00755 | 0.00723 | 0.08670 | 2.74884 | 0.12588 | 0.00000 | 0.89487 | 0.00980 | 0.03680 | 0.00000 | 0.00717 |
| **Reg2** | 0.01212 | 0.00822 | 0.01429 | 0.01055 | 0.01019 | 0.00713 | 0.00768 | 0.01618 | 0.01235 | 0.00884 | 0.01517 | 0.00784 | 0.29842 | 0.01269 | 0.01023 | 0.00758 |
| **Rel** | 0.00804 | 0.00745 | 1.22964 | 0.41949 | 0.00782 | 0.00552 | 0.02678 | 0.48423 | 0.00504 | 0.15090 | 0.00862 | 0.00501 | 0.42532 | 0.00943 | 0.00000 | 0.00677 |
| **Rela** | 1.26078 | 0.02581 | 3.05250 | 4.41542 | 0.03407 | 0.04978 | 3.32078 | 5.41616 | 2.17114 | 2.92897 | 2.06617 | 0.25699 | 1.51791 | 0.03298 | 0.78437 | 2.75245 |
| **Relb** | 0.05283 | 0.00572 | 0.60259 | 3.37766 | 0.07942 | 0.33797 | 2.40678 | 0.00000 | 12.53452 | 8.55725 | 3.63133 | 0.41532 | 0.00000 | 0.00000 | 1.18395 | 0.01978 |
| **Rell1** | 1.67226 | 0.29712 | 3.43671 | 9.49241 | 0.01362 | 4.80745 | 4.23520 | 1.96344 | 3.39241 | 0.00000 | 0.31631 | 0.87337 | 2.01441 | 1.83001 | 1.85079 | 0.47799 |
| **Reln** | 0.01220 | 0.00772 | 0.00588 | 1.97642 | 0.00995 | 0.00632 | 0.00716 | 0.00989 | 0.00000 | 0.00134 | 0.00000 | 0.03517 | 0.01771 | 1.27092 | 1.77994 | 0.00721 |
| **Relt** | 0.00000 | 0.00557 | 1.14319 | 0.01221 | 2.59679 | 0.00730 | 0.00666 | 2.04420 | 0.12347 | 0.79826 | 0.37786 | 0.00652 | 0.50104 | 0.00750 | 0.30881 | 0.36637 |
| **Rem1** | 0.00631 | 0.00732 | 0.02033 | 0.01088 | 0.11946 | 0.00786 | 0.00713 | 0.45028 | 0.01179 | 0.00765 | 0.01055 | 0.00656 | 0.01990 | 0.01434 | 0.00346 | 0.00731 |
| **Renbp** | 10.92456 | 9.70067 | 23.54499 | 6.62360 | 2.02011 | 12.06010 | 10.61373 | 19.53862 | 4.62419 | 9.47849 | 10.97606 | 4.36958 | 17.79614 | 0.01049 | 0.00000 | 14.98666 |
| **Repin1** | 0.58186 | 0.00700 | 0.07175 | 0.53236 | 0.00669 | 0.00511 | 0.00704 | 0.00884 | 0.00089 | 0.25918 | 0.01047 | 0.00593 | 0.01056 | 0.00634 | 0.51061 | 0.20063 |
| **Reps1** | 8.04600 | 7.34197 | 7.20578 | 20.07375 | 5.79700 | 8.13995 | 3.46307 | 4.30192 | 6.84811 | 5.87847 | 4.69448 | 11.85576 | 3.04560 | 0.00000 | 12.50569 | 5.42497 |
| **Reps2** | 0.00000 | 1.99099 | 1.55442 | 0.00397 | 0.00882 | 0.03409 | 0.00558 | 0.09386 | 1.88559 | 2.26863 | 1.45476 | 0.01717 | 0.84542 | 4.54657 | 0.76335 | 0.01945 |
| **Rer1** | 5.79351 | 4.49041 | 4.00862 | 8.61161 | 6.88171 | 6.69420 | 2.69719 | 8.83152 | 9.01842 | 7.87490 | 10.24324 | 11.52959 | 8.90028 | 1.45123 | 2.12424 | 5.88411 |
| **Rere** | 7.35746 | 4.37538 | 3.01405 | 4.41359 | 4.94574 | 2.62128 | 0.00000 | 4.74473 | 6.89682 | 12.97387 | 3.15969 | 3.18830 | 2.40223 | 1.00304 | 6.92551 | 5.71019 |
| **Resp18** | 0.01089 | 0.00797 | 0.01176 | 0.84353 | 0.00875 | 0.00674 | 0.00750 | 0.01462 | 0.00000 | 0.22836 | 0.01175 | 0.00831 | 0.24050 | 0.00761 | 0.11174 | 0.17961 |

| | | | | | | | | | | | | | | | | |
|---|---|---|---|---|---|---|---|---|---|---|---|---|---|---|---|---|
| **Rest** | 1.49748 | 0.00595 | 0.00000 | 6.46495 | 0.63132 | 0.00969 | 0.15615 | 0.53997 | 2.32769 | 5.04803 | 0.34108 | 0.04548 | 2.00554 | 5.78557 | 2.84110 | 0.00345 |
| **Ret** | 0.01773 | 0.00808 | 0.01181 | 0.15667 | 0.00894 | 0.00659 | 0.00748 | 0.01431 | 0.00641 | 0.01489 | 0.01224 | 0.00819 | 0.02531 | 0.01166 | 0.00802 | 0.00731 |
| **Retsat** | 9.70344 | 0.00000 | 4.33935 | 0.11480 | 1.83312 | 1.31396 | 2.67227 | 0.18365 | 0.17206 | 2.84362 | 0.88010 | 0.07153 | 2.29262 | 0.96455 | 0.00000 | 0.22654 |
| **Rev1** | 0.00000 | 0.00649 | 0.00000 | 0.84102 | 0.74257 | 1.13621 | 2.49053 | 0.22465 | 1.41656 | 1.10203 | 0.00000 | 0.06120 | 0.43682 | 0.67023 | 4.26746 | 0.00652 |
| **Rev3l** | 1.80021 | 0.00341 | 0.00000 | 3.90197 | 2.54460 | 4.11389 | 1.08148 | 1.06573 | 1.54529 | 2.77133 | 0.52717 | 1.71705 | 0.13345 | 2.32663 | 5.50236 | 1.04572 |
| **Rex2** | 1.11208 | 3.11099 | 1.07366 | 9.52845 | 0.61005 | 0.04961 | 0.87725 | 2.57246 | 6.32622 | 0.25904 | 3.73453 | 11.61413 | 1.27411 | 19.27462 | 1.17851 | 0.21038 |
| **Rexo1** | 4.93635 | 0.80908 | 1.42079 | 5.40622 | 0.05137 | 0.01445 | 1.23937 | 5.19573 | 6.14000 | 9.68335 | 3.02815 | 0.69011 | 4.66739 | 0.03211 | 0.16330 | 2.33110 |
| **Rexo2** | 11.06596 | 2.88768 | 8.26980 | 2.68209 | 6.26382 | 0.07387 | 0.85923 | 2.59533 | 1.63250 | 2.91272 | 1.14927 | 2.10240 | 4.10713 | 0.09676 | 0.10424 | 2.75628 |
| **Rexo4** | 4.77625 | 3.38906 | 7.02651 | 13.12059 | 0.14168 | 5.61501 | 4.54727 | 5.92866 | 16.71986 | 11.39463 | 7.04733 | 8.65820 | 6.61546 | 3.13318 | 9.06606 | 5.72330 |
| **Rfc1** | 14.53253 | 9.48762 | 11.28778 | 6.65051 | 4.36425 | 3.32524 | 5.69919 | 7.64318 | 7.71841 | 19.48204 | 8.42477 | 10.91272 | 6.53017 | 24.82494 | 20.68215 | 5.11007 |
| **Rfc2** | 6.40289 | 3.94181 | 11.92153 | 1.40271 | 9.00030 | 7.74936 | 6.55448 | 14.56950 | 4.07258 | 8.29881 | 5.34321 | 3.35850 | 12.78287 | 0.55788 | 2.54372 | 10.16101 |
| **Rfc3** | 9.20994 | 3.82516 | 11.19522 | 2.37441 | 3.78630 | 4.02104 | 5.19850 | 9.38044 | 3.45123 | 10.49870 | 8.78455 | 1.84799 | 7.15143 | 2.77921 | 4.00661 | 7.05234 |
| **Rfc4** | 10.40541 | 10.72606 | 11.55414 | 0.57318 | 7.30786 | 3.53794 | 11.78126 | 7.08868 | 1.06526 | 10.33130 | 9.33098 | 3.92727 | 12.90271 | 3.80569 | 6.34870 | 6.50843 |
| **Rfc5** | 13.75795 | 7.64535 | 6.12701 | 7.27710 | 11.79885 | 5.50549 | 6.67402 | 5.17735 | 4.60398 | 6.12325 | 6.67585 | 3.07967 | 6.32543 | 5.99188 | 4.46173 | 4.33317 |
| **Rfesd** | 2.70314 | 0.00353 | 0.35920 | 0.01490 | 0.02125 | 0.65790 | 4.00554 | 0.10848 | 0.01048 | 0.38260 | 0.00456 | 0.61648 | 0.19958 | 0.01250 | 0.00000 | 0.00410 |
| **Rffl** | 0.00000 | 2.39321 | 4.51355 | 4.15651 | 3.49185 | 1.80172 | 4.58985 | 12.72011 | 7.91039 | 7.29965 | 5.40599 | 7.42290 | 1.21205 | 9.93368 | 5.62869 | 0.16230 |
| **Rfk** | 10.01261 | 5.08241 | 13.37287 | 9.69419 | 0.84014 | 8.33047 | 3.32330 | 10.87445 | 12.74518 | 11.14030 | 14.48047 | 5.22977 | 13.57519 | 12.74602 | 3.23563 | 5.26429 |
| **Rfng** | 3.18178 | 4.03794 | 2.97795 | 5.31838 | 0.02363 | 0.59094 | 2.97675 | 14.34016 | 7.22270 | 10.46627 | 3.77065 | 1.89461 | 10.03489 | 0.05282 | 3.67679 | 0.15356 |
| **Rfpl4** | 0.01204 | 0.00782 | 0.00030 | 2.63207 | 0.00941 | 0.00639 | 0.00692 | 0.00000 | 0.77510 | 0.00000 | 0.26565 | 0.00000 | 0.00966 | 2.22322 | 1.06541 | 0.00666 |
| **Rfpl4b** | 0.01247 | 0.00794 | 0.01024 | 9.96345 | 0.00907 | 0.00504 | 0.00714 | 0.01001 | 9.12837 | 0.00000 | 0.00000 | 0.00000 | 0.01307 | 0.04733 | 0.01290 | 0.00501 |
| **Rft1** | 1.48335 | 0.00561 | 0.32457 | 0.24886 | 0.06293 | 0.03526 | 0.22955 | 1.61803 | 0.56149 | 0.04820 | 1.27437 | 0.06133 | 0.62290 | 0.18249 | 2.29381 | 0.21574 |
| **Rftn1** | 0.78340 | 0.00679 | 0.00499 | 0.60369 | 0.00700 | 0.00764 | 0.00713 | 0.01196 | 0.11351 | 0.01251 | 0.02640 | 0.02708 | 0.21433 | 0.01791 | 0.06412 | 0.00671 |
| **Rftn2** | 0.00748 | 0.00741 | 0.02810 | 0.00033 | 0.00834 | 0.00881 | 0.00716 | 3.31030 | 0.27285 | 3.40854 | 0.86936 | 0.72455 | 0.00000 | 1.05465 | 0.88955 | 0.00713 |
| **Rfwd2** | 7.01824 | 3.65293 | 6.27220 | 22.69927 | 1.79918 | 0.00709 | 4.01843 | 9.09259 | 17.15440 | 13.26721 | 5.67932 | 6.48507 | 4.33357 | 19.51409 | 22.58197 | 0.70769 |
| **Rfwd3** | 10.53633 | 4.90762 | 7.97866 | 16.26112 | 5.79189 | 5.23382 | 6.40893 | 7.54425 | 17.25819 | 12.00452 | 8.36573 | 6.15236 | 4.23504 | 26.45007 | 16.34474 | 4.25946 |
| **Rfx1** | 0.00978 | 0.10864 | 0.37801 | 2.66962 | 0.02650 | 2.93278 | 0.22547 | 3.27406 | 4.09858 | 3.93749 | 1.64667 | 0.79688 | 2.68554 | 0.00407 | 0.00000 | 0.01176 |
| **Rfx2** | 9.20294 | 1.09581 | 2.43527 | 1.88741 | 0.73276 | 5.33357 | 6.46536 | 1.41657 | 1.57228 | 0.36285 | 0.00000 | 0.19189 | 0.31534 | 0.00085 | 1.78154 | 1.29378 |
| **Rfx3** | 0.00838 | 0.33187 | 0.02881 | 0.01713 | 0.01736 | 0.00668 | 0.01746 | 0.01537 | 0.02463 | 0.01534 | 0.01190 | 0.01717 | 0.02106 | 0.15698 | 0.02783 | 0.02180 |
| **Rfx4** | 1.04540 | 4.33305 | 1.18508 | 0.28631 | 0.00445 | 1.35228 | 0.00483 | 0.26307 | 0.00451 | 0.00194 | 0.01013 | 0.00541 | 0.34903 | 0.00582 | 0.00000 | 0.00369 |
| **Rfx5** | 0.12986 | 0.00742 | 0.06466 | 1.15156 | 0.00772 | 0.00558 | 0.00701 | 0.01044 | 0.52291 | 0.04797 | 0.00529 | 0.03166 | 0.01909 | 0.17338 | 0.87479 | 0.00725 |
| **Rfx7** | 1.22532 | 0.00420 | 1.85097 | 3.44164 | 1.89679 | 0.00355 | 0.04712 | 0.00417 | 0.64899 | 0.09533 | 0.00000 | 3.84556 | 0.05857 | 0.00049 | 3.23311 | 0.01192 |
| **Rfxank** | 2.68360 | 0.07992 | 4.90310 | 1.17913 | 0.30842 | 0.09723 | 0.72613 | 0.89634 | 2.36792 | 2.36046 | 6.31269 | 0.56545 | 4.71194 | 1.17941 | 2.51597 | 0.32724 |
| **Rfxap** | 1.64095 | 0.01563 | 0.06527 | 9.89340 | 1.70140 | 0.00243 | 2.38212 | 0.00000 | 12.68525 | 5.24478 | 2.32641 | 0.06319 | 0.03856 | 0.58259 | 0.26361 | 0.01745 |
| **Rgcc** | 0.35185 | 0.00810 | 0.19190 | 0.23113 | 0.13973 | 0.94318 | 0.00737 | 0.01982 | 0.95554 | 0.00383 | 0.00422 | 0.01631 | 0.01637 | 0.02399 | 0.06177 | 0.12060 |
| **Rgl1** | 0.06364 | 0.00757 | 0.22450 | 0.40874 | 0.00746 | 0.00882 | 0.00662 | 0.00000 | 0.00401 | 0.63168 | 0.99699 | 0.00720 | 0.00008 | 0.42726 | 0.64679 | 0.01424 |
| **Rgl2** | 2.18634 | 0.00448 | 0.00000 | 2.33070 | 0.02122 | 0.97980 | 0.00503 | 1.34441 | 1.71063 | 1.02353 | 0.92188 | 0.39310 | 0.10164 | 0.00000 | 0.00000 | 0.14487 |
| **Rgmb** | 0.00435 | 0.00714 | 0.28135 | 1.75918 | 0.29395 | 0.00852 | 0.00703 | 0.11712 | 3.24188 | 0.00000 | 0.00000 | 0.00000 | 0.11643 | 0.46393 | 0.14067 | 0.00691 |
| **Rgn** | 0.00000 | 0.02976 | 6.38342 | 4.22345 | 0.00000 | 0.00866 | 0.03763 | 0.04341 | 2.51905 | 4.97322 | 1.21810 | 0.19428 | 2.10797 | 11.83906 | 9.15302 | 0.01015 |
| **Rgp1** | 3.01386 | 2.15165 | 2.59367 | 3.28048 | 0.00900 | 1.66339 | 0.00284 | 1.59375 | 4.01176 | 0.87121 | 2.38752 | 0.24078 | 0.00000 | 0.01240 | 0.82956 | 1.21031 |
| **Rgs10** | 0.00845 | 0.00795 | 0.26285 | 0.20721 | 0.00913 | 0.00916 | 0.00739 | 0.27701 | 0.36067 | 0.09850 | 0.00762 | 0.00616 | 0.00557 | 0.01143 | 0.00000 | 0.00760 |
| **Rgs11** | 0.00056 | 0.00601 | 2.71793 | 1.02180 | 0.00508 | 0.00632 | 0.11654 | 8.51179 | 8.67261 | 3.39418 | 4.33458 | 1.35422 | 0.17612 | 0.28784 | 1.26162 | 0.00478 |
| **Rgs12** | 0.01768 | 0.00807 | 0.01014 | 0.63077 | 0.00913 | 0.00648 | 0.00725 | 0.01092 | 0.00000 | 0.17569 | 0.01051 | 0.01538 | 0.01265 | 0.16108 | 0.44546 | 0.00735 |
| **Rgs14** | 0.00885 | 0.01761 | 0.04016 | 0.22525 | 0.00937 | 0.00760 | 0.00701 | 1.05409 | 0.00518 | 0.00347 | 0.00593 | 0.00555 | 0.14481 | 0.01269 | 0.00000 | 0.00719 |
| **Rgs16** | 0.01365 | 0.00814 | 0.01181 | 0.09373 | 0.00932 | 0.00659 | 0.00751 | 0.01335 | 0.82779 | 0.00862 | 0.00745 | 0.01763 | 0.01288 | 0.00892 | 0.05264 | 0.00734 |
| **Rgs17** | 0.02462 | 0.00761 | 0.01120 | 2.37877 | 0.00859 | 0.01248 | 0.03737 | 0.06573 | 0.16565 | 0.64225 | 0.03794 | 0.00419 | 0.00915 | 0.19065 | 0.25929 | 0.01422 |
| **Rgs19** | 3.22536 | 0.00465 | 0.02988 | 1.14357 | 0.01254 | 0.00597 | 0.00516 | 0.60799 | 0.18155 | 0.64753 | 0.71967 | 0.19238 | 0.42008 | 0.00334 | 1.09006 | 0.02538 |
| **Rgs2** | 0.01216 | 0.00744 | 0.00847 | 0.66827 | 0.00907 | 0.00652 | 0.00706 | 0.02198 | 0.80494 | 1.15381 | 0.32300 | 0.00983 | 0.00825 | 0.00709 | 0.31484 | 0.00725 |
| **Rgs20** | 0.02306 | 0.01787 | 0.01967 | 0.01781 | 0.87873 | 0.04174 | 0.00753 | 0.02292 | 0.06140 | 0.04445 | 0.04114 | 0.02822 | 0.01602 | 0.06953 | 0.03616 | 0.02133 |
| **Rgs3** | 0.44452 | 0.00735 | 0.38777 | 0.41429 | 0.03306 | 0.00828 | 0.00573 | 0.17526 | 0.68292 | 0.00000 | 1.20451 | 0.02564 | 0.60502 | 0.00838 | 0.01222 | 4.35273 |
| **Rgs7bp** | 0.01208 | 0.01807 | 0.05115 | 0.07948 | 0.01849 | 0.02542 | 0.01751 | 0.01617 | 0.03285 | 0.04365 | 0.02314 | 0.01733 | 0.02407 | 0.25433 | 0.06045 | 0.02892 |
| **Rhag** | 0.01769 | 0.00774 | 0.01017 | 0.03994 | 0.00883 | 0.00666 | 0.00727 | 0.00968 | 0.10407 | 0.10908 | 0.06535 | 0.00637 | 0.03850 | 0.11763 | 0.04020 | 0.01512 |
| **Rhbdd1** | 0.18992 | 0.00667 | 2.44575 | 1.15895 | 0.01210 | 0.00741 | 0.89739 | 0.41023 | 0.84503 | 1.76377 | 0.77043 | 2.95501 | 2.81069 | 0.63388 | 0.95281 | 0.39625 |
| **Rhbdd2** | 3.27567 | 0.13824 | 2.55720 | 13.49081 | 4.30855 | 0.01180 | 0.06434 | 0.06822 | 10.32693 | 0.88992 | 0.17942 | 0.47542 | 0.94153 | 1.81349 | 2.93602 | 0.03589 |
| **Rhbdd3** | 2.19470 | 0.70701 | 0.63093 | 1.27344 | 0.74223 | 0.04747 | 2.19668 | 1.83522 | 0.04187 | 0.13299 | 0.34846 | 0.11082 | 2.31250 | 0.00297 | 0.00000 | 2.76621 |
| **Rhbdf1** | 0.00613 | 0.00732 | 0.00000 | 0.54204 | 0.00986 | 0.19672 | 0.34640 | 1.00691 | 0.00332 | 0.53159 | 1.10870 | 0.01676 | 0.00000 | 0.04952 | 0.00000 | 0.02919 |
| **Rhbdf2** | 0.04382 | 0.09411 | 0.08680 | 0.22990 | 0.02287 | 0.00870 | 0.00792 | 0.41289 | 0.00813 | 0.00713 | 0.09332 | 0.00732 | 0.00000 | 0.01096 | 0.00000 | 4.17250 |
| **Rhbdl1** | 0.51738 | 0.00825 | 0.00525 | 0.08478 | 0.01771 | 0.00710 | 0.00712 | 0.01138 | 0.02197 | 0.01467 | 0.03021 | 0.00782 | 0.01178 | 0.01177 | 0.00881 | 0.00708 |
| **Rhbdl2** | 1.97601 | 0.01419 | 0.55437 | 0.05733 | 2.60078 | 2.56455 | 1.19449 | 2.74385 | 0.14601 | 2.61604 | 0.68264 | 0.13788 | 0.66308 | 0.55053 | 0.85085 | 0.02564 |
| **Rhbdl3** | 0.14649 | 0.00779 | 0.13710 | 0.85332 | 0.00727 | 0.00852 | 0.01844 | 0.01804 | 1.83434 | 0.76301 | 0.00000 | 0.07715 | 0.01047 | 1.30159 | 1.99408 | 0.00661 |
| **Rhbg** | 0.00943 | 0.00808 | 0.00661 | 0.29247 | 0.00904 | 0.00698 | 0.00724 | 0.02016 | 0.78880 | 0.18315 | 0.68974 | 0.00243 | 0.73527 | 0.04179 | 0.00000 | 0.00716 |
| **Rhcg** | 0.01438 | 0.00764 | 0.14556 | 0.74241 | 0.00869 | 0.00591 | 0.01736 | 0.25820 | 0.17587 | 3.48279 | 1.74033 | 0.45744 | 0.00001 | 0.03768 | 2.15082 | 0.00707 |
| **Rhd** | 0.43003 | 0.00724 | 0.06203 | 2.66735 | 0.00701 | 0.00538 | 0.00688 | 0.00950 | 0.00000 | 0.65187 | 0.04606 | 0.01590 | 0.01131 | 0.08758 | 0.90801 | 0.02070 |

| | | | | | | | | | | | | | | | | |
|---|---|---|---|---|---|---|---|---|---|---|---|---|---|---|---|---|
| **Rheb** | 12.57880 | 5.29259 | 4.24517 | 8.36273 | 11.59553 | 6.45280 | 4.81307 | 4.02524 | 12.57176 | 11.48149 | 6.96794 | 11.13085 | 5.36726 | 5.22742 | 3.02517 | 3.47213 |
| **Rhebl1** | 6.62969 | 1.53025 | 4.76385 | 5.90231 | 5.39668 | 5.16967 | 1.20586 | 9.33700 | 4.01817 | 6.44664 | 1.59360 | 0.94987 | 9.68968 | 1.03564 | 1.18986 | 0.00000 |
| **Rhno1** | 0.54235 | 2.62863 | 5.82744 | 9.42162 | 6.48788 | 0.44437 | 1.22203 | 3.36658 | 7.09545 | 6.71045 | 3.02187 | 0.43279 | 4.95703 | 0.03556 | 2.40301 | 1.13212 |
| **Rhoa** | 14.13302 | 7.35435 | 11.91891 | 15.03276 | 7.16164 | 12.58179 | 9.90359 | 17.68112 | 19.45996 | 14.09799 | 18.04644 | 10.68622 | 14.79266 | 12.09721 | 12.54722 | 7.02678 |
| **Rhob** | 1.85467 | 0.00464 | 0.51875 | 0.49072 | 0.09892 | 1.19760 | 1.68436 | 1.66160 | 0.51524 | 0.66559 | 0.97885 | 0.05356 | 0.22149 | 0.03896 | 0.34622 | 1.62542 |
| **Rhobtb1** | 0.01325 | 0.00807 | 0.00230 | 1.07393 | 0.00896 | 0.00718 | 0.00689 | 0.00000 | 1.33035 | 0.07945 | 1.42136 | 0.07344 | 0.00000 | 0.00252 | 0.81143 | 0.00663 |
| **Rhobtb2** | 0.41596 | 0.00554 | 1.78351 | 1.77527 | 2.06325 | 2.21406 | 0.98753 | 0.17060 | 1.71643 | 3.71222 | 1.60774 | 0.57523 | 0.17245 | 0.00000 | 1.26459 | 0.01197 |
| **Rhoc** | 7.32317 | 1.75798 | 2.02400 | 0.09777 | 3.22982 | 3.19654 | 2.43930 | 0.83261 | 0.00632 | 1.62797 | 0.06982 | 0.04454 | 1.05481 | 0.00545 | 0.00000 | 5.29484 |
| **Rhod** | 1.02029 | 0.00725 | 1.34987 | 0.39319 | 1.03067 | 0.00448 | 0.01780 | 0.01221 | 0.39624 | 0.00000 | 0.03676 | 0.00535 | 0.01794 | 0.05015 | 0.00000 | 1.08014 |
| **Rhof** | 0.00000 | 0.00735 | 1.71296 | 2.79609 | 0.00756 | 0.00793 | 0.00446 | 0.00000 | 1.67536 | 0.54918 | 0.84094 | 0.05745 | 3.79424 | 0.00000 | 0.57557 | 0.00471 |
| **Rhog** | 1.16284 | 0.05879 | 0.64965 | 0.00000 | 5.12080 | 2.74553 | 0.34544 | 6.39397 | 0.00873 | 6.16370 | 6.42312 | 0.05498 | 3.88329 | 0.00000 | 0.07721 | 7.02230 |
| **Rhoh** | 0.04947 | 0.01947 | 0.11201 | 0.34711 | 0.01063 | 0.01217 | 0.01779 | 0.00622 | 1.87618 | 0.00000 | 0.40724 | 0.06720 | 0.05211 | 0.09892 | 0.33551 | 0.05789 |
| **Rhoq** | 0.01620 | 0.00820 | 0.59754 | 0.03351 | 0.00780 | 0.00581 | 0.00761 | 0.01282 | 0.01225 | 0.40907 | 0.01078 | 0.05964 | 0.01637 | 0.01345 | 0.00208 | 0.00710 |
| **Rhot1** | 6.78433 | 0.11195 | 5.31803 | 12.09172 | 9.81330 | 3.69510 | 0.01786 | 7.45709 | 9.20317 | 8.09102 | 3.99827 | 2.51468 | 3.69332 | 12.56650 | 9.68323 | 1.61062 |
| **Rhot2** | 0.54738 | 2.62494 | 2.68855 | 1.33103 | 8.26779 | 7.49347 | 2.18264 | 5.34400 | 2.16111 | 2.02594 | 0.60975 | 0.09894 | 3.77248 | 0.00000 | 1.28599 | 3.10516 |
| **Rhou** | 1.95168 | 0.00370 | 0.00741 | 0.46030 | 3.57366 | 0.00436 | 0.00569 | 0.13510 | 0.95102 | 1.78967 | 0.00000 | 0.00000 | 0.08641 | 0.00419 | 0.38117 | 2.27408 |
| **Rhov** | 0.01444 | 0.01826 | 0.81409 | 0.00759 | 0.00780 | 0.00585 | 0.00708 | 0.15298 | 0.02231 | 0.45512 | 0.30415 | 0.01692 | 0.00752 | 0.00858 | 0.01697 | 0.00699 |
| **Rhox13** | 0.01242 | 0.00754 | 0.00793 | 0.20995 | 0.00886 | 0.00679 | 0.00706 | 0.49709 | 0.00773 | 1.03986 | 0.01123 | 0.01520 | 0.00549 | 0.01844 | 0.00000 | 0.00705 |
| **Rhox3c** | 0.01369 | 0.00776 | 0.01134 | 0.01031 | 0.00938 | 0.00666 | 0.00742 | 0.01199 | 0.01226 | 0.00833 | 0.01188 | 0.00714 | 0.01297 | 0.12551 | 0.08652 | 0.00744 |
| **Rhox3e** | 0.01374 | 0.00786 | 0.01201 | 0.00996 | 0.00936 | 0.00661 | 0.00747 | 0.01285 | 0.01230 | 0.00917 | 0.01250 | 0.00748 | 0.01280 | 0.04462 | 0.10823 | 0.00742 |
| **Rhox3g** | 0.01371 | 0.00775 | 0.01165 | 0.00937 | 0.00941 | 0.00658 | 0.00742 | 0.01208 | 0.01208 | 0.00557 | 0.01183 | 0.00691 | 0.01294 | 0.07803 | 0.22181 | 0.00743 |
| **Rhox3h** | 0.01372 | 0.00778 | 0.01154 | 0.01029 | 0.00937 | 0.00664 | 0.00744 | 0.01224 | 0.01219 | 0.00833 | 0.01202 | 0.00727 | 0.01292 | 0.09280 | 0.09848 | 0.00744 |
| **Rhox5** | 0.07958 | 0.01519 | 8.26278 | 0.01144 | 0.00000 | 0.01072 | 0.01482 | 5.74815 | 0.01515 | 0.00000 | 0.00000 | 0.00828 | 1.24721 | 0.02302 | 0.00000 | 7.45665 |
| **Rhox6** | 0.01341 | 0.00769 | 0.06556 | 0.01052 | 0.00919 | 0.00746 | 0.00729 | 0.01102 | 0.01264 | 0.01123 | 0.01145 | 0.00667 | 0.01191 | 0.24851 | 0.00234 | 0.00741 |
| **Rhox7a** | 0.01351 | 0.00785 | 0.01210 | 0.54829 | 0.00928 | 0.00649 | 0.00746 | 0.01769 | 0.22858 | 0.02920 | 0.01085 | 0.00682 | 0.01198 | 0.00527 | 0.18875 | 0.00715 |
| **Rhox8** | 0.02651 | 0.00761 | 0.05605 | 0.14855 | 0.00871 | 0.01293 | 0.00735 | 0.01087 | 0.20908 | 0.04210 | 0.04031 | 0.03569 | 0.02322 | 0.09870 | 0.12501 | 0.00736 |
| **Rhox9** | 0.01389 | 0.00761 | 0.39732 | 0.04379 | 0.00858 | 0.00724 | 0.00722 | 0.01118 | 0.01241 | 0.06888 | 0.01244 | 0.02594 | 0.00972 | 0.04511 | 0.00000 | 0.00726 |
| **Rhpn2** | 5.88115 | 0.00000 | 0.00000 | 23.95390 | 3.31147 | 1.40919 | 0.00000 | 1.78834 | 12.17596 | 2.66787 | 2.70401 | 0.48411 | 0.00000 | 5.74727 | 16.06127 | 1.90709 |
| **Ribc1** | 0.96326 | 0.00511 | 1.31932 | 2.99769 | 0.01105 | 0.86452 | 1.43391 | 1.89253 | 1.30049 | 2.87650 | 1.98255 | 0.31342 | 0.00000 | 0.00000 | 0.44015 | 2.84201 |
| **Ric1** | 1.98974 | 0.21091 | 0.41940 | 6.83123 | 0.00345 | 0.00278 | 0.00564 | 1.10339 | 1.57561 | 0.19054 | 0.03466 | 0.03740 | 0.27931 | 0.01920 | 3.71063 | 0.00345 |
| **Ric3** | 0.00000 | 0.00731 | 0.00753 | 2.33878 | 0.00849 | 0.00711 | 0.01777 | 0.63328 | 1.60232 | 0.65985 | 0.14807 | 0.40202 | 0.11542 | 0.00656 | 0.22376 | 1.24110 |
| **Ric8a** | 12.03743 | 0.57173 | 3.60033 | 1.45441 | 6.89062 | 5.73584 | 2.82956 | 3.08509 | 0.22297 | 7.38393 | 0.23338 | 5.60363 | 3.80450 | 0.55525 | 2.05607 | 4.33605 |
| **Ric8b** | 0.20257 | 0.00833 | 0.08136 | 0.62170 | 0.01132 | 1.66527 | 0.12635 | 1.39129 | 0.16940 | 1.82243 | 0.04302 | 0.06313 | 0.00000 | 0.37390 | 2.05071 | 0.00619 |
| **Rictor** | 0.44952 | 0.00395 | 0.34268 | 5.88775 | 0.02401 | 0.00485 | 2.57284 | 0.00000 | 3.23145 | 3.09775 | 2.39437 | 7.96400 | 0.20167 | 6.11713 | 3.11091 | 0.01971 |
| **Rida** | 3.57923 | 4.04502 | 1.10636 | 0.47933 | 0.08553 | 0.00784 | 0.85551 | 0.00000 | 0.40995 | 0.97819 | 0.05919 | 0.00873 | 0.05323 | 0.02129 | 0.90351 | 0.26497 |
| **Rif1** | 7.11168 | 8.17440 | 5.14964 | 19.10099 | 0.06353 | 2.80060 | 5.70459 | 6.98971 | 22.99621 | 15.37657 | 8.82139 | 16.03970 | 3.29017 | 39.77588 | 28.99739 | 3.24716 |
| **Rilp** | 0.00906 | 0.00813 | 0.01272 | 0.38477 | 0.01046 | 0.00716 | 0.00724 | 0.09674 | 0.00369 | 0.00305 | 0.01444 | 0.00750 | 0.73606 | 0.01052 | 0.07186 | 0.00728 |
| **Rilpl1** | 0.00866 | 0.00787 | 0.05484 | 2.96394 | 0.00983 | 0.00577 | 0.00725 | 0.27319 | 1.35410 | 0.02442 | 0.74671 | 0.01305 | 0.02137 | 0.00000 | 0.44642 | 0.01418 |
| **Rilpl2** | 0.01431 | 0.00818 | 0.01297 | 5.53159 | 0.91168 | 0.00686 | 0.00709 | 0.14666 | 11.35557 | 2.31990 | 0.39240 | 0.28561 | 0.92976 | 0.64993 | 0.41890 | 0.02717 |
| **Rimbp2** | 0.04687 | 0.00773 | 0.04776 | 0.00491 | 0.00883 | 0.05010 | 0.02753 | 0.01483 | 0.07831 | 0.08136 | 0.22882 | 0.01475 | 0.04405 | 0.03348 | 0.18202 | 0.01470 |
| **Rimbp3** | 0.01185 | 0.00779 | 0.01984 | 0.59905 | 0.01027 | 0.00665 | 0.00754 | 0.01334 | 0.39822 | 0.23694 | 0.01061 | 0.00477 | 0.25786 | 0.00571 | 0.47660 | 0.00765 |
| **Rimkla** | 1.14181 | 0.02105 | 3.85019 | 7.98974 | 8.80969 | 0.01200 | 0.01292 | 0.00000 | 4.46524 | 0.34562 | 0.51836 | 6.06230 | 2.47889 | 0.00000 | 4.31925 | 0.36591 |
| **Rimklb** | 0.05891 | 0.00653 | 0.00000 | 10.01721 | 0.00750 | 0.00392 | 0.00439 | 1.86856 | 8.48132 | 3.75567 | 5.42438 | 0.33779 | 0.00000 | 4.20071 | 8.28146 | 0.01113 |
| **Rims2** | 0.06098 | 0.00772 | 0.26438 | 0.08794 | 0.00769 | 0.00565 | 0.00729 | 0.01349 | 0.01055 | 0.01522 | 0.01324 | 0.00774 | 0.01007 | 0.01254 | 0.00409 | 0.00711 |
| **Rims3** | 0.02402 | 0.00764 | 0.00119 | 0.58477 | 0.63365 | 0.00816 | 0.02641 | 0.33542 | 0.02529 | 0.82651 | 2.26207 | 0.03356 | 1.24575 | 0.00148 | 0.50527 | 0.00680 |
| **Rin1** | 2.25561 | 0.03507 | 0.61828 | 3.57432 | 0.25371 | 0.06539 | 0.05306 | 1.02689 | 6.19597 | 0.00000 | 5.60163 | 0.28156 | 0.15297 | 1.55305 | 0.09013 | 0.21228 |
| **Rin2** | 0.67748 | 0.00846 | 0.50744 | 1.73039 | 0.01648 | 0.00489 | 0.00751 | 0.37932 | 0.00000 | 0.97679 | 0.00000 | 0.01996 | 0.12688 | 2.14528 | 0.87942 | 0.00649 |
| **Rin3** | 0.01075 | 0.00795 | 0.01300 | 0.13168 | 0.01084 | 1.91202 | 0.00635 | 0.00893 | 0.00689 | 0.00121 | 0.01035 | 0.01702 | 1.92916 | 0.00745 | 0.18831 | 0.00586 |
| **Ring1** | 1.44790 | 0.07198 | 0.94443 | 0.01005 | 0.00752 | 0.74504 | 0.27431 | 2.39783 | 0.30446 | 3.32662 | 3.84715 | 0.05270 | 3.21949 | 0.21469 | 0.25447 | 0.03352 |
| **Rinl** | 3.14171 | 0.01021 | 1.67552 | 1.98695 | 4.42499 | 0.00244 | 0.04207 | 2.18235 | 1.49181 | 2.16646 | 2.29898 | 3.17652 | 1.69024 | 0.05971 | 1.82047 | 0.00178 |
| **Rint1** | 2.95341 | 0.12371 | 2.26482 | 1.08107 | 0.03329 | 0.62651 | 1.74671 | 1.11523 | 1.92509 | 0.40503 | 2.75453 | 0.33860 | 3.25517 | 0.66653 | 1.56077 | 0.80662 |
| **Riok1** | 2.66829 | 0.16723 | 4.58761 | 7.95616 | 3.86322 | 3.37845 | 3.52585 | 6.47815 | 5.61432 | 7.94623 | 2.90577 | 4.16327 | 1.76662 | 6.96892 | 5.97261 | 2.87346 |
| **Riok2** | 10.04087 | 8.91980 | 5.75023 | 6.15323 | 8.43904 | 0.22383 | 5.95767 | 11.82745 | 9.60562 | 9.75652 | 5.64050 | 10.84588 | 8.34673 | 10.13993 | 16.37072 | 2.14979 |
| **Riok3** | 2.03191 | 8.95373 | 8.09779 | 9.30662 | 3.23033 | 6.90973 | 7.83846 | 11.59257 | 9.53452 | 22.81681 | 17.12734 | 9.98437 | 13.32692 | 12.81600 | 12.81373 | 7.79717 |
| **Ripk1** | 1.46159 | 0.00379 | 0.12466 | 1.46780 | 3.18396 | 0.00738 | 2.39260 | 1.13650 | 5.32497 | 0.29621 | 0.22489 | 0.11090 | 0.33725 | 1.49229 | 1.61453 | 0.00558 |
| **Ripk2** | 0.01301 | 0.00781 | 0.01128 | 2.84890 | 0.00953 | 0.00646 | 0.00732 | 0.01066 | 0.03008 | 0.77840 | 0.02851 | 0.02339 | 0.02192 | 0.00235 | 1.16159 | 0.00727 |
| **Ripk3** | 0.01759 | 0.01763 | 0.00972 | 0.00702 | 0.00887 | 0.00663 | 0.00727 | 0.01079 | 0.01074 | 0.21090 | 0.01001 | 0.06975 | 0.01283 | 0.00836 | 0.62292 | 0.00774 |
| **Ripk4** | 2.53238 | 3.27395 | 1.59054 | 2.51057 | 0.00383 | 0.04284 | 0.00347 | 0.01807 | 0.79064 | 0.84708 | 0.83517 | 0.05258 | 0.00000 | 0.00000 | 0.29743 | 0.00239 |
| **Ripply3** | 2.37111 | 0.05715 | 0.24699 | 1.68261 | 0.07456 | 0.06287 | 0.17525 | 0.00000 | 2.49944 | 0.03081 | 4.39385 | 0.17579 | 0.20334 | 1.03567 | 0.24811 | 0.08789 |
| **Rit1** | 3.85967 | 0.14128 | 0.37169 | 3.93473 | 0.05975 | 0.02735 | 0.27678 | 2.86932 | 3.63759 | 3.52357 | 0.75029 | 0.13584 | 4.61682 | 0.00000 | 1.51638 | 4.49241 |
| **Rita1** | 1.27573 | 0.00716 | 0.85569 | 2.89603 | 0.01496 | 1.37707 | 0.00586 | 1.63912 | 2.64297 | 2.09019 | 0.46739 | 0.08217 | 0.00000 | 0.12642 | 0.00081 | 0.01976 |
| **Rlbp1** | 0.00932 | 0.00832 | 0.04133 | 0.28518 | 0.02039 | 0.00854 | 0.00748 | 0.01196 | 0.42980 | 0.33747 | 0.00000 | 0.02045 | 0.08198 | 0.93676 | 1.02880 | 0.00755 |

| | | | | | | | | | | | | | | | | |
|---|---|---|---|---|---|---|---|---|---|---|---|---|---|---|---|---|
| **Rlf** | 1.23265 | 0.23879 | 2.42224 | 9.07865 | 1.84310 | 0.00266 | 3.10432 | 2.25262 | 15.15359 | 3.48771 | 5.71326 | 4.17996 | 0.00000 | 6.85988 | 10.35040 | 0.28669 |
| **Rlim** | 8.02968 | 11.13740 | 6.85939 | 16.16068 | 7.92525 | 0.58400 | 5.22194 | 10.86208 | 22.61167 | 8.67588 | 6.08727 | 9.72480 | 8.63683 | 22.60002 | 13.18133 | 9.79801 |
| **Rln3** | 0.00960 | 0.00809 | 0.00856 | 0.20810 | 0.00943 | 0.00681 | 0.00769 | 0.22835 | 0.00647 | 0.16229 | 0.05991 | 0.00643 | 0.17448 | 0.00627 | 0.09439 | 0.00760 |
| **Rltpr** | 0.02287 | 0.00752 | 0.13417 | 0.07199 | 0.00862 | 1.63396 | 0.00662 | 0.11873 | 0.01997 | 0.23337 | 0.83527 | 0.00240 | 0.00000 | 0.00504 | 0.13449 | 0.00658 |
| **Rmdn1** | 0.02312 | 0.03010 | 0.31228 | 0.00840 | 0.00730 | 0.00878 | 0.00773 | 0.07467 | 0.01190 | 0.01278 | 0.08274 | 0.19205 | 0.00569 | 0.01200 | 0.00540 | 0.53201 |
| **Rmdn2** | 0.01166 | 0.00811 | 0.00558 | 0.00293 | 0.00971 | 0.00647 | 0.00714 | 0.00851 | 0.73900 | 0.00000 | 0.00000 | 0.00000 | 0.01360 | 2.50692 | 0.26104 | 0.00728 |
| **Rmdn3** | 9.83264 | 0.51003 | 5.41895 | 6.62258 | 7.81305 | 5.24936 | 6.87393 | 10.25072 | 0.81535 | 6.56896 | 2.82281 | 9.98314 | 7.38477 | 2.78361 | 6.56236 | 2.86308 |
| **Rmi1** | 2.07860 | 0.05859 | 1.75359 | 7.75312 | 0.01241 | 6.37294 | 0.00338 | 1.97964 | 4.40004 | 1.41843 | 1.95720 | 0.07670 | 1.29254 | 2.93783 | 6.02173 | 0.48784 |
| **Rmi2** | 1.81192 | 0.20483 | 4.12136 | 9.13496 | 0.83126 | 0.39289 | 0.56434 | 0.31563 | 13.71789 | 1.06691 | 15.07772 | 1.15358 | 1.44792 | 5.40395 | 1.68012 | 0.42879 |
| **Rmnd5a** | 2.94271 | 2.59297 | 6.43138 | 20.43266 | 2.86122 | 3.99119 | 1.78091 | 5.65195 | 21.83973 | 9.69100 | 11.01594 | 7.19356 | 9.98452 | 14.37407 | 18.64616 | 1.53370 |
| **Rmnd5b** | 3.48648 | 2.29372 | 2.89790 | 14.81752 | 4.15475 | 6.21527 | 4.98975 | 14.36931 | 15.35221 | 28.30590 | 15.02096 | 6.93972 | 7.35720 | 4.22862 | 10.20196 | 7.44309 |
| **Rn4.5s** | 0.03130 | 0.04098 | 0.12701 | 0.36297 | 0.01032 | 0.00645 | 0.00739 | 0.02623 | 0.73177 | 0.01400 | 0.53747 | 0.03533 | 0.03066 | 0.04225 | 0.06742 | 0.85887 |
| **Rn45s** | 18.77779 | 11.81886 | 2.17697 | 1.73848 | 12.25054 | 9.99766 | 4.51507 | 4.27854 | 2.31797 | 18.91715 | 0.34992 | 1.27664 | 16.77083 | 2.98146 | 1.15453 | 7.70213 |
| **Rnase4** | 0.03314 | 0.00762 | 0.01050 | 0.00793 | 0.00827 | 0.00669 | 0.00729 | 0.01001 | 0.01191 | 0.71419 | 0.01224 | 0.02484 | 0.01162 | 0.01184 | 0.00844 | 0.00761 |
| **Rnaseh1** | 3.05893 | 0.00000 | 2.52499 | 3.28773 | 5.54356 | 1.78131 | 3.94597 | 5.91440 | 5.32062 | 2.75294 | 3.35539 | 1.58340 | 1.93153 | 0.00000 | 6.52868 | 3.35850 |
| **Rnaseh2a** | 9.42084 | 3.25496 | 8.37058 | 2.15673 | 4.16014 | 4.96641 | 6.46620 | 7.28921 | 2.25196 | 5.02629 | 6.72813 | 0.20513 | 4.59524 | 0.00000 | 1.12050 | 6.49164 |
| **Rnaseh2b** | 19.74283 | 4.12847 | 8.05685 | 0.76884 | 6.71462 | 3.56916 | 7.12065 | 14.02621 | 0.00000 | 12.16551 | 5.89966 | 4.88666 | 14.98560 | 0.32193 | 2.36978 | 5.12684 |
| **Rnaseh2c** | 3.97952 | 0.00235 | 0.44713 | 0.71191 | 1.28238 | 0.02214 | 0.04183 | 0.07762 | 0.98289 | 2.83948 | 1.06841 | 0.00526 | 4.49376 | 0.00395 | 0.00000 | 4.01676 |
| **Rnasek** | 13.78372 | 2.98878 | 16.51102 | 3.43871 | 8.35922 | 3.35626 | 4.66278 | 6.44480 | 3.99716 | 11.87618 | 12.82767 | 0.79461 | 13.27258 | 0.00000 | 1.00253 | 4.55460 |
| **Rnasel** | 0.76729 | 0.00703 | 0.54027 | 0.04941 | 0.02208 | 0.13572 | 0.06720 | 0.00207 | 0.11045 | 0.25613 | 0.18273 | 0.03329 | 0.09294 | 3.04560 | 0.00000 | 0.04125 |
| **Rnaset2b** | 0.87922 | 1.75514 | 1.86794 | 0.85057 | 2.82263 | 2.37364 | 1.75769 | 2.75388 | 0.03137 | 3.83427 | 6.68497 | 0.28880 | 4.38259 | 1.03246 | 0.53930 | 0.39148 |
| **Rnd1** | 0.00000 | 3.38886 | 6.61966 | 0.52768 | 0.09770 | 0.01885 | 5.32765 | 4.76649 | 1.25262 | 1.09574 | 1.26754 | 0.03657 | 1.47430 | 0.38281 | 1.34234 | 0.05167 |
| **Rnd2** | 5.11403 | 0.00000 | 3.36603 | 0.00000 | 2.74623 | 0.00850 | 1.40145 | 4.96906 | 0.00421 | 6.63865 | 0.00000 | 0.13267 | 4.62010 | 0.00957 | 0.47174 | 0.02737 |
| **Rnd3** | 8.89957 | 10.13051 | 6.12506 | 0.18857 | 11.93362 | 0.00300 | 4.35127 | 1.32343 | 1.12955 | 0.00000 | 0.62467 | 0.01610 | 0.00000 | 0.00614 | 0.81743 | 3.56260 |
| **Rnf10** | 17.51347 | 9.68145 | 19.05753 | 10.31746 | 14.09611 | 9.67833 | 12.78871 | 17.68515 | 14.15911 | 14.98246 | 9.93457 | 1.60389 | 16.96934 | 5.28445 | 17.13396 | 6.31452 |
| **Rnf103** | 0.65681 | 0.00688 | 0.00000 | 2.91355 | 0.02095 | 0.31337 | 1.46545 | 0.00000 | 0.55740 | 0.98892 | 1.65107 | 0.01827 | 0.10440 | 1.92782 | 0.56542 | 0.57789 |
| **Rnf11** | 2.56564 | 3.48331 | 8.23387 | 20.83991 | 5.60941 | 3.78669 | 4.07884 | 5.56644 | 22.49200 | 10.23175 | 7.66601 | 21.47242 | 4.78286 | 21.36062 | 17.09597 | 1.05034 |
| **Rnf111** | 3.10172 | 3.29817 | 5.33673 | 11.31596 | 4.99336 | 0.28911 | 5.72004 | 4.24402 | 9.81040 | 7.50439 | 5.61048 | 13.21108 | 1.86626 | 15.24590 | 14.22670 | 2.38002 |
| **Rnf113a1** | 2.90849 | 0.00429 | 0.86947 | 0.00968 | 0.99228 | 0.51631 | 0.00367 | 1.23979 | 0.00985 | 0.03649 | 0.00000 | 0.07674 | 3.43458 | 0.01556 | 0.15921 | 0.02452 |
| **Rnf113a2** | 4.89951 | 3.96552 | 6.63687 | 4.26834 | 2.11686 | 1.15921 | 5.33332 | 8.00598 | 8.89279 | 6.76664 | 1.81822 | 0.59619 | 4.75131 | 0.02321 | 4.47006 | 3.54581 |
| **Rnf114** | 6.94551 | 6.59133 | 7.30616 | 13.46647 | 10.04187 | 4.56181 | 0.20652 | 10.05963 | 6.76550 | 12.63668 | 9.88662 | 3.64376 | 9.11984 | 0.55830 | 10.42299 | 5.31857 |
| **Rnf115** | 5.91411 | 14.03928 | 14.00565 | 3.41103 | 1.27127 | 5.15493 | 9.93384 | 10.71169 | 7.86896 | 9.99334 | 11.02852 | 5.98248 | 7.88275 | 21.37952 | 8.49285 | 3.22953 |
| **Rnf121** | 3.62895 | 0.88129 | 5.44796 | 0.00000 | 3.46588 | 3.27710 | 0.01176 | 2.34381 | 0.54808 | 1.34301 | 2.65348 | 0.05452 | 1.08723 | 7.10645 | 1.55118 | 0.38990 |
| **Rnf122** | 0.00401 | 0.02843 | 0.02065 | 4.91109 | 0.01046 | 0.00723 | 0.02736 | 0.55865 | 4.08940 | 0.15445 | 1.11118 | 0.13401 | 0.01657 | 3.77280 | 6.16662 | 0.00632 |
| **Rnf123** | 2.59359 | 0.00441 | 1.60427 | 0.31197 | 0.01411 | 0.00337 | 0.64532 | 0.10839 | 0.25220 | 0.36833 | 0.00723 | 0.06719 | 1.52240 | 0.00968 | 0.00000 | 0.01105 |
| **Rnf125** | 3.33333 | 0.00000 | 0.78926 | 14.62090 | 2.39675 | 0.00000 | 0.00000 | 1.55280 | 15.13276 | 6.39759 | 4.47060 | 7.46784 | 0.23727 | 7.86963 | 4.41922 | 5.78398 |
| **Rnf126** | 0.33594 | 0.00681 | 0.01701 | 1.67123 | 0.78313 | 0.07625 | 0.00651 | 0.00000 | 1.71560 | 5.90036 | 0.94770 | 0.00000 | 0.45613 | 0.00279 | 0.00000 | 0.36406 |
| **Rnf128** | 2.06005 | 3.96896 | 0.00000 | 1.84256 | 5.58428 | 0.59615 | 0.00484 | 0.18409 | 0.35534 | 0.52558 | 0.00471 | 0.11038 | 0.58578 | 0.00139 | 0.00000 | 0.06084 |
| **Rnf13** | 1.52907 | 9.10904 | 5.98577 | 2.02581 | 1.63072 | 3.76696 | 3.16791 | 5.94560 | 0.44418 | 3.61091 | 2.32171 | 3.51683 | 4.21493 | 11.10185 | 4.73327 | 3.14375 |
| **Rnf130** | 11.06276 | 5.01301 | 1.24432 | 0.14594 | 3.83283 | 0.00562 | 7.71118 | 0.55279 | 0.70800 | 1.12587 | 2.01617 | 0.27088 | 1.58848 | 0.14867 | 0.99979 | 2.77757 |
| **Rnf135** | 1.47837 | 0.00423 | 1.30493 | 1.26670 | 0.01184 | 2.46792 | 2.09466 | 1.70695 | 1.35393 | 0.57030 | 1.35445 | 0.00576 | 2.56473 | 0.22674 | 0.79693 | 0.00297 |
| **Rnf138** | 3.43412 | 4.04057 | 2.28512 | 4.18090 | 5.43383 | 1.83659 | 0.74081 | 4.67723 | 3.08472 | 1.26431 | 4.70566 | 16.22473 | 5.87155 | 4.45137 | 0.00000 | 1.27126 |
| **Rnf138rt1** | 0.01324 | 0.00796 | 0.01170 | 0.27097 | 0.00920 | 0.00630 | 0.00728 | 0.01025 | 2.10144 | 0.52633 | 0.00000 | 0.00242 | 0.01200 | 0.00941 | 0.01161 | 0.00729 |
| **Rnf139** | 1.12763 | 0.08259 | 1.28512 | 4.48933 | 0.04930 | 1.14578 | 0.11046 | 3.06836 | 2.89631 | 2.45238 | 4.24390 | 0.91679 | 0.42442 | 7.05754 | 2.97881 | 0.12995 |
| **Rnf14** | 4.44458 | 2.63292 | 3.69377 | 23.04866 | 2.05349 | 8.08459 | 0.41130 | 3.15355 | 14.71226 | 14.77204 | 4.95714 | 5.54223 | 5.38565 | 6.22447 | 15.51337 | 0.26808 |
| **Rnf141** | 0.01828 | 0.00765 | 0.06067 | 0.06895 | 0.00849 | 0.00827 | 0.00696 | 0.14816 | 0.38817 | 0.00000 | 0.00000 | 0.02137 | 0.18589 | 2.40234 | 2.99409 | 0.00697 |
| **Rnf145** | 0.00000 | 0.00760 | 0.91637 | 1.77729 | 0.09259 | 0.00550 | 0.00639 | 0.09431 | 0.00049 | 1.50272 | 0.78364 | 0.19749 | 0.67580 | 1.29136 | 3.37940 | 0.79353 |
| **Rnf146** | 1.24898 | 0.00580 | 0.03950 | 0.84070 | 0.01384 | 0.00438 | 0.00596 | 0.00000 | 0.00000 | 0.92930 | 0.99248 | 0.80890 | 0.00059 | 0.01468 | 0.92623 | 0.07509 |
| **Rnf149** | 10.75448 | 8.15049 | 12.19342 | 1.76217 | 12.75246 | 3.09241 | 8.39929 | 13.48232 | 5.41333 | 10.30661 | 8.71491 | 2.70746 | 6.92623 | 6.60117 | 6.37522 | 0.18688 |
| **Rnf150** | 0.05770 | 0.00787 | 0.03732 | 0.09177 | 0.00917 | 0.00643 | 0.00753 | 0.01812 | 0.22283 | 0.01670 | 0.06602 | 0.01697 | 0.02288 | 0.18005 | 0.01744 | 0.00733 |
| **Rnf152** | 0.03959 | 0.00801 | 0.02617 | 0.02577 | 0.00972 | 0.01321 | 0.01788 | 0.01755 | 0.02243 | 0.01878 | 0.02830 | 0.00787 | 0.05022 | 0.03171 | 0.02225 | 0.01472 |
| **Rnf157** | 0.00706 | 1.74313 | 0.00374 | 0.00000 | 0.00812 | 0.16832 | 0.00655 | 0.00908 | 0.15645 | 0.70077 | 0.00924 | 0.04323 | 0.00955 | 0.00515 | 0.39675 | 0.00664 |
| **Rnf165** | 0.03105 | 0.02863 | 0.01622 | 0.56244 | 0.00936 | 0.04214 | 0.00738 | 0.01141 | 0.00394 | 0.16868 | 0.01191 | 0.01653 | 0.02010 | 0.00611 | 0.22987 | 0.02126 |
| **Rnf166** | 1.06835 | 0.00466 | 1.45254 | 1.06862 | 3.05299 | 0.30488 | 0.80867 | 0.00000 | 0.04571 | 0.82692 | 1.21920 | 0.18049 | 0.00000 | 0.04178 | 0.00000 | 0.38611 |
| **Rnf167** | 15.16049 | 6.65428 | 15.72157 | 12.15065 | 4.38279 | 12.37576 | 12.78929 | 18.54417 | 10.12705 | 15.40415 | 9.43803 | 4.70800 | 17.33223 | 0.12130 | 7.85633 | 6.90095 |
| **Rnf168** | 1.48269 | 0.05797 | 1.06482 | 8.85557 | 0.48154 | 0.34423 | 0.10624 | 0.18920 | 7.81666 | 2.08922 | 8.26612 | 3.92430 | 1.82693 | 8.02354 | 5.41041 | 0.98511 |
| **Rnf169** | 0.00000 | 0.16328 | 2.02181 | 1.67586 | 0.18914 | 0.00529 | 3.12899 | 0.57712 | 1.94120 | 0.31525 | 0.46755 | 0.09398 | 0.32896 | 0.01810 | 1.05137 | 0.04265 |
| **Rnf17** | 0.08847 | 0.00787 | 0.48505 | 1.62640 | 0.00712 | 1.39410 | 0.00772 | 0.13129 | 1.26045 | 0.92220 | 0.05006 | 0.10545 | 0.00000 | 0.00633 | 1.90357 | 0.00633 |
| **Rnf170** | 0.55460 | 1.22015 | 0.00000 | 1.64630 | 0.00666 | 0.00674 | 0.01741 | 0.01730 | 1.27868 | 0.14451 | 0.41071 | 0.17447 | 0.14778 | 1.98470 | 6.07955 | 0.00553 |
| **Rnf181** | 15.09539 | 11.32347 | 16.38030 | 4.56526 | 8.15869 | 10.25092 | 9.89410 | 16.39623 | 7.98893 | 6.73653 | 9.26969 | 3.69565 | 8.66130 | 1.41160 | 8.56041 | 3.26646 |
| **Rnf182** | 0.01262 | 0.00723 | 0.00853 | 0.00464 | 0.00940 | 0.00676 | 0.60234 | 0.01000 | 0.21505 | 0.00808 | 0.01047 | 0.00660 | 0.01115 | 0.01958 | 0.09261 | 0.00733 |
| **Rnf185** | 2.23925 | 7.14340 | 3.37611 | 9.82033 | 1.94434 | 4.22953 | 6.35993 | 4.63877 | 5.72599 | 3.82832 | 2.72006 | 0.52823 | 2.69002 | 2.39281 | 9.87339 | 1.35388 |

| | | | | | | | | | | | | | | | | |
|---|---|---|---|---|---|---|---|---|---|---|---|---|---|---|---|---|
| **Rnf187** | 5.69129 | 2.61101 | 7.19829 | 12.02570 | 12.82570 | 0.80436 | 3.57369 | 4.34665 | 4.53221 | 5.62037 | 2.68689 | 0.34859 | 0.73028 | 0.00000 | 1.73843 | 7.70544 |
| **Rnf19a** | 1.61282 | 0.00374 | 2.40987 | 4.82445 | 0.00369 | 0.12963 | 0.00361 | 0.00000 | 4.67996 | 0.79469 | 1.69580 | 7.74331 | 0.93087 | 0.00000 | 3.16268 | 1.38685 |
| **Rnf19b** | 9.79404 | 2.75883 | 5.10375 | 17.36262 | 9.99588 | 7.94802 | 6.56808 | 14.71569 | 16.04844 | 17.40526 | 9.43762 | 10.41301 | 6.02762 | 0.16009 | 0.00000 | 3.62687 |
| **Rnf2** | 3.50389 | 2.30769 | 1.85272 | 17.87289 | 5.07161 | 2.95171 | 3.59521 | 9.65674 | 11.41992 | 15.02903 | 6.10067 | 14.60908 | 5.33650 | 21.46090 | 16.94502 | 3.36794 |
| **Rnf20** | 4.79886 | 3.26518 | 1.39380 | 6.99279 | 1.35338 | 0.00000 | 0.39525 | 5.92517 | 13.32389 | 6.18712 | 2.46896 | 4.74756 | 1.47769 | 14.45544 | 21.50428 | 2.86965 |
| **Rnf207** | 0.27696 | 0.01658 | 0.55644 | 0.00926 | 0.00619 | 0.00835 | 0.00675 | 0.41149 | 0.01172 | 0.00224 | 0.00974 | 0.00563 | 0.40971 | 0.00714 | 0.23294 | 0.01351 |
| **Rnf208** | 0.01285 | 0.00754 | 0.16685 | 0.00584 | 0.00995 | 0.29279 | 0.00618 | 0.76091 | 0.02109 | 0.51963 | 1.30567 | 0.00000 | 0.92076 | 0.01535 | 0.01661 | 0.00696 |
| **Rnf214** | 3.51270 | 2.52479 | 2.53279 | 7.47155 | 4.01984 | 2.94894 | 0.13794 | 2.88428 | 6.42761 | 5.51608 | 4.45130 | 0.17056 | 1.85231 | 6.46260 | 7.26929 | 1.15770 |
| **Rnf215** | 0.74757 | 0.01660 | 0.81249 | 1.54481 | 0.00525 | 0.00601 | 0.00657 | 1.14662 | 0.74583 | 0.00000 | 0.01847 | 1.08412 | 0.05092 | 0.13976 | 0.48758 | 0.01356 |
| **Rnf216** | 6.51423 | 0.39708 | 5.39856 | 15.90369 | 2.79611 | 0.00000 | 5.20307 | 11.46264 | 18.28157 | 22.95998 | 17.41218 | 6.76367 | 1.94188 | 2.76140 | 13.28178 | 3.65427 |
| **Rnf219** | 0.01208 | 0.00793 | 0.02483 | 0.06644 | 0.01896 | 0.01213 | 0.00729 | 0.01912 | 0.05447 | 0.01090 | 0.02478 | 0.01481 | 0.01179 | 0.60354 | 0.00150 | 0.00747 |
| **Rnf220** | 7.21697 | 5.55962 | 2.09024 | 3.98000 | 0.61314 | 4.11382 | 2.79752 | 3.85881 | 6.36352 | 9.75614 | 1.90629 | 0.60417 | 5.51797 | 0.00000 | 1.28432 | 3.74312 |
| **Rnf223** | 0.00846 | 0.00820 | 0.64074 | 0.42936 | 0.00853 | 0.85261 | 0.00764 | 0.07801 | 0.04365 | 0.37427 | 0.12848 | 0.05460 | 0.35541 | 0.00577 | 0.10646 | 0.00717 |
| **Rnf225** | 0.01161 | 0.00771 | 0.08586 | 0.76423 | 0.01099 | 0.00804 | 0.00706 | 0.73744 | 0.58686 | 0.02624 | 0.05535 | 0.00903 | 0.00602 | 0.00732 | 2.91289 | 0.05727 |
| **Rnf24** | 0.12388 | 0.09333 | 0.00000 | 3.30243 | 0.01598 | 0.09746 | 0.00634 | 1.57983 | 3.77629 | 5.64153 | 0.11552 | 1.38759 | 1.00684 | 8.92493 | 9.02234 | 0.00543 |
| **Rnf25** | 0.00686 | 3.82559 | 2.17291 | 1.06575 | 0.15818 | 3.88194 | 0.00405 | 0.00000 | 1.65426 | 4.82387 | 2.36481 | 0.10340 | 1.48434 | 0.01619 | 2.05095 | 0.00977 |
| **Rnf26** | 4.41637 | 3.31421 | 3.88219 | 11.81217 | 3.13851 | 7.49546 | 4.47368 | 5.68396 | 4.82615 | 3.02709 | 1.88580 | 0.25639 | 7.18441 | 0.00000 | 2.54860 | 2.27284 |
| **Rnf31** | 1.96847 | 0.07371 | 3.59060 | 3.15551 | 4.89085 | 2.92746 | 5.43780 | 5.50711 | 3.42572 | 6.20809 | 5.45004 | 0.02630 | 5.84192 | 0.02236 | 1.87114 | 3.18149 |
| **Rnf32** | 0.02276 | 0.00835 | 0.82327 | 0.00795 | 0.00960 | 0.00700 | 0.00748 | 0.01328 | 0.01144 | 0.08493 | 0.01222 | 0.00656 | 0.23832 | 0.00613 | 0.36496 | 0.02220 |
| **Rnf34** | 5.54485 | 0.15459 | 6.82068 | 13.45493 | 0.04502 | 5.02465 | 4.57868 | 3.90495 | 7.37474 | 11.53569 | 9.51002 | 0.66154 | 4.81464 | 7.50168 | 11.36641 | 2.00148 |
| **Rnf38** | 4.16048 | 1.72839 | 1.91034 | 20.44684 | 0.74413 | 0.00000 | 5.32033 | 7.20188 | 8.81368 | 9.87018 | 5.17784 | 5.64883 | 2.45538 | 7.74204 | 5.89667 | 0.58772 |
| **Rnf39** | 0.21744 | 0.06101 | 0.09496 | 0.00910 | 0.01213 | 0.00550 | 0.00700 | 0.00942 | 0.01206 | 0.33069 | 0.01437 | 0.00714 | 0.01032 | 0.01951 | 0.00459 | 0.00694 |
| **Rnf4** | 15.13056 | 12.39338 | 16.45245 | 25.33088 | 13.15464 | 12.42078 | 9.39154 | 23.55126 | 30.04169 | 23.32704 | 17.78228 | 19.36618 | 9.08463 | 29.71857 | 22.35423 | 5.31690 |
| **Rnf40** | 0.47915 | 0.17220 | 2.23711 | 8.29015 | 4.61437 | 0.38047 | 2.17124 | 7.91437 | 6.75130 | 10.06060 | 1.26612 | 7.17485 | 3.37425 | 0.00000 | 9.07321 | 2.52954 |
| **Rnf41** | 3.11616 | 0.16707 | 3.76898 | 1.65122 | 0.94049 | 0.63809 | 3.74049 | 5.48645 | 4.10435 | 6.27825 | 0.85697 | 0.48393 | 1.51739 | 6.20950 | 2.71196 | 2.27428 |
| **Rnf43** | 0.02201 | 0.00864 | 0.01025 | 0.95655 | 0.00867 | 0.00670 | 0.00798 | 0.06225 | 0.27995 | 0.01269 | 0.01068 | 0.00863 | 0.01013 | 0.01216 | 0.00462 | 0.00716 |
| **Rnf44** | 5.00687 | 1.39179 | 1.89392 | 9.72137 | 4.26215 | 0.14588 | 0.12526 | 5.25803 | 11.40900 | 2.43743 | 3.21584 | 5.07104 | 2.76995 | 3.22600 | 5.54917 | 1.22570 |
| **Rnf5** | 4.79063 | 0.93815 | 0.00000 | 0.18647 | 4.61879 | 4.85965 | 0.92429 | 1.64797 | 0.05737 | 0.95336 | 1.47112 | 0.07525 | 1.09356 | 0.00629 | 0.53712 | 0.60257 |
| **Rnf6** | 0.96398 | 0.42793 | 1.11546 | 7.38820 | 1.33203 | 0.02055 | 0.57749 | 2.19136 | 6.96893 | 7.75924 | 2.72490 | 1.50644 | 1.59734 | 6.30409 | 13.62568 | 0.01000 |
| **Rnf7** | 19.24300 | 13.46021 | 20.60732 | 13.37335 | 14.61509 | 9.89580 | 11.87370 | 34.60391 | 16.56211 | 31.05161 | 28.17570 | 13.49956 | 27.36101 | 20.05654 | 15.23406 | 7.54691 |
| **Rnf8** | 4.43873 | 0.36728 | 3.57146 | 8.64520 | 3.11413 | 2.60868 | 0.02037 | 2.76574 | 13.14242 | 8.44222 | 8.20217 | 1.02706 | 1.96451 | 0.45340 | 3.85877 | 1.92450 |
| **Rnft1** | 3.71871 | 1.01047 | 3.77635 | 0.93176 | 3.08966 | 0.02312 | 2.48581 | 4.61748 | 2.05131 | 3.65825 | 3.73665 | 6.89185 | 2.35461 | 9.58526 | 2.25108 | 1.01662 |
| **Rnft2** | 0.19486 | 0.00812 | 0.07722 | 0.20771 | 0.01011 | 0.00561 | 0.00747 | 0.01389 | 0.27228 | 0.01208 | 0.01119 | 0.01852 | 0.01139 | 0.01263 | 0.02942 | 0.00715 |
| **Rngtt** | 2.77256 | 3.46416 | 3.84805 | 5.13150 | 2.79337 | 0.00322 | 4.24681 | 3.62484 | 12.41931 | 8.51457 | 2.09863 | 1.10696 | 1.50759 | 0.00000 | 10.19463 | 0.07173 |
| **Rnh1** | 1.37243 | 0.00001 | 3.76468 | 1.31277 | 6.25215 | 7.53950 | 0.00149 | 7.13314 | 0.13659 | 5.21130 | 2.91242 | 0.02509 | 4.27023 | 0.01132 | 0.74837 | 4.13043 |
| **Rnmt** | 10.81410 | 10.80896 | 11.03553 | 10.44666 | 10.94421 | 3.57403 | 10.23236 | 15.84891 | 10.81270 | 16.07424 | 9.00868 | 18.09775 | 6.41912 | 21.87984 | 15.68377 | 3.71829 |
| **Rnmtl1** | 5.41603 | 1.43100 | 2.09355 | 0.23203 | 0.00000 | 4.64092 | 4.79976 | 8.79041 | 0.50203 | 13.05310 | 4.37413 | 1.19831 | 5.00174 | 0.32807 | 3.40019 | 2.73511 |
| **Rnpc3** | 2.00943 | 0.01431 | 0.09458 | 4.84470 | 0.95779 | 0.01280 | 0.31200 | 0.97939 | 3.25104 | 1.33575 | 1.12973 | 1.67311 | 0.91564 | 4.96365 | 7.60674 | 0.05618 |
| **Rnpep** | 5.71817 | 1.90757 | 2.68772 | 6.71231 | 5.62542 | 7.54947 | 4.90047 | 10.56676 | 1.63185 | 14.85391 | 7.65796 | 2.28555 | 9.69057 | 0.01499 | 7.87059 | 5.91758 |
| **Rnpepl1** | 2.12722 | 0.00441 | 0.11272 | 1.55942 | 0.39751 | 0.09159 | 0.03326 | 3.88309 | 0.15756 | 4.46756 | 1.74167 | 0.07878 | 3.87517 | 0.04088 | 0.11445 | 0.01009 |
| **Rnps1** | 25.36876 | 8.72356 | 15.02258 | 17.31296 | 17.23606 | 14.52323 | 11.98270 | 28.67004 | 18.84979 | 29.38441 | 15.64210 | 8.09675 | 21.05313 | 3.26259 | 6.04967 | 17.28134 |
| **Rnu73b** | 0.06429 | 0.00792 | 0.05084 | 0.01081 | 0.00792 | 0.00589 | 0.00758 | 0.01535 | 0.01222 | 0.01358 | 0.01252 | 0.00820 | 0.01182 | 0.01385 | 0.03709 | 0.00723 |
| **Robo3** | 0.01477 | 0.00787 | 0.23689 | 0.34503 | 0.00867 | 0.00649 | 0.00724 | 0.01082 | 0.00674 | 0.26108 | 0.01360 | 0.00730 | 0.00961 | 0.01306 | 0.00070 | 0.00696 |
| **Robo4** | 0.01374 | 0.00769 | 0.01034 | 0.07095 | 0.00899 | 0.00659 | 0.00720 | 0.00363 | 0.00824 | 0.53617 | 0.23562 | 0.00594 | 0.00897 | 0.02399 | 0.00432 | 0.00704 |
| **Rock1** | 0.41498 | 4.62242 | 2.67353 | 15.12549 | 0.87893 | 0.93686 | 1.54299 | 3.39346 | 8.34386 | 4.30864 | 0.95269 | 6.38348 | 0.18709 | 19.60806 | 19.48349 | 0.00000 |
| **Rock2** | 4.43160 | 6.57335 | 1.83180 | 1.83603 | 0.06713 | 3.62505 | 0.32675 | 3.14567 | 4.18591 | 2.30587 | 1.51263 | 10.35386 | 1.37356 | 8.18291 | 4.25395 | 0.25024 |
| **Rogdi** | 21.05201 | 6.31762 | 16.07757 | 1.74057 | 15.71170 | 12.98405 | 13.19121 | 17.14215 | 3.56528 | 7.05584 | 7.47844 | 4.63939 | 22.80879 | 0.00523 | 0.44609 | 15.81919 |
| **Rom1** | 0.01908 | 0.00499 | 0.63836 | 0.72185 | 4.30298 | 0.00541 | 0.00699 | 0.00390 | 0.04539 | 3.04916 | 0.00878 | 0.00000 | 0.00843 | 0.00613 | 0.07099 | 0.01395 |
| **Romo1** | 3.81334 | 0.00202 | 1.30854 | 4.67972 | 1.09809 | 0.53023 | 0.34212 | 0.30306 | 4.22104 | 1.77808 | 3.04634 | 0.19401 | 0.66753 | 0.11887 | 0.00000 | 1.49168 |
| **Ropn1l** | 0.01327 | 0.00760 | 0.01152 | 0.42364 | 0.00919 | 0.00649 | 0.00722 | 0.01018 | 0.00557 | 0.76131 | 0.01245 | 0.00509 | 0.01154 | 0.10122 | 0.00000 | 0.00742 |
| **Ror1** | 0.00871 | 0.00662 | 0.00755 | 0.00446 | 0.00920 | 0.00691 | 2.10935 | 0.00980 | 0.01113 | 1.99571 | 0.01231 | 0.00000 | 0.01225 | 0.01885 | 0.00482 | 0.00711 |
| **Rora** | 0.01386 | 0.00774 | 0.00974 | 0.94660 | 0.00923 | 0.00648 | 0.00729 | 0.00668 | 0.00000 | 0.00000 | 0.17248 | 0.02641 | 0.01022 | 0.00908 | 0.43752 | 0.01436 |
| **Rp2** | 0.16687 | 6.89049 | 2.29100 | 0.03150 | 5.19472 | 0.77604 | 0.76028 | 0.55058 | 0.14992 | 0.41568 | 0.16047 | 0.02939 | 0.14417 | 0.08190 | 0.07455 | 0.16328 |
| **Rp9** | 2.69300 | 0.01207 | 2.58485 | 0.37724 | 4.29610 | 0.04848 | 1.25185 | 1.87160 | 2.81424 | 2.32043 | 0.91543 | 0.04369 | 2.30622 | 0.00725 | 0.20592 | 0.37691 |
| **Rpa1** | 5.13703 | 5.96719 | 8.69893 | 1.54690 | 7.01593 | 7.80640 | 11.10938 | 7.10827 | 2.23432 | 2.46881 | 4.51058 | 1.63439 | 11.91807 | 7.91771 | 6.08267 | 7.88122 |
| **Rpa2** | 17.31079 | 10.54822 | 18.48467 | 7.71041 | 16.63668 | 8.03939 | 11.82790 | 21.54941 | 16.53014 | 22.20378 | 5.15863 | 8.00914 | 15.90172 | 6.53071 | 12.98381 | 13.65959 |
| **Rpa3** | 4.01580 | 1.07390 | 0.51721 | 2.58711 | 4.10364 | 2.98775 | 0.78698 | 1.49968 | 3.48282 | 0.31397 | 4.47971 | 1.25519 | 1.70054 | 3.89641 | 1.61794 | 0.57746 |
| **Rpain** | 5.90736 | 0.35625 | 1.42240 | 1.46093 | 2.38557 | 0.00000 | 0.05447 | 1.82866 | 3.15867 | 1.48824 | 1.21873 | 0.45378 | 1.30651 | 2.21897 | 3.51523 | 0.00000 |
| **Rpap1** | 7.98013 | 5.33939 | 5.50363 | 2.37693 | 1.98520 | 0.43826 | 4.24404 | 7.05839 | 1.94999 | 6.15890 | 0.55856 | 0.40305 | 4.43020 | 0.00000 | 2.22955 | 5.46508 |
| **Rpap2** | 3.04185 | 5.62227 | 2.28378 | 4.18266 | 0.00726 | 0.01052 | 0.00000 | 2.32886 | 0.00000 | 3.37153 | 2.04590 | 0.80707 | 2.47546 | 1.69079 | 5.96001 | 4.90303 |
| **Rpap3** | 1.56845 | 6.17496 | 4.82960 | 10.58738 | 8.58622 | 3.82914 | 6.61138 | 12.58183 | 15.55598 | 21.74225 | 7.71363 | 1.81834 | 6.74269 | 0.18011 | 7.72255 | 7.25305 |
| **Rpe** | 6.79152 | 3.40116 | 3.03556 | 5.72015 | 10.70181 | 2.07475 | 2.06013 | 5.52870 | 0.07873 | 1.53261 | 0.94762 | 5.45480 | 3.04382 | 1.02516 | 3.73162 | 3.58583 |

| | | | | | | | | | | | | | | | | |
|---|---|---|---|---|---|---|---|---|---|---|---|---|---|---|---|---|
| **Rpf1** | 3.32896 | 0.30348 | 0.00000 | 7.46306 | 4.66783 | 2.09651 | 2.06629 | 7.26467 | 17.36900 | 8.23539 | 8.71379 | 2.95770 | 7.63717 | 3.64148 | 5.99546 | 0.03972 |
| **Rpf2** | 11.43286 | 12.45741 | 15.56447 | 8.37577 | 10.64672 | 4.78495 | 10.62891 | 14.05914 | 10.04205 | 18.50350 | 14.19069 | 18.53569 | 12.16215 | 28.27645 | 18.57486 | 2.11300 |
| **Rpgr** | 1.82076 | 0.00333 | 1.69551 | 0.23351 | 2.01768 | 0.00449 | 0.96224 | 1.14451 | 0.02067 | 0.41434 | 0.91362 | 1.85294 | 0.00000 | 4.64180 | 0.57949 | 0.00442 |
| **Rpgrip1** | 4.38713 | 5.58858 | 4.88128 | 2.32121 | 4.19770 | 3.35725 | 0.00000 | 4.00289 | 2.52858 | 0.00028 | 0.60659 | 0.84459 | 1.56838 | 0.40160 | 2.74257 | 0.00000 |
| **Rpgrip1l** | 0.03165 | 0.00545 | 0.44242 | 1.23012 | 3.89761 | 0.00532 | 0.36927 | 0.00215 | 0.30636 | 1.06695 | 0.15157 | 0.00000 | 0.00730 | 0.04435 | 2.70247 | 0.00655 |
| **Rph3a** | 0.01252 | 0.00787 | 0.01423 | 8.18782 | 0.00970 | 0.00565 | 0.02841 | 0.01361 | 2.49357 | 0.02545 | 0.16609 | 0.03016 | 0.03984 | 0.00000 | 1.11532 | 0.00586 |
| **Rph3al** | 0.01057 | 0.00790 | 0.11020 | 3.13855 | 0.00869 | 0.00801 | 0.00698 | 0.00722 | 4.23724 | 0.82110 | 0.00000 | 0.00000 | 0.01049 | 0.62404 | 0.06433 | 0.00643 |
| **Rpia** | 2.81781 | 0.20876 | 1.06983 | 11.55988 | 0.03230 | 1.44425 | 0.01024 | 0.10392 | 10.77376 | 3.81146 | 10.61038 | 0.44097 | 0.18821 | 2.64162 | 3.88064 | 0.02861 |
| **Rpl10** | 56.39229 | 25.34710 | 47.12568 | 23.56221 | 32.17831 | 19.22968 | 22.86764 | 49.92595 | 33.24468 | 53.98515 | 47.37198 | 17.99976 | 40.57314 | 33.70440 | 33.09986 | 27.92049 |
| **Rpl10a** | 44.07168 | 19.92956 | 35.08232 | 21.54433 | 25.87766 | 14.94284 | 16.37864 | 34.43526 | 30.21578 | 45.14377 | 41.67917 | 11.83913 | 38.12561 | 6.58627 | 7.81346 | 23.41253 |
| **Rpl10l** | 18.36421 | 5.19294 | 16.63835 | 13.50626 | 11.74239 | 3.36813 | 13.08156 | 17.90983 | 20.96050 | 18.07741 | 9.96387 | 3.62465 | 4.60941 | 2.09188 | 0.54051 | 4.99147 |
| **Rpl11** | 42.81344 | 12.48596 | 33.89514 | 18.62303 | 24.93907 | 14.18120 | 14.77638 | 26.93133 | 33.79518 | 41.03706 | 45.33372 | 9.70810 | 33.53628 | 11.36995 | 18.24704 | 21.44458 |
| **Rpl12** | 47.66475 | 18.02101 | 38.48663 | 19.83606 | 31.83050 | 17.37431 | 20.03357 | 36.38478 | 27.88563 | 50.57517 | 49.94743 | 10.42275 | 42.04813 | 5.10244 | 3.57760 | 26.42615 |
| **Rpl13** | 32.40552 | 7.22214 | 18.72186 | 26.17981 | 28.18897 | 8.54898 | 8.34870 | 5.39766 | 35.67174 | 56.86102 | 51.11073 | 5.96846 | 42.67582 | 2.62477 | 2.95645 | 25.66645 |
| **Rpl13a** | 54.87439 | 25.20902 | 40.49657 | 19.19367 | 32.22886 | 18.42793 | 25.48776 | 48.81722 | 29.14967 | 52.41622 | 49.75691 | 14.02966 | 53.14055 | 6.96853 | 12.33411 | 27.80345 |
| **Rpl14** | 48.41006 | 25.28296 | 37.82468 | 5.34850 | 27.75954 | 17.42841 | 23.52731 | 41.86665 | 17.55907 | 46.12438 | 42.47763 | 12.96566 | 46.29365 | 1.08221 | 14.46331 | 26.42637 |
| **Rpl14.ps1** | 41.44711 | 21.60593 | 31.54994 | 2.52357 | 22.89562 | 14.54804 | 20.09279 | 35.78852 | 10.34000 | 38.92091 | 33.43968 | 10.25211 | 40.27006 | 0.50253 | 10.88156 | 22.82061 |
| **Rpl15** | 51.99669 | 25.88714 | 40.52279 | 17.07793 | 32.17696 | 16.98049 | 21.27159 | 41.95132 | 29.49322 | 54.44267 | 41.73590 | 16.09634 | 43.31581 | 15.39703 | 23.66734 | 26.65890 |
| **Rpl17** | 46.12658 | 18.41051 | 35.36862 | 19.44943 | 25.72539 | 17.14336 | 16.41543 | 33.41088 | 33.13516 | 44.99051 | 48.17330 | 13.15473 | 39.86443 | 33.40673 | 23.98792 | 22.67509 |
| **Rpl18** | 48.79905 | 15.69507 | 31.23408 | 13.53425 | 26.94157 | 16.37746 | 19.37800 | 40.62643 | 21.44432 | 46.54914 | 31.64017 | 8.54672 | 45.85450 | 0.29691 | 4.35595 | 28.96739 |
| **Rpl18a** | 40.77765 | 11.50764 | 31.25246 | 21.40625 | 23.98762 | 10.49318 | 11.80522 | 22.93448 | 30.69600 | 42.23577 | 40.83570 | 8.15949 | 34.39852 | 0.93331 | 14.94386 | 21.36075 |
| **Rpl19** | 45.59271 | 25.00533 | 36.76067 | 17.21596 | 26.20640 | 15.68611 | 23.61248 | 41.82394 | 29.48984 | 44.07339 | 43.45202 | 11.03188 | 39.47200 | 13.19805 | 25.35239 | 24.90323 |
| **Rpl21** | 38.22179 | 9.52901 | 29.46566 | 22.15717 | 22.05097 | 12.55478 | 11.79487 | 23.01020 | 31.20729 | 35.22112 | 49.88887 | 8.67821 | 37.73224 | 26.14490 | 17.64104 | 20.77219 |
| **Rpl22** | 26.21980 | 8.22125 | 23.58606 | 18.86603 | 20.32963 | 6.63584 | 8.93244 | 8.10573 | 23.84039 | 21.73698 | 43.60307 | 6.40775 | 23.80757 | 15.11764 | 13.79062 | 11.98111 |
| **Rpl22l1** | 6.44159 | 0.58093 | 4.37719 | 2.09707 | 6.83099 | 0.44968 | 0.37815 | 0.78671 | 4.82425 | 1.78924 | 21.11736 | 0.98247 | 4.11016 | 3.97517 | 1.92203 | 3.34106 |
| **Rpl23** | 40.05437 | 11.61808 | 30.72212 | 27.02544 | 23.77803 | 11.39898 | 13.39842 | 20.06543 | 40.65040 | 36.36169 | 51.65954 | 9.25351 | 34.94491 | 31.93446 | 23.60315 | 20.56056 |
| **Rpl23a** | 46.22605 | 9.36676 | 32.73039 | 15.65764 | 28.63448 | 14.39114 | 13.76436 | 23.06879 | 31.26280 | 39.12571 | 50.22455 | 11.44373 | 38.40773 | 17.28328 | 23.17492 | 23.03043 |
| **Rpl24** | 39.76697 | 9.99023 | 29.68442 | 17.27594 | 28.08588 | 13.35432 | 14.48754 | 26.73385 | 34.59194 | 38.12831 | 45.08154 | 10.18049 | 35.91547 | 29.51117 | 22.26723 | 20.56922 |
| **Rpl26** | 0.04349 | 0.00819 | 0.02970 | 0.01010 | 0.01364 | 0.00716 | 0.00720 | 0.04848 | 0.01228 | 0.03052 | 0.02509 | 0.00743 | 0.05334 | 0.01158 | 0.00962 | 0.02968 |
| **Rpl27** | 35.62753 | 4.41006 | 22.75085 | 5.60657 | 24.76048 | 8.73487 | 7.63030 | 13.54116 | 21.26025 | 31.63701 | 34.55531 | 4.11203 | 27.49192 | 6.71099 | 13.27628 | 19.10434 |
| **Rpl27a** | 44.92191 | 5.47343 | 19.53765 | 16.25468 | 25.23264 | 13.53510 | 11.68190 | 21.41683 | 30.10122 | 40.74072 | 44.20044 | 11.02554 | 41.30969 | 11.96708 | 0.69792 | 23.61638 |
| **Rpl28** | 30.76414 | 4.75940 | 24.82058 | 17.80140 | 19.43364 | 8.59311 | 7.45348 | 14.43274 | 26.18695 | 28.63575 | 41.09234 | 5.40192 | 22.21055 | 1.14061 | 11.11039 | 14.93692 |
| **Rpl29** | 43.44276 | 17.03516 | 33.48058 | 11.90607 | 24.69252 | 14.96028 | 19.39399 | 38.24963 | 19.07014 | 45.16279 | 35.56997 | 10.93083 | 36.05765 | 0.35810 | 5.97308 | 22.46373 |
| **Rpl3** | 55.69209 | 33.84708 | 48.53004 | 8.57812 | 32.12741 | 23.79206 | 30.31331 | 56.84811 | 13.47412 | 48.67827 | 44.91039 | 14.36436 | 48.21735 | 33.61075 | 36.14778 | 31.08419 |
| **Rpl30** | 0.72975 | 0.06023 | 0.37498 | 0.07282 | 0.43905 | 0.07085 | 0.20190 | 0.39609 | 0.00746 | 1.25223 | 0.81339 | 0.05069 | 0.96659 | 0.32127 | 0.19504 | 0.30499 |
| **Rpl31** | 25.78697 | 2.78538 | 19.55817 | 18.23700 | 19.56521 | 7.07176 | 3.88047 | 5.98636 | 28.24634 | 19.55031 | 40.32574 | 5.05203 | 17.65069 | 13.90155 | 9.34678 | 13.82478 |
| **Rpl31.ps12** | 11.51094 | 0.67434 | 8.17484 | 10.88992 | 8.87111 | 2.60883 | 1.21503 | 1.02263 | 19.45504 | 6.51914 | 29.51434 | 1.56099 | 6.65810 | 6.90885 | 2.83256 | 5.74849 |
| **Rpl32** | 31.49358 | 5.28164 | 25.82147 | 16.84961 | 22.79869 | 9.32045 | 7.68578 | 15.20261 | 28.68149 | 31.42109 | 46.75477 | 8.87319 | 25.03752 | 17.31613 | 12.38160 | 14.99220 |
| **Rpl32l** | 5.09213 | 0.37060 | 4.40808 | 2.57138 | 3.73938 | 0.96971 | 0.67737 | 1.16603 | 6.05171 | 4.56428 | 16.45742 | 1.54824 | 3.27954 | 2.39135 | 0.85921 | 2.47126 |
| **Rpl34** | 1.21723 | 0.04809 | 0.33623 | 0.00007 | 1.50478 | 0.69274 | 0.21807 | 1.82167 | 0.02181 | 2.75691 | 0.79531 | 0.09039 | 1.78086 | 0.02353 | 0.00000 | 0.54122 |
| **Rpl35** | 31.18292 | 3.74362 | 23.47077 | 23.75369 | 18.84497 | 6.61887 | 5.70452 | 7.59714 | 31.94719 | 20.73852 | 43.23149 | 5.37566 | 20.56871 | 10.94993 | 12.59432 | 15.86331 |
| **Rpl35a** | 30.79395 | 4.23564 | 25.16446 | 9.85316 | 17.83149 | 8.18394 | 4.86672 | 7.10591 | 23.10625 | 20.51805 | 42.92301 | 4.71061 | 22.61482 | 19.36075 | 13.09672 | 13.27725 |
| **Rpl36** | 20.95136 | 0.72817 | 13.80733 | 5.75005 | 11.80774 | 3.30879 | 2.56099 | 4.43423 | 6.18869 | 14.99163 | 25.02737 | 2.12783 | 12.37710 | 0.04264 | 0.00000 | 9.91469 |
| **Rpl36a** | 31.41912 | 4.48213 | 24.10293 | 15.75633 | 17.52621 | 6.04094 | 4.35705 | 7.78643 | 25.85737 | 17.62876 | 43.92762 | 4.90234 | 23.54848 | 23.69109 | 14.29251 | 14.06805 |
| **Rpl36al** | 19.10721 | 3.54581 | 17.99667 | 20.27564 | 17.62591 | 4.62370 | 5.37991 | 7.70772 | 34.16130 | 19.15165 | 36.09831 | 3.70803 | 17.18544 | 14.62093 | 12.51230 | 10.49355 |
| **Rpl37** | 16.55040 | 1.67452 | 11.83224 | 17.10642 | 7.78863 | 2.26507 | 2.08341 | 1.68399 | 28.63767 | 6.83492 | 40.82183 | 2.91750 | 13.67627 | 9.44228 | 4.85993 | 9.56356 |
| **Rpl37a** | 25.85232 | 2.30201 | 23.63715 | 17.82786 | 15.17908 | 6.85819 | 4.02461 | 6.89524 | 31.31878 | 16.81086 | 45.94579 | 4.44696 | 19.73069 | 12.08093 | 10.38924 | 12.60051 |
| **Rpl37rt** | 16.04504 | 1.07496 | 11.40754 | 13.30000 | 7.96766 | 2.26896 | 1.21682 | 1.55292 | 23.51445 | 6.41482 | 35.09280 | 2.63834 | 12.83555 | 6.99793 | 3.87486 | 7.78804 |
| **Rpl38** | 20.74999 | 1.07532 | 18.34782 | 11.88951 | 13.41528 | 3.84423 | 2.50011 | 2.16551 | 20.36776 | 7.25075 | 40.62131 | 3.34304 | 15.30455 | 4.91391 | 4.48657 | 10.92233 |
| **Rpl39** | 18.64713 | 2.30724 | 21.38730 | 22.10278 | 8.57373 | 2.43975 | 2.41312 | 2.05339 | 36.99405 | 6.74291 | 44.43904 | 3.98439 | 9.84287 | 25.67581 | 7.21211 | 8.49458 |
| **Rpl39l** | 10.59423 | 0.48335 | 7.89816 | 15.34933 | 9.53541 | 2.84826 | 1.20031 | 5.27638 | 17.56065 | 18.02913 | 23.30840 | 4.28834 | 5.23923 | 3.97412 | 8.07328 | 5.17286 |
| **Rpl3l** | 0.01575 | 0.00783 | 0.11640 | 3.45546 | 0.00876 | 0.00522 | 0.00711 | 0.00530 | 3.09353 | 0.59341 | 0.14272 | 0.01219 | 0.01644 | 0.02196 | 0.03375 | 0.00642 |
| **Rpl4** | 57.69016 | 35.52770 | 53.67924 | 22.21727 | 40.07886 | 27.12442 | 31.70429 | 57.99330 | 34.84694 | 57.61885 | 51.20661 | 24.83230 | 57.86425 | 46.33316 | 36.78439 | 34.44688 |
| **Rpl41** | 38.47675 | 7.29061 | 32.12366 | 34.27549 | 22.98535 | 11.36955 | 10.49820 | 15.89625 | 44.11145 | 35.58262 | 56.01326 | 11.21038 | 33.00141 | 13.50441 | 22.65489 | 20.01723 |
| **Rpl5** | 55.36581 | 35.22930 | 46.74310 | 15.53736 | 35.09672 | 22.56595 | 29.80586 | 51.44518 | 29.12593 | 53.39155 | 47.55184 | 29.91132 | 52.61747 | 43.88649 | 32.00230 | 31.36368 |
| **Rpl6** | 59.04766 | 33.13335 | 48.64463 | 20.66653 | 34.83314 | 23.40642 | 29.54469 | 54.20245 | 37.25490 | 56.19147 | 48.92832 | 28.29675 | 54.10858 | 27.52041 | 31.89842 | 32.17949 |
| **Rpl7** | 54.90794 | 31.07733 | 46.00635 | 8.97087 | 34.31307 | 21.97006 | 26.42881 | 48.72366 | 20.77462 | 53.91167 | 47.72062 | 25.08496 | 50.68596 | 41.10287 | 29.59194 | 31.15689 |
| **Rpl7a** | 50.34188 | 26.92920 | 35.68840 | 10.19712 | 32.16818 | 18.70751 | 25.25506 | 49.40376 | 19.03230 | 37.93012 | 39.13758 | 17.30354 | 40.52517 | 29.83678 | 27.58409 | 26.83024 |
| **Rpl7l1** | 21.54052 | 11.55737 | 18.77265 | 10.20266 | 7.98596 | 8.61961 | 6.03153 | 18.37021 | 11.94002 | 22.85885 | 15.55987 | 8.66387 | 15.23378 | 23.47865 | 19.95355 | 9.97854 |
| **Rpl8** | 52.31952 | 22.64636 | 43.74388 | 15.84964 | 32.40290 | 20.32288 | 27.22552 | 53.92377 | 31.31653 | 52.19004 | 44.59924 | 16.34434 | 52.39677 | 1.41483 | 5.98694 | 29.58456 |
| **Rpl9** | 47.59877 | 24.06562 | 36.75141 | 18.00676 | 26.60463 | 17.36024 | 21.27558 | 41.67098 | 32.01892 | 46.10495 | 45.53208 | 14.21391 | 42.98811 | 37.33142 | 27.30318 | 25.61288 |
| **Rplp0** | 60.47692 | 29.01398 | 50.44765 | 11.49874 | 36.03426 | 27.93663 | 34.65845 | 63.56815 | 20.36617 | 68.51879 | 28.36555 | 30.82331 | 57.87487 | 0.68234 | 3.29696 | 34.00652 |

| | | | | | | | | | | | | | | | | |
|---|---|---|---|---|---|---|---|---|---|---|---|---|---|---|---|---|
| **Rplp1** | 48.25422 | 3.10809 | 28.04833 | 3.22403 | 30.47893 | 15.13431 | 13.64825 | 27.70672 | 11.01993 | 54.77957 | 21.73389 | 10.34979 | 40.21499 | 0.33129 | 0.00000 | 22.77742 |
| **Rplp2** | 30.74171 | 0.87726 | 11.50436 | 8.88093 | 19.87954 | 7.69793 | 2.66730 | 10.36839 | 17.92859 | 26.43293 | 26.63771 | 2.91513 | 25.79266 | 0.00000 | 0.00000 | 17.58936 |
| **Rpn1** | 10.74524 | 2.49449 | 19.80406 | 1.11387 | 4.60896 | 4.77538 | 8.48004 | 15.06274 | 1.88704 | 7.28294 | 8.16018 | 0.57664 | 17.67970 | 5.16947 | 3.91181 | 8.56943 |
| **Rpn2** | 13.64224 | 8.66583 | 21.15380 | 7.44861 | 6.66886 | 9.46138 | 12.54238 | 19.50686 | 3.74760 | 12.62558 | 15.33307 | 12.98187 | 17.69648 | 14.10539 | 12.02508 | 10.66314 |
| **Rpp14** | 6.82620 | 7.83146 | 14.33549 | 7.27634 | 7.71671 | 6.59521 | 3.61090 | 10.92847 | 10.64482 | 7.99564 | 3.83960 | 1.45631 | 4.71261 | 12.40891 | 12.53907 | 8.87978 |
| **Rpp21** | 4.46850 | 0.96757 | 4.31806 | 0.80645 | 2.50095 | 1.60830 | 1.38363 | 1.45180 | 2.13275 | 6.12615 | 15.36552 | 0.35792 | 6.32048 | 0.20668 | 0.22745 | 2.87337 |
| **Rpp25** | 2.02713 | 3.45086 | 0.65819 | 5.10983 | 5.05410 | 0.03030 | 0.04353 | 0.41258 | 7.38198 | 3.12985 | 0.00000 | 0.04199 | 1.50849 | 0.21394 | 0.13938 | 0.43939 |
| **Rpp25l** | 13.51592 | 4.21221 | 1.42372 | 0.03188 | 4.38688 | 2.62390 | 0.62978 | 5.43579 | 0.03254 | 6.60961 | 4.35439 | 0.04822 | 4.39364 | 0.00000 | 0.00347 | 2.35120 |
| **Rpp30** | 12.55827 | 8.55236 | 8.87122 | 3.42224 | 8.57554 | 3.63208 | 3.31886 | 6.58670 | 2.95402 | 11.27595 | 8.31961 | 5.82947 | 13.99422 | 12.24805 | 10.37095 | 2.48546 |
| **Rpp38** | 2.62269 | 3.13098 | 5.71097 | 2.84119 | 2.24797 | 2.86035 | 3.83938 | 6.17857 | 3.39893 | 5.22559 | 5.05064 | 0.91952 | 6.23568 | 2.01916 | 5.35589 | 2.80873 |
| **Rpp40** | 11.53587 | 10.56139 | 6.65587 | 7.78316 | 5.60257 | 0.06159 | 5.79741 | 4.65431 | 11.60382 | 12.75583 | 4.72633 | 7.37583 | 3.11247 | 0.36069 | 3.08637 | 6.25764 |
| **Rprd1a** | 5.72995 | 3.69975 | 7.51500 | 10.32955 | 2.68438 | 4.05611 | 0.78905 | 4.18785 | 17.95804 | 14.97130 | 3.07652 | 0.81584 | 0.01095 | 11.03499 | 14.35242 | 0.04386 |
| **Rprd1b** | 2.41790 | 0.48068 | 8.17596 | 8.32345 | 0.59006 | 1.39719 | 3.89747 | 4.68159 | 9.39955 | 10.34529 | 5.80255 | 11.56371 | 2.25417 | 11.07201 | 14.47279 | 2.17447 |
| **Rprd2** | 1.71894 | 0.25817 | 0.80082 | 5.52686 | 5.01347 | 0.79429 | 3.61689 | 0.10597 | 5.34212 | 1.64582 | 1.11456 | 0.08419 | 0.95613 | 5.65066 | 2.42838 | 0.27346 |
| **Rps10** | 37.87946 | 1.02665 | 6.16587 | 9.03066 | 28.25664 | 14.02523 | 10.25066 | 28.27651 | 20.80133 | 44.51883 | 28.84025 | 3.14028 | 36.01355 | 0.00000 | 0.00000 | 21.98048 |
| **Rps11** | 42.40802 | 13.35196 | 38.12006 | 26.21324 | 28.14950 | 14.63915 | 13.84415 | 27.68061 | 36.76489 | 38.21281 | 52.96609 | 10.84297 | 36.84308 | 27.58112 | 21.49900 | 20.30758 |
| **Rps12** | 46.34409 | 11.78526 | 38.53835 | 23.02692 | 31.80974 | 13.48065 | 16.12978 | 21.31040 | 35.24752 | 41.56772 | 55.44936 | 10.41276 | 36.97724 | 7.56316 | 14.66631 | 21.49239 |
| **Rps13** | 34.02828 | 5.28761 | 24.74879 | 23.55675 | 25.94038 | 12.49315 | 9.97727 | 17.07853 | 39.52525 | 41.43880 | 45.01727 | 10.18069 | 28.53785 | 7.48531 | 4.30978 | 17.70785 |
| **Rps14** | 39.12364 | 9.33044 | 31.74027 | 12.37238 | 27.62842 | 13.98508 | 13.89069 | 18.72966 | 14.34739 | 39.24021 | 33.62666 | 6.53226 | 30.05826 | 8.88424 | 19.89172 | 19.64335 |
| **Rps15** | 30.46119 | 4.52263 | 24.62302 | 19.40176 | 19.34397 | 6.26124 | 8.08615 | 6.38947 | 25.31970 | 26.33569 | 40.80652 | 4.01081 | 24.34705 | 1.30765 | 7.47204 | 14.63829 |
| **Rps15a** | 22.72149 | 3.03170 | 15.54166 | 14.49390 | 15.41537 | 5.35919 | 4.44084 | 5.22406 | 22.71079 | 14.28024 | 38.35431 | 3.15416 | 20.63403 | 21.06135 | 10.18729 | 12.78717 |
| **Rps15a.ps4** | 8.94933 | 0.74078 | 4.90518 | 5.07182 | 6.03291 | 1.59503 | 1.06664 | 0.74481 | 9.48005 | 3.52721 | 20.65601 | 0.76632 | 8.71969 | 9.52002 | 2.47301 | 5.18296 |
| **Rps15a.ps6** | 7.63027 | 0.63781 | 4.54458 | 3.95208 | 4.95195 | 1.34737 | 1.17627 | 0.62847 | 8.46270 | 3.53714 | 18.86411 | 0.54371 | 7.03742 | 7.41168 | 2.38473 | 4.78357 |
| **Rps16** | 45.56061 | 9.26368 | 35.34560 | 19.01236 | 28.98068 | 14.03790 | 16.07968 | 31.83424 | 29.97649 | 46.96161 | 44.80437 | 9.32179 | 44.62588 | 6.79168 | 4.97952 | 24.65890 |
| **Rps17** | 34.96614 | 5.43335 | 26.14296 | 22.60396 | 23.71054 | 11.14327 | 7.85142 | 9.54997 | 35.30675 | 28.63415 | 46.36787 | 7.00280 | 27.00416 | 22.12157 | 14.77220 | 16.39562 |
| **Rps18** | 43.02437 | 10.48952 | 34.61860 | 13.01418 | 29.68998 | 15.04140 | 15.02598 | 24.06520 | 31.13860 | 39.51270 | 49.05431 | 6.08326 | 36.00230 | 3.94854 | 20.44768 | 23.03219 |
| **Rps19** | 35.47692 | 6.60045 | 28.83747 | 20.20996 | 25.04022 | 11.18656 | 11.43689 | 17.74831 | 32.70299 | 37.48319 | 44.51617 | 7.89802 | 26.85044 | 10.74231 | 15.78045 | 16.22499 |
| **Rps19.ps3** | 12.22100 | 1.25895 | 8.75133 | 3.73247 | 8.14336 | 2.96173 | 2.70114 | 4.26693 | 8.23125 | 13.00903 | 17.24552 | 1.11548 | 8.23955 | 2.63555 | 3.77241 | 5.11827 |
| **Rps19bp1** | 4.71803 | 0.22157 | 4.39735 | 15.12841 | 5.30108 | 0.10666 | 0.70801 | 0.99718 | 22.63234 | 20.23055 | 7.79458 | 0.47855 | 6.28053 | 0.08204 | 0.86558 | 5.52507 |
| **Rps2** | 46.06439 | 13.92464 | 34.47682 | 25.28809 | 31.90419 | 16.52993 | 13.83161 | 21.09074 | 35.85172 | 58.02785 | 49.81486 | 13.63346 | 36.36485 | 7.86401 | 8.93067 | 23.59864 |
| **Rps20** | 40.28641 | 8.20857 | 30.21532 | 20.05618 | 33.47848 | 13.33699 | 12.91800 | 19.06612 | 35.19187 | 38.44709 | 46.71352 | 9.57711 | 31.91537 | 6.56684 | 0.88483 | 18.39629 |
| **Rps21** | 26.93869 | 2.41387 | 23.61579 | 20.78151 | 14.89181 | 5.50068 | 3.66999 | 4.15249 | 32.66982 | 12.91379 | 44.15305 | 4.92286 | 16.15810 | 2.87924 | 7.13720 | 13.92243 |
| **Rps23** | 27.83827 | 3.82823 | 20.57614 | 11.76715 | 17.88502 | 7.41708 | 5.10714 | 8.90077 | 22.43085 | 27.07217 | 37.59000 | 3.65004 | 19.17385 | 5.90335 | 9.54962 | 12.28717 |
| **Rps24** | 37.04254 | 8.98971 | 31.58723 | 27.30776 | 27.15610 | 12.09350 | 11.62054 | 16.96028 | 40.26907 | 34.39891 | 53.78693 | 12.22576 | 31.53523 | 34.63638 | 16.93522 | 19.69940 |
| **Rps25** | 30.35602 | 5.41568 | 25.25975 | 25.10084 | 20.99634 | 9.49013 | 7.61643 | 11.06822 | 36.42599 | 29.74579 | 52.44047 | 8.93108 | 23.12477 | 25.20776 | 17.52130 | 14.70437 |
| **Rps26** | 31.93888 | 5.50727 | 26.06710 | 20.84745 | 20.53411 | 8.80897 | 8.75369 | 12.77351 | 33.56282 | 24.35260 | 45.20318 | 6.04264 | 24.18794 | 1.97890 | 8.88421 | 14.65736 |
| **Rps27** | 0.01427 | 0.00812 | 0.01090 | 0.04113 | 0.01162 | 0.38081 | 0.50271 | 0.19199 | 0.22087 | 0.56343 | 0.13758 | 0.01604 | 0.10147 | 0.01036 | 0.00876 | 0.00746 |
| **Rps27a** | 47.13072 | 15.37281 | 33.17273 | 17.76522 | 27.86875 | 15.69076 | 15.35363 | 26.18435 | 31.86178 | 38.65336 | 50.66716 | 11.47650 | 44.61352 | 34.10506 | 23.52263 | 26.99338 |
| **Rps27l** | 19.38024 | 2.25118 | 17.19886 | 22.73987 | 19.72946 | 5.76057 | 5.45670 | 2.31572 | 22.43288 | 15.79545 | 26.92013 | 3.16819 | 11.21400 | 1.65691 | 5.00319 | 6.82748 |
| **Rps28** | 16.96411 | 2.49829 | 17.50807 | 14.29733 | 10.99885 | 4.67757 | 1.84952 | 1.88171 | 19.95603 | 5.35422 | 40.32074 | 1.95959 | 8.86340 | 1.89435 | 3.90615 | 9.04816 |
| **Rps29** | 24.26727 | 2.88755 | 29.51753 | 16.56859 | 15.87437 | 6.28154 | 5.34123 | 3.38282 | 31.22202 | 10.74187 | 50.19418 | 5.42469 | 15.32870 | 1.38956 | 4.64496 | 10.30244 |
| **Rps3** | 52.29451 | 30.82599 | 47.39495 | 16.53742 | 30.16809 | 20.65821 | 29.11066 | 54.57999 | 28.79665 | 48.27501 | 47.44906 | 19.93606 | 50.69021 | 35.34832 | 27.66199 | 29.74111 |
| **Rps3a1** | 55.83867 | 32.16381 | 47.12934 | 21.75300 | 33.86821 | 25.05535 | 27.96657 | 51.05144 | 34.60428 | 56.65377 | 48.75696 | 28.37872 | 51.06235 | 39.36257 | 31.25215 | 29.97240 |
| **Rps4l** | 25.46925 | 4.42645 | 9.46797 | 8.82688 | 17.67599 | 13.24048 | 6.55652 | 20.84833 | 9.69589 | 30.97714 | 18.63686 | 3.00558 | 28.39868 | 0.15332 | 0.00000 | 17.84375 |
| **Rps4x** | 54.48419 | 28.40845 | 49.11351 | 15.70108 | 32.81011 | 21.89345 | 23.07163 | 52.08949 | 30.63907 | 53.87877 | 43.91887 | 31.72086 | 44.86554 | 46.00397 | 35.23612 | 30.81298 |
| **Rps5** | 51.87492 | 25.57976 | 40.86451 | 16.89911 | 31.33902 | 19.46886 | 23.52266 | 44.82443 | 27.62107 | 52.56560 | 41.78331 | 16.79081 | 45.49048 | 4.54544 | 23.58569 | 26.58876 |
| **Rps6** | 55.27339 | 30.69813 | 47.41896 | 20.58629 | 34.73037 | 21.63329 | 28.74396 | 52.89107 | 36.09875 | 52.89555 | 48.19088 | 22.87480 | 51.74806 | 37.84796 | 32.12693 | 30.58135 |
| **Rps6ka1** | 15.56251 | 8.28722 | 12.72112 | 3.94075 | 7.30826 | 4.81250 | 3.35738 | 13.19191 | 0.16534 | 5.70435 | 6.86602 | 0.19778 | 9.93382 | 0.00000 | 0.79287 | 7.78965 |
| **Rps6ka2** | 0.00843 | 0.00643 | 1.02280 | 1.50855 | 2.77304 | 0.00569 | 0.00729 | 0.01961 | 1.31280 | 0.87406 | 0.00000 | 0.00000 | 0.09877 | 0.00369 | 0.07590 | 3.55322 |
| **Rps6ka3** | 0.00180 | 0.04719 | 0.75881 | 2.94425 | 9.88189 | 0.00577 | 0.00715 | 0.45308 | 0.25308 | 0.16307 | 0.00000 | 0.05701 | 0.08714 | 3.07198 | 1.12071 | 1.11277 |
| **Rps6ka4** | 1.93625 | 0.00575 | 0.38210 | 0.21865 | 0.00843 | 0.00683 | 0.02460 | 0.00000 | 0.00972 | 2.28365 | 0.60468 | 0.18320 | 1.08591 | 0.02637 | 0.17776 | 0.00470 |
| **Rps6ka5** | 0.00000 | 0.26042 | 0.81967 | 1.03916 | 0.01555 | 0.00652 | 1.30115 | 0.50585 | 2.22339 | 1.33131 | 1.60579 | 5.46606 | 0.01569 | 0.44846 | 1.39455 | 0.01295 |
| **Rps6ka6** | 0.13907 | 4.58174 | 0.00000 | 11.04865 | 0.01619 | 0.00295 | 0.01423 | 0.00869 | 4.13860 | 1.92637 | 0.00000 | 8.45963 | 0.70866 | 3.61104 | 7.59168 | 0.05487 |
| **Rps6kb1** | 2.31019 | 5.06839 | 2.81878 | 7.25504 | 0.01524 | 2.33567 | 2.64517 | 4.41217 | 5.21246 | 1.48174 | 3.92662 | 1.00878 | 1.66690 | 11.59096 | 7.92013 | 0.08707 |
| **Rps6kb2** | 6.25346 | 0.01866 | 7.46211 | 1.28834 | 0.01615 | 2.10452 | 3.95449 | 8.60883 | 3.53736 | 4.07879 | 2.55529 | 0.14480 | 6.84766 | 0.02691 | 1.85094 | 1.29575 |
| **Rps6kc1** | 0.47504 | 0.12694 | 1.71996 | 0.01099 | 3.81244 | 0.00675 | 0.05714 | 1.21228 | 0.00672 | 1.63019 | 0.00000 | 0.47095 | 0.01784 | 0.13674 | 1.06871 | 0.04490 |
| **Rps6kl1** | 0.01864 | 0.00777 | 0.00713 | 0.03189 | 0.01027 | 0.00685 | 0.00720 | 0.00182 | 0.04914 | 0.00000 | 0.33006 | 0.14782 | 0.02156 | 0.00536 | 0.16845 | 0.00719 |
| **Rps7** | 47.62655 | 20.31579 | 33.82870 | 23.45782 | 29.74005 | 17.57712 | 19.95731 | 41.73652 | 38.80833 | 51.13531 | 49.98524 | 13.89134 | 45.38334 | 17.80195 | 24.25326 | 26.73789 |
| **Rps8** | 55.69313 | 26.89096 | 43.69448 | 28.65863 | 35.08879 | 22.25823 | 24.70592 | 47.98915 | 41.15292 | 55.81979 | 55.17760 | 16.79200 | 50.76532 | 19.02965 | 30.06417 | 29.49452 |
| **Rps9** | 51.28528 | 25.02089 | 38.00783 | 22.30178 | 29.90668 | 19.67446 | 24.15271 | 47.99314 | 35.87209 | 46.79516 | 45.76040 | 11.69560 | 47.46741 | 13.21290 | 26.53278 | 27.88327 |
| **Rpsa** | 57.90127 | 31.25148 | 51.37004 | 16.97447 | 37.41616 | 24.04455 | 29.59609 | 52.21396 | 23.09926 | 59.20330 | 39.90988 | 25.12526 | 52.13303 | 1.90148 | 17.56080 | 32.25717 |
| **Rptor** | 5.22587 | 2.91381 | 2.48003 | 2.45238 | 4.73027 | 1.01944 | 0.00000 | 7.50371 | 2.31761 | 2.78476 | 1.25078 | 0.00000 | 2.51830 | 0.00000 | 0.54871 | 2.01222 |

| | | | | | | | | | | | | | | | | |
|---|---|---|---|---|---|---|---|---|---|---|---|---|---|---|---|---|
| **Rpusd1** | 1.40062 | 0.00583 | 3.00077 | 3.48095 | 0.01754 | 5.48872 | 0.53694 | 2.06161 | 2.94042 | 2.93154 | 0.50453 | 0.02420 | 0.82086 | 0.00000 | 0.33481 | 3.29061 |
| **Rpusd2** | 2.50634 | 0.11295 | 3.59603 | 1.24049 | 0.00679 | 0.03644 | 0.02500 | 1.86368 | 4.27925 | 1.92418 | 2.82191 | 0.38756 | 0.33635 | 0.68669 | 3.25831 | 1.84987 |
| **Rpusd3** | 10.93361 | 0.92420 | 4.08654 | 2.39989 | 0.00000 | 0.00102 | 0.00000 | 4.11837 | 2.47174 | 5.25761 | 1.62851 | 0.16009 | 4.15451 | 0.22881 | 0.01650 | 0.00591 |
| **Rpusd4** | 1.14171 | 2.26322 | 6.32264 | 1.40120 | 3.41174 | 3.09795 | 0.05243 | 2.28674 | 1.05440 | 7.68234 | 3.19479 | 0.38427 | 2.76121 | 0.58781 | 7.62770 | 6.69340 |
| **Rrad** | 0.00679 | 0.01705 | 0.00764 | 0.00366 | 0.00962 | 0.00677 | 1.83812 | 0.04966 | 0.24498 | 0.79811 | 0.00990 | 0.00353 | 0.00921 | 0.01186 | 0.00680 | 0.00757 |
| **Rraga** | 1.92141 | 4.96380 | 5.67444 | 0.86519 | 3.46541 | 0.14653 | 0.04386 | 1.87986 | 0.58055 | 0.00000 | 0.69875 | 0.01796 | 2.23992 | 0.00450 | 0.56069 | 2.54230 |
| **Rragc** | 4.53347 | 4.07171 | 5.96061 | 26.63034 | 3.65111 | 5.70228 | 5.75773 | 3.86840 | 12.82967 | 4.19368 | 2.88855 | 2.63137 | 3.32448 | 0.13984 | 16.35211 | 1.56101 |
| **Rragd** | 1.67427 | 0.16627 | 3.78217 | 0.05459 | 0.13212 | 14.68104 | 6.08984 | 20.41201 | 0.30452 | 12.76003 | 17.65160 | 7.14597 | 12.33348 | 16.25904 | 0.00000 | 0.76465 |
| **Rras** | 1.10223 | 0.00635 | 0.75902 | 0.10092 | 0.00495 | 0.00884 | 0.00663 | 0.00000 | 1.48631 | 0.81428 | 1.01897 | 0.05660 | 0.10777 | 0.00965 | 0.00000 | 0.03804 |
| **Rras2** | 8.92776 | 1.59551 | 4.85283 | 6.04207 | 5.05007 | 0.43793 | 2.06798 | 6.06938 | 7.57219 | 14.21218 | 6.50229 | 14.27950 | 1.70431 | 7.73173 | 6.96242 | 1.36265 |
| **Rrbp1** | 11.64011 | 2.33441 | 13.59231 | 9.30374 | 5.59053 | 4.90833 | 10.40924 | 11.26438 | 11.43399 | 2.44069 | 1.67631 | 1.57932 | 7.26560 | 0.88302 | 3.68662 | 11.61992 |
| **Rreb1** | 2.40539 | 0.05178 | 1.34926 | 8.49651 | 2.44704 | 0.00282 | 2.13868 | 12.16786 | 14.46483 | 8.41751 | 9.08293 | 1.68766 | 2.55207 | 5.60919 | 3.65292 | 4.21791 |
| **Rrm1** | 17.13318 | 17.25571 | 20.61288 | 5.61805 | 19.26392 | 11.24486 | 15.22501 | 18.16024 | 13.94357 | 19.01053 | 9.52542 | 8.68655 | 17.71916 | 14.71378 | 16.34151 | 4.96734 |
| **Rrm2** | 15.56491 | 11.91265 | 15.74089 | 8.79100 | 8.42094 | 9.56510 | 12.59168 | 21.09725 | 21.76338 | 18.85654 | 25.45651 | 7.99061 | 21.10496 | 19.65545 | 20.52852 | 11.00967 |
| **Rrm2b** | 0.12536 | 0.00783 | 0.00000 | 11.64033 | 0.00746 | 0.00420 | 0.00654 | 0.00000 | 4.44041 | 0.00000 | 0.99051 | 0.05046 | 0.00933 | 4.69690 | 10.30804 | 0.00436 |
| **Rrn3** | 15.61615 | 7.84077 | 12.19503 | 14.66061 | 12.76804 | 6.52350 | 5.12319 | 16.02398 | 18.21696 | 20.69396 | 11.38965 | 5.40810 | 8.84395 | 19.71247 | 18.21831 | 11.12198 |
| **Rrnad1** | 1.71885 | 1.50457 | 3.43179 | 2.58805 | 0.03303 | 4.39496 | 0.64256 | 4.06511 | 0.65251 | 3.47224 | 0.53136 | 2.95037 | 2.56049 | 0.40705 | 4.22740 | 2.54811 |
| **Rrp1** | 16.50829 | 7.41238 | 16.63416 | 9.27248 | 5.52961 | 10.71655 | 8.06785 | 25.81364 | 8.36969 | 16.01342 | 16.17170 | 1.30658 | 16.48969 | 0.92939 | 7.63962 | 8.69841 |
| **Rrp12** | 9.76586 | 5.16877 | 7.22486 | 11.75502 | 3.69438 | 0.07344 | 7.47821 | 12.30036 | 12.07904 | 11.60406 | 6.27160 | 0.41845 | 5.09541 | 2.00501 | 9.07672 | 0.06061 |
| **Rrp15** | 13.05541 | 0.11021 | 0.00000 | 1.27241 | 9.81142 | 2.91821 | 2.20961 | 21.77495 | 4.14517 | 27.37177 | 4.52134 | 2.01541 | 13.65705 | 0.00000 | 0.00000 | 4.05743 |
| **Rrp1b** | 13.58911 | 6.80806 | 3.54031 | 1.19201 | 4.07941 | 3.60566 | 2.06784 | 12.85900 | 1.49260 | 6.91590 | 2.78149 | 0.27687 | 2.90769 | 0.35759 | 5.54002 | 2.80310 |
| **Rrp36** | 7.82953 | 7.46040 | 8.85970 | 2.57305 | 5.37222 | 4.93666 | 4.34318 | 5.59939 | 4.56111 | 8.99481 | 3.72274 | 0.52195 | 5.98912 | 0.01211 | 1.51924 | 2.98259 |
| **Rrp7a** | 6.52308 | 1.71080 | 1.89618 | 7.09378 | 7.33952 | 4.38740 | 1.67161 | 3.59321 | 13.77529 | 4.40764 | 2.07042 | 0.16540 | 6.97414 | 1.50275 | 2.10526 | 6.25143 |
| **Rrp8** | 6.14314 | 0.90780 | 5.27001 | 4.48982 | 0.05571 | 7.14490 | 0.28056 | 13.01283 | 6.93045 | 8.86745 | 4.33803 | 0.56146 | 5.19925 | 3.12171 | 8.67622 | 6.36357 |
| **Rrp9** | 8.19601 | 2.78427 | 4.67731 | 11.73656 | 7.00713 | 5.18775 | 1.81129 | 9.10342 | 16.55892 | 20.79114 | 7.73365 | 0.62521 | 8.75070 | 0.00000 | 0.00000 | 7.45140 |
| **Rrs1** | 4.28673 | 1.58208 | 7.18889 | 12.08802 | 4.30606 | 3.68413 | 4.16207 | 12.27367 | 15.60443 | 5.88543 | 4.31251 | 1.33785 | 6.48259 | 11.12920 | 7.90539 | 8.21599 |
| **Rs1** | 0.00942 | 0.00491 | 0.00415 | 0.02333 | 0.00859 | 0.00500 | 0.00606 | 1.03052 | 0.02625 | 0.00000 | 0.00000 | 7.91504 | 0.00551 | 0.55218 | 0.00000 | 0.00685 |
| **Rsad1** | 0.99992 | 0.04551 | 0.09070 | 0.60317 | 0.00585 | 3.46766 | 0.01634 | 0.00000 | 0.19059 | 0.44075 | 0.24175 | 0.04877 | 0.00000 | 0.29225 | 0.48052 | 0.01277 |
| **Rsad2** | 0.01187 | 0.00768 | 0.01222 | 0.00514 | 0.00849 | 0.00653 | 0.00717 | 0.00921 | 0.01088 | 4.18541 | 0.00966 | 0.00000 | 0.01074 | 0.01239 | 0.00628 | 0.00769 |
| **Rsbn1** | 0.01861 | 0.00819 | 0.31430 | 0.55410 | 0.00876 | 0.00617 | 0.00827 | 0.38914 | 1.06071 | 0.45646 | 0.00000 | 0.01681 | 0.03888 | 0.01532 | 0.75751 | 0.02981 |
| **Rsbn1l** | 0.14643 | 2.35691 | 1.56264 | 5.52774 | 0.07871 | 2.96135 | 1.20800 | 0.70577 | 7.52742 | 0.40167 | 1.15734 | 0.37289 | 1.27672 | 2.97451 | 3.38574 | 0.20732 |
| **Rsf1** | 1.00445 | 0.01603 | 1.34974 | 2.72974 | 0.01228 | 0.00714 | 0.98184 | 1.60822 | 6.22534 | 4.34115 | 2.74470 | 1.30229 | 0.00000 | 12.44948 | 2.49937 | 0.00413 |
| **Rsl1** | 0.29027 | 0.00774 | 0.27351 | 0.00450 | 0.01306 | 0.00558 | 0.00688 | 0.00000 | 0.01015 | 0.06023 | 0.36057 | 0.06164 | 0.07998 | 0.26560 | 0.42327 | 0.41141 |
| **Rsl1d1** | 23.17774 | 22.21800 | 23.74538 | 22.71445 | 20.00987 | 10.94423 | 11.39712 | 26.22850 | 26.14506 | 36.49535 | 19.76967 | 21.27166 | 18.00802 | 29.48752 | 28.30711 | 11.94462 |
| **Rsl24d1** | 13.86100 | 10.58951 | 13.09895 | 11.67536 | 8.99021 | 0.21038 | 8.85076 | 11.67530 | 18.42193 | 15.27651 | 11.52481 | 0.83922 | 9.25410 | 32.76909 | 17.36132 | 3.41567 |
| **Rslcan18** | 0.01159 | 0.00695 | 0.30487 | 0.21800 | 0.00971 | 0.00610 | 1.61220 | 0.02511 | 0.00548 | 0.01016 | 0.01342 | 0.00624 | 0.02118 | 0.01139 | 0.00409 | 0.00740 |
| **Rsph1** | 1.30934 | 0.01602 | 0.52279 | 2.99241 | 0.00585 | 0.06742 | 1.95218 | 0.39538 | 4.56930 | 0.43526 | 0.70917 | 0.03078 | 0.06829 | 0.20991 | 0.88112 | 0.24427 |
| **Rsph3a** | 0.01733 | 0.04056 | 0.00669 | 0.17930 | 0.00929 | 0.35918 | 0.00743 | 0.01864 | 0.01087 | 0.20060 | 0.01204 | 0.00744 | 0.01512 | 0.00792 | 0.06710 | 0.00740 |
| **Rsph3b** | 0.01770 | 0.05102 | 0.00548 | 0.19034 | 0.00918 | 0.36717 | 0.00744 | 0.01247 | 0.01730 | 0.22967 | 0.01126 | 0.04914 | 0.01550 | 0.00815 | 0.08347 | 0.00739 |
| **Rsph9** | 0.01152 | 0.00842 | 0.03764 | 0.00465 | 0.01808 | 0.00690 | 0.00774 | 0.01721 | 0.43243 | 0.89380 | 0.00738 | 0.04356 | 0.01080 | 0.01034 | 0.03243 | 0.00721 |
| **Rspo1** | 0.01289 | 0.00780 | 0.01226 | 0.00640 | 0.00896 | 0.00646 | 0.00725 | 0.00995 | 0.01159 | 1.61473 | 0.01092 | 0.00349 | 0.01074 | 0.01221 | 0.00663 | 0.00733 |
| **Rspo2** | 0.01357 | 0.00767 | 0.00526 | 8.41383 | 0.00930 | 0.01795 | 0.00696 | 0.00840 | 1.82566 | 0.00000 | 0.04170 | 0.18463 | 0.01233 | 0.00000 | 2.68447 | 0.00573 |
| **Rspry1** | 2.98779 | 0.00464 | 0.52310 | 2.28031 | 0.63636 | 0.00209 | 0.09642 | 0.43706 | 3.57027 | 1.09861 | 0.23810 | 0.21696 | 0.41671 | 2.48588 | 3.74425 | 0.00336 |
| **Rsrc1** | 6.14591 | 8.16062 | 1.93849 | 0.05318 | 0.19625 | 6.11423 | 3.61176 | 3.16643 | 1.24140 | 2.01575 | 0.17408 | 0.15531 | 3.14612 | 0.81003 | 1.21924 | 1.62435 |
| **Rsrc2** | 6.65870 | 10.03183 | 6.49961 | 15.75709 | 8.37133 | 9.34640 | 10.58066 | 12.47674 | 24.51896 | 15.84448 | 10.70046 | 14.29799 | 8.48616 | 18.56811 | 13.87887 | 6.91044 |
| **Rsrp1** | 4.65918 | 0.04231 | 4.48551 | 9.48748 | 0.03171 | 3.71650 | 5.97353 | 4.64678 | 23.85785 | 5.22203 | 9.90317 | 3.98505 | 2.05685 | 1.43543 | 10.57665 | 1.80217 |
| **Rsu1** | 4.85686 | 1.61978 | 2.24406 | 0.67985 | 4.56396 | 0.91760 | 0.23330 | 0.00000 | 0.43100 | 2.87563 | 0.41120 | 0.05266 | 2.86719 | 0.02314 | 0.28239 | 0.12627 |
| **Rtbdn** | 0.00725 | 0.00832 | 0.00313 | 0.68244 | 0.00919 | 0.00797 | 0.00753 | 0.60122 | 1.53793 | 0.65287 | 1.20615 | 0.03101 | 0.19214 | 0.00460 | 0.01604 | 0.00723 |
| **Rtca** | 4.71361 | 4.18921 | 1.98657 | 4.18702 | 6.58759 | 2.95911 | 3.05733 | 0.41789 | 3.22828 | 6.72505 | 4.95302 | 5.63465 | 5.24618 | 1.15493 | 1.06734 | 5.39345 |
| **Rtcb** | 12.88960 | 11.46624 | 10.68243 | 1.82187 | 10.58281 | 6.91649 | 9.52824 | 10.10171 | 2.09693 | 9.13072 | 3.77107 | 6.08856 | 11.15530 | 7.46133 | 1.61736 | 4.52618 |
| **Rtel1** | 8.05439 | 3.45018 | 9.86514 | 9.90913 | 0.98151 | 0.00000 | 9.75331 | 13.64564 | 2.81967 | 7.87364 | 7.45185 | 2.05383 | 6.41538 | 8.96067 | 10.73431 | 1.04744 |
| **Rtf1** | 6.12621 | 4.10209 | 4.32706 | 4.84967 | 0.11054 | 0.35671 | 5.52935 | 6.21299 | 5.48068 | 8.13529 | 3.57504 | 0.93477 | 2.28641 | 3.41996 | 4.64901 | 1.75970 |
| **Rtfdc1** | 12.27396 | 5.20837 | 11.50258 | 11.81612 | 9.05558 | 5.79530 | 5.53550 | 11.88616 | 18.93679 | 19.61864 | 7.01052 | 2.08380 | 9.24472 | 7.34552 | 14.95830 | 1.04277 |
| **Rtkn** | 3.17737 | 3.25677 | 0.38859 | 0.00000 | 2.23890 | 0.02410 | 3.20034 | 0.84186 | 0.00882 | 0.50848 | 0.27262 | 0.00451 | 1.12314 | 0.00863 | 0.00875 | 0.00225 |
| **Rtkn2** | 0.95415 | 0.19848 | 0.00000 | 0.40288 | 0.06418 | 0.00660 | 0.01625 | 0.01259 | 0.63679 | 0.23297 | 0.16657 | 0.02939 | 0.08082 | 0.38461 | 0.87005 | 0.00614 |
| **Rtn1** | 0.01067 | 0.00792 | 0.00727 | 0.29362 | 0.00824 | 0.66340 | 0.00725 | 0.01026 | 0.00674 | 2.30634 | 0.01047 | 0.01339 | 0.00669 | 0.00728 | 0.06873 | 0.49433 |
| **Rtn2** | 0.00226 | 0.00732 | 0.00000 | 3.58972 | 0.00827 | 0.00651 | 0.00744 | 0.73763 | 8.44430 | 12.86182 | 0.63491 | 2.99981 | 0.18824 | 0.01259 | 0.12123 | 0.00602 |
| **Rtn3** | 6.24994 | 3.06652 | 0.59567 | 1.34780 | 2.18719 | 8.10511 | 0.00000 | 1.02685 | 1.41642 | 5.42981 | 0.71045 | 0.07624 | 5.38943 | 0.02044 | 1.05450 | 5.44842 |
| **Rtn4** | 16.90988 | 16.44885 | 24.11354 | 17.97263 | 12.53068 | 12.72212 | 11.65083 | 22.56639 | 31.43956 | 22.81112 | 30.71438 | 15.24887 | 24.33991 | 35.55701 | 9.44107 | 10.62062 |
| **Rtn4ip1** | 0.64341 | 0.00624 | 2.68790 | 1.95154 | 0.01072 | 1.53756 | 2.66731 | 0.53536 | 0.40256 | 0.68517 | 0.52553 | 0.04407 | 1.26423 | 0.00314 | 1.23528 | 0.01903 |
| **Rtn4r** | 0.00247 | 0.00698 | 0.01514 | 0.00550 | 0.00849 | 0.00833 | 0.30124 | 2.10532 | 0.40154 | 4.76256 | 0.94942 | 0.13649 | 0.02171 | 0.14860 | 0.00000 | 0.00721 |
| **Rtn4rl1** | 0.01029 | 0.00732 | 0.09017 | 0.43083 | 0.00949 | 0.00553 | 0.00679 | 0.51426 | 1.41673 | 0.33214 | 1.10055 | 0.04115 | 0.00000 | 0.00162 | 0.59702 | 0.00707 |

| | | | | | | | | | | | | | | | | |
|---|---|---|---|---|---|---|---|---|---|---|---|---|---|---|---|---|
| **Rtp3** | 1.77877 | 0.52140 | 3.32208 | 0.49021 | 1.13109 | 0.04519 | 1.32162 | 1.57441 | 0.57257 | 1.90514 | 1.95221 | 2.74257 | 0.67797 | 2.23866 | 0.77860 | 0.49517 |
| **Rttn** | 0.02532 | 0.10947 | 0.09483 | 1.99927 | 0.00902 | 0.00771 | 0.00706 | 0.11558 | 0.30610 | 0.28630 | 0.22578 | 0.04263 | 0.00924 | 0.07854 | 0.16183 | 0.00724 |
| **Rubcn** | 0.89411 | 0.01694 | 3.35223 | 5.61950 | 0.00083 | 0.00517 | 0.11690 | 1.38320 | 1.07237 | 0.79175 | 0.26810 | 0.03708 | 1.87833 | 0.00000 | 3.55536 | 1.96445 |
| **Rufy1** | 2.13083 | 0.05595 | 0.27612 | 2.85848 | 0.01209 | 0.00892 | 1.25227 | 2.54333 | 2.47306 | 3.30514 | 1.93113 | 0.04459 | 0.00000 | 0.00000 | 0.64038 | 0.12311 |
| **Rufy2** | 0.05121 | 0.00807 | 0.02830 | 2.78879 | 0.00949 | 0.00486 | 0.00723 | 0.00001 | 7.47062 | 0.15760 | 0.18409 | 0.06497 | 0.07643 | 0.05175 | 1.10784 | 0.02015 |
| **Rufy3** | 3.90181 | 0.52249 | 2.66503 | 1.76418 | 0.07010 | 1.66148 | 3.72987 | 0.92566 | 0.99879 | 1.66040 | 3.76355 | 0.38561 | 1.18550 | 1.56371 | 1.26745 | 0.61734 |
| **Rufy4** | 0.00929 | 0.00769 | 0.41151 | 0.39902 | 0.00902 | 0.00559 | 0.00730 | 1.08059 | 0.00564 | 0.00418 | 0.00711 | 0.00583 | 0.00349 | 0.01188 | 0.00000 | 0.00714 |
| **Rundc1** | 0.46728 | 0.16834 | 1.18336 | 1.97700 | 0.01297 | 0.00519 | 2.47514 | 1.89798 | 4.15488 | 5.03854 | 2.63338 | 0.59358 | 2.50365 | 1.45568 | 1.92508 | 0.00408 |
| **Rundc3a** | 0.99982 | 0.00396 | 0.00000 | 1.47453 | 2.52830 | 0.00598 | 0.93056 | 3.81257 | 0.00000 | 2.91036 | 2.36519 | 1.25113 | 3.37290 | 0.02411 | 0.13925 | 0.53008 |
| **Rundc3b** | 0.01255 | 0.00787 | 0.00967 | 2.54079 | 0.00908 | 0.00622 | 0.00758 | 0.01682 | 0.50207 | 0.98505 | 0.03006 | 0.02328 | 0.01109 | 0.00280 | 1.45086 | 0.00693 |
| **Runx1** | 0.84357 | 0.00667 | 0.19334 | 0.01006 | 0.00695 | 0.02211 | 0.01648 | 0.16081 | 0.01230 | 0.08059 | 0.01184 | 0.01410 | 0.85373 | 0.16636 | 0.00000 | 0.72335 |
| **Runx1t1** | 1.30141 | 0.37694 | 0.02102 | 1.55826 | 0.09453 | 0.00459 | 0.01791 | 0.06322 | 0.64887 | 3.29321 | 0.02428 | 0.08047 | 0.43493 | 3.91602 | 0.05983 | 0.02059 |
| **Runx2** | 0.01337 | 0.01852 | 0.01164 | 0.16641 | 0.00923 | 0.00676 | 0.00755 | 0.01298 | 0.04319 | 0.08191 | 0.00952 | 0.00741 | 0.01327 | 0.00548 | 0.53567 | 0.00739 |
| **Rusc1** | 0.00663 | 0.00359 | 0.23218 | 4.05481 | 0.00688 | 0.48245 | 0.00000 | 13.45571 | 5.47154 | 20.79282 | 18.04258 | 1.57835 | 1.44018 | 0.06086 | 4.19975 | 0.00228 |
| **Rusc2** | 0.00890 | 0.00738 | 0.04151 | 1.71420 | 0.00885 | 0.00870 | 0.00655 | 1.89744 | 0.59816 | 0.13796 | 0.42853 | 0.03747 | 0.96547 | 0.05570 | 0.00000 | 0.00690 |
| **Ruvbl1** | 24.76023 | 9.09349 | 14.49619 | 12.57649 | 17.55558 | 8.95458 | 12.23495 | 20.07217 | 10.66946 | 22.28094 | 9.67592 | 9.39928 | 14.36022 | 5.23765 | 12.20534 | 8.13831 |
| **Ruvbl2** | 21.66589 | 7.48587 | 18.91150 | 3.55143 | 17.26772 | 13.60840 | 14.07665 | 30.09574 | 5.45092 | 25.54854 | 19.07055 | 2.88896 | 21.47747 | 0.17318 | 1.19147 | 11.89997 |
| **Rwdd1** | 7.97699 | 2.80371 | 13.08502 | 13.84489 | 2.32382 | 0.09790 | 3.78131 | 9.92439 | 22.38031 | 29.18537 | 25.22225 | 14.09107 | 11.12870 | 14.82897 | 9.36857 | 3.31584 |
| **Rwdd2a** | 0.00724 | 0.00835 | 0.30547 | 2.81044 | 0.00946 | 0.15873 | 0.04327 | 0.73949 | 5.38690 | 1.74940 | 0.00000 | 0.05602 | 0.10080 | 0.02455 | 0.78815 | 0.00663 |
| **Rwdd2b** | 1.73963 | 0.00295 | 1.92520 | 0.97609 | 5.63256 | 0.28406 | 0.23861 | 0.37103 | 0.90070 | 5.57624 | 0.70420 | 0.11662 | 2.07382 | 1.16755 | 1.50641 | 1.03266 |
| **Rwdd3** | 1.07237 | 0.01417 | 1.12402 | 0.00172 | 0.00397 | 0.00352 | 0.00522 | 0.57299 | 0.33167 | 0.05729 | 0.61897 | 6.32405 | 0.00559 | 0.01211 | 0.07511 | 0.01311 |
| **Rwdd4a** | 2.01303 | 4.13655 | 0.32222 | 11.56325 | 1.13927 | 4.07210 | 0.21345 | 0.00000 | 15.53489 | 16.12948 | 8.80415 | 2.76477 | 8.60908 | 7.13285 | 7.42267 | 4.32746 |
| **Rxra** | 1.06936 | 0.01492 | 3.39637 | 1.49298 | 0.06602 | 0.97809 | 0.77391 | 12.64905 | 1.88295 | 6.69073 | 5.75363 | 0.32142 | 9.12701 | 0.31292 | 1.30040 | 0.73365 |
| **Rxrb** | 4.32818 | 3.44312 | 0.92704 | 0.43687 | 6.51264 | 8.45771 | 4.37664 | 7.91155 | 1.43455 | 6.75090 | 1.69969 | 0.11582 | 4.03118 | 0.00000 | 0.32004 | 2.20600 |
| **Rxrg** | 0.78491 | 0.01307 | 0.01482 | 0.00734 | 4.24423 | 0.14277 | 2.26770 | 0.01373 | 0.01019 | 0.01835 | 0.02428 | 0.00116 | 0.03969 | 0.01290 | 0.03273 | 0.00627 |
| **Rybp** | 7.44677 | 9.33430 | 6.43233 | 14.71179 | 3.16229 | 4.33296 | 4.20846 | 9.81101 | 15.29020 | 15.59582 | 11.39403 | 19.84173 | 5.23032 | 27.60166 | 16.54086 | 8.29125 |
| **Ryk** | 4.15439 | 7.41044 | 9.79285 | 11.11151 | 6.03042 | 0.74616 | 5.62131 | 3.58764 | 2.18273 | 3.96190 | 1.72907 | 0.52103 | 7.40365 | 1.36109 | 9.77986 | 5.55089 |
| **Ryr1** | 0.00000 | 0.05849 | 1.26870 | 0.38942 | 0.00671 | 0.00881 | 2.39335 | 0.15652 | 1.01856 | 0.67138 | 2.06380 | 0.20153 | 1.72859 | 0.05634 | 0.46303 | 0.00530 |
| **Ryr2** | 0.41943 | 0.01695 | 0.05186 | 0.02234 | 0.00721 | 0.01790 | 0.01704 | 2.08264 | 0.00869 | 0.36530 | 0.32077 | 0.00905 | 0.03823 | 0.21470 | 0.00000 | 0.00700 |
| **Ryr3** | 0.01063 | 0.00771 | 1.54263 | 0.03402 | 0.00685 | 0.01387 | 0.00693 | 0.00275 | 0.68819 | 0.00000 | 0.16952 | 0.06823 | 0.00591 | 1.58550 | 0.41359 | 0.00650 |
| **S100a1** | 0.01347 | 0.00801 | 0.01216 | 0.17047 | 0.00912 | 0.00670 | 0.00742 | 0.01169 | 0.00894 | 0.54240 | 0.01282 | 0.00718 | 0.01205 | 0.01093 | 0.00703 | 0.00740 |
| **S100a10** | 4.33955 | 0.11736 | 1.83029 | 0.00000 | 0.09750 | 3.74019 | 0.14649 | 0.00000 | 0.01078 | 2.87678 | 1.14908 | 0.02349 | 4.16061 | 0.01034 | 0.00000 | 10.04290 |
| **S100a11** | 0.10635 | 0.37302 | 2.08816 | 0.10518 | 1.82898 | 6.24771 | 0.63030 | 0.13675 | 0.13552 | 3.54126 | 4.05497 | 0.02469 | 7.30213 | 1.73003 | 0.00000 | 4.18045 |
| **S100a13** | 0.01289 | 0.00805 | 0.00416 | 0.96397 | 0.00905 | 0.00706 | 0.00694 | 0.00000 | 1.21802 | 0.18783 | 0.59786 | 0.09754 | 0.00671 | 0.56089 | 0.00000 | 0.00684 |
| **S100a14** | 0.01557 | 0.00756 | 0.23165 | 3.97146 | 0.00823 | 5.30690 | 0.00621 | 0.00642 | 0.03795 | 0.04116 | 0.02973 | 0.05463 | 1.00171 | 0.01242 | 0.02364 | 0.00414 |
| **S100a16** | 0.01228 | 0.00796 | 0.01134 | 0.00108 | 0.00865 | 2.05974 | 0.00727 | 0.01013 | 0.69352 | 2.98410 | 0.00325 | 0.01083 | 0.01156 | 0.01258 | 0.00653 | 0.00666 |
| **S100a3** | 0.01189 | 0.00812 | 0.01274 | 0.00428 | 0.01001 | 0.00734 | 0.00765 | 0.01370 | 0.40155 | 0.36424 | 0.03913 | 0.04876 | 0.28814 | 0.00993 | 0.22725 | 0.00766 |
| **S100a4** | 0.01361 | 0.00816 | 0.01062 | 0.01023 | 0.00922 | 0.00699 | 0.00767 | 0.01626 | 0.01210 | 0.11023 | 0.01140 | 0.09108 | 0.01298 | 0.01188 | 0.01025 | 0.00742 |
| **S100a6** | 0.01275 | 0.00760 | 0.48739 | 0.38456 | 0.00795 | 0.00607 | 0.00719 | 0.00976 | 0.00578 | 2.46912 | 0.01018 | 0.08984 | 0.00918 | 0.01103 | 0.00000 | 0.00709 |
| **S100g** | 0.01362 | 0.00789 | 0.01219 | 0.00909 | 0.00917 | 0.00664 | 0.00737 | 0.01144 | 0.01214 | 0.35326 | 0.01272 | 0.00680 | 0.01224 | 0.01143 | 0.01004 | 0.00763 |
| **S100pbp** | 1.11005 | 0.09752 | 1.83123 | 2.94589 | 0.02755 | 2.49979 | 0.02534 | 2.52110 | 5.05304 | 2.40991 | 3.28139 | 0.05688 | 0.00000 | 1.70975 | 1.44041 | 0.01714 |
| **S1pr1** | 0.01212 | 0.00855 | 0.01229 | 0.01061 | 0.00994 | 0.00706 | 0.00818 | 0.05451 | 0.01229 | 0.01386 | 0.01357 | 0.01964 | 0.02831 | 0.01323 | 0.00954 | 0.00763 |
| **S1pr2** | 2.97601 | 0.00000 | 1.61392 | 0.51154 | 2.26747 | 4.31127 | 0.00000 | 6.43391 | 0.31571 | 5.25777 | 5.99775 | 2.32225 | 1.53704 | 3.92155 | 0.82040 | 0.00726 |
| **S1pr3** | 0.01379 | 0.00786 | 0.01181 | 0.27983 | 0.00936 | 0.00665 | 0.00750 | 0.01327 | 0.07087 | 0.01291 | 0.01148 | 0.01800 | 0.01280 | 0.01763 | 0.03132 | 0.00737 |
| **S1pr4** | 0.66944 | 0.00678 | 0.10437 | 0.03643 | 0.00660 | 0.01308 | 0.02692 | 0.01677 | 0.12049 | 0.06700 | 0.44204 | 0.07072 | 0.28347 | 0.22190 | 0.05680 | 0.02770 |
| **S1pr5** | 0.00813 | 0.00693 | 0.40215 | 1.50646 | 0.00895 | 0.00894 | 0.00556 | 1.09183 | 0.10872 | 0.31272 | 3.38974 | 0.00000 | 0.24869 | 0.00000 | 0.97784 | 0.00644 |
| **Saa1** | 0.01370 | 0.00797 | 0.01192 | 0.36238 | 0.00924 | 0.00658 | 0.00748 | 0.01296 | 0.13462 | 0.01695 | 0.01136 | 0.01796 | 0.02637 | 0.00742 | 0.06912 | 0.00726 |
| **Saa2** | 0.01386 | 0.00817 | 0.00981 | 0.24783 | 0.00910 | 0.00675 | 0.00752 | 0.00687 | 0.09159 | 0.02895 | 0.29990 | 0.00689 | 0.00899 | 0.00796 | 0.07489 | 0.00715 |
| **Saa3** | 0.01385 | 0.01852 | 0.25157 | 0.16771 | 0.00870 | 0.00565 | 0.00720 | 0.01097 | 0.06292 | 0.00732 | 0.01275 | 0.03875 | 0.26861 | 0.01259 | 0.00112 | 0.00701 |
| **Saal1** | 9.02717 | 6.62255 | 5.72989 | 0.00000 | 6.54767 | 0.00000 | 7.79142 | 7.20470 | 0.00454 | 5.25061 | 5.89054 | 1.69973 | 2.65391 | 2.82683 | 1.04012 | 1.15025 |
| **Sac3d1** | 0.78700 | 0.04548 | 0.00000 | 17.53244 | 3.29225 | 2.03082 | 0.03270 | 1.39769 | 23.24957 | 2.63163 | 8.10594 | 0.38928 | 0.00000 | 0.45739 | 0.00549 | 0.02938 |
| **Sacm1l** | 2.51218 | 5.58516 | 2.63271 | 2.30012 | 3.21265 | 4.75688 | 3.02923 | 0.83241 | 2.00221 | 1.87668 | 1.38533 | 5.37269 | 0.38310 | 2.06174 | 3.08122 | 2.08239 |
| **Sae1** | 15.08080 | 9.37384 | 14.39463 | 5.27847 | 18.14734 | 7.10485 | 9.45180 | 21.25069 | 7.98703 | 18.08916 | 12.25605 | 4.03943 | 17.34930 | 15.52757 | 12.79207 | 8.60745 |
| **Safb** | 7.19590 | 1.59183 | 1.17530 | 2.41286 | 6.78679 | 5.57053 | 0.87296 | 16.30929 | 1.85774 | 14.12908 | 2.70102 | 0.91447 | 11.71558 | 0.01024 | 0.17954 | 5.73688 |
| **Safb2** | 5.38950 | 0.11198 | 0.99020 | 2.70336 | 0.02191 | 0.03164 | 3.03025 | 6.76132 | 3.40558 | 3.98460 | 1.79458 | 0.84679 | 5.41480 | 0.09474 | 0.00000 | 2.98390 |
| **Sag** | 0.01676 | 0.00786 | 0.00626 | 0.00460 | 0.00872 | 0.00718 | 0.00710 | 0.00561 | 0.01113 | 0.02491 | 1.12926 | 0.00220 | 0.13778 | 0.00538 | 0.02437 | 0.00732 |
| **Sall1** | 2.59871 | 5.46186 | 0.61460 | 0.00000 | 0.89208 | 0.01153 | 4.62706 | 6.25201 | 0.01973 | 2.78624 | 3.21970 | 2.23717 | 2.20637 | 0.59220 | 0.00000 | 1.10097 |
| **Sall2** | 0.01360 | 0.00791 | 0.01185 | 0.28298 | 0.00931 | 0.00655 | 0.00751 | 0.01342 | 1.10068 | 0.00767 | 0.00480 | 0.01593 | 0.01293 | 0.00782 | 0.05868 | 0.00728 |
| **Sall4** | 18.01229 | 10.93658 | 17.74552 | 7.59035 | 20.07476 | 9.19304 | 13.02130 | 20.16124 | 10.61100 | 27.35814 | 13.66522 | 18.12302 | 19.72811 | 21.19635 | 14.20560 | 17.21497 |
| **Samd1** | 1.19445 | 0.96398 | 0.73169 | 0.16529 | 0.17502 | 0.31496 | 0.56937 | 1.21445 | 0.34170 | 5.28161 | 1.15660 | 0.86619 | 0.23452 | 0.12791 | 0.87951 | 0.04821 |
| **Samd10** | 0.48865 | 0.00789 | 0.00314 | 1.88542 | 0.00752 | 0.02231 | 0.00713 | 0.01763 | 0.00000 | 0.00771 | 0.01480 | 0.00783 | 0.45745 | 0.01095 | 0.00330 | 0.00655 |
| **Samd4b** | 10.55782 | 5.12228 | 6.69655 | 4.57212 | 9.40338 | 5.33041 | 2.76257 | 14.28682 | 3.84050 | 9.23301 | 5.61299 | 1.16820 | 8.24860 | 0.67416 | 0.87544 | 5.47351 |

| | | | | | | | | | | | | | | | | |
|---|---|---|---|---|---|---|---|---|---|---|---|---|---|---|---|---|
| **Samd7** | 0.01421 | 0.00796 | 0.02289 | 1.67485 | 0.01002 | 0.00642 | 0.00749 | 0.01318 | 0.00000 | 0.00527 | 0.01368 | 0.00652 | 0.05124 | 0.00369 | 0.68446 | 0.00731 |
| **Samd8** | 2.38712 | 0.16022 | 0.80092 | 5.07216 | 3.78166 | 1.45719 | 3.09792 | 1.00872 | 4.50767 | 0.00000 | 1.37014 | 0.95528 | 1.02128 | 0.70650 | 2.44884 | 0.11782 |
| **Samhd1** | 1.17508 | 0.02818 | 0.00000 | 4.67937 | 0.00484 | 0.00391 | 0.01736 | 0.00000 | 5.95675 | 5.18160 | 1.89233 | 3.16844 | 0.03875 | 1.99794 | 2.99992 | 3.55950 |
| **Samm50** | 6.74006 | 4.59243 | 1.35741 | 1.77988 | 0.06769 | 0.06581 | 0.52247 | 3.61102 | 2.52384 | 2.88494 | 3.16424 | 0.82438 | 1.82375 | 0.01746 | 0.00000 | 8.52430 |
| **Sap130** | 5.13517 | 0.27476 | 4.35162 | 5.70918 | 0.00000 | 0.31314 | 1.97770 | 3.52000 | 13.08323 | 2.11122 | 4.94751 | 0.77326 | 1.83847 | 1.68302 | 3.79549 | 1.12818 |
| **Sap18** | 1.52962 | 1.41024 | 3.15327 | 5.09782 | 4.06843 | 1.38310 | 0.97140 | 0.29420 | 6.44178 | 0.16038 | 7.12938 | 2.91816 | 0.96650 | 4.90626 | 1.77789 | 1.43843 |
| **Sap30** | 0.66189 | 0.04847 | 0.02090 | 0.54859 | 0.00698 | 0.00548 | 0.29394 | 1.41235 | 0.97312 | 0.88501 | 0.20332 | 0.07662 | 0.04431 | 1.84750 | 0.69499 | 0.20357 |
| **Sap30bp** | 8.15840 | 6.80932 | 4.24445 | 9.09877 | 8.11552 | 0.02406 | 2.14912 | 11.38001 | 12.93515 | 15.91203 | 6.63633 | 5.33166 | 7.57139 | 0.04662 | 7.13470 | 6.72762 |
| **Sap30l** | 3.02189 | 0.34637 | 0.66118 | 21.42928 | 4.50790 | 0.81738 | 0.77650 | 1.46325 | 20.41291 | 8.21113 | 15.57274 | 0.59684 | 1.96590 | 1.32294 | 0.89363 | 0.11126 |
| **Sapcd1** | 0.00000 | 0.01761 | 0.01033 | 0.01026 | 0.00742 | 0.00640 | 0.00717 | 0.09800 | 0.35098 | 0.00000 | 0.02272 | 0.02692 | 0.00207 | 0.01727 | 0.01035 | 1.90545 |
| **Sapcd2** | 0.49390 | 0.00157 | 2.49719 | 10.02831 | 0.02150 | 0.03551 | 1.55191 | 12.70286 | 13.57162 | 19.09210 | 13.86915 | 6.72860 | 9.48600 | 2.28116 | 7.43872 | 0.01918 |
| **Sar1a** | 14.26011 | 5.66587 | 10.27786 | 10.98917 | 9.93091 | 9.88954 | 6.33954 | 14.13375 | 12.07423 | 20.62262 | 13.46006 | 8.52931 | 10.88768 | 7.98459 | 7.68943 | 8.29501 |
| **Sar1b** | 8.90237 | 11.12362 | 16.35452 | 5.26964 | 7.19569 | 6.27942 | 10.04683 | 17.29263 | 3.14101 | 17.66675 | 14.56558 | 10.45119 | 13.19453 | 11.69832 | 6.65512 | 6.92031 |
| **Saraf** | 1.80951 | 6.17443 | 9.03088 | 5.25741 | 2.87858 | 4.21677 | 6.73643 | 8.06497 | 2.31653 | 12.01243 | 8.12840 | 4.17792 | 4.93477 | 0.24677 | 3.84545 | 0.02230 |
| **Sardh** | 0.00464 | 0.00778 | 0.33946 | 3.93938 | 0.00852 | 0.00543 | 0.01802 | 2.25426 | 1.22123 | 3.05639 | 5.27564 | 0.16514 | 0.03370 | 0.00000 | 0.17524 | 0.00582 |
| **Sarm1** | 1.57458 | 0.34268 | 0.31606 | 0.44508 | 0.03633 | 0.00613 | 0.00488 | 0.72307 | 0.78240 | 1.90106 | 0.36410 | 0.15775 | 1.55953 | 2.36748 | 0.81882 | 0.01248 |
| **Sarnp** | 16.77746 | 12.15149 | 14.12970 | 4.53921 | 15.07914 | 8.32662 | 11.12288 | 11.35787 | 7.14268 | 7.24239 | 8.64574 | 1.58714 | 11.98609 | 11.15088 | 7.68931 | 3.53955 |
| **Sars** | 10.48887 | 9.22431 | 11.57551 | 3.02281 | 7.09261 | 0.60682 | 8.34662 | 12.07519 | 1.60925 | 5.25091 | 6.36378 | 0.58930 | 12.54116 | 0.07599 | 0.97330 | 11.78145 |
| **Sars2** | 8.52916 | 5.77522 | 1.82436 | 1.98591 | 5.46418 | 1.32485 | 1.55587 | 9.70624 | 1.23296 | 1.67961 | 3.27620 | 0.25409 | 3.20521 | 0.00000 | 0.17784 | 5.11282 |
| **Sart1** | 7.53981 | 1.49300 | 6.47290 | 5.54248 | 6.24954 | 4.45400 | 2.29155 | 13.60466 | 6.42409 | 11.92134 | 9.00839 | 1.91162 | 11.17482 | 0.47399 | 2.75381 | 6.43719 |
| **Sart3** | 11.49693 | 5.38364 | 3.12379 | 7.04400 | 3.93112 | 3.53094 | 0.57118 | 8.02525 | 8.26471 | 11.83379 | 6.22830 | 0.83403 | 9.61600 | 1.13384 | 0.95489 | 10.11582 |
| **Sash1** | 0.74058 | 0.00578 | 0.00000 | 1.71792 | 1.65477 | 0.15146 | 0.31064 | 0.00596 | 1.57953 | 0.02192 | 0.00000 | 0.02944 | 0.25836 | 0.00203 | 0.76821 | 0.00618 |
| **Sass6** | 0.97932 | 1.21120 | 2.73031 | 2.05044 | 0.02525 | 0.02778 | 0.06781 | 1.29089 | 1.98473 | 0.69794 | 2.92931 | 0.45710 | 1.03659 | 1.85441 | 2.63123 | 0.19965 |
| **Sat1** | 30.50434 | 19.09387 | 41.86576 | 2.47508 | 20.26879 | 8.86042 | 19.54353 | 29.18834 | 4.18277 | 27.13839 | 15.60738 | 11.34576 | 16.77473 | 41.73670 | 14.58505 | 1.57792 |
| **Sat2** | 0.00557 | 0.00624 | 0.41446 | 4.52591 | 2.22151 | 0.00499 | 0.00659 | 0.00514 | 0.91012 | 2.41260 | 0.00000 | 0.12538 | 0.39912 | 0.80944 | 6.86043 | 0.00594 |
| **Satb1** | 7.01582 | 6.46040 | 4.98960 | 2.48660 | 3.72594 | 0.44948 | 1.99606 | 7.84042 | 1.36180 | 11.77100 | 2.91901 | 17.80073 | 7.37038 | 17.45792 | 2.39592 | 3.12292 |
| **Satb2** | 2.01691 | 6.77276 | 1.00030 | 0.62803 | 5.90901 | 0.60852 | 0.91060 | 4.24813 | 1.08797 | 3.42925 | 0.78621 | 3.65324 | 5.70070 | 4.71286 | 0.00000 | 6.41822 |
| **Sav1** | 1.32208 | 0.78262 | 0.31452 | 3.61813 | 2.68521 | 0.00911 | 3.35326 | 2.75177 | 2.15032 | 0.99274 | 3.22584 | 0.82306 | 2.22652 | 4.24636 | 2.97236 | 0.05303 |
| **Saxo1** | 4.53907 | 1.22158 | 2.92925 | 0.00000 | 0.02395 | 1.43821 | 0.00244 | 0.00183 | 0.41271 | 2.02173 | 0.00000 | 0.00000 | 0.84920 | 2.20090 | 0.00000 | 0.25081 |
| **Saxo1os** | 0.01369 | 0.00810 | 0.01207 | 0.00461 | 0.00933 | 0.00665 | 0.00755 | 0.01387 | 0.42526 | 0.01305 | 0.00962 | 0.00751 | 0.01284 | 0.01345 | 0.00757 | 0.00740 |
| **Saxo2** | 0.01402 | 0.00784 | 0.04414 | 0.03354 | 0.00947 | 0.00593 | 0.00716 | 0.00925 | 0.00991 | 0.00626 | 0.00761 | 0.03875 | 0.01248 | 1.37588 | 0.00269 | 0.00740 |
| **Saysd1** | 0.83200 | 0.00606 | 0.72698 | 1.02092 | 0.02237 | 0.02115 | 0.00554 | 0.74560 | 0.51770 | 1.92891 | 0.47841 | 0.05405 | 1.26032 | 0.42741 | 0.00000 | 0.01952 |
| **Sbds** | 8.33264 | 6.89990 | 9.87995 | 4.58153 | 9.27308 | 4.68389 | 6.96070 | 11.12607 | 2.60277 | 6.08736 | 4.90527 | 3.57138 | 4.74403 | 1.85616 | 11.01321 | 1.90950 |
| **Sbf1** | 0.79509 | 0.02825 | 0.00000 | 0.65325 | 0.08972 | 1.51677 | 0.06768 | 2.99248 | 0.89460 | 2.81745 | 0.00000 | 0.04923 | 1.55907 | 0.13660 | 0.00000 | 1.45018 |
| **Sbf2** | 0.03459 | 0.00500 | 0.00000 | 4.04585 | 2.27178 | 0.00608 | 2.21917 | 0.00000 | 1.07105 | 0.00000 | 0.21541 | 0.02383 | 0.02603 | 0.01900 | 2.05775 | 0.01298 |
| **Sbk1** | 0.00000 | 1.30738 | 1.25545 | 5.20062 | 0.01368 | 0.00803 | 1.16841 | 1.67221 | 2.46703 | 1.09224 | 0.34732 | 0.00000 | 0.76918 | 0.00000 | 1.31815 | 0.57370 |
| **Sbk3** | 0.01338 | 0.00806 | 0.01200 | 0.00814 | 0.00904 | 0.00686 | 0.00750 | 0.01210 | 0.01182 | 0.65624 | 0.01138 | 0.00764 | 0.01211 | 0.01180 | 0.01069 | 0.00739 |
| **Sbno1** | 6.63146 | 5.01217 | 11.03049 | 13.07278 | 5.41759 | 5.95900 | 4.30521 | 9.29082 | 18.13289 | 14.85733 | 5.14698 | 5.77022 | 5.79625 | 25.20897 | 19.81699 | 7.17992 |
| **Sbno2** | 3.34905 | 0.49747 | 0.36089 | 0.50534 | 0.04128 | 2.20858 | 3.03489 | 8.66185 | 0.95769 | 4.43249 | 1.06908 | 0.25572 | 2.90546 | 0.00000 | 0.00000 | 2.32982 |
| **Sbp** | 0.79056 | 0.00648 | 0.25142 | 0.00389 | 0.00603 | 0.00571 | 0.00539 | 0.74432 | 0.48317 | 0.13560 | 0.68277 | 0.05878 | 1.64215 | 0.00417 | 0.37040 | 0.00553 |
| **Sbpl** | 0.45243 | 0.00700 | 0.06828 | 0.00483 | 0.00705 | 0.00554 | 0.00631 | 0.48293 | 0.16427 | 0.12047 | 0.21074 | 0.01419 | 0.86812 | 0.00621 | 0.10553 | 0.00639 |
| **Sbsn** | 0.00000 | 0.00674 | 1.89906 | 0.31633 | 1.19018 | 0.00848 | 0.00713 | 0.22895 | 0.41200 | 0.18187 | 0.05239 | 0.03271 | 0.01588 | 0.00894 | 0.00000 | 0.73746 |
| **Sbspon** | 0.05279 | 0.00763 | 0.05462 | 0.24902 | 0.00809 | 0.00595 | 0.00727 | 0.01012 | 0.00541 | 0.82375 | 0.01175 | 0.00602 | 0.01149 | 0.01075 | 0.00540 | 0.00742 |
| **Sc5d** | 0.83691 | 0.03844 | 0.00000 | 0.36647 | 0.01487 | 4.58854 | 1.47904 | 1.19134 | 0.49058 | 0.06855 | 0.94897 | 0.03154 | 0.00000 | 0.05949 | 0.36229 | 0.00485 |
| **Scaf1** | 0.12548 | 0.00729 | 0.02237 | 1.31000 | 0.01618 | 0.00717 | 0.02692 | 2.05787 | 1.12941 | 0.79785 | 0.19454 | 0.08955 | 1.11902 | 0.00534 | 0.03122 | 0.61600 |
| **Scaf11** | 4.56731 | 4.11607 | 1.11226 | 9.59714 | 5.60651 | 5.21287 | 1.87896 | 6.87779 | 13.94966 | 10.20453 | 6.22777 | 8.29558 | 4.56827 | 8.36977 | 3.91330 | 2.63416 |
| **Scaf4** | 2.97263 | 0.10706 | 2.00846 | 2.51000 | 2.79470 | 2.02690 | 0.16806 | 2.67133 | 3.04608 | 3.94280 | 0.05833 | 0.24943 | 1.43670 | 0.32096 | 2.50018 | 2.83833 |
| **Scaf8** | 6.63472 | 4.10077 | 5.79858 | 6.01840 | 0.09103 | 0.00493 | 3.08707 | 6.79771 | 7.26321 | 6.78516 | 5.17760 | 0.44063 | 4.42620 | 8.54682 | 8.33378 | 4.11680 |
| **Scai** | 0.00428 | 0.00722 | 0.25934 | 1.67848 | 0.00782 | 0.00507 | 0.25697 | 0.00000 | 1.87373 | 0.00000 | 1.38708 | 1.24324 | 0.13654 | 4.70826 | 1.03175 | 0.59192 |
| **Scamp1** | 5.04786 | 2.88444 | 6.85242 | 8.19130 | 4.23256 | 0.31997 | 3.84804 | 2.77504 | 9.11759 | 3.04961 | 7.80882 | 1.51694 | 3.85021 | 9.48004 | 3.61794 | 1.84860 |
| **Scamp2** | 4.76552 | 4.99795 | 10.47253 | 0.47756 | 0.08632 | 3.04473 | 6.87758 | 11.08281 | 1.89992 | 7.66188 | 5.72763 | 0.70930 | 7.54174 | 1.49412 | 4.85004 | 2.55437 |
| **Scamp3** | 0.40011 | 0.09900 | 0.40720 | 0.00843 | 0.02376 | 0.32203 | 0.48527 | 0.64318 | 0.01143 | 0.44692 | 0.00740 | 0.02429 | 1.08360 | 0.01425 | 0.00095 | 1.23337 |
| **Scamp4** | 11.57898 | 8.33134 | 14.00190 | 1.74131 | 2.65264 | 10.38285 | 4.60625 | 14.57756 | 0.87062 | 14.80029 | 8.78510 | 1.53449 | 13.57124 | 0.02304 | 2.96706 | 9.56124 |
| **Scamp5** | 0.58197 | 2.08952 | 0.33761 | 2.60248 | 0.00506 | 5.13218 | 1.33185 | 2.96798 | 1.55051 | 4.58506 | 3.17440 | 0.28065 | 3.22920 | 0.25751 | 2.50534 | 3.10704 |
| **Scand1** | 0.07753 | 0.00767 | 0.01536 | 0.26196 | 0.02673 | 0.00616 | 0.00736 | 0.00000 | 0.32718 | 0.17061 | 0.47474 | 0.04644 | 0.01363 | 0.00931 | 0.00000 | 0.00709 |
| **Scap** | 0.25728 | 0.01428 | 3.54300 | 9.03315 | 0.00946 | 0.25930 | 2.77781 | 9.13989 | 9.48426 | 9.58859 | 6.28513 | 0.78940 | 4.14605 | 1.26170 | 5.62882 | 1.16186 |
| **Scaper** | 0.00000 | 1.68800 | 0.04977 | 0.00000 | 0.26301 | 0.01198 | 0.55710 | 0.23383 | 0.00731 | 2.26653 | 0.00000 | 0.05441 | 0.14269 | 1.27417 | 2.27294 | 0.00666 |
| **Scarb1** | 2.70943 | 1.53136 | 0.00000 | 1.16286 | 2.94536 | 3.64744 | 0.00442 | 1.49945 | 0.04043 | 2.96742 | 0.00000 | 0.13810 | 0.00000 | 0.02363 | 0.36490 | 2.54567 |
| **Scarb2** | 8.26677 | 3.00211 | 9.34436 | 0.16091 | 7.28890 | 1.92031 | 7.20596 | 6.12095 | 0.01048 | 2.46566 | 3.71168 | 0.53927 | 4.96054 | 0.03830 | 1.51871 | 4.77248 |
| **Scarf1** | 0.01346 | 0.00800 | 0.01077 | 1.12106 | 0.00927 | 0.00644 | 0.00734 | 0.01119 | 0.32461 | 0.29017 | 0.00722 | 0.02807 | 0.01248 | 0.00420 | 0.87858 | 0.00719 |
| **Sccpdh** | 1.44181 | 0.07047 | 0.00000 | 0.60319 | 0.03870 | 0.00621 | 0.00616 | 0.01949 | 0.06110 | 0.30663 | 0.01192 | 0.01580 | 0.35588 | 0.00676 | 0.45265 | 7.05919 |
| **Scd1** | 9.52144 | 8.81575 | 20.38382 | 12.15108 | 3.26330 | 6.52726 | 11.00838 | 11.43321 | 8.71431 | 1.33327 | 11.39081 | 1.91024 | 9.52222 | 2.66425 | 10.39273 | 3.71024 |

| | | | | | | | | | | | | | | | | |
|---|---|---|---|---|---|---|---|---|---|---|---|---|---|---|---|---|
| **Scd2** | 13.93845 | 15.48247 | 25.19920 | 1.67924 | 8.65406 | 14.31569 | 13.98846 | 10.65874 | 0.07661 | 3.51914 | 5.25029 | 0.16203 | 10.22216 | 0.00143 | 0.00000 | 2.76049 |
| **Scd3** | 1.75166 | 0.23669 | 3.68863 | 6.12412 | 0.19913 | 1.57550 | 0.47269 | 1.78884 | 5.44826 | 0.06885 | 1.32231 | 0.04331 | 2.47990 | 0.33940 | 0.38829 | 1.22205 |
| **Scd4** | 1.03766 | 0.44182 | 0.41289 | 0.52405 | 0.77532 | 0.05006 | 0.08853 | 0.11477 | 0.37821 | 0.18206 | 0.16108 | 0.46462 | 0.43685 | 0.12186 | 0.19321 | 0.91864 |
| **Scel** | 0.01339 | 0.00771 | 0.01161 | 0.64583 | 0.00906 | 0.00639 | 0.00736 | 0.01051 | 0.00000 | 0.66976 | 0.00975 | 0.00524 | 0.01079 | 0.01170 | 0.38044 | 0.00700 |
| **Scfd1** | 7.73429 | 3.88722 | 0.41325 | 2.24030 | 4.41490 | 6.16815 | 4.92064 | 4.53830 | 2.13105 | 1.64729 | 4.35502 | 4.45524 | 2.54441 | 5.25227 | 7.79452 | 1.27368 |
| **Scfd2** | 0.00000 | 2.95861 | 1.96833 | 1.83730 | 0.01833 | 2.86703 | 2.82257 | 3.64434 | 4.70416 | 3.67586 | 2.87572 | 4.17447 | 0.66085 | 0.02239 | 2.24542 | 0.00279 |
| **Scg3** | 0.01325 | 0.00764 | 0.00784 | 0.78314 | 0.00886 | 0.00648 | 0.00715 | 0.00000 | 0.01670 | 1.10070 | 0.42294 | 0.01138 | 0.00756 | 0.01051 | 0.73508 | 0.00686 |
| **Scgb2b23.ps** | 0.01469 | 0.00796 | 0.02352 | 0.05296 | 0.00902 | 0.00764 | 0.00733 | 0.05727 | 0.06817 | 0.01641 | 0.01220 | 0.00770 | 0.01015 | 0.00992 | 0.02961 | 0.00743 |
| **Scimp** | 0.01377 | 0.00784 | 0.01158 | 0.00494 | 0.00933 | 0.00663 | 0.00747 | 0.01287 | 0.25167 | 0.00930 | 0.01038 | 0.04648 | 0.01282 | 0.00717 | 0.08851 | 0.00739 |
| **Scin** | 0.02030 | 0.00789 | 0.01090 | 0.51319 | 0.00901 | 0.00676 | 0.00746 | 0.01164 | 0.02580 | 0.02151 | 0.02516 | 0.00735 | 0.01543 | 0.16029 | 0.00335 | 0.00734 |
| **Sclt1** | 0.15446 | 0.00751 | 0.25564 | 0.44287 | 0.00727 | 0.00544 | 0.00728 | 0.00897 | 0.19483 | 0.00000 | 0.00000 | 0.00306 | 0.01034 | 0.93055 | 0.83062 | 0.00697 |
| **Scly** | 0.01202 | 0.00817 | 0.08418 | 0.96673 | 0.00907 | 0.00859 | 0.00773 | 0.02632 | 0.44188 | 0.01007 | 0.00968 | 0.01762 | 0.01789 | 0.00691 | 0.05076 | 0.00715 |
| **Scmh1** | 2.50508 | 0.07226 | 2.94643 | 2.31516 | 5.57959 | 0.00942 | 0.00532 | 1.02713 | 1.08260 | 2.98809 | 0.54557 | 0.08274 | 0.82287 | 0.00155 | 1.06149 | 0.01752 |
| **Scml2** | 0.01336 | 0.00767 | 0.00952 | 9.98824 | 0.00943 | 0.00532 | 0.00692 | 0.00955 | 0.36721 | 0.00193 | 0.02348 | 0.02168 | 0.01249 | 0.00000 | 2.73574 | 0.00555 |
| **Scml4** | 0.07795 | 0.01890 | 0.08287 | 0.67403 | 0.02884 | 0.01325 | 0.00743 | 0.00932 | 1.04925 | 0.02301 | 0.21848 | 0.00439 | 0.05955 | 0.24144 | 0.05329 | 0.04345 |
| **Scn10a** | 0.01373 | 0.00774 | 0.01205 | 0.19723 | 0.00943 | 0.00651 | 0.00742 | 0.01222 | 0.00577 | 0.00588 | 0.01122 | 0.00675 | 0.01282 | 0.02314 | 0.36859 | 0.00736 |
| **Scn11a** | 0.01328 | 0.00792 | 0.01174 | 0.00554 | 0.00926 | 0.00636 | 0.00725 | 0.01053 | 0.01070 | 0.62945 | 0.01068 | 0.00515 | 0.01819 | 0.00592 | 0.57108 | 0.00743 |
| **Scn2b** | 0.23688 | 0.04355 | 0.22979 | 0.14317 | 0.05549 | 0.06410 | 0.04839 | 0.00501 | 0.34666 | 0.00034 | 1.98987 | 0.13433 | 0.96337 | 0.31700 | 0.88872 | 0.13236 |
| **Scn3a** | 0.01143 | 0.00785 | 0.01171 | 0.00730 | 0.00871 | 0.00653 | 0.00743 | 0.01217 | 0.01122 | 0.00958 | 0.01086 | 0.00682 | 0.01116 | 0.01167 | 0.60607 | 0.29773 |
| **Scn8a** | 0.01287 | 0.00821 | 0.07803 | 0.84179 | 0.00933 | 0.00572 | 0.00751 | 0.01281 | 0.00000 | 0.08961 | 0.01290 | 0.01855 | 0.01158 | 0.00560 | 0.11669 | 0.00729 |
| **Scnm1** | 5.62274 | 1.94626 | 5.07974 | 2.95602 | 1.69428 | 5.53327 | 3.15055 | 3.29665 | 14.70228 | 7.40671 | 9.20165 | 1.86544 | 7.03524 | 2.35368 | 5.30541 | 2.68168 |
| **Sco1** | 0.70064 | 0.07425 | 0.05098 | 0.32204 | 1.76471 | 0.01121 | 1.50999 | 0.00000 | 2.59107 | 1.04417 | 0.67384 | 7.09333 | 2.10632 | 2.04336 | 1.75611 | 0.10612 |
| **Scoc** | 0.75596 | 4.66506 | 1.83834 | 17.11599 | 1.90011 | 0.00339 | 2.74379 | 2.27360 | 9.46753 | 4.95507 | 2.85583 | 0.10436 | 0.00000 | 8.71962 | 5.35169 | 1.12492 |
| **Scp2** | 19.81343 | 8.30410 | 17.35644 | 1.72859 | 3.01926 | 4.91603 | 9.03865 | 10.58775 | 5.12321 | 11.23955 | 6.49352 | 6.49654 | 9.94079 | 4.39699 | 4.27791 | 7.15888 |
| **Scpep1** | 7.40959 | 6.82955 | 19.54714 | 4.29537 | 3.06967 | 9.35683 | 7.10847 | 18.48516 | 8.74662 | 20.67357 | 12.16151 | 7.71670 | 14.35650 | 22.17472 | 17.15164 | 5.13786 |
| **Scpep1os** | 0.19058 | 0.06081 | 0.29801 | 1.78116 | 0.14297 | 0.02189 | 0.05766 | 0.03693 | 2.01354 | 0.07050 | 1.48020 | 0.20706 | 0.11628 | 0.72941 | 0.15701 | 0.05077 |
| **Scrib** | 6.25845 | 0.01100 | 0.00000 | 1.55499 | 0.11607 | 1.07302 | 0.01768 | 3.46570 | 2.41173 | 2.41421 | 1.83137 | 0.03958 | 7.57758 | 0.01505 | 0.00000 | 7.13785 |
| **Scrn2** | 0.00000 | 0.01676 | 1.91789 | 0.00181 | 0.03171 | 0.87556 | 2.78610 | 3.41028 | 0.00846 | 2.42744 | 0.12874 | 0.03564 | 1.16275 | 0.01350 | 0.00000 | 0.25507 |
| **Scrn3** | 1.29186 | 0.00691 | 0.85213 | 1.36295 | 0.01266 | 0.00997 | 0.00619 | 0.32982 | 1.14222 | 0.54771 | 1.47821 | 1.55773 | 0.04756 | 3.10834 | 0.77669 | 0.00552 |
| **Sct** | 0.58724 | 0.01640 | 0.41243 | 0.18287 | 0.00578 | 0.00739 | 0.00613 | 0.56697 | 0.55924 | 0.12995 | 0.62601 | 0.02053 | 0.62905 | 0.00607 | 0.00000 | 0.91012 |
| **Scube1** | 0.04918 | 0.01798 | 0.03085 | 0.60338 | 0.10155 | 0.00649 | 0.01767 | 0.00087 | 0.34626 | 0.01734 | 0.37041 | 0.02512 | 0.01904 | 0.16605 | 0.03138 | 0.01439 |
| **Scyl1** | 5.06265 | 0.81722 | 4.15412 | 3.09207 | 2.64163 | 5.50740 | 2.79726 | 15.57459 | 3.27406 | 9.91772 | 6.99590 | 0.15808 | 7.71485 | 0.00000 | 1.69492 | 4.34141 |
| **Scyl2** | 2.02782 | 3.78970 | 0.60363 | 2.09406 | 2.85653 | 0.00557 | 1.37745 | 2.00839 | 3.56733 | 3.47866 | 1.40968 | 0.13682 | 0.75423 | 1.02512 | 3.32125 | 1.14740 |
| **Scyl3** | 5.04048 | 0.00237 | 1.06949 | 0.72267 | 0.10755 | 0.01193 | 0.00226 | 2.71394 | 0.21408 | 1.40824 | 0.37670 | 0.93037 | 2.09610 | 0.40278 | 1.78835 | 0.06559 |
| **Sdad1** | 5.50723 | 5.02157 | 7.53700 | 6.59102 | 3.84802 | 0.01289 | 6.06628 | 2.02039 | 4.71814 | 6.22603 | 5.03404 | 0.65915 | 3.88619 | 3.21488 | 5.93721 | 3.73941 |
| **Sdc1** | 0.72899 | 0.00448 | 2.95563 | 8.05160 | 0.04277 | 0.00371 | 4.69024 | 0.21071 | 5.21614 | 0.15267 | 0.00001 | 0.16061 | 1.55656 | 0.01144 | 2.59113 | 0.01777 |
| **Sdc2** | 0.01369 | 0.00811 | 0.01187 | 0.25734 | 0.00926 | 0.00664 | 0.00746 | 0.01300 | 0.55952 | 0.04818 | 0.00891 | 0.01697 | 0.01316 | 0.01113 | 0.00423 | 0.00763 |
| **Sdc3** | 0.01287 | 0.00782 | 0.01032 | 0.23332 | 0.01854 | 0.00654 | 0.00721 | 0.00817 | 0.33416 | 0.62026 | 0.07849 | 0.07347 | 0.01088 | 0.01032 | 0.00105 | 0.00732 |
| **Sdc4** | 14.65918 | 9.37762 | 14.85165 | 0.00000 | 10.34674 | 10.58011 | 10.83777 | 3.51082 | 0.47176 | 0.00000 | 1.51843 | 0.07407 | 8.74413 | 0.00644 | 0.01334 | 10.43772 |
| **Sdcbp** | 2.07543 | 9.81466 | 4.00364 | 2.02472 | 3.32226 | 5.28246 | 6.94234 | 0.79596 | 0.00000 | 3.92956 | 0.85602 | 7.92030 | 4.51892 | 7.45424 | 4.00331 | 1.16249 |
| **Sdcbp2** | 0.00000 | 5.01678 | 3.53610 | 0.95909 | 3.39432 | 0.00796 | 0.01512 | 0.00301 | 1.79403 | 0.02151 | 0.00000 | 0.04479 | 0.02126 | 0.00164 | 0.76290 | 0.01753 |
| **Sdccag3** | 2.46195 | 2.29316 | 1.72061 | 1.10946 | 0.02068 | 0.01331 | 3.37293 | 2.39006 | 1.42817 | 5.26990 | 0.63491 | 0.07397 | 2.62359 | 2.61729 | 4.34445 | 4.45688 |
| **Sdccag8** | 0.62440 | 0.61947 | 1.00609 | 1.29309 | 0.00491 | 0.00582 | 0.07571 | 0.00000 | 0.00000 | 0.19908 | 0.45305 | 0.01343 | 0.10779 | 0.00288 | 0.00000 | 0.00547 |
| **Sde2** | 5.67711 | 3.93219 | 2.18272 | 3.28480 | 1.09204 | 5.33726 | 3.21602 | 6.95104 | 6.20833 | 4.36117 | 3.94956 | 4.76755 | 2.80071 | 2.69629 | 14.08202 | 2.53373 |
| **Sdf2** | 6.77374 | 7.85148 | 6.74644 | 3.54559 | 6.05987 | 3.68398 | 5.38085 | 8.29499 | 5.48627 | 2.18161 | 3.63959 | 2.05373 | 2.60032 | 5.72431 | 6.27802 | 1.76032 |
| **Sdf2l1** | 0.07510 | 0.00793 | 0.04057 | 0.60348 | 0.00839 | 0.01712 | 0.00746 | 0.01108 | 1.97147 | 0.08333 | 0.06555 | 0.02452 | 0.47560 | 0.23610 | 0.13132 | 0.00725 |
| **Sdf4** | 12.14286 | 7.64909 | 17.90556 | 12.23847 | 2.47009 | 11.89124 | 7.18216 | 12.46274 | 17.67400 | 9.86303 | 9.71513 | 6.49076 | 13.40960 | 2.62818 | 9.20690 | 4.23805 |
| **Sdha** | 15.75870 | 8.79619 | 15.82811 | 4.53577 | 9.07988 | 4.28974 | 10.74232 | 13.71114 | 1.65168 | 10.03403 | 13.79833 | 4.02404 | 8.72375 | 5.17112 | 10.63029 | 10.17020 |
| **Sdhaf1** | 0.46115 | 0.06545 | 0.01153 | 1.14424 | 0.03743 | 0.00601 | 0.01715 | 0.02495 | 0.04323 | 0.30533 | 0.04282 | 0.00636 | 0.39352 | 0.00426 | 0.09806 | 0.00678 |
| **Sdhaf2** | 5.86887 | 8.06495 | 4.99076 | 0.82074 | 2.06203 | 4.27863 | 5.25253 | 4.54142 | 0.80626 | 5.39596 | 2.26814 | 5.01237 | 3.74038 | 0.64345 | 2.11751 | 3.90721 |
| **Sdhaf3** | 1.86919 | 0.00530 | 2.09021 | 0.00404 | 0.01080 | 0.05674 | 0.52199 | 1.43163 | 0.25389 | 0.30786 | 0.16715 | 1.82104 | 0.00000 | 0.00503 | 0.30031 | 0.86819 |
| **Sdhaf4** | 1.76809 | 1.75486 | 1.99957 | 0.00380 | 1.52356 | 0.11546 | 1.13742 | 0.18374 | 0.50008 | 2.37820 | 0.85998 | 0.06639 | 1.93383 | 1.53070 | 0.11739 | 0.14628 |
| **Sdhb** | 20.87134 | 7.65753 | 12.69820 | 1.40627 | 12.01957 | 10.30620 | 2.82464 | 10.29435 | 1.93884 | 6.83263 | 7.20262 | 1.95861 | 10.84260 | 0.56598 | 0.17149 | 8.05424 |
| **Sdhc** | 16.59103 | 11.03242 | 15.18648 | 1.24910 | 12.51845 | 8.31360 | 13.63862 | 8.66808 | 2.54535 | 5.26474 | 3.57151 | 2.27721 | 9.71442 | 1.23063 | 2.37939 | 9.24090 |
| **Sdhd** | 7.27707 | 7.52478 | 5.27272 | 0.48469 | 8.31355 | 2.02681 | 2.06967 | 1.50953 | 3.44691 | 0.00000 | 0.74035 | 0.15583 | 0.57554 | 0.66433 | 3.25648 | 0.88904 |
| **Sdk1** | 0.03224 | 0.00806 | 0.11868 | 0.01878 | 0.01428 | 0.00660 | 0.00742 | 1.12593 | 0.98480 | 0.21622 | 0.17027 | 0.01383 | 0.23745 | 0.01208 | 0.02015 | 0.00768 |
| **Sdk2** | 0.00946 | 0.00755 | 0.00819 | 0.01093 | 0.00964 | 0.00687 | 0.00713 | 0.52587 | 0.01172 | 0.02429 | 0.01024 | 0.00684 | 0.00568 | 0.01692 | 0.00176 | 0.00728 |
| **Sdr39u1** | 0.09594 | 0.00772 | 0.25958 | 0.00701 | 0.00742 | 0.00613 | 0.00641 | 0.86688 | 0.01079 | 0.49362 | 0.00482 | 0.04392 | 2.01669 | 0.00511 | 0.32713 | 0.00656 |
| **Sdsl** | 0.01311 | 0.00770 | 0.01183 | 0.01329 | 0.00888 | 0.00645 | 0.00721 | 0.00969 | 0.05781 | 1.27298 | 0.01350 | 0.01591 | 0.01030 | 0.02145 | 0.00373 | 0.00708 |
| **Sebox** | 0.01360 | 0.00776 | 0.00712 | 4.09736 | 0.00940 | 0.00610 | 0.00718 | 0.01061 | 0.40115 | 0.00534 | 0.00999 | 0.18249 | 0.01353 | 0.00000 | 2.32788 | 0.00666 |
| **Sec1** | 0.00000 | 4.15823 | 5.84324 | 8.73710 | 0.00860 | 0.00397 | 0.09314 | 1.52361 | 8.24208 | 2.79234 | 0.12975 | 0.00000 | 3.06338 | 0.38026 | 0.00000 | 0.00000 |
| **Sec11a** | 7.50377 | 6.79131 | 7.41827 | 7.92226 | 0.04092 | 5.65746 | 8.18623 | 10.73179 | 4.88184 | 13.68774 | 11.56545 | 13.60625 | 10.21211 | 6.08942 | 7.75237 | 2.96544 |

| | | | | | | | | | | | | | | | | |
|---|---|---|---|---|---|---|---|---|---|---|---|---|---|---|---|---|
| **Sec11c** | 2.28933 | 0.04412 | 0.00000 | 2.00784 | 1.58194 | 7.20545 | 0.27572 | 2.72440 | 0.00000 | 3.90313 | 3.05486 | 0.59376 | 4.69939 | 0.37706 | 0.16218 | 1.92360 |
| **Sec13** | 16.32979 | 15.56788 | 16.21357 | 5.09792 | 14.34798 | 6.05744 | 13.32920 | 19.42599 | 2.00955 | 18.12639 | 12.96369 | 4.21137 | 16.53200 | 9.03155 | 9.93446 | 14.70108 |
| **Sec14l1** | 5.65735 | 0.01136 | 2.46920 | 1.38472 | 0.17917 | 0.00739 | 1.93841 | 0.00000 | 0.24923 | 2.86453 | 0.00000 | 0.72595 | 0.07822 | 0.09191 | 0.00000 | 0.34940 |
| **Sec14l2** | 0.00000 | 0.01953 | 0.00819 | 0.42595 | 0.00950 | 0.00737 | 0.00807 | 0.14081 | 0.42563 | 0.74068 | 0.34210 | 0.06189 | 0.16078 | 0.00422 | 0.20632 | 1.61448 |
| **Sec14l3** | 0.02651 | 0.00774 | 0.01037 | 0.68952 | 0.00845 | 0.00663 | 0.01753 | 0.00987 | 0.44144 | 0.75786 | 0.02467 | 0.00549 | 0.01127 | 0.07462 | 0.01573 | 0.00713 |
| **Sec14l4** | 0.03920 | 0.00804 | 0.00000 | 0.00925 | 0.00904 | 0.16432 | 0.00686 | 0.07553 | 0.02302 | 0.83287 | 0.22805 | 0.02203 | 0.80251 | 0.01833 | 0.31057 | 0.00698 |
| **Sec16a** | 5.67612 | 6.03677 | 4.62865 | 15.00545 | 0.06238 | 2.89174 | 7.87449 | 5.63047 | 10.55012 | 2.43241 | 3.38873 | 2.27809 | 2.98949 | 4.78240 | 10.65344 | 0.48492 |
| **Sec16b** | 1.69907 | 0.00599 | 0.00000 | 4.28864 | 0.00491 | 0.00618 | 0.00590 | 0.01136 | 1.92032 | 1.71431 | 0.54770 | 0.41964 | 0.01568 | 0.00000 | 0.95245 | 0.00493 |
| **Sec22a** | 1.69800 | 0.80679 | 5.51710 | 3.39498 | 0.02169 | 3.60959 | 3.09530 | 0.06206 | 0.33858 | 2.51545 | 2.87090 | 0.39229 | 3.84156 | 0.00000 | 1.51875 | 0.08070 |
| **Sec22b** | 8.55141 | 5.68766 | 4.14341 | 1.24826 | 5.17098 | 2.17934 | 6.14107 | 5.78674 | 0.86107 | 6.19430 | 3.87432 | 1.24555 | 6.17850 | 2.28245 | 1.16898 | 6.15297 |
| **Sec22c** | 0.01682 | 0.00607 | 0.23524 | 0.66724 | 0.32502 | 0.00832 | 0.00609 | 0.00000 | 0.37616 | 0.00000 | 0.75783 | 3.25777 | 0.42037 | 0.04744 | 1.51795 | 0.00673 |
| **Sec23a** | 0.93232 | 1.41631 | 3.39236 | 1.70581 | 6.85066 | 1.17113 | 0.43617 | 5.89663 | 1.42391 | 8.33607 | 2.81220 | 4.79802 | 2.19378 | 10.67873 | 9.13613 | 3.07284 |
| **Sec23b** | 6.87628 | 11.68081 | 7.73898 | 2.84795 | 6.86635 | 1.80787 | 6.21122 | 9.45173 | 3.26020 | 9.23991 | 4.34760 | 1.80690 | 1.68234 | 1.75565 | 6.69765 | 3.55328 |
| **Sec23ip** | 3.32190 | 0.61819 | 6.70573 | 4.24383 | 0.06994 | 1.34018 | 0.62087 | 3.02351 | 9.71553 | 3.71468 | 5.27760 | 8.22290 | 2.33570 | 7.60655 | 6.18594 | 0.55674 |
| **Sec24a** | 0.19953 | 1.93832 | 2.63550 | 11.85025 | 0.66055 | 0.01160 | 2.61025 | 2.62919 | 10.00341 | 2.91883 | 1.45981 | 0.46479 | 1.62785 | 6.55419 | 5.22035 | 0.54190 |
| **Sec24b** | 4.28506 | 3.32355 | 6.43132 | 1.48687 | 4.82731 | 3.39454 | 4.81849 | 5.65691 | 2.03377 | 6.09675 | 3.42089 | 2.87588 | 3.56087 | 0.89246 | 2.45283 | 1.73613 |
| **Sec24c** | 8.93560 | 8.81922 | 12.39048 | 7.28776 | 4.36741 | 3.44654 | 1.01065 | 7.68536 | 2.29445 | 9.07310 | 6.93583 | 1.90702 | 9.57823 | 7.63502 | 7.72285 | 6.87947 |
| **Sec24d** | 1.81715 | 0.00533 | 0.00000 | 0.54439 | 0.01150 | 0.00556 | 1.86113 | 0.00235 | 0.99243 | 0.71044 | 0.00000 | 0.00700 | 0.01650 | 1.02959 | 0.15068 | 0.30103 |
| **Sec31a** | 10.37909 | 4.10956 | 8.49796 | 8.45965 | 9.93292 | 9.50755 | 5.86242 | 13.16603 | 7.93445 | 11.32264 | 7.75552 | 5.78453 | 7.55472 | 0.69709 | 1.45838 | 6.76753 |
| **Sec31b** | 0.00534 | 1.18131 | 0.00850 | 0.23484 | 0.00863 | 0.00715 | 0.00712 | 0.91024 | 0.40966 | 0.00237 | 0.00900 | 0.00570 | 0.19063 | 0.00765 | 0.04017 | 0.00701 |
| **Sec61a1** | 10.83968 | 10.26548 | 16.12499 | 6.62733 | 8.26771 | 6.19491 | 9.84615 | 12.93334 | 5.09693 | 11.65762 | 5.72016 | 0.79591 | 15.85869 | 0.04491 | 1.75203 | 10.92619 |
| **Sec61a2** | 2.17963 | 0.00444 | 1.43154 | 4.95256 | 0.46437 | 0.00535 | 0.90044 | 0.61025 | 0.89457 | 1.35168 | 0.36786 | 0.34645 | 1.27374 | 0.85326 | 1.20181 | 0.07637 |
| **Sec61b** | 9.69928 | 0.05719 | 1.55063 | 4.13971 | 6.55053 | 2.04256 | 0.68394 | 2.75597 | 2.39729 | 7.30348 | 6.04624 | 0.15099 | 6.72298 | 0.00000 | 0.00000 | 2.58858 |
| **Sec61g** | 3.24987 | 0.57772 | 4.48628 | 4.96073 | 3.50943 | 2.15333 | 0.57235 | 0.38376 | 4.03306 | 0.74744 | 8.55081 | 0.21028 | 1.73549 | 2.30766 | 0.58926 | 3.10923 |
| **Sec62** | 8.14203 | 4.37822 | 8.08876 | 5.13571 | 6.00520 | 9.77349 | 7.50842 | 13.06764 | 5.81319 | 9.37265 | 6.57819 | 13.04723 | 9.55051 | 13.19974 | 12.22386 | 5.46136 |
| **Sec63** | 1.92557 | 3.30491 | 7.28918 | 9.70656 | 2.51918 | 0.00147 | 3.87221 | 2.55798 | 10.86139 | 5.49755 | 3.49764 | 4.72256 | 0.99627 | 12.98757 | 8.20095 | 1.56027 |
| **Secisbp2** | 4.36627 | 4.07766 | 3.62220 | 2.40401 | 0.01285 | 3.41665 | 3.67266 | 4.60840 | 0.63264 | 3.74733 | 2.99552 | 0.18294 | 3.32838 | 0.71962 | 4.65296 | 1.73972 |
| **Secisbp2l** | 0.12997 | 0.00675 | 0.00000 | 7.84661 | 2.02841 | 0.06522 | 0.00719 | 0.00930 | 3.64206 | 0.04854 | 0.51850 | 0.01379 | 0.04059 | 0.00000 | 2.49698 | 0.08278 |
| **Sectm1a** | 0.01758 | 0.00835 | 0.01096 | 0.30201 | 0.00891 | 0.00658 | 0.00758 | 0.01469 | 0.44479 | 0.12436 | 0.00912 | 0.00662 | 0.01177 | 0.11134 | 0.00000 | 0.00708 |
| **Seh1l** | 11.35250 | 9.78950 | 6.09395 | 11.80755 | 7.38963 | 8.14081 | 9.95265 | 8.61624 | 7.55357 | 12.25915 | 8.01295 | 11.47602 | 5.50827 | 18.17777 | 16.43298 | 7.31704 |
| **Sel1l** | 1.91950 | 0.00307 | 3.01027 | 4.06392 | 0.08351 | 3.15734 | 3.61074 | 0.81212 | 4.98932 | 1.72905 | 0.54973 | 1.26590 | 2.02559 | 1.11619 | 2.53163 | 0.02397 |
| **Selenbp1** | 0.22807 | 0.00816 | 0.00217 | 0.08715 | 0.02007 | 0.00817 | 0.00680 | 0.27042 | 0.36093 | 2.39689 | 0.49065 | 0.02557 | 0.43513 | 0.00531 | 0.48822 | 0.00747 |
| **Selenbp2** | 0.09512 | 0.00817 | 0.00464 | 0.34782 | 0.01668 | 0.00734 | 0.00686 | 0.08956 | 0.37242 | 0.93328 | 0.12833 | 0.08105 | 0.50132 | 0.02471 | 0.25953 | 0.00732 |
| **Selk** | 10.96344 | 3.92702 | 6.20375 | 13.29901 | 6.02854 | 7.13276 | 2.91515 | 4.83867 | 17.94126 | 7.86359 | 6.48004 | 6.32807 | 8.10976 | 4.60108 | 7.20782 | 3.29428 |
| **Selm** | 1.42878 | 0.00595 | 0.00000 | 0.00618 | 0.00547 | 0.01558 | 0.23068 | 0.17629 | 0.01124 | 1.64621 | 0.01026 | 0.03413 | 0.00680 | 0.01668 | 0.00520 | 0.00583 |
| **Selo** | 0.02612 | 0.00749 | 0.00000 | 0.08484 | 0.00943 | 0.75081 | 0.29921 | 0.00857 | 0.03123 | 0.22870 | 0.12826 | 0.05361 | 0.20928 | 0.28482 | 0.05418 | 0.01510 |
| **Selplg** | 0.18904 | 0.07132 | 0.16987 | 0.44291 | 0.11062 | 0.01244 | 0.07948 | 0.07779 | 0.35237 | 0.22491 | 0.33445 | 0.04315 | 0.19140 | 0.29647 | 0.10590 | 0.04381 |
| **Selt** | 24.25431 | 17.69409 | 18.32915 | 19.44967 | 17.71042 | 12.54959 | 12.53413 | 21.40879 | 24.61180 | 27.32852 | 23.03014 | 31.17078 | 15.88447 | 38.62587 | 27.69308 | 8.72739 |
| **Sema3b** | 0.71556 | 0.00765 | 0.00000 | 0.00000 | 0.01678 | 6.35521 | 0.00596 | 0.00487 | 0.47427 | 0.00000 | 0.00475 | 0.00668 | 0.52890 | 0.01720 | 0.00622 | 0.00417 |
| **Sema3c** | 0.05052 | 0.00758 | 0.02287 | 0.01830 | 0.00803 | 0.00775 | 0.00734 | 0.01106 | 0.01111 | 0.22019 | 0.01265 | 0.01651 | 0.01242 | 0.01135 | 0.00984 | 0.00765 |
| **Sema3e** | 0.16553 | 0.02824 | 0.33736 | 1.03630 | 0.40960 | 0.00671 | 0.06753 | 0.05714 | 1.38207 | 0.16617 | 1.18856 | 0.13161 | 0.56259 | 0.60405 | 0.04432 | 0.12247 |
| **Sema4a** | 0.37358 | 0.00677 | 0.00000 | 0.72341 | 0.00736 | 1.06575 | 0.00615 | 0.99414 | 1.15292 | 1.17388 | 2.15082 | 0.08343 | 0.00000 | 0.00000 | 0.00000 | 0.00636 |
| **Sema4b** | 1.73058 | 0.05778 | 1.21030 | 0.03955 | 0.02057 | 0.23143 | 0.06680 | 0.08233 | 0.36154 | 0.02665 | 3.39802 | 0.04645 | 0.91673 | 0.12169 | 0.01283 | 0.27535 |
| **Sema4c** | 0.01358 | 0.00788 | 0.00946 | 0.02667 | 0.00951 | 0.00696 | 0.00746 | 0.01035 | 0.26430 | 0.01548 | 0.39093 | 0.01540 | 0.01335 | 0.01355 | 0.02323 | 0.00743 |
| **Sema4d** | 2.11178 | 2.37268 | 3.29399 | 2.08272 | 0.08588 | 2.16769 | 2.88391 | 0.03244 | 2.19531 | 0.38601 | 1.41615 | 0.06313 | 0.00000 | 0.00000 | 0.10165 | 0.01653 |
| **Sema4g** | 0.00000 | 0.00475 | 2.63911 | 0.00000 | 1.41756 | 0.01755 | 3.04550 | 0.55660 | 0.33302 | 3.47717 | 2.21738 | 0.46186 | 0.49611 | 0.01735 | 0.09086 | 0.54800 |
| **Sema5a** | 1.41196 | 0.00584 | 1.50246 | 3.12852 | 0.00404 | 0.00369 | 0.04614 | 0.00263 | 1.29231 | 0.32152 | 0.01931 | 0.00000 | 1.12009 | 0.00000 | 1.05593 | 0.01175 |
| **Sema5b** | 0.00804 | 0.00773 | 0.00559 | 0.01458 | 0.00917 | 0.00709 | 0.00641 | 0.19708 | 0.48170 | 0.00000 | 0.43714 | 0.00319 | 1.11796 | 0.07934 | 0.00000 | 0.00658 |
| **Sema6a** | 0.03646 | 0.00776 | 0.57834 | 0.33447 | 0.00710 | 0.00914 | 0.00733 | 0.01210 | 0.00582 | 0.01468 | 0.01240 | 0.00782 | 0.00892 | 0.01199 | 0.00023 | 0.00703 |
| **Sema6b** | 0.01386 | 0.00784 | 0.01031 | 0.01529 | 0.00905 | 0.00674 | 0.00734 | 0.00533 | 0.12058 | 0.18965 | 0.25177 | 0.00569 | 0.00998 | 0.00869 | 0.01383 | 0.00769 |
| **Sema6c** | 0.01338 | 0.00801 | 0.01253 | 1.54939 | 0.00943 | 0.00654 | 0.01801 | 0.01438 | 0.59688 | 0.01005 | 0.01037 | 0.00670 | 0.02583 | 0.00721 | 0.02610 | 0.02190 |
| **Sema6d** | 0.02518 | 0.01815 | 0.02932 | 0.25197 | 0.00987 | 0.01229 | 0.00737 | 0.02613 | 0.00575 | 0.09196 | 0.02907 | 0.00645 | 0.05325 | 0.03804 | 0.20485 | 0.00770 |
| **Sema7a** | 0.17179 | 0.00814 | 0.00518 | 0.00511 | 0.01687 | 0.00733 | 0.00699 | 0.07351 | 0.06385 | 0.04376 | 0.31384 | 0.01695 | 0.00681 | 0.01140 | 0.01482 | 0.00720 |
| **Senp1** | 1.90330 | 2.20421 | 2.14664 | 3.69680 | 0.70256 | 1.25523 | 0.94313 | 4.65541 | 2.35755 | 5.58886 | 2.36259 | 1.83042 | 3.72625 | 1.24652 | 2.35205 | 0.02909 |
| **Senp2** | 3.97941 | 3.30134 | 0.79727 | 4.07353 | 4.82587 | 4.69174 | 2.92155 | 1.80684 | 7.25490 | 3.52390 | 4.47522 | 4.66299 | 1.19502 | 12.00595 | 5.96588 | 0.39809 |
| **Senp3** | 1.63724 | 4.01278 | 0.00000 | 13.03740 | 0.94607 | 1.69725 | 0.25184 | 13.67323 | 14.45961 | 7.06699 | 7.07042 | 12.36752 | 2.47797 | 14.44952 | 13.56706 | 3.55411 |
| **Senp5** | 2.35915 | 5.95503 | 2.54312 | 3.69469 | 2.97639 | 2.60442 | 2.73069 | 1.43562 | 6.74559 | 5.69891 | 1.19278 | 9.16796 | 2.36448 | 7.59060 | 8.19931 | 0.06288 |
| **Senp6** | 5.84072 | 4.63168 | 1.64273 | 15.23939 | 5.00889 | 5.88550 | 3.48601 | 9.64370 | 14.65328 | 19.27545 | 4.54124 | 16.16357 | 0.71355 | 6.66518 | 15.72564 | 1.68934 |
| **Senp7** | 0.99290 | 0.04493 | 0.05160 | 4.41383 | 0.01416 | 0.01427 | 0.04581 | 0.03564 | 0.40712 | 0.78419 | 0.28443 | 2.09645 | 0.11746 | 1.54681 | 1.29853 | 0.01999 |
| **Senp8** | 0.00000 | 0.01458 | 2.89279 | 0.60788 | 3.75983 | 0.00656 | 0.00618 | 2.33059 | 0.00000 | 0.00000 | 0.00000 | 0.32394 | 1.25184 | 0.08351 | 0.48446 | 0.00533 |
| **X15.Sep** | 26.69823 | 12.46636 | 15.17399 | 3.84069 | 14.08576 | 13.38067 | 12.28285 | 18.35238 | 0.55419 | 14.06328 | 10.31329 | 1.86301 | 18.94167 | 4.31326 | 4.26543 | 14.25703 |
| **Sephs1** | 6.58891 | 4.50594 | 5.81607 | 6.79813 | 13.77145 | 4.09964 | 5.80926 | 16.11430 | 5.69604 | 5.79238 | 8.34319 | 11.72971 | 6.15730 | 4.64905 | 5.79191 | 5.22946 |

| | | | | | | | | | | | | | | | | |
|---|---|---|---|---|---|---|---|---|---|---|---|---|---|---|---|---|
| **Sephs2** | 2.84323 | 5.38754 | 5.59306 | 13.36151 | 3.90127 | 0.42472 | 5.77759 | 2.32223 | 4.49282 | 1.43248 | 1.61589 | 0.10194 | 4.14081 | 0.06731 | 7.72076 | 3.39102 |
| **Sepn1** | 0.00210 | 2.43878 | 0.00000 | 0.16986 | 0.00801 | 0.00728 | 0.00602 | 1.00102 | 0.12015 | 0.00000 | 0.28228 | 0.00403 | 2.33936 | 0.01337 | 0.01173 | 0.01978 |
| **Sepp1** | 0.02387 | 0.00809 | 0.01017 | 0.00561 | 0.02090 | 0.00650 | 0.00720 | 0.00920 | 0.01124 | 2.44678 | 0.03170 | 0.01108 | 0.00736 | 0.01214 | 0.00617 | 1.71788 |
| **Sepsecs** | 0.01625 | 2.07631 | 1.63156 | 0.00000 | 3.80686 | 0.00549 | 0.00616 | 0.53770 | 0.08785 | 0.66442 | 0.08076 | 0.02000 | 0.78494 | 0.16639 | 0.00000 | 0.13686 |
| **X1.Sep** | 0.09464 | 0.00821 | 0.56476 | 0.00484 | 0.01480 | 0.00719 | 0.00701 | 0.00649 | 0.16356 | 0.24339 | 1.03796 | 0.01163 | 0.28500 | 0.00970 | 0.00333 | 0.00733 |
| **X10.Sep** | 1.25849 | 0.25049 | 0.00000 | 0.03330 | 1.41840 | 0.09108 | 0.00553 | 0.00000 | 0.34432 | 0.00000 | 0.69994 | 0.03695 | 0.30627 | 2.07273 | 1.67150 | 0.05085 |
| **X11.Sep** | 14.41891 | 7.12722 | 6.33173 | 1.57942 | 9.61884 | 1.14872 | 0.02352 | 0.18089 | 0.59491 | 0.75533 | 1.80607 | 0.40124 | 2.06050 | 1.70943 | 0.91542 | 1.40588 |
| **X12.Sep** | 3.23086 | 0.06341 | 3.11324 | 0.00763 | 4.56735 | 2.85509 | 0.44329 | 1.34249 | 0.00792 | 0.42009 | 0.52087 | 0.04139 | 1.93411 | 0.00893 | 0.19902 | 0.02371 |
| **X2.Sep** | 15.60177 | 7.54975 | 12.89661 | 3.55084 | 4.74551 | 4.37459 | 11.17548 | 9.51914 | 8.79137 | 15.30820 | 10.18695 | 10.54661 | 4.49773 | 21.91075 | 11.00089 | 5.61446 |
| **X3.Sep** | 0.00000 | 0.00324 | 0.00000 | 1.14243 | 2.84264 | 0.00464 | 0.04416 | 2.76733 | 1.50471 | 5.19215 | 2.79020 | 6.43693 | 2.37871 | 0.14736 | 6.59837 | 0.00497 |
| **X4.Sep** | 0.01121 | 0.00787 | 0.00999 | 0.30733 | 0.02828 | 0.00679 | 0.39785 | 0.01127 | 0.17452 | 0.06609 | 0.01112 | 0.00721 | 0.03123 | 0.01048 | 0.00547 | 0.00716 |
| **X6.Sep** | 0.02863 | 0.00795 | 0.03787 | 0.83924 | 0.01198 | 5.52828 | 0.00680 | 0.00920 | 0.00000 | 0.00433 | 0.00859 | 0.00792 | 0.04443 | 0.01579 | 0.01085 | 0.00523 |
| **X7.Sep** | 2.27993 | 7.92763 | 1.17657 | 4.01774 | 0.71576 | 1.67657 | 7.54352 | 6.74608 | 5.55568 | 0.00000 | 1.95322 | 1.46791 | 2.47841 | 6.98381 | 2.64010 | 0.72925 |
| **X8.Sep** | 2.77050 | 0.03484 | 3.19342 | 0.00000 | 0.01447 | 0.67077 | 4.68422 | 6.95974 | 0.00678 | 2.31183 | 3.39462 | 0.08668 | 5.06348 | 0.58188 | 0.00000 | 0.00000 |
| **X9.Sep** | 2.19958 | 0.00305 | 0.00000 | 0.00436 | 3.72186 | 2.62168 | 0.00515 | 1.43149 | 0.00879 | 0.32842 | 0.00000 | 0.05099 | 0.00000 | 0.22532 | 0.19570 | 4.56280 |
| **Sepw1** | 6.59866 | 2.68764 | 3.97035 | 2.02598 | 1.47487 | 2.21812 | 2.03568 | 5.62297 | 3.57232 | 12.20123 | 15.93476 | 1.17754 | 10.36047 | 0.45015 | 2.32774 | 4.09917 |
| **Serac1** | 3.36725 | 0.00321 | 1.39667 | 0.67667 | 0.37142 | 0.03408 | 0.00360 | 1.69549 | 0.26060 | 0.94886 | 1.28239 | 0.48102 | 0.94343 | 0.01881 | 3.51698 | 0.00259 |
| **Serbp1** | 27.82124 | 15.25467 | 16.84253 | 15.97065 | 20.20739 | 8.47934 | 9.53074 | 24.29408 | 30.41974 | 37.71342 | 32.40316 | 19.57135 | 24.48901 | 19.81027 | 10.87812 | 14.45365 |
| **Serf1** | 1.72784 | 0.01629 | 0.00302 | 0.82468 | 0.04481 | 0.00685 | 0.00498 | 0.15519 | 1.88986 | 0.83788 | 2.19461 | 0.00000 | 0.00000 | 0.01635 | 0.00000 | 0.00501 |
| **Serf2** | 19.37123 | 4.15245 | 18.44595 | 26.72153 | 16.01750 | 5.87885 | 8.00192 | 9.39058 | 20.18873 | 27.93736 | 25.70302 | 4.60762 | 24.68410 | 3.59871 | 9.80533 | 13.51918 |
| **Sergef** | 3.35413 | 0.01266 | 1.11531 | 0.00608 | 1.03263 | 0.00218 | 3.22037 | 1.13595 | 0.01013 | 0.53178 | 0.00431 | 0.00000 | 1.02489 | 0.01052 | 0.01716 | 0.00328 |
| **Serhl** | 0.35537 | 0.00793 | 1.04949 | 0.00000 | 0.02844 | 0.18925 | 0.00647 | 1.89632 | 0.24705 | 2.26535 | 0.15527 | 0.15879 | 1.77996 | 0.58671 | 1.67361 | 0.05654 |
| **Serinc1** | 0.73919 | 0.00643 | 8.07192 | 5.29355 | 2.46372 | 3.28913 | 6.39466 | 13.62556 | 8.84815 | 9.73992 | 10.66907 | 10.62637 | 6.07340 | 21.72289 | 8.76055 | 2.89221 |
| **Serinc2** | 1.87287 | 0.01340 | 2.14508 | 0.05748 | 0.78016 | 0.01525 | 2.86556 | 2.10919 | 0.36672 | 0.84698 | 0.11104 | 0.01003 | 4.67278 | 0.01708 | 0.00501 | 2.03465 |
| **Serinc3** | 10.78977 | 10.38757 | 16.10004 | 13.05489 | 0.96198 | 13.18230 | 16.50792 | 25.64739 | 19.61342 | 12.97497 | 23.31831 | 11.26555 | 18.47815 | 26.96273 | 14.80024 | 9.62552 |
| **Serinc5** | 0.02557 | 0.00750 | 0.00404 | 2.48928 | 0.01233 | 0.00662 | 2.02137 | 0.03994 | 0.39945 | 0.31833 | 0.02836 | 0.07970 | 0.00949 | 0.01910 | 0.98572 | 0.00717 |
| **Serp1** | 6.01243 | 6.75424 | 8.90640 | 17.35883 | 5.31707 | 3.67046 | 1.68449 | 4.13741 | 20.30926 | 11.63199 | 6.48953 | 3.45274 | 4.19322 | 6.29996 | 13.61743 | 3.35387 |
| **Serpina1a** | 0.01526 | 0.01895 | 0.02231 | 0.04072 | 0.02711 | 0.00697 | 0.04942 | 0.01614 | 0.02103 | 0.38596 | 0.01232 | 0.00683 | 0.02044 | 0.03557 | 0.01156 | 0.01499 |
| **Serpina1b** | 0.01371 | 0.00793 | 0.01226 | 0.00990 | 0.00931 | 0.00663 | 0.00743 | 0.01244 | 0.01228 | 0.16169 | 0.01300 | 0.00736 | 0.01266 | 0.01149 | 0.01014 | 0.00758 |
| **Serpina1c** | 0.01364 | 0.00794 | 0.01225 | 0.04893 | 0.00926 | 0.00663 | 0.00740 | 0.01181 | 0.00928 | 0.29161 | 0.01219 | 0.00707 | 0.01233 | 0.01126 | 0.00888 | 0.00755 |
| **Serpina1d** | 0.01366 | 0.00795 | 0.01225 | 0.03392 | 0.00927 | 0.00664 | 0.00741 | 0.01201 | 0.01003 | 0.25200 | 0.01229 | 0.00719 | 0.01242 | 0.01134 | 0.00935 | 0.00755 |
| **Serpina1e** | 0.01195 | 0.00775 | 0.01227 | 0.01394 | 0.00858 | 0.00651 | 0.00722 | 0.00952 | 0.01046 | 3.86947 | 0.00940 | 0.00000 | 0.01057 | 0.01245 | 0.00578 | 0.00745 |
| **Serpina3m** | 2.95689 | 0.01522 | 0.00341 | 0.01063 | 0.00256 | 0.00564 | 0.00504 | 0.01193 | 0.01236 | 0.01492 | 0.01209 | 0.02611 | 0.01087 | 0.01212 | 0.01038 | 0.00419 |
| **Serpinb1a** | 6.66806 | 3.74260 | 0.13301 | 0.00000 | 6.28136 | 0.00000 | 7.64481 | 0.00000 | 0.70563 | 0.02087 | 0.00545 | 0.00029 | 0.03877 | 0.00462 | 0.20987 | 2.81777 |
| **Serpinb1b** | 0.06556 | 0.05999 | 0.01962 | 0.01025 | 0.08470 | 0.00623 | 0.08783 | 0.01226 | 0.01223 | 0.02834 | 0.01333 | 0.00753 | 0.01193 | 0.01049 | 0.01735 | 0.00739 |
| **Serpinb1c** | 0.17997 | 0.06310 | 0.14307 | 0.00963 | 0.10483 | 0.14555 | 0.17064 | 0.01144 | 0.09064 | 0.01304 | 0.01204 | 0.00774 | 0.01709 | 0.01008 | 0.04370 | 0.04364 |
| **Serpinb6a** | 21.09314 | 16.04655 | 16.92778 | 0.00000 | 15.81477 | 14.33137 | 10.56000 | 4.25228 | 0.05026 | 3.22445 | 13.07479 | 0.31185 | 22.17938 | 8.27584 | 0.00000 | 14.80073 |
| **Serpinb6b** | 0.10349 | 0.06964 | 0.11798 | 0.00846 | 3.32879 | 0.30929 | 0.09043 | 0.01177 | 0.01080 | 0.01280 | 0.01174 | 0.00427 | 1.09458 | 0.03917 | 0.02343 | 0.02059 |
| **Serpinb6c** | 4.15378 | 0.13162 | 4.09571 | 0.01627 | 0.19967 | 10.56438 | 4.98963 | 0.00000 | 0.01234 | 0.00000 | 0.00595 | 0.00000 | 8.66593 | 0.05888 | 0.00000 | 1.22602 |
| **Serpinb6d** | 0.09130 | 0.01921 | 0.17355 | 0.01052 | 0.00970 | 0.09929 | 0.12245 | 0.01860 | 0.01237 | 0.02380 | 0.01284 | 0.00748 | 0.18467 | 0.01279 | 0.00691 | 0.01473 |
| **Serpinb6e** | 0.04652 | 0.00827 | 0.05621 | 0.01056 | 0.01017 | 0.09423 | 0.03922 | 0.01756 | 0.01235 | 0.01189 | 0.01307 | 0.00797 | 0.08829 | 0.01283 | 0.00960 | 0.02221 |
| **Serpinb9** | 0.01680 | 0.01879 | 0.11920 | 0.00983 | 0.00849 | 0.01804 | 0.00743 | 0.00532 | 0.01216 | 0.01589 | 0.45238 | 0.00629 | 0.01467 | 0.01143 | 0.00211 | 0.00719 |
| **Serpinb9b** | 2.65036 | 3.28934 | 1.84107 | 0.00706 | 0.06841 | 7.77101 | 0.65444 | 0.31555 | 0.01259 | 0.00000 | 0.00558 | 0.00238 | 4.40045 | 0.01500 | 0.00000 | 0.95312 |
| **Serpinc1** | 0.01221 | 0.00796 | 0.02777 | 3.19498 | 0.01160 | 0.09752 | 0.00676 | 0.12722 | 2.38210 | 1.06769 | 1.79107 | 0.06783 | 0.09605 | 0.20656 | 1.91530 | 0.00658 |
| **Serpind1** | 0.01374 | 0.00793 | 0.01075 | 0.00447 | 0.00927 | 0.00665 | 0.00742 | 0.01175 | 0.86587 | 0.02448 | 0.01933 | 0.06938 | 0.01250 | 0.01650 | 0.00247 | 0.00731 |
| **Serpine1** | 0.01392 | 0.00748 | 0.00804 | 0.00983 | 0.00929 | 0.00684 | 0.00718 | 0.01002 | 0.01210 | 0.00619 | 0.02370 | 0.52602 | 0.01296 | 0.00789 | 0.04025 | 0.00747 |
| **Serpine2** | 2.43646 | 1.01875 | 3.39790 | 2.07097 | 10.87463 | 0.00853 | 3.50079 | 1.75560 | 2.16215 | 2.19079 | 0.04328 | 0.07187 | 5.50345 | 0.00000 | 1.82040 | 2.70290 |
| **Serpine3** | 0.02087 | 0.00821 | 0.00635 | 0.10702 | 0.00917 | 0.08766 | 0.00765 | 0.01005 | 0.32649 | 0.01933 | 0.13359 | 0.00721 | 0.01969 | 0.04592 | 0.22584 | 0.00737 |
| **Serpinf1** | 0.01341 | 0.00781 | 0.01034 | 0.20452 | 0.00917 | 0.00653 | 0.00719 | 0.01014 | 0.00770 | 0.78613 | 0.01176 | 0.11435 | 0.01143 | 0.00701 | 0.12008 | 0.00738 |
| **Serpinf2** | 0.01311 | 0.00791 | 0.01173 | 0.16265 | 0.00915 | 0.00644 | 0.00726 | 0.01019 | 0.47822 | 0.95993 | 0.00750 | 0.00413 | 0.01145 | 0.02450 | 0.24376 | 0.00741 |
| **Serping1** | 0.51963 | 2.09988 | 0.20398 | 1.61075 | 0.01533 | 0.00502 | 5.64235 | 3.50436 | 1.33238 | 5.93979 | 3.15098 | 0.36512 | 2.14845 | 0.69820 | 3.86110 | 0.02632 |
| **Serpinh1** | 12.40814 | 1.14405 | 2.24813 | 0.00353 | 3.31305 | 0.00369 | 0.00000 | 0.00000 | 0.01047 | 0.73792 | 0.00626 | 0.17177 | 0.00530 | 0.00984 | 0.00000 | 6.95314 |
| **Serpini1** | 0.00666 | 0.00825 | 0.00730 | 3.06411 | 0.00880 | 0.00554 | 0.00756 | 0.11703 | 9.42718 | 0.57118 | 0.00000 | 0.23445 | 0.93015 | 0.00000 | 0.76168 | 0.00573 |
| **Sertad1** | 3.35200 | 0.02171 | 0.82651 | 9.70793 | 0.02824 | 2.80194 | 1.73943 | 12.59337 | 20.36123 | 10.89446 | 7.92136 | 0.36423 | 2.31665 | 0.21066 | 0.00563 | 0.06317 |
| **Sertad2** | 3.33743 | 0.00397 | 0.97139 | 5.49856 | 0.13761 | 0.00471 | 0.00367 | 0.00000 | 2.21687 | 1.29863 | 0.69703 | 0.12030 | 0.82470 | 3.60960 | 4.85069 | 0.03035 |
| **Sertad3** | 0.01179 | 0.02866 | 1.88471 | 0.13853 | 0.01623 | 0.08118 | 0.00734 | 0.01215 | 0.62847 | 0.00888 | 0.00856 | 0.01539 | 0.01805 | 0.01170 | 0.00000 | 0.01427 |
| **Sesn1** | 1.51009 | 0.33890 | 6.22034 | 0.57296 | 3.66599 | 3.51230 | 3.51402 | 1.61351 | 3.05663 | 7.38985 | 4.12859 | 7.57086 | 2.49734 | 0.03243 | 4.11808 | 0.65187 |
| **Sesn2** | 0.75606 | 0.00440 | 1.25346 | 0.70258 | 1.37579 | 1.12709 | 2.11367 | 4.74504 | 2.58814 | 3.42003 | 0.24520 | 0.25693 | 0.92720 | 0.04696 | 3.51732 | 1.59664 |
| **Sesn3** | 0.00422 | 4.36154 | 0.87029 | 1.03925 | 0.51100 | 0.79081 | 0.00569 | 0.00000 | 1.93658 | 3.81501 | 0.29129 | 0.07336 | 0.02151 | 0.01961 | 1.64467 | 0.00488 |
| **Sestd1** | 2.20358 | 0.00484 | 0.12654 | 4.69878 | 0.00399 | 1.95730 | 0.00465 | 0.00000 | 0.31309 | 1.68457 | 0.04071 | 0.06515 | 0.82872 | 0.00000 | 5.04006 | 0.00303 |
| **Set** | 37.27685 | 21.32263 | 31.85118 | 10.01278 | 26.58469 | 13.35532 | 20.17765 | 43.47195 | 16.55789 | 50.92397 | 28.25864 | 23.81188 | 34.62755 | 44.35580 | 30.72098 | 22.45903 |
| **Setd1a** | 5.22531 | 2.54486 | 0.75476 | 10.55603 | 3.51928 | 3.12652 | 1.71466 | 5.79177 | 8.54359 | 12.75395 | 1.14887 | 1.54676 | 3.70795 | 0.02918 | 1.99492 | 0.28369 |

| | | | | | | | | | | | | | | | | |
|---|---|---|---|---|---|---|---|---|---|---|---|---|---|---|---|---|
| **Setd1b** | 1.10595 | 0.22972 | 0.95125 | 2.19917 | 0.36872 | 0.39084 | 0.28184 | 0.32424 | 2.06696 | 3.20783 | 3.32536 | 0.16149 | 1.08067 | 0.39088 | 0.14248 | 2.19459 |
| **Setd2** | 0.97267 | 5.80060 | 4.42814 | 19.29069 | 6.54770 | 6.67623 | 4.15209 | 19.73466 | 23.86807 | 22.05934 | 17.67479 | 23.25551 | 3.10564 | 25.68742 | 21.77674 | 0.83511 |
| **Setd3** | 13.40954 | 6.51769 | 7.39604 | 6.35817 | 6.76285 | 7.86320 | 9.19790 | 9.15151 | 13.23827 | 9.95187 | 4.31057 | 1.93368 | 10.55846 | 2.25133 | 5.60349 | 3.75733 |
| **Setd4** | 0.53994 | 0.20304 | 2.51010 | 0.85264 | 0.01039 | 0.01353 | 4.73936 | 8.54732 | 0.80894 | 2.59480 | 1.19452 | 0.10116 | 0.91881 | 0.05149 | 0.52583 | 0.04293 |
| **Setd5** | 0.03648 | 2.60475 | 2.25354 | 0.43684 | 0.51839 | 0.57713 | 1.11625 | 0.16833 | 0.14953 | 0.50172 | 0.17738 | 0.06976 | 1.41340 | 0.00531 | 0.62011 | 0.06742 |
| **Setd6** | 2.72741 | 2.03026 | 4.27632 | 2.10342 | 0.04835 | 0.00981 | 1.85869 | 1.79007 | 1.39110 | 2.08276 | 3.49549 | 0.95392 | 4.22003 | 0.00000 | 1.75621 | 1.20441 |
| **Setd7** | 0.45090 | 0.00752 | 0.00466 | 0.73857 | 0.01487 | 0.00704 | 0.00677 | 0.01110 | 0.00081 | 0.02851 | 0.01224 | 0.04881 | 0.44073 | 0.01135 | 0.00465 | 0.00630 |
| **Setdb1** | 3.21950 | 7.34352 | 2.30655 | 5.42257 | 5.01045 | 8.20248 | 6.19254 | 4.26535 | 10.19954 | 2.86252 | 2.56835 | 8.77680 | 0.93417 | 12.74273 | 11.46319 | 3.13372 |
| **Setdb2** | 0.01257 | 0.00730 | 0.00624 | 1.65571 | 0.00910 | 0.00610 | 0.00697 | 0.00437 | 4.01502 | 0.00000 | 0.10096 | 0.75132 | 0.01120 | 0.44510 | 0.27514 | 0.00666 |
| **Setmar** | 0.56860 | 0.00862 | 1.80374 | 0.00074 | 0.01389 | 0.00851 | 0.00749 | 0.65867 | 0.54770 | 0.00000 | 0.01310 | 0.01434 | 0.00000 | 0.14041 | 1.11455 | 0.00608 |
| **Setx** | 19.07965 | 10.87587 | 10.68718 | 13.44329 | 9.48835 | 1.02832 | 10.53141 | 8.01073 | 15.48655 | 20.67629 | 8.29389 | 17.78878 | 7.25501 | 26.54717 | 23.19519 | 11.13944 |
| **Sez6l** | 0.01337 | 0.00795 | 0.01157 | 1.73901 | 0.00930 | 0.00627 | 0.00737 | 0.01178 | 2.24787 | 0.11975 | 0.00000 | 0.00305 | 0.01283 | 0.03950 | 0.01246 | 0.00703 |
| **Sez6l2** | 0.00134 | 0.00680 | 4.66849 | 1.22171 | 0.00411 | 4.07118 | 1.07981 | 1.95161 | 0.52874 | 0.90528 | 0.12489 | 0.45761 | 0.05556 | 0.00299 | 0.61273 | 0.01218 |
| **Sf1** | 13.93194 | 3.58927 | 0.94814 | 7.58285 | 10.35293 | 1.72115 | 3.97219 | 7.66555 | 16.28879 | 18.70342 | 9.65935 | 7.60291 | 10.51931 | 5.12249 | 2.01839 | 7.16457 |
| **Sf3a1** | 14.26528 | 9.13362 | 9.73798 | 11.55739 | 9.94010 | 3.57645 | 4.15277 | 12.97424 | 17.80579 | 13.25741 | 9.26283 | 1.35665 | 15.03976 | 5.22599 | 2.41870 | 8.57473 |
| **Sf3a2** | 2.72402 | 0.00384 | 0.03981 | 0.42211 | 0.52359 | 0.01301 | 0.01375 | 1.27940 | 0.45741 | 1.22948 | 1.72629 | 0.09982 | 1.29989 | 0.02019 | 0.00000 | 0.39752 |
| **Sf3a3** | 16.20005 | 11.59267 | 10.47177 | 8.81668 | 13.55073 | 10.12012 | 5.65786 | 15.61051 | 8.99902 | 18.65211 | 16.33724 | 2.49659 | 8.63799 | 11.86474 | 11.04658 | 8.79392 |
| **Sf3b1** | 17.66743 | 13.75431 | 14.11071 | 18.27595 | 9.09818 | 8.90459 | 13.72185 | 15.79424 | 31.02923 | 25.21825 | 17.77882 | 21.03464 | 11.21566 | 33.33161 | 25.97849 | 9.79676 |
| **Sf3b2** | 22.04233 | 14.90972 | 12.79356 | 10.57307 | 15.86303 | 9.65844 | 13.85502 | 21.15001 | 8.30473 | 26.31067 | 14.93619 | 13.32326 | 18.99038 | 6.08489 | 14.34327 | 10.31822 |
| **Sf3b3** | 19.85037 | 14.85181 | 17.19421 | 4.27581 | 11.13535 | 0.67045 | 7.93303 | 19.24916 | 6.67386 | 26.81066 | 15.60445 | 12.90900 | 12.94585 | 21.74222 | 15.88352 | 8.44848 |
| **Sf3b4** | 8.49839 | 0.26943 | 0.00000 | 4.21451 | 8.09719 | 4.96392 | 2.65544 | 22.46318 | 5.27972 | 25.82507 | 12.52085 | 1.10124 | 15.03546 | 0.56682 | 0.00000 | 4.82029 |
| **Sf3b5** | 18.08149 | 0.04094 | 0.00000 | 2.98857 | 8.93923 | 5.92792 | 3.98143 | 15.24868 | 13.29003 | 16.00810 | 2.41301 | 0.43399 | 15.72700 | 0.01334 | 0.00000 | 6.90626 |
| **Sf3b6** | 16.14413 | 11.36680 | 15.35917 | 7.60358 | 12.39812 | 4.59136 | 7.49461 | 13.92732 | 13.71074 | 17.51451 | 20.34555 | 3.67268 | 13.36952 | 12.74646 | 11.65577 | 2.75439 |
| **Sfi1** | 3.71216 | 1.13813 | 9.96609 | 24.17377 | 0.00000 | 0.21664 | 8.75851 | 2.64761 | 31.53836 | 5.27391 | 1.94002 | 0.28777 | 8.66877 | 2.91117 | 2.61536 | 3.20838 |
| **Sfmbt1** | 2.77972 | 0.00436 | 0.05202 | 13.57389 | 0.00447 | 0.00178 | 0.29366 | 0.00000 | 6.56529 | 0.50993 | 0.05150 | 0.16735 | 0.01050 | 0.80151 | 8.10087 | 0.01637 |
| **Sfmbt2** | 5.94870 | 1.57812 | 1.10329 | 0.00000 | 0.01233 | 0.00877 | 0.98632 | 0.00000 | 0.41032 | 0.27290 | 0.00000 | 8.68529 | 3.19665 | 4.83708 | 0.00000 | 0.71598 |
| **Sfn** | 10.43001 | 7.11981 | 13.99070 | 0.63477 | 3.79694 | 5.36512 | 12.07073 | 33.63961 | 5.03605 | 36.21126 | 15.56057 | 5.50971 | 18.17981 | 1.92110 | 0.00000 | 5.94292 |
| **Sfpq** | 15.37959 | 14.26241 | 12.11930 | 5.99739 | 13.13841 | 8.73731 | 10.81373 | 21.17343 | 11.52763 | 17.59934 | 17.40832 | 3.96578 | 10.89395 | 24.88013 | 16.13813 | 8.20326 |
| **Sfr1** | 15.89712 | 13.06334 | 12.24629 | 5.60245 | 5.41115 | 6.26290 | 12.06997 | 18.54392 | 9.94922 | 12.17522 | 8.15688 | 12.27715 | 13.64171 | 5.89320 | 11.39860 | 3.94494 |
| **Sfrp1** | 2.87088 | 4.29736 | 4.12077 | 0.47633 | 0.02681 | 0.00493 | 0.03443 | 0.90878 | 0.28564 | 0.00934 | 0.00000 | 0.04768 | 0.17779 | 0.00578 | 0.56980 | 2.19090 |
| **Sfrp5** | 0.01033 | 0.00771 | 0.01113 | 0.01009 | 0.01038 | 0.00712 | 0.00748 | 0.17945 | 0.01205 | 0.00779 | 0.01344 | 0.00683 | 0.03925 | 0.00857 | 0.11619 | 0.00767 |
| **Sfswap** | 2.52340 | 0.03553 | 0.14644 | 0.26474 | 3.44601 | 5.52257 | 1.15627 | 6.79823 | 2.69319 | 7.25765 | 0.62612 | 0.18172 | 1.67472 | 0.17112 | 0.75539 | 4.24359 |
| **Sft2d1** | 8.84253 | 1.78328 | 3.66184 | 2.33607 | 2.61779 | 7.29989 | 2.57305 | 9.35640 | 1.75035 | 7.32493 | 3.70737 | 1.52162 | 4.55025 | 0.00000 | 0.00000 | 4.02105 |
| **Sft2d2** | 0.04094 | 0.00701 | 0.52756 | 0.00000 | 1.90895 | 4.89220 | 0.00664 | 0.00938 | 0.06805 | 0.64493 | 0.00657 | 0.00312 | 0.43031 | 0.01493 | 0.01708 | 1.47684 |
| **Sft2d3** | 0.00792 | 0.00871 | 0.83294 | 2.18179 | 0.00824 | 0.00684 | 0.00780 | 0.17476 | 0.10041 | 0.00803 | 0.00914 | 0.02030 | 0.43553 | 0.06633 | 0.00000 | 0.01377 |
| **Sfta2** | 0.01286 | 0.00792 | 0.05466 | 0.00996 | 0.00913 | 0.00844 | 0.00746 | 0.01333 | 0.01241 | 0.02428 | 0.01218 | 0.00766 | 0.01176 | 0.00716 | 0.08633 | 0.00732 |
| **Sfxn1** | 0.48256 | 5.22761 | 0.97579 | 0.11443 | 0.19781 | 0.00539 | 0.00279 | 2.29024 | 0.01254 | 7.52434 | 1.59736 | 0.17520 | 4.02441 | 3.23497 | 1.44992 | 0.01702 |
| **Sfxn2** | 1.58594 | 2.12504 | 3.64943 | 2.13474 | 0.01647 | 0.03170 | 0.13831 | 1.13660 | 2.79192 | 2.34306 | 0.93643 | 2.97296 | 3.73205 | 1.68468 | 0.81720 | 5.18516 |
| **Sfxn3** | 0.03243 | 0.00761 | 0.80885 | 0.00917 | 0.00684 | 0.00875 | 0.01721 | 0.01178 | 0.06131 | 0.59789 | 0.04257 | 0.02183 | 0.03725 | 0.65339 | 0.07095 | 0.01386 |
| **Sfxn4** | 0.96644 | 0.03931 | 2.13658 | 5.90202 | 0.27399 | 0.02050 | 0.06979 | 0.34163 | 8.98821 | 0.16023 | 5.30562 | 0.52400 | 0.06055 | 5.00582 | 2.52065 | 0.12494 |
| **Sfxn5** | 0.48738 | 0.00687 | 0.59669 | 0.01016 | 0.00609 | 0.00518 | 0.00688 | 0.00903 | 0.01215 | 0.14192 | 0.01242 | 0.00648 | 0.00939 | 0.01302 | 0.00392 | 0.00691 |
| **Sgcb** | 0.01208 | 0.00769 | 0.01139 | 0.00692 | 0.00898 | 0.00654 | 0.00720 | 0.03488 | 0.01157 | 1.10350 | 0.02588 | 0.00513 | 0.00953 | 0.01766 | 0.00517 | 0.00726 |
| **Sgce** | 0.00983 | 0.00882 | 0.01032 | 0.00000 | 0.00920 | 0.00636 | 0.00792 | 0.08555 | 0.46283 | 0.61709 | 0.94984 | 0.27398 | 0.00004 | 0.00504 | 1.52064 | 0.00704 |
| **Sgf29** | 13.19294 | 3.08813 | 5.38412 | 2.54463 | 4.06967 | 8.56730 | 5.20668 | 14.63100 | 5.15692 | 12.59770 | 5.32975 | 1.05908 | 3.87886 | 0.64273 | 4.58115 | 3.54958 |
| **Sgk1** | 7.99420 | 7.62573 | 9.77253 | 12.65505 | 6.36297 | 5.97006 | 5.01752 | 19.06583 | 15.19609 | 26.37131 | 12.38080 | 22.60028 | 13.43928 | 6.90733 | 3.09198 | 5.05245 |
| **Sgk2** | 0.11206 | 0.00832 | 0.66207 | 0.01092 | 0.01526 | 0.00915 | 0.00722 | 0.01152 | 0.01224 | 0.01460 | 0.01172 | 0.00752 | 0.00875 | 0.01518 | 0.00111 | 0.00722 |
| **Sgk3** | 1.91266 | 0.10374 | 1.97091 | 3.85982 | 5.09629 | 2.06494 | 0.02735 | 11.37469 | 6.43628 | 14.28195 | 14.11728 | 20.65055 | 1.57334 | 21.78642 | 11.88801 | 1.63732 |
| **Sgms1** | 1.34787 | 0.93399 | 0.34136 | 3.85885 | 0.00487 | 0.00400 | 0.00509 | 0.00000 | 0.66616 | 0.00000 | 0.21597 | 0.89840 | 1.36917 | 0.00000 | 0.08322 | 0.02501 |
| **Sgms2** | 0.02649 | 0.00812 | 0.08749 | 4.46579 | 0.00919 | 0.02739 | 0.02776 | 0.02416 | 6.19576 | 0.40997 | 0.30044 | 0.00000 | 0.05179 | 0.10291 | 0.00000 | 0.00657 |
| **Sgol1** | 3.60822 | 4.52366 | 4.22994 | 7.63658 | 3.74698 | 4.56799 | 2.97079 | 7.82193 | 7.57044 | 4.98968 | 2.82282 | 7.82615 | 1.27028 | 6.93134 | 8.23046 | 2.25551 |
| **Sgol2a** | 3.93675 | 0.89018 | 0.24855 | 4.91373 | 0.26647 | 0.50754 | 3.64966 | 1.39068 | 11.44376 | 1.06401 | 2.75090 | 0.18395 | 1.32240 | 9.47160 | 7.08248 | 0.71457 |
| **Sgol2b** | 0.01958 | 0.01823 | 0.01183 | 0.08660 | 0.22491 | 0.01882 | 0.32380 | 0.02762 | 0.11098 | 0.01382 | 0.01024 | 0.00647 | 0.01049 | 0.16281 | 0.29127 | 0.00729 |
| **Sgpl1** | 6.25863 | 0.28795 | 6.57654 | 2.14353 | 2.84096 | 2.07246 | 12.02984 | 0.00000 | 1.62768 | 3.14856 | 3.44866 | 0.43809 | 6.89711 | 0.00113 | 1.39878 | 6.02128 |
| **Sgpp1** | 0.49464 | 8.21433 | 3.19888 | 7.14973 | 3.50065 | 9.69924 | 8.09005 | 5.27210 | 13.83922 | 19.23126 | 12.11362 | 20.82504 | 2.28807 | 21.79853 | 20.76113 | 0.16251 |
| **Sgpp2** | 0.17084 | 0.03774 | 0.36587 | 0.75610 | 0.04353 | 0.02434 | 0.11869 | 0.10763 | 0.30679 | 0.22536 | 0.56963 | 0.07534 | 0.23086 | 0.26994 | 0.18743 | 0.06378 |
| **Sgsh** | 0.01940 | 0.00743 | 0.00798 | 1.01296 | 0.00880 | 0.00633 | 0.00710 | 0.20793 | 3.21842 | 0.02704 | 0.00000 | 0.00902 | 0.00723 | 0.00134 | 0.30997 | 0.00700 |
| **Sgsm1** | 2.25553 | 0.21385 | 0.70882 | 2.02465 | 0.09514 | 0.01551 | 0.06659 | 1.95734 | 1.14166 | 1.36843 | 0.59991 | 0.06395 | 0.00000 | 0.21368 | 0.18762 | 0.01841 |
| **Sgsm2** | 0.02259 | 0.00778 | 0.03113 | 2.54056 | 0.01127 | 1.50253 | 0.07436 | 0.82162 | 0.00000 | 0.86933 | 0.97780 | 0.00596 | 0.08854 | 0.40136 | 0.34922 | 0.29083 |
| **Sgsm3** | 3.44374 | 0.03190 | 5.07013 | 7.06223 | 3.60362 | 2.03617 | 2.48483 | 6.65015 | 7.75546 | 3.89075 | 1.78663 | 0.76920 | 5.10413 | 0.03461 | 3.15830 | 4.26546 |
| **Sgta** | 20.54196 | 2.00331 | 7.94309 | 9.22059 | 6.20149 | 10.68408 | 10.77952 | 23.65027 | 11.91329 | 25.19382 | 11.34923 | 2.40550 | 15.76937 | 0.67154 | 0.11494 | 12.57865 |
| **Sgtb** | 0.01296 | 0.01806 | 0.01680 | 0.00349 | 0.00928 | 0.00619 | 0.42481 | 0.00916 | 0.29120 | 0.05017 | 0.02051 | 0.03215 | 0.01111 | 0.43295 | 0.31069 | 0.01460 |
| **Sh2b1** | 6.56097 | 0.03005 | 1.14349 | 5.17931 | 1.30959 | 3.69846 | 4.68119 | 1.03407 | 2.31224 | 3.61035 | 1.86589 | 0.11214 | 4.05634 | 0.65088 | 0.45658 | 0.72339 |

| | | | | | | | | | | | | | | | | |
|---|---|---|---|---|---|---|---|---|---|---|---|---|---|---|---|---|
| **Sh2b2** | 1.11273 | 0.00628 | 0.00369 | 1.81669 | 0.00734 | 0.00622 | 0.38542 | 0.00000 | 5.30189 | 3.41694 | 0.70519 | 0.00000 | 0.00600 | 0.05289 | 0.00000 | 0.01297 |
| **Sh2b3** | 2.04675 | 2.85990 | 1.03846 | 4.66276 | 6.31334 | 6.69003 | 4.48373 | 6.35270 | 8.81717 | 4.38790 | 3.41308 | 0.18847 | 6.68732 | 0.04054 | 3.18033 | 2.15865 |
| **Sh2d3c** | 1.16215 | 0.00609 | 0.23872 | 11.87539 | 0.00523 | 0.00167 | 0.00531 | 0.00000 | 15.25658 | 5.71053 | 1.64079 | 1.14881 | 0.00000 | 0.00530 | 0.05066 | 0.00272 |
| **Sh2d4a** | 0.00792 | 0.00835 | 1.13309 | 0.02306 | 0.00792 | 0.00947 | 0.00804 | 0.25945 | 0.07587 | 0.02419 | 0.00877 | 0.00779 | 0.00028 | 0.02186 | 0.01547 | 0.00698 |
| **Sh2d4b** | 0.02294 | 0.88291 | 0.00589 | 0.05667 | 0.00787 | 0.00677 | 0.00666 | 0.00918 | 0.42381 | 0.00273 | 0.00959 | 0.00673 | 0.38567 | 0.01064 | 0.00000 | 0.00660 |
| **Sh2d5** | 0.01289 | 0.00781 | 0.00689 | 0.40582 | 0.00867 | 0.00715 | 0.00694 | 0.00000 | 0.19473 | 1.72056 | 0.81272 | 0.03099 | 0.00631 | 0.00469 | 0.06772 | 0.00738 |
| **Sh2d7** | 0.01295 | 0.00792 | 0.05060 | 0.01047 | 0.00908 | 0.27528 | 0.00736 | 0.01263 | 0.01240 | 0.01338 | 0.01223 | 0.00785 | 0.00899 | 0.01248 | 0.00985 | 0.00724 |
| **Sh3bgr** | 0.00975 | 0.00767 | 0.01168 | 0.41206 | 0.00835 | 0.00648 | 0.00740 | 0.01822 | 0.00338 | 0.00546 | 0.01150 | 0.00780 | 0.00935 | 0.02093 | 0.06483 | 0.57567 |
| **Sh3bgrl** | 0.01845 | 0.00783 | 0.00864 | 0.58255 | 0.01047 | 0.01888 | 0.01765 | 0.01035 | 0.78275 | 0.12863 | 0.01313 | 0.02403 | 0.01192 | 0.01042 | 0.95129 | 0.00712 |
| **Sh3bgrl2** | 5.27674 | 0.06436 | 0.95423 | 0.64024 | 2.86028 | 2.81996 | 1.93811 | 1.74809 | 1.06107 | 0.07452 | 0.00000 | 0.01213 | 1.79719 | 0.00271 | 0.78071 | 0.26584 |
| **Sh3bgrl3** | 22.99560 | 6.72572 | 13.58613 | 1.01025 | 16.49934 | 1.03659 | 6.69844 | 5.74158 | 1.47364 | 4.79326 | 8.23763 | 0.17367 | 6.73622 | 0.20268 | 1.11716 | 6.84268 |
| **Sh3bp1** | 5.91138 | 0.74241 | 0.00000 | 0.00000 | 0.95242 | 4.17566 | 0.11888 | 2.89712 | 0.03111 | 0.00000 | 0.99206 | 0.00781 | 3.20704 | 0.00930 | 0.00000 | 2.49673 |
| **Sh3bp2** | 0.01059 | 0.00801 | 0.00544 | 0.33750 | 0.00934 | 0.00706 | 0.17906 | 0.04220 | 0.03665 | 0.35741 | 0.99127 | 0.09688 | 0.00365 | 0.18044 | 0.25402 | 0.00691 |
| **Sh3bp4** | 1.01754 | 0.34926 | 0.86161 | 0.94867 | 3.17834 | 0.01560 | 2.58949 | 0.58438 | 0.11105 | 1.42488 | 0.00604 | 0.00000 | 0.00264 | 0.00729 | 0.00000 | 0.00561 |
| **Sh3bp5** | 0.01708 | 0.04227 | 0.02196 | 0.00393 | 0.01882 | 0.00633 | 0.24343 | 0.01089 | 0.13640 | 0.09100 | 1.07042 | 0.00161 | 0.01101 | 0.86994 | 0.00000 | 0.25139 |
| **Sh3bp5l** | 4.39531 | 1.16055 | 2.34261 | 3.86011 | 0.00000 | 3.02520 | 2.33521 | 2.51592 | 1.66567 | 4.34636 | 3.79581 | 1.10268 | 1.76468 | 0.98905 | 3.84617 | 2.62812 |
| **Sh3d19** | 3.53942 | 0.96752 | 0.00000 | 3.21578 | 1.17075 | 5.63444 | 0.88031 | 3.70271 | 0.87623 | 2.30327 | 0.62368 | 0.06627 | 0.88251 | 0.11426 | 3.64865 | 3.85746 |
| **Sh3d21** | 0.50459 | 1.79791 | 4.08780 | 0.18446 | 0.01598 | 6.71939 | 0.00870 | 11.21520 | 0.02043 | 2.18453 | 5.74563 | 0.11942 | 8.94172 | 0.00576 | 1.09613 | 0.01281 |
| **Sh3gl1** | 3.87266 | 1.52614 | 1.80973 | 2.39316 | 4.88885 | 0.08712 | 1.92488 | 4.43693 | 1.36625 | 0.76669 | 0.42578 | 0.44756 | 1.75720 | 0.00150 | 2.13444 | 0.93643 |
| **Sh3gl2** | 0.01158 | 0.00797 | 0.02556 | 0.01550 | 0.01828 | 0.02011 | 0.02822 | 0.01611 | 0.04770 | 0.98086 | 0.00983 | 0.00598 | 0.01962 | 0.06050 | 0.05378 | 0.00730 |
| **Sh3gl3** | 1.62079 | 0.00690 | 0.00000 | 0.01063 | 0.04395 | 0.01317 | 0.00612 | 0.00859 | 0.01239 | 0.00529 | 0.01214 | 0.01744 | 0.00635 | 0.01331 | 0.01041 | 2.70505 |
| **Sh3glb1** | 3.35640 | 3.45193 | 3.60855 | 4.37364 | 5.66024 | 5.66260 | 4.39519 | 4.27372 | 2.08320 | 6.85853 | 7.06791 | 4.84068 | 7.37695 | 1.99087 | 3.07039 | 4.39281 |
| **Sh3glb2** | 6.16080 | 0.00000 | 0.00524 | 1.01866 | 1.64503 | 0.00186 | 0.00178 | 0.06905 | 0.03285 | 3.19095 | 1.46622 | 0.20299 | 0.18963 | 0.15761 | 0.00000 | 0.00826 |
| **Sh3kbp1** | 0.01162 | 0.00820 | 0.00477 | 4.22297 | 0.00915 | 0.00672 | 0.00784 | 0.16541 | 1.02400 | 0.15688 | 0.35384 | 0.06520 | 0.00589 | 0.00000 | 0.67073 | 0.00690 |
| **Sh3pxd2a** | 2.68486 | 0.02305 | 2.93518 | 5.50779 | 0.00000 | 0.00555 | 0.00347 | 0.51013 | 2.84593 | 3.24704 | 2.68506 | 0.10036 | 0.00000 | 0.03383 | 0.66945 | 1.01592 |
| **Sh3pxd2b** | 0.08331 | 0.00813 | 0.00654 | 0.00538 | 0.01543 | 0.00717 | 0.00709 | 0.00000 | 0.01108 | 0.44658 | 0.44530 | 0.04398 | 0.00854 | 0.01171 | 0.01056 | 0.00774 |
| **Sh3rf1** | 0.05034 | 0.03808 | 0.00379 | 5.82547 | 0.00778 | 0.00620 | 0.02670 | 0.00000 | 0.33801 | 0.02092 | 0.41817 | 0.04982 | 0.08469 | 0.50692 | 0.82741 | 0.93686 |
| **Sh3rf3** | 0.42862 | 0.00696 | 0.00000 | 5.56544 | 0.00693 | 0.00598 | 0.00647 | 0.32636 | 0.80893 | 0.07432 | 0.21675 | 0.00000 | 0.00549 | 0.00000 | 1.95832 | 0.01309 |
| **Sh3tc1** | 0.42957 | 0.24136 | 0.49904 | 0.51239 | 0.03040 | 2.61833 | 1.73197 | 5.90140 | 0.00448 | 0.37967 | 0.98999 | 0.00453 | 2.27706 | 0.00016 | 0.00000 | 4.49866 |
| **Sh3tc2** | 0.89188 | 0.00751 | 0.51252 | 0.29762 | 0.01607 | 3.12095 | 0.00582 | 0.25763 | 0.08333 | 0.00154 | 0.00629 | 0.00504 | 1.31451 | 0.00719 | 0.51977 | 0.01187 |
| **Sh3yl1** | 0.73686 | 0.00580 | 1.24898 | 6.67975 | 0.02126 | 0.65883 | 0.36448 | 1.41029 | 2.08856 | 1.24938 | 1.83625 | 1.36893 | 2.44698 | 2.15846 | 0.86907 | 0.00290 |
| **Sharpin** | 5.05435 | 0.07477 | 1.12708 | 0.00000 | 0.00741 | 0.02036 | 3.91236 | 1.27549 | 1.42642 | 2.62289 | 4.10312 | 0.23608 | 4.36984 | 0.01852 | 0.29817 | 3.80120 |
| **Shb** | 0.02362 | 0.00767 | 0.42500 | 4.19755 | 0.00784 | 0.00498 | 0.00685 | 0.00000 | 2.77321 | 0.86717 | 0.91254 | 0.00000 | 0.01049 | 0.00000 | 0.00000 | 0.00627 |
| **Shbg** | 0.00001 | 0.00849 | 0.00000 | 0.24679 | 0.00971 | 0.00636 | 0.00734 | 0.37654 | 7.09710 | 2.12774 | 2.26450 | 0.02814 | 0.01479 | 2.03755 | 1.42161 | 0.00684 |
| **Shc1** | 7.44669 | 4.55437 | 3.87427 | 3.27734 | 6.34234 | 0.34217 | 6.81214 | 6.50530 | 2.68682 | 5.25524 | 0.53905 | 0.04530 | 2.71802 | 1.87180 | 3.77696 | 6.00446 |
| **Shc3** | 0.87165 | 0.13327 | 0.00000 | 0.00477 | 0.02288 | 0.01342 | 2.53493 | 5.33357 | 0.02430 | 2.12440 | 2.12775 | 0.20434 | 0.00001 | 0.04645 | 0.60622 | 0.01312 |
| **Shcbp1** | 4.95236 | 9.37190 | 6.22125 | 0.39018 | 1.70756 | 4.93558 | 7.40819 | 5.12325 | 0.30537 | 4.09444 | 3.09728 | 11.05135 | 2.11711 | 5.47506 | 5.28686 | 3.64269 |
| **Shcbp1l** | 0.01222 | 0.00784 | 0.00000 | 1.09743 | 0.00907 | 1.31446 | 0.00665 | 0.00000 | 0.40342 | 0.13577 | 0.81046 | 0.05641 | 0.00000 | 1.29697 | 0.00000 | 0.00673 |
| **Shf** | 1.40051 | 0.00599 | 0.00692 | 0.34336 | 0.01357 | 0.00567 | 0.00637 | 0.00824 | 0.00555 | 0.01056 | 0.01237 | 0.02608 | 0.01833 | 0.00638 | 0.11350 | 0.00578 |
| **Shfm1** | 8.35984 | 0.56441 | 7.25478 | 4.92604 | 6.99711 | 1.08177 | 0.68473 | 0.80385 | 7.34207 | 2.44840 | 15.77199 | 1.07032 | 2.65096 | 0.40740 | 1.91008 | 3.23257 |
| **Shisa5** | 8.25164 | 2.13084 | 0.00000 | 1.59549 | 4.70941 | 3.06544 | 2.85261 | 1.12473 | 0.11090 | 1.78766 | 0.40458 | 0.05324 | 0.00000 | 0.00000 | 0.00000 | 4.54861 |
| **Shisa7** | 1.62001 | 0.00545 | 1.16323 | 1.01431 | 0.01114 | 0.01275 | 0.02453 | 1.31843 | 0.71866 | 2.56273 | 1.32014 | 0.32868 | 1.35099 | 0.60585 | 0.72105 | 0.00406 |
| **Shisa8** | 0.01221 | 0.00787 | 0.01032 | 1.44757 | 0.00878 | 0.00586 | 0.00716 | 0.00686 | 7.00507 | 1.56867 | 0.00000 | 0.00000 | 0.01125 | 0.00813 | 0.00489 | 0.00680 |
| **Shkbp1** | 8.51906 | 0.00000 | 0.26103 | 0.31291 | 9.88870 | 0.00389 | 1.62748 | 3.41273 | 1.34868 | 1.73120 | 0.15696 | 0.06748 | 2.61998 | 0.03198 | 0.00000 | 4.63659 |
| **Shmt1** | 9.90913 | 13.83798 | 11.80168 | 17.89121 | 11.26242 | 7.19560 | 7.33156 | 7.56344 | 24.13640 | 7.37146 | 7.07008 | 1.23621 | 9.18253 | 5.54530 | 12.22595 | 2.08552 |
| **Shmt2** | 11.00654 | 2.07124 | 7.18916 | 7.70168 | 9.08331 | 3.32039 | 8.45939 | 3.67845 | 2.97315 | 3.73020 | 0.95515 | 0.32588 | 10.21137 | 0.00000 | 0.00000 | 3.03346 |
| **Shoc2** | 4.06636 | 0.19832 | 2.22791 | 4.69140 | 0.74656 | 1.79458 | 1.27880 | 0.00000 | 6.55559 | 1.54181 | 0.64617 | 6.71558 | 1.03844 | 2.04731 | 2.43394 | 0.10567 |
| **Shox2** | 0.01405 | 0.00785 | 0.00829 | 0.00479 | 0.00912 | 0.00671 | 0.00729 | 0.00257 | 0.01093 | 0.00000 | 0.34056 | 0.06568 | 0.00863 | 0.00551 | 0.27192 | 0.00716 |
| **Shprh** | 0.28493 | 2.16671 | 2.34719 | 1.40041 | 0.01086 | 0.00537 | 2.51436 | 0.68376 | 1.56423 | 1.06192 | 1.05653 | 6.57419 | 0.00000 | 4.38973 | 5.50540 | 0.17583 |
| **Shq1** | 2.01712 | 1.66999 | 3.48113 | 0.10669 | 5.98418 | 0.00425 | 1.88116 | 2.18389 | 2.11748 | 3.57296 | 1.06955 | 0.18720 | 4.59457 | 0.00176 | 1.42413 | 0.00826 |
| **Shroom1** | 0.01374 | 0.00806 | 0.01173 | 0.08080 | 0.00930 | 0.00665 | 0.00750 | 0.01336 | 0.35589 | 0.01287 | 0.01128 | 0.02852 | 0.01283 | 0.01263 | 0.00546 | 0.00739 |
| **Shroom2** | 0.00494 | 2.73295 | 1.03136 | 10.45399 | 0.01855 | 0.00685 | 0.00625 | 0.47738 | 5.75874 | 1.76030 | 0.00000 | 0.00000 | 1.24579 | 4.38778 | 2.88906 | 1.26590 |
| **Shroom3** | 5.23015 | 6.29681 | 7.27081 | 1.71216 | 1.81016 | 1.01563 | 12.03641 | 6.89251 | 0.00001 | 0.46378 | 2.86922 | 0.13890 | 5.73654 | 0.00000 | 0.31121 | 4.98037 |
| **Shroom4** | 0.01323 | 0.00776 | 0.01905 | 6.85933 | 0.00987 | 0.01885 | 0.01739 | 0.02475 | 0.46881 | 0.60603 | 0.00000 | 0.01661 | 0.01157 | 0.77271 | 2.88940 | 0.27657 |
| **Shtn1** | 0.01202 | 0.00764 | 0.00000 | 1.61919 | 0.00948 | 1.18934 | 0.00713 | 0.14141 | 0.66200 | 0.06537 | 0.16158 | 0.01145 | 0.00000 | 0.00315 | 2.18031 | 0.00649 |
| **Siae** | 0.45677 | 0.00595 | 0.37466 | 0.12244 | 0.00578 | 0.01643 | 1.49530 | 0.00000 | 0.18240 | 0.94644 | 0.24277 | 0.00000 | 0.02693 | 0.01166 | 0.39265 | 2.01200 |
| **Siah1a** | 1.17946 | 0.01398 | 0.00000 | 3.10563 | 0.00962 | 2.60591 | 0.05405 | 3.34198 | 3.44202 | 7.41531 | 4.66773 | 10.74976 | 0.85647 | 9.69188 | 12.19998 | 0.00240 |
| **Siah1b** | 2.55892 | 0.39295 | 0.73120 | 12.69450 | 0.12475 | 0.42754 | 0.01279 | 0.72560 | 9.53253 | 10.81180 | 2.76496 | 9.08803 | 1.69961 | 3.29234 | 8.07636 | 0.26214 |
| **Siah2** | 1.39410 | 0.00559 | 0.55843 | 29.08310 | 0.00339 | 0.00369 | 0.00443 | 0.60939 | 14.57466 | 4.39535 | 0.86592 | 0.13226 | 0.00000 | 3.51873 | 15.26723 | 2.24672 |
| **Sidt2** | 2.44634 | 3.87785 | 9.76355 | 2.89482 | 2.00686 | 9.38701 | 10.73209 | 22.48499 | 3.23845 | 13.59362 | 12.31942 | 1.27419 | 12.67407 | 0.99974 | 4.53101 | 6.87978 |
| **Sigirr** | 0.68268 | 0.00555 | 0.00000 | 1.26217 | 0.02846 | 4.48929 | 2.30655 | 4.77684 | 0.88874 | 1.35750 | 2.19413 | 0.01106 | 0.00000 | 0.00000 | 0.00000 | 1.55142 |
| **Siglecg** | 0.00546 | 0.00754 | 0.03629 | 0.00758 | 0.01077 | 0.00782 | 0.00740 | 0.77649 | 0.01111 | 0.74862 | 0.00495 | 0.07257 | 0.05611 | 0.67190 | 0.00000 | 0.01544 |

| | | | | | | | | | | | | | | | | |
|---|---|---|---|---|---|---|---|---|---|---|---|---|---|---|---|---|
| **Sigmar1** | 2.56438 | 1.06106 | 1.99240 | 1.87395 | 0.00000 | 0.09347 | 1.73469 | 0.78324 | 0.66755 | 4.60734 | 0.71937 | 0.42725 | 2.37029 | 0.29644 | 0.34466 | 0.93189 |
| **Sik1** | 6.05965 | 0.07296 | 0.57589 | 0.32172 | 3.17745 | 0.00000 | 0.31353 | 0.54834 | 1.30510 | 0.33845 | 1.00382 | 0.07120 | 0.44958 | 0.00355 | 0.08471 | 2.57176 |
| **Sik2** | 0.01236 | 0.00815 | 0.25592 | 0.18421 | 0.00864 | 0.00952 | 0.00739 | 0.01208 | 0.00591 | 0.21424 | 0.01070 | 0.00790 | 0.01055 | 0.05149 | 0.00697 | 0.00723 |
| **Sik3** | 0.92278 | 3.07407 | 1.87213 | 0.92189 | 2.74041 | 0.01373 | 0.00247 | 1.09609 | 1.35713 | 4.32977 | 7.94763 | 0.63044 | 0.78946 | 2.63611 | 0.66088 | 0.01683 |
| **Sike1** | 0.75814 | 0.00414 | 1.11084 | 2.84747 | 2.99978 | 1.95787 | 0.56624 | 1.44479 | 2.64020 | 6.81098 | 8.23476 | 0.79768 | 1.58296 | 2.40743 | 2.90242 | 0.03269 |
| **Sil1** | 0.45644 | 0.00513 | 1.05167 | 0.70346 | 0.02264 | 0.27078 | 3.61108 | 1.96379 | 3.14586 | 0.15478 | 0.05037 | 0.09246 | 2.10970 | 0.00451 | 0.66856 | 0.00505 |
| **Simc1** | 2.13368 | 0.06841 | 1.62754 | 1.84567 | 3.16545 | 0.22926 | 3.60051 | 3.84160 | 4.41711 | 2.13329 | 1.65393 | 0.80184 | 0.49101 | 5.07714 | 3.51321 | 0.02505 |
| **Sin3a** | 2.63018 | 4.48494 | 4.12582 | 8.17334 | 4.95888 | 0.00765 | 3.33898 | 0.71916 | 10.11647 | 7.80947 | 2.56522 | 1.92735 | 1.94026 | 5.45821 | 9.86943 | 4.07558 |
| **Sin3b** | 18.66208 | 2.12348 | 7.60579 | 5.87240 | 12.59653 | 6.89610 | 5.45427 | 13.31971 | 3.86708 | 8.10010 | 8.41223 | 5.70281 | 18.75832 | 1.62452 | 0.83263 | 11.62400 |
| **Sipa1** | 4.44646 | 0.00181 | 0.72499 | 1.13618 | 0.89111 | 3.02828 | 0.23302 | 0.79573 | 0.13076 | 2.90034 | 2.72850 | 0.02326 | 0.11654 | 0.10437 | 0.01252 | 0.02940 |
| **Sipa1l1** | 0.39253 | 2.61834 | 0.26578 | 6.08625 | 0.01821 | 1.14078 | 0.12427 | 0.76162 | 2.19496 | 0.72226 | 0.05043 | 0.05174 | 1.87692 | 0.00000 | 1.67220 | 1.20439 |
| **Sipa1l2** | 2.30517 | 4.01523 | 3.54811 | 6.91912 | 4.29707 | 6.45089 | 1.68047 | 2.17080 | 3.43425 | 1.15861 | 2.56851 | 0.03151 | 3.29614 | 0.00000 | 1.19289 | 0.01788 |
| **Sipa1l3** | 0.19752 | 0.53708 | 0.13781 | 3.69884 | 0.00871 | 0.00540 | 0.41980 | 0.01102 | 0.19424 | 1.43690 | 1.34813 | 0.07870 | 0.00000 | 0.16660 | 0.07329 | 0.11078 |
| **Sirpa** | 0.95986 | 0.00771 | 0.00415 | 0.10895 | 0.01679 | 0.00709 | 0.00676 | 0.01050 | 0.00734 | 0.01527 | 0.01192 | 0.00750 | 0.01172 | 0.01344 | 0.01255 | 0.00661 |
| **Sirt1** | 4.15234 | 4.16799 | 2.76003 | 15.89396 | 0.00083 | 1.00648 | 5.32459 | 9.62606 | 20.43202 | 11.50650 | 6.11297 | 18.43337 | 2.30693 | 19.82423 | 21.04513 | 0.01337 |
| **Sirt2** | 3.86942 | 2.99469 | 2.91853 | 0.41617 | 3.54310 | 5.68430 | 3.81751 | 3.13629 | 2.49535 | 3.30550 | 1.54595 | 2.55742 | 3.21917 | 0.29190 | 0.35270 | 3.41790 |
| **Sirt3** | 2.63463 | 0.00339 | 0.15178 | 1.69054 | 3.73614 | 3.30581 | 0.00517 | 0.00000 | 0.00000 | 0.02415 | 0.01613 | 0.04031 | 0.00402 | 0.02905 | 0.02617 | 0.00301 |
| **Sirt4** | 0.00971 | 0.00882 | 0.10555 | 0.41568 | 0.00906 | 0.00583 | 0.00834 | 0.09843 | 0.02074 | 0.92215 | 0.00924 | 0.05958 | 0.00798 | 0.00886 | 0.15354 | 0.00726 |
| **Sirt5** | 0.01444 | 0.03053 | 0.63931 | 0.61703 | 0.02446 | 0.00538 | 0.00694 | 2.14061 | 2.41366 | 1.69585 | 1.14323 | 0.03331 | 0.12612 | 0.00582 | 0.00000 | 0.01372 |
| **Sirt6** | 1.59575 | 0.04563 | 2.38747 | 1.33957 | 1.60959 | 0.12332 | 0.03232 | 8.10681 | 0.82399 | 10.22272 | 3.01921 | 0.37724 | 4.68567 | 0.00305 | 0.00000 | 0.22938 |
| **Sirt7** | 0.77053 | 0.40644 | 2.02110 | 7.08534 | 0.09978 | 3.78197 | 0.01390 | 7.19810 | 11.19250 | 9.14191 | 4.65646 | 0.81267 | 1.71666 | 0.10484 | 0.00000 | 2.75245 |
| **Siva1** | 5.63055 | 2.12117 | 3.79078 | 23.49830 | 5.95198 | 6.40130 | 2.34961 | 2.66992 | 16.32872 | 14.55272 | 11.08145 | 0.79475 | 10.10297 | 0.00000 | 0.17024 | 8.37236 |
| **Six1** | 3.20762 | 0.00563 | 0.61127 | 0.56164 | 0.00536 | 0.00222 | 0.01758 | 0.44743 | 0.00009 | 0.00674 | 0.00716 | 0.01907 | 0.00306 | 0.00627 | 0.39249 | 0.00367 |
| **Six2** | 0.03110 | 0.00807 | 0.02377 | 0.25555 | 0.00838 | 0.00776 | 0.00762 | 0.01794 | 0.41106 | 0.01842 | 0.02175 | 0.00727 | 0.02244 | 0.02898 | 0.00788 | 0.00743 |
| **Six4** | 0.24198 | 0.00685 | 0.00000 | 0.85736 | 0.00789 | 0.00829 | 0.00682 | 0.01799 | 0.00000 | 0.00000 | 0.00000 | 1.44067 | 0.36652 | 0.00605 | 0.37816 | 0.00691 |
| **Six5** | 0.01248 | 0.00757 | 0.67431 | 0.49757 | 0.00726 | 0.00570 | 0.00699 | 0.00681 | 0.40289 | 0.55864 | 0.07756 | 0.00356 | 0.01496 | 0.16479 | 0.05943 | 0.47452 |
| **Ska1** | 10.59411 | 2.04683 | 4.22035 | 16.11649 | 5.60235 | 3.04025 | 3.48547 | 9.21355 | 18.83785 | 13.84955 | 4.48524 | 2.25654 | 6.99558 | 1.32869 | 4.84013 | 3.73852 |
| **Ska2** | 2.52875 | 3.66515 | 2.74585 | 6.54160 | 2.91196 | 4.19696 | 0.13465 | 1.72164 | 11.22930 | 1.52712 | 1.67654 | 0.14317 | 0.82951 | 7.46631 | 3.44859 | 2.08409 |
| **Ska3** | 4.83877 | 2.17176 | 2.78460 | 10.35318 | 4.03995 | 0.02941 | 4.98140 | 3.59784 | 11.64717 | 7.07045 | 4.21874 | 5.80132 | 2.44517 | 2.00159 | 8.87084 | 0.36845 |
| **Skap1** | 0.01347 | 0.00766 | 0.01123 | 0.92035 | 0.00932 | 0.00638 | 0.00725 | 0.01066 | 0.26851 | 0.40011 | 0.00945 | 0.03204 | 0.01208 | 0.03412 | 0.36955 | 0.00729 |
| **Skap2** | 3.15610 | 1.49531 | 0.00000 | 10.32915 | 0.84091 | 0.08127 | 3.22629 | 1.81773 | 9.72195 | 2.55009 | 0.43677 | 0.00000 | 1.72019 | 0.00000 | 6.43031 | 0.08039 |
| **Ski** | 7.81182 | 0.69696 | 2.76626 | 9.73998 | 1.17011 | 0.18659 | 4.97023 | 5.92622 | 7.08607 | 11.09646 | 6.01731 | 10.44980 | 4.61550 | 0.22019 | 0.23373 | 4.67885 |
| **Skida1** | 0.68956 | 0.01812 | 0.43679 | 0.13453 | 0.00583 | 0.00580 | 0.09851 | 0.41146 | 0.00677 | 0.54713 | 0.00694 | 0.01600 | 0.00396 | 0.00614 | 0.18680 | 0.00631 |
| **Skil** | 0.86586 | 0.00648 | 0.00000 | 1.00045 | 0.15039 | 0.08391 | 0.00667 | 0.04218 | 0.08747 | 0.22327 | 0.06843 | 0.02087 | 0.35872 | 0.00337 | 1.00539 | 0.00656 |
| **Skint2** | 0.01648 | 0.00777 | 0.21989 | 0.32774 | 0.00883 | 0.00566 | 0.00735 | 0.01273 | 0.25452 | 0.01201 | 0.01155 | 0.00711 | 0.01029 | 0.00827 | 0.03910 | 0.00718 |
| **Skint6** | 0.01374 | 0.00786 | 0.01190 | 0.16073 | 0.00935 | 0.00664 | 0.00752 | 0.01340 | 0.28819 | 0.01147 | 0.01002 | 0.00741 | 0.01284 | 0.01876 | 0.03832 | 0.00737 |
| **Skiv2l** | 0.19041 | 2.19019 | 3.79503 | 1.96840 | 3.20619 | 3.03977 | 6.70086 | 2.33714 | 1.88912 | 1.43296 | 2.77443 | 0.07700 | 4.37166 | 0.00000 | 0.00000 | 0.66074 |
| **Skiv2l2** | 9.88126 | 8.79556 | 7.08308 | 7.65765 | 6.91081 | 4.06284 | 3.42736 | 16.26049 | 13.80057 | 25.61089 | 17.18304 | 14.23376 | 3.99071 | 33.86312 | 23.23805 | 6.05953 |
| **Skp1a** | 15.85875 | 11.19249 | 16.27415 | 8.82721 | 10.98242 | 10.26773 | 4.35858 | 3.60198 | 7.94160 | 7.30467 | 9.93564 | 3.27332 | 12.14759 | 9.57311 | 8.68612 | 4.00869 |
| **Skp2** | 2.51669 | 4.29245 | 1.46797 | 6.94845 | 2.50925 | 2.49020 | 0.43673 | 6.57376 | 11.35156 | 8.35014 | 6.01774 | 14.26451 | 1.97307 | 0.44452 | 5.99961 | 0.78286 |
| **Sla2** | 0.01379 | 0.00788 | 0.01199 | 0.09469 | 0.00933 | 0.00659 | 0.00747 | 0.01298 | 0.01076 | 0.01008 | 0.01344 | 0.01786 | 0.01269 | 0.00843 | 0.13118 | 0.00737 |
| **Slain1** | 0.92327 | 0.00735 | 0.00000 | 30.32293 | 0.01647 | 0.00063 | 0.00398 | 0.00000 | 14.44281 | 3.28501 | 1.78371 | 0.80972 | 1.80859 | 4.40272 | 10.10193 | 0.00000 |
| **Slain2** | 6.05858 | 4.28697 | 1.77690 | 18.58977 | 4.19796 | 1.97090 | 3.16448 | 5.72387 | 8.01324 | 2.30989 | 3.92670 | 2.12725 | 2.20090 | 5.42045 | 11.51377 | 1.35052 |
| **Slamf7** | 0.01035 | 0.00794 | 0.00660 | 0.12875 | 0.00948 | 0.00676 | 0.00731 | 0.03083 | 1.46604 | 0.34706 | 0.00000 | 0.00000 | 0.01187 | 1.64293 | 0.11007 | 0.00727 |
| **Slamf8** | 0.04554 | 0.01857 | 0.04555 | 0.30162 | 0.04100 | 0.00623 | 0.00739 | 0.00856 | 0.40244 | 0.05215 | 0.09876 | 0.00612 | 0.02003 | 0.08694 | 0.04553 | 0.00742 |
| **Slamf9** | 0.08350 | 0.00491 | 0.37260 | 0.34901 | 2.56515 | 0.00600 | 3.02374 | 0.02055 | 0.36569 | 0.00000 | 0.08928 | 0.03232 | 0.32188 | 0.01040 | 0.00000 | 0.01439 |
| **Slbp** | 11.57152 | 7.25749 | 8.04204 | 25.16237 | 8.23152 | 8.35560 | 4.53759 | 22.88130 | 19.06899 | 22.34316 | 13.36462 | 13.75058 | 20.48151 | 16.32710 | 16.91191 | 11.54817 |
| **Slc10a1** | 0.02659 | 0.00771 | 0.03028 | 0.12842 | 0.03089 | 0.01258 | 0.02836 | 0.02287 | 0.03852 | 0.14727 | 0.37814 | 0.00433 | 0.06537 | 0.18096 | 0.20891 | 0.00749 |
| **Slc10a2** | 0.02933 | 0.00787 | 0.01155 | 0.01036 | 0.00946 | 0.03080 | 0.00743 | 0.01154 | 0.08291 | 0.00848 | 0.02681 | 0.05916 | 0.01853 | 0.08118 | 0.03715 | 0.01469 |
| **Slc10a3** | 0.00000 | 3.33396 | 1.45374 | 5.19439 | 0.00960 | 2.33157 | 0.02785 | 0.00000 | 5.20736 | 2.96971 | 0.98570 | 0.14886 | 0.00000 | 1.31025 | 5.89543 | 2.23708 |
| **Slc10a4** | 0.01346 | 0.00770 | 0.01022 | 0.04189 | 0.00943 | 0.00671 | 0.00734 | 0.01103 | 0.01041 | 0.00832 | 0.01072 | 0.00629 | 0.01312 | 0.34055 | 0.04853 | 0.00745 |
| **Slc10a6** | 0.00000 | 0.00732 | 1.28568 | 5.17220 | 0.00703 | 0.00447 | 0.00626 | 1.33160 | 6.71269 | 0.00000 | 0.00000 | 2.45920 | 1.87596 | 0.47854 | 0.92235 | 0.01227 |
| **Slc10a7** | 4.49808 | 0.02482 | 1.97648 | 0.33542 | 0.08096 | 0.01411 | 1.44443 | 1.52054 | 1.19921 | 2.02101 | 0.22235 | 0.00000 | 1.21838 | 5.80094 | 2.24284 | 0.00000 |
| **Slc11a1** | 0.13137 | 0.03139 | 0.25495 | 1.23054 | 0.01273 | 0.02300 | 0.02813 | 0.02163 | 1.58742 | 0.04551 | 0.23689 | 0.13533 | 0.33562 | 0.41968 | 0.14453 | 0.05124 |
| **Slc11a2** | 3.04988 | 2.32576 | 9.54088 | 2.38188 | 3.13139 | 0.01594 | 5.06871 | 14.48894 | 6.67124 | 14.33642 | 9.22173 | 2.75521 | 5.04874 | 5.52195 | 3.44716 | 5.09590 |
| **Slc12a2** | 0.92309 | 0.13214 | 2.37972 | 18.23627 | 3.67546 | 0.03803 | 3.13330 | 9.31638 | 13.51057 | 24.97139 | 12.42695 | 19.41025 | 1.60828 | 37.23053 | 30.53241 | 0.03201 |
| **Slc12a4** | 1.66604 | 0.00359 | 1.48189 | 0.00585 | 0.00962 | 0.91210 | 0.00000 | 14.00205 | 0.27890 | 0.11179 | 3.54927 | 0.23098 | 6.65673 | 0.00000 | 0.24775 | 0.01636 |
| **Slc12a5** | 0.01078 | 0.00856 | 0.05901 | 0.11883 | 0.00929 | 0.00853 | 0.00790 | 0.06907 | 0.00965 | 0.08648 | 0.06394 | 0.03012 | 0.00892 | 0.00767 | 0.09378 | 0.00749 |
| **Slc12a6** | 2.79028 | 0.00324 | 0.59257 | 1.21481 | 0.00146 | 0.00435 | 1.24131 | 1.08630 | 1.03224 | 1.49428 | 1.40316 | 1.11446 | 0.00004 | 1.77758 | 1.34500 | 0.13525 |
| **Slc12a7** | 7.91604 | 1.21481 | 10.00232 | 0.00000 | 3.40877 | 0.88324 | 6.59876 | 21.05624 | 2.12090 | 4.68394 | 9.14606 | 3.27997 | 6.06453 | 1.49650 | 4.92560 | 2.17866 |
| **Slc12a8** | 0.07564 | 0.02871 | 0.03804 | 0.65518 | 0.01907 | 0.05435 | 0.01762 | 2.53983 | 0.26751 | 0.24380 | 0.27514 | 0.03268 | 0.00979 | 0.40049 | 0.75206 | 0.01479 |
| **Slc12a9** | 0.00794 | 0.00677 | 2.31484 | 0.61556 | 0.00728 | 0.00539 | 0.85556 | 2.21039 | 0.32479 | 0.17746 | 0.46265 | 0.04908 | 1.72383 | 0.07919 | 0.01942 | 0.00536 |

| | | | | | | | | | | | | | | | | |
|---|---|---|---|---|---|---|---|---|---|---|---|---|---|---|---|---|
| **Slc13a2** | 0.53235 | 2.75891 | 2.09526 | 0.76397 | 0.01033 | 4.36024 | 0.56997 | 9.66596 | 1.03586 | 6.02764 | 6.15736 | 1.02820 | 3.10722 | 0.69180 | 9.39809 | 1.33917 |
| **Slc13a2os** | 0.01078 | 0.00759 | 0.08465 | 0.37929 | 0.00913 | 0.00881 | 0.19391 | 0.11654 | 0.31426 | 0.03794 | 0.88129 | 0.04272 | 0.01065 | 0.13990 | 0.24868 | 0.00711 |
| **Slc13a3** | 0.01355 | 0.00780 | 0.01124 | 0.20014 | 0.00915 | 0.00651 | 0.00728 | 0.01012 | 0.39804 | 0.29078 | 0.02148 | 0.00484 | 0.01247 | 0.00885 | 0.57827 | 0.00775 |
| **Slc13a5** | 2.53323 | 0.00160 | 0.18492 | 0.00665 | 2.63667 | 0.01481 | 3.41423 | 0.00244 | 0.01014 | 0.00000 | 0.01225 | 0.02168 | 0.00545 | 0.00925 | 0.20811 | 8.00681 |
| **Slc14a1** | 0.01300 | 0.00790 | 0.01210 | 0.00682 | 0.00890 | 0.00666 | 0.00735 | 0.01062 | 0.01160 | 1.28407 | 0.01107 | 0.00500 | 0.01126 | 0.01199 | 0.00844 | 0.01477 |
| **Slc15a1** | 0.05878 | 0.01813 | 0.08375 | 0.24785 | 0.00830 | 0.01971 | 0.04761 | 0.01804 | 0.11453 | 0.26155 | 0.21950 | 0.05476 | 0.11495 | 0.12533 | 0.08774 | 0.00700 |
| **Slc15a2** | 0.05363 | 0.00839 | 0.12725 | 0.29570 | 0.05335 | 3.85284 | 0.00671 | 0.02713 | 0.00708 | 0.19492 | 0.00841 | 0.01537 | 1.28780 | 0.10298 | 0.19664 | 1.83357 |
| **Slc15a4** | 2.08069 | 2.91605 | 3.71307 | 2.64273 | 0.00533 | 0.00330 | 0.00225 | 5.95760 | 2.26110 | 1.05294 | 0.99456 | 0.44179 | 3.29142 | 1.42619 | 0.91167 | 2.19110 |
| **Slc16a1** | 5.84428 | 5.25798 | 1.64713 | 1.31515 | 0.02571 | 5.76763 | 4.44609 | 1.19767 | 1.36549 | 1.90815 | 0.42502 | 0.13831 | 5.31184 | 6.52304 | 5.76469 | 2.99915 |
| **Slc16a10** | 2.90869 | 4.70220 | 0.00060 | 7.19521 | 3.47755 | 0.00001 | 1.67241 | 0.00000 | 11.65514 | 2.14845 | 1.19338 | 5.14112 | 2.07045 | 0.32965 | 3.21880 | 0.00599 |
| **Slc16a11** | 0.01361 | 0.00809 | 0.01214 | 0.04889 | 0.00920 | 0.00689 | 0.00759 | 0.01358 | 0.00924 | 0.21942 | 0.01144 | 0.00852 | 0.01284 | 0.00935 | 0.02688 | 0.00741 |
| **Slc16a13** | 1.44611 | 0.00536 | 4.10234 | 0.43367 | 0.00000 | 2.63868 | 0.00514 | 0.36014 | 0.11363 | 0.33912 | 0.11107 | 0.01787 | 0.65452 | 0.01380 | 0.41136 | 0.01819 |
| **Slc16a14** | 0.01371 | 0.00794 | 0.01132 | 0.00588 | 0.00934 | 0.00670 | 0.00744 | 0.01236 | 0.12961 | 0.01109 | 0.01110 | 0.02787 | 0.01298 | 0.09447 | 0.01417 | 0.00745 |
| **Slc16a3** | 0.01352 | 0.00807 | 0.00941 | 0.54316 | 0.00909 | 0.00682 | 0.00730 | 0.00168 | 0.34080 | 0.86150 | 0.31262 | 0.06451 | 0.01224 | 0.00631 | 0.01882 | 0.00708 |
| **Slc16a4** | 0.01345 | 0.00800 | 0.01010 | 0.02558 | 0.00930 | 0.00675 | 0.00745 | 0.01273 | 0.02428 | 0.01102 | 0.01057 | 0.02857 | 0.02690 | 0.23884 | 0.02411 | 0.00741 |
| **Slc16a5** | 1.00328 | 0.02388 | 1.56932 | 0.00000 | 0.00317 | 0.00602 | 2.48685 | 2.78971 | 0.48854 | 1.98537 | 1.90958 | 0.02862 | 1.91995 | 1.29289 | 2.45348 | 0.00440 |
| **Slc16a6** | 0.73751 | 0.00632 | 1.47729 | 0.48693 | 0.02986 | 0.00625 | 0.88244 | 2.89008 | 1.86372 | 6.15410 | 2.86491 | 0.15792 | 1.78037 | 1.10089 | 8.31932 | 0.00463 |
| **Slc17a5** | 0.03737 | 0.01865 | 0.38096 | 0.90616 | 0.02554 | 1.78884 | 0.07291 | 1.71162 | 0.46861 | 2.27490 | 2.11771 | 0.15222 | 0.12327 | 0.81988 | 2.20702 | 0.68653 |
| **Slc17a7** | 0.01716 | 0.00824 | 0.07817 | 0.18540 | 0.01037 | 0.00866 | 0.00743 | 0.01322 | 0.04565 | 0.01893 | 0.01164 | 0.00793 | 0.01134 | 0.01155 | 0.00603 | 0.01465 |
| **Slc17a8** | 0.08348 | 0.06189 | 0.28716 | 0.03658 | 0.01959 | 0.04296 | 0.06939 | 0.03490 | 0.42197 | 0.06564 | 0.40856 | 0.01336 | 0.19587 | 0.29933 | 0.12115 | 0.02166 |
| **Slc17a9** | 0.05014 | 0.00779 | 0.00960 | 0.17355 | 0.00801 | 0.00669 | 0.00728 | 0.01174 | 1.11237 | 0.34703 | 0.00620 | 0.00458 | 0.01239 | 0.01205 | 0.00764 | 0.03033 |
| **Slc18a2** | 0.00674 | 0.00696 | 0.03927 | 1.23742 | 0.00923 | 0.00517 | 0.00691 | 0.00634 | 0.43946 | 0.50670 | 0.14409 | 3.05034 | 0.01032 | 3.60929 | 1.48210 | 0.00686 |
| **Slc18b1** | 1.18148 | 0.88796 | 0.43046 | 1.21536 | 0.02164 | 0.00446 | 0.04537 | 0.05440 | 1.56549 | 1.35505 | 0.06985 | 0.09487 | 0.44683 | 0.89037 | 0.79974 | 0.00509 |
| **Slc19a1** | 0.19462 | 0.00671 | 2.05150 | 0.64359 | 0.01340 | 0.31817 | 0.02569 | 1.38768 | 0.29890 | 0.26254 | 0.71269 | 0.14400 | 1.52214 | 0.00255 | 0.26603 | 0.02805 |
| **Slc19a2** | 0.00000 | 0.19134 | 0.28073 | 0.46107 | 0.00858 | 0.00604 | 1.01702 | 0.80692 | 0.00460 | 0.00000 | 0.12675 | 0.01204 | 0.18165 | 1.45040 | 0.14895 | 0.00681 |
| **Slc19a3** | 0.00000 | 0.00288 | 0.00000 | 6.53175 | 0.00832 | 0.00219 | 0.00462 | 0.18384 | 7.04833 | 0.73980 | 0.46741 | 15.74693 | 0.00473 | 12.42622 | 15.25484 | 0.00420 |
| **Slc1a1** | 9.04200 | 0.88974 | 13.30167 | 0.62257 | 4.42761 | 5.30673 | 9.33808 | 16.55149 | 7.10229 | 9.79872 | 10.52492 | 3.21248 | 13.74210 | 7.35727 | 3.84504 | 2.87729 |
| **Slc1a3** | 3.46416 | 0.01378 | 1.66242 | 0.37217 | 0.00000 | 0.01311 | 0.00347 | 0.00000 | 0.03907 | 1.73138 | 0.72962 | 0.04686 | 0.52346 | 2.94790 | 1.97533 | 2.01413 |
| **Slc1a4** | 1.66512 | 2.69399 | 0.00000 | 1.02047 | 1.60718 | 0.00457 | 0.00486 | 0.00000 | 6.73076 | 0.26023 | 0.08982 | 0.00000 | 0.02959 | 0.01206 | 0.07035 | 0.00431 |
| **Slc1a5** | 4.03630 | 0.00000 | 2.21853 | 11.56104 | 0.00376 | 0.05535 | 2.76011 | 0.72236 | 15.71357 | 2.02049 | 4.35375 | 0.09617 | 0.60973 | 0.77181 | 1.65670 | 3.17859 |
| **Slc1a7** | 0.13627 | 0.02894 | 0.12375 | 0.60903 | 0.02697 | 0.03801 | 0.01768 | 0.03258 | 0.70266 | 0.03457 | 0.27561 | 0.07554 | 0.03625 | 0.31872 | 0.02957 | 0.04339 |
| **Slc20a1** | 11.15525 | 12.24829 | 24.28589 | 3.75674 | 1.42950 | 10.69822 | 13.44670 | 19.25609 | 2.00750 | 8.65975 | 8.48100 | 3.09226 | 28.39835 | 8.37517 | 13.11196 | 6.82677 |
| **Slc20a2** | 1.30160 | 3.39694 | 0.40544 | 0.50942 | 0.00488 | 0.00428 | 0.88322 | 0.23009 | 1.61615 | 1.44062 | 0.01813 | 0.00883 | 2.95159 | 0.00867 | 0.29249 | 0.00275 |
| **Slc22a13** | 0.02545 | 0.01767 | 0.39540 | 0.00334 | 0.01628 | 0.00582 | 0.00732 | 5.62833 | 0.01247 | 0.97715 | 0.00000 | 0.43017 | 0.42291 | 0.25988 | 1.05066 | 0.00714 |
| **Slc22a13b.ps** | 0.59939 | 0.00586 | 0.18429 | 0.00000 | 0.01433 | 0.00537 | 0.00651 | 7.94437 | 0.00624 | 1.63118 | 1.15678 | 0.77708 | 0.41875 | 0.09513 | 1.77657 | 0.00641 |
| **Slc22a15** | 0.09546 | 0.00771 | 0.04978 | 0.26742 | 0.07312 | 0.01967 | 0.03984 | 0.03404 | 0.36517 | 0.04624 | 0.70582 | 0.04361 | 0.32687 | 0.34257 | 0.11252 | 0.02968 |
| **Slc22a16** | 0.20138 | 0.00754 | 0.03836 | 0.07030 | 0.00756 | 0.00579 | 0.00719 | 0.01801 | 0.01156 | 0.01358 | 0.01322 | 0.06721 | 0.01170 | 0.01177 | 0.00739 | 0.00708 |
| **Slc22a17** | 0.83099 | 0.00581 | 0.00000 | 0.42747 | 0.00619 | 2.05386 | 1.73144 | 0.35688 | 1.45169 | 1.34679 | 0.00000 | 0.00997 | 0.00000 | 0.00986 | 0.00186 | 0.00529 |
| **Slc22a20** | 0.00409 | 0.00743 | 0.00000 | 2.73231 | 0.00926 | 0.00646 | 0.00657 | 0.65792 | 0.09433 | 2.15692 | 1.54492 | 0.03791 | 0.02697 | 0.71528 | 2.08985 | 0.00653 |
| **Slc22a21** | 0.00000 | 0.86884 | 0.01746 | 0.22947 | 0.00758 | 0.10247 | 0.00668 | 0.00836 | 1.46873 | 0.00520 | 0.00000 | 0.00439 | 0.47505 | 0.00510 | 0.15764 | 0.72806 |
| **Slc22a23** | 0.00000 | 0.00745 | 0.73877 | 3.36946 | 0.00472 | 7.84688 | 0.00534 | 0.00442 | 0.37002 | 0.00000 | 0.00518 | 0.01439 | 1.72089 | 0.02408 | 0.45218 | 5.50111 |
| **Slc22a28** | 0.01375 | 0.00823 | 0.01224 | 1.66690 | 0.00935 | 0.00646 | 0.00744 | 0.01341 | 0.10342 | 0.01058 | 0.01221 | 0.00822 | 0.01267 | 0.00552 | 0.09557 | 0.00710 |
| **Slc22a5** | 0.00000 | 0.52742 | 0.04262 | 0.00000 | 0.01668 | 5.31184 | 0.03616 | 0.26484 | 0.50939 | 0.33509 | 0.00517 | 0.00231 | 0.59995 | 0.00583 | 1.27631 | 0.87621 |
| **Slc22a7** | 0.93649 | 0.00573 | 0.00267 | 0.00962 | 1.39932 | 0.00747 | 0.00678 | 0.00714 | 0.01105 | 0.00622 | 0.01169 | 0.00581 | 0.01053 | 0.01776 | 0.13255 | 0.00628 |
| **Slc23a1** | 0.64992 | 0.05011 | 0.45307 | 4.08164 | 0.16037 | 0.06810 | 0.16830 | 0.00000 | 6.45646 | 0.82429 | 5.53510 | 0.30651 | 0.14395 | 2.38048 | 0.22062 | 0.10381 |
| **Slc23a2** | 4.60360 | 0.00000 | 3.31323 | 7.44204 | 3.35559 | 2.47226 | 6.49256 | 3.82341 | 8.05857 | 3.04646 | 3.29730 | 0.93604 | 5.05859 | 1.99937 | 5.94777 | 0.16959 |
| **Slc23a3** | 0.00979 | 0.00723 | 0.00398 | 0.54294 | 0.00931 | 0.00723 | 0.43047 | 0.02299 | 0.13637 | 0.04372 | 0.84803 | 0.00187 | 0.31019 | 0.00443 | 0.28734 | 0.00705 |
| **Slc24a3** | 0.01356 | 0.00801 | 0.01194 | 1.36269 | 0.00911 | 0.00655 | 0.00727 | 0.01089 | 0.05466 | 0.43647 | 0.01284 | 0.00640 | 0.01229 | 0.03273 | 0.00779 | 0.00754 |
| **Slc24a5** | 5.03237 | 3.44261 | 1.33888 | 0.10204 | 3.62208 | 0.44658 | 5.78601 | 1.40906 | 0.00445 | 0.51542 | 0.00000 | 0.00253 | 0.11144 | 2.47980 | 0.36732 | 3.13463 |
| **Slc25a1** | 4.16353 | 0.01209 | 0.14419 | 0.02601 | 3.23627 | 3.38225 | 1.33978 | 0.03300 | 0.23693 | 1.92483 | 0.00540 | 0.00590 | 0.01627 | 0.01467 | 0.01921 | 2.44714 |
| **Slc25a10** | 1.62512 | 0.04265 | 1.87508 | 0.68123 | 7.39477 | 1.26935 | 0.08359 | 5.90085 | 2.50889 | 0.87322 | 3.26791 | 0.12833 | 7.18255 | 0.00093 | 0.41193 | 5.31481 |
| **Slc25a11** | 13.62530 | 5.47750 | 8.22924 | 3.45481 | 5.43003 | 8.32687 | 5.82218 | 5.34206 | 4.07241 | 4.80986 | 7.33267 | 0.35614 | 12.55258 | 1.91789 | 4.64389 | 4.20980 |
| **Slc25a12** | 2.56318 | 1.00148 | 0.00000 | 1.04002 | 0.00680 | 1.30272 | 0.00257 | 4.63023 | 4.92444 | 13.64763 | 4.79176 | 4.01525 | 0.00000 | 1.91699 | 7.28230 | 0.41370 |
| **Slc25a13** | 12.78845 | 11.07130 | 6.25318 | 3.58521 | 15.55427 | 0.55057 | 5.88946 | 4.91441 | 4.60882 | 4.02308 | 2.25140 | 0.32295 | 5.17184 | 4.28171 | 6.99669 | 3.99084 |
| **Slc25a14** | 2.42037 | 0.00447 | 4.18382 | 3.17544 | 0.00000 | 0.01275 | 0.01256 | 0.91744 | 2.88385 | 2.07746 | 0.52335 | 0.17187 | 2.92822 | 4.56079 | 0.96737 | 1.86133 |
| **Slc25a15** | 2.94065 | 0.02271 | 3.72932 | 9.03370 | 2.63556 | 0.09246 | 2.63723 | 1.18387 | 2.97492 | 4.15768 | 2.07646 | 2.81306 | 1.42716 | 6.15309 | 5.58866 | 0.09666 |
| **Slc25a16** | 1.96972 | 1.94759 | 2.24848 | 4.04138 | 0.45873 | 1.67451 | 2.39010 | 4.71606 | 6.19189 | 0.31330 | 1.15727 | 8.33516 | 0.91645 | 4.10518 | 10.05748 | 0.17726 |
| **Slc25a17** | 9.49232 | 6.20590 | 4.83479 | 3.89081 | 2.90647 | 7.49841 | 2.32155 | 10.39616 | 4.83478 | 8.43593 | 5.18178 | 7.33118 | 5.86623 | 12.63097 | 6.92454 | 7.23427 |
| **Slc25a18** | 0.00834 | 0.00815 | 0.00531 | 0.20874 | 0.00943 | 0.00768 | 0.00788 | 0.26680 | 0.00545 | 1.17733 | 0.59949 | 0.01203 | 0.00582 | 0.05787 | 0.00000 | 0.00762 |
| **Slc25a19** | 0.40583 | 0.00375 | 3.13097 | 2.00537 | 0.01241 | 3.17086 | 3.55378 | 6.00182 | 1.85828 | 7.39144 | 8.09776 | 0.69121 | 0.70262 | 0.00000 | 1.44871 | 0.02405 |
| **Slc25a2** | 0.01374 | 0.00814 | 0.01210 | 0.01052 | 0.00932 | 0.00666 | 0.00752 | 0.01349 | 0.01240 | 0.01446 | 0.01308 | 0.00795 | 0.01276 | 0.03357 | 0.00919 | 0.00742 |
| **Slc25a20** | 6.52713 | 9.58364 | 6.53960 | 0.46933 | 8.74150 | 6.12084 | 0.59644 | 12.56277 | 2.50798 | 14.09398 | 5.92657 | 0.96337 | 5.51325 | 9.54043 | 9.31366 | 0.25397 |

| | | | | | | | | | | | | | | | | |
|---|---|---|---|---|---|---|---|---|---|---|---|---|---|---|---|---|
| **Slc25a21** | 0.00399 | 0.00743 | 1.99683 | 0.23690 | 0.00803 | 0.00735 | 0.68122 | 0.39552 | 0.43706 | 0.12148 | 0.04076 | 0.00000 | 0.35390 | 1.79497 | 0.00000 | 0.00640 |
| **Slc25a22** | 1.23600 | 0.00338 | 0.57445 | 1.04401 | 0.38639 | 1.46470 | 1.61923 | 6.71782 | 0.78107 | 3.00768 | 3.88275 | 4.33469 | 0.00000 | 0.00000 | 0.94675 | 3.23146 |
| **Slc25a23** | 0.00719 | 0.00726 | 0.01352 | 2.21308 | 0.00941 | 0.13666 | 0.00675 | 1.09406 | 1.35266 | 1.77292 | 1.26067 | 0.00912 | 0.03670 | 0.00000 | 0.24991 | 0.00710 |
| **Slc25a24** | 0.01889 | 0.00796 | 0.00398 | 0.00657 | 0.01006 | 0.40867 | 0.00741 | 0.01127 | 0.01146 | 0.92583 | 0.00893 | 0.09858 | 0.00870 | 0.00823 | 0.12529 | 0.00723 |
| **Slc25a25** | 0.03444 | 3.02671 | 0.05511 | 11.62684 | 1.10771 | 3.71922 | 2.71814 | 6.21773 | 17.82438 | 14.76635 | 7.02827 | 2.65153 | 0.68289 | 0.27869 | 11.99196 | 3.26627 |
| **Slc25a26** | 0.42397 | 0.05032 | 3.13530 | 0.60278 | 0.33095 | 0.20004 | 2.37820 | 3.24152 | 0.90710 | 2.15012 | 1.27409 | 0.02631 | 3.16731 | 0.16879 | 0.71144 | 1.79443 |
| **Slc25a27** | 0.01330 | 0.00802 | 0.01727 | 0.00456 | 0.00924 | 0.00727 | 0.00741 | 0.01164 | 0.69882 | 0.04817 | 0.01981 | 0.05908 | 0.01226 | 0.03836 | 0.02218 | 0.00725 |
| **Slc25a28** | 3.68953 | 1.86039 | 2.02540 | 13.08907 | 3.33835 | 2.72426 | 2.33445 | 6.25261 | 18.27925 | 9.48569 | 5.79844 | 1.42764 | 3.84507 | 0.21381 | 7.15307 | 0.56165 |
| **Slc25a3** | 28.48048 | 14.65892 | 19.70895 | 17.58003 | 22.41449 | 12.92860 | 10.25288 | 21.19512 | 21.16133 | 40.99678 | 35.23487 | 16.65774 | 37.03753 | 17.73246 | 9.82592 | 20.11475 |
| **Slc25a30** | 0.00896 | 0.00755 | 0.00751 | 0.00552 | 0.01025 | 0.00663 | 0.00720 | 0.25357 | 0.02558 | 0.01850 | 0.00522 | 0.02206 | 0.00876 | 0.47171 | 0.80555 | 0.00753 |
| **Slc25a31** | 0.01423 | 0.00801 | 0.09735 | 1.47716 | 0.00889 | 0.00549 | 0.05796 | 0.00632 | 2.48489 | 0.34910 | 0.40962 | 0.11544 | 0.00833 | 0.04389 | 0.83359 | 0.00685 |
| **Slc25a32** | 1.35706 | 0.00411 | 0.26049 | 3.33097 | 0.00497 | 0.00292 | 0.09873 | 0.32637 | 4.95087 | 0.48576 | 0.38495 | 6.27768 | 0.23911 | 0.81312 | 3.78403 | 0.73343 |
| **Slc25a33** | 1.64013 | 0.00408 | 0.39496 | 2.59931 | 0.00507 | 0.00294 | 0.67947 | 0.00000 | 2.16023 | 1.49232 | 0.74157 | 5.23187 | 0.09396 | 0.63469 | 3.18158 | 0.01889 |
| **Slc25a35** | 0.99204 | 2.15306 | 0.97170 | 0.07659 | 0.01412 | 0.01281 | 0.00565 | 0.11977 | 0.17208 | 0.00000 | 0.13426 | 0.04298 | 0.00292 | 0.01034 | 0.00000 | 0.00505 |
| **Slc25a36** | 0.24813 | 0.01344 | 0.76026 | 7.77735 | 3.04679 | 0.75014 | 0.11830 | 2.60259 | 13.60117 | 3.72265 | 5.32915 | 6.14884 | 0.17974 | 9.82848 | 9.45750 | 0.00286 |
| **Slc25a37** | 0.00587 | 0.00813 | 0.01277 | 0.00519 | 0.00781 | 0.00669 | 0.00772 | 0.02164 | 0.01095 | 3.86015 | 0.01009 | 0.00000 | 0.01462 | 0.01325 | 0.00565 | 1.53806 |
| **Slc25a38** | 2.31739 | 5.05282 | 2.59242 | 12.05114 | 2.61918 | 1.01302 | 0.02139 | 4.11617 | 11.11308 | 4.57202 | 2.90680 | 0.38014 | 4.38790 | 0.04855 | 2.39061 | 0.05466 |
| **Slc25a39** | 23.86676 | 7.61122 | 12.04953 | 6.81498 | 16.01375 | 8.66469 | 5.77891 | 7.18495 | 3.27252 | 7.49503 | 7.90900 | 1.70517 | 16.01318 | 0.00000 | 0.21270 | 11.28887 |
| **Slc25a4** | 20.47483 | 10.66222 | 18.16130 | 4.40602 | 16.04445 | 18.16460 | 11.58572 | 19.54522 | 0.24468 | 24.40659 | 17.09891 | 8.64756 | 30.29835 | 4.84840 | 1.21676 | 12.71395 |
| **Slc25a40** | 0.00782 | 0.12962 | 0.00000 | 1.33331 | 1.75119 | 0.00653 | 0.02547 | 0.09652 | 0.00000 | 0.00000 | 0.35854 | 0.27465 | 1.73537 | 1.88780 | 0.65107 | 0.02786 |
| **Slc25a41** | 1.71540 | 0.00479 | 0.96927 | 0.46314 | 0.01450 | 0.00376 | 0.00289 | 2.96426 | 0.26555 | 5.02645 | 5.16467 | 0.52617 | 1.74921 | 0.00000 | 3.64010 | 0.01092 |
| **Slc25a42** | 0.05468 | 0.02922 | 0.16758 | 0.64533 | 0.03178 | 0.02159 | 0.02701 | 0.14072 | 0.48581 | 0.37636 | 1.01111 | 0.29534 | 0.18814 | 0.62612 | 0.58907 | 0.05753 |
| **Slc25a43** | 0.01208 | 0.00783 | 0.51339 | 0.63548 | 0.00821 | 0.00946 | 0.00735 | 0.01282 | 0.00006 | 0.35928 | 0.01112 | 0.00705 | 0.00952 | 0.00901 | 0.17028 | 0.00710 |
| **Slc25a44** | 3.08635 | 7.59299 | 11.15778 | 16.24871 | 4.23236 | 5.96170 | 9.11389 | 10.49861 | 24.24931 | 19.33288 | 12.12297 | 11.78098 | 12.08267 | 15.70392 | 18.61854 | 1.93975 |
| **Slc25a45** | 0.05349 | 2.23887 | 0.00926 | 0.00000 | 0.00744 | 0.00811 | 0.00731 | 0.99126 | 0.00985 | 0.14651 | 0.33268 | 1.37347 | 0.00375 | 0.00554 | 0.00000 | 0.00600 |
| **Slc25a46** | 2.17141 | 0.00004 | 0.95897 | 2.68093 | 5.35122 | 3.05445 | 0.22258 | 0.32493 | 3.64356 | 1.46988 | 2.91462 | 3.67986 | 0.84202 | 6.94716 | 1.44799 | 1.42329 |
| **Slc25a47** | 0.01507 | 0.00802 | 0.02888 | 0.35016 | 0.00923 | 0.00603 | 0.00746 | 0.01353 | 0.84837 | 0.01114 | 0.00864 | 0.00657 | 0.01247 | 0.01111 | 0.00659 | 0.00728 |
| **Slc25a48** | 0.02185 | 0.00809 | 0.01129 | 0.00895 | 0.00872 | 0.00688 | 0.00758 | 0.01484 | 0.03445 | 0.04516 | 0.01208 | 0.02013 | 0.01311 | 0.01188 | 0.01134 | 0.00738 |
| **Slc25a5** | 51.35551 | 33.33523 | 51.22561 | 7.55333 | 36.41250 | 20.50083 | 29.55287 | 40.79161 | 11.32962 | 28.18457 | 30.32690 | 12.60873 | 44.01130 | 13.72761 | 19.22692 | 25.36837 |
| **Slc25a51** | 5.18487 | 2.30672 | 1.38378 | 8.08274 | 4.78738 | 0.46286 | 2.19136 | 1.22338 | 8.15173 | 9.50516 | 3.59831 | 4.99537 | 1.37509 | 8.92981 | 4.15104 | 0.04763 |
| **Slc25a53** | 0.00482 | 0.81346 | 0.29865 | 0.03545 | 0.00734 | 0.00905 | 0.00682 | 0.00329 | 0.00993 | 0.21552 | 0.20733 | 0.00348 | 0.00900 | 0.29279 | 0.66545 | 0.00719 |
| **Slc26a1** | 0.01321 | 0.00816 | 0.01635 | 2.75165 | 0.00921 | 0.00595 | 0.00751 | 0.01303 | 1.92924 | 0.07130 | 0.00000 | 0.00438 | 0.01222 | 0.01212 | 0.01098 | 0.00663 |
| **Slc26a10** | 0.01076 | 0.00759 | 0.00964 | 0.01822 | 0.00964 | 0.00672 | 0.00720 | 0.14965 | 0.01246 | 0.01506 | 0.02893 | 0.00710 | 0.00766 | 0.01352 | 0.02398 | 0.00741 |
| **Slc26a11** | 0.00542 | 0.00617 | 1.48404 | 0.00000 | 0.00803 | 0.00627 | 1.51288 | 3.25629 | 1.16844 | 2.80848 | 3.67175 | 0.32073 | 0.12919 | 0.76488 | 1.88754 | 0.01989 |
| **Slc26a2** | 1.08596 | 0.01824 | 1.14033 | 4.50789 | 0.01464 | 0.00360 | 0.00492 | 0.33993 | 1.38214 | 0.58921 | 0.93681 | 1.88149 | 1.44714 | 0.00000 | 0.26671 | 0.00403 |
| **Slc26a4** | 0.01285 | 0.00748 | 0.00922 | 0.06746 | 0.00903 | 0.00614 | 0.00724 | 0.00894 | 2.77792 | 0.61109 | 0.00000 | 0.01506 | 0.01229 | 0.00418 | 1.67235 | 0.00759 |
| **Slc26a5** | 0.00524 | 0.16290 | 0.13964 | 0.83964 | 0.00924 | 0.00957 | 0.00799 | 0.96611 | 0.00000 | 0.00000 | 0.57573 | 0.04451 | 0.33776 | 0.00455 | 0.00000 | 0.00713 |
| **Slc26a6** | 1.68720 | 0.00444 | 0.47056 | 1.38783 | 0.01344 | 0.00778 | 2.63788 | 0.27084 | 0.09649 | 0.29099 | 0.00916 | 0.00162 | 0.51407 | 0.00273 | 0.69923 | 0.00504 |
| **Slc26a8** | 0.18921 | 0.00720 | 0.02430 | 0.15144 | 0.00726 | 0.00768 | 0.00689 | 0.03902 | 0.81820 | 1.03768 | 0.00547 | 0.00293 | 0.12486 | 0.04200 | 0.32553 | 0.00665 |
| **Slc26a9** | 0.01376 | 0.00773 | 0.01174 | 0.34198 | 0.00936 | 0.00657 | 0.00747 | 0.01270 | 0.30821 | 0.00696 | 0.00948 | 0.00704 | 0.01284 | 0.02015 | 0.14803 | 0.00732 |
| **Slc27a1** | 0.09483 | 0.01816 | 1.19740 | 0.31364 | 0.04424 | 0.02346 | 0.05726 | 1.06822 | 0.78468 | 1.46805 | 1.74126 | 0.02640 | 1.10629 | 0.85625 | 0.38179 | 0.05812 |
| **Slc27a2** | 0.01121 | 0.00767 | 0.00498 | 0.05416 | 0.00967 | 0.00662 | 0.00702 | 0.17347 | 0.01142 | 0.07305 | 0.07886 | 0.00000 | 0.00718 | 1.97660 | 0.00000 | 0.00739 |
| **Slc27a3** | 1.23380 | 0.38607 | 0.00506 | 0.13591 | 0.00666 | 0.00582 | 0.00597 | 0.41278 | 0.00664 | 0.10940 | 0.30139 | 0.13420 | 1.30583 | 0.00944 | 0.00631 | 0.00512 |
| **Slc27a4** | 3.18431 | 0.00001 | 2.24848 | 4.40166 | 0.05087 | 4.92065 | 2.79037 | 5.60819 | 5.83421 | 1.01473 | 3.45042 | 0.35715 | 7.78934 | 0.75368 | 5.13362 | 4.42638 |
| **Slc27a5** | 0.00636 | 0.00795 | 0.71035 | 7.24897 | 0.00798 | 0.00468 | 0.00647 | 0.00000 | 5.36949 | 2.52826 | 1.57644 | 0.12941 | 0.61506 | 1.30516 | 3.08089 | 0.00529 |
| **Slc28a1** | 1.14860 | 0.00657 | 0.00360 | 0.62440 | 0.00564 | 0.00690 | 0.00646 | 0.01094 | 0.42078 | 0.04607 | 0.00969 | 0.00568 | 0.01205 | 0.00530 | 0.12747 | 0.00625 |
| **Slc28a2** | 0.01343 | 0.00781 | 0.01215 | 0.14970 | 0.00903 | 0.00646 | 0.00726 | 0.01028 | 0.00731 | 0.71105 | 0.01397 | 0.00673 | 0.01070 | 0.01866 | 0.00403 | 0.00708 |
| **Slc28a3** | 0.85150 | 2.68077 | 5.81199 | 0.23997 | 0.47724 | 0.49212 | 3.71924 | 0.41308 | 1.56500 | 0.89750 | 12.99405 | 0.45640 | 5.91364 | 5.84811 | 0.00000 | 0.42808 |
| **Slc29a1** | 2.41935 | 0.03667 | 2.16219 | 0.00000 | 0.02066 | 9.49454 | 0.21048 | 2.43588 | 0.00904 | 1.92371 | 0.56093 | 0.03356 | 1.85036 | 0.00300 | 0.52100 | 10.27612 |
| **Slc29a2** | 0.57107 | 0.00584 | 3.37583 | 10.47853 | 0.00654 | 0.02239 | 0.00246 | 3.32287 | 9.72384 | 2.97015 | 2.52179 | 0.28158 | 4.23468 | 0.65382 | 1.23639 | 6.78815 |
| **Slc29a3** | 0.45796 | 0.04734 | 2.75997 | 1.12306 | 0.00186 | 0.64565 | 0.00341 | 4.33269 | 0.74653 | 6.23014 | 4.29809 | 5.88586 | 1.78183 | 0.13745 | 1.39881 | 0.00342 |
| **Slc29a4** | 0.13712 | 0.00631 | 0.10523 | 0.00257 | 0.08980 | 0.00687 | 0.00685 | 0.70366 | 0.80055 | 0.01503 | 0.00000 | 3.17659 | 0.08480 | 0.01026 | 0.00000 | 0.00691 |
| **Slc2a1** | 10.68913 | 1.13704 | 2.81128 | 2.03841 | 2.33687 | 4.97916 | 0.46423 | 2.90309 | 0.09375 | 0.00000 | 0.00000 | 0.17600 | 5.42775 | 0.00298 | 0.00000 | 2.24621 |
| **Slc2a12** | 0.02665 | 0.00768 | 0.01610 | 0.59813 | 0.00857 | 0.00633 | 0.00752 | 0.01256 | 0.41547 | 0.01923 | 0.00946 | 0.00688 | 0.01293 | 0.01452 | 0.12078 | 0.00742 |
| **Slc2a3** | 5.85002 | 2.11244 | 7.30002 | 2.13128 | 0.05105 | 19.23979 | 2.97948 | 5.38541 | 3.38027 | 6.62587 | 21.88468 | 6.28882 | 21.93768 | 4.77679 | 3.60700 | 10.95462 |
| **Slc2a4** | 0.00000 | 0.00633 | 1.74963 | 0.02275 | 0.00709 | 3.41224 | 3.26337 | 0.16591 | 1.49337 | 0.00000 | 0.77949 | 0.09137 | 0.92590 | 0.10541 | 0.71501 | 1.31355 |
| **Slc2a4rg.ps** | 0.02178 | 0.00798 | 0.45101 | 0.01063 | 0.00938 | 0.00571 | 0.00721 | 0.01109 | 0.01232 | 0.01859 | 0.01241 | 0.00741 | 0.14146 | 0.03581 | 0.00046 | 0.02195 |
| **Slc2a7** | 0.53347 | 0.00734 | 0.09222 | 0.00546 | 0.00656 | 0.00839 | 0.00698 | 0.01350 | 0.14345 | 0.01829 | 0.01097 | 0.00729 | 0.01097 | 0.01278 | 0.00816 | 0.00665 |
| **Slc2a8** | 0.90235 | 2.53895 | 9.78040 | 0.30335 | 3.53743 | 5.34866 | 5.98657 | 11.51245 | 1.37339 | 1.90367 | 11.10435 | 0.13682 | 11.86185 | 0.08156 | 0.11689 | 0.40356 |
| **Slc2a9** | 0.01032 | 0.00786 | 0.01651 | 0.00505 | 0.00881 | 0.00705 | 0.00650 | 0.00849 | 0.03870 | 0.19625 | 0.01053 | 0.00520 | 1.30045 | 0.00590 | 0.62243 | 0.01485 |
| **Slc30a1** | 4.41323 | 0.00000 | 4.68783 | 12.36941 | 3.01749 | 4.95898 | 3.99018 | 2.07793 | 19.68249 | 1.17587 | 0.53221 | 0.00000 | 3.87030 | 0.11909 | 0.00000 | 6.38926 |
| **Slc30a2** | 6.84763 | 0.00000 | 4.45876 | 0.56992 | 0.00000 | 0.23988 | 3.55524 | 1.48847 | 4.97712 | 0.32003 | 0.00000 | 0.03604 | 4.40232 | 0.00000 | 0.95584 | 1.86218 |

| | | | | | | | | | | | | | | | | |
|---|---|---|---|---|---|---|---|---|---|---|---|---|---|---|---|---|
| **Slc30a3** | 1.22338 | 0.01936 | 0.00332 | 4.14863 | 0.00660 | 0.00693 | 0.00689 | 0.10367 | 0.54400 | 0.77123 | 0.10941 | 0.01713 | 0.00734 | 0.00919 | 0.00000 | 0.01217 |
| **Slc30a4** | 4.78717 | 8.84553 | 1.59980 | 0.00000 | 4.34283 | 2.37978 | 5.45523 | 5.58112 | 1.75459 | 0.00000 | 5.99891 | 8.22228 | 1.73363 | 7.93171 | 3.18440 | 0.60491 |
| **Slc30a5** | 2.69109 | 2.49323 | 3.81077 | 4.37863 | 2.58054 | 3.68520 | 3.27925 | 5.43715 | 7.43444 | 1.80096 | 1.68559 | 2.92431 | 2.66109 | 7.10069 | 9.03993 | 0.22346 |
| **Slc30a6** | 2.51493 | 2.63074 | 6.66958 | 2.98871 | 1.03137 | 0.17923 | 2.92648 | 3.28683 | 3.72615 | 3.16504 | 1.37851 | 1.01181 | 0.29401 | 3.67919 | 3.96962 | 1.88023 |
| **Slc30a7** | 3.06325 | 0.02247 | 2.68067 | 0.14159 | 0.76870 | 0.01758 | 3.48445 | 1.00017 | 0.49911 | 3.39354 | 2.36483 | 2.72697 | 0.85117 | 3.53250 | 0.77852 | 0.82661 |
| **Slc30a9** | 3.16300 | 4.19391 | 2.27273 | 0.00986 | 2.73170 | 2.45373 | 1.73877 | 0.11328 | 0.16156 | 0.04131 | 0.10530 | 4.32791 | 0.67175 | 0.00439 | 0.00000 | 3.97408 |
| **Slc31a1** | 3.61527 | 1.90881 | 7.09698 | 5.04624 | 0.00000 | 2.70422 | 5.18957 | 4.42002 | 3.08923 | 2.65265 | 2.06802 | 1.79394 | 5.22772 | 5.25649 | 7.77905 | 3.12645 |
| **Slc31a2** | 0.85350 | 0.00652 | 0.00000 | 0.87144 | 0.00689 | 0.02724 | 0.00671 | 0.00794 | 0.00000 | 0.08572 | 0.03571 | 0.02492 | 0.04406 | 0.01139 | 0.22152 | 0.01429 |
| **Slc33a1** | 0.65870 | 1.43960 | 2.13493 | 2.25339 | 0.01482 | 0.22526 | 0.00429 | 0.23538 | 2.76277 | 0.00000 | 0.35350 | 3.24288 | 2.13974 | 2.79959 | 2.24000 | 0.93299 |
| **Slc34a2** | 1.49845 | 8.78398 | 10.17186 | 25.87150 | 0.09360 | 0.18265 | 5.49896 | 16.21256 | 31.37365 | 11.22793 | 17.77938 | 15.00130 | 9.01934 | 23.39600 | 17.91477 | 4.28893 |
| **Slc35a1** | 1.80794 | 0.03578 | 0.99902 | 0.68937 | 4.31489 | 0.00385 | 0.18453 | 0.00464 | 0.02087 | 1.65175 | 0.01648 | 0.04785 | 0.90208 | 0.00673 | 0.87947 | 0.00487 |
| **Slc35a2** | 3.45598 | 0.08466 | 3.89309 | 1.57716 | 1.70389 | 1.74655 | 0.57845 | 2.22627 | 0.39948 | 3.92633 | 1.04526 | 0.06344 | 1.49162 | 0.03386 | 3.18986 | 0.41641 |
| **Slc35a3** | 0.85948 | 0.52048 | 0.63396 | 1.34599 | 0.02582 | 0.01089 | 1.96042 | 0.79151 | 0.00000 | 2.53761 | 1.89307 | 0.77391 | 0.00000 | 8.48767 | 5.79413 | 0.00535 |
| **Slc35a4** | 13.81203 | 4.94916 | 6.43206 | 4.77943 | 0.10502 | 6.94141 | 3.61306 | 8.90971 | 2.46711 | 6.31428 | 2.95012 | 0.59812 | 10.20719 | 0.21658 | 2.41354 | 4.00363 |
| **Slc35a5** | 0.33283 | 1.42493 | 0.00000 | 1.12100 | 0.70720 | 0.02910 | 0.00620 | 0.24049 | 1.74551 | 0.03212 | 0.00000 | 0.06000 | 0.02786 | 5.94008 | 0.00651 | 0.00608 |
| **Slc35b1** | 7.31552 | 8.79108 | 12.70079 | 3.23183 | 5.47734 | 10.85335 | 9.27010 | 9.54199 | 4.37711 | 8.05947 | 4.22217 | 2.62243 | 15.85113 | 3.91620 | 2.17796 | 8.05778 |
| **Slc35b2** | 4.63430 | 0.01034 | 5.20780 | 3.59608 | 3.47102 | 2.16844 | 1.14787 | 1.97802 | 1.11488 | 3.88817 | 1.33517 | 0.08932 | 2.81798 | 0.01266 | 2.93680 | 2.40562 |
| **Slc35b3** | 0.65813 | 0.29175 | 0.94585 | 0.00901 | 0.00543 | 2.22951 | 0.22738 | 0.34192 | 0.03737 | 0.51435 | 0.00517 | 0.06478 | 1.22924 | 0.01139 | 0.42456 | 0.00521 |
| **Slc35b4** | 2.47510 | 2.60623 | 1.25981 | 3.57269 | 5.34455 | 2.27534 | 0.00283 | 3.06291 | 0.46876 | 5.10355 | 0.54141 | 3.69359 | 1.73634 | 2.36486 | 4.67429 | 2.85393 |
| **Slc35c1** | 1.40855 | 0.00541 | 0.28472 | 0.38140 | 0.01604 | 1.89705 | 1.43379 | 1.31368 | 0.39772 | 0.41447 | 0.00000 | 0.02183 | 0.32696 | 0.00350 | 0.46973 | 0.00527 |
| **Slc35c2** | 4.33054 | 2.04724 | 8.72716 | 1.52786 | 3.78523 | 0.01397 | 0.82404 | 6.58685 | 1.47898 | 5.17282 | 3.60952 | 0.46666 | 7.25977 | 0.00000 | 0.45344 | 0.38633 |
| **Slc35d1** | 0.01762 | 0.00759 | 0.31152 | 0.01654 | 0.00800 | 0.00743 | 0.03741 | 0.04498 | 0.01254 | 0.00000 | 0.02899 | 0.00635 | 0.79314 | 0.04208 | 0.10427 | 0.18972 |
| **Slc35d2** | 0.05422 | 0.01786 | 0.05706 | 0.95660 | 0.09120 | 0.00645 | 0.00705 | 0.00150 | 0.86183 | 0.02261 | 0.48707 | 0.03104 | 0.02256 | 0.26533 | 0.02673 | 0.07968 |
| **Slc35e1** | 4.61435 | 2.04075 | 8.63429 | 2.09224 | 4.04452 | 6.14932 | 3.57049 | 2.41940 | 4.15732 | 5.82635 | 1.56381 | 4.12399 | 6.37538 | 3.22439 | 6.30132 | 6.11654 |
| **Slc35e2** | 3.63392 | 0.05519 | 4.15297 | 3.57317 | 0.27211 | 2.74119 | 0.34092 | 0.18458 | 6.09348 | 1.48992 | 5.39880 | 0.52954 | 3.49899 | 8.51016 | 5.18176 | 1.03384 |
| **Slc35e3** | 1.02149 | 0.00636 | 1.73889 | 9.72301 | 0.00277 | 2.44620 | 0.01531 | 3.29048 | 9.76917 | 0.44019 | 1.60462 | 0.10854 | 2.27850 | 0.79949 | 2.23306 | 0.02270 |
| **Slc35e4** | 1.15648 | 0.00499 | 1.89168 | 5.45459 | 0.06085 | 0.00420 | 0.00000 | 9.84693 | 6.94964 | 12.48624 | 2.02829 | 0.74513 | 8.52181 | 0.01474 | 1.25935 | 3.52935 |
| **Slc35f1** | 0.02562 | 0.01835 | 0.02679 | 0.08046 | 0.00866 | 0.00683 | 0.00752 | 0.01098 | 0.02922 | 0.01259 | 0.03720 | 0.00717 | 0.02487 | 0.16600 | 0.03946 | 0.00738 |
| **Slc35f2** | 3.18597 | 7.05826 | 11.53610 | 8.03222 | 0.41268 | 6.05873 | 7.08729 | 11.47693 | 17.96013 | 12.69413 | 17.84579 | 2.48736 | 11.35221 | 10.42724 | 10.52456 | 0.08569 |
| **Slc35f4** | 1.97826 | 0.00623 | 0.00568 | 0.00479 | 0.00819 | 0.00550 | 0.00593 | 0.00853 | 0.26487 | 0.01364 | 0.01013 | 0.01688 | 0.01770 | 0.01318 | 0.00837 | 0.00538 |
| **Slc35f5** | 2.27920 | 1.76036 | 1.45559 | 0.32439 | 4.49439 | 1.85211 | 0.01468 | 0.02820 | 2.41562 | 0.31663 | 0.00000 | 5.13770 | 0.14744 | 0.00254 | 2.04269 | 0.17170 |
| **Slc35f6** | 10.41839 | 0.12055 | 5.59143 | 0.72080 | 0.03412 | 0.89153 | 2.47094 | 5.18117 | 0.86592 | 3.59744 | 6.41475 | 0.10018 | 2.29122 | 4.21180 | 0.45864 | 4.31314 |
| **Slc35g1** | 5.78981 | 3.20254 | 7.78246 | 4.24237 | 5.87463 | 3.48509 | 4.18673 | 6.13164 | 7.11695 | 12.39495 | 10.30332 | 7.19047 | 7.48953 | 20.88618 | 11.89190 | 4.60257 |
| **Slc36a1** | 1.88059 | 0.00571 | 0.12831 | 1.85122 | 0.00478 | 0.06992 | 0.95259 | 0.30074 | 0.96736 | 0.75713 | 0.28136 | 0.03591 | 0.39848 | 0.00608 | 0.30718 | 0.00455 |
| **Slc36a1os** | 0.37977 | 0.57069 | 1.90090 | 5.35103 | 0.12358 | 0.67706 | 0.87718 | 3.13267 | 6.64038 | 2.76871 | 2.80762 | 3.66708 | 0.51445 | 1.79164 | 1.57349 | 0.70739 |
| **Slc36a3** | 0.01045 | 0.00796 | 0.00974 | 0.01063 | 0.00864 | 0.03660 | 0.00732 | 0.01203 | 0.01239 | 0.00754 | 0.01255 | 0.00807 | 0.28431 | 0.01302 | 0.01040 | 0.27185 |
| **Slc37a1** | 5.70518 | 0.00000 | 1.92914 | 0.32243 | 0.22664 | 1.83961 | 0.01199 | 8.79331 | 4.73001 | 6.34607 | 1.18413 | 0.11492 | 1.38112 | 0.29365 | 3.92926 | 0.07789 |
| **Slc37a2** | 4.21821 | 0.16208 | 4.04555 | 0.00923 | 0.01711 | 1.41367 | 5.63226 | 0.00000 | 0.01179 | 0.00000 | 0.00753 | 0.01613 | 1.56713 | 0.02323 | 0.00000 | 3.69287 |
| **Slc37a3** | 5.31861 | 6.85174 | 3.27535 | 2.45487 | 4.80460 | 0.00019 | 5.08650 | 3.93338 | 5.40350 | 1.21197 | 2.74792 | 0.24286 | 3.47685 | 4.78238 | 6.94832 | 1.22184 |
| **Slc37a4** | 0.12001 | 0.00593 | 1.65608 | 0.72067 | 2.37931 | 0.09935 | 0.00656 | 1.91018 | 0.44638 | 1.34438 | 1.05460 | 0.12734 | 0.00663 | 0.37524 | 2.74679 | 0.02150 |
| **Slc38a1** | 0.00000 | 0.00807 | 0.40674 | 0.00565 | 0.00816 | 6.47677 | 0.00592 | 0.00655 | 0.01313 | 0.28766 | 0.00000 | 0.01306 | 2.00801 | 0.72472 | 0.00000 | 0.92086 |
| **Slc38a10** | 3.85792 | 0.00000 | 2.74227 | 0.91991 | 1.96585 | 0.00807 | 3.77192 | 6.68111 | 1.64614 | 4.87749 | 1.72307 | 0.32505 | 2.07773 | 0.23575 | 1.22661 | 3.66286 |
| **Slc38a2** | 1.23553 | 0.00603 | 1.06370 | 0.50462 | 0.01924 | 0.00436 | 0.00632 | 0.63214 | 0.86391 | 1.02676 | 0.00000 | 0.04266 | 0.00000 | 0.00674 | 0.45662 | 3.51539 |
| **Slc38a3** | 0.01379 | 0.00783 | 0.01198 | 0.22473 | 0.00936 | 0.00664 | 0.00751 | 0.01333 | 0.00531 | 0.01255 | 0.01180 | 0.00786 | 0.01277 | 0.01879 | 0.05199 | 0.00738 |
| **Slc38a4** | 8.39445 | 3.94566 | 5.33955 | 0.05049 | 3.87271 | 9.69529 | 8.71892 | 0.90568 | 0.01540 | 0.13557 | 1.25629 | 0.00003 | 6.61475 | 6.03224 | 0.17744 | 4.38369 |
| **Slc38a6** | 0.12654 | 0.00525 | 2.32815 | 1.38176 | 0.05464 | 4.86456 | 0.09496 | 0.36004 | 0.36000 | 1.78037 | 2.34897 | 6.86733 | 1.16048 | 0.65290 | 0.19523 | 0.01767 |
| **Slc38a7** | 6.53651 | 0.03851 | 1.88528 | 2.88966 | 3.39703 | 3.98023 | 4.72517 | 2.44223 | 0.91327 | 4.78809 | 1.77209 | 0.55601 | 6.28845 | 0.27020 | 1.24271 | 3.70964 |
| **Slc38a8** | 0.53286 | 0.13141 | 0.74887 | 1.36590 | 0.00551 | 0.00482 | 0.00654 | 0.00000 | 0.70490 | 0.61495 | 0.59043 | 0.07527 | 0.04902 | 0.00597 | 0.50649 | 0.00590 |
| **Slc38a9** | 0.00463 | 0.00871 | 0.62421 | 0.00000 | 0.00727 | 0.00578 | 0.00808 | 0.04966 | 0.00872 | 1.58149 | 0.00480 | 0.00000 | 0.00743 | 0.31812 | 1.52907 | 0.68203 |
| **Slc39a1** | 2.30693 | 0.79330 | 3.36126 | 4.57044 | 0.08864 | 0.94533 | 0.00000 | 1.19357 | 7.33022 | 5.11010 | 6.34348 | 0.25542 | 2.84588 | 6.75943 | 1.92701 | 0.08643 |
| **Slc39a10** | 1.51492 | 0.12204 | 1.30676 | 10.62887 | 3.41048 | 0.00287 | 3.13193 | 0.91556 | 2.96050 | 5.33115 | 2.69516 | 4.28384 | 3.02642 | 5.76646 | 10.03767 | 0.26539 |
| **Slc39a11** | 0.70596 | 0.30398 | 0.83908 | 0.26847 | 0.00513 | 0.03893 | 0.00578 | 1.13677 | 0.09440 | 1.33858 | 0.26005 | 0.07925 | 1.38868 | 0.00614 | 0.47492 | 0.01295 |
| **Slc39a13** | 0.62785 | 0.02957 | 1.05592 | 0.04222 | 0.01097 | 0.00514 | 0.00670 | 0.15180 | 0.00965 | 0.12584 | 0.28751 | 0.58613 | 0.00000 | 0.00956 | 0.03091 | 2.24151 |
| **Slc39a14** | 8.65498 | 7.23057 | 3.78119 | 0.00001 | 3.46209 | 4.19941 | 5.34493 | 5.03639 | 3.42209 | 0.97264 | 4.24386 | 5.76952 | 4.88103 | 2.45912 | 1.36303 | 0.19536 |
| **Slc39a2** | 0.00807 | 0.00579 | 1.25590 | 0.00000 | 0.01828 | 1.52071 | 1.83044 | 3.77710 | 0.06453 | 4.43497 | 9.67914 | 0.14503 | 1.72862 | 0.03327 | 0.84067 | 0.79939 |
| **Slc39a3** | 0.00377 | 1.76983 | 0.07890 | 0.39215 | 0.61319 | 0.00562 | 0.01510 | 0.02387 | 0.28178 | 1.27742 | 0.35902 | 0.06322 | 2.77473 | 0.00465 | 0.00000 | 0.01268 |
| **Slc39a4** | 4.41806 | 1.44223 | 13.41672 | 12.15550 | 3.18696 | 8.37818 | 10.89247 | 22.71581 | 15.32773 | 15.61244 | 21.73816 | 1.98956 | 12.98717 | 0.00000 | 0.00000 | 5.59378 |
| **Slc39a5** | 0.04719 | 0.00772 | 0.02356 | 0.18687 | 0.00824 | 0.00616 | 0.01778 | 0.02581 | 0.04041 | 0.01970 | 0.02638 | 0.00760 | 0.01150 | 0.06280 | 0.00804 | 0.00741 |
| **Slc39a6** | 0.37362 | 0.01434 | 1.53123 | 12.05757 | 0.00403 | 0.00341 | 6.05819 | 1.39782 | 7.61439 | 3.09924 | 3.74504 | 0.14791 | 2.97796 | 0.12288 | 10.15592 | 1.71455 |
| **Slc39a7** | 7.93326 | 4.72073 | 6.95173 | 1.40471 | 6.30884 | 6.05181 | 3.00357 | 13.27189 | 1.93844 | 6.46893 | 2.69800 | 0.62772 | 11.94633 | 0.30643 | 1.17701 | 9.03132 |
| **Slc39a9** | 6.31008 | 0.04011 | 3.53868 | 3.70673 | 0.03342 | 5.05509 | 3.44555 | 2.70775 | 5.14201 | 6.54020 | 2.75256 | 4.47916 | 2.08777 | 3.62334 | 8.31980 | 3.08512 |
| **Slc3a1** | 0.01161 | 0.00753 | 0.01016 | 0.01014 | 0.00958 | 0.00662 | 0.00724 | 0.08768 | 0.01149 | 0.00680 | 0.01165 | 0.00701 | 0.00907 | 0.01659 | 0.22534 | 0.00737 |

| | | | | | | | | | | | | | | | | |
|---|---|---|---|---|---|---|---|---|---|---|---|---|---|---|---|---|
| Slc3a2 | 14.07692 | 5.06222 | 11.55572 | 6.63901 | 6.08518 | 2.34166 | 4.12925 | 16.33748 | 8.65420 | 9.46546 | 20.07798 | 3.07191 | 17.03041 | 1.03714 | 1.88630 | 10.22116 |
| Slc41a1 | 1.94789 | 3.09009 | 2.80660 | 3.29278 | 0.00000 | 0.00317 | 2.59502 | 0.00000 | 0.54886 | 1.00476 | 0.11814 | 0.03806 | 0.92695 | 0.00000 | 0.49976 | 0.94102 |
| Slc41a2 | 0.01068 | 0.00757 | 1.50214 | 1.01074 | 0.01608 | 0.00794 | 0.38793 | 0.00888 | 0.89601 | 0.08840 | 0.00000 | 0.01614 | 0.00767 | 0.09060 | 0.42099 | 0.00657 |
| Slc41a3 | 0.19699 | 0.00756 | 0.00672 | 0.14250 | 0.00748 | 0.00691 | 0.01726 | 0.01146 | 0.00950 | 0.56551 | 0.01277 | 0.00520 | 0.01384 | 0.00545 | 0.36914 | 0.00752 |
| Slc43a1 | 0.72707 | 0.04939 | 0.00850 | 0.19578 | 0.00649 | 0.00620 | 0.00685 | 0.01235 | 0.00878 | 0.01513 | 0.01375 | 0.00755 | 0.01178 | 0.01099 | 0.00692 | 0.01368 |
| Slc43a2 | 0.04837 | 0.00798 | 0.01048 | 2.04001 | 0.01999 | 0.01439 | 0.02765 | 0.00000 | 0.59125 | 0.00000 | 0.85940 | 0.00000 | 0.01946 | 0.54171 | 0.60696 | 0.02121 |
| Slc43a3 | 3.86640 | 11.67797 | 16.60971 | 4.76195 | 7.59947 | 4.45919 | 13.73507 | 17.82434 | 1.68974 | 8.43250 | 13.36979 | 1.41324 | 11.75736 | 3.33421 | 8.05818 | 2.31130 |
| Slc44a1 | 0.01498 | 0.00750 | 0.00000 | 0.34994 | 0.08428 | 0.44937 | 0.00736 | 0.00521 | 0.91451 | 0.13840 | 0.17643 | 0.01805 | 0.06248 | 1.29604 | 0.88737 | 0.03784 |
| Slc44a2 | 0.33870 | 3.06819 | 0.04477 | 5.85769 | 0.00702 | 7.98149 | 0.00274 | 0.88224 | 8.62407 | 0.98009 | 3.81262 | 0.04280 | 1.81837 | 0.94107 | 6.62790 | 0.00000 |
| Slc44a3 | 2.86875 | 0.00425 | 0.98149 | 0.91328 | 0.94619 | 1.02492 | 0.00500 | 1.17725 | 0.47397 | 0.30986 | 0.06044 | 0.02063 | 0.00000 | 0.00248 | 0.33288 | 0.00331 |
| Slc44a4 | 0.00000 | 1.95041 | 1.13430 | 0.51640 | 0.01501 | 4.42730 | 0.00432 | 1.30380 | 2.22935 | 0.44809 | 1.12042 | 0.09832 | 3.04658 | 0.00000 | 1.09887 | 0.01075 |
| Slc45a3 | 0.12437 | 0.00696 | 0.19166 | 27.26970 | 0.13118 | 0.00175 | 0.00495 | 2.77047 | 14.04443 | 2.02122 | 3.80032 | 0.38299 | 0.53159 | 6.30631 | 12.63631 | 1.19125 |
| Slc45a4 | 7.98834 | 0.89420 | 5.09584 | 0.00000 | 4.01021 | 2.60688 | 5.48570 | 13.56745 | 0.69950 | 7.92177 | 6.02103 | 4.67864 | 4.20972 | 0.05668 | 0.10358 | 5.75757 |
| Slc46a1 | 6.32999 | 5.70905 | 16.23980 | 0.00000 | 0.05771 | 11.39374 | 8.75324 | 13.80502 | 0.90844 | 11.05492 | 7.19694 | 0.70374 | 16.70948 | 0.17508 | 0.70775 | 4.99445 |
| Slc46a3 | 1.26151 | 0.30288 | 0.00000 | 0.13496 | 0.01927 | 0.00724 | 0.00607 | 0.01358 | 0.00714 | 1.12681 | 0.00907 | 0.02134 | 0.00780 | 0.27764 | 0.00000 | 1.33411 |
| Slc48a1 | 1.75479 | 0.00565 | 0.69526 | 1.17802 | 0.00702 | 0.00698 | 0.00571 | 0.00254 | 0.00000 | 3.12164 | 0.00986 | 0.01524 | 0.01250 | 0.00648 | 0.35501 | 0.00547 |
| Slc4a1 | 0.00708 | 1.02037 | 0.00741 | 0.39689 | 0.00796 | 0.00671 | 0.00687 | 0.00939 | 0.00541 | 0.60047 | 0.01214 | 0.00560 | 0.01137 | 0.01121 | 0.14160 | 0.00717 |
| Slc4a11 | 2.16505 | 0.00244 | 5.46301 | 4.73350 | 0.00986 | 7.67846 | 0.13460 | 6.87778 | 8.95372 | 8.32273 | 8.43476 | 0.50751 | 13.31454 | 1.20495 | 4.85328 | 2.80960 |
| Slc4a1ap | 1.19649 | 1.93646 | 2.92008 | 5.15966 | 0.02594 | 0.00578 | 2.40899 | 6.53128 | 4.29887 | 8.44681 | 5.01440 | 0.37496 | 1.77635 | 9.97751 | 11.70044 | 2.06083 |
| Slc4a2 | 5.29246 | 0.00000 | 1.10409 | 0.05149 | 3.35293 | 13.82867 | 4.85219 | 5.68341 | 0.43984 | 0.00000 | 1.40714 | 0.03317 | 3.45939 | 0.00322 | 0.00000 | 0.01571 |
| Slc4a3 | 0.00457 | 0.00741 | 0.85341 | 2.58208 | 0.01880 | 0.00888 | 0.15655 | 1.79953 | 1.39955 | 0.00000 | 0.46866 | 0.13759 | 0.02293 | 0.01793 | 0.00552 | 0.02105 |
| Slc4a5 | 0.00742 | 2.75192 | 0.00000 | 0.84230 | 0.36403 | 0.00668 | 0.00617 | 0.01400 | 0.00000 | 0.00000 | 0.10321 | 0.01464 | 0.01604 | 0.00338 | 0.32917 | 0.00597 |
| Slc4a7 | 1.50830 | 0.00437 | 0.00000 | 1.94308 | 0.01300 | 0.00432 | 1.71896 | 0.00000 | 4.25279 | 0.13296 | 1.45995 | 0.87490 | 0.00000 | 3.26092 | 0.41881 | 1.01074 |
| Slc4a8 | 0.01193 | 0.00736 | 1.03520 | 1.69035 | 0.00738 | 0.00903 | 0.00690 | 0.64192 | 0.03982 | 0.45231 | 0.00634 | 0.01307 | 0.00106 | 0.00885 | 0.00000 | 0.00679 |
| Slc50a1 | 0.31915 | 0.00611 | 1.03285 | 0.00735 | 0.00529 | 0.00899 | 1.26279 | 0.00658 | 0.01173 | 0.76156 | 0.01092 | 0.01265 | 0.00846 | 0.01425 | 0.00000 | 0.00693 |
| Slc51a | 2.34298 | 0.02598 | 3.08862 | 1.46638 | 0.00000 | 0.00523 | 0.00375 | 3.38116 | 1.44287 | 0.42315 | 0.02418 | 0.01654 | 2.94042 | 0.00719 | 0.00000 | 0.00936 |
| Slc51b | 0.01232 | 0.03984 | 0.01195 | 0.26772 | 0.01856 | 0.00662 | 0.00749 | 0.01344 | 0.21947 | 0.01324 | 0.01157 | 0.00748 | 0.01258 | 0.00927 | 0.02067 | 0.00732 |
| Slc52a2 | 0.09982 | 0.21075 | 0.94390 | 2.31409 | 0.30699 | 2.64695 | 0.01698 | 2.47537 | 7.08645 | 1.45158 | 0.36945 | 0.00000 | 0.00000 | 0.00000 | 0.13688 | 0.00531 |
| Slc52a3 | 0.08834 | 0.00686 | 0.00000 | 0.00276 | 0.00745 | 0.78946 | 0.97905 | 0.00000 | 0.02372 | 1.32405 | 0.94973 | 0.00000 | 1.05582 | 0.01108 | 0.00000 | 1.28557 |
| Slc5a11 | 0.00000 | 1.48090 | 9.59608 | 0.74561 | 0.03769 | 8.32636 | 0.41680 | 8.72445 | 2.60941 | 0.81700 | 0.99176 | 0.37842 | 20.63404 | 2.20482 | 2.63691 | 0.01407 |
| Slc5a12 | 0.01077 | 0.00786 | 0.01037 | 0.17415 | 0.07870 | 0.01295 | 0.00747 | 0.01228 | 0.01596 | 0.01508 | 0.02908 | 0.00809 | 0.03840 | 0.01161 | 0.05380 | 0.00721 |
| Slc5a2 | 0.83748 | 0.00649 | 0.13871 | 0.01062 | 0.05698 | 0.00484 | 0.00662 | 0.02843 | 0.01230 | 0.01853 | 0.01289 | 0.00690 | 0.01589 | 0.01249 | 0.00792 | 0.00638 |
| Slc5a3 | 0.19167 | 0.00438 | 4.60380 | 4.93416 | 0.00770 | 3.18055 | 0.04381 | 0.50860 | 6.18983 | 6.01140 | 2.99275 | 7.90827 | 1.80418 | 17.17724 | 4.12746 | 0.18745 |
| Slc5a4b | 0.55476 | 0.00707 | 0.77367 | 1.52152 | 0.01315 | 0.00576 | 1.96385 | 0.36015 | 2.36156 | 0.97906 | 1.38386 | 0.04501 | 0.03374 | 0.00174 | 0.38933 | 0.57569 |
| Slc5a5 | 1.42854 | 0.01703 | 1.12844 | 0.23320 | 0.00398 | 0.00408 | 0.00592 | 1.15347 | 0.00571 | 1.15679 | 0.00000 | 0.03069 | 0.00000 | 0.32660 | 0.00000 | 2.77092 |
| Slc5a6 | 3.23928 | 5.35052 | 13.71450 | 19.77251 | 4.43003 | 11.08787 | 15.24118 | 39.68842 | 20.52794 | 38.96249 | 33.47520 | 18.37576 | 25.00615 | 20.10858 | 34.94643 | 4.95341 |
| Slc5a8 | 0.01224 | 0.00785 | 0.00002 | 0.00455 | 0.00935 | 0.00728 | 0.00693 | 0.00000 | 0.10398 | 0.00000 | 0.40960 | 0.00000 | 0.00870 | 2.09591 | 0.09077 | 0.00744 |
| Slc6a12 | 0.01910 | 0.01884 | 0.00611 | 0.00965 | 0.01031 | 1.64499 | 0.00728 | 0.01123 | 0.01264 | 0.00835 | 0.01063 | 0.00808 | 0.00461 | 0.01265 | 0.01016 | 0.01391 |
| Slc6a13 | 0.03059 | 0.01767 | 1.56918 | 0.00949 | 0.00819 | 1.38065 | 0.90912 | 0.00665 | 0.01213 | 0.00000 | 0.01035 | 0.00556 | 1.33064 | 0.01680 | 0.04368 | 0.00560 |
| Slc6a14 | 0.02161 | 0.00787 | 0.00002 | 0.07530 | 0.01007 | 3.34781 | 0.37705 | 0.11137 | 0.37342 | 0.00000 | 0.00000 | 0.37621 | 0.00000 | 0.01816 | 0.81015 | 0.01303 |
| Slc6a15 | 0.84947 | 0.00596 | 0.00000 | 8.60085 | 0.97585 | 0.29816 | 0.00568 | 0.89113 | 3.55125 | 1.89756 | 0.34418 | 0.00000 | 0.00000 | 6.79105 | 4.78156 | 2.35644 |
| Slc6a16 | 0.01375 | 0.00815 | 0.01229 | 0.46317 | 0.00930 | 0.00660 | 0.00752 | 0.01396 | 0.03023 | 0.01527 | 0.01379 | 0.00824 | 0.01266 | 0.01112 | 0.00528 | 0.00733 |
| Slc6a18 | 0.05292 | 0.01865 | 0.12321 | 0.04680 | 0.00829 | 0.02102 | 0.09298 | 0.05934 | 0.08286 | 0.08863 | 0.00736 | 0.02709 | 0.02844 | 0.59538 | 0.04287 | 0.01442 |
| Slc6a2 | 0.03849 | 0.00755 | 0.18163 | 0.11792 | 0.00799 | 0.01185 | 0.03769 | 0.01317 | 0.20225 | 0.04493 | 0.54684 | 0.02339 | 0.06011 | 0.26535 | 0.07851 | 0.01419 |
| Slc6a20b | 0.04947 | 0.03098 | 0.05426 | 0.09192 | 0.02134 | 0.01900 | 0.01862 | 0.04491 | 0.04631 | 0.06909 | 0.05606 | 0.01867 | 0.07201 | 0.07094 | 0.06408 | 0.00721 |
| Slc6a4 | 0.60108 | 0.00722 | 0.00005 | 0.63472 | 0.01649 | 0.00858 | 0.00656 | 0.72844 | 0.19315 | 0.08981 | 0.00569 | 0.00375 | 0.00650 | 0.00698 | 0.44818 | 0.00724 |
| Slc6a6 | 7.93505 | 4.33077 | 6.27454 | 1.30708 | 0.00000 | 4.74114 | 6.19970 | 18.36432 | 1.56223 | 11.61470 | 13.27528 | 5.16351 | 4.91168 | 11.57225 | 9.93981 | 1.72654 |
| Slc6a7 | 0.01372 | 0.00784 | 0.01181 | 2.12709 | 0.00925 | 0.00638 | 0.00739 | 0.01163 | 0.33862 | 0.03858 | 0.04202 | 0.00627 | 0.01207 | 0.00165 | 0.25839 | 0.00706 |
| Slc6a8 | 23.45436 | 10.50889 | 26.00626 | 0.32157 | 10.12671 | 16.12178 | 17.42612 | 33.60075 | 1.61389 | 22.53026 | 17.65633 | 6.96910 | 24.88163 | 11.93410 | 22.70141 | 12.48262 |
| Slc6a9 | 0.00950 | 0.00776 | 0.00876 | 2.32921 | 0.00920 | 0.00612 | 0.00720 | 0.14112 | 4.35625 | 0.67902 | 0.00000 | 0.00678 | 0.00791 | 0.03036 | 0.56305 | 0.00674 |
| Slc7a1 | 1.61294 | 2.04532 | 0.65991 | 0.87077 | 0.01553 | 0.00405 | 0.00536 | 0.01864 | 0.01935 | 0.00509 | 0.01221 | 0.01618 | 0.00802 | 0.00453 | 0.16519 | 0.01883 |
| Slc7a11 | 6.15882 | 2.42097 | 4.05581 | 1.04166 | 2.73756 | 1.19209 | 1.21824 | 2.45033 | 0.39240 | 1.87112 | 0.13121 | 0.47254 | 2.76396 | 0.34787 | 0.85015 | 3.30029 |
| Slc7a14 | 1.48736 | 0.19424 | 1.78323 | 9.33911 | 0.85133 | 0.14415 | 0.27204 | 0.00000 | 13.37988 | 1.11535 | 10.98321 | 1.34076 | 1.05478 | 7.97061 | 7.09005 | 0.32741 |
| Slc7a2 | 0.88373 | 0.09018 | 2.38539 | 1.10441 | 0.38197 | 0.25820 | 0.19256 | 0.52167 | 0.52499 | 1.18820 | 6.56008 | 0.97490 | 3.92361 | 0.75393 | 1.47953 | 0.39187 |
| Slc7a3 | 2.68470 | 0.00477 | 0.00344 | 0.01042 | 0.00297 | 0.00579 | 0.00515 | 0.02218 | 0.01250 | 0.02269 | 0.01217 | 0.02469 | 0.01054 | 0.00960 | 0.02463 | 0.01168 |
| Slc7a4 | 0.01384 | 0.00789 | 0.01113 | 0.00874 | 0.00920 | 0.00671 | 0.00739 | 0.01088 | 0.01207 | 0.07875 | 0.06213 | 0.03643 | 0.01210 | 0.01122 | 0.00927 | 0.00773 |
| Slc7a5 | 6.06703 | 0.20676 | 1.16440 | 5.52914 | 5.03784 | 0.00605 | 3.93913 | 1.35199 | 7.55782 | 6.00209 | 4.99887 | 0.83006 | 4.84637 | 1.59039 | 2.01219 | 1.98240 |
| Slc7a6 | 3.95286 | 9.49993 | 12.42867 | 0.60179 | 4.81332 | 8.29377 | 11.26999 | 10.05910 | 1.56505 | 6.28179 | 6.28158 | 4.18221 | 13.90708 | 9.96959 | 10.28084 | 10.98285 |
| Slc7a6os | 4.78259 | 1.66596 | 0.75879 | 12.68594 | 4.74322 | 0.49771 | 1.42572 | 1.49949 | 16.85775 | 8.49844 | 3.53765 | 0.12718 | 1.00698 | 2.81155 | 1.76811 | 2.52375 |
| Slc7a7 | 1.65363 | 0.00704 | 0.59747 | 0.05280 | 0.01796 | 3.42039 | 0.04065 | 1.42280 | 0.01018 | 0.81566 | 0.46428 | 0.03265 | 1.59144 | 0.54205 | 0.49862 | 2.80230 |
| Slc7a8 | 1.54100 | 0.00688 | 1.07597 | 0.54883 | 0.00315 | 0.00499 | 0.01785 | 0.34448 | 0.00215 | 0.11279 | 0.00646 | 0.00556 | 0.00112 | 0.00527 | 0.68365 | 0.00493 |
| Slc7a9 | 0.01292 | 0.00816 | 0.01359 | 0.09839 | 0.00998 | 0.00736 | 0.00783 | 0.01681 | 0.01034 | 0.09033 | 0.01430 | 0.00861 | 0.12443 | 0.01398 | 0.02629 | 0.00770 |

| | | | | | | | | | | | | | | | | |
|---|---|---|---|---|---|---|---|---|---|---|---|---|---|---|---|---|
| **Slc8b1** | 1.36209 | 0.29882 | 5.30391 | 1.58402 | 2.11100 | 2.34361 | 0.41309 | 9.36407 | 1.25595 | 9.44804 | 8.30379 | 0.80986 | 3.36385 | 2.48069 | 5.65694 | 4.68366 |
| **Slc9a1** | 2.76072 | 0.00287 | 1.64388 | 3.46076 | 0.01594 | 2.64976 | 1.69667 | 3.05081 | 6.61302 | 0.66988 | 0.80389 | 0.19108 | 2.20983 | 0.00000 | 0.46182 | 1.81455 |
| **Slc9a3r1** | 8.98421 | 0.11817 | 0.00000 | 3.62307 | 3.10096 | 6.46224 | 0.00000 | 0.00000 | 1.12284 | 1.30632 | 0.09302 | 0.02683 | 4.36917 | 0.00000 | 0.30284 | 7.36010 |
| **Slc9a3r2** | 0.02104 | 0.00529 | 0.02559 | 0.12431 | 2.68059 | 0.00664 | 2.36162 | 0.47592 | 0.00643 | 0.37195 | 0.00980 | 0.01183 | 0.00517 | 0.01190 | 0.00000 | 0.00753 |
| **Slc9a5** | 1.54297 | 0.00629 | 0.00098 | 0.21043 | 0.02613 | 0.00645 | 0.00612 | 0.00711 | 0.01974 | 0.11610 | 0.01175 | 0.00607 | 0.00698 | 0.01074 | 0.00809 | 2.64751 |
| **Slc9a6** | 0.00000 | 3.55100 | 0.78644 | 0.70782 | 0.03235 | 0.90953 | 3.46548 | 0.01376 | 0.00696 | 1.27113 | 0.00000 | 4.29028 | 2.51247 | 0.00321 | 0.00000 | 0.01794 |
| **Slc9a8** | 0.75506 | 0.18002 | 4.33269 | 0.05110 | 1.81707 | 0.01467 | 1.80234 | 4.32751 | 0.19888 | 0.76956 | 1.47148 | 0.00766 | 1.71508 | 0.45069 | 1.13412 | 0.85372 |
| **Slc9b1** | 1.42556 | 0.00529 | 5.84083 | 0.01943 | 0.00000 | 0.12741 | 0.01433 | 1.52716 | 1.14921 | 0.58176 | 2.33975 | 0.06924 | 0.24684 | 2.74594 | 1.42213 | 0.01765 |
| **Slc9b2** | 0.01284 | 0.00767 | 0.13361 | 0.00913 | 0.00887 | 3.13289 | 0.00696 | 0.00990 | 0.01280 | 0.00139 | 0.00828 | 0.08509 | 0.00255 | 0.01354 | 0.00621 | 0.00614 |
| **Slco1a5** | 0.00966 | 0.00758 | 0.00881 | 0.82637 | 0.00963 | 0.00668 | 0.00712 | 0.32713 | 0.55368 | 0.00475 | 0.00762 | 0.00574 | 0.00633 | 0.04410 | 0.00202 | 0.00727 |
| **Slco2a1** | 0.00000 | 0.00822 | 0.03810 | 0.00754 | 0.00779 | 6.25479 | 0.06291 | 0.18242 | 0.01296 | 0.05111 | 0.00627 | 0.02824 | 1.48271 | 0.04926 | 0.00269 | 2.61753 |
| **Slco2b1** | 0.02183 | 0.00775 | 0.00005 | 0.00848 | 0.47535 | 2.15483 | 0.00716 | 0.00984 | 0.01250 | 0.11323 | 0.00982 | 0.00689 | 0.01081 | 0.01141 | 0.00989 | 0.78219 |
| **Slco3a1** | 0.01009 | 0.00839 | 0.41145 | 5.25992 | 0.00838 | 0.00508 | 0.00725 | 0.00954 | 0.64790 | 1.20668 | 0.00142 | 0.04146 | 0.00897 | 0.62612 | 1.47472 | 0.00599 |
| **Slco4a1** | 0.01323 | 0.00788 | 0.01112 | 0.62673 | 0.00922 | 0.00633 | 0.00753 | 0.01286 | 3.72350 | 0.00428 | 0.00000 | 0.00000 | 0.01309 | 0.01294 | 0.03909 | 0.00701 |
| **Slco4c1** | 0.36393 | 0.00774 | 0.38760 | 7.58957 | 0.01586 | 0.00499 | 0.00584 | 0.90675 | 1.30855 | 3.86589 | 2.30363 | 0.00000 | 0.16009 | 12.76378 | 0.72760 | 0.00535 |
| **Slco5a1** | 0.00988 | 0.00765 | 0.00706 | 1.49245 | 0.00963 | 0.00665 | 0.00702 | 0.93718 | 0.04603 | 0.00193 | 0.02060 | 0.00566 | 0.00524 | 0.00475 | 0.34262 | 0.00698 |
| **Slf1** | 0.37035 | 0.00705 | 0.00212 | 0.18882 | 0.02558 | 0.00763 | 0.00732 | 0.03753 | 0.15187 | 0.48919 | 0.00053 | 0.42744 | 0.17866 | 0.03627 | 0.88780 | 0.00697 |
| **Slfn2** | 0.01157 | 0.09052 | 0.01015 | 0.10963 | 0.00876 | 0.00649 | 0.06820 | 0.01060 | 0.02579 | 0.42345 | 0.02026 | 0.04361 | 0.02289 | 0.00508 | 0.31059 | 0.01401 |
| **Slfn4** | 0.00818 | 0.37316 | 0.00959 | 0.04573 | 0.00821 | 0.00661 | 0.00721 | 0.01047 | 0.03775 | 0.01509 | 0.02849 | 0.00738 | 0.01718 | 0.01998 | 0.03075 | 0.15542 |
| **Slfn5** | 1.96190 | 0.27834 | 1.44508 | 9.21390 | 0.68623 | 0.25596 | 0.15407 | 0.00000 | 15.13841 | 0.00000 | 11.15047 | 1.07429 | 0.70860 | 5.97474 | 1.09956 | 0.28391 |
| **Slfn9** | 0.01891 | 0.00779 | 0.04705 | 0.01770 | 0.00856 | 0.00843 | 0.00735 | 0.01584 | 0.01255 | 0.04088 | 0.01147 | 0.00761 | 0.05178 | 0.10263 | 0.03419 | 0.00724 |
| **Slirp** | 5.42423 | 0.32426 | 2.12547 | 1.22224 | 2.17533 | 1.44874 | 0.31728 | 0.00000 | 1.33031 | 0.43289 | 9.77298 | 0.12746 | 0.11602 | 1.97252 | 0.64145 | 0.65541 |
| **Slit3** | 0.01364 | 0.00772 | 0.01134 | 0.10988 | 0.00927 | 0.00656 | 0.00733 | 0.01122 | 0.01086 | 0.29678 | 0.01257 | 0.05348 | 0.01191 | 0.04105 | 0.01696 | 0.00734 |
| **Slk** | 7.03491 | 3.18550 | 4.21826 | 1.33040 | 2.20208 | 0.97136 | 1.18862 | 2.25374 | 0.79771 | 0.61875 | 1.00617 | 0.14397 | 1.34413 | 0.61102 | 1.29432 | 2.54592 |
| **Slmap** | 2.53952 | 5.50170 | 1.74246 | 11.35577 | 5.34194 | 1.06519 | 3.64151 | 6.12581 | 10.18014 | 8.25837 | 9.83722 | 12.53960 | 3.93221 | 22.79907 | 15.06088 | 1.37463 |
| **Slmo1** | 0.00979 | 0.00789 | 0.01025 | 0.08877 | 0.00964 | 0.02438 | 0.01794 | 0.16407 | 2.92800 | 0.57724 | 0.66476 | 0.00000 | 0.21947 | 0.00607 | 0.00000 | 0.01473 |
| **Slmo2** | 6.02073 | 4.82703 | 8.39181 | 6.43754 | 3.76133 | 5.02050 | 6.13281 | 6.27088 | 1.98968 | 3.98749 | 3.48519 | 0.40995 | 8.51608 | 1.79447 | 1.53933 | 6.57305 |
| **Sln** | 0.01390 | 0.00810 | 0.00800 | 0.00460 | 0.00906 | 0.00686 | 0.00754 | 0.00260 | 0.01116 | 0.00000 | 0.67222 | 0.00481 | 0.00619 | 0.00528 | 0.12426 | 0.00707 |
| **Sltm** | 4.72425 | 4.54171 | 3.22380 | 6.95775 | 4.93143 | 0.70051 | 2.49348 | 8.73905 | 7.69324 | 19.37173 | 4.22858 | 12.60641 | 3.95470 | 16.97696 | 20.51333 | 2.41518 |
| **Slu7** | 5.65274 | 4.73906 | 12.04438 | 8.36240 | 1.42513 | 0.00294 | 2.86485 | 9.11319 | 16.43552 | 11.33054 | 8.84323 | 10.03974 | 3.18154 | 21.19319 | 18.45257 | 3.23606 |
| **Slx1b** | 5.66649 | 0.50848 | 7.02906 | 1.20853 | 1.12817 | 3.63534 | 1.38923 | 5.08347 | 6.08531 | 4.55102 | 17.17459 | 1.74561 | 9.16158 | 3.11107 | 2.42446 | 1.30070 |
| **Slx4** | 1.11892 | 0.03283 | 2.03898 | 3.30268 | 0.00199 | 3.11649 | 5.91202 | 3.88347 | 2.81215 | 3.93187 | 0.54712 | 4.40001 | 0.00000 | 0.00000 | 0.68773 | 0.79468 |
| **Slx4ip** | 0.02249 | 0.10179 | 0.00045 | 11.15924 | 0.00831 | 0.01792 | 0.06765 | 0.05441 | 6.80745 | 0.66182 | 0.26399 | 0.03417 | 0.01326 | 2.14401 | 4.46241 | 0.02624 |
| **Smad1** | 0.00856 | 0.00631 | 0.03425 | 0.39974 | 0.00963 | 0.00593 | 2.84136 | 0.00981 | 0.00535 | 0.27705 | 0.01305 | 0.01436 | 0.01034 | 0.00895 | 0.03712 | 0.00756 |
| **Smad2** | 7.39012 | 3.80278 | 4.23106 | 6.87582 | 8.18691 | 1.58582 | 5.41205 | 4.35312 | 3.95474 | 1.42720 | 3.82180 | 3.89527 | 3.91166 | 5.43073 | 4.33122 | 2.62554 |
| **Smad3** | 2.73921 | 0.00579 | 1.56843 | 10.08547 | 0.21195 | 0.00814 | 0.06447 | 1.42566 | 1.09700 | 1.08073 | 0.86562 | 0.06239 | 0.07060 | 1.89290 | 2.67059 | 0.00225 |
| **Smad4** | 3.74432 | 3.29584 | 4.08678 | 6.68852 | 8.70359 | 7.23003 | 0.97243 | 5.58045 | 2.09826 | 8.62211 | 2.61259 | 3.31669 | 1.29147 | 2.02357 | 4.08506 | 0.00001 |
| **Smad5** | 3.51575 | 0.00000 | 0.00000 | 0.58091 | 3.13498 | 0.00293 | 4.32516 | 0.00000 | 1.27405 | 1.67734 | 0.15688 | 0.01603 | 0.48674 | 0.00339 | 1.42636 | 0.06060 |
| **Smad7** | 0.02811 | 0.00693 | 0.13207 | 6.40271 | 0.15770 | 0.02441 | 0.01587 | 0.32085 | 12.04233 | 0.37293 | 2.80785 | 0.00004 | 0.07129 | 0.13850 | 0.68896 | 0.01911 |
| **Smad9** | 0.10248 | 0.00782 | 0.18539 | 0.11561 | 0.00850 | 0.00586 | 0.00732 | 0.01526 | 0.04845 | 0.02345 | 0.29829 | 0.00608 | 0.03976 | 0.07666 | 0.03915 | 0.00708 |
| **Smagp** | 1.83360 | 2.82576 | 1.44979 | 11.80796 | 2.49556 | 2.95607 | 0.80696 | 0.11703 | 4.21578 | 0.11580 | 3.71186 | 1.13918 | 0.00196 | 0.00000 | 0.04757 | 0.42346 |
| **Smap1** | 2.66229 | 3.18148 | 6.10875 | 0.00000 | 6.17525 | 4.06431 | 3.72119 | 5.70030 | 1.04118 | 3.33414 | 0.04930 | 0.13170 | 3.73688 | 0.31256 | 1.71585 | 1.84562 |
| **Smap2** | 6.89564 | 0.00000 | 4.88857 | 2.60359 | 7.63647 | 2.18961 | 6.49329 | 10.00833 | 6.53349 | 12.67694 | 10.78140 | 11.18534 | 2.56145 | 4.88268 | 4.40610 | 3.35176 |
| **Smarca2** | 4.03566 | 1.56911 | 2.39838 | 5.87653 | 0.04869 | 0.00765 | 3.63351 | 0.00000 | 1.65471 | 4.64195 | 0.06131 | 0.31889 | 0.02957 | 1.52073 | 2.66551 | 1.77237 |
| **Smarca4** | 12.12003 | 2.53680 | 1.21473 | 11.77925 | 11.84046 | 4.12446 | 5.43403 | 29.87753 | 15.55961 | 27.72758 | 17.30619 | 2.32074 | 16.11375 | 1.29051 | 5.73525 | 4.87495 |
| **Smarca5** | 9.59505 | 13.13456 | 8.18789 | 14.21919 | 8.78861 | 3.05048 | 8.46423 | 15.05500 | 20.19317 | 15.02343 | 11.82847 | 11.38689 | 7.28008 | 34.64324 | 21.97731 | 3.22640 |
| **Smarca5.ps** | 3.01560 | 4.46765 | 2.78104 | 7.70214 | 3.31001 | 0.94010 | 3.08950 | 5.52418 | 7.97182 | 5.23578 | 2.10828 | 4.78424 | 2.67834 | 17.92037 | 11.97056 | 1.02518 |
| **Smarcad1** | 4.27700 | 3.58985 | 4.06827 | 5.41555 | 1.92504 | 0.01015 | 1.78781 | 2.56537 | 8.14274 | 3.82036 | 7.70393 | 12.52345 | 2.32890 | 14.10109 | 7.02755 | 1.35873 |
| **Smarcal1** | 4.60527 | 3.04227 | 1.80501 | 1.54567 | 6.13143 | 0.00000 | 3.02525 | 4.35583 | 0.37581 | 3.58480 | 0.90665 | 0.07366 | 2.23166 | 0.00000 | 0.61168 | 6.14186 |
| **Smarcb1** | 12.54390 | 3.01653 | 8.47492 | 7.58013 | 6.18012 | 4.08912 | 4.82043 | 3.78624 | 5.09868 | 10.24469 | 8.86361 | 0.49457 | 10.20221 | 1.54952 | 5.34842 | 5.56616 |
| **Smarcc1** | 9.56340 | 0.01898 | 6.16832 | 9.16021 | 10.16192 | 4.39432 | 3.45528 | 7.46519 | 8.08483 | 14.62616 | 4.27585 | 9.32727 | 7.10016 | 11.56168 | 6.61398 | 3.79703 |
| **Smarcc2** | 2.26150 | 3.73456 | 0.00000 | 1.39328 | 0.00582 | 0.81507 | 0.01417 | 1.29351 | 2.59145 | 2.93118 | 0.87443 | 0.09230 | 0.00003 | 0.00000 | 0.61966 | 2.83402 |
| **Smarcd1** | 0.96691 | 0.26340 | 0.60206 | 4.51610 | 2.60160 | 0.00490 | 0.04448 | 1.43633 | 3.29596 | 2.68587 | 0.25421 | 0.14754 | 2.29035 | 0.00000 | 1.07182 | 2.86706 |
| **Smarcd2** | 10.56557 | 4.75018 | 9.32151 | 10.00535 | 2.52965 | 0.00000 | 4.77965 | 20.64609 | 16.19516 | 15.04597 | 14.72180 | 4.86466 | 12.56529 | 1.03076 | 13.93797 | 4.05541 |
| **Smarcd3** | 0.00784 | 0.00736 | 0.59710 | 0.24913 | 0.02111 | 0.00568 | 0.18942 | 1.71501 | 0.02587 | 0.24668 | 0.23090 | 0.14176 | 1.53966 | 0.02360 | 0.00000 | 0.00659 |
| **Smarce1** | 9.71591 | 4.85418 | 10.02039 | 4.38315 | 5.94977 | 4.26217 | 7.59604 | 15.59096 | 4.11598 | 14.00086 | 7.82493 | 1.57465 | 5.17718 | 4.20426 | 13.02386 | 4.67440 |
| **Smc1a** | 8.76681 | 9.29539 | 7.84599 | 5.75771 | 10.19718 | 3.99490 | 5.05607 | 11.63959 | 4.17881 | 10.54470 | 4.63058 | 2.92057 | 7.22242 | 22.37320 | 11.57512 | 7.07442 |
| **Smc1b** | 0.56027 | 0.00611 | 0.00000 | 0.98753 | 0.01487 | 0.00590 | 1.25439 | 0.00677 | 1.38215 | 0.53477 | 0.00000 | 0.00000 | 0.01028 | 0.00591 | 2.21028 | 0.00628 |
| **Smc2** | 8.40135 | 6.48115 | 5.12178 | 2.26943 | 0.55094 | 3.33575 | 3.66194 | 1.51238 | 4.16434 | 5.88466 | 2.28113 | 15.52212 | 2.15852 | 13.79675 | 8.34886 | 4.01578 |
| **Smc3** | 7.41715 | 6.85270 | 7.27392 | 6.01860 | 9.62224 | 5.33156 | 4.45662 | 10.71626 | 14.35701 | 14.87397 | 7.14800 | 12.62506 | 4.35929 | 22.80762 | 18.50830 | 1.96436 |
| **Smc4** | 9.01865 | 13.54423 | 4.06751 | 8.71182 | 0.15580 | 4.60569 | 4.28778 | 8.32755 | 16.29640 | 17.93592 | 8.04596 | 17.94207 | 4.89112 | 31.69823 | 17.60305 | 3.14823 |
| **Smc5** | 5.18136 | 2.86673 | 4.27847 | 3.16914 | 7.76457 | 3.77367 | 9.32973 | 5.41515 | 1.69540 | 11.89020 | 6.86274 | 9.61454 | 3.96240 | 14.79746 | 8.07478 | 0.05105 |

| | | | | | | | | | | | | | | | | |
|---|---|---|---|---|---|---|---|---|---|---|---|---|---|---|---|---|
| **Smc6** | 10.76358 | 10.28917 | 6.05535 | 15.49667 | 7.54230 | 5.55917 | 5.32159 | 6.55114 | 14.88277 | 9.62304 | 6.26950 | 17.33196 | 3.26711 | 15.85102 | 16.22491 | 2.05014 |
| **Smchd1** | 1.35032 | 3.01025 | 1.22929 | 8.75540 | 4.17177 | 5.68690 | 6.05972 | 5.27555 | 7.92675 | 16.50291 | 3.91714 | 6.28170 | 0.00000 | 18.91303 | 16.18930 | 2.15693 |
| **Smco1** | 0.34495 | 0.10267 | 0.72211 | 0.68742 | 0.19412 | 0.09210 | 0.25227 | 0.07979 | 1.01898 | 0.40909 | 3.44765 | 0.17336 | 0.43872 | 1.07519 | 0.42034 | 0.19874 |
| **Smco2** | 0.01246 | 0.00737 | 0.44332 | 0.24951 | 0.00872 | 0.00559 | 0.16377 | 0.28335 | 0.45695 | 0.07590 | 0.42315 | 0.00332 | 0.00157 | 0.00785 | 0.02404 | 0.00683 |
| **Smco4** | 0.19302 | 0.02919 | 0.56402 | 2.91049 | 0.00800 | 0.00514 | 0.00703 | 0.31101 | 1.52187 | 0.48612 | 0.05203 | 0.00027 | 0.00309 | 0.17117 | 0.03241 | 0.17958 |
| **Smcr8** | 1.73918 | 4.00549 | 4.66394 | 4.01232 | 0.00000 | 0.46929 | 2.85142 | 0.73615 | 7.10824 | 3.31695 | 0.46740 | 4.59348 | 2.09650 | 1.12700 | 8.26009 | 0.04288 |
| **Smdt1** | 6.34786 | 0.01589 | 2.95417 | 5.66582 | 8.14797 | 2.02259 | 0.35511 | 0.89151 | 3.11220 | 8.66987 | 11.52544 | 0.20131 | 13.43337 | 0.72998 | 0.00000 | 2.99901 |
| **Smek1** | 4.94233 | 4.79133 | 2.71310 | 4.70663 | 2.49860 | 0.64192 | 2.45690 | 3.20270 | 16.24179 | 6.34523 | 2.78207 | 0.52762 | 1.29774 | 16.22775 | 11.67788 | 3.50397 |
| **Smek2** | 2.73937 | 6.23935 | 2.99574 | 8.50418 | 2.39250 | 0.00000 | 2.62853 | 1.62536 | 5.03903 | 3.03041 | 1.79929 | 13.11604 | 0.76126 | 14.37482 | 9.90109 | 0.23348 |
| **Smg1** | 7.93477 | 6.57027 | 6.46643 | 7.97811 | 0.68201 | 5.03834 | 3.16684 | 4.19809 | 12.39250 | 7.95961 | 4.67577 | 14.10776 | 0.77610 | 17.06103 | 9.91668 | 3.85970 |
| **Smg5** | 6.09999 | 1.10469 | 8.06671 | 11.06067 | 0.56402 | 2.93059 | 2.14509 | 9.90342 | 14.43869 | 16.18921 | 12.94048 | 1.12757 | 11.74713 | 0.50971 | 4.06133 | 7.79212 |
| **Smg6** | 1.15196 | 1.14543 | 1.88838 | 2.80785 | 0.01453 | 2.28231 | 0.00521 | 3.13348 | 2.92522 | 1.79297 | 1.78039 | 0.20460 | 0.00000 | 0.97324 | 4.44999 | 0.70358 |
| **Smg7** | 4.77669 | 0.26311 | 7.16839 | 3.76846 | 3.03054 | 0.01961 | 4.89403 | 7.23540 | 8.43630 | 7.55438 | 9.86984 | 1.62800 | 3.68179 | 12.21374 | 5.05219 | 1.20134 |
| **Smg8** | 1.58477 | 4.68152 | 4.82404 | 9.26729 | 2.46769 | 2.60247 | 2.26570 | 7.83867 | 11.83407 | 10.59731 | 3.60343 | 7.64299 | 1.93186 | 9.44827 | 14.07773 | 2.38389 |
| **Smg9** | 2.33676 | 0.09347 | 1.87823 | 1.05189 | 0.00000 | 0.00446 | 1.40316 | 3.80420 | 0.09850 | 4.31609 | 0.77035 | 0.24321 | 4.69237 | 0.04624 | 0.05677 | 1.16334 |
| **Smim1** | 0.72850 | 0.00616 | 0.10263 | 0.20053 | 0.00672 | 0.32984 | 0.01515 | 0.51291 | 0.00552 | 0.00000 | 2.90466 | 0.03706 | 1.08991 | 0.00417 | 0.00000 | 0.00531 |
| **Smim10l1** | 0.01130 | 0.00843 | 0.01095 | 0.19521 | 0.00985 | 0.00724 | 0.00833 | 0.08852 | 0.00805 | 0.14317 | 0.01142 | 0.00879 | 0.02351 | 0.01467 | 0.25795 | 0.00757 |
| **Smim10l2a** | 0.01296 | 0.00788 | 0.09756 | 0.05270 | 0.00897 | 0.00719 | 0.00709 | 0.00998 | 0.01073 | 0.00281 | 0.01240 | 0.00724 | 0.19628 | 0.00768 | 0.51956 | 0.00710 |
| **Smim11** | 1.94085 | 0.07770 | 0.65344 | 3.36018 | 1.73703 | 0.22052 | 0.05401 | 0.03932 | 1.18450 | 0.49790 | 1.38311 | 0.04882 | 2.00639 | 0.09978 | 0.00000 | 0.00357 |
| **Smim12** | 9.24915 | 3.51971 | 4.06905 | 0.90769 | 0.03228 | 1.34782 | 0.43054 | 1.35452 | 3.21352 | 2.44143 | 0.33005 | 0.06928 | 4.22306 | 0.00000 | 0.00000 | 0.17620 |
| **Smim13** | 1.89642 | 2.31079 | 3.35510 | 8.02523 | 2.22802 | 3.23538 | 4.08181 | 6.24737 | 11.50489 | 12.93925 | 14.24204 | 8.14634 | 2.66351 | 21.94879 | 11.92471 | 0.03158 |
| **Smim14** | 0.00000 | 0.00742 | 1.52696 | 13.79575 | 0.05373 | 0.00780 | 0.00626 | 0.56541 | 3.50607 | 1.07713 | 1.10210 | 0.00504 | 0.26513 | 0.02929 | 2.38808 | 0.46949 |
| **Smim15** | 0.98414 | 2.58905 | 1.07164 | 2.87165 | 2.23243 | 2.06340 | 0.96799 | 3.19400 | 6.90972 | 0.82749 | 0.40935 | 1.68964 | 2.21339 | 4.89039 | 4.29563 | 0.03004 |
| **Smim19** | 0.00000 | 0.01571 | 1.35815 | 0.18446 | 2.87099 | 0.00529 | 1.12607 | 0.28927 | 1.80646 | 0.49975 | 0.99646 | 0.02420 | 0.16078 | 0.00302 | 0.20431 | 0.09419 |
| **Smim20** | 3.54708 | 0.02065 | 3.81100 | 1.08995 | 0.02243 | 5.36054 | 5.53884 | 1.12467 | 0.05197 | 5.05571 | 4.51911 | 0.75404 | 7.19243 | 1.04763 | 0.00000 | 2.58435 |
| **Smim22** | 0.99162 | 0.00672 | 0.51485 | 0.00451 | 0.04860 | 0.00638 | 0.00628 | 0.00336 | 0.01094 | 0.00447 | 0.97303 | 0.00229 | 0.22757 | 0.12241 | 0.00000 | 0.00591 |
| **Smim24** | 0.01057 | 0.00815 | 0.00784 | 0.01409 | 0.00927 | 0.00718 | 0.00755 | 0.46948 | 0.12326 | 0.39011 | 0.00648 | 0.06977 | 0.00633 | 0.00700 | 0.11901 | 0.00737 |
| **Smim4** | 0.92591 | 0.00645 | 0.24091 | 3.08112 | 0.00590 | 0.00577 | 0.15629 | 0.05592 | 1.31958 | 0.53261 | 1.27128 | 0.08592 | 0.35332 | 0.03624 | 0.05347 | 0.03452 |
| **Smim5** | 0.01355 | 0.00781 | 0.01222 | 0.00858 | 0.00917 | 0.00657 | 0.00730 | 0.01083 | 0.01209 | 0.49675 | 0.01277 | 0.00618 | 0.01201 | 0.01154 | 0.00904 | 0.00765 |
| **Smim6** | 0.79248 | 0.00625 | 0.00000 | 0.98970 | 0.00611 | 0.00685 | 0.42157 | 0.73206 | 0.00000 | 0.00000 | 0.52409 | 0.00000 | 0.00332 | 1.17974 | 0.29768 | 0.00609 |
| **Smim7** | 11.49602 | 3.39305 | 12.19181 | 9.73412 | 11.05752 | 2.52626 | 9.06117 | 7.09549 | 11.89514 | 12.53622 | 10.04236 | 2.81375 | 6.68853 | 11.46628 | 12.33447 | 1.77604 |
| **Smim8** | 3.10620 | 0.20928 | 2.16507 | 1.67445 | 1.79661 | 2.06004 | 1.01275 | 0.34084 | 1.20488 | 0.81615 | 0.43337 | 0.16760 | 1.83271 | 0.92282 | 0.06089 | 0.02404 |
| **Smim9** | 0.01563 | 0.00788 | 0.04554 | 0.12190 | 0.00916 | 0.00586 | 0.00742 | 0.01214 | 0.60191 | 0.04055 | 0.00835 | 0.01588 | 0.01164 | 0.02395 | 0.10025 | 0.00714 |
| **Smkr.ps** | 0.01375 | 0.00783 | 0.01237 | 0.00963 | 0.00935 | 0.00658 | 0.00749 | 0.01327 | 0.01235 | 0.01017 | 0.01281 | 0.00742 | 0.01265 | 0.00670 | 0.14729 | 0.00738 |
| **Smn1** | 6.75009 | 5.08972 | 4.70110 | 3.38040 | 1.57422 | 2.46946 | 5.99512 | 11.63015 | 10.58270 | 14.69669 | 13.81838 | 7.33997 | 9.90212 | 4.61776 | 2.66726 | 3.68874 |
| **Smndc1** | 4.33747 | 9.05678 | 6.01183 | 13.14652 | 7.97625 | 0.03737 | 5.40598 | 8.75855 | 15.54453 | 7.79223 | 5.85382 | 9.55604 | 6.37933 | 15.00529 | 8.98168 | 0.99074 |
| **Smo** | 0.01311 | 0.00806 | 0.01297 | 0.00725 | 0.00949 | 0.00693 | 0.00749 | 0.01192 | 0.01171 | 0.91774 | 0.01280 | 0.01608 | 0.02412 | 0.01001 | 0.02327 | 0.00773 |
| **Smoc1** | 0.62671 | 2.89111 | 3.00797 | 1.28791 | 4.65495 | 0.00564 | 0.00527 | 1.32844 | 0.19743 | 0.00000 | 0.43532 | 0.00313 | 0.00000 | 0.00521 | 0.71953 | 3.09831 |
| **Smoc2** | 0.00913 | 0.00841 | 0.00830 | 0.07071 | 0.00923 | 0.00679 | 0.00818 | 0.15046 | 0.31932 | 0.98907 | 0.01657 | 0.03786 | 0.00759 | 0.25990 | 0.35145 | 0.00742 |
| **Smox** | 0.24604 | 0.00621 | 0.31907 | 2.27912 | 0.00727 | 0.00890 | 2.89460 | 0.01692 | 0.46613 | 1.55102 | 0.00867 | 0.01959 | 0.50519 | 0.00587 | 0.02489 | 0.00638 |
| **Smpd1** | 5.34498 | 4.81338 | 14.71304 | 1.20621 | 4.80292 | 21.05435 | 11.98115 | 17.85922 | 11.03443 | 9.43031 | 9.38960 | 9.53099 | 27.79135 | 0.00000 | 0.35605 | 8.93693 |
| **Smpd2** | 3.13670 | 0.03245 | 0.85667 | 0.12178 | 0.02553 | 0.01056 | 1.50018 | 4.37681 | 0.07989 | 1.67885 | 2.44116 | 1.76956 | 4.68406 | 0.06263 | 0.02792 | 0.00836 |
| **Smpd3** | 0.43593 | 0.02666 | 0.78801 | 1.02525 | 1.50187 | 0.00617 | 0.38998 | 0.00260 | 0.04847 | 0.00002 | 0.15660 | 0.01256 | 0.10528 | 0.00502 | 0.49570 | 0.00628 |
| **Smpd4** | 8.17337 | 6.09285 | 9.33501 | 11.97028 | 4.69033 | 5.34174 | 7.81250 | 12.53650 | 15.00675 | 16.50189 | 7.91049 | 5.70085 | 11.08503 | 0.65128 | 14.48029 | 4.77833 |
| **Smpdl3a** | 0.00000 | 4.15592 | 4.62589 | 0.00000 | 0.01666 | 5.07393 | 0.99819 | 4.31187 | 0.03821 | 5.92473 | 10.67935 | 2.61381 | 9.47647 | 8.41822 | 0.95417 | 0.00000 |
| **Smpdl3b** | 6.74035 | 2.86479 | 2.42566 | 0.51520 | 0.61212 | 0.04462 | 1.18458 | 0.37218 | 1.11709 | 3.15593 | 2.40938 | 0.21148 | 2.39169 | 0.73847 | 0.16109 | 2.62527 |
| **Sms** | 26.05641 | 11.58957 | 17.62071 | 15.00789 | 22.52465 | 4.38891 | 8.63986 | 31.31398 | 9.18412 | 33.85008 | 23.48831 | 18.80704 | 25.01550 | 34.67378 | 24.15817 | 14.69920 |
| **Smtn** | 2.70117 | 0.00362 | 0.00000 | 0.08230 | 0.98052 | 4.19612 | 0.00353 | 4.54949 | 0.93675 | 2.02861 | 0.00000 | 0.05146 | 2.44801 | 0.01455 | 0.00000 | 2.38077 |
| **Smtnl1** | 1.60879 | 0.09698 | 0.00880 | 0.01055 | 0.00498 | 0.00524 | 0.00619 | 0.00709 | 0.01239 | 0.01478 | 0.01277 | 0.00688 | 0.01127 | 0.01168 | 0.00954 | 0.02003 |
| **Smtnl2** | 0.38683 | 0.05972 | 0.74175 | 0.23559 | 0.42671 | 0.02740 | 0.11883 | 0.04393 | 0.55397 | 0.14328 | 1.83005 | 0.22884 | 0.69288 | 0.76502 | 0.27662 | 2.14437 |
| **Smu1** | 21.32691 | 13.90312 | 18.32950 | 9.43215 | 12.70060 | 7.97140 | 9.57669 | 14.62788 | 19.60211 | 19.40844 | 11.41385 | 12.59675 | 14.72658 | 17.00163 | 14.21135 | 10.30039 |
| **Smug1** | 4.83011 | 1.59737 | 4.60717 | 2.42806 | 1.20399 | 3.92254 | 0.01098 | 1.20674 | 0.01906 | 2.62296 | 2.55263 | 0.19511 | 2.51473 | 2.05441 | 2.23316 | 0.00000 |
| **Smurf1** | 0.06956 | 5.61296 | 1.34528 | 5.74617 | 0.01615 | 0.81074 | 5.05105 | 3.04563 | 1.61333 | 3.62520 | 0.82869 | 0.47523 | 4.60198 | 0.43230 | 0.66206 | 1.61839 |
| **Smurf2** | 4.10107 | 1.78316 | 0.00000 | 12.52399 | 6.54334 | 6.94624 | 1.45587 | 2.68271 | 9.63905 | 11.14868 | 4.68825 | 12.38142 | 0.13877 | 0.96445 | 16.75489 | 0.38577 |
| **Smyd1** | 0.01772 | 0.00762 | 0.12958 | 0.14947 | 0.01182 | 2.04451 | 0.01640 | 0.26453 | 0.02003 | 0.00000 | 0.00000 | 0.00940 | 1.08545 | 1.77953 | 0.00000 | 0.01329 |
| **Smyd2** | 1.46591 | 0.13091 | 0.00000 | 0.00000 | 0.00504 | 2.68002 | 0.00000 | 2.28813 | 0.03532 | 2.76156 | 5.12520 | 0.06421 | 7.29907 | 0.90402 | 0.00000 | 0.00000 |
| **Smyd3** | 6.31712 | 3.77664 | 0.43008 | 0.29195 | 2.15185 | 5.32529 | 0.00000 | 1.35324 | 1.95357 | 2.22603 | 0.42927 | 2.17263 | 2.22285 | 0.42940 | 0.25649 | 2.18009 |
| **Smyd4** | 0.51930 | 0.11880 | 1.04188 | 0.34592 | 0.03367 | 1.84011 | 0.05727 | 0.04400 | 0.64124 | 0.18603 | 0.34287 | 0.05811 | 1.74199 | 1.39062 | 0.02352 | 0.04898 |
| **Smyd5** | 14.41232 | 11.15750 | 13.82227 | 7.13015 | 15.29153 | 3.84635 | 9.80949 | 27.41988 | 9.14194 | 26.00145 | 16.10631 | 2.94690 | 16.47292 | 3.04660 | 12.51297 | 6.20547 |
| **Snai1** | 3.63861 | 1.69599 | 1.52187 | 8.77419 | 0.21242 | 0.00000 | 0.00000 | 0.93441 | 18.98450 | 1.10961 | 5.34109 | 0.00000 | 1.03453 | 0.01357 | 1.60011 | 2.46819 |
| **Snai2** | 0.02345 | 0.00746 | 0.02060 | 0.35644 | 0.00842 | 0.02215 | 0.00721 | 0.00965 | 0.00647 | 1.13603 | 0.02492 | 0.00346 | 0.01231 | 0.00489 | 0.27984 | 0.00769 |
| **Snai3** | 0.00000 | 0.00747 | 0.00857 | 1.37630 | 0.00798 | 0.00653 | 0.00709 | 0.55782 | 0.28299 | 0.00386 | 0.02397 | 0.01616 | 0.00000 | 0.00746 | 0.02648 | 1.67109 |

| | | | | | | | | | | | | | | | | |
|---|---|---|---|---|---|---|---|---|---|---|---|---|---|---|---|---|
| **Snap23** | 3.39275 | 7.46909 | 0.70431 | 10.07877 | 5.14444 | 5.75671 | 3.27168 | 4.10337 | 15.77220 | 7.37149 | 4.68789 | 5.34928 | 3.75138 | 14.00288 | 17.40209 | 3.46740 |
| **Snap29** | 0.00000 | 1.84572 | 2.80860 | 7.17466 | 7.48073 | 0.14768 | 0.03434 | 6.92575 | 4.90113 | 2.56568 | 4.27788 | 3.23035 | 1.86096 | 0.98646 | 1.18926 | 0.08922 |
| **Snap47** | 6.40587 | 0.41643 | 2.16331 | 3.78400 | 3.17065 | 0.02076 | 0.51889 | 1.85501 | 2.19023 | 2.13232 | 0.71972 | 0.98564 | 7.05045 | 0.21030 | 1.21607 | 2.98347 |
| **Snap91** | 0.00599 | 0.00766 | 0.18720 | 0.90888 | 0.00838 | 0.01069 | 0.00740 | 0.06745 | 1.80583 | 1.19318 | 0.57822 | 3.35413 | 0.34141 | 0.52714 | 2.53192 | 0.01372 |
| **Snapc1** | 0.69637 | 3.29638 | 1.89425 | 5.65116 | 0.00328 | 0.00942 | 0.02448 | 3.30751 | 11.04025 | 2.98023 | 2.61813 | 0.00000 | 0.38115 | 0.00037 | 2.26569 | 0.14673 |
| **Snapc2** | 6.25994 | 2.36505 | 3.48872 | 17.79538 | 4.70173 | 0.94099 | 0.78736 | 7.08398 | 19.99851 | 10.75651 | 2.63427 | 0.27202 | 2.54802 | 0.00000 | 1.76350 | 4.27338 |
| **Snapc3** | 2.38218 | 1.04612 | 1.76115 | 8.65845 | 7.48689 | 0.00101 | 0.00000 | 3.16869 | 15.30542 | 18.24959 | 5.94912 | 1.54466 | 4.06852 | 2.02931 | 6.53510 | 1.92178 |
| **Snapc4** | 0.91126 | 2.13140 | 1.36582 | 9.64640 | 5.29082 | 2.62690 | 4.64358 | 5.53632 | 16.80757 | 5.46806 | 4.85966 | 1.09771 | 4.69599 | 0.07917 | 2.51863 | 3.62231 |
| **Snapc5** | 4.30247 | 5.67602 | 7.32547 | 8.12507 | 3.92586 | 0.15966 | 0.10781 | 4.38226 | 11.73935 | 5.97846 | 3.22101 | 0.63465 | 4.26157 | 3.13134 | 5.89552 | 1.00639 |
| **Snapin** | 1.99816 | 0.35695 | 4.28878 | 3.00367 | 3.61245 | 0.03169 | 0.02358 | 1.31171 | 0.00000 | 2.93332 | 2.54956 | 1.13826 | 4.03629 | 0.17367 | 2.45239 | 2.64735 |
| **Snca** | 0.01149 | 0.00766 | 0.01228 | 0.00493 | 0.00842 | 0.00648 | 0.00717 | 0.00910 | 0.01073 | 5.55903 | 0.00891 | 0.00000 | 0.01056 | 0.01264 | 0.00550 | 0.00758 |
| **Sncb** | 0.02623 | 0.00798 | 0.01125 | 0.01052 | 0.08847 | 0.00699 | 0.00763 | 0.01627 | 0.01229 | 0.01314 | 0.01407 | 0.00784 | 0.02435 | 0.01235 | 0.01060 | 0.46203 |
| **Sncg** | 0.00888 | 0.00732 | 0.00823 | 0.00753 | 0.08749 | 0.00676 | 0.61945 | 0.00983 | 0.01199 | 0.70550 | 0.01162 | 0.00523 | 0.01043 | 0.01163 | 0.00812 | 0.01431 |
| **Snd1** | 3.06591 | 0.00414 | 0.00000 | 2.43703 | 0.95548 | 2.35766 | 0.04916 | 1.12697 | 1.72380 | 2.16111 | 1.51476 | 0.04824 | 0.05689 | 0.10045 | 2.75676 | 0.76614 |
| **Snf8** | 13.09185 | 6.35017 | 10.89824 | 6.00137 | 8.19147 | 6.24534 | 4.82493 | 14.82536 | 7.35966 | 18.65031 | 14.37092 | 1.48625 | 14.33295 | 2.76664 | 13.42997 | 6.41592 |
| **Snhg1** | 3.69073 | 4.23626 | 6.65382 | 12.34038 | 1.02446 | 1.32115 | 3.25379 | 4.44562 | 20.40786 | 8.25481 | 21.70997 | 2.76135 | 8.30504 | 9.23038 | 4.76425 | 3.82439 |
| **Snhg10** | 0.03056 | 0.00769 | 0.02166 | 1.20359 | 0.00770 | 0.00744 | 0.02754 | 0.01392 | 0.21217 | 0.00000 | 0.17451 | 0.00614 | 0.00768 | 0.00472 | 0.16341 | 0.68530 |
| **Snhg11** | 4.64177 | 1.42736 | 3.02658 | 2.65211 | 0.13679 | 0.21925 | 0.82732 | 0.16735 | 3.65833 | 0.36987 | 4.28496 | 0.18087 | 3.76205 | 3.81622 | 1.56136 | 0.08041 |
| **Snhg12** | 1.80187 | 2.34225 | 3.93210 | 4.90841 | 0.01316 | 0.17861 | 3.41347 | 3.44111 | 10.56238 | 6.41372 | 9.91940 | 0.95257 | 3.63489 | 0.15417 | 4.31958 | 3.33776 |
| **Snhg14** | 0.02374 | 0.00423 | 0.06395 | 0.00669 | 0.00777 | 0.00710 | 0.00384 | 0.98487 | 0.89668 | 1.19278 | 2.22498 | 10.35243 | 1.47458 | 27.30599 | 0.37773 | 0.01225 |
| **Snhg15** | 0.43253 | 0.01601 | 1.57793 | 7.20967 | 0.01283 | 0.01050 | 0.05271 | 1.82276 | 11.33947 | 3.66706 | 6.33872 | 0.84889 | 1.76517 | 0.95683 | 0.01373 | 2.35689 |
| **Snhg17** | 4.71845 | 0.27718 | 1.88204 | 3.84834 | 0.09432 | 2.56577 | 1.23057 | 5.25219 | 7.04526 | 8.64322 | 7.07924 | 0.49747 | 3.90566 | 1.21284 | 4.59342 | 2.12282 |
| **Snhg18** | 0.03378 | 0.00782 | 0.47705 | 0.67659 | 0.01170 | 0.00551 | 0.00686 | 0.23470 | 1.60262 | 0.12499 | 0.00000 | 0.06419 | 0.48024 | 0.01147 | 0.31868 | 0.05410 |
| **Snhg20** | 0.42448 | 0.00866 | 0.41156 | 2.54741 | 0.01461 | 0.00591 | 0.03945 | 0.03580 | 2.28657 | 0.84987 | 0.78578 | 0.00916 | 0.03576 | 0.01775 | 0.01696 | 1.31088 |
| **Snhg3** | 0.98772 | 0.04862 | 5.01184 | 9.53750 | 1.21881 | 0.00941 | 0.98956 | 0.21276 | 10.49712 | 1.29848 | 2.97807 | 0.54433 | 1.91565 | 0.44738 | 1.69882 | 0.49423 |
| **Snhg4** | 2.96831 | 0.01404 | 1.70280 | 1.06320 | 0.02130 | 1.50919 | 0.80774 | 0.02147 | 2.47850 | 3.43934 | 0.57087 | 1.07901 | 0.06223 | 0.00394 | 1.86458 | 1.49201 |
| **Snhg5** | 5.95163 | 7.46124 | 9.66726 | 4.61454 | 5.40940 | 5.65225 | 5.61689 | 17.59374 | 8.54884 | 18.40314 | 16.93845 | 6.17387 | 9.30130 | 9.35321 | 12.18702 | 9.44738 |
| **Snhg6** | 0.40600 | 0.01740 | 0.05433 | 1.33385 | 0.50061 | 0.07625 | 0.03820 | 0.10925 | 1.71423 | 0.14124 | 1.00855 | 0.13699 | 0.05544 | 0.06474 | 0.29722 | 0.00648 |
| **Snhg7** | 0.36075 | 1.42950 | 3.00132 | 0.83787 | 0.00182 | 1.14435 | 1.06810 | 1.21267 | 3.89741 | 2.13352 | 0.00000 | 0.05493 | 2.15314 | 0.95831 | 0.62202 | 1.77877 |
| **Snhg8** | 0.62911 | 0.08133 | 1.08762 | 2.49329 | 0.13813 | 0.00554 | 0.03421 | 0.34870 | 5.65043 | 0.66366 | 3.84741 | 0.11759 | 0.25413 | 0.03552 | 0.41655 | 0.01960 |
| **Snhg9** | 0.02108 | 0.00788 | 0.12177 | 1.37327 | 0.00866 | 0.03023 | 0.00731 | 0.02535 | 1.20315 | 0.02349 | 0.00001 | 0.00549 | 0.05051 | 0.09033 | 0.00000 | 0.00689 |
| **Snip1** | 0.67696 | 6.14669 | 4.25691 | 7.79431 | 2.59008 | 0.00338 | 1.30979 | 3.37284 | 11.53024 | 3.15850 | 3.43716 | 2.79777 | 1.23867 | 5.04256 | 7.70567 | 1.23491 |
| **Snn** | 1.89720 | 0.00576 | 0.83693 | 1.09812 | 0.00296 | 0.00398 | 0.00597 | 1.03745 | 0.79429 | 0.12118 | 0.35070 | 0.22018 | 0.00931 | 0.00287 | 0.75605 | 0.96593 |
| **Snora15** | 0.01215 | 0.00769 | 0.08114 | 0.01090 | 0.00898 | 0.00857 | 0.00728 | 0.01261 | 0.01203 | 0.00866 | 0.01196 | 0.00751 | 0.12625 | 0.01639 | 0.06685 | 0.00720 |
| **Snora16a** | 0.01135 | 0.00774 | 0.00894 | 0.00552 | 0.00939 | 0.00700 | 0.00731 | 0.24845 | 0.11936 | 0.04674 | 0.00943 | 0.00739 | 0.00707 | 0.00740 | 0.05910 | 0.00738 |
| **Snora21** | 0.11140 | 0.01881 | 0.00152 | 0.00580 | 0.01476 | 0.00774 | 0.00688 | 0.16426 | 0.03913 | 0.55489 | 0.27136 | 0.03223 | 0.26275 | 0.52115 | 0.32239 | 0.00722 |
| **Snora23** | 0.01704 | 0.00772 | 0.00890 | 0.00593 | 0.04436 | 0.00696 | 0.00735 | 0.05070 | 0.09382 | 0.08915 | 0.00979 | 0.10235 | 0.01824 | 0.09126 | 0.00193 | 0.00726 |
| **Snora28** | 0.19739 | 0.00697 | 0.06136 | 0.00740 | 0.00776 | 0.00713 | 0.51013 | 0.03645 | 0.04178 | 0.00688 | 0.01028 | 0.00525 | 0.04895 | 0.42366 | 0.05818 | 0.00715 |
| **Snora2b** | 0.01356 | 0.00787 | 0.01147 | 0.77218 | 0.00927 | 0.00635 | 0.00741 | 0.01167 | 1.60944 | 0.00285 | 0.01555 | 0.00452 | 0.01249 | 0.00534 | 0.30038 | 0.00706 |
| **Snora3** | 0.90985 | 0.56008 | 0.00000 | 0.01697 | 0.82982 | 0.00832 | 0.35590 | 0.07966 | 0.01039 | 0.14134 | 0.02651 | 0.00522 | 0.37447 | 0.02146 | 0.01213 | 0.00591 |
| **Snora31** | 0.41384 | 0.00740 | 0.09013 | 0.44982 | 0.00698 | 0.00543 | 0.00667 | 0.05426 | 0.47102 | 0.49540 | 0.83854 | 0.06369 | 0.01213 | 0.01556 | 0.32728 | 0.30813 |
| **Snora33** | 0.16793 | 0.79750 | 0.89337 | 0.33305 | 0.00575 | 0.02411 | 0.63339 | 0.00440 | 0.70256 | 0.57244 | 0.34519 | 0.11872 | 0.12809 | 0.70839 | 0.15597 | 0.22950 |
| **Snora34** | 0.01045 | 0.00827 | 0.13890 | 0.00829 | 0.04643 | 0.00591 | 0.00774 | 0.01682 | 0.01161 | 0.05103 | 0.02334 | 0.00702 | 0.09745 | 0.09545 | 0.31245 | 0.00734 |
| **Snora41** | 0.00395 | 0.28462 | 0.14110 | 0.00284 | 0.00869 | 0.01898 | 0.26326 | 0.21298 | 0.00978 | 0.74652 | 0.37576 | 0.32942 | 0.06958 | 0.02738 | 0.48386 | 0.00738 |
| **Snora43** | 0.00585 | 0.52236 | 0.66157 | 0.00867 | 0.00696 | 0.00862 | 0.29918 | 0.07057 | 0.01217 | 0.26103 | 0.01078 | 0.00700 | 0.26977 | 0.01020 | 0.02216 | 0.00676 |
| **Snora44** | 0.01578 | 0.40082 | 0.26446 | 0.06480 | 0.00754 | 0.00930 | 0.53460 | 0.02053 | 0.05568 | 0.02588 | 0.00971 | 0.00780 | 0.23317 | 0.00832 | 0.22451 | 0.59315 |
| **Snora47** | 0.01360 | 0.00817 | 0.01191 | 0.13995 | 0.00929 | 0.00660 | 0.00754 | 0.01386 | 0.97722 | 0.01086 | 0.00686 | 0.00671 | 0.01294 | 0.01278 | 0.00445 | 0.00734 |
| **Snora52** | 0.01378 | 0.00777 | 0.01199 | 0.00933 | 0.00938 | 0.00655 | 0.00744 | 0.01241 | 0.01167 | 0.00757 | 0.01219 | 0.00728 | 0.01273 | 0.00969 | 0.28471 | 0.00737 |
| **Snora5c** | 0.12260 | 0.00763 | 0.00842 | 0.01076 | 0.00811 | 0.00732 | 0.08202 | 0.01381 | 0.01223 | 0.01393 | 0.01410 | 0.00779 | 0.03300 | 0.01422 | 0.01557 | 0.00746 |
| **Snora61** | 0.22916 | 0.46511 | 0.41629 | 0.05309 | 0.02280 | 0.00675 | 0.58380 | 0.20036 | 0.00816 | 0.37116 | 0.00735 | 0.05953 | 0.50530 | 0.10644 | 0.46871 | 0.56469 |
| **Snora62** | 0.01527 | 0.00812 | 0.07942 | 0.01026 | 0.00917 | 0.00586 | 0.00783 | 0.03177 | 0.01237 | 0.01260 | 0.01249 | 0.00806 | 0.01053 | 0.00911 | 0.06344 | 0.00732 |
| **Snora64** | 0.28897 | 0.37348 | 0.64304 | 0.00301 | 0.60228 | 0.00704 | 0.01672 | 0.51093 | 0.03380 | 2.22910 | 0.00610 | 0.02845 | 0.08140 | 0.01262 | 0.23784 | 0.13688 |
| **Snora65** | 0.65639 | 0.37586 | 1.48464 | 0.00259 | 0.01733 | 0.17318 | 0.78858 | 0.63240 | 0.01919 | 0.86756 | 0.12771 | 0.22929 | 0.58635 | 0.52046 | 0.49190 | 0.44882 |
| **Snora68** | 0.63817 | 0.31949 | 0.90121 | 0.33385 | 0.00459 | 0.75205 | 1.10268 | 0.69752 | 0.06896 | 0.97892 | 0.31904 | 0.02112 | 0.44390 | 0.10074 | 0.36082 | 1.27010 |
| **Snora69** | 0.01040 | 0.00775 | 0.01022 | 0.21914 | 0.00909 | 0.00645 | 0.00704 | 0.07223 | 0.20988 | 0.84231 | 0.01098 | 0.00465 | 0.20684 | 0.03146 | 0.28969 | 0.00707 |
| **Snora70** | 0.26397 | 0.00646 | 1.34788 | 0.00088 | 0.00478 | 0.00729 | 1.47318 | 0.51861 | 0.01806 | 0.39494 | 0.50454 | 0.03438 | 0.99541 | 1.01423 | 0.78524 | 0.68447 |
| **Snora74a** | 0.01014 | 0.00821 | 0.01193 | 0.09729 | 0.00833 | 0.00672 | 0.00755 | 0.01406 | 0.02126 | 0.04761 | 0.01124 | 0.00930 | 0.01040 | 0.00764 | 0.10402 | 0.54487 |
| **Snora75** | 0.01265 | 0.00755 | 0.01284 | 0.02257 | 0.00931 | 0.00653 | 0.40013 | 0.01051 | 0.09058 | 0.08821 | 0.01199 | 0.00664 | 0.01182 | 0.02520 | 0.04282 | 0.00771 |
| **Snora78** | 0.01772 | 0.00774 | 0.01929 | 0.01042 | 0.00894 | 0.00628 | 0.00725 | 0.03506 | 0.01227 | 0.05323 | 0.01310 | 0.01676 | 0.01775 | 0.01138 | 0.01034 | 0.00764 |
| **Snora7a** | 0.00913 | 0.00847 | 0.20869 | 0.06404 | 0.00927 | 0.00653 | 0.00777 | 0.17716 | 0.01079 | 0.06324 | 0.00987 | 0.08451 | 0.00669 | 0.01844 | 0.15117 | 0.00732 |
| **Snora81** | 0.01335 | 0.00848 | 0.02072 | 0.01665 | 0.00944 | 0.00843 | 0.10752 | 0.10305 | 0.01005 | 0.05321 | 0.00661 | 0.00494 | 0.06949 | 0.97051 | 0.34631 | 0.10017 |
| **Snord100** | 0.26383 | 1.96459 | 0.82407 | 0.23267 | 0.00572 | 0.00774 | 1.53048 | 0.36254 | 0.00648 | 0.52117 | 0.12398 | 0.00180 | 0.21979 | 0.15583 | 0.68290 | 0.19020 |

| | | | | | | | | | | | | | | | | |
|---|---|---|---|---|---|---|---|---|---|---|---|---|---|---|---|---|
| **Snord104** | 0.01143 | 0.00793 | 0.06264 | 0.73925 | 0.13221 | 0.00586 | 0.00750 | 0.01222 | 0.53253 | 0.01263 | 0.06021 | 0.02841 | 0.01376 | 0.01109 | 0.00019 | 0.00716 |
| **Snord12** | 0.04643 | 0.47676 | 0.50685 | 0.06235 | 0.01324 | 0.00597 | 0.05017 | 0.47539 | 0.12518 | 0.65353 | 0.05101 | 0.01523 | 0.04028 | 0.00581 | 0.39375 | 0.17202 |
| **Snord15a** | 0.01121 | 0.00861 | 0.01129 | 0.01045 | 0.00965 | 0.00684 | 0.00805 | 0.07290 | 0.01210 | 0.01450 | 0.01285 | 0.00873 | 0.16357 | 0.01249 | 0.03442 | 0.00748 |
| **Snord15b** | 0.03881 | 0.00762 | 0.01013 | 0.06224 | 0.00800 | 0.00653 | 0.00712 | 0.65662 | 0.12435 | 0.49137 | 0.19626 | 0.30039 | 0.22587 | 0.47320 | 0.00000 | 0.17787 |
| **Snord16a** | 0.36242 | 0.73086 | 0.16296 | 0.00238 | 0.25702 | 0.74215 | 0.33589 | 0.46336 | 0.03980 | 0.49309 | 0.00000 | 0.00000 | 0.41061 | 0.80261 | 0.40094 | 0.57218 |
| **Snord17** | 0.00940 | 0.00789 | 0.01167 | 0.00949 | 0.01015 | 0.00666 | 0.01744 | 0.21806 | 0.01190 | 0.00000 | 0.01090 | 0.00595 | 0.03980 | 0.20071 | 0.19520 | 0.00757 |
| **Snord1b** | 0.01747 | 0.00869 | 0.01020 | 0.01028 | 0.00891 | 0.00678 | 0.00807 | 0.08128 | 0.01213 | 0.01282 | 0.01224 | 0.00922 | 0.00937 | 0.01105 | 0.03877 | 0.00734 |
| **Snord2** | 0.01183 | 0.00837 | 0.01096 | 0.00846 | 0.00956 | 0.00676 | 0.00751 | 0.06456 | 0.02651 | 0.02566 | 0.01244 | 0.00790 | 0.03769 | 0.04227 | 0.08204 | 0.00750 |
| **Snord22** | 0.05232 | 0.09819 | 0.89324 | 0.00147 | 0.65140 | 0.00670 | 0.76736 | 0.42252 | 0.50929 | 1.47775 | 0.32089 | 0.36678 | 2.00010 | 0.41769 | 0.26266 | 0.47048 |
| **Snord32a** | 0.08962 | 0.06165 | 0.12528 | 0.01056 | 0.01626 | 0.00569 | 0.02896 | 0.28262 | 0.01207 | 0.02213 | 0.01111 | 0.00768 | 0.33099 | 0.01227 | 0.00995 | 0.00706 |
| **Snord33** | 0.49361 | 0.16922 | 0.70476 | 0.00447 | 0.01580 | 0.01101 | 0.21439 | 0.69661 | 0.09019 | 0.33964 | 0.13987 | 0.02583 | 0.29363 | 0.02244 | 0.05807 | 0.14482 |
| **Snord34** | 0.26970 | 0.06687 | 0.75638 | 0.00636 | 0.01046 | 0.00570 | 0.14104 | 0.63023 | 0.01088 | 0.15868 | 0.08885 | 0.03677 | 0.48911 | 0.02798 | 0.25147 | 0.09325 |
| **Snord35a** | 0.38807 | 0.01861 | 1.06127 | 0.01478 | 0.00529 | 0.00538 | 0.40100 | 0.86988 | 0.04313 | 0.60720 | 0.41752 | 0.13107 | 0.24604 | 0.04203 | 1.03000 | 0.11506 |
| **Snord35b** | 0.01719 | 0.00767 | 0.24091 | 0.00965 | 0.00869 | 0.00571 | 0.17647 | 0.41190 | 0.01164 | 0.12215 | 0.01032 | 0.00631 | 0.02574 | 0.00838 | 0.06406 | 0.00750 |
| **Snord38a** | 0.03683 | 0.00815 | 0.04154 | 0.02538 | 0.00823 | 0.00598 | 0.00761 | 0.01482 | 0.06231 | 0.01379 | 0.03617 | 0.00826 | 0.01161 | 0.01238 | 0.00855 | 0.00728 |
| **Snord42a** | 0.16993 | 0.00790 | 0.12150 | 0.00810 | 0.03463 | 0.00852 | 0.00695 | 0.13729 | 0.03404 | 0.09553 | 0.01014 | 0.00608 | 0.08230 | 0.00762 | 0.19083 | 0.00725 |
| **Snord42b** | 0.08905 | 0.02857 | 0.02792 | 0.00942 | 0.01622 | 0.00584 | 0.00732 | 0.32061 | 0.01183 | 0.05555 | 0.00968 | 0.06815 | 0.12717 | 0.00667 | 0.15416 | 0.00713 |
| **Snord45b** | 0.01341 | 0.00772 | 0.01119 | 0.00965 | 0.00935 | 0.00659 | 0.07838 | 0.01704 | 0.01177 | 0.01776 | 0.01295 | 0.00761 | 0.01180 | 0.01154 | 0.19770 | 0.00729 |
| **Snord47** | 0.35078 | 0.34149 | 0.67581 | 0.01338 | 0.00843 | 0.01251 | 0.18050 | 0.55100 | 0.46941 | 0.62165 | 0.61153 | 0.92206 | 0.24267 | 0.55714 | 0.93997 | 0.21871 |
| **Snord49a** | 0.01425 | 0.00801 | 0.00784 | 0.00999 | 0.00968 | 0.45682 | 0.00761 | 0.01411 | 0.01258 | 0.01140 | 0.01320 | 0.00778 | 0.01730 | 0.00906 | 0.03840 | 0.00735 |
| **Snord49b** | 0.01325 | 0.00795 | 0.00919 | 0.00671 | 0.01095 | 0.00731 | 0.00742 | 0.13375 | 0.01149 | 0.09221 | 0.17800 | 0.00707 | 0.01752 | 0.01099 | 0.02820 | 0.15567 |
| **Snord4a** | 0.05437 | 0.01792 | 0.00190 | 0.02765 | 0.00970 | 0.01482 | 0.13790 | 0.27114 | 0.01191 | 0.09394 | 0.19922 | 0.04480 | 0.01535 | 0.00646 | 0.33812 | 0.00740 |
| **Snord52** | 0.01465 | 0.39137 | 0.63818 | 0.00482 | 0.01604 | 0.59001 | 0.10768 | 0.19684 | 0.02557 | 0.41159 | 0.05466 | 0.01670 | 0.05927 | 0.25692 | 0.15259 | 0.00688 |
| **Snord53** | 0.01245 | 0.00842 | 0.01053 | 0.01004 | 0.00939 | 0.00673 | 0.15979 | 0.03394 | 0.01220 | 0.01222 | 0.01262 | 0.00872 | 0.01089 | 0.00941 | 0.06224 | 0.00737 |
| **Snord55** | 0.01269 | 0.00796 | 0.01332 | 0.00894 | 0.00978 | 0.00693 | 0.00749 | 0.01889 | 0.01210 | 0.38465 | 0.01426 | 0.00661 | 0.10952 | 0.01209 | 0.00907 | 0.00777 |
| **Snord57** | 0.00985 | 0.18573 | 0.02618 | 0.00819 | 0.00785 | 0.00706 | 0.09621 | 0.19432 | 0.02217 | 0.24584 | 0.01189 | 0.00664 | 0.41837 | 0.01641 | 0.12460 | 0.54529 |
| **Snord58b** | 0.01592 | 0.00783 | 0.00346 | 0.00878 | 0.00918 | 0.42216 | 0.12184 | 0.06835 | 0.01213 | 0.20900 | 0.00995 | 0.02690 | 0.00569 | 0.06675 | 0.12238 | 0.00710 |
| **Snord65** | 0.01312 | 0.00821 | 0.34596 | 0.11004 | 0.00909 | 0.00743 | 0.00776 | 0.01892 | 0.24409 | 0.33998 | 0.11771 | 0.00589 | 0.02431 | 0.00783 | 0.15560 | 0.00757 |
| **Snord68** | 0.01348 | 0.00818 | 0.01298 | 0.01023 | 0.00976 | 0.00695 | 0.00765 | 0.01487 | 0.01225 | 0.05986 | 0.01381 | 0.01864 | 0.02683 | 0.01087 | 0.01765 | 0.01520 |
| **Snord69** | 0.04896 | 0.00764 | 0.17623 | 0.00896 | 0.00808 | 0.00768 | 0.03803 | 0.01061 | 0.01201 | 0.21907 | 0.01310 | 0.00587 | 0.02296 | 0.24829 | 0.08976 | 0.13477 |
| **Snord72** | 0.10943 | 0.22393 | 0.04017 | 0.35980 | 0.01610 | 0.00851 | 0.63100 | 0.15223 | 0.15180 | 0.60813 | 0.41317 | 0.02220 | 0.00502 | 0.02770 | 0.45116 | 0.02161 |
| **Snord73a** | 0.01219 | 0.00848 | 0.11982 | 0.00930 | 0.00905 | 0.43217 | 0.01927 | 0.07961 | 0.01244 | 0.13868 | 0.00994 | 0.00823 | 0.10565 | 0.00798 | 0.05982 | 0.00696 |
| **Snord8** | 0.01370 | 0.00807 | 0.01153 | 0.00964 | 0.00929 | 0.00659 | 0.00748 | 0.01199 | 0.01204 | 0.05569 | 0.01111 | 0.00705 | 0.01219 | 0.08029 | 0.17012 | 0.00722 |
| **Snord83b** | 0.37253 | 0.32815 | 0.19191 | 0.00417 | 0.00777 | 0.00629 | 0.16902 | 0.01926 | 0.08457 | 0.28195 | 0.20260 | 0.01700 | 0.01154 | 0.09653 | 0.13081 | 0.00666 |
| **Snord87** | 0.02027 | 0.00799 | 0.01122 | 0.01847 | 0.00885 | 0.00668 | 0.01813 | 0.02179 | 0.01109 | 0.01192 | 0.01243 | 0.01867 | 0.01206 | 0.00890 | 0.07285 | 0.00737 |
| **Snord89** | 0.01332 | 0.00846 | 0.01195 | 0.01022 | 0.00934 | 0.00667 | 0.00780 | 0.02319 | 0.01218 | 0.01361 | 0.01293 | 0.00886 | 0.01205 | 0.01036 | 0.04198 | 0.00740 |
| **Snord92** | 0.01316 | 0.00794 | 0.01273 | 0.01069 | 0.00973 | 0.00687 | 0.00756 | 0.02602 | 0.01228 | 0.01408 | 0.01380 | 0.00775 | 0.02138 | 0.01263 | 0.02086 | 0.00757 |
| **Snord95** | 0.01939 | 0.00741 | 0.14735 | 0.00713 | 0.00864 | 0.04012 | 0.32381 | 0.18062 | 0.01152 | 0.08270 | 0.10565 | 0.00626 | 0.05555 | 0.02776 | 0.09222 | 0.00749 |
| **Snord96a** | 0.59254 | 0.19849 | 0.65972 | 0.00457 | 0.00593 | 0.87734 | 0.16136 | 0.46865 | 0.13698 | 0.68173 | 0.00000 | 0.32328 | 0.67516 | 0.46857 | 0.10535 | 0.25234 |
| **Snord98** | 0.01684 | 0.00782 | 0.01084 | 0.01046 | 0.00908 | 0.00669 | 0.00735 | 0.02857 | 0.01225 | 0.01035 | 0.01224 | 0.00747 | 0.01185 | 0.10570 | 0.04041 | 0.00741 |
| **Snord99** | 0.22489 | 0.03744 | 0.16894 | 0.07343 | 0.01534 | 0.00701 | 0.34415 | 0.16478 | 0.01087 | 0.23417 | 0.14150 | 0.06693 | 0.05277 | 0.00718 | 0.26911 | 0.49737 |
| **Snph** | 0.01366 | 0.00814 | 0.01190 | 0.68595 | 0.00929 | 0.00656 | 0.00750 | 0.01368 | 0.71131 | 0.01105 | 0.00858 | 0.01758 | 0.01287 | 0.01068 | 0.00545 | 0.00726 |
| **Snrk** | 1.46299 | 0.22071 | 0.83570 | 0.54474 | 0.03303 | 0.00437 | 1.89959 | 0.32626 | 0.00446 | 0.63461 | 1.18792 | 0.02224 | 0.75427 | 0.04509 | 0.06753 | 0.30400 |
| **Snrnp200** | 11.01088 | 13.39104 | 8.04095 | 4.87316 | 7.17774 | 5.56899 | 4.97626 | 15.51758 | 10.81627 | 16.29636 | 10.16208 | 8.76431 | 5.39103 | 16.64890 | 11.57384 | 9.39625 |
| **Snrnp25** | 6.49491 | 0.03768 | 9.48372 | 9.28216 | 1.56446 | 0.00000 | 2.00910 | 5.25407 | 16.91817 | 10.41173 | 10.46154 | 2.58926 | 9.69721 | 1.26614 | 2.89203 | 3.08954 |
| **Snrnp27** | 12.85198 | 6.77992 | 5.43496 | 5.48889 | 11.87432 | 4.87816 | 2.02689 | 3.27538 | 4.74651 | 6.88779 | 10.69724 | 3.06418 | 1.53737 | 7.07234 | 8.91251 | 2.27571 |
| **Snrnp35** | 1.38253 | 1.67609 | 0.56181 | 3.18846 | 0.00511 | 0.00953 | 0.35312 | 0.85695 | 9.17066 | 0.73272 | 2.67673 | 0.09252 | 1.09248 | 0.98365 | 2.51643 | 0.00292 |
| **Snrnp40** | 12.82194 | 7.32622 | 9.22450 | 2.98519 | 8.08088 | 1.37250 | 7.81969 | 10.00458 | 3.72539 | 16.51432 | 5.88946 | 2.77869 | 12.11027 | 1.68296 | 6.18366 | 9.30094 |
| **Snrnp48** | 1.44175 | 3.77967 | 4.29032 | 3.65005 | 2.84781 | 0.00405 | 1.42147 | 4.85485 | 15.31004 | 5.98586 | 3.98135 | 0.95377 | 2.82712 | 1.12863 | 3.35203 | 1.30827 |
| **Snrnp70** | 3.22316 | 0.53472 | 0.95623 | 1.66852 | 1.24196 | 2.46214 | 0.68722 | 2.69653 | 2.26567 | 2.53077 | 2.51261 | 0.16478 | 2.21982 | 1.33545 | 0.71477 | 2.52314 |
| **Snrpa** | 19.03404 | 0.22325 | 0.00000 | 5.61496 | 14.31958 | 5.34172 | 0.92291 | 10.06244 | 15.26476 | 25.08327 | 9.69757 | 0.62639 | 19.28678 | 0.03563 | 0.64071 | 10.22533 |
| **Snrpa1** | 14.31124 | 8.11910 | 16.09162 | 14.86582 | 1.43288 | 5.58962 | 4.57801 | 10.99394 | 13.39895 | 20.88059 | 14.08093 | 7.51689 | 15.59100 | 0.78656 | 3.94551 | 6.08435 |
| **Snrpb** | 33.41202 | 1.08936 | 9.89425 | 6.67464 | 19.23335 | 12.61766 | 12.73913 | 38.19345 | 13.89852 | 35.46178 | 16.27755 | 2.38771 | 29.25992 | 1.19621 | 0.00000 | 18.14332 |
| **Snrpb2** | 20.78980 | 10.84338 | 11.65274 | 2.89751 | 8.31403 | 4.97897 | 11.47675 | 19.86804 | 1.62688 | 14.43495 | 11.53913 | 4.42652 | 10.70320 | 22.74447 | 10.95378 | 8.84580 |
| **Snrpc** | 20.89805 | 0.30552 | 0.38079 | 13.63592 | 11.00182 | 6.59668 | 5.76244 | 22.08856 | 23.25924 | 33.67394 | 19.12015 | 4.89324 | 20.76953 | 1.14898 | 0.00000 | 6.88112 |
| **Snrpd1** | 14.41171 | 5.35579 | 13.40713 | 19.61995 | 14.58223 | 2.66283 | 3.89069 | 7.99250 | 27.24841 | 10.79668 | 22.08621 | 2.57978 | 16.97997 | 11.54267 | 9.14783 | 7.71085 |
| **Snrpd2** | 20.05110 | 1.59391 | 12.59384 | 9.26873 | 7.91991 | 2.28097 | 3.88813 | 3.40246 | 18.77277 | 14.19343 | 18.81779 | 1.64667 | 11.22605 | 0.12342 | 4.33710 | 8.43986 |
| **Snrpd3** | 4.87943 | 1.96863 | 4.45831 | 6.56808 | 9.79173 | 5.86420 | 0.77700 | 5.66691 | 9.63384 | 6.57385 | 21.71005 | 0.77732 | 9.75935 | 8.54095 | 4.85674 | 7.72656 |
| **Snrpe** | 12.91586 | 1.75622 | 10.62092 | 13.77545 | 10.96820 | 2.25174 | 2.15094 | 2.02864 | 22.25333 | 10.54945 | 29.48556 | 2.15662 | 12.07245 | 12.98650 | 9.37705 | 7.31996 |
| **Snrpf** | 1.72119 | 0.02570 | 0.16198 | 0.66318 | 0.68587 | 0.02463 | 0.01526 | 0.00000 | 0.85293 | 1.71098 | 1.21448 | 0.05244 | 0.76956 | 0.27946 | 0.00000 | 0.61478 |
| **Snrpg** | 13.55386 | 0.78851 | 12.62167 | 24.56939 | 11.99380 | 2.53790 | 1.66862 | 0.42847 | 23.07568 | 6.77697 | 26.57575 | 1.99824 | 5.37155 | 6.08656 | 5.48023 | 3.71633 |
| **Snta1** | 1.04854 | 0.00691 | 0.00000 | 2.58532 | 0.01653 | 0.01329 | 0.17721 | 1.20217 | 3.64701 | 3.40431 | 1.52420 | 0.04917 | 3.10442 | 0.03137 | 0.40476 | 0.00439 |

| | | | | | | | | | | | | | | | | |
|---|---|---|---|---|---|---|---|---|---|---|---|---|---|---|---|---|
| **Sntb2** | 0.18946 | 3.11718 | 0.66904 | 3.03822 | 0.07416 | 0.00445 | 0.00532 | 0.00803 | 6.08533 | 1.27976 | 0.53658 | 0.85902 | 0.47115 | 3.99234 | 1.30546 | 0.03367 |
| **Sntg2** | 0.01372 | 0.00827 | 0.01225 | 0.93095 | 0.00932 | 0.00657 | 0.00751 | 0.01400 | 0.18646 | 0.01398 | 0.01133 | 0.00813 | 0.01273 | 0.01090 | 0.00352 | 0.00725 |
| **Snupn** | 4.99878 | 1.22067 | 4.48436 | 17.12046 | 0.09530 | 0.00400 | 0.02828 | 5.87579 | 18.86595 | 19.88353 | 9.56882 | 6.90121 | 0.24507 | 10.62809 | 16.72858 | 2.00472 |
| **Snw1** | 4.36042 | 4.23123 | 4.41564 | 5.91810 | 5.49038 | 0.29259 | 1.51809 | 0.05323 | 5.00076 | 2.45094 | 5.17259 | 0.28864 | 1.74724 | 8.95936 | 1.83307 | 2.46809 |
| **Snx1** | 15.83888 | 10.11512 | 20.90756 | 2.13224 | 3.92069 | 14.02341 | 11.19630 | 23.29217 | 6.21966 | 15.12646 | 19.79216 | 6.95798 | 15.87113 | 0.32297 | 5.12310 | 4.17916 |
| **Snx10** | 9.39901 | 4.41956 | 3.42716 | 1.20081 | 4.46994 | 2.37209 | 2.01854 | 2.44028 | 1.11471 | 3.81653 | 1.52440 | 0.03245 | 0.90832 | 4.12243 | 3.31231 | 3.38454 |
| **Snx11** | 0.56842 | 0.02077 | 0.09521 | 1.62567 | 0.01770 | 0.00787 | 0.03923 | 2.62353 | 1.62810 | 6.85807 | 0.16235 | 0.35238 | 1.10912 | 0.00861 | 0.00000 | 0.01372 |
| **Snx12** | 6.05674 | 2.63650 | 3.30607 | 4.74523 | 1.40542 | 0.80497 | 3.18058 | 4.21355 | 5.08806 | 4.88189 | 2.27444 | 0.19012 | 2.61701 | 3.25306 | 4.09549 | 1.87102 |
| **Snx13** | 2.41692 | 3.20246 | 0.83176 | 2.79766 | 0.01436 | 2.28599 | 1.57203 | 0.07851 | 0.44425 | 0.44399 | 0.28811 | 7.37266 | 1.26621 | 0.57361 | 2.92467 | 1.21174 |
| **Snx14** | 3.68069 | 0.90772 | 0.00000 | 2.06617 | 0.00000 | 0.00681 | 2.25098 | 3.95862 | 3.56442 | 5.33822 | 1.84765 | 7.11681 | 0.14892 | 5.36541 | 0.47583 | 0.84803 |
| **Snx15** | 7.95807 | 0.02940 | 0.97655 | 6.31455 | 0.13634 | 2.43055 | 2.76857 | 2.44200 | 1.44958 | 3.86018 | 1.83504 | 0.15653 | 2.36120 | 0.07121 | 0.78388 | 0.03946 |
| **Snx16** | 4.50982 | 0.03977 | 2.20925 | 2.88708 | 0.30773 | 0.01405 | 2.40756 | 4.52890 | 3.63435 | 12.29003 | 7.09435 | 9.94685 | 4.12503 | 14.67234 | 6.77102 | 0.01930 |
| **Snx17** | 7.38635 | 4.91635 | 7.48896 | 0.40392 | 4.52676 | 0.00617 | 2.11841 | 2.95683 | 5.59519 | 3.28269 | 0.31869 | 0.10904 | 4.26059 | 0.76214 | 0.60651 | 7.02290 |
| **Snx18** | 2.46611 | 2.11909 | 2.71098 | 10.59293 | 3.99724 | 2.87252 | 0.00156 | 3.13843 | 8.30057 | 5.59593 | 4.75100 | 0.13020 | 0.67392 | 5.02444 | 5.75781 | 2.07837 |
| **Snx19** | 0.01167 | 0.00794 | 0.16838 | 0.29469 | 0.01133 | 0.00572 | 0.00728 | 0.41782 | 0.00433 | 0.32806 | 0.09212 | 0.02202 | 0.00651 | 0.33150 | 0.96642 | 0.00774 |
| **Snx2** | 24.56614 | 26.52733 | 26.99609 | 2.48174 | 29.35023 | 22.52202 | 17.95807 | 22.86652 | 1.51726 | 14.81541 | 13.37062 | 9.33017 | 25.08960 | 15.71994 | 12.24734 | 24.74433 |
| **Snx21** | 1.04394 | 0.00746 | 0.00712 | 0.65932 | 0.01588 | 0.00574 | 0.00651 | 0.01543 | 0.25102 | 0.33506 | 0.00934 | 0.04689 | 0.05067 | 0.01306 | 0.03274 | 0.96968 |
| **Snx22** | 0.01333 | 0.00764 | 0.00000 | 1.40224 | 0.01046 | 0.95030 | 0.02602 | 0.27532 | 0.02992 | 0.95945 | 2.06311 | 0.03138 | 0.04156 | 0.00407 | 1.51827 | 0.00680 |
| **Snx24** | 0.00898 | 0.00801 | 0.07578 | 0.00730 | 0.00870 | 0.00866 | 0.00709 | 0.01239 | 0.01182 | 0.99918 | 0.01069 | 0.01485 | 0.67576 | 0.01229 | 0.00713 | 0.00708 |
| **Snx25** | 0.96673 | 0.00568 | 0.40326 | 0.01055 | 0.00593 | 0.03706 | 1.01664 | 0.00697 | 0.01181 | 0.00392 | 0.01295 | 0.01484 | 0.05021 | 0.05718 | 0.09103 | 0.00649 |
| **Snx27** | 12.09718 | 5.22874 | 9.56729 | 1.52744 | 9.67460 | 5.25052 | 9.28975 | 10.53471 | 4.27238 | 12.19443 | 10.81789 | 2.75736 | 5.76841 | 12.87862 | 7.93874 | 8.58549 |
| **Snx29** | 0.24722 | 0.00736 | 0.00614 | 0.80914 | 0.00744 | 0.00671 | 0.00749 | 0.01636 | 0.00000 | 0.94156 | 0.01093 | 0.00457 | 0.01096 | 0.00604 | 0.31890 | 0.00701 |
| **Snx3** | 18.18539 | 7.56283 | 17.37136 | 11.85656 | 19.49654 | 10.10393 | 6.90661 | 15.66380 | 23.40425 | 30.82849 | 28.78547 | 19.04892 | 22.84595 | 3.29286 | 7.29903 | 11.79091 |
| **Snx30** | 1.68298 | 3.00164 | 1.91408 | 1.51308 | 0.05091 | 6.32330 | 4.24224 | 5.09593 | 1.53331 | 9.71570 | 6.55335 | 1.54471 | 4.33128 | 11.47370 | 5.32699 | 2.11674 |
| **Snx32** | 0.01387 | 0.00832 | 0.00957 | 0.00558 | 0.00913 | 0.00679 | 0.00753 | 0.00697 | 0.01136 | 0.00472 | 0.32055 | 0.01832 | 0.00881 | 0.00833 | 0.03283 | 0.00722 |
| **Snx33** | 0.01377 | 0.00811 | 0.01207 | 0.00900 | 0.00930 | 0.00661 | 0.00743 | 0.01253 | 0.01172 | 0.02484 | 0.01284 | 0.00776 | 0.01302 | 0.00686 | 0.22275 | 0.00762 |
| **Snx4** | 12.33674 | 6.51357 | 4.75423 | 5.67881 | 6.10569 | 6.44581 | 7.64888 | 9.02134 | 8.50345 | 11.04120 | 5.21111 | 2.73639 | 2.69859 | 15.75653 | 7.37739 | 1.00658 |
| **Snx5** | 24.87195 | 17.31219 | 15.28996 | 9.54005 | 14.85740 | 12.54803 | 13.07827 | 23.09103 | 14.51410 | 28.46612 | 32.28735 | 27.46321 | 19.44193 | 27.02326 | 21.73206 | 9.40565 |
| **Snx6** | 17.77539 | 11.80979 | 15.16205 | 10.12599 | 9.57811 | 10.05851 | 11.97061 | 17.01265 | 10.59614 | 25.20532 | 14.24459 | 13.39180 | 16.02325 | 7.70542 | 15.45887 | 7.78958 |
| **Snx7** | 0.01182 | 0.00765 | 0.03106 | 0.00491 | 0.00899 | 0.30706 | 0.01748 | 0.01223 | 0.40483 | 0.22083 | 0.00790 | 0.01679 | 0.19440 | 0.03025 | 0.22572 | 0.00685 |
| **Snx8** | 3.68280 | 1.79829 | 4.97372 | 3.50899 | 0.00874 | 0.00794 | 0.00000 | 4.68639 | 2.54149 | 5.88215 | 9.47106 | 0.35814 | 2.61738 | 0.04029 | 1.23108 | 2.91190 |
| **Snx9** | 0.95035 | 3.03716 | 2.17916 | 0.85897 | 3.30702 | 1.09215 | 0.72663 | 0.82790 | 0.05560 | 0.63044 | 0.30587 | 0.05479 | 1.69244 | 0.12356 | 2.84541 | 3.77148 |
| **Soat1** | 0.82859 | 2.62782 | 2.54523 | 2.43934 | 0.05089 | 0.01267 | 1.78779 | 0.00000 | 1.43495 | 3.04525 | 3.92706 | 0.16745 | 1.34906 | 7.51589 | 5.76370 | 1.66442 |
| **Soat2** | 0.03266 | 0.02866 | 0.02983 | 0.92028 | 0.11684 | 0.02994 | 0.03628 | 0.59026 | 3.16548 | 0.68864 | 2.55115 | 0.16808 | 0.21721 | 1.25851 | 1.90606 | 0.04980 |
| **Socs1** | 0.01760 | 0.00786 | 0.09457 | 1.22650 | 0.00834 | 0.00864 | 0.04677 | 0.10393 | 0.93339 | 0.01229 | 0.79920 | 0.01147 | 0.24761 | 0.02099 | 0.00000 | 0.14096 |
| **Socs2** | 7.34753 | 0.57577 | 0.87763 | 4.64700 | 10.87658 | 0.03508 | 0.08840 | 0.13354 | 2.32964 | 3.35061 | 0.10172 | 0.15461 | 2.33582 | 0.00000 | 0.36925 | 0.13702 |
| **Socs3** | 4.07729 | 0.16842 | 0.00000 | 3.76900 | 8.16709 | 0.00000 | 4.50284 | 9.37033 | 15.34440 | 13.16892 | 9.72735 | 1.14943 | 0.00000 | 0.95620 | 10.34198 | 2.02940 |
| **Socs4** | 2.15483 | 0.00444 | 0.00000 | 3.13661 | 0.00515 | 0.00432 | 0.00539 | 0.35848 | 2.56923 | 0.51975 | 0.45383 | 5.47841 | 0.20701 | 2.05554 | 1.35674 | 1.67681 |
| **Socs5** | 0.38397 | 0.00738 | 0.09984 | 2.59226 | 0.00923 | 0.00845 | 0.00701 | 0.00833 | 1.48600 | 0.46770 | 0.02471 | 0.01431 | 0.01138 | 0.00265 | 0.58433 | 0.00650 |
| **Socs6** | 7.29926 | 1.43139 | 0.00000 | 3.51919 | 6.16121 | 0.00000 | 0.23750 | 4.33020 | 2.25457 | 3.68552 | 0.00000 | 0.33766 | 0.62186 | 0.11632 | 5.12228 | 1.38468 |
| **Socs7** | 0.58092 | 0.00409 | 0.83973 | 10.66954 | 2.25152 | 1.08752 | 2.60565 | 0.00000 | 11.83527 | 6.79009 | 2.42130 | 0.14423 | 0.03047 | 9.08833 | 11.13216 | 0.14996 |
| **Sod1** | 28.27008 | 4.94256 | 19.86451 | 11.44098 | 23.03016 | 7.96176 | 5.42870 | 10.63731 | 8.72010 | 10.33783 | 14.09304 | 1.62710 | 17.45070 | 0.75046 | 0.25721 | 9.01343 |
| **Sod2** | 17.09965 | 9.40303 | 14.79682 | 15.12294 | 15.38460 | 10.49009 | 9.92147 | 9.34213 | 19.25982 | 16.12095 | 19.22152 | 15.34770 | 23.31210 | 10.86732 | 4.76625 | 15.36154 |
| **Soga1** | 0.64509 | 0.02126 | 0.03729 | 0.85307 | 0.03603 | 0.01345 | 0.04082 | 0.20479 | 1.21693 | 0.00000 | 1.40927 | 0.15949 | 0.01451 | 0.04099 | 0.31625 | 0.01377 |
| **Sohlh1** | 0.00883 | 0.00866 | 0.00638 | 0.14690 | 0.00939 | 0.00719 | 0.00821 | 0.43659 | 0.00939 | 0.49200 | 0.23689 | 0.02927 | 0.00464 | 0.01030 | 0.00000 | 0.00706 |
| **Son** | 11.24609 | 5.35054 | 5.11621 | 14.57666 | 7.05967 | 2.50532 | 7.39604 | 7.72453 | 22.11678 | 15.32078 | 14.80181 | 17.14739 | 8.70451 | 20.56813 | 15.67823 | 4.54134 |
| **Sorbs1** | 3.14874 | 2.86051 | 6.27606 | 13.80186 | 0.03768 | 4.24037 | 0.72407 | 6.45963 | 5.88098 | 10.43949 | 5.26540 | 6.59539 | 3.82221 | 5.40318 | 8.83831 | 0.00000 |
| **Sorbs2** | 0.01373 | 0.00788 | 0.01118 | 1.95850 | 0.00923 | 0.00670 | 0.00750 | 0.01235 | 0.09010 | 0.10002 | 0.03950 | 0.01971 | 0.01258 | 0.01116 | 0.15492 | 0.00707 |
| **Sorbs3** | 0.01426 | 0.00660 | 0.16695 | 0.02721 | 0.00888 | 0.03504 | 2.11633 | 0.08936 | 0.50599 | 0.12272 | 0.75639 | 0.00094 | 0.00884 | 0.00512 | 0.08183 | 0.00712 |
| **Sorcs2** | 0.01241 | 0.00803 | 0.01183 | 0.12154 | 0.00894 | 0.00662 | 0.00749 | 0.01342 | 0.00710 | 0.01203 | 0.01211 | 0.02894 | 0.01150 | 0.01186 | 0.00788 | 0.17232 |
| **Sorcs3** | 0.01372 | 0.00796 | 0.01238 | 0.11716 | 0.00940 | 0.00655 | 0.00743 | 0.01289 | 0.00996 | 0.09186 | 0.01379 | 0.00745 | 0.01264 | 0.01101 | 0.00757 | 0.00746 |
| **Sord** | 20.08670 | 25.94641 | 33.79241 | 11.74386 | 24.21257 | 25.29549 | 22.11388 | 41.07154 | 15.59977 | 42.87077 | 39.25278 | 19.41987 | 40.77818 | 40.05382 | 31.77186 | 18.78723 |
| **Sorl1** | 0.00760 | 0.00853 | 0.00773 | 1.43673 | 0.00908 | 0.00646 | 0.00784 | 0.37162 | 0.18339 | 0.99281 | 0.00422 | 0.09530 | 0.58093 | 0.02114 | 0.44830 | 0.00656 |
| **Sort1** | 0.02110 | 0.00774 | 0.01044 | 0.94953 | 0.00857 | 0.00645 | 0.00729 | 0.01114 | 0.00000 | 1.08906 | 0.00862 | 0.00349 | 0.01827 | 0.00588 | 0.97899 | 0.00728 |
| **Sos1** | 0.98971 | 0.00000 | 0.55303 | 10.63113 | 1.55511 | 2.53136 | 0.01563 | 10.19134 | 19.14413 | 12.84249 | 11.82159 | 19.14030 | 5.46413 | 20.57952 | 19.04313 | 0.77333 |
| **Sos2** | 2.62126 | 2.83767 | 0.00000 | 1.99441 | 0.02292 | 1.57746 | 2.43654 | 3.14887 | 0.18531 | 0.81041 | 1.86068 | 11.65868 | 0.89532 | 2.04426 | 1.96974 | 0.61590 |
| **Sowaha** | 0.01125 | 0.00768 | 0.00828 | 4.48177 | 0.00920 | 0.00606 | 0.00689 | 0.15690 | 2.06516 | 0.00000 | 0.00000 | 0.05625 | 0.11823 | 0.26708 | 0.00000 | 0.00629 |
| **Sowahb** | 1.10881 | 0.00688 | 0.00238 | 0.01023 | 0.03644 | 0.00763 | 0.01595 | 0.79855 | 0.02238 | 0.00865 | 0.00815 | 0.00508 | 0.00553 | 0.01402 | 0.00326 | 0.00646 |
| **Sowahc** | 0.17023 | 0.00666 | 1.97594 | 1.15455 | 0.00387 | 0.00694 | 0.00676 | 1.27697 | 0.00000 | 1.88025 | 0.07326 | 1.89416 | 0.00000 | 0.50748 | 1.62483 | 0.10455 |
| **Sox11** | 0.01317 | 0.00749 | 0.00932 | 0.21843 | 0.00928 | 0.00633 | 0.00725 | 0.01427 | 0.00612 | 0.20549 | 0.01479 | 0.02916 | 0.01259 | 0.00772 | 2.36371 | 0.00776 |
| **Sox12** | 1.43123 | 0.00440 | 0.00000 | 0.17630 | 0.83517 | 0.00497 | 3.91391 | 2.57661 | 0.56014 | 3.04680 | 0.15959 | 0.12362 | 1.29426 | 0.34141 | 1.24572 | 0.93570 |
| **Sox13** | 4.97339 | 1.70111 | 4.85744 | 5.97550 | 0.30956 | 3.67217 | 4.21727 | 3.36495 | 4.74749 | 5.70504 | 5.75355 | 1.56771 | 10.22650 | 0.00000 | 1.17335 | 6.01720 |

| | | | | | | | | | | | | | | | | |
|---|---|---|---|---|---|---|---|---|---|---|---|---|---|---|---|---|
| **Sox15** | 6.09701 | 0.94010 | 0.89841 | 0.14585 | 2.03991 | 6.08321 | 2.74754 | 22.91585 | 6.53560 | 33.54810 | 12.91232 | 2.00248 | 10.37550 | 0.15502 | 1.31474 | 1.82367 |
| **Sox17** | 6.76537 | 0.00000 | 0.00000 | 0.00000 | 5.29114 | 0.00007 | 0.00161 | 0.00000 | 0.00998 | 1.31257 | 0.00980 | 0.00000 | 0.01671 | 0.01091 | 0.01264 | 4.60382 |
| **Sox2** | 13.88850 | 8.24872 | 0.00000 | 0.41138 | 2.93616 | 0.00000 | 8.29069 | 0.62117 | 0.32373 | 0.00000 | 0.13411 | 0.03440 | 0.01158 | 0.01364 | 0.12162 | 0.04551 |
| **Sox21** | 3.38709 | 3.62649 | 0.00000 | 0.00000 | 0.00911 | 0.76758 | 1.71751 | 8.63116 | 0.02249 | 10.78204 | 3.28878 | 11.71746 | 0.60038 | 5.23853 | 3.35049 | 0.00000 |
| **Sox2ot** | 0.01501 | 0.00783 | 0.02672 | 0.11488 | 0.01811 | 0.00742 | 0.02776 | 0.02269 | 0.02772 | 0.01875 | 0.19884 | 0.01657 | 0.02568 | 0.14333 | 0.06586 | 0.05830 |
| **Sox30** | 0.28106 | 0.00756 | 0.00618 | 0.18001 | 0.00735 | 0.00697 | 0.00733 | 0.01154 | 0.23094 | 0.00773 | 0.01095 | 0.00775 | 0.01248 | 0.02804 | 0.12154 | 0.00703 |
| **Sox4** | 0.01360 | 0.00800 | 0.01180 | 0.37066 | 0.00925 | 0.00649 | 0.00740 | 0.01196 | 0.90444 | 0.11799 | 0.00736 | 0.00667 | 0.01211 | 0.03907 | 0.00651 | 0.00712 |
| **Sox6** | 0.00764 | 0.00842 | 1.07492 | 0.01935 | 0.00812 | 0.00944 | 0.00808 | 0.36986 | 0.01071 | 0.00845 | 0.00845 | 0.00761 | 0.00001 | 0.01699 | 0.00000 | 0.00705 |
| **Sox7** | 0.01654 | 0.00836 | 0.30167 | 0.01992 | 0.00842 | 0.00569 | 0.00762 | 0.01336 | 0.04372 | 0.02355 | 0.02811 | 0.00848 | 0.00976 | 0.03624 | 0.01798 | 0.01441 |
| **Sp1** | 3.75156 | 9.25567 | 5.31820 | 8.01600 | 5.06854 | 5.50099 | 7.96419 | 8.08974 | 9.37462 | 13.33855 | 5.09118 | 3.45423 | 4.44709 | 23.03167 | 13.78619 | 5.07921 |
| **Sp100** | 0.22656 | 0.03965 | 0.09134 | 0.51180 | 0.00787 | 0.02771 | 0.01758 | 0.08326 | 0.97956 | 0.36760 | 0.51013 | 0.11412 | 0.24348 | 0.72367 | 0.50106 | 0.04381 |
| **Sp110** | 0.00000 | 2.70603 | 14.55283 | 29.03194 | 0.17022 | 4.20735 | 9.37866 | 41.04313 | 31.40897 | 53.01300 | 32.94491 | 25.42797 | 21.60966 | 36.99385 | 43.91001 | 2.34916 |
| **Sp140** | 2.94279 | 0.29761 | 1.33128 | 15.96341 | 1.91818 | 1.02427 | 0.37976 | 6.07663 | 14.43697 | 15.16494 | 5.54456 | 2.76120 | 1.75383 | 16.99182 | 17.55358 | 0.32799 |
| **Sp2** | 2.53314 | 0.02615 | 0.03647 | 2.08639 | 2.39695 | 0.00589 | 0.76780 | 2.86921 | 2.58256 | 7.30232 | 1.55142 | 0.32000 | 0.57559 | 0.68390 | 0.41184 | 1.15077 |
| **Sp3** | 0.69653 | 3.07342 | 0.35961 | 4.25258 | 0.33377 | 0.54265 | 0.08779 | 0.23590 | 4.41408 | 0.00000 | 1.73355 | 0.10128 | 0.42901 | 20.05002 | 6.63894 | 0.40742 |
| **Sp3os** | 0.66406 | 0.00627 | 0.72923 | 18.04575 | 0.04454 | 0.01580 | 0.08406 | 0.00545 | 10.16254 | 1.58269 | 8.21019 | 0.30983 | 0.38440 | 3.03001 | 1.41598 | 0.00850 |
| **Sp4** | 0.00000 | 2.88795 | 0.89202 | 1.71279 | 2.12751 | 0.65825 | 0.00613 | 0.02681 | 1.90834 | 0.23151 | 0.00000 | 0.00000 | 0.00476 | 4.08691 | 0.00000 | 0.00547 |
| **Sp5** | 0.01371 | 0.00817 | 0.01225 | 0.85686 | 0.00930 | 0.00656 | 0.00750 | 0.01396 | 0.23485 | 0.01375 | 0.01236 | 0.00785 | 0.01271 | 0.01048 | 0.00540 | 0.00726 |
| **Sp9** | 1.98385 | 0.00570 | 0.00349 | 0.07230 | 0.00480 | 0.00647 | 0.00591 | 0.01255 | 0.13662 | 0.01449 | 0.01088 | 0.01687 | 0.01107 | 0.01226 | 0.00798 | 0.00527 |
| **Spa17** | 1.64021 | 0.00614 | 0.00402 | 0.98545 | 0.00473 | 0.00592 | 0.00547 | 0.15830 | 0.24860 | 0.03218 | 0.03873 | 0.01596 | 0.78265 | 0.06354 | 0.00002 | 0.00493 |
| **Spaca6** | 0.07300 | 0.00726 | 0.00489 | 0.00789 | 0.00779 | 0.00747 | 0.00713 | 0.26832 | 0.01144 | 0.11571 | 0.10892 | 0.01759 | 0.01416 | 0.01962 | 0.00896 | 0.00707 |
| **Spag1** | 0.63812 | 0.03286 | 0.00000 | 3.44667 | 0.01516 | 0.21494 | 0.02884 | 0.03472 | 4.97454 | 2.52280 | 0.40253 | 1.08647 | 0.00535 | 0.05094 | 5.44449 | 0.00575 |
| **Spag16** | 0.01304 | 0.00815 | 0.02335 | 0.09132 | 0.00934 | 0.00686 | 0.00744 | 0.01852 | 0.00782 | 0.01302 | 0.01238 | 0.00816 | 0.01214 | 0.00765 | 0.12095 | 0.00743 |
| **Spag17** | 0.01361 | 0.00775 | 0.01207 | 0.00897 | 0.00911 | 0.00664 | 0.00732 | 0.01089 | 0.01198 | 0.43002 | 0.01280 | 0.00629 | 0.01228 | 0.01273 | 0.01600 | 0.00780 |
| **Spag4** | 0.01186 | 0.00725 | 0.00000 | 0.23442 | 0.02038 | 0.01467 | 0.30265 | 0.82335 | 0.06820 | 0.00000 | 1.23259 | 0.05387 | 1.34489 | 0.00335 | 0.26561 | 0.00640 |
| **Spag5** | 3.85309 | 4.00838 | 0.35063 | 6.60317 | 1.03802 | 4.00804 | 1.77722 | 3.50792 | 6.32749 | 1.78686 | 4.36959 | 1.55714 | 1.33263 | 5.04783 | 7.92532 | 1.37413 |
| **Spag6l** | 0.44439 | 0.00756 | 0.00267 | 0.00601 | 0.00627 | 0.00691 | 0.00695 | 0.01687 | 0.01069 | 0.48818 | 0.01000 | 0.00620 | 0.01522 | 0.00608 | 0.53439 | 1.66594 |
| **Spag7** | 20.01880 | 13.36344 | 16.22074 | 1.73540 | 11.01565 | 7.41473 | 9.97718 | 18.85709 | 2.44806 | 16.27370 | 15.87155 | 4.43770 | 20.66354 | 1.37934 | 10.30385 | 7.95507 |
| **Spag9** | 5.34979 | 9.03231 | 2.21493 | 3.90653 | 5.77681 | 0.36435 | 0.86803 | 1.85997 | 7.89945 | 2.68943 | 4.15716 | 13.85303 | 2.42738 | 15.25836 | 7.23110 | 1.91616 |
| **Spam1** | 0.01374 | 0.00754 | 0.01122 | 0.52261 | 0.00945 | 0.00641 | 0.00734 | 0.01105 | 0.00307 | 0.00459 | 0.01052 | 0.00596 | 0.01292 | 0.01896 | 0.80709 | 0.00729 |
| **Sparc** | 11.60000 | 3.71946 | 0.92324 | 1.97487 | 8.35873 | 0.00653 | 0.00000 | 0.04007 | 0.40391 | 3.57470 | 1.09810 | 0.58867 | 0.00000 | 0.00000 | 1.21212 | 18.68813 |
| **Sparcl1** | 0.01153 | 0.00780 | 0.00650 | 0.00665 | 0.00913 | 0.00672 | 0.00715 | 0.00948 | 0.01151 | 1.01121 | 0.00602 | 0.00000 | 0.01175 | 1.58313 | 0.00000 | 0.00775 |
| **Spast** | 5.65848 | 5.15546 | 1.13887 | 18.07977 | 0.02148 | 3.97114 | 5.45043 | 3.98211 | 15.68451 | 8.28204 | 4.53354 | 3.84673 | 1.35377 | 6.99195 | 13.39234 | 0.02921 |
| **Spata1** | 0.01091 | 0.00810 | 0.17488 | 0.09302 | 0.02756 | 0.00574 | 0.00759 | 0.02370 | 0.00781 | 0.01462 | 0.01186 | 0.00834 | 0.01007 | 0.01157 | 0.01022 | 0.00727 |
| **Spata13** | 0.00171 | 0.00752 | 1.05834 | 0.36907 | 0.00824 | 0.00784 | 0.00462 | 0.00572 | 0.06976 | 0.31194 | 0.04379 | 0.00247 | 4.50560 | 0.00510 | 0.13737 | 0.00502 |
| **Spata17** | 0.02476 | 0.00799 | 0.20616 | 0.00495 | 0.00836 | 0.00569 | 0.00747 | 0.00481 | 0.07390 | 0.00000 | 0.48514 | 0.00588 | 0.00678 | 0.00656 | 0.06072 | 0.00703 |
| **Spata2** | 6.96970 | 5.04800 | 6.68051 | 17.83850 | 0.30976 | 2.82784 | 8.01943 | 6.52491 | 19.15846 | 8.48003 | 8.34964 | 1.76294 | 6.01328 | 3.95892 | 8.33065 | 2.39322 |
| **Spata21** | 0.01411 | 0.01755 | 0.00568 | 4.09747 | 0.00812 | 0.00573 | 0.00697 | 0.00799 | 3.26458 | 2.19746 | 0.00000 | 0.31706 | 0.04273 | 0.06352 | 1.88223 | 0.41176 |
| **Spata22** | 0.01182 | 0.00704 | 0.02306 | 0.01019 | 0.00943 | 0.00672 | 1.26495 | 0.00963 | 0.01230 | 0.01159 | 0.01324 | 0.00673 | 0.01050 | 0.01230 | 0.00966 | 0.00732 |
| **Spata24** | 0.36878 | 0.85417 | 0.44234 | 0.64069 | 0.04280 | 1.80215 | 1.12580 | 0.03847 | 0.12087 | 0.99613 | 1.02465 | 0.00706 | 0.07249 | 0.00532 | 0.47425 | 0.01316 |
| **Spata25** | 0.01370 | 0.00791 | 0.01236 | 0.01854 | 0.00941 | 0.00655 | 0.00741 | 0.01240 | 0.01125 | 0.14913 | 0.01275 | 0.00721 | 0.01252 | 0.01151 | 0.00913 | 0.00748 |
| **Spata2l** | 0.01648 | 0.00801 | 0.17793 | 0.65462 | 0.00892 | 0.00568 | 0.00732 | 0.01232 | 0.05511 | 0.01362 | 0.01240 | 0.01786 | 0.01063 | 0.03286 | 0.00000 | 0.00718 |
| **Spata33** | 0.01374 | 0.00820 | 0.01228 | 0.61869 | 0.00930 | 0.00659 | 0.00751 | 0.01395 | 0.07064 | 0.01484 | 0.01324 | 0.00825 | 0.01268 | 0.01093 | 0.00598 | 0.00731 |
| **Spata5** | 4.03223 | 0.02986 | 4.91111 | 8.74700 | 2.69119 | 0.00000 | 2.71539 | 2.59769 | 20.04546 | 8.30003 | 8.18565 | 4.86686 | 2.26092 | 2.77301 | 8.99833 | 5.19409 |
| **Spata5l1** | 0.01096 | 0.00770 | 0.39573 | 0.00973 | 0.00874 | 0.00755 | 0.00715 | 0.01633 | 0.01216 | 0.18802 | 0.01193 | 0.00648 | 0.24629 | 0.01046 | 0.05864 | 0.00722 |
| **Spata6** | 0.01215 | 0.00774 | 0.02610 | 1.05402 | 0.01944 | 0.00781 | 0.00748 | 0.01302 | 0.34358 | 0.11048 | 0.01127 | 0.00549 | 0.03386 | 0.00321 | 0.33064 | 0.00782 |
| **Spata7** | 4.36862 | 6.82127 | 1.94618 | 1.13164 | 0.00739 | 0.01038 | 1.97657 | 3.94731 | 0.45156 | 6.13779 | 1.78968 | 4.32604 | 1.10889 | 0.01869 | 3.88678 | 0.01338 |
| **Spats1** | 0.07656 | 0.00731 | 0.18243 | 1.05402 | 0.00916 | 0.00539 | 0.00645 | 0.00094 | 0.46451 | 0.81819 | 0.24560 | 0.02094 | 0.84343 | 0.00514 | 0.68859 | 0.00651 |
| **Spats2** | 3.12521 | 5.76172 | 2.26498 | 0.18196 | 0.49111 | 0.73989 | 2.13808 | 0.51164 | 0.10073 | 0.23982 | 0.00000 | 0.06077 | 0.14380 | 0.13108 | 0.64287 | 1.97746 |
| **Spats2l** | 0.01277 | 0.00805 | 0.00722 | 0.00827 | 0.00960 | 0.00668 | 0.00720 | 0.00969 | 0.01111 | 0.00407 | 0.00674 | 0.00378 | 0.01365 | 1.01660 | 0.40252 | 0.00753 |
| **Spc24** | 15.50906 | 0.87108 | 3.79980 | 0.89753 | 7.52302 | 4.57673 | 0.97970 | 4.78182 | 1.62074 | 7.41920 | 11.00514 | 0.40324 | 6.97377 | 0.47669 | 0.83918 | 3.77643 |
| **Spc25** | 8.79978 | 6.60469 | 6.49740 | 10.18857 | 2.98342 | 6.24428 | 1.26791 | 4.72157 | 15.64150 | 5.28777 | 4.98348 | 16.69730 | 3.21039 | 2.04477 | 7.48246 | 4.81581 |
| **Spcs1** | 15.74126 | 5.57244 | 10.53281 | 5.59052 | 9.47907 | 4.48107 | 2.61724 | 7.83731 | 6.23610 | 12.24618 | 10.13721 | 2.58463 | 11.66063 | 8.88727 | 5.64046 | 5.41214 |
| **Spcs2** | 7.08009 | 7.21561 | 10.27479 | 4.45253 | 6.83615 | 8.67327 | 4.15857 | 4.97613 | 1.17653 | 10.75748 | 8.24470 | 8.07695 | 7.60240 | 0.57012 | 2.69505 | 8.01265 |
| **Spcs3** | 5.70359 | 4.41268 | 7.82534 | 0.00000 | 10.98031 | 6.17601 | 4.00711 | 8.31059 | 0.49997 | 0.83116 | 2.90576 | 1.24348 | 1.89795 | 4.07009 | 1.89975 | 1.62812 |
| **Spdef** | 0.01348 | 0.00787 | 0.01180 | 0.00879 | 0.00909 | 0.00658 | 0.00732 | 0.01076 | 0.98199 | 0.40854 | 0.00643 | 0.00471 | 0.01244 | 0.01197 | 0.00706 | 0.00774 |
| **Spdl1** | 3.60395 | 5.74502 | 1.74253 | 9.94676 | 7.11117 | 4.79555 | 3.59346 | 6.21306 | 15.29814 | 11.52250 | 3.73503 | 4.38811 | 1.41076 | 1.42526 | 12.61693 | 1.27262 |
| **Spdya** | 0.00763 | 0.00872 | 0.02131 | 0.01489 | 0.00984 | 0.00626 | 0.00819 | 0.06341 | 0.00925 | 0.31970 | 0.00000 | 0.07886 | 0.01007 | 1.66152 | 0.67198 | 0.00738 |
| **Spdye4a** | 0.01384 | 0.00814 | 0.01081 | 0.18291 | 0.00928 | 0.00679 | 0.00746 | 0.01274 | 0.00819 | 0.01981 | 0.01271 | 0.08685 | 0.01307 | 0.00841 | 0.06161 | 0.00739 |
| **Spdye4b** | 0.01338 | 0.00819 | 0.01114 | 1.65517 | 0.00917 | 0.00654 | 0.00748 | 0.01232 | 1.42341 | 0.20680 | 0.00000 | 0.00557 | 0.01324 | 0.00218 | 0.52723 | 0.00706 |
| **Specc1** | 1.60045 | 0.00628 | 0.16573 | 0.00802 | 0.00455 | 0.00754 | 0.01819 | 0.29716 | 1.11641 | 1.47709 | 0.88575 | 0.00620 | 0.03493 | 0.86352 | 0.00000 | 0.07573 |
| **Specc1l** | 1.35135 | 0.00800 | 0.95833 | 8.08360 | 0.10191 | 0.00366 | 0.00619 | 0.74625 | 5.35631 | 1.07999 | 1.51103 | 0.06705 | 0.38731 | 0.58320 | 5.04195 | 0.73048 |

| | | | | | | | | | | | | | | | | |
|---|---|---|---|---|---|---|---|---|---|---|---|---|---|---|---|---|
| **Speer1** | 0.01795 | 0.00793 | 0.04247 | 0.11180 | 0.00869 | 0.00752 | 0.00744 | 0.00768 | 0.13180 | 0.00245 | 0.30846 | 0.00623 | 0.00873 | 0.00647 | 0.05416 | 0.00717 |
| **Speer2** | 0.05256 | 0.00781 | 0.15790 | 0.01069 | 0.00768 | 0.00845 | 0.00733 | 0.01411 | 0.01232 | 0.01514 | 0.01252 | 0.00777 | 0.01071 | 0.01279 | 0.00803 | 0.00720 |
| **Speer3** | 0.10920 | 0.00768 | 0.28597 | 0.01754 | 0.00718 | 0.00749 | 0.00725 | 0.01382 | 0.02452 | 0.01494 | 0.01252 | 0.00765 | 0.00997 | 0.01326 | 0.00563 | 0.00706 |
| **Speer4b** | 0.01773 | 0.00816 | 0.02775 | 0.11078 | 0.01060 | 0.01531 | 0.00735 | 0.01171 | 0.06545 | 0.00800 | 0.01143 | 0.00740 | 0.01213 | 0.00738 | 0.37707 | 0.00735 |
| **Speer4c** | 0.01212 | 0.00802 | 0.09593 | 0.01858 | 0.00924 | 0.00590 | 0.00758 | 0.03031 | 0.01092 | 0.04466 | 0.01169 | 0.00814 | 0.01086 | 0.00861 | 0.12045 | 0.00749 |
| **Speer4cos** | 0.01363 | 0.00795 | 0.01192 | 1.92421 | 0.00925 | 0.00663 | 0.00753 | 0.01383 | 0.48614 | 0.06750 | 0.00861 | 0.00740 | 0.01327 | 0.00380 | 0.15910 | 0.00707 |
| **Speer4d** | 0.01006 | 0.00789 | 0.08860 | 0.06462 | 0.00948 | 0.01160 | 0.00744 | 0.11818 | 0.04163 | 0.07412 | 0.01110 | 0.01771 | 0.00795 | 0.00786 | 0.13985 | 0.00750 |
| **Speer4e** | 0.01118 | 0.00796 | 0.16421 | 0.14372 | 0.00917 | 0.00602 | 0.00754 | 0.05153 | 0.13961 | 0.08832 | 0.00941 | 0.00754 | 0.00960 | 0.00723 | 0.33870 | 0.00740 |
| **Speer5.ps1** | 0.04074 | 0.00826 | 0.19045 | 1.38742 | 0.02905 | 0.10539 | 0.00711 | 0.05720 | 0.60093 | 0.03954 | 0.01779 | 0.00605 | 0.00730 | 0.00611 | 0.08664 | 0.00697 |
| **Speer6.ps1** | 0.04236 | 0.00778 | 0.21097 | 0.01063 | 0.00775 | 0.00774 | 0.00732 | 0.01385 | 0.01238 | 0.01487 | 0.01250 | 0.00766 | 0.01035 | 0.01179 | 0.01191 | 0.00719 |
| **Speer8.ps1** | 0.01287 | 0.00799 | 0.06807 | 0.03062 | 0.00927 | 0.00582 | 0.00762 | 0.01345 | 0.36924 | 0.00759 | 0.00950 | 0.00718 | 0.01158 | 0.00777 | 0.16384 | 0.00730 |
| **Spef1** | 0.01310 | 0.00771 | 0.01082 | 0.00530 | 0.00889 | 0.00655 | 0.00718 | 0.00546 | 0.12482 | 1.41323 | 0.15321 | 0.01276 | 0.01601 | 0.01276 | 0.00314 | 0.00733 |
| **Speg** | 0.06954 | 0.00768 | 0.00425 | 0.48518 | 0.03144 | 0.00757 | 0.42961 | 0.00000 | 0.44216 | 1.36209 | 0.63414 | 0.02784 | 0.00692 | 0.00671 | 0.00000 | 0.00746 |
| **Spen** | 3.71208 | 0.02370 | 1.86769 | 1.38104 | 0.01202 | 0.00985 | 2.96560 | 1.49721 | 0.63093 | 5.96677 | 2.87255 | 1.07119 | 0.51834 | 0.87318 | 0.06583 | 0.00901 |
| **Spesp1** | 0.71309 | 0.00702 | 0.00354 | 1.19392 | 0.00642 | 0.00681 | 0.00673 | 0.00928 | 3.36280 | 0.25132 | 0.00000 | 0.00844 | 0.01202 | 0.00423 | 0.25354 | 0.00632 |
| **Spg11** | 4.63868 | 5.57559 | 2.47906 | 0.36499 | 2.72871 | 0.79007 | 0.33401 | 1.94611 | 1.00230 | 6.61224 | 2.59971 | 3.69390 | 0.71322 | 1.76216 | 1.74681 | 0.85899 |
| **Spg20** | 4.04861 | 0.00698 | 4.38081 | 8.51497 | 9.03138 | 2.92356 | 1.82440 | 4.44564 | 0.49627 | 0.00730 | 0.02143 | 0.08236 | 2.66248 | 0.00000 | 1.82911 | 0.46449 |
| **Spg21** | 7.06698 | 6.77075 | 6.78050 | 0.08780 | 4.32727 | 2.70800 | 2.44894 | 2.01508 | 0.27150 | 4.60338 | 3.98497 | 2.13105 | 2.99451 | 0.00000 | 0.85952 | 4.60990 |
| **Spg7** | 3.73405 | 0.03409 | 6.22000 | 6.59471 | 2.70091 | 0.00829 | 3.48028 | 1.83895 | 6.53970 | 1.40202 | 1.20955 | 0.11993 | 6.25765 | 0.03530 | 0.81499 | 0.40545 |
| **Sphk1** | 0.01398 | 0.00733 | 0.00000 | 0.00000 | 0.01099 | 1.79578 | 0.00651 | 0.79635 | 0.00989 | 0.33340 | 1.18209 | 0.03245 | 0.00000 | 0.00283 | 0.45852 | 0.00666 |
| **Sphk2** | 1.27560 | 1.14182 | 1.21740 | 4.57728 | 3.94656 | 5.99437 | 2.50791 | 3.74707 | 6.81748 | 3.67648 | 1.40415 | 0.04343 | 4.78324 | 0.44067 | 0.00000 | 2.28711 |
| **Spi1** | 0.06351 | 0.00813 | 0.09504 | 0.00498 | 0.01229 | 0.07374 | 0.02752 | 0.02665 | 0.12724 | 1.23503 | 0.19616 | 0.00354 | 0.30670 | 0.01534 | 0.00025 | 0.01455 |
| **Spib** | 0.03069 | 0.00831 | 0.05316 | 0.02101 | 0.02080 | 0.00648 | 0.00751 | 0.01063 | 0.24930 | 0.25096 | 0.09247 | 0.02688 | 0.02016 | 0.13674 | 0.01109 | 0.02141 |
| **Spic** | 12.33137 | 2.66598 | 16.02293 | 9.37350 | 12.85489 | 1.17411 | 14.07931 | 6.52731 | 28.66398 | 29.42267 | 30.43805 | 13.38952 | 0.00000 | 33.71868 | 24.55448 | 1.39510 |
| **Spice1** | 3.24526 | 4.04084 | 0.61261 | 0.00000 | 1.30671 | 0.00600 | 3.77159 | 1.01366 | 0.00663 | 4.42081 | 0.19509 | 0.15532 | 0.11141 | 0.00753 | 1.22457 | 0.01728 |
| **Spidr** | 1.72331 | 0.00664 | 0.00000 | 0.43036 | 0.01259 | 0.00459 | 0.10222 | 3.58054 | 0.42423 | 3.08100 | 0.18278 | 0.43595 | 0.56935 | 0.88411 | 2.21666 | 2.41485 |
| **Spin1** | 9.64883 | 7.19653 | 5.84919 | 37.79245 | 9.84828 | 8.33801 | 7.29559 | 7.61751 | 15.28744 | 15.40772 | 10.47095 | 8.14171 | 6.43021 | 17.17405 | 16.91756 | 4.32052 |
| **Spin2c** | 0.00705 | 0.00898 | 0.34461 | 0.34171 | 0.01035 | 0.01239 | 0.00854 | 0.76490 | 0.96524 | 0.00199 | 0.00000 | 0.00644 | 0.22772 | 0.01530 | 0.00191 | 0.00738 |
| **Spin2d** | 0.60470 | 0.00754 | 0.01438 | 1.42068 | 0.00673 | 0.07497 | 0.00747 | 0.05011 | 2.63656 | 0.12856 | 0.00005 | 0.02366 | 0.01700 | 0.07750 | 0.50924 | 0.00628 |
| **Spink2** | 1.97721 | 0.00287 | 0.00000 | 1.34781 | 4.04227 | 0.00382 | 0.00352 | 0.51977 | 2.29845 | 6.13415 | 5.16035 | 0.16849 | 0.07955 | 3.26617 | 1.65560 | 1.71725 |
| **Spink4** | 0.01355 | 0.00812 | 0.01050 | 0.08743 | 0.00921 | 0.00675 | 0.00768 | 0.01501 | 0.08574 | 0.00736 | 0.17077 | 0.01784 | 0.01010 | 0.00745 | 0.06638 | 0.00730 |
| **Spint1** | 9.36405 | 0.59720 | 4.06673 | 5.83871 | 6.35156 | 9.09981 | 5.94339 | 10.64867 | 1.27488 | 8.23014 | 5.10670 | 2.23761 | 9.74201 | 0.57067 | 5.36145 | 3.96129 |
| **Spint2** | 4.94731 | 0.05807 | 3.73798 | 0.92150 | 0.45706 | 1.24342 | 6.07825 | 7.15637 | 1.02567 | 10.47486 | 8.99585 | 0.47957 | 4.15342 | 4.93976 | 5.03119 | 3.40202 |
| **Spint5** | 0.00514 | 0.00730 | 0.09397 | 0.01008 | 0.88207 | 0.00667 | 0.00724 | 0.01178 | 0.01192 | 0.00641 | 0.01255 | 0.00695 | 0.36278 | 0.01347 | 0.00883 | 0.41529 |
| **Spire1** | 1.01354 | 2.49998 | 1.24016 | 14.50147 | 0.00385 | 0.00392 | 0.27269 | 0.02463 | 8.23459 | 4.77431 | 0.52865 | 0.26107 | 1.34044 | 0.18538 | 13.85368 | 0.05072 |
| **Spire2** | 0.00923 | 0.00777 | 0.00521 | 0.00485 | 0.00971 | 0.00718 | 0.00738 | 0.49569 | 0.01050 | 0.00000 | 0.27569 | 0.04624 | 0.00466 | 0.00547 | 0.35784 | 0.00728 |
| **Spn** | 0.01381 | 0.00802 | 0.01056 | 1.66799 | 0.00911 | 0.00648 | 0.00739 | 0.00807 | 0.32034 | 0.06074 | 0.21273 | 0.00688 | 0.00944 | 0.00958 | 0.01041 | 0.00684 |
| **Spns1** | 15.80233 | 1.97039 | 7.29369 | 10.36509 | 4.37139 | 9.36129 | 11.39890 | 33.10924 | 11.36832 | 32.92505 | 18.22865 | 2.37079 | 25.13361 | 0.00000 | 0.00000 | 11.05230 |
| **Spns2** | 0.19572 | 0.14636 | 0.60660 | 2.50474 | 0.47679 | 0.06591 | 0.00618 | 1.67616 | 2.62101 | 1.88284 | 2.52272 | 0.02583 | 0.05507 | 0.70988 | 2.91594 | 0.35359 |
| **Spns3** | 0.00906 | 0.00791 | 0.01258 | 0.00908 | 0.01043 | 0.00715 | 0.00743 | 0.10228 | 0.01185 | 0.00166 | 0.01498 | 0.00674 | 0.71428 | 0.00804 | 0.25502 | 0.00734 |
| **Spock1** | 0.10239 | 0.00812 | 0.00856 | 0.20301 | 0.00986 | 0.00688 | 0.00732 | 0.01258 | 0.29274 | 0.13362 | 0.01227 | 0.00681 | 0.01285 | 0.01079 | 0.00602 | 0.00767 |
| **Spock2** | 0.03334 | 0.00837 | 0.04788 | 0.00340 | 0.00915 | 0.00583 | 0.00798 | 0.33242 | 0.44752 | 0.02826 | 0.45637 | 0.01763 | 0.00374 | 0.03782 | 0.37278 | 0.00699 |
| **Spock3** | 0.01374 | 0.00776 | 0.01220 | 0.46357 | 0.00927 | 0.00659 | 0.00742 | 0.01243 | 0.04363 | 0.05739 | 0.01221 | 0.00673 | 0.01310 | 0.00484 | 0.24691 | 0.00763 |
| **Spon1** | 0.02025 | 0.00777 | 0.25704 | 0.48117 | 0.00877 | 0.00601 | 0.00738 | 0.00964 | 1.47718 | 0.27403 | 0.00000 | 0.00000 | 0.01934 | 1.53772 | 0.00000 | 0.00723 |
| **Spon2** | 0.02228 | 0.00847 | 0.05750 | 0.01061 | 0.01168 | 0.00590 | 0.00764 | 0.01392 | 0.01237 | 0.01949 | 0.01290 | 0.00841 | 0.01179 | 0.01253 | 0.00951 | 0.00747 |
| **Spop** | 2.07617 | 0.76556 | 5.39623 | 2.20163 | 3.26704 | 4.72609 | 2.41032 | 8.46821 | 1.18305 | 4.18461 | 4.14602 | 1.04559 | 1.44958 | 8.72376 | 12.83419 | 0.31024 |
| **Spopl** | 0.70975 | 0.00659 | 0.00000 | 0.61321 | 0.01678 | 0.02855 | 3.35977 | 0.03646 | 0.03592 | 0.88231 | 0.07563 | 0.02306 | 0.02971 | 0.00627 | 2.09144 | 0.00662 |
| **Spp1** | 24.08850 | 16.75114 | 7.08745 | 0.00732 | 18.34373 | 0.56715 | 3.25182 | 0.00000 | 0.00581 | 0.00000 | 0.00000 | 0.29404 | 3.39878 | 0.00443 | 0.00000 | 14.79645 |
| **Sppl2a** | 1.33352 | 1.14465 | 0.68956 | 1.80759 | 5.89312 | 0.63340 | 0.09614 | 1.76054 | 0.15795 | 0.76881 | 1.51373 | 0.60383 | 2.57535 | 2.68983 | 0.82694 | 3.77879 |
| **Sppl2b** | 0.56381 | 0.00714 | 0.00000 | 0.00357 | 1.19822 | 0.01346 | 0.05029 | 0.45264 | 0.22539 | 0.12096 | 0.08018 | 0.02588 | 3.89710 | 0.00787 | 0.05194 | 3.22688 |
| **Sppl3** | 1.91776 | 0.63527 | 5.48050 | 1.99735 | 1.11874 | 5.26496 | 2.91041 | 15.22501 | 4.52816 | 18.83812 | 5.78023 | 3.22524 | 3.41884 | 2.06578 | 9.15981 | 1.40173 |
| **Spr** | 7.90023 | 2.49781 | 2.14027 | 0.30085 | 2.25618 | 0.03550 | 0.07716 | 0.93758 | 0.24459 | 10.13890 | 1.03781 | 0.16231 | 5.62666 | 0.33456 | 0.00000 | 4.11827 |
| **Spred1** | 0.59836 | 1.95454 | 0.27996 | 2.77090 | 1.00982 | 0.20382 | 0.00660 | 0.01204 | 2.19229 | 6.78723 | 0.00000 | 0.04354 | 0.13950 | 12.39062 | 1.66125 | 0.00531 |
| **Spred2** | 0.01241 | 0.00796 | 0.01852 | 0.43777 | 0.00960 | 0.00775 | 0.00761 | 0.01381 | 0.56183 | 0.06118 | 0.02320 | 0.01865 | 0.16865 | 0.01423 | 0.10535 | 0.00744 |
| **Spred3** | 0.02161 | 0.00842 | 0.00030 | 0.34193 | 0.00872 | 0.08915 | 0.00813 | 0.25136 | 0.00689 | 0.16845 | 0.02749 | 0.04299 | 0.00527 | 0.00755 | 0.13090 | 0.00698 |
| **Sprtn** | 0.02099 | 0.15852 | 0.06776 | 1.24475 | 0.01179 | 0.00772 | 0.46358 | 0.66169 | 2.50283 | 0.00000 | 0.01950 | 0.00860 | 0.43800 | 0.22692 | 0.36390 | 0.00661 |
| **Spry1** | 0.03810 | 0.49416 | 0.54558 | 1.37620 | 0.00645 | 0.00888 | 0.49729 | 0.48672 | 0.42252 | 0.39797 | 0.01594 | 0.03378 | 0.13496 | 0.00284 | 0.10693 | 1.38901 |
| **Spry2** | 2.04935 | 0.38907 | 0.81430 | 2.82853 | 0.63747 | 0.00384 | 0.09245 | 0.96310 | 4.80872 | 0.00000 | 2.34996 | 1.29035 | 0.61891 | 9.45922 | 0.48265 | 1.46159 |
| **Spry4** | 2.40818 | 0.00460 | 0.00000 | 18.97164 | 0.00338 | 0.00092 | 0.00447 | 0.20873 | 11.49589 | 2.52951 | 0.28065 | 0.04753 | 0.00247 | 0.00000 | 4.94503 | 1.04033 |
| **Spryd3** | 1.28635 | 2.72197 | 1.88970 | 1.40509 | 0.44613 | 0.01165 | 0.00417 | 3.61305 | 1.04661 | 2.07343 | 0.33442 | 0.06112 | 1.40981 | 0.05273 | 1.69228 | 0.69894 |
| **Spryd4** | 5.99674 | 4.09364 | 1.84415 | 0.73642 | 5.01777 | 7.54526 | 5.23035 | 3.15723 | 0.24223 | 1.01856 | 1.34496 | 0.23491 | 0.23755 | 0.13132 | 4.59160 | 0.08454 |
| **Spryd7** | 5.19471 | 4.08774 | 1.33481 | 7.19850 | 1.63978 | 1.48560 | 1.57014 | 2.33686 | 5.15794 | 5.98150 | 5.42029 | 4.79449 | 2.02040 | 3.00226 | 1.46389 | 0.49157 |

| | | | | | | | | | | | | | | | | |
|---|---|---|---|---|---|---|---|---|---|---|---|---|---|---|---|---|
| **Spsb1** | 0.00075 | 0.00705 | 0.00754 | 0.01030 | 0.00930 | 0.01353 | 1.26771 | 0.53503 | 0.01158 | 0.06605 | 0.00881 | 0.00661 | 0.00558 | 0.01305 | 0.00458 | 0.34178 |
| **Spsb2** | 3.69568 | 0.00473 | 0.00000 | 0.28231 | 0.01882 | 1.60386 | 0.00503 | 0.44047 | 0.82776 | 0.53077 | 0.13148 | 0.00173 | 0.00000 | 0.01202 | 0.00001 | 0.01076 |
| **Spsb3** | 2.21166 | 3.30269 | 2.61198 | 5.21692 | 0.06910 | 3.38381 | 3.75010 | 6.25110 | 5.77469 | 7.94787 | 0.63330 | 0.47442 | 5.15098 | 0.00000 | 1.34364 | 5.06712 |
| **Spsb4** | 4.35391 | 0.01275 | 0.00000 | 3.35358 | 0.54865 | 6.71158 | 2.05766 | 0.12065 | 6.60974 | 3.44248 | 2.23824 | 0.46348 | 2.69451 | 2.67847 | 2.50450 | 1.86750 |
| **Sptan1** | 8.71538 | 5.39600 | 9.86002 | 12.08003 | 1.95482 | 10.06165 | 3.30970 | 16.12785 | 6.87566 | 18.84666 | 13.93822 | 20.63214 | 9.08298 | 6.21659 | 12.25705 | 2.43128 |
| **Sptb** | 0.00863 | 0.00741 | 0.00654 | 0.05251 | 0.00891 | 0.00687 | 0.00609 | 0.29493 | 0.19030 | 0.75065 | 0.18459 | 0.02957 | 1.73104 | 0.04117 | 0.01111 | 0.00668 |
| **Sptbn1** | 3.85161 | 0.15382 | 2.33432 | 4.37826 | 9.20474 | 0.97886 | 1.78331 | 0.08478 | 1.36246 | 1.25329 | 2.05654 | 0.09654 | 1.70295 | 0.00000 | 1.54324 | 2.98587 |
| **Sptbn2** | 0.65151 | 0.00000 | 2.06097 | 6.01059 | 0.02710 | 0.04387 | 3.27240 | 16.83039 | 2.23529 | 14.90093 | 6.51795 | 8.69092 | 6.90193 | 0.40292 | 4.82827 | 0.00713 |
| **Sptbn4** | 0.92003 | 0.00769 | 0.00371 | 0.19182 | 0.30797 | 0.00718 | 0.00668 | 0.00599 | 0.06943 | 0.01206 | 0.16257 | 0.01673 | 0.12544 | 0.01024 | 0.00241 | 0.00650 |
| **Sptlc1** | 6.43248 | 4.76905 | 5.39010 | 3.41410 | 0.18967 | 3.22996 | 5.07143 | 8.63564 | 2.77491 | 2.38470 | 4.75683 | 2.77081 | 3.08372 | 2.38481 | 9.19049 | 2.79503 |
| **Sptlc2** | 6.12089 | 3.66243 | 6.56381 | 6.61993 | 6.25393 | 8.10923 | 10.00133 | 5.93229 | 12.35231 | 5.60468 | 2.96628 | 3.33293 | 5.33236 | 5.94402 | 9.52379 | 2.84029 |
| **Sptssa** | 11.49406 | 9.02566 | 10.31863 | 5.91891 | 11.77313 | 8.48926 | 7.80430 | 8.51734 | 3.35111 | 11.71224 | 5.73662 | 2.97042 | 13.08586 | 3.22031 | 9.35250 | 5.88751 |
| **Spty2d1** | 2.31306 | 0.47934 | 2.89263 | 4.66735 | 2.39879 | 0.00119 | 2.33273 | 8.01793 | 14.60741 | 8.57157 | 9.89633 | 7.90773 | 1.62996 | 23.64932 | 13.71272 | 0.80745 |
| **Spz1** | 0.01240 | 0.00789 | 0.00822 | 4.81716 | 0.00896 | 0.00522 | 0.00716 | 0.00724 | 7.22366 | 0.45929 | 0.00000 | 0.00919 | 0.01766 | 0.00807 | 1.58888 | 0.00584 |
| **Sqle** | 4.69335 | 3.53035 | 8.77619 | 14.03931 | 0.19738 | 0.66304 | 3.73982 | 7.13211 | 8.83635 | 19.15246 | 7.79247 | 7.40693 | 0.72207 | 8.46433 | 11.76804 | 2.76735 |
| **Sqstm1** | 18.12215 | 10.96082 | 23.44913 | 0.82216 | 20.69766 | 4.18828 | 5.89534 | 9.25362 | 4.09602 | 9.01450 | 7.82998 | 1.07703 | 15.20893 | 1.01996 | 2.09748 | 3.91601 |
| **Sra1** | 15.45597 | 4.16505 | 10.66942 | 2.14703 | 6.24652 | 6.05021 | 1.71305 | 3.51801 | 3.62236 | 6.38954 | 7.88128 | 0.22171 | 7.26855 | 1.53982 | 2.34545 | 5.77050 |
| **Srbd1** | 2.87007 | 0.00296 | 1.59543 | 2.83738 | 0.01328 | 0.00155 | 0.58749 | 0.81172 | 5.33489 | 3.63479 | 1.59383 | 3.78884 | 0.90880 | 1.74939 | 4.30788 | 0.56001 |
| **Src** | 0.74033 | 0.00558 | 1.90938 | 0.43882 | 0.00916 | 2.32190 | 2.84769 | 0.26385 | 0.38513 | 1.10395 | 0.00423 | 0.00000 | 0.00000 | 0.00741 | 0.02900 | 2.83858 |
| **Srcap** | 0.34077 | 0.86160 | 0.08009 | 3.10945 | 0.01584 | 1.79470 | 0.83932 | 2.75858 | 4.50756 | 1.36515 | 0.31705 | 0.04638 | 0.08207 | 2.83929 | 3.01345 | 0.08363 |
| **Srcin1** | 0.01345 | 0.00790 | 0.00916 | 0.00532 | 0.00916 | 0.00679 | 0.01769 | 0.01419 | 0.02527 | 0.01476 | 0.34488 | 0.01604 | 0.00817 | 0.02544 | 0.01991 | 0.00716 |
| **Srd5a1** | 0.01259 | 0.01830 | 0.00703 | 1.25032 | 0.00908 | 1.05848 | 0.00709 | 0.34071 | 0.00000 | 0.55025 | 0.00992 | 0.04619 | 0.00000 | 0.00592 | 0.07847 | 0.00700 |
| **Srd5a3** | 0.00189 | 0.00672 | 1.18358 | 2.40385 | 0.00938 | 0.52652 | 1.94418 | 0.75054 | 0.64753 | 0.42531 | 0.61428 | 0.09113 | 0.00000 | 0.00298 | 2.17993 | 0.02032 |
| **Srebf1** | 2.12496 | 0.02406 | 0.34708 | 0.39375 | 2.77352 | 0.23550 | 0.08446 | 2.85790 | 0.00000 | 0.50921 | 1.52899 | 0.06024 | 2.23200 | 0.01456 | 0.22424 | 2.68132 |
| **Srebf2** | 3.22952 | 1.26979 | 2.70472 | 0.16900 | 0.09696 | 0.07802 | 4.17287 | 0.57600 | 0.25612 | 0.36484 | 0.47366 | 0.01773 | 0.00000 | 0.33554 | 0.00000 | 0.16915 |
| **Srek1** | 1.39308 | 0.32516 | 2.30976 | 9.16092 | 7.09944 | 0.00307 | 2.94642 | 6.33233 | 7.11588 | 8.58946 | 4.05688 | 9.44621 | 3.12992 | 9.55146 | 6.82844 | 1.73788 |
| **Srek1ip1** | 2.32207 | 1.29770 | 1.29776 | 0.91241 | 0.02994 | 5.37972 | 0.02382 | 0.67774 | 0.03545 | 0.09474 | 0.00000 | 0.02099 | 0.51735 | 0.19665 | 4.03912 | 2.01103 |
| **Srf** | 1.45320 | 0.01377 | 0.97042 | 4.18303 | 2.58977 | 0.61096 | 2.26302 | 2.19304 | 4.03229 | 1.65456 | 1.07544 | 0.18464 | 2.31093 | 2.05841 | 1.29306 | 0.63973 |
| **Srfbp1** | 6.55018 | 8.21921 | 5.10143 | 2.13413 | 10.09920 | 0.00000 | 6.43801 | 13.04982 | 7.37843 | 12.73540 | 6.92788 | 1.97160 | 6.77961 | 2.91492 | 10.65779 | 6.50404 |
| **Srgap1** | 0.28913 | 0.00853 | 0.52876 | 0.88559 | 0.01924 | 0.00621 | 0.00742 | 0.01265 | 0.58649 | 0.00164 | 0.00937 | 0.02877 | 0.23260 | 0.01040 | 0.25747 | 0.01487 |
| **Srgap2** | 0.38238 | 0.95755 | 0.00000 | 3.03145 | 0.01492 | 2.00491 | 2.53726 | 1.09170 | 1.73405 | 0.00000 | 0.00000 | 0.08967 | 0.00000 | 2.28110 | 0.00000 | 2.08473 |
| **Srgap3** | 0.02496 | 0.00790 | 0.77154 | 0.00740 | 0.00840 | 0.01126 | 0.00740 | 0.01706 | 0.03817 | 0.00322 | 0.01035 | 0.05825 | 0.00811 | 0.02398 | 0.22260 | 0.00703 |
| **Srgn** | 0.01097 | 0.01896 | 0.00000 | 10.98971 | 0.00599 | 0.01266 | 0.08665 | 0.61103 | 0.00000 | 0.08416 | 0.65053 | 0.02720 | 0.04988 | 0.00000 | 0.72493 | 5.48566 |
| **Sri** | 1.44116 | 1.67045 | 1.79566 | 0.67163 | 5.30020 | 3.23539 | 0.06598 | 0.00012 | 0.83402 | 0.45437 | 0.11418 | 0.01897 | 2.82147 | 0.00362 | 0.00000 | 0.00956 |
| **Srl** | 0.87692 | 0.00691 | 0.00000 | 2.15544 | 0.01141 | 2.12592 | 0.07509 | 1.13385 | 6.83889 | 0.00000 | 0.18632 | 0.04235 | 0.00000 | 2.69283 | 1.61502 | 0.00478 |
| **Srm** | 8.69804 | 3.18027 | 10.73224 | 6.38051 | 11.80080 | 2.53510 | 4.63228 | 12.18985 | 5.80720 | 15.89517 | 8.43192 | 4.59923 | 10.34063 | 0.10988 | 2.81751 | 7.55553 |
| **Srp14** | 17.30698 | 13.57775 | 20.60556 | 9.89558 | 16.75299 | 7.32424 | 10.34256 | 13.26430 | 15.58129 | 16.02645 | 15.90495 | 4.50120 | 12.77305 | 3.67223 | 8.88513 | 3.29202 |
| **Srp19** | 13.05315 | 5.09897 | 3.74398 | 4.42348 | 11.15548 | 5.37924 | 2.65813 | 6.34784 | 9.37817 | 11.13352 | 5.97658 | 2.42788 | 9.96225 | 4.77488 | 0.52687 | 5.53569 |
| **Srp54a** | 5.69202 | 3.68379 | 6.01574 | 9.89393 | 4.68623 | 3.88970 | 3.14327 | 6.53386 | 3.77663 | 8.94892 | 3.34391 | 11.56341 | 6.07435 | 3.94091 | 6.09543 | 3.66509 |
| **Srp54b** | 5.06070 | 2.87122 | 4.49056 | 3.42895 | 4.09341 | 3.77634 | 3.07053 | 6.69480 | 2.74437 | 6.91168 | 3.62573 | 11.52770 | 4.86773 | 1.50070 | 5.62619 | 2.40578 |
| **Srp68** | 15.17989 | 13.24093 | 10.37100 | 4.81932 | 7.45680 | 4.27945 | 10.35970 | 9.83201 | 3.83097 | 8.37692 | 7.58135 | 3.47391 | 9.34963 | 3.48462 | 2.81854 | 9.12224 |
| **Srp72** | 10.27966 | 4.79275 | 3.62674 | 5.84951 | 10.54569 | 6.80737 | 8.45310 | 3.81942 | 5.17295 | 8.83596 | 8.59425 | 6.08978 | 4.65529 | 11.38171 | 10.43308 | 4.82606 |
| **Srp9** | 13.64229 | 2.98361 | 12.99532 | 19.99936 | 7.79238 | 6.77955 | 2.71525 | 7.82346 | 22.49701 | 33.94109 | 21.16948 | 6.32359 | 21.00549 | 6.47487 | 3.32049 | 9.12871 |
| **Srpk1** | 8.20261 | 5.46768 | 11.10300 | 7.93875 | 9.61291 | 6.14741 | 9.49373 | 24.64454 | 13.79717 | 24.63901 | 13.03503 | 2.11517 | 8.74648 | 15.22710 | 19.06399 | 1.74415 |
| **Srpk2** | 2.36873 | 4.86473 | 4.23128 | 7.35313 | 0.75547 | 1.44701 | 2.84630 | 0.00000 | 4.77474 | 1.74789 | 1.29315 | 6.47720 | 1.33639 | 7.34944 | 5.47465 | 1.02419 |
| **Srpk3** | 0.06142 | 0.00824 | 0.15157 | 0.98856 | 0.01524 | 0.00878 | 0.00731 | 0.01235 | 0.00000 | 0.01138 | 0.01349 | 0.00726 | 0.01203 | 0.00545 | 0.09197 | 0.00731 |
| **Srpr** | 8.59209 | 5.89145 | 6.16445 | 10.91062 | 3.74902 | 4.52015 | 7.31488 | 16.66902 | 14.22168 | 6.81655 | 7.53084 | 2.46941 | 6.96495 | 12.90894 | 21.83866 | 6.32078 |
| **Srprb** | 6.96095 | 4.21017 | 14.20757 | 6.11332 | 6.27082 | 7.43112 | 5.65955 | 13.80001 | 9.24714 | 15.45376 | 11.68563 | 4.58863 | 10.38846 | 0.94056 | 6.67519 | 4.90342 |
| **Srr** | 0.00000 | 3.84196 | 2.68479 | 0.80262 | 3.44785 | 0.46166 | 3.59989 | 2.31096 | 2.04495 | 0.95172 | 0.12265 | 0.05229 | 0.96637 | 3.15019 | 2.39600 | 0.00382 |
| **Srrd** | 0.75684 | 0.07777 | 0.40677 | 0.00153 | 0.04489 | 1.18964 | 0.00733 | 0.06029 | 0.30800 | 1.57560 | 0.16445 | 0.43271 | 0.01812 | 0.00707 | 0.39336 | 0.02823 |
| **Srrm1** | 12.63024 | 10.27184 | 9.62500 | 5.15974 | 3.18042 | 2.67829 | 9.79932 | 11.06629 | 8.38103 | 15.04074 | 6.53968 | 6.46572 | 7.37872 | 7.25748 | 11.13396 | 4.23231 |
| **Srrm2** | 4.65506 | 0.30237 | 2.99842 | 8.96997 | 1.40363 | 0.12524 | 1.81494 | 1.10684 | 11.54057 | 7.37294 | 5.52674 | 0.31687 | 3.57262 | 7.74296 | 5.82283 | 3.64908 |
| **Srrt** | 21.26117 | 8.62942 | 17.30972 | 10.38338 | 9.70658 | 0.19425 | 10.85278 | 18.37618 | 4.67228 | 16.02864 | 8.70899 | 5.06129 | 11.58041 | 0.22500 | 1.71535 | 7.59780 |
| **Srsf1** | 15.88860 | 12.66412 | 12.83871 | 14.06221 | 14.25313 | 4.48847 | 10.48833 | 22.35245 | 21.64224 | 22.11608 | 18.84749 | 17.46715 | 11.52061 | 42.24989 | 27.65495 | 5.53047 |
| **Srsf10** | 10.08187 | 9.52233 | 7.98929 | 14.79323 | 6.37906 | 2.33216 | 6.79854 | 7.23530 | 18.89226 | 10.91650 | 8.19729 | 14.21183 | 8.30157 | 10.43899 | 8.95846 | 3.48746 |
| **Srsf11** | 10.73049 | 11.26486 | 14.52551 | 10.72910 | 12.48744 | 6.96895 | 11.20101 | 11.07018 | 12.10293 | 14.11334 | 6.38482 | 18.60714 | 8.27332 | 11.66637 | 15.67067 | 7.30178 |
| **Srsf2** | 16.96824 | 12.51960 | 15.42698 | 14.94390 | 16.02228 | 7.35425 | 11.34423 | 18.47235 | 27.07425 | 32.65735 | 29.61659 | 18.37526 | 17.04992 | 18.86389 | 21.83562 | 7.41885 |
| **Srsf3** | 30.55552 | 23.28598 | 24.84537 | 9.84493 | 22.35602 | 16.00975 | 15.37390 | 33.04829 | 25.48591 | 37.86262 | 27.08390 | 18.62105 | 28.34743 | 35.57148 | 30.48103 | 17.86420 |
| **Srsf4** | 0.01973 | 0.00770 | 0.01779 | 0.01095 | 0.00832 | 0.00633 | 0.32053 | 0.72658 | 0.01147 | 0.05025 | 0.00838 | 0.00615 | 0.17198 | 0.01650 | 0.00826 | 0.38372 |
| **Srsf5** | 10.60409 | 10.42613 | 8.72243 | 17.30238 | 4.71404 | 6.17992 | 4.66293 | 13.53513 | 27.93095 | 5.35167 | 20.75714 | 6.90313 | 8.16453 | 15.72326 | 10.11772 | 5.32607 |
| **Srsf6** | 20.20525 | 10.90763 | 11.35201 | 9.35784 | 17.73907 | 9.52513 | 5.09420 | 16.02584 | 16.80756 | 28.41356 | 18.96930 | 17.65478 | 12.35978 | 21.34590 | 16.91927 | 5.42582 |
| **Srsf7** | 25.36392 | 19.74903 | 20.46267 | 13.53868 | 16.30666 | 10.90864 | 6.38149 | 29.84213 | 26.50793 | 37.67807 | 26.19329 | 19.21400 | 25.18897 | 15.94356 | 22.96259 | 14.73403 |

| | | | | | | | | | | | | | | | | |
|---|---|---|---|---|---|---|---|---|---|---|---|---|---|---|---|---|
| **Srsf9** | 1.73611 | 2.12730 | 0.20666 | 2.13642 | 1.43914 | 0.04323 | 0.22755 | 1.40223 | 3.25158 | 2.96635 | 7.04087 | 0.22160 | 0.35082 | 1.76723 | 1.63963 | 0.21705 |
| **Srxn1** | 0.04160 | 3.28977 | 3.68338 | 0.43439 | 9.06743 | 0.12359 | 2.26231 | 0.00000 | 0.00000 | 0.25884 | 0.06288 | 0.05682 | 0.14569 | 0.00998 | 0.01235 | 3.32358 |
| **Ss18** | 9.80954 | 11.15848 | 12.25676 | 2.56943 | 8.04932 | 8.62328 | 6.37673 | 17.61765 | 1.72061 | 10.79586 | 9.39399 | 8.05201 | 12.42324 | 11.49389 | 14.33960 | 4.63912 |
| **Ss18l1** | 0.00531 | 0.00642 | 0.71007 | 1.04076 | 0.00856 | 1.90947 | 0.00559 | 0.01068 | 0.89956 | 0.00000 | 0.40890 | 5.67153 | 2.17922 | 0.00294 | 0.36800 | 0.01961 |
| **Ssb** | 15.07401 | 15.41597 | 12.97185 | 3.23225 | 13.94499 | 9.63727 | 10.04115 | 14.20401 | 3.71994 | 7.67253 | 4.16263 | 11.16613 | 10.44575 | 17.51608 | 8.57353 | 6.59582 |
| **Ssbp1** | 12.51278 | 4.57640 | 3.31716 | 2.96416 | 5.62250 | 1.84929 | 1.66363 | 5.23752 | 6.28066 | 1.12960 | 5.15012 | 1.65278 | 2.48275 | 4.33278 | 5.05663 | 5.01142 |
| **Ssbp2** | 0.15391 | 2.53733 | 0.00000 | 5.83270 | 0.00727 | 0.01141 | 0.00584 | 0.34443 | 2.43992 | 3.17716 | 0.56981 | 0.02495 | 0.00365 | 0.07802 | 3.61802 | 0.00485 |
| **Ssbp3** | 1.40891 | 3.45384 | 3.24733 | 8.59922 | 4.79843 | 6.03696 | 1.79746 | 3.79252 | 3.35519 | 3.07203 | 2.68059 | 0.12092 | 2.75524 | 0.10708 | 0.49965 | 1.08083 |
| **Ssbp4** | 0.04936 | 0.00766 | 0.08331 | 0.08573 | 0.45982 | 0.00601 | 0.01734 | 0.03890 | 0.01069 | 0.03241 | 0.03201 | 0.00678 | 0.95130 | 0.01357 | 0.01038 | 0.02155 |
| **Ssc4d** | 2.70913 | 1.03192 | 0.05166 | 5.18675 | 0.02021 | 0.00286 | 1.85093 | 0.03586 | 5.34912 | 2.50054 | 0.10897 | 5.80718 | 0.34338 | 0.09405 | 0.00000 | 0.00278 |
| **Ssfa2** | 0.59065 | 0.00649 | 0.25194 | 1.16991 | 0.00656 | 0.00839 | 0.00655 | 0.00452 | 1.37721 | 0.54737 | 0.18269 | 0.28840 | 0.00985 | 0.02577 | 4.74250 | 0.00659 |
| **Ssh1** | 0.06775 | 0.00715 | 0.00302 | 0.00931 | 0.66894 | 0.00783 | 0.43353 | 0.00908 | 0.01163 | 0.02309 | 0.01203 | 0.01932 | 0.00913 | 0.00829 | 0.07995 | 0.73697 |
| **Ssh2** | 0.00000 | 0.00538 | 0.00000 | 1.77222 | 0.00717 | 0.00513 | 0.39009 | 0.52956 | 0.00000 | 1.86220 | 0.40334 | 5.37819 | 0.00000 | 0.00673 | 2.48571 | 2.10230 |
| **Ssh3** | 0.28171 | 0.00800 | 0.09141 | 0.41672 | 0.02921 | 0.05174 | 0.00704 | 0.00919 | 0.65839 | 0.58791 | 0.00557 | 0.04340 | 0.01004 | 0.02392 | 0.06325 | 0.00687 |
| **Ssmem1** | 0.01120 | 0.00822 | 0.04603 | 0.64011 | 0.00997 | 0.00639 | 0.00776 | 0.02098 | 0.86206 | 0.43744 | 0.15279 | 0.00207 | 0.16816 | 0.00460 | 1.08109 | 0.00796 |
| **Ssna1** | 10.00315 | 7.11700 | 12.06952 | 3.78256 | 3.29640 | 0.21150 | 4.25172 | 9.85374 | 4.91722 | 14.75248 | 3.79364 | 3.13421 | 6.98986 | 0.62092 | 2.73648 | 5.05157 |
| **Sspn** | 0.01354 | 0.00779 | 0.01069 | 1.80235 | 0.00919 | 0.00635 | 0.00734 | 0.01047 | 1.03244 | 0.06121 | 0.03654 | 0.00445 | 0.01934 | 0.00332 | 0.69979 | 0.00716 |
| **Sspo** | 0.15513 | 0.00793 | 0.00862 | 0.01058 | 0.00795 | 0.00725 | 0.00756 | 0.01731 | 0.01234 | 0.01470 | 0.01407 | 0.00794 | 0.03015 | 0.01227 | 0.01059 | 0.00737 |
| **Ssr1** | 8.30861 | 7.29391 | 13.58461 | 3.23203 | 10.13916 | 8.24972 | 11.49496 | 12.84643 | 3.46677 | 8.00454 | 7.26341 | 2.38521 | 15.62590 | 10.44398 | 7.77980 | 8.37569 |
| **Ssr2** | 18.86182 | 8.65514 | 24.37167 | 7.97411 | 11.56132 | 11.30110 | 10.93168 | 17.62544 | 10.60883 | 17.23422 | 9.53959 | 6.99979 | 22.58230 | 12.10114 | 16.32052 | 11.67108 |
| **Ssr3** | 11.69850 | 8.04508 | 16.43353 | 10.12143 | 10.76578 | 6.77317 | 10.53514 | 11.56747 | 10.20470 | 7.39471 | 7.46217 | 3.67612 | 12.25924 | 10.91296 | 14.05923 | 7.47702 |
| **Ssr4** | 20.21711 | 5.37126 | 12.43012 | 5.71752 | 10.95835 | 4.29412 | 4.91353 | 15.92562 | 6.89338 | 18.54549 | 14.77117 | 1.72151 | 13.81859 | 8.00209 | 7.97343 | 12.57016 |
| **Ssrp1** | 23.43677 | 13.01799 | 18.37362 | 11.24239 | 16.92786 | 8.74433 | 13.39327 | 20.23407 | 7.01458 | 12.22457 | 10.48942 | 11.09768 | 15.40248 | 6.00756 | 16.98150 | 13.94753 |
| **Sssca1** | 14.53498 | 0.23918 | 0.00000 | 0.24532 | 6.85508 | 1.97011 | 2.03896 | 12.47278 | 0.81688 | 19.68705 | 3.92779 | 0.61447 | 17.44820 | 0.00000 | 0.00000 | 2.75441 |
| **Ssu72** | 11.48044 | 2.96764 | 12.20050 | 15.99238 | 11.28010 | 6.04149 | 3.94396 | 13.49577 | 8.14902 | 22.65824 | 10.75908 | 8.75998 | 12.66543 | 2.57558 | 6.01538 | 6.99775 |
| **Ssx2ip** | 2.89403 | 3.83763 | 1.57450 | 8.93367 | 1.16692 | 0.00225 | 2.77417 | 4.34889 | 9.44177 | 4.67731 | 4.04025 | 0.83854 | 1.41529 | 1.75652 | 6.29265 | 0.24891 |
| **Ssxa1** | 0.01646 | 0.00796 | 0.20187 | 0.00987 | 0.00884 | 0.00567 | 0.00739 | 0.01264 | 0.46015 | 0.02651 | 0.00931 | 0.00705 | 0.01019 | 0.01230 | 0.00923 | 0.00713 |
| **St13** | 20.61892 | 12.25910 | 22.00651 | 12.51881 | 15.48437 | 10.91495 | 10.73352 | 18.84458 | 12.65354 | 11.91425 | 9.29963 | 2.94521 | 18.70405 | 2.99904 | 9.44983 | 10.21084 |
| **St14** | 1.20477 | 0.40892 | 2.88115 | 0.00000 | 0.02505 | 5.36298 | 4.46681 | 1.04472 | 0.00946 | 1.54523 | 0.75062 | 0.00372 | 4.27582 | 0.00791 | 0.01897 | 0.28175 |
| **St3gal1** | 0.01408 | 0.00811 | 0.02407 | 0.26156 | 0.00926 | 0.00619 | 0.01826 | 0.01535 | 0.32313 | 0.02306 | 0.07428 | 0.00746 | 0.01138 | 0.05164 | 0.01471 | 0.00736 |
| **St3gal2** | 0.88614 | 0.00668 | 0.45053 | 1.94810 | 0.02648 | 0.00445 | 0.00580 | 0.76540 | 1.21105 | 2.88543 | 1.24378 | 0.06281 | 0.00311 | 0.03839 | 2.61268 | 0.00566 |
| **St3gal3** | 2.68988 | 0.00397 | 0.07050 | 1.39180 | 2.31800 | 0.00271 | 0.00517 | 0.00000 | 0.85030 | 0.15748 | 0.00699 | 0.01199 | 0.26643 | 0.03107 | 0.00663 | 0.01206 |
| **St3gal4** | 3.34094 | 0.62825 | 0.00000 | 0.00711 | 0.00238 | 0.00556 | 0.00467 | 0.00819 | 0.01206 | 0.68116 | 0.01148 | 0.01331 | 0.00843 | 0.00999 | 0.00329 | 1.38949 |
| **St3gal5** | 0.01491 | 0.00786 | 0.08864 | 0.52321 | 0.01750 | 0.00848 | 0.02769 | 0.01861 | 0.33587 | 1.04089 | 0.00994 | 0.01302 | 0.01171 | 0.02540 | 0.03347 | 0.00764 |
| **St3gal6** | 1.59131 | 0.00457 | 1.23387 | 4.73342 | 0.01225 | 2.90900 | 0.00309 | 0.76678 | 3.32405 | 1.57368 | 0.14568 | 3.53960 | 3.45879 | 2.38549 | 1.74588 | 3.20426 |
| **St5** | 0.51864 | 0.27520 | 3.16059 | 0.42607 | 2.66756 | 0.00685 | 0.00525 | 2.14629 | 0.40472 | 2.46891 | 0.80698 | 0.07057 | 1.44368 | 0.89491 | 1.51571 | 0.01228 |
| **St6gal1** | 0.40192 | 0.00815 | 0.00466 | 1.71418 | 0.01764 | 0.00710 | 0.00717 | 0.01024 | 0.35925 | 0.06113 | 0.03854 | 0.00699 | 0.01235 | 0.00433 | 0.35677 | 0.00691 |
| **St6galnac2** | 6.98841 | 3.22847 | 8.16742 | 0.50592 | 1.73612 | 2.72184 | 3.70734 | 4.73203 | 0.15105 | 4.21306 | 4.11588 | 0.00000 | 1.62807 | 0.74427 | 2.89370 | 2.09185 |
| **St6galnac4** | 0.01748 | 0.00775 | 0.01143 | 0.92818 | 0.00890 | 0.00658 | 0.01742 | 0.01699 | 0.10091 | 0.19931 | 0.01292 | 0.00664 | 0.01242 | 0.00837 | 0.10202 | 0.00762 |
| **St6galnac5** | 0.01374 | 0.00804 | 0.01215 | 0.01062 | 0.00927 | 0.00664 | 0.00752 | 0.01353 | 0.01235 | 0.01505 | 0.01297 | 0.00812 | 0.01973 | 0.01236 | 0.01063 | 0.00739 |
| **St6galnac6** | 0.01122 | 0.00773 | 0.01157 | 0.40908 | 0.06215 | 0.00657 | 0.00745 | 0.01202 | 0.02317 | 0.18346 | 0.01220 | 0.00715 | 0.01163 | 0.01456 | 0.03152 | 0.00721 |
| **St7** | 0.02128 | 0.00839 | 0.01118 | 1.05448 | 0.00870 | 0.00659 | 0.00763 | 0.01895 | 0.30560 | 0.01138 | 0.04004 | 0.00807 | 0.01163 | 0.01074 | 0.00213 | 0.02890 |
| **St7l** | 0.02147 | 2.55718 | 0.76655 | 0.45459 | 0.00654 | 0.05788 | 0.03559 | 1.49750 | 0.18370 | 5.41296 | 1.36946 | 3.87583 | 0.07498 | 0.67996 | 1.72895 | 0.01952 |
| **St8sia1** | 0.04976 | 0.00758 | 0.04015 | 0.06146 | 0.02696 | 0.01270 | 0.03772 | 0.01529 | 0.05378 | 0.05019 | 0.04902 | 0.00683 | 0.01870 | 0.17577 | 0.18749 | 0.01454 |
| **Stab1** | 0.01360 | 0.00827 | 0.01303 | 0.09455 | 0.00967 | 0.00684 | 0.00764 | 0.01489 | 0.09338 | 0.01270 | 0.01431 | 0.00821 | 0.02291 | 0.00957 | 0.04434 | 0.00754 |
| **Stac2** | 2.64955 | 1.24269 | 5.67010 | 1.16700 | 5.01268 | 6.20197 | 0.68542 | 9.11365 | 1.40219 | 9.94339 | 0.26646 | 0.84583 | 2.75918 | 0.00000 | 0.99082 | 0.02040 |
| **Stag1** | 5.10806 | 8.77565 | 2.92680 | 4.48271 | 5.16007 | 3.06105 | 2.72295 | 1.20536 | 3.25509 | 1.09758 | 1.78691 | 8.14203 | 1.39511 | 1.34865 | 2.93282 | 2.58892 |
| **Stag2** | 0.56465 | 0.48041 | 4.46092 | 10.04890 | 0.01733 | 0.00231 | 0.01619 | 10.04047 | 10.35266 | 12.16605 | 17.74248 | 16.77024 | 2.33446 | 25.72361 | 19.03613 | 0.31398 |
| **Stag3** | 0.00000 | 2.14673 | 3.02830 | 3.21308 | 0.00204 | 0.02430 | 1.65893 | 6.42719 | 0.95971 | 1.78398 | 1.94759 | 0.89960 | 6.01917 | 4.36444 | 0.28619 | 0.01609 |
| **Stam** | 3.88682 | 5.00311 | 2.75461 | 1.38313 | 3.06325 | 0.00788 | 3.75374 | 1.95320 | 2.15265 | 2.51519 | 2.84969 | 10.29974 | 1.60589 | 5.56073 | 1.62243 | 2.68552 |
| **Stam2** | 2.55182 | 5.40996 | 4.30347 | 8.62215 | 5.39822 | 2.76939 | 5.91528 | 11.38676 | 4.77344 | 12.56266 | 7.37104 | 10.74409 | 6.36485 | 3.69309 | 11.68951 | 4.09246 |
| **Stambp** | 1.74195 | 0.00527 | 1.74276 | 0.91224 | 0.01844 | 3.38067 | 0.00423 | 0.00000 | 1.50160 | 0.22043 | 0.65615 | 0.04981 | 2.02220 | 0.00201 | 2.12397 | 0.72042 |
| **Stambpl1** | 2.77817 | 0.00211 | 0.12402 | 0.00799 | 5.10999 | 0.00275 | 0.11743 | 0.00000 | 0.01018 | 0.01974 | 0.01142 | 0.02322 | 0.00973 | 0.01079 | 0.02244 | 0.00422 |
| **Stamos** | 0.22158 | 0.00822 | 0.13960 | 0.00406 | 0.82947 | 1.42954 | 0.00726 | 0.00464 | 0.01169 | 0.00000 | 0.17428 | 0.01771 | 0.01043 | 0.01201 | 0.00530 | 0.00661 |
| **Stap2** | 0.01054 | 0.00790 | 0.01110 | 0.00438 | 0.10349 | 0.00653 | 0.00742 | 0.01079 | 0.57597 | 0.55507 | 0.00694 | 0.00544 | 0.01071 | 0.01204 | 0.00715 | 0.00703 |
| **Star** | 0.81014 | 0.00756 | 0.20716 | 0.71822 | 0.01425 | 0.00517 | 0.00700 | 0.00823 | 0.09927 | 3.52663 | 0.01668 | 0.12784 | 0.03650 | 0.00675 | 0.39782 | 0.00676 |
| **Stard10** | 12.76776 | 0.38509 | 11.46411 | 0.00000 | 4.57250 | 10.60147 | 2.05104 | 15.34630 | 0.00140 | 1.07618 | 0.31907 | 0.70774 | 23.46595 | 0.00320 | 0.00000 | 7.91068 |
| **Stard13** | 0.01373 | 0.00790 | 0.01149 | 1.07890 | 0.00930 | 0.00645 | 0.00741 | 0.01143 | 0.46195 | 0.00249 | 0.04979 | 0.01605 | 0.01190 | 0.00955 | 0.21135 | 0.00714 |
| **Stard3** | 2.15788 | 0.17060 | 1.14811 | 0.24880 | 2.70394 | 3.52550 | 2.40126 | 0.61042 | 0.89798 | 2.77820 | 0.91781 | 0.12065 | 1.86509 | 0.17729 | 0.25489 | 1.43302 |
| **Stard3nl** | 7.98583 | 2.53604 | 4.80503 | 0.86865 | 4.24346 | 9.94221 | 5.53793 | 1.61976 | 0.00958 | 6.06241 | 0.91240 | 0.95946 | 7.29802 | 2.60494 | 0.15929 | 3.91065 |
| **Stard4** | 0.03095 | 0.00835 | 0.00000 | 1.16565 | 0.03731 | 0.30159 | 0.00705 | 1.85731 | 0.06173 | 0.00000 | 0.43068 | 0.03300 | 1.70595 | 0.00286 | 0.13153 | 0.00619 |
| **Stard5** | 1.47689 | 0.02507 | 0.00000 | 0.00357 | 0.00504 | 0.00777 | 0.07553 | 0.00348 | 0.83891 | 1.03822 | 0.00000 | 0.69455 | 0.22168 | 0.00841 | 0.25684 | 0.00578 |

| | | | | | | | | | | | | | | | | |
|---|---|---|---|---|---|---|---|---|---|---|---|---|---|---|---|---|
| **Stard7** | 5.96811 | 6.77162 | 6.38647 | 7.01199 | 4.93743 | 6.61668 | 2.83797 | 1.71443 | 3.56535 | 2.38800 | 0.00000 | 9.01951 | 2.87886 | 0.00000 | 6.22034 | 2.62628 |
| **Stat1** | 1.67698 | 0.01738 | 0.34605 | 0.38130 | 0.19553 | 0.02868 | 1.27529 | 0.01280 | 0.33121 | 0.44296 | 0.95494 | 0.09640 | 0.17089 | 0.29331 | 0.53395 | 0.07777 |
| **Stat2** | 0.03494 | 0.01752 | 1.78475 | 7.32345 | 0.00510 | 0.03318 | 0.00431 | 3.18774 | 9.33004 | 3.92551 | 6.50334 | 0.94676 | 0.00000 | 2.82941 | 4.37659 | 0.00373 |
| **Stat3** | 12.74012 | 13.08654 | 12.73397 | 3.31704 | 9.30657 | 5.89818 | 11.28318 | 8.57471 | 3.95658 | 2.59502 | 2.45855 | 4.93482 | 3.57635 | 0.09477 | 2.81566 | 5.44249 |
| **Stat4** | 0.01378 | 0.00791 | 0.01207 | 0.87188 | 0.00934 | 0.00655 | 0.00748 | 0.01314 | 0.00000 | 0.02103 | 0.02513 | 0.00769 | 0.01233 | 0.01147 | 0.04214 | 0.00723 |
| **Stat5a** | 0.12145 | 0.00782 | 1.73600 | 1.80076 | 1.03198 | 0.03529 | 0.00680 | 0.84016 | 1.09737 | 0.44350 | 0.03203 | 0.03108 | 1.66647 | 0.25760 | 0.59798 | 0.01400 |
| **Stat5b** | 0.02093 | 0.00774 | 1.09625 | 0.00705 | 0.00724 | 0.00602 | 0.00743 | 0.01725 | 0.01159 | 1.24638 | 0.01122 | 0.01396 | 0.03008 | 0.01435 | 0.00000 | 0.14164 |
| **Stat6** | 0.99482 | 0.35666 | 0.00000 | 0.56540 | 0.00597 | 10.05403 | 1.67010 | 0.77361 | 0.00062 | 0.97096 | 0.00000 | 0.02564 | 6.40392 | 0.40368 | 0.34169 | 0.00710 |
| **Stau1** | 5.50497 | 8.45527 | 3.03763 | 13.57460 | 10.82510 | 5.89651 | 1.50898 | 4.23531 | 18.20057 | 4.67583 | 3.90527 | 2.34379 | 0.46490 | 4.97417 | 15.48281 | 3.01570 |
| **Stau2** | 0.00570 | 0.00723 | 1.54721 | 0.00658 | 0.00757 | 0.00905 | 2.16382 | 0.04849 | 0.01166 | 0.42631 | 0.00880 | 0.00473 | 0.00404 | 0.02494 | 0.17819 | 0.00648 |
| **Stbd1** | 0.00956 | 0.00768 | 0.00000 | 2.07132 | 0.00938 | 0.00693 | 0.00670 | 0.00000 | 0.53837 | 0.50864 | 0.64926 | 0.06628 | 0.01809 | 2.37354 | 2.52023 | 0.00673 |
| **Stc2** | 0.01378 | 0.00824 | 0.00951 | 0.37506 | 0.00912 | 0.00677 | 0.00757 | 0.00775 | 0.26812 | 0.00460 | 0.34829 | 0.00642 | 0.00859 | 0.00802 | 0.00000 | 0.00714 |
| **Steap1** | 0.01966 | 0.00771 | 0.20471 | 0.26057 | 0.00955 | 0.00571 | 0.00731 | 0.52169 | 0.09179 | 0.60442 | 0.00761 | 0.00403 | 0.01057 | 0.24404 | 0.00000 | 0.01540 |
| **Steap2** | 0.80889 | 0.00625 | 0.00294 | 2.98159 | 0.12408 | 0.00689 | 0.01629 | 0.00000 | 0.00000 | 1.64335 | 0.36424 | 0.01917 | 0.01530 | 0.08667 | 0.26978 | 0.01344 |
| **Steap3** | 1.39212 | 0.01207 | 1.13584 | 0.29408 | 0.01989 | 0.15884 | 4.97743 | 9.18037 | 0.25471 | 4.56561 | 3.79304 | 0.10389 | 7.98872 | 1.84376 | 0.45666 | 0.98640 |
| **Stil** | 4.26056 | 0.33318 | 0.00053 | 3.81728 | 0.00000 | 0.00083 | 1.66879 | 1.02671 | 6.30724 | 2.64243 | 1.06356 | 0.65969 | 0.59997 | 0.01968 | 3.26858 | 1.85398 |
| **Stim1** | 1.42921 | 0.00460 | 0.57028 | 0.52887 | 0.00934 | 0.43665 | 3.41096 | 1.53193 | 3.07319 | 3.01502 | 0.00000 | 1.88697 | 0.67393 | 0.62689 | 1.41523 | 0.10390 |
| **Stim2** | 0.53517 | 1.34320 | 0.00577 | 0.70265 | 0.11636 | 0.00605 | 0.00649 | 0.00876 | 0.94855 | 0.00072 | 0.00727 | 0.00364 | 0.04257 | 0.01112 | 0.98963 | 1.42694 |
| **Stip1** | 31.13605 | 27.92799 | 46.09564 | 9.56724 | 20.34358 | 19.20425 | 24.65711 | 49.94402 | 12.34746 | 44.45212 | 28.24688 | 13.98262 | 40.24100 | 20.37498 | 29.09081 | 13.41607 |
| **Stk10** | 0.00600 | 0.00643 | 0.52944 | 6.78895 | 1.68954 | 0.00481 | 0.01669 | 0.31752 | 3.48318 | 5.59995 | 1.31290 | 0.30236 | 0.00000 | 0.00000 | 1.76106 | 0.00550 |
| **Stk11** | 9.11609 | 4.18318 | 1.32222 | 1.11072 | 1.24505 | 9.55948 | 7.62954 | 19.31012 | 1.67070 | 17.26426 | 2.32325 | 0.66692 | 7.58584 | 0.00000 | 0.49381 | 7.82608 |
| **Stk11ip** | 0.33672 | 0.00547 | 2.11278 | 3.98341 | 0.00352 | 0.85094 | 3.61747 | 1.51845 | 2.43484 | 5.70449 | 1.85583 | 0.18474 | 0.97100 | 0.55778 | 1.23421 | 0.00418 |
| **Stk16** | 9.52268 | 6.60212 | 14.09536 | 5.74106 | 1.04961 | 5.84164 | 3.58678 | 6.68244 | 10.51712 | 14.64427 | 7.66094 | 7.62882 | 13.67604 | 4.03546 | 3.75854 | 2.16828 |
| **Stk17b** | 0.00000 | 2.73449 | 0.58812 | 1.59655 | 0.25821 | 0.00624 | 0.65927 | 2.35088 | 1.87908 | 1.56483 | 2.40462 | 0.01703 | 0.81551 | 2.85895 | 2.46177 | 0.01871 |
| **Stk19** | 3.68044 | 0.04250 | 0.34536 | 6.61386 | 4.75069 | 0.01344 | 0.73843 | 4.34171 | 12.70274 | 8.77691 | 3.25056 | 0.80175 | 2.58303 | 0.00208 | 0.00000 | 4.30391 |
| **Stk24** | 8.06634 | 5.50516 | 6.14886 | 7.01040 | 5.80656 | 7.90879 | 5.84364 | 21.10807 | 3.61312 | 11.55048 | 13.54485 | 6.83669 | 13.53827 | 7.65918 | 1.56677 | 3.59997 |
| **Stk25** | 3.92245 | 2.13638 | 2.70427 | 0.00000 | 2.02412 | 6.35297 | 4.59859 | 3.34156 | 0.31171 | 5.10882 | 0.39938 | 0.33921 | 7.51364 | 4.20486 | 5.60769 | 5.37763 |
| **Stk3** | 2.18588 | 0.40609 | 0.66109 | 5.66576 | 4.11747 | 0.00260 | 0.03558 | 1.36115 | 1.61176 | 0.75505 | 0.68260 | 0.02282 | 0.00000 | 1.77834 | 1.12593 | 0.04104 |
| **Stk31** | 3.97137 | 7.45965 | 2.52295 | 1.80986 | 3.00207 | 0.01101 | 3.70067 | 2.53854 | 3.13919 | 3.74653 | 1.23932 | 1.91852 | 0.00000 | 4.14052 | 3.55646 | 0.01300 |
| **Stk35** | 2.05765 | 0.20331 | 0.00000 | 19.92890 | 0.02030 | 0.00000 | 0.70002 | 5.02751 | 24.21704 | 7.57959 | 9.03531 | 9.74435 | 0.00000 | 4.88974 | 5.00193 | 0.04756 |
| **Stk36** | 0.13997 | 0.00738 | 0.00336 | 4.65144 | 0.01610 | 0.01234 | 0.00618 | 0.01490 | 0.99279 | 0.73002 | 0.00000 | 0.00772 | 1.56219 | 0.00000 | 1.93341 | 0.00604 |
| **Stk38** | 14.16761 | 2.87782 | 3.18131 | 8.21254 | 5.60696 | 1.10745 | 4.16680 | 5.81395 | 7.82270 | 14.54487 | 9.38285 | 12.43538 | 5.52905 | 7.49231 | 12.94785 | 1.45855 |
| **Stk38l** | 5.90531 | 4.17132 | 4.09669 | 5.22614 | 0.00000 | 0.00460 | 2.93913 | 1.15937 | 6.57555 | 6.35556 | 1.62787 | 0.38633 | 0.67001 | 4.13818 | 8.66271 | 0.01029 |
| **Stk39** | 0.01345 | 0.00722 | 0.00670 | 0.28694 | 0.00902 | 0.00685 | 0.00711 | 0.01006 | 0.23948 | 0.55260 | 0.00000 | 0.85146 | 0.01242 | 0.01184 | 0.28105 | 0.00752 |
| **Stk4** | 2.45010 | 0.07954 | 2.29867 | 3.09585 | 0.01560 | 7.78088 | 2.88774 | 7.09122 | 4.34516 | 8.77613 | 5.50522 | 10.30220 | 5.30779 | 2.91595 | 2.22416 | 1.48576 |
| **Stk40** | 7.95277 | 8.16494 | 6.60787 | 7.72957 | 3.30259 | 1.87057 | 2.63568 | 5.66683 | 7.70361 | 5.20172 | 4.92749 | 0.29596 | 4.82085 | 0.00000 | 1.57949 | 2.45488 |
| **Stkld1** | 0.01368 | 0.00814 | 0.01186 | 0.08419 | 0.00933 | 0.00666 | 0.00752 | 0.01335 | 0.42576 | 0.01163 | 0.00940 | 0.00731 | 0.01294 | 0.03904 | 0.00651 | 0.00740 |
| **Stmn1** | 11.57871 | 9.25654 | 8.97247 | 4.11256 | 3.17672 | 3.30815 | 5.85736 | 9.66732 | 11.91270 | 1.59446 | 3.77185 | 1.98440 | 5.67403 | 1.54040 | 3.34816 | 7.00004 |
| **Stmn1.rs1** | 0.08217 | 0.04236 | 0.04455 | 0.04438 | 0.02701 | 0.00792 | 0.01781 | 0.03381 | 0.10124 | 0.02059 | 0.04940 | 0.03769 | 0.03853 | 0.01090 | 0.02284 | 0.03684 |
| **Stmn3** | 9.06854 | 0.37917 | 0.22097 | 7.54272 | 0.60002 | 3.47363 | 1.87762 | 18.77431 | 6.19218 | 28.10147 | 12.99940 | 3.18075 | 19.10489 | 0.00000 | 0.00000 | 1.67212 |
| **Stom** | 0.01701 | 0.00791 | 0.00706 | 1.01563 | 0.02065 | 0.00684 | 0.00740 | 0.00263 | 0.00000 | 0.07174 | 0.40306 | 0.03844 | 0.00891 | 0.01028 | 0.62486 | 0.00727 |
| **Stoml1** | 0.84569 | 1.05981 | 1.57967 | 0.40688 | 0.00430 | 0.00491 | 3.08460 | 0.00000 | 0.09914 | 0.66756 | 0.95598 | 0.01520 | 1.41244 | 0.00510 | 0.00000 | 0.06589 |
| **Stoml2** | 11.58282 | 0.09327 | 4.68783 | 0.48876 | 5.64185 | 0.21534 | 4.50373 | 9.46258 | 0.76085 | 16.30237 | 7.08711 | 7.22599 | 15.01134 | 0.03041 | 6.23234 | 11.97550 |
| **Stoml3** | 0.02051 | 0.00805 | 0.01164 | 0.01034 | 0.00878 | 0.00688 | 0.00763 | 0.01405 | 0.01220 | 0.07850 | 0.01230 | 0.00889 | 0.01303 | 0.01171 | 0.01195 | 0.00739 |
| **Ston2** | 0.06966 | 0.09177 | 0.14068 | 6.74364 | 0.01657 | 0.06220 | 0.16902 | 0.01146 | 2.87301 | 0.58669 | 1.55895 | 0.14218 | 0.28939 | 0.43669 | 1.63581 | 0.01994 |
| **Stox1** | 0.01033 | 0.00745 | 0.00758 | 0.21660 | 0.00929 | 0.00634 | 0.00686 | 0.00923 | 0.29918 | 0.00000 | 0.00008 | 0.00307 | 0.50351 | 0.72253 | 0.53394 | 0.00685 |
| **Stox2** | 0.00719 | 0.00718 | 0.00449 | 0.44222 | 1.47833 | 0.00769 | 0.00747 | 0.01170 | 0.00000 | 0.01343 | 0.01177 | 0.00690 | 0.01007 | 0.01130 | 0.00599 | 0.00716 |
| **Stra13** | 5.09907 | 1.43080 | 9.16669 | 7.09046 | 2.03933 | 1.02781 | 1.07801 | 2.21192 | 1.61731 | 4.05911 | 8.54521 | 1.21830 | 3.66015 | 1.17098 | 1.07472 | 0.51737 |
| **Stra6** | 0.00790 | 0.00748 | 1.53942 | 0.16257 | 0.00723 | 0.00912 | 0.00698 | 0.90020 | 0.00907 | 0.12184 | 0.00504 | 0.00440 | 0.01748 | 0.01018 | 0.00000 | 0.00659 |
| **Stra8** | 0.00748 | 0.39374 | 0.27448 | 0.14660 | 0.00771 | 0.00909 | 0.00711 | 0.01021 | 0.00959 | 0.01266 | 0.01335 | 0.01724 | 0.01078 | 0.00924 | 0.00000 | 0.00710 |
| **Strada** | 3.84863 | 3.53582 | 5.50527 | 2.02401 | 5.66491 | 3.88228 | 0.07841 | 13.86725 | 0.02457 | 9.19284 | 5.82063 | 0.29557 | 7.98569 | 0.47800 | 0.70717 | 4.14529 |
| **Stradb** | 0.35396 | 0.04094 | 0.88729 | 13.05695 | 1.36486 | 3.11914 | 0.00613 | 0.48262 | 6.76472 | 1.49900 | 0.00000 | 0.34164 | 1.64217 | 0.01949 | 2.68869 | 2.04906 |
| **Strap** | 13.30452 | 7.45199 | 11.23275 | 16.22739 | 13.44580 | 9.39921 | 10.08748 | 16.67621 | 18.47165 | 30.25174 | 21.89218 | 21.82211 | 15.22534 | 23.86081 | 21.12545 | 6.58405 |
| **Strbp** | 9.88166 | 3.71424 | 3.88264 | 12.31477 | 7.46841 | 3.42959 | 10.51404 | 3.02720 | 8.05764 | 6.53868 | 5.54644 | 9.73471 | 4.40188 | 0.99673 | 13.71265 | 1.29555 |
| **Strc** | 0.00975 | 0.00832 | 0.01335 | 0.00818 | 0.00936 | 0.00792 | 0.00797 | 0.57396 | 0.01077 | 0.26866 | 0.00656 | 0.00780 | 0.00611 | 0.00822 | 0.31054 | 0.00739 |
| **Strip1** | 2.91529 | 6.06070 | 2.66029 | 9.56928 | 3.71851 | 2.39380 | 1.43132 | 7.14379 | 13.30399 | 7.83628 | 6.05620 | 0.45910 | 3.93343 | 0.08091 | 5.21115 | 4.95499 |
| **Strip2** | 0.00591 | 2.12495 | 0.00000 | 0.66060 | 0.00799 | 0.02059 | 0.00652 | 0.00000 | 2.08200 | 0.01665 | 0.45684 | 0.00220 | 0.02088 | 0.00663 | 0.04567 | 0.00629 |
| **Strn** | 0.00035 | 0.00725 | 0.00000 | 4.94415 | 0.32877 | 0.00623 | 0.00717 | 0.82075 | 3.85604 | 0.82026 | 0.32458 | 0.02225 | 0.08902 | 0.79361 | 3.86059 | 0.00683 |
| **Strn3** | 1.73899 | 5.81546 | 4.49279 | 6.03631 | 0.88644 | 3.51742 | 3.78298 | 8.31159 | 7.21763 | 12.12239 | 5.34370 | 15.55605 | 0.73305 | 25.12323 | 14.19110 | 1.46736 |
| **Strn4** | 7.96826 | 2.36209 | 1.81385 | 2.13312 | 2.58299 | 7.06492 | 1.72102 | 3.39123 | 1.67690 | 5.30560 | 4.71967 | 0.65717 | 5.61341 | 0.68906 | 0.37754 | 4.58028 |
| **Stt3a** | 8.30684 | 4.47078 | 10.52170 | 1.50564 | 5.46610 | 5.56933 | 6.69623 | 4.49891 | 2.16196 | 5.05108 | 4.72477 | 0.08115 | 6.13748 | 19.54477 | 8.25166 | 3.51711 |
| **Stt3b** | 6.86345 | 7.57517 | 13.57143 | 21.37654 | 4.07243 | 7.79369 | 6.49418 | 10.31163 | 15.04763 | 15.96596 | 6.99934 | 17.40993 | 9.56714 | 13.49603 | 19.76269 | 2.57457 |

| | | | | | | | | | | | | | | | | |
|---|---|---|---|---|---|---|---|---|---|---|---|---|---|---|---|---|
| **Stub1** | 3.60214 | 1.26002 | 2.30202 | 0.00000 | 1.75499 | 2.80177 | 0.41273 | 1.37697 | 0.00868 | 3.25722 | 0.00000 | 0.02641 | 5.54677 | 0.00552 | 0.06656 | 2.82616 |
| **Stx11** | 0.01358 | 0.00830 | 0.00729 | 0.92835 | 0.00906 | 0.00701 | 0.00759 | 0.00434 | 0.01310 | 0.00465 | 0.82762 | 0.00478 | 0.03814 | 0.00448 | 0.00000 | 0.00688 |
| **Stx12** | 5.55309 | 5.84558 | 3.55886 | 6.24138 | 4.95716 | 7.24665 | 0.25898 | 7.14759 | 2.66262 | 5.43959 | 2.63949 | 4.93604 | 4.03681 | 8.78375 | 9.01487 | 5.33834 |
| **Stx16** | 5.43930 | 2.12505 | 0.20048 | 0.68623 | 3.04831 | 3.42315 | 0.00000 | 1.37857 | 0.82005 | 0.00000 | 1.01205 | 0.02447 | 2.55760 | 0.12927 | 1.51838 | 0.34012 |
| **Stx17** | 2.08254 | 4.55006 | 4.32317 | 1.72021 | 0.00567 | 2.03960 | 6.62292 | 1.37073 | 1.56289 | 8.00706 | 6.58822 | 7.16484 | 2.15961 | 2.53251 | 7.59683 | 3.14793 |
| **Stx18** | 8.33716 | 0.49713 | 2.44067 | 0.46203 | 9.34537 | 4.62687 | 5.68853 | 4.99749 | 0.00000 | 2.95984 | 1.25641 | 0.20917 | 3.04640 | 0.00067 | 0.11872 | 1.30775 |
| **Stx1a** | 0.00000 | 0.00651 | 0.43947 | 2.00119 | 0.01848 | 1.89585 | 1.32775 | 5.27335 | 3.51051 | 5.24279 | 2.50142 | 0.34372 | 1.44218 | 0.00000 | 0.49824 | 0.00494 |
| **Stx1b** | 0.01380 | 0.00810 | 0.01156 | 0.02645 | 0.00925 | 0.00662 | 0.00750 | 0.01270 | 0.01286 | 0.06584 | 0.01230 | 0.04854 | 0.01208 | 0.01004 | 0.01949 | 0.00721 |
| **Stx2** | 1.62088 | 0.00629 | 0.13475 | 0.60938 | 0.00754 | 1.99210 | 0.00588 | 0.00000 | 0.82600 | 0.40298 | 0.73889 | 0.09032 | 0.00000 | 0.00487 | 0.00000 | 0.00464 |
| **Stx3** | 3.33716 | 7.33322 | 1.08093 | 0.60844 | 4.15821 | 6.12436 | 3.87166 | 1.06854 | 0.01194 | 6.09754 | 3.03596 | 6.48099 | 6.82427 | 2.87168 | 1.52986 | 5.17359 |
| **Stx4a** | 3.95586 | 0.01950 | 8.98294 | 0.06247 | 3.86875 | 3.50924 | 1.87801 | 7.56263 | 0.01965 | 2.03788 | 4.50590 | 0.10724 | 4.65123 | 0.00795 | 1.22171 | 5.87144 |
| **Stx5a** | 4.33076 | 0.63520 | 2.70078 | 11.02989 | 2.12117 | 6.02152 | 0.32699 | 3.53516 | 14.60073 | 4.09782 | 2.70020 | 2.72985 | 3.88354 | 2.90047 | 4.98418 | 5.40243 |
| **Stx6** | 0.02876 | 0.00775 | 0.03707 | 0.51984 | 0.00829 | 0.00594 | 0.01742 | 0.04069 | 0.00300 | 2.35747 | 0.01081 | 0.08115 | 0.75580 | 0.01191 | 0.19484 | 0.00681 |
| **Stx7** | 4.46287 | 3.58895 | 4.60304 | 3.72497 | 6.11027 | 0.09840 | 4.42458 | 7.85054 | 0.00003 | 7.37313 | 5.07541 | 0.76692 | 14.13039 | 6.46780 | 1.43133 | 1.99008 |
| **Stx8** | 9.16416 | 4.74593 | 4.34655 | 0.84964 | 0.46669 | 5.25909 | 2.15327 | 1.35127 | 0.00000 | 3.46786 | 0.11908 | 0.12115 | 2.86406 | 3.03079 | 0.00000 | 3.61870 |
| **Stxbp1** | 2.84371 | 0.00601 | 1.21119 | 4.28818 | 0.02902 | 0.00807 | 0.00541 | 0.87586 | 1.68129 | 1.28823 | 1.39788 | 0.03492 | 0.00000 | 0.47735 | 0.29970 | 0.61542 |
| **Stxbp2** | 7.95207 | 1.47734 | 3.39523 | 2.59627 | 0.87533 | 2.57300 | 2.59782 | 7.86358 | 3.34807 | 4.11448 | 2.98312 | 0.34399 | 7.09789 | 1.21550 | 0.00000 | 7.34932 |
| **Stxbp3** | 0.81311 | 1.06921 | 0.19017 | 0.56670 | 0.00582 | 3.02636 | 0.00495 | 1.84624 | 1.41390 | 0.85086 | 0.87625 | 0.01892 | 0.96727 | 2.99139 | 2.00115 | 2.21829 |
| **Stxbp3.ps** | 0.02859 | 0.00763 | 0.06867 | 0.77519 | 0.00955 | 0.00631 | 0.00706 | 0.11553 | 0.63590 | 0.00928 | 0.08917 | 0.35614 | 0.06955 | 0.51565 | 0.09501 | 0.29854 |
| **Stxbp4** | 0.68530 | 0.05939 | 1.19441 | 0.49593 | 0.40207 | 0.01683 | 0.12721 | 0.06559 | 0.73012 | 0.14389 | 1.79326 | 0.18587 | 0.64309 | 0.92481 | 0.20114 | 0.11491 |
| **Stxbp5** | 2.04876 | 0.01227 | 2.22727 | 12.98719 | 2.99811 | 0.39677 | 1.40633 | 1.65844 | 5.68303 | 1.67235 | 0.76691 | 0.31842 | 0.03890 | 0.00000 | 7.48942 | 0.00140 |
| **Stxbp5l** | 0.32612 | 0.01909 | 0.03524 | 1.78123 | 0.03053 | 0.00638 | 0.01683 | 0.17654 | 1.10935 | 0.77882 | 0.02985 | 0.01551 | 0.02955 | 1.16741 | 1.93988 | 0.00655 |
| **Stxbp6** | 0.57009 | 0.24543 | 0.06564 | 0.96277 | 0.04101 | 0.00567 | 1.04292 | 0.03877 | 0.40253 | 0.20480 | 0.34851 | 0.00302 | 2.23278 | 0.06140 | 0.00000 | 0.00591 |
| **Styk1** | 0.01348 | 0.00801 | 0.01172 | 0.21485 | 0.00914 | 0.00683 | 0.00752 | 0.01232 | 0.09444 | 0.43842 | 0.00970 | 0.00782 | 0.01248 | 0.00897 | 0.19071 | 0.00736 |
| **Styx** | 0.64525 | 1.85227 | 1.82656 | 3.95716 | 0.16881 | 0.29383 | 0.01441 | 0.75016 | 3.25495 | 3.12869 | 0.76170 | 5.67442 | 0.80907 | 7.66474 | 2.98743 | 0.15001 |
| **Styxl1** | 0.01356 | 0.00759 | 0.01207 | 0.35458 | 0.00929 | 0.00647 | 0.00728 | 0.01077 | 0.02403 | 0.46788 | 0.01272 | 0.00603 | 0.01188 | 0.01597 | 0.03359 | 0.00740 |
| **Sub1** | 20.60555 | 10.92824 | 17.47854 | 18.13117 | 22.58033 | 6.91969 | 6.73923 | 7.40987 | 26.58252 | 14.09512 | 19.61164 | 3.41444 | 17.84982 | 11.38021 | 12.63234 | 11.25667 |
| **Sucla2** | 12.62049 | 6.96139 | 13.15656 | 5.03831 | 10.52810 | 6.82330 | 10.73630 | 14.90978 | 9.06402 | 10.21074 | 15.65635 | 5.43835 | 13.04203 | 16.14984 | 3.06802 | 5.97648 |
| **Suclg1** | 25.08013 | 17.07513 | 22.57754 | 0.27615 | 20.79571 | 13.61282 | 14.99320 | 19.28993 | 0.78370 | 11.78232 | 8.41526 | 9.76636 | 17.04798 | 1.43633 | 1.57659 | 18.66775 |
| **Suclg2** | 16.20479 | 6.53211 | 6.73989 | 4.21445 | 16.81007 | 9.73820 | 5.56994 | 11.26401 | 1.63780 | 8.21630 | 6.80067 | 0.32471 | 10.83248 | 7.87616 | 5.07033 | 6.80782 |
| **Suco** | 2.95787 | 0.83251 | 4.36482 | 2.54475 | 0.56508 | 0.00849 | 2.49629 | 1.73194 | 4.25407 | 0.96966 | 0.19610 | 8.92224 | 0.54468 | 7.69055 | 1.72240 | 0.03800 |
| **Suds3** | 4.49762 | 4.40210 | 5.87199 | 14.03657 | 4.35177 | 1.05091 | 6.12092 | 9.61089 | 8.54092 | 24.42585 | 11.72325 | 7.98287 | 4.84137 | 0.15682 | 8.41728 | 0.01287 |
| **Sufu** | 1.76650 | 0.00388 | 3.19708 | 5.35935 | 0.00747 | 0.00416 | 1.45256 | 0.00000 | 3.09223 | 0.84141 | 1.33079 | 1.02979 | 0.13828 | 0.00000 | 0.40371 | 0.00291 |
| **Sugct** | 1.42343 | 0.00666 | 0.00381 | 3.37902 | 0.00515 | 0.00641 | 0.00622 | 0.01184 | 1.82838 | 0.00678 | 0.00000 | 0.00426 | 0.01152 | 0.01010 | 0.00328 | 0.00509 |
| **Sugp1** | 3.18999 | 0.85354 | 1.71672 | 5.74367 | 6.56008 | 0.94465 | 3.00818 | 7.83227 | 11.18877 | 6.46598 | 3.70929 | 0.22256 | 4.64808 | 0.00000 | 1.19602 | 7.31596 |
| **Sugp2** | 0.00677 | 0.34393 | 0.94534 | 0.84178 | 0.00653 | 0.00619 | 0.05600 | 0.28074 | 0.81096 | 1.38374 | 0.61889 | 0.00000 | 0.02522 | 0.02202 | 0.31215 | 0.13788 |
| **Sugt1** | 29.62468 | 14.99716 | 30.67735 | 18.69956 | 19.05329 | 16.86278 | 17.18831 | 39.32702 | 29.86734 | 48.59246 | 44.66553 | 27.88981 | 35.29895 | 39.55650 | 31.46265 | 13.51998 |
| **Sulf2** | 9.21078 | 3.31322 | 6.49925 | 2.16029 | 6.33371 | 0.00467 | 5.69313 | 1.49050 | 0.77553 | 2.18264 | 0.97637 | 0.00000 | 3.31580 | 0.00000 | 0.52366 | 0.60644 |
| **Sult1a1** | 0.01372 | 0.00817 | 0.01218 | 0.10330 | 0.00928 | 0.00676 | 0.00758 | 0.01434 | 0.01056 | 0.03039 | 0.01343 | 0.00876 | 0.01298 | 0.00998 | 0.02604 | 0.00740 |
| **Sult2a6** | 0.01378 | 0.00812 | 0.01192 | 0.32662 | 0.00935 | 0.00666 | 0.00750 | 0.01351 | 0.00428 | 0.01479 | 0.01228 | 0.02901 | 0.01276 | 0.01126 | 0.00554 | 0.00737 |
| **Sult2b1** | 0.02121 | 0.00669 | 0.75869 | 3.87718 | 0.03275 | 0.00490 | 2.02635 | 0.98882 | 2.75608 | 2.92746 | 1.14193 | 1.37339 | 0.71966 | 0.00000 | 0.07973 | 0.01256 |
| **Sult4a1** | 0.00000 | 0.00663 | 0.06137 | 0.14339 | 0.01007 | 0.00847 | 3.30420 | 0.55642 | 0.00618 | 0.86899 | 1.63078 | 0.04783 | 1.23914 | 0.00773 | 0.20931 | 0.03821 |
| **Sult5a1** | 2.26636 | 0.00387 | 6.27131 | 1.38257 | 0.00000 | 0.01374 | 0.00405 | 2.05790 | 2.55287 | 1.58290 | 2.02537 | 0.06084 | 0.00000 | 0.00000 | 2.14379 | 0.03110 |
| **Sult6b1** | 0.43634 | 0.00686 | 0.38409 | 0.00281 | 0.00668 | 0.00536 | 0.85306 | 0.00000 | 0.00979 | 1.35726 | 0.97428 | 0.00000 | 0.01354 | 0.00544 | 0.59519 | 0.00676 |
| **Sumf1** | 3.76775 | 0.01627 | 0.02949 | 1.54576 | 0.01081 | 0.00235 | 0.59796 | 0.98353 | 1.29801 | 2.61247 | 0.40424 | 0.02061 | 0.26139 | 0.77521 | 2.09292 | 0.00305 |
| **Sumf2** | 5.34672 | 3.07270 | 1.86733 | 0.22350 | 5.89903 | 0.00182 | 3.97371 | 1.58972 | 0.00000 | 0.85993 | 0.00000 | 0.00000 | 1.82415 | 0.00585 | 0.27296 | 0.28126 |
| **Sumo1** | 28.03855 | 16.89612 | 25.65544 | 14.74738 | 11.21279 | 12.51526 | 18.00149 | 28.59006 | 27.32226 | 31.53432 | 28.72076 | 10.82309 | 25.86920 | 35.54880 | 26.10772 | 10.29030 |
| **Sumo2** | 25.27734 | 14.56541 | 29.56287 | 11.91457 | 19.27695 | 14.99724 | 16.16578 | 37.35493 | 16.79771 | 49.31684 | 35.29728 | 27.00941 | 30.67227 | 33.04728 | 23.54743 | 14.20921 |
| **Sumo3** | 6.95008 | 1.04090 | 6.09196 | 5.76780 | 0.64119 | 5.45166 | 1.29724 | 7.96073 | 6.01161 | 14.54303 | 6.58594 | 2.39481 | 7.04905 | 13.12124 | 8.53324 | 6.57808 |
| **Sun1** | 3.04952 | 4.30740 | 10.84816 | 10.60055 | 3.48848 | 5.90689 | 9.56329 | 25.92061 | 17.16294 | 21.11060 | 20.44850 | 8.23256 | 10.63377 | 23.03135 | 23.64815 | 2.07449 |
| **Sun2** | 0.01228 | 0.00788 | 0.04963 | 0.72621 | 0.00931 | 0.00676 | 0.00735 | 0.02725 | 0.19820 | 0.00758 | 0.01052 | 0.01747 | 0.01110 | 0.00670 | 0.28173 | 0.00725 |
| **Suox** | 5.82671 | 0.03047 | 6.86659 | 0.00000 | 1.91662 | 0.01016 | 2.88242 | 3.99377 | 0.00683 | 1.07072 | 0.00000 | 0.03332 | 0.86459 | 0.19371 | 0.31999 | 5.12601 |
| **Supt16** | 0.28007 | 2.52705 | 0.77624 | 0.49927 | 1.05909 | 0.00566 | 0.82829 | 1.69025 | 0.74517 | 1.23655 | 2.03031 | 0.04035 | 0.98748 | 4.00653 | 1.46471 | 0.01178 |
| **Supt20** | 0.31131 | 2.71289 | 2.31792 | 0.99979 | 4.73617 | 3.47450 | 4.12346 | 1.88846 | 1.04218 | 2.29020 | 0.37750 | 0.14377 | 0.96428 | 0.00157 | 4.17820 | 0.31804 |
| **Supt3** | 0.03032 | 0.00769 | 2.49383 | 1.03688 | 0.03230 | 0.42237 | 0.00655 | 1.56273 | 0.83785 | 7.06477 | 1.70040 | 0.14944 | 0.13186 | 0.00073 | 0.84798 | 0.30469 |
| **Supt4a** | 8.95854 | 8.17480 | 11.12843 | 14.54328 | 9.69644 | 1.94946 | 4.28917 | 9.25482 | 22.64123 | 18.66699 | 14.76463 | 3.78195 | 10.50233 | 9.68118 | 13.69020 | 2.06954 |
| **Supt4b** | 3.38873 | 3.39942 | 4.45329 | 5.05899 | 3.49415 | 0.57454 | 1.50919 | 3.28458 | 8.42534 | 8.46623 | 5.09968 | 1.27997 | 4.41827 | 3.76920 | 6.04327 | 0.65980 |
| **Supt5** | 12.68947 | 7.37004 | 11.10681 | 17.00054 | 6.64004 | 5.43775 | 7.54809 | 20.88745 | 19.58742 | 29.91591 | 25.03348 | 7.77390 | 15.50897 | 3.74953 | 13.75725 | 6.24115 |
| **Supt6** | 11.86770 | 10.26953 | 7.45958 | 6.46234 | 5.72848 | 3.43066 | 1.47916 | 13.42796 | 6.58547 | 17.19987 | 5.98222 | 0.62138 | 4.62409 | 0.74810 | 3.81963 | 8.70743 |
| **Supt7l** | 2.10106 | 4.63389 | 0.43681 | 5.58222 | 0.04801 | 1.00671 | 0.04057 | 8.79033 | 8.04835 | 5.09176 | 3.87078 | 0.44818 | 2.93190 | 8.98096 | 7.00027 | 0.08541 |
| **Supv3l1** | 4.79474 | 3.37987 | 4.79317 | 7.18366 | 2.74660 | 5.90355 | 4.08127 | 8.17345 | 6.25171 | 10.90591 | 5.19620 | 1.10871 | 7.00319 | 1.66368 | 12.55142 | 4.64136 |
| **Surf1** | 5.95158 | 6.20158 | 9.26783 | 1.83462 | 0.05186 | 2.58085 | 3.26122 | 8.69362 | 0.61226 | 8.60287 | 8.39623 | 0.16724 | 6.05878 | 0.00000 | 0.84003 | 2.87874 |

| | | | | | | | | | | | | | | | | |
|---|---|---|---|---|---|---|---|---|---|---|---|---|---|---|---|---|
| **Surf2** | 11.15691 | 8.58833 | 14.86003 | 2.14396 | 5.37682 | 7.36814 | 7.51215 | 15.49361 | 2.04528 | 14.96990 | 3.81525 | 1.65679 | 7.29027 | 5.90051 | 11.75321 | 0.50785 |
| **Surf4** | 13.24795 | 3.54839 | 16.54885 | 9.32472 | 3.43425 | 10.76819 | 5.30123 | 7.18619 | 6.36066 | 4.19583 | 6.42456 | 0.56879 | 8.43292 | 1.19977 | 6.29528 | 10.67806 |
| **Surf6** | 0.46119 | 1.94175 | 1.23619 | 12.27389 | 0.05733 | 0.00324 | 0.04348 | 3.12479 | 15.67859 | 4.14264 | 9.05858 | 0.81904 | 0.26821 | 1.59773 | 6.55043 | 6.13734 |
| **Susd1** | 0.01386 | 0.00787 | 0.01046 | 0.25158 | 0.00930 | 0.00648 | 0.00731 | 0.00694 | 0.00722 | 0.00080 | 0.17066 | 0.00600 | 0.01043 | 0.00551 | 0.53290 | 0.00731 |
| **Susd2** | 0.08535 | 0.00729 | 1.24078 | 0.20481 | 0.00394 | 2.98230 | 0.00605 | 0.00604 | 0.00839 | 0.01617 | 0.00837 | 0.01693 | 1.34825 | 0.01154 | 0.00000 | 4.69113 |
| **Susd3** | 0.49289 | 0.00752 | 0.03343 | 5.46969 | 0.01616 | 0.00780 | 0.00551 | 0.82158 | 2.45870 | 3.13387 | 2.85025 | 0.00871 | 0.00000 | 0.00000 | 0.10700 | 0.00559 |
| **Susd4** | 0.01299 | 0.00822 | 0.05275 | 0.01064 | 0.00927 | 0.00594 | 0.00767 | 0.01436 | 0.01236 | 0.01524 | 0.01288 | 0.00852 | 0.01184 | 0.01269 | 0.00978 | 0.00738 |
| **Susd6** | 1.58873 | 3.75896 | 0.91145 | 3.21071 | 1.94298 | 0.88973 | 1.64884 | 2.88354 | 7.17812 | 0.07147 | 2.14736 | 0.08108 | 0.30115 | 4.46358 | 4.58246 | 0.03825 |
| **Suv39h1** | 8.54297 | 2.73911 | 4.56790 | 9.08707 | 4.84988 | 0.00087 | 3.70146 | 5.18817 | 12.94536 | 4.90293 | 0.66746 | 0.35852 | 5.01299 | 7.03335 | 7.80729 | 6.14928 |
| **Suv39h2** | 1.38062 | 0.00511 | 1.68883 | 7.27713 | 0.01114 | 0.34213 | 0.00459 | 0.00000 | 3.23323 | 1.65203 | 0.98773 | 1.62772 | 1.20833 | 2.18321 | 4.25644 | 0.11263 |
| **Suv420h1** | 5.20062 | 4.73561 | 1.04728 | 13.65311 | 1.41679 | 1.69311 | 2.90868 | 2.65943 | 14.41495 | 10.45782 | 6.21654 | 8.19022 | 2.91391 | 16.28930 | 8.18171 | 7.98564 |
| **Suv420h2** | 3.84984 | 0.00000 | 0.73681 | 0.85444 | 2.49677 | 0.25480 | 1.53230 | 0.69391 | 2.01956 | 0.51104 | 0.44407 | 0.05199 | 0.42721 | 0.04517 | 0.00000 | 1.67830 |
| **Suz12** | 14.69032 | 13.18005 | 9.88205 | 23.05114 | 12.63302 | 7.41894 | 14.96696 | 17.64045 | 22.11553 | 25.05034 | 18.85427 | 25.29555 | 8.46242 | 32.09200 | 27.96091 | 2.46485 |
| **Sva** | 0.01370 | 0.00829 | 0.01221 | 1.80280 | 0.00931 | 0.00648 | 0.00747 | 0.01394 | 0.31602 | 0.01279 | 0.01115 | 0.00793 | 0.01273 | 0.01027 | 0.00289 | 0.00710 |
| **Svbp** | 0.05432 | 0.00593 | 0.55491 | 0.00000 | 2.58361 | 0.41193 | 0.03625 | 0.34989 | 0.00987 | 2.21521 | 0.00712 | 0.00751 | 0.64211 | 0.01559 | 0.00000 | 0.00660 |
| **Svil** | 0.05173 | 0.00722 | 1.50143 | 0.01042 | 0.00531 | 0.05442 | 0.01716 | 0.09719 | 0.01240 | 0.01514 | 0.01028 | 0.01599 | 0.00243 | 0.00880 | 0.03309 | 0.00672 |
| **Svip** | 1.06188 | 0.00594 | 0.17871 | 3.73017 | 0.57218 | 0.00549 | 0.00586 | 0.36008 | 0.45726 | 1.53249 | 0.48586 | 0.01351 | 0.00241 | 4.63306 | 1.44406 | 0.15631 |
| **Svop** | 0.01286 | 0.00783 | 0.01168 | 1.14398 | 0.00880 | 0.00629 | 0.00715 | 0.00955 | 0.00000 | 1.64726 | 0.01346 | 0.01558 | 0.01014 | 0.00860 | 0.27183 | 0.00688 |
| **Svopl** | 0.01513 | 0.02983 | 0.01900 | 0.14597 | 0.03599 | 0.00610 | 0.00744 | 0.00863 | 1.03503 | 0.67751 | 0.12141 | 0.02401 | 0.01906 | 0.05300 | 0.28205 | 0.01433 |
| **Swap70** | 0.90981 | 0.57263 | 6.31437 | 0.40791 | 5.16435 | 3.57229 | 2.49500 | 3.50639 | 0.98257 | 2.69747 | 0.50501 | 0.15320 | 3.60722 | 0.21710 | 0.93560 | 1.00435 |
| **Swi5** | 9.80537 | 9.13211 | 8.40927 | 12.08361 | 8.68980 | 3.80370 | 2.67539 | 2.56812 | 4.95197 | 3.99428 | 5.62703 | 0.09619 | 6.70497 | 1.41732 | 1.67715 | 9.55786 |
| **Swsap1** | 0.64742 | 0.01703 | 1.91351 | 0.42646 | 1.92939 | 0.00833 | 0.00642 | 0.36939 | 1.33242 | 0.00000 | 0.48016 | 0.11342 | 1.69209 | 0.00655 | 1.24796 | 0.00510 |
| **Swt1** | 3.44250 | 1.32168 | 3.37505 | 12.63856 | 1.59361 | 0.90188 | 3.48070 | 7.01691 | 11.90654 | 12.17377 | 11.81534 | 13.18111 | 2.95441 | 21.92937 | 16.93356 | 2.31399 |
| **Syap1** | 2.79757 | 0.06409 | 7.39503 | 12.75703 | 11.59377 | 2.25356 | 3.48755 | 7.45879 | 15.60127 | 22.08848 | 13.73745 | 13.55340 | 9.31113 | 22.24227 | 18.64923 | 0.77434 |
| **Sybu** | 0.01379 | 0.00779 | 0.01195 | 0.38093 | 0.00934 | 0.00661 | 0.00749 | 0.01321 | 0.00475 | 0.01201 | 0.01249 | 0.00779 | 0.01275 | 0.01890 | 0.06329 | 0.00735 |
| **Syce1** | 0.01287 | 0.00783 | 0.01690 | 0.00562 | 0.00919 | 0.00615 | 0.00738 | 0.01249 | 0.08152 | 0.00480 | 0.02771 | 0.00728 | 0.13144 | 0.00667 | 0.19158 | 0.00717 |
| **Syce1l** | 0.36490 | 0.00664 | 1.19782 | 0.00473 | 0.00518 | 0.00619 | 0.09420 | 0.05908 | 0.11236 | 0.86753 | 0.74261 | 0.09073 | 1.96464 | 0.05037 | 0.04390 | 0.00561 |
| **Syce2** | 8.47012 | 1.05952 | 11.22365 | 17.32637 | 3.31674 | 6.19263 | 7.24287 | 12.61946 | 19.02489 | 11.78437 | 16.18883 | 0.86042 | 14.00476 | 1.25622 | 0.31220 | 4.79558 |
| **Syce3** | 0.02022 | 0.00812 | 0.20019 | 0.76772 | 0.00972 | 0.00615 | 0.00729 | 0.00457 | 0.97342 | 0.08623 | 0.27764 | 0.01403 | 0.00805 | 0.02601 | 0.00000 | 0.04241 |
| **Sycn** | 1.44901 | 0.43054 | 0.95222 | 0.00000 | 1.92126 | 0.00468 | 0.71665 | 0.35902 | 0.03012 | 3.56893 | 2.45060 | 1.22126 | 1.02591 | 0.14567 | 0.55524 | 0.05700 |
| **Sycp1** | 0.04776 | 0.00644 | 1.01938 | 6.18840 | 0.00700 | 0.01226 | 0.79762 | 0.02452 | 0.27683 | 0.78898 | 0.03450 | 3.32807 | 1.02378 | 14.72278 | 3.35239 | 0.00522 |
| **Sycp1.ps1** | 0.01088 | 0.00824 | 0.04099 | 0.07128 | 0.00936 | 0.00776 | 0.01849 | 0.02136 | 0.02030 | 0.06131 | 0.00807 | 0.01622 | 0.07733 | 0.63268 | 0.24353 | 0.00752 |
| **Sycp2** | 0.00767 | 0.00738 | 0.00506 | 5.03119 | 0.01001 | 0.00565 | 0.62739 | 0.04851 | 2.05829 | 0.00000 | 0.00009 | 0.03930 | 0.00940 | 3.40805 | 0.48782 | 0.00616 |
| **Sycp2l** | 0.01369 | 0.00809 | 0.01208 | 0.00493 | 0.00931 | 0.00665 | 0.00755 | 0.01390 | 0.39895 | 0.01318 | 0.00980 | 0.00757 | 0.01283 | 0.01338 | 0.00772 | 0.00740 |
| **Sycp3** | 0.35078 | 4.39930 | 2.55168 | 1.02760 | 1.40599 | 0.01346 | 2.05601 | 0.58848 | 3.46062 | 0.72617 | 0.58927 | 0.49928 | 2.96148 | 1.41698 | 3.84637 | 0.05307 |
| **Syde1** | 0.01381 | 0.00812 | 0.00947 | 0.17330 | 0.00905 | 0.00666 | 0.00744 | 0.00504 | 0.66941 | 0.14426 | 0.27266 | 0.02657 | 0.00891 | 0.00882 | 0.00000 | 0.00702 |
| **Syde2** | 0.66813 | 0.09718 | 0.84591 | 1.42930 | 0.01245 | 0.00627 | 0.00581 | 0.52363 | 3.38920 | 0.00000 | 2.25044 | 3.16230 | 0.04514 | 0.08438 | 0.00000 | 0.01317 |
| **Syf2** | 8.10613 | 5.11416 | 5.84796 | 13.36814 | 8.28173 | 5.59180 | 5.93025 | 17.85813 | 16.07566 | 24.22457 | 13.48105 | 4.85421 | 16.26610 | 2.26652 | 7.81908 | 9.84734 |
| **Sympk** | 0.43826 | 0.00740 | 0.17157 | 0.43287 | 0.05327 | 0.00699 | 0.07342 | 0.00813 | 0.47194 | 0.86947 | 0.49124 | 0.02429 | 0.21354 | 0.00845 | 0.00000 | 0.15985 |
| **Syn1** | 0.06348 | 0.00782 | 0.14788 | 0.25605 | 0.00764 | 0.00552 | 0.00734 | 0.01145 | 0.40923 | 0.00300 | 0.04787 | 0.24158 | 0.06703 | 0.01072 | 0.00000 | 0.00706 |
| **Syn2** | 0.01109 | 0.00865 | 0.00911 | 0.00815 | 0.00950 | 0.00671 | 0.00813 | 0.11643 | 0.01121 | 0.01062 | 0.01025 | 0.07091 | 0.00829 | 0.00802 | 0.41254 | 0.00729 |
| **Syna** | 0.01196 | 0.00769 | 0.41661 | 1.78742 | 0.00843 | 0.00873 | 0.00718 | 0.01145 | 1.08211 | 0.47959 | 0.00250 | 0.00384 | 0.00965 | 0.00945 | 0.03746 | 0.00692 |
| **Sync** | 0.00525 | 0.00771 | 0.10291 | 0.02380 | 0.01857 | 0.00874 | 0.00677 | 0.99503 | 0.01132 | 0.00000 | 0.00445 | 0.01311 | 1.10728 | 0.12994 | 0.35523 | 0.00681 |
| **Syncrip** | 12.63692 | 13.29420 | 13.83256 | 14.76114 | 12.29139 | 6.00696 | 10.69388 | 23.76448 | 22.65351 | 22.59553 | 18.12507 | 12.69485 | 14.82572 | 28.60259 | 17.09121 | 4.56978 |
| **Syne1** | 0.03704 | 0.00812 | 0.05543 | 0.11192 | 0.00884 | 0.00613 | 0.00758 | 0.04875 | 0.24303 | 0.04095 | 0.03686 | 0.01824 | 0.05096 | 0.21730 | 0.04041 | 0.00725 |
| **Syne2** | 0.00000 | 1.37309 | 1.84665 | 7.14504 | 0.04801 | 0.88099 | 1.67621 | 4.79128 | 8.17067 | 13.47161 | 5.36477 | 8.39579 | 0.89436 | 1.25733 | 5.35924 | 0.00218 |
| **Syne3** | 0.00842 | 0.00725 | 0.00438 | 0.00148 | 0.00912 | 0.00672 | 0.00705 | 1.09211 | 0.59547 | 1.25622 | 0.13717 | 0.13821 | 0.00477 | 0.02311 | 1.06057 | 0.02201 |
| **Syngr1** | 8.77829 | 1.64294 | 6.35205 | 2.73454 | 6.22356 | 3.42849 | 5.74041 | 10.67095 | 2.19375 | 13.40536 | 16.11064 | 2.83218 | 13.90804 | 2.12695 | 2.17790 | 0.18561 |
| **Syngr2** | 8.54249 | 0.88845 | 10.09763 | 7.12353 | 0.12837 | 12.53783 | 4.14027 | 8.63296 | 2.01534 | 6.35716 | 4.35625 | 0.59528 | 10.84484 | 0.00410 | 4.23818 | 6.18660 |
| **Syngr3** | 0.01301 | 0.36428 | 0.02306 | 7.68005 | 0.00856 | 0.06209 | 0.00670 | 0.00000 | 9.90722 | 0.38504 | 1.34562 | 0.08590 | 0.05619 | 0.00000 | 0.67084 | 0.01291 |
| **Syngr4** | 0.57528 | 0.01979 | 0.01219 | 0.28880 | 0.01599 | 0.00595 | 0.00695 | 0.01044 | 0.00727 | 0.02122 | 0.01449 | 0.01914 | 0.01168 | 0.01006 | 0.05350 | 0.00685 |
| **Synj1** | 1.71812 | 2.45844 | 1.03155 | 9.13361 | 1.56728 | 0.55743 | 0.01502 | 1.20580 | 8.59515 | 3.32111 | 1.41857 | 1.17011 | 1.30495 | 7.96111 | 8.27931 | 6.58898 |
| **Synj2** | 0.01319 | 0.00791 | 0.01048 | 0.00950 | 0.00917 | 0.00695 | 0.19093 | 0.01176 | 0.01207 | 0.24342 | 0.01201 | 0.00818 | 0.01971 | 0.01150 | 0.01166 | 0.00743 |
| **Synj2bp** | 1.74581 | 0.83634 | 0.73088 | 1.19761 | 2.37193 | 4.64733 | 5.06434 | 0.17870 | 1.00603 | 0.00000 | 1.63189 | 6.91417 | 0.29621 | 3.25153 | 2.98505 | 0.10445 |
| **Synm** | 0.01370 | 0.00770 | 0.01218 | 0.97650 | 0.00923 | 0.00652 | 0.00734 | 0.01166 | 0.00000 | 0.15771 | 0.01223 | 0.00626 | 0.01285 | 0.00372 | 0.28038 | 0.00762 |
| **Synpo2l** | 0.00000 | 0.00733 | 0.88989 | 0.03353 | 0.00653 | 0.00972 | 0.00762 | 2.34436 | 0.00959 | 0.00000 | 0.03244 | 0.00247 | 0.06393 | 0.01534 | 0.00000 | 1.43735 |
| **Synpr** | 0.00617 | 0.66441 | 0.02154 | 0.32177 | 0.00819 | 0.00591 | 0.00702 | 0.06896 | 0.00684 | 0.00205 | 0.00607 | 0.58581 | 0.00927 | 0.00491 | 0.59031 | 0.00703 |
| **Synrg** | 5.48120 | 2.87229 | 0.62857 | 5.04354 | 4.68690 | 5.64221 | 0.07021 | 1.17479 | 7.41315 | 2.26230 | 6.18584 | 2.27208 | 2.19155 | 3.17078 | 4.01197 | 0.35315 |
| **Syp** | 0.00965 | 0.00864 | 0.34102 | 1.05592 | 0.00945 | 0.00797 | 0.00766 | 0.03274 | 0.45595 | 0.10749 | 1.22940 | 0.00234 | 0.42941 | 0.00943 | 0.24485 | 0.00706 |
| **Sypl** | 5.74318 | 6.53205 | 9.51161 | 4.56799 | 4.37705 | 4.32091 | 10.45337 | 1.85852 | 3.59575 | 2.90937 | 2.73698 | 10.36332 | 6.02702 | 4.94000 | 5.58959 | 1.23893 |
| **Sys1** | 5.26720 | 0.28989 | 3.24962 | 15.23874 | 7.62767 | 0.00718 | 0.18834 | 5.31275 | 11.43417 | 11.66343 | 11.37304 | 0.62315 | 2.12787 | 0.12773 | 2.78689 | 2.08532 |
| **Syt11** | 0.05987 | 0.00761 | 0.16535 | 8.11457 | 0.02674 | 0.01619 | 0.03604 | 0.00000 | 7.76938 | 0.00000 | 0.88619 | 0.02024 | 1.43719 | 0.55302 | 2.45984 | 0.02527 |

| | | | | | | | | | | | | | | | | |
|---|---|---|---|---|---|---|---|---|---|---|---|---|---|---|---|---|
| **Syt12** | 0.06474 | 0.00812 | 0.10620 | 0.13724 | 0.04177 | 0.01261 | 0.05049 | 0.02689 | 0.23283 | 0.05237 | 0.47706 | 0.01575 | 0.07670 | 0.11896 | 0.13457 | 0.02169 |
| **Syt14** | 0.01511 | 0.00812 | 0.01415 | 0.01858 | 0.00984 | 0.00713 | 0.00761 | 0.01645 | 0.01131 | 0.00793 | 0.01465 | 0.00756 | 0.40448 | 0.00905 | 0.04272 | 0.00749 |
| **Syt15** | 0.28872 | 0.10861 | 1.03290 | 0.47842 | 0.06385 | 0.02356 | 0.42610 | 0.00000 | 0.63751 | 0.17859 | 2.31683 | 0.13031 | 0.53971 | 1.57139 | 0.43420 | 0.07025 |
| **Syt16** | 0.01252 | 0.00768 | 0.00721 | 0.89949 | 0.00952 | 0.00666 | 0.00723 | 0.00969 | 0.03773 | 0.01854 | 0.00715 | 0.00319 | 0.01305 | 1.35821 | 0.00000 | 0.00716 |
| **Syt17** | 0.01293 | 0.00801 | 0.04760 | 0.01063 | 0.00918 | 0.00789 | 0.00748 | 0.01361 | 0.01234 | 0.01529 | 0.01286 | 0.00800 | 0.01195 | 0.01244 | 0.00999 | 0.00739 |
| **Syt2** | 0.01375 | 0.00812 | 0.01118 | 0.01002 | 0.02084 | 0.00681 | 0.00718 | 0.01221 | 0.01190 | 0.19470 | 0.01148 | 0.00808 | 0.64798 | 0.04130 | 0.03171 | 0.00714 |
| **Syt5** | 0.00680 | 0.00809 | 0.00570 | 1.19159 | 0.00894 | 0.00726 | 0.00637 | 0.00000 | 0.59724 | 0.29528 | 0.71900 | 0.09090 | 1.68335 | 0.00424 | 0.04124 | 0.00634 |
| **Syt7** | 0.40880 | 0.00805 | 0.00198 | 0.01526 | 0.01737 | 0.00723 | 0.00632 | 0.00777 | 0.13681 | 0.13784 | 0.01016 | 0.18497 | 0.90358 | 0.04377 | 0.05821 | 1.03614 |
| **Syt9** | 12.10687 | 0.00000 | 3.14763 | 0.00000 | 2.17339 | 0.00000 | 2.31689 | 0.00000 | 1.16625 | 0.00000 | 0.00000 | 0.03178 | 0.01204 | 0.01732 | 0.00000 | 0.31956 |
| **Sytl1** | 0.01493 | 0.00815 | 0.47774 | 0.10809 | 0.00933 | 0.00639 | 0.00766 | 0.01618 | 0.00729 | 0.04403 | 0.01306 | 0.00818 | 0.17668 | 0.01253 | 0.00103 | 0.00751 |
| **Sytl2** | 0.01370 | 0.00741 | 0.00593 | 1.74226 | 0.00939 | 0.00633 | 0.00704 | 0.00899 | 0.56166 | 0.11461 | 0.02813 | 0.59161 | 0.01264 | 0.00369 | 1.13327 | 0.00711 |
| **Sytl3** | 1.43263 | 0.00660 | 0.00226 | 0.00424 | 0.01502 | 0.00771 | 0.00610 | 0.30911 | 0.00983 | 3.56655 | 1.11489 | 0.06957 | 1.02662 | 0.06304 | 0.17621 | 0.02766 |
| **Sytl4** | 0.01307 | 0.00781 | 0.01570 | 1.38001 | 0.00902 | 0.00728 | 0.00725 | 0.00661 | 0.60985 | 0.44073 | 0.13732 | 0.00445 | 0.01097 | 0.09398 | 0.34678 | 0.00726 |
| **Syvn1** | 1.80860 | 0.00370 | 1.79913 | 0.78639 | 1.09344 | 3.97178 | 1.88713 | 2.78753 | 1.81927 | 2.55973 | 0.19191 | 0.29636 | 3.79506 | 0.00847 | 0.00000 | 3.24518 |
| **Szrd1** | 10.83328 | 3.39184 | 11.78156 | 1.60255 | 5.49919 | 2.03141 | 8.31750 | 7.93241 | 0.62073 | 9.50765 | 4.45305 | 0.46852 | 9.58695 | 0.01674 | 0.19631 | 5.24398 |
| **Szt2** | 2.43024 | 0.23908 | 2.32315 | 0.77530 | 0.00699 | 0.00560 | 1.24010 | 4.04433 | 2.04255 | 1.08554 | 0.84034 | 0.08454 | 1.42436 | 0.02716 | 0.35133 | 2.16469 |
| **T2** | 0.02986 | 0.00772 | 0.06378 | 1.30740 | 0.04035 | 0.01229 | 0.00685 | 0.00000 | 2.31303 | 0.01831 | 1.31517 | 0.00000 | 0.01891 | 0.32491 | 0.00000 | 0.02212 |
| **Tab1** | 0.45703 | 0.02958 | 2.91231 | 1.77935 | 0.74525 | 2.61699 | 0.00585 | 0.00000 | 0.67361 | 1.41503 | 0.76578 | 0.13077 | 0.63786 | 0.02262 | 0.00000 | 0.01198 |
| **Tab2** | 3.44661 | 4.51453 | 1.50343 | 10.59492 | 5.56563 | 3.64413 | 3.99160 | 3.89921 | 7.29919 | 3.85297 | 5.58541 | 6.09100 | 3.00247 | 10.05221 | 11.86290 | 1.95051 |
| **Tab3** | 0.92436 | 0.95177 | 0.93530 | 7.18135 | 0.00896 | 1.48175 | 0.00482 | 2.12402 | 2.76311 | 0.93010 | 1.13894 | 1.10308 | 1.30706 | 2.87737 | 3.43340 | 0.00334 |
| **Tac2** | 0.01616 | 0.00815 | 0.08161 | 2.75185 | 0.01024 | 0.00826 | 0.00713 | 0.01242 | 0.40372 | 0.00658 | 0.00964 | 0.01766 | 0.20669 | 0.01274 | 0.00318 | 0.00670 |
| **Tacc1** | 2.53021 | 0.01620 | 0.39988 | 5.00655 | 0.01039 | 4.71009 | 0.00386 | 1.70558 | 3.62650 | 4.29256 | 1.58120 | 0.08965 | 1.58018 | 1.00425 | 2.57810 | 1.57386 |
| **Tacc2** | 6.22662 | 2.59781 | 6.72946 | 7.50401 | 3.05336 | 1.16709 | 0.62716 | 4.00105 | 1.11459 | 1.28411 | 0.71860 | 0.09703 | 6.37443 | 0.00000 | 2.76933 | 2.22453 |
| **Tacc3** | 9.10600 | 8.31177 | 8.76029 | 13.20514 | 4.17001 | 4.38938 | 6.91315 | 18.72776 | 15.75006 | 8.02301 | 9.55055 | 1.48185 | 7.99669 | 6.29535 | 16.67369 | 9.88948 |
| **Taco1** | 1.42335 | 0.14153 | 2.05391 | 1.60375 | 4.47552 | 0.00665 | 0.00596 | 2.54837 | 1.38483 | 0.01037 | 0.02922 | 0.01248 | 0.46328 | 0.00440 | 0.41678 | 3.33048 |
| **Tacr3** | 0.01353 | 0.00738 | 0.00767 | 0.87824 | 0.00916 | 0.00653 | 0.00710 | 0.00908 | 2.53428 | 0.00000 | 0.00000 | 0.71710 | 0.01350 | 0.00919 | 0.00000 | 0.00706 |
| **Tacstd2** | 0.12187 | 0.00824 | 0.02718 | 0.00818 | 0.06661 | 4.96735 | 0.00580 | 0.00565 | 0.01297 | 0.00000 | 0.00889 | 0.01681 | 1.68577 | 0.01512 | 0.00919 | 5.87785 |
| **Tada1** | 5.15109 | 4.58639 | 5.57006 | 9.35257 | 1.41618 | 5.46643 | 3.58382 | 5.95569 | 9.27800 | 8.91893 | 7.21407 | 13.40517 | 4.18727 | 10.42491 | 5.26132 | 5.38704 |
| **Tada2a** | 8.09536 | 11.24479 | 0.00000 | 1.97927 | 9.28340 | 7.09991 | 4.13518 | 9.31869 | 6.64683 | 13.70138 | 4.93955 | 7.30645 | 5.13492 | 6.29704 | 17.54097 | 3.92345 |
| **Tada2b** | 0.95860 | 4.58984 | 0.46361 | 3.42621 | 3.16326 | 3.62511 | 1.53705 | 0.20713 | 0.77286 | 1.93028 | 0.00000 | 0.15988 | 0.88722 | 0.38629 | 1.51415 | 1.23973 |
| **Tada3** | 1.62140 | 2.93784 | 6.78119 | 1.50168 | 0.15239 | 5.75699 | 1.60719 | 12.52752 | 1.56492 | 8.44869 | 5.01316 | 0.28247 | 7.58941 | 0.00000 | 0.14184 | 1.03645 |
| **Taf1** | 3.70611 | 4.91094 | 5.98894 | 7.80406 | 3.77197 | 0.06041 | 5.19093 | 3.91251 | 14.83723 | 9.09692 | 4.59116 | 11.41989 | 3.21860 | 23.37012 | 13.43020 | 0.07171 |
| **Taf10** | 2.53786 | 1.09252 | 0.70646 | 19.96436 | 2.15866 | 0.10504 | 0.04277 | 0.03866 | 20.85100 | 1.78646 | 12.70647 | 0.12923 | 1.06222 | 0.49748 | 0.00000 | 1.68625 |
| **Taf11** | 8.45850 | 5.04065 | 6.99562 | 7.93341 | 0.79766 | 10.50441 | 8.19185 | 12.62715 | 6.53261 | 13.08726 | 3.86939 | 0.66865 | 11.03381 | 3.00541 | 4.45906 | 9.06246 |
| **Taf12** | 3.90306 | 4.80707 | 9.27558 | 4.46662 | 6.42443 | 5.75951 | 3.78309 | 6.04445 | 8.95551 | 8.29594 | 4.53993 | 1.54018 | 5.96094 | 5.86367 | 8.72995 | 5.17596 |
| **Taf13** | 4.88432 | 5.57660 | 9.53059 | 10.32415 | 0.17382 | 0.58012 | 4.03644 | 9.97133 | 11.79776 | 12.18874 | 10.65020 | 7.18560 | 9.40064 | 21.88603 | 13.35834 | 1.45506 |
| **Taf15** | 0.08306 | 0.00794 | 0.00193 | 0.00335 | 0.17461 | 0.00774 | 0.04599 | 0.32688 | 0.17704 | 0.00000 | 0.48239 | 0.02030 | 0.00949 | 0.19638 | 0.34260 | 0.00718 |
| **Taf1a** | 5.11109 | 0.00000 | 3.25962 | 7.25015 | 0.03862 | 0.01181 | 0.00946 | 4.29166 | 14.79404 | 6.33941 | 5.21083 | 1.54508 | 2.83570 | 1.07444 | 8.85768 | 3.87132 |
| **Taf1b** | 3.12605 | 4.07711 | 3.34691 | 8.52280 | 2.96568 | 1.20878 | 4.29406 | 5.76721 | 11.53351 | 6.81789 | 4.25351 | 4.61652 | 2.59172 | 7.57715 | 8.62933 | 2.00913 |
| **Taf1c** | 5.78292 | 0.42813 | 2.19212 | 0.00000 | 0.00995 | 0.89400 | 1.20731 | 12.77074 | 0.33471 | 10.76709 | 0.28174 | 0.43320 | 4.55650 | 0.03327 | 0.73352 | 0.01301 |
| **Taf1d** | 9.27386 | 6.95310 | 0.80452 | 22.07859 | 3.68334 | 2.80938 | 2.36614 | 14.31387 | 28.42208 | 12.76761 | 14.72885 | 14.92337 | 7.73778 | 26.76399 | 19.21948 | 7.44963 |
| **Taf2** | 3.89985 | 5.36439 | 0.57485 | 6.71366 | 8.69668 | 3.28870 | 4.58054 | 5.75252 | 3.46060 | 4.17410 | 3.53873 | 6.12415 | 1.30954 | 11.85788 | 12.21711 | 0.01092 |
| **Taf3** | 2.24681 | 0.23658 | 0.44492 | 8.08391 | 2.38713 | 0.08415 | 0.90301 | 0.30683 | 4.23337 | 3.72410 | 0.84067 | 1.67663 | 1.95750 | 2.33133 | 9.12537 | 0.22605 |
| **Taf4** | 1.59148 | 2.86021 | 0.60448 | 9.55611 | 5.41144 | 4.27611 | 2.10062 | 2.52165 | 4.74798 | 3.02149 | 1.12379 | 3.62983 | 2.15778 | 1.09704 | 1.95275 | 1.23153 |
| **Taf4b** | 3.71673 | 5.08924 | 0.30644 | 14.74858 | 0.30908 | 0.08210 | 3.18307 | 7.80487 | 11.39190 | 6.37420 | 3.79997 | 10.78622 | 1.07439 | 23.98401 | 16.58747 | 2.57352 |
| **Taf5** | 1.25485 | 3.71878 | 0.00000 | 10.76857 | 0.00555 | 0.17080 | 0.16142 | 1.79971 | 20.23698 | 5.32076 | 1.21241 | 2.18029 | 1.69235 | 5.55252 | 3.52798 | 0.02184 |
| **Taf5l** | 2.09427 | 1.42881 | 0.00000 | 12.63700 | 1.09343 | 3.39794 | 1.24703 | 7.38520 | 15.09822 | 1.70725 | 7.95737 | 1.85824 | 1.08199 | 5.81590 | 11.40470 | 0.40301 |
| **Taf6** | 6.75195 | 6.10098 | 2.19393 | 8.74259 | 9.34551 | 3.85394 | 4.83535 | 8.30615 | 7.82340 | 12.88477 | 6.88677 | 1.14916 | 3.56818 | 0.64365 | 5.82403 | 2.86730 |
| **Taf6l** | 0.04879 | 0.00762 | 0.00691 | 0.00427 | 0.01008 | 0.00698 | 0.00717 | 0.39498 | 0.33449 | 0.38830 | 0.00623 | 0.00403 | 0.10589 | 0.00685 | 0.34777 | 0.01510 |
| **Taf7** | 13.29978 | 3.11014 | 4.85025 | 0.00000 | 10.14962 | 5.23763 | 4.87492 | 13.17300 | 0.05111 | 10.53227 | 0.47575 | 0.92009 | 1.83141 | 9.80617 | 14.24441 | 4.91438 |
| **Taf8** | 2.22295 | 2.84968 | 4.04697 | 5.22104 | 0.00976 | 0.34376 | 0.48777 | 4.42940 | 4.49109 | 4.60114 | 1.94402 | 0.33508 | 1.02177 | 1.94433 | 7.26105 | 0.00825 |
| **Taf9** | 14.32137 | 11.19377 | 10.29627 | 29.30986 | 10.05063 | 11.01721 | 8.49576 | 16.20186 | 34.97665 | 36.43979 | 18.11245 | 22.64191 | 12.71832 | 27.78646 | 25.53880 | 8.38867 |
| **Taf9b** | 0.01358 | 0.00778 | 0.00994 | 0.78216 | 0.00932 | 0.00625 | 0.00727 | 0.00902 | 0.93779 | 0.00641 | 0.04484 | 0.01462 | 0.01183 | 0.01381 | 1.21994 | 0.00702 |
| **Tagap** | 0.24716 | 0.56921 | 1.27492 | 1.38124 | 0.01596 | 0.01283 | 0.75748 | 1.73215 | 0.92994 | 3.45427 | 0.11499 | 1.60342 | 0.02595 | 0.60094 | 1.48811 | 0.00595 |
| **Tagap1** | 0.29043 | 0.93783 | 1.50372 | 2.51628 | 0.01741 | 0.00540 | 1.03937 | 2.32788 | 1.89133 | 4.88797 | 0.37372 | 2.91920 | 0.00000 | 0.95433 | 1.93997 | 0.01988 |
| **Tagln** | 0.00000 | 0.00769 | 1.16153 | 0.16363 | 0.00785 | 0.00684 | 0.00686 | 0.85754 | 0.24709 | 0.85535 | 0.00000 | 1.55537 | 0.75069 | 0.74989 | 0.00000 | 0.00636 |
| **Tagln2** | 22.29405 | 14.60050 | 20.95337 | 7.93520 | 4.68465 | 20.28594 | 15.39509 | 9.65631 | 0.39849 | 13.69488 | 13.19013 | 2.33505 | 23.12959 | 1.30223 | 4.80292 | 11.03558 |
| **Taldo1** | 25.18253 | 13.42959 | 31.36260 | 10.54190 | 20.15202 | 7.51914 | 17.94823 | 33.86748 | 18.46494 | 25.69154 | 21.98981 | 3.92918 | 29.85850 | 0.45763 | 9.98404 | 13.38806 |
| **Tamm41** | 2.00118 | 0.06353 | 1.95092 | 0.34632 | 0.00000 | 0.02576 | 0.23267 | 1.33132 | 0.02523 | 1.52914 | 0.40703 | 0.02114 | 3.31523 | 0.00515 | 0.27829 | 0.00322 |
| **Tanc1** | 3.83466 | 0.81347 | 0.53907 | 3.92140 | 0.01620 | 1.32597 | 3.02019 | 0.91123 | 4.23656 | 2.14115 | 1.32887 | 0.85550 | 4.81114 | 0.43302 | 1.67164 | 1.29013 |
| **Tanc2** | 0.01342 | 0.00766 | 0.00254 | 2.93823 | 0.00925 | 0.00554 | 0.13451 | 0.00696 | 3.18298 | 4.93430 | 0.14105 | 0.18943 | 0.01579 | 0.94548 | 9.85602 | 0.02079 |
| **Tango2** | 7.02118 | 0.16129 | 4.91845 | 3.71225 | 6.86124 | 0.01203 | 3.95605 | 4.61490 | 3.28572 | 4.66374 | 2.24989 | 1.31307 | 3.90663 | 0.01258 | 3.91864 | 4.29913 |

| | | | | | | | | | | | | | | | | |
|---|---|---|---|---|---|---|---|---|---|---|---|---|---|---|---|---|
| **Tango6** | 3.94932 | 3.55789 | 4.79714 | 1.35045 | 5.21479 | 0.01023 | 3.38024 | 1.25964 | 5.77208 | 1.90776 | 1.45856 | 1.70724 | 2.08189 | 0.58661 | 2.83609 | 2.19572 |
| **Tank** | 0.00000 | 1.97305 | 0.28754 | 1.89376 | 0.00676 | 0.00791 | 0.13739 | 0.88935 | 0.00000 | 1.57175 | 0.00000 | 0.91089 | 0.55510 | 1.87988 | 3.61205 | 1.05587 |
| **Taok1** | 3.90310 | 7.89256 | 7.71792 | 15.81840 | 1.39998 | 4.49557 | 1.04190 | 8.67538 | 15.10836 | 19.76110 | 12.50028 | 17.73054 | 0.59273 | 28.84480 | 20.46276 | 1.30592 |
| **Taok2** | 5.33296 | 0.08975 | 3.47068 | 6.57424 | 3.76210 | 0.12005 | 5.07653 | 8.12426 | 1.42278 | 7.74316 | 5.81616 | 1.45303 | 2.29858 | 0.00153 | 0.89283 | 2.22012 |
| **Taok3** | 0.01558 | 0.00771 | 0.19681 | 0.52673 | 0.00906 | 0.00872 | 0.00737 | 0.17657 | 0.00377 | 0.93659 | 0.51206 | 0.01251 | 0.11436 | 0.01251 | 0.23966 | 0.01535 |
| **Tap1** | 0.00626 | 0.00770 | 1.07457 | 0.18340 | 0.03253 | 0.00585 | 0.07965 | 0.77116 | 0.00572 | 0.01228 | 0.31233 | 0.00354 | 0.79187 | 0.00799 | 0.00000 | 0.02274 |
| **Tapbp** | 4.20384 | 0.00270 | 0.00000 | 1.70631 | 0.00000 | 0.42519 | 0.08266 | 0.85969 | 0.24041 | 1.73234 | 1.27158 | 0.08238 | 0.01764 | 0.28446 | 0.26092 | 1.22568 |
| **Tapbpl** | 1.11817 | 0.03746 | 0.93193 | 6.27926 | 0.80593 | 0.05021 | 0.12532 | 0.00000 | 9.79216 | 0.00000 | 8.50625 | 0.50988 | 0.24271 | 3.76285 | 0.38555 | 0.17380 |
| **Tapt1** | 0.08944 | 0.04795 | 0.42496 | 1.07483 | 0.78377 | 0.00519 | 0.00666 | 1.21706 | 0.14405 | 2.13339 | 0.25272 | 2.48104 | 0.18741 | 1.71147 | 2.88951 | 0.00648 |
| **Tarbp1** | 0.02300 | 0.00757 | 0.00568 | 0.50068 | 0.00856 | 0.00698 | 0.00721 | 0.42081 | 0.81681 | 0.02277 | 0.00000 | 0.07730 | 0.00634 | 0.27060 | 0.34737 | 0.01449 |
| **Tarbp2** | 15.96291 | 4.89790 | 13.57971 | 2.97277 | 7.91067 | 6.40416 | 8.13707 | 17.61188 | 10.02076 | 19.20409 | 8.54201 | 2.57366 | 17.94033 | 1.01841 | 4.16350 | 3.97706 |
| **Tardbp** | 22.94958 | 12.68739 | 20.17212 | 7.62621 | 18.34650 | 14.00498 | 13.96493 | 29.22431 | 10.58975 | 23.71798 | 22.46877 | 7.40080 | 18.48139 | 26.47208 | 26.30955 | 9.38154 |
| **Tarm1** | 0.00976 | 0.00787 | 0.00753 | 0.16697 | 0.00940 | 0.00719 | 0.00761 | 0.98999 | 0.18087 | 0.26571 | 0.00445 | 0.05180 | 0.00578 | 0.00683 | 0.07350 | 0.00734 |
| **Tars** | 18.22777 | 7.11176 | 7.93307 | 1.64423 | 13.62080 | 8.11317 | 12.28672 | 17.38376 | 0.41909 | 11.00806 | 3.66274 | 1.55807 | 11.79089 | 0.00000 | 0.00000 | 9.87562 |
| **Tars2** | 12.00225 | 11.38168 | 5.67234 | 2.44504 | 4.68912 | 7.49236 | 7.13816 | 11.82380 | 6.43863 | 3.22640 | 9.74588 | 0.34449 | 3.48328 | 0.03952 | 1.28528 | 6.09420 |
| **Tarsl2** | 3.52045 | 0.00000 | 0.63397 | 0.58431 | 3.88949 | 0.00191 | 2.85385 | 2.05051 | 0.37063 | 0.00000 | 1.58275 | 0.00000 | 0.98833 | 0.00325 | 0.74742 | 0.00226 |
| **Tas1r1** | 0.00000 | 0.00711 | 3.44517 | 0.51802 | 0.01099 | 0.00632 | 0.00677 | 1.21437 | 1.15360 | 0.00000 | 0.00000 | 0.04432 | 0.00000 | 0.00677 | 0.16909 | 1.68478 |
| **Tas1r3** | 0.01373 | 0.00822 | 0.01110 | 0.76319 | 0.00921 | 0.00660 | 0.00745 | 0.01099 | 0.47129 | 0.01491 | 0.09400 | 0.00678 | 0.01142 | 0.03277 | 0.00000 | 0.00734 |
| **Tasp1** | 1.26635 | 0.00598 | 0.37927 | 2.68789 | 0.00531 | 0.00498 | 0.02578 | 2.11310 | 4.50446 | 2.99906 | 3.54309 | 0.34069 | 1.17653 | 10.06814 | 9.17516 | 1.67975 |
| **Tat** | 1.10665 | 5.75396 | 0.49050 | 0.00000 | 0.00607 | 0.00479 | 0.56075 | 1.67778 | 0.71007 | 0.64529 | 0.15897 | 0.01404 | 0.01088 | 0.58697 | 1.22461 | 0.01759 |
| **Tatdn1** | 2.37051 | 1.43623 | 1.58082 | 0.73180 | 0.00000 | 0.04114 | 0.48954 | 2.44106 | 0.66782 | 4.94753 | 2.93862 | 0.76850 | 1.43643 | 3.27368 | 1.48898 | 0.75650 |
| **Tatdn2** | 0.02046 | 0.00632 | 0.00433 | 5.94409 | 0.01646 | 0.00743 | 1.65059 | 0.57643 | 7.92111 | 1.70938 | 2.54650 | 0.04533 | 0.00000 | 1.91643 | 1.47957 | 0.94591 |
| **Tatdn3** | 2.70031 | 0.00239 | 0.01181 | 2.73158 | 3.80104 | 0.00330 | 0.03402 | 1.33142 | 1.04719 | 3.36847 | 2.94372 | 0.03810 | 0.03367 | 3.91269 | 2.71155 | 0.01808 |
| **Tax1bp1** | 10.86814 | 11.38488 | 10.90082 | 6.65759 | 12.64609 | 7.69106 | 8.15026 | 18.96114 | 14.59917 | 22.74116 | 15.76223 | 20.70700 | 4.95528 | 23.41003 | 17.62345 | 5.46262 |
| **Tax1bp3** | 15.20919 | 2.88747 | 19.14558 | 3.22194 | 13.12771 | 7.74664 | 12.84619 | 15.18521 | 0.12195 | 5.98649 | 7.11980 | 1.10084 | 14.04066 | 0.59070 | 0.85528 | 4.14982 |
| **Taz** | 7.25820 | 5.15802 | 5.55936 | 2.93771 | 0.03379 | 2.31028 | 0.00000 | 0.61252 | 0.18388 | 2.33258 | 3.38070 | 0.16307 | 4.83161 | 0.00000 | 1.19308 | 3.68874 |
| **Tbc1d1** | 3.43920 | 2.03210 | 1.22040 | 5.14420 | 5.69955 | 0.89907 | 0.17029 | 1.20501 | 7.52295 | 4.09875 | 3.85673 | 0.21120 | 2.68352 | 4.09532 | 3.93389 | 4.78810 |
| **Tbc1d10a** | 2.81877 | 0.69504 | 0.79845 | 2.30852 | 0.81482 | 0.06400 | 0.08315 | 1.32684 | 1.90953 | 1.94713 | 0.23347 | 0.15810 | 1.92845 | 0.00097 | 0.01623 | 1.75770 |
| **Tbc1d10b** | 0.93876 | 0.01378 | 2.48683 | 6.45246 | 0.07322 | 5.27404 | 2.82683 | 11.87510 | 8.71225 | 6.97914 | 9.09893 | 0.75676 | 8.93837 | 1.24578 | 4.08007 | 6.49271 |
| **Tbc1d10c** | 0.00960 | 0.02892 | 0.58294 | 0.69589 | 0.00841 | 0.00922 | 0.00726 | 0.04126 | 2.39876 | 0.68559 | 0.06837 | 0.07237 | 0.01710 | 0.00306 | 0.13872 | 0.00711 |
| **Tbc1d12** | 0.13240 | 0.02696 | 0.23762 | 1.74257 | 2.52652 | 0.63856 | 2.22117 | 0.01492 | 0.00000 | 0.87700 | 0.00000 | 0.19628 | 0.13340 | 0.58385 | 0.47695 | 0.00638 |
| **Tbc1d13** | 8.76872 | 5.12827 | 2.86647 | 9.98164 | 7.90073 | 1.57620 | 1.61770 | 3.48452 | 10.79138 | 7.72898 | 3.39051 | 0.33879 | 4.13584 | 0.71994 | 5.18930 | 6.54913 |
| **Tbc1d14** | 1.51484 | 2.38722 | 1.06438 | 6.61960 | 5.54143 | 2.41490 | 2.44432 | 2.19617 | 1.36908 | 3.65968 | 2.78352 | 7.06647 | 1.75637 | 1.47586 | 4.87767 | 0.79219 |
| **Tbc1d15** | 2.50350 | 7.41383 | 5.79173 | 9.38593 | 7.04101 | 8.73132 | 5.40888 | 14.49762 | 12.46608 | 6.89876 | 5.41280 | 10.03657 | 5.27271 | 13.77010 | 15.28576 | 4.21281 |
| **Tbc1d16** | 0.32492 | 0.00717 | 0.00530 | 0.19019 | 0.00721 | 0.00713 | 0.00722 | 0.00753 | 0.00595 | 0.12133 | 0.11473 | 0.03576 | 0.01101 | 0.00540 | 0.17793 | 0.00704 |
| **Tbc1d17** | 2.52588 | 0.78079 | 2.30987 | 0.00000 | 9.86244 | 0.70797 | 0.02417 | 11.11786 | 0.47599 | 3.88585 | 2.26035 | 0.17174 | 1.79126 | 0.01746 | 0.60011 | 2.99602 |
| **Tbc1d19** | 2.67861 | 2.55115 | 0.00000 | 1.88154 | 0.00506 | 1.62430 | 1.55235 | 0.00000 | 0.00000 | 0.00000 | 0.39075 | 0.01389 | 0.00778 | 3.23987 | 0.13525 | 0.76381 |
| **Tbc1d2** | 0.00000 | 2.25404 | 1.09581 | 2.03996 | 0.01492 | 6.00024 | 2.60768 | 0.01160 | 0.52450 | 0.00000 | 0.00741 | 0.00177 | 2.78577 | 0.23408 | 0.00000 | 0.00233 |
| **Tbc1d20** | 0.76180 | 0.03951 | 0.35366 | 2.84623 | 0.00682 | 0.80004 | 1.25604 | 3.08992 | 1.06214 | 3.36089 | 1.40858 | 0.15764 | 0.39847 | 1.05343 | 3.12148 | 0.49759 |
| **Tbc1d22a** | 4.26850 | 1.94780 | 2.86282 | 2.70335 | 0.01209 | 0.00224 | 4.76444 | 2.61640 | 2.14270 | 5.56608 | 5.52191 | 0.42780 | 2.33564 | 0.13719 | 2.30513 | 0.00624 |
| **Tbc1d22b** | 0.09612 | 1.41325 | 3.39769 | 3.33715 | 1.98722 | 0.48913 | 0.02692 | 1.31323 | 1.82544 | 1.38656 | 1.47210 | 0.00000 | 0.05625 | 0.00000 | 0.39362 | 1.15639 |
| **Tbc1d23** | 1.39758 | 0.02352 | 3.70896 | 9.57549 | 4.38766 | 1.29515 | 2.82234 | 5.09944 | 8.14637 | 4.07816 | 5.06746 | 3.08320 | 6.81935 | 9.23633 | 4.33703 | 0.16279 |
| **Tbc1d24** | 1.01186 | 0.02699 | 0.80509 | 1.98742 | 0.17146 | 0.01045 | 0.82160 | 0.16778 | 1.85939 | 0.05584 | 4.60496 | 0.32970 | 0.12917 | 1.34613 | 0.36319 | 0.04867 |
| **Tbc1d25** | 4.24599 | 0.01082 | 1.54269 | 3.45935 | 0.34331 | 4.12178 | 2.64542 | 4.60853 | 4.01074 | 8.19012 | 2.87240 | 0.84223 | 4.68613 | 0.00000 | 8.02349 | 2.42860 |
| **Tbc1d2b** | 0.00444 | 0.00856 | 0.35203 | 2.28028 | 0.01932 | 0.00907 | 0.00771 | 0.07172 | 0.38829 | 1.07358 | 0.01650 | 0.02784 | 0.01648 | 0.02047 | 0.31649 | 2.28360 |
| **Tbc1d31** | 1.99992 | 0.01259 | 2.98304 | 0.28332 | 0.05669 | 0.00549 | 1.38259 | 1.66283 | 0.00388 | 1.26788 | 3.67650 | 4.70831 | 0.53077 | 0.04413 | 0.82760 | 4.22235 |
| **Tbc1d32** | 0.41682 | 0.00720 | 0.26717 | 0.00437 | 0.00679 | 0.00521 | 0.00702 | 0.01004 | 0.34861 | 0.00659 | 0.00967 | 0.00697 | 0.00994 | 0.00755 | 0.19838 | 0.00669 |
| **Tbc1d5** | 0.00000 | 2.87187 | 0.33816 | 0.43669 | 3.25857 | 0.00736 | 0.32599 | 0.01405 | 0.00000 | 0.14571 | 0.00986 | 0.00423 | 0.04158 | 0.00527 | 0.15124 | 0.89163 |
| **Tbc1d7** | 3.34951 | 4.34277 | 2.10537 | 0.44611 | 2.18013 | 0.00277 | 0.04183 | 0.96405 | 0.18584 | 0.25741 | 0.19739 | 0.03601 | 1.29082 | 0.00719 | 0.00000 | 0.01652 |
| **Tbc1d8** | 0.01838 | 0.05794 | 0.00053 | 3.54418 | 0.22821 | 0.00695 | 0.00648 | 1.03220 | 0.98921 | 0.56113 | 0.01371 | 0.01707 | 1.32106 | 0.00000 | 1.78759 | 0.00632 |
| **Tbc1d8b** | 0.01527 | 0.00783 | 0.01917 | 0.03982 | 0.05332 | 0.00630 | 0.00747 | 0.01136 | 0.04961 | 0.03339 | 0.04721 | 0.03842 | 0.01169 | 0.09691 | 0.02654 | 0.00736 |
| **Tbc1d9** | 0.00000 | 0.01509 | 2.43334 | 7.20913 | 1.07772 | 0.03536 | 0.00300 | 3.35548 | 9.50676 | 12.84967 | 7.30667 | 3.49731 | 0.88808 | 8.91980 | 11.03040 | 0.00986 |
| **Tbc1d9b** | 6.75233 | 0.89497 | 1.94977 | 2.16363 | 1.20108 | 0.00000 | 4.27157 | 0.08828 | 1.46932 | 1.25319 | 1.29017 | 0.06100 | 2.76753 | 0.28201 | 0.87848 | 1.29982 |
| **Tbca** | 8.84634 | 0.37706 | 3.57259 | 5.70249 | 4.22682 | 3.24728 | 1.57595 | 0.59242 | 6.82201 | 3.68413 | 4.75181 | 0.18853 | 2.52535 | 1.56027 | 1.39089 | 5.30846 |
| **Tbcb** | 8.59719 | 1.84028 | 9.51647 | 7.83642 | 8.51846 | 9.81402 | 8.28331 | 5.69647 | 11.74060 | 13.72320 | 6.37936 | 0.30141 | 8.64517 | 0.84448 | 0.89781 | 9.20915 |
| **Tbcc** | 4.82522 | 0.00209 | 0.00000 | 7.60623 | 0.01894 | 0.90178 | 0.01197 | 2.11690 | 24.60631 | 4.76111 | 2.92032 | 0.07105 | 0.20538 | 0.05520 | 0.12456 | 4.89875 |
| **Tbccd1** | 2.91894 | 5.51955 | 4.52279 | 0.45204 | 0.01206 | 3.34476 | 4.66181 | 4.26121 | 0.63244 | 2.29051 | 2.06624 | 0.61529 | 0.99034 | 2.38616 | 3.78468 | 0.77127 |
| **Tbcd** | 2.36819 | 3.17667 | 0.81664 | 0.13108 | 0.02117 | 0.00340 | 0.00430 | 0.06181 | 0.01802 | 0.74808 | 0.00732 | 0.07656 | 0.02446 | 0.80207 | 0.00000 | 0.01862 |
| **Tbce** | 2.91805 | 0.31735 | 0.00000 | 0.71988 | 2.21229 | 0.12818 | 2.97244 | 2.33628 | 0.71602 | 2.65125 | 0.51957 | 6.49228 | 2.12947 | 8.22945 | 3.61870 | 1.03150 |
| **Tbcel** | 0.26168 | 0.00850 | 0.00624 | 0.00992 | 0.01678 | 0.00697 | 0.00746 | 0.01415 | 0.01189 | 0.12475 | 0.01103 | 0.00755 | 0.01148 | 0.01257 | 0.12576 | 0.00715 |
| **Tbck** | 0.94059 | 2.96181 | 1.73006 | 0.64721 | 0.07835 | 0.01649 | 0.02783 | 1.95675 | 0.48072 | 1.85384 | 1.15329 | 0.11584 | 1.29735 | 0.77663 | 1.10614 | 0.06092 |
| **Tbk1** | 0.00000 | 0.64651 | 0.44394 | 0.46688 | 0.13344 | 1.32164 | 0.07712 | 2.08238 | 3.83025 | 0.00000 | 2.10830 | 0.30663 | 0.14089 | 0.79328 | 1.27769 | 0.78231 |

| | | | | | | | | | | | | | | | | |
|---|---|---|---|---|---|---|---|---|---|---|---|---|---|---|---|---|
| **Tbkbp1** | 1.56157 | 0.28330 | 0.00267 | 0.00750 | 0.00571 | 0.00729 | 0.00616 | 0.00489 | 0.01202 | 0.44487 | 0.04409 | 0.01653 | 0.01626 | 0.02773 | 0.00356 | 0.00578 |
| **Tbl1x** | 0.96049 | 0.00692 | 0.00000 | 2.89055 | 0.00584 | 0.00521 | 0.00523 | 0.00481 | 1.24661 | 0.24772 | 0.00000 | 0.00000 | 1.83050 | 2.82082 | 0.23719 | 0.00458 |
| **Tbl1xr1** | 7.08307 | 0.29213 | 6.51281 | 1.84999 | 3.57052 | 2.59858 | 3.65877 | 4.91889 | 0.49877 | 3.15039 | 4.71175 | 0.51565 | 4.59229 | 1.80190 | 1.59560 | 2.14282 |
| **Tbl2** | 0.00000 | 1.29446 | 8.48200 | 4.94450 | 0.00000 | 0.01896 | 3.94416 | 6.74039 | 5.31531 | 7.68922 | 4.85360 | 0.43671 | 4.05878 | 4.79613 | 7.98451 | 4.21489 |
| **Tbl3** | 9.23152 | 5.41295 | 6.13209 | 0.89144 | 8.04223 | 3.66005 | 3.76195 | 19.46271 | 2.99166 | 31.52197 | 3.42587 | 7.89727 | 12.95206 | 0.00000 | 0.30361 | 5.09957 |
| **Tbp** | 0.59229 | 11.14975 | 7.51323 | 8.15838 | 3.93193 | 0.70687 | 7.09272 | 4.14095 | 6.85447 | 6.96425 | 1.57113 | 1.83162 | 2.69495 | 5.19579 | 17.56649 | 0.72793 |
| **Tbpl1** | 5.38366 | 3.07629 | 6.04953 | 5.02596 | 5.76326 | 6.33283 | 5.16091 | 2.51472 | 4.01916 | 3.74714 | 1.29911 | 0.10933 | 3.44871 | 2.96396 | 1.30374 | 2.21545 |
| **Tbrg1** | 18.93045 | 5.77529 | 14.69464 | 11.58958 | 15.85578 | 13.70545 | 7.16723 | 17.50367 | 12.12085 | 23.32797 | 21.01048 | 13.15734 | 21.23456 | 11.37702 | 7.81719 | 17.23905 |
| **Tbrg3** | 0.12316 | 0.01845 | 0.86177 | 0.05438 | 0.05752 | 0.03031 | 0.01725 | 0.03800 | 0.29867 | 0.10254 | 1.07294 | 0.02860 | 0.48842 | 0.50459 | 0.05903 | 0.05085 |
| **Tbrg4** | 9.46353 | 4.29777 | 5.25598 | 10.38063 | 4.27579 | 4.42248 | 2.49188 | 14.69095 | 11.52525 | 11.19919 | 9.22649 | 1.64719 | 9.00131 | 0.50314 | 2.04161 | 3.39016 |
| **Tbx15** | 2.82366 | 0.00000 | 0.50290 | 0.00569 | 4.11898 | 0.00250 | 4.41208 | 0.00000 | 0.00990 | 0.29770 | 0.01235 | 0.01008 | 0.47879 | 0.02307 | 0.00000 | 0.00355 |
| **Tbx18** | 0.03223 | 0.01777 | 0.01992 | 0.02139 | 0.00840 | 0.02007 | 0.04779 | 0.02828 | 0.05018 | 0.04091 | 0.06330 | 0.02554 | 0.01958 | 0.33802 | 0.15233 | 0.00730 |
| **Tbx19** | 0.01371 | 0.00783 | 0.01138 | 5.09341 | 0.00928 | 0.00639 | 0.00737 | 0.01273 | 0.21985 | 0.04112 | 0.02135 | 0.00760 | 0.01307 | 0.03869 | 0.19026 | 0.00655 |
| **Tbx2** | 0.01272 | 0.00801 | 0.01198 | 0.11948 | 0.00898 | 0.00649 | 0.00726 | 0.01096 | 0.01009 | 0.92138 | 0.01224 | 0.00602 | 0.10560 | 0.00776 | 0.06637 | 0.00725 |
| **Tbx20** | 2.17632 | 2.00016 | 2.31738 | 0.20114 | 1.56899 | 0.95147 | 2.27906 | 5.75289 | 0.01535 | 1.67000 | 4.65502 | 0.06145 | 3.64405 | 2.32572 | 0.00000 | 0.14820 |
| **Tbx21** | 0.01377 | 0.00788 | 0.00858 | 0.31149 | 0.00891 | 0.00692 | 0.00718 | 0.00000 | 0.30799 | 0.43912 | 0.50964 | 0.00365 | 0.00812 | 0.02502 | 0.00000 | 0.00749 |
| **Tbx3** | 10.52955 | 4.64400 | 4.78475 | 5.48186 | 5.50404 | 2.89853 | 4.19977 | 13.65285 | 9.56669 | 11.25057 | 18.09526 | 0.58977 | 8.50608 | 9.28043 | 1.39552 | 11.99015 |
| **Tbx4** | 0.01367 | 0.00815 | 0.01203 | 1.88735 | 0.00931 | 0.00645 | 0.00747 | 0.01377 | 0.63335 | 0.01082 | 0.00915 | 0.00718 | 0.01282 | 0.01211 | 0.00468 | 0.00706 |
| **Tbx6** | 0.00628 | 0.29118 | 0.69491 | 0.02138 | 0.00713 | 0.00596 | 0.00706 | 0.00926 | 0.27419 | 0.00288 | 0.01084 | 0.00772 | 0.47016 | 0.01235 | 0.01021 | 0.00661 |
| **Tbxa2r** | 0.01325 | 0.00832 | 0.01136 | 7.45558 | 0.00921 | 0.00588 | 0.00732 | 0.01373 | 3.26114 | 0.01027 | 0.00000 | 0.00271 | 0.01333 | 0.00867 | 0.00426 | 0.00594 |
| **Tc2n** | 0.31751 | 0.02830 | 0.45702 | 2.45782 | 0.20188 | 0.05592 | 0.05672 | 0.00000 | 4.46021 | 0.00000 | 4.81756 | 0.24808 | 0.15961 | 3.35927 | 0.59496 | 0.22112 |
| **Tcaim** | 0.16108 | 0.00654 | 0.10692 | 2.11223 | 0.00764 | 2.47117 | 1.43433 | 0.26735 | 1.89490 | 0.47863 | 0.45789 | 0.01747 | 2.04957 | 2.28829 | 0.81363 | 0.01991 |
| **Tcam1** | 1.76138 | 1.59782 | 2.71806 | 0.21984 | 0.02464 | 0.00404 | 0.02453 | 2.51242 | 1.93723 | 2.48855 | 1.48390 | 1.42317 | 0.01747 | 2.65407 | 2.24079 | 0.00336 |
| **Tcea1** | 13.34897 | 9.61059 | 8.42341 | 17.92148 | 8.35168 | 9.81645 | 4.77005 | 12.78952 | 23.41044 | 25.48372 | 13.87663 | 21.19019 | 7.72662 | 24.75722 | 23.84441 | 3.28295 |
| **Tcea3** | 1.14163 | 0.00436 | 3.89985 | 0.28094 | 0.00000 | 0.00556 | 1.62498 | 2.02239 | 2.49327 | 0.00000 | 1.22553 | 0.01547 | 1.79847 | 0.01809 | 0.00000 | 0.01073 |
| **Tceal8** | 0.01071 | 0.00775 | 0.00468 | 0.01235 | 0.00911 | 0.00636 | 0.00711 | 0.79929 | 0.01727 | 0.97795 | 0.00000 | 0.03261 | 0.00541 | 2.31282 | 0.89517 | 0.00717 |
| **Tceanc** | 0.03103 | 0.00821 | 0.56587 | 0.55963 | 0.02714 | 0.00561 | 0.01827 | 0.01925 | 0.44836 | 0.38549 | 0.24462 | 0.01804 | 0.12985 | 0.01002 | 0.14787 | 0.08308 |
| **Tceanc2** | 2.71704 | 4.66091 | 3.11836 | 6.42097 | 3.71337 | 4.29562 | 2.69998 | 2.66800 | 6.43182 | 4.01957 | 0.47624 | 0.18891 | 1.94077 | 0.00000 | 0.00000 | 0.34361 |
| **Tceb1** | 14.81665 | 14.76084 | 20.01746 | 20.29491 | 12.09464 | 11.81577 | 11.89021 | 20.68737 | 35.68511 | 23.21819 | 42.09115 | 24.93894 | 18.57112 | 40.29883 | 20.32187 | 5.28539 |
| **Tceb2** | 9.90437 | 0.51724 | 7.92980 | 12.80791 | 9.89463 | 4.11634 | 0.73735 | 1.39892 | 20.06217 | 8.19750 | 17.58327 | 1.12354 | 7.58822 | 0.02873 | 0.05791 | 8.40940 |
| **Tceb3** | 4.23549 | 2.72655 | 5.58393 | 18.00981 | 8.25261 | 5.77467 | 3.23433 | 13.17859 | 25.64789 | 20.87866 | 7.69776 | 17.71672 | 6.15622 | 20.55921 | 20.13586 | 1.10990 |
| **Tcerg1** | 5.68338 | 4.59600 | 6.71855 | 8.74891 | 4.83549 | 2.36949 | 6.50468 | 8.12705 | 14.03072 | 11.62381 | 7.72138 | 16.52958 | 3.49203 | 20.34746 | 13.91632 | 2.60850 |
| **Tcerg1l** | 0.01353 | 0.00762 | 0.01121 | 0.72155 | 0.00931 | 0.00628 | 0.00736 | 0.01111 | 1.60916 | 0.00342 | 0.00000 | 0.00349 | 0.01291 | 0.00695 | 0.66395 | 0.00704 |
| **Tcf12** | 0.30362 | 1.21632 | 0.16512 | 4.91420 | 2.77979 | 0.01028 | 0.49945 | 0.00000 | 4.39047 | 0.61111 | 0.07612 | 0.23231 | 0.95914 | 4.36309 | 2.55264 | 0.33226 |
| **Tcf15** | 0.01386 | 0.09395 | 0.16337 | 0.23567 | 0.41833 | 0.13956 | 0.00678 | 0.00681 | 2.09229 | 0.16617 | 1.27510 | 0.00674 | 0.02672 | 0.01558 | 0.00000 | 0.00682 |
| **Tcf19** | 1.79362 | 0.00486 | 1.96319 | 2.02651 | 0.00000 | 0.03418 | 0.31146 | 1.04364 | 3.92922 | 1.46638 | 0.00000 | 0.20600 | 0.00289 | 0.00000 | 0.84231 | 0.00415 |
| **Tcf20** | 0.99169 | 0.18667 | 0.92917 | 5.84087 | 0.06226 | 2.68152 | 0.48898 | 0.39643 | 0.33244 | 0.39737 | 0.02961 | 0.09796 | 0.75675 | 0.42123 | 2.48931 | 0.13140 |
| **Tcf23** | 6.01195 | 0.00000 | 2.11795 | 0.00000 | 5.42711 | 1.62001 | 0.70694 | 0.00000 | 0.01653 | 0.07337 | 1.39510 | 2.20889 | 0.04125 | 1.38163 | 1.34902 | 0.00632 |
| **Tcf24** | 0.00111 | 0.00467 | 0.00000 | 0.68811 | 6.12664 | 0.00935 | 0.34829 | 3.38680 | 0.68720 | 0.93372 | 0.16808 | 0.07193 | 0.00228 | 0.00656 | 0.00000 | 0.00693 |
| **Tcf25** | 13.01597 | 2.38811 | 12.98892 | 11.75460 | 7.84447 | 11.07907 | 6.78203 | 21.18333 | 6.60005 | 22.98704 | 13.48221 | 5.63846 | 13.30004 | 3.64434 | 16.69627 | 9.15536 |
| **Tcf3** | 4.70223 | 0.02051 | 0.00000 | 3.98279 | 3.25874 | 1.97484 | 0.30632 | 4.62954 | 1.62431 | 4.38487 | 0.76149 | 0.11733 | 2.00859 | 0.16634 | 0.55906 | 9.36578 |
| **Tcf4** | 0.99809 | 0.01672 | 0.00158 | 0.01320 | 0.00598 | 0.00691 | 0.02641 | 0.00827 | 0.13241 | 0.68980 | 0.00826 | 0.02267 | 0.01150 | 0.00869 | 0.68181 | 0.00630 |
| **Tcf7** | 0.03864 | 0.00627 | 0.00719 | 4.20639 | 3.28756 | 0.00633 | 0.00772 | 0.10380 | 1.00351 | 0.69389 | 0.02940 | 0.06143 | 0.00751 | 0.00000 | 0.31492 | 0.00649 |
| **Tcf7l1** | 3.37584 | 0.00442 | 0.29630 | 0.58748 | 0.00452 | 0.00538 | 0.00434 | 0.00000 | 0.55596 | 0.27026 | 0.02458 | 0.05014 | 0.25734 | 0.00974 | 0.00000 | 1.78552 |
| **Tcf7l2** | 1.81064 | 0.00586 | 0.00107 | 0.00620 | 0.00449 | 0.00690 | 0.18397 | 0.00674 | 0.01166 | 0.46387 | 0.13995 | 0.01375 | 0.30559 | 0.25454 | 0.00000 | 1.00758 |
| **Tcfl5** | 0.00000 | 0.00573 | 1.70801 | 1.20939 | 0.00589 | 5.22855 | 3.60737 | 1.21694 | 2.49734 | 4.56514 | 0.81327 | 0.20336 | 3.66955 | 0.00000 | 1.47647 | 2.15556 |
| **Tchh** | 0.01126 | 0.00754 | 0.00848 | 3.05701 | 0.00846 | 0.00500 | 0.00706 | 0.00000 | 14.02741 | 2.92250 | 0.00000 | 0.05727 | 0.01069 | 0.01655 | 0.37650 | 0.00586 |
| **Tchp** | 1.44787 | 0.05560 | 3.58880 | 11.84119 | 0.00731 | 0.00277 | 3.52789 | 1.70166 | 2.16023 | 1.46644 | 0.77287 | 0.08285 | 1.59830 | 0.73657 | 0.74914 | 0.00140 |
| **Tcirg1** | 0.12416 | 0.00703 | 0.47970 | 3.79659 | 0.04105 | 0.00832 | 0.03606 | 3.00012 | 2.74303 | 5.07467 | 2.60893 | 0.12250 | 2.96462 | 0.00000 | 0.05052 | 0.00457 |
| **Tcl1** | 6.33328 | 0.00000 | 0.07465 | 22.19317 | 5.31975 | 0.00000 | 0.00000 | 0.64745 | 8.48148 | 1.25087 | 0.93917 | 0.13815 | 0.00000 | 0.07481 | 2.46835 | 2.97052 |
| **Tcl1b1** | 0.01263 | 0.00784 | 0.00475 | 12.39456 | 0.00921 | 0.00506 | 0.00689 | 0.00815 | 2.66452 | 0.02440 | 0.21062 | 0.02808 | 0.01053 | 0.00000 | 4.15489 | 0.00506 |
| **Tcl1b2** | 0.01362 | 0.00787 | 0.00962 | 5.23137 | 0.00937 | 0.00614 | 0.00715 | 0.00927 | 0.42537 | 0.00420 | 0.06294 | 0.02585 | 0.01219 | 0.12588 | 0.59114 | 0.00651 |
| **Tcl1b3** | 0.01351 | 0.00783 | 0.00958 | 1.95731 | 0.00950 | 0.00619 | 0.00722 | 0.00947 | 0.01895 | 0.00000 | 0.04228 | 0.01322 | 0.01248 | 0.08216 | 1.62283 | 0.00709 |
| **Tcl1b4** | 0.01286 | 0.00777 | 0.00972 | 2.99590 | 0.00944 | 0.00612 | 0.00714 | 0.02691 | 0.82745 | 0.06875 | 0.06096 | 0.03244 | 0.01100 | 0.00004 | 0.88991 | 0.00689 |
| **Tcl1b5** | 0.01758 | 0.00799 | 0.00975 | 3.73587 | 0.00898 | 0.00630 | 0.00732 | 0.01119 | 0.11221 | 0.03174 | 0.03434 | 0.02837 | 0.01290 | 0.00169 | 1.38820 | 0.00673 |
| **Tcn2** | 5.25597 | 0.03685 | 3.86833 | 0.37414 | 4.32985 | 2.75868 | 6.41600 | 2.09196 | 0.12786 | 0.82532 | 2.80437 | 0.00461 | 2.80459 | 0.00614 | 1.39464 | 0.10288 |
| **Tcof1** | 11.98768 | 7.82807 | 4.60961 | 10.17599 | 4.58310 | 0.25068 | 4.36001 | 13.72426 | 9.25643 | 12.45847 | 6.37731 | 1.17174 | 10.45138 | 0.72392 | 4.75442 | 1.83839 |
| **Tcp1** | 29.34629 | 20.99129 | 28.24864 | 6.74984 | 23.62786 | 11.91095 | 16.49170 | 30.60053 | 10.01432 | 31.41923 | 14.38465 | 16.66366 | 25.67853 | 23.56531 | 22.24501 | 19.58950 |
| **Tcp10a** | 0.01269 | 0.00800 | 0.08963 | 0.00487 | 0.00906 | 0.00879 | 0.00743 | 0.01328 | 0.30937 | 0.01359 | 0.01006 | 0.00745 | 0.01142 | 0.01344 | 0.00703 | 0.00735 |
| **Tcp10b** | 0.01269 | 0.00799 | 0.09553 | 0.00511 | 0.00905 | 0.00882 | 0.00743 | 0.01327 | 0.23106 | 0.01400 | 0.01061 | 0.00756 | 0.01133 | 0.01328 | 0.00739 | 0.00735 |
| **Tcp10c** | 0.01287 | 0.00801 | 0.04832 | 0.00513 | 0.00917 | 0.00837 | 0.00746 | 0.01348 | 0.21364 | 0.01413 | 0.01090 | 0.00764 | 0.01200 | 0.01306 | 0.00825 | 0.00739 |
| **Tcp11** | 1.29794 | 0.00693 | 0.00289 | 0.95442 | 0.00526 | 0.00731 | 0.00720 | 0.30103 | 0.15760 | 0.61789 | 0.28042 | 0.01579 | 0.00476 | 0.00730 | 0.00000 | 0.00550 |

| | | | | | | | | | | | | | | | | |
|---|---|---|---|---|---|---|---|---|---|---|---|---|---|---|---|---|
| **Tcp11l1** | 0.01427 | 0.00767 | 0.02138 | 0.43483 | 0.00908 | 0.12805 | 0.00714 | 0.00812 | 0.99395 | 0.12583 | 0.06014 | 0.16639 | 0.01002 | 0.00628 | 0.49828 | 0.00718 |
| **Tcp11l2** | 0.00250 | 0.00755 | 0.17179 | 0.03102 | 0.14799 | 0.00569 | 0.41068 | 0.03219 | 0.02427 | 1.13629 | 0.13075 | 0.21033 | 0.01544 | 3.94575 | 1.71320 | 0.01421 |
| **Tcstv1** | 0.00856 | 0.00871 | 0.24314 | 7.99479 | 0.00916 | 0.00459 | 0.00799 | 0.06806 | 7.18220 | 0.05028 | 0.00000 | 0.01906 | 0.20714 | 0.01119 | 0.23089 | 0.00567 |
| **Tcstv3** | 0.01461 | 0.00803 | 0.00026 | 8.75093 | 0.00919 | 0.30776 | 0.00737 | 0.44179 | 14.23789 | 0.41286 | 0.49921 | 0.31411 | 0.13326 | 0.25155 | 0.66839 | 0.00517 |
| **Tcta** | 3.20653 | 1.79389 | 2.65655 | 0.23839 | 0.00000 | 0.00412 | 0.04307 | 0.77866 | 1.07205 | 1.93939 | 0.01653 | 0.00550 | 0.00766 | 0.04650 | 0.09748 | 0.00207 |
| **Tcte2** | 0.01154 | 0.00773 | 1.67989 | 1.79472 | 0.00622 | 0.00899 | 0.00703 | 0.00836 | 0.30771 | 0.51555 | 0.00744 | 0.00456 | 0.03548 | 0.00761 | 0.00000 | 0.00619 |
| **Tctex1d2** | 0.63505 | 0.00720 | 0.00000 | 1.55972 | 0.01362 | 2.23159 | 0.00643 | 0.89514 | 0.01284 | 0.04246 | 0.00078 | 0.03470 | 0.04770 | 0.00294 | 0.52533 | 0.00595 |
| **Tctn1** | 0.01067 | 0.00883 | 0.15203 | 0.27296 | 0.01106 | 1.49633 | 0.00774 | 0.41845 | 0.10725 | 0.00000 | 0.41173 | 0.04168 | 0.84074 | 0.01145 | 0.01512 | 0.00631 |
| **Tctn2** | 1.88962 | 0.00275 | 2.29412 | 1.62020 | 3.64635 | 0.00971 | 1.39011 | 0.65075 | 0.22836 | 0.93285 | 1.72732 | 0.03463 | 2.32807 | 0.00074 | 0.76066 | 0.19980 |
| **Tctn3** | 0.13190 | 0.00732 | 3.40425 | 0.38242 | 0.01018 | 0.00619 | 0.09171 | 2.66037 | 0.36914 | 1.74205 | 1.50105 | 0.42795 | 3.99842 | 0.35195 | 2.87838 | 0.00376 |
| **Tdgf1** | 23.79751 | 13.44864 | 4.27279 | 0.00000 | 16.34820 | 0.00000 | 13.35434 | 3.11107 | 0.10780 | 14.50521 | 16.73482 | 1.45174 | 0.00000 | 21.13076 | 7.63808 | 6.95691 |
| **Tdh** | 8.65132 | 8.74622 | 13.03021 | 4.96645 | 5.94103 | 0.00000 | 18.46784 | 16.34453 | 11.95272 | 6.63398 | 10.18637 | 1.10001 | 1.42999 | 0.11570 | 0.48345 | 1.70358 |
| **Tdp1** | 3.73321 | 7.60147 | 1.63316 | 1.05203 | 3.14452 | 0.18258 | 0.00000 | 0.54360 | 0.00000 | 0.08558 | 0.63108 | 0.00000 | 0.00000 | 0.00000 | 0.39167 | 3.02440 |
| **Tdp2** | 6.21841 | 3.40048 | 1.34433 | 10.75929 | 5.45023 | 1.81719 | 2.59965 | 2.64459 | 13.40647 | 10.32547 | 3.16535 | 3.64670 | 1.97947 | 8.69376 | 9.32657 | 2.82951 |
| **Tdpoz1** | 0.00562 | 0.00809 | 0.00460 | 9.50976 | 0.00890 | 0.00424 | 0.00731 | 0.32403 | 8.67711 | 1.66776 | 0.00000 | 0.84124 | 0.82612 | 0.18187 | 5.69515 | 0.00461 |
| **Tdpoz2** | 0.00641 | 0.00777 | 1.23906 | 0.76417 | 0.00668 | 0.00557 | 0.00711 | 0.00382 | 1.31601 | 3.38232 | 0.14232 | 0.04281 | 0.03206 | 0.74261 | 5.64522 | 0.00654 |
| **Tdpoz3** | 0.01241 | 0.00781 | 0.01147 | 0.07064 | 0.00917 | 0.00635 | 0.00723 | 0.01190 | 0.00826 | 0.01668 | 0.01055 | 0.01698 | 0.28389 | 0.00680 | 0.28809 | 0.00705 |
| **Tdpoz4** | 0.01780 | 0.00820 | 0.00651 | 0.33266 | 0.01078 | 0.00668 | 0.00745 | 0.01397 | 0.04046 | 0.06601 | 0.00616 | 0.00351 | 0.01291 | 0.96199 | 0.49522 | 0.00748 |
| **Tdrd1** | 0.01288 | 0.00780 | 0.00978 | 4.27848 | 0.00939 | 0.00594 | 0.00728 | 0.01070 | 0.28507 | 0.55111 | 0.00523 | 0.02937 | 0.01500 | 0.02545 | 2.54302 | 0.00649 |
| **Tdrd12** | 0.00569 | 0.05822 | 1.85344 | 1.42128 | 0.00661 | 0.00554 | 4.30167 | 0.01351 | 0.00000 | 0.08442 | 0.01937 | 0.00301 | 0.00599 | 0.00305 | 0.42283 | 0.01388 |
| **Tdrd3** | 0.00000 | 9.25276 | 3.12756 | 1.59018 | 7.24451 | 0.01164 | 2.20424 | 5.06115 | 0.78521 | 1.47926 | 1.34730 | 0.03049 | 0.00000 | 1.10646 | 1.69878 | 2.43916 |
| **Tdrd5** | 0.02252 | 0.00806 | 0.03418 | 1.80591 | 0.01961 | 0.00569 | 0.00757 | 0.26986 | 1.19137 | 0.41176 | 0.55994 | 0.05046 | 0.00388 | 0.16942 | 1.34175 | 0.00705 |
| **Tdrd7** | 0.00973 | 0.00848 | 0.00852 | 0.00918 | 0.00960 | 0.00701 | 0.00826 | 0.24599 | 0.01178 | 0.04882 | 0.00867 | 0.00809 | 0.00713 | 0.14167 | 0.22051 | 0.00745 |
| **Tdrd9** | 0.18369 | 0.00778 | 1.10289 | 2.60898 | 0.04972 | 0.00498 | 0.00589 | 2.29025 | 2.76195 | 1.71576 | 3.16666 | 0.26315 | 0.00000 | 0.37487 | 1.72731 | 0.43868 |
| **Tdrkh** | 0.01813 | 0.00741 | 0.04600 | 0.14076 | 0.45217 | 0.00938 | 0.00722 | 0.00000 | 0.00650 | 0.47676 | 0.55793 | 0.21527 | 0.25276 | 1.22561 | 0.34511 | 0.00744 |
| **Tdrp** | 9.17242 | 12.60843 | 9.38974 | 3.36274 | 10.38029 | 15.72792 | 3.35604 | 19.91646 | 0.00000 | 0.35188 | 9.54610 | 4.87687 | 22.54950 | 6.60052 | 0.00000 | 9.39861 |
| **Tead1** | 9.59788 | 7.60586 | 7.65468 | 1.27280 | 2.47393 | 0.00000 | 3.02848 | 4.75557 | 4.18449 | 3.73955 | 2.13450 | 5.13803 | 4.28003 | 5.22997 | 3.23913 | 5.89723 |
| **Tead2** | 2.96400 | 0.59604 | 0.00000 | 2.15451 | 0.03582 | 4.32508 | 0.32072 | 0.04596 | 1.60360 | 0.55510 | 0.37621 | 0.07530 | 3.62294 | 0.00109 | 0.45697 | 2.04811 |
| **Tead3** | 0.07805 | 0.00787 | 0.01791 | 0.05950 | 0.01012 | 0.02255 | 0.02764 | 0.28445 | 0.43462 | 0.25733 | 0.69864 | 0.06317 | 0.07864 | 0.21217 | 0.04225 | 0.05884 |
| **Tead4** | 6.19897 | 0.00000 | 5.94236 | 6.60734 | 5.36869 | 4.33757 | 6.18328 | 17.81064 | 9.10372 | 22.68863 | 20.38800 | 6.51313 | 11.72325 | 3.47247 | 8.73496 | 2.73493 |
| **Tec** | 0.01373 | 0.00816 | 0.01225 | 0.22230 | 0.00934 | 0.00665 | 0.00752 | 0.01384 | 0.10851 | 0.01830 | 0.01182 | 0.00799 | 0.01283 | 0.01139 | 0.00795 | 0.00744 |
| **Tecpr1** | 0.69203 | 0.00548 | 0.64565 | 0.00323 | 0.01437 | 0.00763 | 1.45986 | 2.63457 | 0.10725 | 0.00000 | 0.46258 | 0.00597 | 1.78675 | 0.00375 | 0.23088 | 0.07776 |
| **Tecpr2** | 0.49320 | 2.42131 | 0.86509 | 0.68442 | 0.09843 | 2.84229 | 0.04494 | 1.11507 | 2.20174 | 1.16996 | 2.32996 | 0.11735 | 0.20126 | 0.67124 | 1.72705 | 2.94600 |
| **Tecr** | 13.91857 | 2.16820 | 16.17170 | 0.83692 | 9.09193 | 6.30912 | 6.11086 | 9.04309 | 0.78701 | 5.78232 | 7.76543 | 4.41152 | 11.78388 | 1.00604 | 2.22499 | 9.41703 |
| **Teddm1b** | 0.53616 | 0.09367 | 0.80494 | 4.72264 | 0.22021 | 0.07149 | 0.09839 | 0.00000 | 8.42007 | 0.00007 | 5.57120 | 0.44410 | 0.17530 | 2.27106 | 0.31789 | 0.17536 |
| **Teddm3** | 0.01136 | 0.00759 | 0.87797 | 0.01099 | 0.00667 | 0.00559 | 0.00725 | 0.01200 | 0.01222 | 0.00553 | 0.01166 | 0.01759 | 0.00684 | 0.01625 | 0.00000 | 0.83086 |
| **Tef** | 0.00000 | 0.00517 | 0.06077 | 8.24553 | 0.00715 | 0.00426 | 3.96845 | 1.54969 | 9.63624 | 4.19719 | 0.27327 | 0.07450 | 0.00000 | 1.10385 | 0.00000 | 1.93502 |
| **Tefm** | 3.94488 | 5.52515 | 7.80586 | 0.70743 | 7.45583 | 1.84488 | 0.90576 | 3.62123 | 5.08591 | 9.04174 | 5.52289 | 0.91208 | 3.59014 | 8.86831 | 8.97684 | 0.03069 |
| **Tek** | 0.01334 | 0.00756 | 0.01108 | 1.64672 | 0.00908 | 0.00633 | 0.00720 | 0.01013 | 0.39667 | 0.55525 | 0.00797 | 0.01270 | 0.01219 | 0.00339 | 0.77721 | 0.00750 |
| **Tekt1** | 0.18472 | 0.00729 | 0.00430 | 0.21749 | 0.00731 | 0.00765 | 0.00659 | 0.00000 | 0.00678 | 0.49263 | 1.51813 | 0.00000 | 0.00000 | 0.00369 | 0.09496 | 0.00707 |
| **Tekt2** | 0.00664 | 0.00823 | 0.00000 | 0.46984 | 0.00797 | 0.00592 | 0.00753 | 0.06098 | 3.21611 | 2.29331 | 0.91800 | 0.56384 | 0.00000 | 0.25205 | 3.70354 | 0.67145 |
| **Telo2** | 5.47877 | 0.00000 | 2.79398 | 0.51813 | 2.19535 | 1.71690 | 3.98935 | 3.04743 | 2.60055 | 2.93775 | 0.72584 | 0.02265 | 3.63922 | 0.00000 | 1.06187 | 0.02736 |
| **Ten1** | 1.79268 | 1.20216 | 4.17526 | 1.38770 | 6.61293 | 2.34117 | 0.96306 | 6.32763 | 2.47912 | 3.44030 | 3.82725 | 0.17674 | 7.05498 | 0.69145 | 0.00102 | 2.61816 |
| **Tenm3** | 0.01767 | 0.00800 | 0.01001 | 0.27194 | 0.00897 | 0.00673 | 0.17876 | 0.01167 | 0.34695 | 0.01107 | 0.00993 | 0.02770 | 0.01220 | 0.03385 | 0.00209 | 0.00733 |
| **Tenm4** | 0.02138 | 0.00837 | 0.01157 | 0.00602 | 0.01093 | 0.00649 | 0.00732 | 0.01003 | 0.01140 | 1.42333 | 0.01025 | 0.00450 | 0.01037 | 0.00701 | 0.11043 | 0.00720 |
| **Tep1** | 1.34937 | 2.68714 | 5.80598 | 1.52646 | 0.00371 | 0.01150 | 2.17094 | 4.65652 | 0.77922 | 6.29063 | 5.99555 | 0.17067 | 1.16087 | 0.40898 | 4.50816 | 0.60995 |
| **Terb1** | 0.01342 | 0.00785 | 0.01070 | 0.27956 | 0.00912 | 0.00670 | 0.00748 | 0.01079 | 0.76919 | 0.35368 | 0.03706 | 0.00563 | 0.01229 | 0.00900 | 0.56457 | 0.00730 |
| **Terb2** | 2.68984 | 6.88925 | 1.31713 | 2.65984 | 0.00000 | 0.00248 | 3.58051 | 0.00000 | 2.98085 | 3.65365 | 1.35652 | 4.20736 | 3.06295 | 6.15051 | 1.17933 | 0.00604 |
| **Terf1** | 6.94296 | 4.94300 | 2.37253 | 7.06259 | 5.74657 | 3.73975 | 1.94411 | 12.06426 | 13.43103 | 7.86947 | 8.79361 | 10.33095 | 3.21293 | 7.37541 | 4.09977 | 5.80449 |
| **Terf2** | 2.86512 | 2.20323 | 4.37063 | 2.08357 | 0.01547 | 0.00247 | 0.00000 | 2.57313 | 1.58883 | 4.97638 | 1.37798 | 4.24085 | 4.96567 | 0.24131 | 3.00723 | 0.29070 |
| **Terf2ip** | 2.13050 | 3.07540 | 4.55553 | 3.55123 | 10.27081 | 0.02779 | 0.06579 | 1.10737 | 4.45987 | 3.49051 | 1.02733 | 7.47734 | 1.71282 | 1.74871 | 1.13875 | 1.76350 |
| **Tert** | 0.30248 | 0.00680 | 0.15105 | 0.00607 | 0.02205 | 0.00551 | 0.79461 | 0.01314 | 0.01108 | 0.20880 | 0.01176 | 0.00437 | 0.49674 | 0.00622 | 0.62481 | 0.00687 |
| **Tes** | 15.73486 | 4.13783 | 8.60510 | 1.89593 | 17.31016 | 14.91500 | 13.09239 | 4.02513 | 2.14449 | 3.52659 | 3.02264 | 0.18834 | 8.88537 | 0.96660 | 0.00000 | 9.06306 |
| **Tesc** | 0.18415 | 0.00633 | 0.00000 | 21.02586 | 0.02446 | 2.86885 | 0.89759 | 1.65514 | 8.63285 | 1.98547 | 3.34171 | 1.16947 | 1.58090 | 0.00000 | 0.21885 | 0.00799 |
| **Tescl** | 0.50996 | 0.00809 | 0.00754 | 0.01063 | 0.01630 | 0.00643 | 0.00673 | 0.00946 | 0.01239 | 0.00811 | 0.01255 | 0.00749 | 0.52792 | 0.01286 | 0.01058 | 0.00671 |
| **Tesk1** | 1.11789 | 0.00719 | 0.00000 | 2.38069 | 0.00835 | 1.67535 | 0.00516 | 0.34186 | 0.39789 | 0.18491 | 0.05827 | 0.02490 | 2.82179 | 0.00801 | 0.00000 | 0.82177 |
| **Tesk2** | 0.00000 | 0.01504 | 1.64875 | 3.09826 | 2.23739 | 0.00664 | 0.47488 | 1.19270 | 7.01377 | 3.99307 | 0.42644 | 3.79016 | 0.00000 | 1.46135 | 7.55249 | 3.60715 |
| **Tet1** | 16.28453 | 6.67562 | 4.57148 | 2.96835 | 9.28537 | 0.00000 | 5.99863 | 3.61716 | 0.87211 | 11.16076 | 2.22611 | 6.33897 | 0.96117 | 21.99464 | 17.37010 | 3.00186 |
| **Tet2** | 2.61558 | 1.57436 | 2.96518 | 0.40148 | 4.95425 | 0.00357 | 4.08057 | 2.13981 | 0.18802 | 0.24689 | 0.00000 | 0.00000 | 4.62137 | 0.55616 | 0.00000 | 3.47206 |
| **Tet3** | 0.11447 | 0.00672 | 0.00000 | 3.95652 | 0.00750 | 0.87435 | 0.39627 | 0.00000 | 1.30810 | 0.25228 | 0.51627 | 2.52045 | 0.12059 | 0.00000 | 1.17820 | 0.00595 |
| **Tex10** | 9.99645 | 3.46511 | 5.27967 | 1.99395 | 14.33775 | 4.93476 | 5.05138 | 6.74010 | 2.67374 | 7.56214 | 2.95882 | 10.47816 | 4.17316 | 8.56502 | 9.90636 | 2.84305 |
| **Tex101** | 0.00848 | 0.00882 | 0.00853 | 2.20838 | 0.00915 | 0.00670 | 0.00822 | 0.45695 | 0.00000 | 1.26803 | 0.00812 | 0.00511 | 0.00562 | 0.00962 | 0.00000 | 0.00677 |

| | | | | | | | | | | | | | | | | |
|---|---|---|---|---|---|---|---|---|---|---|---|---|---|---|---|---|
| **Tex11** | 0.01271 | 0.00766 | 0.00285 | 5.83530 | 0.00930 | 0.00489 | 0.00668 | 0.00870 | 5.13377 | 0.58722 | 0.00000 | 0.00000 | 0.01211 | 7.29714 | 3.46183 | 0.00597 |
| **Tex12** | 0.05662 | 0.75397 | 0.08349 | 0.80106 | 0.00788 | 0.00562 | 0.76219 | 0.51183 | 1.52934 | 0.13738 | 1.44444 | 0.41822 | 0.46692 | 1.09414 | 0.23076 | 0.00610 |
| **Tex13** | 0.01379 | 0.00797 | 0.01211 | 2.26863 | 0.00933 | 0.00639 | 0.00739 | 0.01298 | 0.00000 | 0.00979 | 0.01354 | 0.00795 | 0.01265 | 0.00689 | 0.14300 | 0.00700 |
| **Tex14** | 1.61349 | 0.00712 | 0.15638 | 1.25337 | 0.01073 | 0.00420 | 0.00692 | 0.69612 | 1.16370 | 0.59676 | 0.00000 | 0.08770 | 0.01089 | 0.16681 | 1.92364 | 0.00567 |
| **Tex15** | 0.00587 | 0.00500 | 0.00402 | 5.62998 | 0.00853 | 0.00379 | 0.27672 | 0.01029 | 4.61986 | 2.31124 | 0.00000 | 7.63047 | 0.01067 | 0.00000 | 6.55958 | 0.00563 |
| **Tex19.1** | 10.38671 | 18.82098 | 20.75186 | 4.85375 | 14.15944 | 6.10321 | 12.21328 | 17.86655 | 7.48757 | 11.03185 | 3.97661 | 0.60899 | 20.02395 | 9.87658 | 4.88776 | 3.54227 |
| **Tex19.2** | 9.79317 | 11.50054 | 9.44611 | 1.84393 | 6.92085 | 4.58510 | 0.57798 | 8.69862 | 0.00000 | 3.81888 | 2.40993 | 0.10807 | 7.83079 | 0.00000 | 0.90436 | 3.02938 |
| **Tex2** | 2.32989 | 0.82674 | 4.69695 | 3.25633 | 0.09280 | 4.17618 | 0.76657 | 8.83357 | 5.20714 | 6.25577 | 5.11296 | 0.41064 | 4.66992 | 12.32141 | 7.68541 | 2.41198 |
| **Tex21** | 0.01292 | 0.00792 | 0.01351 | 0.01091 | 0.01014 | 0.00712 | 0.00771 | 0.01530 | 0.01188 | 0.00733 | 0.01411 | 0.01760 | 0.14645 | 0.01999 | 0.08044 | 0.00766 |
| **Tex26** | 0.01375 | 0.00780 | 0.01229 | 0.33932 | 0.00940 | 0.00652 | 0.00745 | 0.01272 | 0.00421 | 0.00816 | 0.01172 | 0.00716 | 0.01269 | 0.00571 | 0.27109 | 0.00731 |
| **Tex261** | 3.49903 | 2.55611 | 8.01845 | 3.29098 | 0.79640 | 3.16089 | 4.98213 | 4.25763 | 1.25322 | 2.70320 | 2.98606 | 0.19688 | 0.47854 | 0.00741 | 1.42350 | 0.69630 |
| **Tex264** | 11.93644 | 3.01597 | 10.94154 | 6.85434 | 0.88673 | 7.22412 | 10.16061 | 12.63895 | 6.71526 | 9.54027 | 7.90542 | 0.56718 | 10.39659 | 0.40602 | 0.00000 | 7.74867 |
| **Tex30** | 9.40027 | 3.51513 | 4.53758 | 11.90029 | 2.19761 | 0.16555 | 0.00000 | 4.48722 | 13.55147 | 5.66507 | 3.72543 | 0.36240 | 3.54818 | 2.35678 | 4.47204 | 3.71972 |
| **Tex37** | 0.01323 | 0.00785 | 0.01168 | 0.00869 | 0.00908 | 0.00666 | 0.00759 | 0.01614 | 0.01166 | 0.56001 | 0.01199 | 0.00641 | 0.01161 | 0.01850 | 0.05943 | 0.00763 |
| **Tex38** | 0.00994 | 0.00813 | 0.00784 | 0.00482 | 0.00950 | 0.00692 | 0.00783 | 0.75939 | 0.59827 | 0.00426 | 0.00000 | 0.00619 | 0.00555 | 0.01860 | 0.00000 | 0.00732 |
| **Tex40** | 0.01429 | 0.00786 | 0.00276 | 0.58376 | 0.01762 | 0.13896 | 0.06626 | 0.00838 | 0.00375 | 0.07977 | 0.22599 | 0.03885 | 3.15113 | 0.00828 | 0.12916 | 0.00579 |
| **Tex43** | 0.01329 | 0.00792 | 0.01151 | 0.00603 | 0.00921 | 0.00647 | 0.00739 | 0.01223 | 1.13987 | 0.02432 | 0.00197 | 0.01570 | 0.07856 | 0.01095 | 0.01364 | 0.00729 |
| **Tex9** | 2.66838 | 0.05221 | 0.97690 | 1.76888 | 0.02257 | 2.47956 | 1.30284 | 0.17746 | 0.20004 | 5.73381 | 2.44391 | 0.40574 | 0.48165 | 4.04236 | 1.68964 | 2.35268 |
| **Tfam** | 15.48072 | 3.98753 | 7.86041 | 3.54746 | 9.37834 | 5.87842 | 3.63540 | 7.27677 | 1.73163 | 11.89143 | 5.90485 | 7.79477 | 6.07904 | 1.35287 | 7.22891 | 4.86225 |
| **Tfap2a** | 0.02160 | 0.00770 | 0.01545 | 0.14415 | 0.00865 | 1.82112 | 0.01743 | 0.01070 | 0.09070 | 0.15543 | 0.00872 | 0.01739 | 0.04027 | 0.04621 | 0.08572 | 0.00662 |
| **Tfap2c** | 4.20170 | 3.53649 | 7.90694 | 2.54341 | 6.45273 | 8.02457 | 8.25070 | 18.21673 | 5.24204 | 19.89645 | 9.68826 | 1.01579 | 11.03817 | 10.74159 | 7.35936 | 0.29743 |
| **Tfap2e** | 0.00791 | 0.01805 | 0.00331 | 4.01004 | 0.00965 | 0.00681 | 0.00732 | 1.17483 | 4.37533 | 0.45176 | 0.43206 | 0.05683 | 0.00976 | 0.04240 | 0.73006 | 0.00632 |
| **Tfap4** | 0.18915 | 0.00827 | 0.14723 | 0.05728 | 0.20134 | 0.00569 | 0.00768 | 0.86270 | 0.16820 | 2.71446 | 0.36793 | 0.11620 | 0.29111 | 0.08950 | 0.37631 | 0.61898 |
| **Tfb1m** | 7.24447 | 0.47572 | 1.70537 | 0.82264 | 1.83190 | 5.64431 | 1.75575 | 2.21545 | 0.13727 | 0.95621 | 0.58811 | 6.16156 | 1.31747 | 2.25850 | 1.16937 | 0.85600 |
| **Tfb2m** | 2.39429 | 0.00568 | 1.24810 | 10.95158 | 6.57284 | 1.89293 | 2.25806 | 9.98195 | 21.79933 | 16.50987 | 7.84909 | 10.28926 | 4.95723 | 11.29322 | 11.13419 | 0.47673 |
| **Tfcp2** | 0.34085 | 0.01663 | 0.00000 | 2.52955 | 0.00615 | 0.00532 | 0.94491 | 1.78857 | 3.87942 | 5.68838 | 1.16693 | 0.44181 | 0.84195 | 9.43898 | 8.78321 | 2.56686 |
| **Tfcp2l1** | 20.56760 | 10.94982 | 27.29720 | 0.78987 | 10.42162 | 13.58894 | 19.26803 | 21.67702 | 2.53075 | 0.00000 | 19.91225 | 0.19528 | 21.69384 | 11.15631 | 0.00000 | 10.66261 |
| **Tfdp1** | 9.46659 | 9.22720 | 7.98407 | 7.54296 | 5.53164 | 8.37470 | 5.90253 | 6.57334 | 3.11248 | 14.72724 | 5.50737 | 16.78181 | 12.20265 | 11.13365 | 12.17267 | 3.01575 |
| **Tfdp2** | 2.15850 | 0.00000 | 0.80947 | 4.90873 | 0.75271 | 0.00570 | 4.69865 | 0.92892 | 3.31312 | 4.72509 | 3.61486 | 1.85123 | 1.56845 | 1.40457 | 5.96053 | 0.81978 |
| **Tfe3** | 6.39668 | 1.39988 | 5.26322 | 4.61256 | 5.16403 | 8.06193 | 2.76808 | 8.53011 | 1.42585 | 3.91002 | 0.57216 | 0.21532 | 3.13475 | 0.01060 | 3.79310 | 3.17604 |
| **Tfeb** | 5.07818 | 0.65652 | 0.00000 | 0.59109 | 0.77441 | 5.77379 | 3.71803 | 12.29066 | 1.98209 | 16.39263 | 5.18042 | 0.43922 | 10.17907 | 0.00792 | 0.00000 | 4.64080 |
| **Tfg** | 9.91666 | 8.52594 | 8.72440 | 15.76123 | 6.78057 | 3.84770 | 6.05826 | 6.88271 | 15.79631 | 22.20549 | 8.52873 | 18.06150 | 13.23899 | 1.21498 | 12.46417 | 8.02317 |
| **Tfip11** | 2.79427 | 1.89607 | 3.86256 | 6.61127 | 3.92663 | 0.82798 | 5.22801 | 9.18267 | 5.61087 | 9.97103 | 2.23948 | 2.50157 | 7.52124 | 0.14488 | 2.48790 | 1.46349 |
| **Tfpi** | 4.23158 | 0.72309 | 1.03395 | 5.55137 | 6.31896 | 0.00128 | 0.00304 | 0.00000 | 2.51764 | 0.98200 | 0.02337 | 0.00000 | 0.00572 | 0.00756 | 0.60072 | 11.52365 |
| **Tfpt** | 3.70285 | 0.01274 | 0.00000 | 2.89404 | 0.01566 | 4.11984 | 0.00000 | 6.62752 | 7.79804 | 8.92547 | 2.06524 | 0.12887 | 7.35133 | 0.00000 | 0.00000 | 4.50091 |
| **Tfr2** | 0.01839 | 0.00805 | 0.00733 | 0.24846 | 0.00856 | 0.00715 | 0.00731 | 0.00142 | 0.00655 | 0.05047 | 0.72206 | 0.00580 | 0.47514 | 0.00690 | 0.00000 | 0.00693 |
| **Tfrc** | 1.45229 | 0.12970 | 8.81041 | 21.80145 | 0.02246 | 0.00827 | 5.61263 | 15.48305 | 23.98107 | 24.22055 | 30.70104 | 20.31294 | 7.20679 | 44.28933 | 24.64181 | 1.09859 |
| **Tgds** | 3.69630 | 0.60291 | 0.88825 | 3.86128 | 0.01948 | 0.00175 | 0.01516 | 1.64584 | 0.00000 | 0.79928 | 0.03258 | 0.03117 | 0.17936 | 1.28675 | 4.18227 | 0.76463 |
| **Tgfa** | 0.01356 | 0.00810 | 0.01212 | 0.14506 | 0.00924 | 0.00666 | 0.00758 | 0.01618 | 0.00968 | 0.12340 | 0.01413 | 0.00752 | 0.01260 | 0.01096 | 0.00829 | 0.00772 |
| **Tgfb1i1** | 0.01293 | 0.00765 | 0.01097 | 0.00630 | 0.00880 | 0.00657 | 0.00718 | 0.00955 | 0.01135 | 1.69648 | 0.01151 | 0.06369 | 0.01062 | 0.01665 | 0.00321 | 0.00738 |
| **Tgfb2** | 0.00912 | 0.00835 | 0.00645 | 6.25892 | 0.01007 | 0.00586 | 0.00807 | 0.42254 | 2.90718 | 0.00000 | 0.00000 | 0.15585 | 0.01382 | 0.00000 | 2.38608 | 0.00614 |
| **Tgfb3** | 0.01343 | 0.00762 | 0.00971 | 1.54414 | 0.00915 | 0.00657 | 0.00720 | 0.01031 | 0.44562 | 0.71448 | 0.00734 | 0.16868 | 0.01179 | 0.01221 | 0.01187 | 0.00714 |
| **Tgfbr1** | 2.13500 | 2.07279 | 0.17117 | 4.03908 | 0.01169 | 0.00251 | 3.97065 | 1.25697 | 10.12152 | 0.00000 | 0.21930 | 4.44019 | 2.08452 | 0.18528 | 2.69931 | 0.00153 |
| **Tgfbr2** | 0.01978 | 0.00807 | 0.01318 | 0.01057 | 0.00954 | 0.00715 | 0.00771 | 0.01545 | 0.01234 | 0.00987 | 0.01482 | 0.02826 | 0.18821 | 0.01251 | 0.00987 | 0.00761 |
| **Tgfbr3** | 0.01347 | 0.00787 | 0.01168 | 0.62781 | 0.00920 | 0.00658 | 0.00736 | 0.01087 | 0.27782 | 0.47563 | 0.01970 | 0.00603 | 0.01203 | 0.00726 | 0.24485 | 0.00735 |
| **Tgfbr3l** | 0.01269 | 0.00833 | 0.01365 | 0.00985 | 0.01015 | 0.00714 | 0.00769 | 0.01555 | 0.01198 | 0.00806 | 0.01490 | 0.02997 | 0.18696 | 0.00889 | 0.09479 | 0.00765 |
| **Tgfbrap1** | 2.51095 | 1.55270 | 0.00036 | 5.69535 | 1.91989 | 0.00171 | 0.00285 | 0.88102 | 2.51465 | 6.49203 | 2.72370 | 6.37674 | 0.04520 | 2.85559 | 2.94933 | 0.00215 |
| **Tgif1** | 5.05263 | 2.71750 | 4.90781 | 0.16826 | 12.14547 | 0.05297 | 1.70807 | 1.59642 | 0.27371 | 1.97234 | 0.28051 | 0.77571 | 0.67625 | 2.37596 | 2.54104 | 2.15679 |
| **Tgif2** | 5.29169 | 2.32349 | 3.95721 | 8.02011 | 0.93710 | 4.63829 | 3.56995 | 4.76193 | 1.00705 | 7.54944 | 6.29664 | 0.33192 | 0.62953 | 1.32443 | 2.90095 | 4.78461 |
| **Tgm2** | 0.01278 | 0.00799 | 0.62892 | 0.00421 | 0.00801 | 0.00562 | 0.00616 | 0.00000 | 0.01070 | 0.69599 | 3.31874 | 0.00000 | 0.00000 | 0.02969 | 0.02247 | 0.00662 |
| **Tgm4** | 0.00408 | 0.08721 | 0.08694 | 0.98239 | 0.00909 | 0.00556 | 0.30410 | 0.63894 | 2.31286 | 1.88296 | 1.09743 | 2.14117 | 0.04246 | 0.13395 | 2.01132 | 0.00676 |
| **Tgoln1** | 2.30194 | 3.10872 | 6.74227 | 6.55903 | 2.43927 | 0.00996 | 3.08120 | 3.01304 | 11.26021 | 8.60004 | 3.02565 | 1.71131 | 0.92763 | 14.72940 | 13.07706 | 2.76538 |
| **Tgs1** | 3.32497 | 0.11059 | 4.02108 | 8.62537 | 0.88719 | 0.80859 | 5.66547 | 0.77488 | 14.91857 | 5.38607 | 3.06124 | 3.82754 | 1.70798 | 8.09893 | 6.94055 | 0.01947 |
| **Th** | 0.01304 | 0.00790 | 0.00962 | 8.74085 | 0.00908 | 0.00539 | 0.00707 | 0.00845 | 5.98860 | 0.00000 | 0.04750 | 0.04752 | 0.01184 | 0.00000 | 0.20882 | 0.00537 |
| **Thada** | 3.77345 | 1.57926 | 1.22776 | 0.26833 | 0.00000 | 0.59608 | 1.55477 | 1.97558 | 0.57830 | 0.46176 | 1.61474 | 0.00612 | 1.18670 | 1.60975 | 2.31613 | 0.00000 |
| **Thap1** | 0.00000 | 1.59051 | 0.88321 | 8.39888 | 0.00810 | 0.03073 | 1.37624 | 0.24309 | 11.91879 | 0.85026 | 2.35247 | 0.25948 | 0.28486 | 2.08315 | 5.73305 | 0.00378 |
| **Thap11** | 5.90800 | 0.54688 | 1.42148 | 17.50917 | 1.94207 | 1.24233 | 1.08041 | 6.85107 | 20.90551 | 14.67906 | 6.26522 | 1.67058 | 7.70820 | 0.03962 | 9.11169 | 0.06461 |
| **Thap2** | 0.24874 | 0.00841 | 1.26124 | 0.78529 | 0.03406 | 2.69923 | 0.03781 | 0.00309 | 1.14588 | 1.34887 | 0.15678 | 0.01084 | 0.24571 | 0.07451 | 0.21303 | 0.02014 |
| **Thap3** | 1.65986 | 0.49140 | 0.68499 | 1.16561 | 1.75877 | 0.12565 | 0.01392 | 1.50114 | 1.76668 | 1.70750 | 2.65294 | 0.02773 | 2.28111 | 0.00000 | 0.00000 | 0.01903 |
| **Thap4** | 8.84576 | 4.82307 | 8.55027 | 5.00004 | 5.58338 | 4.55493 | 0.20235 | 5.14277 | 5.85292 | 11.69501 | 7.65735 | 0.38963 | 6.96282 | 0.89338 | 4.90163 | 6.01339 |
| **Thap6** | 0.34359 | 0.00657 | 1.46213 | 1.96244 | 0.43284 | 0.11326 | 0.14273 | 0.92075 | 1.25654 | 1.35404 | 0.49198 | 0.16883 | 0.93042 | 3.07352 | 1.50460 | 0.07076 |
| **Thap7** | 5.20229 | 0.00000 | 0.00000 | 2.11535 | 3.48172 | 0.95248 | 0.62367 | 6.53836 | 0.38844 | 6.71107 | 0.50343 | 0.08252 | 5.73635 | 0.00015 | 0.00000 | 0.02063 |

| | | | | | | | | | | | | | | | | |
|---|---|---|---|---|---|---|---|---|---|---|---|---|---|---|---|---|
| **Thbd** | 0.79620 | 0.00686 | 0.00267 | 0.15326 | 0.00621 | 0.00773 | 0.00684 | 0.34238 | 0.00878 | 0.01028 | 0.01131 | 0.00729 | 0.00624 | 0.00878 | 0.06971 | 0.00645 |
| **Thbs1** | 0.01217 | 0.00706 | 0.00629 | 0.00373 | 0.00884 | 0.00678 | 0.00709 | 0.01040 | 0.45905 | 0.02860 | 0.00000 | 1.63919 | 0.01266 | 0.01435 | 0.10930 | 0.17415 |
| **Thbs3** | 0.02590 | 0.00792 | 0.01137 | 0.55135 | 0.00844 | 0.00663 | 0.00727 | 0.01139 | 0.00321 | 0.47823 | 0.01335 | 0.00640 | 0.01213 | 0.01000 | 0.00498 | 0.02294 |
| **Theg** | 0.01380 | 0.00811 | 0.01046 | 0.70348 | 0.00919 | 0.00663 | 0.00746 | 0.00857 | 0.26781 | 0.00035 | 0.18931 | 0.00659 | 0.01029 | 0.01023 | 0.18703 | 0.00720 |
| **Thegl** | 0.01227 | 0.00794 | 0.02773 | 0.58277 | 0.00909 | 0.00583 | 0.00694 | 0.01085 | 0.18420 | 0.00107 | 0.02931 | 0.00756 | 0.63983 | 0.00861 | 0.13468 | 0.00674 |
| **Them4** | 3.77498 | 2.10162 | 1.99007 | 2.34557 | 1.65094 | 0.00001 | 0.01294 | 5.95345 | 3.84880 | 7.43748 | 2.63268 | 0.76806 | 0.12894 | 6.05784 | 2.43480 | 0.00754 |
| **Them5** | 0.12848 | 0.00834 | 0.71086 | 0.01096 | 0.01041 | 0.00550 | 0.00747 | 0.01319 | 0.01225 | 0.01378 | 0.01161 | 0.01942 | 0.01518 | 0.01584 | 0.00024 | 0.00704 |
| **Themis** | 0.01941 | 0.00764 | 0.05379 | 0.40128 | 0.01790 | 0.00629 | 0.01769 | 0.02193 | 0.13757 | 0.04066 | 0.10478 | 0.00522 | 0.05420 | 0.34569 | 0.26385 | 0.00720 |
| **Themis2** | 0.01074 | 0.00726 | 0.00756 | 0.00219 | 0.00904 | 0.00660 | 0.59290 | 0.01498 | 0.03742 | 1.74110 | 0.00815 | 0.00000 | 0.01066 | 0.00581 | 1.08841 | 0.00743 |
| **Themis3** | 0.00923 | 0.00726 | 0.00733 | 0.04268 | 0.00959 | 0.00680 | 0.00706 | 1.76708 | 0.01053 | 0.00199 | 0.00000 | 0.00425 | 0.02525 | 0.01619 | 0.00000 | 0.00725 |
| **Thg1l** | 1.28897 | 0.16818 | 0.00000 | 0.08114 | 0.02178 | 0.04240 | 0.00594 | 0.12097 | 1.46357 | 2.64269 | 0.58809 | 0.02512 | 0.00418 | 2.53439 | 0.62783 | 0.47853 |
| **Thnsl1** | 0.00524 | 0.00778 | 1.50318 | 0.82059 | 0.00738 | 0.10148 | 0.00652 | 0.00798 | 0.00000 | 0.00000 | 0.00000 | 0.01750 | 0.89598 | 2.16607 | 0.78340 | 0.00610 |
| **Thoc1** | 2.47816 | 7.81138 | 5.09417 | 5.78424 | 6.37178 | 5.20283 | 7.13552 | 9.06725 | 8.60692 | 15.44588 | 7.48756 | 17.99664 | 6.55166 | 18.20814 | 12.89277 | 5.56727 |
| **Thoc2** | 1.37957 | 4.73600 | 3.27327 | 2.83712 | 2.86501 | 0.00328 | 3.11881 | 3.63469 | 3.91667 | 7.50302 | 4.10260 | 5.47970 | 0.00000 | 28.54959 | 9.51187 | 1.84527 |
| **Thoc3** | 3.09493 | 0.91998 | 6.43597 | 6.59736 | 6.67274 | 2.26520 | 1.42750 | 6.81403 | 5.32700 | 14.12705 | 9.43105 | 0.60457 | 4.95870 | 9.65020 | 8.61178 | 1.20531 |
| **Thoc5** | 7.98163 | 5.58777 | 4.69210 | 8.61811 | 4.66545 | 5.59676 | 0.00000 | 8.33470 | 9.96219 | 11.03965 | 4.65243 | 2.61722 | 6.31895 | 2.26612 | 10.68022 | 0.12458 |
| **Thoc6** | 15.06422 | 8.63029 | 8.12653 | 0.87518 | 5.81926 | 0.60751 | 7.28007 | 9.11000 | 2.74033 | 10.34224 | 5.57728 | 1.08404 | 12.41873 | 0.00000 | 0.27545 | 7.15281 |
| **Thoc7** | 15.08386 | 11.08014 | 8.50006 | 9.09472 | 8.71443 | 8.89454 | 7.38569 | 10.46752 | 7.16854 | 17.90981 | 15.68227 | 5.49508 | 11.87531 | 9.08786 | 3.78221 | 10.74731 |
| **Thop1** | 9.41266 | 0.00441 | 0.00000 | 0.05279 | 5.15832 | 2.67151 | 0.01395 | 8.21479 | 0.80082 | 5.94350 | 0.31717 | 0.11797 | 11.85531 | 0.00677 | 0.00000 | 3.99629 |
| **Thra** | 4.09386 | 0.00303 | 0.96809 | 0.39752 | 1.89641 | 0.75178 | 0.00448 | 0.71450 | 0.00314 | 3.68934 | 0.01736 | 0.07958 | 0.95900 | 0.00499 | 0.86944 | 0.00193 |
| **Thrap3** | 11.60878 | 3.93685 | 4.44811 | 4.40484 | 7.33411 | 2.54426 | 6.57511 | 10.21940 | 10.00783 | 4.45290 | 6.84728 | 0.59552 | 7.21087 | 3.19800 | 1.34531 | 5.76914 |
| **Thrsp** | 0.01046 | 0.00768 | 0.00934 | 0.00744 | 0.00922 | 0.01320 | 0.00718 | 0.11602 | 0.01165 | 0.93316 | 0.01169 | 0.00482 | 0.00831 | 0.01193 | 0.00748 | 0.00775 |
| **Thsd1** | 0.01368 | 0.00816 | 0.01230 | 0.55520 | 0.00929 | 0.00659 | 0.00751 | 0.01394 | 0.00434 | 0.01528 | 0.01398 | 0.00829 | 0.01259 | 0.01102 | 0.00494 | 0.01463 |
| **Thsd4** | 0.01335 | 0.00798 | 0.00969 | 0.31569 | 0.00946 | 0.00667 | 0.00731 | 0.01066 | 0.17641 | 0.00431 | 0.00879 | 0.00553 | 0.01327 | 0.42088 | 0.10538 | 0.00741 |
| **Thsd7a** | 0.01378 | 0.00815 | 0.01200 | 0.82303 | 0.00935 | 0.00656 | 0.00746 | 0.01323 | 0.00000 | 0.01146 | 0.01284 | 0.01908 | 0.01270 | 0.00796 | 0.08138 | 0.00726 |
| **Thsd7b** | 0.01365 | 0.00826 | 0.01208 | 0.63765 | 0.00932 | 0.00657 | 0.00752 | 0.01391 | 0.63396 | 0.01185 | 0.00886 | 0.00736 | 0.01285 | 0.01144 | 0.00338 | 0.00728 |
| **Thtpa** | 0.37215 | 0.00705 | 0.00370 | 0.40013 | 0.00719 | 0.00661 | 0.00710 | 0.01397 | 0.00484 | 0.00573 | 0.01075 | 0.00540 | 0.01207 | 0.00566 | 1.07072 | 0.00684 |
| **Thumpd1** | 4.12327 | 8.51024 | 8.36532 | 4.25697 | 4.77305 | 4.14897 | 3.72324 | 7.63834 | 8.20251 | 9.31784 | 9.28467 | 7.88970 | 10.50197 | 12.91126 | 12.55047 | 3.97388 |
| **Thumpd2** | 0.01042 | 0.00816 | 0.02002 | 0.01049 | 0.01008 | 0.00685 | 0.00759 | 0.01558 | 0.01238 | 0.02229 | 0.01517 | 0.01748 | 0.44706 | 0.01204 | 0.00953 | 0.09191 |
| **Thumpd3** | 5.89375 | 9.88155 | 6.12330 | 2.65599 | 3.93905 | 5.30854 | 4.97832 | 2.96512 | 1.49745 | 1.81977 | 3.79475 | 4.00162 | 3.65974 | 3.70852 | 3.92380 | 1.19360 |
| **Thy1** | 0.06460 | 0.00809 | 0.00813 | 1.27471 | 0.01517 | 0.00662 | 0.00726 | 0.01052 | 0.00000 | 0.00524 | 0.02512 | 0.05457 | 0.01939 | 0.00359 | 0.36812 | 0.00725 |
| **Thyn1** | 4.27223 | 1.15259 | 4.63673 | 0.01253 | 0.62526 | 2.15718 | 1.08659 | 2.40742 | 0.00557 | 1.23102 | 0.75955 | 2.29633 | 4.08506 | 2.27755 | 1.11085 | 0.26873 |
| **Tia1** | 0.85329 | 0.00640 | 0.67888 | 0.00365 | 0.01511 | 0.00471 | 1.82421 | 0.00611 | 0.00994 | 1.14978 | 0.00936 | 0.00212 | 0.01485 | 0.01735 | 0.70006 | 0.02108 |
| **Tial1** | 11.40736 | 4.14668 | 4.71834 | 11.06592 | 7.04418 | 4.99677 | 2.92101 | 6.14289 | 11.01733 | 13.41833 | 8.96595 | 10.82934 | 3.33805 | 9.83993 | 11.32822 | 1.68959 |
| **Tiam1** | 0.00000 | 0.00700 | 0.00000 | 8.19751 | 0.00877 | 0.01848 | 0.28909 | 2.39903 | 10.88739 | 3.06479 | 1.88794 | 0.70325 | 0.00000 | 1.67702 | 0.20421 | 0.06278 |
| **Tiam2** | 3.11470 | 0.00342 | 0.00000 | 5.94550 | 0.51372 | 0.00444 | 0.67191 | 0.69536 | 2.60897 | 1.79663 | 0.00000 | 0.30265 | 0.18500 | 1.56687 | 1.91465 | 0.33188 |
| **Ticam1** | 0.00953 | 0.01905 | 1.04272 | 4.86964 | 0.00906 | 1.52011 | 0.00698 | 2.16488 | 3.38392 | 1.73911 | 0.27251 | 0.03839 | 0.00000 | 0.40636 | 0.54609 | 0.05733 |
| **Ticam2** | 0.01741 | 0.00744 | 0.09307 | 2.10493 | 0.00872 | 0.00560 | 0.00708 | 0.05710 | 0.00000 | 0.97749 | 0.00874 | 0.02379 | 0.00872 | 0.02550 | 1.00757 | 0.00676 |
| **Ticrr** | 8.16943 | 5.03909 | 10.47966 | 8.01769 | 9.95055 | 9.23649 | 4.51452 | 11.13931 | 16.13488 | 13.35246 | 8.76599 | 10.10514 | 10.02857 | 10.12956 | 16.96428 | 5.06972 |
| **Tie1** | 0.00000 | 0.00610 | 3.33143 | 0.44340 | 0.01853 | 0.00798 | 0.00503 | 6.75487 | 0.50197 | 0.29567 | 0.40149 | 0.00000 | 2.98759 | 0.00596 | 0.00000 | 1.23692 |
| **Tifab** | 0.84467 | 0.00684 | 0.39425 | 0.00582 | 0.00606 | 0.00950 | 0.00705 | 0.00477 | 0.01138 | 0.21883 | 0.29037 | 0.00603 | 0.13665 | 0.03020 | 0.00000 | 0.03707 |
| **Tigar** | 19.22388 | 13.16636 | 18.49051 | 14.88422 | 7.50090 | 9.60232 | 13.42894 | 29.65489 | 20.72232 | 26.58457 | 20.72049 | 9.13865 | 20.45262 | 11.12662 | 17.64389 | 6.35345 |
| **Tigd2** | 0.13146 | 0.00850 | 1.58400 | 0.63591 | 0.01764 | 0.00818 | 0.00730 | 0.01034 | 0.68827 | 0.37970 | 0.00613 | 0.00362 | 0.27777 | 0.00707 | 0.39089 | 0.00685 |
| **Tigd3** | 0.08677 | 0.01820 | 0.22732 | 0.77909 | 0.02249 | 0.01253 | 0.02670 | 0.02240 | 2.56809 | 0.00004 | 1.00135 | 0.03022 | 0.13409 | 0.35677 | 1.78050 | 0.03661 |
| **Tigd5** | 0.02895 | 0.00717 | 0.12851 | 0.34846 | 0.00833 | 0.00567 | 0.95033 | 0.01485 | 0.00509 | 0.01056 | 0.01320 | 0.01680 | 0.00982 | 0.01140 | 0.00283 | 0.07169 |
| **Tigit** | 0.01899 | 0.00747 | 0.02661 | 0.51432 | 0.00842 | 0.00820 | 0.00686 | 0.15061 | 0.34611 | 0.91620 | 0.62761 | 0.31598 | 0.03940 | 0.22775 | 1.14971 | 0.00700 |
| **Timd2** | 7.69983 | 16.24231 | 44.83319 | 16.25745 | 5.23831 | 14.15656 | 23.77898 | 49.57338 | 28.51201 | 45.82025 | 52.05875 | 29.53882 | 46.33890 | 53.60286 | 35.21746 | 7.65540 |
| **Timd4** | 0.01302 | 0.00755 | 0.00803 | 1.22898 | 0.00950 | 0.00668 | 0.00720 | 0.00980 | 0.00000 | 0.00817 | 0.00935 | 0.00464 | 0.01330 | 0.86307 | 0.00000 | 0.00725 |
| **Timeless** | 3.35173 | 4.03051 | 5.46445 | 0.77360 | 0.07709 | 0.07313 | 4.30340 | 2.76929 | 2.85337 | 2.05523 | 1.56413 | 0.35520 | 2.66577 | 0.79909 | 3.25328 | 2.50293 |
| **Timm10** | 12.33536 | 2.64762 | 12.50896 | 11.93396 | 5.92369 | 1.92543 | 3.85473 | 8.84383 | 15.00758 | 7.97048 | 13.51791 | 1.26572 | 5.58345 | 1.53151 | 5.64553 | 5.10392 |
| **Timm10b** | 0.85842 | 0.01582 | 0.12898 | 0.23194 | 1.33113 | 0.88171 | 0.06814 | 0.44778 | 0.87166 | 4.17296 | 0.46079 | 0.02086 | 0.10171 | 0.00783 | 0.00000 | 1.21295 |
| **Timm13** | 18.37880 | 1.05887 | 2.96102 | 4.13097 | 14.99714 | 4.06466 | 5.50238 | 7.51788 | 5.35792 | 15.78282 | 13.49102 | 1.01926 | 17.23214 | 0.00000 | 0.00000 | 6.41694 |
| **Timm17a** | 20.84860 | 11.70058 | 19.68368 | 13.48859 | 9.96446 | 6.48737 | 9.06501 | 27.22806 | 20.75099 | 35.92462 | 32.70411 | 16.86049 | 24.08371 | 27.95025 | 27.04780 | 10.49601 |
| **Timm17b** | 12.83512 | 4.11299 | 14.91698 | 7.07942 | 7.32368 | 7.35058 | 7.13954 | 18.86323 | 5.51133 | 14.79505 | 8.56850 | 0.52717 | 10.57574 | 0.60811 | 2.17022 | 6.33609 |
| **Timm21** | 0.21674 | 0.23376 | 0.00000 | 1.52087 | 0.07299 | 0.01465 | 0.00693 | 0.01163 | 1.06522 | 1.30361 | 0.05154 | 0.00000 | 0.01046 | 0.25298 | 0.35401 | 0.00709 |
| **Timm22** | 14.01791 | 8.80978 | 12.23478 | 1.56160 | 8.37762 | 3.47581 | 6.76012 | 21.76756 | 7.98171 | 23.58227 | 18.38293 | 3.29903 | 13.39743 | 10.32497 | 10.39095 | 4.27622 |
| **Timm23** | 35.77719 | 7.77901 | 25.62901 | 21.29182 | 23.91568 | 18.91699 | 17.28345 | 39.26098 | 18.43443 | 46.67749 | 33.76330 | 20.27791 | 35.18317 | 20.40931 | 12.05435 | 17.19025 |
| **Timm44** | 15.66420 | 8.36208 | 9.28301 | 3.95224 | 5.70583 | 7.50218 | 2.47963 | 16.07667 | 3.62033 | 15.82782 | 6.09759 | 5.79536 | 8.49874 | 0.08641 | 3.94299 | 6.45678 |
| **Timm50** | 10.15819 | 0.96223 | 9.12408 | 0.42412 | 2.71934 | 5.05097 | 4.26411 | 7.47398 | 0.71117 | 16.90838 | 1.47516 | 2.41788 | 17.99700 | 0.00000 | 0.00000 | 1.65860 |
| **Timm8a1** | 12.61580 | 3.65402 | 15.07905 | 16.94514 | 11.06687 | 5.45250 | 5.28884 | 25.35293 | 12.45521 | 42.15066 | 20.59399 | 3.83935 | 16.06024 | 23.16920 | 15.14300 | 3.37218 |
| **Timm8a2** | 1.44302 | 1.82061 | 6.03271 | 2.61946 | 8.48114 | 0.01724 | 1.92139 | 8.18301 | 2.33914 | 6.57379 | 4.42702 | 0.38337 | 1.82412 | 1.20401 | 1.03107 | 0.05206 |
| **Timm8b** | 5.79289 | 0.24570 | 2.71715 | 14.29539 | 5.30181 | 0.97172 | 0.60692 | 0.19751 | 4.84307 | 5.06738 | 4.76669 | 0.27516 | 0.50770 | 0.10229 | 0.79590 | 1.48612 |

| | | | | | | | | | | | | | | | | |
|---|---|---|---|---|---|---|---|---|---|---|---|---|---|---|---|---|
| **Timm9** | 8.43712 | 7.26666 | 9.39498 | 2.84064 | 0.39183 | 4.58143 | 3.29819 | 7.64890 | 3.97479 | 10.09090 | 4.38901 | 4.68548 | 5.13462 | 0.72402 | 3.86330 | 3.05585 |
| **Timmdc1** | 0.17302 | 0.00707 | 1.33902 | 1.54054 | 0.03198 | 0.63836 | 0.12578 | 1.04195 | 2.26416 | 1.49709 | 2.10355 | 0.40678 | 0.36118 | 0.75590 | 0.37558 | 0.02058 |
| **Timp1** | 5.49043 | 0.00000 | 0.00211 | 0.57370 | 5.89348 | 0.00939 | 0.00240 | 0.67154 | 1.49317 | 1.27560 | 0.20454 | 0.00000 | 0.00000 | 0.27782 | 0.53760 | 5.49928 |
| **Timp2** | 0.01302 | 0.00785 | 0.01070 | 0.24285 | 0.00885 | 0.00657 | 0.00722 | 0.00984 | 0.00726 | 1.51506 | 0.01103 | 0.09098 | 0.01076 | 0.01112 | 0.00000 | 0.00736 |
| **Timp3** | 1.50157 | 0.00476 | 0.00000 | 0.00515 | 0.00499 | 0.00815 | 2.13098 | 0.00397 | 0.01152 | 1.35904 | 0.00986 | 0.05212 | 0.01001 | 0.01116 | 0.01198 | 0.00552 |
| **Tinagl1** | 3.10520 | 0.54464 | 1.01495 | 0.00000 | 1.66318 | 8.60115 | 1.88703 | 4.75611 | 0.00581 | 0.11525 | 2.64904 | 0.00000 | 7.77589 | 0.01107 | 0.00000 | 2.81079 |
| **Tinf2** | 9.09249 | 6.28024 | 5.82384 | 18.70374 | 11.26454 | 1.58102 | 2.33572 | 6.32823 | 12.40835 | 12.96945 | 8.31032 | 3.97281 | 10.51632 | 3.65035 | 5.57588 | 3.10374 |
| **Tiparp** | 3.53849 | 6.08336 | 1.81323 | 14.65905 | 2.77119 | 0.00256 | 0.00000 | 0.59760 | 8.64179 | 5.28690 | 2.19443 | 1.24728 | 0.93836 | 7.44442 | 10.71696 | 0.11174 |
| **Tipin** | 18.80798 | 14.71639 | 14.24400 | 16.80909 | 8.98495 | 4.84314 | 6.22559 | 15.14200 | 23.17123 | 22.60610 | 15.76000 | 17.27189 | 14.76657 | 22.47537 | 19.67141 | 11.46320 |
| **Tiprl** | 6.94087 | 7.31423 | 6.51683 | 11.82985 | 11.21681 | 5.77213 | 6.09522 | 6.98439 | 20.89950 | 14.67544 | 6.74849 | 2.88742 | 5.54721 | 6.03202 | 11.13677 | 3.05073 |
| **Tirap** | 0.07262 | 0.05322 | 0.46882 | 1.10432 | 0.78206 | 0.09624 | 0.23124 | 1.09293 | 2.95518 | 1.03008 | 1.06005 | 0.11719 | 0.30377 | 0.80724 | 0.61048 | 0.14754 |
| **Tjap1** | 2.03332 | 0.00441 | 1.38574 | 11.09120 | 0.18862 | 2.23298 | 0.05329 | 4.22638 | 4.58359 | 4.97651 | 3.84248 | 0.59538 | 3.70121 | 0.04645 | 0.00000 | 1.58164 |
| **Tjp1** | 6.90924 | 3.83140 | 2.69459 | 5.74880 | 3.69902 | 2.07889 | 1.39694 | 5.84112 | 4.00732 | 6.68241 | 3.39922 | 7.25054 | 5.75398 | 16.33237 | 2.94386 | 2.39707 |
| **Tjp2** | 7.59997 | 0.13651 | 0.84270 | 5.19670 | 3.58853 | 10.85544 | 2.96121 | 6.09078 | 5.47049 | 2.94195 | 1.48811 | 0.09335 | 5.69699 | 2.08071 | 1.97679 | 4.12047 |
| **Tjp3** | 0.25143 | 0.00756 | 0.00000 | 0.94963 | 0.00659 | 1.40480 | 0.00649 | 0.49564 | 0.80692 | 0.42976 | 0.04411 | 0.01348 | 0.62590 | 0.01656 | 0.10613 | 1.59024 |
| **Tk1** | 15.09587 | 13.08986 | 24.28526 | 5.30366 | 7.11473 | 8.78649 | 9.04980 | 26.94328 | 4.56049 | 20.09852 | 15.73600 | 10.47115 | 14.82658 | 2.27197 | 14.06234 | 13.50092 |
| **Tk2** | 1.37924 | 0.00684 | 0.00679 | 2.50673 | 0.02297 | 0.00443 | 0.00583 | 1.47370 | 5.82234 | 1.18502 | 0.68095 | 0.92073 | 1.20125 | 0.02135 | 0.59248 | 0.01173 |
| **Tkfc** | 0.04933 | 0.00791 | 0.00417 | 0.11952 | 0.14593 | 0.00733 | 0.05718 | 0.00343 | 0.59767 | 0.00000 | 0.18724 | 0.00414 | 0.28522 | 0.51043 | 0.00000 | 0.00710 |
| **Tkt** | 3.40690 | 5.98698 | 2.82038 | 0.14040 | 7.23337 | 4.28429 | 6.99004 | 16.20667 | 5.86071 | 14.55972 | 10.49603 | 6.21167 | 2.76192 | 0.24012 | 6.12633 | 3.58860 |
| **Tktl1** | 0.00000 | 10.92737 | 4.94855 | 7.18830 | 0.01459 | 0.00424 | 2.53469 | 1.89094 | 5.10188 | 9.39172 | 5.30581 | 0.98711 | 1.17723 | 12.19507 | 7.46343 | 0.00534 |
| **Tktl2** | 0.01806 | 0.00835 | 0.01138 | 1.61698 | 0.01033 | 0.00668 | 0.00744 | 0.01215 | 0.01222 | 0.40788 | 0.01121 | 0.00813 | 0.01250 | 0.00334 | 0.35748 | 0.00718 |
| **Tlcd1** | 2.13706 | 0.00967 | 4.81778 | 0.75319 | 5.32444 | 0.00360 | 1.82712 | 0.37278 | 0.81745 | 4.96123 | 1.59758 | 7.15485 | 1.86322 | 0.03984 | 0.98113 | 0.00928 |
| **Tlcd2** | 0.08881 | 0.00769 | 0.53670 | 8.76006 | 0.01007 | 0.06669 | 0.05608 | 0.07195 | 7.44666 | 3.85152 | 0.00000 | 0.17749 | 0.23642 | 7.11025 | 15.21355 | 0.00472 |
| **Tldc1** | 0.78687 | 0.00464 | 0.36297 | 0.00992 | 0.00639 | 0.00497 | 2.91674 | 2.08980 | 0.01029 | 0.13117 | 0.00000 | 0.08985 | 0.18017 | 0.01352 | 0.00000 | 0.01433 |
| **Tle1** | 3.79957 | 0.01911 | 1.06493 | 3.75299 | 6.06792 | 3.93927 | 3.63240 | 0.02165 | 1.20475 | 0.12624 | 0.03422 | 1.37562 | 0.24659 | 0.16553 | 0.47443 | 1.60676 |
| **Tle2** | 0.01334 | 0.00793 | 0.02608 | 0.34363 | 0.01845 | 0.00692 | 0.00749 | 0.01316 | 0.00651 | 0.14210 | 0.02514 | 0.01961 | 0.01921 | 0.00827 | 0.04100 | 0.00731 |
| **Tle3** | 0.19479 | 0.36839 | 1.09961 | 0.88783 | 0.01188 | 2.24645 | 1.04548 | 0.43510 | 0.55544 | 0.87514 | 2.43719 | 0.03050 | 0.81919 | 0.08516 | 0.75985 | 2.63209 |
| **Tle4** | 4.19597 | 9.98487 | 6.55538 | 8.44018 | 6.19482 | 2.22814 | 5.66558 | 10.45341 | 15.05675 | 5.44330 | 4.53528 | 17.20364 | 1.86445 | 14.08786 | 7.83483 | 2.60714 |
| **Tle6** | 1.25840 | 0.00607 | 2.21747 | 14.95852 | 0.80419 | 0.00456 | 0.06935 | 2.04280 | 4.17578 | 6.76345 | 2.90596 | 0.34590 | 2.26130 | 0.00000 | 1.54560 | 2.94109 |
| **Tlk1** | 3.12283 | 4.69384 | 0.78670 | 4.79876 | 4.45138 | 3.07845 | 4.17081 | 1.99100 | 1.37124 | 4.52919 | 1.92466 | 6.37842 | 1.36202 | 5.82521 | 5.16642 | 1.22409 |
| **Tlk2** | 0.00000 | 3.20709 | 3.32546 | 14.83822 | 3.79610 | 0.03140 | 2.71557 | 6.48055 | 12.75035 | 11.35216 | 1.53235 | 7.52461 | 5.76859 | 16.58882 | 14.58244 | 4.62721 |
| **Tll1** | 0.02528 | 0.00816 | 0.01060 | 0.34845 | 0.00985 | 0.00678 | 0.00767 | 0.01602 | 0.01926 | 0.01041 | 0.05240 | 0.03897 | 0.02162 | 0.00869 | 0.04027 | 0.00737 |
| **Tln1** | 4.25494 | 0.34158 | 2.45995 | 0.82241 | 7.48993 | 1.19348 | 6.03975 | 5.13432 | 1.39970 | 2.02200 | 0.82629 | 0.37051 | 6.00385 | 0.00029 | 0.01720 | 5.73245 |
| **Tln2** | 0.17051 | 1.65814 | 0.10782 | 1.84681 | 10.96971 | 0.66824 | 0.18955 | 2.50745 | 0.56104 | 1.40233 | 0.16497 | 0.08901 | 0.25171 | 0.22247 | 0.06043 | 0.03015 |
| **Tlr3** | 0.01383 | 0.00792 | 0.01095 | 1.59373 | 0.00927 | 0.00655 | 0.00737 | 0.01211 | 0.27642 | 0.01022 | 0.02539 | 0.07651 | 0.01254 | 0.00891 | 0.01208 | 0.00713 |
| **Tm2d1** | 1.17529 | 0.07469 | 2.80243 | 2.14698 | 2.37152 | 2.32847 | 0.09467 | 0.80426 | 0.62558 | 0.28427 | 2.66871 | 5.02788 | 0.24890 | 1.02026 | 0.00000 | 0.50513 |
| **Tm2d2** | 7.76592 | 1.81631 | 1.60909 | 2.10890 | 4.07323 | 5.64955 | 2.93009 | 4.48970 | 2.70691 | 1.52544 | 0.16295 | 0.23975 | 1.86837 | 0.00300 | 0.08007 | 5.30499 |
| **Tm2d3** | 6.98655 | 0.03716 | 7.17155 | 17.20235 | 1.82894 | 3.11924 | 3.39204 | 9.20849 | 4.25860 | 9.28839 | 4.51357 | 0.13127 | 8.19985 | 0.01535 | 0.17135 | 4.98910 |
| **Tm4sf1** | 0.01241 | 0.00788 | 0.01237 | 0.53414 | 0.00876 | 0.00645 | 0.00720 | 0.00971 | 0.00331 | 2.62950 | 0.01112 | 0.00000 | 0.01079 | 0.01034 | 0.00335 | 0.00738 |
| **Tm4sf5** | 0.28812 | 0.00737 | 0.00177 | 0.73234 | 0.01522 | 0.00801 | 0.04668 | 0.24020 | 0.00446 | 0.00001 | 0.28285 | 0.00561 | 0.27727 | 0.00467 | 0.40537 | 0.00670 |
| **Tm6sf1** | 0.00319 | 0.39203 | 1.37511 | 0.00635 | 0.00586 | 0.00905 | 0.53616 | 0.00792 | 0.01184 | 0.89390 | 0.01045 | 0.02267 | 0.00760 | 0.00882 | 0.01001 | 0.00682 |
| **Tm6sf2** | 0.01279 | 0.00785 | 0.00891 | 0.00581 | 0.00923 | 0.00685 | 0.00726 | 0.03154 | 0.01144 | 0.00576 | 0.33741 | 0.01637 | 0.00780 | 0.01054 | 0.00450 | 0.00722 |
| **Tm7sf2** | 0.29668 | 0.00669 | 0.02629 | 0.72613 | 0.15429 | 0.00749 | 0.00626 | 0.00000 | 0.15542 | 0.87489 | 0.90905 | 0.00000 | 0.62539 | 0.00414 | 0.00000 | 0.00650 |
| **Tm7sf3** | 1.63728 | 2.24278 | 4.99465 | 3.44765 | 0.00446 | 1.56544 | 7.42828 | 0.84463 | 2.01659 | 0.00000 | 1.55642 | 0.03128 | 2.59620 | 0.76298 | 1.72553 | 3.95370 |
| **Tm9sf1** | 0.58336 | 4.23900 | 2.99544 | 1.73522 | 0.01050 | 4.18313 | 1.73833 | 2.02983 | 0.98389 | 0.82646 | 0.73675 | 0.37943 | 1.34894 | 0.01586 | 4.33962 | 1.19774 |
| **Tm9sf2** | 4.69361 | 6.76805 | 6.15974 | 9.40007 | 1.70293 | 1.94300 | 5.59620 | 4.98267 | 7.31833 | 5.25708 | 2.62362 | 21.07542 | 4.98917 | 18.91359 | 3.07286 | 4.59231 |
| **Tm9sf3** | 7.55311 | 5.08388 | 7.18441 | 1.04528 | 6.75319 | 2.29713 | 8.28910 | 6.81917 | 2.06176 | 6.23610 | 3.67536 | 2.69682 | 7.44526 | 9.96981 | 10.04037 | 1.90921 |
| **Tm9sf4** | 7.09667 | 2.54474 | 4.80198 | 2.62899 | 0.76488 | 0.81276 | 6.80989 | 6.01487 | 2.34563 | 2.23531 | 3.35131 | 1.90031 | 5.81725 | 5.54999 | 1.66783 | 5.68901 |
| **Tma16** | 6.35950 | 6.45139 | 3.35935 | 3.01377 | 2.03159 | 4.87114 | 1.38207 | 12.92316 | 5.71899 | 11.63136 | 8.02462 | 3.66916 | 5.06511 | 5.20271 | 8.16095 | 1.53472 |
| **Tma7** | 18.78701 | 4.68998 | 18.20560 | 16.44828 | 7.47122 | 6.80190 | 5.41664 | 11.13027 | 19.46748 | 21.31233 | 35.47263 | 4.71249 | 17.08672 | 15.95060 | 13.96716 | 8.67359 |
| **Tmbim1** | 10.46554 | 0.11552 | 8.29387 | 14.51177 | 3.85422 | 11.74033 | 6.38712 | 11.56288 | 8.51206 | 4.44597 | 8.97529 | 0.84873 | 12.80473 | 1.40661 | 4.23427 | 1.74522 |
| **Tmbim4** | 10.56054 | 1.17998 | 0.32274 | 0.00000 | 6.58991 | 5.38828 | 0.06350 | 4.35606 | 0.01414 | 4.15473 | 4.76100 | 0.11655 | 7.03633 | 0.94515 | 0.00000 | 3.48102 |
| **Tmbim6** | 13.46305 | 6.80557 | 12.90696 | 8.05996 | 10.51418 | 8.42302 | 10.80472 | 12.28134 | 11.33623 | 9.69572 | 7.27138 | 2.59051 | 15.08515 | 0.26182 | 0.00000 | 9.02387 |
| **Tmc2** | 0.10540 | 0.09211 | 0.10350 | 0.28146 | 0.01028 | 0.01246 | 0.03764 | 0.04705 | 0.28519 | 0.03398 | 0.29938 | 0.02077 | 0.19623 | 0.99392 | 0.06033 | 0.05174 |
| **Tmc4** | 0.00757 | 0.00718 | 0.00850 | 2.38296 | 0.00922 | 0.00600 | 0.00695 | 1.97398 | 2.77085 | 0.66000 | 0.46580 | 0.70413 | 0.00373 | 0.00000 | 0.82002 | 0.00656 |
| **Tmc5** | 0.01343 | 0.00800 | 0.01202 | 0.44956 | 0.00904 | 0.00672 | 0.00746 | 0.01171 | 0.00503 | 0.62324 | 0.01193 | 0.00724 | 0.01173 | 0.01228 | 0.01201 | 0.00724 |
| **Tmc6** | 4.20014 | 0.41311 | 5.47697 | 2.28306 | 0.00694 | 0.02696 | 6.24982 | 9.10101 | 0.10860 | 3.04771 | 2.71197 | 0.35412 | 4.73396 | 0.02280 | 0.05119 | 1.61200 |
| **Tmc7** | 0.60183 | 0.00715 | 0.00484 | 0.01803 | 0.00673 | 0.00701 | 0.00707 | 0.01174 | 0.01101 | 0.01025 | 0.01251 | 0.00762 | 0.02769 | 0.00762 | 0.12410 | 0.00669 |
| **Tmc8** | 0.04010 | 0.00768 | 0.02081 | 0.99732 | 0.00894 | 0.01309 | 0.01699 | 0.00753 | 2.26928 | 0.85410 | 0.22231 | 0.00000 | 0.01026 | 0.02478 | 0.04036 | 0.00706 |
| **Tmcc1** | 0.83639 | 0.00595 | 0.10188 | 5.28235 | 0.01414 | 0.00763 | 1.23222 | 0.56675 | 1.13225 | 2.83883 | 0.00000 | 0.50345 | 0.01887 | 0.00040 | 5.30621 | 0.01928 |
| **Tmcc2** | 0.01295 | 0.00800 | 0.01085 | 5.89896 | 0.00931 | 0.00589 | 0.00759 | 0.02336 | 1.82099 | 0.00711 | 0.00000 | 0.00340 | 0.01225 | 0.00000 | 0.65929 | 0.00628 |
| **Tmcc3** | 0.00582 | 0.00769 | 0.00000 | 7.65511 | 0.00996 | 0.01676 | 0.00658 | 0.00000 | 0.27435 | 0.54047 | 0.54532 | 0.08952 | 0.00696 | 4.67812 | 8.04577 | 0.00561 |

| | | | | | | | | | | | | | | | | |
|---|---|---|---|---|---|---|---|---|---|---|---|---|---|---|---|---|
| **Tmco1** | 10.53650 | 9.56027 | 19.48593 | 11.68576 | 3.65958 | 12.22208 | 12.85804 | 19.39369 | 9.36785 | 15.96034 | 9.61897 | 10.47056 | 11.68196 | 7.25697 | 4.93105 | 6.73488 |
| **Tmco2** | 0.27198 | 0.00773 | 0.05198 | 0.02371 | 0.00698 | 0.00840 | 0.00719 | 0.61003 | 0.82673 | 0.86666 | 0.00000 | 0.02366 | 0.00532 | 0.02220 | 0.61828 | 0.00668 |
| **Tmco3** | 1.86426 | 0.59605 | 3.33842 | 2.85511 | 0.50292 | 3.60163 | 0.11436 | 1.05930 | 2.51043 | 2.18395 | 2.75145 | 0.46437 | 1.56375 | 0.87857 | 1.63733 | 0.07509 |
| **Tmco4** | 1.13410 | 0.42618 | 3.28422 | 1.41062 | 0.00000 | 2.08883 | 0.68701 | 4.40385 | 0.00000 | 2.04293 | 1.15647 | 0.00533 | 1.01629 | 0.00000 | 0.00002 | 0.00315 |
| **Tmco6** | 0.01367 | 0.00798 | 0.01215 | 0.01057 | 0.00930 | 0.00666 | 0.00752 | 0.01359 | 0.01242 | 0.01451 | 0.01293 | 0.00801 | 0.01269 | 0.03058 | 0.00819 | 0.01475 |
| **Tmed1** | 5.08245 | 1.22106 | 5.81327 | 0.09830 | 4.05346 | 0.02744 | 2.79799 | 11.21615 | 0.01891 | 7.38345 | 2.13626 | 0.26770 | 5.39539 | 0.05724 | 1.19983 | 3.74908 |
| **Tmed10** | 16.68131 | 8.37313 | 21.12364 | 6.76844 | 13.45670 | 7.31039 | 14.87531 | 19.12536 | 9.53727 | 22.91435 | 13.40366 | 10.58574 | 19.57213 | 12.35842 | 10.18492 | 10.77345 |
| **Tmed2** | 9.37222 | 6.41361 | 9.36080 | 6.47371 | 6.81247 | 8.95320 | 8.98028 | 13.96636 | 7.21152 | 14.74979 | 7.81965 | 4.81340 | 12.50185 | 15.90823 | 11.45399 | 5.54744 |
| **Tmed3** | 0.01152 | 0.00701 | 0.00497 | 0.21363 | 0.01847 | 0.13502 | 1.14160 | 0.05392 | 0.00634 | 3.14831 | 0.55780 | 0.02282 | 0.24761 | 0.00414 | 0.22868 | 0.61965 |
| **Tmed4** | 4.12435 | 3.33596 | 4.40739 | 1.99528 | 0.15425 | 6.12875 | 4.34231 | 3.96223 | 1.23302 | 8.74398 | 1.84919 | 1.96122 | 2.43846 | 7.05133 | 8.94135 | 0.01043 |
| **Tmed5** | 0.88921 | 3.84974 | 3.58022 | 10.63497 | 8.67848 | 0.56733 | 2.03263 | 10.75273 | 14.24411 | 14.35394 | 6.41943 | 13.66450 | 5.29051 | 23.91752 | 8.20327 | 1.42056 |
| **Tmed6** | 0.09719 | 0.00713 | 0.06995 | 13.04524 | 0.00760 | 0.01529 | 0.00468 | 0.00000 | 2.94241 | 3.61601 | 2.65080 | 0.13291 | 1.32775 | 2.85653 | 3.63245 | 0.00367 |
| **Tmed7** | 1.59795 | 2.74027 | 1.87587 | 0.79571 | 6.64271 | 4.88615 | 0.00000 | 1.58919 | 1.02356 | 0.48335 | 2.85390 | 12.71951 | 1.26819 | 12.61179 | 4.42748 | 0.80975 |
| **Tmed8** | 3.90131 | 0.00000 | 0.04215 | 3.60903 | 0.00317 | 0.00697 | 2.25770 | 0.15058 | 0.63751 | 3.77793 | 0.32650 | 7.57880 | 1.79538 | 0.70620 | 0.54751 | 0.04598 |
| **Tmed9** | 1.16955 | 0.04342 | 4.24789 | 11.92526 | 7.19590 | 2.77675 | 0.23593 | 1.98073 | 14.44663 | 3.72248 | 5.52982 | 0.13263 | 1.72045 | 1.40748 | 3.02804 | 1.11506 |
| **Tmeff1** | 2.83152 | 0.39788 | 1.42829 | 0.00261 | 0.25868 | 0.01018 | 2.43219 | 1.54149 | 0.01788 | 0.00000 | 0.22135 | 0.09646 | 0.61457 | 0.02736 | 0.38163 | 0.13599 |
| **Tmem101** | 4.04956 | 3.24488 | 4.32849 | 0.63557 | 0.01894 | 0.01322 | 7.91466 | 4.75016 | 1.08472 | 2.12239 | 3.40441 | 0.02146 | 3.97895 | 0.02694 | 1.14916 | 2.67311 |
| **Tmem104** | 1.46187 | 0.00565 | 0.02377 | 0.19082 | 0.02039 | 0.01056 | 0.18219 | 1.78629 | 0.97878 | 0.35216 | 0.28665 | 0.00850 | 0.89310 | 0.00981 | 0.00000 | 2.83786 |
| **Tmem106a** | 1.68516 | 0.16465 | 1.15479 | 2.31113 | 2.44390 | 0.00354 | 0.02573 | 0.89362 | 1.00204 | 0.65738 | 3.00792 | 0.09148 | 0.00000 | 0.00000 | 0.25325 | 0.01189 |
| **Tmem106b** | 2.76773 | 2.92156 | 0.00000 | 0.00000 | 0.07736 | 2.12601 | 1.43514 | 0.42083 | 0.01023 | 0.83249 | 1.42093 | 0.03284 | 1.11711 | 0.06497 | 1.48359 | 0.03215 |
| **Tmem106c** | 0.00000 | 1.81920 | 0.00000 | 1.82119 | 0.00791 | 2.71445 | 4.58982 | 0.00529 | 0.06139 | 0.32434 | 0.24307 | 0.03965 | 1.65536 | 0.01669 | 0.00000 | 0.00433 |
| **Tmem107** | 1.90173 | 0.01697 | 0.13868 | 1.49940 | 0.00739 | 0.04532 | 1.91974 | 0.14493 | 0.10419 | 0.68512 | 0.66872 | 0.02080 | 0.93416 | 0.18896 | 0.37300 | 0.00464 |
| **Tmem108** | 0.01367 | 0.00783 | 0.01247 | 2.09935 | 0.00938 | 0.00633 | 0.00730 | 0.01187 | 0.02111 | 0.21178 | 0.01264 | 0.00676 | 0.01226 | 0.00313 | 0.10379 | 0.00708 |
| **Tmem109** | 2.65517 | 1.84568 | 0.12944 | 4.79148 | 0.02037 | 1.15121 | 2.85876 | 1.62503 | 0.56692 | 3.05183 | 0.01819 | 0.02164 | 4.06025 | 0.29047 | 0.00000 | 2.89058 |
| **Tmem11** | 14.64193 | 7.62498 | 13.03871 | 12.63348 | 5.59366 | 2.92641 | 5.87358 | 14.20055 | 16.58783 | 13.83082 | 9.61296 | 0.98168 | 8.70135 | 2.02466 | 4.15527 | 6.26633 |
| **Tmem110** | 0.43648 | 0.04020 | 1.48354 | 0.53223 | 0.01214 | 2.17583 | 0.00656 | 1.67305 | 0.45823 | 0.40498 | 0.36363 | 0.02756 | 0.11983 | 0.00606 | 0.09503 | 3.69837 |
| **Tmem115** | 5.13250 | 2.22193 | 5.44613 | 0.48331 | 0.00000 | 5.42414 | 1.40895 | 1.59355 | 1.04732 | 2.48998 | 0.03816 | 0.09930 | 0.40277 | 0.00451 | 0.13460 | 2.92965 |
| **Tmem116** | 0.12403 | 0.00674 | 0.51125 | 0.24382 | 0.01540 | 0.00530 | 0.00611 | 0.93469 | 0.62941 | 0.00000 | 1.08464 | 0.03365 | 0.60994 | 0.03420 | 0.06312 | 1.44768 |
| **Tmem117** | 0.01775 | 0.00816 | 0.01122 | 1.36881 | 0.00895 | 0.00650 | 0.00744 | 0.01277 | 0.91200 | 0.00474 | 0.03469 | 0.00623 | 0.01231 | 0.02983 | 0.03864 | 0.00710 |
| **Tmem120a** | 4.90719 | 0.02244 | 3.20162 | 7.03572 | 0.82004 | 4.61052 | 0.55017 | 4.82490 | 9.27725 | 18.67257 | 4.13080 | 0.42292 | 11.36882 | 0.07713 | 4.22136 | 0.70841 |
| **Tmem120b** | 1.33301 | 0.01620 | 0.00000 | 0.93526 | 0.00539 | 2.12022 | 0.01610 | 2.65847 | 1.91087 | 3.71543 | 1.71616 | 0.05800 | 0.00000 | 0.00000 | 0.00000 | 0.02665 |
| **Tmem123** | 1.13186 | 0.01587 | 0.51302 | 2.14973 | 0.00467 | 0.00341 | 0.00476 | 1.29889 | 1.98895 | 3.96589 | 1.82740 | 5.30653 | 0.97189 | 2.40303 | 2.89602 | 2.01876 |
| **Tmem125** | 0.00000 | 0.00717 | 0.00535 | 0.00912 | 0.01543 | 4.50223 | 0.00626 | 0.83776 | 0.01190 | 0.00000 | 0.00357 | 0.00524 | 0.73955 | 0.01864 | 0.00000 | 3.91915 |
| **Tmem126a** | 7.40635 | 0.37336 | 1.31468 | 0.01120 | 3.11625 | 2.61157 | 0.07313 | 0.00000 | 0.00984 | 0.57975 | 1.46621 | 0.02731 | 0.83849 | 0.52064 | 0.44610 | 2.85324 |
| **Tmem126b** | 3.62937 | 2.06655 | 3.33350 | 0.02547 | 0.24056 | 0.05426 | 0.00321 | 0.31071 | 0.04339 | 0.00000 | 0.57909 | 0.10392 | 0.05008 | 0.00742 | 0.00000 | 0.91803 |
| **Tmem127** | 1.36144 | 0.12368 | 4.45755 | 0.24991 | 0.00000 | 0.01867 | 1.50645 | 1.63908 | 0.77141 | 0.10440 | 1.66961 | 0.05185 | 2.55724 | 0.00739 | 0.73302 | 3.37873 |
| **Tmem128** | 1.05553 | 1.28343 | 1.10334 | 0.50318 | 1.24735 | 0.00512 | 0.46564 | 4.46344 | 1.52176 | 1.36410 | 2.02350 | 3.92444 | 1.96834 | 3.06844 | 4.46548 | 3.28809 |
| **Tmem129** | 2.37317 | 3.07325 | 0.46564 | 4.57349 | 0.02009 | 0.25982 | 0.21488 | 1.40618 | 3.01912 | 3.24536 | 0.52818 | 0.88770 | 1.20896 | 1.03745 | 1.25817 | 0.00166 |
| **Tmem130** | 0.00880 | 0.00782 | 0.01792 | 1.78172 | 0.00952 | 0.01527 | 0.00704 | 0.31785 | 0.73020 | 0.17773 | 0.29205 | 0.01347 | 0.00455 | 0.02709 | 0.00000 | 0.00678 |
| **Tmem131** | 8.88760 | 2.47071 | 0.32784 | 2.13946 | 1.98765 | 0.57612 | 5.81213 | 7.45207 | 1.13178 | 1.44162 | 0.90895 | 2.87207 | 5.98696 | 0.00296 | 0.38377 | 4.99926 |
| **Tmem132b** | 0.01366 | 0.00782 | 0.00945 | 3.04536 | 0.00924 | 0.00636 | 0.00731 | 0.00835 | 0.16800 | 0.02289 | 0.10770 | 0.01650 | 0.01170 | 0.00383 | 1.32548 | 0.00684 |
| **Tmem132c** | 0.01341 | 0.00784 | 0.01157 | 2.21633 | 0.00912 | 0.00624 | 0.00733 | 0.01272 | 1.14406 | 0.01911 | 0.02215 | 0.00515 | 0.05056 | 0.00245 | 0.08301 | 0.00705 |
| **Tmem134** | 2.41880 | 0.16250 | 0.35640 | 6.72918 | 1.96885 | 4.48644 | 0.10722 | 0.40293 | 0.66319 | 1.26672 | 1.93100 | 0.03592 | 0.34410 | 0.00000 | 0.34660 | 4.66639 |
| **Tmem135** | 1.68409 | 5.21260 | 0.00000 | 3.54569 | 0.00792 | 0.36266 | 0.01545 | 0.03257 | 0.00000 | 0.74713 | 0.00000 | 0.00000 | 0.60528 | 3.47862 | 0.44304 | 1.52367 |
| **Tmem136** | 0.02324 | 0.00804 | 0.05545 | 0.59864 | 0.00861 | 0.00575 | 0.00748 | 0.01164 | 0.54581 | 0.00313 | 0.00753 | 0.02827 | 0.01193 | 0.03597 | 0.56769 | 0.00718 |
| **Tmem138** | 1.45669 | 2.56871 | 1.14426 | 2.47385 | 0.02126 | 0.01629 | 2.42586 | 2.15265 | 2.38806 | 1.29393 | 2.41537 | 0.39636 | 0.83652 | 0.28302 | 3.08579 | 0.00345 |
| **Tmem139** | 0.01287 | 0.00779 | 2.76613 | 0.00481 | 0.00462 | 0.01361 | 0.00717 | 0.00419 | 0.01086 | 0.00000 | 0.98113 | 0.00200 | 0.05018 | 0.01137 | 0.00000 | 0.01309 |
| **Tmem141** | 4.34760 | 0.26110 | 6.07276 | 0.21427 | 0.28464 | 1.03814 | 0.76305 | 3.70670 | 0.00681 | 3.30678 | 0.55757 | 0.06509 | 2.20360 | 0.00425 | 0.76061 | 1.22908 |
| **Tmem143** | 0.01451 | 0.04106 | 0.27439 | 1.47176 | 0.00874 | 0.00552 | 0.00743 | 0.01569 | 0.18476 | 0.06060 | 0.01207 | 0.00758 | 0.01000 | 0.00778 | 0.02014 | 0.00702 |
| **Tmem144** | 0.23609 | 0.02589 | 2.57454 | 0.00000 | 1.29600 | 0.00662 | 1.53745 | 4.44647 | 0.00626 | 0.00000 | 0.34433 | 2.27529 | 2.32245 | 0.01034 | 0.22481 | 0.03302 |
| **Tmem145** | 0.01319 | 0.00765 | 0.00625 | 0.01416 | 0.00903 | 0.00728 | 0.00701 | 0.00602 | 0.20768 | 0.39049 | 0.88829 | 0.00217 | 0.00535 | 0.10416 | 0.00000 | 0.00729 |
| **Tmem147** | 12.93073 | 1.66765 | 8.03228 | 0.00000 | 5.32553 | 5.93072 | 6.36293 | 7.75086 | 0.00069 | 0.95742 | 4.90246 | 0.11614 | 14.19208 | 0.00000 | 0.21054 | 8.64355 |
| **Tmem14a** | 0.02128 | 0.00824 | 0.09576 | 0.40558 | 0.00973 | 0.00867 | 0.00740 | 0.01294 | 0.00710 | 0.06017 | 0.01177 | 0.00777 | 0.01077 | 0.01127 | 0.00589 | 0.00712 |
| **Tmem14c** | 7.36263 | 6.91886 | 5.04639 | 6.96731 | 0.05970 | 4.64215 | 4.86514 | 6.54586 | 3.08349 | 8.39020 | 4.72561 | 0.38065 | 5.90801 | 1.12577 | 1.94162 | 7.91318 |
| **Tmem150a** | 2.59260 | 0.03081 | 8.05981 | 0.34140 | 0.03032 | 2.85690 | 3.97095 | 2.59693 | 0.05388 | 0.00000 | 0.23550 | 0.00704 | 2.47946 | 0.00502 | 0.00000 | 5.85990 |
| **Tmem150c** | 0.00869 | 0.00799 | 0.00713 | 0.70880 | 0.00979 | 0.00653 | 0.00718 | 0.23600 | 1.10127 | 0.15256 | 0.00941 | 0.00209 | 0.00711 | 0.48569 | 0.75033 | 0.00725 |
| **Tmem158** | 0.01046 | 0.00857 | 0.01050 | 0.00661 | 0.01845 | 0.00671 | 0.04168 | 0.06339 | 1.14528 | 0.58469 | 0.00000 | 0.00549 | 0.00958 | 0.01354 | 0.00628 | 0.00738 |
| **Tmem159** | 10.60912 | 6.28806 | 20.09409 | 1.74594 | 0.71557 | 9.73822 | 10.68831 | 18.58847 | 8.33442 | 18.17812 | 14.92353 | 6.15548 | 17.06649 | 1.64307 | 6.53625 | 3.31561 |
| **Tmem160** | 0.70090 | 0.00846 | 0.01109 | 0.31418 | 0.01741 | 0.00765 | 0.00718 | 0.01351 | 0.79514 | 0.18595 | 0.14574 | 0.04062 | 0.01100 | 0.00975 | 0.00000 | 0.00668 |
| **Tmem161a** | 1.75347 | 0.06926 | 0.06272 | 0.00412 | 0.04851 | 0.00595 | 0.13327 | 0.64789 | 0.81982 | 0.84322 | 1.08773 | 0.08758 | 0.17218 | 0.44578 | 0.01992 | 0.01390 |
| **Tmem161b** | 0.03515 | 0.03650 | 0.00000 | 0.21707 | 0.00825 | 0.01212 | 2.17978 | 0.10230 | 1.41730 | 0.00000 | 0.00000 | 0.11635 | 0.01631 | 3.64566 | 0.61618 | 0.00674 |
| **Tmem164** | 1.13465 | 0.07922 | 0.00000 | 4.36413 | 0.00571 | 0.08032 | 0.00601 | 0.00397 | 2.92430 | 2.59005 | 0.00000 | 0.01077 | 0.01080 | 3.60895 | 1.11225 | 0.01919 |

| | | | | | | | | | | | | | | | | |
|---|---|---|---|---|---|---|---|---|---|---|---|---|---|---|---|---|
| **Tmem165** | 0.74548 | 3.60937 | 2.01430 | 2.67498 | 2.26783 | 0.38101 | 0.01395 | 0.07765 | 2.58781 | 0.94718 | 0.21109 | 0.01933 | 2.19529 | 0.00000 | 1.97649 | 1.71512 |
| **Tmem167** | 9.76982 | 4.50799 | 7.49684 | 8.13523 | 7.45901 | 6.20851 | 7.24782 | 4.34233 | 8.81200 | 11.12217 | 9.76234 | 0.90268 | 9.40693 | 9.07551 | 8.29243 | 1.53243 |
| **Tmem167b** | 1.92909 | 2.14073 | 4.24651 | 1.48277 | 6.64840 | 0.00379 | 0.26986 | 1.99371 | 0.53439 | 2.67179 | 2.39802 | 0.18909 | 0.37352 | 2.33020 | 4.53442 | 2.54148 |
| **Tmem168** | 3.42098 | 0.00221 | 0.44351 | 6.42267 | 0.00000 | 2.04935 | 1.10779 | 2.19096 | 0.67936 | 0.42030 | 0.96632 | 2.66203 | 2.66479 | 0.21121 | 3.85781 | 1.60437 |
| **Tmem169** | 0.01223 | 0.00749 | 0.00624 | 1.48157 | 0.00975 | 0.00646 | 0.00711 | 0.00928 | 0.00000 | 0.00285 | 0.00429 | 0.00000 | 0.01375 | 1.68872 | 0.42235 | 0.00723 |
| **Tmem17** | 1.28661 | 0.00562 | 1.02798 | 0.00500 | 0.01235 | 0.00762 | 1.61074 | 0.42924 | 0.01077 | 0.35629 | 0.19567 | 0.04928 | 2.69015 | 0.02225 | 0.00000 | 0.01143 |
| **Tmem170** | 1.50348 | 0.07279 | 0.81959 | 4.82658 | 1.70795 | 0.00956 | 0.00602 | 0.51648 | 1.04523 | 0.00000 | 0.79509 | 0.00000 | 0.40830 | 1.86876 | 3.38992 | 0.05594 |
| **Tmem170b** | 0.21021 | 0.00705 | 0.02816 | 0.39218 | 0.00791 | 0.00629 | 1.23372 | 0.00000 | 0.00568 | 0.07359 | 0.66281 | 0.01363 | 0.04882 | 0.01126 | 0.02851 | 0.01421 |
| **Tmem171** | 0.00000 | 0.99356 | 1.96154 | 0.50767 | 0.00411 | 0.36553 | 1.08504 | 0.03834 | 0.26653 | 0.00155 | 0.00961 | 0.00503 | 0.00498 | 0.00729 | 0.00000 | 1.58639 |
| **Tmem173** | 0.01312 | 0.00779 | 0.00989 | 0.01027 | 0.00915 | 0.00635 | 0.00746 | 0.01068 | 5.29214 | 0.00000 | 0.00000 | 0.07189 | 0.01342 | 0.02061 | 0.01198 | 0.00703 |
| **Tmem175** | 5.20604 | 0.11501 | 1.81022 | 0.47912 | 2.96113 | 0.01139 | 0.51508 | 0.11321 | 0.37998 | 1.02387 | 0.89967 | 0.00000 | 2.75911 | 0.00438 | 0.82756 | 0.22909 |
| **Tmem176b** | 0.01178 | 0.00785 | 0.01219 | 0.00539 | 0.00848 | 0.00657 | 0.00729 | 0.00977 | 0.08974 | 4.36563 | 0.00829 | 0.00000 | 0.01045 | 0.01258 | 0.00634 | 0.00731 |
| **Tmem177** | 0.00000 | 2.68720 | 0.98112 | 3.95842 | 0.00536 | 0.19547 | 0.00599 | 2.25136 | 1.68687 | 1.02572 | 0.49738 | 0.02331 | 0.00000 | 0.00000 | 1.49011 | 2.24226 |
| **Tmem179b** | 2.13148 | 0.56539 | 5.03205 | 2.21729 | 0.18971 | 2.94778 | 0.03552 | 0.83786 | 0.05078 | 0.00000 | 4.09506 | 0.06984 | 2.24624 | 0.00000 | 0.67345 | 1.90599 |
| **Tmem18** | 0.81672 | 0.04124 | 1.86019 | 0.21991 | 0.00201 | 0.00483 | 0.09713 | 0.36141 | 0.00478 | 0.00000 | 0.57544 | 0.04629 | 2.45486 | 0.50627 | 0.68766 | 4.08564 |
| **Tmem181a** | 0.04466 | 0.00797 | 0.00661 | 0.36641 | 0.00913 | 0.03901 | 0.27315 | 0.39572 | 0.00608 | 0.89747 | 0.24435 | 0.01331 | 0.42321 | 0.02080 | 0.08635 | 0.01458 |
| **Tmem181b.p** | 0.47806 | 0.11665 | 0.94119 | 0.50493 | 0.04620 | 1.56083 | 0.46209 | 1.61101 | 1.02354 | 0.98496 | 0.37901 | 1.18147 | 1.38459 | 0.65362 | 0.83470 | 0.52506 |
| **Tmem181c.p** | 0.00940 | 0.12509 | 0.03367 | 0.68157 | 0.00850 | 0.36576 | 0.70869 | 0.08800 | 0.27737 | 0.48314 | 0.64474 | 0.02667 | 1.03098 | 0.14218 | 0.21629 | 0.06346 |
| **Tmem183a** | 13.19160 | 7.17766 | 7.12271 | 7.76913 | 8.25147 | 4.72560 | 6.62959 | 6.16337 | 3.06413 | 9.78270 | 4.67627 | 4.17036 | 5.71677 | 10.58316 | 8.12180 | 5.09860 |
| **Tmem184a** | 0.01774 | 0.00737 | 0.06650 | 0.07386 | 0.00811 | 0.00873 | 0.00706 | 1.67726 | 0.02827 | 1.76014 | 0.75228 | 0.05767 | 0.00000 | 0.00982 | 0.00000 | 0.00691 |
| **Tmem184b** | 2.49831 | 0.00524 | 0.69506 | 1.15306 | 0.01752 | 2.78281 | 0.12164 | 0.38116 | 0.86070 | 0.17959 | 0.70910 | 0.00041 | 0.13303 | 0.01007 | 0.02628 | 0.14576 |
| **Tmem184c** | 0.00000 | 0.02325 | 2.75005 | 4.52990 | 3.33658 | 3.53147 | 1.55839 | 9.08909 | 12.24296 | 2.82838 | 1.36809 | 5.03403 | 1.33609 | 3.87430 | 12.51337 | 0.00966 |
| **Tmem185a** | 3.98549 | 0.02094 | 0.00000 | 2.37120 | 4.78463 | 2.70176 | 3.36212 | 1.86164 | 0.51237 | 0.64290 | 0.46797 | 0.11727 | 1.45307 | 0.00299 | 7.40819 | 3.10881 |
| **Tmem185b** | 1.76601 | 1.53475 | 4.17094 | 2.23213 | 2.92754 | 0.00558 | 5.80745 | 8.36829 | 0.13938 | 2.41078 | 2.68650 | 0.02266 | 1.76884 | 4.86621 | 3.85368 | 0.10201 |
| **Tmem186** | 1.46134 | 0.00498 | 1.60957 | 0.49602 | 0.00213 | 0.02398 | 0.00478 | 2.20379 | 2.93303 | 2.39815 | 2.34666 | 0.31194 | 0.28829 | 0.31231 | 3.22060 | 0.01171 |
| **Tmem189** | 0.69781 | 0.10655 | 0.64847 | 4.79898 | 1.70841 | 0.30869 | 1.84003 | 2.92148 | 5.49065 | 4.59087 | 4.46446 | 0.19683 | 2.68651 | 0.98905 | 1.35585 | 0.24115 |
| **Tmem19** | 2.25454 | 2.55931 | 4.35140 | 1.03980 | 0.00495 | 0.00473 | 3.59345 | 1.32411 | 1.47292 | 3.19928 | 0.26385 | 0.07340 | 1.58296 | 1.62286 | 2.72479 | 0.00000 |
| **Tmem190** | 0.01371 | 0.00805 | 0.01240 | 0.65696 | 0.00941 | 0.00650 | 0.00739 | 0.01266 | 0.00012 | 0.11858 | 0.01427 | 0.00752 | 0.01257 | 0.01012 | 0.00407 | 0.00738 |
| **Tmem191c** | 6.02210 | 0.01002 | 0.46551 | 1.01989 | 0.05068 | 1.88416 | 0.65631 | 9.67355 | 2.19785 | 3.50451 | 5.02105 | 0.00000 | 9.81536 | 0.00000 | 0.00000 | 2.30805 |
| **Tmem192** | 7.56689 | 0.70111 | 6.59476 | 5.03631 | 7.91162 | 3.71383 | 4.45077 | 3.35508 | 1.82952 | 9.85609 | 8.39166 | 0.73909 | 4.07263 | 0.04414 | 0.41964 | 0.46823 |
| **Tmem198** | 1.09017 | 0.00660 | 0.45053 | 0.00657 | 0.00566 | 0.00859 | 0.05843 | 0.08230 | 0.08005 | 0.00681 | 0.01305 | 0.01590 | 0.21453 | 0.01509 | 0.00180 | 0.00624 |
| **Tmem198b** | 0.01561 | 0.07210 | 0.01291 | 0.00936 | 0.00936 | 0.00719 | 0.01814 | 0.01389 | 0.01217 | 0.24913 | 0.01488 | 0.00682 | 0.09466 | 0.01077 | 0.00954 | 0.00802 |
| **Tmem199** | 7.73416 | 8.65256 | 10.29340 | 4.97308 | 9.50855 | 5.62557 | 3.44816 | 9.75586 | 3.84898 | 3.23229 | 4.31185 | 0.70190 | 3.48096 | 2.05218 | 4.96505 | 5.37612 |
| **Tmem2** | 0.00081 | 0.02736 | 0.03374 | 0.02524 | 0.00943 | 0.04537 | 2.26940 | 0.21342 | 0.81023 | 1.93852 | 0.32277 | 0.00000 | 0.39531 | 0.12804 | 2.48245 | 0.01381 |
| **Tmem201** | 1.66837 | 0.94813 | 2.14657 | 6.90699 | 1.41912 | 0.00368 | 2.09827 | 6.11064 | 8.96128 | 13.61028 | 3.53259 | 4.22128 | 3.22980 | 0.49292 | 3.66340 | 3.95282 |
| **Tmem203** | 0.05601 | 0.00747 | 0.28893 | 0.89033 | 0.00768 | 0.00666 | 0.00713 | 0.03853 | 0.55758 | 1.67107 | 0.00888 | 0.03905 | 0.24327 | 0.00501 | 0.35557 | 0.50708 |
| **Tmem205** | 2.14117 | 0.28119 | 2.86440 | 0.02641 | 0.00000 | 0.00396 | 1.45110 | 0.03071 | 0.01188 | 0.83143 | 0.01190 | 0.01089 | 0.17165 | 0.01110 | 0.00068 | 2.56305 |
| **Tmem206** | 1.23100 | 0.09891 | 0.90365 | 6.44230 | 0.44450 | 0.08842 | 2.55146 | 0.40246 | 9.13819 | 0.67228 | 8.54803 | 0.76535 | 0.37466 | 2.72385 | 0.41577 | 0.31744 |
| **Tmem208** | 11.85376 | 7.34207 | 8.40275 | 1.89781 | 4.11916 | 1.25110 | 2.77436 | 9.22294 | 4.23474 | 7.70093 | 7.94538 | 1.27132 | 10.14922 | 1.69841 | 3.08080 | 7.06569 |
| **Tmem209** | 6.73968 | 4.39077 | 6.56018 | 5.20662 | 5.51264 | 3.92685 | 3.03824 | 9.02007 | 4.02338 | 3.49857 | 5.47988 | 14.50401 | 5.79391 | 17.15627 | 10.00013 | 2.28829 |
| **Tmem213** | 0.01851 | 0.00828 | 0.01193 | 0.01050 | 0.01054 | 0.00663 | 0.00755 | 0.01351 | 0.01242 | 0.03830 | 0.01243 | 0.00802 | 0.01223 | 0.01170 | 0.01074 | 0.00731 |
| **Tmem214** | 1.30834 | 4.08749 | 10.74349 | 3.80799 | 7.91469 | 0.04092 | 6.25607 | 1.76871 | 0.00000 | 1.18383 | 1.02963 | 0.11744 | 4.54242 | 0.02871 | 0.50832 | 6.06373 |
| **Tmem215** | 0.01359 | 0.00812 | 0.01174 | 0.01084 | 0.00930 | 0.00647 | 0.00746 | 0.01258 | 1.28159 | 0.00488 | 0.00001 | 0.00603 | 0.01293 | 0.00741 | 0.23257 | 0.00724 |
| **Tmem216** | 0.92636 | 0.08999 | 4.69339 | 2.21843 | 3.35331 | 0.01125 | 1.97341 | 1.53962 | 4.51757 | 4.66762 | 1.64549 | 1.01294 | 1.03713 | 2.69199 | 6.31407 | 0.01035 |
| **Tmem218** | 0.01156 | 0.00784 | 0.61892 | 0.92122 | 0.00784 | 0.00564 | 0.00741 | 0.00990 | 0.00000 | 1.57417 | 0.02669 | 0.01476 | 0.00905 | 0.00455 | 0.19555 | 0.00703 |
| **Tmem219** | 2.62821 | 0.00302 | 1.28862 | 0.01711 | 0.00001 | 0.00292 | 1.55713 | 3.22331 | 0.30704 | 0.81304 | 0.53269 | 0.00696 | 0.81229 | 1.34839 | 0.00000 | 0.00375 |
| **Tmem222** | 11.33751 | 3.78125 | 9.66262 | 2.48948 | 3.21968 | 5.12933 | 7.25467 | 11.01594 | 2.51269 | 4.35129 | 3.97259 | 1.20932 | 10.22921 | 0.03404 | 1.56827 | 8.89663 |
| **Tmem223** | 5.09321 | 1.02113 | 2.17461 | 4.63598 | 2.34944 | 0.35229 | 2.07089 | 2.29169 | 6.29672 | 8.16439 | 7.15077 | 0.41861 | 1.65407 | 0.35729 | 0.00000 | 0.78069 |
| **Tmem229b** | 0.09561 | 0.00282 | 4.16697 | 9.92770 | 0.00860 | 3.44309 | 0.00000 | 3.16372 | 8.60885 | 25.50658 | 13.07682 | 7.76898 | 3.47554 | 11.57784 | 21.77287 | 0.00000 |
| **Tmem230** | 4.84842 | 2.38549 | 7.07673 | 1.10683 | 0.00000 | 3.84843 | 2.74608 | 1.38410 | 0.86555 | 3.15494 | 0.66975 | 0.11287 | 3.03825 | 0.00000 | 1.45909 | 2.82185 |
| **Tmem231** | 9.40908 | 4.07845 | 9.51687 | 7.19445 | 5.15064 | 5.48840 | 6.36683 | 8.84856 | 9.05143 | 7.73542 | 7.55675 | 2.50811 | 12.80932 | 5.82686 | 6.06056 | 5.48641 |
| **Tmem232** | 0.01353 | 0.00762 | 0.01217 | 1.61569 | 0.00908 | 0.00632 | 0.00723 | 0.01016 | 0.00000 | 0.52964 | 0.02689 | 0.00626 | 0.01077 | 0.00672 | 0.15040 | 0.00684 |
| **Tmem234** | 4.32485 | 0.89694 | 1.81230 | 0.70795 | 4.80066 | 1.81255 | 0.49182 | 2.53718 | 2.17143 | 3.24052 | 0.97858 | 0.09474 | 5.22944 | 0.17850 | 0.26968 | 2.02276 |
| **Tmem235** | 0.01372 | 0.00805 | 0.01220 | 0.00508 | 0.00935 | 0.00662 | 0.00751 | 0.01341 | 0.15819 | 0.01083 | 0.01149 | 0.00733 | 0.01277 | 0.02236 | 0.06647 | 0.00740 |
| **Tmem237** | 0.00116 | 0.00804 | 1.08712 | 0.01181 | 0.00823 | 0.00535 | 0.00724 | 0.41550 | 0.95698 | 1.30301 | 0.07238 | 1.19171 | 0.40324 | 1.68334 | 0.46098 | 0.00656 |
| **Tmem241** | 0.30960 | 0.00784 | 0.00717 | 0.79280 | 0.00760 | 0.00729 | 0.00741 | 0.01715 | 0.00000 | 0.01452 | 0.01503 | 0.00795 | 0.03297 | 0.01173 | 0.00574 | 0.00708 |
| **Tmem242** | 3.94468 | 5.00979 | 2.16391 | 0.31732 | 3.09840 | 2.60283 | 6.71582 | 7.86484 | 0.00000 | 5.58699 | 1.39254 | 0.09159 | 1.52136 | 0.29841 | 0.74166 | 0.87540 |
| **Tmem243** | 0.06566 | 0.00882 | 0.00801 | 0.03519 | 0.02359 | 0.00726 | 0.00810 | 0.04685 | 0.01232 | 0.10878 | 0.01195 | 0.02287 | 0.01113 | 0.01241 | 0.01069 | 0.00744 |
| **Tmem245** | 1.61445 | 0.38260 | 0.39769 | 1.08498 | 2.42491 | 0.01596 | 1.56420 | 1.23959 | 0.36127 | 0.37498 | 2.43012 | 0.52351 | 0.61594 | 0.78206 | 2.07011 | 0.03986 |
| **Tmem246** | 0.04248 | 0.00866 | 0.01025 | 0.86577 | 0.01354 | 0.00689 | 0.00743 | 0.01161 | 0.18838 | 0.57745 | 0.01143 | 0.00778 | 0.01218 | 0.01014 | 0.00462 | 0.00735 |
| **Tmem248** | 5.38828 | 0.47952 | 1.36135 | 5.07530 | 0.80249 | 4.24861 | 0.77028 | 4.89965 | 2.69462 | 7.11921 | 4.00779 | 0.51435 | 4.94711 | 3.46170 | 2.34949 | 3.11995 |
| **Tmem25** | 0.01375 | 0.00812 | 0.01230 | 0.26475 | 0.00932 | 0.00663 | 0.00753 | 0.01396 | 0.00648 | 0.01546 | 0.01342 | 0.00821 | 0.01268 | 0.01136 | 0.00681 | 0.00737 |

| | | | | | | | | | | | | | | | | |
|---|---|---|---|---|---|---|---|---|---|---|---|---|---|---|---|---|
| **Tmem251** | 3.84871 | 2.69696 | 3.78661 | 9.36221 | 2.82154 | 1.49195 | 1.61103 | 1.20287 | 3.20784 | 2.78458 | 0.07229 | 0.21844 | 0.39782 | 0.15654 | 5.26578 | 0.09799 |
| **Tmem253** | 1.45271 | 0.00607 | 0.00357 | 0.00518 | 0.00520 | 0.00679 | 0.00606 | 0.00000 | 0.01119 | 0.20411 | 0.65962 | 0.00295 | 0.01380 | 0.01174 | 0.00000 | 0.01297 |
| **Tmem254a** | 0.01328 | 0.00819 | 0.01387 | 0.01056 | 0.01002 | 0.00708 | 0.00776 | 0.01627 | 0.01233 | 0.01345 | 0.01479 | 0.00797 | 0.07014 | 0.01236 | 0.01043 | 0.00769 |
| **Tmem255b** | 0.01373 | 0.00815 | 0.01226 | 0.29522 | 0.00932 | 0.00663 | 0.00753 | 0.01395 | 0.09575 | 0.01491 | 0.01265 | 0.00810 | 0.01271 | 0.01148 | 0.00581 | 0.00736 |
| **Tmem256** | 3.29708 | 0.00353 | 1.38537 | 0.00310 | 0.12740 | 0.42950 | 0.20590 | 0.00000 | 0.73261 | 0.33917 | 1.90287 | 0.00260 | 1.12148 | 0.03449 | 0.06089 | 1.77347 |
| **Tmem258** | 4.97269 | 0.34275 | 2.49893 | 3.88209 | 5.78572 | 0.20376 | 0.25029 | 0.00000 | 1.83492 | 0.77335 | 8.06986 | 0.17496 | 1.01058 | 1.38912 | 0.98891 | 3.22400 |
| **Tmem259** | 0.54570 | 0.00759 | 0.00919 | 0.00887 | 0.01059 | 0.69809 | 0.00687 | 0.01049 | 0.01175 | 2.10606 | 0.01012 | 0.02266 | 0.53152 | 0.01557 | 0.10008 | 0.09835 |
| **Tmem260** | 1.99608 | 0.00414 | 0.56482 | 0.01024 | 0.00303 | 0.00607 | 1.92936 | 0.00202 | 0.01217 | 0.00675 | 0.01216 | 0.01391 | 0.61742 | 0.01539 | 0.00116 | 0.30776 |
| **Tmem261** | 1.94878 | 0.15469 | 1.77881 | 0.64824 | 2.60058 | 2.11162 | 0.98178 | 1.41541 | 0.59803 | 1.12714 | 0.49099 | 0.09945 | 4.27426 | 0.02890 | 0.00000 | 2.23501 |
| **Tmem263** | 2.54852 | 2.93019 | 10.18329 | 4.73907 | 1.88630 | 5.03305 | 0.45993 | 2.74435 | 1.47217 | 1.18466 | 2.60482 | 5.41743 | 2.77970 | 8.23054 | 0.73884 | 1.33821 |
| **Tmem265** | 0.62693 | 0.00590 | 1.05371 | 2.11302 | 1.17963 | 2.18650 | 0.09279 | 1.52038 | 0.26354 | 2.00263 | 2.30609 | 0.95551 | 1.32214 | 0.10437 | 1.17398 | 0.41914 |
| **Tmem266** | 0.01241 | 0.00791 | 0.64149 | 0.15602 | 0.00804 | 0.00928 | 0.00734 | 0.01134 | 0.20479 | 0.01089 | 0.06615 | 0.00695 | 0.00843 | 0.01159 | 0.00000 | 0.00708 |
| **Tmem267** | 0.00000 | 0.00329 | 0.03811 | 6.38642 | 0.00805 | 0.00321 | 0.00439 | 1.64129 | 0.38248 | 8.43650 | 2.09303 | 12.30581 | 0.00000 | 10.08249 | 8.82890 | 0.00468 |
| **Tmem268** | 0.51736 | 0.00699 | 0.02585 | 0.09493 | 0.00596 | 0.00817 | 0.00566 | 0.00000 | 0.01051 | 0.16647 | 2.10554 | 0.01504 | 0.39138 | 0.13805 | 0.18925 | 2.05273 |
| **Tmem29** | 5.24932 | 0.08755 | 0.81367 | 5.29984 | 7.44499 | 0.00000 | 0.00932 | 0.00000 | 5.56418 | 3.13181 | 5.55592 | 4.77070 | 1.92462 | 4.65551 | 4.10330 | 0.00515 |
| **Tmem30a** | 1.87303 | 2.90537 | 2.11415 | 4.36337 | 3.39336 | 3.23761 | 3.57403 | 4.44052 | 2.73005 | 3.06427 | 4.27799 | 12.21219 | 1.68969 | 18.49220 | 5.19020 | 0.15373 |
| **Tmem30b** | 0.00000 | 0.05759 | 2.94413 | 0.00652 | 0.00644 | 0.15570 | 0.56556 | 0.11381 | 0.40714 | 1.26708 | 0.02813 | 0.00000 | 1.67345 | 0.46707 | 0.00000 | 0.01316 |
| **Tmem33** | 5.40832 | 2.09987 | 8.42874 | 2.77701 | 2.78016 | 7.37917 | 9.21301 | 8.19849 | 5.25281 | 7.94944 | 6.13236 | 1.57537 | 7.32684 | 11.49673 | 2.78645 | 4.78831 |
| **Tmem35b** | 2.34560 | 2.48331 | 7.78325 | 2.12846 | 0.00000 | 3.80460 | 2.91817 | 0.00915 | 3.04039 | 0.00000 | 0.06325 | 0.01871 | 0.06645 | 0.81283 | 1.09365 | 0.00000 |
| **Tmem37** | 0.80958 | 0.01418 | 1.97403 | 0.06265 | 0.05229 | 0.00415 | 0.00000 | 3.89813 | 9.77474 | 20.68621 | 13.05528 | 5.17776 | 0.94157 | 0.00000 | 1.47240 | 0.00219 |
| **Tmem38a** | 0.00321 | 0.00746 | 0.02800 | 1.64597 | 0.01055 | 0.00810 | 0.00685 | 0.44405 | 0.54013 | 1.09763 | 1.08097 | 0.05098 | 0.00913 | 1.48810 | 0.00000 | 0.00734 |
| **Tmem38b** | 2.87349 | 0.05604 | 1.32130 | 1.14400 | 1.69588 | 5.85592 | 0.00000 | 4.74324 | 0.24902 | 1.25182 | 0.52532 | 0.09444 | 7.91100 | 6.54076 | 0.29784 | 0.42245 |
| **Tmem39a** | 10.45365 | 3.57310 | 7.17119 | 10.38495 | 4.21655 | 4.36070 | 7.32766 | 9.57125 | 9.88056 | 2.97698 | 6.14328 | 6.60710 | 2.61932 | 9.29785 | 5.07718 | 2.19821 |
| **Tmem39b** | 2.50083 | 2.11110 | 1.57645 | 2.08675 | 0.00000 | 1.28486 | 0.00324 | 2.73467 | 3.32759 | 4.20027 | 1.14287 | 0.03138 | 1.68003 | 0.02339 | 0.33373 | 0.00914 |
| **Tmem40** | 0.01088 | 0.00776 | 1.96349 | 0.01087 | 0.00673 | 0.00977 | 0.00709 | 0.01281 | 0.01212 | 0.00000 | 0.01311 | 0.02619 | 0.79207 | 0.01677 | 0.00000 | 0.00661 |
| **Tmem41a** | 1.50695 | 0.00000 | 0.86926 | 0.01791 | 5.81904 | 0.00503 | 4.77890 | 1.20580 | 0.00388 | 1.79536 | 0.04454 | 0.02903 | 0.27400 | 0.00481 | 0.44085 | 0.06376 |
| **Tmem41b** | 26.97940 | 21.37212 | 33.81916 | 27.31395 | 23.92749 | 13.02151 | 18.56925 | 43.73572 | 32.59040 | 43.16851 | 38.10380 | 28.11069 | 33.67297 | 39.61822 | 25.63787 | 9.60017 |
| **Tmem42** | 0.40542 | 0.63829 | 0.00240 | 1.21003 | 0.01781 | 0.00698 | 0.00653 | 0.03002 | 1.47351 | 0.00714 | 0.28557 | 0.01242 | 0.00815 | 0.00660 | 0.16900 | 0.00659 |
| **Tmem43** | 6.37290 | 3.34128 | 2.37301 | 1.77035 | 0.00561 | 3.66873 | 7.27883 | 4.21185 | 2.28831 | 2.39598 | 1.91903 | 0.22291 | 0.00000 | 0.00348 | 0.55726 | 0.00000 |
| **Tmem44** | 0.03629 | 0.00784 | 0.00454 | 0.49547 | 0.06948 | 0.00762 | 0.03729 | 0.00000 | 0.70061 | 0.39202 | 1.41198 | 0.01992 | 0.00000 | 0.00925 | 0.06093 | 0.00716 |
| **Tmem45a** | 0.58683 | 1.06448 | 0.00000 | 0.00695 | 0.03491 | 0.00799 | 0.00631 | 0.00697 | 0.01215 | 0.60067 | 0.01207 | 0.03241 | 0.01127 | 0.01026 | 0.00000 | 0.01495 |
| **Tmem45b** | 0.00000 | 0.08029 | 2.45447 | 0.09667 | 0.03150 | 8.58696 | 0.07490 | 1.63158 | 0.25308 | 0.00000 | 0.03641 | 0.07867 | 4.08382 | 0.47053 | 0.80660 | 1.36839 |
| **Tmem47** | 0.90261 | 0.00670 | 0.00000 | 0.19624 | 0.01686 | 0.01256 | 0.00656 | 0.01673 | 0.64060 | 0.00000 | 0.27082 | 3.19244 | 0.00845 | 0.01532 | 0.10457 | 0.00618 |
| **Tmem5** | 0.42276 | 2.85970 | 2.64591 | 2.84742 | 0.00277 | 0.01884 | 0.01538 | 0.58585 | 2.63014 | 1.86872 | 0.82519 | 0.92715 | 1.13605 | 0.05451 | 1.25421 | 0.01069 |
| **Tmem50a** | 8.63851 | 2.57036 | 10.53124 | 0.66257 | 1.06493 | 3.90184 | 2.43788 | 5.13289 | 0.56423 | 2.43666 | 4.73181 | 0.62805 | 1.56507 | 0.63943 | 4.65199 | 7.04947 |
| **Tmem50b** | 1.09166 | 0.00585 | 6.84213 | 16.48985 | 0.11118 | 8.36064 | 5.00120 | 18.15108 | 22.06692 | 31.71339 | 20.12546 | 16.37490 | 6.18263 | 9.20380 | 20.84974 | 5.96510 |
| **Tmem51** | 2.27099 | 0.00548 | 0.00307 | 0.00445 | 0.00361 | 0.00673 | 0.00559 | 0.00000 | 0.37490 | 0.32372 | 0.35606 | 0.00470 | 0.00932 | 0.01052 | 0.03212 | 0.00483 |
| **Tmem52** | 0.01814 | 0.00684 | 0.36959 | 0.08898 | 0.00765 | 0.00617 | 0.00591 | 2.61632 | 0.00665 | 1.72187 | 0.97370 | 0.02259 | 1.38123 | 0.01082 | 0.00000 | 0.00615 |
| **Tmem53** | 0.08739 | 0.00773 | 0.00902 | 0.56863 | 0.00788 | 0.00688 | 0.00757 | 0.01350 | 0.33357 | 0.02989 | 0.01077 | 0.00766 | 0.01294 | 0.00873 | 0.02060 | 0.00719 |
| **Tmem54** | 0.02676 | 0.00826 | 0.00915 | 0.14618 | 0.21545 | 0.00678 | 0.00741 | 0.01203 | 0.00949 | 0.01274 | 0.01344 | 0.00799 | 0.08301 | 0.01178 | 0.00834 | 0.00735 |
| **Tmem55a** | 2.61896 | 3.54437 | 7.85043 | 0.00000 | 2.49752 | 8.62204 | 2.89934 | 5.89230 | 0.51138 | 1.95927 | 9.67409 | 3.60888 | 6.99642 | 9.82692 | 3.89224 | 1.23427 |
| **Tmem55b** | 6.37033 | 5.91012 | 5.76675 | 3.83757 | 0.01830 | 0.66364 | 2.84171 | 4.40113 | 6.20763 | 3.45558 | 5.86416 | 0.21500 | 4.48103 | 6.37863 | 5.37184 | 0.29042 |
| **Tmem56** | 0.00000 | 0.00000 | 0.00000 | 7.47620 | 0.00730 | 0.00178 | 1.88637 | 5.56066 | 4.33816 | 23.86402 | 10.25679 | 18.54841 | 0.00000 | 28.14718 | 14.98889 | 0.00190 |
| **Tmem57** | 6.90906 | 3.98942 | 3.52412 | 6.05451 | 7.84767 | 2.59787 | 2.36054 | 7.25231 | 5.74265 | 1.24845 | 1.38418 | 1.31554 | 1.31725 | 7.15252 | 4.91621 | 0.89947 |
| **Tmem59** | 11.83358 | 5.19014 | 21.67212 | 10.44446 | 11.87526 | 15.52406 | 13.81600 | 27.66122 | 11.21763 | 25.28196 | 22.48170 | 19.36819 | 28.90118 | 16.96611 | 8.21090 | 11.80338 |
| **Tmem60** | 2.68646 | 0.36815 | 7.04576 | 8.69127 | 1.85718 | 3.89449 | 3.37644 | 0.74400 | 5.02593 | 4.73472 | 2.14427 | 6.31786 | 4.37791 | 1.56669 | 1.41804 | 0.03947 |
| **Tmem62** | 1.18282 | 5.93087 | 5.78987 | 0.45983 | 0.03696 | 12.08388 | 4.83774 | 6.02473 | 0.46727 | 2.90725 | 4.37961 | 0.15028 | 6.44211 | 10.22334 | 5.65072 | 1.91575 |
| **Tmem63a** | 0.00000 | 0.00528 | 0.30542 | 0.15908 | 0.02764 | 0.00504 | 2.27834 | 0.89259 | 0.35776 | 0.23073 | 0.33468 | 4.09590 | 0.45998 | 0.07339 | 1.27613 | 0.00682 |
| **Tmem63b** | 0.81273 | 0.00457 | 1.29681 | 0.08627 | 0.29730 | 0.32561 | 3.58542 | 0.00075 | 0.14209 | 0.00000 | 0.48005 | 0.06093 | 0.07337 | 0.00609 | 0.22798 | 0.01260 |
| **Tmem64** | 2.67069 | 6.28067 | 2.98028 | 0.16897 | 2.26651 | 4.37497 | 7.33644 | 12.03957 | 0.63632 | 3.24793 | 6.59968 | 15.73274 | 4.92421 | 6.76300 | 2.95852 | 0.00000 |
| **Tmem65** | 4.12154 | 1.68199 | 0.92599 | 7.30121 | 7.28889 | 3.56498 | 7.22534 | 7.39404 | 4.86432 | 4.24846 | 6.13279 | 2.51944 | 2.42259 | 8.51259 | 2.57969 | 2.23683 |
| **Tmem67** | 1.42002 | 0.67875 | 0.00000 | 0.81887 | 0.00506 | 0.00541 | 0.00572 | 0.76456 | 0.05715 | 0.25340 | 0.00000 | 0.02494 | 0.00579 | 1.28684 | 0.69420 | 0.00566 |
| **Tmem68** | 0.79991 | 0.42618 | 1.46265 | 7.32359 | 0.01288 | 0.00644 | 0.00471 | 0.47705 | 3.60863 | 0.75923 | 2.25424 | 2.92899 | 0.26035 | 6.60537 | 7.98098 | 0.01824 |
| **Tmem69** | 1.32848 | 0.10934 | 1.46239 | 7.89849 | 0.58832 | 0.31676 | 0.26912 | 0.00000 | 11.91610 | 0.57504 | 12.77265 | 3.39901 | 1.70475 | 4.19183 | 2.44874 | 1.40818 |
| **Tmem70** | 10.81601 | 3.13127 | 6.15701 | 6.84556 | 9.16972 | 2.98935 | 0.62338 | 3.32011 | 6.91550 | 13.33620 | 4.59395 | 5.63978 | 2.73151 | 6.53201 | 4.11806 | 0.38171 |
| **Tmem72** | 0.01370 | 0.00770 | 0.01151 | 0.68814 | 0.00930 | 0.00639 | 0.00732 | 0.01098 | 0.00000 | 0.09867 | 0.01167 | 0.00662 | 0.01206 | 0.01140 | 0.70150 | 0.00709 |
| **Tmem79** | 2.26416 | 0.02688 | 0.02629 | 10.40305 | 0.00352 | 1.30257 | 0.00000 | 2.40990 | 2.76623 | 3.81721 | 4.69241 | 0.03766 | 4.21957 | 0.04401 | 0.97012 | 0.45985 |
| **Tmem8** | 6.10320 | 3.26054 | 4.05298 | 4.60443 | 3.07127 | 0.57865 | 7.61464 | 5.50945 | 2.62118 | 3.89149 | 5.20796 | 0.33528 | 1.27186 | 0.00000 | 2.21837 | 5.12238 |
| **Tmem80** | 1.82133 | 0.00649 | 1.12674 | 0.01030 | 0.00283 | 0.08768 | 0.07400 | 0.25277 | 0.01202 | 0.14811 | 0.00792 | 0.00612 | 0.32080 | 0.01554 | 0.00000 | 0.00458 |
| **Tmem81** | 0.10594 | 0.00740 | 1.10533 | 0.00577 | 0.00533 | 0.00916 | 0.00703 | 0.01054 | 0.01096 | 0.65796 | 0.00879 | 0.04914 | 0.09146 | 0.00585 | 0.55301 | 1.50163 |
| **Tmem82** | 1.33853 | 0.14099 | 0.80866 | 0.24447 | 0.06378 | 0.09604 | 0.59479 | 0.71125 | 0.33918 | 0.04054 | 0.90973 | 0.10728 | 1.96437 | 0.35875 | 0.17072 | 0.09688 |
| **Tmem86a** | 0.72078 | 0.00731 | 1.22868 | 1.02259 | 0.01026 | 0.00493 | 0.09044 | 0.00141 | 1.15462 | 0.85074 | 0.39374 | 0.00000 | 0.99951 | 0.00501 | 0.14942 | 2.38645 |

| | | | | | | | | | | | | | | | | |
|---|---|---|---|---|---|---|---|---|---|---|---|---|---|---|---|---|
| **Tmem86b** | 0.58734 | 0.00712 | 0.24647 | 0.07572 | 0.00626 | 0.00505 | 0.00670 | 0.00279 | 0.02215 | 0.00996 | 0.25567 | 0.00651 | 0.88890 | 0.03486 | 0.00000 | 0.00599 |
| **Tmem87a** | 0.00000 | 3.25947 | 1.02400 | 0.86121 | 0.89326 | 0.01192 | 2.73302 | 0.71188 | 1.44755 | 0.51773 | 3.98586 | 0.35497 | 0.15224 | 2.49576 | 4.81019 | 0.56400 |
| **Tmem87b** | 2.34558 | 0.00000 | 2.24227 | 7.49044 | 0.01324 | 0.56428 | 4.02505 | 4.26841 | 5.29781 | 7.16062 | 3.43330 | 5.92763 | 1.18865 | 3.81658 | 7.17369 | 0.31759 |
| **Tmem88** | 0.07917 | 0.47970 | 0.00000 | 0.00797 | 2.68966 | 0.92841 | 0.00696 | 0.00513 | 0.01156 | 0.06595 | 0.01092 | 0.00440 | 0.01749 | 0.00762 | 0.01466 | 0.00671 |
| **Tmem88b** | 0.04310 | 0.00792 | 0.05780 | 0.51877 | 0.02710 | 0.01386 | 0.01800 | 0.01493 | 0.43091 | 0.01416 | 0.30065 | 0.03674 | 0.01697 | 0.24332 | 0.00635 | 0.01439 |
| **Tmem8b** | 1.38996 | 0.01751 | 0.01771 | 0.01451 | 0.02137 | 0.00706 | 0.00643 | 0.02700 | 0.43935 | 0.14107 | 0.18727 | 0.00480 | 0.02341 | 0.03166 | 0.01762 | 0.00608 |
| **Tmem9** | 7.79883 | 4.35536 | 5.94347 | 0.31670 | 3.75324 | 6.71488 | 8.55461 | 0.00000 | 0.01705 | 0.44936 | 0.28055 | 0.05644 | 12.48159 | 0.00287 | 0.15670 | 4.79513 |
| **Tmem91** | 0.01375 | 0.00807 | 0.01222 | 0.10667 | 0.00933 | 0.00665 | 0.00753 | 0.01358 | 0.00747 | 0.01371 | 0.02454 | 0.00795 | 0.01241 | 0.00952 | 0.02918 | 0.00738 |
| **Tmem92** | 0.00000 | 0.01856 | 0.95988 | 18.46970 | 0.02300 | 0.17996 | 0.80985 | 2.52415 | 13.01425 | 1.34585 | 0.44738 | 0.14302 | 1.43468 | 5.66540 | 2.60638 | 0.02528 |
| **Tmem94** | 0.60740 | 0.00686 | 0.14068 | 0.22218 | 0.00639 | 0.08958 | 0.00692 | 0.01027 | 0.11712 | 0.28765 | 0.01137 | 0.02506 | 0.01028 | 0.01132 | 0.02080 | 0.00697 |
| **Tmem97** | 12.48253 | 15.29364 | 23.47420 | 8.28805 | 9.61193 | 4.64604 | 15.48092 | 26.57665 | 9.47413 | 14.07271 | 10.47801 | 2.81332 | 17.62422 | 3.27793 | 8.63793 | 9.84840 |
| **Tmem98** | 0.01344 | 0.00786 | 0.01156 | 0.00856 | 0.00921 | 0.01289 | 0.00735 | 0.01119 | 0.01210 | 0.50951 | 0.01233 | 0.00651 | 0.01177 | 0.01174 | 0.00865 | 0.02214 |
| **Tmem9b** | 1.75055 | 1.66560 | 1.45406 | 3.46446 | 0.04261 | 3.37762 | 3.53913 | 1.64082 | 0.35866 | 9.02999 | 6.06798 | 1.17421 | 4.51978 | 0.00000 | 2.23850 | 0.57618 |
| **Tmf1** | 0.78377 | 0.03599 | 1.46373 | 2.67074 | 0.00360 | 0.00600 | 3.34761 | 1.15959 | 1.56615 | 0.83158 | 0.80240 | 0.08862 | 0.24305 | 1.06670 | 5.26798 | 1.27377 |
| **Tmigd1** | 0.00000 | 0.00803 | 0.00000 | 0.60536 | 0.02051 | 0.00740 | 0.00718 | 0.94279 | 0.04389 | 1.88276 | 1.45119 | 0.06463 | 0.00589 | 3.09254 | 0.00000 | 0.00735 |
| **Tmod1** | 0.01208 | 0.00820 | 0.01103 | 0.01032 | 0.00946 | 0.00668 | 0.00760 | 0.06276 | 0.01209 | 0.01240 | 0.01269 | 0.00838 | 0.01000 | 0.01077 | 0.04835 | 0.00739 |
| **Tmod2** | 1.70195 | 0.00471 | 0.81156 | 1.13871 | 0.00308 | 0.00698 | 2.01725 | 0.01828 | 0.74730 | 0.00000 | 0.49226 | 0.04888 | 0.03973 | 1.92071 | 0.75502 | 0.38636 |
| **Tmod3** | 5.40999 | 5.32522 | 6.77606 | 2.46311 | 5.60188 | 4.59045 | 7.95848 | 3.48082 | 3.07623 | 2.30081 | 3.48310 | 3.12960 | 3.58385 | 8.96151 | 6.20053 | 0.16902 |
| **Tmod4** | 0.01303 | 0.00808 | 0.01406 | 0.08667 | 0.01009 | 0.00706 | 0.00775 | 0.01627 | 0.28559 | 0.00902 | 0.01210 | 0.00714 | 0.10611 | 0.00882 | 0.05430 | 0.00764 |
| **Tmpo** | 14.01434 | 7.87313 | 3.60676 | 9.80751 | 9.24490 | 8.56181 | 6.00493 | 10.70885 | 12.08711 | 11.85570 | 3.38603 | 9.85346 | 6.61094 | 19.99643 | 12.83078 | 1.89608 |
| **Tmppe** | 3.47273 | 3.06857 | 9.51275 | 5.44271 | 1.25822 | 3.62470 | 4.34725 | 5.55989 | 7.32193 | 11.30071 | 22.23624 | 1.79361 | 10.13518 | 12.73830 | 15.84435 | 2.07210 |
| **Tmprss11a** | 0.01315 | 0.00750 | 0.00674 | 9.08257 | 0.00939 | 0.00527 | 0.00686 | 0.00436 | 2.24548 | 0.03020 | 0.14692 | 0.01524 | 0.01101 | 0.04276 | 2.59796 | 0.00573 |
| **Tmprss12** | 0.01032 | 0.00844 | 0.00731 | 0.31860 | 0.00949 | 0.00699 | 0.00813 | 0.17249 | 0.13194 | 0.00861 | 0.15586 | 0.03106 | 0.02913 | 0.00522 | 0.45154 | 0.00727 |
| **Tmprss2** | 0.02169 | 0.01797 | 0.08652 | 0.00718 | 0.09098 | 10.66736 | 0.00643 | 0.00762 | 0.01328 | 0.00000 | 0.00763 | 0.00773 | 0.02204 | 0.01563 | 0.00793 | 0.88802 |
| **Tmprss9** | 0.01307 | 0.00809 | 0.01225 | 0.04982 | 0.00932 | 0.12702 | 0.00780 | 0.02334 | 0.23469 | 0.26407 | 0.01069 | 0.02693 | 0.01019 | 0.00689 | 0.12938 | 0.00745 |
| **Tmsb10** | 6.75691 | 0.77996 | 7.41750 | 7.68658 | 2.68776 | 8.44262 | 2.27837 | 1.08124 | 10.70744 | 7.54436 | 10.47525 | 1.37109 | 11.45314 | 0.00000 | 0.00000 | 9.29508 |
| **Tmsb15a** | 0.01360 | 0.00787 | 0.01167 | 0.23484 | 0.00927 | 0.00658 | 0.00752 | 0.01352 | 1.21256 | 0.00855 | 0.00366 | 0.00591 | 0.01308 | 0.01874 | 0.02055 | 0.00731 |
| **Tmsb15b1** | 0.01363 | 0.00814 | 0.01101 | 1.11446 | 0.00933 | 0.00657 | 0.00733 | 0.01136 | 0.01877 | 0.01566 | 0.01270 | 0.00736 | 0.02002 | 0.16616 | 0.00507 | 0.00725 |
| **Tmsb4x** | 0.00000 | 0.33642 | 0.05780 | 0.73095 | 5.63783 | 1.49441 | 0.00690 | 0.34585 | 0.83210 | 16.72120 | 0.24086 | 0.00000 | 0.14863 | 0.00229 | 0.64607 | 3.50345 |
| **Tmtc1** | 0.22857 | 1.07350 | 0.36508 | 6.40634 | 0.00675 | 0.00480 | 0.00662 | 0.19070 | 1.28490 | 0.63320 | 0.02836 | 0.04460 | 0.01915 | 0.06001 | 3.95866 | 0.00575 |
| **Tmtc2** | 0.00910 | 0.00765 | 0.00952 | 0.01546 | 0.01668 | 0.00655 | 0.00739 | 0.00782 | 0.11233 | 0.00000 | 0.19011 | 0.02703 | 0.00736 | 0.01768 | 0.11687 | 0.88009 |
| **Tmtc3** | 0.55168 | 2.78325 | 1.74766 | 1.96249 | 3.55077 | 0.00590 | 3.46776 | 0.98806 | 2.74950 | 1.62614 | 0.60258 | 1.96767 | 0.01042 | 4.28808 | 6.63974 | 0.63138 |
| **Tmtc4** | 0.89046 | 0.00486 | 0.37857 | 0.79203 | 2.85777 | 1.56845 | 0.00620 | 2.69016 | 0.01557 | 0.00000 | 0.94546 | 0.00000 | 0.00000 | 0.00153 | 0.67512 | 0.15135 |
| **Tmub1** | 0.01606 | 0.00787 | 0.05075 | 0.00491 | 0.01063 | 0.00809 | 0.00715 | 0.05104 | 0.28081 | 0.00344 | 0.31325 | 0.06376 | 0.00797 | 0.01162 | 0.00132 | 0.00733 |
| **Tmub2** | 6.01117 | 0.00800 | 4.61422 | 1.07717 | 0.23652 | 0.02656 | 3.74424 | 6.10931 | 2.30412 | 5.62235 | 2.80102 | 0.41920 | 6.71168 | 0.09120 | 3.27650 | 3.32369 |
| **Tmx1** | 0.01761 | 3.20667 | 3.28003 | 2.59757 | 1.12226 | 0.00545 | 0.18660 | 1.18772 | 0.69026 | 0.00688 | 1.81337 | 0.06584 | 0.22481 | 0.11473 | 1.91137 | 0.14369 |
| **Tmx2** | 7.12003 | 7.61626 | 14.50905 | 0.46022 | 4.46131 | 5.69032 | 8.52306 | 6.08298 | 0.95318 | 8.95550 | 4.71790 | 4.53558 | 13.66072 | 11.49033 | 7.65220 | 1.23990 |
| **Tmx3** | 0.13653 | 0.02538 | 0.45158 | 1.56401 | 0.17446 | 0.42674 | 0.00580 | 1.90751 | 1.27992 | 0.08751 | 0.66610 | 6.43429 | 0.48280 | 5.84367 | 1.66449 | 2.36206 |
| **Tmx4** | 2.34624 | 0.03063 | 5.66958 | 0.31617 | 2.67796 | 3.16919 | 5.46880 | 0.13607 | 0.83942 | 0.00000 | 1.50921 | 0.14404 | 1.99108 | 1.12585 | 0.00000 | 0.38869 |
| **Tnc** | 0.01383 | 0.00784 | 0.01102 | 0.02649 | 0.00923 | 0.00674 | 0.00736 | 0.01168 | 0.01046 | 0.11697 | 0.01241 | 0.07384 | 0.01297 | 0.01143 | 0.00892 | 0.00776 |
| **Tnfaip1** | 6.62455 | 9.78476 | 9.39227 | 7.08375 | 8.59352 | 1.14996 | 3.09462 | 5.10207 | 5.03778 | 5.82842 | 2.99101 | 1.11344 | 7.45062 | 3.13509 | 5.76989 | 2.07298 |
| **Tnfaip2** | 0.01311 | 0.00786 | 0.01189 | 0.00689 | 0.00889 | 0.00665 | 0.00731 | 0.01035 | 0.01163 | 1.22924 | 0.01121 | 0.01495 | 0.01126 | 0.01190 | 0.00817 | 0.00752 |
| **Tnfaip3** | 0.01372 | 0.00825 | 0.01161 | 0.51120 | 0.00934 | 0.00672 | 0.00752 | 0.01308 | 0.00005 | 0.02910 | 0.01069 | 0.02072 | 0.01325 | 0.00573 | 0.34997 | 0.00733 |
| **Tnfaip6** | 0.02154 | 0.01881 | 0.04358 | 0.03992 | 0.00991 | 0.00653 | 0.01776 | 0.00788 | 0.75442 | 0.08130 | 0.18536 | 0.01466 | 0.08397 | 0.22259 | 0.01271 | 0.00723 |
| **Tnfaip8** | 1.61466 | 0.00318 | 0.00000 | 2.46151 | 3.79061 | 0.79391 | 1.20935 | 0.00725 | 0.80583 | 0.36650 | 0.01750 | 0.01470 | 0.60448 | 0.02820 | 2.36194 | 0.27825 |
| **Tnfaip8l2** | 0.71032 | 0.00794 | 0.00000 | 4.39971 | 0.01710 | 0.00463 | 0.20841 | 0.00380 | 7.34581 | 1.08662 | 1.01262 | 0.24339 | 0.00969 | 0.18646 | 2.98589 | 0.00568 |
| **Tnfrsf10b** | 2.22769 | 0.11168 | 2.55951 | 0.74492 | 0.29582 | 0.19772 | 1.63888 | 0.05718 | 1.79265 | 0.49562 | 5.12651 | 0.18167 | 1.88322 | 2.43728 | 1.41164 | 0.85349 |
| **Tnfrsf11a** | 0.02191 | 0.00748 | 0.02098 | 0.24588 | 0.00841 | 0.00791 | 0.00703 | 0.05595 | 0.24006 | 0.42385 | 0.24722 | 0.11396 | 0.03231 | 1.14875 | 0.77181 | 0.00703 |
| **Tnfrsf12a** | 0.72319 | 0.00703 | 0.33106 | 3.71134 | 0.03112 | 5.95256 | 0.00414 | 1.91458 | 4.24122 | 2.05436 | 1.76823 | 0.08760 | 2.86124 | 0.01612 | 0.56486 | 0.00234 |
| **Tnfrsf13c** | 0.02183 | 0.00822 | 0.00876 | 0.20400 | 0.00856 | 0.00690 | 0.00757 | 0.00869 | 1.28702 | 0.00824 | 0.33355 | 0.00510 | 0.00912 | 0.00857 | 0.00000 | 0.00714 |
| **Tnfrsf17** | 0.00194 | 0.00678 | 0.00000 | 0.52528 | 0.00937 | 0.00611 | 0.00675 | 2.89791 | 0.50915 | 1.26769 | 0.45694 | 2.09053 | 0.00338 | 1.00649 | 1.56090 | 0.00700 |
| **Tnfrsf19** | 0.50595 | 0.00696 | 0.00443 | 0.27960 | 0.00673 | 0.00682 | 0.00672 | 0.00860 | 0.00474 | 0.78005 | 0.22571 | 0.00417 | 0.02702 | 0.00817 | 0.24387 | 0.00638 |
| **Tnfrsf1a** | 0.00808 | 0.00748 | 0.03152 | 0.18842 | 0.00991 | 0.14790 | 0.00728 | 0.92349 | 0.00664 | 0.37792 | 0.11155 | 0.02354 | 0.00811 | 0.01278 | 0.10190 | 0.00770 |
| **Tnfrsf1b** | 0.78240 | 0.00662 | 0.00395 | 0.00880 | 0.00646 | 0.00700 | 0.00691 | 0.01462 | 0.01202 | 0.01072 | 0.01206 | 0.00653 | 0.01365 | 0.00608 | 0.29767 | 0.00653 |
| **Tnfrsf21** | 0.00000 | 0.54883 | 0.53910 | 0.01537 | 0.00754 | 0.00594 | 0.01665 | 0.48812 | 0.00869 | 0.61632 | 0.37798 | 0.00000 | 0.00385 | 2.61643 | 0.79619 | 0.00721 |
| **Tnfrsf22** | 0.00943 | 0.32365 | 0.04680 | 14.18669 | 2.30457 | 0.00313 | 0.64146 | 3.35253 | 10.04162 | 14.56847 | 0.41458 | 1.62363 | 0.36152 | 28.99621 | 41.63737 | 0.00993 |
| **Tnfrsf23** | 0.28951 | 0.00756 | 0.00362 | 0.16163 | 0.17908 | 0.00737 | 0.00782 | 0.03184 | 0.00604 | 0.16585 | 0.02437 | 0.04150 | 0.01074 | 0.00554 | 0.48569 | 0.00704 |
| **Tnfrsf25** | 0.00797 | 0.00875 | 0.05837 | 0.00903 | 0.00972 | 0.00586 | 0.00844 | 0.63124 | 0.02233 | 0.32716 | 0.00753 | 0.01922 | 0.00553 | 0.01628 | 0.00000 | 0.00741 |
| **Tnfrsf26** | 0.28770 | 0.04061 | 0.02650 | 0.60839 | 0.00725 | 0.00852 | 0.00725 | 0.01303 | 0.02247 | 0.34173 | 0.04897 | 0.00798 | 0.01537 | 0.00936 | 0.00662 | 0.00696 |
| **Tnfrsf4** | 0.01370 | 0.00820 | 0.00794 | 0.01769 | 0.00902 | 0.00698 | 0.00761 | 0.00510 | 0.21807 | 0.00319 | 0.64043 | 0.00524 | 0.00643 | 0.01146 | 0.00000 | 0.01421 |
| **Tnfrsf8** | 0.02265 | 0.00760 | 0.00761 | 0.01085 | 0.00854 | 0.00696 | 0.00712 | 0.30537 | 0.01203 | 0.00985 | 0.01116 | 0.00699 | 0.01384 | 0.01330 | 0.00578 | 0.00731 |
| **Tnfrsf9** | 0.00708 | 0.00765 | 0.06770 | 0.00000 | 0.01877 | 4.97305 | 0.00708 | 0.82310 | 0.78887 | 0.65600 | 0.00000 | 0.38130 | 0.00000 | 0.00426 | 0.43215 | 0.01309 |

| | | | | | | | | | | | | | | | | |
|---|---|---|---|---|---|---|---|---|---|---|---|---|---|---|---|---|
| **Tnfsf10** | 0.45552 | 0.20731 | 0.53112 | 3.26146 | 0.33826 | 0.03531 | 0.34688 | 0.05200 | 6.38010 | 0.00000 | 4.22979 | 0.41000 | 0.18520 | 2.57837 | 0.20022 | 0.16460 |
| **Tnfsf12** | 0.00737 | 0.00782 | 0.00536 | 3.34583 | 0.00921 | 0.00682 | 0.00712 | 0.59759 | 1.79008 | 1.16064 | 0.27446 | 0.00862 | 0.00450 | 0.27468 | 0.05067 | 0.00662 |
| **Tnfsf13b** | 0.00000 | 0.02175 | 1.41569 | 6.78529 | 0.02164 | 0.00300 | 0.61807 | 9.53281 | 11.65329 | 23.57372 | 19.54913 | 10.03201 | 6.06780 | 22.14257 | 13.77221 | 0.00000 |
| **Tnik** | 0.01374 | 0.00816 | 0.01230 | 0.27970 | 0.00935 | 0.00664 | 0.00753 | 0.01399 | 0.03221 | 0.01522 | 0.01230 | 0.00820 | 0.01271 | 0.01142 | 0.00645 | 0.00737 |
| **Tnip1** | 2.96383 | 1.26596 | 5.05264 | 12.42800 | 0.00001 | 3.08619 | 6.15775 | 4.52330 | 10.36572 | 15.37240 | 9.09503 | 5.69195 | 4.88840 | 0.87850 | 6.50177 | 1.28049 |
| **Tnip2** | 0.22558 | 0.00804 | 0.00000 | 3.42194 | 0.01320 | 0.01439 | 0.00657 | 0.00000 | 2.33239 | 2.81892 | 0.88881 | 0.07291 | 0.12425 | 0.88459 | 0.13327 | 0.21200 |
| **Tnk1** | 0.38013 | 0.00730 | 0.09208 | 0.00982 | 0.00658 | 0.00847 | 0.00704 | 0.01310 | 0.01235 | 0.00783 | 0.01208 | 0.00724 | 0.01544 | 0.00711 | 0.12389 | 0.62935 |
| **Tnk2** | 2.44595 | 0.00476 | 0.26049 | 4.00520 | 0.01278 | 0.35465 | 0.00373 | 1.33092 | 2.52556 | 1.92890 | 1.43922 | 0.00000 | 1.44608 | 0.00000 | 0.73831 | 2.08276 |
| **Tnks** | 0.06032 | 0.00802 | 0.00000 | 2.14448 | 0.01505 | 0.00631 | 0.95706 | 0.71856 | 2.34396 | 0.70412 | 1.13989 | 0.03295 | 0.00000 | 1.41819 | 1.59706 | 0.00654 |
| **Tnks1bp1** | 0.00001 | 0.16442 | 0.39119 | 1.34507 | 0.00718 | 0.00669 | 0.36701 | 1.25227 | 0.50303 | 0.51964 | 0.82081 | 0.13313 | 1.28198 | 0.00180 | 0.10828 | 1.88008 |
| **Tnks2** | 8.95623 | 6.28639 | 7.56139 | 11.37234 | 1.93719 | 4.74366 | 6.92812 | 13.45429 | 11.23182 | 17.57842 | 8.33593 | 17.59404 | 4.18974 | 32.63060 | 20.96929 | 2.88848 |
| **Tnnc2** | 0.01559 | 0.00851 | 0.00668 | 0.00468 | 0.01014 | 0.00664 | 0.00784 | 0.04064 | 3.32445 | 0.68525 | 0.29170 | 0.03168 | 0.01357 | 0.01095 | 0.38785 | 0.00741 |
| **Tnni3** | 1.10144 | 0.00718 | 0.03473 | 0.34072 | 0.01491 | 0.00608 | 0.01636 | 0.02252 | 0.00662 | 0.01608 | 0.01411 | 0.00700 | 0.01395 | 0.01193 | 0.01325 | 1.06644 |
| **Tnnt1** | 0.55310 | 0.00748 | 0.05484 | 0.11291 | 0.02623 | 0.00531 | 0.00619 | 0.00666 | 0.02125 | 0.31251 | 0.01214 | 0.00550 | 1.01087 | 0.01080 | 0.00746 | 0.00688 |
| **Tnpo1** | 1.81774 | 0.64654 | 3.30116 | 9.30429 | 5.22036 | 0.31856 | 0.00000 | 2.63935 | 4.49499 | 7.02534 | 10.72094 | 0.54808 | 2.97397 | 8.39128 | 8.72495 | 0.23254 |
| **Tnpo2** | 0.35634 | 1.07049 | 1.57103 | 3.94510 | 0.00475 | 0.00621 | 0.10406 | 0.19654 | 5.10231 | 0.05842 | 8.26615 | 0.17556 | 0.17630 | 7.79729 | 0.91014 | 0.17555 |
| **Tnpo3** | 3.10739 | 4.13586 | 3.04682 | 10.77532 | 3.76823 | 4.17501 | 2.90424 | 4.74819 | 16.49422 | 8.54987 | 6.06742 | 0.61150 | 5.23375 | 16.62290 | 11.32433 | 0.13072 |
| **Tnrc18** | 4.19153 | 3.36954 | 2.95935 | 0.03383 | 2.71667 | 5.67618 | 4.47860 | 13.46871 | 0.11160 | 8.58788 | 8.50447 | 0.39575 | 1.62622 | 1.93316 | 1.69908 | 1.58377 |
| **Tnrc6a** | 3.99288 | 3.85764 | 5.92438 | 0.72689 | 0.02404 | 1.05709 | 1.88762 | 2.81596 | 1.74679 | 6.20446 | 5.10471 | 1.49272 | 2.14136 | 0.82802 | 4.00127 | 0.01864 |
| **Tnrc6b** | 2.28268 | 0.01779 | 0.72483 | 2.78194 | 0.11292 | 0.01316 | 0.00592 | 0.11200 | 1.67788 | 2.07999 | 0.13878 | 0.02133 | 0.00647 | 1.76554 | 0.76683 | 0.00410 |
| **Tnrc6c** | 2.09912 | 0.00347 | 0.00659 | 2.98132 | 3.02755 | 0.02135 | 0.00524 | 0.00520 | 2.27900 | 5.01247 | 0.76336 | 0.00000 | 0.02023 | 0.21800 | 0.00000 | 0.01160 |
| **Tns1** | 4.32679 | 0.00000 | 2.58179 | 0.00000 | 4.70288 | 0.00237 | 0.00267 | 1.05299 | 0.01858 | 0.15639 | 1.12957 | 0.13847 | 0.87240 | 0.01169 | 0.00429 | 0.02464 |
| **Tns2** | 0.01380 | 0.00810 | 0.01083 | 0.47470 | 0.00928 | 0.01332 | 0.00750 | 0.01316 | 0.03477 | 0.10561 | 0.01149 | 0.04044 | 0.01289 | 0.00988 | 0.00859 | 0.00742 |
| **Tns3** | 1.96810 | 4.02981 | 0.56803 | 0.45558 | 2.18265 | 2.03628 | 0.90089 | 1.72496 | 0.10238 | 0.03030 | 5.45994 | 0.14718 | 0.14167 | 0.37541 | 0.54328 | 1.82875 |
| **Tns4** | 0.10791 | 0.02565 | 3.19993 | 0.23018 | 0.04635 | 0.09181 | 2.47062 | 0.42950 | 1.50703 | 0.33228 | 1.09428 | 0.38898 | 0.07225 | 0.18402 | 1.04543 | 0.05778 |
| **Tob1** | 0.08919 | 0.00786 | 0.01278 | 0.01024 | 0.18639 | 0.00671 | 0.00760 | 0.01447 | 0.01166 | 0.03505 | 0.01173 | 0.02773 | 0.03469 | 0.01526 | 0.15917 | 0.00728 |
| **Tob2** | 0.54681 | 0.00631 | 0.00000 | 13.27024 | 0.00717 | 0.00439 | 0.45617 | 0.85979 | 16.76522 | 0.55666 | 1.45670 | 0.05610 | 0.00000 | 0.59344 | 0.76597 | 0.00313 |
| **Toe1** | 5.46337 | 6.99249 | 11.64656 | 1.68271 | 5.85733 | 2.98660 | 7.05005 | 12.36594 | 1.19665 | 9.28083 | 4.01390 | 1.47307 | 9.32581 | 4.96808 | 7.30222 | 5.41544 |
| **Tollip** | 6.89139 | 6.07998 | 10.76109 | 6.61439 | 11.52587 | 6.53241 | 6.36613 | 15.90081 | 3.88508 | 11.96147 | 9.37270 | 1.44884 | 5.06413 | 10.35392 | 5.58351 | 0.47953 |
| **Tom1** | 5.58416 | 5.36904 | 4.68498 | 0.98105 | 4.99777 | 1.44977 | 7.16652 | 11.43007 | 1.76623 | 6.86661 | 4.80154 | 0.63588 | 7.58015 | 0.45280 | 1.74739 | 3.55009 |
| **Tom1l1** | 0.01830 | 0.00717 | 2.37453 | 0.27019 | 0.02450 | 0.00767 | 0.41581 | 1.88609 | 0.16013 | 0.82479 | 0.78635 | 0.25140 | 0.82099 | 1.04923 | 0.56114 | 0.01358 |
| **Tom1l2** | 0.57581 | 1.14067 | 0.56985 | 6.80195 | 2.74689 | 1.99501 | 2.15556 | 5.48073 | 5.49304 | 9.42740 | 5.70476 | 0.16765 | 1.72915 | 0.18527 | 1.36338 | 0.00924 |
| **Tomm20** | 23.20016 | 10.75169 | 20.43145 | 26.79969 | 16.27402 | 8.43960 | 7.39608 | 13.48240 | 31.62974 | 47.40720 | 41.74661 | 19.58008 | 23.68365 | 21.60457 | 13.69748 | 6.69190 |
| **Tomm20l** | 0.01475 | 0.00770 | 0.07597 | 0.02102 | 0.00882 | 0.00601 | 0.00706 | 0.04194 | 0.01275 | 0.37194 | 0.89061 | 0.08404 | 0.00430 | 0.00556 | 0.05301 | 0.00735 |
| **Tomm22** | 24.67694 | 18.86439 | 25.81776 | 16.32523 | 13.98737 | 14.69396 | 13.16393 | 32.50783 | 23.10229 | 41.94664 | 21.51713 | 8.64099 | 24.58640 | 11.66362 | 11.82572 | 6.89341 |
| **Tomm34** | 4.65222 | 3.62779 | 0.56824 | 6.35671 | 2.82606 | 0.01378 | 4.08555 | 0.86994 | 9.03742 | 4.81688 | 0.35713 | 0.41858 | 1.92275 | 0.20600 | 1.65466 | 3.90103 |
| **Tomm40** | 15.25009 | 0.52499 | 4.30661 | 5.04876 | 10.54448 | 6.47205 | 3.33501 | 16.97906 | 6.34977 | 17.78326 | 8.52321 | 0.91356 | 17.07345 | 0.00000 | 0.00000 | 9.05945 |
| **Tomm40l** | 3.84001 | 0.00067 | 6.05240 | 0.24252 | 1.85696 | 0.00234 | 0.32564 | 0.00000 | 0.48781 | 0.94680 | 0.00402 | 0.02273 | 0.93621 | 0.01011 | 0.00000 | 2.99511 |
| **Tomm5** | 22.65873 | 8.66265 | 21.52810 | 10.59215 | 18.52668 | 6.94101 | 9.60139 | 20.89247 | 27.33068 | 35.62637 | 27.25288 | 8.31174 | 23.45242 | 12.28978 | 18.45244 | 5.74059 |
| **Tomm6** | 6.83603 | 0.43001 | 6.66597 | 6.83322 | 7.53255 | 2.49865 | 1.15420 | 1.71443 | 12.25300 | 6.79759 | 10.81599 | 1.35302 | 3.89256 | 1.36837 | 0.00000 | 1.92627 |
| **Tomm7** | 13.08383 | 0.82133 | 11.19487 | 10.93757 | 7.05119 | 3.95867 | 2.22585 | 1.32622 | 8.46098 | 4.16732 | 25.56344 | 1.66724 | 10.93333 | 4.97892 | 5.42249 | 7.60349 |
| **Tomm70a** | 12.96356 | 12.68450 | 6.57248 | 3.64506 | 10.47540 | 7.14644 | 4.96071 | 12.26922 | 3.40608 | 21.94570 | 12.08901 | 15.75530 | 8.80459 | 24.21176 | 17.83204 | 6.89144 |
| **Tomt** | 0.65047 | 0.00685 | 0.63908 | 0.03943 | 0.00550 | 0.15469 | 0.00621 | 0.00902 | 0.07527 | 0.06160 | 0.00470 | 0.56292 | 1.06269 | 0.00954 | 0.15474 | 0.00595 |
| **Tonsl** | 4.24885 | 0.34343 | 2.74713 | 3.06668 | 5.71992 | 0.01450 | 3.58952 | 4.46680 | 3.37044 | 4.50043 | 5.44927 | 0.42523 | 3.57483 | 0.01956 | 0.00000 | 0.43127 |
| **Top1** | 5.41638 | 6.93750 | 8.44306 | 11.16202 | 5.24677 | 0.30824 | 8.06918 | 14.48992 | 18.92578 | 11.81066 | 9.40844 | 4.74143 | 6.44963 | 30.03867 | 24.08837 | 3.04798 |
| **Top1mt** | 0.01349 | 0.00753 | 0.00000 | 0.47391 | 0.00859 | 0.00858 | 0.01526 | 1.71029 | 0.00478 | 6.37457 | 2.16245 | 0.26937 | 3.30825 | 0.00655 | 0.02945 | 0.02054 |
| **Top2a** | 14.56855 | 19.23211 | 10.90596 | 12.03767 | 0.92430 | 0.17124 | 10.36247 | 16.03366 | 20.43350 | 21.30471 | 15.40402 | 9.01491 | 5.63402 | 28.50303 | 21.96676 | 3.33929 |
| **Top2b** | 1.50959 | 4.16934 | 1.95408 | 0.85638 | 0.36929 | 1.03107 | 0.18782 | 0.02660 | 0.64267 | 1.58840 | 0.01229 | 0.02631 | 0.04059 | 0.00362 | 0.47306 | 1.99318 |
| **Top3a** | 3.26230 | 3.34845 | 2.01170 | 3.20034 | 0.00753 | 0.21571 | 1.47382 | 1.63976 | 3.92980 | 2.76389 | 1.40671 | 0.04892 | 0.29171 | 0.00000 | 1.73632 | 0.00000 |
| **Top3b** | 3.86614 | 0.30583 | 1.60266 | 0.11443 | 2.10223 | 2.46618 | 5.28161 | 5.03713 | 0.06915 | 1.32224 | 0.55110 | 0.01922 | 2.19844 | 1.02544 | 0.00000 | 3.89762 |
| **Topaz1** | 0.29127 | 0.35518 | 0.00291 | 0.14857 | 0.01739 | 0.00695 | 0.00689 | 0.00881 | 0.01607 | 0.01073 | 0.01112 | 0.00591 | 0.01191 | 0.01630 | 0.62964 | 0.00717 |
| **Topbp1** | 6.68938 | 9.66866 | 6.30618 | 11.03654 | 6.34513 | 5.75041 | 7.58409 | 14.53616 | 13.47814 | 19.64931 | 6.74349 | 19.03760 | 6.78034 | 13.79114 | 15.88586 | 3.26367 |
| **Topors** | 3.27596 | 12.14664 | 4.70286 | 9.74484 | 5.07082 | 0.01883 | 5.84328 | 12.03855 | 15.16660 | 21.43217 | 2.43384 | 8.99029 | 2.61761 | 9.70207 | 19.63938 | 1.03975 |
| **Toporsos** | 0.02582 | 0.01901 | 0.12539 | 0.13291 | 0.00819 | 0.00616 | 0.00766 | 0.00703 | 0.19247 | 0.01514 | 0.33261 | 0.00677 | 0.00804 | 0.00946 | 0.00000 | 0.00705 |
| **Tor1a** | 3.46145 | 5.18311 | 3.38901 | 8.18454 | 4.05911 | 3.07632 | 2.39438 | 4.91973 | 14.27915 | 5.67783 | 3.29393 | 0.84348 | 1.34405 | 1.04798 | 5.65856 | 2.76381 |
| **Tor1aip1** | 1.41419 | 0.95909 | 2.84957 | 11.53776 | 3.00655 | 0.00227 | 0.00000 | 6.22512 | 18.02800 | 9.62010 | 7.47481 | 0.40137 | 3.14910 | 8.42389 | 8.35331 | 1.30720 |
| **Tor1aip2** | 13.20862 | 8.41028 | 11.87025 | 8.61427 | 9.55179 | 4.31324 | 9.37605 | 13.86912 | 10.08951 | 10.27265 | 10.65809 | 8.50212 | 7.51358 | 17.33254 | 8.74738 | 6.47754 |
| **Tor1b** | 5.61778 | 3.18236 | 2.59037 | 10.35783 | 0.00988 | 0.89937 | 4.40892 | 8.75671 | 20.23796 | 17.60750 | 11.42010 | 8.03400 | 1.97093 | 18.02644 | 20.31780 | 1.74808 |
| **Tor2a** | 5.46927 | 0.14430 | 4.87431 | 11.39449 | 0.68338 | 0.28422 | 1.67944 | 10.03479 | 7.34072 | 17.28059 | 11.33214 | 3.47991 | 11.44720 | 0.13242 | 1.43179 | 0.00000 |
| **Tor3a** | 0.02650 | 0.00803 | 0.62489 | 6.07007 | 0.01177 | 0.00498 | 0.00602 | 0.84292 | 0.55393 | 1.68380 | 2.35145 | 0.02064 | 0.00000 | 4.53873 | 0.94563 | 0.01294 |
| **Tor4a** | 2.02256 | 0.14991 | 1.88443 | 7.66382 | 0.31254 | 0.54976 | 1.33138 | 0.86776 | 14.89223 | 0.38801 | 9.90622 | 0.72170 | 2.03227 | 4.78006 | 0.69177 | 0.29522 |
| **Tox3** | 0.01376 | 0.00785 | 0.01241 | 0.54010 | 0.00934 | 0.00654 | 0.00748 | 0.01342 | 0.00401 | 0.01093 | 0.01320 | 0.00749 | 0.01264 | 0.00595 | 0.11848 | 0.00730 |

| | | | | | | | | | | | | | | | | |
|---|---|---|---|---|---|---|---|---|---|---|---|---|---|---|---|---|
| **Tox4** | 9.62486 | 9.27713 | 6.81537 | 10.50979 | 10.06904 | 1.05974 | 6.00586 | 4.84662 | 17.05040 | 7.47902 | 9.46124 | 3.20990 | 7.01932 | 11.60265 | 13.69837 | 6.39406 |
| **Tpbg** | 0.11654 | 0.00760 | 0.00687 | 0.01068 | 0.00725 | 0.00679 | 0.00742 | 0.01198 | 0.01238 | 0.00603 | 0.01234 | 0.03965 | 0.00836 | 0.01301 | 0.00980 | 1.03147 |
| **Tpbgl** | 1.34601 | 0.00673 | 0.00361 | 0.00624 | 0.01351 | 0.00677 | 0.00606 | 0.00121 | 0.01159 | 0.80223 | 0.14379 | 0.04980 | 0.01029 | 0.01151 | 0.00358 | 0.00622 |
| **Tpcn1** | 0.46298 | 0.00740 | 0.00347 | 0.19733 | 0.03133 | 0.00793 | 0.00661 | 0.44608 | 0.36864 | 0.40950 | 0.75686 | 0.03864 | 0.00609 | 0.00544 | 0.00000 | 0.01466 |
| **Tpcn2** | 0.00822 | 0.00550 | 1.99435 | 0.00471 | 0.00679 | 0.09408 | 3.30071 | 0.00000 | 0.01140 | 0.00000 | 0.46017 | 0.01062 | 0.39966 | 0.01178 | 0.00000 | 0.01341 |
| **Tpd52** | 16.66767 | 13.37290 | 17.95440 | 15.37258 | 7.66822 | 11.99142 | 12.92482 | 26.56354 | 2.09806 | 28.26113 | 25.88623 | 15.37577 | 16.06987 | 12.32553 | 11.38217 | 7.83417 |
| **Tpd52l1** | 0.01191 | 0.00792 | 0.01212 | 0.63584 | 0.00852 | 0.00646 | 0.00722 | 0.00923 | 0.16554 | 3.81846 | 0.02593 | 0.00000 | 0.01048 | 0.00857 | 0.01954 | 0.01475 |
| **Tpd52l2** | 17.23354 | 8.14435 | 20.23679 | 15.81101 | 12.30563 | 2.95925 | 12.87377 | 20.16628 | 12.70652 | 24.03709 | 18.99478 | 12.20775 | 16.40543 | 7.94791 | 12.98640 | 6.28726 |
| **Tpgs1** | 1.75031 | 0.00609 | 0.14813 | 1.20723 | 0.23424 | 0.02167 | 0.01446 | 0.01562 | 1.40561 | 3.95465 | 0.01298 | 0.00000 | 3.26290 | 0.01043 | 0.00177 | 1.08087 |
| **Tpgs2** | 0.64448 | 0.06994 | 0.66209 | 6.44754 | 0.52673 | 0.01046 | 0.24962 | 0.05966 | 2.76610 | 0.38672 | 2.82315 | 6.53044 | 0.37207 | 0.26293 | 0.69666 | 0.01169 |
| **Tph2** | 0.01379 | 0.00774 | 0.01207 | 1.13982 | 0.00938 | 0.00645 | 0.00741 | 0.01250 | 0.00000 | 0.00795 | 0.01238 | 0.00723 | 0.01270 | 0.00736 | 0.26627 | 0.00718 |
| **Tpi1** | 10.12793 | 12.10085 | 7.29244 | 1.72819 | 0.54023 | 10.62590 | 3.33103 | 4.40379 | 3.91570 | 4.94422 | 2.32135 | 0.06873 | 6.51547 | 0.00000 | 1.08912 | 11.04482 |
| **Tpk1** | 0.66284 | 0.00703 | 0.81145 | 1.51145 | 0.00791 | 2.00401 | 0.00526 | 0.00550 | 0.00000 | 0.00000 | 0.01084 | 0.02543 | 2.25115 | 0.01172 | 0.00000 | 0.02588 |
| **Tpm1** | 10.03913 | 6.06014 | 7.72143 | 0.00000 | 9.83128 | 15.43120 | 4.75481 | 9.74990 | 0.75733 | 6.21206 | 7.14536 | 0.75013 | 17.77626 | 1.25663 | 0.00000 | 11.72076 |
| **Tpm2** | 0.00000 | 0.00728 | 0.64546 | 0.00454 | 0.00621 | 0.00568 | 0.00623 | 0.00736 | 0.21159 | 3.88756 | 0.00668 | 0.09688 | 1.40572 | 0.01374 | 0.00000 | 2.66490 |
| **Tpm3** | 25.03828 | 15.92810 | 23.78610 | 13.83790 | 19.21710 | 17.58599 | 17.14658 | 32.63919 | 18.59970 | 30.95555 | 18.34831 | 7.56568 | 24.62546 | 18.57527 | 23.47423 | 14.84525 |
| **Tpm4** | 16.55948 | 13.41164 | 10.58675 | 0.12107 | 20.67840 | 17.90168 | 11.51183 | 16.72896 | 0.19104 | 11.60961 | 8.69057 | 12.79232 | 18.86400 | 9.65346 | 0.92347 | 14.54430 |
| **Tpmt** | 5.57202 | 1.00012 | 0.63647 | 1.85298 | 0.04350 | 0.04268 | 0.03373 | 0.08751 | 3.56211 | 1.75169 | 2.45490 | 0.13734 | 2.38052 | 1.00910 | 1.47558 | 0.01307 |
| **Tpo** | 0.01246 | 0.00776 | 0.01073 | 0.42188 | 0.00897 | 0.00604 | 0.00732 | 0.01195 | 4.07306 | 0.00424 | 0.00000 | 0.00000 | 0.15344 | 0.00533 | 0.08987 | 0.00678 |
| **Tpp1** | 2.15228 | 0.02299 | 4.13200 | 0.00000 | 3.39332 | 5.48314 | 1.31531 | 0.76043 | 0.00955 | 1.24946 | 0.00000 | 0.02146 | 1.31145 | 0.01091 | 0.01468 | 0.01589 |
| **Tpp2** | 2.92969 | 7.43202 | 5.94284 | 4.98246 | 6.03314 | 2.99823 | 5.41243 | 5.79478 | 7.67854 | 3.95125 | 7.68152 | 10.73891 | 3.29852 | 14.14664 | 6.98916 | 0.02953 |
| **Tppp** | 0.01365 | 0.00775 | 0.01190 | 0.15572 | 0.00933 | 0.00651 | 0.00745 | 0.01239 | 0.56926 | 0.01321 | 0.00884 | 0.00590 | 0.01303 | 0.00512 | 0.41232 | 0.00734 |
| **Tppp3** | 7.44189 | 0.00000 | 0.57391 | 0.05870 | 0.01797 | 0.00000 | 0.00000 | 1.72336 | 0.72307 | 3.48588 | 0.59354 | 0.02030 | 1.76910 | 0.00704 | 0.00000 | 0.02015 |
| **Tpr** | 13.40536 | 10.86384 | 10.14694 | 16.56285 | 8.22999 | 10.73370 | 7.87835 | 14.17364 | 25.86339 | 26.04210 | 10.30715 | 16.67410 | 7.59838 | 18.11571 | 21.32856 | 3.72691 |
| **Tpra1** | 2.93516 | 0.00363 | 0.29621 | 1.00408 | 0.27513 | 0.87424 | 0.93237 | 2.70361 | 0.97261 | 0.00000 | 0.06817 | 0.03431 | 0.09340 | 0.00831 | 0.00000 | 0.04751 |
| **Tprgl** | 6.29444 | 4.28480 | 2.73488 | 13.62014 | 5.02727 | 6.04391 | 0.63634 | 5.49541 | 1.29557 | 8.73576 | 6.24990 | 0.57083 | 5.24578 | 0.08467 | 7.14990 | 2.05559 |
| **Tprkb** | 8.27944 | 1.58607 | 9.77868 | 2.14690 | 2.87993 | 2.40111 | 4.16754 | 5.21711 | 2.94430 | 6.71897 | 3.38089 | 1.29691 | 4.67979 | 8.08251 | 5.52187 | 1.61316 |
| **Tprn** | 2.83791 | 0.01174 | 2.36118 | 5.18851 | 0.03437 | 5.10513 | 0.18276 | 10.30591 | 11.53016 | 13.95369 | 12.49082 | 2.24441 | 6.95401 | 2.26863 | 16.55679 | 5.48748 |
| **Tpst1** | 5.02856 | 1.96667 | 3.30095 | 1.18968 | 1.02125 | 0.09871 | 0.01186 | 0.93662 | 0.22864 | 0.83013 | 1.02147 | 0.05366 | 0.00000 | 0.01935 | 2.33275 | 0.01464 |
| **Tpst2** | 2.84283 | 0.04557 | 0.85660 | 0.00000 | 2.32399 | 0.17947 | 0.44087 | 2.08979 | 1.09051 | 1.62126 | 0.43480 | 0.04526 | 3.37900 | 0.02246 | 0.23707 | 4.53546 |
| **Tpt1** | 37.37383 | 17.28283 | 23.79299 | 27.15174 | 28.80407 | 13.37109 | 14.93130 | 17.15157 | 35.51287 | 59.20414 | 51.76850 | 15.15357 | 43.01443 | 20.10316 | 9.80601 | 25.37017 |
| **Tpx2** | 10.47344 | 13.17282 | 9.81908 | 17.30055 | 7.27296 | 3.12333 | 4.69116 | 19.79957 | 27.88404 | 21.28146 | 13.91916 | 19.68093 | 5.53061 | 22.76673 | 23.51291 | 4.46428 |
| **Tra2a** | 19.78231 | 17.10749 | 17.88240 | 14.34844 | 7.41307 | 13.40145 | 10.67932 | 16.05696 | 27.43603 | 19.53911 | 19.55284 | 22.86392 | 16.68950 | 27.13234 | 12.48498 | 13.30367 |
| **Tra2b** | 13.45145 | 17.59768 | 16.59795 | 16.39687 | 13.06951 | 4.83679 | 13.28584 | 20.61874 | 30.14553 | 20.89007 | 20.88513 | 19.00211 | 16.33449 | 27.44567 | 15.48693 | 6.63287 |
| **Trabd** | 1.31410 | 0.11028 | 0.00000 | 0.50362 | 0.02142 | 2.50247 | 2.34407 | 1.40192 | 0.81082 | 0.75511 | 0.01151 | 0.01904 | 2.95581 | 0.00818 | 0.01184 | 0.24394 |
| **Trabd2b** | 0.01877 | 0.01856 | 0.01926 | 1.22317 | 0.01059 | 0.00610 | 0.00726 | 0.01038 | 0.97132 | 0.24647 | 0.01856 | 0.03257 | 0.01435 | 0.08188 | 0.00000 | 0.02224 |
| **Tradd** | 1.78330 | 0.02815 | 0.40347 | 0.00561 | 0.00687 | 0.00489 | 0.01656 | 1.94427 | 0.00996 | 0.00000 | 0.31932 | 0.02118 | 0.15447 | 0.00713 | 0.01545 | 1.67986 |
| **Traf1** | 0.74137 | 0.00710 | 0.00812 | 0.05608 | 0.01524 | 0.00647 | 0.00704 | 0.01422 | 0.00900 | 0.01665 | 0.02850 | 0.00705 | 0.01870 | 0.00710 | 0.12190 | 0.00665 |
| **Traf2** | 2.76081 | 0.10341 | 5.96498 | 7.03949 | 0.01543 | 0.40270 | 3.69975 | 7.16652 | 5.72755 | 8.35735 | 0.20764 | 0.88269 | 3.38340 | 0.07658 | 3.83103 | 0.00000 |
| **Traf3** | 5.08883 | 0.00000 | 2.24236 | 5.99411 | 2.00864 | 0.00000 | 0.00219 | 0.74846 | 3.84667 | 3.09541 | 0.86781 | 2.06289 | 0.09990 | 0.01330 | 1.03450 | 0.00000 |
| **Traf3ip1** | 1.83821 | 0.00504 | 0.17515 | 0.89056 | 0.00406 | 0.00779 | 0.48842 | 3.72167 | 0.00000 | 0.81839 | 1.12480 | 0.07971 | 0.03204 | 0.84484 | 0.90582 | 0.00517 |
| **Traf3ip2** | 0.60457 | 0.00813 | 0.06701 | 0.66193 | 0.01683 | 0.00505 | 1.12092 | 0.18062 | 0.00000 | 0.73392 | 0.01153 | 0.01945 | 0.03793 | 0.01204 | 0.06579 | 0.00644 |
| **Traf4** | 3.30831 | 0.00000 | 7.21912 | 4.11982 | 4.15002 | 0.89741 | 3.38806 | 10.39596 | 0.65423 | 5.65492 | 3.19260 | 0.29621 | 4.02096 | 0.17759 | 2.32969 | 4.06734 |
| **Traf5** | 0.00875 | 0.00734 | 0.00773 | 0.10232 | 0.00961 | 0.00679 | 0.00707 | 1.31712 | 0.43968 | 0.52696 | 0.00000 | 0.01127 | 0.00542 | 0.01112 | 0.00000 | 0.00742 |
| **Traf6** | 0.00484 | 0.00801 | 0.27420 | 1.09620 | 0.00655 | 0.00557 | 0.00739 | 0.01261 | 0.00000 | 0.00000 | 0.01092 | 0.01979 | 0.00584 | 0.03556 | 0.43895 | 3.13486 |
| **Traf7** | 8.86997 | 0.58143 | 11.65678 | 8.54148 | 7.77544 | 9.65104 | 10.19209 | 10.53427 | 6.25007 | 11.41768 | 2.33593 | 0.58325 | 8.82727 | 0.38770 | 8.73210 | 7.27377 |
| **Trafd1** | 1.30498 | 0.00601 | 2.69675 | 5.59673 | 1.08863 | 3.03830 | 0.00415 | 1.77531 | 1.19931 | 1.78042 | 3.84876 | 0.48185 | 0.25755 | 1.64586 | 0.13913 | 0.77141 |
| **Traip** | 3.97107 | 3.51633 | 5.47702 | 9.75546 | 2.44566 | 2.92194 | 3.37134 | 4.27804 | 13.22207 | 5.05861 | 6.54954 | 1.01146 | 2.65503 | 3.82976 | 6.15644 | 0.00000 |
| **Trak1** | 3.64554 | 1.87141 | 1.12688 | 9.65140 | 1.57899 | 1.11240 | 0.60145 | 4.40391 | 1.72126 | 5.71878 | 5.25606 | 0.32978 | 0.00000 | 1.50801 | 3.91544 | 1.33410 |
| **Trak2** | 1.04677 | 6.00317 | 6.17554 | 10.92558 | 2.59399 | 2.43159 | 3.37460 | 0.44034 | 10.20132 | 5.52591 | 2.25910 | 0.57348 | 2.15356 | 1.59678 | 5.60021 | 0.01900 |
| **Tram1** | 5.75636 | 0.14273 | 8.91560 | 1.65692 | 3.98496 | 1.74884 | 0.35466 | 5.24021 | 0.24762 | 0.90704 | 3.06976 | 0.20365 | 1.96222 | 3.87644 | 0.00000 | 0.15303 |
| **Tram2** | 0.01483 | 0.00822 | 0.05390 | 0.33408 | 0.00902 | 0.00580 | 0.00747 | 0.04627 | 0.85009 | 0.00988 | 0.00688 | 0.01817 | 0.01029 | 0.01220 | 0.00170 | 0.00726 |
| **Trank1** | 0.01390 | 0.00807 | 0.00943 | 0.25150 | 0.00914 | 0.00675 | 0.00746 | 0.00448 | 0.24078 | 0.10927 | 0.32859 | 0.02632 | 0.00931 | 0.01134 | 0.00282 | 0.00707 |
| **Trap1** | 12.27032 | 5.07492 | 7.82682 | 3.39702 | 4.43842 | 6.64626 | 7.58213 | 11.47068 | 3.22772 | 5.64515 | 5.15278 | 0.86618 | 9.01237 | 0.06060 | 2.43836 | 8.34004 |
| **Trap1a** | 5.05910 | 6.80501 | 3.25146 | 1.09773 | 5.83639 | 11.03977 | 3.75482 | 0.44652 | 5.76201 | 0.58086 | 2.83206 | 0.04829 | 8.33037 | 0.00000 | 0.94255 | 8.42768 |
| **Trappc1** | 9.74754 | 6.14091 | 5.34189 | 0.88894 | 7.26432 | 1.68388 | 3.18684 | 3.17981 | 0.00441 | 2.84940 | 2.99764 | 0.36863 | 5.33284 | 1.75235 | 0.17409 | 2.68314 |
| **Trappc10** | 1.79569 | 0.00458 | 0.58212 | 8.92465 | 1.50379 | 0.00351 | 0.00377 | 1.34925 | 5.90708 | 3.48900 | 2.66732 | 0.36649 | 1.00390 | 2.76345 | 7.36069 | 0.00257 |
| **Trappc11** | 2.57210 | 0.15095 | 3.86365 | 1.90516 | 0.60751 | 4.22610 | 2.20244 | 2.69576 | 0.99186 | 3.45318 | 3.02022 | 7.82173 | 4.27322 | 2.10533 | 5.59483 | 1.87844 |
| **Trappc12** | 0.92284 | 4.08169 | 2.53946 | 3.01355 | 0.00411 | 0.01663 | 3.11939 | 4.66001 | 2.93120 | 0.81165 | 0.21325 | 0.20356 | 3.90330 | 0.00000 | 2.12216 | 2.13259 |
| **Trappc13** | 10.06028 | 4.05468 | 7.33821 | 1.31186 | 11.86757 | 3.09673 | 6.46935 | 7.28595 | 4.07208 | 9.20088 | 8.57867 | 3.97815 | 5.84752 | 9.17204 | 8.18267 | 3.66974 |
| **Trappc2** | 1.62220 | 0.02578 | 0.55289 | 2.78015 | 0.44563 | 0.44922 | 0.00469 | 0.00000 | 0.05212 | 0.00000 | 0.89841 | 0.01641 | 1.14365 | 0.29471 | 1.14772 | 2.23875 |
| **Trappc2l** | 7.10616 | 1.05429 | 5.15517 | 0.29146 | 4.46215 | 3.56937 | 2.12064 | 5.99974 | 4.36672 | 2.81443 | 10.77651 | 0.66191 | 6.49886 | 2.89070 | 3.30349 | 3.19201 |

| | | | | | | | | | | | | | | | | |
|---|---|---|---|---|---|---|---|---|---|---|---|---|---|---|---|---|
| **Trappc3** | 0.39237 | 4.07368 | 2.57944 | 0.34884 | 4.66332 | 0.09678 | 0.04317 | 4.60309 | 0.21807 | 2.86139 | 4.98254 | 0.08304 | 2.30719 | 0.53988 | 0.63103 | 0.81191 |
| **Trappc4** | 9.07123 | 4.20811 | 10.48433 | 3.87082 | 6.62271 | 7.87787 | 2.61247 | 9.63982 | 7.53590 | 15.19824 | 9.01799 | 5.73696 | 12.63554 | 3.84292 | 4.34329 | 2.90507 |
| **Trappc5** | 8.47478 | 0.00000 | 3.36016 | 1.32340 | 3.22997 | 0.02144 | 0.05930 | 4.43253 | 0.00000 | 4.19299 | 0.90264 | 0.02589 | 3.67406 | 0.00489 | 0.32388 | 2.61310 |
| **Trappc6a** | 4.09043 | 0.01171 | 0.28130 | 4.54795 | 2.64817 | 0.00245 | 0.19729 | 1.97753 | 1.35147 | 0.91878 | 0.20303 | 0.13674 | 3.15380 | 0.00523 | 0.00000 | 2.35288 |
| **Trappc6b** | 4.56635 | 0.00917 | 2.51336 | 5.48778 | 0.02051 | 3.48862 | 1.62200 | 2.03197 | 3.61717 | 4.47490 | 4.98674 | 5.68106 | 2.48515 | 0.63883 | 4.61925 | 3.81761 |
| **Trappc8** | 4.27220 | 5.98998 | 4.63012 | 6.44559 | 1.20730 | 5.23089 | 4.64756 | 6.83209 | 7.82186 | 0.12025 | 4.44952 | 17.58892 | 2.32008 | 15.33319 | 12.51419 | 0.87542 |
| **Trappc9** | 1.61160 | 0.28117 | 1.18284 | 26.78313 | 0.76147 | 2.40848 | 2.57689 | 1.55492 | 22.22185 | 6.11979 | 2.22053 | 1.95046 | 0.42129 | 9.41242 | 14.03981 | 0.40333 |
| **Trdmt1** | 0.38328 | 3.04068 | 0.51745 | 0.02238 | 0.20387 | 0.01107 | 1.10970 | 0.00808 | 0.04226 | 0.00000 | 0.15841 | 0.00000 | 0.07330 | 3.93060 | 1.84648 | 0.01292 |
| **Trerf1** | 0.01358 | 0.00809 | 0.01168 | 0.38528 | 0.00919 | 0.00658 | 0.00740 | 0.01218 | 1.20255 | 0.12257 | 0.00415 | 0.01543 | 0.01318 | 0.01052 | 0.00358 | 0.00765 |
| **Trex1** | 4.47200 | 3.54140 | 5.46493 | 0.01677 | 0.10657 | 1.08557 | 0.69101 | 1.10926 | 0.00954 | 0.89301 | 0.03450 | 0.09058 | 0.14081 | 0.00988 | 0.00382 | 0.25984 |
| **Trf** | 0.01327 | 0.00791 | 0.01211 | 0.07048 | 0.00898 | 0.00669 | 0.00733 | 0.01071 | 0.00831 | 0.92760 | 0.01145 | 0.00571 | 0.01173 | 0.01135 | 0.00778 | 0.00762 |
| **Trhr2** | 0.01110 | 0.00784 | 0.59989 | 0.01093 | 0.00813 | 0.00923 | 0.00717 | 0.01117 | 0.01227 | 0.00809 | 0.01174 | 0.00748 | 0.25510 | 0.01561 | 0.00130 | 0.00696 |
| **Triap1** | 10.35683 | 8.73234 | 11.92199 | 2.39260 | 1.31842 | 9.53608 | 9.59191 | 12.22203 | 2.00151 | 12.37748 | 3.66273 | 1.51356 | 10.57081 | 2.45456 | 8.55015 | 4.79594 |
| **Trib1** | 2.87099 | 3.64238 | 2.38331 | 1.05961 | 0.02906 | 1.14992 | 0.00000 | 7.00532 | 0.30964 | 1.73094 | 5.65120 | 1.00441 | 6.31101 | 6.57157 | 0.39915 | 2.11485 |
| **Trib2** | 0.01342 | 0.00791 | 0.01031 | 2.72771 | 0.00898 | 0.00665 | 0.00734 | 0.00611 | 0.05902 | 0.78726 | 0.22995 | 0.00553 | 0.00997 | 0.00903 | 0.02622 | 0.00688 |
| **Trib3** | 0.23418 | 0.37060 | 0.00000 | 9.90402 | 0.00766 | 0.00533 | 0.00600 | 9.68935 | 4.90941 | 0.78660 | 1.23606 | 0.12913 | 0.00923 | 0.13131 | 1.53878 | 0.01965 |
| **Trim11** | 5.28481 | 2.48737 | 1.39470 | 3.18672 | 3.18740 | 0.98670 | 3.22270 | 1.24514 | 5.39941 | 0.00000 | 0.67160 | 0.13564 | 2.45462 | 0.00000 | 0.32415 | 0.25691 |
| **Trim12c** | 0.01229 | 0.00808 | 0.10110 | 0.10704 | 0.01818 | 0.07685 | 0.02932 | 0.07349 | 0.28093 | 0.30092 | 0.09294 | 0.06272 | 0.05926 | 0.06120 | 0.38775 | 0.10227 |
| **Trim13** | 5.57459 | 0.00000 | 6.26639 | 12.87884 | 1.37453 | 4.16321 | 7.03195 | 3.76280 | 15.59684 | 5.39822 | 3.91222 | 0.93885 | 1.31016 | 12.72240 | 12.30371 | 0.07129 |
| **Trim14** | 0.18649 | 0.01522 | 0.12941 | 0.21836 | 3.17912 | 0.01277 | 1.68527 | 0.06484 | 0.34142 | 0.39322 | 1.97898 | 0.02015 | 0.07034 | 0.10558 | 0.32686 | 0.02043 |
| **Trim16** | 0.05406 | 0.00779 | 0.00943 | 0.00803 | 0.00797 | 0.00681 | 0.00731 | 0.01270 | 0.01191 | 0.67721 | 0.01232 | 0.02498 | 0.01188 | 0.01167 | 0.00875 | 0.00767 |
| **Trim17** | 0.01137 | 0.00796 | 0.15605 | 0.03586 | 0.00900 | 0.00903 | 0.00728 | 0.01222 | 0.01253 | 0.00904 | 0.01324 | 0.00771 | 0.26048 | 0.01302 | 0.00719 | 0.00719 |
| **Trim2** | 0.03598 | 0.00765 | 0.01573 | 0.03259 | 0.01834 | 3.75068 | 0.00719 | 0.01484 | 0.05320 | 0.32263 | 0.01249 | 0.01524 | 0.02308 | 0.21649 | 0.40211 | 0.03575 |
| **Trim23** | 0.10704 | 1.42846 | 0.26582 | 0.26027 | 0.15899 | 0.17595 | 0.13518 | 0.88175 | 0.00549 | 1.28050 | 0.00370 | 0.00000 | 0.16530 | 0.01010 | 1.48452 | 0.08474 |
| **Trim24** | 11.24817 | 2.50003 | 5.85373 | 8.45523 | 4.88270 | 7.33150 | 3.26967 | 17.41537 | 13.25772 | 17.42398 | 12.05772 | 17.28753 | 6.05034 | 12.13643 | 12.04707 | 6.86073 |
| **Trim25** | 5.71584 | 4.33708 | 7.59398 | 3.44905 | 11.50018 | 2.42023 | 2.80850 | 5.04493 | 4.59356 | 4.57237 | 4.87721 | 5.48448 | 2.60371 | 2.69099 | 1.69793 | 7.94812 |
| **Trim26** | 1.98650 | 0.01554 | 2.87562 | 2.11332 | 0.01033 | 2.26389 | 0.01363 | 0.00000 | 0.80265 | 1.03113 | 1.14796 | 0.01140 | 2.17795 | 0.00083 | 0.53640 | 3.87865 |
| **Trim27** | 10.32046 | 5.59750 | 11.39550 | 6.78494 | 9.14373 | 3.85044 | 5.06157 | 5.70408 | 11.78300 | 15.05263 | 5.91684 | 4.62920 | 7.79542 | 2.61798 | 8.16775 | 5.17328 |
| **Trim28** | 39.20524 | 23.77249 | 28.82337 | 21.13725 | 27.86708 | 9.65792 | 21.99133 | 38.14875 | 30.97825 | 38.34564 | 32.27486 | 14.07646 | 34.63085 | 7.85692 | 16.36563 | 24.19702 |
| **Trim3** | 0.03202 | 0.00668 | 0.57208 | 2.23900 | 0.02836 | 0.00629 | 1.02068 | 1.62693 | 0.00000 | 0.00000 | 0.44417 | 0.03641 | 0.37593 | 0.00060 | 0.07986 | 0.00656 |
| **Trim31** | 0.01275 | 0.00789 | 0.59093 | 0.33970 | 0.00816 | 0.00866 | 0.00730 | 0.01229 | 0.00452 | 0.01431 | 0.01141 | 0.01765 | 0.00892 | 0.01144 | 0.00000 | 0.00709 |
| **Trim32** | 1.36745 | 0.00438 | 0.11362 | 1.39679 | 2.93146 | 0.73966 | 0.00521 | 0.87648 | 0.44329 | 0.12354 | 0.00839 | 0.08611 | 1.67578 | 0.00464 | 0.65592 | 2.46298 |
| **Trim33** | 5.12728 | 8.09215 | 4.37781 | 14.11760 | 9.46919 | 2.28177 | 7.36213 | 8.63140 | 19.09705 | 17.60313 | 7.12965 | 17.14297 | 1.28972 | 23.72310 | 19.30463 | 1.70196 |
| **Trim34a** | 0.01767 | 0.00821 | 0.01173 | 0.43905 | 0.00895 | 0.00660 | 0.00751 | 0.01425 | 0.67474 | 0.01169 | 0.00877 | 0.00728 | 0.01279 | 0.01172 | 0.00366 | 0.01458 |
| **Trim35** | 0.83229 | 0.00752 | 0.00419 | 0.00852 | 0.01801 | 0.00698 | 0.00656 | 0.00710 | 0.08405 | 0.60419 | 0.01141 | 0.04284 | 0.01144 | 0.01200 | 0.00628 | 0.00684 |
| **Trim36** | 0.06473 | 0.06481 | 1.17525 | 2.99916 | 0.03854 | 0.04212 | 0.06252 | 0.23893 | 1.69149 | 0.14528 | 0.51719 | 0.12875 | 0.10490 | 0.39515 | 1.31049 | 0.07867 |
| **Trim37** | 0.53472 | 4.72213 | 1.76695 | 6.07990 | 2.65691 | 0.02020 | 2.98642 | 3.08310 | 3.28791 | 5.74588 | 3.55552 | 12.54061 | 1.24538 | 10.64879 | 5.79787 | 1.03791 |
| **Trim38** | 3.11503 | 1.77257 | 10.81866 | 0.29032 | 5.33718 | 0.01532 | 0.23994 | 2.70638 | 0.00000 | 0.00000 | 0.40143 | 0.03304 | 0.04141 | 0.00289 | 0.00000 | 0.00000 |
| **Trim39** | 3.28608 | 1.73482 | 2.47405 | 2.57825 | 0.03442 | 2.48585 | 0.61056 | 5.41960 | 2.28923 | 5.15584 | 6.50341 | 0.24854 | 6.69620 | 14.29636 | 8.61963 | 2.62406 |
| **Trim40** | 0.01316 | 0.00801 | 0.45033 | 0.01048 | 0.00858 | 0.00568 | 0.00722 | 0.00000 | 0.00877 | 0.75650 | 0.60480 | 0.00000 | 0.00581 | 2.72262 | 0.66183 | 0.00705 |
| **Trim41** | 4.46136 | 3.57603 | 3.66757 | 11.32504 | 4.21996 | 6.90782 | 7.00729 | 4.94428 | 8.67593 | 1.34182 | 3.14176 | 0.35640 | 2.69292 | 0.83630 | 0.12336 | 2.34507 |
| **Trim43a** | 16.77860 | 21.63907 | 28.34253 | 14.93651 | 10.70714 | 8.46085 | 15.07984 | 35.02231 | 23.80447 | 48.36180 | 36.41618 | 22.16537 | 20.94998 | 53.52126 | 38.12878 | 3.89226 |
| **Trim43b** | 9.67831 | 15.74476 | 23.12216 | 18.62555 | 5.18737 | 4.56527 | 10.74278 | 29.79263 | 29.22294 | 47.16138 | 40.37176 | 20.92684 | 16.70211 | 55.23356 | 40.14847 | 1.97437 |
| **Trim43c** | 6.14753 | 10.37520 | 16.65973 | 14.31384 | 2.59336 | 2.47186 | 6.06649 | 18.90243 | 22.33667 | 35.70045 | 33.44706 | 17.88708 | 8.65070 | 42.16029 | 30.69043 | 1.40092 |
| **Trim44** | 6.79032 | 2.33013 | 0.85526 | 10.90438 | 0.03966 | 0.00000 | 5.06542 | 0.00000 | 4.81381 | 4.12183 | 0.36195 | 10.90587 | 0.12706 | 11.30405 | 8.18481 | 3.53010 |
| **Trim45** | 1.26374 | 0.00750 | 0.00384 | 0.15662 | 0.01676 | 0.00690 | 0.00650 | 0.00995 | 0.00641 | 0.01540 | 0.01195 | 0.00738 | 0.01794 | 0.01169 | 0.00836 | 0.00627 |
| **Trim47** | 1.64759 | 1.54114 | 4.09430 | 0.14479 | 0.00000 | 0.00468 | 1.65737 | 1.29192 | 0.00621 | 0.04879 | 0.00000 | 0.00852 | 0.26169 | 0.00521 | 0.00000 | 0.02515 |
| **Trim5** | 0.04294 | 0.00770 | 0.01452 | 0.00889 | 0.00815 | 0.01450 | 0.01747 | 0.02191 | 0.02697 | 0.04060 | 0.01165 | 0.00731 | 0.02621 | 0.08756 | 0.04348 | 0.00730 |
| **Trim52** | 0.01520 | 0.00767 | 0.03477 | 0.27732 | 0.00893 | 0.00719 | 0.00714 | 0.17693 | 0.52107 | 0.11192 | 0.12303 | 0.14650 | 0.00974 | 0.21752 | 0.20454 | 0.00702 |
| **Trim54** | 1.23051 | 0.00654 | 0.00286 | 0.03177 | 0.00603 | 0.00762 | 0.00686 | 0.95242 | 0.01173 | 1.12683 | 0.08946 | 0.01285 | 0.09985 | 0.01314 | 0.00000 | 0.00596 |
| **Trim56** | 0.01422 | 0.00758 | 0.01972 | 0.00906 | 0.00909 | 0.00627 | 0.00725 | 0.01590 | 0.01182 | 0.45093 | 0.01228 | 0.01558 | 0.01845 | 0.01433 | 0.03508 | 0.01531 |
| **Trim59** | 2.60015 | 4.65015 | 0.17161 | 7.62046 | 0.24248 | 0.47873 | 0.01191 | 0.33028 | 10.87590 | 0.00000 | 1.73395 | 3.17874 | 0.91055 | 3.22496 | 3.88206 | 0.18679 |
| **Trim6** | 4.33391 | 0.86730 | 3.07939 | 2.56209 | 4.15977 | 0.04436 | 3.38904 | 5.18617 | 6.56229 | 6.32115 | 5.83956 | 11.66714 | 4.72216 | 13.16112 | 13.06659 | 2.76973 |
| **Trim60** | 0.01701 | 0.00765 | 0.00876 | 3.15112 | 0.00909 | 0.00590 | 0.00712 | 0.00999 | 1.04494 | 0.39615 | 0.00000 | 0.00000 | 0.01234 | 0.11189 | 2.03930 | 0.00685 |
| **Trim61** | 0.00000 | 0.00684 | 0.42431 | 21.90125 | 0.00898 | 0.00259 | 0.00580 | 0.00000 | 5.02353 | 0.00000 | 0.46400 | 3.86881 | 0.00693 | 4.37683 | 18.22265 | 0.00279 |
| **Trim62** | 0.01281 | 0.00774 | 0.01133 | 0.00633 | 0.00905 | 0.00645 | 0.00747 | 0.01173 | 0.83076 | 0.02791 | 0.00481 | 0.00542 | 0.01172 | 0.01422 | 0.39313 | 0.10616 |
| **Trim63** | 0.01413 | 0.00821 | 0.02038 | 0.00579 | 0.00938 | 0.00620 | 0.00757 | 0.01308 | 2.97939 | 0.00308 | 0.00000 | 0.00291 | 0.01662 | 0.05693 | 0.00066 | 0.00732 |
| **Trim65** | 0.01336 | 0.00761 | 0.00912 | 0.00484 | 0.00934 | 0.00680 | 0.00727 | 0.00946 | 0.27801 | 0.02411 | 0.03404 | 0.00521 | 0.01288 | 0.46263 | 0.00000 | 0.00764 |
| **Trim66** | 0.11325 | 0.00746 | 0.00000 | 0.01396 | 0.00752 | 0.00717 | 0.00693 | 0.70549 | 0.16327 | 1.37191 | 0.23013 | 0.03620 | 0.00488 | 1.97608 | 0.25744 | 0.00713 |
| **Trim67** | 0.01619 | 0.00739 | 0.00490 | 0.27828 | 0.01003 | 0.00656 | 0.00695 | 0.01530 | 0.00685 | 0.05391 | 0.00000 | 2.18498 | 0.21014 | 0.06786 | 0.56423 | 0.00712 |
| **Trim68** | 0.01375 | 0.00833 | 0.01224 | 0.35522 | 0.00932 | 0.00655 | 0.00752 | 0.01312 | 0.00392 | 0.04094 | 0.01133 | 0.00825 | 0.01221 | 0.00657 | 0.13606 | 0.00718 |
| **Trim69** | 0.01629 | 0.00800 | 0.12066 | 0.47752 | 0.00901 | 0.00571 | 0.00738 | 0.01306 | 0.43017 | 0.01301 | 0.01115 | 0.00736 | 0.01107 | 0.01158 | 0.00192 | 0.00721 |

| | | | | | | | | | | | | | | | | |
|---|---|---|---|---|---|---|---|---|---|---|---|---|---|---|---|---|
| **Trim7** | 0.41323 | 0.01737 | 0.59116 | 0.05418 | 0.05850 | 0.20527 | 0.74188 | 8.11043 | 0.05488 | 6.22971 | 0.00000 | 0.79288 | 2.96485 | 3.66838 | 7.37030 | 0.10777 |
| **Trim71** | 0.01501 | 0.00743 | 0.05414 | 5.26287 | 0.00930 | 0.00521 | 0.00701 | 0.00944 | 1.87695 | 0.00000 | 0.00000 | 0.00000 | 0.01228 | 0.31517 | 0.62383 | 0.00632 |
| **Trim75** | 0.01371 | 0.00767 | 0.01122 | 1.43861 | 0.00943 | 0.00631 | 0.00730 | 0.01109 | 0.15544 | 0.00454 | 0.00962 | 0.01577 | 0.01284 | 0.00695 | 0.84938 | 0.00709 |
| **Trim8** | 0.34746 | 0.10104 | 2.22297 | 1.53771 | 1.22849 | 0.60325 | 0.00630 | 0.64223 | 1.17874 | 1.45067 | 0.48413 | 0.01466 | 1.13266 | 0.05640 | 0.00000 | 1.75263 |
| **Triml1** | 2.23200 | 2.68816 | 1.12069 | 0.00000 | 0.00624 | 0.00651 | 0.00439 | 0.00559 | 0.34288 | 0.00103 | 0.00869 | 0.00535 | 0.43505 | 0.00836 | 0.30927 | 0.90607 |
| **Triml2** | 7.09597 | 0.00983 | 5.59305 | 1.04447 | 0.02294 | 6.49924 | 0.00000 | 2.96127 | 2.49542 | 8.63578 | 11.59468 | 0.18953 | 7.84632 | 8.92554 | 2.01066 | 3.65640 |
| **Trio** | 0.05796 | 0.00797 | 0.11836 | 1.19023 | 0.81055 | 0.00786 | 0.00819 | 0.33010 | 0.00000 | 1.32000 | 0.01036 | 0.02452 | 0.12688 | 0.01007 | 0.00000 | 0.07600 |
| **Triobp** | 3.08886 | 0.00363 | 0.00000 | 0.58077 | 0.28222 | 0.55851 | 0.00453 | 4.29502 | 0.25097 | 4.24769 | 0.96852 | 0.09251 | 0.27481 | 0.00310 | 0.00000 | 2.66744 |
| **Trip10** | 0.27543 | 0.00732 | 0.00000 | 0.00717 | 0.71036 | 0.12191 | 0.00768 | 0.21051 | 0.01140 | 0.04255 | 0.05386 | 0.05841 | 0.13143 | 0.00647 | 0.19324 | 0.03638 |
| **Trip11** | 0.20933 | 0.00742 | 0.00360 | 1.89314 | 0.00755 | 0.00586 | 0.00734 | 0.01030 | 0.96512 | 0.01764 | 0.00000 | 0.07574 | 0.01364 | 0.00268 | 5.41328 | 0.00672 |
| **Trip12** | 18.03460 | 12.79363 | 15.35392 | 26.45884 | 15.81772 | 10.06616 | 10.98180 | 21.13386 | 25.79967 | 21.90829 | 21.25797 | 17.83376 | 11.41326 | 34.93138 | 27.22983 | 9.66607 |
| **Trip13** | 10.25380 | 8.73387 | 13.06375 | 7.69573 | 8.21889 | 1.93346 | 7.38193 | 13.51013 | 10.44573 | 23.61755 | 15.66553 | 14.31359 | 11.54397 | 22.70626 | 19.88656 | 1.43149 |
| **Trip4** | 3.68211 | 5.31335 | 3.02213 | 4.26627 | 3.16323 | 0.03748 | 3.88481 | 2.87752 | 5.91653 | 5.18363 | 5.18497 | 1.41571 | 2.92679 | 9.40519 | 7.11107 | 3.06132 |
| **Trip6** | 1.35544 | 1.11425 | 0.20283 | 0.16194 | 0.00519 | 0.00427 | 1.34642 | 0.00000 | 0.51145 | 0.25440 | 0.68269 | 0.00146 | 0.00000 | 0.00479 | 0.02224 | 4.36837 |
| **Trit1** | 6.27084 | 4.94236 | 1.16546 | 0.68204 | 2.98408 | 1.62718 | 2.36520 | 3.40771 | 0.27488 | 3.38453 | 1.48935 | 0.13074 | 3.49854 | 1.58684 | 0.62357 | 0.15945 |
| **Trmo** | 0.41354 | 0.67426 | 0.01423 | 0.00000 | 0.09421 | 0.00558 | 1.40247 | 1.17758 | 0.47133 | 2.45386 | 1.37506 | 0.07159 | 0.44329 | 0.52818 | 2.03527 | 0.00609 |
| **Trmt1** | 15.82970 | 2.64218 | 6.25619 | 1.09120 | 1.37938 | 3.45257 | 9.83302 | 11.83201 | 1.40343 | 4.13700 | 4.84793 | 0.57239 | 16.05271 | 0.00000 | 0.00000 | 7.25471 |
| **Trmt10a** | 1.51410 | 7.86591 | 4.69578 | 1.08779 | 7.40231 | 4.13307 | 3.85403 | 4.11098 | 3.66779 | 4.82246 | 3.49088 | 2.10946 | 2.13042 | 1.50689 | 5.99568 | 0.04056 |
| **Trmt10b** | 5.05518 | 2.61435 | 4.89579 | 1.10236 | 0.00000 | 0.45071 | 3.12191 | 1.38979 | 0.00000 | 2.39649 | 0.99753 | 0.00000 | 3.86816 | 5.11916 | 0.75744 | 0.00000 |
| **Trmt10c** | 2.46948 | 4.82039 | 1.80857 | 2.08795 | 2.78838 | 0.03821 | 0.03267 | 4.37230 | 4.77214 | 3.06623 | 1.86749 | 1.25149 | 0.95518 | 7.76800 | 7.47076 | 3.11032 |
| **Trmt11** | 3.17874 | 0.15777 | 3.00921 | 2.34866 | 5.17473 | 0.00253 | 4.00351 | 3.98534 | 2.82982 | 2.50304 | 2.89024 | 8.16512 | 0.62298 | 6.61214 | 2.78233 | 0.92502 |
| **Trmt112** | 12.00872 | 0.07292 | 2.88528 | 10.10359 | 10.23187 | 4.29162 | 3.44673 | 9.55571 | 10.15823 | 13.20281 | 11.04803 | 0.59495 | 13.36038 | 0.34626 | 0.00000 | 3.41014 |
| **Trmt12** | 0.41320 | 3.04726 | 1.09226 | 3.14018 | 0.01908 | 0.00441 | 2.93072 | 1.28851 | 10.33075 | 0.41261 | 0.78421 | 0.09914 | 1.44518 | 0.01796 | 2.44685 | 1.36328 |
| **Trmt13** | 0.13527 | 1.56052 | 0.12796 | 0.00938 | 0.11211 | 0.74361 | 0.00682 | 0.65752 | 0.00861 | 0.38547 | 0.00000 | 0.82879 | 0.18451 | 0.37597 | 2.31925 | 0.00648 |
| **Trmt1l** | 5.32275 | 1.23567 | 5.44557 | 1.71594 | 3.69836 | 3.72480 | 2.30384 | 12.75171 | 0.56197 | 7.41293 | 3.69464 | 8.55472 | 3.11589 | 9.55696 | 4.25412 | 1.49438 |
| **Trmt2a** | 7.11369 | 5.81973 | 6.37790 | 1.45267 | 6.54487 | 6.37856 | 4.35784 | 12.29334 | 1.56271 | 6.84464 | 2.44305 | 4.46947 | 6.80371 | 0.46928 | 5.33945 | 1.99223 |
| **Trmt2b** | 0.00000 | 0.00851 | 0.70824 | 0.95604 | 0.00883 | 0.03553 | 0.00785 | 0.04262 | 2.23018 | 2.24958 | 0.00000 | 0.20566 | 0.24564 | 3.04284 | 0.91401 | 0.15730 |
| **Trmt44** | 1.13887 | 2.04084 | 1.49099 | 0.30609 | 0.04441 | 0.18346 | 0.00563 | 1.26678 | 0.82252 | 0.41483 | 1.14920 | 0.07879 | 0.28768 | 0.00334 | 0.62487 | 1.02850 |
| **Trmt5** | 3.31289 | 3.87381 | 4.66917 | 1.49906 | 3.63227 | 5.50575 | 0.00768 | 4.41441 | 1.17448 | 9.84161 | 3.72821 | 9.66751 | 4.33230 | 3.22579 | 7.49916 | 1.35629 |
| **Trmt6** | 6.03626 | 11.16730 | 10.97230 | 8.33344 | 6.44544 | 4.74236 | 7.90702 | 15.21411 | 6.34134 | 11.12780 | 7.87947 | 5.88629 | 11.09262 | 9.33830 | 13.01041 | 8.54294 |
| **Trmt61a** | 2.04128 | 1.54138 | 1.22173 | 7.05353 | 4.38129 | 0.00264 | 0.00266 | 2.17199 | 8.77531 | 8.16035 | 3.74698 | 0.45473 | 1.39227 | 0.13045 | 3.09703 | 0.06623 |
| **Trmt61b** | 0.00000 | 0.01676 | 0.18492 | 1.19997 | 1.75004 | 0.00530 | 1.02109 | 2.37591 | 2.02058 | 9.72461 | 1.07081 | 1.96053 | 1.44429 | 1.32334 | 1.32655 | 0.02051 |
| **Trmu** | 1.17919 | 0.00736 | 0.44488 | 0.08084 | 0.01189 | 0.98899 | 0.01818 | 0.12029 | 0.01130 | 0.00000 | 0.62133 | 0.02259 | 0.08662 | 0.00491 | 0.15871 | 5.11567 |
| **Trnau1ap** | 4.25339 | 3.78137 | 1.41116 | 8.14974 | 6.29068 | 0.23498 | 0.10077 | 0.26665 | 7.86240 | 4.07949 | 6.62100 | 0.51871 | 2.88736 | 0.77471 | 1.87258 | 0.55177 |
| **Trnp1** | 0.00957 | 0.00710 | 0.00000 | 6.59597 | 0.01277 | 0.00913 | 1.49279 | 1.19751 | 1.86662 | 0.03088 | 1.90020 | 0.02192 | 0.01874 | 0.00000 | 0.12799 | 0.00593 |
| **Trnt1** | 4.64070 | 4.38259 | 2.11344 | 5.85512 | 1.40111 | 2.94140 | 3.12591 | 3.71845 | 10.06906 | 3.97926 | 5.48902 | 7.83629 | 2.40256 | 11.06848 | 10.66236 | 2.35107 |
| **Tro** | 0.01265 | 0.01894 | 0.00949 | 0.00715 | 0.00907 | 0.01351 | 0.00743 | 0.02131 | 0.30564 | 0.39622 | 0.06828 | 0.00699 | 0.02436 | 0.10142 | 0.01006 | 0.00733 |
| **Troap** | 1.35836 | 0.00574 | 0.15218 | 1.68428 | 0.36444 | 0.45665 | 0.51290 | 1.63195 | 4.83415 | 2.37152 | 0.86499 | 0.04250 | 0.26894 | 0.46370 | 0.09832 | 4.83351 |
| **Trove2** | 0.13816 | 0.01812 | 0.70911 | 1.72586 | 0.00965 | 0.04979 | 0.01552 | 0.67488 | 0.28559 | 0.29179 | 3.34126 | 0.70468 | 0.16356 | 5.67562 | 2.11914 | 0.04273 |
| **Trp53** | 11.33375 | 6.68507 | 0.84450 | 0.15094 | 6.74295 | 0.01091 | 3.57393 | 0.83252 | 0.02052 | 0.00003 | 0.28718 | 0.10850 | 3.47139 | 3.05116 | 3.88331 | 1.51326 |
| **Trp53bp1** | 3.66446 | 2.28310 | 4.83668 | 3.88972 | 6.63283 | 2.67873 | 3.25817 | 6.62772 | 3.29106 | 11.68826 | 1.07202 | 8.77913 | 1.13052 | 6.45656 | 14.22681 | 0.83936 |
| **Trp53bp2** | 0.16405 | 0.22253 | 1.04603 | 6.24356 | 0.04586 | 5.09142 | 2.33287 | 1.35517 | 0.59923 | 1.90852 | 0.00000 | 0.02412 | 11.01479 | 0.02004 | 0.02592 | 0.01440 |
| **Trp53i11** | 0.63587 | 0.00578 | 1.39482 | 0.00000 | 0.02074 | 3.64950 | 2.63669 | 0.65576 | 0.03011 | 0.13680 | 1.98162 | 0.00000 | 2.61775 | 0.01626 | 0.00000 | 0.01785 |
| **Trp53i13** | 0.01530 | 0.00801 | 0.01213 | 0.01678 | 0.01865 | 0.00676 | 0.00762 | 0.01910 | 0.07486 | 0.00919 | 0.02603 | 0.00761 | 0.22274 | 0.00690 | 0.13012 | 0.00730 |
| **Trp53inp1** | 2.08995 | 0.00295 | 0.12063 | 6.55763 | 3.52071 | 0.50343 | 0.60904 | 1.56032 | 4.50539 | 0.67947 | 2.27316 | 0.65361 | 0.77155 | 12.27476 | 2.78243 | 0.00998 |
| **Trp53inp2** | 1.23387 | 1.22824 | 3.21605 | 7.89909 | 0.00784 | 4.64760 | 3.02001 | 2.42995 | 3.80093 | 3.16585 | 3.39393 | 0.31986 | 3.52388 | 4.17282 | 5.84924 | 1.75002 |
| **Trp53rka** | 0.51989 | 0.06761 | 2.88633 | 1.64811 | 0.00001 | 0.04086 | 3.17663 | 0.19786 | 2.24476 | 0.26583 | 0.91471 | 0.21428 | 0.42738 | 0.01921 | 1.62645 | 2.00358 |
| **Trp53rkb** | 0.00000 | 0.02531 | 1.27115 | 2.04437 | 2.44647 | 0.00746 | 0.57201 | 0.93751 | 2.40946 | 1.49446 | 1.17426 | 1.66215 | 0.00000 | 0.03994 | 0.50142 | 0.05054 |
| **Trp73** | 0.00000 | 1.87368 | 0.00625 | 1.09966 | 0.00785 | 0.00627 | 2.88659 | 0.03326 | 0.49443 | 0.46221 | 0.00458 | 0.01124 | 0.00950 | 0.84084 | 0.00000 | 0.00657 |
| **Trpa1** | 0.01012 | 0.00869 | 0.00864 | 0.21657 | 0.00950 | 0.00692 | 0.00823 | 0.21783 | 0.00527 | 0.14647 | 0.02434 | 0.00920 | 0.00970 | 0.00739 | 0.30116 | 0.00726 |
| **Trpc1** | 0.00632 | 0.02834 | 0.09664 | 0.09452 | 0.10294 | 0.00718 | 0.01782 | 0.07554 | 1.81618 | 0.08183 | 0.12472 | 0.02273 | 0.09838 | 0.07239 | 0.28443 | 0.01428 |
| **Trpc4ap** | 3.39464 | 1.62682 | 4.66695 | 1.75864 | 3.58709 | 4.18428 | 2.30780 | 4.78716 | 2.92845 | 10.20888 | 3.24380 | 0.47504 | 3.31250 | 0.00000 | 1.01307 | 3.31212 |
| **Trpc5** | 0.07696 | 0.04953 | 0.00000 | 0.52417 | 0.01946 | 1.26710 | 0.01682 | 0.00000 | 0.74960 | 0.00000 | 0.44017 | 0.03388 | 0.02190 | 2.27668 | 0.11702 | 0.02100 |
| **Trpc5os** | 0.00533 | 0.00837 | 0.53477 | 0.00318 | 0.00842 | 0.44777 | 0.00711 | 0.17849 | 0.00967 | 0.00000 | 1.44347 | 0.80273 | 1.13002 | 0.00856 | 0.14681 | 0.00602 |
| **Trpc6** | 1.77693 | 2.07601 | 7.09346 | 6.50619 | 0.88400 | 3.02292 | 3.14909 | 12.23431 | 14.68476 | 11.84921 | 12.36950 | 12.95775 | 2.04889 | 9.90743 | 6.06410 | 1.42694 |
| **Trpd52l3** | 0.01380 | 0.00789 | 0.01173 | 0.01044 | 0.00933 | 0.00663 | 0.00745 | 0.01284 | 0.01238 | 0.03464 | 0.01338 | 0.03832 | 0.01252 | 0.01249 | 0.01421 | 0.00735 |
| **Trpm3** | 0.01674 | 0.04052 | 0.01471 | 0.07784 | 0.01035 | 0.01262 | 0.01768 | 0.02066 | 0.04630 | 0.02605 | 0.02653 | 0.00772 | 0.01869 | 0.05632 | 0.02275 | 0.00743 |
| **Trpm4** | 0.47421 | 0.01761 | 0.35018 | 0.15490 | 0.01568 | 1.68446 | 0.00676 | 0.90752 | 0.00849 | 0.38505 | 0.19189 | 0.01355 | 0.30906 | 0.00889 | 0.00000 | 0.01422 |
| **Trpm6** | 0.21149 | 0.00730 | 1.06302 | 0.00987 | 0.01381 | 0.00930 | 0.00722 | 0.01025 | 0.01203 | 0.02262 | 0.04137 | 0.00755 | 0.03311 | 0.01544 | 0.00005 | 0.00670 |
| **Trpm7** | 6.09846 | 9.19382 | 3.76376 | 7.74015 | 5.59940 | 5.21036 | 5.39504 | 1.30186 | 13.07179 | 1.63898 | 1.66003 | 14.49660 | 0.55637 | 11.01370 | 12.50061 | 2.11363 |
| **Trps1** | 0.26470 | 0.00715 | 0.00540 | 2.25475 | 0.01574 | 0.00671 | 0.00713 | 0.01027 | 0.00000 | 0.00386 | 0.03956 | 0.01651 | 0.01189 | 0.01368 | 0.40335 | 0.01385 |
| **Trpt1** | 0.00741 | 0.01944 | 0.00009 | 2.11074 | 0.03118 | 4.28176 | 0.00610 | 1.66250 | 1.34645 | 0.69204 | 1.20043 | 0.13992 | 1.77651 | 0.00638 | 0.39066 | 0.00479 |

| | | | | | | | | | | | | | | | | |
|---|---|---|---|---|---|---|---|---|---|---|---|---|---|---|---|---|
| **Trpv1** | 1.91000 | 0.00570 | 0.00377 | 0.01060 | 0.00452 | 0.00665 | 0.00606 | 0.01467 | 0.01235 | 0.01499 | 0.01313 | 0.00666 | 0.02561 | 0.01230 | 0.01086 | 0.00543 |
| **Trpv2** | 0.01355 | 0.00818 | 0.01127 | 0.86615 | 0.00926 | 0.00665 | 0.00750 | 0.01247 | 0.23321 | 0.10434 | 0.00811 | 0.00735 | 0.01334 | 0.00438 | 0.82638 | 0.00726 |
| **Trpv4** | 0.11767 | 0.00785 | 0.00733 | 0.01752 | 0.00765 | 1.77683 | 0.00726 | 0.01270 | 0.01273 | 0.00816 | 0.01044 | 0.00826 | 0.00413 | 0.01281 | 0.01004 | 0.00642 |
| **Trpv5** | 0.13454 | 0.00802 | 0.00000 | 0.01801 | 0.00788 | 1.48114 | 0.00805 | 0.04044 | 0.01320 | 0.02194 | 0.00917 | 0.00795 | 0.00841 | 0.17427 | 0.00375 | 0.01391 |
| **Trpv6** | 0.16697 | 0.00753 | 0.02108 | 0.01062 | 0.00753 | 0.00783 | 0.00743 | 0.01263 | 0.01235 | 0.01535 | 0.01293 | 0.00793 | 0.01225 | 0.01241 | 0.01051 | 0.00719 |
| **Trrap** | 3.25102 | 2.73572 | 2.71382 | 2.61171 | 2.27120 | 0.00335 | 1.94221 | 2.41008 | 3.15105 | 1.99871 | 0.62460 | 0.48193 | 0.08179 | 0.07625 | 2.89820 | 0.72334 |
| **Trub1** | 3.73816 | 1.03827 | 7.90222 | 0.05675 | 3.03913 | 2.25574 | 1.14337 | 8.07081 | 0.94295 | 0.80219 | 2.71567 | 0.26694 | 2.54082 | 0.69782 | 0.46463 | 1.17023 |
| **Trub2** | 1.70466 | 0.41582 | 5.55608 | 13.08456 | 0.10501 | 4.01732 | 5.95385 | 6.98368 | 17.91368 | 6.27250 | 3.25134 | 0.20794 | 7.67641 | 1.54962 | 5.86311 | 4.58055 |
| **Tsacc** | 0.93091 | 0.10699 | 0.34177 | 0.47281 | 0.23794 | 0.00524 | 0.04674 | 0.10131 | 0.38280 | 0.14382 | 1.25215 | 0.04911 | 0.15428 | 0.00798 | 0.36167 | 0.00582 |
| **Tsc1** | 0.92995 | 2.56060 | 4.68081 | 6.76431 | 0.46688 | 0.08257 | 0.30171 | 2.31160 | 7.17589 | 3.91785 | 2.61934 | 0.48893 | 0.00000 | 0.33381 | 3.03680 | 0.00164 |
| **Tsc2** | 6.49033 | 0.00437 | 5.70529 | 1.01441 | 3.65626 | 10.12212 | 4.79006 | 7.99885 | 2.94843 | 6.46915 | 7.54259 | 2.86922 | 6.38475 | 0.80021 | 3.54390 | 2.30752 |
| **Tsc22d1** | 6.40147 | 0.01441 | 2.35260 | 10.32270 | 11.14506 | 3.88491 | 1.69552 | 4.93098 | 9.22169 | 9.35462 | 1.31579 | 0.18023 | 2.07445 | 0.00000 | 1.77743 | 14.30894 |
| **Tsc22d2** | 0.14726 | 1.26281 | 2.53448 | 21.56155 | 2.88539 | 0.12011 | 0.04349 | 1.77435 | 21.15319 | 6.02135 | 3.34860 | 5.54382 | 0.57669 | 8.65970 | 18.30552 | 0.00000 |
| **Tsc22d3** | 0.01357 | 0.00773 | 0.01168 | 0.32509 | 0.00916 | 0.00671 | 0.00745 | 0.01188 | 0.19144 | 0.34186 | 0.01048 | 0.00692 | 0.01253 | 0.01340 | 0.14722 | 0.00741 |
| **Tsc22d4** | 4.10685 | 0.01255 | 0.00000 | 11.10624 | 0.01030 | 2.64407 | 0.10286 | 3.52543 | 13.02288 | 8.94352 | 9.65522 | 0.15854 | 3.81455 | 0.35278 | 1.67366 | 0.60655 |
| **Tsen15** | 6.88334 | 0.87409 | 1.37436 | 3.95753 | 4.76959 | 2.23650 | 0.25008 | 1.37594 | 3.13775 | 3.15652 | 1.10521 | 0.06383 | 5.33389 | 1.42893 | 2.84988 | 0.94211 |
| **Tsen2** | 3.41601 | 3.92574 | 4.14495 | 2.38959 | 3.91111 | 1.54524 | 0.00000 | 7.58359 | 5.08980 | 8.25254 | 4.96369 | 1.04102 | 1.41844 | 0.00000 | 3.15144 | 0.04235 |
| **Tsen34** | 7.32526 | 2.95018 | 4.08546 | 1.66495 | 10.09270 | 4.19677 | 3.82554 | 5.62683 | 4.73247 | 12.05154 | 10.05421 | 3.46507 | 2.44231 | 0.09888 | 1.10208 | 0.69612 |
| **Tsen54** | 5.03121 | 0.42980 | 0.00000 | 1.91920 | 0.00974 | 2.85508 | 0.01343 | 0.83307 | 1.81523 | 2.84038 | 1.40255 | 0.13114 | 1.35482 | 0.00000 | 1.21196 | 0.42961 |
| **Tsfm** | 8.23815 | 0.06277 | 2.17091 | 0.05981 | 0.94887 | 0.00920 | 0.32799 | 3.11818 | 0.02043 | 5.05177 | 0.39138 | 0.09275 | 3.84090 | 0.00784 | 0.00000 | 5.08763 |
| **Tsg101** | 12.25159 | 7.04484 | 5.86954 | 4.81564 | 5.78638 | 0.02024 | 3.55884 | 7.19424 | 7.60772 | 8.44578 | 3.70069 | 2.55001 | 11.54379 | 6.46023 | 6.42497 | 5.17886 |
| **Tsga10** | 0.01332 | 0.00760 | 0.00673 | 0.94506 | 0.00958 | 0.02506 | 0.00725 | 0.01087 | 0.00000 | 0.05552 | 0.00747 | 0.00414 | 0.01285 | 0.59569 | 0.53753 | 0.00731 |
| **Tsga8** | 0.01290 | 0.00774 | 0.01218 | 0.02395 | 0.00888 | 0.00648 | 0.00721 | 0.00972 | 0.05079 | 1.62873 | 0.01137 | 0.00354 | 0.01075 | 0.01423 | 0.00559 | 0.00739 |
| **Tshz1** | 0.13259 | 0.00779 | 0.11659 | 0.15425 | 0.01626 | 1.08238 | 0.00690 | 0.01363 | 0.19667 | 0.14203 | 0.01517 | 0.38779 | 0.00519 | 0.16123 | 0.36298 | 0.16397 |
| **Tshz2** | 0.01340 | 0.00755 | 0.00909 | 1.66763 | 0.00902 | 0.00626 | 0.00716 | 0.00463 | 2.07321 | 0.33130 | 0.16159 | 0.00938 | 0.01082 | 0.00112 | 0.72779 | 0.00734 |
| **Tsix** | 0.01161 | 0.00803 | 0.05804 | 0.06991 | 0.00892 | 0.00589 | 0.00759 | 0.01328 | 0.01118 | 0.07398 | 0.01488 | 0.00869 | 0.01203 | 0.02103 | 0.02351 | 0.16948 |
| **Tsks** | 0.01214 | 0.00775 | 0.00758 | 0.85505 | 0.00891 | 0.00661 | 0.00706 | 0.02204 | 2.57357 | 0.64710 | 0.50618 | 0.00000 | 0.00667 | 0.00577 | 0.02652 | 0.00690 |
| **Tsku** | 0.00815 | 0.00787 | 0.07001 | 0.00876 | 0.00775 | 0.00847 | 0.00737 | 0.01170 | 0.01216 | 0.01043 | 0.07146 | 0.02809 | 0.01464 | 0.01211 | 0.00783 | 0.86310 |
| **Tslp** | 0.01301 | 0.00760 | 0.00959 | 0.92398 | 0.00956 | 0.00604 | 0.00724 | 0.01019 | 0.16500 | 0.20462 | 0.00581 | 0.01794 | 0.01270 | 0.00442 | 3.40643 | 0.00730 |
| **Tsn** | 18.85863 | 12.72587 | 18.32821 | 2.60509 | 9.61967 | 6.75176 | 11.17517 | 8.51850 | 6.04962 | 17.04275 | 7.14101 | 5.81421 | 13.63805 | 8.39772 | 9.12253 | 8.76429 |
| **Tsnax** | 4.43772 | 5.99055 | 4.96164 | 3.03891 | 4.67188 | 3.81093 | 6.93443 | 2.01491 | 6.16828 | 4.96851 | 4.97947 | 7.85356 | 3.57845 | 1.85174 | 6.91552 | 2.93811 |
| **Tsnaxip1** | 0.01218 | 0.00712 | 0.06449 | 2.45759 | 0.00932 | 0.00550 | 0.00641 | 0.62507 | 0.00000 | 0.00000 | 1.49857 | 0.03245 | 0.00000 | 0.00000 | 0.64754 | 0.00642 |
| **Tspan1** | 0.91497 | 3.36753 | 1.74839 | 0.00000 | 0.05489 | 1.13787 | 2.90130 | 6.07407 | 0.04002 | 1.55348 | 3.61620 | 0.56408 | 6.15523 | 3.42630 | 0.00000 | 5.32128 |
| **Tspan10** | 0.00495 | 0.00782 | 1.63343 | 0.00353 | 0.00825 | 0.00684 | 1.52270 | 0.36727 | 0.49716 | 0.00490 | 0.00450 | 0.00587 | 0.00000 | 0.01839 | 0.00000 | 0.00676 |
| **Tspan13** | 0.01437 | 0.00731 | 0.00738 | 0.39289 | 0.01073 | 0.00720 | 0.00718 | 0.01143 | 0.13092 | 2.28103 | 0.00000 | 1.83693 | 0.18186 | 0.00973 | 0.00000 | 0.00768 |
| **Tspan14** | 1.57068 | 0.31060 | 2.83176 | 0.92832 | 0.01545 | 3.11887 | 2.17916 | 3.76702 | 1.69467 | 1.81544 | 2.09676 | 0.84095 | 0.00000 | 0.74694 | 0.44863 | 2.75449 |
| **Tspan15** | 0.70117 | 0.01894 | 0.00000 | 0.24353 | 0.02334 | 2.74057 | 0.02957 | 0.83357 | 0.23414 | 0.26765 | 0.53564 | 0.04720 | 0.14017 | 0.23960 | 0.01043 | 0.01249 |
| **Tspan17** | 3.43269 | 0.00263 | 1.80606 | 0.47359 | 1.81963 | 0.00340 | 0.00397 | 3.60348 | 2.40415 | 1.46084 | 1.73339 | 0.01059 | 0.55747 | 0.04530 | 0.00000 | 6.22700 |
| **Tspan3** | 4.55674 | 0.10965 | 9.99444 | 0.50938 | 0.29540 | 0.09048 | 2.49781 | 2.09226 | 0.00523 | 1.22797 | 0.00000 | 0.10137 | 2.50862 | 0.00536 | 0.00000 | 0.28831 |
| **Tspan31** | 8.71798 | 5.41538 | 7.96237 | 0.67918 | 5.89122 | 4.47073 | 4.96700 | 1.28255 | 0.54434 | 6.17973 | 1.74280 | 0.51029 | 7.06078 | 5.24620 | 7.27597 | 6.68944 |
| **Tspan33** | 0.02405 | 0.00829 | 0.00211 | 1.03502 | 0.01241 | 1.91128 | 0.00750 | 0.18901 | 0.20695 | 0.30679 | 0.00802 | 0.00699 | 0.14900 | 0.00421 | 0.17980 | 0.00694 |
| **Tspan4** | 1.42801 | 0.60049 | 0.41272 | 0.51905 | 0.02922 | 0.00766 | 0.00395 | 4.99214 | 0.00000 | 18.23863 | 1.00960 | 0.82716 | 3.11775 | 0.42018 | 3.68477 | 0.06217 |
| **Tspan5** | 0.01321 | 0.00769 | 0.00945 | 6.58346 | 0.00896 | 0.00590 | 0.00696 | 0.00779 | 1.72993 | 0.71512 | 0.05220 | 0.06810 | 0.01155 | 0.00000 | 0.54522 | 0.00656 |
| **Tspan6** | 0.01915 | 0.00778 | 0.05228 | 0.25421 | 0.00855 | 0.00844 | 0.00720 | 0.01006 | 0.00473 | 0.45850 | 0.01064 | 0.00550 | 0.01182 | 0.00549 | 0.49110 | 0.00773 |
| **Tspan7** | 0.01236 | 0.00794 | 0.69743 | 0.00930 | 0.00798 | 0.00568 | 0.00741 | 0.01072 | 0.01205 | 0.36297 | 0.01189 | 0.00659 | 0.00911 | 0.01410 | 0.00132 | 0.00751 |
| **Tspan8** | 0.00000 | 0.01775 | 0.30288 | 0.58548 | 0.22381 | 20.75750 | 0.22238 | 0.00000 | 1.27826 | 2.61940 | 0.00000 | 0.01481 | 3.19639 | 0.05167 | 0.40817 | 7.62256 |
| **Tspan9** | 1.11751 | 0.00557 | 0.05327 | 1.62078 | 0.00560 | 0.00781 | 0.92116 | 0.36611 | 0.00000 | 1.01112 | 1.25655 | 2.56054 | 0.79420 | 0.00404 | 0.00000 | 0.00515 |
| **Tspo** | 0.13849 | 0.00798 | 0.04386 | 0.08006 | 0.00863 | 0.00716 | 0.00738 | 0.01293 | 0.01076 | 1.42264 | 0.01360 | 0.00376 | 0.18959 | 0.01139 | 0.00547 | 0.00733 |
| **Tspyl1** | 6.97522 | 2.86143 | 5.38515 | 3.20036 | 2.36085 | 0.16769 | 1.46707 | 0.07652 | 3.44214 | 5.20297 | 0.30948 | 0.11982 | 1.68376 | 2.37797 | 4.68809 | 1.17615 |
| **Tspyl2** | 0.00046 | 0.00836 | 0.03986 | 3.67437 | 0.01294 | 0.00622 | 1.55146 | 0.62445 | 0.32116 | 0.00268 | 0.00479 | 0.07657 | 0.54592 | 0.00520 | 1.50706 | 0.00695 |
| **Tspyl3** | 0.00792 | 0.00839 | 0.00915 | 0.01066 | 0.01092 | 0.00742 | 0.00839 | 0.58325 | 0.01176 | 0.00000 | 0.00837 | 0.00586 | 0.05551 | 0.41273 | 0.00000 | 0.00783 |
| **Tspyl4** | 0.01227 | 0.00763 | 0.01110 | 1.67823 | 0.00944 | 0.00643 | 0.00723 | 0.05564 | 0.06823 | 0.00798 | 0.01151 | 0.01642 | 0.01019 | 0.00337 | 0.20773 | 0.00707 |
| **Tsr1** | 4.38854 | 5.79360 | 5.37919 | 3.74353 | 4.28039 | 2.04280 | 4.20439 | 9.73652 | 6.59615 | 10.48837 | 6.78230 | 8.61796 | 3.34519 | 14.83665 | 11.11870 | 3.48816 |
| **Tsr2** | 8.81928 | 10.78427 | 16.88010 | 0.00000 | 6.63306 | 1.55756 | 7.41873 | 17.04914 | 0.02909 | 20.30513 | 2.83763 | 2.73898 | 8.56519 | 15.17845 | 14.59911 | 0.00000 |
| **Tsr3** | 3.74529 | 0.01232 | 1.73119 | 4.60307 | 0.23129 | 0.59287 | 1.81025 | 1.39066 | 6.17773 | 6.38984 | 7.85034 | 0.35685 | 7.00074 | 0.00000 | 1.04092 | 4.96852 |
| **Tssc1** | 7.21411 | 1.80217 | 8.76863 | 1.94868 | 5.14869 | 0.00388 | 4.46124 | 7.67021 | 4.80522 | 6.81884 | 2.92816 | 0.26354 | 5.25249 | 0.00000 | 0.92047 | 0.27780 |
| **Tssc4** | 0.09730 | 2.09579 | 3.72775 | 5.68895 | 2.32655 | 0.27438 | 0.00293 | 3.18093 | 5.14559 | 6.56904 | 0.85917 | 0.15047 | 5.10172 | 0.00000 | 0.45732 | 0.06641 |
| **Tst** | 3.52320 | 0.36732 | 0.05360 | 0.00755 | 0.01366 | 0.00576 | 0.00430 | 0.00000 | 0.01200 | 0.67211 | 0.01123 | 0.00403 | 0.51095 | 0.01015 | 0.00511 | 0.00339 |
| **Tsta3** | 10.57997 | 6.74986 | 3.55820 | 0.91105 | 5.39853 | 0.00000 | 2.58345 | 3.87727 | 0.31782 | 2.59467 | 1.65667 | 0.01543 | 6.48685 | 0.01747 | 1.02637 | 6.13656 |
| **Tstd1** | 0.00777 | 0.15593 | 1.10087 | 0.14706 | 0.00669 | 1.83992 | 0.00629 | 0.81123 | 2.90671 | 0.05832 | 1.71190 | 0.06483 | 0.44595 | 0.01744 | 0.15759 | 0.82517 |
| **Tstd2** | 0.07682 | 0.00746 | 0.38566 | 0.00688 | 0.01195 | 2.91186 | 1.68896 | 0.21326 | 0.01136 | 0.51565 | 0.05382 | 0.02519 | 0.17733 | 0.00895 | 0.63161 | 0.01342 |
| **Tstd3** | 3.79315 | 0.00325 | 1.24438 | 0.55388 | 0.15866 | 1.17785 | 0.04429 | 0.00000 | 0.23170 | 0.31708 | 1.49306 | 0.15135 | 0.74256 | 1.98196 | 0.35327 | 2.01523 |

| | | | | | | | | | | | | | | | | |
|---|---|---|---|---|---|---|---|---|---|---|---|---|---|---|---|---|
| **Tsx** | 1.05011 | 0.00767 | 0.47521 | 0.54175 | 0.01542 | 0.00518 | 0.00655 | 0.00000 | 0.83727 | 0.31040 | 0.53404 | 0.06687 | 0.20051 | 0.53425 | 1.32208 | 0.02210 |
| **Ttbk1** | 0.00977 | 0.00769 | 0.00878 | 0.04597 | 0.00960 | 0.00673 | 0.00715 | 0.31557 | 0.61110 | 0.00702 | 0.00647 | 0.00606 | 0.01035 | 0.01444 | 0.00150 | 0.00733 |
| **Ttbk2** | 2.10873 | 0.00000 | 0.34762 | 2.84335 | 3.62692 | 0.00854 | 2.23729 | 0.91090 | 3.46343 | 1.20994 | 1.19961 | 4.38720 | 1.68711 | 0.32736 | 5.36917 | 0.05469 |
| **Ttc1** | 20.71489 | 7.21319 | 22.39054 | 20.67739 | 12.98133 | 4.06058 | 9.87998 | 16.86422 | 21.94468 | 25.00299 | 13.62172 | 20.12555 | 14.30943 | 22.68198 | 17.87649 | 5.48934 |
| **Ttc12** | 0.01233 | 0.00779 | 0.92242 | 0.23955 | 0.00764 | 0.39928 | 0.00721 | 0.01151 | 0.00530 | 0.01071 | 0.01043 | 0.00714 | 0.00661 | 0.01063 | 0.00857 | 0.00683 |
| **Ttc13** | 2.72274 | 2.95259 | 3.38149 | 0.40822 | 5.04104 | 2.05416 | 2.52616 | 9.03077 | 0.76890 | 5.07735 | 5.82522 | 1.74481 | 6.28631 | 0.31739 | 2.42797 | 2.91605 |
| **Ttc14** | 0.35157 | 0.20851 | 2.66464 | 9.38294 | 2.28598 | 0.63003 | 1.00691 | 0.00022 | 8.33134 | 2.71292 | 1.90104 | 12.51445 | 0.13408 | 0.58182 | 9.32395 | 0.00974 |
| **Ttc16** | 0.01368 | 0.00791 | 0.01164 | 2.45145 | 0.00921 | 0.00631 | 0.00740 | 0.01193 | 0.78419 | 0.01943 | 0.06139 | 0.00628 | 0.01139 | 0.00484 | 0.06013 | 0.00680 |
| **Ttc17** | 1.91156 | 0.24580 | 0.83116 | 0.33583 | 0.00255 | 1.76648 | 4.37774 | 2.64000 | 0.45598 | 1.29026 | 0.00000 | 0.30125 | 1.13047 | 3.24138 | 1.87131 | 1.25250 |
| **Ttc19** | 2.67860 | 2.67374 | 2.66888 | 2.89061 | 3.87386 | 1.84900 | 0.54022 | 0.12385 | 2.81349 | 3.93354 | 1.39704 | 2.24458 | 1.22887 | 4.26825 | 2.94953 | 3.89978 |
| **Ttc21a** | 0.56037 | 0.00727 | 0.42048 | 0.31698 | 0.00614 | 0.00501 | 0.00676 | 0.00000 | 0.00729 | 0.15969 | 0.65830 | 0.00350 | 0.00485 | 0.01392 | 0.03858 | 0.00611 |
| **Ttc23** | 0.00880 | 0.00733 | 0.00857 | 0.35564 | 0.31924 | 0.00689 | 0.00726 | 0.01905 | 0.28920 | 1.11856 | 0.01984 | 0.00376 | 0.01068 | 0.01304 | 0.04847 | 0.00750 |
| **Ttc23l** | 0.01372 | 0.00811 | 0.01220 | 0.16261 | 0.00930 | 0.00663 | 0.00753 | 0.01389 | 0.19458 | 0.01438 | 0.01276 | 0.00790 | 0.01272 | 0.01193 | 0.00636 | 0.00738 |
| **Ttc25** | 1.26545 | 0.68397 | 0.00957 | 0.05041 | 0.00520 | 0.00543 | 0.00639 | 2.44628 | 0.04711 | 0.18995 | 0.34177 | 0.00149 | 0.00401 | 0.05339 | 0.24392 | 0.02008 |
| **Ttc26** | 0.11869 | 0.01597 | 0.29576 | 0.62833 | 2.84949 | 0.01318 | 0.07857 | 0.00000 | 0.87287 | 0.72656 | 1.24392 | 0.03611 | 0.50709 | 0.93585 | 1.78020 | 0.05102 |
| **Ttc27** | 4.83340 | 7.72894 | 4.10265 | 8.77441 | 4.15782 | 3.24807 | 3.97148 | 12.35967 | 5.39067 | 12.34254 | 8.52131 | 9.11894 | 4.97377 | 6.08002 | 8.36916 | 2.32703 |
| **Ttc28** | 0.35496 | 0.00838 | 0.00419 | 0.53024 | 0.02018 | 0.00754 | 0.00729 | 0.01047 | 1.57607 | 0.77228 | 0.00000 | 0.07212 | 0.10094 | 0.00635 | 0.55810 | 0.00746 |
| **Ttc3** | 2.21648 | 0.00471 | 0.00000 | 0.74339 | 0.91660 | 0.29699 | 0.12739 | 0.00558 | 1.21065 | 3.52780 | 0.03556 | 0.25506 | 0.83117 | 2.56732 | 6.39066 | 0.04946 |
| **Ttc30a1** | 0.63768 | 0.00546 | 0.00000 | 0.00000 | 0.00652 | 0.00631 | 1.89563 | 0.09528 | 0.00843 | 1.71165 | 0.00699 | 0.01556 | 0.00836 | 0.01179 | 1.69695 | 0.00685 |
| **Ttc30a2** | 0.01251 | 0.00798 | 0.01061 | 0.00909 | 0.00948 | 0.00677 | 0.09761 | 0.03179 | 0.01196 | 0.15226 | 0.01304 | 0.01609 | 0.01353 | 0.07421 | 0.10562 | 0.00779 |
| **Ttc30b** | 0.00769 | 0.00789 | 0.00644 | 1.50059 | 0.02059 | 0.00654 | 0.12758 | 0.23762 | 0.87517 | 0.18426 | 0.00000 | 0.02644 | 0.02856 | 1.47966 | 1.07219 | 0.00747 |
| **Ttc32** | 2.48458 | 0.00344 | 0.04505 | 1.79962 | 3.15396 | 0.00355 | 0.00534 | 0.02137 | 0.00000 | 0.39671 | 0.01410 | 0.01980 | 0.47357 | 0.00372 | 0.07647 | 0.17904 |
| **Ttc33** | 4.19983 | 5.80262 | 3.69159 | 3.30076 | 5.02910 | 0.01238 | 3.58681 | 4.69900 | 1.86783 | 3.67155 | 4.00381 | 0.96305 | 4.21942 | 5.83277 | 9.89184 | 3.39309 |
| **Ttc34** | 0.01375 | 0.00816 | 0.01231 | 0.52272 | 0.00931 | 0.00660 | 0.00751 | 0.01394 | 0.00455 | 0.01543 | 0.01393 | 0.00828 | 0.01266 | 0.01104 | 0.00509 | 0.00733 |
| **Ttc37** | 0.50326 | 0.92745 | 0.80151 | 0.12439 | 3.43385 | 0.00489 | 0.07314 | 0.81736 | 0.02223 | 0.23579 | 0.00000 | 0.82971 | 0.00811 | 0.00680 | 1.68601 | 0.00621 |
| **Ttc38** | 0.26738 | 0.03915 | 0.82748 | 1.13583 | 0.02145 | 0.02919 | 0.13557 | 0.05126 | 0.48416 | 0.98463 | 0.33744 | 0.08512 | 0.19348 | 0.48199 | 0.11309 | 0.02871 |
| **Ttc39a** | 0.00000 | 0.00767 | 0.00547 | 0.13403 | 0.00961 | 0.00732 | 0.00583 | 1.32809 | 0.54144 | 0.13092 | 0.38363 | 0.00000 | 1.88608 | 2.28262 | 0.47042 | 0.00681 |
| **Ttc39b** | 2.67321 | 0.20747 | 2.33621 | 0.78207 | 0.02966 | 0.04564 | 2.45137 | 2.38836 | 0.47966 | 0.90854 | 3.40016 | 7.10146 | 0.94735 | 3.31444 | 0.67121 | 1.58801 |
| **Ttc39c** | 0.94621 | 2.57224 | 0.00000 | 1.49873 | 2.30302 | 0.01313 | 0.00571 | 0.00443 | 0.00000 | 0.00189 | 0.01401 | 0.00265 | 0.36693 | 0.00351 | 0.84769 | 0.00512 |
| **Ttc4** | 2.85994 | 12.77977 | 7.77844 | 3.03363 | 2.83369 | 7.66167 | 4.54341 | 7.99910 | 1.10186 | 7.74753 | 5.50609 | 0.94902 | 5.17827 | 1.80632 | 7.54273 | 4.55714 |
| **Ttc41** | 2.57190 | 2.58744 | 0.21052 | 0.19389 | 0.00528 | 0.00893 | 0.00405 | 0.00000 | 0.00567 | 0.00000 | 0.01105 | 0.01516 | 0.89004 | 0.02286 | 0.26324 | 0.00317 |
| **Ttc5** | 3.39291 | 0.18719 | 0.24134 | 0.08784 | 0.05362 | 0.00445 | 0.04725 | 1.20058 | 0.35388 | 3.22948 | 1.07574 | 0.01570 | 0.51290 | 0.17591 | 1.58855 | 3.18608 |
| **Ttc6** | 0.01111 | 0.00748 | 0.00650 | 1.64677 | 0.00938 | 0.00654 | 0.00703 | 0.47218 | 0.00000 | 0.09592 | 0.01690 | 0.03104 | 0.00727 | 0.00191 | 1.02559 | 0.00746 |
| **Ttc7** | 0.04706 | 0.00815 | 0.20227 | 0.20475 | 0.00809 | 0.00896 | 0.00818 | 0.26263 | 0.45759 | 0.12023 | 0.00609 | 0.01733 | 0.00529 | 0.10183 | 0.01589 | 0.94193 |
| **Ttc7b** | 5.14197 | 0.46717 | 0.00000 | 0.71362 | 8.42937 | 6.66023 | 2.10176 | 13.67804 | 0.25313 | 4.71887 | 7.34310 | 1.07884 | 5.14135 | 2.30493 | 1.46924 | 1.37502 |
| **Ttc8** | 0.02561 | 0.00851 | 0.08247 | 0.74392 | 0.00958 | 0.00578 | 0.00762 | 0.01472 | 0.00000 | 0.01516 | 0.01276 | 0.00882 | 0.01138 | 0.01119 | 0.00288 | 0.00723 |
| **Ttc9** | 0.01657 | 0.00797 | 0.24941 | 2.08694 | 0.00877 | 0.00550 | 0.00727 | 0.01293 | 0.00000 | 0.01419 | 0.01340 | 0.00791 | 0.01652 | 0.00985 | 0.00000 | 0.00688 |
| **Ttc9c** | 3.08573 | 0.04502 | 1.08068 | 4.12213 | 0.11210 | 1.64716 | 0.11726 | 1.36260 | 3.68595 | 2.33583 | 2.15962 | 0.16496 | 1.98535 | 2.46586 | 2.29174 | 0.32472 |
| **Ttf1** | 1.59217 | 0.03405 | 2.49695 | 11.52861 | 0.71947 | 0.11155 | 0.84386 | 4.30964 | 16.45997 | 2.45061 | 9.24444 | 3.67690 | 2.09691 | 9.74001 | 8.08669 | 0.36323 |
| **Ttf2** | 3.90535 | 0.00228 | 3.25786 | 0.62198 | 0.01621 | 0.01706 | 2.15590 | 3.47803 | 0.14110 | 0.69180 | 1.25032 | 0.13422 | 0.88072 | 0.00064 | 1.11331 | 1.01735 |
| **Tti1** | 2.95585 | 0.19118 | 4.43234 | 4.93309 | 0.03907 | 0.01609 | 3.45228 | 4.22540 | 1.57236 | 6.11514 | 3.24069 | 0.69049 | 3.01316 | 0.00686 | 7.57971 | 1.45324 |
| **Tti2** | 2.69076 | 0.07098 | 0.70836 | 1.40508 | 0.01518 | 3.33359 | 1.92082 | 3.75144 | 3.41956 | 0.78813 | 2.20733 | 0.82393 | 0.48270 | 0.40432 | 2.40357 | 0.40938 |
| **Ttk** | 2.72870 | 9.43298 | 3.48741 | 5.96505 | 8.30488 | 3.18531 | 0.00948 | 4.24038 | 2.46263 | 4.15285 | 3.13743 | 4.49226 | 1.54485 | 5.62185 | 6.29795 | 0.02455 |
| **Ttl** | 2.87488 | 0.39659 | 5.95811 | 1.33312 | 0.15793 | 2.76452 | 3.10561 | 9.78221 | 2.48865 | 8.81268 | 7.14816 | 2.71684 | 6.96409 | 2.35164 | 1.93069 | 2.58183 |
| **Ttll1** | 1.39314 | 4.34668 | 4.83421 | 1.56003 | 1.91450 | 1.66598 | 0.01317 | 4.98150 | 2.39331 | 2.48018 | 0.00000 | 0.21059 | 3.04426 | 0.07051 | 0.80376 | 0.00000 |
| **Ttll10** | 0.00970 | 0.00720 | 0.10913 | 0.00585 | 0.00914 | 0.00581 | 0.00721 | 2.81334 | 0.06786 | 1.12837 | 0.13982 | 0.01694 | 0.01313 | 0.01375 | 0.03687 | 0.00750 |
| **Ttll11** | 0.01287 | 0.00774 | 0.01068 | 2.62984 | 0.00915 | 0.00634 | 0.00721 | 0.01079 | 0.07275 | 0.01703 | 0.00901 | 0.02819 | 0.18835 | 0.00214 | 0.77866 | 0.00680 |
| **Ttll12** | 7.26231 | 3.79234 | 5.73665 | 1.59462 | 7.08537 | 0.11405 | 5.54713 | 3.97247 | 4.64011 | 3.32039 | 4.55050 | 0.86209 | 4.43039 | 0.00000 | 1.24630 | 2.26820 |
| **Ttll13** | 0.35155 | 0.00787 | 0.00025 | 0.94634 | 0.00644 | 0.00725 | 0.00618 | 0.00896 | 2.65756 | 0.21365 | 0.00000 | 0.02427 | 1.47512 | 0.24501 | 0.29375 | 1.55521 |
| **Ttll2** | 0.01364 | 0.00811 | 0.01197 | 0.06621 | 0.00927 | 0.00661 | 0.00754 | 0.01385 | 0.74208 | 0.01178 | 0.00882 | 0.00705 | 0.01288 | 0.01314 | 0.00549 | 0.00736 |
| **Ttll3** | 0.02864 | 0.00670 | 0.91383 | 0.37052 | 0.02308 | 0.00864 | 0.92974 | 2.44262 | 0.67403 | 0.11727 | 1.39632 | 0.05355 | 1.07566 | 0.03320 | 0.00000 | 0.01385 |
| **Ttll4** | 4.74816 | 5.85389 | 0.54654 | 3.50024 | 0.02111 | 1.38961 | 0.00972 | 0.04762 | 3.07430 | 2.77158 | 0.24128 | 7.67181 | 2.24034 | 2.01715 | 6.07578 | 0.00000 |
| **Ttll5** | 2.99828 | 2.76752 | 2.40100 | 1.32383 | 0.01433 | 0.85724 | 0.00248 | 0.12601 | 1.49471 | 1.13367 | 1.32352 | 0.81430 | 1.56229 | 1.73963 | 3.54407 | 2.33773 |
| **Ttll6** | 0.58660 | 1.57289 | 6.80520 | 0.07682 | 0.00343 | 0.00584 | 0.84548 | 0.47812 | 0.66863 | 3.07500 | 0.98945 | 3.20288 | 1.35101 | 0.25113 | 0.19151 | 0.01681 |
| **Ttll9** | 0.10888 | 0.00783 | 0.00374 | 1.67509 | 0.00772 | 0.00707 | 0.00741 | 0.18187 | 0.84543 | 1.04408 | 0.00000 | 0.05292 | 0.00739 | 0.00188 | 0.43324 | 0.00690 |
| **Ttn** | 0.00670 | 0.00753 | 0.00000 | 0.15005 | 0.00977 | 0.00607 | 1.05664 | 0.01179 | 0.00548 | 0.84660 | 0.26617 | 0.02846 | 0.02193 | 6.75416 | 2.98488 | 0.00719 |
| **Ttpal** | 0.79479 | 2.57962 | 2.98580 | 1.15113 | 0.00000 | 4.19748 | 2.69605 | 2.12419 | 0.87136 | 4.31929 | 1.34372 | 0.05353 | 0.88736 | 4.31557 | 2.97637 | 0.98223 |
| **Ttyh1** | 0.01649 | 0.00762 | 0.59792 | 0.00409 | 0.00818 | 0.13895 | 0.00726 | 0.01093 | 1.12399 | 0.00918 | 0.01462 | 0.01519 | 0.00821 | 0.03184 | 0.00000 | 0.00701 |
| **Ttyh2** | 4.13290 | 1.57740 | 4.63658 | 0.99166 | 0.02543 | 3.36833 | 2.78030 | 2.79335 | 8.34618 | 3.32368 | 9.14510 | 0.30309 | 3.54125 | 4.05855 | 2.92914 | 1.94951 |
| **Ttyh3** | 1.60958 | 0.00592 | 0.10192 | 2.53405 | 0.00459 | 0.03293 | 0.04530 | 0.03333 | 3.50575 | 0.46385 | 0.30327 | 0.03737 | 0.55968 | 0.01684 | 0.13167 | 0.08935 |
| **Tub** | 1.32789 | 0.00639 | 1.91347 | 5.21860 | 0.02593 | 3.23861 | 0.00460 | 0.55292 | 0.05648 | 0.25906 | 3.42625 | 0.30029 | 0.00000 | 1.47557 | 4.99476 | 0.01066 |
| **Tuba1a** | 30.69051 | 14.80676 | 25.48957 | 2.85610 | 19.18803 | 10.13920 | 9.28740 | 17.63684 | 9.67197 | 17.22204 | 10.62857 | 3.55982 | 17.60294 | 3.18115 | 7.90602 | 15.12389 |

| | | | | | | | | | | | | | | | | |
|---|---|---|---|---|---|---|---|---|---|---|---|---|---|---|---|---|
| **Tuba1b** | 35.32724 | 17.71941 | 30.04707 | 2.73734 | 22.00586 | 11.58894 | 11.94233 | 22.19590 | 3.45859 | 23.49188 | 17.33644 | 5.44102 | 23.18100 | 4.89861 | 9.94793 | 19.14579 |
| **Tuba1c** | 38.48560 | 20.49451 | 33.07958 | 5.53625 | 23.39205 | 13.42082 | 13.23671 | 24.67884 | 8.10679 | 22.77991 | 14.96025 | 5.52374 | 23.08340 | 4.60216 | 10.94415 | 18.81166 |
| **Tuba3a** | 0.00000 | 0.00654 | 0.76126 | 1.93743 | 0.00737 | 0.00652 | 1.42898 | 2.07620 | 4.10273 | 1.61438 | 1.90023 | 0.20834 | 1.03456 | 0.00000 | 0.43257 | 0.45219 |
| **Tuba3b** | 0.00000 | 0.00388 | 3.05135 | 1.97478 | 0.00250 | 0.00528 | 6.07413 | 1.55495 | 5.74340 | 5.40941 | 2.00218 | 0.44180 | 0.04519 | 0.01203 | 1.05182 | 2.43068 |
| **Tuba4a** | 16.69796 | 7.96680 | 18.49394 | 5.09598 | 13.87690 | 4.76761 | 6.72632 | 18.23182 | 16.47301 | 2.32266 | 7.88372 | 0.31376 | 20.89631 | 0.16054 | 2.84217 | 5.74384 |
| **Tuba8** | 0.01180 | 0.00778 | 0.90612 | 0.00821 | 0.00761 | 0.00929 | 0.00727 | 0.01761 | 0.04598 | 0.02200 | 0.01088 | 0.00737 | 0.01478 | 0.00961 | 0.02519 | 0.02151 |
| **Tubb2a** | 0.50507 | 0.01816 | 0.52721 | 4.48899 | 0.13051 | 0.00535 | 0.03561 | 0.00000 | 0.00000 | 0.97239 | 1.22961 | 0.02487 | 1.31697 | 0.00000 | 0.00000 | 0.00561 |
| **Tubb2b** | 0.04688 | 0.00813 | 0.04683 | 0.39767 | 0.01320 | 0.00590 | 0.00728 | 0.00931 | 0.00396 | 0.16056 | 0.07424 | 0.00674 | 0.06862 | 0.01313 | 0.01787 | 0.00773 |
| **Tubb3** | 1.86200 | 0.00503 | 0.93519 | 5.48215 | 0.01502 | 0.00853 | 0.00395 | 3.63245 | 8.40081 | 2.65899 | 1.42400 | 0.06355 | 1.80337 | 0.00000 | 0.19046 | 0.01052 |
| **Tubb4b** | 27.47115 | 18.48870 | 33.35389 | 22.44909 | 23.95929 | 9.24209 | 11.73084 | 36.26298 | 36.15741 | 41.62648 | 30.64614 | 12.75792 | 29.75012 | 4.60990 | 23.78590 | 15.97733 |
| **Tubb5** | 37.02337 | 15.65525 | 21.08924 | 9.13493 | 17.86115 | 11.42639 | 13.35534 | 2.30349 | 20.27049 | 19.08311 | 11.87499 | 2.56096 | 17.33669 | 4.93847 | 4.96458 | 23.52900 |
| **Tubb6** | 1.03235 | 0.00619 | 0.52211 | 6.98114 | 0.41192 | 3.73947 | 0.51829 | 3.72348 | 2.06980 | 7.26695 | 1.44614 | 0.71721 | 7.80054 | 0.57026 | 0.76722 | 0.10917 |
| **Tubd1** | 0.83494 | 6.09618 | 0.00000 | 0.92815 | 0.00689 | 3.74003 | 0.00216 | 3.03393 | 0.01727 | 0.55849 | 1.05177 | 6.25926 | 1.73636 | 2.85026 | 0.41418 | 0.00897 |
| **Tube1** | 3.22569 | 0.00297 | 2.92567 | 1.11665 | 0.76218 | 1.85826 | 0.31270 | 0.00263 | 0.00000 | 1.02848 | 0.95557 | 0.48770 | 0.24250 | 0.03548 | 0.63516 | 0.03663 |
| **Tubg1** | 4.07741 | 5.55328 | 3.50671 | 4.93259 | 5.38590 | 3.19213 | 2.60204 | 8.79123 | 7.94494 | 11.28772 | 7.28344 | 0.62726 | 8.20771 | 0.06090 | 3.75503 | 6.04031 |
| **Tubg2** | 0.19032 | 0.43505 | 0.25696 | 12.29578 | 1.02947 | 0.34173 | 2.10895 | 2.13176 | 2.02317 | 1.77420 | 0.57190 | 0.11996 | 1.18823 | 0.00610 | 2.49592 | 1.18538 |
| **Tubgcp2** | 6.93602 | 1.98071 | 9.71716 | 2.93558 | 6.83486 | 2.91253 | 1.47559 | 4.33721 | 3.67654 | 2.51524 | 2.44684 | 0.14139 | 3.45838 | 0.91278 | 3.88662 | 2.81801 |
| **Tubgcp3** | 3.53787 | 1.29825 | 4.57438 | 1.62743 | 0.02370 | 2.26857 | 2.60487 | 6.92571 | 3.26621 | 11.39712 | 5.31338 | 2.45476 | 3.42762 | 0.34738 | 5.41454 | 2.45191 |
| **Tubgcp4** | 0.32815 | 4.16908 | 0.24172 | 0.32869 | 0.64381 | 1.82510 | 2.34595 | 0.44473 | 0.00494 | 0.20085 | 0.00529 | 0.02077 | 0.35491 | 0.00419 | 1.41391 | 2.09699 |
| **Tubgcp5** | 0.46538 | 6.79585 | 1.41481 | 1.15746 | 0.00534 | 0.72283 | 0.88361 | 3.04847 | 1.20665 | 0.00000 | 0.69200 | 7.92053 | 1.21579 | 1.31939 | 6.20817 | 0.00173 |
| **Tubgcp6** | 2.58796 | 1.74242 | 0.74132 | 1.94672 | 1.33548 | 1.18505 | 0.69801 | 0.66663 | 0.88294 | 1.99753 | 1.61431 | 0.48317 | 1.02113 | 0.27258 | 2.03665 | 0.02336 |
| **Tufm** | 18.29593 | 13.25009 | 18.56748 | 4.82954 | 18.63951 | 11.77330 | 11.10285 | 26.75659 | 6.28590 | 28.46354 | 10.90174 | 6.46606 | 23.38976 | 3.90199 | 12.22732 | 9.37148 |
| **Tuft1** | 4.76745 | 0.00000 | 5.68425 | 5.40285 | 0.05683 | 2.03754 | 0.00000 | 8.76010 | 13.65425 | 6.16072 | 6.51969 | 0.09092 | 5.20110 | 0.91276 | 3.68497 | 0.85817 |
| **Tug1** | 3.47491 | 0.81907 | 5.04176 | 4.19286 | 2.01400 | 3.32541 | 3.69945 | 4.63270 | 4.25514 | 6.12662 | 1.11490 | 17.85781 | 0.56154 | 13.64305 | 8.18428 | 3.04903 |
| **Tulp1** | 0.00379 | 0.00676 | 0.00829 | 0.08533 | 0.00836 | 0.00814 | 0.00712 | 6.62837 | 0.35254 | 4.26964 | 0.61116 | 0.07952 | 0.00347 | 0.00077 | 0.36414 | 0.00724 |
| **Tulp2** | 0.01510 | 0.00736 | 0.88298 | 0.78740 | 0.01725 | 0.01097 | 0.00699 | 0.00878 | 0.22649 | 0.67666 | 0.00892 | 0.00355 | 0.00847 | 0.00394 | 0.43870 | 0.00691 |
| **Tulp3** | 1.45936 | 1.10999 | 1.25711 | 9.41106 | 0.00290 | 1.01999 | 0.52006 | 0.00000 | 7.55437 | 1.46540 | 0.98904 | 0.51853 | 0.00000 | 2.93512 | 3.71819 | 0.57349 |
| **Tulp4** | 2.58528 | 0.00000 | 1.07304 | 7.07680 | 0.10140 | 0.42123 | 2.91416 | 0.09494 | 1.69263 | 6.31988 | 1.85357 | 9.07776 | 0.00000 | 3.51652 | 5.39765 | 0.00193 |
| **Tusc2** | 4.44782 | 4.23796 | 5.42639 | 0.60724 | 0.00736 | 3.46443 | 2.25213 | 1.07639 | 0.15893 | 1.27806 | 1.14927 | 0.27176 | 5.16155 | 0.04546 | 0.19629 | 0.01728 |
| **Tusc3** | 0.77885 | 0.76039 | 0.00000 | 0.61142 | 0.01579 | 3.79886 | 0.76292 | 0.00000 | 0.00000 | 1.48988 | 0.77326 | 1.36025 | 0.00000 | 0.00965 | 1.04778 | 0.44015 |
| **Tut1** | 4.11003 | 3.48201 | 1.60276 | 4.07309 | 2.34823 | 0.00199 | 4.87289 | 4.97575 | 6.04837 | 8.32556 | 5.19297 | 0.54484 | 4.53126 | 0.03219 | 4.09520 | 4.56822 |
| **Tvp23a** | 0.01152 | 0.00720 | 0.03307 | 0.01061 | 0.00934 | 0.00813 | 0.62698 | 0.00993 | 0.01180 | 0.00885 | 0.01212 | 0.00683 | 0.01091 | 0.01897 | 0.11144 | 0.00734 |
| **Tvp23b** | 2.41138 | 0.14428 | 2.83044 | 1.17281 | 0.05340 | 3.78936 | 4.52109 | 1.75706 | 0.39619 | 1.31853 | 1.70760 | 0.99688 | 5.36431 | 0.00845 | 4.64260 | 5.10878 |
| **Twf1** | 6.70824 | 8.55739 | 7.30684 | 4.16088 | 10.12755 | 2.66892 | 8.05545 | 4.69134 | 5.89280 | 8.78543 | 7.31093 | 10.38849 | 3.83834 | 6.84220 | 7.09409 | 5.03690 |
| **Twf2** | 4.61224 | 0.07377 | 0.30604 | 0.49405 | 0.00761 | 0.73400 | 0.99307 | 2.66937 | 0.34719 | 1.42252 | 0.63490 | 0.00921 | 1.72739 | 0.00420 | 0.00000 | 3.01926 |
| **Twistnb** | 7.85703 | 6.64535 | 2.42149 | 5.16415 | 1.10067 | 0.00202 | 4.04828 | 4.55791 | 6.37220 | 8.02415 | 3.42556 | 2.15082 | 3.56342 | 14.01610 | 6.58607 | 1.23218 |
| **Twsg1** | 0.79326 | 0.00727 | 0.00000 | 0.36076 | 0.01610 | 2.99880 | 0.00612 | 0.00000 | 0.98369 | 0.34056 | 0.84192 | 0.10979 | 0.03307 | 0.00920 | 0.36199 | 0.00541 |
| **Txk** | 0.00543 | 0.00825 | 0.55246 | 0.01851 | 0.00963 | 0.00556 | 0.00722 | 1.04493 | 0.34412 | 0.32788 | 0.00000 | 0.05147 | 0.00298 | 1.21002 | 0.45025 | 0.00717 |
| **Txlna** | 11.35365 | 1.86325 | 8.66426 | 7.80053 | 9.15805 | 5.30073 | 8.47240 | 15.27824 | 13.83013 | 16.08320 | 6.69463 | 3.46301 | 12.74052 | 0.48145 | 12.71190 | 7.63256 |
| **Txlng** | 4.49368 | 5.18600 | 4.30453 | 7.24864 | 1.22443 | 1.61457 | 0.00000 | 6.68827 | 13.18449 | 14.64851 | 5.28507 | 12.54395 | 1.89141 | 21.15580 | 19.87058 | 2.69693 |
| **Txn1** | 25.27350 | 3.50065 | 13.52185 | 0.76698 | 19.25616 | 7.04376 | 2.55308 | 0.64923 | 0.85805 | 1.96029 | 7.19620 | 0.65530 | 11.86322 | 0.95091 | 1.35123 | 12.15563 |
| **Txn2** | 16.90629 | 11.52677 | 14.59779 | 8.94427 | 12.33956 | 13.47861 | 12.64474 | 20.69100 | 5.10923 | 18.28419 | 7.72447 | 7.91916 | 24.32634 | 0.00000 | 3.39907 | 9.02997 |
| **Txndc11** | 5.89088 | 0.05733 | 0.04689 | 3.53373 | 0.23301 | 0.00000 | 0.10602 | 0.97697 | 2.77764 | 1.93798 | 0.48138 | 0.05780 | 2.27449 | 0.00924 | 0.07198 | 0.39290 |
| **Txndc12** | 15.42410 | 3.06946 | 13.60319 | 2.12486 | 7.84834 | 9.35631 | 4.08563 | 5.49646 | 1.32839 | 1.11142 | 2.15168 | 0.15996 | 7.69601 | 1.26588 | 0.11886 | 8.30194 |
| **Txndc15** | 2.59671 | 0.08312 | 0.97823 | 5.12136 | 0.02719 | 0.16005 | 2.65254 | 6.18945 | 3.81024 | 6.98031 | 2.71554 | 1.03899 | 1.52838 | 7.16499 | 0.69432 | 2.64353 |
| **Txndc16** | 0.01490 | 0.01855 | 1.33123 | 4.56659 | 0.03382 | 4.52390 | 0.00575 | 0.00000 | 0.77924 | 0.02434 | 0.29788 | 0.18971 | 0.99202 | 4.24520 | 3.60679 | 0.00386 |
| **Txndc17** | 12.04630 | 4.97928 | 8.53865 | 4.06938 | 4.59553 | 7.32885 | 7.77863 | 3.62807 | 2.27195 | 6.20404 | 3.63460 | 0.33741 | 8.65999 | 1.03428 | 2.01338 | 11.15156 |
| **Txndc2** | 0.01383 | 0.00739 | 0.00801 | 3.54641 | 0.00924 | 0.00637 | 0.00705 | 0.01023 | 0.48825 | 0.03001 | 0.00585 | 0.50428 | 0.01271 | 0.00062 | 0.26854 | 0.00658 |
| **Txndc5** | 1.96439 | 0.00407 | 0.99171 | 0.14802 | 2.39960 | 0.03498 | 0.00426 | 1.21913 | 0.10816 | 0.03749 | 0.03030 | 0.03719 | 3.15486 | 0.00551 | 1.32496 | 3.37038 |
| **Txndc9** | 21.15899 | 11.24860 | 19.49098 | 10.05431 | 8.34800 | 7.50967 | 6.90857 | 13.95257 | 8.35654 | 13.88296 | 11.94882 | 8.93294 | 12.63756 | 11.18939 | 12.23932 | 7.98998 |
| **Txnip** | 1.02031 | 0.00642 | 0.00000 | 2.02677 | 0.00546 | 0.00480 | 0.00626 | 0.00122 | 3.23986 | 4.91798 | 0.04611 | 0.00000 | 0.00867 | 1.16761 | 2.86086 | 0.95828 |
| **Txnl1** | 22.86981 | 12.27879 | 8.64496 | 9.74138 | 14.09636 | 9.79061 | 9.54897 | 16.65432 | 11.78889 | 17.69478 | 16.44949 | 13.98993 | 23.68060 | 7.75496 | 5.62298 | 9.69982 |
| **Txnl4a** | 20.57224 | 8.59591 | 16.52814 | 13.36208 | 13.50513 | 5.95757 | 9.39723 | 18.31517 | 13.57690 | 28.39131 | 19.63695 | 3.67301 | 17.49317 | 3.61639 | 9.02016 | 12.80313 |
| **Txnl4b** | 1.11454 | 0.00775 | 0.13816 | 0.80930 | 0.01612 | 1.27706 | 0.00630 | 0.83593 | 0.46714 | 0.72493 | 0.31758 | 0.01104 | 1.22151 | 0.00671 | 0.00000 | 2.83830 |
| **Txnrd1** | 5.54838 | 5.36470 | 5.52749 | 1.00664 | 8.54987 | 2.18503 | 6.77401 | 1.83540 | 0.30850 | 6.61594 | 2.49713 | 0.28460 | 3.93314 | 0.51424 | 3.68727 | 0.54915 |
| **Txnrd2** | 4.54945 | 2.94629 | 3.89091 | 0.00000 | 1.01074 | 0.00177 | 6.54030 | 5.82311 | 0.00931 | 2.51346 | 5.18973 | 0.14065 | 4.12183 | 0.00000 | 1.18051 | 2.22545 |
| **Txnrd3** | 0.01498 | 0.00784 | 0.18445 | 0.20345 | 0.00983 | 0.00612 | 0.00742 | 0.01227 | 0.24248 | 0.49773 | 0.01210 | 0.00541 | 0.12080 | 0.01238 | 0.00365 | 0.00765 |
| **Tyk2** | 3.12677 | 2.17741 | 8.14068 | 5.13666 | 0.00976 | 0.23539 | 1.06461 | 12.32627 | 4.17517 | 8.48666 | 11.02465 | 0.38503 | 1.62508 | 4.17919 | 5.47090 | 0.29214 |
| **Tymp** | 0.01834 | 0.00814 | 0.21807 | 0.16077 | 0.00822 | 0.00584 | 0.00757 | 0.01170 | 0.00886 | 0.25427 | 0.01237 | 0.00815 | 0.06380 | 0.01151 | 0.00399 | 0.07284 |
| **Tyms** | 15.76092 | 3.89746 | 8.42981 | 2.07918 | 12.67317 | 3.72893 | 2.24876 | 17.84934 | 9.13153 | 13.93113 | 16.08016 | 4.53219 | 13.72236 | 4.91705 | 2.75726 | 7.55218 |
| **Tyms.ps** | 1.58956 | 0.26273 | 0.61019 | 0.12618 | 1.39731 | 0.26089 | 0.19377 | 1.92716 | 0.98591 | 1.09227 | 2.19724 | 0.44703 | 1.57536 | 0.33177 | 0.12941 | 0.51613 |
| **Tyr** | 0.71308 | 0.07096 | 0.59738 | 3.82057 | 0.26458 | 0.07602 | 0.14630 | 0.00000 | 5.51687 | 0.10490 | 5.06690 | 0.33206 | 0.25770 | 2.45711 | 0.24601 | 0.12083 |

| | | | | | | | | | | | | | | | | |
|---|---|---|---|---|---|---|---|---|---|---|---|---|---|---|---|---|
| **Tyro3** | 2.69203 | 1.39307 | 4.35364 | 2.09232 | 0.07985 | 4.35304 | 5.75550 | 4.17156 | 1.35711 | 3.88360 | 0.37182 | 0.26023 | 0.90728 | 0.00325 | 0.49103 | 7.42755 |
| **Tysnd1** | 0.69374 | 0.02773 | 0.97174 | 3.21064 | 0.01388 | 0.01135 | 0.13686 | 0.00000 | 3.54817 | 3.59326 | 1.03626 | 0.16891 | 1.82029 | 0.00000 | 0.06074 | 0.60938 |
| **Tyw1** | 5.05344 | 2.19811 | 7.90569 | 2.45988 | 0.01216 | 0.01384 | 5.74232 | 5.12111 | 4.28939 | 4.69264 | 1.97477 | 0.27383 | 5.70151 | 0.12913 | 2.69863 | 1.39254 |
| **Tyw3** | 1.08362 | 0.32447 | 3.06491 | 0.16071 | 3.09848 | 0.01364 | 3.15895 | 1.97503 | 1.40654 | 7.06135 | 2.80992 | 0.18617 | 0.77442 | 0.03156 | 0.04219 | 0.00393 |
| **Tyw5** | 3.69017 | 1.36956 | 4.40611 | 3.37023 | 1.35598 | 3.55602 | 2.59004 | 3.71353 | 2.64563 | 3.45968 | 1.43784 | 9.71473 | 2.67202 | 6.23479 | 1.02207 | 0.26660 |
| **U2af1** | 11.59478 | 11.17216 | 8.36241 | 6.77774 | 2.28014 | 6.37092 | 5.58863 | 8.78159 | 11.93152 | 9.25319 | 9.11125 | 0.50509 | 7.21061 | 0.42228 | 1.59413 | 7.57812 |
| **U2af1l4** | 6.50499 | 4.96771 | 9.16996 | 3.36891 | 4.94869 | 5.13191 | 4.70606 | 3.45344 | 3.28753 | 4.62892 | 5.55377 | 0.34064 | 5.33450 | 0.08225 | 0.05076 | 1.99734 |
| **U2af2** | 3.71147 | 0.07438 | 4.43091 | 5.72357 | 2.03265 | 0.54924 | 0.82281 | 0.20020 | 12.27329 | 4.79341 | 14.92559 | 0.55680 | 1.54911 | 1.84672 | 2.09587 | 0.88209 |
| **U2surp** | 9.00558 | 12.57714 | 6.44484 | 4.66003 | 8.41430 | 5.50252 | 7.68050 | 7.42418 | 12.25697 | 16.00607 | 4.65498 | 9.84678 | 3.37969 | 18.56680 | 13.56739 | 2.42908 |
| **Uaca** | 5.72136 | 0.01262 | 0.00000 | 0.12374 | 0.12100 | 0.00261 | 0.00293 | 1.30191 | 0.00587 | 2.80245 | 0.48795 | 1.22907 | 0.00348 | 0.04706 | 0.56680 | 0.00000 |
| **Uap1** | 18.17450 | 14.31306 | 12.69278 | 3.82966 | 11.59338 | 7.23537 | 9.43250 | 19.36619 | 5.98340 | 24.88375 | 12.57920 | 6.74762 | 11.44697 | 29.68843 | 19.63575 | 10.05501 |
| **Uap1l1** | 9.09692 | 0.01559 | 3.55552 | 7.02550 | 5.33386 | 0.70262 | 0.02095 | 5.76116 | 3.03410 | 12.47558 | 11.17319 | 1.03483 | 11.39408 | 0.02073 | 2.67975 | 2.09327 |
| **Uba1** | 32.92222 | 18.48567 | 27.82487 | 19.16548 | 21.68020 | 10.81386 | 17.94733 | 37.18267 | 15.93761 | 30.66299 | 17.37752 | 15.89788 | 20.77910 | 12.93539 | 16.32386 | 16.09385 |
| **Uba1y** | 0.08646 | 0.07307 | 0.03954 | 0.00894 | 0.00770 | 0.00708 | 0.56664 | 0.00000 | 0.08623 | 3.04389 | 8.55465 | 6.84639 | 0.00000 | 0.07278 | 0.00000 | 0.02035 |
| **Uba2** | 21.77712 | 15.18997 | 14.03037 | 14.43614 | 12.44036 | 9.34699 | 10.87564 | 14.78593 | 15.56844 | 25.70029 | 12.71541 | 12.97307 | 13.99663 | 21.49138 | 21.30803 | 8.01428 |
| **Uba3** | 1.51500 | 8.60288 | 0.00000 | 5.27353 | 2.11698 | 4.03429 | 1.55350 | 6.82393 | 4.27771 | 0.57649 | 3.81349 | 5.52853 | 1.20739 | 2.31678 | 5.57622 | 0.46646 |
| **Uba5** | 10.42489 | 7.56460 | 5.97376 | 2.71332 | 8.45544 | 0.03218 | 7.42504 | 5.56391 | 9.13372 | 12.34444 | 12.41462 | 13.47609 | 3.97923 | 3.40515 | 4.33202 | 5.35307 |
| **Uba52** | 34.76432 | 5.89039 | 30.46335 | 23.81168 | 22.66853 | 10.50640 | 9.31848 | 14.91140 | 33.97331 | 36.21937 | 45.56433 | 8.52615 | 28.45576 | 9.66808 | 17.66878 | 17.64478 |
| **Uba6** | 0.19490 | 0.00489 | 0.13924 | 2.27103 | 0.01733 | 0.06770 | 1.12802 | 1.34374 | 4.11320 | 1.42094 | 0.00000 | 6.77236 | 0.25144 | 2.73144 | 3.55936 | 1.51235 |
| **Ubac1** | 9.71131 | 8.10357 | 5.16997 | 2.11063 | 9.07141 | 6.53147 | 1.42810 | 6.22040 | 4.04269 | 2.17316 | 7.28790 | 4.16035 | 3.90302 | 3.55503 | 2.88382 | 8.77593 |
| **Ubac2** | 1.88084 | 0.11515 | 4.17245 | 1.99168 | 0.30243 | 6.42966 | 1.32665 | 8.04648 | 3.46029 | 7.52978 | 3.25886 | 0.13877 | 2.82713 | 0.76009 | 0.45054 | 4.33812 |
| **Ubald1** | 2.88580 | 0.00288 | 0.00000 | 4.66478 | 0.98653 | 3.77253 | 0.06073 | 6.09830 | 5.41551 | 2.56235 | 5.41336 | 0.18823 | 5.51518 | 0.06333 | 0.16952 | 1.21271 |
| **Ubap1** | 6.02644 | 6.57534 | 5.14505 | 14.84902 | 6.26342 | 5.40167 | 3.73365 | 6.16878 | 17.88221 | 13.74688 | 6.65087 | 0.67213 | 4.11969 | 0.12062 | 5.04970 | 5.49358 |
| **Ubap2** | 7.78542 | 4.58298 | 3.53752 | 13.88239 | 1.47543 | 1.27880 | 3.06370 | 11.93043 | 11.07490 | 16.98118 | 11.16017 | 2.15384 | 16.42195 | 1.31249 | 4.55760 | 6.35427 |
| **Ubap2l** | 10.99431 | 8.59320 | 4.85980 | 9.12883 | 4.00082 | 5.37511 | 6.29136 | 9.46610 | 9.26221 | 12.96918 | 7.33616 | 11.84143 | 9.30475 | 0.23933 | 4.88265 | 3.03701 |
| **Ubash3a** | 0.67423 | 0.00741 | 1.86648 | 0.01038 | 0.01482 | 0.00644 | 0.00683 | 0.00986 | 0.01216 | 0.12180 | 0.01260 | 0.01458 | 0.08969 | 0.01385 | 0.00000 | 0.00665 |
| **Ubash3b** | 2.07008 | 0.08455 | 0.00000 | 0.00965 | 0.33970 | 0.15960 | 3.17942 | 0.00494 | 1.21386 | 1.21514 | 0.00000 | 0.00000 | 0.20541 | 0.00418 | 1.66921 | 0.00509 |
| **Ubb** | 41.31867 | 26.79546 | 41.49578 | 8.60394 | 28.13567 | 14.39126 | 23.68519 | 30.49540 | 16.92337 | 27.87058 | 13.58625 | 14.41410 | 36.20126 | 6.02173 | 21.31746 | 22.04468 |
| **Ubc** | 23.00502 | 15.45213 | 22.20243 | 21.82958 | 7.84904 | 8.64862 | 13.78171 | 24.00278 | 31.09283 | 29.58618 | 18.25954 | 7.65774 | 27.46167 | 13.18714 | 21.19189 | 16.67982 |
| **Ube2a** | 18.88958 | 1.14696 | 2.27430 | 16.39705 | 13.77393 | 5.36294 | 3.32090 | 9.28092 | 16.66734 | 13.04596 | 8.75263 | 7.38599 | 7.73708 | 3.57985 | 7.81243 | 14.73617 |
| **Ube2b** | 11.00463 | 5.02961 | 10.83050 | 12.72344 | 7.18932 | 9.75435 | 2.22677 | 10.30454 | 7.63581 | 19.02492 | 11.06746 | 9.82738 | 8.31015 | 6.02123 | 11.14651 | 1.99836 |
| **Ube2c** | 21.33940 | 3.63067 | 14.10927 | 12.59406 | 15.70810 | 8.63660 | 5.50075 | 37.55279 | 20.56300 | 45.92139 | 32.37789 | 11.66421 | 28.01401 | 12.11152 | 8.45848 | 7.66810 |
| **Ube2cbp** | 3.00909 | 0.02129 | 3.15941 | 0.00000 | 4.28050 | 0.00533 | 2.84043 | 4.62529 | 0.82570 | 3.15809 | 1.97725 | 0.09210 | 0.00000 | 0.00232 | 1.89359 | 0.01674 |
| **Ube2d.ps** | 0.37923 | 0.00477 | 2.67908 | 0.18472 | 1.74731 | 0.00582 | 0.00505 | 0.00000 | 0.24342 | 0.52011 | 0.76527 | 3.90968 | 1.44317 | 0.00387 | 0.66663 | 0.92173 |
| **Ube2d1** | 1.61389 | 0.00624 | 0.00000 | 0.00335 | 0.04525 | 0.00588 | 0.00576 | 0.00000 | 0.01010 | 0.66167 | 0.66140 | 0.00000 | 0.04847 | 1.12847 | 0.38901 | 1.28062 |
| **Ube2d2a** | 6.65648 | 5.90731 | 6.95095 | 35.12108 | 1.97726 | 4.37155 | 7.25934 | 5.84458 | 30.25369 | 9.99961 | 8.77486 | 16.55534 | 6.74943 | 17.15335 | 20.68931 | 5.85053 |
| **Ube2d3** | 20.45187 | 11.04878 | 14.26480 | 23.75946 | 17.32841 | 11.57185 | 8.51253 | 17.32607 | 27.31945 | 24.87372 | 17.84312 | 13.15973 | 19.34056 | 17.36244 | 11.48864 | 7.40553 |
| **Ube2dnl1** | 0.48578 | 0.00677 | 0.40536 | 0.68575 | 0.00633 | 0.00500 | 0.00684 | 0.00313 | 0.45292 | 0.07511 | 0.18235 | 0.03267 | 0.00789 | 0.00619 | 0.46289 | 0.06857 |
| **Ube2dnl2** | 0.03653 | 0.00777 | 0.03892 | 0.34015 | 0.00837 | 0.00593 | 0.00749 | 0.01240 | 0.18967 | 0.02724 | 0.02244 | 0.01758 | 0.01155 | 0.01123 | 0.02519 | 0.01428 |
| **Ube2e1** | 3.86705 | 3.36343 | 4.69089 | 0.35540 | 2.05538 | 6.70428 | 2.40082 | 6.81515 | 0.23843 | 8.80056 | 0.00000 | 4.77106 | 8.90660 | 10.05050 | 6.24957 | 4.97668 |
| **Ube2e2** | 0.00958 | 0.00744 | 0.00988 | 0.13764 | 0.02816 | 0.00657 | 0.00720 | 0.10766 | 0.00943 | 0.01766 | 0.02688 | 0.01622 | 0.01252 | 0.00640 | 0.24666 | 0.00724 |
| **Ube2e3** | 6.45386 | 2.98073 | 6.49729 | 12.39962 | 8.22858 | 5.25873 | 3.77033 | 4.68796 | 12.43772 | 25.19722 | 11.42463 | 16.63174 | 7.15002 | 24.63703 | 18.35162 | 3.00997 |
| **Ube2f** | 5.14652 | 1.50272 | 9.68657 | 2.99419 | 5.42626 | 6.08221 | 0.13263 | 4.70511 | 0.02894 | 6.19782 | 6.32415 | 0.48417 | 8.29698 | 4.76614 | 0.47412 | 2.76513 |
| **Ube2g1** | 0.82073 | 0.06635 | 0.72274 | 4.08571 | 4.31608 | 4.12763 | 0.00602 | 6.90344 | 4.73760 | 7.48548 | 0.30194 | 0.47025 | 0.00000 | 6.57105 | 3.57080 | 1.33516 |
| **Ube2g2** | 3.31159 | 2.95449 | 1.23479 | 6.88278 | 0.25502 | 1.55232 | 0.01209 | 1.37942 | 1.51947 | 7.17844 | 2.29586 | 0.20236 | 4.97105 | 0.02297 | 1.51365 | 2.58802 |
| **Ube2h** | 4.50115 | 2.80590 | 9.27263 | 7.00143 | 2.48440 | 11.67523 | 6.03895 | 13.79340 | 8.56934 | 6.97070 | 5.89756 | 1.13436 | 16.28914 | 10.52327 | 2.18269 | 4.16615 |
| **Ube2i** | 22.03156 | 6.69444 | 15.55131 | 13.79389 | 14.67975 | 7.66009 | 6.91250 | 16.39577 | 13.20551 | 18.87822 | 13.71120 | 9.41276 | 17.00038 | 5.44894 | 6.39245 | 13.51551 |
| **Ube2j1** | 2.92421 | 0.22115 | 0.77765 | 1.00079 | 5.01959 | 0.79843 | 2.77895 | 1.14775 | 3.54343 | 4.56734 | 4.60438 | 6.43640 | 1.27935 | 0.19257 | 2.58905 | 0.00002 |
| **Ube2j2** | 0.33232 | 0.04040 | 0.23490 | 2.85478 | 0.27187 | 0.06801 | 0.12144 | 0.22991 | 3.58977 | 1.06555 | 2.87781 | 0.12166 | 0.38027 | 0.88327 | 0.31800 | 0.25898 |
| **Ube2k** | 22.05419 | 12.53112 | 14.16979 | 18.05591 | 12.78646 | 9.82432 | 8.53774 | 13.96821 | 16.22896 | 15.62443 | 10.09407 | 9.16780 | 13.37379 | 11.70783 | 15.93524 | 8.27988 |
| **Ube2l3** | 27.23614 | 12.34374 | 23.86719 | 10.60384 | 18.87015 | 8.95887 | 9.91965 | 19.75152 | 15.03500 | 30.31391 | 28.50756 | 6.55733 | 17.94860 | 13.51377 | 15.86917 | 11.02992 |
| **Ube2l6** | 19.67672 | 5.19781 | 3.49746 | 2.06720 | 9.05189 | 1.87107 | 1.18660 | 1.99746 | 1.31315 | 2.44762 | 0.82209 | 2.11882 | 2.59581 | 0.76904 | 0.20806 | 18.81608 |
| **Ube2m** | 8.30782 | 1.83018 | 10.16179 | 0.27833 | 8.01431 | 2.71836 | 0.09942 | 4.97458 | 0.40869 | 3.49428 | 2.31435 | 0.34147 | 3.77057 | 0.00860 | 0.02625 | 3.70008 |
| **Ube2n** | 15.50215 | 10.00800 | 12.69291 | 14.57199 | 11.84522 | 10.18585 | 9.99755 | 9.68525 | 14.39223 | 10.98048 | 6.70209 | 2.50612 | 11.08433 | 13.59157 | 10.70009 | 5.69412 |
| **Ube2o** | 12.10691 | 3.21281 | 6.28550 | 20.98365 | 5.96254 | 4.06968 | 0.40943 | 12.14398 | 24.63858 | 19.35100 | 4.26283 | 2.48167 | 6.79226 | 0.00000 | 2.29996 | 0.00000 |
| **Ube2q1** | 7.77334 | 3.26629 | 5.77493 | 0.46883 | 6.16238 | 6.82869 | 3.00427 | 4.68906 | 0.92595 | 3.21709 | 2.62609 | 0.84734 | 5.36496 | 0.15332 | 1.87364 | 3.49353 |
| **Ube2q2** | 2.02422 | 5.76790 | 7.20838 | 3.10667 | 0.47611 | 1.52628 | 8.31593 | 6.50204 | 0.59508 | 3.93018 | 3.82267 | 1.66557 | 3.99693 | 9.01290 | 2.48001 | 0.15574 |
| **Ube2r2** | 4.75477 | 4.41478 | 0.44069 | 4.47130 | 6.44199 | 4.39791 | 0.16459 | 1.53073 | 5.08619 | 4.72763 | 1.50917 | 1.14282 | 1.68446 | 2.15619 | 1.54858 | 0.96138 |
| **Ube2s** | 12.36197 | 4.69228 | 13.66835 | 19.35108 | 12.25643 | 5.62966 | 6.30305 | 16.52010 | 18.77742 | 33.10367 | 30.05407 | 7.65964 | 10.81534 | 0.50261 | 2.41899 | 2.75576 |
| **Ube2t** | 2.02947 | 1.55051 | 3.89953 | 4.30140 | 3.91720 | 0.05946 | 2.47914 | 1.64766 | 3.73275 | 2.85462 | 2.83848 | 1.87004 | 1.63399 | 3.86190 | 2.42307 | 1.38545 |
| **Ube2u** | 0.15628 | 0.00720 | 0.03208 | 0.00402 | 0.00799 | 0.00612 | 0.03884 | 1.37616 | 0.49960 | 0.35832 | 0.21633 | 0.02775 | 0.01258 | 0.63439 | 0.12012 | 0.00710 |
| **Ube2v1** | 23.24726 | 9.29486 | 17.80250 | 11.04167 | 13.28857 | 12.68848 | 11.30376 | 22.42506 | 9.01820 | 30.76009 | 14.53953 | 2.04468 | 18.59557 | 0.41307 | 0.96406 | 13.74750 |

| | | | | | | | | | | | | | | | | |
|---|---|---|---|---|---|---|---|---|---|---|---|---|---|---|---|---|
| **Ube2v2** | 10.76576 | 13.35768 | 12.45636 | 9.57878 | 9.46715 | 6.23960 | 7.02420 | 7.50892 | 9.53937 | 11.84574 | 8.66446 | 13.34867 | 5.03132 | 18.98973 | 9.84411 | 5.01284 |
| **Ube2w** | 2.47701 | 5.73573 | 0.59114 | 5.84517 | 0.01483 | 4.59801 | 3.00406 | 1.24195 | 5.61750 | 0.77405 | 1.72766 | 0.03201 | 2.90503 | 0.00000 | 2.29982 | 0.00541 |
| **Ube2z** | 14.06242 | 11.08517 | 12.24004 | 13.37947 | 10.04552 | 8.26785 | 5.92673 | 12.71878 | 15.94803 | 19.51733 | 15.62064 | 2.37878 | 5.82000 | 13.50625 | 10.56460 | 6.06227 |
| **Ube3a** | 2.58966 | 4.70674 | 4.94966 | 4.50695 | 0.26242 | 0.00308 | 2.94811 | 0.97645 | 3.72587 | 1.12674 | 1.61669 | 0.05031 | 3.51159 | 17.41910 | 0.00000 | 0.01295 |
| **Ube3b** | 7.35164 | 3.55378 | 7.21881 | 2.60116 | 4.97347 | 2.90473 | 7.80479 | 7.64330 | 2.37225 | 4.99280 | 1.72428 | 0.81233 | 2.37818 | 0.04393 | 1.15755 | 3.48094 |
| **Ube3c** | 7.15228 | 7.18434 | 4.84068 | 9.40020 | 8.45895 | 6.69922 | 6.83560 | 3.60987 | 5.01045 | 10.37264 | 3.53454 | 1.64311 | 4.12713 | 13.98278 | 11.09457 | 2.94192 |
| **Ube4a** | 0.00000 | 0.21219 | 3.24899 | 0.53101 | 0.01955 | 0.00671 | 0.50681 | 1.12461 | 2.13161 | 0.63908 | 2.62434 | 0.18703 | 3.38011 | 7.39730 | 0.00000 | 0.89422 |
| **Ube4b** | 10.45644 | 0.87750 | 1.50838 | 20.65543 | 4.89518 | 6.33093 | 3.12457 | 7.87998 | 14.85192 | 12.92357 | 4.86873 | 1.01536 | 2.28946 | 1.59948 | 7.96836 | 2.18640 |
| **Ubfd1** | 7.86012 | 9.59090 | 12.95949 | 14.83605 | 4.51149 | 5.63824 | 6.38831 | 7.78335 | 11.59011 | 16.45001 | 13.35886 | 5.80427 | 6.20709 | 10.21083 | 14.41376 | 0.87659 |
| **Ubiad1** | 1.21814 | 0.00383 | 3.92034 | 4.77422 | 0.02063 | 2.91396 | 3.69627 | 4.12654 | 7.40961 | 2.98341 | 0.25114 | 0.20287 | 4.75617 | 2.83183 | 5.89386 | 1.57823 |
| **Ubl3** | 3.37276 | 4.49297 | 2.84841 | 2.03018 | 6.34132 | 8.40601 | 4.91757 | 8.07911 | 4.06912 | 6.43218 | 1.97034 | 7.13161 | 4.50800 | 3.06835 | 0.00000 | 0.50715 |
| **Ubl4a** | 0.22860 | 4.67345 | 6.34801 | 0.42453 | 3.57311 | 0.02679 | 2.09234 | 1.12709 | 1.41467 | 4.80555 | 0.62220 | 0.23912 | 2.62387 | 1.89966 | 3.47522 | 3.62278 |
| **Ubl5** | 7.18220 | 0.71649 | 4.79441 | 22.13320 | 6.36432 | 1.01723 | 1.15740 | 1.58165 | 26.12696 | 7.40911 | 24.77641 | 2.26636 | 5.96999 | 10.05708 | 9.08908 | 2.90165 |
| **Ubl7** | 8.77958 | 3.84409 | 9.35476 | 10.57366 | 4.57392 | 5.03358 | 3.88117 | 15.76938 | 12.60733 | 27.10130 | 13.34039 | 4.62212 | 9.83917 | 0.03132 | 0.62944 | 5.45936 |
| **Ublcp1** | 9.27452 | 5.74393 | 3.33379 | 2.56150 | 8.21266 | 5.18349 | 0.42977 | 6.31376 | 2.13579 | 3.80727 | 2.16719 | 1.31346 | 0.73804 | 5.35667 | 13.82153 | 0.60415 |
| **Ubn1** | 2.19492 | 0.02497 | 5.79189 | 15.02323 | 0.00000 | 2.04873 | 1.71129 | 5.29012 | 15.45119 | 4.35109 | 4.82064 | 0.51237 | 2.56297 | 2.70221 | 8.36145 | 5.61861 |
| **Ubn2** | 1.87437 | 2.32759 | 2.27872 | 9.04493 | 0.13768 | 1.96546 | 1.93765 | 1.80942 | 9.11601 | 4.25881 | 1.61685 | 6.94405 | 0.45447 | 7.70245 | 11.22799 | 0.89699 |
| **Ubox5** | 0.72501 | 1.96882 | 8.21224 | 1.86163 | 0.02006 | 0.00496 | 2.01363 | 1.16146 | 5.05156 | 3.55668 | 1.91148 | 0.98231 | 1.87650 | 0.09458 | 0.40679 | 1.88016 |
| **Ubp1** | 3.49725 | 1.66724 | 7.96662 | 17.24554 | 10.94421 | 4.09422 | 4.22024 | 2.15583 | 13.55249 | 4.95708 | 5.18537 | 8.57894 | 5.51579 | 3.00255 | 11.08380 | 2.92202 |
| **Ubqln1** | 14.50099 | 10.86609 | 14.31902 | 17.23414 | 12.84067 | 8.08684 | 7.94184 | 16.89910 | 11.37959 | 15.44612 | 11.68598 | 2.67118 | 13.16579 | 18.00580 | 11.55776 | 6.53672 |
| **Ubqln2** | 1.30156 | 1.91187 | 0.89999 | 7.12922 | 0.02104 | 0.10354 | 0.32498 | 0.44046 | 9.23016 | 0.00000 | 1.63549 | 0.21052 | 4.07267 | 0.14545 | 0.00000 | 0.04270 |
| **Ubqln4** | 6.86235 | 2.43127 | 5.41771 | 3.33847 | 10.37919 | 2.35477 | 3.52103 | 4.53116 | 0.32090 | 0.73962 | 1.03258 | 2.07833 | 3.30595 | 0.00000 | 1.26635 | 6.74401 |
| **Ubr1** | 2.69209 | 3.36530 | 0.61032 | 0.52348 | 0.26403 | 0.07366 | 0.26507 | 1.74263 | 0.00000 | 0.00000 | 0.99747 | 0.52977 | 1.55241 | 0.87683 | 0.71332 | 0.27906 |
| **Ubr2** | 1.14317 | 1.73470 | 1.63024 | 1.61233 | 1.28709 | 1.70231 | 1.81098 | 2.33270 | 4.94520 | 2.84355 | 0.36468 | 0.06234 | 1.36824 | 2.62775 | 1.47759 | 1.06173 |
| **Ubr3** | 4.51787 | 10.23252 | 10.68362 | 6.86631 | 3.35488 | 6.94400 | 6.62604 | 9.35274 | 10.05223 | 3.41218 | 7.79504 | 12.63855 | 6.20855 | 11.22673 | 5.17067 | 2.52741 |
| **Ubr4** | 18.03532 | 10.50869 | 15.65983 | 3.77406 | 13.85912 | 8.47685 | 10.59778 | 13.65199 | 5.26222 | 8.95800 | 11.81162 | 6.31970 | 16.57306 | 0.68189 | 3.26705 | 5.92098 |
| **Ubr5** | 8.07525 | 6.32951 | 8.52010 | 16.30398 | 7.12226 | 1.91480 | 8.24710 | 7.50547 | 21.15263 | 28.24373 | 14.66309 | 24.67446 | 6.10818 | 29.13148 | 21.85914 | 1.62485 |
| **Ubr7** | 3.66377 | 3.03703 | 4.72422 | 4.89053 | 5.18804 | 3.09548 | 4.79060 | 9.40295 | 4.75430 | 6.90120 | 4.52692 | 1.61043 | 0.23952 | 0.00000 | 2.43693 | 5.61597 |
| **Ubtd1** | 0.00114 | 0.52973 | 0.92407 | 1.58320 | 0.01543 | 3.13470 | 1.92688 | 6.81243 | 1.13699 | 11.49872 | 2.26777 | 0.56887 | 1.51166 | 0.00000 | 0.00000 | 0.02593 |
| **Ubtd2** | 0.10793 | 0.00555 | 1.16266 | 2.70340 | 0.95564 | 0.08197 | 1.68366 | 1.91883 | 7.98478 | 4.53730 | 1.11572 | 2.69133 | 0.00867 | 6.22168 | 5.65119 | 0.00535 |
| **Ubtf** | 21.31266 | 9.21412 | 15.39286 | 4.91044 | 11.11447 | 1.58707 | 8.33452 | 25.80949 | 10.22785 | 35.45930 | 13.50975 | 2.60156 | 8.97609 | 9.43700 | 14.81723 | 4.90389 |
| **Ubtfl1** | 0.00000 | 0.01570 | 0.03902 | 1.80235 | 0.00851 | 0.04642 | 0.00581 | 2.62339 | 0.28257 | 1.58391 | 0.00000 | 7.49293 | 0.18401 | 19.52670 | 16.00834 | 0.02062 |
| **Ubxn1** | 26.47990 | 13.21930 | 21.28344 | 14.05246 | 13.58416 | 13.85767 | 18.25733 | 36.85130 | 13.89095 | 40.28578 | 35.35043 | 5.05929 | 34.82592 | 0.84057 | 12.66517 | 15.80716 |
| **Ubxn11** | 0.25546 | 0.00769 | 0.00201 | 1.26276 | 0.01231 | 0.00801 | 0.00683 | 1.01090 | 1.97060 | 1.12944 | 0.00000 | 0.06555 | 0.00569 | 0.02097 | 0.11653 | 0.00717 |
| **Ubxn2a** | 7.12470 | 0.42015 | 2.69844 | 6.08027 | 2.23215 | 3.47677 | 1.01245 | 4.14179 | 4.03020 | 8.83437 | 3.63824 | 1.24963 | 2.59617 | 9.66156 | 10.93393 | 0.14528 |
| **Ubxn2b** | 2.64803 | 0.11265 | 1.89299 | 0.01238 | 0.00000 | 0.00510 | 2.63674 | 0.18101 | 0.64734 | 0.48412 | 1.46417 | 0.02567 | 0.00000 | 0.22943 | 1.10109 | 3.90790 |
| **Ubxn4** | 12.27472 | 3.78304 | 10.22696 | 16.81671 | 0.57549 | 2.46688 | 9.29628 | 10.46527 | 19.59293 | 11.03238 | 7.95783 | 1.62968 | 10.89619 | 5.14561 | 10.11428 | 1.78809 |
| **Ubxn6** | 11.16393 | 0.06832 | 0.00000 | 4.78521 | 0.79362 | 7.02026 | 1.28881 | 14.13291 | 7.05459 | 12.49826 | 6.68315 | 1.36301 | 8.94267 | 0.15068 | 0.06392 | 3.10290 |
| **Ubxn7** | 0.46863 | 0.00689 | 0.05700 | 6.97150 | 0.06898 | 0.00500 | 1.26736 | 3.76777 | 7.36063 | 4.37333 | 2.56936 | 0.36447 | 1.55228 | 1.73341 | 4.00557 | 0.01155 |
| **Ubxn8** | 8.57500 | 6.89100 | 13.68273 | 7.42486 | 5.33403 | 4.56507 | 10.07591 | 12.89222 | 7.96083 | 13.80192 | 14.81345 | 7.26105 | 11.33896 | 10.34763 | 7.71077 | 1.76271 |
| **Uchl1** | 21.65132 | 6.80631 | 22.47139 | 16.41568 | 0.50636 | 2.73164 | 16.10900 | 16.62032 | 8.90304 | 6.51196 | 6.89625 | 2.67684 | 11.72072 | 0.22399 | 0.00000 | 0.08966 |
| **Uchl3** | 10.80776 | 5.17611 | 5.45497 | 8.31766 | 7.98953 | 2.44598 | 4.05559 | 4.16246 | 11.09844 | 10.76542 | 8.17944 | 5.87425 | 10.21697 | 2.44672 | 1.85559 | 4.96982 |
| **Uchl4** | 2.33983 | 1.17063 | 0.92573 | 0.93482 | 1.76728 | 0.47363 | 0.64574 | 0.61577 | 1.41863 | 2.54345 | 1.48176 | 1.11190 | 1.97938 | 0.33206 | 0.46488 | 0.85540 |
| **Uchl5** | 7.10416 | 7.90705 | 8.71484 | 1.28174 | 8.61161 | 1.58793 | 8.45223 | 10.25189 | 2.63590 | 9.81817 | 6.81686 | 5.58277 | 8.10660 | 4.35692 | 3.65258 | 7.13915 |
| **Uck1** | 1.92015 | 0.17408 | 3.00371 | 0.49792 | 0.31369 | 3.79426 | 3.54112 | 0.95698 | 0.07710 | 3.31203 | 0.81433 | 0.10269 | 0.80142 | 0.00476 | 0.72070 | 0.32921 |
| **Uck2** | 13.64388 | 6.30694 | 14.87555 | 25.53908 | 8.41133 | 0.23957 | 9.50549 | 14.98764 | 28.12668 | 9.95400 | 11.48786 | 1.45545 | 5.18583 | 0.46478 | 1.03920 | 5.03501 |
| **Uckl1** | 1.56329 | 0.20649 | 2.27010 | 0.86195 | 0.00437 | 0.01369 | 0.00540 | 0.00322 | 0.01664 | 0.06624 | 0.01963 | 0.05221 | 0.56282 | 0.00449 | 0.29731 | 0.12945 |
| **Ucn2** | 0.01499 | 0.02972 | 0.85641 | 0.02618 | 0.00812 | 0.00819 | 0.00760 | 0.01517 | 0.01224 | 0.01425 | 0.01400 | 0.00731 | 0.10653 | 0.01461 | 0.00000 | 0.00741 |
| **Ucp2** | 0.29829 | 0.00595 | 0.07813 | 0.27146 | 0.71725 | 0.00858 | 1.95502 | 0.00000 | 0.14858 | 7.89347 | 0.20774 | 0.04060 | 0.00512 | 0.01264 | 0.00684 | 2.66308 |
| **Uevld** | 0.83672 | 3.62544 | 0.65201 | 3.25855 | 4.40843 | 5.35308 | 0.02570 | 0.00000 | 3.01214 | 0.94594 | 0.92881 | 0.05921 | 0.03888 | 0.06839 | 3.09023 | 0.00212 |
| **Ufc1** | 14.67517 | 11.69058 | 12.56979 | 2.87167 | 6.84096 | 6.51104 | 11.52484 | 17.62301 | 3.83968 | 19.94636 | 8.64852 | 3.58745 | 16.31367 | 3.38298 | 7.05943 | 9.32178 |
| **Ufd1l** | 18.78660 | 12.89539 | 11.64595 | 6.31689 | 10.81857 | 7.20591 | 8.36841 | 13.68600 | 10.02057 | 9.68100 | 12.83698 | 1.16273 | 8.47858 | 12.79566 | 12.37933 | 4.33647 |
| **Ufl1** | 5.62730 | 0.00000 | 1.94201 | 0.00000 | 4.43650 | 0.00000 | 0.08347 | 0.94132 | 0.04152 | 0.00000 | 1.06670 | 0.01784 | 0.13648 | 0.27529 | 1.15226 | 0.69021 |
| **Ufm1** | 8.47041 | 8.20235 | 10.43985 | 7.52163 | 1.96840 | 4.13138 | 1.46010 | 2.91571 | 6.50782 | 10.42954 | 10.99261 | 2.03317 | 3.39556 | 6.20920 | 10.34017 | 2.77758 |
| **Ufsp1** | 0.08485 | 0.00878 | 0.40111 | 0.00427 | 0.01716 | 0.06714 | 0.02916 | 0.79457 | 0.90524 | 0.21732 | 0.00000 | 0.00373 | 0.11625 | 0.03537 | 0.00003 | 0.01617 |
| **Ufsp2** | 7.59576 | 8.21916 | 6.02445 | 0.04073 | 1.56256 | 0.97675 | 7.33893 | 1.70843 | 0.98955 | 1.75938 | 3.43621 | 1.72695 | 2.63480 | 7.15807 | 1.69169 | 3.15099 |
| **Ugcg** | 8.25458 | 0.27994 | 7.34784 | 10.41812 | 5.83202 | 3.66343 | 9.74174 | 15.80501 | 10.19996 | 20.57112 | 19.38886 | 21.05026 | 3.65365 | 32.02703 | 17.95363 | 2.87078 |
| **Ugdh** | 6.47261 | 6.11595 | 3.85685 | 0.39640 | 2.86078 | 1.88547 | 9.15367 | 1.20452 | 1.50444 | 5.54852 | 3.49495 | 4.15253 | 1.59855 | 3.43569 | 7.13249 | 3.41712 |
| **Uggt1** | 3.89429 | 0.01301 | 8.66793 | 3.60338 | 3.79467 | 0.06765 | 7.08699 | 10.65398 | 5.35955 | 15.52224 | 7.81191 | 9.36132 | 4.89328 | 12.03492 | 15.09284 | 0.23126 |
| **Uggt2** | 0.01268 | 0.00787 | 0.01120 | 0.19586 | 0.01883 | 0.00690 | 0.00768 | 0.01505 | 0.03460 | 0.20447 | 0.01038 | 0.01928 | 0.01552 | 0.01159 | 0.50740 | 0.00743 |
| **Ugp2** | 5.42514 | 4.17923 | 2.93141 | 0.63090 | 7.30437 | 10.86027 | 4.56187 | 1.73288 | 0.19349 | 6.66072 | 0.97242 | 0.89055 | 3.18033 | 4.66970 | 0.54318 | 2.00078 |
| **Ugt2b36** | 0.01177 | 0.00730 | 0.00138 | 9.25015 | 0.00868 | 0.00541 | 0.00657 | 0.00000 | 6.63639 | 1.30218 | 0.64228 | 0.07611 | 0.00619 | 0.00000 | 1.68362 | 0.00503 |

| | | | | | | | | | | | | | | | | |
|---|---|---|---|---|---|---|---|---|---|---|---|---|---|---|---|---|
| **Ugt8a** | 0.01355 | 0.00774 | 0.01046 | 1.00820 | 0.00948 | 0.00622 | 0.00727 | 0.01050 | 0.08478 | 0.00390 | 0.00878 | 0.01457 | 0.01293 | 0.00845 | 1.75169 | 0.00716 |
| **Uhmk1** | 1.93695 | 0.00539 | 0.17157 | 8.20333 | 0.01346 | 0.00286 | 0.00510 | 0.44612 | 4.65385 | 2.62440 | 2.23899 | 0.17482 | 0.00003 | 5.78990 | 4.43317 | 0.00334 |
| **Uhrf1** | 20.77281 | 10.97532 | 5.60430 | 4.80499 | 9.18533 | 3.05488 | 7.16803 | 0.80621 | 2.43879 | 1.36948 | 1.26902 | 0.07435 | 0.67093 | 0.01582 | 0.69050 | 4.43570 |
| **Uhrf1bp1** | 0.03812 | 0.90836 | 5.49370 | 8.97226 | 4.29009 | 2.19880 | 8.50063 | 2.90097 | 9.09545 | 4.17133 | 0.26948 | 0.89851 | 3.70198 | 0.00000 | 5.49612 | 0.42391 |
| **Uhrf1bp1l** | 4.13061 | 0.04049 | 3.29884 | 4.66999 | 2.94426 | 0.00776 | 0.20492 | 1.97910 | 5.04266 | 2.58781 | 6.00153 | 5.17135 | 0.80133 | 9.66386 | 6.32812 | 0.09479 |
| **Uhrf2** | 2.59346 | 1.62223 | 0.00000 | 8.22995 | 2.73900 | 0.00367 | 0.00425 | 0.00000 | 0.00000 | 0.00004 | 0.02971 | 0.01230 | 0.02616 | 0.00000 | 0.42349 | 0.88820 |
| **Uimc1** | 0.96270 | 0.00384 | 0.19190 | 2.02728 | 2.61472 | 3.21023 | 1.60634 | 2.52911 | 3.44135 | 0.72836 | 1.13716 | 0.86347 | 0.51804 | 0.44510 | 3.51743 | 0.02552 |
| **Ulbp1** | 0.01244 | 0.00810 | 0.30438 | 0.01051 | 0.00878 | 0.00567 | 0.00723 | 0.00956 | 0.01249 | 0.00425 | 0.01024 | 0.00646 | 0.20960 | 0.36735 | 0.00000 | 0.00709 |
| **Ulk1** | 16.00561 | 7.71252 | 12.86832 | 13.70364 | 17.41485 | 3.01155 | 9.72272 | 28.36083 | 16.21557 | 28.13493 | 16.38731 | 12.20003 | 9.92935 | 9.70830 | 24.73780 | 4.22608 |
| **Ulk2** | 1.63626 | 0.00548 | 0.00000 | 5.94646 | 0.17297 | 0.00919 | 2.24477 | 0.39018 | 2.75101 | 4.11592 | 0.10929 | 0.49160 | 0.05255 | 5.90134 | 6.25506 | 0.00422 |
| **Ulk3** | 0.09808 | 0.00801 | 0.60325 | 0.13948 | 0.01521 | 0.14254 | 0.01918 | 0.37260 | 0.07926 | 0.58953 | 0.00619 | 0.04630 | 0.00611 | 0.08764 | 0.44737 | 0.02218 |
| **Ulk4** | 0.28497 | 0.08305 | 0.00347 | 0.98051 | 0.00784 | 0.00846 | 0.00729 | 0.53827 | 0.20312 | 0.00225 | 0.01142 | 0.00628 | 0.31691 | 0.01127 | 0.03593 | 0.00694 |
| **Umad1** | 0.79177 | 0.00790 | 0.43844 | 0.09485 | 0.01278 | 0.00474 | 0.00744 | 0.51434 | 1.62784 | 0.57017 | 0.01196 | 0.06500 | 0.00408 | 0.00515 | 0.19601 | 0.00674 |
| **Umodl1** | 0.01235 | 0.00748 | 0.00736 | 4.99231 | 0.00870 | 0.00607 | 0.00695 | 0.00570 | 0.27057 | 2.32471 | 0.25840 | 0.07676 | 0.00843 | 0.00000 | 0.97584 | 0.00616 |
| **Umps** | 13.87154 | 10.74108 | 14.26181 | 6.69598 | 6.29412 | 6.75861 | 8.97860 | 16.50953 | 11.97481 | 15.85682 | 7.73733 | 9.03482 | 10.67633 | 8.98909 | 14.99088 | 8.00505 |
| **Unc119** | 0.86697 | 0.02644 | 0.00608 | 1.13518 | 0.02599 | 0.00539 | 0.35927 | 0.18161 | 2.08657 | 1.47496 | 1.92051 | 0.14961 | 3.08157 | 0.00000 | 0.44350 | 2.03492 |
| **Unc119b** | 3.23542 | 0.03312 | 2.85659 | 4.03738 | 4.12329 | 5.67762 | 0.20196 | 1.21719 | 13.46836 | 11.05039 | 0.73725 | 0.54903 | 3.37791 | 0.29774 | 6.19859 | 4.91724 |
| **Unc13a** | 0.01340 | 0.00744 | 0.49767 | 0.19349 | 0.00873 | 0.00560 | 0.00705 | 0.10273 | 0.00869 | 0.12334 | 0.01239 | 0.01522 | 0.00614 | 0.00697 | 0.04899 | 0.00715 |
| **Unc13b** | 0.00000 | 0.00772 | 0.78160 | 10.16862 | 0.56363 | 0.00423 | 0.00717 | 0.85339 | 5.48212 | 5.71335 | 1.28416 | 0.20533 | 0.00001 | 3.26606 | 4.26984 | 0.00461 |
| **Unc13c** | 0.00000 | 0.00764 | 0.00000 | 16.57055 | 0.01004 | 0.00430 | 0.00643 | 0.68246 | 0.00000 | 2.27914 | 0.40207 | 0.21067 | 0.00483 | 3.37836 | 18.69343 | 0.00442 |
| **Unc13d** | 0.00988 | 0.00769 | 0.98478 | 0.57068 | 0.00753 | 0.00970 | 0.00693 | 0.20589 | 0.00091 | 0.12536 | 0.00957 | 0.00690 | 0.34426 | 0.00982 | 0.00000 | 0.00681 |
| **Unc45a** | 1.91159 | 0.38584 | 0.96014 | 2.38458 | 0.38942 | 6.96211 | 4.48135 | 6.99109 | 0.83669 | 3.15276 | 4.70714 | 0.19550 | 8.69774 | 0.00000 | 0.00000 | 6.83146 |
| **Unc45b** | 0.00941 | 0.00589 | 0.00000 | 0.45257 | 0.00823 | 0.00583 | 0.00564 | 0.00000 | 0.36393 | 2.36003 | 3.43381 | 5.43654 | 0.00000 | 0.00193 | 0.29661 | 0.01348 |
| **Unc50** | 2.80151 | 3.11267 | 0.14268 | 15.50507 | 1.03545 | 4.02111 | 2.84268 | 4.37689 | 7.59159 | 1.33304 | 1.36421 | 5.34905 | 2.32015 | 0.45563 | 6.06471 | 1.71236 |
| **Unc5b** | 0.04021 | 0.00812 | 0.00721 | 0.78790 | 0.01039 | 0.00655 | 0.00687 | 0.00113 | 0.03582 | 0.00000 | 0.32310 | 0.02609 | 0.62416 | 0.00459 | 0.28925 | 1.14748 |
| **Unc5c** | 0.01329 | 0.00763 | 0.00906 | 4.61722 | 0.00949 | 0.00582 | 0.00728 | 0.01219 | 0.79912 | 0.01700 | 0.02104 | 0.03971 | 0.01275 | 0.00000 | 2.24324 | 0.00659 |
| **Unc5cl** | 0.01377 | 0.00785 | 0.01164 | 0.00952 | 0.00931 | 0.00672 | 0.00751 | 0.01318 | 0.01220 | 0.01760 | 0.01198 | 0.03862 | 0.01301 | 0.00635 | 0.16835 | 0.00741 |
| **Unc80** | 0.01253 | 0.00761 | 0.01105 | 0.09748 | 0.00950 | 0.00663 | 0.00732 | 0.04721 | 0.02140 | 0.00947 | 0.01144 | 0.00735 | 0.01070 | 0.01891 | 0.11103 | 0.00738 |
| **Unc93a** | 1.08081 | 1.75226 | 0.00000 | 0.04439 | 0.02749 | 4.10159 | 0.01717 | 0.08638 | 0.02894 | 0.32060 | 0.43186 | 0.00425 | 1.03498 | 0.05791 | 0.40828 | 0.01799 |
| **Unc93b1** | 0.01265 | 0.00796 | 0.01285 | 0.00606 | 0.00908 | 0.00671 | 0.00741 | 0.01076 | 0.01142 | 1.93551 | 0.01095 | 0.00297 | 0.01543 | 0.01185 | 0.00863 | 0.00744 |
| **Uncx** | 0.01533 | 0.02897 | 0.00836 | 0.45144 | 0.00877 | 0.00689 | 0.00660 | 0.00000 | 0.43473 | 0.59541 | 1.78534 | 0.00000 | 0.23307 | 0.00694 | 0.00000 | 0.01408 |
| **Ung** | 10.00779 | 2.45920 | 2.48477 | 7.45244 | 4.46374 | 3.27612 | 5.66999 | 7.89266 | 7.24078 | 6.23202 | 8.37322 | 0.76574 | 5.52592 | 2.73525 | 6.66176 | 5.62738 |
| **Unk** | 2.53946 | 1.16769 | 1.97173 | 4.33034 | 0.03900 | 2.09547 | 0.00268 | 2.47929 | 4.99748 | 3.55112 | 0.76768 | 0.28237 | 3.84124 | 0.00000 | 0.56164 | 1.53924 |
| **Unkl** | 2.28960 | 4.10928 | 5.85361 | 4.66165 | 6.49782 | 4.86204 | 0.32676 | 8.48031 | 4.24857 | 7.61452 | 2.99963 | 0.95293 | 4.66388 | 2.58362 | 6.51922 | 1.66698 |
| **Upb1** | 0.01269 | 0.00789 | 0.01189 | 0.14853 | 0.01865 | 0.00653 | 0.00742 | 0.01783 | 0.20981 | 0.04252 | 0.01167 | 0.00742 | 0.01831 | 0.01005 | 0.05067 | 0.00729 |
| **Upf1** | 2.07832 | 0.01477 | 0.02254 | 1.43966 | 0.58692 | 0.00560 | 0.01292 | 1.00350 | 5.00855 | 2.05083 | 0.73055 | 0.01905 | 4.11959 | 0.00000 | 0.03694 | 0.08312 |
| **Upf2** | 6.28231 | 9.25246 | 4.61891 | 8.08744 | 1.07859 | 6.73285 | 4.96738 | 3.02452 | 7.45576 | 11.49065 | 2.23128 | 1.17603 | 4.62507 | 7.21741 | 6.37217 | 5.42026 |
| **Upf3a** | 5.20757 | 0.00000 | 2.90009 | 0.65599 | 0.77907 | 2.37904 | 1.27897 | 5.30633 | 0.51118 | 5.36534 | 3.72650 | 0.27507 | 1.73653 | 0.50807 | 2.95696 | 4.92299 |
| **Upf3b** | 3.32879 | 1.97080 | 9.29513 | 2.50673 | 3.22122 | 6.46419 | 2.90405 | 8.90003 | 1.69732 | 12.32415 | 2.51031 | 9.25783 | 0.16214 | 13.21231 | 11.29117 | 2.46626 |
| **Upk1a** | 1.18086 | 0.00619 | 1.60899 | 1.26881 | 0.00280 | 0.00661 | 0.00611 | 0.00000 | 1.54783 | 0.69846 | 0.34268 | 0.01048 | 0.00572 | 0.00286 | 0.26201 | 0.00519 |
| **Upk3a** | 0.00227 | 0.00704 | 0.76432 | 0.94712 | 0.01062 | 0.00577 | 2.30469 | 1.10520 | 0.53016 | 0.10406 | 2.62965 | 0.15559 | 0.00000 | 0.05079 | 0.36848 | 0.00672 |
| **Upk3b** | 0.80353 | 0.00811 | 0.55382 | 0.31028 | 0.04458 | 1.57599 | 0.00662 | 0.00751 | 0.00550 | 1.06553 | 0.01057 | 0.04264 | 0.46827 | 0.10889 | 0.00000 | 0.00649 |
| **Upk3bl** | 0.02213 | 0.00859 | 0.60873 | 0.34422 | 0.01055 | 0.00570 | 0.00769 | 0.00803 | 0.00503 | 0.00850 | 0.33930 | 0.00741 | 0.00671 | 0.00826 | 0.00000 | 0.00704 |
| **Upp1** | 29.93329 | 11.10887 | 6.52914 | 1.81200 | 21.74418 | 0.00000 | 6.98413 | 0.00000 | 2.62274 | 0.00000 | 0.00000 | 0.20043 | 0.17046 | 0.00000 | 0.91361 | 6.59884 |
| **Uqcc1** | 3.61160 | 11.15605 | 9.02450 | 1.73842 | 2.32019 | 9.26608 | 8.94359 | 13.16377 | 4.89087 | 9.53018 | 9.86602 | 1.33629 | 6.66090 | 19.74965 | 6.96688 | 5.15921 |
| **Uqcc2** | 2.52294 | 0.05718 | 2.05122 | 3.26017 | 1.72065 | 0.78088 | 0.57120 | 0.08140 | 0.14669 | 1.28083 | 3.72247 | 0.09381 | 2.06461 | 0.00000 | 0.00000 | 2.64973 |
| **Uqcc3** | 6.41158 | 1.46617 | 5.94545 | 7.98286 | 3.19449 | 1.59064 | 7.13225 | 5.44939 | 23.26546 | 12.38197 | 14.05045 | 2.69041 | 5.45719 | 0.08011 | 5.71722 | 5.42777 |
| **Uqcr10** | 13.70979 | 0.83891 | 10.85351 | 6.26541 | 8.54086 | 3.43722 | 0.63444 | 0.89054 | 11.46458 | 8.51080 | 17.36582 | 1.02361 | 8.32939 | 0.44004 | 3.20290 | 7.02024 |
| **Uqcr11** | 5.64863 | 0.04131 | 3.01308 | 2.14679 | 3.26440 | 1.78897 | 0.55487 | 1.40879 | 0.87108 | 1.81155 | 6.56308 | 0.09614 | 4.63468 | 0.00000 | 0.00000 | 5.21421 |
| **Uqcrb** | 10.25998 | 2.54802 | 9.00032 | 2.69654 | 6.24503 | 1.86698 | 2.36887 | 1.77086 | 3.84473 | 4.50664 | 26.41167 | 1.78545 | 8.86033 | 7.49573 | 2.03659 | 6.47279 |
| **Uqcrc1** | 26.68377 | 6.98336 | 13.64690 | 2.57038 | 16.78178 | 17.20351 | 7.97507 | 23.98747 | 3.85421 | 24.77523 | 8.32719 | 2.72068 | 28.38133 | 0.01577 | 0.00000 | 16.14843 |
| **Uqcrc2** | 23.50923 | 15.56903 | 20.70395 | 5.05719 | 15.05964 | 9.95232 | 12.34601 | 19.15080 | 2.24567 | 16.10675 | 10.20842 | 10.22902 | 14.92833 | 18.62791 | 10.61426 | 14.86065 |
| **Uqcrfs1** | 15.00424 | 3.45674 | 6.75451 | 9.44031 | 12.82564 | 6.71547 | 0.08937 | 1.60596 | 15.41547 | 15.60516 | 12.38958 | 5.81017 | 7.22116 | 2.44992 | 3.13448 | 4.03185 |
| **Uqcrh** | 10.66051 | 0.52240 | 9.26424 | 4.39241 | 9.13193 | 2.25287 | 2.35627 | 1.51279 | 3.39990 | 9.42823 | 22.11306 | 1.39176 | 7.13824 | 3.89379 | 5.41973 | 2.52967 |
| **Uqcrq** | 10.44704 | 0.31040 | 2.38483 | 0.94504 | 11.07335 | 3.51664 | 1.34511 | 1.97501 | 1.97766 | 6.23365 | 4.13518 | 0.25655 | 5.80722 | 0.00000 | 0.00000 | 2.54318 |
| **Urah** | 1.83156 | 0.17225 | 0.00000 | 0.36604 | 0.00661 | 0.00618 | 0.00562 | 0.00350 | 0.00626 | 1.34239 | 0.01272 | 0.00261 | 0.00656 | 0.00852 | 0.04359 | 2.93742 |
| **Urb1** | 0.88561 | 0.06097 | 4.12533 | 6.19502 | 1.94740 | 0.01234 | 5.17814 | 9.02509 | 8.22115 | 13.03237 | 3.71808 | 1.17251 | 1.37796 | 0.26862 | 4.86085 | 0.79132 |
| **Urb2** | 1.64708 | 0.06191 | 1.59705 | 4.28265 | 0.01512 | 0.00329 | 0.25567 | 1.73708 | 2.79123 | 2.25316 | 0.92232 | 0.42396 | 0.04865 | 3.16979 | 2.92709 | 0.01848 |
| **Urgcp** | 6.50975 | 0.46546 | 0.92322 | 4.44624 | 0.00000 | 1.57889 | 0.58411 | 4.79142 | 0.50154 | 5.90704 | 1.53843 | 0.20808 | 0.46648 | 0.36678 | 0.91353 | 4.23082 |
| **Uri1** | 11.89451 | 2.93925 | 1.68382 | 2.52061 | 12.35969 | 2.87626 | 5.32814 | 5.34007 | 2.75705 | 5.79945 | 3.62689 | 0.67982 | 1.54236 | 0.99229 | 2.73226 | 0.15496 |
| **Urm1** | 0.86859 | 1.87460 | 0.92167 | 3.97225 | 0.07791 | 0.29904 | 0.83666 | 3.18948 | 5.90683 | 9.91597 | 9.85502 | 1.89598 | 1.14362 | 1.80650 | 0.56862 | 0.00919 |
| **Urod** | 8.82590 | 10.02880 | 9.07912 | 5.87339 | 3.89257 | 0.87659 | 3.66185 | 10.77005 | 5.31686 | 12.03971 | 7.66273 | 1.11998 | 10.75380 | 3.29077 | 10.72428 | 2.57753 |

| | | | | | | | | | | | | | | | | |
|---|---|---|---|---|---|---|---|---|---|---|---|---|---|---|---|---|
| **Uros** | 0.42237 | 0.03547 | 0.03984 | 1.13387 | 0.15012 | 2.88369 | 0.00639 | 0.89764 | 1.90350 | 0.00000 | 0.20933 | 0.02732 | 0.09901 | 0.00316 | 1.00610 | 0.00520 |
| **Usb1** | 3.64317 | 1.35159 | 2.08263 | 0.72890 | 2.53652 | 0.10158 | 0.06378 | 5.52862 | 0.35615 | 3.75671 | 1.04595 | 0.06825 | 2.29595 | 0.56111 | 0.08234 | 0.02905 |
| **Use1** | 9.39704 | 0.66879 | 6.89939 | 3.11170 | 5.84471 | 1.69695 | 2.40343 | 6.34751 | 4.46317 | 8.55154 | 4.46800 | 1.66570 | 8.73261 | 0.02886 | 1.76697 | 3.79014 |
| **Usf1** | 2.50827 | 3.78739 | 0.15099 | 0.12223 | 0.02295 | 0.00589 | 1.11707 | 1.66759 | 0.62601 | 1.85949 | 0.00000 | 0.09123 | 0.03115 | 0.04117 | 2.68142 | 0.00314 |
| **Usf2** | 6.51937 | 2.08190 | 3.56654 | 2.66988 | 3.02654 | 4.65497 | 2.46947 | 2.85460 | 0.99223 | 1.09000 | 1.98872 | 0.02754 | 3.45705 | 0.01475 | 0.00000 | 0.27027 |
| **Usf3** | 0.92608 | 0.00429 | 0.00000 | 1.76891 | 0.00593 | 2.70726 | 1.33568 | 0.00000 | 3.87212 | 3.69604 | 1.95400 | 3.81624 | 0.00000 | 5.92052 | 5.23044 | 0.00396 |
| **Ush2a** | 0.02183 | 0.00781 | 0.01134 | 0.23757 | 0.00879 | 0.00657 | 0.00746 | 0.01356 | 0.70410 | 0.00676 | 0.00747 | 0.01620 | 0.01279 | 0.00610 | 0.13905 | 0.00724 |
| **Usmg5** | 6.93598 | 0.22427 | 5.51571 | 11.48171 | 3.85681 | 1.32399 | 0.41595 | 0.00000 | 15.39635 | 1.76975 | 16.77010 | 2.50467 | 2.28660 | 2.88525 | 1.85035 | 1.90571 |
| **Uso1** | 4.36498 | 10.15935 | 8.24415 | 8.28926 | 8.99716 | 8.19930 | 6.29214 | 5.03815 | 0.88693 | 9.52050 | 5.14487 | 2.85538 | 3.29670 | 8.01072 | 9.57858 | 3.18625 |
| **Usp1** | 6.28912 | 5.99477 | 7.98945 | 19.27684 | 7.13922 | 5.76971 | 10.14574 | 16.54180 | 22.36317 | 31.53633 | 8.99910 | 31.96323 | 11.05406 | 37.02895 | 27.95514 | 3.04998 |
| **Usp10** | 13.88163 | 1.53700 | 2.72364 | 10.43628 | 3.61241 | 6.02464 | 4.58778 | 26.99063 | 12.69806 | 34.52548 | 13.28107 | 3.31851 | 9.50766 | 4.40766 | 3.53070 | 6.76705 |
| **Usp11** | 0.00488 | 0.00523 | 0.64571 | 0.08842 | 0.00671 | 0.00444 | 0.93168 | 0.02462 | 1.99812 | 0.00000 | 0.36635 | 6.32415 | 0.00207 | 0.00310 | 0.79457 | 1.17263 |
| **Usp12** | 4.63153 | 1.51131 | 1.54424 | 5.40517 | 0.22181 | 3.72909 | 3.57095 | 3.71901 | 5.99264 | 7.70231 | 2.16217 | 3.30978 | 2.79627 | 0.28733 | 3.91250 | 1.74944 |
| **Usp13** | 0.01057 | 0.00734 | 0.00815 | 0.07482 | 0.00970 | 0.00677 | 0.00711 | 0.35651 | 0.00798 | 0.09321 | 0.00880 | 0.07836 | 0.00634 | 0.01283 | 0.10746 | 0.00744 |
| **Usp14** | 17.47281 | 15.37460 | 16.86807 | 2.78464 | 17.99197 | 12.14047 | 6.86574 | 19.61521 | 4.62735 | 18.55044 | 16.01911 | 19.51651 | 15.61294 | 20.98397 | 14.53988 | 4.39875 |
| **Usp15** | 6.50542 | 7.88997 | 9.53949 | 12.59789 | 8.58772 | 2.89494 | 9.27151 | 13.74287 | 1.57806 | 6.28024 | 8.66332 | 6.01206 | 1.31029 | 12.68487 | 14.00632 | 2.25258 |
| **Usp16** | 4.60718 | 7.24565 | 7.45715 | 19.47336 | 12.31924 | 2.17178 | 4.41148 | 16.44937 | 20.18377 | 23.98849 | 13.72222 | 17.80073 | 6.19370 | 30.10402 | 27.16435 | 3.94420 |
| **Usp17la** | 0.03673 | 0.01875 | 0.06474 | 2.11899 | 0.00890 | 0.00560 | 0.00736 | 0.01682 | 6.00239 | 0.03277 | 0.00000 | 0.00000 | 0.03841 | 0.17013 | 0.08100 | 0.05198 |
| **Usp17lb** | 0.01252 | 0.00780 | 0.01028 | 1.39505 | 0.00930 | 0.00613 | 0.00725 | 0.03209 | 2.66483 | 0.00350 | 0.00000 | 0.00010 | 0.01134 | 0.00516 | 0.69997 | 0.00683 |
| **Usp17lc** | 0.01349 | 0.00775 | 0.01197 | 0.19696 | 0.00934 | 0.00649 | 0.00741 | 0.01729 | 0.49792 | 0.00606 | 0.00937 | 0.00630 | 0.01236 | 0.00596 | 0.25281 | 0.00728 |
| **Usp17ld** | 0.01368 | 0.00805 | 0.01209 | 0.17783 | 0.00931 | 0.00660 | 0.00753 | 0.01377 | 0.52332 | 0.01131 | 0.00956 | 0.00720 | 0.01281 | 0.01060 | 0.03000 | 0.00735 |
| **Usp17le** | 0.01360 | 0.00809 | 0.01176 | 0.26633 | 0.00927 | 0.00658 | 0.00753 | 0.01346 | 1.08820 | 0.00896 | 0.01843 | 0.00622 | 0.01265 | 0.01176 | 0.01487 | 0.00730 |
| **Usp19** | 4.88737 | 0.03132 | 0.00000 | 19.05065 | 1.83996 | 3.54967 | 2.68137 | 4.51529 | 15.14573 | 8.13224 | 5.78823 | 0.34085 | 2.40772 | 1.03242 | 0.11026 | 4.71943 |
| **Usp2** | 0.88700 | 0.00725 | 1.38975 | 0.64134 | 0.01450 | 0.02148 | 0.00528 | 0.07733 | 1.40336 | 0.00000 | 1.67360 | 0.00000 | 1.14958 | 0.02437 | 0.59719 | 0.01235 |
| **Usp20** | 0.15593 | 1.86118 | 0.14440 | 0.74510 | 0.00732 | 2.10982 | 2.17881 | 0.88527 | 2.24583 | 1.32125 | 0.21274 | 0.01630 | 0.00000 | 0.09589 | 0.48398 | 0.01261 |
| **Usp21** | 1.20530 | 0.01427 | 2.66326 | 2.47097 | 0.03807 | 0.04175 | 3.21540 | 0.97914 | 0.99136 | 3.14644 | 0.74161 | 0.08218 | 0.34072 | 1.04030 | 1.69617 | 4.63332 |
| **Usp22** | 5.38835 | 5.42372 | 4.87141 | 12.91756 | 2.66487 | 4.39213 | 0.23671 | 11.57814 | 13.67858 | 11.93952 | 12.40277 | 11.92800 | 8.11992 | 16.31333 | 14.59961 | 4.39788 |
| **Usp24** | 4.99962 | 5.78387 | 5.87662 | 5.71657 | 3.11372 | 8.36968 | 10.32310 | 6.05981 | 1.82810 | 4.95128 | 5.45293 | 6.04614 | 5.57116 | 1.32416 | 6.55878 | 4.80754 |
| **Usp25** | 0.98799 | 0.00514 | 3.21528 | 0.49660 | 2.86142 | 0.99050 | 0.00602 | 2.05899 | 0.20370 | 3.22098 | 0.65470 | 0.35861 | 0.75475 | 0.08624 | 5.37716 | 3.46203 |
| **Usp26** | 0.01340 | 0.00813 | 0.01131 | 2.01271 | 0.00919 | 0.00630 | 0.00747 | 0.01291 | 2.42816 | 0.00335 | 0.03252 | 0.00369 | 0.01227 | 0.00882 | 0.00753 | 0.00688 |
| **Usp27x** | 0.01200 | 0.00823 | 0.01806 | 0.95484 | 0.00921 | 0.00712 | 0.00731 | 0.01179 | 0.51981 | 0.00456 | 0.03496 | 0.00726 | 0.27952 | 0.01120 | 0.00208 | 0.00700 |
| **Usp28** | 4.11923 | 3.52936 | 1.55080 | 5.56905 | 1.77201 | 5.28206 | 2.29593 | 2.84250 | 1.67991 | 2.30646 | 2.02218 | 0.30110 | 0.00000 | 0.02997 | 9.40559 | 2.22539 |
| **Usp29** | 0.00000 | 0.00630 | 0.24126 | 12.92473 | 0.01729 | 2.06758 | 0.00206 | 0.49323 | 16.60388 | 11.21856 | 9.99173 | 0.90944 | 0.91579 | 5.55724 | 9.60890 | 0.02280 |
| **Usp3** | 4.53757 | 4.04266 | 1.90677 | 5.31873 | 5.10615 | 2.97838 | 3.65519 | 0.13421 | 6.95390 | 1.49406 | 2.34486 | 0.24344 | 2.33653 | 1.95238 | 1.83419 | 5.80687 |
| **Usp30** | 1.28651 | 0.28078 | 2.64917 | 0.85122 | 0.23134 | 0.00580 | 0.00471 | 2.72680 | 0.75561 | 4.15134 | 1.33364 | 0.83646 | 1.19086 | 0.00094 | 1.79450 | 0.00395 |
| **Usp31** | 1.60100 | 0.54038 | 2.56099 | 5.47341 | 0.00357 | 0.64975 | 1.96325 | 2.10845 | 3.63848 | 0.27725 | 0.22383 | 1.52562 | 0.71009 | 1.88972 | 2.50293 | 0.85386 |
| **Usp32** | 4.48072 | 0.00000 | 1.27846 | 5.01302 | 0.00931 | 1.75802 | 2.91544 | 2.18411 | 8.31237 | 2.87429 | 1.87816 | 0.74657 | 1.32448 | 1.61928 | 4.11287 | 2.74466 |
| **Usp33** | 4.77384 | 4.68786 | 4.82494 | 16.99166 | 4.85094 | 2.32643 | 4.47533 | 10.01474 | 19.89523 | 12.92710 | 14.20261 | 18.86671 | 4.35346 | 28.36974 | 23.50631 | 0.00000 |
| **Usp34** | 2.60619 | 2.59217 | 0.34972 | 8.08032 | 0.16305 | 4.81049 | 0.20180 | 1.28271 | 7.38796 | 4.95802 | 0.00000 | 0.39408 | 1.17651 | 20.30432 | 14.77626 | 0.80556 |
| **Usp35** | 0.01132 | 0.00859 | 0.01045 | 0.03058 | 0.00947 | 0.00677 | 0.00815 | 0.09877 | 0.02870 | 0.01387 | 0.01226 | 0.00885 | 0.00884 | 0.01373 | 0.00836 | 0.00738 |
| **Usp36** | 1.40087 | 3.49041 | 5.46200 | 5.06408 | 0.05670 | 0.00515 | 3.33564 | 4.56124 | 6.02966 | 4.67270 | 2.31067 | 0.50099 | 1.45795 | 0.00000 | 8.88849 | 0.30904 |
| **Usp37** | 3.18940 | 5.20376 | 7.49043 | 9.74808 | 1.49056 | 5.00910 | 3.78972 | 9.49571 | 14.33274 | 16.02702 | 8.12566 | 16.61289 | 6.76897 | 21.44708 | 16.85558 | 4.15772 |
| **Usp38** | 0.26981 | 5.55949 | 2.19222 | 15.06449 | 6.10729 | 0.00776 | 4.00903 | 6.90229 | 19.66469 | 10.92516 | 7.37810 | 14.49729 | 2.22361 | 11.52004 | 19.52520 | 1.83183 |
| **Usp39** | 9.54969 | 2.95087 | 7.01185 | 3.70454 | 7.71902 | 5.79971 | 4.12184 | 7.62055 | 5.81713 | 13.80025 | 6.20465 | 4.30635 | 5.38153 | 11.10224 | 10.82931 | 5.04585 |
| **Usp4** | 4.68212 | 3.46915 | 3.13681 | 4.32301 | 4.40118 | 4.64842 | 5.50821 | 5.10468 | 5.86466 | 5.51008 | 2.26485 | 7.73145 | 6.31992 | 3.28328 | 7.33080 | 1.89860 |
| **Usp40** | 0.03929 | 0.08520 | 1.96869 | 0.03372 | 0.69723 | 0.00552 | 1.60930 | 0.13973 | 1.80465 | 0.08488 | 0.75859 | 0.40461 | 0.88424 | 0.00451 | 1.40622 | 0.00604 |
| **Usp42** | 3.04231 | 1.72874 | 2.49343 | 5.19952 | 1.62507 | 3.23640 | 1.38403 | 2.95137 | 3.95091 | 4.59724 | 2.95206 | 0.50181 | 2.60191 | 0.00000 | 7.40950 | 0.02738 |
| **Usp43** | 5.70331 | 1.40428 | 4.56238 | 3.50718 | 2.74804 | 0.00000 | 2.90177 | 5.25614 | 0.00000 | 1.34409 | 3.30413 | 2.04219 | 3.28969 | 0.00004 | 0.34212 | 0.01778 |
| **Usp44** | 0.90574 | 0.00655 | 0.00373 | 0.00905 | 0.00605 | 0.00715 | 0.00656 | 0.00903 | 0.01163 | 0.29597 | 0.02394 | 0.00561 | 0.01194 | 0.01380 | 0.11761 | 0.00675 |
| **Usp45** | 5.40422 | 0.07317 | 2.10876 | 2.01076 | 2.04373 | 2.97136 | 0.06345 | 2.63529 | 0.16444 | 3.38953 | 1.89515 | 1.56107 | 1.58488 | 0.78147 | 3.26529 | 0.01957 |
| **Usp46** | 1.10803 | 0.00689 | 1.18951 | 3.02674 | 0.05230 | 3.12768 | 0.96462 | 1.21184 | 0.09836 | 0.00000 | 0.00000 | 0.44705 | 1.40875 | 0.05420 | 0.86482 | 0.55368 |
| **Usp47** | 3.34916 | 1.72261 | 2.37727 | 10.38420 | 2.68259 | 0.00089 | 0.00320 | 1.46160 | 2.36103 | 1.20704 | 0.45732 | 0.38541 | 1.98800 | 2.08214 | 5.79064 | 0.63129 |
| **Usp48** | 18.60645 | 13.07451 | 26.39624 | 10.07130 | 10.97968 | 9.90420 | 15.93315 | 28.07265 | 12.60388 | 35.55014 | 23.65931 | 21.37225 | 19.52540 | 21.45999 | 16.49567 | 11.78503 |
| **Usp49** | 1.55281 | 0.00568 | 0.03845 | 4.72325 | 0.00494 | 0.00344 | 0.00524 | 0.37304 | 7.10499 | 1.99061 | 2.07768 | 0.01553 | 0.00000 | 0.00675 | 1.88852 | 0.00426 |
| **Usp5** | 12.80629 | 11.55765 | 10.55431 | 4.19668 | 11.80816 | 6.18623 | 9.36735 | 14.69119 | 4.30327 | 11.28762 | 6.42540 | 0.77428 | 11.08504 | 0.11058 | 5.12602 | 7.12107 |
| **Usp50** | 0.01363 | 0.00772 | 0.01028 | 2.48321 | 0.00914 | 0.00633 | 0.00717 | 0.00831 | 0.41287 | 0.26800 | 0.07413 | 0.02399 | 0.01178 | 0.00308 | 0.61157 | 0.00734 |
| **Usp53** | 0.00000 | 0.60669 | 0.00000 | 6.08599 | 0.01636 | 0.20052 | 0.00638 | 1.47152 | 3.94074 | 0.35324 | 1.14380 | 0.00000 | 0.00000 | 3.14785 | 2.08322 | 0.01266 |
| **Usp54** | 1.26435 | 0.00001 | 2.66989 | 4.54570 | 3.72924 | 1.59035 | 1.18547 | 3.26684 | 5.33293 | 3.24679 | 3.46671 | 7.09320 | 1.37867 | 10.34238 | 7.62696 | 2.31268 |
| **Usp6nl** | 0.49397 | 0.00862 | 0.01597 | 0.59314 | 0.02068 | 0.02248 | 0.00724 | 0.01771 | 0.04507 | 0.00589 | 0.10219 | 0.00734 | 0.29023 | 0.01044 | 0.00720 | 0.00717 |
| **Usp7** | 13.55354 | 10.00008 | 6.31698 | 13.33328 | 6.60327 | 8.19798 | 7.99796 | 16.61339 | 14.22447 | 15.20108 | 13.39950 | 7.69910 | 7.67763 | 16.76295 | 14.38765 | 5.53211 |
| **Usp8** | 8.65433 | 3.61835 | 2.70032 | 3.79211 | 11.05164 | 0.83871 | 4.73712 | 8.09419 | 3.19828 | 6.53788 | 1.72657 | 7.68343 | 3.58403 | 1.73878 | 2.25399 | 1.16648 |
| **Usp9x** | 7.41533 | 10.73534 | 9.66094 | 15.88451 | 9.88977 | 2.13648 | 1.59027 | 2.73456 | 12.65027 | 8.53734 | 3.17434 | 7.65701 | 2.19747 | 23.28956 | 18.46312 | 2.12346 |

| | | | | | | | | | | | | | | | | |
|---|---|---|---|---|---|---|---|---|---|---|---|---|---|---|---|---|
| **Uspl1** | 0.17812 | 1.15149 | 6.69581 | 16.66372 | 7.07186 | 1.14614 | 2.50083 | 5.28431 | 18.28655 | 10.54220 | 3.17489 | 8.31060 | 3.51650 | 3.41935 | 9.78770 | 1.56258 |
| **Utf1** | 5.01821 | 0.23760 | 0.97025 | 0.10260 | 4.93496 | 0.02661 | 0.35893 | 0.00000 | 0.37066 | 5.78307 | 2.38047 | 0.19710 | 6.23947 | 0.01299 | 0.00000 | 7.98794 |
| **Utp11l** | 11.69761 | 7.11910 | 11.24170 | 8.92953 | 11.79243 | 6.90296 | 5.06236 | 12.37210 | 15.19095 | 15.57645 | 14.56170 | 7.22017 | 7.13184 | 5.50007 | 11.99508 | 4.79418 |
| **Utp14a** | 5.32722 | 3.42976 | 5.06890 | 2.24811 | 3.65263 | 0.05622 | 2.57780 | 12.51962 | 2.93870 | 7.08544 | 2.38083 | 0.30174 | 4.73880 | 15.72784 | 14.48277 | 0.21641 |
| **Utp14b** | 0.65660 | 0.01627 | 1.87373 | 20.49096 | 0.00290 | 1.13379 | 0.00416 | 6.49799 | 15.10635 | 2.34914 | 3.15641 | 0.09403 | 0.51317 | 5.70751 | 10.94843 | 0.00738 |
| **Utp15** | 9.64039 | 8.19613 | 7.73847 | 10.25892 | 0.22083 | 6.87444 | 1.76862 | 11.07643 | 13.28911 | 8.23812 | 3.49891 | 4.63910 | 2.34031 | 16.79851 | 16.26257 | 2.34668 |
| **Utp18** | 12.15136 | 6.21130 | 9.54765 | 4.84100 | 8.14360 | 3.64807 | 8.21592 | 8.55731 | 11.31036 | 14.36949 | 7.47924 | 10.92253 | 6.92010 | 17.49543 | 10.75089 | 1.10625 |
| **Utp20** | 2.44148 | 5.88391 | 4.71918 | 4.23881 | 1.02427 | 0.01083 | 2.63748 | 7.88930 | 8.42017 | 4.62940 | 3.96604 | 2.61036 | 3.53749 | 9.81310 | 12.19147 | 1.36865 |
| **Utp23** | 4.60344 | 4.37226 | 9.72242 | 24.26761 | 0.38998 | 0.27954 | 2.92269 | 5.09618 | 23.95581 | 11.95490 | 9.80158 | 6.97729 | 5.06392 | 12.52370 | 15.13273 | 0.51541 |
| **Utp3** | 13.83402 | 4.06921 | 6.85217 | 15.29168 | 4.01833 | 5.40633 | 8.84116 | 15.67613 | 27.80883 | 27.09771 | 12.73310 | 10.20770 | 8.08907 | 15.80770 | 18.12040 | 3.00655 |
| **Utp6** | 6.58077 | 1.77097 | 5.87626 | 7.11236 | 3.59006 | 0.62319 | 2.31987 | 7.65194 | 12.02048 | 10.31683 | 6.78051 | 3.62789 | 3.63504 | 10.55272 | 9.45239 | 2.34719 |
| **Utrn** | 0.00000 | 0.00786 | 1.07685 | 2.85101 | 0.01053 | 0.68697 | 0.18759 | 0.00000 | 3.00652 | 0.00000 | 0.69664 | 0.23269 | 1.88993 | 3.02126 | 0.24120 | 0.03385 |
| **Uts2r** | 0.01257 | 0.00776 | 0.00574 | 0.16182 | 0.00904 | 0.00757 | 0.51163 | 0.00000 | 0.17279 | 0.41815 | 0.49025 | 0.00533 | 0.00826 | 0.00733 | 0.01430 | 0.00722 |
| **Uty** | 0.01462 | 0.00822 | 0.01635 | 1.07422 | 0.00963 | 0.00664 | 0.02925 | 0.00587 | 0.00000 | 0.00879 | 0.46660 | 0.01655 | 0.02069 | 0.00582 | 0.00000 | 0.00735 |
| **Uvrag** | 2.20300 | 1.41051 | 0.52136 | 4.57207 | 3.45320 | 2.03190 | 0.00499 | 0.67686 | 0.72045 | 2.40464 | 1.55931 | 0.45038 | 0.29326 | 0.22388 | 0.56673 | 2.07387 |
| **Uvssa** | 0.47521 | 2.51289 | 0.01701 | 0.27471 | 0.00686 | 0.01496 | 0.03865 | 0.02389 | 0.18452 | 0.00000 | 1.11335 | 0.03115 | 0.02685 | 0.27434 | 0.89532 | 0.00549 |
| **Uxs1** | 0.00848 | 0.01861 | 0.03437 | 0.00867 | 0.01988 | 0.01729 | 0.00740 | 0.01130 | 0.09116 | 0.00605 | 0.00151 | 0.00000 | 0.02653 | 2.17311 | 0.39805 | 0.01511 |
| **Uxt** | 2.30603 | 2.26968 | 3.04476 | 7.19773 | 4.41985 | 0.12277 | 0.71518 | 3.04214 | 2.02438 | 7.24417 | 12.01067 | 0.39213 | 2.82861 | 1.96097 | 4.23473 | 1.42401 |
| **Vac14** | 10.01103 | 2.30778 | 5.78638 | 7.58691 | 3.32509 | 4.49671 | 6.70703 | 5.75819 | 4.50758 | 6.59820 | 1.74011 | 2.76001 | 5.59419 | 0.12744 | 7.90229 | 6.57433 |
| **Vamp1** | 0.01244 | 0.00762 | 0.01102 | 0.20378 | 0.00919 | 0.00659 | 0.03776 | 0.03400 | 0.66286 | 0.38210 | 0.00840 | 0.00472 | 0.01121 | 0.00658 | 0.17318 | 0.00769 |
| **Vamp2** | 0.03464 | 0.00746 | 0.03723 | 0.13293 | 0.00879 | 0.00667 | 0.01754 | 0.12735 | 0.00967 | 0.83719 | 0.16838 | 0.02174 | 0.03861 | 0.00616 | 0.27836 | 0.01525 |
| **Vamp3** | 8.55688 | 6.32167 | 14.84398 | 0.78523 | 7.43788 | 6.23437 | 6.03442 | 12.12767 | 1.02304 | 12.79491 | 7.08420 | 3.31607 | 13.33562 | 10.08129 | 5.03664 | 3.90657 |
| **Vamp4** | 1.36511 | 0.00579 | 0.41395 | 2.46106 | 1.05200 | 0.00965 | 0.00626 | 0.11641 | 1.61626 | 0.31630 | 0.01324 | 0.04179 | 0.00000 | 0.00211 | 1.84383 | 1.99398 |
| **Vamp5** | 0.01672 | 0.00800 | 0.02372 | 0.00741 | 0.00865 | 0.00755 | 0.00742 | 0.01266 | 0.01186 | 0.85461 | 0.02282 | 0.01557 | 0.01064 | 0.01225 | 0.00764 | 0.00729 |
| **Vamp8** | 6.47204 | 2.79224 | 4.66866 | 1.17549 | 0.24191 | 7.77593 | 2.61464 | 9.34370 | 9.59446 | 16.66238 | 14.99450 | 1.50405 | 8.44237 | 4.89394 | 6.05603 | 5.18813 |
| **Vangl1** | 4.96640 | 0.45640 | 2.41373 | 0.00000 | 0.69943 | 0.16637 | 2.15258 | 3.08937 | 0.00769 | 0.62956 | 1.09733 | 0.03769 | 1.34622 | 0.00936 | 0.00000 | 4.74552 |
| **Vangl2** | 0.01714 | 0.01816 | 0.04226 | 0.01852 | 0.00877 | 0.01228 | 0.01726 | 0.01752 | 0.01108 | 0.06070 | 0.03917 | 0.02631 | 0.72226 | 0.01230 | 0.03628 | 0.00704 |
| **Vapa** | 12.63478 | 9.24972 | 10.29475 | 9.83997 | 15.88462 | 10.08924 | 8.55030 | 13.54395 | 10.13373 | 19.20845 | 18.98249 | 2.44214 | 9.96658 | 20.98320 | 9.74495 | 5.57348 |
| **Vapb** | 0.02613 | 0.01686 | 0.41215 | 0.51564 | 0.00725 | 0.53940 | 1.06386 | 0.00999 | 0.12289 | 1.86792 | 1.03728 | 0.06713 | 0.27089 | 0.00658 | 0.00000 | 1.85815 |
| **Vars** | 8.74479 | 0.00740 | 0.22392 | 0.12956 | 2.89454 | 0.10554 | 0.02168 | 0.38597 | 0.22065 | 6.34677 | 0.67099 | 0.05579 | 17.54846 | 0.00662 | 0.00000 | 14.65137 |
| **Vars2** | 3.77692 | 0.00361 | 0.52607 | 0.29565 | 0.00000 | 0.06818 | 0.32526 | 1.53789 | 0.25240 | 2.56238 | 0.12545 | 0.14477 | 0.50258 | 0.04694 | 0.06186 | 2.89234 |
| **Vash1** | 0.01360 | 0.00757 | 0.01181 | 0.07692 | 0.00909 | 0.00662 | 0.00730 | 0.01064 | 0.08287 | 0.43071 | 0.01239 | 0.00586 | 0.01230 | 0.01400 | 0.09729 | 0.00780 |
| **Vasn** | 0.00000 | 0.00564 | 0.00000 | 1.50950 | 2.93038 | 0.00756 | 0.00542 | 1.86199 | 1.17302 | 0.34435 | 1.07550 | 0.07913 | 2.19604 | 0.69412 | 0.07048 | 0.00569 |
| **Vasp** | 3.68238 | 1.44015 | 3.37543 | 0.91114 | 0.35834 | 4.84842 | 1.30007 | 6.69679 | 2.34951 | 1.62826 | 2.82405 | 0.00000 | 3.92649 | 0.62630 | 0.39882 | 3.07339 |
| **Vat1** | 0.09745 | 0.29845 | 0.12121 | 0.90854 | 0.00836 | 0.10563 | 0.29493 | 0.32664 | 2.93541 | 1.72056 | 4.56912 | 0.16450 | 0.10329 | 0.48829 | 0.64007 | 0.45725 |
| **Vat1l** | 1.84719 | 0.00595 | 0.00380 | 0.01723 | 0.00553 | 0.00631 | 0.00590 | 0.00833 | 0.01264 | 0.11895 | 0.01275 | 0.00602 | 0.01143 | 0.01150 | 0.00967 | 0.01279 |
| **Vav1** | 0.01967 | 0.05850 | 1.54015 | 0.27679 | 0.06100 | 0.04621 | 0.01744 | 1.01541 | 0.26364 | 0.96244 | 0.61663 | 0.00741 | 0.09495 | 0.20342 | 0.00131 | 0.00661 |
| **Vav2** | 2.39282 | 1.61246 | 6.96233 | 1.29273 | 0.03523 | 0.18874 | 0.08249 | 5.93005 | 0.39428 | 4.31987 | 4.10572 | 1.32319 | 3.70350 | 0.16146 | 2.24763 | 2.46930 |
| **Vax2os** | 0.01510 | 0.00792 | 0.02850 | 0.05294 | 0.00917 | 0.00606 | 0.00743 | 0.01359 | 0.01207 | 0.01475 | 0.01367 | 0.00796 | 0.02495 | 0.01204 | 0.00944 | 0.01457 |
| **Vbp1** | 7.08606 | 11.75924 | 9.99701 | 4.31393 | 12.56937 | 5.94216 | 3.74630 | 6.11393 | 8.96556 | 8.85733 | 10.93321 | 2.31880 | 10.12037 | 25.93489 | 11.46918 | 4.00888 |
| **Vcl** | 2.89647 | 4.10921 | 3.04646 | 2.95862 | 0.70120 | 2.35578 | 4.64077 | 1.35836 | 0.55427 | 2.74558 | 1.86297 | 11.54265 | 2.37561 | 7.88046 | 3.54220 | 0.04179 |
| **Vcp** | 33.48171 | 19.80414 | 24.93029 | 12.05293 | 25.90559 | 12.62809 | 18.33345 | 33.78979 | 23.29165 | 25.39378 | 12.60399 | 3.53748 | 27.16943 | 1.52584 | 6.37508 | 15.06722 |
| **Vcpip1** | 1.83247 | 3.10020 | 1.65825 | 4.12718 | 0.15415 | 0.00190 | 2.67097 | 0.80674 | 12.23957 | 2.16285 | 2.29108 | 3.24806 | 0.57676 | 5.98367 | 8.24702 | 0.00000 |
| **Vcpkmt** | 1.21892 | 0.37904 | 0.11227 | 0.10506 | 0.38986 | 0.00536 | 0.31341 | 0.00000 | 0.04147 | 1.16501 | 0.62361 | 0.87747 | 1.30111 | 0.62835 | 0.13373 | 1.16939 |
| **Vdac1** | 14.90675 | 8.72915 | 4.80180 | 7.01111 | 13.94849 | 13.86306 | 5.69176 | 15.02036 | 6.72518 | 10.03164 | 14.72673 | 0.75959 | 12.85434 | 15.09782 | 9.00955 | 13.21736 |
| **Vdac2** | 27.95803 | 9.70941 | 18.66451 | 3.51506 | 15.34538 | 15.63617 | 8.01209 | 19.85024 | 3.70097 | 16.96701 | 7.34400 | 3.41954 | 23.02888 | 8.60320 | 3.33482 | 13.19275 |
| **Vdac3** | 19.55725 | 11.52754 | 21.76150 | 6.57346 | 9.67436 | 12.97479 | 12.66638 | 14.58476 | 2.18956 | 10.72348 | 13.14720 | 8.31908 | 12.83218 | 12.66882 | 10.66537 | 12.97760 |
| **Vdr** | 0.01104 | 0.00840 | 0.00285 | 0.00615 | 0.00946 | 1.79857 | 0.00818 | 0.33379 | 0.01090 | 0.25497 | 0.00508 | 0.01827 | 0.00000 | 0.01566 | 0.50667 | 0.00646 |
| **Vegfa** | 0.01743 | 0.01919 | 0.01731 | 0.37190 | 0.01042 | 0.00727 | 0.01763 | 0.01144 | 0.19363 | 0.00929 | 0.02583 | 0.04696 | 0.03614 | 0.02736 | 0.02747 | 0.01475 |
| **Vegfb** | 0.47524 | 0.01763 | 0.00035 | 0.22742 | 2.43324 | 0.00811 | 0.00680 | 0.00000 | 0.44863 | 0.34834 | 0.18979 | 0.01378 | 0.16790 | 0.00945 | 0.00000 | 2.44387 |
| **Veph1** | 0.02049 | 0.02956 | 0.08177 | 0.09300 | 0.00981 | 0.01415 | 0.07005 | 0.05847 | 0.42245 | 0.12866 | 0.16063 | 0.03553 | 0.18333 | 0.30939 | 0.04231 | 0.04410 |
| **Vezf1** | 0.62445 | 0.18369 | 2.68230 | 3.96097 | 1.79472 | 0.00759 | 1.51734 | 1.43857 | 0.75944 | 1.35523 | 2.69650 | 0.06729 | 0.11005 | 0.14154 | 4.32051 | 0.01225 |
| **Vezt** | 0.00000 | 0.00577 | 2.41087 | 0.34185 | 0.01962 | 4.76196 | 3.40740 | 0.00000 | 0.27103 | 1.83203 | 0.30156 | 0.03175 | 0.90978 | 0.66064 | 0.41673 | 2.48093 |
| **Vgf** | 0.01384 | 0.00782 | 0.00953 | 0.26087 | 0.00905 | 0.00678 | 0.00723 | 0.00472 | 0.22976 | 0.30156 | 0.21603 | 0.05321 | 0.01018 | 0.00904 | 0.00000 | 0.00760 |
| **Vgll1** | 0.00876 | 0.00813 | 0.01361 | 0.09558 | 0.00981 | 0.00683 | 0.00705 | 0.01309 | 0.01090 | 0.00547 | 0.01527 | 0.00737 | 1.19552 | 0.01254 | 0.00856 | 0.00694 |
| **Vgll3** | 0.01577 | 0.00803 | 0.00588 | 0.01042 | 0.01040 | 0.32975 | 0.00693 | 0.00998 | 0.01246 | 0.00953 | 0.01188 | 0.03810 | 0.79014 | 0.01251 | 0.00958 | 0.01406 |
| **Vgll4** | 0.00967 | 0.00736 | 0.03147 | 0.00884 | 0.00923 | 0.00836 | 0.00706 | 1.15700 | 0.01087 | 0.57664 | 0.00527 | 0.00337 | 0.00553 | 0.01413 | 0.00000 | 0.00766 |
| **Vhl** | 1.41117 | 0.69141 | 4.16451 | 13.93219 | 0.02256 | 7.74563 | 5.35406 | 9.00262 | 14.39951 | 18.25583 | 10.76932 | 4.91377 | 9.01622 | 15.41065 | 17.17522 | 2.98450 |
| **Vil1** | 0.01153 | 0.00778 | 0.00527 | 9.57641 | 0.00915 | 0.00477 | 0.00689 | 0.01132 | 6.08542 | 0.42082 | 0.00000 | 0.09089 | 0.01217 | 0.32705 | 4.18654 | 0.00549 |
| **Vill** | 0.54623 | 0.00642 | 1.82111 | 0.00092 | 0.01224 | 2.38189 | 0.00446 | 0.95826 | 0.01004 | 0.55730 | 2.80813 | 0.00000 | 2.66608 | 0.00597 | 0.00337 | 0.00388 |
| **Vim** | 1.90329 | 0.00513 | 0.00148 | 0.08780 | 0.00389 | 0.00656 | 0.00562 | 0.00224 | 0.01095 | 6.12185 | 0.00577 | 0.27086 | 0.00875 | 0.00762 | 0.03788 | 1.15696 |
| **Vimp** | 9.66453 | 5.08339 | 17.34833 | 22.10606 | 7.11099 | 5.25831 | 10.84972 | 13.38914 | 27.02000 | 22.95288 | 13.60566 | 14.56099 | 13.02062 | 18.81776 | 13.34630 | 10.34918 |

| | | | | | | | | | | | | | | | | |
|---|---|---|---|---|---|---|---|---|---|---|---|---|---|---|---|---|
| **Vipas39** | 5.41040 | 7.42056 | 7.04476 | 2.86787 | 1.00113 | 10.62103 | 3.48326 | 6.50555 | 1.15149 | 4.04976 | 1.20745 | 2.32882 | 1.49317 | 2.73364 | 2.53106 | 1.84773 |
| **Vkorc1** | 7.33460 | 0.28935 | 3.74385 | 4.41582 | 4.35788 | 3.35925 | 4.37566 | 9.32471 | 9.51507 | 6.71910 | 6.07204 | 0.31810 | 5.95882 | 0.51971 | 0.00000 | 4.43599 |
| **Vkorc1l1** | 6.30735 | 2.45270 | 3.36330 | 9.89678 | 11.96445 | 2.75008 | 4.00069 | 3.00016 | 10.66870 | 8.43578 | 4.76600 | 1.71615 | 2.97279 | 10.80463 | 8.42445 | 0.41249 |
| **Vldlr** | 0.01374 | 0.00807 | 0.01121 | 0.27205 | 0.00922 | 0.00691 | 0.00755 | 0.01348 | 0.00744 | 0.10180 | 0.01259 | 0.05506 | 0.01313 | 0.01172 | 0.00947 | 0.00738 |
| **Vma21** | 0.13301 | 1.86844 | 0.90512 | 3.99373 | 4.21546 | 1.56289 | 0.40362 | 1.13000 | 1.04011 | 2.28970 | 2.24319 | 0.25576 | 2.03708 | 0.28510 | 2.07204 | 0.11842 |
| **Vmac** | 0.00000 | 2.30054 | 0.00589 | 0.29252 | 0.00945 | 0.00627 | 1.63077 | 0.20363 | 0.46193 | 1.18543 | 0.00451 | 0.01640 | 1.59121 | 0.00418 | 1.43190 | 0.03636 |
| **Vmn1r132** | 0.01371 | 0.00785 | 0.01182 | 0.42749 | 0.00930 | 0.00661 | 0.00748 | 0.01306 | 0.57292 | 0.02339 | 0.00870 | 0.00662 | 0.01311 | 0.01509 | 0.04777 | 0.00750 |
| **Vmn1r148** | 0.01374 | 0.00789 | 0.01200 | 0.09155 | 0.00928 | 0.00667 | 0.00747 | 0.01296 | 0.27459 | 0.05219 | 0.01097 | 0.00704 | 0.01309 | 0.01401 | 0.01665 | 0.00768 |
| **Vmn1r32** | 0.01381 | 0.00763 | 0.01166 | 0.01723 | 0.00931 | 0.00662 | 0.00741 | 0.01192 | 0.01174 | 0.02556 | 0.01205 | 0.00688 | 0.01313 | 0.01498 | 0.30152 | 0.00764 |
| **Vmn1r51** | 0.01338 | 0.00821 | 0.01151 | 1.98449 | 0.00937 | 0.00627 | 0.00741 | 0.01281 | 2.48888 | 0.03781 | 0.00000 | 0.01372 | 0.01324 | 0.00954 | 0.00280 | 0.00700 |
| **Vmn1r58** | 0.02629 | 0.03841 | 0.23980 | 0.45728 | 0.06336 | 0.23415 | 0.30455 | 0.33205 | 0.69015 | 0.38149 | 0.02099 | 0.11636 | 0.07838 | 1.43756 | 0.40002 | 0.04296 |
| **Vmn1r65** | 0.08289 | 0.00835 | 0.18185 | 1.28174 | 0.03161 | 0.07758 | 0.15670 | 0.03021 | 1.51939 | 0.01428 | 0.12203 | 0.03842 | 0.14804 | 0.08218 | 0.30180 | 0.25554 |
| **Vmn2r.ps129** | 0.61631 | 0.06893 | 0.83520 | 5.29330 | 0.21826 | 0.11113 | 0.06581 | 0.00000 | 6.59975 | 0.00000 | 4.76375 | 0.33227 | 0.15603 | 2.97395 | 0.28665 | 0.12595 |
| **Vmn2r100** | 0.01375 | 0.00810 | 0.01202 | 0.73528 | 0.00929 | 0.00658 | 0.00751 | 0.01344 | 0.09400 | 0.01369 | 0.02870 | 0.00806 | 0.01233 | 0.01249 | 0.00622 | 0.00728 |
| **Vmn2r12** | 0.11702 | 0.00761 | 0.07478 | 0.00631 | 0.00785 | 0.00588 | 0.00752 | 0.03455 | 0.34369 | 0.02293 | 0.00961 | 0.00740 | 0.01423 | 0.08256 | 0.02136 | 0.00722 |
| **Vmn2r121** | 0.13841 | 0.00745 | 0.11721 | 0.02584 | 0.00755 | 0.00563 | 0.02808 | 0.03182 | 0.01235 | 0.03961 | 0.01233 | 0.00758 | 0.01017 | 0.09517 | 0.01201 | 0.00719 |
| **Vmn2r28** | 0.01372 | 0.00787 | 0.01226 | 0.58880 | 0.00933 | 0.00654 | 0.00749 | 0.01350 | 0.28616 | 0.00988 | 0.01065 | 0.00712 | 0.01271 | 0.00687 | 0.09582 | 0.00727 |
| **Vmn2r29** | 0.01660 | 0.01844 | 0.01130 | 0.00597 | 0.00879 | 0.00685 | 0.00748 | 0.02310 | 0.06551 | 0.49355 | 0.01013 | 0.00709 | 0.01180 | 0.00599 | 0.20819 | 0.00737 |
| **Vmn2r55** | 0.01290 | 0.00790 | 0.02415 | 0.01003 | 0.00949 | 0.00624 | 0.00748 | 0.01135 | 0.01233 | 0.01864 | 0.02763 | 0.02733 | 0.03271 | 0.22933 | 0.00673 | 0.00752 |
| **Vmn2r62** | 0.00919 | 0.00772 | 0.00800 | 0.00921 | 0.00930 | 0.00651 | 0.00685 | 0.00876 | 0.02440 | 0.00000 | 0.01994 | 0.00459 | 0.70756 | 0.81533 | 0.00299 | 0.00684 |
| **Vmn2r94** | 0.01364 | 0.00788 | 0.01214 | 0.22207 | 0.00929 | 0.00654 | 0.00751 | 0.01356 | 0.77992 | 0.00864 | 0.00877 | 0.00627 | 0.01278 | 0.00701 | 0.07363 | 0.00730 |
| **Vmn2r98** | 0.01377 | 0.00807 | 0.01170 | 0.76144 | 0.00925 | 0.00658 | 0.00750 | 0.01267 | 0.11023 | 0.01221 | 0.05788 | 0.00787 | 0.01176 | 0.01266 | 0.00638 | 0.00725 |
| **Vmp1** | 7.69927 | 8.59185 | 17.70844 | 10.52507 | 6.11859 | 10.67829 | 16.72880 | 27.44367 | 9.27398 | 24.74379 | 21.46199 | 18.10541 | 16.35051 | 25.06457 | 25.66959 | 7.05948 |
| **Vnn1** | 2.24068 | 9.96187 | 22.48368 | 0.00000 | 0.26559 | 6.26023 | 4.20328 | 16.52501 | 0.71868 | 16.59168 | 8.09819 | 1.25215 | 7.88578 | 30.69496 | 3.05384 | 0.98334 |
| **Vprbp** | 1.38480 | 1.28236 | 0.76313 | 3.82307 | 0.80074 | 0.12534 | 0.30700 | 3.08368 | 2.19722 | 0.00000 | 3.65034 | 9.14561 | 0.01279 | 4.54788 | 10.19700 | 1.08872 |
| **Vpreb3** | 0.00744 | 0.00690 | 0.62120 | 0.65909 | 0.00777 | 0.00670 | 0.46446 | 1.99162 | 0.00000 | 5.00097 | 5.48633 | 0.19876 | 0.00000 | 0.95712 | 0.88103 | 0.01360 |
| **Vps11** | 4.86647 | 7.36009 | 8.02222 | 2.23359 | 6.18647 | 5.26221 | 2.87538 | 8.35267 | 1.67060 | 4.24893 | 5.98934 | 0.54746 | 4.12367 | 0.39678 | 3.97540 | 0.09512 |
| **Vps13a** | 0.05989 | 0.00626 | 0.03933 | 6.87902 | 0.00784 | 0.00472 | 0.00547 | 0.10043 | 0.00000 | 1.70129 | 0.72090 | 2.84917 | 1.22051 | 2.13666 | 4.76817 | 0.25183 |
| **Vps13b** | 3.78097 | 4.08563 | 1.98481 | 4.34094 | 0.00849 | 0.01827 | 0.00000 | 2.49009 | 0.42238 | 2.54203 | 1.11257 | 3.95653 | 0.84703 | 4.70907 | 2.46950 | 4.54505 |
| **Vps13c** | 1.81956 | 3.67702 | 2.96407 | 0.27927 | 0.06254 | 0.00482 | 1.07846 | 1.34714 | 0.00645 | 0.00000 | 0.03232 | 0.29497 | 0.83640 | 0.00411 | 0.39687 | 2.08619 |
| **Vps13d** | 2.65851 | 0.00000 | 5.44100 | 3.87707 | 5.34903 | 0.40933 | 2.05788 | 5.77170 | 5.59282 | 7.59573 | 3.42780 | 2.64988 | 2.92014 | 1.77049 | 7.02815 | 1.36618 |
| **Vps16** | 9.50496 | 0.37218 | 6.11222 | 3.75055 | 4.31785 | 5.04266 | 7.06389 | 8.54147 | 2.19297 | 10.80661 | 9.90651 | 1.23640 | 7.76625 | 8.40479 | 10.55023 | 4.76751 |
| **Vps18** | 2.92337 | 0.14887 | 0.34939 | 3.78466 | 3.97181 | 0.00255 | 0.45944 | 2.00253 | 2.85545 | 4.59384 | 0.97522 | 0.69177 | 2.68077 | 0.00000 | 0.48545 | 0.06045 |
| **Vps25** | 16.87617 | 12.39957 | 16.90812 | 3.98023 | 8.31546 | 7.24998 | 5.61512 | 4.80649 | 2.30096 | 7.98750 | 6.44702 | 4.00407 | 9.77425 | 0.02956 | 1.11449 | 9.29999 |
| **Vps26a** | 12.96641 | 6.18627 | 11.75274 | 8.96270 | 3.71462 | 5.80360 | 8.19456 | 14.48923 | 8.31195 | 10.48152 | 8.07637 | 4.44759 | 8.66800 | 1.76645 | 8.97529 | 3.78790 |
| **Vps26b** | 3.40521 | 2.84025 | 2.97548 | 1.62551 | 4.93133 | 4.48278 | 1.60652 | 3.64028 | 5.80211 | 1.85735 | 2.04619 | 0.15439 | 2.45679 | 1.74568 | 1.48779 | 0.59571 |
| **Vps28** | 8.12547 | 4.57718 | 10.58178 | 1.32598 | 0.02685 | 0.89419 | 6.38364 | 7.64476 | 1.85263 | 4.76499 | 4.45013 | 0.07583 | 6.93239 | 0.00000 | 0.80906 | 4.19810 |
| **Vps29** | 15.26561 | 8.90672 | 17.43072 | 5.23286 | 11.82414 | 6.02190 | 6.09954 | 6.11940 | 5.04666 | 2.29715 | 1.77274 | 0.18661 | 12.48886 | 0.00000 | 0.00000 | 4.39611 |
| **Vps33a** | 6.19951 | 7.59665 | 5.45499 | 4.32856 | 6.19776 | 7.87464 | 1.91208 | 11.53412 | 7.54836 | 6.30866 | 5.16485 | 7.35178 | 3.83466 | 3.22149 | 7.12442 | 4.35220 |
| **Vps33b** | 5.90011 | 0.62910 | 3.28649 | 3.15539 | 3.39479 | 0.00990 | 2.47499 | 7.25378 | 3.16283 | 1.61960 | 5.31137 | 0.15328 | 1.26644 | 3.59099 | 0.00000 | 0.17581 |
| **Vps35** | 7.36155 | 15.23149 | 15.35732 | 8.03191 | 14.52544 | 9.93667 | 7.62207 | 12.15051 | 6.10975 | 11.11768 | 6.67468 | 18.63231 | 7.77478 | 17.87659 | 12.64557 | 7.57119 |
| **Vps36** | 4.47300 | 1.77517 | 1.06720 | 0.00514 | 0.00000 | 4.80399 | 3.00335 | 0.65651 | 0.01175 | 1.73909 | 0.67082 | 0.34487 | 5.31774 | 0.86320 | 0.00000 | 1.83215 |
| **Vps37a** | 2.68955 | 0.21118 | 1.60469 | 4.91554 | 1.70144 | 1.48237 | 1.53754 | 3.44862 | 0.45655 | 3.13521 | 2.16465 | 0.51126 | 1.66161 | 12.25513 | 2.09921 | 0.82009 |
| **Vps37b** | 3.64125 | 0.16124 | 0.39404 | 4.95442 | 0.68150 | 0.69460 | 1.24096 | 12.23056 | 7.71026 | 13.22690 | 5.37551 | 0.51305 | 5.02824 | 0.00000 | 0.00000 | 3.28591 |
| **Vps37c** | 0.37033 | 0.00686 | 0.65474 | 0.27027 | 0.04402 | 0.00535 | 0.86123 | 1.21623 | 0.00572 | 0.41911 | 0.50276 | 0.03651 | 2.61552 | 0.00626 | 0.12153 | 0.58860 |
| **Vps37d** | 0.07866 | 0.00806 | 0.02195 | 0.30244 | 0.00864 | 0.00648 | 0.00723 | 0.00000 | 0.44910 | 2.00609 | 0.74859 | 0.05042 | 0.03330 | 0.00592 | 0.00000 | 0.01494 |
| **Vps39** | 3.65948 | 6.25768 | 5.90959 | 1.01525 | 0.05198 | 1.63194 | 1.04869 | 3.08125 | 0.39343 | 3.01295 | 1.78621 | 1.67040 | 0.46766 | 0.43388 | 1.22640 | 0.20804 |
| **Vps41** | 11.06627 | 5.94995 | 4.77979 | 0.77262 | 0.05732 | 7.87743 | 6.93290 | 5.59869 | 1.97147 | 6.71210 | 9.37245 | 7.06712 | 4.84512 | 3.10707 | 3.35830 | 4.76687 |
| **Vps45** | 2.82765 | 0.71302 | 4.97743 | 0.00168 | 0.00689 | 3.16642 | 0.25097 | 1.23278 | 0.61471 | 1.49644 | 0.60722 | 0.02979 | 2.91236 | 0.04717 | 0.00000 | 0.00000 |
| **Vps4a** | 7.64643 | 4.29966 | 5.93826 | 5.02605 | 10.37220 | 0.01873 | 0.95231 | 9.84540 | 10.49573 | 8.56376 | 4.13602 | 9.07377 | 3.77462 | 7.67303 | 6.32273 | 7.12216 |
| **Vps4b** | 5.07553 | 7.42896 | 2.56897 | 1.92052 | 3.98073 | 2.62545 | 3.74484 | 5.74183 | 3.95600 | 6.18324 | 4.04947 | 18.52034 | 3.26982 | 0.99423 | 11.97405 | 1.49009 |
| **Vps50** | 2.59238 | 3.66962 | 3.29853 | 2.31107 | 0.26118 | 1.73189 | 6.81951 | 3.51176 | 0.15701 | 2.84835 | 5.44656 | 11.77967 | 2.87498 | 5.39639 | 1.35231 | 0.08675 |
| **Vps51** | 4.40425 | 0.05749 | 2.18467 | 1.58635 | 1.24451 | 0.43814 | 0.09881 | 5.78924 | 0.47332 | 4.16890 | 2.01509 | 0.17059 | 8.09424 | 0.00000 | 0.00000 | 1.34502 |
| **Vps52** | 8.04303 | 3.73276 | 3.94354 | 0.62149 | 1.37414 | 1.79543 | 4.73829 | 6.63123 | 0.59570 | 4.67801 | 2.57726 | 0.40196 | 8.87905 | 0.20060 | 0.17252 | 2.52172 |
| **Vps53** | 5.73641 | 4.45499 | 2.15294 | 0.00000 | 0.02090 | 4.13204 | 1.56353 | 2.67841 | 0.47039 | 5.48402 | 3.29287 | 0.54459 | 2.30955 | 0.02585 | 0.94983 | 1.85896 |
| **Vps54** | 4.41497 | 8.54906 | 8.14771 | 2.36701 | 6.98034 | 5.80205 | 0.06777 | 7.71037 | 0.37349 | 4.20253 | 3.77874 | 0.31360 | 4.73444 | 4.73261 | 2.95971 | 0.41897 |
| **Vps72** | 7.62669 | 0.05958 | 0.00000 | 0.37185 | 2.18410 | 2.26454 | 0.31871 | 7.79845 | 0.72256 | 11.08933 | 1.58934 | 0.37428 | 6.12274 | 0.26505 | 0.00000 | 4.20172 |
| **Vps8** | 4.98893 | 2.96137 | 0.00000 | 1.77831 | 0.01505 | 2.60385 | 1.69132 | 7.72943 | 2.96570 | 4.83488 | 2.98900 | 2.41567 | 7.76486 | 0.54185 | 0.72482 | 0.01074 |
| **Vps9d1** | 1.63021 | 0.38687 | 1.57163 | 1.23812 | 0.00232 | 0.00506 | 2.23964 | 0.90433 | 2.18802 | 1.36201 | 0.67353 | 0.00848 | 0.09947 | 0.00000 | 0.55866 | 1.12442 |
| **Vrk1** | 15.06639 | 6.75508 | 8.81700 | 1.99376 | 4.52923 | 4.94322 | 4.85013 | 13.88008 | 4.31963 | 11.96364 | 11.05184 | 4.85613 | 9.31046 | 3.61201 | 13.86660 | 3.04180 |
| **Vrk2** | 0.66409 | 0.00715 | 0.73270 | 0.00534 | 0.03023 | 0.00505 | 0.00641 | 1.59635 | 0.00947 | 2.22269 | 0.24642 | 0.04790 | 0.38182 | 0.02341 | 0.00000 | 0.00620 |
| **Vrk3** | 0.00000 | 3.13920 | 1.60056 | 0.94093 | 0.01225 | 4.05669 | 2.50914 | 5.39242 | 1.27308 | 3.61624 | 4.90066 | 0.12546 | 6.12689 | 1.25383 | 2.77118 | 3.62987 |

| | | | | | | | | | | | | | | | | |
|---|---|---|---|---|---|---|---|---|---|---|---|---|---|---|---|---|
| **Vrtn** | 0.01374 | 0.00799 | 0.01228 | 1.01235 | 0.00936 | 0.00650 | 0.00746 | 0.01324 | 0.13018 | 0.00944 | 0.01148 | 0.00756 | 0.01268 | 0.00548 | 0.14090 | 0.00720 |
| **Vsig10** | 0.02704 | 0.00782 | 0.30414 | 0.14644 | 0.03657 | 0.02261 | 0.02713 | 0.33904 | 0.40590 | 0.50918 | 0.07038 | 0.06656 | 0.03476 | 0.07612 | 0.27952 | 0.02954 |
| **Vsig10l** | 0.11344 | 0.01794 | 1.12765 | 0.25312 | 0.00552 | 0.01365 | 0.00578 | 1.83203 | 3.65621 | 0.00000 | 1.17288 | 0.04137 | 1.54449 | 0.00000 | 0.03198 | 0.00539 |
| **Vsir** | 0.01089 | 0.00777 | 0.00982 | 0.00991 | 0.00955 | 0.00687 | 0.00733 | 0.13769 | 0.01205 | 0.03323 | 0.01137 | 0.01833 | 0.00831 | 0.00708 | 0.10547 | 0.00740 |
| **Vstm5** | 3.80534 | 1.12635 | 0.01163 | 0.00245 | 0.00935 | 0.00234 | 0.00382 | 0.00000 | 0.27609 | 0.00466 | 0.01024 | 0.00455 | 0.01002 | 0.01274 | 0.00000 | 0.30947 |
| **Vsx2** | 0.01290 | 0.01911 | 0.00911 | 0.86646 | 0.00896 | 0.00641 | 0.00751 | 0.00889 | 4.37287 | 0.00000 | 0.19193 | 0.00000 | 0.01004 | 0.00938 | 0.00339 | 0.00683 |
| **Vta1** | 7.09697 | 6.05029 | 10.73813 | 6.08068 | 10.34250 | 2.47670 | 1.14960 | 3.72902 | 2.92746 | 3.49214 | 1.73488 | 0.89710 | 5.79749 | 0.04621 | 0.63873 | 4.03271 |
| **Vtcn1** | 0.00000 | 0.00827 | 2.90029 | 0.00203 | 0.02520 | 0.74284 | 0.80104 | 1.91199 | 0.00879 | 1.43949 | 0.01855 | 0.00663 | 4.08266 | 0.00575 | 0.78668 | 0.00453 |
| **Vti1a** | 1.51798 | 0.49084 | 0.02092 | 0.20800 | 0.03105 | 1.38383 | 0.01501 | 0.08395 | 0.00816 | 0.63257 | 0.04765 | 0.05079 | 1.55697 | 0.00571 | 0.05127 | 2.57632 |
| **Vti1b** | 4.65745 | 6.41373 | 6.41898 | 3.18620 | 0.21281 | 3.26063 | 3.45300 | 1.74305 | 8.25784 | 7.51960 | 7.47822 | 0.68757 | 10.37300 | 1.56028 | 0.29703 | 5.09016 |
| **Vwa2** | 0.01370 | 0.00788 | 0.01201 | 0.47216 | 0.00937 | 0.00651 | 0.00734 | 0.01134 | 0.00417 | 0.22774 | 0.02602 | 0.01656 | 0.01211 | 0.01147 | 0.00615 | 0.00739 |
| **Vwa3a** | 0.01049 | 0.00776 | 0.00673 | 1.19081 | 0.00877 | 0.00642 | 0.00710 | 0.00878 | 0.47118 | 2.81540 | 0.00000 | 0.04516 | 0.01047 | 1.36558 | 0.18847 | 0.00702 |
| **Vwa5b1** | 0.00415 | 0.00818 | 0.00000 | 0.00282 | 0.00932 | 0.00819 | 0.00658 | 2.46876 | 0.00813 | 0.63847 | 1.23125 | 0.01632 | 1.42018 | 0.00959 | 0.65070 | 0.00647 |
| **Vwa5b2** | 0.01282 | 0.00776 | 0.01156 | 1.05188 | 0.00878 | 0.00643 | 0.00726 | 0.01005 | 0.17092 | 1.74812 | 0.00945 | 0.04101 | 0.01048 | 0.00521 | 0.05443 | 0.00706 |
| **Vwa7** | 0.10821 | 0.01756 | 0.02205 | 0.08919 | 0.00767 | 0.01477 | 0.01753 | 0.00689 | 0.37527 | 0.00787 | 0.27397 | 0.00515 | 0.03048 | 0.22659 | 0.04746 | 0.00710 |
| **Vwa8** | 2.65429 | 0.00429 | 0.00000 | 0.60117 | 0.20099 | 3.46171 | 0.00466 | 0.93980 | 0.57873 | 0.35217 | 0.57388 | 0.02757 | 0.00000 | 0.00588 | 0.46397 | 0.20496 |
| **Vwa9** | 3.70921 | 1.20806 | 2.48604 | 7.80372 | 2.44998 | 2.09668 | 3.79305 | 3.08488 | 0.86272 | 4.64898 | 0.98728 | 0.20194 | 3.94625 | 0.00000 | 0.75111 | 0.80478 |
| **Vwc2** | 0.08449 | 0.00850 | 0.00898 | 0.56277 | 0.01570 | 0.00682 | 0.00728 | 0.01074 | 0.21220 | 0.28368 | 0.01112 | 0.00655 | 0.01254 | 0.01064 | 0.00565 | 0.00785 |
| **Vwce** | 0.00411 | 0.00710 | 0.00463 | 0.20249 | 0.83147 | 0.00746 | 0.00690 | 0.14404 | 0.01450 | 0.00461 | 0.01240 | 0.00650 | 0.53198 | 0.01251 | 0.00472 | 0.00687 |
| **Vwf** | 0.01353 | 0.00794 | 0.01026 | 0.01623 | 0.00930 | 0.00672 | 0.00729 | 0.01069 | 0.01207 | 0.06799 | 0.01190 | 0.00664 | 0.01329 | 0.25480 | 0.16147 | 0.00777 |
| **Wac** | 5.66267 | 0.64502 | 3.80745 | 12.38329 | 6.77683 | 4.32037 | 6.81771 | 12.23830 | 19.34352 | 13.57328 | 11.88158 | 6.36986 | 4.69833 | 31.38372 | 15.18271 | 1.38397 |
| **Wapl** | 7.32920 | 9.76667 | 7.25657 | 21.66765 | 10.60596 | 5.55099 | 6.13020 | 9.98185 | 21.98576 | 12.85895 | 7.09927 | 14.32497 | 5.90298 | 26.13944 | 20.36478 | 2.10200 |
| **Wars** | 15.23226 | 8.03715 | 10.51586 | 1.42894 | 9.43719 | 0.07435 | 8.65120 | 13.63779 | 3.09140 | 9.94366 | 5.52115 | 1.32993 | 6.99841 | 0.00000 | 2.40655 | 7.68678 |
| **Wars2** | 3.00352 | 5.07063 | 2.58390 | 0.56077 | 0.38982 | 1.96727 | 8.15922 | 5.20295 | 1.71291 | 8.48694 | 4.94317 | 3.54360 | 0.68158 | 4.70752 | 3.06402 | 0.00604 |
| **Was** | 0.63840 | 0.03865 | 0.19382 | 0.46035 | 0.06044 | 0.00596 | 0.04873 | 0.05585 | 1.04246 | 0.82628 | 0.79328 | 0.03986 | 0.17822 | 0.32721 | 0.62865 | 0.08356 |
| **Wasf1** | 1.39533 | 0.00617 | 0.04605 | 4.05389 | 0.06282 | 0.00542 | 0.00642 | 1.01533 | 3.63229 | 0.72978 | 0.01749 | 0.00553 | 0.04005 | 0.00750 | 0.66853 | 0.20424 |
| **Wasf2** | 1.35406 | 1.38932 | 2.16837 | 4.42860 | 0.60563 | 5.11998 | 0.70434 | 3.53962 | 4.93833 | 2.99183 | 5.56562 | 0.16569 | 3.36182 | 0.18919 | 0.78095 | 0.92194 |
| **Wash1** | 2.74226 | 1.36627 | 0.94151 | 4.08682 | 0.03801 | 0.04439 | 0.40259 | 0.85634 | 3.47755 | 2.71193 | 0.72141 | 0.14390 | 0.54941 | 0.19868 | 0.09212 | 0.23508 |
| **Wasl** | 6.20104 | 2.03741 | 0.55029 | 16.06420 | 1.22944 | 4.99821 | 3.03402 | 6.14043 | 9.94550 | 7.00805 | 4.17050 | 11.59848 | 3.91782 | 9.02715 | 8.49415 | 3.89045 |
| **Wbp1** | 3.63857 | 0.76141 | 7.54364 | 0.73027 | 3.27363 | 3.25972 | 6.47001 | 2.93924 | 0.00000 | 4.82160 | 0.69737 | 0.00997 | 3.70817 | 0.00258 | 0.00000 | 4.60407 |
| **Wbp11** | 11.38293 | 8.82391 | 7.53638 | 13.21200 | 7.84757 | 5.81456 | 2.47070 | 9.27668 | 15.40320 | 13.70323 | 9.32510 | 9.44410 | 9.35716 | 1.80048 | 3.18694 | 6.96050 |
| **Wbp1l** | 8.70434 | 1.79010 | 6.76684 | 7.32815 | 5.77799 | 9.34735 | 6.05828 | 6.17523 | 6.29587 | 9.09250 | 4.91851 | 0.49342 | 2.62429 | 0.29882 | 6.19984 | 4.07347 |
| **Wbp2** | 8.56900 | 0.50825 | 3.27166 | 3.56340 | 5.66480 | 6.15340 | 0.38574 | 7.86533 | 3.11477 | 5.36780 | 8.60127 | 0.96974 | 10.18222 | 0.00000 | 0.74806 | 6.12422 |
| **Wbp2nl** | 0.00920 | 0.00667 | 0.00595 | 0.00283 | 0.00932 | 0.01878 | 2.30230 | 0.01120 | 0.33921 | 0.97410 | 0.00850 | 0.02381 | 0.00938 | 0.15995 | 0.40141 | 0.00694 |
| **Wbp4** | 8.01533 | 0.92253 | 4.02824 | 5.45922 | 0.89881 | 0.00000 | 1.64136 | 3.58468 | 3.07244 | 10.28965 | 3.32900 | 2.74648 | 2.95988 | 4.41950 | 3.18764 | 0.11334 |
| **Wbp5** | 0.00533 | 0.03063 | 0.03104 | 0.00462 | 0.03163 | 0.00650 | 0.00591 | 0.04611 | 0.01094 | 3.27358 | 0.00864 | 0.00000 | 3.68373 | 0.00591 | 0.20831 | 0.18936 |
| **Wbscr16** | 2.28022 | 3.24793 | 5.50753 | 2.19462 | 7.77236 | 3.62638 | 0.83649 | 1.89586 | 3.81362 | 5.62683 | 4.72448 | 0.36683 | 2.32994 | 0.00000 | 0.86844 | 0.17493 |
| **Wbscr17** | 0.01369 | 0.00787 | 0.01173 | 0.22546 | 0.00927 | 0.00653 | 0.00752 | 0.01245 | 0.60161 | 0.07020 | 0.00681 | 0.00633 | 0.01218 | 0.01475 | 0.13542 | 0.00713 |
| **Wbscr22** | 13.00315 | 10.58636 | 13.74383 | 1.85049 | 7.20598 | 3.64761 | 7.39543 | 9.02786 | 6.15532 | 16.62394 | 8.49253 | 1.20899 | 12.39153 | 0.00000 | 1.69268 | 4.02884 |
| **Wbscr25** | 0.01371 | 0.00813 | 0.01225 | 0.59050 | 0.00929 | 0.00658 | 0.00751 | 0.01396 | 0.23005 | 0.01394 | 0.01249 | 0.00782 | 0.01271 | 0.01065 | 0.00643 | 0.00730 |
| **Wbscr27** | 1.54758 | 0.34294 | 1.55188 | 2.93748 | 5.62754 | 1.00531 | 0.13739 | 3.07009 | 0.73993 | 2.75584 | 0.53154 | 0.07047 | 1.15573 | 0.00000 | 0.54606 | 1.66947 |
| **Wdfy1** | 6.02802 | 4.84875 | 7.35993 | 0.42241 | 7.00815 | 2.74253 | 4.98972 | 7.27539 | 0.17325 | 8.46392 | 5.77435 | 11.33485 | 8.13845 | 18.08168 | 4.24970 | 5.35573 |
| **Wdfy2** | 0.00411 | 0.00750 | 0.61734 | 2.03931 | 0.00711 | 0.00819 | 0.00706 | 0.08021 | 4.16543 | 0.21010 | 0.00000 | 0.48587 | 0.02754 | 0.01738 | 0.46739 | 0.55826 |
| **Wdfy3** | 3.77697 | 4.29633 | 0.00000 | 3.28129 | 0.17631 | 0.00000 | 0.05402 | 0.00000 | 1.94816 | 6.42768 | 4.92487 | 8.13711 | 0.00000 | 0.82060 | 3.78644 | 0.00000 |
| **Wdfy4** | 0.01141 | 0.00804 | 0.01420 | 0.00731 | 0.00984 | 0.00723 | 0.00748 | 0.01255 | 0.01171 | 0.96770 | 0.01406 | 0.00507 | 0.33784 | 0.01168 | 0.00888 | 0.00798 |
| **Wdhd1** | 4.89300 | 3.73506 | 2.11978 | 4.31528 | 0.00557 | 0.00709 | 5.97115 | 2.69523 | 4.00496 | 7.35658 | 2.23222 | 1.95415 | 4.71986 | 7.97427 | 6.59176 | 4.06594 |
| **Wdpcp** | 0.01637 | 0.00749 | 0.29854 | 0.00835 | 0.00855 | 0.00570 | 0.00712 | 0.00917 | 0.01184 | 0.52039 | 0.02917 | 0.00557 | 0.01024 | 0.01342 | 0.01244 | 0.00764 |
| **Wdr1** | 13.69344 | 8.02215 | 8.02800 | 3.77126 | 5.29636 | 9.43094 | 8.15451 | 5.03259 | 1.29546 | 3.33902 | 3.64464 | 0.25701 | 7.51598 | 0.01022 | 2.98230 | 6.42874 |
| **Wdr11** | 2.35589 | 3.97473 | 4.60590 | 1.41131 | 6.85976 | 4.02406 | 1.11926 | 2.31659 | 2.25793 | 3.12923 | 3.21451 | 1.30341 | 5.60895 | 0.68018 | 1.42190 | 1.61357 |
| **Wdr12** | 9.08196 | 14.51725 | 14.75770 | 4.64821 | 10.97188 | 2.58556 | 11.96021 | 13.59870 | 7.11040 | 19.28296 | 11.68442 | 10.21147 | 10.99249 | 17.30318 | 12.00273 | 3.89494 |
| **Wdr13** | 0.00000 | 0.00606 | 0.85082 | 1.44784 | 2.54553 | 0.00597 | 0.00731 | 2.23779 | 0.00000 | 1.60129 | 0.60627 | 0.28924 | 0.00000 | 0.00253 | 1.79495 | 2.47837 |
| **Wdr17** | 0.01048 | 0.00743 | 0.00801 | 0.17455 | 0.00968 | 0.00677 | 0.00714 | 0.25958 | 0.14390 | 0.00291 | 0.00738 | 0.00544 | 0.00677 | 0.36774 | 0.03529 | 0.00743 |
| **Wdr18** | 15.98256 | 5.71802 | 10.17086 | 4.83431 | 11.03759 | 7.11522 | 5.78391 | 25.38580 | 8.53962 | 24.57956 | 13.25468 | 3.06837 | 18.67797 | 3.29795 | 2.11855 | 3.61776 |
| **Wdr19** | 0.98207 | 0.00661 | 0.00078 | 2.61062 | 0.03808 | 0.00656 | 0.00628 | 0.00407 | 0.18601 | 0.13463 | 0.16492 | 0.02333 | 0.00771 | 0.00099 | 0.60909 | 1.20767 |
| **Wdr20** | 2.38264 | 4.03949 | 0.85832 | 12.06125 | 0.16878 | 2.77646 | 0.03203 | 0.91775 | 6.74358 | 1.20051 | 1.16276 | 0.31719 | 0.53974 | 3.21288 | 8.72166 | 1.57608 |
| **Wdr20rt** | 2.44544 | 0.40434 | 0.07555 | 0.29251 | 0.00350 | 0.51632 | 0.24995 | 0.13293 | 0.07202 | 0.12407 | 0.00502 | 0.02081 | 0.03647 | 0.27280 | 1.35431 | 0.09958 |
| **Wdr24** | 0.03852 | 2.28773 | 1.16262 | 1.65068 | 0.04236 | 0.00849 | 2.61100 | 3.82271 | 0.54598 | 6.42345 | 2.64613 | 0.78937 | 2.42329 | 0.00000 | 1.00048 | 4.73244 |
| **Wdr25** | 2.22980 | 0.00534 | 0.34744 | 1.37838 | 0.03026 | 0.00862 | 0.00508 | 0.07588 | 2.17592 | 0.32019 | 0.47522 | 0.02675 | 0.01730 | 0.00277 | 0.04480 | 0.00488 |
| **Wdr26** | 10.09006 | 3.16509 | 9.90167 | 19.24239 | 10.95108 | 9.09212 | 4.76917 | 10.75956 | 17.05365 | 15.35807 | 8.84308 | 11.07084 | 6.99450 | 21.99763 | 21.14699 | 5.29512 |
| **Wdr27** | 0.00928 | 0.00757 | 0.00956 | 0.18530 | 0.00906 | 0.00667 | 0.00717 | 0.18581 | 0.08544 | 1.49442 | 0.01010 | 0.00382 | 0.00669 | 0.01231 | 0.06693 | 0.00711 |
| **Wdr3** | 10.28550 | 12.65172 | 7.78037 | 6.19216 | 9.43307 | 4.00165 | 5.31570 | 8.73555 | 8.99748 | 13.91692 | 9.31151 | 11.84941 | 7.33825 | 12.34548 | 14.11506 | 1.07215 |
| **Wdr31** | 0.68417 | 0.00723 | 0.01351 | 0.23587 | 0.00690 | 0.00559 | 0.02890 | 0.26411 | 0.00789 | 0.00000 | 1.76538 | 0.00000 | 0.00000 | 0.00271 | 0.41052 | 0.04758 |

| | | | | | | | | | | | | | | | | |
|---|---|---|---|---|---|---|---|---|---|---|---|---|---|---|---|---|
| **Wdr33** | 14.11742 | 1.40802 | 8.03174 | 9.35340 | 0.79765 | 0.05880 | 2.81840 | 9.86042 | 15.14360 | 10.96183 | 6.63434 | 5.97540 | 3.74832 | 1.56152 | 6.58917 | 6.30664 |
| **Wdr34** | 3.18769 | 0.16808 | 1.77086 | 1.90277 | 0.53331 | 1.61012 | 1.15253 | 3.28060 | 3.77518 | 4.05287 | 7.06315 | 0.86035 | 0.57898 | 0.50235 | 1.05263 | 0.39275 |
| **Wdr35** | 0.00675 | 4.71080 | 0.00334 | 0.00000 | 0.06243 | 0.00625 | 0.01555 | 0.00865 | 0.28663 | 0.61171 | 0.00851 | 0.01447 | 0.01052 | 0.01097 | 0.16674 | 0.00555 |
| **Wdr36** | 6.01575 | 11.10279 | 3.69420 | 13.84085 | 5.75494 | 4.67921 | 7.52972 | 13.32161 | 20.23265 | 24.16402 | 13.40450 | 15.39390 | 6.44866 | 19.16990 | 21.51941 | 5.89753 |
| **Wdr37** | 0.95867 | 1.57141 | 2.90267 | 7.47591 | 0.00986 | 0.00309 | 0.22492 | 3.08351 | 9.74628 | 1.08512 | 1.78357 | 12.79847 | 0.31042 | 3.93647 | 4.78702 | 0.04419 |
| **Wdr4** | 3.77189 | 2.20816 | 2.08580 | 3.91366 | 6.94260 | 3.66200 | 5.19542 | 10.93268 | 5.85935 | 14.37685 | 6.79972 | 0.63798 | 9.67299 | 0.60704 | 3.42187 | 4.34529 |
| **Wdr41** | 0.01266 | 0.00769 | 0.01017 | 0.25822 | 0.00997 | 0.00722 | 0.00731 | 0.00000 | 0.00583 | 1.00031 | 0.53627 | 0.03694 | 0.13232 | 0.00412 | 0.23990 | 0.00767 |
| **Wdr43** | 11.32913 | 9.94396 | 9.23624 | 21.64172 | 12.60697 | 3.08339 | 6.84538 | 23.24727 | 30.13908 | 39.89680 | 27.43487 | 26.73600 | 6.77452 | 21.92938 | 34.72556 | 6.13199 |
| **Wdr44** | 0.00670 | 0.00804 | 0.02126 | 6.21899 | 0.02398 | 0.52949 | 0.01714 | 1.81644 | 12.12309 | 0.97492 | 0.57942 | 0.00000 | 0.00000 | 6.74112 | 1.27821 | 0.07699 |
| **Wdr45** | 7.92449 | 8.90085 | 22.77547 | 8.53311 | 4.59706 | 4.66664 | 10.82038 | 31.07744 | 18.61287 | 27.68353 | 19.50212 | 13.84412 | 15.89168 | 6.59966 | 17.83889 | 9.49725 |
| **Wdr45b** | 5.69244 | 1.05895 | 4.33680 | 12.34473 | 3.44807 | 9.54816 | 5.81202 | 25.66679 | 9.32562 | 34.68019 | 15.03781 | 17.67020 | 12.24945 | 14.01422 | 9.62066 | 5.05910 |
| **Wdr46** | 11.00223 | 4.30937 | 6.02057 | 6.38663 | 5.93784 | 0.89353 | 6.10162 | 15.86048 | 7.27663 | 19.56225 | 13.25043 | 1.13497 | 9.51276 | 2.93896 | 4.08460 | 5.12549 |
| **Wdr47** | 0.00000 | 0.00690 | 0.84894 | 9.90438 | 0.00742 | 3.51028 | 0.01722 | 0.55509 | 9.52157 | 3.48776 | 2.65372 | 3.87803 | 0.10824 | 9.40910 | 4.49124 | 0.01731 |
| **Wdr48** | 1.95992 | 4.59578 | 0.00000 | 0.89135 | 7.90696 | 3.31538 | 2.24208 | 4.82208 | 0.68594 | 4.78607 | 2.11555 | 0.32474 | 1.35463 | 6.20038 | 0.83870 | 0.00000 |
| **Wdr5** | 12.90657 | 6.15451 | 18.49264 | 17.98642 | 6.79750 | 7.37256 | 12.91383 | 23.52245 | 15.23205 | 16.47097 | 20.58036 | 6.47856 | 19.98414 | 12.89874 | 13.55367 | 3.07959 |
| **Wdr53** | 1.92782 | 5.16038 | 5.92288 | 2.97238 | 2.79256 | 3.34259 | 5.32849 | 7.11235 | 7.02475 | 5.11646 | 3.41097 | 5.53415 | 4.86513 | 5.46901 | 3.32679 | 0.00290 |
| **Wdr54** | 0.00602 | 0.00763 | 0.42741 | 0.00591 | 0.00916 | 0.00585 | 0.22130 | 0.97221 | 0.01039 | 1.05110 | 0.15109 | 0.00216 | 0.00565 | 0.01262 | 0.00147 | 0.00778 |
| **Wdr55** | 3.59481 | 4.04384 | 5.49086 | 11.25824 | 5.49321 | 0.63572 | 2.54596 | 5.04550 | 13.76300 | 9.88948 | 1.92445 | 0.31735 | 2.97314 | 0.49692 | 4.48731 | 6.10587 |
| **Wdr59** | 1.53727 | 6.08718 | 4.89489 | 1.08930 | 0.51109 | 0.63592 | 0.45862 | 0.35401 | 3.19306 | 2.79731 | 1.21701 | 0.92984 | 0.18670 | 0.25951 | 1.43554 | 1.07284 |
| **Wdr5b** | 0.01863 | 0.03945 | 0.06941 | 0.09193 | 0.00882 | 0.00576 | 0.00760 | 0.02103 | 0.56980 | 0.00456 | 0.00521 | 0.05498 | 0.01128 | 0.00782 | 1.80869 | 0.00729 |
| **Wdr6** | 5.61415 | 3.26215 | 1.09078 | 0.05389 | 4.22754 | 4.80696 | 0.00000 | 3.92742 | 0.55317 | 2.84390 | 0.74418 | 0.11930 | 1.69240 | 0.35085 | 1.01751 | 3.50018 |
| **Wdr60** | 0.01166 | 0.00753 | 0.46871 | 0.00392 | 0.00833 | 0.00739 | 0.00695 | 0.00867 | 0.79939 | 0.43950 | 0.00027 | 0.05980 | 0.27948 | 0.05207 | 0.40688 | 0.00719 |
| **Wdr61** | 17.38502 | 10.60656 | 13.87863 | 0.73570 | 13.91571 | 6.83419 | 10.18993 | 10.34151 | 0.93919 | 16.03141 | 8.01280 | 10.08033 | 9.39280 | 9.70269 | 9.12003 | 3.97883 |
| **Wdr62** | 3.90296 | 2.37388 | 1.75200 | 5.89319 | 0.01545 | 0.00190 | 0.17926 | 2.47655 | 5.86380 | 4.77412 | 2.91722 | 0.17596 | 0.00000 | 0.00663 | 3.61899 | 0.22042 |
| **Wdr63** | 0.01332 | 0.00768 | 0.01134 | 0.00452 | 0.00900 | 0.00660 | 0.00725 | 0.01003 | 0.64770 | 0.79989 | 0.00723 | 0.03197 | 0.01186 | 0.01161 | 0.01872 | 0.00774 |
| **Wdr7** | 0.52923 | 3.93715 | 0.03650 | 0.98484 | 0.01713 | 0.00873 | 2.27851 | 1.33560 | 0.38638 | 0.42068 | 1.01941 | 0.19633 | 0.01331 | 0.90084 | 3.28729 | 0.02081 |
| **Wdr70** | 0.54179 | 5.05746 | 0.67248 | 0.94513 | 4.33626 | 0.23969 | 0.76828 | 5.06034 | 1.04479 | 2.91028 | 1.71436 | 0.07675 | 1.62103 | 0.63359 | 2.17681 | 3.02325 |
| **Wdr72** | 0.00087 | 0.00821 | 0.01317 | 0.00000 | 0.01163 | 3.45122 | 1.75773 | 1.81251 | 0.00798 | 0.30080 | 0.69261 | 0.07581 | 0.25556 | 1.57183 | 1.50616 | 0.02024 |
| **Wdr73** | 7.96994 | 5.21078 | 12.84978 | 9.21132 | 0.78643 | 4.26024 | 8.83531 | 13.78564 | 7.61219 | 16.42849 | 6.62439 | 1.30205 | 4.31094 | 1.85998 | 5.57521 | 6.23459 |
| **Wdr74** | 15.45037 | 9.93875 | 17.17263 | 7.74018 | 7.95666 | 7.56702 | 8.52819 | 26.14885 | 14.08416 | 24.39049 | 19.35962 | 3.75482 | 16.46937 | 3.13498 | 7.73802 | 10.54347 |
| **Wdr75** | 7.04771 | 11.25138 | 9.63460 | 7.54847 | 8.17130 | 0.04144 | 8.25762 | 16.08996 | 13.96281 | 13.26029 | 12.77741 | 12.67011 | 5.78091 | 24.29910 | 19.13701 | 5.35385 |
| **Wdr76** | 0.78825 | 1.54446 | 1.28402 | 0.55786 | 0.01899 | 0.01634 | 2.37318 | 0.00000 | 1.38901 | 0.43712 | 1.21230 | 0.03320 | 0.12168 | 2.44661 | 0.20578 | 0.18963 |
| **Wdr77** | 19.08112 | 8.15611 | 13.60880 | 15.67377 | 18.92500 | 5.01268 | 8.25762 | 22.80237 | 16.31593 | 27.99715 | 13.13249 | 15.20910 | 12.23202 | 23.33032 | 21.41074 | 3.56492 |
| **Wdr78** | 0.02211 | 0.00793 | 0.11126 | 0.29943 | 0.01013 | 0.00574 | 0.00728 | 0.01189 | 0.00731 | 0.07607 | 0.01418 | 0.01692 | 0.01136 | 0.01065 | 0.00398 | 0.00750 |
| **Wdr8** | 4.07907 | 0.14539 | 3.53479 | 0.00000 | 3.20367 | 0.40857 | 2.28524 | 2.64195 | 0.00628 | 0.87075 | 0.10150 | 0.01833 | 5.39195 | 0.01593 | 0.22622 | 1.18078 |
| **Wdr81** | 0.21897 | 1.19100 | 3.94911 | 7.04827 | 0.00000 | 0.11455 | 7.85340 | 7.00860 | 4.67533 | 11.37652 | 3.85277 | 0.28976 | 5.03459 | 0.00000 | 3.62865 | 0.01539 |
| **Wdr82** | 8.35508 | 9.21712 | 8.58803 | 10.22128 | 5.88574 | 5.18234 | 9.82188 | 5.84733 | 6.18745 | 7.42580 | 2.56340 | 12.64856 | 8.77208 | 14.40581 | 6.66899 | 5.93729 |
| **Wdr83** | 3.81198 | 0.00000 | 3.12772 | 0.44920 | 3.29272 | 3.71260 | 2.65038 | 0.98641 | 0.00000 | 2.37575 | 1.33321 | 0.07813 | 6.17270 | 0.00301 | 0.01390 | 1.46726 |
| **Wdr83os** | 5.05259 | 1.16201 | 6.60927 | 1.30459 | 2.27748 | 3.52410 | 4.40810 | 4.43087 | 1.06340 | 3.61328 | 8.25620 | 0.87223 | 9.48575 | 1.73455 | 2.09958 | 3.43179 |
| **Wdr89** | 0.02101 | 0.00759 | 0.43611 | 1.07469 | 0.00912 | 0.04680 | 0.43301 | 1.68925 | 1.35541 | 0.94416 | 0.32226 | 0.13710 | 1.24803 | 0.42987 | 3.53911 | 0.23867 |
| **Wdr90** | 0.93359 | 0.32752 | 0.40365 | 0.20610 | 0.28756 | 2.51067 | 1.17040 | 6.35393 | 0.22015 | 2.75437 | 3.88790 | 0.09622 | 2.33289 | 0.67154 | 1.51672 | 0.16462 |
| **Wdr91** | 3.80723 | 0.02963 | 6.99173 | 1.57639 | 2.39228 | 0.00212 | 0.98007 | 12.45484 | 1.57661 | 5.82185 | 8.05541 | 1.80924 | 3.06813 | 0.04371 | 1.25651 | 0.01227 |
| **Wdr92** | 4.21507 | 5.39326 | 2.66447 | 9.29859 | 5.42601 | 1.90497 | 0.95020 | 2.08383 | 10.60058 | 3.53140 | 2.92135 | 11.03625 | 1.04059 | 5.49448 | 5.75440 | 0.00994 |
| **Wdr93** | 0.00462 | 0.00810 | 1.32530 | 1.00194 | 0.00702 | 0.00640 | 0.00690 | 0.27365 | 0.94015 | 0.43255 | 0.21576 | 0.37246 | 1.55850 | 0.02234 | 1.27784 | 0.00616 |
| **Wdr95** | 2.34538 | 0.05706 | 1.90623 | 1.13035 | 0.00411 | 1.56831 | 2.04124 | 0.00000 | 0.00000 | 1.00263 | 0.42323 | 0.12189 | 1.45609 | 0.72964 | 0.00000 | 0.18283 |
| **Wdsub1** | 1.55647 | 0.00503 | 0.25451 | 0.58347 | 0.00459 | 0.00584 | 0.96230 | 2.24792 | 0.71928 | 3.68716 | 0.00000 | 0.00000 | 1.70808 | 3.30436 | 0.81375 | 0.00473 |
| **Wdtc1** | 5.06998 | 0.90858 | 2.35631 | 11.72293 | 4.24721 | 4.82477 | 5.09296 | 17.84770 | 10.72764 | 26.64138 | 16.01988 | 1.85655 | 10.16283 | 0.92539 | 4.08500 | 5.53583 |
| **Wdyhv1** | 4.80584 | 0.04171 | 0.00000 | 3.15783 | 0.00000 | 5.56060 | 2.73829 | 1.34949 | 3.34960 | 1.91293 | 1.75931 | 0.29140 | 0.00000 | 1.57661 | 2.73516 | 0.06645 |
| **Wee1** | 10.11288 | 5.62209 | 6.88656 | 5.91561 | 0.78675 | 1.81146 | 6.04826 | 6.45070 | 2.77658 | 6.77055 | 3.35375 | 10.78393 | 3.38554 | 6.26525 | 10.44312 | 2.61503 |
| **Wee2** | 0.01257 | 0.00802 | 0.00672 | 4.29290 | 0.00927 | 0.00589 | 0.00751 | 0.01501 | 2.26090 | 0.23106 | 0.28744 | 0.00000 | 0.00900 | 0.01623 | 1.55785 | 0.00651 |
| **Wfdc15a** | 0.80012 | 0.12829 | 0.20025 | 1.08901 | 1.41190 | 0.00670 | 0.00632 | 0.05200 | 0.00000 | 0.82058 | 0.75048 | 0.06887 | 0.12857 | 0.00690 | 0.43539 | 0.02839 |
| **Wfdc15b** | 0.01302 | 0.00796 | 0.00997 | 0.00784 | 0.00912 | 0.00679 | 0.00730 | 0.01044 | 0.01202 | 0.67350 | 0.01047 | 0.00508 | 0.01200 | 0.32825 | 0.00000 | 0.00755 |
| **Wfdc18** | 0.01369 | 0.00803 | 0.01221 | 0.00986 | 0.00924 | 0.00669 | 0.00749 | 0.01274 | 0.01225 | 0.17303 | 0.01248 | 0.00774 | 0.01248 | 0.01148 | 0.01071 | 0.00745 |
| **Wfdc2** | 2.08417 | 0.02502 | 0.15457 | 0.74817 | 0.01030 | 0.27575 | 0.00509 | 0.00000 | 0.00000 | 3.32379 | 0.00914 | 0.05320 | 0.31879 | 0.01445 | 0.01194 | 4.36961 |
| **Wfdc3** | 0.00640 | 0.00760 | 0.00821 | 1.77314 | 0.00897 | 0.00620 | 0.00608 | 0.00001 | 0.46457 | 0.00000 | 0.33177 | 0.00250 | 1.99381 | 0.00289 | 0.97649 | 0.00595 |
| **Wfikkn1** | 0.02350 | 0.00741 | 0.00222 | 0.22956 | 0.00981 | 0.10307 | 0.82847 | 0.05552 | 0.49759 | 0.28977 | 0.00729 | 0.00668 | 0.00872 | 0.01466 | 0.02191 | 0.00723 |
| **Wfikkn2** | 1.06756 | 0.00751 | 0.01609 | 0.18568 | 0.01691 | 0.01309 | 0.00662 | 0.00885 | 0.03557 | 0.03217 | 0.01296 | 0.00718 | 0.04824 | 0.06975 | 0.02061 | 0.01371 |
| **Wfs1** | 0.01784 | 0.00689 | 1.80948 | 0.46964 | 0.00534 | 0.00650 | 0.03541 | 1.88527 | 1.34293 | 0.02043 | 1.15340 | 0.01157 | 2.16570 | 0.01790 | 0.00000 | 0.07545 |
| **Whamm** | 5.87241 | 3.95440 | 7.94433 | 6.19445 | 6.55481 | 1.05398 | 6.48445 | 8.11332 | 1.36146 | 0.00000 | 5.97711 | 0.02305 | 2.62468 | 0.00000 | 0.00000 | 1.94572 |
| **Whrn** | 2.33452 | 0.07671 | 0.12826 | 0.66374 | 0.00461 | 1.67216 | 0.61587 | 3.25412 | 0.00000 | 0.11091 | 2.99301 | 0.00354 | 0.77223 | 0.01427 | 0.00000 | 0.33184 |
| **Whsc1** | 0.15714 | 0.01838 | 1.48987 | 5.66451 | 2.61242 | 5.63754 | 0.00530 | 0.22065 | 2.54652 | 2.50729 | 0.20098 | 0.46550 | 3.22082 | 0.20032 | 4.20663 | 5.66323 |
| **Whsc1l1** | 2.13788 | 0.02699 | 0.59891 | 3.49791 | 2.09375 | 1.66097 | 0.09657 | 0.85163 | 3.92253 | 0.93712 | 1.20316 | 0.09458 | 2.14244 | 0.23105 | 5.28489 | 0.16660 |

| | | | | | | | | | | | | | | | | |
|---|---|---|---|---|---|---|---|---|---|---|---|---|---|---|---|---|
| **Wif1** | 0.01371 | 0.00805 | 0.01151 | 0.00951 | 0.00917 | 0.00659 | 0.00745 | 0.01170 | 0.01232 | 0.25642 | 0.01136 | 0.04745 | 0.01145 | 0.01140 | 0.00901 | 0.00714 |
| **Wipf1** | 1.62984 | 0.00368 | 0.22717 | 1.75562 | 0.01245 | 5.30857 | 0.00000 | 1.16224 | 0.00000 | 5.90533 | 6.26941 | 4.64668 | 4.51723 | 7.43263 | 9.15454 | 0.67171 |
| **Wipf2** | 2.43169 | 0.93287 | 3.43198 | 7.74766 | 0.13048 | 0.03386 | 0.05367 | 1.20983 | 7.35798 | 7.24290 | 4.49645 | 3.36807 | 1.37734 | 4.91053 | 6.44130 | 0.02830 |
| **Wipi1** | 0.00772 | 0.00746 | 0.00689 | 0.70124 | 0.00860 | 0.00671 | 0.00603 | 0.00000 | 0.22538 | 2.38498 | 0.59583 | 0.03440 | 1.74654 | 0.01928 | 0.24944 | 0.00650 |
| **Wipi2** | 6.46749 | 3.07258 | 8.17424 | 16.30013 | 5.09552 | 12.30304 | 6.52447 | 20.74910 | 23.57497 | 26.88611 | 20.27631 | 3.42395 | 8.97148 | 7.83057 | 16.01583 | 7.47809 |
| **Wisp1** | 0.01391 | 0.00777 | 0.00960 | 0.00483 | 0.00933 | 0.00680 | 0.00733 | 0.01132 | 0.24217 | 0.00939 | 0.00935 | 0.20896 | 0.01316 | 0.01285 | 0.00386 | 0.00746 |
| **Wisp3** | 0.01376 | 0.00797 | 0.01216 | 0.01082 | 0.00932 | 0.00666 | 0.00753 | 0.01376 | 0.01224 | 0.01490 | 0.01291 | 0.00805 | 0.01272 | 0.01427 | 0.01563 | 0.00741 |
| **Wiz** | 3.50838 | 3.67368 | 3.61181 | 4.80215 | 0.28332 | 0.08610 | 5.61514 | 7.15907 | 3.00316 | 9.15032 | 5.82349 | 1.51994 | 5.77197 | 0.17485 | 0.81127 | 3.14776 |
| **Wls** | 0.01359 | 0.00785 | 0.01140 | 0.00435 | 0.00918 | 0.00662 | 0.00732 | 0.01090 | 0.67653 | 0.32500 | 0.00765 | 0.03420 | 0.01253 | 0.01237 | 0.00600 | 0.00772 |
| **Wnk1** | 17.25412 | 8.70532 | 9.59559 | 21.30552 | 7.67695 | 7.20245 | 10.47091 | 14.65340 | 24.51806 | 29.61093 | 19.22791 | 21.23090 | 7.77341 | 33.35778 | 24.93130 | 5.98877 |
| **Wnk2** | 0.00000 | 0.00764 | 0.00849 | 2.79536 | 0.01734 | 4.05001 | 0.00651 | 0.85508 | 0.36777 | 0.75040 | 0.57394 | 0.02014 | 0.47680 | 0.00308 | 0.00000 | 3.22645 |
| **Wnk3** | 1.64180 | 5.39385 | 0.61823 | 1.37158 | 0.01391 | 0.00380 | 4.09114 | 1.75149 | 0.46067 | 7.86875 | 0.73898 | 0.12471 | 1.68184 | 18.86335 | 7.12405 | 0.00159 |
| **Wnk4** | 0.05895 | 0.00737 | 0.30583 | 2.02015 | 0.01544 | 0.00517 | 0.00606 | 0.02439 | 1.29862 | 0.08078 | 0.01929 | 0.03870 | 1.49186 | 0.21738 | 0.00000 | 2.94606 |
| **Wnt10b** | 0.00733 | 0.00718 | 0.04223 | 0.00463 | 0.00854 | 0.00650 | 0.00670 | 1.01208 | 0.00949 | 3.84292 | 1.44808 | 0.01951 | 0.00000 | 0.02399 | 0.05722 | 0.00687 |
| **Wnt2b** | 0.01302 | 0.00792 | 0.01052 | 0.00720 | 0.00931 | 0.00670 | 0.00734 | 0.02923 | 0.01182 | 0.00945 | 0.15061 | 0.00738 | 0.00963 | 0.01117 | 0.00866 | 0.00732 |
| **Wnt3a** | 0.00000 | 0.00744 | 0.65248 | 0.00865 | 0.00644 | 4.64435 | 0.00605 | 0.00645 | 0.01278 | 0.00000 | 0.00853 | 0.00641 | 1.31352 | 0.01839 | 0.00000 | 2.05374 |
| **Wnt4** | 0.13539 | 0.08357 | 0.42554 | 0.24911 | 0.09349 | 0.05828 | 0.09054 | 0.07043 | 0.62928 | 0.81770 | 1.08150 | 0.24385 | 0.33870 | 0.57837 | 0.30044 | 1.67656 |
| **Wnt6** | 0.01405 | 0.00756 | 0.05422 | 0.01036 | 0.43830 | 0.02240 | 0.00743 | 0.01257 | 0.01214 | 0.00839 | 0.01248 | 0.00763 | 0.01534 | 0.01307 | 0.00971 | 0.59137 |
| **Wnt7a** | 0.01360 | 0.00786 | 0.01213 | 0.05358 | 0.00936 | 0.00652 | 0.00732 | 0.01124 | 0.08790 | 0.36206 | 0.01159 | 0.01606 | 0.01207 | 0.01149 | 0.00656 | 0.00746 |
| **Wnt7b** | 0.00000 | 0.00720 | 0.01965 | 0.00879 | 0.05606 | 6.05350 | 0.00521 | 1.35850 | 0.01155 | 0.00000 | 0.00000 | 0.00307 | 2.84993 | 0.01885 | 0.00000 | 2.36393 |
| **Wrap53** | 15.58888 | 4.59596 | 5.69311 | 4.95366 | 6.54937 | 0.00000 | 5.57705 | 4.32117 | 4.32826 | 8.96495 | 3.64943 | 0.27402 | 5.50480 | 0.11191 | 2.87620 | 3.25529 |
| **Wrb** | 1.80401 | 2.22877 | 4.61018 | 0.48967 | 0.00478 | 2.16595 | 2.95120 | 3.50551 | 0.24050 | 5.24871 | 2.26120 | 0.67830 | 2.86417 | 2.14371 | 1.11173 | 0.00728 |
| **Wrn** | 0.47870 | 0.00716 | 0.14931 | 1.90338 | 0.00724 | 0.00534 | 0.00714 | 0.00822 | 0.07101 | 0.00000 | 0.03238 | 0.01374 | 0.11861 | 0.39189 | 0.79119 | 0.00662 |
| **Wrnip1** | 11.64798 | 5.23220 | 5.84499 | 4.24359 | 0.61528 | 0.00231 | 2.45391 | 4.36203 | 1.59960 | 2.34999 | 1.58962 | 1.15556 | 3.28173 | 1.63183 | 1.42134 | 4.64836 |
| **Wsb1** | 7.37739 | 11.12997 | 4.56868 | 13.53387 | 6.78581 | 7.65431 | 5.13711 | 22.90270 | 21.88363 | 25.96975 | 21.45732 | 9.66884 | 5.75623 | 35.10077 | 29.54985 | 4.86669 |
| **Wsb2** | 4.55095 | 3.30724 | 0.96708 | 1.15799 | 5.51667 | 7.44126 | 0.07072 | 4.39383 | 0.44113 | 5.60196 | 7.66383 | 0.43310 | 1.66222 | 3.23324 | 2.24731 | 0.31885 |
| **Wscd2** | 0.00000 | 0.00818 | 0.00612 | 0.00570 | 0.00766 | 0.00686 | 0.00776 | 0.47726 | 0.01079 | 0.00000 | 0.38010 | 0.00588 | 0.47362 | 0.01440 | 0.00000 | 1.28365 |
| **Wt1** | 0.01369 | 0.00797 | 0.01218 | 0.00983 | 0.00924 | 0.00671 | 0.00746 | 0.01257 | 0.01221 | 0.17867 | 0.01283 | 0.00758 | 0.01275 | 0.01147 | 0.01076 | 0.00760 |
| **Wtap** | 13.17828 | 15.02679 | 24.24764 | 20.89243 | 9.00618 | 14.83978 | 18.31431 | 30.39478 | 26.29264 | 35.50340 | 39.23853 | 16.86160 | 26.79099 | 44.32542 | 35.00927 | 5.64141 |
| **Wtip** | 1.90521 | 0.00385 | 0.00000 | 1.20675 | 3.09925 | 0.38864 | 0.31134 | 1.13360 | 0.66527 | 2.83862 | 0.06041 | 0.10611 | 2.26515 | 0.04732 | 0.00000 | 0.01918 |
| **Wwc1** | 0.00000 | 0.00719 | 0.68743 | 0.00715 | 0.00644 | 0.01894 | 0.44431 | 0.90746 | 0.01056 | 1.05556 | 0.00443 | 0.01175 | 1.40045 | 0.01493 | 0.07602 | 1.33804 |
| **Wwc2** | 0.00000 | 0.59092 | 4.36852 | 7.79199 | 0.99435 | 3.09862 | 6.94251 | 15.84313 | 5.22099 | 4.51341 | 4.96528 | 0.20464 | 6.96344 | 0.58775 | 3.72842 | 0.86154 |
| **Wwox** | 0.11821 | 0.00860 | 0.64266 | 0.32102 | 0.31772 | 0.00563 | 0.00742 | 0.00272 | 0.05205 | 0.10907 | 0.34987 | 0.00644 | 0.01174 | 0.00491 | 0.31961 | 2.09531 |
| **Wwp1** | 0.32619 | 6.77466 | 0.56417 | 4.79288 | 2.39361 | 0.97435 | 2.20007 | 1.33963 | 1.00429 | 0.88014 | 0.48587 | 1.87174 | 0.07306 | 4.45608 | 1.96815 | 1.46560 |
| **Wwp2** | 3.36791 | 6.14442 | 4.58792 | 7.79276 | 4.81019 | 4.43388 | 1.48773 | 10.29128 | 5.96905 | 12.42015 | 12.74628 | 2.47983 | 4.06516 | 1.13803 | 9.71688 | 3.92655 |
| **Wwtr1** | 2.80342 | 6.37305 | 6.90279 | 5.11867 | 2.66576 | 2.57928 | 3.29856 | 3.23636 | 10.09652 | 7.99572 | 9.20639 | 4.65900 | 0.94930 | 10.61052 | 2.40451 | 1.15656 |
| **Xab2** | 10.91606 | 5.35178 | 11.83967 | 3.56546 | 5.89643 | 7.17459 | 9.40824 | 15.23833 | 4.97184 | 9.27725 | 5.66055 | 0.86179 | 13.03936 | 0.13687 | 3.92601 | 0.38223 |
| **Xaf1** | 0.42658 | 0.08857 | 0.87392 | 3.47559 | 1.94835 | 0.11848 | 0.01381 | 0.18127 | 4.86125 | 0.03396 | 7.39128 | 0.27946 | 0.10451 | 1.37684 | 0.33959 | 0.14667 |
| **Xbp1** | 6.30131 | 4.19853 | 9.12835 | 0.14634 | 3.06592 | 4.04806 | 4.61166 | 5.19662 | 0.06634 | 0.00054 | 1.48489 | 0.16490 | 4.73713 | 0.00408 | 0.32732 | 6.20410 |
| **Xcr1** | 0.03496 | 0.03215 | 1.40686 | 0.50963 | 0.09051 | 0.01145 | 0.00818 | 0.70272 | 0.22331 | 0.03505 | 0.16259 | 0.01561 | 0.07507 | 0.71049 | 0.09003 | 0.00671 |
| **Xdh** | 0.01368 | 0.00801 | 0.00942 | 0.88482 | 0.00971 | 0.00750 | 0.00764 | 0.00215 | 0.00000 | 0.18590 | 0.49470 | 0.01731 | 0.03997 | 0.01078 | 0.21531 | 0.00745 |
| **Xiap** | 3.32254 | 6.53679 | 6.40066 | 5.90231 | 0.40985 | 0.00707 | 3.21798 | 1.62563 | 2.89571 | 3.94954 | 3.97893 | 14.60349 | 1.34073 | 23.77778 | 9.30318 | 1.91887 |
| **Xist** | 10.86946 | 0.00000 | 12.63464 | 0.15632 | 0.76803 | 0.00000 | 0.01514 | 9.71999 | 0.21341 | 8.24615 | 0.00133 | 0.02063 | 3.35083 | 13.94413 | 6.71352 | 12.35011 |
| **Xk** | 0.59188 | 0.00587 | 0.23455 | 0.01087 | 0.01707 | 0.00782 | 0.00592 | 0.00588 | 0.00928 | 0.00000 | 0.00000 | 4.69275 | 0.11982 | 0.01632 | 0.07136 | 0.00660 |
| **Xkr8** | 4.81592 | 0.97595 | 6.18443 | 0.15540 | 4.24852 | 3.55417 | 3.02639 | 4.23505 | 0.35116 | 0.00000 | 0.31770 | 0.16644 | 0.59211 | 0.28639 | 0.70423 | 0.01969 |
| **Xkr9** | 0.00534 | 0.00887 | 0.00196 | 0.00000 | 0.00938 | 0.00619 | 0.00802 | 0.17312 | 0.00747 | 2.02748 | 0.15742 | 0.00000 | 0.00597 | 2.39119 | 3.16330 | 0.00701 |
| **Xkrx** | 0.05052 | 0.06053 | 0.02517 | 0.03606 | 0.01028 | 0.01484 | 0.00703 | 0.00794 | 0.06972 | 0.01573 | 0.20494 | 0.04642 | 0.13879 | 2.62462 | 0.01546 | 0.01475 |
| **Xlr** | 0.01105 | 0.00701 | 0.00662 | 0.14931 | 0.00930 | 0.00653 | 0.00703 | 0.06356 | 0.02586 | 0.51384 | 0.00000 | 1.15843 | 0.01049 | 0.05067 | 0.45268 | 0.00715 |
| **Xlr3a** | 2.21724 | 3.06858 | 5.82266 | 0.70114 | 1.13124 | 0.00508 | 2.69951 | 0.00000 | 1.25897 | 0.64851 | 1.22849 | 0.05474 | 3.48932 | 0.17689 | 1.76744 | 1.63989 |
| **Xlr3b** | 2.78751 | 3.42699 | 4.01472 | 0.47257 | 1.01516 | 0.02058 | 3.43516 | 0.00000 | 1.36385 | 0.59399 | 1.84359 | 0.06446 | 1.93209 | 0.55053 | 1.18369 | 0.97307 |
| **Xlr3c** | 2.16521 | 3.41435 | 5.14136 | 0.28781 | 1.21571 | 0.01323 | 3.36666 | 0.00000 | 1.54976 | 0.89517 | 1.11684 | 0.07990 | 1.16312 | 0.48260 | 0.74481 | 0.52989 |
| **Xlr4a** | 2.33453 | 1.54108 | 0.29200 | 1.36460 | 0.94797 | 0.19560 | 2.61386 | 0.19433 | 1.35245 | 0.20352 | 1.59673 | 0.84767 | 0.87170 | 0.04388 | 1.04826 | 0.03151 |
| **Xlr4b** | 3.33718 | 4.59409 | 0.96006 | 2.14253 | 5.52319 | 1.24192 | 3.84618 | 1.10736 | 2.53527 | 0.62877 | 2.44066 | 5.51044 | 3.20707 | 0.14900 | 2.67618 | 0.05513 |
| **Xlr4c** | 1.19642 | 1.86681 | 0.52375 | 1.89554 | 2.14436 | 0.15068 | 2.25343 | 0.30959 | 1.13671 | 0.27117 | 1.74473 | 2.09388 | 0.98319 | 0.05110 | 1.55931 | 0.03928 |
| **Xlr5a** | 1.89123 | 0.12192 | 0.00000 | 1.39457 | 0.06276 | 0.00411 | 3.72518 | 0.00000 | 0.00000 | 1.70180 | 0.91580 | 0.59951 | 1.95322 | 4.69232 | 1.98102 | 0.11447 |
| **Xlr5b** | 1.44801 | 0.04785 | 0.00000 | 1.71662 | 0.05378 | 0.00525 | 3.81365 | 0.00000 | 0.00673 | 1.61709 | 0.54006 | 0.38094 | 1.68708 | 5.13598 | 1.66269 | 0.06364 |
| **Xlr5c** | 0.53384 | 0.01782 | 0.00000 | 0.00853 | 0.04703 | 0.00661 | 1.44323 | 0.01460 | 0.01086 | 0.48814 | 0.04411 | 0.07710 | 0.57572 | 1.62902 | 0.60210 | 0.04382 |
| **Xndc1** | 1.91264 | 0.50553 | 0.78377 | 2.89456 | 3.02612 | 0.28388 | 0.16361 | 1.29224 | 3.39780 | 1.74226 | 0.23189 | 0.03848 | 0.71911 | 0.00000 | 0.22203 | 0.62909 |
| **Xpa** | 0.98006 | 0.01608 | 0.65496 | 2.00239 | 0.08018 | 0.00741 | 0.00526 | 0.00000 | 3.45726 | 0.53277 | 2.24297 | 0.02240 | 0.55601 | 0.73902 | 0.29786 | 0.01219 |
| **Xpc** | 4.20657 | 0.21439 | 0.00000 | 0.22711 | 0.01986 | 5.70816 | 2.51519 | 4.36468 | 0.00715 | 1.28089 | 0.27925 | 0.00750 | 3.53396 | 0.34775 | 0.32125 | 1.83215 |
| **Xpnpep1** | 17.43324 | 11.39617 | 17.47637 | 13.50683 | 12.16220 | 9.19482 | 9.28252 | 16.51320 | 5.92096 | 18.05658 | 9.33643 | 4.54555 | 14.99016 | 0.32509 | 9.05761 | 14.19669 |
| **Xpnpep2** | 0.00553 | 0.00845 | 0.07262 | 0.00658 | 0.00969 | 0.00886 | 0.00814 | 0.27888 | 0.01102 | 0.79684 | 0.00351 | 0.08294 | 0.00850 | 0.78406 | 0.22868 | 0.00741 |

| | | | | | | | | | | | | | | | | |
|---|---|---|---|---|---|---|---|---|---|---|---|---|---|---|---|---|
| **Xpnpep3** | 1.03706 | 0.13817 | 0.94716 | 0.70751 | 0.09886 | 0.04498 | 0.15103 | 1.28655 | 1.59029 | 1.02255 | 1.40882 | 0.11271 | 1.54289 | 1.08780 | 2.57736 | 0.02737 |
| **Xpo1** | 8.96860 | 15.15451 | 12.88664 | 18.69015 | 8.76915 | 2.85810 | 10.52597 | 21.64222 | 26.39281 | 21.96540 | 20.84072 | 30.66844 | 12.32719 | 42.78398 | 24.98445 | 1.92681 |
| **Xpo4** | 2.78377 | 2.14210 | 0.29700 | 1.81218 | 0.04986 | 0.00146 | 0.85197 | 2.72042 | 1.07792 | 4.78500 | 1.59787 | 7.73561 | 0.28979 | 6.63903 | 4.32169 | 0.00239 |
| **Xpo5** | 4.98937 | 8.00669 | 1.95928 | 0.47294 | 1.80006 | 4.49418 | 1.84879 | 8.29522 | 0.49219 | 6.62390 | 5.11380 | 0.16920 | 7.61378 | 0.31814 | 7.39163 | 1.23443 |
| **Xpo6** | 8.43276 | 1.90530 | 4.59391 | 8.45863 | 7.64722 | 9.23298 | 6.95624 | 13.53225 | 2.50324 | 12.01419 | 6.96823 | 1.57366 | 5.37211 | 0.57619 | 4.53493 | 2.19654 |
| **Xpo7** | 9.09970 | 0.09720 | 8.52564 | 5.60871 | 2.56953 | 3.17285 | 7.30297 | 4.08940 | 3.50335 | 6.13967 | 7.14429 | 4.60034 | 6.83232 | 7.08213 | 4.96864 | 6.65113 |
| **Xpot** | 12.68805 | 14.29571 | 6.21904 | 6.06918 | 10.78040 | 5.03318 | 10.34370 | 19.86728 | 10.84885 | 20.26230 | 13.84544 | 19.19388 | 10.89108 | 23.26692 | 15.54374 | 4.29023 |
| **Xpr1** | 4.27257 | 0.00000 | 2.19527 | 7.75621 | 3.13024 | 1.38240 | 4.58721 | 9.43452 | 8.46910 | 18.75617 | 12.38844 | 10.97260 | 2.42605 | 20.91856 | 9.78392 | 0.03168 |
| **Xrcc1** | 8.34691 | 4.93880 | 9.88349 | 0.88909 | 8.71709 | 7.94230 | 7.35845 | 15.54961 | 0.50286 | 11.37004 | 10.69682 | 0.60606 | 15.01121 | 0.03099 | 1.56354 | 5.95347 |
| **Xrcc2** | 3.25769 | 0.03575 | 2.02957 | 2.19295 | 0.01995 | 0.05894 | 2.04606 | 0.21909 | 0.97649 | 0.37308 | 1.61645 | 0.01112 | 0.35094 | 2.17448 | 2.85172 | 0.29339 |
| **Xrcc3** | 0.00000 | 1.43644 | 2.18204 | 0.21174 | 2.98309 | 2.89450 | 1.72651 | 2.31855 | 0.06324 | 0.97831 | 0.48722 | 0.04028 | 2.31734 | 0.00334 | 0.25452 | 0.01170 |
| **Xrcc4** | 2.66850 | 3.15221 | 1.96281 | 1.13741 | 1.26687 | 4.55025 | 0.00275 | 0.04557 | 0.00000 | 1.48859 | 0.79239 | 0.06701 | 2.00547 | 2.45194 | 1.01635 | 0.00000 |
| **Xrcc5** | 9.50452 | 4.53901 | 1.97807 | 1.06291 | 4.82089 | 1.89651 | 4.95036 | 0.81834 | 1.12831 | 0.60037 | 0.31799 | 0.08043 | 1.85780 | 0.00001 | 0.51536 | 4.22024 |
| **Xrcc6** | 9.40542 | 10.30213 | 8.96233 | 0.08602 | 6.56427 | 3.62996 | 0.63438 | 5.45736 | 0.02813 | 5.08064 | 4.88645 | 0.15230 | 2.11390 | 4.18610 | 2.23839 | 0.06071 |
| **Xrcc6bp1** | 0.01638 | 0.00695 | 0.52890 | 0.55467 | 0.84793 | 0.00594 | 0.00691 | 0.10591 | 3.12769 | 4.08738 | 0.32608 | 0.01599 | 0.11398 | 0.00519 | 0.84179 | 0.02087 |
| **Xrn1** | 0.04782 | 0.00644 | 0.75068 | 5.40351 | 2.40225 | 0.02970 | 0.06999 | 2.27829 | 2.39481 | 2.50323 | 1.45676 | 0.21623 | 0.86898 | 1.03920 | 2.13339 | 0.52972 |
| **Xrn2** | 12.20326 | 13.90390 | 9.25497 | 6.20508 | 12.53708 | 4.95095 | 9.57756 | 13.17262 | 8.95908 | 9.69775 | 5.21492 | 14.55033 | 5.46774 | 12.83091 | 12.99265 | 1.65841 |
| **Xxylt1** | 0.01970 | 1.71547 | 0.78919 | 0.05565 | 0.01514 | 0.87476 | 0.00642 | 0.58175 | 0.16934 | 0.04140 | 0.00570 | 0.01376 | 0.00000 | 0.00456 | 0.59163 | 0.02048 |
| **Xylb** | 3.51286 | 0.88268 | 2.02191 | 6.61878 | 3.49401 | 5.80571 | 0.00000 | 7.84566 | 10.26332 | 10.98399 | 11.28777 | 4.76129 | 4.50482 | 7.94178 | 13.69883 | 2.92007 |
| **Xylt2** | 4.29799 | 0.18751 | 2.89931 | 0.61335 | 0.55821 | 0.00000 | 0.00296 | 0.00000 | 0.78706 | 0.08185 | 0.56586 | 0.03781 | 0.80359 | 0.00325 | 0.20408 | 3.93410 |
| **Yae1d1** | 6.08712 | 3.49611 | 6.44687 | 3.17757 | 1.95754 | 0.00551 | 4.48004 | 2.99423 | 2.78551 | 12.18861 | 5.09309 | 7.66408 | 4.27117 | 12.04874 | 9.79318 | 1.81823 |
| **Yaf2** | 0.00000 | 0.01756 | 0.00608 | 1.62959 | 2.09853 | 0.07631 | 1.56045 | 0.07591 | 2.45457 | 1.36473 | 0.04011 | 0.34413 | 1.43043 | 0.00643 | 0.57358 | 0.00652 |
| **Yap1** | 8.11746 | 3.75396 | 4.65827 | 16.36212 | 7.07682 | 5.44220 | 5.86090 | 7.57036 | 17.01288 | 10.02970 | 9.72085 | 3.28503 | 4.70504 | 19.58402 | 15.17922 | 2.19675 |
| **Yars** | 11.74250 | 14.13881 | 10.13206 | 2.70342 | 13.20968 | 5.75017 | 12.18006 | 26.11046 | 7.42878 | 17.45783 | 14.05664 | 7.81407 | 18.16727 | 2.30999 | 11.79251 | 10.99669 |
| **Yars2** | 4.32826 | 2.13692 | 0.44406 | 2.98994 | 0.01119 | 0.29974 | 0.01119 | 2.62097 | 1.64305 | 3.51186 | 4.56541 | 1.30659 | 0.89509 | 1.68611 | 0.57525 | 0.00725 |
| **Ybey** | 0.52641 | 0.01781 | 3.86509 | 1.86067 | 0.23824 | 0.01401 | 0.08609 | 1.22534 | 3.55503 | 1.85932 | 1.13560 | 0.62865 | 0.65262 | 3.35590 | 0.24200 | 0.07901 |
| **Ybx1** | 26.61496 | 21.42595 | 23.92640 | 20.63809 | 20.65508 | 10.57981 | 20.73938 | 27.48930 | 26.60297 | 38.59717 | 36.64362 | 22.84051 | 26.44386 | 24.40133 | 28.32156 | 11.77531 |
| **Ybx2** | 0.75574 | 0.00698 | 0.04389 | 5.29491 | 0.68483 | 0.77190 | 0.04667 | 0.85893 | 3.56702 | 4.30859 | 1.93980 | 0.04361 | 3.32933 | 0.00000 | 0.00000 | 1.24091 |
| **Ybx3** | 25.41934 | 13.71647 | 21.83728 | 8.29984 | 10.43362 | 14.97109 | 12.81294 | 25.01677 | 3.16783 | 25.93286 | 27.41792 | 12.00710 | 21.46764 | 3.77061 | 5.68312 | 12.55633 |
| **Ydjc** | 0.08587 | 0.00760 | 0.75371 | 0.77705 | 0.00762 | 0.00828 | 0.00655 | 0.02601 | 2.72361 | 0.48147 | 0.88201 | 0.01460 | 0.61445 | 0.01378 | 0.00000 | 0.03616 |
| **Yeats2** | 5.02294 | 6.69082 | 4.30282 | 5.12511 | 3.23261 | 2.86363 | 0.98977 | 2.93803 | 3.88512 | 6.17094 | 1.62640 | 3.32425 | 2.14130 | 4.78067 | 3.00563 | 7.90742 |
| **Yeats4** | 10.98856 | 0.62939 | 2.05175 | 4.95506 | 1.75636 | 5.07442 | 0.15475 | 2.88353 | 5.43872 | 13.77545 | 4.20645 | 0.56237 | 5.07120 | 0.50310 | 0.00000 | 0.60928 |
| **Yes1** | 0.72085 | 1.94140 | 0.00000 | 2.87130 | 0.49556 | 0.04223 | 0.00522 | 0.68463 | 0.58908 | 0.00000 | 0.50329 | 1.46642 | 0.02955 | 9.64892 | 3.12653 | 0.17901 |
| **Yif1a** | 13.79390 | 9.89392 | 12.00228 | 1.99119 | 2.78884 | 5.16823 | 7.21347 | 10.62072 | 2.37504 | 7.35553 | 3.76093 | 0.17658 | 10.24811 | 0.00381 | 1.72378 | 6.08770 |
| **Yif1b** | 11.54658 | 0.20811 | 2.10462 | 0.06444 | 7.74522 | 9.87544 | 4.12882 | 15.92379 | 0.02067 | 11.82118 | 0.00000 | 0.45191 | 16.91143 | 0.00386 | 0.00000 | 13.77930 |
| **Yipf1** | 4.59620 | 0.10260 | 8.28810 | 8.86599 | 0.02636 | 0.16525 | 0.09763 | 5.79824 | 14.83861 | 7.77917 | 10.38725 | 1.91704 | 6.70163 | 12.92279 | 5.50590 | 2.06755 |
| **Yipf2** | 0.81181 | 1.20680 | 2.90605 | 0.09894 | 1.13188 | 3.78675 | 0.18001 | 1.62669 | 0.00816 | 0.08283 | 0.00000 | 0.03837 | 2.84069 | 0.00370 | 0.49115 | 0.77487 |
| **Yipf3** | 3.97096 | 0.04164 | 4.45294 | 5.49322 | 1.57944 | 0.36182 | 1.96907 | 9.58114 | 5.47064 | 6.72022 | 3.96581 | 0.39468 | 10.00558 | 0.00846 | 0.41707 | 7.35770 |
| **Yipf4** | 0.05218 | 3.04745 | 0.77556 | 3.42007 | 0.01229 | 0.00504 | 0.26870 | 2.97044 | 10.77910 | 0.65837 | 3.20632 | 2.60665 | 2.68150 | 12.48439 | 5.35906 | 0.04837 |
| **Yipf5** | 4.66864 | 5.51579 | 10.75629 | 1.27171 | 6.42422 | 8.77947 | 6.03602 | 9.67856 | 2.47456 | 14.05566 | 4.88003 | 7.85511 | 3.69538 | 9.62049 | 10.27909 | 6.63433 |
| **Yipf6** | 2.49500 | 2.85153 | 4.06698 | 4.78256 | 5.31772 | 2.60912 | 0.47226 | 2.41966 | 0.77652 | 4.52342 | 1.89699 | 2.96717 | 4.16628 | 0.64376 | 8.31747 | 1.50611 |
| **Ykt6** | 6.37358 | 0.24025 | 10.10936 | 2.51853 | 7.45237 | 8.90132 | 1.75209 | 8.32348 | 2.50019 | 9.34929 | 2.69781 | 0.22256 | 8.48215 | 3.32541 | 2.13662 | 5.06226 |
| **Ylpm1** | 2.85323 | 0.80345 | 3.45509 | 8.01377 | 0.29447 | 3.64901 | 1.36595 | 3.15098 | 8.38908 | 8.90963 | 1.55154 | 0.24897 | 1.77004 | 6.08163 | 8.95153 | 0.56726 |
| **Yme1l1** | 7.67297 | 10.39134 | 7.38352 | 12.63436 | 7.03749 | 8.12802 | 7.89208 | 12.06327 | 13.24133 | 15.56631 | 10.99723 | 16.08800 | 3.98490 | 30.66757 | 22.78608 | 1.41608 |
| **Yod1** | 0.02187 | 0.00810 | 0.01177 | 0.84781 | 0.00876 | 0.00660 | 0.00748 | 0.01449 | 0.00000 | 0.01516 | 0.01410 | 0.00816 | 0.01264 | 0.01019 | 0.00672 | 0.00723 |
| **Ypel1** | 0.01312 | 0.00777 | 0.00383 | 0.56468 | 0.00947 | 0.45713 | 0.00720 | 0.05148 | 0.02198 | 0.00539 | 0.00971 | 0.00651 | 0.01932 | 0.29088 | 0.19516 | 0.00710 |
| **Ypel2** | 2.73644 | 4.00050 | 6.34931 | 5.25737 | 0.72462 | 3.78676 | 6.73882 | 11.98468 | 14.88157 | 18.70272 | 16.00574 | 12.64774 | 7.40386 | 17.24877 | 19.60055 | 0.00381 |
| **Ypel3** | 0.33184 | 1.32686 | 0.95473 | 8.23905 | 5.32647 | 0.00401 | 0.71164 | 2.56860 | 3.83959 | 8.58613 | 4.33990 | 1.04488 | 0.93233 | 0.01405 | 0.00000 | 0.00326 |
| **Ypel5** | 9.73970 | 5.79374 | 6.22243 | 11.49048 | 1.84300 | 1.89011 | 5.85613 | 11.17798 | 20.96352 | 16.76156 | 11.51958 | 6.01597 | 6.89937 | 29.43041 | 9.38872 | 1.08959 |
| **Yrdc** | 4.44740 | 0.77190 | 1.58536 | 11.19031 | 0.10505 | 3.26112 | 0.00000 | 8.21693 | 16.63500 | 13.16770 | 12.39950 | 3.14803 | 7.10413 | 6.26447 | 7.40111 | 0.36821 |
| **Ythdc1** | 4.59069 | 7.79074 | 3.63162 | 9.35286 | 5.58651 | 2.83437 | 3.22904 | 7.55737 | 12.18879 | 6.50196 | 9.20162 | 9.64321 | 6.04526 | 22.33681 | 5.90581 | 1.74080 |
| **Ythdc2** | 0.00629 | 0.42110 | 0.04974 | 0.35090 | 0.11075 | 0.02389 | 0.05531 | 0.00675 | 0.02251 | 0.06056 | 0.16575 | 1.73045 | 0.38119 | 0.53342 | 0.77402 | 0.01478 |
| **Ythdf1** | 2.81920 | 9.43340 | 8.52854 | 12.35072 | 7.25691 | 3.82672 | 7.38114 | 10.46574 | 13.73138 | 12.87302 | 7.11441 | 11.94351 | 7.48622 | 13.43440 | 11.39887 | 6.93814 |
| **Ythdf2** | 2.81476 | 0.34228 | 3.67360 | 16.88121 | 2.22263 | 0.79226 | 2.02510 | 3.54953 | 26.00521 | 12.57179 | 17.23607 | 5.29506 | 4.02391 | 18.70896 | 14.99371 | 3.60451 |
| **Ythdf3** | 2.01826 | 2.56227 | 0.75611 | 11.99279 | 0.02759 | 1.60394 | 0.02296 | 2.24584 | 4.02186 | 0.00678 | 0.43480 | 3.00247 | 2.26106 | 9.12690 | 4.52953 | 0.09968 |
| **Ywhab** | 18.10444 | 9.59173 | 17.61249 | 6.34563 | 14.23578 | 10.43959 | 13.43803 | 16.08867 | 15.14136 | 23.09935 | 15.89561 | 13.07174 | 13.39460 | 20.02515 | 11.48720 | 8.40637 |
| **Ywhae** | 30.29820 | 21.49190 | 26.57377 | 12.69016 | 23.55772 | 15.20102 | 19.05076 | 29.65757 | 18.06140 | 30.01166 | 27.07122 | 9.75668 | 26.90059 | 25.50757 | 20.59361 | 17.38760 |
| **Ywhag** | 22.85141 | 6.00133 | 14.41270 | 8.01760 | 16.71604 | 12.06354 | 11.57272 | 17.33608 | 10.02990 | 19.00725 | 16.50986 | 1.19588 | 16.72768 | 5.30675 | 2.48101 | 10.13460 |
| **Ywhah** | 16.87321 | 12.25281 | 9.81422 | 8.43947 | 14.67650 | 4.74786 | 7.34082 | 8.44941 | 9.54449 | 15.45743 | 11.39056 | 17.20569 | 15.69342 | 18.23364 | 9.30923 | 6.78586 |
| **Ywhaq** | 26.24520 | 15.37971 | 21.79873 | 19.86192 | 23.81502 | 20.89057 | 15.97252 | 41.60473 | 32.04327 | 42.61154 | 31.84482 | 29.54403 | 36.91431 | 24.80617 | 15.69692 | 14.92476 |
| **Ywhaz** | 25.11537 | 14.01401 | 22.80035 | 5.60726 | 16.91533 | 9.49063 | 14.23652 | 13.81818 | 2.74355 | 12.24012 | 14.61180 | 9.78514 | 16.81946 | 11.02910 | 4.02901 | 12.28983 |
| **Yy1** | 8.37822 | 8.61084 | 4.84570 | 17.36541 | 5.90453 | 6.31421 | 5.93976 | 8.96482 | 16.53585 | 17.48624 | 13.06404 | 18.00400 | 5.11724 | 17.06150 | 16.03733 | 0.33560 |

| | | | | | | | | | | | | | | | | |
|---|---|---|---|---|---|---|---|---|---|---|---|---|---|---|---|---|
| **Yy2** | 0.00656 | 0.00745 | 0.95179 | 0.62855 | 0.00725 | 0.28421 | 0.00679 | 1.82103 | 0.76512 | 2.05819 | 0.08904 | 0.70579 | 0.68461 | 5.65795 | 11.12189 | 0.00620 |
| **Zadh2** | 0.00380 | 0.45259 | 0.79842 | 0.00365 | 0.84501 | 1.88124 | 0.00742 | 0.00988 | 0.06840 | 0.00245 | 0.01084 | 0.00688 | 0.02651 | 0.01492 | 0.00000 | 0.00637 |
| **Zak** | 0.22821 | 1.68113 | 0.38257 | 0.23281 | 0.02615 | 2.42762 | 0.03937 | 0.03049 | 0.08863 | 0.55923 | 0.21164 | 0.06366 | 0.19432 | 0.15644 | 0.23453 | 0.02795 |
| **Zan** | 0.47481 | 0.00721 | 0.34029 | 0.00595 | 0.00662 | 0.00526 | 0.01658 | 0.40345 | 0.01064 | 0.59004 | 0.06764 | 0.00000 | 0.07420 | 1.01674 | 0.19020 | 0.00695 |
| **Zap70** | 0.13261 | 0.00760 | 0.01501 | 0.60980 | 0.00747 | 0.00709 | 0.00636 | 0.00000 | 0.01035 | 4.15273 | 0.38570 | 0.10703 | 1.33836 | 0.00710 | 0.20253 | 0.00643 |
| **Zar1** | 0.01147 | 0.00782 | 0.00757 | 4.01818 | 0.00842 | 0.00626 | 0.00715 | 0.00174 | 1.85970 | 0.00000 | 0.35145 | 0.05224 | 0.00753 | 0.01371 | 0.20965 | 0.35807 |
| **Zar1l** | 0.00760 | 0.00859 | 0.00601 | 5.51976 | 0.00920 | 0.00592 | 0.00782 | 0.20711 | 9.19900 | 0.00000 | 0.36425 | 0.00000 | 0.00484 | 0.17198 | 0.00921 | 0.00562 |
| **Zbed3** | 0.02834 | 0.00755 | 0.00000 | 12.37913 | 0.00844 | 0.00537 | 0.00657 | 0.00000 | 6.57506 | 0.60997 | 0.56477 | 0.01316 | 0.03031 | 0.32632 | 2.08058 | 0.00499 |
| **Zbed4** | 7.31106 | 3.85953 | 5.35426 | 8.24800 | 0.01818 | 4.36352 | 2.92803 | 6.14949 | 7.23822 | 9.49932 | 3.44973 | 2.25947 | 3.12873 | 19.19522 | 17.02282 | 1.69257 |
| **Zbed5** | 9.89820 | 0.14032 | 1.38476 | 8.12776 | 8.39197 | 0.95151 | 0.15182 | 0.89205 | 13.08732 | 19.71867 | 9.39033 | 0.34088 | 13.33695 | 0.57457 | 0.14933 | 5.86891 |
| **Zbed6** | 0.00690 | 0.00696 | 0.00010 | 1.77971 | 1.03903 | 0.00640 | 0.00714 | 0.00707 | 2.61146 | 0.00000 | 0.00000 | 0.00817 | 0.01620 | 0.01463 | 2.00169 | 0.02047 |
| **Zbp1** | 0.01378 | 0.00796 | 0.01093 | 0.00577 | 0.00920 | 0.00670 | 0.00752 | 0.01056 | 0.13646 | 0.00872 | 0.12693 | 0.00732 | 0.01789 | 0.01497 | 0.01972 | 0.00732 |
| **Zbtb1** | 0.02669 | 0.32838 | 0.00601 | 1.43314 | 2.32308 | 0.00802 | 0.00683 | 0.71738 | 1.19269 | 0.07126 | 0.89483 | 0.16397 | 0.06858 | 0.02271 | 3.47703 | 0.00669 |
| **Zbtb10** | 0.50845 | 0.63538 | 3.00213 | 19.90006 | 5.71995 | 0.28984 | 0.00000 | 9.56300 | 21.76095 | 11.92742 | 15.36190 | 22.91199 | 3.11647 | 32.28067 | 27.60521 | 2.13631 |
| **Zbtb11** | 0.13428 | 2.85189 | 0.00000 | 9.51686 | 0.11148 | 0.00390 | 0.19147 | 0.91789 | 13.50413 | 0.31470 | 4.86618 | 5.39742 | 0.17610 | 9.09061 | 10.95277 | 0.00997 |
| **Zbtb12** | 0.01757 | 0.00778 | 0.05363 | 0.14017 | 0.00875 | 0.00579 | 0.00708 | 0.02705 | 0.00815 | 0.03257 | 0.76979 | 0.02467 | 0.41399 | 0.00691 | 0.00000 | 0.00700 |
| **Zbtb14** | 3.79849 | 1.96919 | 2.23754 | 0.40483 | 0.00697 | 0.61243 | 0.01218 | 2.90771 | 2.27004 | 2.66598 | 1.42458 | 0.37524 | 2.02392 | 0.00101 | 5.55765 | 0.00000 |
| **Zbtb16** | 0.01253 | 0.00791 | 0.25886 | 0.00570 | 0.00873 | 0.01811 | 0.00736 | 0.01280 | 0.16841 | 0.01800 | 0.01075 | 0.00744 | 0.01007 | 0.01362 | 0.00521 | 0.00728 |
| **Zbtb17** | 7.93182 | 2.94141 | 1.93278 | 5.65572 | 2.66196 | 6.87544 | 4.53020 | 14.19881 | 8.21268 | 20.97898 | 7.20654 | 3.91846 | 8.22987 | 0.00000 | 2.23983 | 2.64532 |
| **Zbtb18** | 2.40612 | 0.00516 | 0.77493 | 0.86536 | 0.28580 | 1.47332 | 0.00507 | 0.00000 | 0.52117 | 0.44264 | 0.14689 | 0.32362 | 0.13995 | 0.00443 | 1.75797 | 0.00370 |
| **Zbtb2** | 0.11128 | 3.85694 | 2.82623 | 1.80992 | 0.01008 | 0.00563 | 0.00461 | 3.61785 | 5.27824 | 2.46029 | 0.00000 | 0.08290 | 2.42486 | 2.63784 | 2.04882 | 2.23548 |
| **Zbtb20** | 0.21463 | 0.00852 | 0.00607 | 0.00973 | 0.01673 | 0.00729 | 0.00691 | 0.01483 | 0.01209 | 0.20781 | 0.01140 | 0.02009 | 0.72346 | 0.01158 | 0.01171 | 0.00707 |
| **Zbtb21** | 0.30052 | 0.00642 | 0.88965 | 6.08433 | 1.39697 | 0.44936 | 0.04709 | 2.97377 | 5.18136 | 0.15258 | 1.63265 | 0.07015 | 1.09102 | 1.28406 | 0.85752 | 0.00427 |
| **Zbtb22** | 4.45133 | 0.89998 | 0.00000 | 3.53810 | 2.16273 | 6.35540 | 1.44339 | 2.58460 | 0.18214 | 2.63169 | 1.05936 | 0.17413 | 2.32604 | 0.00000 | 0.76357 | 2.55188 |
| **Zbtb24** | 2.34220 | 3.55561 | 0.26945 | 2.46449 | 0.01449 | 2.67458 | 0.02204 | 0.99753 | 5.07161 | 5.02409 | 3.20348 | 0.35698 | 1.67773 | 0.03181 | 2.32483 | 0.01525 |
| **Zbtb25** | 1.13989 | 0.08413 | 1.02930 | 0.42579 | 2.34840 | 0.00448 | 0.00654 | 0.00620 | 0.79108 | 0.24803 | 0.00000 | 0.01546 | 0.07023 | 0.28695 | 0.00000 | 2.69552 |
| **Zbtb26** | 0.01056 | 0.00758 | 0.67995 | 0.14102 | 0.00759 | 0.00799 | 0.00713 | 0.00052 | 0.47944 | 0.00000 | 0.31913 | 0.03144 | 0.00798 | 0.02963 | 1.67720 | 0.22681 |
| **Zbtb3** | 0.28656 | 0.00644 | 0.62689 | 0.22533 | 0.00884 | 0.00639 | 1.78469 | 0.78848 | 0.69494 | 0.22248 | 0.00000 | 0.01782 | 0.01720 | 0.00536 | 1.95277 | 0.02247 |
| **Zbtb32** | 0.49502 | 0.00715 | 0.00000 | 0.95307 | 0.00698 | 0.13204 | 0.00685 | 0.04168 | 0.26715 | 0.01190 | 0.02959 | 0.00717 | 0.11166 | 0.01056 | 0.00239 | 0.00650 |
| **Zbtb33** | 0.00833 | 0.00843 | 0.00813 | 0.68092 | 0.00959 | 0.00666 | 0.00799 | 0.25857 | 0.24071 | 0.00666 | 0.00616 | 0.03904 | 0.36464 | 0.29095 | 0.18501 | 0.00700 |
| **Zbtb34** | 2.11704 | 0.01349 | 2.33585 | 6.47938 | 0.00830 | 8.38163 | 2.10838 | 3.46016 | 7.30970 | 8.86019 | 5.42938 | 3.83851 | 0.21985 | 0.71237 | 6.11752 | 0.00000 |
| **Zbtb37** | 0.00000 | 0.00736 | 0.00000 | 3.92764 | 0.01000 | 0.00506 | 0.00673 | 0.61942 | 3.42042 | 1.24591 | 0.26682 | 1.39127 | 0.00470 | 7.65457 | 5.45564 | 0.00626 |
| **Zbtb39** | 1.14136 | 0.00607 | 0.38803 | 3.26144 | 0.00556 | 0.00550 | 1.46880 | 0.08349 | 0.48946 | 0.94066 | 1.50831 | 0.11622 | 0.05124 | 0.45071 | 0.20616 | 0.47798 |
| **Zbtb4** | 0.01169 | 0.00763 | 0.00682 | 0.04095 | 0.00914 | 0.00668 | 0.00724 | 0.00922 | 0.58718 | 0.94799 | 0.00000 | 0.00973 | 0.01120 | 1.38487 | 0.00000 | 0.00717 |
| **Zbtb40** | 1.39578 | 0.00431 | 2.04571 | 0.18141 | 0.04701 | 0.00640 | 2.63149 | 5.02164 | 1.07470 | 5.07186 | 2.19072 | 0.88033 | 1.93978 | 0.02567 | 0.27258 | 0.01150 |
| **Zbtb41** | 1.31287 | 0.06900 | 0.60801 | 6.90008 | 0.01294 | 0.01350 | 0.02793 | 0.40073 | 3.33570 | 1.98707 | 0.00000 | 6.76939 | 0.89278 | 4.15932 | 6.07293 | 0.00362 |
| **Zbtb42** | 0.01777 | 0.00784 | 0.01161 | 0.09658 | 0.00879 | 0.00665 | 0.00729 | 0.01108 | 0.06084 | 0.35501 | 0.01335 | 0.01604 | 0.01976 | 0.01094 | 0.00759 | 0.00775 |
| **Zbtb43** | 1.08162 | 5.34816 | 3.86556 | 9.59225 | 3.73255 | 3.05598 | 6.76964 | 7.84967 | 12.40401 | 8.53935 | 2.69761 | 0.88430 | 2.44483 | 1.44396 | 8.84951 | 0.03303 |
| **Zbtb44** | 4.89638 | 0.17752 | 3.84201 | 9.82528 | 3.90303 | 4.66744 | 6.29013 | 3.21719 | 11.83035 | 4.81339 | 4.04410 | 4.24801 | 1.57476 | 14.73025 | 8.23976 | 0.41466 |
| **Zbtb45** | 0.79722 | 0.00522 | 1.59744 | 2.54007 | 0.01375 | 0.27192 | 1.51062 | 0.67554 | 2.77460 | 1.34416 | 1.84126 | 0.14453 | 1.11284 | 0.00000 | 0.33438 | 0.01211 |
| **Zbtb46** | 0.00927 | 0.00851 | 0.00010 | 0.99381 | 0.01083 | 0.01377 | 0.00730 | 0.85562 | 0.29754 | 0.38581 | 0.25257 | 0.13619 | 1.34830 | 0.23375 | 0.88253 | 0.01406 |
| **Zbtb48** | 1.58709 | 0.09012 | 2.44060 | 4.25456 | 0.00957 | 1.25745 | 0.00383 | 2.65440 | 0.58716 | 1.32799 | 1.43690 | 0.90292 | 2.14465 | 0.01135 | 0.46374 | 2.01551 |
| **Zbtb49** | 0.01008 | 0.00761 | 0.00735 | 0.49434 | 0.00941 | 0.00687 | 0.00705 | 0.37946 | 0.55352 | 0.00000 | 0.11694 | 0.01494 | 0.07169 | 0.00936 | 0.00000 | 0.01437 |
| **Zbtb5** | 0.00000 | 0.08962 | 2.10518 | 6.74631 | 0.01346 | 3.07579 | 0.00474 | 0.12535 | 8.47631 | 4.62359 | 0.40151 | 6.51578 | 1.83454 | 1.34951 | 4.84462 | 0.00250 |
| **Zbtb6** | 1.13267 | 0.01544 | 0.60718 | 3.89220 | 0.02099 | 0.03159 | 0.05354 | 0.02812 | 2.15308 | 0.35443 | 2.27430 | 0.08929 | 2.48503 | 5.26317 | 3.32174 | 0.01811 |
| **Zbtb7a** | 0.08525 | 0.00703 | 0.21332 | 1.79459 | 1.05461 | 0.01515 | 0.01637 | 0.01747 | 3.23812 | 2.07223 | 2.60997 | 0.06096 | 0.49964 | 0.01011 | 0.79125 | 0.01292 |
| **Zbtb7b** | 3.39350 | 0.45776 | 1.50329 | 1.53306 | 3.51258 | 2.09931 | 2.50347 | 7.57495 | 0.67441 | 7.56846 | 6.85699 | 0.13200 | 9.00160 | 0.06353 | 0.26414 | 6.19625 |
| **Zbtb7c** | 0.02280 | 0.00875 | 0.00912 | 0.58950 | 0.00917 | 0.00789 | 0.00810 | 0.14535 | 0.30632 | 0.51281 | 0.18830 | 0.00684 | 0.40195 | 0.00903 | 0.00000 | 0.00742 |
| **Zbtb8a** | 0.23344 | 0.00748 | 0.93108 | 0.84862 | 0.00957 | 0.00851 | 0.01532 | 0.40079 | 0.55650 | 3.40391 | 1.95646 | 0.14733 | 1.21028 | 0.20454 | 5.35752 | 0.00594 |
| **Zbtb8b** | 0.01388 | 0.01707 | 2.00511 | 2.51454 | 0.01601 | 0.00886 | 1.81828 | 1.98607 | 2.25807 | 2.98257 | 2.73180 | 0.02989 | 0.52958 | 2.25691 | 2.14717 | 0.00547 |
| **Zbtb8os** | 5.24485 | 4.25930 | 4.42899 | 1.90396 | 3.42032 | 1.49525 | 2.26361 | 1.03709 | 1.09901 | 2.37152 | 2.98941 | 0.21839 | 4.04544 | 0.13737 | 0.89126 | 0.20746 |
| **Zbtb9** | 1.28162 | 0.00553 | 0.70500 | 3.13478 | 0.01397 | 4.05479 | 0.00475 | 0.83205 | 8.97671 | 2.62116 | 0.88827 | 2.47166 | 1.32260 | 0.68020 | 0.90421 | 0.02458 |
| **Zc2hc1a** | 1.34729 | 0.00638 | 0.10700 | 0.22753 | 0.36243 | 0.46902 | 0.00598 | 0.00835 | 1.42928 | 0.81818 | 0.00000 | 0.05812 | 0.00782 | 0.01086 | 0.00063 | 0.01338 |
| **Zc2hc1c** | 0.72730 | 0.00549 | 0.56692 | 0.00383 | 0.00537 | 1.51454 | 0.00614 | 0.00567 | 0.05588 | 0.10134 | 0.00000 | 3.56543 | 0.00933 | 0.00641 | 0.11378 | 0.00533 |
| **Zc3h10** | 0.01200 | 0.00811 | 0.30442 | 0.45641 | 0.01088 | 3.04159 | 0.06956 | 2.24293 | 0.90902 | 1.47157 | 0.69417 | 0.07919 | 3.96245 | 1.35185 | 0.11964 | 0.04233 |
| **Zc3h11a** | 6.69023 | 6.62474 | 11.65460 | 11.56489 | 9.32640 | 3.89220 | 10.51551 | 6.53527 | 19.33605 | 13.90834 | 7.03257 | 9.88787 | 6.49733 | 14.25039 | 11.06512 | 3.84323 |
| **Zc3h12a** | 0.36565 | 0.02732 | 0.07060 | 5.32181 | 0.04590 | 0.48919 | 0.42518 | 1.06747 | 6.84271 | 3.49707 | 4.53293 | 0.76314 | 0.02984 | 1.04056 | 0.13528 | 0.09286 |
| **Zc3h12c** | 0.00000 | 0.00556 | 0.00000 | 1.73631 | 0.00864 | 0.03843 | 0.00552 | 0.94015 | 2.02651 | 1.08531 | 1.90898 | 6.34628 | 0.00000 | 8.83130 | 6.68190 | 0.01998 |
| **Zc3h13** | 2.85373 | 0.62234 | 6.54036 | 2.96322 | 6.39690 | 0.79741 | 0.02202 | 5.13777 | 7.24865 | 7.23945 | 3.85162 | 1.37441 | 3.95340 | 2.76842 | 9.12336 | 2.06461 |
| **Zc3h14** | 5.40900 | 2.57882 | 5.22082 | 4.94244 | 2.72788 | 5.53387 | 6.59211 | 3.89927 | 0.77855 | 11.96829 | 4.79086 | 8.74950 | 4.33740 | 11.51831 | 8.80800 | 2.93706 |
| **Zc3h15** | 13.41959 | 8.61189 | 12.08063 | 19.25092 | 13.73035 | 8.53855 | 4.92199 | 12.57350 | 18.28482 | 26.80264 | 13.36803 | 17.72145 | 12.12075 | 21.51281 | 19.31849 | 6.17932 |
| **Zc3h18** | 2.45982 | 0.93054 | 1.29908 | 12.02869 | 2.69320 | 1.82597 | 1.35025 | 4.03556 | 15.63914 | 5.99927 | 4.79630 | 2.74118 | 2.08168 | 3.13591 | 2.68466 | 1.64386 |

| | | | | | | | | | | | | | | | | |
|---|---|---|---|---|---|---|---|---|---|---|---|---|---|---|---|---|
| **Zc3h3** | 0.92712 | 0.36444 | 1.15413 | 6.21197 | 0.01715 | 0.00516 | 0.89741 | 1.28313 | 3.80497 | 4.79761 | 1.75544 | 2.18550 | 1.15314 | 0.04136 | 0.09357 | 0.39773 |
| **Zc3h4** | 3.71472 | 0.41182 | 1.76516 | 4.27805 | 0.00000 | 1.44608 | 1.01796 | 0.45467 | 1.88091 | 5.36767 | 0.46826 | 0.21483 | 1.81610 | 1.28829 | 5.54516 | 0.05273 |
| **Zc3h6** | 0.03029 | 0.00761 | 0.00860 | 2.57602 | 0.00833 | 0.00639 | 0.00726 | 0.00644 | 1.13311 | 0.00000 | 0.22756 | 0.00388 | 0.00950 | 0.01088 | 0.27418 | 0.00661 |
| **Zc3h7a** | 2.18046 | 1.62408 | 5.20890 | 2.75525 | 5.19857 | 1.96527 | 0.00252 | 1.40407 | 0.83940 | 3.28823 | 2.83171 | 0.54847 | 1.17958 | 4.52024 | 2.08087 | 0.00000 |
| **Zc3h7b** | 0.06958 | 0.00767 | 0.00426 | 2.44895 | 0.00846 | 0.00747 | 0.00688 | 0.28381 | 0.48757 | 0.94525 | 0.42250 | 0.03049 | 0.00466 | 0.00456 | 0.01974 | 0.00699 |
| **Zc3h8** | 0.92689 | 2.98930 | 3.48241 | 4.02365 | 2.56159 | 2.95475 | 3.71030 | 5.20965 | 3.08912 | 6.99176 | 2.77608 | 3.80215 | 1.93052 | 2.67048 | 5.84314 | 0.72952 |
| **Zc3hav1** | 11.53830 | 0.00000 | 0.90304 | 3.01725 | 0.21588 | 0.00000 | 1.99483 | 0.00000 | 3.81914 | 0.19221 | 0.14005 | 0.22511 | 0.02506 | 0.00328 | 0.77078 | 0.18463 |
| **Zc3hav1l** | 0.39171 | 0.04299 | 0.42596 | 0.38795 | 0.00887 | 0.84329 | 0.00769 | 0.20558 | 0.66275 | 0.00000 | 0.00000 | 0.02908 | 0.08782 | 0.06730 | 0.57328 | 0.09899 |
| **Zc3hc1** | 16.42618 | 10.43431 | 9.89638 | 11.34910 | 9.63419 | 1.37357 | 8.55853 | 25.75808 | 20.52719 | 13.82054 | 19.31951 | 2.41385 | 16.58307 | 2.21657 | 13.58594 | 4.35925 |
| **Zcchc10** | 3.56298 | 3.56136 | 4.11475 | 9.45313 | 4.12161 | 0.00744 | 0.66971 | 9.20351 | 12.93000 | 5.47723 | 2.67929 | 6.31477 | 7.99687 | 8.23439 | 8.36202 | 0.00000 |
| **Zcchc11** | 0.11351 | 3.50286 | 3.63520 | 0.81976 | 1.37322 | 0.00726 | 0.05008 | 1.76308 | 0.00000 | 1.93012 | 0.45901 | 0.18617 | 0.00000 | 0.09707 | 1.87616 | 2.64215 |
| **Zcchc13** | 0.01350 | 0.00777 | 0.01179 | 0.24562 | 0.00926 | 0.00640 | 0.00747 | 0.01269 | 1.81793 | 0.00511 | 0.00000 | 0.00411 | 0.01290 | 0.00539 | 0.19756 | 0.00716 |
| **Zcchc14** | 1.96111 | 0.00416 | 0.49081 | 5.05540 | 0.00338 | 1.66128 | 2.11391 | 0.00000 | 0.67765 | 1.73470 | 0.02090 | 0.07271 | 0.78595 | 0.00000 | 3.49823 | 0.03824 |
| **Zcchc16** | 0.01359 | 0.00817 | 0.01181 | 0.01909 | 0.00928 | 0.00652 | 0.00749 | 0.01308 | 1.19335 | 0.00608 | 0.00591 | 0.00631 | 0.01284 | 0.00798 | 0.11459 | 0.00729 |
| **Zcchc17** | 13.81288 | 0.11693 | 5.37381 | 1.92734 | 6.61244 | 0.35169 | 2.89098 | 6.38072 | 4.55643 | 6.26671 | 0.00000 | 0.22728 | 5.52773 | 0.55285 | 2.16441 | 5.25833 |
| **Zcchc2** | 0.60762 | 1.00366 | 0.96135 | 7.53997 | 0.05660 | 0.65530 | 0.00640 | 0.83665 | 2.84841 | 2.91996 | 0.00000 | 0.07731 | 0.42338 | 0.03606 | 2.02518 | 0.37217 |
| **Zcchc3** | 2.82579 | 0.05654 | 5.54326 | 4.09760 | 3.58674 | 0.00853 | 3.47116 | 5.59763 | 2.05963 | 4.89995 | 4.45187 | 0.13701 | 0.03458 | 0.61651 | 0.89776 | 0.64733 |
| **Zcchc4** | 0.62672 | 3.26285 | 1.04263 | 0.88929 | 0.04845 | 1.60658 | 4.88331 | 0.58132 | 1.87101 | 5.17121 | 2.08629 | 0.20368 | 1.78906 | 0.95649 | 5.37996 | 0.03973 |
| **Zcchc6** | 0.15242 | 1.57857 | 0.49186 | 1.51973 | 0.03186 | 1.39008 | 1.85420 | 2.38580 | 0.78600 | 1.30522 | 0.05016 | 0.09488 | 0.13040 | 1.24102 | 3.85155 | 0.29458 |
| **Zcchc7** | 2.79204 | 2.47143 | 0.41421 | 4.99148 | 0.01305 | 0.00125 | 0.00215 | 5.61021 | 8.01371 | 3.06643 | 2.08950 | 1.50442 | 1.32402 | 7.58011 | 3.42673 | 1.04262 |
| **Zcchc8** | 8.40718 | 8.20651 | 5.72110 | 3.04498 | 5.20288 | 4.11283 | 3.79206 | 4.24288 | 6.44968 | 5.82754 | 5.70335 | 4.53213 | 4.31560 | 5.31632 | 4.04026 | 5.36984 |
| **Zcchc9** | 2.07410 | 4.23630 | 2.88434 | 3.53094 | 6.10399 | 2.68531 | 1.77518 | 0.99938 | 4.24405 | 2.15862 | 3.17543 | 1.02010 | 2.42409 | 5.07144 | 3.55042 | 2.70597 |
| **Zcrb1** | 2.84244 | 1.07742 | 1.37270 | 10.95051 | 5.05777 | 1.91343 | 0.07040 | 1.56708 | 16.43900 | 12.84497 | 10.95202 | 2.22857 | 5.99201 | 1.70948 | 0.53806 | 1.14111 |
| **Zcwpw1** | 1.71533 | 0.00587 | 2.24614 | 0.10283 | 0.00238 | 0.00526 | 0.08780 | 0.83419 | 0.55386 | 1.82320 | 0.21113 | 0.18609 | 1.23112 | 4.58381 | 4.04641 | 0.02702 |
| **Zdbf2** | 0.36058 | 0.00628 | 1.52020 | 0.02958 | 0.00447 | 0.00507 | 2.55751 | 0.24260 | 1.01531 | 0.00000 | 0.00000 | 0.03017 | 0.61887 | 0.01638 | 0.10210 | 0.02662 |
| **Zdhhc11** | 0.01375 | 0.00812 | 0.01230 | 0.29389 | 0.00931 | 0.00662 | 0.00752 | 0.01394 | 0.00695 | 0.01549 | 0.01388 | 0.00821 | 0.01266 | 0.01130 | 0.00654 | 0.00736 |
| **Zdhhc12** | 10.28729 | 1.06492 | 0.82286 | 1.17966 | 5.72019 | 6.60378 | 1.82687 | 5.42227 | 3.68458 | 1.23461 | 1.15267 | 0.02759 | 2.63343 | 0.03254 | 0.00000 | 4.55393 |
| **Zdhhc13** | 0.52097 | 0.00514 | 1.38640 | 1.73199 | 0.00439 | 0.00559 | 2.21695 | 1.12177 | 0.00000 | 1.75818 | 0.02986 | 0.01317 | 1.07578 | 1.18511 | 0.75132 | 0.28903 |
| **Zdhhc14** | 0.01660 | 0.00752 | 0.55634 | 0.17773 | 0.00825 | 0.00560 | 0.00724 | 0.01147 | 0.00915 | 0.00807 | 0.01281 | 0.00685 | 0.00892 | 0.00702 | 0.17231 | 0.00704 |
| **Zdhhc15** | 0.00690 | 2.52596 | 0.30183 | 0.01592 | 0.00762 | 0.00563 | 0.00627 | 0.00841 | 0.01141 | 0.00768 | 0.01263 | 0.00712 | 0.00956 | 0.00997 | 0.00000 | 0.00623 |
| **Zdhhc16** | 4.30489 | 2.43746 | 2.85155 | 0.06809 | 3.65469 | 4.58501 | 1.66083 | 1.90067 | 0.04170 | 0.99692 | 0.00000 | 3.02360 | 4.24154 | 0.27646 | 0.81806 | 4.10474 |
| **Zdhhc17** | 2.73270 | 2.15013 | 0.87269 | 0.58239 | 5.59487 | 0.00444 | 4.77970 | 0.69584 | 2.21277 | 1.44789 | 0.74024 | 0.00023 | 0.17454 | 5.74657 | 2.01247 | 0.00270 |
| **Zdhhc18** | 1.33442 | 0.00475 | 0.68951 | 3.93447 | 0.45841 | 1.13459 | 0.02628 | 7.84560 | 1.53276 | 7.26261 | 2.52339 | 0.30719 | 0.00000 | 0.01071 | 0.03251 | 0.17871 |
| **Zdhhc19** | 0.01331 | 0.00830 | 0.01196 | 0.01062 | 0.00933 | 0.00669 | 0.00782 | 0.02358 | 0.01234 | 0.01526 | 0.01281 | 0.00851 | 0.01206 | 0.01285 | 0.01044 | 0.00741 |
| **Zdhhc2** | 0.02970 | 0.00787 | 0.00654 | 0.01024 | 0.01038 | 0.00743 | 0.00724 | 0.36798 | 0.01159 | 0.10136 | 0.02157 | 0.01841 | 0.12648 | 0.01394 | 0.02176 | 0.00737 |
| **Zdhhc20** | 3.39182 | 4.62128 | 1.80278 | 3.69608 | 2.07182 | 5.43648 | 6.32482 | 4.97094 | 9.53701 | 9.41357 | 8.99317 | 15.83647 | 3.97006 | 24.66195 | 8.10596 | 2.90794 |
| **Zdhhc21** | 0.01891 | 0.00728 | 0.86510 | 0.01147 | 0.00742 | 0.01907 | 0.00727 | 0.58165 | 0.01150 | 0.00000 | 0.00995 | 0.01543 | 0.02145 | 0.02385 | 0.00842 | 0.15637 |
| **Zdhhc24** | 1.89429 | 0.03432 | 0.00000 | 0.45482 | 1.21874 | 0.01233 | 2.04041 | 0.00642 | 0.83192 | 0.04069 | 0.07121 | 0.01860 | 0.40789 | 0.02081 | 0.11020 | 1.19672 |
| **Zdhhc3** | 6.51300 | 5.35584 | 5.88172 | 2.24470 | 0.02326 | 8.34444 | 3.32389 | 8.50559 | 3.02773 | 3.68516 | 6.35052 | 7.59178 | 5.44435 | 3.04805 | 3.52122 | 1.67347 |
| **Zdhhc4** | 3.43399 | 2.69484 | 7.07277 | 0.03967 | 3.39206 | 6.95913 | 2.32813 | 6.61531 | 2.34901 | 2.57136 | 2.48070 | 0.03322 | 6.95758 | 0.48737 | 0.74745 | 3.87221 |
| **Zdhhc5** | 2.16606 | 3.02724 | 3.15815 | 13.44648 | 1.80901 | 2.26531 | 1.75653 | 3.72664 | 7.22915 | 2.98927 | 3.90830 | 0.60991 | 2.80990 | 6.40366 | 4.48040 | 0.78211 |
| **Zdhhc6** | 9.45009 | 2.04565 | 6.01451 | 3.19864 | 4.26361 | 2.28958 | 1.23655 | 4.23982 | 2.23060 | 1.45166 | 2.86394 | 0.46101 | 4.81196 | 3.44626 | 4.34804 | 1.75844 |
| **Zdhhc7** | 5.12564 | 2.42432 | 2.64343 | 11.39265 | 2.89366 | 0.70411 | 0.00529 | 10.40995 | 13.38374 | 21.91152 | 11.62356 | 5.01809 | 3.29896 | 0.04991 | 4.44987 | 0.00000 |
| **Zdhhc8** | 0.13364 | 0.00785 | 0.00767 | 2.44496 | 0.01598 | 0.00657 | 0.00728 | 0.00996 | 0.84885 | 0.46457 | 0.00649 | 0.00534 | 0.01104 | 0.03277 | 0.00330 | 0.00650 |
| **Zdhhc9** | 0.51448 | 0.00696 | 0.00028 | 0.02478 | 0.00673 | 0.00747 | 0.71414 | 0.01337 | 0.01163 | 1.93794 | 0.10562 | 0.02198 | 1.21591 | 0.02441 | 0.49652 | 0.21536 |
| **Zeb1** | 0.64427 | 0.00768 | 0.00092 | 0.59635 | 0.01779 | 0.01441 | 0.00681 | 0.00868 | 0.71366 | 0.00225 | 0.00766 | 0.06491 | 0.01193 | 0.00485 | 0.40031 | 0.00674 |
| **Zeb2** | 0.01384 | 0.00795 | 0.01118 | 0.01062 | 0.00932 | 0.00671 | 0.00745 | 0.01270 | 0.01231 | 0.01356 | 0.01264 | 0.06869 | 0.01284 | 0.01206 | 0.01041 | 0.00743 |
| **Zer1** | 1.68213 | 0.00535 | 2.79279 | 1.00744 | 0.05069 | 0.05658 | 0.06478 | 0.79460 | 0.15678 | 4.25074 | 0.48324 | 1.43121 | 2.03753 | 0.00508 | 0.16754 | 0.02576 |
| **Zfa.ps** | 0.01735 | 0.08039 | 0.00866 | 0.33340 | 0.00871 | 0.00654 | 0.00719 | 0.04074 | 0.10614 | 0.15324 | 0.12281 | 0.80520 | 0.01152 | 0.18568 | 0.20303 | 0.00707 |
| **Zfand1** | 5.99350 | 5.32140 | 2.69243 | 0.25078 | 8.48840 | 0.01153 | 3.53844 | 1.89587 | 0.32512 | 4.30315 | 1.49437 | 4.13262 | 3.11055 | 2.25965 | 0.55026 | 0.00341 |
| **Zfand2a** | 4.61596 | 11.35130 | 10.56544 | 14.73770 | 2.03113 | 4.67274 | 7.70568 | 17.54691 | 21.33964 | 19.92297 | 14.73574 | 3.32829 | 7.03141 | 14.41454 | 16.50056 | 0.79428 |
| **Zfand2b** | 2.51511 | 3.10729 | 5.63363 | 0.66132 | 8.09811 | 1.33240 | 1.79547 | 9.81912 | 0.98198 | 6.08310 | 2.02899 | 0.15644 | 8.70525 | 0.02260 | 0.22474 | 5.60638 |
| **Zfand3** | 2.93956 | 0.52653 | 4.95011 | 3.71458 | 0.83317 | 1.14772 | 0.28450 | 3.69839 | 2.98002 | 7.74270 | 4.10263 | 0.23594 | 5.21326 | 0.04626 | 0.00000 | 2.83111 |
| **Zfand4** | 0.00983 | 0.00773 | 0.00888 | 0.03346 | 0.00959 | 0.00674 | 0.00722 | 0.29319 | 0.10747 | 0.12563 | 0.00975 | 0.00654 | 0.01324 | 0.01333 | 0.01099 | 0.00714 |
| **Zfand5** | 3.08363 | 7.59838 | 4.79887 | 27.66062 | 5.53555 | 2.85440 | 4.55695 | 5.05568 | 37.70890 | 20.62776 | 26.52206 | 27.43569 | 7.01454 | 27.29904 | 23.60026 | 0.26402 |
| **Zfand6** | 1.15042 | 3.95074 | 6.30104 | 7.51152 | 1.66451 | 1.13946 | 5.78647 | 4.84050 | 11.42035 | 10.48135 | 2.74033 | 0.67320 | 1.79310 | 6.32351 | 4.34827 | 0.30832 |
| **Zfat** | 0.10291 | 0.00864 | 0.67519 | 0.00747 | 0.01592 | 0.00551 | 0.00727 | 0.11353 | 0.54513 | 0.00507 | 0.00771 | 0.00626 | 0.00644 | 0.00995 | 0.44966 | 0.00722 |
| **Zfc3h1** | 1.57619 | 0.49511 | 4.51577 | 7.02278 | 0.00000 | 0.33843 | 1.63914 | 0.00000 | 8.01336 | 0.72541 | 0.77996 | 7.96134 | 0.26744 | 3.87371 | 12.32507 | 0.91063 |
| **Zfhx2** | 0.14285 | 0.00694 | 0.05789 | 0.46790 | 1.92755 | 0.00609 | 0.00768 | 0.01538 | 0.03651 | 0.00928 | 0.01435 | 0.00587 | 0.04226 | 0.00798 | 0.05503 | 0.01468 |
| **Zfhx3** | 0.02325 | 0.00717 | 0.20396 | 1.62972 | 0.02377 | 0.01497 | 1.82091 | 0.04553 | 1.00004 | 0.37879 | 0.01838 | 0.03300 | 0.10488 | 0.49216 | 0.00000 | 0.76911 |
| **Zfp1** | 5.93837 | 0.03927 | 2.78616 | 5.64234 | 1.63446 | 0.00000 | 4.11922 | 2.00247 | 3.66666 | 3.66289 | 0.66105 | 2.57080 | 1.53704 | 3.55490 | 2.95671 | 0.02002 |
| **Zfp101** | 0.88305 | 0.00783 | 2.06918 | 1.04448 | 0.02683 | 0.00483 | 0.00681 | 1.29042 | 4.63830 | 0.00000 | 0.28833 | 0.06194 | 0.16781 | 1.16458 | 5.61468 | 0.00564 |

| | | | | | | | | | | | | | | | | |
|---|---|---|---|---|---|---|---|---|---|---|---|---|---|---|---|---|
| **Zfp105** | 5.10582 | 0.00000 | 1.18777 | 0.00000 | 0.00000 | 0.00378 | 5.40668 | 0.87272 | 0.43898 | 0.76836 | 1.59045 | 0.08392 | 3.83443 | 0.02817 | 2.28299 | 3.78340 |
| **Zfp106** | 17.45894 | 10.28567 | 8.69573 | 6.69925 | 9.07103 | 8.55581 | 7.77167 | 11.84066 | 7.46915 | 22.14075 | 10.90060 | 10.11370 | 11.89731 | 17.99305 | 21.97345 | 5.43381 |
| **Zfp108** | 0.08918 | 0.00774 | 0.05796 | 3.85017 | 0.00923 | 0.00616 | 0.00728 | 0.02002 | 0.83487 | 0.21108 | 0.00645 | 0.00414 | 0.51554 | 0.00000 | 0.78752 | 0.00679 |
| **Zfp109** | 0.03670 | 0.00725 | 0.00250 | 0.01739 | 2.26878 | 0.01455 | 0.00797 | 0.09951 | 0.00921 | 0.00556 | 0.01009 | 0.00604 | 0.01655 | 0.08912 | 0.13268 | 0.00721 |
| **Zfp11** | 2.09112 | 0.04035 | 0.06413 | 1.22717 | 0.05895 | 0.02633 | 0.01644 | 0.09009 | 2.28403 | 0.00000 | 0.34220 | 0.03935 | 0.23948 | 0.65290 | 0.86082 | 0.06236 |
| **Zfp110** | 4.63808 | 8.13584 | 2.98282 | 1.44914 | 0.65312 | 4.06609 | 5.45595 | 5.70441 | 2.09300 | 6.80767 | 4.40741 | 1.60135 | 10.34368 | 8.58413 | 2.53322 | 2.31712 |
| **Zfp111** | 1.52077 | 0.00680 | 0.00000 | 0.13969 | 0.02993 | 0.42636 | 0.01833 | 0.09127 | 0.10717 | 0.46140 | 0.34661 | 0.06594 | 0.06563 | 0.41652 | 0.31731 | 0.01911 |
| **Zfp112** | 0.01519 | 0.00793 | 0.02863 | 0.01062 | 0.00923 | 0.00610 | 0.00745 | 0.01359 | 0.01234 | 0.01507 | 0.01307 | 0.00796 | 0.01873 | 0.01234 | 0.01032 | 0.00735 |
| **Zfp113** | 1.03911 | 0.17475 | 0.88367 | 8.15800 | 0.42545 | 0.13825 | 3.24847 | 0.09656 | 13.30890 | 1.66788 | 6.53527 | 0.37140 | 0.59154 | 8.61830 | 4.47945 | 0.20073 |
| **Zfp114** | 0.04795 | 0.00791 | 0.01030 | 0.01875 | 0.00874 | 0.00680 | 0.00751 | 0.01391 | 0.01116 | 0.02324 | 0.01261 | 0.00791 | 0.01289 | 0.00968 | 0.04389 | 0.00745 |
| **Zfp119a** | 0.03941 | 0.09136 | 0.00827 | 0.06046 | 0.01741 | 0.02200 | 0.04752 | 0.01996 | 0.01124 | 0.00000 | 0.49017 | 0.02580 | 0.73391 | 0.09272 | 0.22994 | 0.04203 |
| **Zfp119b** | 0.09443 | 0.47490 | 0.11827 | 0.83056 | 0.02812 | 0.02425 | 0.16430 | 0.11526 | 0.49866 | 0.16398 | 0.09006 | 0.17115 | 0.14859 | 0.48341 | 0.30520 | 0.01428 |
| **Zfp12** | 0.28960 | 4.51832 | 0.59120 | 2.58637 | 0.11631 | 0.13778 | 0.22743 | 0.05091 | 4.49776 | 0.27367 | 3.80816 | 0.22917 | 0.34838 | 3.64920 | 3.66231 | 0.11342 |
| **Zfp120** | 0.09642 | 0.10247 | 0.85289 | 2.35033 | 0.10391 | 0.01101 | 2.54845 | 0.76167 | 2.41672 | 2.04381 | 0.22330 | 0.38304 | 0.57936 | 0.53680 | 4.80973 | 0.01982 |
| **Zfp13** | 2.24964 | 0.00416 | 0.02507 | 0.06465 | 2.00004 | 0.00545 | 0.05528 | 0.02526 | 0.02069 | 0.41350 | 0.42216 | 0.01878 | 0.00959 | 0.00625 | 0.07608 | 2.05815 |
| **Zfp131** | 0.28703 | 5.41975 | 5.39704 | 11.42205 | 2.13416 | 0.32489 | 0.60311 | 7.10904 | 16.42243 | 12.32696 | 4.22886 | 17.45169 | 1.43126 | 25.26449 | 20.10509 | 1.28163 |
| **Zfp133.ps** | 0.56543 | 0.00774 | 0.00343 | 0.00851 | 0.01807 | 0.00697 | 0.00692 | 0.00909 | 0.01096 | 0.00603 | 0.01093 | 0.01670 | 0.01217 | 0.01394 | 0.50818 | 0.00702 |
| **Zfp14** | 0.32768 | 0.00830 | 0.01001 | 0.01030 | 0.01515 | 0.00619 | 0.00712 | 0.01003 | 0.02726 | 0.14482 | 0.01122 | 0.00637 | 0.01227 | 0.00604 | 0.60519 | 0.00732 |
| **Zfp141** | 0.50426 | 3.47162 | 2.64851 | 1.75309 | 3.03554 | 0.12623 | 0.54188 | 0.60293 | 1.88450 | 3.97313 | 3.06610 | 5.08783 | 0.83122 | 13.11021 | 4.35160 | 0.26191 |
| **Zfp142** | 0.03303 | 2.22073 | 1.04862 | 6.80224 | 0.50396 | 0.53506 | 0.02442 | 3.77360 | 4.65684 | 4.30603 | 1.57803 | 0.27658 | 3.58520 | 0.00270 | 2.16886 | 2.97622 |
| **Zfp143** | 1.67317 | 6.50682 | 2.22041 | 10.21888 | 0.03433 | 1.62428 | 4.43384 | 5.74873 | 11.92563 | 13.99112 | 8.86758 | 11.87960 | 4.58317 | 23.50173 | 18.91281 | 0.54477 |
| **Zfp146** | 0.01606 | 0.00776 | 0.09987 | 0.01084 | 0.00908 | 0.00577 | 0.00737 | 0.01263 | 0.01225 | 0.02218 | 0.01245 | 0.02750 | 0.01764 | 0.01409 | 0.01385 | 0.00728 |
| **Zfp148** | 1.95003 | 1.17568 | 2.13616 | 3.59061 | 3.01699 | 0.00924 | 3.88882 | 5.48353 | 6.51319 | 8.34335 | 5.11919 | 5.14458 | 0.68643 | 15.54737 | 6.80418 | 0.90036 |
| **Zfp157** | 0.05525 | 3.70613 | 0.41136 | 0.00000 | 1.08041 | 0.22951 | 0.95963 | 0.21191 | 0.01000 | 0.40504 | 0.72129 | 0.01315 | 0.67072 | 0.01558 | 0.00000 | 0.01217 |
| **Zfp160** | 0.28172 | 3.48556 | 1.51886 | 0.00000 | 0.01418 | 0.29957 | 0.00508 | 0.56166 | 0.07169 | 1.60635 | 0.38498 | 3.28065 | 0.00000 | 5.79715 | 2.32128 | 0.85239 |
| **Zfp174** | 0.01256 | 0.00806 | 0.16363 | 0.84204 | 0.00920 | 0.00863 | 0.02778 | 0.01070 | 0.76723 | 0.04374 | 0.04095 | 0.00568 | 0.01624 | 0.00810 | 0.41851 | 0.00718 |
| **Zfp180** | 1.61440 | 0.62898 | 2.31624 | 0.00000 | 0.12036 | 1.31592 | 0.00162 | 3.96609 | 1.00561 | 11.78525 | 3.43359 | 13.13442 | 1.60385 | 5.18910 | 13.76790 | 1.98497 |
| **Zfp182** | 0.06899 | 0.00830 | 2.05988 | 2.33765 | 0.01073 | 0.01400 | 0.04824 | 0.01009 | 0.00000 | 0.00000 | 0.00000 | 0.00000 | 1.13304 | 2.94084 | 0.02878 | 0.00557 |
| **Zfp184** | 0.19990 | 0.35924 | 0.02881 | 0.72331 | 0.01520 | 0.00861 | 0.01829 | 2.03554 | 0.00000 | 0.54185 | 0.00000 | 0.02385 | 0.02426 | 0.98708 | 0.07308 | 1.37201 |
| **Zfp185** | 0.01049 | 0.00732 | 0.00240 | 0.20969 | 0.00956 | 3.45565 | 0.00675 | 1.09922 | 0.16637 | 0.00001 | 0.00000 | 0.00540 | 0.20974 | 0.01551 | 0.00000 | 0.00580 |
| **Zfp189** | 0.02167 | 0.00793 | 0.73366 | 0.01654 | 0.00743 | 0.00558 | 0.00744 | 0.01216 | 0.69361 | 0.00805 | 0.00699 | 0.03826 | 0.00870 | 0.05215 | 0.00000 | 0.00703 |
| **Zfp2** | 0.03988 | 0.00821 | 0.00000 | 0.12602 | 0.01353 | 0.39145 | 0.00712 | 0.33950 | 0.00675 | 0.04462 | 0.01045 | 0.00674 | 0.15196 | 0.00805 | 0.05172 | 0.00769 |
| **Zfp202** | 0.05208 | 0.02905 | 0.14221 | 0.86880 | 0.00838 | 0.01214 | 0.03855 | 0.00151 | 1.46747 | 0.32546 | 0.72123 | 0.08785 | 0.04341 | 0.11951 | 0.53410 | 0.08444 |
| **Zfp207** | 15.33610 | 16.67044 | 15.74173 | 6.21907 | 12.99829 | 9.44762 | 14.31779 | 23.16423 | 21.24609 | 32.41257 | 13.72138 | 9.24004 | 10.23621 | 27.18055 | 26.95493 | 7.79103 |
| **Zfp212** | 0.91340 | 2.68699 | 4.67058 | 4.02593 | 0.01510 | 2.74740 | 3.06730 | 0.97234 | 3.04274 | 3.22435 | 2.13762 | 0.15814 | 2.47000 | 0.00000 | 2.40538 | 0.95714 |
| **Zfp213** | 0.01064 | 2.10076 | 1.22965 | 0.30760 | 0.00577 | 0.01108 | 0.00658 | 0.89653 | 0.96920 | 0.07513 | 0.00000 | 0.08373 | 0.35815 | 0.01899 | 0.05454 | 0.00611 |
| **Zfp217** | 5.00537 | 1.79288 | 1.40705 | 13.63659 | 0.53340 | 2.41583 | 0.13958 | 3.87901 | 16.30449 | 7.39630 | 8.04347 | 0.36335 | 1.58183 | 14.23157 | 7.46744 | 1.38414 |
| **Zfp219** | 1.31572 | 0.03141 | 0.32852 | 0.07737 | 0.01244 | 0.02255 | 0.21491 | 0.21708 | 0.01085 | 0.90691 | 0.57054 | 0.02117 | 3.24353 | 0.00898 | 0.00000 | 0.83090 |
| **Zfp229** | 0.93717 | 0.00346 | 0.06837 | 4.56861 | 0.01731 | 0.01591 | 3.06825 | 0.02713 | 3.99436 | 0.29341 | 0.24685 | 8.34494 | 0.71522 | 4.48166 | 4.13579 | 0.59757 |
| **Zfp235** | 1.54777 | 0.00644 | 0.00000 | 0.74071 | 0.04322 | 2.44575 | 0.92540 | 0.00472 | 0.49354 | 0.01495 | 0.06356 | 0.01285 | 0.07616 | 0.08492 | 0.58498 | 0.32868 |
| **Zfp236** | 0.00000 | 0.00658 | 0.22291 | 3.51567 | 1.66589 | 2.08140 | 0.72939 | 1.26951 | 2.92539 | 1.23608 | 1.32135 | 0.08456 | 0.06807 | 3.96552 | 2.84264 | 0.25771 |
| **Zfp239** | 2.59968 | 0.00000 | 3.66495 | 3.85715 | 6.09438 | 0.05432 | 3.62869 | 3.44256 | 6.23384 | 7.21786 | 3.87757 | 0.59206 | 2.82557 | 2.79400 | 10.89422 | 1.39959 |
| **Zfp24** | 1.37533 | 2.06672 | 2.15536 | 1.65786 | 0.01789 | 2.16550 | 0.34485 | 0.57216 | 1.20185 | 3.17701 | 0.49218 | 0.31235 | 0.14395 | 2.81499 | 2.66772 | 3.11925 |
| **Zfp248** | 0.91392 | 0.01758 | 0.17558 | 0.26724 | 0.00592 | 0.01048 | 0.02805 | 0.01366 | 0.01431 | 0.05269 | 0.02330 | 0.01818 | 0.01027 | 0.05615 | 0.01808 | 0.00633 |
| **Zfp251** | 0.77665 | 0.00666 | 2.28868 | 0.54757 | 0.03151 | 0.00816 | 1.16834 | 2.00504 | 0.83202 | 2.05357 | 0.31980 | 0.10121 | 0.33552 | 0.75191 | 2.49067 | 1.19803 |
| **Zfp26** | 0.15332 | 0.00582 | 1.00136 | 0.19100 | 0.00611 | 0.02306 | 2.31361 | 0.60684 | 0.88539 | 0.12322 | 1.41549 | 0.06482 | 0.93870 | 2.39264 | 0.23009 | 1.00672 |
| **Zfp260** | 2.78081 | 0.00413 | 1.57966 | 1.06387 | 0.02595 | 0.87545 | 0.01480 | 0.26255 | 0.00000 | 1.29029 | 0.02091 | 0.03501 | 0.55690 | 0.00709 | 2.53012 | 0.01715 |
| **Zfp263** | 0.63422 | 0.02493 | 0.00000 | 7.61019 | 3.15043 | 0.01103 | 1.14026 | 1.85007 | 6.06729 | 0.00000 | 2.02207 | 0.06769 | 0.00515 | 0.00000 | 2.41916 | 0.01139 |
| **Zfp266** | 0.26172 | 0.00793 | 0.00760 | 0.00463 | 0.00795 | 0.00748 | 0.00740 | 0.01780 | 0.90468 | 0.00334 | 0.00840 | 0.00628 | 0.37551 | 0.01546 | 0.00561 | 0.00710 |
| **Zfp27** | 0.04838 | 2.06760 | 0.15479 | 0.32056 | 0.01432 | 0.00597 | 4.07100 | 0.00000 | 0.94821 | 0.00000 | 0.68677 | 0.01639 | 0.02652 | 0.03268 | 0.21405 | 0.00626 |
| **Zfp273** | 0.00000 | 1.78918 | 0.34887 | 0.01555 | 0.00750 | 0.00856 | 0.03528 | 1.26971 | 0.11160 | 0.15206 | 1.03655 | 0.00000 | 0.00000 | 4.76157 | 0.00000 | 0.04879 |
| **Zfp275** | 2.00788 | 0.01599 | 0.28774 | 0.22138 | 0.01146 | 1.75948 | 0.00572 | 0.04153 | 0.00827 | 0.63562 | 0.00871 | 0.01428 | 0.00000 | 0.00547 | 0.13033 | 1.14091 |
| **Zfp276** | 1.74487 | 0.07006 | 1.01718 | 0.61740 | 0.01134 | 0.00387 | 0.30799 | 1.37119 | 3.47597 | 0.68426 | 0.04085 | 0.11482 | 0.90750 | 0.00657 | 0.53055 | 0.08093 |
| **Zfp277** | 4.33115 | 3.99585 | 3.49829 | 1.90323 | 4.49727 | 4.52245 | 0.45309 | 2.86717 | 0.89585 | 6.95892 | 1.43919 | 9.78023 | 2.72354 | 3.63697 | 1.60951 | 0.20211 |
| **Zfp28** | 0.00381 | 0.00822 | 0.00000 | 0.00000 | 0.00923 | 0.21031 | 0.00782 | 0.69016 | 0.35832 | 3.13736 | 0.31225 | 0.06953 | 0.08108 | 1.03543 | 1.26036 | 0.00694 |
| **Zfp280b** | 6.28790 | 2.80053 | 14.40094 | 22.02368 | 7.27906 | 2.09603 | 8.44366 | 9.61211 | 22.68420 | 19.84960 | 8.40555 | 11.33015 | 10.58754 | 2.25585 | 13.99145 | 3.50224 |
| **Zfp280c** | 0.01885 | 0.00798 | 0.63554 | 0.26029 | 0.00917 | 0.00891 | 0.00691 | 0.00779 | 0.02469 | 0.00000 | 0.00000 | 0.00000 | 0.00901 | 2.49802 | 1.28621 | 0.00702 |
| **Zfp280d** | 0.00001 | 5.71968 | 0.98623 | 2.05943 | 3.24774 | 1.97480 | 1.90800 | 2.06131 | 1.63282 | 3.49155 | 0.30285 | 9.29837 | 2.32832 | 14.14063 | 6.63786 | 2.55206 |
| **Zfp281** | 3.08695 | 5.05858 | 8.75915 | 11.44245 | 0.02564 | 6.11997 | 2.84689 | 8.15505 | 8.46195 | 9.10396 | 8.69888 | 10.67596 | 2.74905 | 13.92583 | 10.82898 | 2.84203 |
| **Zfp282** | 3.67577 | 0.78084 | 0.13036 | 2.66080 | 0.00452 | 0.08751 | 0.00310 | 0.54368 | 1.54715 | 0.42052 | 0.30172 | 0.09515 | 1.67365 | 0.38821 | 0.99810 | 0.07657 |
| **Zfp287** | 0.04157 | 0.00768 | 0.05030 | 0.15496 | 0.00816 | 0.01893 | 0.02765 | 0.01746 | 0.13177 | 0.39212 | 0.15912 | 0.00534 | 0.03638 | 0.13243 | 0.02861 | 0.00699 |
| **Zfp292** | 2.03435 | 5.66530 | 1.64631 | 5.36399 | 0.01628 | 0.00323 | 2.89071 | 2.94924 | 2.32614 | 3.33105 | 1.78662 | 3.02546 | 0.64514 | 10.77208 | 10.86349 | 0.12504 |

| | | | | | | | | | | | | | | | | |
|---|---|---|---|---|---|---|---|---|---|---|---|---|---|---|---|---|
| **Zfp296** | 15.55813 | 0.80457 | 0.00000 | 11.98580 | 17.09067 | 3.80633 | 5.71516 | 22.25154 | 22.28458 | 28.83298 | 13.68280 | 1.13508 | 0.76756 | 0.00000 | 0.73525 | 4.47207 |
| **Zfp3** | 0.02454 | 0.00778 | 0.00246 | 0.70447 | 0.00829 | 0.00739 | 0.00668 | 0.09517 | 0.43427 | 0.52562 | 1.24609 | 0.00000 | 0.00000 | 0.05747 | 0.66579 | 0.00696 |
| **Zfp30** | 0.01194 | 0.01896 | 0.01240 | 1.49349 | 0.02023 | 0.01392 | 0.00763 | 0.01475 | 0.09700 | 0.01097 | 0.01365 | 0.00726 | 0.12460 | 0.07615 | 0.01067 | 0.00750 |
| **Zfp300** | 0.09957 | 0.05022 | 0.10486 | 0.13133 | 0.00775 | 0.10868 | 0.01816 | 0.10459 | 0.29544 | 0.17615 | 0.04542 | 0.01560 | 0.07941 | 0.41863 | 0.11482 | 0.01434 |
| **Zfp316** | 0.63316 | 0.30761 | 1.24234 | 2.21488 | 0.39380 | 0.17371 | 0.62995 | 1.00814 | 4.21755 | 0.87083 | 5.10646 | 0.49197 | 0.78786 | 2.38854 | 0.94455 | 0.11358 |
| **Zfp317** | 1.28006 | 2.86564 | 1.33614 | 1.44090 | 3.04276 | 0.02314 | 1.12524 | 0.51760 | 1.02311 | 0.68623 | 0.73867 | 0.07588 | 0.88198 | 2.51740 | 2.36181 | 0.03338 |
| **Zfp318** | 0.41217 | 0.01839 | 0.17208 | 5.04273 | 0.02005 | 0.02336 | 0.00683 | 0.43220 | 2.51060 | 0.08824 | 0.05091 | 0.10014 | 0.04228 | 0.06486 | 6.31393 | 1.23361 |
| **Zfp319** | 0.46751 | 0.00723 | 1.57090 | 1.49537 | 0.00417 | 0.18332 | 0.00703 | 0.05439 | 0.37881 | 0.00000 | 0.14559 | 0.00533 | 0.67064 | 0.01095 | 0.10011 | 0.00544 |
| **Zfp322a** | 1.90251 | 1.40176 | 2.10838 | 3.86897 | 0.54902 | 0.00614 | 0.01585 | 1.02182 | 2.22878 | 0.78474 | 1.61831 | 0.08921 | 0.75927 | 0.10869 | 2.55474 | 2.57909 |
| **Zfp324** | 0.00782 | 0.00745 | 0.01280 | 0.05050 | 0.00955 | 0.00694 | 0.71121 | 0.02335 | 1.38028 | 0.80373 | 0.00000 | 0.00191 | 0.13665 | 0.00429 | 0.28120 | 0.19597 |
| **Zfp326** | 0.00000 | 1.65226 | 0.59243 | 0.05126 | 1.77850 | 0.01822 | 1.85853 | 1.08579 | 0.32815 | 0.22412 | 0.12482 | 0.00000 | 0.01960 | 1.15796 | 0.00000 | 0.70176 |
| **Zfp329** | 0.00415 | 0.00786 | 0.01788 | 1.43860 | 0.01026 | 0.00788 | 0.00711 | 0.98646 | 0.77227 | 0.77993 | 1.04224 | 0.02954 | 0.04169 | 1.24256 | 2.85541 | 0.02167 |
| **Zfp330** | 8.52714 | 9.28554 | 10.65724 | 6.27483 | 1.14650 | 2.04232 | 6.23279 | 7.52779 | 13.43712 | 5.87879 | 6.42376 | 4.07313 | 8.78980 | 4.40047 | 7.68250 | 7.01416 |
| **Zfp334** | 0.01479 | 0.00759 | 0.01968 | 0.03530 | 0.00934 | 0.00623 | 0.00738 | 0.01146 | 0.01971 | 0.00567 | 0.02782 | 0.00705 | 0.01226 | 0.01933 | 0.27525 | 0.00736 |
| **Zfp335** | 0.99302 | 0.00348 | 0.79392 | 7.76201 | 0.00463 | 0.42916 | 0.76043 | 7.92145 | 11.24740 | 11.66205 | 3.61749 | 4.35042 | 4.11604 | 0.03255 | 0.00000 | 0.37798 |
| **Zfp341** | 1.00661 | 0.00618 | 0.08384 | 3.33188 | 0.01148 | 0.01208 | 1.69856 | 0.02276 | 0.91442 | 1.44997 | 0.00001 | 0.03749 | 3.12049 | 0.00001 | 0.55957 | 0.00459 |
| **Zfp345** | 1.96925 | 0.08334 | 1.91763 | 0.50122 | 0.15602 | 0.18834 | 2.99949 | 2.10797 | 0.51026 | 0.58937 | 5.78489 | 0.20540 | 1.51556 | 9.27075 | 2.74978 | 0.09750 |
| **Zfp346** | 0.00000 | 0.86535 | 0.38246 | 5.74339 | 1.47357 | 0.00479 | 2.90076 | 2.06920 | 6.04273 | 0.56899 | 0.35584 | 0.03705 | 0.04122 | 0.00000 | 1.57146 | 2.35596 |
| **Zfp35** | 4.90186 | 5.18706 | 1.95124 | 1.22175 | 5.21910 | 2.43358 | 1.27333 | 4.74333 | 1.08571 | 1.88213 | 1.37504 | 1.29084 | 2.28959 | 9.54924 | 2.10239 | 0.01131 |
| **Zfp352** | 0.01240 | 0.00808 | 0.01961 | 7.76993 | 0.00905 | 0.01259 | 0.00728 | 0.01128 | 6.09007 | 0.00000 | 0.04073 | 0.00000 | 0.01171 | 0.01288 | 0.00565 | 0.00558 |
| **Zfp354a** | 0.00582 | 2.61618 | 1.68434 | 0.00661 | 0.00534 | 0.00701 | 0.00715 | 0.13998 | 0.01227 | 0.35579 | 0.01010 | 0.00589 | 0.01076 | 0.01510 | 0.02921 | 0.00595 |
| **Zfp354b** | 0.00887 | 0.23507 | 0.06213 | 0.00787 | 0.00842 | 0.00852 | 0.00733 | 0.01308 | 0.01077 | 0.01459 | 0.01017 | 0.00618 | 0.01143 | 0.01708 | 0.62399 | 0.00727 |
| **Zfp354c** | 0.01602 | 0.00776 | 0.49839 | 0.00810 | 0.00829 | 0.00559 | 0.00714 | 0.01017 | 0.01125 | 0.00314 | 0.01123 | 0.05954 | 0.00903 | 0.00717 | 0.41427 | 0.05704 |
| **Zfp358** | 0.22307 | 0.00849 | 0.00692 | 0.00978 | 0.01733 | 0.00717 | 0.00732 | 0.01152 | 0.01216 | 0.18921 | 0.01248 | 0.00784 | 0.01261 | 0.01147 | 0.01147 | 0.00759 |
| **Zfp36** | 0.57509 | 1.50863 | 0.33147 | 4.94239 | 0.51634 | 0.67999 | 0.49146 | 4.96152 | 9.62766 | 4.44135 | 5.38964 | 0.83353 | 0.99447 | 0.84658 | 1.67934 | 0.69655 |
| **Zfp362** | 0.13890 | 0.00749 | 0.00335 | 0.12592 | 0.79686 | 0.00773 | 0.00660 | 0.16441 | 0.00877 | 0.00362 | 0.01223 | 0.00625 | 0.77926 | 0.01303 | 0.00531 | 0.00675 |
| **Zfp367** | 0.00000 | 0.52089 | 0.02701 | 9.55471 | 0.05595 | 1.04806 | 0.08194 | 0.51223 | 3.85965 | 0.56353 | 0.39711 | 1.65241 | 0.38238 | 6.88977 | 5.66964 | 0.01175 |
| **Zfp369** | 0.06676 | 0.08465 | 0.01584 | 0.34444 | 0.16803 | 0.04536 | 0.12842 | 0.08574 | 0.42679 | 0.11545 | 0.00683 | 0.01783 | 0.11172 | 0.14720 | 0.08234 | 0.02880 |
| **Zfp36l1** | 5.62830 | 0.00987 | 1.90222 | 0.00618 | 3.19867 | 0.00000 | 1.59436 | 0.00000 | 0.01047 | 0.36285 | 0.00886 | 0.02936 | 0.36908 | 0.03317 | 0.00000 | 2.41715 |
| **Zfp36l2** | 6.36336 | 4.58650 | 0.00000 | 1.69646 | 2.65259 | 10.37974 | 1.99754 | 0.14647 | 3.31220 | 3.60844 | 11.59036 | 0.13437 | 2.31429 | 0.06772 | 1.18127 | 2.82791 |
| **Zfp37** | 0.32751 | 0.00769 | 0.00610 | 0.18695 | 0.00725 | 0.00706 | 0.00718 | 0.01976 | 0.01606 | 0.01488 | 0.01352 | 0.00798 | 0.01198 | 0.01106 | 0.01328 | 0.00694 |
| **Zfp382** | 0.01382 | 0.00817 | 0.01111 | 0.86911 | 0.00922 | 0.00662 | 0.00748 | 0.01099 | 0.00000 | 0.01052 | 0.12580 | 0.01815 | 0.01073 | 0.00896 | 0.00260 | 0.00719 |
| **Zfp383** | 0.00557 | 0.34252 | 0.68268 | 0.21241 | 1.50060 | 0.00663 | 0.00788 | 0.50109 | 0.01794 | 0.11017 | 0.31733 | 0.00446 | 0.00001 | 0.00645 | 0.12230 | 0.00701 |
| **Zfp384** | 0.10825 | 0.18517 | 0.00525 | 0.06699 | 0.03389 | 0.00769 | 0.00737 | 0.01262 | 0.71023 | 0.70876 | 0.06290 | 0.10119 | 0.34778 | 0.05635 | 0.21123 | 0.00755 |
| **Zfp385a** | 0.01217 | 0.00734 | 0.13767 | 1.44305 | 0.35131 | 0.00590 | 0.00655 | 0.17065 | 1.36522 | 0.44684 | 1.77492 | 0.01522 | 0.00000 | 0.00053 | 0.00000 | 0.07009 |
| **Zfp385c** | 0.04585 | 0.00790 | 0.16215 | 0.95721 | 0.05263 | 0.00619 | 0.08871 | 0.04058 | 2.17607 | 0.01581 | 0.66550 | 0.00941 | 0.12486 | 0.18351 | 0.33829 | 0.01406 |
| **Zfp386** | 0.70349 | 3.10900 | 0.81214 | 8.39096 | 0.15258 | 0.00411 | 2.62519 | 0.32650 | 0.00000 | 3.48519 | 2.64940 | 0.08166 | 2.66431 | 5.27211 | 3.84393 | 0.00964 |
| **Zfp39** | 0.00805 | 0.00810 | 0.03005 | 0.66898 | 0.00779 | 0.01180 | 0.00744 | 0.01456 | 0.00485 | 0.35049 | 0.01371 | 0.00839 | 0.05240 | 0.01143 | 0.03552 | 1.06732 |
| **Zfp395** | 1.07136 | 0.05766 | 0.49807 | 0.00563 | 0.03040 | 0.43990 | 0.14743 | 0.94664 | 0.02087 | 1.30419 | 0.00000 | 0.13513 | 0.55421 | 0.15898 | 0.04255 | 0.45291 |
| **Zfp397** | 1.34977 | 0.03226 | 0.01267 | 0.32657 | 0.00539 | 0.01202 | 2.77539 | 0.11706 | 2.54399 | 0.00000 | 0.16696 | 3.19641 | 0.49672 | 1.37765 | 1.20821 | 0.40745 |
| **Zfp398** | 0.01902 | 0.00751 | 0.03806 | 1.84434 | 0.02655 | 0.01227 | 0.00719 | 1.20544 | 0.32804 | 0.00255 | 0.01861 | 0.00472 | 0.00546 | 0.00486 | 0.08016 | 0.00688 |
| **Zfp40** | 0.01251 | 0.01766 | 0.01039 | 0.24745 | 0.01848 | 0.01307 | 0.00727 | 0.01569 | 0.00507 | 0.38180 | 0.01139 | 0.01538 | 0.01199 | 0.04541 | 0.09679 | 0.00774 |
| **Zfp407** | 1.86671 | 1.55054 | 0.69866 | 2.33968 | 0.00306 | 0.04207 | 1.85402 | 0.97336 | 4.23967 | 2.56281 | 0.71627 | 5.86112 | 1.18086 | 1.69468 | 6.87617 | 0.00261 |
| **Zfp408** | 0.17765 | 0.00824 | 0.67186 | 2.13342 | 0.05717 | 0.00520 | 0.08232 | 0.23203 | 4.23449 | 0.51256 | 0.53824 | 0.06263 | 1.30539 | 0.15250 | 1.03426 | 1.68505 |
| **Zfp41** | 0.01202 | 0.00790 | 0.04301 | 0.12833 | 0.03063 | 0.00643 | 0.00743 | 0.01837 | 0.00694 | 0.45735 | 0.01402 | 0.00598 | 0.22934 | 0.01093 | 0.00535 | 0.01545 |
| **Zfp410** | 2.43963 | 4.37852 | 6.82697 | 11.33630 | 4.31078 | 0.02480 | 3.07985 | 9.25516 | 11.71819 | 9.39392 | 3.59958 | 9.57170 | 4.19050 | 3.77364 | 11.91995 | 2.79071 |
| **Zfp414** | 8.82670 | 1.58808 | 4.95335 | 0.46042 | 0.28775 | 0.00000 | 3.25090 | 3.39579 | 0.00072 | 5.58705 | 0.84257 | 0.11599 | 3.05692 | 0.00076 | 0.55522 | 0.74194 |
| **Zfp418** | 0.75529 | 0.00679 | 0.02523 | 0.00626 | 0.00699 | 0.00524 | 0.00662 | 0.00908 | 0.01043 | 0.00482 | 0.01028 | 0.03497 | 0.01173 | 0.00771 | 0.88877 | 0.00646 |
| **Zfp42** | 15.92187 | 13.57883 | 18.69323 | 4.99793 | 9.53081 | 10.13787 | 11.08916 | 12.97130 | 9.47807 | 4.57346 | 7.17885 | 0.71131 | 9.60091 | 3.38534 | 2.02296 | 4.77905 |
| **Zfp420** | 0.00000 | 2.79316 | 1.90216 | 1.26649 | 0.01715 | 0.00776 | 0.00676 | 0.56691 | 2.25770 | 0.25618 | 0.10820 | 0.02508 | 0.38755 | 0.00291 | 3.52509 | 1.11354 |
| **Zfp422** | 0.13471 | 0.01930 | 0.29670 | 0.20072 | 0.07524 | 0.19039 | 0.02983 | 0.13931 | 0.29809 | 0.92367 | 1.00532 | 0.03947 | 0.05240 | 0.84520 | 0.20953 | 0.04507 |
| **Zfp423** | 0.01356 | 0.00814 | 0.01188 | 0.63059 | 0.00915 | 0.00677 | 0.00755 | 0.01333 | 0.20552 | 0.19621 | 0.02503 | 0.00826 | 0.01936 | 0.00837 | 0.02102 | 0.00726 |
| **Zfp426** | 1.47101 | 0.00435 | 1.34944 | 1.09074 | 2.47136 | 0.00487 | 0.00522 | 1.09540 | 1.23514 | 1.46994 | 0.81698 | 0.20283 | 1.05626 | 0.01901 | 0.25130 | 0.04213 |
| **Zfp428** | 0.49374 | 0.01875 | 0.02495 | 1.67301 | 0.01228 | 0.00534 | 0.00643 | 0.00865 | 0.77236 | 4.34795 | 0.08290 | 0.08311 | 0.50119 | 0.02332 | 0.00000 | 0.01391 |
| **Zfp429** | 1.31972 | 0.00696 | 0.00000 | 0.04571 | 0.01633 | 0.00571 | 0.00568 | 0.00343 | 0.11993 | 0.00000 | 0.01944 | 0.01042 | 0.74419 | 1.42487 | 0.17526 | 3.18069 |
| **Zfp433** | 0.05412 | 0.51295 | 2.50784 | 2.51481 | 0.15876 | 0.24540 | 0.14294 | 1.81194 | 7.17438 | 2.97700 | 4.14208 | 7.49114 | 1.11127 | 13.95449 | 2.75412 | 0.00334 |
| **Zfp438** | 1.14684 | 0.70099 | 0.61752 | 0.15956 | 0.01320 | 2.66750 | 0.00513 | 3.11465 | 0.58645 | 0.19260 | 1.17135 | 0.18521 | 1.06394 | 1.34277 | 0.22592 | 0.00388 |
| **Zfp442** | 0.03395 | 0.13844 | 0.50131 | 4.84271 | 0.09613 | 0.12848 | 0.15291 | 0.23078 | 2.53581 | 3.59503 | 0.17675 | 0.64029 | 0.35081 | 6.12293 | 13.70812 | 0.12671 |
| **Zfp444** | 0.44765 | 0.02863 | 0.22844 | 1.72041 | 0.89354 | 0.15139 | 0.07852 | 0.00000 | 3.36405 | 1.06375 | 1.78252 | 0.01066 | 0.22632 | 2.07919 | 0.54980 | 0.79631 |
| **Zfp445** | 2.31709 | 3.10730 | 2.29908 | 3.31702 | 2.75758 | 0.08882 | 3.03982 | 3.70001 | 7.03636 | 1.42290 | 4.23644 | 6.62879 | 1.02786 | 7.85666 | 5.40454 | 1.63375 |
| **Zfp446** | 0.89731 | 0.04828 | 0.52389 | 4.32644 | 0.01463 | 4.29695 | 1.65245 | 3.84526 | 3.18786 | 2.19818 | 0.33237 | 0.05456 | 2.24237 | 0.11526 | 0.81812 | 0.03814 |
| **Zfp449** | 0.01340 | 0.00755 | 0.01013 | 0.15987 | 0.00924 | 0.00632 | 0.00728 | 0.00970 | 1.38630 | 0.10604 | 0.01005 | 0.00030 | 0.01329 | 0.00886 | 1.41556 | 0.00769 |

| | | | | | | | | | | | | | | | | |
|---|---|---|---|---|---|---|---|---|---|---|---|---|---|---|---|---|
| **Zfp451** | 2.81289 | 0.14703 | 3.13778 | 6.77878 | 4.77338 | 0.00227 | 0.76680 | 2.90447 | 4.86787 | 3.13602 | 3.34667 | 2.80591 | 0.87768 | 1.30933 | 1.90371 | 0.82888 |
| **Zfp454** | 0.01332 | 0.00774 | 0.01869 | 0.34950 | 0.00949 | 0.00719 | 0.00744 | 0.01261 | 0.00648 | 0.00710 | 0.01340 | 0.01648 | 0.02632 | 0.00552 | 0.31877 | 0.00738 |
| **Zfp455** | 0.03216 | 0.00827 | 0.15086 | 0.00579 | 0.01144 | 0.00599 | 0.00741 | 0.00549 | 0.01119 | 0.09621 | 0.25684 | 0.09042 | 0.01182 | 0.00799 | 0.05976 | 0.05917 |
| **Zfp456** | 0.14365 | 0.00837 | 0.17719 | 0.00954 | 0.01431 | 0.00594 | 0.00741 | 0.01272 | 0.01172 | 0.05322 | 0.01155 | 0.07799 | 0.09526 | 0.00785 | 0.16080 | 0.21636 |
| **Zfp457** | 0.01697 | 0.01870 | 0.01577 | 0.11810 | 0.00897 | 0.00634 | 0.00742 | 0.01648 | 0.19271 | 0.01679 | 0.06042 | 0.00692 | 0.01151 | 0.06372 | 0.04403 | 0.00736 |
| **Zfp458** | 0.14360 | 0.03033 | 0.00000 | 0.00720 | 0.01687 | 0.00663 | 0.00677 | 0.01083 | 0.02643 | 0.29182 | 0.00973 | 0.00000 | 0.17973 | 3.13734 | 0.00000 | 0.00736 |
| **Zfp459** | 4.19446 | 1.97958 | 1.06682 | 0.38734 | 0.89718 | 0.01282 | 0.07325 | 0.02011 | 0.43240 | 0.46343 | 1.01893 | 0.06828 | 0.18619 | 0.28586 | 0.34508 | 1.54376 |
| **Zfp46** | 0.01213 | 0.00690 | 0.54530 | 3.49434 | 0.00856 | 0.00516 | 1.29019 | 0.00799 | 1.15083 | 0.00325 | 0.01237 | 0.01252 | 0.01455 | 0.00089 | 1.10910 | 0.00625 |
| **Zfp462** | 3.35494 | 0.18260 | 0.00000 | 0.30766 | 0.01362 | 0.00411 | 0.00465 | 0.00000 | 0.00618 | 0.01093 | 0.00610 | 0.00539 | 0.04378 | 2.86225 | 0.00000 | 0.01884 |
| **Zfp467** | 0.01267 | 0.00756 | 0.01213 | 0.00609 | 0.00896 | 0.00647 | 0.00721 | 0.01525 | 0.01164 | 1.35956 | 0.01130 | 0.00331 | 0.02394 | 0.00588 | 0.13770 | 0.00762 |
| **Zfp472** | 2.27599 | 6.61264 | 4.13603 | 1.54450 | 0.01504 | 0.00430 | 2.13979 | 4.46175 | 0.32051 | 4.74585 | 1.88894 | 0.18690 | 1.75117 | 0.16305 | 8.97014 | 0.47919 |
| **Zfp473** | 0.53769 | 0.01768 | 0.00000 | 2.00299 | 0.00655 | 0.00715 | 0.07769 | 0.73927 | 0.18143 | 0.20699 | 0.47818 | 0.02423 | 1.03329 | 0.00217 | 1.08254 | 0.00583 |
| **Zfp488** | 0.38553 | 0.26304 | 0.73624 | 0.65175 | 0.06290 | 0.20826 | 0.59572 | 0.89410 | 1.15178 | 0.51449 | 0.09540 | 0.93281 | 0.32221 | 4.38212 | 0.82033 | 0.12616 |
| **Zfp493** | 0.02170 | 0.00797 | 0.13589 | 0.00475 | 0.00914 | 0.00845 | 0.00702 | 0.04590 | 0.01098 | 1.13961 | 0.69723 | 0.00000 | 0.02203 | 0.03774 | 0.03781 | 0.00702 |
| **Zfp503** | 0.01348 | 0.00810 | 0.01207 | 1.78136 | 0.00910 | 0.00673 | 0.00746 | 0.01248 | 0.00000 | 0.48789 | 0.01144 | 0.00804 | 0.01234 | 0.00965 | 0.01187 | 0.00710 |
| **Zfp507** | 0.74447 | 2.33099 | 1.91569 | 1.69870 | 0.02078 | 0.26674 | 0.45547 | 1.20124 | 2.38796 | 0.93758 | 0.00000 | 0.03342 | 2.07249 | 0.14420 | 2.00571 | 0.95718 |
| **Zfp51** | 0.00000 | 6.40544 | 0.76266 | 3.90609 | 0.00771 | 0.04133 | 0.02191 | 1.43078 | 5.95782 | 5.36926 | 4.43955 | 9.74390 | 0.81477 | 8.86842 | 8.64923 | 0.78096 |
| **Zfp511** | 4.52068 | 0.08008 | 1.56315 | 1.52283 | 5.61865 | 0.00659 | 0.01079 | 0.73445 | 0.92540 | 6.40388 | 2.98890 | 1.34145 | 3.27566 | 0.56679 | 0.00000 | 2.00643 |
| **Zfp512** | 6.38201 | 0.00000 | 0.67656 | 0.38801 | 3.99676 | 2.28713 | 1.48745 | 0.00000 | 0.59183 | 1.70965 | 1.06683 | 0.63557 | 2.47262 | 0.00803 | 0.05427 | 1.90961 |
| **Zfp512b** | 2.85713 | 0.00461 | 0.00000 | 3.17498 | 0.01283 | 0.00290 | 0.00432 | 2.66965 | 1.80432 | 0.60485 | 0.00000 | 0.21680 | 1.48285 | 4.34553 | 1.75366 | 0.03319 |
| **Zfp513** | 0.28345 | 2.09727 | 1.73684 | 18.92481 | 0.18586 | 0.01078 | 0.73525 | 2.57790 | 14.39975 | 7.83051 | 4.86694 | 1.61285 | 5.70347 | 0.38264 | 1.64226 | 4.02191 |
| **Zfp516** | 0.03302 | 0.00802 | 0.05800 | 0.19804 | 0.00732 | 0.00834 | 0.00728 | 0.01241 | 0.00583 | 0.16754 | 0.01088 | 0.00732 | 0.03798 | 0.01063 | 0.00640 | 1.14714 |
| **Zfp518a** | 0.07086 | 1.12665 | 0.20923 | 5.51232 | 3.77163 | 0.01142 | 6.10219 | 0.30360 | 1.83449 | 2.55996 | 0.95703 | 6.46888 | 0.54931 | 8.11942 | 14.48059 | 0.53806 |
| **Zfp518b** | 2.17755 | 0.02575 | 1.80150 | 3.56034 | 0.11961 | 0.07512 | 0.08483 | 2.05025 | 1.49447 | 1.18717 | 1.10819 | 0.05375 | 0.44962 | 3.05134 | 1.54944 | 2.63411 |
| **Zfp52** | 0.28013 | 0.00863 | 0.00000 | 0.56099 | 0.01897 | 0.00601 | 0.00727 | 0.01692 | 3.45539 | 0.57756 | 0.00000 | 0.00000 | 0.29946 | 10.75244 | 0.24328 | 0.01539 |
| **Zfp523** | 0.27533 | 0.00791 | 0.00648 | 0.93986 | 0.00722 | 0.00685 | 0.00717 | 0.01219 | 0.04226 | 0.36426 | 0.01181 | 0.01750 | 0.01102 | 0.00972 | 0.00336 | 0.02748 |
| **Zfp524** | 3.80690 | 0.00308 | 0.00000 | 0.21287 | 0.00280 | 3.25496 | 0.00209 | 1.97488 | 0.00520 | 0.64311 | 0.31643 | 0.05218 | 3.18371 | 0.01250 | 0.08845 | 0.00000 |
| **Zfp526** | 0.00743 | 1.41691 | 0.00684 | 0.39165 | 0.00867 | 0.02721 | 0.00699 | 0.01138 | 0.00606 | 0.00412 | 0.01611 | 0.01623 | 0.16707 | 0.00865 | 0.20540 | 0.00701 |
| **Zfp53** | 1.46448 | 0.92323 | 0.00000 | 1.46509 | 0.00688 | 0.65142 | 0.00528 | 0.00949 | 0.20473 | 0.50348 | 0.55944 | 0.00979 | 0.61527 | 3.50632 | 0.33919 | 0.01191 |
| **Zfp532** | 0.30769 | 1.21397 | 0.00000 | 4.53276 | 0.01160 | 2.07426 | 1.25912 | 2.88697 | 0.00000 | 3.52400 | 1.34724 | 0.20017 | 0.95517 | 0.01124 | 1.18520 | 0.23884 |
| **Zfp534** | 0.45202 | 3.88847 | 0.68602 | 0.31421 | 1.40577 | 0.31365 | 0.93375 | 1.79470 | 0.33483 | 1.49014 | 1.32383 | 1.69584 | 0.42401 | 12.09267 | 1.62458 | 0.55892 |
| **Zfp54** | 0.36163 | 0.00736 | 0.00000 | 4.94752 | 0.00744 | 0.00624 | 0.00664 | 0.54415 | 7.84408 | 0.67529 | 0.52923 | 0.00000 | 0.96133 | 0.00000 | 1.59733 | 0.00543 |
| **Zfp541** | 0.00727 | 0.00684 | 0.00000 | 0.95795 | 0.00907 | 0.00714 | 1.98146 | 0.00000 | 0.86500 | 1.66009 | 0.60451 | 0.00545 | 0.15155 | 0.00803 | 1.20688 | 0.00671 |
| **Zfp553** | 1.77377 | 3.45074 | 2.00799 | 3.54672 | 0.01940 | 4.17184 | 0.42262 | 5.81019 | 5.39656 | 0.86343 | 0.99388 | 0.00000 | 2.07110 | 0.22269 | 2.66357 | 0.77753 |
| **Zfp558** | 0.07745 | 0.00894 | 0.02764 | 0.87730 | 0.03453 | 0.00710 | 0.04256 | 0.05922 | 0.03400 | 0.29839 | 0.00000 | 0.07271 | 0.14161 | 2.39468 | 1.25552 | 0.01541 |
| **Zfp560** | 0.91792 | 0.00750 | 0.17204 | 3.72727 | 0.01665 | 0.00398 | 0.00651 | 0.00000 | 3.74251 | 0.37050 | 0.28380 | 0.00992 | 0.00805 | 0.04706 | 0.00000 | 0.01262 |
| **Zfp563** | 0.16588 | 3.22271 | 0.50135 | 0.18122 | 0.01136 | 0.00902 | 0.00575 | 0.00867 | 0.07274 | 0.01019 | 0.01288 | 0.00580 | 0.46702 | 0.01006 | 0.75268 | 0.07861 |
| **Zfp566** | 1.35051 | 2.82434 | 7.76271 | 1.87331 | 0.00000 | 0.00354 | 5.33947 | 3.43068 | 3.95085 | 3.22439 | 1.72925 | 4.44219 | 0.03576 | 3.01363 | 1.46747 | 0.78704 |
| **Zfp568** | 5.34594 | 5.97853 | 11.02321 | 9.46879 | 2.12060 | 1.62666 | 0.01795 | 11.54179 | 5.66860 | 15.38729 | 11.24634 | 12.22692 | 12.23946 | 19.71283 | 18.41113 | 6.77709 |
| **Zfp57** | 6.74082 | 3.31787 | 1.03624 | 3.71586 | 0.01480 | 1.22271 | 2.37277 | 1.23635 | 6.57885 | 1.65178 | 0.34811 | 0.12153 | 0.21956 | 1.50654 | 0.61637 | 2.21481 |
| **Zfp572** | 0.01375 | 0.00790 | 0.01198 | 0.35392 | 0.00938 | 0.00646 | 0.00738 | 0.01192 | 0.00656 | 0.00660 | 0.01315 | 0.00712 | 0.01270 | 0.00606 | 0.47071 | 0.00730 |
| **Zfp574** | 3.17615 | 0.46518 | 0.59800 | 6.36840 | 0.00000 | 2.48224 | 2.23521 | 1.23570 | 9.69424 | 10.62767 | 0.77278 | 0.59805 | 5.06283 | 0.00000 | 0.00000 | 2.24391 |
| **Zfp575** | 0.78559 | 0.85909 | 3.19664 | 3.09390 | 0.49712 | 1.48800 | 1.55758 | 7.29463 | 6.42426 | 7.60862 | 5.89371 | 8.91149 | 1.06487 | 4.39561 | 3.40766 | 0.67706 |
| **Zfp579** | 0.62963 | 0.00761 | 0.00000 | 0.23627 | 0.01638 | 0.35583 | 0.00658 | 0.80775 | 0.02295 | 0.60679 | 0.42565 | 0.07656 | 0.03265 | 0.00695 | 0.00000 | 0.88614 |
| **Zfp58** | 0.37873 | 0.02011 | 0.00822 | 0.05414 | 0.03994 | 0.00780 | 0.00697 | 0.00000 | 0.01756 | 0.07522 | 0.27059 | 0.00932 | 0.03084 | 2.00238 | 0.00000 | 0.00736 |
| **Zfp580** | 0.01344 | 0.00835 | 0.01728 | 1.04406 | 0.00930 | 0.00635 | 0.00758 | 0.01443 | 0.21532 | 0.01977 | 0.01194 | 0.00851 | 0.01274 | 0.01067 | 0.00348 | 0.00723 |
| **Zfp583** | 0.02935 | 0.49016 | 0.10891 | 1.48699 | 0.00842 | 0.00529 | 0.80000 | 0.01198 | 1.34365 | 0.35389 | 0.00000 | 4.10945 | 0.06590 | 0.26771 | 0.47506 | 0.01377 |
| **Zfp59** | 1.26040 | 0.00550 | 1.00494 | 0.00000 | 0.01263 | 0.00555 | 2.77632 | 0.14198 | 0.01286 | 0.19458 | 0.39550 | 0.07660 | 0.04752 | 0.00691 | 3.43684 | 0.03389 |
| **Zfp592** | 5.53627 | 1.62812 | 0.64445 | 12.04118 | 0.03765 | 6.29947 | 2.80204 | 6.00999 | 5.89703 | 3.05504 | 4.84816 | 2.22004 | 4.08647 | 1.80883 | 4.37067 | 0.02284 |
| **Zfp593** | 1.23027 | 0.00554 | 0.84470 | 1.87118 | 1.10249 | 3.37247 | 0.21334 | 1.56793 | 3.83229 | 2.63383 | 4.82990 | 0.04777 | 2.30730 | 0.00000 | 0.78308 | 1.73399 |
| **Zfp595** | 0.00493 | 0.11890 | 1.05651 | 0.00940 | 0.40898 | 0.00897 | 0.01707 | 0.00905 | 0.01203 | 0.25312 | 0.01004 | 0.00579 | 0.31409 | 0.01340 | 0.00001 | 0.00645 |
| **Zfp597** | 0.00000 | 0.12248 | 0.47637 | 3.48286 | 0.01261 | 1.17529 | 0.00724 | 0.31937 | 0.26858 | 0.07964 | 0.93095 | 3.92386 | 0.00000 | 7.72159 | 3.30140 | 0.00588 |
| **Zfp598** | 6.79505 | 2.37748 | 4.35164 | 13.37508 | 0.46237 | 1.46590 | 6.99551 | 10.46068 | 11.72474 | 15.41680 | 10.45604 | 8.25196 | 6.61754 | 0.00000 | 1.76144 | 7.02130 |
| **Zfp599** | 0.00000 | 0.32052 | 2.64669 | 5.89776 | 1.41119 | 0.38808 | 0.03614 | 1.44657 | 12.80728 | 0.00000 | 2.42750 | 0.00000 | 0.08406 | 20.81708 | 4.19266 | 0.08705 |
| **Zfp60** | 1.28706 | 0.00641 | 0.00000 | 0.47554 | 0.01111 | 0.65495 | 0.52066 | 0.17454 | 0.02086 | 0.00000 | 0.00000 | 0.39031 | 1.47061 | 3.00612 | 0.93708 | 0.01217 |
| **Zfp600** | 2.05274 | 4.31939 | 1.84751 | 13.42297 | 1.78931 | 0.18170 | 1.34098 | 4.54043 | 10.61317 | 0.62679 | 5.56131 | 16.66920 | 2.55348 | 26.23579 | 2.47782 | 0.74310 |
| **Zfp605** | 0.16201 | 0.00739 | 0.75191 | 2.56168 | 0.12645 | 0.01589 | 0.03712 | 0.03470 | 3.38983 | 0.03769 | 1.35265 | 0.06691 | 0.72678 | 1.13029 | 1.70968 | 0.12015 |
| **Zfp606** | 0.00615 | 0.00605 | 0.13477 | 0.03835 | 3.00135 | 0.00610 | 0.00732 | 0.02482 | 0.00941 | 0.72341 | 0.01385 | 0.00232 | 0.02128 | 0.01154 | 0.00649 | 0.03125 |
| **Zfp607a** | 0.42038 | 0.12220 | 1.05582 | 3.92430 | 0.14289 | 0.03595 | 0.10700 | 0.33991 | 5.96280 | 0.12002 | 2.96284 | 0.14441 | 0.23676 | 2.19652 | 0.18120 | 0.15071 |
| **Zfp607b** | 0.12227 | 0.01716 | 0.21711 | 0.52270 | 0.06272 | 0.04464 | 0.08546 | 0.02979 | 0.62277 | 0.05416 | 1.03337 | 3.53625 | 0.32352 | 2.53592 | 0.13328 | 0.11735 |
| **Zfp609** | 0.91663 | 0.00724 | 0.00248 | 1.07705 | 0.00600 | 0.00644 | 0.05127 | 0.30792 | 1.21719 | 1.07701 | 0.00000 | 0.03252 | 0.00796 | 0.08157 | 0.62056 | 0.23045 |
| **Zfp61** | 2.07714 | 0.00348 | 1.51934 | 0.46608 | 3.11952 | 2.21098 | 0.95260 | 1.57893 | 1.88981 | 1.05576 | 0.31261 | 0.32927 | 0.08060 | 0.00388 | 1.70831 | 2.70862 |

| | | | | | | | | | | | | | | | | |
|---|---|---|---|---|---|---|---|---|---|---|---|---|---|---|---|---|
| **Zfp612** | 0.02349 | 0.01680 | 1.94329 | 0.59576 | 0.01056 | 0.00598 | 1.69890 | 1.65175 | 0.00000 | 0.38863 | 0.00000 | 0.09357 | 0.30394 | 0.00311 | 2.56433 | 0.00666 |
| **Zfp616** | 0.01374 | 0.00801 | 0.01178 | 0.23308 | 0.00930 | 0.00665 | 0.00749 | 0.01260 | 0.06931 | 0.01286 | 0.03827 | 0.00775 | 0.01222 | 0.04570 | 0.00259 | 0.00737 |
| **Zfp617** | 2.90231 | 0.08814 | 1.02006 | 5.08022 | 3.66959 | 2.53363 | 0.83779 | 1.51812 | 4.89030 | 2.98584 | 2.51721 | 4.43895 | 3.63426 | 1.55047 | 5.16671 | 0.55891 |
| **Zfp619** | 0.01167 | 0.00872 | 0.01060 | 0.12080 | 0.00916 | 0.00664 | 0.00782 | 0.03388 | 0.06901 | 0.00225 | 0.69288 | 0.02855 | 0.13781 | 0.04254 | 0.00000 | 0.00699 |
| **Zfp62** | 1.21988 | 0.02688 | 0.62042 | 0.63085 | 0.01162 | 0.00327 | 0.19685 | 0.54751 | 8.12150 | 0.00000 | 1.32305 | 1.03315 | 0.10533 | 0.00000 | 1.17773 | 0.00495 |
| **Zfp622** | 8.12923 | 9.41359 | 8.47782 | 24.52247 | 5.47937 | 0.00369 | 4.85785 | 7.10395 | 37.01403 | 23.28514 | 17.66124 | 6.74140 | 8.94512 | 20.17287 | 10.42593 | 8.08925 |
| **Zfp623** | 0.69717 | 0.00538 | 1.09815 | 8.34144 | 0.18629 | 0.00325 | 2.45204 | 0.00865 | 7.69913 | 3.19806 | 0.94348 | 2.06362 | 2.36089 | 0.00716 | 5.33285 | 1.59409 |
| **Zfp626** | 0.49903 | 0.08970 | 1.30168 | 4.56639 | 0.31624 | 0.09910 | 0.16570 | 0.48747 | 5.52728 | 0.43755 | 7.74604 | 0.37692 | 0.40658 | 4.42752 | 1.21332 | 0.34464 |
| **Zfp628** | 0.92290 | 0.00733 | 0.22724 | 0.85188 | 0.00565 | 0.00730 | 0.00723 | 0.22315 | 1.64116 | 1.59822 | 0.40032 | 0.11619 | 0.01818 | 0.00592 | 0.01314 | 0.05799 |
| **Zfp629** | 0.01040 | 0.00894 | 0.00957 | 0.07986 | 0.00932 | 0.00674 | 0.00821 | 0.14164 | 0.01074 | 0.38228 | 0.04582 | 0.00803 | 0.01047 | 0.00913 | 0.04541 | 0.00710 |
| **Zfp637** | 2.08884 | 2.30639 | 4.20121 | 1.42923 | 0.05654 | 2.35466 | 0.01411 | 4.37047 | 2.06200 | 5.35565 | 0.50691 | 0.44449 | 0.91479 | 0.82756 | 2.20986 | 2.57779 |
| **Zfp638** | 8.02419 | 6.67407 | 2.71314 | 10.13011 | 1.47207 | 3.74176 | 4.31461 | 3.35748 | 5.79753 | 12.45384 | 2.86978 | 15.16537 | 2.61963 | 30.30975 | 15.47719 | 0.00000 |
| **Zfp639** | 3.32569 | 3.27253 | 2.58904 | 9.08907 | 6.28910 | 0.00890 | 0.00170 | 2.29726 | 8.53595 | 7.29666 | 1.82660 | 0.15606 | 0.97154 | 1.98488 | 11.35431 | 0.68953 |
| **Zfp64** | 4.03630 | 0.07746 | 1.70598 | 0.38481 | 3.23899 | 0.01017 | 4.50923 | 1.76305 | 0.00001 | 0.92165 | 0.42921 | 0.17550 | 4.04212 | 0.00559 | 0.00000 | 3.86942 |
| **Zfp644** | 0.13456 | 0.00520 | 1.46555 | 11.30681 | 0.00541 | 0.00500 | 0.00392 | 0.00000 | 13.32982 | 2.49264 | 4.44464 | 4.71164 | 0.00000 | 11.54878 | 10.66343 | 0.00257 |
| **Zfp646** | 2.94509 | 2.18524 | 2.16439 | 0.00000 | 0.00654 | 0.01039 | 1.05238 | 1.25826 | 0.01045 | 3.74275 | 0.00000 | 0.02770 | 2.13982 | 0.00409 | 0.25356 | 2.87354 |
| **Zfp647** | 0.76618 | 0.00729 | 0.00240 | 0.46411 | 0.01972 | 0.00832 | 0.00674 | 1.01826 | 0.45587 | 0.00000 | 0.02224 | 0.03228 | 0.02555 | 0.01147 | 0.09877 | 0.00695 |
| **Zfp65** | 0.10066 | 1.80088 | 0.56969 | 0.41521 | 0.84927 | 0.05117 | 0.20404 | 0.88244 | 1.26669 | 0.73567 | 0.31620 | 1.11666 | 0.14803 | 1.58111 | 0.93394 | 0.02716 |
| **Zfp651** | 2.43402 | 0.00491 | 0.00349 | 0.10529 | 0.00350 | 0.00584 | 0.00526 | 0.00000 | 0.00727 | 1.77065 | 0.01164 | 0.00003 | 0.02268 | 0.04831 | 0.00000 | 0.00455 |
| **Zfp652** | 0.49250 | 0.00675 | 0.00000 | 1.11985 | 0.80556 | 0.02383 | 0.00701 | 0.10896 | 1.36220 | 1.11048 | 0.39205 | 0.01730 | 0.12021 | 0.00722 | 0.01661 | 0.01423 |
| **Zfp653** | 1.69213 | 0.03512 | 1.03894 | 3.07204 | 0.00276 | 0.00560 | 0.00280 | 2.51570 | 0.83096 | 3.72904 | 1.69711 | 1.69419 | 3.82822 | 0.00000 | 0.73789 | 0.01728 |
| **Zfp654** | 0.81306 | 2.67304 | 2.12636 | 8.52449 | 2.62806 | 0.87883 | 1.01262 | 1.81523 | 15.26217 | 2.28426 | 3.33219 | 4.77206 | 0.89369 | 16.77626 | 0.79407 | 0.25407 |
| **Zfp655** | 3.97295 | 7.00444 | 6.53195 | 18.72925 | 5.31347 | 7.50333 | 7.49246 | 11.34418 | 13.68494 | 14.08397 | 14.09983 | 6.19261 | 2.66006 | 15.17785 | 20.39710 | 0.00875 |
| **Zfp658** | 0.00554 | 0.00755 | 0.00756 | 0.01313 | 0.00951 | 0.16233 | 0.00681 | 1.88693 | 0.04770 | 0.21543 | 0.01362 | 0.02895 | 1.48008 | 0.00407 | 0.79624 | 0.00684 |
| **Zfp661** | 0.02888 | 0.00766 | 0.01895 | 0.00898 | 0.16264 | 0.00679 | 0.00759 | 0.01505 | 0.01150 | 0.00354 | 0.01410 | 0.01655 | 0.13576 | 0.00985 | 0.32021 | 0.00753 |
| **Zfp664** | 9.39552 | 0.00095 | 0.00000 | 18.47382 | 0.27727 | 5.15627 | 6.70872 | 3.89830 | 22.52224 | 3.00402 | 1.69268 | 0.90159 | 2.65034 | 0.26027 | 6.80024 | 0.07820 |
| **Zfp667** | 0.01544 | 0.00823 | 0.00248 | 0.17340 | 0.01124 | 2.85674 | 0.09348 | 0.28000 | 0.74768 | 0.00225 | 0.00002 | 0.00514 | 0.32303 | 0.00875 | 0.77085 | 0.00616 |
| **Zfp668** | 1.78081 | 0.09978 | 0.55817 | 4.13724 | 0.02444 | 0.00582 | 1.75928 | 2.06687 | 3.86125 | 2.45816 | 1.19387 | 0.23128 | 3.30061 | 0.00412 | 0.06166 | 0.00260 |
| **Zfp672** | 1.18004 | 0.00703 | 0.21862 | 5.24888 | 0.01543 | 0.00922 | 0.01535 | 0.02758 | 2.96647 | 2.69730 | 1.65524 | 0.15526 | 0.51572 | 0.33637 | 2.24166 | 0.03451 |
| **Zfp677** | 0.00964 | 0.00742 | 0.00528 | 1.78200 | 0.00929 | 0.00611 | 0.00666 | 0.00756 | 0.41594 | 0.00000 | 0.00818 | 0.51205 | 0.52163 | 1.00564 | 1.12663 | 0.00655 |
| **Zfp68** | 0.24670 | 2.48490 | 0.00000 | 6.73350 | 0.00654 | 0.01545 | 2.23406 | 0.00000 | 2.09454 | 0.98413 | 0.78244 | 1.00549 | 0.03250 | 2.27244 | 6.11986 | 1.70709 |
| **Zfp687** | 1.46414 | 2.19698 | 0.00000 | 0.37404 | 0.02051 | 0.98792 | 0.01712 | 0.28210 | 0.13190 | 0.13046 | 0.00735 | 0.02914 | 0.52763 | 0.00481 | 0.02522 | 0.01152 |
| **Zfp688** | 2.56381 | 0.00524 | 0.77970 | 0.17819 | 0.44101 | 0.08321 | 0.01587 | 0.31529 | 0.12786 | 0.87969 | 0.39723 | 0.01231 | 0.24427 | 0.00815 | 0.00000 | 0.00437 |
| **Zfp689** | 0.14151 | 0.00807 | 0.01647 | 0.68858 | 0.00755 | 0.00600 | 0.00736 | 0.02475 | 1.21634 | 0.01893 | 0.03709 | 0.00610 | 0.01700 | 0.01065 | 0.00567 | 0.03507 |
| **Zfp69** | 0.67263 | 0.00828 | 0.00512 | 0.49864 | 0.01770 | 0.00759 | 0.00710 | 0.02479 | 0.13174 | 0.03320 | 0.01184 | 0.00807 | 0.02571 | 0.01074 | 0.00528 | 0.00706 |
| **Zfp691** | 0.34310 | 0.03670 | 0.46401 | 0.00695 | 0.02234 | 0.00862 | 0.01670 | 0.03555 | 0.11146 | 1.09464 | 0.03401 | 0.02986 | 0.19268 | 0.05922 | 0.03096 | 0.00711 |
| **Zfp692** | 1.18693 | 1.48134 | 3.45770 | 0.14716 | 0.04420 | 7.23711 | 2.58901 | 2.28280 | 0.14166 | 1.37080 | 1.47998 | 0.06467 | 4.08869 | 0.88996 | 0.09870 | 0.91555 |
| **Zfp7** | 1.49739 | 2.91334 | 1.91848 | 5.78275 | 1.25006 | 8.17567 | 3.16414 | 5.16673 | 8.55005 | 5.72060 | 3.79365 | 0.04766 | 1.57511 | 0.00000 | 7.28476 | 2.50685 |
| **Zfp703** | 0.00889 | 0.00759 | 0.00000 | 2.78771 | 0.01024 | 3.42866 | 0.00698 | 1.04968 | 3.15958 | 0.54264 | 0.35759 | 0.06709 | 0.00000 | 0.00170 | 0.00000 | 0.00622 |
| **Zfp704** | 9.57368 | 1.87891 | 4.07849 | 1.46632 | 3.52515 | 3.84690 | 5.22182 | 3.29403 | 0.16849 | 3.07643 | 2.10539 | 0.00000 | 2.81647 | 1.79300 | 1.84471 | 3.33506 |
| **Zfp706** | 18.72925 | 9.86586 | 19.79636 | 29.76128 | 17.67294 | 6.93452 | 10.68781 | 25.98648 | 32.24713 | 33.92664 | 32.48673 | 17.15071 | 18.17919 | 23.23408 | 15.74545 | 7.77305 |
| **Zfp707** | 1.96042 | 0.01574 | 0.32209 | 23.65119 | 0.15820 | 4.66520 | 0.45263 | 2.56003 | 30.47604 | 6.64092 | 5.88090 | 0.14786 | 3.80694 | 0.59448 | 0.11919 | 1.58862 |
| **Zfp708** | 0.32234 | 0.00698 | 0.43264 | 0.87831 | 0.00671 | 0.00639 | 0.00707 | 0.44388 | 2.15785 | 0.61176 | 1.06061 | 1.64986 | 0.00000 | 0.05946 | 1.58796 | 0.00629 |
| **Zfp709** | 0.26414 | 0.07682 | 0.00419 | 1.36860 | 0.00729 | 0.00680 | 0.00719 | 0.00946 | 0.46235 | 0.25007 | 0.00627 | 0.00498 | 0.01747 | 0.04428 | 0.48721 | 0.00653 |
| **Zfp710** | 4.74562 | 0.00000 | 0.00000 | 3.41175 | 4.59738 | 3.76871 | 0.00000 | 4.16418 | 9.77464 | 6.69393 | 4.24256 | 0.72696 | 1.91194 | 0.21465 | 1.69227 | 0.83329 |
| **Zfp712** | 0.01356 | 0.00753 | 0.01104 | 0.01135 | 0.00944 | 0.00640 | 0.01740 | 0.01069 | 0.01098 | 0.19398 | 0.01002 | 0.01419 | 0.01253 | 0.01096 | 0.86519 | 0.00748 |
| **Zfp715** | 0.05960 | 0.00603 | 0.33121 | 1.28095 | 0.01519 | 0.07579 | 2.55568 | 1.07610 | 2.80731 | 0.06840 | 0.13908 | 2.55417 | 1.32831 | 0.82801 | 1.16227 | 0.54440 |
| **Zfp719** | 0.11229 | 1.64372 | 0.02498 | 1.37747 | 0.01400 | 0.00465 | 0.00608 | 0.01697 | 0.20670 | 0.00000 | 0.06130 | 6.16864 | 0.20063 | 0.00379 | 2.88755 | 0.00578 |
| **Zfp72** | 0.00264 | 1.00601 | 0.00000 | 0.00000 | 0.00798 | 2.21790 | 0.00635 | 0.19816 | 0.01907 | 0.13410 | 0.38237 | 1.65806 | 0.00000 | 1.40534 | 0.18152 | 0.00579 |
| **Zfp729a** | 0.04640 | 3.28946 | 0.37798 | 0.19363 | 0.01469 | 0.00589 | 1.84537 | 1.49037 | 2.59447 | 0.00414 | 2.32323 | 0.74964 | 0.04306 | 5.71300 | 3.65758 | 0.24758 |
| **Zfp729b** | 0.61480 | 0.02570 | 2.98384 | 1.63376 | 0.06921 | 0.00453 | 1.06041 | 3.06082 | 2.06861 | 3.29102 | 4.72047 | 7.46423 | 0.00000 | 3.45503 | 4.46688 | 0.00394 |
| **Zfp738** | 0.22817 | 0.21591 | 0.25198 | 0.13012 | 0.05397 | 0.00688 | 1.83074 | 0.01389 | 0.81537 | 0.00000 | 0.89807 | 0.06788 | 0.92308 | 2.55368 | 1.67824 | 0.01337 |
| **Zfp74** | 0.00498 | 2.08064 | 0.41578 | 1.07248 | 0.82567 | 0.00567 | 1.32722 | 0.03818 | 1.35074 | 1.70910 | 0.00000 | 0.00000 | 0.01340 | 1.42400 | 0.94744 | 0.01357 |
| **Zfp740** | 5.51585 | 3.63484 | 5.40651 | 2.71120 | 0.90548 | 3.36580 | 7.80067 | 6.27478 | 1.08649 | 10.96044 | 2.30754 | 0.65262 | 1.77176 | 5.46995 | 9.50226 | 3.99541 |
| **Zfp746** | 0.93496 | 0.00513 | 3.25083 | 1.37410 | 0.00841 | 1.44153 | 1.76790 | 3.89824 | 2.98840 | 2.87123 | 1.00396 | 0.03464 | 0.60814 | 0.76775 | 0.30765 | 1.88208 |
| **Zfp747** | 0.68555 | 1.66520 | 0.42985 | 0.01242 | 0.00553 | 0.93634 | 0.00583 | 1.00900 | 0.03838 | 0.00753 | 0.00460 | 0.00442 | 0.73895 | 0.01228 | 0.25764 | 1.06120 |
| **Zfp748** | 0.63714 | 6.27107 | 0.68188 | 0.51199 | 0.00605 | 0.00492 | 0.20757 | 1.09963 | 1.20455 | 0.94893 | 1.12600 | 0.29717 | 0.19343 | 0.01222 | 2.40078 | 0.00363 |
| **Zfp750** | 0.01346 | 0.00821 | 0.01187 | 1.47123 | 0.00928 | 0.00646 | 0.00745 | 0.01795 | 0.64181 | 0.02443 | 0.02023 | 0.00691 | 0.01189 | 0.01023 | 0.00466 | 0.00701 |
| **Zfp758** | 0.09155 | 0.05751 | 0.00000 | 2.15531 | 0.07664 | 0.09571 | 0.08559 | 0.19386 | 2.39593 | 1.11244 | 0.00000 | 0.03724 | 0.01260 | 3.54219 | 1.56158 | 0.03747 |
| **Zfp759** | 0.08557 | 0.00883 | 0.00000 | 0.00223 | 0.03183 | 0.17822 | 0.00773 | 0.45405 | 0.04883 | 0.07526 | 0.26388 | 0.36285 | 0.21263 | 1.00142 | 0.91680 | 0.00711 |
| **Zfp760** | 0.05691 | 0.00785 | 0.03016 | 1.01969 | 0.01273 | 0.00583 | 0.00714 | 0.01019 | 0.03133 | 0.20266 | 0.01050 | 0.00532 | 0.01137 | 0.00463 | 0.65805 | 0.16293 |
| **Zfp763** | 0.13452 | 0.00697 | 0.29508 | 0.00428 | 0.30340 | 0.00545 | 0.55549 | 0.55760 | 0.00904 | 0.15220 | 0.26096 | 0.15270 | 0.00292 | 0.00898 | 1.49323 | 0.00671 |

| | | | | | | | | | | | | | | | | |
|---|---|---|---|---|---|---|---|---|---|---|---|---|---|---|---|---|
| **Zfp764** | 0.70464 | 0.62847 | 0.05955 | 0.00847 | 0.00677 | 0.14793 | 0.00644 | 0.07199 | 0.01188 | 0.00168 | 0.01234 | 0.00656 | 0.34373 | 0.00968 | 0.24963 | 0.11217 |
| **Zfp768** | 0.01557 | 0.29304 | 0.00126 | 0.13321 | 0.00989 | 0.35624 | 0.00694 | 1.11130 | 0.14854 | 0.00331 | 0.00549 | 0.00512 | 0.00000 | 0.01066 | 0.00000 | 0.01424 |
| **Zfp770** | 0.20140 | 0.00758 | 0.11312 | 0.46905 | 0.01442 | 0.00536 | 0.46771 | 0.78100 | 1.31138 | 0.00000 | 0.25137 | 0.09505 | 0.00000 | 1.70541 | 2.79028 | 0.95785 |
| **Zfp771** | 0.08504 | 0.00869 | 0.00858 | 0.09737 | 0.01587 | 0.00695 | 0.00748 | 0.01259 | 0.08044 | 0.06484 | 0.01238 | 0.02968 | 0.02884 | 0.01109 | 0.00951 | 0.00738 |
| **Zfp772** | 0.00156 | 5.52163 | 0.95342 | 0.00000 | 0.01166 | 0.01113 | 0.11677 | 1.23290 | 1.55429 | 1.20219 | 0.11678 | 1.92929 | 0.90241 | 3.47079 | 1.23176 | 0.02756 |
| **Zfp773** | 0.02098 | 0.00792 | 0.07343 | 0.34200 | 0.34589 | 0.00575 | 0.00740 | 0.00969 | 0.23652 | 0.05403 | 0.06177 | 0.28380 | 0.01151 | 0.02285 | 0.05460 | 0.00730 |
| **Zfp775** | 0.00005 | 0.00651 | 0.41797 | 17.94924 | 0.01589 | 0.00261 | 0.00281 | 0.00000 | 18.99951 | 14.63409 | 6.86301 | 2.15176 | 1.65475 | 4.33903 | 3.89828 | 0.00000 |
| **Zfp777** | 3.00808 | 0.01382 | 0.07342 | 3.08105 | 3.15636 | 0.21209 | 0.00204 | 1.82526 | 3.82920 | 7.65941 | 1.14642 | 0.05823 | 4.22571 | 0.00000 | 0.27727 | 3.00620 |
| **Zfp78** | 1.02823 | 0.00589 | 2.01527 | 0.48812 | 1.48854 | 0.00499 | 0.01704 | 0.00337 | 0.00000 | 0.19575 | 0.75310 | 0.07927 | 0.20448 | 0.02011 | 1.71219 | 0.00544 |
| **Zfp780b** | 0.16750 | 0.04836 | 0.77600 | 1.67928 | 0.00739 | 0.00663 | 0.67804 | 0.05896 | 1.72623 | 0.03975 | 0.98102 | 0.06679 | 0.24836 | 1.17134 | 2.53251 | 0.06368 |
| **Zfp781** | 0.80844 | 0.12334 | 0.46185 | 3.79590 | 0.22605 | 0.35781 | 0.30579 | 1.13840 | 4.93393 | 2.23692 | 4.03737 | 0.39586 | 1.85141 | 5.54448 | 4.77836 | 0.00354 |
| **Zfp783** | 0.01167 | 0.00653 | 0.03301 | 0.66556 | 0.00872 | 0.00788 | 0.00662 | 0.00156 | 0.51228 | 0.00000 | 0.23808 | 2.89990 | 0.00949 | 0.00647 | 0.00000 | 0.00689 |
| **Zfp784** | 0.01265 | 0.00774 | 0.00794 | 0.67960 | 0.00925 | 0.05857 | 0.06723 | 0.01069 | 0.05808 | 0.38425 | 0.01046 | 0.01559 | 0.22027 | 0.00444 | 0.35439 | 0.00715 |
| **Zfp786** | 0.01828 | 0.00770 | 0.01091 | 0.00938 | 0.01072 | 0.00656 | 0.00735 | 0.02125 | 0.01116 | 0.00575 | 0.01135 | 0.00683 | 0.01215 | 0.01959 | 0.39506 | 0.00743 |
| **Zfp787** | 1.56778 | 0.00615 | 0.00000 | 0.41999 | 0.00865 | 1.06231 | 0.01633 | 0.00269 | 1.86376 | 1.36932 | 0.04931 | 0.00000 | 0.01273 | 0.00747 | 0.15792 | 0.01278 |
| **Zfp788** | 0.00000 | 0.01586 | 1.30838 | 1.53789 | 0.00694 | 0.00493 | 1.59895 | 1.57126 | 0.96942 | 0.48901 | 1.81309 | 5.48085 | 1.92643 | 4.28663 | 1.20726 | 0.43147 |
| **Zfp790** | 0.01542 | 0.03703 | 3.11717 | 0.67401 | 3.30459 | 0.00729 | 0.02634 | 0.00400 | 3.16419 | 0.41074 | 0.82135 | 0.19759 | 0.05507 | 1.98418 | 2.76789 | 0.00560 |
| **Zfp791** | 0.01403 | 0.00738 | 1.94920 | 7.61736 | 0.01840 | 0.22883 | 0.01685 | 2.36300 | 8.40092 | 0.00000 | 2.10288 | 0.00000 | 0.00000 | 1.27250 | 2.81711 | 0.00433 |
| **Zfp799** | 0.50404 | 0.14180 | 1.08384 | 0.42606 | 0.06062 | 0.11228 | 0.18761 | 0.23788 | 0.93399 | 0.81344 | 1.95461 | 0.16399 | 0.37909 | 0.85710 | 0.39913 | 0.12574 |
| **Zfp800** | 0.03643 | 0.38642 | 0.14856 | 0.66763 | 0.00774 | 0.06128 | 0.00698 | 1.25461 | 0.76135 | 0.23723 | 0.48138 | 0.99702 | 0.00524 | 0.00421 | 3.68079 | 0.00677 |
| **Zfp808** | 1.89642 | 5.73284 | 2.79513 | 4.45849 | 1.15789 | 0.75209 | 3.56925 | 1.74096 | 3.53234 | 2.35055 | 2.22124 | 7.25391 | 1.61303 | 14.32675 | 0.71389 | 0.57314 |
| **Zfp809** | 0.33690 | 0.61752 | 7.38502 | 8.40969 | 0.07992 | 0.87667 | 1.72897 | 3.88695 | 5.70694 | 1.34033 | 0.43339 | 0.13437 | 1.90792 | 3.81923 | 2.58456 | 0.20586 |
| **Zfp81** | 0.86497 | 0.26963 | 0.37648 | 0.93960 | 0.01626 | 0.00510 | 0.00739 | 0.29399 | 0.41349 | 2.24612 | 0.00668 | 0.02488 | 0.04611 | 0.04823 | 1.75904 | 0.00631 |
| **Zfp810** | 1.26715 | 2.95718 | 0.57790 | 2.80112 | 0.00627 | 0.02819 | 0.00523 | 0.68394 | 4.18274 | 0.00000 | 0.70300 | 0.00674 | 0.03699 | 1.23120 | 1.39200 | 0.01858 |
| **Zfp811** | 0.01301 | 0.00805 | 0.03539 | 0.31318 | 0.00921 | 0.00750 | 0.02810 | 0.01316 | 0.00665 | 0.01842 | 0.01370 | 0.00804 | 0.01218 | 0.01156 | 0.01432 | 0.00741 |
| **Zfp819** | 2.34712 | 0.26021 | 0.31293 | 1.21311 | 0.00292 | 0.00968 | 0.00523 | 0.01258 | 1.65597 | 0.92601 | 0.06262 | 0.11420 | 0.00726 | 0.00553 | 0.00000 | 0.67921 |
| **Zfp82** | 1.74014 | 0.68911 | 0.00000 | 1.89751 | 0.00773 | 1.45384 | 0.00574 | 0.01378 | 3.03293 | 1.46222 | 0.00000 | 0.14261 | 0.12931 | 0.04855 | 3.92260 | 0.00471 |
| **Zfp820** | 0.58069 | 2.17271 | 0.00000 | 4.33915 | 0.03233 | 0.00000 | 0.24145 | 5.24370 | 5.31302 | 1.20818 | 6.84397 | 20.06443 | 0.00000 | 21.53110 | 2.64549 | 0.01677 |
| **Zfp821** | 0.36556 | 0.47478 | 0.89278 | 6.24972 | 0.01275 | 0.06471 | 0.00174 | 12.17323 | 10.94362 | 8.03499 | 4.51301 | 1.61125 | 5.28632 | 0.00000 | 0.72961 | 0.01620 |
| **Zfp825** | 2.38849 | 6.79279 | 6.92763 | 0.57827 | 4.37216 | 2.37487 | 1.98891 | 4.74277 | 0.79126 | 4.50152 | 7.77327 | 0.71859 | 0.78429 | 9.97202 | 6.69631 | 0.48741 |
| **Zfp827** | 0.01354 | 0.00817 | 0.01164 | 0.01642 | 0.00913 | 0.00689 | 0.00752 | 0.01256 | 0.01255 | 0.35801 | 0.01208 | 0.01976 | 0.01255 | 0.03262 | 0.00914 | 0.00741 |
| **Zfp830** | 0.40535 | 4.18757 | 1.57274 | 6.71187 | 2.28187 | 0.06940 | 2.55720 | 5.67733 | 8.34713 | 2.02829 | 1.15155 | 2.10279 | 3.53896 | 1.33278 | 6.97719 | 0.55318 |
| **Zfp839** | 1.34982 | 0.10648 | 1.25261 | 2.18881 | 0.00532 | 0.34381 | 0.10361 | 1.06566 | 1.80506 | 0.12843 | 0.78950 | 0.08230 | 0.00000 | 0.08187 | 2.60063 | 1.06032 |
| **Zfp84** | 0.26360 | 0.90700 | 2.28449 | 1.94259 | 1.98605 | 0.00433 | 0.08279 | 2.06513 | 2.67834 | 1.73128 | 1.22890 | 4.73546 | 0.00000 | 2.26729 | 1.71800 | 0.32471 |
| **Zfp846** | 0.78833 | 5.48370 | 0.14577 | 0.30802 | 0.01354 | 2.25989 | 0.00440 | 1.09620 | 0.75389 | 0.87418 | 1.95997 | 0.08234 | 0.00000 | 0.02998 | 2.87052 | 0.00318 |
| **Zfp85** | 0.14438 | 0.29094 | 0.47491 | 0.00243 | 0.02946 | 0.00576 | 0.05753 | 0.51004 | 0.46254 | 0.52161 | 0.32784 | 0.00000 | 0.00252 | 2.79555 | 0.00000 | 0.00726 |
| **Zfp850** | 0.61804 | 0.04235 | 2.48140 | 0.64922 | 0.62936 | 0.06209 | 0.00619 | 0.23725 | 1.31640 | 0.45890 | 2.21464 | 0.07824 | 0.95250 | 0.00263 | 5.06220 | 1.36944 |
| **Zfp862.ps** | 0.05064 | 0.04159 | 0.00000 | 0.01811 | 0.01449 | 0.03277 | 0.61017 | 0.31933 | 0.01950 | 0.02066 | 0.00864 | 0.06214 | 0.00592 | 0.03102 | 0.33809 | 0.03626 |
| **Zfp865** | 0.84855 | 0.00521 | 0.55954 | 2.65693 | 2.66717 | 0.00602 | 0.00643 | 0.12734 | 0.00000 | 3.77763 | 0.19398 | 0.11838 | 0.05981 | 0.00204 | 0.13127 | 0.00594 |
| **Zfp866** | 0.05673 | 0.77983 | 0.42911 | 1.04244 | 0.01660 | 1.40404 | 0.83708 | 1.13626 | 1.80549 | 1.44789 | 0.79454 | 0.13261 | 0.02336 | 0.56117 | 0.97898 | 0.71654 |
| **Zfp867** | 0.01750 | 0.00778 | 0.00668 | 1.62335 | 0.00891 | 0.00682 | 0.00714 | 0.00000 | 0.27512 | 0.59753 | 0.66302 | 0.00000 | 0.01190 | 0.00261 | 1.02770 | 0.00724 |
| **Zfp868** | 1.97025 | 0.31645 | 0.34807 | 6.33485 | 0.01664 | 0.00211 | 2.66332 | 0.33421 | 6.60678 | 1.65594 | 1.42182 | 2.14280 | 4.80234 | 2.57925 | 4.76380 | 1.20108 |
| **Zfp869** | 2.42689 | 9.11086 | 1.51256 | 3.47722 | 0.00899 | 0.00332 | 5.34881 | 5.19161 | 3.43957 | 5.14016 | 2.51095 | 6.64962 | 3.32284 | 2.15028 | 3.94732 | 0.25368 |
| **Zfp87** | 0.00000 | 7.05570 | 2.45008 | 0.00000 | 0.00380 | 0.00690 | 2.12321 | 2.55540 | 0.90323 | 0.00000 | 0.63487 | 0.02197 | 2.52096 | 0.02194 | 2.24043 | 0.92337 |
| **Zfp870** | 0.79170 | 0.01643 | 1.65442 | 2.33256 | 0.09263 | 0.04468 | 0.00445 | 2.02654 | 3.57143 | 0.37075 | 4.31984 | 0.27499 | 0.05671 | 1.45803 | 0.59549 | 2.14971 |
| **Zfp871** | 0.51282 | 2.57956 | 0.12205 | 0.00000 | 2.05188 | 4.70192 | 1.89851 | 0.23638 | 2.63054 | 3.10738 | 2.00802 | 0.01219 | 3.38270 | 0.09592 | 3.47682 | 0.01721 |
| **Zfp872** | 0.01821 | 0.00793 | 0.03476 | 0.01572 | 0.01069 | 0.00615 | 0.00746 | 0.13845 | 0.38654 | 0.03199 | 0.00856 | 0.00665 | 0.01154 | 0.08971 | 0.01104 | 0.00764 |
| **Zfp873** | 0.14264 | 0.00709 | 0.90901 | 0.81059 | 1.31235 | 0.00918 | 0.00709 | 0.27352 | 1.09415 | 0.82606 | 0.10395 | 0.15448 | 0.05159 | 5.98392 | 3.41917 | 0.01405 |
| **Zfp874a** | 2.48908 | 1.53296 | 1.06959 | 0.52480 | 6.16059 | 0.73868 | 0.98828 | 0.53383 | 0.79499 | 1.37184 | 0.76048 | 5.33805 | 0.26028 | 11.32603 | 6.17001 | 0.00264 |
| **Zfp874b** | 1.38187 | 1.46938 | 0.86093 | 2.01482 | 0.31710 | 0.21309 | 1.18799 | 0.56920 | 3.07150 | 0.75721 | 1.84171 | 0.39612 | 0.19061 | 4.48636 | 1.86738 | 0.06317 |
| **Zfp882** | 0.38106 | 2.73417 | 0.37235 | 2.20729 | 0.07785 | 0.12882 | 1.10751 | 0.02820 | 0.99931 | 0.10234 | 1.18764 | 0.11812 | 0.30480 | 0.82393 | 1.30525 | 0.32357 |
| **Zfp9** | 0.15032 | 0.09965 | 0.11328 | 2.21330 | 0.11655 | 0.07950 | 0.04932 | 0.16715 | 0.83413 | 1.20255 | 0.63373 | 0.02613 | 0.11865 | 1.84914 | 0.12013 | 2.39561 |
| **Zfp90** | 0.01385 | 0.00768 | 0.01127 | 0.01071 | 0.00936 | 0.00665 | 0.00744 | 0.01233 | 0.01174 | 0.00879 | 0.01192 | 0.03735 | 0.01284 | 0.01919 | 0.13610 | 0.00741 |
| **Zfp91** | 5.44083 | 4.90555 | 5.28640 | 9.66187 | 0.34431 | 3.71516 | 4.84076 | 5.27455 | 6.57281 | 6.84956 | 4.49053 | 4.04421 | 5.19098 | 10.53706 | 6.55702 | 1.79032 |
| **Zfp93** | 1.00838 | 0.00636 | 0.38059 | 0.54272 | 0.00560 | 0.00452 | 0.00635 | 0.00514 | 1.01601 | 0.62494 | 0.01659 | 0.00261 | 0.01062 | 0.00434 | 0.23867 | 0.01398 |
| **Zfp930** | 3.16651 | 5.86625 | 1.98783 | 1.37806 | 0.98558 | 0.08227 | 0.04314 | 3.09638 | 1.55376 | 1.07316 | 1.71218 | 3.36712 | 2.55123 | 0.73775 | 4.30794 | 2.24389 |
| **Zfp931** | 0.15234 | 0.23381 | 0.42101 | 0.46071 | 0.01833 | 0.46061 | 0.10179 | 0.06363 | 0.06153 | 0.11916 | 0.29098 | 0.02645 | 0.16109 | 0.15652 | 0.09348 | 0.21346 |
| **Zfp932** | 2.11986 | 6.86111 | 3.83164 | 3.86188 | 1.35636 | 3.90111 | 3.44128 | 12.24865 | 1.51656 | 20.83669 | 10.33419 | 19.88527 | 3.57156 | 31.06814 | 14.57878 | 0.36504 |
| **Zfp933** | 0.20726 | 0.06974 | 1.18836 | 0.18744 | 0.43446 | 0.05288 | 0.16955 | 0.01687 | 1.00610 | 1.11878 | 1.89876 | 0.04647 | 0.28785 | 2.32369 | 0.36203 | 0.22612 |
| **Zfp934** | 1.44541 | 2.98206 | 2.40127 | 3.00462 | 0.48484 | 0.55662 | 2.00888 | 0.95116 | 1.05740 | 1.73010 | 1.33591 | 7.26694 | 1.41324 | 7.87529 | 0.81793 | 0.31120 |
| **Zfp935** | 0.57448 | 0.67549 | 1.58386 | 6.01909 | 0.40860 | 3.44232 | 0.36665 | 2.23197 | 7.32621 | 0.81231 | 5.85176 | 3.33123 | 1.02487 | 7.48008 | 2.56706 | 0.02373 |
| **Zfp936** | 3.09119 | 3.10119 | 3.18984 | 14.14190 | 2.18243 | 0.00405 | 2.02086 | 3.18986 | 22.48098 | 2.24726 | 7.05600 | 11.35570 | 1.38646 | 17.73509 | 10.03046 | 0.42316 |

| | | | | | | | | | | | | | | | | |
|---|---|---|---|---|---|---|---|---|---|---|---|---|---|---|---|---|
| **Zfp937** | 0.14667 | 0.08166 | 1.24778 | 3.14227 | 0.59909 | 0.08269 | 0.22306 | 0.22938 | 1.29115 | 1.92205 | 0.29273 | 0.05951 | 0.32498 | 0.83382 | 0.44118 | 0.12876 |
| **Zfp938** | 0.00987 | 0.06220 | 0.03014 | 0.00968 | 0.01824 | 1.71718 | 0.03825 | 0.03290 | 0.79422 | 0.05465 | 0.00000 | 0.14944 | 0.01017 | 0.03058 | 0.63115 | 0.02110 |
| **Zfp939** | 0.00000 | 0.00681 | 0.84558 | 0.88875 | 0.00984 | 2.48252 | 1.43213 | 2.09545 | 1.89780 | 0.00000 | 2.93406 | 0.09076 | 0.82011 | 4.44144 | 4.48444 | 0.00521 |
| **Zfp94** | 0.01150 | 0.00717 | 0.01252 | 0.26951 | 0.01075 | 0.00736 | 1.45336 | 0.00000 | 0.00543 | 0.34069 | 0.46353 | 0.02277 | 0.61156 | 0.00685 | 0.00000 | 0.01372 |
| **Zfp940** | 0.23532 | 0.00773 | 0.00407 | 0.00906 | 0.00787 | 0.00689 | 0.00696 | 0.01148 | 0.01193 | 0.13219 | 0.01146 | 0.08720 | 0.12260 | 0.00707 | 0.13656 | 1.18365 |
| **Zfp941** | 0.30808 | 0.00720 | 0.10250 | 0.16660 | 0.00779 | 0.00591 | 0.01742 | 0.03896 | 0.93419 | 0.20670 | 0.48434 | 0.03359 | 0.13372 | 0.32510 | 0.09645 | 0.02077 |
| **Zfp942** | 0.64056 | 1.59932 | 1.01247 | 6.00874 | 0.10975 | 2.54028 | 2.05760 | 2.43434 | 2.68788 | 0.00035 | 1.70244 | 1.81470 | 0.00000 | 3.13996 | 3.02503 | 0.98484 |
| **Zfp943** | 0.60806 | 0.00600 | 0.05623 | 1.30276 | 0.02770 | 0.00787 | 0.48087 | 0.45008 | 0.87438 | 0.23495 | 0.00000 | 2.80346 | 0.02585 | 0.14631 | 0.00000 | 0.01396 |
| **Zfp944** | 0.87184 | 0.61374 | 0.00000 | 7.11358 | 0.02914 | 0.51328 | 0.03535 | 0.61674 | 1.45047 | 1.51646 | 0.76889 | 11.20918 | 0.06352 | 1.30646 | 2.41789 | 1.88249 |
| **Zfp945** | 0.01519 | 0.04302 | 0.00907 | 0.01335 | 0.01054 | 0.00715 | 0.05037 | 0.05347 | 0.01160 | 0.34675 | 0.02238 | 0.03237 | 0.02499 | 0.14936 | 0.28062 | 0.01494 |
| **Zfp946** | 0.34380 | 0.03952 | 0.00000 | 1.22724 | 0.00732 | 0.06180 | 0.01771 | 0.03455 | 0.45459 | 0.07903 | 0.03230 | 0.01555 | 0.01169 | 0.25307 | 0.01823 | 0.01397 |
| **Zfp947** | 0.11371 | 0.01634 | 0.12173 | 1.42420 | 0.01877 | 0.52230 | 0.04928 | 2.80574 | 2.57090 | 0.00000 | 1.14381 | 5.22315 | 0.76960 | 6.88145 | 1.32233 | 0.03486 |
| **Zfp948** | 0.23000 | 0.00620 | 0.00000 | 8.26143 | 0.01734 | 0.86876 | 0.00588 | 0.00757 | 5.69079 | 0.16460 | 0.38466 | 6.81719 | 0.07078 | 0.96296 | 4.82928 | 0.53126 |
| **Zfp949** | 0.00000 | 0.06240 | 1.61415 | 2.92010 | 0.02810 | 2.89699 | 0.10860 | 0.56210 | 1.64826 | 2.91312 | 2.13699 | 0.73432 | 0.57372 | 8.02793 | 4.47386 | 0.01927 |
| **Zfp950** | 0.88984 | 3.06414 | 1.78391 | 6.04131 | 0.14197 | 0.36032 | 0.30005 | 2.80582 | 4.71114 | 3.32099 | 2.08347 | 0.36160 | 0.14606 | 3.33429 | 4.00939 | 0.20574 |
| **Zfp951** | 1.05691 | 9.05500 | 6.13364 | 4.41679 | 0.08674 | 6.25329 | 2.72630 | 5.97556 | 9.15947 | 2.84526 | 4.02275 | 12.29885 | 1.84422 | 4.61063 | 6.68847 | 1.36766 |
| **Zfp952** | 0.00000 | 1.30064 | 0.31114 | 0.91660 | 0.01779 | 0.00608 | 0.00703 | 0.00341 | 0.02956 | 1.34168 | 0.45439 | 0.31007 | 0.20360 | 2.46853 | 2.09030 | 0.00669 |
| **Zfp953** | 0.12347 | 0.00754 | 0.19516 | 0.00227 | 0.00983 | 1.00298 | 0.00668 | 1.07430 | 0.00950 | 0.19123 | 0.45428 | 0.01162 | 0.39285 | 0.02090 | 0.51450 | 0.19581 |
| **Zfp954** | 1.89954 | 5.60941 | 5.43204 | 3.33246 | 1.76111 | 1.05043 | 0.00000 | 4.59802 | 11.12163 | 6.64753 | 4.23765 | 0.28657 | 3.56437 | 2.22550 | 4.76096 | 0.63851 |
| **Zfp955a** | 2.14101 | 0.52188 | 0.00000 | 1.26957 | 1.95658 | 0.27909 | 0.00544 | 0.55306 | 0.08457 | 0.72710 | 0.86204 | 2.22719 | 0.01241 | 5.85660 | 3.71420 | 0.04825 |
| **Zfp955b** | 0.95601 | 0.04775 | 0.91460 | 0.90372 | 0.02166 | 2.13749 | 0.95773 | 1.16465 | 1.45228 | 1.11678 | 1.11251 | 0.06112 | 0.00000 | 6.46162 | 3.59605 | 0.04076 |
| **Zfp956** | 0.87576 | 0.00383 | 1.50657 | 3.02964 | 0.00335 | 0.01547 | 4.56768 | 0.76167 | 3.54128 | 0.05940 | 0.02461 | 0.04978 | 3.58012 | 0.00000 | 0.52317 | 0.81983 |
| **Zfp957** | 0.01689 | 0.00779 | 0.00781 | 0.31227 | 0.00919 | 0.00665 | 0.00722 | 0.01943 | 0.31758 | 0.00000 | 0.00555 | 0.00405 | 0.01283 | 0.65701 | 0.30572 | 0.00738 |
| **Zfp958** | 2.84672 | 2.69974 | 2.34866 | 6.79037 | 0.32810 | 0.01892 | 0.07380 | 5.48005 | 11.82842 | 2.50983 | 5.36931 | 1.42262 | 1.38622 | 12.03019 | 5.45405 | 2.44870 |
| **Zfp959** | 0.84764 | 2.93602 | 0.03968 | 0.46967 | 0.04288 | 0.02777 | 0.03570 | 0.04285 | 0.07892 | 0.49444 | 0.29514 | 0.37002 | 0.40349 | 0.30204 | 1.11342 | 0.06211 |
| **Zfp960** | 1.59622 | 8.01843 | 4.33948 | 1.37436 | 2.95365 | 0.40701 | 4.50206 | 6.27987 | 6.75584 | 7.37981 | 11.28816 | 17.18723 | 4.55984 | 22.54727 | 4.60421 | 0.36260 |
| **Zfp961** | 4.90230 | 4.70868 | 2.34058 | 9.22237 | 5.29223 | 0.00616 | 3.81564 | 5.13761 | 8.65506 | 4.10848 | 3.45442 | 4.69168 | 4.65071 | 12.46993 | 9.72569 | 4.55497 |
| **Zfp963** | 0.01185 | 0.02756 | 1.72939 | 0.29476 | 0.00605 | 0.00906 | 0.00681 | 0.00075 | 0.00719 | 0.00000 | 0.27700 | 0.38690 | 0.00509 | 0.00728 | 0.00000 | 0.00651 |
| **Zfp97** | 1.19407 | 6.39279 | 3.49237 | 0.96601 | 1.98621 | 0.26615 | 2.85942 | 4.76880 | 4.16862 | 5.19380 | 8.54917 | 15.85942 | 3.63090 | 18.26772 | 2.74844 | 0.21889 |
| **Zfp970** | 4.54974 | 3.09642 | 5.72268 | 22.26424 | 1.89323 | 1.71616 | 5.08677 | 4.18384 | 27.44199 | 13.59798 | 3.85065 | 3.06258 | 2.45821 | 20.46160 | 23.74445 | 2.48819 |
| **Zfp971** | 0.45312 | 1.12837 | 1.56166 | 0.16055 | 0.05862 | 1.51617 | 0.62914 | 0.30374 | 0.22505 | 0.75288 | 0.18159 | 0.31148 | 0.73705 | 3.42850 | 3.57491 | 0.42026 |
| **Zfp972** | 0.15750 | 0.04702 | 0.99950 | 9.76471 | 0.00723 | 0.19487 | 0.02558 | 0.00000 | 8.35638 | 0.97116 | 1.47934 | 4.24001 | 0.05187 | 0.79412 | 1.12288 | 0.01124 |
| **Zfp973** | 0.01089 | 0.08076 | 0.01277 | 0.01744 | 0.01845 | 0.14697 | 0.03827 | 0.01036 | 0.02202 | 0.02277 | 0.03877 | 0.00670 | 0.02149 | 0.27870 | 0.01361 | 0.00741 |
| **Zfp974** | 0.00000 | 0.02964 | 0.00519 | 0.94547 | 0.00958 | 0.00569 | 0.00663 | 0.00578 | 1.25871 | 1.85547 | 0.00000 | 0.04933 | 0.61000 | 6.30515 | 5.37600 | 0.00659 |
| **Zfp975** | 1.46563 | 1.55945 | 0.72545 | 3.61682 | 0.10560 | 0.03597 | 0.85179 | 0.22022 | 2.32789 | 0.88317 | 1.53176 | 0.72364 | 1.21821 | 2.97206 | 1.20082 | 0.21527 |
| **Zfp976** | 0.32205 | 0.65319 | 0.34887 | 0.53158 | 0.04155 | 0.03539 | 0.14461 | 0.37615 | 0.23569 | 0.49695 | 0.43094 | 0.26831 | 0.34517 | 2.02558 | 0.95755 | 0.02821 |
| **Zfp979** | 3.51440 | 0.62531 | 0.91259 | 11.74534 | 1.36331 | 0.01766 | 2.44268 | 1.84809 | 5.73950 | 2.04727 | 5.86015 | 6.55819 | 2.14769 | 15.27811 | 3.42070 | 0.97594 |
| **Zfp980** | 1.42362 | 3.91914 | 1.42194 | 14.64353 | 1.04941 | 0.06287 | 1.23435 | 3.92292 | 8.73346 | 0.65615 | 4.94561 | 14.58697 | 1.59785 | 23.68474 | 1.98496 | 0.45161 |
| **Zfp981** | 1.18283 | 4.68593 | 1.21424 | 10.05376 | 1.03912 | 0.11778 | 1.36154 | 3.57164 | 10.94616 | 0.68390 | 4.62725 | 19.94041 | 1.68929 | 19.90480 | 1.22895 | 0.36651 |
| **Zfp982** | 0.72302 | 0.75179 | 0.22837 | 2.60639 | 0.33099 | 0.57361 | 0.73705 | 0.40441 | 0.00000 | 1.83895 | 0.99800 | 0.47606 | 0.39034 | 12.00191 | 3.22119 | 0.46622 |
| **Zfp983** | 0.16050 | 0.09260 | 0.86102 | 1.15138 | 0.00632 | 2.68948 | 0.00700 | 0.00939 | 0.44733 | 0.30643 | 0.06020 | 0.16075 | 0.02558 | 0.00595 | 1.61147 | 0.30717 |
| **Zfp984** | 3.02425 | 6.89086 | 4.11748 | 6.23468 | 2.83435 | 1.39661 | 4.31213 | 7.48133 | 8.67017 | 4.74453 | 4.80566 | 19.41368 | 4.69901 | 24.84367 | 3.29679 | 2.05230 |
| **Zfp985** | 3.50015 | 8.83049 | 4.80832 | 2.91936 | 3.05919 | 1.70579 | 3.07118 | 3.86195 | 5.17516 | 1.85283 | 6.33733 | 8.14753 | 4.02619 | 16.54634 | 0.28024 | 1.78985 |
| **Zfp987** | 2.21892 | 5.66977 | 1.88491 | 8.29178 | 2.88838 | 0.45956 | 1.25489 | 3.32697 | 9.55365 | 2.61024 | 3.60412 | 13.65637 | 1.12505 | 20.22586 | 1.31616 | 1.84793 |
| **Zfp991** | 1.20187 | 0.23439 | 0.44499 | 0.00310 | 0.37212 | 0.06682 | 0.32948 | 0.10130 | 0.24176 | 0.20544 | 0.67901 | 0.00709 | 0.98148 | 1.35091 | 0.00000 | 0.89252 |
| **Zfp992** | 1.54643 | 0.32151 | 1.09698 | 2.06225 | 0.11210 | 0.28129 | 2.01559 | 0.29188 | 2.60190 | 0.24925 | 0.54094 | 1.31383 | 1.25304 | 0.87033 | 0.31498 | 0.15226 |
| **Zfp994** | 1.38088 | 0.98645 | 1.73005 | 2.95923 | 0.21854 | 0.28343 | 0.97599 | 0.45163 | 3.24592 | 0.76684 | 1.04731 | 1.10378 | 0.49818 | 3.33129 | 0.59940 | 0.72159 |
| **Zfp995** | 0.22756 | 0.10466 | 0.52436 | 0.12438 | 0.04432 | 0.03377 | 0.05252 | 0.40048 | 0.00714 | 0.04908 | 0.02871 | 0.01493 | 0.04298 | 1.05561 | 0.11328 | 0.02887 |
| **Zfpl1** | 1.50785 | 0.01716 | 0.16869 | 1.15745 | 0.13046 | 0.02465 | 0.03245 | 0.15922 | 1.35588 | 7.38115 | 0.90237 | 0.04737 | 7.67891 | 0.00545 | 0.00000 | 5.30160 |
| **Zfpm1** | 0.14329 | 0.90022 | 0.00902 | 0.01191 | 0.01257 | 0.00784 | 0.00678 | 0.03206 | 0.04321 | 0.31219 | 0.01115 | 0.01607 | 0.20670 | 0.01026 | 0.00001 | 0.07807 |
| **Zfr** | 6.94438 | 9.21858 | 5.96887 | 9.55912 | 9.89659 | 6.53968 | 5.56254 | 6.99600 | 8.80239 | 5.08501 | 2.97888 | 11.09397 | 2.23089 | 9.16288 | 4.26293 | 0.01697 |
| **Zfr2** | 0.01369 | 0.00803 | 0.01200 | 0.13932 | 0.00931 | 0.00663 | 0.00753 | 0.01372 | 0.45966 | 0.01214 | 0.00929 | 0.01752 | 0.01284 | 0.01056 | 0.01510 | 0.00737 |
| **Zfx** | 2.03970 | 0.70336 | 1.89128 | 11.88009 | 2.05614 | 3.71761 | 1.12231 | 2.44874 | 4.62996 | 7.49151 | 1.53214 | 7.93294 | 0.86666 | 10.14918 | 7.11722 | 0.18177 |
| **Zfyve1** | 1.10912 | 0.82046 | 2.09595 | 1.91562 | 1.43948 | 3.61788 | 0.03338 | 5.53349 | 1.30241 | 1.03170 | 3.97512 | 0.08558 | 5.33241 | 0.00000 | 0.00000 | 0.00806 |
| **Zfyve16** | 0.20696 | 0.85135 | 1.47213 | 0.01277 | 0.03771 | 0.00576 | 0.76315 | 0.14298 | 0.89507 | 1.49572 | 1.29456 | 0.00000 | 0.19603 | 2.23604 | 1.04911 | 0.00604 |
| **Zfyve19** | 0.00572 | 0.26888 | 4.79444 | 0.39027 | 10.97100 | 3.14838 | 0.00685 | 3.48177 | 0.23919 | 5.37415 | 0.07612 | 0.32036 | 0.11900 | 0.30923 | 1.66387 | 2.71936 |
| **Zfyve21** | 1.54226 | 0.31368 | 0.83568 | 3.39491 | 3.82178 | 0.06920 | 0.47094 | 1.83431 | 4.91626 | 0.26000 | 1.57769 | 0.06869 | 5.84808 | 0.00000 | 0.00000 | 0.97444 |
| **Zfyve26** | 6.71247 | 9.14647 | 6.99577 | 7.65721 | 0.00000 | 0.20670 | 5.71130 | 4.26149 | 5.76928 | 5.63424 | 6.19928 | 3.52149 | 3.56443 | 2.81671 | 5.94189 | 0.83043 |
| **Zfyve27** | 0.60660 | 0.26628 | 6.09177 | 2.18476 | 0.01436 | 0.03581 | 0.26993 | 1.01906 | 3.39236 | 4.05752 | 1.21959 | 0.21358 | 1.20971 | 2.56106 | 1.16798 | 0.06123 |
| **Zfyve9** | 0.01465 | 0.00757 | 0.02808 | 2.54988 | 0.00875 | 0.00800 | 0.00671 | 0.00000 | 0.84570 | 0.56699 | 0.86151 | 0.21691 | 0.00573 | 1.59356 | 1.61317 | 0.00643 |
| **Zg16** | 0.02090 | 0.00803 | 0.01123 | 0.00918 | 0.00865 | 0.00677 | 0.00744 | 0.02388 | 0.01212 | 0.34099 | 0.01161 | 0.00772 | 0.01869 | 0.01151 | 0.01057 | 0.00724 |
| **Zglp1** | 0.02466 | 0.00771 | 0.37394 | 0.00617 | 0.00783 | 0.00888 | 0.00709 | 0.00542 | 0.01170 | 0.30653 | 0.20966 | 0.00526 | 0.01499 | 0.03027 | 0.00020 | 0.00719 |

| | | | | | | | | | | | | | | | | |
|---|---|---|---|---|---|---|---|---|---|---|---|---|---|---|---|---|
| **Zgpat** | 1.57464 | 0.00522 | 0.18393 | 4.48899 | 1.18453 | 0.00482 | 0.65738 | 3.66147 | 6.59437 | 2.63567 | 1.14913 | 0.20431 | 1.39782 | 0.00000 | 1.62338 | 3.64922 |
| **Zgrf1** | 0.22894 | 3.14241 | 0.92469 | 2.60764 | 0.02284 | 0.39657 | 2.24297 | 0.32547 | 0.52301 | 1.36684 | 0.82830 | 0.22839 | 1.12404 | 1.73926 | 2.42278 | 0.00397 |
| **Zhx1** | 0.02988 | 0.00813 | 0.01084 | 0.81666 | 0.00834 | 0.00686 | 0.00748 | 0.01344 | 0.26751 | 0.41393 | 0.01038 | 0.00796 | 0.01249 | 0.01098 | 0.00647 | 0.00718 |
| **Zhx3** | 0.05632 | 0.01775 | 0.00991 | 2.94546 | 0.00763 | 0.00763 | 0.01744 | 1.87936 | 4.54914 | 0.78484 | 0.56743 | 0.24928 | 0.00262 | 0.17245 | 1.58218 | 0.01339 |
| **Zic3** | 0.02896 | 0.00802 | 0.01556 | 0.91852 | 0.00708 | 0.00687 | 0.00728 | 0.01373 | 1.04294 | 0.00000 | 0.00597 | 0.00643 | 0.00680 | 0.03184 | 0.00001 | 2.00250 |
| **Zik1** | 1.68322 | 0.00526 | 0.99635 | 0.01001 | 1.32881 | 0.00378 | 0.00635 | 0.00642 | 0.01162 | 0.03867 | 0.01038 | 0.01653 | 0.00770 | 0.01535 | 0.00000 | 0.00500 |
| **Zkscan1** | 1.38143 | 0.00458 | 1.81567 | 4.27467 | 1.05280 | 0.00395 | 0.89935 | 0.00000 | 3.16718 | 4.68808 | 2.29573 | 1.34677 | 1.24532 | 7.06852 | 2.68490 | 3.11146 |
| **Zkscan14** | 1.88756 | 0.35511 | 1.85245 | 4.05585 | 0.03274 | 0.26828 | 0.00485 | 2.05917 | 6.46353 | 1.55318 | 0.18818 | 0.05988 | 1.06660 | 0.00000 | 3.15434 | 0.00294 |
| **Zkscan16** | 0.01378 | 0.00783 | 0.01211 | 0.01046 | 0.00934 | 0.00662 | 0.00750 | 0.01327 | 0.01208 | 0.01120 | 0.01262 | 0.00771 | 0.01271 | 0.01174 | 0.09731 | 0.00740 |
| **Zkscan17** | 1.59647 | 0.00638 | 1.56559 | 3.36696 | 0.09194 | 0.02982 | 0.01567 | 0.94694 | 3.28751 | 3.37886 | 0.48436 | 0.02167 | 0.37493 | 0.02746 | 1.60371 | 8.04623 |
| **Zkscan3** | 2.75540 | 0.00481 | 1.39109 | 0.67100 | 0.00000 | 2.29760 | 0.04613 | 0.00000 | 0.00000 | 1.19012 | 0.41524 | 0.05462 | 0.01384 | 0.00510 | 0.49674 | 0.46796 |
| **Zkscan4** | 0.04685 | 0.00816 | 0.00585 | 0.79589 | 0.00821 | 0.06667 | 0.01789 | 0.01475 | 0.04388 | 0.01445 | 0.01256 | 0.00838 | 0.01162 | 0.01091 | 0.00412 | 0.00713 |
| **Zkscan5** | 0.00113 | 0.00615 | 0.00854 | 3.85429 | 2.98568 | 0.01160 | 0.35897 | 2.99081 | 4.31680 | 1.99156 | 0.00000 | 0.00000 | 0.04245 | 0.47561 | 1.13769 | 0.02778 |
| **Zkscan6** | 2.45337 | 0.27431 | 1.57977 | 1.03687 | 0.01712 | 0.00616 | 0.00449 | 2.34808 | 0.79257 | 0.72958 | 1.31263 | 0.27065 | 0.00000 | 0.14025 | 0.39133 | 0.01069 |
| **Zkscan7** | 0.01376 | 0.00776 | 0.01223 | 0.00907 | 0.00937 | 0.00654 | 0.00745 | 0.01267 | 0.01203 | 0.00812 | 0.01245 | 0.00713 | 0.01268 | 0.00674 | 0.26561 | 0.00737 |
| **Zkscan8** | 0.01317 | 0.01876 | 1.22941 | 0.43370 | 0.03107 | 0.00691 | 0.27428 | 0.00341 | 0.00492 | 0.20172 | 0.33185 | 0.00539 | 0.06676 | 0.01187 | 0.23415 | 0.00707 |
| **Zmat2** | 8.66372 | 3.64305 | 2.23943 | 3.17090 | 1.75555 | 4.95373 | 0.17875 | 3.16790 | 2.77727 | 1.08821 | 3.73487 | 0.26991 | 5.26044 | 6.21766 | 4.64014 | 2.34758 |
| **Zmat3** | 4.79966 | 1.27338 | 0.61491 | 0.54285 | 1.12793 | 0.04921 | 4.12807 | 0.00000 | 0.48084 | 0.76349 | 0.86354 | 0.15101 | 0.57139 | 1.95687 | 0.50951 | 0.01750 |
| **Zmat4** | 0.01360 | 0.00786 | 0.01058 | 2.23079 | 0.00932 | 0.00622 | 0.00731 | 0.01104 | 1.19670 | 0.01064 | 0.00000 | 0.03517 | 0.01306 | 0.00325 | 0.76527 | 0.00690 |
| **Zmat5** | 13.61866 | 6.55487 | 12.37432 | 2.95217 | 1.00030 | 1.51702 | 4.13857 | 11.40585 | 5.32553 | 12.03333 | 8.19557 | 2.02727 | 8.59358 | 0.00000 | 0.00000 | 2.39640 |
| **Zmiz1** | 17.28424 | 5.45641 | 14.12105 | 5.84623 | 5.18018 | 7.09442 | 9.10187 | 14.65963 | 9.97728 | 25.80689 | 15.32502 | 14.90393 | 7.23977 | 6.24020 | 13.57996 | 3.72883 |
| **Zmiz2** | 3.72920 | 0.07608 | 0.33070 | 7.63371 | 0.02966 | 4.34256 | 1.51137 | 9.52516 | 2.84724 | 6.67234 | 4.35233 | 0.40781 | 8.15169 | 0.03290 | 0.22098 | 3.72627 |
| **Zmpste24** | 10.80074 | 11.99182 | 24.39601 | 8.93078 | 7.81683 | 13.89767 | 15.91123 | 33.07769 | 19.38244 | 34.12761 | 32.47319 | 24.65604 | 26.05055 | 40.54811 | 32.74909 | 6.56556 |
| **Zmym1** | 5.58697 | 3.65552 | 2.21886 | 9.68366 | 5.52460 | 4.47148 | 0.65295 | 5.55313 | 13.73592 | 4.59679 | 8.05096 | 10.52130 | 4.68480 | 1.17566 | 1.13953 | 5.16854 |
| **Zmym2** | 3.67063 | 5.88560 | 0.00000 | 6.90866 | 1.31918 | 4.87456 | 2.34234 | 2.10492 | 11.62174 | 3.28159 | 0.95496 | 10.76433 | 1.03757 | 12.01991 | 7.13932 | 0.30167 |
| **Zmym3** | 0.00000 | 0.00127 | 2.06886 | 2.56936 | 6.75502 | 0.00487 | 1.39377 | 9.03954 | 2.80582 | 10.48981 | 6.97312 | 0.69676 | 0.40226 | 0.94125 | 12.81901 | 2.31662 |
| **Zmym4** | 4.20442 | 3.18038 | 1.60294 | 1.39478 | 4.03540 | 1.41068 | 5.30541 | 1.95754 | 3.02601 | 6.80459 | 1.86546 | 0.44972 | 1.29185 | 4.52491 | 4.34989 | 0.01341 |
| **Zmym5** | 2.85245 | 0.79658 | 0.58993 | 2.16840 | 0.66581 | 0.06984 | 0.08365 | 1.34828 | 4.66337 | 0.18820 | 0.56934 | 4.47941 | 0.63460 | 4.70310 | 1.62212 | 0.95277 |
| **Zmym6** | 0.08709 | 0.76306 | 1.72059 | 0.80451 | 0.03232 | 0.01882 | 0.06619 | 0.00000 | 2.77878 | 0.40216 | 1.04614 | 4.40835 | 0.56007 | 0.94393 | 0.01794 | 0.04025 |
| **Zmynd11** | 0.00248 | 0.00782 | 1.28416 | 1.89039 | 0.00793 | 0.00764 | 1.77907 | 0.07106 | 0.00000 | 1.59836 | 0.02179 | 0.00000 | 0.03258 | 0.01548 | 1.41403 | 0.00650 |
| **Zmynd12** | 0.01361 | 0.00794 | 0.01232 | 0.00705 | 0.00931 | 0.00672 | 0.00743 | 0.01202 | 0.40768 | 0.22394 | 0.01068 | 0.00623 | 0.01544 | 0.01184 | 0.00825 | 0.00786 |
| **Zmynd19** | 1.32681 | 3.35312 | 2.03357 | 18.10222 | 3.39992 | 2.72346 | 0.01904 | 3.48416 | 16.01823 | 8.54134 | 6.53672 | 1.01504 | 3.46796 | 17.25740 | 14.28735 | 6.45453 |
| **Zmynd8** | 5.61718 | 1.79563 | 2.83207 | 7.36169 | 1.66449 | 0.06839 | 3.26193 | 3.68172 | 9.55828 | 6.82695 | 11.62399 | 0.58180 | 1.10700 | 2.60397 | 3.62389 | 3.62264 |
| **Znf41.ps** | 1.67958 | 0.09306 | 1.37927 | 1.13771 | 0.06185 | 0.11192 | 2.19161 | 0.33812 | 1.37792 | 0.46446 | 0.43839 | 0.89189 | 1.38692 | 0.43804 | 0.70710 | 0.20005 |
| **Znfx1** | 1.17935 | 2.76590 | 0.92282 | 3.23812 | 0.00431 | 0.00423 | 0.01356 | 0.00000 | 1.40972 | 0.50930 | 0.67897 | 0.59959 | 2.61675 | 0.00000 | 0.32892 | 0.15151 |
| **Znhit1** | 7.77145 | 5.28459 | 2.44926 | 1.63979 | 3.13046 | 1.31053 | 3.22211 | 6.54524 | 4.05706 | 10.59850 | 9.98109 | 1.13455 | 11.65374 | 0.14246 | 0.33524 | 2.83759 |
| **Znhit2** | 3.55146 | 0.01440 | 0.17854 | 8.06514 | 0.00442 | 0.02145 | 0.02353 | 1.47937 | 16.95590 | 2.37541 | 1.47994 | 0.00000 | 0.42527 | 0.00011 | 0.00000 | 0.02993 |
| **Znhit3** | 13.64967 | 8.04454 | 10.11953 | 18.44113 | 7.56387 | 5.85274 | 3.20315 | 15.52400 | 26.36405 | 27.47743 | 12.48659 | 15.71231 | 9.60486 | 25.32223 | 20.70427 | 2.21861 |
| **Znhit6** | 10.79640 | 9.73608 | 10.40953 | 16.71280 | 5.40286 | 6.64486 | 8.01604 | 8.03177 | 27.15969 | 13.50527 | 10.81045 | 9.71012 | 13.34861 | 8.00177 | 9.76572 | 7.88481 |
| **Znrd1** | 12.75372 | 5.32128 | 9.30710 | 10.51463 | 6.85369 | 5.13994 | 2.52390 | 9.24310 | 18.90971 | 19.93182 | 17.38922 | 1.61437 | 10.31180 | 5.91946 | 14.61747 | 3.12214 |
| **Znrd1as** | 4.27718 | 2.30292 | 0.91931 | 1.23064 | 0.01763 | 0.37494 | 0.00293 | 0.02053 | 0.58780 | 0.66557 | 0.00000 | 2.84088 | 0.03179 | 0.05705 | 0.19632 | 0.00000 |
| **Znrf1** | 1.65614 | 0.16972 | 0.80502 | 6.21728 | 0.01925 | 0.05345 | 0.14728 | 4.98415 | 3.29009 | 8.16755 | 4.49064 | 0.41022 | 2.48997 | 0.49885 | 0.98987 | 0.39597 |
| **Znrf2** | 10.56921 | 10.30158 | 1.17054 | 12.13393 | 7.61331 | 6.39436 | 9.13926 | 5.14613 | 18.10547 | 12.92863 | 11.55643 | 8.43055 | 6.21142 | 19.71293 | 12.57602 | 2.78765 |
| **Znrf3** | 0.55672 | 0.00604 | 0.00000 | 1.31651 | 1.07616 | 0.00505 | 2.03753 | 1.30124 | 4.88297 | 2.62412 | 1.43645 | 0.04573 | 0.81622 | 4.20263 | 1.94620 | 0.45441 |
| **Zp1** | 0.00467 | 0.00769 | 0.88674 | 5.09721 | 0.00784 | 0.00498 | 0.00718 | 2.06334 | 2.02591 | 1.63114 | 1.03674 | 0.10740 | 0.00000 | 0.00961 | 2.76882 | 0.00603 |
| **Zp2** | 0.91333 | 0.00509 | 1.12573 | 4.92999 | 0.00368 | 0.00619 | 0.00514 | 0.23877 | 6.76766 | 0.00000 | 1.50067 | 4.82065 | 0.00000 | 0.65173 | 6.46010 | 0.95310 |
| **Zp3** | 0.04481 | 0.01935 | 0.00049 | 15.68493 | 0.23891 | 0.00212 | 0.00507 | 3.91220 | 21.39124 | 7.69418 | 3.85787 | 1.98928 | 0.49951 | 0.21809 | 5.31620 | 0.00239 |
| **Zpbp2** | 0.78457 | 0.00696 | 0.00000 | 0.00629 | 0.00656 | 0.58897 | 0.00672 | 0.00729 | 0.01138 | 0.01214 | 0.07942 | 0.00713 | 0.37519 | 0.00779 | 0.29113 | 0.00612 |
| **Zpr1** | 10.09518 | 3.97194 | 8.12163 | 12.22728 | 8.36678 | 4.72866 | 3.39533 | 11.58917 | 17.40261 | 24.45021 | 11.34592 | 8.57308 | 10.91356 | 5.29588 | 16.66215 | 4.99062 |
| **Zranb1** | 1.83982 | 0.01501 | 2.27805 | 3.47891 | 0.01015 | 0.01586 | 1.06530 | 0.55617 | 3.18596 | 2.77760 | 0.11017 | 3.29402 | 0.43222 | 5.84088 | 7.90307 | 0.09250 |
| **Zranb2** | 3.02740 | 0.07665 | 6.55822 | 7.97506 | 8.60110 | 2.36309 | 2.85682 | 11.03598 | 6.16206 | 11.79415 | 6.64957 | 17.56093 | 9.59173 | 11.98574 | 15.61255 | 1.45025 |
| **Zranb3** | 1.57925 | 3.66466 | 1.78568 | 0.57199 | 1.42418 | 0.00378 | 2.80392 | 0.95331 | 0.41466 | 2.50584 | 0.92071 | 0.50740 | 2.32755 | 0.00519 | 3.59607 | 0.00952 |
| **Zrsr1** | 0.77860 | 1.84528 | 0.35829 | 2.71871 | 0.00650 | 0.00411 | 1.54064 | 2.47769 | 5.75574 | 0.00000 | 0.58044 | 0.00000 | 0.78449 | 3.08124 | 4.98298 | 0.72771 |
| **Zrsr2** | 5.50701 | 0.08540 | 3.15036 | 1.04911 | 4.14720 | 0.00000 | 0.00000 | 2.70754 | 7.09894 | 5.13517 | 5.05693 | 0.23240 | 5.47184 | 4.35685 | 6.58437 | 2.56387 |
| **Zscan10** | 2.84801 | 0.37804 | 0.00000 | 2.26161 | 2.18755 | 0.00318 | 1.00373 | 0.00951 | 7.38450 | 1.76941 | 0.00000 | 2.04471 | 0.02764 | 0.01714 | 0.03821 | 1.58430 |
| **Zscan12** | 3.00490 | 2.26178 | 4.07911 | 1.22854 | 0.01384 | 0.00367 | 0.01471 | 0.11229 | 3.07844 | 2.20826 | 1.43889 | 0.00000 | 0.05440 | 0.00000 | 3.44509 | 2.90137 |
| **Zscan18** | 0.01767 | 0.00716 | 0.07901 | 0.69560 | 0.31786 | 0.07493 | 0.00522 | 0.00000 | 0.69839 | 0.72633 | 1.75024 | 0.00000 | 2.28008 | 1.28521 | 1.05371 | 0.07223 |
| **Zscan20** | 1.66325 | 0.00624 | 0.20674 | 0.06586 | 0.00505 | 0.00463 | 0.00619 | 0.00000 | 0.35889 | 0.00000 | 0.70587 | 0.15441 | 0.07087 | 0.00462 | 0.28156 | 0.01323 |
| **Zscan21** | 0.03734 | 4.84330 | 11.37274 | 8.88070 | 6.59051 | 5.30504 | 6.47421 | 9.37439 | 17.29848 | 6.86597 | 3.52794 | 1.39561 | 5.04091 | 3.45568 | 4.43113 | 2.62226 |
| **Zscan22** | 0.13916 | 0.00714 | 2.29459 | 0.25722 | 0.00491 | 0.00787 | 0.00675 | 1.01011 | 0.00615 | 0.66909 | 0.00000 | 0.00635 | 0.00000 | 1.49995 | 0.58651 | 0.00612 |
| **Zscan25** | 0.25925 | 0.00766 | 0.00393 | 0.03377 | 0.00787 | 0.00873 | 0.03015 | 0.40095 | 0.00972 | 0.20435 | 0.01137 | 0.01669 | 0.29732 | 0.00956 | 0.02295 | 0.00763 |
| **Zscan26** | 0.08497 | 0.02676 | 4.87088 | 2.53591 | 1.43475 | 0.06588 | 1.27180 | 0.55013 | 4.21172 | 1.20879 | 2.96476 | 0.18234 | 0.53604 | 0.19912 | 2.16551 | 1.65263 |

| | | | | | | | | | | | | | | | | |
|---|---|---|---|---|---|---|---|---|---|---|---|---|---|---|---|---|
| **Zscan29** | 1.04469 | 1.60743 | 0.18390 | 6.42087 | 1.07976 | 0.00374 | 2.97348 | 2.27167 | 5.05023 | 1.40187 | 0.14210 | 0.00000 | 1.92808 | 1.49732 | 1.60398 | 0.01104 |
| **Zscan4a** | 0.01213 | 0.00755 | 0.00813 | 8.51409 | 0.00936 | 0.00575 | 0.00696 | 0.01365 | 3.87897 | 0.00000 | 0.00000 | 0.01668 | 0.01326 | 0.80348 | 0.00000 | 0.00568 |
| **Zscan4b** | 0.01314 | 0.00772 | 0.00969 | 4.29709 | 0.00933 | 0.00629 | 0.00723 | 0.01036 | 2.26081 | 0.00000 | 0.00000 | 0.00243 | 0.01384 | 0.36127 | 0.00000 | 0.00662 |
| **Zscan4c** | 0.01227 | 0.00760 | 0.00845 | 10.16463 | 0.00924 | 0.00546 | 0.00695 | 0.00770 | 6.06646 | 0.00000 | 0.00000 | 0.00014 | 0.01391 | 0.66255 | 0.00000 | 0.00528 |
| **Zscan4d** | 0.01122 | 0.00751 | 0.00614 | 8.68432 | 0.00946 | 0.00546 | 0.00685 | 0.00655 | 4.73302 | 0.00000 | 0.00000 | 0.00000 | 0.01375 | 2.25530 | 0.00000 | 0.00551 |
| **Zscan4f** | 0.01232 | 0.00765 | 0.00836 | 10.38295 | 0.00925 | 0.00549 | 0.00697 | 0.00806 | 5.67030 | 0.00000 | 0.00000 | 0.00000 | 0.01404 | 0.63300 | 0.10877 | 0.00528 |
| **Zscan5b** | 0.00000 | 0.00520 | 1.13031 | 15.16311 | 0.00689 | 0.00749 | 2.68675 | 2.73721 | 22.88745 | 7.21023 | 6.11595 | 1.46822 | 1.01033 | 19.81150 | 22.62357 | 0.00799 |
| **Zswim1** | 6.47758 | 0.92853 | 3.90113 | 12.88710 | 0.02501 | 4.11055 | 2.24650 | 8.82728 | 6.84578 | 10.90825 | 4.59856 | 4.41982 | 6.25583 | 0.29776 | 0.75643 | 4.39162 |
| **Zswim3** | 1.23501 | 6.94780 | 1.63744 | 4.00154 | 0.00142 | 4.07318 | 0.00349 | 5.28011 | 8.44121 | 6.27905 | 0.85278 | 0.55952 | 0.00000 | 0.58482 | 5.96915 | 2.77465 |
| **Zswim4** | 0.02656 | 0.00798 | 0.00478 | 5.25805 | 0.02227 | 0.00593 | 0.01687 | 0.00000 | 3.60107 | 1.83694 | 0.35391 | 0.00934 | 0.79568 | 0.00000 | 1.82876 | 0.00596 |
| **Zswim5** | 0.01093 | 0.00806 | 0.01218 | 0.58901 | 0.00909 | 0.00685 | 0.00761 | 0.01361 | 0.00366 | 0.02264 | 0.01161 | 0.01837 | 0.02792 | 0.00496 | 0.95922 | 0.32971 |
| **Zswim6** | 0.00448 | 0.00626 | 0.00000 | 0.73650 | 0.14091 | 0.00628 | 0.00645 | 0.06269 | 1.11143 | 0.02996 | 0.43775 | 3.61773 | 0.00498 | 0.00269 | 0.38902 | 0.48665 |
| **Zswim7** | 1.84702 | 0.32569 | 0.43316 | 1.37528 | 4.45629 | 0.01089 | 0.00423 | 0.32801 | 0.98871 | 0.16512 | 0.01402 | 0.00452 | 2.25032 | 0.36872 | 0.00000 | 0.72484 |
| **Zswim8** | 3.43047 | 0.00299 | 0.38479 | 0.32867 | 0.00969 | 0.00500 | 1.39154 | 0.07118 | 0.03327 | 0.40723 | 0.82111 | 0.06041 | 0.00000 | 0.00561 | 0.08764 | 3.86009 |
| **Zufsp** | 0.86950 | 6.09042 | 2.01023 | 0.28248 | 2.35422 | 0.15088 | 1.02954 | 1.98350 | 0.90356 | 2.33144 | 3.05453 | 3.41900 | 0.61312 | 2.73692 | 6.13674 | 0.00988 |
| **Zw10** | 10.18890 | 0.70010 | 5.87500 | 2.34211 | 0.20905 | 0.33059 | 2.95036 | 4.32056 | 1.94575 | 1.60392 | 3.23257 | 0.80506 | 6.43386 | 1.37506 | 5.71302 | 5.65670 |
| **Zwilch** | 5.35721 | 9.89667 | 7.27157 | 1.91024 | 1.24199 | 3.22191 | 5.35504 | 3.86852 | 3.44064 | 3.23130 | 3.33374 | 5.29143 | 3.37142 | 8.05842 | 5.55950 | 4.00085 |
| **Zwint** | 14.57251 | 14.45369 | 13.51828 | 3.50178 | 15.66633 | 5.98120 | 8.02079 | 12.18583 | 2.26864 | 4.91629 | 6.43270 | 7.75023 | 9.53759 | 12.45215 | 3.55341 | 3.10712 |
| **Zxda** | 0.10570 | 0.00806 | 0.06989 | 0.12304 | 0.03672 | 0.01323 | 0.04934 | 0.03903 | 0.14793 | 0.22201 | 0.15329 | 0.06093 | 0.10233 | 0.22064 | 0.14296 | 0.03701 |
| **Zxdc** | 0.00000 | 0.00477 | 5.21470 | 2.27207 | 1.84775 | 0.00691 | 1.79141 | 0.80148 | 1.42016 | 0.63228 | 1.25613 | 0.02687 | 0.04126 | 0.00000 | 0.22650 | 0.02769 |
| **Zyg11a** | 3.22584 | 2.77405 | 2.18195 | 0.52779 | 0.00000 | 0.95077 | 2.36157 | 2.41748 | 5.33248 | 6.75216 | 3.40903 | 2.65158 | 0.54148 | 0.87076 | 10.54962 | 0.01471 |
| **Zyg11b** | 4.28223 | 1.15581 | 2.54603 | 6.46039 | 2.91875 | 0.36916 | 2.22757 | 0.96295 | 7.77340 | 2.49823 | 4.84069 | 0.20367 | 0.97694 | 4.53515 | 4.44858 | 0.53453 |
| **Zyx** | 7.77280 | 2.01248 | 0.19286 | 0.41094 | 5.07324 | 7.39151 | 0.97728 | 4.72891 | 0.00000 | 1.14484 | 4.14072 | 0.23640 | 3.75139 | 0.00322 | 0.02011 | 9.19855 |
| **Zzef1** | 1.31047 | 1.68389 | 1.45635 | 3.10288 | 2.11730 | 4.24406 | 1.73209 | 0.77455 | 6.09185 | 5.44218 | 1.34775 | 0.57576 | 1.99952 | 0.66243 | 6.73690 | 0.02285 |
| **Zzz3** | 4.70793 | 0.06356 | 3.15980 | 12.36814 | 4.49668 | 4.10289 | 6.26845 | 5.66940 | 12.86815 | 11.94895 | 10.05666 | 18.03282 | 3.62709 | 25.79878 | 18.87905 | 0.80381 |

### Supplementary Table S9

#### Mouse female NMF coefficient matrix

| Gene | Cluster 1 | Cluster 2 | Cluster 3 | Cluster 4 | Cluster 5 | Cluster 6 | Cluster 7 | Cluster 8 | Cluster 9 | Cluster 10 | Cluster 11 | Cluster 12 | Cluster 13 | Cluster 14 |
|---|---|---|---|---|---|---|---|---|---|---|---|---|---|---|
| A4galt | 0.02671 | 0.02023 | 0.02768 | 0.07404 | 0.05681 | 0.06383 | 0.03403 | 0.03975 | 0.07820 | 0.00835 | 0.02867 | 0.02492 | 0.24193 | 0.21522 |
| A630020A06 | 0.22306 | 0.00410 | 0.05225 | 0.02955 | 0.46592 | 0.08625 | 0.06806 | 0.03596 | 1.00538 | 0.00734 | 0.01013 | 0.05604 | 1.43993 | 0.33161 |
| AA388235 | 0.58477 | 0.18071 | 0.36327 | 0.00000 | 0.87568 | 0.25290 | 0.01771 | 0.08457 | 0.24461 | 0.00540 | 0.07358 | 0.27103 | 0.94976 | 0.40729 |
| AA414768 | 0.02918 | 0.01993 | 0.00000 | 1.19582 | 0.09711 | 2.94784 | 0.00386 | 0.00192 | 1.70546 | 0.00000 | 2.93727 | 0.03276 | 1.79708 | 1.53086 |
| AA415398 | 0.57501 | 0.00400 | 0.01226 | 0.01864 | 0.00000 | 0.00794 | 0.00418 | 0.01332 | 0.00000 | 0.00728 | 0.00700 | 0.00326 | 3.48687 | 0.88779 |
| AA465934 | 0.02026 | 0.00420 | 0.00968 | 0.01419 | 0.00864 | 0.01717 | 0.00430 | 0.00717 | 0.10571 | 0.00627 | 0.00590 | 0.12245 | 0.13067 | 0.00000 |
| AA467197 | 2.46522 | 7.90902 | 1.29057 | 7.14523 | 1.10621 | 12.71531 | 7.80431 | 27.79439 | 0.76571 | 13.34619 | 2.94699 | 16.46669 | 0.01666 | 0.72899 |
| AA536875 | 0.07114 | 0.02063 | 0.02437 | 0.05419 | 0.03410 | 0.02847 | 0.00429 | 0.04098 | 0.05471 | 0.00817 | 0.01617 | 0.02193 | 0.02451 | 0.16186 |
| AA543186 | 0.02394 | 0.00443 | 0.00721 | 0.04491 | 0.07940 | 0.02458 | 0.00423 | 0.01273 | 0.52695 | 0.00781 | 0.01111 | 0.02528 | 0.12545 | 0.00221 |
| AA543401 | 0.00241 | 0.00402 | 0.01472 | 0.01220 | 0.00580 | 0.01191 | 0.00411 | 0.00828 | 0.01557 | 0.00591 | 0.00917 | 0.00732 | 2.71778 | 0.02835 |
| AA545190 | 0.00766 | 0.00445 | 0.01534 | 0.00920 | 0.26633 | 0.01268 | 0.00494 | 0.01016 | 0.05129 | 0.00708 | 0.00829 | 0.00866 | 0.04038 | 0.00000 |
| AA619741 | 0.00288 | 0.00388 | 0.01434 | 0.01310 | 0.00196 | 0.01157 | 0.00403 | 0.00792 | 0.00931 | 0.00657 | 0.00810 | 0.00650 | 3.72293 | 0.34118 |
| AA792892 | 1.53653 | 0.12445 | 0.74223 | 0.63335 | 2.07875 | 0.17990 | 0.15659 | 0.07930 | 4.74125 | 0.42909 | 0.14632 | 0.03598 | 15.74528 | 9.60641 |
| AA986860 | 0.01342 | 0.00437 | 0.09518 | 0.15535 | 0.00740 | 0.00000 | 0.44093 | 1.85168 | 0.00969 | 0.52054 | 0.00000 | 0.00929 | 0.05817 | 0.00613 |
| Aaas | 5.58775 | 2.59928 | 19.10015 | 9.99800 | 8.59039 | 12.01458 | 3.01745 | 6.89007 | 5.12666 | 0.13673 | 9.60349 | 10.25610 | 1.08476 | 8.55212 |
| Aacs | 6.49651 | 8.93253 | 20.60611 | 5.99260 | 17.44201 | 2.78760 | 2.67114 | 0.00000 | 2.33515 | 0.00000 | 5.02097 | 7.88310 | 0.12528 | 12.74749 |
| Aaed1 | 0.34271 | 0.00392 | 0.04879 | 0.00000 | 0.47414 | 0.48616 | 0.00420 | 0.00526 | 0.00997 | 0.02552 | 0.00386 | 0.12000 | 0.41005 | 0.00486 |
| Aagab | 0.28991 | 3.49580 | 1.53382 | 0.70588 | 2.19115 | 5.97870 | 0.36686 | 3.70375 | 1.56192 | 0.07660 | 0.00000 | 2.61641 | 0.15422 | 3.21707 |
| Aak1 | 3.82788 | 0.03977 | 3.60861 | 0.49906 | 1.72302 | 4.57110 | 0.01284 | 0.00000 | 4.74654 | 0.00759 | 3.43381 | 2.60553 | 0.61654 | 3.65485 |
| Aamdc | 0.40960 | 2.47127 | 0.90465 | 2.63319 | 0.58722 | 2.55778 | 0.96696 | 1.74009 | 1.02237 | 0.01254 | 0.01572 | 1.08885 | 2.05677 | 0.16296 |
| Aamp | 7.85578 | 5.18847 | 26.80973 | 16.95174 | 16.93221 | 21.38049 | 4.00541 | 10.94021 | 0.66919 | 4.33682 | 15.82868 | 12.58253 | 2.63859 | 11.14904 |
| Aar2 | 0.83066 | 4.21632 | 5.16210 | 1.55621 | 6.56962 | 6.38709 | 1.08792 | 3.23824 | 1.70370 | 0.06745 | 4.22548 | 2.79717 | 2.10619 | 5.57450 |
| Aard | 0.00517 | 0.01395 | 0.00792 | 0.00000 | 0.15277 | 0.00801 | 2.50047 | 0.24514 | 0.00909 | 0.06868 | 0.00048 | 0.35625 | 0.01138 | 0.02310 |
| Aars | 5.97242 | 7.45721 | 15.21016 | 11.65795 | 14.59462 | 14.54929 | 5.67057 | 10.38614 | 0.25842 | 9.41860 | 17.07980 | 13.14116 | 8.98571 | 10.80659 |
| Aars2 | 0.16862 | 0.03317 | 3.14786 | 2.68546 | 1.46146 | 0.00000 | 0.00274 | 1.87695 | 0.01704 | 0.01066 | 0.50719 | 0.92727 | 0.00957 | 0.00000 |
| Aarsd1 | 4.35783 | 0.09238 | 13.52642 | 9.78923 | 5.08518 | 6.70807 | 1.82604 | 3.56863 | 0.00000 | 13.83458 | 4.39038 | 8.44352 | 0.56050 | 1.65085 |
| Aasdh | 1.15857 | 0.02410 | 2.07996 | 0.99097 | 0.34146 | 0.54268 | 0.69527 | 0.57301 | 0.28482 | 0.03038 | 0.55770 | 2.59969 | 1.18392 | 3.63525 |
| Aasdhppt | 5.65505 | 0.02705 | 2.70880 | 6.18646 | 2.94218 | 7.58300 | 2.58465 | 2.66054 | 2.14747 | 0.03081 | 4.41378 | 4.24381 | 4.12279 | 1.80710 |
| Aass | 0.00000 | 0.00441 | 0.03384 | 0.00959 | 0.08253 | 0.00779 | 0.00424 | 0.01016 | 0.19082 | 0.00617 | 0.00683 | 0.00858 | 0.29411 | 0.43456 |
| Aatf | 0.75478 | 0.05064 | 10.19342 | 4.99105 | 9.66429 | 6.24495 | 1.43217 | 2.15894 | 7.97351 | 0.02854 | 2.85166 | 6.16723 | 0.94442 | 14.98064 |
| AB124611 | 0.00708 | 0.00437 | 0.23742 | 0.00632 | 0.00443 | 0.00187 | 0.00400 | 0.01097 | 0.19798 | 0.00725 | 0.00587 | 0.00769 | 0.00301 | 0.67357 |
| Abat | 0.27423 | 0.00454 | 1.31636 | 0.41433 | 2.84337 | 1.11445 | 1.27646 | 1.94233 | 4.59965 | 0.01101 | 0.00000 | 0.08375 | 0.10488 | 0.86478 |
| Abca1 | 0.29405 | 0.00444 | 0.19000 | 0.00350 | 0.03785 | 1.03353 | 0.00404 | 0.00497 | 0.00851 | 0.00806 | 0.00959 | 0.00368 | 0.40546 | 0.08954 |
| Abca13 | 0.02863 | 0.01285 | 0.13013 | 0.40283 | 0.23102 | 0.21547 | 0.00472 | 0.03334 | 0.00566 | 0.00498 | 0.00528 | 0.00695 | 2.07718 | 1.40167 |
| Abca16 | 0.00399 | 0.00437 | 0.07915 | 0.04756 | 0.16899 | 0.05460 | 0.00419 | 0.00616 | 0.01129 | 0.00725 | 0.00621 | 0.01031 | 0.00628 | 0.11021 |
| Abca17 | 1.44925 | 0.12297 | 1.02333 | 1.14549 | 2.69625 | 0.55126 | 0.12916 | 0.02855 | 0.08289 | 0.01157 | 0.06642 | 0.00000 | 0.37827 | 2.07827 |
| Abca2 | 0.02944 | 0.01961 | 0.87502 | 2.41006 | 0.07219 | 1.81244 | 0.00460 | 0.60093 | 0.00590 | 0.00849 | 0.24390 | 0.00388 | 0.46571 | 0.00000 |
| Abca3 | 0.00000 | 2.61340 | 4.63334 | 3.59276 | 0.00000 | 6.13536 | 2.54871 | 6.99310 | 0.03221 | 0.08691 | 2.69911 | 0.00000 | 0.00000 | 1.18332 |
| Abca4 | 0.00860 | 0.00457 | 0.17064 | 0.00704 | 0.00539 | 0.00958 | 0.00436 | 0.01486 | 0.00651 | 0.00952 | 0.00642 | 0.00840 | 0.02655 | 0.44930 |
| Abca5 | 0.67619 | 0.00424 | 0.12710 | 0.00210 | 0.00000 | 0.33309 | 0.00411 | 0.00121 | 0.01841 | 0.00086 | 1.86811 | 0.00719 | 0.27707 | 0.17604 |
| Abca7 | 0.00269 | 0.01117 | 0.30150 | 0.47690 | 0.00000 | 0.00000 | 0.00419 | 1.24516 | 0.10194 | 0.00302 | 0.67079 | 0.00441 | 0.00343 | 0.00000 |
| Abca8b | 0.04528 | 0.00444 | 0.02230 | 0.01878 | 0.01258 | 0.01116 | 0.00435 | 0.01292 | 0.11326 | 0.00700 | 0.00765 | 0.01248 | 0.01759 | 0.00284 |
| Abcb10 | 1.21916 | 0.02871 | 3.25934 | 1.26412 | 1.47413 | 1.71819 | 3.22252 | 1.53980 | 2.75640 | 0.16127 | 1.64442 | 0.18479 | 1.58564 | 6.53190 |
| Abcb11 | 0.00317 | 0.00413 | 0.02215 | 0.01071 | 0.00000 | 0.01012 | 0.00415 | 0.00826 | 0.00961 | 0.00613 | 0.00753 | 0.00779 | 1.33895 | 0.63609 |
| Abcb1a | 1.85229 | 0.00377 | 1.26990 | 0.01276 | 0.00000 | 0.68189 | 0.00955 | 0.01456 | 0.34568 | 0.00185 | 1.28905 | 0.00105 | 0.00415 | 0.80654 |
| Abcb1b | 0.25027 | 0.00382 | 0.95496 | 0.37791 | 0.86457 | 2.61043 | 0.00371 | 0.00733 | 0.19734 | 0.00713 | 1.82102 | 0.00000 | 1.18360 | 0.00654 |
| Abcb4 | 0.06033 | 0.00459 | 0.07034 | 0.02628 | 0.02029 | 0.05864 | 0.00449 | 0.01202 | 0.01173 | 0.00699 | 0.05610 | 0.00645 | 0.00695 | 0.12038 |
| Abcb5 | 2.09112 | 0.00328 | 1.37104 | 0.42378 | 0.63465 | 0.00000 | 0.00374 | 0.00316 | 0.07349 | 0.00301 | 0.00461 | 0.00154 | 4.14100 | 3.70039 |
| Abcb6 | 5.53206 | 0.03684 | 9.87924 | 10.27421 | 0.37299 | 11.09004 | 0.03542 | 4.03292 | 1.16214 | 0.10058 | 5.02016 | 5.58457 | 0.00000 | 3.13445 |
| Abcb7 | 5.91389 | 0.04398 | 3.63963 | 0.51115 | 1.78653 | 4.45195 | 0.02978 | 0.11992 | 9.21342 | 0.28856 | 3.39360 | 3.67224 | 2.03552 | 1.64789 |
| Abcb8 | 0.00000 | 0.04054 | 3.41737 | 8.28445 | 0.82421 | 1.45842 | 1.66208 | 4.15903 | 0.01224 | 0.05574 | 4.59364 | 1.83543 | 0.17490 | 0.00000 |
| Abcb9 | 0.00240 | 0.00443 | 0.28929 | 0.03222 | 0.53509 | 0.00355 | 0.00427 | 0.00969 | 0.01032 | 0.00571 | 0.00648 | 0.04849 | 0.19917 | 0.00234 |

| | | | | | | | | | | | | | | |
|---|---|---|---|---|---|---|---|---|---|---|---|---|---|---|
| **Abcc1** | 0.59969 | 0.01792 | 0.77426 | 1.84212 | 0.00000 | 2.33116 | 0.45684 | 0.24138 | 0.21209 | 3.93642 | 1.93175 | 1.79417 | 0.04498 | 1.44648 |
| **Abcc10** | 0.01743 | 4.32608 | 3.26951 | 1.82466 | 1.98836 | 0.13876 | 0.70229 | 0.07406 | 0.08207 | 0.07128 | 0.96771 | 0.25129 | 0.60296 | 0.02139 |
| **Abcc2** | 0.41163 | 0.00444 | 0.14510 | 0.00000 | 0.08267 | 0.00103 | 0.00401 | 0.01202 | 1.52642 | 0.00811 | 0.00409 | 0.00643 | 0.00215 | 0.39990 |
| **Abcc3** | 0.00000 | 0.01214 | 2.47187 | 1.07843 | 0.77518 | 0.00000 | 0.00886 | 0.00000 | 0.20808 | 0.00225 | 0.00564 | 0.46073 | 0.00454 | 4.75019 |
| **Abcc4** | 0.00000 | 0.03744 | 2.24526 | 2.26027 | 0.00000 | 0.71145 | 3.17280 | 0.18370 | 1.17150 | 0.06735 | 0.00000 | 1.12328 | 0.00000 | 0.00000 |
| **Abcc5** | 4.24578 | 0.07592 | 9.32860 | 1.14508 | 5.54996 | 9.30002 | 1.37264 | 2.87623 | 2.45287 | 11.91181 | 0.00000 | 6.16176 | 3.46676 | 7.19794 |
| **Abcc6** | 0.00576 | 0.00439 | 0.01402 | 0.01222 | 0.00909 | 0.01265 | 0.00433 | 0.00980 | 0.01061 | 0.00644 | 0.00893 | 0.00972 | 0.13994 | 0.08239 |
| **Abcc8** | 0.05788 | 0.00361 | 0.00000 | 0.02206 | 0.02426 | 0.10952 | 0.00390 | 0.00164 | 0.00600 | 0.01304 | 0.52571 | 0.50337 | 0.02254 | 0.09074 |
| **Abcc9** | 0.18922 | 0.07898 | 0.07407 | 0.47254 | 0.42515 | 0.42746 | 0.08635 | 0.18432 | 0.42676 | 0.31390 | 0.05785 | 0.08691 | 0.40230 | 0.62056 |
| **Abcd1** | 2.18511 | 6.37724 | 1.49256 | 0.00000 | 1.73533 | 0.59693 | 2.36544 | 2.30462 | 0.00000 | 0.10206 | 2.16037 | 3.19004 | 1.57564 | 1.92464 |
| **Abcd2** | 0.08016 | 0.04762 | 0.00630 | 0.13747 | 0.08834 | 0.10998 | 0.01019 | 0.01717 | 0.11571 | 0.00921 | 0.01001 | 0.05455 | 0.21317 | 0.23957 |
| **Abcd3** | 2.09627 | 0.01578 | 3.77935 | 2.69450 | 1.42982 | 2.19583 | 2.94122 | 0.82649 | 1.66003 | 0.13224 | 4.67918 | 5.30477 | 0.00000 | 0.75817 |
| **Abcd4** | 0.06749 | 0.99114 | 1.15858 | 6.13381 | 0.77289 | 3.77415 | 4.23546 | 2.36219 | 0.23266 | 0.26471 | 4.85555 | 0.55091 | 0.44604 | 1.75892 |
| **Abce1** | 15.42425 | 0.26371 | 17.39901 | 8.71131 | 17.29020 | 12.95380 | 0.76548 | 3.51673 | 27.75565 | 0.75063 | 8.45382 | 10.85920 | 6.40480 | 13.03335 |
| **Abcf1** | 8.48650 | 7.72635 | 11.73236 | 18.87768 | 24.93957 | 15.26989 | 3.11248 | 8.80470 | 2.49271 | 0.34825 | 4.07956 | 10.89514 | 10.60723 | 22.42880 |
| **Abcf2** | 13.27308 | 3.24516 | 26.74905 | 16.88657 | 22.11772 | 23.14231 | 2.66384 | 6.53959 | 0.93978 | 0.32918 | 15.73466 | 9.49213 | 12.31555 | 23.03665 |
| **Abcf3** | 1.46438 | 0.05318 | 9.29963 | 10.51705 | 8.34893 | 8.66195 | 2.75765 | 5.46373 | 10.51765 | 1.63981 | 8.26908 | 3.03558 | 4.57581 | 11.04645 |
| **Abcg1** | 0.00607 | 0.01908 | 0.08206 | 0.00573 | 0.02401 | 0.00698 | 0.00381 | 0.01195 | 0.01887 | 0.00640 | 0.00377 | 2.33272 | 0.00754 | 0.08154 |
| **Abcg2** | 0.02521 | 1.68637 | 0.00000 | 0.00000 | 0.00748 | 2.18292 | 0.03045 | 0.01531 | 0.00775 | 0.00232 | 1.72972 | 1.02500 | 0.02573 | 0.00000 |
| **Abcg4** | 1.53144 | 0.02332 | 1.80512 | 1.44278 | 0.81844 | 0.00000 | 0.00390 | 1.19959 | 0.03291 | 0.00959 | 0.00000 | 0.38290 | 0.68428 | 0.55660 |
| **Abhd1** | 0.00811 | 0.00431 | 0.02100 | 0.12155 | 0.01247 | 0.01056 | 0.00421 | 0.00914 | 0.01184 | 0.00671 | 0.00772 | 0.00929 | 0.00683 | 0.01149 |
| **Abhd10** | 0.72423 | 0.04484 | 0.18408 | 0.94990 | 1.34961 | 0.00000 | 1.18147 | 1.27673 | 2.95456 | 0.02476 | 0.00000 | 1.75771 | 0.23386 | 0.45064 |
| **Abhd11** | 5.34706 | 4.28513 | 12.60245 | 5.23804 | 4.55784 | 8.34951 | 4.23683 | 9.23408 | 0.36682 | 0.32555 | 5.03923 | 6.56177 | 0.13870 | 0.33776 |
| **Abhd11os** | 1.68351 | 0.00482 | 0.07452 | 0.87024 | 0.66618 | 0.26443 | 0.01103 | 1.09747 | 0.00599 | 0.04276 | 0.06471 | 0.07121 | 0.03465 | 0.12671 |
| **Abhd12** | 0.00000 | 0.03183 | 1.34669 | 2.76062 | 0.13336 | 10.28190 | 0.00652 | 3.09590 | 0.08109 | 11.11004 | 4.31593 | 2.65819 | 1.82927 | 0.13510 |
| **Abhd12b** | 0.43840 | 0.00484 | 1.40393 | 0.00000 | 0.24867 | 0.17191 | 0.00385 | 0.00223 | 0.57105 | 0.00423 | 0.00414 | 0.00422 | 0.00000 | 0.85853 |
| **Abhd13** | 4.42829 | 0.01778 | 3.15713 | 1.33428 | 3.13869 | 3.86705 | 1.65419 | 5.03896 | 3.13912 | 0.06823 | 0.00000 | 0.42864 | 3.05405 | 6.51680 |
| **Abhd14a** | 0.28628 | 0.08816 | 0.54319 | 1.28336 | 0.34840 | 1.38561 | 3.51982 | 6.54650 | 0.00000 | 0.04140 | 0.49433 | 1.05823 | 0.00372 | 0.33035 |
| **Abhd14b** | 0.38953 | 0.02820 | 0.00000 | 0.00871 | 1.00856 | 0.36649 | 0.00415 | 3.64718 | 0.00384 | 0.00839 | 0.00164 | 0.00302 | 0.47959 | 0.40954 |
| **Abhd15** | 0.03009 | 0.00456 | 0.01574 | 0.07262 | 0.44238 | 0.09943 | 0.01815 | 0.05744 | 0.06797 | 0.01570 | 0.00741 | 0.02745 | 0.15918 | 0.13816 |
| **Abhd16a** | 1.17161 | 0.06934 | 2.05712 | 1.70751 | 0.32927 | 0.86827 | 0.03564 | 5.37121 | 0.00145 | 0.01529 | 0.00000 | 2.93392 | 0.26176 | 0.80384 |
| **Abhd17a** | 0.26926 | 0.08564 | 3.39340 | 7.52862 | 12.11342 | 0.00000 | 0.01363 | 5.03500 | 0.00271 | 0.00000 | 2.13359 | 1.98899 | 0.03134 | 0.00000 |
| **Abhd17b** | 0.56416 | 1.43918 | 0.90527 | 0.34411 | 0.39475 | 0.52833 | 0.50293 | 0.74144 | 0.22295 | 0.00799 | 0.86305 | 0.52831 | 4.29781 | 0.00000 |
| **Abhd17c** | 2.37039 | 0.07891 | 4.75696 | 3.42949 | 0.80993 | 0.00000 | 2.82955 | 5.44043 | 1.72629 | 0.03678 | 2.73071 | 0.54903 | 3.53908 | 2.44163 |
| **Abhd18** | 1.44597 | 0.00357 | 0.26680 | 0.00000 | 0.65736 | 0.17838 | 0.00926 | 0.85955 | 1.53803 | 0.00013 | 1.83429 | 0.03686 | 0.23289 | 0.07625 |
| **Abhd2** | 0.00300 | 0.00339 | 0.01935 | 1.37118 | 0.00000 | 0.00000 | 0.00915 | 0.77906 | 0.08607 | 0.01058 | 0.00000 | 0.69529 | 0.33537 | 0.95315 |
| **Abhd3** | 1.06762 | 0.00406 | 0.73899 | 0.00000 | 0.00000 | 0.00000 | 0.00454 | 0.09890 | 1.97003 | 0.00220 | 0.83928 | 0.00202 | 0.37092 | 0.94340 |
| **Abhd4** | 0.52223 | 0.04496 | 2.78891 | 2.46465 | 0.11219 | 4.84987 | 0.03636 | 7.17722 | 1.23757 | 0.00868 | 2.80435 | 6.36715 | 0.73851 | 0.55220 |
| **Abhd5** | 3.02897 | 0.01453 | 1.85546 | 1.86075 | 1.58842 | 4.27894 | 0.78178 | 2.08867 | 0.00000 | 0.04157 | 5.88612 | 4.32783 | 0.05606 | 1.77185 |
| **Abhd6** | 9.05240 | 0.02215 | 16.33625 | 6.00149 | 6.02552 | 4.25432 | 2.16946 | 0.00000 | 15.83163 | 0.04295 | 0.00000 | 0.00000 | 9.56609 | 19.70527 |
| **Abhd8** | 0.00301 | 0.00318 | 0.01381 | 0.53261 | 0.39817 | 0.00842 | 0.00409 | 0.55018 | 0.02252 | 0.00606 | 0.01204 | 1.05486 | 0.00523 | 0.00303 |
| **Abi1** | 10.87295 | 6.36095 | 8.15117 | 2.47348 | 6.65561 | 7.25448 | 5.12756 | 2.59733 | 10.14182 | 0.02999 | 10.86886 | 5.11816 | 7.29768 | 6.28954 |
| **Abi2** | 0.99629 | 1.06543 | 1.15641 | 1.28412 | 0.06120 | 2.03712 | 1.51679 | 0.01930 | 0.25445 | 0.08124 | 0.61185 | 0.71321 | 0.18389 | 3.06171 |
| **Abi3bp** | 0.01889 | 0.00464 | 0.06019 | 0.00000 | 0.00708 | 0.04896 | 0.00412 | 0.00896 | 0.08016 | 0.00520 | 0.20399 | 0.01775 | 0.09979 | 0.00690 |
| **Abl1** | 5.48137 | 0.08478 | 6.47270 | 1.64109 | 6.66216 | 3.87786 | 0.07966 | 0.76019 | 0.02157 | 0.01604 | 0.26188 | 4.48282 | 2.65775 | 1.44505 |
| **Abl2** | 5.11188 | 0.04533 | 1.65504 | 1.06087 | 5.18694 | 2.25463 | 0.03193 | 1.57232 | 6.29904 | 0.03487 | 2.41023 | 2.37487 | 8.58320 | 10.87600 |
| **Ablim1** | 0.13896 | 0.03948 | 2.12990 | 0.00481 | 0.62135 | 2.57425 | 0.00799 | 1.54760 | 0.17908 | 0.03614 | 0.02248 | 2.60704 | 0.79065 | 1.00108 |
| **Ablim2** | 0.00411 | 0.00482 | 0.13211 | 0.08417 | 0.02173 | 1.05268 | 0.01520 | 0.56562 | 0.00712 | 0.01799 | 0.25929 | 0.03000 | 0.65113 | 0.00000 |
| **Ablim3** | 0.00755 | 0.00439 | 0.09640 | 0.00565 | 0.00130 | 0.00762 | 0.00434 | 0.01107 | 0.01000 | 0.00776 | 0.00684 | 0.00935 | 0.00603 | 0.50459 |
| **Abo** | 0.00000 | 0.00446 | 0.34307 | 1.30577 | 0.15645 | 0.00000 | 1.45181 | 0.00372 | 0.00644 | 0.03738 | 0.00181 | 0.00451 | 0.00366 | 0.14872 |
| **Abr** | 0.20235 | 0.13984 | 4.21694 | 1.36606 | 3.92965 | 3.97119 | 0.68499 | 6.20935 | 0.04543 | 0.07607 | 0.00000 | 2.24337 | 0.38603 | 2.19606 |
| **Abracl** | 0.36396 | 0.06733 | 0.00000 | 0.00000 | 0.07511 | 0.00000 | 0.04599 | 2.07539 | 0.08356 | 10.16659 | 0.24201 | 1.06636 | 0.00798 | 0.86702 |
| **Abt1** | 1.95468 | 0.02347 | 7.69938 | 1.25456 | 2.85754 | 2.33025 | 1.52383 | 1.44358 | 12.19336 | 0.05074 | 6.19400 | 0.90475 | 5.79556 | 25.55782 |
| **Abtb1** | 0.16397 | 0.07358 | 3.62627 | 3.03342 | 1.16889 | 0.62892 | 0.01455 | 2.20056 | 0.00836 | 0.01117 | 2.62743 | 3.26864 | 0.06267 | 0.00000 |
| **Abtb2** | 0.31157 | 0.00986 | 1.33453 | 0.01629 | 0.30324 | 2.32772 | 0.00306 | 0.06219 | 0.06841 | 0.01916 | 0.00000 | 1.90003 | 2.71946 | 0.00000 |

| | | | | | | | | | | | | | | |
|---|---|---|---|---|---|---|---|---|---|---|---|---|---|---|
| **Acaa1a** | 0.00000 | 0.03340 | 1.41853 | 4.29821 | 0.40695 | 3.47896 | 1.85460 | 2.48860 | 0.00000 | 2.16498 | 0.46922 | 4.16858 | 0.03943 | 0.12933 |
| **Acaa1b** | 0.00000 | 0.01886 | 0.61969 | 1.98317 | 0.13501 | 1.25767 | 0.81016 | 0.94125 | 0.00421 | 0.72265 | 0.19569 | 2.45351 | 0.04369 | 0.04295 |
| **Acaa2** | 0.00000 | 8.88551 | 3.82258 | 7.33785 | 0.76486 | 10.46230 | 6.91881 | 13.65138 | 0.00000 | 17.90667 | 8.46090 | 13.08083 | 0.00000 | 0.14223 |
| **Acaca** | 1.76509 | 0.02208 | 5.83538 | 2.54004 | 1.46047 | 3.37296 | 0.00137 | 1.69243 | 0.02407 | 0.00000 | 2.18946 | 4.15803 | 1.42152 | 2.92031 |
| **Acacb** | 1.47943 | 0.00198 | 8.18387 | 0.02895 | 6.74824 | 2.02018 | 0.00195 | 0.14627 | 0.32581 | 0.02723 | 0.18492 | 2.29536 | 0.75510 | 2.46263 |
| **Acad10** | 0.12465 | 0.01851 | 1.01553 | 0.00000 | 0.13930 | 1.95422 | 0.07732 | 1.66394 | 0.00796 | 0.00961 | 0.49981 | 1.75974 | 0.50841 | 0.42632 |
| **Acad11** | 3.41623 | 0.00737 | 3.62926 | 0.55400 | 4.38819 | 0.72487 | 3.08924 | 0.36209 | 3.30280 | 0.07796 | 1.04300 | 4.41928 | 3.02002 | 1.61077 |
| **Acad12** | 0.00785 | 0.04052 | 0.88110 | 0.26051 | 0.00000 | 1.64767 | 0.39626 | 0.76899 | 0.00218 | 0.01547 | 2.07226 | 1.38175 | 0.03896 | 0.06600 |
| **Acad8** | 5.73719 | 0.08602 | 13.50625 | 11.86057 | 4.16728 | 6.46437 | 3.80532 | 3.26625 | 6.89030 | 0.14563 | 8.30161 | 4.72665 | 6.20969 | 5.51279 |
| **Acad9** | 3.82197 | 0.74848 | 5.14858 | 4.85270 | 2.62961 | 4.93216 | 3.04890 | 3.21584 | 1.86995 | 0.08194 | 0.01414 | 5.87415 | 0.83592 | 4.83840 |
| **Acadl** | 19.28450 | 1.03959 | 21.49580 | 17.90940 | 14.05618 | 21.16297 | 7.63493 | 15.16223 | 15.97096 | 9.92633 | 10.47110 | 14.49806 | 1.12113 | 12.86665 |
| **Acadm** | 16.59129 | 4.95578 | 27.94025 | 15.87038 | 20.48246 | 21.92681 | 7.93413 | 12.74883 | 24.53441 | 12.18669 | 15.63664 | 9.57446 | 1.93396 | 24.58260 |
| **Acads** | 0.69867 | 2.81709 | 5.33662 | 2.34937 | 8.21336 | 1.24866 | 0.44014 | 0.80729 | 0.00285 | 0.93750 | 2.02068 | 0.80626 | 0.00000 | 2.11361 |
| **Acadsb** | 0.15996 | 0.01005 | 0.00000 | 0.00000 | 0.50868 | 1.77135 | 0.00352 | 0.02888 | 0.00836 | 0.00401 | 0.68474 | 1.46185 | 0.53142 | 0.92277 |
| **Acadvl** | 0.00000 | 0.15360 | 5.83340 | 11.25387 | 1.58883 | 12.08617 | 2.16282 | 9.17327 | 1.42975 | 0.16410 | 13.23174 | 12.24504 | 0.00000 | 1.16987 |
| **Acan** | 0.05738 | 0.00430 | 0.01224 | 0.08156 | 0.03423 | 0.01206 | 0.00441 | 0.00884 | 0.00810 | 0.00637 | 0.00745 | 0.01149 | 0.00462 | 0.04213 |
| **Acap1** | 0.01140 | 0.00445 | 0.07238 | 0.01098 | 0.17102 | 0.01103 | 0.00425 | 0.01313 | 0.00986 | 0.00811 | 0.00838 | 0.00887 | 0.00772 | 0.01625 |
| **Acap2** | 0.71963 | 0.02011 | 0.72608 | 0.26163 | 0.27640 | 1.48275 | 0.00264 | 0.44090 | 0.92475 | 0.01966 | 0.00000 | 4.24884 | 0.00400 | 0.00000 |
| **Acap3** | 0.77096 | 8.22226 | 3.19729 | 3.12663 | 0.78938 | 0.44566 | 0.02466 | 3.31300 | 0.00000 | 0.03170 | 0.00000 | 0.80859 | 0.82821 | 0.00000 |
| **Acat1** | 0.00000 | 0.01681 | 4.40990 | 4.07150 | 2.51738 | 2.15512 | 1.54442 | 3.26146 | 5.89585 | 0.00582 | 5.41420 | 2.60611 | 1.81490 | 8.77428 |
| **Acat2** | 4.47420 | 0.13946 | 4.58956 | 1.93424 | 6.32927 | 9.40980 | 4.14822 | 5.64084 | 5.29879 | 0.16103 | 4.58745 | 6.19030 | 3.05343 | 3.44613 |
| **Acat3** | 0.04808 | 0.01100 | 0.54227 | 0.00000 | 0.35744 | 0.88618 | 1.21169 | 0.23731 | 0.30093 | 0.00821 | 1.52510 | 0.69099 | 0.36535 | 0.36175 |
| **Acbd3** | 6.28896 | 0.02346 | 4.64493 | 0.00000 | 4.95937 | 3.93857 | 3.54778 | 0.33834 | 10.53441 | 0.07494 | 7.94188 | 1.75211 | 2.82262 | 6.52649 |
| **Acbd4** | 0.70084 | 0.03924 | 1.49638 | 0.80655 | 1.08316 | 1.73965 | 0.01047 | 0.78337 | 0.00721 | 0.00578 | 0.94566 | 0.29654 | 0.00249 | 1.14188 |
| **Acbd5** | 6.03828 | 1.89326 | 5.79861 | 2.19364 | 6.33876 | 2.05437 | 0.00000 | 0.02769 | 15.22578 | 0.00000 | 1.64781 | 2.27099 | 6.35873 | 7.59988 |
| **Acbd6** | 0.82672 | 0.04996 | 2.50734 | 5.44635 | 5.86745 | 1.51386 | 2.79584 | 1.92783 | 0.41533 | 0.21056 | 1.55131 | 3.24355 | 2.66572 | 0.48844 |
| **Acbd7** | 0.00742 | 0.00445 | 0.01523 | 0.00000 | 0.11224 | 0.18098 | 0.00411 | 0.00573 | 0.01230 | 0.01317 | 0.00767 | 0.01089 | 0.15414 | 0.00045 |
| **Accs** | 0.01542 | 0.00388 | 0.84371 | 0.54130 | 1.47093 | 0.19484 | 0.00352 | 0.00000 | 0.09359 | 0.00154 | 1.55039 | 0.00480 | 0.19011 | 0.10577 |
| **Accsl** | 0.35439 | 0.00287 | 0.46091 | 0.00000 | 0.76337 | 0.00000 | 0.00365 | 0.29997 | 0.06886 | 0.00630 | 0.00416 | 0.00000 | 8.31023 | 5.72888 |
| **Acd** | 4.63471 | 5.41020 | 9.21519 | 13.36347 | 7.05253 | 2.57738 | 0.02164 | 7.57197 | 0.37361 | 0.09183 | 5.15651 | 3.19837 | 2.40451 | 2.26146 |
| **Ace** | 0.01178 | 0.00476 | 0.00293 | 0.10119 | 0.28447 | 0.01223 | 0.12942 | 0.04367 | 0.01081 | 0.03476 | 0.03346 | 0.00712 | 0.01145 | 0.05660 |
| **Ace2** | 0.01165 | 0.00443 | 0.01117 | 0.01177 | 0.04210 | 0.01190 | 0.00463 | 0.01013 | 0.01794 | 0.00654 | 0.00812 | 0.00955 | 0.02440 | 0.26594 |
| **Acer1** | 0.00000 | 2.37826 | 0.75807 | 0.26977 | 0.00000 | 0.50218 | 3.40495 | 5.90514 | 0.39676 | 0.26705 | 0.00000 | 0.83804 | 0.98442 | 1.21398 |
| **Acer2** | 7.79478 | 0.16882 | 5.54224 | 7.87111 | 10.75242 | 5.65619 | 2.71706 | 4.69086 | 9.59813 | 8.02941 | 1.78261 | 4.72456 | 2.15828 | 8.52473 |
| **Acer3** | 0.77296 | 0.27332 | 2.41920 | 3.08661 | 1.32201 | 0.26714 | 1.96195 | 0.77194 | 4.00041 | 0.07347 | 1.24582 | 1.94889 | 0.17631 | 0.87362 |
| **Acin1** | 0.00000 | 0.08031 | 3.40118 | 2.99805 | 8.53640 | 4.18054 | 0.75868 | 4.13430 | 0.97723 | 0.07028 | 0.31523 | 4.69398 | 0.00000 | 4.47100 |
| **Ackr4** | 0.01030 | 0.00444 | 0.00890 | 0.01671 | 0.18692 | 0.01228 | 0.00419 | 0.01002 | 0.00919 | 0.00794 | 0.00903 | 0.00837 | 0.01013 | 0.06137 |
| **Acly** | 4.42622 | 0.75590 | 18.37811 | 4.83775 | 9.96352 | 11.53543 | 4.83699 | 6.19486 | 8.91270 | 0.14490 | 8.07956 | 12.38405 | 2.63207 | 7.75384 |
| **Acmsd** | 0.08482 | 0.07881 | 0.04768 | 0.03925 | 0.08442 | 0.07651 | 0.00420 | 0.01373 | 0.28330 | 0.04506 | 0.01777 | 0.03914 | 0.57439 | 0.16176 |
| **Acnat1** | 0.10241 | 0.06938 | 0.09656 | 0.09551 | 0.21289 | 0.04373 | 0.00449 | 0.04634 | 0.93521 | 0.02011 | 0.08438 | 0.09008 | 0.15682 | 1.84002 |
| **Aco1** | 1.73704 | 0.06874 | 5.57010 | 8.10690 | 0.41463 | 3.44612 | 1.65220 | 0.16047 | 1.33507 | 10.59932 | 3.90958 | 11.19331 | 0.47904 | 3.01695 |
| **Aco2** | 8.07093 | 7.44635 | 20.44811 | 16.70974 | 5.86855 | 21.68620 | 5.20004 | 12.45634 | 0.76729 | 3.01916 | 14.41423 | 13.79295 | 0.42094 | 9.02231 |
| **Acot1** | 0.00469 | 0.00384 | 0.57819 | 0.08281 | 0.45806 | 0.00201 | 0.00394 | 0.00223 | 0.00842 | 0.00422 | 0.12421 | 0.30076 | 0.03283 | 0.05238 |
| **Acot10** | 1.12165 | 0.01832 | 0.88455 | 0.36814 | 0.51237 | 1.17629 | 0.62968 | 0.48913 | 1.31847 | 0.02263 | 0.72367 | 0.83070 | 0.18749 | 0.52388 |
| **Acot11** | 0.28014 | 0.01048 | 1.40801 | 1.50946 | 1.50064 | 0.14649 | 0.00334 | 0.14571 | 1.90975 | 0.00195 | 0.87412 | 0.13441 | 0.31558 | 0.78525 |
| **Acot13** | 2.63744 | 0.01768 | 0.85089 | 0.61608 | 3.60414 | 1.89793 | 0.00911 | 0.07580 | 0.37118 | 10.61165 | 0.02235 | 1.06446 | 2.15225 | 1.94382 |
| **Acot2** | 0.00522 | 0.00424 | 0.10709 | 0.05019 | 0.13988 | 1.37509 | 0.14258 | 1.64038 | 0.00652 | 0.02759 | 0.00082 | 0.15217 | 0.01508 | 0.03003 |
| **Acot3** | 0.01513 | 0.00437 | 0.26362 | 0.00000 | 0.07876 | 0.00615 | 0.00414 | 0.01132 | 0.00794 | 0.00732 | 0.00677 | 0.01066 | 0.00793 | 0.09372 |
| **Acot5** | 0.00662 | 0.00436 | 0.07899 | 0.01420 | 0.17935 | 0.00945 | 0.00440 | 0.01056 | 0.00918 | 0.02757 | 0.00752 | 0.00848 | 0.00790 | 0.01469 |
| **Acot7** | 6.90073 | 0.05013 | 9.97546 | 8.59300 | 4.48820 | 10.27681 | 0.02083 | 1.94697 | 0.48543 | 0.04840 | 3.27253 | 0.00000 | 3.60758 | 1.69375 |
| **Acot8** | 1.17280 | 0.07540 | 1.14333 | 4.48751 | 3.95309 | 5.68805 | 1.83476 | 5.00579 | 0.01358 | 0.11766 | 3.66406 | 2.30805 | 1.13930 | 0.33817 |
| **Acot9** | 7.69693 | 0.02034 | 7.78767 | 3.18045 | 4.11754 | 8.05868 | 3.84790 | 3.71380 | 7.01839 | 0.09504 | 4.77904 | 6.63690 | 0.78793 | 4.98695 |
| **Acox1** | 2.90448 | 0.56630 | 5.94242 | 3.72331 | 1.66169 | 3.37574 | 0.39163 | 4.65052 | 0.15937 | 0.03065 | 9.11302 | 5.50142 | 6.02342 | 3.29954 |
| **Acox3** | 0.01017 | 0.00354 | 0.04859 | 0.00000 | 0.00000 | 0.00000 | 0.74321 | 0.02320 | 0.00182 | 0.02416 | 0.87797 | 0.01871 | 2.12726 | 0.12934 |
| **Acoxl** | 0.61889 | 0.00412 | 0.48923 | 0.12822 | 1.27970 | 0.00000 | 0.00385 | 0.00609 | 0.00751 | 0.00623 | 0.00798 | 0.01388 | 0.06268 | 0.24419 |

| | | | | | | | | | | | | | | |
|---|---|---|---|---|---|---|---|---|---|---|---|---|---|---|
| **Acp1** | 5.46147 | 1.78920 | 13.81964 | 16.29740 | 11.74942 | 18.22068 | 4.40045 | 11.11015 | 7.26118 | 0.41807 | 8.47865 | 15.14717 | 0.00000 | 8.41663 |
| **Acp2** | 0.18217 | 0.05357 | 1.10143 | 2.04501 | 0.94231 | 1.74474 | 0.00333 | 0.27969 | 0.21748 | 0.00000 | 1.66591 | 3.13289 | 0.08799 | 2.10315 |
| **Acp5** | 0.00000 | 0.05883 | 11.17206 | 10.48274 | 5.38465 | 6.17840 | 0.00931 | 3.75852 | 0.16613 | 8.83117 | 2.18956 | 6.00652 | 0.00000 | 0.93368 |
| **Acp6** | 0.59268 | 0.01923 | 1.69514 | 1.96131 | 0.08914 | 1.79684 | 2.83555 | 2.36962 | 0.40660 | 0.05880 | 0.33072 | 0.65025 | 0.00012 | 0.24077 |
| **Acp7** | 0.02112 | 0.01220 | 0.02438 | 0.00862 | 0.22366 | 0.01239 | 0.00425 | 0.00887 | 0.00881 | 0.00594 | 0.00741 | 0.04851 | 0.01183 | 0.11966 |
| **Acpp** | 4.24853 | 0.02515 | 5.91856 | 0.24783 | 4.03077 | 2.52140 | 0.19706 | 0.33479 | 6.85349 | 0.02567 | 3.42683 | 2.89668 | 3.05733 | 11.27781 |
| **Acrbp** | 0.00000 | 0.01680 | 1.42736 | 0.00000 | 0.29783 | 1.89543 | 0.02933 | 0.00000 | 0.51232 | 12.98307 | 0.26032 | 0.60110 | 0.00373 | 0.00410 |
| **Acsbg1** | 2.07856 | 0.01929 | 8.01238 | 2.67811 | 1.53221 | 3.77714 | 0.95011 | 0.59387 | 0.00000 | 0.04404 | 2.56789 | 2.29890 | 0.00000 | 0.03460 |
| **Acsbg2** | 2.92861 | 0.00370 | 2.71302 | 1.17899 | 3.19137 | 0.42472 | 0.00339 | 0.00000 | 0.52075 | 0.02467 | 0.00000 | 0.00275 | 0.00000 | 2.27198 |
| **Acsf2** | 0.82850 | 0.83540 | 1.10953 | 2.39847 | 0.64002 | 1.35966 | 0.01634 | 2.66801 | 0.01263 | 0.02086 | 0.00000 | 0.27935 | 0.00000 | 0.54548 |
| **Acsf3** | 0.43767 | 0.02002 | 0.00000 | 1.19061 | 0.01696 | 0.79811 | 0.00349 | 2.68921 | 0.00424 | 0.01079 | 4.92517 | 1.70938 | 0.00684 | 0.00000 |
| **Acsl1** | 0.52106 | 0.00335 | 0.10076 | 1.08674 | 0.50851 | 1.60893 | 0.00329 | 0.12104 | 1.46232 | 0.00000 | 2.41959 | 0.40551 | 0.89010 | 1.47497 |
| **Acsl3** | 1.37022 | 0.09579 | 6.91112 | 1.90961 | 5.57564 | 8.12466 | 2.23466 | 1.93673 | 18.02800 | 0.03623 | 5.16998 | 8.48190 | 9.21496 | 10.25511 |
| **Acsl4** | 25.77437 | 0.00000 | 11.48557 | 0.00000 | 15.95908 | 4.39869 | 0.00143 | 0.00000 | 35.26430 | 0.00000 | 4.74972 | 0.61127 | 7.13105 | 22.13160 |
| **Acsl5** | 0.00970 | 0.02664 | 0.00820 | 0.00000 | 0.01889 | 0.23284 | 0.00359 | 0.02999 | 0.01287 | 0.00526 | 0.00097 | 1.58187 | 0.00706 | 0.01063 |
| **Acsl6** | 0.00837 | 0.03639 | 0.20103 | 0.00710 | 0.22596 | 3.12390 | 0.00373 | 2.94183 | 0.00418 | 0.02097 | 0.03085 | 0.00000 | 0.00912 | 0.63592 |
| **Acsm2** | 0.02567 | 0.00472 | 0.01984 | 0.03042 | 0.03292 | 0.03399 | 0.00430 | 0.01977 | 0.09638 | 0.00701 | 0.01521 | 0.01048 | 0.15579 | 0.02552 |
| **Acsm3** | 0.05082 | 0.00439 | 0.21991 | 1.01295 | 0.00000 | 2.08224 | 0.59881 | 0.28527 | 0.12089 | 0.01345 | 0.03752 | 0.00000 | 0.00295 | 1.05534 |
| **Acss1** | 0.42241 | 6.25246 | 1.00991 | 0.91468 | 0.00000 | 4.36597 | 0.89992 | 2.68439 | 0.95775 | 0.03708 | 0.00000 | 2.77377 | 0.00000 | 0.99385 |
| **Acss2** | 0.72580 | 0.01553 | 1.45779 | 1.85716 | 1.70977 | 3.12866 | 0.37088 | 0.00000 | 0.91625 | 0.02069 | 0.00000 | 3.61230 | 2.01973 | 1.03019 |
| **Acta1** | 0.00899 | 1.15165 | 0.89697 | 1.42914 | 0.15562 | 1.84566 | 0.77824 | 2.20298 | 0.00625 | 1.01671 | 1.34119 | 0.84795 | 0.00569 | 0.06408 |
| **Acta2** | 1.85106 | 1.12012 | 0.58884 | 1.42382 | 1.54459 | 2.50290 | 0.87825 | 1.99721 | 0.03157 | 0.64108 | 1.45790 | 1.02756 | 0.00001 | 0.01990 |
| **Actb** | 4.70062 | 14.96883 | 10.33005 | 27.36457 | 15.01016 | 43.70960 | 10.04883 | 38.88575 | 18.17081 | 21.81934 | 11.02979 | 19.67170 | 4.57336 | 12.92872 |
| **Actg1** | 18.98980 | 23.77610 | 50.79859 | 62.78916 | 30.03686 | 66.24350 | 21.03315 | 50.59086 | 13.20551 | 32.71191 | 43.12775 | 44.67875 | 10.26227 | 14.83077 |
| **Actg2** | 0.71789 | 0.00386 | 0.00221 | 0.00354 | 0.46260 | 0.01122 | 0.00424 | 0.00819 | 0.01008 | 0.00703 | 0.00570 | 0.68666 | 0.00357 | 0.00378 |
| **Actl6a** | 9.76577 | 9.42627 | 15.50627 | 11.35715 | 13.19809 | 10.49827 | 0.32516 | 6.52339 | 6.14645 | 0.05165 | 6.37258 | 10.99066 | 5.12748 | 14.81767 |
| **Actn1** | 1.99258 | 0.01958 | 0.07528 | 1.36610 | 0.68426 | 1.76310 | 0.08436 | 1.24938 | 0.04454 | 0.02538 | 0.07964 | 1.44992 | 1.63224 | 0.00000 |
| **Actn2** | 11.57529 | 0.04633 | 25.73089 | 20.45203 | 12.87059 | 5.28377 | 3.39420 | 0.29740 | 2.03829 | 0.15128 | 1.25008 | 0.00869 | 2.00792 | 9.50190 |
| **Actn4** | 6.47605 | 12.83003 | 33.53818 | 16.95047 | 12.47880 | 28.50184 | 8.34258 | 25.17064 | 0.00000 | 18.22562 | 20.53501 | 9.18050 | 0.33148 | 4.45775 |
| **Actr10** | 10.64699 | 0.21446 | 12.21744 | 8.86105 | 10.40081 | 11.40470 | 3.10089 | 2.06385 | 8.91954 | 0.30240 | 7.52484 | 5.51477 | 4.86838 | 10.42285 |
| **Actr1a** | 7.84921 | 0.11032 | 11.84212 | 4.73071 | 8.49160 | 10.50349 | 3.14866 | 8.74317 | 3.45791 | 0.19701 | 7.10323 | 7.89595 | 3.87781 | 4.02732 |
| **Actr1b** | 3.28927 | 0.09546 | 8.05864 | 4.71714 | 1.55260 | 4.38334 | 0.06978 | 7.14943 | 0.47186 | 0.05874 | 6.21443 | 6.22268 | 0.00000 | 3.20727 |
| **Actr2** | 9.59642 | 0.64956 | 6.53821 | 6.22205 | 4.15407 | 10.77962 | 2.33198 | 5.08081 | 9.78786 | 0.21610 | 3.01389 | 4.42841 | 3.80805 | 7.70169 |
| **Actr3** | 9.19565 | 11.17990 | 16.79422 | 14.98164 | 8.31082 | 25.52959 | 5.05032 | 19.40188 | 14.30378 | 9.75782 | 21.78369 | 11.92074 | 6.27341 | 10.08937 |
| **Actr3b** | 0.00707 | 0.01065 | 0.23965 | 0.92406 | 0.00000 | 0.52166 | 1.93360 | 0.00266 | 0.00724 | 0.03736 | 0.03250 | 0.23628 | 0.01858 | 0.75500 |
| **Actr5** | 1.79933 | 0.03415 | 7.09495 | 1.06672 | 13.30648 | 0.17251 | 2.42659 | 0.58937 | 0.00000 | 0.01734 | 0.22373 | 2.18175 | 3.53866 | 1.01807 |
| **Actr6** | 1.87707 | 0.00220 | 1.63808 | 0.25908 | 0.16487 | 3.85397 | 0.00155 | 0.03361 | 12.25119 | 0.00996 | 4.05887 | 0.18969 | 3.46625 | 9.08142 |
| **Actr8** | 4.93183 | 0.09622 | 6.79517 | 2.68328 | 3.83481 | 2.55518 | 0.00000 | 4.06825 | 4.78011 | 0.67033 | 5.79882 | 3.86822 | 0.00000 | 3.58377 |
| **Acvr1** | 0.14855 | 0.00379 | 0.73227 | 0.04001 | 0.00000 | 0.05906 | 0.00336 | 0.01042 | 3.11105 | 0.00567 | 0.09247 | 0.12311 | 0.00295 | 1.56375 |
| **Acvr1b** | 0.00983 | 4.02726 | 0.00144 | 0.24759 | 0.03146 | 0.30199 | 1.65454 | 0.03628 | 0.00934 | 0.03327 | 0.00197 | 0.04260 | 0.00986 | 0.00653 |
| **Acvr2a** | 0.49178 | 0.01078 | 2.13762 | 0.17322 | 0.38734 | 2.65431 | 1.85319 | 0.07260 | 1.96337 | 0.03030 | 3.12274 | 0.72120 | 1.15778 | 1.71971 |
| **Acvr2b** | 0.30642 | 0.04466 | 1.68023 | 1.17712 | 0.64253 | 3.22917 | 0.23678 | 3.36912 | 0.41991 | 0.03739 | 0.11772 | 1.98668 | 0.20727 | 0.54605 |
| **Acvrl1** | 0.01934 | 0.00446 | 0.11937 | 0.00442 | 0.00851 | 0.00888 | 0.00426 | 0.01145 | 0.01288 | 0.00667 | 0.00776 | 0.00935 | 0.03714 | 0.05803 |
| **Acy1** | 0.52835 | 0.03505 | 0.54989 | 2.67903 | 0.10062 | 0.10637 | 0.25339 | 4.09499 | 0.17453 | 0.01723 | 0.00000 | 0.27310 | 0.42272 | 1.69952 |
| **Acy3** | 0.04111 | 0.00405 | 0.54777 | 1.38232 | 0.03717 | 0.96744 | 0.00392 | 0.00191 | 0.00348 | 0.01248 | 0.03713 | 0.00464 | 1.62909 | 0.00000 |
| **Acyp1** | 0.67819 | 0.01210 | 0.00489 | 0.05795 | 1.50001 | 0.86279 | 0.01500 | 1.11201 | 0.12403 | 0.01833 | 0.19437 | 0.39469 | 0.58573 | 2.14566 |
| **Acyp2** | 0.09091 | 0.01328 | 0.05562 | 0.00000 | 0.38680 | 0.19531 | 0.00400 | 0.02701 | 0.18514 | 0.00773 | 0.00969 | 0.26084 | 0.03479 | 0.06518 |
| **Ada** | 0.03071 | 0.00443 | 0.24847 | 0.01296 | 0.04789 | 1.15483 | 0.00403 | 0.00390 | 0.00348 | 0.00716 | 0.00453 | 0.00331 | 0.77920 | 0.10609 |
| **Adad2** | 0.01248 | 0.13884 | 0.78071 | 1.02431 | 0.00543 | 0.00000 | 0.00982 | 0.61753 | 0.00657 | 0.01525 | 0.89520 | 0.01219 | 0.14294 | 0.00000 |
| **Adal** | 1.65982 | 0.00268 | 0.74321 | 1.00059 | 1.12496 | 1.62718 | 0.00339 | 0.72487 | 1.48536 | 0.01109 | 0.00000 | 2.15947 | 0.00000 | 0.58488 |
| **Adam10** | 3.74308 | 0.38186 | 3.08807 | 1.84530 | 2.93328 | 4.60676 | 0.00482 | 0.95946 | 18.69429 | 0.00000 | 3.52106 | 5.45649 | 3.03448 | 8.09757 |
| **Adam12** | 0.06045 | 0.00448 | 0.00326 | 0.00709 | 0.67814 | 0.01330 | 0.00499 | 0.01047 | 0.01336 | 0.00643 | 0.00859 | 0.00767 | 0.01781 | 0.01549 |
| **Adam15** | 0.16284 | 0.05695 | 1.44531 | 2.64536 | 0.51597 | 0.88518 | 0.76349 | 2.37097 | 0.00425 | 0.00746 | 0.55945 | 2.07629 | 0.01644 | 0.00000 |
| **Adam17** | 10.14045 | 0.08766 | 9.79392 | 2.55695 | 8.63909 | 2.79379 | 0.00525 | 3.93190 | 9.05291 | 0.05187 | 2.36217 | 2.41579 | 8.67119 | 12.93409 |
| **Adam19** | 0.00538 | 0.03674 | 0.20297 | 0.38030 | 0.20913 | 2.52037 | 1.92285 | 2.47850 | 0.01508 | 5.58750 | 1.00404 | 7.37180 | 0.98061 | 0.97865 |

| | | | | | | | | | | | | | | |
|---|---|---|---|---|---|---|---|---|---|---|---|---|---|---|
| **Adam1a** | 0.52133 | 0.00399 | 0.02406 | 0.32520 | 0.21492 | 0.00000 | 0.00920 | 0.00559 | 0.40866 | 0.00611 | 0.01083 | 0.01371 | 0.25194 | 0.11103 |
| **Adam2** | 0.00000 | 0.00400 | 0.20527 | 0.07217 | 0.00194 | 0.00000 | 0.00428 | 0.00686 | 0.17562 | 0.00602 | 0.07460 | 0.09270 | 0.00830 | 0.61022 |
| **Adam22** | 0.01165 | 0.00436 | 0.00952 | 0.09902 | 0.00000 | 0.00538 | 0.00401 | 0.00782 | 0.14132 | 0.00568 | 0.01350 | 0.02495 | 0.05095 | 0.25736 |
| **Adam28** | 0.12300 | 0.08013 | 0.05367 | 0.32686 | 0.12713 | 0.29165 | 0.05921 | 0.09965 | 0.43770 | 0.10236 | 0.02263 | 0.09716 | 0.41726 | 0.29603 |
| **Adam33** | 0.00439 | 0.00432 | 0.01380 | 0.01222 | 0.07533 | 0.01311 | 0.00436 | 0.01066 | 0.00890 | 0.00635 | 0.00896 | 0.00943 | 0.30547 | 0.00392 |
| **Adam4** | 0.07546 | 0.00406 | 0.00872 | 0.14897 | 0.01567 | 0.00722 | 0.00393 | 0.00599 | 0.00854 | 0.00655 | 0.00682 | 0.01295 | 0.62908 | 0.05049 |
| **Adam8** | 0.31200 | 0.00386 | 0.00660 | 0.00000 | 0.14376 | 0.15582 | 0.00401 | 0.16967 | 0.00862 | 0.00548 | 0.00413 | 0.00812 | 0.14065 | 0.00071 |
| **Adam9** | 1.38907 | 0.04048 | 0.55825 | 0.61208 | 0.00000 | 1.65450 | 0.55618 | 0.06874 | 1.27413 | 0.01182 | 0.37684 | 1.75350 | 0.47594 | 0.52761 |
| **Adamts10** | 0.00000 | 0.00412 | 0.85232 | 1.40298 | 0.58549 | 0.00000 | 0.00353 | 0.00158 | 0.01252 | 0.00326 | 0.00868 | 0.52702 | 0.00154 | 0.00000 |
| **Adamts12** | 0.20119 | 0.00459 | 0.01277 | 0.01339 | 0.00892 | 0.01001 | 0.00430 | 0.01446 | 0.01671 | 0.00678 | 0.00730 | 0.00980 | 0.01401 | 0.01063 |
| **Adamts13** | 9.00925 | 0.08752 | 2.22754 | 0.69828 | 5.73575 | 3.53453 | 0.80438 | 0.95952 | 3.37083 | 2.88681 | 1.26434 | 0.98847 | 4.28550 | 1.84387 |
| **Adamts14** | 0.11112 | 0.00429 | 0.31746 | 0.00000 | 0.62626 | 0.00008 | 0.00396 | 0.01302 | 0.02399 | 0.00635 | 0.00542 | 0.00703 | 0.00165 | 0.98472 |
| **Adamts15** | 0.00661 | 0.00436 | 0.01157 | 0.00109 | 0.01781 | 0.36273 | 0.00418 | 0.00458 | 0.01110 | 0.00544 | 0.00765 | 0.01077 | 0.13897 | 0.01065 |
| **Adamts17** | 0.01445 | 0.00438 | 0.01389 | 0.03415 | 0.03992 | 0.01958 | 0.00432 | 0.01477 | 0.01776 | 0.01624 | 0.01059 | 0.01219 | 0.03106 | 0.01441 |
| **Adamts2** | 1.22816 | 0.06583 | 0.00000 | 0.38931 | 1.53918 | 0.10341 | 0.14770 | 0.00881 | 4.39870 | 0.06783 | 0.06181 | 0.07434 | 9.84169 | 6.89584 |
| **Adamts4** | 0.40659 | 0.02098 | 0.00000 | 0.07095 | 0.03733 | 0.29617 | 0.04709 | 0.04158 | 0.39199 | 0.01636 | 0.01787 | 0.03669 | 1.47111 | 0.65763 |
| **Adamts5** | 0.07339 | 0.00395 | 0.01226 | 0.00000 | 0.08165 | 0.00963 | 0.00386 | 0.39441 | 0.01035 | 0.00667 | 0.00659 | 0.01017 | 0.00550 | 0.01108 |
| **Adamts7** | 0.01051 | 0.00383 | 0.32111 | 0.09288 | 0.00055 | 0.18857 | 0.00412 | 0.03720 | 0.01210 | 0.02049 | 0.36696 | 0.52163 | 0.00650 | 0.00415 |
| **Adamtsl1** | 0.00295 | 0.00426 | 0.00000 | 0.01151 | 0.81622 | 0.01190 | 0.00414 | 0.00789 | 0.00986 | 0.00606 | 0.00819 | 0.00690 | 0.84471 | 0.51319 |
| **Adamtsl4** | 0.18142 | 0.00411 | 1.46402 | 0.19517 | 1.21239 | 0.00000 | 0.00369 | 0.00529 | 0.05313 | 0.00328 | 0.09040 | 0.00825 | 0.10705 | 0.70280 |
| **Adamtsl5** | 0.82752 | 0.00446 | 0.61470 | 1.74198 | 0.89452 | 0.00000 | 0.00398 | 0.00873 | 0.34354 | 0.00614 | 0.00240 | 0.00767 | 0.00000 | 0.00000 |
| **Adap1** | 0.56264 | 0.00414 | 0.68074 | 0.64719 | 0.62478 | 0.96993 | 0.00417 | 0.45509 | 0.00306 | 0.02871 | 0.87896 | 0.00000 | 0.01627 | 0.14147 |
| **Adap2** | 0.00000 | 0.00419 | 0.22125 | 0.22394 | 0.22134 | 1.67712 | 0.00405 | 0.22289 | 0.54695 | 0.00718 | 0.16026 | 0.14579 | 0.00358 | 0.47318 |
| **Adar** | 0.15064 | 0.02725 | 3.53173 | 4.71945 | 0.53133 | 2.10998 | 0.00299 | 0.21472 | 2.01593 | 0.01342 | 0.98301 | 1.13256 | 0.72381 | 2.54985 |
| **Adarb1** | 0.29015 | 0.05051 | 0.52950 | 1.41942 | 0.47949 | 1.24438 | 0.08049 | 0.43196 | 0.32460 | 0.23069 | 0.03197 | 0.22784 | 1.13832 | 1.93281 |
| **Adarb2** | 3.42607 | 2.03625 | 1.81133 | 3.65508 | 2.38069 | 3.42510 | 1.42067 | 1.26705 | 2.16512 | 2.54341 | 1.46650 | 5.59550 | 4.80614 | 1.79411 |
| **Adat1** | 0.26672 | 0.00411 | 1.47031 | 0.50794 | 0.14332 | 0.46651 | 0.00442 | 0.21817 | 0.72030 | 0.01218 | 0.00000 | 0.84370 | 0.00948 | 2.07067 |
| **Adat2** | 0.47501 | 0.22041 | 0.04397 | 1.76925 | 0.73866 | 0.67899 | 0.02267 | 0.55913 | 0.00611 | 8.42103 | 0.54400 | 2.62811 | 0.00872 | 0.58276 |
| **Adat3** | 0.46113 | 0.01757 | 0.03080 | 2.02567 | 1.97774 | 0.00000 | 0.00386 | 0.00930 | 0.00481 | 0.01252 | 0.40112 | 0.49530 | 0.74354 | 0.00000 |
| **Adck1** | 2.34313 | 0.00859 | 2.51287 | 3.62437 | 1.79651 | 2.77857 | 0.00780 | 0.00000 | 1.06158 | 3.96650 | 2.04102 | 2.64268 | 0.00182 | 3.99632 |
| **Adck2** | 0.00338 | 0.01241 | 0.17589 | 0.02238 | 0.00000 | 0.00000 | 0.02292 | 1.66368 | 0.00915 | 0.02187 | 0.86280 | 0.28386 | 0.11139 | 0.88581 |
| **Adck5** | 1.17437 | 3.19555 | 3.57930 | 1.87875 | 2.34809 | 2.11030 | 0.01219 | 5.17764 | 0.00000 | 0.03187 | 0.33895 | 6.83316 | 0.00000 | 2.12572 |
| **Adcy1** | 0.00917 | 0.01219 | 0.07095 | 0.00682 | 0.00645 | 0.00644 | 0.00403 | 0.00942 | 0.56230 | 0.01063 | 0.00373 | 0.12973 | 0.00824 | 0.00000 |
| **Adcy10** | 0.00370 | 0.01547 | 0.05061 | 0.20150 | 0.02508 | 0.17331 | 0.00407 | 0.11615 | 0.19962 | 0.01106 | 0.00527 | 0.00955 | 0.00347 | 0.57640 |
| **Adcy3** | 0.13240 | 0.00485 | 0.22024 | 0.00519 | 0.00556 | 0.00000 | 0.00444 | 0.56080 | 0.02371 | 0.00400 | 0.87706 | 0.00564 | 0.00872 | 0.05996 |
| **Adcy5** | 0.01381 | 0.00436 | 0.00755 | 0.01063 | 0.23078 | 0.01325 | 0.00413 | 0.00912 | 0.01055 | 0.00669 | 0.00840 | 0.00836 | 0.27016 | 0.21082 |
| **Adcy6** | 0.86804 | 0.00435 | 0.29020 | 0.00759 | 0.23788 | 0.00000 | 0.00457 | 0.14362 | 2.24612 | 0.02094 | 0.00402 | 0.00334 | 0.00422 | 1.03776 |
| **Adcy7** | 0.01318 | 0.00429 | 0.01233 | 0.01208 | 0.00532 | 0.01056 | 0.00431 | 0.00925 | 0.01016 | 0.00612 | 0.00751 | 0.00877 | 0.32402 | 0.75500 |
| **Adcy9** | 1.11391 | 0.03511 | 2.42442 | 1.71284 | 2.97235 | 2.41572 | 0.00833 | 1.36222 | 1.23723 | 0.01177 | 2.01737 | 2.20585 | 0.45575 | 7.14718 |
| **Adcyap1r1** | 0.00000 | 0.00453 | 0.13884 | 0.00000 | 0.00528 | 0.29718 | 0.00392 | 0.01241 | 0.02120 | 0.00379 | 0.54643 | 0.49079 | 0.23428 | 0.00032 |
| **Add1** | 0.52337 | 0.07149 | 5.32043 | 7.92491 | 6.29392 | 8.33561 | 1.93533 | 2.25549 | 0.00000 | 1.58706 | 8.47173 | 8.41404 | 2.83679 | 1.47778 |
| **Add2** | 0.00138 | 0.02159 | 0.07531 | 0.00000 | 0.28423 | 0.00914 | 0.00385 | 0.01487 | 0.01125 | 0.00641 | 0.00454 | 0.38935 | 0.35453 | 0.00235 |
| **Add3** | 0.00000 | 0.01687 | 4.40256 | 0.81798 | 0.47643 | 1.04023 | 0.00265 | 0.01956 | 0.00223 | 0.00000 | 3.85651 | 2.47641 | 0.57873 | 0.00000 |
| **Adgra2** | 0.03481 | 0.00397 | 0.09921 | 0.03715 | 0.30083 | 0.03934 | 0.00402 | 0.06184 | 0.00190 | 0.00668 | 0.04944 | 0.00993 | 1.76958 | 0.39804 |
| **Adgra3** | 0.21570 | 0.00352 | 0.21164 | 0.00000 | 0.19524 | 0.59995 | 0.00400 | 1.83536 | 0.01039 | 0.00486 | 0.00000 | 0.81766 | 0.07414 | 0.86923 |
| **Adgre5** | 0.00000 | 0.05452 | 10.57075 | 6.37419 | 0.43059 | 9.12526 | 0.13510 | 4.36981 | 0.01497 | 0.05272 | 3.43494 | 6.33393 | 0.00000 | 0.00000 |
| **Adgrf3** | 0.10742 | 0.00406 | 0.71771 | 0.44150 | 0.78371 | 0.00000 | 0.00375 | 0.00484 | 0.18429 | 0.00470 | 0.01015 | 0.01361 | 0.00000 | 0.63487 |
| **Adgrg1** | 0.00000 | 0.00403 | 0.53074 | 1.63970 | 1.23128 | 0.30779 | 0.00389 | 0.00083 | 0.01907 | 0.01013 | 0.00788 | 0.00625 | 0.64596 | 0.02475 |
| **Adgrg2** | 0.00791 | 0.00451 | 0.05025 | 0.00000 | 0.47532 | 0.02840 | 0.00500 | 0.00933 | 0.02837 | 0.00689 | 0.00740 | 0.00910 | 0.00495 | 0.00194 |
| **Adgrg3** | 0.00290 | 0.00451 | 0.07189 | 0.00310 | 0.46923 | 0.01101 | 0.00478 | 0.01023 | 0.01079 | 0.00803 | 0.00730 | 0.00797 | 0.00233 | 0.30218 |
| **Adgrg6** | 0.00488 | 0.02797 | 0.01224 | 0.00809 | 0.01039 | 0.01538 | 0.00355 | 0.00812 | 0.01083 | 0.00662 | 0.00365 | 2.30625 | 0.00566 | 0.02162 |
| **Adgrl1** | 0.06189 | 0.02642 | 0.09002 | 0.29411 | 0.43231 | 1.29314 | 2.21821 | 0.64004 | 0.27201 | 0.06786 | 0.00000 | 1.75911 | 0.73385 | 0.19473 |
| **Adgrl2** | 0.01887 | 0.01169 | 0.72931 | 0.02479 | 0.00000 | 0.83813 | 0.00362 | 1.58891 | 0.00534 | 0.00983 | 0.00384 | 0.00417 | 0.00066 | 0.59419 |
| **Adh1** | 0.00497 | 11.98133 | 0.00335 | 0.00000 | 0.01389 | 0.01135 | 1.24073 | 24.00476 | 0.00896 | 0.80171 | 0.00000 | 0.68848 | 0.00403 | 0.00710 |
| **Adh5** | 4.15272 | 4.73608 | 8.59225 | 10.48666 | 4.33200 | 14.31077 | 5.93625 | 12.89653 | 4.84902 | 8.72097 | 9.65846 | 14.93239 | 2.66486 | 6.20969 |

| | | | | | | | | | | | | | | |
|---|---|---|---|---|---|---|---|---|---|---|---|---|---|---|
| **Adhfe1** | 0.46748 | 0.00450 | 0.28505 | 0.00000 | 0.00000 | 0.00054 | 0.00403 | 0.01991 | 0.00948 | 0.00652 | 0.02094 | 0.36159 | 0.00726 | 0.02424 |
| **Adi1** | 3.94288 | 0.32508 | 6.00650 | 13.50270 | 8.38174 | 5.35298 | 2.20015 | 4.22934 | 3.69098 | 8.71573 | 4.79509 | 5.62019 | 2.83899 | 1.73196 |
| **Adipor1** | 8.85195 | 0.19140 | 13.68496 | 8.49583 | 7.48724 | 11.64179 | 4.04367 | 8.41718 | 4.83498 | 0.11956 | 4.84222 | 10.24716 | 6.49025 | 9.48376 |
| **Adipor2** | 2.22568 | 0.16945 | 4.59455 | 0.00004 | 2.46884 | 7.72598 | 0.28076 | 9.36691 | 3.76002 | 0.10049 | 6.26072 | 7.35557 | 3.82196 | 4.70894 |
| **Adk** | 0.27725 | 0.05138 | 2.16095 | 0.00000 | 0.90429 | 2.40834 | 2.55783 | 5.25545 | 1.96390 | 0.05038 | 4.10709 | 1.20286 | 2.35686 | 2.17180 |
| **Adm** | 0.04234 | 0.01080 | 0.26756 | 0.27972 | 0.16461 | 0.00000 | 0.00382 | 0.00318 | 0.00760 | 0.00544 | 0.00723 | 0.75471 | 0.16911 | 0.00000 |
| **Adm2** | 0.02184 | 0.00442 | 0.01480 | 0.01196 | 0.01115 | 0.01262 | 0.00427 | 0.01026 | 0.01190 | 0.00652 | 0.00857 | 0.00983 | 0.06741 | 0.03762 |
| **Adnp2** | 0.93463 | 0.02364 | 1.70574 | 0.48520 | 0.77290 | 3.28989 | 1.50709 | 0.72032 | 0.89631 | 0.00988 | 5.18361 | 3.52800 | 0.00000 | 2.46451 |
| **Ado** | 2.62182 | 0.01850 | 0.63331 | 1.36513 | 1.05769 | 0.89160 | 0.06186 | 0.24848 | 1.54181 | 0.00964 | 0.69030 | 1.62557 | 0.00000 | 0.41775 |
| **Adora1** | 0.00305 | 0.03870 | 0.00000 | 0.86154 | 0.08873 | 0.50198 | 0.00400 | 0.84757 | 0.00711 | 0.00605 | 0.00000 | 1.83865 | 0.00566 | 0.07122 |
| **Adpgk** | 5.65115 | 4.96695 | 3.11399 | 2.62729 | 1.15539 | 4.41667 | 2.35967 | 0.02187 | 0.56875 | 0.07412 | 7.18683 | 3.49489 | 1.08983 | 4.28434 |
| **Adprh** | 6.84324 | 1.05196 | 7.83048 | 13.00637 | 6.81244 | 8.70980 | 0.89262 | 10.29472 | 0.42550 | 0.17658 | 6.70737 | 12.64079 | 0.22189 | 1.16830 |
| **Adprhl1** | 0.39887 | 0.00403 | 0.48996 | 0.02609 | 0.00000 | 0.00000 | 0.00393 | 0.00962 | 0.00880 | 0.01272 | 1.09467 | 0.00376 | 0.00400 | 0.00227 |
| **Adprhl2** | 0.99577 | 0.03355 | 6.04764 | 2.97979 | 3.92655 | 5.82982 | 0.98137 | 4.12751 | 0.00000 | 0.01670 | 4.15449 | 0.82121 | 0.00000 | 1.20391 |
| **Adprm** | 0.00000 | 0.00489 | 0.77344 | 1.84586 | 0.30439 | 0.32519 | 3.10196 | 0.61451 | 0.10542 | 0.04588 | 0.56999 | 0.97153 | 0.17663 | 0.71433 |
| **Adrb3** | 0.03112 | 0.01887 | 0.82355 | 1.93245 | 0.28527 | 0.62352 | 0.00349 | 0.00000 | 0.00448 | 0.01625 | 1.07438 | 0.03098 | 0.00000 | 0.26916 |
| **Adrbk1** | 0.83034 | 4.07239 | 10.94567 | 5.10751 | 12.15405 | 1.58939 | 1.92108 | 3.09125 | 0.19852 | 0.09139 | 2.37435 | 7.38957 | 0.38510 | 4.72656 |
| **Adrm1** | 3.60948 | 2.09531 | 15.50751 | 39.53402 | 33.89941 | 3.48496 | 6.47008 | 10.23953 | 0.29397 | 7.51254 | 2.03704 | 14.22311 | 9.65015 | 0.57331 |
| **Adsl** | 10.02218 | 1.44348 | 15.15918 | 6.00466 | 11.94800 | 9.34455 | 5.16883 | 0.54147 | 2.52035 | 8.33558 | 10.42941 | 5.89966 | 7.03495 | 9.82201 |
| **Adss** | 4.89863 | 4.12367 | 8.69198 | 8.65253 | 10.69698 | 13.05669 | 2.49169 | 1.64114 | 3.19685 | 0.10452 | 8.21375 | 5.69296 | 5.70520 | 7.27335 |
| **Adssl1** | 1.38198 | 0.00369 | 0.82206 | 0.00000 | 0.00000 | 0.10820 | 0.00375 | 0.00000 | 0.00485 | 0.00898 | 0.00323 | 0.29709 | 0.00536 | 0.45258 |
| **Adtrp** | 17.07417 | 0.02126 | 13.75241 | 9.30936 | 8.02837 | 6.57438 | 1.58187 | 0.00000 | 15.45969 | 0.07347 | 2.55547 | 0.97184 | 3.82128 | 9.00789 |
| **Aebp1** | 0.16944 | 0.00447 | 0.00000 | 0.00000 | 0.82020 | 0.43286 | 0.00417 | 0.00527 | 0.00389 | 0.00613 | 0.00389 | 0.02516 | 0.01077 | 0.24623 |
| **Aebp2** | 17.33134 | 2.14426 | 15.29438 | 8.41760 | 14.70704 | 12.41037 | 2.50644 | 4.79009 | 17.46022 | 0.14527 | 5.04621 | 10.75610 | 6.57966 | 16.61404 |
| **Aen** | 3.85357 | 0.05581 | 10.90351 | 12.40957 | 5.44245 | 7.20141 | 0.12029 | 7.32954 | 4.18934 | 0.06186 | 3.01120 | 5.02568 | 3.79259 | 2.94097 |
| **Aes** | 0.11057 | 0.03238 | 1.77330 | 1.47750 | 1.86787 | 0.26069 | 1.18849 | 0.76542 | 0.01465 | 7.09107 | 0.82783 | 1.65439 | 2.70205 | 0.00000 |
| **AF067061** | 0.12933 | 0.00421 | 0.03886 | 0.07636 | 0.04365 | 0.16221 | 0.17004 | 0.00818 | 0.15560 | 0.00623 | 0.00658 | 0.00314 | 3.12011 | 0.20099 |
| **AF067063** | 9.69151 | 0.00055 | 1.75578 | 1.66193 | 8.97213 | 2.73297 | 0.00177 | 0.00000 | 9.35532 | 0.03240 | 2.26803 | 0.00000 | 14.52794 | 13.41229 |
| **AF357399** | 0.00824 | 0.00439 | 0.10327 | 0.17289 | 0.00986 | 0.09716 | 0.00412 | 0.00425 | 0.01608 | 0.00533 | 0.00648 | 0.01002 | 0.00950 | 0.24244 |
| **AF366264** | 0.00634 | 0.00406 | 0.03812 | 0.00112 | 0.00424 | 0.13861 | 0.00408 | 0.03321 | 0.14776 | 0.00757 | 0.00565 | 0.00936 | 0.00819 | 0.18925 |
| **Afap1** | 0.09977 | 0.03639 | 0.00001 | 0.40962 | 0.01456 | 0.05312 | 0.03250 | 0.02684 | 0.13908 | 0.00634 | 0.01461 | 0.11089 | 1.18757 | 0.12482 |
| **Afap1l1** | 0.00000 | 0.00466 | 0.69548 | 1.50508 | 0.20533 | 0.30682 | 0.00403 | 0.31606 | 0.02042 | 0.00557 | 0.10220 | 0.00324 | 0.00070 | 0.47895 |
| **Afap1l2** | 0.00350 | 0.00421 | 0.01302 | 0.01252 | 0.00464 | 0.01219 | 0.00416 | 0.00851 | 0.02541 | 0.00625 | 0.00861 | 0.00816 | 1.04403 | 0.11073 |
| **Aff1** | 3.60037 | 0.04987 | 3.41802 | 0.00000 | 6.32790 | 0.85032 | 0.05621 | 1.40938 | 4.46846 | 0.04359 | 2.47096 | 2.85016 | 11.02898 | 7.21432 |
| **Aff3** | 0.89755 | 0.00408 | 0.01855 | 0.61711 | 0.28770 | 0.00000 | 0.00417 | 0.01330 | 0.61826 | 0.00555 | 0.00385 | 0.00473 | 0.08871 | 0.37334 |
| **Aff4** | 5.75296 | 0.06196 | 3.70669 | 2.67355 | 7.19990 | 4.60969 | 3.02120 | 2.42293 | 15.58205 | 0.15107 | 1.65192 | 5.13523 | 6.26356 | 6.78675 |
| **Afg3l1** | 4.80528 | 0.06079 | 6.97637 | 10.43890 | 3.20253 | 5.72533 | 1.90518 | 4.36832 | 0.56326 | 0.05738 | 3.95947 | 6.14093 | 0.03177 | 3.62215 |
| **Afg3l2** | 1.22768 | 0.07520 | 5.01459 | 4.64756 | 3.53213 | 6.47438 | 2.39923 | 5.78866 | 0.03402 | 0.07743 | 6.08551 | 3.59311 | 3.37214 | 0.53108 |
| **Afmid** | 11.06658 | 3.71435 | 5.42250 | 4.68592 | 8.53393 | 6.89165 | 0.01302 | 2.45690 | 6.10347 | 0.03545 | 9.94195 | 0.03500 | 3.04739 | 11.26095 |
| **Aftph** | 8.16701 | 0.13077 | 3.42050 | 0.90727 | 2.89426 | 1.18799 | 0.00835 | 1.17633 | 4.95968 | 0.02787 | 4.32146 | 1.02409 | 4.22641 | 6.35493 |
| **Aga** | 0.48010 | 0.00467 | 0.38463 | 0.00000 | 0.10755 | 0.00139 | 0.00403 | 0.01846 | 0.00778 | 0.00699 | 0.00517 | 0.05587 | 0.00377 | 0.00672 |
| **Agap1** | 0.07062 | 0.00308 | 1.10646 | 0.00000 | 0.42402 | 0.97682 | 0.31922 | 1.53954 | 0.13395 | 0.02163 | 1.20282 | 1.03211 | 0.71742 | 0.11586 |
| **Agap3** | 0.00000 | 0.09128 | 3.66422 | 2.89466 | 2.12773 | 3.53664 | 0.16760 | 5.76269 | 0.00078 | 2.74369 | 0.00000 | 3.54790 | 0.35675 | 0.00000 |
| **Agbl2** | 0.40299 | 0.00410 | 0.29306 | 0.00000 | 0.66218 | 0.00004 | 0.00412 | 0.00908 | 0.00247 | 0.00647 | 0.00567 | 0.00727 | 1.43513 | 0.63461 |
| **Agbl3** | 0.00751 | 0.00438 | 0.01474 | 0.01208 | 0.01467 | 0.01595 | 0.00431 | 0.00900 | 0.01310 | 0.00646 | 0.00890 | 0.00963 | 0.21897 | 0.00849 |
| **Agbl5** | 0.12228 | 0.00410 | 0.34039 | 0.20669 | 0.25960 | 0.00511 | 0.00393 | 0.02306 | 0.03227 | 0.00973 | 0.00529 | 0.00523 | 0.02766 | 0.01314 |
| **Ager** | 0.00343 | 0.00384 | 0.00159 | 1.00336 | 0.00664 | 0.05919 | 0.00407 | 0.44001 | 0.01494 | 0.01098 | 0.00114 | 0.01034 | 0.00703 | 0.00584 |
| **Agfg1** | 5.06341 | 0.01324 | 5.64610 | 2.86247 | 3.43716 | 12.07949 | 2.65466 | 0.21015 | 6.07661 | 5.82971 | 8.03337 | 3.68039 | 5.50962 | 2.17676 |
| **Agfg2** | 0.15766 | 0.48093 | 2.37541 | 0.38474 | 0.64672 | 1.06048 | 0.21318 | 0.72181 | 1.61577 | 0.00966 | 0.16458 | 0.03004 | 2.35584 | 2.06476 |
| **Aggf1** | 3.86645 | 0.07676 | 12.53189 | 2.74495 | 15.79432 | 9.23304 | 7.13359 | 2.05255 | 15.37027 | 0.17114 | 4.84040 | 4.51834 | 8.13222 | 21.64239 |
| **Agk** | 0.35286 | 0.00369 | 1.17426 | 0.00379 | 0.28472 | 0.14335 | 0.02112 | 0.00000 | 0.04610 | 0.00506 | 0.02456 | 0.47113 | 0.00475 | 2.50319 |
| **Agl** | 1.35948 | 0.15805 | 0.78860 | 1.45107 | 0.21569 | 3.09679 | 0.34893 | 0.33767 | 1.36172 | 0.78234 | 0.02443 | 2.38313 | 8.25289 | 3.53260 |
| **Agmat** | 0.66582 | 0.00402 | 0.61317 | 0.86322 | 1.89019 | 0.71415 | 0.00358 | 0.00615 | 0.67639 | 0.00629 | 0.26600 | 0.00432 | 0.60474 | 1.94187 |
| **Ago1** | 0.41294 | 0.00970 | 4.43191 | 1.15844 | 4.15887 | 0.00000 | 0.00150 | 1.07333 | 5.27555 | 0.00557 | 1.66561 | 1.99870 | 1.11428 | 3.12861 |
| **Ago2** | 10.13482 | 0.05570 | 13.25780 | 9.80992 | 13.93507 | 5.27333 | 0.00439 | 3.55569 | 2.96324 | 0.02326 | 7.70085 | 3.57552 | 7.25942 | 15.38421 |

| | | | | | | | | | | | | | | |
|---|---|---|---|---|---|---|---|---|---|---|---|---|---|---|
| **Ago3** | 0.82874 | 0.00320 | 1.14422 | 0.00000 | 0.29287 | 0.54302 | 0.00304 | 0.00000 | 3.65132 | 0.00896 | 0.00000 | 0.55738 | 1.57979 | 1.42819 |
| **Ago4** | 2.57799 | 0.08726 | 1.35035 | 0.27917 | 5.41971 | 0.00000 | 0.17625 | 1.11599 | 13.24645 | 0.03103 | 0.00000 | 0.00150 | 0.52250 | 5.53406 |
| **Agpat1** | 0.04733 | 0.00456 | 0.08914 | 1.05934 | 0.00081 | 0.24022 | 0.01089 | 0.30974 | 0.00655 | 0.00641 | 0.00000 | 0.33228 | 0.38831 | 0.19915 |
| **Agpat2** | 13.50249 | 1.21381 | 17.81175 | 18.95442 | 11.36866 | 17.59006 | 6.33815 | 8.56347 | 0.40494 | 10.89033 | 8.41305 | 11.90666 | 1.20880 | 8.27470 |
| **Agpat3** | 0.14986 | 0.02465 | 0.51084 | 1.37122 | 0.54483 | 1.75354 | 0.32537 | 0.35607 | 0.00309 | 0.02254 | 0.00000 | 0.00000 | 0.00926 | 0.22524 |
| **Agpat4** | 0.04757 | 0.05130 | 0.00000 | 1.02189 | 0.57166 | 3.21658 | 2.51127 | 2.20653 | 0.41083 | 0.51460 | 0.03393 | 2.77615 | 0.00232 | 0.08344 |
| **Agpat5** | 17.57649 | 0.05472 | 9.67934 | 6.46684 | 10.04451 | 2.46757 | 0.17214 | 1.79655 | 12.25447 | 0.22833 | 2.62255 | 3.19104 | 11.50601 | 13.16233 |
| **Agpat9** | 3.82326 | 0.09286 | 11.42406 | 2.42179 | 5.28008 | 2.19189 | 0.38281 | 1.68205 | 25.83098 | 0.01147 | 4.59643 | 2.36220 | 5.68891 | 13.56927 |
| **Agps** | 7.57992 | 1.71669 | 2.33323 | 5.24462 | 7.83748 | 7.19604 | 0.05731 | 1.13915 | 13.47258 | 0.09943 | 8.22119 | 4.71658 | 2.81442 | 6.16251 |
| **Agrn** | 1.09022 | 3.62021 | 0.80797 | 1.29256 | 0.16559 | 2.21540 | 0.00269 | 2.48173 | 0.03512 | 0.01766 | 2.57158 | 2.12759 | 1.23591 | 0.00000 |
| **Agrp** | 0.19236 | 0.01137 | 0.53684 | 1.75477 | 1.07266 | 2.28262 | 0.01649 | 2.33124 | 0.26582 | 0.00647 | 0.30625 | 0.01409 | 0.00135 | 1.85363 |
| **Agtpbp1** | 0.00000 | 0.00307 | 0.00000 | 1.13548 | 1.22114 | 0.21345 | 0.00351 | 0.00935 | 1.71125 | 0.00505 | 0.00000 | 1.15166 | 1.17295 | 2.52383 |
| **Agtr1a** | 0.05604 | 0.01489 | 0.03376 | 0.00000 | 0.00633 | 0.00825 | 0.00405 | 0.04171 | 0.00968 | 0.00643 | 0.00468 | 0.35254 | 0.60673 | 0.00879 |
| **Agtrap** | 1.13415 | 0.24923 | 5.30116 | 8.01896 | 3.40555 | 7.34372 | 3.47595 | 7.39343 | 0.05691 | 0.22369 | 1.90105 | 14.44050 | 2.16955 | 1.12766 |
| **Ahctf1** | 15.98094 | 0.09444 | 12.92886 | 4.51963 | 16.96160 | 6.75979 | 3.77675 | 4.68580 | 5.31322 | 0.06454 | 8.38170 | 4.50570 | 18.25492 | 13.21716 |
| **Ahcy** | 3.30932 | 0.17471 | 13.63384 | 12.65048 | 4.38446 | 12.87547 | 1.83853 | 5.15162 | 0.90571 | 0.12006 | 10.89422 | 15.54996 | 0.08056 | 3.79886 |
| **Ahcyl1** | 4.98909 | 5.05208 | 4.40180 | 2.57294 | 3.17795 | 8.73332 | 0.00827 | 0.25733 | 10.81093 | 8.84650 | 9.45010 | 7.20571 | 1.29662 | 3.92068 |
| **Ahcyl2** | 1.17658 | 0.02879 | 2.54472 | 0.00000 | 1.75887 | 3.19738 | 0.00670 | 3.64481 | 1.74210 | 0.00000 | 4.97417 | 2.99791 | 3.09032 | 1.57254 |
| **Ahdc1** | 0.02231 | 0.01309 | 0.00000 | 0.08803 | 0.19936 | 0.23836 | 0.06057 | 0.13919 | 0.06014 | 0.00725 | 0.00154 | 0.54685 | 0.24952 | 0.00000 |
| **Ahi1** | 0.03669 | 0.00438 | 0.00953 | 0.15642 | 0.13688 | 0.00809 | 0.00427 | 0.01095 | 0.01023 | 0.00670 | 0.01394 | 0.00983 | 0.26448 | 0.05514 |
| **Ahnak** | 0.88768 | 0.39182 | 0.00000 | 0.45538 | 0.89636 | 3.53891 | 1.25683 | 8.80712 | 0.02228 | 2.41035 | 0.00000 | 0.21075 | 0.89883 | 1.30414 |
| **Ahr** | 0.21570 | 0.00505 | 0.00943 | 0.04689 | 0.01551 | 0.01887 | 0.00465 | 0.02140 | 0.01538 | 0.00650 | 0.00626 | 0.01595 | 0.30796 | 0.50806 |
| **Ahrr** | 0.04667 | 0.00437 | 0.01419 | 0.01168 | 0.00946 | 0.01192 | 0.00424 | 0.00983 | 0.00607 | 0.00695 | 0.00814 | 0.00921 | 0.21924 | 0.11312 |
| **Ahsa1** | 11.09873 | 9.59127 | 19.26335 | 24.52129 | 9.11546 | 29.61750 | 6.03364 | 14.59588 | 5.18206 | 14.70575 | 19.22547 | 19.06933 | 10.79083 | 8.41836 |
| **Ahsa2** | 0.07873 | 0.02694 | 0.83649 | 0.99973 | 0.37423 | 1.37583 | 0.91692 | 3.12873 | 0.19426 | 0.04039 | 0.00000 | 0.79872 | 0.00808 | 0.11785 |
| **Ahsg** | 0.00000 | 0.00353 | 0.66990 | 0.20941 | 0.00000 | 0.26456 | 0.00372 | 0.00000 | 0.01051 | 0.00965 | 0.00183 | 0.73889 | 0.00384 | 0.00976 |
| **AI314180** | 8.90964 | 0.00775 | 5.87248 | 5.19291 | 5.85718 | 8.40695 | 0.00636 | 0.86300 | 9.35203 | 0.03871 | 2.01277 | 4.19760 | 3.54139 | 6.55436 |
| **AI314278** | 0.62242 | 0.05041 | 0.15565 | 0.04361 | 0.36817 | 0.03790 | 0.05529 | 0.05570 | 0.50107 | 0.00567 | 0.02343 | 0.00929 | 0.45425 | 0.34664 |
| **AI413582** | 0.52294 | 0.01126 | 0.21901 | 1.67982 | 1.41261 | 2.52932 | 0.71809 | 2.41231 | 0.00147 | 0.01921 | 0.00000 | 1.19192 | 0.04522 | 0.42395 |
| **AI427809** | 0.00481 | 0.00385 | 0.00200 | 0.22873 | 0.00724 | 0.06071 | 0.00450 | 0.06132 | 0.01074 | 0.00970 | 0.00483 | 0.93556 | 0.22699 | 0.00000 |
| **AI450353** | 0.00744 | 0.00565 | 0.11684 | 0.00691 | 0.00639 | 0.00266 | 0.00430 | 0.18156 | 0.00666 | 0.00733 | 0.00783 | 0.05511 | 0.03160 | 0.24857 |
| **AI464131** | 3.37965 | 6.19927 | 1.48805 | 3.08633 | 5.43335 | 2.49939 | 3.93676 | 3.55552 | 12.84666 | 8.06380 | 0.04710 | 1.14776 | 3.76627 | 12.82753 |
| **AI467606** | 0.00472 | 0.00405 | 0.08925 | 0.02542 | 0.79721 | 0.00990 | 0.00433 | 0.00604 | 0.01531 | 0.00653 | 0.00561 | 0.21095 | 0.00348 | 0.00790 |
| **AI506816** | 1.67209 | 0.00332 | 0.08908 | 0.00000 | 0.53492 | 0.09370 | 0.00335 | 0.23954 | 2.92500 | 0.00556 | 0.06536 | 0.34038 | 0.51532 | 3.17867 |
| **AI507597** | 0.00572 | 0.00407 | 0.06824 | 0.23540 | 0.00684 | 0.00600 | 0.00409 | 0.00531 | 0.01197 | 0.00561 | 0.00805 | 0.05543 | 0.00642 | 0.01234 |
| **AI597479** | 1.70798 | 1.58621 | 2.76683 | 0.00001 | 4.42878 | 4.83780 | 2.75328 | 0.06690 | 8.26634 | 2.07285 | 2.60048 | 2.48309 | 2.05010 | 5.88709 |
| **AI661453** | 0.25649 | 0.01169 | 0.24235 | 0.76900 | 2.00617 | 0.76878 | 0.00971 | 0.72597 | 1.31594 | 0.01691 | 0.05867 | 0.00386 | 7.06965 | 1.69717 |
| **AI662270** | 0.00000 | 0.04207 | 0.00000 | 0.00000 | 1.46441 | 7.74244 | 0.55536 | 1.50919 | 0.01593 | 0.04707 | 2.95985 | 3.59715 | 0.00000 | 1.64923 |
| **AI837181** | 0.84138 | 0.03408 | 2.03942 | 5.26626 | 0.51169 | 1.79748 | 1.40818 | 4.25407 | 0.08334 | 0.06227 | 1.78561 | 2.90547 | 0.29864 | 0.00000 |
| **AI846148** | 1.75081 | 1.14282 | 4.18279 | 1.42328 | 3.47887 | 1.92034 | 2.98257 | 1.90238 | 1.60435 | 0.08495 | 1.75223 | 2.73202 | 3.40927 | 7.14709 |
| **AI854703** | 0.00184 | 0.00381 | 0.00000 | 1.41656 | 0.53309 | 0.00000 | 0.00378 | 0.00573 | 0.03402 | 0.00600 | 0.01131 | 0.01153 | 1.20204 | 1.04978 |
| **AI987944** | 0.17126 | 0.00324 | 0.03923 | 0.17806 | 0.03084 | 1.17384 | 0.09288 | 0.27785 | 0.39457 | 0.02798 | 1.93113 | 0.00000 | 1.33471 | 0.43468 |
| **Aicda** | 0.00232 | 0.00430 | 0.01647 | 0.01197 | 0.00167 | 0.01175 | 0.00429 | 0.00866 | 0.00634 | 0.00685 | 0.00753 | 0.00857 | 0.36281 | 0.68689 |
| **Aida** | 0.00820 | 6.99445 | 0.31158 | 0.22768 | 0.67108 | 3.62451 | 0.05769 | 1.07201 | 1.35137 | 0.02018 | 0.00000 | 3.84431 | 3.84142 | 0.56726 |
| **Aif1** | 0.01430 | 0.00430 | 0.01007 | 0.03200 | 0.05916 | 0.01118 | 0.00406 | 0.00863 | 0.02341 | 0.01288 | 0.00797 | 0.00781 | 0.33533 | 0.00169 |
| **Aif1l** | 0.24447 | 0.01432 | 0.00416 | 0.23546 | 1.77045 | 0.00264 | 0.01191 | 0.29962 | 0.02815 | 0.00756 | 0.00345 | 0.30672 | 0.91297 | 0.00000 |
| **Aifm1** | 1.98322 | 2.42146 | 5.86727 | 0.32366 | 9.11927 | 11.33664 | 4.23476 | 4.31667 | 9.57335 | 0.09409 | 6.47365 | 8.69252 | 3.48503 | 14.61304 |
| **Aifm2** | 0.02330 | 0.00411 | 0.00666 | 0.44399 | 0.00000 | 0.71293 | 0.00412 | 0.03360 | 0.02019 | 0.03182 | 3.35500 | 0.00000 | 0.19238 | 0.00000 |
| **Aifm3** | 2.64913 | 0.00344 | 1.16543 | 0.06272 | 1.40709 | 0.00000 | 0.00375 | 0.00000 | 0.70889 | 0.00502 | 0.00264 | 0.30234 | 0.64103 | 1.00811 |
| **Aig1** | 0.33342 | 0.00434 | 0.01353 | 0.00844 | 0.00426 | 0.02462 | 0.00428 | 0.00810 | 0.01091 | 0.00626 | 0.00667 | 0.00808 | 0.32835 | 0.00491 |
| **Aim1** | 0.08984 | 0.00414 | 4.61560 | 2.04075 | 0.12082 | 3.65970 | 3.19578 | 1.42363 | 4.64356 | 0.11643 | 0.75087 | 0.00000 | 0.14447 | 0.48303 |
| **Aim1l** | 0.00000 | 5.66566 | 2.64446 | 1.43557 | 1.49304 | 3.27993 | 3.51387 | 7.57384 | 0.00630 | 0.14846 | 4.45392 | 2.15776 | 0.56710 | 0.09161 |
| **Aim2** | 0.57854 | 0.01351 | 0.00000 | 0.25818 | 0.00000 | 0.04114 | 0.02165 | 0.01773 | 0.46808 | 0.01329 | 0.00350 | 0.10237 | 0.04400 | 0.00000 |
| **Aimp1** | 18.13716 | 0.48983 | 19.88620 | 17.97274 | 11.30808 | 17.90476 | 6.54456 | 11.16899 | 4.73232 | 17.30206 | 13.39787 | 18.34824 | 0.00000 | 11.13467 |
| **Aimp2** | 13.06464 | 8.87667 | 20.12217 | 15.21390 | 11.12243 | 15.81176 | 6.17345 | 10.26520 | 1.29175 | 15.21809 | 11.14999 | 8.51775 | 6.61332 | 11.11934 |

| | | | | | | | | | | | | | | |
|---|---|---|---|---|---|---|---|---|---|---|---|---|---|---|
| **Aip** | 2.44691 | 0.01656 | 14.67841 | 6.78140 | 10.51966 | 4.13426 | 2.07178 | 2.43572 | 0.00000 | 0.05328 | 4.64585 | 1.30182 | 0.77541 | 1.50195 |
| **Aire** | 0.01150 | 0.00470 | 0.01317 | 0.00000 | 0.01417 | 0.01243 | 0.15431 | 0.00938 | 0.01280 | 0.02024 | 0.00507 | 0.22231 | 0.00912 | 0.01177 |
| **Airn** | 0.00684 | 0.00439 | 0.01398 | 0.01729 | 0.01381 | 0.01339 | 0.01647 | 0.01035 | 0.01706 | 0.00681 | 0.00886 | 0.00924 | 0.13662 | 0.01266 |
| **Ajuba** | 4.17786 | 0.04493 | 7.13725 | 4.94781 | 5.42013 | 3.42280 | 4.56900 | 0.00000 | 5.46448 | 0.17155 | 1.64138 | 0.59470 | 3.54605 | 8.02723 |
| **AK010878** | 2.46020 | 2.80910 | 2.39983 | 3.54276 | 2.89211 | 3.13288 | 0.70526 | 0.33681 | 1.59303 | 1.93349 | 1.23929 | 2.57083 | 6.95722 | 1.07108 |
| **Ak1** | 0.06564 | 0.00434 | 0.00966 | 0.09875 | 0.03798 | 0.01115 | 0.00431 | 0.00896 | 0.01066 | 0.00586 | 0.00699 | 0.02407 | 0.00602 | 0.00997 |
| **Ak2** | 2.89272 | 0.39431 | 9.61848 | 13.84090 | 8.43302 | 15.99326 | 6.71418 | 16.36117 | 1.41109 | 0.49318 | 10.49325 | 11.68831 | 0.95773 | 3.57779 |
| **Ak3** | 0.02436 | 0.00468 | 0.00508 | 0.00000 | 0.49966 | 0.02716 | 0.00461 | 1.35144 | 0.00927 | 0.00575 | 0.00631 | 0.00809 | 0.00424 | 0.01552 |
| **Ak4** | 16.13431 | 0.40087 | 27.13872 | 23.65325 | 15.03796 | 29.38498 | 6.30446 | 12.05889 | 17.87018 | 15.04174 | 20.35151 | 23.90904 | 0.03569 | 8.40085 |
| **Ak6** | 12.04360 | 0.04277 | 7.79299 | 6.42751 | 12.52169 | 10.37188 | 2.88821 | 1.57153 | 7.18323 | 10.03150 | 4.07930 | 6.17911 | 10.35378 | 9.41637 |
| **Ak7** | 8.21496 | 4.95354 | 6.43404 | 0.77888 | 6.28992 | 8.57158 | 0.00021 | 1.46621 | 3.18660 | 4.19422 | 3.89029 | 0.18796 | 2.54074 | 4.09752 |
| **Ak8** | 0.34381 | 0.00445 | 1.92588 | 0.14627 | 0.18965 | 0.03687 | 0.00375 | 0.00003 | 0.00814 | 0.00252 | 0.00544 | 0.00524 | 0.40061 | 0.03996 |
| **Akap1** | 5.61498 | 0.80969 | 18.17808 | 5.64868 | 15.78869 | 9.17081 | 3.09742 | 10.14995 | 2.14861 | 7.07773 | 5.10302 | 8.37210 | 2.97261 | 30.11464 |
| **Akap10** | 0.23530 | 0.00406 | 0.06845 | 0.02172 | 0.25571 | 0.02302 | 0.00440 | 0.01013 | 0.10691 | 0.00598 | 0.33672 | 0.10145 | 1.09695 | 1.50699 |
| **Akap11** | 0.00000 | 0.00333 | 0.07306 | 0.00000 | 0.47217 | 0.47583 | 0.00374 | 0.00485 | 1.34364 | 0.00511 | 0.00000 | 1.18044 | 0.71526 | 1.85277 |
| **Akap12** | 0.00000 | 1.35199 | 10.44089 | 7.91996 | 2.99638 | 6.96706 | 1.10410 | 3.89274 | 10.34903 | 0.08912 | 9.03908 | 11.34354 | 0.97675 | 9.21551 |
| **Akap13** | 0.19189 | 0.01018 | 0.57084 | 0.00000 | 1.09549 | 0.32271 | 0.00863 | 1.08538 | 0.23008 | 0.01516 | 0.00000 | 0.88927 | 1.23991 | 0.66484 |
| **Akap14** | 0.15123 | 0.00447 | 0.24997 | 0.00000 | 0.95158 | 0.00444 | 0.00409 | 0.01092 | 0.01196 | 0.00626 | 0.00589 | 0.00708 | 0.00163 | 0.00523 |
| **Akap17b** | 0.21422 | 0.01258 | 0.00000 | 0.00000 | 1.98524 | 0.16764 | 0.95669 | 0.42116 | 4.56305 | 0.02356 | 0.00000 | 0.19641 | 1.22803 | 5.82420 |
| **Akap5** | 0.11977 | 0.02439 | 0.01252 | 0.01944 | 0.08788 | 0.02098 | 0.01045 | 0.58219 | 0.55232 | 0.02849 | 0.01679 | 0.00602 | 0.82795 | 0.00000 |
| **Akap7** | 0.33218 | 0.00347 | 0.45377 | 0.00000 | 1.04907 | 0.61896 | 0.00321 | 0.00000 | 4.12288 | 0.00586 | 0.00000 | 0.51778 | 0.65876 | 2.66261 |
| **Akap8** | 20.77573 | 0.16849 | 18.27401 | 10.50725 | 27.70686 | 8.35749 | 4.50212 | 4.44918 | 26.22312 | 0.14790 | 5.34986 | 9.32021 | 18.67086 | 26.10610 |
| **Akap8l** | 0.00000 | 0.02734 | 1.92476 | 5.07383 | 4.13295 | 0.00000 | 0.00897 | 1.13355 | 0.07208 | 0.02305 | 0.11208 | 2.93397 | 0.00000 | 0.26017 |
| **Akap9** | 0.64657 | 0.00334 | 0.06734 | 0.63840 | 0.22037 | 0.54192 | 0.00856 | 0.37271 | 1.68578 | 0.00406 | 0.37011 | 0.44921 | 0.70882 | 1.11915 |
| **Akip1** | 0.34662 | 4.51786 | 1.18100 | 6.56398 | 1.28894 | 3.62513 | 3.22160 | 1.62473 | 0.15110 | 6.05636 | 2.45980 | 4.23825 | 0.72007 | 0.00000 |
| **Akirin1** | 14.40594 | 0.06513 | 6.66292 | 5.84232 | 16.01043 | 7.89217 | 2.32862 | 3.73749 | 6.90037 | 0.05852 | 4.73551 | 1.97749 | 5.33931 | 13.22541 |
| **Akirin2** | 6.76858 | 0.03825 | 5.32433 | 2.25728 | 9.37835 | 3.40458 | 0.10647 | 0.90805 | 9.12212 | 0.00000 | 6.00444 | 3.91613 | 11.79646 | 1.53958 |
| **Akna** | 0.01285 | 0.00432 | 0.46704 | 0.11905 | 0.00037 | 0.01624 | 0.00410 | 0.00634 | 0.01973 | 0.00659 | 0.01147 | 0.00891 | 0.00605 | 0.01330 |
| **Akr1a1** | 17.60171 | 11.32717 | 25.15877 | 20.72275 | 25.97696 | 23.50001 | 9.17186 | 25.38572 | 8.65100 | 18.49345 | 17.41940 | 20.15193 | 7.28355 | 11.87290 |
| **Akr1b10** | 0.00000 | 0.11455 | 0.28053 | 0.48253 | 0.00000 | 0.76922 | 0.92042 | 1.61867 | 0.01223 | 0.49165 | 0.34980 | 0.04842 | 0.01108 | 0.00001 |
| **Akr1b7** | 0.00000 | 0.00485 | 0.19591 | 0.00000 | 0.35047 | 0.02330 | 0.02078 | 0.06234 | 0.00818 | 0.01277 | 0.01156 | 0.24746 | 1.17314 | 0.01330 |
| **Akr1b8** | 0.00000 | 12.12091 | 22.38070 | 21.53424 | 0.00000 | 38.98158 | 15.20769 | 37.43832 | 2.70040 | 16.29456 | 20.21092 | 8.43947 | 0.00000 | 0.00000 |
| **Akr1c13** | 0.00825 | 0.01371 | 0.01055 | 0.00009 | 0.36072 | 0.01352 | 0.00452 | 0.01551 | 0.01826 | 0.00630 | 0.00455 | 0.41351 | 0.00471 | 0.00742 |
| **Akr1c14** | 0.01328 | 0.00469 | 0.10457 | 0.14345 | 0.00670 | 0.74794 | 0.00424 | 0.00472 | 0.01120 | 0.00666 | 0.05064 | 0.00461 | 0.00997 | 0.00476 |
| **Akr1c21** | 15.29382 | 0.03845 | 3.78016 | 0.00000 | 0.00000 | 0.01375 | 0.00226 | 0.18671 | 2.01680 | 0.00537 | 0.43364 | 5.18594 | 0.03763 | 1.01755 |
| **Akr1cl** | 0.01197 | 0.00450 | 0.00301 | 0.01215 | 0.79296 | 0.01127 | 0.00437 | 0.01066 | 0.00639 | 0.00629 | 0.00794 | 0.00734 | 0.00441 | 0.40115 |
| **Akr1d1** | 0.01455 | 0.00439 | 0.01769 | 0.01990 | 0.01674 | 0.03373 | 0.01027 | 0.00986 | 0.05449 | 0.00763 | 0.01103 | 0.01883 | 0.22817 | 0.00532 |
| **Akr1e1** | 0.00000 | 0.07385 | 0.52771 | 0.00000 | 0.15079 | 1.82131 | 0.01466 | 0.04105 | 0.09738 | 11.82010 | 0.15091 | 2.76580 | 0.00395 | 0.55719 |
| **Akr7a5** | 0.00319 | 0.06227 | 0.13453 | 0.50171 | 0.02635 | 0.05793 | 0.00342 | 0.83243 | 0.01166 | 0.03199 | 0.00000 | 1.76434 | 0.48870 | 0.02020 |
| **Akt1** | 1.80038 | 2.71825 | 12.34148 | 4.15332 | 9.40245 | 5.21497 | 2.99127 | 12.23823 | 0.00000 | 0.15336 | 1.60715 | 5.25273 | 2.33692 | 0.86812 |
| **Akt1s1** | 0.29860 | 0.08290 | 11.71348 | 8.18223 | 18.49160 | 0.00000 | 0.14407 | 7.39007 | 0.03598 | 0.08053 | 0.68344 | 3.79233 | 0.78741 | 0.13956 |
| **Akt2** | 3.67942 | 0.11876 | 13.53924 | 7.03303 | 8.25580 | 11.13955 | 4.08240 | 1.90245 | 0.02758 | 0.12913 | 8.96471 | 12.51289 | 4.54786 | 7.09094 |
| **Akt3** | 0.73514 | 0.00337 | 2.35038 | 1.66741 | 0.05027 | 4.42725 | 0.05966 | 0.53640 | 1.36503 | 0.01075 | 2.26290 | 0.63178 | 1.76064 | 0.76576 |
| **Aktip** | 0.30603 | 0.02707 | 0.54460 | 1.62169 | 0.89995 | 0.65521 | 0.00410 | 1.55300 | 0.28731 | 0.04136 | 1.53227 | 1.02995 | 1.18890 | 2.15703 |
| **Alad** | 2.05368 | 12.55359 | 15.18375 | 11.71858 | 5.29623 | 15.89874 | 4.13859 | 12.50740 | 0.82363 | 0.18069 | 6.39883 | 8.38317 | 0.01697 | 2.88366 |
| **Alas1** | 6.15185 | 0.28051 | 3.58852 | 0.22929 | 0.75980 | 1.28421 | 0.79849 | 2.94043 | 0.00000 | 0.02837 | 2.44837 | 1.03603 | 4.58198 | 2.93404 |
| **Alas2** | 0.39400 | 0.00430 | 0.00000 | 0.00000 | 6.85094 | 1.42277 | 0.00417 | 0.00194 | 0.00772 | 0.01879 | 1.06278 | 0.00422 | 0.00000 | 0.00000 |
| **Aldh16a1** | 1.09360 | 0.07689 | 9.07867 | 11.94791 | 3.22223 | 0.59854 | 2.81238 | 10.55303 | 0.00000 | 0.23441 | 5.39603 | 3.81266 | 1.17514 | 0.22481 |
| **Aldh18a1** | 0.00000 | 0.15815 | 0.31788 | 0.00000 | 1.52114 | 1.59333 | 0.02826 | 4.97721 | 0.05098 | 0.03003 | 0.62607 | 5.33114 | 0.80691 | 0.83226 |
| **Aldh1a1** | 0.01219 | 0.00448 | 0.00528 | 0.00991 | 0.45364 | 0.01377 | 0.00498 | 0.01006 | 0.02248 | 0.00704 | 0.00879 | 0.00827 | 0.00325 | 0.01823 |
| **Aldh1b1** | 0.00399 | 0.00394 | 0.14736 | 0.16417 | 0.00000 | 0.98733 | 0.00498 | 0.41454 | 0.00919 | 0.02508 | 1.63938 | 0.10364 | 0.00636 | 0.00462 |
| **Aldh1l1** | 0.00000 | 7.01942 | 0.90079 | 1.21342 | 0.06551 | 2.10532 | 0.00219 | 0.66967 | 0.36406 | 0.03235 | 0.00000 | 0.03631 | 0.36649 | 0.14188 |
| **Aldh1l2** | 0.05833 | 0.00282 | 0.00000 | 0.00000 | 0.12294 | 0.38041 | 0.00343 | 0.42450 | 0.25575 | 0.00560 | 0.00000 | 1.56066 | 0.15346 | 0.00000 |
| **Aldh2** | 0.69293 | 0.03992 | 0.07327 | 2.23180 | 1.23056 | 1.48245 | 0.69346 | 1.64991 | 0.00037 | 0.01058 | 0.47862 | 0.68189 | 1.38896 | 0.10879 |
| **Aldh3a2** | 0.01890 | 0.00331 | 0.26187 | 0.85779 | 0.59328 | 1.25618 | 0.00394 | 1.12670 | 0.54289 | 0.00013 | 2.19490 | 0.67256 | 1.02902 | 0.00000 |

| | | | | | | | | | | | | | | |
|---|---|---|---|---|---|---|---|---|---|---|---|---|---|---|
| **Aldh3b1** | 0.00913 | 0.00446 | 0.00893 | 0.00708 | 0.19081 | 0.01108 | 0.00481 | 0.05946 | 0.01757 | 0.00638 | 0.00790 | 0.02341 | 0.00760 | 0.06344 |
| **Aldh3b2** | 0.01544 | 0.02918 | 0.00986 | 0.00000 | 0.01177 | 0.05356 | 0.18456 | 2.76056 | 0.01085 | 0.02118 | 0.00319 | 0.04656 | 0.00690 | 0.01237 |
| **Aldh3b3** | 0.01645 | 0.00408 | 0.01048 | 0.04336 | 0.02281 | 0.00702 | 0.00391 | 0.21568 | 0.01320 | 0.01048 | 0.00757 | 0.00781 | 0.01631 | 0.01231 |
| **Aldh4a1** | 0.00000 | 0.06099 | 1.25102 | 3.70319 | 0.13675 | 3.77555 | 2.57382 | 5.34235 | 0.07656 | 0.05244 | 0.00000 | 3.90411 | 0.65383 | 2.51835 |
| **Aldh5a1** | 0.00604 | 0.02688 | 0.42742 | 0.00264 | 0.00119 | 0.00459 | 0.00408 | 0.00559 | 0.00660 | 0.00463 | 0.28276 | 1.33352 | 0.16958 | 0.31124 |
| **Aldh6a1** | 1.19138 | 0.03425 | 1.62695 | 2.61916 | 2.30164 | 7.35792 | 1.90407 | 2.22481 | 0.15323 | 0.04675 | 1.80013 | 2.15531 | 0.48944 | 2.29715 |
| **Aldh7a1** | 0.14867 | 0.00214 | 1.62100 | 2.19917 | 2.27943 | 1.30962 | 0.00366 | 0.50979 | 0.50693 | 0.05824 | 2.66309 | 1.89400 | 0.00000 | 2.30569 |
| **Aldh9a1** | 1.34693 | 0.16162 | 4.24616 | 4.50131 | 3.19621 | 7.31683 | 1.51738 | 5.03169 | 1.30562 | 0.12464 | 7.81400 | 11.96379 | 1.86266 | 5.62564 |
| **Aldoa** | 11.47272 | 8.27287 | 21.21829 | 28.82008 | 8.54630 | 37.40776 | 9.46299 | 30.53222 | 1.39759 | 24.57466 | 24.31722 | 24.38902 | 3.91919 | 5.78998 |
| **Aldoart1** | 0.15992 | 0.26910 | 0.63388 | 1.10634 | 0.09154 | 2.30118 | 0.50223 | 2.07896 | 0.02374 | 2.27105 | 1.67473 | 1.17164 | 0.00537 | 0.09980 |
| **Aldoart2** | 0.02373 | 0.00432 | 0.01998 | 0.04400 | 0.01391 | 0.07937 | 0.00423 | 0.05626 | 0.01188 | 0.06460 | 0.07699 | 0.04886 | 0.00676 | 0.01629 |
| **Aldoc** | 0.00000 | 0.01113 | 0.52583 | 1.26204 | 1.72192 | 0.18305 | 0.00347 | 0.00000 | 0.00874 | 0.00865 | 1.91311 | 0.00795 | 0.00000 | 2.06962 |
| **Alg1** | 1.45539 | 0.00938 | 2.49996 | 2.15620 | 0.58156 | 3.45733 | 0.22932 | 0.00000 | 0.00194 | 0.00585 | 0.29290 | 2.15040 | 0.00000 | 0.71154 |
| **Alg10b** | 0.27931 | 0.01142 | 0.32367 | 0.00000 | 0.59958 | 1.27641 | 0.00347 | 0.04890 | 0.31233 | 0.00484 | 0.00000 | 1.04578 | 1.97716 | 0.00000 |
| **Alg11** | 0.72927 | 0.02217 | 0.63245 | 1.26764 | 2.69016 | 2.52865 | 0.00310 | 0.06495 | 5.39354 | 0.02638 | 4.19247 | 0.37375 | 0.00000 | 0.54806 |
| **Alg12** | 0.70307 | 0.01977 | 0.87305 | 2.88512 | 1.14410 | 1.56036 | 1.50099 | 0.15034 | 0.54356 | 0.06133 | 0.38155 | 0.69744 | 0.47241 | 1.55502 |
| **Alg13** | 21.51885 | 0.09218 | 6.91513 | 3.69514 | 8.00963 | 6.35132 | 2.16593 | 2.84217 | 12.68019 | 14.42590 | 1.33229 | 8.25932 | 9.98224 | 3.54568 |
| **Alg14** | 5.33894 | 0.01289 | 3.99451 | 1.02748 | 1.26427 | 3.02575 | 0.00057 | 1.53506 | 1.52657 | 0.03320 | 5.73425 | 5.49212 | 0.00000 | 2.12992 |
| **Alg2** | 1.44767 | 0.00307 | 2.06605 | 1.36946 | 2.71932 | 2.23284 | 1.58396 | 0.59974 | 0.06156 | 0.03636 | 0.00000 | 1.35193 | 1.28272 | 5.13017 |
| **Alg3** | 4.91282 | 0.04533 | 5.03079 | 5.40861 | 1.85232 | 8.33865 | 0.03531 | 0.03917 | 0.00551 | 0.03140 | 4.52314 | 3.57094 | 0.52778 | 1.84106 |
| **Alg5** | 5.74695 | 0.03840 | 3.19392 | 6.69913 | 0.31737 | 4.78857 | 2.48552 | 3.18345 | 4.93789 | 0.07194 | 7.18810 | 6.59986 | 0.00000 | 1.02026 |
| **Alg6** | 1.79514 | 3.07667 | 0.30542 | 0.00000 | 0.52846 | 0.57864 | 0.00278 | 0.39341 | 0.88286 | 0.01187 | 0.05843 | 0.52034 | 0.04218 | 0.37802 |
| **Alg8** | 1.55541 | 0.04799 | 1.72464 | 1.60895 | 0.05252 | 2.76151 | 0.94394 | 2.39874 | 1.95629 | 0.03900 | 1.60209 | 0.77324 | 0.00000 | 2.34634 |
| **Alg9** | 1.06974 | 0.02782 | 1.70869 | 3.29419 | 0.50439 | 4.72555 | 3.14428 | 1.08526 | 0.61173 | 0.03834 | 3.38342 | 2.42344 | 0.64215 | 0.00000 |
| **Alkbh1** | 1.66147 | 0.01505 | 3.44854 | 1.85246 | 3.98724 | 3.42772 | 0.18189 | 0.05955 | 1.93896 | 0.16380 | 2.36128 | 1.87156 | 3.65871 | 8.01184 |
| **Alkbh2** | 1.27262 | 0.00935 | 1.86814 | 1.64189 | 5.11353 | 1.02525 | 0.00296 | 0.00000 | 0.00000 | 0.01663 | 0.01427 | 2.06238 | 1.10036 | 0.72318 |
| **Alkbh3** | 7.02662 | 9.08879 | 3.25322 | 9.38182 | 2.54628 | 6.20093 | 0.00545 | 2.24767 | 1.70461 | 0.02588 | 6.80327 | 3.30895 | 1.56698 | 3.55592 |
| **Alkbh3os1** | 0.00560 | 0.00416 | 0.13053 | 0.17927 | 0.38155 | 0.05925 | 0.00455 | 0.00346 | 0.02610 | 0.00644 | 0.06162 | 0.03352 | 0.00583 | 0.00212 |
| **Alkbh4** | 0.50788 | 0.01791 | 4.69419 | 2.63872 | 4.05483 | 2.68774 | 0.14964 | 3.25511 | 0.00000 | 0.00000 | 2.36658 | 2.16267 | 2.96301 | 0.00000 |
| **Alkbh5** | 1.23937 | 0.08506 | 4.62385 | 1.22343 | 3.29247 | 3.76281 | 3.63259 | 3.99182 | 2.77625 | 0.08361 | 1.74717 | 1.74000 | 5.48924 | 6.81169 |
| **Alkbh6** | 0.00000 | 0.01035 | 0.22628 | 2.91247 | 1.83474 | 0.00000 | 0.06051 | 0.28889 | 0.02547 | 0.02475 | 0.00843 | 1.54673 | 0.01825 | 0.00000 |
| **Alkbh7** | 0.00000 | 0.06145 | 0.95357 | 5.37647 | 2.06538 | 0.55920 | 0.02576 | 4.91780 | 0.00094 | 3.61406 | 0.00000 | 4.89549 | 1.32020 | 0.19009 |
| **Alkbh8** | 0.00000 | 0.04058 | 0.51549 | 0.06772 | 0.83268 | 0.69872 | 0.00416 | 0.33243 | 0.01288 | 0.00319 | 0.44759 | 0.21925 | 0.68426 | 1.03435 |
| **Allc** | 1.68842 | 0.01069 | 1.22323 | 2.92636 | 2.58098 | 0.00000 | 0.00339 | 0.00000 | 0.00621 | 0.00411 | 0.02637 | 0.74188 | 0.00000 | 3.77193 |
| **Alms1** | 0.00000 | 0.01066 | 1.44592 | 0.08845 | 0.15435 | 0.01902 | 0.00340 | 0.00000 | 1.01615 | 0.00000 | 0.78972 | 1.37396 | 0.90503 | 2.76298 |
| **Alms1.ps2** | 0.03971 | 0.01080 | 0.68355 | 0.14668 | 0.06762 | 0.00000 | 0.00349 | 0.00000 | 0.57133 | 0.00230 | 0.50387 | 1.04715 | 0.01812 | 0.86711 |
| **Alox12** | 1.89398 | 0.04750 | 0.96179 | 3.71917 | 3.63457 | 3.84516 | 0.02349 | 0.54429 | 1.54557 | 0.00458 | 0.60266 | 1.52383 | 5.24672 | 0.22356 |
| **Alox12e** | 0.02236 | 0.00455 | 0.07637 | 0.00000 | 1.10349 | 0.16281 | 0.00456 | 0.00735 | 0.02212 | 0.00605 | 0.00468 | 0.00591 | 0.00000 | 0.28805 |
| **Alox15** | 0.00789 | 0.00448 | 0.00523 | 0.01075 | 0.60510 | 0.01353 | 0.00456 | 0.01080 | 0.01284 | 0.00654 | 0.00894 | 0.00805 | 0.00515 | 0.00728 |
| **Alox5ap** | 0.00822 | 0.00471 | 0.15235 | 0.00525 | 0.05505 | 0.00891 | 0.00430 | 0.01532 | 0.00980 | 0.00724 | 0.00756 | 0.00892 | 0.00688 | 0.01415 |
| **Alox8** | 0.14486 | 0.04415 | 0.09594 | 0.09198 | 0.03739 | 0.09237 | 0.01062 | 0.04218 | 0.49772 | 0.03165 | 0.02953 | 0.03540 | 0.44281 | 0.23996 |
| **Aloxe3** | 0.40488 | 0.00373 | 2.01540 | 0.17491 | 1.80799 | 0.00000 | 0.00359 | 0.00307 | 0.51525 | 0.00390 | 0.00595 | 0.00701 | 1.59692 | 2.61992 |
| **Alpi** | 0.05525 | 0.00424 | 0.32198 | 0.41137 | 0.23503 | 0.03785 | 0.00395 | 0.00425 | 0.01218 | 0.00787 | 0.01570 | 0.00794 | 0.01281 | 0.00905 |
| **Alpk3** | 0.01171 | 0.00421 | 0.00079 | 0.27971 | 0.06934 | 0.34741 | 0.00436 | 0.00368 | 0.03292 | 0.00604 | 0.01546 | 0.02726 | 0.00567 | 0.00000 |
| **Alpl** | 0.27006 | 1.31490 | 3.10984 | 6.98866 | 2.02184 | 1.45154 | 0.77245 | 9.45291 | 0.06244 | 0.09085 | 0.00000 | 13.96180 | 0.66958 | 0.00000 |
| **Alppl2** | 21.66814 | 3.46017 | 56.66862 | 60.54444 | 43.81117 | 27.28374 | 2.99927 | 2.89595 | 11.00825 | 1.13992 | 18.08525 | 0.00000 | 15.60524 | 12.39858 |
| **Als2** | 8.20124 | 0.05908 | 10.95420 | 4.43398 | 10.20926 | 2.60950 | 0.50796 | 1.36588 | 14.10545 | 0.26574 | 4.66861 | 2.89190 | 9.79974 | 15.03654 |
| **Als2cl** | 0.13103 | 0.00432 | 0.12288 | 0.04394 | 0.03110 | 0.03434 | 0.00426 | 0.01774 | 0.01455 | 0.00759 | 0.01226 | 0.03598 | 0.08903 | 0.10133 |
| **Als2cr11** | 0.00220 | 0.00445 | 0.03455 | 0.00804 | 0.24529 | 0.00855 | 0.00436 | 0.01229 | 0.37905 | 0.00782 | 0.00636 | 0.00832 | 0.00357 | 0.21313 |
| **Alx4** | 2.02677 | 0.01150 | 0.46049 | 0.10994 | 1.64709 | 0.05054 | 0.02184 | 0.03308 | 1.20511 | 0.00525 | 0.01101 | 0.05860 | 0.96476 | 1.48809 |
| **Alyref** | 0.00000 | 0.11748 | 2.83310 | 4.98360 | 7.57744 | 2.76112 | 1.47489 | 2.76824 | 3.55900 | 0.81648 | 1.13087 | 1.25040 | 2.54242 | 9.49383 |
| **Alyref2** | 0.00000 | 0.00473 | 0.04571 | 0.17075 | 0.23324 | 0.07728 | 0.05313 | 0.16656 | 0.16829 | 0.03062 | 0.03864 | 0.06004 | 0.90104 | 0.21781 |
| **Amacr** | 0.08302 | 0.00333 | 0.01423 | 0.14500 | 0.01765 | 0.01681 | 1.23952 | 0.00312 | 0.00692 | 0.05676 | 0.00154 | 0.98323 | 0.37564 | 0.01141 |
| **Ambra1** | 2.49426 | 0.02673 | 1.76754 | 2.96008 | 1.64333 | 1.78808 | 0.01072 | 0.35759 | 0.26587 | 0.01390 | 3.09978 | 1.23472 | 4.11712 | 2.89306 |
| **Amdhd2** | 1.39006 | 2.49490 | 6.48785 | 12.45653 | 1.39827 | 3.67519 | 0.03009 | 6.06687 | 0.00000 | 2.83964 | 5.37835 | 2.45699 | 0.09347 | 0.00000 |

| | | | | | | | | | | | | | | |
|---|---|---|---|---|---|---|---|---|---|---|---|---|---|---|
| **Amer1** | 0.00309 | 0.00435 | 0.09559 | 0.00001 | 0.01129 | 0.00642 | 0.00382 | 0.02223 | 0.05394 | 0.00959 | 0.00426 | 0.77691 | 0.02810 | 0.09835 |
| **Amfr** | 0.02511 | 0.04968 | 0.00000 | 0.00000 | 0.58488 | 2.00205 | 0.00951 | 1.25848 | 0.02038 | 0.00665 | 0.00000 | 2.23025 | 1.07877 | 2.15024 |
| **Amhr2** | 2.03929 | 0.01422 | 10.39544 | 5.64396 | 11.75878 | 0.66920 | 1.68502 | 0.00000 | 0.01879 | 0.10796 | 2.00841 | 3.81392 | 4.47473 | 1.88752 |
| **Amigo2** | 0.00000 | 5.13969 | 0.14547 | 0.30174 | 0.75932 | 0.00000 | 0.00274 | 0.00893 | 0.00323 | 0.00621 | 0.00389 | 0.00000 | 0.83646 | 0.00812 |
| **Amigo3** | 0.85761 | 0.00429 | 0.76580 | 0.00000 | 1.53262 | 0.37400 | 0.00493 | 0.82646 | 0.00320 | 0.00444 | 0.00622 | 0.00127 | 0.73495 | 0.10312 |
| **Ammecr1l** | 9.88140 | 7.80458 | 12.84407 | 5.78932 | 16.56104 | 7.83097 | 0.00996 | 0.69823 | 24.54048 | 0.00000 | 0.00000 | 0.63105 | 13.73935 | 20.71016 |
| **Amn** | 0.00300 | 0.00404 | 0.00150 | 1.29783 | 1.34950 | 0.05283 | 0.00396 | 0.00455 | 0.01425 | 0.00596 | 0.01094 | 0.00715 | 0.29997 | 0.00984 |
| **Amn1** | 0.80701 | 0.02335 | 3.08330 | 2.77337 | 1.70313 | 5.75340 | 1.28331 | 2.25644 | 4.91189 | 0.01290 | 7.37267 | 2.05854 | 1.31995 | 1.75336 |
| **Amot** | 8.16361 | 0.07423 | 10.88555 | 3.26349 | 1.61038 | 8.60557 | 0.02667 | 3.65175 | 7.41744 | 7.25718 | 2.84550 | 6.81270 | 0.58761 | 0.00000 |
| **Amotl1** | 0.30769 | 0.00509 | 0.00000 | 0.29053 | 0.16674 | 0.08029 | 0.02442 | 0.16800 | 0.04046 | 0.01333 | 0.00364 | 0.00841 | 0.12854 | 0.00801 |
| **Amotl2** | 0.82848 | 0.06902 | 12.67834 | 0.18599 | 5.68668 | 18.81031 | 3.89783 | 5.06172 | 4.58814 | 13.40988 | 2.58441 | 0.00000 | 0.00000 | 3.05582 |
| **Ampd1** | 0.00246 | 0.00408 | 0.00818 | 0.01729 | 0.20411 | 0.01273 | 0.00405 | 0.00806 | 0.00826 | 0.00640 | 0.00905 | 0.00681 | 2.23690 | 0.00650 |
| **Ampd2** | 0.55094 | 0.01165 | 0.66227 | 1.88476 | 0.08673 | 2.52979 | 0.01566 | 1.60712 | 0.01667 | 0.01285 | 0.00000 | 0.23457 | 0.09167 | 0.00000 |
| **Ampd3** | 0.02612 | 0.00407 | 0.02942 | 0.00707 | 0.48132 | 0.01104 | 0.00425 | 0.00812 | 0.03939 | 0.00619 | 0.00893 | 0.00719 | 2.66762 | 0.08190 |
| **Amph** | 0.00532 | 0.01355 | 0.37998 | 0.00000 | 0.00307 | 0.00000 | 0.01078 | 0.17376 | 0.00778 | 0.00662 | 0.00343 | 0.25265 | 0.00279 | 0.11688 |
| **Amt** | 0.00331 | 0.01995 | 0.00810 | 0.00000 | 0.04132 | 0.00276 | 0.00475 | 1.04212 | 0.07924 | 0.00633 | 0.00585 | 0.02601 | 0.01526 | 0.37135 |
| **Amy1** | 0.00226 | 0.00447 | 0.07158 | 0.00000 | 0.04169 | 0.14020 | 0.00437 | 0.00543 | 0.06345 | 0.00612 | 0.00717 | 0.00931 | 0.01122 | 0.47559 |
| **Amz1** | 0.00242 | 0.00409 | 0.00291 | 0.92568 | 0.42854 | 0.00000 | 0.00402 | 0.00850 | 0.00623 | 0.00523 | 0.01122 | 0.00568 | 0.85995 | 0.00000 |
| **Amz2** | 3.08018 | 0.03993 | 2.56626 | 1.79889 | 2.16643 | 2.34794 | 0.00113 | 2.94537 | 0.00000 | 0.03702 | 3.69848 | 5.61300 | 0.59322 | 4.86330 |
| **Anapc1** | 4.84475 | 0.02314 | 10.33796 | 4.97946 | 6.22262 | 6.18555 | 2.63392 | 3.46115 | 6.31732 | 0.00000 | 10.75709 | 9.34030 | 2.61979 | 8.87225 |
| **Anapc10** | 14.93192 | 0.05732 | 9.12579 | 7.03566 | 15.31056 | 13.86085 | 3.02334 | 1.35726 | 18.12660 | 0.44154 | 6.26873 | 5.22727 | 2.65071 | 10.40314 |
| **Anapc11** | 4.02445 | 1.35020 | 2.11831 | 4.77587 | 10.29228 | 1.89068 | 0.52605 | 2.45738 | 0.04587 | 1.95148 | 2.27470 | 3.22277 | 2.89529 | 1.38951 |
| **Anapc13** | 1.27601 | 0.11595 | 0.08206 | 0.00000 | 1.00156 | 2.27983 | 0.91483 | 3.12158 | 0.20537 | 4.29702 | 0.02357 | 2.73888 | 1.30735 | 0.34700 |
| **Anapc15** | 12.67556 | 0.12610 | 8.12035 | 12.06263 | 13.65655 | 13.57814 | 5.05389 | 6.26988 | 6.12197 | 9.49657 | 3.81732 | 8.09337 | 11.72633 | 10.31467 |
| **Anapc16** | 3.19594 | 0.00309 | 0.67993 | 0.56005 | 1.40548 | 0.73221 | 0.01467 | 0.44846 | 1.25522 | 0.05107 | 0.15527 | 0.40518 | 3.56953 | 2.38320 |
| **Anapc2** | 7.15172 | 0.25484 | 13.73909 | 8.14284 | 10.98958 | 5.52128 | 4.49818 | 10.34809 | 4.44654 | 1.32719 | 11.17829 | 9.39294 | 2.62635 | 5.79691 |
| **Anapc4** | 7.43539 | 0.04077 | 7.94066 | 1.28086 | 6.83950 | 2.27568 | 2.94481 | 4.98118 | 9.01907 | 0.14647 | 8.26614 | 5.20467 | 3.79027 | 12.33059 |
| **Anapc5** | 18.41103 | 7.36363 | 28.10484 | 28.46204 | 20.86685 | 16.13025 | 2.14053 | 15.40589 | 12.85137 | 2.40427 | 12.15519 | 15.80054 | 7.03947 | 24.74960 |
| **Anapc7** | 10.52892 | 7.69663 | 11.63263 | 11.66372 | 17.93685 | 6.06824 | 0.13432 | 1.12430 | 1.17201 | 0.02990 | 6.99088 | 3.10173 | 8.31796 | 18.30212 |
| **Angel1** | 0.00000 | 0.02255 | 0.13145 | 0.32142 | 0.07611 | 2.50294 | 0.00383 | 0.24236 | 0.09223 | 0.01314 | 0.42598 | 0.36943 | 1.03057 | 0.53270 |
| **Angel2** | 0.15310 | 0.02296 | 1.54403 | 0.00000 | 1.10456 | 2.23502 | 0.00216 | 2.04087 | 1.19740 | 0.01058 | 4.82623 | 2.71824 | 6.33993 | 8.01351 |
| **Angpt2** | 0.08126 | 0.00427 | 0.00203 | 0.01196 | 0.03307 | 0.00803 | 0.00417 | 0.00974 | 0.88568 | 0.00657 | 0.00773 | 0.04480 | 0.20560 | 0.15359 |
| **Angptl1** | 0.00752 | 0.00443 | 0.04373 | 0.01303 | 0.01914 | 0.01000 | 0.00424 | 0.01022 | 0.05569 | 0.00709 | 0.00772 | 0.01059 | 0.00884 | 0.10604 |
| **Angptl2** | 0.22911 | 0.00372 | 0.07113 | 0.00256 | 0.00000 | 0.00000 | 0.00413 | 0.00470 | 0.01507 | 0.00599 | 0.33210 | 0.44818 | 0.11557 | 0.45579 |
| **Angptl4** | 0.65041 | 0.00137 | 2.81665 | 3.09575 | 2.87407 | 0.11699 | 0.01048 | 0.31641 | 0.34355 | 0.00693 | 3.24123 | 2.87992 | 0.00667 | 4.36486 |
| **Angptl6** | 0.05495 | 0.00389 | 0.07744 | 0.07259 | 0.31720 | 0.44176 | 0.00396 | 0.00048 | 0.01015 | 0.00636 | 0.57073 | 0.13754 | 0.01355 | 0.05413 |
| **Ank** | 1.17597 | 1.81661 | 3.88144 | 0.03729 | 0.95636 | 13.83809 | 6.95279 | 15.79268 | 0.00000 | 0.48011 | 4.27042 | 0.11885 | 0.74715 | 2.08552 |
| **Ank2** | 0.15331 | 0.01008 | 0.07170 | 0.00255 | 0.01060 | 0.00960 | 0.00374 | 0.01327 | 0.02792 | 0.00695 | 0.12295 | 1.91663 | 0.01201 | 0.02128 |
| **Ank3** | 0.01437 | 0.01426 | 0.01109 | 0.12605 | 0.01482 | 0.06442 | 0.00417 | 0.02043 | 0.02964 | 0.01086 | 0.00725 | 0.00922 | 0.01780 | 0.05014 |
| **Ankar** | 0.13924 | 0.00448 | 0.04065 | 0.00768 | 0.00000 | 0.00476 | 0.00430 | 0.01596 | 1.13759 | 0.00679 | 0.00486 | 0.00818 | 0.01432 | 0.05600 |
| **Ankdd1b** | 1.21559 | 0.00487 | 0.15038 | 0.00000 | 0.37668 | 1.01111 | 0.07824 | 0.16353 | 2.50583 | 0.02243 | 0.00000 | 0.57149 | 0.66334 | 0.00000 |
| **Ankfy1** | 5.66219 | 0.06164 | 6.49103 | 3.46209 | 5.00554 | 6.05797 | 2.09711 | 2.76907 | 3.18607 | 0.17376 | 4.90922 | 2.43141 | 4.31052 | 6.33129 |
| **Ankhd1** | 3.93423 | 0.03337 | 3.03756 | 1.96281 | 3.49091 | 1.54168 | 0.00079 | 0.98915 | 10.50041 | 0.01184 | 2.44790 | 3.02585 | 0.78240 | 7.48899 |
| **Ankib1** | 4.95451 | 0.01552 | 2.95108 | 0.00809 | 4.44089 | 3.80672 | 0.00185 | 1.26219 | 5.27862 | 0.00000 | 2.07491 | 3.05701 | 2.26202 | 5.02046 |
| **Ankle1** | 0.27627 | 0.01116 | 1.61609 | 2.64272 | 0.02607 | 0.58323 | 0.00966 | 0.83053 | 0.02151 | 0.03642 | 1.45413 | 0.15849 | 0.47524 | 0.34533 |
| **Ankle2** | 8.80714 | 0.00690 | 7.00260 | 0.71855 | 5.86560 | 4.41754 | 0.49461 | 2.16533 | 10.78751 | 0.04200 | 0.00000 | 4.23220 | 9.68100 | 10.72939 |
| **Ankmy2** | 4.61572 | 0.02385 | 6.04588 | 3.33192 | 7.20409 | 4.87535 | 0.00000 | 2.39305 | 11.81214 | 0.03760 | 0.48990 | 2.40345 | 1.72538 | 7.33810 |
| **Ankra2** | 2.08373 | 0.00359 | 0.40699 | 0.00000 | 0.18913 | 1.02986 | 0.00374 | 0.07461 | 1.39816 | 0.00414 | 0.32814 | 0.00779 | 2.47654 | 0.00000 |
| **Ankrd1** | 0.01170 | 0.00436 | 0.00444 | 0.00857 | 0.01128 | 0.59255 | 0.00418 | 0.00289 | 0.01145 | 0.00331 | 1.65256 | 0.00758 | 0.00958 | 0.00019 |
| **Ankrd10** | 23.79270 | 0.10367 | 27.22911 | 10.54151 | 35.70686 | 4.61074 | 4.39471 | 1.20004 | 11.55413 | 0.09607 | 0.00000 | 7.89300 | 17.94924 | 31.87345 |
| **Ankrd11** | 0.31950 | 0.84491 | 6.79111 | 0.43801 | 1.98402 | 2.24732 | 1.39884 | 1.33138 | 1.16542 | 0.01350 | 4.33822 | 3.96741 | 0.00000 | 1.81728 |
| **Ankrd12** | 0.42156 | 0.00306 | 0.64418 | 1.09673 | 2.60713 | 1.06019 | 0.00290 | 0.13132 | 7.19846 | 0.02902 | 1.31054 | 0.35986 | 1.04178 | 4.54057 |
| **Ankrd13a** | 6.92067 | 7.53944 | 13.07172 | 6.90264 | 13.75568 | 10.00495 | 6.39505 | 9.22888 | 0.16561 | 0.36705 | 9.83864 | 6.41348 | 3.73055 | 5.47823 |
| **Ankrd13b** | 0.04209 | 0.00447 | 0.13778 | 0.21818 | 0.24898 | 0.03249 | 0.00482 | 0.05355 | 0.01819 | 0.00958 | 0.00797 | 0.09444 | 0.12706 | 0.00553 |
| **Ankrd13c** | 11.89847 | 0.00000 | 8.35668 | 0.00000 | 10.09724 | 0.00000 | 1.48352 | 4.32380 | 14.32695 | 0.04418 | 2.10246 | 0.74773 | 11.38238 | 12.40224 |

| | | | | | | | | | | | | | | |
|---|---|---|---|---|---|---|---|---|---|---|---|---|---|---|
| **Ankrd13d** | 0.00000 | 0.00509 | 0.48887 | 0.02369 | 1.54028 | 0.03357 | 0.00395 | 0.01747 | 0.01268 | 0.00884 | 0.00673 | 0.00447 | 0.00000 | 0.30573 |
| **Ankrd16** | 1.79601 | 0.02569 | 2.23026 | 0.96394 | 0.29799 | 1.03282 | 0.00298 | 0.46650 | 1.58664 | 0.02393 | 0.08294 | 2.07667 | 0.25349 | 0.11200 |
| **Ankrd17** | 14.64185 | 0.04935 | 14.80125 | 4.61889 | 17.91816 | 13.15903 | 3.21015 | 2.41169 | 21.76037 | 10.44783 | 10.52146 | 5.75888 | 10.19823 | 16.01178 |
| **Ankrd22** | 0.01630 | 0.00435 | 0.01480 | 0.02517 | 0.01050 | 0.01368 | 0.00426 | 0.00879 | 0.01041 | 0.00660 | 0.00842 | 0.00855 | 0.23663 | 0.00528 |
| **Ankrd23** | 0.04124 | 0.00442 | 0.62567 | 0.57181 | 0.72174 | 0.78334 | 0.00368 | 0.00204 | 0.25275 | 0.00557 | 0.00240 | 0.00490 | 1.40235 | 0.13205 |
| **Ankrd24** | 0.22692 | 0.00391 | 0.00404 | 0.00000 | 0.11430 | 0.15370 | 0.00373 | 0.09910 | 0.01141 | 0.01022 | 0.14972 | 0.07652 | 0.00461 | 0.00398 |
| **Ankrd26** | 0.12229 | 0.00521 | 0.01094 | 0.00000 | 0.19873 | 0.03179 | 0.00471 | 0.02996 | 0.15385 | 0.00759 | 0.00574 | 0.03425 | 0.01166 | 1.74104 |
| **Ankrd27** | 0.80709 | 0.01676 | 1.67916 | 3.63569 | 0.06340 | 1.83137 | 0.01961 | 0.21713 | 0.56330 | 9.03052 | 1.70725 | 1.17199 | 0.12819 | 1.27160 |
| **Ankrd28** | 1.00592 | 1.46355 | 1.03606 | 1.83672 | 1.04811 | 1.59355 | 0.16724 | 0.33841 | 6.86550 | 0.04751 | 1.98156 | 1.20569 | 6.10228 | 1.94163 |
| **Ankrd33b** | 0.00000 | 0.00950 | 0.16193 | 0.00000 | 0.83251 | 0.00000 | 0.00279 | 0.81378 | 0.00722 | 0.00603 | 0.00000 | 3.03435 | 0.00385 | 0.11011 |
| **Ankrd35** | 0.04608 | 0.00355 | 0.00677 | 0.00000 | 0.54133 | 0.02761 | 0.00459 | 0.21785 | 0.02892 | 0.00688 | 0.00348 | 0.40750 | 5.28664 | 0.03458 |
| **Ankrd36** | 0.00473 | 0.00411 | 0.02748 | 0.20625 | 0.27495 | 0.00000 | 0.00392 | 0.00742 | 0.01284 | 0.00632 | 0.11232 | 0.02002 | 0.10735 | 0.01410 |
| **Ankrd37** | 0.66004 | 0.00415 | 0.14233 | 0.31820 | 0.44012 | 0.50796 | 0.00395 | 0.00402 | 0.01668 | 0.00984 | 0.00446 | 0.00564 | 0.77213 | 0.00036 |
| **Ankrd39** | 0.10721 | 0.00318 | 1.84251 | 0.97152 | 0.91464 | 0.75775 | 0.00923 | 0.02104 | 0.50212 | 0.01551 | 0.00000 | 1.30263 | 0.21888 | 0.00000 |
| **Ankrd40** | 8.30198 | 5.59338 | 5.05653 | 0.00000 | 6.53210 | 5.63924 | 3.87288 | 2.89573 | 5.95429 | 0.12143 | 2.61448 | 4.53254 | 4.41521 | 9.04951 |
| **Ankrd42** | 1.98612 | 0.01124 | 3.24022 | 0.00000 | 2.71157 | 1.07854 | 0.00322 | 0.00427 | 3.91048 | 0.00000 | 0.00000 | 0.82252 | 2.62497 | 3.54027 |
| **Ankrd44** | 0.54115 | 0.00279 | 0.47680 | 0.65280 | 2.49304 | 3.08754 | 0.00285 | 0.00000 | 4.68293 | 0.01671 | 0.00000 | 0.39232 | 4.91833 | 4.04044 |
| **Ankrd45** | 0.52836 | 0.00389 | 0.20795 | 0.32305 | 0.56020 | 0.13231 | 0.00375 | 0.00408 | 0.13338 | 0.01125 | 0.00642 | 0.52283 | 0.01075 | 1.76691 |
| **Ankrd46** | 0.66502 | 0.00099 | 1.62113 | 0.03213 | 2.13261 | 4.31284 | 2.28935 | 1.62324 | 1.90094 | 0.04082 | 3.85480 | 2.58441 | 1.58012 | 2.98028 |
| **Ankrd49** | 1.89957 | 0.00162 | 1.48898 | 0.87862 | 2.26778 | 1.43606 | 0.00270 | 0.86023 | 5.95875 | 0.01387 | 1.18462 | 3.30716 | 0.00000 | 2.25437 |
| **Ankrd50** | 0.00000 | 0.01210 | 0.15900 | 0.00000 | 0.06894 | 0.89287 | 2.52848 | 1.08612 | 0.00261 | 0.04285 | 0.10384 | 0.68381 | 2.13361 | 0.83198 |
| **Ankrd52** | 0.93191 | 0.01108 | 0.74651 | 0.64809 | 0.29597 | 0.12866 | 0.00358 | 0.29669 | 0.10759 | 0.01499 | 0.01186 | 0.00377 | 1.17762 | 0.47926 |
| **Ankrd54** | 1.74814 | 8.00088 | 0.24803 | 2.82474 | 1.63518 | 2.56924 | 1.94962 | 0.29592 | 0.00000 | 7.83790 | 0.50671 | 2.03455 | 1.94264 | 1.38823 |
| **Ankrd6** | 0.96095 | 0.01162 | 1.13430 | 0.26245 | 1.99052 | 2.61657 | 0.87346 | 0.02254 | 3.55028 | 0.02527 | 4.22462 | 0.01050 | 3.86798 | 5.86187 |
| **Ankrd9** | 0.00623 | 0.02985 | 0.01569 | 0.85325 | 1.83106 | 0.11628 | 0.18675 | 1.10100 | 0.00738 | 0.09293 | 0.00000 | 0.55610 | 0.06970 | 0.00000 |
| **Anks1** | 0.07573 | 0.05670 | 3.17705 | 0.61382 | 1.45507 | 3.11551 | 0.55649 | 2.10989 | 0.00000 | 0.00000 | 5.17690 | 2.14605 | 2.19006 | 1.94931 |
| **Anks1b** | 0.44127 | 0.00423 | 0.12994 | 0.00438 | 0.00000 | 0.00322 | 0.00416 | 0.00992 | 0.03459 | 0.00937 | 0.00533 | 0.17662 | 0.03293 | 0.40199 |
| **Anks3** | 0.81395 | 0.02652 | 2.70139 | 2.16648 | 2.42743 | 1.12616 | 0.97830 | 1.88933 | 0.00712 | 0.04892 | 2.42714 | 0.58428 | 1.06900 | 0.40322 |
| **Anks4b** | 0.00353 | 0.00411 | 0.00857 | 0.00000 | 0.00733 | 0.63679 | 0.00389 | 0.00348 | 0.02808 | 0.00560 | 0.08291 | 0.00802 | 0.71258 | 0.00393 |
| **Ankzf1** | 0.21775 | 0.01180 | 1.55124 | 1.56231 | 0.46445 | 1.05613 | 2.22866 | 0.66025 | 0.00000 | 0.03454 | 0.00000 | 0.90829 | 1.80244 | 1.51023 |
| **Anln** | 2.13613 | 3.73653 | 7.56238 | 0.81426 | 5.71015 | 5.24668 | 0.73701 | 1.30388 | 19.10470 | 0.55165 | 7.87591 | 2.07690 | 13.10728 | 9.19051 |
| **Ano1** | 0.00731 | 0.00453 | 0.02556 | 0.00969 | 0.37116 | 0.01258 | 0.00424 | 0.01072 | 0.00841 | 0.00631 | 0.00879 | 0.00815 | 0.00502 | 0.02214 |
| **Ano10** | 0.02140 | 0.03229 | 0.00000 | 0.05876 | 0.00000 | 1.95592 | 1.48902 | 0.03106 | 0.07943 | 0.02711 | 0.18090 | 0.00000 | 0.27275 | 0.04725 |
| **Ano2** | 0.03143 | 0.00452 | 0.42841 | 0.00000 | 0.02332 | 0.17643 | 0.00425 | 0.00457 | 0.00860 | 0.00650 | 0.00569 | 0.00836 | 0.03060 | 0.02011 |
| **Ano6** | 1.02281 | 0.11274 | 2.68848 | 2.66221 | 1.85403 | 4.68472 | 0.00221 | 6.59936 | 0.70403 | 0.02318 | 1.56222 | 0.19080 | 0.00000 | 3.50251 |
| **Ano7** | 0.00397 | 0.00439 | 0.19600 | 0.01098 | 0.37321 | 0.00696 | 0.00468 | 0.01023 | 0.01186 | 0.00759 | 0.00673 | 0.00717 | 0.09866 | 0.04269 |
| **Ano9** | 4.20848 | 0.00301 | 10.25205 | 2.53924 | 1.27342 | 0.00000 | 1.88713 | 0.00000 | 2.67941 | 0.08418 | 0.01846 | 0.67066 | 0.00000 | 2.68698 |
| **Anp32a** | 0.97275 | 0.12073 | 0.60234 | 0.79515 | 0.70587 | 2.49085 | 0.01952 | 3.83815 | 0.00979 | 0.01833 | 1.92255 | 0.00004 | 0.01928 | 1.00982 |
| **Anp32b** | 24.62402 | 10.84421 | 22.59538 | 19.12976 | 22.25506 | 24.10675 | 8.30796 | 8.84627 | 21.16757 | 12.89855 | 16.70092 | 16.78302 | 15.02764 | 19.60735 |
| **Anp32e** | 8.01133 | 0.21038 | 9.62057 | 7.63070 | 13.43812 | 17.92630 | 4.57856 | 7.08634 | 15.70446 | 3.84093 | 16.39560 | 8.87340 | 14.08528 | 12.26468 |
| **Anpep** | 0.84040 | 0.00373 | 0.67067 | 0.00000 | 0.65783 | 0.04889 | 0.00395 | 0.00278 | 0.93548 | 0.00484 | 0.00133 | 0.30840 | 0.09163 | 0.92388 |
| **Antxr1** | 0.19742 | 0.00400 | 0.32859 | 0.17400 | 0.67369 | 0.19785 | 0.00379 | 0.00211 | 0.01341 | 0.00541 | 0.00870 | 0.00586 | 1.41052 | 0.14667 |
| **Antxr2** | 0.00750 | 0.00425 | 0.00893 | 0.01200 | 0.13619 | 0.01148 | 0.00437 | 0.01025 | 0.01056 | 0.00590 | 0.00832 | 0.00854 | 0.57251 | 0.33580 |
| **Anxa1** | 0.14026 | 0.00452 | 0.00638 | 0.00992 | 0.40765 | 0.01155 | 0.00437 | 0.01174 | 0.01004 | 0.00618 | 0.00769 | 0.00742 | 0.00435 | 0.00812 |
| **Anxa11** | 1.60891 | 0.48080 | 1.20124 | 0.91096 | 5.50897 | 0.00000 | 0.92549 | 1.21309 | 0.42037 | 0.01950 | 0.80332 | 1.08675 | 1.22513 | 1.68514 |
| **Anxa2** | 0.00000 | 8.77299 | 0.00000 | 3.49746 | 1.30182 | 17.55314 | 7.59917 | 23.47982 | 2.28068 | 17.85037 | 4.44348 | 0.55128 | 0.00000 | 0.00000 |
| **Anxa3** | 0.33715 | 0.02871 | 0.40699 | 1.13349 | 0.39109 | 9.59450 | 0.00367 | 3.01846 | 0.00432 | 0.03041 | 0.00000 | 0.00000 | 0.00351 | 0.01472 |
| **Anxa4** | 0.20813 | 0.00356 | 2.16070 | 3.22594 | 1.55632 | 0.44617 | 0.00298 | 0.00000 | 1.65850 | 0.00154 | 2.54509 | 0.36402 | 0.00000 | 2.01894 |
| **Anxa5** | 0.06013 | 0.01395 | 0.35842 | 0.00000 | 1.51527 | 1.05818 | 0.00375 | 1.23688 | 0.00661 | 0.00037 | 1.68560 | 0.58989 | 0.00188 | 0.00726 |
| **Anxa6** | 0.00000 | 4.21146 | 6.23056 | 1.07339 | 0.10764 | 3.41258 | 6.47483 | 9.73846 | 0.00000 | 16.95544 | 5.11414 | 2.03564 | 0.45111 | 0.99549 |
| **Anxa7** | 7.69465 | 6.31515 | 16.53118 | 15.99525 | 19.30159 | 21.11239 | 7.51998 | 5.22913 | 7.21823 | 12.60024 | 11.99561 | 8.75882 | 3.23991 | 22.49737 |
| **Anxa8** | 0.07467 | 0.01396 | 0.84413 | 1.45191 | 0.27373 | 2.18649 | 0.00426 | 0.00751 | 0.01427 | 0.00434 | 0.47056 | 0.00000 | 0.00177 | 0.02437 |
| **Anxa9** | 0.67308 | 5.36940 | 2.77485 | 2.43252 | 0.00000 | 2.85075 | 4.19312 | 3.43093 | 0.10664 | 0.05829 | 4.62308 | 0.00000 | 0.36873 | 0.35929 |
| **Aoc1** | 0.00388 | 0.00406 | 0.00209 | 0.50285 | 0.60021 | 0.00002 | 0.00431 | 0.00648 | 0.03517 | 0.00600 | 0.01140 | 0.01577 | 0.00320 | 0.00000 |
| **Aoc2** | 0.85799 | 0.01128 | 1.18464 | 0.18050 | 0.23017 | 0.58108 | 0.00356 | 0.00000 | 0.19747 | 1.00161 | 0.00134 | 0.00000 | 0.21523 | 1.17388 |

| | | | | | | | | | | | | | | |
|---|---|---|---|---|---|---|---|---|---|---|---|---|---|---|
| **Aoc3** | 0.00495 | 0.01264 | 0.05937 | 0.00000 | 0.23957 | 0.07122 | 0.00414 | 0.00824 | 0.01087 | 0.00634 | 0.00725 | 0.00955 | 0.30099 | 0.05308 |
| **Aox2** | 0.00758 | 0.00478 | 0.02056 | 0.00338 | 0.00810 | 0.20583 | 0.00426 | 0.01274 | 0.00770 | 0.01025 | 0.00664 | 0.00982 | 0.00391 | 0.14253 |
| **Aox4** | 1.32535 | 0.00396 | 0.00000 | 0.00000 | 0.06377 | 0.80888 | 0.00378 | 0.00140 | 0.00519 | 0.00400 | 0.44552 | 0.00663 | 0.60979 | 0.14748 |
| **Ap1ar** | 1.54561 | 0.00858 | 2.68272 | 0.46896 | 3.59229 | 3.71866 | 0.17299 | 1.93579 | 4.10304 | 0.02224 | 1.78266 | 2.52587 | 1.82312 | 10.15440 |
| **Ap1b1** | 6.96829 | 0.11246 | 9.81940 | 6.54567 | 6.72795 | 10.66421 | 0.48562 | 5.96604 | 0.21695 | 3.23471 | 3.71761 | 4.07792 | 2.76094 | 5.23413 |
| **Ap1g1** | 6.65644 | 2.00384 | 12.81233 | 8.39152 | 10.92134 | 11.26002 | 0.96339 | 5.90968 | 14.46780 | 5.11365 | 12.36218 | 8.18706 | 9.98173 | 11.77219 |
| **Ap1g2** | 0.00251 | 0.00448 | 0.04042 | 0.93830 | 0.00670 | 0.14234 | 0.00415 | 0.00502 | 0.02178 | 0.01246 | 0.00503 | 0.00700 | 0.00703 | 0.00328 |
| **Ap1m1** | 3.83516 | 0.05985 | 6.84370 | 0.00000 | 8.17265 | 1.83791 | 0.99341 | 2.76264 | 0.10817 | 4.88269 | 0.57548 | 4.31667 | 3.39203 | 3.06078 |
| **Ap1m2** | 3.53056 | 5.07060 | 12.08733 | 14.83606 | 6.03602 | 9.63871 | 4.92836 | 8.04675 | 0.69854 | 14.98868 | 7.46376 | 2.96197 | 1.57274 | 12.66956 |
| **Ap1s1** | 6.09840 | 4.30435 | 20.82690 | 20.67092 | 19.92313 | 11.29113 | 1.74336 | 17.04093 | 0.16695 | 3.84794 | 7.70169 | 4.22128 | 1.97265 | 6.68339 |
| **Ap1s2** | 0.60387 | 0.00426 | 0.06708 | 0.00277 | 0.14941 | 0.09225 | 0.00417 | 0.00538 | 0.00824 | 0.00590 | 0.01529 | 0.00912 | 0.35178 | 0.05960 |
| **Ap1s3** | 3.44264 | 0.03283 | 0.34652 | 0.39165 | 2.79146 | 0.55210 | 0.01490 | 1.14113 | 11.63670 | 0.00535 | 2.71924 | 0.22765 | 0.39527 | 3.07664 |
| **Ap2a1** | 0.08011 | 3.66067 | 3.80413 | 3.73626 | 1.66786 | 0.30950 | 0.23992 | 0.35059 | 0.00492 | 6.74846 | 0.97306 | 3.42340 | 0.00718 | 0.00000 |
| **Ap2a2** | 5.24546 | 8.04586 | 19.74813 | 14.67698 | 9.74578 | 15.20652 | 3.31528 | 19.44341 | 1.83180 | 0.45493 | 7.29816 | 2.64986 | 1.11651 | 2.18816 |
| **Ap2b1** | 2.71193 | 0.00073 | 7.33345 | 6.98093 | 3.40893 | 8.59908 | 0.01850 | 2.27884 | 0.79615 | 0.07353 | 5.52912 | 1.62264 | 2.06204 | 6.09466 |
| **Ap2m1** | 19.28087 | 7.07391 | 27.05056 | 22.54594 | 14.21960 | 28.10607 | 7.58371 | 19.84971 | 11.38257 | 18.03521 | 21.09325 | 12.96169 | 3.30748 | 20.34706 |
| **Ap2s1** | 0.00000 | 5.88996 | 6.40505 | 18.68244 | 10.16272 | 0.00000 | 5.52175 | 14.38267 | 0.00000 | 4.37119 | 0.00083 | 8.93813 | 0.01352 | 0.00000 |
| **Ap3b1** | 2.84583 | 9.78261 | 12.32697 | 5.91480 | 6.32657 | 12.23703 | 4.23797 | 2.63957 | 4.12281 | 0.07079 | 8.95123 | 9.74626 | 0.94117 | 3.06335 |
| **Ap3b2** | 0.12598 | 0.00403 | 2.96881 | 0.89039 | 1.15909 | 0.08762 | 0.00363 | 0.00000 | 0.02107 | 0.01851 | 0.41876 | 0.00802 | 0.00000 | 0.36234 |
| **Ap3d1** | 6.17888 | 11.36416 | 15.09265 | 10.25915 | 11.44046 | 12.42382 | 3.21131 | 6.09618 | 9.62783 | 0.20211 | 8.16090 | 11.41313 | 5.25568 | 14.09906 |
| **Ap3m1** | 8.77863 | 0.02970 | 6.97369 | 2.27023 | 4.85132 | 8.21677 | 0.28840 | 2.78068 | 8.83119 | 0.03907 | 4.30475 | 2.41642 | 1.72034 | 3.51455 |
| **Ap3m2** | 0.01729 | 0.00446 | 0.06938 | 0.00000 | 0.45499 | 0.01127 | 0.00405 | 0.01146 | 0.63542 | 0.00853 | 0.00535 | 0.00822 | 0.00577 | 0.00000 |
| **Ap3s1** | 7.48980 | 0.00918 | 1.67077 | 0.64620 | 1.98730 | 2.25030 | 0.02010 | 2.01140 | 1.29369 | 0.00648 | 3.16232 | 1.11574 | 1.12076 | 1.49533 |
| **Ap3s2** | 4.78658 | 0.10861 | 8.78132 | 3.31608 | 7.49669 | 6.38936 | 0.33532 | 6.48916 | 1.94365 | 5.59837 | 3.12470 | 5.42459 | 1.01024 | 3.70810 |
| **Ap4b1** | 12.16253 | 0.00649 | 6.74310 | 2.08214 | 10.62513 | 2.57202 | 0.07999 | 1.28609 | 5.50228 | 0.00000 | 9.90611 | 1.99297 | 5.85956 | 15.08519 |
| **Ap4e1** | 1.62140 | 0.02730 | 0.91031 | 0.41696 | 1.20521 | 1.97234 | 1.19245 | 0.18798 | 1.79431 | 0.01783 | 4.40452 | 0.18221 | 3.96322 | 1.35877 |
| **Ap4m1** | 0.63016 | 0.01416 | 2.11572 | 0.35921 | 1.32008 | 0.00000 | 0.00335 | 0.79673 | 0.95897 | 0.01160 | 3.38461 | 0.00291 | 0.00072 | 3.85945 |
| **Ap4s1** | 11.50060 | 0.00870 | 6.79890 | 8.50900 | 13.05729 | 4.71447 | 0.01876 | 1.06105 | 7.52501 | 0.00000 | 0.00000 | 0.32101 | 5.28613 | 11.13273 |
| **Ap5b1** | 0.03681 | 0.00386 | 0.03923 | 0.68990 | 0.07022 | 0.00000 | 0.00374 | 0.36479 | 0.01029 | 0.00511 | 0.02285 | 0.01235 | 0.00881 | 0.01383 |
| **Ap5m1** | 2.07900 | 0.00224 | 0.68220 | 1.15175 | 0.61303 | 0.17411 | 0.01483 | 0.00693 | 2.22196 | 0.02064 | 0.95135 | 2.19233 | 1.38396 | 2.94042 |
| **Ap5s1** | 0.03920 | 0.02713 | 1.12282 | 1.89351 | 0.85498 | 2.38068 | 0.40035 | 3.87010 | 0.00000 | 0.04558 | 0.00000 | 0.35396 | 0.42399 | 1.63086 |
| **Ap5z1** | 0.74344 | 0.03476 | 3.50269 | 7.54929 | 3.60035 | 1.10937 | 0.77844 | 2.78883 | 0.85723 | 0.08153 | 1.67904 | 2.06706 | 2.94233 | 0.49114 |
| **Apaf1** | 4.68675 | 1.30452 | 3.43209 | 0.00000 | 12.45572 | 3.82641 | 0.87332 | 4.76722 | 7.88031 | 0.06230 | 5.89204 | 1.84325 | 5.25221 | 11.56860 |
| **Apba1** | 0.01275 | 0.00444 | 0.01380 | 0.01324 | 0.01088 | 0.01272 | 0.00433 | 0.01032 | 0.00957 | 0.00692 | 0.00877 | 0.01000 | 0.00722 | 0.07208 |
| **Apba2** | 0.00858 | 0.00466 | 0.01312 | 0.02171 | 0.04535 | 0.01660 | 0.01624 | 0.01786 | 0.10674 | 0.00905 | 0.01267 | 0.01230 | 0.00828 | 0.03114 |
| **Apba3** | 0.00000 | 0.01196 | 1.13941 | 1.94295 | 0.89017 | 0.00000 | 0.00419 | 0.54676 | 0.01526 | 0.01793 | 0.02530 | 0.71366 | 0.54386 | 0.09022 |
| **Apbb1** | 0.06100 | 0.06233 | 1.21392 | 3.27482 | 0.07227 | 2.73912 | 0.21316 | 3.68911 | 0.00000 | 7.31816 | 0.00000 | 2.63111 | 0.37182 | 0.41541 |
| **Apbb1ip** | 0.05569 | 0.00380 | 0.00000 | 0.70359 | 0.00000 | 0.36385 | 0.00370 | 0.00127 | 0.05486 | 0.00392 | 0.26196 | 0.09915 | 0.00427 | 0.00000 |
| **Apbb2** | 0.59111 | 0.00272 | 0.45115 | 0.00000 | 1.17906 | 1.97972 | 0.00399 | 0.48165 | 1.53942 | 0.02769 | 0.04950 | 1.96897 | 1.59070 | 2.38966 |
| **Apbb3** | 0.00003 | 0.02857 | 0.07985 | 1.82620 | 0.09851 | 0.02713 | 0.01721 | 1.20768 | 0.01178 | 0.00446 | 0.31325 | 0.01199 | 0.00774 | 0.00027 |
| **Apc** | 3.02725 | 0.00852 | 3.25892 | 2.95133 | 5.36350 | 6.74049 | 0.70310 | 0.89327 | 11.20246 | 0.07202 | 0.15182 | 2.23742 | 6.59158 | 12.65553 |
| **Apc2** | 0.00231 | 0.00414 | 0.01841 | 0.16762 | 0.00376 | 0.02916 | 0.00410 | 0.00691 | 0.00523 | 0.00676 | 0.00571 | 0.00615 | 0.55565 | 0.21449 |
| **Apcdd1** | 0.05425 | 0.01008 | 0.00000 | 1.01070 | 0.03522 | 0.06442 | 0.00372 | 0.00264 | 0.02057 | 0.00462 | 0.01070 | 1.19081 | 0.00945 | 0.12275 |
| **Apeh** | 0.85916 | 0.05505 | 1.39126 | 1.78432 | 0.00000 | 2.87215 | 0.86570 | 6.91286 | 0.00000 | 0.08547 | 2.48482 | 5.31219 | 0.45504 | 0.00823 |
| **Apela** | 0.00295 | 0.00311 | 0.00000 | 1.08414 | 0.00118 | 0.41371 | 0.00361 | 0.00000 | 0.01317 | 0.01007 | 1.86989 | 0.55658 | 0.00705 | 0.00000 |
| **Apex1** | 18.03530 | 8.07802 | 31.04808 | 30.07954 | 24.13035 | 14.52792 | 6.34119 | 9.78808 | 15.45216 | 0.57668 | 18.39712 | 17.00455 | 9.02295 | 22.94779 |
| **Apex2** | 0.77444 | 0.02659 | 1.14365 | 2.77571 | 1.76035 | 3.16695 | 3.82768 | 3.72038 | 2.24752 | 0.11668 | 1.81376 | 1.72135 | 1.95865 | 2.73347 |
| **Aph1a** | 0.00000 | 0.02512 | 1.42555 | 3.02488 | 2.12057 | 2.33604 | 0.00923 | 2.05500 | 0.28415 | 0.01690 | 0.37471 | 2.24226 | 1.60360 | 0.76268 |
| **Aph1b** | 0.00791 | 0.00461 | 0.00552 | 0.01099 | 0.10022 | 0.48770 | 0.01219 | 0.13866 | 0.01879 | 0.00676 | 0.00676 | 0.00530 | 0.17333 | 0.04216 |
| **Aph1c** | 0.01249 | 0.00459 | 0.00805 | 0.01697 | 0.01447 | 1.13419 | 0.01920 | 0.43620 | 0.05040 | 0.00756 | 0.00651 | 0.00633 | 0.08378 | 0.00385 |
| **Api5** | 11.47462 | 0.10117 | 18.81515 | 13.22577 | 14.91519 | 16.05728 | 3.92512 | 5.41463 | 18.11773 | 0.18247 | 11.90785 | 10.62996 | 0.66040 | 5.35597 |
| **Apip** | 4.43876 | 0.00832 | 1.39186 | 0.00000 | 1.81597 | 3.80183 | 2.47242 | 2.69859 | 0.34760 | 0.06860 | 3.33170 | 2.88570 | 0.53174 | 1.23450 |
| **Apitd1** | 3.04194 | 0.08550 | 4.73595 | 4.79752 | 5.84582 | 3.61930 | 3.42015 | 2.88223 | 0.06477 | 0.13896 | 0.17219 | 2.06953 | 6.78907 | 5.29703 |
| **Aplf** | 1.35918 | 0.25693 | 0.85115 | 2.06668 | 2.43171 | 1.45128 | 0.01487 | 0.09627 | 1.04056 | 0.84300 | 1.02776 | 0.88698 | 1.90401 | 0.91807 |
| **Aplp1** | 0.02578 | 0.00502 | 0.00817 | 0.04832 | 0.12098 | 0.02334 | 0.00435 | 0.01668 | 0.01663 | 0.01815 | 0.00835 | 0.00697 | 0.01572 | 0.01238 |

| | | | | | | | | | | | | | | |
|---|---|---|---|---|---|---|---|---|---|---|---|---|---|---|
| **Aplp2** | 3.91520 | 0.09646 | 4.72851 | 8.23821 | 4.57648 | 10.60245 | 4.62961 | 8.37055 | 7.07352 | 0.08260 | 8.28898 | 12.96173 | 15.06425 | 3.39221 |
| **Apmap** | 1.78657 | 0.03669 | 2.01405 | 5.19155 | 0.82223 | 4.32209 | 0.01943 | 4.13144 | 2.54177 | 0.02480 | 3.71629 | 3.56608 | 1.52614 | 2.80209 |
| **Apoa1** | 0.00000 | 0.03469 | 7.86209 | 1.08001 | 0.00000 | 0.17482 | 0.52739 | 4.04804 | 0.00493 | 0.01855 | 0.02529 | 0.00566 | 0.00933 | 0.00000 |
| **Apoa1bp** | 0.25989 | 0.25914 | 1.23499 | 4.65626 | 1.13713 | 0.53369 | 0.46443 | 2.69221 | 0.00000 | 6.08387 | 1.33585 | 4.62587 | 2.02530 | 1.12992 |
| **Apoa4** | 0.00000 | 0.00402 | 1.03950 | 0.94030 | 0.06526 | 0.12214 | 0.00405 | 0.00000 | 0.01863 | 0.00302 | 0.00991 | 0.01420 | 0.00312 | 0.00000 |
| **Apoa5** | 0.00000 | 0.00407 | 0.25437 | 0.09736 | 0.00000 | 0.00740 | 0.00420 | 0.01098 | 0.00875 | 0.00570 | 0.00640 | 0.00579 | 1.14638 | 0.00000 |
| **Apobec1** | 0.00000 | 0.00475 | 0.12437 | 0.00000 | 0.19966 | 0.09209 | 0.00444 | 0.00770 | 0.03655 | 0.00553 | 0.00386 | 0.12253 | 0.00326 | 1.41602 |
| **Apobec2** | 0.00137 | 0.01533 | 0.23260 | 0.00000 | 0.00635 | 0.00193 | 0.00374 | 0.14866 | 0.01160 | 0.01879 | 0.00272 | 0.77337 | 0.00579 | 0.01543 |
| **Apobec3** | 3.61563 | 0.00296 | 2.62566 | 2.06026 | 2.27807 | 0.00000 | 0.00788 | 0.00000 | 5.43589 | 0.01124 | 2.10907 | 0.02093 | 0.90134 | 3.44570 |
| **Apobr** | 0.01081 | 0.02575 | 0.23881 | 0.25273 | 0.00000 | 0.73007 | 1.84929 | 2.16443 | 0.00693 | 0.05189 | 1.02365 | 1.21926 | 0.04876 | 0.01990 |
| **Apoc1** | 0.98913 | 0.17604 | 0.02510 | 2.36512 | 0.94755 | 2.23457 | 0.73171 | 5.86673 | 0.10303 | 0.21080 | 0.00000 | 5.86398 | 1.13470 | 0.00000 |
| **Apoc2** | 0.08979 | 0.00437 | 0.04230 | 0.02453 | 0.25394 | 0.01070 | 0.00470 | 0.01095 | 0.01222 | 0.00870 | 0.00715 | 0.06586 | 0.02419 | 0.11248 |
| **Apod** | 0.02900 | 0.00364 | 0.00762 | 0.04777 | 0.01911 | 0.03283 | 0.03035 | 0.00967 | 0.02238 | 0.00817 | 0.00565 | 0.90150 | 0.02289 | 0.14547 |
| **Apoe** | 0.07927 | 0.04267 | 0.30774 | 3.21307 | 0.95964 | 0.02983 | 0.00937 | 0.05004 | 0.00804 | 0.02037 | 0.04938 | 1.79341 | 1.29122 | 0.00000 |
| **Apof** | 0.03399 | 0.01320 | 0.02807 | 0.07208 | 0.05509 | 0.04289 | 0.01035 | 0.01465 | 0.13846 | 0.01157 | 0.01242 | 0.01154 | 0.09522 | 0.06151 |
| **Apoh** | 0.00486 | 0.00435 | 0.00967 | 0.00609 | 0.04652 | 0.03108 | 0.00431 | 0.00822 | 0.09057 | 0.00615 | 0.07430 | 0.00786 | 0.19783 | 0.01438 |
| **Apol10a** | 0.00982 | 0.00444 | 0.03371 | 0.01474 | 0.02783 | 0.01244 | 0.00433 | 0.01088 | 0.01082 | 0.00678 | 0.00863 | 0.00953 | 0.00789 | 0.01241 |
| **Apol6** | 0.04087 | 0.02151 | 0.01376 | 0.05038 | 0.04558 | 0.07365 | 0.02283 | 0.03396 | 0.08518 | 0.03768 | 0.00748 | 0.02791 | 0.11438 | 0.15573 |
| **Apol7a** | 0.01660 | 0.00430 | 0.05339 | 0.00776 | 0.01441 | 0.02748 | 0.07948 | 0.00556 | 0.01354 | 0.00608 | 0.01325 | 0.00933 | 0.00567 | 0.08179 |
| **Apol7b** | 0.03333 | 0.01201 | 1.03228 | 0.00000 | 0.24946 | 0.42077 | 1.75843 | 0.01047 | 0.10417 | 0.03491 | 0.03416 | 0.00868 | 3.35609 | 1.62961 |
| **Apol7c** | 0.01635 | 0.00436 | 0.02901 | 0.01158 | 0.01320 | 0.01166 | 0.05133 | 0.00809 | 0.01212 | 0.00674 | 0.00874 | 0.01038 | 0.00551 | 0.06400 |
| **Apol7e** | 0.00000 | 0.01182 | 0.62410 | 0.00000 | 0.21173 | 0.34887 | 1.05555 | 0.05983 | 0.22369 | 0.01090 | 0.02299 | 0.00193 | 3.40698 | 0.84613 |
| **Apom** | 0.45586 | 0.00297 | 0.29184 | 0.31663 | 1.24547 | 0.28075 | 0.00351 | 0.00164 | 0.01342 | 0.00456 | 0.01611 | 0.39482 | 6.66708 | 0.63437 |
| **Apon** | 0.00000 | 0.00577 | 0.56573 | 0.00000 | 0.00000 | 0.04434 | 0.00423 | 0.30235 | 0.22752 | 0.00865 | 0.01328 | 0.00826 | 0.07427 | 0.59279 |
| **Apoo** | 2.92166 | 0.27992 | 1.10683 | 0.65177 | 2.34159 | 2.18948 | 1.49557 | 4.10612 | 1.59420 | 0.05575 | 1.56487 | 2.04146 | 0.96142 | 1.17060 |
| **Apool** | 1.52323 | 0.22823 | 1.99304 | 2.50872 | 1.20093 | 2.40225 | 0.00684 | 5.72118 | 3.22657 | 2.51026 | 5.40902 | 2.74407 | 0.22899 | 0.70219 |
| **Apopt1** | 1.80930 | 0.10559 | 0.37844 | 1.51015 | 0.73094 | 0.84181 | 0.10582 | 0.76134 | 1.59814 | 0.41494 | 0.06164 | 1.75121 | 2.18804 | 1.27431 |
| **App** | 0.84936 | 0.01213 | 0.28204 | 0.65694 | 0.22904 | 0.00000 | 0.00408 | 1.39804 | 0.00894 | 0.01009 | 0.00015 | 0.05861 | 0.54485 | 0.03269 |
| **Appbp2** | 3.46229 | 0.02936 | 9.33091 | 0.00000 | 9.44205 | 2.82588 | 0.06909 | 0.03446 | 11.83148 | 0.00512 | 6.15949 | 6.15721 | 3.29166 | 14.99351 |
| **Appbp2os** | 0.00784 | 0.00496 | 0.01338 | 0.00000 | 0.01195 | 0.12599 | 0.00417 | 0.00824 | 0.01014 | 0.02506 | 0.16806 | 0.08676 | 0.00734 | 0.02927 |
| **Appl1** | 7.58868 | 0.00766 | 4.03084 | 1.83912 | 4.02703 | 4.86224 | 0.00781 | 0.05841 | 5.09329 | 0.01368 | 3.35322 | 4.67545 | 1.00254 | 3.97850 |
| **Appl2** | 0.00451 | 0.02067 | 0.04594 | 0.27728 | 0.54792 | 0.54798 | 3.41940 | 0.12755 | 0.00287 | 0.09159 | 1.23568 | 1.87246 | 0.46401 | 0.16005 |
| **Aprt** | 21.14973 | 9.01579 | 29.91254 | 36.87130 | 32.91543 | 23.06065 | 4.40711 | 15.94693 | 0.60422 | 20.86455 | 13.52382 | 9.72580 | 8.76930 | 21.56926 |
| **Aptx** | 4.50549 | 0.03617 | 6.89821 | 6.76063 | 5.08761 | 3.79482 | 0.42966 | 0.00000 | 3.39586 | 7.43634 | 3.48437 | 4.38386 | 1.55581 | 7.25055 |
| **Aqp11** | 3.63247 | 0.00372 | 2.19672 | 0.01925 | 2.12127 | 0.00000 | 0.00378 | 0.00446 | 0.12000 | 0.00360 | 0.00380 | 0.00135 | 0.52614 | 3.02197 |
| **Aqp3** | 1.00071 | 0.03640 | 1.24431 | 4.38495 | 0.00000 | 3.73274 | 0.02828 | 1.40919 | 0.84004 | 1.04603 | 3.23077 | 0.37507 | 0.14558 | 2.50663 |
| **Aqp7** | 0.03982 | 0.02935 | 0.07725 | 0.14759 | 0.02254 | 0.11325 | 0.01606 | 0.04631 | 0.23913 | 0.04288 | 0.01622 | 0.05277 | 0.12293 | 0.26738 |
| **Aqp8** | 0.00133 | 0.67314 | 0.07727 | 0.00000 | 0.00216 | 0.01108 | 2.92415 | 0.10697 | 0.00584 | 0.06440 | 0.00000 | 16.29780 | 0.00487 | 0.13707 |
| **Aqp9** | 10.16720 | 0.03659 | 11.37414 | 11.60851 | 9.23910 | 9.38304 | 7.84375 | 5.92295 | 19.13657 | 0.25993 | 6.46907 | 0.00000 | 6.33574 | 21.22659 |
| **Aqr** | 7.30759 | 0.03786 | 6.97129 | 2.70095 | 7.36222 | 5.58533 | 0.00661 | 3.03283 | 1.89507 | 0.05224 | 3.67336 | 4.97610 | 5.97220 | 5.40358 |
| **Araf** | 3.46079 | 0.06565 | 12.00126 | 8.49817 | 4.76025 | 10.52149 | 1.66602 | 4.80362 | 8.13041 | 0.08505 | 0.92890 | 3.76085 | 1.11353 | 6.78661 |
| **Arap1** | 0.52300 | 0.01114 | 3.62190 | 1.23391 | 2.33290 | 1.56397 | 1.38267 | 0.33787 | 0.11586 | 0.11257 | 0.03674 | 0.73281 | 0.10920 | 4.45193 |
| **Arap2** | 0.36048 | 0.00285 | 0.63076 | 0.23734 | 2.22302 | 0.21166 | 0.00328 | 0.49359 | 2.69814 | 0.00017 | 1.81583 | 0.79413 | 4.03558 | 6.16614 |
| **Arcn1** | 7.17121 | 0.11531 | 5.97701 | 2.01220 | 6.93310 | 7.95094 | 3.20918 | 7.00406 | 7.82329 | 0.20497 | 8.30215 | 6.88144 | 9.24420 | 8.86558 |
| **Arel1** | 2.94013 | 0.02375 | 3.28009 | 0.39459 | 3.37043 | 4.82422 | 2.77509 | 0.15102 | 3.82218 | 0.12737 | 1.41556 | 3.32088 | 3.51322 | 4.80534 |
| **Arf1** | 8.74098 | 8.91733 | 35.79721 | 29.75333 | 22.85582 | 30.40708 | 7.65751 | 28.65772 | 0.29245 | 17.85057 | 19.92212 | 21.68758 | 5.11660 | 16.78276 |
| **Arf2** | 2.23546 | 5.93213 | 3.65734 | 5.49698 | 5.33012 | 10.74699 | 2.54954 | 2.81940 | 1.42315 | 0.14156 | 2.88401 | 0.02689 | 4.32582 | 7.37718 |
| **Arf3** | 4.67810 | 0.10160 | 10.96966 | 5.73054 | 7.05255 | 12.41265 | 0.05488 | 7.23175 | 4.94347 | 0.08771 | 9.08091 | 10.88137 | 1.41314 | 10.53749 |
| **Arf4** | 6.42918 | 0.18048 | 16.35357 | 11.23947 | 8.13052 | 19.19515 | 5.14953 | 8.60517 | 9.76194 | 0.28745 | 10.50161 | 15.85069 | 3.63052 | 4.36250 |
| **Arf5** | 0.00000 | 0.14466 | 1.22383 | 2.47813 | 0.38254 | 0.00000 | 0.06589 | 3.34374 | 0.00849 | 0.03687 | 0.15804 | 2.01895 | 0.05606 | 0.02174 |
| **Arf6** | 1.15526 | 0.06273 | 8.52718 | 5.76497 | 8.89798 | 6.59660 | 2.97905 | 6.26056 | 6.36740 | 0.12206 | 3.65942 | 3.94398 | 13.03666 | 2.06113 |
| **Arfgap1** | 3.91376 | 0.17051 | 8.29058 | 12.91826 | 4.92118 | 7.14488 | 5.04240 | 5.62010 | 0.01145 | 0.15182 | 3.95320 | 6.29681 | 0.00000 | 5.90386 |
| **Arfgap2** | 0.00000 | 0.00347 | 1.40016 | 0.91013 | 0.30890 | 0.27702 | 0.00464 | 0.70887 | 0.06146 | 0.00131 | 3.17744 | 0.07450 | 0.00000 | 0.65201 |
| **Arfgap3** | 0.02295 | 0.01264 | 0.00000 | 1.08807 | 0.22407 | 0.21170 | 0.60982 | 0.62051 | 0.02871 | 0.02249 | 0.00000 | 0.05862 | 0.11805 | 0.33058 |
| **Arfgef1** | 0.80558 | 0.43590 | 1.15532 | 0.55712 | 2.13574 | 3.95907 | 2.91289 | 4.53069 | 8.06130 | 0.01273 | 2.65492 | 5.08753 | 2.56125 | 8.68095 |

| | | | | | | | | | | | | | | |
|---|---|---|---|---|---|---|---|---|---|---|---|---|---|---|
| **Arfgef2** | 3.93572 | 0.17230 | 5.09089 | 2.35359 | 5.56821 | 6.22508 | 5.28513 | 3.24798 | 0.31283 | 0.12920 | 5.93404 | 2.81091 | 3.52246 | 3.76512 |
| **Arfgef3** | 0.00817 | 0.00442 | 0.01701 | 0.01191 | 0.00663 | 0.01219 | 0.00430 | 0.00997 | 0.01571 | 0.00678 | 0.00816 | 0.01384 | 0.02149 | 0.18680 |
| **Arfip1** | 3.49135 | 0.03169 | 3.66281 | 1.48236 | 4.59713 | 4.58673 | 1.11085 | 4.37793 | 8.28556 | 0.06516 | 1.61349 | 1.99345 | 1.77526 | 3.24391 |
| **Arfip2** | 6.02652 | 0.06251 | 3.86526 | 4.18243 | 2.61266 | 4.53479 | 1.90209 | 7.76819 | 0.74109 | 0.04915 | 0.13581 | 3.30465 | 0.25930 | 0.23610 |
| **Arfrp1** | 5.21310 | 0.09325 | 6.81479 | 11.29747 | 9.56464 | 4.31453 | 0.16757 | 4.53298 | 1.02949 | 0.04708 | 3.26840 | 3.64633 | 7.78428 | 8.23603 |
| **Arg1** | 0.28099 | 0.00345 | 3.88339 | 0.00000 | 1.70936 | 0.00000 | 0.65653 | 0.00000 | 2.70401 | 0.00000 | 0.00260 | 0.00017 | 1.90999 | 7.91050 |
| **Arg2** | 0.14011 | 0.00372 | 1.57315 | 0.05256 | 0.35650 | 1.67835 | 0.00357 | 0.06169 | 0.96957 | 0.01542 | 0.44105 | 0.00000 | 2.12538 | 1.28347 |
| **Arglu1** | 3.22968 | 0.60439 | 6.55061 | 5.12895 | 4.58235 | 5.43182 | 1.85063 | 4.43833 | 8.50281 | 4.30001 | 1.88364 | 4.52020 | 20.04309 | 3.40406 |
| **Arhgap1** | 15.70822 | 8.22128 | 15.27767 | 5.86109 | 16.51974 | 21.32713 | 3.22041 | 8.72306 | 8.63619 | 0.28549 | 9.10743 | 4.01417 | 5.32990 | 14.92439 |
| **Arhgap10** | 0.78879 | 0.00397 | 0.83522 | 0.66689 | 2.20950 | 0.04499 | 0.00380 | 0.20346 | 0.15600 | 0.00589 | 0.08623 | 0.10949 | 0.03699 | 2.55253 |
| **Arhgap11a** | 4.71887 | 0.02141 | 4.36590 | 3.48968 | 7.83044 | 5.21177 | 2.36096 | 4.69480 | 9.33149 | 0.08802 | 1.93376 | 4.86100 | 19.45016 | 16.91922 |
| **Arhgap12** | 0.22884 | 0.01030 | 1.03523 | 0.00000 | 1.62946 | 1.73237 | 0.00183 | 0.23038 | 8.55154 | 0.00000 | 5.34337 | 1.50268 | 6.79774 | 1.22856 |
| **Arhgap15** | 0.01291 | 0.00444 | 0.01552 | 0.01281 | 0.04758 | 0.01277 | 0.00425 | 0.01068 | 0.02119 | 0.00668 | 0.00865 | 0.00978 | 0.00881 | 0.01055 |
| **Arhgap17** | 2.84295 | 0.06128 | 1.47125 | 2.31080 | 0.30481 | 0.13728 | 1.22671 | 2.28770 | 0.00998 | 0.04489 | 0.92922 | 2.38357 | 0.00000 | 1.08938 |
| **Arhgap18** | 3.32160 | 0.00320 | 12.03227 | 2.27099 | 8.39103 | 2.77378 | 0.00098 | 0.00000 | 12.43117 | 0.00000 | 0.15338 | 0.39161 | 0.00000 | 4.84110 |
| **Arhgap19** | 2.48735 | 0.01696 | 3.02958 | 0.36451 | 0.51061 | 2.31997 | 0.03564 | 0.89234 | 3.13963 | 0.03208 | 3.64838 | 2.06160 | 1.82803 | 9.28961 |
| **Arhgap20** | 0.00784 | 0.00452 | 0.00262 | 0.01522 | 0.07920 | 0.00791 | 0.00437 | 0.00988 | 1.16782 | 0.00723 | 0.00581 | 0.00921 | 0.00275 | 0.00000 |
| **Arhgap20os** | 0.38265 | 0.00454 | 0.04455 | 0.11071 | 0.20966 | 0.01911 | 0.00436 | 0.03044 | 0.83129 | 0.00686 | 0.00676 | 0.08369 | 1.31088 | 0.00000 |
| **Arhgap21** | 11.92813 | 0.04205 | 16.72447 | 4.17074 | 15.93667 | 4.09560 | 3.14901 | 4.70101 | 2.70855 | 2.42329 | 3.05220 | 5.11739 | 6.26854 | 2.86112 |
| **Arhgap22** | 0.04047 | 0.03277 | 0.00715 | 0.00000 | 0.00488 | 0.02103 | 2.38218 | 4.31620 | 0.01234 | 5.97014 | 0.06375 | 0.01619 | 0.05321 | 0.00172 |
| **Arhgap23** | 0.05327 | 0.00377 | 0.13707 | 0.00000 | 0.96859 | 0.31786 | 0.00400 | 0.00160 | 0.00984 | 0.00387 | 0.34914 | 0.57386 | 0.46860 | 0.00578 |
| **Arhgap26** | 0.01565 | 0.01060 | 0.51343 | 0.00000 | 0.02955 | 0.07243 | 0.00330 | 1.07242 | 0.04937 | 0.01738 | 0.00000 | 0.37253 | 0.18160 | 0.08990 |
| **Arhgap27** | 0.00980 | 0.00442 | 0.01314 | 0.01937 | 0.04763 | 0.01288 | 0.00424 | 0.01029 | 0.01063 | 0.00682 | 0.00862 | 0.01003 | 0.00701 | 0.01239 |
| **Arhgap28** | 0.70522 | 0.04800 | 0.02493 | 0.00000 | 0.00000 | 0.00000 | 0.00409 | 0.65241 | 0.07143 | 0.00631 | 0.00000 | 0.61664 | 0.07004 | 2.58133 |
| **Arhgap29** | 3.48429 | 0.00094 | 4.17500 | 0.51508 | 5.02948 | 4.03740 | 1.72283 | 0.11705 | 6.40705 | 0.08127 | 5.52013 | 0.88813 | 8.59369 | 10.47441 |
| **Arhgap30** | 0.00004 | 0.01952 | 0.26900 | 0.00863 | 0.24822 | 0.00579 | 0.02477 | 0.00868 | 0.13879 | 0.00614 | 0.00928 | 0.36618 | 0.90705 | 0.02381 |
| **Arhgap31** | 0.06519 | 0.00384 | 0.01270 | 0.00000 | 0.00951 | 0.00437 | 0.01417 | 1.05118 | 0.01157 | 0.00565 | 0.00646 | 0.01429 | 0.00529 | 0.01203 |
| **Arhgap32** | 0.00000 | 0.00390 | 2.18940 | 2.05756 | 2.71204 | 0.44276 | 0.00334 | 0.24001 | 1.38679 | 0.00128 | 0.00000 | 0.02554 | 0.00000 | 2.12968 |
| **Arhgap33** | 0.00667 | 0.00417 | 0.10021 | 0.07107 | 1.22908 | 0.00399 | 0.00410 | 0.00853 | 0.01267 | 0.00593 | 0.00577 | 0.01067 | 0.36587 | 0.37704 |
| **Arhgap35** | 3.04827 | 0.03027 | 7.66298 | 5.87494 | 4.89780 | 6.82800 | 0.43687 | 2.35661 | 4.02053 | 0.07561 | 5.27224 | 5.40822 | 2.74303 | 7.22338 |
| **Arhgap39** | 0.08345 | 0.00399 | 0.16548 | 0.07924 | 0.27229 | 0.36438 | 0.00418 | 0.06323 | 0.00765 | 0.01145 | 0.00463 | 0.00909 | 1.36282 | 0.03494 |
| **Arhgap4** | 0.00232 | 0.00548 | 0.00885 | 0.00000 | 0.00714 | 0.03655 | 0.00417 | 0.10233 | 0.01092 | 0.00621 | 0.00379 | 0.34157 | 0.23906 | 0.50849 |
| **Arhgap42** | 4.36512 | 0.02724 | 2.74761 | 1.84146 | 4.54130 | 1.38176 | 0.08374 | 0.25475 | 3.58100 | 0.00849 | 0.00000 | 0.75838 | 0.61130 | 1.46934 |
| **Arhgap44** | 0.19141 | 0.00375 | 0.01363 | 0.00000 | 0.90234 | 1.02922 | 0.77885 | 0.02923 | 0.00972 | 0.01278 | 0.13536 | 0.47746 | 0.46937 | 0.04968 |
| **Arhgap5** | 0.79858 | 0.01769 | 1.86192 | 0.00000 | 0.13992 | 1.53174 | 0.04631 | 0.07749 | 5.85790 | 0.01000 | 2.49797 | 0.45456 | 0.80484 | 0.76362 |
| **Arhgap8** | 0.53340 | 0.06417 | 4.31511 | 1.39436 | 3.52731 | 5.06530 | 3.59691 | 10.48197 | 0.13499 | 0.04394 | 1.61451 | 0.13209 | 1.70469 | 0.29588 |
| **Arhgap9** | 0.18953 | 0.07982 | 1.81754 | 0.66280 | 0.17849 | 0.70809 | 0.56492 | 7.81846 | 0.64952 | 0.03287 | 2.65134 | 1.66747 | 0.00000 | 0.00000 |
| **Arhgdia** | 2.86512 | 4.84079 | 19.44067 | 20.84084 | 18.89765 | 12.01258 | 5.89391 | 21.00322 | 0.00000 | 13.83591 | 12.36225 | 12.97061 | 1.39840 | 0.98761 |
| **Arhgdib** | 0.00837 | 0.08245 | 0.30572 | 0.29658 | 0.46010 | 0.18062 | 0.00398 | 5.04955 | 0.00982 | 0.03987 | 0.00163 | 0.01447 | 0.00522 | 0.00313 |
| **Arhgdig** | 4.27537 | 0.01867 | 2.91124 | 3.28755 | 1.89276 | 3.37118 | 0.06089 | 0.67265 | 0.43321 | 0.02061 | 2.68347 | 0.08139 | 4.15714 | 1.41041 |
| **Arhgef1** | 1.25284 | 0.59446 | 4.60074 | 5.61391 | 0.88304 | 0.77623 | 0.01483 | 3.16639 | 0.00333 | 0.01673 | 1.49142 | 0.92461 | 0.02881 | 0.66241 |
| **Arhgef10** | 0.01354 | 0.00383 | 0.14566 | 0.01954 | 0.27588 | 0.00176 | 0.00422 | 0.65574 | 0.17528 | 0.01636 | 0.00485 | 0.00553 | 0.00277 | 0.02119 |
| **Arhgef10l** | 0.03014 | 0.00444 | 0.00829 | 0.00857 | 0.27483 | 0.01328 | 0.00423 | 0.01007 | 0.00935 | 0.00627 | 0.00896 | 0.01344 | 0.00539 | 0.05256 |
| **Arhgef11** | 0.01540 | 0.01248 | 0.00522 | 0.00000 | 0.24853 | 0.01093 | 0.00524 | 0.33120 | 0.01100 | 0.00499 | 0.07686 | 0.21952 | 0.34752 | 0.01863 |
| **Arhgef12** | 5.89946 | 0.07651 | 4.49361 | 1.62044 | 7.74497 | 3.28331 | 1.75675 | 1.56021 | 7.98730 | 0.03901 | 5.65548 | 3.12696 | 7.85968 | 11.19114 |
| **Arhgef15** | 0.00691 | 0.01387 | 0.01216 | 0.00808 | 0.01319 | 0.01433 | 0.00405 | 0.01741 | 0.01332 | 0.00681 | 0.01125 | 0.23780 | 0.27227 | 0.03285 |
| **Arhgef16** | 5.34234 | 0.02548 | 17.18242 | 3.40577 | 14.75726 | 2.11078 | 5.97891 | 3.08394 | 0.20988 | 0.21556 | 1.17560 | 0.00000 | 2.56497 | 0.57955 |
| **Arhgef17** | 0.16540 | 0.00456 | 0.22080 | 0.00101 | 0.77910 | 0.35862 | 0.00571 | 0.40338 | 0.01837 | 0.00512 | 0.02191 | 0.00953 | 0.01308 | 0.16348 |
| **Arhgef18** | 0.00000 | 0.01688 | 0.71027 | 0.42844 | 3.02982 | 1.89265 | 1.68422 | 2.72542 | 0.01045 | 0.08785 | 0.00000 | 2.77569 | 0.62354 | 0.45942 |
| **Arhgef19** | 0.08279 | 0.02375 | 3.12303 | 2.09044 | 2.25320 | 0.00000 | 0.00244 | 0.00000 | 0.15915 | 0.00552 | 0.59508 | 3.08034 | 0.00000 | 0.34317 |
| **Arhgef2** | 0.05183 | 0.04337 | 0.18620 | 1.40947 | 0.25123 | 0.48020 | 0.00364 | 0.01062 | 0.01465 | 0.01042 | 0.00000 | 1.30637 | 0.56653 | 0.07122 |
| **Arhgef26** | 1.53863 | 2.97781 | 2.53011 | 0.52137 | 0.58162 | 3.73095 | 0.00936 | 0.49244 | 0.61085 | 0.01929 | 3.29408 | 0.00000 | 0.86408 | 0.25310 |
| **Arhgef28** | 0.02191 | 0.00444 | 0.00754 | 0.01435 | 0.27546 | 0.01335 | 0.00452 | 0.00950 | 0.00930 | 0.00658 | 0.00857 | 0.00830 | 0.16845 | 0.00411 |
| **Arhgef3** | 6.25362 | 0.02034 | 11.38628 | 0.00000 | 12.21642 | 0.61787 | 0.46935 | 2.78273 | 20.41923 | 0.00000 | 4.36389 | 3.55569 | 1.97425 | 14.69569 |
| **Arhgef39** | 1.02926 | 3.19059 | 9.21854 | 6.61732 | 0.94458 | 4.90484 | 0.04242 | 4.46590 | 0.11504 | 0.04379 | 4.29348 | 1.16188 | 4.55553 | 0.24915 |

| | | | | | | | | | | | | | | |
|---|---|---|---|---|---|---|---|---|---|---|---|---|---|---|
| **Arhgef4** | 0.00000 | 0.01164 | 0.93238 | 0.53137 | 0.34630 | 0.00000 | 0.00357 | 1.36966 | 0.00000 | 0.00123 | 1.17425 | 0.72371 | 2.76203 | 0.22187 |
| **Arhgef40** | 0.03689 | 0.00450 | 0.06663 | 0.01650 | 0.10591 | 0.01252 | 0.00474 | 0.02347 | 0.02967 | 0.00794 | 0.00822 | 0.00736 | 0.03205 | 0.20585 |
| **Arhgef5** | 0.01366 | 8.65066 | 0.36048 | 0.00000 | 0.28049 | 0.97962 | 2.95561 | 0.11096 | 0.00787 | 0.05346 | 0.00298 | 0.63116 | 0.01443 | 0.00203 |
| **Arhgef7** | 1.43368 | 0.01793 | 1.24369 | 2.19103 | 2.39405 | 0.73250 | 0.64109 | 0.21919 | 0.52676 | 0.04193 | 0.00000 | 1.51034 | 2.42700 | 1.32384 |
| **Arid1a** | 18.60120 | 2.81244 | 23.09085 | 6.02600 | 27.27459 | 15.25043 | 8.26789 | 9.35763 | 14.23121 | 0.70507 | 8.00107 | 6.90964 | 10.93447 | 21.32529 |
| **Arid1b** | 0.89130 | 0.00080 | 2.87077 | 0.67228 | 1.94683 | 0.70862 | 2.89283 | 0.00000 | 5.06513 | 0.04788 | 3.47746 | 3.33079 | 2.44606 | 0.84533 |
| **Arid2** | 1.86962 | 0.01590 | 7.17802 | 2.47118 | 6.07295 | 6.33310 | 0.00077 | 0.67441 | 3.22043 | 0.00000 | 3.42756 | 5.77941 | 4.61989 | 5.16680 |
| **Arid3a** | 0.00430 | 0.00343 | 0.00638 | 0.00000 | 0.01063 | 0.13094 | 0.00383 | 0.47807 | 0.02036 | 0.00555 | 0.00000 | 0.83714 | 0.01830 | 0.00830 |
| **Arid3b** | 1.39781 | 0.04159 | 4.89378 | 3.25861 | 3.43559 | 2.80359 | 0.00866 | 3.51286 | 0.20678 | 0.01110 | 1.09191 | 1.85055 | 0.21154 | 2.60055 |
| **Arid3c** | 0.00486 | 0.00404 | 0.01273 | 0.60092 | 1.01988 | 0.28873 | 0.00411 | 0.02868 | 0.01328 | 0.00518 | 0.00647 | 0.01175 | 0.39513 | 0.04238 |
| **Arid4a** | 0.73575 | 1.47558 | 0.08428 | 0.32065 | 0.53276 | 0.49066 | 0.00371 | 0.01139 | 0.11804 | 0.00562 | 0.11262 | 0.05650 | 1.24176 | 2.86657 |
| **Arid4b** | 1.12885 | 0.00161 | 2.13437 | 0.00000 | 1.59338 | 1.82514 | 0.92005 | 0.32835 | 7.41673 | 0.02279 | 0.96026 | 2.22144 | 4.06890 | 0.76665 |
| **Arid5a** | 0.00576 | 0.00482 | 0.00537 | 0.00000 | 0.00632 | 0.37325 | 0.00407 | 0.37980 | 0.03514 | 0.00819 | 0.00253 | 0.10561 | 3.37452 | 0.00316 |
| **Arid5b** | 0.02881 | 0.01231 | 0.00000 | 0.04616 | 0.00000 | 0.00000 | 0.00382 | 0.09345 | 0.53268 | 0.02542 | 0.46458 | 1.85073 | 0.00421 | 0.39675 |
| **Arih1** | 9.53316 | 0.04608 | 8.14825 | 1.97859 | 4.98679 | 9.54699 | 1.47703 | 1.89826 | 8.33847 | 3.94390 | 3.11480 | 4.03818 | 18.66682 | 10.20899 |
| **Arih2** | 30.14291 | 1.16508 | 42.10900 | 16.23330 | 34.46896 | 27.73705 | 11.20591 | 7.33139 | 11.33426 | 0.99897 | 8.30684 | 20.11859 | 10.53826 | 35.17650 |
| **Arl1** | 1.03002 | 0.10331 | 9.37866 | 6.35940 | 3.86479 | 3.71766 | 4.26904 | 6.52558 | 2.38174 | 0.18048 | 0.94511 | 8.50797 | 2.18287 | 0.58299 |
| **Arl10** | 0.00942 | 0.00421 | 0.00621 | 0.39699 | 0.08284 | 0.00709 | 0.00424 | 0.00731 | 0.04098 | 0.00614 | 0.00651 | 0.00751 | 0.00719 | 0.00301 |
| **Arl13b** | 0.03371 | 0.00795 | 1.30390 | 0.00000 | 0.14668 | 0.26476 | 0.87392 | 0.53817 | 0.00272 | 9.40445 | 1.59913 | 1.19726 | 0.53597 | 0.67352 |
| **Arl14ep** | 1.36081 | 0.05790 | 1.19096 | 0.44890 | 3.17294 | 1.96599 | 0.03551 | 2.47343 | 2.37506 | 0.01254 | 2.06300 | 3.75083 | 6.96916 | 4.92623 |
| **Arl14epl** | 0.96731 | 0.03976 | 0.82196 | 3.67223 | 2.09503 | 7.52935 | 1.35708 | 3.01638 | 0.28894 | 0.05712 | 6.13773 | 1.56826 | 0.00000 | 2.27449 |
| **Arl15** | 0.93234 | 0.06693 | 0.55552 | 1.01557 | 0.08145 | 0.72753 | 0.00933 | 0.59769 | 0.08136 | 0.01874 | 0.13540 | 0.14001 | 1.49230 | 1.04033 |
| **Arl16** | 0.31352 | 4.70378 | 1.05366 | 0.49173 | 0.39904 | 0.61927 | 1.96851 | 0.10279 | 0.52580 | 0.02185 | 1.24591 | 0.00000 | 1.42163 | 0.27432 |
| **Arl2** | 1.54905 | 0.12570 | 1.51009 | 4.08054 | 0.00000 | 6.00637 | 2.11101 | 4.16349 | 0.00000 | 0.06798 | 4.05728 | 6.70359 | 0.22933 | 0.00000 |
| **Arl2bp** | 0.10375 | 0.01093 | 0.00085 | 0.28566 | 0.00000 | 0.00419 | 0.00404 | 0.00502 | 0.07894 | 0.00611 | 0.00463 | 0.51958 | 2.13671 | 1.10912 |
| **Arl3** | 1.02362 | 0.00944 | 0.43250 | 0.46736 | 2.53004 | 1.41276 | 0.00861 | 0.38630 | 0.00361 | 0.03210 | 2.03722 | 2.25908 | 2.13795 | 0.69628 |
| **Arl4a** | 0.53770 | 0.00347 | 0.56224 | 0.71763 | 1.20857 | 0.58573 | 0.05494 | 0.42367 | 3.28600 | 0.04358 | 0.00000 | 1.06736 | 10.18832 | 0.00000 |
| **Arl4c** | 0.01023 | 0.00414 | 0.00884 | 0.00000 | 0.24502 | 0.02800 | 0.00461 | 0.00782 | 0.01338 | 0.00587 | 0.00483 | 0.37313 | 0.00542 | 0.00887 |
| **Arl4d** | 0.00000 | 0.09349 | 0.48366 | 0.65318 | 0.43221 | 0.00000 | 0.00349 | 2.17861 | 0.00522 | 0.02907 | 1.13112 | 1.03547 | 0.54143 | 0.06700 |
| **Arl5a** | 0.50195 | 0.02034 | 1.39183 | 0.00000 | 4.01590 | 7.32330 | 1.78402 | 5.29554 | 1.48812 | 0.10634 | 0.14880 | 7.82310 | 2.69195 | 1.65615 |
| **Arl5b** | 0.78092 | 0.01452 | 0.22521 | 0.00865 | 0.04876 | 0.00000 | 0.01036 | 0.89267 | 0.94932 | 0.00835 | 1.34501 | 0.00477 | 0.98842 | 0.12273 |
| **Arl5c** | 0.05686 | 0.00429 | 0.01536 | 0.02149 | 0.03717 | 0.01070 | 0.00407 | 0.00898 | 0.02380 | 0.00912 | 0.00770 | 0.00776 | 0.46101 | 0.01202 |
| **Arl6** | 0.26476 | 0.00327 | 1.40924 | 0.77918 | 0.19209 | 0.53115 | 0.00336 | 0.00000 | 0.01025 | 0.00368 | 0.00000 | 0.84965 | 0.33452 | 1.02268 |
| **Arl6ip1** | 15.94868 | 9.07773 | 21.13447 | 8.05229 | 13.88609 | 19.21059 | 4.70121 | 15.18855 | 20.42658 | 12.04653 | 13.40719 | 18.04039 | 3.16335 | 20.23507 |
| **Arl6ip4** | 4.22024 | 0.11107 | 6.90061 | 9.67712 | 9.75092 | 2.46389 | 2.90788 | 5.48313 | 0.36111 | 0.13794 | 2.46036 | 6.57016 | 3.90762 | 1.10205 |
| **Arl6ip5** | 0.03980 | 0.00387 | 0.00209 | 0.24058 | 0.00442 | 0.19963 | 0.14944 | 0.10507 | 0.01038 | 0.01026 | 0.38226 | 0.12765 | 0.00495 | 0.02013 |
| **Arl6ip6** | 5.04638 | 0.00933 | 0.69517 | 1.53754 | 2.47022 | 2.18776 | 0.77061 | 1.46476 | 5.16513 | 0.03226 | 0.24669 | 2.42359 | 0.82822 | 0.94763 |
| **Arl8a** | 1.10279 | 1.14653 | 0.11190 | 0.18797 | 0.08738 | 0.96786 | 0.13003 | 0.64111 | 0.03674 | 0.11018 | 0.13239 | 0.89529 | 1.38402 | 0.82415 |
| **Arl8b** | 19.83820 | 0.06015 | 16.55437 | 12.26841 | 19.91598 | 13.00976 | 1.94686 | 7.69340 | 22.26008 | 4.50148 | 9.67235 | 6.93039 | 4.70130 | 14.84287 |
| **Armc1** | 5.72935 | 0.01154 | 3.65031 | 4.68411 | 5.14652 | 7.49462 | 0.96618 | 5.86749 | 2.30162 | 0.05027 | 7.28158 | 6.48908 | 4.58450 | 3.75063 |
| **Armc10** | 1.53679 | 0.05777 | 1.44292 | 3.61295 | 5.16570 | 1.80653 | 0.04146 | 2.81766 | 1.50807 | 0.00878 | 4.76020 | 1.48478 | 0.62106 | 2.16435 |
| **Armc12** | 0.01843 | 0.00424 | 0.01410 | 0.02528 | 0.03394 | 0.01385 | 0.00408 | 0.00767 | 0.02173 | 0.00631 | 0.00815 | 0.00998 | 0.60459 | 0.00268 |
| **Armc2** | 0.11189 | 0.00423 | 0.15554 | 0.01796 | 0.35154 | 0.01815 | 0.00994 | 0.00663 | 0.00715 | 0.01113 | 0.00606 | 0.00952 | 0.49636 | 0.01703 |
| **Armc5** | 0.24786 | 0.01045 | 1.14970 | 2.75185 | 2.76415 | 0.15274 | 0.00334 | 0.00596 | 0.00262 | 0.01835 | 0.01781 | 1.20798 | 0.53033 | 0.00000 |
| **Armc6** | 1.59881 | 0.03305 | 8.35392 | 2.40148 | 5.62526 | 4.62264 | 3.31897 | 1.62852 | 0.01122 | 0.00000 | 2.97363 | 1.83901 | 0.85993 | 1.01427 |
| **Armc7** | 1.87026 | 0.03902 | 2.30009 | 0.00000 | 3.85394 | 2.28718 | 0.65249 | 0.80804 | 0.20771 | 0.01990 | 0.00000 | 0.86137 | 9.69680 | 0.89340 |
| **Armc8** | 1.90503 | 0.03143 | 4.74800 | 3.41579 | 4.73363 | 5.12502 | 0.00146 | 3.07861 | 3.19066 | 0.03347 | 4.52873 | 2.51394 | 0.00000 | 3.30464 |
| **Armc9** | 0.53647 | 0.01077 | 1.27516 | 0.00000 | 0.47140 | 0.66311 | 0.06312 | 0.00000 | 0.00091 | 0.00294 | 0.00000 | 0.66095 | 0.96919 | 0.08696 |
| **Armcx4** | 0.02534 | 0.10736 | 0.00000 | 0.35888 | 0.45212 | 0.21818 | 0.07867 | 0.15046 | 2.53128 | 0.05802 | 0.00364 | 0.60296 | 0.41149 | 0.00000 |
| **Armcx5** | 0.82175 | 0.00306 | 0.32916 | 0.77344 | 2.08927 | 1.41275 | 0.00331 | 1.01384 | 2.92551 | 0.00554 | 0.00000 | 0.85539 | 2.59349 | 2.73249 |
| **Armt1** | 0.33156 | 0.02803 | 0.16038 | 0.52948 | 0.27403 | 3.28105 | 4.01304 | 2.76661 | 0.01633 | 0.20678 | 0.18319 | 1.54823 | 0.29491 | 0.00213 |
| **Arnt** | 4.01868 | 0.00000 | 4.48271 | 2.64131 | 5.24019 | 0.37692 | 0.15897 | 0.05960 | 3.41902 | 4.95639 | 1.18412 | 2.87222 | 1.28927 | 1.87734 |
| **Arntl** | 0.49818 | 0.00066 | 0.62409 | 0.18432 | 0.21367 | 1.70165 | 0.00245 | 0.00000 | 0.59417 | 0.00211 | 3.51581 | 3.08676 | 2.18149 | 2.20149 |
| **Arntl2** | 0.00489 | 0.00445 | 0.01113 | 0.02070 | 0.05691 | 0.04915 | 0.00421 | 0.00657 | 0.02119 | 0.00605 | 0.00826 | 0.00973 | 0.02701 | 0.55411 |
| **Arpc1a** | 8.41021 | 0.23312 | 16.89997 | 11.27119 | 16.25199 | 13.71461 | 7.32433 | 10.48825 | 1.89565 | 0.38097 | 9.78695 | 9.43036 | 4.84281 | 8.86292 |

| | | | | | | | | | | | | | | |
|---|---|---|---|---|---|---|---|---|---|---|---|---|---|---|
| **Arpc1b** | 4.61137 | 4.12986 | 22.11235 | 17.15630 | 18.19101 | 15.74518 | 2.85997 | 23.35338 | 0.21515 | 6.08344 | 11.46487 | 16.10946 | 2.09075 | 3.93130 |
| **Arpc2** | 14.97877 | 1.73406 | 19.64830 | 13.26081 | 16.25518 | 24.87596 | 10.69048 | 20.21857 | 9.19542 | 15.50292 | 11.24811 | 14.80241 | 10.75560 | 9.32161 |
| **Arpc3** | 5.84796 | 5.57347 | 1.40499 | 1.73658 | 3.33546 | 1.92627 | 2.44669 | 5.42395 | 5.79911 | 0.18918 | 6.76111 | 1.50070 | 1.96767 | 3.36611 |
| **Arpc4** | 1.41635 | 9.55736 | 16.88434 | 18.51328 | 10.19721 | 5.83932 | 4.64522 | 21.67174 | 0.00000 | 4.31491 | 10.53360 | 4.70113 | 0.00000 | 0.84240 |
| **Arpc5** | 8.80750 | 12.43558 | 18.16431 | 12.50637 | 7.40447 | 23.68561 | 8.15002 | 26.39993 | 3.67697 | 23.25815 | 14.42495 | 9.57369 | 1.71169 | 2.28153 |
| **Arpc5l** | 6.58254 | 0.09745 | 8.59190 | 10.21025 | 12.03384 | 4.61397 | 4.49512 | 5.31410 | 3.01423 | 0.24203 | 1.66941 | 6.00757 | 6.43419 | 1.76163 |
| **Arpin** | 0.00000 | 0.00354 | 0.20357 | 0.00000 | 0.25008 | 0.84571 | 0.00363 | 0.00246 | 0.00133 | 0.00627 | 0.00000 | 0.55242 | 1.32134 | 0.06661 |
| **Arpp19** | 3.54419 | 0.10619 | 0.88882 | 1.29674 | 4.23596 | 12.51258 | 1.87266 | 8.30490 | 4.14421 | 3.80027 | 2.19985 | 5.70753 | 2.83995 | 3.12010 |
| **Arrb1** | 0.21885 | 0.00187 | 1.64794 | 3.20132 | 0.08382 | 1.10448 | 0.01867 | 0.00600 | 0.00067 | 0.01221 | 0.17729 | 2.96352 | 0.00000 | 1.68955 |
| **Arrb2** | 0.35874 | 0.00237 | 1.14384 | 1.20873 | 1.77041 | 0.17321 | 0.79862 | 0.00000 | 0.00325 | 0.02336 | 1.08321 | 1.60037 | 2.19475 | 0.15749 |
| **Arrdc1** | 2.34079 | 6.76625 | 6.26544 | 1.24280 | 11.20161 | 0.28819 | 0.00708 | 1.49324 | 0.31649 | 0.00812 | 0.00000 | 0.00000 | 3.96467 | 7.92548 |
| **Arrdc2** | 1.21715 | 0.01016 | 1.94329 | 0.77786 | 3.56168 | 0.00000 | 0.00356 | 0.15850 | 0.22212 | 0.02382 | 1.00947 | 0.00000 | 6.86707 | 0.26363 |
| **Arrdc3** | 0.02282 | 2.24165 | 0.19350 | 0.00070 | 0.05929 | 0.00000 | 0.00301 | 1.43487 | 0.00424 | 0.00190 | 1.18682 | 0.00339 | 0.00958 | 0.64176 |
| **Arrdc4** | 0.02993 | 0.00394 | 0.23599 | 0.00000 | 0.00143 | 0.37810 | 0.00416 | 0.34361 | 0.03431 | 0.00487 | 0.00398 | 0.00601 | 0.03281 | 0.07442 |
| **Arrdc5** | 0.04319 | 0.00436 | 0.17828 | 0.00000 | 0.17561 | 0.00563 | 0.00423 | 0.01284 | 0.00621 | 0.00679 | 0.00664 | 0.01063 | 0.00351 | 0.17593 |
| **Arsa** | 0.02197 | 0.00399 | 0.01449 | 0.68082 | 0.04983 | 0.56485 | 0.00972 | 0.00000 | 0.01454 | 0.01261 | 1.36743 | 0.01767 | 0.00753 | 0.00000 |
| **Arsb** | 0.13044 | 0.00483 | 0.21469 | 0.00000 | 0.00000 | 0.16552 | 0.01058 | 0.00494 | 1.60772 | 0.00577 | 0.00417 | 0.01294 | 1.02791 | 0.15280 |
| **Arsg** | 0.01110 | 0.00456 | 0.04934 | 0.01020 | 0.00488 | 0.00949 | 0.00417 | 0.01199 | 0.29659 | 0.00717 | 0.00749 | 0.00917 | 0.03066 | 0.00328 |
| **Arsk** | 0.00038 | 0.00450 | 0.00997 | 0.02373 | 0.00227 | 0.15217 | 0.00402 | 0.00561 | 0.25834 | 0.00534 | 0.00985 | 0.04659 | 3.73056 | 0.00000 |
| **Art2a.ps** | 0.01266 | 0.00444 | 0.01471 | 0.01309 | 0.01338 | 0.01300 | 0.00428 | 0.01033 | 0.01224 | 0.00693 | 0.00873 | 0.01001 | 0.00959 | 0.01954 |
| **Art3** | 0.30943 | 0.00393 | 1.60609 | 0.00000 | 1.43600 | 0.00000 | 0.00334 | 0.00364 | 2.71887 | 0.00363 | 0.00386 | 0.00904 | 0.49311 | 2.55414 |
| **Art5** | 0.03437 | 0.00409 | 0.00000 | 0.80651 | 0.00000 | 0.00000 | 0.00387 | 0.00527 | 1.02356 | 0.00587 | 0.00226 | 0.01290 | 0.00374 | 0.87626 |
| **Artn** | 0.00756 | 0.00446 | 0.08543 | 0.00000 | 0.33909 | 0.01088 | 0.00416 | 0.01089 | 0.01159 | 0.00697 | 0.00771 | 0.01024 | 0.00533 | 0.00836 |
| **Arv1** | 1.26900 | 0.00293 | 1.58099 | 2.83622 | 0.47409 | 1.93467 | 0.96402 | 0.00117 | 0.34450 | 0.01394 | 0.16438 | 1.00728 | 1.96597 | 1.15980 |
| **Arvcf** | 2.78792 | 3.33831 | 13.33639 | 13.90993 | 3.91900 | 3.85895 | 3.88662 | 0.00000 | 0.23984 | 0.11088 | 3.22177 | 3.26897 | 0.59726 | 1.46217 |
| **Arx** | 0.02718 | 0.00444 | 0.01431 | 0.01357 | 0.01308 | 0.01282 | 0.00432 | 0.01023 | 0.01196 | 0.00708 | 0.00855 | 0.00973 | 0.03959 | 0.00690 |
| **Arxes1** | 0.00609 | 0.00441 | 0.00731 | 0.00000 | 0.17726 | 0.19099 | 0.00412 | 0.00670 | 0.05322 | 0.00730 | 0.00724 | 0.01046 | 0.00600 | 0.06397 |
| **Arxes2** | 0.00599 | 0.00445 | 0.00000 | 0.00885 | 0.85521 | 0.06293 | 0.00412 | 0.00765 | 0.51752 | 0.00692 | 0.00734 | 0.00821 | 0.00607 | 0.19381 |
| **As3mt** | 0.00000 | 0.03628 | 9.28533 | 4.41238 | 0.00000 | 7.50642 | 0.11633 | 2.70461 | 0.00000 | 0.00000 | 9.41706 | 6.55897 | 0.23437 | 0.00000 |
| **Asah1** | 6.38755 | 6.71190 | 10.28123 | 9.67109 | 6.30228 | 10.13393 | 1.63551 | 2.20401 | 9.23962 | 0.10023 | 5.06454 | 8.91963 | 1.49335 | 14.51115 |
| **Asah2** | 0.01707 | 0.05753 | 0.01418 | 0.01265 | 0.01476 | 0.01276 | 0.00420 | 0.00782 | 0.01889 | 0.00664 | 0.00873 | 0.00916 | 0.01683 | 0.01383 |
| **Asap1** | 1.20420 | 0.00320 | 0.23864 | 0.42561 | 0.79109 | 1.58302 | 0.00457 | 0.10862 | 0.00347 | 0.00119 | 1.86955 | 0.86871 | 0.03981 | 1.56557 |
| **Asap2** | 0.38607 | 0.01908 | 1.15451 | 0.50341 | 0.39512 | 0.87079 | 0.02110 | 0.76990 | 0.72443 | 0.02142 | 0.45671 | 0.03750 | 0.00000 | 0.12788 |
| **Asap3** | 0.00000 | 0.01982 | 2.27413 | 3.13112 | 0.00000 | 2.31834 | 0.01078 | 0.04091 | 0.21858 | 0.09962 | 0.01932 | 0.45638 | 0.10779 | 0.36870 |
| **Asb1** | 1.05857 | 0.00367 | 0.00000 | 0.00000 | 0.18917 | 0.85439 | 0.02812 | 0.01779 | 0.00354 | 0.00593 | 0.04115 | 0.86507 | 0.04599 | 0.47478 |
| **Asb10** | 0.01004 | 0.00422 | 0.01270 | 0.00437 | 0.01338 | 0.11153 | 0.09273 | 0.00416 | 0.01102 | 0.01338 | 0.00835 | 0.01171 | 0.00744 | 0.01101 |
| **Asb13** | 3.10118 | 0.03776 | 5.92883 | 3.51906 | 3.53171 | 6.03519 | 0.32941 | 3.15946 | 2.26292 | 8.66063 | 2.45786 | 3.16493 | 7.75547 | 0.49544 |
| **Asb14** | 0.00422 | 0.00443 | 0.05953 | 0.00801 | 0.00928 | 0.01345 | 0.00426 | 0.00963 | 0.01198 | 0.00667 | 0.00845 | 0.00915 | 0.28175 | 0.00379 |
| **Asb16** | 0.00076 | 0.00406 | 0.07092 | 0.62613 | 0.42545 | 0.02430 | 0.00963 | 0.00376 | 0.01848 | 0.00495 | 0.00596 | 0.03667 | 0.01120 | 0.00778 |
| **Asb17** | 0.11014 | 0.00428 | 0.04941 | 0.05891 | 0.00945 | 0.32385 | 0.00403 | 0.13026 | 0.01985 | 0.01483 | 0.00517 | 0.00377 | 0.00918 | 0.05037 |
| **Asb17os** | 0.16394 | 0.00562 | 0.04195 | 0.39835 | 0.02155 | 0.69210 | 0.00441 | 0.27264 | 0.07819 | 0.00714 | 0.10894 | 0.00285 | 0.01585 | 0.06601 |
| **Asb3** | 0.01868 | 0.00413 | 1.43718 | 1.36489 | 0.00000 | 0.09663 | 0.00362 | 0.00000 | 0.96602 | 0.00365 | 0.00386 | 0.01558 | 0.29420 | 0.00000 |
| **Asb4** | 0.01546 | 0.00456 | 0.13821 | 0.00000 | 0.10596 | 0.00000 | 0.00378 | 0.01932 | 0.80806 | 0.00659 | 0.00276 | 0.18673 | 0.26644 | 1.04395 |
| **Asb6** | 2.48214 | 2.26267 | 7.96198 | 0.00000 | 1.13528 | 0.64733 | 0.85611 | 1.61907 | 0.03478 | 0.00000 | 1.31808 | 1.00955 | 0.02532 | 1.97015 |
| **Asb7** | 0.00000 | 0.00401 | 0.40368 | 0.98012 | 0.00000 | 0.84678 | 0.00407 | 1.59483 | 0.15244 | 0.01148 | 0.00188 | 0.02437 | 0.96902 | 0.29189 |
| **Asb8** | 1.42623 | 0.06034 | 1.86185 | 2.25329 | 2.38753 | 4.41241 | 0.86822 | 4.39623 | 2.42998 | 0.03806 | 2.14376 | 0.52913 | 2.10097 | 1.07288 |
| **Ascc1** | 0.49210 | 0.07130 | 3.66752 | 1.05249 | 2.16741 | 0.76179 | 0.01322 | 3.75216 | 0.64293 | 0.01603 | 3.00838 | 1.67436 | 1.32458 | 2.03851 |
| **Ascc2** | 2.90698 | 0.01376 | 7.40849 | 3.98257 | 6.87792 | 2.07626 | 2.00512 | 2.91071 | 0.92577 | 1.92143 | 5.60256 | 4.63251 | 4.91804 | 8.38841 |
| **Ascc3** | 8.45259 | 0.00223 | 4.58548 | 3.78570 | 6.87543 | 0.73967 | 0.00093 | 0.33866 | 14.39290 | 2.05593 | 0.91082 | 0.94274 | 3.38481 | 3.03747 |
| **Asf1a** | 5.78811 | 0.02381 | 1.22820 | 0.70905 | 2.44815 | 5.51831 | 1.76416 | 0.04906 | 1.57015 | 11.85326 | 0.37633 | 1.00168 | 4.70866 | 0.00000 |
| **Asf1b** | 2.58471 | 0.07426 | 7.54128 | 1.37899 | 5.04458 | 15.73333 | 3.52220 | 3.56755 | 2.76916 | 0.18357 | 8.45335 | 7.24373 | 7.85960 | 8.67040 |
| **Asgr2** | 0.00989 | 0.00486 | 0.06224 | 0.06138 | 0.28695 | 0.01927 | 0.00500 | 0.11046 | 0.05225 | 0.00686 | 0.02280 | 0.00622 | 0.00316 | 0.21205 |
| **Ash1l** | 0.31631 | 0.00402 | 1.32537 | 0.20994 | 0.76883 | 0.51048 | 0.00324 | 0.31136 | 2.88186 | 0.01259 | 0.23133 | 0.94620 | 4.48988 | 1.33913 |
| **Ash2l** | 5.84798 | 9.17213 | 17.61158 | 15.34449 | 9.41822 | 26.02244 | 6.71501 | 14.22673 | 11.65849 | 16.59025 | 17.48798 | 12.55705 | 5.98558 | 12.91806 |
| **Asic1** | 0.11300 | 0.00473 | 0.23286 | 0.00000 | 0.55613 | 0.36787 | 0.00401 | 0.01162 | 0.01101 | 0.00835 | 0.00381 | 0.04349 | 0.23960 | 1.75956 |

| | | | | | | | | | | | | | | |
|---|---|---|---|---|---|---|---|---|---|---|---|---|---|---|
| **Asic3** | 0.00000 | 0.00520 | 0.67105 | 1.09195 | 0.00000 | 1.12230 | 1.86271 | 0.84732 | 0.00970 | 0.02409 | 0.55708 | 0.00000 | 0.00510 | 0.00000 |
| **Asl** | 1.05803 | 5.94766 | 2.35319 | 0.00073 | 3.27756 | 4.82900 | 0.25249 | 4.08144 | 0.08127 | 0.10160 | 2.81702 | 2.43541 | 0.81666 | 2.27400 |
| **Asna1** | 10.46577 | 9.23400 | 12.29981 | 17.58124 | 13.57103 | 16.59526 | 2.84986 | 7.54632 | 0.80373 | 0.24704 | 11.47706 | 6.77036 | 1.19115 | 8.25449 |
| **Asns** | 0.67563 | 5.21545 | 0.85193 | 0.00000 | 0.93357 | 10.26143 | 0.04731 | 0.41373 | 0.00000 | 0.12331 | 7.62974 | 26.22068 | 0.88487 | 1.10878 |
| **Asnsd1** | 5.11202 | 0.02534 | 6.24373 | 8.54413 | 9.81433 | 6.22255 | 1.33017 | 1.90231 | 4.21401 | 0.09054 | 7.12759 | 6.66633 | 1.91228 | 6.22922 |
| **Aspa** | 0.02199 | 0.00248 | 0.03697 | 0.00000 | 0.01592 | 0.04098 | 0.00952 | 0.37798 | 0.01320 | 0.01043 | 0.00000 | 2.07480 | 0.00599 | 0.00850 |
| **Aspg** | 2.95993 | 0.00314 | 5.00384 | 2.12101 | 7.77413 | 0.00000 | 0.00797 | 2.24746 | 0.49120 | 0.01114 | 0.00000 | 0.00000 | 4.03448 | 3.43910 |
| **Asph** | 0.57030 | 0.00161 | 0.91605 | 0.00000 | 0.32341 | 1.64213 | 0.00314 | 0.00000 | 0.97536 | 0.00393 | 1.86906 | 2.37666 | 2.45850 | 1.16880 |
| **Asphd1** | 0.00548 | 0.00358 | 0.00292 | 0.39488 | 0.00785 | 0.00000 | 0.00369 | 0.00267 | 0.01795 | 0.00534 | 0.09833 | 0.47059 | 0.00776 | 0.00591 |
| **Asphd2** | 0.00499 | 0.00439 | 0.01381 | 0.01239 | 0.11946 | 0.01206 | 0.00439 | 0.01173 | 0.00797 | 0.00664 | 0.00823 | 0.00949 | 0.00620 | 0.20759 |
| **Aspm** | 2.84527 | 0.55529 | 3.83424 | 1.26050 | 1.34564 | 0.16364 | 1.95197 | 0.00000 | 4.85238 | 0.01894 | 3.58363 | 0.30894 | 9.84445 | 4.54441 |
| **Aspn** | 0.01909 | 0.01130 | 0.01495 | 0.00000 | 0.03179 | 0.03096 | 0.00433 | 0.06462 | 0.02433 | 0.00539 | 0.00411 | 0.22523 | 0.05890 | 0.02557 |
| **Aspscr1** | 2.07763 | 0.19754 | 6.01677 | 3.96913 | 7.83757 | 6.32418 | 1.39920 | 4.50038 | 0.00000 | 13.58375 | 4.34254 | 3.57054 | 1.41838 | 0.82602 |
| **Asrgl1** | 0.01177 | 0.00420 | 0.10104 | 0.03924 | 0.00658 | 0.61512 | 0.00395 | 0.00629 | 0.01114 | 0.01971 | 0.00000 | 0.61095 | 0.15627 | 0.04557 |
| **Ass1** | 0.01199 | 0.01157 | 0.02247 | 0.04340 | 0.01605 | 0.05636 | 0.00397 | 0.11455 | 0.01255 | 0.00826 | 0.05756 | 0.00719 | 0.00903 | 0.00941 |
| **Aste1** | 0.48308 | 0.00275 | 2.81609 | 1.34107 | 4.37461 | 0.85040 | 1.01382 | 0.87662 | 3.36340 | 0.02671 | 0.00000 | 0.95654 | 4.93724 | 1.56354 |
| **Astl** | 1.13748 | 0.00373 | 0.00000 | 0.01220 | 0.81233 | 0.00277 | 0.00400 | 0.00746 | 0.00137 | 0.00675 | 0.00545 | 0.00203 | 2.77009 | 2.99806 |
| **Asun** | 8.12657 | 0.77160 | 8.21997 | 2.06839 | 9.91456 | 3.62978 | 0.51954 | 2.21523 | 9.45377 | 0.00950 | 2.81365 | 6.15888 | 3.56451 | 10.54804 |
| **Asxl1** | 1.20909 | 0.01582 | 4.63256 | 0.71357 | 5.58497 | 2.39428 | 0.65353 | 1.35685 | 5.95743 | 0.04087 | 2.57456 | 1.97126 | 9.74451 | 4.14926 |
| **Asxl2** | 3.44270 | 0.03877 | 3.87243 | 2.88349 | 4.06181 | 0.37976 | 2.16068 | 0.09048 | 8.66456 | 0.08363 | 3.01134 | 1.31106 | 6.97060 | 7.13027 |
| **Asz1** | 1.88836 | 0.00310 | 1.90964 | 0.76919 | 3.32720 | 2.66305 | 0.01083 | 0.58166 | 3.14321 | 0.02696 | 3.52340 | 0.00933 | 0.00000 | 2.64904 |
| **Atad1** | 4.11148 | 0.00059 | 4.77288 | 1.71485 | 1.65327 | 6.58029 | 2.65192 | 0.28608 | 11.13980 | 0.04301 | 0.54206 | 3.54614 | 3.40509 | 3.30722 |
| **Atad2** | 2.95014 | 0.07155 | 1.91327 | 3.13143 | 5.85166 | 2.41117 | 1.98973 | 3.93585 | 7.37340 | 0.03762 | 2.07703 | 4.08615 | 13.09802 | 6.77396 |
| **Atad2b** | 10.89349 | 0.06597 | 10.58111 | 1.39030 | 14.33924 | 5.55233 | 1.56435 | 4.21096 | 22.28113 | 0.03832 | 4.95965 | 2.82709 | 6.37836 | 11.78636 |
| **Atad3a** | 6.10280 | 0.32687 | 15.82453 | 11.34913 | 7.36926 | 15.59120 | 4.06519 | 12.99293 | 0.58396 | 0.26285 | 12.14452 | 12.07183 | 3.52303 | 7.73550 |
| **Atad5** | 1.46648 | 0.00255 | 2.26021 | 0.00000 | 1.85262 | 1.33355 | 1.06791 | 0.00000 | 11.16900 | 0.00772 | 1.74038 | 0.57988 | 2.93568 | 1.44632 |
| **Atat1** | 0.00852 | 0.06551 | 0.70894 | 0.00000 | 0.66443 | 0.04362 | 0.00377 | 0.75577 | 0.04845 | 0.00536 | 0.00000 | 0.80873 | 1.41912 | 0.07583 |
| **Atcay** | 0.00725 | 0.00440 | 0.01089 | 0.01389 | 0.00477 | 0.01101 | 0.00432 | 0.01005 | 0.00905 | 0.00692 | 0.00757 | 0.00956 | 0.01341 | 0.64184 |
| **Ate1** | 4.27288 | 0.01694 | 2.99396 | 1.46033 | 2.15303 | 4.00073 | 1.40661 | 0.97610 | 0.55107 | 0.03858 | 4.43999 | 2.70783 | 2.60571 | 2.28779 |
| **Atf1** | 11.09618 | 0.26638 | 11.59008 | 5.67658 | 14.48388 | 5.86460 | 4.26572 | 0.81993 | 19.16362 | 4.23706 | 5.10385 | 3.31918 | 12.18925 | 18.39297 |
| **Atf2** | 3.38321 | 0.00778 | 3.89943 | 0.54245 | 5.68443 | 0.66033 | 1.41838 | 0.00000 | 18.08545 | 0.04591 | 1.33540 | 1.95261 | 7.67070 | 7.68135 |
| **Atf4** | 15.56236 | 9.44121 | 25.52427 | 21.46927 | 27.18238 | 15.42406 | 10.15639 | 26.01063 | 3.99526 | 3.92945 | 10.89068 | 17.49347 | 14.11482 | 7.63648 |
| **Atf5** | 0.19669 | 0.00399 | 0.33977 | 0.00000 | 1.21349 | 0.42947 | 0.00386 | 0.00407 | 0.01282 | 0.01398 | 0.00554 | 0.68767 | 0.32760 | 0.03438 |
| **Atf6** | 5.23356 | 0.01977 | 3.91660 | 0.34796 | 2.62002 | 3.24356 | 1.10729 | 0.38241 | 4.00699 | 0.04052 | 2.59900 | 1.58476 | 0.28257 | 6.29669 |
| **Atf6b** | 0.52674 | 0.04111 | 5.24783 | 4.59312 | 2.92978 | 1.75260 | 1.77089 | 0.90419 | 0.05490 | 0.07225 | 0.02162 | 2.33373 | 2.68878 | 0.05045 |
| **Atf7** | 1.12264 | 0.02162 | 1.60099 | 0.00000 | 0.24508 | 1.46688 | 0.00376 | 0.10502 | 0.02186 | 0.02656 | 1.68647 | 0.78465 | 1.00789 | 3.32063 |
| **Atf7ip** | 1.74639 | 0.01746 | 5.22734 | 0.70027 | 5.87824 | 0.38513 | 3.31884 | 0.22291 | 0.09879 | 0.07052 | 0.67671 | 2.00612 | 7.00179 | 0.00000 |
| **Atf7ip2** | 2.46097 | 0.00195 | 2.70269 | 0.00000 | 6.48121 | 0.00000 | 2.57388 | 0.13739 | 10.47639 | 0.07171 | 1.02098 | 0.00845 | 10.04796 | 5.57936 |
| **Atg10** | 0.00007 | 0.00412 | 0.53724 | 0.24164 | 1.43714 | 0.37367 | 0.00908 | 0.00084 | 1.17226 | 0.01004 | 0.00193 | 0.00905 | 0.26480 | 0.94466 |
| **Atg101** | 3.62049 | 0.18319 | 14.68795 | 9.96501 | 10.61921 | 6.45172 | 0.01835 | 5.30247 | 0.62711 | 0.06434 | 2.74047 | 4.08290 | 3.95585 | 4.46824 |
| **Atg12** | 5.56202 | 2.46409 | 9.14264 | 7.12452 | 12.81776 | 8.06974 | 2.28940 | 2.87600 | 9.50511 | 0.04617 | 5.75917 | 1.79336 | 6.23620 | 11.78057 |
| **Atg13** | 7.07270 | 0.22561 | 15.44671 | 5.03790 | 15.90839 | 8.84589 | 5.30193 | 11.47689 | 5.26415 | 0.35750 | 11.84416 | 14.36410 | 7.32971 | 13.67461 |
| **Atg14** | 8.41147 | 0.06190 | 25.66798 | 11.07318 | 24.55517 | 9.52005 | 3.79479 | 2.41684 | 11.45185 | 0.22241 | 8.20786 | 4.98575 | 13.23956 | 19.53043 |
| **Atg16l1** | 4.66395 | 0.02551 | 5.68634 | 2.11852 | 9.23239 | 5.42898 | 0.03353 | 0.33917 | 0.13386 | 0.00000 | 3.50471 | 2.48383 | 6.04627 | 12.51556 |
| **Atg16l2** | 0.63759 | 0.22057 | 12.20343 | 13.70651 | 5.55749 | 9.71107 | 1.60459 | 7.08546 | 0.74065 | 14.02379 | 8.06972 | 3.76710 | 0.00000 | 5.54157 |
| **Atg2a** | 7.41091 | 10.63718 | 22.56864 | 17.92487 | 18.82326 | 15.21620 | 4.25994 | 3.52539 | 0.08165 | 14.86654 | 9.57517 | 11.22886 | 3.92081 | 7.31545 |
| **Atg2b** | 4.71644 | 0.09451 | 9.33638 | 3.45974 | 7.37527 | 6.81072 | 0.01000 | 5.06049 | 6.31614 | 0.03383 | 2.27947 | 6.52889 | 3.59000 | 9.81865 |
| **Atg3** | 10.65357 | 0.00907 | 4.33472 | 3.45558 | 18.13415 | 4.51328 | 2.83586 | 0.26156 | 14.64430 | 0.08361 | 2.05292 | 1.66138 | 8.34128 | 8.09278 |
| **Atg4a** | 0.72862 | 0.00235 | 0.16449 | 0.00000 | 0.33033 | 1.37080 | 0.00915 | 0.00000 | 0.83714 | 0.00110 | 3.24620 | 0.98019 | 0.00177 | 0.69038 |
| **Atg4b** | 15.68554 | 2.96392 | 34.40898 | 24.85646 | 21.34854 | 21.28214 | 4.48729 | 9.41631 | 7.98656 | 10.17835 | 22.96291 | 14.27528 | 5.49858 | 22.90814 |
| **Atg4c** | 0.08743 | 0.01264 | 0.00575 | 0.03322 | 0.08834 | 0.03472 | 0.01033 | 0.40380 | 0.12874 | 0.00742 | 1.17873 | 0.00949 | 0.33374 | 0.02876 |
| **Atg4d** | 0.13675 | 0.08618 | 7.72467 | 5.68238 | 5.51329 | 2.01112 | 1.73742 | 4.41201 | 0.00000 | 0.05131 | 0.00000 | 0.92916 | 0.04321 | 1.85858 |
| **Atg5** | 3.17590 | 0.04487 | 8.21161 | 1.43546 | 3.63255 | 6.44688 | 3.10110 | 2.48366 | 1.14046 | 0.14745 | 4.48630 | 3.24164 | 3.80924 | 5.85813 |
| **Atg7** | 5.66991 | 0.05952 | 16.19348 | 14.53105 | 15.42664 | 5.50770 | 0.02341 | 1.80655 | 0.31364 | 0.66582 | 8.11032 | 0.96545 | 1.04290 | 7.27062 |
| **Atg9a** | 10.94938 | 0.10585 | 20.55382 | 18.58718 | 20.54066 | 5.55752 | 2.16219 | 2.70985 | 3.49656 | 0.01781 | 7.28341 | 0.86936 | 11.18941 | 11.49257 |

| | | | | | | | | | | | | | | |
|---|---|---|---|---|---|---|---|---|---|---|---|---|---|---|
| **Atg9b** | 0.12398 | 0.00338 | 0.04212 | 0.62779 | 0.00000 | 0.12029 | 0.04174 | 0.00766 | 0.21504 | 0.00482 | 0.00527 | 0.80695 | 0.00431 | 0.00000 |
| **Atic** | 0.07284 | 0.04636 | 4.71387 | 3.21885 | 2.23876 | 3.73232 | 0.01327 | 1.07068 | 2.01898 | 0.00273 | 2.94520 | 3.57471 | 0.01471 | 2.95821 |
| **Atl1** | 0.15321 | 0.00447 | 0.50635 | 0.00000 | 0.94578 | 0.05392 | 0.00390 | 0.00345 | 1.60904 | 0.00522 | 0.02758 | 0.00438 | 0.01426 | 2.42256 |
| **Atl2** | 3.79266 | 0.07046 | 6.84218 | 1.14719 | 0.57709 | 3.04451 | 1.58962 | 0.94838 | 7.70086 | 0.00748 | 2.24838 | 3.01664 | 4.53499 | 4.98963 |
| **Atl3** | 0.63301 | 0.02280 | 0.64693 | 0.00000 | 0.15398 | 1.18216 | 0.00131 | 0.89087 | 5.95362 | 0.00672 | 1.58220 | 6.21775 | 5.24398 | 6.47096 |
| **Atm** | 8.20344 | 0.05557 | 8.31177 | 1.87046 | 9.86540 | 5.45054 | 0.00000 | 1.29457 | 16.73953 | 0.03466 | 7.47764 | 3.57407 | 3.87232 | 10.68182 |
| **Atmin** | 6.94941 | 0.14492 | 7.93479 | 6.26870 | 5.18255 | 8.91217 | 2.29391 | 5.95252 | 7.27332 | 0.08566 | 0.00000 | 6.51315 | 6.43810 | 8.69119 |
| **Atn1** | 2.46057 | 2.73375 | 6.14816 | 3.32205 | 6.53689 | 2.64405 | 0.37699 | 3.59661 | 1.18541 | 2.30116 | 2.70763 | 3.97472 | 4.65459 | 2.16033 |
| **Atox1** | 4.34561 | 0.14867 | 0.00000 | 1.20826 | 6.64915 | 10.92976 | 2.76376 | 7.30749 | 0.59379 | 2.70880 | 0.18386 | 8.09571 | 3.43243 | 2.06718 |
| **Atp10a** | 0.00864 | 0.00451 | 0.00543 | 0.01599 | 0.04543 | 0.00955 | 0.00427 | 0.01072 | 0.30013 | 0.00666 | 0.00711 | 0.00927 | 0.01688 | 0.05081 |
| **Atp10b** | 0.01231 | 0.01167 | 0.16731 | 0.00116 | 0.00000 | 0.00978 | 0.00416 | 0.00861 | 0.01047 | 0.00661 | 0.00954 | 0.01285 | 0.79847 | 0.50004 |
| **Atp10d** | 1.21149 | 0.00381 | 0.24790 | 0.93896 | 0.00000 | 0.30916 | 0.00368 | 0.00156 | 0.00742 | 0.01221 | 0.00616 | 0.00739 | 0.06026 | 2.96278 |
| **Atp11a** | 0.25462 | 0.01133 | 1.33315 | 1.89586 | 1.64479 | 0.45137 | 0.00331 | 2.42087 | 0.34337 | 0.00209 | 0.68415 | 0.60301 | 3.15925 | 6.16437 |
| **Atp11b** | 3.66419 | 0.17818 | 2.15864 | 0.53101 | 0.81311 | 0.09552 | 0.00126 | 0.01061 | 9.34236 | 6.02118 | 0.51206 | 0.40379 | 1.06610 | 3.98584 |
| **Atp11c** | 2.60238 | 0.02539 | 2.14877 | 1.79254 | 0.65679 | 1.35768 | 0.00205 | 0.09992 | 12.24375 | 0.00272 | 1.79934 | 0.49222 | 0.50295 | 0.29551 |
| **Atp12a** | 0.00294 | 5.72532 | 0.00111 | 0.00847 | 0.00258 | 2.04332 | 6.05224 | 1.37413 | 0.01077 | 0.21850 | 1.64076 | 0.00000 | 0.00639 | 0.00000 |
| **Atp13a1** | 6.45015 | 0.10740 | 15.79886 | 7.92471 | 4.52925 | 6.34065 | 4.94014 | 7.54520 | 0.30085 | 0.16744 | 4.78726 | 10.07514 | 1.40836 | 2.51032 |
| **Atp13a2** | 4.21008 | 0.03967 | 2.07868 | 1.70812 | 1.26736 | 1.71236 | 0.02836 | 0.91038 | 1.85268 | 0.00264 | 0.04748 | 1.80336 | 3.88099 | 0.36859 |
| **Atp13a3** | 19.46634 | 0.59878 | 14.58626 | 4.03564 | 15.34115 | 13.95818 | 1.96754 | 0.32132 | 39.05156 | 10.80321 | 10.17341 | 11.02156 | 14.66631 | 20.37711 |
| **Atp13a5** | 0.00520 | 0.00449 | 0.00963 | 0.01489 | 0.09551 | 0.01119 | 0.00431 | 0.01065 | 0.01715 | 0.00630 | 0.00797 | 0.00882 | 0.00560 | 0.38694 |
| **Atp1a1** | 0.21641 | 8.31039 | 20.50577 | 19.13858 | 11.01991 | 17.15648 | 1.90091 | 15.90874 | 3.66632 | 2.17507 | 9.46709 | 9.72647 | 1.57219 | 14.13193 |
| **Atp1a2** | 0.03779 | 0.01427 | 0.02049 | 0.00000 | 0.02077 | 0.02425 | 0.05450 | 0.07429 | 0.10695 | 0.00673 | 0.01204 | 0.10238 | 0.23506 | 0.07956 |
| **Atp1a3** | 0.00756 | 0.00377 | 0.00810 | 0.11012 | 0.06039 | 0.00773 | 0.00380 | 0.00414 | 0.00727 | 0.00979 | 0.01297 | 0.41527 | 0.32237 | 0.12861 |
| **Atp1b1** | 0.00000 | 1.98378 | 3.60665 | 2.56069 | 0.74786 | 8.56884 | 4.22863 | 16.35614 | 1.67950 | 5.96516 | 5.09130 | 4.81909 | 4.34027 | 1.49528 |
| **Atp1b2** | 0.13158 | 0.00358 | 0.05024 | 0.00008 | 1.03855 | 0.00005 | 0.00407 | 0.00584 | 1.34868 | 0.00788 | 0.00000 | 0.89286 | 0.88885 | 1.15574 |
| **Atp1b3** | 20.35915 | 6.28874 | 8.24273 | 0.00005 | 16.07034 | 8.25488 | 2.29380 | 7.86582 | 13.56844 | 0.17015 | 2.43271 | 7.46823 | 12.37549 | 19.29940 |
| **Atp2a1** | 0.00893 | 0.00455 | 0.07084 | 0.05561 | 0.42910 | 0.01844 | 0.00423 | 0.00892 | 0.00752 | 0.00974 | 0.00672 | 0.00848 | 0.00624 | 0.06018 |
| **Atp2a2** | 8.26919 | 0.09670 | 13.52246 | 11.48719 | 7.75343 | 22.93914 | 7.15379 | 4.42731 | 14.61898 | 0.35682 | 10.43645 | 10.19998 | 10.56152 | 12.76837 |
| **Atp2a3** | 0.01535 | 0.00428 | 0.00158 | 0.82397 | 0.00565 | 0.52206 | 0.00422 | 0.00685 | 0.01290 | 0.01497 | 0.00255 | 0.00667 | 0.01963 | 0.00000 |
| **Atp2b1** | 0.78632 | 1.18129 | 1.86244 | 1.73612 | 1.09776 | 3.32409 | 0.01450 | 1.52633 | 7.72497 | 0.00956 | 2.69192 | 3.37828 | 1.02350 | 2.72232 |
| **Atp2b2** | 0.00000 | 0.00460 | 0.22479 | 0.71721 | 0.00000 | 0.20665 | 0.78424 | 0.05421 | 0.00684 | 0.02420 | 0.02242 | 0.00054 | 0.03283 | 0.01729 |
| **Atp2b3** | 0.00767 | 0.00435 | 0.27477 | 0.12234 | 0.03057 | 0.00372 | 0.00427 | 0.00941 | 0.00724 | 0.00583 | 0.00948 | 0.00723 | 0.00415 | 0.05636 |
| **Atp2b4** | 0.00000 | 0.00371 | 0.46628 | 1.61060 | 2.32821 | 0.69980 | 0.00347 | 0.00000 | 0.00463 | 0.00346 | 0.84735 | 0.45334 | 0.34013 | 2.91313 |
| **Atp2c1** | 2.30155 | 0.02900 | 1.55768 | 0.40179 | 0.86478 | 1.64688 | 0.00121 | 2.59813 | 3.63517 | 0.00352 | 1.75185 | 4.77141 | 2.23304 | 4.61934 |
| **Atp2c2** | 0.00423 | 0.00410 | 0.01750 | 0.25416 | 0.01790 | 0.00423 | 0.00392 | 0.00816 | 0.03021 | 0.00620 | 0.00426 | 0.51021 | 0.00526 | 0.07782 |
| **Atp4b** | 0.14395 | 0.00408 | 0.06071 | 3.25966 | 0.00000 | 3.81336 | 0.01495 | 0.00392 | 0.00875 | 0.01952 | 0.00000 | 0.00289 | 1.66963 | 0.00000 |
| **Atp5a1** | 26.16538 | 9.60750 | 39.45969 | 40.72036 | 27.90939 | 36.79292 | 15.30763 | 31.88990 | 15.47284 | 21.59802 | 24.46322 | 30.08779 | 10.55500 | 26.91582 |
| **Atp5b** | 29.43395 | 14.01265 | 38.51120 | 46.54665 | 34.97862 | 37.64440 | 12.14303 | 29.84878 | 20.05238 | 22.29209 | 21.32940 | 31.46483 | 19.91574 | 26.02991 |
| **Atp5c1** | 21.60419 | 2.76612 | 30.76177 | 23.85177 | 26.52926 | 29.05048 | 10.92983 | 26.32570 | 6.52215 | 22.19646 | 18.64234 | 24.58325 | 6.94867 | 17.17788 |
| **Atp5d** | 3.32625 | 5.23082 | 3.86283 | 36.99335 | 14.60498 | 7.85175 | 5.18950 | 10.38122 | 0.00000 | 13.34486 | 2.53691 | 10.10782 | 0.60956 | 0.35633 |
| **Atp5e** | 2.10599 | 2.65597 | 0.05135 | 0.81261 | 3.45619 | 5.51926 | 3.42028 | 10.59307 | 0.00000 | 12.96411 | 0.13194 | 6.30203 | 0.59579 | 0.83332 |
| **Atp5f1** | 27.69887 | 9.09341 | 36.36766 | 33.19474 | 30.51002 | 28.73402 | 10.79555 | 30.49051 | 14.22004 | 18.20909 | 19.96215 | 25.37332 | 11.45062 | 12.13624 |
| **Atp5g1** | 18.87172 | 10.20195 | 8.29675 | 21.14285 | 24.41211 | 27.66623 | 9.33036 | 25.47716 | 3.14538 | 19.31894 | 3.95017 | 22.07417 | 19.97228 | 15.14125 |
| **Atp5g2** | 5.05113 | 9.73621 | 4.19854 | 13.40291 | 10.02152 | 20.49041 | 6.80937 | 19.51385 | 1.50704 | 15.17638 | 5.65948 | 16.98698 | 1.56540 | 6.50931 |
| **Atp5g3** | 2.09179 | 11.83566 | 5.58002 | 18.50587 | 20.89388 | 16.28047 | 9.63383 | 26.02774 | 2.28571 | 23.05895 | 5.26429 | 26.92466 | 4.28786 | 1.61726 |
| **Atp5h** | 11.03592 | 10.69127 | 1.33878 | 15.71235 | 16.59975 | 18.76231 | 6.44638 | 21.92130 | 8.40307 | 19.41229 | 3.61876 | 17.71558 | 7.44493 | 9.75397 |
| **Atp5j** | 7.38480 | 7.97632 | 0.10026 | 3.17932 | 15.63938 | 16.46057 | 5.68021 | 11.66896 | 2.65846 | 11.67414 | 1.05849 | 14.77681 | 10.98494 | 3.14471 |
| **Atp5j2** | 4.41903 | 3.27676 | 0.17830 | 1.00081 | 5.80233 | 6.63785 | 2.27669 | 8.45601 | 2.37466 | 8.99481 | 0.29228 | 7.66685 | 4.60806 | 2.97667 |
| **Atp5k** | 2.56220 | 1.21224 | 0.08325 | 2.42319 | 2.41073 | 5.79376 | 0.95592 | 6.68542 | 1.07680 | 13.15312 | 0.01058 | 4.43030 | 5.56682 | 1.55580 |
| **Atp5l** | 5.06995 | 3.92316 | 0.00000 | 7.29656 | 9.15661 | 12.46018 | 4.16015 | 17.82728 | 4.43053 | 15.58092 | 0.95799 | 11.88437 | 2.64727 | 3.43396 |
| **Atp5o** | 13.79931 | 7.26274 | 10.37773 | 24.79502 | 18.75104 | 22.45791 | 9.58479 | 24.32153 | 3.79302 | 22.08343 | 11.78405 | 21.88457 | 1.81895 | 12.66610 |
| **Atp5s** | 1.10886 | 0.00319 | 0.84829 | 2.29838 | 0.56293 | 0.10963 | 1.50033 | 0.00000 | 0.90173 | 0.06438 | 0.94750 | 0.14803 | 3.80034 | 0.00000 |
| **Atp5sl** | 3.24274 | 0.96553 | 9.00662 | 4.65679 | 7.35674 | 3.48665 | 0.01447 | 3.36581 | 0.39359 | 0.02213 | 2.54508 | 4.86566 | 2.27167 | 0.41091 |
| **Atp6ap1** | 3.04483 | 4.16201 | 9.92321 | 4.86452 | 8.19334 | 6.69276 | 5.84561 | 3.17370 | 12.75549 | 0.16999 | 5.60203 | 6.72908 | 2.64433 | 9.53210 |
| **Atp6ap1l** | 3.56100 | 0.03374 | 1.79426 | 1.83148 | 0.35829 | 0.50615 | 1.64719 | 3.08168 | 2.76574 | 0.06733 | 1.80210 | 0.17142 | 1.15991 | 2.89407 |

| | | | | | | | | | | | | | | |
|---|---|---|---|---|---|---|---|---|---|---|---|---|---|---|
| **Atp6ap2** | 12.04740 | 0.06092 | 19.09399 | 12.16121 | 5.13447 | 19.77256 | 0.03886 | 4.18850 | 21.59008 | 9.93088 | 9.01602 | 5.53675 | 1.44093 | 13.08735 |
| **Atp6v0a1** | 2.63683 | 0.00157 | 5.28422 | 3.98938 | 2.71687 | 2.48000 | 0.00814 | 0.00000 | 0.02330 | 0.00000 | 1.24365 | 2.52502 | 0.00000 | 0.95558 |
| **Atp6v0a2** | 1.80658 | 0.01789 | 1.56403 | 2.56894 | 1.64996 | 4.69039 | 0.00245 | 1.64825 | 0.59519 | 0.02049 | 0.00000 | 4.28418 | 1.66915 | 3.45903 |
| **Atp6v0a4** | 0.65623 | 0.02708 | 1.66542 | 1.13198 | 0.48509 | 2.16928 | 0.05025 | 1.54929 | 0.35553 | 0.00977 | 0.00000 | 0.00000 | 0.19012 | 1.75557 |
| **Atp6v0b** | 9.97392 | 9.92439 | 16.94503 | 26.59884 | 8.45468 | 19.59760 | 7.82664 | 16.34023 | 0.00000 | 0.79041 | 6.82597 | 9.29713 | 1.27222 | 4.60850 |
| **Atp6v0d1** | 19.06570 | 6.75806 | 14.02470 | 16.98612 | 12.18469 | 14.57804 | 2.36604 | 9.40462 | 4.36555 | 9.60030 | 11.86110 | 3.94997 | 6.88956 | 12.74300 |
| **Atp6v0e** | 10.14421 | 5.85802 | 13.09446 | 19.98909 | 20.48541 | 21.22137 | 5.38312 | 16.67717 | 5.88004 | 21.15277 | 7.81673 | 12.57347 | 8.82304 | 17.16719 |
| **Atp6v0e2** | 0.00300 | 0.00416 | 0.00636 | 0.62667 | 0.01615 | 0.00613 | 0.00424 | 0.00666 | 0.03207 | 0.00523 | 0.00607 | 0.00751 | 0.10241 | 0.00000 |
| **Atp6v1a** | 6.59319 | 0.19169 | 8.29432 | 2.13847 | 4.45122 | 17.33393 | 1.49888 | 3.96691 | 12.67169 | 7.85400 | 10.96970 | 7.22013 | 10.66265 | 10.99117 |
| **Atp6v1b2** | 3.56033 | 5.07061 | 13.48230 | 10.20313 | 9.16442 | 10.62611 | 1.84137 | 9.66684 | 5.68505 | 0.22524 | 8.83172 | 9.70188 | 2.78772 | 7.57446 |
| **Atp6v1c1** | 11.39714 | 4.01288 | 18.24023 | 14.62647 | 9.96526 | 12.93494 | 4.55364 | 4.06914 | 4.75999 | 0.21996 | 11.52471 | 14.36002 | 3.61061 | 9.16772 |
| **Atp6v1c2** | 0.07884 | 0.01240 | 0.48683 | 0.00000 | 2.18260 | 0.04008 | 0.00401 | 0.00409 | 0.00685 | 0.00545 | 0.00550 | 0.00510 | 0.04018 | 1.16992 |
| **Atp6v1d** | 10.89608 | 7.39239 | 15.40453 | 9.41828 | 19.75517 | 16.76911 | 7.62859 | 9.07045 | 8.43406 | 13.71084 | 9.85879 | 7.11654 | 5.48869 | 16.37763 |
| **Atp6v1e1** | 17.93709 | 5.42341 | 26.13838 | 21.90427 | 16.84859 | 24.17174 | 7.77815 | 10.68754 | 4.09418 | 13.90639 | 18.08467 | 15.14496 | 5.22092 | 19.70647 |
| **Atp6v1f** | 3.03363 | 0.13424 | 0.24153 | 2.87768 | 3.49594 | 8.97377 | 3.79358 | 5.81589 | 0.02524 | 1.34844 | 1.73951 | 5.44274 | 0.00000 | 0.47892 |
| **Atp6v1g1** | 5.43060 | 5.97438 | 6.63167 | 14.96099 | 7.21456 | 19.04271 | 2.93934 | 16.62414 | 0.20411 | 16.57802 | 10.96668 | 13.81560 | 0.37933 | 4.35596 |
| **Atp6v1g2** | 0.03344 | 0.00433 | 0.38803 | 0.00081 | 0.00362 | 0.00535 | 0.00421 | 0.01142 | 0.01071 | 0.00671 | 0.00632 | 0.00920 | 0.07304 | 0.03674 |
| **Atp6v1h** | 7.32339 | 0.05525 | 1.85628 | 4.28270 | 1.45768 | 9.23381 | 0.24722 | 5.63365 | 3.70143 | 0.00000 | 7.84847 | 5.42039 | 0.04436 | 4.46797 |
| **Atp7a** | 0.46792 | 0.83080 | 0.39984 | 0.00000 | 0.62461 | 0.73963 | 0.00743 | 0.00257 | 9.21587 | 0.01259 | 0.00000 | 0.52203 | 2.25521 | 6.99114 |
| **Atp8a1** | 1.84748 | 0.00937 | 0.99150 | 0.76866 | 0.44792 | 0.40633 | 0.01298 | 0.00000 | 2.59316 | 0.01476 | 1.46531 | 1.84494 | 0.07505 | 0.00000 |
| **Atp8b1** | 0.00000 | 0.00168 | 0.30876 | 0.00000 | 0.16357 | 0.68196 | 0.00175 | 0.52320 | 2.11943 | 5.89042 | 0.00000 | 1.82585 | 0.31607 | 0.00000 |
| **Atp8b2** | 0.42303 | 0.01098 | 0.63629 | 0.06330 | 0.34418 | 0.00000 | 0.00901 | 0.00184 | 0.71594 | 0.00729 | 0.00160 | 0.35884 | 0.00344 | 0.00000 |
| **Atp8b4** | 0.34620 | 0.00438 | 2.36038 | 0.00000 | 2.55687 | 0.00000 | 0.00328 | 0.00496 | 4.09951 | 0.00725 | 0.00195 | 0.00566 | 0.00000 | 0.21294 |
| **Atp8b5** | 0.00000 | 0.09133 | 0.01371 | 0.00354 | 0.22546 | 0.03190 | 0.00402 | 0.00526 | 0.65021 | 0.00595 | 0.00667 | 0.00645 | 0.00350 | 0.48636 |
| **Atp9a** | 0.57497 | 0.04319 | 5.03261 | 2.65394 | 6.63989 | 0.16848 | 0.00254 | 2.66719 | 0.03887 | 0.02177 | 2.94931 | 1.12796 | 0.22132 | 0.27090 |
| **Atp9b** | 1.13098 | 0.06508 | 0.52148 | 1.00631 | 0.47917 | 3.91691 | 2.58283 | 1.83719 | 0.00000 | 0.03768 | 0.00000 | 2.13082 | 0.75229 | 0.86428 |
| **Atpaf1** | 5.87943 | 0.04694 | 3.55708 | 1.82645 | 5.32328 | 3.32092 | 0.73196 | 4.71790 | 4.89003 | 0.05603 | 4.64669 | 7.82008 | 1.43507 | 2.74108 |
| **Atpaf2** | 2.53710 | 0.06632 | 7.69454 | 7.92924 | 6.30031 | 1.85366 | 0.01813 | 4.00315 | 0.65544 | 0.07865 | 1.80602 | 2.77104 | 1.57362 | 5.79723 |
| **Atpif1** | 1.58950 | 1.66184 | 0.00000 | 2.95023 | 3.77758 | 6.24035 | 2.19798 | 10.30771 | 0.85287 | 11.19263 | 0.51739 | 12.16434 | 8.13930 | 1.55195 |
| **Atr** | 2.91644 | 0.00866 | 0.00000 | 0.99622 | 0.75605 | 1.08379 | 0.00927 | 0.47576 | 1.01007 | 0.00269 | 7.03874 | 2.08726 | 1.55472 | 1.29165 |
| **Atraid** | 4.71210 | 0.03669 | 6.09935 | 1.67273 | 9.99984 | 5.69507 | 4.04613 | 7.38250 | 0.86095 | 0.18729 | 2.50289 | 4.28173 | 4.61489 | 9.31043 |
| **Atrip** | 0.35944 | 0.06341 | 2.74454 | 1.65843 | 1.15082 | 1.31670 | 3.02230 | 0.70920 | 0.05442 | 0.07754 | 1.86408 | 0.47515 | 0.00000 | 1.40145 |
| **Atrn** | 2.50862 | 0.00345 | 0.08701 | 1.81207 | 1.64239 | 2.50701 | 0.00390 | 0.00734 | 1.42269 | 0.00594 | 0.00000 | 1.07403 | 1.84898 | 1.37370 |
| **Atrnl1** | 0.00891 | 0.00427 | 0.00212 | 0.00000 | 0.28696 | 0.42196 | 0.00366 | 0.25567 | 0.00623 | 0.01304 | 0.00365 | 0.04636 | 0.09425 | 0.07226 |
| **Atrx** | 3.70884 | 0.01635 | 7.64680 | 0.95516 | 6.72505 | 1.91882 | 0.15944 | 2.28540 | 25.43067 | 0.03608 | 3.06960 | 2.41458 | 1.24512 | 7.55932 |
| **Atxn1** | 0.07421 | 1.19538 | 0.94832 | 0.07246 | 0.16325 | 0.03812 | 0.00358 | 0.43807 | 0.00880 | 0.01828 | 0.00730 | 0.21459 | 0.00798 | 1.43308 |
| **Atxn10** | 7.29040 | 0.88138 | 7.94301 | 7.49058 | 4.36959 | 4.52663 | 5.61501 | 15.11200 | 1.84290 | 0.17953 | 2.75946 | 8.58313 | 0.00000 | 2.00083 |
| **Atxn1l** | 0.84212 | 0.01869 | 2.20577 | 1.76796 | 1.08618 | 1.42758 | 0.00278 | 0.96024 | 3.39930 | 0.03593 | 1.44530 | 2.32339 | 3.73026 | 3.80801 |
| **Atxn2** | 6.23270 | 0.01428 | 9.70879 | 0.00000 | 4.92687 | 3.38908 | 1.41185 | 0.43053 | 1.22535 | 5.73796 | 6.17087 | 4.34425 | 2.77477 | 11.10392 |
| **Atxn2l** | 0.42286 | 0.02145 | 0.34833 | 0.62081 | 2.76515 | 1.29232 | 0.02261 | 0.26006 | 0.19412 | 0.02330 | 0.23784 | 0.53024 | 5.01670 | 1.79820 |
| **Atxn3** | 3.74488 | 0.00000 | 5.73425 | 0.82966 | 6.09418 | 1.27467 | 1.85202 | 0.00000 | 13.00273 | 2.41956 | 4.15662 | 1.81383 | 7.51152 | 6.19265 |
| **Atxn7** | 5.61619 | 0.01616 | 1.57346 | 0.41946 | 0.78706 | 2.18759 | 0.08402 | 0.01525 | 8.69621 | 0.00000 | 1.34318 | 0.90455 | 4.89294 | 7.70853 |
| **Atxn7l1** | 0.14703 | 0.01643 | 0.00000 | 0.32593 | 0.00000 | 4.42036 | 0.00301 | 0.61981 | 0.01154 | 0.03323 | 3.79382 | 4.39930 | 0.00519 | 0.00000 |
| **Atxn7l2** | 0.00306 | 0.02250 | 0.34125 | 0.49660 | 0.00098 | 0.00000 | 0.01030 | 0.02520 | 0.01189 | 0.00745 | 0.00559 | 0.08962 | 0.00268 | 0.37388 |
| **Atxn7l3** | 1.31831 | 0.90226 | 5.10974 | 2.88040 | 9.89857 | 1.36321 | 0.03383 | 0.47013 | 0.05354 | 0.60590 | 0.38931 | 3.03369 | 0.44929 | 5.28418 |
| **Atxn7l3b** | 1.27333 | 1.45790 | 2.02442 | 5.52591 | 4.43055 | 3.75188 | 0.18533 | 2.04401 | 6.65029 | 0.05009 | 0.40713 | 3.54583 | 0.85942 | 2.13895 |
| **AU015228** | 3.33728 | 0.00273 | 0.16424 | 0.00827 | 0.04281 | 0.00000 | 0.00356 | 0.01204 | 0.02823 | 0.00570 | 0.09574 | 0.00382 | 14.82480 | 0.20545 |
| **AU015836** | 1.04850 | 0.01192 | 0.63986 | 0.00000 | 0.72656 | 0.84735 | 0.00471 | 0.51764 | 1.17911 | 0.01801 | 0.29923 | 0.00000 | 1.44380 | 1.43176 |
| **AU016765** | 0.00872 | 0.00449 | 0.01195 | 0.01770 | 0.00534 | 0.01220 | 0.00425 | 0.01023 | 0.00809 | 0.00920 | 0.00747 | 0.01194 | 0.03601 | 0.39580 |
| **AU018091** | 17.23882 | 2.02228 | 14.65840 | 13.65109 | 12.67854 | 26.15107 | 2.12381 | 14.56106 | 32.69103 | 0.32894 | 5.99048 | 5.01544 | 7.77736 | 15.72711 |
| **AU019823** | 4.73850 | 0.02679 | 1.43983 | 1.52987 | 1.51343 | 1.92826 | 0.96054 | 0.94665 | 5.05219 | 0.02930 | 1.41637 | 0.95161 | 3.30009 | 2.62042 |
| **AU019990** | 0.01486 | 0.00425 | 0.39039 | 0.00000 | 0.00000 | 0.01194 | 0.00399 | 0.00819 | 0.83034 | 0.00620 | 0.00476 | 0.00592 | 1.42485 | 0.38377 |
| **AU021092** | 0.39023 | 0.05842 | 0.72958 | 2.14635 | 0.00000 | 4.19796 | 0.16372 | 9.88983 | 0.00000 | 0.04736 | 0.57646 | 0.00000 | 0.00002 | 0.31398 |
| **AU022252** | 1.14119 | 0.00103 | 1.67457 | 5.39877 | 3.27393 | 1.53856 | 0.39731 | 1.80537 | 0.00000 | 0.06668 | 0.05969 | 4.34399 | 2.30080 | 0.12833 |
| **AU022751** | 0.07506 | 0.00221 | 2.26194 | 0.00109 | 0.03800 | 0.76048 | 1.28776 | 0.00000 | 0.02325 | 0.02474 | 3.01699 | 1.19900 | 2.57209 | 0.17554 |

| | | | | | | | | | | | | | | |
|---|---|---|---|---|---|---|---|---|---|---|---|---|---|---|
| **AU022754** | 0.01575 | 0.00420 | 0.02880 | 0.06123 | 0.02034 | 0.03267 | 0.00426 | 0.00591 | 0.12339 | 0.00626 | 0.01566 | 0.02498 | 0.41979 | 0.01274 |
| **AU023762** | 0.12174 | 0.22091 | 0.02139 | 0.08670 | 0.04460 | 0.10563 | 0.06292 | 0.06505 | 0.46502 | 0.00779 | 0.02376 | 0.05007 | 0.40323 | 0.29563 |
| **AU040320** | 0.00000 | 0.02829 | 1.99074 | 1.57468 | 2.94781 | 0.85198 | 0.00388 | 1.18502 | 2.84583 | 0.00021 | 0.67341 | 0.65258 | 2.09767 | 3.27546 |
| **AU041133** | 0.00000 | 0.00383 | 0.20150 | 0.00000 | 0.66609 | 0.34269 | 0.00361 | 0.05909 | 2.62012 | 0.02001 | 0.68709 | 0.00920 | 0.00000 | 0.50732 |
| **Auh** | 0.88583 | 0.00405 | 0.50318 | 0.34339 | 1.01535 | 0.50623 | 0.01644 | 0.21332 | 0.00191 | 0.00074 | 2.05351 | 0.92401 | 1.28067 | 0.06812 |
| **Aunip** | 0.97109 | 0.04120 | 0.34968 | 0.16369 | 3.04948 | 1.60125 | 0.01636 | 0.78057 | 0.00000 | 0.04287 | 0.01316 | 0.51874 | 5.20635 | 1.34704 |
| **Aup1** | 0.48899 | 0.08383 | 9.59389 | 16.60476 | 6.77054 | 5.76491 | 1.47394 | 7.41445 | 0.01455 | 0.02492 | 4.23037 | 7.16274 | 0.05030 | 0.79619 |
| **Aurka** | 14.02427 | 8.74625 | 33.17316 | 14.00630 | 26.37425 | 20.49818 | 6.68733 | 14.72154 | 21.54395 | 0.49981 | 15.54869 | 7.41836 | 11.75329 | 28.78244 |
| **Aurkaip1** | 7.19812 | 2.14367 | 9.92121 | 14.71271 | 11.22579 | 8.46366 | 5.37741 | 9.17522 | 0.35660 | 17.62484 | 5.66594 | 8.20996 | 4.38913 | 2.80366 |
| **Aurkb** | 5.05554 | 6.30392 | 8.89137 | 1.41515 | 8.24506 | 2.73819 | 0.13496 | 5.29209 | 0.68559 | 0.04210 | 8.21816 | 4.64253 | 3.91028 | 11.51881 |
| **Aurkc** | 8.70663 | 0.00144 | 13.98785 | 6.68628 | 13.64198 | 0.21728 | 0.00000 | 0.00000 | 7.08048 | 0.00000 | 1.31005 | 0.52317 | 5.35447 | 14.35084 |
| **Auts2** | 0.35889 | 0.00315 | 3.58574 | 0.84770 | 2.85430 | 0.40323 | 0.00825 | 2.86461 | 3.46634 | 0.01385 | 0.00000 | 0.72419 | 1.61892 | 4.77266 |
| **AV051173** | 0.02658 | 0.01248 | 0.28674 | 0.23270 | 0.16126 | 0.01078 | 0.00476 | 0.43475 | 0.00785 | 0.00564 | 0.02918 | 0.01371 | 0.00207 | 0.11074 |
| **AV320801** | 0.01724 | 0.00396 | 0.01188 | 0.00000 | 0.01188 | 0.03214 | 1.37288 | 0.00286 | 0.01010 | 0.00413 | 0.00740 | 0.01130 | 0.02238 | 0.30085 |
| **Aven** | 2.28100 | 0.01531 | 3.98600 | 3.05195 | 8.80856 | 6.38356 | 0.11424 | 1.45011 | 2.70829 | 0.01138 | 4.21986 | 3.02158 | 1.20988 | 5.94229 |
| **Avl9** | 7.54099 | 0.00110 | 7.15857 | 0.00000 | 12.54952 | 1.51542 | 0.00000 | 1.53909 | 14.77899 | 0.01710 | 0.00000 | 0.00000 | 13.59552 | 12.64173 |
| **Avpi1** | 0.38180 | 3.92476 | 3.17692 | 11.89144 | 4.24848 | 3.70696 | 3.16672 | 6.14939 | 0.07188 | 9.55006 | 1.26141 | 5.41139 | 0.00000 | 0.05466 |
| **AW011738** | 0.00543 | 0.00437 | 0.01459 | 0.01175 | 0.02745 | 0.01346 | 0.00421 | 0.00943 | 0.01233 | 0.00649 | 0.00909 | 0.00944 | 0.24860 | 0.00263 |
| **AW146154** | 0.29667 | 0.00366 | 0.27614 | 0.06720 | 0.32310 | 1.35647 | 0.18440 | 0.29084 | 1.73401 | 0.17901 | 1.17334 | 0.06869 | 0.67659 | 0.06473 |
| **AW209491** | 2.80219 | 0.01023 | 3.19944 | 0.17349 | 3.25095 | 0.12022 | 0.00719 | 2.56714 | 6.70435 | 0.01026 | 0.85675 | 0.00262 | 3.55413 | 8.22193 |
| **AW495222** | 0.08632 | 0.00461 | 0.01526 | 0.00617 | 0.17625 | 0.01886 | 0.00426 | 0.01039 | 0.13641 | 0.00605 | 0.00694 | 0.00830 | 0.00445 | 0.11137 |
| **AW549542** | 0.15025 | 0.00492 | 0.22001 | 0.00242 | 0.00000 | 0.02441 | 0.00428 | 0.02029 | 0.01098 | 0.01287 | 0.01028 | 0.00827 | 0.04093 | 0.00593 |
| **AW549877** | 0.93813 | 0.01217 | 0.35664 | 1.79447 | 0.71777 | 2.51854 | 1.23696 | 1.61671 | 1.08255 | 0.03353 | 0.00000 | 0.70484 | 0.00696 | 0.00000 |
| **AW554918** | 0.00000 | 0.00385 | 0.06284 | 0.16498 | 0.59448 | 0.39798 | 0.00386 | 0.02279 | 0.00923 | 0.00606 | 0.01156 | 0.13833 | 1.42562 | 1.31375 |
| **AW822252** | 0.00842 | 0.00402 | 0.01333 | 0.00000 | 0.04863 | 0.02718 | 0.00418 | 0.02851 | 0.01024 | 0.00692 | 0.00568 | 0.12721 | 0.00850 | 0.01255 |
| **Awat2** | 0.00848 | 0.00460 | 0.00102 | 0.40763 | 0.06130 | 0.18539 | 0.00452 | 0.00184 | 0.02359 | 0.00411 | 0.56547 | 0.01591 | 0.00556 | 0.00000 |
| **Axin1** | 2.78764 | 0.06033 | 7.05576 | 2.34220 | 7.14872 | 8.14637 | 0.00568 | 3.84387 | 2.22026 | 0.02353 | 1.80847 | 7.29190 | 6.99799 | 14.30781 |
| **Axin2** | 0.00000 | 0.00293 | 0.05970 | 1.08400 | 0.02644 | 0.00000 | 0.22833 | 0.00365 | 0.04495 | 0.01149 | 0.00866 | 1.50493 | 1.05869 | 0.00000 |
| **Axl** | 0.23459 | 0.00448 | 0.00293 | 0.01019 | 0.28517 | 0.00929 | 0.00431 | 0.01206 | 0.14459 | 0.00650 | 0.00639 | 0.00739 | 0.00296 | 0.00000 |
| **AY074887** | 0.05283 | 0.00429 | 0.20751 | 0.23658 | 0.15934 | 0.17405 | 0.03031 | 0.12537 | 0.18000 | 0.11732 | 0.23298 | 0.15700 | 0.20205 | 0.29198 |
| **AY358078** | 0.22108 | 0.07312 | 0.01712 | 0.03558 | 0.02581 | 0.10045 | 0.02812 | 0.05413 | 0.02230 | 0.02217 | 0.01751 | 0.25576 | 0.26972 | 0.07448 |
| **AY512915** | 0.00000 | 0.00429 | 0.00347 | 0.01802 | 0.05169 | 0.00898 | 0.00417 | 0.00835 | 0.08105 | 0.00599 | 0.00717 | 0.00757 | 0.24513 | 1.41667 |
| **AY512931** | 1.14021 | 0.00438 | 0.48821 | 0.07108 | 0.48824 | 0.21401 | 0.11325 | 0.07506 | 0.79698 | 0.00754 | 0.00578 | 0.33822 | 2.24449 | 0.99622 |
| **AY761184** | 0.09625 | 0.00398 | 0.00734 | 0.11174 | 0.01239 | 0.07270 | 0.00414 | 0.00645 | 0.13639 | 0.00854 | 0.00808 | 0.07995 | 0.96945 | 0.00000 |
| **AY761185** | 0.01903 | 0.00440 | 0.00990 | 0.03424 | 0.06847 | 0.01118 | 0.00408 | 0.00926 | 0.01197 | 0.01298 | 0.00809 | 0.00829 | 0.01547 | 0.00763 |
| **Azi2** | 4.65547 | 0.03776 | 1.45103 | 1.91897 | 2.21544 | 4.47626 | 3.27716 | 1.37650 | 3.73278 | 0.08999 | 0.93310 | 1.57622 | 1.45397 | 1.02244 |
| **Azin1** | 22.23518 | 3.57011 | 20.80092 | 9.28595 | 21.16213 | 18.43631 | 4.24026 | 4.95914 | 36.66361 | 12.24107 | 15.53846 | 7.21711 | 12.80079 | 11.44512 |
| **Azin2** | 0.02238 | 0.00524 | 0.13779 | 0.00000 | 0.01951 | 0.11766 | 0.00988 | 0.00983 | 0.05491 | 0.01125 | 0.00466 | 0.10821 | 0.04536 | 0.00041 |
| **B2m** | 4.91557 | 0.05056 | 1.40583 | 4.38424 | 3.78985 | 7.37110 | 5.10654 | 5.33019 | 2.40538 | 0.17055 | 2.88630 | 0.85117 | 4.34247 | 3.69853 |
| **B3galnt1** | 0.00484 | 0.01455 | 0.04365 | 0.05514 | 0.00009 | 0.01344 | 0.01036 | 0.04474 | 0.04478 | 0.00703 | 0.03129 | 0.60903 | 0.50155 | 0.16178 |
| **B3galnt2** | 1.04846 | 0.00339 | 0.14563 | 0.00000 | 0.03970 | 0.86959 | 0.05440 | 0.00033 | 3.38875 | 0.01189 | 0.34114 | 0.42738 | 2.68367 | 0.23382 |
| **B3galt4** | 0.42160 | 0.00410 | 0.85050 | 0.00569 | 0.12730 | 0.47301 | 0.00384 | 0.00000 | 0.01747 | 0.00222 | 0.84682 | 0.00638 | 0.29736 | 0.53876 |
| **B3galt5** | 0.00750 | 0.00497 | 0.01545 | 0.01079 | 0.01618 | 0.01056 | 0.00443 | 0.02893 | 0.01030 | 0.00728 | 0.00920 | 0.00896 | 0.35709 | 0.00712 |
| **B3galt6** | 3.80679 | 0.00268 | 0.93056 | 0.00000 | 0.55670 | 0.73802 | 2.94833 | 0.00000 | 0.20432 | 0.11530 | 1.12538 | 0.69141 | 0.44556 | 0.35107 |
| **B3gat2** | 0.01078 | 0.00449 | 0.12048 | 0.00375 | 0.01120 | 0.01247 | 0.00485 | 0.03948 | 0.01177 | 0.00674 | 0.00841 | 0.00950 | 0.00726 | 0.01181 |
| **B3gat3** | 0.21190 | 0.08003 | 2.43746 | 1.12155 | 1.28531 | 3.91295 | 0.15891 | 4.80365 | 0.41995 | 10.63510 | 3.35149 | 2.86429 | 1.37086 | 1.38497 |
| **B3glct** | 0.00000 | 0.03041 | 1.75869 | 2.74974 | 1.30246 | 0.09200 | 0.03083 | 0.02635 | 0.06393 | 0.01179 | 2.90077 | 3.20116 | 0.00000 | 1.60742 |
| **B3gnt2** | 0.04963 | 0.00334 | 1.23843 | 0.09838 | 0.32871 | 0.00000 | 2.55115 | 0.00000 | 2.04794 | 0.05175 | 0.00000 | 0.36094 | 2.17074 | 0.73576 |
| **B3gnt3** | 0.01277 | 0.01748 | 0.03384 | 0.62935 | 0.12991 | 0.00000 | 0.00370 | 0.00291 | 0.00721 | 0.00555 | 0.00433 | 0.49080 | 0.08433 | 0.28964 |
| **B3gnt4** | 0.01830 | 0.00432 | 0.36337 | 0.00000 | 0.00636 | 0.24969 | 0.00408 | 0.00389 | 0.01024 | 0.00806 | 0.00625 | 0.03916 | 0.01166 | 0.03424 |
| **B3gnt5** | 0.00464 | 2.83484 | 0.00000 | 0.00000 | 0.00450 | 1.28123 | 3.53034 | 9.58177 | 0.00802 | 0.04258 | 1.49308 | 0.83220 | 0.01365 | 0.00000 |
| **B3gnt6** | 0.00000 | 0.00268 | 0.28965 | 0.21912 | 0.00000 | 0.00000 | 0.00359 | 0.00287 | 0.01326 | 0.00539 | 0.00652 | 1.97205 | 0.00546 | 0.00751 |
| **B3gnt7** | 0.02512 | 0.08738 | 0.24199 | 0.00000 | 0.89714 | 0.00000 | 0.00312 | 0.92798 | 0.23209 | 0.00519 | 0.00000 | 4.74287 | 0.38768 | 0.02148 |
| **B3gnt8** | 0.03174 | 0.02073 | 0.63866 | 0.09992 | 1.18781 | 0.06991 | 0.00384 | 0.00484 | 0.01240 | 0.01203 | 0.00968 | 0.55790 | 0.00000 | 0.00187 |
| **B3gntl1** | 0.31818 | 0.00432 | 0.34939 | 0.09925 | 0.09059 | 0.27090 | 0.00473 | 0.29762 | 0.46277 | 0.00662 | 0.00346 | 0.00280 | 0.12279 | 0.83327 |

| | | | | | | | | | | | | | | |
|---|---|---|---|---|---|---|---|---|---|---|---|---|---|---|
| **B4galnt1** | 0.13685 | 0.02131 | 4.02465 | 3.10128 | 0.71326 | 0.85509 | 0.86535 | 0.00662 | 0.75607 | 0.07225 | 0.96525 | 1.35082 | 0.25998 | 0.98871 |
| **B4galnt2** | 1.25210 | 0.49451 | 1.08140 | 0.00000 | 0.86255 | 0.81960 | 0.04147 | 0.03776 | 0.97768 | 0.07574 | 0.01809 | 0.47886 | 3.96673 | 1.87749 |
| **B4galnt3** | 0.00795 | 0.00495 | 0.22660 | 0.00000 | 0.12738 | 0.00324 | 0.00423 | 0.03119 | 0.16039 | 0.00838 | 0.00635 | 0.05052 | 0.00374 | 0.01224 |
| **B4galnt4** | 0.02144 | 0.00412 | 0.00272 | 0.41432 | 0.36553 | 0.00506 | 0.00427 | 0.00602 | 0.01251 | 0.00663 | 0.00748 | 0.01137 | 0.02520 | 0.01887 |
| **B4galt1** | 0.60794 | 0.06495 | 3.06926 | 2.42443 | 0.57057 | 5.55178 | 0.00143 | 1.86096 | 0.16488 | 2.63113 | 0.42407 | 5.31631 | 0.00489 | 1.10571 |
| **B4galt2** | 0.00000 | 0.01968 | 1.15225 | 0.00000 | 0.37337 | 0.00000 | 0.00335 | 0.61776 | 0.00225 | 0.00359 | 0.00176 | 0.26351 | 0.88951 | 0.68348 |
| **B4galt3** | 4.02171 | 8.47633 | 11.26129 | 13.63952 | 15.77158 | 3.58950 | 0.26831 | 9.50928 | 0.52404 | 0.15261 | 4.73342 | 2.10561 | 5.62405 | 3.47237 |
| **B4galt4** | 0.01141 | 0.00384 | 0.01313 | 0.00000 | 0.05048 | 0.00403 | 0.00361 | 0.72685 | 0.01188 | 0.00545 | 0.00875 | 0.01078 | 0.00769 | 0.04163 |
| **B4galt5** | 0.04439 | 0.05383 | 1.20828 | 0.68113 | 0.00000 | 9.26507 | 1.58646 | 3.39884 | 0.00000 | 0.03296 | 0.13038 | 6.30281 | 1.57577 | 0.00000 |
| **B4galt6** | 0.92380 | 0.28436 | 0.95513 | 3.19641 | 0.00000 | 2.05452 | 0.19632 | 3.88961 | 0.85543 | 0.08746 | 4.14582 | 5.61386 | 3.28884 | 1.28576 |
| **B4galt7** | 0.47070 | 0.12809 | 0.44758 | 1.79113 | 0.42451 | 1.75350 | 0.21748 | 0.52865 | 0.93549 | 1.73568 | 0.03792 | 0.74360 | 4.01528 | 1.81159 |
| **B4gat1** | 0.93325 | 0.04524 | 0.29628 | 0.38534 | 0.09810 | 0.50021 | 0.00313 | 1.67462 | 0.01109 | 0.01243 | 1.58092 | 0.94858 | 0.01636 | 0.64003 |
| **B9d1** | 0.13536 | 0.02751 | 0.50534 | 0.17151 | 0.15927 | 1.22879 | 0.00348 | 1.20877 | 0.00000 | 0.00717 | 0.02892 | 3.60275 | 0.95134 | 1.82680 |
| **B9d2** | 0.00542 | 0.01922 | 0.10135 | 0.00000 | 0.12770 | 0.01458 | 0.00407 | 0.00879 | 0.01607 | 0.00927 | 0.00370 | 0.85733 | 0.05501 | 0.00719 |
| **Baat** | 0.30749 | 0.04023 | 0.00407 | 0.00458 | 0.01871 | 0.00748 | 0.00402 | 0.01514 | 0.20104 | 0.00594 | 0.00355 | 0.40456 | 0.16146 | 0.00000 |
| **Babam1** | 2.11587 | 3.35780 | 4.60207 | 7.30982 | 2.22559 | 0.91941 | 0.30162 | 2.79420 | 0.03856 | 11.80158 | 1.85046 | 5.16795 | 0.00000 | 0.59212 |
| **Bace1** | 2.52084 | 4.75944 | 5.44164 | 1.76170 | 3.68141 | 8.40655 | 0.69949 | 3.60529 | 5.64287 | 0.02988 | 3.03218 | 3.26509 | 5.38751 | 8.55679 |
| **Bace2** | 0.22131 | 0.00477 | 0.00922 | 0.00895 | 0.16984 | 0.01057 | 0.00473 | 0.01568 | 0.01052 | 0.00682 | 0.00725 | 0.01079 | 0.00414 | 0.01008 |
| **Bach1** | 4.30551 | 0.01408 | 8.37358 | 7.44102 | 8.69122 | 3.28784 | 2.06384 | 2.37066 | 26.11975 | 0.34843 | 7.73909 | 3.16215 | 4.89201 | 8.88655 |
| **Bach2** | 0.07536 | 0.01278 | 0.01930 | 0.14941 | 0.03898 | 0.13543 | 0.04054 | 0.04813 | 0.08419 | 0.03650 | 0.01628 | 0.02238 | 0.13012 | 0.17978 |
| **Bad** | 2.02357 | 7.58389 | 8.98239 | 8.76820 | 10.26992 | 5.74035 | 2.61340 | 13.99184 | 0.01927 | 1.54683 | 6.36438 | 5.51594 | 0.34888 | 5.66850 |
| **Bag1** | 18.47511 | 0.07685 | 10.51720 | 11.54059 | 14.76954 | 12.27595 | 2.79522 | 0.87312 | 5.05099 | 13.59197 | 6.61070 | 8.06432 | 13.55439 | 4.42882 |
| **Bag2** | 0.04494 | 0.02035 | 0.04551 | 0.53321 | 0.46499 | 0.44980 | 0.00419 | 1.65494 | 0.00735 | 0.01165 | 0.00346 | 0.21828 | 0.02906 | 0.00000 |
| **Bag3** | 3.43132 | 7.36345 | 5.96064 | 19.40281 | 2.73509 | 10.32336 | 5.22235 | 8.10052 | 0.73074 | 4.39682 | 9.84988 | 1.39351 | 10.49830 | 6.50025 |
| **Bag4** | 2.48941 | 0.00851 | 6.38600 | 3.17363 | 3.14385 | 6.53838 | 2.91985 | 2.23743 | 2.37189 | 0.08844 | 4.13190 | 3.30111 | 3.85884 | 4.81868 |
| **Bag5** | 4.23818 | 0.32645 | 7.79782 | 2.40300 | 10.84423 | 6.12254 | 3.86037 | 6.81579 | 3.50319 | 0.17830 | 2.87346 | 4.20350 | 15.18392 | 10.05494 |
| **Bag6** | 5.67330 | 0.23668 | 17.55376 | 15.96914 | 17.82865 | 4.24607 | 2.81171 | 11.82028 | 1.16483 | 0.23282 | 5.54802 | 7.37315 | 5.08823 | 3.72917 |
| **Bahcc1** | 0.02130 | 0.00433 | 0.01727 | 0.00000 | 0.01352 | 0.14484 | 0.00428 | 0.00534 | 0.02243 | 0.00574 | 0.01403 | 0.03895 | 0.00700 | 0.00659 |
| **Bahd1** | 0.00000 | 0.00368 | 1.20337 | 1.58612 | 0.10810 | 1.70175 | 1.31628 | 0.07932 | 0.41369 | 0.02015 | 0.00000 | 0.55943 | 3.35442 | 0.63615 |
| **Baiap2** | 0.20303 | 0.12321 | 19.15090 | 8.08718 | 7.20132 | 0.93667 | 0.01018 | 0.53429 | 0.00000 | 0.01151 | 3.30662 | 8.16410 | 1.01126 | 1.57445 |
| **Baiap2l1** | 0.87798 | 0.04039 | 1.57203 | 0.35367 | 0.86953 | 0.70879 | 1.21598 | 0.38852 | 0.01294 | 0.02581 | 0.03415 | 0.00000 | 1.12294 | 1.48732 |
| **Baiap2l2** | 0.10783 | 0.00443 | 0.01243 | 0.01261 | 0.05690 | 0.01691 | 0.00420 | 0.00977 | 0.01843 | 0.00656 | 0.00823 | 0.00969 | 0.02382 | 0.00597 |
| **Baiap3** | 0.01012 | 0.00438 | 0.01468 | 0.01090 | 0.01400 | 0.01332 | 0.00434 | 0.00932 | 0.01164 | 0.01422 | 0.00853 | 0.02133 | 0.01642 | 0.01005 |
| **Bak1** | 0.07010 | 0.00446 | 0.01197 | 0.01745 | 0.06633 | 0.01985 | 0.00428 | 0.00978 | 0.01054 | 0.00707 | 0.00770 | 0.00866 | 0.00690 | 0.00958 |
| **Bambi** | 1.74444 | 0.00202 | 0.64678 | 0.00000 | 0.18772 | 0.07923 | 0.76718 | 0.00004 | 0.00157 | 0.06564 | 0.76111 | 2.61023 | 1.91566 | 0.00289 |
| **Bambi.ps1** | 0.36148 | 0.00487 | 0.01104 | 0.02682 | 0.00948 | 0.02004 | 0.00413 | 0.02383 | 0.11200 | 0.00790 | 0.00662 | 0.01959 | 0.03196 | 0.02002 |
| **Banf1** | 11.19752 | 11.28036 | 9.20782 | 20.16727 | 17.80699 | 19.08087 | 6.95200 | 15.29792 | 0.38400 | 18.16809 | 7.14000 | 16.96672 | 3.42714 | 11.00428 |
| **Banp** | 0.07720 | 0.02665 | 4.22645 | 5.85046 | 7.37994 | 1.69112 | 0.02613 | 3.82250 | 0.00037 | 0.02042 | 4.11432 | 0.33666 | 3.51920 | 0.00000 |
| **Bap1** | 5.89032 | 0.07666 | 10.55585 | 8.25995 | 6.96191 | 3.76499 | 3.53142 | 5.44270 | 1.04586 | 0.17546 | 5.48704 | 6.01100 | 5.23313 | 2.46656 |
| **Bard1** | 5.38734 | 0.06558 | 0.28748 | 0.00000 | 3.15727 | 0.16997 | 1.69852 | 1.42558 | 1.58486 | 0.02689 | 0.04884 | 2.34696 | 7.11208 | 1.11791 |
| **Barx2** | 0.02269 | 0.00436 | 0.02772 | 0.04280 | 0.15734 | 0.00507 | 0.00425 | 0.00926 | 0.00964 | 0.01351 | 0.26563 | 0.01142 | 0.00548 | 0.01169 |
| **Basp1** | 0.43912 | 4.35101 | 0.19265 | 0.59504 | 0.00000 | 1.50829 | 0.26476 | 0.28932 | 0.00236 | 0.01393 | 0.00000 | 1.58407 | 0.29390 | 0.19971 |
| **Batf** | 0.71309 | 0.00437 | 0.03349 | 0.01035 | 0.00199 | 0.00741 | 0.00427 | 0.01182 | 0.01028 | 0.00649 | 0.00669 | 0.00876 | 0.00634 | 0.01700 |
| **Batf3** | 0.00663 | 0.00446 | 0.16385 | 0.00000 | 0.23440 | 0.00879 | 0.00416 | 0.01105 | 0.00825 | 0.00670 | 0.00730 | 0.00931 | 0.01031 | 0.04009 |
| **Bax** | 1.30657 | 0.06553 | 0.74644 | 5.73288 | 3.52536 | 2.34103 | 0.02007 | 0.10073 | 0.31126 | 3.68367 | 0.00000 | 3.86193 | 1.49372 | 0.76513 |
| **Baz1a** | 18.91331 | 7.01087 | 19.37741 | 5.20857 | 26.78861 | 15.59176 | 4.77622 | 6.17185 | 33.20219 | 0.16992 | 14.33464 | 4.19535 | 8.39966 | 23.80627 |
| **Baz1b** | 7.27957 | 0.04327 | 11.21669 | 5.57171 | 8.96502 | 7.90396 | 1.01593 | 2.30582 | 2.27921 | 0.06583 | 6.64376 | 4.61186 | 7.65758 | 10.70032 |
| **Baz2a** | 14.09088 | 8.36071 | 15.91115 | 9.58174 | 18.61448 | 9.74929 | 3.91717 | 4.99755 | 11.95567 | 1.21629 | 11.29212 | 10.50471 | 12.04066 | 23.67524 |
| **Baz2b** | 1.61433 | 0.00000 | 0.45324 | 1.22085 | 1.77773 | 0.00000 | 1.08444 | 0.00268 | 5.85668 | 0.03504 | 2.21484 | 4.20165 | 2.47943 | 3.58840 |
| **BB283400** | 0.04407 | 0.00451 | 0.23754 | 0.01813 | 0.48350 | 0.02929 | 0.01534 | 0.01055 | 0.25828 | 0.01505 | 0.00574 | 0.02144 | 0.08314 | 0.12472 |
| **BB287469** | 0.00000 | 0.00392 | 0.01982 | 0.04777 | 0.01503 | 0.03969 | 0.02729 | 0.00949 | 0.94605 | 0.00686 | 0.02867 | 0.01356 | 3.53408 | 0.80946 |
| **BB557941** | 0.10535 | 0.00392 | 0.02851 | 0.00515 | 0.00738 | 0.63187 | 0.00399 | 0.00455 | 0.02691 | 0.01212 | 0.00460 | 0.00533 | 3.42406 | 0.00125 |
| **Bbc3** | 0.07358 | 0.00383 | 0.06918 | 0.68459 | 0.40473 | 0.06711 | 0.00443 | 0.62066 | 0.05445 | 0.00668 | 0.00000 | 0.33686 | 0.21038 | 0.03609 |
| **Bbip1** | 1.73777 | 0.00763 | 1.19572 | 2.62879 | 2.40423 | 5.01109 | 1.17158 | 2.74769 | 2.77681 | 6.25244 | 0.61181 | 4.05014 | 3.12594 | 0.00000 |
| **Bbof1** | 0.90708 | 0.00396 | 0.04265 | 0.00000 | 0.00000 | 0.16070 | 0.00440 | 0.00666 | 0.63045 | 0.00546 | 0.00293 | 0.27179 | 1.01343 | 0.00000 |

| | | | | | | | | | | | | | | |
|---|---|---|---|---|---|---|---|---|---|---|---|---|---|---|
| **Bbs1** | 0.00787 | 0.00389 | 0.37454 | 0.05073 | 0.09452 | 0.04913 | 0.01781 | 0.02398 | 0.05176 | 0.01410 | 0.01184 | 0.25875 | 2.68859 | 0.07140 |
| **Bbs10** | 0.42266 | 0.00476 | 0.00000 | 0.01174 | 0.00000 | 0.65168 | 0.00397 | 0.01940 | 0.51151 | 0.00887 | 0.00473 | 0.00388 | 0.91535 | 0.52231 |
| **Bbs12** | 0.05237 | 0.00413 | 0.00848 | 0.00124 | 0.00241 | 0.05880 | 1.48868 | 0.00289 | 0.03430 | 0.71262 | 0.00004 | 0.00365 | 0.40502 | 0.07330 |
| **Bbs2** | 2.97092 | 0.00228 | 0.65117 | 1.30274 | 3.64588 | 1.56729 | 0.00328 | 0.00216 | 0.57320 | 0.00248 | 0.95371 | 0.70371 | 6.21988 | 1.87434 |
| **Bbs4** | 0.09287 | 0.00250 | 0.79454 | 0.00000 | 1.22644 | 1.80553 | 1.70519 | 0.39627 | 2.63779 | 0.08430 | 5.73001 | 0.33375 | 2.86705 | 7.28174 |
| **Bbs5** | 7.57373 | 0.01074 | 4.14566 | 1.06230 | 2.05278 | 2.36102 | 1.28898 | 0.50395 | 3.55230 | 0.04538 | 4.61618 | 1.13832 | 0.59549 | 6.21172 |
| **Bbs7** | 0.01071 | 0.00443 | 0.00685 | 0.01017 | 0.31686 | 0.01345 | 0.00502 | 0.00971 | 0.01863 | 0.00671 | 0.00897 | 0.00850 | 0.11075 | 0.01927 |
| **Bbs9** | 0.00000 | 0.01340 | 0.07857 | 0.16467 | 0.00000 | 0.01016 | 0.00437 | 0.00791 | 0.56710 | 0.00563 | 0.01027 | 0.16812 | 0.50593 | 0.69750 |
| **Bbx** | 2.11873 | 0.00357 | 4.03986 | 0.10052 | 3.00779 | 0.22946 | 0.01302 | 0.23557 | 17.51215 | 0.00993 | 0.62625 | 0.15008 | 0.00000 | 5.18281 |
| **BC002163** | 1.89670 | 1.84402 | 0.00000 | 0.37735 | 2.07044 | 3.92372 | 1.94410 | 3.34061 | 0.06298 | 5.70756 | 0.06239 | 3.83798 | 0.64970 | 1.12126 |
| **BC003331** | 5.91443 | 0.01689 | 1.94496 | 2.32629 | 4.13538 | 7.03626 | 0.68700 | 0.55005 | 10.00116 | 0.01414 | 5.39417 | 3.46699 | 0.00000 | 4.00506 |
| **BC003965** | 4.10002 | 3.76810 | 4.55194 | 10.30759 | 9.36479 | 3.07796 | 4.15283 | 6.85848 | 1.19516 | 9.06049 | 2.10170 | 5.61991 | 5.80086 | 2.59091 |
| **BC004004** | 4.02414 | 0.09553 | 2.61967 | 3.06281 | 2.16406 | 2.34422 | 5.22161 | 4.16333 | 0.27088 | 0.30393 | 1.25186 | 5.28566 | 1.68518 | 3.15521 |
| **BC005537** | 6.28877 | 1.17690 | 8.77745 | 1.35189 | 3.59756 | 9.72518 | 1.65487 | 3.22148 | 11.46483 | 0.01847 | 8.81670 | 6.56957 | 3.97156 | 3.87847 |
| **BC005561** | 0.00000 | 0.00392 | 0.36355 | 0.26139 | 1.04079 | 0.82345 | 0.00193 | 0.00000 | 14.23148 | 0.01110 | 0.00089 | 0.06327 | 0.00000 | 1.30279 |
| **BC005624** | 2.14701 | 0.05602 | 5.98324 | 5.17060 | 4.64535 | 8.26541 | 0.00000 | 6.72819 | 3.32834 | 0.21648 | 7.25510 | 4.97571 | 2.24706 | 6.57750 |
| **BC006965** | 0.01658 | 0.00462 | 0.01170 | 0.00925 | 0.01571 | 0.01400 | 0.00416 | 0.01178 | 0.06634 | 0.00667 | 0.01202 | 0.02522 | 0.02993 | 0.36263 |
| **BC017158** | 0.00000 | 0.02448 | 0.69947 | 0.46891 | 0.00000 | 0.29757 | 0.00991 | 0.58670 | 0.03506 | 0.02636 | 0.00000 | 1.69315 | 0.00310 | 0.00547 |
| **BC017643** | 0.00000 | 0.00154 | 2.35435 | 0.38800 | 0.82637 | 3.48549 | 2.97590 | 4.12493 | 2.51657 | 0.12125 | 1.51602 | 2.13008 | 5.13289 | 3.48143 |
| **BC018473** | 0.06505 | 0.00446 | 0.01655 | 0.01351 | 0.01353 | 0.07419 | 0.00448 | 0.01645 | 0.18361 | 0.06957 | 0.01094 | 0.05650 | 0.59717 | 0.07792 |
| **BC020402** | 0.00598 | 0.00466 | 0.03242 | 0.13429 | 0.02216 | 0.05569 | 0.01928 | 0.49554 | 0.01337 | 0.00658 | 0.04845 | 0.00961 | 0.00591 | 0.02564 |
| **BC021614** | 0.16078 | 0.01242 | 0.13612 | 0.00000 | 0.77649 | 0.66959 | 0.00395 | 0.00517 | 0.48048 | 0.00676 | 0.02647 | 0.00675 | 1.80870 | 1.26918 |
| **BC021767** | 1.14636 | 0.70210 | 2.64611 | 0.81789 | 1.11824 | 1.45445 | 0.20984 | 1.40314 | 0.98917 | 0.02530 | 0.38030 | 0.74320 | 0.23793 | 0.48139 |
| **BC021891** | 0.00443 | 0.00401 | 0.04237 | 0.05214 | 0.14544 | 0.00167 | 0.00362 | 0.00529 | 2.11686 | 0.02965 | 0.00160 | 0.11428 | 0.57416 | 0.00000 |
| **BC022687** | 1.50826 | 0.01118 | 0.98248 | 0.00000 | 0.61332 | 0.53577 | 0.00406 | 1.50420 | 0.04059 | 0.00406 | 0.00000 | 0.77112 | 0.21762 | 1.34008 |
| **BC024386** | 0.11226 | 0.08586 | 0.00000 | 0.14920 | 0.07941 | 0.44374 | 0.08093 | 0.06082 | 0.27425 | 0.06863 | 0.00736 | 0.15020 | 1.60787 | 0.23940 |
| **BC024978** | 0.56807 | 0.01219 | 0.37565 | 0.00516 | 0.45562 | 0.08569 | 0.12732 | 0.12398 | 0.14821 | 0.00765 | 0.01207 | 0.41846 | 0.93113 | 0.31173 |
| **BC025920** | 0.00000 | 0.00415 | 0.05129 | 0.26117 | 0.92700 | 0.01185 | 0.00435 | 0.00484 | 0.41293 | 0.00593 | 0.00433 | 0.00680 | 0.67253 | 0.08681 |
| **BC026585** | 0.24441 | 0.02176 | 0.30900 | 1.05450 | 0.81319 | 0.20256 | 0.00388 | 2.86817 | 0.01410 | 0.00000 | 3.53210 | 0.02849 | 0.13946 | 0.25018 |
| **BC028528** | 0.43825 | 0.00362 | 0.22183 | 0.17626 | 0.08145 | 0.05256 | 0.00364 | 0.00866 | 0.75940 | 0.00578 | 0.08852 | 0.43054 | 0.43888 | 0.33618 |
| **BC029214** | 0.02277 | 4.48800 | 0.30456 | 0.03209 | 0.52477 | 0.12705 | 0.00349 | 2.64506 | 0.01161 | 0.01633 | 1.15016 | 0.00000 | 0.01702 | 0.00000 |
| **BC029722** | 0.02019 | 0.00447 | 0.09577 | 0.02926 | 0.00853 | 0.00855 | 0.00430 | 0.01075 | 0.01189 | 0.00626 | 0.01640 | 0.01235 | 0.00732 | 0.02970 |
| **BC030336** | 1.34953 | 0.00352 | 0.55538 | 0.00000 | 1.33933 | 1.14503 | 0.00335 | 0.74799 | 5.04438 | 0.00391 | 0.77444 | 0.38240 | 1.73166 | 1.52878 |
| **BC030867** | 8.27068 | 0.08673 | 14.21482 | 3.65000 | 14.62511 | 3.54764 | 0.01577 | 5.41739 | 3.22834 | 0.06053 | 7.73262 | 6.58348 | 6.91288 | 13.19927 |
| **BC031181** | 11.23350 | 7.84380 | 16.45115 | 15.97869 | 13.23252 | 20.02478 | 6.84709 | 11.84665 | 2.36414 | 8.37180 | 12.02357 | 6.98439 | 5.46329 | 5.74592 |
| **BC031361** | 0.01209 | 0.00418 | 0.01178 | 0.01814 | 0.00740 | 0.01165 | 0.00418 | 0.00884 | 0.01006 | 0.00678 | 0.01116 | 0.00801 | 1.41487 | 0.16618 |
| **BC037034** | 0.02533 | 0.02543 | 0.77842 | 0.32585 | 1.82313 | 0.38771 | 0.00342 | 0.87358 | 0.01370 | 1.49737 | 0.00000 | 2.39608 | 0.00000 | 0.04168 |
| **BC037704** | 0.59919 | 0.00404 | 0.12034 | 0.35639 | 0.16012 | 0.28827 | 0.00393 | 0.01811 | 0.00851 | 0.00757 | 0.00382 | 0.00372 | 0.48996 | 0.00000 |
| **BC039771** | 0.05930 | 0.02919 | 0.01459 | 0.39058 | 0.05951 | 0.44120 | 0.02301 | 0.13696 | 0.38555 | 0.12156 | 0.01534 | 0.03038 | 0.18074 | 0.47907 |
| **BC046251** | 0.01189 | 0.00429 | 0.01781 | 0.01807 | 0.01359 | 0.02025 | 0.00990 | 0.01693 | 0.09185 | 0.00802 | 0.01568 | 0.00910 | 0.07245 | 0.01414 |
| **BC048403** | 0.12685 | 0.00383 | 0.84953 | 0.35354 | 0.12808 | 0.00000 | 0.00407 | 0.33534 | 0.03171 | 0.00400 | 0.32743 | 0.09002 | 0.00887 | 0.18187 |
| **BC048507** | 0.10578 | 0.18835 | 0.05355 | 0.03830 | 0.31115 | 0.24344 | 0.34692 | 0.44513 | 0.02257 | 0.56513 | 0.03081 | 0.42521 | 0.51741 | 0.00000 |
| **BC048546** | 0.01041 | 0.00410 | 0.00000 | 0.00000 | 2.09912 | 0.02063 | 0.00469 | 0.00596 | 0.01215 | 0.00588 | 0.29103 | 0.12298 | 0.00010 | 0.00929 |
| **BC048679** | 0.58522 | 3.18352 | 2.61253 | 10.41990 | 0.00000 | 2.53589 | 1.34324 | 2.93213 | 0.00780 | 0.09131 | 0.28296 | 1.18703 | 0.00126 | 0.00000 |
| **BC049715** | 0.01859 | 0.03548 | 0.07833 | 0.02492 | 0.27774 | 0.03764 | 0.01716 | 0.02546 | 0.07812 | 0.00725 | 0.00818 | 0.01519 | 0.01404 | 0.04420 |
| **BC051019** | 0.21145 | 0.00358 | 1.52533 | 0.35177 | 2.21841 | 1.22864 | 0.00372 | 0.00000 | 0.13272 | 0.00918 | 0.00000 | 0.49449 | 0.05375 | 0.62911 |
| **BC051142** | 0.72361 | 0.00474 | 1.00248 | 0.00000 | 0.00000 | 0.43370 | 0.00380 | 0.00132 | 0.27426 | 0.00434 | 0.16956 | 0.09364 | 0.00012 | 0.19389 |
| **BC051226** | 0.09742 | 0.00472 | 0.40222 | 0.01470 | 0.21881 | 0.07767 | 0.00430 | 0.00882 | 0.00860 | 0.00658 | 0.07837 | 0.00535 | 0.01534 | 0.01989 |
| **BC051537** | 0.05617 | 0.00439 | 0.13471 | 0.00885 | 0.00000 | 0.08024 | 0.00408 | 0.00834 | 0.21012 | 0.01102 | 0.00566 | 0.00702 | 0.00842 | 0.37092 |
| **BC051665** | 0.25639 | 5.24395 | 0.08522 | 0.34255 | 0.00000 | 0.00000 | 0.00241 | 1.68125 | 0.93539 | 0.00630 | 0.00177 | 0.11461 | 0.00334 | 0.19554 |
| **BC052040** | 6.13070 | 0.01344 | 7.58740 | 4.17983 | 6.29201 | 0.45076 | 0.00650 | 1.02395 | 1.56963 | 0.00000 | 7.30385 | 2.79946 | 0.00000 | 7.70775 |
| **BC052688** | 10.05798 | 0.61136 | 13.24409 | 15.42038 | 13.45542 | 23.12276 | 5.02159 | 2.95938 | 24.20088 | 9.61722 | 15.05462 | 7.16646 | 3.58398 | 7.96168 |
| **BC053393** | 0.00000 | 7.30927 | 0.00000 | 0.00000 | 0.00000 | 5.15193 | 2.78757 | 19.47017 | 0.18897 | 11.99498 | 14.20687 | 3.34580 | 0.43823 | 0.10022 |
| **BC055324** | 7.66136 | 8.24091 | 7.96570 | 3.90633 | 11.69756 | 4.13185 | 0.01537 | 0.55106 | 8.60335 | 0.00000 | 2.43250 | 4.76134 | 0.34317 | 6.82146 |
| **BC064078** | 0.00436 | 0.00455 | 0.02847 | 0.08258 | 0.03719 | 0.02181 | 0.05515 | 0.01825 | 0.00505 | 0.00926 | 0.00770 | 0.00690 | 0.15290 | 0.48933 |

| | | | | | | | | | | | | | | |
|---|---|---|---|---|---|---|---|---|---|---|---|---|---|---|
| **BC065397** | 1.29810 | 0.03643 | 0.01073 | 0.00000 | 0.00000 | 0.63247 | 0.00347 | 0.01029 | 0.51670 | 0.00859 | 3.28765 | 1.58757 | 0.03881 | 0.06702 |
| **BC068281** | 0.02591 | 0.00507 | 0.17819 | 0.00000 | 0.01679 | 0.39276 | 0.01674 | 0.11811 | 0.03335 | 0.01092 | 0.00709 | 0.35208 | 0.20642 | 0.20406 |
| **BC080695** | 0.01131 | 0.00418 | 0.01730 | 0.01197 | 0.00844 | 0.01208 | 0.00411 | 0.00851 | 0.04798 | 0.00628 | 0.00855 | 0.00790 | 1.37396 | 0.00000 |
| **BC100530** | 0.01724 | 0.01983 | 0.01127 | 0.01530 | 0.01329 | 0.05758 | 0.02832 | 0.15494 | 0.01277 | 0.23662 | 0.00428 | 0.00690 | 0.00928 | 0.01313 |
| **BC147527** | 0.27029 | 0.00416 | 0.63751 | 0.46553 | 0.19778 | 0.27401 | 0.00381 | 0.00258 | 0.08886 | 0.00568 | 0.00369 | 0.00332 | 2.04413 | 0.26667 |
| **Bcam** | 0.00911 | 0.00453 | 0.02577 | 0.02806 | 0.01337 | 0.01527 | 0.00446 | 0.01747 | 0.00800 | 0.00842 | 0.01009 | 0.00833 | 0.05054 | 0.03038 |
| **Bcan** | 0.06200 | 0.00416 | 0.40260 | 0.18812 | 0.51501 | 0.00000 | 0.00399 | 0.00815 | 0.11498 | 0.00669 | 0.00520 | 0.01172 | 0.01812 | 0.01903 |
| **Bcap29** | 5.47770 | 0.01013 | 1.39588 | 2.38283 | 2.78881 | 4.19782 | 0.03168 | 0.08913 | 9.97564 | 0.00645 | 1.77862 | 1.02310 | 0.71057 | 1.41721 |
| **Bcap31** | 19.33215 | 7.78195 | 20.92723 | 19.18117 | 13.64677 | 23.71195 | 8.97385 | 20.87744 | 20.73280 | 11.71770 | 11.88589 | 18.18238 | 5.68132 | 23.84938 |
| **Bcar1** | 0.13874 | 0.00314 | 0.56181 | 0.62447 | 1.32234 | 0.75402 | 0.01515 | 1.46150 | 0.00225 | 0.00571 | 0.00000 | 1.81280 | 0.45996 | 0.00000 |
| **Bcar3** | 0.00521 | 0.02546 | 0.00000 | 0.02262 | 0.00000 | 0.01153 | 0.00384 | 0.00550 | 0.41294 | 0.00641 | 0.00235 | 0.60959 | 0.06856 | 0.26821 |
| **Bcas1** | 0.00731 | 0.00429 | 0.01531 | 0.01169 | 0.00852 | 0.01280 | 0.00421 | 0.00898 | 0.01145 | 0.00613 | 0.00891 | 0.00886 | 0.57210 | 0.03117 |
| **Bcas2** | 14.69788 | 7.63430 | 20.75791 | 22.24627 | 19.14061 | 19.06015 | 4.06540 | 14.18731 | 3.06566 | 10.43778 | 15.21107 | 13.16693 | 5.00068 | 10.86270 |
| **Bcas3** | 5.31534 | 0.07799 | 18.42644 | 13.06124 | 14.70278 | 7.32229 | 0.45779 | 2.14945 | 10.29409 | 0.08333 | 6.62817 | 9.41182 | 3.28200 | 14.02610 |
| **Bcas3os1** | 0.01167 | 0.01961 | 0.06662 | 0.11974 | 0.13390 | 0.06209 | 0.01078 | 0.03700 | 0.90275 | 0.00718 | 0.04770 | 0.02329 | 0.09589 | 0.02921 |
| **Bcat1** | 16.15148 | 0.62883 | 33.13851 | 28.17254 | 27.67606 | 22.14737 | 6.64928 | 0.45053 | 22.85983 | 0.40506 | 16.82181 | 9.15584 | 2.22605 | 27.57008 |
| **Bcat2** | 2.62177 | 2.60005 | 5.89272 | 10.50349 | 2.84378 | 10.40149 | 3.86803 | 13.00800 | 0.44060 | 0.33889 | 2.62068 | 7.32896 | 0.00000 | 5.83237 |
| **Bccip** | 0.00000 | 0.05819 | 8.85444 | 0.65397 | 1.64979 | 6.73533 | 4.11707 | 3.88958 | 6.20587 | 0.14952 | 6.68094 | 4.90167 | 0.00000 | 9.08570 |
| **Bcdin3d** | 1.36586 | 0.01937 | 0.99983 | 2.07651 | 0.32633 | 2.36676 | 0.00347 | 0.04574 | 0.00000 | 0.02997 | 1.65791 | 0.54170 | 0.90105 | 0.96036 |
| **Bckdha** | 1.01085 | 0.02176 | 11.13438 | 11.02286 | 9.25141 | 2.30209 | 0.00000 | 1.60884 | 0.00000 | 0.01409 | 5.71620 | 2.70602 | 0.24891 | 2.09621 |
| **Bckdhb** | 5.75538 | 5.79295 | 6.64593 | 1.72812 | 6.32262 | 4.95057 | 0.75107 | 0.27045 | 7.38630 | 0.03626 | 2.05760 | 3.99070 | 2.04013 | 4.16196 |
| **Bckdk** | 0.29263 | 0.20055 | 8.64602 | 11.31540 | 6.49868 | 4.11076 | 0.07172 | 3.49323 | 0.32772 | 0.11819 | 4.98567 | 6.71192 | 1.03401 | 5.91504 |
| **Bcl10** | 3.01715 | 0.03785 | 4.03033 | 0.00000 | 2.66478 | 4.75451 | 2.02913 | 3.25087 | 0.00000 | 2.96222 | 4.56221 | 3.80063 | 0.73727 | 3.89121 |
| **Bcl11b** | 0.00033 | 0.03239 | 0.25667 | 0.05501 | 0.45044 | 0.84424 | 0.00405 | 0.08506 | 0.35985 | 0.02028 | 0.00000 | 0.33388 | 0.00319 | 0.00000 |
| **Bcl2** | 0.42616 | 0.00431 | 0.01041 | 0.00945 | 0.05437 | 0.01185 | 0.00431 | 0.01605 | 0.02040 | 0.00715 | 0.01012 | 0.00846 | 0.00478 | 0.02356 |
| **Bcl2l1** | 0.47873 | 1.77326 | 5.40704 | 2.64276 | 4.38420 | 7.54603 | 0.10537 | 2.95220 | 0.00400 | 0.13868 | 0.45371 | 1.93321 | 1.10254 | 1.06051 |
| **Bcl2l10** | 0.32008 | 0.00361 | 0.00000 | 0.12287 | 0.69792 | 0.01405 | 0.07380 | 0.00427 | 0.03589 | 0.00778 | 0.00581 | 0.00712 | 4.58616 | 0.41569 |
| **Bcl2l11** | 0.91087 | 0.02774 | 3.54881 | 1.30270 | 4.08304 | 0.84930 | 2.36063 | 1.95088 | 1.30539 | 0.08380 | 0.06769 | 0.19655 | 2.88278 | 0.33787 |
| **Bcl2l12** | 0.00000 | 0.02122 | 0.61422 | 0.46199 | 0.66031 | 0.12663 | 0.00369 | 1.48658 | 0.01122 | 0.01702 | 0.10873 | 0.91602 | 0.48334 | 0.00000 |
| **Bcl2l13** | 9.49894 | 0.02972 | 8.16081 | 2.52448 | 10.93068 | 3.10136 | 4.50999 | 3.25977 | 10.73623 | 0.06635 | 3.25019 | 1.91935 | 10.15146 | 15.43830 |
| **Bcl2l14** | 4.79521 | 0.01460 | 14.95848 | 2.37909 | 16.96442 | 4.26901 | 1.56609 | 0.00000 | 6.35714 | 0.00000 | 7.52831 | 3.53755 | 4.92118 | 12.65479 |
| **Bcl2l15** | 0.00286 | 0.00360 | 0.04506 | 0.14125 | 0.19085 | 0.14456 | 0.00404 | 0.24814 | 0.01571 | 0.00609 | 0.00048 | 0.38783 | 0.22202 | 0.04176 |
| **Bcl2l2** | 0.42657 | 0.00365 | 0.62725 | 0.19688 | 0.03814 | 0.17387 | 0.02771 | 0.22559 | 2.97760 | 0.00765 | 0.00351 | 0.19252 | 1.55647 | 2.04200 |
| **Bcl3** | 0.67298 | 0.02582 | 1.18189 | 0.00000 | 0.97915 | 0.59058 | 0.82696 | 0.90086 | 0.00319 | 0.06098 | 0.52138 | 3.34384 | 0.45675 | 0.04583 |
| **Bcl7a** | 1.56483 | 0.00337 | 2.87229 | 0.00000 | 6.95296 | 1.26350 | 0.09783 | 0.49601 | 0.30404 | 0.03852 | 0.03618 | 0.53097 | 1.06044 | 3.38173 |
| **Bcl7b** | 1.55147 | 0.01103 | 4.80788 | 3.38492 | 8.26650 | 2.25004 | 2.63091 | 0.14768 | 0.01402 | 0.12397 | 2.00537 | 3.03891 | 0.91090 | 3.04451 |
| **Bcl7c** | 0.37474 | 0.04136 | 0.09580 | 0.19135 | 2.16859 | 0.12052 | 0.00391 | 0.41215 | 0.00000 | 0.02632 | 0.12484 | 1.27852 | 1.82211 | 0.88279 |
| **Bcl9** | 0.05737 | 0.00425 | 0.07679 | 0.14043 | 0.07026 | 0.02559 | 0.00444 | 0.01723 | 0.03021 | 0.00764 | 0.01325 | 0.00524 | 0.96100 | 0.04398 |
| **Bcl9l** | 0.02217 | 0.04049 | 0.34865 | 0.58091 | 0.00000 | 0.02910 | 0.00400 | 1.61127 | 0.01248 | 0.00639 | 0.00000 | 0.25503 | 0.00608 | 0.00421 |
| **Bclaf1** | 10.33871 | 5.68390 | 20.79040 | 10.86105 | 16.51475 | 14.80095 | 2.57036 | 3.49228 | 27.05712 | 10.54721 | 7.79471 | 8.80298 | 16.36625 | 28.05461 |
| **Bco1** | 0.13853 | 0.00514 | 0.06359 | 0.13946 | 0.42289 | 0.01547 | 0.00444 | 0.03155 | 0.08841 | 0.00783 | 0.02286 | 0.01400 | 0.02767 | 0.65753 |
| **Bco2** | 0.00340 | 0.00438 | 0.00548 | 0.17637 | 0.01010 | 0.01682 | 0.00402 | 0.00568 | 0.01331 | 0.00695 | 0.00603 | 0.01480 | 0.00464 | 0.42790 |
| **Bcor** | 0.00000 | 0.02874 | 1.85164 | 0.40654 | 2.15612 | 3.99148 | 1.70216 | 0.00528 | 0.34647 | 0.01737 | 3.25885 | 3.65246 | 7.03526 | 1.64351 |
| **Bcorl1** | 0.00000 | 0.04736 | 0.95558 | 0.75897 | 0.14395 | 0.04241 | 0.00370 | 2.37085 | 0.05186 | 0.01516 | 0.07890 | 0.07363 | 0.44535 | 0.00904 |
| **Bcr** | 0.50048 | 0.05514 | 1.38628 | 0.82349 | 0.00000 | 3.48888 | 0.00386 | 0.84456 | 0.12474 | 0.01677 | 2.26837 | 0.79234 | 1.93676 | 1.58863 |
| **Bcs1l** | 2.03079 | 0.05132 | 4.43735 | 0.66263 | 2.76366 | 2.57023 | 0.01002 | 5.10220 | 3.44337 | 14.35514 | 2.95795 | 3.69189 | 0.66544 | 2.39044 |
| **Bdh1** | 1.78975 | 0.02517 | 9.24016 | 6.08077 | 5.37261 | 1.62873 | 0.66116 | 3.10208 | 0.81819 | 0.04645 | 3.10613 | 2.54186 | 2.11812 | 2.19843 |
| **Bdh2** | 0.00311 | 0.00499 | 0.00892 | 0.06073 | 0.01289 | 0.02500 | 0.00412 | 0.01770 | 0.00653 | 0.01025 | 0.00828 | 0.00644 | 0.32758 | 0.17358 |
| **Bdp1** | 4.10943 | 0.00968 | 3.38415 | 0.00000 | 6.07348 | 2.52062 | 1.20556 | 0.53169 | 12.91696 | 0.03561 | 1.94431 | 1.68550 | 7.04013 | 5.74342 |
| **Becn1** | 1.40764 | 7.61157 | 6.80783 | 8.27250 | 5.64830 | 5.65134 | 2.68980 | 2.18660 | 8.10941 | 0.07141 | 5.38600 | 6.18524 | 2.20119 | 10.48665 |
| **Becn2** | 14.28233 | 0.04107 | 19.25060 | 15.96038 | 11.46034 | 11.03385 | 4.02098 | 1.08424 | 15.03320 | 0.09649 | 6.25183 | 0.23687 | 8.27607 | 19.16715 |
| **Bend3** | 8.12778 | 0.01693 | 3.93417 | 0.25783 | 3.42986 | 1.60394 | 0.26244 | 0.08538 | 10.82558 | 0.60538 | 2.10116 | 1.07110 | 7.06768 | 8.80483 |
| **Bend4** | 0.73121 | 6.46917 | 3.66047 | 0.77416 | 5.95150 | 3.74308 | 0.30100 | 3.34541 | 0.63540 | 0.07970 | 0.00000 | 0.07906 | 0.00000 | 3.58339 |
| **Bend5** | 0.01785 | 0.01190 | 0.01879 | 0.00000 | 0.01242 | 0.01784 | 0.00420 | 0.01586 | 0.01020 | 0.00746 | 0.00358 | 0.90273 | 0.00561 | 0.03436 |
| **Bend6** | 0.00000 | 0.00502 | 0.00000 | 0.12587 | 0.00000 | 0.02233 | 0.00387 | 0.05751 | 0.64312 | 0.01389 | 0.00573 | 0.00633 | 0.02405 | 0.69452 |

| | | | | | | | | | | | | | | |
|---|---|---|---|---|---|---|---|---|---|---|---|---|---|---|
| **Bend7** | 0.20947 | 0.01874 | 0.04235 | 0.02770 | 0.00311 | 0.28382 | 0.00368 | 0.00563 | 0.00661 | 0.01907 | 0.10156 | 1.12041 | 0.54745 | 0.00000 |
| **Best2** | 0.07980 | 0.00402 | 0.23593 | 0.32322 | 0.24702 | 0.03171 | 0.00376 | 0.00275 | 0.00528 | 0.00573 | 0.00758 | 0.01137 | 0.58820 | 0.13200 |
| **Bet1** | 4.23148 | 7.45521 | 0.60221 | 2.67047 | 2.18412 | 3.13791 | 5.13620 | 0.44525 | 3.02553 | 0.16572 | 0.00000 | 2.96843 | 1.27252 | 4.05326 |
| **Bet1l** | 0.16943 | 0.04372 | 5.16994 | 3.36811 | 5.71898 | 2.47517 | 4.21835 | 1.92206 | 0.80602 | 0.08256 | 3.67919 | 1.07709 | 1.77475 | 7.39873 |
| **Bex1** | 2.28986 | 0.00405 | 0.52202 | 0.15176 | 0.29991 | 0.02706 | 0.00405 | 0.00533 | 0.01019 | 0.00515 | 0.01114 | 0.00371 | 0.61279 | 0.06458 |
| **Bex2** | 0.20319 | 0.04573 | 0.04429 | 0.06632 | 0.07775 | 0.19135 | 0.01003 | 0.01235 | 0.01352 | 0.00633 | 0.00459 | 0.04385 | 0.02443 | 0.03838 |
| **Bex4** | 0.00902 | 0.00438 | 0.54511 | 0.77898 | 0.25361 | 0.13441 | 0.00391 | 0.00307 | 0.02539 | 0.00540 | 0.00376 | 0.00476 | 0.06332 | 0.01055 |
| **Bex6** | 8.05976 | 0.00167 | 3.98342 | 1.73311 | 11.98284 | 0.00000 | 0.00157 | 0.00000 | 10.56692 | 0.00000 | 0.35736 | 0.00000 | 6.15320 | 14.07804 |
| **Bfar** | 2.97764 | 0.02341 | 5.14633 | 1.35097 | 3.50288 | 8.94670 | 1.67094 | 0.37609 | 6.41137 | 0.04351 | 3.40504 | 3.75242 | 3.39853 | 11.08646 |
| **Bfsp1** | 0.30494 | 0.00384 | 0.00698 | 0.00176 | 0.00387 | 2.05267 | 0.00365 | 0.01065 | 0.00362 | 0.01883 | 0.00310 | 0.00449 | 1.92909 | 0.00351 |
| **Bfsp2** | 0.43239 | 0.00389 | 0.22810 | 0.00000 | 0.05028 | 0.00469 | 0.01026 | 0.02094 | 0.00840 | 0.00634 | 0.00562 | 0.35598 | 0.00426 | 0.00595 |
| **Bglap** | 0.04758 | 0.00527 | 0.01290 | 0.02407 | 0.00960 | 0.06147 | 0.00445 | 0.09200 | 0.01272 | 0.00917 | 0.06565 | 0.01094 | 0.10661 | 0.00223 |
| **Bglap2** | 0.06100 | 0.00535 | 0.01476 | 0.04644 | 0.00663 | 0.09802 | 0.00441 | 0.16293 | 0.03002 | 0.00843 | 0.12033 | 0.02704 | 0.06893 | 0.00589 |
| **Bglap3** | 0.08094 | 0.00533 | 0.22258 | 0.90230 | 0.00000 | 0.19098 | 0.05445 | 0.34364 | 0.04236 | 0.00716 | 0.16599 | 0.06166 | 0.09450 | 0.00000 |
| **Bgn** | 1.46060 | 0.00513 | 0.00000 | 0.23311 | 1.21656 | 0.02044 | 0.00447 | 0.04155 | 0.01004 | 0.01028 | 0.00748 | 0.00252 | 0.00249 | 0.00217 |
| **Bhlha15** | 0.00874 | 0.00443 | 0.48810 | 0.16201 | 0.00454 | 0.00000 | 0.00442 | 0.04170 | 0.04666 | 0.01380 | 0.01088 | 0.01200 | 0.01075 | 0.00907 |
| **Bhlhb9** | 2.99770 | 0.01666 | 0.18108 | 0.41736 | 3.12118 | 0.39063 | 0.00347 | 0.42540 | 1.29969 | 0.00695 | 0.00000 | 1.17621 | 0.00964 | 2.58866 |
| **Bhlhe40** | 0.19889 | 0.00368 | 0.09318 | 0.57522 | 0.07799 | 0.00000 | 0.00376 | 0.01491 | 0.03386 | 0.00480 | 1.81734 | 0.03783 | 0.02281 | 0.79811 |
| **Bhlhe41** | 0.35580 | 0.03231 | 0.00000 | 0.12600 | 0.25264 | 0.23478 | 0.00506 | 0.02320 | 0.01731 | 0.01114 | 0.00434 | 0.00864 | 0.28742 | 0.13046 |
| **Bhmt** | 35.61915 | 1.03128 | 64.46531 | 67.93085 | 51.78433 | 44.62928 | 10.07573 | 10.70538 | 25.94076 | 16.96481 | 41.75128 | 30.50785 | 5.19654 | 41.38039 |
| **Bhmt2** | 5.23801 | 0.01720 | 17.67584 | 11.61581 | 12.51853 | 8.06255 | 0.10684 | 0.13930 | 2.68668 | 0.18188 | 3.71773 | 0.90141 | 1.48183 | 9.56631 |
| **Bicc1** | 0.02540 | 0.00432 | 0.01125 | 0.01335 | 0.00860 | 0.00967 | 0.00420 | 0.00943 | 0.08823 | 0.00684 | 0.03020 | 0.01163 | 0.30014 | 0.00000 |
| **Bicd2** | 0.83629 | 0.00322 | 0.26358 | 0.14387 | 1.11648 | 0.52771 | 0.11856 | 0.75389 | 0.10237 | 0.00686 | 0.00000 | 0.74692 | 5.40003 | 2.61689 |
| **Bid** | 6.74622 | 0.04396 | 8.90290 | 5.01629 | 9.80317 | 1.53112 | 1.03265 | 0.34052 | 3.79100 | 0.00848 | 4.90963 | 1.33850 | 4.60879 | 7.24165 |
| **Bik** | 0.01023 | 0.00407 | 0.01669 | 0.00239 | 0.10866 | 0.00955 | 0.01496 | 2.02485 | 0.01178 | 0.09941 | 0.01802 | 0.01045 | 0.01357 | 0.05629 |
| **Bin1** | 0.00000 | 0.00282 | 0.34788 | 1.13529 | 0.07802 | 0.00000 | 1.03296 | 0.00118 | 0.00968 | 0.03410 | 0.00366 | 1.67368 | 0.00586 | 0.00325 |
| **Bin2** | 0.71679 | 4.32102 | 10.45644 | 8.88827 | 5.01277 | 0.63415 | 3.39825 | 11.41670 | 0.01239 | 0.17810 | 2.03304 | 1.52118 | 0.08501 | 0.66171 |
| **Bin3** | 0.86357 | 0.00203 | 0.00000 | 1.85798 | 0.00000 | 1.08247 | 0.00314 | 0.00852 | 0.05981 | 0.00000 | 4.18366 | 0.87412 | 1.02253 | 1.88485 |
| **Birc2** | 1.59419 | 0.03717 | 2.07760 | 0.83073 | 1.39348 | 1.73744 | 0.00343 | 0.03385 | 4.06127 | 0.00000 | 1.00063 | 1.35545 | 2.39371 | 0.91012 |
| **Birc3** | 4.45311 | 0.00273 | 6.02914 | 0.04704 | 6.54896 | 1.30138 | 0.00194 | 0.20270 | 10.50160 | 0.00075 | 0.30731 | 0.01318 | 4.90221 | 8.81150 |
| **Birc5** | 13.99933 | 10.46435 | 20.18729 | 24.10587 | 18.01247 | 23.39564 | 8.55921 | 15.42828 | 2.99035 | 18.24763 | 16.30544 | 15.90886 | 11.00598 | 19.68702 |
| **Birc6** | 0.81668 | 0.01070 | 1.32163 | 0.77272 | 1.08375 | 2.01242 | 0.00967 | 0.61743 | 8.74259 | 0.00443 | 0.13271 | 0.87381 | 4.74346 | 5.34994 |
| **Bivm** | 0.00554 | 0.00386 | 0.12152 | 0.00000 | 0.06418 | 0.00696 | 0.00399 | 0.00848 | 0.01010 | 0.00642 | 0.00367 | 1.00949 | 0.00762 | 0.01689 |
| **Blcap** | 0.02984 | 0.03022 | 2.23120 | 1.18015 | 0.45304 | 2.89711 | 3.09160 | 1.40610 | 3.05713 | 10.50294 | 2.55188 | 3.10355 | 1.16110 | 3.59442 |
| **Blm** | 8.45296 | 0.37497 | 5.92629 | 2.07984 | 8.17867 | 4.46352 | 0.00000 | 2.30544 | 6.94020 | 0.05650 | 4.12556 | 3.51369 | 3.90560 | 8.18282 |
| **Blmh** | 1.83718 | 0.06495 | 4.30722 | 3.62036 | 5.27544 | 6.28548 | 4.52448 | 2.09301 | 0.78129 | 0.29004 | 8.85294 | 7.31765 | 2.22542 | 5.59862 |
| **Blnk** | 0.00295 | 4.14677 | 0.00516 | 0.00000 | 0.02041 | 0.00373 | 0.00187 | 5.58219 | 0.01108 | 0.25247 | 0.00000 | 0.71833 | 0.04628 | 0.00311 |
| **Bloc1s1** | 2.29782 | 7.36882 | 0.00000 | 1.31029 | 2.31718 | 4.11252 | 1.93447 | 1.51832 | 0.00000 | 8.01468 | 0.08499 | 4.02693 | 0.00000 | 1.14416 |
| **Bloc1s2** | 1.64178 | 2.75493 | 5.69189 | 11.44377 | 5.81014 | 6.04056 | 5.18290 | 4.85187 | 1.42111 | 4.83257 | 4.70813 | 4.21751 | 3.52188 | 0.81209 |
| **Bloc1s3** | 0.50267 | 0.05644 | 0.16537 | 0.52708 | 0.29705 | 0.63408 | 0.07021 | 0.18514 | 0.96909 | 0.22583 | 0.01511 | 0.22234 | 0.93018 | 0.64906 |
| **Bloc1s4** | 0.36924 | 0.01112 | 0.01088 | 0.96151 | 1.29430 | 0.29737 | 0.00379 | 0.02446 | 0.13188 | 0.00624 | 0.51822 | 1.08634 | 1.57452 | 0.03107 |
| **Bloc1s5** | 0.84912 | 0.01656 | 3.40428 | 2.41637 | 2.89537 | 5.21195 | 1.72608 | 2.07088 | 0.00476 | 0.05949 | 1.63048 | 4.59878 | 0.00000 | 0.99978 |
| **Bloc1s6** | 2.25470 | 0.05875 | 1.27678 | 3.05770 | 3.69098 | 4.99401 | 0.03338 | 0.88821 | 0.04899 | 0.05234 | 4.95980 | 2.55241 | 0.24270 | 2.31745 |
| **Blvra** | 1.65944 | 6.93580 | 0.46038 | 0.00000 | 0.00000 | 1.64944 | 3.01098 | 8.49487 | 1.21726 | 0.05645 | 0.00000 | 0.75682 | 1.02857 | 0.84259 |
| **Blvrb** | 0.45365 | 0.06555 | 0.25814 | 1.38038 | 2.44872 | 1.18157 | 2.04903 | 1.13668 | 0.00251 | 0.14223 | 1.59248 | 3.28799 | 0.00811 | 0.10828 |
| **Blzf1** | 6.49800 | 0.04174 | 3.03527 | 1.68245 | 1.35279 | 1.23916 | 0.27256 | 0.57214 | 5.41410 | 0.00000 | 6.34404 | 1.54542 | 1.37626 | 3.41208 |
| **Bmf** | 0.51837 | 2.50194 | 1.00125 | 0.00000 | 0.94187 | 1.62811 | 0.00274 | 0.06617 | 0.00147 | 0.02084 | 0.00000 | 1.53937 | 0.83214 | 0.03977 |
| **Bmi1** | 2.88890 | 0.37966 | 1.12380 | 2.87974 | 2.57463 | 2.43293 | 0.00649 | 0.76826 | 16.30923 | 0.01685 | 0.03544 | 1.85888 | 12.65201 | 10.71213 |
| **Bmp1** | 0.22402 | 0.00342 | 0.00000 | 0.88768 | 0.26772 | 0.04362 | 0.00360 | 0.00000 | 0.28375 | 0.00184 | 1.52053 | 0.22820 | 0.00240 | 0.00000 |
| **Bmp15** | 0.60437 | 0.00355 | 0.00000 | 0.03161 | 0.42658 | 0.37487 | 0.00386 | 0.02135 | 1.00529 | 0.00635 | 0.00313 | 0.00358 | 4.35104 | 0.00000 |
| **Bmp2k** | 0.03816 | 0.00389 | 0.01170 | 0.39226 | 0.05231 | 0.00000 | 0.00364 | 0.00400 | 0.00876 | 0.01137 | 0.01314 | 0.46442 | 0.57381 | 0.15030 |
| **Bmp4** | 0.01743 | 0.07363 | 0.02502 | 0.02473 | 0.01854 | 0.02084 | 0.00918 | 0.06447 | 0.01639 | 0.00646 | 0.00417 | 4.64307 | 0.00719 | 0.02172 |
| **Bmp5** | 0.00000 | 0.00412 | 0.08120 | 0.19356 | 0.00749 | 0.00741 | 0.00401 | 0.00761 | 0.09788 | 0.00559 | 0.01219 | 0.00590 | 1.50193 | 0.02344 |
| **Bmp6** | 0.00000 | 0.01913 | 0.59654 | 0.00000 | 0.69950 | 0.00623 | 0.00363 | 0.00475 | 0.00554 | 0.00469 | 0.08180 | 0.83920 | 3.09754 | 2.06428 |
| **Bmp7** | 0.01264 | 0.00404 | 0.01228 | 0.40878 | 0.00945 | 0.00324 | 0.00393 | 0.00621 | 0.01931 | 0.00633 | 0.01686 | 0.01463 | 0.22333 | 0.00000 |

| | | | | | | | | | | | | | | |
|---|---|---|---|---|---|---|---|---|---|---|---|---|---|---|
| **Bmp8a** | 0.01631 | 0.01160 | 0.01259 | 0.00000 | 0.00848 | 0.01420 | 2.11247 | 0.48280 | 0.01096 | 0.06754 | 0.00482 | 0.01638 | 0.00664 | 0.00992 |
| **Bmp8b** | 0.02096 | 0.00389 | 0.21142 | 0.10369 | 0.00887 | 0.24560 | 0.00368 | 0.28111 | 0.05898 | 0.04345 | 0.00357 | 0.00235 | 0.04157 | 0.00000 |
| **Bmper** | 0.03928 | 0.01467 | 0.12785 | 0.00000 | 0.01161 | 0.00410 | 0.00412 | 0.05149 | 0.01048 | 0.00706 | 0.00490 | 0.19391 | 0.00583 | 0.01124 |
| **Bmpr1a** | 2.96057 | 0.07819 | 2.32816 | 0.17097 | 2.55419 | 4.00918 | 3.05702 | 0.76049 | 10.19218 | 0.06289 | 2.31423 | 3.96556 | 3.05744 | 1.79999 |
| **Bmpr1b** | 0.00599 | 0.00427 | 0.01055 | 0.01351 | 0.05475 | 0.01202 | 0.00443 | 0.00840 | 0.01422 | 0.00613 | 0.00863 | 0.00822 | 0.87393 | 0.17884 |
| **Bmpr2** | 1.81924 | 0.00247 | 1.00531 | 0.00000 | 1.42608 | 1.45533 | 3.01162 | 0.57616 | 4.91187 | 0.06346 | 1.27397 | 1.82098 | 3.48737 | 0.00000 |
| **Bms1** | 9.18515 | 5.86945 | 12.57934 | 4.03083 | 10.29986 | 6.67251 | 3.63526 | 2.99689 | 3.55007 | 0.06679 | 7.87576 | 5.89621 | 10.21257 | 15.37476 |
| **Bmt2** | 0.41414 | 0.00357 | 0.05918 | 0.00000 | 0.14624 | 1.03658 | 0.01109 | 0.99688 | 0.04123 | 0.01501 | 0.83320 | 0.52410 | 0.55423 | 0.14274 |
| **Bmyc** | 0.00425 | 0.06318 | 0.01505 | 11.34004 | 0.53994 | 1.13049 | 1.82320 | 3.84372 | 0.01125 | 7.76850 | 0.77081 | 0.24293 | 0.00169 | 0.00000 |
| **Bnc1** | 0.00802 | 0.00406 | 0.00341 | 0.01446 | 0.10945 | 0.00874 | 0.00415 | 0.00716 | 0.00428 | 0.00589 | 0.00679 | 0.00603 | 1.24639 | 2.45285 |
| **Bnc2** | 0.00000 | 0.00427 | 0.10105 | 0.00404 | 0.13519 | 0.00498 | 0.00428 | 0.01065 | 0.09167 | 0.00661 | 0.00645 | 0.00740 | 0.58125 | 1.00737 |
| **Bnip1** | 1.60458 | 0.02547 | 4.26787 | 1.29761 | 3.50635 | 4.24587 | 3.18134 | 4.95760 | 0.00315 | 0.24714 | 4.79753 | 3.25726 | 0.73321 | 2.07962 |
| **Bnip2** | 1.53204 | 0.00363 | 0.00000 | 0.09699 | 2.22957 | 0.50556 | 0.00321 | 0.01954 | 6.55549 | 0.01784 | 0.00000 | 1.11983 | 0.40079 | 5.26138 |
| **Bnip3** | 23.30596 | 0.31425 | 3.89463 | 1.61866 | 6.00542 | 2.69970 | 0.24987 | 2.90309 | 16.75951 | 0.03748 | 4.50434 | 0.00000 | 10.95407 | 20.06608 |
| **Bnip3l** | 23.64945 | 0.30930 | 32.41987 | 24.21731 | 33.33882 | 22.44524 | 0.41150 | 9.18021 | 24.88898 | 0.16196 | 12.34734 | 7.40907 | 8.48462 | 23.96718 |
| **Bnipl** | 0.00688 | 0.00488 | 0.32311 | 0.01152 | 0.62039 | 0.02615 | 0.00456 | 0.06252 | 0.02309 | 0.01163 | 0.00855 | 0.00513 | 0.36302 | 0.05161 |
| **Boc** | 0.17339 | 0.01152 | 0.16276 | 0.00000 | 0.10774 | 0.20313 | 0.00403 | 0.00381 | 0.01067 | 0.01393 | 0.00181 | 1.16230 | 0.05317 | 0.02437 |
| **Bod1** | 10.59286 | 0.17450 | 7.49285 | 6.72500 | 12.38177 | 4.92180 | 4.33158 | 2.27571 | 9.07343 | 0.12343 | 6.23869 | 6.30149 | 7.33760 | 16.09281 |
| **Bod1l** | 0.82559 | 0.00119 | 1.56892 | 0.62830 | 3.72180 | 2.88363 | 0.54738 | 0.43995 | 27.07298 | 0.00401 | 1.30159 | 1.65791 | 4.78506 | 15.63592 |
| **Bok** | 4.78793 | 8.23523 | 9.92611 | 13.63797 | 4.77773 | 1.93912 | 3.71966 | 10.06494 | 0.02125 | 13.68335 | 0.75654 | 2.10501 | 0.00000 | 0.65357 |
| **Bola1** | 0.00000 | 0.04072 | 1.47846 | 12.83635 | 1.72255 | 0.00000 | 0.88379 | 2.91353 | 0.00000 | 0.01563 | 0.06698 | 2.04410 | 1.35419 | 0.00000 |
| **Bola2** | 0.72083 | 2.82021 | 0.09943 | 2.84799 | 4.95979 | 1.67791 | 0.81402 | 5.08758 | 0.03057 | 6.04446 | 0.37918 | 4.08109 | 3.70074 | 0.65721 |
| **Bola3** | 0.03092 | 0.00382 | 0.14774 | 0.13242 | 0.39662 | 0.56971 | 0.00378 | 0.11291 | 0.02166 | 0.02021 | 0.00000 | 0.37142 | 0.00424 | 0.06803 |
| **Bop1** | 3.62301 | 0.05647 | 4.76213 | 7.98757 | 15.22027 | 1.21126 | 0.33879 | 3.03788 | 0.19872 | 0.05212 | 3.16392 | 4.77652 | 10.19416 | 0.88315 |
| **Bora** | 5.16957 | 4.43964 | 7.05117 | 2.45279 | 7.81787 | 1.43635 | 2.25520 | 0.37980 | 4.40189 | 0.04884 | 7.47145 | 1.63129 | 1.62858 | 8.53880 |
| **Borcs5** | 0.20337 | 0.12692 | 1.49320 | 1.38519 | 3.77721 | 0.90532 | 2.83654 | 0.00265 | 0.02678 | 0.02969 | 0.27061 | 0.45546 | 0.85806 | 1.14093 |
| **Borcs6** | 0.56772 | 0.00402 | 0.18424 | 0.54640 | 2.33309 | 0.09418 | 0.00470 | 0.44517 | 0.08550 | 0.00702 | 0.00065 | 0.00151 | 2.78528 | 0.74193 |
| **Borcs7** | 0.43157 | 0.01754 | 0.06635 | 1.79803 | 1.01382 | 1.32581 | 0.43704 | 0.90850 | 0.24075 | 0.09164 | 0.18854 | 1.80577 | 0.73846 | 0.35742 |
| **Borcs8** | 2.69494 | 0.00794 | 1.66342 | 0.00000 | 3.99978 | 2.71953 | 0.12665 | 2.33551 | 0.15545 | 6.19078 | 0.18913 | 3.51160 | 3.70485 | 0.54449 |
| **Bpgm** | 0.95869 | 0.00340 | 0.00000 | 0.73853 | 1.76097 | 1.18526 | 0.00398 | 1.00997 | 1.45672 | 0.00717 | 0.00000 | 0.31357 | 4.71893 | 2.77933 |
| **Bphl** | 0.00291 | 0.18465 | 0.00000 | 0.00484 | 0.31333 | 0.82134 | 1.80862 | 0.00000 | 0.01174 | 0.07742 | 1.68236 | 0.00000 | 0.00527 | 0.00174 |
| **Bpifb9a** | 0.02503 | 0.00443 | 0.07252 | 0.00544 | 0.03791 | 0.00891 | 0.00428 | 0.01168 | 0.14504 | 0.00699 | 0.00698 | 0.01022 | 0.00496 | 0.00063 |
| **Bpifb9b** | 0.02259 | 0.00442 | 0.12455 | 0.00282 | 0.02213 | 0.00752 | 0.00423 | 0.01179 | 0.18211 | 0.00725 | 0.00664 | 0.01015 | 0.00486 | 0.00098 |
| **Bpifc** | 0.00553 | 0.00434 | 0.01407 | 0.01226 | 0.02332 | 0.01266 | 0.00428 | 0.00990 | 0.01173 | 0.00682 | 0.00886 | 0.00944 | 0.29626 | 0.04007 |
| **Bpnt1** | 0.70616 | 2.82076 | 3.57293 | 2.91261 | 1.79523 | 3.21768 | 1.52578 | 0.92032 | 2.96654 | 0.13578 | 3.49057 | 0.30585 | 0.65348 | 5.07662 |
| **Bptf** | 3.26943 | 1.42888 | 6.47080 | 0.99704 | 17.03199 | 8.31849 | 0.00980 | 5.80329 | 12.22962 | 0.04889 | 3.12184 | 4.21193 | 5.89868 | 17.50913 |
| **Braf** | 2.53691 | 0.01932 | 2.68950 | 1.94769 | 3.36014 | 0.75891 | 1.47270 | 1.83368 | 7.30247 | 0.03910 | 3.38403 | 2.15463 | 0.69425 | 4.84907 |
| **Brap** | 1.32220 | 0.34365 | 5.67952 | 3.03350 | 8.02107 | 0.87383 | 3.23972 | 5.02404 | 0.06134 | 0.03872 | 0.14528 | 0.61759 | 3.79112 | 2.03847 |
| **Brat1** | 0.00000 | 0.00456 | 1.86555 | 2.20108 | 0.33803 | 0.83861 | 0.00385 | 1.50822 | 0.00234 | 0.00373 | 0.00000 | 0.15269 | 1.81050 | 0.86394 |
| **Brca1** | 1.62439 | 0.02259 | 3.69674 | 2.20221 | 3.66392 | 3.47933 | 0.50882 | 2.47792 | 2.29475 | 0.01059 | 4.44838 | 4.40603 | 2.27126 | 5.09814 |
| **Brca2** | 3.15560 | 0.30243 | 7.80707 | 4.27483 | 15.91755 | 1.40676 | 0.07607 | 0.00000 | 25.65428 | 0.00347 | 0.00000 | 3.18492 | 12.11558 | 21.14812 |
| **Brcc3** | 2.42736 | 0.07071 | 1.36676 | 2.04090 | 2.05613 | 6.14450 | 0.00871 | 0.23916 | 3.00501 | 0.02782 | 2.80182 | 0.26822 | 0.00000 | 0.00000 |
| **Brd1** | 9.36027 | 0.25850 | 11.06106 | 6.61144 | 10.43546 | 6.31952 | 4.98627 | 7.59204 | 0.55579 | 0.05783 | 7.05385 | 2.78517 | 10.97100 | 15.60193 |
| **Brd2** | 5.45887 | 2.55627 | 17.34600 | 8.26033 | 15.29375 | 12.63130 | 2.82322 | 6.01308 | 10.24599 | 2.69598 | 10.16448 | 3.62366 | 10.67051 | 21.42835 |
| **Brd3** | 2.39814 | 0.05026 | 16.80140 | 4.80850 | 14.37959 | 8.92304 | 1.77968 | 5.57140 | 3.26734 | 0.48064 | 5.41743 | 6.41269 | 3.67663 | 12.40941 |
| **Brd4** | 5.20196 | 1.85610 | 13.54021 | 5.10189 | 15.32373 | 7.88641 | 2.66835 | 3.81360 | 1.69443 | 0.11785 | 5.82577 | 4.75264 | 6.44601 | 14.77524 |
| **Brd7** | 9.51361 | 2.73308 | 5.77507 | 4.91644 | 8.79924 | 8.69521 | 5.05723 | 3.13558 | 9.36241 | 0.15558 | 9.21664 | 6.90054 | 5.49400 | 12.77244 |
| **Brd8** | 13.48396 | 0.25856 | 11.17415 | 4.63132 | 9.76682 | 7.10430 | 4.49334 | 0.24908 | 11.04461 | 0.85720 | 2.38937 | 2.97390 | 13.20166 | 13.01968 |
| **Brd9** | 11.03703 | 2.90047 | 20.89936 | 4.77200 | 17.45418 | 7.87824 | 3.11886 | 6.25350 | 13.80454 | 9.54204 | 10.98628 | 6.06610 | 8.28566 | 18.42727 |
| **Brdt** | 0.30605 | 0.00348 | 0.81077 | 0.46062 | 0.44357 | 0.00000 | 0.00349 | 0.01124 | 0.25437 | 0.01209 | 0.00441 | 0.01350 | 6.95116 | 3.22365 |
| **Bre** | 0.11404 | 0.01618 | 3.97697 | 6.64298 | 2.55223 | 4.35521 | 1.39497 | 3.88375 | 0.24493 | 0.09973 | 2.63627 | 4.47364 | 0.00000 | 0.19013 |
| **Brf1** | 2.16235 | 0.04091 | 2.63213 | 1.47299 | 2.41973 | 0.48552 | 2.91045 | 0.84223 | 0.00000 | 0.01285 | 0.55486 | 3.28319 | 3.56712 | 1.70912 |
| **Brf2** | 0.16135 | 0.01890 | 1.20663 | 1.61275 | 3.26082 | 1.00013 | 0.01459 | 0.63442 | 0.00000 | 0.00893 | 0.63045 | 0.22869 | 4.10593 | 1.35227 |
| **Bri3** | 0.62418 | 2.24366 | 0.11261 | 1.31743 | 1.15290 | 0.45621 | 1.73134 | 0.48713 | 0.09270 | 3.53106 | 0.02680 | 2.17279 | 10.78373 | 0.16427 |
| **Bri3bp** | 15.80202 | 0.19349 | 16.39564 | 6.28877 | 15.13079 | 2.67651 | 0.33790 | 2.40667 | 3.78684 | 0.15276 | 0.00000 | 6.06524 | 8.67343 | 11.09509 |

| | | | | | | | | | | | | | | |
|---|---|---|---|---|---|---|---|---|---|---|---|---|---|---|
| **Brip1** | 1.05099 | 0.02383 | 0.13916 | 0.58761 | 2.08555 | 1.32819 | 0.12099 | 0.07168 | 1.43269 | 0.03309 | 1.21853 | 1.62518 | 6.13665 | 3.66954 |
| **Brip1os** | 1.93456 | 0.12355 | 0.92388 | 2.06179 | 0.09882 | 5.24005 | 0.34464 | 0.92893 | 2.61737 | 0.56222 | 0.48987 | 2.57562 | 10.30430 | 2.07595 |
| **Brix1** | 20.79096 | 0.40596 | 27.00762 | 14.16395 | 27.65529 | 16.91726 | 0.30245 | 6.54604 | 16.38564 | 0.19873 | 13.35476 | 12.52043 | 14.51301 | 19.47003 |
| **Brk1** | 15.19225 | 10.07782 | 16.61142 | 17.19902 | 14.58823 | 22.92010 | 8.73071 | 19.84795 | 9.12479 | 14.89903 | 15.61220 | 9.12423 | 3.75109 | 15.97111 |
| **Brms1** | 4.66150 | 0.13541 | 15.64423 | 6.77965 | 11.48348 | 8.35646 | 0.79356 | 6.94073 | 0.29627 | 0.10632 | 5.10492 | 6.02020 | 5.38682 | 7.72042 |
| **Brms1l** | 0.81274 | 0.00188 | 3.44518 | 1.40829 | 3.03095 | 1.02892 | 1.25629 | 0.00000 | 3.30311 | 0.01767 | 2.41539 | 0.45334 | 8.57285 | 7.23350 |
| **Brox** | 2.42843 | 0.01572 | 3.52571 | 0.83075 | 2.89619 | 4.38682 | 3.12067 | 1.73941 | 0.78392 | 0.06682 | 2.53157 | 3.56373 | 0.91250 | 6.36238 |
| **Brpf1** | 1.42520 | 5.86746 | 4.46367 | 1.72748 | 7.61938 | 4.93544 | 0.90938 | 3.40390 | 0.80796 | 0.06081 | 2.09422 | 3.51506 | 6.50763 | 6.20202 |
| **Brpf3** | 4.05248 | 0.01687 | 2.53898 | 2.36831 | 3.67411 | 0.85947 | 0.22262 | 0.11937 | 0.47283 | 0.02388 | 0.00000 | 2.95213 | 7.26428 | 1.10116 |
| **Brsk1** | 0.01444 | 0.00404 | 0.23293 | 0.00000 | 0.00802 | 0.00632 | 0.00429 | 0.00700 | 0.01258 | 0.00890 | 0.00493 | 0.26716 | 0.10564 | 0.08831 |
| **Brsk2** | 0.04783 | 0.00403 | 0.01548 | 0.47194 | 0.26286 | 0.14251 | 0.00404 | 0.03334 | 0.00856 | 0.00596 | 0.00614 | 0.00875 | 0.15553 | 0.03183 |
| **Brwd1** | 3.64902 | 0.17459 | 9.90756 | 2.37151 | 11.14823 | 4.97458 | 2.47598 | 0.18179 | 12.87579 | 0.04800 | 3.60945 | 1.94633 | 2.87629 | 13.63412 |
| **Brwd3** | 7.64056 | 0.00159 | 0.46964 | 1.82578 | 6.19262 | 1.04121 | 0.00114 | 0.00000 | 16.04203 | 0.00757 | 1.12989 | 0.40699 | 4.46063 | 5.68131 |
| **Bscl2** | 0.00530 | 0.00222 | 1.03276 | 0.49715 | 0.81566 | 2.67876 | 4.02331 | 5.15464 | 0.00000 | 0.14468 | 0.48706 | 2.46372 | 0.51392 | 1.03115 |
| **Bsdc1** | 2.71664 | 0.03918 | 5.74476 | 2.61096 | 6.06892 | 2.90552 | 1.62442 | 2.89094 | 2.88400 | 0.07609 | 5.11470 | 5.34243 | 2.03970 | 8.92830 |
| **Bsg** | 5.27557 | 1.27408 | 10.17419 | 26.80756 | 10.28915 | 13.49530 | 2.99054 | 7.27845 | 0.12051 | 13.23381 | 7.54534 | 12.05747 | 0.48372 | 0.66498 |
| **Bsn** | 0.00724 | 0.00448 | 0.02576 | 0.00587 | 0.64869 | 0.01319 | 0.00418 | 0.01011 | 0.01275 | 0.00717 | 0.00836 | 0.00933 | 0.01152 | 0.00831 |
| **Bspry** | 3.68319 | 0.04102 | 4.87801 | 2.77336 | 5.73933 | 0.31918 | 0.75585 | 3.01219 | 1.39956 | 0.00415 | 0.00000 | 0.75736 | 10.21339 | 6.03539 |
| **Bst2** | 0.00625 | 0.01190 | 0.22895 | 0.37340 | 0.42628 | 0.56729 | 0.22235 | 1.46994 | 0.00610 | 0.01960 | 0.45132 | 0.00213 | 0.00479 | 0.09309 |
| **Btaf1** | 5.97951 | 0.02940 | 3.13409 | 1.25440 | 6.52589 | 5.03124 | 3.45235 | 0.21611 | 6.23627 | 0.09614 | 4.87538 | 5.16707 | 10.10899 | 13.13338 |
| **Btbd1** | 5.98958 | 2.86397 | 4.69681 | 2.40284 | 2.55535 | 4.09661 | 1.55040 | 3.97555 | 4.78920 | 0.09355 | 6.10007 | 4.02385 | 1.77872 | 5.00168 |
| **Btbd10** | 7.11497 | 4.43352 | 4.28429 | 2.57421 | 5.25678 | 2.91241 | 1.44008 | 0.03296 | 4.99415 | 0.01661 | 2.11908 | 4.28401 | 1.74573 | 8.20439 |
| **Btbd16** | 0.00000 | 0.05597 | 0.09901 | 0.02964 | 0.09824 | 0.03759 | 0.01113 | 0.03581 | 0.60521 | 0.11931 | 0.01779 | 0.04398 | 1.12329 | 0.39975 |
| **Btbd18** | 10.88675 | 0.02131 | 11.47963 | 5.85869 | 18.53613 | 0.00000 | 0.00388 | 0.00000 | 15.69722 | 11.24832 | 6.81160 | 0.00000 | 17.56604 | 20.95353 |
| **Btbd19** | 0.14356 | 0.09211 | 0.56260 | 0.65291 | 1.30799 | 0.57079 | 0.07588 | 0.33443 | 1.65061 | 0.06783 | 0.00918 | 1.65895 | 0.97835 | 0.65228 |
| **Btbd2** | 0.01361 | 0.02290 | 0.02913 | 0.78324 | 0.87018 | 0.40562 | 0.00426 | 1.05324 | 0.01094 | 0.00667 | 0.00000 | 0.97162 | 1.52951 | 0.29028 |
| **Btbd3** | 0.19543 | 0.01152 | 0.00000 | 0.00000 | 0.00000 | 0.40358 | 0.00359 | 0.49369 | 0.02334 | 0.01084 | 1.62222 | 0.21264 | 0.01504 | 0.86699 |
| **Btbd6** | 0.40577 | 0.06264 | 2.24338 | 0.94517 | 1.09315 | 0.09943 | 0.00321 | 1.84842 | 0.00167 | 0.01531 | 1.64548 | 0.71879 | 1.86801 | 0.19774 |
| **Btbd7** | 0.94553 | 0.00299 | 0.21444 | 0.43677 | 1.28349 | 0.48612 | 0.00381 | 0.00212 | 0.87070 | 0.00295 | 0.92620 | 0.50512 | 0.79961 | 3.25945 |
| **Btbd8** | 0.01808 | 0.00447 | 0.02637 | 0.00005 | 0.01210 | 0.01412 | 0.00519 | 0.43443 | 0.01150 | 0.00629 | 0.00733 | 0.01102 | 0.00899 | 0.01536 |
| **Btbd9** | 0.00000 | 0.00460 | 1.39639 | 0.78512 | 0.89141 | 0.89446 | 0.00377 | 1.02475 | 1.23313 | 0.00395 | 0.15942 | 0.00180 | 0.49548 | 0.00000 |
| **Btd** | 0.00000 | 0.00538 | 0.47641 | 0.62817 | 0.00000 | 0.12003 | 0.00413 | 0.03495 | 0.13768 | 0.00889 | 0.00715 | 0.00523 | 0.24071 | 1.42325 |
| **Btf3** | 31.78787 | 13.41205 | 33.99219 | 40.50973 | 32.88862 | 37.48932 | 11.64195 | 33.71996 | 19.06947 | 22.85712 | 22.55387 | 29.73856 | 12.07862 | 25.49901 |
| **Btf3l4** | 12.44566 | 0.26412 | 12.18734 | 11.58889 | 12.04056 | 10.25819 | 5.39128 | 11.82230 | 17.67477 | 0.36054 | 7.18578 | 10.87733 | 5.15326 | 9.42080 |
| **Btg1** | 13.02649 | 6.01860 | 15.24627 | 10.05393 | 18.91871 | 11.03207 | 2.40281 | 6.98897 | 29.02453 | 0.14354 | 1.70487 | 6.92705 | 15.50957 | 13.51775 |
| **Btg2** | 0.52412 | 0.02455 | 0.26898 | 0.00000 | 0.82297 | 0.28436 | 0.02584 | 0.87725 | 0.00447 | 0.03498 | 0.00000 | 1.10279 | 1.35217 | 1.44573 |
| **Btg3** | 0.39240 | 0.00426 | 0.08497 | 0.01303 | 0.28928 | 0.13604 | 0.18071 | 0.00801 | 0.07632 | 0.00480 | 1.15900 | 0.52728 | 1.16247 | 0.00000 |
| **Btg4** | 0.33835 | 0.00334 | 0.00000 | 0.08557 | 2.21519 | 0.00000 | 0.00408 | 0.00576 | 0.33932 | 0.00589 | 0.00384 | 0.00006 | 5.46953 | 5.16578 |
| **Btk** | 1.66163 | 0.00381 | 0.00000 | 0.38832 | 0.00000 | 0.00000 | 0.00359 | 0.00441 | 1.51646 | 0.00598 | 0.00971 | 0.01930 | 0.32937 | 0.35208 |
| **Btn2a2** | 0.00066 | 0.00355 | 0.00811 | 0.01505 | 0.00000 | 0.01728 | 0.00438 | 0.01200 | 0.04938 | 0.00611 | 0.00909 | 1.23115 | 0.09016 | 0.30626 |
| **Btnl1** | 0.16762 | 0.00442 | 0.07007 | 0.00210 | 0.03293 | 0.03141 | 0.00437 | 0.01030 | 0.14962 | 0.00694 | 0.00562 | 0.00908 | 0.00444 | 0.00000 |
| **Btrc** | 12.18293 | 0.58335 | 33.32613 | 16.67537 | 19.03116 | 19.03317 | 7.62925 | 7.52253 | 1.28808 | 1.76460 | 19.48574 | 15.26837 | 2.63590 | 15.73892 |
| **Bub1** | 11.53566 | 0.01836 | 5.27489 | 1.82241 | 9.69023 | 4.91073 | 0.02609 | 6.00950 | 14.24934 | 0.01679 | 5.37590 | 4.76003 | 4.65619 | 10.92355 |
| **Bub1b** | 5.73367 | 4.64121 | 9.64110 | 2.71667 | 6.55564 | 4.31797 | 3.16952 | 5.84694 | 2.73358 | 0.06026 | 5.98358 | 5.17593 | 5.48826 | 7.59436 |
| **Bub3** | 9.53563 | 0.47535 | 19.72289 | 13.86481 | 15.61001 | 18.56502 | 6.94442 | 13.19708 | 1.40485 | 3.01713 | 16.84096 | 14.94865 | 4.22719 | 19.13075 |
| **Bud13** | 0.83688 | 0.03134 | 3.79808 | 0.00000 | 1.53115 | 2.03108 | 3.13140 | 0.12007 | 4.54499 | 2.44984 | 1.30195 | 3.33218 | 0.06134 | 9.08601 |
| **Bud31** | 10.53017 | 0.18957 | 10.39806 | 11.80040 | 12.20868 | 10.98172 | 6.13060 | 11.28877 | 7.34932 | 0.37545 | 9.70452 | 8.49068 | 5.98427 | 12.81783 |
| **Bves** | 0.00773 | 0.00429 | 0.00556 | 0.19452 | 0.01001 | 0.08252 | 0.00429 | 0.00747 | 0.01272 | 0.00687 | 0.00610 | 0.00949 | 0.00701 | 0.00925 |
| **Bvht** | 0.00645 | 0.00408 | 0.02483 | 0.17439 | 0.00579 | 0.05060 | 0.00390 | 0.00340 | 0.02559 | 0.01067 | 0.00834 | 0.32171 | 0.05701 | 0.02342 |
| **Bysl** | 5.95124 | 0.17975 | 21.83073 | 15.65538 | 19.62365 | 13.66487 | 6.16456 | 3.01745 | 1.61857 | 0.32319 | 6.00584 | 7.82483 | 5.53109 | 8.50758 |
| **Bzw1** | 20.69222 | 1.52726 | 28.70126 | 17.72307 | 24.12680 | 23.24353 | 4.94236 | 8.57047 | 41.18684 | 18.42312 | 20.50979 | 15.56269 | 14.09741 | 29.14257 |
| **Bzw2** | 6.43238 | 4.38259 | 14.85967 | 11.72228 | 18.58272 | 10.26043 | 3.02727 | 3.96226 | 10.44723 | 0.13227 | 10.58243 | 10.73844 | 6.23554 | 14.01131 |
| **C1d** | 6.18423 | 0.06988 | 11.02653 | 15.37719 | 14.07593 | 13.11041 | 5.46563 | 7.47309 | 20.05741 | 0.25615 | 14.30087 | 5.75760 | 4.60229 | 13.65803 |
| **C1galt1** | 0.06010 | 0.00440 | 0.05135 | 0.00000 | 0.00578 | 0.02953 | 0.00506 | 0.08900 | 0.05199 | 0.00632 | 0.00836 | 0.02675 | 0.08002 | 0.03535 |
| **C1galt1c1** | 1.34929 | 0.00367 | 2.00555 | 0.00000 | 0.31002 | 2.03964 | 0.02296 | 1.05759 | 1.33596 | 0.01823 | 0.00000 | 0.41829 | 0.00000 | 2.69770 |

| | | | | | | | | | | | | | | |
|---|---|---|---|---|---|---|---|---|---|---|---|---|---|---|
| **C1qa** | 0.00000 | 0.00411 | 0.87050 | 0.78225 | 0.17552 | 0.00000 | 0.00389 | 0.00645 | 0.00939 | 0.00505 | 0.00449 | 0.01263 | 0.00278 | 0.02148 |
| **C1qb** | 0.00886 | 0.00448 | 0.01080 | 0.01252 | 0.14216 | 0.01332 | 0.00462 | 0.01045 | 0.01407 | 0.00676 | 0.00886 | 0.00913 | 0.00546 | 0.01445 |
| **C1qbp** | 31.16655 | 7.15026 | 16.28799 | 17.58236 | 34.29075 | 19.93267 | 8.31246 | 3.98717 | 12.45961 | 11.04784 | 10.26596 | 16.87912 | 18.85481 | 14.54852 |
| **C1ql1** | 0.38771 | 0.00404 | 0.29074 | 0.00000 | 1.16136 | 0.00000 | 0.00399 | 0.00865 | 0.00078 | 0.00654 | 0.00624 | 0.00528 | 1.66219 | 0.95367 |
| **C1ql4** | 0.02001 | 0.01179 | 0.85523 | 0.65082 | 0.00000 | 0.11202 | 0.00366 | 0.00049 | 0.01299 | 0.00581 | 0.11690 | 0.83137 | 0.01590 | 0.00000 |
| **C1qtnf2** | 0.01079 | 0.00447 | 0.01446 | 0.01252 | 0.01904 | 0.01787 | 0.00432 | 0.00983 | 0.01127 | 0.00693 | 0.01082 | 0.01066 | 0.00763 | 0.01213 |
| **C1qtnf4** | 0.00412 | 0.00467 | 0.00000 | 0.29292 | 1.50551 | 0.01163 | 0.00438 | 0.02064 | 0.01325 | 0.00737 | 0.00854 | 0.00620 | 0.01020 | 0.00000 |
| **C1ra** | 0.33961 | 0.00435 | 0.00000 | 0.06740 | 0.37214 | 0.04905 | 0.03048 | 0.00575 | 0.04941 | 0.00979 | 0.03718 | 0.00516 | 0.01148 | 0.02469 |
| **C1rb** | 0.01370 | 0.00443 | 0.00000 | 0.11881 | 0.22694 | 0.05112 | 0.00417 | 0.00589 | 0.01695 | 0.00692 | 0.11658 | 0.00503 | 0.01137 | 0.00276 |
| **C1rl** | 0.02790 | 0.02330 | 0.00933 | 0.00000 | 0.18986 | 0.02914 | 0.00396 | 0.02448 | 0.03250 | 0.06263 | 0.00394 | 0.60720 | 0.04730 | 0.02074 |
| **C1s1** | 0.35608 | 0.00439 | 0.19746 | 0.00000 | 0.09380 | 0.01268 | 0.00402 | 0.00857 | 0.27096 | 0.00755 | 0.00865 | 0.00820 | 0.04772 | 0.53704 |
| **C1s2** | 0.01054 | 0.00431 | 0.17379 | 0.00000 | 0.49795 | 0.06139 | 0.00383 | 0.00569 | 1.44455 | 0.00569 | 0.00580 | 0.00572 | 0.44435 | 0.00000 |
| **C2** | 2.57871 | 0.01258 | 8.03216 | 6.75856 | 2.70856 | 3.81371 | 0.00146 | 1.38484 | 8.66726 | 0.01027 | 0.00000 | 0.00000 | 0.76190 | 2.33827 |
| **C2cd2** | 0.00000 | 0.00310 | 0.85000 | 1.47292 | 0.71350 | 0.00000 | 0.04425 | 0.14686 | 0.06377 | 0.06359 | 0.27588 | 1.77004 | 0.39887 | 1.57721 |
| **C2cd2l** | 6.60306 | 4.45079 | 8.77451 | 1.65065 | 9.00673 | 1.05415 | 1.90784 | 2.76863 | 2.43034 | 0.01516 | 0.00000 | 1.39962 | 11.37290 | 12.25896 |
| **C2cd3** | 0.46903 | 1.44316 | 0.47046 | 0.17689 | 2.25075 | 1.44548 | 2.04278 | 0.03012 | 0.31499 | 0.06459 | 0.00000 | 1.26663 | 5.86950 | 3.67524 |
| **C2cd4a** | 0.00873 | 0.00451 | 0.01454 | 0.00000 | 0.01437 | 0.18362 | 0.00420 | 0.00635 | 0.01098 | 0.01432 | 0.01271 | 0.01849 | 0.02848 | 0.00817 |
| **C2cd4b** | 0.00451 | 0.00430 | 0.01481 | 0.01245 | 0.01059 | 0.01337 | 0.00421 | 0.00883 | 0.00844 | 0.00677 | 0.00905 | 0.00892 | 0.54126 | 0.00841 |
| **C2cd5** | 0.35593 | 0.00123 | 1.76863 | 0.00000 | 0.00000 | 1.41539 | 0.00344 | 0.32409 | 1.20807 | 0.00000 | 3.74672 | 1.66280 | 6.94054 | 5.30982 |
| **C3** | 0.00000 | 0.00438 | 0.10845 | 0.11144 | 1.76761 | 0.00423 | 0.00455 | 0.00811 | 0.00805 | 0.00608 | 0.00597 | 0.00559 | 0.00000 | 0.43646 |
| **C4a** | 0.00000 | 0.00433 | 0.35538 | 0.32141 | 0.09507 | 0.75426 | 0.00389 | 0.00000 | 0.02497 | 0.00283 | 0.76896 | 0.00456 | 0.21923 | 0.00000 |
| **C4b** | 0.00397 | 0.01272 | 0.11301 | 0.02440 | 0.04225 | 0.16854 | 0.00427 | 0.00534 | 0.03533 | 0.00534 | 0.13187 | 0.00454 | 0.11567 | 0.00000 |
| **C5ar2** | 0.02173 | 0.00450 | 0.01484 | 0.02792 | 0.04715 | 0.06082 | 0.01012 | 0.06442 | 0.07406 | 0.01770 | 0.01094 | 0.01091 | 0.12489 | 0.10783 |
| **C6** | 0.06528 | 0.00446 | 0.02379 | 0.03641 | 0.07460 | 0.01991 | 0.06185 | 0.02543 | 0.19488 | 0.00600 | 0.02368 | 0.01607 | 0.00372 | 0.02872 |
| **C7** | 0.00960 | 0.00446 | 0.04350 | 0.00704 | 0.08451 | 0.01255 | 0.00422 | 0.01073 | 0.01505 | 0.00751 | 0.00818 | 0.00967 | 0.00729 | 0.01212 |
| **C77080** | 3.66333 | 7.11839 | 10.94648 | 2.10898 | 9.74021 | 10.64382 | 3.83833 | 7.40006 | 0.68015 | 6.26837 | 4.12553 | 3.41538 | 2.94143 | 8.16186 |
| **C77370** | 0.75666 | 0.00950 | 1.70838 | 1.86424 | 0.64374 | 0.90186 | 0.06221 | 0.41998 | 3.32546 | 0.01673 | 1.18887 | 3.03872 | 0.00000 | 0.00000 |
| **C86187** | 0.03996 | 0.00352 | 0.34054 | 0.00000 | 0.36068 | 0.00762 | 0.00372 | 0.00575 | 0.22745 | 0.00656 | 0.00751 | 0.00530 | 6.28220 | 1.18373 |
| **C87198** | 0.00470 | 0.00451 | 0.01539 | 0.00669 | 0.02657 | 0.03318 | 0.01620 | 0.03354 | 0.08389 | 0.00611 | 0.00698 | 0.02378 | 0.68125 | 0.22404 |
| **C87414** | 0.49304 | 0.00375 | 0.01738 | 0.28176 | 0.40572 | 0.03500 | 0.03945 | 0.00218 | 4.47193 | 0.01910 | 0.08292 | 0.02212 | 2.30570 | 1.04644 |
| **C87436** | 2.42855 | 0.04409 | 2.28130 | 0.44653 | 2.13297 | 2.52379 | 0.00909 | 1.15530 | 0.80762 | 0.01042 | 0.23853 | 1.51354 | 3.06968 | 6.24568 |
| **C87499** | 0.67449 | 0.00342 | 0.08637 | 0.01190 | 0.00000 | 0.00182 | 0.00401 | 0.00903 | 0.05069 | 0.00665 | 0.02077 | 0.00510 | 7.34903 | 0.69962 |
| **C87977** | 0.58213 | 0.00362 | 3.43454 | 0.00000 | 0.79915 | 0.27198 | 0.00277 | 0.00000 | 3.84176 | 0.00000 | 0.02026 | 0.00103 | 4.05502 | 1.70563 |
| **C8b** | 0.00533 | 0.00424 | 0.00779 | 0.01903 | 0.15384 | 0.17403 | 0.00423 | 0.00813 | 0.00807 | 0.00540 | 0.00840 | 0.00839 | 0.22573 | 0.02792 |
| **Caap1** | 7.13803 | 0.00314 | 0.25470 | 0.00000 | 0.00000 | 0.22660 | 0.01587 | 0.03295 | 2.57713 | 0.00588 | 0.01256 | 0.20563 | 2.40640 | 5.96092 |
| **Cab39** | 14.78610 | 7.45398 | 21.44800 | 7.16360 | 19.61747 | 15.41794 | 6.10382 | 4.77624 | 27.92499 | 0.21295 | 15.21429 | 6.51537 | 18.27823 | 21.74849 |
| **Cab39l** | 1.48664 | 0.10440 | 7.28884 | 4.66128 | 7.58465 | 6.39703 | 0.01545 | 7.09076 | 11.65583 | 0.01090 | 5.95737 | 4.63704 | 2.43392 | 7.35886 |
| **Cabin1** | 3.09821 | 0.02847 | 3.35073 | 3.20433 | 6.87636 | 2.79573 | 3.10898 | 0.43100 | 0.34908 | 0.12077 | 2.51758 | 2.54812 | 5.18809 | 4.20313 |
| **Cables1** | 0.00000 | 0.05291 | 0.34386 | 0.00000 | 0.01211 | 0.09231 | 0.01615 | 0.72058 | 0.00749 | 0.00731 | 0.00000 | 0.52274 | 0.89927 | 0.62019 |
| **Cables2** | 3.66005 | 0.01595 | 2.83941 | 1.69906 | 3.74052 | 2.49112 | 2.33963 | 1.93001 | 0.00000 | 0.02843 | 4.65455 | 1.94095 | 1.10482 | 4.36301 |
| **Cabp4** | 0.01039 | 0.00444 | 0.01460 | 0.01326 | 0.01387 | 0.01322 | 0.00429 | 0.01025 | 0.01167 | 0.00701 | 0.00884 | 0.01003 | 0.01628 | 0.01015 |
| **Cabp5** | 0.00589 | 0.01208 | 0.02182 | 0.01796 | 0.01913 | 0.47490 | 0.01047 | 0.00532 | 0.01079 | 0.00619 | 0.00229 | 0.20579 | 0.00874 | 0.00942 |
| **Cabp7** | 0.00698 | 0.00440 | 0.02888 | 0.06740 | 0.02014 | 0.01209 | 0.00432 | 0.00869 | 0.01075 | 0.00614 | 0.00785 | 0.00801 | 0.05952 | 0.00586 |
| **Cabyr** | 0.00389 | 0.00412 | 0.01081 | 0.21244 | 0.00601 | 0.00537 | 0.00396 | 0.00607 | 0.06466 | 0.00585 | 0.01051 | 0.01624 | 0.19750 | 0.00000 |
| **Cacfd1** | 0.64853 | 0.00298 | 0.00000 | 1.49779 | 0.21879 | 0.79214 | 0.00337 | 0.00000 | 0.00779 | 0.00613 | 2.73969 | 0.51969 | 0.26469 | 0.68603 |
| **Cachd1** | 0.41551 | 0.01211 | 0.00597 | 0.00376 | 0.00000 | 0.00505 | 0.00405 | 0.01889 | 0.00779 | 0.00575 | 0.00413 | 1.53686 | 0.00844 | 1.06990 |
| **Cacna1a** | 0.00957 | 0.00494 | 0.07421 | 0.21641 | 0.57410 | 0.12821 | 0.00434 | 0.16328 | 0.02411 | 0.00958 | 0.00347 | 0.04361 | 0.69748 | 0.01024 |
| **Cacna1b** | 0.00266 | 0.12038 | 0.00000 | 0.00572 | 0.83371 | 0.00000 | 0.00404 | 0.00609 | 0.00698 | 0.00368 | 0.72748 | 0.00221 | 1.36380 | 0.04591 |
| **Cacna1d** | 0.00632 | 0.00474 | 0.06721 | 0.17298 | 0.06036 | 0.09271 | 0.00405 | 0.00412 | 0.06927 | 0.00335 | 0.62962 | 0.05216 | 0.00586 | 0.05156 |
| **Cacna1h** | 0.01998 | 0.00419 | 0.16957 | 0.00000 | 0.17317 | 0.00710 | 0.00403 | 0.00959 | 0.01916 | 0.00622 | 0.00710 | 0.00864 | 1.08674 | 0.02669 |
| **Cacna2d1** | 0.10326 | 0.00416 | 0.01572 | 0.00180 | 0.15221 | 0.01177 | 0.00437 | 0.00805 | 0.09804 | 0.00594 | 0.00550 | 0.22895 | 0.06900 | 0.08998 |
| **Cacna2d3** | 0.03448 | 0.00443 | 0.01072 | 0.01147 | 0.23433 | 0.01108 | 0.00412 | 0.01010 | 0.03620 | 0.00807 | 0.00734 | 0.00809 | 0.00858 | 0.36454 |
| **Cacnb1** | 0.14108 | 0.00300 | 1.87428 | 2.40280 | 2.54258 | 0.13877 | 0.00316 | 0.00000 | 0.00527 | 0.00770 | 0.26531 | 1.24273 | 0.14994 | 0.00000 |
| **Cacnb2** | 0.00643 | 0.00439 | 0.03207 | 0.00974 | 0.00844 | 0.01181 | 0.00423 | 0.00998 | 0.05116 | 0.00669 | 0.00818 | 0.00949 | 0.17827 | 0.00983 |
| **Cacnb3** | 0.22789 | 0.13615 | 5.72637 | 9.40298 | 1.55606 | 0.00000 | 0.21952 | 10.72931 | 0.00000 | 0.07701 | 1.39854 | 0.34231 | 0.76343 | 0.00000 |

| | | | | | | | | | | | | | | |
|---|---|---|---|---|---|---|---|---|---|---|---|---|---|---|
| **Cactin** | 0.99325 | 0.14358 | 7.12456 | 6.90628 | 10.58788 | 1.64855 | 2.50879 | 4.65370 | 0.00000 | 0.08739 | 2.54103 | 2.51647 | 5.65945 | 0.00000 |
| **Cacul1** | 10.22083 | 0.01941 | 11.76781 | 7.86848 | 4.35788 | 13.97003 | 3.43787 | 3.11210 | 6.39462 | 0.12571 | 4.81162 | 7.72544 | 0.19788 | 2.15129 |
| **Cacybp** | 17.36604 | 5.29189 | 6.80830 | 17.71641 | 12.78051 | 17.73127 | 2.97753 | 2.25944 | 4.33057 | 0.11740 | 6.18454 | 15.35277 | 7.09533 | 2.56702 |
| **Cad** | 9.94848 | 0.92059 | 23.86256 | 21.70695 | 22.97181 | 9.91828 | 1.13807 | 7.41723 | 0.10550 | 0.17807 | 9.56219 | 10.18418 | 4.73543 | 6.53934 |
| **Cadm1** | 0.87356 | 0.02021 | 5.04584 | 2.34125 | 0.71911 | 3.39345 | 0.00222 | 0.18280 | 0.73362 | 0.01436 | 8.20071 | 1.99274 | 2.69367 | 0.00000 |
| **Cadm4** | 0.28637 | 0.00417 | 0.00000 | 0.26374 | 0.00000 | 0.73227 | 0.00488 | 0.04724 | 0.20335 | 0.02158 | 0.02297 | 0.08681 | 1.97229 | 0.33690 |
| **Cadps2** | 0.00430 | 0.00452 | 0.01445 | 0.00921 | 0.00783 | 0.01256 | 0.00413 | 0.01157 | 0.01853 | 0.00615 | 0.00843 | 0.02178 | 0.77858 | 0.01484 |
| **Cage1** | 0.00706 | 0.00434 | 0.01416 | 0.00832 | 0.01155 | 0.02532 | 0.01026 | 0.01025 | 0.01259 | 0.00635 | 0.03217 | 0.00889 | 0.09639 | 0.02208 |
| **Calb1** | 0.00786 | 0.00446 | 0.00546 | 0.01091 | 0.61559 | 0.01339 | 0.00445 | 0.01120 | 0.01045 | 0.00645 | 0.00891 | 0.00819 | 0.00585 | 0.00874 |
| **Calb2** | 0.01751 | 0.00433 | 0.01257 | 0.01976 | 0.09998 | 0.01124 | 0.00423 | 0.01035 | 0.00984 | 0.00813 | 0.00818 | 0.00842 | 0.20675 | 0.07385 |
| **Calcoco1** | 2.24490 | 0.00960 | 1.20653 | 0.35761 | 0.79751 | 1.23055 | 0.37718 | 0.29915 | 0.71803 | 0.00299 | 2.32797 | 1.69735 | 0.10538 | 0.00000 |
| **Calcoco2** | 28.01357 | 18.67725 | 59.47764 | 66.34898 | 40.28755 | 65.29891 | 17.93993 | 42.21612 | 40.19989 | 28.99418 | 42.65848 | 43.79996 | 16.94490 | 32.15015 |
| **Calcrl** | 0.00271 | 0.00436 | 0.00216 | 0.01310 | 0.00196 | 0.01633 | 0.00402 | 0.00817 | 1.10350 | 0.00667 | 0.00949 | 0.00874 | 0.29508 | 0.00000 |
| **Cald1** | 0.64025 | 0.00447 | 0.00401 | 0.00833 | 0.41893 | 0.00898 | 0.00443 | 0.01248 | 0.00914 | 0.00632 | 0.00710 | 0.00619 | 0.00364 | 0.00545 |
| **Calm1** | 4.36382 | 11.12354 | 19.69478 | 27.99442 | 15.54572 | 26.30793 | 10.04483 | 28.30385 | 6.50849 | 14.60290 | 11.06185 | 19.75006 | 8.46693 | 8.27639 |
| **Calm2** | 13.41427 | 11.54304 | 25.50253 | 17.85314 | 15.87363 | 28.06604 | 9.23575 | 27.88036 | 17.10046 | 15.89362 | 20.72637 | 23.53942 | 11.31096 | 16.26497 |
| **Calm3** | 1.31522 | 4.32037 | 0.69534 | 3.55465 | 4.17911 | 2.42659 | 0.20912 | 6.90360 | 0.03537 | 12.24380 | 0.78875 | 6.96315 | 2.88826 | 0.93756 |
| **Calm4** | 0.21016 | 0.00451 | 0.74339 | 0.00000 | 0.49795 | 0.23597 | 0.00393 | 0.00353 | 0.00713 | 0.00634 | 0.00396 | 0.00830 | 0.00119 | 1.01267 |
| **Calm5** | 0.34844 | 0.00466 | 1.50049 | 0.00000 | 0.83464 | 0.44237 | 0.00384 | 0.00203 | 0.00577 | 0.00556 | 0.00310 | 0.00596 | 0.00000 | 1.88063 |
| **Calml4** | 0.79370 | 0.00414 | 0.00000 | 0.36013 | 0.95037 | 0.37137 | 0.00403 | 0.00412 | 0.00638 | 0.01484 | 0.00300 | 0.00724 | 0.08437 | 1.50759 |
| **Caln1** | 0.00782 | 0.00477 | 0.00782 | 0.00778 | 0.24461 | 0.05412 | 0.00425 | 0.00804 | 0.00761 | 0.00690 | 0.00833 | 0.00870 | 0.04747 | 0.01401 |
| **Calr** | 6.25095 | 3.21860 | 5.39572 | 29.87701 | 4.31550 | 25.25949 | 7.49330 | 13.62252 | 6.60668 | 6.86601 | 6.71356 | 13.72575 | 10.68007 | 5.78594 |
| **Calr3** | 0.00161 | 0.00450 | 0.26054 | 0.00000 | 0.00000 | 0.05109 | 0.00408 | 0.00583 | 0.33340 | 0.00619 | 0.00539 | 0.00716 | 0.00795 | 1.15685 |
| **Calr4** | 0.29326 | 0.00379 | 0.78332 | 0.39462 | 0.12994 | 0.69545 | 0.00320 | 0.00000 | 2.29034 | 0.02056 | 0.04664 | 0.06230 | 0.92036 | 0.54819 |
| **Calu** | 8.11190 | 3.30657 | 6.08560 | 9.36273 | 2.68166 | 12.34009 | 0.33777 | 5.66668 | 15.27601 | 1.07858 | 10.21973 | 6.68733 | 3.32408 | 12.98979 |
| **Camk1** | 0.00000 | 4.26220 | 3.24489 | 4.70746 | 0.22205 | 0.78372 | 3.42179 | 7.10932 | 0.00000 | 2.57592 | 0.53495 | 0.39860 | 0.00974 | 0.11505 |
| **Camk1d** | 0.00664 | 0.00428 | 0.01255 | 0.01310 | 0.02635 | 0.01208 | 0.00418 | 0.00881 | 0.01237 | 0.00649 | 0.01215 | 0.00837 | 0.62952 | 0.07057 |
| **Camk2a** | 0.02342 | 0.05194 | 0.02139 | 0.07750 | 0.03590 | 0.02195 | 0.02217 | 0.03514 | 0.11972 | 0.00655 | 0.01399 | 0.04494 | 0.00642 | 0.04876 |
| **Camk2b** | 0.24337 | 0.04394 | 0.02199 | 0.56282 | 0.73122 | 0.37011 | 0.00369 | 2.17372 | 0.04459 | 0.00576 | 0.03341 | 0.11935 | 0.00385 | 0.09882 |
| **Camk2d** | 0.54259 | 0.00160 | 1.61575 | 1.56079 | 0.30667 | 2.23259 | 0.67259 | 0.22162 | 0.96505 | 0.00000 | 3.05788 | 2.59102 | 0.00000 | 0.00000 |
| **Camk2g** | 0.23431 | 0.05234 | 6.55635 | 2.65247 | 5.14577 | 2.32680 | 0.03255 | 1.33229 | 0.91393 | 0.01717 | 4.96838 | 4.23909 | 3.92067 | 5.38567 |
| **Camk2n2** | 0.72741 | 0.00425 | 0.04871 | 0.06642 | 0.00000 | 0.00000 | 0.00423 | 0.01400 | 0.08859 | 0.00745 | 0.17424 | 0.00639 | 0.37597 | 0.06415 |
| **Camk4** | 0.18810 | 0.05162 | 0.06703 | 0.31705 | 0.13661 | 0.17641 | 0.01057 | 0.50792 | 0.32889 | 0.15850 | 0.04030 | 0.01858 | 0.51959 | 0.31764 |
| **Camkk2** | 0.02811 | 0.00415 | 0.01800 | 0.07044 | 0.33270 | 0.00906 | 0.00411 | 0.00671 | 0.00534 | 0.00624 | 0.00697 | 0.01275 | 0.67555 | 0.28286 |
| **Camkmt** | 0.76140 | 0.00306 | 0.67698 | 0.06617 | 0.17912 | 0.18157 | 0.00886 | 0.45181 | 0.00457 | 0.02201 | 0.64860 | 0.85989 | 0.11411 | 0.00000 |
| **Caml** | 5.10351 | 1.26882 | 6.79946 | 4.80301 | 11.50430 | 6.57493 | 1.48101 | 2.32952 | 1.05067 | 0.03738 | 2.13395 | 7.39099 | 8.38889 | 3.10104 |
| **Camp** | 0.23534 | 0.00444 | 0.00000 | 0.01353 | 0.62631 | 0.00940 | 0.00427 | 0.00996 | 0.02362 | 0.00795 | 0.00678 | 0.00723 | 0.02271 | 0.29676 |
| **Camsap1** | 6.07182 | 0.05377 | 8.27624 | 2.92038 | 7.10980 | 6.30341 | 5.80507 | 1.30674 | 2.93810 | 0.22786 | 6.73710 | 5.02670 | 6.31632 | 13.09747 |
| **Camsap2** | 4.52634 | 0.00122 | 1.12007 | 0.00000 | 1.85116 | 2.49884 | 2.21104 | 0.74437 | 3.13961 | 0.03392 | 0.62087 | 2.77401 | 5.01947 | 5.04519 |
| **Camsap3** | 0.00000 | 0.07940 | 0.13560 | 4.35231 | 0.05991 | 1.71421 | 2.66179 | 5.49292 | 0.01385 | 0.08354 | 0.00000 | 1.50491 | 0.18657 | 0.14581 |
| **Camta1** | 0.01173 | 0.00456 | 0.04226 | 0.03380 | 0.22607 | 0.16376 | 0.00517 | 0.23883 | 0.02540 | 0.00803 | 0.05068 | 0.15672 | 0.93724 | 0.03857 |
| **Camta2** | 0.00000 | 0.02762 | 0.10002 | 2.03075 | 0.59753 | 0.01062 | 0.01069 | 2.29159 | 0.00638 | 0.02587 | 0.00000 | 0.00646 | 0.08164 | 0.00000 |
| **Cand1** | 29.75030 | 4.94763 | 36.11988 | 20.57931 | 37.95929 | 21.03717 | 5.47630 | 3.32110 | 32.29681 | 1.09418 | 21.57751 | 12.77450 | 17.25044 | 31.03034 |
| **Cand2** | 0.00484 | 0.05000 | 0.03676 | 0.11126 | 0.24147 | 0.11222 | 0.04574 | 0.05299 | 0.24622 | 0.03410 | 0.01418 | 0.45392 | 2.28289 | 0.56047 |
| **Cant1** | 0.30464 | 0.03189 | 2.32502 | 2.20625 | 1.69103 | 2.50873 | 0.79604 | 2.42183 | 0.01598 | 0.08345 | 0.13251 | 2.85731 | 1.70750 | 0.52604 |
| **Canx** | 11.54439 | 8.80419 | 12.39127 | 11.91525 | 11.23921 | 27.03691 | 4.82923 | 8.65223 | 31.83394 | 6.90491 | 11.81344 | 11.13705 | 2.88146 | 14.51992 |
| **Cap1** | 0.11455 | 0.01060 | 0.44415 | 0.98575 | 1.49251 | 0.00000 | 0.65256 | 3.11046 | 0.56045 | 0.03270 | 1.51373 | 0.51923 | 0.00000 | 3.09685 |
| **Capg** | 0.00000 | 0.22784 | 1.33939 | 0.53682 | 0.42925 | 0.46441 | 2.32159 | 16.53394 | 0.00000 | 4.21799 | 0.00000 | 4.03627 | 0.00000 | 0.20379 |
| **Capn1** | 0.36215 | 0.02809 | 0.10265 | 1.37962 | 1.17791 | 0.11214 | 0.02263 | 0.95762 | 0.01278 | 0.00556 | 0.62873 | 0.78010 | 1.77142 | 0.61269 |
| **Capn10** | 0.29685 | 0.03336 | 3.35669 | 3.06451 | 1.67728 | 1.95202 | 0.04903 | 1.29984 | 0.00467 | 0.02061 | 3.90924 | 3.60969 | 1.45135 | 1.34910 |
| **Capn11** | 0.02889 | 0.00391 | 0.25379 | 0.17346 | 1.67297 | 0.00000 | 0.00402 | 0.00487 | 0.00973 | 0.00613 | 0.00554 | 0.20157 | 0.34403 | 0.19774 |
| **Capn15** | 0.00000 | 0.02963 | 1.74777 | 0.90288 | 0.56181 | 0.24229 | 0.46597 | 0.71535 | 0.00795 | 0.00844 | 0.00000 | 0.89423 | 0.00000 | 0.28979 |
| **Capn2** | 0.08450 | 0.07521 | 0.23960 | 0.76910 | 0.00000 | 3.87602 | 0.00296 | 6.98238 | 0.00755 | 0.04277 | 0.05856 | 0.00000 | 0.00661 | 0.00000 |
| **Capn5** | 0.00000 | 0.01905 | 0.24336 | 1.91114 | 0.00000 | 3.08206 | 0.00312 | 2.29317 | 0.00819 | 0.02943 | 1.83037 | 1.09607 | 0.00616 | 0.00000 |
| **Capn6** | 0.00352 | 0.01259 | 0.22060 | 0.44042 | 0.00167 | 0.09622 | 0.00397 | 0.00676 | 0.01273 | 0.00589 | 0.00718 | 0.85498 | 0.00625 | 0.00724 |

| | | | | | | | | | | | | | | |
|---|---|---|---|---|---|---|---|---|---|---|---|---|---|---|
| **Capn7** | 2.94739 | 0.01143 | 0.96108 | 0.92531 | 0.15064 | 1.77518 | 0.01099 | 1.14217 | 3.29435 | 0.01606 | 0.08246 | 0.85054 | 0.13841 | 0.05311 |
| **Capns1** | 0.08516 | 5.52451 | 5.28265 | 8.02113 | 1.31871 | 22.11331 | 6.86948 | 23.08019 | 0.16677 | 13.60208 | 10.14528 | 16.43256 | 1.77359 | 0.47073 |
| **Caprin1** | 13.11002 | 6.79527 | 21.13367 | 11.37897 | 17.89584 | 20.88578 | 6.43254 | 12.55547 | 24.32610 | 0.37247 | 16.04750 | 16.43104 | 9.03221 | 15.36853 |
| **Caprin2** | 1.57715 | 0.06937 | 0.36299 | 1.60231 | 2.20626 | 0.52431 | 2.31278 | 0.00000 | 2.85826 | 0.04955 | 4.51704 | 5.18707 | 4.77254 | 4.21185 |
| **Capsl** | 0.00709 | 0.00441 | 0.01414 | 0.01225 | 0.07228 | 0.01495 | 0.00439 | 0.01100 | 0.00561 | 0.00652 | 0.00844 | 0.00988 | 0.00592 | 0.13274 |
| **Capza1** | 11.27195 | 3.96032 | 11.65182 | 6.48822 | 9.69039 | 18.68251 | 3.77692 | 7.28581 | 20.37930 | 0.21331 | 11.05941 | 7.64041 | 3.17000 | 5.96174 |
| **Capza2** | 9.52880 | 0.66055 | 18.74021 | 9.29240 | 15.47473 | 16.08896 | 1.59132 | 9.91086 | 25.25543 | 0.09442 | 13.70614 | 6.21570 | 2.49125 | 10.87247 |
| **Capzb** | 5.45679 | 4.81053 | 17.93318 | 14.66072 | 9.06976 | 17.63852 | 6.21865 | 24.70224 | 0.15398 | 6.48586 | 7.58048 | 14.90880 | 0.00000 | 2.35537 |
| **Car2** | 0.01171 | 0.00445 | 0.00881 | 0.01833 | 0.29751 | 0.01336 | 0.00426 | 0.01011 | 0.01158 | 0.00659 | 0.00857 | 0.00806 | 0.00479 | 0.01323 |
| **Car4** | 0.46591 | 2.74675 | 5.37567 | 1.00265 | 0.26079 | 1.70634 | 0.00241 | 0.35226 | 0.81865 | 0.00000 | 0.01611 | 0.00795 | 0.00000 | 1.06035 |
| **Car5a** | 0.03941 | 0.01871 | 0.06031 | 0.00033 | 0.05453 | 0.19969 | 0.00976 | 0.15152 | 0.05528 | 0.02662 | 0.00757 | 0.21141 | 0.13717 | 0.10832 |
| **Car5b** | 0.53946 | 0.11665 | 1.02976 | 0.56647 | 0.65049 | 1.10342 | 0.08558 | 0.24168 | 0.83237 | 0.09093 | 0.59250 | 0.29670 | 1.31360 | 0.60446 |
| **Car6** | 0.00804 | 0.00416 | 0.00942 | 0.13817 | 0.00517 | 0.00766 | 0.00399 | 0.00636 | 0.02081 | 0.00629 | 0.00664 | 0.01472 | 0.20762 | 0.10729 |
| **Car8** | 0.06449 | 0.03663 | 0.00651 | 0.13057 | 0.03893 | 0.20079 | 0.04652 | 0.04289 | 0.03925 | 0.02590 | 0.00535 | 0.03212 | 0.22109 | 0.12545 |
| **Car9** | 0.00000 | 0.00178 | 0.31328 | 0.04150 | 0.20484 | 0.86687 | 0.00298 | 0.00000 | 0.03188 | 0.00247 | 1.32391 | 2.59648 | 0.91552 | 0.00000 |
| **Card10** | 0.82767 | 0.00131 | 0.65384 | 0.44772 | 0.53717 | 0.44460 | 2.22865 | 0.00000 | 0.00511 | 0.09143 | 1.04600 | 3.40514 | 0.23552 | 0.00000 |
| **Card11** | 0.02127 | 0.00425 | 0.80405 | 0.02366 | 0.01301 | 0.00275 | 0.00390 | 0.01268 | 0.00923 | 0.02188 | 0.00590 | 0.00639 | 0.04343 | 0.00697 |
| **Card14** | 0.04405 | 0.00451 | 0.25023 | 0.00000 | 0.65051 | 0.02352 | 0.00418 | 0.00894 | 0.00639 | 0.00644 | 0.00615 | 0.00783 | 0.00249 | 0.09081 |
| **Card19** | 0.01761 | 0.00418 | 0.01056 | 0.00000 | 0.14402 | 0.01306 | 0.01044 | 0.26396 | 0.02099 | 0.01806 | 0.00766 | 0.01467 | 0.00542 | 0.00566 |
| **Card6** | 1.19010 | 0.00282 | 0.16468 | 0.00000 | 2.62118 | 0.00000 | 0.00318 | 0.18040 | 2.14808 | 0.00508 | 0.00000 | 0.71109 | 6.61093 | 2.01081 |
| **Card9** | 0.00587 | 0.00411 | 0.01832 | 0.32617 | 0.00864 | 0.00543 | 0.00395 | 0.00595 | 0.01200 | 0.00569 | 0.00634 | 0.01888 | 0.01016 | 0.01395 |
| **Carf** | 0.80706 | 0.09654 | 0.54581 | 0.11554 | 0.10257 | 0.18522 | 0.24498 | 0.12422 | 1.93846 | 0.03707 | 0.03719 | 0.10256 | 0.78283 | 0.47343 |
| **Carhsp1** | 0.97868 | 0.09650 | 4.70393 | 4.39506 | 1.64389 | 5.51179 | 3.30013 | 7.76836 | 1.99840 | 6.80076 | 2.64349 | 5.14229 | 6.85590 | 5.86863 |
| **Carkd** | 1.09322 | 10.23183 | 5.93825 | 6.98005 | 1.90119 | 3.41534 | 0.54914 | 4.47127 | 0.00000 | 0.71762 | 6.99746 | 6.19804 | 0.00000 | 1.13936 |
| **Carm1** | 0.00000 | 0.04847 | 2.95938 | 3.07044 | 1.50097 | 0.80036 | 0.41446 | 1.98944 | 0.00095 | 0.02877 | 2.20611 | 1.67416 | 0.09233 | 0.07595 |
| **Carnmt1** | 6.12953 | 0.01031 | 2.46124 | 6.39720 | 2.67559 | 6.64701 | 1.50151 | 1.87212 | 2.05800 | 0.10487 | 1.06049 | 0.56072 | 3.83722 | 2.32147 |
| **Carns1** | 0.00000 | 0.00389 | 0.85202 | 0.00000 | 1.44319 | 0.00000 | 0.00913 | 0.49314 | 0.72155 | 0.00883 | 0.00401 | 0.00559 | 0.75925 | 0.00000 |
| **Cars** | 4.08399 | 0.10475 | 6.07879 | 5.78452 | 5.15278 | 4.77211 | 1.06329 | 4.61998 | 0.01837 | 0.09466 | 10.75573 | 5.45791 | 0.25342 | 4.70988 |
| **Cars2** | 4.27576 | 0.00225 | 5.62771 | 2.03122 | 5.29298 | 3.38715 | 0.01353 | 0.52928 | 0.05080 | 0.00000 | 2.45800 | 0.72974 | 0.22428 | 3.07762 |
| **Casc1** | 0.00961 | 0.00439 | 0.00784 | 0.01156 | 0.27594 | 0.01259 | 0.00449 | 0.01047 | 0.03763 | 0.00609 | 0.00858 | 0.00850 | 0.23478 | 0.00986 |
| **Casc3** | 7.99509 | 1.39626 | 10.00606 | 5.04416 | 12.20158 | 5.86883 | 2.63700 | 3.44566 | 2.93410 | 0.07018 | 4.23449 | 4.58824 | 7.43552 | 10.18546 |
| **Casc4** | 6.86259 | 0.22583 | 5.21916 | 3.46199 | 2.23141 | 9.87458 | 1.68289 | 6.02850 | 13.52467 | 1.90433 | 2.83634 | 4.19433 | 17.37416 | 6.63443 |
| **Casc5** | 0.54386 | 0.00366 | 0.49997 | 0.24683 | 0.66517 | 0.23911 | 0.00911 | 0.07209 | 6.82454 | 0.00575 | 0.50859 | 0.00551 | 0.60255 | 2.03967 |
| **Casd1** | 0.89967 | 0.00300 | 0.00000 | 0.35992 | 0.00000 | 1.46264 | 0.00318 | 1.36350 | 0.01648 | 0.00902 | 0.00000 | 1.15398 | 0.01170 | 0.00000 |
| **Cask** | 0.77422 | 0.00406 | 0.94120 | 0.00000 | 0.18035 | 0.12353 | 0.00388 | 0.00001 | 0.00279 | 0.00295 | 0.16796 | 0.28673 | 2.09988 | 2.21910 |
| **Caskin1** | 2.68885 | 0.07684 | 5.23283 | 2.96296 | 7.43916 | 0.84304 | 0.24403 | 5.41209 | 0.01856 | 1.93074 | 1.70056 | 4.14936 | 1.07353 | 1.51791 |
| **Caskin2** | 0.22147 | 7.75969 | 4.64908 | 1.68133 | 0.40635 | 0.82763 | 1.96995 | 0.22444 | 0.00000 | 0.09144 | 1.12141 | 2.26099 | 1.39608 | 0.28844 |
| **Casp2** | 1.02203 | 0.01160 | 0.59928 | 1.21955 | 0.15158 | 1.83274 | 0.03010 | 0.66389 | 2.63810 | 0.00505 | 0.51572 | 2.65551 | 2.05313 | 0.42129 |
| **Casp3** | 3.52544 | 0.00000 | 8.49057 | 1.11492 | 7.26729 | 2.00447 | 0.00000 | 0.00000 | 9.95601 | 4.94029 | 4.88108 | 3.95338 | 7.55254 | 12.13054 |
| **Casp6** | 6.81331 | 0.22580 | 19.13333 | 10.80353 | 10.19600 | 10.45661 | 5.03289 | 3.44950 | 3.22799 | 0.28899 | 7.89398 | 9.86607 | 0.30039 | 11.57806 |
| **Casp7** | 0.24890 | 0.01748 | 0.77127 | 0.99752 | 0.73245 | 0.68795 | 0.00908 | 2.25578 | 0.01511 | 0.00390 | 2.94041 | 2.47854 | 0.07779 | 0.05187 |
| **Casp8ap2** | 1.46083 | 0.00193 | 2.38370 | 0.00770 | 3.82583 | 2.92796 | 3.00941 | 1.48318 | 21.70298 | 0.04086 | 1.49826 | 0.46596 | 4.18749 | 7.72563 |
| **Casp9** | 1.22753 | 0.00820 | 1.33718 | 1.58428 | 0.98624 | 2.79893 | 0.00220 | 1.29885 | 0.03609 | 0.03230 | 2.53524 | 4.23335 | 0.56563 | 0.04370 |
| **Casq1** | 0.00877 | 0.00449 | 0.01006 | 0.01225 | 0.17412 | 0.01830 | 0.00452 | 0.00973 | 0.01103 | 0.00653 | 0.00842 | 0.00909 | 0.02831 | 0.00823 |
| **Casq2** | 0.02451 | 0.00434 | 0.01171 | 0.01241 | 0.10091 | 0.01307 | 0.00440 | 0.01076 | 0.01131 | 0.00629 | 0.00800 | 0.02502 | 0.00517 | 0.03896 |
| **Cass4** | 0.00000 | 0.00450 | 0.13955 | 0.00000 | 1.75680 | 0.00000 | 0.00433 | 0.01487 | 0.04084 | 0.00358 | 0.55880 | 0.02086 | 0.43378 | 0.72837 |
| **Cast** | 0.34494 | 0.05405 | 1.26114 | 1.95685 | 0.45845 | 1.03370 | 4.26749 | 0.13189 | 2.02349 | 0.13582 | 2.13363 | 1.37414 | 1.03870 | 1.13053 |
| **Casz1** | 0.01610 | 0.00431 | 0.25092 | 0.01214 | 0.18142 | 0.53850 | 0.00418 | 0.01014 | 0.00776 | 0.01408 | 0.00407 | 0.08683 | 0.05572 | 0.07842 |
| **Cat** | 0.00000 | 0.04105 | 5.98858 | 5.16897 | 0.00000 | 6.08715 | 5.82799 | 1.14354 | 0.00776 | 0.19506 | 9.16507 | 5.60704 | 0.00000 | 0.00000 |
| **Catip** | 0.00939 | 0.00442 | 0.01678 | 0.01205 | 0.01172 | 0.01299 | 0.00437 | 0.01008 | 0.01105 | 0.00694 | 0.00875 | 0.01386 | 0.00833 | 0.04835 |
| **Catsper2** | 0.66342 | 0.00405 | 0.01745 | 0.26432 | 0.23800 | 0.37383 | 0.00395 | 0.00013 | 0.07053 | 0.00959 | 0.81074 | 0.00646 | 0.41126 | 0.00000 |
| **Catsperd** | 0.00000 | 0.00385 | 0.00000 | 0.25582 | 1.54399 | 0.00000 | 2.66604 | 0.00272 | 2.09821 | 0.03187 | 0.00040 | 0.00566 | 1.04528 | 2.05055 |
| **Catsperg1** | 1.57036 | 0.48780 | 0.46076 | 0.00000 | 0.00000 | 0.07000 | 0.00378 | 0.00109 | 0.00479 | 0.00407 | 0.00505 | 0.19074 | 1.21160 | 0.28092 |
| **Catsperg2** | 0.51101 | 0.09846 | 0.11255 | 0.01214 | 0.00000 | 0.03669 | 0.00418 | 0.00406 | 0.00625 | 0.00530 | 0.00675 | 0.05460 | 0.28908 | 0.16666 |
| **Cav1** | 0.24192 | 0.00422 | 0.00891 | 0.07535 | 0.00365 | 0.00490 | 0.00422 | 0.00923 | 0.01763 | 0.00598 | 0.01063 | 0.01257 | 0.00333 | 0.86592 |

| | | | | | | | | | | | | | | |
|---|---|---|---|---|---|---|---|---|---|---|---|---|---|---|
| **Cav2** | 0.38453 | 0.00441 | 0.00000 | 0.01661 | 0.08566 | 0.00600 | 0.00397 | 0.00940 | 1.08815 | 0.00904 | 0.00491 | 0.00723 | 0.01048 | 0.34154 |
| **Cbarp** | 0.62150 | 0.01768 | 1.75172 | 1.37415 | 0.22734 | 0.44535 | 0.00317 | 1.23080 | 0.00299 | 0.06597 | 0.05073 | 0.76760 | 0.00000 | 0.04277 |
| **Cbfa2t2** | 3.22084 | 0.10380 | 2.03374 | 3.68918 | 5.92900 | 3.21550 | 0.02488 | 1.48087 | 7.67930 | 0.03393 | 0.21990 | 5.03139 | 7.73547 | 5.73581 |
| **Cbfa2t3** | 0.00000 | 0.00415 | 0.69323 | 0.76005 | 0.07760 | 0.40205 | 0.00455 | 0.05025 | 0.01623 | 0.00843 | 0.00086 | 0.45774 | 0.02067 | 0.00000 |
| **Cbfb** | 3.39318 | 0.04848 | 2.66812 | 0.75998 | 2.11404 | 4.13454 | 0.00710 | 1.12804 | 9.20380 | 0.28773 | 2.92884 | 5.64073 | 6.08026 | 5.82628 |
| **Cbl** | 2.77664 | 0.00919 | 2.14483 | 0.00000 | 2.49699 | 2.09625 | 0.00699 | 0.00000 | 13.48665 | 0.01488 | 0.03067 | 1.42499 | 4.16281 | 4.74083 |
| **Cblb** | 0.04422 | 0.01145 | 0.02232 | 0.48085 | 0.00282 | 0.89470 | 0.00378 | 0.09460 | 0.01190 | 0.00682 | 0.00000 | 1.50118 | 0.00816 | 0.00000 |
| **Cblc** | 0.00437 | 0.01215 | 0.07107 | 1.21681 | 0.59792 | 0.04532 | 0.00425 | 0.45051 | 0.01457 | 0.01908 | 0.00167 | 0.01396 | 0.01257 | 0.09994 |
| **Cbll1** | 5.01630 | 0.02679 | 4.76414 | 2.39812 | 6.41096 | 3.18308 | 1.27436 | 0.73555 | 12.36595 | 0.06428 | 3.01130 | 0.87100 | 12.53325 | 6.44110 |
| **Cbln1** | 0.43937 | 0.00358 | 0.24472 | 0.43368 | 0.29304 | 0.07976 | 0.00382 | 0.00308 | 0.00926 | 0.00568 | 0.00269 | 0.90402 | 0.07913 | 0.26305 |
| **Cbln3** | 0.46973 | 0.16057 | 0.31875 | 0.76536 | 0.58519 | 1.74608 | 0.26260 | 0.38540 | 0.42886 | 0.42493 | 0.03558 | 0.33285 | 1.17674 | 1.07271 |
| **Cbr1** | 4.83058 | 0.11172 | 17.98798 | 19.79830 | 8.68529 | 3.33235 | 0.86189 | 6.32047 | 1.16848 | 0.04225 | 5.92956 | 8.40090 | 1.55188 | 1.52755 |
| **Cbr2** | 0.00000 | 0.01734 | 7.64057 | 9.52230 | 6.55016 | 6.36280 | 0.00230 | 0.04020 | 0.02122 | 0.05659 | 5.34794 | 1.05029 | 0.45710 | 1.52578 |
| **Cbr3** | 0.01055 | 0.00424 | 0.39081 | 0.30904 | 0.00563 | 0.00000 | 0.00414 | 0.01443 | 0.01217 | 0.00805 | 0.01268 | 0.02015 | 0.00916 | 0.00655 |
| **Cbr4** | 1.15421 | 0.03897 | 0.37433 | 0.18537 | 0.13980 | 2.30093 | 0.00865 | 3.87693 | 0.00226 | 0.01567 | 3.16417 | 0.91977 | 0.00407 | 0.76461 |
| **Cbs** | 0.82145 | 4.02242 | 0.71911 | 0.26943 | 1.11478 | 2.78585 | 5.22755 | 16.52454 | 0.00428 | 0.33589 | 1.95509 | 0.00000 | 0.06837 | 0.05788 |
| **Cbwd1** | 9.51453 | 1.48166 | 2.44885 | 0.00000 | 1.61540 | 0.45692 | 0.18514 | 0.00984 | 14.52501 | 0.00221 | 3.68636 | 0.42831 | 3.47904 | 7.80458 |
| **Cbx1** | 2.48831 | 2.88087 | 1.67206 | 2.40569 | 1.01463 | 2.78979 | 1.15057 | 2.23640 | 0.79676 | 0.01015 | 1.93309 | 3.21292 | 0.56377 | 0.17295 |
| **Cbx2** | 0.00050 | 0.00315 | 1.37308 | 0.00000 | 0.00000 | 1.39603 | 0.00359 | 0.00000 | 0.00000 | 0.02016 | 0.00000 | 0.96088 | 4.17485 | 5.49722 |
| **Cbx3** | 19.78685 | 0.07287 | 3.44867 | 4.01157 | 9.94507 | 5.99923 | 2.04653 | 5.67526 | 23.77075 | 0.18920 | 4.17396 | 7.35103 | 11.40786 | 7.40724 |
| **Cbx4** | 0.34182 | 0.00444 | 0.01068 | 0.00765 | 0.02551 | 0.00538 | 0.00579 | 0.21672 | 0.01882 | 0.00981 | 0.00642 | 0.01131 | 0.45072 | 0.00328 |
| **Cbx5** | 10.09996 | 0.10384 | 14.39360 | 5.05526 | 14.77993 | 12.75878 | 3.29441 | 8.13420 | 19.65591 | 8.11238 | 10.19392 | 10.02979 | 6.01744 | 13.56413 |
| **Cbx6** | 0.01800 | 0.01947 | 0.00000 | 0.13042 | 0.02291 | 0.01055 | 0.00357 | 0.00591 | 0.15598 | 0.00639 | 0.00961 | 1.49856 | 0.07596 | 0.09675 |
| **Cbx7** | 1.14226 | 0.06992 | 8.67045 | 5.34935 | 2.41204 | 4.32210 | 1.63421 | 3.20600 | 3.17913 | 0.07163 | 4.97478 | 4.79917 | 2.77248 | 5.02440 |
| **Cbx8** | 0.09355 | 0.00392 | 0.03905 | 0.00000 | 0.10579 | 0.00000 | 0.00475 | 0.11262 | 0.00618 | 0.00356 | 0.96003 | 0.18407 | 1.25726 | 0.04538 |
| **Cby1** | 0.03914 | 0.02500 | 1.13722 | 0.00681 | 0.78318 | 1.85103 | 0.00814 | 4.10960 | 0.22403 | 0.01937 | 2.83674 | 1.55302 | 2.13168 | 1.04862 |
| **Cc2d1a** | 0.00000 | 0.01818 | 1.60269 | 5.13974 | 2.43229 | 0.59930 | 0.04573 | 0.70254 | 0.00199 | 12.89376 | 1.18659 | 0.55058 | 0.12791 | 0.00000 |
| **Cc2d1b** | 0.79836 | 0.02388 | 1.71651 | 7.88003 | 0.33662 | 6.39912 | 0.30468 | 3.97692 | 0.00000 | 0.06136 | 0.00000 | 3.42263 | 0.70092 | 0.60375 |
| **Ccar1** | 13.19388 | 3.61851 | 13.04959 | 5.93250 | 15.30547 | 8.44527 | 3.79248 | 1.14426 | 29.85292 | 12.51296 | 9.78289 | 4.05575 | 8.86204 | 20.32554 |
| **Ccar2** | 0.06060 | 0.01365 | 2.21894 | 3.39708 | 0.29501 | 0.15787 | 0.00400 | 1.16193 | 0.06074 | 0.02408 | 0.35281 | 0.03423 | 0.34778 | 0.71755 |
| **Ccbe1** | 0.03704 | 0.00435 | 0.01453 | 0.01190 | 0.00956 | 0.01326 | 0.00426 | 0.00943 | 0.00982 | 0.00672 | 0.00842 | 0.00914 | 0.30931 | 0.00506 |
| **Ccdc106** | 0.00962 | 0.00433 | 0.02517 | 0.04146 | 0.01141 | 0.00935 | 0.00422 | 0.00846 | 0.01146 | 0.00618 | 0.01911 | 0.01300 | 0.05916 | 0.00759 |
| **Ccdc107** | 0.04614 | 0.03452 | 0.06643 | 4.66646 | 0.80203 | 0.00000 | 0.00428 | 1.34107 | 0.01065 | 0.02501 | 0.01590 | 0.25665 | 1.83250 | 0.00000 |
| **Ccdc109b** | 0.00445 | 0.00436 | 0.16048 | 0.00000 | 0.23079 | 0.00824 | 0.00407 | 0.00998 | 0.00707 | 0.00688 | 0.00796 | 0.00924 | 0.31656 | 0.01723 |
| **Ccdc112** | 0.01119 | 0.00453 | 0.02509 | 0.00000 | 0.16399 | 1.65499 | 0.00386 | 0.00739 | 0.00683 | 0.00577 | 0.01625 | 0.11783 | 0.22501 | 0.04992 |
| **Ccdc114** | 0.00000 | 0.00376 | 0.51812 | 0.00000 | 0.94441 | 0.00020 | 0.00396 | 0.00335 | 0.01710 | 0.00507 | 0.00395 | 0.66272 | 0.01418 | 0.00734 |
| **Ccdc115** | 10.59130 | 0.04640 | 15.19931 | 13.05185 | 12.11026 | 12.25806 | 4.09082 | 2.87161 | 3.07954 | 0.16435 | 6.75185 | 3.69583 | 5.61564 | 4.87330 |
| **Ccdc116** | 0.52543 | 0.00401 | 0.09887 | 0.03505 | 0.80444 | 0.00601 | 0.00453 | 0.00693 | 0.04550 | 0.00565 | 0.00808 | 0.01030 | 1.32963 | 0.00000 |
| **Ccdc117** | 23.03054 | 4.15402 | 22.98182 | 2.17587 | 28.18043 | 9.77231 | 2.97253 | 3.74298 | 24.67966 | 0.11195 | 6.57389 | 6.80946 | 15.75745 | 27.26406 |
| **Ccdc12** | 3.74768 | 5.71511 | 5.25944 | 9.55173 | 10.53282 | 7.37508 | 4.17673 | 12.04090 | 0.15383 | 8.79414 | 2.66841 | 5.85843 | 3.92825 | 4.30160 |
| **Ccdc120** | 1.48805 | 4.94123 | 0.40407 | 0.00000 | 0.73511 | 0.00000 | 0.00352 | 0.88204 | 0.00970 | 0.00471 | 0.00419 | 0.02748 | 1.25641 | 0.49101 |
| **Ccdc124** | 0.98526 | 1.81109 | 29.90494 | 28.12253 | 26.67816 | 0.00000 | 5.44655 | 17.58507 | 0.00000 | 16.70219 | 7.27180 | 9.15432 | 2.08353 | 0.00000 |
| **Ccdc125** | 0.35929 | 0.00397 | 0.00000 | 0.02895 | 0.78772 | 0.04978 | 0.00420 | 0.01379 | 1.49274 | 0.00673 | 0.00652 | 0.03041 | 3.73708 | 1.15934 |
| **Ccdc126** | 2.48510 | 0.05273 | 2.96955 | 0.50185 | 0.90686 | 2.68682 | 0.46616 | 0.54767 | 1.37968 | 0.02655 | 2.22474 | 2.28542 | 5.48652 | 1.55711 |
| **Ccdc127** | 2.64944 | 6.37789 | 3.90467 | 3.50733 | 2.85606 | 8.18427 | 6.36469 | 4.10267 | 1.54266 | 0.23676 | 3.79141 | 4.33033 | 0.16732 | 6.21381 |
| **Ccdc13** | 0.00274 | 0.00364 | 0.00000 | 2.35579 | 0.00000 | 1.94732 | 0.00351 | 0.00000 | 0.01297 | 0.00000 | 2.85456 | 0.00724 | 0.00637 | 0.00000 |
| **Ccdc130** | 1.70743 | 4.00862 | 5.12366 | 5.52197 | 2.57728 | 4.67453 | 0.01565 | 1.39929 | 0.03191 | 0.03931 | 1.21760 | 2.06568 | 1.01854 | 0.59217 |
| **Ccdc134** | 1.34768 | 0.03906 | 1.90221 | 2.47918 | 1.60342 | 3.11769 | 1.53887 | 0.02198 | 0.19200 | 0.04206 | 3.32531 | 1.19565 | 1.75022 | 0.81106 |
| **Ccdc136** | 0.00103 | 0.00451 | 0.00419 | 0.06674 | 0.00000 | 0.01628 | 0.00461 | 0.03373 | 0.02733 | 0.00832 | 0.00896 | 0.00552 | 0.37701 | 1.06591 |
| **Ccdc137** | 8.17364 | 0.02179 | 10.06204 | 1.62819 | 12.95359 | 8.14065 | 2.67886 | 2.59173 | 9.34174 | 0.06894 | 4.74981 | 1.27973 | 14.10199 | 12.02838 |
| **Ccdc138** | 1.38476 | 0.00342 | 0.87605 | 0.00000 | 0.06470 | 1.13380 | 0.00338 | 0.00000 | 2.63830 | 0.00785 | 0.00000 | 0.53003 | 0.00059 | 0.00000 |
| **Ccdc14** | 0.39019 | 1.20369 | 1.16594 | 0.70778 | 0.91395 | 0.82011 | 0.14489 | 0.00000 | 1.06423 | 0.01086 | 0.07572 | 0.72507 | 0.77777 | 1.81571 |
| **Ccdc141** | 0.00704 | 0.00441 | 0.00884 | 0.02293 | 0.15046 | 0.01174 | 0.00424 | 0.00894 | 0.00725 | 0.00653 | 0.00830 | 0.01075 | 0.07227 | 0.08337 |
| **Ccdc142** | 0.01427 | 0.00461 | 0.01242 | 0.10537 | 0.01068 | 0.01708 | 0.00456 | 0.02404 | 0.01192 | 0.00805 | 0.01154 | 0.01451 | 0.00667 | 0.01075 |
| **Ccdc146** | 0.00830 | 0.03852 | 0.14060 | 0.00301 | 0.07666 | 0.00885 | 0.00422 | 0.01633 | 0.01108 | 0.00658 | 0.00710 | 0.01572 | 0.00683 | 0.01295 |

| | | | | | | | | | | | | | | |
|---|---|---|---|---|---|---|---|---|---|---|---|---|---|---|
| **Ccdc148** | 0.01405 | 0.00479 | 0.01889 | 0.03470 | 0.02353 | 0.03496 | 0.00451 | 0.02712 | 0.10321 | 0.01493 | 0.00764 | 0.00983 | 0.15722 | 0.05087 |
| **Ccdc15** | 0.43211 | 0.01188 | 0.06127 | 0.18228 | 0.14963 | 0.00000 | 0.00498 | 0.59251 | 0.00689 | 0.00000 | 1.94743 | 0.00298 | 0.24896 | 0.09057 |
| **Ccdc150** | 0.26911 | 0.02090 | 0.45612 | 0.00000 | 0.29698 | 0.00000 | 0.00354 | 1.48650 | 1.45919 | 0.00559 | 0.00000 | 0.48097 | 0.00000 | 0.00000 |
| **Ccdc152** | 0.83699 | 0.00400 | 0.00000 | 0.43910 | 0.18981 | 0.00000 | 0.00362 | 0.00494 | 0.91259 | 0.01282 | 0.00277 | 0.01282 | 0.00198 | 0.08572 |
| **Ccdc153** | 0.00427 | 0.00428 | 0.00000 | 0.02986 | 1.07338 | 0.06053 | 0.01016 | 0.01134 | 0.01392 | 0.00653 | 0.00959 | 0.00900 | 0.03586 | 0.25933 |
| **Ccdc154** | 0.28770 | 0.00421 | 0.23399 | 0.00000 | 0.23519 | 0.02132 | 0.00421 | 0.00888 | 0.01830 | 0.00697 | 0.00837 | 0.00853 | 0.50336 | 0.19236 |
| **Ccdc155** | 0.51299 | 0.00368 | 1.18498 | 0.22114 | 1.33589 | 0.69020 | 0.00357 | 0.00000 | 0.00652 | 1.08543 | 0.00000 | 0.00000 | 0.01043 | 0.71377 |
| **Ccdc157** | 0.00095 | 0.00439 | 0.00464 | 0.07589 | 0.09804 | 0.26843 | 0.00448 | 0.00642 | 0.05746 | 0.00519 | 0.00571 | 0.00544 | 3.01826 | 0.01376 |
| **Ccdc158** | 0.01284 | 0.00434 | 0.02136 | 0.10101 | 0.01023 | 0.00741 | 0.00420 | 0.00838 | 0.03528 | 0.00659 | 0.00675 | 0.01285 | 0.32314 | 0.01585 |
| **Ccdc160** | 0.12403 | 0.38251 | 0.09016 | 0.01770 | 0.00000 | 0.01239 | 0.00389 | 0.00802 | 0.71581 | 0.00681 | 0.00626 | 0.03047 | 0.01292 | 0.03417 |
| **Ccdc163** | 6.31424 | 0.00818 | 6.46175 | 4.63204 | 6.81238 | 2.77308 | 2.53988 | 0.01841 | 4.63548 | 0.02747 | 2.49073 | 1.61778 | 3.15811 | 3.45910 |
| **Ccdc166** | 0.11159 | 0.00379 | 0.28839 | 0.88504 | 0.85941 | 0.00000 | 0.00356 | 0.66336 | 0.00870 | 0.00452 | 0.00251 | 0.00607 | 0.17577 | 0.00000 |
| **Ccdc167** | 0.00000 | 0.00469 | 0.13724 | 0.12870 | 0.04420 | 0.00000 | 0.00406 | 0.01541 | 1.42893 | 0.01000 | 0.00448 | 0.02582 | 0.05762 | 0.17754 |
| **Ccdc169** | 0.00644 | 0.15717 | 0.01139 | 0.01016 | 0.00995 | 0.01580 | 0.00415 | 0.00714 | 0.02354 | 0.00702 | 0.00519 | 0.32552 | 0.01055 | 0.04956 |
| **Ccdc17** | 0.00109 | 0.00379 | 0.53693 | 0.37528 | 0.14743 | 0.00000 | 0.00386 | 0.00157 | 0.43025 | 0.00579 | 0.00816 | 0.29348 | 0.00419 | 0.00371 |
| **Ccdc170** | 0.00000 | 0.00417 | 0.19738 | 0.25894 | 0.00000 | 0.03537 | 0.00386 | 0.00380 | 0.28156 | 0.00538 | 0.00520 | 0.00833 | 0.36494 | 0.01345 |
| **Ccdc171** | 0.00000 | 0.00402 | 0.23105 | 0.18924 | 0.00000 | 0.00000 | 0.00382 | 0.00574 | 0.53369 | 0.00634 | 0.00397 | 0.01161 | 0.01284 | 4.28176 |
| **Ccdc172** | 0.57301 | 0.00446 | 0.03842 | 0.00672 | 0.35873 | 0.00703 | 0.00421 | 0.01172 | 0.00512 | 0.00728 | 0.00688 | 0.00630 | 0.00360 | 0.17355 |
| **Ccdc173** | 0.57976 | 0.00423 | 0.38748 | 0.44226 | 0.90780 | 0.21812 | 0.00484 | 0.16901 | 4.35027 | 0.00472 | 0.00376 | 0.00230 | 0.66226 | 1.50242 |
| **Ccdc174** | 3.11031 | 0.03457 | 7.27869 | 6.40995 | 6.18134 | 3.22805 | 3.11910 | 0.37495 | 1.04975 | 0.03890 | 1.68972 | 4.34571 | 2.98763 | 5.96441 |
| **Ccdc177** | 0.38997 | 0.00438 | 0.00776 | 0.01194 | 0.16264 | 0.00958 | 0.00445 | 0.01280 | 0.00955 | 0.00726 | 0.00710 | 0.00758 | 0.00784 | 0.00664 |
| **Ccdc18** | 0.00000 | 0.24612 | 0.00000 | 0.04337 | 0.29437 | 0.02177 | 0.00381 | 0.00760 | 3.46265 | 0.00729 | 0.00201 | 0.18500 | 0.33540 | 0.00000 |
| **Ccdc181** | 0.01542 | 0.00381 | 0.00381 | 0.00000 | 0.20234 | 0.00572 | 0.00514 | 0.84425 | 0.00753 | 0.01035 | 0.00000 | 0.90232 | 0.06270 | 1.09926 |
| **Ccdc183** | 0.07668 | 0.00407 | 0.10904 | 0.00000 | 0.04580 | 0.01558 | 0.00437 | 0.00748 | 0.00846 | 0.01002 | 0.00633 | 0.22443 | 0.07532 | 0.00511 |
| **Ccdc184** | 0.16705 | 0.00392 | 0.01194 | 0.00844 | 0.00724 | 0.06811 | 0.00386 | 0.13974 | 0.00975 | 0.02687 | 0.00509 | 0.00686 | 0.03471 | 0.00291 |
| **Ccdc186** | 0.01032 | 0.00410 | 0.01526 | 0.23581 | 0.06183 | 0.00000 | 0.00440 | 0.28393 | 1.06033 | 0.00571 | 0.00949 | 0.00893 | 0.66003 | 2.42709 |
| **Ccdc189** | 0.94976 | 0.01044 | 0.52641 | 0.00000 | 1.09095 | 0.71536 | 0.00387 | 0.00444 | 0.32321 | 0.01262 | 0.00000 | 1.68501 | 0.07004 | 1.13006 |
| **Ccdc191** | 0.00000 | 0.00261 | 0.72359 | 0.56785 | 0.25361 | 0.00000 | 0.00309 | 0.28677 | 0.00493 | 0.00000 | 5.08475 | 0.51534 | 0.25802 | 0.18597 |
| **Ccdc22** | 0.08145 | 0.00993 | 3.92813 | 5.95202 | 0.00000 | 4.54317 | 1.26759 | 0.86681 | 0.00000 | 0.01264 | 2.61117 | 1.94812 | 0.00000 | 0.01146 |
| **Ccdc24** | 0.01306 | 0.00444 | 0.01257 | 0.01096 | 0.09200 | 0.01343 | 0.00420 | 0.01009 | 0.01251 | 0.00645 | 0.00907 | 0.00944 | 0.06139 | 0.00537 |
| **Ccdc25** | 8.50357 | 0.00000 | 8.39061 | 3.37352 | 11.31141 | 7.40080 | 1.59467 | 1.83743 | 15.54457 | 5.69816 | 4.66767 | 2.63736 | 8.15804 | 14.85823 |
| **Ccdc28a** | 1.53959 | 0.07086 | 3.53108 | 1.06344 | 2.13370 | 2.97113 | 0.01400 | 1.93970 | 0.08199 | 0.03848 | 1.41839 | 2.40375 | 1.65846 | 5.72601 |
| **Ccdc28b** | 0.68875 | 0.03089 | 0.20145 | 0.43163 | 0.02105 | 0.12908 | 0.01054 | 2.53154 | 0.13985 | 0.68116 | 0.00000 | 0.20712 | 0.48477 | 0.00000 |
| **Ccdc30** | 0.02500 | 0.00261 | 0.00000 | 0.13051 | 0.41561 | 0.14445 | 0.00324 | 0.00000 | 0.05291 | 0.00995 | 0.55554 | 1.60011 | 0.19275 | 0.26920 |
| **Ccdc32** | 0.70946 | 0.05507 | 1.03218 | 0.47122 | 1.67160 | 2.48720 | 4.14499 | 5.88373 | 1.02239 | 0.08225 | 2.60430 | 0.00000 | 1.80191 | 2.40827 |
| **Ccdc34** | 0.04740 | 0.03191 | 0.85388 | 3.10922 | 1.78624 | 2.07651 | 0.00253 | 1.43424 | 0.40981 | 0.01350 | 2.15202 | 3.98588 | 0.45772 | 0.71043 |
| **Ccdc36** | 0.11706 | 0.01210 | 0.35704 | 0.02661 | 0.58228 | 0.32584 | 0.03621 | 0.07680 | 2.58711 | 0.03484 | 0.04242 | 0.06813 | 2.87177 | 0.03588 |
| **Ccdc38** | 0.12538 | 0.00426 | 0.12592 | 0.02695 | 0.05350 | 0.01750 | 0.00396 | 0.01460 | 0.01649 | 0.01095 | 0.00779 | 0.00876 | 0.12377 | 0.09645 |
| **Ccdc39** | 0.01023 | 0.00419 | 0.06386 | 0.08094 | 0.07826 | 0.00535 | 0.15339 | 0.00493 | 0.01044 | 0.01112 | 0.01738 | 0.01327 | 0.01271 | 0.03012 |
| **Ccdc40** | 0.13802 | 0.00393 | 1.99636 | 2.43227 | 2.10136 | 0.99414 | 0.00359 | 0.00000 | 0.00713 | 0.00647 | 0.82421 | 0.02016 | 0.20020 | 1.57226 |
| **Ccdc42** | 0.00267 | 0.01297 | 0.00690 | 0.00000 | 0.00655 | 0.15921 | 0.08700 | 0.77700 | 0.01111 | 2.42019 | 0.00000 | 0.01289 | 0.09904 | 0.00000 |
| **Ccdc43** | 5.64513 | 0.06718 | 12.47790 | 8.91797 | 5.07328 | 3.19716 | 0.11244 | 4.45955 | 12.72038 | 0.03941 | 11.36450 | 2.98024 | 0.00000 | 1.52437 |
| **Ccdc47** | 6.49567 | 0.03841 | 7.20429 | 2.96092 | 1.08857 | 6.48145 | 0.32226 | 3.95488 | 6.78907 | 0.04825 | 5.56513 | 4.46709 | 1.05738 | 8.68418 |
| **Ccdc50** | 2.58340 | 0.27029 | 0.25209 | 1.48777 | 0.63081 | 4.20709 | 1.32386 | 2.08244 | 0.82836 | 0.04328 | 0.01406 | 3.50865 | 2.38072 | 2.36109 |
| **Ccdc51** | 1.14999 | 0.00280 | 3.85223 | 0.59899 | 4.08399 | 1.39459 | 3.88389 | 0.00000 | 0.00000 | 0.21061 | 1.88076 | 1.37138 | 0.10941 | 2.64475 |
| **Ccdc53** | 4.97345 | 0.09143 | 3.89153 | 1.59595 | 2.51573 | 9.28499 | 3.87395 | 3.40663 | 2.97901 | 10.44536 | 6.48069 | 7.04227 | 0.00000 | 3.22180 |
| **Ccdc57** | 1.44120 | 0.00459 | 1.71291 | 0.35668 | 3.79515 | 0.00000 | 0.00381 | 0.12398 | 2.85494 | 0.00899 | 0.00399 | 0.06261 | 0.79946 | 2.08568 |
| **Ccdc58** | 6.43353 | 6.39532 | 2.44969 | 7.15433 | 10.07355 | 8.59515 | 6.72874 | 4.93078 | 5.52726 | 8.82983 | 2.83250 | 8.02617 | 2.68119 | 7.47200 |
| **Ccdc59** | 8.12466 | 0.08464 | 8.21035 | 7.90409 | 11.52459 | 6.42090 | 1.32212 | 1.61819 | 3.18029 | 12.45811 | 5.20022 | 7.08641 | 8.49360 | 5.23245 |
| **Ccdc6** | 3.30065 | 3.65246 | 2.97851 | 0.79163 | 6.24257 | 5.67564 | 0.00612 | 3.04180 | 12.50182 | 0.02914 | 3.94905 | 1.50124 | 12.90831 | 27.26527 |
| **Ccdc60** | 0.04276 | 0.00438 | 0.21802 | 0.03605 | 0.00000 | 0.06965 | 0.00409 | 0.01170 | 0.00463 | 0.00939 | 0.00644 | 0.00583 | 0.59735 | 0.56955 |
| **Ccdc61** | 0.71111 | 0.02156 | 0.10541 | 1.74354 | 1.82609 | 0.39080 | 2.03971 | 1.30127 | 0.02477 | 0.08540 | 0.00449 | 0.78140 | 0.53640 | 0.02508 |
| **Ccdc62** | 0.00394 | 0.00426 | 0.00196 | 0.00000 | 0.35066 | 0.45816 | 0.00400 | 0.00441 | 0.01877 | 0.00508 | 0.02553 | 0.32094 | 0.23332 | 0.00233 |
| **Ccdc63** | 0.29686 | 0.00405 | 0.55014 | 0.00000 | 1.42516 | 0.08235 | 0.00407 | 0.00194 | 0.05852 | 0.00471 | 0.08165 | 0.00586 | 2.09180 | 0.91007 |
| **Ccdc64** | 0.00000 | 0.00430 | 0.12512 | 0.50353 | 0.41176 | 0.21102 | 0.00471 | 0.19474 | 0.01075 | 0.00518 | 0.03920 | 0.00786 | 0.00536 | 0.03582 |

| | | | | | | | | | | | | | | |
|---|---|---|---|---|---|---|---|---|---|---|---|---|---|---|
| **Ccdc64b** | 0.04225 | 0.00437 | 0.48484 | 0.00000 | 0.62228 | 0.04479 | 0.00411 | 0.00637 | 0.00690 | 0.00597 | 0.00572 | 0.00723 | 0.18440 | 0.00474 |
| **Ccdc65** | 0.56916 | 0.00437 | 1.15142 | 0.00410 | 0.03776 | 0.00399 | 0.00381 | 0.00693 | 1.44807 | 0.00615 | 0.00508 | 0.00702 | 0.00000 | 0.26991 |
| **Ccdc66** | 0.00000 | 0.02551 | 2.04101 | 0.00000 | 2.06340 | 3.29002 | 4.47569 | 0.01412 | 1.91124 | 0.12916 | 1.49278 | 2.63931 | 0.63004 | 1.53061 |
| **Ccdc68** | 0.00462 | 0.02362 | 0.08435 | 0.00000 | 0.00162 | 0.11834 | 0.00397 | 0.56897 | 1.10809 | 0.02477 | 0.00438 | 0.00620 | 0.00489 | 0.00000 |
| **Ccdc69** | 0.03166 | 0.00416 | 0.00458 | 0.00937 | 0.45714 | 0.01128 | 0.00409 | 0.00802 | 0.01335 | 0.00617 | 0.00788 | 0.00692 | 1.31333 | 0.53842 |
| **Ccdc71** | 1.07444 | 0.00348 | 0.59333 | 2.96617 | 1.05553 | 1.17889 | 0.00330 | 0.85754 | 3.18481 | 0.00551 | 2.66060 | 0.21814 | 0.01174 | 0.43718 |
| **Ccdc71l** | 0.26044 | 0.00435 | 0.07910 | 0.00552 | 0.13780 | 0.00874 | 0.00427 | 0.01125 | 0.01641 | 0.00670 | 0.00650 | 0.00746 | 0.31941 | 0.03948 |
| **Ccdc73** | 1.70792 | 0.00364 | 0.07603 | 0.00000 | 0.12203 | 0.21404 | 0.00358 | 0.64811 | 0.20018 | 0.00950 | 0.00000 | 0.50555 | 0.03560 | 1.01398 |
| **Ccdc74a** | 0.04921 | 0.00420 | 0.12809 | 0.25316 | 0.70421 | 0.02944 | 0.00427 | 0.00391 | 0.01516 | 0.00766 | 0.00518 | 0.00960 | 0.07770 | 0.00297 |
| **Ccdc77** | 3.22841 | 0.06158 | 4.66357 | 4.23976 | 4.76374 | 5.51367 | 1.80619 | 2.03547 | 4.24045 | 0.02470 | 10.65527 | 1.54454 | 4.19287 | 7.64328 |
| **Ccdc78** | 0.22802 | 0.03000 | 0.02221 | 0.29582 | 0.22483 | 0.27906 | 0.04711 | 0.05791 | 0.24020 | 0.08421 | 0.01037 | 0.05327 | 0.20363 | 0.54996 |
| **Ccdc7b** | 0.14328 | 0.00437 | 0.00166 | 0.04561 | 0.08370 | 0.01276 | 0.06602 | 0.14349 | 0.29752 | 0.00948 | 0.00551 | 0.00839 | 0.01001 | 0.00000 |
| **Ccdc80** | 0.52240 | 0.00436 | 0.01155 | 0.00975 | 0.03923 | 0.00941 | 0.00439 | 0.01304 | 0.00911 | 0.00647 | 0.00695 | 0.00842 | 0.00503 | 0.00783 |
| **Ccdc81** | 0.00485 | 0.00457 | 0.09321 | 0.17375 | 0.00540 | 0.01071 | 0.00474 | 0.03610 | 0.01068 | 0.00911 | 0.00924 | 0.00919 | 0.00627 | 0.05114 |
| **Ccdc82** | 7.44103 | 1.21367 | 3.14103 | 3.75013 | 33.29994 | 2.55259 | 0.68500 | 1.51278 | 36.31431 | 0.14211 | 1.22028 | 1.99297 | 22.75188 | 57.47667 |
| **Ccdc84** | 1.74196 | 0.02078 | 1.92865 | 1.26930 | 1.72349 | 0.17203 | 1.24139 | 0.71308 | 0.64982 | 0.01978 | 0.11049 | 0.31953 | 0.69087 | 0.09899 |
| **Ccdc85a** | 0.02538 | 0.00442 | 0.01095 | 0.03805 | 0.07781 | 0.01351 | 0.00444 | 0.00889 | 0.01642 | 0.01033 | 0.01506 | 0.00889 | 0.03362 | 0.00669 |
| **Ccdc85b** | 0.03610 | 0.00419 | 0.00564 | 0.05027 | 0.22595 | 0.15927 | 0.00442 | 0.00786 | 0.02674 | 0.00663 | 0.00715 | 0.04065 | 0.95158 | 0.12095 |
| **Ccdc86** | 6.37002 | 2.84839 | 20.95317 | 12.53615 | 17.93635 | 0.00000 | 6.51871 | 5.87919 | 1.01807 | 1.96641 | 3.00040 | 6.60331 | 10.59771 | 3.14294 |
| **Ccdc87** | 0.00742 | 0.00470 | 0.00287 | 0.00953 | 0.06275 | 0.03043 | 0.00477 | 0.00897 | 0.65312 | 0.00688 | 0.00711 | 0.00930 | 0.00431 | 0.00204 |
| **Ccdc88a** | 0.00000 | 0.00365 | 0.03815 | 0.00000 | 1.52051 | 0.12244 | 0.00375 | 0.00382 | 1.71447 | 0.00568 | 0.00000 | 0.98839 | 0.02124 | 1.35391 |
| **Ccdc88b** | 0.00449 | 0.00433 | 0.00564 | 0.01142 | 0.57089 | 0.01303 | 0.00422 | 0.00959 | 0.00731 | 0.00686 | 0.00909 | 0.00786 | 0.40807 | 0.00685 |
| **Ccdc88c** | 0.00000 | 0.02085 | 3.75402 | 0.73013 | 0.62086 | 0.49882 | 0.00322 | 0.25966 | 0.04471 | 0.00000 | 0.79662 | 0.83144 | 0.07072 | 0.70465 |
| **Ccdc89** | 0.00669 | 0.00367 | 0.01102 | 0.00542 | 0.01139 | 0.01300 | 0.00415 | 0.00643 | 0.01524 | 0.00684 | 0.00412 | 0.83124 | 0.01439 | 0.02231 |
| **Ccdc9** | 2.83880 | 0.12362 | 7.48292 | 4.59600 | 8.09412 | 3.96385 | 0.04832 | 3.14449 | 0.32512 | 0.07356 | 2.31497 | 4.22490 | 4.57429 | 3.19038 |
| **Ccdc90b** | 9.73031 | 0.06640 | 4.34407 | 4.62264 | 4.31004 | 3.46160 | 2.36403 | 0.70588 | 0.03686 | 0.06674 | 10.06417 | 4.85549 | 1.30976 | 6.12043 |
| **Ccdc91** | 1.68589 | 0.05810 | 1.24455 | 2.15735 | 1.85416 | 2.38453 | 0.01472 | 3.61317 | 0.08216 | 0.09114 | 0.04830 | 1.93191 | 0.90737 | 3.63410 |
| **Ccdc92** | 0.96646 | 0.01058 | 2.52476 | 0.30259 | 0.50785 | 1.87879 | 2.06648 | 0.10710 | 0.15103 | 0.05496 | 1.49735 | 0.92525 | 2.22298 | 0.00000 |
| **Ccdc92b** | 4.28824 | 0.00139 | 7.21578 | 0.36863 | 4.73375 | 0.74497 | 0.00182 | 0.00000 | 2.36540 | 0.38273 | 3.52242 | 1.18913 | 2.14749 | 2.27164 |
| **Ccdc93** | 9.11562 | 7.14105 | 10.23555 | 1.27018 | 8.22028 | 9.75701 | 3.08021 | 7.30477 | 9.17150 | 0.12684 | 3.28285 | 6.72998 | 0.00000 | 4.05701 |
| **Ccdc94** | 0.09025 | 0.01514 | 8.75763 | 1.88941 | 7.91725 | 1.68860 | 2.82307 | 2.57047 | 0.00000 | 12.67732 | 2.58696 | 1.79417 | 0.05812 | 0.54912 |
| **Ccdc96** | 0.00190 | 0.00547 | 0.25816 | 0.00000 | 0.11260 | 0.05188 | 0.00430 | 0.11436 | 0.01513 | 0.00867 | 0.01031 | 0.00731 | 0.35385 | 0.00150 |
| **Ccdc97** | 0.83579 | 0.06303 | 3.43407 | 3.15291 | 2.84697 | 0.84050 | 2.21136 | 2.00946 | 0.30330 | 0.11537 | 1.05310 | 2.49044 | 0.61442 | 0.66647 |
| **Cchcr1** | 0.54111 | 0.09393 | 1.94288 | 1.11412 | 4.57618 | 0.00000 | 0.00346 | 0.56179 | 0.00689 | 0.00663 | 0.00000 | 1.53039 | 0.57110 | 0.28838 |
| **Cck** | 0.01722 | 0.00402 | 0.00438 | 0.66632 | 0.00573 | 0.05677 | 0.00386 | 0.00337 | 0.01123 | 0.00624 | 0.02276 | 0.01492 | 0.00384 | 0.03737 |
| **Ccl22** | 0.00698 | 0.00439 | 0.41892 | 0.00000 | 0.55599 | 0.00122 | 0.00419 | 0.01076 | 0.00701 | 0.00781 | 0.00700 | 0.00784 | 0.00950 | 0.36286 |
| **Ccl25** | 0.80858 | 0.02489 | 1.13932 | 0.00000 | 0.60838 | 0.99154 | 0.00301 | 0.00000 | 1.34018 | 0.00970 | 0.06627 | 2.07103 | 0.00149 | 0.11429 |
| **Ccl27a** | 0.01247 | 0.00428 | 0.00820 | 0.04332 | 0.06855 | 0.07580 | 0.00412 | 0.00599 | 0.01006 | 0.00775 | 0.13315 | 0.01763 | 0.00791 | 0.01139 |
| **Ccl28** | 0.01570 | 0.00457 | 0.01413 | 0.06386 | 0.02340 | 0.00819 | 0.00972 | 0.02889 | 0.06983 | 0.03804 | 0.01274 | 0.00882 | 0.13440 | 0.00327 |
| **Ccl9** | 0.02849 | 0.03597 | 0.79951 | 0.47240 | 0.00000 | 0.39208 | 1.22611 | 0.01079 | 0.16044 | 0.01674 | 0.00761 | 0.02658 | 0.15247 | 0.12509 |
| **Ccm2** | 0.15617 | 0.04898 | 2.39726 | 0.00000 | 1.41692 | 3.86285 | 2.51985 | 3.36674 | 0.00000 | 7.85817 | 0.00000 | 2.72835 | 0.00000 | 0.61422 |
| **Ccm2l** | 0.00266 | 0.00426 | 1.74124 | 0.08913 | 0.62731 | 0.00000 | 0.00366 | 0.00650 | 0.01218 | 0.00423 | 0.00682 | 0.01182 | 0.00000 | 0.00334 |
| **Ccna2** | 28.82889 | 3.27868 | 20.92202 | 8.41645 | 30.74957 | 14.14535 | 2.17634 | 6.77649 | 23.18940 | 0.09199 | 5.07398 | 8.77893 | 25.03056 | 27.80775 |
| **Ccnb1** | 34.79082 | 0.39625 | 37.01099 | 21.66819 | 43.21370 | 20.99841 | 5.78002 | 15.28701 | 42.86658 | 9.22119 | 18.10978 | 14.60996 | 22.75415 | 41.12436 |
| **Ccnb1ip1** | 5.03875 | 0.00000 | 7.18639 | 4.39125 | 4.98495 | 5.21106 | 1.73549 | 0.00000 | 6.84087 | 8.70907 | 4.34061 | 1.55008 | 0.58296 | 8.80135 |
| **Ccnb2** | 12.90965 | 3.82265 | 9.23175 | 11.22519 | 14.53224 | 13.02484 | 5.34509 | 7.61286 | 3.48553 | 15.93849 | 6.30371 | 10.86648 | 3.94135 | 7.67784 |
| **Ccnb3** | 0.00544 | 0.00428 | 0.26624 | 0.00000 | 0.23245 | 0.00152 | 0.00396 | 0.00959 | 0.28969 | 0.00718 | 0.00560 | 0.00677 | 0.39660 | 0.63125 |
| **Ccnc** | 0.67300 | 0.09103 | 2.69438 | 3.27114 | 3.23870 | 1.65847 | 0.50764 | 3.19331 | 8.02700 | 0.00000 | 7.65546 | 1.61835 | 4.78585 | 7.60261 |
| **Ccnd1** | 0.39763 | 0.00443 | 0.77730 | 3.31979 | 2.73211 | 0.24487 | 0.00381 | 0.00234 | 0.01171 | 0.01519 | 0.00623 | 0.00288 | 0.00000 | 1.52565 |
| **Ccnd2** | 0.52656 | 0.00436 | 0.00000 | 0.18940 | 0.54927 | 0.00000 | 0.00481 | 0.02943 | 0.27994 | 0.00818 | 0.00466 | 0.01427 | 0.00209 | 0.69206 |
| **Ccnd3** | 7.95638 | 0.31434 | 19.09327 | 15.99488 | 6.53466 | 11.94707 | 5.55232 | 7.62471 | 0.34364 | 0.22129 | 4.15605 | 13.51328 | 0.51011 | 1.54397 |
| **Ccndbp1** | 7.53304 | 0.07374 | 12.76333 | 6.18687 | 6.80066 | 9.81288 | 1.12645 | 8.04136 | 3.71906 | 0.08394 | 2.53135 | 6.17938 | 9.58309 | 6.53662 |
| **Ccne1** | 21.73976 | 0.04874 | 26.31639 | 10.93925 | 26.09104 | 13.03012 | 3.48959 | 0.41707 | 13.08645 | 0.24928 | 5.10122 | 3.17013 | 23.07222 | 26.42146 |
| **Ccne2** | 0.00317 | 0.00426 | 0.45866 | 0.10370 | 0.26859 | 0.01114 | 0.01138 | 0.20135 | 2.14968 | 0.00557 | 0.00298 | 0.05492 | 0.33015 | 0.37190 |
| **Ccnf** | 5.73792 | 0.08211 | 12.25707 | 4.28953 | 10.91620 | 8.70258 | 4.83683 | 8.83595 | 1.91577 | 0.19362 | 7.93975 | 5.05687 | 8.81128 | 10.95793 |

| | | | | | | | | | | | | | | |
|---|---|---|---|---|---|---|---|---|---|---|---|---|---|---|
| **Ccng1** | 2.36125 | 0.51144 | 4.03205 | 3.82004 | 0.00000 | 9.28305 | 1.17486 | 4.66561 | 0.94396 | 0.07008 | 4.39847 | 9.31826 | 0.11091 | 0.00000 |
| **Ccng2** | 0.01556 | 0.00389 | 0.00350 | 0.00291 | 0.48444 | 0.01636 | 0.00429 | 0.00709 | 0.00266 | 0.00825 | 0.00562 | 0.20737 | 2.27430 | 0.06018 |
| **Ccnh** | 12.30947 | 0.06556 | 7.17184 | 8.03517 | 11.24597 | 3.38062 | 0.01105 | 7.74081 | 2.44950 | 0.00968 | 3.97358 | 6.56038 | 2.62188 | 2.94594 |
| **Ccni** | 1.36692 | 3.65487 | 1.08530 | 2.52769 | 1.91922 | 3.98128 | 0.02408 | 4.48115 | 1.76964 | 0.05003 | 0.61356 | 8.70089 | 0.94800 | 2.04253 |
| **Ccnj** | 0.21013 | 0.00393 | 0.23088 | 0.63177 | 0.00000 | 0.17410 | 0.00367 | 0.00233 | 0.09572 | 0.00586 | 0.00455 | 0.01165 | 1.17431 | 0.92048 |
| **Ccnjl** | 0.00870 | 0.00442 | 0.01475 | 0.01305 | 0.01271 | 0.01355 | 0.00428 | 0.00985 | 0.01133 | 0.00694 | 0.00886 | 0.00985 | 0.08815 | 0.00469 |
| **Ccnk** | 2.99065 | 0.07638 | 1.32633 | 1.71316 | 4.47670 | 3.59709 | 2.51537 | 1.73209 | 0.73464 | 0.11453 | 0.31567 | 4.51465 | 9.32792 | 0.24284 |
| **Ccnl1** | 6.50419 | 6.34713 | 8.03321 | 4.20273 | 10.50723 | 3.12406 | 3.38855 | 1.26549 | 6.84394 | 5.44343 | 0.11226 | 3.44904 | 5.66021 | 10.91776 |
| **Ccnl2** | 8.37699 | 3.01393 | 11.12345 | 11.75140 | 15.02156 | 10.76904 | 4.84539 | 5.95252 | 2.64115 | 0.21070 | 6.09075 | 4.87479 | 12.91639 | 9.76156 |
| **Ccno** | 0.16272 | 0.00404 | 0.07208 | 0.00450 | 0.29462 | 0.00648 | 0.00406 | 0.00888 | 0.00210 | 0.00690 | 0.00647 | 0.00673 | 2.44130 | 0.61618 |
| **Ccnt1** | 1.87091 | 0.01109 | 1.19662 | 0.03357 | 3.77708 | 3.59346 | 0.00429 | 0.56116 | 1.24402 | 0.10511 | 2.03303 | 0.26385 | 2.49331 | 8.73858 |
| **Ccnt2** | 2.17932 | 0.03696 | 1.96752 | 0.74339 | 1.09990 | 1.09288 | 0.01381 | 0.42327 | 3.36242 | 0.00000 | 2.14286 | 1.40869 | 5.60683 | 2.72404 |
| **Ccny** | 1.49540 | 0.03499 | 1.84992 | 0.00000 | 1.53274 | 3.20466 | 2.43219 | 2.21399 | 0.01207 | 0.10359 | 0.30571 | 1.65012 | 1.23186 | 0.00000 |
| **Ccnyl1** | 0.26429 | 0.00736 | 0.56839 | 1.75457 | 0.90503 | 3.40157 | 1.56743 | 0.02417 | 0.00000 | 0.00280 | 0.33790 | 4.35316 | 6.56749 | 3.49966 |
| **Ccp110** | 2.56332 | 0.03188 | 3.05023 | 1.15083 | 3.01877 | 1.77839 | 0.12015 | 1.70908 | 8.97590 | 0.03206 | 2.81698 | 2.58731 | 1.67709 | 5.22096 |
| **Ccpg1** | 13.42511 | 0.01190 | 18.12420 | 7.49854 | 8.06719 | 8.47001 | 2.52073 | 1.02747 | 14.32572 | 0.04777 | 5.13686 | 3.72334 | 10.04899 | 16.85489 |
| **Ccpg1os** | 1.25546 | 3.87845 | 0.02381 | 0.47700 | 1.17032 | 2.06167 | 1.59968 | 2.47405 | 1.40432 | 0.06526 | 3.32551 | 1.23635 | 1.44121 | 2.22108 |
| **Ccr4** | 0.01172 | 0.01312 | 0.02864 | 0.00000 | 1.26717 | 0.00710 | 0.01050 | 0.03468 | 2.03877 | 0.00803 | 0.00383 | 0.01481 | 0.05815 | 0.22864 |
| **Ccr8** | 0.32728 | 0.00443 | 0.00480 | 0.01196 | 0.17262 | 0.00848 | 0.00424 | 0.01091 | 0.01158 | 0.00656 | 0.00693 | 0.00717 | 0.00461 | 0.38070 |
| **Ccr9** | 0.00999 | 0.00480 | 0.01445 | 0.00646 | 0.01484 | 0.06706 | 0.00438 | 0.00810 | 0.02367 | 0.00649 | 0.00814 | 0.01040 | 0.01285 | 0.01111 |
| **Ccs** | 4.13878 | 4.32610 | 6.79333 | 2.25025 | 3.14417 | 5.57252 | 0.01135 | 4.78662 | 0.30414 | 0.05355 | 7.35261 | 1.90177 | 1.55814 | 2.35104 |
| **Ccsap** | 0.45909 | 0.00369 | 0.19189 | 0.00000 | 0.00000 | 1.37825 | 0.00453 | 0.84560 | 0.00088 | 0.03353 | 0.03354 | 0.69091 | 1.18973 | 0.59142 |
| **Ccser1** | 0.00080 | 0.01427 | 0.01550 | 0.00000 | 0.00211 | 0.00509 | 1.70240 | 0.01791 | 0.05199 | 0.03732 | 0.00649 | 0.01080 | 0.47572 | 0.13200 |
| **Ccser2** | 0.39065 | 3.21401 | 1.45398 | 1.15970 | 1.98984 | 1.21101 | 0.00197 | 0.13370 | 3.68810 | 0.00276 | 0.94257 | 1.89493 | 5.58515 | 4.43246 |
| **Cct2** | 30.76764 | 8.85710 | 36.63458 | 28.46656 | 30.46346 | 32.80446 | 9.91301 | 17.98154 | 33.38652 | 9.70347 | 25.13675 | 26.03260 | 9.17641 | 26.72057 |
| **Cct3** | 23.76289 | 12.48368 | 39.96822 | 29.26512 | 28.54144 | 35.38835 | 10.11157 | 22.76321 | 28.93934 | 18.77251 | 26.04891 | 27.96517 | 14.20297 | 30.05671 |
| **Cct4** | 24.82130 | 7.93833 | 35.13110 | 27.55225 | 33.78520 | 21.23923 | 7.13444 | 14.40070 | 25.27149 | 17.43138 | 14.86673 | 23.75430 | 14.56216 | 16.67740 |
| **Cct5** | 24.78507 | 14.61368 | 40.51863 | 37.11411 | 28.97596 | 33.72908 | 10.21463 | 25.85020 | 30.16536 | 5.02149 | 26.65943 | 27.64375 | 1.84317 | 29.69492 |
| **Cct6a** | 19.58767 | 10.30342 | 35.65354 | 31.20497 | 26.31311 | 29.86708 | 6.96494 | 16.45939 | 17.75449 | 13.39845 | 23.61764 | 25.93777 | 8.77371 | 26.86222 |
| **Cct6b** | 0.00000 | 0.03871 | 0.96402 | 0.03431 | 1.63189 | 1.48956 | 0.01059 | 1.32426 | 0.66959 | 0.04757 | 0.00063 | 0.00000 | 0.96945 | 0.57200 |
| **Cct7** | 26.58431 | 13.63136 | 45.47102 | 34.56640 | 32.63968 | 36.24341 | 12.48595 | 18.98163 | 12.09456 | 11.35049 | 25.98867 | 28.32559 | 3.82733 | 35.06015 |
| **Cct8** | 29.79066 | 9.82802 | 35.13935 | 31.65429 | 27.27873 | 28.45811 | 8.54941 | 18.72654 | 22.26170 | 16.15094 | 21.94862 | 24.30834 | 15.86311 | 17.37168 |
| **Cct8l1** | 0.05381 | 0.00447 | 0.00015 | 0.00847 | 0.84282 | 0.01062 | 0.00421 | 0.01003 | 0.06787 | 0.00617 | 0.00761 | 0.00707 | 0.06991 | 0.26353 |
| **Ccz1** | 13.48872 | 0.11474 | 9.21404 | 1.09403 | 15.16416 | 8.39149 | 0.01461 | 5.28831 | 7.38469 | 0.21083 | 7.72626 | 8.24879 | 3.13162 | 18.94538 |
| **Cd109** | 0.07549 | 0.00216 | 0.26992 | 0.03100 | 0.00000 | 0.00093 | 0.00324 | 0.01380 | 0.00282 | 0.00679 | 0.00582 | 2.83789 | 1.34415 | 0.03984 |
| **Cd14** | 0.00776 | 0.00441 | 0.01284 | 0.01236 | 0.08936 | 0.01286 | 0.00440 | 0.01139 | 0.00952 | 0.00652 | 0.00890 | 0.00979 | 0.00930 | 0.04316 |
| **Cd151** | 1.96839 | 0.05874 | 9.22245 | 9.57081 | 5.47461 | 12.75268 | 6.39043 | 9.45191 | 0.41938 | 0.27331 | 3.93776 | 7.88848 | 0.00000 | 4.83546 |
| **Cd160** | 0.00488 | 0.01531 | 0.01301 | 0.00747 | 0.00944 | 0.07304 | 0.00449 | 0.54312 | 0.02069 | 0.01657 | 0.00663 | 0.01314 | 0.20250 | 0.01374 |
| **Cd163** | 0.01322 | 0.00472 | 0.01805 | 0.04166 | 0.02785 | 0.01194 | 0.00448 | 0.02105 | 0.05507 | 0.00704 | 0.01063 | 0.02568 | 0.04276 | 0.09753 |
| **Cd164** | 14.36816 | 0.04497 | 11.31645 | 10.37887 | 8.63828 | 16.60908 | 1.29145 | 2.23563 | 16.29013 | 7.09700 | 10.52723 | 8.45297 | 7.60347 | 4.70797 |
| **Cd164l2** | 3.15657 | 0.00397 | 0.76175 | 0.92764 | 2.91612 | 0.05627 | 0.00378 | 0.00167 | 0.17682 | 0.00469 | 0.00315 | 0.00697 | 2.56864 | 0.95141 |
| **Cd177** | 0.00428 | 0.00416 | 0.13330 | 0.23171 | 0.35939 | 0.00001 | 0.00385 | 0.00739 | 0.00709 | 0.00725 | 0.00613 | 0.01254 | 0.09376 | 0.07403 |
| **Cd19** | 0.21638 | 0.00444 | 0.17316 | 0.00000 | 1.01280 | 0.00277 | 0.00415 | 0.01210 | 0.01332 | 0.00589 | 0.00598 | 0.00726 | 0.00541 | 0.18431 |
| **Cd1d1** | 0.18740 | 0.00453 | 0.24050 | 0.00006 | 0.00000 | 0.56647 | 0.00404 | 0.00424 | 0.02424 | 0.00548 | 0.00411 | 0.00691 | 0.11411 | 0.03343 |
| **Cd1d2** | 0.04819 | 0.00453 | 0.06112 | 0.00239 | 0.00887 | 0.14848 | 0.00421 | 0.00638 | 0.02363 | 0.00624 | 0.00595 | 0.00930 | 0.01461 | 0.04847 |
| **Cd2** | 0.00908 | 0.00424 | 0.13763 | 0.00000 | 0.01390 | 0.07648 | 0.00425 | 0.00578 | 0.01059 | 0.00675 | 0.00540 | 0.06443 | 0.00861 | 0.01392 |
| **Cd200** | 0.04484 | 0.01360 | 0.15090 | 0.00000 | 0.22579 | 0.00431 | 0.00392 | 0.01640 | 0.01156 | 0.00567 | 0.00575 | 0.06433 | 0.00628 | 0.39300 |
| **Cd209c** | 32.32726 | 0.01781 | 13.96105 | 1.21202 | 17.60931 | 8.19105 | 0.00661 | 0.42603 | 15.91818 | 0.01547 | 0.20886 | 0.21847 | 11.57463 | 7.90228 |
| **Cd209e** | 0.34644 | 0.00430 | 0.00696 | 0.00000 | 0.00583 | 0.17304 | 0.00480 | 0.02320 | 0.00922 | 0.00707 | 0.00408 | 0.65319 | 0.00521 | 0.00398 |
| **Cd209g** | 0.00346 | 0.00429 | 0.00889 | 0.00791 | 0.07679 | 0.01252 | 0.00438 | 0.00918 | 0.14795 | 0.00608 | 0.00621 | 0.07017 | 0.27253 | 0.00000 |
| **Cd22** | 0.27722 | 0.01003 | 0.12680 | 0.46233 | 0.28125 | 0.00000 | 0.04362 | 0.01042 | 0.49256 | 0.02129 | 0.49831 | 2.35763 | 0.32754 | 1.09025 |
| **Cd24a** | 0.21499 | 0.37534 | 1.72473 | 2.23299 | 4.56474 | 0.00000 | 2.76802 | 0.00000 | 3.17303 | 0.03573 | 0.89183 | 2.84852 | 0.26570 | 0.31550 |
| **Cd274** | 0.02285 | 0.01287 | 0.02120 | 0.03056 | 0.01396 | 0.02449 | 0.01644 | 0.01925 | 0.20314 | 0.00653 | 0.02221 | 0.02700 | 0.04691 | 0.07887 |
| **Cd276** | 0.05995 | 0.00425 | 0.03735 | 0.12505 | 0.01091 | 0.00664 | 0.00404 | 0.00741 | 0.01378 | 0.00743 | 0.00662 | 0.01386 | 0.00652 | 0.00913 |
| **Cd28** | 0.00700 | 0.00499 | 0.60451 | 0.00363 | 0.00323 | 0.37278 | 0.00408 | 0.00730 | 0.01065 | 0.01637 | 0.00738 | 0.00502 | 0.00730 | 0.00589 |

| | | | | | | | | | | | | | | |
|---|---|---|---|---|---|---|---|---|---|---|---|---|---|---|
| **Cd2ap** | 7.06009 | 1.58292 | 8.75377 | 4.65682 | 9.76929 | 10.96630 | 3.72123 | 10.22568 | 16.55334 | 0.12400 | 11.32126 | 10.41956 | 4.27060 | 10.26519 |
| **Cd2bp2** | 10.71160 | 4.14194 | 13.78382 | 12.45843 | 15.00917 | 7.03718 | 3.68045 | 8.77562 | 2.45429 | 0.21519 | 6.02395 | 3.59751 | 4.97595 | 9.79746 |
| **Cd300a** | 0.14965 | 0.08423 | 1.41239 | 0.58174 | 1.31578 | 0.45150 | 0.09405 | 0.07256 | 1.22098 | 0.24199 | 0.01142 | 0.26335 | 2.56152 | 1.61836 |
| **Cd300c2** | 0.01028 | 0.00432 | 0.06921 | 0.01078 | 0.07396 | 0.01145 | 0.00442 | 0.00962 | 0.02217 | 0.00730 | 0.00676 | 0.04014 | 0.00413 | 0.03733 |
| **Cd300ld** | 0.01098 | 0.00429 | 0.00557 | 0.00122 | 0.06533 | 0.83883 | 0.00390 | 0.00446 | 0.00670 | 0.00706 | 0.00605 | 0.00910 | 0.13654 | 0.05661 |
| **Cd300lf** | 0.01399 | 0.00440 | 0.01147 | 0.02654 | 0.00000 | 0.07893 | 0.00415 | 0.06738 | 0.07126 | 0.01677 | 0.00656 | 0.00982 | 0.00362 | 0.46264 |
| **Cd300lg** | 0.00753 | 0.00446 | 0.00000 | 0.03097 | 0.55148 | 0.01139 | 0.00410 | 0.00877 | 0.54655 | 0.00714 | 0.00598 | 0.00723 | 0.00360 | 0.00000 |
| **Cd320** | 16.20718 | 0.03482 | 23.58688 | 13.48926 | 23.71202 | 2.44778 | 0.01450 | 4.45852 | 6.38403 | 0.04133 | 0.00000 | 3.46097 | 12.45296 | 11.94263 |
| **Cd33** | 0.05666 | 0.02896 | 0.02955 | 0.02428 | 0.02768 | 0.04334 | 0.00449 | 0.04297 | 0.10195 | 0.00696 | 0.01496 | 0.02014 | 0.03211 | 0.05722 |
| **Cd36** | 0.01566 | 0.00445 | 0.01460 | 0.01297 | 0.01412 | 0.01294 | 0.00431 | 0.01033 | 0.01192 | 0.00689 | 0.00866 | 0.00998 | 0.00875 | 0.02690 |
| **Cd38** | 0.32262 | 0.02632 | 0.47939 | 0.08746 | 0.96847 | 0.32761 | 0.06612 | 0.06420 | 2.80416 | 0.02713 | 0.12102 | 1.96454 | 7.16354 | 4.47688 |
| **Cd3d** | 0.29071 | 0.21555 | 0.07588 | 0.00945 | 0.09508 | 0.06084 | 0.02212 | 0.01811 | 0.23154 | 0.01918 | 0.01784 | 0.01480 | 0.12112 | 0.13898 |
| **Cd3eap** | 1.28783 | 0.02317 | 4.36442 | 5.13786 | 2.78905 | 3.75009 | 1.50898 | 1.46524 | 0.86995 | 0.03712 | 5.15150 | 1.76559 | 6.45370 | 4.14331 |
| **Cd40** | 1.01054 | 0.00425 | 0.00909 | 0.00000 | 0.00361 | 0.30068 | 0.00416 | 0.00457 | 0.00721 | 0.01076 | 0.00501 | 0.02383 | 0.05362 | 0.00446 |
| **Cd44** | 0.20305 | 0.00443 | 0.01349 | 0.01058 | 0.00882 | 0.01121 | 0.00428 | 0.01121 | 0.00978 | 0.00662 | 0.00742 | 0.00916 | 0.02360 | 0.01014 |
| **Cd47** | 0.03199 | 0.01057 | 0.01010 | 0.01920 | 0.00589 | 0.00460 | 0.00348 | 1.26470 | 0.01285 | 0.01098 | 0.00553 | 0.02150 | 1.17666 | 0.02524 |
| **Cd52** | 0.09751 | 0.00422 | 0.07464 | 0.02926 | 0.43621 | 0.00557 | 0.00415 | 0.00870 | 0.00626 | 0.00688 | 0.00695 | 0.00822 | 1.26916 | 0.22768 |
| **Cd55** | 0.00000 | 0.01038 | 0.36007 | 0.24536 | 0.00000 | 0.30966 | 0.01422 | 1.40433 | 1.00160 | 0.00826 | 0.98116 | 0.06177 | 1.19685 | 0.49392 |
| **Cd55b** | 0.00911 | 0.00387 | 0.07072 | 0.05216 | 0.00707 | 0.01809 | 0.00390 | 0.22809 | 0.01448 | 0.01055 | 0.22871 | 0.02502 | 0.02307 | 0.04321 |
| **Cd59a** | 4.14152 | 0.00305 | 0.98497 | 1.63733 | 0.80710 | 1.43981 | 0.00404 | 1.17327 | 1.78051 | 0.02685 | 2.72675 | 0.31529 | 0.65266 | 3.97714 |
| **Cd59b** | 0.21584 | 0.00432 | 0.01716 | 0.14077 | 0.03865 | 0.07592 | 0.01180 | 0.07549 | 0.06644 | 0.01412 | 0.55884 | 0.00412 | 0.82322 | 0.07838 |
| **Cd6** | 0.33210 | 0.00436 | 0.40201 | 0.00000 | 0.17684 | 0.00200 | 0.00490 | 0.01152 | 0.01421 | 0.00626 | 0.00583 | 0.00800 | 0.00384 | 0.13019 |
| **Cd63** | 33.61560 | 12.02651 | 36.67297 | 49.15786 | 29.38143 | 48.41932 | 14.43335 | 33.33836 | 26.12985 | 16.86354 | 23.20065 | 15.98380 | 13.77173 | 33.81541 |
| **Cd68** | 1.35710 | 0.00345 | 1.46624 | 0.35261 | 0.14433 | 1.52113 | 0.00923 | 0.42585 | 0.45016 | 0.00521 | 0.51468 | 0.91206 | 0.00000 | 0.40622 |
| **Cd7** | 0.00000 | 0.00374 | 3.75418 | 3.45524 | 0.19597 | 0.89192 | 0.00309 | 0.00000 | 0.39095 | 0.01057 | 0.00000 | 0.23606 | 0.00000 | 0.25262 |
| **Cd74** | 0.00000 | 0.00435 | 0.14433 | 0.76434 | 1.87617 | 0.00000 | 0.00450 | 0.01007 | 0.01803 | 0.00649 | 0.01852 | 0.00825 | 0.15740 | 0.00000 |
| **Cd79a** | 0.00945 | 0.00433 | 0.34487 | 0.00000 | 0.16999 | 0.10774 | 0.00953 | 0.00350 | 0.00562 | 0.00771 | 0.00544 | 0.00774 | 0.16502 | 0.70129 |
| **Cd79b** | 0.00000 | 0.00411 | 0.44333 | 0.42423 | 0.21581 | 1.65003 | 0.00956 | 0.01171 | 0.01011 | 0.00623 | 0.00160 | 0.00575 | 0.00632 | 0.00000 |
| **Cd81** | 0.11855 | 0.00396 | 0.00106 | 0.14375 | 1.13224 | 0.00989 | 0.00440 | 0.00476 | 0.01096 | 0.00638 | 0.01084 | 0.18432 | 0.01364 | 0.00318 |
| **Cd82** | 0.00000 | 0.25419 | 0.77524 | 0.66595 | 1.35378 | 0.00000 | 0.00186 | 7.15753 | 0.03741 | 4.22008 | 0.00000 | 0.07552 | 0.00674 | 0.22035 |
| **Cd84** | 27.72530 | 4.95993 | 39.09457 | 21.38673 | 28.45795 | 30.19293 | 5.51737 | 9.36181 | 42.65561 | 8.68232 | 23.30229 | 27.44402 | 1.00419 | 28.18967 |
| **Cd8a** | 0.61075 | 0.01240 | 0.09829 | 0.09739 | 0.38489 | 0.05873 | 0.02050 | 0.11772 | 0.26101 | 0.00689 | 0.02092 | 0.02677 | 0.00880 | 0.11337 |
| **Cd8b1** | 0.10601 | 0.01151 | 0.04810 | 0.07959 | 0.08741 | 0.15572 | 0.02229 | 0.04511 | 0.08904 | 0.02948 | 0.06218 | 0.09303 | 0.12999 | 0.19295 |
| **Cd9** | 18.92871 | 9.41755 | 37.78354 | 35.77728 | 22.27845 | 33.90517 | 11.64533 | 29.04922 | 27.23107 | 1.50166 | 21.49591 | 21.33181 | 0.77426 | 32.67124 |
| **Cd93** | 0.06850 | 0.03554 | 0.01106 | 0.09713 | 0.05399 | 0.12947 | 0.01018 | 0.04169 | 0.14809 | 0.05829 | 0.01463 | 0.03443 | 0.22587 | 0.21439 |
| **Cd99l2** | 0.10244 | 0.01231 | 0.54756 | 1.06027 | 0.94690 | 0.72156 | 0.00514 | 0.69999 | 0.61991 | 0.02625 | 0.00000 | 0.06053 | 0.00000 | 0.89720 |
| **Cda** | 0.00313 | 0.00440 | 0.03716 | 0.05823 | 0.00873 | 0.01223 | 0.00423 | 0.01932 | 0.00985 | 0.00806 | 0.00738 | 0.00687 | 0.48453 | 0.00297 |
| **Cdadc1** | 6.56059 | 0.03502 | 2.12185 | 0.89497 | 1.34513 | 1.47670 | 4.17479 | 0.00000 | 0.85754 | 0.18358 | 0.65936 | 1.70533 | 2.60394 | 1.02946 |
| **Cdan1** | 10.01762 | 0.04265 | 16.28890 | 4.03664 | 15.20731 | 4.39720 | 5.57295 | 3.89880 | 4.60158 | 0.17197 | 7.69252 | 2.48821 | 11.02874 | 10.11709 |
| **Cdc123** | 2.21608 | 0.37540 | 2.03058 | 2.71492 | 0.20336 | 7.49187 | 2.44997 | 4.72784 | 2.19918 | 0.18883 | 1.89882 | 8.33002 | 4.56543 | 1.67363 |
| **Cdc14a** | 0.12643 | 0.00402 | 0.16235 | 0.00000 | 0.13424 | 0.00000 | 0.96798 | 0.66977 | 0.00655 | 0.02399 | 0.03952 | 0.66780 | 1.02317 | 0.61296 |
| **Cdc14b** | 11.34227 | 0.10302 | 12.41007 | 4.05955 | 13.72921 | 7.69778 | 1.50251 | 1.00155 | 20.23309 | 0.10832 | 13.25717 | 3.28127 | 11.54318 | 18.88088 |
| **Cdc16** | 9.36226 | 0.05176 | 7.95271 | 5.31224 | 10.81419 | 5.39947 | 0.07844 | 2.46155 | 7.77698 | 11.64921 | 6.47752 | 4.06850 | 3.21573 | 8.54732 |
| **Cdc20** | 11.13311 | 7.88016 | 30.60717 | 19.55744 | 29.20595 | 15.24055 | 7.19273 | 15.46476 | 16.88880 | 0.39844 | 6.80983 | 10.10667 | 12.48164 | 29.37689 |
| **Cdc23** | 4.08249 | 0.04707 | 4.67571 | 1.38643 | 3.66882 | 3.14717 | 0.62052 | 1.59570 | 4.14047 | 0.00000 | 7.74192 | 3.04384 | 0.01420 | 4.25971 |
| **Cdc25a** | 6.73701 | 0.03285 | 7.90386 | 5.05298 | 10.09996 | 8.31429 | 2.82236 | 2.16163 | 11.40180 | 16.95521 | 0.56797 | 7.27429 | 6.87172 | 17.55282 |
| **Cdc25b** | 6.86460 | 0.04075 | 2.93849 | 1.36283 | 6.13307 | 3.53705 | 0.00280 | 3.51387 | 3.69680 | 0.01441 | 0.97916 | 0.00000 | 0.13520 | 6.35536 |
| **Cdc25c** | 5.84566 | 0.15044 | 3.97620 | 5.10439 | 6.75563 | 5.79730 | 1.34246 | 3.60356 | 0.24659 | 0.05580 | 7.44016 | 3.81901 | 8.73229 | 11.63184 |
| **Cdc26** | 0.77787 | 1.66208 | 2.54739 | 2.39569 | 1.66785 | 3.46239 | 2.02058 | 4.80083 | 0.64977 | 0.03426 | 5.00600 | 2.90218 | 0.40052 | 2.93846 |
| **Cdc27** | 8.28572 | 3.97314 | 10.77580 | 5.08397 | 13.61222 | 5.15542 | 1.13797 | 2.92788 | 24.35194 | 0.04074 | 8.41020 | 2.55277 | 11.53151 | 13.98209 |
| **Cdc34** | 4.04696 | 8.83293 | 28.98489 | 30.27810 | 20.94726 | 18.75836 | 8.77481 | 18.78808 | 0.00000 | 20.57805 | 7.50805 | 15.79743 | 8.32239 | 0.00000 |
| **Cdc37** | 6.70087 | 4.82519 | 27.93204 | 22.75459 | 22.76158 | 16.32162 | 5.73947 | 13.58539 | 0.01132 | 2.62529 | 13.16247 | 13.44145 | 1.60637 | 8.57878 |
| **Cdc37l1** | 2.76246 | 3.55021 | 0.97233 | 0.00000 | 0.50675 | 2.45792 | 0.00148 | 1.59020 | 4.92908 | 0.00347 | 0.00000 | 1.01546 | 6.23036 | 1.89397 |
| **Cdc40** | 6.58172 | 0.03627 | 5.46516 | 0.00000 | 8.68718 | 3.22025 | 1.34111 | 0.35145 | 10.06862 | 0.00833 | 4.22900 | 3.56263 | 7.24908 | 13.25798 |
| **Cdc42** | 14.73072 | 3.64368 | 23.40070 | 22.94716 | 23.31476 | 30.75471 | 8.19696 | 23.73674 | 23.73559 | 18.25539 | 21.44242 | 19.50453 | 10.30755 | 17.19799 |

| | | | | | | | | | | | | | | |
|---|---|---|---|---|---|---|---|---|---|---|---|---|---|---|
| **Cdc42bpa** | 0.46324 | 0.01086 | 0.41872 | 0.22487 | 0.70371 | 0.15861 | 0.00400 | 0.19098 | 0.03881 | 0.00451 | 0.00369 | 0.20269 | 0.08554 | 0.09421 |
| **Cdc42bpb** | 0.00000 | 0.14352 | 5.25365 | 2.87025 | 9.99231 | 0.00000 | 0.01315 | 5.51395 | 0.00000 | 5.27651 | 1.74273 | 4.27306 | 0.34320 | 0.00000 |
| **Cdc42bpg** | 0.44902 | 0.49672 | 0.00000 | 0.24152 | 0.18354 | 0.76117 | 0.00399 | 0.28474 | 1.14496 | 0.04626 | 0.02989 | 0.26647 | 1.44685 | 0.29390 |
| **Cdc42ep1** | 1.72821 | 0.07990 | 5.50979 | 4.00236 | 0.80376 | 0.31313 | 0.10845 | 2.83494 | 0.02125 | 0.00000 | 2.67838 | 1.10302 | 0.02310 | 0.70250 |
| **Cdc42ep2** | 0.09420 | 0.01149 | 3.08005 | 0.43041 | 3.43827 | 3.63606 | 0.01139 | 0.10827 | 0.00000 | 0.02849 | 1.66433 | 0.10522 | 2.73026 | 2.78098 |
| **Cdc42ep3** | 1.57539 | 0.01054 | 3.17669 | 4.63406 | 0.45373 | 0.25904 | 1.94557 | 0.38709 | 0.00643 | 0.01639 | 4.68337 | 1.21575 | 0.00000 | 0.49002 |
| **Cdc42ep4** | 1.34852 | 0.01201 | 7.14706 | 0.46031 | 6.41391 | 2.85492 | 0.13734 | 4.60802 | 0.10325 | 0.06233 | 0.27256 | 0.00000 | 1.93318 | 6.53669 |
| **Cdc42ep5** | 0.00000 | 0.51793 | 1.48101 | 7.87003 | 2.76886 | 0.00000 | 1.48063 | 10.83797 | 0.00146 | 0.12081 | 0.00000 | 0.07413 | 0.02716 | 0.00000 |
| **Cdc42se1** | 0.56457 | 0.14524 | 3.66541 | 5.08328 | 4.28134 | 0.59987 | 2.23194 | 8.84252 | 4.05925 | 0.95647 | 2.13014 | 2.87583 | 7.12853 | 2.02883 |
| **Cdc42se2** | 2.28705 | 0.06548 | 2.01967 | 0.00000 | 4.71019 | 7.04615 | 1.42037 | 3.00643 | 6.66813 | 0.13200 | 5.86698 | 3.39068 | 8.46833 | 19.70163 |
| **Cdc45** | 13.99235 | 0.06498 | 17.31578 | 4.40410 | 16.74466 | 13.79257 | 3.12083 | 3.01040 | 4.69743 | 0.04337 | 5.97730 | 6.64683 | 12.88959 | 23.49438 |
| **Cdc5l** | 19.30197 | 4.46733 | 30.74455 | 17.06699 | 23.58939 | 20.72619 | 9.09699 | 6.07357 | 25.23051 | 0.64261 | 16.02078 | 17.41397 | 8.91157 | 20.93424 |
| **Cdc6** | 15.64716 | 3.08892 | 15.35303 | 6.43892 | 17.18559 | 11.83867 | 4.83269 | 1.62592 | 20.87609 | 0.18493 | 10.44139 | 4.54663 | 9.05242 | 21.30294 |
| **Cdc7** | 4.91751 | 0.03590 | 3.63505 | 5.79976 | 3.94638 | 4.11092 | 2.77499 | 0.12360 | 6.70730 | 0.92940 | 1.99259 | 3.09339 | 0.00000 | 4.94335 |
| **Cdc73** | 5.66725 | 0.03607 | 5.63744 | 0.76261 | 8.94658 | 8.19394 | 3.82022 | 0.98123 | 11.73172 | 0.04030 | 5.32775 | 6.62587 | 4.13695 | 6.03990 |
| **Cdca2** | 0.45720 | 0.10892 | 4.94101 | 1.65793 | 3.63970 | 2.81519 | 0.26899 | 5.02320 | 0.81696 | 0.01745 | 6.98565 | 3.02514 | 1.96990 | 3.59217 |
| **Cdca3** | 13.01398 | 12.70021 | 24.90156 | 17.62036 | 20.09090 | 19.89468 | 8.07065 | 21.36915 | 4.01468 | 19.82732 | 10.22570 | 18.46393 | 6.46700 | 7.11077 |
| **Cdca4** | 9.71407 | 0.09654 | 7.09747 | 2.59637 | 12.81502 | 6.15773 | 4.09473 | 2.84569 | 5.83421 | 0.13249 | 1.00811 | 2.10301 | 6.89129 | 7.82892 |
| **Cdca5** | 9.98678 | 9.39351 | 18.07712 | 11.72486 | 16.12681 | 8.34023 | 4.53646 | 3.12941 | 6.73056 | 0.28168 | 5.29712 | 5.79198 | 16.65934 | 16.59396 |
| **Cdca7** | 3.23753 | 3.77897 | 9.36895 | 2.74364 | 6.81955 | 5.87174 | 4.06264 | 0.00000 | 1.99525 | 0.27531 | 5.17353 | 4.83590 | 0.17336 | 10.88064 |
| **Cdca7l** | 0.02460 | 0.00438 | 0.02867 | 0.00762 | 0.06434 | 0.01607 | 0.00432 | 0.00878 | 0.00761 | 0.00601 | 0.00743 | 0.00913 | 0.16689 | 0.50877 |
| **Cdca8** | 12.71482 | 0.67200 | 24.71483 | 16.27961 | 25.08877 | 12.02572 | 7.84375 | 16.30037 | 1.82943 | 13.86613 | 8.75703 | 13.27398 | 16.24377 | 16.00269 |
| **Cdcp1** | 0.00671 | 0.00407 | 0.01201 | 0.02327 | 0.00352 | 0.00657 | 0.21117 | 0.04956 | 0.12501 | 0.00588 | 0.00782 | 0.01162 | 0.00811 | 0.24638 |
| **Cdh1** | 7.79165 | 8.27848 | 16.30755 | 14.96821 | 9.28682 | 25.84013 | 6.40511 | 20.84594 | 13.78568 | 6.68639 | 6.86504 | 12.38861 | 3.15808 | 9.10499 |
| **Cdh11** | 0.33301 | 0.00377 | 0.73059 | 0.21332 | 0.22272 | 0.84894 | 0.05016 | 0.01517 | 0.22098 | 1.64129 | 0.59366 | 0.66148 | 0.15780 | 0.17060 |
| **Cdh15** | 0.01220 | 0.00437 | 0.10942 | 0.05117 | 0.01326 | 0.00878 | 0.00402 | 0.00953 | 0.01084 | 0.01412 | 0.00776 | 0.00668 | 0.51367 | 0.00098 |
| **Cdh16** | 0.13675 | 0.00462 | 0.01055 | 0.08120 | 0.02460 | 0.07525 | 0.02224 | 0.00891 | 0.07559 | 0.00804 | 0.01752 | 0.05986 | 0.10690 | 0.03076 |
| **Cdh17** | 0.00999 | 0.00442 | 0.00332 | 0.01559 | 0.04239 | 0.00932 | 0.00428 | 0.01132 | 0.52943 | 0.00690 | 0.00663 | 0.00975 | 0.00454 | 0.01097 |
| **Cdh2** | 0.06361 | 0.00329 | 0.00000 | 0.06289 | 0.00000 | 0.00380 | 0.00363 | 0.00434 | 0.46702 | 0.00648 | 0.00175 | 1.15815 | 0.01093 | 0.00913 |
| **Cdh23** | 0.14769 | 0.00464 | 0.50898 | 0.00000 | 0.75210 | 0.03126 | 0.00418 | 0.01185 | 0.11401 | 0.00631 | 0.00597 | 0.00599 | 0.00000 | 0.06310 |
| **Cdh26** | 0.04774 | 0.00442 | 0.00612 | 0.00874 | 0.34471 | 0.01406 | 0.00417 | 0.00993 | 0.01341 | 0.00764 | 0.00842 | 0.00832 | 0.10593 | 0.00484 |
| **Cdh3** | 0.00616 | 0.00422 | 0.16969 | 0.22947 | 0.00091 | 0.00000 | 0.00465 | 0.01243 | 0.00840 | 0.00424 | 0.43794 | 0.00416 | 0.00777 | 0.23265 |
| **Cdh4** | 0.00547 | 0.00432 | 0.01064 | 0.01284 | 0.91172 | 0.01736 | 0.00455 | 0.03310 | 0.00773 | 0.01986 | 0.00637 | 0.42690 | 0.28288 | 0.21401 |
| **Cdip1** | 2.82061 | 0.07693 | 4.91265 | 0.92083 | 4.16591 | 6.87873 | 0.48437 | 0.82259 | 5.22727 | 0.06725 | 2.83556 | 2.76072 | 6.75867 | 5.14365 |
| **Cdipt** | 16.78552 | 3.88790 | 30.76100 | 24.39334 | 24.30818 | 23.76136 | 7.17978 | 18.35719 | 1.78212 | 0.69783 | 11.41850 | 14.42031 | 10.26411 | 17.69381 |
| **Cdk1** | 26.07427 | 7.22886 | 28.25421 | 13.59942 | 20.68089 | 18.67174 | 4.02901 | 10.91928 | 18.20336 | 15.54391 | 9.92883 | 13.81677 | 17.22939 | 24.94450 |
| **Cdk10** | 0.00000 | 0.73115 | 3.81851 | 7.82473 | 1.98686 | 1.88528 | 0.00000 | 7.27523 | 0.09998 | 5.02001 | 3.99918 | 2.10648 | 0.70819 | 0.67890 |
| **Cdk11b** | 7.38706 | 0.16648 | 17.78233 | 13.97180 | 17.45191 | 10.75778 | 5.34220 | 5.56373 | 0.00000 | 0.17998 | 13.90912 | 6.05491 | 1.32478 | 8.40918 |
| **Cdk12** | 0.91132 | 0.03913 | 1.52939 | 1.94590 | 2.33990 | 0.88321 | 0.00205 | 1.33400 | 10.19934 | 0.01398 | 1.96755 | 0.71908 | 7.06359 | 7.25238 |
| **Cdk13** | 0.55390 | 0.00263 | 1.11860 | 0.00000 | 1.26576 | 2.01512 | 0.36715 | 0.00000 | 1.10183 | 0.00286 | 0.17881 | 1.12369 | 5.08269 | 0.65302 |
| **Cdk14** | 0.00107 | 0.02418 | 0.30626 | 0.09147 | 0.05108 | 0.28416 | 0.00865 | 2.56096 | 1.62954 | 0.00409 | 2.34765 | 0.84592 | 0.01949 | 0.00000 |
| **Cdk16** | 4.54157 | 0.55682 | 16.67242 | 5.70873 | 11.33031 | 10.25711 | 3.50464 | 4.23451 | 2.19133 | 0.10413 | 4.33488 | 9.51621 | 3.12751 | 11.60050 |
| **Cdk17** | 0.08314 | 0.07734 | 0.78456 | 0.22116 | 2.16570 | 1.83636 | 0.00243 | 0.59323 | 2.19948 | 0.02906 | 2.04864 | 3.40849 | 0.22235 | 1.15472 |
| **Cdk18** | 1.48970 | 0.01588 | 4.56653 | 5.38908 | 1.17727 | 2.22999 | 0.03012 | 4.18608 | 0.00000 | 0.04005 | 3.97582 | 2.87801 | 0.00000 | 1.01492 |
| **Cdk19** | 2.77167 | 0.02478 | 2.08280 | 0.00000 | 2.09180 | 1.30865 | 0.00218 | 0.30920 | 6.05262 | 0.00185 | 0.00527 | 2.80631 | 3.51600 | 2.88380 |
| **Cdk2** | 0.22950 | 2.70527 | 4.00935 | 2.97763 | 6.12168 | 2.82484 | 5.45617 | 7.70962 | 11.73389 | 0.18647 | 5.23068 | 3.21636 | 1.83970 | 12.24682 |
| **Cdk20** | 0.00000 | 0.01134 | 0.18468 | 0.46809 | 0.23237 | 0.18643 | 0.00941 | 0.00000 | 0.00000 | 0.02059 | 1.80611 | 0.01518 | 3.37360 | 1.85712 |
| **Cdk2ap1** | 24.37519 | 0.11973 | 43.06096 | 26.16916 | 39.37219 | 14.44510 | 1.98117 | 7.50479 | 30.13049 | 7.36660 | 5.62167 | 4.32216 | 17.21414 | 35.67305 |
| **Cdk2ap2** | 2.78282 | 3.96986 | 3.46725 | 6.44325 | 7.83790 | 2.59700 | 1.64667 | 4.21026 | 0.28835 | 0.16586 | 1.36590 | 3.53131 | 6.21875 | 0.10424 |
| **Cdk3.ps** | 0.00999 | 0.00466 | 0.03410 | 0.11607 | 0.00372 | 0.23447 | 0.00424 | 0.00671 | 0.18754 | 0.00759 | 0.00620 | 0.00754 | 0.00766 | 0.00000 |
| **Cdk4** | 11.96318 | 6.12601 | 27.97443 | 30.82476 | 23.10983 | 20.70975 | 7.05324 | 15.96667 | 1.95056 | 2.71252 | 15.82031 | 19.44649 | 5.52982 | 10.78289 |
| **Cdk5** | 0.15925 | 0.08761 | 3.20306 | 2.83004 | 0.00000 | 7.99081 | 3.42145 | 6.48472 | 0.00000 | 1.30521 | 6.34928 | 3.80388 | 0.01442 | 0.00000 |
| **Cdk5r1** | 0.77623 | 0.00360 | 0.11379 | 0.00000 | 0.58399 | 0.66804 | 0.03095 | 0.00168 | 0.00394 | 0.00557 | 0.18134 | 0.17979 | 1.96843 | 0.17760 |
| **Cdk5r2** | 0.01286 | 0.00429 | 0.00985 | 0.11394 | 0.01668 | 0.01013 | 0.00410 | 0.00782 | 0.01248 | 0.00988 | 0.00794 | 0.00840 | 0.07647 | 0.00244 |
| **Cdk5rap1** | 0.92259 | 0.01564 | 5.44694 | 0.00000 | 3.29705 | 4.07690 | 1.08757 | 0.75356 | 0.64989 | 0.06137 | 6.01679 | 3.44614 | 2.06625 | 5.65932 |

| | | | | | | | | | | | | | | |
|---|---|---|---|---|---|---|---|---|---|---|---|---|---|---|
| **Cdk5rap2** | 1.38226 | 0.18702 | 1.92905 | 0.48198 | 0.34335 | 1.97332 | 0.00265 | 1.16150 | 2.24401 | 0.02005 | 2.63493 | 0.87015 | 0.00000 | 0.67641 |
| **Cdk5rap3** | 3.16929 | 3.17688 | 6.44192 | 8.27717 | 3.97354 | 11.68885 | 0.64018 | 1.14027 | 0.12834 | 0.09519 | 8.02853 | 7.44732 | 1.34735 | 3.78108 |
| **Cdk6** | 0.04708 | 0.00428 | 0.05075 | 0.41975 | 0.19470 | 1.51528 | 0.06822 | 3.36549 | 0.86806 | 0.06001 | 0.02057 | 0.03092 | 0.27354 | 0.18065 |
| **Cdk7** | 16.59961 | 0.07258 | 15.43634 | 1.51018 | 12.47092 | 9.69778 | 2.08800 | 2.45676 | 11.52746 | 0.05159 | 8.68083 | 5.83876 | 9.59429 | 20.84612 |
| **Cdk8** | 9.51168 | 0.03275 | 9.67718 | 3.07228 | 15.40678 | 7.22816 | 0.00000 | 1.05335 | 1.33448 | 0.98336 | 9.04588 | 5.23699 | 5.99518 | 23.15650 |
| **Cdk9** | 8.47302 | 0.16618 | 9.96423 | 8.45316 | 15.41216 | 11.79937 | 6.55514 | 5.62917 | 0.69828 | 0.22361 | 5.45791 | 7.05709 | 11.82703 | 17.86035 |
| **Cdkal1** | 0.27369 | 0.03481 | 0.42738 | 0.57520 | 0.55535 | 2.24844 | 0.04315 | 0.20121 | 0.98119 | 0.33613 | 0.00000 | 0.85279 | 4.01683 | 0.95181 |
| **Cdkl2** | 1.54601 | 0.02489 | 1.07814 | 0.78850 | 2.20837 | 1.83855 | 0.00893 | 0.94485 | 11.56524 | 0.05006 | 0.00000 | 1.79530 | 2.36191 | 5.00530 |
| **Cdkl3** | 0.39202 | 0.00399 | 0.09868 | 0.28893 | 0.00000 | 0.00000 | 0.00348 | 0.00500 | 2.69385 | 0.00595 | 0.00877 | 0.01669 | 0.10932 | 0.31772 |
| **Cdkl4** | 0.87693 | 0.00382 | 0.92625 | 0.12194 | 0.83864 | 0.00000 | 0.00379 | 0.00572 | 0.00556 | 0.00514 | 0.00544 | 0.00698 | 2.32247 | 1.18205 |
| **Cdkl5** | 0.00838 | 0.00443 | 0.01245 | 0.01327 | 0.00697 | 0.01214 | 0.00433 | 0.01013 | 0.01108 | 0.00683 | 0.00857 | 0.00990 | 0.01893 | 0.20195 |
| **Cdkn1a** | 0.15444 | 0.01731 | 3.45430 | 7.30166 | 7.00007 | 1.33650 | 0.01612 | 0.13877 | 1.46526 | 0.82593 | 0.00000 | 2.65732 | 0.84899 | 3.03025 |
| **Cdkn1b** | 0.02260 | 0.00439 | 0.01306 | 0.00661 | 0.01102 | 0.02366 | 0.00470 | 0.02195 | 0.01289 | 0.01497 | 0.03614 | 0.02501 | 0.00781 | 0.05664 |
| **Cdkn1c** | 0.01248 | 0.00381 | 0.65772 | 1.07858 | 2.89250 | 0.00000 | 0.00371 | 0.00420 | 0.00576 | 0.00524 | 0.00719 | 0.01014 | 1.84281 | 0.24976 |
| **Cdkn2aip** | 0.01645 | 0.03366 | 2.46413 | 0.00000 | 4.29909 | 2.31012 | 2.54153 | 0.34300 | 4.76374 | 0.11888 | 0.60718 | 0.53735 | 1.59622 | 5.69951 |
| **Cdkn2aipnl** | 3.57277 | 0.00071 | 6.01344 | 9.13479 | 4.92035 | 6.93168 | 3.05850 | 0.15315 | 1.89241 | 7.83483 | 0.18334 | 2.15947 | 1.20825 | 1.10755 |
| **Cdkn2b** | 0.02246 | 0.00419 | 0.01036 | 0.07258 | 0.03887 | 0.00908 | 0.00411 | 0.00695 | 0.01203 | 0.01288 | 0.00872 | 0.04980 | 0.02421 | 0.00659 |
| **Cdkn2c** | 0.49290 | 0.01124 | 0.22044 | 0.26435 | 1.16621 | 2.00914 | 0.07688 | 0.54259 | 0.00665 | 0.01831 | 0.00418 | 0.02834 | 0.00488 | 0.00000 |
| **Cdkn2d** | 0.83319 | 3.49510 | 3.88481 | 0.00000 | 5.72378 | 0.73300 | 2.81361 | 7.27043 | 0.00000 | 0.05767 | 1.87741 | 1.24835 | 7.45957 | 1.12785 |
| **Cdkn3** | 0.42439 | 0.00461 | 0.00000 | 0.08148 | 0.57612 | 0.22699 | 0.00439 | 0.73484 | 0.04376 | 0.05660 | 0.34360 | 0.00909 | 1.56187 | 0.68551 |
| **Cdnf** | 0.05953 | 0.06158 | 0.04128 | 0.00000 | 0.12520 | 0.05657 | 0.02808 | 0.00844 | 0.12465 | 2.41308 | 0.00906 | 0.12259 | 0.12174 | 0.20355 |
| **Cdo1** | 0.00483 | 0.02019 | 0.00664 | 0.00539 | 0.24181 | 0.00654 | 0.00508 | 0.62458 | 0.03706 | 0.00550 | 0.00827 | 0.00685 | 1.75948 | 0.02029 |
| **Cdon** | 0.25375 | 0.00439 | 0.11115 | 0.00000 | 0.01369 | 0.40129 | 0.01146 | 0.04852 | 0.16017 | 0.01030 | 1.19396 | 0.03363 | 0.00460 | 0.00000 |
| **Cdpf1** | 0.50583 | 0.05022 | 2.82179 | 7.65341 | 2.12289 | 7.64944 | 4.22825 | 4.70083 | 0.00000 | 0.40393 | 7.12556 | 1.94515 | 0.13418 | 1.72556 |
| **Cdr2** | 0.94587 | 0.37572 | 1.07131 | 2.12412 | 0.36513 | 0.41574 | 0.00330 | 0.00000 | 0.00062 | 0.00242 | 0.12422 | 0.55576 | 10.22462 | 2.58798 |
| **Cdr2l** | 0.00000 | 0.00235 | 0.30690 | 0.21248 | 0.44879 | 0.04109 | 0.00418 | 0.00364 | 0.00301 | 0.00576 | 0.00325 | 2.32164 | 1.60588 | 0.33241 |
| **Cds1** | 3.78699 | 0.02353 | 6.00088 | 5.41435 | 2.10897 | 0.63549 | 0.00208 | 1.86457 | 0.02719 | 0.00834 | 0.05850 | 1.49706 | 1.78141 | 1.75742 |
| **Cds2** | 5.07321 | 0.19183 | 15.70865 | 21.38508 | 16.48208 | 13.55123 | 1.81864 | 4.20996 | 6.36047 | 0.35092 | 11.27019 | 10.24174 | 1.03880 | 19.18500 |
| **Cdsn** | 0.76610 | 0.01038 | 0.96624 | 0.00000 | 0.98928 | 0.00000 | 0.55544 | 0.18756 | 0.13122 | 0.00611 | 0.05826 | 1.15366 | 1.02826 | 0.00000 |
| **Cdt1** | 9.12368 | 0.32816 | 22.88543 | 18.90428 | 31.85472 | 4.73843 | 6.18757 | 7.87746 | 0.33892 | 0.32923 | 3.75115 | 13.62418 | 12.48718 | 1.44756 |
| **Cdv3** | 19.19386 | 10.25717 | 25.76299 | 27.07956 | 22.83948 | 33.66341 | 9.85864 | 16.89369 | 28.87423 | 1.35461 | 14.87252 | 20.17024 | 18.70156 | 28.89888 |
| **Cdx1** | 0.00360 | 3.14293 | 0.01001 | 0.00372 | 0.00657 | 0.00149 | 0.04011 | 1.68290 | 0.01410 | 1.02707 | 0.00252 | 0.00000 | 0.00540 | 0.00692 |
| **Cdx2** | 0.00000 | 3.82494 | 0.91692 | 3.10762 | 0.13103 | 4.94986 | 2.17303 | 4.08295 | 0.09748 | 0.10004 | 4.44119 | 0.00000 | 0.00198 | 0.00000 |
| **Cdyl** | 0.06542 | 0.01069 | 0.10891 | 0.13966 | 2.08982 | 0.90288 | 0.01092 | 0.06875 | 0.02927 | 0.02004 | 0.64401 | 1.03725 | 4.20370 | 3.80073 |
| **Cdyl2** | 0.42023 | 0.01673 | 4.35416 | 0.94756 | 6.13167 | 4.04353 | 0.00768 | 0.28484 | 2.52856 | 0.28536 | 6.52801 | 2.15886 | 0.00000 | 1.43023 |
| **Ceacam1** | 0.15174 | 0.01010 | 0.17369 | 0.00000 | 0.07414 | 0.00041 | 0.00369 | 0.01109 | 0.04498 | 0.00000 | 2.36660 | 0.74747 | 0.50708 | 0.03203 |
| **Ceacam10** | 1.72500 | 0.05053 | 2.69283 | 2.29221 | 2.72051 | 5.47591 | 0.00747 | 0.19038 | 0.66615 | 1.44303 | 7.05170 | 7.69526 | 0.00000 | 0.00000 |
| **Ceacam18** | 0.02340 | 0.00438 | 0.00745 | 0.04658 | 0.04169 | 0.01016 | 0.00413 | 0.00884 | 0.01804 | 0.01392 | 0.00718 | 0.00812 | 0.00833 | 0.20576 |
| **Ceacam19** | 0.21883 | 0.00417 | 0.13704 | 0.22280 | 0.09300 | 0.06657 | 0.00431 | 0.03827 | 0.03603 | 0.00673 | 0.08222 | 0.16062 | 0.00706 | 0.07681 |
| **Ceacam2** | 0.05272 | 0.00402 | 0.12200 | 0.00000 | 0.06855 | 0.00702 | 0.00438 | 0.01551 | 0.03525 | 0.00590 | 0.27079 | 0.18007 | 0.33942 | 0.01405 |
| **Ceacam20** | 0.16475 | 0.00454 | 0.23235 | 0.01724 | 0.03231 | 1.38097 | 0.01616 | 0.00025 | 0.03008 | 0.00397 | 0.86394 | 0.00303 | 0.23293 | 0.00000 |
| **Ceacam5** | 0.00388 | 0.00429 | 0.01507 | 0.01228 | 0.02355 | 0.01308 | 0.00431 | 0.00977 | 0.00926 | 0.00631 | 0.00909 | 0.00921 | 0.49534 | 0.00508 |
| **Cebpa** | 2.61773 | 4.63009 | 3.73713 | 1.00383 | 5.03818 | 0.00000 | 0.23520 | 3.11081 | 1.23892 | 0.00655 | 0.00000 | 0.00000 | 0.18703 | 2.63353 |
| **Cebpb** | 0.03233 | 0.00448 | 0.01134 | 0.02577 | 0.09218 | 0.03333 | 0.00448 | 0.00901 | 0.02525 | 0.00758 | 0.00718 | 0.06756 | 0.13630 | 0.00425 |
| **Cebpd** | 0.13615 | 0.00441 | 0.00642 | 0.00810 | 0.36627 | 0.01244 | 0.00427 | 0.01009 | 0.01096 | 0.00626 | 0.00768 | 0.00746 | 0.25188 | 0.00238 |
| **Cebpg** | 7.25117 | 0.01037 | 4.90660 | 1.12235 | 8.89196 | 0.73080 | 0.00150 | 1.39071 | 14.09494 | 0.00414 | 0.08997 | 0.34186 | 3.06735 | 7.88969 |
| **Cebpz** | 10.73968 | 0.46721 | 14.82339 | 4.78187 | 11.79528 | 6.24024 | 3.69738 | 0.55638 | 13.71974 | 0.10576 | 1.09636 | 3.98216 | 4.42920 | 20.17253 |
| **Cebpzos** | 0.87068 | 0.34704 | 0.00000 | 2.37350 | 1.90065 | 1.73863 | 0.68851 | 2.78770 | 0.42937 | 3.40612 | 0.19964 | 1.42769 | 0.01493 | 0.84306 |
| **Cecr2** | 10.32572 | 0.73805 | 11.53073 | 5.99081 | 19.92479 | 2.85988 | 2.06132 | 6.62668 | 25.01676 | 0.06364 | 5.65572 | 5.77530 | 14.60146 | 20.46160 |
| **Cecr5** | 0.01338 | 0.00394 | 0.04589 | 0.00000 | 0.01728 | 0.01681 | 0.00393 | 0.19473 | 0.00790 | 0.00723 | 0.01058 | 0.01026 | 0.00553 | 0.04911 |
| **Cela1** | 0.99115 | 0.01145 | 0.40884 | 0.43745 | 0.11010 | 0.66163 | 0.00418 | 0.00288 | 0.00435 | 0.00535 | 0.00000 | 0.48035 | 0.16650 | 1.10238 |
| **Cela2a** | 0.03462 | 0.00364 | 0.05459 | 1.72690 | 0.00238 | 0.15011 | 0.00386 | 0.00161 | 0.01300 | 0.01110 | 0.31558 | 0.44159 | 0.00757 | 0.00000 |
| **Cela3b** | 0.00788 | 0.00447 | 0.00582 | 0.00980 | 0.54455 | 0.01359 | 0.00424 | 0.01042 | 0.01315 | 0.00642 | 0.00920 | 0.00818 | 0.00545 | 0.01102 |
| **Celf1** | 6.75620 | 0.06730 | 16.49560 | 11.43897 | 15.97182 | 21.17705 | 2.77712 | 3.12274 | 13.23841 | 10.15335 | 12.89911 | 10.84132 | 8.85118 | 12.13144 |
| **Celf2** | 0.30059 | 0.01149 | 0.00000 | 1.30078 | 0.97498 | 2.80267 | 0.00416 | 2.10895 | 1.32416 | 0.00606 | 0.00000 | 0.83851 | 0.15462 | 0.53988 |

| | | | | | | | | | | | | | | |
|---|---|---|---|---|---|---|---|---|---|---|---|---|---|---|
| **Celf3** | 0.00768 | 0.00469 | 0.16959 | 0.00053 | 0.00601 | 0.02262 | 0.00430 | 0.08053 | 0.00930 | 0.01016 | 0.01676 | 0.00912 | 0.00387 | 0.16429 |
| **Celf5** | 0.01426 | 0.00436 | 0.01114 | 0.01305 | 0.01722 | 0.04363 | 0.00398 | 0.14383 | 0.05057 | 0.00661 | 0.00971 | 0.01890 | 0.00511 | 0.04244 |
| **Celsr1** | 0.30633 | 0.00448 | 0.03974 | 0.01631 | 0.40258 | 0.68959 | 0.00457 | 0.86106 | 0.09656 | 0.00552 | 0.67921 | 0.00409 | 0.98015 | 0.00000 |
| **Celsr3** | 0.01559 | 0.00388 | 0.24302 | 0.00538 | 0.01741 | 0.01163 | 0.00404 | 0.00491 | 0.01124 | 0.01716 | 0.00459 | 0.37762 | 0.01580 | 0.06795 |
| **Cend1** | 0.28826 | 0.00431 | 0.01049 | 0.35978 | 0.00676 | 0.03496 | 0.00413 | 0.00549 | 0.01039 | 0.00573 | 0.00540 | 0.01173 | 0.00598 | 0.00247 |
| **Cenpa** | 15.72237 | 7.69414 | 15.72378 | 31.00799 | 19.12922 | 9.42246 | 4.99252 | 10.42995 | 2.42419 | 0.40168 | 1.45678 | 9.06433 | 17.68350 | 2.54451 |
| **Cenpb** | 0.57827 | 0.02523 | 0.07224 | 1.08147 | 0.42156 | 0.02230 | 0.06160 | 3.16944 | 0.01096 | 0.01199 | 0.01129 | 1.61939 | 0.00258 | 0.18711 |
| **Cenpc1** | 0.41340 | 0.02243 | 1.00909 | 0.18713 | 0.45563 | 1.18815 | 1.06545 | 0.45992 | 4.15826 | 0.03002 | 0.46392 | 0.00544 | 2.77724 | 1.79789 |
| **Cenpe** | 16.55852 | 0.07899 | 9.93765 | 2.32278 | 23.96318 | 3.82250 | 1.10320 | 4.70279 | 27.16228 | 0.07178 | 6.36229 | 4.88516 | 12.00748 | 22.95610 |
| **Cenpf** | 1.84229 | 0.12933 | 3.40792 | 0.00000 | 3.01796 | 3.48228 | 0.14670 | 2.75971 | 2.33790 | 0.18716 | 3.06291 | 4.36016 | 8.78524 | 1.00779 |
| **Cenph** | 4.44526 | 0.02239 | 8.99208 | 3.29541 | 7.53568 | 5.72233 | 0.47993 | 2.71709 | 1.68457 | 1.04626 | 3.40870 | 4.51486 | 7.62909 | 1.53945 |
| **Cenpi** | 6.53148 | 0.00841 | 1.61119 | 1.97302 | 4.56858 | 1.47795 | 0.67589 | 0.90294 | 12.16076 | 0.00000 | 1.49292 | 1.89936 | 0.04132 | 1.97635 |
| **Cenpj** | 0.30292 | 0.00299 | 0.77342 | 0.29263 | 0.32066 | 1.24958 | 0.00330 | 0.00000 | 0.57143 | 0.00000 | 3.25201 | 0.00000 | 3.88786 | 2.99310 |
| **Cenpk** | 4.60158 | 0.00309 | 1.17692 | 2.04556 | 1.54192 | 2.94998 | 1.18430 | 0.00747 | 3.25495 | 0.06050 | 0.27063 | 0.20179 | 0.57099 | 0.00000 |
| **Cenpl** | 3.17602 | 0.03919 | 3.64939 | 0.00000 | 3.92462 | 5.05340 | 2.35116 | 0.96690 | 0.69514 | 0.02620 | 2.34982 | 2.39596 | 5.39305 | 6.70664 |
| **Cenpm** | 1.62820 | 0.02302 | 4.59703 | 9.67928 | 4.07978 | 5.62165 | 0.03368 | 0.09309 | 0.00000 | 0.00981 | 2.76996 | 4.83591 | 0.66535 | 0.11502 |
| **Cenpn** | 4.26740 | 5.39291 | 10.40157 | 1.46378 | 9.62222 | 8.77384 | 0.14006 | 3.90602 | 7.35055 | 0.05191 | 12.95359 | 6.46676 | 3.59117 | 13.68315 |
| **Cenpo** | 1.02971 | 0.00985 | 2.67824 | 2.31173 | 1.77756 | 3.56705 | 3.44991 | 0.64249 | 0.01263 | 0.00360 | 3.15920 | 2.12841 | 0.05133 | 2.42965 |
| **Cenpp** | 5.01367 | 0.03713 | 3.65342 | 0.35272 | 3.86862 | 1.35413 | 0.01325 | 4.21698 | 0.01135 | 0.01146 | 4.13305 | 3.11960 | 0.76782 | 3.82473 |
| **Cenpq** | 3.17721 | 2.94036 | 2.05557 | 1.56073 | 4.28841 | 1.36822 | 0.00200 | 0.00000 | 3.27622 | 0.00019 | 2.28681 | 0.71474 | 4.46902 | 7.96116 |
| **Cenpt** | 3.57259 | 0.06809 | 10.10313 | 6.60641 | 8.00951 | 5.02383 | 0.09528 | 3.73055 | 1.74277 | 0.05911 | 0.67225 | 7.87509 | 5.78681 | 8.95624 |
| **Cenpu** | 1.21080 | 0.08958 | 1.25190 | 0.86468 | 2.38459 | 2.93751 | 0.00318 | 0.06378 | 0.56929 | 0.02168 | 0.34394 | 2.28375 | 0.03375 | 2.85642 |
| **Cenpv** | 0.16434 | 0.09555 | 0.00000 | 0.11022 | 0.55557 | 0.03705 | 0.12127 | 0.00717 | 0.01060 | 0.01644 | 0.00552 | 0.80324 | 0.67400 | 0.00000 |
| **Cenpw** | 0.91445 | 1.60879 | 0.41155 | 1.14521 | 4.04883 | 2.15486 | 1.75754 | 3.17832 | 1.75658 | 0.09857 | 0.00000 | 2.86158 | 4.79987 | 0.00000 |
| **Cep104** | 4.48251 | 0.01798 | 2.71698 | 1.01941 | 2.27761 | 1.53183 | 0.00327 | 1.56662 | 0.55808 | 0.01962 | 0.00000 | 0.70352 | 1.49500 | 2.15668 |
| **Cep120** | 4.56107 | 0.00097 | 4.76177 | 1.04848 | 3.27117 | 4.73074 | 1.93481 | 0.91900 | 5.89684 | 0.03807 | 4.61975 | 1.09685 | 3.20563 | 10.78310 |
| **Cep126** | 0.39652 | 0.00406 | 0.18913 | 0.22511 | 0.06317 | 0.18129 | 0.00400 | 0.01387 | 2.71639 | 0.00592 | 0.00886 | 0.12246 | 0.00000 | 0.00000 |
| **Cep128** | 0.00438 | 0.00380 | 0.02723 | 0.00000 | 0.00501 | 0.45931 | 0.00354 | 1.56984 | 0.01857 | 0.00499 | 0.00184 | 0.12275 | 0.94443 | 0.05215 |
| **Cep131** | 0.03437 | 0.01156 | 1.30406 | 2.91084 | 1.19550 | 0.00000 | 0.00842 | 1.79101 | 0.03026 | 0.00819 | 0.01238 | 0.08869 | 0.21542 | 0.00000 |
| **Cep135** | 0.71388 | 0.00364 | 0.25860 | 0.04572 | 0.15963 | 0.47492 | 0.00380 | 0.00000 | 1.58966 | 0.00152 | 1.49610 | 0.01280 | 0.72172 | 1.27707 |
| **Cep152** | 0.01967 | 0.01020 | 1.20237 | 1.50030 | 0.61082 | 2.27637 | 0.00351 | 0.00000 | 0.00178 | 0.01625 | 0.00000 | 1.00234 | 0.25528 | 0.94472 |
| **Cep162** | 0.28828 | 0.01823 | 0.38712 | 0.84143 | 0.15522 | 0.82224 | 0.00264 | 2.57195 | 1.63839 | 0.03344 | 0.76570 | 0.40077 | 0.00870 | 1.60506 |
| **Cep164** | 0.68629 | 0.05437 | 3.29583 | 4.12524 | 1.84766 | 1.80433 | 0.01933 | 3.05820 | 0.19019 | 0.25766 | 4.38883 | 3.02928 | 0.00000 | 2.54343 |
| **Cep170** | 0.65551 | 0.00980 | 1.73985 | 0.00000 | 2.31627 | 0.00000 | 0.00222 | 0.00160 | 6.73342 | 0.00000 | 1.64159 | 1.09237 | 4.99287 | 0.89604 |
| **Cep170b** | 0.11422 | 0.01929 | 0.99226 | 4.05499 | 0.52760 | 1.43278 | 0.01026 | 3.49651 | 0.28815 | 0.02613 | 0.00001 | 0.17986 | 2.81129 | 0.00104 |
| **Cep19** | 0.00000 | 0.02479 | 1.20055 | 0.75219 | 3.05948 | 1.15789 | 0.43942 | 0.05483 | 0.01315 | 0.02124 | 0.00000 | 3.43888 | 0.56066 | 2.35206 |
| **Cep192** | 5.09824 | 0.01666 | 4.13662 | 0.00000 | 3.50219 | 2.50150 | 0.00092 | 2.32001 | 8.05151 | 1.59926 | 2.75528 | 1.51733 | 5.84751 | 11.54027 |
| **Cep250** | 1.08356 | 0.01712 | 1.69791 | 0.59298 | 2.83051 | 2.41174 | 0.14129 | 1.48397 | 0.13488 | 0.01849 | 0.70526 | 3.07259 | 1.15591 | 2.18160 |
| **Cep290** | 0.32863 | 0.00376 | 0.09627 | 0.10485 | 0.85077 | 0.89758 | 0.02411 | 0.99078 | 6.42818 | 0.00417 | 0.28255 | 0.01311 | 0.35302 | 0.00000 |
| **Cep295** | 1.00257 | 0.02856 | 2.13569 | 0.73401 | 1.70192 | 1.21292 | 1.51551 | 0.98636 | 4.64123 | 0.07465 | 1.66168 | 1.73740 | 2.06355 | 1.25658 |
| **Cep350** | 0.44166 | 0.00342 | 0.21992 | 0.10870 | 0.17781 | 0.76031 | 0.00372 | 0.24393 | 0.01053 | 0.00543 | 2.16020 | 1.49673 | 0.55280 | 0.54411 |
| **Cep41** | 0.32638 | 0.00392 | 2.03233 | 0.27514 | 1.60833 | 1.88400 | 0.00336 | 0.00000 | 1.38563 | 0.01510 | 0.22469 | 0.08155 | 0.26396 | 0.68689 |
| **Cep44** | 1.79925 | 0.00344 | 0.53104 | 0.00000 | 0.00000 | 0.34250 | 0.01020 | 0.16149 | 1.21075 | 0.00365 | 0.53773 | 0.51287 | 0.00166 | 1.78615 |
| **Cep55** | 0.17316 | 0.02148 | 0.39175 | 0.32977 | 0.00000 | 0.58277 | 0.00327 | 0.83566 | 3.02306 | 0.31726 | 2.95276 | 0.04243 | 0.43385 | 2.06928 |
| **Cep57** | 1.74776 | 0.04572 | 2.47906 | 2.40530 | 1.92178 | 3.39029 | 1.01518 | 1.96688 | 1.65308 | 0.07241 | 3.60385 | 5.31762 | 4.24328 | 1.81284 |
| **Cep57l1** | 5.04020 | 0.02106 | 4.36503 | 4.20151 | 5.88889 | 7.15418 | 3.24883 | 0.11427 | 4.76992 | 0.11838 | 2.84681 | 3.90633 | 0.00000 | 5.07092 |
| **Cep63** | 6.25455 | 0.02424 | 5.90778 | 3.33424 | 8.77204 | 3.03367 | 0.02643 | 2.04036 | 8.70725 | 0.00000 | 2.63571 | 2.64130 | 4.42966 | 12.17049 |
| **Cep68** | 0.54793 | 0.01958 | 0.07312 | 0.00000 | 1.59433 | 0.00000 | 0.00414 | 1.18143 | 0.22958 | 0.01435 | 1.68954 | 1.24535 | 0.80991 | 1.20433 |
| **Cep70** | 1.60793 | 0.09243 | 1.37332 | 0.63826 | 1.67332 | 1.79512 | 3.81339 | 0.67243 | 4.66760 | 0.40265 | 0.76431 | 1.89871 | 2.37235 | 3.03783 |
| **Cep72** | 0.85875 | 0.00350 | 0.99999 | 2.03288 | 0.54504 | 0.85042 | 0.03479 | 0.05992 | 1.38182 | 0.01140 | 0.00000 | 0.51609 | 0.00000 | 1.78124 |
| **Cep76** | 0.18165 | 0.00135 | 3.43838 | 1.58446 | 2.39221 | 2.03894 | 0.00148 | 0.48399 | 7.11524 | 0.03265 | 1.33232 | 2.62753 | 2.84921 | 8.80453 |
| **Cep78** | 0.13578 | 5.04995 | 0.57272 | 0.57621 | 1.24264 | 1.07495 | 0.00255 | 0.01178 | 0.40082 | 0.01599 | 0.84441 | 2.40906 | 0.23433 | 2.62954 |
| **Cep83** | 0.38250 | 0.00146 | 1.33861 | 0.48536 | 1.62270 | 2.33532 | 3.44733 | 0.27956 | 1.36196 | 0.07734 | 2.09039 | 2.60670 | 1.13696 | 2.64941 |
| **Cep83os** | 0.01819 | 0.00420 | 0.03064 | 0.08423 | 0.00554 | 0.01557 | 0.00403 | 0.00603 | 0.00915 | 0.00658 | 0.02339 | 0.01327 | 0.22604 | 0.12708 |
| **Cep85** | 1.53036 | 0.02343 | 4.32019 | 2.52963 | 4.80152 | 1.63881 | 0.00200 | 0.59379 | 1.09166 | 0.00000 | 3.13957 | 2.22281 | 1.17537 | 6.68360 |

| | | | | | | | | | | | | | | |
|---|---|---|---|---|---|---|---|---|---|---|---|---|---|---|
| **Cep85l** | 0.02034 | 0.00441 | 0.00595 | 0.01314 | 0.00000 | 0.00895 | 0.00417 | 0.01005 | 0.16254 | 0.00648 | 0.00698 | 0.00938 | 0.04893 | 0.50977 |
| **Cep89** | 1.22670 | 0.02540 | 1.77831 | 1.09642 | 3.26941 | 0.56662 | 2.02967 | 0.57165 | 2.40649 | 0.04245 | 1.36680 | 3.61044 | 2.09764 | 4.59709 |
| **Cep95** | 1.50242 | 0.00351 | 1.80579 | 0.00000 | 0.13948 | 0.90794 | 0.00418 | 0.02211 | 1.83470 | 0.00743 | 0.62749 | 0.71273 | 0.18915 | 1.28604 |
| **Cep97** | 0.15845 | 0.01279 | 0.21409 | 0.87494 | 0.41172 | 0.66096 | 0.06310 | 0.03062 | 0.32121 | 0.02735 | 0.40884 | 0.78956 | 2.98978 | 1.46922 |
| **Cept1** | 1.54260 | 0.01017 | 4.83022 | 1.96048 | 0.00000 | 3.89014 | 2.93472 | 3.62928 | 6.49838 | 0.03514 | 2.47050 | 1.12285 | 1.07614 | 0.00000 |
| **Cercam** | 0.04005 | 0.16787 | 0.05749 | 0.06184 | 0.12582 | 0.22645 | 0.04249 | 0.03255 | 0.13437 | 0.00749 | 0.01068 | 0.05463 | 0.96157 | 0.07316 |
| **Cerk** | 0.06261 | 0.00373 | 0.81181 | 1.99051 | 0.34245 | 0.00000 | 0.00366 | 0.80703 | 0.14471 | 0.00875 | 2.16598 | 0.00719 | 0.00000 | 0.00000 |
| **Cers2** | 11.11617 | 4.96900 | 21.33464 | 4.69669 | 9.94884 | 8.59213 | 3.86074 | 7.59508 | 18.83749 | 0.20637 | 10.15952 | 6.05304 | 5.93222 | 24.55101 |
| **Cers4** | 0.33904 | 0.00387 | 0.34873 | 2.00451 | 0.03244 | 0.00000 | 0.00446 | 0.01483 | 0.00838 | 0.00576 | 0.28977 | 0.01072 | 0.00589 | 1.10845 |
| **Cers5** | 8.84138 | 4.44729 | 6.93005 | 7.75630 | 10.14251 | 6.02540 | 3.22083 | 3.32473 | 11.62656 | 0.11762 | 4.13590 | 0.34658 | 4.62252 | 11.17929 |
| **Cers6** | 0.00717 | 0.00308 | 0.83602 | 0.28192 | 0.00000 | 0.90798 | 0.00441 | 0.51327 | 0.00341 | 0.01072 | 0.68243 | 1.58703 | 1.06985 | 0.04984 |
| **Ces1a** | 0.00947 | 0.00445 | 0.25940 | 0.00000 | 0.12622 | 0.00668 | 0.00440 | 0.01103 | 0.01046 | 0.00680 | 0.00668 | 0.00984 | 0.00616 | 0.01399 |
| **Ces1d** | 0.00795 | 0.00451 | 0.00741 | 0.01225 | 0.38111 | 0.01334 | 0.00448 | 0.01074 | 0.01206 | 0.00645 | 0.00892 | 0.00825 | 0.00558 | 0.00938 |
| **Ces1f** | 0.00852 | 0.00445 | 0.00728 | 0.01063 | 0.34496 | 0.01369 | 0.00477 | 0.01064 | 0.01635 | 0.00676 | 0.00894 | 0.00867 | 0.00563 | 0.00633 |
| **Cetn2** | 0.85853 | 0.00888 | 0.94773 | 0.32945 | 1.56165 | 3.47316 | 0.00174 | 3.31863 | 3.07967 | 0.00074 | 5.98735 | 1.41432 | 0.60779 | 0.81664 |
| **Cetn3** | 4.48491 | 3.98954 | 1.90511 | 3.23486 | 3.67334 | 8.39818 | 1.58115 | 4.79523 | 4.85626 | 0.16188 | 8.65564 | 4.16245 | 0.12106 | 2.69657 |
| **Cetn4** | 0.04961 | 0.61102 | 0.01389 | 0.01381 | 0.01163 | 0.01233 | 0.00391 | 0.00402 | 0.01148 | 0.00596 | 0.00793 | 0.00496 | 0.00587 | 0.01004 |
| **Cfap100** | 0.00603 | 0.00435 | 0.05488 | 0.00000 | 0.54681 | 0.10933 | 0.00483 | 0.01070 | 0.16147 | 0.00619 | 0.00659 | 0.00680 | 0.00184 | 0.00000 |
| **Cfap126** | 0.00000 | 0.00387 | 0.15941 | 0.09903 | 0.02938 | 0.02809 | 0.00387 | 0.00336 | 0.01020 | 0.00543 | 0.01315 | 0.22927 | 0.17858 | 0.51218 |
| **Cfap157** | 0.00000 | 0.01173 | 0.17383 | 0.33520 | 1.83008 | 0.10435 | 0.00377 | 0.00273 | 1.10256 | 0.00660 | 0.00191 | 0.00369 | 1.31874 | 0.62652 |
| **Cfap161** | 0.01229 | 0.00456 | 0.72588 | 0.00000 | 0.12677 | 0.36359 | 0.00420 | 0.00369 | 0.01576 | 0.00617 | 0.00442 | 0.00933 | 0.00292 | 0.22730 |
| **Cfap20** | 5.40072 | 0.01913 | 2.60217 | 5.19985 | 5.34765 | 1.51539 | 0.10197 | 0.50673 | 1.25347 | 0.02485 | 1.01784 | 2.59504 | 8.81515 | 0.04972 |
| **Cfap206** | 0.44088 | 0.00408 | 0.26020 | 1.20635 | 0.11085 | 0.00000 | 0.00408 | 0.00774 | 0.00858 | 0.01052 | 0.00441 | 0.00545 | 0.23915 | 0.00000 |
| **Cfap221** | 0.01456 | 0.00442 | 0.01386 | 0.01239 | 0.01061 | 0.01275 | 0.00428 | 0.01015 | 0.01534 | 0.00661 | 0.00875 | 0.00990 | 0.06300 | 0.04953 |
| **Cfap36** | 0.93435 | 0.02276 | 0.56036 | 0.00000 | 0.53972 | 0.40875 | 0.00359 | 0.44431 | 0.00689 | 0.00562 | 0.44473 | 1.85089 | 1.03450 | 0.03468 |
| **Cfap43** | 0.82008 | 0.01076 | 0.54951 | 0.08308 | 0.81479 | 0.79634 | 0.00344 | 0.02111 | 0.04835 | 0.00000 | 4.93152 | 0.24820 | 0.12142 | 1.33528 |
| **Cfap45** | 0.00608 | 0.00389 | 0.00397 | 0.00000 | 0.00698 | 0.82862 | 0.00380 | 0.36939 | 0.00719 | 0.00695 | 0.03842 | 0.12091 | 0.13250 | 0.13847 |
| **Cfap46** | 0.00469 | 0.00429 | 0.15877 | 0.10926 | 0.03254 | 0.01061 | 0.00412 | 0.00788 | 0.06667 | 0.00678 | 0.03080 | 0.01114 | 0.01548 | 0.04478 |
| **Cfap53** | 0.00232 | 0.00374 | 0.01142 | 0.00000 | 0.00732 | 0.03730 | 0.00397 | 0.59930 | 0.01892 | 0.00513 | 0.00589 | 0.01149 | 0.98119 | 0.11986 |
| **Cfap57** | 0.00307 | 0.17628 | 0.00000 | 0.04817 | 0.11652 | 0.98020 | 0.13360 | 1.15897 | 0.00614 | 2.13129 | 0.00000 | 0.00000 | 0.16066 | 0.04322 |
| **Cfap61** | 0.00000 | 0.00446 | 0.21374 | 0.00204 | 0.14041 | 0.10082 | 0.00982 | 0.00678 | 0.02176 | 0.03576 | 0.00587 | 0.01057 | 0.01905 | 1.58571 |
| **Cfap70** | 0.51102 | 0.00428 | 0.00000 | 0.00881 | 0.44083 | 0.00596 | 0.00424 | 0.00959 | 0.00650 | 0.00724 | 0.00597 | 0.00545 | 0.14749 | 1.62420 |
| **Cfap74** | 5.12477 | 0.08933 | 2.79415 | 1.35324 | 5.31991 | 2.73575 | 0.65011 | 1.22668 | 3.04111 | 0.17872 | 0.77206 | 0.81324 | 6.57990 | 2.08465 |
| **Cfap77** | 0.00825 | 0.00431 | 0.01193 | 0.13118 | 0.01061 | 0.01158 | 0.00428 | 0.00751 | 0.01224 | 0.00618 | 0.00955 | 0.00865 | 0.00685 | 0.01090 |
| **Cfap97** | 1.05516 | 0.01709 | 0.10645 | 0.83092 | 0.48490 | 1.45037 | 3.00172 | 2.58967 | 0.01913 | 0.01827 | 1.61495 | 0.00878 | 3.94872 | 2.80070 |
| **Cfb** | 1.07762 | 0.00370 | 0.53068 | 0.27284 | 0.58302 | 0.98490 | 0.00376 | 0.40679 | 0.38751 | 0.01705 | 0.48536 | 0.00610 | 1.09048 | 0.03606 |
| **Cfc1** | 0.00298 | 0.00433 | 0.00512 | 0.03978 | 0.00695 | 0.38953 | 0.34283 | 1.21952 | 0.01259 | 2.10662 | 0.00000 | 0.00000 | 0.00740 | 0.00171 |
| **Cfd** | 0.00775 | 0.00452 | 0.08195 | 0.00390 | 0.39638 | 0.01012 | 0.00433 | 0.01217 | 0.00867 | 0.00654 | 0.00803 | 0.00837 | 0.01082 | 0.01581 |
| **Cfdp1** | 23.99434 | 9.19244 | 28.60302 | 21.09082 | 28.57417 | 21.80176 | 8.68873 | 11.26192 | 28.35838 | 13.68477 | 16.36649 | 14.83153 | 11.60576 | 25.58383 |
| **Cfh** | 0.02049 | 0.00436 | 0.01164 | 0.01149 | 0.06990 | 0.01296 | 0.00474 | 0.00957 | 0.02288 | 0.00674 | 0.00851 | 0.00913 | 0.24909 | 0.02372 |
| **Cfi** | 0.00306 | 0.00411 | 0.00972 | 0.18254 | 0.13786 | 0.00424 | 0.00439 | 0.00608 | 0.10243 | 0.00556 | 0.01818 | 0.01557 | 0.26951 | 0.00000 |
| **Cfl1** | 10.07566 | 15.46663 | 36.76745 | 37.63290 | 34.08462 | 35.27488 | 7.76793 | 33.94807 | 2.29724 | 25.20051 | 22.45289 | 25.41917 | 11.63078 | 17.34905 |
| **Cfl2** | 9.57091 | 2.65185 | 7.07269 | 9.28815 | 11.14878 | 9.45754 | 2.81192 | 3.91662 | 12.65386 | 0.11146 | 4.41894 | 4.42566 | 6.32235 | 11.06588 |
| **Cflar** | 4.73778 | 0.50717 | 0.00000 | 2.28827 | 2.47646 | 7.63596 | 1.36044 | 1.63878 | 7.81561 | 3.30051 | 0.16927 | 2.59150 | 23.16620 | 11.59250 |
| **Cfp** | 0.11673 | 0.04516 | 0.66040 | 0.27748 | 0.62331 | 1.13108 | 0.00492 | 0.50464 | 0.01949 | 0.00996 | 0.00000 | 0.10153 | 0.00550 | 0.04788 |
| **Cftr** | 1.07111 | 0.00435 | 0.00664 | 0.00961 | 0.12078 | 0.00755 | 0.00421 | 0.01221 | 0.00940 | 0.00693 | 0.00704 | 0.00663 | 0.00865 | 0.02310 |
| **Cga** | 0.11787 | 0.00388 | 0.01298 | 0.04483 | 0.00459 | 0.01380 | 0.57195 | 0.09702 | 0.01021 | 0.02959 | 0.00995 | 0.01215 | 0.54135 | 0.00255 |
| **Cggbp1** | 8.69369 | 3.14103 | 6.63311 | 3.34131 | 7.57741 | 6.76386 | 4.66100 | 2.00776 | 17.78750 | 0.06982 | 7.02390 | 5.45354 | 18.53821 | 1.27242 |
| **Cgn** | 1.31912 | 2.12742 | 6.13610 | 3.70026 | 2.17302 | 5.11370 | 0.87289 | 6.12126 | 3.75185 | 0.08798 | 2.04331 | 0.00000 | 2.46297 | 1.53049 |
| **Cgnl1** | 0.00000 | 2.39837 | 0.92005 | 0.00000 | 0.00000 | 2.67126 | 1.36019 | 11.31840 | 1.71988 | 0.06721 | 0.00000 | 1.65115 | 0.00936 | 2.68959 |
| **Cgref1** | 4.39870 | 0.00333 | 0.74356 | 1.15877 | 2.37903 | 0.53296 | 0.00362 | 0.15595 | 1.28037 | 0.01427 | 0.00000 | 0.00000 | 0.01906 | 3.06918 |
| **Cgrrf1** | 0.00997 | 0.02681 | 0.00000 | 0.76022 | 0.00505 | 0.27739 | 0.21900 | 2.24056 | 0.01421 | 0.03007 | 0.01162 | 0.00314 | 0.00749 | 0.00379 |
| **Chac1** | 0.01489 | 0.02000 | 0.01826 | 0.01140 | 0.10413 | 0.01256 | 0.00420 | 0.00905 | 0.01197 | 0.00610 | 0.00887 | 0.00868 | 0.14273 | 0.01486 |
| **Chac2** | 0.00097 | 0.00390 | 0.03877 | 0.12409 | 0.07364 | 0.27705 | 0.00382 | 0.00235 | 0.03398 | 0.00516 | 0.09245 | 0.10253 | 0.56033 | 0.00138 |
| **Chadl** | 0.06083 | 0.00439 | 0.02614 | 0.00609 | 0.30458 | 0.01024 | 0.00430 | 0.01020 | 0.01101 | 0.00631 | 0.02324 | 0.00923 | 0.22320 | 0.01697 |

| | | | | | | | | | | | | | | |
|---|---|---|---|---|---|---|---|---|---|---|---|---|---|---|
| **Chaf1a** | 7.28586 | 0.08004 | 6.80194 | 8.42585 | 6.87943 | 12.58092 | 0.03764 | 5.28393 | 0.92170 | 0.09469 | 3.20195 | 4.05202 | 6.20315 | 10.44324 |
| **Chaf1b** | 5.47981 | 0.07009 | 14.25559 | 8.44295 | 11.58060 | 5.92254 | 2.52162 | 0.10877 | 2.25286 | 0.04821 | 2.70790 | 4.38374 | 2.76613 | 6.92967 |
| **Champ1** | 8.48798 | 3.87105 | 11.85634 | 3.39534 | 15.90083 | 4.94580 | 2.81074 | 3.85128 | 10.02894 | 0.71534 | 2.59265 | 5.41217 | 6.90958 | 11.73465 |
| **Chchd1** | 4.04328 | 5.61744 | 0.04830 | 3.07395 | 6.17110 | 3.96278 | 3.00160 | 4.85638 | 3.06253 | 13.53180 | 0.82876 | 5.31800 | 4.57823 | 1.21188 |
| **Chchd10** | 0.00787 | 0.03380 | 0.00000 | 0.49252 | 3.95155 | 0.51367 | 0.38977 | 1.96953 | 0.01753 | 3.32598 | 0.00000 | 1.93846 | 1.53413 | 0.00000 |
| **Chchd2** | 4.52213 | 0.44057 | 0.45654 | 39.34828 | 36.52376 | 10.60640 | 1.44180 | 8.42630 | 2.12666 | 6.00818 | 0.07613 | 14.04542 | 20.26139 | 0.00000 |
| **Chchd3** | 3.52744 | 7.41093 | 8.34554 | 8.42997 | 2.86017 | 10.83809 | 4.79682 | 7.75616 | 0.00000 | 0.14608 | 3.25644 | 8.81261 | 0.20074 | 3.30187 |
| **Chchd4** | 6.58677 | 0.22309 | 7.03308 | 13.19145 | 4.30665 | 17.36453 | 5.63418 | 7.43609 | 0.64967 | 0.32554 | 5.90828 | 13.48309 | 1.13220 | 5.93352 |
| **Chchd5** | 1.61316 | 2.33396 | 7.72534 | 10.35198 | 5.17537 | 10.53146 | 5.80526 | 5.54962 | 0.38524 | 20.81669 | 7.61170 | 5.82361 | 0.46083 | 6.17734 |
| **Chchd6** | 0.19126 | 0.00778 | 0.50918 | 0.98836 | 1.31395 | 0.90119 | 0.14866 | 2.42964 | 0.00000 | 0.00509 | 2.07124 | 4.39557 | 0.53202 | 0.00000 |
| **Chchd7** | 0.01189 | 0.02999 | 0.00000 | 2.53292 | 0.67824 | 0.18270 | 1.04361 | 0.62160 | 0.02166 | 0.29116 | 0.13703 | 1.50890 | 0.08238 | 0.09067 |
| **Chd1** | 5.11247 | 0.03631 | 8.06445 | 0.00000 | 5.44380 | 5.42686 | 0.71585 | 1.69263 | 14.28179 | 0.02315 | 6.35919 | 5.74059 | 9.65868 | 12.51935 |
| **Chd1l** | 0.07445 | 0.01164 | 1.27490 | 1.21862 | 0.91980 | 1.57054 | 0.00300 | 0.00000 | 2.80356 | 0.01943 | 0.07212 | 0.57327 | 0.05124 | 0.36825 |
| **Chd2** | 0.04734 | 0.01003 | 0.78978 | 0.36090 | 0.22936 | 0.84486 | 0.00331 | 0.00000 | 0.13567 | 0.01620 | 0.49087 | 0.86242 | 0.57858 | 2.48561 |
| **Chd3** | 0.18669 | 0.01289 | 0.37080 | 0.29132 | 0.73975 | 0.00000 | 0.00521 | 0.93284 | 0.08794 | 0.00732 | 0.00000 | 0.45286 | 0.38315 | 0.00000 |
| **Chd3os** | 0.01296 | 0.00446 | 0.36341 | 0.52320 | 0.34700 | 0.09066 | 0.00412 | 0.00375 | 0.08629 | 0.00628 | 0.00417 | 0.00624 | 0.20567 | 0.01818 |
| **Chd4** | 1.19085 | 5.48707 | 16.67242 | 11.54805 | 21.86329 | 4.47364 | 3.21729 | 12.33682 | 0.49421 | 0.08867 | 8.33891 | 11.53646 | 1.74192 | 0.99385 |
| **Chd5** | 0.00353 | 0.00406 | 0.00768 | 0.48228 | 0.34536 | 0.00442 | 0.00394 | 0.00593 | 0.00783 | 0.00567 | 0.00666 | 0.01818 | 0.00938 | 0.03994 |
| **Chd6** | 1.88333 | 0.00370 | 2.06612 | 0.48093 | 2.79354 | 1.12684 | 0.00973 | 1.19721 | 2.61912 | 0.02627 | 0.00000 | 0.46020 | 0.78728 | 2.09368 |
| **Chd7** | 0.72221 | 3.34799 | 1.67287 | 3.71451 | 9.00754 | 1.09431 | 0.00106 | 0.44646 | 6.46262 | 0.00070 | 2.88141 | 2.83270 | 4.12828 | 8.58778 |
| **Chd8** | 0.61116 | 0.02483 | 1.91630 | 1.04137 | 3.88814 | 2.16361 | 0.00224 | 2.70695 | 3.41857 | 0.01926 | 1.04849 | 1.61066 | 6.76403 | 5.63660 |
| **Chd9** | 0.34280 | 0.00397 | 0.34966 | 0.00000 | 1.20795 | 0.00339 | 0.00452 | 0.03537 | 0.11878 | 0.00343 | 0.45610 | 0.51207 | 2.13183 | 0.05785 |
| **Chdh** | 0.00171 | 0.01136 | 0.26378 | 0.55857 | 0.35648 | 0.05138 | 0.34755 | 1.23199 | 1.14045 | 0.35099 | 0.16276 | 0.06962 | 0.01645 | 0.21642 |
| **Chek1** | 7.37704 | 0.02611 | 4.18361 | 0.31370 | 3.70650 | 4.39456 | 0.95884 | 0.00007 | 8.39281 | 0.03348 | 7.73568 | 4.58851 | 2.45292 | 5.11632 |
| **Chek2** | 17.59102 | 0.03665 | 14.84087 | 1.25967 | 13.73533 | 0.55045 | 0.94086 | 0.86284 | 19.10551 | 0.04625 | 6.31107 | 2.22353 | 7.22415 | 22.37297 |
| **Cherp** | 12.96529 | 0.19319 | 15.32380 | 13.96627 | 21.12192 | 6.74770 | 1.39080 | 7.87827 | 6.12076 | 0.08598 | 5.66745 | 6.69131 | 11.10375 | 10.29548 |
| **Chfr** | 4.50400 | 4.96552 | 3.39792 | 0.75428 | 3.03074 | 6.35894 | 3.71637 | 0.93893 | 0.84026 | 0.58079 | 7.74566 | 4.42880 | 2.00176 | 2.70359 |
| **Chga** | 0.00000 | 3.31182 | 2.22349 | 0.00000 | 0.30247 | 0.00000 | 0.00118 | 2.82976 | 0.00355 | 0.02276 | 0.27695 | 4.17993 | 0.00000 | 0.15164 |
| **Chgb** | 0.00848 | 0.00413 | 0.15435 | 0.27030 | 0.00379 | 0.00000 | 0.00400 | 0.00791 | 0.01452 | 0.00580 | 0.01525 | 0.01502 | 0.00635 | 0.00822 |
| **Chic1** | 2.32221 | 0.04349 | 0.91379 | 0.89264 | 4.67823 | 0.00000 | 0.03212 | 0.09674 | 13.86162 | 0.03902 | 0.01202 | 0.00798 | 0.37153 | 3.27015 |
| **Chic2** | 3.88955 | 0.02346 | 0.65304 | 0.00000 | 3.85892 | 1.60660 | 0.00318 | 1.64353 | 0.06823 | 0.00407 | 0.31239 | 2.43610 | 1.68403 | 0.63604 |
| **Chid1** | 0.14062 | 0.08209 | 3.46540 | 4.32243 | 0.29703 | 11.42352 | 0.77192 | 3.99250 | 0.16857 | 0.05718 | 4.15651 | 5.66702 | 0.35512 | 4.10431 |
| **Chil1** | 0.00351 | 0.00423 | 0.18526 | 0.00000 | 0.00888 | 0.44856 | 0.00381 | 0.00437 | 0.01036 | 0.00596 | 0.00084 | 0.43920 | 0.00780 | 0.00800 |
| **Chil5** | 0.00309 | 0.00401 | 0.01212 | 0.00772 | 0.00439 | 0.01434 | 0.00401 | 0.06122 | 0.00760 | 0.00724 | 0.00778 | 0.00872 | 0.26248 | 0.31781 |
| **Chit1** | 0.00299 | 0.00379 | 0.00165 | 0.01910 | 0.08554 | 4.21166 | 3.80321 | 0.04757 | 0.00550 | 0.17657 | 0.09357 | 0.00000 | 0.81931 | 0.00000 |
| **Chka** | 32.46770 | 0.20381 | 36.72298 | 4.75210 | 43.00595 | 3.92875 | 7.92307 | 14.24129 | 6.52604 | 0.45393 | 1.78955 | 2.60710 | 15.50196 | 30.90136 |
| **Chkb** | 0.00935 | 0.00409 | 0.04821 | 0.16480 | 0.08190 | 0.04236 | 0.01036 | 0.20940 | 0.03100 | 0.01367 | 0.07074 | 0.01798 | 0.20095 | 0.16604 |
| **Chm** | 1.76362 | 0.00291 | 0.92801 | 0.70555 | 0.81027 | 0.25224 | 0.00239 | 0.00000 | 8.01913 | 0.00835 | 0.43473 | 0.86378 | 0.00000 | 0.29395 |
| **Chml** | 0.80242 | 0.00426 | 0.00000 | 0.00000 | 0.00000 | 0.01070 | 0.13262 | 0.00516 | 1.04356 | 0.00947 | 0.39269 | 0.51457 | 0.00906 | 0.00000 |
| **Chmp1a** | 5.52584 | 7.45729 | 6.10318 | 7.44770 | 6.28245 | 6.58613 | 2.84472 | 7.71293 | 1.83575 | 0.13934 | 6.23425 | 4.83259 | 2.97364 | 5.01701 |
| **Chmp1b** | 4.65531 | 0.01128 | 6.40638 | 1.61303 | 3.25699 | 3.35579 | 1.23082 | 1.50425 | 0.95156 | 0.03106 | 2.49124 | 0.51425 | 1.49585 | 2.96863 |
| **Chmp2a** | 7.75676 | 0.93588 | 7.05329 | 12.58281 | 9.70553 | 13.20972 | 6.90223 | 15.61668 | 4.39491 | 7.03151 | 13.23420 | 11.61587 | 6.22898 | 12.75617 |
| **Chmp2b** | 0.43611 | 0.09635 | 0.05121 | 3.71641 | 0.00000 | 2.96872 | 0.08628 | 6.67067 | 0.12262 | 10.48554 | 0.03639 | 1.55364 | 0.09500 | 0.00000 |
| **Chmp3** | 7.52973 | 0.44733 | 6.80035 | 6.12087 | 3.26608 | 10.54620 | 0.03595 | 4.81731 | 6.37773 | 2.59607 | 4.21072 | 5.36098 | 0.00000 | 6.93343 |
| **Chmp4b** | 1.31393 | 4.02098 | 0.00000 | 0.09392 | 1.47134 | 2.76581 | 0.62432 | 3.49873 | 0.60645 | 3.57611 | 0.00000 | 1.13028 | 5.22452 | 0.27402 |
| **Chmp4c** | 1.29487 | 0.05930 | 3.83875 | 5.31069 | 2.26067 | 0.29800 | 1.56745 | 0.65609 | 0.00950 | 0.01229 | 0.86106 | 6.57721 | 0.00000 | 0.66661 |
| **Chmp5** | 0.88429 | 0.02168 | 4.10466 | 2.92952 | 2.82481 | 4.56117 | 0.03526 | 4.92128 | 5.65259 | 0.02931 | 7.49894 | 5.18973 | 2.20700 | 6.16978 |
| **Chmp6** | 0.46289 | 0.06952 | 3.76954 | 7.56814 | 3.24947 | 3.43443 | 3.02004 | 2.05532 | 0.32702 | 0.23250 | 0.00000 | 4.19449 | 0.73076 | 0.00000 |
| **Chmp7** | 3.01921 | 5.63298 | 9.34003 | 5.90309 | 7.11311 | 8.07112 | 5.48054 | 3.39714 | 0.00000 | 6.84779 | 8.51053 | 7.66112 | 2.50891 | 5.43687 |
| **Chn1** | 0.27038 | 0.00312 | 0.27130 | 1.12597 | 0.73591 | 0.16131 | 0.00337 | 0.00000 | 0.98101 | 0.00933 | 0.45232 | 0.56199 | 1.67998 | 1.12979 |
| **Chn2** | 0.00000 | 0.00451 | 0.41677 | 0.39126 | 0.16490 | 0.49394 | 0.00407 | 0.00011 | 0.04422 | 0.00207 | 1.18334 | 0.00453 | 0.00299 | 0.04657 |
| **Chordc1** | 7.67044 | 0.06257 | 7.43680 | 4.19483 | 6.55466 | 10.79746 | 0.00000 | 2.50548 | 7.42944 | 0.02765 | 6.37075 | 3.41432 | 0.45776 | 5.12326 |
| **Chp1** | 9.59153 | 2.88264 | 16.25354 | 9.50419 | 18.95886 | 18.06903 | 2.43631 | 6.22094 | 18.34654 | 11.78770 | 9.39505 | 4.91289 | 8.59819 | 14.51778 |
| **Chpf** | 0.11371 | 0.07517 | 2.87664 | 2.16269 | 0.52216 | 0.20236 | 2.24302 | 1.76556 | 0.01451 | 0.05748 | 0.01217 | 5.76982 | 0.00000 | 0.00000 |
| **Chpf2** | 0.03404 | 0.02741 | 0.75492 | 1.09627 | 0.00000 | 1.30498 | 0.42393 | 1.12262 | 0.00398 | 0.00354 | 0.00000 | 1.51278 | 0.15811 | 0.00000 |

| | | | | | | | | | | | | | | |
|---|---|---|---|---|---|---|---|---|---|---|---|---|---|---|
| **Chpt1** | 10.80536 | 0.11912 | 5.24793 | 2.49415 | 4.15938 | 4.31248 | 0.16667 | 2.50686 | 14.81600 | 0.00000 | 2.75042 | 2.37711 | 2.47058 | 3.87377 |
| **Chrac1** | 4.71244 | 2.55205 | 3.67623 | 2.80406 | 7.25730 | 5.43043 | 1.37061 | 0.63789 | 0.14538 | 0.03684 | 2.99345 | 5.50750 | 0.82628 | 3.16345 |
| **Chrna1** | 1.37325 | 0.18763 | 0.00000 | 0.40782 | 0.28808 | 2.58211 | 0.42272 | 0.64798 | 1.62564 | 1.51161 | 0.02404 | 0.57557 | 10.89825 | 0.77116 |
| **Chrna10** | 0.03859 | 0.01323 | 0.34545 | 0.36870 | 0.11459 | 0.00000 | 0.00515 | 0.38533 | 0.01652 | 0.00574 | 0.00355 | 0.01237 | 0.08389 | 0.89266 |
| **Chrna5** | 0.41477 | 0.00447 | 0.16036 | 0.00000 | 0.00000 | 0.40133 | 0.00384 | 0.00384 | 0.79623 | 0.00803 | 0.00418 | 0.00437 | 0.40776 | 0.03110 |
| **Chrna9** | 1.35118 | 0.00402 | 0.74278 | 0.07388 | 0.76565 | 0.00000 | 0.00378 | 0.00379 | 0.64232 | 0.00572 | 0.00241 | 0.13779 | 0.68912 | 1.85925 |
| **Chrnb1** | 0.01145 | 0.00428 | 0.24338 | 1.01785 | 0.29649 | 0.11843 | 0.00366 | 0.00210 | 2.21010 | 0.00647 | 0.17936 | 0.01291 | 0.00227 | 1.35612 |
| **Chrnb4** | 0.00000 | 0.03348 | 0.00962 | 0.00000 | 0.00553 | 0.34109 | 0.51388 | 0.59098 | 0.22468 | 0.01503 | 0.00000 | 1.26739 | 1.42264 | 0.51335 |
| **Chrnd** | 0.04065 | 0.00445 | 0.10412 | 0.00751 | 0.17805 | 0.00738 | 0.00432 | 0.01098 | 0.01413 | 0.00834 | 0.00669 | 0.00818 | 0.00509 | 0.24164 |
| **Chrne** | 0.28822 | 0.00432 | 0.33646 | 0.00452 | 0.03208 | 0.02974 | 0.00400 | 0.00660 | 0.01314 | 0.00762 | 0.00672 | 0.00829 | 0.28696 | 0.07102 |
| **Chst10** | 0.01273 | 0.00444 | 0.01435 | 0.01373 | 0.01537 | 0.01512 | 0.00429 | 0.00995 | 0.01192 | 0.00732 | 0.01031 | 0.00986 | 0.00916 | 0.01140 |
| **Chst11** | 0.25035 | 0.00389 | 0.00505 | 0.00000 | 0.31151 | 0.01091 | 0.00389 | 0.33349 | 0.02051 | 0.00636 | 0.00531 | 0.00704 | 0.44282 | 0.00945 |
| **Chst12** | 1.84066 | 0.03774 | 3.98853 | 0.73775 | 1.32683 | 2.49780 | 0.40049 | 1.04269 | 0.01302 | 0.02206 | 0.00000 | 0.84634 | 0.44739 | 0.58683 |
| **Chst14** | 0.03397 | 0.05392 | 0.01450 | 0.00750 | 0.01318 | 0.10002 | 0.00424 | 0.00533 | 0.01138 | 0.00599 | 0.00861 | 0.01000 | 0.00680 | 0.00924 |
| **Chst2** | 0.03001 | 0.00452 | 0.01820 | 0.00924 | 0.00903 | 0.01424 | 0.01040 | 0.01192 | 0.01002 | 0.00680 | 0.01199 | 0.01270 | 0.13303 | 0.09129 |
| **Chst3** | 0.00003 | 0.01170 | 0.54921 | 0.06642 | 0.49130 | 2.85585 | 0.00427 | 0.75029 | 0.02547 | 0.04749 | 0.01094 | 0.00000 | 0.04336 | 0.93302 |
| **Chst5** | 0.24190 | 0.00431 | 0.03552 | 0.01384 | 0.02370 | 0.01117 | 0.00428 | 0.00953 | 0.00684 | 0.00600 | 0.00710 | 0.00666 | 0.91988 | 0.00800 |
| **Chst7** | 0.00903 | 0.00434 | 0.01080 | 0.05873 | 0.01240 | 0.01058 | 0.00416 | 0.00792 | 0.01229 | 0.00627 | 0.00755 | 0.01428 | 0.01560 | 0.00971 |
| **Chst8** | 0.00360 | 0.00420 | 0.01473 | 0.19094 | 0.00753 | 0.02470 | 0.00416 | 0.00507 | 0.01264 | 0.00558 | 0.00731 | 0.00679 | 0.39164 | 0.00000 |
| **Chsy1** | 0.06347 | 0.00393 | 0.00000 | 0.00000 | 0.07290 | 0.43360 | 0.00394 | 0.00424 | 0.02524 | 0.00686 | 0.00319 | 0.18295 | 2.07502 | 0.86013 |
| **Chtf18** | 2.20209 | 0.07969 | 8.77665 | 6.74379 | 6.86897 | 5.12248 | 0.05362 | 1.81938 | 0.00000 | 0.02087 | 2.04585 | 2.24486 | 0.44771 | 1.28090 |
| **Chtf8** | 0.78996 | 0.03320 | 2.58897 | 3.50966 | 3.67388 | 2.64343 | 1.71986 | 2.32282 | 0.41529 | 0.05954 | 1.22364 | 2.71090 | 1.94739 | 2.30626 |
| **Chtop** | 7.10155 | 8.56696 | 8.63051 | 10.50424 | 14.73685 | 14.16812 | 5.99496 | 6.56328 | 7.53920 | 0.21889 | 3.06765 | 9.25905 | 12.10312 | 7.57480 |
| **Chuk** | 1.09326 | 0.00303 | 2.78607 | 0.00001 | 1.23125 | 2.91815 | 0.49616 | 0.39689 | 3.78800 | 0.02796 | 0.34455 | 0.93594 | 3.14711 | 4.14742 |
| **Churc1** | 2.49994 | 1.60725 | 0.00000 | 2.30078 | 2.90881 | 3.45276 | 1.51089 | 1.83871 | 5.25482 | 8.81473 | 0.83367 | 2.53843 | 12.88216 | 4.05657 |
| **Ciao1** | 6.22794 | 0.09221 | 13.81489 | 8.02348 | 6.51100 | 6.42957 | 2.07007 | 2.70905 | 1.58303 | 7.84591 | 12.62751 | 5.14601 | 2.41120 | 6.00255 |
| **Ciapin1** | 4.11309 | 0.06836 | 7.00435 | 9.95153 | 3.23092 | 10.22692 | 4.28238 | 4.98023 | 0.21707 | 0.20486 | 8.08219 | 6.97368 | 0.00926 | 5.03470 |
| **Ciart** | 0.37863 | 0.01189 | 1.51134 | 1.11812 | 1.35010 | 0.03710 | 0.00375 | 0.00171 | 0.42148 | 0.00181 | 0.70328 | 0.00280 | 0.22400 | 4.23553 |
| **Cib1** | 2.01821 | 10.57227 | 5.83762 | 10.31960 | 7.86399 | 4.41933 | 3.08426 | 5.59790 | 0.00000 | 0.49816 | 5.36658 | 1.90512 | 0.04841 | 0.00000 |
| **Cib2** | 0.01184 | 0.00400 | 0.00559 | 0.04546 | 0.00898 | 0.78007 | 0.00408 | 0.45794 | 0.01199 | 0.00640 | 0.34154 | 0.00129 | 0.00793 | 0.00385 |
| **Cic** | 0.42851 | 3.34847 | 0.75576 | 6.22568 | 1.79860 | 0.46474 | 1.99085 | 6.19592 | 0.93453 | 0.74093 | 0.00000 | 3.28726 | 3.60421 | 1.08889 |
| **Cidea** | 0.00000 | 0.71080 | 0.25260 | 0.74010 | 0.17398 | 2.53404 | 1.29068 | 0.00157 | 0.56687 | 0.06816 | 2.21285 | 0.00000 | 0.01297 | 1.50974 |
| **Cideb** | 0.43639 | 0.00444 | 0.10760 | 0.04046 | 0.09656 | 0.04167 | 0.00450 | 0.00722 | 0.01689 | 0.00656 | 0.00570 | 0.00584 | 0.19223 | 0.03058 |
| **Cidec** | 0.01114 | 0.00457 | 0.01222 | 0.24408 | 0.01649 | 0.44509 | 0.00429 | 0.00954 | 0.01152 | 0.01647 | 0.03610 | 0.00317 | 0.09762 | 0.00142 |
| **Ciita** | 0.00000 | 0.00398 | 0.50767 | 0.29037 | 1.48774 | 0.00000 | 0.00407 | 0.00637 | 0.00891 | 0.00638 | 0.01159 | 0.01426 | 1.54126 | 0.09200 |
| **Cilp** | 2.02082 | 0.00375 | 3.49449 | 0.61049 | 3.05709 | 2.27257 | 0.00345 | 0.00000 | 0.01583 | 0.00000 | 1.80610 | 0.00001 | 0.27379 | 2.11838 |
| **Cilp2** | 0.13086 | 0.00398 | 0.01141 | 0.11044 | 0.95574 | 0.28821 | 0.00416 | 0.23507 | 0.00745 | 0.00826 | 0.00306 | 0.00201 | 0.13103 | 0.00000 |
| **Cinp** | 3.28572 | 0.04200 | 2.73281 | 2.35540 | 3.63157 | 3.65646 | 0.02029 | 0.74876 | 1.84073 | 0.04952 | 2.17864 | 2.07654 | 3.29575 | 3.62945 |
| **Cipc** | 0.39787 | 1.10729 | 0.20581 | 0.44343 | 0.08589 | 0.00000 | 1.18538 | 3.67349 | 0.12773 | 0.05784 | 2.51399 | 0.00006 | 0.99367 | 0.60206 |
| **Cir1** | 5.24058 | 5.67842 | 5.85328 | 2.82556 | 4.36451 | 1.49073 | 4.20647 | 1.88561 | 6.36620 | 0.21209 | 6.59220 | 3.51592 | 7.01817 | 8.04057 |
| **Cirbp** | 0.03679 | 0.11903 | 1.38370 | 4.24137 | 0.88152 | 0.26565 | 1.32556 | 5.60597 | 0.00621 | 0.05203 | 1.75678 | 3.21126 | 0.64362 | 1.42994 |
| **Cirh1a** | 18.84548 | 5.54372 | 24.02689 | 10.34396 | 27.25753 | 11.57992 | 4.85335 | 4.33113 | 14.00846 | 0.13545 | 10.91382 | 9.19837 | 11.32535 | 23.98926 |
| **Cisd1** | 1.79853 | 3.74802 | 0.15973 | 9.94307 | 4.39249 | 5.58243 | 1.67573 | 4.21321 | 1.08104 | 0.15909 | 0.04274 | 6.00768 | 0.27221 | 1.15949 |
| **Cisd2** | 2.20860 | 0.02327 | 1.86568 | 5.78793 | 1.69985 | 1.54060 | 0.71771 | 2.50145 | 1.67045 | 0.01708 | 1.66761 | 3.44683 | 1.95531 | 0.96901 |
| **Cisd3** | 0.26947 | 0.03835 | 0.11462 | 0.00634 | 0.12894 | 0.49797 | 0.01586 | 0.01387 | 0.00511 | 0.00714 | 0.16026 | 0.83520 | 0.06446 | 0.00716 |
| **Cish** | 1.83229 | 4.81175 | 6.48363 | 0.72542 | 12.80364 | 8.55639 | 7.66308 | 5.34199 | 2.00599 | 0.42904 | 0.02053 | 0.00000 | 0.16137 | 3.91505 |
| **Cit** | 0.18622 | 0.03490 | 2.07495 | 0.58555 | 3.81223 | 0.34610 | 0.94896 | 1.25664 | 1.09733 | 0.02211 | 0.01485 | 0.90246 | 0.00000 | 1.40506 |
| **Cited1** | 27.94452 | 0.05273 | 34.96181 | 34.19566 | 41.67152 | 11.98417 | 0.91549 | 0.00000 | 1.52046 | 0.09993 | 7.23007 | 3.18976 | 4.14346 | 13.32482 |
| **Cited2** | 0.57888 | 0.03054 | 0.74641 | 0.26889 | 0.00000 | 0.10678 | 0.00387 | 1.14250 | 0.00199 | 0.02977 | 0.00000 | 1.01465 | 3.07291 | 0.71071 |
| **Cited4** | 0.00983 | 3.55073 | 0.00963 | 0.17136 | 0.05047 | 0.00000 | 0.00335 | 0.21529 | 0.01330 | 0.02484 | 0.05252 | 0.00506 | 0.05802 | 0.00000 |
| **Ciz1** | 0.55689 | 0.08425 | 0.16578 | 0.94360 | 0.75050 | 1.27414 | 3.48204 | 1.64580 | 0.00000 | 11.74925 | 4.96385 | 2.12917 | 1.27187 | 1.88953 |
| **Ckap2** | 4.41941 | 5.13002 | 7.21124 | 0.52594 | 4.90999 | 3.93795 | 0.45823 | 1.60832 | 0.51207 | 13.74342 | 8.16805 | 2.93531 | 10.41826 | 13.22241 |
| **Ckap2l** | 0.00000 | 0.11666 | 2.52910 | 0.52626 | 1.97724 | 4.45626 | 1.43902 | 7.43301 | 2.91813 | 0.08409 | 10.37256 | 4.52863 | 0.71791 | 3.90668 |
| **Ckap4** | 0.36928 | 0.20104 | 1.33523 | 0.00000 | 0.01984 | 0.68215 | 0.00170 | 0.00000 | 0.00924 | 0.02686 | 0.00000 | 7.42859 | 0.00000 | 0.31534 |
| **Ckap5** | 5.29071 | 0.04197 | 3.23698 | 1.49683 | 6.59894 | 5.14057 | 0.01105 | 0.14844 | 7.65350 | 6.46981 | 7.83137 | 4.27452 | 4.00068 | 10.16922 |

| | | | | | | | | | | | | | | |
|---|---|---|---|---|---|---|---|---|---|---|---|---|---|---|
| **Ckb** | 15.38504 | 9.19004 | 35.06030 | 47.82909 | 30.67122 | 7.05227 | 6.03641 | 24.27701 | 0.00000 | 11.45178 | 6.99477 | 26.08570 | 1.06821 | 0.78881 |
| **Cklf** | 0.00000 | 0.49291 | 0.09376 | 0.27515 | 0.12954 | 0.52157 | 2.01057 | 0.63624 | 0.33759 | 0.07749 | 0.00000 | 1.06717 | 0.77227 | 0.00000 |
| **Ckm** | 0.00681 | 0.00391 | 0.17473 | 0.12993 | 0.01164 | 0.00013 | 0.00389 | 0.13333 | 0.01049 | 0.00807 | 0.09213 | 0.03848 | 0.00665 | 0.02432 |
| **Ckmt1** | 7.58652 | 0.36250 | 14.96108 | 20.81179 | 9.82018 | 16.92570 | 4.27963 | 10.64089 | 4.48074 | 12.85454 | 13.08611 | 9.69675 | 0.00000 | 5.46794 |
| **Cks1b** | 6.76423 | 5.84496 | 7.37845 | 11.39367 | 12.20421 | 18.74620 | 7.62220 | 7.64928 | 3.26007 | 9.84761 | 4.73926 | 9.21704 | 6.64263 | 9.88057 |
| **Cks1brt** | 0.40427 | 0.53446 | 0.29680 | 0.78369 | 0.71786 | 1.58627 | 0.79179 | 0.39687 | 0.25731 | 0.76771 | 0.30692 | 0.64493 | 0.61641 | 0.59576 |
| **Cks2** | 25.96903 | 4.83249 | 7.07316 | 15.48837 | 38.78464 | 21.02663 | 9.05383 | 12.30567 | 12.12391 | 14.06418 | 5.44882 | 13.36613 | 24.55913 | 28.09602 |
| **Clasp1** | 0.00000 | 0.02740 | 1.01599 | 1.61454 | 1.21188 | 0.22584 | 0.32395 | 1.03669 | 0.03321 | 0.02388 | 0.37396 | 2.31668 | 0.48725 | 0.00454 |
| **Clasp2** | 0.10842 | 0.33449 | 0.12449 | 0.50223 | 0.57741 | 0.07715 | 0.00374 | 0.00430 | 0.66715 | 0.01313 | 0.01047 | 0.11628 | 1.20467 | 2.59596 |
| **Clasrp** | 0.16653 | 0.00367 | 0.16286 | 4.56181 | 0.52421 | 0.48733 | 0.06772 | 1.83521 | 0.00570 | 0.02329 | 0.19351 | 1.42721 | 0.66093 | 0.31445 |
| **Clca3a1** | 0.05525 | 0.03606 | 0.08675 | 0.12362 | 0.04559 | 0.17101 | 0.03446 | 0.12458 | 0.44705 | 0.15250 | 0.15681 | 0.11511 | 0.40854 | 0.22082 |
| **Clca3a2** | 0.00784 | 0.25617 | 0.01911 | 0.06781 | 0.05107 | 0.03935 | 0.01028 | 0.01005 | 0.11021 | 0.02294 | 0.02065 | 0.01555 | 0.47482 | 0.10161 |
| **Clca4c.ps** | 0.07823 | 0.00443 | 0.02762 | 0.01809 | 0.02890 | 0.02564 | 0.00430 | 0.00883 | 0.01572 | 0.00673 | 0.00787 | 0.01193 | 0.02066 | 0.05518 |
| **Clcc1** | 2.04264 | 0.02384 | 2.97983 | 0.26467 | 1.52034 | 3.21857 | 0.18715 | 0.27786 | 1.46835 | 0.00589 | 0.51410 | 0.93977 | 0.50700 | 3.26905 |
| **Clcn2** | 1.01886 | 0.01982 | 3.41835 | 1.02931 | 1.28375 | 0.33417 | 0.00391 | 0.46002 | 0.00438 | 0.00709 | 0.78752 | 1.51991 | 0.59697 | 1.87140 |
| **Clcn3** | 6.58332 | 0.00976 | 3.42104 | 1.30876 | 2.17694 | 9.13929 | 1.64495 | 1.79948 | 6.48170 | 0.05881 | 0.73567 | 1.03157 | 0.42101 | 2.45212 |
| **Clcn4** | 3.29524 | 0.03297 | 3.10017 | 2.13001 | 1.70065 | 4.58660 | 0.00215 | 2.11507 | 7.08217 | 0.02167 | 0.07037 | 2.51888 | 0.00000 | 1.41145 |
| **Clcn5** | 0.06383 | 0.00384 | 0.28436 | 0.04023 | 0.97100 | 0.69155 | 0.00344 | 1.53456 | 1.50134 | 0.00682 | 0.00215 | 0.00000 | 0.52043 | 0.00000 |
| **Clcn6** | 1.79249 | 3.38943 | 1.75892 | 1.26035 | 1.05360 | 0.81018 | 1.15667 | 0.05267 | 0.01392 | 0.05941 | 1.49718 | 1.48187 | 1.17458 | 3.24943 |
| **Clcn7** | 2.03758 | 0.03187 | 7.70400 | 6.16191 | 1.56777 | 5.50624 | 0.01408 | 4.16154 | 0.25366 | 0.01914 | 0.00000 | 4.41189 | 0.31082 | 2.75796 |
| **Clcnka** | 1.93182 | 0.00354 | 5.47862 | 1.70321 | 1.90796 | 0.54420 | 0.00294 | 0.00000 | 0.02505 | 0.00000 | 1.98573 | 0.00172 | 0.04951 | 1.29586 |
| **Clcnkb** | 0.00000 | 0.00379 | 0.24867 | 1.06558 | 0.22526 | 0.48252 | 0.00326 | 2.38107 | 0.01600 | 0.00451 | 0.39830 | 0.00109 | 0.00440 | 0.00000 |
| **Cldn10** | 0.01189 | 0.00430 | 0.01062 | 0.01058 | 0.13255 | 0.01293 | 0.00415 | 0.00901 | 0.01113 | 0.00626 | 0.00916 | 0.00856 | 0.54857 | 0.04460 |
| **Cldn12** | 1.35679 | 0.00913 | 0.58728 | 0.44244 | 0.00000 | 1.81546 | 0.39102 | 0.00767 | 3.46557 | 5.96457 | 3.27801 | 1.04889 | 1.49904 | 0.47334 |
| **Cldn15** | 0.01579 | 0.00430 | 0.25580 | 0.00526 | 0.00279 | 0.00543 | 0.00416 | 0.01028 | 0.01051 | 0.00636 | 0.00680 | 0.00802 | 0.29373 | 0.09185 |
| **Cldn19** | 0.01014 | 0.00374 | 0.00269 | 0.16394 | 0.00736 | 0.00000 | 0.00386 | 0.00762 | 0.01338 | 0.00129 | 1.90294 | 0.00812 | 0.00703 | 0.00345 |
| **Cldn22** | 0.01304 | 0.01522 | 0.35183 | 0.00000 | 0.00600 | 0.00000 | 0.00433 | 0.14772 | 0.01096 | 0.01011 | 0.00706 | 0.01240 | 0.00576 | 0.01583 |
| **Cldn3** | 0.00476 | 3.28492 | 0.00397 | 0.04231 | 1.13539 | 0.00631 | 0.17893 | 4.21939 | 0.01425 | 0.02803 | 0.00436 | 0.01804 | 0.01720 | 0.00015 |
| **Cldn34b1** | 0.00941 | 0.00466 | 0.16404 | 0.00027 | 0.02197 | 0.03170 | 0.00426 | 0.00886 | 0.01072 | 0.00647 | 0.00864 | 0.01000 | 0.00749 | 0.01183 |
| **Cldn34b2** | 0.01266 | 0.00479 | 0.02711 | 0.00554 | 0.06202 | 0.07956 | 0.00429 | 0.00760 | 0.00993 | 0.00695 | 0.00839 | 0.00944 | 0.01131 | 0.01629 |
| **Cldn34c1** | 0.06379 | 0.00416 | 0.02045 | 0.02292 | 0.03403 | 0.01689 | 0.00419 | 0.01959 | 0.07202 | 0.00886 | 0.02365 | 0.13251 | 0.04625 | 0.02893 |
| **Cldn34d** | 0.01092 | 0.00445 | 0.19986 | 0.00000 | 0.01124 | 0.00000 | 0.00489 | 0.15851 | 0.87235 | 0.00586 | 0.00504 | 0.00664 | 0.00410 | 0.00049 |
| **Cldn4** | 0.98851 | 16.14407 | 28.20544 | 18.95108 | 13.45595 | 24.45262 | 11.86742 | 37.18392 | 4.51715 | 12.91006 | 5.13588 | 1.46753 | 0.00000 | 2.03954 |
| **Cldn5** | 0.14123 | 0.00441 | 0.00744 | 0.00906 | 0.23089 | 0.01142 | 0.00431 | 0.01212 | 0.00592 | 0.00624 | 0.00781 | 0.00833 | 0.00645 | 0.08423 |
| **Cldn6** | 0.00000 | 6.16231 | 5.50560 | 4.76631 | 6.30252 | 17.89442 | 5.18126 | 17.62404 | 2.27044 | 2.00789 | 2.66178 | 0.00000 | 0.36332 | 10.85578 |
| **Cldn7** | 5.52106 | 4.84195 | 12.58692 | 19.86643 | 5.71337 | 20.67015 | 8.77614 | 20.29511 | 0.00000 | 1.01120 | 3.67768 | 2.04668 | 0.00000 | 4.66466 |
| **Cldn9** | 0.03834 | 0.00391 | 0.37052 | 0.49448 | 0.14065 | 2.31508 | 0.35108 | 0.10317 | 0.00514 | 0.00815 | 0.00000 | 0.45796 | 0.55186 | 0.00000 |
| **Cldnd1** | 3.12814 | 0.00806 | 2.04126 | 0.52722 | 0.34381 | 4.70907 | 0.00818 | 0.01504 | 1.03690 | 0.02568 | 2.15180 | 2.87001 | 4.28557 | 1.89337 |
| **Clec10a** | 0.03415 | 0.01176 | 1.47729 | 1.42719 | 0.43946 | 4.36585 | 0.88886 | 0.00000 | 0.28063 | 0.02480 | 1.44489 | 0.00000 | 3.23160 | 0.70865 |
| **Clec11a** | 0.01499 | 0.00444 | 0.02781 | 0.07674 | 0.01179 | 0.01341 | 0.00442 | 0.03118 | 0.01248 | 0.00767 | 0.00918 | 0.00838 | 0.00677 | 0.01092 |
| **Clec16a** | 0.16742 | 2.00415 | 0.85451 | 0.45584 | 0.81115 | 1.41471 | 0.00361 | 0.98262 | 0.00047 | 0.04370 | 2.80457 | 0.00000 | 0.64672 | 0.06198 |
| **Clec2e** | 0.01386 | 0.00443 | 0.01968 | 0.01341 | 0.01529 | 0.01590 | 0.00431 | 0.00966 | 0.01117 | 0.00662 | 0.00841 | 0.00957 | 0.08155 | 0.00533 |
| **Clec2f** | 0.00350 | 0.00316 | 0.08279 | 0.75866 | 0.26242 | 0.09626 | 0.00358 | 0.00206 | 0.01866 | 0.00543 | 0.05178 | 0.99319 | 0.00571 | 0.00000 |
| **Clec2g** | 0.06140 | 0.00429 | 0.07263 | 0.00934 | 0.03580 | 0.35338 | 0.00432 | 0.00918 | 0.10175 | 0.01099 | 0.00538 | 0.00861 | 0.61950 | 1.25384 |
| **Clec2h** | 0.01502 | 0.00442 | 0.01585 | 0.01860 | 0.02460 | 0.02962 | 0.00454 | 0.01779 | 0.04036 | 0.00663 | 0.00965 | 0.01341 | 0.03738 | 0.06213 |
| **Clec2i** | 0.02940 | 0.00437 | 0.03041 | 0.01140 | 0.00000 | 0.01659 | 0.00417 | 0.00881 | 0.10021 | 0.00594 | 0.04881 | 0.00735 | 0.15591 | 0.28117 |
| **Clec2l** | 0.90667 | 0.00445 | 0.05020 | 0.00164 | 0.00264 | 0.21689 | 0.00411 | 0.00668 | 0.00817 | 0.00742 | 0.00445 | 0.00782 | 0.00546 | 0.00236 |
| **Clec3b** | 0.01769 | 0.00454 | 0.02555 | 0.02393 | 0.01182 | 0.02472 | 0.00446 | 0.01370 | 0.00885 | 0.00823 | 0.00826 | 0.01393 | 0.00456 | 0.10405 |
| **Clec4a1** | 0.13006 | 0.00516 | 0.11781 | 0.09141 | 0.12553 | 0.27888 | 0.00414 | 0.01842 | 0.01436 | 0.02486 | 0.00570 | 0.06609 | 0.85469 | 0.00000 |
| **Clgn** | 0.00611 | 0.00416 | 0.08752 | 0.09399 | 0.00977 | 0.00676 | 0.00402 | 0.01046 | 0.00729 | 0.00776 | 0.00639 | 0.01842 | 0.10664 | 0.04523 |
| **Clhc1** | 0.00000 | 0.00392 | 0.14715 | 0.58982 | 0.54595 | 0.00000 | 0.00357 | 0.00468 | 1.64176 | 0.00541 | 0.00427 | 0.01226 | 0.79887 | 0.12002 |
| **Clic1** | 0.69530 | 0.07872 | 0.03181 | 0.27135 | 0.62016 | 0.75819 | 0.00254 | 1.73780 | 0.00243 | 0.03613 | 0.00000 | 4.77631 | 0.00399 | 0.45852 |
| **Clic3** | 0.00464 | 0.01920 | 0.02036 | 2.36191 | 0.02614 | 6.34366 | 0.03426 | 0.69711 | 0.01098 | 0.03362 | 0.09840 | 0.00000 | 0.01336 | 0.00000 |
| **Clic4** | 3.73934 | 0.94508 | 9.77753 | 4.73246 | 2.84911 | 26.81160 | 5.81001 | 18.74684 | 4.07658 | 12.88741 | 9.84161 | 8.71508 | 7.65378 | 0.71699 |
| **Clint1** | 4.04105 | 0.20998 | 4.85598 | 0.00000 | 3.81693 | 7.10285 | 0.06333 | 0.32730 | 2.54388 | 0.03747 | 3.86609 | 5.77493 | 1.62686 | 6.56718 |

| | | | | | | | | | | | | | | |
|---|---|---|---|---|---|---|---|---|---|---|---|---|---|---|
| **Clip1** | 3.28386 | 0.00989 | 3.08559 | 0.70332 | 5.79493 | 0.99297 | 0.00308 | 1.29344 | 1.02932 | 0.00279 | 1.17702 | 1.07307 | 1.73237 | 8.07916 |
| **Clip2** | 0.06273 | 0.00384 | 0.01771 | 0.31062 | 0.00499 | 0.00449 | 0.00398 | 0.00435 | 0.03722 | 0.00592 | 0.00524 | 0.17423 | 0.09933 | 0.01794 |
| **Clip3** | 0.29171 | 0.00449 | 0.00939 | 0.01019 | 0.11152 | 0.01084 | 0.00436 | 0.01157 | 0.01014 | 0.00673 | 0.00720 | 0.00798 | 0.00417 | 0.01199 |
| **Clip4** | 0.00157 | 0.00410 | 0.00477 | 0.30631 | 0.00860 | 0.00828 | 0.00423 | 0.00633 | 0.00694 | 0.00542 | 0.00602 | 0.00668 | 0.35186 | 0.63767 |
| **Clk1** | 14.60147 | 0.05453 | 10.40916 | 2.24724 | 12.19017 | 5.01360 | 5.53604 | 5.11952 | 22.33869 | 0.23806 | 3.94694 | 8.08589 | 8.72984 | 15.31681 |
| **Clk2** | 2.75857 | 0.01416 | 1.36106 | 2.82180 | 2.45794 | 0.82967 | 0.02086 | 0.96198 | 0.30190 | 9.19209 | 2.68826 | 3.39663 | 2.15543 | 0.80128 |
| **Clk3** | 11.02213 | 6.95417 | 9.50439 | 3.64257 | 4.02754 | 9.63532 | 4.58165 | 4.55722 | 0.04457 | 0.23567 | 2.75290 | 4.92269 | 3.65692 | 0.40977 |
| **Clk4** | 0.57990 | 5.87412 | 1.31975 | 2.36025 | 3.38613 | 1.47634 | 0.00197 | 0.00000 | 0.88250 | 0.00000 | 1.36611 | 0.37843 | 2.85792 | 5.95841 |
| **Clmn** | 0.49348 | 0.02009 | 0.48108 | 1.15414 | 0.29328 | 2.53020 | 0.42901 | 4.00276 | 0.65064 | 0.12658 | 0.00731 | 0.75790 | 1.19236 | 1.73160 |
| **Clmp** | 0.00000 | 0.00392 | 1.51094 | 0.33547 | 1.84758 | 0.60873 | 0.00915 | 0.00000 | 0.11952 | 0.00970 | 0.27432 | 0.00390 | 1.44160 | 0.93480 |
| **Cln3** | 5.40649 | 4.76799 | 16.77448 | 9.49059 | 8.03152 | 6.02565 | 3.33668 | 6.36801 | 2.30120 | 0.20703 | 3.48619 | 2.36783 | 1.43931 | 9.01376 |
| **Cln5** | 1.45253 | 5.70998 | 0.63849 | 1.70762 | 0.00000 | 1.85116 | 0.00584 | 1.17716 | 4.35424 | 0.02100 | 3.39176 | 4.73047 | 0.00000 | 2.38480 |
| **Cln6** | 0.40985 | 0.01792 | 0.37350 | 0.00000 | 0.00035 | 0.77024 | 0.00958 | 4.37033 | 0.00470 | 0.00789 | 1.60796 | 0.16977 | 0.00233 | 0.33597 |
| **Cln8** | 7.23629 | 3.03338 | 15.37155 | 12.31180 | 4.68455 | 9.43890 | 6.68106 | 7.08286 | 3.36642 | 0.38586 | 7.15675 | 4.71848 | 1.55953 | 8.34226 |
| **Clns1a** | 20.51132 | 0.30914 | 26.63859 | 16.79177 | 30.54884 | 8.93399 | 0.18890 | 9.16453 | 10.73854 | 0.14307 | 4.27733 | 14.56142 | 8.46678 | 11.98974 |
| **Clock** | 0.10509 | 0.00995 | 0.00000 | 1.16345 | 0.21448 | 2.00273 | 0.00323 | 0.00000 | 1.07731 | 0.00253 | 1.14635 | 0.35199 | 1.70334 | 2.74230 |
| **Clp1** | 8.84625 | 0.06147 | 7.80826 | 2.39320 | 8.06629 | 7.09447 | 2.51140 | 4.88138 | 8.63782 | 0.08374 | 3.13132 | 0.93657 | 11.90955 | 11.80800 |
| **Clpb** | 3.03823 | 0.05248 | 6.57199 | 6.51419 | 1.10590 | 4.55500 | 0.01830 | 5.78069 | 0.52584 | 14.97725 | 3.16126 | 3.78513 | 0.00000 | 1.78282 |
| **Clpp** | 10.81492 | 8.13558 | 20.68860 | 22.10116 | 16.34428 | 19.00608 | 6.12406 | 5.97389 | 0.00000 | 2.85647 | 8.86026 | 15.39368 | 6.96088 | 1.97012 |
| **Clps** | 0.03001 | 0.00436 | 0.01055 | 0.01650 | 0.06385 | 0.01228 | 0.00446 | 0.00882 | 0.00580 | 0.00620 | 0.00817 | 0.00841 | 0.36865 | 0.17199 |
| **Clptm1** | 2.32648 | 3.65270 | 16.04840 | 14.09097 | 10.67547 | 0.00000 | 3.49279 | 12.45834 | 1.91563 | 8.61863 | 1.39425 | 9.59919 | 2.16126 | 0.04283 |
| **Clptm1l** | 7.36794 | 0.03216 | 13.06936 | 6.35010 | 6.89640 | 13.09129 | 5.40453 | 2.58877 | 5.40904 | 8.55241 | 6.50961 | 6.27026 | 1.77403 | 11.11378 |
| **Clpx** | 28.99084 | 4.67581 | 29.81112 | 7.30313 | 36.61324 | 11.66443 | 5.72593 | 6.11011 | 32.63954 | 0.11830 | 9.93789 | 8.59691 | 15.67851 | 28.32644 |
| **Clrn1** | 0.02043 | 0.00445 | 0.02908 | 0.01346 | 0.03104 | 0.03367 | 0.00436 | 0.00904 | 0.08161 | 0.00690 | 0.01784 | 0.03133 | 0.00732 | 0.03914 |
| **Clspn** | 5.87114 | 0.00987 | 3.86470 | 0.81519 | 3.75472 | 4.83741 | 0.03317 | 0.24889 | 5.89023 | 0.00000 | 4.48815 | 2.10561 | 7.73405 | 6.67450 |
| **Clstn1** | 1.06770 | 7.32194 | 0.11629 | 8.54005 | 1.89956 | 5.16559 | 0.00498 | 4.68672 | 0.06089 | 7.56478 | 2.26276 | 5.31107 | 0.38113 | 0.53373 |
| **Clstn3** | 0.00000 | 0.01169 | 1.84893 | 0.76096 | 1.21736 | 0.00000 | 0.00392 | 0.74244 | 0.00764 | 0.01176 | 0.80525 | 1.58645 | 0.00000 | 0.56332 |
| **Clta** | 10.25051 | 2.60283 | 2.85211 | 31.80526 | 27.50299 | 14.72839 | 3.02750 | 7.07896 | 2.76397 | 11.75897 | 1.84798 | 12.17905 | 9.41419 | 5.87104 |
| **Cltb** | 3.01241 | 0.01818 | 1.53355 | 1.48943 | 1.82217 | 2.90588 | 2.18106 | 2.67151 | 0.82042 | 0.22801 | 0.00000 | 1.47332 | 5.02579 | 0.50086 |
| **Cltc** | 1.28689 | 2.98926 | 13.96366 | 8.20584 | 10.37407 | 16.63808 | 4.81289 | 11.24722 | 15.36256 | 11.53958 | 17.23767 | 13.20129 | 5.84905 | 8.71562 |
| **Clu** | 0.01619 | 7.13819 | 0.12877 | 0.00000 | 1.24496 | 0.00000 | 0.00229 | 0.00589 | 0.37105 | 0.00523 | 0.00258 | 0.00652 | 0.03362 | 0.20245 |
| **Cluap1** | 3.85502 | 0.02345 | 4.63327 | 2.77152 | 2.21487 | 1.19256 | 4.40107 | 3.53223 | 1.22873 | 10.10084 | 0.96526 | 1.71665 | 0.15902 | 3.91235 |
| **Cluh** | 3.27299 | 2.54049 | 18.76064 | 15.00793 | 7.17004 | 13.16292 | 1.88260 | 7.79542 | 6.16256 | 0.21649 | 8.20397 | 9.66424 | 2.56823 | 10.49292 |
| **Clvs2** | 0.00522 | 0.00436 | 0.04225 | 0.00662 | 0.28586 | 0.01218 | 0.00448 | 0.00905 | 0.01746 | 0.00610 | 0.00905 | 0.00818 | 0.63632 | 0.00726 |
| **Clybl** | 0.89726 | 1.10113 | 1.90807 | 3.83032 | 2.26284 | 1.79338 | 3.34817 | 0.03893 | 2.17224 | 0.13260 | 0.00000 | 0.78078 | 3.98954 | 2.62357 |
| **Cmah** | 9.08730 | 1.18967 | 5.43703 | 4.25101 | 5.00420 | 4.39365 | 1.71853 | 0.92118 | 18.33981 | 1.64339 | 4.17240 | 5.80423 | 8.87082 | 8.96159 |
| **Cmas** | 1.22507 | 0.01444 | 1.09025 | 1.95666 | 0.91124 | 3.87115 | 1.26155 | 3.13824 | 0.00419 | 0.06046 | 0.91633 | 6.37665 | 2.17915 | 0.45365 |
| **Cmbl** | 0.00000 | 0.06724 | 2.03622 | 2.28246 | 0.56412 | 5.39701 | 2.41332 | 2.43130 | 0.11370 | 13.98741 | 4.59535 | 2.62349 | 0.00000 | 0.18852 |
| **Cmc1** | 0.17481 | 0.11018 | 0.09943 | 0.00000 | 0.09089 | 0.82373 | 1.00969 | 0.86117 | 0.41152 | 0.91240 | 0.66076 | 1.91682 | 1.22664 | 1.20456 |
| **Cmc2** | 2.87076 | 0.13385 | 1.06412 | 7.31732 | 6.25691 | 5.72123 | 3.28296 | 6.60386 | 0.21139 | 0.24101 | 0.91274 | 5.79865 | 0.67145 | 4.06029 |
| **Cmc4** | 0.49737 | 0.00446 | 0.08981 | 0.00000 | 0.60512 | 0.54441 | 0.00411 | 0.00588 | 0.00525 | 0.00714 | 0.04217 | 0.00921 | 0.00316 | 0.25036 |
| **Cmip** | 1.25680 | 0.12321 | 9.77844 | 3.62029 | 1.46395 | 2.69153 | 2.12350 | 7.39047 | 0.92377 | 6.81219 | 0.91607 | 0.72933 | 0.00000 | 1.76671 |
| **Cmklr1** | 0.01239 | 0.00443 | 0.00429 | 0.24453 | 0.13634 | 0.42814 | 0.00368 | 0.70290 | 0.01376 | 0.02210 | 0.00217 | 0.05077 | 0.00547 | 0.01513 |
| **Cmpk1** | 4.30733 | 0.02293 | 3.52126 | 2.14576 | 3.25339 | 4.45560 | 1.30430 | 2.94151 | 4.12565 | 0.00000 | 4.89401 | 4.25108 | 1.78236 | 3.89846 |
| **Cmpk2** | 0.00053 | 0.00433 | 0.01947 | 0.01133 | 0.09540 | 0.00923 | 0.00433 | 0.01112 | 0.00531 | 0.00639 | 0.00707 | 0.00909 | 0.02822 | 1.26056 |
| **Cmss1** | 5.17603 | 0.08099 | 7.13463 | 4.22043 | 3.78040 | 3.44250 | 0.13614 | 1.40293 | 0.07386 | 0.01549 | 5.10484 | 4.72816 | 2.98350 | 5.25079 |
| **Cmtm3** | 0.01581 | 0.00409 | 0.00553 | 0.42173 | 0.04114 | 0.01317 | 0.00427 | 0.00455 | 0.01807 | 0.00618 | 0.00603 | 0.01640 | 0.00637 | 0.00513 |
| **Cmtm4** | 4.12776 | 0.08897 | 6.91657 | 1.52395 | 7.00157 | 9.69114 | 0.01937 | 3.69731 | 6.87058 | 0.07413 | 4.39291 | 2.82665 | 7.23037 | 6.46403 |
| **Cmtm6** | 4.71323 | 6.81349 | 10.19735 | 7.05791 | 5.15128 | 10.93888 | 3.32968 | 4.06523 | 3.98448 | 12.82854 | 7.75164 | 7.25376 | 4.17068 | 7.04654 |
| **Cmtm7** | 6.83014 | 0.82996 | 5.15528 | 9.90435 | 8.76532 | 13.21260 | 4.13962 | 8.97917 | 1.79906 | 11.72297 | 2.61553 | 8.67338 | 0.00000 | 0.86507 |
| **Cmtm8** | 0.01796 | 0.00391 | 0.00463 | 0.11329 | 0.01336 | 0.07564 | 0.00399 | 0.57589 | 0.00986 | 0.00904 | 0.00437 | 0.00526 | 0.00863 | 0.12876 |
| **Cmtr1** | 0.01052 | 0.00456 | 0.04139 | 0.02629 | 0.02369 | 0.01128 | 0.00403 | 0.02051 | 0.01528 | 0.00697 | 0.03880 | 0.02962 | 0.00495 | 0.03157 |
| **Cmtr2** | 0.04476 | 0.01162 | 0.00000 | 1.07939 | 0.43569 | 0.90439 | 0.00377 | 0.04018 | 0.06631 | 0.00287 | 0.83002 | 0.00438 | 0.46042 | 0.63492 |
| **Cmya5** | 0.00397 | 0.00426 | 0.00842 | 0.01011 | 0.22323 | 0.01420 | 0.00411 | 0.00841 | 0.00697 | 0.00735 | 0.00929 | 0.00827 | 0.81240 | 0.01986 |
| **Cnbd2** | 7.42431 | 0.03431 | 4.35346 | 5.54423 | 7.21427 | 4.75256 | 2.76758 | 1.59210 | 4.60618 | 0.25382 | 0.15453 | 2.23249 | 6.25541 | 5.25802 |

| | | | | | | | | | | | | | | |
|---|---|---|---|---|---|---|---|---|---|---|---|---|---|---|
| **Cnbp** | 31.79297 | 10.42201 | 45.59480 | 42.47744 | 42.54003 | 35.11465 | 12.45317 | 21.56176 | 29.15407 | 19.61888 | 27.49030 | 23.97194 | 16.50267 | 36.79741 |
| **Cndp1** | 0.00364 | 0.00424 | 0.00000 | 0.00000 | 0.24295 | 0.62182 | 0.06588 | 0.00177 | 0.01372 | 0.00901 | 0.54592 | 0.00580 | 0.26511 | 0.00000 |
| **Cndp2** | 8.74440 | 10.68282 | 28.73760 | 31.78650 | 13.40850 | 30.44534 | 11.14506 | 27.77084 | 8.19599 | 19.70952 | 23.70931 | 10.97337 | 1.99167 | 14.33463 |
| **Cnep1r1** | 5.59400 | 0.00978 | 1.40447 | 1.13698 | 1.84389 | 2.97709 | 0.12558 | 0.82393 | 5.71912 | 0.00000 | 1.59974 | 0.00000 | 6.94972 | 5.41933 |
| **Cnga1** | 0.01102 | 0.00444 | 0.01339 | 0.01343 | 0.00915 | 0.01256 | 0.00435 | 0.01021 | 0.01090 | 0.00694 | 0.00868 | 0.00995 | 0.00928 | 0.10820 |
| **Cnga3** | 0.00398 | 0.09582 | 0.03190 | 0.06924 | 0.00870 | 0.03372 | 0.00411 | 0.02335 | 0.05843 | 0.08197 | 0.01046 | 0.01928 | 0.06070 | 0.75731 |
| **Cnga4** | 0.11170 | 0.03589 | 0.00966 | 0.04333 | 0.00110 | 0.00145 | 0.00449 | 0.58042 | 0.00643 | 0.00903 | 0.00679 | 0.01952 | 0.00855 | 0.57266 |
| **Cngb1** | 0.00071 | 0.00416 | 0.00938 | 0.01264 | 0.03507 | 0.01012 | 0.00406 | 0.00811 | 0.04580 | 0.00585 | 0.00781 | 0.00731 | 1.16696 | 0.67547 |
| **Cnih1** | 8.35735 | 6.52985 | 8.01303 | 8.29443 | 7.26619 | 14.03066 | 3.75856 | 6.86866 | 5.00006 | 1.20280 | 4.76357 | 11.65540 | 6.15108 | 3.45924 |
| **Cnih2** | 0.16433 | 0.01500 | 0.10544 | 0.00912 | 0.00203 | 0.00318 | 0.00452 | 0.22172 | 0.01080 | 0.01488 | 0.00366 | 0.33144 | 0.03802 | 0.00372 |
| **Cnih4** | 4.78740 | 0.26776 | 0.37687 | 4.47017 | 4.75129 | 8.02382 | 3.58186 | 2.96185 | 1.87304 | 0.38056 | 1.94006 | 6.15038 | 2.98262 | 4.64218 |
| **Cnksr1** | 0.00000 | 0.01306 | 0.70646 | 0.00000 | 0.54055 | 0.72708 | 0.12198 | 4.10413 | 0.01063 | 0.03584 | 0.00177 | 0.03669 | 0.03511 | 0.72569 |
| **Cnksr3** | 0.27231 | 0.01208 | 0.43161 | 0.36841 | 0.30749 | 0.83383 | 0.00523 | 0.04649 | 0.00494 | 0.01315 | 0.00000 | 0.24596 | 1.73771 | 0.19355 |
| **Cnn1** | 0.06518 | 0.00454 | 0.00160 | 0.01342 | 0.18688 | 0.00811 | 0.00447 | 0.01041 | 0.73144 | 0.00699 | 0.00572 | 0.00815 | 0.01115 | 0.00000 |
| **Cnn2** | 0.83703 | 9.62849 | 1.49993 | 3.02785 | 3.26209 | 8.14244 | 8.44435 | 19.45492 | 0.00000 | 11.97413 | 2.68096 | 0.00000 | 0.00000 | 0.01508 |
| **Cnn3** | 8.80570 | 8.68049 | 14.04036 | 20.96130 | 4.58229 | 27.09744 | 7.88751 | 14.19311 | 5.02871 | 14.69555 | 17.96037 | 11.96358 | 3.93645 | 2.81209 |
| **Cnnm1** | 0.37702 | 0.00417 | 0.00156 | 0.35905 | 0.24185 | 0.00508 | 0.00436 | 0.00681 | 0.01632 | 0.00684 | 0.00640 | 0.00683 | 0.00273 | 0.15415 |
| **Cnnm2** | 1.84432 | 0.03823 | 7.42014 | 2.24964 | 1.32259 | 3.22393 | 1.76660 | 2.37919 | 0.00000 | 4.26707 | 0.81887 | 4.29638 | 5.49011 | 5.75769 |
| **Cnnm3** | 0.15125 | 0.86431 | 2.38653 | 0.91845 | 2.49966 | 0.52857 | 2.42844 | 1.82274 | 2.41306 | 0.07372 | 0.00000 | 1.17030 | 3.37382 | 2.22166 |
| **Cnnm4** | 9.37105 | 0.03213 | 10.50934 | 1.39468 | 5.67950 | 4.86447 | 0.27295 | 3.78352 | 5.33533 | 0.02559 | 2.99276 | 0.52308 | 0.49911 | 10.54888 |
| **Cnot1** | 20.60540 | 6.66214 | 28.45664 | 9.70408 | 27.92054 | 13.84552 | 5.34923 | 7.08035 | 26.36091 | 0.13749 | 12.03912 | 17.13297 | 8.52775 | 25.90252 |
| **Cnot10** | 4.22120 | 0.08589 | 10.79810 | 4.29349 | 7.44341 | 7.43994 | 0.14829 | 0.53272 | 1.26264 | 0.05238 | 9.23683 | 9.15913 | 3.16323 | 5.84523 |
| **Cnot11** | 5.05785 | 0.01030 | 3.54043 | 1.28855 | 2.39804 | 1.62697 | 0.01010 | 0.45737 | 1.45555 | 0.02821 | 0.58434 | 2.61617 | 2.11252 | 1.96600 |
| **Cnot2** | 10.16205 | 2.26589 | 12.15414 | 2.68229 | 14.14578 | 9.08959 | 3.51163 | 6.38654 | 22.39276 | 0.13068 | 11.76585 | 5.49272 | 3.83094 | 10.61857 |
| **Cnot3** | 0.39287 | 0.02542 | 4.09626 | 2.05940 | 1.35250 | 0.34299 | 1.62560 | 2.92106 | 0.00243 | 1.97080 | 0.04434 | 1.39740 | 0.07509 | 0.26088 |
| **Cnot4** | 1.96456 | 0.04524 | 4.16467 | 3.40309 | 3.40717 | 2.83874 | 0.10049 | 0.52697 | 2.59621 | 0.01427 | 0.31856 | 2.85685 | 2.67493 | 6.27884 |
| **Cnot6** | 4.60334 | 0.02078 | 4.14779 | 0.00000 | 3.38303 | 7.55396 | 1.71543 | 2.55730 | 10.25491 | 0.00234 | 7.53447 | 4.53454 | 4.50680 | 6.25616 |
| **Cnot6l** | 5.92786 | 0.00230 | 1.39489 | 0.00000 | 1.47745 | 3.17762 | 1.31001 | 0.95013 | 0.89393 | 0.03343 | 0.23179 | 1.06701 | 3.86950 | 1.57951 |
| **Cnot7** | 8.24520 | 0.02274 | 3.07440 | 0.00000 | 5.78093 | 3.21300 | 0.00108 | 0.25237 | 11.78755 | 0.00813 | 0.00000 | 2.58350 | 7.46490 | 4.07272 |
| **Cnot8** | 11.55975 | 0.07891 | 15.14310 | 10.68587 | 19.28332 | 11.31532 | 1.87149 | 4.91334 | 7.85412 | 0.09400 | 5.45779 | 7.96910 | 13.10036 | 22.26124 |
| **Cnot9** | 4.93580 | 4.22936 | 11.30207 | 8.73098 | 9.73680 | 11.76648 | 4.47201 | 3.99513 | 9.70403 | 0.11338 | 9.29103 | 6.69213 | 1.89525 | 11.95873 |
| **Cnp** | 1.44376 | 0.00249 | 0.93624 | 1.27255 | 0.41829 | 0.88653 | 0.00334 | 0.83501 | 0.00265 | 0.02136 | 0.00000 | 1.64958 | 0.00315 | 1.77053 |
| **Cnppd1** | 9.94276 | 0.01456 | 7.75199 | 4.74719 | 14.78349 | 5.46385 | 0.03053 | 1.79472 | 0.16804 | 0.04229 | 6.86430 | 1.51181 | 17.42155 | 6.87699 |
| **Cnpy1** | 0.05168 | 0.00383 | 0.00000 | 0.45972 | 0.00730 | 0.00000 | 0.00366 | 0.00395 | 1.15237 | 0.01081 | 0.00801 | 0.07516 | 0.63429 | 0.00000 |
| **Cnpy2** | 2.06828 | 8.15097 | 3.15343 | 5.66958 | 0.99451 | 6.35423 | 3.96826 | 10.11409 | 0.01042 | 0.16172 | 5.77195 | 13.46325 | 0.00000 | 0.88268 |
| **Cnpy3** | 2.30399 | 0.04249 | 2.75325 | 2.61400 | 1.82884 | 2.18245 | 2.21358 | 2.96377 | 0.00000 | 4.57237 | 2.27894 | 1.82750 | 1.19688 | 1.17289 |
| **Cnpy4** | 0.33562 | 0.01065 | 0.90836 | 0.00000 | 0.75137 | 1.05987 | 0.00256 | 1.52445 | 5.59015 | 0.00988 | 0.00000 | 0.69360 | 2.44937 | 3.12510 |
| **Cnr2** | 0.16204 | 0.04320 | 0.06990 | 0.05671 | 0.10642 | 0.12479 | 0.04729 | 0.02796 | 0.62637 | 0.02296 | 0.05290 | 0.02669 | 0.42249 | 0.12263 |
| **Cnrip1** | 0.91511 | 0.00429 | 0.13211 | 0.05686 | 0.06698 | 0.00414 | 0.00430 | 0.01024 | 0.02071 | 0.00615 | 0.00613 | 0.00604 | 0.00541 | 0.13776 |
| **Cnst** | 1.13734 | 0.03204 | 1.93035 | 1.28139 | 5.33455 | 3.36352 | 1.18143 | 3.97504 | 2.77149 | 0.01027 | 3.12290 | 0.00000 | 3.79118 | 4.73075 |
| **Cntd1** | 0.00682 | 0.63578 | 0.00629 | 0.19976 | 0.00934 | 0.00000 | 0.00466 | 0.02178 | 0.01248 | 0.00575 | 0.10717 | 0.00521 | 0.00666 | 0.00948 |
| **Cntf** | 0.00888 | 0.24548 | 0.04207 | 0.05231 | 0.00415 | 0.02904 | 0.00387 | 0.00366 | 0.23580 | 0.00577 | 0.01366 | 0.00989 | 0.00626 | 0.00360 |
| **Cntln** | 0.00000 | 0.00396 | 0.22589 | 0.00000 | 0.64504 | 0.19904 | 0.00400 | 0.00198 | 1.40027 | 0.00540 | 0.00355 | 0.00279 | 1.41482 | 1.69536 |
| **Cntn1** | 0.00516 | 0.00443 | 0.01260 | 0.01389 | 0.02353 | 0.01176 | 0.00418 | 0.00992 | 0.05762 | 0.00707 | 0.00783 | 0.00952 | 0.01097 | 0.25914 |
| **Cntn2** | 0.02901 | 0.06942 | 0.01892 | 0.13984 | 0.19423 | 0.10821 | 0.02985 | 0.36423 | 0.21715 | 0.04083 | 0.01581 | 0.07455 | 0.58553 | 0.25566 |
| **Cntn4** | 0.00098 | 0.00438 | 0.00342 | 0.01404 | 0.02287 | 0.00911 | 0.00412 | 0.00923 | 0.55328 | 0.00657 | 0.00720 | 0.00888 | 0.32218 | 0.00000 |
| **Cntn6** | 0.00636 | 0.00441 | 0.01412 | 0.01234 | 0.00947 | 0.01265 | 0.00435 | 0.00995 | 0.01062 | 0.00648 | 0.00896 | 0.00982 | 0.10020 | 0.07589 |
| **Cntnap1** | 0.00313 | 0.00428 | 0.01525 | 0.01126 | 0.07584 | 0.01384 | 0.00415 | 0.00859 | 0.01113 | 0.00657 | 0.00897 | 0.00854 | 0.68926 | 0.00928 |
| **Cntrl** | 0.53523 | 0.00413 | 0.28495 | 0.00000 | 0.00000 | 0.71934 | 0.00405 | 0.00412 | 0.18133 | 0.00994 | 0.00678 | 0.24684 | 1.07311 | 0.54983 |
| **Cntrob** | 0.74861 | 0.01455 | 0.08940 | 1.05997 | 0.12027 | 1.32374 | 0.00410 | 0.47942 | 0.06308 | 0.01231 | 2.03932 | 0.00000 | 0.69618 | 0.25880 |
| **Coa3** | 3.38777 | 0.72364 | 1.01615 | 10.25198 | 3.17850 | 8.73921 | 3.44766 | 5.97501 | 0.17237 | 0.18147 | 3.07601 | 7.54540 | 0.49381 | 2.70020 |
| **Coa4** | 1.65590 | 0.06590 | 1.52788 | 6.52685 | 2.58582 | 5.33894 | 1.62332 | 5.07339 | 0.00400 | 0.16899 | 8.40319 | 4.33647 | 0.00000 | 0.58956 |
| **Coa5** | 3.57713 | 0.17162 | 7.06631 | 9.45115 | 4.54009 | 13.63002 | 1.05823 | 9.82343 | 2.21545 | 0.10431 | 12.73440 | 6.85960 | 0.80409 | 3.63616 |
| **Coa6** | 0.00000 | 4.38213 | 1.04335 | 4.17202 | 3.53074 | 0.00000 | 0.96319 | 2.99848 | 0.07851 | 6.16675 | 0.00000 | 2.65925 | 0.45629 | 0.10651 |
| **Coa7** | 1.66833 | 0.11362 | 6.87677 | 10.19693 | 10.58994 | 5.96576 | 1.63391 | 4.24724 | 3.37455 | 0.09467 | 8.95318 | 3.97580 | 1.23784 | 10.28721 |

| | | | | | | | | | | | | | | |
|---|---|---|---|---|---|---|---|---|---|---|---|---|---|---|
| **Coasy** | 0.18784 | 1.10046 | 5.21913 | 7.37301 | 0.80037 | 4.20393 | 4.04147 | 2.60520 | 0.02363 | 0.10683 | 5.66729 | 5.11271 | 0.08420 | 0.62205 |
| **Cobl** | 0.00886 | 0.00428 | 0.01261 | 0.01268 | 0.05277 | 0.01194 | 0.00436 | 0.01021 | 0.01185 | 0.00602 | 0.00798 | 0.00892 | 0.40091 | 0.31140 |
| **Cobll1** | 0.02793 | 0.00454 | 0.03900 | 0.00615 | 0.17276 | 0.00897 | 0.00413 | 0.01151 | 0.00557 | 0.00649 | 0.00754 | 0.01099 | 0.09050 | 0.60896 |
| **Coch** | 0.00141 | 0.00429 | 0.02852 | 0.01083 | 0.02370 | 0.00994 | 0.00436 | 0.00864 | 0.01217 | 0.00607 | 0.00744 | 0.00800 | 0.61523 | 0.73666 |
| **Cog1** | 0.50011 | 0.18031 | 3.14560 | 1.81179 | 2.63751 | 2.10942 | 1.59851 | 1.82492 | 0.88803 | 0.01232 | 1.29710 | 0.58978 | 0.00000 | 2.28026 |
| **Cog2** | 2.45264 | 2.20783 | 5.39021 | 5.03629 | 5.22704 | 4.05339 | 0.02519 | 0.68552 | 0.27900 | 1.02996 | 5.60852 | 2.59929 | 0.00000 | 4.96043 |
| **Cog3** | 3.02882 | 0.00233 | 1.93627 | 0.00000 | 1.15395 | 1.95530 | 0.00315 | 0.00000 | 0.78324 | 0.00319 | 0.03621 | 1.51475 | 0.39721 | 7.06323 |
| **Cog4** | 6.94034 | 0.08850 | 8.96447 | 4.10164 | 10.21457 | 10.49860 | 0.04633 | 6.58379 | 7.64766 | 1.74993 | 3.76832 | 7.03930 | 10.85075 | 14.57290 |
| **Cog5** | 0.00000 | 0.00282 | 1.92645 | 0.44174 | 0.75532 | 1.91890 | 0.00280 | 0.13228 | 2.09345 | 0.03250 | 2.43115 | 0.53313 | 0.00000 | 1.10645 |
| **Cog6** | 1.34153 | 7.50805 | 0.47923 | 1.63073 | 1.45972 | 1.99221 | 0.06208 | 0.78477 | 0.00232 | 0.01081 | 2.03236 | 0.00331 | 0.14671 | 1.42080 |
| **Cog7** | 3.51675 | 1.91050 | 3.84967 | 4.69454 | 1.58920 | 2.92391 | 0.02394 | 4.55123 | 0.01202 | 0.02627 | 3.63384 | 3.82107 | 1.76549 | 0.00000 |
| **Cog8** | 0.15732 | 0.02801 | 1.10810 | 1.23954 | 2.75757 | 0.00000 | 0.00973 | 0.35078 | 0.01181 | 0.01147 | 0.03158 | 1.97679 | 0.00000 | 0.00342 |
| **Coil** | 5.46582 | 0.07230 | 6.29267 | 6.24659 | 7.90738 | 16.00500 | 5.74050 | 2.74946 | 0.00000 | 0.44905 | 12.03164 | 3.19896 | 0.78129 | 0.90756 |
| **Col10a1** | 0.49197 | 0.00415 | 0.07321 | 0.12952 | 0.11419 | 0.21642 | 0.00406 | 0.01523 | 0.01209 | 0.00724 | 0.00454 | 0.00454 | 0.16203 | 0.01632 |
| **Col11a2** | 0.30646 | 0.00458 | 0.09120 | 0.01922 | 0.00000 | 0.05171 | 0.00422 | 0.00623 | 0.00660 | 0.00587 | 0.00992 | 0.00914 | 0.36630 | 0.00545 |
| **Col12a1** | 1.80040 | 0.03511 | 0.01178 | 0.00947 | 0.00222 | 0.00684 | 0.00419 | 0.01069 | 0.00794 | 0.00658 | 0.00684 | 0.00611 | 0.00409 | 0.00307 |
| **Col13a1** | 0.00421 | 0.00444 | 0.01806 | 0.01279 | 0.06236 | 0.01327 | 0.00444 | 0.01222 | 0.00811 | 0.00669 | 0.00783 | 0.05201 | 0.28778 | 0.00731 |
| **Col16a1** | 0.08320 | 0.00443 | 0.08204 | 0.00516 | 0.01942 | 0.00958 | 0.00438 | 0.01139 | 0.01144 | 0.00694 | 0.00717 | 0.01033 | 0.00546 | 0.01031 |
| **Col18a1** | 0.07588 | 0.00433 | 0.01598 | 0.00906 | 0.07784 | 0.02191 | 0.00434 | 0.00940 | 0.00852 | 0.00631 | 0.02379 | 0.00847 | 0.23940 | 0.00926 |
| **Col19a1** | 0.00338 | 0.00431 | 0.01466 | 0.01220 | 0.00855 | 0.01299 | 0.00422 | 0.00897 | 0.01973 | 0.00631 | 0.00896 | 0.00899 | 0.52353 | 0.00826 |
| **Col1a1** | 3.88729 | 0.00410 | 0.01108 | 0.00922 | 0.01237 | 0.00519 | 0.00434 | 0.01401 | 0.00638 | 0.00654 | 0.00694 | 0.00479 | 0.00511 | 0.00664 |
| **Col1a2** | 2.03159 | 0.00423 | 0.00864 | 0.00905 | 0.08145 | 0.00649 | 0.00439 | 0.01400 | 0.00644 | 0.00681 | 0.00687 | 0.00602 | 0.00418 | 0.01164 |
| **Col20a1** | 0.19129 | 0.00446 | 0.06742 | 0.16330 | 0.10650 | 0.11238 | 0.00514 | 0.05531 | 0.01491 | 0.00653 | 0.01000 | 0.01210 | 0.01024 | 0.01243 |
| **Col22a1** | 0.00572 | 0.00435 | 0.01321 | 0.01155 | 0.03946 | 0.01335 | 0.00470 | 0.00927 | 0.01979 | 0.00661 | 0.00899 | 0.00932 | 0.30357 | 0.00268 |
| **Col23a1** | 0.00289 | 0.00425 | 0.06243 | 0.00435 | 0.18306 | 0.00967 | 0.00435 | 0.01065 | 0.00461 | 0.00743 | 0.00651 | 0.04181 | 0.00608 | 0.40788 |
| **Col24a1** | 0.01036 | 0.00445 | 0.01218 | 0.01369 | 0.01067 | 0.01197 | 0.00426 | 0.01038 | 0.08745 | 0.00711 | 0.00814 | 0.00999 | 0.01064 | 0.00638 |
| **Col26a1** | 0.00172 | 0.01793 | 0.05413 | 0.09385 | 0.31769 | 0.00000 | 1.76202 | 2.02999 | 0.01999 | 0.03324 | 0.08232 | 0.00781 | 0.00969 | 0.04268 |
| **Col2a1** | 0.01632 | 0.00442 | 0.00188 | 0.01756 | 0.00271 | 0.00781 | 0.00407 | 0.00961 | 0.83938 | 0.00744 | 0.00616 | 0.00916 | 0.13432 | 0.00000 |
| **Col3a1** | 0.37690 | 0.00441 | 0.01304 | 0.01012 | 0.00796 | 0.01014 | 0.00430 | 0.01191 | 0.00978 | 0.00658 | 0.00709 | 0.00882 | 0.00488 | 0.00543 |
| **Col4a1** | 0.73719 | 0.31189 | 3.47989 | 3.68866 | 0.00000 | 1.71697 | 0.01224 | 1.36522 | 0.16346 | 0.03056 | 0.23332 | 14.43853 | 0.00000 | 0.04009 |
| **Col4a2** | 0.68758 | 0.57051 | 5.53438 | 4.08516 | 0.00000 | 2.95153 | 0.45417 | 2.80370 | 0.00000 | 0.67410 | 3.95427 | 20.00957 | 0.00000 | 0.20272 |
| **Col4a3bp** | 1.89196 | 0.03124 | 1.09954 | 0.00002 | 2.04216 | 1.93513 | 0.49396 | 4.39746 | 5.80084 | 0.01688 | 3.17218 | 2.72184 | 6.78561 | 5.59656 |
| **Col5a1** | 1.03949 | 0.00430 | 0.01225 | 0.00760 | 0.00353 | 0.00990 | 0.00431 | 0.01124 | 0.00887 | 0.00649 | 0.00871 | 0.01829 | 0.00432 | 0.00362 |
| **Col5a2** | 1.07459 | 0.00436 | 0.14746 | 0.00338 | 0.00000 | 0.00427 | 0.00416 | 0.01332 | 0.00865 | 0.00661 | 0.00643 | 0.00733 | 0.00366 | 0.00370 |
| **Col5a3** | 0.43828 | 0.00408 | 1.54079 | 2.97292 | 1.41361 | 0.00000 | 0.00351 | 0.00956 | 0.18521 | 0.00384 | 0.01842 | 0.34008 | 0.49992 | 1.70748 |
| **Col6a1** | 0.22449 | 0.00450 | 0.01265 | 0.01119 | 0.13587 | 0.01040 | 0.00432 | 0.01191 | 0.00977 | 0.00628 | 0.00730 | 0.00796 | 0.00425 | 0.01535 |
| **Col6a2** | 0.14257 | 0.00440 | 0.01114 | 0.00984 | 0.15796 | 0.01162 | 0.00444 | 0.01239 | 0.00954 | 0.00652 | 0.00755 | 0.00870 | 0.00541 | 0.01101 |
| **Col6a3** | 1.05432 | 0.00428 | 0.00610 | 0.00974 | 0.09880 | 0.00787 | 0.00420 | 0.01091 | 0.01195 | 0.00673 | 0.00645 | 0.00601 | 0.35371 | 0.21152 |
| **Col6a4** | 0.00000 | 0.01149 | 4.76890 | 0.44400 | 6.55919 | 0.00000 | 0.00318 | 0.00302 | 0.32809 | 0.00502 | 0.00363 | 0.00782 | 0.00000 | 0.19288 |
| **Col6a5** | 0.00000 | 0.00440 | 0.21561 | 0.70504 | 0.00000 | 0.00000 | 0.00416 | 0.01135 | 0.01047 | 0.01367 | 0.01024 | 0.00584 | 0.70707 | 0.00000 |
| **Col6a6** | 0.00566 | 0.00446 | 0.02832 | 0.00533 | 0.48387 | 0.01153 | 0.00414 | 0.01024 | 0.00764 | 0.00655 | 0.00803 | 0.00807 | 0.00607 | 0.38626 |
| **Col7a1** | 0.11083 | 0.01249 | 0.34967 | 2.60184 | 0.00000 | 0.20362 | 0.00378 | 0.10025 | 0.01236 | 0.00587 | 0.35317 | 0.56618 | 0.00711 | 0.00484 |
| **Col8a2** | 0.01846 | 0.00440 | 0.00000 | 0.01830 | 0.10174 | 0.00490 | 0.00410 | 0.01031 | 0.57296 | 0.00733 | 0.00570 | 0.00701 | 0.13857 | 1.58446 |
| **Col9a2** | 0.10729 | 0.00393 | 0.94019 | 3.99346 | 0.11289 | 0.13335 | 0.00370 | 0.00000 | 0.00976 | 0.01065 | 0.01881 | 0.01760 | 0.00124 | 1.34716 |
| **Col9a3** | 0.00647 | 0.00436 | 0.01467 | 0.01306 | 0.07429 | 0.01269 | 0.00419 | 0.00919 | 0.00999 | 0.00651 | 0.00894 | 0.00882 | 0.30637 | 0.01896 |
| **Colec12** | 1.16563 | 0.01808 | 0.00000 | 0.58491 | 4.74833 | 0.18592 | 0.17696 | 0.31728 | 7.47047 | 0.01174 | 0.38571 | 0.31573 | 9.03438 | 15.07865 |
| **Colgalt1** | 6.37845 | 5.27821 | 2.09581 | 2.64118 | 2.15990 | 2.41663 | 0.00000 | 3.17116 | 3.02537 | 0.04140 | 2.67970 | 8.97404 | 2.97309 | 5.83339 |
| **Colgalt2** | 0.00000 | 0.00390 | 0.00915 | 0.00637 | 0.00136 | 0.01153 | 0.00386 | 0.07615 | 0.21775 | 0.00688 | 0.00769 | 0.00735 | 1.15033 | 0.01650 |
| **Colq** | 0.00436 | 0.01151 | 0.09399 | 0.30262 | 0.02323 | 0.00000 | 0.00394 | 0.00735 | 0.02166 | 0.00598 | 0.10874 | 0.01340 | 0.01646 | 0.06105 |
| **Commd1** | 1.41463 | 0.14767 | 1.66882 | 3.53212 | 1.96534 | 3.80124 | 3.14562 | 5.09385 | 0.00425 | 8.24861 | 0.00000 | 5.32661 | 0.00000 | 0.66810 |
| **Commd10** | 1.76584 | 0.07345 | 3.16068 | 4.67638 | 5.49653 | 6.91923 | 2.99992 | 8.85191 | 3.61384 | 2.91654 | 6.11674 | 5.80351 | 0.53845 | 5.31096 |
| **Commd2** | 7.96297 | 0.11976 | 11.94818 | 4.80494 | 11.94420 | 11.47949 | 3.77937 | 4.69878 | 2.07934 | 0.12690 | 11.04654 | 8.57246 | 3.24882 | 9.12817 |
| **Commd3** | 2.93533 | 0.08771 | 0.47941 | 3.75801 | 0.32213 | 6.52431 | 5.17877 | 11.88858 | 0.00000 | 0.26059 | 5.08110 | 8.94486 | 1.02055 | 0.00000 |
| **Commd4** | 3.07021 | 6.00264 | 3.00205 | 9.60955 | 3.16821 | 4.07166 | 3.55241 | 12.31877 | 0.00000 | 1.02061 | 5.21882 | 4.43947 | 0.90783 | 1.41930 |
| **Commd5** | 3.46267 | 8.83598 | 17.35352 | 17.35641 | 17.61646 | 2.11739 | 1.56147 | 9.30240 | 0.08235 | 0.07830 | 4.82713 | 7.54298 | 5.48647 | 0.65138 |

| | | | | | | | | | | | | | | |
|---|---|---|---|---|---|---|---|---|---|---|---|---|---|---|
| **Commd6** | 1.48177 | 0.04190 | 0.98247 | 1.50311 | 3.93191 | 2.77914 | 2.02213 | 2.84009 | 2.46424 | 0.10458 | 2.39895 | 4.20185 | 0.30768 | 2.05731 |
| **Commd7** | 5.74580 | 0.08259 | 4.48967 | 1.80976 | 9.01027 | 5.02065 | 0.53748 | 2.68384 | 1.66324 | 0.12608 | 0.00000 | 2.76631 | 8.44613 | 3.48283 |
| **Commd8** | 3.97118 | 0.05693 | 3.50798 | 8.69877 | 3.49773 | 5.79660 | 4.02447 | 2.29694 | 0.75792 | 0.09118 | 8.20638 | 2.47867 | 0.03301 | 3.68385 |
| **Commd9** | 2.86420 | 0.08467 | 5.97950 | 6.68838 | 4.27012 | 10.76711 | 4.84903 | 2.02958 | 0.03193 | 0.24753 | 0.00000 | 5.90924 | 0.00000 | 1.10540 |
| **Comp** | 0.02168 | 0.00457 | 0.00000 | 0.45258 | 0.50071 | 0.00474 | 0.00424 | 0.02103 | 0.00588 | 0.01460 | 0.00660 | 0.00650 | 0.00489 | 1.29354 |
| **Comt** | 2.40542 | 1.22802 | 4.56608 | 1.48378 | 0.91555 | 4.22551 | 3.79123 | 1.09035 | 3.55623 | 0.70026 | 3.59192 | 3.38179 | 12.21881 | 4.68643 |
| **Comtd1** | 0.01454 | 0.03872 | 0.30887 | 0.35891 | 0.34103 | 0.00000 | 0.00454 | 1.64058 | 0.00804 | 0.00548 | 0.07083 | 0.38392 | 0.25150 | 0.00000 |
| **Copa** | 8.38720 | 0.19514 | 6.37383 | 3.61797 | 9.53803 | 12.88761 | 2.07882 | 5.49518 | 7.33096 | 0.13244 | 5.84703 | 9.73760 | 3.44905 | 6.59271 |
| **Copb1** | 3.34157 | 0.03720 | 2.58058 | 0.00000 | 0.78802 | 5.02362 | 3.08492 | 1.40005 | 3.45937 | 0.07301 | 7.95740 | 2.60213 | 2.59145 | 2.43673 |
| **Copb2** | 10.99780 | 0.12538 | 13.74562 | 9.36861 | 11.73518 | 15.04869 | 3.34507 | 4.08602 | 14.83621 | 0.20944 | 13.97069 | 12.51263 | 1.95685 | 14.18590 |
| **Cope** | 6.72407 | 0.46831 | 19.79415 | 18.45199 | 16.60991 | 16.09085 | 3.24649 | 16.58923 | 0.00000 | 0.26819 | 11.09395 | 11.72645 | 0.55344 | 1.88142 |
| **Copg1** | 4.36430 | 7.25155 | 11.16694 | 8.27482 | 6.17431 | 11.72652 | 2.09607 | 4.73275 | 6.49412 | 1.14065 | 9.62552 | 10.69836 | 2.51074 | 8.44213 |
| **Copg2** | 0.03516 | 0.01802 | 0.00000 | 0.65196 | 0.00000 | 0.17856 | 0.00313 | 3.05182 | 0.94632 | 0.02783 | 0.47361 | 0.00967 | 1.25286 | 0.84408 |
| **Coprs** | 1.49937 | 0.01064 | 0.05648 | 7.78584 | 4.78234 | 1.20939 | 0.30757 | 0.26352 | 0.97088 | 0.03896 | 0.04021 | 1.23926 | 0.78868 | 0.00000 |
| **Cops2** | 12.18845 | 0.05990 | 13.20826 | 7.80638 | 12.94013 | 14.19298 | 4.75223 | 4.62889 | 23.35396 | 0.19173 | 12.88322 | 10.73774 | 4.95167 | 11.73393 |
| **Cops3** | 4.96199 | 9.79429 | 14.60400 | 3.95311 | 7.67099 | 14.58402 | 3.89922 | 8.19021 | 5.13443 | 8.05394 | 11.84953 | 10.94964 | 1.28932 | 13.31418 |
| **Cops4** | 18.22391 | 0.89743 | 24.99026 | 15.66984 | 24.23988 | 18.83304 | 6.45804 | 10.62871 | 7.15784 | 0.39116 | 15.20015 | 14.69794 | 7.12758 | 20.51316 |
| **Cops5** | 15.35468 | 7.20127 | 18.93680 | 17.70476 | 15.56408 | 23.17727 | 5.99214 | 18.43502 | 14.39100 | 9.32119 | 22.74073 | 15.41917 | 0.26574 | 12.77086 |
| **Cops6** | 16.56940 | 0.22568 | 11.99425 | 15.33725 | 17.04013 | 11.96494 | 2.43140 | 9.97784 | 1.64873 | 0.20485 | 5.90633 | 9.42931 | 12.50177 | 9.77782 |
| **Cops7a** | 13.05111 | 0.27810 | 25.84118 | 17.39709 | 14.22337 | 13.19506 | 0.55671 | 8.27516 | 8.51149 | 4.08258 | 17.63740 | 13.14148 | 10.90937 | 13.12364 |
| **Cops7b** | 0.37180 | 0.03643 | 7.31221 | 3.41751 | 8.99416 | 1.66095 | 0.00154 | 3.26064 | 0.74287 | 1.11453 | 4.02297 | 1.07574 | 2.07904 | 1.43612 |
| **Cops8** | 8.38834 | 0.22127 | 17.82037 | 6.13873 | 9.22518 | 13.27317 | 4.03560 | 12.47434 | 6.14337 | 0.18673 | 8.53443 | 10.61522 | 3.91120 | 11.22943 |
| **Copz1** | 12.71441 | 0.40672 | 22.03838 | 17.24505 | 14.75096 | 22.32102 | 5.79177 | 14.85933 | 14.33526 | 8.59164 | 15.63811 | 16.70923 | 2.29101 | 11.55202 |
| **Copz2** | 0.43144 | 0.00420 | 0.01054 | 0.11360 | 0.05790 | 0.01805 | 0.00399 | 0.00646 | 0.01168 | 0.00855 | 0.00643 | 0.00628 | 0.72402 | 0.10279 |
| **Coq10a** | 0.50646 | 0.00416 | 0.21872 | 0.28065 | 0.00000 | 0.65986 | 0.00391 | 0.00556 | 0.00574 | 0.01694 | 0.00337 | 0.00184 | 1.83137 | 0.56020 |
| **Coq10b** | 2.01561 | 0.01690 | 3.45080 | 0.49703 | 1.69455 | 1.52883 | 0.94880 | 4.65342 | 1.18279 | 0.06767 | 3.36493 | 2.14585 | 0.47579 | 0.65795 |
| **Coq2** | 0.74510 | 6.43437 | 1.87184 | 0.25026 | 0.93612 | 1.67289 | 0.02735 | 3.12938 | 0.01850 | 0.03607 | 1.77392 | 3.61738 | 1.38509 | 0.94989 |
| **Coq3** | 5.68950 | 0.09464 | 3.31616 | 2.67615 | 6.38118 | 5.91286 | 3.67431 | 2.22122 | 2.61364 | 12.08150 | 6.87886 | 3.70103 | 2.26432 | 8.32204 |
| **Coq4** | 2.25612 | 7.08961 | 5.29045 | 6.04777 | 8.43222 | 1.21403 | 2.03924 | 3.22423 | 0.14473 | 0.07960 | 0.00000 | 3.37091 | 4.43623 | 2.68294 |
| **Coq5** | 0.73871 | 0.05001 | 7.14678 | 8.52976 | 2.39960 | 4.05760 | 3.64880 | 2.47024 | 0.73760 | 0.08766 | 7.11395 | 4.92115 | 1.56486 | 4.04841 |
| **Coq6** | 3.77775 | 1.51813 | 10.92563 | 8.97924 | 6.40863 | 4.01709 | 1.26329 | 5.05418 | 0.16546 | 0.02011 | 8.45547 | 3.44243 | 3.12531 | 1.53089 |
| **Coq7** | 5.86270 | 0.10343 | 7.91684 | 9.24525 | 6.71434 | 6.87030 | 2.27752 | 10.40342 | 0.11702 | 7.12081 | 7.63282 | 6.10036 | 1.41014 | 2.29640 |
| **Coq8a** | 0.01885 | 0.00383 | 1.40005 | 0.94985 | 0.18170 | 0.00000 | 0.00353 | 0.07924 | 0.28432 | 0.00128 | 0.46506 | 0.00321 | 0.00000 | 0.00000 |
| **Coq8b** | 1.87843 | 0.14295 | 3.52718 | 0.69134 | 2.42932 | 0.75675 | 2.51300 | 3.00996 | 0.16562 | 0.00659 | 0.01804 | 1.07095 | 0.33135 | 0.63462 |
| **Coq9** | 4.34938 | 6.64030 | 6.63555 | 3.70753 | 3.93160 | 5.86884 | 0.00000 | 3.08739 | 0.30453 | 0.58912 | 2.60761 | 3.34239 | 0.36273 | 5.64570 |
| **Coro1a** | 0.56441 | 0.00415 | 2.14201 | 0.00000 | 1.27709 | 0.00098 | 0.00379 | 0.01686 | 0.01258 | 0.00344 | 0.00454 | 0.00688 | 0.00000 | 0.09290 |
| **Coro1b** | 3.53543 | 9.62524 | 4.85017 | 8.79835 | 3.59481 | 11.47859 | 5.77859 | 14.57065 | 0.00000 | 0.31883 | 5.97221 | 4.34157 | 0.30736 | 0.94816 |
| **Coro1c** | 3.78300 | 8.99921 | 21.28444 | 13.43800 | 9.30431 | 23.97478 | 5.09922 | 16.42070 | 0.45445 | 0.42230 | 9.56803 | 10.82614 | 0.68834 | 3.24965 |
| **Coro2a** | 2.41879 | 0.00383 | 2.00185 | 0.71332 | 0.59839 | 0.25048 | 0.00367 | 0.01445 | 1.00815 | 0.00000 | 1.12778 | 0.00609 | 0.35799 | 0.14284 |
| **Coro2b** | 0.02057 | 0.02712 | 0.01052 | 0.02172 | 0.09183 | 0.01372 | 0.00401 | 0.00802 | 0.05861 | 0.00661 | 0.00549 | 0.39340 | 0.00877 | 0.07109 |
| **Coro7** | 0.00000 | 0.08693 | 0.42843 | 1.06813 | 0.32299 | 2.05258 | 0.04658 | 2.22485 | 0.00087 | 0.00926 | 3.48306 | 1.95371 | 0.00216 | 0.30668 |
| **Cort** | 0.02484 | 0.00443 | 0.02721 | 0.00873 | 0.14745 | 0.01281 | 0.00440 | 0.00989 | 0.01256 | 0.00680 | 0.00896 | 0.01152 | 0.10437 | 0.01352 |
| **Cotl1** | 0.03127 | 0.02205 | 2.33165 | 2.83209 | 0.62701 | 0.00000 | 0.00994 | 0.49810 | 0.00000 | 0.00346 | 0.00000 | 2.13399 | 1.30705 | 0.10074 |
| **Cox10** | 3.02111 | 0.06936 | 8.85077 | 2.48944 | 6.05961 | 6.01649 | 1.62627 | 8.21731 | 2.72445 | 6.93467 | 0.49903 | 3.23905 | 1.10280 | 2.56212 |
| **Cox11** | 1.71785 | 0.02703 | 0.16066 | 2.06723 | 3.99632 | 0.62514 | 0.01141 | 0.76746 | 0.00684 | 0.00780 | 0.01011 | 1.37634 | 0.80460 | 0.67690 |
| **Cox14** | 0.22077 | 0.09061 | 0.22831 | 6.12765 | 4.43867 | 1.80558 | 1.67210 | 9.13778 | 0.00245 | 8.69267 | 0.00000 | 5.44615 | 2.87442 | 0.00000 |
| **Cox15** | 0.98091 | 0.03382 | 2.84410 | 2.16772 | 0.87766 | 1.97843 | 0.00208 | 2.65029 | 3.66655 | 0.02093 | 2.37908 | 1.49884 | 0.00000 | 2.56883 |
| **Cox16** | 0.35974 | 0.01101 | 0.19334 | 0.04705 | 1.10299 | 0.63514 | 0.29556 | 1.40945 | 0.00445 | 0.03783 | 0.08795 | 1.06156 | 0.03430 | 0.16111 |
| **Cox17** | 1.73821 | 0.19365 | 0.03825 | 1.74619 | 2.42561 | 4.23997 | 0.77192 | 5.55029 | 0.00477 | 5.70446 | 0.35487 | 4.09832 | 0.46057 | 0.41967 |
| **Cox18** | 4.78615 | 0.63334 | 3.53981 | 4.63635 | 3.63446 | 5.20011 | 0.02027 | 3.26728 | 0.11344 | 0.25929 | 4.09998 | 4.24868 | 0.82303 | 1.94775 |
| **Cox19** | 2.48449 | 0.09437 | 1.03895 | 4.27870 | 4.17441 | 5.17802 | 0.00908 | 4.64628 | 0.00000 | 0.08098 | 0.00000 | 3.61732 | 0.25662 | 2.22579 |
| **Cox20** | 2.05922 | 0.02567 | 0.71262 | 0.54712 | 0.13345 | 2.58423 | 2.35628 | 3.08747 | 2.53741 | 3.29156 | 0.43087 | 3.03787 | 0.18617 | 0.00000 |
| **Cox4i1** | 11.52955 | 11.37444 | 8.19467 | 36.12144 | 22.44794 | 25.50347 | 9.86294 | 29.95517 | 1.82075 | 22.12433 | 8.42531 | 24.88469 | 6.79644 | 8.56123 |
| **Cox4i2** | 0.78937 | 0.00388 | 0.23696 | 0.01645 | 1.32437 | 0.00000 | 0.00411 | 0.00708 | 0.12055 | 0.00703 | 0.00531 | 0.09135 | 1.34349 | 1.03623 |
| **Cox5a** | 0.52123 | 2.34021 | 0.00000 | 10.45073 | 20.08949 | 17.42268 | 2.15678 | 10.48905 | 1.51418 | 10.49397 | 3.24529 | 7.87671 | 13.59129 | 2.18893 |

| | | | | | | | | | | | | | | |
|---|---|---|---|---|---|---|---|---|---|---|---|---|---|---|
| **Cox5b** | 0.04675 | 0.48281 | 0.00000 | 5.26436 | 6.74578 | 3.65918 | 1.55939 | 8.51141 | 0.20866 | 5.97170 | 0.00000 | 5.90909 | 2.29192 | 0.00000 |
| **Cox6a1** | 1.33212 | 4.55700 | 0.26064 | 10.38183 | 16.72322 | 6.38738 | 3.16966 | 14.61396 | 0.51009 | 13.21711 | 0.00000 | 16.66280 | 10.05427 | 0.16915 |
| **Cox6b1** | 4.27785 | 4.74561 | 0.00000 | 6.23619 | 7.37408 | 13.49241 | 3.43231 | 19.80615 | 3.48659 | 14.37241 | 1.02142 | 12.19786 | 6.74125 | 4.41110 |
| **Cox6b2** | 0.22690 | 0.00335 | 0.02992 | 0.14711 | 0.13718 | 0.05420 | 0.00383 | 0.02069 | 0.00420 | 0.01788 | 0.01039 | 1.07480 | 0.94313 | 1.53844 |
| **Cox6c** | 7.59807 | 3.05478 | 0.00000 | 4.06994 | 7.28645 | 14.82983 | 4.62090 | 14.60150 | 2.68867 | 16.18173 | 0.32584 | 12.57157 | 10.66077 | 2.16332 |
| **Cox7a1** | 0.12949 | 0.00412 | 0.00000 | 0.20575 | 0.53468 | 0.19696 | 0.00391 | 0.00478 | 0.01624 | 0.05663 | 0.00458 | 0.07260 | 0.10586 | 0.14353 |
| **Cox7a2** | 4.85994 | 0.35347 | 0.00000 | 1.08513 | 7.03081 | 10.19562 | 3.32779 | 10.27280 | 2.06973 | 13.37588 | 0.00000 | 8.36313 | 9.34739 | 3.26932 |
| **Cox7a2l** | 10.76479 | 4.54585 | 24.85720 | 28.54723 | 22.37853 | 15.49451 | 6.73887 | 22.74515 | 1.85855 | 10.60822 | 19.27638 | 11.02603 | 3.35673 | 16.98018 |
| **Cox7b** | 1.83167 | 3.23388 | 0.00000 | 0.32841 | 1.68991 | 8.57647 | 3.06636 | 10.73274 | 1.79151 | 8.02891 | 1.94825 | 9.27296 | 5.56824 | 1.37119 |
| **Cox7b2** | 0.05031 | 0.00437 | 0.01698 | 0.01183 | 0.01206 | 0.01279 | 0.00425 | 0.00996 | 0.01073 | 0.00679 | 0.00815 | 0.00926 | 0.19863 | 0.00708 |
| **Cox7c** | 3.34092 | 2.03549 | 0.16342 | 1.96463 | 1.77342 | 5.67928 | 1.13522 | 5.54765 | 0.67837 | 3.37087 | 0.18898 | 4.86449 | 3.95638 | 0.95583 |
| **Cox8a** | 0.11206 | 2.03192 | 0.00000 | 7.79869 | 15.45740 | 4.65302 | 1.13756 | 9.28246 | 0.74154 | 4.44572 | 0.00000 | 10.46905 | 6.18470 | 0.00000 |
| **Cox8c** | 0.00684 | 0.00445 | 0.01781 | 0.00771 | 0.35848 | 0.01315 | 0.00418 | 0.00995 | 0.00692 | 0.00779 | 0.00863 | 0.00847 | 0.07482 | 0.09682 |
| **Cp** | 0.02636 | 0.00445 | 0.02230 | 0.01091 | 0.01898 | 0.01454 | 0.00425 | 0.01003 | 0.04267 | 0.00694 | 0.00866 | 0.00971 | 0.01440 | 0.02283 |
| **Cpa1** | 0.39809 | 0.01041 | 3.19020 | 0.00000 | 0.00000 | 0.00000 | 0.81001 | 0.00713 | 1.57128 | 0.02054 | 0.00348 | 0.00000 | 6.94059 | 3.89876 |
| **Cpa2** | 0.00460 | 0.00461 | 0.00156 | 0.44669 | 0.00468 | 0.00000 | 0.00387 | 0.01910 | 0.00699 | 0.00923 | 1.01933 | 0.00597 | 0.81067 | 0.01668 |
| **Cpa4** | 0.00252 | 0.00437 | 0.00941 | 0.01349 | 0.00121 | 0.01174 | 0.00429 | 0.00921 | 0.01717 | 0.00625 | 0.00739 | 0.00923 | 0.09568 | 0.74705 |
| **Cpb2** | 0.00847 | 0.00451 | 0.01226 | 0.01429 | 0.11563 | 0.01291 | 0.00432 | 0.01079 | 0.01153 | 0.00642 | 0.00882 | 0.00905 | 0.00832 | 0.01434 |
| **Cpd** | 0.02323 | 0.00291 | 0.42251 | 0.71614 | 0.50120 | 2.76079 | 2.10522 | 0.59962 | 0.72224 | 0.00000 | 1.70317 | 0.68761 | 2.14487 | 2.96450 |
| **Cpe** | 0.07116 | 0.00241 | 0.09193 | 0.00000 | 0.82987 | 0.00508 | 0.00363 | 0.01676 | 0.00800 | 0.01271 | 0.00464 | 2.56547 | 0.39123 | 0.01734 |
| **Cpeb1** | 0.56841 | 0.00386 | 0.11277 | 0.41430 | 1.21259 | 0.62742 | 0.00391 | 0.00376 | 0.12807 | 0.00577 | 0.00181 | 0.00000 | 2.29416 | 1.57095 |
| **Cpeb2** | 3.89941 | 0.00871 | 2.38830 | 0.07590 | 0.76124 | 2.01508 | 0.12285 | 0.19141 | 9.48054 | 5.02257 | 1.24087 | 1.95848 | 1.12687 | 3.06279 |
| **Cpeb3** | 1.38833 | 0.00396 | 2.01709 | 0.35409 | 1.96272 | 1.10016 | 0.00901 | 0.25161 | 7.61344 | 0.02344 | 0.02221 | 0.01753 | 2.48961 | 4.66105 |
| **Cpeb4** | 0.21983 | 0.00325 | 0.10076 | 0.19362 | 0.00000 | 0.15888 | 1.87620 | 0.00000 | 6.54426 | 0.00282 | 1.05205 | 0.01141 | 4.61884 | 0.00000 |
| **Cplx1** | 0.00477 | 0.00430 | 0.19093 | 0.00000 | 0.00632 | 0.02660 | 0.00418 | 0.00570 | 0.01550 | 0.02444 | 0.25637 | 0.00731 | 0.13051 | 0.00247 |
| **Cplx2** | 0.02534 | 0.00418 | 0.01093 | 0.02552 | 0.07899 | 0.06944 | 0.00402 | 0.03547 | 0.27467 | 0.20660 | 0.01010 | 0.01884 | 0.13369 | 0.14315 |
| **Cpm** | 0.00000 | 3.10876 | 2.57526 | 1.22906 | 0.42068 | 2.05547 | 0.00300 | 0.00000 | 0.51671 | 0.00813 | 0.87517 | 0.18416 | 2.19842 | 2.33808 |
| **Cpn1** | 1.49720 | 6.41474 | 5.45233 | 6.16673 | 1.82691 | 13.37234 | 0.07311 | 9.81373 | 0.36287 | 0.96949 | 9.33414 | 4.71024 | 2.11063 | 1.67660 |
| **Cpne1** | 5.21236 | 0.00927 | 5.72459 | 0.00000 | 2.42724 | 1.12954 | 0.70615 | 0.05404 | 6.61003 | 0.00759 | 0.00000 | 2.53120 | 4.43283 | 7.40744 |
| **Cpne2** | 0.01721 | 0.00438 | 0.26682 | 0.00000 | 0.00551 | 0.23364 | 0.00424 | 0.00473 | 0.01353 | 0.00609 | 0.00502 | 0.03188 | 0.03383 | 0.02274 |
| **Cpne3** | 0.94367 | 4.61215 | 0.91667 | 1.79149 | 0.00000 | 6.72924 | 0.01984 | 2.78039 | 0.06146 | 0.01282 | 1.42572 | 1.83076 | 0.52085 | 0.06630 |
| **Cpne5** | 0.06338 | 0.00410 | 0.67167 | 0.28472 | 0.46043 | 0.25330 | 0.00386 | 0.00141 | 0.00305 | 0.00738 | 0.00586 | 0.00514 | 0.95932 | 0.01803 |
| **Cpne7** | 1.28985 | 0.01203 | 0.89424 | 0.00000 | 1.83391 | 0.44872 | 0.00412 | 0.53279 | 0.47430 | 0.02062 | 0.64851 | 1.06432 | 0.60858 | 0.69563 |
| **Cpne8** | 7.74918 | 0.35888 | 1.81611 | 1.37795 | 5.42072 | 5.89780 | 1.45777 | 3.54744 | 2.97752 | 1.74829 | 4.59948 | 2.27353 | 5.98508 | 3.04321 |
| **Cpne9** | 0.96072 | 0.00392 | 0.71507 | 0.60413 | 0.32437 | 0.01794 | 0.00964 | 0.00757 | 0.35343 | 0.00505 | 0.02918 | 0.02680 | 0.52881 | 0.75772 |
| **Cpox** | 0.42361 | 0.05819 | 0.32417 | 0.55521 | 0.95172 | 1.72734 | 0.04109 | 0.02192 | 0.80840 | 0.00440 | 0.77940 | 0.64301 | 0.06123 | 0.18526 |
| **Cpped1** | 0.12443 | 0.02641 | 0.45781 | 0.00000 | 0.31053 | 0.85525 | 0.00395 | 0.90197 | 0.00073 | 0.00761 | 0.00000 | 1.94224 | 1.10502 | 1.07067 |
| **Cpsf1** | 3.46123 | 0.06194 | 11.64868 | 11.94620 | 6.72430 | 6.59689 | 2.03906 | 7.25303 | 0.23246 | 0.07215 | 5.75459 | 3.12217 | 1.57469 | 4.06690 |
| **Cpsf2** | 4.36822 | 0.01602 | 7.12921 | 4.59745 | 5.84985 | 6.99211 | 2.29080 | 0.76025 | 8.02412 | 0.08638 | 4.13140 | 5.12943 | 0.55880 | 8.18124 |
| **Cpsf3** | 10.79324 | 2.93595 | 19.16502 | 16.67017 | 15.57693 | 13.89984 | 3.74090 | 9.37509 | 13.22148 | 7.99529 | 15.51739 | 8.48823 | 1.52985 | 15.98736 |
| **Cpsf3l** | 4.07373 | 7.48717 | 14.38795 | 17.71791 | 11.25934 | 9.74309 | 2.89310 | 0.09234 | 4.17845 | 0.09203 | 9.46041 | 5.86722 | 4.12123 | 14.55590 |
| **Cpsf4** | 0.00000 | 0.91815 | 6.54720 | 1.22440 | 6.80211 | 5.62940 | 1.08379 | 3.41228 | 0.00214 | 0.07490 | 5.29908 | 4.24335 | 2.94371 | 1.49438 |
| **Cpsf4l** | 0.67033 | 0.00386 | 0.50252 | 0.00000 | 0.37519 | 0.00000 | 0.00398 | 0.00774 | 0.00467 | 0.00299 | 0.64450 | 0.00680 | 1.02225 | 0.17928 |
| **Cpsf6** | 10.63302 | 0.05357 | 13.27800 | 11.21704 | 7.96771 | 6.38156 | 0.61745 | 2.56626 | 9.80804 | 0.02202 | 7.94112 | 8.19589 | 11.98239 | 1.33198 |
| **Cpsf7** | 1.48359 | 0.19236 | 4.62941 | 2.10832 | 2.76774 | 4.37207 | 2.50855 | 1.36071 | 1.03395 | 0.09840 | 2.37817 | 3.91881 | 0.22646 | 6.83150 |
| **Cpt1a** | 3.76243 | 0.06801 | 13.21466 | 3.95740 | 6.37225 | 3.68419 | 0.22631 | 0.00007 | 4.62938 | 0.00687 | 2.49966 | 4.04331 | 1.91784 | 9.35295 |
| **Cpt1c** | 0.24620 | 0.00405 | 0.00000 | 0.29999 | 0.85468 | 0.12812 | 0.00399 | 0.00398 | 0.11461 | 0.00581 | 0.00363 | 0.00364 | 0.88828 | 0.55857 |
| **Cpt2** | 0.36188 | 0.00858 | 0.00000 | 0.00000 | 0.31625 | 1.68214 | 1.93692 | 1.74436 | 0.45552 | 0.07889 | 1.29596 | 3.36546 | 0.86809 | 0.13407 |
| **Cptp** | 1.76015 | 0.00833 | 5.75607 | 0.00000 | 0.97297 | 4.79665 | 2.06035 | 4.74081 | 1.79542 | 0.08687 | 0.00000 | 2.30467 | 8.55448 | 1.06801 |
| **Cpxm1** | 0.00000 | 0.02603 | 0.74755 | 4.92899 | 0.18272 | 1.37655 | 0.19205 | 0.05045 | 0.00243 | 0.01173 | 0.01748 | 1.64599 | 0.07243 | 0.58591 |
| **Cpxm2** | 0.77900 | 0.00452 | 0.00931 | 0.09753 | 0.00000 | 0.01185 | 0.00462 | 0.05741 | 0.00973 | 0.00875 | 0.00969 | 0.00505 | 0.00509 | 0.00326 |
| **Cr1l** | 1.15300 | 0.04605 | 0.62966 | 1.67114 | 1.86572 | 4.70697 | 2.90543 | 0.00000 | 1.70202 | 0.02250 | 3.94989 | 2.99627 | 0.00000 | 0.38546 |
| **Crabp2** | 0.00634 | 0.00466 | 0.00000 | 0.02671 | 0.04952 | 0.00438 | 0.00413 | 0.01289 | 1.63131 | 0.00766 | 0.00577 | 0.00965 | 0.00720 | 0.37292 |
| **Cracr2a** | 0.00682 | 0.00481 | 0.00320 | 0.21883 | 0.00867 | 0.32856 | 0.00417 | 0.00860 | 0.01219 | 0.01452 | 0.00557 | 0.00467 | 0.00759 | 0.00722 |
| **Cracr2b** | 0.13181 | 0.09627 | 0.88399 | 0.99384 | 1.31250 | 0.15490 | 0.05874 | 1.14042 | 0.10480 | 0.53687 | 0.02522 | 0.05331 | 0.73480 | 0.21878 |

| | | | | | | | | | | | | | | |
|---|---|---|---|---|---|---|---|---|---|---|---|---|---|---|
| **Cradd** | 0.00000 | 0.01239 | 0.53609 | 0.43617 | 0.00000 | 0.21319 | 1.32079 | 0.00517 | 0.10458 | 0.01130 | 0.03148 | 0.02698 | 0.00092 | 0.10874 |
| **Cramp1l** | 4.51359 | 0.06544 | 3.60062 | 0.37515 | 3.50395 | 4.59707 | 0.37711 | 2.93690 | 4.97074 | 0.03409 | 3.89104 | 0.98147 | 5.25482 | 5.93165 |
| **Crat** | 0.21725 | 0.02596 | 0.57445 | 1.60160 | 0.34743 | 2.15517 | 0.10801 | 3.64090 | 0.07978 | 0.04689 | 1.92506 | 1.09873 | 0.33078 | 0.32564 |
| **Crb3** | 7.88207 | 6.59280 | 17.41184 | 5.68325 | 28.25509 | 0.80854 | 4.16183 | 9.35379 | 0.01820 | 10.36760 | 1.88160 | 0.00000 | 7.86186 | 1.62433 |
| **Crbn** | 2.50270 | 0.00894 | 3.13678 | 2.53207 | 4.06578 | 4.67035 | 0.01842 | 2.17285 | 9.21023 | 0.00000 | 3.45110 | 2.52675 | 0.54377 | 2.37940 |
| **Crcp** | 2.43796 | 0.03124 | 6.28531 | 7.71546 | 9.92343 | 7.36239 | 3.97143 | 2.06718 | 4.99047 | 10.69219 | 1.23158 | 3.07624 | 5.98696 | 10.99083 |
| **Creb1** | 1.50807 | 0.00883 | 2.43376 | 0.66628 | 1.40555 | 3.41229 | 0.22219 | 1.17506 | 2.28556 | 0.00000 | 4.36646 | 1.64339 | 3.03803 | 2.13433 |
| **Creb3** | 4.27623 | 0.02966 | 7.68516 | 2.40841 | 5.30174 | 6.46127 | 1.17585 | 7.37225 | 2.09793 | 0.69523 | 5.48377 | 2.79252 | 4.56178 | 6.30191 |
| **Creb3l1** | 0.07893 | 0.00438 | 0.35566 | 0.20595 | 0.30945 | 0.12092 | 0.00488 | 0.42449 | 0.00874 | 0.00597 | 0.00769 | 0.01136 | 0.07276 | 0.00000 |
| **Creb3l2** | 0.04399 | 1.50938 | 0.01000 | 0.01780 | 0.01056 | 0.00527 | 0.00297 | 0.00652 | 0.01072 | 0.00612 | 0.00281 | 2.64397 | 0.16363 | 0.01433 |
| **Creb3l3** | 0.09752 | 0.01561 | 0.76728 | 0.00000 | 0.42053 | 0.00000 | 0.00421 | 0.39601 | 0.08773 | 0.00637 | 0.00337 | 0.00637 | 0.02586 | 0.53997 |
| **Creb3l4** | 0.00000 | 0.00310 | 0.10729 | 0.00000 | 0.33533 | 0.00000 | 0.00367 | 0.43545 | 0.00823 | 0.00009 | 1.91465 | 0.00326 | 2.49891 | 0.01239 |
| **Crebbp** | 0.00000 | 0.00445 | 1.63257 | 1.76864 | 1.17498 | 0.50118 | 0.00348 | 0.18795 | 3.90732 | 0.00470 | 0.00000 | 0.00416 | 1.01614 | 9.70356 |
| **Crebl2** | 1.51457 | 0.00384 | 0.44892 | 0.00000 | 0.20572 | 0.00000 | 0.00407 | 0.00457 | 0.15452 | 0.00618 | 0.09262 | 0.13441 | 0.43739 | 0.78608 |
| **Crebrf** | 10.66045 | 0.00183 | 2.45358 | 0.41267 | 10.97631 | 1.33137 | 0.00171 | 0.00000 | 13.63413 | 0.00860 | 0.12038 | 0.00000 | 7.65075 | 8.66471 |
| **Crebzf** | 0.74618 | 0.00349 | 0.40507 | 0.98184 | 0.35206 | 0.15112 | 0.00321 | 0.00000 | 3.29577 | 0.01157 | 0.08931 | 0.19498 | 1.44030 | 0.56152 |
| **Creg1** | 3.79858 | 0.06186 | 5.51669 | 5.65645 | 8.67399 | 2.25165 | 0.01189 | 0.64278 | 3.42927 | 0.03040 | 2.28351 | 9.57516 | 0.04833 | 3.08922 |
| **Creld1** | 0.05203 | 0.02105 | 0.00000 | 0.12176 | 0.70103 | 0.00000 | 0.00968 | 2.41565 | 0.00692 | 0.06931 | 0.74005 | 0.06222 | 0.04207 | 0.15878 |
| **Creld2** | 9.52170 | 0.07473 | 7.42044 | 10.81800 | 3.68963 | 10.75909 | 5.57798 | 3.06092 | 2.01484 | 0.21000 | 2.46633 | 2.83040 | 1.95825 | 3.13582 |
| **Crem** | 0.23430 | 0.00383 | 0.90259 | 2.20908 | 0.04313 | 0.87553 | 0.00390 | 0.00721 | 0.01048 | 0.01651 | 0.01421 | 0.08013 | 1.20247 | 0.06670 |
| **Crim1** | 0.08771 | 0.00465 | 0.06474 | 0.00268 | 0.00783 | 0.03728 | 0.00434 | 0.00862 | 0.00914 | 0.00645 | 0.00685 | 0.01020 | 0.12036 | 0.01131 |
| **Crip1** | 0.00000 | 2.93825 | 0.00000 | 0.58902 | 0.72880 | 1.41381 | 3.74833 | 11.24049 | 0.09321 | 8.53284 | 0.00000 | 0.12071 | 0.00527 | 0.28312 |
| **Crip2** | 1.17859 | 13.60996 | 36.58809 | 39.78084 | 18.15823 | 25.05853 | 8.34338 | 28.77500 | 0.06111 | 19.21058 | 12.06305 | 11.03976 | 0.00807 | 0.00000 |
| **Cript** | 8.72768 | 3.20254 | 7.86068 | 10.89989 | 18.34689 | 11.40393 | 5.69199 | 4.45318 | 12.98505 | 0.22257 | 3.10663 | 7.64097 | 12.70738 | 8.16630 |
| **Crispld2** | 0.78456 | 0.00402 | 1.35159 | 0.00568 | 0.00000 | 0.00316 | 0.00377 | 0.00698 | 0.00312 | 0.00450 | 0.00596 | 0.00735 | 1.01136 | 0.08191 |
| **Crk** | 7.89551 | 0.03675 | 1.19111 | 0.98367 | 7.65455 | 6.26979 | 1.61423 | 4.16057 | 17.88200 | 0.03917 | 3.85378 | 7.81555 | 5.46469 | 10.15329 |
| **Crkl** | 3.62708 | 0.01647 | 3.95214 | 0.86027 | 4.44894 | 4.83373 | 5.31298 | 2.56421 | 9.46984 | 0.19243 | 2.28750 | 2.02959 | 1.75839 | 6.52747 |
| **Crlf1** | 0.10381 | 0.00440 | 0.02208 | 0.00895 | 0.01092 | 0.01451 | 0.02820 | 0.00935 | 0.00825 | 0.00644 | 0.00754 | 0.00972 | 0.00533 | 0.02666 |
| **Crlf2** | 0.03772 | 0.03233 | 0.00326 | 0.26440 | 0.35885 | 0.00338 | 0.00451 | 0.15312 | 0.03625 | 0.01087 | 0.00865 | 0.10747 | 0.20865 | 0.00000 |
| **Crlf3** | 8.20517 | 0.05740 | 8.14465 | 1.37374 | 9.11928 | 4.00763 | 0.37253 | 2.45792 | 7.43796 | 0.04016 | 3.74340 | 3.23515 | 5.65822 | 8.45124 |
| **Crls1** | 8.06574 | 0.02420 | 2.22226 | 1.22574 | 4.07800 | 0.14942 | 0.02029 | 2.62460 | 4.29722 | 0.00941 | 0.30551 | 1.07863 | 3.53780 | 4.97551 |
| **Crmp1** | 0.01804 | 0.00411 | 0.00387 | 0.33326 | 0.00172 | 0.00843 | 0.00430 | 0.01408 | 0.01205 | 0.00596 | 0.00986 | 0.00762 | 0.00469 | 0.67020 |
| **Crnkl1** | 15.83768 | 0.02887 | 24.53775 | 2.06707 | 32.28051 | 7.53681 | 4.11707 | 0.00000 | 23.31295 | 0.05662 | 7.28704 | 1.51460 | 13.43903 | 35.85298 |
| **Crocc** | 0.11419 | 0.00461 | 0.30927 | 0.29881 | 1.23242 | 0.07470 | 0.00462 | 0.80780 | 0.00521 | 0.10341 | 0.00000 | 0.45061 | 0.17844 | 0.03710 |
| **Crot** | 1.59725 | 0.02565 | 2.47397 | 1.48394 | 2.05484 | 3.24242 | 1.24741 | 1.58029 | 2.43899 | 0.03090 | 3.03063 | 7.61475 | 0.36735 | 2.58100 |
| **Crtac1** | 0.00674 | 0.00449 | 0.02130 | 0.01605 | 0.07215 | 0.01275 | 0.00433 | 0.01060 | 0.00589 | 0.00625 | 0.00899 | 0.00889 | 0.00515 | 0.11932 |
| **Crtam** | 0.00288 | 0.00425 | 0.00000 | 0.18005 | 0.42151 | 0.00000 | 0.00385 | 0.00717 | 0.54093 | 0.00658 | 0.00461 | 0.01351 | 0.00462 | 0.00000 |
| **Crtap** | 8.86263 | 0.84853 | 7.77732 | 0.00000 | 6.79019 | 7.94945 | 2.29331 | 6.32186 | 2.87213 | 0.05658 | 3.67255 | 8.54250 | 3.90220 | 4.47256 |
| **Crtc1** | 0.00000 | 0.02791 | 0.50565 | 1.10588 | 0.76582 | 0.12042 | 0.00377 | 0.25405 | 0.00976 | 0.00446 | 0.01184 | 1.69060 | 0.01781 | 0.00000 |
| **Crtc2** | 0.22428 | 0.00382 | 0.13807 | 0.15185 | 0.00000 | 0.09272 | 0.15146 | 0.08034 | 0.39764 | 0.00622 | 0.02684 | 0.23527 | 0.89707 | 0.65810 |
| **Crtc3** | 0.03484 | 0.04111 | 0.16802 | 0.42795 | 0.08663 | 0.86096 | 0.01125 | 1.03843 | 0.18684 | 0.00662 | 0.01149 | 0.49764 | 2.20083 | 0.00000 |
| **Crx** | 1.63801 | 0.01178 | 0.02806 | 0.06374 | 5.22748 | 0.02212 | 0.00983 | 0.02944 | 1.02930 | 0.01357 | 0.03471 | 0.01476 | 1.71431 | 4.13849 |
| **Crxos** | 28.97269 | 11.83277 | 17.95809 | 29.06479 | 29.55277 | 32.40530 | 11.22276 | 27.79822 | 20.46202 | 20.36593 | 16.20943 | 19.91951 | 22.46512 | 20.94615 |
| **Cry1** | 11.74241 | 0.02105 | 9.28534 | 0.00000 | 8.20206 | 3.15196 | 0.00000 | 1.89691 | 3.97851 | 0.00000 | 2.60088 | 3.82119 | 7.80334 | 10.06436 |
| **Cry2** | 1.62198 | 0.01333 | 2.20467 | 0.09006 | 2.31900 | 1.43175 | 0.01564 | 0.31392 | 0.13255 | 0.02468 | 1.25668 | 0.04312 | 0.74498 | 2.21051 |
| **Cryab** | 0.00000 | 0.17171 | 0.61005 | 0.75220 | 0.00000 | 5.80183 | 4.95977 | 11.94964 | 0.08316 | 0.14912 | 0.46460 | 0.00000 | 0.00657 | 0.00000 |
| **Cryba1** | 0.00000 | 0.00430 | 0.00000 | 0.00916 | 0.78117 | 0.01001 | 0.00413 | 0.00877 | 0.32214 | 0.00712 | 0.00729 | 0.00718 | 0.91714 | 0.00000 |
| **Crybg3** | 0.09653 | 0.00394 | 0.60837 | 0.33084 | 0.71359 | 0.42590 | 0.00381 | 0.14386 | 0.96072 | 0.00638 | 0.00000 | 0.56728 | 2.71174 | 0.67312 |
| **Cryga** | 0.04127 | 0.04508 | 0.01395 | 0.00734 | 0.01297 | 0.01587 | 0.00430 | 0.01172 | 0.01118 | 0.00707 | 0.00703 | 0.05535 | 0.00647 | 0.01130 |
| **Crygc** | 0.00657 | 0.00502 | 0.01914 | 0.00000 | 0.01325 | 0.00873 | 0.00438 | 0.11345 | 0.01214 | 0.01793 | 0.00369 | 0.53295 | 0.00772 | 0.01143 |
| **Crygd** | 0.00450 | 4.76109 | 0.01312 | 0.00279 | 0.02373 | 0.01277 | 0.00817 | 0.00585 | 0.00560 | 0.02023 | 0.00000 | 3.27064 | 0.00368 | 0.04641 |
| **Cryge** | 0.06452 | 1.32428 | 0.02722 | 0.00000 | 0.06787 | 0.03578 | 0.00385 | 0.31030 | 0.00457 | 0.00691 | 0.00000 | 1.11375 | 0.02524 | 0.16056 |
| **Crygf** | 0.02385 | 0.83731 | 0.02461 | 0.00000 | 0.03841 | 0.02325 | 0.00419 | 0.05399 | 0.00658 | 0.00734 | 0.00453 | 0.27909 | 0.00506 | 0.05584 |
| **Crygs** | 0.21218 | 0.00332 | 0.08235 | 0.27559 | 3.76418 | 0.00000 | 0.00303 | 0.00198 | 0.63652 | 3.67596 | 0.08605 | 0.00000 | 0.00343 | 0.65650 |
| **Cryl1** | 0.00000 | 0.01085 | 0.00000 | 0.49745 | 0.85964 | 0.89969 | 0.00371 | 0.02159 | 0.25991 | 0.00582 | 0.00441 | 0.17268 | 0.36614 | 0.37360 |

| | | | | | | | | | | | | | | |
|---|---|---|---|---|---|---|---|---|---|---|---|---|---|---|
| **Crym** | 0.04960 | 0.00355 | 0.00753 | 0.01635 | 0.10081 | 3.04568 | 2.70772 | 0.01665 | 0.01442 | 0.07534 | 3.38943 | 0.00000 | 0.01341 | 0.04204 |
| **Cryz** | 0.04880 | 0.12844 | 0.79756 | 4.16620 | 0.00000 | 4.86423 | 0.02497 | 5.52265 | 0.00000 | 0.46741 | 2.49843 | 6.29882 | 0.00371 | 0.55532 |
| **Cryzl1** | 2.27411 | 0.05636 | 2.51846 | 0.00000 | 0.74254 | 2.74055 | 3.60704 | 2.00698 | 0.01959 | 0.08252 | 1.90851 | 0.81991 | 0.01074 | 1.01963 |
| **Cs** | 11.86104 | 7.78880 | 25.35103 | 18.44304 | 12.21940 | 20.78795 | 7.50002 | 12.06301 | 8.07130 | 0.42321 | 15.81835 | 12.67251 | 10.79812 | 15.39750 |
| **Csad** | 1.18313 | 0.02504 | 2.01145 | 0.25259 | 0.02623 | 1.19928 | 0.00872 | 0.00000 | 0.12324 | 0.01976 | 0.00000 | 1.68543 | 0.67870 | 1.13549 |
| **Csdc2** | 0.64584 | 0.00413 | 3.33437 | 0.86288 | 1.68027 | 1.29255 | 0.00335 | 0.00000 | 0.00135 | 0.00781 | 0.00000 | 0.00875 | 0.22601 | 1.71529 |
| **Csde1** | 17.00056 | 3.82844 | 21.84571 | 12.56316 | 30.12670 | 15.28683 | 4.26788 | 6.90727 | 40.54780 | 0.23545 | 15.58341 | 12.24284 | 14.04948 | 35.03164 |
| **Cse1l** | 21.05756 | 4.05345 | 13.37382 | 5.47128 | 21.43084 | 11.30999 | 0.66758 | 3.85314 | 15.08420 | 13.32399 | 10.45502 | 11.27058 | 8.73024 | 22.76705 |
| **Csf1** | 0.45114 | 0.00440 | 0.01209 | 0.00966 | 0.00639 | 0.00950 | 0.00446 | 0.01166 | 0.01076 | 0.00640 | 0.00710 | 0.00853 | 0.02641 | 0.03829 |
| **Csf2ra** | 0.80276 | 0.01754 | 0.89356 | 1.66350 | 1.79987 | 0.08718 | 0.00354 | 0.54064 | 0.01742 | 0.00370 | 0.27572 | 1.00494 | 1.14087 | 0.00000 |
| **Csf2rb** | 0.01774 | 0.00438 | 0.00540 | 0.05139 | 0.05077 | 0.03260 | 0.01666 | 0.01091 | 0.11399 | 0.00869 | 0.02006 | 0.01184 | 0.17650 | 0.07074 |
| **Csf2rb2** | 1.43430 | 0.02165 | 0.31720 | 0.03706 | 0.69109 | 0.08092 | 0.06707 | 0.03713 | 0.93489 | 0.04022 | 0.01242 | 0.01478 | 0.85587 | 0.50065 |
| **Csf3r** | 0.00000 | 0.77591 | 5.04173 | 5.73584 | 1.20822 | 12.42182 | 6.41511 | 12.83755 | 3.62550 | 14.23759 | 9.88410 | 4.24382 | 0.28310 | 2.12043 |
| **Csgalnact2** | 0.00338 | 0.00415 | 0.39522 | 0.48404 | 0.61211 | 0.27954 | 0.00325 | 0.09817 | 3.84597 | 0.01110 | 0.00940 | 0.01960 | 0.00000 | 0.80190 |
| **Csk** | 5.47316 | 0.26941 | 9.26783 | 6.06981 | 8.12960 | 7.50889 | 0.05560 | 8.39941 | 0.00000 | 0.03001 | 3.90486 | 8.22540 | 1.50052 | 0.00000 |
| **Csl** | 0.24463 | 0.79491 | 1.72863 | 1.47874 | 0.75020 | 1.19853 | 0.46597 | 0.75112 | 0.28340 | 0.01384 | 1.07612 | 0.73571 | 0.62488 | 0.78459 |
| **Csnk1a1** | 1.59790 | 0.02899 | 9.47142 | 4.79885 | 8.40671 | 4.36804 | 0.15900 | 2.93179 | 7.29469 | 0.00787 | 7.99910 | 5.20876 | 16.31294 | 7.26190 |
| **Csnk1d** | 2.70919 | 0.12807 | 6.13326 | 7.19151 | 7.45188 | 6.12105 | 3.17517 | 2.09109 | 1.57245 | 0.15259 | 1.86623 | 6.18456 | 1.99230 | 5.82933 |
| **Csnk1e** | 14.45990 | 0.06353 | 21.87088 | 9.20019 | 25.38722 | 4.58147 | 0.68141 | 7.30604 | 8.01881 | 5.81661 | 6.51815 | 5.43643 | 16.17431 | 15.40600 |
| **Csnk1g1** | 1.14480 | 0.01058 | 3.18646 | 1.31283 | 6.35703 | 4.93732 | 0.00885 | 0.98202 | 10.49809 | 0.00335 | 1.61911 | 1.67339 | 4.67520 | 6.29886 |
| **Csnk1g2** | 0.00000 | 4.90610 | 6.11656 | 7.72311 | 5.62036 | 0.00000 | 1.78065 | 10.75063 | 0.00000 | 3.84179 | 0.00000 | 7.11294 | 0.03578 | 0.00218 |
| **Csnk1g3** | 9.78109 | 0.03305 | 6.17892 | 1.33119 | 7.84202 | 3.68740 | 1.57003 | 1.81328 | 12.57737 | 0.05404 | 0.04431 | 1.22891 | 9.72570 | 11.06666 |
| **Csnk2a1** | 8.36179 | 5.48417 | 8.52182 | 2.37357 | 10.98504 | 11.90347 | 0.14110 | 1.10198 | 14.44955 | 2.86680 | 4.13395 | 4.73024 | 6.44339 | 9.46867 |
| **Csnk2a2** | 8.60383 | 0.01017 | 6.93785 | 2.50985 | 5.66912 | 5.41948 | 2.06499 | 2.39434 | 4.69758 | 0.03525 | 2.82817 | 1.05565 | 4.51175 | 8.04830 |
| **Csnk2b** | 16.80996 | 10.61353 | 24.30440 | 22.74439 | 22.23995 | 18.06884 | 5.82340 | 13.56660 | 6.50345 | 8.45690 | 16.90260 | 13.84484 | 11.50385 | 13.76467 |
| **Cspg5** | 0.13147 | 0.00432 | 0.00000 | 0.63513 | 0.28142 | 0.33010 | 0.00387 | 0.00571 | 0.00574 | 0.01940 | 0.00353 | 0.01097 | 0.53083 | 0.00000 |
| **Cspp1** | 0.23795 | 0.00299 | 1.76764 | 0.52674 | 1.34067 | 2.15367 | 1.61218 | 0.62984 | 2.29343 | 0.01917 | 3.78853 | 1.33402 | 0.12098 | 1.11897 |
| **Csrnp1** | 4.03941 | 0.78788 | 5.89028 | 2.33890 | 10.24760 | 8.55216 | 3.07329 | 9.02052 | 0.46601 | 0.10999 | 3.93702 | 2.40967 | 7.89461 | 10.84954 |
| **Csrnp2** | 7.38811 | 0.00235 | 3.60313 | 1.86790 | 2.43073 | 0.00000 | 1.44959 | 0.18101 | 4.29997 | 0.04416 | 0.15589 | 0.00000 | 8.85698 | 13.32334 |
| **Csrnp3** | 0.02149 | 0.01243 | 0.01509 | 0.04180 | 0.21452 | 0.07322 | 0.01539 | 0.02773 | 0.04779 | 0.07144 | 0.00637 | 0.02139 | 0.76258 | 0.94568 |
| **Csrp1** | 0.50957 | 4.44233 | 7.29446 | 3.87625 | 7.11194 | 2.61574 | 2.18119 | 10.39821 | 0.00000 | 12.75156 | 3.57101 | 0.00000 | 0.00000 | 0.40369 |
| **Csrp2** | 0.15968 | 5.33610 | 1.74145 | 6.87890 | 1.04652 | 2.69286 | 0.17602 | 8.19555 | 0.00000 | 19.77439 | 1.72190 | 2.78912 | 0.12375 | 0.54251 |
| **Csrp2bp** | 5.64477 | 0.07390 | 7.06363 | 5.59490 | 9.79403 | 5.57633 | 0.95049 | 2.17561 | 2.57580 | 0.03770 | 3.84315 | 3.23159 | 4.52987 | 11.51290 |
| **Cst13** | 0.23885 | 0.00403 | 0.01038 | 0.00938 | 0.00677 | 0.18260 | 0.00417 | 0.03952 | 0.00946 | 0.00852 | 0.00507 | 0.00698 | 0.03849 | 0.00161 |
| **Cst3** | 0.61014 | 0.31227 | 1.57122 | 12.30648 | 3.46617 | 1.13039 | 4.55839 | 8.55741 | 0.00000 | 0.29668 | 1.61658 | 7.59380 | 0.28736 | 0.00000 |
| **Cst6** | 0.00758 | 0.00430 | 0.00706 | 0.09083 | 0.29410 | 0.00585 | 0.00401 | 0.40633 | 0.01028 | 0.00778 | 0.02795 | 0.02485 | 0.00688 | 0.02743 |
| **Cst7** | 0.98074 | 0.00362 | 2.46713 | 2.01944 | 1.83791 | 0.33380 | 0.01200 | 0.11078 | 2.03679 | 0.00912 | 0.00000 | 0.00000 | 7.09202 | 1.23262 |
| **Cst8** | 0.00869 | 0.00448 | 0.00518 | 0.00982 | 0.49115 | 0.01390 | 0.00498 | 0.01006 | 0.02281 | 0.00689 | 0.00894 | 0.00822 | 0.00427 | 0.00317 |
| **Csta1** | 0.00006 | 2.39581 | 0.00000 | 0.36971 | 0.09651 | 2.16189 | 2.22151 | 8.65806 | 0.10578 | 10.15386 | 0.19745 | 0.24329 | 0.77089 | 0.00000 |
| **Cstb** | 6.61249 | 4.66835 | 0.10753 | 4.97094 | 7.61754 | 16.22135 | 7.05975 | 17.64203 | 1.69798 | 20.22252 | 1.01068 | 9.55863 | 4.90506 | 2.72297 |
| **Cstf1** | 5.47838 | 0.33155 | 10.81594 | 10.75181 | 11.73878 | 6.28091 | 2.69071 | 3.79807 | 0.00000 | 0.17407 | 11.15385 | 4.57627 | 6.35589 | 6.92741 |
| **Cstf2** | 4.91427 | 1.01914 | 7.27221 | 3.25791 | 7.57702 | 3.97931 | 0.01686 | 5.36933 | 1.97542 | 0.00470 | 2.97693 | 6.06801 | 0.49811 | 3.14492 |
| **Cstf2t** | 5.90257 | 0.00186 | 4.68690 | 0.62450 | 3.87959 | 1.03644 | 0.00149 | 0.19986 | 15.20035 | 0.01264 | 0.28699 | 1.03538 | 9.81763 | 3.16021 |
| **Cstf3** | 2.43025 | 0.03092 | 5.55254 | 3.47158 | 6.38227 | 4.37588 | 0.58417 | 1.86357 | 2.90508 | 0.03431 | 9.55329 | 3.38621 | 5.71417 | 1.30560 |
| **Ctbp1** | 0.08659 | 0.01503 | 5.44034 | 2.92170 | 5.00102 | 2.22066 | 3.34253 | 6.59008 | 0.01821 | 0.02502 | 2.39634 | 4.89970 | 0.00000 | 0.00796 |
| **Ctbp2** | 1.78650 | 0.15351 | 8.95942 | 2.96761 | 2.45256 | 10.72628 | 1.34373 | 9.36726 | 0.59775 | 0.09756 | 8.76966 | 9.52063 | 0.92131 | 2.03547 |
| **Ctbs** | 0.32812 | 0.01226 | 0.32508 | 0.01132 | 0.10478 | 0.41666 | 0.00382 | 0.96527 | 0.00503 | 0.00592 | 1.24167 | 0.01875 | 0.66037 | 0.00000 |
| **Ctc1** | 4.44332 | 0.06570 | 7.96958 | 0.96120 | 6.57793 | 5.87304 | 4.20855 | 3.53272 | 2.98857 | 0.13432 | 5.85354 | 7.52607 | 3.67137 | 5.14440 |
| **Ctcf** | 10.92156 | 4.20436 | 16.70618 | 7.98987 | 16.26621 | 8.59759 | 1.49906 | 2.82465 | 5.64774 | 0.08677 | 10.14165 | 6.17542 | 10.61878 | 10.75805 |
| **Ctcfl** | 0.07513 | 0.03651 | 0.01536 | 0.33454 | 0.06839 | 0.23308 | 0.08088 | 0.09463 | 0.15618 | 0.08388 | 0.01089 | 0.06819 | 0.30190 | 0.18774 |
| **Ctdnep1** | 2.34733 | 0.08439 | 4.14178 | 3.41711 | 4.89529 | 8.78663 | 0.06939 | 5.97725 | 0.08126 | 3.01075 | 0.45191 | 5.99842 | 5.22225 | 3.49129 |
| **Ctdp1** | 1.53917 | 0.08892 | 11.98755 | 3.45161 | 8.37614 | 3.99070 | 0.22512 | 2.72752 | 0.00000 | 0.03396 | 3.38030 | 2.48892 | 0.84798 | 0.34279 |
| **Ctdsp1** | 0.23711 | 0.06575 | 0.61541 | 0.00000 | 0.04086 | 0.01804 | 0.00257 | 3.04962 | 0.12563 | 0.01722 | 0.50439 | 2.97279 | 0.97234 | 0.36557 |
| **Ctdsp2** | 1.24764 | 0.14982 | 3.12842 | 0.40890 | 3.59702 | 7.89551 | 5.05098 | 8.50236 | 1.23720 | 0.20319 | 0.00000 | 9.79159 | 1.95767 | 6.05386 |
| **Ctdspl** | 0.00867 | 0.02706 | 0.64788 | 0.09183 | 0.48019 | 1.04179 | 0.00860 | 2.50131 | 0.00868 | 0.03491 | 0.00168 | 0.01357 | 0.19868 | 0.00000 |

| | | | | | | | | | | | | | | |
|---|---|---|---|---|---|---|---|---|---|---|---|---|---|---|
| **Ctdspl2** | 2.57215 | 0.56352 | 5.56462 | 2.01796 | 7.05853 | 0.92606 | 1.17100 | 1.63396 | 14.48222 | 0.01075 | 0.30844 | 2.89945 | 6.63567 | 3.52493 |
| **Ctf1** | 0.00689 | 0.00430 | 0.01995 | 0.00991 | 0.01137 | 0.01317 | 0.07922 | 0.00667 | 0.01140 | 0.00651 | 0.00879 | 0.00995 | 0.13236 | 0.00379 |
| **Ctf2** | 0.01015 | 0.00485 | 0.04327 | 0.00213 | 0.01276 | 0.01587 | 0.00415 | 0.01604 | 0.01243 | 0.00692 | 0.00610 | 0.09241 | 0.00764 | 0.01134 |
| **Ctgf** | 1.28299 | 0.00394 | 0.02162 | 0.11698 | 0.00000 | 0.02097 | 0.37017 | 0.02666 | 0.00872 | 0.01295 | 0.03240 | 0.03013 | 0.00486 | 0.00000 |
| **Cth** | 20.08398 | 0.08650 | 30.30411 | 14.28358 | 23.60732 | 10.11827 | 7.59063 | 2.89022 | 28.00663 | 0.26378 | 14.31942 | 3.87237 | 9.90593 | 27.61663 |
| **Cthrc1** | 6.79168 | 0.01104 | 10.83641 | 2.31133 | 5.41484 | 0.12395 | 0.00729 | 0.00000 | 2.41419 | 0.00000 | 0.00000 | 0.00595 | 0.00000 | 1.43894 |
| **Ctif** | 0.01996 | 0.00388 | 0.02411 | 0.06666 | 0.01330 | 0.57011 | 0.00415 | 0.03758 | 0.00880 | 0.00549 | 0.00978 | 0.04167 | 0.25074 | 0.00239 |
| **Ctla2a** | 0.01171 | 0.09569 | 0.03569 | 0.08299 | 0.05129 | 0.06017 | 0.02361 | 0.08803 | 0.38429 | 0.00722 | 0.05429 | 0.06717 | 0.03607 | 0.18869 |
| **Ctnna1** | 9.67328 | 7.44352 | 23.57961 | 15.16372 | 12.54981 | 23.62279 | 9.14912 | 14.60598 | 6.74778 | 0.92677 | 19.72895 | 11.59099 | 7.21495 | 12.97877 |
| **Ctnnal1** | 4.41547 | 3.01904 | 7.27846 | 0.00000 | 7.79378 | 6.87794 | 5.24965 | 0.00000 | 11.28542 | 0.27234 | 2.63820 | 2.28667 | 8.00302 | 5.58155 |
| **Ctnnb1** | 9.75392 | 0.15016 | 17.15100 | 9.09804 | 14.75159 | 15.99270 | 10.48974 | 11.73124 | 9.45995 | 12.18140 | 12.21377 | 13.86241 | 10.97822 | 8.58674 |
| **Ctnnbip1** | 1.19971 | 0.01901 | 1.18287 | 1.36203 | 3.08107 | 0.96326 | 0.01033 | 2.07932 | 0.00552 | 0.01001 | 0.12000 | 2.35724 | 4.66033 | 2.20032 |
| **Ctnnbl1** | 5.13037 | 1.83652 | 9.36712 | 9.86419 | 11.31460 | 7.25679 | 3.53649 | 1.10065 | 7.23447 | 0.14249 | 1.83143 | 6.72020 | 5.92220 | 14.09917 |
| **Ctnnd1** | 8.45986 | 6.87912 | 10.23767 | 3.80915 | 10.49804 | 10.87745 | 4.63132 | 9.30569 | 16.01199 | 1.54810 | 10.75725 | 7.27921 | 5.49639 | 12.44332 |
| **Ctns** | 0.00000 | 0.00433 | 0.70884 | 0.78689 | 0.64191 | 0.16005 | 0.00900 | 1.78940 | 0.00702 | 0.01564 | 0.00741 | 0.15623 | 0.00000 | 1.41950 |
| **Ctps** | 5.98060 | 0.39878 | 15.19549 | 10.12989 | 13.02328 | 10.03482 | 2.64486 | 5.84863 | 10.41750 | 0.09749 | 11.85854 | 6.64343 | 6.82499 | 17.67398 |
| **Ctps2** | 0.63981 | 0.04216 | 1.56576 | 0.00000 | 4.24257 | 1.81046 | 0.00215 | 3.46190 | 3.31717 | 0.02244 | 0.78189 | 0.96737 | 1.88786 | 3.66024 |
| **Ctr9** | 1.66663 | 0.57770 | 5.53064 | 6.23850 | 6.24136 | 3.74428 | 0.87324 | 0.00000 | 1.88966 | 1.53266 | 1.39007 | 3.63366 | 2.44864 | 7.32638 |
| **Ctrb1** | 0.01085 | 0.00425 | 0.00668 | 0.05460 | 0.26763 | 0.01090 | 0.00410 | 0.00772 | 0.01160 | 0.00691 | 0.00796 | 0.01032 | 0.30701 | 0.03191 |
| **Ctrc** | 0.02409 | 0.00398 | 0.00857 | 0.00330 | 0.13293 | 0.01320 | 0.00428 | 0.00733 | 0.00996 | 0.00679 | 0.00486 | 0.52196 | 0.00944 | 0.25674 |
| **Ctrcos** | 0.06093 | 0.02701 | 0.01463 | 0.04018 | 0.02125 | 0.07519 | 0.00981 | 0.00674 | 0.14837 | 0.02507 | 0.01110 | 0.01925 | 0.35890 | 0.18599 |
| **Ctrl** | 1.43669 | 0.01254 | 1.09753 | 0.62421 | 2.74839 | 0.08648 | 0.00493 | 0.66014 | 0.05978 | 0.00513 | 0.00865 | 0.00170 | 0.06182 | 2.18808 |
| **Cts8** | 0.03472 | 0.00473 | 0.02457 | 0.09085 | 0.04479 | 0.07313 | 0.01658 | 0.04865 | 0.08721 | 0.00868 | 0.01420 | 0.02567 | 0.03683 | 0.07951 |
| **Ctsa** | 19.73351 | 6.24082 | 39.21820 | 41.19696 | 24.68123 | 20.27996 | 7.52442 | 14.20564 | 9.05592 | 11.77015 | 13.90844 | 16.08754 | 7.32653 | 36.21204 |
| **Ctsb** | 16.71644 | 0.25911 | 21.15008 | 19.88248 | 22.65631 | 9.89196 | 1.27437 | 0.98704 | 21.75469 | 0.16115 | 6.10418 | 23.06886 | 17.62918 | 33.76311 |
| **Ctsc** | 24.52210 | 5.45812 | 28.73689 | 15.21460 | 25.69003 | 9.13254 | 0.04655 | 6.31999 | 37.01645 | 0.04299 | 7.64453 | 6.21740 | 13.73949 | 31.33559 |
| **Ctsd** | 7.38851 | 9.29954 | 25.78757 | 34.88665 | 14.86304 | 29.97729 | 5.65304 | 14.15426 | 10.81013 | 13.54812 | 18.22664 | 17.68442 | 0.51386 | 17.14778 |
| **Ctsf** | 0.02814 | 0.00430 | 0.29248 | 0.01921 | 0.59659 | 0.00466 | 0.00408 | 0.00847 | 0.01907 | 0.00747 | 0.00562 | 0.03442 | 0.00347 | 0.00482 |
| **Ctsh** | 2.52705 | 0.02533 | 6.38863 | 3.26304 | 9.18978 | 0.25937 | 2.85877 | 5.00865 | 3.37136 | 4.86947 | 0.07393 | 0.00000 | 3.51154 | 8.50840 |
| **Ctsk** | 0.02786 | 0.00428 | 0.00629 | 0.00824 | 0.26071 | 0.02409 | 0.00403 | 0.00607 | 0.00803 | 0.00623 | 0.64699 | 0.00475 | 0.01888 | 0.85943 |
| **Ctsl** | 37.65742 | 9.92064 | 36.48883 | 34.14527 | 35.00889 | 25.64694 | 10.04647 | 17.48704 | 49.54234 | 1.30249 | 18.60038 | 23.58824 | 24.40924 | 55.72194 |
| **Ctss** | 0.00985 | 0.00450 | 0.00466 | 0.01180 | 0.57488 | 0.01350 | 0.00434 | 0.00988 | 0.01110 | 0.00690 | 0.00894 | 0.00798 | 0.00459 | 0.00744 |
| **Ctsz** | 14.48154 | 0.05521 | 17.17638 | 20.76053 | 15.12951 | 2.62304 | 0.02806 | 0.64801 | 5.82690 | 0.04554 | 2.19407 | 1.28561 | 4.55140 | 13.97635 |
| **Cttn** | 10.24852 | 0.26900 | 23.19208 | 15.44842 | 17.80352 | 20.94532 | 5.76470 | 15.60171 | 1.94657 | 0.31323 | 15.62362 | 8.63954 | 5.43845 | 16.78636 |
| **Cttnbp2nl** | 0.77046 | 0.23981 | 1.37793 | 0.66137 | 1.18274 | 1.79647 | 1.84754 | 0.46772 | 1.07364 | 0.03337 | 0.00000 | 1.02480 | 4.75971 | 2.44573 |
| **Ctu1** | 2.21564 | 0.00176 | 2.97631 | 2.57453 | 6.26146 | 1.82866 | 0.00242 | 0.00000 | 1.49588 | 0.00418 | 0.00000 | 2.25615 | 1.87785 | 2.86313 |
| **Ctu2** | 1.02362 | 0.91464 | 8.87129 | 11.26822 | 9.64429 | 1.24523 | 0.49974 | 2.30028 | 0.00000 | 0.01649 | 0.00000 | 3.48161 | 0.54817 | 0.00000 |
| **Ctxn1** | 1.04853 | 0.00428 | 0.40436 | 0.96338 | 0.99845 | 0.10371 | 0.00413 | 0.72903 | 0.00518 | 0.01687 | 0.00000 | 0.68399 | 0.39539 | 0.14336 |
| **Cubn** | 0.00377 | 0.01196 | 0.08329 | 0.92579 | 0.07639 | 0.00000 | 0.00377 | 0.00430 | 0.46574 | 0.00528 | 0.05987 | 0.78357 | 0.33721 | 0.00000 |
| **Cuedc1** | 0.02968 | 0.00449 | 0.01013 | 0.01215 | 0.15662 | 0.01277 | 0.00448 | 0.01056 | 0.01337 | 0.00651 | 0.00851 | 0.00905 | 0.00711 | 0.01467 |
| **Cuedc2** | 0.49237 | 0.16614 | 1.80216 | 2.34066 | 2.38439 | 0.56532 | 3.42516 | 7.95406 | 0.00000 | 0.16300 | 1.13772 | 6.24618 | 1.14607 | 1.08597 |
| **Cul1** | 16.02639 | 5.91489 | 12.49753 | 4.14517 | 17.09707 | 16.96898 | 5.05887 | 1.01130 | 18.98227 | 0.09071 | 14.16978 | 6.50427 | 13.22016 | 19.30647 |
| **Cul2** | 4.61259 | 6.39295 | 10.55722 | 4.57273 | 8.49584 | 6.40183 | 1.28424 | 3.69879 | 6.28390 | 0.03799 | 14.96331 | 8.23853 | 0.32340 | 14.02022 |
| **Cul3** | 11.72281 | 1.12759 | 11.06598 | 2.30726 | 12.09474 | 12.02583 | 1.69689 | 5.52403 | 23.47808 | 0.13591 | 7.40381 | 10.44360 | 17.86918 | 12.45766 |
| **Cul4a** | 4.12995 | 0.04404 | 2.60930 | 0.50855 | 3.63202 | 5.19912 | 0.00130 | 2.45676 | 0.52603 | 0.02408 | 5.33643 | 5.33951 | 1.15761 | 5.67009 |
| **Cul4b** | 1.89608 | 1.05629 | 6.56559 | 5.41211 | 7.90678 | 17.78417 | 1.35008 | 3.92146 | 21.41587 | 0.07307 | 13.18704 | 8.00874 | 10.98765 | 3.80376 |
| **Cul5** | 15.87225 | 0.03134 | 24.58195 | 6.97605 | 24.26491 | 13.69947 | 3.66126 | 0.54306 | 31.25886 | 0.34938 | 17.67498 | 7.92018 | 3.72839 | 16.60392 |
| **Cul7** | 0.81586 | 0.08337 | 0.56186 | 0.00000 | 0.22006 | 0.00000 | 0.01319 | 0.00000 | 0.00303 | 12.23789 | 2.00536 | 2.68353 | 0.05468 | 0.00000 |
| **Cul9** | 0.03125 | 0.00439 | 0.07726 | 0.00551 | 0.01506 | 0.01196 | 0.00419 | 0.00995 | 0.01169 | 0.00648 | 0.00730 | 0.00987 | 0.19620 | 0.16688 |
| **Cuta** | 2.19691 | 0.19837 | 3.90755 | 14.69520 | 8.27920 | 9.96953 | 2.61233 | 9.23354 | 0.00087 | 0.26440 | 1.32112 | 8.61529 | 0.52084 | 1.53467 |
| **Cutc** | 4.58080 | 0.02441 | 1.72624 | 1.34895 | 2.02374 | 3.30620 | 0.00784 | 0.00433 | 2.66768 | 0.01059 | 0.05098 | 3.69279 | 0.58250 | 1.46471 |
| **Cux1** | 0.82490 | 0.02451 | 3.11311 | 2.18188 | 2.06162 | 1.45727 | 0.00904 | 0.94892 | 0.43159 | 0.02258 | 0.18115 | 2.86943 | 0.00000 | 1.28864 |
| **Cux2** | 2.83903 | 0.01105 | 6.60625 | 3.21155 | 3.79726 | 0.59769 | 0.00260 | 0.00000 | 0.89568 | 0.00000 | 0.16168 | 0.00245 | 0.50552 | 1.69966 |
| **Cuzd1** | 0.00429 | 0.00411 | 0.01822 | 0.29687 | 0.00343 | 0.00362 | 0.00411 | 0.00650 | 0.00891 | 0.00564 | 0.01989 | 0.01731 | 0.00324 | 0.24465 |
| **Cwc15** | 19.61431 | 0.24713 | 15.45767 | 16.32670 | 21.05520 | 16.85877 | 4.59098 | 11.18944 | 14.01790 | 11.36413 | 12.09092 | 11.84118 | 15.17979 | 19.31363 |

| | | | | | | | | | | | | | | |
|---|---|---|---|---|---|---|---|---|---|---|---|---|---|---|
| **Cwc22** | 8.53266 | 0.08890 | 11.16388 | 2.96953 | 13.08330 | 8.03792 | 3.77103 | 2.15326 | 19.72689 | 0.10993 | 9.06059 | 8.47069 | 8.48866 | 12.46113 |
| **Cwc25** | 1.82971 | 2.04853 | 5.40913 | 2.93266 | 6.79446 | 4.18916 | 1.41845 | 0.41770 | 0.33819 | 0.12329 | 0.00000 | 2.07384 | 2.91639 | 9.20136 |
| **Cwc27** | 2.28575 | 0.03894 | 4.93343 | 4.88662 | 1.10817 | 2.57889 | 1.17583 | 2.00955 | 1.98927 | 0.06405 | 4.54257 | 5.80389 | 0.00000 | 3.57559 |
| **Cwf19l1** | 1.33340 | 3.11259 | 8.57604 | 2.09928 | 4.14322 | 5.18714 | 0.12711 | 0.00000 | 2.63685 | 0.03249 | 4.04474 | 1.99177 | 0.70441 | 3.16250 |
| **Cwf19l2** | 0.73488 | 0.03030 | 1.16787 | 0.14635 | 0.00000 | 0.40877 | 0.00999 | 0.62560 | 3.37050 | 0.00000 | 2.33799 | 1.16062 | 0.80383 | 0.96682 |
| **Cwh43** | 0.00650 | 0.00443 | 0.05347 | 0.03187 | 0.90924 | 0.01294 | 0.00427 | 0.00802 | 0.01270 | 0.00659 | 0.01320 | 0.01117 | 0.00518 | 0.02881 |
| **Cx3cl1** | 0.02203 | 0.00429 | 0.01062 | 0.10436 | 0.01014 | 0.00890 | 0.00411 | 0.00722 | 0.00967 | 0.00663 | 0.00720 | 0.01476 | 0.00592 | 0.02875 |
| **Cx3cr1** | 0.01650 | 0.02924 | 0.01877 | 0.01998 | 0.06725 | 0.03282 | 0.01090 | 0.01677 | 0.07124 | 0.00677 | 0.02505 | 0.02106 | 0.00945 | 0.11287 |
| **Cxadr** | 6.35808 | 0.07263 | 6.52583 | 4.06183 | 6.76407 | 4.13920 | 3.11571 | 4.12615 | 19.24823 | 3.02702 | 0.90848 | 1.49944 | 9.53844 | 9.90335 |
| **Cxcl10** | 0.00222 | 0.00447 | 0.32823 | 0.10799 | 0.00000 | 0.48403 | 0.00431 | 0.00335 | 0.00624 | 0.00490 | 0.00536 | 0.00471 | 0.01328 | 0.16915 |
| **Cxcl11** | 0.01374 | 0.00445 | 0.01419 | 0.02324 | 0.00992 | 0.02484 | 0.00424 | 0.00816 | 0.01326 | 0.00827 | 0.00862 | 0.00888 | 0.00884 | 0.17166 |
| **Cxcl12** | 0.08303 | 0.00437 | 0.01020 | 0.04296 | 0.02749 | 0.00966 | 0.00408 | 0.00925 | 0.01216 | 0.01599 | 0.00749 | 0.00813 | 0.01895 | 0.00553 |
| **Cxcl16** | 0.00000 | 0.00334 | 0.00000 | 1.69488 | 0.00000 | 0.00000 | 0.33246 | 0.54775 | 0.47776 | 0.03753 | 0.00000 | 0.00396 | 4.94950 | 0.04159 |
| **Cxcl9** | 0.00242 | 0.00407 | 0.08359 | 0.01103 | 0.04098 | 0.01017 | 0.00421 | 0.00939 | 0.01709 | 0.00679 | 0.00889 | 0.00778 | 2.20634 | 0.01313 |
| **Cxcr2** | 1.29296 | 0.09852 | 0.00000 | 0.61925 | 0.31356 | 2.10236 | 0.29262 | 0.38245 | 2.34507 | 0.63363 | 0.02963 | 0.70235 | 10.01585 | 1.29473 |
| **Cxx1a** | 0.08157 | 0.00433 | 0.02091 | 0.01560 | 0.10420 | 0.01346 | 0.00431 | 0.01155 | 0.01185 | 0.00662 | 0.00849 | 0.01509 | 0.00558 | 0.00759 |
| **Cxx1b** | 0.04347 | 0.00443 | 0.01950 | 0.00986 | 0.08323 | 0.01271 | 0.00433 | 0.01041 | 0.01240 | 0.00687 | 0.00845 | 0.01223 | 0.00652 | 0.00997 |
| **Cxx1c** | 0.02516 | 0.00435 | 0.03344 | 0.01033 | 0.04450 | 0.01401 | 0.00433 | 0.01209 | 0.01214 | 0.00681 | 0.00802 | 0.01532 | 0.00681 | 0.01029 |
| **Cxxc1** | 4.00386 | 1.06174 | 12.15351 | 13.16128 | 6.60983 | 5.05093 | 2.60838 | 5.80008 | 0.09644 | 12.63644 | 4.85951 | 5.86760 | 2.09045 | 2.61453 |
| **Cxxc5** | 0.10845 | 0.01049 | 0.00000 | 0.00000 | 0.42140 | 0.51178 | 0.10757 | 3.66067 | 0.00379 | 0.18345 | 0.83582 | 1.40973 | 0.14244 | 0.18821 |
| **Cyb561** | 0.62918 | 5.89272 | 0.00000 | 0.56151 | 0.32155 | 1.72604 | 0.00249 | 1.47307 | 0.00078 | 0.00963 | 1.06591 | 0.45091 | 0.00296 | 0.43233 |
| **Cyb561a3** | 3.72966 | 0.11484 | 6.74086 | 1.87931 | 3.38626 | 3.13480 | 4.20123 | 2.94389 | 1.34470 | 0.30010 | 1.62794 | 1.25835 | 2.11605 | 5.16427 |
| **Cyb561d1** | 0.17550 | 1.65178 | 0.00610 | 0.03296 | 0.00526 | 0.00000 | 0.00347 | 0.00536 | 0.00912 | 0.00587 | 0.95230 | 0.00004 | 0.04298 | 0.00231 |
| **Cyb561d2** | 1.22908 | 0.06232 | 2.73633 | 3.03761 | 1.96645 | 4.88191 | 3.52982 | 3.92149 | 0.02087 | 0.07709 | 2.63016 | 1.51136 | 0.03765 | 1.44892 |
| **Cyb5a** | 0.67544 | 0.08461 | 1.28631 | 10.35712 | 5.92261 | 8.36003 | 5.19366 | 8.28763 | 0.41329 | 0.28793 | 4.74799 | 8.21636 | 0.80709 | 2.41215 |
| **Cyb5b** | 1.67492 | 1.42125 | 2.58153 | 0.01882 | 7.51304 | 9.01617 | 0.00141 | 0.45347 | 5.87189 | 0.05077 | 0.43032 | 3.74652 | 3.08635 | 4.33787 |
| **Cyb5d1** | 0.01297 | 0.00430 | 0.02169 | 0.00740 | 0.34858 | 0.05406 | 0.00421 | 0.00768 | 0.00931 | 0.00740 | 0.06869 | 0.02472 | 0.00292 | 0.36783 |
| **Cyb5d2** | 0.05125 | 0.06028 | 0.62937 | 0.49658 | 0.61157 | 1.20202 | 0.24612 | 0.84909 | 0.78510 | 0.06086 | 0.82444 | 1.20311 | 0.09058 | 2.21613 |
| **Cyb5r1** | 2.49591 | 1.10895 | 6.01879 | 8.52136 | 0.00000 | 6.28562 | 4.90681 | 5.84474 | 0.37171 | 0.17817 | 8.18289 | 9.00901 | 1.27231 | 0.14402 |
| **Cyb5r2** | 0.66729 | 0.00401 | 0.08009 | 0.43692 | 0.25699 | 0.00000 | 0.00401 | 0.00613 | 0.06376 | 0.01954 | 0.09009 | 0.02266 | 0.18694 | 0.17812 |
| **Cyb5r3** | 0.00000 | 3.56537 | 3.14866 | 1.48716 | 2.54293 | 10.68993 | 5.53166 | 10.79623 | 0.00000 | 0.42818 | 7.28137 | 13.93015 | 0.93219 | 0.16079 |
| **Cyb5r4** | 11.97410 | 0.03975 | 5.22551 | 6.69673 | 6.47824 | 4.01352 | 0.00746 | 1.83775 | 11.12081 | 0.02995 | 0.27451 | 3.39651 | 2.64789 | 5.54573 |
| **Cyb5rl** | 0.00000 | 0.00359 | 0.24618 | 0.95715 | 0.14740 | 0.66897 | 0.00338 | 0.78708 | 0.01179 | 0.00670 | 1.06374 | 0.18117 | 0.01033 | 0.41154 |
| **Cyba** | 0.00000 | 0.01085 | 0.91263 | 0.33942 | 1.24498 | 0.00000 | 0.00322 | 0.31740 | 0.00361 | 8.95697 | 0.00271 | 0.00000 | 0.00000 | 0.37526 |
| **Cybb** | 0.01049 | 0.02034 | 0.07806 | 0.00000 | 0.13026 | 0.00724 | 0.00401 | 0.01013 | 1.32009 | 0.00928 | 0.00521 | 0.00804 | 0.00217 | 0.00000 |
| **Cybrd1** | 0.06276 | 0.01236 | 0.04431 | 0.12027 | 0.13997 | 0.19819 | 0.01711 | 0.07733 | 0.08931 | 0.01764 | 0.01502 | 0.01744 | 0.36795 | 0.26107 |
| **Cyc1** | 8.99762 | 3.49003 | 11.33691 | 15.13757 | 12.95078 | 13.13935 | 5.13818 | 14.81691 | 4.41847 | 12.92633 | 3.91853 | 14.30068 | 1.01185 | 2.97494 |
| **Cycs** | 10.16731 | 5.05841 | 15.80527 | 16.51607 | 17.72027 | 23.18279 | 6.07952 | 22.41971 | 16.67939 | 17.56600 | 13.34669 | 14.34634 | 11.00684 | 13.85299 |
| **Cyct** | 0.72050 | 3.98311 | 0.00000 | 0.47469 | 0.57179 | 0.90088 | 0.00354 | 0.24314 | 0.43163 | 0.45368 | 0.27280 | 0.00760 | 0.29562 | 0.00000 |
| **Cyfip1** | 5.76060 | 1.21718 | 3.17203 | 2.80528 | 4.41598 | 6.80640 | 3.13023 | 0.92300 | 5.92498 | 0.11738 | 4.10076 | 1.75135 | 2.43586 | 5.99420 |
| **Cyfip2** | 0.11749 | 0.00392 | 0.30329 | 0.51044 | 0.56438 | 0.00001 | 0.00378 | 0.00578 | 0.04238 | 0.00496 | 0.02244 | 0.01354 | 1.09351 | 0.08860 |
| **Cyhr1** | 0.92894 | 0.02115 | 0.63152 | 5.85292 | 0.05565 | 5.39400 | 5.23580 | 0.69129 | 0.00699 | 10.38119 | 2.91325 | 2.55306 | 2.20307 | 0.64318 |
| **Cyld** | 1.11927 | 0.20051 | 0.09579 | 0.28481 | 0.35255 | 1.37595 | 0.14975 | 0.27157 | 2.09843 | 0.12461 | 0.11681 | 0.23221 | 3.45168 | 1.40846 |
| **Cyp11a1** | 0.07152 | 0.00450 | 0.00000 | 0.00607 | 2.12274 | 0.01240 | 0.00439 | 0.01121 | 0.00824 | 0.00662 | 0.00798 | 0.00678 | 0.00000 | 0.09007 |
| **Cyp17a1** | 0.00000 | 0.00426 | 0.51518 | 0.90014 | 0.00000 | 0.44452 | 0.00375 | 0.00177 | 0.16389 | 0.01171 | 0.00294 | 0.00512 | 0.39920 | 0.92115 |
| **Cyp1a2** | 0.00249 | 0.00435 | 0.27040 | 0.00782 | 0.13850 | 0.00446 | 0.00400 | 0.01029 | 0.01927 | 0.00897 | 0.00660 | 0.00767 | 0.18961 | 0.06730 |
| **Cyp1b1** | 0.01987 | 0.00444 | 0.00185 | 0.00972 | 0.40801 | 0.01125 | 0.00497 | 0.01001 | 0.18643 | 0.00659 | 0.00762 | 0.00801 | 0.11719 | 0.00299 |
| **Cyp20a1** | 1.64891 | 0.06233 | 4.17089 | 3.15119 | 1.32515 | 3.11973 | 0.17674 | 0.81413 | 0.47903 | 0.33530 | 0.03023 | 1.82703 | 1.24757 | 3.44149 |
| **Cyp26a1** | 0.00792 | 0.00377 | 0.01749 | 0.00000 | 0.01300 | 0.01681 | 0.00397 | 0.07636 | 0.01183 | 0.00795 | 0.00394 | 0.32892 | 0.00756 | 0.01153 |
| **Cyp26b1** | 0.01151 | 0.00449 | 0.12052 | 0.00000 | 0.49721 | 0.00970 | 0.00412 | 0.01049 | 0.01469 | 0.00893 | 0.00718 | 0.00894 | 0.00403 | 0.00619 |
| **Cyp2ab1** | 0.02162 | 0.00444 | 0.01057 | 0.00690 | 0.07614 | 0.04318 | 0.00425 | 0.00742 | 0.08540 | 0.00620 | 0.00790 | 0.00964 | 0.08660 | 0.00000 |
| **Cyp2b10** | 0.00253 | 0.00455 | 0.38770 | 0.00000 | 0.56820 | 0.31979 | 0.00413 | 0.00411 | 0.01035 | 0.00518 | 0.00584 | 0.01141 | 0.02700 | 0.07943 |
| **Cyp2b13** | 0.01042 | 0.00438 | 0.06307 | 0.02369 | 0.03972 | 0.03654 | 0.00419 | 0.00624 | 0.01298 | 0.00582 | 0.00765 | 0.01155 | 0.30548 | 0.05774 |
| **Cyp2b23** | 0.01191 | 0.00437 | 0.10000 | 0.00000 | 0.02202 | 0.01489 | 0.00449 | 0.01483 | 0.01210 | 0.00679 | 0.00741 | 0.02760 | 0.01358 | 0.02314 |
| **Cyp2c44** | 0.00000 | 0.00358 | 0.08914 | 0.57066 | 0.03988 | 0.54195 | 0.00965 | 0.69167 | 1.18305 | 0.01880 | 0.90503 | 0.00000 | 0.58019 | 0.28566 |

| | | | | | | | | | | | | | | |
|---|---|---|---|---|---|---|---|---|---|---|---|---|---|---|
| **Cyp2c55** | 0.00886 | 0.00442 | 0.03438 | 0.00000 | 0.01333 | 0.35607 | 0.00414 | 0.00634 | 0.01096 | 0.00657 | 0.00543 | 0.01096 | 0.00802 | 0.00892 |
| **Cyp2d10** | 0.01831 | 0.00443 | 0.08098 | 0.00503 | 0.02780 | 0.01304 | 0.00419 | 0.01044 | 0.01954 | 0.00662 | 0.01111 | 0.00971 | 0.01464 | 0.01924 |
| **Cyp2d12** | 0.00943 | 0.00443 | 0.17114 | 0.00000 | 0.12572 | 0.01474 | 0.00411 | 0.00953 | 0.01560 | 0.00638 | 0.01582 | 0.00997 | 0.01669 | 0.01037 |
| **Cyp2d22** | 0.00930 | 0.00418 | 0.03424 | 0.12123 | 0.03387 | 0.00642 | 0.00425 | 0.00626 | 0.01630 | 0.00561 | 0.01685 | 0.03494 | 0.05355 | 0.02508 |
| **Cyp2d40** | 0.00551 | 0.00429 | 0.05660 | 0.00955 | 0.01411 | 0.01258 | 0.00417 | 0.00882 | 0.01812 | 0.00609 | 0.00950 | 0.00953 | 0.50027 | 0.01501 |
| **Cyp2d9** | 0.01373 | 0.00451 | 1.88268 | 0.00000 | 1.12418 | 0.00921 | 0.00364 | 0.00433 | 0.20923 | 0.00191 | 0.19857 | 0.00548 | 0.03873 | 0.00000 |
| **Cyp2e1** | 0.59973 | 0.04429 | 0.02113 | 0.28879 | 0.04919 | 0.80912 | 0.19888 | 0.12908 | 0.78612 | 0.39337 | 0.01652 | 0.22609 | 4.41875 | 0.47493 |
| **Cyp2f2** | 0.00000 | 0.00402 | 0.97625 | 0.00000 | 0.81360 | 0.05321 | 0.00428 | 0.00058 | 0.02448 | 0.00502 | 0.00428 | 0.17116 | 0.00969 | 0.51980 |
| **Cyp2j13** | 0.18500 | 0.00458 | 0.08723 | 0.01343 | 0.06198 | 0.08492 | 0.01058 | 0.03896 | 0.24542 | 0.00647 | 0.02106 | 0.01394 | 0.05196 | 0.14914 |
| **Cyp2j6** | 0.01655 | 0.00441 | 0.03076 | 0.01366 | 0.00152 | 0.00990 | 0.00431 | 0.01002 | 0.01274 | 0.00662 | 0.00725 | 0.00911 | 0.00670 | 0.54440 |
| **Cyp2s1** | 16.28454 | 1.46611 | 26.30565 | 17.31493 | 5.83952 | 29.73242 | 8.83274 | 19.42019 | 4.62217 | 14.73123 | 10.56254 | 5.16627 | 2.59417 | 9.45452 |
| **Cyp2t4** | 0.00000 | 0.00429 | 0.66147 | 0.79426 | 0.09529 | 0.84663 | 0.00380 | 0.00000 | 0.15548 | 0.00404 | 0.00377 | 0.00584 | 0.00387 | 0.00747 |
| **Cyp39a1** | 0.00000 | 0.00413 | 0.00000 | 0.01423 | 0.00000 | 0.00574 | 0.00392 | 0.00774 | 0.79097 | 0.00649 | 0.00562 | 0.00646 | 1.53917 | 0.79993 |
| **Cyp46a1** | 0.19055 | 0.00444 | 0.00961 | 0.00652 | 0.00590 | 0.05710 | 0.00425 | 0.00555 | 0.00846 | 0.00994 | 0.22435 | 0.02828 | 0.19486 | 0.00098 |
| **Cyp4f13** | 0.00240 | 0.00371 | 0.00822 | 1.84014 | 0.01898 | 0.00000 | 0.00959 | 0.00978 | 0.01581 | 0.00504 | 0.00611 | 0.35637 | 0.00681 | 0.00428 |
| **Cyp4f16** | 0.00298 | 0.03644 | 0.00000 | 0.09689 | 0.00533 | 0.22740 | 0.00411 | 0.02523 | 0.02335 | 0.00546 | 0.53935 | 1.86564 | 0.01124 | 0.00402 |
| **Cyp4f17** | 0.01931 | 0.00422 | 0.20906 | 0.17532 | 0.01305 | 0.00273 | 0.00412 | 0.00861 | 0.01184 | 0.00864 | 0.01383 | 0.00848 | 0.01022 | 0.00809 |
| **Cyp4f37** | 0.00963 | 0.00432 | 0.01243 | 0.00998 | 0.01343 | 0.01266 | 0.00431 | 0.00982 | 0.01165 | 0.00689 | 0.08239 | 0.13519 | 0.00752 | 0.01128 |
| **Cyp4f39** | 0.00000 | 0.01972 | 0.57654 | 4.28506 | 0.00000 | 2.12532 | 0.12583 | 3.35684 | 0.00693 | 0.00591 | 0.00000 | 0.00000 | 0.00934 | 0.05749 |
| **Cyp4f41.ps** | 0.00411 | 0.00523 | 0.10012 | 0.23335 | 0.04875 | 0.00000 | 0.00442 | 0.09982 | 0.01115 | 0.00692 | 0.00797 | 0.02294 | 0.00608 | 0.00902 |
| **Cyp4v3** | 0.01581 | 0.00435 | 0.14645 | 0.00326 | 0.01975 | 0.00622 | 0.00425 | 0.01239 | 0.00868 | 0.00689 | 0.00674 | 0.01033 | 0.02198 | 0.31940 |
| **Cyp51** | 3.72020 | 0.00854 | 1.03935 | 0.00000 | 0.00000 | 8.88836 | 0.01298 | 1.18809 | 1.61415 | 0.04530 | 1.45357 | 2.19526 | 0.04135 | 1.51776 |
| **Cypt3** | 0.00501 | 0.00429 | 0.01412 | 0.01682 | 0.01174 | 0.01271 | 0.00417 | 0.00872 | 0.01000 | 0.00780 | 0.00886 | 0.00846 | 0.55065 | 0.00356 |
| **Cyr61** | 0.75193 | 0.32533 | 0.51170 | 0.73681 | 0.00000 | 0.43076 | 0.03114 | 0.09360 | 1.13115 | 0.19392 | 0.68955 | 0.00000 | 0.74581 | 2.95225 |
| **Cys1** | 0.00593 | 0.00454 | 0.09689 | 0.01287 | 0.19201 | 0.01084 | 0.00473 | 0.01354 | 0.01842 | 0.00758 | 0.00786 | 0.00772 | 0.00353 | 0.08788 |
| **Cysrt1** | 0.01233 | 0.00433 | 0.15904 | 0.00312 | 0.01016 | 0.00966 | 0.00416 | 0.01445 | 0.01148 | 0.00772 | 0.00799 | 0.01020 | 0.00710 | 0.01171 |
| **Cystm1** | 1.97675 | 5.67800 | 6.85368 | 10.41123 | 7.06926 | 9.89809 | 3.11805 | 13.30695 | 0.00000 | 10.63626 | 2.50750 | 0.61256 | 0.00000 | 1.37872 |
| **Cyth1** | 0.14148 | 0.00436 | 0.05513 | 0.22574 | 0.15542 | 0.00780 | 0.00491 | 0.01421 | 0.01143 | 0.01068 | 0.00766 | 0.00797 | 0.33033 | 0.00000 |
| **Cyth2** | 7.80858 | 0.06523 | 17.54020 | 8.25968 | 14.28544 | 7.36191 | 0.41045 | 1.94385 | 2.72663 | 0.06902 | 3.41685 | 4.63301 | 4.62798 | 15.88497 |
| **Cyth3** | 1.71167 | 0.01136 | 0.65473 | 0.00000 | 0.49280 | 2.43044 | 0.00332 | 0.61024 | 0.04312 | 0.01804 | 0.01265 | 0.65046 | 1.20781 | 1.29479 |
| **D10Jhu81e** | 0.00000 | 0.04504 | 0.26181 | 0.78825 | 0.00226 | 0.16470 | 0.01026 | 3.25321 | 0.01207 | 0.04924 | 1.35194 | 0.03515 | 0.00680 | 0.00000 |
| **D10Wsu102e** | 3.25259 | 0.03548 | 9.53281 | 7.50058 | 9.02829 | 7.34754 | 4.06294 | 3.21854 | 8.86132 | 7.69052 | 7.06951 | 4.56130 | 4.01939 | 12.33010 |
| **D11Wsu47e** | 0.84660 | 0.16541 | 2.65859 | 1.84095 | 3.41398 | 1.60180 | 0.00303 | 1.00532 | 1.18589 | 0.00006 | 2.25885 | 0.20156 | 2.06587 | 2.81128 |
| **D16Ertd472e** | 0.56957 | 0.04982 | 0.28221 | 2.55587 | 4.72699 | 1.01725 | 4.76816 | 1.30497 | 3.19053 | 0.12980 | 2.39831 | 1.75884 | 12.40509 | 3.63938 |
| **D16Ertd519e** | 0.01039 | 0.00443 | 0.01296 | 0.01498 | 0.01003 | 0.01245 | 0.00426 | 0.01008 | 0.00720 | 0.00730 | 0.00864 | 0.00971 | 0.01781 | 0.13389 |
| **D17Ertd648e** | 5.46772 | 0.02521 | 2.42885 | 1.56518 | 6.27427 | 1.13216 | 0.66513 | 0.23812 | 10.26530 | 0.69735 | 0.30874 | 0.11502 | 24.22172 | 17.47690 |
| **D17H6S53E** | 0.17518 | 0.06648 | 0.03137 | 0.54540 | 0.72919 | 0.63813 | 0.10618 | 0.55686 | 0.33369 | 0.22040 | 0.06248 | 0.27905 | 1.37546 | 0.42748 |
| **D17H6S56E.5** | 3.68955 | 0.11312 | 4.18804 | 2.47540 | 4.03541 | 4.36895 | 0.19360 | 1.18451 | 2.93547 | 0.08170 | 3.93703 | 3.37491 | 5.32581 | 5.31329 |
| **D17Wsu92e** | 4.03565 | 0.08750 | 8.55879 | 6.05357 | 6.96446 | 11.33114 | 1.55640 | 5.87310 | 2.59129 | 10.71533 | 2.41357 | 4.11948 | 9.87984 | 8.37858 |
| **D1Ertd622e** | 0.63979 | 0.00388 | 0.56460 | 0.96357 | 0.44027 | 1.02738 | 0.00339 | 0.00937 | 1.17892 | 0.00524 | 0.07796 | 0.25472 | 4.87203 | 1.45283 |
| **D1Pas1** | 0.55257 | 0.00372 | 0.25746 | 0.22010 | 1.02275 | 0.10069 | 0.00369 | 0.00049 | 0.00718 | 0.00378 | 0.27328 | 0.00822 | 1.90750 | 0.52093 |
| **D2hgdh** | 0.00001 | 0.01311 | 4.02828 | 2.61137 | 0.44534 | 1.43682 | 0.00315 | 0.46472 | 0.14605 | 0.03193 | 0.95756 | 0.88486 | 0.30500 | 0.53942 |
| **D2Wsu81e** | 1.11256 | 0.04306 | 6.32344 | 7.13579 | 4.94721 | 6.67905 | 0.02085 | 0.25637 | 0.00000 | 0.05310 | 2.76189 | 4.48089 | 2.24448 | 4.99525 |
| **D3Ertd254e** | 2.69773 | 0.01848 | 1.30981 | 1.35967 | 2.93401 | 0.63451 | 0.08550 | 0.01767 | 1.59833 | 0.12718 | 2.20789 | 2.16453 | 5.76905 | 0.67308 |
| **D3Ertd751e** | 0.02875 | 0.00357 | 0.00000 | 0.02913 | 0.00000 | 0.46697 | 0.00367 | 0.01334 | 0.95722 | 0.00513 | 0.00271 | 0.55885 | 0.00648 | 0.04520 |
| **D4Ertd617e** | 0.00790 | 0.00421 | 0.01515 | 0.00796 | 0.01206 | 0.02707 | 0.00416 | 0.01420 | 0.01394 | 0.00596 | 0.00850 | 0.02504 | 0.10083 | 0.00902 |
| **D5Ertd577e** | 4.24253 | 0.00156 | 1.29959 | 0.91753 | 3.34293 | 0.57628 | 2.23622 | 0.00000 | 8.03188 | 6.32846 | 0.93243 | 0.08414 | 4.23250 | 2.35742 |
| **D5Ertd579e** | 0.58074 | 0.02262 | 0.95559 | 1.62046 | 0.53308 | 2.32852 | 0.00344 | 0.74010 | 0.00327 | 0.03857 | 1.88104 | 2.03744 | 0.00656 | 1.17687 |
| **D5Ertd605e** | 0.94934 | 0.15006 | 0.24921 | 0.00000 | 1.81148 | 0.72818 | 0.05069 | 3.93449 | 1.57784 | 0.02600 | 0.02855 | 0.00000 | 0.48100 | 1.02854 |
| **D5Ertd615e** | 0.10829 | 0.28086 | 0.10004 | 0.22180 | 0.21296 | 0.14455 | 0.07591 | 0.12553 | 1.24589 | 0.07789 | 0.09058 | 0.28964 | 0.24853 | 0.31785 |
| **D6Ertd474e** | 0.00000 | 0.00389 | 0.00000 | 0.01470 | 0.00000 | 0.00152 | 0.00397 | 0.00711 | 0.96517 | 0.00640 | 0.00492 | 0.00417 | 3.06077 | 2.97656 |
| **D6Ertd527e** | 0.03418 | 0.00398 | 0.47879 | 0.00000 | 0.13628 | 0.01129 | 0.00382 | 0.00409 | 0.08461 | 0.00417 | 0.30467 | 0.00516 | 1.14149 | 0.86894 |
| **D6Wsu163e** | 7.51218 | 0.10685 | 4.16718 | 0.58009 | 3.90191 | 2.85648 | 0.00000 | 0.00000 | 6.56000 | 11.06882 | 5.89264 | 1.04650 | 2.28794 | 10.10631 |
| **D7Ertd143e** | 0.43749 | 0.69205 | 2.86750 | 1.81936 | 0.71875 | 0.78484 | 0.10458 | 0.37097 | 0.48234 | 0.15826 | 0.01636 | 3.78476 | 1.50886 | 0.25458 |
| **D7Ertd443e** | 0.06113 | 0.00413 | 0.02398 | 0.00255 | 1.31301 | 0.00001 | 0.00363 | 0.00755 | 3.64553 | 0.01475 | 0.00214 | 0.00334 | 0.00000 | 8.63856 |

| | | | | | | | | | | | | | | |
|---|---|---|---|---|---|---|---|---|---|---|---|---|---|---|
| **D8Ertd738e** | 4.54127 | 6.37009 | 3.06618 | 13.37285 | 22.24532 | 8.80362 | 5.44965 | 13.75718 | 0.21239 | 15.60217 | 2.45709 | 8.61103 | 12.24401 | 0.56805 |
| **Daam1** | 0.46800 | 0.02645 | 1.86386 | 0.23696 | 0.24355 | 1.62921 | 0.00992 | 1.56326 | 0.00106 | 0.02848 | 4.03235 | 2.16650 | 2.82453 | 1.35111 |
| **Dab1** | 0.00694 | 0.00454 | 0.02649 | 0.01036 | 0.01082 | 0.00848 | 0.00491 | 0.05768 | 0.01099 | 0.00644 | 0.00882 | 0.00948 | 0.19515 | 0.00331 |
| **Dab2** | 3.74818 | 1.01237 | 18.73003 | 7.74123 | 1.11846 | 43.35698 | 9.39724 | 17.70411 | 2.77516 | 2.19660 | 12.82169 | 1.05683 | 1.60041 | 2.26109 |
| **Dab2ip** | 0.02926 | 0.03574 | 1.11543 | 1.83411 | 0.22205 | 5.24789 | 0.26717 | 2.63442 | 0.00484 | 1.76395 | 0.78717 | 2.32660 | 0.16631 | 0.00000 |
| **Dact1** | 0.00964 | 0.00389 | 0.49046 | 0.00000 | 0.19538 | 0.00000 | 0.00417 | 0.36639 | 0.01934 | 0.00662 | 0.00518 | 0.00695 | 0.00457 | 0.00487 |
| **Dact3** | 0.01522 | 0.00411 | 0.00547 | 0.06800 | 0.02542 | 0.42744 | 0.00388 | 0.00343 | 0.00953 | 0.00735 | 0.03959 | 0.00755 | 0.80253 | 0.00000 |
| **Dad1** | 6.38805 | 0.62155 | 1.19039 | 8.21225 | 7.67149 | 12.23204 | 3.76892 | 6.71312 | 2.22107 | 11.15915 | 1.12622 | 8.61049 | 2.49669 | 4.22076 |
| **Dag1** | 0.97646 | 8.68117 | 6.21086 | 3.15638 | 2.18939 | 8.74953 | 2.64992 | 14.77299 | 2.23094 | 0.23451 | 5.24463 | 7.12746 | 5.00820 | 1.02517 |
| **Dagla** | 0.01203 | 0.00437 | 0.01424 | 0.01536 | 0.01220 | 0.01334 | 0.00424 | 0.00974 | 0.00983 | 0.00689 | 0.00869 | 0.00927 | 0.20001 | 0.00553 |
| **Daglb** | 0.20209 | 0.01324 | 0.90370 | 0.00000 | 0.11646 | 0.00000 | 0.00385 | 0.01359 | 0.16125 | 0.00533 | 0.00708 | 0.20934 | 0.00703 | 1.18373 |
| **Dalrd3** | 2.84172 | 0.02461 | 10.21141 | 9.09081 | 7.10553 | 1.46941 | 0.07107 | 2.90310 | 0.37629 | 0.03877 | 0.84702 | 2.99723 | 1.28353 | 0.80279 |
| **Dancr** | 0.80563 | 0.02186 | 0.00000 | 0.57149 | 3.34603 | 0.15309 | 0.01748 | 1.48807 | 0.10168 | 0.00588 | 0.78474 | 1.60691 | 2.75519 | 0.18284 |
| **Dand5** | 0.03424 | 0.00408 | 0.00000 | 0.12158 | 0.01458 | 0.28865 | 0.00431 | 1.03914 | 0.01564 | 0.00824 | 0.00000 | 0.26537 | 0.50063 | 0.27550 |
| **Dap** | 2.19703 | 6.13866 | 17.95918 | 22.23158 | 0.88463 | 7.38838 | 9.56635 | 16.68090 | 0.00000 | 0.99009 | 3.57429 | 10.34385 | 0.00000 | 0.00292 |
| **Dap3** | 3.61024 | 4.66422 | 10.12473 | 12.97775 | 7.95721 | 16.32063 | 3.08692 | 11.63464 | 1.48317 | 0.28658 | 10.06252 | 14.51656 | 1.35041 | 5.74341 |
| **Dapk1** | 0.00000 | 0.01133 | 0.86762 | 0.79727 | 1.70513 | 0.65339 | 0.02325 | 3.74258 | 0.09092 | 0.00821 | 0.88166 | 0.00101 | 0.00000 | 1.19686 |
| **Dapk2** | 0.00732 | 0.00444 | 0.00877 | 0.00874 | 0.04708 | 0.39241 | 0.00416 | 0.00678 | 0.01108 | 0.01552 | 0.00601 | 0.00739 | 0.00925 | 0.01006 |
| **Dapk3** | 0.04963 | 0.00534 | 0.17242 | 0.06220 | 0.11126 | 0.23906 | 0.00467 | 0.11142 | 0.00999 | 0.00846 | 0.04866 | 0.01064 | 0.01354 | 0.11830 |
| **Dapp1** | 0.00351 | 0.02748 | 0.00017 | 0.00001 | 0.57900 | 0.71715 | 1.24782 | 2.02800 | 0.01508 | 0.01987 | 0.00319 | 0.01411 | 0.00589 | 0.00131 |
| **Dars** | 10.88255 | 5.81399 | 13.56332 | 10.68002 | 11.83774 | 12.57515 | 2.70410 | 6.05442 | 13.79803 | 0.10366 | 16.87220 | 13.67119 | 2.46384 | 18.77239 |
| **Dars2** | 2.41254 | 0.03355 | 4.09416 | 3.62070 | 3.65136 | 2.24818 | 0.01682 | 2.32052 | 2.83399 | 0.02005 | 6.96577 | 7.52784 | 1.17112 | 4.97865 |
| **Daw1** | 0.00649 | 0.00440 | 0.01354 | 0.01228 | 0.00735 | 0.01227 | 0.00432 | 0.00987 | 0.01928 | 0.00641 | 0.00869 | 0.00971 | 0.12185 | 0.11616 |
| **Daxx** | 1.66817 | 0.14554 | 7.37020 | 2.18647 | 6.17407 | 5.85696 | 0.10720 | 4.61568 | 0.00000 | 11.12303 | 0.05548 | 4.34166 | 2.14571 | 2.67356 |
| **Dazap1** | 0.04149 | 0.57795 | 0.06692 | 6.48205 | 3.21292 | 0.14608 | 1.22833 | 2.01715 | 0.00284 | 0.10724 | 0.00000 | 1.48048 | 0.57698 | 0.00000 |
| **Dazap2** | 11.60163 | 0.95343 | 19.21812 | 10.05848 | 17.23749 | 15.11902 | 5.89687 | 11.68293 | 14.53587 | 12.22700 | 7.55751 | 5.68248 | 5.98650 | 13.35160 |
| **Dazl** | 2.50573 | 0.00386 | 1.49222 | 0.00000 | 1.21406 | 0.00000 | 0.00316 | 0.00343 | 4.77653 | 0.00172 | 0.94356 | 0.00387 | 0.50672 | 1.27095 |
| **Dbf4** | 7.48580 | 0.13447 | 7.21303 | 2.15006 | 12.16938 | 6.89251 | 3.60095 | 1.39165 | 14.77829 | 0.96306 | 4.42972 | 5.70033 | 13.38824 | 24.65671 |
| **Dbhos** | 12.60006 | 0.01032 | 3.43737 | 0.40266 | 4.09339 | 1.92747 | 0.00937 | 0.45617 | 5.69367 | 0.00611 | 0.06662 | 0.02593 | 8.08996 | 1.37634 |
| **Dbi** | 0.42945 | 2.66111 | 0.00000 | 0.15420 | 0.97947 | 2.67539 | 0.58826 | 4.12551 | 2.13963 | 11.71220 | 0.09752 | 6.45439 | 2.78993 | 0.31287 |
| **Dbil5** | 0.01571 | 0.00404 | 0.00878 | 0.00827 | 0.24760 | 0.01665 | 1.00464 | 0.00924 | 0.01096 | 0.05459 | 0.00613 | 0.00776 | 0.00906 | 0.01187 |
| **Dbn1** | 0.11355 | 0.00429 | 0.00732 | 0.15216 | 0.00676 | 0.01252 | 0.00431 | 0.19144 | 0.01170 | 0.00814 | 0.00816 | 0.00598 | 0.00559 | 0.00611 |
| **Dbndd1** | 0.02551 | 0.00413 | 0.22899 | 0.00218 | 0.00000 | 0.00860 | 0.00406 | 0.00826 | 0.01137 | 0.00707 | 0.00606 | 0.00756 | 0.98554 | 0.71049 |
| **Dbndd2** | 0.14502 | 2.20316 | 0.28991 | 0.22448 | 0.50531 | 0.45973 | 1.11341 | 0.99740 | 0.03032 | 0.02305 | 0.00154 | 0.00000 | 2.60652 | 0.00000 |
| **Dbnl** | 5.78885 | 2.84172 | 15.74871 | 12.72500 | 9.83188 | 5.29569 | 0.03837 | 9.35015 | 1.97250 | 0.06574 | 7.27500 | 7.18711 | 3.31152 | 6.52452 |
| **Dbp** | 0.05652 | 0.00422 | 0.10143 | 0.00000 | 0.04656 | 0.01221 | 0.00418 | 0.00888 | 0.01160 | 0.00712 | 0.00605 | 0.08761 | 0.00528 | 0.00885 |
| **Dbr1** | 2.31222 | 0.00982 | 3.17684 | 3.32319 | 4.67080 | 2.85189 | 1.69595 | 0.92507 | 4.29803 | 0.04330 | 0.45620 | 2.26152 | 3.96421 | 2.72440 |
| **Dbt** | 4.81896 | 0.02576 | 2.91701 | 2.24939 | 1.74063 | 3.56562 | 0.26271 | 2.20233 | 12.38342 | 0.01613 | 3.19303 | 1.72012 | 2.13833 | 5.70475 |
| **Dcaf10** | 6.99217 | 0.04803 | 3.58516 | 1.71164 | 5.05621 | 3.08977 | 0.29272 | 0.74058 | 8.94707 | 0.08689 | 1.90859 | 0.44945 | 8.10461 | 9.59166 |
| **Dcaf11** | 3.12272 | 1.94727 | 8.15521 | 1.83170 | 6.93484 | 7.19261 | 2.87335 | 6.90628 | 4.87663 | 0.13332 | 7.02215 | 1.50606 | 6.45735 | 7.60253 |
| **Dcaf12** | 4.98929 | 0.08076 | 7.15086 | 6.53590 | 7.11130 | 11.27009 | 3.29145 | 4.20291 | 5.67235 | 0.16716 | 10.24607 | 5.37747 | 10.98249 | 7.38302 |
| **Dcaf12l1** | 1.26422 | 0.03011 | 1.66858 | 0.53884 | 0.28152 | 3.10652 | 0.00295 | 0.02001 | 1.22214 | 0.02568 | 0.00000 | 4.43664 | 0.34188 | 0.00000 |
| **Dcaf13** | 19.30401 | 0.06510 | 22.27239 | 11.60294 | 22.75621 | 17.36180 | 5.43582 | 4.13615 | 27.20309 | 0.25061 | 12.31641 | 8.28828 | 6.47417 | 22.71654 |
| **Dcaf15** | 1.30498 | 7.08850 | 9.08211 | 7.90549 | 6.12214 | 4.61072 | 0.00988 | 5.46453 | 0.00000 | 0.04934 | 6.27113 | 4.65230 | 0.00004 | 0.45235 |
| **Dcaf17** | 3.75053 | 0.06962 | 2.76084 | 0.66404 | 0.88992 | 2.03308 | 0.19642 | 2.74579 | 5.42654 | 0.40159 | 4.30011 | 1.13963 | 4.09696 | 4.20238 |
| **Dcaf4** | 8.87300 | 7.05007 | 23.67391 | 8.04654 | 25.94701 | 7.27485 | 3.98355 | 0.96804 | 0.30353 | 0.19905 | 10.81698 | 6.60672 | 3.43138 | 11.05100 |
| **Dcaf5** | 0.71145 | 0.02694 | 3.79868 | 2.05564 | 3.61793 | 0.78671 | 0.00206 | 2.67195 | 2.38188 | 0.03466 | 2.11612 | 0.96695 | 7.84056 | 5.73483 |
| **Dcaf6** | 0.21503 | 0.00454 | 0.19304 | 0.00419 | 0.01942 | 0.01100 | 0.00411 | 0.00990 | 0.23199 | 0.00595 | 0.00542 | 0.00793 | 0.36271 | 1.97694 |
| **Dcaf7** | 8.07666 | 0.08948 | 4.67787 | 2.46770 | 5.34228 | 9.00410 | 4.04173 | 1.69183 | 1.17525 | 0.13202 | 5.09111 | 4.44082 | 8.40495 | 4.48659 |
| **Dcaf8** | 3.46693 | 0.07734 | 3.63166 | 3.04805 | 3.86637 | 4.50124 | 0.16122 | 5.35349 | 0.95097 | 0.02994 | 4.35406 | 3.46560 | 4.65798 | 5.52403 |
| **Dcakd** | 0.00000 | 7.62444 | 2.10242 | 2.28804 | 0.87696 | 2.23889 | 0.17825 | 4.16098 | 0.00000 | 1.88164 | 1.41068 | 5.06626 | 1.70280 | 0.00000 |
| **Dcbld1** | 3.62950 | 0.07356 | 10.78171 | 3.15611 | 3.98695 | 5.06475 | 4.03757 | 3.14273 | 2.81207 | 11.86353 | 7.80078 | 8.04445 | 0.40028 | 8.49090 |
| **Dcbld2** | 0.48167 | 0.00230 | 2.11390 | 0.24115 | 0.24025 | 2.33407 | 0.00274 | 0.00000 | 1.76067 | 0.00000 | 3.55104 | 0.22778 | 3.84147 | 1.33283 |
| **Dcc** | 0.00000 | 0.05898 | 0.00000 | 0.02395 | 0.59708 | 0.00493 | 0.00388 | 0.02974 | 0.36137 | 0.02006 | 0.00811 | 0.00849 | 0.78410 | 1.52201 |
| **Dcdc2a** | 0.01166 | 0.01983 | 0.02757 | 0.01945 | 0.01214 | 0.04431 | 0.01012 | 0.00608 | 0.02994 | 0.00589 | 0.00946 | 0.01589 | 0.93963 | 0.09748 |

| | | | | | | | | | | | | | | |
|---|---|---|---|---|---|---|---|---|---|---|---|---|---|---|
| **Dcdc2b** | 0.72711 | 0.00364 | 0.36977 | 0.76381 | 0.36975 | 0.23063 | 0.00403 | 0.19416 | 0.18918 | 0.00652 | 0.01142 | 0.66683 | 0.32009 | 0.26862 |
| **Dcdc2c** | 24.42146 | 0.34960 | 2.33830 | 2.19734 | 30.33875 | 2.31915 | 2.41873 | 0.78761 | 38.84451 | 0.08604 | 1.04998 | 1.09912 | 39.74908 | 47.49741 |
| **Dck** | 6.73947 | 0.00739 | 3.67466 | 0.00000 | 2.21421 | 4.64493 | 1.03672 | 1.46076 | 5.56532 | 0.06091 | 6.30889 | 1.73280 | 8.00001 | 4.97487 |
| **Dclk2** | 0.17621 | 0.00345 | 0.53905 | 0.58189 | 1.02948 | 0.21119 | 0.00339 | 0.00000 | 0.33061 | 0.14592 | 0.00669 | 0.52270 | 1.45236 | 0.61247 |
| **Dclk3** | 0.02219 | 0.00443 | 0.01191 | 0.01279 | 0.00534 | 0.01167 | 0.00430 | 0.01034 | 0.02017 | 0.00667 | 0.00810 | 0.00983 | 0.01687 | 0.20844 |
| **Dclre1a** | 0.83797 | 0.00374 | 0.69456 | 0.00000 | 0.28076 | 0.84527 | 0.00489 | 0.29216 | 1.47173 | 0.00507 | 0.00519 | 0.71402 | 0.95106 | 1.23576 |
| **Dclre1b** | 3.53061 | 0.13350 | 1.98039 | 1.02540 | 2.61536 | 0.75965 | 0.10493 | 1.47447 | 3.06706 | 0.27136 | 1.99050 | 1.70495 | 4.69713 | 5.54776 |
| **Dclre1c** | 2.80183 | 0.09878 | 2.16395 | 3.17489 | 2.31397 | 4.08235 | 2.43113 | 0.51648 | 6.65620 | 1.66345 | 0.04898 | 1.82808 | 14.56100 | 4.07376 |
| **Dcn** | 0.00790 | 0.00445 | 0.00314 | 0.00876 | 1.10863 | 0.01346 | 0.00437 | 0.01102 | 0.01047 | 0.00655 | 0.00896 | 0.00782 | 0.00303 | 0.01637 |
| **Dcp1a** | 0.37539 | 0.04207 | 3.02746 | 4.26606 | 2.48474 | 7.00779 | 0.25540 | 2.46829 | 0.16357 | 0.10634 | 5.88850 | 0.93712 | 4.10456 | 4.34464 |
| **Dcp1b** | 0.00381 | 0.00381 | 0.04204 | 0.00000 | 0.00669 | 0.48611 | 1.93606 | 0.00758 | 0.00875 | 0.03391 | 0.00298 | 0.07440 | 0.25453 | 0.00017 |
| **Dcp2** | 1.45128 | 0.03479 | 1.85376 | 2.14095 | 4.39898 | 4.26534 | 1.88428 | 0.83837 | 0.22091 | 0.07398 | 3.48100 | 4.53306 | 4.99036 | 6.65815 |
| **Dcps** | 0.84330 | 0.04179 | 0.00000 | 1.77341 | 0.87274 | 0.33016 | 0.01425 | 3.86762 | 0.00812 | 0.01752 | 4.39735 | 3.00967 | 1.12218 | 4.19913 |
| **Dcst1** | 0.01050 | 0.03477 | 0.00873 | 0.04784 | 0.02043 | 0.00983 | 0.00368 | 0.16830 | 0.01249 | 0.01039 | 0.00295 | 0.46608 | 0.01114 | 0.00968 |
| **Dctd** | 12.96315 | 5.31812 | 14.12099 | 11.02057 | 15.34657 | 12.51975 | 5.24365 | 5.68200 | 16.17779 | 0.22929 | 6.20300 | 8.94151 | 3.20228 | 20.11945 |
| **Dctn1** | 1.60828 | 0.11365 | 2.33673 | 6.87880 | 4.60285 | 1.77682 | 0.11866 | 6.25689 | 0.01111 | 0.04590 | 3.14742 | 7.32951 | 0.00000 | 0.00000 |
| **Dctn2** | 2.14241 | 8.82179 | 13.02702 | 12.83705 | 5.00634 | 6.12672 | 4.67183 | 10.90801 | 0.21929 | 13.92383 | 8.47392 | 9.96047 | 0.15677 | 2.70345 |
| **Dctn3** | 3.06215 | 0.43665 | 4.27316 | 7.63338 | 4.36174 | 8.44532 | 4.48383 | 8.09071 | 0.04613 | 1.28061 | 2.29381 | 8.95607 | 0.38405 | 0.56777 |
| **Dctn4** | 2.69133 | 0.00222 | 2.23333 | 3.88621 | 4.90831 | 1.03247 | 0.56866 | 3.29351 | 7.71717 | 0.05110 | 1.97608 | 0.84953 | 2.08991 | 3.52091 |
| **Dctn5** | 2.89402 | 0.09408 | 7.27034 | 6.02162 | 6.82948 | 7.00066 | 0.00000 | 4.88192 | 10.97431 | 0.09137 | 6.57911 | 4.64703 | 0.77219 | 10.74908 |
| **Dctn6** | 9.08031 | 0.02777 | 4.43156 | 3.54872 | 6.98863 | 5.55597 | 0.29988 | 2.76995 | 0.44570 | 0.03468 | 1.98392 | 6.83132 | 0.63528 | 2.05743 |
| **Dctpp1** | 2.06297 | 4.14207 | 0.64885 | 10.79580 | 7.28453 | 4.92969 | 1.07727 | 8.40630 | 0.47854 | 6.47832 | 0.35123 | 8.55439 | 1.67522 | 0.47152 |
| **Dcun1d1** | 0.42549 | 0.03194 | 2.16319 | 0.39360 | 1.78952 | 6.34084 | 2.43568 | 0.12305 | 6.01863 | 0.04341 | 1.74498 | 2.84338 | 1.08090 | 4.93090 |
| **Dcun1d2** | 4.85111 | 0.15815 | 1.16705 | 0.02968 | 2.97487 | 3.46388 | 0.03403 | 1.40780 | 1.22519 | 0.01978 | 3.90832 | 1.23163 | 2.63239 | 2.45459 |
| **Dcun1d3** | 3.10681 | 0.02769 | 3.20788 | 0.83424 | 2.91819 | 0.68451 | 2.34776 | 0.08677 | 8.46856 | 0.06718 | 0.00000 | 1.63363 | 7.39033 | 14.50108 |
| **Dcun1d4** | 0.56074 | 0.00375 | 0.14520 | 0.88250 | 0.24534 | 1.21142 | 0.30883 | 0.02344 | 0.40340 | 0.00425 | 0.04309 | 0.39018 | 0.06664 | 0.45312 |
| **Dcun1d5** | 14.96679 | 0.02904 | 3.51835 | 6.73392 | 10.66968 | 6.90255 | 3.21405 | 5.94673 | 6.16340 | 0.19016 | 2.84482 | 5.14157 | 9.50462 | 0.00000 |
| **Dcx** | 0.04741 | 0.00430 | 0.02093 | 0.02463 | 0.07777 | 0.03582 | 0.02222 | 0.03589 | 0.04895 | 0.01314 | 0.01943 | 0.01316 | 0.03873 | 0.05826 |
| **Dcxr** | 0.19976 | 0.01994 | 0.00000 | 2.62893 | 1.71738 | 0.54857 | 0.06051 | 0.83774 | 0.00614 | 0.01874 | 0.17457 | 3.05526 | 0.01142 | 0.00000 |
| **Dda1** | 3.16241 | 0.07515 | 4.10972 | 8.30228 | 8.73750 | 0.00000 | 2.51970 | 5.63597 | 0.97593 | 0.12372 | 2.48030 | 6.77860 | 4.82182 | 1.71763 |
| **Ddah1** | 5.95531 | 0.81560 | 11.25460 | 3.09772 | 9.14351 | 15.41175 | 3.96068 | 20.14624 | 10.47063 | 0.29903 | 8.18029 | 5.19061 | 0.00000 | 2.26857 |
| **Ddah2** | 0.09626 | 0.07629 | 4.35384 | 4.39880 | 4.54764 | 0.00000 | 0.01605 | 1.53169 | 0.00962 | 0.00432 | 0.00000 | 3.96621 | 0.00151 | 0.00000 |
| **Ddb1** | 8.24062 | 11.39032 | 16.29454 | 19.53526 | 20.51533 | 23.99786 | 2.26312 | 15.52319 | 26.20904 | 0.88489 | 11.20728 | 15.29770 | 7.33702 | 23.38203 |
| **Ddb2** | 0.26412 | 0.01254 | 3.21623 | 2.62501 | 1.40991 | 1.19621 | 0.00390 | 0.54053 | 1.34004 | 0.00000 | 1.35726 | 0.21313 | 0.00000 | 2.99110 |
| **Ddc** | 0.00587 | 0.00273 | 0.09810 | 0.00000 | 0.01344 | 0.18311 | 0.00382 | 0.00000 | 0.17375 | 0.00000 | 3.74675 | 0.91989 | 0.27281 | 0.00000 |
| **Ddhd1** | 0.00000 | 0.02410 | 1.70102 | 2.13081 | 1.07932 | 1.80976 | 0.00249 | 0.22266 | 0.51024 | 0.02516 | 0.00000 | 2.74823 | 0.86988 | 1.48273 |
| **Ddhd2** | 2.07106 | 0.24731 | 3.88786 | 6.61309 | 5.09849 | 3.02144 | 2.48666 | 1.31004 | 4.43081 | 0.17784 | 3.27510 | 3.90792 | 2.56800 | 8.43243 |
| **Ddi2** | 0.01405 | 0.01122 | 1.02200 | 0.09377 | 0.17063 | 0.11225 | 0.14130 | 0.03577 | 0.57199 | 0.00292 | 0.28363 | 0.23487 | 0.04781 | 0.16351 |
| **Ddias** | 15.10030 | 0.00000 | 7.19177 | 2.74275 | 12.97009 | 5.81411 | 0.00000 | 1.90138 | 24.91752 | 0.01165 | 0.20227 | 1.25417 | 13.27138 | 10.75348 |
| **Ddit3** | 0.00000 | 3.68102 | 0.59834 | 2.20431 | 0.89470 | 0.34762 | 0.04474 | 4.20236 | 0.00147 | 13.21261 | 0.57491 | 0.00001 | 0.49949 | 0.51689 |
| **Ddit4** | 0.17207 | 0.16772 | 0.07029 | 2.36822 | 3.08919 | 0.00000 | 0.81171 | 0.00111 | 0.47553 | 0.04485 | 0.26077 | 0.51596 | 3.83979 | 0.77540 |
| **Ddit4l** | 0.67430 | 0.00236 | 0.00000 | 1.98640 | 0.13508 | 2.84628 | 1.05859 | 0.00427 | 0.11133 | 0.01643 | 0.00000 | 0.85809 | 9.19019 | 0.19414 |
| **Ddo** | 0.03700 | 0.04667 | 0.00550 | 0.08407 | 0.03829 | 0.06071 | 0.13456 | 0.75877 | 0.03603 | 0.00840 | 0.00462 | 0.37779 | 0.08036 | 0.05913 |
| **Ddost** | 3.66051 | 8.75260 | 7.77722 | 9.12452 | 5.63218 | 10.49749 | 1.80222 | 5.45968 | 3.09541 | 2.93774 | 5.02019 | 9.45552 | 1.18798 | 5.54157 |
| **Ddr1** | 0.04433 | 0.01026 | 4.52228 | 1.98355 | 2.81874 | 0.27606 | 0.00297 | 0.01536 | 0.02022 | 0.00435 | 0.00140 | 0.45742 | 2.27539 | 2.77599 |
| **Ddr2** | 0.34388 | 0.00331 | 4.12030 | 1.18983 | 2.80291 | 0.18094 | 0.00117 | 0.00000 | 13.01858 | 0.01144 | 0.00000 | 1.06581 | 3.85780 | 1.04137 |
| **Ddrgk1** | 3.62630 | 0.10400 | 6.25808 | 15.10286 | 2.15810 | 8.13391 | 1.83278 | 6.17934 | 0.00000 | 0.09514 | 7.65348 | 5.08594 | 0.08125 | 0.22047 |
| **Ddt** | 0.07798 | 0.14747 | 0.00000 | 2.08556 | 1.49839 | 1.16435 | 0.00634 | 6.09505 | 0.00000 | 6.52275 | 1.75116 | 6.68841 | 1.94397 | 0.85942 |
| **Ddx1** | 13.70857 | 0.09201 | 8.94740 | 5.46250 | 7.99433 | 11.01343 | 3.67583 | 5.16286 | 14.34634 | 0.19190 | 11.96486 | 10.43930 | 1.94621 | 6.50711 |
| **Ddx10** | 8.71942 | 0.02338 | 5.86477 | 0.00000 | 7.42992 | 6.11349 | 1.39921 | 0.84139 | 16.97503 | 0.00000 | 9.21421 | 3.88068 | 6.44752 | 11.79492 |
| **Ddx11** | 0.02591 | 0.01047 | 1.12012 | 1.37433 | 0.00000 | 1.02123 | 0.03269 | 0.08999 | 0.08259 | 0.01395 | 0.75982 | 1.10369 | 0.83310 | 0.20772 |
| **Ddx17** | 19.41943 | 5.54906 | 22.29807 | 7.76374 | 21.26880 | 22.09262 | 4.88029 | 11.06388 | 22.52018 | 0.34669 | 13.01504 | 16.34700 | 6.64840 | 21.38221 |
| **Ddx18** | 13.40936 | 0.23118 | 27.22314 | 10.44180 | 22.57743 | 18.05403 | 6.24487 | 3.15406 | 22.50553 | 0.25391 | 14.72508 | 15.20792 | 9.60410 | 29.27890 |
| **Ddx19a** | 1.52326 | 0.01191 | 2.02191 | 0.56932 | 2.18464 | 0.59852 | 0.36509 | 0.32585 | 0.39943 | 0.34039 | 0.01812 | 0.99974 | 3.07554 | 1.18564 |
| **Ddx19b** | 4.81237 | 0.02937 | 8.54688 | 2.62136 | 6.91632 | 3.63866 | 1.84537 | 0.23560 | 7.12000 | 11.36135 | 1.09857 | 2.89178 | 0.90708 | 8.56068 |

| | | | | | | | | | | | | | | |
|---|---|---|---|---|---|---|---|---|---|---|---|---|---|---|
| **Ddx20** | 7.03169 | 0.14102 | 9.73093 | 4.36506 | 10.19933 | 3.89842 | 1.26960 | 0.14155 | 11.42879 | 0.04446 | 8.56243 | 4.28433 | 11.39127 | 11.51801 |
| **Ddx21** | 20.36191 | 0.15801 | 28.72445 | 10.87347 | 27.06794 | 17.33240 | 6.26634 | 5.17038 | 21.48496 | 0.30449 | 15.10343 | 10.85611 | 12.55443 | 26.26837 |
| **Ddx23** | 0.00033 | 0.02625 | 0.00000 | 0.46298 | 0.16595 | 0.00000 | 0.00388 | 1.07546 | 0.30782 | 0.00542 | 0.00000 | 1.38099 | 0.00319 | 0.07236 |
| **Ddx24** | 6.78649 | 4.17449 | 10.90201 | 5.81190 | 14.25083 | 8.93994 | 2.83130 | 3.36045 | 8.52456 | 0.18518 | 7.58581 | 8.62471 | 10.52499 | 15.79449 |
| **Ddx25** | 0.00970 | 0.00429 | 0.03272 | 0.20138 | 0.00615 | 0.00911 | 0.00432 | 0.00742 | 0.02027 | 0.00559 | 0.00664 | 0.00784 | 0.01449 | 0.05584 |
| **Ddx26b** | 0.06952 | 0.00380 | 0.06353 | 0.00000 | 0.51589 | 0.13832 | 0.00375 | 0.00234 | 1.77035 | 0.00503 | 0.00071 | 0.23276 | 0.21498 | 1.80866 |
| **Ddx27** | 13.99737 | 0.32699 | 18.89827 | 11.69042 | 10.02426 | 11.70095 | 5.32119 | 10.09403 | 3.30773 | 0.26858 | 8.00215 | 10.12893 | 10.23722 | 9.37471 |
| **Ddx28** | 2.16946 | 0.03173 | 5.96860 | 6.50144 | 6.36599 | 2.22073 | 2.39358 | 0.00000 | 1.21891 | 0.03209 | 3.66838 | 2.01745 | 5.22140 | 5.18475 |
| **Ddx31** | 4.13268 | 0.00169 | 9.80012 | 2.05314 | 8.43256 | 2.83975 | 0.91900 | 2.37957 | 3.14973 | 0.00000 | 0.49681 | 1.29394 | 4.45151 | 8.82385 |
| **Ddx39** | 17.49406 | 9.70620 | 33.31361 | 30.56152 | 21.58413 | 25.43818 | 9.05499 | 18.88635 | 9.86589 | 14.40776 | 21.30215 | 16.01215 | 16.87062 | 19.81803 |
| **Ddx39b** | 9.29451 | 9.71640 | 23.93463 | 16.73573 | 14.28608 | 17.73188 | 5.40333 | 15.73451 | 6.72137 | 0.41165 | 12.42862 | 15.36586 | 4.85748 | 16.25062 |
| **Ddx3x** | 24.88999 | 1.20423 | 33.63767 | 17.83358 | 27.75832 | 26.81402 | 6.51737 | 4.73833 | 43.64861 | 1.14194 | 15.73604 | 9.01363 | 16.03920 | 17.99938 |
| **Ddx3y** | 0.00917 | 0.00235 | 0.00000 | 2.91240 | 0.26207 | 0.00000 | 0.68746 | 0.00000 | 0.00349 | 0.00275 | 4.85065 | 0.49367 | 0.35148 | 0.53320 |
| **Ddx4** | 0.00000 | 0.00408 | 0.00504 | 0.00000 | 0.41973 | 0.28476 | 0.00369 | 0.00316 | 2.20256 | 0.00568 | 0.00187 | 0.15173 | 0.00594 | 1.71010 |
| **Ddx41** | 9.33384 | 0.15515 | 13.63631 | 11.01646 | 11.26170 | 5.43528 | 3.40537 | 7.41283 | 0.20601 | 0.17640 | 3.44477 | 10.73747 | 3.48246 | 7.44880 |
| **Ddx42** | 0.95835 | 8.43764 | 6.93271 | 7.22773 | 4.60059 | 9.63049 | 2.67097 | 1.61538 | 5.08233 | 0.07861 | 5.90705 | 6.48907 | 3.37104 | 9.11890 |
| **Ddx43** | 0.00990 | 0.00357 | 0.32698 | 0.65869 | 0.44550 | 0.00000 | 0.00372 | 1.19187 | 0.82915 | 0.00999 | 0.08141 | 0.18251 | 1.05445 | 0.15642 |
| **Ddx46** | 12.90553 | 3.74360 | 14.10944 | 5.68155 | 14.97386 | 6.25801 | 3.70873 | 2.94180 | 13.11871 | 0.96220 | 6.19282 | 8.87894 | 10.05975 | 17.56540 |
| **Ddx47** | 7.70162 | 5.57050 | 15.90828 | 4.20328 | 9.86279 | 21.51276 | 4.45046 | 2.26572 | 17.46243 | 12.64448 | 15.05244 | 8.87485 | 11.15290 | 20.45390 |
| **Ddx49** | 1.39320 | 3.45370 | 14.35594 | 10.89645 | 12.45967 | 0.00000 | 1.31598 | 2.85373 | 0.01274 | 0.00000 | 6.76687 | 1.81958 | 0.06907 | 7.03234 |
| **Ddx5** | 35.40206 | 8.15031 | 36.64285 | 30.18381 | 39.74850 | 26.81568 | 10.94096 | 16.87892 | 37.82330 | 6.25322 | 23.23269 | 20.75780 | 19.17524 | 29.26569 |
| **Ddx50** | 4.89695 | 0.08060 | 4.91246 | 0.00000 | 1.91016 | 4.38217 | 1.74779 | 4.44339 | 2.18650 | 0.97414 | 3.51013 | 3.85370 | 0.75916 | 4.59727 |
| **Ddx51** | 4.29939 | 0.01304 | 7.06875 | 10.27606 | 4.73601 | 4.77791 | 0.00651 | 0.00000 | 3.04335 | 0.02409 | 0.23797 | 5.54852 | 7.70115 | 4.15696 |
| **Ddx52** | 7.34513 | 2.88642 | 11.06551 | 8.72330 | 14.47235 | 7.04490 | 4.78069 | 2.95092 | 16.50332 | 0.17790 | 4.17916 | 6.69267 | 11.10785 | 20.09326 |
| **Ddx54** | 1.45770 | 0.10468 | 15.35039 | 15.70904 | 20.74861 | 0.00002 | 0.37792 | 7.73647 | 0.08492 | 0.04047 | 0.25563 | 5.91799 | 7.59840 | 0.80833 |
| **Ddx55** | 4.68017 | 1.38350 | 9.18972 | 4.20742 | 6.80381 | 3.31658 | 2.02639 | 1.22700 | 2.94098 | 0.04464 | 2.73279 | 0.65475 | 3.74890 | 4.06197 |
| **Ddx56** | 12.49398 | 9.54342 | 22.12703 | 19.31549 | 14.97023 | 14.12105 | 5.87141 | 9.03871 | 5.36204 | 11.30965 | 12.88465 | 12.81507 | 6.75195 | 15.10907 |
| **Ddx58** | 0.05654 | 0.00447 | 0.08192 | 0.08471 | 0.43598 | 0.04251 | 0.00354 | 0.00946 | 2.53630 | 0.00693 | 0.00271 | 0.32040 | 0.33235 | 0.00000 |
| **Ddx59** | 0.46780 | 0.00263 | 1.25125 | 0.00000 | 1.64827 | 0.68136 | 0.00298 | 2.24800 | 0.00000 | 0.00000 | 2.82405 | 0.73855 | 1.60071 | 1.38667 |
| **Ddx6** | 0.65939 | 0.02505 | 2.81898 | 0.89124 | 2.57028 | 6.13037 | 0.02621 | 1.20016 | 6.17193 | 3.09250 | 0.00000 | 3.25648 | 3.11334 | 3.25339 |
| **Ddx60** | 0.75051 | 0.00395 | 0.39394 | 0.13851 | 0.40312 | 0.00000 | 0.00345 | 0.00479 | 3.93069 | 0.00553 | 0.00333 | 0.01098 | 0.04400 | 2.54101 |
| **Deaf1** | 3.10329 | 0.02749 | 8.32691 | 1.62109 | 3.91172 | 4.04882 | 0.00808 | 2.42313 | 0.32548 | 0.00825 | 0.00000 | 3.95115 | 1.34859 | 4.09759 |
| **Deb1** | 1.35021 | 0.03446 | 0.00000 | 1.69785 | 3.19902 | 3.16317 | 0.76442 | 2.36991 | 1.37089 | 0.11841 | 0.12111 | 2.43055 | 7.33441 | 0.80668 |
| **Decr1** | 0.00000 | 0.04398 | 1.21095 | 2.18394 | 0.14635 | 3.14749 | 2.82962 | 1.01512 | 0.00444 | 0.08537 | 0.00000 | 2.55348 | 0.00416 | 2.14092 |
| **Decr2** | 0.38272 | 0.07941 | 1.70372 | 2.22213 | 0.20679 | 1.41640 | 0.02089 | 0.62548 | 5.75380 | 0.01925 | 1.31722 | 0.78406 | 2.33733 | 3.32921 |
| **Dedd** | 0.45967 | 0.01057 | 2.34797 | 1.02726 | 4.89552 | 4.32379 | 0.00917 | 1.37661 | 5.55987 | 0.01793 | 2.83941 | 0.35732 | 5.66159 | 4.10151 |
| **Dedd2** | 2.82859 | 0.01908 | 2.64831 | 1.51460 | 9.55546 | 0.61750 | 3.05335 | 3.24515 | 0.11507 | 0.04833 | 0.70341 | 0.03645 | 3.04215 | 1.13310 |
| **Def6** | 0.21025 | 0.00311 | 1.26881 | 0.00003 | 0.10585 | 0.11445 | 0.00327 | 0.02229 | 0.00498 | 0.00465 | 0.32538 | 0.97170 | 0.16086 | 0.03823 |
| **Def8** | 5.25195 | 0.04061 | 14.00072 | 15.78145 | 14.83283 | 8.02616 | 1.55438 | 0.00000 | 17.28313 | 0.11431 | 2.57724 | 0.50777 | 3.42401 | 14.93771 |
| **Defa29** | 0.11950 | 0.00392 | 0.00622 | 0.09641 | 0.00874 | 0.01514 | 0.00409 | 0.00662 | 0.17817 | 0.00762 | 0.00547 | 0.10169 | 0.97259 | 0.00000 |
| **Defb23** | 0.01087 | 0.00436 | 0.01480 | 0.01228 | 0.01107 | 0.01343 | 0.00424 | 0.00943 | 0.00945 | 0.00657 | 0.00898 | 0.00948 | 0.26220 | 0.00607 |
| **Defb30** | 0.02785 | 0.00440 | 0.01187 | 0.01688 | 0.04439 | 0.01181 | 0.00419 | 0.00975 | 0.01093 | 0.00826 | 0.00827 | 0.00878 | 0.12934 | 0.07959 |
| **Defb42** | 0.60875 | 0.00430 | 0.04476 | 0.18484 | 1.05175 | 0.01551 | 0.00443 | 0.00639 | 0.00740 | 0.00639 | 0.00540 | 0.00535 | 0.56884 | 1.30212 |
| **Degs1** | 5.96743 | 0.10568 | 8.41761 | 5.32381 | 2.70000 | 3.49314 | 2.75205 | 7.34492 | 2.66290 | 0.08000 | 1.08956 | 7.58308 | 7.35337 | 2.88528 |
| **Degs2** | 0.29420 | 0.00373 | 0.03300 | 0.19425 | 0.00000 | 0.46321 | 0.00401 | 0.00386 | 0.02318 | 0.00452 | 0.13727 | 0.32697 | 0.01247 | 0.15982 |
| **Dek** | 4.02031 | 0.07601 | 3.61232 | 0.85513 | 8.85432 | 4.45573 | 1.62232 | 0.81711 | 12.90604 | 0.03321 | 3.89930 | 4.75426 | 0.44563 | 11.82964 |
| **Dennd1a** | 0.42002 | 0.02406 | 1.45145 | 0.00000 | 0.14507 | 0.00000 | 0.95796 | 2.40383 | 0.02056 | 6.21253 | 0.56996 | 2.08418 | 0.19342 | 0.04145 |
| **Dennd1b** | 1.04124 | 0.01908 | 2.15173 | 1.00878 | 0.69617 | 1.50922 | 0.01029 | 0.98702 | 0.01732 | 2.61010 | 0.02345 | 1.11292 | 0.15574 | 1.27824 |
| **Dennd1c** | 1.00127 | 0.00439 | 0.75282 | 0.97764 | 0.68330 | 0.05140 | 0.00415 | 0.01048 | 0.02284 | 0.00603 | 0.00540 | 0.00586 | 0.04982 | 0.37982 |
| **Dennd2a** | 1.12556 | 0.02852 | 0.69499 | 1.27499 | 1.83800 | 0.00000 | 0.00413 | 1.00072 | 0.28165 | 1.02657 | 0.00000 | 0.10274 | 5.73870 | 1.85986 |
| **Dennd2c** | 0.01613 | 0.00908 | 0.04133 | 0.00602 | 0.00285 | 0.00000 | 0.00336 | 0.00000 | 0.01013 | 0.00418 | 3.17701 | 3.01642 | 0.34389 | 0.16751 |
| **Dennd3** | 0.00000 | 6.10833 | 0.53229 | 0.22599 | 1.95638 | 0.00000 | 0.82704 | 0.03683 | 0.03789 | 0.03707 | 0.01533 | 0.00000 | 2.10979 | 0.65001 |
| **Dennd4a** | 2.50092 | 0.00485 | 0.98214 | 0.11509 | 0.33360 | 0.00000 | 0.00370 | 0.37493 | 3.67423 | 0.00664 | 0.00576 | 0.00182 | 1.36375 | 1.71931 |
| **Dennd4b** | 0.03252 | 0.00370 | 0.43252 | 0.00000 | 0.00000 | 0.05775 | 0.00379 | 0.01048 | 0.00817 | 0.00646 | 0.00526 | 0.36863 | 0.86162 | 0.78057 |
| **Dennd4c** | 10.16180 | 0.10164 | 6.19568 | 0.45983 | 1.11586 | 0.87819 | 0.39819 | 0.00007 | 9.49370 | 0.01466 | 1.54091 | 0.38349 | 2.23632 | 0.64024 |

| | | | | | | | | | | | | | | |
|---|---|---|---|---|---|---|---|---|---|---|---|---|---|---|
| **Dennd5a** | 2.99616 | 2.18906 | 5.41822 | 1.59956 | 10.02847 | 1.78675 | 3.98035 | 4.02168 | 0.65000 | 0.07359 | 2.68754 | 7.21715 | 3.55934 | 2.93192 |
| **Dennd5b** | 4.26812 | 0.09033 | 2.39281 | 2.35931 | 5.00220 | 3.52732 | 0.74553 | 0.37578 | 13.30047 | 1.03939 | 0.22154 | 0.81763 | 9.35702 | 2.11796 |
| **Dennd6a** | 0.36261 | 0.16487 | 0.54269 | 1.04226 | 0.12624 | 3.28685 | 0.11033 | 0.95814 | 1.29271 | 0.26083 | 0.53908 | 3.03112 | 3.16693 | 2.22790 |
| **Dennd6b** | 0.86306 | 0.00328 | 1.33060 | 2.42108 | 0.41590 | 0.88476 | 0.01520 | 0.01155 | 0.00327 | 0.00539 | 1.66761 | 0.44466 | 0.73926 | 0.34823 |
| **Denr** | 9.22516 | 0.24898 | 15.21591 | 14.27954 | 16.36270 | 9.60262 | 2.16358 | 11.11643 | 6.07387 | 0.19737 | 6.19109 | 12.37978 | 7.46102 | 4.93594 |
| **Depdc1a** | 0.05382 | 0.00413 | 0.23013 | 0.65761 | 0.00361 | 0.00000 | 0.00508 | 0.05127 | 0.01348 | 0.00378 | 0.61054 | 0.32543 | 0.00508 | 0.13434 |
| **Depdc1b** | 0.00000 | 0.00350 | 0.67693 | 0.22916 | 0.00000 | 0.00000 | 0.00384 | 0.00000 | 0.00677 | 0.00413 | 0.20474 | 0.43950 | 0.26657 | 0.92535 |
| **Depdc5** | 1.07657 | 0.08751 | 1.29007 | 0.84723 | 0.85318 | 0.69939 | 0.02735 | 1.06848 | 0.91483 | 0.80492 | 0.44031 | 0.90687 | 3.67681 | 2.43631 |
| **Depdc7** | 13.62225 | 0.03516 | 8.46368 | 5.02216 | 18.64922 | 9.76574 | 7.29659 | 0.70305 | 13.48774 | 0.27840 | 4.72910 | 2.67613 | 13.34577 | 15.41839 |
| **Deptor** | 2.83133 | 0.10677 | 6.68297 | 0.12653 | 6.97657 | 1.91543 | 0.01668 | 0.37492 | 14.79928 | 0.01458 | 3.94458 | 7.26746 | 2.24478 | 6.29543 |
| **Dera** | 4.53626 | 0.06857 | 2.81814 | 3.02360 | 0.97918 | 2.85043 | 0.01858 | 7.91204 | 0.00000 | 0.01503 | 7.17310 | 3.89886 | 0.06192 | 1.10616 |
| **Derl1** | 5.07263 | 0.11807 | 4.20188 | 7.42262 | 2.75099 | 10.65145 | 0.87392 | 8.42595 | 1.59924 | 0.09306 | 6.08241 | 6.56895 | 3.28884 | 5.73204 |
| **Derl2** | 4.34769 | 7.09630 | 5.34475 | 7.94081 | 1.74645 | 10.32940 | 5.12424 | 3.99728 | 3.15843 | 0.16801 | 8.46435 | 6.81257 | 0.00000 | 3.40244 |
| **Derl3** | 0.00000 | 0.04333 | 0.55215 | 0.00000 | 0.27522 | 0.03300 | 0.00316 | 0.00293 | 0.02215 | 0.00996 | 0.01064 | 2.07445 | 1.82836 | 0.02147 |
| **Des** | 0.01015 | 0.00429 | 0.01202 | 0.00486 | 0.12395 | 0.01751 | 0.00445 | 0.00821 | 0.00996 | 0.00628 | 0.00658 | 0.13743 | 0.00608 | 0.01599 |
| **Desi1** | 7.35391 | 11.22291 | 3.17451 | 4.52653 | 1.69453 | 2.83851 | 0.15617 | 0.32937 | 1.06192 | 0.02062 | 1.24793 | 4.46152 | 1.14666 | 2.67499 |
| **Desi2** | 0.44884 | 0.00965 | 0.84955 | 1.26878 | 0.11766 | 1.92990 | 0.00280 | 1.42774 | 3.03801 | 0.00421 | 0.31024 | 2.09790 | 3.99844 | 2.94331 |
| **Det1** | 4.05829 | 0.00979 | 3.02789 | 0.95919 | 3.69905 | 1.16312 | 0.10412 | 2.25475 | 0.21467 | 0.01052 | 0.00000 | 1.21080 | 1.02704 | 3.41165 |
| **Dexi** | 0.19042 | 0.00449 | 0.02947 | 0.00000 | 0.00689 | 0.06547 | 0.42878 | 0.01686 | 0.01872 | 0.01712 | 0.01183 | 0.36382 | 0.47531 | 0.00223 |
| **Dffa** | 6.60365 | 0.04406 | 3.53951 | 3.41109 | 5.05255 | 5.13490 | 1.15132 | 0.33038 | 1.83265 | 0.03948 | 2.33941 | 1.46668 | 3.03704 | 2.95399 |
| **Dffb** | 4.05360 | 0.01746 | 5.18937 | 2.95954 | 7.96051 | 1.67553 | 2.11215 | 1.11539 | 0.92030 | 0.05145 | 3.20296 | 2.26299 | 0.38010 | 3.97225 |
| **Dfna5** | 0.02959 | 0.00479 | 0.01040 | 0.00545 | 0.01276 | 0.03961 | 0.00506 | 0.43485 | 0.00947 | 0.00724 | 0.00594 | 0.01140 | 0.18100 | 0.35258 |
| **Dgat1** | 0.31627 | 0.09043 | 2.56775 | 7.02774 | 0.10438 | 3.29492 | 1.60526 | 5.34795 | 0.00636 | 0.06154 | 2.47321 | 1.53498 | 0.26396 | 0.44131 |
| **Dgat2** | 3.54281 | 0.06302 | 14.88672 | 3.38898 | 11.16316 | 0.18168 | 0.00716 | 0.15798 | 0.64358 | 0.00000 | 0.40457 | 2.88043 | 0.37152 | 14.11112 |
| **Dgcr14** | 3.39015 | 0.06536 | 7.13059 | 5.35079 | 7.89378 | 3.25770 | 2.83032 | 5.09067 | 1.06788 | 0.08396 | 6.28581 | 1.43210 | 4.58265 | 3.89684 |
| **Dgcr2** | 1.99609 | 0.26003 | 4.71493 | 2.92052 | 3.69264 | 6.87512 | 2.45591 | 4.52104 | 4.52906 | 0.07887 | 6.23184 | 4.97921 | 1.72885 | 6.34666 |
| **Dgcr6** | 1.01144 | 0.01677 | 1.18331 | 4.39670 | 0.83202 | 1.73190 | 0.02018 | 2.87209 | 0.00681 | 0.04434 | 0.02103 | 2.28863 | 0.00000 | 0.13862 |
| **Dgcr8** | 3.95946 | 0.02496 | 6.35597 | 1.32628 | 2.02800 | 3.27249 | 1.52376 | 0.30865 | 3.43425 | 0.01404 | 0.00000 | 4.63959 | 3.75821 | 5.50947 |
| **Dgka** | 3.28510 | 0.68438 | 8.89630 | 1.73681 | 3.02923 | 3.54135 | 3.23950 | 2.02922 | 4.06739 | 15.59930 | 4.80887 | 3.35957 | 1.39455 | 1.12870 |
| **Dgkd** | 2.37353 | 0.02275 | 2.52757 | 1.27328 | 2.98360 | 2.89457 | 0.37204 | 2.69788 | 0.92674 | 12.17408 | 1.80514 | 1.66615 | 5.29021 | 1.50210 |
| **Dgke** | 0.26497 | 0.02629 | 2.04784 | 1.25802 | 3.70274 | 3.18484 | 1.00908 | 0.27073 | 6.09583 | 0.02214 | 0.00000 | 0.54115 | 10.27852 | 9.28927 |
| **Dgkeos** | 0.02603 | 0.00443 | 0.01929 | 0.01081 | 0.30167 | 0.00373 | 0.00416 | 0.01021 | 0.51493 | 0.00660 | 0.00560 | 0.00715 | 0.13101 | 1.34625 |
| **Dgkh** | 0.01287 | 0.00382 | 0.01136 | 0.01903 | 0.01925 | 0.01324 | 0.00411 | 0.00551 | 0.01128 | 0.00671 | 0.00441 | 0.51975 | 0.00844 | 0.02382 |
| **Dgkk** | 0.01416 | 0.00458 | 0.00617 | 0.00564 | 0.00347 | 0.01440 | 0.00381 | 0.02915 | 0.15086 | 0.00856 | 0.00334 | 0.49310 | 0.73549 | 0.00000 |
| **Dgkq** | 0.10363 | 0.00375 | 0.24317 | 0.98425 | 0.00000 | 0.27014 | 0.00356 | 0.58205 | 0.13338 | 0.00446 | 0.00000 | 0.38530 | 0.00470 | 0.88362 |
| **Dgkz** | 0.57259 | 0.03034 | 8.95954 | 4.65092 | 1.80981 | 2.46882 | 2.42830 | 2.65605 | 0.38204 | 0.05234 | 3.54833 | 3.97950 | 0.00000 | 4.73661 |
| **Dguok** | 0.99194 | 0.03650 | 1.82062 | 2.90719 | 1.45918 | 1.70213 | 0.00312 | 1.60129 | 0.01555 | 0.03473 | 0.40819 | 2.21955 | 0.40391 | 0.00000 |
| **Dhcr24** | 0.00000 | 6.96048 | 5.12916 | 3.49949 | 1.59903 | 7.22725 | 0.00782 | 0.03566 | 1.39586 | 0.05728 | 1.27461 | 1.38243 | 0.05339 | 5.61108 |
| **Dhcr7** | 0.53981 | 0.00444 | 0.69091 | 0.99833 | 1.03536 | 1.79433 | 0.30545 | 0.72580 | 0.24563 | 0.03252 | 0.01755 | 0.02090 | 0.76393 | 1.14801 |
| **Dhdds** | 2.47330 | 0.04866 | 6.57040 | 0.44237 | 7.26784 | 5.09957 | 0.00744 | 1.89850 | 0.26734 | 0.02334 | 3.43844 | 3.28901 | 3.92536 | 3.50969 |
| **Dhdh** | 0.00793 | 0.00415 | 0.28160 | 0.04319 | 0.16361 | 0.00313 | 0.00432 | 0.00858 | 0.00813 | 0.00553 | 0.00659 | 0.00873 | 0.96356 | 0.09176 |
| **Dhfr** | 5.01228 | 0.08208 | 2.54376 | 6.10019 | 3.45185 | 7.03213 | 0.03444 | 6.80910 | 8.84915 | 0.03174 | 4.20941 | 7.36059 | 2.26199 | 3.92792 |
| **Dhh** | 0.29647 | 0.00390 | 0.00263 | 0.33708 | 0.23047 | 0.01523 | 0.00432 | 0.27841 | 0.01298 | 0.00906 | 0.00466 | 0.00451 | 0.00842 | 0.00439 |
| **Dhodh** | 2.79148 | 0.05286 | 6.29272 | 6.87533 | 6.10372 | 3.84902 | 6.19845 | 5.41744 | 0.00000 | 0.18809 | 7.43310 | 6.19190 | 1.00617 | 0.95130 |
| **Dhps** | 3.45474 | 0.01641 | 0.96167 | 11.84731 | 5.51080 | 5.21255 | 4.06108 | 0.26602 | 0.82876 | 5.76249 | 2.39366 | 2.75202 | 0.71084 | 3.06058 |
| **Dhrs1** | 2.39419 | 0.03301 | 2.84674 | 2.78788 | 0.77911 | 2.04869 | 2.75356 | 1.49029 | 0.31695 | 0.05935 | 2.50534 | 1.50357 | 1.05866 | 0.03644 |
| **Dhrs11** | 0.13346 | 2.73741 | 0.28860 | 2.44690 | 0.00000 | 1.20962 | 0.12425 | 1.63198 | 0.10503 | 0.04851 | 0.00000 | 0.32727 | 0.60954 | 0.14558 |
| **Dhrs13** | 1.90640 | 0.05078 | 2.84757 | 3.85018 | 1.52104 | 2.76273 | 0.00337 | 1.70672 | 0.00368 | 0.02191 | 1.88230 | 0.45442 | 0.15818 | 1.92940 |
| **Dhrs2** | 0.02585 | 0.00442 | 0.01119 | 0.01274 | 0.00418 | 0.01138 | 0.00429 | 0.01033 | 0.01753 | 0.00665 | 0.00794 | 0.00978 | 0.01500 | 0.29478 |
| **Dhrs3** | 0.00419 | 0.01537 | 0.00000 | 0.00000 | 0.48424 | 0.00000 | 0.00436 | 0.39415 | 0.01153 | 0.00373 | 1.14049 | 0.56778 | 0.13189 | 0.00177 |
| **Dhrs4** | 0.00362 | 0.05791 | 0.07877 | 0.00000 | 0.05806 | 0.16475 | 0.04291 | 0.00000 | 0.01074 | 0.01684 | 1.46958 | 1.15202 | 0.01772 | 0.00777 |
| **Dhrs7** | 0.00000 | 8.41932 | 1.73944 | 2.77494 | 0.00000 | 2.96823 | 0.03250 | 7.00878 | 1.55355 | 8.09523 | 2.65691 | 2.11784 | 0.00000 | 0.62974 |
| **Dhrs7b** | 3.23900 | 6.42045 | 10.89622 | 5.76844 | 8.41327 | 11.55658 | 3.66009 | 5.51179 | 4.50328 | 0.13370 | 0.38408 | 7.94622 | 1.55645 | 12.73035 |
| **Dhtkd1** | 0.72301 | 0.00338 | 0.89770 | 0.12918 | 0.75274 | 0.67512 | 2.86011 | 0.00000 | 3.53679 | 0.02286 | 0.00000 | 0.47696 | 0.00000 | 1.07104 |
| **Dhx15** | 16.40226 | 0.15679 | 21.15130 | 6.78255 | 21.96198 | 16.83118 | 4.41344 | 4.26269 | 34.11214 | 0.19101 | 13.26450 | 12.74081 | 8.01244 | 17.31610 |

| | | | | | | | | | | | | | | |
|---|---|---|---|---|---|---|---|---|---|---|---|---|---|---|
| **Dhx16** | 10.86534 | 0.76136 | 18.65932 | 6.87294 | 11.92995 | 14.23146 | 0.02638 | 6.69325 | 1.71014 | 0.23862 | 13.51174 | 10.77405 | 3.15061 | 8.05552 |
| **Dhx29** | 3.54095 | 0.03443 | 5.34121 | 3.46439 | 5.54666 | 4.00677 | 0.18005 | 1.05178 | 5.83272 | 0.04729 | 1.33749 | 3.49738 | 1.66251 | 7.07345 |
| **Dhx30** | 13.42424 | 0.19812 | 11.87694 | 0.00000 | 16.05851 | 5.70802 | 2.21943 | 5.19941 | 0.40421 | 0.03319 | 4.60423 | 4.18248 | 5.69077 | 17.27438 |
| **Dhx32** | 6.65774 | 0.59027 | 9.01603 | 5.46848 | 2.58891 | 8.58225 | 4.63077 | 2.87265 | 4.54866 | 0.13077 | 5.16449 | 6.01353 | 2.46416 | 6.02517 |
| **Dhx33** | 8.93240 | 0.44462 | 11.28401 | 4.39369 | 12.55088 | 6.59229 | 1.54536 | 1.37433 | 10.63325 | 2.39970 | 1.43619 | 1.23643 | 5.06497 | 10.66351 |
| **Dhx34** | 0.00000 | 0.06735 | 2.98300 | 5.78995 | 2.56713 | 0.13289 | 2.40234 | 2.25554 | 0.00317 | 0.04481 | 0.54492 | 1.91254 | 0.01246 | 0.00000 |
| **Dhx35** | 1.62726 | 0.02018 | 1.73804 | 2.54410 | 2.56326 | 1.72967 | 0.01085 | 1.07999 | 0.00460 | 0.00860 | 1.47216 | 0.22265 | 0.00737 | 0.41864 |
| **Dhx36** | 1.76012 | 0.01551 | 2.05262 | 0.52011 | 4.06076 | 3.55429 | 0.13029 | 0.95640 | 13.04762 | 0.03499 | 2.41550 | 2.31549 | 5.60280 | 0.87305 |
| **Dhx37** | 1.30691 | 0.04892 | 6.71623 | 1.12963 | 1.79285 | 1.01370 | 0.00305 | 1.25352 | 0.38379 | 0.05006 | 0.15921 | 0.58421 | 1.85272 | 2.62624 |
| **Dhx38** | 2.02998 | 3.48424 | 7.10173 | 1.70688 | 5.92643 | 2.86198 | 0.37047 | 3.04374 | 0.01635 | 1.62074 | 5.51443 | 3.27086 | 2.12771 | 0.38515 |
| **Dhx40** | 10.28952 | 0.03812 | 9.15863 | 0.89828 | 11.13184 | 4.27916 | 3.40243 | 0.79270 | 12.79219 | 0.02188 | 4.78837 | 5.00927 | 5.58451 | 8.06001 |
| **Dhx57** | 0.06603 | 0.01001 | 5.19666 | 0.83580 | 7.35524 | 0.37542 | 0.32705 | 0.08301 | 2.44646 | 0.02129 | 0.14346 | 1.38502 | 1.17357 | 0.00000 |
| **Dhx58** | 0.86738 | 0.00392 | 0.40583 | 0.33027 | 0.14434 | 0.08209 | 0.00387 | 0.00252 | 0.13209 | 0.00673 | 0.00508 | 0.06220 | 1.47509 | 0.32285 |
| **Dhx8** | 5.89100 | 5.71179 | 13.16852 | 3.77366 | 10.08126 | 5.85008 | 4.93804 | 1.02052 | 3.97466 | 0.21653 | 10.09589 | 2.79977 | 6.89644 | 11.22367 |
| **Dhx9** | 2.57341 | 0.03862 | 0.00000 | 1.68659 | 2.43610 | 4.31570 | 0.49995 | 1.39740 | 6.58590 | 3.80670 | 0.00000 | 3.57738 | 14.55619 | 3.66166 |
| **Diablo** | 5.28534 | 0.78486 | 13.20942 | 10.45964 | 12.32372 | 16.63298 | 6.71029 | 1.57087 | 6.80849 | 14.52295 | 11.74253 | 8.09084 | 5.01591 | 9.45838 |
| **Diaph1** | 1.06756 | 1.21025 | 5.20498 | 3.05578 | 3.62725 | 4.73973 | 1.14573 | 4.05732 | 0.49453 | 0.18140 | 3.37998 | 4.22248 | 1.97142 | 1.94438 |
| **Diaph2** | 0.04888 | 0.02864 | 0.69645 | 0.00000 | 0.40465 | 2.91762 | 0.06644 | 0.05548 | 0.70999 | 0.05085 | 0.04432 | 0.95132 | 0.28693 | 0.14297 |
| **Diaph3** | 1.19201 | 0.01851 | 1.00213 | 0.00000 | 3.28750 | 2.11218 | 0.14178 | 1.19124 | 5.21581 | 0.00158 | 0.81940 | 1.14893 | 2.26796 | 3.97541 |
| **Dicer1** | 2.43049 | 0.04178 | 6.00197 | 8.69489 | 7.90845 | 6.63226 | 3.02621 | 0.69731 | 3.22195 | 0.06933 | 0.85872 | 4.63995 | 4.25307 | 12.72474 |
| **Dido1** | 5.37550 | 0.07077 | 5.80491 | 0.74195 | 7.65405 | 2.77584 | 4.93270 | 1.29832 | 6.03954 | 0.14081 | 0.43165 | 5.28641 | 7.93491 | 6.09567 |
| **Diexf** | 0.94343 | 0.04127 | 3.87309 | 1.05461 | 7.43053 | 3.67113 | 2.43689 | 0.00000 | 5.30698 | 2.29765 | 2.53934 | 1.24346 | 1.19080 | 10.79508 |
| **Dimt1** | 11.38828 | 0.46470 | 8.11426 | 0.88415 | 2.96640 | 1.87524 | 0.12330 | 0.25379 | 6.66380 | 0.44340 | 6.44416 | 1.59556 | 11.28677 | 2.39064 |
| **Dio1** | 0.12683 | 0.00428 | 1.12049 | 2.28761 | 2.47016 | 0.46546 | 0.00380 | 0.00000 | 0.41564 | 0.08204 | 0.50031 | 0.00669 | 0.04318 | 0.00000 |
| **Dio3** | 0.01128 | 0.00406 | 0.00588 | 0.34975 | 0.17445 | 0.00480 | 0.00383 | 0.01219 | 0.01631 | 0.00565 | 0.01317 | 0.01042 | 0.12714 | 0.00524 |
| **Dio3os** | 0.00947 | 0.00415 | 0.05508 | 0.14561 | 0.00610 | 0.00544 | 0.00406 | 0.00659 | 0.01485 | 0.00586 | 0.01646 | 0.01508 | 0.22394 | 0.00438 |
| **Dip2a** | 0.28592 | 0.01774 | 0.00000 | 0.00013 | 0.11514 | 0.10106 | 0.00879 | 1.44683 | 0.00503 | 0.00712 | 1.52258 | 0.00649 | 0.38649 | 1.38313 |
| **Dip2b** | 2.23910 | 5.44901 | 3.17184 | 1.00631 | 1.45197 | 5.81274 | 0.76071 | 1.60592 | 1.65241 | 0.05100 | 5.19008 | 2.36076 | 2.13485 | 1.82859 |
| **Dip2c** | 0.00000 | 0.01009 | 0.00000 | 0.00000 | 0.93769 | 1.58628 | 0.00923 | 0.00842 | 1.65615 | 0.00522 | 0.00000 | 1.70097 | 0.00787 | 0.84529 |
| **Diras2** | 0.00341 | 0.00410 | 0.03311 | 0.16909 | 0.44906 | 0.12445 | 0.00387 | 0.00318 | 0.01129 | 0.00642 | 0.00567 | 0.01243 | 0.50730 | 0.02154 |
| **Dirc2** | 17.84084 | 0.06582 | 24.04057 | 3.07032 | 20.57106 | 7.24669 | 0.25616 | 8.26278 | 9.41708 | 0.03406 | 9.47868 | 5.12644 | 3.74524 | 26.78722 |
| **Dis3** | 4.82014 | 0.00942 | 5.28013 | 0.46601 | 6.94534 | 0.30510 | 4.20707 | 0.30596 | 9.62888 | 0.04006 | 3.81962 | 1.29507 | 5.82021 | 13.57390 |
| **Dis3l** | 0.18502 | 0.01090 | 0.39179 | 1.39779 | 2.67515 | 1.10921 | 0.00379 | 1.79220 | 1.48290 | 0.00610 | 2.05564 | 0.00000 | 8.63154 | 3.88260 |
| **Dis3l2** | 0.00000 | 0.01437 | 0.15538 | 0.23327 | 0.07985 | 0.80564 | 0.00407 | 0.99220 | 0.00602 | 0.02867 | 0.00000 | 0.29325 | 0.00509 | 0.39018 |
| **Disp1** | 0.04427 | 0.00370 | 0.00000 | 0.00255 | 0.06352 | 0.05433 | 0.00393 | 0.04289 | 0.02698 | 0.00390 | 0.82732 | 0.00429 | 0.61935 | 0.55173 |
| **Dixdc1** | 0.01593 | 0.00449 | 0.15054 | 0.00000 | 0.14981 | 0.01984 | 0.00447 | 0.00952 | 0.01085 | 0.00654 | 0.00663 | 0.01014 | 0.00538 | 0.01140 |
| **Dkc1** | 12.64657 | 0.47025 | 22.18042 | 13.08105 | 22.03565 | 19.37973 | 2.85126 | 5.88149 | 33.78861 | 9.88555 | 15.73863 | 15.90908 | 2.70223 | 30.35673 |
| **Dkk1** | 0.32117 | 0.02300 | 0.00636 | 0.00000 | 0.14107 | 0.00000 | 3.39031 | 0.43226 | 0.06446 | 0.16625 | 0.00000 | 4.21166 | 0.05750 | 0.03469 |
| **Dkk3** | 0.08400 | 0.00449 | 0.01969 | 0.00768 | 0.38701 | 0.01155 | 0.00489 | 0.01083 | 0.02015 | 0.00690 | 0.00755 | 0.00796 | 0.00432 | 0.00557 |
| **Dkkl1** | 0.25777 | 0.17394 | 0.63720 | 3.93230 | 1.02930 | 4.10561 | 4.98388 | 13.52526 | 0.00008 | 12.30491 | 2.59690 | 0.00000 | 0.84719 | 0.31601 |
| **Dlat** | 2.67760 | 0.06640 | 6.01126 | 7.56502 | 3.11814 | 2.14661 | 0.01157 | 4.88077 | 2.38786 | 0.01995 | 4.51046 | 8.48580 | 0.22287 | 3.78825 |
| **Dlc1** | 0.07034 | 1.09282 | 2.67931 | 1.64403 | 0.00000 | 4.00274 | 3.09276 | 2.23391 | 0.33170 | 0.10561 | 7.89636 | 2.79696 | 1.40037 | 0.00000 |
| **Dld** | 9.04675 | 0.06435 | 9.04560 | 2.07750 | 3.34823 | 13.82017 | 0.85056 | 6.06376 | 6.76681 | 0.03973 | 12.19106 | 9.68834 | 3.02624 | 7.37505 |
| **Dlec1** | 0.00728 | 0.00440 | 0.01289 | 0.06333 | 0.01453 | 0.01200 | 0.00425 | 0.00899 | 0.00819 | 0.00676 | 0.00893 | 0.00988 | 0.00381 | 0.08521 |
| **Dleu2** | 0.00152 | 0.02343 | 0.00000 | 0.00000 | 0.03814 | 0.12784 | 0.00455 | 0.46930 | 0.11797 | 0.00542 | 1.02600 | 0.01343 | 0.63159 | 0.00000 |
| **Dlg1** | 1.20082 | 0.00300 | 1.89238 | 0.00000 | 0.37783 | 2.17578 | 0.26938 | 0.21836 | 1.35456 | 0.00224 | 0.19022 | 1.36309 | 1.20532 | 1.11586 |
| **Dlg3** | 1.00120 | 6.71945 | 2.98378 | 1.06714 | 4.15706 | 3.67460 | 3.26336 | 1.31513 | 2.21638 | 0.11407 | 0.09358 | 0.00000 | 0.57336 | 1.09063 |
| **Dlg5** | 3.12113 | 0.01155 | 0.86321 | 1.10103 | 3.80622 | 0.75768 | 0.00204 | 0.00000 | 10.36879 | 0.00000 | 1.89832 | 0.48548 | 2.58446 | 8.97116 |
| **Dlgap1** | 0.24842 | 0.01757 | 0.48885 | 0.00000 | 0.24473 | 0.00000 | 0.00347 | 0.24510 | 0.21123 | 0.00515 | 0.01354 | 0.33365 | 0.01410 | 1.12327 |
| **Dlgap4** | 0.18092 | 7.56397 | 9.33537 | 8.49988 | 7.54874 | 1.31138 | 4.23567 | 11.78598 | 0.00000 | 0.13534 | 1.01922 | 3.14311 | 0.67808 | 0.00000 |
| **Dlgap5** | 3.97057 | 0.10947 | 6.16255 | 3.05044 | 5.81138 | 7.51286 | 5.19955 | 2.14044 | 4.60245 | 6.95572 | 9.35985 | 3.71516 | 4.36495 | 7.84747 |
| **Dlk2** | 0.05376 | 0.00445 | 0.10779 | 0.00487 | 0.32883 | 0.01957 | 0.00432 | 0.00813 | 0.02217 | 0.00587 | 0.13174 | 0.00603 | 0.05517 | 0.02885 |
| **Dll3** | 0.24464 | 0.00366 | 0.00000 | 0.23425 | 0.92205 | 0.33219 | 0.00399 | 0.00279 | 0.00756 | 0.01913 | 0.04506 | 0.14167 | 1.04010 | 0.10838 |
| **Dlst** | 1.74268 | 3.46039 | 4.22466 | 8.30917 | 0.25607 | 8.83113 | 3.93089 | 7.44759 | 1.63790 | 0.18776 | 5.81284 | 6.35676 | 2.67304 | 2.30764 |
| **Dlx2** | 0.00853 | 0.00441 | 0.04950 | 0.01642 | 0.01077 | 0.01063 | 0.00428 | 0.00992 | 0.01058 | 0.00656 | 0.00826 | 0.00982 | 0.04667 | 0.02242 |

| | | | | | | | | | | | | | | |
|---|---|---|---|---|---|---|---|---|---|---|---|---|---|---|
| **Dlx3** | 0.00308 | 0.00421 | 0.01730 | 0.01702 | 0.14092 | 0.00819 | 0.00445 | 0.00813 | 0.02865 | 0.00792 | 0.00696 | 0.00658 | 0.68151 | 1.27278 |
| **Dmap1** | 0.00000 | 0.13205 | 11.80472 | 11.24320 | 14.77230 | 0.00000 | 1.90165 | 7.65306 | 0.00055 | 0.07001 | 0.61598 | 6.21688 | 0.03573 | 0.00000 |
| **Dmbx1** | 1.12005 | 0.01129 | 2.89364 | 0.00000 | 0.67029 | 0.00000 | 0.00355 | 0.00429 | 0.02400 | 0.00151 | 0.00875 | 0.00534 | 0.74372 | 1.95034 |
| **Dmc1** | 1.29915 | 2.11225 | 1.34245 | 2.63728 | 0.08387 | 4.84276 | 2.98485 | 0.79967 | 0.18980 | 0.07534 | 4.45424 | 2.37276 | 0.22545 | 0.46164 |
| **Dmd** | 1.39614 | 0.00332 | 1.70816 | 0.45501 | 0.21818 | 0.43034 | 0.00300 | 0.90847 | 2.15306 | 0.00059 | 1.05073 | 0.44227 | 0.00010 | 0.04140 |
| **Dmgdh** | 6.93734 | 0.00755 | 12.47239 | 10.88960 | 9.88755 | 8.35961 | 1.87695 | 0.00000 | 7.01573 | 0.11939 | 6.83593 | 2.07091 | 1.23700 | 11.25010 |
| **Dmkn** | 0.00408 | 0.05612 | 0.00516 | 0.00028 | 0.01044 | 1.62921 | 0.00952 | 3.94405 | 0.00993 | 0.03998 | 0.35049 | 0.00908 | 0.00883 | 0.00527 |
| **Dmpk** | 0.06305 | 0.00397 | 0.02260 | 0.54194 | 0.07407 | 0.03866 | 0.00369 | 0.00329 | 0.00590 | 0.00608 | 0.00569 | 0.00917 | 0.91682 | 0.00000 |
| **Dmrt1** | 0.00000 | 0.00391 | 0.10880 | 1.45229 | 0.23791 | 0.00000 | 0.00389 | 0.00486 | 0.00620 | 0.00526 | 0.01221 | 0.07283 | 0.01252 | 0.21175 |
| **Dmrta2** | 0.00927 | 0.00507 | 0.01717 | 0.02098 | 0.03001 | 0.01546 | 0.00456 | 0.13002 | 0.01261 | 0.00777 | 0.03936 | 0.00941 | 0.00743 | 0.01193 |
| **Dmrtb1** | 5.02169 | 0.00263 | 1.54526 | 0.86163 | 5.13170 | 0.00000 | 0.00336 | 0.00014 | 0.05254 | 0.00416 | 0.00347 | 0.00000 | 9.12293 | 2.47895 |
| **Dmtf1** | 9.63774 | 0.00000 | 9.73813 | 1.29778 | 6.90555 | 4.36710 | 2.36045 | 0.00153 | 24.24289 | 0.01153 | 6.19552 | 3.34107 | 2.74236 | 8.29748 |
| **Dmtn** | 2.24245 | 0.00800 | 4.17866 | 0.91978 | 1.24151 | 1.40385 | 0.00851 | 0.02985 | 0.31757 | 0.02964 | 2.94766 | 3.26197 | 0.49496 | 1.81011 |
| **Dmwd** | 1.02385 | 4.53322 | 4.09120 | 5.96784 | 4.22329 | 4.30330 | 5.27373 | 3.55376 | 0.07863 | 0.27409 | 2.30671 | 4.99285 | 4.51006 | 0.77347 |
| **Dmxl1** | 1.11865 | 0.00364 | 0.36762 | 0.00000 | 0.00000 | 0.19552 | 0.00439 | 0.24583 | 2.88602 | 0.00107 | 1.44548 | 0.09721 | 0.85167 | 1.55426 |
| **Dmxl2** | 1.05040 | 0.01602 | 1.30088 | 0.44104 | 3.61362 | 2.36897 | 1.17042 | 0.00000 | 4.10523 | 0.01112 | 1.20022 | 2.28079 | 1.58035 | 11.20366 |
| **Dna2** | 2.82377 | 0.01954 | 4.77588 | 2.29533 | 6.12705 | 1.28940 | 0.00257 | 0.14056 | 0.00502 | 0.01986 | 2.66295 | 1.99365 | 6.31263 | 3.67610 |
| **Dnaaf1** | 2.50436 | 0.00356 | 1.90459 | 1.10837 | 3.38943 | 1.72707 | 0.00903 | 0.78266 | 0.04607 | 0.00737 | 0.10594 | 0.00000 | 0.13338 | 1.18845 |
| **Dnaaf2** | 4.25339 | 0.08153 | 11.68106 | 1.27280 | 8.45504 | 4.24401 | 0.42050 | 5.90281 | 3.32217 | 0.03659 | 2.33963 | 0.60689 | 4.68620 | 6.33880 |
| **Dnaaf3** | 0.01094 | 0.00428 | 0.00838 | 0.03974 | 0.00998 | 0.00933 | 0.00427 | 0.00887 | 0.00989 | 0.00509 | 0.51570 | 0.00429 | 0.02669 | 0.04897 |
| **Dnaaf5** | 4.39118 | 0.01789 | 2.70506 | 2.99035 | 0.53358 | 1.88240 | 0.60231 | 0.04370 | 1.51500 | 0.00000 | 2.75004 | 1.26197 | 5.06904 | 3.43762 |
| **Dnah1** | 0.01542 | 0.00428 | 0.01256 | 0.09591 | 0.00707 | 0.00928 | 0.00410 | 0.00721 | 0.01010 | 0.00743 | 0.00703 | 0.01433 | 0.00435 | 0.12339 |
| **Dnah10** | 0.16018 | 0.00441 | 0.16984 | 0.00000 | 0.48252 | 0.00405 | 0.00431 | 0.01255 | 0.00917 | 0.00713 | 0.00632 | 0.00822 | 0.06239 | 0.15964 |
| **Dnah11** | 0.00447 | 0.00482 | 0.12793 | 0.00000 | 0.05759 | 0.06763 | 0.00414 | 0.00739 | 1.00220 | 0.00720 | 0.00675 | 0.00864 | 0.00351 | 0.04494 |
| **Dnah17** | 0.74143 | 0.00414 | 0.92284 | 0.00404 | 0.06414 | 0.00000 | 0.00391 | 0.00859 | 0.12471 | 0.00761 | 0.25375 | 0.00455 | 0.11989 | 0.99789 |
| **Dnah2** | 0.19837 | 0.02481 | 1.85893 | 0.00000 | 1.04340 | 1.06379 | 0.00306 | 0.00000 | 0.00749 | 0.00257 | 0.00000 | 1.80915 | 0.70429 | 0.63201 |
| **Dnah5** | 0.00119 | 0.00443 | 0.00770 | 0.00776 | 0.81998 | 0.01066 | 0.00427 | 0.01023 | 0.03214 | 0.00614 | 0.00808 | 0.00744 | 0.03408 | 0.60168 |
| **Dnah6** | 0.07847 | 0.00433 | 0.11385 | 0.14747 | 0.00000 | 0.33269 | 0.00412 | 0.00343 | 0.00570 | 0.00592 | 0.09994 | 0.00819 | 0.00539 | 0.38122 |
| **Dnah7a** | 0.01944 | 0.00444 | 0.19025 | 0.00681 | 0.00656 | 0.00974 | 0.00408 | 0.00836 | 0.49883 | 0.00621 | 0.00650 | 0.01378 | 0.16714 | 0.35624 |
| **Dnah7b** | 0.10589 | 0.00423 | 0.01999 | 0.01876 | 0.06153 | 0.00867 | 0.00414 | 0.00739 | 0.10511 | 0.00586 | 0.00842 | 0.01690 | 0.24805 | 1.60950 |
| **Dnah7c** | 0.03682 | 0.00443 | 0.05757 | 0.10114 | 0.02991 | 0.03901 | 0.00405 | 0.00466 | 0.06917 | 0.00564 | 0.01038 | 0.02800 | 0.07713 | 0.51796 |
| **Dnah8** | 0.02448 | 0.03342 | 1.00858 | 0.00000 | 0.98995 | 0.00000 | 0.00388 | 0.02286 | 0.00904 | 0.00461 | 0.00393 | 0.14988 | 0.01537 | 0.23692 |
| **Dnah9** | 0.00278 | 0.00405 | 0.00854 | 0.00897 | 0.08418 | 0.02198 | 0.00424 | 0.00729 | 0.00212 | 0.00575 | 0.03269 | 0.00690 | 1.52356 | 0.71295 |
| **Dnaic1** | 0.54489 | 0.00406 | 0.41882 | 0.45190 | 0.18718 | 0.00000 | 0.00396 | 0.00573 | 0.00715 | 0.01883 | 0.16943 | 0.01128 | 0.04572 | 0.00000 |
| **Dnaic2** | 0.00576 | 0.00408 | 0.09622 | 0.41400 | 0.27207 | 0.17527 | 0.00384 | 0.00452 | 0.00946 | 0.00660 | 0.00362 | 0.00374 | 1.32469 | 0.00000 |
| **Dnaja1** | 45.34266 | 9.44479 | 54.50144 | 53.69751 | 40.14529 | 42.46526 | 12.15564 | 17.73929 | 36.04573 | 7.93829 | 25.48232 | 25.08448 | 14.43143 | 27.42183 |
| **Dnaja2** | 21.36746 | 1.82304 | 24.20691 | 13.49376 | 26.15451 | 21.62005 | 5.59547 | 6.56988 | 18.93818 | 8.12601 | 12.00288 | 13.47967 | 16.44745 | 25.61853 |
| **Dnaja3** | 4.00522 | 5.60657 | 8.40896 | 5.30332 | 6.96027 | 5.68863 | 5.50488 | 4.62391 | 7.56426 | 0.46594 | 8.38193 | 6.42270 | 0.24816 | 8.12251 |
| **Dnaja4** | 0.52081 | 0.00344 | 0.00000 | 0.34831 | 0.89289 | 0.07814 | 0.00378 | 0.00285 | 0.00264 | 0.00528 | 0.00745 | 0.46742 | 1.04398 | 0.59156 |
| **Dnajb1** | 6.44619 | 8.73059 | 10.64918 | 10.43683 | 13.16229 | 14.40546 | 3.45290 | 4.59173 | 2.15378 | 1.12454 | 7.14186 | 4.82830 | 5.99581 | 6.35154 |
| **Dnajb11** | 2.57447 | 0.13178 | 11.20153 | 12.10670 | 5.63711 | 14.03320 | 4.73249 | 3.11050 | 13.01257 | 0.25051 | 5.03631 | 5.88823 | 4.25367 | 11.01907 |
| **Dnajb12** | 0.04762 | 0.00449 | 0.32566 | 0.70034 | 0.87672 | 0.19253 | 0.00413 | 0.52367 | 0.30910 | 0.00672 | 0.00660 | 0.01450 | 1.02585 | 0.95721 |
| **Dnajb13** | 0.56244 | 0.00442 | 0.00991 | 0.01370 | 0.07571 | 0.00806 | 0.00431 | 0.01167 | 0.01261 | 0.00686 | 0.00682 | 0.00709 | 0.07503 | 0.10835 |
| **Dnajb14** | 9.05515 | 0.00145 | 1.14934 | 0.42110 | 3.77725 | 0.00000 | 0.00269 | 1.10226 | 5.14947 | 0.00401 | 0.07181 | 0.84043 | 7.87887 | 11.49153 |
| **Dnajb2** | 1.50286 | 0.00394 | 2.24988 | 6.56662 | 1.97448 | 5.45336 | 0.75062 | 0.86860 | 0.13736 | 0.03479 | 1.34269 | 0.73511 | 0.00697 | 2.92472 |
| **Dnajb3** | 0.00196 | 0.00434 | 0.02539 | 1.76179 | 0.04294 | 0.01175 | 0.00401 | 0.01917 | 0.00704 | 0.01552 | 0.00528 | 0.03904 | 0.34433 | 0.01141 |
| **Dnajb4** | 0.07068 | 0.02304 | 0.00632 | 0.00733 | 0.00313 | 0.02744 | 0.02388 | 0.04403 | 0.00954 | 0.00566 | 0.19911 | 0.42376 | 0.32711 | 0.53020 |
| **Dnajb6** | 31.51381 | 1.76935 | 29.26269 | 28.61142 | 41.64512 | 22.16079 | 5.45547 | 14.92556 | 29.67559 | 15.27990 | 14.20396 | 13.96774 | 15.96205 | 25.10545 |
| **Dnajb9** | 22.03481 | 0.01154 | 9.09722 | 8.87434 | 13.02249 | 1.21103 | 2.92868 | 0.00000 | 29.94178 | 0.02865 | 1.80814 | 1.60047 | 19.43799 | 20.74185 |
| **Dnajc1** | 3.04030 | 0.00923 | 1.65593 | 2.36716 | 1.31088 | 3.10130 | 0.00299 | 0.21259 | 1.44874 | 0.02094 | 0.52747 | 1.18722 | 3.51960 | 2.17474 |
| **Dnajc10** | 3.78788 | 0.02120 | 4.98162 | 0.96379 | 1.15415 | 5.65794 | 1.22861 | 0.42325 | 1.33914 | 0.00473 | 3.45154 | 7.21366 | 0.19573 | 3.27247 |
| **Dnajc11** | 5.12027 | 9.69139 | 15.23650 | 12.27432 | 10.52936 | 17.35322 | 3.37946 | 12.01762 | 0.80852 | 12.15607 | 14.69172 | 9.61586 | 1.30841 | 11.62011 |
| **Dnajc12** | 0.79585 | 0.00277 | 5.01513 | 0.00000 | 3.95506 | 0.00000 | 0.00359 | 0.91360 | 0.09591 | 0.00466 | 2.80763 | 0.59843 | 4.32609 | 2.24930 |
| **Dnajc13** | 5.12225 | 0.04135 | 9.58660 | 2.81307 | 6.87451 | 10.48717 | 3.89673 | 3.10917 | 11.45103 | 0.01555 | 9.42723 | 5.18018 | 2.57466 | 11.21696 |
| **Dnajc14** | 0.12493 | 0.14903 | 5.14188 | 2.82148 | 3.35423 | 3.16293 | 2.95297 | 2.14510 | 1.50683 | 0.09399 | 1.07760 | 2.95800 | 0.65275 | 4.83929 |

| | | | | | | | | | | | | | | |
|---|---|---|---|---|---|---|---|---|---|---|---|---|---|---|
| **Dnajc15** | 10.64209 | 6.17367 | 5.89990 | 7.77685 | 15.31260 | 3.50322 | 3.56309 | 4.91249 | 1.37464 | 8.12271 | 2.86561 | 5.30491 | 3.70776 | 3.12195 |
| **Dnajc16** | 0.40367 | 0.00390 | 1.33821 | 1.60168 | 1.23550 | 0.41498 | 0.00465 | 0.67461 | 0.00270 | 0.00297 | 0.01385 | 0.85526 | 0.47878 | 0.65050 |
| **Dnajc17** | 3.60512 | 0.05356 | 4.54498 | 4.03544 | 4.86247 | 3.10691 | 2.13958 | 2.45784 | 0.00000 | 0.09092 | 0.94674 | 3.94867 | 0.15990 | 1.78646 |
| **Dnajc18** | 0.01899 | 0.01463 | 0.03268 | 0.00000 | 0.00000 | 0.00000 | 0.00416 | 0.28490 | 0.00689 | 0.02118 | 1.36886 | 0.56744 | 0.16971 | 0.86268 |
| **Dnajc19** | 0.19626 | 1.16973 | 0.33016 | 3.49450 | 3.96530 | 1.34952 | 2.74668 | 11.09946 | 0.05003 | 0.39879 | 0.03190 | 7.32268 | 0.57207 | 0.24612 |
| **Dnajc2** | 23.38921 | 2.33289 | 17.36219 | 8.67114 | 25.97856 | 16.92586 | 5.58964 | 2.31925 | 26.55572 | 15.25552 | 10.36187 | 9.04008 | 2.89583 | 15.20077 |
| **Dnajc21** | 10.67763 | 0.06985 | 24.04460 | 5.69674 | 30.41965 | 11.28247 | 2.78154 | 3.93580 | 29.37724 | 0.88922 | 9.56301 | 7.10761 | 5.88057 | 34.38573 |
| **Dnajc22** | 0.43094 | 0.04668 | 0.74984 | 0.00000 | 0.04498 | 0.01819 | 0.20885 | 0.00124 | 0.26352 | 0.00638 | 0.00677 | 2.81019 | 0.04530 | 0.00000 |
| **Dnajc24** | 2.33216 | 0.03610 | 3.22165 | 1.90747 | 5.98163 | 1.46349 | 0.00950 | 2.03147 | 0.01468 | 0.01848 | 0.03965 | 3.16457 | 0.03240 | 0.43471 |
| **Dnajc25** | 0.62354 | 0.02138 | 1.16584 | 1.18322 | 1.65275 | 0.15788 | 0.00339 | 2.20218 | 2.24696 | 0.01311 | 1.04506 | 0.15765 | 2.98011 | 0.00000 |
| **Dnajc27** | 0.21700 | 0.05876 | 2.01967 | 0.00000 | 0.41882 | 1.55218 | 0.00323 | 0.04281 | 0.00000 | 0.01540 | 1.59889 | 2.06817 | 0.00000 | 2.93839 |
| **Dnajc28** | 0.02181 | 0.00436 | 0.17756 | 0.01113 | 0.09329 | 0.00306 | 0.00402 | 0.01095 | 0.00574 | 0.00805 | 0.00644 | 0.00735 | 0.00397 | 0.77225 |
| **Dnajc3** | 6.67298 | 0.03215 | 2.73367 | 0.00000 | 6.12559 | 8.01467 | 0.02612 | 1.64968 | 11.26572 | 0.02652 | 2.05062 | 2.29400 | 4.03787 | 12.56575 |
| **Dnajc30** | 0.65966 | 4.22002 | 2.51264 | 3.43555 | 2.04214 | 0.55776 | 0.00263 | 0.49658 | 0.29084 | 0.03342 | 0.01590 | 0.47296 | 1.36121 | 0.00000 |
| **Dnajc4** | 0.21654 | 0.01129 | 0.35360 | 1.95080 | 1.46089 | 1.94279 | 0.01757 | 1.63807 | 0.01462 | 0.02854 | 0.22520 | 1.88949 | 2.05059 | 0.68033 |
| **Dnajc5** | 8.27021 | 5.39371 | 3.04451 | 3.44653 | 9.22134 | 10.53945 | 0.03701 | 3.47731 | 4.62482 | 0.05137 | 0.97837 | 4.12921 | 3.98203 | 7.58406 |
| **Dnajc5g** | 1.11558 | 0.00405 | 0.77460 | 0.00000 | 2.64745 | 0.00000 | 0.00366 | 0.00529 | 2.08283 | 0.00564 | 0.00108 | 0.00401 | 0.59614 | 0.98323 |
| **Dnajc6** | 0.00000 | 0.00284 | 0.25580 | 0.03744 | 0.01926 | 0.00000 | 0.00357 | 0.00000 | 0.16925 | 0.00000 | 1.96224 | 0.93706 | 0.26826 | 0.39808 |
| **Dnajc7** | 10.24450 | 5.73547 | 9.98955 | 8.24932 | 6.28274 | 5.62816 | 1.89703 | 3.15670 | 15.86040 | 1.54811 | 5.28942 | 3.15681 | 5.37859 | 11.10628 |
| **Dnajc8** | 10.71533 | 4.27478 | 18.19382 | 13.99106 | 14.42038 | 20.02069 | 1.81791 | 9.72429 | 14.75471 | 7.60100 | 13.67814 | 15.07967 | 5.92158 | 15.31221 |
| **Dnajc9** | 1.99292 | 0.02788 | 0.00239 | 1.85615 | 2.21141 | 1.40684 | 0.00362 | 1.14106 | 0.60181 | 0.05191 | 0.04393 | 0.89770 | 2.57797 | 2.06977 |
| **Dnal1** | 0.11507 | 0.00372 | 0.22663 | 0.16401 | 0.77652 | 0.00000 | 0.00983 | 0.23942 | 2.17491 | 0.01329 | 0.35395 | 0.24922 | 0.00196 | 0.74270 |
| **Dnal4** | 1.70583 | 0.15468 | 3.16266 | 2.46796 | 2.26865 | 2.44451 | 3.79959 | 4.45722 | 0.00746 | 7.18517 | 0.06754 | 4.24353 | 2.61986 | 1.76486 |
| **Dnase1** | 0.03703 | 0.00451 | 0.14832 | 0.00000 | 0.00667 | 0.22728 | 0.00409 | 0.00561 | 0.12494 | 0.00596 | 0.00520 | 0.00785 | 0.00597 | 0.08467 |
| **Dnase1l1** | 0.08528 | 0.00420 | 0.13809 | 0.21312 | 0.05455 | 0.00000 | 0.00458 | 0.01082 | 0.30803 | 0.00810 | 0.00486 | 0.00667 | 0.00509 | 0.35598 |
| **Dnase1l2** | 0.61695 | 0.02088 | 0.06087 | 0.97328 | 0.33019 | 0.03050 | 1.67868 | 0.02581 | 0.01339 | 0.04051 | 0.00175 | 0.00087 | 0.11736 | 0.02241 |
| **Dnase1l3** | 0.00508 | 0.00423 | 0.02145 | 0.01586 | 0.00870 | 0.01551 | 0.00416 | 0.00823 | 0.01139 | 0.00683 | 0.00910 | 0.00790 | 0.85158 | 0.00717 |
| **Dnase2a** | 0.04828 | 0.03490 | 0.56351 | 1.52474 | 0.35255 | 0.87046 | 0.03292 | 1.17460 | 0.14256 | 0.11197 | 0.00070 | 0.29418 | 0.12578 | 1.33346 |
| **Dnd1** | 0.11350 | 0.01381 | 0.10106 | 0.01387 | 0.43501 | 0.11566 | 0.01761 | 1.18061 | 0.05749 | 0.04628 | 0.13140 | 0.02991 | 0.61570 | 0.07709 |
| **Dner** | 0.00422 | 0.00444 | 0.20744 | 0.00183 | 0.06078 | 0.00703 | 0.00425 | 0.01352 | 0.00823 | 0.00683 | 0.00717 | 0.01254 | 0.11866 | 0.00609 |
| **Dnlz** | 0.11519 | 0.05242 | 0.26157 | 0.88341 | 0.19256 | 0.99728 | 0.26274 | 0.02261 | 0.01164 | 1.04641 | 0.14624 | 0.34138 | 0.33201 | 0.05780 |
| **Dnm1** | 0.23245 | 0.00439 | 0.01313 | 0.00939 | 0.00751 | 0.01159 | 0.00428 | 0.01066 | 0.00983 | 0.00666 | 0.00737 | 0.01674 | 0.01070 | 0.01791 |
| **Dnm1l** | 3.85070 | 4.10908 | 4.14598 | 1.33453 | 4.40837 | 9.81169 | 1.42826 | 2.06484 | 4.89509 | 0.07252 | 9.72619 | 6.24411 | 3.59868 | 6.08256 |
| **Dnm2** | 0.49474 | 0.08134 | 10.15505 | 6.95752 | 5.33300 | 1.04618 | 0.11729 | 9.90218 | 0.00000 | 8.62019 | 2.24974 | 5.25451 | 0.89875 | 0.00000 |
| **Dnm3** | 0.19507 | 0.00444 | 0.11017 | 0.00245 | 0.21210 | 0.00715 | 0.00409 | 0.01151 | 0.01185 | 0.00672 | 0.00806 | 0.00874 | 0.00565 | 0.01020 |
| **Dnmbp** | 0.05489 | 0.24550 | 0.00000 | 0.26241 | 0.03880 | 2.25647 | 0.44840 | 3.34587 | 0.00303 | 0.01085 | 3.84723 | 0.00000 | 1.76680 | 0.00000 |
| **Dnmt1** | 0.83733 | 0.05900 | 8.45837 | 9.83806 | 8.44523 | 2.71107 | 0.01398 | 1.51856 | 0.07170 | 2.76318 | 4.47883 | 4.18822 | 5.45228 | 0.32248 |
| **Dnmt3a** | 1.91257 | 0.07837 | 6.27185 | 0.77486 | 3.77648 | 9.21245 | 0.00069 | 1.41535 | 3.78121 | 0.04338 | 4.78013 | 4.47578 | 3.20795 | 8.20553 |
| **Dnmt3b** | 4.90270 | 6.29867 | 10.63691 | 2.91884 | 7.27255 | 9.52172 | 6.33687 | 13.67314 | 1.38654 | 15.24178 | 5.45562 | 7.01238 | 0.78757 | 10.38496 |
| **Dnmt3l** | 5.33023 | 5.29976 | 11.11741 | 8.83769 | 5.17644 | 20.86753 | 7.86234 | 18.29721 | 0.11083 | 0.48435 | 2.49594 | 9.07054 | 0.90275 | 7.04243 |
| **Dnpep** | 6.81483 | 0.28144 | 16.63127 | 14.49434 | 9.15233 | 18.86804 | 6.02290 | 11.88413 | 0.24487 | 1.09052 | 6.92982 | 10.16534 | 5.87883 | 2.62672 |
| **Dnph1** | 0.95704 | 0.04240 | 4.03996 | 6.43065 | 1.94073 | 0.66286 | 2.67464 | 4.61064 | 0.19364 | 2.81016 | 0.06760 | 1.07588 | 0.54858 | 0.23280 |
| **Dnttip1** | 3.30224 | 0.05506 | 2.53022 | 1.58568 | 4.12314 | 3.25252 | 0.01398 | 1.44073 | 0.03873 | 0.00565 | 2.21201 | 3.12991 | 0.61573 | 2.43121 |
| **Dnttip2** | 18.23263 | 0.09915 | 19.97677 | 12.84548 | 22.53113 | 11.98099 | 4.77483 | 0.78341 | 23.07733 | 9.94849 | 9.15807 | 9.72135 | 7.93423 | 26.74339 |
| **Doc2a** | 2.48369 | 0.14976 | 4.98479 | 3.51901 | 2.49822 | 4.31832 | 0.51285 | 2.42013 | 0.00000 | 0.09963 | 3.85197 | 4.46387 | 3.59025 | 0.32849 |
| **Doc2b** | 0.11137 | 0.00443 | 0.01295 | 0.01780 | 0.00939 | 0.01551 | 0.00424 | 0.00922 | 0.01058 | 0.01000 | 0.00726 | 0.00900 | 0.02672 | 0.00554 |
| **Doc2g** | 0.00986 | 0.00439 | 0.31490 | 0.00203 | 0.00792 | 0.00958 | 0.00420 | 0.01026 | 0.01162 | 0.00730 | 0.00721 | 0.00985 | 0.00739 | 0.01083 |
| **Dock1** | 0.12348 | 0.00346 | 0.34399 | 0.00000 | 0.38951 | 1.25548 | 1.20286 | 0.25399 | 0.32048 | 0.01428 | 0.08087 | 1.65109 | 0.00142 | 0.00000 |
| **Dock10** | 0.00401 | 0.01444 | 0.10746 | 0.00000 | 0.02851 | 0.02370 | 0.00416 | 0.38686 | 0.00929 | 0.00790 | 0.01789 | 0.29441 | 0.05355 | 0.01244 |
| **Dock11** | 0.00000 | 0.01330 | 0.13808 | 0.00000 | 0.07893 | 0.52532 | 0.00381 | 0.00481 | 0.00407 | 0.01016 | 0.00078 | 0.32855 | 1.31260 | 0.85673 |
| **Dock3** | 0.05610 | 0.00419 | 0.02147 | 0.15925 | 0.40803 | 0.00199 | 0.00395 | 0.00695 | 0.25970 | 0.00621 | 0.00497 | 0.01205 | 0.27399 | 0.08563 |
| **Dock4** | 0.01862 | 0.00416 | 0.36582 | 0.21600 | 0.70519 | 0.65621 | 0.00384 | 0.00732 | 1.11847 | 0.01589 | 0.01395 | 0.00342 | 0.17608 | 0.00982 |
| **Dock5** | 0.90342 | 0.05993 | 2.29803 | 2.69915 | 2.77241 | 0.69251 | 0.01053 | 0.22201 | 2.95458 | 0.01395 | 0.00000 | 3.42555 | 1.28250 | 2.82868 |
| **Dock6** | 0.74788 | 0.01337 | 4.55502 | 3.45673 | 1.68099 | 1.58070 | 0.00354 | 0.00003 | 0.02124 | 0.01865 | 0.00000 | 0.39602 | 0.66874 | 0.83294 |
| **Dock7** | 0.03130 | 0.00409 | 0.60364 | 0.00000 | 0.92957 | 0.37080 | 0.00468 | 0.18306 | 0.73732 | 0.00484 | 0.00000 | 0.64872 | 1.26086 | 0.98695 |

| | | | | | | | | | | | | | | |
|---|---|---|---|---|---|---|---|---|---|---|---|---|---|---|
| **Dock8** | 2.30632 | 0.00411 | 0.00000 | 0.02727 | 0.00000 | 0.52149 | 0.00408 | 0.00593 | 0.27144 | 0.00626 | 0.00577 | 0.03530 | 0.23996 | 0.00000 |
| **Dock9** | 2.62540 | 0.00867 | 4.94686 | 0.00000 | 3.71238 | 4.21011 | 0.00241 | 0.36456 | 2.29872 | 0.00356 | 0.00000 | 4.66317 | 1.99389 | 2.08498 |
| **Dohh** | 0.00000 | 0.09455 | 3.03097 | 11.48996 | 9.50759 | 0.00000 | 0.57752 | 3.11045 | 0.00000 | 0.02226 | 0.00846 | 4.47498 | 0.48595 | 0.00000 |
| **Dok1** | 0.00000 | 4.46974 | 1.47770 | 0.80367 | 0.38747 | 2.46822 | 1.15532 | 4.28854 | 0.00295 | 0.66327 | 0.56123 | 1.06332 | 0.01170 | 0.00000 |
| **Dok2** | 0.00000 | 0.14761 | 3.48488 | 6.09414 | 0.00000 | 6.69935 | 0.00225 | 0.38352 | 0.03645 | 0.01785 | 2.44656 | 2.67467 | 0.00000 | 0.00000 |
| **Dok3** | 0.01215 | 0.00461 | 0.02746 | 0.31320 | 0.08762 | 0.00000 | 0.04407 | 0.60264 | 0.01034 | 0.00554 | 0.00924 | 0.01055 | 0.07636 | 0.00000 |
| **Dok4** | 0.03615 | 0.05192 | 0.16745 | 2.01231 | 0.08740 | 1.07675 | 0.00890 | 6.22154 | 0.00311 | 0.04252 | 0.00006 | 0.73596 | 0.85894 | 0.59299 |
| **Dok5** | 0.01250 | 0.01359 | 0.05407 | 0.43252 | 0.13579 | 0.00000 | 0.01137 | 0.71515 | 0.45403 | 0.03072 | 0.00577 | 0.00947 | 0.02351 | 0.00000 |
| **Dolk** | 2.96488 | 0.01507 | 2.53721 | 2.34671 | 2.36151 | 0.74273 | 0.01869 | 1.69150 | 1.07373 | 0.00000 | 2.76654 | 2.68589 | 4.02896 | 4.10488 |
| **Dolpp1** | 0.93066 | 5.16762 | 6.59310 | 7.10553 | 4.85597 | 5.16001 | 0.23064 | 2.79348 | 0.10077 | 0.06851 | 0.63235 | 4.98716 | 0.22860 | 0.91116 |
| **Donson** | 0.95443 | 0.01793 | 2.06565 | 0.59045 | 3.08700 | 1.31017 | 0.40627 | 1.59959 | 2.49395 | 0.40796 | 1.87103 | 1.29830 | 4.86733 | 2.62820 |
| **Dopey1** | 1.16682 | 0.31316 | 0.36590 | 0.15591 | 0.12622 | 0.14312 | 0.00356 | 0.16775 | 0.93862 | 0.01550 | 1.35945 | 0.00296 | 0.72542 | 1.52929 |
| **Dopey2** | 0.05088 | 0.01167 | 3.53348 | 0.00000 | 0.70605 | 0.00000 | 0.00410 | 0.02545 | 0.39052 | 0.00000 | 0.13829 | 0.49786 | 0.31604 | 1.09121 |
| **Dot1l** | 0.00710 | 0.00285 | 2.26297 | 3.32015 | 1.40126 | 3.00115 | 0.31533 | 0.00001 | 0.18625 | 0.00649 | 1.15117 | 1.16557 | 0.09950 | 1.65513 |
| **Dpagt1** | 1.45922 | 0.01797 | 4.62296 | 4.81157 | 1.91245 | 0.66236 | 2.98055 | 1.27666 | 1.61139 | 0.05040 | 1.43093 | 4.29528 | 5.04292 | 2.47182 |
| **Dpcd** | 4.30885 | 0.02297 | 1.18676 | 1.06494 | 1.85460 | 2.10754 | 0.00000 | 3.87497 | 2.35294 | 9.58337 | 4.61718 | 2.01469 | 0.80974 | 4.15082 |
| **Dpcr1** | 0.01348 | 0.00473 | 0.01503 | 0.01637 | 0.00757 | 0.02433 | 0.00462 | 0.04016 | 0.03012 | 0.00660 | 0.00836 | 0.01048 | 0.28950 | 0.27387 |
| **Dpep1** | 0.45122 | 0.00366 | 0.67312 | 0.00000 | 0.08643 | 0.00000 | 0.00390 | 0.70092 | 0.00447 | 0.00350 | 0.00356 | 0.01009 | 1.61222 | 0.06497 |
| **Dpep2** | 0.39536 | 0.00345 | 0.64143 | 0.00000 | 0.87760 | 0.00000 | 0.00363 | 0.00560 | 2.35180 | 0.00560 | 0.00431 | 0.00102 | 5.45057 | 3.93332 |
| **Dpep3** | 2.13161 | 0.02866 | 1.80533 | 1.70247 | 0.78863 | 0.35083 | 1.70050 | 2.31915 | 0.83503 | 5.61591 | 4.48242 | 0.00121 | 1.04323 | 0.48597 |
| **Dpf1** | 0.00356 | 0.00424 | 0.06134 | 0.00109 | 0.00750 | 0.03143 | 0.00420 | 0.00714 | 0.07889 | 0.00694 | 0.00491 | 0.12449 | 0.23118 | 0.00830 |
| **Dpf2** | 11.32231 | 8.32105 | 21.80878 | 11.04486 | 21.63093 | 12.93541 | 3.63322 | 7.39777 | 5.03135 | 0.26418 | 13.23626 | 7.26581 | 6.42202 | 18.89207 |
| **Dpf3** | 0.83677 | 0.00345 | 6.74315 | 1.42465 | 3.65851 | 2.48361 | 0.00245 | 0.93673 | 0.20700 | 0.00716 | 0.00000 | 0.02829 | 2.77601 | 2.09147 |
| **Dph1** | 0.00000 | 5.54070 | 5.52511 | 3.08210 | 4.49798 | 0.74980 | 0.14359 | 1.63394 | 0.00000 | 0.00000 | 2.21398 | 2.12752 | 0.00000 | 0.03262 |
| **Dph2** | 3.37276 | 0.02344 | 4.45938 | 7.29787 | 6.73120 | 5.78971 | 1.02383 | 1.82066 | 1.12604 | 0.02110 | 0.29244 | 2.10092 | 4.17410 | 5.43031 |
| **Dph3** | 8.80728 | 0.07955 | 2.25834 | 14.28136 | 14.06034 | 8.09302 | 2.20389 | 3.89579 | 4.40984 | 0.18178 | 1.87984 | 4.92158 | 7.15960 | 6.20629 |
| **Dph5** | 5.89727 | 0.08448 | 4.40976 | 6.93901 | 4.35824 | 5.34183 | 0.02639 | 3.21667 | 9.49098 | 0.06617 | 2.08082 | 4.54143 | 1.45220 | 4.96727 |
| **Dph6** | 2.39869 | 0.05590 | 3.57173 | 1.09578 | 2.61128 | 1.83915 | 0.36690 | 1.27576 | 0.36879 | 0.09841 | 5.28198 | 2.12978 | 1.25439 | 3.25344 |
| **Dph7** | 0.08923 | 1.53323 | 0.00000 | 0.97659 | 0.90912 | 0.00000 | 0.01011 | 0.04303 | 3.57516 | 0.00000 | 2.71156 | 0.46400 | 0.72820 | 4.83232 |
| **Dpm1** | 0.10878 | 0.02129 | 0.39114 | 0.79950 | 0.14236 | 0.47761 | 1.39896 | 0.25138 | 0.13377 | 0.04215 | 0.95891 | 0.13286 | 1.11236 | 0.08288 |
| **Dpm2** | 6.29216 | 4.52011 | 4.76599 | 12.64740 | 4.82028 | 11.19119 | 2.72888 | 6.19406 | 0.98066 | 1.59592 | 3.67576 | 8.83703 | 2.11432 | 3.49068 |
| **Dpm3** | 0.15838 | 0.58672 | 0.00000 | 0.40676 | 0.97585 | 1.77524 | 0.29661 | 2.46040 | 0.00699 | 0.23626 | 0.00861 | 2.03693 | 0.07924 | 0.00704 |
| **Dpp10** | 0.00719 | 0.00440 | 0.01375 | 0.01281 | 0.03855 | 0.01256 | 0.00441 | 0.01127 | 0.00913 | 0.00651 | 0.00881 | 0.01005 | 0.02757 | 0.07516 |
| **Dpp3** | 1.59836 | 0.34588 | 3.50236 | 10.21040 | 5.33819 | 1.54890 | 0.03099 | 5.76423 | 0.24118 | 0.03425 | 0.00000 | 7.11440 | 0.16245 | 0.39689 |
| **Dpp4** | 0.65808 | 0.01474 | 0.00000 | 0.00000 | 0.49006 | 0.48846 | 0.01203 | 0.57433 | 0.02859 | 0.00854 | 0.00000 | 0.78852 | 0.00507 | 0.00000 |
| **Dpp7** | 0.30984 | 0.13673 | 11.78815 | 22.73613 | 2.11748 | 13.42646 | 3.46034 | 5.11437 | 0.00000 | 10.83605 | 2.10294 | 4.39417 | 0.00000 | 1.36844 |
| **Dpp8** | 0.09750 | 0.01772 | 0.73473 | 0.00000 | 0.85834 | 0.76671 | 0.00304 | 0.00781 | 0.22860 | 0.00544 | 0.00000 | 2.46653 | 1.83909 | 1.82141 |
| **Dpp9** | 0.16235 | 9.64079 | 1.98522 | 0.00000 | 2.12981 | 2.85986 | 0.65900 | 2.51051 | 0.00351 | 0.03311 | 0.00313 | 2.05106 | 2.33337 | 2.37506 |
| **Dppa1** | 0.00111 | 2.29071 | 0.00000 | 0.00000 | 0.00000 | 3.50068 | 7.08335 | 15.86603 | 0.03755 | 11.11913 | 1.64038 | 0.14293 | 0.03911 | 0.00000 |
| **Dppa2** | 23.17207 | 1.35682 | 29.21711 | 15.87708 | 25.05505 | 18.35483 | 2.76405 | 4.75295 | 36.23738 | 0.92091 | 28.18016 | 18.55014 | 13.79646 | 39.42001 |
| **Dppa3** | 40.27824 | 11.46976 | 40.40558 | 52.14249 | 51.87218 | 52.27226 | 14.92892 | 25.94624 | 36.47353 | 21.77077 | 32.81862 | 31.14290 | 26.86850 | 44.79915 |
| **Dppa4** | 22.08545 | 0.41921 | 28.70712 | 19.35464 | 27.22427 | 14.50085 | 5.75490 | 11.06032 | 19.72774 | 0.29130 | 14.30337 | 15.44133 | 13.64399 | 27.12789 |
| **Dppa5a** | 24.62866 | 13.09239 | 6.51344 | 25.34852 | 26.93618 | 39.07486 | 10.99954 | 34.25431 | 15.50996 | 31.82535 | 10.48010 | 34.26712 | 23.90926 | 17.18409 |
| **Dpy19l1** | 0.51503 | 0.01000 | 0.03491 | 0.00000 | 0.00566 | 1.68885 | 0.00355 | 0.39381 | 0.01206 | 0.01233 | 0.37402 | 2.19653 | 0.03331 | 0.18882 |
| **Dpy19l2** | 0.00909 | 0.00408 | 0.24465 | 0.05979 | 0.00410 | 0.00000 | 0.00431 | 0.01170 | 0.01473 | 0.00275 | 1.08078 | 0.00352 | 0.01069 | 0.01116 |
| **Dpy19l3** | 0.05731 | 0.00395 | 0.28885 | 0.43234 | 0.08923 | 0.39382 | 0.00386 | 0.00168 | 0.01436 | 0.00392 | 0.13869 | 0.01223 | 0.51705 | 0.00000 |
| **Dpy19l4** | 0.00000 | 0.00950 | 0.66290 | 0.00000 | 0.00000 | 0.00000 | 0.00725 | 0.00000 | 0.10192 | 7.42901 | 0.00239 | 0.54034 | 2.37750 | 0.28868 |
| **Dpy30** | 5.13446 | 0.53620 | 1.98314 | 11.77979 | 12.85549 | 11.37826 | 4.20822 | 8.89281 | 0.47170 | 8.33615 | 3.71821 | 14.21532 | 9.35608 | 6.05649 |
| **Dpys** | 0.01177 | 0.00446 | 0.00688 | 0.00000 | 0.47099 | 0.00430 | 0.00505 | 0.44893 | 0.01111 | 0.00539 | 0.00913 | 0.00859 | 0.03694 | 0.16133 |
| **Dpysl2** | 0.02259 | 0.00459 | 0.00000 | 0.68785 | 0.16003 | 0.14628 | 0.00400 | 0.00726 | 1.14790 | 0.00764 | 0.00321 | 0.00591 | 0.00134 | 0.00000 |
| **Dpysl3** | 0.18355 | 0.00416 | 0.08722 | 0.00349 | 0.00000 | 0.00170 | 0.00414 | 0.01013 | 0.84345 | 0.00725 | 0.00480 | 0.00599 | 1.05895 | 0.90912 |
| **Dpysl4** | 0.01277 | 0.01429 | 3.06169 | 1.13878 | 0.00000 | 0.00000 | 0.00373 | 0.09214 | 0.09123 | 0.00816 | 0.72414 | 0.66108 | 0.00412 | 0.08571 |
| **Dpysl5** | 0.00218 | 0.00448 | 0.66302 | 0.02218 | 0.00085 | 0.61706 | 0.00415 | 0.00462 | 0.00772 | 0.02348 | 0.00542 | 0.00439 | 0.16344 | 0.00170 |
| **Dqx1** | 0.09684 | 0.09752 | 0.27400 | 0.22703 | 0.07870 | 0.20530 | 0.08205 | 0.40099 | 0.17068 | 0.05160 | 0.08499 | 0.70753 | 0.49082 | 0.39452 |
| **Dr1** | 0.99217 | 0.02231 | 3.37856 | 0.00000 | 4.03840 | 4.08404 | 2.65097 | 0.92712 | 8.95897 | 6.77731 | 0.13829 | 2.41489 | 9.68508 | 3.37709 |

| | | | | | | | | | | | | | | |
|---|---|---|---|---|---|---|---|---|---|---|---|---|---|---|
| **Dram1** | 0.00000 | 0.01148 | 0.40572 | 0.16445 | 0.19804 | 0.01476 | 0.00366 | 0.00000 | 0.00967 | 0.00562 | 2.33007 | 0.41003 | 0.02228 | 4.49026 |
| **Dram2** | 1.31323 | 0.03156 | 3.15248 | 2.56102 | 1.13586 | 4.57928 | 4.43572 | 1.12526 | 0.34822 | 0.21737 | 2.65734 | 4.05319 | 0.43878 | 0.70566 |
| **Drap1** | 0.81354 | 0.06983 | 0.07391 | 0.32463 | 0.76405 | 1.37454 | 2.23442 | 2.05275 | 0.09277 | 11.59859 | 0.15984 | 3.01992 | 0.06079 | 0.76682 |
| **Draxin** | 0.02298 | 0.00445 | 0.01276 | 0.01902 | 0.01514 | 0.01361 | 0.00458 | 0.02213 | 0.01932 | 0.00852 | 0.00823 | 0.00799 | 0.23883 | 0.01645 |
| **Drc1** | 0.00000 | 0.00329 | 0.71309 | 0.00000 | 2.69258 | 0.40467 | 0.00293 | 0.00000 | 4.24193 | 0.01980 | 0.34564 | 0.23000 | 0.45797 | 6.54288 |
| **Drc7** | 0.06317 | 0.00430 | 0.01674 | 0.00500 | 0.17798 | 0.14604 | 0.00431 | 0.00529 | 0.08107 | 0.00587 | 0.00659 | 0.00901 | 0.48869 | 0.04017 |
| **Drd3** | 0.14837 | 0.00359 | 0.09513 | 0.64717 | 0.45614 | 0.06295 | 0.00351 | 0.00000 | 0.16506 | 0.00000 | 1.75442 | 0.00804 | 0.11963 | 0.39811 |
| **Drg1** | 19.55227 | 2.98143 | 21.38732 | 15.77269 | 23.06719 | 15.21915 | 3.49719 | 14.39143 | 19.00863 | 0.24908 | 13.70352 | 15.37452 | 6.48916 | 19.34011 |
| **Drg2** | 13.02894 | 0.16303 | 30.50633 | 18.55539 | 25.65159 | 13.35052 | 3.64559 | 5.20493 | 14.40253 | 0.10466 | 13.96282 | 6.95351 | 4.40552 | 23.73364 |
| **Drosha** | 4.80799 | 3.35036 | 1.00548 | 8.55230 | 6.14128 | 4.86198 | 1.32122 | 1.74069 | 6.14088 | 12.87482 | 2.98360 | 2.29042 | 2.94002 | 6.59658 |
| **Dsc2** | 0.00427 | 0.01068 | 0.00859 | 0.00000 | 0.01006 | 0.04683 | 0.56974 | 1.55141 | 0.00647 | 0.02247 | 0.00109 | 0.14833 | 0.12749 | 0.04836 |
| **Dscam** | 0.00658 | 0.00438 | 0.01455 | 0.01620 | 0.01124 | 0.01289 | 0.00427 | 0.00932 | 0.01235 | 0.00651 | 0.00871 | 0.01027 | 0.15523 | 0.01497 |
| **Dscc1** | 8.74588 | 0.02184 | 4.93215 | 8.13637 | 12.44927 | 0.71616 | 0.01193 | 0.00000 | 6.39533 | 0.00956 | 0.00000 | 3.19665 | 3.45373 | 7.34273 |
| **Dscr3** | 0.46212 | 0.01428 | 5.76186 | 3.63436 | 2.71726 | 6.80460 | 2.85040 | 4.20969 | 0.00000 | 0.15848 | 1.09275 | 6.71708 | 0.78644 | 1.04246 |
| **Dse** | 0.03811 | 3.46407 | 0.11485 | 0.00000 | 0.06103 | 0.00734 | 0.00295 | 0.01881 | 0.00772 | 0.00525 | 0.00000 | 0.73726 | 0.19701 | 0.05059 |
| **Dsel** | 0.04004 | 0.00394 | 0.00898 | 0.01184 | 0.00943 | 0.03271 | 0.00417 | 0.10583 | 0.13589 | 0.00697 | 0.00615 | 0.00890 | 0.01038 | 0.01928 |
| **Dsg1b** | 0.07859 | 0.03670 | 0.00678 | 0.20236 | 0.06834 | 0.18239 | 0.02917 | 0.02254 | 0.11385 | 0.02736 | 0.01036 | 0.07691 | 0.28749 | 0.15759 |
| **Dsg2** | 0.00000 | 0.06247 | 0.41696 | 2.50013 | 0.00000 | 6.88278 | 4.84291 | 5.17819 | 0.13612 | 0.13819 | 4.66280 | 0.79574 | 0.00061 | 0.66576 |
| **Dsn1** | 1.28551 | 0.05376 | 5.62060 | 6.46803 | 8.16649 | 1.65588 | 0.01177 | 3.70784 | 14.36627 | 0.00735 | 4.20918 | 0.35842 | 6.21775 | 15.80394 |
| **Dsp** | 0.07829 | 0.07944 | 2.83293 | 0.71434 | 0.00000 | 11.49172 | 2.65805 | 8.39645 | 0.00000 | 0.16691 | 3.88154 | 0.00000 | 0.55251 | 0.00000 |
| **Dst** | 1.39975 | 0.00224 | 2.98970 | 1.13941 | 1.84981 | 1.92072 | 0.64954 | 1.46864 | 0.50800 | 0.06258 | 1.63452 | 1.91909 | 3.40231 | 2.14712 |
| **Dstn** | 14.06460 | 10.84041 | 9.94909 | 9.50345 | 11.38975 | 29.02008 | 10.09171 | 25.60802 | 12.14081 | 10.68800 | 19.09009 | 14.66516 | 6.41717 | 14.28294 |
| **Dstyk** | 0.00000 | 0.00362 | 0.00000 | 0.28996 | 0.22339 | 0.58354 | 0.01409 | 0.83645 | 2.18634 | 0.00611 | 0.00211 | 0.00142 | 0.77968 | 2.16344 |
| **Dtd1** | 1.64553 | 0.01620 | 3.88777 | 0.80628 | 1.39193 | 0.00000 | 0.01269 | 0.99274 | 0.01930 | 6.37975 | 2.07214 | 1.45440 | 0.25319 | 1.88322 |
| **Dtd2** | 0.00000 | 0.00267 | 0.74680 | 0.81289 | 0.00000 | 1.23421 | 1.18663 | 0.00913 | 0.00822 | 4.19680 | 0.25817 | 0.68778 | 0.10099 | 0.11220 |
| **Dtl** | 14.33257 | 0.03631 | 11.85520 | 2.88749 | 20.42992 | 4.87366 | 0.05104 | 0.15584 | 21.85115 | 0.06101 | 10.50229 | 5.58650 | 11.95984 | 17.57801 |
| **Dtnb** | 3.87329 | 0.00261 | 3.94029 | 4.45051 | 2.40323 | 0.80403 | 0.00346 | 0.42508 | 1.57432 | 0.00000 | 0.00000 | 1.21702 | 1.84422 | 3.36179 |
| **Dtnbp1** | 2.09474 | 0.00197 | 1.26623 | 4.20677 | 4.41156 | 2.45410 | 0.00311 | 0.01753 | 2.76922 | 0.00373 | 2.75955 | 1.98561 | 0.77658 | 2.42176 |
| **Dtwd1** | 1.07064 | 0.03098 | 0.33771 | 0.32477 | 1.33954 | 0.82954 | 0.08791 | 0.00000 | 0.01717 | 0.00000 | 4.01001 | 3.92981 | 0.00000 | 1.79224 |
| **Dtwd2** | 0.75098 | 0.00351 | 0.60459 | 0.52897 | 1.75618 | 0.60527 | 0.00341 | 0.00445 | 3.49679 | 0.02375 | 0.13135 | 0.56165 | 2.16430 | 4.34301 |
| **Dtx2** | 0.15588 | 0.01975 | 7.58944 | 4.93728 | 11.71447 | 0.07851 | 0.05638 | 1.99033 | 0.00000 | 0.01146 | 0.01964 | 0.07780 | 4.73312 | 0.00000 |
| **Dtx3** | 0.00558 | 0.01129 | 0.00923 | 0.00000 | 0.22272 | 0.01524 | 0.04119 | 0.00643 | 0.01185 | 0.00598 | 0.00425 | 0.55691 | 0.40227 | 0.00666 |
| **Dtx3l** | 0.01791 | 0.00453 | 0.01656 | 0.11615 | 0.42287 | 0.10938 | 0.03613 | 0.09595 | 0.08603 | 0.04212 | 0.03015 | 0.05087 | 0.54875 | 1.04407 |
| **Dtx4** | 0.10718 | 0.00450 | 0.00640 | 0.00384 | 0.03152 | 0.16316 | 0.00415 | 0.00506 | 0.01511 | 0.00568 | 0.00620 | 0.00726 | 1.17223 | 0.33040 |
| **Dtymk** | 6.83844 | 1.12986 | 9.96405 | 16.28684 | 15.47866 | 9.52031 | 4.54915 | 8.51026 | 0.95754 | 8.94086 | 0.48990 | 10.76847 | 0.83843 | 1.97385 |
| **Dubr** | 0.04341 | 0.00445 | 0.01024 | 0.00333 | 0.01275 | 0.06244 | 0.00424 | 0.00693 | 0.01501 | 0.00549 | 0.29098 | 0.00687 | 0.00638 | 0.00687 |
| **Duox1** | 0.00625 | 0.01210 | 0.01187 | 0.00477 | 0.01093 | 0.01369 | 0.00415 | 0.00840 | 0.01147 | 0.00663 | 0.00450 | 0.55417 | 0.09470 | 0.01986 |
| **Duoxa2** | 0.00071 | 0.00312 | 0.05419 | 0.52993 | 0.00476 | 0.01881 | 0.00382 | 0.04580 | 0.01168 | 0.00493 | 0.00266 | 1.14184 | 0.07980 | 0.00000 |
| **Dus1l** | 6.06023 | 6.11699 | 11.77353 | 10.39331 | 8.39702 | 6.52125 | 2.57164 | 6.31993 | 0.64872 | 0.06697 | 1.45996 | 6.64178 | 1.76971 | 5.07655 |
| **Dus2** | 1.95950 | 0.01471 | 3.33829 | 1.33183 | 1.41668 | 5.41060 | 0.00811 | 0.02353 | 0.48875 | 0.02597 | 0.27102 | 4.35587 | 2.18293 | 2.84214 |
| **Dus3l** | 4.04947 | 5.89336 | 14.67935 | 9.09441 | 8.30615 | 0.65527 | 0.02887 | 4.43747 | 0.00448 | 0.01873 | 2.89173 | 6.98648 | 2.44460 | 0.58290 |
| **Dus4l** | 0.30829 | 0.00446 | 0.23370 | 0.00428 | 0.00000 | 0.00000 | 0.00388 | 0.01446 | 0.03296 | 0.00427 | 1.29970 | 0.02440 | 0.01749 | 0.00006 |
| **Dusp1** | 0.02910 | 0.04784 | 0.28012 | 0.00000 | 0.05850 | 1.65773 | 2.72491 | 3.00924 | 0.02346 | 0.01112 | 0.00000 | 1.87480 | 4.02459 | 0.02738 |
| **Dusp10** | 2.06291 | 0.01918 | 0.85812 | 0.82574 | 3.87995 | 2.75218 | 0.00312 | 2.00142 | 2.01455 | 0.19684 | 2.45637 | 0.30300 | 2.95758 | 7.14831 |
| **Dusp11** | 7.16666 | 0.06919 | 11.20543 | 10.34741 | 6.88563 | 6.61853 | 4.70679 | 3.88122 | 8.77024 | 0.20858 | 6.04535 | 6.91941 | 1.65926 | 2.13900 |
| **Dusp12** | 11.11700 | 0.03766 | 8.08296 | 7.95164 | 15.72895 | 4.21951 | 2.52635 | 1.42277 | 5.58525 | 11.31765 | 1.79341 | 3.86559 | 1.59534 | 4.20093 |
| **Dusp14** | 5.85572 | 2.02166 | 16.93951 | 9.32527 | 7.59227 | 25.62130 | 8.28038 | 12.29353 | 4.57501 | 1.15447 | 11.57619 | 0.00019 | 16.44207 | 9.88952 |
| **Dusp16** | 3.09663 | 0.00280 | 0.10770 | 0.20418 | 1.11837 | 1.12217 | 1.12213 | 1.65758 | 4.50545 | 1.05717 | 1.05098 | 0.21328 | 4.39658 | 6.90611 |
| **Dusp18** | 1.00753 | 0.00379 | 0.41515 | 0.00000 | 1.25399 | 0.79402 | 0.79667 | 0.47233 | 0.44473 | 0.02971 | 0.36878 | 0.61953 | 0.02837 | 0.41344 |
| **Dusp19** | 0.07781 | 0.03668 | 0.87176 | 0.64260 | 1.05455 | 1.38466 | 0.00392 | 1.61724 | 0.64179 | 0.74766 | 1.48998 | 0.31819 | 0.00000 | 0.88893 |
| **Dusp2** | 0.67926 | 0.00428 | 0.13143 | 0.00000 | 0.87230 | 0.00052 | 0.00401 | 0.01084 | 0.29600 | 0.00892 | 0.00478 | 0.00521 | 0.61959 | 0.15893 |
| **Dusp22** | 0.96717 | 0.00376 | 0.29036 | 0.70309 | 0.45830 | 0.14938 | 0.04418 | 0.14638 | 0.11677 | 0.04508 | 0.01226 | 0.04398 | 3.81582 | 0.81940 |
| **Dusp23** | 0.04298 | 0.00393 | 0.00593 | 0.19604 | 0.01560 | 0.02560 | 0.00392 | 0.04256 | 0.01102 | 0.00652 | 0.00545 | 0.04240 | 0.10602 | 0.00000 |
| **Dusp26** | 0.10677 | 0.00411 | 3.29996 | 0.16550 | 4.33692 | 0.00000 | 0.00322 | 0.00553 | 0.78416 | 0.00899 | 0.00120 | 0.01299 | 0.10370 | 1.07294 |
| **Dusp28** | 0.01477 | 0.00542 | 0.04223 | 0.00000 | 0.27746 | 0.00346 | 0.00434 | 0.16155 | 0.01210 | 0.00779 | 0.00784 | 0.00925 | 0.08248 | 0.15389 |

| | | | | | | | | | | | | | | |
|---|---|---|---|---|---|---|---|---|---|---|---|---|---|---|
| **Dusp3** | 0.01686 | 0.02068 | 1.09109 | 1.67919 | 0.78059 | 2.35116 | 3.89000 | 1.53004 | 0.07557 | 0.12552 | 0.02492 | 0.00000 | 0.27139 | 0.71458 |
| **Dusp4** | 0.77364 | 0.31516 | 0.19540 | 6.38569 | 0.00000 | 10.11379 | 1.10873 | 3.33929 | 0.00177 | 0.02563 | 5.61616 | 4.37174 | 0.26467 | 0.00000 |
| **Dusp5** | 0.03102 | 0.00458 | 0.45061 | 0.00000 | 0.05245 | 0.00379 | 0.00428 | 0.02095 | 0.01010 | 0.00674 | 0.00557 | 0.08367 | 0.00396 | 0.08875 |
| **Dusp6** | 0.00259 | 0.05176 | 0.05305 | 0.00000 | 0.18453 | 0.04153 | 2.17481 | 3.20042 | 0.01326 | 0.05941 | 0.01112 | 0.02460 | 0.36972 | 1.33140 |
| **Dusp7** | 0.00000 | 0.00332 | 0.65031 | 0.17447 | 0.36292 | 0.20157 | 0.00349 | 0.01108 | 0.07445 | 0.00420 | 0.07758 | 0.65955 | 1.55228 | 0.65915 |
| **Dusp8** | 0.14779 | 0.00445 | 0.01416 | 0.00425 | 0.08334 | 0.00609 | 0.59355 | 0.01553 | 0.06394 | 0.02043 | 0.00588 | 0.01471 | 0.01589 | 0.00002 |
| **Dusp9** | 0.00000 | 0.00465 | 1.55393 | 1.88890 | 0.39962 | 0.92947 | 0.00401 | 0.08502 | 0.30947 | 0.00553 | 0.83700 | 0.05608 | 0.00437 | 0.40723 |
| **Dut** | 0.70401 | 3.35748 | 0.08843 | 19.95159 | 9.67499 | 10.37935 | 2.90501 | 1.28896 | 2.66579 | 10.84022 | 0.77203 | 9.39735 | 10.43529 | 0.44674 |
| **Dvl1** | 0.00000 | 0.01819 | 3.16857 | 6.75540 | 1.68356 | 0.84260 | 2.03777 | 1.59376 | 0.00282 | 0.06026 | 0.02119 | 0.83945 | 0.81483 | 1.51456 |
| **Dvl2** | 0.01162 | 0.01085 | 0.33673 | 2.76496 | 1.31944 | 0.52364 | 0.03525 | 0.64729 | 0.00209 | 0.02748 | 0.73075 | 1.03974 | 1.34530 | 0.05150 |
| **Dvl3** | 0.00000 | 0.00458 | 0.92370 | 0.92663 | 0.46260 | 0.16918 | 0.00457 | 0.37246 | 0.87552 | 0.01015 | 0.00000 | 0.19604 | 0.01089 | 0.42352 |
| **Dxo** | 2.27875 | 0.05489 | 3.26647 | 6.97164 | 1.36264 | 0.56466 | 0.02445 | 2.77976 | 0.05375 | 0.06496 | 1.98055 | 4.87678 | 5.33983 | 0.00000 |
| **Dydc1** | 0.00462 | 0.00425 | 0.02893 | 0.07577 | 0.03832 | 0.01016 | 0.00419 | 0.00755 | 0.00804 | 0.00597 | 0.00685 | 0.00781 | 0.25029 | 0.27488 |
| **Dydc2** | 0.48029 | 0.00401 | 0.14473 | 0.31452 | 0.28977 | 0.00000 | 0.00396 | 0.01050 | 0.70961 | 0.00620 | 0.00335 | 0.00536 | 0.05832 | 1.65785 |
| **Dym** | 5.23895 | 0.05279 | 7.68845 | 3.50839 | 3.05883 | 3.50560 | 0.06609 | 3.84504 | 0.00000 | 0.01768 | 0.52650 | 5.16061 | 2.92972 | 2.43749 |
| **Dynap** | 0.40515 | 0.00442 | 0.34484 | 0.00000 | 0.48217 | 0.00000 | 0.00405 | 0.00977 | 1.59933 | 0.00716 | 0.00323 | 0.00530 | 0.00153 | 0.64307 |
| **Dync1h1** | 3.89220 | 8.56219 | 12.10356 | 12.15740 | 2.29948 | 14.09221 | 2.68243 | 7.78529 | 0.09298 | 0.23730 | 9.46261 | 12.83263 | 1.79771 | 3.27001 |
| **Dync1i1** | 0.00712 | 0.00435 | 0.00707 | 0.05752 | 0.35748 | 0.01170 | 0.00474 | 0.01168 | 0.02174 | 0.00720 | 0.00745 | 0.01065 | 0.00760 | 0.02786 |
| **Dync1i2** | 6.40222 | 0.02730 | 4.53544 | 1.76095 | 3.99616 | 6.62137 | 0.96806 | 1.15143 | 4.53258 | 0.00554 | 9.22932 | 4.12919 | 2.31870 | 8.34123 |
| **Dync1li1** | 1.73359 | 0.01662 | 3.29950 | 1.77099 | 3.90380 | 3.17448 | 0.53418 | 0.36882 | 2.34304 | 0.00588 | 0.84319 | 1.96544 | 0.04635 | 3.51855 |
| **Dync1li2** | 3.10028 | 0.01635 | 5.16394 | 2.45233 | 5.05568 | 6.34629 | 0.00135 | 1.15063 | 7.24735 | 0.03532 | 5.47366 | 1.54433 | 0.42384 | 6.10839 |
| **Dync2h1** | 1.21541 | 0.05166 | 1.39702 | 0.00000 | 1.36017 | 0.00000 | 0.00271 | 0.10339 | 1.83528 | 0.01365 | 1.88411 | 2.80005 | 0.00000 | 1.67683 |
| **Dync2li1** | 0.00668 | 0.00384 | 0.01104 | 0.23072 | 0.29654 | 0.00378 | 0.00398 | 0.00377 | 0.01015 | 0.00523 | 0.01602 | 0.31030 | 0.00454 | 0.01472 |
| **Dynll1** | 12.07520 | 8.50017 | 5.61065 | 13.64476 | 28.61440 | 20.46721 | 12.64717 | 23.25984 | 2.64079 | 19.99351 | 4.75268 | 24.59668 | 20.52535 | 4.37998 |
| **Dynll2** | 0.11374 | 0.00283 | 1.44035 | 1.52515 | 7.90537 | 1.52386 | 0.00921 | 1.18946 | 2.89989 | 0.01584 | 0.17820 | 1.38175 | 4.09416 | 1.59161 |
| **Dynlrb1** | 3.91810 | 0.17770 | 0.75647 | 7.55810 | 11.44473 | 10.15264 | 4.55271 | 9.42964 | 2.30380 | 10.55753 | 2.59930 | 7.73857 | 4.38396 | 3.88089 |
| **Dynlrb2** | 0.00880 | 0.00447 | 0.00595 | 0.00976 | 0.46679 | 0.01425 | 0.00426 | 0.01027 | 0.01521 | 0.00724 | 0.00899 | 0.00839 | 0.00489 | 0.00595 |
| **Dynlt1a** | 0.71289 | 1.39453 | 0.65496 | 2.54150 | 0.96001 | 1.42304 | 5.25255 | 1.95640 | 2.20998 | 5.41687 | 2.19160 | 5.56696 | 2.18455 | 2.00518 |
| **Dynlt1b** | 0.62893 | 1.47917 | 0.43551 | 2.45545 | 0.81381 | 1.69960 | 4.00949 | 2.02186 | 1.14477 | 6.48648 | 2.35781 | 5.92671 | 0.17434 | 1.72568 |
| **Dynlt1f** | 0.01971 | 0.00488 | 0.02895 | 0.10325 | 0.03360 | 0.02961 | 0.45760 | 0.02620 | 0.01011 | 0.01865 | 0.16714 | 0.01545 | 0.00908 | 0.00944 |
| **Dynlt3** | 2.28531 | 1.21302 | 0.10291 | 2.76126 | 1.37641 | 4.81184 | 2.81367 | 1.97109 | 4.06761 | 0.04582 | 4.16477 | 3.60578 | 0.77666 | 0.49567 |
| **Dyrk1a** | 6.54128 | 0.03881 | 8.27765 | 1.40313 | 15.16647 | 6.07949 | 1.27386 | 3.88440 | 2.21969 | 0.02829 | 6.11363 | 2.94936 | 9.75078 | 14.15340 |
| **Dyrk1b** | 0.01385 | 0.00428 | 0.06639 | 0.28290 | 0.03194 | 0.01895 | 0.00421 | 0.00596 | 0.00992 | 0.00597 | 0.00589 | 0.00743 | 0.05798 | 0.00238 |
| **Dyrk3** | 3.06479 | 0.05698 | 5.24225 | 4.50227 | 2.77874 | 4.17306 | 1.21515 | 1.89070 | 1.87751 | 0.07596 | 5.03034 | 1.15835 | 5.58906 | 6.16631 |
| **Dyrk4** | 0.68399 | 0.00476 | 0.14218 | 0.04773 | 0.37123 | 0.00000 | 0.00424 | 0.02035 | 0.06382 | 0.00739 | 0.00505 | 0.01026 | 0.03612 | 0.97062 |
| **Dysf** | 3.34623 | 0.01900 | 5.52864 | 1.48345 | 3.78814 | 0.50505 | 0.00221 | 3.44670 | 5.23184 | 0.00000 | 0.00000 | 0.38962 | 1.14733 | 4.23675 |
| **Dyx1c1** | 0.52210 | 0.00396 | 0.11730 | 0.00856 | 0.02318 | 0.00000 | 0.00392 | 0.00631 | 1.03375 | 0.00621 | 0.86795 | 0.04458 | 0.00637 | 0.13855 |
| **Dzank1** | 0.00000 | 0.02138 | 0.34630 | 0.05128 | 0.26145 | 0.05279 | 0.01091 | 0.02145 | 2.09925 | 0.00790 | 0.02469 | 0.01114 | 1.41176 | 1.96151 |
| **Dzip1** | 0.02775 | 0.00410 | 0.00000 | 0.41403 | 0.49727 | 0.00000 | 0.00490 | 0.32400 | 1.24131 | 0.00678 | 0.00000 | 0.38552 | 0.00000 | 0.00000 |
| **Dzip3** | 0.13470 | 0.01081 | 0.00000 | 0.00000 | 0.18141 | 0.75178 | 0.00335 | 0.00398 | 1.31184 | 0.00655 | 0.00000 | 1.65526 | 0.93545 | 1.52984 |
| **E2f1** | 1.33723 | 4.98000 | 3.20840 | 1.82481 | 7.27551 | 4.46934 | 0.23640 | 0.20703 | 0.10399 | 0.02564 | 1.05231 | 2.69500 | 7.13750 | 10.30150 |
| **E2f2** | 0.85467 | 0.00422 | 0.77462 | 0.64274 | 0.49480 | 0.00000 | 0.00438 | 0.06714 | 0.76950 | 0.00614 | 0.00167 | 0.15956 | 2.33096 | 1.63370 |
| **E2f3** | 0.03444 | 0.00463 | 0.61963 | 0.10841 | 0.06334 | 0.26645 | 0.00487 | 0.26457 | 0.05016 | 0.01928 | 0.18629 | 0.14963 | 0.04674 | 0.16036 |
| **E2f4** | 5.90119 | 1.00391 | 9.45395 | 9.77493 | 7.38732 | 7.67788 | 0.45382 | 2.88859 | 1.13489 | 0.06885 | 3.17507 | 5.96369 | 6.63573 | 10.32456 |
| **E2f5** | 1.15908 | 0.01894 | 0.14371 | 0.00000 | 1.71938 | 0.29391 | 0.00331 | 0.00266 | 3.47415 | 0.00439 | 0.30835 | 1.68567 | 4.31449 | 2.73622 |
| **E2f6** | 0.69191 | 0.01680 | 3.59784 | 0.00000 | 0.51304 | 5.61243 | 0.00212 | 0.00494 | 3.42628 | 0.00000 | 1.11676 | 3.91159 | 0.00000 | 2.88794 |
| **E2f7** | 0.00000 | 0.00392 | 0.20934 | 0.35947 | 0.04020 | 0.48688 | 0.00409 | 0.00288 | 0.00251 | 0.01596 | 0.08806 | 0.10557 | 0.51913 | 0.31579 |
| **E2f8** | 0.28185 | 0.01214 | 0.23047 | 0.28267 | 0.83915 | 2.06914 | 0.00381 | 3.64346 | 0.00450 | 0.00523 | 0.77383 | 0.00000 | 0.09546 | 0.96078 |
| **E4f1** | 2.37648 | 7.49125 | 9.02281 | 7.22262 | 5.40064 | 2.35152 | 3.28188 | 1.19869 | 0.22523 | 0.07810 | 1.09687 | 0.62887 | 4.97845 | 0.66148 |
| **Eaf1** | 3.71228 | 0.02940 | 8.50256 | 2.19616 | 3.37380 | 2.86472 | 2.20201 | 1.75404 | 5.00999 | 0.05346 | 3.36540 | 2.83578 | 5.13852 | 5.85654 |
| **Eaf2** | 0.56821 | 0.00444 | 0.84366 | 0.69534 | 1.07757 | 0.12736 | 0.12101 | 0.05366 | 1.32908 | 0.00476 | 0.22983 | 0.04889 | 1.86992 | 0.31451 |
| **Eapp** | 7.79674 | 0.16203 | 9.55995 | 10.82577 | 7.46862 | 4.68796 | 2.71848 | 4.85248 | 6.32343 | 12.59180 | 12.08015 | 5.61964 | 3.54564 | 6.53111 |
| **Ears2** | 0.10630 | 0.09701 | 3.10300 | 5.85845 | 2.90728 | 5.07485 | 0.11417 | 0.74861 | 0.70658 | 0.19095 | 0.31380 | 3.71626 | 6.16870 | 2.92501 |
| **Ebag9** | 1.16215 | 0.01100 | 0.45316 | 0.00000 | 0.02917 | 1.15749 | 2.49295 | 1.89961 | 0.03716 | 0.10988 | 0.64370 | 1.23778 | 0.78253 | 0.89884 |
| **Ebf3** | 0.00116 | 0.00427 | 0.00966 | 0.01300 | 0.00526 | 0.01216 | 0.00446 | 0.01053 | 0.00838 | 0.01354 | 0.00778 | 0.04021 | 0.44633 | 0.71267 |

| | | | | | | | | | | | | | | |
|---|---|---|---|---|---|---|---|---|---|---|---|---|---|---|
| **Ebf4** | 0.01130 | 0.00405 | 0.02500 | 0.10168 | 0.01063 | 0.01704 | 0.00404 | 0.09942 | 0.01151 | 0.00751 | 0.00605 | 0.00667 | 0.00669 | 0.01123 |
| **Ebna1bp2** | 17.66995 | 5.82524 | 26.11268 | 13.16706 | 21.91006 | 19.65981 | 5.88461 | 5.96977 | 7.49501 | 0.64330 | 15.42056 | 12.28597 | 18.89518 | 20.36611 |
| **Ebp** | 5.12787 | 6.76497 | 9.48908 | 6.46753 | 6.10657 | 13.88671 | 0.00924 | 6.47186 | 1.87673 | 1.29766 | 4.07939 | 7.05261 | 1.30814 | 13.86803 |
| **Ebpl** | 0.32111 | 0.01100 | 0.00000 | 0.85907 | 0.79079 | 0.40300 | 0.74206 | 0.01586 | 0.66945 | 0.03066 | 0.00306 | 0.72808 | 0.08757 | 0.16025 |
| **Ecd** | 7.39651 | 0.12694 | 10.21262 | 3.62771 | 9.61641 | 6.23929 | 0.19989 | 3.88890 | 6.30477 | 14.81577 | 8.74301 | 4.64957 | 0.70793 | 9.68981 |
| **Ece1** | 0.08834 | 0.00435 | 0.07688 | 0.00567 | 0.23711 | 0.01904 | 0.00442 | 0.01531 | 0.23888 | 0.01698 | 0.00783 | 0.09311 | 0.01305 | 0.44651 |
| **Ece2** | 3.63788 | 0.02506 | 2.91941 | 1.50458 | 7.47602 | 1.95967 | 0.01589 | 1.55647 | 2.30826 | 0.01926 | 0.00003 | 1.06719 | 3.31941 | 4.04823 |
| **Ecel1** | 0.00473 | 0.00401 | 0.00872 | 1.14923 | 0.00517 | 0.00000 | 0.00403 | 0.00694 | 0.02642 | 0.00528 | 0.01793 | 0.01187 | 0.00633 | 0.00380 |
| **Ech1** | 0.32152 | 1.59125 | 0.59792 | 3.51473 | 0.90317 | 9.61246 | 3.83037 | 7.35548 | 0.06853 | 1.92682 | 9.05175 | 5.45385 | 0.00000 | 0.00000 |
| **Echdc1** | 0.00210 | 0.00434 | 0.24090 | 0.05143 | 0.00000 | 0.17360 | 0.00420 | 0.00248 | 0.00978 | 0.00556 | 0.26744 | 0.02063 | 0.00466 | 0.41593 |
| **Echdc2** | 14.32656 | 1.28448 | 31.31656 | 30.17370 | 21.33100 | 18.29276 | 9.02605 | 23.64283 | 2.31791 | 9.54385 | 10.18866 | 18.87216 | 5.68076 | 2.68904 |
| **Echdc3** | 0.00105 | 0.00468 | 0.11973 | 0.00000 | 0.01849 | 0.12980 | 0.02534 | 0.30104 | 0.00702 | 0.00691 | 0.00585 | 0.01036 | 1.07124 | 0.01376 |
| **Echs1** | 0.00000 | 0.52402 | 2.83959 | 11.60735 | 0.79475 | 8.15443 | 4.20566 | 8.58655 | 0.00000 | 0.24209 | 10.88950 | 10.93598 | 0.75607 | 2.33489 |
| **Eci1** | 5.25690 | 0.00903 | 2.14562 | 4.71215 | 7.76417 | 1.33384 | 0.16182 | 2.06793 | 3.14169 | 0.04906 | 2.17823 | 0.34662 | 6.57258 | 3.80664 |
| **Eci2** | 2.95804 | 0.10099 | 4.31411 | 6.07714 | 2.84221 | 1.79036 | 5.30899 | 3.43172 | 1.55539 | 0.07360 | 5.72135 | 4.68867 | 0.64153 | 3.56969 |
| **Eci3** | 0.02855 | 0.00443 | 0.07711 | 0.11323 | 0.04379 | 0.00000 | 0.13732 | 0.05024 | 0.02626 | 0.01650 | 0.42034 | 0.02883 | 0.00836 | 0.07672 |
| **Ecm1** | 0.14325 | 0.00397 | 1.82679 | 0.00000 | 0.51204 | 0.22816 | 0.00422 | 0.93815 | 0.79632 | 0.00520 | 0.00000 | 1.16153 | 0.29380 | 0.19991 |
| **Ecm2** | 0.06288 | 0.05183 | 0.29863 | 0.12451 | 0.25890 | 0.18350 | 0.03428 | 0.05438 | 0.33876 | 0.21359 | 0.01404 | 0.05479 | 0.55483 | 0.29775 |
| **Ecscr** | 0.01100 | 0.00427 | 0.28231 | 0.00093 | 0.01030 | 0.00648 | 0.00432 | 0.01025 | 0.01474 | 0.00622 | 0.00794 | 0.00887 | 0.41112 | 0.09425 |
| **Ecsit** | 6.10868 | 0.04883 | 10.02311 | 4.42573 | 12.41152 | 7.12638 | 2.11537 | 3.56038 | 2.01443 | 0.05789 | 4.42292 | 2.83479 | 2.60614 | 14.06634 |
| **Ect2** | 0.56617 | 3.94733 | 7.75218 | 1.11058 | 10.75840 | 4.31330 | 2.90900 | 0.93687 | 25.29610 | 0.05300 | 8.64280 | 1.47580 | 9.54921 | 12.47794 |
| **Eda** | 5.36448 | 0.00260 | 5.31550 | 0.82678 | 6.97797 | 1.48121 | 0.01017 | 0.59208 | 11.45414 | 0.00026 | 0.15433 | 0.17770 | 5.41787 | 11.30639 |
| **Eda2r** | 0.69553 | 0.00338 | 0.82785 | 0.00000 | 0.00000 | 0.04278 | 0.01000 | 0.00266 | 0.00743 | 0.01638 | 1.43748 | 0.52808 | 0.02318 | 0.00000 |
| **Edar** | 0.00659 | 0.00425 | 0.00945 | 0.13350 | 0.00866 | 0.00813 | 0.00404 | 0.00683 | 0.01977 | 0.00667 | 0.00682 | 0.01514 | 0.01924 | 0.04269 |
| **Edaradd** | 0.04061 | 0.00437 | 0.23798 | 0.00620 | 0.21433 | 0.00323 | 0.00424 | 0.01050 | 0.02785 | 0.00902 | 0.00609 | 0.00691 | 0.00760 | 1.57887 |
| **Edc3** | 0.56317 | 0.05543 | 4.19502 | 2.90786 | 4.68799 | 3.44578 | 1.63010 | 1.67539 | 0.15824 | 0.06691 | 0.02366 | 5.64036 | 0.78950 | 5.48663 |
| **Edc4** | 4.14297 | 0.03624 | 10.87168 | 4.67687 | 7.63535 | 5.11665 | 4.10273 | 2.68158 | 0.26987 | 0.14337 | 1.65469 | 9.27533 | 0.85244 | 2.49739 |
| **Edem1** | 1.26654 | 0.03136 | 1.25951 | 0.63019 | 1.19506 | 3.45858 | 0.00188 | 3.06964 | 1.45430 | 0.01180 | 4.35405 | 2.76883 | 0.54098 | 2.14426 |
| **Edem2** | 0.04341 | 0.05144 | 4.09820 | 4.69125 | 4.82433 | 0.17314 | 0.00163 | 3.21623 | 0.00250 | 0.00541 | 0.15407 | 4.41124 | 0.09375 | 0.00000 |
| **Edem3** | 1.00406 | 0.00357 | 1.58438 | 0.59606 | 0.00000 | 1.75478 | 0.00340 | 0.11233 | 4.86584 | 0.01108 | 0.21317 | 0.30317 | 1.41992 | 1.69937 |
| **Edf1** | 3.39370 | 1.48047 | 1.48781 | 20.35555 | 21.02663 | 7.04094 | 4.67744 | 10.40284 | 0.02618 | 6.31146 | 0.43981 | 6.35500 | 2.14680 | 1.31265 |
| **Edn2** | 0.00841 | 0.00456 | 0.21662 | 0.03215 | 0.00725 | 0.00998 | 0.00458 | 0.02509 | 0.01154 | 0.00902 | 0.00925 | 0.01529 | 0.00675 | 0.01116 |
| **Edn3** | 0.00430 | 0.00447 | 0.00679 | 0.01225 | 0.26859 | 0.01202 | 0.00435 | 0.00983 | 0.00666 | 0.00632 | 0.00816 | 0.00835 | 0.00607 | 0.43264 |
| **Ednra** | 0.01136 | 0.00422 | 0.00997 | 0.21459 | 0.00922 | 0.00933 | 0.00415 | 0.00676 | 0.01335 | 0.00582 | 0.00901 | 0.01388 | 0.00673 | 0.00952 |
| **Ednrb** | 0.02046 | 0.02242 | 0.01993 | 0.00000 | 0.02000 | 0.03267 | 0.00411 | 0.02742 | 0.09231 | 0.00623 | 0.00724 | 0.12412 | 0.02276 | 0.02984 |
| **Edrf1** | 0.74312 | 0.01930 | 1.71176 | 0.81913 | 0.12442 | 1.11348 | 0.02449 | 0.61444 | 2.71127 | 0.00212 | 0.19559 | 1.11547 | 1.20817 | 3.37257 |
| **Eea1** | 2.39734 | 0.00878 | 0.31605 | 0.16185 | 0.00000 | 1.21879 | 0.00212 | 1.48754 | 6.26015 | 0.00000 | 3.73494 | 0.00384 | 4.88119 | 4.51975 |
| **Eed** | 10.84938 | 0.58176 | 18.18823 | 7.04583 | 12.14654 | 15.02934 | 5.96525 | 3.88890 | 13.72006 | 0.22807 | 14.50979 | 15.88951 | 6.70882 | 14.39265 |
| **Eef1a1** | 47.96195 | 22.76397 | 68.31038 | 69.14379 | 54.16653 | 70.21443 | 20.11206 | 54.70809 | 56.91273 | 34.88571 | 44.53726 | 52.44065 | 16.58097 | 53.19021 |
| **Eef1akmt1** | 10.50481 | 0.04671 | 9.87006 | 7.33867 | 15.03327 | 9.30994 | 2.51983 | 2.19579 | 5.57793 | 0.11770 | 5.09618 | 3.51106 | 5.55615 | 11.25696 |
| **Eef1b2** | 34.83509 | 12.66953 | 37.42719 | 46.31012 | 39.34091 | 42.25375 | 15.36255 | 33.78005 | 9.04695 | 29.80839 | 26.69878 | 35.84125 | 10.21284 | 23.89412 |
| **Eef1d** | 21.43500 | 0.51302 | 17.14686 | 23.72361 | 13.96607 | 23.26521 | 4.75681 | 14.42129 | 7.23944 | 9.04234 | 13.98467 | 16.05389 | 11.22057 | 13.40661 |
| **Eef1e1** | 15.07504 | 0.24271 | 21.88988 | 15.16665 | 19.89841 | 21.59548 | 8.12661 | 11.87171 | 18.24087 | 18.13358 | 11.26245 | 12.53937 | 2.96338 | 19.71625 |
| **Eef1g** | 28.18252 | 17.86119 | 53.89182 | 51.14669 | 38.15166 | 45.46603 | 14.88831 | 38.60668 | 28.51286 | 24.80362 | 31.11943 | 36.17222 | 4.48183 | 31.97074 |
| **Eef2** | 25.64403 | 12.08087 | 47.43850 | 47.49759 | 37.27740 | 36.82684 | 9.40250 | 29.15356 | 3.83234 | 19.93387 | 27.83816 | 29.68705 | 12.07162 | 35.38013 |
| **Eef2k** | 0.05972 | 0.01815 | 0.30035 | 0.36206 | 0.40252 | 0.84680 | 0.00363 | 0.05763 | 0.66950 | 0.00673 | 0.01353 | 0.17842 | 0.00564 | 1.06048 |
| **Eef2kmt** | 0.44933 | 0.01674 | 2.59470 | 2.76975 | 0.37303 | 3.34566 | 0.00225 | 0.00820 | 0.15871 | 0.05472 | 2.48152 | 2.97180 | 0.33745 | 0.16239 |
| **Eefsec** | 0.53211 | 0.01570 | 0.76346 | 0.69326 | 2.95808 | 1.81418 | 2.06689 | 2.56944 | 0.00000 | 0.07545 | 0.56349 | 4.00975 | 0.00000 | 0.06977 |
| **Eepd1** | 0.23415 | 0.02072 | 0.67763 | 1.03058 | 0.15697 | 0.00000 | 0.00370 | 2.60778 | 0.10771 | 0.01199 | 0.00062 | 0.00530 | 2.06841 | 0.00000 |
| **Efcab10** | 0.00863 | 0.01587 | 0.01341 | 0.00000 | 0.01240 | 0.01188 | 0.00420 | 0.38871 | 0.01189 | 0.01097 | 0.00526 | 0.17986 | 0.00774 | 0.01158 |
| **Efcab11** | 0.00293 | 0.00377 | 0.00000 | 0.27396 | 0.00316 | 0.10120 | 0.00491 | 0.14140 | 0.00664 | 0.00456 | 0.77084 | 0.67483 | 0.48026 | 0.11832 |
| **Efcab12** | 0.00000 | 0.01180 | 0.07305 | 0.55824 | 0.44098 | 0.29084 | 0.00359 | 0.00302 | 0.87234 | 0.00576 | 0.00000 | 0.65254 | 0.00894 | 0.00000 |
| **Efcab14** | 1.47789 | 0.00798 | 0.47133 | 0.75873 | 0.00000 | 2.58070 | 0.15895 | 0.00000 | 0.64605 | 0.01168 | 2.30396 | 3.78181 | 0.36457 | 4.71973 |
| **Efcab2** | 0.00118 | 0.00406 | 0.00569 | 0.00000 | 0.00981 | 0.41426 | 0.00384 | 0.50194 | 0.09556 | 0.01861 | 0.00000 | 0.78711 | 0.16668 | 0.03559 |
| **Efcab3** | 0.00001 | 0.00432 | 0.02399 | 0.03342 | 0.77477 | 0.00138 | 0.00404 | 0.00743 | 1.54619 | 0.01699 | 0.00431 | 0.00921 | 0.11131 | 0.61783 |

| | | | | | | | | | | | | | | |
|---|---|---|---|---|---|---|---|---|---|---|---|---|---|---|
| **Efcab5** | 0.00000 | 0.01361 | 0.36681 | 0.36167 | 0.11237 | 1.48524 | 0.00387 | 0.70401 | 0.49415 | 0.01373 | 0.00229 | 0.00000 | 1.69015 | 0.00000 |
| **Efcab6** | 0.01636 | 0.00433 | 0.00689 | 0.01250 | 0.28752 | 0.01321 | 0.00411 | 0.00882 | 0.00967 | 0.00696 | 0.00888 | 0.00801 | 0.47130 | 0.00291 |
| **Efcab7** | 0.03820 | 0.00448 | 0.03666 | 0.00383 | 0.01670 | 0.04387 | 0.00424 | 0.40784 | 0.01622 | 0.01047 | 0.17185 | 0.00639 | 0.13037 | 0.01382 |
| **Efcc1** | 0.06491 | 0.00444 | 0.02835 | 0.02653 | 0.03610 | 0.22918 | 0.02384 | 0.02694 | 0.37083 | 0.03822 | 0.00439 | 0.01689 | 2.65720 | 0.08683 |
| **Efemp1** | 0.00283 | 0.13565 | 0.00000 | 0.26537 | 0.92723 | 0.03089 | 2.44711 | 0.46956 | 0.01808 | 0.03700 | 0.00218 | 0.00133 | 0.00248 | 0.00000 |
| **Efemp2** | 0.10419 | 0.03645 | 0.00489 | 0.06042 | 0.00825 | 0.00756 | 0.00372 | 0.01759 | 0.01113 | 0.00704 | 0.00331 | 1.42195 | 0.09386 | 0.25659 |
| **Efhb** | 1.78367 | 0.00351 | 1.02567 | 0.16031 | 1.52861 | 0.00000 | 0.00359 | 0.23663 | 0.76470 | 0.00330 | 0.03452 | 0.00420 | 0.43199 | 4.48020 |
| **Efhc2** | 1.11851 | 0.00437 | 0.25125 | 0.00000 | 0.39884 | 0.34246 | 0.00508 | 0.01159 | 0.16805 | 0.00634 | 0.00388 | 0.00346 | 0.27139 | 0.00000 |
| **Efhd1** | 3.07844 | 0.01159 | 3.50754 | 2.40670 | 5.32825 | 2.22652 | 0.00296 | 0.00000 | 0.85250 | 0.03471 | 0.00000 | 0.53626 | 1.14025 | 2.43491 |
| **Efhd2** | 1.55274 | 6.27230 | 4.44846 | 0.95146 | 2.50591 | 13.47144 | 5.35631 | 15.98491 | 0.00000 | 11.91060 | 4.63540 | 0.32874 | 0.00000 | 0.00902 |
| **Efl1** | 2.86979 | 0.00909 | 1.01305 | 2.00116 | 0.97614 | 4.14302 | 0.06338 | 0.02095 | 0.04099 | 0.04593 | 2.73510 | 1.69728 | 0.31946 | 3.43404 |
| **Efna1** | 1.86200 | 0.01823 | 6.54443 | 0.00000 | 3.30946 | 0.00000 | 0.00727 | 5.43062 | 0.76893 | 0.01012 | 0.42083 | 0.90049 | 0.26565 | 3.65468 |
| **Efna2** | 1.11865 | 0.01905 | 2.40232 | 1.80661 | 2.48714 | 1.13578 | 0.29911 | 1.90874 | 0.01269 | 0.02528 | 0.15440 | 0.69936 | 0.06966 | 0.00000 |
| **Efna3** | 0.00998 | 0.00403 | 0.03366 | 0.30282 | 0.00244 | 0.01922 | 0.00413 | 0.31180 | 0.00708 | 0.01736 | 0.00105 | 0.46593 | 0.00308 | 0.28417 |
| **Efna4** | 0.00329 | 0.00357 | 0.00000 | 0.33626 | 0.06908 | 0.48935 | 0.00399 | 0.69984 | 0.03917 | 0.00608 | 0.08245 | 1.27215 | 0.01339 | 0.00974 |
| **Efna5** | 0.07727 | 0.01239 | 0.00351 | 0.00492 | 0.14908 | 0.15005 | 0.00994 | 0.03921 | 0.00854 | 0.00765 | 0.02393 | 0.01324 | 0.00855 | 0.60423 |
| **Efnb1** | 0.10662 | 0.60742 | 0.42202 | 0.28102 | 0.37101 | 0.00000 | 0.00424 | 1.07421 | 0.00856 | 0.01955 | 0.00298 | 0.00217 | 0.04730 | 0.00000 |
| **Efnb2** | 1.04842 | 1.13846 | 1.17382 | 0.44186 | 0.19362 | 6.71680 | 1.48706 | 2.46908 | 5.89748 | 6.15583 | 0.97748 | 0.17987 | 4.19430 | 1.59900 |
| **Efnb3** | 0.00412 | 0.01944 | 0.00317 | 0.11566 | 0.17046 | 0.00831 | 0.00998 | 0.66335 | 0.00573 | 0.01450 | 0.00000 | 1.88164 | 0.30806 | 0.15429 |
| **Efr3a** | 2.18137 | 0.02362 | 0.28093 | 0.00000 | 1.19570 | 4.07755 | 0.00876 | 1.61677 | 2.67301 | 0.00031 | 4.23883 | 3.88150 | 2.24463 | 1.20248 |
| **Efr3b** | 0.45745 | 0.01130 | 0.00000 | 0.12269 | 0.36179 | 0.32929 | 0.03388 | 0.31981 | 0.83149 | 0.17560 | 0.01634 | 0.02244 | 0.67378 | 1.22606 |
| **Efs** | 0.00000 | 0.00310 | 0.15036 | 0.00112 | 0.00636 | 0.00000 | 0.00299 | 0.00198 | 0.01058 | 4.38625 | 0.00349 | 0.00361 | 0.00639 | 0.00589 |
| **Eftud2** | 9.55931 | 7.07627 | 19.74543 | 13.13488 | 17.63547 | 13.38786 | 1.69039 | 6.67208 | 2.47873 | 0.15625 | 14.81047 | 7.91643 | 0.96513 | 17.11425 |
| **Egf** | 0.13006 | 0.00452 | 0.10698 | 0.35240 | 0.00000 | 0.31637 | 0.00374 | 0.43363 | 0.00775 | 0.01001 | 0.00462 | 0.06710 | 0.08332 | 1.06904 |
| **Egfl7** | 0.06731 | 0.04021 | 0.30080 | 1.29263 | 0.59164 | 1.03029 | 0.00193 | 1.90134 | 0.00248 | 0.01579 | 5.09549 | 3.28132 | 0.44662 | 0.14849 |
| **Egfl8** | 0.20226 | 0.00410 | 0.63978 | 0.21772 | 0.55257 | 0.00000 | 0.00989 | 0.01003 | 0.00997 | 0.02023 | 0.01128 | 0.01155 | 0.00493 | 0.00314 |
| **Egflam** | 0.00631 | 0.00458 | 0.01038 | 0.00277 | 0.07279 | 0.01351 | 0.00443 | 0.01381 | 0.01466 | 0.00642 | 0.00433 | 0.58048 | 0.00690 | 0.01025 |
| **Egfr** | 0.06620 | 0.00417 | 0.01362 | 0.00781 | 0.00549 | 0.02230 | 0.00423 | 0.00853 | 0.02361 | 0.00599 | 0.00776 | 0.01565 | 0.85572 | 0.03598 |
| **Egln1** | 8.57450 | 1.29270 | 17.20636 | 9.09698 | 6.93041 | 16.12410 | 7.00259 | 16.36687 | 2.36890 | 12.53806 | 9.56536 | 7.84934 | 2.81287 | 7.26302 |
| **Egln2** | 3.37458 | 0.10427 | 8.52307 | 4.31798 | 5.47655 | 3.17066 | 0.94389 | 4.86027 | 0.00000 | 10.42344 | 2.61146 | 2.87116 | 2.84178 | 0.32936 |
| **Egln3** | 0.01378 | 0.00394 | 0.06359 | 1.61164 | 0.62415 | 0.00000 | 0.00377 | 0.00574 | 0.01474 | 0.00598 | 0.01003 | 0.01118 | 0.53861 | 0.00000 |
| **Egr1** | 0.08768 | 1.95839 | 0.03090 | 0.00000 | 0.01036 | 0.00714 | 0.00382 | 0.03396 | 0.01123 | 0.00664 | 0.00629 | 0.06577 | 0.01912 | 0.07306 |
| **Egr3** | 0.00797 | 0.00448 | 0.00322 | 0.00804 | 1.07343 | 0.01381 | 0.00421 | 0.01031 | 0.01441 | 0.00646 | 0.00917 | 0.00770 | 0.00333 | 0.00460 |
| **Ehbp1** | 0.00000 | 0.00459 | 0.28723 | 0.00000 | 0.11888 | 1.05971 | 0.36083 | 0.00511 | 0.00072 | 0.03466 | 0.03771 | 0.25927 | 2.30430 | 0.17588 |
| **Ehbp1l1** | 0.18857 | 0.04442 | 0.33736 | 0.08357 | 0.16940 | 0.27473 | 0.02271 | 0.12182 | 0.43754 | 0.46000 | 0.07900 | 0.15362 | 0.63509 | 0.20992 |
| **Ehd1** | 9.56007 | 0.75038 | 13.68178 | 7.86927 | 10.57007 | 15.20114 | 0.11858 | 6.91022 | 0.19421 | 3.82069 | 7.43517 | 4.96340 | 0.00000 | 4.12248 |
| **Ehd2** | 0.11288 | 0.00421 | 0.09465 | 0.22371 | 0.35385 | 0.52314 | 0.00413 | 0.00412 | 0.00408 | 0.01427 | 0.00362 | 0.00179 | 0.03052 | 2.38882 |
| **Ehd4** | 6.42304 | 0.05093 | 2.84657 | 4.33602 | 1.17373 | 9.12148 | 0.21483 | 7.99436 | 3.19121 | 3.57520 | 3.89160 | 4.10389 | 0.78623 | 0.84621 |
| **Ehf** | 0.01430 | 0.00422 | 0.00385 | 0.01091 | 0.23716 | 0.01102 | 0.00412 | 0.00813 | 0.03008 | 0.00639 | 0.00752 | 0.00741 | 0.80316 | 0.70928 |
| **Ehmt1** | 2.00848 | 0.02117 | 6.52455 | 0.00000 | 8.67314 | 2.31969 | 0.00669 | 1.96956 | 1.78267 | 0.00000 | 1.95018 | 6.32976 | 4.24603 | 7.37972 |
| **Ehmt2** | 0.00000 | 7.06386 | 16.38332 | 12.75904 | 15.97424 | 0.73910 | 7.86753 | 11.79415 | 0.00000 | 0.48791 | 0.80173 | 5.02024 | 0.72911 | 0.61175 |
| **Ei24** | 14.95362 | 2.54111 | 12.44136 | 16.33345 | 17.66093 | 6.25178 | 1.20187 | 3.35681 | 23.60324 | 0.02959 | 2.18509 | 6.25653 | 15.84765 | 28.37316 |
| **Eid1** | 0.00879 | 0.00485 | 0.31855 | 0.00000 | 1.00565 | 0.01891 | 0.00447 | 0.00996 | 0.02276 | 0.00557 | 0.00642 | 0.00979 | 0.02061 | 0.06642 |
| **Eid2** | 1.21235 | 0.01612 | 0.26032 | 0.96445 | 0.66637 | 0.43542 | 0.14067 | 0.00768 | 0.01744 | 0.01181 | 0.00000 | 1.00076 | 8.68373 | 0.61804 |
| **Eid2b** | 1.18750 | 0.00388 | 0.03211 | 0.60008 | 1.22687 | 0.22866 | 0.00405 | 0.02159 | 0.01436 | 0.00645 | 0.00460 | 0.00296 | 0.32821 | 0.62870 |
| **Eid3** | 0.01131 | 0.00461 | 0.01770 | 0.02195 | 0.01265 | 0.65128 | 0.00419 | 0.00677 | 0.01286 | 0.00651 | 0.07427 | 0.00375 | 0.01000 | 0.00802 |
| **Eif1** | 22.63417 | 6.67050 | 11.75333 | 22.77803 | 36.65594 | 17.23175 | 6.91750 | 17.40302 | 9.76094 | 12.01974 | 9.47314 | 17.67772 | 26.68533 | 14.87363 |
| **Eif1a** | 19.53067 | 6.13151 | 20.74320 | 13.62794 | 24.09014 | 23.47948 | 8.16470 | 7.79203 | 28.32931 | 0.45223 | 13.68712 | 7.37291 | 11.07473 | 18.39162 |
| **Eif1ad** | 9.61254 | 3.21982 | 17.78337 | 16.08050 | 12.27805 | 12.75664 | 4.07011 | 8.51052 | 3.50485 | 7.91247 | 5.08722 | 10.02846 | 11.61191 | 4.85436 |
| **Eif1ax** | 5.87073 | 0.21426 | 10.94729 | 22.84246 | 17.47389 | 7.92718 | 4.85018 | 6.30548 | 2.15948 | 14.62813 | 0.54682 | 9.79738 | 4.71539 | 1.46985 |
| **Eif1b** | 5.27705 | 0.01535 | 2.75403 | 2.36765 | 5.27055 | 3.07674 | 0.10026 | 3.99490 | 0.96402 | 0.11815 | 0.64898 | 5.01078 | 4.00641 | 2.48729 |
| **Eif2a** | 11.39125 | 2.96176 | 18.24424 | 6.29124 | 16.40921 | 11.77271 | 0.66295 | 5.18065 | 25.17667 | 0.08025 | 10.29203 | 8.20411 | 3.43572 | 16.09357 |
| **Eif2ak1** | 4.11048 | 6.44958 | 2.00248 | 1.91073 | 2.67824 | 1.23785 | 0.01816 | 2.63021 | 0.62234 | 0.01629 | 0.08995 | 1.55534 | 1.17984 | 5.95871 |
| **Eif2ak2** | 0.02778 | 0.02768 | 0.01721 | 0.02786 | 0.20943 | 0.06522 | 0.14053 | 0.00938 | 1.85406 | 0.01033 | 0.00000 | 0.50726 | 0.09440 | 0.01190 |
| **Eif2ak3** | 0.31874 | 0.02206 | 1.76887 | 0.00000 | 0.38752 | 0.40708 | 0.00395 | 0.28684 | 0.04113 | 0.00055 | 0.99024 | 0.51766 | 1.00656 | 6.00061 |

| | | | | | | | | | | | | | | |
|---|---|---|---|---|---|---|---|---|---|---|---|---|---|---|
| **Eif2ak4** | 0.00000 | 0.00884 | 0.37092 | 1.76083 | 0.45744 | 2.05256 | 2.52074 | 1.83099 | 0.06182 | 0.05090 | 2.81116 | 1.55263 | 3.24124 | 0.00000 |
| **Eif2b1** | 15.46981 | 8.08398 | 27.10810 | 8.99760 | 23.84285 | 16.18868 | 3.14471 | 5.72740 | 5.52788 | 0.13217 | 15.28627 | 8.06770 | 8.01722 | 23.35686 |
| **Eif2b2** | 12.28437 | 7.18026 | 11.05395 | 10.67468 | 12.22807 | 10.64913 | 3.30074 | 7.16720 | 0.39572 | 0.08944 | 6.15255 | 5.02979 | 4.11011 | 3.07243 |
| **Eif2b3** | 10.37385 | 0.04493 | 18.65412 | 13.32699 | 15.07014 | 12.12129 | 0.90324 | 3.18277 | 13.65800 | 0.92662 | 7.59518 | 2.92209 | 5.64737 | 17.36258 |
| **Eif2b4** | 6.54102 | 1.00595 | 22.41380 | 15.49388 | 15.83025 | 9.90202 | 2.74542 | 9.50505 | 0.84057 | 0.22298 | 6.37896 | 8.13369 | 11.72825 | 5.82389 |
| **Eif2b5** | 11.94305 | 2.60468 | 21.29016 | 15.78621 | 18.44708 | 19.07177 | 5.23356 | 11.99373 | 14.13818 | 0.35176 | 9.32088 | 9.10313 | 5.50434 | 17.79692 |
| **Eif2d** | 7.53928 | 0.48047 | 15.77364 | 16.81980 | 11.30004 | 17.20052 | 6.23339 | 4.68007 | 3.72608 | 0.25881 | 16.60481 | 9.89382 | 0.62647 | 14.27952 |
| **Eif2s1** | 19.50435 | 6.39211 | 26.59730 | 15.61872 | 26.14872 | 26.33345 | 8.46774 | 12.05863 | 28.77782 | 13.79560 | 19.00473 | 18.20279 | 11.44545 | 25.87694 |
| **Eif2s2** | 37.90805 | 11.07871 | 47.41522 | 36.65905 | 43.56405 | 43.34912 | 14.74840 | 26.61136 | 41.20074 | 12.18740 | 36.16296 | 36.03124 | 18.28543 | 37.15688 |
| **Eif2s3x** | 13.97774 | 2.00136 | 29.49921 | 20.01222 | 28.24382 | 15.84968 | 5.24587 | 9.55505 | 34.81664 | 16.28909 | 11.67922 | 12.04409 | 4.85060 | 25.50432 |
| **Eif2s3y** | 0.00035 | 0.01769 | 0.00000 | 5.63782 | 0.07615 | 0.00000 | 5.57258 | 0.15437 | 0.00000 | 0.05641 | 3.64848 | 2.07097 | 2.80719 | 2.49795 |
| **Eif3a** | 25.11091 | 6.48380 | 28.26983 | 18.17960 | 34.05664 | 24.06037 | 7.38007 | 8.82198 | 40.15899 | 9.86373 | 18.27268 | 13.33493 | 16.62328 | 32.85635 |
| **Eif3b** | 17.19145 | 3.22087 | 37.31061 | 25.96333 | 25.55669 | 25.34295 | 7.94477 | 13.56883 | 0.19198 | 7.20160 | 17.96190 | 18.00391 | 11.46468 | 21.00380 |
| **Eif3c** | 28.54043 | 6.96307 | 43.17483 | 26.70460 | 36.30377 | 28.18296 | 9.37961 | 15.57348 | 3.51343 | 0.88844 | 25.16920 | 18.81029 | 18.65035 | 26.18822 |
| **Eif3d** | 16.08477 | 3.17315 | 37.57952 | 25.60044 | 25.09407 | 23.48354 | 7.78643 | 19.45167 | 14.39041 | 12.73702 | 23.56308 | 18.10115 | 3.19759 | 25.65283 |
| **Eif3e** | 18.15172 | 7.44635 | 30.34003 | 18.15177 | 25.61244 | 26.82441 | 11.26578 | 19.30705 | 34.09378 | 7.82986 | 19.73842 | 16.84294 | 5.02196 | 27.03212 |
| **Eif3f** | 24.70208 | 6.38692 | 16.62611 | 12.12945 | 19.87544 | 7.26288 | 3.43154 | 12.52384 | 7.39165 | 7.76386 | 8.90830 | 11.96586 | 15.76217 | 6.69705 |
| **Eif3g** | 15.70700 | 7.40058 | 41.64047 | 36.86371 | 36.58980 | 16.94326 | 4.66660 | 24.54004 | 0.06536 | 20.52351 | 19.17344 | 19.51127 | 7.69023 | 1.01303 |
| **Eif3h** | 30.30434 | 9.69639 | 39.31392 | 37.48987 | 35.73339 | 30.67333 | 9.32009 | 22.74266 | 11.51240 | 1.60918 | 18.31711 | 20.62940 | 21.45714 | 15.73497 |
| **Eif3i** | 23.30148 | 11.05551 | 37.78194 | 24.20639 | 26.83891 | 29.11698 | 7.74309 | 17.74894 | 18.23109 | 17.85083 | 21.29286 | 20.77664 | 6.54933 | 24.75844 |
| **Eif3k** | 17.35337 | 3.77004 | 15.73582 | 30.48654 | 19.54356 | 17.47864 | 9.05464 | 19.11561 | 4.99502 | 16.47696 | 9.10239 | 12.55839 | 3.63640 | 13.27137 |
| **Eif3l** | 15.75604 | 8.83116 | 33.45027 | 25.53786 | 28.54575 | 24.67558 | 7.25021 | 14.96481 | 5.13766 | 0.60942 | 18.84337 | 16.81274 | 3.79591 | 26.18685 |
| **Eif3m** | 22.66599 | 2.55926 | 33.10790 | 23.91291 | 28.37577 | 27.50611 | 5.76816 | 13.77387 | 21.85686 | 12.89778 | 14.53604 | 21.81484 | 6.65901 | 23.78339 |
| **Eif4a1** | 20.38007 | 14.61806 | 43.89443 | 37.89933 | 35.34590 | 38.82918 | 9.41180 | 27.59530 | 30.81218 | 19.59752 | 30.55117 | 25.49950 | 12.97770 | 36.49137 |
| **Eif4a2** | 24.44262 | 9.34881 | 28.70246 | 30.87429 | 18.61005 | 34.56916 | 11.64110 | 12.46215 | 28.53844 | 2.01683 | 23.82643 | 14.05211 | 8.02943 | 24.66823 |
| **Eif4a3** | 24.05914 | 12.19536 | 38.37112 | 28.66852 | 32.56430 | 24.34566 | 6.13184 | 14.54879 | 5.62777 | 0.53979 | 18.57533 | 17.05136 | 13.97448 | 22.98996 |
| **Eif4b** | 13.81041 | 7.41790 | 24.51299 | 22.11380 | 24.49306 | 24.79678 | 6.52126 | 16.99505 | 22.29039 | 9.33863 | 16.21916 | 18.77703 | 9.68823 | 26.70225 |
| **Eif4e** | 35.74120 | 8.06765 | 35.20548 | 18.32474 | 41.75072 | 21.42872 | 7.21630 | 7.34735 | 45.12747 | 0.53710 | 14.18140 | 13.37256 | 22.95581 | 43.31015 |
| **Eif4e1b** | 0.05567 | 0.00401 | 0.37779 | 0.53193 | 0.73010 | 0.00000 | 0.00387 | 0.00465 | 0.02887 | 0.00520 | 0.01142 | 0.11187 | 1.47413 | 0.02488 |
| **Eif4e2** | 15.72796 | 0.51223 | 16.96901 | 15.83193 | 17.03328 | 12.97509 | 5.05013 | 13.38964 | 10.59853 | 0.51463 | 6.62908 | 13.03160 | 6.99969 | 17.54909 |
| **Eif4e3** | 0.67101 | 0.01861 | 0.00000 | 0.01730 | 0.00000 | 0.01484 | 0.00377 | 0.05965 | 0.63916 | 0.00743 | 0.01875 | 0.01251 | 3.58199 | 1.79110 |
| **Eif4ebp1** | 4.11379 | 0.36200 | 4.97804 | 32.09318 | 6.31680 | 8.74697 | 2.77219 | 4.00537 | 0.00000 | 9.22900 | 3.36960 | 15.31079 | 4.16266 | 0.00000 |
| **Eif4ebp2** | 0.47363 | 0.00350 | 0.31494 | 0.00000 | 4.07268 | 1.36382 | 0.00327 | 1.31381 | 1.47079 | 0.00592 | 0.00000 | 0.82824 | 0.83585 | 2.06734 |
| **Eif4ebp3** | 3.57862 | 0.00302 | 1.50983 | 0.80642 | 1.98537 | 2.00409 | 0.00314 | 0.98867 | 1.55910 | 0.00252 | 0.64876 | 1.35504 | 1.38817 | 0.77497 |
| **Eif4enif1** | 9.04629 | 0.05861 | 8.52264 | 3.23345 | 3.58495 | 7.96821 | 0.76090 | 3.07949 | 0.95562 | 0.06424 | 6.89762 | 5.21784 | 11.90085 | 11.61400 |
| **Eif4g1** | 0.00000 | 4.09243 | 21.65276 | 25.61259 | 22.02383 | 1.19098 | 5.04156 | 12.72041 | 3.65265 | 0.34692 | 2.22917 | 11.35974 | 0.55252 | 2.01585 |
| **Eif4g2** | 30.59462 | 5.05484 | 36.28965 | 22.10750 | 36.06082 | 31.69641 | 9.39369 | 16.49462 | 47.86801 | 9.95180 | 27.36709 | 21.67258 | 24.66177 | 27.87071 |
| **Eif4g3** | 14.41681 | 3.50196 | 9.25923 | 2.38379 | 6.39829 | 7.64681 | 0.01416 | 3.87645 | 8.80502 | 10.17484 | 6.32732 | 6.71388 | 1.73190 | 9.84510 |
| **Eif4h** | 24.74265 | 7.45590 | 46.15526 | 21.99680 | 42.79043 | 25.18112 | 7.74150 | 14.62127 | 4.66377 | 4.87494 | 14.98050 | 17.92104 | 12.28482 | 16.50157 |
| **Eif5** | 41.45883 | 3.40475 | 34.96608 | 20.94846 | 40.03565 | 28.83339 | 7.30164 | 8.46510 | 44.66244 | 6.21870 | 22.08517 | 13.78124 | 18.72773 | 26.53540 |
| **Eif5a** | 18.26557 | 14.03157 | 42.35802 | 49.86818 | 46.42645 | 47.04269 | 11.27798 | 36.51534 | 9.63866 | 18.09871 | 25.79490 | 31.81428 | 12.84094 | 29.42385 |
| **Eif5a2** | 0.57585 | 0.00400 | 0.00000 | 0.01660 | 0.00902 | 0.52581 | 0.00405 | 0.00947 | 0.32845 | 0.00660 | 0.02978 | 0.00176 | 0.85040 | 1.65923 |
| **Eif5b** | 18.74942 | 0.12730 | 19.27158 | 13.17147 | 28.91038 | 13.21027 | 1.79855 | 5.70654 | 32.55601 | 5.62837 | 13.84623 | 10.25211 | 12.22438 | 26.61391 |
| **Eif6** | 18.18736 | 0.44123 | 26.40226 | 25.92767 | 34.26142 | 12.55417 | 6.09871 | 14.11520 | 2.00669 | 3.79274 | 7.90890 | 13.20253 | 14.05616 | 6.76975 |
| **Elac1** | 1.45739 | 0.00389 | 0.54032 | 0.00000 | 0.18622 | 1.72596 | 4.23221 | 0.01340 | 0.01534 | 0.10873 | 0.34503 | 0.49696 | 0.21361 | 0.71926 |
| **Elac2** | 3.61433 | 0.05825 | 5.37403 | 3.56343 | 5.93415 | 4.49119 | 1.41275 | 0.64051 | 0.26990 | 12.11673 | 4.77627 | 3.44287 | 0.85398 | 5.84323 |
| **Elavl1** | 12.15682 | 4.40260 | 9.87887 | 4.90017 | 10.95654 | 17.45141 | 1.43116 | 4.25147 | 6.69458 | 1.86147 | 3.32839 | 9.04411 | 1.53425 | 3.53264 |
| **Elavl2** | 1.01337 | 0.01090 | 0.00000 | 0.00000 | 0.51837 | 0.11430 | 0.00366 | 0.00256 | 1.68617 | 0.01891 | 0.00250 | 0.06289 | 2.76416 | 2.32604 |
| **Elavl3** | 1.45517 | 0.00393 | 3.33263 | 1.59200 | 0.73548 | 0.11408 | 0.00390 | 0.03102 | 0.19011 | 0.02977 | 0.23875 | 0.20762 | 0.97297 | 2.56854 |
| **Eldr** | 0.02237 | 0.00411 | 0.11751 | 0.16471 | 0.01152 | 0.00453 | 0.00404 | 0.00589 | 0.01210 | 0.00887 | 0.01643 | 0.04951 | 0.01200 | 0.00775 |
| **Elf1** | 1.78581 | 0.03388 | 0.00373 | 0.00000 | 0.44314 | 0.65077 | 0.12528 | 0.05688 | 1.91521 | 0.02633 | 0.08760 | 0.88696 | 11.48326 | 0.45743 |
| **Elf2** | 1.13333 | 0.00945 | 2.13762 | 0.43725 | 1.28290 | 2.06657 | 3.37478 | 0.67605 | 1.59020 | 0.03926 | 0.41273 | 2.77097 | 2.82956 | 2.82288 |
| **Elf3** | 0.00000 | 11.86034 | 23.44034 | 21.47635 | 0.69600 | 42.57879 | 11.24832 | 30.73094 | 0.88631 | 12.89541 | 19.63681 | 10.99939 | 0.00000 | 1.36282 |
| **Elf4** | 2.08424 | 1.62355 | 0.15676 | 0.00000 | 1.00817 | 1.81685 | 2.81583 | 2.45926 | 3.21395 | 0.07228 | 0.00000 | 0.06957 | 0.00000 | 2.59135 |
| **Elf5** | 0.01018 | 4.02352 | 0.01549 | 0.00822 | 0.01229 | 0.00762 | 0.00869 | 0.05611 | 0.01163 | 0.00590 | 0.00804 | 0.00399 | 0.00664 | 0.01144 |

| | | | | | | | | | | | | | | |
|---|---|---|---|---|---|---|---|---|---|---|---|---|---|---|
| **Elk1** | 0.01218 | 0.02007 | 0.84716 | 0.52569 | 1.37018 | 0.84357 | 0.00463 | 0.43705 | 0.82120 | 0.00410 | 0.00184 | 0.75342 | 2.30666 | 2.06109 |
| **Elk3** | 0.03296 | 0.03478 | 0.10588 | 0.01848 | 0.00322 | 0.00815 | 0.00411 | 0.00692 | 0.00925 | 0.00578 | 0.00931 | 0.65357 | 0.00662 | 0.13585 |
| **Elk4** | 0.00000 | 0.00374 | 0.58438 | 0.00000 | 0.32302 | 0.00000 | 0.00341 | 0.94505 | 0.04002 | 0.00964 | 0.00336 | 0.00463 | 0.80449 | 2.67507 |
| **Ell** | 2.19590 | 4.27113 | 3.71516 | 1.63004 | 3.27150 | 2.07096 | 3.58845 | 2.29723 | 0.12204 | 0.11088 | 2.48492 | 0.60531 | 3.38531 | 2.12796 |
| **Ell2** | 14.15834 | 2.77143 | 6.85168 | 1.09604 | 18.01160 | 1.17579 | 3.30816 | 3.44128 | 22.50180 | 0.09157 | 4.47405 | 4.07800 | 13.92190 | 20.48870 |
| **Ell3** | 5.65584 | 0.00340 | 6.00258 | 2.46091 | 8.60616 | 2.18828 | 0.01463 | 1.14808 | 0.91328 | 0.00000 | 1.72961 | 0.16067 | 0.24898 | 10.91963 |
| **Elmo2** | 1.14995 | 0.03880 | 0.41801 | 1.06970 | 2.49203 | 2.22405 | 0.00289 | 0.46240 | 0.04436 | 0.01175 | 1.88497 | 2.56059 | 0.00000 | 3.11237 |
| **Elmo3** | 0.00000 | 0.03674 | 0.61189 | 0.00298 | 0.54368 | 0.31270 | 0.00326 | 2.25967 | 0.02202 | 0.00595 | 1.45357 | 0.98985 | 0.03675 | 0.14120 |
| **Elmod1** | 0.00850 | 0.00426 | 0.01447 | 0.00328 | 0.03531 | 0.32321 | 0.00425 | 0.01772 | 0.01636 | 0.00602 | 0.00711 | 0.00938 | 0.00832 | 0.00731 |
| **Elmod2** | 5.07559 | 0.01753 | 0.63919 | 0.00000 | 1.47739 | 0.56435 | 0.00225 | 0.08026 | 5.56133 | 7.99441 | 0.00000 | 2.32386 | 3.54400 | 4.72946 |
| **Elmod3** | 0.47282 | 0.02485 | 0.47675 | 0.52821 | 0.00000 | 0.39307 | 1.14162 | 2.53168 | 0.00431 | 0.02300 | 0.22685 | 0.95651 | 1.57380 | 0.34153 |
| **Elmsan1** | 0.07098 | 0.00382 | 0.64421 | 0.36792 | 1.76886 | 0.33136 | 0.00391 | 0.29675 | 0.10125 | 0.04062 | 0.00001 | 0.22070 | 0.08643 | 0.53779 |
| **Eln** | 0.00641 | 0.00475 | 0.00000 | 0.08349 | 0.07632 | 0.44636 | 0.00445 | 0.00635 | 0.00377 | 0.00652 | 0.00798 | 0.00409 | 0.21111 | 0.18473 |
| **Elof1** | 17.40564 | 4.15338 | 13.67525 | 8.96161 | 22.49615 | 12.63418 | 1.62812 | 9.13981 | 3.78327 | 4.88013 | 5.24918 | 6.82841 | 8.74729 | 17.86193 |
| **Elovl1** | 1.25436 | 1.12343 | 1.46548 | 1.41060 | 0.46729 | 7.68944 | 3.80239 | 7.48102 | 0.06758 | 16.56761 | 0.00000 | 4.77371 | 0.14462 | 0.47305 |
| **Elovl2** | 0.00879 | 0.00436 | 0.00751 | 0.02220 | 0.00416 | 0.00954 | 0.00423 | 0.00934 | 0.00957 | 0.00925 | 0.00719 | 0.00863 | 0.02451 | 0.97405 |
| **Elovl3** | 0.03133 | 0.00450 | 0.00452 | 0.00657 | 0.92148 | 0.01201 | 0.00422 | 0.01044 | 0.07872 | 0.00689 | 0.00764 | 0.00747 | 0.01693 | 0.03894 |
| **Elovl5** | 1.58157 | 0.08279 | 2.30824 | 3.11440 | 0.21114 | 10.80120 | 0.01349 | 2.71798 | 2.61224 | 0.08646 | 9.74018 | 4.38372 | 0.01611 | 0.00000 |
| **Elovl6** | 7.50660 | 9.26994 | 23.65605 | 8.01825 | 14.36213 | 18.03321 | 3.87952 | 10.28749 | 31.91917 | 14.58022 | 14.93154 | 14.27822 | 8.52498 | 26.32633 |
| **Elovl7** | 0.11856 | 0.11684 | 0.00000 | 6.12768 | 0.00000 | 8.41912 | 0.03808 | 2.67574 | 0.13601 | 0.05091 | 3.62025 | 8.86104 | 0.49763 | 0.00000 |
| **Elp2** | 8.69636 | 0.07059 | 7.25298 | 8.88387 | 5.43524 | 8.14924 | 0.69600 | 2.74776 | 13.02560 | 0.07247 | 8.35397 | 7.49479 | 0.65342 | 8.24822 |
| **Elp3** | 4.49582 | 0.06357 | 10.59574 | 10.35907 | 6.83272 | 13.50702 | 2.36483 | 7.35797 | 2.60284 | 0.02860 | 18.47409 | 7.48311 | 0.00000 | 4.92936 |
| **Elp4** | 3.79393 | 0.01048 | 0.81956 | 3.86051 | 1.25148 | 2.00965 | 0.00319 | 0.01195 | 0.38495 | 0.02428 | 0.24358 | 1.33005 | 0.00100 | 0.40761 |
| **Elp5** | 8.33609 | 0.18233 | 15.49272 | 5.99818 | 13.00167 | 13.90578 | 3.22751 | 4.82737 | 11.09234 | 8.14864 | 9.92419 | 11.45440 | 14.88260 | 7.09772 |
| **Elp6** | 1.67094 | 0.04104 | 13.38190 | 3.86375 | 9.25517 | 3.79569 | 2.98887 | 0.74902 | 0.08224 | 4.65248 | 3.75173 | 1.97957 | 3.45508 | 1.10740 |
| **Emb** | 0.00000 | 0.01237 | 0.33897 | 0.57516 | 1.07897 | 0.00000 | 0.00981 | 0.01644 | 2.50015 | 0.01311 | 0.00000 | 2.16898 | 0.23784 | 0.00000 |
| **Emc1** | 5.52606 | 0.22397 | 6.38746 | 8.59472 | 5.42164 | 4.54498 | 0.17536 | 2.28089 | 15.49091 | 0.03652 | 3.01372 | 3.65313 | 4.45773 | 9.26753 |
| **Emc10** | 1.25001 | 0.06866 | 1.21344 | 12.69988 | 3.24205 | 6.31600 | 1.04262 | 1.43092 | 0.00269 | 0.12922 | 1.23526 | 7.47516 | 0.18083 | 0.78884 |
| **Emc2** | 6.88293 | 0.10530 | 5.67535 | 4.91721 | 5.59276 | 7.03921 | 3.25716 | 4.47858 | 9.83298 | 0.10899 | 8.29135 | 6.96992 | 1.85095 | 4.86749 |
| **Emc3** | 18.52962 | 4.22656 | 26.70265 | 13.19342 | 21.47709 | 10.94401 | 5.67586 | 4.52581 | 19.36671 | 0.10951 | 5.26979 | 6.03748 | 14.14544 | 23.78622 |
| **Emc4** | 10.88724 | 0.19699 | 16.64303 | 21.77084 | 17.19828 | 11.48261 | 4.33004 | 7.22206 | 6.65985 | 10.27575 | 6.94186 | 10.20094 | 4.60364 | 4.25235 |
| **Emc6** | 4.59224 | 0.02714 | 1.91969 | 9.47363 | 8.02993 | 5.52304 | 1.14163 | 2.45700 | 1.34982 | 2.72251 | 3.31156 | 5.22858 | 1.51283 | 3.61182 |
| **Emc7** | 12.32778 | 0.11319 | 13.15844 | 4.00020 | 6.06366 | 13.46929 | 2.72341 | 6.13102 | 2.20019 | 0.12711 | 1.73825 | 10.43487 | 6.90350 | 8.21973 |
| **Emc8** | 3.28777 | 1.48917 | 2.10894 | 10.10902 | 7.00290 | 4.63770 | 2.71821 | 2.21193 | 1.28193 | 0.15032 | 1.45692 | 3.80650 | 6.39018 | 1.13588 |
| **Emc9** | 0.03018 | 0.01856 | 0.35918 | 2.24160 | 0.21180 | 0.11999 | 2.20460 | 0.97216 | 0.07077 | 0.03133 | 0.46062 | 0.66683 | 0.38330 | 0.00000 |
| **Emd** | 0.58763 | 0.02621 | 0.81641 | 0.88710 | 0.38752 | 1.45371 | 0.00301 | 1.27465 | 0.45118 | 0.01515 | 1.21015 | 1.36118 | 0.71702 | 1.44440 |
| **Eme1** | 3.85716 | 0.08650 | 6.54439 | 2.27786 | 4.84553 | 6.95949 | 2.86833 | 0.42397 | 0.76130 | 0.10372 | 3.76450 | 3.28711 | 1.28068 | 9.28573 |
| **Eme2** | 0.00103 | 0.00534 | 0.05505 | 1.71847 | 0.01411 | 0.74032 | 0.00450 | 0.47345 | 0.01113 | 0.02411 | 0.11443 | 0.00265 | 0.00829 | 0.00422 |
| **Emg1** | 7.87646 | 5.98748 | 24.65222 | 17.97322 | 19.38507 | 17.91866 | 6.12061 | 9.22234 | 6.57315 | 0.54308 | 11.24965 | 9.54871 | 0.92152 | 25.59476 |
| **Emid1** | 0.01481 | 0.00422 | 0.15061 | 0.00000 | 0.01076 | 0.15122 | 0.00422 | 0.00411 | 0.01004 | 0.00514 | 0.00647 | 0.00969 | 0.62262 | 0.00677 |
| **Emilin1** | 0.02869 | 0.01256 | 0.21827 | 0.00000 | 0.00861 | 0.00000 | 0.00492 | 0.80047 | 0.01170 | 0.00515 | 0.35436 | 0.00879 | 0.13155 | 0.00635 |
| **Emilin2** | 0.03186 | 0.00435 | 0.00545 | 0.01649 | 0.16446 | 0.00891 | 0.00443 | 0.00888 | 0.19204 | 0.00617 | 0.00797 | 0.00868 | 0.44672 | 0.14745 |
| **Eml1** | 0.04864 | 0.00377 | 0.11212 | 0.10016 | 0.20879 | 0.00000 | 0.00411 | 0.00280 | 0.00826 | 0.00466 | 0.36534 | 0.33396 | 0.06301 | 0.18634 |
| **Eml2** | 0.00000 | 0.01138 | 0.20146 | 1.23177 | 0.35379 | 0.40796 | 0.14413 | 1.33001 | 0.00860 | 0.01481 | 1.80010 | 0.00253 | 0.00540 | 0.02615 |
| **Eml3** | 1.17520 | 0.07938 | 1.93185 | 1.00174 | 2.00291 | 0.39115 | 0.83774 | 3.63200 | 0.07241 | 0.06311 | 0.97010 | 3.39945 | 3.16760 | 1.83320 |
| **Eml4** | 1.34940 | 5.05335 | 1.00870 | 0.27154 | 0.62325 | 1.32617 | 1.30275 | 0.75950 | 0.08175 | 0.02247 | 2.74995 | 3.47809 | 2.08114 | 0.97121 |
| **Eml5** | 0.11885 | 0.00402 | 0.00000 | 0.00000 | 0.08994 | 0.44837 | 0.00474 | 0.03308 | 0.81206 | 0.00537 | 0.00000 | 0.38066 | 1.39684 | 1.68161 |
| **Eml6** | 0.00000 | 0.00402 | 0.03258 | 0.67391 | 0.53085 | 0.05040 | 0.00385 | 0.00291 | 0.12311 | 0.00531 | 0.16843 | 0.01475 | 0.01062 | 0.06181 |
| **Emp1** | 0.18565 | 0.00450 | 0.01027 | 0.01139 | 0.12402 | 0.01107 | 0.00433 | 0.01179 | 0.00968 | 0.00624 | 0.00748 | 0.00817 | 0.00496 | 0.00873 |
| **Emp2** | 0.18865 | 0.34613 | 1.20363 | 0.18298 | 0.01543 | 7.05790 | 2.24831 | 10.87060 | 0.02912 | 17.52931 | 0.00000 | 1.23303 | 1.12276 | 0.96725 |
| **Emp3** | 0.05844 | 0.00444 | 0.01091 | 0.01693 | 0.13527 | 0.01129 | 0.00439 | 0.00995 | 0.00574 | 0.00780 | 0.00799 | 0.00830 | 0.06053 | 0.13559 |
| **Emsy** | 0.44338 | 0.00978 | 2.42064 | 0.46061 | 1.43226 | 0.97591 | 0.00776 | 2.92856 | 2.41717 | 0.03258 | 5.48955 | 0.72827 | 0.74336 | 2.33941 |
| **Emx2** | 0.01805 | 0.00442 | 0.01375 | 0.01187 | 0.03034 | 0.01358 | 0.00425 | 0.01015 | 0.01266 | 0.00723 | 0.00868 | 0.00987 | 0.06710 | 0.00497 |
| **Emx2os** | 0.01160 | 0.00442 | 0.01327 | 0.02681 | 0.08478 | 0.01489 | 0.01028 | 0.01324 | 0.04776 | 0.00717 | 0.01300 | 0.01262 | 0.07131 | 0.05986 |
| **En2** | 0.00619 | 0.00405 | 0.01321 | 0.15735 | 0.03248 | 0.14774 | 0.00406 | 0.00397 | 0.00946 | 0.00636 | 0.01291 | 0.05673 | 0.07666 | 0.00181 |

| | | | | | | | | | | | | | | |
|---|---|---|---|---|---|---|---|---|---|---|---|---|---|---|
| **Enah** | 14.61765 | 0.17078 | 23.99040 | 20.63439 | 18.09042 | 25.10747 | 5.96061 | 8.64387 | 22.58221 | 1.72945 | 17.91918 | 17.12285 | 1.72761 | 15.75652 |
| **Enc1** | 0.24396 | 0.00168 | 1.70502 | 4.46756 | 0.20789 | 0.48400 | 0.00244 | 0.08753 | 1.52609 | 0.01027 | 3.32746 | 1.80256 | 2.23483 | 2.70453 |
| **Endod1** | 0.02244 | 0.05575 | 0.80413 | 0.00000 | 0.20780 | 3.46774 | 0.56268 | 3.52280 | 0.00238 | 0.05833 | 0.00000 | 1.32464 | 0.87296 | 1.29077 |
| **Endog** | 0.36154 | 0.01732 | 0.01869 | 0.52462 | 1.85542 | 0.00000 | 0.30511 | 0.25302 | 0.00775 | 0.01950 | 0.04104 | 0.22935 | 0.39878 | 0.41555 |
| **Endov** | 0.88814 | 0.22788 | 1.98526 | 1.72208 | 1.35058 | 1.20386 | 0.00252 | 2.76207 | 1.23299 | 0.02581 | 0.37766 | 1.40796 | 1.58090 | 0.24987 |
| **Eng** | 0.03512 | 0.00410 | 0.97193 | 1.15447 | 2.90339 | 0.00000 | 0.00391 | 0.01086 | 0.00573 | 0.01416 | 0.00893 | 0.01227 | 0.00000 | 0.16668 |
| **Engase** | 0.01903 | 0.00380 | 0.25474 | 0.17506 | 0.02259 | 0.19050 | 0.01150 | 0.39985 | 0.01144 | 0.01813 | 0.13507 | 1.66859 | 0.00888 | 0.01092 |
| **Enho** | 0.00915 | 0.00518 | 1.44382 | 0.18001 | 0.12302 | 0.00000 | 0.00415 | 0.07025 | 0.02123 | 0.01989 | 0.00735 | 0.00831 | 0.02371 | 0.00663 |
| **Enkd1** | 0.05534 | 0.02411 | 2.52613 | 2.90238 | 0.95774 | 0.14124 | 0.01953 | 5.68213 | 0.02300 | 0.00625 | 0.00000 | 2.22797 | 0.25678 | 0.00000 |
| **Eno1** | 0.01519 | 0.00428 | 0.05957 | 0.11458 | 0.00738 | 0.25564 | 0.06748 | 0.20588 | 0.01293 | 0.00485 | 0.46473 | 0.23804 | 0.00451 | 0.02877 |
| **Eno1b** | 0.32250 | 3.91720 | 2.11945 | 4.46294 | 0.94489 | 10.12374 | 2.57254 | 7.66212 | 0.00000 | 3.12677 | 7.92535 | 6.43451 | 0.31248 | 0.75101 |
| **Eno2** | 0.15271 | 0.00524 | 0.14553 | 0.28060 | 0.00000 | 0.16523 | 0.00393 | 0.40227 | 0.01023 | 0.01007 | 0.00000 | 0.76057 | 0.04897 | 0.00581 |
| **Eno3** | 0.73197 | 0.00347 | 1.27147 | 1.69846 | 1.69913 | 0.36682 | 0.00325 | 1.78414 | 0.61275 | 0.01135 | 1.25695 | 0.91257 | 0.00000 | 0.61291 |
| **Eno4** | 0.00424 | 0.01294 | 0.00265 | 0.02646 | 0.00970 | 0.00016 | 0.00400 | 0.01225 | 0.01981 | 0.02471 | 0.84303 | 0.62150 | 0.20309 | 0.00000 |
| **Enoph1** | 7.23344 | 5.38491 | 17.11643 | 5.37789 | 11.66271 | 11.21724 | 3.27430 | 4.80736 | 0.38369 | 15.66781 | 4.80386 | 4.98115 | 2.51430 | 3.74687 |
| **Enox1** | 1.07173 | 0.00391 | 1.15260 | 0.00000 | 1.49817 | 0.00000 | 0.00355 | 0.00416 | 1.08985 | 0.00645 | 0.00706 | 0.00621 | 1.12879 | 1.28923 |
| **Enox2** | 0.01381 | 0.00444 | 0.01273 | 0.01357 | 0.01826 | 0.01260 | 0.00426 | 0.01017 | 0.01217 | 0.00690 | 0.00859 | 0.00982 | 0.01153 | 0.09162 |
| **Enpep** | 0.00000 | 0.82602 | 0.16456 | 1.27269 | 0.00000 | 6.17422 | 3.32747 | 9.76584 | 0.01108 | 7.86706 | 7.75983 | 2.73897 | 0.00412 | 0.00000 |
| **Enpp1** | 0.01473 | 0.04133 | 0.00000 | 0.34487 | 0.00593 | 0.75560 | 0.00888 | 5.17110 | 0.01663 | 0.01033 | 0.01052 | 0.07349 | 0.00832 | 0.00087 |
| **Enpp2** | 0.00270 | 0.00458 | 0.02082 | 0.01077 | 0.02079 | 0.01180 | 0.00448 | 0.01519 | 0.02009 | 0.00670 | 0.00896 | 0.00870 | 0.67560 | 0.00715 |
| **Enpp3** | 2.82083 | 0.00216 | 2.04344 | 0.00000 | 1.70297 | 0.00000 | 0.00681 | 0.00000 | 11.41832 | 0.00158 | 0.00000 | 0.90732 | 2.03857 | 10.63881 |
| **Enpp4** | 0.05717 | 0.00449 | 0.31601 | 0.16623 | 0.23061 | 0.68108 | 0.00378 | 0.00170 | 1.05452 | 0.00520 | 0.24437 | 0.00440 | 0.00238 | 0.58569 |
| **Enpp5** | 0.00006 | 0.00467 | 0.00000 | 0.01612 | 0.00000 | 0.00764 | 0.00441 | 0.01459 | 0.45415 | 0.00691 | 0.00634 | 0.00797 | 0.35876 | 0.57173 |
| **Ensa** | 1.13531 | 0.13987 | 1.17040 | 0.15169 | 0.00000 | 0.74972 | 0.00269 | 2.99891 | 3.12110 | 0.02847 | 0.91133 | 0.00196 | 3.55575 | 1.98236 |
| **Enthd2** | 0.37337 | 1.51291 | 5.49556 | 3.86347 | 1.65688 | 3.48582 | 0.01684 | 0.65993 | 0.06918 | 1.05066 | 0.47652 | 2.51677 | 1.21184 | 1.23750 |
| **Entpd1** | 0.00844 | 0.08735 | 0.08895 | 0.00000 | 0.00667 | 0.00059 | 0.00793 | 8.72554 | 0.00665 | 0.04118 | 0.00218 | 0.09669 | 0.00503 | 0.05229 |
| **Entpd2** | 1.29684 | 0.01796 | 5.28051 | 4.65386 | 1.81224 | 0.80314 | 0.00280 | 2.40409 | 0.00328 | 0.01123 | 1.59911 | 0.00000 | 0.72744 | 0.66160 |
| **Entpd3** | 0.69835 | 0.00331 | 6.63748 | 0.00000 | 6.98567 | 0.00000 | 0.00196 | 0.00198 | 6.26726 | 0.00180 | 0.00000 | 0.00393 | 0.99251 | 9.11970 |
| **Entpd4** | 0.02296 | 0.00444 | 0.01627 | 0.01294 | 0.01190 | 0.01241 | 0.00427 | 0.01049 | 0.01800 | 0.00676 | 0.00844 | 0.00991 | 0.01276 | 0.02397 |
| **Entpd5** | 0.86574 | 0.00143 | 0.76270 | 0.31139 | 0.87794 | 0.93743 | 0.00303 | 0.00000 | 0.88655 | 0.00389 | 0.00000 | 2.81220 | 2.97792 | 4.31996 |
| **Entpd6** | 0.92935 | 0.07185 | 1.12967 | 1.64876 | 0.94126 | 5.96687 | 0.04407 | 4.98330 | 0.01785 | 0.05495 | 2.84359 | 4.89542 | 1.20334 | 1.54107 |
| **Entpd7** | 0.19427 | 0.00416 | 0.88092 | 0.67733 | 0.05903 | 0.78660 | 0.01067 | 0.33721 | 0.21880 | 0.00960 | 0.00274 | 0.01047 | 0.02163 | 1.38234 |
| **Eny2** | 2.14249 | 0.13230 | 0.02595 | 0.74765 | 2.85605 | 3.32699 | 2.86072 | 5.91056 | 2.14402 | 7.74517 | 0.06870 | 5.89794 | 9.88773 | 1.58656 |
| **Eogt** | 2.14853 | 0.00413 | 1.30649 | 0.88174 | 0.25594 | 0.95542 | 0.00335 | 0.97501 | 4.33947 | 0.01117 | 0.00000 | 0.00158 | 0.00000 | 0.04527 |
| **Eomes** | 0.00000 | 10.44499 | 7.35773 | 1.82166 | 0.00000 | 9.07062 | 4.61937 | 20.44326 | 0.57038 | 0.41989 | 1.32806 | 4.91408 | 0.38893 | 1.79282 |
| **Ep300** | 0.03692 | 0.00303 | 0.00000 | 0.00000 | 0.66081 | 0.19169 | 0.00328 | 0.06320 | 2.14031 | 0.00488 | 0.24627 | 0.41571 | 7.32792 | 1.95232 |
| **Ep400** | 10.30719 | 0.04445 | 11.68112 | 7.95637 | 9.85293 | 6.73210 | 2.57808 | 2.30090 | 2.86264 | 11.96428 | 5.12684 | 8.13969 | 6.57770 | 10.59734 |
| **Epas1** | 7.76892 | 0.09901 | 21.41673 | 12.46620 | 6.56061 | 7.12834 | 1.09692 | 9.50411 | 0.34632 | 5.39905 | 7.21008 | 1.80669 | 0.44871 | 0.15095 |
| **Epb41** | 0.08395 | 0.05112 | 3.54978 | 0.00000 | 2.45738 | 6.45149 | 2.27830 | 7.84561 | 0.35389 | 3.17942 | 6.47090 | 3.56498 | 2.82658 | 8.05098 |
| **Epb41l1** | 0.00000 | 0.05145 | 2.12385 | 3.40135 | 0.00000 | 6.57173 | 0.17514 | 4.78763 | 0.00770 | 1.07826 | 1.25085 | 0.35619 | 0.00000 | 0.01294 |
| **Epb41l2** | 2.57195 | 0.38300 | 5.66243 | 4.41257 | 0.65530 | 7.26646 | 1.66171 | 3.75360 | 2.17605 | 0.08653 | 5.85428 | 5.91876 | 1.09361 | 3.99313 |
| **Epb41l3** | 0.10887 | 0.00415 | 0.00809 | 0.19705 | 0.00000 | 0.00026 | 0.00391 | 0.00740 | 0.15251 | 0.00576 | 0.01899 | 0.02355 | 0.17523 | 0.17114 |
| **Epb41l4a** | 0.49670 | 0.00420 | 0.16319 | 0.00000 | 0.13805 | 0.11998 | 0.01255 | 0.22513 | 0.00825 | 0.00976 | 0.00797 | 1.03489 | 0.00867 | 0.33413 |
| **Epb41l4aos** | 0.14508 | 0.00419 | 0.00000 | 1.31468 | 1.47147 | 0.50367 | 0.00433 | 0.42209 | 0.03840 | 0.04187 | 0.00000 | 0.25468 | 0.40203 | 0.29142 |
| **Epb41l4b** | 5.18446 | 0.04559 | 8.18268 | 6.63738 | 3.89617 | 3.27300 | 4.67377 | 6.19990 | 2.36145 | 13.65406 | 4.16887 | 9.24672 | 1.76711 | 5.69571 |
| **Epb41l5** | 0.57229 | 0.00272 | 4.86192 | 0.49437 | 0.00000 | 5.54282 | 0.19384 | 1.40047 | 1.17539 | 0.04573 | 2.89332 | 0.87065 | 1.99051 | 1.76758 |
| **Epb42** | 0.04324 | 0.16828 | 0.02383 | 0.08391 | 0.04072 | 0.05834 | 0.00984 | 0.00650 | 0.09738 | 0.00877 | 0.02157 | 0.01559 | 0.09020 | 0.13106 |
| **Epc1** | 0.40636 | 0.02974 | 0.99637 | 1.43090 | 0.93160 | 3.03588 | 0.19526 | 1.74497 | 1.63167 | 0.79919 | 1.76933 | 0.97797 | 1.97078 | 4.05965 |
| **Epc2** | 6.67316 | 0.00121 | 4.25141 | 1.54991 | 5.84362 | 2.95848 | 0.00727 | 0.68835 | 5.45779 | 0.03259 | 1.82483 | 0.89221 | 5.80952 | 14.79237 |
| **Epcam** | 0.41327 | 8.21450 | 6.39939 | 14.58615 | 2.31847 | 24.79731 | 11.33983 | 20.83212 | 3.22999 | 5.95050 | 15.57694 | 5.33739 | 2.68506 | 3.81729 |
| **Epg5** | 1.67728 | 0.09709 | 5.42363 | 2.01423 | 1.51099 | 5.76502 | 3.26563 | 3.78564 | 2.99616 | 0.10886 | 1.09141 | 0.83263 | 1.79289 | 1.98734 |
| **Epha1** | 0.01615 | 0.00430 | 0.00909 | 0.01227 | 0.01118 | 0.00000 | 0.01153 | 0.05950 | 0.01163 | 0.02117 | 0.35845 | 0.00896 | 0.01111 | 0.03337 |
| **Epha10** | 0.23398 | 6.81076 | 0.23740 | 0.82779 | 1.96168 | 1.01193 | 0.00748 | 2.13634 | 0.00000 | 0.01402 | 0.00000 | 0.00000 | 1.28451 | 1.69879 |
| **Epha2** | 0.11013 | 0.01518 | 1.14098 | 0.02008 | 0.00000 | 0.00000 | 0.05691 | 0.25690 | 0.00765 | 0.03058 | 0.00395 | 0.19379 | 0.00279 | 0.13212 |
| **Epha3** | 0.01984 | 0.00437 | 0.01468 | 0.01246 | 0.01080 | 0.01300 | 0.00427 | 0.00958 | 0.01113 | 0.00664 | 0.00878 | 0.00938 | 0.24432 | 0.00620 |

| | | | | | | | | | | | | | | |
|---|---|---|---|---|---|---|---|---|---|---|---|---|---|---|
| **Epha4** | 4.56155 | 0.02492 | 3.45286 | 1.06575 | 0.00000 | 0.00082 | 0.00000 | 0.00000 | 5.78821 | 0.00000 | 1.68159 | 9.97187 | 2.35805 | 0.46282 |
| **Epha7** | 0.00000 | 0.00444 | 0.28992 | 0.30919 | 0.00040 | 0.00000 | 0.00427 | 0.00939 | 0.01129 | 0.00582 | 0.03310 | 0.06663 | 0.01640 | 0.14624 |
| **Ephb2** | 0.60582 | 0.01819 | 0.09301 | 0.00000 | 0.30271 | 0.69289 | 0.03063 | 2.06757 | 0.46796 | 0.04227 | 0.00000 | 2.07968 | 0.39639 | 0.34540 |
| **Ephb3** | 0.00000 | 0.08019 | 0.56640 | 7.00865 | 0.45059 | 4.61729 | 2.32682 | 5.89155 | 0.00000 | 0.06256 | 3.05985 | 8.07449 | 0.90272 | 0.12117 |
| **Ephb4** | 0.44961 | 0.01535 | 3.50409 | 0.39034 | 0.82232 | 0.42204 | 0.00740 | 0.53171 | 0.03617 | 0.01758 | 0.03793 | 3.94338 | 0.27928 | 1.47364 |
| **Ephb6** | 0.00000 | 0.01760 | 0.30988 | 0.35544 | 0.00000 | 0.23205 | 3.08826 | 0.49040 | 0.00313 | 0.07494 | 0.00000 | 3.87430 | 0.00619 | 0.09613 |
| **Ephx1** | 0.00000 | 0.00827 | 2.17021 | 2.25241 | 0.05623 | 1.45681 | 0.00290 | 0.00000 | 0.00905 | 0.01378 | 2.70022 | 3.33538 | 0.01545 | 0.00000 |
| **Ephx2** | 0.35774 | 0.06520 | 0.00000 | 0.00000 | 0.56163 | 0.00635 | 0.01011 | 0.05505 | 0.00442 | 0.01417 | 0.00316 | 5.00655 | 0.02494 | 0.41371 |
| **Ephx3** | 0.00490 | 0.00445 | 0.04557 | 0.00191 | 0.00813 | 0.02903 | 0.00434 | 0.00737 | 0.00676 | 0.00737 | 0.00885 | 0.07229 | 0.00382 | 0.60203 |
| **Ephx4** | 0.00000 | 0.00389 | 0.25076 | 0.82040 | 0.00000 | 0.15403 | 0.00928 | 0.00000 | 0.01542 | 0.00865 | 0.48658 | 0.01273 | 0.00697 | 0.00000 |
| **Epm2a** | 0.00227 | 0.00446 | 0.17683 | 0.00356 | 0.11257 | 0.00492 | 0.00419 | 0.01180 | 0.01851 | 0.00702 | 0.00667 | 0.00871 | 0.03447 | 0.34257 |
| **Epm2aip1** | 0.54650 | 0.00361 | 0.43169 | 1.63559 | 1.43381 | 1.52555 | 0.77524 | 0.27401 | 3.86134 | 0.01876 | 1.77298 | 0.67524 | 5.22096 | 1.02440 |
| **Epn1** | 0.52749 | 0.00365 | 0.08342 | 0.61479 | 0.90778 | 0.19815 | 1.15156 | 0.29962 | 0.06674 | 0.21812 | 0.02004 | 0.51265 | 2.31181 | 0.13064 |
| **Epn2** | 9.86224 | 0.01778 | 9.03558 | 0.00000 | 10.40529 | 0.60373 | 2.43572 | 3.37091 | 2.61757 | 0.07086 | 3.72546 | 0.03232 | 8.72477 | 12.61852 |
| **Epn3** | 0.06688 | 0.03223 | 0.02013 | 0.00611 | 0.01156 | 0.01676 | 0.06922 | 1.32245 | 0.01132 | 0.01275 | 0.00547 | 0.00798 | 0.04159 | 0.00486 |
| **Eprs** | 16.42323 | 0.32412 | 17.92121 | 10.35810 | 16.22242 | 19.02308 | 0.17271 | 10.75592 | 21.63308 | 0.24014 | 18.38742 | 14.53193 | 5.07544 | 18.65699 |
| **Eps15** | 2.71587 | 0.00675 | 11.88458 | 9.27548 | 2.55627 | 8.59681 | 1.41096 | 0.00000 | 8.43998 | 0.08181 | 7.83176 | 3.44125 | 1.17381 | 3.73299 |
| **Eps15l1** | 2.76210 | 0.00812 | 5.19276 | 1.39432 | 3.59244 | 3.39708 | 4.20867 | 0.60392 | 0.94418 | 9.09174 | 1.39983 | 3.31711 | 2.80393 | 3.90175 |
| **Eps8** | 0.00000 | 0.02282 | 0.03351 | 0.02942 | 0.00000 | 0.00000 | 0.00369 | 0.08194 | 2.04208 | 0.00856 | 0.00000 | 0.63630 | 2.44225 | 2.85675 |
| **Eps8l1** | 0.00357 | 0.00381 | 0.00916 | 0.94480 | 0.06173 | 0.08342 | 0.00975 | 0.00350 | 0.01111 | 0.00633 | 0.00366 | 0.31061 | 0.01013 | 0.00866 |
| **Eps8l2** | 0.40891 | 0.01166 | 3.12523 | 2.90347 | 0.01417 | 1.37180 | 0.01856 | 0.42825 | 0.00212 | 0.02207 | 0.65977 | 0.00000 | 0.01172 | 0.00000 |
| **Ept1** | 0.00000 | 0.01530 | 3.95009 | 2.72011 | 1.60879 | 5.43297 | 0.00720 | 0.12179 | 12.78832 | 0.00000 | 7.17056 | 2.79922 | 0.00000 | 10.70231 |
| **Eqtn** | 0.00922 | 0.00443 | 0.17847 | 0.00000 | 0.09740 | 0.00857 | 0.00476 | 0.01125 | 0.01956 | 0.00753 | 0.00729 | 0.00994 | 0.00606 | 0.00672 |
| **Eral1** | 0.74828 | 0.03525 | 2.62646 | 1.60134 | 1.16974 | 2.14152 | 0.24000 | 0.20816 | 2.14986 | 0.02372 | 1.83691 | 0.45956 | 0.04252 | 4.88759 |
| **Erap1** | 0.52197 | 0.00444 | 0.44034 | 0.00000 | 0.07704 | 0.12068 | 0.00391 | 0.00210 | 1.91962 | 0.00440 | 0.00353 | 0.05820 | 0.48660 | 1.01229 |
| **Eras** | 0.01899 | 0.00534 | 0.77980 | 2.12164 | 0.00000 | 0.00000 | 0.00450 | 0.51280 | 0.01196 | 0.00684 | 0.00417 | 0.00565 | 0.00042 | 0.25433 |
| **Erbb2** | 0.00000 | 0.01899 | 0.91212 | 1.79554 | 0.00000 | 0.96795 | 0.61702 | 7.38435 | 0.04161 | 0.07075 | 1.83352 | 0.00000 | 0.00268 | 0.00000 |
| **Erbb3** | 0.00000 | 0.02833 | 0.06591 | 1.24730 | 0.00600 | 0.76996 | 0.02190 | 0.82194 | 0.08072 | 0.01962 | 0.45274 | 0.19325 | 0.00557 | 0.06188 |
| **Erbin** | 3.54157 | 4.83913 | 0.57438 | 0.00000 | 1.92888 | 3.48073 | 0.19021 | 1.80843 | 4.97922 | 0.00335 | 2.69274 | 0.25242 | 0.55428 | 4.41630 |
| **Erc1** | 1.41759 | 0.01755 | 1.42286 | 0.00002 | 1.27852 | 1.64341 | 0.01447 | 0.00000 | 1.21586 | 0.01488 | 1.72831 | 0.80456 | 1.25832 | 1.03111 |
| **Erc2** | 0.00571 | 0.00424 | 0.01444 | 0.01089 | 0.00407 | 0.01212 | 0.00420 | 0.00839 | 0.07083 | 0.00614 | 0.00802 | 0.02562 | 0.52744 | 0.06231 |
| **Ercc1** | 1.95903 | 0.06192 | 5.40407 | 6.59749 | 3.96966 | 1.32168 | 0.09222 | 3.16253 | 0.09468 | 0.00704 | 3.78731 | 0.64274 | 2.01295 | 4.48987 |
| **Ercc2** | 0.13836 | 0.03434 | 2.76274 | 1.58855 | 3.04359 | 1.37173 | 0.00331 | 1.29801 | 0.00000 | 0.00757 | 0.88073 | 1.36089 | 1.20735 | 0.20067 |
| **Ercc3** | 6.40302 | 0.01638 | 8.72340 | 2.00637 | 5.19732 | 3.84072 | 1.67883 | 1.85429 | 0.65999 | 0.01102 | 5.31122 | 3.21168 | 1.65833 | 10.12250 |
| **Ercc4** | 0.05796 | 0.03053 | 3.31041 | 1.97974 | 0.98005 | 1.13983 | 1.57140 | 2.87426 | 0.15065 | 4.19420 | 0.03943 | 1.12111 | 2.36373 | 0.01186 |
| **Ercc5** | 0.34647 | 0.00285 | 0.82850 | 1.17401 | 0.56509 | 0.69143 | 0.00869 | 0.34942 | 0.01655 | 0.00466 | 0.10110 | 1.46356 | 0.59296 | 1.49385 |
| **Ercc6** | 1.58272 | 0.06851 | 2.55738 | 0.75817 | 3.19434 | 1.29264 | 0.09979 | 0.28761 | 3.87789 | 0.11366 | 0.24310 | 0.02127 | 4.71611 | 7.52106 |
| **Ercc6l** | 1.71666 | 0.01078 | 2.00801 | 0.66107 | 6.16851 | 3.20165 | 0.00802 | 0.96789 | 11.24441 | 0.03084 | 2.38028 | 0.74546 | 6.50539 | 7.13272 |
| **Ercc6l2** | 1.24259 | 0.00250 | 1.01777 | 0.58671 | 2.19183 | 1.10262 | 0.00243 | 0.00000 | 5.21555 | 0.00000 | 3.90100 | 0.15128 | 1.22099 | 0.39732 |
| **Ercc8** | 2.71063 | 0.02979 | 5.28251 | 4.04436 | 5.39917 | 6.85413 | 3.03858 | 0.70486 | 0.83984 | 0.02973 | 0.77808 | 1.82878 | 0.00000 | 6.77237 |
| **Erdr1** | 1.07695 | 2.07179 | 1.05169 | 3.50533 | 9.38617 | 1.84923 | 1.01439 | 3.15478 | 0.09371 | 0.65219 | 0.04669 | 1.68478 | 0.53496 | 0.29023 |
| **Erf** | 1.88540 | 0.29698 | 0.00000 | 0.00000 | 1.23185 | 0.98981 | 1.06608 | 9.06030 | 0.00001 | 0.36296 | 0.03839 | 0.64194 | 1.40882 | 0.00000 |
| **Erg** | 0.00009 | 0.00442 | 0.00000 | 0.01310 | 0.28043 | 0.00834 | 0.00459 | 0.00940 | 0.91862 | 0.00672 | 0.00619 | 0.00778 | 0.32898 | 0.00000 |
| **Ergic1** | 0.91634 | 0.02859 | 3.21921 | 0.79188 | 0.24366 | 1.93055 | 0.00857 | 2.04170 | 0.39409 | 0.00360 | 1.40644 | 0.50784 | 0.50485 | 0.20054 |
| **Ergic2** | 4.50805 | 0.03627 | 7.36487 | 2.50895 | 4.41983 | 5.16887 | 2.74931 | 2.11487 | 11.93851 | 0.04047 | 6.89545 | 4.96362 | 0.33817 | 9.18377 |
| **Ergic3** | 9.38572 | 0.25354 | 16.02007 | 21.11670 | 7.97316 | 14.49282 | 3.09147 | 11.79441 | 3.98725 | 0.20163 | 11.28952 | 11.95778 | 4.76276 | 7.93747 |
| **Erh** | 18.27391 | 10.61123 | 12.52353 | 25.41221 | 26.15414 | 25.48561 | 6.88475 | 17.46224 | 12.46120 | 15.33884 | 9.14474 | 18.45993 | 13.17967 | 17.47086 |
| **Eri1** | 10.94549 | 0.01366 | 5.46009 | 0.94716 | 8.30355 | 6.93516 | 0.03072 | 0.01745 | 10.32883 | 0.01247 | 4.16106 | 2.87093 | 7.83095 | 9.00584 |
| **Eri2** | 1.11339 | 0.00444 | 2.45917 | 1.95922 | 1.71107 | 1.35495 | 0.11951 | 0.19052 | 4.68824 | 0.02417 | 0.09675 | 0.32563 | 0.05666 | 1.75173 |
| **Eri3** | 2.08822 | 0.02174 | 2.01807 | 4.41037 | 2.95333 | 5.56161 | 3.37796 | 0.91168 | 0.00000 | 0.05524 | 5.18713 | 7.91960 | 0.07902 | 0.30616 |
| **Erich1** | 1.02991 | 0.13092 | 2.10811 | 0.60321 | 0.63147 | 4.18480 | 0.04385 | 0.00007 | 1.62696 | 0.54850 | 0.90117 | 1.54909 | 0.05208 | 2.43100 |
| **Erich5** | 0.26713 | 0.12992 | 0.00000 | 0.89537 | 0.60467 | 0.69739 | 0.19301 | 0.34716 | 0.25568 | 0.28816 | 0.01833 | 0.23056 | 2.10900 | 0.70347 |
| **Erlec1** | 3.02449 | 0.00142 | 0.17397 | 0.09453 | 0.61644 | 0.73425 | 0.00295 | 0.23382 | 0.10984 | 0.01421 | 1.12366 | 3.03702 | 0.68537 | 0.00000 |
| **Erlin1** | 2.94057 | 0.06105 | 2.68907 | 1.87016 | 2.01206 | 0.40096 | 1.41187 | 3.70062 | 10.11424 | 0.01054 | 1.46791 | 1.25553 | 1.80949 | 7.08560 |
| **Erlin2** | 3.66425 | 0.00743 | 4.38344 | 6.80274 | 5.19941 | 3.62448 | 0.00653 | 4.41429 | 8.06951 | 0.07953 | 4.35358 | 3.30730 | 0.00000 | 8.99941 |

| | | | | | | | | | | | | | | |
|---|---|---|---|---|---|---|---|---|---|---|---|---|---|---|
| **Ermap** | 0.01226 | 0.01190 | 0.02411 | 0.02391 | 0.43872 | 0.03830 | 0.00481 | 0.01234 | 0.06927 | 0.00675 | 0.01010 | 0.00881 | 0.41371 | 0.03538 |
| **Ermard** | 1.09709 | 0.01180 | 0.47045 | 1.19885 | 0.00000 | 0.70891 | 0.64449 | 0.08778 | 3.11996 | 0.02585 | 2.09870 | 2.01188 | 0.96863 | 0.17866 |
| **Ermn** | 0.01865 | 0.02652 | 0.03132 | 0.05012 | 0.01109 | 0.03102 | 0.01644 | 0.05881 | 0.04478 | 0.00698 | 0.00873 | 0.00836 | 0.03831 | 0.03829 |
| **Ermp1** | 8.33362 | 1.09578 | 12.85995 | 3.60305 | 4.22317 | 19.31173 | 2.66646 | 5.81011 | 14.93397 | 0.04115 | 6.71149 | 3.66072 | 0.54115 | 12.31668 |
| **Ern1** | 0.66127 | 0.00382 | 0.89494 | 0.00000 | 0.85036 | 0.00000 | 0.00389 | 0.00854 | 0.00238 | 0.00408 | 0.00502 | 0.00511 | 2.37992 | 1.35908 |
| **Ero1l** | 0.82897 | 0.00803 | 0.85837 | 0.38359 | 1.47078 | 1.42502 | 0.00281 | 0.71061 | 1.83273 | 0.01231 | 2.76116 | 3.81189 | 1.04268 | 2.50472 |
| **Ero1lb** | 0.14351 | 0.00363 | 0.04448 | 0.55154 | 0.28908 | 0.00000 | 0.61531 | 0.00173 | 0.51934 | 0.01528 | 0.13896 | 0.01547 | 1.84851 | 2.08126 |
| **Erp29** | 6.52994 | 0.23817 | 11.81401 | 14.94992 | 8.12324 | 10.68219 | 5.60499 | 12.63332 | 0.52087 | 4.33170 | 3.92465 | 11.08039 | 2.49097 | 5.42184 |
| **Erp44** | 6.83631 | 5.25822 | 7.55111 | 4.89593 | 4.41623 | 6.27331 | 5.92037 | 0.04972 | 2.91180 | 0.20787 | 4.06044 | 5.58103 | 1.98197 | 3.15961 |
| **Errfi1** | 0.75678 | 5.03446 | 0.44918 | 0.00000 | 1.23668 | 0.99242 | 3.91220 | 8.70239 | 0.52479 | 0.02842 | 0.00000 | 1.17666 | 8.03019 | 3.60759 |
| **Esam** | 0.00671 | 0.00383 | 0.82596 | 0.68063 | 0.31059 | 0.00000 | 0.00391 | 0.00282 | 0.00603 | 0.00850 | 0.00404 | 0.46329 | 0.00942 | 0.26983 |
| **Esco1** | 3.71804 | 0.00755 | 2.60029 | 0.11705 | 2.29120 | 3.45130 | 0.00000 | 0.00000 | 17.41617 | 0.00286 | 1.80523 | 2.01938 | 12.71714 | 3.66718 |
| **Esco2** | 11.68390 | 0.01046 | 13.57350 | 4.56392 | 13.75962 | 4.25198 | 0.33592 | 1.67602 | 25.39690 | 0.00000 | 9.13679 | 3.92327 | 11.21803 | 14.00198 |
| **Esd** | 7.42018 | 0.13873 | 3.27588 | 3.16756 | 1.36929 | 6.06343 | 0.56763 | 2.27720 | 2.49503 | 1.41978 | 5.05299 | 5.42270 | 0.19728 | 3.01124 |
| **Esf1** | 8.44505 | 0.00637 | 4.09355 | 0.00000 | 2.83304 | 3.03817 | 0.01640 | 1.09613 | 23.87199 | 0.01134 | 6.63003 | 2.32931 | 7.41848 | 9.48339 |
| **Esp24** | 0.06655 | 0.00432 | 0.02198 | 0.02635 | 0.01586 | 0.03387 | 0.00425 | 0.00780 | 0.04318 | 0.00641 | 0.00881 | 0.00731 | 1.25567 | 0.07111 |
| **Espl1** | 0.41386 | 0.22957 | 13.54851 | 12.39622 | 20.76864 | 0.00000 | 1.85186 | 4.69810 | 0.04255 | 0.04152 | 0.07909 | 5.01874 | 8.16117 | 0.00000 |
| **Esr1** | 0.10856 | 0.00453 | 0.00506 | 0.00912 | 0.71732 | 0.01219 | 0.00433 | 0.01125 | 0.00910 | 0.00642 | 0.00769 | 0.00711 | 0.01758 | 0.01957 |
| **Esr2** | 0.03253 | 0.00463 | 0.01172 | 0.03713 | 0.03083 | 0.05066 | 0.00431 | 0.01402 | 0.27027 | 0.03462 | 0.00778 | 0.01933 | 0.83177 | 0.02612 |
| **Esrp1** | 5.17239 | 0.80793 | 1.66442 | 3.08319 | 5.46284 | 2.85705 | 4.29193 | 4.05239 | 5.94689 | 2.57060 | 0.00000 | 1.60899 | 3.88754 | 6.41866 |
| **Esrp2** | 0.55906 | 4.20459 | 9.83682 | 4.07044 | 7.58946 | 5.46523 | 0.43001 | 3.11206 | 2.71686 | 0.04089 | 5.03489 | 0.44131 | 0.00000 | 8.20759 |
| **Esrra** | 2.07118 | 9.94292 | 3.77110 | 5.69490 | 11.12665 | 2.77879 | 1.89254 | 0.92614 | 0.00000 | 7.11670 | 0.00000 | 3.16168 | 1.94081 | 3.45081 |
| **Esrrb** | 14.71691 | 0.10200 | 20.94734 | 8.35824 | 18.43047 | 2.70546 | 0.02108 | 0.00000 | 7.70558 | 1.39035 | 9.96309 | 13.18656 | 3.74184 | 7.75077 |
| **Esrrg** | 0.00292 | 0.00434 | 0.01155 | 0.01271 | 0.00324 | 0.01231 | 0.00424 | 0.00914 | 0.02437 | 0.00617 | 0.00809 | 0.00915 | 0.27331 | 0.27197 |
| **Esyt1** | 0.88348 | 0.08389 | 4.90816 | 2.47989 | 0.61486 | 3.98739 | 3.11739 | 5.73664 | 0.00000 | 0.05474 | 0.00306 | 0.00000 | 4.72048 | 3.83285 |
| **Esyt2** | 5.57501 | 0.01715 | 2.57771 | 0.00000 | 1.60195 | 3.21676 | 1.35072 | 2.98653 | 13.49751 | 0.01175 | 1.02014 | 2.69230 | 1.23469 | 3.29253 |
| **Esyt3** | 1.19854 | 0.17609 | 0.04321 | 0.46082 | 0.20765 | 2.21096 | 1.69709 | 0.23064 | 1.36206 | 0.03331 | 1.37344 | 1.23098 | 0.81528 | 1.66981 |
| **Etaa1** | 6.48864 | 0.00169 | 3.71694 | 0.00000 | 9.29946 | 2.18090 | 0.57109 | 0.73216 | 11.83082 | 0.03830 | 0.00000 | 0.68284 | 10.20398 | 5.32067 |
| **Etaa1os** | 0.00673 | 0.00437 | 0.01857 | 0.01161 | 0.03536 | 0.01254 | 0.00432 | 0.01050 | 0.00668 | 0.00689 | 0.00873 | 0.00978 | 0.15615 | 0.03982 |
| **Etd** | 0.00462 | 0.00439 | 0.01179 | 0.01281 | 0.00685 | 0.01150 | 0.00421 | 0.00968 | 0.11487 | 0.00649 | 0.00818 | 0.00949 | 0.21936 | 0.00000 |
| **Etf1** | 22.58805 | 6.21301 | 25.70697 | 19.45754 | 25.21125 | 27.04010 | 1.92170 | 12.91091 | 27.59667 | 6.72510 | 18.51684 | 11.99429 | 15.66409 | 24.79416 |
| **Etfa** | 13.00964 | 7.49827 | 15.71354 | 11.88521 | 13.55557 | 15.23494 | 7.10709 | 10.50665 | 6.08348 | 0.37202 | 8.26008 | 14.19098 | 4.36472 | 6.49711 |
| **Etfb** | 6.08774 | 8.20895 | 9.59622 | 19.38575 | 6.89739 | 24.20996 | 7.25393 | 21.87579 | 0.11704 | 18.96173 | 11.55090 | 14.80177 | 0.00000 | 4.17887 |
| **Etfdh** | 0.26837 | 0.00286 | 1.25964 | 0.00000 | 0.29354 | 2.10627 | 0.00962 | 0.79402 | 4.61690 | 0.33211 | 1.22375 | 1.41166 | 0.00000 | 0.49923 |
| **Ethe1** | 0.20498 | 0.03133 | 1.79493 | 7.77842 | 2.30057 | 6.22501 | 0.02117 | 1.58020 | 0.00200 | 0.02913 | 0.00000 | 1.21148 | 1.16689 | 0.00000 |
| **Etl4** | 0.19526 | 0.02634 | 2.62683 | 1.43369 | 0.86804 | 3.70930 | 2.03918 | 1.77786 | 3.57753 | 0.08193 | 3.66100 | 2.63892 | 0.00000 | 2.19106 |
| **Etnk1** | 9.83267 | 0.10475 | 4.80763 | 0.00000 | 14.51828 | 6.56452 | 3.76624 | 2.75701 | 12.41726 | 0.07397 | 0.00000 | 4.34164 | 2.59013 | 22.90638 |
| **Etnk2** | 0.00265 | 0.01473 | 0.36855 | 0.00000 | 0.00695 | 0.20129 | 0.00396 | 0.00852 | 0.01003 | 0.00556 | 0.00209 | 0.53280 | 0.00884 | 0.01033 |
| **Etohd2** | 0.00304 | 0.00530 | 0.05068 | 0.05492 | 0.47918 | 0.57699 | 0.01067 | 0.02464 | 0.00608 | 0.00997 | 0.01168 | 0.00134 | 0.41610 | 0.05256 |
| **Ets2** | 0.00582 | 3.50249 | 0.01315 | 0.00000 | 0.59176 | 0.45028 | 0.02965 | 0.89225 | 0.03040 | 0.00595 | 0.00000 | 0.50767 | 1.09998 | 0.12873 |
| **Etv1** | 0.00408 | 0.00375 | 0.00903 | 0.00728 | 0.00388 | 0.01056 | 0.00400 | 0.00638 | 0.01111 | 0.01492 | 0.00380 | 0.92205 | 0.01654 | 0.52114 |
| **Etv3** | 0.25795 | 0.00384 | 0.09145 | 0.02731 | 0.50576 | 0.14388 | 0.00967 | 0.29804 | 0.11909 | 0.01267 | 0.00417 | 0.04758 | 1.00088 | 3.00701 |
| **Etv4** | 0.01361 | 0.02216 | 0.07114 | 0.00000 | 0.01278 | 0.01005 | 0.00388 | 0.01909 | 0.01128 | 0.01355 | 0.00445 | 0.60984 | 0.00688 | 0.01113 |
| **Etv5** | 0.19750 | 0.10583 | 4.86518 | 0.78067 | 2.98557 | 0.77931 | 0.00437 | 0.00000 | 0.87432 | 0.05001 | 7.48736 | 12.28449 | 4.17358 | 0.87548 |
| **Etv6** | 0.20200 | 0.00200 | 5.24885 | 0.00000 | 6.87182 | 0.00000 | 0.00119 | 0.00000 | 8.85262 | 0.00000 | 0.31058 | 1.83708 | 2.23054 | 6.17750 |
| **EU599041** | 0.35162 | 0.01377 | 0.19111 | 0.78785 | 0.24045 | 0.32430 | 0.01633 | 0.27337 | 2.97263 | 0.54351 | 0.03109 | 0.64444 | 0.43057 | 0.46883 |
| **Eva1a** | 0.00000 | 6.48697 | 9.92134 | 4.85091 | 9.40203 | 3.42000 | 0.66898 | 5.96539 | 2.41784 | 0.04451 | 7.02159 | 6.64569 | 2.24939 | 1.10146 |
| **Eva1b** | 0.01399 | 0.00381 | 0.01018 | 0.06092 | 0.01514 | 0.02145 | 0.00390 | 0.60565 | 0.01527 | 0.01158 | 0.00557 | 0.08230 | 0.01796 | 0.01090 |
| **Evi5** | 12.62209 | 0.01917 | 10.60752 | 2.22721 | 10.89219 | 11.68966 | 0.00441 | 0.74131 | 18.35326 | 0.43692 | 3.67536 | 6.02427 | 10.17744 | 18.11162 |
| **Evi5l** | 0.00262 | 0.00305 | 0.00347 | 1.74548 | 0.39603 | 0.00000 | 0.00352 | 0.00309 | 0.00767 | 0.00498 | 0.00649 | 1.04931 | 0.43019 | 0.00000 |
| **Evl** | 0.89090 | 0.00317 | 0.12749 | 0.49268 | 0.10322 | 0.04410 | 0.00921 | 0.00947 | 0.14077 | 0.00511 | 0.00263 | 1.58878 | 0.01648 | 0.64839 |
| **Evpl** | 0.00000 | 0.02444 | 0.23684 | 1.75297 | 0.04027 | 0.00000 | 0.00442 | 0.42594 | 0.00915 | 0.00542 | 0.35422 | 0.00311 | 0.03804 | 0.00000 |
| **Evx1** | 0.01223 | 0.00442 | 0.02048 | 0.00838 | 0.02000 | 0.16014 | 0.00419 | 0.00560 | 0.01192 | 0.00850 | 0.00736 | 0.00870 | 0.08851 | 0.00188 |
| **Evx1os** | 0.00785 | 0.00426 | 0.01302 | 0.12818 | 0.01080 | 0.00846 | 0.00407 | 0.00694 | 0.01226 | 0.00676 | 0.00702 | 0.01482 | 0.00688 | 0.01341 |
| **Ewsr1** | 4.49012 | 5.32114 | 14.69544 | 20.92695 | 12.26099 | 6.76428 | 6.73444 | 12.16670 | 2.18317 | 14.32358 | 12.78601 | 10.59660 | 7.20695 | 5.16581 |

| | | | | | | | | | | | | | | |
|---|---|---|---|---|---|---|---|---|---|---|---|---|---|---|
| **Exd1** | 0.81273 | 0.00315 | 1.07200 | 0.00000 | 4.79296 | 0.00000 | 0.00332 | 0.00329 | 2.17540 | 0.00387 | 1.53476 | 0.00069 | 2.26428 | 5.26983 |
| **Exd2** | 3.74419 | 0.01503 | 2.78602 | 1.32053 | 6.31259 | 1.27428 | 0.00222 | 0.00000 | 2.06726 | 0.00000 | 0.00000 | 2.96230 | 5.47258 | 7.45469 |
| **Exo1** | 8.81775 | 0.03922 | 8.83418 | 3.73815 | 12.96298 | 4.33785 | 0.00021 | 1.23536 | 2.15848 | 0.00000 | 7.45612 | 4.01465 | 6.23259 | 7.13845 |
| **Exo5** | 0.44487 | 0.01120 | 3.48194 | 0.10937 | 3.73081 | 2.72967 | 3.24660 | 0.74063 | 0.17199 | 0.13098 | 1.72505 | 1.20076 | 0.12905 | 3.53904 |
| **Exoc1** | 4.58751 | 4.32718 | 4.66673 | 1.63762 | 3.93302 | 2.66812 | 0.00850 | 0.15437 | 1.25460 | 0.00000 | 5.62343 | 2.06881 | 0.82905 | 9.84974 |
| **Exoc2** | 1.26405 | 0.00274 | 1.59985 | 0.10581 | 2.10331 | 3.28426 | 0.00338 | 0.00000 | 0.52639 | 0.00706 | 0.32529 | 1.32515 | 1.16588 | 2.85667 |
| **Exoc3** | 11.26480 | 0.02174 | 6.43405 | 4.23672 | 5.31369 | 5.91788 | 0.01301 | 1.54251 | 3.83651 | 0.00000 | 6.74273 | 3.60612 | 1.35963 | 6.04160 |
| **Exoc3l** | 0.00525 | 0.00412 | 0.05697 | 0.31983 | 0.16678 | 0.00374 | 0.00392 | 0.00643 | 0.01267 | 0.00573 | 0.00970 | 0.01603 | 0.00597 | 0.00597 |
| **Exoc3l4** | 0.02097 | 0.00427 | 4.26286 | 0.05756 | 0.00000 | 0.02579 | 0.00354 | 0.00866 | 0.01240 | 0.01561 | 0.11076 | 0.00437 | 0.00000 | 0.01081 |
| **Exoc4** | 0.09199 | 0.01725 | 0.51995 | 0.00000 | 0.68454 | 1.40323 | 0.00828 | 0.09969 | 0.76607 | 0.01429 | 1.65834 | 2.31585 | 0.00474 | 0.54813 |
| **Exoc5** | 9.02932 | 0.00000 | 3.99397 | 0.89077 | 4.53519 | 3.77513 | 0.06393 | 1.74972 | 12.11384 | 0.01651 | 2.67226 | 4.88355 | 1.46992 | 2.73737 |
| **Exoc6** | 0.56158 | 0.00745 | 3.36075 | 0.03414 | 1.73500 | 0.00000 | 0.10967 | 0.15099 | 2.17555 | 8.77146 | 6.70803 | 0.83247 | 0.63780 | 2.83012 |
| **Exoc6b** | 0.40987 | 0.00282 | 0.62533 | 0.00000 | 0.23924 | 0.64148 | 0.00303 | 0.41027 | 0.73744 | 0.00401 | 0.06384 | 1.41391 | 0.00000 | 0.00000 |
| **Exoc7** | 0.89257 | 0.04523 | 0.69300 | 1.34745 | 1.17811 | 0.91892 | 2.93911 | 1.14362 | 0.00000 | 0.05202 | 3.22470 | 2.40227 | 0.68403 | 0.77257 |
| **Exoc8** | 1.71897 | 0.04296 | 1.51538 | 0.52077 | 2.45576 | 5.05805 | 0.08303 | 1.29274 | 4.97682 | 1.01498 | 2.24903 | 2.38010 | 4.77104 | 0.77452 |
| **Exog** | 1.82124 | 0.01139 | 2.39340 | 0.24659 | 0.50151 | 2.51866 | 1.05761 | 0.00000 | 0.05135 | 0.02512 | 1.21749 | 0.87459 | 0.01044 | 3.30821 |
| **Exosc1** | 9.19579 | 2.11302 | 10.50497 | 4.77579 | 10.10541 | 10.95501 | 3.00136 | 2.16355 | 3.31765 | 0.03922 | 4.61082 | 7.37893 | 3.33661 | 13.59923 |
| **Exosc10** | 10.53065 | 0.15380 | 11.77621 | 6.30249 | 13.73028 | 3.92540 | 3.14177 | 2.74813 | 0.66002 | 6.18226 | 7.43772 | 3.93903 | 10.01321 | 13.07844 |
| **Exosc2** | 4.38510 | 0.19397 | 8.44412 | 2.34350 | 2.90546 | 8.65296 | 1.20344 | 2.14174 | 8.10691 | 1.32505 | 6.98773 | 6.29473 | 11.58554 | 7.59323 |
| **Exosc3** | 4.24127 | 0.01678 | 0.68682 | 7.89331 | 12.87058 | 2.16693 | 1.31731 | 0.00765 | 4.09886 | 0.06896 | 0.05555 | 0.66283 | 7.05495 | 1.99308 |
| **Exosc4** | 0.23633 | 3.43207 | 4.70388 | 12.41091 | 7.86948 | 0.63011 | 0.81406 | 4.43895 | 0.01126 | 7.81938 | 0.18548 | 5.57917 | 0.39087 | 0.62590 |
| **Exosc5** | 0.08629 | 3.90929 | 12.82884 | 15.15785 | 13.00824 | 0.48438 | 0.26756 | 6.69848 | 0.00000 | 0.06362 | 5.75525 | 6.17217 | 0.07354 | 0.00000 |
| **Exosc6** | 1.04758 | 0.03153 | 0.00047 | 2.93699 | 2.73129 | 0.27366 | 0.00406 | 0.28525 | 0.00478 | 0.07889 | 0.00821 | 0.22506 | 6.87542 | 0.05677 |
| **Exosc7** | 13.64541 | 5.62666 | 17.53176 | 12.60096 | 13.62115 | 15.67547 | 3.88396 | 3.99128 | 6.02467 | 6.77159 | 14.74387 | 9.84409 | 1.28732 | 12.21217 |
| **Exosc8** | 11.73245 | 0.25189 | 14.71410 | 11.31361 | 14.67771 | 13.07192 | 6.19643 | 11.05034 | 8.83806 | 0.68191 | 14.85573 | 14.96648 | 3.22415 | 11.95704 |
| **Exosc9** | 8.26857 | 0.04163 | 13.75844 | 6.35209 | 6.31931 | 12.19288 | 4.77969 | 1.31588 | 10.30465 | 0.16729 | 8.31830 | 4.54378 | 0.22249 | 6.39106 |
| **Exph5** | 0.00000 | 0.00460 | 0.45021 | 1.09466 | 0.00000 | 0.99749 | 0.00414 | 0.09749 | 0.00906 | 0.02933 | 0.14230 | 0.48780 | 0.32887 | 0.00000 |
| **Ext1** | 1.02914 | 0.00990 | 0.84533 | 0.00000 | 4.58708 | 0.92256 | 0.00326 | 0.00691 | 0.02360 | 0.00136 | 0.99544 | 2.17437 | 2.24220 | 4.79807 |
| **Ext2** | 0.20720 | 0.01594 | 2.01323 | 0.72738 | 1.03147 | 1.36391 | 0.00243 | 2.14531 | 0.00269 | 0.04414 | 0.77095 | 3.08463 | 0.00000 | 0.02057 |
| **Extl1** | 0.00000 | 0.02956 | 0.44859 | 0.73435 | 0.26213 | 2.47520 | 1.12489 | 3.23314 | 0.00043 | 0.02098 | 0.00602 | 0.00000 | 1.19695 | 0.09048 |
| **Extl2** | 0.01194 | 0.00376 | 0.01288 | 0.03034 | 0.00290 | 0.00911 | 0.00412 | 0.00476 | 0.00780 | 0.00687 | 0.00751 | 0.52573 | 0.00379 | 0.52607 |
| **Extl3** | 0.00000 | 0.01284 | 1.38423 | 1.83538 | 1.02083 | 2.26707 | 0.87178 | 1.36691 | 0.68466 | 0.03166 | 0.00000 | 0.99961 | 0.82059 | 1.10702 |
| **Eya1** | 0.07438 | 0.00439 | 0.01293 | 0.01395 | 0.00681 | 0.01479 | 0.00420 | 0.01388 | 0.00833 | 0.00906 | 0.00759 | 0.01144 | 0.32144 | 0.04882 |
| **Eya2** | 0.00753 | 0.00442 | 0.01364 | 0.01239 | 0.02795 | 0.01307 | 0.00453 | 0.00997 | 0.02471 | 0.00669 | 0.00877 | 0.00972 | 0.10398 | 0.00350 |
| **Eya3** | 2.45991 | 0.01494 | 7.23101 | 1.88997 | 6.17923 | 3.49766 | 2.16082 | 2.39062 | 2.11501 | 0.00713 | 4.96324 | 3.82521 | 1.83864 | 9.38921 |
| **Ezh1** | 1.01120 | 0.01009 | 0.94366 | 1.20028 | 3.28401 | 0.84803 | 0.00259 | 1.15721 | 1.84847 | 0.77592 | 3.33878 | 1.57039 | 5.25335 | 7.86095 |
| **Ezh2** | 19.64459 | 0.15042 | 19.34840 | 9.49822 | 17.42838 | 9.72389 | 3.86281 | 3.74704 | 19.17524 | 0.10448 | 16.05963 | 17.05302 | 12.43529 | 22.16096 |
| **Ezr** | 0.00000 | 0.19441 | 3.07243 | 9.64387 | 0.00000 | 0.88399 | 4.53298 | 9.39776 | 0.00000 | 0.26490 | 12.17194 | 16.35102 | 0.44589 | 0.00000 |
| **F10** | 0.02271 | 0.00477 | 0.02572 | 0.07531 | 0.04043 | 0.05222 | 0.04167 | 0.05541 | 0.11873 | 0.03058 | 0.00706 | 0.04776 | 0.32252 | 0.03888 |
| **F11r** | 9.69160 | 9.19148 | 22.71213 | 21.29809 | 12.35985 | 28.06814 | 9.30310 | 19.91173 | 25.55795 | 0.83295 | 15.62030 | 3.10311 | 2.48268 | 26.54946 |
| **F12** | 0.02638 | 0.00435 | 0.63423 | 0.00502 | 0.00367 | 0.00550 | 0.00410 | 0.00968 | 0.01174 | 0.00667 | 0.00691 | 0.01003 | 0.00500 | 0.00957 |
| **F13a1** | 0.01011 | 0.00444 | 0.01454 | 0.01322 | 0.01359 | 0.01314 | 0.00429 | 0.01026 | 0.01281 | 0.00697 | 0.00882 | 0.01003 | 0.01837 | 0.01188 |
| **F2** | 1.68347 | 0.02139 | 1.68397 | 0.00000 | 0.41790 | 0.42977 | 0.00358 | 0.00065 | 0.00030 | 0.00333 | 0.00186 | 0.24021 | 0.93146 | 0.41242 |
| **F2r** | 0.47603 | 0.00461 | 0.00771 | 0.00014 | 0.15191 | 0.00750 | 0.00482 | 0.09319 | 0.00820 | 0.00814 | 0.00631 | 0.05803 | 0.01906 | 0.01503 |
| **F2rl1** | 0.00000 | 2.11796 | 0.05432 | 0.20482 | 0.00098 | 0.38748 | 0.18232 | 2.39845 | 0.08081 | 0.01211 | 0.00000 | 2.24491 | 0.00340 | 0.00645 |
| **F3** | 0.18581 | 0.00429 | 0.01121 | 0.07419 | 0.00611 | 0.00826 | 0.00414 | 0.00937 | 0.01293 | 0.00640 | 0.01027 | 0.01110 | 0.00452 | 0.01139 |
| **F8** | 0.01730 | 0.00449 | 0.08688 | 0.00000 | 0.01147 | 0.22493 | 0.00421 | 0.00538 | 0.01886 | 0.00587 | 0.00927 | 0.01016 | 0.00701 | 0.00682 |
| **F8a** | 0.22838 | 0.00407 | 0.01003 | 0.53513 | 1.36508 | 0.02286 | 0.00405 | 0.00569 | 0.01223 | 0.01210 | 0.01069 | 0.01118 | 0.00587 | 0.00301 |
| **Fa2h** | 0.00000 | 0.00414 | 0.00814 | 0.01303 | 0.00477 | 0.01129 | 0.00438 | 0.00815 | 0.12269 | 0.00613 | 0.00797 | 0.00749 | 1.71335 | 0.13222 |
| **Faah** | 0.00000 | 3.73676 | 1.61187 | 1.41423 | 0.00000 | 1.26368 | 0.09737 | 3.05447 | 0.00000 | 3.94026 | 0.00000 | 1.80997 | 0.00000 | 0.42446 |
| **Faap100** | 0.00000 | 0.01642 | 1.73230 | 0.61115 | 1.02515 | 0.13626 | 0.00258 | 2.56924 | 0.00942 | 0.00864 | 3.78164 | 0.93555 | 0.00000 | 0.01302 |
| **Faap20** | 2.66449 | 4.89317 | 1.42843 | 4.72975 | 0.88914 | 0.76505 | 2.96062 | 4.48145 | 0.01021 | 0.09498 | 0.00000 | 0.49558 | 0.25263 | 0.13690 |
| **Faap24** | 0.69387 | 0.01918 | 6.20808 | 3.51278 | 8.70063 | 2.88201 | 0.00279 | 0.38937 | 0.08953 | 0.01993 | 1.26048 | 2.50768 | 2.87717 | 1.33847 |
| **Fabp3** | 7.17150 | 15.14617 | 0.86181 | 14.85194 | 2.72250 | 38.15852 | 12.14641 | 40.56993 | 0.04418 | 27.74823 | 8.63120 | 2.68659 | 1.83587 | 0.00000 |
| **Fabp4** | 4.30337 | 0.00307 | 0.00000 | 0.23498 | 1.11359 | 0.00000 | 0.00345 | 0.00348 | 1.66604 | 0.01313 | 0.00194 | 0.00452 | 4.23136 | 2.88582 |

| | | | | | | | | | | | | | | |
|---|---|---|---|---|---|---|---|---|---|---|---|---|---|---|
| **Fabp5** | 0.18825 | 0.98472 | 0.00000 | 6.32051 | 0.15567 | 13.06195 | 2.77417 | 1.39128 | 0.00000 | 1.50096 | 8.56867 | 27.56577 | 0.45028 | 0.73694 |
| **Fabp6** | 0.01069 | 0.00450 | 0.01337 | 0.01078 | 0.01134 | 0.00786 | 0.00498 | 0.06413 | 0.06438 | 0.00669 | 0.00850 | 0.00889 | 0.00694 | 0.00353 |
| **Fabp9** | 0.00456 | 0.00395 | 0.00000 | 0.05509 | 0.21970 | 0.41311 | 0.00442 | 0.00368 | 0.01587 | 0.00833 | 0.00384 | 0.06643 | 0.48112 | 0.00000 |
| **Fadd** | 1.48166 | 0.07217 | 1.12143 | 1.07717 | 3.64027 | 1.33985 | 0.07806 | 1.72612 | 0.54685 | 0.06431 | 0.01254 | 0.38116 | 1.76046 | 1.60185 |
| **Fads1** | 0.33007 | 0.02590 | 1.26752 | 0.00000 | 0.73992 | 1.48338 | 0.00243 | 0.00000 | 3.99462 | 0.00648 | 0.00000 | 2.44704 | 0.05509 | 1.42822 |
| **Fads2** | 0.02031 | 0.00390 | 0.02030 | 0.00000 | 0.08176 | 0.01393 | 0.00381 | 0.15507 | 0.00596 | 0.00722 | 0.00617 | 0.06952 | 0.29870 | 0.04215 |
| **Fads3** | 0.17616 | 0.00324 | 1.75581 | 0.20181 | 0.00000 | 0.31337 | 0.00331 | 0.00000 | 0.05097 | 0.00380 | 0.00001 | 0.85034 | 0.15513 | 1.02630 |
| **Faf1** | 11.81194 | 0.14894 | 13.69591 | 7.02553 | 13.11459 | 9.24461 | 3.25597 | 6.98381 | 10.57566 | 0.14766 | 9.97977 | 9.57633 | 10.12785 | 12.46230 |
| **Faf2** | 3.42216 | 0.06299 | 5.08482 | 1.87351 | 3.38643 | 3.82955 | 0.02976 | 0.94447 | 5.25116 | 6.38138 | 6.90625 | 2.59800 | 4.76519 | 14.16866 |
| **Fah** | 0.00284 | 0.00421 | 0.46475 | 0.02083 | 0.11088 | 0.00000 | 0.00397 | 0.00967 | 0.00507 | 0.00669 | 0.00587 | 0.01010 | 0.14564 | 1.16799 |
| **Fahd1** | 0.00182 | 0.05116 | 0.05873 | 0.00000 | 0.00713 | 0.00000 | 0.02051 | 3.16355 | 0.01074 | 0.01963 | 0.10621 | 2.28686 | 0.01420 | 0.00848 |
| **Fahd2a** | 0.56593 | 0.00392 | 0.57834 | 0.86729 | 0.02192 | 0.09063 | 0.00412 | 0.27416 | 0.71844 | 0.00580 | 0.00599 | 0.24336 | 0.01052 | 0.00000 |
| **Faim** | 2.01293 | 0.03961 | 0.37872 | 2.63958 | 2.02570 | 1.86556 | 0.01000 | 1.29504 | 0.95477 | 0.00456 | 1.50975 | 0.19155 | 1.30586 | 0.08089 |
| **Faiml** | 1.67040 | 0.01169 | 0.26297 | 1.54903 | 0.83547 | 0.20777 | 0.00459 | 0.49198 | 0.90424 | 0.01225 | 0.29199 | 0.63327 | 0.41604 | 2.27558 |
| **Fam101a** | 0.08765 | 0.00444 | 0.01402 | 0.01138 | 0.01158 | 0.01219 | 0.00427 | 0.01096 | 0.01105 | 0.00673 | 0.00794 | 0.00971 | 0.00624 | 0.00940 |
| **Fam101b** | 16.95545 | 5.48786 | 15.02011 | 3.21795 | 12.46492 | 11.02011 | 6.40566 | 13.48694 | 4.87513 | 5.34994 | 7.71249 | 1.96480 | 0.23125 | 8.04791 |
| **Fam102a** | 0.05492 | 1.43823 | 12.55450 | 2.64537 | 7.91626 | 6.06807 | 4.96355 | 3.59072 | 1.59990 | 0.12576 | 1.18973 | 6.45544 | 6.19420 | 6.54797 |
| **Fam102b** | 0.00658 | 0.00421 | 0.10026 | 0.32919 | 0.19656 | 0.00000 | 0.00400 | 0.00695 | 1.28270 | 0.00571 | 0.00320 | 0.00631 | 0.06968 | 0.57470 |
| **Fam103a1** | 14.62601 | 0.01711 | 13.12832 | 7.91957 | 19.09130 | 12.91963 | 2.70993 | 5.34059 | 18.04254 | 0.12613 | 7.68692 | 6.13996 | 13.59848 | 17.49153 |
| **Fam104a** | 2.93882 | 0.00819 | 1.66967 | 4.76292 | 12.53845 | 3.24158 | 0.29640 | 2.94598 | 0.13424 | 1.56649 | 0.73880 | 3.13164 | 5.72048 | 2.92471 |
| **Fam105a** | 0.00000 | 0.00433 | 0.50030 | 0.00000 | 0.21077 | 0.00163 | 0.00397 | 0.01041 | 0.00751 | 0.00752 | 0.00650 | 0.00862 | 0.01042 | 0.57432 |
| **Fam107a** | 0.00434 | 0.00428 | 0.00551 | 0.01193 | 0.43403 | 0.01390 | 0.00488 | 0.00852 | 0.01795 | 0.00630 | 0.00877 | 0.01207 | 0.52910 | 0.00495 |
| **Fam107b** | 0.57537 | 0.03372 | 1.86303 | 3.64845 | 0.00000 | 4.80336 | 1.94962 | 5.13900 | 0.74894 | 0.08644 | 0.00000 | 1.71695 | 1.38817 | 0.00000 |
| **Fam109a** | 0.00000 | 0.01710 | 1.19489 | 0.50335 | 5.59082 | 1.18971 | 3.42240 | 4.53294 | 0.00227 | 0.21135 | 0.00000 | 0.42277 | 10.07116 | 0.54637 |
| **Fam110a** | 0.02277 | 0.04463 | 0.00821 | 0.08637 | 0.01735 | 0.00282 | 0.00429 | 1.32197 | 0.01321 | 0.01755 | 0.00594 | 0.00674 | 0.01817 | 0.00668 |
| **Fam110b** | 0.25153 | 0.00565 | 0.00478 | 0.10784 | 0.00456 | 0.40521 | 0.01091 | 0.27035 | 0.00607 | 0.00920 | 0.00496 | 0.00607 | 0.10001 | 0.01889 |
| **Fam110c** | 0.00535 | 0.00485 | 0.01281 | 0.00307 | 0.01082 | 0.01304 | 0.00408 | 0.01708 | 0.02681 | 0.00634 | 0.00509 | 0.24299 | 0.14640 | 0.00696 |
| **Fam111a** | 0.02117 | 0.00552 | 0.01488 | 0.00000 | 0.00928 | 0.00396 | 0.00441 | 0.28756 | 0.03944 | 0.00826 | 0.00833 | 0.01190 | 0.15458 | 0.00205 |
| **Fam114a1** | 0.04755 | 0.00447 | 0.00631 | 0.00982 | 0.34866 | 0.01282 | 0.00472 | 0.01083 | 0.02280 | 0.00654 | 0.00830 | 0.00836 | 0.00443 | 0.00424 |
| **Fam114a2** | 5.15672 | 0.01197 | 5.79572 | 0.00000 | 2.91703 | 6.20117 | 2.01937 | 2.75807 | 4.48803 | 7.82267 | 7.59369 | 5.52953 | 0.91507 | 7.13827 |
| **Fam117a** | 0.00000 | 0.02526 | 0.32295 | 4.13466 | 0.35539 | 12.15348 | 0.21701 | 4.27868 | 0.07236 | 0.04416 | 1.71696 | 1.15631 | 1.47075 | 0.92091 |
| **Fam117b** | 2.64213 | 0.06613 | 7.12293 | 0.76192 | 1.62724 | 2.77739 | 0.00100 | 0.00000 | 0.14856 | 0.01069 | 0.65388 | 5.74941 | 8.89441 | 3.73922 |
| **Fam118a** | 8.38847 | 0.00635 | 10.60261 | 5.73610 | 14.67280 | 0.32921 | 0.00000 | 0.02515 | 15.02301 | 0.01074 | 2.57454 | 2.64686 | 9.16071 | 15.30350 |
| **Fam118b** | 7.61908 | 0.02766 | 6.66059 | 4.60474 | 10.87158 | 6.10741 | 0.63055 | 5.04963 | 7.54258 | 1.84805 | 1.26905 | 4.61141 | 1.62991 | 9.52230 |
| **Fam120a** | 14.45693 | 0.08139 | 12.19480 | 4.07234 | 11.33626 | 12.53988 | 5.37189 | 5.11615 | 21.88734 | 3.17739 | 8.13848 | 6.90957 | 3.12138 | 6.69153 |
| **Fam120aos** | 0.18145 | 0.00533 | 0.08717 | 0.06021 | 0.18804 | 0.00000 | 0.00455 | 0.46582 | 0.74021 | 0.00880 | 0.00330 | 0.01386 | 0.05540 | 0.23447 |
| **Fam120b** | 1.88307 | 0.00232 | 1.68733 | 0.09866 | 0.47939 | 0.78028 | 0.00226 | 0.36603 | 0.05120 | 7.69777 | 0.00000 | 0.00000 | 1.36492 | 1.08165 |
| **Fam120c** | 0.01251 | 0.00458 | 0.00300 | 0.41035 | 0.40558 | 0.00867 | 0.00403 | 0.00923 | 0.02571 | 0.00665 | 0.00975 | 0.00771 | 0.00242 | 0.06731 |
| **Fam122a** | 1.26038 | 0.00371 | 0.34706 | 0.47212 | 0.55910 | 0.20502 | 2.36673 | 0.00463 | 0.03540 | 0.06379 | 0.00629 | 0.98260 | 0.79600 | 0.15674 |
| **Fam122b** | 1.95097 | 0.01228 | 0.62061 | 0.27807 | 0.14210 | 3.23411 | 0.28715 | 0.75499 | 4.95372 | 0.00814 | 2.27132 | 0.00000 | 0.40123 | 0.78510 |
| **Fam124a** | 0.02090 | 0.01190 | 0.00355 | 0.01133 | 0.01183 | 0.00000 | 0.02657 | 0.02122 | 0.05052 | 0.02452 | 1.03233 | 0.01387 | 0.06136 | 0.02546 |
| **Fam124b** | 0.00921 | 0.00444 | 0.00224 | 0.01650 | 0.00372 | 0.00726 | 0.00405 | 0.00995 | 1.20206 | 0.00750 | 0.01255 | 0.01114 | 0.00886 | 0.00263 |
| **Fam126a** | 0.00000 | 0.00310 | 1.11023 | 0.60878 | 0.00000 | 1.54435 | 1.86225 | 0.03830 | 0.55225 | 0.02922 | 0.00000 | 1.92021 | 0.63908 | 1.16568 |
| **Fam126b** | 2.94509 | 0.63834 | 0.81492 | 0.00000 | 3.26123 | 0.95087 | 2.20535 | 0.36494 | 14.39187 | 0.00378 | 2.04638 | 0.00000 | 4.05681 | 2.37861 |
| **Fam129a** | 1.57279 | 4.76315 | 5.85530 | 7.14291 | 0.00000 | 6.32248 | 0.51265 | 6.31263 | 2.14586 | 0.11735 | 4.91460 | 4.06680 | 0.10846 | 0.00000 |
| **Fam129b** | 0.56283 | 4.60951 | 0.28426 | 1.64982 | 0.51620 | 4.47791 | 1.88426 | 0.23102 | 0.00815 | 0.08838 | 0.00009 | 1.34658 | 0.02025 | 0.00000 |
| **Fam129c** | 0.01539 | 0.00597 | 0.01120 | 0.06042 | 0.00918 | 0.12836 | 0.00456 | 0.28106 | 0.04310 | 0.00821 | 0.00707 | 0.01340 | 0.55172 | 0.00000 |
| **Fam131a** | 0.00000 | 0.00409 | 0.25721 | 0.27009 | 0.36302 | 0.00000 | 0.00417 | 0.00974 | 0.05442 | 0.00514 | 0.00537 | 0.00539 | 1.32116 | 0.02745 |
| **Fam131c** | 0.07651 | 0.00426 | 0.02527 | 0.26270 | 0.37059 | 0.00710 | 0.00411 | 0.00839 | 0.01450 | 0.00812 | 0.00604 | 0.00658 | 0.18878 | 0.00000 |
| **Fam132a** | 1.61665 | 0.01671 | 4.34427 | 6.69061 | 2.18641 | 2.41512 | 1.88012 | 2.52743 | 0.07401 | 0.02162 | 2.20344 | 0.32794 | 2.26092 | 0.00000 |
| **Fam132b** | 0.13399 | 5.68487 | 0.22438 | 1.68119 | 0.30524 | 0.91346 | 0.00325 | 0.69496 | 0.00524 | 0.01191 | 0.10940 | 0.10977 | 0.10512 | 0.59980 |
| **Fam133b** | 2.60904 | 2.59933 | 2.24022 | 1.26453 | 2.12272 | 5.04329 | 2.12198 | 1.67718 | 3.00748 | 0.09614 | 1.20328 | 4.39040 | 1.99829 | 3.68590 |
| **Fam134a** | 10.77830 | 5.97373 | 11.76161 | 8.94927 | 11.14085 | 5.94895 | 0.00732 | 0.67100 | 3.67427 | 0.01462 | 2.78829 | 1.25673 | 7.91596 | 8.26575 |
| **Fam134b** | 0.00703 | 0.00449 | 0.00276 | 0.00965 | 0.56941 | 0.01228 | 0.00506 | 0.01022 | 0.10879 | 0.00675 | 0.00828 | 0.00792 | 0.00563 | 0.02078 |
| **Fam134c** | 1.76360 | 2.59435 | 3.52115 | 1.71493 | 3.63024 | 0.43636 | 0.03866 | 0.59347 | 5.75673 | 0.01002 | 2.32558 | 0.57809 | 4.72107 | 3.25765 |

| | | | | | | | | | | | | | | |
|---|---|---|---|---|---|---|---|---|---|---|---|---|---|---|
| **Fam135a** | 0.35481 | 0.00297 | 0.26037 | 0.27314 | 0.26692 | 0.22238 | 1.54640 | 0.00000 | 2.18661 | 0.00988 | 1.54788 | 0.53255 | 0.03319 | 1.95939 |
| **Fam136a** | 1.71846 | 4.33954 | 7.70166 | 15.59482 | 4.07218 | 12.21578 | 3.72251 | 5.79635 | 0.32903 | 0.23079 | 8.61126 | 9.86642 | 0.00000 | 2.71517 |
| **Fam13b** | 1.70339 | 0.00380 | 1.88731 | 0.41472 | 0.92556 | 0.00000 | 0.00325 | 0.00391 | 0.73891 | 0.00000 | 2.59455 | 0.37059 | 1.33670 | 3.09583 |
| **Fam13c** | 0.00752 | 0.00423 | 0.05212 | 0.00752 | 0.00000 | 0.00944 | 0.00416 | 0.00920 | 0.03561 | 0.00628 | 0.00738 | 0.00888 | 0.74353 | 0.28933 |
| **Fam149b** | 0.56721 | 0.03036 | 0.34349 | 0.00000 | 0.80047 | 0.39460 | 0.00293 | 1.23619 | 0.20762 | 0.00925 | 1.47441 | 1.67408 | 1.02183 | 1.78739 |
| **Fam151a** | 0.00000 | 0.07071 | 5.79549 | 2.24671 | 0.04740 | 4.63955 | 0.00222 | 1.36123 | 0.00000 | 0.00000 | 1.36290 | 2.60058 | 0.00000 | 2.43698 |
| **Fam159a** | 0.01340 | 0.00419 | 0.08983 | 0.00885 | 0.18918 | 0.00663 | 0.00421 | 0.00899 | 0.00910 | 0.00608 | 0.09970 | 0.01511 | 0.53485 | 0.01583 |
| **Fam159b** | 0.42438 | 0.04032 | 0.01156 | 0.52049 | 0.00000 | 0.36399 | 0.00340 | 0.02343 | 1.17665 | 0.01591 | 0.05093 | 2.65258 | 0.27918 | 0.44504 |
| **Fam160a1** | 0.07274 | 0.00439 | 0.90175 | 0.08984 | 0.64300 | 1.37205 | 0.00474 | 1.10218 | 0.20401 | 0.00358 | 0.15283 | 0.00000 | 1.41307 | 0.96683 |
| **Fam160a2** | 1.10391 | 0.05913 | 0.80706 | 3.63617 | 1.07659 | 2.09519 | 0.00939 | 1.85046 | 0.01067 | 0.02939 | 2.40930 | 1.63536 | 0.00911 | 0.09299 |
| **Fam160b1** | 0.71763 | 0.00982 | 1.35740 | 0.00000 | 1.20208 | 2.46330 | 2.09137 | 0.00000 | 0.02623 | 0.04346 | 0.00000 | 2.47815 | 1.72843 | 0.03124 |
| **Fam160b2** | 2.33873 | 6.56163 | 6.03717 | 7.24802 | 4.93269 | 1.53223 | 1.85410 | 2.29515 | 0.76513 | 0.11834 | 1.27647 | 4.50333 | 2.30641 | 1.43311 |
| **Fam161a** | 1.13156 | 0.00400 | 0.46889 | 2.09832 | 1.21354 | 0.00000 | 0.00435 | 0.70139 | 0.05073 | 0.00384 | 1.08905 | 0.01052 | 1.88571 | 1.03572 |
| **Fam161b** | 2.89865 | 0.01383 | 2.30120 | 0.25720 | 2.90730 | 0.68421 | 0.01063 | 0.31272 | 0.46791 | 0.02105 | 1.24571 | 0.20834 | 1.48502 | 1.91178 |
| **Fam162a** | 0.00445 | 0.02148 | 0.00280 | 0.00000 | 0.00330 | 0.83988 | 0.00333 | 0.29734 | 0.00343 | 0.03510 | 0.00000 | 2.79085 | 0.28292 | 0.39579 |
| **Fam163a** | 0.00655 | 0.01134 | 0.01224 | 0.00380 | 0.01172 | 0.01415 | 0.00410 | 0.01346 | 0.01111 | 0.00640 | 0.00494 | 0.42160 | 0.00828 | 0.03750 |
| **Fam166a** | 0.02316 | 0.00444 | 0.01741 | 0.05251 | 0.04097 | 0.01165 | 0.00419 | 0.01253 | 0.01370 | 0.01296 | 0.00840 | 0.00757 | 0.06242 | 0.01788 |
| **Fam167a** | 0.00241 | 0.01361 | 0.01273 | 0.00228 | 0.00845 | 0.01335 | 0.00400 | 0.01555 | 0.00931 | 0.00609 | 0.00473 | 0.40119 | 0.81548 | 0.00403 |
| **Fam167b** | 0.01081 | 0.00443 | 0.00604 | 0.00915 | 0.50315 | 0.01431 | 0.00421 | 0.01006 | 0.01206 | 0.00739 | 0.00892 | 0.00836 | 0.07992 | 0.00288 |
| **Fam168a** | 4.08164 | 0.03347 | 6.63675 | 1.67132 | 4.48728 | 1.41327 | 0.00830 | 0.53079 | 7.61599 | 0.00000 | 2.67200 | 2.73208 | 3.36800 | 6.76585 |
| **Fam168b** | 4.30350 | 0.04403 | 1.40266 | 4.64767 | 5.47282 | 7.08244 | 2.22571 | 3.11358 | 1.68278 | 0.13003 | 3.06052 | 4.28619 | 6.38227 | 7.41566 |
| **Fam169a** | 1.48275 | 0.00979 | 0.89272 | 0.00000 | 0.43013 | 0.12285 | 1.57980 | 2.61476 | 2.09809 | 0.02140 | 0.56229 | 1.49017 | 2.14343 | 4.82447 |
| **Fam169b** | 0.46738 | 0.02755 | 2.48985 | 0.87219 | 3.67140 | 0.23569 | 0.07799 | 0.18823 | 2.73408 | 0.03534 | 0.19867 | 0.15842 | 6.38889 | 7.21789 |
| **Fam171a1** | 0.00807 | 0.00404 | 0.03496 | 0.66405 | 0.01102 | 0.01066 | 0.00391 | 0.00743 | 0.01306 | 0.00551 | 0.01503 | 0.02352 | 0.49714 | 0.00000 |
| **Fam171a2** | 0.07108 | 0.00419 | 0.00817 | 0.01356 | 0.14144 | 0.02229 | 0.00452 | 0.02262 | 0.01560 | 0.01166 | 0.00826 | 0.00762 | 0.03946 | 0.00451 |
| **Fam172a** | 0.00000 | 0.00278 | 0.71117 | 0.00000 | 0.02448 | 1.66387 | 2.39156 | 0.00000 | 1.68729 | 0.09190 | 0.32150 | 1.73940 | 0.64278 | 0.00000 |
| **Fam173a** | 0.10766 | 0.05668 | 1.62702 | 5.29304 | 0.59887 | 0.00000 | 3.57457 | 2.00593 | 0.00492 | 7.16120 | 0.40903 | 1.14829 | 0.07384 | 0.01588 |
| **Fam173b** | 1.90825 | 0.01864 | 3.17438 | 1.09885 | 1.58400 | 5.24054 | 0.02700 | 2.04999 | 0.00000 | 0.01232 | 4.36596 | 0.22755 | 0.64914 | 0.20711 |
| **Fam174a** | 0.64849 | 0.00434 | 0.01165 | 0.10116 | 0.00000 | 0.45373 | 0.00529 | 0.08638 | 0.03089 | 0.00551 | 0.01743 | 0.09808 | 0.09920 | 0.22295 |
| **Fam174b** | 0.00595 | 0.00441 | 0.00578 | 0.00000 | 0.19314 | 1.64681 | 0.00411 | 0.00483 | 0.00729 | 0.00556 | 0.01122 | 0.00781 | 0.01829 | 0.01132 |
| **Fam175a** | 4.34776 | 0.03103 | 3.45679 | 2.90232 | 4.00834 | 0.71014 | 2.54966 | 0.12103 | 1.60775 | 0.09854 | 2.88255 | 1.86152 | 1.68168 | 3.90418 |
| **Fam175b** | 2.42253 | 3.07906 | 2.94993 | 0.40663 | 6.95724 | 3.80194 | 2.71136 | 1.09262 | 4.46101 | 10.04426 | 2.54008 | 1.01789 | 2.66088 | 9.25418 |
| **Fam178a** | 2.08322 | 0.02033 | 1.12406 | 2.95588 | 4.58082 | 3.14702 | 3.26143 | 0.04243 | 8.57781 | 0.02565 | 5.40424 | 4.47032 | 5.52260 | 9.90416 |
| **Fam178b** | 0.81533 | 0.00380 | 0.46272 | 0.79869 | 0.45145 | 0.08256 | 0.00353 | 0.00133 | 0.46521 | 0.00505 | 0.00409 | 0.00614 | 2.01359 | 0.72420 |
| **Fam179b** | 0.57584 | 0.00883 | 2.14079 | 0.00000 | 1.58448 | 1.40661 | 0.34723 | 0.18572 | 6.53268 | 0.01665 | 0.00000 | 3.37589 | 4.36034 | 4.69627 |
| **Fam183b** | 0.00960 | 0.00448 | 0.01203 | 0.01270 | 0.09209 | 0.01334 | 0.00462 | 0.01037 | 0.01458 | 0.00677 | 0.00884 | 0.00942 | 0.00645 | 0.01003 |
| **Fam184a** | 0.00111 | 0.00414 | 0.01306 | 0.01290 | 0.00130 | 0.01090 | 0.00414 | 0.00780 | 0.00592 | 0.00621 | 0.00741 | 0.00716 | 1.21738 | 0.92289 |
| **Fam184b** | 0.00385 | 0.00427 | 0.06419 | 0.18690 | 0.00358 | 0.00835 | 0.00433 | 0.00747 | 0.00992 | 0.00602 | 0.00667 | 0.00781 | 0.00349 | 0.15124 |
| **Fam185a** | 2.29341 | 0.01749 | 0.50710 | 0.19616 | 0.09453 | 0.85079 | 0.69986 | 0.00100 | 0.03316 | 0.01018 | 0.01166 | 0.64831 | 0.00882 | 0.13452 |
| **Fam187b** | 0.02413 | 0.02803 | 0.01451 | 0.00774 | 0.01830 | 0.01227 | 0.00502 | 0.31572 | 0.01562 | 0.00654 | 0.00968 | 0.00995 | 0.00972 | 0.01299 |
| **Fam188a** | 1.39502 | 0.02022 | 1.21518 | 0.70317 | 2.58860 | 1.80379 | 0.13580 | 1.00380 | 0.15807 | 0.01973 | 0.31854 | 0.94001 | 1.55374 | 1.25770 |
| **Fam188b** | 0.00370 | 0.00495 | 0.07398 | 0.03580 | 0.16337 | 0.00983 | 0.00418 | 0.02650 | 0.00794 | 0.00659 | 0.00684 | 0.05604 | 0.00289 | 0.10818 |
| **Fam189a2** | 0.01405 | 3.03090 | 0.63941 | 0.00000 | 0.28712 | 0.00000 | 0.00232 | 0.79157 | 0.12864 | 0.00580 | 0.00000 | 3.15535 | 0.05696 | 0.03848 |
| **Fam189b** | 0.00000 | 0.00325 | 0.92768 | 0.77993 | 0.04200 | 0.05639 | 0.86445 | 0.00565 | 0.16119 | 0.01668 | 0.00358 | 0.89131 | 1.00398 | 0.00000 |
| **Fam192a** | 8.55536 | 0.06241 | 10.50288 | 2.94423 | 7.62611 | 12.22450 | 2.59197 | 3.25670 | 0.18549 | 0.14163 | 5.71761 | 8.52944 | 7.00944 | 6.82321 |
| **Fam193a** | 0.71879 | 0.01295 | 1.11728 | 0.65979 | 1.86785 | 1.78152 | 0.00416 | 1.70871 | 1.26873 | 0.03950 | 0.60719 | 0.57894 | 0.29861 | 2.36638 |
| **Fam193b** | 0.01806 | 3.77212 | 2.78871 | 2.92833 | 1.52761 | 1.39040 | 0.00149 | 4.27192 | 0.01375 | 0.00652 | 0.09244 | 3.14530 | 0.03067 | 0.00000 |
| **Fam195a** | 2.57492 | 0.01280 | 0.72223 | 18.58747 | 17.03756 | 0.45733 | 0.04140 | 0.20050 | 0.00706 | 0.04140 | 0.04200 | 1.07115 | 0.00000 | 1.52131 |
| **Fam195b** | 0.00275 | 0.25019 | 0.00000 | 0.46692 | 0.54745 | 0.30296 | 0.00330 | 2.05211 | 0.02070 | 0.00534 | 0.00000 | 1.67568 | 0.27253 | 0.00000 |
| **Fam196a** | 0.00534 | 0.00408 | 0.01266 | 0.23781 | 0.02166 | 0.00640 | 0.00393 | 0.00571 | 0.01440 | 0.00638 | 0.01325 | 0.01608 | 0.29156 | 0.00000 |
| **Fam198b** | 0.11918 | 0.03774 | 0.05101 | 0.00000 | 0.03201 | 0.96320 | 0.05885 | 2.28660 | 0.41602 | 0.04932 | 0.00000 | 1.14565 | 0.09249 | 0.06501 |
| **Fam199x** | 0.00000 | 0.01212 | 0.00000 | 0.00000 | 0.51357 | 0.07997 | 0.00541 | 0.23992 | 0.65178 | 0.00615 | 0.00563 | 0.00303 | 3.02031 | 1.22964 |
| **Fam19a4** | 0.00681 | 0.00419 | 0.24224 | 0.00000 | 0.00194 | 0.00622 | 0.14814 | 0.00535 | 0.00845 | 0.01374 | 0.00680 | 0.00984 | 0.17982 | 0.12666 |
| **Fam19a5** | 0.01097 | 0.00420 | 0.00417 | 0.35008 | 0.00312 | 0.19985 | 1.15047 | 0.15447 | 0.01338 | 0.01214 | 0.43262 | 0.01464 | 0.00611 | 0.00667 |
| **Fam204a** | 6.60723 | 0.00185 | 1.30649 | 0.61886 | 3.19490 | 2.50812 | 0.91875 | 0.00000 | 2.26669 | 0.04269 | 1.91321 | 1.25964 | 2.71681 | 1.59047 |

| | | | | | | | | | | | | | | |
|---|---|---|---|---|---|---|---|---|---|---|---|---|---|---|
| **Fam206a** | 1.42676 | 1.42714 | 2.85054 | 1.87523 | 1.52266 | 4.67957 | 1.40024 | 0.36632 | 0.69417 | 0.10265 | 0.10760 | 5.63896 | 0.16165 | 3.15702 |
| **Fam207a** | 1.24270 | 0.03251 | 3.18177 | 2.11666 | 7.99611 | 3.50698 | 1.83031 | 0.85283 | 1.22397 | 0.04019 | 1.73078 | 0.73705 | 2.11143 | 2.28745 |
| **Fam208a** | 4.90072 | 0.03708 | 1.93772 | 0.55720 | 4.01203 | 2.93777 | 2.66576 | 0.57520 | 11.81046 | 0.44478 | 1.78927 | 1.63654 | 4.01821 | 2.59669 |
| **Fam208b** | 3.42478 | 0.01063 | 4.05451 | 2.31358 | 4.27882 | 2.97371 | 0.00876 | 1.25192 | 12.63372 | 0.00307 | 0.16225 | 1.53590 | 5.76733 | 5.01427 |
| **Fam209** | 0.06511 | 0.00454 | 0.03620 | 0.00705 | 0.30432 | 0.06423 | 0.00413 | 0.00610 | 0.02034 | 0.00787 | 0.00658 | 0.00648 | 0.80737 | 0.16303 |
| **Fam20b** | 1.00187 | 3.29289 | 2.12920 | 0.49350 | 0.98156 | 3.62708 | 0.00777 | 1.05212 | 4.59421 | 0.01243 | 2.53116 | 1.58470 | 1.91245 | 4.72934 |
| **Fam21** | 13.09206 | 4.53126 | 12.48242 | 8.37468 | 12.34008 | 4.05033 | 0.65595 | 0.26963 | 4.75990 | 0.08620 | 7.51466 | 5.75993 | 7.92159 | 16.46159 |
| **Fam210a** | 4.55266 | 0.44224 | 3.18032 | 1.05011 | 2.10117 | 4.77457 | 0.51854 | 0.36431 | 9.54240 | 0.02859 | 0.28598 | 1.46715 | 8.11218 | 5.61478 |
| **Fam210b** | 0.41343 | 0.00348 | 0.04239 | 0.77366 | 1.75462 | 0.03173 | 0.22175 | 1.69045 | 0.92903 | 0.01903 | 0.00000 | 0.61757 | 0.19835 | 0.14822 |
| **Fam212a** | 0.98932 | 0.00334 | 0.19556 | 0.13123 | 3.37072 | 0.15680 | 0.00368 | 0.00227 | 0.00505 | 0.00518 | 0.00756 | 0.00444 | 5.40104 | 2.01158 |
| **Fam212b** | 0.00000 | 0.01747 | 0.12468 | 0.59166 | 0.00626 | 0.00000 | 0.02024 | 0.00275 | 0.04435 | 0.01124 | 0.00394 | 1.08328 | 0.00502 | 0.01455 |
| **Fam213a** | 0.00000 | 6.95440 | 1.52461 | 2.49422 | 0.00000 | 8.09444 | 7.95318 | 16.11820 | 0.11171 | 0.43949 | 4.99845 | 0.00000 | 0.41823 | 0.00000 |
| **Fam213b** | 0.39857 | 0.00422 | 0.00000 | 0.31698 | 0.00119 | 0.24128 | 0.02739 | 1.37825 | 0.00884 | 0.02306 | 0.35341 | 0.00119 | 0.01271 | 0.00000 |
| **Fam214a** | 6.62598 | 0.00175 | 4.83709 | 0.00000 | 7.95409 | 0.35938 | 0.00212 | 0.00000 | 5.06097 | 0.00000 | 0.84987 | 0.00530 | 8.55848 | 13.37457 |
| **Fam214b** | 5.88192 | 0.00679 | 10.36598 | 0.37195 | 3.76059 | 0.00000 | 2.99485 | 1.22119 | 2.79495 | 9.84332 | 2.03714 | 1.69328 | 7.84344 | 9.67629 |
| **Fam216a** | 2.41132 | 0.00287 | 0.36588 | 0.35112 | 2.42753 | 1.84196 | 0.00380 | 0.00658 | 0.00000 | 0.01260 | 0.06716 | 0.83863 | 3.88747 | 0.12819 |
| **Fam217a** | 0.00094 | 0.00394 | 0.11608 | 0.00615 | 0.00487 | 0.01045 | 0.00412 | 0.00549 | 0.05974 | 0.00575 | 0.03735 | 0.32705 | 0.32873 | 0.00000 |
| **Fam219a** | 0.00000 | 0.00279 | 0.25499 | 0.24360 | 0.00000 | 0.11021 | 0.00366 | 0.00442 | 0.01383 | 0.00414 | 0.00325 | 1.59984 | 0.04812 | 0.17379 |
| **Fam219b** | 0.12411 | 0.07869 | 0.86939 | 3.74648 | 0.32842 | 0.82525 | 2.15554 | 0.00000 | 0.94029 | 0.10088 | 2.96382 | 1.11300 | 0.38499 | 1.72954 |
| **Fam220a** | 0.44937 | 0.21033 | 1.44365 | 1.51678 | 2.35618 | 2.72973 | 0.02923 | 1.83570 | 0.56288 | 0.00000 | 2.71620 | 1.68493 | 3.96288 | 1.94738 |
| **Fam221a** | 0.10748 | 0.02791 | 2.57582 | 0.00000 | 0.84638 | 1.18140 | 1.99745 | 3.35578 | 0.00088 | 0.03297 | 0.78851 | 0.08195 | 0.00758 | 0.60064 |
| **Fam222a** | 0.00463 | 0.00490 | 0.01502 | 0.04186 | 0.16116 | 0.04537 | 0.00437 | 0.04142 | 0.00877 | 0.00866 | 0.00784 | 0.05914 | 0.62636 | 0.00256 |
| **Fam222b** | 0.00000 | 0.02254 | 1.84908 | 1.55933 | 4.38877 | 1.28378 | 3.68271 | 2.24761 | 0.00000 | 0.12060 | 3.76502 | 3.10051 | 6.32451 | 4.12804 |
| **Fam227a** | 0.07991 | 0.03642 | 0.08035 | 0.63574 | 0.38524 | 0.14111 | 0.06680 | 0.07560 | 0.56575 | 0.18435 | 0.05635 | 0.10495 | 0.73722 | 0.39402 |
| **Fam228b** | 0.91647 | 0.01251 | 0.54160 | 0.33558 | 1.54222 | 0.00000 | 0.00424 | 0.01125 | 0.61331 | 0.00603 | 0.00341 | 0.03367 | 0.08263 | 0.04107 |
| **Fam234a** | 6.99289 | 3.86693 | 18.11699 | 14.82544 | 11.79408 | 12.56275 | 1.22838 | 12.12726 | 0.56978 | 0.18594 | 7.07654 | 3.79858 | 2.71402 | 9.22154 |
| **Fam234b** | 0.00891 | 0.00446 | 0.00926 | 0.01321 | 0.16725 | 0.01840 | 0.00416 | 0.00946 | 0.01047 | 0.00728 | 0.00843 | 0.00893 | 0.00613 | 0.01039 |
| **Fam25c** | 0.00239 | 0.17814 | 0.00878 | 0.00000 | 0.00925 | 0.05900 | 0.09907 | 0.69545 | 0.00912 | 0.02285 | 0.00087 | 5.27722 | 0.00433 | 0.04050 |
| **Fam26f** | 0.00000 | 0.00411 | 0.08940 | 0.02763 | 0.75225 | 0.00360 | 0.00435 | 0.00729 | 0.11954 | 0.00644 | 0.00911 | 0.00880 | 1.37475 | 0.28323 |
| **Fam32a** | 10.22722 | 4.47658 | 17.88430 | 11.94082 | 13.78118 | 17.15283 | 3.34701 | 9.49850 | 3.35518 | 1.22172 | 13.58719 | 11.05739 | 4.98233 | 16.72815 |
| **Fam35a** | 0.00000 | 0.02574 | 0.46608 | 0.00000 | 0.10995 | 0.90837 | 0.00404 | 0.09621 | 0.03992 | 0.00743 | 0.76702 | 1.67825 | 1.24709 | 0.93869 |
| **Fam3a** | 1.03340 | 0.03293 | 1.44006 | 1.52600 | 1.93911 | 2.61936 | 0.00294 | 0.00000 | 0.00313 | 0.02151 | 1.87171 | 2.15275 | 0.84510 | 2.64689 |
| **Fam3b** | 0.00700 | 0.01221 | 0.00451 | 0.56088 | 0.00853 | 0.01202 | 0.00367 | 0.00417 | 0.01928 | 0.00626 | 0.00675 | 0.49267 | 0.04029 | 0.06418 |
| **Fam3c** | 1.10464 | 0.03307 | 2.46197 | 2.83333 | 2.14110 | 1.59070 | 1.97515 | 1.67652 | 2.19755 | 0.02839 | 1.55209 | 4.74224 | 1.49718 | 1.72606 |
| **Fam43a** | 0.01150 | 0.00435 | 0.01147 | 0.01121 | 0.12105 | 0.01307 | 0.00467 | 0.01013 | 0.01434 | 0.00635 | 0.00892 | 0.00912 | 0.27332 | 0.00733 |
| **Fam43b** | 0.00181 | 0.00402 | 0.01408 | 0.01312 | 0.01166 | 0.01231 | 0.00404 | 0.00812 | 0.01451 | 0.00646 | 0.00923 | 0.00723 | 2.77493 | 0.00139 |
| **Fam45a** | 5.80027 | 0.06443 | 10.34572 | 7.02783 | 7.50153 | 7.76343 | 1.25786 | 2.37187 | 21.94744 | 1.92145 | 6.09715 | 0.86862 | 3.04658 | 12.84991 |
| **Fam46a** | 0.00064 | 0.00450 | 0.01391 | 0.00000 | 0.24457 | 0.04296 | 0.00415 | 0.00871 | 0.01631 | 0.00586 | 0.00603 | 0.04012 | 0.05516 | 1.07858 |
| **Fam46b** | 0.00000 | 0.01155 | 0.32224 | 0.23765 | 0.66136 | 0.00000 | 0.00510 | 0.51943 | 0.00013 | 0.00513 | 0.00000 | 0.58277 | 3.95460 | 0.00906 |
| **Fam46c** | 0.55191 | 0.02464 | 15.26240 | 6.76085 | 2.02898 | 1.09773 | 2.19880 | 0.23387 | 0.02162 | 0.00000 | 1.81421 | 3.30100 | 0.00418 | 0.72999 |
| **Fam47e** | 0.01070 | 0.00445 | 0.01368 | 0.01271 | 0.03057 | 0.01333 | 0.00453 | 0.01024 | 0.01431 | 0.00702 | 0.00883 | 0.00994 | 0.00707 | 0.01043 |
| **Fam49a** | 0.28625 | 0.00419 | 0.00000 | 0.00090 | 0.00000 | 0.34761 | 0.00400 | 0.00429 | 0.05981 | 0.00687 | 0.00460 | 0.00788 | 0.71381 | 0.83494 |
| **Fam49b** | 3.31145 | 0.00000 | 2.64202 | 1.78036 | 1.72114 | 2.84128 | 1.17750 | 0.70394 | 0.28863 | 0.02853 | 2.96467 | 4.26406 | 0.00000 | 3.86184 |
| **Fam50a** | 13.44765 | 3.92437 | 7.89191 | 14.77646 | 13.40763 | 9.92587 | 0.02812 | 3.33293 | 4.47930 | 2.14669 | 4.08634 | 8.43373 | 2.43894 | 7.89945 |
| **Fam53a** | 1.62530 | 0.02527 | 0.58734 | 1.37845 | 2.03176 | 1.37558 | 1.06896 | 0.32301 | 0.00000 | 0.01990 | 0.86887 | 1.65822 | 1.12762 | 0.00000 |
| **Fam53b** | 0.00356 | 0.02501 | 0.05364 | 0.46607 | 0.81820 | 0.19658 | 0.00336 | 0.00204 | 0.00968 | 1.41135 | 0.16926 | 1.39881 | 0.01256 | 0.00000 |
| **Fam53c** | 1.29187 | 0.01838 | 4.49096 | 2.35831 | 2.79218 | 2.33727 | 0.13471 | 4.25601 | 0.03190 | 0.04274 | 3.03982 | 0.65383 | 3.16800 | 5.11379 |
| **Fam57a** | 0.00569 | 0.00400 | 0.06565 | 0.04229 | 0.19751 | 0.11615 | 0.00385 | 0.00592 | 0.05039 | 0.00610 | 0.00546 | 0.74358 | 0.00267 | 0.32058 |
| **Fam57b** | 0.86873 | 0.01313 | 6.81852 | 2.93406 | 11.14468 | 0.00000 | 0.04793 | 1.48015 | 0.00286 | 0.02990 | 0.77019 | 0.06673 | 0.01790 | 0.21848 |
| **Fam58b** | 4.47400 | 0.00201 | 4.65376 | 7.32287 | 6.64229 | 3.46168 | 0.16995 | 0.06970 | 0.10476 | 0.03551 | 2.29285 | 1.82028 | 5.06887 | 0.81783 |
| **Fam60a** | 0.30484 | 0.02430 | 6.24592 | 3.97850 | 3.96517 | 3.78401 | 0.81964 | 4.23250 | 14.79706 | 0.05221 | 5.12639 | 3.38199 | 0.80152 | 7.18531 |
| **Fam63a** | 10.22372 | 6.94906 | 14.11348 | 5.83412 | 13.44987 | 13.71035 | 5.85210 | 12.24003 | 4.43567 | 12.71158 | 7.58106 | 5.56510 | 11.00554 | 10.11152 |
| **Fam63b** | 0.59569 | 0.00396 | 0.00000 | 0.00000 | 0.23098 | 0.42798 | 0.00442 | 0.01410 | 1.01826 | 0.00563 | 0.00384 | 0.00681 | 0.42721 | 1.36593 |
| **Fam65a** | 0.90328 | 4.03666 | 6.72977 | 4.75884 | 7.93588 | 6.27297 | 4.05790 | 9.64553 | 0.17115 | 9.45721 | 1.53590 | 0.32029 | 0.82990 | 2.50829 |
| **Fam65b** | 0.09064 | 0.02877 | 0.01541 | 0.09018 | 0.05542 | 0.50933 | 0.02288 | 0.03997 | 1.13961 | 0.03727 | 0.00719 | 0.05276 | 0.85135 | 0.02536 |

| | | | | | | | | | | | | | | |
|---|---|---|---|---|---|---|---|---|---|---|---|---|---|---|
| **Fam65c** | 0.01227 | 0.00444 | 1.42781 | 0.00000 | 3.43290 | 0.00114 | 0.00386 | 0.00712 | 0.01055 | 0.00422 | 0.00448 | 0.00645 | 0.00000 | 0.03227 |
| **Fam69a** | 1.71106 | 0.00762 | 2.23601 | 0.67316 | 1.67533 | 7.21551 | 1.85572 | 1.48410 | 2.97226 | 0.00000 | 8.92616 | 2.61266 | 0.46625 | 2.37217 |
| **Fam69b** | 0.01331 | 0.00456 | 0.00884 | 0.01451 | 0.50067 | 0.00540 | 0.00540 | 0.32604 | 0.00436 | 0.00603 | 0.00782 | 0.00808 | 0.29203 | 0.26436 |
| **Fam72a** | 0.80039 | 0.06213 | 0.61700 | 0.56013 | 0.17237 | 1.41519 | 0.00866 | 4.76499 | 0.00821 | 0.01478 | 5.55206 | 2.15458 | 4.80796 | 0.00000 |
| **Fam73a** | 2.20840 | 0.02710 | 2.79106 | 0.87748 | 2.11096 | 0.00000 | 0.00343 | 1.26431 | 2.01369 | 0.01202 | 1.58373 | 0.45444 | 1.21450 | 4.33336 |
| **Fam73b** | 0.95656 | 0.01264 | 0.35734 | 0.49288 | 0.08912 | 2.19980 | 0.04628 | 0.68188 | 0.00429 | 0.02991 | 0.12607 | 0.00000 | 0.07423 | 0.59237 |
| **Fam76a** | 7.67269 | 0.04086 | 9.13209 | 5.69443 | 14.47959 | 5.60625 | 0.01159 | 4.79874 | 10.97573 | 0.13362 | 5.53881 | 1.15702 | 2.97203 | 15.07198 |
| **Fam76b** | 3.65043 | 0.01344 | 1.07898 | 0.67167 | 0.82176 | 1.39623 | 0.00301 | 0.08119 | 6.51316 | 0.02488 | 2.53315 | 0.27903 | 2.80952 | 0.64135 |
| **Fam78a** | 0.00799 | 0.00437 | 0.15405 | 0.02429 | 0.35463 | 0.00446 | 0.00429 | 0.00971 | 0.00771 | 0.00837 | 0.00691 | 0.00791 | 0.01366 | 0.28747 |
| **Fam83a** | 0.06055 | 1.04255 | 0.01292 | 0.60395 | 0.02978 | 0.08886 | 0.02287 | 0.12411 | 0.26685 | 0.01781 | 0.03239 | 0.06667 | 0.65799 | 0.05342 |
| **Fam83b** | 0.00902 | 0.00407 | 0.39484 | 0.01450 | 0.23236 | 0.00000 | 0.00513 | 0.34234 | 0.01276 | 0.00136 | 1.28085 | 0.00303 | 0.01452 | 0.00612 |
| **Fam83d** | 0.17026 | 0.04143 | 2.66910 | 0.66878 | 0.00000 | 1.33341 | 2.03777 | 2.13268 | 0.00000 | 0.03248 | 3.51834 | 2.12648 | 4.41109 | 0.00000 |
| **Fam83e** | 0.00553 | 0.00454 | 0.00797 | 0.10250 | 0.00418 | 0.01175 | 0.00431 | 0.01240 | 0.02123 | 0.00655 | 0.00642 | 0.01061 | 0.00333 | 0.21116 |
| **Fam83g** | 0.12518 | 0.00430 | 1.54614 | 0.72806 | 1.51182 | 1.54401 | 1.70928 | 0.06000 | 0.00306 | 0.09182 | 0.00000 | 0.00000 | 0.04305 | 0.45500 |
| **Fam83h** | 0.79839 | 0.00291 | 1.00380 | 0.94620 | 0.24830 | 2.69825 | 1.13298 | 1.66661 | 0.02113 | 0.03806 | 0.33205 | 1.30851 | 0.00090 | 0.00000 |
| **Fam84a** | 0.03118 | 0.00449 | 0.01118 | 0.01225 | 0.12629 | 0.01277 | 0.00448 | 0.01088 | 0.01262 | 0.00648 | 0.00855 | 0.00905 | 0.00578 | 0.00992 |
| **Fam84b** | 0.00000 | 0.00326 | 0.00000 | 0.15160 | 0.00000 | 5.30724 | 2.84227 | 2.54047 | 1.46377 | 0.03411 | 2.53663 | 0.01029 | 1.35321 | 2.03032 |
| **Fam89a** | 0.01271 | 0.00440 | 0.04920 | 0.00785 | 0.32470 | 0.05895 | 0.00434 | 0.00700 | 0.01845 | 0.01076 | 0.00621 | 0.04402 | 0.02404 | 0.00546 |
| **Fam89b** | 2.66770 | 0.02153 | 1.04794 | 0.01213 | 0.18847 | 0.41994 | 0.00389 | 0.21870 | 0.01645 | 0.02625 | 0.00134 | 1.47371 | 1.55739 | 1.69577 |
| **Fam8a1** | 0.33659 | 0.00300 | 0.08758 | 0.37829 | 0.00000 | 0.00000 | 0.00888 | 0.00272 | 0.33830 | 5.15868 | 0.00293 | 0.00152 | 0.00523 | 0.00000 |
| **Fam90a1a** | 0.01196 | 0.00443 | 0.01418 | 0.01529 | 0.01417 | 0.01329 | 0.00426 | 0.00998 | 0.01242 | 0.00765 | 0.00867 | 0.00976 | 0.03105 | 0.00817 |
| **Fam91a1** | 14.76893 | 0.64098 | 7.35191 | 2.46768 | 11.75955 | 6.40763 | 1.30411 | 3.56900 | 14.69167 | 0.15991 | 8.69959 | 2.94125 | 8.07503 | 13.18846 |
| **Fam92a** | 1.98620 | 0.00363 | 0.05577 | 0.23064 | 0.35318 | 0.45410 | 0.00555 | 0.01147 | 0.99059 | 0.00535 | 0.00480 | 0.72435 | 0.00070 | 0.00000 |
| **Fam92b** | 0.00961 | 0.00443 | 0.06953 | 0.00439 | 0.08135 | 0.01213 | 0.00467 | 0.01115 | 0.01665 | 0.00700 | 0.00802 | 0.00970 | 0.00679 | 0.00852 |
| **Fam96a** | 15.18062 | 6.99196 | 26.36851 | 21.05606 | 15.13478 | 31.55801 | 8.53033 | 25.94630 | 5.75050 | 10.15072 | 17.20329 | 14.88474 | 1.87005 | 17.45809 |
| **Fam96b** | 2.56789 | 0.05312 | 0.93056 | 6.01907 | 4.87451 | 8.24117 | 3.17238 | 4.81902 | 0.00000 | 0.20031 | 2.30520 | 6.92275 | 2.85771 | 0.31116 |
| **Fam98a** | 10.48084 | 6.65235 | 8.75081 | 9.09808 | 8.06021 | 5.79726 | 0.05680 | 4.03843 | 9.76523 | 0.00974 | 4.80558 | 3.12859 | 3.75248 | 9.10652 |
| **Fam98b** | 1.76366 | 0.03325 | 0.00000 | 3.15223 | 4.26778 | 2.85972 | 0.56697 | 2.02691 | 4.76499 | 0.05510 | 0.17337 | 2.85931 | 6.08623 | 2.23807 |
| **Fam98c** | 0.38209 | 0.01327 | 0.29326 | 0.58108 | 0.23592 | 0.18751 | 0.00981 | 0.32414 | 0.01528 | 0.00642 | 0.00000 | 0.95702 | 0.05485 | 0.00965 |
| **Fan1** | 0.01932 | 0.00450 | 0.02172 | 0.02605 | 0.01353 | 0.00369 | 0.00502 | 0.41657 | 0.00835 | 0.01009 | 0.00787 | 0.01052 | 0.01719 | 0.15152 |
| **Fanca** | 0.09716 | 0.37416 | 1.67356 | 0.86880 | 0.69875 | 1.09145 | 1.20168 | 0.22452 | 0.13387 | 0.00100 | 0.84408 | 0.00000 | 0.35317 | 0.45089 |
| **Fancb** | 1.32274 | 0.00348 | 2.13021 | 0.34347 | 1.02508 | 0.13476 | 0.00294 | 0.00000 | 2.27767 | 0.00000 | 2.84433 | 0.01717 | 0.01909 | 0.28902 |
| **Fancc** | 0.56243 | 0.01746 | 2.38052 | 0.23818 | 3.47687 | 0.47301 | 0.00811 | 1.97237 | 3.85259 | 0.01939 | 2.27648 | 0.13113 | 1.57128 | 2.19343 |
| **Fancd2** | 1.88487 | 0.05231 | 2.16384 | 2.93272 | 2.30171 | 1.13222 | 2.15643 | 1.06791 | 3.60191 | 0.10571 | 1.17703 | 4.69497 | 0.92895 | 3.45579 |
| **Fancd2os** | 0.92946 | 0.00386 | 0.31759 | 0.96085 | 1.96746 | 2.85472 | 0.13400 | 0.19859 | 2.62085 | 0.01477 | 0.06515 | 0.14324 | 0.70739 | 3.03855 |
| **Fance** | 2.93687 | 0.03228 | 2.11362 | 2.65155 | 2.68547 | 1.36660 | 0.03191 | 4.71387 | 0.01198 | 0.05155 | 1.63787 | 2.06293 | 2.64997 | 0.76234 |
| **Fancf** | 0.02687 | 0.00450 | 0.02318 | 0.00400 | 0.00928 | 0.05007 | 0.00412 | 0.00835 | 0.01150 | 0.00768 | 0.00567 | 0.04852 | 0.43324 | 0.00407 |
| **Fancg** | 0.91704 | 0.00260 | 1.02184 | 1.19362 | 1.32513 | 1.67936 | 0.01440 | 2.02045 | 1.13704 | 0.04809 | 0.00000 | 2.18043 | 0.83810 | 1.14628 |
| **Fanci** | 1.09048 | 0.00282 | 2.28581 | 1.61726 | 2.38709 | 2.66931 | 0.01046 | 0.04023 | 4.27622 | 0.00681 | 3.63863 | 0.60138 | 0.00000 | 2.28846 |
| **Fancl** | 1.90566 | 0.03016 | 2.61285 | 2.54554 | 1.92982 | 2.23446 | 0.01498 | 1.10635 | 3.45391 | 0.01156 | 4.03870 | 1.29286 | 0.20817 | 1.40000 |
| **Fancm** | 7.35830 | 0.00731 | 2.00633 | 1.71803 | 5.76020 | 2.41869 | 0.00669 | 0.71891 | 10.38999 | 0.01080 | 0.10394 | 4.10186 | 6.40625 | 10.61085 |
| **Fank1** | 0.00000 | 0.04507 | 1.05145 | 1.72799 | 0.42625 | 4.55744 | 0.00319 | 1.26412 | 0.00000 | 0.02967 | 0.15808 | 2.61107 | 0.00580 | 1.78070 |
| **Far1** | 1.27790 | 0.00254 | 0.41934 | 0.51090 | 0.53874 | 3.45468 | 0.95221 | 0.28307 | 1.04506 | 0.02131 | 2.36675 | 1.77310 | 1.36557 | 0.62277 |
| **Far2** | 0.45296 | 0.00388 | 0.35315 | 0.27621 | 0.00000 | 0.00002 | 0.00399 | 0.00320 | 0.08444 | 0.00557 | 0.01635 | 0.27896 | 0.01014 | 0.01317 |
| **Farp1** | 0.55833 | 0.02596 | 0.10268 | 0.00000 | 0.83015 | 0.93969 | 0.02697 | 5.02254 | 0.20286 | 0.02074 | 0.43326 | 0.01148 | 1.90046 | 0.11343 |
| **Farp2** | 0.00000 | 0.00335 | 1.24214 | 0.02154 | 1.23592 | 0.71364 | 1.35539 | 0.00480 | 0.54487 | 0.00518 | 1.97715 | 0.07489 | 0.00000 | 1.42743 |
| **Fars2** | 0.02598 | 0.02381 | 2.22721 | 1.50070 | 1.03835 | 0.00000 | 0.00862 | 4.09967 | 0.00664 | 0.02151 | 2.22872 | 2.32396 | 0.00000 | 2.32019 |
| **Farsa** | 4.80798 | 1.20013 | 14.79371 | 10.99727 | 9.16375 | 9.11970 | 2.35100 | 5.46128 | 4.20433 | 0.17207 | 1.48838 | 3.36480 | 2.22735 | 6.79208 |
| **Farsb** | 19.04106 | 6.42181 | 21.85622 | 15.35465 | 16.71416 | 21.03556 | 7.02993 | 12.08469 | 15.00323 | 0.32866 | 14.25803 | 17.29946 | 3.17257 | 15.20758 |
| **Fasn** | 8.47862 | 6.66804 | 21.80009 | 13.66254 | 16.35178 | 17.11916 | 1.94710 | 5.27144 | 0.00000 | 0.14799 | 9.70974 | 9.42493 | 1.24155 | 1.77412 |
| **Fastk** | 0.07092 | 0.09480 | 5.46916 | 7.89275 | 0.65504 | 4.47576 | 0.04968 | 5.87381 | 0.00000 | 0.10971 | 1.90679 | 6.65141 | 0.00000 | 0.88664 |
| **Fastkd1** | 2.46292 | 0.00309 | 0.05061 | 0.18534 | 0.00000 | 0.25222 | 0.00234 | 0.00187 | 9.40948 | 0.01114 | 1.00306 | 0.00312 | 3.41644 | 3.28718 |
| **Fastkd2** | 4.89765 | 0.06216 | 3.03978 | 3.55707 | 3.94005 | 1.45355 | 2.83889 | 2.62858 | 5.59897 | 0.04765 | 5.07096 | 3.96312 | 0.95891 | 5.40253 |
| **Fastkd3** | 2.16445 | 0.08475 | 6.41682 | 1.20353 | 6.02335 | 5.06472 | 2.51480 | 1.37017 | 3.51308 | 5.32317 | 0.30041 | 3.18653 | 6.71920 | 5.38373 |
| **Fastkd5** | 5.82594 | 0.00234 | 2.43733 | 0.45645 | 0.73157 | 5.38797 | 2.90131 | 0.03875 | 4.22545 | 0.07930 | 2.13036 | 0.36153 | 2.62886 | 1.92583 |

| | | | | | | | | | | | | | | |
|---|---|---|---|---|---|---|---|---|---|---|---|---|---|---|
| **Fat1** | 0.94742 | 0.01782 | 2.00778 | 2.46617 | 0.70143 | 2.57257 | 0.00260 | 0.06821 | 0.35851 | 0.01636 | 2.40391 | 2.95115 | 0.41192 | 2.01057 |
| **Fau** | 15.58971 | 13.98256 | 7.37148 | 32.20955 | 34.97134 | 26.65064 | 8.32712 | 26.73138 | 0.23809 | 23.86379 | 5.23645 | 23.24279 | 11.87341 | 8.93070 |
| **Faxc** | 0.85759 | 0.00447 | 0.25868 | 0.00000 | 0.79155 | 0.19449 | 0.00412 | 0.00329 | 1.09639 | 0.00480 | 0.00324 | 0.00580 | 1.02372 | 0.83843 |
| **Fbf1** | 0.09520 | 0.00336 | 0.11048 | 0.76887 | 0.05329 | 0.83802 | 0.23276 | 0.28588 | 0.00648 | 0.05971 | 1.29095 | 0.28106 | 0.21993 | 0.16541 |
| **Fbl** | 18.17761 | 5.36864 | 21.93926 | 29.96844 | 27.88259 | 25.35500 | 4.62056 | 15.02130 | 9.56128 | 0.54453 | 15.14643 | 15.82526 | 15.12212 | 22.79582 |
| **Fblim1** | 7.28575 | 0.10584 | 16.80616 | 5.57875 | 7.13747 | 8.19161 | 0.00000 | 2.43410 | 6.56879 | 0.03802 | 4.46344 | 6.04889 | 3.21453 | 4.01943 |
| **Fbln1** | 0.00000 | 0.06057 | 0.15664 | 0.78564 | 0.51017 | 0.87424 | 0.01418 | 4.37615 | 0.02664 | 4.16114 | 0.23839 | 0.80496 | 0.04331 | 0.04565 |
| **Fbln2** | 0.46486 | 0.00437 | 0.00392 | 0.01200 | 0.21169 | 0.01404 | 0.00420 | 0.00970 | 0.00917 | 0.00752 | 0.00679 | 0.00669 | 0.13525 | 0.15505 |
| **Fbln5** | 0.10210 | 0.00428 | 0.01002 | 0.05278 | 0.00759 | 0.03026 | 0.00438 | 0.36539 | 0.01128 | 0.01898 | 0.00735 | 0.00590 | 0.00558 | 0.00703 |
| **Fbln7** | 0.05845 | 0.00449 | 0.27464 | 0.00000 | 1.58188 | 0.00218 | 0.00382 | 0.01049 | 1.47180 | 0.00675 | 0.00288 | 0.00587 | 0.00000 | 0.00000 |
| **Fbn1** | 0.64825 | 0.00416 | 0.00801 | 0.00693 | 0.00545 | 0.00656 | 0.00419 | 0.01065 | 0.01565 | 0.00701 | 0.30822 | 0.01717 | 0.00401 | 0.00700 |
| **Fbp1** | 0.00341 | 0.01786 | 0.00750 | 0.31732 | 0.05234 | 0.74008 | 0.01046 | 0.29372 | 0.12573 | 0.02207 | 0.03752 | 3.67083 | 0.02206 | 0.00000 |
| **Fbp2** | 8.02470 | 5.69115 | 30.31863 | 30.14155 | 8.77340 | 29.92727 | 9.53691 | 19.44838 | 6.29859 | 15.37304 | 22.33141 | 25.02388 | 0.00000 | 2.71426 |
| **Fbrs** | 1.62162 | 0.00196 | 3.68972 | 6.07946 | 2.39988 | 1.79128 | 3.00844 | 5.22858 | 0.60200 | 0.03939 | 2.49489 | 1.49083 | 4.07326 | 0.60810 |
| **Fbrsl1** | 6.88639 | 0.04708 | 9.80041 | 6.88685 | 15.32746 | 2.68136 | 3.73449 | 1.94816 | 1.13270 | 0.03076 | 2.77080 | 4.26445 | 18.72760 | 7.91659 |
| **Fbxl12** | 0.00000 | 0.04200 | 1.26137 | 1.24765 | 0.00755 | 0.84776 | 2.93663 | 1.78574 | 0.01929 | 0.05955 | 0.00358 | 0.11188 | 3.54081 | 0.00000 |
| **Fbxl12os** | 0.01773 | 0.38168 | 0.06274 | 0.02371 | 0.02922 | 0.03407 | 0.01053 | 0.09645 | 0.52278 | 0.00742 | 0.04121 | 0.07798 | 0.52486 | 0.11789 |
| **Fbxl13** | 0.07788 | 0.00425 | 0.41302 | 0.00000 | 0.33768 | 0.00944 | 0.00393 | 0.00682 | 0.01391 | 0.00512 | 0.09611 | 0.00725 | 0.15218 | 0.93433 |
| **Fbxl14** | 1.68680 | 0.01039 | 0.28610 | 1.51153 | 1.26488 | 1.86264 | 0.00332 | 0.00178 | 2.54774 | 0.04550 | 0.00000 | 0.51224 | 2.00393 | 1.67963 |
| **Fbxl15** | 0.00449 | 0.00412 | 0.01032 | 0.15930 | 1.44789 | 0.01193 | 0.00445 | 0.00651 | 0.01491 | 0.00559 | 0.00767 | 0.16780 | 0.00215 | 0.00210 |
| **Fbxl16** | 1.62575 | 0.00335 | 0.34095 | 0.00000 | 0.64190 | 0.01222 | 0.88734 | 0.40443 | 0.02789 | 0.01428 | 1.44022 | 0.45101 | 0.27240 | 0.00000 |
| **Fbxl17** | 0.27130 | 0.06602 | 1.52275 | 0.56906 | 0.78985 | 0.91293 | 0.78842 | 0.78887 | 0.94398 | 0.05495 | 0.45764 | 1.12928 | 1.10434 | 1.16046 |
| **Fbxl18** | 0.05437 | 0.04016 | 2.42683 | 0.85147 | 2.27983 | 0.40763 | 0.00389 | 1.56506 | 0.00129 | 0.00929 | 0.35063 | 0.56740 | 0.57165 | 0.61949 |
| **Fbxl19** | 0.21134 | 0.02392 | 0.75101 | 0.74685 | 0.42623 | 0.41456 | 0.05433 | 4.02719 | 0.22509 | 0.04520 | 0.00000 | 3.33959 | 0.88259 | 0.20221 |
| **Fbxl2** | 0.37012 | 0.00489 | 0.22617 | 0.00761 | 0.09608 | 0.10989 | 0.00426 | 0.01225 | 0.52452 | 0.00636 | 0.00536 | 0.00448 | 0.54362 | 0.00000 |
| **Fbxl20** | 10.08735 | 0.00000 | 27.73590 | 7.61050 | 17.81607 | 0.00000 | 0.00000 | 0.00000 | 11.10440 | 1.97893 | 8.23610 | 8.98685 | 3.40671 | 25.38673 |
| **Fbxl21** | 0.00737 | 0.00397 | 0.00002 | 0.82032 | 0.00478 | 0.04404 | 0.00386 | 0.00239 | 0.00608 | 0.01022 | 0.13456 | 0.01532 | 0.00703 | 0.19149 |
| **Fbxl22** | 0.00975 | 0.00447 | 0.01157 | 0.01670 | 0.09423 | 0.01344 | 0.00420 | 0.01078 | 0.04643 | 0.00716 | 0.00860 | 0.00890 | 0.00570 | 0.03409 |
| **Fbxl3** | 1.71274 | 0.10176 | 1.00336 | 1.03962 | 0.88046 | 0.86144 | 0.19585 | 0.63088 | 5.30770 | 0.21420 | 2.46282 | 0.22004 | 6.52174 | 5.28082 |
| **Fbxl4** | 0.13507 | 0.00384 | 0.00271 | 0.00000 | 0.16414 | 0.71270 | 1.30231 | 0.01854 | 0.00160 | 0.03437 | 0.01080 | 0.70487 | 0.80210 | 0.30149 |
| **Fbxl5** | 3.62134 | 0.00680 | 4.25815 | 0.39054 | 3.52855 | 0.73552 | 1.86969 | 0.63505 | 5.22838 | 0.01598 | 1.41589 | 6.27779 | 6.54140 | 8.06830 |
| **Fbxl6** | 0.00000 | 0.07575 | 7.18703 | 5.75463 | 3.17855 | 2.76880 | 0.04696 | 2.28004 | 0.17640 | 0.00000 | 1.11747 | 3.40349 | 1.03422 | 4.82603 |
| **Fbxl7** | 0.00947 | 0.00440 | 0.01486 | 0.01243 | 0.01174 | 0.01338 | 0.00429 | 0.00968 | 0.01117 | 0.00669 | 0.00890 | 0.00969 | 0.16002 | 0.00346 |
| **Fbxl8** | 0.00267 | 0.00370 | 0.00860 | 0.31705 | 0.05199 | 0.01283 | 0.01022 | 0.01019 | 0.00947 | 0.00505 | 0.01185 | 0.36393 | 0.36567 | 0.00000 |
| **Fbxo10** | 0.08545 | 0.00450 | 0.47593 | 0.00000 | 0.20043 | 0.00000 | 0.00374 | 0.01191 | 0.00610 | 0.00586 | 0.00352 | 0.48464 | 0.14351 | 1.69765 |
| **Fbxo11** | 1.15200 | 5.54429 | 3.06800 | 1.44493 | 5.77549 | 3.56237 | 2.19497 | 1.76098 | 21.78891 | 0.08319 | 6.42570 | 2.63997 | 10.79254 | 4.37092 |
| **Fbxo15** | 48.18741 | 13.33257 | 60.38772 | 46.68916 | 59.96784 | 34.25324 | 12.51261 | 24.77110 | 63.15211 | 9.07781 | 33.80934 | 22.53824 | 21.34331 | 54.01160 |
| **Fbxo16** | 1.51411 | 0.00507 | 2.97532 | 0.36587 | 0.45615 | 1.68769 | 1.76401 | 0.36643 | 0.00005 | 0.14170 | 0.00044 | 0.00000 | 1.65128 | 0.00000 |
| **Fbxo17** | 0.00000 | 0.01772 | 0.96290 | 0.00000 | 0.46925 | 0.33620 | 1.75966 | 1.68285 | 0.05485 | 0.01649 | 0.66094 | 1.36238 | 2.51470 | 0.27279 |
| **Fbxo18** | 2.27021 | 0.03658 | 0.10936 | 3.58000 | 1.59500 | 2.46426 | 1.08652 | 3.54868 | 1.52486 | 0.02421 | 4.59043 | 3.15184 | 1.98878 | 3.83841 |
| **Fbxo2** | 0.08502 | 0.00980 | 0.08741 | 0.00000 | 0.02819 | 0.00593 | 0.00322 | 1.01727 | 0.00808 | 0.00899 | 1.32056 | 0.83665 | 0.30193 | 0.01404 |
| **Fbxo21** | 0.00000 | 0.08084 | 4.52939 | 2.62525 | 0.99126 | 2.44939 | 0.07872 | 2.83637 | 0.14732 | 0.01035 | 6.43321 | 7.25920 | 0.00000 | 0.80902 |
| **Fbxo22** | 3.44000 | 4.92823 | 10.65511 | 3.96336 | 11.83563 | 3.80242 | 0.02042 | 5.27577 | 2.43742 | 12.25674 | 6.03489 | 6.26798 | 1.68582 | 6.68478 |
| **Fbxo25** | 0.04951 | 0.02753 | 1.38691 | 1.42100 | 0.04373 | 1.30727 | 0.00342 | 0.00000 | 0.00365 | 0.01071 | 0.00414 | 1.10364 | 0.00206 | 0.91751 |
| **Fbxo27** | 0.75786 | 0.01935 | 0.50800 | 0.00000 | 0.00000 | 1.57749 | 0.00418 | 0.71295 | 0.02517 | 0.00449 | 0.20514 | 0.17988 | 0.06929 | 0.00000 |
| **Fbxo28** | 1.98352 | 0.01039 | 2.51592 | 0.00000 | 2.07129 | 3.20222 | 2.76084 | 0.54065 | 0.17525 | 0.13163 | 4.34032 | 3.52783 | 4.11857 | 2.72087 |
| **Fbxo3** | 0.00000 | 0.03575 | 2.86990 | 3.86755 | 0.70526 | 3.92486 | 0.00104 | 1.08696 | 0.00000 | 1.35354 | 10.46592 | 5.72806 | 0.21372 | 0.00000 |
| **Fbxo30** | 2.02769 | 0.41141 | 1.25424 | 1.71859 | 2.43509 | 2.25922 | 0.94924 | 0.61185 | 5.97351 | 0.01808 | 2.20586 | 2.07668 | 2.72510 | 0.66434 |
| **Fbxo31** | 1.35994 | 2.53789 | 2.86448 | 12.55815 | 17.93470 | 0.56251 | 1.31802 | 0.87961 | 0.00242 | 0.08567 | 0.61345 | 4.62058 | 0.62838 | 0.09947 |
| **Fbxo32** | 0.16359 | 0.02005 | 0.01184 | 0.04189 | 0.09325 | 0.16312 | 0.04986 | 0.08222 | 0.30686 | 0.07410 | 0.00254 | 0.62077 | 1.34982 | 0.29356 |
| **Fbxo33** | 9.06929 | 0.02355 | 5.73783 | 0.00000 | 11.88285 | 1.97127 | 1.89632 | 2.62711 | 16.61772 | 0.07725 | 1.00857 | 1.05521 | 17.17481 | 14.58943 |
| **Fbxo34** | 8.53792 | 1.64111 | 7.72535 | 0.98118 | 12.14959 | 0.00000 | 0.70323 | 0.63599 | 2.54378 | 0.00000 | 0.00000 | 1.31702 | 10.16093 | 10.73012 |
| **Fbxo36** | 0.24356 | 0.00393 | 0.03028 | 0.34585 | 1.20354 | 0.52347 | 0.00374 | 0.00337 | 1.39174 | 0.00638 | 0.00658 | 0.01196 | 4.23777 | 0.22344 |
| **Fbxo38** | 9.40832 | 0.04715 | 4.60449 | 2.67888 | 4.15639 | 5.59844 | 2.80092 | 1.55418 | 1.57249 | 0.09539 | 4.75375 | 4.46675 | 2.59692 | 2.64860 |
| **Fbxo4** | 0.01393 | 0.00446 | 0.05339 | 0.01033 | 0.01193 | 0.01460 | 0.00431 | 0.01067 | 0.01228 | 0.00706 | 0.00808 | 0.00943 | 0.01799 | 0.01085 |

| | | | | | | | | | | | | | | |
|---|---|---|---|---|---|---|---|---|---|---|---|---|---|---|
| **Fbxo41** | 0.14957 | 0.01271 | 0.00000 | 0.11909 | 0.31207 | 0.70791 | 0.00432 | 0.01343 | 0.00444 | 0.03193 | 0.00382 | 0.00095 | 0.00536 | 0.35165 |
| **Fbxo42** | 1.46812 | 0.04321 | 6.39872 | 0.90931 | 8.71357 | 4.50449 | 5.79437 | 3.19349 | 0.26609 | 0.32267 | 5.37498 | 1.15330 | 4.87979 | 5.08750 |
| **Fbxo43** | 0.00000 | 0.06161 | 0.00000 | 0.41484 | 0.00000 | 0.16989 | 0.04036 | 0.00895 | 1.22377 | 0.00639 | 0.00837 | 0.01031 | 2.12991 | 2.54351 |
| **Fbxo44** | 0.00000 | 0.00422 | 0.26070 | 1.29210 | 0.59708 | 0.20674 | 0.00406 | 0.01231 | 0.00664 | 0.00633 | 0.00308 | 0.00362 | 0.07776 | 0.41176 |
| **Fbxo45** | 4.46071 | 0.02489 | 1.25840 | 0.37053 | 7.79398 | 2.10864 | 1.34500 | 3.33712 | 9.14342 | 0.08517 | 0.10943 | 1.27823 | 6.86884 | 7.54199 |
| **Fbxo46** | 0.01268 | 0.00433 | 0.00000 | 0.27748 | 0.26669 | 0.45252 | 0.00430 | 0.20175 | 0.10481 | 0.02558 | 0.00206 | 0.12419 | 1.01397 | 0.39302 |
| **Fbxo47** | 0.38931 | 0.00407 | 0.18098 | 0.00000 | 0.46091 | 0.00000 | 0.00403 | 0.00842 | 0.15115 | 0.00608 | 0.00520 | 0.00489 | 1.06906 | 2.24809 |
| **Fbxo5** | 6.47794 | 8.62105 | 11.42821 | 2.74465 | 9.46097 | 7.07203 | 0.01518 | 5.75460 | 7.41949 | 0.03536 | 4.10657 | 5.79297 | 6.50375 | 10.13214 |
| **Fbxo6** | 5.39832 | 0.68069 | 18.54408 | 10.14532 | 14.52068 | 10.19292 | 5.34960 | 6.39567 | 0.32206 | 0.34360 | 8.28380 | 11.71412 | 6.43281 | 11.03920 |
| **Fbxo7** | 0.09545 | 0.43533 | 0.42238 | 0.00000 | 0.59827 | 0.73036 | 0.00304 | 0.00853 | 0.01165 | 0.00206 | 0.79670 | 1.97406 | 1.29692 | 0.62889 |
| **Fbxo8** | 3.81717 | 0.05557 | 4.89348 | 2.09685 | 5.14652 | 3.05447 | 2.39513 | 2.15979 | 3.21851 | 0.16083 | 0.00000 | 1.58970 | 3.16234 | 3.45179 |
| **Fbxo9** | 3.34195 | 0.03538 | 4.36760 | 3.13302 | 4.05185 | 1.99686 | 2.20114 | 2.61687 | 1.56465 | 0.23029 | 4.02934 | 1.96694 | 3.41051 | 6.46815 |
| **Fbxw11** | 2.30793 | 0.03622 | 5.26441 | 3.09385 | 5.42057 | 11.88146 | 2.59609 | 1.29061 | 4.01322 | 0.07796 | 4.73034 | 7.52032 | 11.17461 | 4.31182 |
| **Fbxw13** | 0.57289 | 0.00426 | 0.01216 | 0.01133 | 0.00000 | 0.02546 | 0.00409 | 0.00695 | 0.16316 | 0.00795 | 0.00573 | 0.00636 | 0.63638 | 0.14408 |
| **Fbxw14** | 0.21822 | 0.00394 | 0.21690 | 0.00000 | 0.99229 | 0.03570 | 0.00395 | 0.00875 | 1.05853 | 0.00604 | 0.00368 | 0.00218 | 4.86160 | 1.20159 |
| **Fbxw15** | 0.28769 | 0.00348 | 0.49275 | 0.09545 | 1.14609 | 0.00247 | 0.00381 | 0.00498 | 0.78291 | 0.00565 | 0.00406 | 0.00436 | 6.22852 | 0.36153 |
| **Fbxw16** | 0.00000 | 0.00394 | 0.11493 | 0.00000 | 0.77616 | 0.00000 | 0.00382 | 0.00810 | 2.38285 | 0.00780 | 0.00309 | 0.00445 | 3.39708 | 1.17692 |
| **Fbxw17** | 0.00695 | 0.00478 | 0.01004 | 0.06039 | 0.01112 | 0.01215 | 0.00980 | 0.01955 | 0.01189 | 0.01918 | 0.00480 | 0.24805 | 0.00626 | 0.01374 |
| **Fbxw18** | 0.83162 | 0.00370 | 0.20125 | 0.00266 | 0.34027 | 0.00192 | 0.00385 | 0.00934 | 0.02354 | 0.00790 | 0.00575 | 0.00417 | 5.37589 | 0.20449 |
| **Fbxw19** | 0.16727 | 0.00391 | 0.39381 | 0.00000 | 0.82063 | 0.26523 | 0.00350 | 0.00122 | 2.39840 | 0.00494 | 0.00285 | 0.00150 | 3.64995 | 1.38096 |
| **Fbxw2** | 4.78670 | 0.06860 | 4.51092 | 1.77753 | 7.64814 | 4.61818 | 2.27312 | 3.27690 | 4.88097 | 0.04544 | 0.86935 | 3.86433 | 4.77496 | 9.41840 |
| **Fbxw20** | 1.35525 | 0.00370 | 0.07620 | 0.14411 | 0.24894 | 0.01049 | 0.00391 | 0.02885 | 0.23488 | 0.00647 | 0.00417 | 0.01095 | 1.25831 | 0.83646 |
| **Fbxw21** | 0.66163 | 0.00417 | 0.07390 | 0.00000 | 0.00000 | 0.29546 | 0.00396 | 0.00340 | 0.14144 | 0.00538 | 0.00487 | 0.00564 | 0.40039 | 1.20956 |
| **Fbxw22** | 0.02284 | 0.00405 | 0.01790 | 0.01032 | 0.18592 | 0.01203 | 0.00423 | 0.00690 | 0.11763 | 0.00602 | 0.00699 | 0.00609 | 2.20392 | 0.92132 |
| **Fbxw24** | 1.22166 | 0.00372 | 0.38589 | 0.33116 | 1.46784 | 0.00000 | 0.00323 | 0.01033 | 3.86285 | 0.01114 | 0.00000 | 0.11333 | 5.62588 | 4.67365 |
| **Fbxw26** | 0.03779 | 0.00406 | 0.01027 | 0.00000 | 0.08649 | 0.40807 | 0.00387 | 0.00403 | 0.08607 | 0.00576 | 0.00479 | 0.00810 | 1.22113 | 0.25618 |
| **Fbxw27** | 0.70001 | 0.00397 | 0.00000 | 0.05970 | 0.00000 | 0.00305 | 0.00378 | 0.00694 | 1.88747 | 0.00606 | 0.00389 | 0.00419 | 2.03686 | 0.16930 |
| **Fbxw28** | 0.39097 | 0.00386 | 0.22024 | 0.00000 | 0.53951 | 0.03879 | 0.00376 | 0.01418 | 0.95797 | 0.00686 | 0.00375 | 0.00187 | 3.64514 | 1.77097 |
| **Fbxw4** | 0.41775 | 0.00417 | 0.26420 | 0.17163 | 0.38310 | 0.00000 | 0.00411 | 0.00857 | 0.01751 | 0.00688 | 0.00554 | 0.00513 | 0.35226 | 0.21608 |
| **Fbxw5** | 0.06527 | 0.04414 | 0.30095 | 2.15752 | 0.07455 | 0.11162 | 0.00320 | 3.21597 | 0.00324 | 0.00000 | 2.33322 | 0.44377 | 1.02555 | 0.26843 |
| **Fbxw7** | 0.00411 | 0.01771 | 0.11767 | 0.33164 | 0.00000 | 0.58771 | 0.00346 | 1.16196 | 0.00817 | 0.00502 | 0.27865 | 3.48348 | 1.20695 | 0.02634 |
| **Fbxw8** | 1.03762 | 0.06800 | 5.12581 | 3.45368 | 3.67841 | 5.53241 | 0.79155 | 2.42450 | 0.00000 | 0.11981 | 4.75518 | 4.57702 | 0.00052 | 1.41026 |
| **Fbxw9** | 3.21560 | 0.03555 | 5.14224 | 6.45709 | 5.21156 | 2.08480 | 1.85903 | 1.41041 | 0.02020 | 0.05343 | 0.27228 | 4.88408 | 5.12427 | 3.16734 |
| **Fcer1g** | 0.00798 | 0.00423 | 0.01294 | 0.00507 | 0.03665 | 0.01557 | 0.00448 | 0.00783 | 0.01010 | 0.00640 | 0.00626 | 0.13709 | 0.21090 | 0.00406 |
| **Fcer2a** | 0.10897 | 0.03661 | 0.02864 | 0.33570 | 0.15322 | 0.29236 | 0.05953 | 0.18020 | 0.19212 | 0.09418 | 0.02643 | 0.05060 | 0.48687 | 0.30243 |
| **Fcf1** | 8.99116 | 0.11193 | 4.06555 | 8.16084 | 21.23474 | 9.54270 | 3.84803 | 3.89494 | 13.14941 | 0.24421 | 3.48922 | 7.08860 | 7.89344 | 15.03814 |
| **Fcgbp** | 0.00000 | 0.00508 | 0.43911 | 0.34553 | 0.14814 | 0.18463 | 0.00384 | 0.00696 | 0.21006 | 0.01477 | 0.00483 | 0.00306 | 0.49737 | 1.22300 |
| **Fcgr1** | 0.58874 | 0.00458 | 0.02889 | 0.01724 | 0.00000 | 0.01101 | 0.00407 | 0.01415 | 0.01256 | 0.00771 | 0.00689 | 0.00634 | 0.50353 | 0.18628 |
| **Fcgr2b** | 0.90864 | 0.00371 | 3.93320 | 0.39814 | 3.34479 | 0.00000 | 0.00260 | 0.05687 | 4.96045 | 0.00477 | 0.45638 | 0.00430 | 0.00000 | 6.24645 |
| **Fcgr3** | 13.84071 | 0.00691 | 23.20098 | 3.64800 | 23.46005 | 0.39535 | 0.33688 | 0.30292 | 24.01222 | 0.00000 | 3.79537 | 0.00534 | 11.71554 | 26.13202 |
| **Fcgrt** | 1.01263 | 0.01252 | 2.07613 | 1.33088 | 1.11355 | 1.45868 | 0.00386 | 0.00000 | 0.00207 | 0.00173 | 0.58213 | 0.00015 | 1.39718 | 0.67330 |
| **Fcho1** | 0.00473 | 0.00404 | 0.00888 | 0.40087 | 0.14852 | 0.00656 | 0.00410 | 0.00588 | 0.01048 | 0.00534 | 0.00667 | 0.03170 | 0.06532 | 0.00000 |
| **Fcho2** | 1.23284 | 0.00960 | 0.65909 | 0.56153 | 0.58394 | 0.65644 | 0.11430 | 0.02162 | 9.41717 | 0.00994 | 1.33272 | 0.80769 | 2.48356 | 2.62006 |
| **Fchsd1** | 0.00912 | 0.01456 | 0.01591 | 0.11153 | 0.24893 | 0.00425 | 0.00475 | 0.33044 | 0.01093 | 0.00703 | 0.00662 | 0.01319 | 0.00471 | 0.04340 |
| **Fchsd2** | 0.54760 | 0.00364 | 0.94364 | 0.00000 | 0.40721 | 0.24124 | 0.00396 | 0.00106 | 4.47400 | 0.00366 | 0.01416 | 0.17350 | 3.98705 | 1.58894 |
| **Fcrl5** | 0.22856 | 0.00399 | 0.10133 | 0.06226 | 0.03427 | 0.23046 | 0.00408 | 0.11622 | 0.05786 | 0.00652 | 0.02702 | 0.00911 | 0.11977 | 0.06444 |
| **Fcrl6** | 0.01729 | 0.00409 | 0.02825 | 0.05520 | 0.02810 | 0.04824 | 0.00409 | 0.03895 | 0.03424 | 0.00629 | 0.01487 | 0.00851 | 0.00748 | 0.04984 |
| **Fdft1** | 25.00517 | 0.16140 | 28.40290 | 0.50911 | 23.27280 | 16.87218 | 0.00611 | 1.26769 | 15.10415 | 4.34932 | 3.26821 | 8.87845 | 13.49444 | 22.84121 |
| **Fdps** | 19.85327 | 0.77622 | 29.47941 | 15.06571 | 17.09544 | 23.40593 | 3.02088 | 9.68285 | 13.11869 | 13.10381 | 16.71518 | 10.94442 | 4.79243 | 16.74250 |
| **Fdx1** | 1.70531 | 0.00397 | 0.65737 | 0.61718 | 2.27003 | 1.45417 | 0.28957 | 0.33111 | 2.57981 | 0.06870 | 0.09918 | 0.54652 | 0.22043 | 0.98534 |
| **Fdx1l** | 0.00451 | 0.00404 | 0.00602 | 0.50928 | 0.32002 | 0.01583 | 0.01574 | 0.07917 | 0.01249 | 0.00516 | 0.00956 | 0.05153 | 0.00525 | 0.00461 |
| **Fdxacb1** | 1.04515 | 0.00409 | 1.87367 | 1.63295 | 1.60861 | 1.20869 | 0.76098 | 0.35205 | 0.01781 | 0.03593 | 0.29340 | 0.43280 | 0.00000 | 1.85462 |
| **Fdxr** | 0.08480 | 6.54274 | 1.14097 | 2.65602 | 0.73312 | 1.30947 | 0.00801 | 0.00000 | 0.00969 | 0.00362 | 2.17624 | 5.33908 | 0.00086 | 0.00000 |
| **Fech** | 1.94927 | 0.02797 | 2.22135 | 0.83755 | 3.18161 | 1.54364 | 3.97123 | 0.46049 | 2.20585 | 0.12031 | 2.26409 | 1.28375 | 0.00000 | 3.87469 |
| **Fem1a** | 0.00000 | 0.02845 | 0.20598 | 1.32809 | 0.50974 | 0.83673 | 0.15350 | 1.86910 | 0.23075 | 0.03976 | 0.00000 | 0.75867 | 4.12726 | 1.53884 |

| | | | | | | | | | | | | | | |
|---|---|---|---|---|---|---|---|---|---|---|---|---|---|---|
| **Fem1b** | 6.36593 | 4.36585 | 5.04208 | 4.09075 | 7.87274 | 6.50582 | 0.29352 | 4.10224 | 15.16024 | 1.69740 | 0.00000 | 6.77735 | 10.10464 | 11.37070 |
| **Fem1c** | 2.81532 | 0.01033 | 3.93552 | 0.93527 | 2.00948 | 0.00000 | 1.79961 | 1.60777 | 4.64456 | 0.04726 | 0.00000 | 0.01147 | 10.72753 | 2.58742 |
| **Fen1** | 3.12446 | 7.58267 | 9.54760 | 9.80654 | 5.40034 | 8.07430 | 2.25341 | 3.99440 | 4.02112 | 0.14732 | 6.21747 | 7.43677 | 4.37452 | 7.41203 |
| **Fendrr** | 0.11245 | 0.05847 | 0.29538 | 0.00000 | 0.26963 | 0.00975 | 0.00438 | 0.01350 | 0.03515 | 0.00987 | 0.00938 | 0.02765 | 0.42391 | 0.02233 |
| **Fer** | 0.82413 | 0.00285 | 0.39631 | 1.22008 | 0.04624 | 0.38292 | 3.33112 | 0.81498 | 0.02098 | 0.11975 | 2.57311 | 0.64163 | 0.03713 | 1.01400 |
| **Fer1l4** | 0.01005 | 0.00444 | 0.01411 | 0.01335 | 0.01238 | 0.01321 | 0.00429 | 0.01023 | 0.00781 | 0.00695 | 0.00894 | 0.01003 | 0.00588 | 0.04776 |
| **Fer1l5** | 0.00237 | 0.00411 | 0.12210 | 0.22251 | 0.03805 | 0.00325 | 0.00398 | 0.00658 | 0.00856 | 0.00512 | 0.02930 | 0.01365 | 0.16096 | 0.06226 |
| **Fermt2** | 3.89292 | 0.02478 | 7.01948 | 4.41259 | 7.07138 | 2.48603 | 1.72521 | 1.49190 | 9.34999 | 0.07755 | 2.02348 | 2.21018 | 1.71653 | 11.66198 |
| **Fermt3** | 1.76577 | 0.00386 | 0.74034 | 1.29261 | 1.07047 | 1.04217 | 0.00401 | 0.24632 | 0.18329 | 0.00449 | 0.03592 | 0.00000 | 0.00000 | 0.56539 |
| **Fes** | 0.00784 | 0.01241 | 0.20924 | 0.61612 | 0.13493 | 0.01928 | 0.00501 | 0.28970 | 0.00765 | 0.00675 | 0.00309 | 0.00432 | 0.22538 | 0.00776 |
| **Fetub** | 0.57155 | 0.00407 | 0.31263 | 0.17246 | 0.00000 | 0.00542 | 0.00957 | 0.00557 | 0.04037 | 0.01148 | 0.00558 | 0.01082 | 0.28720 | 0.37578 |
| **Fez1** | 0.38955 | 0.00398 | 1.25042 | 0.21382 | 3.26189 | 0.00000 | 0.00347 | 0.00629 | 2.49078 | 0.00562 | 0.00000 | 0.00347 | 0.00000 | 2.92857 |
| **Fez2** | 0.00000 | 0.10583 | 8.16639 | 9.43304 | 4.36828 | 4.24918 | 5.21423 | 3.17792 | 0.98075 | 0.40040 | 4.75427 | 1.70639 | 2.20989 | 1.40151 |
| **Ffar2** | 0.09239 | 0.00403 | 0.04916 | 0.01619 | 0.02273 | 0.01393 | 0.00411 | 0.00606 | 0.05629 | 0.01048 | 0.01078 | 0.18795 | 0.17276 | 0.01482 |
| **Ffar3** | 0.01092 | 0.00380 | 0.01541 | 0.00000 | 0.00569 | 0.57822 | 0.00393 | 0.00489 | 0.00840 | 0.00564 | 0.00007 | 0.81898 | 0.01725 | 0.19442 |
| **Fgd1** | 0.14209 | 0.00270 | 0.72748 | 0.03730 | 0.18577 | 0.30954 | 0.00357 | 0.00000 | 0.00324 | 0.00417 | 0.01176 | 1.92266 | 0.16768 | 0.92225 |
| **Fgd2** | 0.01106 | 0.00437 | 0.61128 | 0.00000 | 0.34475 | 0.00337 | 0.00398 | 0.01027 | 0.01428 | 0.00782 | 0.00572 | 0.00892 | 0.00509 | 0.02803 |
| **Fgd3** | 0.19235 | 0.00384 | 0.00402 | 0.04014 | 0.00683 | 0.65339 | 0.00404 | 0.21259 | 0.01090 | 0.01068 | 0.00329 | 0.00262 | 0.11955 | 0.04512 |
| **Fgd4** | 2.04537 | 0.00382 | 1.04061 | 0.08132 | 0.04172 | 0.29772 | 0.85806 | 0.00569 | 1.08454 | 0.01051 | 0.01749 | 0.33978 | 0.13104 | 0.02426 |
| **Fgd6** | 0.00255 | 0.00423 | 0.04260 | 0.00000 | 0.08817 | 1.82525 | 2.77616 | 0.14813 | 0.13760 | 0.04266 | 0.01545 | 0.37452 | 0.23352 | 2.28416 |
| **Fgf1** | 2.00164 | 0.00403 | 0.62425 | 0.00000 | 1.82267 | 0.47592 | 0.00399 | 0.00127 | 1.27267 | 0.00430 | 0.00210 | 0.00324 | 1.61427 | 0.00000 |
| **Fgf10** | 1.95829 | 0.00000 | 0.01213 | 0.00000 | 0.00000 | 0.03855 | 0.00203 | 0.04146 | 0.03145 | 0.00468 | 0.35629 | 7.85076 | 0.24842 | 0.02235 |
| **Fgf11** | 0.00256 | 0.00358 | 0.00916 | 0.03964 | 0.01640 | 0.01855 | 0.01037 | 0.00491 | 0.02586 | 0.00644 | 0.01137 | 0.82931 | 0.27321 | 0.00000 |
| **Fgf13** | 0.50019 | 0.05229 | 0.43424 | 0.36309 | 0.18397 | 0.10735 | 0.03588 | 0.09161 | 0.11820 | 0.14264 | 0.00971 | 0.02771 | 0.00194 | 0.00000 |
| **Fgf17** | 0.00546 | 0.00437 | 0.01499 | 0.01237 | 0.01098 | 0.01345 | 0.00426 | 0.00937 | 0.01110 | 0.00663 | 0.00900 | 0.00948 | 0.26188 | 0.00280 |
| **Fgf18** | 0.00868 | 0.00445 | 0.00829 | 0.00948 | 0.32288 | 0.01267 | 0.00417 | 0.01012 | 0.00987 | 0.00618 | 0.00904 | 0.00859 | 0.03442 | 0.14156 |
| **Fgf3** | 0.34682 | 0.00398 | 1.87071 | 0.60551 | 0.64434 | 0.00000 | 0.00367 | 0.00420 | 0.08046 | 0.00473 | 0.00757 | 0.00643 | 0.83626 | 0.02510 |
| **Fgf4** | 0.34434 | 0.00207 | 1.63266 | 0.65400 | 4.42322 | 0.00000 | 0.00259 | 0.00000 | 3.26644 | 0.00250 | 0.00000 | 1.45913 | 3.78841 | 7.04678 |
| **Fgf5** | 0.00276 | 0.01227 | 0.00360 | 0.01213 | 0.19933 | 0.00982 | 0.00411 | 0.00922 | 0.08131 | 0.00640 | 0.00734 | 0.00799 | 0.01203 | 0.85316 |
| **Fgf7** | 0.01024 | 0.00394 | 0.05251 | 0.00738 | 0.00731 | 0.01514 | 0.00395 | 0.06303 | 0.01318 | 0.00759 | 0.00889 | 0.00784 | 0.95348 | 0.00216 |
| **Fgf9** | 0.09738 | 0.00452 | 0.01779 | 0.01411 | 0.00639 | 0.00907 | 0.00446 | 0.02260 | 0.06206 | 0.00777 | 0.01081 | 0.00721 | 0.51876 | 0.01922 |
| **Fgfbp1** | 0.00000 | 0.01946 | 0.46182 | 0.05516 | 0.18756 | 5.59478 | 0.00430 | 1.02241 | 0.01198 | 0.03050 | 0.00223 | 0.00000 | 0.00463 | 0.00000 |
| **Fgfbp3** | 0.01208 | 0.00443 | 0.11394 | 0.02229 | 0.12806 | 0.00762 | 0.00431 | 0.00943 | 0.01096 | 0.01337 | 0.00737 | 0.01722 | 0.00594 | 0.01185 |
| **Fgfr1** | 2.34532 | 0.10862 | 11.08865 | 0.00000 | 5.71852 | 2.72858 | 2.67639 | 7.67311 | 0.00338 | 0.11506 | 4.93934 | 11.78631 | 1.09659 | 1.15708 |
| **Fgfr1op** | 5.23966 | 0.03615 | 4.36623 | 2.66449 | 8.34582 | 3.69199 | 2.95119 | 1.75042 | 9.41116 | 0.97572 | 2.29235 | 0.14307 | 3.40983 | 4.63687 |
| **Fgfr1op2** | 2.72271 | 0.04953 | 4.01172 | 6.41597 | 7.57001 | 3.49464 | 1.35455 | 0.95323 | 8.18657 | 0.03530 | 4.67443 | 5.56582 | 0.76317 | 0.00000 |
| **Fgfr2** | 3.55705 | 0.10726 | 2.41910 | 0.20280 | 3.46773 | 4.74859 | 0.43520 | 2.54088 | 11.74869 | 0.02678 | 1.71547 | 4.83830 | 0.00000 | 2.75732 |
| **Fgfr3** | 0.20944 | 0.00385 | 1.06499 | 0.87237 | 1.78942 | 0.16942 | 0.00388 | 0.00000 | 0.18607 | 0.01092 | 0.09792 | 0.26260 | 0.97529 | 1.03712 |
| **Fgfr4** | 0.22533 | 0.00985 | 0.19656 | 1.20750 | 1.41518 | 2.17127 | 0.00933 | 0.32206 | 0.28924 | 0.01278 | 1.11531 | 0.81116 | 2.83162 | 2.68517 |
| **Fgfrl1** | 0.19329 | 3.08231 | 0.17779 | 0.10355 | 0.09549 | 0.00000 | 0.38559 | 0.00309 | 0.00077 | 0.01122 | 0.00546 | 0.00318 | 1.83466 | 0.05038 |
| **Fgl1** | 0.00907 | 0.00396 | 0.01142 | 0.00000 | 0.02692 | 0.15333 | 0.00375 | 0.43448 | 0.01044 | 0.00752 | 0.00566 | 0.01077 | 0.00673 | 0.01076 |
| **Fgl2** | 0.01832 | 0.00450 | 0.01068 | 0.01297 | 0.13326 | 0.01297 | 0.00452 | 0.01015 | 0.01159 | 0.00667 | 0.00865 | 0.00929 | 0.00503 | 0.01937 |
| **Fgr** | 1.41802 | 0.00344 | 2.08323 | 0.00000 | 4.53331 | 0.00000 | 0.00308 | 0.00238 | 2.20541 | 0.00260 | 0.00000 | 0.00099 | 3.11238 | 3.37394 |
| **Fh1** | 1.66397 | 0.17923 | 3.37151 | 4.53611 | 0.33602 | 5.02955 | 5.09339 | 5.80614 | 1.59291 | 12.31871 | 3.19942 | 10.91502 | 1.11777 | 1.23962 |
| **Fhad1** | 0.00963 | 0.00444 | 0.01090 | 0.01416 | 0.00819 | 0.01911 | 0.00425 | 0.00879 | 0.01708 | 0.00751 | 0.00780 | 0.00967 | 0.01095 | 0.26121 |
| **Fhl1** | 0.08913 | 0.00468 | 0.00408 | 0.00310 | 0.74077 | 0.01439 | 0.00473 | 0.01857 | 0.01847 | 0.00638 | 0.01039 | 0.03018 | 0.00310 | 0.00351 |
| **Fhl2** | 0.00000 | 7.96000 | 0.48568 | 2.25907 | 0.00065 | 3.54029 | 3.76099 | 7.22297 | 0.00712 | 0.17793 | 0.00000 | 4.60610 | 0.00552 | 0.00000 |
| **Fhl3** | 0.07801 | 0.01264 | 0.00611 | 0.15164 | 0.56477 | 0.00360 | 0.00497 | 0.16786 | 0.16886 | 0.01011 | 0.00514 | 0.00432 | 1.01493 | 0.13752 |
| **Fhl4** | 0.00798 | 0.00413 | 0.00328 | 0.14201 | 0.00951 | 0.42704 | 0.00391 | 0.00331 | 0.01124 | 0.00867 | 0.17312 | 0.02269 | 0.00789 | 0.00628 |
| **Fhod1** | 0.08767 | 0.08007 | 2.24501 | 0.00000 | 0.00050 | 0.18738 | 0.01396 | 3.01020 | 0.00809 | 0.04376 | 0.00000 | 8.62683 | 0.07963 | 0.04307 |
| **Fhod3** | 0.98053 | 0.00283 | 0.75672 | 0.00000 | 1.85404 | 0.08462 | 0.00332 | 0.00000 | 0.91825 | 0.00436 | 0.00032 | 0.71837 | 3.17255 | 4.17128 |
| **Fibcd1** | 0.00599 | 0.00429 | 0.01475 | 0.01189 | 0.00934 | 0.01304 | 0.00421 | 0.00892 | 0.00970 | 0.00630 | 0.00937 | 0.00899 | 0.57836 | 0.01349 |
| **Fibp** | 1.01381 | 0.91398 | 4.54951 | 9.04698 | 1.41753 | 4.16340 | 0.02344 | 8.62891 | 0.00000 | 0.05392 | 2.83910 | 6.92979 | 0.24102 | 2.49335 |
| **Ficd** | 7.74511 | 0.00311 | 7.26733 | 1.86108 | 7.00859 | 0.69096 | 0.00216 | 0.38198 | 6.56006 | 0.01178 | 0.21313 | 0.00000 | 5.26014 | 18.81305 |
| **Fig4** | 7.03378 | 0.01087 | 12.64042 | 3.48716 | 13.83193 | 3.61594 | 0.02475 | 0.56676 | 2.17925 | 0.00000 | 4.49366 | 0.13456 | 1.00891 | 1.16651 |

| | | | | | | | | | | | | | | |
|---|---|---|---|---|---|---|---|---|---|---|---|---|---|---|
| **Fignl1** | 4.04111 | 1.43758 | 3.68863 | 2.12275 | 2.81204 | 4.18935 | 1.46242 | 0.00000 | 8.89596 | 0.02458 | 8.07627 | 3.55518 | 1.62674 | 8.59291 |
| **Filip1l** | 0.30841 | 0.01248 | 0.10850 | 0.00000 | 0.48633 | 0.00000 | 2.48312 | 0.12366 | 0.32171 | 0.05259 | 0.00266 | 0.00276 | 0.27462 | 0.54024 |
| **Fip1l1** | 4.30378 | 0.78963 | 6.96368 | 4.62428 | 5.80840 | 3.80420 | 1.35499 | 7.38133 | 0.16412 | 0.10274 | 7.41220 | 8.10685 | 2.58737 | 1.95843 |
| **Firre** | 0.55522 | 0.02370 | 1.12226 | 2.19721 | 0.45568 | 0.26726 | 0.65631 | 0.56021 | 0.00901 | 0.02506 | 0.00000 | 0.32891 | 0.00879 | 0.01535 |
| **Fis1** | 3.85060 | 7.77887 | 2.88238 | 12.49365 | 6.21908 | 7.99446 | 2.66311 | 8.16312 | 0.00000 | 16.22254 | 9.05623 | 4.37625 | 1.33048 | 3.12504 |
| **Fitm2** | 0.00240 | 0.00379 | 0.07755 | 0.35486 | 0.48223 | 0.45914 | 0.00382 | 0.00343 | 0.00349 | 0.01142 | 0.28077 | 0.23117 | 0.38147 | 0.33500 |
| **Fiz1** | 7.40243 | 0.06245 | 6.17306 | 7.36667 | 17.08512 | 1.29349 | 4.99134 | 1.54968 | 2.73183 | 0.11817 | 0.17388 | 2.47648 | 16.92898 | 3.41386 |
| **Fjx1** | 0.16617 | 0.02245 | 0.01118 | 0.00582 | 0.00808 | 0.01099 | 0.00417 | 0.02392 | 0.01067 | 0.00619 | 0.00460 | 0.41402 | 0.00515 | 0.00771 |
| **Fkbp10** | 0.91723 | 0.00402 | 2.21692 | 0.12349 | 6.06681 | 0.00000 | 0.00377 | 0.00440 | 0.28192 | 0.00319 | 0.00246 | 0.00301 | 0.08945 | 2.73179 |
| **Fkbp11** | 1.26603 | 0.03189 | 0.67274 | 11.02282 | 1.78524 | 4.83613 | 0.00848 | 0.03653 | 0.00722 | 0.05353 | 1.59757 | 1.64681 | 0.00000 | 0.71412 |
| **Fkbp14** | 0.00966 | 0.00401 | 0.01030 | 0.00000 | 0.23093 | 0.00886 | 1.08993 | 0.00405 | 0.01072 | 0.01125 | 0.00793 | 0.01127 | 0.01051 | 0.01077 |
| **Fkbp15** | 0.07330 | 0.05688 | 1.88578 | 1.04615 | 0.82930 | 3.78035 | 1.08495 | 2.08467 | 0.00000 | 0.03446 | 2.95562 | 3.78684 | 1.13846 | 1.87506 |
| **Fkbp1a** | 4.75715 | 1.25522 | 9.78453 | 2.67852 | 7.68597 | 7.97113 | 4.25468 | 7.51071 | 6.53950 | 10.93749 | 5.41712 | 6.08926 | 10.09896 | 3.16711 |
| **Fkbp1b** | 2.03335 | 0.00355 | 0.06176 | 1.46245 | 1.15768 | 0.00000 | 0.00392 | 0.00550 | 0.00000 | 0.00642 | 0.01309 | 0.02031 | 2.47184 | 0.50854 |
| **Fkbp2** | 2.56753 | 4.05964 | 0.19496 | 5.91886 | 3.65440 | 4.31817 | 0.44875 | 1.86390 | 0.09863 | 0.26139 | 0.50495 | 2.82992 | 0.78032 | 2.24461 |
| **Fkbp3** | 9.64833 | 2.95378 | 10.14805 | 9.74429 | 12.17812 | 15.47124 | 4.59111 | 12.64928 | 1.33830 | 7.54877 | 11.19973 | 12.44564 | 0.44955 | 3.10369 |
| **Fkbp4** | 17.01076 | 10.07569 | 23.04699 | 28.01412 | 18.25191 | 31.27519 | 6.65106 | 19.58060 | 3.09453 | 6.62205 | 17.45902 | 15.03207 | 10.89870 | 16.38720 |
| **Fkbp5** | 0.97480 | 0.59275 | 0.91256 | 1.94565 | 0.92550 | 5.98735 | 0.00818 | 1.41881 | 1.86520 | 0.03133 | 2.44374 | 5.23953 | 6.09755 | 9.57883 |
| **Fkbp6** | 6.95254 | 0.12410 | 17.57400 | 11.72121 | 10.32949 | 17.95850 | 0.07006 | 7.20396 | 4.83280 | 0.14876 | 9.86814 | 2.50064 | 5.73334 | 6.86752 |
| **Fkbp8** | 1.27553 | 11.19091 | 13.99811 | 17.45202 | 11.20704 | 8.72840 | 2.27182 | 12.96510 | 0.02900 | 3.24214 | 5.74883 | 9.38884 | 1.07592 | 0.00000 |
| **Fkbp9** | 0.29417 | 0.10606 | 2.65548 | 0.89601 | 2.85428 | 3.14005 | 3.16294 | 2.60900 | 1.89421 | 0.13771 | 0.00000 | 6.19875 | 0.00000 | 1.22584 |
| **Fkbpl** | 1.04488 | 0.01853 | 1.72281 | 0.76834 | 3.68919 | 1.25349 | 0.00943 | 0.12657 | 0.00109 | 0.00891 | 0.64388 | 0.60580 | 0.98274 | 0.41234 |
| **Fkrp** | 0.23972 | 0.01065 | 1.32644 | 0.43779 | 0.43144 | 0.94162 | 0.00353 | 0.08961 | 0.00868 | 0.01051 | 0.00000 | 1.17452 | 0.00394 | 1.22069 |
| **Fktn** | 0.02529 | 0.03099 | 0.32807 | 0.58821 | 0.37403 | 1.46924 | 0.01448 | 0.83322 | 0.64843 | 0.04978 | 0.25124 | 1.71656 | 0.23475 | 0.93821 |
| **Flad1** | 0.83728 | 0.05157 | 1.56639 | 5.33452 | 3.34473 | 5.14780 | 2.55733 | 5.03620 | 0.74130 | 0.88065 | 4.52567 | 1.15762 | 0.00000 | 3.80369 |
| **Flcn** | 0.66261 | 0.19051 | 2.07993 | 4.99160 | 1.09482 | 4.78800 | 0.51399 | 0.01609 | 1.73636 | 11.24192 | 0.43554 | 0.98746 | 5.22658 | 2.15020 |
| **Flii** | 1.14501 | 0.04773 | 11.46078 | 1.76466 | 2.33351 | 7.78694 | 3.31927 | 5.74595 | 0.00000 | 10.14316 | 8.16642 | 0.93747 | 0.02206 | 1.46622 |
| **Flna** | 2.61034 | 0.21621 | 6.62861 | 2.58975 | 1.52933 | 0.00000 | 0.19907 | 5.63647 | 1.46164 | 0.00788 | 0.00000 | 0.00739 | 3.03963 | 2.33136 |
| **Flnb** | 7.61672 | 4.28754 | 14.28753 | 7.17014 | 4.18221 | 16.53134 | 4.66920 | 12.25270 | 5.50747 | 0.21104 | 13.54508 | 9.12616 | 2.48154 | 6.38664 |
| **Flot1** | 0.13152 | 0.00270 | 1.28526 | 3.44937 | 2.90879 | 0.46268 | 0.00352 | 0.88113 | 0.01955 | 0.01591 | 0.10462 | 1.94615 | 0.56698 | 0.30561 |
| **Flot2** | 0.03034 | 6.02356 | 1.22199 | 2.50534 | 1.74312 | 6.26546 | 2.62108 | 6.01420 | 0.00000 | 0.07771 | 0.00000 | 0.00000 | 0.00523 | 0.58253 |
| **Flrt1** | 0.00000 | 0.01900 | 0.49571 | 0.27967 | 0.00146 | 0.48002 | 0.00479 | 0.01234 | 0.02966 | 0.00543 | 0.00000 | 1.03186 | 0.00765 | 0.00000 |
| **Flrt2** | 0.09510 | 0.00436 | 0.01412 | 0.01073 | 0.00852 | 0.01233 | 0.00429 | 0.00989 | 0.00962 | 0.00659 | 0.00793 | 0.01241 | 0.19392 | 0.00734 |
| **Flrt3** | 0.00945 | 0.01408 | 0.01558 | 0.00000 | 0.01081 | 0.00957 | 0.00423 | 0.07244 | 0.01552 | 0.00731 | 0.00360 | 0.90710 | 0.21342 | 0.00926 |
| **Flt1** | 0.03848 | 0.00422 | 0.05521 | 0.00838 | 0.22488 | 0.01191 | 0.00435 | 0.01000 | 0.00646 | 0.00755 | 0.00747 | 0.00868 | 0.73626 | 0.01588 |
| **Flt3** | 0.00771 | 0.00418 | 0.01689 | 0.14474 | 0.00835 | 0.18257 | 0.00400 | 0.00335 | 0.01120 | 0.00504 | 0.02630 | 0.01360 | 0.12196 | 0.00000 |
| **Flt3l** | 0.04980 | 0.00314 | 3.04411 | 2.06782 | 2.62589 | 0.00000 | 0.00300 | 0.00000 | 0.41278 | 0.00595 | 0.00769 | 0.78229 | 1.60286 | 0.00000 |
| **Flywch1** | 0.01323 | 0.00438 | 0.03458 | 0.49905 | 0.02459 | 0.01631 | 0.00418 | 0.00720 | 0.01524 | 0.00621 | 0.00860 | 0.09978 | 0.59408 | 0.00000 |
| **Flywch2** | 0.48440 | 0.00489 | 0.02705 | 0.02792 | 0.54725 | 0.40627 | 0.00415 | 0.01656 | 0.00842 | 0.01786 | 0.00514 | 0.00172 | 0.60097 | 0.10843 |
| **Fmc1** | 0.03532 | 0.04242 | 0.01085 | 0.14388 | 0.59328 | 0.64869 | 0.69361 | 1.64614 | 0.01886 | 0.99035 | 0.00000 | 0.65151 | 0.38094 | 0.00881 |
| **Fmn1** | 0.00864 | 0.00395 | 0.11559 | 0.00000 | 0.03939 | 0.00000 | 0.00365 | 0.42723 | 0.02759 | 0.00558 | 0.74550 | 0.00447 | 0.40015 | 0.25972 |
| **Fmn2** | 0.00000 | 0.00439 | 0.06809 | 0.00456 | 0.01481 | 0.00498 | 0.00403 | 0.01036 | 0.84130 | 0.00701 | 0.00562 | 0.00800 | 0.22610 | 0.32138 |
| **Fmnl1** | 0.00787 | 0.01170 | 0.01590 | 0.00000 | 0.03233 | 0.00773 | 0.11281 | 0.64157 | 0.01179 | 0.00698 | 0.00746 | 0.01199 | 0.03904 | 0.00873 |
| **Fmnl2** | 0.00000 | 0.01020 | 1.77770 | 0.77258 | 0.43448 | 6.21543 | 1.74542 | 1.90285 | 0.00340 | 0.01229 | 3.62833 | 0.70830 | 0.28127 | 0.34052 |
| **Fmnl3** | 0.01580 | 0.00302 | 0.00000 | 0.00107 | 0.16614 | 2.34488 | 0.59190 | 2.31418 | 0.33315 | 0.03884 | 0.00591 | 0.75917 | 7.48108 | 0.67610 |
| **Fmo1** | 6.64996 | 0.45173 | 2.42895 | 2.95019 | 7.08703 | 4.04983 | 0.83827 | 2.20055 | 13.23437 | 4.27750 | 2.00588 | 1.57539 | 2.89364 | 9.45991 |
| **Fmo5** | 0.09842 | 0.00445 | 0.00836 | 0.05847 | 0.00886 | 0.03055 | 0.00442 | 0.03466 | 0.02720 | 0.01227 | 0.00974 | 0.00672 | 0.17636 | 0.06914 |
| **Fmr1** | 5.00262 | 0.27498 | 5.22006 | 4.57345 | 4.08548 | 6.89160 | 0.81370 | 0.62545 | 17.26698 | 0.01417 | 3.93504 | 4.90681 | 4.80893 | 2.73871 |
| **Fmr1nb** | 6.13499 | 0.25821 | 6.78072 | 12.59448 | 8.92393 | 10.98077 | 6.27819 | 10.61193 | 8.66897 | 0.43239 | 3.01815 | 5.02324 | 0.76302 | 9.56636 |
| **Fn1** | 3.48510 | 0.13250 | 2.33643 | 2.51473 | 1.56425 | 2.20022 | 0.00000 | 0.01515 | 0.00000 | 0.01996 | 0.27526 | 12.06466 | 0.00000 | 0.24514 |
| **Fnbp1** | 0.32930 | 0.04243 | 2.21848 | 2.46894 | 2.55700 | 0.68572 | 0.00230 | 1.00728 | 1.90556 | 0.01295 | 0.85492 | 5.08667 | 1.80295 | 2.53304 |
| **Fnbp1l** | 0.18313 | 0.01004 | 0.00000 | 0.40688 | 0.00000 | 3.04985 | 1.23160 | 0.85873 | 1.45434 | 5.34300 | 2.58481 | 0.29743 | 2.35457 | 0.00000 |
| **Fnbp4** | 8.78525 | 0.18249 | 10.25064 | 5.77746 | 12.95018 | 4.11258 | 4.16359 | 1.48719 | 6.20145 | 0.17150 | 5.83395 | 4.68270 | 9.61015 | 16.41129 |
| **Fndc10** | 0.00072 | 0.00400 | 0.00000 | 0.39958 | 0.00000 | 0.03996 | 0.00487 | 0.73416 | 0.05614 | 0.00654 | 0.00070 | 0.63936 | 0.02808 | 0.25623 |
| **Fndc3a** | 0.55222 | 0.00885 | 0.12026 | 0.00000 | 0.18175 | 1.19418 | 3.03499 | 0.00303 | 2.09871 | 0.01113 | 1.55118 | 3.79135 | 1.27305 | 0.00000 |

| | | | | | | | | | | | | | | |
|---|---|---|---|---|---|---|---|---|---|---|---|---|---|---|
| **Fndc3b** | 0.10610 | 0.02518 | 0.35763 | 0.00000 | 0.99798 | 0.82948 | 1.34994 | 1.34021 | 0.55579 | 0.01873 | 0.00000 | 0.54683 | 4.02501 | 0.00000 |
| **Fndc4** | 0.08360 | 0.02145 | 0.43519 | 0.19456 | 0.00000 | 0.49035 | 0.00439 | 0.14153 | 0.19170 | 0.02407 | 0.01101 | 0.56221 | 0.00681 | 0.01120 |
| **Fndc5** | 0.00947 | 0.00533 | 0.00995 | 0.00649 | 0.26519 | 0.00723 | 0.00456 | 0.09844 | 0.01853 | 0.01513 | 0.00907 | 0.00833 | 0.05802 | 0.01158 |
| **Fndc7** | 0.00729 | 0.00485 | 0.00344 | 0.07445 | 0.27388 | 0.11144 | 0.00450 | 0.00673 | 0.01192 | 0.00648 | 0.00644 | 0.00560 | 0.00667 | 0.00702 |
| **Fndc8** | 0.01681 | 0.00494 | 0.03620 | 0.00000 | 0.01378 | 0.10460 | 0.00447 | 0.00853 | 0.01025 | 0.00644 | 0.00618 | 0.04777 | 0.00855 | 0.42780 |
| **Fnip1** | 0.20193 | 0.00212 | 0.07181 | 0.77893 | 0.95718 | 1.72008 | 0.00265 | 1.47845 | 2.70364 | 0.37637 | 3.28490 | 0.69797 | 3.22254 | 0.84022 |
| **Fnip2** | 0.17078 | 0.02685 | 0.39414 | 1.18637 | 0.30096 | 0.24442 | 0.08624 | 0.06109 | 3.19900 | 0.05221 | 0.09454 | 1.27136 | 1.10666 | 2.13656 |
| **Fnta** | 10.01935 | 0.02550 | 5.81177 | 1.58762 | 9.28185 | 7.61809 | 1.80494 | 2.74681 | 8.00815 | 0.05446 | 1.10304 | 7.77365 | 3.33001 | 4.97281 |
| **Fntb** | 0.74390 | 0.06939 | 4.23619 | 3.45338 | 5.04046 | 1.15986 | 3.63857 | 5.02860 | 0.00021 | 0.08537 | 6.37781 | 3.13574 | 1.73299 | 2.22615 |
| **Focad** | 0.12675 | 0.01636 | 0.84083 | 2.56117 | 0.89143 | 1.47357 | 0.00785 | 0.00000 | 0.18722 | 0.02031 | 2.19015 | 4.15636 | 1.26165 | 0.06004 |
| **Folr1** | 19.77609 | 8.14135 | 35.50709 | 29.83635 | 28.53058 | 30.84973 | 2.16930 | 17.86790 | 6.50429 | 13.44812 | 14.57178 | 3.15337 | 9.43840 | 28.93159 |
| **Folr2** | 0.23811 | 0.00351 | 0.77271 | 0.52071 | 0.48796 | 0.98779 | 2.07287 | 0.75847 | 0.01177 | 0.09064 | 0.77542 | 0.06159 | 0.01104 | 0.84050 |
| **Fopnl** | 3.41612 | 0.06867 | 4.08438 | 4.47874 | 7.69400 | 6.25112 | 3.34266 | 5.08920 | 5.94166 | 0.09304 | 5.84430 | 6.43022 | 1.22114 | 5.93884 |
| **Fos** | 0.00280 | 12.46285 | 0.00420 | 0.00000 | 0.00839 | 0.18682 | 0.00000 | 6.68941 | 0.03183 | 0.01448 | 0.00000 | 1.64514 | 0.12169 | 0.00000 |
| **Fosb** | 0.00580 | 0.02121 | 0.01918 | 0.15018 | 0.00917 | 0.01092 | 0.00472 | 0.05256 | 0.01303 | 0.00772 | 0.00818 | 0.01082 | 0.14710 | 0.00000 |
| **Fosl2** | 0.07718 | 0.03597 | 0.00298 | 0.00000 | 0.03401 | 0.13560 | 3.21393 | 1.26875 | 0.01273 | 0.13996 | 0.00000 | 0.24467 | 0.05747 | 0.12444 |
| **Foxa1** | 8.35807 | 0.00311 | 5.87229 | 0.00000 | 14.63027 | 0.00000 | 0.00068 | 0.00067 | 17.36851 | 0.00000 | 0.00000 | 0.00000 | 0.00000 | 7.60450 |
| **Foxa3** | 0.02669 | 0.00398 | 0.01236 | 0.01495 | 0.01216 | 0.06165 | 0.00419 | 0.06469 | 0.01852 | 0.00720 | 0.00627 | 0.00975 | 0.01751 | 0.00605 |
| **Foxc1** | 0.30712 | 0.00372 | 0.41098 | 0.06000 | 0.00000 | 0.46247 | 0.00380 | 0.00247 | 0.00457 | 0.00764 | 0.00340 | 0.16341 | 0.94674 | 0.00000 |
| **Foxc2** | 0.01811 | 0.00407 | 0.25469 | 0.00000 | 2.48636 | 0.00000 | 2.26764 | 0.00150 | 0.00833 | 0.18461 | 0.14868 | 0.00212 | 0.02212 | 0.00000 |
| **Foxe1** | 0.01609 | 0.00424 | 0.16068 | 0.21736 | 0.36266 | 0.00000 | 0.00401 | 0.00946 | 0.01060 | 0.00680 | 0.00856 | 0.00880 | 0.00952 | 0.01769 |
| **Foxf1** | 0.00000 | 0.00390 | 0.41483 | 0.00000 | 0.23478 | 0.00223 | 0.00414 | 0.00865 | 0.01500 | 0.00671 | 0.00690 | 0.00775 | 3.02243 | 0.37570 |
| **Foxf2** | 0.01516 | 0.00999 | 0.04518 | 0.49450 | 0.01014 | 0.00000 | 0.00334 | 0.00295 | 0.01518 | 0.00885 | 0.00834 | 1.41931 | 0.01384 | 0.00644 |
| **Foxg1** | 0.03054 | 0.00438 | 0.02278 | 0.01022 | 0.01520 | 0.02573 | 0.01018 | 0.00809 | 0.10488 | 0.00795 | 0.00960 | 0.01341 | 0.08981 | 0.00239 |
| **Foxh1** | 0.00000 | 0.20938 | 0.34072 | 0.94244 | 0.21342 | 0.43283 | 0.01847 | 0.06057 | 0.01186 | 0.03518 | 0.00000 | 7.05366 | 0.07376 | 0.15583 |
| **Foxi3** | 0.12388 | 0.00445 | 0.02930 | 0.00611 | 0.31693 | 0.01135 | 0.00424 | 0.01054 | 0.00822 | 0.00648 | 0.00776 | 0.00785 | 0.13140 | 0.00365 |
| **Foxj1** | 0.00880 | 0.00446 | 0.00853 | 0.01058 | 0.26684 | 0.01373 | 0.00448 | 0.01032 | 0.01668 | 0.00669 | 0.00904 | 0.00881 | 0.00542 | 0.00640 |
| **Foxj2** | 0.02326 | 0.02013 | 0.00476 | 0.37950 | 0.01514 | 0.85531 | 0.01692 | 2.08755 | 0.00778 | 0.01988 | 0.48528 | 0.00375 | 0.69576 | 0.00000 |
| **Foxj3** | 2.13026 | 0.96724 | 3.26936 | 0.40900 | 3.04975 | 6.03388 | 4.12120 | 0.16755 | 6.78649 | 0.09701 | 0.80201 | 1.44069 | 23.79182 | 9.33282 |
| **Foxk1** | 0.06809 | 0.00928 | 0.10053 | 0.18648 | 0.21502 | 1.44246 | 0.13586 | 2.69712 | 0.72015 | 2.22610 | 1.89487 | 1.86708 | 0.53574 | 0.00000 |
| **Foxk2** | 0.00000 | 0.01865 | 1.78871 | 0.37001 | 2.08826 | 1.57308 | 1.59549 | 0.85138 | 0.80123 | 0.01962 | 1.51794 | 2.47873 | 1.80691 | 13.54481 |
| **Foxl1** | 0.04995 | 0.02111 | 0.02349 | 0.06626 | 0.03848 | 0.05885 | 0.01617 | 0.00928 | 0.05268 | 0.03265 | 0.01173 | 0.01751 | 0.11829 | 0.04560 |
| **Foxm1** | 1.28951 | 0.06594 | 2.40556 | 1.04871 | 2.14199 | 3.77262 | 1.93446 | 2.73834 | 0.52141 | 0.97834 | 1.93377 | 4.33670 | 9.69710 | 5.00171 |
| **Foxn1** | 0.00754 | 0.00421 | 0.00834 | 0.15829 | 0.14423 | 0.00966 | 0.00415 | 0.01521 | 0.00961 | 0.00637 | 0.00753 | 0.00661 | 0.00787 | 0.03718 |
| **Foxn2** | 2.94647 | 0.00165 | 2.49021 | 3.00793 | 2.64637 | 3.66207 | 1.03411 | 1.49070 | 10.90400 | 0.00403 | 2.22796 | 1.84222 | 7.09775 | 2.08799 |
| **Foxn3** | 0.06325 | 0.00392 | 0.42566 | 0.74006 | 0.68457 | 0.18810 | 0.00372 | 0.05094 | 3.09751 | 0.00588 | 0.00070 | 0.00281 | 0.01162 | 2.40839 |
| **Foxo1** | 0.07349 | 0.07590 | 1.30095 | 1.26328 | 4.39341 | 5.84967 | 6.56637 | 3.35409 | 4.36813 | 0.67687 | 5.36927 | 2.04297 | 8.20942 | 2.75389 |
| **Foxo3** | 0.28616 | 6.37949 | 3.09155 | 2.11994 | 2.70690 | 0.12719 | 0.00876 | 0.72161 | 0.02701 | 0.00000 | 1.39169 | 0.80063 | 0.00843 | 0.61824 |
| **Foxo4** | 0.04945 | 0.00490 | 0.03034 | 0.02630 | 0.03822 | 0.01670 | 0.00430 | 0.01643 | 0.00755 | 0.00746 | 0.00696 | 0.06271 | 0.01243 | 0.07779 |
| **Foxo6** | 0.00932 | 0.00546 | 0.02266 | 0.00870 | 0.01029 | 0.00498 | 0.74242 | 0.12662 | 0.01042 | 0.00899 | 0.00845 | 0.00742 | 0.01095 | 0.01417 |
| **Foxp1** | 0.05678 | 0.00440 | 0.00594 | 0.01412 | 0.28873 | 0.01182 | 0.00458 | 0.01106 | 0.03528 | 0.00702 | 0.00789 | 0.00798 | 0.07670 | 0.02599 |
| **Foxp4** | 0.04512 | 0.01950 | 0.72374 | 1.52610 | 2.36917 | 0.00000 | 0.00334 | 1.14776 | 0.01600 | 0.02090 | 0.00000 | 0.34634 | 0.02686 | 0.00000 |
| **Foxq1** | 0.15456 | 0.00475 | 0.01651 | 0.01375 | 0.00996 | 0.01481 | 0.00410 | 0.01279 | 0.01016 | 0.01018 | 0.00653 | 0.02980 | 0.03552 | 0.00479 |
| **Foxr1** | 2.78876 | 0.04226 | 2.54001 | 2.41723 | 5.64502 | 1.50797 | 0.04440 | 1.96358 | 0.75600 | 0.10641 | 0.19792 | 1.94435 | 15.99429 | 6.63658 |
| **Foxred1** | 0.79076 | 0.03519 | 1.72451 | 3.76461 | 0.65724 | 3.62677 | 0.00882 | 2.24710 | 2.12055 | 0.05150 | 3.83851 | 2.96265 | 0.00000 | 4.02625 |
| **Foxred2** | 0.00000 | 4.58102 | 1.26840 | 0.00000 | 0.35719 | 0.64760 | 0.89138 | 1.15791 | 0.10719 | 0.02830 | 0.00314 | 0.23132 | 0.00000 | 0.79347 |
| **Fpgs** | 0.62642 | 0.09291 | 2.89847 | 5.14121 | 13.26274 | 0.94061 | 2.99803 | 3.31645 | 0.10409 | 0.10047 | 0.31713 | 2.30400 | 6.05047 | 4.42914 |
| **Fpgt** | 0.87465 | 0.00456 | 0.00000 | 0.00000 | 0.00000 | 0.43753 | 0.00340 | 0.00583 | 3.82904 | 0.00579 | 0.00168 | 0.08350 | 0.17228 | 0.18053 |
| **Fra10ac1** | 2.59936 | 0.00222 | 0.38410 | 2.86235 | 1.82384 | 1.93219 | 3.13202 | 0.46621 | 2.72642 | 0.09015 | 1.72455 | 1.67219 | 0.01268 | 1.00710 |
| **Frat1** | 0.03024 | 0.01511 | 0.22985 | 2.85407 | 0.96804 | 0.00000 | 0.01036 | 0.37648 | 0.18018 | 0.02834 | 0.92737 | 0.00237 | 2.36908 | 0.34920 |
| **Frat2** | 0.72012 | 0.08091 | 0.98497 | 0.48134 | 0.42434 | 0.39354 | 0.00413 | 0.86550 | 0.09532 | 0.04046 | 0.39122 | 0.00000 | 5.03172 | 3.13068 |
| **Frem2** | 0.81358 | 0.00421 | 0.34751 | 0.00005 | 0.12058 | 0.10178 | 0.00404 | 0.00427 | 0.06330 | 0.00551 | 0.44499 | 0.00460 | 0.14834 | 1.46109 |
| **Frg1** | 8.96421 | 0.12098 | 9.20634 | 10.65983 | 6.96325 | 9.23652 | 2.05639 | 5.58735 | 4.66891 | 0.22793 | 5.16085 | 4.35764 | 8.10817 | 7.15853 |
| **Frk** | 0.34748 | 0.00409 | 0.28021 | 0.00000 | 1.01460 | 1.96817 | 0.00908 | 2.96740 | 1.80443 | 0.00755 | 0.00158 | 0.00000 | 0.02033 | 0.62143 |
| **Frmd3** | 0.01579 | 0.02230 | 0.01096 | 0.01617 | 0.01324 | 0.02288 | 0.00431 | 0.03782 | 0.08000 | 0.01482 | 0.01866 | 0.01773 | 0.28180 | 0.01640 |

| | | | | | | | | | | | | | | |
|---|---|---|---|---|---|---|---|---|---|---|---|---|---|---|
| **Frmd4a** | 0.06985 | 0.01335 | 0.38578 | 0.09066 | 0.00912 | 0.00000 | 0.00386 | 0.01492 | 0.00814 | 0.00732 | 0.00435 | 0.19364 | 0.04952 | 0.67287 |
| **Frmd4b** | 1.15186 | 0.83656 | 5.12467 | 3.49944 | 1.06588 | 13.51329 | 9.30970 | 2.74766 | 0.00000 | 0.25801 | 7.15723 | 0.00000 | 0.00841 | 2.54245 |
| **Frmd6** | 1.36443 | 0.00762 | 2.01771 | 0.40567 | 0.99222 | 3.49876 | 0.00756 | 0.00000 | 0.01124 | 0.00852 | 1.79453 | 4.15094 | 0.60316 | 1.00394 |
| **Frmd8** | 5.23989 | 0.05838 | 15.53723 | 11.28090 | 6.82657 | 13.40107 | 2.52977 | 9.34989 | 0.90609 | 0.22301 | 7.51763 | 3.97438 | 1.08229 | 3.21402 |
| **Frmpd1** | 1.96300 | 0.02806 | 7.34937 | 0.00000 | 4.66144 | 7.01790 | 5.00061 | 2.42645 | 0.16777 | 3.22969 | 1.10140 | 0.44001 | 1.06185 | 0.37952 |
| **Frmpd3** | 0.01635 | 0.00437 | 0.01395 | 0.03459 | 0.02010 | 0.01111 | 0.00413 | 0.00937 | 0.01808 | 0.01186 | 0.00791 | 0.00853 | 0.06206 | 0.00516 |
| **Frrs1** | 5.08672 | 4.50049 | 11.50355 | 8.86768 | 4.21693 | 8.96864 | 0.03132 | 6.38526 | 16.57618 | 0.07756 | 9.42923 | 5.39753 | 5.61177 | 16.50400 |
| **Frrs1l** | 0.00375 | 0.01051 | 0.16886 | 0.00000 | 0.38015 | 1.09155 | 0.00460 | 0.33269 | 0.05518 | 0.00292 | 2.40728 | 1.55294 | 0.00403 | 1.47967 |
| **Frs2** | 1.74971 | 0.01791 | 1.15772 | 0.00000 | 0.61698 | 1.45782 | 0.03453 | 2.64031 | 1.14542 | 0.00000 | 2.18336 | 1.92119 | 1.13989 | 4.41766 |
| **Fry** | 0.22184 | 0.00409 | 0.63666 | 0.00000 | 0.33115 | 1.78261 | 0.01036 | 0.38553 | 0.72219 | 0.00464 | 0.02568 | 0.00000 | 0.00000 | 0.36087 |
| **Fryl** | 0.00000 | 0.00254 | 1.34118 | 0.00000 | 1.61061 | 3.52395 | 0.00946 | 0.27418 | 1.54132 | 0.00887 | 3.04916 | 1.09838 | 1.70171 | 2.85557 |
| **Fsbp** | 0.26126 | 0.00443 | 0.68320 | 0.02961 | 0.22344 | 0.15117 | 0.79140 | 0.00107 | 0.84117 | 0.03194 | 0.04030 | 0.00270 | 0.00375 | 0.64457 |
| **Fscn1** | 0.87782 | 0.06700 | 0.11374 | 0.62230 | 8.54287 | 0.00000 | 0.00329 | 1.45759 | 0.00397 | 0.01675 | 0.06842 | 3.23761 | 5.29752 | 0.80926 |
| **Fscn2** | 0.00784 | 0.00448 | 0.22742 | 0.00009 | 0.00700 | 0.39362 | 0.00412 | 0.00516 | 0.00982 | 0.00729 | 0.00530 | 0.00796 | 0.00858 | 0.00942 |
| **Fsd1** | 0.00678 | 0.00432 | 2.00238 | 0.00408 | 0.00935 | 0.00715 | 0.00387 | 0.00530 | 0.00524 | 0.00244 | 0.01465 | 0.00887 | 0.53238 | 0.09177 |
| **Fsd1l** | 0.47993 | 0.01231 | 0.21675 | 0.53046 | 1.09363 | 1.32476 | 0.00460 | 0.09275 | 1.07367 | 0.06387 | 0.01792 | 0.11441 | 0.01109 | 0.31790 |
| **Fsd2** | 0.08925 | 0.01211 | 0.48595 | 0.00000 | 0.01438 | 0.00000 | 0.00522 | 1.03426 | 0.00616 | 0.00454 | 0.00000 | 0.71558 | 0.14516 | 0.76214 |
| **Fsip1** | 0.65782 | 0.00429 | 0.24118 | 0.08142 | 0.33034 | 0.03612 | 0.00401 | 0.00939 | 0.37179 | 0.00845 | 0.00440 | 0.00454 | 0.00307 | 0.04910 |
| **Fstl1** | 0.41970 | 0.00436 | 0.00853 | 0.01004 | 0.19464 | 0.01008 | 0.00433 | 0.01132 | 0.00615 | 0.00651 | 0.00705 | 0.00691 | 0.34932 | 0.00701 |
| **Fstl3** | 0.00000 | 0.00419 | 1.72038 | 1.27222 | 1.04026 | 0.14708 | 0.00391 | 0.23842 | 0.00729 | 0.00325 | 0.06683 | 0.16873 | 0.00000 | 0.07878 |
| **Fth1** | 7.31519 | 7.26430 | 1.90490 | 43.09632 | 28.26229 | 16.33188 | 2.11225 | 9.51829 | 2.93290 | 15.01206 | 5.93462 | 17.38030 | 17.91626 | 0.57517 |
| **Fthl17.ps1** | 2.58897 | 0.00411 | 0.66100 | 0.88868 | 1.49106 | 1.55551 | 0.48652 | 0.00174 | 0.24305 | 0.01632 | 0.13475 | 0.00000 | 0.03255 | 1.56940 |
| **Fthl17a** | 0.21223 | 0.00342 | 0.04216 | 0.00273 | 0.00316 | 0.38230 | 0.00862 | 0.00318 | 0.01004 | 3.20121 | 0.01262 | 0.00000 | 0.00585 | 0.00000 |
| **Fthl17b** | 2.91533 | 0.00408 | 0.54708 | 0.87090 | 1.25135 | 1.44682 | 0.22392 | 0.00382 | 0.13575 | 0.01075 | 0.18067 | 0.00328 | 0.10016 | 1.32252 |
| **Fthl17d** | 2.75602 | 0.00393 | 0.39092 | 0.89286 | 1.40109 | 1.06212 | 0.16102 | 0.00167 | 0.25306 | 0.00484 | 0.11560 | 0.00442 | 0.01762 | 1.44846 |
| **Fthl17e** | 2.64101 | 0.00401 | 0.56381 | 1.46074 | 2.49910 | 1.15467 | 0.14195 | 0.00093 | 0.44531 | 0.01060 | 0.26440 | 0.00002 | 0.08789 | 2.00337 |
| **Fthl17f** | 2.51217 | 0.00401 | 1.21648 | 1.69404 | 3.06483 | 1.67427 | 0.20615 | 0.12959 | 0.27693 | 0.01531 | 0.72545 | 0.00000 | 0.04018 | 2.06421 |
| **Ftl1** | 20.67158 | 17.53146 | 52.59872 | 63.81708 | 46.11853 | 55.03775 | 15.80344 | 45.85979 | 0.00000 | 27.84157 | 36.96205 | 37.26834 | 4.69446 | 22.04555 |
| **Fto** | 1.29971 | 0.00085 | 2.38632 | 0.42324 | 0.22499 | 1.29541 | 0.00623 | 0.95076 | 0.31105 | 12.25050 | 1.40181 | 1.01792 | 0.01495 | 1.41258 |
| **Ftsj1** | 5.21830 | 0.69960 | 13.72076 | 7.08449 | 8.63981 | 8.80132 | 2.75116 | 4.37965 | 2.97747 | 0.09806 | 2.93728 | 5.61206 | 3.08398 | 12.13279 |
| **Ftsj3** | 10.32261 | 7.57251 | 20.38950 | 14.32638 | 13.53336 | 10.09129 | 4.09145 | 5.10835 | 21.87854 | 0.19853 | 12.46696 | 8.14205 | 2.03879 | 16.83515 |
| **Ftx** | 0.03617 | 0.00376 | 0.32602 | 0.00000 | 0.01241 | 0.04082 | 0.00378 | 0.59478 | 0.21258 | 0.00999 | 0.00000 | 0.37610 | 0.02559 | 0.50885 |
| **Fubp1** | 8.82054 | 0.04942 | 4.98317 | 1.17995 | 6.17134 | 8.91783 | 0.06606 | 4.50879 | 7.06877 | 0.01451 | 10.07481 | 9.67547 | 5.99489 | 6.05136 |
| **Fubp3** | 2.87819 | 2.18389 | 3.15831 | 1.78366 | 1.17732 | 3.80225 | 0.02503 | 1.90048 | 3.81150 | 0.02761 | 5.77294 | 2.84614 | 2.83024 | 3.10243 |
| **Fuca1** | 0.56334 | 1.25109 | 1.03216 | 1.10715 | 1.61895 | 2.66856 | 2.01052 | 6.97271 | 0.99610 | 0.07236 | 2.10796 | 4.25917 | 1.59959 | 0.00000 |
| **Fuca2** | 1.45088 | 0.06715 | 0.00000 | 3.51744 | 0.13656 | 6.06606 | 1.87325 | 5.46764 | 0.28121 | 0.14744 | 3.46630 | 3.52133 | 0.03295 | 1.16225 |
| **Fuk** | 0.38977 | 0.00288 | 2.33034 | 3.12709 | 1.04435 | 0.00000 | 0.00938 | 0.36474 | 0.07664 | 1.80059 | 0.00000 | 1.86660 | 0.95112 | 0.53177 |
| **Fundc1** | 0.71619 | 0.03186 | 2.18917 | 1.42677 | 1.98063 | 8.30725 | 2.92800 | 0.09536 | 3.80796 | 0.10469 | 0.00000 | 5.24125 | 1.93840 | 4.00694 |
| **Fundc2** | 0.36359 | 0.00451 | 0.44304 | 0.41944 | 0.84604 | 0.00000 | 0.00439 | 0.47860 | 1.06236 | 0.01548 | 0.00000 | 0.40499 | 0.02968 | 0.78505 |
| **Fuom** | 0.00229 | 0.00385 | 0.00743 | 2.12756 | 0.38968 | 0.32302 | 0.00920 | 0.52524 | 0.02124 | 0.01545 | 0.37849 | 0.16325 | 0.00326 | 0.15800 |
| **Furin** | 0.16265 | 4.29366 | 0.48402 | 1.10404 | 1.33669 | 4.26566 | 1.33287 | 6.38826 | 0.28601 | 0.10720 | 0.10304 | 2.26534 | 0.31763 | 1.21591 |
| **Fus** | 1.65280 | 0.58575 | 0.10114 | 2.14926 | 1.89465 | 3.29484 | 0.77264 | 2.69434 | 0.97927 | 0.33491 | 0.29216 | 2.12877 | 13.11686 | 0.87452 |
| **Fut10** | 0.17152 | 0.00368 | 0.00662 | 1.58349 | 0.05538 | 0.00000 | 0.00374 | 0.00476 | 0.00322 | 0.00504 | 0.25356 | 0.03198 | 0.73271 | 1.37125 |
| **Fut11** | 0.58047 | 0.00460 | 0.08071 | 0.10960 | 0.00000 | 0.00257 | 0.00450 | 0.03177 | 0.02911 | 0.01891 | 0.00768 | 0.00804 | 0.42405 | 0.75621 |
| **Fut2** | 0.13310 | 0.00427 | 0.04087 | 0.12074 | 0.12401 | 0.06977 | 0.00408 | 0.00478 | 0.01619 | 0.00649 | 0.03152 | 0.07939 | 0.00432 | 0.31991 |
| **Fut4** | 0.01632 | 0.00444 | 0.12906 | 0.01558 | 0.00000 | 0.12380 | 0.01088 | 0.00843 | 0.01580 | 0.00624 | 0.00551 | 0.00628 | 0.04487 | 0.65111 |
| **Fut8** | 0.85543 | 0.11549 | 1.31621 | 1.51267 | 2.41587 | 0.89093 | 0.27844 | 0.29693 | 3.37164 | 0.00666 | 1.32284 | 5.66627 | 1.51511 | 3.47784 |
| **Fut9** | 0.04929 | 0.00502 | 0.04800 | 0.00000 | 0.32085 | 0.17086 | 0.00432 | 0.00700 | 1.42491 | 0.00570 | 0.00501 | 0.00564 | 0.00722 | 0.00000 |
| **Fuz** | 0.00215 | 0.02971 | 0.09223 | 0.38484 | 0.00550 | 0.00357 | 0.00396 | 0.01503 | 0.01305 | 0.00611 | 0.00512 | 0.31185 | 0.00659 | 0.00849 |
| **Fv1** | 0.03141 | 0.00409 | 0.00676 | 0.04855 | 0.01347 | 0.10289 | 0.00430 | 0.00501 | 0.00985 | 0.00782 | 0.00506 | 0.13293 | 0.00711 | 0.29375 |
| **Fxn** | 0.09782 | 0.01064 | 0.07496 | 0.46286 | 1.60537 | 1.00377 | 1.13035 | 0.04651 | 0.09225 | 0.11560 | 0.00142 | 0.33200 | 1.05330 | 0.12526 |
| **Fxr1** | 12.31282 | 0.10227 | 8.25045 | 10.58610 | 7.66214 | 9.68967 | 5.18050 | 0.55714 | 10.69382 | 0.87143 | 11.78235 | 11.51010 | 11.53965 | 8.51943 |
| **Fxr2** | 7.23954 | 0.17381 | 6.88870 | 5.76614 | 10.44261 | 10.82113 | 4.12133 | 6.65429 | 1.52744 | 0.28759 | 2.69591 | 7.44813 | 6.32886 | 6.40395 |
| **Fxyd3** | 0.00995 | 0.00444 | 0.01345 | 0.01308 | 0.04817 | 0.01323 | 0.00426 | 0.01047 | 0.01163 | 0.00695 | 0.00887 | 0.00983 | 0.00723 | 0.01113 |
| **Fxyd4** | 0.65656 | 0.05797 | 0.00000 | 1.55856 | 1.17948 | 1.02744 | 1.43084 | 5.35327 | 0.00677 | 0.21313 | 0.05906 | 0.62475 | 0.00565 | 0.20455 |

| | | | | | | | | | | | | | | |
|---|---|---|---|---|---|---|---|---|---|---|---|---|---|---|
| **Fxyd5** | 0.02908 | 0.00479 | 0.01737 | 0.08571 | 0.05513 | 0.10561 | 0.00480 | 0.03008 | 0.01501 | 0.00778 | 0.00837 | 0.03194 | 0.01419 | 0.06781 |
| **Fxyd6** | 0.00268 | 1.71374 | 0.00000 | 0.00000 | 0.19554 | 2.99742 | 0.02336 | 10.63483 | 0.02796 | 0.02620 | 0.55264 | 0.13540 | 0.54907 | 0.00000 |
| **Fyb** | 0.04065 | 0.00399 | 0.00812 | 0.08625 | 0.01270 | 0.03027 | 0.00398 | 0.00761 | 0.01180 | 0.00872 | 0.02120 | 0.16786 | 0.02274 | 0.00699 |
| **Fyco1** | 1.84310 | 0.01861 | 1.26298 | 1.03697 | 2.79510 | 1.10901 | 0.00944 | 1.87199 | 0.04751 | 0.00000 | 2.34421 | 0.65723 | 0.90156 | 4.45761 |
| **Fyn** | 0.01993 | 0.00395 | 0.00000 | 0.00000 | 0.38059 | 0.35343 | 0.00382 | 0.00947 | 0.35841 | 0.00527 | 0.00562 | 0.00385 | 1.17417 | 1.34734 |
| **Fyttd1** | 2.19749 | 1.60320 | 3.45487 | 6.01099 | 4.44983 | 5.33437 | 1.85444 | 1.85260 | 10.24357 | 0.08333 | 0.13721 | 3.97942 | 2.26590 | 3.36924 |
| **Fzd2** | 0.03170 | 0.00444 | 0.01325 | 0.00980 | 0.14331 | 0.01310 | 0.00419 | 0.01026 | 0.01228 | 0.00649 | 0.00866 | 0.00900 | 0.06108 | 0.00484 |
| **Fzd3** | 0.58062 | 0.01219 | 2.17634 | 0.00000 | 2.43000 | 1.08092 | 0.00917 | 1.31993 | 4.99174 | 0.02497 | 0.00081 | 0.01441 | 3.61814 | 0.94485 |
| **Fzd4** | 0.00323 | 0.00433 | 0.08256 | 0.00858 | 0.00000 | 0.00884 | 0.00411 | 0.00906 | 0.45326 | 0.00793 | 0.00632 | 0.00799 | 0.49606 | 0.09722 |
| **Fzd5** | 0.00000 | 0.00325 | 0.01816 | 1.20878 | 0.23188 | 0.30966 | 0.86695 | 0.15571 | 2.18942 | 0.11784 | 0.00143 | 1.32664 | 0.02392 | 1.43721 |
| **Fzd6** | 0.00833 | 0.00455 | 0.01279 | 0.02260 | 0.04927 | 0.01545 | 0.00429 | 0.01307 | 0.01523 | 0.00724 | 0.00889 | 0.00839 | 0.27265 | 0.00210 |
| **Fzd7** | 0.00192 | 0.00359 | 0.05950 | 0.28736 | 0.01920 | 0.00856 | 0.00401 | 0.00323 | 0.02476 | 0.00502 | 0.01363 | 0.31828 | 2.93649 | 0.00000 |
| **Fzd8** | 0.01356 | 0.00429 | 0.00752 | 0.10001 | 0.13171 | 0.01741 | 0.00419 | 0.00649 | 0.00906 | 0.00600 | 0.00738 | 0.00685 | 0.31140 | 0.00548 |
| **Fzd9** | 0.00000 | 0.00422 | 0.38361 | 1.48511 | 0.17891 | 0.00000 | 0.00396 | 0.01172 | 0.00552 | 0.00400 | 0.78239 | 0.07530 | 0.76836 | 0.00000 |
| **Fzr1** | 2.00432 | 9.80905 | 8.97386 | 4.32319 | 4.57430 | 9.07457 | 1.78087 | 10.59543 | 0.00800 | 0.15663 | 5.82300 | 3.74898 | 1.31381 | 0.25256 |
| **G2e3** | 0.16048 | 0.46600 | 0.33108 | 0.00000 | 1.10349 | 0.62072 | 0.13143 | 0.65315 | 5.61920 | 0.01645 | 0.00352 | 0.00484 | 7.58518 | 0.00000 |
| **G3bp1** | 9.56089 | 7.39759 | 26.49070 | 27.31833 | 25.84411 | 18.89101 | 7.28379 | 12.37183 | 8.84998 | 0.64950 | 17.07344 | 18.01590 | 13.64524 | 16.37942 |
| **G3bp2** | 24.27265 | 3.07389 | 29.97103 | 26.92853 | 29.75016 | 39.00615 | 8.23250 | 16.71949 | 35.14004 | 13.82573 | 21.96616 | 15.99912 | 16.57837 | 24.72640 |
| **G6pc** | 0.00381 | 0.00431 | 0.01508 | 0.01228 | 0.00963 | 0.01344 | 0.00423 | 0.00894 | 0.01080 | 0.00652 | 0.00907 | 0.00904 | 0.48802 | 0.00233 |
| **G6pc3** | 1.14186 | 0.12370 | 5.22404 | 1.70631 | 3.06552 | 0.22625 | 0.00217 | 3.79115 | 0.25255 | 0.00549 | 2.13514 | 1.23072 | 1.20814 | 2.83028 |
| **G6pd2** | 0.00714 | 0.00416 | 0.05899 | 0.07262 | 0.03545 | 0.15662 | 0.00419 | 0.00704 | 0.04149 | 0.00598 | 0.13645 | 0.04634 | 0.00564 | 0.15228 |
| **G6pdx** | 1.74544 | 0.03548 | 3.21533 | 3.96934 | 3.76116 | 4.28143 | 0.00710 | 0.63854 | 2.89979 | 0.04249 | 4.40806 | 2.72542 | 1.96422 | 4.72209 |
| **Gaa** | 10.78170 | 0.92235 | 29.81729 | 27.18613 | 15.50217 | 14.43162 | 0.72649 | 6.13762 | 0.21728 | 4.67665 | 9.78768 | 8.17565 | 0.67144 | 8.01694 |
| **Gab1** | 4.10947 | 1.53739 | 6.49438 | 2.73609 | 5.56438 | 5.27779 | 2.29114 | 3.63540 | 3.39827 | 0.08115 | 4.40110 | 1.16828 | 0.35917 | 5.58993 |
| **Gab2** | 0.00000 | 0.01864 | 1.50279 | 1.52003 | 0.00000 | 1.34278 | 0.00316 | 2.49288 | 1.26934 | 0.01671 | 0.34123 | 1.64499 | 0.05229 | 0.33265 |
| **Gab3** | 0.00561 | 0.00391 | 0.00404 | 0.33744 | 0.00000 | 0.00000 | 1.51306 | 0.01315 | 0.12385 | 0.04588 | 0.00622 | 0.01101 | 0.22412 | 0.09238 |
| **Gabarap** | 3.95730 | 11.00650 | 3.23048 | 7.24200 | 10.78874 | 2.83033 | 2.49950 | 11.33201 | 2.14056 | 1.33625 | 3.12538 | 2.95944 | 7.14750 | 3.52602 |
| **Gabarapl1** | 0.00000 | 0.03559 | 0.33399 | 2.97829 | 0.29301 | 11.41581 | 0.26503 | 0.02846 | 0.88694 | 0.05093 | 3.09860 | 6.77640 | 0.00126 | 0.00165 |
| **Gabarapl2** | 35.27932 | 15.68373 | 41.21419 | 47.31974 | 38.12544 | 48.47078 | 16.30148 | 38.58853 | 23.01227 | 22.33842 | 25.46417 | 36.28539 | 15.61210 | 28.46722 |
| **Gabbr1** | 0.00283 | 0.01867 | 0.01732 | 0.02537 | 0.22908 | 0.01559 | 0.00413 | 0.00751 | 0.01235 | 0.00606 | 0.00708 | 1.04516 | 0.79654 | 0.01093 |
| **Gabpa** | 17.11003 | 0.06909 | 13.42711 | 3.93835 | 18.53704 | 7.55368 | 0.05655 | 3.39242 | 19.83708 | 0.00000 | 10.97117 | 6.28375 | 15.44590 | 23.31701 |
| **Gabpb1** | 9.01909 | 0.26832 | 4.37471 | 0.00000 | 3.26393 | 5.76139 | 1.13566 | 0.00000 | 13.49720 | 0.05721 | 1.60340 | 2.13467 | 5.86091 | 4.47147 |
| **Gabpb2** | 2.08560 | 3.82763 | 2.47315 | 1.04127 | 1.34707 | 1.92460 | 0.96591 | 0.16912 | 16.05578 | 0.10548 | 1.00518 | 1.27486 | 4.31371 | 8.45690 |
| **Gabra3** | 0.00847 | 0.00459 | 0.00782 | 0.01922 | 0.24803 | 0.01242 | 0.00417 | 0.01242 | 0.01567 | 0.00644 | 0.00967 | 0.00822 | 0.21131 | 0.05685 |
| **Gabrb3** | 0.13948 | 0.00429 | 0.01203 | 0.01898 | 0.00447 | 0.01573 | 0.00412 | 0.00778 | 0.02889 | 0.01041 | 0.00684 | 0.00764 | 0.50406 | 0.00000 |
| **Gabrd** | 2.83648 | 0.01193 | 2.96062 | 1.55423 | 1.23974 | 0.04455 | 0.00328 | 0.01068 | 0.16606 | 0.00119 | 0.14082 | 0.03553 | 0.00000 | 0.64288 |
| **Gabre** | 0.00507 | 0.96935 | 0.10386 | 0.00235 | 0.01742 | 0.01075 | 0.00370 | 0.00321 | 0.01295 | 0.00638 | 0.00710 | 0.01018 | 0.16256 | 0.03770 |
| **Gabrp** | 0.00913 | 0.00409 | 0.00288 | 0.33144 | 0.00653 | 0.00000 | 0.00385 | 0.01110 | 0.15452 | 0.00721 | 0.01368 | 0.01590 | 0.00444 | 0.01837 |
| **Gad2** | 0.09268 | 0.05234 | 0.05091 | 0.19591 | 0.06796 | 0.15483 | 0.02885 | 0.11958 | 0.25043 | 0.08545 | 0.03636 | 0.20872 | 0.25708 | 0.28037 |
| **Gadd45a** | 0.01195 | 0.00400 | 0.01932 | 1.44805 | 0.24580 | 0.00000 | 0.01098 | 0.00582 | 0.01492 | 0.00530 | 0.04394 | 0.04030 | 0.02671 | 0.00000 |
| **Gadd45b** | 0.00000 | 0.00394 | 1.27674 | 6.89767 | 1.55952 | 0.59878 | 1.24130 | 0.53090 | 0.00000 | 0.04832 | 0.07906 | 0.00123 | 1.24820 | 0.00000 |
| **Gadd45g** | 0.01757 | 0.02994 | 1.84361 | 0.00000 | 0.00112 | 0.00000 | 0.00332 | 1.41704 | 0.00716 | 0.02198 | 0.00000 | 1.65108 | 0.06093 | 0.00767 |
| **Gadd45gip1** | 1.21588 | 0.06398 | 0.10714 | 10.99813 | 14.91045 | 0.37230 | 0.03276 | 1.79083 | 0.11160 | 0.05006 | 0.26003 | 2.19934 | 6.92908 | 0.38707 |
| **Gak** | 4.63856 | 0.15141 | 9.50006 | 11.07473 | 4.21170 | 9.85332 | 3.43003 | 14.85249 | 1.74732 | 0.20904 | 4.93983 | 3.83069 | 4.56858 | 3.86290 |
| **Gal** | 0.12594 | 0.00444 | 0.02605 | 0.01740 | 0.00786 | 0.01514 | 0.00422 | 0.00917 | 0.00879 | 0.01020 | 0.00720 | 0.00871 | 0.04090 | 0.01063 |
| **Gal3st1** | 0.03706 | 0.00386 | 0.21115 | 0.00000 | 0.02239 | 0.01806 | 0.00396 | 0.39652 | 0.04001 | 0.00632 | 0.00442 | 0.01293 | 0.06402 | 0.02432 |
| **Gal3st2** | 0.00000 | 0.00414 | 0.08293 | 0.10844 | 0.02284 | 5.37888 | 0.00386 | 0.00300 | 0.00590 | 0.00518 | 0.24084 | 0.00369 | 1.59067 | 0.00000 |
| **Gal3st3** | 0.06537 | 0.05562 | 0.36609 | 0.81614 | 0.31544 | 0.14375 | 0.00841 | 0.06275 | 0.43036 | 4.02064 | 0.00762 | 0.02268 | 0.26439 | 0.41108 |
| **Galc** | 0.43390 | 0.00388 | 0.07559 | 0.09491 | 0.17227 | 0.00323 | 0.00404 | 0.00525 | 0.06153 | 0.00643 | 0.00470 | 0.20556 | 0.36467 | 0.05641 |
| **Gale** | 0.07436 | 6.94711 | 1.42588 | 4.89741 | 0.07666 | 3.29331 | 0.52958 | 7.75424 | 0.00000 | 0.07491 | 6.71863 | 3.04722 | 0.38419 | 0.00000 |
| **Galk1** | 10.74426 | 1.21565 | 22.93320 | 27.72995 | 23.51452 | 4.46325 | 3.95189 | 15.48165 | 0.00000 | 2.17655 | 5.21746 | 8.81792 | 0.00000 | 1.60589 |
| **Galk2** | 0.00000 | 0.02427 | 3.05125 | 5.28081 | 3.17946 | 0.33376 | 0.02510 | 4.13294 | 0.45310 | 0.32841 | 5.03414 | 1.05533 | 0.00603 | 3.51190 |
| **Galm** | 0.73992 | 0.00202 | 0.38495 | 1.42828 | 1.25929 | 0.21351 | 2.33321 | 0.00121 | 0.40827 | 0.03930 | 0.00000 | 2.75946 | 0.33470 | 1.78180 |
| **Galns** | 0.00681 | 0.00437 | 0.45519 | 0.00000 | 0.89239 | 0.79027 | 0.01200 | 0.66267 | 0.00553 | 0.01577 | 0.00112 | 0.20257 | 0.15591 | 0.30705 |
| **Galnt1** | 3.47600 | 0.12617 | 1.98384 | 0.00000 | 0.52901 | 6.27402 | 1.51407 | 3.93213 | 6.95367 | 1.77504 | 3.58156 | 6.10993 | 3.62642 | 2.96196 |

| | | | | | | | | | | | | | | |
|---|---|---|---|---|---|---|---|---|---|---|---|---|---|---|
| **Galnt10** | 0.01923 | 0.03953 | 0.01324 | 0.00000 | 0.05428 | 0.06600 | 0.00402 | 0.00856 | 0.01076 | 0.00533 | 0.00497 | 0.34146 | 0.00693 | 0.01643 |
| **Galnt11** | 4.40555 | 0.06971 | 5.42171 | 3.32155 | 5.44031 | 0.97114 | 0.00831 | 0.48121 | 4.38623 | 0.00001 | 0.63085 | 5.41845 | 3.24536 | 3.84288 |
| **Galnt12** | 0.01395 | 0.00443 | 0.01444 | 0.01311 | 0.01687 | 0.01297 | 0.00431 | 0.01058 | 0.01216 | 0.00694 | 0.00875 | 0.01007 | 0.01067 | 0.01544 |
| **Galnt16** | 2.07832 | 0.00401 | 1.75056 | 0.01489 | 0.93210 | 1.29921 | 0.00367 | 0.00000 | 0.96959 | 0.00353 | 0.12604 | 0.28544 | 0.55676 | 0.50034 |
| **Galnt2** | 0.82744 | 0.05736 | 5.60858 | 3.28608 | 4.45064 | 2.39857 | 1.10784 | 1.05868 | 0.37110 | 0.04518 | 2.50952 | 5.18498 | 0.00000 | 3.35685 |
| **Galnt3** | 1.35665 | 0.00474 | 0.02031 | 0.01315 | 0.07821 | 0.54276 | 0.00429 | 0.01632 | 0.00877 | 0.00782 | 0.00455 | 0.00238 | 0.55617 | 0.17545 |
| **Galnt4** | 0.01289 | 0.00404 | 0.00000 | 0.40626 | 0.21488 | 0.69277 | 0.00395 | 0.00294 | 0.10827 | 0.01801 | 0.00325 | 0.02606 | 0.01503 | 0.03860 |
| **Galnt6** | 0.00000 | 0.01796 | 2.21607 | 3.46809 | 1.52883 | 1.50925 | 0.84499 | 0.00000 | 0.00287 | 0.00647 | 0.47784 | 0.89050 | 0.63531 | 4.88021 |
| **Galnt7** | 0.41755 | 0.00379 | 0.00219 | 0.29608 | 0.02707 | 0.00000 | 0.00377 | 0.00620 | 0.07626 | 0.00566 | 0.01419 | 0.02159 | 1.61416 | 0.02705 |
| **Galnt9** | 0.00000 | 0.00406 | 1.86417 | 0.69845 | 0.44427 | 0.00000 | 0.00360 | 0.00548 | 0.06801 | 0.00405 | 0.00424 | 0.00740 | 0.00000 | 0.71658 |
| **Galntl5** | 0.00638 | 0.00536 | 0.05154 | 0.05568 | 0.00000 | 0.08500 | 0.00410 | 0.40404 | 0.48840 | 0.01017 | 0.29263 | 0.00495 | 0.20980 | 0.72712 |
| **Galntl6** | 0.06676 | 0.00457 | 0.02293 | 0.04227 | 0.02891 | 0.08091 | 0.02828 | 0.04808 | 0.04222 | 0.00807 | 0.00787 | 0.02099 | 0.08075 | 0.12583 |
| **Galt** | 0.54746 | 0.17465 | 2.29887 | 4.03142 | 3.56585 | 0.61218 | 4.37700 | 8.88622 | 0.02781 | 0.10851 | 8.04609 | 3.42922 | 1.72712 | 0.92178 |
| **Gamt** | 0.17639 | 0.00342 | 0.09664 | 0.45907 | 1.98186 | 0.19582 | 0.00387 | 0.00100 | 0.00604 | 0.02125 | 1.25390 | 0.25021 | 0.09030 | 0.09468 |
| **Gan** | 0.46583 | 0.00440 | 0.40023 | 0.96272 | 0.35302 | 0.29692 | 0.00393 | 0.00490 | 0.00633 | 0.02090 | 0.00419 | 0.06638 | 1.12613 | 0.38911 |
| **Ganab** | 1.55101 | 0.18219 | 7.34627 | 7.81838 | 1.19201 | 10.86365 | 1.06543 | 3.64548 | 11.18816 | 0.06679 | 3.98194 | 9.98581 | 2.13116 | 8.05402 |
| **Ganc** | 0.00000 | 0.00226 | 0.36731 | 0.00000 | 0.31154 | 0.16084 | 2.22690 | 0.00000 | 0.01614 | 0.01215 | 3.16969 | 1.14120 | 0.00359 | 0.46754 |
| **Gapdh** | 0.15906 | 12.78196 | 10.12446 | 28.59073 | 2.93240 | 32.43748 | 9.23776 | 37.67900 | 0.13975 | 22.14446 | 19.02299 | 28.75114 | 6.87535 | 8.37720 |
| **Gapdhs** | 0.05787 | 0.00409 | 0.33575 | 0.40614 | 0.05010 | 0.02658 | 0.83661 | 0.00338 | 0.01586 | 0.00585 | 0.00267 | 0.26784 | 0.08376 | 0.00823 |
| **Gapt** | 0.30412 | 0.00392 | 0.01737 | 0.07920 | 1.47094 | 0.02336 | 0.00406 | 0.00456 | 0.07299 | 0.00454 | 0.34883 | 0.00230 | 2.88197 | 0.49938 |
| **Gapvd1** | 3.51292 | 3.79960 | 5.85235 | 0.00000 | 5.51778 | 5.69943 | 1.50144 | 2.48431 | 3.97505 | 0.04210 | 3.23343 | 3.93626 | 3.58479 | 9.21583 |
| **Gar1** | 1.57293 | 0.04684 | 2.10533 | 4.48962 | 2.91204 | 4.68656 | 2.86642 | 1.32995 | 1.08322 | 7.28199 | 0.72729 | 4.81238 | 4.85647 | 8.10872 |
| **Garem** | 0.00000 | 0.00376 | 0.71045 | 0.08973 | 0.16636 | 0.45412 | 0.00363 | 0.05218 | 0.00372 | 0.00187 | 1.51855 | 0.13836 | 1.07612 | 0.13600 |
| **Gareml** | 0.06865 | 0.00402 | 0.49584 | 0.11514 | 0.45912 | 0.00000 | 0.08847 | 0.00522 | 0.00863 | 0.00897 | 0.00610 | 0.00888 | 0.90956 | 0.00699 |
| **Garnl3** | 0.40505 | 0.00322 | 0.91114 | 0.48477 | 0.19363 | 0.11665 | 0.00349 | 0.09230 | 0.00485 | 0.00968 | 1.18379 | 0.50467 | 0.51211 | 0.56083 |
| **Gars** | 16.74435 | 7.82127 | 15.22849 | 10.90528 | 12.47181 | 19.09298 | 2.49484 | 11.57702 | 11.40524 | 13.40630 | 19.13588 | 15.85297 | 1.90786 | 14.02384 |
| **Gart** | 10.15487 | 0.14471 | 20.37103 | 12.25215 | 14.23337 | 10.15472 | 5.34102 | 6.77839 | 1.99445 | 12.31956 | 13.36813 | 9.73811 | 1.22904 | 15.72734 |
| **Gas1** | 0.01172 | 0.00457 | 0.01251 | 0.01032 | 0.09589 | 0.02344 | 0.00452 | 0.01080 | 0.01768 | 0.00643 | 0.01246 | 0.00953 | 0.00749 | 0.00936 |
| **Gas2** | 0.92516 | 0.00365 | 0.08893 | 0.31688 | 0.00000 | 0.01618 | 0.00373 | 0.00000 | 0.01291 | 0.00000 | 2.16700 | 0.01122 | 0.00691 | 0.00000 |
| **Gas2l1** | 0.91637 | 0.07465 | 0.51520 | 1.15325 | 0.75587 | 1.22135 | 0.89066 | 2.14821 | 0.00000 | 0.08780 | 1.36462 | 1.72176 | 1.97780 | 0.00000 |
| **Gas2l3** | 2.37780 | 0.13593 | 1.64372 | 1.06851 | 0.99095 | 3.93491 | 1.15453 | 1.85872 | 5.09151 | 1.44663 | 2.12602 | 0.52151 | 12.43827 | 2.41917 |
| **Gas5** | 22.29700 | 2.19257 | 10.48302 | 14.17219 | 12.51559 | 9.61902 | 5.25809 | 12.07205 | 13.53410 | 4.77933 | 9.33261 | 10.61739 | 12.39764 | 9.93875 |
| **Gas6** | 0.00724 | 0.00437 | 1.98466 | 0.67172 | 0.53050 | 0.27692 | 0.00373 | 0.00000 | 0.00864 | 0.00233 | 0.00396 | 0.00716 | 0.06376 | 0.00000 |
| **Gas8** | 0.00000 | 0.01090 | 1.59472 | 1.79337 | 0.75814 | 0.33936 | 0.00302 | 3.96713 | 0.22635 | 0.00244 | 0.06204 | 0.93693 | 1.58177 | 2.30021 |
| **Gata1** | 2.57773 | 0.04021 | 17.86838 | 2.22189 | 13.96194 | 2.94096 | 1.19284 | 0.72680 | 0.69443 | 0.06827 | 0.33489 | 0.00000 | 8.10378 | 12.55413 |
| **Gata2** | 0.00404 | 0.53567 | 0.00473 | 0.00000 | 0.03029 | 0.00000 | 4.29940 | 12.14770 | 0.01092 | 0.14095 | 0.00227 | 0.11021 | 0.06321 | 0.00171 |
| **Gata3** | 0.00000 | 6.34852 | 2.82862 | 6.21663 | 2.10651 | 24.59489 | 9.81364 | 13.49908 | 0.72627 | 14.23515 | 6.11392 | 0.00000 | 0.19566 | 0.27968 |
| **Gata4** | 3.30325 | 0.06889 | 1.24888 | 0.00000 | 11.11156 | 0.00000 | 0.01892 | 0.43928 | 2.24361 | 0.02927 | 0.00000 | 7.91470 | 10.85929 | 17.88349 |
| **Gata6** | 19.86140 | 0.21938 | 29.19799 | 12.75628 | 22.17436 | 17.00723 | 9.40933 | 7.80513 | 7.33570 | 0.48067 | 17.89015 | 13.11521 | 0.68203 | 11.09938 |
| **Gatad1** | 2.62963 | 5.28571 | 0.05704 | 2.43507 | 0.05475 | 2.06593 | 0.00239 | 1.78414 | 0.00199 | 0.02112 | 2.49663 | 1.57314 | 0.00340 | 1.23025 |
| **Gatad2a** | 10.67506 | 0.19801 | 14.08322 | 10.23914 | 16.76456 | 11.52068 | 5.17086 | 11.42958 | 2.63396 | 2.07011 | 6.49854 | 7.60251 | 15.40623 | 4.57737 |
| **Gatad2b** | 0.30371 | 0.00413 | 0.78379 | 0.00000 | 2.08882 | 0.76591 | 0.00349 | 0.00000 | 4.97421 | 0.00480 | 0.01276 | 0.09950 | 0.54521 | 4.60043 |
| **Gatb** | 0.06805 | 0.05587 | 0.89983 | 0.97048 | 0.31977 | 3.86662 | 0.55357 | 3.56876 | 1.16904 | 0.02420 | 4.79548 | 5.88734 | 0.00000 | 0.48135 |
| **Gatc** | 0.58849 | 0.03462 | 0.75807 | 0.00000 | 1.57777 | 2.32977 | 0.00877 | 2.26545 | 2.36949 | 5.10256 | 0.00000 | 2.17559 | 0.91798 | 1.43322 |
| **Gatm** | 0.01619 | 0.01202 | 0.02975 | 0.04931 | 0.07038 | 0.00422 | 0.00408 | 0.01005 | 0.00673 | 0.01402 | 0.04192 | 0.00702 | 0.00630 | 0.34740 |
| **Gatsl2** | 0.00374 | 0.00427 | 0.01054 | 0.01130 | 0.08811 | 0.01502 | 0.01004 | 0.00820 | 0.05925 | 0.00610 | 0.00870 | 0.00842 | 0.75040 | 0.00000 |
| **Gatsl3** | 6.45554 | 0.47642 | 10.56091 | 9.51712 | 10.89680 | 10.52272 | 1.89467 | 6.45548 | 0.03232 | 0.10857 | 6.29887 | 3.36196 | 1.58552 | 8.84491 |
| **Gba** | 5.02694 | 0.13420 | 14.08863 | 17.77359 | 4.60281 | 15.02375 | 1.72396 | 3.66141 | 6.95154 | 0.08302 | 10.22279 | 9.44388 | 1.03962 | 9.41633 |
| **Gba2** | 0.19665 | 0.02289 | 1.48696 | 0.44895 | 0.57112 | 0.34694 | 0.00364 | 0.94216 | 0.00866 | 0.03172 | 0.00003 | 0.49007 | 2.35865 | 0.04410 |
| **Gbas** | 0.08210 | 0.06997 | 0.21138 | 0.00000 | 0.66930 | 1.13200 | 0.20130 | 1.51443 | 0.00086 | 0.01345 | 0.30443 | 2.23368 | 2.73229 | 0.19494 |
| **Gbe1** | 0.49638 | 0.00388 | 0.28866 | 0.47794 | 0.34576 | 2.35168 | 0.00373 | 0.00700 | 0.63005 | 0.00581 | 0.00000 | 0.56831 | 0.18994 | 0.47160 |
| **Gbf1** | 0.28394 | 0.02353 | 1.77519 | 4.07874 | 1.23706 | 3.94123 | 1.08492 | 2.69203 | 0.12674 | 0.02748 | 2.52360 | 3.95403 | 0.61048 | 1.05820 |
| **Gbp11** | 0.01059 | 0.00432 | 0.01800 | 0.03359 | 0.01291 | 0.01077 | 0.00429 | 0.00788 | 0.03416 | 0.00639 | 0.01053 | 0.06007 | 0.00471 | 0.02806 |
| **Gbp4** | 0.00000 | 0.00396 | 0.00340 | 0.90693 | 1.10631 | 0.00000 | 0.00372 | 0.00469 | 1.29834 | 0.00604 | 0.00612 | 0.01395 | 0.00000 | 0.40929 |
| **Gbp7** | 0.03949 | 0.00436 | 0.01572 | 0.01080 | 0.00827 | 0.01476 | 0.00424 | 0.00924 | 0.01068 | 0.00633 | 0.00878 | 0.00929 | 0.26730 | 0.06261 |

| | | | | | | | | | | | | | | |
|---|---|---|---|---|---|---|---|---|---|---|---|---|---|---|
| **Gbp9** | 0.00000 | 0.00422 | 0.67617 | 0.15542 | 0.31165 | 0.00000 | 0.00377 | 0.00360 | 1.32829 | 0.00819 | 0.00043 | 0.45656 | 0.00000 | 0.01655 |
| **Gbx1** | 0.02193 | 0.00450 | 0.02725 | 0.01062 | 0.07082 | 0.01209 | 0.00450 | 0.01051 | 0.01164 | 0.00653 | 0.00838 | 0.00952 | 0.00893 | 0.02770 |
| **Gbx2** | 0.31609 | 0.00354 | 0.00000 | 1.65562 | 0.23741 | 0.00000 | 0.00931 | 0.00445 | 0.00553 | 0.00256 | 1.20942 | 0.02029 | 0.56887 | 0.00000 |
| **Gc** | 0.84333 | 0.00439 | 0.00000 | 0.06046 | 0.01979 | 0.03936 | 0.00402 | 0.00735 | 1.05297 | 0.02260 | 0.00387 | 0.03749 | 0.18689 | 0.00000 |
| **Gca** | 4.53826 | 0.08252 | 6.67830 | 3.83833 | 2.53135 | 5.39486 | 1.66947 | 5.68174 | 11.66845 | 8.58679 | 3.07253 | 2.81582 | 2.41378 | 3.46284 |
| **Gcat** | 0.14467 | 7.04186 | 3.59386 | 12.06798 | 0.47560 | 3.17721 | 3.88504 | 0.04041 | 0.00529 | 0.15375 | 3.57024 | 5.65866 | 0.01639 | 0.13615 |
| **Gcc1** | 0.35161 | 0.00357 | 1.76042 | 0.86616 | 0.55722 | 1.08248 | 0.12898 | 1.07897 | 0.00559 | 0.00216 | 0.00000 | 1.17136 | 0.03165 | 0.00000 |
| **Gcc2** | 2.58197 | 0.00417 | 1.67165 | 1.66011 | 3.61449 | 2.21851 | 0.00387 | 0.08242 | 0.00569 | 0.00287 | 0.61945 | 0.56860 | 2.36558 | 2.95537 |
| **Gcdh** | 4.59125 | 0.21334 | 7.85157 | 3.52480 | 7.29209 | 3.54197 | 0.00000 | 4.13443 | 4.22195 | 0.01778 | 4.47347 | 7.62719 | 2.09426 | 7.01719 |
| **Gcfc2** | 0.05598 | 0.00417 | 1.00780 | 0.50444 | 1.24834 | 0.08327 | 0.00406 | 0.01717 | 0.76217 | 0.00360 | 0.00349 | 0.00590 | 0.00000 | 0.00000 |
| **Gch1** | 8.83267 | 5.69045 | 2.13417 | 4.74523 | 10.63709 | 7.23123 | 1.55461 | 0.84213 | 2.29956 | 0.05171 | 2.82855 | 1.02080 | 6.20916 | 2.31215 |
| **Gchfr** | 0.01133 | 0.05851 | 0.03520 | 0.36915 | 0.00491 | 0.01188 | 0.00360 | 2.28678 | 0.01126 | 0.05842 | 0.00000 | 0.26651 | 0.00938 | 0.02547 |
| **Gck** | 0.02251 | 0.00445 | 0.00766 | 0.01030 | 0.21158 | 0.01353 | 0.00417 | 0.01011 | 0.01690 | 0.00765 | 0.00866 | 0.00889 | 0.00732 | 0.10281 |
| **Gckr** | 0.00690 | 0.00439 | 0.01045 | 0.04303 | 0.02242 | 0.01269 | 0.00429 | 0.01035 | 0.00534 | 0.00676 | 0.00741 | 0.00839 | 0.04493 | 0.20908 |
| **Gclc** | 2.56785 | 0.03362 | 0.75595 | 1.28765 | 1.82062 | 3.03665 | 0.70591 | 2.43550 | 0.53131 | 0.01839 | 1.36487 | 2.01123 | 6.93751 | 2.36024 |
| **Gclm** | 8.14630 | 0.02005 | 3.66624 | 2.99479 | 2.64087 | 4.75278 | 0.02320 | 6.92395 | 0.00000 | 0.02125 | 3.46469 | 4.61230 | 1.91239 | 3.96682 |
| **Gcm1** | 0.00000 | 0.00436 | 0.00000 | 0.01694 | 0.00000 | 0.00000 | 0.00386 | 0.00890 | 2.03241 | 0.00740 | 0.00419 | 0.00770 | 0.00455 | 4.06587 |
| **Gcm2** | 0.00098 | 0.00401 | 0.04219 | 0.04017 | 0.01912 | 0.00000 | 0.00396 | 0.00926 | 0.01565 | 0.00656 | 0.14177 | 0.00875 | 1.36515 | 0.00077 |
| **Gcn1l1** | 0.61080 | 2.13961 | 13.52088 | 10.83698 | 7.63936 | 2.21269 | 3.90541 | 0.13118 | 0.00000 | 0.05283 | 2.20279 | 9.81585 | 0.08919 | 0.00000 |
| **Gcnt1** | 0.00262 | 2.17701 | 5.26598 | 3.11670 | 2.17494 | 10.34790 | 2.05645 | 6.07650 | 6.16412 | 13.34479 | 1.89883 | 1.88913 | 0.00000 | 1.51435 |
| **Gcnt2** | 1.23409 | 0.02072 | 1.58451 | 0.00000 | 0.86416 | 0.00000 | 0.97587 | 2.09361 | 3.30588 | 0.01636 | 0.00000 | 2.00875 | 2.95487 | 1.66107 |
| **Gcnt3** | 0.00910 | 0.00447 | 0.01191 | 0.04512 | 0.05065 | 0.01073 | 0.00446 | 0.01343 | 0.01050 | 0.00702 | 0.00800 | 0.01087 | 0.00659 | 0.01597 |
| **Gcnt4** | 0.03874 | 0.00389 | 0.00478 | 0.00000 | 0.30706 | 0.00916 | 0.00389 | 1.18472 | 0.00770 | 0.00682 | 0.00533 | 0.01335 | 0.01695 | 0.19549 |
| **Gcnt7** | 0.00879 | 0.00447 | 0.06317 | 0.00000 | 0.00974 | 0.00179 | 0.00522 | 0.27421 | 0.00863 | 0.00667 | 0.00765 | 0.01059 | 0.00554 | 0.08961 |
| **Gcsh** | 38.22182 | 1.56312 | 37.38169 | 33.69458 | 49.27822 | 13.66888 | 3.14587 | 10.51287 | 22.05600 | 3.54678 | 5.44106 | 10.83156 | 23.56134 | 21.96726 |
| **Gda** | 0.00000 | 0.00401 | 0.02004 | 0.15798 | 0.00000 | 0.00024 | 0.00397 | 0.01591 | 0.38160 | 0.01548 | 0.14163 | 0.00519 | 0.99980 | 2.77276 |
| **Gdap1** | 0.00000 | 0.00320 | 0.00000 | 0.01345 | 0.12980 | 0.00000 | 0.00287 | 0.00490 | 7.74842 | 0.00681 | 0.00299 | 0.00099 | 6.07064 | 5.82189 |
| **Gdap10** | 0.00260 | 0.00446 | 0.00000 | 0.23467 | 0.00253 | 0.00650 | 0.00413 | 0.01955 | 1.26258 | 0.00635 | 0.00672 | 0.00772 | 0.00649 | 0.00379 |
| **Gdap2** | 17.32697 | 2.44706 | 19.23810 | 4.01014 | 20.97945 | 14.77325 | 4.80437 | 4.20246 | 20.98454 | 0.22799 | 6.19828 | 3.27067 | 11.25207 | 18.46364 |
| **Gde1** | 13.38577 | 0.56820 | 12.42037 | 8.67351 | 14.17088 | 3.85695 | 0.23020 | 5.11643 | 3.56872 | 0.03603 | 1.64218 | 3.81419 | 12.08055 | 11.76642 |
| **Gdf11** | 0.00504 | 0.00442 | 0.01303 | 0.44316 | 0.00567 | 0.04780 | 0.00400 | 0.00494 | 0.00969 | 0.00773 | 0.33484 | 0.11992 | 0.00542 | 0.02223 |
| **Gdf15** | 0.01363 | 0.00447 | 0.01291 | 0.01323 | 0.06404 | 0.01315 | 0.00437 | 0.01013 | 0.01122 | 0.00668 | 0.00877 | 0.00964 | 0.00643 | 0.01165 |
| **Gdf3** | 0.01340 | 2.32075 | 10.19418 | 6.23592 | 0.46878 | 5.69753 | 2.20852 | 0.12102 | 0.51925 | 0.07116 | 4.29912 | 10.22916 | 1.75413 | 0.45176 |
| **Gdf5** | 0.00501 | 0.00440 | 0.00973 | 0.01066 | 0.13794 | 0.01249 | 0.00417 | 0.00971 | 0.06629 | 0.00623 | 0.00870 | 0.00896 | 0.19204 | 0.00000 |
| **Gdf9** | 3.15542 | 0.00190 | 2.76656 | 0.00000 | 3.99187 | 1.03295 | 0.00295 | 0.11287 | 0.36774 | 0.02107 | 0.00000 | 0.78529 | 8.86541 | 5.46372 |
| **Gdi1** | 1.70945 | 2.96496 | 1.29747 | 1.54545 | 1.20975 | 0.28160 | 0.30900 | 3.23405 | 0.29212 | 0.73917 | 1.71497 | 2.44790 | 0.00000 | 0.97805 |
| **Gdi2** | 27.36714 | 7.99902 | 31.40219 | 22.87560 | 28.10555 | 24.25876 | 7.37973 | 13.34277 | 27.86508 | 0.55613 | 17.21099 | 20.08344 | 11.35837 | 16.77042 |
| **Gdpd1** | 0.39476 | 0.00363 | 1.55126 | 0.19471 | 1.02203 | 0.83543 | 0.00329 | 0.00000 | 1.40215 | 0.00089 | 0.75333 | 0.00946 | 1.00716 | 1.24296 |
| **Gdpd2** | 0.00210 | 0.00452 | 0.39124 | 0.00455 | 0.00000 | 0.30479 | 0.00411 | 0.00379 | 0.01400 | 0.00681 | 0.00551 | 0.00814 | 0.00227 | 0.50767 |
| **Gdpd3** | 0.00427 | 0.00433 | 0.01924 | 0.01129 | 0.00907 | 0.01315 | 0.00423 | 0.00920 | 0.02077 | 0.00654 | 0.00879 | 0.00922 | 0.38759 | 0.00581 |
| **Gdpd5** | 0.01582 | 0.07875 | 0.01274 | 0.00000 | 0.00746 | 0.34708 | 0.01877 | 0.14126 | 0.00706 | 0.00532 | 0.00000 | 3.70380 | 0.00926 | 0.14353 |
| **Gdpgp1** | 0.21105 | 0.06856 | 1.61286 | 0.47084 | 0.36750 | 1.05666 | 0.23895 | 1.28529 | 0.26793 | 0.02029 | 1.51718 | 0.37110 | 1.33655 | 1.39603 |
| **Gemin2** | 4.60007 | 1.21981 | 9.68612 | 7.70558 | 14.50955 | 4.58579 | 3.26529 | 5.67923 | 6.46502 | 6.01630 | 4.97822 | 4.30415 | 2.24411 | 8.66184 |
| **Gemin4** | 5.84023 | 0.05467 | 11.30627 | 4.84992 | 9.28677 | 5.06619 | 2.87405 | 0.25319 | 6.88899 | 0.06414 | 7.03548 | 4.85522 | 3.67714 | 8.09082 |
| **Gemin5** | 15.21786 | 0.17616 | 18.93448 | 11.87696 | 23.39929 | 13.15878 | 3.91134 | 7.49159 | 14.38715 | 0.21636 | 9.29801 | 10.82113 | 10.43729 | 27.08685 |
| **Gemin6** | 6.31333 | 0.03018 | 3.91452 | 0.00000 | 0.10183 | 4.48801 | 4.80199 | 1.91700 | 3.16226 | 0.16890 | 6.77668 | 5.35868 | 2.49151 | 2.98011 |
| **Gemin7** | 0.55216 | 0.51935 | 1.07736 | 2.31668 | 4.31535 | 2.00309 | 0.87661 | 0.62089 | 0.09350 | 0.00820 | 0.91529 | 1.91340 | 2.45228 | 1.17275 |
| **Gemin8** | 2.09533 | 0.04841 | 2.15330 | 1.88312 | 2.31174 | 3.19333 | 0.11052 | 0.26311 | 1.57524 | 0.19819 | 0.04849 | 2.24000 | 0.36838 | 4.59027 |
| **Gen1** | 0.80977 | 0.00395 | 0.63284 | 1.10622 | 0.47623 | 0.08665 | 0.02373 | 0.77055 | 0.69074 | 0.00428 | 0.00000 | 0.28775 | 0.01122 | 0.00000 |
| **Get4** | 2.48717 | 1.26600 | 10.83873 | 11.61615 | 10.98804 | 6.01874 | 2.54971 | 5.77676 | 0.00226 | 0.09424 | 0.40385 | 7.53136 | 0.00405 | 0.01501 |
| **Gfer** | 0.78917 | 3.96786 | 1.33787 | 4.34977 | 5.94787 | 2.52435 | 1.36924 | 1.18672 | 2.55819 | 0.03463 | 0.80642 | 1.87628 | 6.56185 | 0.00000 |
| **Gfi1** | 0.00000 | 0.00537 | 0.36433 | 0.29927 | 0.30058 | 0.02111 | 0.00468 | 0.03908 | 0.01518 | 0.01201 | 0.00732 | 0.00441 | 0.27107 | 0.00000 |
| **Gfi1b** | 0.11920 | 0.00444 | 0.24553 | 0.00173 | 0.16251 | 0.01767 | 0.00405 | 0.00830 | 0.00898 | 0.00954 | 0.00618 | 0.00823 | 0.01836 | 0.00588 |
| **Gfm1** | 5.60437 | 0.04632 | 7.00259 | 0.88945 | 1.55601 | 4.42912 | 0.00708 | 3.08946 | 3.80974 | 0.00000 | 3.63680 | 3.50210 | 3.07192 | 7.49542 |
| **Gfm2** | 2.79310 | 2.93071 | 2.43117 | 2.34895 | 3.36164 | 5.42685 | 0.00189 | 2.23322 | 1.01867 | 0.00765 | 2.32364 | 2.49133 | 1.46916 | 3.98495 |

| | | | | | | | | | | | | | | |
|---|---|---|---|---|---|---|---|---|---|---|---|---|---|---|
| **Gfod1** | 0.00438 | 0.00407 | 0.00000 | 0.25169 | 0.03934 | 0.00000 | 0.00410 | 0.00822 | 0.63051 | 0.00571 | 0.00462 | 0.00583 | 1.32917 | 0.25921 |
| **Gfod2** | 0.56862 | 0.01177 | 0.85333 | 1.04352 | 1.04101 | 0.00000 | 0.00409 | 0.54632 | 0.26486 | 0.00275 | 0.80166 | 1.25920 | 2.43271 | 1.72611 |
| **Gfpt1** | 8.96224 | 0.02767 | 6.62746 | 3.08750 | 10.32362 | 2.81988 | 0.01888 | 1.03362 | 10.72045 | 0.00000 | 5.70949 | 3.53667 | 8.39194 | 12.13824 |
| **Gfpt2** | 0.08153 | 0.00413 | 0.00000 | 0.00000 | 0.33853 | 1.13308 | 0.00374 | 0.00444 | 0.00226 | 0.00829 | 0.00000 | 0.58142 | 0.41205 | 0.38937 |
| **Gfra1** | 0.38474 | 0.00429 | 0.03441 | 0.00000 | 1.24870 | 0.00423 | 0.00433 | 0.01101 | 0.58904 | 0.00725 | 0.00438 | 0.00512 | 1.41552 | 0.00000 |
| **Gfra2** | 0.02408 | 0.01376 | 0.02424 | 0.03221 | 0.02343 | 0.03331 | 0.00439 | 0.05487 | 0.05828 | 0.00888 | 0.02187 | 0.02326 | 0.02289 | 0.06802 |
| **Gfra3** | 1.57819 | 0.00385 | 0.79323 | 0.00000 | 2.94599 | 0.89148 | 0.00363 | 0.00000 | 0.04115 | 0.01599 | 1.95144 | 0.00000 | 1.28857 | 2.46223 |
| **Gfra4** | 0.00634 | 0.00420 | 0.00645 | 0.29987 | 0.11355 | 0.00764 | 0.00395 | 0.00822 | 0.02689 | 0.00947 | 0.00646 | 0.00690 | 0.04642 | 0.07707 |
| **Gga1** | 0.67262 | 5.02541 | 6.60422 | 8.12541 | 4.45272 | 1.30834 | 5.27959 | 7.59766 | 0.00000 | 11.04590 | 3.09965 | 3.81523 | 1.98666 | 0.00000 |
| **Gga2** | 9.70283 | 3.87098 | 12.39220 | 16.80203 | 11.36997 | 15.39738 | 0.47411 | 9.15294 | 5.78546 | 0.16969 | 7.63546 | 11.45178 | 0.73440 | 4.43849 |
| **Gga3** | 1.07087 | 0.11347 | 6.83585 | 5.92374 | 2.16591 | 7.07473 | 0.15807 | 3.45028 | 3.08468 | 0.80306 | 7.59495 | 4.09598 | 3.93185 | 5.67583 |
| **Ggact** | 0.14530 | 0.00406 | 0.11112 | 0.00000 | 0.16303 | 0.02747 | 0.00432 | 0.02063 | 0.21067 | 0.00843 | 0.00713 | 0.15171 | 0.04789 | 0.78654 |
| **Ggct** | 0.81707 | 0.03857 | 0.34793 | 0.00000 | 0.77104 | 0.93714 | 0.01417 | 3.47092 | 0.03559 | 0.01279 | 2.63978 | 2.51804 | 3.42832 | 1.16334 |
| **Ggcx** | 1.13508 | 0.00393 | 0.45690 | 0.36037 | 0.00000 | 0.21534 | 0.00379 | 0.03283 | 0.85641 | 0.01641 | 0.00239 | 0.18430 | 0.44873 | 0.18817 |
| **Ggh** | 0.17060 | 0.00437 | 0.06148 | 0.00527 | 0.00709 | 0.01236 | 0.00436 | 0.01616 | 0.01150 | 0.00617 | 0.00705 | 0.00922 | 0.00440 | 0.01331 |
| **Ggnbp1** | 0.00600 | 0.00506 | 0.03324 | 0.70799 | 0.97855 | 0.06251 | 0.00416 | 0.00862 | 0.01990 | 0.00602 | 0.00627 | 0.04446 | 0.02880 | 0.06802 |
| **Ggnbp2** | 1.25026 | 0.05304 | 0.54536 | 0.84038 | 4.74154 | 2.21245 | 5.20187 | 4.34465 | 5.26958 | 0.12947 | 3.52717 | 5.69306 | 2.44106 | 2.66042 |
| **Ggnbp2os** | 0.00364 | 0.00429 | 0.00744 | 0.08756 | 0.12322 | 0.02055 | 0.00473 | 0.03427 | 0.00712 | 0.00889 | 0.00683 | 0.42169 | 0.05762 | 0.25253 |
| **Ggps1** | 0.50646 | 0.02062 | 1.41197 | 2.47342 | 1.17367 | 3.30936 | 0.06606 | 0.45296 | 1.61145 | 0.10876 | 0.07858 | 1.32698 | 5.77194 | 2.18864 |
| **Ggt1** | 1.24283 | 0.04235 | 5.25346 | 6.05963 | 2.66434 | 6.35353 | 3.55912 | 2.40796 | 0.19037 | 0.10224 | 2.77829 | 7.45994 | 0.00000 | 1.14039 |
| **Ggt5** | 0.00995 | 0.00478 | 0.00940 | 0.04444 | 0.01233 | 0.00763 | 0.00463 | 1.25982 | 0.01197 | 0.00988 | 0.00795 | 0.00662 | 0.00936 | 0.01161 |
| **Ggt6** | 0.07862 | 0.00387 | 0.48705 | 0.49760 | 0.78269 | 0.00000 | 0.00371 | 0.00431 | 0.13712 | 0.00533 | 0.00746 | 0.01898 | 1.03235 | 0.47680 |
| **Ggt7** | 0.02719 | 0.00443 | 0.07549 | 0.00509 | 0.31049 | 0.00931 | 0.00411 | 0.01079 | 0.03107 | 0.00792 | 0.00734 | 0.00818 | 0.06514 | 0.01809 |
| **Ggta1** | 0.00000 | 0.60813 | 0.51851 | 0.83627 | 0.00000 | 2.49331 | 0.00340 | 3.83047 | 0.00629 | 0.02211 | 0.00099 | 0.00000 | 0.32644 | 1.36824 |
| **Ghdc** | 0.29675 | 0.07420 | 1.83138 | 4.26157 | 0.13059 | 5.12450 | 0.03308 | 3.05346 | 0.63608 | 0.08884 | 3.65477 | 2.30326 | 1.30237 | 1.24361 |
| **Ghitm** | 14.52267 | 5.32559 | 20.04919 | 22.05332 | 11.04022 | 29.25281 | 5.05013 | 12.71603 | 14.80098 | 0.52969 | 21.60001 | 18.58473 | 8.39766 | 12.13319 |
| **Ghrh** | 0.01355 | 0.00445 | 0.00920 | 0.02499 | 0.15036 | 0.01178 | 0.00415 | 0.00992 | 0.01425 | 0.00963 | 0.00839 | 0.00812 | 0.01439 | 0.00996 |
| **Ghrl** | 0.01426 | 0.00425 | 0.17944 | 0.01211 | 0.07231 | 0.01232 | 0.00404 | 0.01315 | 0.01034 | 0.01116 | 0.00775 | 0.00858 | 0.13527 | 0.01985 |
| **Gid4** | 4.38724 | 2.45144 | 3.46145 | 2.61858 | 5.93373 | 4.16586 | 0.01268 | 3.59220 | 5.88951 | 0.04581 | 0.10539 | 3.99257 | 2.77575 | 6.21193 |
| **Gid8** | 1.17676 | 6.93334 | 3.73070 | 0.00000 | 3.77166 | 3.05844 | 0.67928 | 4.79928 | 2.87862 | 0.05367 | 2.68172 | 5.71939 | 1.98541 | 1.22217 |
| **Gigyf1** | 0.00820 | 0.00419 | 0.03239 | 1.15512 | 0.82893 | 0.19100 | 0.00406 | 0.67944 | 0.01744 | 0.02465 | 0.02529 | 0.41578 | 0.08586 | 0.08764 |
| **Gigyf2** | 0.79312 | 0.05388 | 5.44715 | 2.15928 | 5.43083 | 2.09018 | 0.00149 | 0.45411 | 5.45182 | 0.00690 | 2.07167 | 2.80389 | 0.71915 | 7.39002 |
| **Gimap1** | 0.23208 | 0.65663 | 0.24336 | 0.15052 | 0.24351 | 0.15077 | 0.00397 | 0.01441 | 0.72439 | 0.00697 | 0.02407 | 0.08391 | 0.80272 | 0.38695 |
| **Gimap3** | 0.00503 | 0.00449 | 0.01304 | 0.01329 | 0.12909 | 0.01184 | 0.00432 | 0.01059 | 0.01335 | 0.00644 | 0.00845 | 0.00878 | 0.06181 | 0.14643 |
| **Gimap8** | 0.04073 | 0.02666 | 0.02273 | 0.02476 | 0.01566 | 0.01404 | 0.01590 | 0.01807 | 0.06589 | 0.00627 | 0.00726 | 0.01097 | 0.00567 | 0.03156 |
| **Gimap9** | 0.00443 | 0.00418 | 0.00186 | 0.35927 | 0.00421 | 0.11184 | 0.01000 | 0.00398 | 0.00755 | 0.01200 | 0.04494 | 0.01850 | 0.00407 | 0.23450 |
| **Gin1** | 4.45590 | 0.00264 | 2.46164 | 0.60526 | 2.31273 | 2.24504 | 0.00910 | 0.67562 | 7.89616 | 0.00467 | 1.67033 | 1.27335 | 0.39276 | 3.70372 |
| **Ginm1** | 0.78133 | 0.06303 | 0.89189 | 1.42783 | 0.70686 | 0.00000 | 2.02936 | 1.95690 | 1.03119 | 0.04034 | 0.03497 | 0.77471 | 0.00000 | 0.00000 |
| **Gins1** | 9.19761 | 0.05750 | 14.72168 | 11.78641 | 15.61573 | 9.07812 | 3.54669 | 5.31600 | 3.51106 | 0.06763 | 10.61335 | 5.64688 | 4.35866 | 7.55362 |
| **Gins2** | 3.04764 | 0.04959 | 6.25129 | 0.00000 | 9.03237 | 2.85021 | 0.02225 | 1.93670 | 0.36993 | 10.35234 | 0.55456 | 10.01197 | 4.27095 | 7.12017 |
| **Gins3** | 1.57151 | 0.00961 | 1.82552 | 3.16692 | 2.27930 | 4.27125 | 2.34476 | 3.52733 | 5.62052 | 0.02397 | 0.39719 | 2.05392 | 7.55143 | 6.24743 |
| **Gins4** | 9.96722 | 0.81723 | 9.13172 | 16.09304 | 13.64490 | 5.38800 | 1.63476 | 3.79664 | 4.74844 | 0.09748 | 4.28813 | 5.49224 | 7.88251 | 8.15912 |
| **Gip** | 0.14800 | 0.00419 | 0.04335 | 0.00287 | 0.01822 | 0.10557 | 0.00413 | 0.00573 | 0.00837 | 0.00866 | 0.00567 | 0.00774 | 0.92754 | 0.00620 |
| **Gipc1** | 0.40392 | 5.57766 | 5.56832 | 7.15424 | 0.11100 | 0.58495 | 1.85673 | 11.99513 | 0.00000 | 0.11245 | 0.84553 | 2.85080 | 0.07981 | 0.00000 |
| **Gipc2** | 0.08771 | 0.03600 | 0.70596 | 0.91050 | 0.00000 | 2.04984 | 0.02224 | 2.94102 | 0.02516 | 0.01661 | 1.23033 | 0.23199 | 1.49674 | 0.00000 |
| **Gipr** | 0.00649 | 0.00488 | 0.09428 | 0.14640 | 0.01476 | 0.02410 | 0.00426 | 0.01895 | 0.01053 | 0.00923 | 0.00713 | 0.02629 | 0.00823 | 0.01212 |
| **Git1** | 0.00000 | 3.33676 | 3.24263 | 5.39941 | 1.58496 | 1.76683 | 0.95705 | 9.41193 | 0.00000 | 0.07065 | 0.44676 | 3.62886 | 0.51112 | 0.00000 |
| **Git2** | 0.49512 | 0.02584 | 0.89572 | 2.34477 | 2.05607 | 5.32842 | 4.16713 | 1.63573 | 0.28301 | 0.08645 | 2.15796 | 1.54966 | 1.27207 | 3.77326 |
| **Gja1** | 10.32093 | 0.18397 | 11.00298 | 1.47520 | 1.89989 | 10.62899 | 3.50470 | 2.72317 | 32.94336 | 1.18744 | 10.46635 | 8.15791 | 4.74209 | 9.87862 |
| **Gja10** | 0.01108 | 0.00441 | 0.00970 | 0.01300 | 0.00239 | 0.01051 | 0.00419 | 0.00993 | 0.15900 | 0.00648 | 0.00759 | 0.00947 | 0.13135 | 0.09054 |
| **Gja3** | 0.00556 | 0.00428 | 0.01462 | 0.01117 | 0.02651 | 0.01285 | 0.00417 | 0.00894 | 0.00994 | 0.00611 | 0.00935 | 0.00891 | 0.61501 | 0.03233 |
| **Gja4** | 0.00000 | 0.00564 | 1.09446 | 0.00000 | 0.10236 | 0.00000 | 0.00435 | 0.55046 | 0.00561 | 0.00535 | 0.00405 | 0.01155 | 0.50098 | 0.65710 |
| **Gjb3** | 3.01614 | 7.01576 | 21.21492 | 15.97217 | 2.28366 | 8.17902 | 2.93470 | 19.78206 | 0.00000 | 14.74814 | 11.51194 | 15.45210 | 0.00000 | 0.78254 |
| **Gjb4** | 0.00829 | 0.00412 | 0.01177 | 0.00442 | 0.01349 | 0.00000 | 0.00406 | 0.01307 | 0.01101 | 0.02016 | 0.51671 | 0.40719 | 0.00702 | 0.02220 |
| **Gjb5** | 0.00000 | 0.26886 | 1.57174 | 0.00000 | 0.00000 | 0.06821 | 3.99125 | 11.25107 | 0.00000 | 0.12450 | 7.30000 | 22.84603 | 0.00000 | 0.08843 |

| | | | | | | | | | | | | | | |
|---|---|---|---|---|---|---|---|---|---|---|---|---|---|---|
| **Gjc1** | 0.00373 | 0.01177 | 0.00606 | 0.01353 | 0.09677 | 0.20083 | 0.00399 | 0.00469 | 0.06059 | 0.02155 | 0.00531 | 0.02034 | 0.46694 | 0.06200 |
| **Gjc3** | 0.13771 | 0.12780 | 0.04324 | 0.28005 | 0.08294 | 0.17051 | 0.05490 | 0.13426 | 0.31060 | 0.01784 | 0.03754 | 0.14339 | 0.52414 | 0.33821 |
| **Gje1** | 0.12613 | 0.07476 | 0.02008 | 0.22652 | 0.19612 | 0.41021 | 0.12431 | 0.10840 | 0.14242 | 0.14000 | 0.00951 | 0.06756 | 0.80304 | 0.23004 |
| **Gk** | 0.00000 | 0.00940 | 0.21166 | 0.00000 | 1.58043 | 3.04980 | 3.85340 | 1.17686 | 5.70893 | 4.18362 | 1.95791 | 0.13234 | 1.76852 | 0.00000 |
| **Gk5** | 0.08498 | 0.00416 | 2.88407 | 0.00000 | 1.07856 | 0.30181 | 0.51357 | 0.04741 | 2.56999 | 0.01966 | 0.33379 | 0.00057 | 0.00000 | 0.73770 |
| **Gkap1** | 4.63043 | 0.04969 | 2.49926 | 0.37108 | 0.88399 | 4.35616 | 4.98248 | 7.22695 | 1.76947 | 0.24824 | 2.00076 | 5.11217 | 1.38838 | 2.06751 |
| **Gla** | 10.26631 | 0.05736 | 12.27284 | 11.26161 | 9.48577 | 16.31929 | 5.93491 | 8.15837 | 12.95832 | 0.38099 | 8.15973 | 3.12432 | 6.64802 | 16.25461 |
| **Glb1** | 5.62397 | 6.86976 | 16.73554 | 8.19356 | 16.31509 | 8.24844 | 3.90176 | 7.66764 | 4.64816 | 7.27205 | 3.47696 | 9.31665 | 3.43859 | 13.07454 |
| **Glb1l** | 0.00000 | 0.00434 | 0.10797 | 0.06417 | 0.18132 | 0.04974 | 0.27247 | 0.00350 | 0.72119 | 0.01679 | 0.08723 | 0.00471 | 0.34669 | 0.31979 |
| **Glcci1** | 0.86432 | 2.26643 | 1.63071 | 0.24805 | 2.88670 | 1.00642 | 0.00912 | 0.78652 | 1.02827 | 0.00156 | 0.67409 | 0.93784 | 6.89003 | 3.36448 |
| **Glce** | 0.26696 | 0.01630 | 1.41520 | 0.00000 | 0.81499 | 1.56475 | 0.00714 | 2.09325 | 5.58063 | 0.00730 | 0.97729 | 4.43317 | 1.09532 | 2.47085 |
| **Gldc** | 31.78362 | 1.51404 | 38.39686 | 22.33085 | 28.65015 | 21.92005 | 3.29695 | 10.80802 | 19.74163 | 0.22481 | 10.60166 | 11.07471 | 12.81451 | 34.52941 |
| **Gle1** | 12.00293 | 0.07345 | 19.27724 | 10.85403 | 15.39213 | 9.29850 | 3.96442 | 6.38591 | 9.76763 | 0.12553 | 6.67949 | 3.92133 | 10.12373 | 19.08534 |
| **Glg1** | 0.63106 | 0.07161 | 6.02069 | 1.40643 | 2.73185 | 5.73424 | 0.00107 | 2.46825 | 0.01279 | 0.44240 | 1.57468 | 5.09667 | 0.08081 | 9.41110 |
| **Gli1** | 0.01619 | 0.00422 | 0.14329 | 0.00000 | 0.08241 | 0.00899 | 0.00425 | 0.00994 | 0.01056 | 0.00623 | 0.32544 | 0.01715 | 0.00935 | 0.01040 |
| **Gli2** | 0.00237 | 0.00425 | 0.46598 | 0.00101 | 0.01440 | 0.00303 | 0.00408 | 0.01051 | 0.03445 | 0.00609 | 0.00702 | 0.00823 | 0.40704 | 0.31708 |
| **Gli3** | 0.00000 | 0.00424 | 0.28690 | 0.00000 | 0.15462 | 0.14659 | 0.00392 | 0.00323 | 0.01556 | 0.00446 | 0.00576 | 0.00582 | 3.26671 | 0.62074 |
| **Glipr1** | 0.01495 | 0.12468 | 0.36751 | 0.00000 | 0.00037 | 0.11559 | 0.00313 | 5.27006 | 0.03206 | 0.09725 | 0.50133 | 0.49868 | 0.00517 | 0.00000 |
| **Glipr2** | 0.00000 | 0.22475 | 5.11897 | 1.95000 | 0.00000 | 1.62318 | 0.02703 | 3.93526 | 0.00000 | 12.62436 | 1.81713 | 7.57872 | 0.00000 | 1.11176 |
| **Glis1** | 0.00307 | 0.00422 | 0.24615 | 0.01939 | 0.03869 | 0.00618 | 0.00428 | 0.00998 | 0.00922 | 0.00603 | 0.00699 | 0.00714 | 0.54555 | 0.09835 |
| **Glis2** | 0.96006 | 0.02146 | 1.61356 | 0.00000 | 5.46716 | 0.22298 | 0.00314 | 0.86298 | 2.90068 | 0.00355 | 0.00000 | 2.96638 | 0.93823 | 6.52245 |
| **Glis3** | 0.16893 | 0.01373 | 0.00286 | 0.00000 | 0.21701 | 0.00862 | 0.00438 | 0.04757 | 0.04364 | 0.00746 | 0.00427 | 0.54667 | 1.06175 | 0.30603 |
| **Glmn** | 3.23749 | 1.97946 | 2.29706 | 1.40466 | 0.48706 | 0.90368 | 0.00236 | 0.66539 | 4.11107 | 0.00000 | 1.16243 | 0.16992 | 0.00000 | 0.76766 |
| **Glmp** | 3.77972 | 2.57905 | 16.37427 | 17.77310 | 10.16029 | 18.25202 | 6.90573 | 9.18946 | 8.75441 | 13.66492 | 9.31342 | 8.47626 | 4.44329 | 17.68233 |
| **Glo1** | 2.66224 | 0.19019 | 1.72639 | 1.69969 | 2.54877 | 7.91847 | 5.55949 | 6.57969 | 0.93521 | 6.72019 | 8.18461 | 11.17000 | 0.98171 | 1.16269 |
| **Glod4** | 3.45021 | 0.11807 | 5.89929 | 7.24734 | 3.66609 | 11.33288 | 2.78900 | 7.86171 | 8.39890 | 8.84841 | 12.31837 | 10.33094 | 4.43888 | 2.79721 |
| **Glod5** | 0.00000 | 0.00421 | 0.00000 | 0.01210 | 0.96057 | 0.00729 | 0.00419 | 0.00823 | 0.47287 | 0.00635 | 0.00586 | 0.00726 | 1.26280 | 0.35256 |
| **Glrb** | 0.00527 | 0.00391 | 0.01481 | 0.11614 | 0.00632 | 0.00000 | 1.22884 | 1.64686 | 0.01156 | 0.04788 | 0.01564 | 0.00696 | 0.00665 | 0.00976 |
| **Glrx** | 0.00000 | 0.26613 | 0.68102 | 0.00000 | 0.13901 | 9.64555 | 7.06372 | 10.51994 | 0.00663 | 13.00273 | 3.72829 | 18.71935 | 0.00289 | 0.52777 |
| **Glrx2** | 2.26974 | 0.22481 | 0.36035 | 2.55309 | 4.34606 | 1.36547 | 0.19474 | 0.55315 | 5.01492 | 0.06342 | 0.00000 | 0.82323 | 8.84450 | 1.28884 |
| **Glrx3** | 9.33395 | 1.31179 | 6.12084 | 9.16567 | 7.12263 | 6.01408 | 0.33597 | 3.95168 | 3.09683 | 0.40573 | 2.02935 | 4.26795 | 2.83646 | 0.90371 |
| **Glrx5** | 1.35418 | 0.02409 | 0.19571 | 1.39246 | 3.44575 | 3.18709 | 0.18215 | 2.70467 | 0.08976 | 0.05597 | 0.35136 | 3.55349 | 0.74500 | 0.00000 |
| **Gls** | 6.71781 | 0.17551 | 7.44620 | 1.34521 | 6.73849 | 6.08066 | 2.42676 | 1.53686 | 13.14479 | 0.09870 | 2.71312 | 2.22866 | 4.67925 | 1.42915 |
| **Gls2** | 1.24232 | 0.01031 | 4.40095 | 0.64338 | 5.38054 | 0.53314 | 0.00228 | 0.42845 | 2.08291 | 0.00769 | 0.10495 | 3.25680 | 2.45617 | 5.49152 |
| **Glt1d1** | 0.18382 | 0.00401 | 0.42492 | 0.77468 | 0.50796 | 0.58666 | 0.00401 | 0.02264 | 0.12909 | 0.00469 | 0.02743 | 0.00305 | 0.20073 | 0.00000 |
| **Glt28d2** | 0.11633 | 0.00410 | 0.02721 | 0.00000 | 0.48870 | 0.11712 | 1.31371 | 0.05215 | 0.13095 | 0.05881 | 0.01373 | 0.05045 | 0.05787 | 0.03142 |
| **Glt8d1** | 2.27731 | 0.01055 | 2.31814 | 1.35846 | 0.19801 | 3.01198 | 1.11487 | 3.22016 | 3.66685 | 0.04451 | 2.83513 | 1.02844 | 2.01506 | 2.64945 |
| **Glt8d2** | 0.00932 | 0.00488 | 0.01130 | 0.02085 | 0.02518 | 0.13016 | 0.00469 | 0.01296 | 0.01231 | 0.00754 | 0.00862 | 0.00732 | 0.00713 | 0.00935 |
| **Gltp** | 2.99705 | 0.08509 | 1.37503 | 1.59094 | 0.65303 | 10.82282 | 4.16373 | 5.07577 | 0.20409 | 0.15168 | 3.41951 | 4.92406 | 0.58278 | 0.05244 |
| **Gltscr1** | 0.25992 | 0.00345 | 1.07114 | 0.11467 | 0.98366 | 0.84834 | 0.24815 | 0.41749 | 0.01080 | 0.01503 | 0.02597 | 0.55032 | 0.29554 | 0.48061 |
| **Gltscr1l** | 4.31970 | 0.00887 | 1.82435 | 1.41974 | 6.11780 | 2.55711 | 0.00883 | 0.89161 | 5.14119 | 0.11134 | 3.67614 | 1.75298 | 6.75980 | 11.08604 |
| **Gltscr2** | 0.00000 | 5.65675 | 13.36558 | 25.94285 | 26.40283 | 0.00000 | 1.71357 | 5.85594 | 0.11849 | 2.86874 | 0.06696 | 7.34143 | 3.66691 | 0.00000 |
| **Glud1** | 9.79887 | 6.87527 | 9.69463 | 13.00004 | 11.84092 | 11.04595 | 6.60948 | 10.09973 | 5.33669 | 2.12663 | 16.57476 | 19.11392 | 6.44203 | 11.60914 |
| **Glul** | 8.75500 | 3.24034 | 20.34833 | 19.16266 | 15.26830 | 19.48768 | 3.82308 | 13.84984 | 10.84760 | 18.67938 | 12.77414 | 12.19449 | 15.56316 | 18.02758 |
| **Glyctk** | 0.01452 | 0.01225 | 0.12830 | 0.53232 | 0.01209 | 0.09959 | 0.00518 | 0.51325 | 0.00819 | 0.01044 | 0.00323 | 0.27527 | 0.41545 | 0.04617 |
| **Glyr1** | 5.46982 | 0.50246 | 9.33621 | 5.81375 | 7.08902 | 12.72242 | 5.80282 | 4.09457 | 10.25824 | 0.35014 | 6.03521 | 8.32347 | 3.91230 | 10.26222 |
| **Gm10012** | 3.19277 | 1.67146 | 0.00000 | 1.10194 | 2.34941 | 6.54474 | 1.02511 | 5.74445 | 0.56218 | 4.92871 | 0.18618 | 5.30180 | 5.69914 | 0.74069 |
| **Gm10033** | 0.03699 | 0.05649 | 0.00000 | 0.64303 | 0.13152 | 0.50234 | 0.05056 | 0.06781 | 2.15621 | 0.03372 | 0.64625 | 0.77411 | 0.75823 | 0.00000 |
| **Gm10046** | 0.00321 | 0.02050 | 0.00996 | 0.00727 | 0.00451 | 0.06834 | 0.00444 | 0.38463 | 0.04254 | 1.21591 | 0.00000 | 0.00489 | 0.00424 | 0.04259 |
| **Gm10069** | 0.87832 | 0.00949 | 2.15638 | 2.48233 | 1.92310 | 2.86890 | 0.97057 | 1.02810 | 2.53949 | 1.00574 | 1.03409 | 2.54678 | 0.33201 | 2.86336 |
| **Gm10190** | 0.10255 | 0.00451 | 0.06365 | 0.28892 | 0.03331 | 0.15285 | 0.03987 | 0.00801 | 0.37044 | 0.02244 | 0.08224 | 0.18180 | 0.68243 | 0.05576 |
| **Gm10319** | 0.00653 | 0.00436 | 0.01459 | 0.01203 | 0.00948 | 0.01275 | 0.00424 | 0.00946 | 0.02357 | 0.00636 | 0.00891 | 0.00945 | 0.28077 | 0.01710 |
| **Gm10324** | 0.39337 | 0.02506 | 4.40223 | 3.48933 | 3.27723 | 3.10103 | 1.42984 | 1.16947 | 12.72916 | 0.03949 | 4.03826 | 3.28106 | 0.63524 | 4.75286 |
| **Gm10334** | 0.00764 | 0.00435 | 0.01369 | 0.07979 | 0.00931 | 0.01139 | 0.00429 | 0.00811 | 0.01797 | 0.00612 | 0.00869 | 0.00921 | 0.01206 | 0.03974 |
| **Gm10336** | 2.97617 | 0.00258 | 0.40883 | 0.67291 | 0.29814 | 1.23119 | 0.00329 | 0.00637 | 0.62406 | 0.04761 | 0.22965 | 1.34480 | 1.79248 | 0.00000 |

| | | | | | | | | | | | | | | |
|---|---|---|---|---|---|---|---|---|---|---|---|---|---|---|
| **Gm10354** | 0.10680 | 0.00435 | 0.05475 | 0.02524 | 0.08769 | 0.02739 | 0.00414 | 0.00695 | 0.06564 | 0.00607 | 0.00925 | 0.01112 | 0.07739 | 0.13855 |
| **Gm10364** | 0.02317 | 0.04456 | 0.08017 | 0.07740 | 0.02308 | 0.24561 | 0.02380 | 0.08395 | 0.01767 | 0.10714 | 0.05594 | 0.07356 | 0.46483 | 0.16494 |
| **Gm10389** | 0.03702 | 0.00468 | 0.04816 | 0.03451 | 0.05491 | 0.04998 | 0.01704 | 0.08630 | 0.24393 | 0.00719 | 0.01759 | 0.05465 | 0.32814 | 0.12076 |
| **Gm10406** | 0.06333 | 0.00450 | 0.01497 | 0.04906 | 0.08859 | 0.02300 | 0.00405 | 0.00741 | 0.02085 | 0.00651 | 0.00753 | 0.03077 | 0.29349 | 0.00597 |
| **Gm10408** | 0.02816 | 0.00418 | 0.01541 | 0.02973 | 0.03620 | 0.02519 | 0.00401 | 0.01940 | 0.08705 | 0.00682 | 0.00781 | 0.00900 | 0.01809 | 0.03711 |
| **Gm10409** | 0.02545 | 0.00452 | 0.00743 | 0.08296 | 0.08310 | 0.01316 | 0.00400 | 0.00797 | 0.04472 | 0.00676 | 0.00806 | 0.03896 | 0.03268 | 0.01118 |
| **Gm10416** | 0.80765 | 0.00289 | 0.56228 | 0.83977 | 0.25210 | 1.13440 | 0.22563 | 0.00000 | 0.30621 | 0.01955 | 1.69561 | 1.30209 | 0.05261 | 0.01875 |
| **Gm10432** | 0.02616 | 0.01233 | 0.02635 | 0.03053 | 0.04268 | 0.02682 | 0.00425 | 0.01189 | 0.10152 | 0.00822 | 0.00874 | 0.03555 | 0.00722 | 0.08009 |
| **Gm10433** | 0.60122 | 0.00421 | 0.21226 | 0.00268 | 0.00000 | 0.00942 | 0.01588 | 0.01080 | 0.01224 | 0.00707 | 0.02275 | 0.00757 | 0.09941 | 0.83666 |
| **Gm10436** | 0.45312 | 0.00268 | 0.00000 | 0.13055 | 2.25015 | 0.00000 | 0.00249 | 0.00367 | 9.86757 | 0.00681 | 0.00058 | 0.00000 | 6.24372 | 12.73366 |
| **Gm10439** | 0.01010 | 0.00444 | 0.02098 | 0.01190 | 0.01166 | 0.01295 | 0.00429 | 0.01032 | 0.01152 | 0.00690 | 0.00912 | 0.00996 | 0.01463 | 0.04463 |
| **Gm10509** | 0.06898 | 0.00441 | 0.04862 | 0.09361 | 0.29975 | 0.11476 | 0.08479 | 0.00407 | 0.49651 | 0.01308 | 0.63835 | 0.00787 | 0.31378 | 0.00000 |
| **Gm10512** | 0.00000 | 0.00425 | 0.00000 | 0.01071 | 0.44612 | 0.01007 | 0.00415 | 0.00861 | 0.29268 | 0.00632 | 0.00795 | 0.00699 | 1.29845 | 0.13888 |
| **Gm10548** | 0.15829 | 0.16317 | 0.02295 | 0.11527 | 0.10778 | 0.15811 | 0.04693 | 0.06065 | 0.02785 | 0.09296 | 0.01334 | 0.08959 | 0.07310 | 0.10386 |
| **Gm10636** | 0.07475 | 0.02934 | 0.02263 | 0.08950 | 0.01791 | 0.07360 | 0.01736 | 0.02808 | 0.13613 | 0.02355 | 0.00694 | 0.02989 | 0.30886 | 0.07546 |
| **Gm10638** | 0.01056 | 0.00434 | 0.01384 | 0.01304 | 0.05909 | 0.01308 | 0.00434 | 0.01067 | 0.00912 | 0.00665 | 0.00878 | 0.00952 | 0.21457 | 0.00437 |
| **Gm10639** | 0.07932 | 0.00413 | 0.87500 | 0.39757 | 0.29320 | 0.00000 | 0.79266 | 0.75201 | 0.32693 | 1.58233 | 0.23840 | 0.00000 | 0.03804 | 0.33962 |
| **Gm10653** | 10.25954 | 1.74352 | 1.12321 | 8.25184 | 17.16050 | 9.35322 | 1.25393 | 3.65454 | 8.32612 | 9.08013 | 1.59505 | 9.42128 | 23.18275 | 13.10612 |
| **Gm10658** | 0.01644 | 0.00420 | 0.00548 | 0.46393 | 0.14093 | 0.00753 | 0.00445 | 0.00646 | 0.01394 | 0.00534 | 0.00644 | 0.00715 | 0.04102 | 0.01978 |
| **Gm10696** | 2.56685 | 0.00267 | 0.60214 | 0.24879 | 2.17749 | 0.00000 | 0.08479 | 0.00000 | 2.71532 | 0.01696 | 0.00000 | 0.72728 | 6.22408 | 5.82798 |
| **Gm10767** | 3.62488 | 0.00917 | 3.04067 | 1.69506 | 4.97289 | 3.17899 | 0.51668 | 0.99420 | 5.84918 | 0.02314 | 2.21911 | 1.45002 | 4.29223 | 5.76370 |
| **Gm10768** | 0.00506 | 0.00438 | 0.01193 | 0.01280 | 0.00749 | 0.01148 | 0.00426 | 0.01011 | 0.06430 | 0.00645 | 0.00816 | 0.00963 | 0.15518 | 0.10886 |
| **Gm10778** | 0.19183 | 0.01076 | 4.29340 | 3.72794 | 3.78439 | 2.70011 | 0.03797 | 0.04502 | 5.54820 | 0.22812 | 1.25978 | 0.42631 | 4.77530 | 0.86787 |
| **Gm10785** | 0.02710 | 0.01244 | 0.02553 | 0.02411 | 0.02320 | 0.02074 | 0.00419 | 0.01036 | 0.15373 | 0.00660 | 0.02530 | 0.00982 | 0.00770 | 0.02047 |
| **Gm10789** | 0.15927 | 0.01266 | 0.10176 | 0.02186 | 0.11692 | 0.04769 | 0.05849 | 0.02818 | 0.54594 | 0.00696 | 0.00538 | 0.03193 | 0.62218 | 0.15656 |
| **Gm10791** | 0.14200 | 0.01215 | 0.04247 | 0.05696 | 0.08639 | 0.05496 | 0.02409 | 0.01721 | 0.24784 | 0.00690 | 0.01370 | 0.03278 | 0.07738 | 0.16663 |
| **Gm10814** | 0.00661 | 0.00443 | 0.00885 | 0.01463 | 0.00258 | 0.01048 | 0.00425 | 0.01048 | 0.11714 | 0.00698 | 0.00752 | 0.00983 | 0.00404 | 0.23169 |
| **Gm10845** | 16.47021 | 2.04864 | 8.38363 | 1.79615 | 13.21436 | 10.05879 | 1.63040 | 2.53696 | 19.44616 | 2.54666 | 4.70304 | 1.43160 | 6.52047 | 12.87114 |
| **Gm10857** | 0.05715 | 0.01402 | 0.18500 | 0.07912 | 0.47046 | 0.20418 | 0.14825 | 0.25017 | 0.61957 | 0.01707 | 1.17958 | 0.01022 | 1.59238 | 2.38730 |
| **Gm10865** | 0.38519 | 0.01230 | 0.79842 | 0.65141 | 0.16475 | 0.26280 | 0.08203 | 0.01213 | 0.76957 | 0.00491 | 0.05156 | 0.06349 | 3.26499 | 1.01126 |
| **Gm10941** | 0.01300 | 0.01212 | 0.02303 | 0.02270 | 0.01180 | 0.01937 | 0.01549 | 0.02350 | 0.03851 | 0.00619 | 0.00892 | 0.02050 | 0.07911 | 0.05338 |
| **Gm1110** | 0.38922 | 0.00419 | 0.73647 | 0.46957 | 1.23926 | 0.00000 | 0.00412 | 0.00549 | 0.71110 | 0.00429 | 0.00248 | 0.00521 | 0.00000 | 0.00000 |
| **Gm11127** | 16.37979 | 0.79000 | 3.83374 | 1.15637 | 8.99803 | 7.60903 | 0.77066 | 1.27804 | 7.25719 | 0.32384 | 2.35937 | 1.06902 | 8.63607 | 3.69022 |
| **Gm11166** | 0.00807 | 0.00378 | 0.02715 | 0.00000 | 0.00523 | 0.04174 | 0.89303 | 0.00272 | 0.00539 | 0.04629 | 0.00223 | 0.23093 | 0.00484 | 0.14473 |
| **Gm1123** | 0.00825 | 0.00445 | 0.01701 | 0.01226 | 0.00608 | 0.00896 | 0.00417 | 0.01055 | 0.50297 | 0.00748 | 0.00679 | 0.01022 | 0.00646 | 0.00019 |
| **Gm11238** | 0.02777 | 0.00418 | 0.01371 | 0.01182 | 0.01868 | 0.02178 | 0.00418 | 0.00758 | 0.00712 | 0.00627 | 0.00795 | 0.00784 | 1.41449 | 0.00329 |
| **Gm1140** | 0.32304 | 0.00411 | 0.02028 | 0.00000 | 0.00262 | 0.07901 | 0.00420 | 0.00919 | 0.03051 | 0.00491 | 0.24346 | 0.05319 | 0.03627 | 0.05572 |
| **Gm11413** | 0.00000 | 0.00421 | 0.50542 | 0.03277 | 0.26057 | 0.00316 | 0.00415 | 0.01188 | 0.02345 | 0.00561 | 0.00713 | 0.00522 | 1.12827 | 0.32425 |
| **Gm11423** | 0.67517 | 0.00362 | 0.00272 | 0.00000 | 0.10922 | 0.38185 | 0.00356 | 0.82984 | 0.00583 | 0.02983 | 0.00271 | 0.23672 | 0.29786 | 0.12114 |
| **Gm11487** | 0.20825 | 0.00403 | 0.00670 | 0.00682 | 0.05836 | 0.03351 | 0.00407 | 0.00687 | 0.23719 | 0.00608 | 0.00587 | 0.00609 | 2.51654 | 0.13546 |
| **Gm11517** | 28.20671 | 0.59742 | 13.57426 | 14.34294 | 27.10169 | 4.79347 | 0.49242 | 1.52850 | 26.49276 | 2.49258 | 1.60000 | 1.84064 | 24.68523 | 30.93802 |
| **Gm11532** | 0.00682 | 0.00439 | 0.01291 | 0.01484 | 0.01600 | 0.01205 | 0.00426 | 0.00994 | 0.01063 | 0.00732 | 0.00854 | 0.00943 | 0.12461 | 0.11814 |
| **Gm11544** | 4.16047 | 0.04399 | 4.66306 | 0.13252 | 4.17469 | 0.14344 | 0.04274 | 0.00514 | 3.39253 | 0.00000 | 0.09197 | 0.00000 | 13.55851 | 10.48459 |
| **Gm11545** | 16.85929 | 5.37468 | 31.79990 | 8.05576 | 18.75695 | 24.97423 | 10.19398 | 7.79738 | 29.08556 | 0.90026 | 15.08766 | 8.64508 | 8.07637 | 23.57672 |
| **Gm11549** | 0.06067 | 0.00415 | 0.02557 | 0.09509 | 0.03815 | 0.07026 | 0.00954 | 0.01361 | 0.04562 | 0.00845 | 0.02278 | 0.01067 | 0.47863 | 0.05470 |
| **Gm11665** | 0.00859 | 0.00442 | 0.11937 | 0.00353 | 0.00763 | 0.00998 | 0.00424 | 0.01114 | 0.02401 | 0.00713 | 0.00754 | 0.00967 | 0.02064 | 0.04081 |
| **Gm11696** | 0.03451 | 0.08523 | 0.01624 | 0.03594 | 0.01617 | 0.12043 | 0.02317 | 0.00701 | 0.23689 | 0.01899 | 0.85523 | 0.05283 | 0.23613 | 0.54918 |
| **Gm11744** | 0.00394 | 0.00458 | 0.03486 | 0.12976 | 0.17256 | 0.13655 | 0.00438 | 0.01228 | 0.00859 | 0.00886 | 0.00627 | 0.00595 | 0.07508 | 0.00315 |
| **Gm11944** | 0.16216 | 0.00480 | 0.12271 | 0.00000 | 0.26767 | 0.02739 | 0.00409 | 0.01503 | 0.34569 | 0.00608 | 0.00450 | 0.08461 | 0.00309 | 0.36131 |
| **Gm11961** | 0.00056 | 0.00414 | 0.00000 | 0.01779 | 0.39263 | 0.04233 | 0.00414 | 0.00457 | 0.00447 | 0.00607 | 0.00673 | 0.00673 | 0.75283 | 1.64587 |
| **Gm11992** | 0.00939 | 0.00444 | 0.01328 | 0.01299 | 0.01711 | 0.01316 | 0.00445 | 0.01017 | 0.00835 | 0.00691 | 0.00888 | 0.00998 | 0.00640 | 0.08199 |
| **Gm12018** | 0.02582 | 0.11472 | 0.02456 | 0.02464 | 0.01446 | 0.02859 | 0.01693 | 0.05243 | 0.41193 | 0.01052 | 0.00594 | 0.01492 | 0.08086 | 0.00000 |
| **Gm12060** | 0.45510 | 0.08798 | 0.14007 | 0.67616 | 0.58620 | 0.62543 | 0.22404 | 0.18202 | 0.52123 | 0.94208 | 0.00902 | 0.47610 | 0.55525 | 0.43862 |
| **Gm12070** | 0.28923 | 9.71841 | 5.80338 | 20.79040 | 1.25267 | 25.21449 | 7.05575 | 31.06790 | 0.11227 | 18.41903 | 13.87445 | 22.69191 | 5.79317 | 5.20970 |
| **Gm12169** | 0.00869 | 0.04984 | 0.12925 | 0.00000 | 0.00269 | 0.58932 | 0.01625 | 1.85371 | 0.01123 | 0.20896 | 0.14695 | 0.05297 | 0.00579 | 0.00784 |

| | | | | | | | | | | | | | | |
|---|---|---|---|---|---|---|---|---|---|---|---|---|---|---|
| **Gm12171** | 0.01580 | 0.00405 | 0.01077 | 0.00447 | 0.00911 | 0.04119 | 0.52013 | 0.09196 | 0.02324 | 0.01103 | 0.29304 | 0.00660 | 0.03832 | 0.00687 |
| **Gm12185** | 0.30035 | 0.05169 | 0.01813 | 0.15458 | 0.63290 | 0.02871 | 0.05561 | 0.08697 | 1.22106 | 0.00666 | 0.11842 | 0.16557 | 1.64017 | 2.54872 |
| **Gm12238** | 0.03030 | 0.00441 | 0.02273 | 0.01516 | 0.00494 | 0.01438 | 0.00417 | 0.00894 | 0.16484 | 0.00870 | 0.00719 | 0.00937 | 0.10321 | 0.01038 |
| **Gm12250** | 0.00000 | 0.00416 | 0.51196 | 0.05945 | 0.49679 | 0.00141 | 0.00394 | 0.01044 | 0.03072 | 0.00571 | 0.00593 | 0.00548 | 0.53498 | 0.20754 |
| **Gm12253** | 0.06299 | 0.01194 | 0.03120 | 0.02487 | 0.01153 | 0.02740 | 0.00416 | 0.00643 | 0.09081 | 0.00792 | 0.01373 | 0.02264 | 0.26746 | 0.03676 |
| **Gm12338** | 2.23486 | 1.39765 | 0.00000 | 1.22884 | 1.60595 | 3.12168 | 0.62464 | 4.52378 | 0.33267 | 2.75521 | 0.08145 | 2.99633 | 2.22919 | 0.76789 |
| **Gm12504** | 11.30344 | 6.53480 | 23.11412 | 26.81554 | 21.13037 | 20.02179 | 5.99017 | 16.80811 | 8.04136 | 10.62449 | 11.60850 | 16.08455 | 1.70300 | 9.11387 |
| **Gm12505** | 0.07289 | 0.00424 | 0.16738 | 0.13962 | 0.11148 | 0.09495 | 0.02173 | 0.05061 | 0.49305 | 0.00635 | 0.16561 | 0.12302 | 0.18915 | 0.12220 |
| **Gm12530** | 0.01198 | 0.00530 | 0.00964 | 0.10970 | 0.00922 | 0.00000 | 0.00441 | 0.04297 | 0.01263 | 0.00680 | 0.14061 | 0.01121 | 0.01122 | 0.03300 |
| **Gm12596** | 0.04313 | 0.00411 | 0.10719 | 0.06985 | 0.43368 | 0.00000 | 0.00382 | 0.00684 | 1.12259 | 0.01554 | 0.00635 | 0.01703 | 0.76396 | 0.90804 |
| **Gm12657** | 11.46411 | 3.09982 | 22.23290 | 19.46419 | 23.21386 | 14.84691 | 4.24828 | 8.26512 | 4.24898 | 11.69956 | 9.51385 | 8.88301 | 8.59153 | 5.77438 |
| **Gm12789** | 0.00000 | 0.00369 | 0.27267 | 0.26252 | 0.14191 | 0.37614 | 0.23039 | 0.01665 | 0.50679 | 0.01300 | 0.04700 | 0.07680 | 3.01550 | 0.68648 |
| **Gm12794** | 0.00000 | 0.00401 | 0.08504 | 0.00000 | 0.02952 | 0.18391 | 0.07515 | 0.00320 | 0.09093 | 0.00496 | 0.05964 | 0.02525 | 1.26770 | 0.05377 |
| **Gm128** | 0.19736 | 0.00421 | 0.53140 | 0.07218 | 0.42068 | 0.00893 | 0.00398 | 0.00546 | 0.00670 | 0.00595 | 0.00661 | 0.00554 | 0.40904 | 0.92049 |
| **Gm12992** | 0.01938 | 0.00528 | 0.00963 | 0.20721 | 0.07720 | 0.02389 | 0.00412 | 0.07112 | 0.07532 | 0.01002 | 0.00492 | 0.41646 | 0.01142 | 0.14080 |
| **Gm13023** | 0.15980 | 0.00306 | 0.43915 | 0.17391 | 1.07451 | 0.00139 | 0.00333 | 0.00414 | 4.71559 | 0.00492 | 0.00275 | 0.00000 | 7.41370 | 4.98619 |
| **Gm13034** | 0.03496 | 0.00515 | 0.10550 | 0.01171 | 0.15190 | 0.10673 | 0.00408 | 0.03169 | 2.08295 | 0.00718 | 0.09755 | 0.01592 | 0.31699 | 0.06617 |
| **Gm13078** | 0.00000 | 0.05618 | 1.33862 | 0.26355 | 0.12456 | 1.09049 | 0.15421 | 0.03814 | 0.84893 | 0.09066 | 0.41333 | 0.10733 | 3.37869 | 0.09358 |
| **Gm13083** | 0.13602 | 0.03506 | 1.98271 | 0.43479 | 0.16182 | 1.78679 | 0.23623 | 0.10431 | 1.42382 | 0.09388 | 0.78923 | 0.12736 | 1.44469 | 0.29740 |
| **Gm13084** | 0.00000 | 0.00379 | 0.15879 | 0.07423 | 0.50032 | 0.00000 | 0.00377 | 0.00568 | 0.22853 | 0.00656 | 0.00922 | 0.00861 | 2.06812 | 2.44344 |
| **Gm13088** | 0.00529 | 0.00440 | 0.01391 | 0.01762 | 0.00900 | 0.01004 | 0.00423 | 0.00969 | 0.09058 | 0.00678 | 0.00728 | 0.00894 | 0.08201 | 0.38970 |
| **Gm13102** | 22.58282 | 6.74298 | 26.44189 | 17.88056 | 17.23788 | 28.04762 | 6.74861 | 3.43240 | 29.83284 | 6.36519 | 12.07275 | 10.86704 | 8.94112 | 16.47775 |
| **Gm13103** | 0.00325 | 0.00434 | 0.01235 | 0.01917 | 0.00000 | 0.01008 | 0.00425 | 0.00907 | 0.03540 | 0.00644 | 0.00712 | 0.00851 | 0.13042 | 0.87401 |
| **Gm13119** | 0.00000 | 0.00414 | 0.04321 | 0.00000 | 0.19846 | 0.02855 | 0.00394 | 0.00620 | 0.62360 | 0.00652 | 0.00506 | 0.00635 | 1.21168 | 0.80974 |
| **Gm13125** | 0.24346 | 0.00440 | 0.00146 | 0.00703 | 0.04750 | 0.03453 | 0.00400 | 0.00723 | 1.24077 | 0.00736 | 0.00444 | 0.00787 | 0.14041 | 0.02748 |
| **Gm13128** | 0.11124 | 0.90428 | 0.24329 | 0.26322 | 0.41596 | 2.71335 | 3.46806 | 0.00831 | 1.29163 | 0.10469 | 0.54663 | 0.07804 | 1.19325 | 1.85411 |
| **Gm13212** | 3.32166 | 0.14596 | 9.21047 | 5.56893 | 8.74394 | 3.85369 | 0.33155 | 0.33628 | 27.94830 | 3.06848 | 3.58517 | 5.01449 | 8.86350 | 0.00000 |
| **Gm13238** | 0.00997 | 0.00481 | 0.04684 | 0.00607 | 0.01874 | 0.06468 | 0.01169 | 0.02447 | 0.02402 | 0.00635 | 0.00800 | 0.00870 | 0.15104 | 0.00465 |
| **Gm13315** | 0.00832 | 0.00447 | 0.01788 | 0.02699 | 0.01492 | 0.01413 | 0.00417 | 0.01174 | 0.01665 | 0.00721 | 0.01031 | 0.01954 | 0.04224 | 0.04087 |
| **Gm13375** | 0.00620 | 0.00425 | 0.07756 | 0.15545 | 0.21299 | 0.02031 | 0.00475 | 0.43848 | 0.01774 | 0.00633 | 0.00858 | 0.16971 | 0.01136 | 0.15230 |
| **Gm13498** | 6.57560 | 2.34831 | 13.15733 | 7.42768 | 5.90442 | 12.54191 | 1.70363 | 3.85679 | 6.85804 | 1.99616 | 6.93950 | 6.52659 | 0.00000 | 7.46646 |
| **Gm13648** | 0.13784 | 0.00522 | 0.55685 | 0.06766 | 0.13017 | 1.30453 | 0.00427 | 0.79013 | 0.89102 | 0.00775 | 0.00287 | 0.00000 | 0.85576 | 0.00000 |
| **Gm13752** | 0.22598 | 0.00442 | 0.06764 | 1.58569 | 0.34293 | 0.00000 | 0.00539 | 0.40662 | 0.00718 | 0.00507 | 0.00330 | 0.00540 | 0.00442 | 0.29134 |
| **Gm13830** | 0.06615 | 0.01182 | 0.01779 | 0.05397 | 0.05298 | 0.07942 | 0.00999 | 0.01617 | 0.02100 | 0.02856 | 0.01081 | 0.00987 | 0.16985 | 0.05317 |
| **Gm13884** | 0.00443 | 0.00443 | 0.10994 | 0.08099 | 0.07322 | 0.15584 | 0.00498 | 0.05246 | 0.00749 | 0.01427 | 0.00713 | 0.00426 | 0.00488 | 0.05822 |
| **Gm13985** | 0.15193 | 0.02137 | 0.02770 | 0.21788 | 0.09690 | 0.16507 | 0.07243 | 0.03503 | 0.15849 | 0.10104 | 0.05095 | 0.13151 | 0.36361 | 0.26875 |
| **Gm13986** | 0.03255 | 0.00440 | 0.01580 | 0.01513 | 0.04788 | 0.01331 | 0.00455 | 0.02083 | 0.08462 | 0.00753 | 0.01500 | 0.01114 | 0.01497 | 0.05910 |
| **Gm14005** | 0.65829 | 1.88507 | 0.06366 | 0.43124 | 0.24378 | 1.63677 | 0.13113 | 0.01244 | 1.47583 | 0.41544 | 0.07408 | 0.31847 | 5.70682 | 0.63819 |
| **Gm14023** | 0.00592 | 0.00409 | 0.00292 | 0.52133 | 0.01441 | 0.50491 | 0.00413 | 0.08993 | 0.01508 | 0.00641 | 0.01262 | 0.01354 | 0.01080 | 0.01321 |
| **Gm14057** | 0.90717 | 0.00488 | 0.70382 | 0.25148 | 0.97465 | 0.28280 | 0.06808 | 0.11865 | 0.02081 | 0.01668 | 0.04139 | 0.00000 | 1.02772 | 0.04482 |
| **Gm14085** | 0.05696 | 0.00443 | 0.02344 | 0.01524 | 0.25338 | 0.01492 | 0.00411 | 0.00897 | 0.01253 | 0.01277 | 0.00719 | 0.00833 | 0.01844 | 0.00497 |
| **Gm14124** | 0.37755 | 0.00369 | 0.28327 | 0.31791 | 0.32464 | 0.11621 | 1.20233 | 0.00018 | 1.80260 | 0.01554 | 0.25632 | 0.34022 | 0.09465 | 0.05759 |
| **Gm14137** | 0.00117 | 0.00442 | 0.00000 | 0.11616 | 0.00000 | 0.00000 | 0.00494 | 0.07045 | 0.17211 | 0.00634 | 0.00783 | 0.01216 | 0.22504 | 0.34627 |
| **Gm14139** | 0.17427 | 0.01332 | 0.31057 | 0.75295 | 0.43348 | 0.36036 | 0.58874 | 0.05096 | 2.82959 | 0.03715 | 1.43468 | 0.66783 | 0.00000 | 0.47303 |
| **Gm14164** | 0.00000 | 0.00444 | 0.13144 | 0.00000 | 0.39005 | 0.00384 | 0.00398 | 0.01010 | 0.67641 | 0.00828 | 0.00550 | 0.00680 | 0.31895 | 0.20438 |
| **Gm14204** | 0.05919 | 0.00445 | 0.04984 | 0.03822 | 0.10557 | 0.04804 | 0.03528 | 0.03553 | 0.20076 | 0.03069 | 0.01810 | 0.02590 | 0.47847 | 0.40303 |
| **Gm14288** | 0.00988 | 0.00434 | 0.06897 | 0.11653 | 0.00813 | 0.16079 | 0.01551 | 0.12324 | 0.04722 | 0.09208 | 0.12976 | 0.02514 | 0.04157 | 0.00000 |
| **Gm14295** | 0.00000 | 3.87684 | 1.30961 | 4.31998 | 0.09405 | 3.85188 | 3.21691 | 3.91230 | 1.06216 | 0.29065 | 5.63845 | 0.58216 | 0.16174 | 0.00000 |
| **Gm14305** | 0.06544 | 0.03648 | 0.04460 | 0.13965 | 0.05685 | 0.05246 | 0.08208 | 0.06985 | 0.50689 | 0.06548 | 0.13449 | 0.07253 | 0.00445 | 0.03623 |
| **Gm14322** | 2.57311 | 1.41039 | 1.46221 | 1.07740 | 1.39328 | 6.58324 | 1.78086 | 2.03110 | 3.40995 | 2.46699 | 2.71790 | 2.56496 | 0.94354 | 1.82591 |
| **Gm14325** | 3.32684 | 0.10658 | 2.69065 | 1.90005 | 3.51922 | 6.66867 | 1.11981 | 0.03633 | 5.52621 | 6.09452 | 3.08477 | 3.67311 | 0.87362 | 3.13847 |
| **Gm14326** | 0.83082 | 0.15616 | 0.47825 | 0.30496 | 0.44795 | 0.78152 | 0.60671 | 0.11146 | 1.38522 | 0.87700 | 1.32044 | 0.86896 | 0.12914 | 0.14903 |
| **Gm14327** | 0.01484 | 0.84140 | 0.01976 | 0.02777 | 0.05791 | 0.01734 | 0.14509 | 0.01128 | 0.01395 | 0.01154 | 0.02255 | 0.00632 | 0.00908 | 0.12225 |
| **Gm14391** | 0.01019 | 0.00526 | 0.02717 | 0.00000 | 0.01177 | 0.12651 | 0.00411 | 0.01047 | 0.03782 | 0.00672 | 0.17203 | 0.07291 | 0.00788 | 0.05791 |
| **Gm14393** | 0.09772 | 0.02186 | 0.08723 | 0.29787 | 0.08416 | 0.20736 | 0.16403 | 0.05250 | 0.26991 | 0.12709 | 0.12718 | 0.21797 | 0.05424 | 0.01677 |

| | | | | | | | | | | | | | | |
|---|---|---|---|---|---|---|---|---|---|---|---|---|---|---|
| **Gm14403** | 0.85409 | 0.37435 | 0.84205 | 1.57627 | 1.46606 | 6.56153 | 0.75541 | 0.08460 | 6.52842 | 6.91000 | 3.54129 | 1.66548 | 1.50979 | 8.38367 |
| **Gm14405** | 0.03843 | 0.00403 | 0.12425 | 0.36398 | 0.41398 | 0.09725 | 0.11291 | 0.03112 | 1.11633 | 0.48329 | 0.07189 | 0.13074 | 0.20931 | 0.09172 |
| **Gm14440** | 0.19653 | 0.51992 | 1.02049 | 0.93983 | 0.28685 | 1.74809 | 0.30469 | 1.56446 | 1.83561 | 0.04634 | 2.02962 | 0.15056 | 0.16249 | 0.00000 |
| **Gm14461** | 0.21665 | 0.02791 | 0.12423 | 0.08796 | 0.17150 | 0.13580 | 0.05872 | 0.05380 | 0.21904 | 0.15390 | 0.08682 | 0.07241 | 0.28962 | 0.29779 |
| **Gm14634** | 0.01069 | 0.00445 | 0.02699 | 0.01080 | 0.01182 | 0.01273 | 0.00427 | 0.01050 | 0.04121 | 0.00712 | 0.00904 | 0.00990 | 0.00643 | 0.00811 |
| **Gm14635** | 0.00373 | 0.00433 | 0.01435 | 0.01218 | 0.00847 | 0.01277 | 0.00422 | 0.00916 | 0.02775 | 0.00631 | 0.00886 | 0.00917 | 0.41871 | 0.00945 |
| **Gm14827** | 0.13987 | 0.00423 | 0.06232 | 0.10637 | 0.02513 | 0.02931 | 0.01590 | 0.00887 | 0.00776 | 0.01698 | 0.00950 | 0.01048 | 0.09871 | 0.07009 |
| **Gm14851** | 0.05253 | 0.00404 | 0.00957 | 0.06250 | 0.00980 | 0.02452 | 0.00420 | 0.00739 | 0.08421 | 0.00796 | 0.00747 | 0.06202 | 0.73380 | 0.00000 |
| **Gm15056** | 0.01008 | 0.00441 | 0.00750 | 0.02694 | 0.17829 | 0.01338 | 0.00412 | 0.00988 | 0.01937 | 0.00954 | 0.00786 | 0.00791 | 0.00897 | 0.08789 |
| **Gm15107** | 0.01390 | 0.00443 | 0.05752 | 0.01068 | 0.01043 | 0.01119 | 0.00427 | 0.01037 | 0.01107 | 0.00689 | 0.00982 | 0.01031 | 0.01364 | 0.03969 |
| **Gm1527** | 0.00630 | 0.00434 | 0.00490 | 0.00965 | 0.53035 | 0.01373 | 0.00418 | 0.00891 | 0.01128 | 0.00663 | 0.00943 | 0.00790 | 0.58644 | 0.01041 |
| **Gm15319** | 0.00586 | 0.00442 | 0.00866 | 0.01444 | 0.00416 | 0.01039 | 0.00420 | 0.00994 | 0.18258 | 0.00698 | 0.00767 | 0.00956 | 0.10580 | 0.05456 |
| **Gm15401** | 0.06044 | 0.03580 | 0.01607 | 0.48997 | 0.07361 | 0.17819 | 0.01586 | 0.07606 | 0.11090 | 0.05714 | 0.02542 | 0.05943 | 0.26758 | 0.17888 |
| **Gm15417** | 0.01671 | 0.00461 | 0.00911 | 0.01803 | 0.01369 | 0.02918 | 0.00464 | 0.04079 | 0.33598 | 0.01542 | 0.00837 | 0.01586 | 0.15112 | 0.10439 |
| **Gm15421** | 0.67502 | 0.22188 | 0.00000 | 2.17028 | 1.78698 | 1.50758 | 2.15182 | 4.45118 | 2.16075 | 2.28685 | 0.16851 | 3.29813 | 2.52289 | 0.45040 |
| **Gm15446** | 0.30580 | 0.01140 | 0.66745 | 1.21738 | 0.33293 | 0.34825 | 0.04183 | 1.56203 | 3.89046 | 0.23100 | 0.08115 | 0.27309 | 0.07012 | 0.01108 |
| **Gm15455** | 1.89134 | 0.01132 | 1.92454 | 1.27684 | 1.33045 | 1.12636 | 0.03694 | 1.19323 | 1.01234 | 0.00783 | 0.69286 | 0.49101 | 2.77931 | 5.92720 |
| **Gm15545** | 0.01657 | 0.00431 | 0.09997 | 0.02756 | 0.00753 | 0.00800 | 0.00422 | 0.00893 | 0.01049 | 0.00605 | 0.01191 | 0.01180 | 0.12561 | 0.04014 |
| **Gm15645** | 12.77482 | 1.96199 | 7.44323 | 4.33938 | 9.44726 | 7.12924 | 1.60306 | 3.81119 | 2.77460 | 1.35033 | 2.48624 | 2.73628 | 10.33261 | 10.72301 |
| **Gm15694** | 0.04589 | 0.04274 | 0.27834 | 0.46066 | 0.80276 | 0.15342 | 0.05699 | 0.05585 | 0.06137 | 0.14024 | 0.00184 | 0.73302 | 0.88038 | 0.67321 |
| **Gm15698** | 21.66067 | 2.12028 | 13.86343 | 13.63488 | 11.92807 | 19.61193 | 7.78649 | 12.27679 | 7.60564 | 19.12142 | 14.06198 | 16.38540 | 10.35596 | 9.63223 |
| **Gm15706** | 0.13928 | 0.00428 | 0.11225 | 0.00506 | 0.00505 | 0.01279 | 0.00426 | 0.00826 | 0.00956 | 0.01174 | 0.00629 | 0.04500 | 0.05654 | 0.00428 |
| **Gm15708** | 0.00743 | 0.00443 | 0.27478 | 0.00137 | 0.06205 | 0.00663 | 0.00413 | 0.01121 | 0.00769 | 0.00688 | 0.00702 | 0.00945 | 0.00513 | 0.03700 |
| **Gm15760** | 0.19927 | 0.02587 | 0.63512 | 0.37489 | 0.10590 | 0.00000 | 0.00988 | 1.40383 | 2.33024 | 0.01181 | 0.53451 | 0.01740 | 0.42032 | 0.79534 |
| **Gm15787** | 0.01070 | 0.00435 | 0.01399 | 0.00720 | 0.01191 | 0.00845 | 0.00429 | 0.01025 | 0.01590 | 0.00630 | 0.10293 | 0.02169 | 0.05242 | 0.02763 |
| **Gm15800** | 3.22628 | 0.04963 | 5.43041 | 0.01832 | 3.17000 | 4.14531 | 2.99639 | 1.93104 | 0.55193 | 0.04034 | 3.51772 | 3.01150 | 4.24101 | 3.26622 |
| **Gm15910** | 0.11216 | 0.02214 | 0.03451 | 0.08560 | 0.10899 | 0.07091 | 0.04706 | 0.03263 | 0.10964 | 0.02043 | 0.00799 | 0.05748 | 0.38347 | 0.07396 |
| **Gm15915** | 0.00603 | 0.00420 | 0.05538 | 0.12757 | 0.00857 | 0.00891 | 0.00408 | 0.00855 | 0.01243 | 0.00651 | 0.00657 | 0.01466 | 0.07305 | 0.00266 |
| **Gm15941** | 0.12608 | 0.00423 | 0.01360 | 0.01234 | 0.00982 | 0.01252 | 0.00414 | 0.00877 | 0.00737 | 0.00684 | 0.00763 | 0.00760 | 0.93153 | 0.00341 |
| **Gm1600** | 0.00882 | 0.00452 | 0.00335 | 0.19158 | 0.08982 | 0.03934 | 0.00453 | 0.01429 | 0.09617 | 0.00922 | 0.01258 | 0.01413 | 0.08701 | 0.01066 |
| **Gm16023** | 0.00386 | 0.00456 | 0.00640 | 0.14829 | 0.03006 | 0.01673 | 0.00451 | 0.02993 | 0.00702 | 0.00951 | 0.00951 | 0.00717 | 0.00361 | 0.32077 |
| **Gm16062** | 0.00707 | 0.00491 | 0.00502 | 0.13489 | 0.00459 | 0.09127 | 0.00421 | 0.01588 | 0.09680 | 0.00999 | 0.00731 | 0.01515 | 0.05745 | 0.02022 |
| **Gm16070** | 0.02928 | 0.00335 | 0.00000 | 0.09253 | 2.72110 | 0.03739 | 0.00375 | 0.01568 | 4.67175 | 0.00640 | 0.01548 | 0.04199 | 3.33166 | 14.00513 |
| **Gm16157** | 0.01151 | 0.00385 | 0.17991 | 0.00000 | 0.15533 | 0.01609 | 0.00408 | 0.00509 | 0.11329 | 0.00645 | 0.00374 | 0.54527 | 0.36756 | 0.00000 |
| **Gm16223** | 0.12490 | 0.00408 | 0.18082 | 0.08255 | 0.38974 | 0.42624 | 0.07550 | 0.07261 | 1.03337 | 0.00533 | 0.13296 | 0.05889 | 0.01657 | 0.44565 |
| **Gm16287** | 0.08950 | 0.09248 | 0.03280 | 0.17710 | 0.17878 | 0.19216 | 0.01645 | 0.23210 | 0.15609 | 0.04771 | 0.00856 | 0.02735 | 0.78668 | 0.23976 |
| **Gm16381** | 0.00000 | 0.00352 | 0.08869 | 0.00988 | 0.00000 | 0.00313 | 0.00364 | 0.01590 | 1.20137 | 0.00727 | 0.00532 | 0.01246 | 6.28016 | 0.00000 |
| **Gm16386** | 0.48943 | 0.00408 | 0.03741 | 0.02120 | 0.08339 | 0.02576 | 0.00444 | 0.00605 | 0.37067 | 0.00580 | 0.04244 | 0.05946 | 0.62785 | 0.00000 |
| **Gm16513** | 11.39550 | 0.01433 | 4.99649 | 3.26098 | 10.41408 | 3.07804 | 1.31824 | 0.00000 | 18.45475 | 0.10920 | 3.99873 | 2.72067 | 6.59141 | 6.05487 |
| **Gm16523** | 0.02858 | 0.02939 | 0.04756 | 0.21198 | 0.20094 | 0.24214 | 0.08440 | 0.10826 | 2.83725 | 0.04836 | 0.06714 | 0.04574 | 0.20862 | 0.03769 |
| **Gm1653** | 0.00727 | 0.00443 | 0.00000 | 0.01564 | 0.15262 | 0.00834 | 0.00428 | 0.01166 | 0.75835 | 0.00691 | 0.00629 | 0.00902 | 0.00363 | 0.00000 |
| **Gm16551** | 0.01050 | 0.00427 | 0.13603 | 0.00252 | 0.00499 | 0.00977 | 0.00421 | 0.00983 | 0.00820 | 0.00635 | 0.00774 | 0.00884 | 0.57725 | 0.02904 |
| **Gm16702** | 2.75662 | 0.61920 | 0.00000 | 0.91085 | 1.61586 | 4.03657 | 0.73268 | 0.87949 | 5.31923 | 1.33179 | 0.11584 | 1.87829 | 18.51239 | 7.06693 |
| **Gm16740** | 0.00984 | 0.00400 | 0.04381 | 0.00000 | 0.01172 | 0.02512 | 0.00967 | 0.07274 | 0.01731 | 0.01579 | 0.00709 | 0.01787 | 0.01257 | 0.01569 |
| **Gm16796** | 0.03479 | 0.00446 | 0.01843 | 0.01603 | 0.18447 | 0.01209 | 0.00433 | 0.01006 | 0.02330 | 0.00653 | 0.00800 | 0.00895 | 0.00442 | 0.01048 |
| **Gm16845** | 0.00417 | 0.00400 | 0.00929 | 0.57829 | 0.63820 | 0.03962 | 0.00437 | 0.00358 | 0.02820 | 0.00505 | 0.03060 | 0.02015 | 0.41528 | 0.00000 |
| **Gm16853** | 0.00357 | 0.00445 | 0.04543 | 0.00553 | 0.02816 | 0.00474 | 0.00534 | 0.12069 | 0.01095 | 0.00622 | 0.00906 | 0.00838 | 0.55160 | 0.00785 |
| **Gm16907** | 0.01083 | 0.01172 | 0.17179 | 0.00000 | 0.11144 | 0.00508 | 0.00401 | 0.00679 | 0.01160 | 0.00668 | 0.00480 | 0.31500 | 0.00836 | 0.31988 |
| **Gm16938** | 0.31107 | 0.02022 | 0.91594 | 2.00912 | 0.60392 | 0.48066 | 0.11231 | 0.92129 | 1.24442 | 0.01141 | 0.10526 | 0.26005 | 3.79268 | 1.09228 |
| **Gm16973** | 0.27024 | 0.02527 | 0.31868 | 0.37394 | 0.11457 | 0.12719 | 0.00430 | 0.10204 | 0.05179 | 0.00774 | 0.00404 | 0.04833 | 2.38644 | 0.13606 |
| **Gm17019** | 0.15033 | 0.00419 | 0.06635 | 0.12281 | 0.19103 | 0.01900 | 0.00394 | 0.00543 | 0.07801 | 0.00562 | 0.01822 | 0.01232 | 0.16626 | 0.09308 |
| **Gm17026** | 0.00395 | 0.00374 | 0.03194 | 0.07212 | 0.00352 | 0.09097 | 0.54193 | 0.11260 | 0.03198 | 0.00505 | 0.00198 | 0.17179 | 0.28153 | 0.05159 |
| **Gm17066** | 0.72154 | 0.01931 | 0.64404 | 1.23617 | 0.47003 | 0.70420 | 0.00295 | 0.07444 | 7.17084 | 0.00551 | 0.00000 | 1.04972 | 0.65983 | 0.09147 |
| **Gm17644** | 0.00000 | 0.15203 | 1.58924 | 1.64637 | 5.49330 | 0.23247 | 0.88599 | 1.16707 | 5.12546 | 0.05021 | 0.03327 | 0.94109 | 0.00000 | 2.09524 |
| **Gm17751** | 0.07808 | 0.00439 | 0.00971 | 0.01840 | 0.00277 | 0.01522 | 0.00423 | 0.00856 | 0.01260 | 0.00934 | 0.00698 | 0.00878 | 0.06853 | 0.30335 |

| | | | | | | | | | | | | | | |
|---|---|---|---|---|---|---|---|---|---|---|---|---|---|---|
| **Gm17762** | 0.00452 | 0.01322 | 0.01546 | 0.50617 | 0.00619 | 0.01610 | 0.01951 | 0.34674 | 0.01334 | 0.00626 | 0.13979 | 0.00565 | 0.00722 | 0.00655 |
| **Gm17801** | 0.05591 | 0.00448 | 0.00684 | 0.10171 | 0.03324 | 0.05905 | 0.00418 | 0.04801 | 0.04704 | 0.01309 | 0.02112 | 0.03114 | 0.27787 | 0.10884 |
| **Gm1821** | 16.43064 | 10.48972 | 26.14403 | 29.09326 | 15.91167 | 34.66512 | 9.09665 | 27.90422 | 6.96765 | 9.14200 | 23.87359 | 25.19546 | 14.82577 | 18.67713 |
| **Gm1943** | 10.25905 | 0.03141 | 9.12025 | 4.09416 | 13.86247 | 5.52238 | 0.31308 | 0.57118 | 2.12666 | 0.02258 | 0.83425 | 4.50908 | 8.47299 | 6.58857 |
| **Gm19434** | 0.25209 | 0.00395 | 0.02493 | 0.15941 | 0.52120 | 0.10725 | 0.05039 | 0.08552 | 0.72690 | 0.00592 | 0.08582 | 0.12884 | 1.07164 | 0.78368 |
| **Gm19510** | 0.00903 | 0.00448 | 0.01549 | 0.00685 | 0.01617 | 0.01445 | 0.03502 | 0.01244 | 0.01184 | 0.01462 | 0.00708 | 0.16760 | 0.01814 | 0.01354 |
| **Gm19557** | 0.00481 | 0.00443 | 0.05591 | 0.00725 | 0.00280 | 0.01016 | 0.00416 | 0.01031 | 0.14372 | 0.00839 | 0.00681 | 0.00932 | 0.00353 | 0.34034 |
| **Gm19583** | 0.00000 | 0.00408 | 0.17441 | 0.49924 | 0.23618 | 0.00000 | 0.00365 | 0.00406 | 0.34199 | 0.00601 | 0.00572 | 0.17917 | 0.00239 | 0.00000 |
| **Gm19589** | 0.01633 | 0.00523 | 0.01482 | 0.02156 | 0.05250 | 0.01678 | 0.00442 | 0.15501 | 0.05716 | 0.00760 | 0.13184 | 0.01399 | 0.31511 | 0.09345 |
| **Gm19619** | 0.01382 | 0.01203 | 0.04271 | 0.04533 | 0.03868 | 0.03625 | 0.01092 | 0.02298 | 0.15194 | 0.01233 | 0.04036 | 0.04104 | 0.00525 | 0.11716 |
| **Gm1965** | 0.00098 | 0.00366 | 0.11146 | 0.18526 | 0.00458 | 0.00424 | 0.00395 | 0.00805 | 0.15728 | 0.00561 | 0.00625 | 0.00446 | 6.14240 | 0.00000 |
| **Gm19705** | 0.00934 | 0.00402 | 0.00839 | 0.03849 | 0.25090 | 0.00946 | 0.00415 | 0.00536 | 0.01532 | 0.00767 | 0.00624 | 0.23719 | 0.15524 | 0.10095 |
| **Gm19710** | 0.01739 | 0.00448 | 0.04102 | 0.00000 | 0.05451 | 0.15188 | 0.00420 | 0.00579 | 0.01788 | 0.00569 | 0.00520 | 0.07386 | 0.02649 | 0.09051 |
| **Gm19757** | 0.58610 | 1.05746 | 1.60607 | 1.38354 | 6.64684 | 3.40750 | 0.42225 | 1.85458 | 13.27659 | 1.00176 | 2.38874 | 0.80035 | 2.09183 | 9.81343 |
| **Gm1976** | 0.04427 | 0.02057 | 0.09428 | 0.18335 | 0.03789 | 0.05871 | 0.01082 | 0.03185 | 0.20046 | 0.00718 | 0.12716 | 0.03589 | 0.55541 | 0.11461 |
| **Gm19784** | 0.00478 | 0.00439 | 0.21050 | 0.00486 | 0.00418 | 0.00613 | 0.00420 | 0.01089 | 0.03187 | 0.00758 | 0.00702 | 0.00889 | 0.10853 | 0.06535 |
| **Gm1995** | 1.53501 | 0.00282 | 1.13102 | 0.02185 | 2.35090 | 0.02609 | 0.00341 | 0.00503 | 0.83771 | 0.00461 | 0.01081 | 0.00628 | 14.96565 | 1.59829 |
| **Gm2011** | 0.01453 | 0.00448 | 0.03041 | 0.01809 | 0.04436 | 0.03737 | 0.00468 | 0.00884 | 0.01647 | 0.00701 | 0.03874 | 0.00724 | 0.01036 | 0.05716 |
| **Gm20110** | 0.06109 | 0.00455 | 0.01704 | 0.00802 | 0.04120 | 0.02361 | 0.00439 | 0.02023 | 0.04616 | 0.00640 | 0.00917 | 0.01456 | 0.00869 | 0.01773 |
| **Gm20125** | 0.01976 | 0.00474 | 0.01801 | 0.00964 | 0.06003 | 0.01129 | 0.00448 | 0.02417 | 0.05721 | 0.00735 | 0.01651 | 0.07029 | 0.00595 | 0.07255 |
| **Gm20139** | 0.01497 | 0.00422 | 0.01514 | 0.05600 | 0.01481 | 0.05052 | 0.00445 | 0.01996 | 0.03408 | 0.00669 | 0.01769 | 0.03304 | 0.06361 | 0.05622 |
| **Gm2016** | 0.12460 | 0.00380 | 0.05536 | 0.03970 | 0.16758 | 0.07853 | 0.02129 | 0.02512 | 2.21274 | 0.00684 | 0.00435 | 0.01804 | 3.80017 | 0.00000 |
| **Gm20172** | 1.62006 | 0.08774 | 0.41706 | 0.04744 | 1.31982 | 0.10179 | 0.05208 | 0.01625 | 1.14112 | 0.00568 | 0.05661 | 0.02072 | 1.16009 | 0.98775 |
| **Gm20187** | 0.05177 | 0.74154 | 0.04148 | 0.02923 | 0.04219 | 0.04523 | 0.00373 | 0.05750 | 0.03606 | 0.00586 | 0.01029 | 0.00901 | 0.07626 | 0.06382 |
| **Gm20199** | 0.00783 | 0.00471 | 0.00000 | 0.24533 | 0.00612 | 0.62024 | 0.00427 | 0.01041 | 0.10489 | 0.00742 | 0.03843 | 0.00315 | 0.02014 | 0.19211 |
| **Gm20219** | 0.00842 | 0.00397 | 0.04313 | 0.14035 | 0.10645 | 0.00815 | 0.00396 | 0.77290 | 0.01508 | 0.00667 | 0.00400 | 0.00522 | 0.10068 | 0.21100 |
| **Gm2022** | 0.23867 | 0.00357 | 0.00967 | 0.05284 | 0.07602 | 0.07347 | 0.02133 | 0.13364 | 0.82791 | 0.00696 | 0.03938 | 0.00712 | 4.45096 | 0.49923 |
| **Gm2027** | 1.02142 | 0.11441 | 0.14319 | 0.40369 | 0.12141 | 0.08254 | 0.10677 | 0.00591 | 1.84337 | 0.01356 | 0.06303 | 0.06579 | 2.24627 | 0.00000 |
| **Gm20300** | 0.42133 | 0.03953 | 0.35420 | 0.09461 | 0.06554 | 0.92615 | 0.13986 | 0.33107 | 0.63558 | 0.02635 | 0.01912 | 0.33375 | 0.00510 | 0.12404 |
| **Gm20324** | 0.50947 | 0.00503 | 0.00000 | 0.04811 | 0.00000 | 0.66118 | 0.98323 | 0.03329 | 0.15692 | 0.02270 | 0.00418 | 0.00565 | 0.00373 | 0.01710 |
| **Gm20337** | 0.66785 | 0.04177 | 0.00000 | 0.35897 | 1.20084 | 0.41300 | 0.02242 | 0.03831 | 0.89543 | 0.05654 | 0.64663 | 0.71787 | 1.57440 | 0.72572 |
| **Gm20356** | 0.00220 | 0.00454 | 0.00869 | 0.01424 | 0.02781 | 0.01491 | 0.00449 | 0.01737 | 0.14274 | 0.00775 | 0.00720 | 0.00896 | 0.01761 | 0.63171 |
| **Gm20362** | 2.96894 | 0.54886 | 3.55546 | 1.65225 | 0.70550 | 10.61896 | 5.40350 | 5.91909 | 2.56284 | 5.73531 | 2.24336 | 13.80833 | 0.06567 | 0.69068 |
| **Gm2042** | 0.18174 | 0.00308 | 0.09493 | 0.04955 | 1.80104 | 0.00000 | 0.00290 | 0.00378 | 7.46034 | 0.00615 | 0.00154 | 0.00068 | 4.28413 | 9.95997 |
| **Gm20594** | 14.07911 | 5.76660 | 19.31895 | 4.62927 | 21.47906 | 5.41764 | 2.19859 | 5.21558 | 39.25353 | 2.66476 | 9.90772 | 4.14222 | 6.33079 | 27.48757 |
| **Gm20597** | 0.02148 | 0.00463 | 0.02323 | 0.02037 | 0.01480 | 0.02144 | 0.00413 | 0.00998 | 0.09249 | 0.00667 | 0.02769 | 0.01993 | 0.04799 | 0.06433 |
| **Gm20605** | 0.00975 | 0.00457 | 0.04689 | 0.25731 | 0.10257 | 0.02059 | 0.00464 | 0.01840 | 0.01337 | 0.00687 | 0.00653 | 0.00809 | 0.00588 | 0.00657 |
| **Gm20611** | 0.01235 | 0.00448 | 0.04875 | 0.01338 | 0.04670 | 0.01051 | 0.00427 | 0.01092 | 0.00996 | 0.00630 | 0.00808 | 0.00892 | 0.01184 | 0.05142 |
| **Gm20750** | 0.00565 | 0.00447 | 0.04099 | 0.00000 | 0.75344 | 0.01939 | 0.00417 | 0.00951 | 0.01475 | 0.00699 | 0.01283 | 0.00829 | 0.00351 | 0.17632 |
| **Gm20752** | 0.05119 | 0.92206 | 0.04434 | 0.10415 | 0.06985 | 0.07542 | 0.24216 | 0.08577 | 0.06350 | 0.02318 | 0.01184 | 0.08723 | 0.19624 | 0.01425 |
| **Gm20759** | 0.00508 | 0.00437 | 0.01505 | 0.01246 | 0.01227 | 0.01331 | 0.00427 | 0.00939 | 0.01087 | 0.00650 | 0.00904 | 0.00938 | 0.29148 | 0.00415 |
| **Gm20767** | 3.60776 | 0.04900 | 1.17377 | 0.92061 | 4.26404 | 0.76574 | 1.17524 | 0.10739 | 7.99397 | 0.30162 | 0.14123 | 0.06452 | 21.32455 | 14.37325 |
| **Gm20939** | 4.07587 | 0.03428 | 6.54476 | 2.26379 | 6.71431 | 2.15855 | 0.92281 | 0.29787 | 14.76897 | 0.12576 | 3.55551 | 2.78325 | 0.16788 | 1.91646 |
| **Gm21119** | 0.00819 | 0.00443 | 0.01219 | 0.01311 | 0.00831 | 0.01166 | 0.00426 | 0.01023 | 0.07579 | 0.00672 | 0.00818 | 0.00984 | 0.05886 | 0.04981 |
| **Gm21190** | 0.04341 | 0.00426 | 0.04839 | 0.05289 | 0.13489 | 0.03862 | 0.00406 | 0.00576 | 0.06729 | 0.00585 | 0.01038 | 0.01002 | 0.30649 | 0.04439 |
| **Gm21283** | 0.00000 | 0.00363 | 0.10799 | 0.17847 | 0.00248 | 0.00000 | 0.00479 | 0.53372 | 0.01597 | 0.00493 | 0.39425 | 1.15310 | 0.00600 | 0.00687 |
| **Gm21284** | 0.10265 | 0.05101 | 0.01463 | 0.18314 | 0.11047 | 0.57105 | 0.05645 | 0.07097 | 0.23249 | 0.01881 | 0.03788 | 0.10668 | 0.61193 | 0.41611 |
| **Gm21319** | 0.00000 | 0.01926 | 0.01075 | 0.10594 | 0.16488 | 0.05208 | 0.01562 | 0.01705 | 0.47844 | 0.01457 | 0.02142 | 0.02270 | 1.69190 | 1.72092 |
| **Gm21637** | 0.24076 | 0.00404 | 0.12246 | 0.00000 | 0.19710 | 0.04511 | 0.00398 | 0.03285 | 0.21053 | 0.00642 | 0.00552 | 0.00564 | 0.40510 | 0.08248 |
| **Gm2381** | 0.61201 | 2.44272 | 0.92674 | 0.56227 | 1.32379 | 0.75665 | 0.00775 | 1.12703 | 4.42018 | 0.62738 | 0.53402 | 0.38096 | 3.43428 | 1.66756 |
| **Gm2518** | 0.06385 | 0.00472 | 0.00965 | 0.06173 | 0.03130 | 0.01660 | 0.00415 | 0.02436 | 0.01116 | 0.01377 | 0.00760 | 0.00829 | 0.00828 | 0.03146 |
| **Gm2560** | 0.00600 | 0.00444 | 0.11145 | 0.00000 | 0.51486 | 0.00653 | 0.01082 | 0.01600 | 0.00897 | 0.02214 | 0.00738 | 0.00844 | 0.01739 | 0.14024 |
| **Gm26705** | 0.03997 | 0.02344 | 0.01970 | 0.01704 | 0.03153 | 0.01743 | 0.00433 | 0.10031 | 0.15622 | 0.00817 | 0.00945 | 0.02261 | 0.04502 | 0.03588 |
| **Gm2694** | 0.03021 | 0.00413 | 0.01316 | 0.19398 | 0.24740 | 0.05896 | 0.00402 | 0.00389 | 0.08041 | 0.00551 | 0.00700 | 0.01311 | 0.86946 | 0.00000 |
| **Gm28042** | 0.01149 | 0.00388 | 0.03813 | 0.00000 | 0.01229 | 0.00682 | 0.00405 | 0.09327 | 0.01269 | 0.00533 | 0.20673 | 0.03474 | 0.00751 | 0.01023 |

| | | | | | | | | | | | | | | |
|---|---|---|---|---|---|---|---|---|---|---|---|---|---|---|
| **Gm2837** | 0.48997 | 0.00382 | 0.19577 | 0.00000 | 0.31508 | 0.03234 | 0.00383 | 0.12919 | 0.35348 | 0.00630 | 0.00462 | 0.00358 | 0.74220 | 0.22713 |
| **Gm2897** | 0.01532 | 0.00457 | 0.01276 | 0.08127 | 0.06266 | 0.01294 | 0.00416 | 0.01140 | 0.01485 | 0.00658 | 0.00909 | 0.03936 | 0.03067 | 0.00998 |
| **Gm29443** | 0.03098 | 0.00454 | 0.02834 | 0.04946 | 0.02988 | 0.06517 | 0.01143 | 0.04517 | 0.18957 | 0.07089 | 0.00601 | 0.01905 | 0.99360 | 0.08941 |
| **Gm29683** | 0.06516 | 0.03906 | 0.02841 | 0.06094 | 0.10266 | 0.05777 | 0.08396 | 0.04541 | 0.30308 | 0.01666 | 0.01341 | 0.02921 | 0.04392 | 0.14879 |
| **Gm29811** | 0.00221 | 0.00411 | 0.01455 | 0.01264 | 0.00620 | 0.01256 | 0.00410 | 0.00829 | 0.01984 | 0.00618 | 0.00923 | 0.00768 | 1.94253 | 0.00419 |
| **Gm29968** | 0.03354 | 0.11250 | 0.03144 | 0.05194 | 0.02824 | 0.06339 | 0.03650 | 0.03199 | 0.45584 | 0.00713 | 0.02083 | 0.01185 | 0.05606 | 0.11699 |
| **Gm2a** | 9.72860 | 10.65253 | 7.32000 | 17.70122 | 11.43286 | 24.21235 | 6.99737 | 33.38020 | 13.99534 | 9.36027 | 6.73117 | 1.26506 | 5.77616 | 15.60283 |
| **Gm3002** | 0.02245 | 0.00467 | 0.00884 | 0.10311 | 0.03814 | 0.01486 | 0.00407 | 0.00944 | 0.04609 | 0.00649 | 0.00664 | 0.04942 | 0.02794 | 0.00647 |
| **Gm30505** | 0.01108 | 0.00441 | 0.06910 | 0.02850 | 0.04487 | 0.02562 | 0.00421 | 0.00818 | 0.00817 | 0.00642 | 0.00724 | 0.01179 | 0.04494 | 0.02535 |
| **Gm30551** | 0.02305 | 0.00484 | 0.01212 | 0.04541 | 0.04031 | 0.03295 | 0.00456 | 0.06879 | 0.03243 | 0.00687 | 0.01649 | 0.01486 | 0.10635 | 0.01686 |
| **Gm3086** | 0.00956 | 0.00449 | 0.00532 | 0.01357 | 0.08196 | 0.01146 | 0.00432 | 0.01452 | 0.35886 | 0.00654 | 0.00689 | 0.00887 | 0.00741 | 0.03726 |
| **Gm3139** | 22.13158 | 1.17383 | 29.90655 | 25.62371 | 25.85882 | 19.21328 | 9.35613 | 2.41305 | 33.77648 | 8.44359 | 17.84418 | 15.81467 | 12.85247 | 28.46250 |
| **Gm3143** | 0.01004 | 0.01160 | 0.24652 | 0.10861 | 0.10705 | 0.00831 | 0.01548 | 0.00455 | 0.03338 | 0.00608 | 0.00621 | 0.10345 | 0.00687 | 0.44629 |
| **Gm3219** | 0.22414 | 0.00370 | 0.04072 | 0.07479 | 1.02317 | 0.16194 | 0.00379 | 0.61779 | 0.00541 | 0.02322 | 0.34657 | 0.82231 | 1.67671 | 0.50445 |
| **Gm32511** | 0.75641 | 0.00487 | 0.00000 | 0.30002 | 0.04602 | 0.19683 | 0.00457 | 0.21939 | 0.05114 | 0.00895 | 0.00409 | 0.00294 | 0.13173 | 0.70412 |
| **Gm3259** | 0.03816 | 0.00414 | 0.00000 | 0.06468 | 0.19461 | 0.00000 | 0.00404 | 0.01443 | 0.84925 | 0.03140 | 0.00793 | 0.00561 | 0.49479 | 0.75732 |
| **Gm3264** | 0.05913 | 0.00462 | 0.01434 | 0.03474 | 0.04252 | 0.01515 | 0.00420 | 0.01006 | 0.07495 | 0.00616 | 0.00670 | 0.02016 | 0.00650 | 0.00844 |
| **Gm3286** | 2.95465 | 0.00053 | 6.44623 | 4.14190 | 6.96177 | 2.78027 | 2.42019 | 0.00000 | 13.25681 | 1.99993 | 3.87587 | 1.12883 | 4.80480 | 8.92709 |
| **Gm3336** | 0.10480 | 0.00434 | 0.30083 | 0.01713 | 0.01949 | 0.00269 | 0.00405 | 0.01113 | 0.00593 | 0.01471 | 0.00624 | 0.00748 | 0.00921 | 0.13374 |
| **Gm3383** | 0.07722 | 0.01232 | 0.00748 | 0.13526 | 0.13111 | 0.01514 | 0.00387 | 0.00666 | 0.08225 | 0.00645 | 0.00663 | 0.14274 | 0.02334 | 0.01956 |
| **Gm3414** | 0.08681 | 0.07410 | 0.10993 | 0.15467 | 0.04602 | 0.08406 | 0.02788 | 0.05452 | 0.05337 | 0.06095 | 0.01253 | 0.00437 | 0.15244 | 0.03301 |
| **Gm3435** | 0.05618 | 0.00425 | 0.23805 | 0.47727 | 0.00000 | 0.16337 | 0.18265 | 0.07613 | 0.57505 | 0.02174 | 0.74648 | 0.72229 | 0.02150 | 0.30669 |
| **Gm3500** | 0.04247 | 0.00441 | 0.01075 | 0.12284 | 0.09784 | 0.00596 | 0.00389 | 0.00729 | 0.06410 | 0.00608 | 0.00891 | 0.09307 | 0.00921 | 0.00960 |
| **Gm3558** | 0.01626 | 0.00470 | 0.01573 | 0.07757 | 0.08272 | 0.02333 | 0.00400 | 0.00891 | 0.02451 | 0.00687 | 0.00618 | 0.09344 | 0.03833 | 0.02002 |
| **Gm3604** | 1.24554 | 0.03942 | 1.11507 | 1.73305 | 0.86157 | 2.34829 | 1.12800 | 0.11115 | 6.43809 | 0.05887 | 0.32718 | 0.84167 | 1.61257 | 0.00000 |
| **Gm3636** | 0.05597 | 0.00470 | 0.01290 | 0.02241 | 0.07243 | 0.02437 | 0.00409 | 0.01000 | 0.06505 | 0.00794 | 0.00654 | 0.03701 | 0.09592 | 0.00714 |
| **Gm364** | 3.18870 | 0.00371 | 1.08512 | 0.71836 | 0.52763 | 0.00000 | 0.00308 | 0.00245 | 5.22961 | 0.00436 | 0.00260 | 0.00476 | 0.00239 | 0.90148 |
| **Gm3696** | 0.03729 | 0.00452 | 0.00893 | 0.06805 | 0.05428 | 0.01314 | 0.00404 | 0.00827 | 0.04088 | 0.00612 | 0.00811 | 0.04041 | 0.00729 | 0.01380 |
| **Gm3776** | 0.32550 | 0.02007 | 1.12695 | 0.25674 | 1.33218 | 0.19651 | 0.22748 | 1.86163 | 1.28740 | 0.20754 | 0.61620 | 0.02998 | 0.00000 | 0.05894 |
| **Gm38407** | 0.16663 | 0.00436 | 0.01340 | 0.01051 | 0.00530 | 0.01174 | 0.00431 | 0.00991 | 0.00612 | 0.00620 | 0.00768 | 0.00845 | 0.30325 | 0.04175 |
| **Gm38415** | 0.02523 | 0.00403 | 0.13600 | 0.04039 | 0.00000 | 0.15539 | 0.01002 | 0.00321 | 0.01565 | 0.00487 | 0.01297 | 0.01023 | 1.48004 | 0.83739 |
| **Gm38416** | 0.01219 | 0.01187 | 0.00941 | 0.01787 | 0.36816 | 0.01306 | 0.00422 | 0.01735 | 0.01734 | 0.00640 | 0.01354 | 0.01116 | 0.00797 | 0.01434 |
| **Gm38426** | 0.00681 | 0.00419 | 0.00000 | 0.10837 | 0.22907 | 0.45355 | 0.00425 | 0.01331 | 0.01037 | 0.01093 | 0.00570 | 0.08732 | 0.00617 | 0.00704 |
| **Gm3893** | 0.00873 | 0.00464 | 0.00883 | 0.14470 | 0.06117 | 0.03770 | 0.00486 | 0.06809 | 0.00952 | 0.00858 | 0.00931 | 0.00567 | 0.02582 | 0.00800 |
| **Gm40893** | 0.03773 | 0.23303 | 0.04024 | 0.00000 | 0.01154 | 0.18665 | 0.00960 | 0.08216 | 0.23269 | 0.02442 | 0.01078 | 0.54255 | 0.19206 | 0.09974 |
| **Gm4224** | 0.00971 | 0.00444 | 0.01657 | 0.01673 | 0.01315 | 0.01648 | 0.00428 | 0.01014 | 0.01142 | 0.00787 | 0.00893 | 0.00929 | 0.07472 | 0.01716 |
| **Gm4262** | 17.12739 | 1.02667 | 4.63395 | 0.98737 | 9.41628 | 9.45454 | 0.82802 | 1.44882 | 10.04848 | 0.44379 | 2.89465 | 1.16841 | 9.70452 | 5.59188 |
| **Gm4285** | 0.03526 | 0.02300 | 0.05570 | 0.10036 | 0.04194 | 0.11351 | 0.03505 | 0.14265 | 1.25957 | 0.01966 | 0.00763 | 0.04338 | 0.09091 | 0.09884 |
| **Gm4349** | 0.50354 | 0.00971 | 0.20741 | 0.33444 | 0.00000 | 0.71641 | 1.00595 | 0.00697 | 1.17666 | 0.00986 | 0.07549 | 1.35564 | 2.25905 | 1.14325 |
| **Gm4371** | 0.00527 | 0.00427 | 0.00641 | 0.00301 | 0.47701 | 0.01445 | 0.00437 | 0.00821 | 0.01184 | 0.00607 | 0.00718 | 0.11497 | 0.00673 | 0.11563 |
| **Gm4425** | 0.19706 | 0.00427 | 0.28412 | 0.11416 | 0.65819 | 0.53486 | 0.16995 | 0.08261 | 1.28992 | 0.02218 | 0.55480 | 0.22811 | 0.05191 | 0.00000 |
| **Gm4432** | 0.00000 | 0.00454 | 0.01139 | 0.06882 | 0.00000 | 0.00883 | 0.00443 | 0.02947 | 0.45509 | 0.00846 | 0.00735 | 0.00913 | 0.32641 | 0.48195 |
| **Gm4532** | 0.00685 | 0.00425 | 0.00785 | 0.05026 | 0.10185 | 0.04693 | 0.00433 | 0.00815 | 0.02273 | 0.00630 | 0.00745 | 0.00707 | 0.29320 | 0.01177 |
| **Gm4541** | 0.01092 | 0.00440 | 0.01530 | 0.02137 | 0.09002 | 0.01547 | 0.00423 | 0.01269 | 0.23485 | 0.00669 | 0.01028 | 0.01018 | 0.01644 | 0.00000 |
| **Gm4566** | 0.00733 | 0.00439 | 0.01486 | 0.01225 | 0.01182 | 0.01319 | 0.00429 | 0.00971 | 0.01070 | 0.00659 | 0.00908 | 0.00972 | 0.16657 | 0.00930 |
| **Gm4567** | 0.04558 | 0.01246 | 0.10251 | 0.04485 | 0.05560 | 0.00980 | 0.00454 | 0.01526 | 0.21254 | 0.00600 | 0.00933 | 0.00832 | 0.04426 | 0.08608 |
| **Gm4598** | 0.00580 | 0.00381 | 0.00991 | 0.00000 | 0.01169 | 0.03194 | 0.00431 | 0.36393 | 0.01212 | 0.00791 | 0.00127 | 0.58424 | 0.00707 | 0.01115 |
| **Gm4737** | 1.38137 | 0.10884 | 11.78440 | 8.67527 | 2.71736 | 9.85706 | 0.87838 | 4.34562 | 0.46447 | 0.05711 | 9.28079 | 12.30514 | 0.00000 | 3.19541 |
| **Gm4791** | 0.01210 | 0.00438 | 0.01680 | 0.01917 | 0.01856 | 0.01331 | 0.01020 | 0.00840 | 0.06577 | 0.00752 | 0.03751 | 0.01690 | 0.00534 | 0.05112 |
| **Gm4827** | 0.00459 | 0.00422 | 0.01147 | 0.02712 | 0.00939 | 0.10492 | 0.00409 | 0.00445 | 0.01163 | 0.00541 | 0.00815 | 0.01194 | 0.42268 | 0.00035 |
| **Gm4832** | 0.03689 | 0.03753 | 0.07503 | 0.26680 | 0.20447 | 0.16685 | 0.02912 | 0.08734 | 0.12447 | 0.10519 | 0.00366 | 0.06689 | 0.76520 | 0.01210 |
| **Gm4841** | 0.15678 | 0.02863 | 0.05574 | 0.08037 | 0.11560 | 0.04626 | 0.00439 | 0.04873 | 0.11007 | 0.01490 | 0.07574 | 0.02502 | 0.03638 | 0.08047 |
| **Gm4850** | 0.00000 | 0.00370 | 0.17101 | 0.00000 | 0.60959 | 0.00000 | 0.00391 | 0.00735 | 0.42566 | 0.00614 | 0.02520 | 0.00650 | 5.03332 | 1.38759 |
| **Gm4858** | 0.03641 | 0.00436 | 0.00477 | 0.01743 | 0.00000 | 0.00585 | 0.01555 | 0.01718 | 0.50687 | 0.00777 | 0.00667 | 0.00735 | 0.14179 | 2.46678 |
| **Gm4871** | 0.00001 | 0.00442 | 0.09120 | 0.00000 | 0.10860 | 0.00874 | 0.00410 | 0.01050 | 0.43926 | 0.00715 | 0.00630 | 0.00847 | 0.16773 | 0.11572 |

| | | | | | | | | | | | | | | |
|---|---|---|---|---|---|---|---|---|---|---|---|---|---|---|
| **Gm4922** | 0.05789 | 0.00460 | 0.04754 | 0.01005 | 0.09581 | 0.08622 | 0.09117 | 0.05059 | 0.07740 | 0.00668 | 0.01213 | 0.06243 | 0.02521 | 0.11723 |
| **Gm4925** | 0.07567 | 0.11872 | 0.08143 | 0.13648 | 0.14244 | 0.17564 | 0.03130 | 0.02109 | 0.10200 | 0.04948 | 0.01252 | 0.04908 | 0.14124 | 0.15991 |
| **Gm4926** | 1.24403 | 6.41675 | 1.14035 | 0.42953 | 0.15042 | 4.77930 | 0.40876 | 14.63484 | 3.38127 | 0.03546 | 1.16525 | 0.10227 | 0.69440 | 0.00000 |
| **Gm4951** | 0.08195 | 0.08728 | 0.01254 | 0.12593 | 0.05318 | 0.14688 | 0.07584 | 0.05737 | 0.12678 | 0.01847 | 0.03139 | 0.04802 | 0.26439 | 0.17993 |
| **Gm4956** | 0.00805 | 0.00436 | 0.01674 | 0.01604 | 0.00607 | 0.00996 | 0.00426 | 0.00986 | 0.02550 | 0.00635 | 0.00870 | 0.01026 | 0.10776 | 0.43981 |
| **Gm4961** | 4.96460 | 0.00346 | 4.42749 | 0.91854 | 3.42000 | 1.26753 | 0.09292 | 0.67272 | 1.35579 | 0.72260 | 0.25550 | 0.08094 | 2.68429 | 2.74913 |
| **Gm4971** | 0.00560 | 0.00490 | 0.03478 | 0.00725 | 0.01624 | 0.01056 | 0.01696 | 0.04422 | 0.03354 | 0.00576 | 0.00566 | 0.14700 | 1.14841 | 0.02287 |
| **Gm4984** | 0.00828 | 0.00438 | 0.00000 | 0.49106 | 0.04778 | 0.24264 | 0.00398 | 0.00463 | 0.02682 | 0.00643 | 0.00689 | 0.00551 | 0.11014 | 0.20117 |
| **Gm5** | 0.04430 | 0.00363 | 0.01093 | 0.01071 | 0.57761 | 0.00725 | 0.00390 | 0.00743 | 0.00000 | 0.00667 | 0.00665 | 0.00425 | 5.97015 | 1.57888 |
| **Gm5039** | 0.01936 | 0.00363 | 0.45683 | 0.07327 | 0.67537 | 0.39214 | 0.16352 | 0.07874 | 2.03663 | 0.01121 | 0.26684 | 0.04157 | 4.09062 | 0.20768 |
| **Gm5065** | 0.01525 | 0.00433 | 0.01865 | 0.02347 | 0.02007 | 0.02193 | 0.05684 | 0.00600 | 0.03903 | 0.00810 | 0.01819 | 0.00996 | 0.15380 | 0.00351 |
| **Gm5069** | 0.00000 | 1.66915 | 0.36399 | 2.05473 | 0.02087 | 3.47248 | 0.75474 | 7.35332 | 0.01118 | 3.61729 | 1.84209 | 3.93771 | 0.15214 | 0.31706 |
| **Gm5071** | 0.00686 | 0.00436 | 0.01755 | 0.01183 | 0.03050 | 0.01220 | 0.00433 | 0.01072 | 0.01094 | 0.00628 | 0.00888 | 0.00975 | 0.18561 | 0.06979 |
| **Gm5072** | 0.01086 | 0.00434 | 0.02849 | 0.01005 | 0.03016 | 0.01148 | 0.00432 | 0.01085 | 0.01078 | 0.00617 | 0.00862 | 0.00942 | 0.28317 | 0.10502 |
| **Gm5088** | 4.26200 | 5.54343 | 18.54412 | 30.05988 | 28.03999 | 2.42445 | 2.39181 | 6.82744 | 2.10636 | 1.10444 | 1.06320 | 7.03626 | 9.67664 | 2.56199 |
| **Gm5113** | 0.64199 | 0.18440 | 0.25559 | 0.31122 | 0.49855 | 0.62558 | 0.04760 | 0.02445 | 0.59830 | 0.02765 | 0.12798 | 0.10136 | 0.42661 | 1.15609 |
| **Gm5114** | 0.00784 | 0.00450 | 0.01264 | 0.03634 | 0.03792 | 0.07422 | 0.00439 | 0.00656 | 0.00809 | 0.00562 | 0.00835 | 0.00770 | 0.00649 | 0.05879 |
| **Gm5127** | 0.00567 | 0.00437 | 0.01000 | 0.00151 | 0.00905 | 0.25958 | 0.00410 | 0.00563 | 0.08386 | 0.00626 | 0.00606 | 0.01087 | 0.12612 | 0.00000 |
| **Gm5129** | 0.00938 | 0.00438 | 0.01867 | 0.04818 | 0.03443 | 0.01110 | 0.00419 | 0.00868 | 0.01125 | 0.00652 | 0.00806 | 0.01084 | 0.00662 | 0.01174 |
| **Gm5134** | 0.00000 | 0.00992 | 2.77291 | 0.00000 | 1.14042 | 1.08550 | 0.00285 | 0.00000 | 1.78833 | 0.00560 | 0.11038 | 0.80616 | 3.16220 | 1.39418 |
| **Gm5141** | 0.15887 | 0.04974 | 0.17378 | 0.20238 | 0.38181 | 0.38249 | 0.54288 | 0.02022 | 2.83082 | 0.10535 | 0.15851 | 0.22461 | 0.76199 | 0.00000 |
| **Gm5148** | 12.39546 | 8.16375 | 3.06085 | 14.75630 | 22.59057 | 19.99153 | 7.86065 | 19.07675 | 4.77560 | 17.55907 | 1.58777 | 18.63517 | 12.00357 | 11.17177 |
| **Gm5176** | 0.83226 | 0.36188 | 1.65421 | 0.41787 | 1.33127 | 0.81436 | 0.40855 | 0.49901 | 1.36284 | 2.43696 | 1.11471 | 0.90990 | 1.73087 | 1.68983 |
| **Gm5177** | 0.01592 | 0.79353 | 0.13184 | 1.11224 | 0.04534 | 1.77403 | 0.36107 | 3.52598 | 0.00665 | 1.71144 | 0.75209 | 2.00099 | 0.49093 | 0.14393 |
| **Gm525** | 0.01749 | 0.00474 | 0.01348 | 0.01218 | 0.00957 | 0.01148 | 0.00437 | 0.01741 | 0.00823 | 0.00738 | 0.00871 | 0.00996 | 0.00679 | 0.11103 |
| **Gm527** | 0.37964 | 0.00383 | 0.02013 | 0.35160 | 0.44893 | 0.02810 | 0.00409 | 0.00407 | 0.00842 | 0.00617 | 0.02696 | 0.25631 | 1.94224 | 0.24194 |
| **Gm53** | 0.00759 | 0.00435 | 0.02077 | 0.04677 | 0.48748 | 0.01284 | 0.00411 | 0.00778 | 0.01124 | 0.00651 | 0.00741 | 0.01392 | 0.00502 | 0.00855 |
| **Gm5415** | 0.00993 | 0.00400 | 0.02108 | 0.00621 | 0.01092 | 0.06930 | 0.00415 | 0.05064 | 0.06581 | 0.00700 | 0.00660 | 0.00959 | 0.03415 | 0.03410 |
| **Gm5431** | 0.04835 | 0.00428 | 0.02168 | 0.01858 | 0.09985 | 0.04021 | 0.15754 | 0.00638 | 0.07034 | 0.00705 | 0.02531 | 0.07293 | 0.04932 | 0.01989 |
| **Gm5434** | 0.14163 | 0.01297 | 0.52893 | 0.45146 | 0.32192 | 0.52671 | 0.20500 | 0.10801 | 0.05792 | 0.00668 | 0.11267 | 0.13475 | 0.02984 | 0.41833 |
| **Gm5458** | 0.25965 | 0.00439 | 0.00960 | 0.06604 | 0.12362 | 0.07242 | 0.00415 | 0.00519 | 0.25771 | 0.00629 | 0.00612 | 0.01089 | 0.30133 | 0.00000 |
| **Gm5475** | 0.00691 | 0.00439 | 0.03769 | 0.00724 | 0.04790 | 0.01112 | 0.00443 | 0.00930 | 0.18324 | 0.00618 | 0.03642 | 0.00771 | 0.40329 | 0.00645 |
| **Gm5485** | 0.40345 | 0.00439 | 0.11330 | 0.00757 | 0.00000 | 0.00475 | 0.00413 | 0.01207 | 0.15243 | 0.00686 | 0.00587 | 0.00830 | 0.00559 | 0.10334 |
| **Gm5512** | 0.06468 | 1.89749 | 0.35078 | 0.03824 | 0.18849 | 1.08279 | 0.48043 | 0.29420 | 0.00409 | 0.02399 | 0.08139 | 0.80265 | 0.31688 | 0.31867 |
| **Gm5523** | 0.00000 | 2.10893 | 0.47425 | 2.78096 | 0.06278 | 4.98470 | 1.17969 | 8.94269 | 0.00000 | 5.07803 | 2.31781 | 5.49253 | 0.42491 | 0.34918 |
| **Gm5532** | 0.00000 | 0.00408 | 0.70683 | 0.31875 | 0.67802 | 0.00000 | 0.00381 | 0.00662 | 0.32963 | 0.00592 | 0.00383 | 0.00634 | 0.30535 | 0.63298 |
| **Gm5547** | 0.00506 | 0.00401 | 0.33116 | 0.14439 | 0.08543 | 0.00000 | 0.00384 | 0.00605 | 0.01006 | 0.00481 | 0.38992 | 0.03811 | 0.00461 | 0.00750 |
| **Gm5577** | 0.02290 | 0.00397 | 0.02686 | 0.34350 | 0.00853 | 0.00969 | 0.05792 | 0.00361 | 0.06274 | 0.00490 | 0.10608 | 0.02070 | 0.44946 | 0.00293 |
| **Gm5607** | 0.19674 | 0.03697 | 0.08324 | 0.21731 | 0.13794 | 0.19173 | 0.07308 | 0.14961 | 0.49164 | 0.26591 | 0.01178 | 0.09506 | 0.58779 | 0.58548 |
| **Gm561** | 0.91459 | 0.01128 | 0.00000 | 0.00000 | 0.26735 | 0.53793 | 0.89056 | 0.44846 | 1.71399 | 0.06995 | 0.00449 | 0.40907 | 3.99516 | 0.47973 |
| **Gm5617** | 0.00228 | 0.01308 | 0.02391 | 0.31377 | 0.13943 | 0.02844 | 0.14994 | 0.33438 | 0.01598 | 0.00672 | 0.00000 | 0.65105 | 0.02360 | 0.01100 |
| **Gm5622** | 0.43629 | 0.31780 | 0.02203 | 0.04729 | 0.00705 | 0.84103 | 0.00377 | 0.00460 | 0.01335 | 0.00596 | 0.01302 | 0.56157 | 0.29306 | 0.45076 |
| **Gm5640** | 0.00504 | 0.01513 | 0.04623 | 0.14432 | 0.04014 | 0.04501 | 0.00423 | 0.10882 | 0.04321 | 0.00868 | 0.00578 | 0.15413 | 0.00490 | 0.02123 |
| **Gm5643** | 0.49011 | 2.45944 | 5.20965 | 7.56310 | 9.33889 | 7.48208 | 2.53834 | 7.23917 | 17.89010 | 2.50956 | 8.87170 | 6.08939 | 3.04534 | 15.09312 |
| **Gm5662** | 0.07692 | 0.01076 | 0.15635 | 0.11887 | 0.30793 | 0.11652 | 0.06761 | 0.04877 | 2.82522 | 0.02692 | 0.06388 | 0.02028 | 5.81313 | 0.13538 |
| **Gm5712** | 0.01065 | 0.00451 | 0.05704 | 0.05879 | 0.02981 | 0.06044 | 0.02194 | 0.02456 | 0.01113 | 0.00644 | 0.03566 | 0.04010 | 0.00920 | 0.03250 |
| **Gm572** | 0.02538 | 0.00443 | 0.01662 | 0.01198 | 0.01236 | 0.01276 | 0.00430 | 0.01040 | 0.01110 | 0.00667 | 0.00869 | 0.00987 | 0.05510 | 0.01255 |
| **Gm5766** | 0.11190 | 0.21840 | 0.37955 | 0.28213 | 0.06727 | 0.25286 | 0.08467 | 0.28340 | 0.23463 | 0.47441 | 0.19387 | 0.23296 | 0.09645 | 0.35135 |
| **Gm5779** | 0.36001 | 0.68835 | 2.37224 | 1.91439 | 1.41355 | 1.06269 | 0.33281 | 1.95443 | 0.00877 | 0.88340 | 0.91588 | 1.11805 | 0.00000 | 0.26589 |
| **Gm5795** | 0.07046 | 0.00450 | 0.02069 | 0.02412 | 0.06634 | 0.01802 | 0.00427 | 0.00921 | 0.05140 | 0.00698 | 0.00709 | 0.01073 | 0.16980 | 0.00632 |
| **Gm5796** | 0.03689 | 0.01255 | 0.01098 | 0.08579 | 0.04738 | 0.00791 | 0.00394 | 0.00821 | 0.04177 | 0.00650 | 0.00849 | 0.06438 | 0.00777 | 0.00534 |
| **Gm5797** | 0.18623 | 0.00428 | 0.02312 | 0.04676 | 0.10544 | 0.03979 | 0.00424 | 0.00558 | 0.19451 | 0.00597 | 0.00586 | 0.00612 | 0.48667 | 0.00000 |
| **Gm5801** | 0.18340 | 0.35091 | 2.44576 | 1.25049 | 3.70024 | 0.07438 | 1.02415 | 1.58434 | 0.00251 | 0.02272 | 0.36272 | 1.20393 | 0.54708 | 0.00000 |
| **Gm5803** | 0.00000 | 1.33208 | 2.54429 | 1.92023 | 0.88205 | 1.45064 | 0.98438 | 2.14473 | 10.52775 | 0.91161 | 3.90272 | 1.82174 | 0.07504 | 8.73824 |
| **Gm5820** | 0.00577 | 0.00444 | 0.01583 | 0.01439 | 0.01293 | 0.01037 | 0.01603 | 0.00959 | 0.04808 | 0.00625 | 0.00763 | 0.01284 | 0.19151 | 0.27836 |

| | | | | | | | | | | | | | | |
|---|---|---|---|---|---|---|---|---|---|---|---|---|---|---|
| **Gm5860** | 1.77183 | 0.63231 | 0.72280 | 1.60851 | 0.94505 | 1.15984 | 0.39408 | 0.64773 | 0.82755 | 0.24613 | 0.58345 | 1.97451 | 2.29528 | 0.46929 |
| **Gm5861** | 0.08756 | 0.00440 | 0.03329 | 0.01497 | 0.05522 | 0.03319 | 0.00437 | 0.00885 | 0.01466 | 0.00672 | 0.00683 | 0.01108 | 0.09727 | 0.01671 |
| **Gm5868** | 0.01066 | 0.00448 | 0.07367 | 0.09272 | 0.00861 | 0.01300 | 0.00448 | 0.01593 | 0.01071 | 0.00753 | 0.00994 | 0.01168 | 0.00635 | 0.01454 |
| **Gm5907** | 0.00108 | 0.01014 | 0.04882 | 0.01320 | 0.11588 | 1.21976 | 0.64646 | 0.00814 | 0.01021 | 4.42353 | 0.04503 | 0.00000 | 0.00759 | 0.00000 |
| **Gm5936** | 0.00510 | 0.00542 | 0.03538 | 0.13861 | 0.04876 | 0.03315 | 0.00425 | 0.13039 | 0.03795 | 0.00845 | 0.00566 | 0.20312 | 0.00488 | 0.02467 |
| **Gm5941** | 0.00330 | 0.00432 | 0.04271 | 0.00296 | 0.02815 | 0.03623 | 0.00423 | 0.00689 | 0.01263 | 0.00550 | 0.05449 | 0.00820 | 0.40435 | 0.00503 |
| **Gm595** | 0.08916 | 0.00422 | 0.00000 | 0.00281 | 0.18727 | 0.00000 | 0.00389 | 0.01083 | 1.04421 | 0.00393 | 0.78189 | 0.00413 | 0.00131 | 0.00000 |
| **Gm6083** | 9.46518 | 5.87782 | 0.48911 | 6.45381 | 8.78865 | 16.59614 | 4.24468 | 15.99188 | 3.96184 | 18.79685 | 2.06778 | 15.60450 | 12.60445 | 4.04252 |
| **Gm6086** | 0.01950 | 0.00439 | 0.03700 | 0.04583 | 0.04705 | 0.62589 | 0.00420 | 0.00480 | 0.03238 | 0.00600 | 0.04695 | 0.00759 | 0.18148 | 0.01914 |
| **Gm6093** | 0.04458 | 0.00447 | 0.02431 | 0.02437 | 0.05336 | 0.01265 | 0.01028 | 0.01027 | 0.08850 | 0.00710 | 0.00743 | 0.00891 | 0.01351 | 0.06962 |
| **Gm6194** | 0.00746 | 0.00422 | 0.10787 | 0.11479 | 0.12180 | 0.08369 | 0.00426 | 0.14888 | 0.01804 | 0.00738 | 0.01789 | 0.01250 | 0.17789 | 0.09731 |
| **Gm6225** | 0.03123 | 0.00454 | 0.01947 | 0.01360 | 0.09041 | 0.03833 | 0.00435 | 0.00998 | 0.01723 | 0.01037 | 0.00713 | 0.01509 | 0.08379 | 0.05178 |
| **Gm6297** | 0.01916 | 0.00459 | 0.03741 | 0.00376 | 0.02654 | 0.04505 | 0.00426 | 0.00777 | 0.01059 | 0.00610 | 0.00862 | 0.00981 | 0.21653 | 0.00227 |
| **Gm6313** | 0.16447 | 0.04483 | 0.11056 | 0.23718 | 0.50308 | 0.23996 | 0.08914 | 0.29102 | 1.50352 | 0.00609 | 0.13417 | 0.13570 | 1.60508 | 3.11845 |
| **Gm6367** | 0.08618 | 0.00411 | 0.00000 | 0.05513 | 0.16602 | 0.00717 | 0.00993 | 0.00633 | 1.25826 | 0.00709 | 0.01802 | 0.00541 | 1.10320 | 0.96127 |
| **Gm6402** | 1.27216 | 0.82356 | 0.26497 | 3.04735 | 2.33604 | 4.01969 | 1.17981 | 3.80671 | 1.79759 | 4.31375 | 0.17562 | 3.01633 | 4.51998 | 1.05654 |
| **Gm6455** | 0.16789 | 0.00435 | 0.03810 | 0.01918 | 0.05992 | 0.04498 | 0.00433 | 0.00780 | 0.01627 | 0.00671 | 0.00639 | 0.01158 | 0.10987 | 0.01201 |
| **Gm6460** | 0.06100 | 0.00437 | 0.03368 | 0.01735 | 0.02931 | 0.04446 | 0.00430 | 0.00832 | 0.01230 | 0.00716 | 0.00654 | 0.01213 | 0.01836 | 0.01819 |
| **Gm6498** | 0.04653 | 1.38719 | 0.30027 | 1.60437 | 0.05444 | 3.01303 | 0.65260 | 5.77328 | 0.13499 | 2.89255 | 1.42167 | 3.01832 | 0.33792 | 0.31604 |
| **Gm6524** | 0.09147 | 0.15589 | 0.13744 | 0.08001 | 0.16433 | 0.11664 | 0.17046 | 0.02781 | 0.18333 | 0.02054 | 0.13001 | 0.04521 | 0.52949 | 0.48650 |
| **Gm6525** | 1.29089 | 0.84064 | 0.00000 | 2.25421 | 1.80154 | 1.80817 | 0.55973 | 2.57267 | 1.46344 | 1.86030 | 0.00664 | 1.70551 | 1.66818 | 0.89248 |
| **Gm6548** | 21.59475 | 12.59878 | 35.85205 | 37.38927 | 22.84586 | 39.58685 | 10.31960 | 30.87977 | 29.55832 | 18.79946 | 24.56724 | 29.14248 | 2.90387 | 26.24517 |
| **Gm6566** | 0.26569 | 0.00523 | 0.14773 | 0.25360 | 0.02008 | 0.01157 | 0.00449 | 0.48187 | 0.67697 | 0.00786 | 0.02472 | 0.02668 | 0.50176 | 0.07298 |
| **Gm6568** | 0.84904 | 0.00292 | 0.79881 | 1.11007 | 0.95186 | 0.13754 | 0.18945 | 0.00109 | 2.37648 | 0.00482 | 0.15010 | 0.09801 | 6.93467 | 0.88508 |
| **Gm6578** | 0.42638 | 0.00404 | 0.14826 | 0.23129 | 0.47434 | 0.26698 | 0.06792 | 0.13903 | 0.03795 | 0.32199 | 0.19874 | 0.28743 | 0.31155 | 0.47199 |
| **Gm6602** | 0.01494 | 0.01294 | 0.02660 | 0.02996 | 0.01509 | 0.01089 | 0.00410 | 0.01398 | 0.13698 | 0.00713 | 0.03266 | 0.05235 | 0.01442 | 0.02534 |
| **Gm6607** | 0.02192 | 0.01286 | 0.01498 | 0.00875 | 0.02003 | 0.05829 | 0.01716 | 0.06452 | 0.01135 | 0.02691 | 0.00583 | 0.03619 | 0.06036 | 0.01498 |
| **Gm6623** | 4.02508 | 0.00349 | 1.57373 | 1.33233 | 4.32572 | 1.82628 | 0.48008 | 0.90770 | 1.75413 | 0.00848 | 1.22159 | 0.29968 | 0.90729 | 4.05676 |
| **Gm6639** | 1.05821 | 0.00371 | 0.12277 | 0.00000 | 0.20232 | 0.04915 | 0.02748 | 0.00363 | 1.73509 | 0.00603 | 0.00000 | 0.40505 | 0.79867 | 0.45194 |
| **Gm6654** | 9.14875 | 4.85401 | 2.01474 | 11.92015 | 12.78047 | 16.40887 | 4.86606 | 18.17036 | 0.26154 | 15.21810 | 1.10381 | 12.62748 | 11.89695 | 6.80946 |
| **Gm6710** | 0.44379 | 0.00267 | 1.09315 | 0.00268 | 0.46680 | 1.59217 | 0.07187 | 0.00000 | 2.84543 | 2.35740 | 0.89459 | 1.02477 | 0.12666 | 1.23698 |
| **Gm6712** | 5.27588 | 2.69931 | 7.93808 | 1.90295 | 4.39734 | 3.32911 | 2.60282 | 0.28436 | 19.02775 | 0.03280 | 0.84070 | 0.50820 | 1.54708 | 0.62268 |
| **Gm6756** | 2.05936 | 1.76525 | 9.12392 | 7.72976 | 4.21933 | 3.62532 | 0.82176 | 2.39648 | 0.00000 | 0.38996 | 3.87179 | 0.57073 | 0.01166 | 1.49366 |
| **Gm6787** | 0.01508 | 0.00456 | 0.00491 | 0.22010 | 0.00921 | 0.03805 | 0.00458 | 0.08602 | 0.18531 | 0.00950 | 0.03203 | 0.01089 | 0.00821 | 0.00000 |
| **Gm6792** | 0.00355 | 0.03491 | 0.01109 | 0.00000 | 0.00924 | 0.01009 | 0.00438 | 0.49033 | 0.01173 | 0.02504 | 0.00220 | 0.24453 | 0.23971 | 0.07698 |
| **Gm6815** | 0.43446 | 0.12444 | 1.06098 | 0.17116 | 0.67628 | 0.90955 | 0.19962 | 0.17551 | 2.17046 | 0.00177 | 1.07444 | 0.35819 | 0.06881 | 3.86288 |
| **Gm6880** | 0.00000 | 0.00403 | 0.44243 | 1.03137 | 0.85162 | 0.30538 | 0.00375 | 0.00112 | 0.31937 | 0.01068 | 0.00721 | 0.00597 | 0.00403 | 1.83285 |
| **Gm6890** | 0.00000 | 0.00406 | 0.19166 | 0.60820 | 0.57138 | 0.20602 | 0.00380 | 0.00245 | 0.36566 | 0.01174 | 0.02445 | 0.00973 | 0.01970 | 1.16248 |
| **Gm6902** | 0.00000 | 0.00376 | 0.46474 | 0.44877 | 0.99949 | 0.01813 | 0.00362 | 0.00242 | 1.95266 | 0.00482 | 0.00204 | 0.00491 | 3.90731 | 0.00000 |
| **Gm6904** | 0.00894 | 0.00461 | 0.04916 | 0.01410 | 0.01969 | 0.14265 | 0.03368 | 0.01538 | 0.01148 | 0.00657 | 0.03287 | 0.04440 | 0.00706 | 0.01034 |
| **Gm694** | 0.14644 | 0.01240 | 0.03373 | 0.04023 | 0.03037 | 0.25537 | 0.01053 | 0.37844 | 0.00980 | 0.01468 | 0.11367 | 0.00473 | 0.02407 | 0.02337 |
| **Gm6981** | 0.02681 | 0.79080 | 0.36855 | 0.87639 | 0.08923 | 2.78128 | 0.46750 | 4.59915 | 0.02200 | 2.58309 | 0.84839 | 2.62599 | 0.43001 | 0.13292 |
| **Gm7008** | 0.00999 | 0.00442 | 0.06732 | 0.00097 | 0.01813 | 0.05641 | 0.00421 | 0.00807 | 0.01209 | 0.00840 | 0.00767 | 0.01067 | 0.00738 | 0.01047 |
| **Gm7030** | 0.00608 | 0.05016 | 0.11019 | 0.16067 | 0.01719 | 0.05896 | 0.02918 | 0.09171 | 0.01062 | 0.05413 | 0.00659 | 0.03882 | 0.00926 | 0.01890 |
| **Gm7056** | 0.04368 | 0.00351 | 0.10005 | 0.00000 | 1.27399 | 0.00000 | 0.00402 | 0.00764 | 0.07884 | 0.00610 | 0.00538 | 0.00256 | 6.14601 | 4.60181 |
| **Gm7102** | 20.64936 | 0.32871 | 1.32502 | 1.72888 | 26.41146 | 1.45694 | 1.56647 | 0.55943 | 35.75709 | 0.12988 | 0.84055 | 0.74599 | 35.43635 | 44.96060 |
| **Gm7104** | 1.00384 | 0.00300 | 0.39188 | 0.00000 | 1.40648 | 0.01169 | 0.00352 | 0.00442 | 0.42195 | 0.00491 | 0.00474 | 0.00300 | 11.63761 | 0.70301 |
| **Gm715** | 0.02193 | 0.00423 | 0.01002 | 0.04678 | 0.02516 | 0.01055 | 0.00405 | 0.00742 | 0.01889 | 0.01655 | 0.00733 | 0.02575 | 0.17112 | 0.00152 |
| **Gm7244** | 0.00270 | 0.00420 | 0.00727 | 0.15673 | 0.01127 | 0.03014 | 0.00422 | 0.00883 | 0.01126 | 0.00661 | 0.02253 | 0.00550 | 0.80261 | 0.00214 |
| **Gm7325** | 0.00317 | 0.01234 | 0.00000 | 1.39318 | 0.00661 | 0.16472 | 0.00371 | 0.05469 | 0.01402 | 0.70918 | 0.00877 | 1.54183 | 0.00708 | 0.00214 |
| **Gm7361** | 0.05948 | 0.00433 | 0.03094 | 0.00753 | 0.03472 | 0.02807 | 0.00415 | 0.01990 | 0.05703 | 0.00650 | 0.01403 | 0.01554 | 0.00716 | 0.00690 |
| **Gm7534** | 0.50361 | 0.00430 | 0.89425 | 0.36948 | 0.63636 | 0.12417 | 0.00369 | 0.00000 | 0.57964 | 0.00409 | 0.00854 | 0.00907 | 1.33519 | 0.00000 |
| **Gm7609** | 0.56661 | 0.02989 | 0.42791 | 0.06029 | 0.05237 | 0.01508 | 0.01608 | 0.02822 | 2.32188 | 0.01055 | 0.00769 | 0.02413 | 1.14689 | 0.68445 |
| **Gm7616** | 0.05091 | 0.00452 | 0.01515 | 0.02338 | 0.01283 | 0.05470 | 0.02268 | 0.02068 | 0.08860 | 0.03344 | 0.02712 | 0.01763 | 0.62918 | 0.03385 |
| **Gm7694** | 0.00601 | 0.00424 | 0.05128 | 0.10994 | 0.00845 | 0.14384 | 0.00399 | 0.00363 | 0.01183 | 0.00593 | 0.00660 | 0.01268 | 0.10720 | 0.00000 |

| | | | | | | | | | | | | | | |
|---|---|---|---|---|---|---|---|---|---|---|---|---|---|---|
| **Gm7788** | 0.02513 | 0.00428 | 0.02405 | 0.01566 | 0.01314 | 0.01658 | 0.00415 | 0.01398 | 0.05281 | 0.00731 | 0.01047 | 0.01428 | 0.03498 | 0.02535 |
| **Gm7827** | 0.00000 | 0.00415 | 0.02674 | 0.01888 | 0.00000 | 0.03615 | 0.00413 | 0.00543 | 0.08826 | 0.00538 | 0.03430 | 0.00569 | 1.02303 | 0.66006 |
| **Gm7849** | 0.00851 | 0.00440 | 0.01617 | 0.01197 | 0.02496 | 0.01279 | 0.00427 | 0.00989 | 0.02300 | 0.00669 | 0.00846 | 0.01280 | 0.09669 | 0.00990 |
| **Gm7904** | 0.01173 | 0.00432 | 0.01458 | 0.00694 | 0.01691 | 0.02215 | 0.01630 | 0.00958 | 0.00906 | 0.00641 | 0.03682 | 0.01597 | 0.05660 | 0.05567 |
| **Gm7977** | 2.44500 | 0.51657 | 5.18275 | 3.21273 | 2.22288 | 1.64434 | 0.24566 | 1.15789 | 0.75210 | 0.85351 | 0.81483 | 0.85863 | 0.00373 | 0.86291 |
| **Gm805** | 0.00561 | 0.00438 | 0.00305 | 0.31594 | 0.00819 | 0.21270 | 0.00400 | 0.00510 | 0.01331 | 0.00838 | 0.00421 | 0.01162 | 0.00833 | 0.02288 |
| **Gm8096** | 1.48361 | 1.26146 | 6.32043 | 5.49837 | 3.38118 | 2.70261 | 0.63247 | 1.39571 | 0.00381 | 0.73336 | 2.48327 | 0.23789 | 0.36817 | 0.88226 |
| **Gm813** | 0.01738 | 0.00405 | 0.01124 | 0.01250 | 0.06973 | 0.01973 | 0.00390 | 0.04007 | 0.00930 | 0.00997 | 0.00787 | 0.00753 | 0.43933 | 0.00220 |
| **Gm8234** | 2.41280 | 0.20668 | 4.44856 | 1.95979 | 4.72423 | 0.84891 | 0.27853 | 0.46632 | 5.45164 | 0.03771 | 0.46423 | 1.34636 | 7.76634 | 9.23521 |
| **Gm8267** | 1.06509 | 0.26858 | 0.01136 | 0.16165 | 0.00280 | 0.74664 | 0.00354 | 0.00965 | 0.01230 | 0.00604 | 0.00931 | 1.18982 | 1.07289 | 0.24453 |
| **Gm8300** | 0.06148 | 0.00347 | 0.18854 | 0.34273 | 0.30994 | 0.30751 | 0.08063 | 0.06613 | 2.05414 | 0.00556 | 0.12635 | 0.04174 | 6.51367 | 1.45773 |
| **Gm8363** | 0.03151 | 0.00411 | 0.06052 | 0.10207 | 0.04692 | 0.09504 | 0.03156 | 0.04866 | 0.01180 | 0.00677 | 0.06624 | 0.05508 | 0.02958 | 0.04596 |
| **Gm839** | 0.02775 | 0.00433 | 0.07943 | 0.00734 | 0.00436 | 0.00867 | 0.00422 | 0.01004 | 0.00936 | 0.00658 | 0.00748 | 0.00874 | 0.33680 | 0.22499 |
| **Gm8579** | 0.40051 | 0.00386 | 0.29133 | 0.00000 | 0.06334 | 0.30152 | 0.00395 | 0.13464 | 0.00988 | 0.03537 | 0.10012 | 0.15077 | 0.00439 | 0.01172 |
| **Gm8580** | 8.05084 | 4.53623 | 4.17339 | 11.44869 | 12.33976 | 10.47800 | 3.86612 | 9.23132 | 0.01198 | 9.97563 | 2.52536 | 10.38906 | 1.14149 | 3.99847 |
| **Gm8615** | 3.28636 | 0.00726 | 4.45570 | 1.60472 | 2.39809 | 2.36325 | 0.09646 | 0.79015 | 3.85761 | 0.00000 | 8.42666 | 2.32348 | 0.80634 | 6.09857 |
| **Gm8624** | 4.65781 | 3.53190 | 0.29211 | 4.41750 | 8.97773 | 9.50105 | 3.14646 | 8.26094 | 1.70550 | 8.69289 | 0.49965 | 8.21810 | 7.46588 | 3.78531 |
| **Gm867** | 0.01473 | 0.00442 | 0.27412 | 0.00000 | 0.00248 | 0.00543 | 0.00412 | 0.01174 | 0.38475 | 0.00764 | 0.00589 | 0.00831 | 0.00829 | 0.01122 |
| **Gm8709** | 0.01049 | 0.02014 | 0.01970 | 0.03628 | 0.01332 | 0.04161 | 0.01055 | 0.06019 | 0.02006 | 0.07031 | 0.02272 | 0.02324 | 0.02615 | 0.01872 |
| **Gm8801** | 2.99270 | 2.07206 | 3.18161 | 0.00000 | 3.11461 | 1.45770 | 1.48587 | 1.70397 | 0.32957 | 0.01236 | 0.87487 | 0.96775 | 1.52218 | 2.63601 |
| **Gm884** | 0.00701 | 0.00448 | 0.01302 | 0.01465 | 0.03839 | 0.01231 | 0.00435 | 0.01062 | 0.00840 | 0.00659 | 0.00861 | 0.00952 | 0.00605 | 0.12657 |
| **Gm8909** | 0.01089 | 0.05771 | 0.50543 | 0.67696 | 0.04071 | 0.31501 | 0.07823 | 0.27988 | 0.02282 | 0.22962 | 0.03952 | 0.37749 | 0.00902 | 0.02890 |
| **Gm8994** | 7.68430 | 3.29066 | 10.53802 | 7.47979 | 9.83036 | 5.55585 | 1.37233 | 2.50156 | 1.03751 | 0.12360 | 4.06166 | 3.46972 | 6.33378 | 4.97077 |
| **Gm9** | 0.00000 | 0.00433 | 0.75639 | 0.03340 | 1.20777 | 0.00000 | 0.00427 | 0.02020 | 1.19353 | 0.00661 | 0.00268 | 0.00695 | 0.01243 | 2.20910 |
| **Gm9079** | 0.04460 | 0.04888 | 0.11608 | 0.04855 | 0.08571 | 0.15389 | 0.67783 | 0.06860 | 0.18092 | 0.01921 | 0.04011 | 0.07332 | 0.05526 | 0.21648 |
| **Gm9125** | 0.58603 | 0.00319 | 0.08712 | 0.03169 | 0.43762 | 0.00000 | 0.00902 | 0.02152 | 0.56273 | 0.01185 | 0.00449 | 0.02821 | 6.57998 | 2.45873 |
| **Gm9159** | 0.09067 | 0.32460 | 0.05761 | 0.04768 | 0.09088 | 0.09329 | 0.06390 | 0.07788 | 0.31143 | 0.00807 | 0.01381 | 0.12155 | 0.15770 | 0.31940 |
| **Gm9199** | 14.33942 | 0.09509 | 6.73538 | 8.13401 | 16.66643 | 1.91337 | 0.07821 | 0.91184 | 5.14335 | 0.49518 | 0.94738 | 1.81744 | 6.37085 | 4.38400 |
| **Gm9268** | 0.01040 | 0.00446 | 0.00804 | 0.08055 | 0.01176 | 0.01134 | 0.00429 | 0.01295 | 0.07616 | 0.00999 | 0.00803 | 0.00816 | 0.05454 | 0.00307 |
| **Gm9376** | 8.02468 | 4.92475 | 1.36187 | 13.82334 | 12.61327 | 14.20196 | 5.79442 | 12.99635 | 7.03576 | 12.01636 | 1.63240 | 5.86333 | 13.48553 | 5.82949 |
| **Gm960** | 0.00459 | 0.00444 | 0.32588 | 0.00000 | 0.52583 | 0.01874 | 0.00420 | 0.00756 | 0.01424 | 0.00528 | 0.01175 | 0.00775 | 0.07881 | 0.01003 |
| **Gm973** | 0.50378 | 0.01854 | 0.90333 | 0.00000 | 0.00000 | 0.39283 | 0.96539 | 1.04813 | 0.86021 | 0.04104 | 1.95019 | 2.06405 | 0.00265 | 0.13461 |
| **Gm9731** | 0.05255 | 0.00416 | 0.05497 | 0.08489 | 0.08926 | 0.06072 | 0.00405 | 0.00529 | 0.02235 | 0.00652 | 0.04764 | 0.01903 | 0.22323 | 0.02228 |
| **Gm9758** | 0.33534 | 0.00408 | 0.03633 | 0.18556 | 0.17315 | 0.01862 | 0.00395 | 0.00676 | 0.02837 | 0.00594 | 0.01488 | 0.01032 | 0.35280 | 0.01929 |
| **Gm9776** | 0.38542 | 0.04268 | 0.01508 | 0.49641 | 0.04001 | 0.47187 | 0.08908 | 0.09965 | 0.58431 | 0.07747 | 0.03664 | 0.14695 | 3.80940 | 0.32635 |
| **Gm9833** | 0.03474 | 0.00356 | 0.64091 | 0.27519 | 0.25781 | 0.30653 | 0.00367 | 0.00040 | 0.38561 | 0.00431 | 0.28356 | 0.57938 | 0.01932 | 1.62867 |
| **Gm9895** | 0.14486 | 0.00467 | 0.03124 | 0.00000 | 0.59305 | 0.10110 | 0.00545 | 0.04951 | 0.00980 | 0.00646 | 0.00567 | 0.00540 | 0.39490 | 0.45297 |
| **Gm9926** | 0.01973 | 0.03024 | 0.01600 | 0.05549 | 0.09979 | 0.07011 | 0.03691 | 0.02002 | 0.10113 | 0.03391 | 0.00666 | 0.03934 | 0.37716 | 0.08986 |
| **Gm9958** | 0.44756 | 0.00421 | 0.09172 | 0.13880 | 0.00000 | 0.00360 | 0.00390 | 0.00752 | 0.09636 | 0.00821 | 0.00540 | 0.01133 | 0.00419 | 0.13767 |
| **Gm996** | 0.02269 | 0.00473 | 0.31977 | 0.06132 | 0.01052 | 0.51122 | 0.00392 | 0.00732 | 0.01009 | 0.03666 | 0.00570 | 0.00304 | 0.02341 | 0.00129 |
| **Gm9992** | 0.00036 | 0.01902 | 0.18147 | 0.00000 | 0.10722 | 0.10486 | 1.01728 | 2.34885 | 0.00995 | 0.02952 | 0.00405 | 0.16974 | 0.01676 | 0.12290 |
| **Gm9994** | 0.00810 | 0.00454 | 0.02383 | 0.00328 | 0.02158 | 0.23479 | 0.00429 | 0.00654 | 0.01054 | 0.00642 | 0.02449 | 0.00965 | 0.00819 | 0.01783 |
| **Gmcl1** | 0.27984 | 0.00237 | 1.62436 | 0.00000 | 0.80476 | 2.04474 | 0.00288 | 0.00000 | 2.11614 | 0.00344 | 2.51045 | 1.24547 | 2.45766 | 0.44962 |
| **Gmds** | 0.26702 | 0.00346 | 0.01576 | 0.00000 | 0.08313 | 0.35786 | 0.00404 | 0.00443 | 0.01377 | 0.00964 | 1.00734 | 0.80797 | 0.46456 | 0.00924 |
| **Gmeb1** | 6.66178 | 0.44169 | 4.92385 | 3.04708 | 4.78541 | 5.21402 | 0.76556 | 2.97205 | 9.30964 | 1.16327 | 1.50779 | 2.57125 | 14.62817 | 3.38292 |
| **Gmeb2** | 1.00688 | 0.01049 | 0.92965 | 0.73651 | 2.67769 | 0.65283 | 0.00874 | 0.27237 | 3.43262 | 0.02492 | 0.21823 | 0.63291 | 2.27998 | 2.52572 |
| **Gmfb** | 3.48691 | 0.04733 | 0.95241 | 2.27332 | 0.42169 | 6.36328 | 3.06757 | 1.68536 | 7.54750 | 15.27854 | 1.04864 | 2.45015 | 2.48195 | 1.69801 |
| **Gmfg** | 0.00196 | 0.01275 | 0.00000 | 0.09904 | 0.01259 | 0.82536 | 1.72813 | 0.46995 | 0.02675 | 0.06751 | 0.22486 | 1.63906 | 0.29041 | 0.00000 |
| **Gmip** | 0.47468 | 0.04867 | 0.13034 | 0.25589 | 0.24743 | 0.87894 | 0.10169 | 0.68793 | 0.08392 | 0.01491 | 0.00000 | 1.08658 | 0.74371 | 0.21636 |
| **Gml2** | 0.00516 | 0.00362 | 0.01103 | 0.00485 | 0.01411 | 0.01571 | 0.00413 | 0.00613 | 0.01225 | 0.00696 | 0.00446 | 0.91156 | 0.00721 | 0.01108 |
| **Gmnn** | 20.14482 | 0.20781 | 19.15879 | 17.94976 | 26.38733 | 19.11534 | 3.43421 | 8.49377 | 8.94281 | 0.34223 | 14.04960 | 14.56298 | 20.21339 | 20.83365 |
| **Gmppa** | 0.47544 | 0.10817 | 3.67981 | 3.78264 | 0.70162 | 2.23107 | 0.34829 | 4.80265 | 0.01309 | 0.06100 | 3.20602 | 4.51271 | 0.00000 | 1.75082 |
| **Gmppb** | 2.56618 | 0.03828 | 4.39752 | 6.13397 | 4.47378 | 5.42634 | 0.00708 | 5.42710 | 0.15647 | 0.06876 | 4.01614 | 1.62962 | 0.00000 | 1.77960 |
| **Gmpr** | 0.10053 | 0.00415 | 2.01241 | 0.51336 | 0.79677 | 0.63537 | 0.00902 | 0.00226 | 0.00933 | 0.01903 | 0.02862 | 0.00611 | 0.00000 | 0.23658 |
| **Gmpr2** | 0.84289 | 0.00223 | 2.22975 | 1.06083 | 0.06595 | 3.14159 | 1.74094 | 0.23133 | 0.00000 | 0.10318 | 2.11745 | 3.11327 | 1.32351 | 0.62651 |

| | | | | | | | | | | | | | | |
|---|---|---|---|---|---|---|---|---|---|---|---|---|---|---|
| **Gmps** | 12.93090 | 0.10623 | 14.67123 | 8.65947 | 15.56211 | 17.41564 | 4.18155 | 0.27609 | 13.88937 | 0.11435 | 8.07144 | 9.86291 | 6.24350 | 12.19636 |
| **Gna11** | 1.18326 | 0.03304 | 10.78049 | 3.14477 | 4.21517 | 10.25002 | 0.01847 | 4.55464 | 0.00000 | 0.08427 | 2.50919 | 3.38620 | 0.44814 | 0.76193 |
| **Gna12** | 0.06152 | 0.01320 | 0.39155 | 0.00000 | 1.15971 | 1.02368 | 0.02374 | 0.44854 | 0.00788 | 0.00656 | 0.08360 | 0.24380 | 0.14354 | 0.00595 |
| **Gna13** | 1.74013 | 6.75197 | 5.72226 | 4.84968 | 2.86504 | 7.20391 | 1.87323 | 2.37563 | 3.96950 | 5.14794 | 7.69472 | 8.55098 | 0.03943 | 6.51046 |
| **Gna14** | 1.33820 | 0.00389 | 0.29184 | 0.07226 | 0.00000 | 0.00110 | 0.00399 | 0.00838 | 0.00470 | 0.00538 | 0.01303 | 0.01087 | 1.14693 | 0.00476 |
| **Gna15** | 0.00000 | 0.01845 | 0.56712 | 0.67211 | 0.28491 | 0.00000 | 0.00400 | 0.45162 | 0.01697 | 0.00633 | 0.06490 | 0.88931 | 0.00302 | 0.00000 |
| **Gnai1** | 0.01347 | 0.00990 | 0.00000 | 0.17883 | 0.15223 | 0.80707 | 3.42342 | 0.22981 | 0.00565 | 0.08183 | 0.00000 | 2.19144 | 0.03849 | 3.33222 |
| **Gnai2** | 0.78150 | 0.41009 | 2.84300 | 3.28845 | 5.02726 | 4.25293 | 1.97594 | 11.31385 | 0.81834 | 0.71200 | 2.10272 | 1.30791 | 0.01224 | 1.31612 |
| **Gnai3** | 10.32554 | 0.60064 | 5.26042 | 5.14773 | 5.60798 | 9.73985 | 3.16115 | 3.15781 | 5.59698 | 9.25579 | 10.23926 | 8.91778 | 7.17826 | 4.67798 |
| **Gnal** | 2.54354 | 0.00245 | 1.38243 | 1.43401 | 2.25795 | 1.35474 | 1.54219 | 1.21638 | 0.85892 | 0.05438 | 1.49136 | 1.81241 | 1.03669 | 1.62474 |
| **Gnaq** | 0.23193 | 0.06165 | 1.47462 | 0.00000 | 2.02289 | 3.05891 | 0.00643 | 4.32072 | 4.53789 | 0.02854 | 1.18937 | 4.63579 | 0.16226 | 4.08890 |
| **Gnas** | 14.92387 | 0.17621 | 10.89194 | 5.61247 | 10.84544 | 7.76963 | 5.69962 | 2.71779 | 3.71715 | 0.20561 | 3.87414 | 7.11417 | 2.62263 | 3.55932 |
| **Gnb1** | 10.79876 | 6.75070 | 12.57591 | 18.99086 | 9.21356 | 20.30353 | 1.66208 | 9.40110 | 10.97300 | 0.23963 | 14.84606 | 11.49467 | 2.93232 | 9.35982 |
| **Gnb1l** | 1.22525 | 0.03238 | 7.35515 | 5.74270 | 8.99207 | 2.29467 | 2.33470 | 4.34308 | 0.05077 | 0.13985 | 1.18941 | 3.01215 | 1.63369 | 1.11610 |
| **Gnb2** | 0.60245 | 0.16061 | 4.28527 | 2.28792 | 4.03079 | 5.84636 | 2.71752 | 5.78522 | 0.00615 | 12.53733 | 0.86757 | 7.75746 | 0.16782 | 0.17212 |
| **Gnb3** | 0.01063 | 0.00443 | 0.07356 | 0.00831 | 0.01261 | 0.01075 | 0.00426 | 0.01072 | 0.01147 | 0.00740 | 0.00795 | 0.01014 | 0.00741 | 0.01213 |
| **Gnb4** | 0.03442 | 0.00554 | 0.13016 | 0.00000 | 0.00000 | 0.00165 | 0.00453 | 0.16482 | 0.01186 | 0.00899 | 0.00613 | 0.00927 | 0.00613 | 1.62964 |
| **Gnb5** | 0.00000 | 0.00438 | 0.28978 | 0.26047 | 0.68751 | 0.01265 | 0.00418 | 0.00734 | 0.00619 | 0.00773 | 0.00514 | 0.00563 | 0.16325 | 0.02373 |
| **Gne** | 0.02850 | 0.00402 | 1.18407 | 1.10685 | 0.16539 | 0.49753 | 0.00369 | 0.00000 | 0.00419 | 0.00393 | 0.00923 | 0.01171 | 0.18596 | 0.11804 |
| **Gng10** | 0.64487 | 0.00848 | 0.00000 | 1.79816 | 0.00443 | 5.49713 | 0.00867 | 0.07718 | 0.00573 | 11.71556 | 1.46914 | 1.59153 | 0.00582 | 0.00000 |
| **Gng12** | 1.19895 | 0.00410 | 0.00000 | 1.96983 | 0.98640 | 0.45165 | 0.07335 | 1.26183 | 0.20301 | 0.03575 | 1.13738 | 0.36220 | 1.27086 | 0.00000 |
| **Gng13** | 0.02302 | 0.00481 | 0.00598 | 0.11318 | 0.01760 | 0.10889 | 0.00425 | 0.00949 | 0.01207 | 0.00703 | 0.01360 | 0.01087 | 0.00792 | 0.01251 |
| **Gng2** | 0.00397 | 5.91314 | 0.02267 | 0.00000 | 0.17572 | 0.88074 | 0.59068 | 14.52277 | 0.00000 | 4.88552 | 0.00000 | 0.68726 | 0.00097 | 0.37945 |
| **Gng3** | 0.00552 | 0.00417 | 0.04500 | 0.01289 | 0.00470 | 0.01207 | 0.00409 | 0.00810 | 0.02413 | 0.00746 | 0.00829 | 0.00764 | 1.41331 | 0.00228 |
| **Gng5** | 1.62036 | 0.20199 | 0.00000 | 2.34529 | 6.33899 | 4.21140 | 2.28063 | 10.26608 | 0.48953 | 7.39351 | 0.05130 | 7.74795 | 3.24818 | 0.00000 |
| **Gng7** | 0.13200 | 0.00427 | 0.35342 | 0.05370 | 0.00000 | 0.00000 | 0.00382 | 0.00774 | 0.68107 | 0.01620 | 0.00422 | 0.00669 | 0.00300 | 0.86614 |
| **Gngt1** | 0.07267 | 0.02418 | 0.01576 | 0.15509 | 0.06349 | 0.19084 | 0.03111 | 0.05787 | 0.28535 | 0.02451 | 0.02456 | 0.01707 | 1.48097 | 0.07050 |
| **Gnl1** | 1.94355 | 0.18257 | 14.88297 | 8.61843 | 10.69571 | 7.57173 | 3.11819 | 10.84649 | 0.15310 | 9.08079 | 5.14651 | 5.33996 | 3.83337 | 3.37308 |
| **Gnl2** | 12.25051 | 0.08786 | 16.61884 | 5.65131 | 10.48120 | 13.03496 | 2.09061 | 4.11320 | 15.81423 | 10.75015 | 10.97271 | 8.63091 | 5.36891 | 18.20076 |
| **Gnl3** | 27.17406 | 0.33446 | 24.85929 | 15.00446 | 25.19438 | 21.10128 | 7.00989 | 11.63560 | 30.13771 | 0.44880 | 17.69963 | 13.39489 | 14.07459 | 28.42889 |
| **Gnl3l** | 9.24649 | 0.30594 | 6.94972 | 5.87540 | 15.21327 | 8.03377 | 0.27767 | 2.86266 | 18.89866 | 0.47098 | 4.31921 | 4.45236 | 4.35856 | 13.29792 |
| **Gnmt** | 0.34205 | 0.00438 | 0.09892 | 0.00383 | 0.10371 | 0.00737 | 0.00433 | 0.01354 | 0.00772 | 0.00672 | 0.00675 | 0.00830 | 0.06322 | 0.00484 |
| **Gnpat** | 1.06750 | 4.02929 | 1.75912 | 3.88939 | 1.30488 | 5.79412 | 2.61795 | 5.27282 | 3.96259 | 0.07181 | 5.27763 | 8.63790 | 1.98936 | 2.66557 |
| **Gnpda1** | 2.39245 | 0.01397 | 4.88764 | 2.15117 | 3.08540 | 3.02590 | 0.14600 | 1.89781 | 5.92496 | 0.04209 | 10.38979 | 2.85687 | 0.66567 | 8.34569 |
| **Gnpda2** | 2.75864 | 0.02510 | 0.53779 | 0.00000 | 0.29878 | 2.31676 | 0.00921 | 0.99324 | 1.06617 | 0.01373 | 0.03032 | 1.82852 | 0.00430 | 1.34329 |
| **Gnpnat1** | 25.39348 | 0.01613 | 17.81175 | 15.49877 | 16.12528 | 7.93774 | 1.58334 | 0.00000 | 30.76920 | 0.09441 | 15.68547 | 3.92574 | 8.24283 | 12.79953 |
| **Gnptab** | 7.52675 | 0.07877 | 11.68005 | 0.78293 | 13.58275 | 10.26227 | 3.98309 | 5.69766 | 21.58353 | 9.99501 | 7.71564 | 8.71188 | 3.94884 | 13.32308 |
| **Gnptg** | 4.29157 | 0.04316 | 8.15728 | 6.62332 | 10.76028 | 3.12741 | 0.00802 | 2.65985 | 2.94392 | 0.03177 | 1.48911 | 1.85392 | 4.29750 | 6.21669 |
| **Gnrh1** | 0.15491 | 0.00438 | 0.00806 | 0.01141 | 0.03623 | 0.01015 | 0.00432 | 0.00998 | 0.01121 | 0.00659 | 0.00711 | 0.00846 | 0.14952 | 0.39468 |
| **Gns** | 1.22795 | 0.02975 | 2.96188 | 9.43480 | 0.37142 | 4.59787 | 3.04238 | 1.12765 | 0.00140 | 0.03018 | 5.37698 | 1.92306 | 0.41253 | 1.13819 |
| **Golga1** | 3.20069 | 0.00241 | 2.79264 | 0.52056 | 2.79261 | 1.73470 | 0.09842 | 0.00000 | 3.26352 | 0.00543 | 0.39098 | 1.14082 | 4.56964 | 2.35448 |
| **Golga2** | 1.94281 | 0.08564 | 4.39841 | 5.11770 | 2.95691 | 3.33268 | 2.75090 | 3.74684 | 0.95021 | 0.04487 | 2.76620 | 1.93570 | 2.54692 | 4.93082 |
| **Golga3** | 1.27317 | 0.85222 | 1.73036 | 0.00000 | 2.44106 | 2.06985 | 3.74396 | 0.50846 | 2.71260 | 0.13641 | 3.00731 | 3.54844 | 0.25074 | 3.16922 |
| **Golga4** | 2.25653 | 1.79031 | 3.50554 | 2.08641 | 2.42357 | 6.07940 | 4.52256 | 5.34586 | 0.45870 | 0.06285 | 3.45792 | 7.77922 | 0.69288 | 3.09516 |
| **Golga5** | 0.96730 | 3.17630 | 3.95970 | 0.77271 | 2.54938 | 1.76600 | 3.84544 | 2.91488 | 2.57586 | 0.15897 | 2.88289 | 1.36888 | 4.49712 | 6.12433 |
| **Golga7** | 8.08326 | 0.56543 | 16.31591 | 9.92888 | 10.56648 | 12.57955 | 2.41169 | 6.47471 | 5.39321 | 0.06101 | 8.92684 | 10.64828 | 4.05295 | 4.15896 |
| **Golga7b** | 0.13520 | 0.00423 | 0.00980 | 0.06597 | 0.07169 | 0.01102 | 0.00407 | 0.01518 | 0.01413 | 0.00745 | 0.00732 | 0.00713 | 0.00563 | 0.00601 |
| **Golgb1** | 4.13615 | 3.31029 | 2.23048 | 1.90387 | 2.46828 | 2.33140 | 0.25221 | 0.59495 | 0.24851 | 0.01752 | 0.19917 | 2.67244 | 1.22152 | 3.76068 |
| **Golm1** | 0.00000 | 0.57389 | 1.47139 | 2.07051 | 0.80795 | 2.29628 | 0.01214 | 2.90017 | 1.01305 | 0.00353 | 4.48157 | 6.21205 | 0.96789 | 0.00000 |
| **Golph3** | 2.92703 | 0.02289 | 4.65236 | 0.00000 | 5.84820 | 5.95081 | 1.75252 | 5.06330 | 8.21995 | 0.08301 | 4.57679 | 4.07314 | 7.43484 | 5.10310 |
| **Golph3l** | 0.74274 | 0.01563 | 0.42788 | 0.95607 | 0.51233 | 0.00000 | 0.00760 | 2.62778 | 2.92365 | 0.01895 | 1.31476 | 3.05650 | 0.61257 | 0.00000 |
| **Golt1a** | 0.94085 | 0.01190 | 0.03337 | 0.02519 | 0.22222 | 0.00572 | 0.00413 | 0.00916 | 0.00621 | 0.00688 | 0.00627 | 0.00471 | 0.25879 | 0.70593 |
| **Golt1b** | 11.38100 | 4.38275 | 3.85971 | 3.35456 | 5.07007 | 9.47583 | 3.25765 | 5.73336 | 9.96453 | 0.08650 | 2.82712 | 5.13489 | 3.61561 | 8.09821 |
| **Gon4l** | 1.79601 | 0.03138 | 5.73729 | 2.36962 | 3.08933 | 4.01806 | 2.09533 | 2.33446 | 1.19894 | 0.06258 | 6.19639 | 2.54522 | 1.57225 | 5.80999 |
| **Gopc** | 0.73925 | 0.02761 | 0.52599 | 0.00000 | 0.44737 | 2.40988 | 0.00397 | 0.22330 | 0.65674 | 0.00479 | 0.03323 | 2.41108 | 0.01065 | 0.36830 |

| | | | | | | | | | | | | | | |
|---|---|---|---|---|---|---|---|---|---|---|---|---|---|---|
| **Gorab** | 0.21158 | 0.00386 | 0.99984 | 0.00000 | 0.83823 | 0.39347 | 2.71484 | 0.62447 | 0.01694 | 0.02714 | 0.02929 | 0.51235 | 1.56716 | 1.62404 |
| **Gorasp1** | 5.68839 | 0.00961 | 14.42105 | 5.18498 | 23.15567 | 2.44191 | 0.00070 | 0.90594 | 0.45316 | 0.00000 | 1.53003 | 2.98473 | 1.79626 | 10.73585 |
| **Gorasp2** | 6.06375 | 7.59877 | 5.41946 | 3.83105 | 6.99663 | 12.24830 | 6.54344 | 4.04288 | 1.61533 | 0.21808 | 6.11694 | 9.53314 | 1.80416 | 3.57558 |
| **Gosr1** | 3.63545 | 6.14013 | 5.89606 | 0.84706 | 3.54278 | 2.12116 | 0.01369 | 1.81950 | 2.75545 | 0.01998 | 4.77007 | 1.24222 | 1.41855 | 7.74350 |
| **Gosr2** | 4.01014 | 0.05437 | 5.27414 | 6.12566 | 7.16340 | 8.74427 | 1.66247 | 6.98859 | 10.74996 | 0.10041 | 4.17151 | 2.68430 | 0.74175 | 11.71566 |
| **Got1** | 3.99298 | 7.00482 | 23.64166 | 22.59100 | 7.40441 | 22.52550 | 7.17343 | 16.33314 | 6.22642 | 1.38626 | 19.38818 | 12.01064 | 4.90117 | 7.33010 |
| **Got1l1** | 0.92575 | 0.02277 | 5.26663 | 3.92298 | 0.80653 | 2.40791 | 0.00174 | 0.00000 | 0.18881 | 0.00000 | 2.45556 | 3.60766 | 0.00000 | 1.57581 |
| **Got2** | 5.28921 | 0.21969 | 5.11060 | 7.71765 | 8.31894 | 7.08356 | 0.02953 | 4.44106 | 4.93280 | 0.04491 | 6.69047 | 6.49457 | 0.76602 | 6.67905 |
| **Gp1ba** | 0.00679 | 0.00442 | 0.09199 | 0.06757 | 0.17697 | 0.06040 | 0.00489 | 0.06904 | 0.01144 | 0.00648 | 0.01201 | 0.00654 | 0.00548 | 0.00962 |
| **Gpa33** | 0.00000 | 5.52282 | 0.27987 | 0.00000 | 0.12954 | 0.00000 | 4.11436 | 3.11443 | 0.01237 | 9.08937 | 0.00000 | 0.59379 | 0.00533 | 0.00366 |
| **Gpaa1** | 2.12378 | 4.33681 | 7.81368 | 6.82474 | 3.01590 | 3.37142 | 0.00497 | 6.33104 | 0.00000 | 2.38939 | 1.29350 | 5.20104 | 0.00000 | 0.07548 |
| **Gpalpp1** | 5.71748 | 0.01763 | 6.34740 | 3.41729 | 3.04207 | 4.44079 | 0.04981 | 2.84602 | 7.85131 | 0.10367 | 3.80451 | 1.49994 | 3.84104 | 4.03818 |
| **Gpam** | 0.12930 | 0.00421 | 0.04032 | 0.04360 | 0.19903 | 0.09158 | 0.00382 | 0.00345 | 0.00434 | 0.02150 | 0.23937 | 0.04933 | 0.32094 | 0.50164 |
| **Gpank1** | 3.19639 | 0.05025 | 9.80415 | 6.87639 | 8.99823 | 3.90304 | 2.39632 | 2.03241 | 0.21260 | 0.15745 | 5.90971 | 4.76489 | 7.72841 | 2.04427 |
| **Gpat2** | 0.85067 | 0.00370 | 1.22850 | 4.33756 | 1.70427 | 0.55496 | 0.00360 | 0.00000 | 0.01206 | 0.00693 | 0.48107 | 0.23521 | 0.00000 | 1.71692 |
| **Gpat4** | 0.00000 | 0.05669 | 2.85935 | 7.60659 | 0.96470 | 6.75089 | 2.38304 | 3.76503 | 0.15125 | 0.09086 | 0.83042 | 6.17303 | 3.73982 | 2.44185 |
| **Gpatch1** | 2.30885 | 0.01038 | 3.25118 | 1.31148 | 2.14154 | 1.16791 | 0.00309 | 2.10437 | 0.39573 | 0.00000 | 0.26594 | 1.28994 | 1.02516 | 1.19105 |
| **Gpatch11** | 2.89614 | 0.02550 | 5.60199 | 2.78788 | 2.10896 | 6.27303 | 0.00282 | 1.83083 | 1.67352 | 0.02422 | 2.52123 | 0.87104 | 0.36890 | 2.38525 |
| **Gpatch2** | 1.27264 | 0.00881 | 2.73993 | 1.53519 | 1.06875 | 1.43426 | 0.00969 | 0.01959 | 0.15959 | 0.00000 | 2.99184 | 2.60195 | 0.87241 | 1.02590 |
| **Gpatch2l** | 0.82396 | 0.01730 | 2.31995 | 2.30965 | 0.56877 | 0.46958 | 0.00272 | 1.44081 | 0.59927 | 0.01726 | 1.29683 | 1.24750 | 2.03775 | 1.25639 |
| **Gpatch3** | 0.58081 | 0.00970 | 0.61477 | 2.18085 | 1.32281 | 0.00000 | 0.00319 | 1.63941 | 0.00026 | 0.02158 | 0.61785 | 1.71279 | 4.96137 | 0.00000 |
| **Gpatch4** | 4.38956 | 0.02421 | 8.46406 | 5.57828 | 4.40511 | 7.57783 | 2.53421 | 0.99489 | 4.33921 | 0.06795 | 4.69234 | 5.11121 | 2.99306 | 9.96487 |
| **Gpatch8** | 4.51052 | 0.00855 | 4.11497 | 0.41080 | 5.22699 | 1.29658 | 0.00860 | 0.92934 | 4.44460 | 0.02197 | 2.27590 | 2.82360 | 4.89572 | 4.39825 |
| **Gpbp1** | 17.24478 | 0.20399 | 13.25805 | 2.75226 | 18.91487 | 7.99089 | 5.98728 | 1.82775 | 28.05925 | 7.34651 | 11.41857 | 8.05418 | 20.12193 | 22.13322 |
| **Gpbp1l1** | 0.00000 | 0.01096 | 1.68281 | 0.53487 | 3.69104 | 0.31522 | 0.00178 | 1.45368 | 11.93413 | 0.00654 | 1.40597 | 0.34840 | 5.88256 | 6.77298 |
| **Gpc2** | 0.01392 | 0.00487 | 0.00470 | 0.13981 | 0.01026 | 0.11793 | 0.00409 | 0.01089 | 0.01523 | 0.01141 | 0.00602 | 0.00607 | 0.06133 | 0.02641 |
| **Gpc3** | 0.01016 | 0.00427 | 0.15197 | 0.00000 | 0.01170 | 0.00896 | 0.00429 | 0.00948 | 0.01159 | 0.00786 | 0.01390 | 0.07496 | 0.00756 | 0.01151 |
| **Gpc4** | 2.49559 | 0.01424 | 4.69452 | 0.53323 | 0.15836 | 0.00000 | 0.00353 | 0.80052 | 0.00721 | 0.00801 | 0.38118 | 0.00776 | 1.37411 | 2.50438 |
| **Gpc6** | 0.05492 | 0.01293 | 0.01409 | 0.04879 | 0.02577 | 0.03956 | 0.00440 | 0.03199 | 0.27511 | 0.06030 | 0.01110 | 0.01000 | 0.09243 | 0.09931 |
| **Gpcpd1** | 20.90004 | 0.03565 | 12.11839 | 7.01392 | 14.87148 | 1.93601 | 5.77040 | 2.65939 | 24.43364 | 0.22910 | 14.60008 | 3.61011 | 12.20242 | 12.03771 |
| **Gpd1** | 0.00000 | 0.00449 | 0.56487 | 0.00000 | 0.88938 | 0.01746 | 0.00406 | 0.00549 | 0.10077 | 0.00549 | 0.00543 | 0.00647 | 0.67571 | 0.67819 |
| **Gpd1l** | 33.29702 | 0.28129 | 53.55065 | 39.44555 | 44.46812 | 38.23771 | 7.72097 | 12.60985 | 40.41475 | 1.28539 | 21.85312 | 6.75024 | 15.05525 | 39.73882 |
| **Gpd2** | 17.98937 | 5.10433 | 21.11327 | 14.96122 | 20.06429 | 16.96249 | 4.59474 | 5.27307 | 30.55707 | 0.30770 | 12.08816 | 8.20064 | 1.19076 | 24.27250 |
| **Gpha2** | 0.00473 | 0.01074 | 0.03165 | 0.25527 | 0.54963 | 1.94066 | 0.00376 | 0.54135 | 0.00964 | 0.01942 | 0.37865 | 0.01576 | 0.24146 | 0.00366 |
| **Gphn** | 1.58108 | 1.85137 | 0.24082 | 0.54433 | 0.00000 | 5.22728 | 0.01692 | 1.32403 | 2.07805 | 0.02912 | 0.16733 | 0.30115 | 0.30773 | 1.05358 |
| **Gpi1** | 3.66913 | 8.28443 | 12.92024 | 15.55942 | 5.66903 | 17.96216 | 4.22834 | 12.87280 | 1.60360 | 0.28246 | 10.51539 | 13.67996 | 2.41678 | 6.66227 |
| **Gpkow** | 7.21847 | 0.02075 | 4.00022 | 1.01103 | 0.40166 | 4.48408 | 0.46842 | 0.09421 | 9.20629 | 0.01292 | 1.89992 | 1.32948 | 7.39439 | 8.14704 |
| **Gpld1** | 0.78608 | 0.00409 | 1.38602 | 0.47752 | 0.35468 | 0.20947 | 0.00409 | 1.41766 | 0.08509 | 0.00165 | 0.54751 | 0.02390 | 0.00263 | 0.90758 |
| **Gpm6a** | 0.00369 | 0.00438 | 0.01013 | 0.01308 | 0.00000 | 0.01105 | 0.00427 | 0.00958 | 0.03049 | 0.00652 | 0.00775 | 0.00943 | 0.10418 | 0.50656 |
| **Gpn1** | 4.98564 | 0.10315 | 6.60806 | 5.82416 | 12.03033 | 10.92137 | 4.48394 | 5.23958 | 6.25032 | 0.16913 | 9.11244 | 10.18764 | 5.45331 | 14.89270 |
| **Gpn2** | 1.35933 | 0.01044 | 1.28099 | 0.44199 | 1.15769 | 0.51697 | 0.04778 | 0.73229 | 0.13221 | 0.38209 | 0.34564 | 0.42455 | 1.59267 | 2.03401 |
| **Gpn3** | 18.08465 | 0.05295 | 12.54720 | 2.24646 | 9.63365 | 7.84952 | 1.96112 | 1.06977 | 0.64313 | 0.04278 | 3.76774 | 1.93635 | 12.17979 | 11.63668 |
| **Gpnmb** | 0.02965 | 0.00435 | 0.01268 | 0.01233 | 0.00703 | 0.01202 | 0.00419 | 0.00959 | 0.06874 | 0.00646 | 0.00799 | 0.00918 | 0.31107 | 0.00000 |
| **Gpr1** | 0.01522 | 0.00444 | 0.01509 | 0.01046 | 0.07034 | 0.01295 | 0.00480 | 0.01031 | 0.02109 | 0.00668 | 0.00873 | 0.00959 | 0.01405 | 0.04377 |
| **Gpr107** | 1.02393 | 0.00130 | 7.31587 | 4.51802 | 2.68250 | 5.69080 | 0.00739 | 0.08983 | 0.04923 | 0.02931 | 0.00000 | 3.40874 | 1.27383 | 4.47756 |
| **Gpr108** | 0.47561 | 0.12752 | 2.40808 | 7.01193 | 0.21558 | 5.33466 | 1.27289 | 3.80542 | 0.03223 | 0.17051 | 2.48651 | 3.87847 | 0.21826 | 0.89003 |
| **Gpr137** | 0.35727 | 0.04324 | 1.33524 | 1.18589 | 0.79531 | 0.98502 | 0.00340 | 2.14562 | 0.00397 | 0.00586 | 0.00000 | 1.02279 | 0.00479 | 0.05031 |
| **Gpr137b** | 16.68234 | 0.11477 | 18.69287 | 18.04650 | 17.14160 | 16.66100 | 4.36342 | 5.50144 | 16.96108 | 2.75950 | 7.73675 | 2.14768 | 4.06422 | 13.03800 |
| **Gpr137b.ps** | 12.53220 | 0.10476 | 15.27084 | 15.98677 | 13.45983 | 12.72089 | 3.68295 | 3.57253 | 10.40916 | 1.61535 | 6.53933 | 2.40539 | 2.16933 | 8.28513 |
| **Gpr143** | 0.45207 | 0.00446 | 0.00950 | 0.00699 | 0.20623 | 0.00730 | 0.00423 | 0.01180 | 0.30388 | 0.00658 | 0.00623 | 0.00689 | 0.00254 | 0.00000 |
| **Gpr146** | 0.12563 | 0.00390 | 0.15229 | 0.00167 | 0.00152 | 0.00571 | 0.00379 | 0.06228 | 0.00873 | 0.00671 | 0.00684 | 0.00677 | 0.88187 | 0.00709 |
| **Gpr155** | 1.69706 | 0.00428 | 0.00773 | 0.00863 | 0.08069 | 0.00680 | 0.00395 | 0.01374 | 0.00929 | 0.00593 | 0.00585 | 0.17973 | 2.29157 | 0.01324 |
| **Gpr156** | 0.00800 | 0.00520 | 0.06493 | 0.13217 | 0.33444 | 0.15672 | 0.00415 | 0.02478 | 0.02239 | 0.00603 | 0.00839 | 0.07072 | 0.42429 | 0.52275 |
| **Gpr157** | 0.84742 | 0.08389 | 0.46813 | 0.62697 | 0.75707 | 0.58870 | 0.19062 | 0.17400 | 1.20477 | 0.03985 | 0.03807 | 0.13450 | 1.84517 | 1.00060 |
| **Gpr160** | 0.12394 | 0.00257 | 0.74453 | 1.68219 | 1.83377 | 2.34801 | 0.00224 | 0.00000 | 7.01726 | 0.01185 | 0.13218 | 1.09713 | 3.56772 | 0.00000 |

| | | | | | | | | | | | | | | |
|---|---|---|---|---|---|---|---|---|---|---|---|---|---|---|
| **Gpr161** | 12.64407 | 0.17291 | 10.65974 | 10.19458 | 11.45895 | 5.74811 | 4.14500 | 7.65960 | 21.11398 | 0.41860 | 8.05989 | 0.98323 | 9.52232 | 13.88256 |
| **Gpr171** | 0.00000 | 0.00392 | 0.21092 | 0.00000 | 0.00000 | 0.66376 | 0.52458 | 0.01932 | 2.33272 | 0.04185 | 0.00727 | 0.00847 | 0.00096 | 0.50837 |
| **Gpr173** | 0.03024 | 0.00429 | 0.04547 | 0.32040 | 0.00818 | 0.02709 | 0.01657 | 0.02916 | 0.07041 | 0.01246 | 0.01436 | 0.02222 | 0.00716 | 0.03349 |
| **Gpr174** | 0.03482 | 0.02926 | 0.03043 | 0.03568 | 0.03457 | 0.06042 | 0.01079 | 0.08917 | 0.05335 | 0.00711 | 0.01603 | 0.02580 | 0.02488 | 0.04239 |
| **Gpr176** | 0.09490 | 0.00436 | 0.01419 | 0.01131 | 0.00811 | 0.01250 | 0.00424 | 0.00984 | 0.00973 | 0.00666 | 0.00785 | 0.00887 | 0.27641 | 0.00242 |
| **Gpr180** | 0.68892 | 0.00355 | 0.83859 | 0.15711 | 0.00000 | 0.63389 | 0.00422 | 0.01103 | 0.09895 | 0.00576 | 0.04135 | 0.76567 | 0.91437 | 2.34657 |
| **Gpr19** | 0.00000 | 0.03439 | 2.73381 | 0.54032 | 0.79208 | 0.82014 | 1.34137 | 0.76960 | 0.00045 | 0.05688 | 0.66163 | 1.34854 | 0.36221 | 0.72309 |
| **Gpr20** | 0.00542 | 0.00391 | 0.00424 | 0.26888 | 0.00819 | 0.01975 | 0.00388 | 1.31983 | 0.01323 | 0.01362 | 0.00278 | 0.00956 | 0.00675 | 0.00920 |
| **Gpr3** | 0.08169 | 0.00400 | 0.00000 | 0.00530 | 0.71911 | 0.00000 | 0.00436 | 0.00891 | 0.06722 | 0.00591 | 0.17988 | 0.00391 | 1.65702 | 1.00098 |
| **Gpr35** | 0.01904 | 0.00451 | 0.01226 | 0.00663 | 0.05814 | 0.16159 | 0.00423 | 0.00600 | 0.01723 | 0.00636 | 0.00756 | 0.00897 | 0.00834 | 0.03632 |
| **Gpr4** | 0.00000 | 5.03063 | 2.23234 | 1.46597 | 0.00000 | 0.37967 | 0.00886 | 3.70963 | 0.01045 | 3.10563 | 0.45443 | 0.00000 | 0.10564 | 0.00000 |
| **Gpr50** | 0.00096 | 0.00360 | 0.00000 | 0.24067 | 0.70793 | 0.04174 | 0.00411 | 0.00383 | 0.00000 | 0.00544 | 0.00876 | 0.00725 | 4.06065 | 1.17479 |
| **Gpr68** | 0.00955 | 0.00452 | 0.02700 | 0.06666 | 0.01342 | 0.07217 | 0.02636 | 0.04885 | 0.04161 | 0.01297 | 0.00594 | 0.01855 | 0.39807 | 0.19390 |
| **Gpr75** | 0.01501 | 0.00446 | 0.01350 | 0.01497 | 0.01024 | 0.01300 | 0.00439 | 0.01129 | 0.01019 | 0.00704 | 0.00894 | 0.00968 | 0.00819 | 0.07939 |
| **Gpr85** | 0.00777 | 0.00448 | 0.23502 | 0.00000 | 0.00789 | 0.29717 | 0.00413 | 0.00525 | 0.01090 | 0.00768 | 0.00859 | 0.00938 | 0.00799 | 0.00754 |
| **Gpr89** | 0.82334 | 0.01275 | 1.74691 | 3.21755 | 1.13261 | 6.43346 | 0.00483 | 1.80356 | 3.37375 | 13.20871 | 4.17487 | 5.02312 | 0.00000 | 0.92188 |
| **Gprasp1** | 1.50468 | 0.84140 | 0.06269 | 0.27469 | 0.96709 | 2.46457 | 0.00331 | 0.94861 | 0.00438 | 0.02267 | 0.00000 | 2.01321 | 1.50903 | 0.75379 |
| **Gprasp2** | 0.00910 | 0.00405 | 0.01229 | 0.02699 | 0.01759 | 0.06028 | 0.00414 | 0.05700 | 0.01241 | 0.00601 | 0.00615 | 0.01505 | 0.00760 | 0.01248 |
| **Gprc5a** | 2.94951 | 3.10533 | 20.31961 | 12.21196 | 6.74010 | 16.87997 | 7.96060 | 7.66933 | 0.85572 | 0.62846 | 9.18082 | 2.05032 | 0.18198 | 9.04999 |
| **Gprc5b** | 0.03285 | 0.00438 | 0.01187 | 0.04485 | 0.02996 | 0.01445 | 0.00421 | 0.00810 | 0.01575 | 0.00913 | 0.00744 | 0.00794 | 0.17030 | 0.01192 |
| **Gprc5c** | 2.43019 | 0.03948 | 9.12433 | 4.73908 | 5.97326 | 3.53874 | 0.00051 | 3.05469 | 6.10159 | 0.02233 | 2.98033 | 1.78881 | 1.52146 | 14.80932 |
| **Gprin1** | 0.09970 | 0.00436 | 0.24009 | 0.00086 | 0.00000 | 0.00413 | 0.00404 | 0.01148 | 0.21093 | 0.00708 | 0.00552 | 0.00863 | 0.02890 | 0.35494 |
| **Gps1** | 9.12250 | 1.62834 | 14.40892 | 7.44625 | 12.69515 | 11.23621 | 3.45397 | 13.13037 | 1.37874 | 0.18529 | 11.94348 | 13.44622 | 11.00018 | 14.58502 |
| **Gps2** | 7.61489 | 0.11717 | 12.02675 | 9.00993 | 8.16093 | 2.51086 | 2.40125 | 3.50866 | 0.39602 | 0.05961 | 3.18391 | 1.83323 | 2.01131 | 2.91507 |
| **Gpsm1** | 0.29797 | 0.01047 | 1.90351 | 0.24900 | 5.89715 | 0.00000 | 0.00351 | 0.62743 | 0.49673 | 0.01167 | 0.18041 | 0.42413 | 0.30211 | 1.03611 |
| **Gpsm2** | 1.46783 | 0.00912 | 4.04532 | 1.82735 | 2.02508 | 3.51428 | 3.27866 | 1.30016 | 1.36446 | 0.07982 | 3.19845 | 0.77311 | 1.67780 | 1.37044 |
| **Gpsm3** | 0.01390 | 0.00397 | 0.03921 | 0.17993 | 0.18342 | 0.04022 | 0.00416 | 0.70771 | 0.01072 | 0.00573 | 0.00345 | 0.00476 | 0.01136 | 0.00777 |
| **Gpt** | 0.00000 | 0.01112 | 0.14438 | 0.93740 | 0.12890 | 0.49139 | 0.00378 | 0.00240 | 0.00889 | 0.00537 | 0.00004 | 0.50100 | 0.30251 | 0.00000 |
| **Gpt2** | 1.98619 | 0.02513 | 1.35317 | 1.80858 | 3.17798 | 0.70648 | 0.00244 | 0.41528 | 4.18001 | 0.00000 | 1.41185 | 4.83747 | 5.13909 | 7.48332 |
| **Gpx1** | 9.66848 | 13.47272 | 27.83092 | 35.04858 | 27.33966 | 10.85073 | 8.30154 | 29.19538 | 0.17256 | 20.92144 | 2.75735 | 22.03628 | 7.29812 | 4.24760 |
| **Gpx2** | 0.00000 | 0.00404 | 2.27926 | 4.14192 | 0.05696 | 2.27763 | 0.34461 | 0.98838 | 0.45602 | 0.04452 | 0.00000 | 0.10309 | 0.00282 | 0.00000 |
| **Gpx2.ps1** | 0.00000 | 0.00425 | 0.62240 | 1.33310 | 0.04057 | 0.31663 | 0.14203 | 0.25432 | 0.14290 | 0.03150 | 0.00489 | 0.02274 | 0.00281 | 0.00000 |
| **Gpx3** | 0.00000 | 0.10381 | 4.41535 | 1.15880 | 4.28736 | 0.00000 | 0.00701 | 0.07984 | 0.00261 | 0.02665 | 0.02897 | 7.81902 | 0.04980 | 1.05266 |
| **Gpx4** | 5.97311 | 10.60361 | 21.83911 | 33.49807 | 20.90548 | 24.88918 | 8.18071 | 24.95800 | 0.00000 | 15.50914 | 16.98582 | 18.48089 | 0.00000 | 0.68153 |
| **Gpx5** | 0.00458 | 0.01285 | 0.21069 | 0.02645 | 0.00458 | 1.82508 | 0.00420 | 0.00597 | 0.01170 | 0.00603 | 0.02111 | 0.00202 | 0.00864 | 0.00427 |
| **Gpx6** | 0.00609 | 0.00429 | 0.00817 | 0.06648 | 0.11399 | 0.00988 | 0.00406 | 0.00746 | 0.01108 | 0.00769 | 0.00759 | 0.01101 | 0.14627 | 0.03760 |
| **Gpx7** | 0.00000 | 0.00184 | 1.33074 | 1.60966 | 0.00000 | 0.00000 | 0.82783 | 0.47867 | 0.05932 | 0.07451 | 2.02719 | 2.46062 | 4.82530 | 0.00000 |
| **Gramd1a** | 0.00432 | 0.00348 | 0.00000 | 0.56473 | 0.00724 | 0.25272 | 0.00384 | 0.00339 | 0.00983 | 0.00505 | 0.00168 | 0.92973 | 0.19943 | 0.00000 |
| **Gramd1b** | 0.01372 | 0.00498 | 0.04336 | 0.00660 | 0.00628 | 0.00648 | 0.00432 | 0.03153 | 0.15903 | 0.00734 | 0.01557 | 0.00878 | 0.26082 | 0.00000 |
| **Gramd1c** | 0.00000 | 0.00446 | 0.00000 | 0.00527 | 1.00527 | 0.02886 | 0.00452 | 0.00845 | 0.86853 | 0.00670 | 0.00513 | 0.00696 | 0.04404 | 0.17973 |
| **Gramd2** | 0.00000 | 0.00362 | 0.54833 | 0.00513 | 0.00000 | 0.00000 | 0.00399 | 0.01076 | 0.00666 | 0.00650 | 2.40553 | 0.00423 | 0.77091 | 0.03406 |
| **Gramd3** | 0.01018 | 0.00483 | 0.00642 | 0.01238 | 0.39646 | 0.01065 | 0.00444 | 0.01820 | 0.00940 | 0.00646 | 0.00835 | 0.00771 | 0.00896 | 0.24281 |
| **Gramd4** | 0.00000 | 0.01995 | 0.87400 | 0.27326 | 0.66349 | 0.26056 | 0.00908 | 0.31071 | 0.13230 | 0.00583 | 0.00000 | 0.83013 | 0.00834 | 0.78702 |
| **Grap** | 0.00767 | 0.00430 | 0.01146 | 0.01638 | 0.24049 | 0.01351 | 0.00429 | 0.01573 | 0.00622 | 0.00750 | 0.00865 | 0.00800 | 0.22657 | 0.05120 |
| **Grasp** | 0.00000 | 0.01136 | 0.02559 | 0.14947 | 0.00229 | 0.13869 | 1.25080 | 0.32270 | 0.21803 | 0.06095 | 0.00000 | 0.41823 | 0.25346 | 0.00000 |
| **Grb10** | 0.22207 | 0.01695 | 0.45397 | 0.16393 | 0.66380 | 0.00000 | 0.00315 | 0.00536 | 2.84913 | 0.00000 | 2.17644 | 0.81452 | 0.74264 | 3.24090 |
| **Grb14** | 0.81395 | 0.00411 | 0.06896 | 0.00549 | 0.34463 | 0.01268 | 0.00414 | 0.02619 | 0.01137 | 0.00599 | 0.00652 | 0.00747 | 0.01319 | 0.03792 |
| **Grb2** | 11.92693 | 2.41887 | 13.49152 | 4.61652 | 18.47180 | 5.53732 | 1.86717 | 2.01850 | 10.20537 | 0.03465 | 5.44714 | 4.71668 | 8.02505 | 11.68755 |
| **Grb7** | 0.00000 | 0.03220 | 7.05122 | 0.03655 | 0.04691 | 0.00000 | 0.00000 | 4.96961 | 0.00000 | 10.08559 | 0.00000 | 0.00000 | 0.00000 | 0.33324 |
| **Grcc10** | 0.06675 | 2.36531 | 0.01102 | 1.04882 | 0.90308 | 0.24539 | 0.51125 | 2.62957 | 0.01071 | 4.62994 | 0.01815 | 1.42464 | 0.05134 | 0.03316 |
| **Greb1** | 0.00716 | 0.01824 | 0.00381 | 1.14932 | 0.09308 | 0.00000 | 0.00956 | 0.00487 | 0.10365 | 0.02896 | 0.00882 | 0.02086 | 0.00910 | 0.03487 |
| **Greb1l** | 3.61610 | 0.01037 | 6.36926 | 1.62494 | 5.33795 | 2.63235 | 0.00104 | 0.00000 | 12.81933 | 0.01085 | 1.19145 | 0.05975 | 0.00000 | 0.53198 |
| **Grhl1** | 0.00000 | 2.54967 | 0.00000 | 0.03328 | 0.37309 | 0.12358 | 1.87456 | 0.75234 | 3.26159 | 0.04758 | 1.19009 | 0.00000 | 0.13757 | 0.00000 |
| **Grhl2** | 0.04531 | 0.01713 | 3.70012 | 4.46724 | 2.45053 | 5.07571 | 0.14263 | 3.09409 | 3.14025 | 0.05730 | 1.62180 | 0.86447 | 6.99445 | 5.32802 |
| **Grhl3** | 1.16243 | 0.00391 | 0.08009 | 0.29000 | 0.37927 | 0.00000 | 0.00426 | 0.00734 | 0.21799 | 0.01234 | 0.01034 | 0.00762 | 1.19052 | 0.84943 |

| | | | | | | | | | | | | | | |
|---|---|---|---|---|---|---|---|---|---|---|---|---|---|---|
| **Grhpr** | 3.40574 | 0.14068 | 5.67959 | 5.98466 | 3.05157 | 2.82120 | 3.17027 | 6.94640 | 0.03589 | 0.28240 | 3.16652 | 6.76685 | 1.33951 | 1.23264 |
| **Gria4** | 0.00924 | 0.00444 | 0.01408 | 0.01343 | 0.01141 | 0.01283 | 0.00434 | 0.01027 | 0.01089 | 0.00695 | 0.00885 | 0.01000 | 0.00828 | 0.05528 |
| **Grid1** | 0.10221 | 0.00450 | 0.01801 | 0.02665 | 0.00737 | 0.04684 | 0.00417 | 0.00609 | 0.02437 | 0.01004 | 0.00647 | 0.00818 | 0.41978 | 0.00156 |
| **Grid2** | 0.00617 | 0.00441 | 0.00095 | 0.08991 | 0.08185 | 0.01006 | 0.00412 | 0.00933 | 0.21546 | 0.00683 | 0.00616 | 0.00775 | 0.00671 | 0.00250 |
| **Grid2ip** | 0.00272 | 0.00415 | 0.27700 | 0.27063 | 0.09656 | 0.00000 | 0.00413 | 0.00709 | 0.05661 | 0.00586 | 0.01256 | 0.01480 | 0.04028 | 0.05927 |
| **Grik1** | 0.58347 | 0.00470 | 0.01201 | 0.01404 | 0.00339 | 0.00747 | 0.00415 | 0.01460 | 0.01568 | 0.00623 | 0.00636 | 0.05134 | 0.00813 | 0.01404 |
| **Grik5** | 0.01607 | 0.00441 | 0.02122 | 0.02091 | 0.01292 | 0.01378 | 0.00423 | 0.00922 | 0.01981 | 0.00665 | 0.01061 | 0.01181 | 0.00725 | 0.01076 |
| **Grin1** | 0.00000 | 0.00407 | 0.56921 | 0.38144 | 1.79794 | 0.00000 | 0.00395 | 0.00900 | 0.02085 | 0.01344 | 0.00905 | 0.01287 | 0.00000 | 0.03755 |
| **Grin2d** | 1.54465 | 0.00462 | 0.99045 | 0.04298 | 2.65872 | 0.16524 | 0.00391 | 0.00395 | 0.00647 | 0.00630 | 0.00400 | 0.00577 | 0.10478 | 0.07702 |
| **Grin3a** | 0.00478 | 0.00391 | 0.06164 | 0.35106 | 0.00382 | 0.50106 | 0.00386 | 0.00260 | 0.02041 | 0.00513 | 0.00567 | 0.15537 | 0.00707 | 0.00561 |
| **Grin3b** | 0.02463 | 0.00421 | 0.00530 | 0.39691 | 0.06019 | 0.12207 | 0.01081 | 0.03164 | 0.00958 | 0.00587 | 0.00479 | 0.00458 | 0.16782 | 0.02594 |
| **Grina** | 1.13254 | 0.02618 | 0.18697 | 0.69431 | 0.99405 | 0.61648 | 0.00311 | 2.14534 | 0.00489 | 0.02093 | 0.08488 | 1.12464 | 0.39961 | 0.15096 |
| **Grip1** | 0.00000 | 0.00419 | 0.12542 | 0.00280 | 0.03548 | 0.00851 | 0.00409 | 0.00948 | 0.00553 | 0.00616 | 0.00812 | 0.00830 | 1.13170 | 0.08037 |
| **Gripap1** | 0.56502 | 0.00362 | 0.78926 | 0.63237 | 1.13257 | 1.19925 | 0.00340 | 0.00000 | 0.62494 | 0.00453 | 0.00000 | 0.61985 | 2.09567 | 2.17245 |
| **Grk1** | 0.06271 | 0.17001 | 0.05451 | 0.04555 | 0.10040 | 0.03428 | 0.01044 | 0.04992 | 0.38226 | 0.00796 | 0.01419 | 0.12653 | 0.30551 | 0.23670 |
| **Grk4** | 0.48778 | 0.14297 | 0.58688 | 0.35911 | 0.46336 | 0.36257 | 0.35501 | 0.27370 | 4.09866 | 0.00518 | 0.27075 | 0.27505 | 2.07849 | 1.06819 |
| **Grk5** | 2.45600 | 0.00324 | 5.87053 | 1.95654 | 3.62188 | 4.59394 | 0.00296 | 0.08894 | 6.23907 | 0.00793 | 0.49617 | 0.00000 | 5.38972 | 4.31987 |
| **Grk6** | 4.80483 | 4.90687 | 9.77463 | 5.70278 | 3.91280 | 4.21333 | 0.26420 | 4.32930 | 0.39113 | 0.08469 | 2.69204 | 5.65273 | 0.67386 | 2.29018 |
| **Grm2** | 0.01273 | 0.00426 | 0.10065 | 0.00782 | 0.00397 | 0.01169 | 0.00415 | 0.00856 | 0.00656 | 0.00761 | 0.00812 | 0.00770 | 1.00137 | 0.04218 |
| **Grm4** | 0.00668 | 0.00403 | 0.00795 | 0.02965 | 0.01120 | 1.55649 | 0.00403 | 0.06441 | 0.00982 | 0.00751 | 0.00435 | 0.00000 | 0.01887 | 0.00575 |
| **Grm8** | 0.00809 | 0.00444 | 0.01304 | 0.01392 | 0.00866 | 0.01252 | 0.00432 | 0.01016 | 0.01012 | 0.00710 | 0.00857 | 0.00991 | 0.00601 | 0.13598 |
| **Grn** | 17.32461 | 0.35336 | 40.63231 | 37.49500 | 20.91162 | 17.25621 | 6.27200 | 7.90188 | 5.13209 | 0.75511 | 7.67284 | 10.92425 | 1.86451 | 14.24269 |
| **Grpel1** | 11.45950 | 7.44049 | 18.84778 | 19.59782 | 18.14547 | 18.53471 | 5.15298 | 17.34554 | 3.84323 | 12.77222 | 12.94485 | 15.71610 | 1.62887 | 9.36373 |
| **Grpel2** | 6.33880 | 0.05206 | 7.14154 | 4.28593 | 4.84303 | 5.41020 | 4.16113 | 2.82606 | 2.33971 | 0.24223 | 4.12608 | 0.76535 | 0.06826 | 3.07422 |
| **Grrp1** | 0.01218 | 0.00442 | 0.01453 | 0.01306 | 0.01454 | 0.01339 | 0.00426 | 0.00994 | 0.01291 | 0.00690 | 0.00877 | 0.00983 | 0.08058 | 0.00520 |
| **Grsf1** | 14.01381 | 0.08758 | 9.47760 | 4.50690 | 13.90228 | 11.81460 | 3.26954 | 1.67333 | 17.47290 | 0.08861 | 6.16714 | 10.81037 | 13.69030 | 17.89735 |
| **Grtp1** | 1.05789 | 0.15721 | 6.21640 | 3.28866 | 2.83345 | 1.84255 | 1.95921 | 7.42316 | 0.13088 | 12.93657 | 2.57175 | 4.58242 | 0.19058 | 0.70168 |
| **Grwd1** | 8.59305 | 0.22223 | 24.57919 | 13.97434 | 21.58833 | 11.46672 | 0.87903 | 8.95663 | 0.63372 | 8.35508 | 8.75408 | 10.34321 | 9.37771 | 17.41902 |
| **Grxcr2** | 0.00319 | 0.00363 | 0.01019 | 0.00774 | 0.00358 | 0.01089 | 0.00411 | 0.00544 | 0.00898 | 0.00732 | 0.00383 | 0.77593 | 0.00471 | 0.50722 |
| **Gsap** | 0.00712 | 0.00384 | 0.20459 | 0.00000 | 0.05496 | 0.00702 | 0.00426 | 0.00668 | 0.00942 | 0.00635 | 0.00555 | 0.54592 | 0.00819 | 0.01184 |
| **Gsc** | 0.00264 | 0.01061 | 0.04966 | 0.38283 | 0.16969 | 0.00865 | 0.00341 | 0.00311 | 0.01431 | 0.00511 | 0.00980 | 1.03608 | 0.00457 | 0.01124 |
| **Gsdma** | 0.81853 | 0.00365 | 0.39559 | 0.40801 | 0.01612 | 0.00000 | 0.09447 | 0.47283 | 0.07867 | 0.01340 | 0.04018 | 0.71553 | 0.02969 | 0.70988 |
| **Gsdma2** | 0.00358 | 0.00479 | 0.06484 | 0.39014 | 0.00754 | 0.01128 | 0.02695 | 0.01323 | 0.11094 | 0.01101 | 0.00686 | 0.04706 | 0.04062 | 0.00000 |
| **Gsdma3** | 0.00744 | 0.00439 | 0.38496 | 0.22408 | 0.12877 | 0.00000 | 1.81607 | 0.33217 | 0.65106 | 0.01243 | 0.08002 | 0.07975 | 0.00367 | 0.00000 |
| **Gsdmc** | 0.01628 | 0.00425 | 0.11574 | 0.10596 | 0.11692 | 0.10965 | 0.00399 | 0.00451 | 0.23471 | 0.00632 | 0.01416 | 0.00804 | 0.24094 | 0.11667 |
| **Gsdmc2** | 0.00000 | 0.22253 | 1.03978 | 0.68526 | 1.28455 | 0.69339 | 0.00333 | 0.00000 | 1.95354 | 0.00396 | 0.00631 | 0.06559 | 1.10610 | 0.54656 |
| **Gsdmc3** | 0.00000 | 0.05401 | 0.27883 | 0.16538 | 0.44132 | 0.25428 | 0.00388 | 0.00203 | 0.94134 | 0.00577 | 0.00629 | 0.08438 | 0.36210 | 0.42282 |
| **Gsdmc4** | 0.00845 | 0.53969 | 0.25707 | 0.27489 | 0.47506 | 0.17229 | 0.00348 | 0.00648 | 1.02550 | 0.00595 | 0.00863 | 0.02410 | 0.04588 | 0.16934 |
| **Gsdmd** | 0.00490 | 0.03377 | 0.00103 | 0.00000 | 0.01082 | 0.46869 | 0.00302 | 2.23264 | 0.00751 | 0.01684 | 0.00000 | 0.90133 | 0.67687 | 0.01289 |
| **Gse1** | 0.84006 | 0.00315 | 2.70282 | 0.00000 | 5.01441 | 0.33626 | 0.00310 | 0.58236 | 0.00348 | 0.00062 | 0.32966 | 0.58859 | 1.14821 | 1.17226 |
| **Gsg1** | 0.05155 | 0.00436 | 0.01093 | 0.04911 | 0.00998 | 0.01199 | 0.00432 | 0.00874 | 0.01067 | 0.00639 | 0.00732 | 0.00815 | 0.05980 | 0.16658 |
| **Gsg1l** | 0.05985 | 0.00429 | 0.07308 | 0.01675 | 0.03339 | 0.08101 | 0.01150 | 0.06329 | 0.03617 | 0.01210 | 0.01446 | 0.02396 | 0.00484 | 0.08877 |
| **Gsg2** | 3.47789 | 0.01046 | 3.68465 | 3.70630 | 3.46196 | 1.07102 | 0.57025 | 2.05255 | 5.08680 | 0.03315 | 2.34717 | 0.07386 | 6.70203 | 5.81899 |
| **Gsk3a** | 0.41560 | 0.01027 | 0.00000 | 0.88870 | 0.86493 | 1.42083 | 0.00388 | 0.27581 | 0.11317 | 0.01757 | 0.13510 | 0.94201 | 3.66003 | 0.00000 |
| **Gsk3b** | 5.08430 | 0.09882 | 5.71313 | 3.59962 | 3.35756 | 2.24062 | 0.75998 | 2.57106 | 6.33336 | 0.01151 | 3.26192 | 0.53838 | 7.15331 | 7.65649 |
| **Gskip** | 2.05129 | 0.00407 | 1.26812 | 0.33758 | 1.95611 | 2.33525 | 0.00900 | 0.01003 | 5.10528 | 0.88753 | 0.13895 | 0.41766 | 1.68572 | 1.62649 |
| **Gsn** | 0.03270 | 5.02151 | 1.65138 | 5.57242 | 0.00000 | 8.31127 | 2.95548 | 4.53257 | 0.01045 | 5.97008 | 2.09196 | 0.00000 | 0.00000 | 0.00000 |
| **Gspt1** | 19.63617 | 8.01268 | 21.17017 | 8.60830 | 20.05644 | 18.04094 | 4.74413 | 5.28068 | 29.01019 | 0.12137 | 15.43926 | 13.80615 | 8.69322 | 12.15518 |
| **Gspt2** | 0.84973 | 0.01080 | 0.16982 | 1.88507 | 1.25388 | 1.22456 | 0.00332 | 0.00081 | 2.43537 | 0.00912 | 0.00101 | 0.01024 | 0.27675 | 0.85308 |
| **Gsr** | 3.22837 | 0.03534 | 6.58286 | 3.01122 | 8.47826 | 3.91366 | 0.90802 | 2.63018 | 1.58350 | 0.05533 | 2.68538 | 6.96360 | 1.08197 | 9.13512 |
| **Gss** | 0.00000 | 0.32603 | 7.41399 | 14.54713 | 0.85326 | 5.60936 | 3.16822 | 13.70544 | 0.02882 | 2.34484 | 9.71602 | 12.30531 | 1.70992 | 0.00000 |
| **Gsta1** | 0.36219 | 0.01178 | 1.25349 | 0.33487 | 1.42951 | 0.22164 | 0.34827 | 1.93384 | 1.24250 | 0.20772 | 0.79179 | 0.04216 | 0.00000 | 0.09254 |
| **Gsta2** | 0.11968 | 0.00411 | 1.09042 | 0.39766 | 0.42628 | 0.04541 | 0.51133 | 0.79689 | 0.68349 | 0.94933 | 0.30231 | 0.00164 | 0.00000 | 0.58087 |
| **Gsta3** | 0.01255 | 0.00425 | 0.01157 | 0.00000 | 0.01560 | 0.10659 | 0.00422 | 0.00605 | 0.01122 | 0.01681 | 0.24992 | 0.02615 | 0.00749 | 0.00889 |
| **Gsta4** | 1.63841 | 12.27011 | 3.35799 | 0.00020 | 0.89761 | 11.20911 | 5.54615 | 22.66556 | 0.39915 | 22.31763 | 6.88849 | 6.31314 | 0.36791 | 1.03501 |

| | | | | | | | | | | | | | | |
|---|---|---|---|---|---|---|---|---|---|---|---|---|---|---|
| **Gstcd** | 0.48122 | 0.00365 | 0.15396 | 0.34720 | 0.51047 | 0.30761 | 1.48506 | 0.16890 | 0.02249 | 0.04799 | 0.49182 | 0.17542 | 1.02813 | 0.15783 |
| **Gstk1** | 0.00478 | 0.00410 | 0.00305 | 0.33601 | 0.00792 | 0.43569 | 0.00412 | 1.97260 | 0.01227 | 0.01595 | 0.00512 | 0.00408 | 0.00766 | 0.00846 |
| **Gstm1** | 0.00788 | 0.07012 | 0.00000 | 0.00000 | 3.93616 | 0.62611 | 0.00459 | 0.71943 | 0.00595 | 0.00561 | 0.00000 | 2.85117 | 0.00000 | 0.12877 |
| **Gstm2** | 0.00273 | 0.07233 | 0.00000 | 0.00000 | 2.67922 | 1.10832 | 0.02187 | 4.53796 | 0.00687 | 0.05993 | 0.00000 | 1.44238 | 0.00000 | 0.00684 |
| **Gstm3** | 0.00000 | 0.00443 | 0.09682 | 0.27386 | 0.35497 | 0.08904 | 0.00484 | 0.18714 | 0.05434 | 0.00779 | 0.01170 | 0.42470 | 0.00358 | 0.01343 |
| **Gstm4** | 0.04290 | 0.00095 | 0.05013 | 0.00000 | 0.00000 | 2.87580 | 4.57464 | 0.03096 | 0.00051 | 0.10154 | 2.63340 | 3.35787 | 0.00307 | 0.46400 |
| **Gstm5** | 0.00708 | 0.00438 | 0.01822 | 0.00908 | 0.23468 | 0.01071 | 0.00445 | 0.00999 | 0.07501 | 0.00637 | 0.00750 | 0.01459 | 0.16793 | 0.18607 |
| **Gstm6** | 0.00426 | 0.01196 | 0.00426 | 0.00000 | 0.00831 | 2.20416 | 0.00401 | 0.00735 | 0.00386 | 0.01279 | 0.10474 | 0.42046 | 0.00536 | 0.09168 |
| **Gstm7** | 0.00000 | 0.00314 | 0.45581 | 0.00000 | 0.35586 | 0.00864 | 0.00358 | 0.00410 | 0.01117 | 0.00589 | 0.00359 | 1.65625 | 0.00344 | 0.01543 |
| **Gsto1** | 8.04504 | 13.77549 | 37.10606 | 41.17380 | 8.23357 | 34.23607 | 16.69214 | 32.88300 | 0.23782 | 21.98064 | 14.45691 | 18.34390 | 0.56855 | 0.00000 |
| **Gstp1** | 2.68586 | 1.58331 | 10.68757 | 24.69538 | 2.38502 | 16.88585 | 6.91289 | 17.36816 | 0.96802 | 14.19754 | 13.93300 | 14.22134 | 0.03153 | 3.13812 |
| **Gstp2** | 1.63683 | 2.54760 | 9.18515 | 22.16701 | 1.83414 | 18.95142 | 6.68722 | 16.84602 | 0.64499 | 14.25229 | 14.13866 | 16.88649 | 0.09649 | 1.88678 |
| **Gstt2** | 0.00348 | 0.01108 | 0.00000 | 0.00000 | 0.90738 | 0.28548 | 0.00392 | 0.00343 | 0.01081 | 0.00541 | 0.00359 | 0.44416 | 0.12841 | 0.34835 |
| **Gstz1** | 2.19302 | 0.13057 | 4.29464 | 4.50633 | 1.58621 | 5.03818 | 2.44065 | 1.36042 | 0.00000 | 0.03939 | 6.12601 | 4.94301 | 0.00000 | 4.13511 |
| **Gt.ROSA.26S** | 1.59074 | 0.04076 | 0.63982 | 0.43773 | 1.21521 | 0.75980 | 0.00353 | 2.18748 | 0.02312 | 0.00687 | 0.00000 | 0.67092 | 1.92702 | 0.41207 |
| **Gtdc1** | 0.46731 | 0.02587 | 1.42353 | 0.00000 | 2.01181 | 1.50687 | 1.70922 | 0.06630 | 1.55800 | 0.01410 | 0.28169 | 0.93816 | 6.35235 | 6.64625 |
| **Gtf2a1** | 2.18033 | 0.18430 | 2.66519 | 1.74388 | 4.31137 | 3.62946 | 0.87876 | 2.40807 | 13.37658 | 0.00000 | 5.66172 | 3.51615 | 9.95879 | 6.92983 |
| **Gtf2a1l** | 0.02400 | 0.00443 | 0.01305 | 0.01324 | 0.00937 | 0.01259 | 0.00433 | 0.01019 | 0.00860 | 0.00720 | 0.00842 | 0.00983 | 0.01554 | 0.11577 |
| **Gtf2a2** | 1.48643 | 0.10271 | 1.28897 | 6.40810 | 2.85415 | 4.99095 | 2.97644 | 5.12601 | 0.64141 | 0.15578 | 3.45202 | 8.29870 | 0.31156 | 4.41941 |
| **Gtf2b** | 3.90120 | 0.06604 | 7.01609 | 6.93410 | 3.62584 | 10.79465 | 5.48413 | 6.34995 | 3.24240 | 0.25943 | 12.22410 | 8.18945 | 6.89855 | 6.63739 |
| **Gtf2e1** | 2.36277 | 0.10038 | 5.55486 | 1.93862 | 3.02134 | 4.60853 | 0.93817 | 2.50981 | 4.30613 | 0.04283 | 8.77169 | 4.32635 | 1.94018 | 5.78929 |
| **Gtf2e2** | 7.09794 | 0.05899 | 7.73179 | 4.04249 | 5.14726 | 9.60651 | 0.02290 | 4.79151 | 7.75893 | 0.03131 | 6.39603 | 7.75631 | 5.28331 | 3.51395 |
| **Gtf2f1** | 4.59761 | 0.29086 | 23.83248 | 18.04271 | 19.01516 | 7.63267 | 6.45710 | 13.57399 | 0.00000 | 1.43626 | 7.30735 | 10.19709 | 5.52055 | 4.22301 |
| **Gtf2f2** | 9.65843 | 1.41934 | 17.15010 | 13.75312 | 12.32660 | 9.47777 | 2.05368 | 6.16379 | 4.46340 | 0.07689 | 5.69685 | 6.95248 | 13.34038 | 6.92153 |
| **Gtf2h1** | 14.00985 | 0.33449 | 9.06153 | 8.63716 | 15.04156 | 8.82975 | 0.02864 | 5.46291 | 9.91028 | 0.04156 | 3.27490 | 3.69884 | 11.22516 | 15.54328 |
| **Gtf2h2** | 16.82216 | 1.47294 | 14.93051 | 7.54806 | 17.16527 | 9.55708 | 0.06605 | 5.70098 | 12.58648 | 9.55641 | 6.05234 | 6.38035 | 6.29741 | 13.15045 |
| **Gtf2h3** | 0.21238 | 0.04712 | 5.43518 | 0.87238 | 5.18428 | 3.51891 | 0.01877 | 0.25474 | 3.75652 | 0.00000 | 5.84987 | 0.58577 | 0.60307 | 13.27335 |
| **Gtf2h4** | 0.00000 | 0.10800 | 8.45172 | 9.35288 | 14.27598 | 0.00000 | 0.87692 | 4.38115 | 0.02034 | 0.03739 | 0.11010 | 4.46278 | 0.00000 | 0.00000 |
| **Gtf2h5** | 7.92280 | 0.60333 | 3.68132 | 10.33093 | 10.02043 | 7.71537 | 0.40585 | 7.51681 | 4.54599 | 3.93215 | 5.26074 | 7.65338 | 2.40667 | 10.63577 |
| **Gtf2i** | 2.06848 | 0.01508 | 6.21486 | 2.07355 | 9.85411 | 3.55914 | 0.53739 | 1.21319 | 0.00000 | 0.01449 | 0.83296 | 3.52504 | 1.66985 | 5.37555 |
| **Gtf2ird1** | 1.16877 | 0.06421 | 7.84563 | 5.35857 | 2.48163 | 2.81385 | 3.37851 | 3.02688 | 0.39471 | 0.09731 | 3.60943 | 5.22636 | 2.84251 | 4.84979 |
| **Gtf2ird2** | 0.08229 | 0.00383 | 0.92283 | 0.00000 | 1.36163 | 0.00000 | 0.00386 | 0.41367 | 0.00681 | 0.00457 | 0.00287 | 0.00679 | 0.96771 | 0.25860 |
| **Gtf3a** | 0.18049 | 0.00423 | 0.26195 | 0.22111 | 1.09065 | 0.43933 | 1.01543 | 0.23509 | 0.01361 | 0.09791 | 0.06725 | 0.00159 | 0.00317 | 0.00021 |
| **Gtf3c1** | 1.88653 | 0.09900 | 6.79066 | 6.54377 | 13.00621 | 2.92704 | 0.03601 | 1.87487 | 0.69416 | 0.08848 | 0.24002 | 3.26196 | 7.90046 | 2.22465 |
| **Gtf3c2** | 6.95238 | 0.06668 | 11.00193 | 7.78096 | 13.26145 | 9.30881 | 5.81614 | 3.10968 | 15.91085 | 0.16324 | 6.13978 | 6.37814 | 10.61719 | 14.00673 |
| **Gtf3c3** | 2.32971 | 3.36905 | 3.03075 | 1.24096 | 2.85239 | 2.20656 | 1.10146 | 0.10073 | 6.86043 | 10.29062 | 6.05159 | 1.85898 | 5.95722 | 6.55341 |
| **Gtf3c4** | 8.26149 | 0.02928 | 5.53871 | 3.70807 | 8.34252 | 5.17020 | 1.87877 | 1.27924 | 5.88765 | 0.11373 | 2.11124 | 2.44380 | 9.51409 | 10.93425 |
| **Gtf3c5** | 5.96359 | 0.03184 | 1.95438 | 1.85598 | 10.08267 | 3.49189 | 0.04143 | 1.22878 | 3.36791 | 0.04761 | 2.79117 | 2.65231 | 9.33635 | 11.61888 |
| **Gtf3c6** | 2.60950 | 0.03840 | 3.07733 | 4.76646 | 4.48027 | 3.10835 | 0.01331 | 3.69058 | 4.89811 | 0.64140 | 5.03480 | 3.73493 | 9.78667 | 5.88090 |
| **Gtpbp1** | 0.74390 | 1.56879 | 2.93420 | 0.39874 | 1.11397 | 2.89251 | 0.00696 | 6.45984 | 0.00464 | 0.01804 | 1.18862 | 2.44042 | 0.22561 | 0.34983 |
| **Gtpbp10** | 2.86770 | 1.59658 | 2.92626 | 0.84825 | 0.78653 | 0.88036 | 0.17062 | 0.00730 | 7.74616 | 0.00505 | 4.14833 | 0.65098 | 1.08764 | 9.41129 |
| **Gtpbp2** | 0.88841 | 0.00444 | 0.01020 | 0.02155 | 0.03276 | 0.00574 | 0.00504 | 0.17597 | 0.01499 | 0.00813 | 0.00664 | 0.01779 | 0.01278 | 0.08845 |
| **Gtpbp3** | 0.00000 | 0.00968 | 1.87210 | 2.68830 | 0.67450 | 0.94987 | 0.07676 | 0.95795 | 0.04941 | 0.01005 | 0.01301 | 2.23558 | 3.10905 | 0.36496 |
| **Gtpbp4** | 13.17652 | 0.44517 | 16.36913 | 6.49822 | 16.84846 | 16.44569 | 3.16688 | 6.96779 | 27.00982 | 0.11235 | 13.53632 | 9.39833 | 10.21938 | 21.19986 |
| **Gtpbp6** | 0.00000 | 0.00377 | 0.67769 | 1.45791 | 0.90002 | 0.00000 | 0.00389 | 0.54686 | 0.01152 | 0.00418 | 0.00207 | 0.51956 | 0.00319 | 0.00000 |
| **Gtpbp8** | 0.23732 | 0.01074 | 1.05473 | 1.23055 | 3.50058 | 1.02479 | 0.00947 | 1.07716 | 2.25510 | 0.00877 | 1.42077 | 1.16074 | 0.00000 | 1.08611 |
| **Gtse1** | 1.24973 | 0.05976 | 6.22894 | 4.78698 | 1.66392 | 8.98693 | 2.61444 | 3.17213 | 0.15936 | 0.16619 | 5.07430 | 5.13129 | 9.35536 | 2.02039 |
| **Gtsf1** | 0.00000 | 0.01034 | 0.51247 | 0.17846 | 0.85126 | 3.15619 | 0.00323 | 0.00000 | 2.18111 | 0.01071 | 0.00000 | 0.40194 | 1.49927 | 0.30641 |
| **Gtsf1l** | 1.76127 | 3.88221 | 1.52722 | 3.87837 | 2.09948 | 1.31225 | 2.40069 | 2.75132 | 0.03115 | 0.12725 | 2.78110 | 5.76784 | 0.74709 | 1.76110 |
| **Guca1a** | 0.44797 | 3.98911 | 24.43579 | 31.10864 | 26.64278 | 1.20185 | 7.96093 | 9.29248 | 0.00000 | 14.74212 | 6.72362 | 2.86631 | 0.87326 | 0.00000 |
| **Guca1b** | 0.45474 | 0.00406 | 0.05305 | 0.39790 | 0.17047 | 0.23413 | 0.00904 | 0.00285 | 0.02618 | 0.00609 | 0.00565 | 0.01329 | 0.06185 | 1.12632 |
| **Gucd1** | 1.47363 | 0.04137 | 3.47028 | 0.92602 | 1.90947 | 4.23806 | 1.01104 | 5.61299 | 2.35892 | 0.04641 | 2.51352 | 2.13293 | 0.00000 | 3.96649 |
| **Gucy1a2** | 0.02577 | 0.09352 | 0.00000 | 0.04420 | 0.00820 | 0.00375 | 0.01684 | 0.06934 | 1.55412 | 0.00768 | 0.02030 | 0.03316 | 0.03560 | 0.02623 |
| **Gucy1b2** | 0.04706 | 0.00435 | 1.93620 | 0.00323 | 0.09590 | 0.00253 | 0.00366 | 0.00802 | 0.08760 | 0.00468 | 0.00512 | 0.00838 | 0.00000 | 0.06230 |
| **Gucy2c** | 0.01073 | 0.00436 | 0.00744 | 0.08156 | 0.04390 | 0.01082 | 0.00457 | 0.00816 | 0.01061 | 0.00584 | 0.00687 | 0.00781 | 0.03475 | 0.39054 |

| | | | | | | | | | | | | | | |
|---|---|---|---|---|---|---|---|---|---|---|---|---|---|---|
| **Gucy2e** | 3.35783 | 0.00284 | 3.67499 | 0.86728 | 1.87454 | 0.34750 | 2.76934 | 0.00000 | 4.39992 | 0.10339 | 1.30182 | 0.00000 | 2.02478 | 7.50719 |
| **Guf1** | 0.96883 | 0.00358 | 0.47835 | 0.79459 | 0.18040 | 0.22110 | 2.26548 | 0.00000 | 1.98631 | 0.02344 | 0.73837 | 0.18111 | 0.00072 | 0.62944 |
| **Guk1** | 4.40918 | 0.10068 | 2.94651 | 9.81098 | 6.44144 | 8.08832 | 0.54935 | 4.29436 | 0.86551 | 0.09449 | 1.93951 | 5.34294 | 0.10076 | 1.87836 |
| **Gulo** | 18.28046 | 0.08477 | 39.17903 | 26.01421 | 28.03797 | 14.93642 | 2.77848 | 0.00000 | 8.93274 | 0.12300 | 19.55195 | 8.70125 | 0.00000 | 31.03431 |
| **Gusb** | 1.74266 | 0.03163 | 2.30439 | 2.00920 | 3.68194 | 1.86709 | 0.00806 | 0.30213 | 0.02194 | 0.00000 | 3.63310 | 2.35530 | 2.17322 | 1.71749 |
| **Gxylt1** | 7.63621 | 0.01492 | 1.82861 | 0.00000 | 4.67260 | 4.10500 | 0.61047 | 0.60947 | 8.29049 | 0.03138 | 0.80269 | 3.51520 | 6.09656 | 4.35469 |
| **Gyg** | 0.85619 | 7.81387 | 0.32059 | 1.22878 | 0.08438 | 2.08472 | 0.00153 | 0.21810 | 2.99567 | 0.00576 | 1.05758 | 1.23237 | 5.05263 | 3.01941 |
| **Gyltl1b** | 0.00000 | 0.00390 | 1.09916 | 0.00000 | 0.21697 | 1.07051 | 0.00954 | 3.26012 | 0.00875 | 0.00871 | 1.30353 | 0.30507 | 0.00292 | 0.00000 |
| **Gys1** | 2.55414 | 0.04434 | 5.65495 | 3.40054 | 2.78680 | 5.94553 | 0.60009 | 1.64130 | 0.00399 | 0.02282 | 1.69181 | 3.11225 | 1.18167 | 1.54461 |
| **Gzf1** | 0.10121 | 0.03351 | 2.13670 | 0.57237 | 1.23332 | 3.39715 | 0.02839 | 1.69500 | 1.00559 | 0.03389 | 3.45786 | 3.21588 | 1.35788 | 3.44791 |
| **Gzmm** | 0.78418 | 0.00429 | 1.07282 | 0.67296 | 0.75966 | 0.17596 | 1.17829 | 0.19629 | 0.02597 | 0.03120 | 0.00074 | 0.00795 | 0.78777 | 0.09953 |
| **H13** | 6.89687 | 0.02131 | 6.22490 | 5.37848 | 6.09535 | 5.13957 | 1.78259 | 1.68198 | 0.18711 | 0.08661 | 1.28919 | 6.74484 | 0.99975 | 1.53736 |
| **H19** | 0.00774 | 0.05841 | 0.01416 | 0.03055 | 0.24282 | 0.01397 | 0.00426 | 0.05167 | 0.01136 | 0.00870 | 0.00913 | 0.02118 | 0.00537 | 0.00793 |
| **H1f0** | 2.44170 | 0.02268 | 0.56529 | 1.60266 | 1.72052 | 0.00000 | 0.00663 | 2.09355 | 2.39597 | 8.05405 | 0.73639 | 1.38710 | 1.05546 | 3.27566 |
| **H1foo** | 0.75898 | 0.00360 | 0.42154 | 0.90400 | 1.68696 | 0.36948 | 0.00889 | 0.00205 | 0.82315 | 0.01902 | 0.00043 | 0.00000 | 5.87541 | 1.27165 |
| **H1fx** | 0.00864 | 0.00445 | 0.01232 | 0.03342 | 0.03427 | 0.00926 | 0.00457 | 0.01484 | 0.01154 | 0.05756 | 0.00815 | 0.01049 | 0.00677 | 0.01281 |
| **H2.Aa** | 0.00661 | 0.00447 | 0.07134 | 0.00582 | 0.53164 | 0.01275 | 0.00463 | 0.01107 | 0.01311 | 0.00668 | 0.00852 | 0.00780 | 0.00307 | 0.05295 |
| **H2.Ab1** | 0.00343 | 0.00426 | 0.00000 | 0.65921 | 0.43425 | 0.04120 | 0.00406 | 0.00261 | 0.01193 | 0.00870 | 0.91858 | 0.00422 | 0.00550 | 0.00000 |
| **H2.Bl** | 0.00000 | 5.74767 | 1.23033 | 2.22134 | 0.07483 | 0.62622 | 3.08467 | 5.04002 | 0.02531 | 5.20319 | 0.00000 | 0.11052 | 0.15451 | 0.10236 |
| **H2.D1** | 0.83647 | 2.29885 | 0.79568 | 1.22055 | 0.53767 | 0.46358 | 0.12344 | 2.29742 | 0.83465 | 0.02734 | 0.01301 | 0.58602 | 0.92897 | 0.22934 |
| **H2.DMa** | 0.08245 | 0.01112 | 1.89830 | 1.22031 | 1.76691 | 0.43900 | 0.00326 | 0.00000 | 0.15417 | 0.00000 | 2.33745 | 0.39917 | 0.18759 | 0.26460 |
| **H2.DMb1** | 0.00455 | 0.00474 | 0.04365 | 1.48549 | 0.00718 | 0.40904 | 0.00416 | 0.00649 | 0.00716 | 0.00906 | 0.00373 | 0.00732 | 0.11797 | 0.35545 |
| **H2.DMb2** | 0.00413 | 0.00493 | 0.02593 | 0.33616 | 0.00214 | 0.06900 | 0.00419 | 0.01346 | 0.03208 | 0.00861 | 0.00806 | 0.00747 | 0.00803 | 0.09976 |
| **H2.Eb1** | 0.00000 | 0.00350 | 1.57096 | 0.00000 | 1.98833 | 0.00000 | 0.00394 | 0.00000 | 0.00408 | 0.01039 | 0.00473 | 0.85889 | 0.65405 | 0.42912 |
| **H2.K1** | 0.00000 | 6.05754 | 12.17703 | 19.59615 | 0.00000 | 14.10270 | 1.41222 | 9.66318 | 2.47735 | 0.21247 | 0.33316 | 3.81953 | 2.24242 | 0.15434 |
| **H2.K2** | 1.41970 | 0.08989 | 0.86567 | 3.05668 | 0.43327 | 0.37686 | 0.01908 | 0.41001 | 0.11539 | 0.05340 | 0.33628 | 4.75440 | 0.15069 | 0.13647 |
| **H2.Ke6** | 0.00903 | 0.00543 | 0.01322 | 0.88884 | 0.03323 | 0.09732 | 0.00434 | 0.23197 | 0.02099 | 0.01116 | 0.00591 | 0.00908 | 0.30415 | 0.00000 |
| **H2.M5** | 0.20705 | 0.00434 | 0.01766 | 0.00000 | 0.00305 | 0.10402 | 0.00427 | 0.00696 | 0.08455 | 0.00693 | 0.00576 | 0.02493 | 0.00885 | 0.04128 |
| **H2.Oa** | 0.01662 | 0.00474 | 0.68282 | 0.75518 | 0.15985 | 2.25155 | 0.00387 | 0.00417 | 0.00884 | 0.01578 | 0.00000 | 0.01303 | 0.02107 | 0.09887 |
| **H2.Q1** | 0.11173 | 0.33184 | 0.75897 | 1.25208 | 0.03689 | 0.24220 | 0.07694 | 0.48516 | 0.02875 | 0.29840 | 0.01515 | 0.17681 | 0.10133 | 0.06549 |
| **H2.Q10** | 1.14445 | 0.24712 | 1.21154 | 0.82140 | 1.44315 | 0.31249 | 0.03544 | 0.37210 | 0.17434 | 0.18899 | 0.01954 | 0.09296 | 5.29099 | 0.74910 |
| **H2.Q2** | 0.00914 | 0.29887 | 0.31506 | 0.80193 | 0.05192 | 0.05965 | 0.04477 | 0.19076 | 0.00870 | 0.05039 | 0.03262 | 0.36324 | 0.03415 | 0.06346 |
| **H2.Q4** | 0.00000 | 0.22396 | 1.33480 | 1.58395 | 0.30390 | 0.03686 | 0.07035 | 0.69886 | 1.50144 | 0.00337 | 0.00150 | 0.46520 | 0.53886 | 1.38462 |
| **H2.Q5** | 0.37202 | 0.02410 | 2.62791 | 2.76643 | 1.76480 | 0.00000 | 0.00879 | 0.00000 | 0.00000 | 0.00150 | 0.05965 | 0.91942 | 2.71731 | 0.05557 |
| **H2.T.ps** | 0.01816 | 0.21579 | 1.02732 | 1.62954 | 0.00525 | 0.62287 | 0.08875 | 1.05640 | 0.06460 | 0.13550 | 0.06325 | 0.29442 | 0.03147 | 0.05454 |
| **H2.T10** | 0.28834 | 0.06964 | 0.09096 | 0.13861 | 0.09394 | 0.14377 | 0.05269 | 0.07496 | 0.18617 | 0.05663 | 0.44455 | 0.22142 | 0.90359 | 1.29975 |
| **H2.T23** | 1.79284 | 0.18871 | 0.24717 | 1.33927 | 0.30396 | 2.91481 | 0.56162 | 0.65128 | 2.31940 | 1.26312 | 0.03890 | 0.80662 | 12.95293 | 1.52567 |
| **H2.T24** | 0.15218 | 0.02945 | 0.02379 | 0.12280 | 0.04760 | 0.44207 | 0.04761 | 0.03461 | 0.44380 | 0.06325 | 0.03875 | 0.10155 | 2.43631 | 1.03356 |
| **H2.T3** | 0.02938 | 0.00412 | 0.00589 | 0.82817 | 0.01081 | 0.04154 | 0.77878 | 0.00633 | 0.09849 | 0.00543 | 0.00950 | 0.02867 | 0.05217 | 0.15886 |
| **H2afj** | 0.28806 | 0.00874 | 0.71655 | 0.97477 | 1.62558 | 1.01861 | 0.02464 | 0.94477 | 0.01343 | 7.85265 | 0.87023 | 1.05493 | 0.00000 | 1.04350 |
| **H2afv** | 2.41215 | 3.41420 | 0.28773 | 7.80292 | 15.15036 | 5.24669 | 1.75645 | 1.03539 | 1.93841 | 0.16833 | 0.25296 | 4.56981 | 9.02343 | 2.54888 |
| **H2afx** | 2.31518 | 0.20472 | 0.47615 | 13.67934 | 18.35621 | 0.88541 | 1.35000 | 4.27630 | 0.78152 | 9.34724 | 0.71681 | 3.49823 | 11.88813 | 2.46704 |
| **H2afy** | 0.03606 | 1.20179 | 6.34847 | 3.95701 | 0.50345 | 13.88479 | 3.34413 | 14.24231 | 1.32573 | 0.33698 | 8.56621 | 15.13571 | 0.00000 | 0.92853 |
| **H2afy2** | 0.02133 | 0.00416 | 0.06792 | 0.26754 | 0.00533 | 0.09538 | 0.00418 | 0.00384 | 0.01179 | 0.00647 | 0.01889 | 0.15941 | 0.01823 | 0.05439 |
| **H2afy3** | 0.01151 | 0.00450 | 0.02257 | 0.06401 | 0.01322 | 0.03099 | 0.00430 | 0.00708 | 0.01304 | 0.00691 | 0.00762 | 0.00848 | 0.01825 | 0.01726 |
| **H2afz** | 26.43857 | 15.16545 | 35.18295 | 41.10631 | 37.91812 | 39.44006 | 13.68054 | 38.63665 | 5.68035 | 28.62797 | 24.77863 | 39.04418 | 24.05446 | 12.80551 |
| **H3f3a** | 24.65500 | 6.98837 | 42.32563 | 37.71800 | 42.66514 | 31.18411 | 9.30297 | 20.05505 | 14.06964 | 20.97284 | 20.77903 | 20.12155 | 20.36137 | 15.56089 |
| **H3f3b** | 32.00608 | 18.86553 | 39.43512 | 36.26184 | 48.21785 | 33.53036 | 11.44822 | 27.08763 | 18.92408 | 15.96409 | 25.87700 | 21.43608 | 27.33515 | 26.00191 |
| **H60b** | 0.26172 | 0.00479 | 0.48453 | 0.00221 | 0.37667 | 0.07047 | 0.00410 | 0.00551 | 0.25091 | 0.00972 | 0.00535 | 0.01012 | 0.00722 | 0.00000 |
| **H6pd** | 0.01049 | 0.00356 | 0.01135 | 0.00000 | 0.08198 | 0.10007 | 0.00416 | 0.00562 | 0.00819 | 0.00560 | 0.00302 | 0.80817 | 0.31381 | 0.00262 |
| **Habp4** | 0.02023 | 0.00482 | 0.05492 | 0.32488 | 0.08101 | 0.17176 | 0.01058 | 0.57580 | 0.00704 | 0.01861 | 0.02342 | 1.20036 | 0.14718 | 0.17331 |
| **Hacd1** | 0.25910 | 0.01106 | 0.00000 | 0.36822 | 0.25530 | 0.63834 | 0.00417 | 0.06366 | 0.00912 | 0.02973 | 0.00213 | 0.01079 | 0.00569 | 0.00000 |
| **Hacd2** | 2.85907 | 0.13407 | 0.61397 | 0.77959 | 0.74518 | 4.31330 | 0.15020 | 2.30359 | 0.18456 | 0.02866 | 0.12673 | 4.08179 | 4.05207 | 2.39422 |
| **Hacd3** | 3.16780 | 0.00000 | 2.82956 | 5.17978 | 4.85633 | 3.24375 | 0.00164 | 0.57424 | 7.78772 | 0.01596 | 2.59104 | 2.94018 | 8.28653 | 11.75946 |
| **Hace1** | 0.80968 | 2.44238 | 0.62187 | 0.00000 | 0.17329 | 0.82734 | 0.00280 | 0.01184 | 1.54868 | 0.00448 | 0.00000 | 1.34210 | 0.28369 | 0.38863 |

| | | | | | | | | | | | | | | |
|---|---|---|---|---|---|---|---|---|---|---|---|---|---|---|
| **Hacl1** | 0.00000 | 0.01046 | 0.39940 | 0.36864 | 0.36219 | 0.22445 | 0.00367 | 0.66773 | 0.04153 | 0.01495 | 1.92081 | 0.00242 | 0.17115 | 2.76945 |
| **Hadh** | 0.50877 | 6.07402 | 17.60995 | 17.04265 | 5.75783 | 15.04737 | 4.29704 | 13.63051 | 0.30624 | 0.36736 | 15.18335 | 10.48367 | 0.17893 | 3.59059 |
| **Hadha** | 2.66281 | 6.35354 | 5.96789 | 12.29291 | 4.54059 | 11.78497 | 4.87552 | 6.24886 | 5.31046 | 0.16359 | 10.47628 | 8.13497 | 1.07193 | 6.66514 |
| **Hadhb** | 11.12322 | 4.70341 | 12.93230 | 6.42774 | 10.13724 | 13.94143 | 2.76256 | 6.63341 | 12.86038 | 0.05656 | 11.23956 | 8.77816 | 1.68328 | 10.83160 |
| **Hagh** | 0.05740 | 0.05884 | 0.26546 | 3.27422 | 0.56679 | 1.00943 | 3.71494 | 1.70552 | 0.00333 | 0.04024 | 0.16817 | 2.12449 | 1.15372 | 0.03659 |
| **Haghl** | 0.00555 | 0.00436 | 0.34385 | 0.17262 | 0.17293 | 0.00000 | 0.00391 | 0.10592 | 0.00938 | 0.00486 | 0.00513 | 0.01475 | 0.00420 | 0.01208 |
| **Hal** | 0.00000 | 0.01589 | 0.11315 | 0.00000 | 0.40501 | 1.01879 | 0.00296 | 1.24085 | 0.00595 | 0.01426 | 0.00000 | 2.80855 | 0.00906 | 0.03601 |
| **Hamp** | 0.00969 | 0.00415 | 0.01291 | 0.00484 | 0.01514 | 0.01422 | 0.00430 | 0.00808 | 0.00941 | 0.00732 | 0.00595 | 0.13999 | 0.14708 | 0.06010 |
| **Hand1** | 14.05799 | 0.00215 | 6.67545 | 7.03521 | 18.92511 | 0.00000 | 0.00249 | 0.00000 | 0.49676 | 0.00499 | 0.00147 | 0.00000 | 0.00000 | 7.96124 |
| **Hap1** | 0.00317 | 0.00389 | 0.08776 | 0.28169 | 0.00559 | 0.02054 | 0.00393 | 0.00375 | 0.01128 | 0.00576 | 0.00855 | 0.35436 | 0.02882 | 0.00462 |
| **Hapln4** | 0.01048 | 0.01353 | 1.49933 | 0.94747 | 1.01056 | 1.45769 | 0.00401 | 1.23775 | 1.24218 | 0.06028 | 0.00000 | 0.00000 | 0.01018 | 0.96069 |
| **Harbi1** | 0.12211 | 0.01935 | 0.16962 | 3.40439 | 1.68151 | 2.03900 | 1.86232 | 1.04727 | 0.02277 | 0.07000 | 4.42770 | 1.52958 | 1.49586 | 0.69590 |
| **Hars** | 12.16132 | 5.10364 | 13.79122 | 9.40182 | 10.92535 | 13.05032 | 0.39551 | 8.64478 | 1.44717 | 0.13317 | 14.00695 | 10.45933 | 5.41921 | 14.96788 |
| **Hars2** | 3.49817 | 0.01642 | 5.57383 | 5.05560 | 6.73959 | 3.97865 | 3.43269 | 3.05336 | 0.93301 | 0.05390 | 0.29662 | 5.07313 | 4.60847 | 4.86158 |
| **Has3** | 0.00000 | 0.00396 | 0.31858 | 0.00000 | 0.89000 | 0.00000 | 0.00434 | 0.00653 | 0.27322 | 0.00671 | 0.00456 | 0.00424 | 0.86863 | 4.18040 |
| **Hat1** | 15.46616 | 2.96322 | 20.27890 | 13.57219 | 13.93769 | 27.35238 | 5.25980 | 12.57706 | 15.16107 | 0.28445 | 14.04961 | 18.02965 | 3.60067 | 14.52132 |
| **Haus1** | 3.79510 | 5.76458 | 3.14628 | 4.10405 | 2.04868 | 8.93179 | 0.01697 | 5.89650 | 2.21442 | 0.10205 | 4.96441 | 5.74346 | 0.82123 | 2.26404 |
| **Haus2** | 9.28588 | 0.12721 | 6.38481 | 6.14768 | 10.43294 | 5.10005 | 1.09808 | 2.62746 | 12.72496 | 0.13246 | 2.74354 | 4.62872 | 3.22613 | 10.39023 |
| **Haus3** | 5.57360 | 0.06115 | 3.94892 | 0.57099 | 7.93752 | 5.49054 | 5.32496 | 1.99587 | 14.34056 | 0.07738 | 4.84833 | 2.75605 | 7.72111 | 5.71895 |
| **Haus4** | 3.60616 | 0.97964 | 8.05407 | 11.59676 | 12.95156 | 6.08904 | 0.72823 | 3.29555 | 0.42908 | 0.87543 | 3.29593 | 2.02194 | 6.73065 | 7.53621 |
| **Haus5** | 3.13921 | 5.58745 | 9.02947 | 6.12714 | 2.97319 | 6.33550 | 2.40525 | 7.21401 | 0.03150 | 0.25347 | 8.59399 | 8.21571 | 0.26117 | 2.68725 |
| **Haus6** | 1.03895 | 0.00000 | 0.72646 | 0.77930 | 3.91998 | 0.47607 | 0.00000 | 0.09685 | 13.79062 | 8.19127 | 4.21791 | 1.17994 | 5.39604 | 8.28435 |
| **Haus7** | 0.83069 | 2.26496 | 1.95410 | 2.55049 | 2.16390 | 3.67904 | 0.00841 | 1.65692 | 0.63416 | 0.01123 | 1.85566 | 1.29498 | 0.79672 | 3.51836 |
| **Haus8** | 5.09656 | 2.54763 | 7.54977 | 8.44040 | 4.06412 | 7.07429 | 3.98169 | 4.47480 | 0.00000 | 0.19553 | 8.69224 | 8.61322 | 0.66562 | 2.11718 |
| **Havcr1** | 5.64252 | 9.16639 | 9.25958 | 4.49515 | 8.26122 | 13.00684 | 6.01023 | 14.30033 | 16.92654 | 9.14712 | 9.14875 | 8.62112 | 0.48885 | 5.49095 |
| **Havcr2** | 0.01113 | 0.00436 | 0.01371 | 0.01492 | 0.25631 | 0.02683 | 0.00406 | 0.02123 | 0.03713 | 0.00589 | 0.01802 | 0.01415 | 0.41302 | 0.09291 |
| **Hax1** | 20.50416 | 0.34974 | 29.67394 | 22.69781 | 27.88371 | 20.51954 | 7.96577 | 7.40541 | 24.09172 | 0.83344 | 15.52385 | 11.24763 | 14.40556 | 23.20045 |
| **Hbegf** | 0.07043 | 0.05522 | 0.11059 | 0.69522 | 0.00000 | 0.00000 | 0.00401 | 1.23516 | 0.00617 | 0.00680 | 0.10718 | 0.09751 | 0.59467 | 0.07735 |
| **Hbp1** | 10.88392 | 0.00616 | 10.46497 | 0.87135 | 15.70362 | 3.55812 | 4.25045 | 1.10975 | 16.13303 | 0.08084 | 5.13431 | 1.69589 | 11.32844 | 18.66319 |
| **Hbs1l** | 5.17353 | 0.07944 | 4.95989 | 4.38558 | 6.02453 | 7.70231 | 1.54623 | 5.96902 | 0.63926 | 0.05186 | 1.89748 | 7.26877 | 1.53828 | 3.30127 |
| **Hccs** | 8.98195 | 0.02340 | 3.60273 | 0.46261 | 6.24631 | 4.97827 | 0.04243 | 1.10407 | 5.12859 | 0.03167 | 2.64810 | 2.91286 | 6.56310 | 5.22046 |
| **Hcfc1** | 8.12119 | 5.49600 | 12.52518 | 7.87166 | 15.83710 | 10.22432 | 0.35018 | 5.84182 | 11.41497 | 13.16316 | 8.23370 | 8.15021 | 4.66966 | 16.32099 |
| **Hcfc1r1** | 0.62507 | 0.10751 | 5.51024 | 2.26166 | 5.76508 | 2.91304 | 3.96856 | 3.25218 | 0.00000 | 0.16431 | 0.90571 | 5.86933 | 0.03092 | 1.07185 |
| **Hcfc2** | 0.01617 | 0.00410 | 0.00109 | 0.13098 | 0.08905 | 0.00000 | 0.02226 | 1.25715 | 0.03182 | 0.01077 | 0.20968 | 0.00388 | 0.00370 | 0.05089 |
| **Hck** | 0.00000 | 0.00324 | 0.73091 | 0.04974 | 0.55972 | 0.00000 | 0.00360 | 0.00074 | 0.01025 | 0.00507 | 0.00330 | 1.13924 | 0.00231 | 0.00899 |
| **Hcn3** | 3.72185 | 0.00274 | 1.37521 | 0.00000 | 1.75552 | 0.00000 | 0.00349 | 0.00000 | 0.04808 | 0.00339 | 0.01422 | 1.24936 | 0.00000 | 0.46106 |
| **Hcrt** | 0.00000 | 0.00405 | 3.44131 | 0.96191 | 5.09354 | 0.00000 | 0.00351 | 0.00618 | 0.01654 | 0.00000 | 0.00336 | 0.00757 | 0.00000 | 0.01598 |
| **Hdac1** | 22.77397 | 5.45537 | 35.34729 | 15.42488 | 29.20116 | 16.05032 | 3.88655 | 5.01534 | 30.38599 | 0.21351 | 19.09042 | 15.62183 | 13.11906 | 30.69065 |
| **Hdac10** | 0.00000 | 0.00374 | 0.11605 | 1.29137 | 0.00000 | 0.00000 | 0.00392 | 0.00095 | 0.00867 | 0.00283 | 1.14106 | 0.15042 | 0.19025 | 0.59331 |
| **Hdac11** | 0.00650 | 0.00440 | 0.01016 | 0.01188 | 0.21203 | 0.01271 | 0.00443 | 0.01171 | 0.00768 | 0.00647 | 0.00885 | 0.00921 | 0.01373 | 0.06965 |
| **Hdac2** | 10.35093 | 1.64239 | 7.36064 | 1.72695 | 11.00418 | 14.09000 | 2.63021 | 4.03991 | 6.91833 | 0.11947 | 9.08134 | 8.83836 | 4.72992 | 5.36709 |
| **Hdac3** | 0.00000 | 0.03775 | 3.18294 | 1.63909 | 0.79608 | 1.12823 | 0.06845 | 1.25170 | 1.23794 | 0.03312 | 1.57642 | 3.62551 | 0.00000 | 0.49810 |
| **Hdac4** | 1.60552 | 0.00293 | 0.11498 | 0.97331 | 0.20661 | 0.63712 | 0.00891 | 0.00790 | 0.07108 | 0.00594 | 0.01367 | 1.35343 | 0.63762 | 0.53514 |
| **Hdac5** | 0.28870 | 2.39122 | 4.08309 | 3.30843 | 7.06563 | 0.63959 | 0.01379 | 0.72846 | 0.00790 | 0.06021 | 1.33830 | 0.07891 | 0.00537 | 0.00000 |
| **Hdac6** | 3.74121 | 0.01839 | 9.48793 | 4.50453 | 5.81405 | 2.39145 | 0.76076 | 2.62970 | 4.05259 | 0.02743 | 2.32802 | 0.47097 | 1.83498 | 10.60908 |
| **Hdac7** | 0.04579 | 0.00413 | 0.00351 | 0.52270 | 0.13415 | 0.00688 | 0.00427 | 0.01021 | 0.01319 | 0.00691 | 0.00615 | 0.00706 | 0.00641 | 0.00683 |
| **Hdac8** | 0.55974 | 0.00406 | 0.19215 | 0.00000 | 0.12447 | 0.00000 | 0.00419 | 0.01206 | 0.00447 | 0.00541 | 0.14497 | 0.00788 | 0.59174 | 0.04902 |
| **Hdac9** | 0.01157 | 0.00427 | 0.01556 | 0.01147 | 0.08196 | 0.01197 | 0.00425 | 0.01864 | 0.01000 | 0.00636 | 0.00887 | 0.00955 | 0.00887 | 0.01291 |
| **Hdc** | 0.44739 | 0.00490 | 0.23433 | 0.55644 | 1.63022 | 0.00000 | 0.00409 | 0.11208 | 0.01068 | 0.01693 | 0.00388 | 0.37402 | 0.71597 | 0.85660 |
| **Hddc2** | 0.73117 | 0.09864 | 0.02586 | 0.00000 | 2.71376 | 2.44019 | 0.00380 | 1.17232 | 0.00217 | 3.85042 | 0.00000 | 2.16053 | 0.24437 | 0.35322 |
| **Hddc3** | 0.62787 | 0.07339 | 0.50025 | 3.46853 | 0.80429 | 1.90756 | 0.04808 | 0.03019 | 0.28386 | 0.02251 | 0.00000 | 0.20647 | 0.00207 | 0.34214 |
| **Hdgf** | 28.58872 | 11.35417 | 46.75177 | 38.50341 | 39.74789 | 34.83147 | 10.00085 | 22.29038 | 32.75619 | 8.95612 | 18.48838 | 23.23897 | 13.73077 | 35.30681 |
| **Hdgfrp2** | 7.09290 | 0.08062 | 9.27200 | 11.99865 | 9.84112 | 8.74716 | 0.15821 | 9.72115 | 0.08006 | 0.06437 | 6.75390 | 6.86471 | 4.30216 | 8.46548 |
| **Hdgfrp3** | 0.01624 | 0.00434 | 0.02623 | 0.14958 | 0.02322 | 0.01842 | 0.00418 | 0.00652 | 0.01136 | 0.00589 | 0.12610 | 0.09250 | 0.01649 | 0.01137 |
| **Hdhd2** | 0.82862 | 0.00069 | 2.02529 | 0.73429 | 0.15353 | 2.13528 | 0.00223 | 0.96437 | 0.23318 | 5.32266 | 3.89856 | 2.25569 | 0.01761 | 1.25307 |

| | | | | | | | | | | | | | | |
|---|---|---|---|---|---|---|---|---|---|---|---|---|---|---|
| **Hdhd3** | 0.00422 | 0.02308 | 0.00000 | 0.18976 | 0.00781 | 0.00000 | 0.00394 | 0.23644 | 0.00741 | 0.01162 | 1.14698 | 0.58486 | 0.02603 | 0.62009 |
| **Hdlbp** | 17.81957 | 1.67350 | 32.22574 | 21.43319 | 28.57311 | 20.47465 | 5.58405 | 12.94374 | 6.41885 | 0.47371 | 18.04282 | 17.96161 | 3.27166 | 16.57472 |
| **Hdx** | 0.00500 | 0.00415 | 0.00648 | 0.10794 | 0.00605 | 0.06140 | 0.00405 | 0.00425 | 0.00588 | 0.00624 | 0.00532 | 0.05241 | 0.00479 | 0.23236 |
| **Heatr1** | 10.61588 | 0.08975 | 17.46656 | 5.78650 | 22.63531 | 9.22921 | 4.05363 | 1.67253 | 7.65713 | 0.08561 | 5.74090 | 6.25380 | 9.72744 | 19.12743 |
| **Heatr3** | 4.15207 | 0.03195 | 7.40897 | 4.10352 | 5.00597 | 2.41896 | 0.00656 | 0.76741 | 14.18896 | 0.00964 | 2.39769 | 2.54824 | 5.78263 | 11.91327 |
| **Heatr5a** | 1.05737 | 3.88211 | 5.11574 | 1.34738 | 1.13779 | 6.45303 | 0.00086 | 0.82415 | 1.00453 | 0.01623 | 4.14411 | 2.73050 | 1.44709 | 2.08387 |
| **Heatr5b** | 1.91700 | 0.01049 | 1.61858 | 1.75140 | 0.80415 | 2.87184 | 0.00407 | 0.75307 | 1.81766 | 0.01027 | 0.14946 | 2.12077 | 0.00000 | 1.16889 |
| **Heatr6** | 1.13546 | 0.04695 | 2.78320 | 0.92832 | 1.23036 | 0.78054 | 0.76064 | 1.74155 | 0.77648 | 0.01838 | 0.20310 | 1.20125 | 0.30274 | 1.70964 |
| **Hebp1** | 0.00396 | 0.15882 | 0.00201 | 0.09359 | 0.20346 | 0.01079 | 0.00930 | 1.24340 | 0.01157 | 0.00714 | 0.00000 | 1.19514 | 0.00521 | 0.01121 |
| **Hebp2** | 0.00762 | 0.00396 | 0.00618 | 0.44702 | 0.00854 | 0.08133 | 0.00385 | 0.00286 | 0.01239 | 0.00480 | 0.02301 | 0.05059 | 0.00602 | 0.01663 |
| **Heca** | 0.65710 | 0.00446 | 0.48276 | 0.00587 | 0.00000 | 0.16554 | 0.05488 | 0.00442 | 0.01644 | 0.02218 | 0.00606 | 0.00732 | 0.02305 | 1.67179 |
| **Hectd1** | 7.82828 | 4.67410 | 7.81062 | 2.49915 | 8.60540 | 10.53245 | 0.03806 | 9.58721 | 13.79841 | 0.09723 | 8.53867 | 7.15800 | 6.61243 | 12.48207 |
| **Hectd2** | 0.01687 | 0.00397 | 0.01039 | 0.00353 | 0.01570 | 0.01163 | 0.00397 | 0.15913 | 0.13712 | 0.00743 | 0.00709 | 0.00989 | 0.00481 | 0.04586 |
| **Hectd3** | 0.69302 | 0.04117 | 1.01466 | 2.27807 | 1.57937 | 2.83290 | 1.08544 | 0.91169 | 0.41889 | 0.03516 | 0.53156 | 0.25890 | 0.69090 | 1.97819 |
| **Hecw2** | 0.00000 | 0.00461 | 0.00000 | 0.17081 | 0.91257 | 0.36167 | 0.00369 | 0.00496 | 3.62068 | 0.00636 | 0.00303 | 0.00355 | 0.00916 | 0.31565 |
| **Heg1** | 0.02429 | 0.00416 | 0.01082 | 0.15646 | 0.01219 | 0.02026 | 0.00402 | 0.00474 | 0.01180 | 0.00652 | 0.00623 | 0.03529 | 0.00662 | 0.00965 |
| **Helb** | 0.14681 | 0.01099 | 0.74281 | 0.72351 | 0.45014 | 0.09478 | 0.00368 | 0.00905 | 0.68857 | 0.02593 | 0.00234 | 0.11193 | 1.96260 | 1.53644 |
| **Hells** | 2.27452 | 0.00915 | 2.74549 | 0.51946 | 3.17890 | 2.66849 | 1.02935 | 0.01083 | 11.23677 | 0.03176 | 0.28075 | 1.29834 | 5.39367 | 1.70749 |
| **Helq** | 0.66650 | 0.65966 | 1.16517 | 0.23397 | 0.00000 | 0.09559 | 0.00301 | 0.00000 | 3.50677 | 0.00405 | 0.00956 | 0.09495 | 1.94634 | 9.22500 |
| **Helz** | 0.00569 | 0.01111 | 0.07460 | 4.26590 | 3.08596 | 0.12135 | 0.01583 | 0.00626 | 0.12666 | 0.01216 | 0.04149 | 0.61314 | 0.00000 | 0.12334 |
| **Hemk1** | 0.54216 | 0.04863 | 1.48321 | 1.78748 | 0.17862 | 1.34276 | 1.53073 | 2.88837 | 0.00651 | 0.07740 | 0.14479 | 1.42576 | 0.00718 | 0.78323 |
| **Henmt1** | 0.02457 | 0.00441 | 0.00641 | 0.01220 | 0.08342 | 0.01216 | 0.00414 | 0.00961 | 0.01159 | 0.00738 | 0.00730 | 0.00893 | 0.00343 | 0.76196 |
| **Hepacam** | 0.01725 | 0.00483 | 0.24226 | 0.00000 | 0.20964 | 0.51611 | 0.00414 | 0.00483 | 0.01382 | 0.01245 | 0.13759 | 0.02279 | 0.00683 | 0.00332 |
| **Herc1** | 1.48965 | 0.02571 | 3.46344 | 0.47857 | 2.17453 | 3.86629 | 0.01650 | 0.37997 | 1.34638 | 0.01649 | 0.42825 | 4.18540 | 0.02445 | 2.04288 |
| **Herc2** | 1.89285 | 0.34154 | 0.72964 | 2.50350 | 2.13815 | 2.95007 | 0.00319 | 0.33609 | 3.32350 | 0.00968 | 1.75279 | 1.19295 | 2.92386 | 4.01632 |
| **Herc3** | 0.06738 | 0.00373 | 0.06761 | 1.24622 | 0.00000 | 0.00000 | 0.00364 | 0.12888 | 0.00603 | 0.01634 | 0.00347 | 0.67032 | 0.05189 | 0.27260 |
| **Herc4** | 12.03625 | 1.82309 | 9.08245 | 3.10996 | 8.34612 | 9.18947 | 4.63392 | 1.79774 | 11.80377 | 0.13666 | 3.76378 | 5.57539 | 5.78221 | 8.84407 |
| **Herpud1** | 0.01120 | 0.00342 | 0.00000 | 3.07036 | 0.02714 | 0.00000 | 1.02259 | 0.00351 | 0.01767 | 0.03904 | 1.72393 | 0.45170 | 0.00570 | 0.00000 |
| **Herpud2** | 3.20264 | 0.01701 | 1.29056 | 0.00000 | 1.85861 | 1.28995 | 0.00268 | 1.16054 | 1.80358 | 0.00094 | 1.05186 | 1.61875 | 3.85567 | 2.36615 |
| **Hes1** | 0.88501 | 0.28537 | 0.00064 | 0.00000 | 4.49548 | 0.11528 | 1.15269 | 0.92035 | 0.10956 | 0.04296 | 0.00000 | 0.56379 | 2.98111 | 1.28898 |
| **Hes6** | 0.08457 | 0.00398 | 0.00000 | 0.24013 | 0.03586 | 0.24683 | 0.00400 | 0.00373 | 0.01192 | 0.01305 | 0.00430 | 0.13457 | 0.00888 | 0.31610 |
| **Hesx1** | 1.26274 | 0.00179 | 0.00000 | 0.03757 | 0.27721 | 0.00000 | 0.00327 | 0.00000 | 0.75203 | 0.01396 | 1.57877 | 2.23702 | 2.01868 | 0.43178 |
| **Hexa** | 0.00000 | 3.79143 | 9.70328 | 22.47671 | 2.05794 | 9.56627 | 7.81542 | 12.32546 | 0.22539 | 2.04691 | 6.06488 | 11.61514 | 0.27661 | 1.09381 |
| **Hexb** | 6.28436 | 5.44444 | 14.27313 | 15.89588 | 11.28909 | 5.34652 | 2.62750 | 3.67906 | 19.49245 | 0.07702 | 6.89324 | 6.94615 | 3.00927 | 15.51662 |
| **Hexdc** | 0.55223 | 0.02748 | 0.54518 | 1.73001 | 1.73562 | 0.18494 | 0.00966 | 1.04753 | 0.00656 | 0.01488 | 2.00500 | 1.48640 | 0.01322 | 0.00000 |
| **Hexim1** | 0.15240 | 2.41731 | 2.96219 | 5.08205 | 4.22782 | 0.00000 | 1.77177 | 3.81849 | 0.08724 | 0.10061 | 0.52934 | 3.00730 | 5.81329 | 0.00000 |
| **Hexim2** | 0.28328 | 0.01883 | 0.82139 | 0.00000 | 1.03932 | 0.45528 | 4.68702 | 1.37003 | 0.00950 | 0.07466 | 1.23221 | 1.41258 | 1.12346 | 0.00000 |
| **Hey1** | 0.16167 | 0.00393 | 0.35057 | 1.28228 | 0.28846 | 0.26837 | 0.00361 | 0.00030 | 0.00638 | 0.00378 | 0.17365 | 0.00998 | 0.45649 | 0.06429 |
| **Heyl** | 0.01020 | 0.00442 | 0.04816 | 0.00597 | 0.01321 | 0.01001 | 0.00449 | 0.01490 | 0.11208 | 0.00713 | 0.00838 | 0.01925 | 0.00784 | 0.12297 |
| **Hfm1** | 0.33088 | 0.00425 | 0.15189 | 0.03225 | 0.04704 | 0.08251 | 0.01061 | 0.00610 | 0.22205 | 0.00619 | 0.01080 | 0.03005 | 0.45326 | 0.23420 |
| **Hgh1** | 1.56933 | 0.05606 | 5.95700 | 3.85193 | 3.39172 | 5.84650 | 0.01989 | 3.74100 | 0.14423 | 0.03315 | 3.53892 | 1.85751 | 0.36689 | 3.06481 |
| **Hgs** | 0.41628 | 1.66695 | 5.63912 | 12.17189 | 10.35087 | 1.37409 | 0.26892 | 7.09230 | 0.00000 | 0.21522 | 0.24271 | 6.28506 | 2.86338 | 0.23823 |
| **Hgsnat** | 0.82453 | 5.52103 | 1.60938 | 1.63303 | 0.51542 | 2.54524 | 0.05276 | 3.88136 | 0.77055 | 0.02953 | 0.00000 | 0.65659 | 0.89608 | 2.93936 |
| **Hhex** | 0.00000 | 0.00295 | 1.69756 | 1.02843 | 0.24394 | 0.49250 | 0.00409 | 0.07546 | 0.00000 | 1.33833 | 0.17803 | 1.14015 | 4.02370 | 0.45951 |
| **Hhipl2** | 0.00000 | 0.00461 | 0.43233 | 0.00000 | 0.58294 | 0.19317 | 0.00402 | 0.00418 | 0.01532 | 0.00501 | 0.00591 | 0.00615 | 0.00031 | 0.40049 |
| **Hibadh** | 1.49875 | 0.03215 | 1.73669 | 0.00000 | 0.44787 | 3.92019 | 2.50662 | 1.18290 | 1.36593 | 0.09853 | 2.00027 | 4.36726 | 0.25837 | 0.14451 |
| **Hibch** | 3.03206 | 0.01338 | 3.19085 | 3.70967 | 4.89579 | 7.10355 | 0.01838 | 4.34178 | 9.05008 | 0.04612 | 10.59537 | 4.33780 | 0.49254 | 4.43462 |
| **Hic1** | 0.07803 | 0.01084 | 0.15904 | 1.39732 | 0.22973 | 0.08891 | 0.00381 | 0.54778 | 0.00607 | 0.03239 | 0.00000 | 0.12146 | 0.16928 | 0.14347 |
| **Hic2** | 0.75776 | 5.73487 | 10.72881 | 2.41462 | 6.00497 | 8.43318 | 7.80303 | 12.95164 | 2.80351 | 0.44584 | 3.42250 | 4.24702 | 11.45962 | 10.27913 |
| **Hif1a** | 0.55590 | 0.00804 | 0.10992 | 0.00000 | 0.06110 | 1.44006 | 0.34367 | 0.76213 | 0.00956 | 0.05205 | 2.90229 | 4.46942 | 0.71395 | 0.00224 |
| **Hif1an** | 1.93617 | 0.03890 | 5.95444 | 4.44611 | 5.21538 | 2.37005 | 0.01782 | 0.00000 | 5.58140 | 0.00078 | 0.00000 | 3.03629 | 1.20922 | 6.80815 |
| **Hif3a** | 0.00000 | 0.00443 | 0.31966 | 0.21770 | 0.20736 | 0.54805 | 0.00412 | 0.00563 | 0.01212 | 0.00682 | 0.00432 | 0.00323 | 0.41741 | 0.26088 |
| **Higd1a** | 1.35936 | 0.09320 | 0.00000 | 0.99664 | 2.83104 | 7.98454 | 3.28428 | 5.58475 | 0.26699 | 8.17569 | 1.90240 | 4.23641 | 7.61286 | 0.44096 |
| **Higd2a** | 2.54451 | 0.12013 | 0.58957 | 5.06979 | 2.06709 | 6.22933 | 0.40689 | 9.30426 | 0.22715 | 0.16946 | 0.99887 | 6.63107 | 2.26506 | 4.37404 |
| **Hilpda** | 0.07864 | 0.06520 | 3.42671 | 4.64991 | 0.97559 | 6.01760 | 4.80601 | 4.33584 | 0.69068 | 0.23217 | 0.24849 | 3.19644 | 1.54068 | 1.39673 |

| | | | | | | | | | | | | | | |
|---|---|---|---|---|---|---|---|---|---|---|---|---|---|---|
| **Hinfp** | 3.60215 | 0.02085 | 4.92106 | 4.65551 | 6.27797 | 0.21601 | 1.43637 | 1.19638 | 0.56515 | 0.03496 | 0.00201 | 0.60953 | 6.57503 | 12.94464 |
| **Hint1** | 3.19000 | 2.56295 | 0.00000 | 4.40394 | 11.76949 | 7.23297 | 3.09295 | 14.64480 | 0.30248 | 8.84490 | 0.03184 | 11.01593 | 6.22556 | 0.21821 |
| **Hint2** | 0.00516 | 0.39512 | 0.00000 | 0.83009 | 0.12039 | 0.44768 | 0.68173 | 0.43152 | 0.00880 | 0.89490 | 0.10028 | 1.16045 | 0.01333 | 0.00814 |
| **Hint3** | 0.36550 | 0.04234 | 0.12649 | 1.18486 | 1.81570 | 1.54687 | 2.11327 | 2.42659 | 0.06293 | 2.75156 | 0.08527 | 2.84717 | 0.28824 | 0.17086 |
| **Hip1** | 5.06941 | 0.57701 | 10.36977 | 8.68980 | 11.95047 | 11.85947 | 5.03514 | 11.47774 | 6.87276 | 0.17134 | 11.11713 | 13.81357 | 0.47422 | 6.19254 |
| **Hip1r** | 0.00000 | 0.00194 | 0.39052 | 2.58314 | 1.20938 | 0.28234 | 1.30563 | 0.00000 | 0.00300 | 4.70349 | 0.91999 | 1.02074 | 1.47566 | 0.00000 |
| **Hipk1** | 8.63902 | 5.32462 | 7.40098 | 3.26244 | 11.54692 | 7.56875 | 2.95213 | 6.13779 | 15.43852 | 3.62166 | 3.69441 | 7.21997 | 16.36319 | 12.82922 |
| **Hipk2** | 0.09421 | 0.00311 | 0.09827 | 0.00000 | 0.69379 | 0.30511 | 1.27487 | 0.00583 | 0.33612 | 0.02118 | 0.67330 | 1.01989 | 0.06250 | 0.00000 |
| **Hipk3** | 18.70103 | 0.33876 | 12.23619 | 3.38241 | 15.15815 | 11.35593 | 0.29105 | 3.09150 | 18.47948 | 0.08546 | 9.93746 | 3.06505 | 14.16223 | 7.30028 |
| **Hipk4** | 0.00938 | 0.00453 | 0.04117 | 0.00660 | 0.15047 | 0.06547 | 0.00425 | 0.00878 | 0.00975 | 0.00717 | 0.00738 | 0.00844 | 0.00619 | 0.01189 |
| **Hira** | 4.37588 | 0.01844 | 6.24168 | 2.01041 | 10.29009 | 1.40900 | 0.00866 | 0.49037 | 5.87123 | 0.00000 | 3.70861 | 1.86836 | 0.47569 | 9.52118 |
| **Hirip3** | 8.01114 | 0.07123 | 8.37276 | 12.74384 | 3.10214 | 10.50352 | 1.92863 | 0.36642 | 0.34371 | 3.66720 | 5.55617 | 5.01990 | 4.30502 | 2.59864 |
| **Hist1h1a** | 0.22317 | 0.07029 | 0.00355 | 0.10700 | 0.00234 | 0.00877 | 0.00364 | 0.59229 | 0.01541 | 0.00877 | 0.00022 | 0.78049 | 0.24130 | 0.05673 |
| **Hist1h1b** | 0.01438 | 0.01905 | 0.02136 | 0.00907 | 0.01797 | 0.04688 | 0.00399 | 0.05037 | 0.01045 | 0.00679 | 0.00695 | 0.00870 | 0.00644 | 0.01508 |
| **Hist1h1c** | 0.01774 | 0.00525 | 0.03814 | 0.22025 | 0.34736 | 0.11485 | 0.00429 | 0.87720 | 0.01005 | 0.00722 | 0.00303 | 0.00945 | 0.01222 | 0.06753 |
| **Hist1h1d** | 0.00511 | 0.00496 | 0.00863 | 0.37688 | 0.01348 | 0.00465 | 0.00424 | 0.05098 | 0.01400 | 0.00711 | 0.01318 | 0.23124 | 0.00683 | 0.00821 |
| **Hist1h2ab** | 0.00767 | 0.02053 | 0.01635 | 0.02995 | 0.25566 | 0.01052 | 0.00447 | 0.03161 | 0.01184 | 0.00705 | 0.01624 | 0.03812 | 0.00554 | 0.00801 |
| **Hist1h2ac** | 0.00886 | 0.00446 | 0.00976 | 0.03122 | 0.12556 | 0.01313 | 0.00448 | 0.01535 | 0.01183 | 0.00686 | 0.00866 | 0.01098 | 0.00637 | 0.01075 |
| **Hist1h2ad** | 0.00730 | 0.07154 | 0.01492 | 0.04184 | 0.26167 | 0.01578 | 0.00447 | 0.01835 | 0.01175 | 0.00735 | 0.00811 | 0.03807 | 0.00492 | 0.01056 |
| **Hist1h2ae** | 0.00850 | 0.01928 | 0.01398 | 0.06065 | 0.07156 | 0.01360 | 0.00417 | 0.01671 | 0.01195 | 0.00602 | 0.01443 | 0.01504 | 0.00621 | 0.00987 |
| **Hist1h2af** | 0.00737 | 0.01271 | 0.01575 | 0.03423 | 0.22262 | 0.01222 | 0.00438 | 0.01687 | 0.01153 | 0.00683 | 0.01036 | 0.02605 | 0.00589 | 0.02595 |
| **Hist1h2ag** | 0.00958 | 0.01223 | 0.01225 | 0.01478 | 0.07589 | 0.01313 | 0.00422 | 0.00919 | 0.01149 | 0.00666 | 0.00876 | 0.01354 | 0.00648 | 0.01057 |
| **Hist1h2ah** | 0.00779 | 0.01219 | 0.01611 | 0.01753 | 0.06430 | 0.01239 | 0.00421 | 0.00903 | 0.01085 | 0.00657 | 0.01025 | 0.01036 | 0.07233 | 0.00487 |
| **Hist1h2ai** | 0.01228 | 0.05187 | 0.00966 | 0.04216 | 0.24430 | 0.01244 | 0.00452 | 0.01481 | 0.01237 | 0.00631 | 0.00851 | 0.01904 | 0.00519 | 0.00766 |
| **Hist1h2al** | 0.04799 | 0.82942 | 0.03770 | 0.07961 | 0.08015 | 0.01785 | 0.00387 | 0.15870 | 0.11784 | 0.00548 | 0.15844 | 0.00526 | 0.03392 | 0.16207 |
| **Hist1h2an** | 0.00932 | 0.01253 | 0.01425 | 0.02155 | 0.06841 | 0.01395 | 0.00430 | 0.01495 | 0.01183 | 0.00700 | 0.00872 | 0.00915 | 0.00656 | 0.01074 |
| **Hist1h2bb** | 0.00765 | 0.00406 | 0.01271 | 0.00000 | 0.01247 | 0.01204 | 0.23946 | 0.00453 | 0.01159 | 0.02097 | 0.00459 | 0.27859 | 0.00693 | 0.01401 |
| **Hist1h2bc** | 0.00194 | 3.67228 | 0.07892 | 0.00000 | 0.29550 | 0.14325 | 0.00315 | 1.25196 | 0.00535 | 0.00698 | 0.00000 | 0.60678 | 1.64329 | 0.00134 |
| **Hist1h2be** | 0.00305 | 0.10405 | 0.01332 | 0.03102 | 0.15695 | 0.00651 | 0.02925 | 0.12568 | 0.00885 | 0.00702 | 0.00918 | 0.07898 | 0.60348 | 0.00306 |
| **Hist1h4c** | 0.00879 | 0.15605 | 0.03237 | 0.01599 | 0.01491 | 0.00369 | 0.00502 | 0.10847 | 0.01330 | 0.05533 | 0.01504 | 0.00722 | 0.01508 | 0.01183 |
| **Hist1h4d** | 0.01633 | 0.00526 | 0.01814 | 0.03023 | 0.01058 | 0.00279 | 0.00444 | 0.13759 | 0.01383 | 0.25488 | 0.01257 | 0.00789 | 0.00607 | 0.01284 |
| **Hist1h4f** | 0.01066 | 0.01450 | 0.01764 | 0.00881 | 0.01576 | 0.00000 | 0.00449 | 0.14337 | 0.01327 | 0.19135 | 0.03515 | 0.01122 | 0.01511 | 0.01046 |
| **Hist1h4h** | 0.01194 | 0.01374 | 0.01310 | 0.01179 | 0.01512 | 0.00854 | 0.00452 | 0.14011 | 0.01147 | 0.10520 | 0.01532 | 0.02881 | 0.00735 | 0.01128 |
| **Hist1h4i** | 0.00423 | 0.01273 | 0.04744 | 0.28779 | 0.02548 | 0.00000 | 0.00420 | 0.21609 | 0.00801 | 2.55237 | 0.16787 | 0.00000 | 0.27179 | 0.00000 |
| **Hist2h2bb** | 0.00668 | 0.00531 | 0.01264 | 0.02201 | 0.05103 | 0.00458 | 0.00453 | 0.20911 | 0.02663 | 0.00820 | 0.00708 | 0.02428 | 0.02276 | 0.04875 |
| **Hist2h2be** | 0.00814 | 0.00417 | 0.01024 | 0.00000 | 0.14399 | 0.09645 | 0.08046 | 0.00546 | 0.01078 | 0.00712 | 0.00715 | 0.01846 | 0.00833 | 0.01269 |
| **Hist3h2a** | 0.00000 | 4.29314 | 0.65943 | 0.69313 | 0.07063 | 0.00000 | 0.00432 | 0.19769 | 0.00908 | 0.00505 | 0.00297 | 0.00272 | 0.00540 | 0.00854 |
| **Hivep1** | 0.00000 | 0.01899 | 1.42419 | 3.35534 | 2.02100 | 0.68237 | 0.13436 | 0.31222 | 0.10904 | 0.02940 | 0.39661 | 2.13296 | 0.76343 | 0.26519 |
| **Hivep2** | 0.02462 | 0.00395 | 0.03427 | 0.00000 | 0.00480 | 0.27762 | 0.00406 | 0.00347 | 0.02110 | 0.00484 | 0.00384 | 0.22361 | 0.22789 | 0.10577 |
| **Hivep3** | 0.03575 | 0.00403 | 0.03932 | 0.27002 | 0.00537 | 0.00256 | 0.00404 | 0.00555 | 0.01098 | 0.00547 | 0.02326 | 0.06237 | 0.02250 | 0.02960 |
| **Hjurp** | 0.00000 | 0.04054 | 0.86465 | 0.00000 | 0.13826 | 0.49571 | 0.02552 | 0.76555 | 1.13055 | 0.00406 | 0.00000 | 0.51519 | 1.83137 | 0.68993 |
| **Hk1** | 1.18107 | 0.01099 | 3.47114 | 3.20504 | 6.86403 | 0.00000 | 0.00322 | 0.00218 | 0.25306 | 0.00179 | 0.01886 | 0.00420 | 0.95435 | 0.97301 |
| **Hk1os** | 0.00978 | 0.00447 | 0.03583 | 0.00475 | 0.29844 | 0.01419 | 0.00419 | 0.01027 | 0.01465 | 0.00779 | 0.00954 | 0.00885 | 0.00790 | 0.01835 |
| **Hk2** | 0.00000 | 3.02312 | 1.25090 | 1.05508 | 0.00000 | 7.35549 | 3.23750 | 6.03643 | 1.47197 | 2.02125 | 7.42935 | 0.33553 | 0.00000 | 0.95731 |
| **Hkdc1** | 2.38731 | 0.00244 | 6.07849 | 7.31697 | 2.68735 | 5.57792 | 0.00890 | 0.36623 | 0.00000 | 0.00000 | 2.21941 | 1.19164 | 0.50812 | 2.03958 |
| **Hlcs** | 0.77135 | 4.25210 | 4.58665 | 0.43329 | 1.85859 | 1.86172 | 0.00266 | 0.66425 | 0.20141 | 0.00000 | 0.00000 | 1.47805 | 0.00000 | 1.88003 |
| **Hlf** | 0.13369 | 0.00429 | 0.56796 | 0.00000 | 0.69815 | 0.00900 | 0.00400 | 0.00718 | 0.00857 | 0.01392 | 0.57711 | 0.00624 | 0.00000 | 0.04793 |
| **Hltf** | 1.60905 | 0.02229 | 1.21338 | 0.21189 | 0.18730 | 1.42706 | 0.83467 | 0.28930 | 1.95847 | 0.06319 | 0.85096 | 2.00671 | 0.68407 | 2.11447 |
| **Hmbox1** | 0.68166 | 0.00365 | 2.04811 | 0.40015 | 3.37013 | 0.11622 | 0.02217 | 0.02000 | 1.20188 | 0.00201 | 0.06523 | 0.00037 | 2.31035 | 4.68608 |
| **Hmbs** | 7.57149 | 6.54600 | 5.25029 | 10.05166 | 14.40706 | 4.62388 | 0.08250 | 5.64714 | 1.06378 | 0.05177 | 6.24983 | 6.11597 | 6.92657 | 6.11751 |
| **Hmces** | 0.33719 | 0.07993 | 0.87118 | 0.77052 | 0.75458 | 1.14870 | 1.20181 | 1.78905 | 0.00000 | 0.04837 | 1.08404 | 8.32621 | 4.31220 | 0.04522 |
| **Hmg20a** | 1.70184 | 0.00859 | 4.27463 | 0.00000 | 2.62811 | 5.88723 | 0.26725 | 1.89436 | 0.66391 | 0.04274 | 6.58845 | 1.76343 | 0.04486 | 4.40966 |
| **Hmg20b** | 0.00000 | 0.10054 | 0.81875 | 6.34358 | 5.52604 | 0.00000 | 0.01011 | 3.48101 | 0.02025 | 0.22343 | 0.00000 | 1.45158 | 0.08387 | 0.00268 |
| **Hmga2** | 23.07024 | 0.19286 | 25.18501 | 12.11132 | 15.14857 | 17.08828 | 2.61719 | 6.79155 | 28.04894 | 10.18818 | 18.16867 | 14.44727 | 1.50780 | 9.06042 |
| **Hmga2.ps1** | 0.02784 | 0.01211 | 0.00374 | 0.13369 | 0.00956 | 0.00443 | 0.00391 | 2.14756 | 0.04184 | 0.04138 | 0.37583 | 0.01906 | 0.18549 | 0.18369 |

| | | | | | | | | | | | | | | |
|---|---|---|---|---|---|---|---|---|---|---|---|---|---|---|
| **Hmgb1** | 15.93383 | 8.11223 | 26.15693 | 15.14231 | 25.15254 | 20.52543 | 5.91165 | 13.22910 | 29.57326 | 20.34882 | 19.56396 | 15.53698 | 13.46415 | 23.35316 |
| **Hmgb1.rs17** | 1.59322 | 0.86003 | 2.85158 | 0.79023 | 2.14781 | 1.39166 | 0.65777 | 0.91930 | 2.51489 | 3.91824 | 2.01343 | 1.68636 | 2.20411 | 3.41281 |
| **Hmgb2** | 8.27602 | 4.12299 | 19.14930 | 5.41189 | 6.37563 | 11.93242 | 6.45996 | 6.68306 | 18.79998 | 14.66873 | 16.77075 | 11.26652 | 6.13200 | 28.19941 |
| **Hmgb3** | 14.70560 | 0.06544 | 16.22443 | 6.63956 | 14.48599 | 12.75989 | 2.41490 | 3.39009 | 18.01834 | 0.12345 | 5.61297 | 6.93710 | 14.05940 | 22.25871 |
| **Hmgcl** | 2.74612 | 0.53151 | 9.78236 | 9.07245 | 5.86686 | 14.32862 | 2.90936 | 8.77144 | 0.00000 | 8.36021 | 12.39763 | 6.66639 | 2.48351 | 6.83845 |
| **Hmgcll1** | 0.00000 | 0.00403 | 0.00000 | 0.01620 | 0.00000 | 0.00648 | 0.00395 | 0.00856 | 1.43170 | 0.00653 | 0.00615 | 0.00701 | 2.68126 | 0.00000 |
| **Hmgcr** | 0.61725 | 0.00102 | 0.00000 | 0.00000 | 0.00000 | 5.15319 | 4.12004 | 0.30689 | 2.08936 | 1.50339 | 0.81087 | 4.47097 | 0.55982 | 1.26302 |
| **Hmgcs1** | 3.50804 | 5.75846 | 5.82245 | 1.68545 | 5.70306 | 13.83952 | 1.04306 | 2.69199 | 1.40472 | 0.05906 | 2.01922 | 4.54313 | 6.10498 | 13.50037 |
| **Hmgcs2** | 0.01931 | 0.00449 | 0.00425 | 0.01528 | 0.61295 | 0.01305 | 0.00423 | 0.00989 | 0.01296 | 0.00724 | 0.00838 | 0.00852 | 0.00323 | 0.01898 |
| **Hmgn1** | 21.76377 | 4.74626 | 8.24950 | 31.48544 | 26.21993 | 13.70463 | 6.10940 | 10.28685 | 9.52453 | 0.68413 | 3.32835 | 15.00449 | 7.05123 | 8.71455 |
| **Hmgn2** | 19.20656 | 0.49388 | 23.56535 | 15.41938 | 26.82146 | 14.20191 | 7.98518 | 15.78609 | 9.50268 | 1.62622 | 11.37599 | 14.62525 | 12.35727 | 9.13346 |
| **Hmgn3** | 0.35260 | 0.00444 | 0.00287 | 0.00584 | 0.44443 | 0.01787 | 0.00435 | 0.00997 | 0.00847 | 0.00717 | 0.00695 | 0.00679 | 0.32297 | 0.00193 |
| **Hmgn5** | 0.61281 | 6.16778 | 1.51995 | 4.57015 | 1.43882 | 8.41071 | 3.45167 | 1.78996 | 6.99910 | 0.15856 | 5.15682 | 11.32264 | 9.48319 | 3.55361 |
| **Hmgxb3** | 1.27166 | 0.02952 | 8.06598 | 3.25658 | 10.58987 | 5.23894 | 4.20642 | 0.00000 | 0.33260 | 0.11877 | 6.07255 | 2.92989 | 9.51346 | 4.31692 |
| **Hmgxb4** | 3.90265 | 0.09407 | 5.35901 | 0.15906 | 5.17047 | 6.81902 | 3.60646 | 1.89856 | 13.65325 | 10.21988 | 4.71560 | 4.76079 | 4.89916 | 15.07951 |
| **Hmha1** | 0.00000 | 0.03705 | 0.43609 | 0.30069 | 0.85716 | 1.57366 | 0.00400 | 0.91174 | 0.00169 | 0.00715 | 0.16719 | 1.12743 | 0.02024 | 0.43440 |
| **Hmmr** | 6.88693 | 4.16497 | 5.01981 | 0.73446 | 5.67706 | 4.19114 | 0.15127 | 1.11908 | 21.80224 | 0.14981 | 2.47808 | 3.17019 | 7.98186 | 20.57244 |
| **Hmox1** | 0.27884 | 0.00402 | 0.00347 | 0.46881 | 0.20169 | 0.00000 | 0.00385 | 0.00668 | 0.01171 | 0.00634 | 0.01329 | 0.02038 | 0.00480 | 0.00000 |
| **Hmox2** | 3.52901 | 0.07182 | 4.28626 | 6.78120 | 4.65829 | 3.78102 | 2.09864 | 8.35769 | 6.06670 | 0.50978 | 3.33337 | 2.98122 | 0.40095 | 6.72860 |
| **Hn1** | 12.95410 | 4.07301 | 17.30366 | 12.07363 | 21.51330 | 13.52643 | 4.70403 | 23.61977 | 0.00000 | 11.92734 | 7.05536 | 4.44824 | 8.84388 | 11.38721 |
| **Hn1l** | 0.85496 | 4.98267 | 0.84589 | 0.99333 | 1.38722 | 7.62714 | 0.01811 | 3.81350 | 1.58861 | 0.02783 | 3.67508 | 8.92506 | 2.69069 | 5.10099 |
| **Hnf1a** | 0.01109 | 0.00413 | 0.01045 | 0.41475 | 0.53121 | 0.02114 | 0.00400 | 0.00672 | 0.01053 | 0.00567 | 0.06250 | 0.00857 | 0.25258 | 0.07117 |
| **Hnf1b** | 0.00000 | 0.07102 | 0.20791 | 0.00000 | 0.01614 | 0.00760 | 0.00361 | 0.01282 | 0.00705 | 0.00667 | 0.00340 | 1.88352 | 0.07286 | 0.04502 |
| **Hnf4a** | 0.00000 | 0.00000 | 0.09196 | 0.44419 | 0.00906 | 0.03300 | 0.00236 | 0.00383 | 0.01049 | 0.01375 | 0.01844 | 6.19057 | 0.00510 | 0.01871 |
| **Hnf4aos** | 0.04248 | 0.00435 | 0.02784 | 0.05482 | 0.00991 | 0.10962 | 0.01583 | 0.00446 | 0.01182 | 0.00542 | 0.01632 | 0.01371 | 0.00737 | 0.00917 |
| **Hnmt** | 0.00143 | 0.00409 | 0.01447 | 0.01223 | 0.00479 | 0.01212 | 0.00412 | 0.00833 | 0.02694 | 0.00601 | 0.00914 | 0.00759 | 2.10802 | 0.01011 |
| **Hnrnpa0** | 5.21220 | 0.21823 | 10.76163 | 7.01739 | 8.47625 | 13.62249 | 5.99533 | 11.96647 | 21.79281 | 0.43408 | 10.16856 | 9.17272 | 5.91100 | 12.03891 |
| **Hnrnpa1** | 0.34710 | 0.01288 | 1.27516 | 0.52623 | 1.52144 | 0.66383 | 0.28990 | 0.67465 | 6.35611 | 0.17659 | 1.53874 | 0.43560 | 0.09494 | 4.63897 |
| **Hnrnpa2b1** | 23.62317 | 2.53141 | 14.84619 | 14.86511 | 13.57743 | 19.61766 | 8.73242 | 13.63235 | 19.89843 | 4.08146 | 10.76898 | 17.03292 | 7.51938 | 15.27246 |
| **Hnrnpa3** | 20.82299 | 4.03427 | 15.91043 | 13.60745 | 20.39398 | 24.62295 | 5.11653 | 13.71867 | 26.63147 | 5.28666 | 12.88973 | 12.96176 | 5.24758 | 11.33063 |
| **Hnrnpab** | 38.35751 | 3.62756 | 36.46861 | 26.33570 | 44.25713 | 32.39834 | 9.91451 | 18.79882 | 22.09844 | 11.47058 | 21.14562 | 20.13674 | 23.65415 | 34.93752 |
| **Hnrnpc** | 26.51595 | 4.81781 | 40.19592 | 35.73460 | 37.93225 | 35.69731 | 12.27250 | 23.95442 | 39.58919 | 15.32777 | 24.96062 | 24.85167 | 16.53417 | 40.05603 |
| **Hnrnpd** | 18.45453 | 5.43263 | 18.97925 | 16.18791 | 20.79382 | 23.03022 | 8.85431 | 8.24004 | 19.72092 | 0.62464 | 15.05786 | 14.47820 | 6.79353 | 18.93808 |
| **Hnrnpdl** | 17.72754 | 1.65684 | 15.33290 | 13.95888 | 27.76221 | 16.31261 | 6.66963 | 8.67224 | 11.79521 | 8.43396 | 4.93722 | 11.22195 | 12.28600 | 14.66987 |
| **Hnrnpf** | 28.38892 | 11.52874 | 43.33727 | 35.28080 | 39.05552 | 38.10525 | 12.28706 | 23.68271 | 18.23433 | 5.64752 | 26.50478 | 24.57433 | 15.80373 | 42.72021 |
| **Hnrnph1** | 18.66474 | 1.19409 | 28.62043 | 12.74889 | 28.48227 | 19.45697 | 9.43324 | 17.49122 | 24.36607 | 0.83563 | 13.15565 | 15.88390 | 1.87426 | 34.34320 |
| **Hnrnph2** | 1.77425 | 0.06871 | 7.39242 | 7.85293 | 8.96064 | 7.18115 | 0.06855 | 3.32408 | 23.91144 | 0.00747 | 2.78321 | 3.61149 | 1.83059 | 14.73274 |
| **Hnrnph3** | 0.05137 | 0.00490 | 0.00000 | 0.92151 | 0.04611 | 0.23268 | 0.00407 | 0.05450 | 0.78097 | 0.00750 | 0.00329 | 0.39639 | 0.01203 | 0.34937 |
| **Hnrnpk** | 14.38751 | 9.43928 | 36.69088 | 24.81202 | 29.72501 | 29.71080 | 9.62801 | 16.92817 | 20.36167 | 0.93140 | 25.82870 | 23.26648 | 10.91636 | 31.44876 |
| **Hnrnpl** | 0.26939 | 0.12552 | 0.14349 | 4.50211 | 0.00000 | 1.76164 | 0.03167 | 1.69257 | 7.97388 | 0.50847 | 0.40875 | 1.19328 | 3.00785 | 9.04858 |
| **Hnrnpll** | 15.67324 | 0.03312 | 8.43350 | 0.68109 | 13.90122 | 5.47193 | 0.85363 | 3.38171 | 6.58979 | 0.02219 | 7.40965 | 6.66515 | 11.05315 | 16.40265 |
| **Hnrnpm** | 10.65415 | 8.38754 | 22.59532 | 23.72343 | 24.75724 | 7.18869 | 6.07310 | 11.28903 | 3.97984 | 13.60805 | 4.91918 | 17.42582 | 11.76031 | 2.23914 |
| **Hnrnpr** | 11.09646 | 0.02975 | 10.57335 | 6.57313 | 14.03800 | 11.40620 | 1.53413 | 5.13916 | 5.25346 | 0.07552 | 8.03152 | 9.06396 | 8.39659 | 12.58334 |
| **Hnrnpu** | 19.76975 | 8.91254 | 26.10233 | 23.27017 | 27.01752 | 25.83176 | 6.72242 | 13.90554 | 32.42443 | 6.34680 | 17.53624 | 19.49894 | 17.40145 | 28.47827 |
| **Hnrnpul1** | 0.05901 | 0.03656 | 0.62576 | 1.02228 | 1.97275 | 0.20517 | 0.72812 | 1.25218 | 0.03477 | 0.03140 | 0.46533 | 1.94178 | 0.00760 | 0.30163 |
| **Hnrnpul2** | 5.70435 | 0.98441 | 2.78118 | 2.99836 | 6.35872 | 7.92367 | 0.26188 | 2.50447 | 8.82385 | 0.09024 | 2.82502 | 4.15178 | 7.21749 | 8.76386 |
| **Hoga1** | 0.49942 | 0.00434 | 0.45921 | 0.00482 | 0.25611 | 0.00402 | 0.00414 | 0.01881 | 0.00770 | 0.00591 | 0.00688 | 0.00595 | 0.00659 | 0.04429 |
| **Homer1** | 0.17905 | 0.00310 | 0.75114 | 0.00000 | 0.41543 | 1.13900 | 0.00438 | 0.33793 | 0.01912 | 0.00000 | 2.01415 | 0.85778 | 1.83571 | 0.22761 |
| **Homer2** | 1.96227 | 0.66526 | 1.74767 | 1.29500 | 2.09024 | 2.40197 | 0.33137 | 2.85459 | 1.38956 | 0.05472 | 1.03571 | 1.16031 | 1.11862 | 0.75426 |
| **Homer3** | 0.13241 | 0.00387 | 0.05690 | 0.25051 | 0.14609 | 0.00000 | 0.93457 | 0.00267 | 0.00701 | 0.03911 | 0.00836 | 0.04930 | 0.13246 | 0.08160 |
| **Homez** | 0.00746 | 0.00389 | 0.00589 | 0.72320 | 0.02097 | 0.02780 | 0.00911 | 0.91015 | 0.01113 | 0.00537 | 0.01899 | 0.01059 | 0.80859 | 0.00000 |
| **Hook1** | 0.00000 | 0.00423 | 0.13670 | 0.00000 | 0.33870 | 0.00230 | 0.00363 | 0.00860 | 3.22483 | 0.00670 | 0.00335 | 0.00544 | 0.81123 | 0.92658 |
| **Hook2** | 0.17686 | 0.07834 | 5.65431 | 11.32719 | 3.78652 | 0.00000 | 1.45638 | 5.25563 | 0.01799 | 12.59619 | 2.03478 | 0.91289 | 0.43573 | 0.00000 |
| **Hook3** | 2.10192 | 0.01018 | 0.16242 | 0.13125 | 0.16434 | 1.33160 | 0.00375 | 1.07112 | 0.09961 | 0.00548 | 0.00000 | 0.61016 | 0.27074 | 2.68156 |
| **Hopx** | 0.01366 | 0.00433 | 0.01309 | 0.01195 | 0.00687 | 0.01242 | 0.00426 | 0.00925 | 0.01235 | 0.00620 | 0.00864 | 0.00912 | 0.35862 | 0.13854 |

| | | | | | | | | | | | | | | |
|---|---|---|---|---|---|---|---|---|---|---|---|---|---|---|
| **Hormad1** | 1.86450 | 0.00159 | 3.12921 | 0.00000 | 2.28163 | 1.35689 | 0.00104 | 0.00000 | 10.88368 | 0.00000 | 2.97911 | 1.06851 | 3.42037 | 5.08088 |
| **Hormad2** | 0.42678 | 0.02898 | 0.44044 | 0.13848 | 0.21090 | 0.14129 | 0.00468 | 0.77217 | 0.12427 | 0.00467 | 0.00311 | 0.63797 | 0.24869 | 0.00000 |
| **Hotairm1** | 0.01490 | 0.00449 | 0.03368 | 0.01485 | 0.01516 | 0.02310 | 0.00486 | 0.06593 | 0.01023 | 0.00788 | 0.00768 | 0.00780 | 0.03212 | 0.02955 |
| **Hoxa1** | 0.00488 | 0.00381 | 0.00422 | 0.00000 | 0.29197 | 0.12036 | 0.00363 | 0.41169 | 0.00683 | 0.00529 | 0.04363 | 0.00611 | 0.01097 | 0.75537 |
| **Hoxa10** | 0.00196 | 0.00417 | 0.00652 | 0.35160 | 0.07542 | 0.00408 | 0.00398 | 0.00654 | 0.02715 | 0.00523 | 0.00570 | 0.01591 | 0.00326 | 0.40697 |
| **Hoxa7** | 0.01027 | 0.00436 | 0.01472 | 0.01226 | 0.01080 | 0.01321 | 0.00424 | 0.00949 | 0.01326 | 0.00657 | 0.00887 | 0.00950 | 0.25843 | 0.00418 |
| **Hoxb13** | 0.00436 | 0.02020 | 0.00117 | 0.00000 | 0.01022 | 0.61297 | 0.00355 | 0.32082 | 0.00913 | 0.02002 | 0.14911 | 1.29062 | 0.01449 | 0.01292 |
| **Hoxb7** | 0.00485 | 0.00412 | 0.00752 | 0.46940 | 0.03161 | 0.00390 | 0.00400 | 0.00622 | 0.01851 | 0.00532 | 0.00954 | 0.01707 | 0.00649 | 0.00855 |
| **Hoxb9** | 0.00000 | 0.00435 | 0.18360 | 0.12080 | 0.50131 | 0.00176 | 0.01073 | 0.02936 | 0.00712 | 0.01199 | 0.00631 | 0.00926 | 0.01542 | 0.19345 |
| **Hoxc13** | 0.00768 | 0.00428 | 0.18716 | 0.00000 | 0.01020 | 0.18314 | 0.00421 | 0.00397 | 0.01032 | 0.00638 | 0.00663 | 0.05713 | 0.00749 | 0.00992 |
| **Hoxd11** | 0.13742 | 0.00449 | 0.01247 | 0.03611 | 0.01741 | 0.01198 | 0.00425 | 0.02909 | 0.02700 | 0.01946 | 0.00683 | 0.00789 | 0.05867 | 0.01417 |
| **Hoxd8** | 0.05590 | 0.00444 | 0.15603 | 0.10209 | 0.30099 | 0.15190 | 0.12290 | 0.23674 | 0.02545 | 0.00531 | 0.02030 | 0.01550 | 0.00477 | 0.00308 |
| **Hoxd9** | 0.00348 | 0.00434 | 0.13347 | 0.13079 | 0.34385 | 0.00559 | 0.00428 | 0.01310 | 0.00949 | 0.00676 | 0.00598 | 0.00983 | 0.00628 | 0.01224 |
| **Hp** | 0.15689 | 0.02086 | 0.10842 | 0.01905 | 0.18768 | 0.00000 | 0.00422 | 0.85244 | 0.00811 | 0.00745 | 0.00609 | 0.00457 | 0.97855 | 0.00481 |
| **Hp1bp3** | 4.50012 | 0.03641 | 2.81274 | 2.50739 | 2.49132 | 4.29785 | 3.63761 | 0.05246 | 1.62317 | 1.62340 | 3.87615 | 1.95890 | 2.78017 | 2.51634 |
| **Hpcal1** | 0.94381 | 0.05590 | 0.77181 | 1.25810 | 0.96300 | 0.78579 | 0.01682 | 2.42640 | 0.07425 | 0.01708 | 0.36028 | 0.76932 | 2.17598 | 0.00000 |
| **Hpd** | 0.00218 | 0.00432 | 0.02010 | 0.01140 | 0.00551 | 0.01147 | 0.00422 | 0.00890 | 0.00784 | 0.00639 | 0.00770 | 0.00860 | 0.37530 | 0.52524 |
| **Hpdl** | 4.15431 | 0.17622 | 11.45268 | 11.10263 | 13.52916 | 1.68177 | 3.87089 | 9.68685 | 0.15991 | 13.63786 | 1.45553 | 1.12573 | 1.85084 | 1.89613 |
| **Hpf1** | 13.97868 | 0.14162 | 9.39289 | 10.00537 | 9.27570 | 11.12333 | 3.63527 | 6.27645 | 3.49312 | 0.26538 | 8.10189 | 4.98938 | 5.73201 | 2.48999 |
| **Hpgds** | 0.09266 | 0.01249 | 0.02160 | 0.21858 | 0.45906 | 0.20986 | 0.03049 | 0.28021 | 0.31104 | 0.39519 | 0.00496 | 0.06399 | 0.71721 | 0.53537 |
| **Hprt** | 35.36902 | 10.01403 | 44.58837 | 37.31445 | 38.03631 | 40.21550 | 11.68867 | 23.34303 | 29.43100 | 14.38370 | 25.07181 | 14.54949 | 9.74250 | 27.44912 |
| **Hps1** | 0.00559 | 0.00374 | 0.00000 | 0.76123 | 0.53744 | 0.00000 | 0.12296 | 0.55126 | 0.00861 | 0.01171 | 0.03014 | 1.06540 | 0.12672 | 0.00000 |
| **Hps3** | 0.00000 | 0.13738 | 1.28831 | 1.32277 | 0.75084 | 1.88593 | 0.05152 | 0.06637 | 1.36837 | 0.00369 | 0.34440 | 1.05680 | 0.25975 | 0.60861 |
| **Hps4** | 0.58433 | 6.44855 | 3.92942 | 1.88735 | 1.47325 | 0.59737 | 0.00767 | 2.11383 | 0.00013 | 0.00418 | 0.46168 | 0.52912 | 0.00000 | 0.22793 |
| **Hps5** | 2.33472 | 0.01766 | 2.07798 | 2.28798 | 1.96889 | 0.61013 | 0.37048 | 0.56196 | 4.00198 | 0.01279 | 2.05979 | 1.20198 | 3.30088 | 2.45619 |
| **Hps6** | 0.52753 | 0.01272 | 2.95560 | 0.74490 | 2.94479 | 0.11690 | 0.00990 | 1.54175 | 0.00397 | 0.01518 | 0.00000 | 0.10361 | 1.33431 | 0.40221 |
| **Hpse** | 0.00000 | 0.00357 | 0.43335 | 1.34873 | 0.11294 | 0.46964 | 0.00369 | 0.00109 | 0.89228 | 0.00545 | 0.00158 | 0.29441 | 0.75330 | 1.24618 |
| **Hr** | 0.00811 | 0.01103 | 0.02658 | 0.00000 | 0.01388 | 0.15730 | 0.00419 | 0.09080 | 0.01159 | 0.00553 | 0.00411 | 0.09458 | 0.00756 | 0.01016 |
| **Hras** | 10.46661 | 2.94732 | 5.10881 | 4.73105 | 5.92972 | 2.52572 | 0.04424 | 2.64878 | 0.04470 | 1.54016 | 0.53427 | 3.52805 | 7.45418 | 1.16655 |
| **Hs1bp3** | 1.66612 | 1.26285 | 3.18082 | 3.83272 | 1.26525 | 8.29554 | 3.74704 | 4.17091 | 2.20617 | 0.20725 | 2.12965 | 5.89053 | 0.00000 | 0.85583 |
| **Hs2st1** | 3.39022 | 3.86584 | 4.30342 | 1.62865 | 1.39487 | 8.16093 | 0.18520 | 0.06875 | 0.03500 | 0.03634 | 3.98414 | 1.70938 | 2.23589 | 3.41051 |
| **Hs3st1** | 0.65490 | 0.00226 | 0.05984 | 0.00000 | 0.02835 | 0.00000 | 0.00301 | 0.24980 | 0.00781 | 0.00169 | 1.11388 | 2.29168 | 0.01295 | 0.67040 |
| **Hs3st3b1** | 0.00000 | 0.00419 | 0.49631 | 0.04406 | 0.00000 | 5.20909 | 1.99889 | 1.72517 | 1.38710 | 0.09537 | 0.00000 | 0.18356 | 0.00221 | 0.10640 |
| **Hs3st5** | 0.23115 | 0.00445 | 0.00961 | 0.00000 | 0.00733 | 0.34837 | 0.00422 | 0.00533 | 0.00918 | 0.00558 | 0.00550 | 0.01079 | 0.00667 | 0.02751 |
| **Hs3st6** | 0.00820 | 0.00424 | 0.01852 | 0.04797 | 0.22284 | 0.03528 | 0.00425 | 0.00586 | 0.00928 | 0.00574 | 0.00754 | 0.01226 | 0.27369 | 0.00372 |
| **Hs6st1** | 2.26358 | 0.03985 | 6.22065 | 0.73985 | 7.96193 | 6.20735 | 0.00065 | 5.84831 | 5.43540 | 0.04573 | 6.84536 | 0.47753 | 2.08223 | 4.71330 |
| **Hs6st2** | 0.03169 | 0.00463 | 0.00897 | 0.02278 | 0.00000 | 0.02172 | 0.00429 | 0.01064 | 0.08426 | 0.00680 | 0.01436 | 0.02066 | 0.06731 | 0.65286 |
| **Hsbp1** | 9.24624 | 0.33918 | 7.58847 | 6.56445 | 4.96974 | 17.36305 | 3.82032 | 14.58993 | 3.53500 | 5.85718 | 5.59518 | 8.19844 | 0.41533 | 4.37100 |
| **Hsbp1l1** | 0.00707 | 0.01617 | 0.04820 | 0.49357 | 0.00000 | 0.11151 | 0.41779 | 2.35731 | 0.11054 | 13.26542 | 0.27344 | 0.00465 | 0.07273 | 0.09195 |
| **Hscb** | 1.47920 | 0.04789 | 0.55341 | 0.00000 | 0.32524 | 1.81841 | 0.00234 | 2.62315 | 0.00884 | 0.02222 | 1.85051 | 5.36144 | 1.76887 | 0.03002 |
| **Hsd11b2** | 0.00590 | 0.02747 | 0.01152 | 0.00000 | 0.01289 | 0.04824 | 0.00416 | 0.00550 | 0.01155 | 0.00646 | 0.00372 | 0.72850 | 0.00753 | 0.01085 |
| **Hsd17b1** | 0.02490 | 0.00344 | 1.35186 | 0.12037 | 0.13303 | 0.00000 | 0.00361 | 0.00000 | 0.01061 | 0.00441 | 0.46447 | 0.56713 | 0.00656 | 0.00485 |
| **Hsd17b10** | 2.59941 | 0.17077 | 3.50613 | 3.38354 | 4.38152 | 3.93242 | 2.14853 | 9.42308 | 1.34786 | 0.10896 | 3.67285 | 4.87002 | 0.53234 | 1.60988 |
| **Hsd17b11** | 0.11350 | 0.00404 | 0.04038 | 0.34766 | 0.01721 | 0.23246 | 0.00397 | 0.00352 | 0.00872 | 0.00852 | 0.00633 | 0.32384 | 0.19559 | 0.01991 |
| **Hsd17b12** | 7.39676 | 0.10542 | 4.13119 | 8.01394 | 1.99408 | 11.80868 | 5.24957 | 2.66495 | 0.17039 | 0.12949 | 6.54582 | 8.72692 | 2.38027 | 2.41704 |
| **Hsd17b13** | 0.00346 | 0.01173 | 0.00889 | 0.03382 | 0.09273 | 0.01814 | 0.01619 | 0.00539 | 0.72665 | 0.01897 | 0.00755 | 0.01813 | 0.96774 | 0.30453 |
| **Hsd17b14** | 11.53851 | 0.01801 | 11.26765 | 7.98090 | 20.10435 | 1.42567 | 0.00158 | 2.12615 | 1.49663 | 0.01261 | 0.00000 | 1.59588 | 5.60241 | 1.13191 |
| **Hsd17b2** | 0.02488 | 0.00420 | 0.03263 | 0.09682 | 0.03398 | 0.00328 | 0.00388 | 0.00677 | 0.42799 | 0.00711 | 0.00505 | 0.01436 | 0.22029 | 0.02951 |
| **Hsd17b4** | 2.03465 | 0.07142 | 7.77681 | 5.00391 | 2.59493 | 3.59582 | 0.01125 | 1.04607 | 4.67263 | 0.01473 | 9.20576 | 6.89879 | 0.04298 | 0.98110 |
| **Hsd17b6** | 0.00000 | 7.34354 | 0.43905 | 0.00000 | 1.37596 | 0.00000 | 2.43574 | 8.31587 | 1.21940 | 13.95825 | 0.53178 | 0.00000 | 3.38153 | 2.24917 |
| **Hsd17b7** | 0.06953 | 0.01089 | 0.50412 | 1.58463 | 0.71513 | 1.86242 | 1.35270 | 0.00176 | 0.01912 | 0.03713 | 0.00000 | 0.57871 | 1.77108 | 0.10001 |
| **Hsd3b1** | 0.06764 | 0.00444 | 0.00007 | 0.02632 | 1.55413 | 0.01003 | 0.00440 | 0.00939 | 0.00997 | 0.00673 | 0.00730 | 0.00863 | 0.00000 | 0.14207 |
| **Hsd3b7** | 0.47971 | 4.65472 | 0.00000 | 0.00161 | 0.00000 | 2.19300 | 0.00232 | 1.68836 | 0.02477 | 0.85151 | 1.43815 | 0.38910 | 0.53655 | 0.21975 |
| **Hsdl1** | 0.06973 | 0.02684 | 1.69664 | 0.35313 | 1.11279 | 1.03547 | 0.00274 | 4.30132 | 0.55462 | 0.03968 | 1.14454 | 1.81052 | 0.00191 | 0.46804 |
| **Hsdl2** | 2.65192 | 2.13160 | 3.30083 | 0.50507 | 0.95711 | 1.95092 | 1.27518 | 0.21989 | 0.00624 | 0.02830 | 1.59137 | 0.10799 | 0.34540 | 0.13009 |

| | | | | | | | | | | | | | | |
|---|---|---|---|---|---|---|---|---|---|---|---|---|---|---|
| **Hsf1** | 2.50509 | 6.12363 | 1.41215 | 3.86000 | 1.48521 | 0.40403 | 0.00133 | 0.61339 | 0.76532 | 2.38305 | 2.53873 | 2.07810 | 8.04362 | 1.70711 |
| **Hsf2** | 1.47096 | 0.00962 | 0.84104 | 1.21412 | 0.65079 | 0.92761 | 2.59458 | 0.89451 | 0.81593 | 0.08769 | 2.69882 | 0.63154 | 0.35978 | 0.41561 |
| **Hsf2bp** | 1.20795 | 0.04718 | 0.28936 | 3.46274 | 0.00000 | 4.03744 | 0.00752 | 1.13177 | 0.43372 | 0.01897 | 4.55427 | 4.02023 | 1.40470 | 2.80547 |
| **Hsf5** | 0.00862 | 0.01358 | 0.00453 | 0.23886 | 0.01879 | 0.32610 | 0.00428 | 0.05060 | 0.01187 | 0.02233 | 0.00377 | 0.00319 | 0.04755 | 0.00000 |
| **Hsh2d** | 0.29039 | 0.00424 | 0.48355 | 0.02035 | 0.00000 | 0.00140 | 0.00991 | 0.00954 | 0.03477 | 0.00739 | 0.00623 | 0.01884 | 0.00416 | 0.44796 |
| **Hsp90aa1** | 45.92840 | 13.64103 | 60.50566 | 59.06492 | 46.68268 | 67.15729 | 16.34200 | 32.26281 | 54.54089 | 25.30687 | 46.27132 | 38.17918 | 17.46236 | 37.41975 |
| **Hsp90ab1** | 46.33073 | 18.65366 | 69.95512 | 59.72962 | 54.61815 | 65.72738 | 16.43250 | 37.16257 | 52.33946 | 25.31952 | 41.13315 | 43.36924 | 17.56812 | 52.13190 |
| **Hsp90b1** | 7.50868 | 4.20464 | 5.10692 | 15.41497 | 3.72140 | 34.55592 | 10.37754 | 9.56840 | 10.92892 | 0.67035 | 9.99924 | 21.47040 | 1.74789 | 8.68514 |
| **Hspa12a** | 0.02627 | 0.00437 | 0.31713 | 0.00003 | 0.03823 | 1.14865 | 0.00487 | 0.59068 | 0.00321 | 0.00579 | 0.02116 | 0.00619 | 0.41494 | 0.57004 |
| **Hspa12b** | 0.00705 | 0.00429 | 0.01073 | 0.11447 | 0.19762 | 0.00873 | 0.00396 | 0.00696 | 0.01138 | 0.00735 | 0.00707 | 0.01229 | 0.00581 | 0.00776 |
| **Hspa13** | 0.01130 | 0.00201 | 0.00000 | 0.50806 | 0.16990 | 0.38958 | 0.75849 | 0.00907 | 0.00722 | 2.94954 | 2.00745 | 1.44116 | 0.00964 | 0.07605 |
| **Hspa14** | 11.23980 | 7.00514 | 15.52889 | 7.46969 | 7.67038 | 13.79746 | 2.95700 | 3.06824 | 9.92630 | 0.16434 | 13.27850 | 8.23478 | 7.59141 | 13.41166 |
| **Hspa1a** | 0.57047 | 0.00349 | 0.52759 | 0.84674 | 0.80187 | 0.17639 | 0.00372 | 0.00184 | 0.03876 | 0.00507 | 0.24740 | 0.08029 | 2.49820 | 0.00000 |
| **Hspa1b** | 0.07773 | 0.00380 | 0.22394 | 0.19646 | 0.16559 | 0.06256 | 0.00395 | 0.00909 | 0.02213 | 0.00664 | 0.00733 | 0.02740 | 2.03459 | 0.01650 |
| **Hspa1l** | 0.01038 | 0.00551 | 0.05545 | 0.08782 | 0.06221 | 0.29629 | 0.00411 | 0.01617 | 0.02173 | 0.00940 | 0.00666 | 0.00396 | 0.00561 | 0.01736 |
| **Hspa2** | 0.05880 | 0.00362 | 1.79963 | 1.55295 | 0.89779 | 1.00319 | 0.00889 | 0.00000 | 0.39399 | 0.00293 | 0.00000 | 0.01662 | 2.98109 | 0.99970 |
| **Hspa4** | 21.24161 | 0.56983 | 18.06316 | 5.51812 | 15.77990 | 13.73377 | 5.95422 | 8.52159 | 15.41179 | 7.78827 | 10.81040 | 15.36536 | 10.87134 | 17.54434 |
| **Hspa4l** | 0.00000 | 0.00373 | 0.00000 | 0.15851 | 1.18237 | 0.32238 | 0.00991 | 0.26959 | 0.16512 | 0.00631 | 0.01138 | 0.00414 | 0.71417 | 1.45251 |
| **Hspa5** | 9.19984 | 9.85283 | 12.06398 | 31.00850 | 3.07756 | 32.73654 | 12.00575 | 14.78741 | 18.42136 | 4.13315 | 16.22114 | 15.55845 | 12.75866 | 17.49559 |
| **Hspa8** | 49.61170 | 18.79130 | 77.83423 | 70.93843 | 63.55854 | 73.64697 | 19.34927 | 41.53337 | 30.95111 | 26.45183 | 41.05164 | 44.37023 | 18.53364 | 40.94309 |
| **Hspa9** | 21.32275 | 7.82767 | 24.87595 | 25.61792 | 21.81830 | 23.63745 | 7.48036 | 10.33532 | 25.00365 | 0.77188 | 17.95743 | 16.39636 | 14.27596 | 18.46350 |
| **Hspb1** | 1.24060 | 0.00372 | 0.00960 | 0.36418 | 3.80546 | 0.20380 | 0.00425 | 0.33101 | 0.00000 | 0.02792 | 0.00164 | 0.00000 | 7.18446 | 0.09417 |
| **Hspb11** | 1.11031 | 0.00317 | 0.03027 | 0.65877 | 1.21662 | 0.35079 | 0.42514 | 0.63222 | 0.55595 | 0.01579 | 0.01127 | 0.70825 | 0.89865 | 0.31161 |
| **Hspb2** | 0.00338 | 0.00462 | 0.02151 | 0.66040 | 0.01697 | 0.00552 | 0.00521 | 0.94998 | 0.01680 | 0.00660 | 0.00699 | 0.01509 | 0.00690 | 0.00596 |
| **Hspb6** | 0.00000 | 0.02634 | 0.08282 | 0.74449 | 0.00558 | 1.07923 | 1.58720 | 0.00358 | 0.00502 | 6.05887 | 0.37138 | 1.06965 | 0.02204 | 0.13264 |
| **Hspb7** | 0.03824 | 0.00422 | 0.01319 | 0.00577 | 0.08692 | 0.00437 | 0.01008 | 0.02903 | 0.05485 | 0.01707 | 0.74455 | 0.00714 | 0.02870 | 0.05138 |
| **Hspb8** | 0.09923 | 0.00952 | 0.00000 | 0.37268 | 0.10892 | 1.26101 | 0.00334 | 0.00263 | 0.00804 | 0.02938 | 0.00000 | 1.67537 | 0.02521 | 0.00000 |
| **Hspb9** | 0.00915 | 0.00440 | 0.13779 | 0.03252 | 0.01132 | 0.00755 | 0.00416 | 0.00965 | 0.01157 | 0.00745 | 0.00767 | 0.01184 | 0.00688 | 0.01135 |
| **Hspbap1** | 2.49104 | 0.01015 | 1.55402 | 1.06159 | 2.14373 | 1.02769 | 0.41159 | 0.21220 | 0.13550 | 0.01478 | 2.73090 | 1.12043 | 1.77535 | 3.00678 |
| **Hspbp1** | 6.16107 | 0.27591 | 22.20964 | 13.48730 | 16.22212 | 8.78514 | 4.73086 | 10.15621 | 0.00000 | 0.30464 | 6.67354 | 4.07292 | 0.97605 | 1.83248 |
| **Hspd1** | 12.94764 | 14.44304 | 24.52482 | 24.17152 | 11.59934 | 32.71479 | 10.42355 | 16.14368 | 18.19576 | 15.05463 | 15.78571 | 23.59521 | 1.51959 | 18.76969 |
| **Hspe1** | 5.06790 | 7.30402 | 0.63927 | 10.15334 | 8.70966 | 22.51494 | 6.00392 | 19.87566 | 1.34104 | 9.18732 | 2.29599 | 17.82541 | 3.15829 | 1.88565 |
| **Hspg2** | 0.60151 | 0.01562 | 0.11766 | 0.00000 | 0.01264 | 0.14693 | 0.00414 | 0.05622 | 0.00736 | 0.00543 | 0.00225 | 0.45624 | 0.02784 | 0.01574 |
| **Hsph1** | 7.14760 | 0.59419 | 16.10491 | 16.88120 | 10.72235 | 36.83016 | 5.54789 | 10.48196 | 10.65204 | 0.51932 | 21.47311 | 23.83154 | 13.48712 | 11.94401 |
| **Htatip2** | 2.48251 | 0.02761 | 1.77648 | 10.53091 | 1.18693 | 11.98470 | 0.34607 | 1.89769 | 0.16535 | 0.07401 | 9.21317 | 4.72574 | 0.12571 | 0.00000 |
| **Htatsf1** | 11.79951 | 0.08632 | 5.03727 | 2.50671 | 10.36609 | 4.43231 | 3.47201 | 2.27651 | 11.83147 | 0.10970 | 4.14030 | 5.21816 | 1.55015 | 11.00907 |
| **Htr5b** | 0.55838 | 0.00422 | 5.18116 | 0.74683 | 0.57234 | 0.12590 | 0.00302 | 0.00000 | 0.00000 | 0.00000 | 0.00469 | 0.00648 | 0.85923 | 1.17282 |
| **Htr7** | 0.00730 | 0.00443 | 0.13205 | 0.00218 | 0.00496 | 0.00708 | 0.00418 | 0.01159 | 0.32376 | 0.00796 | 0.00648 | 0.00901 | 0.03190 | 0.00461 |
| **Htra1** | 0.11433 | 0.03438 | 0.00000 | 0.00000 | 0.36849 | 2.61231 | 0.00656 | 2.47353 | 0.00000 | 11.83563 | 0.00000 | 0.93958 | 1.57851 | 0.62384 |
| **Htra2** | 0.39024 | 0.17581 | 6.03805 | 6.01194 | 0.62116 | 3.36704 | 2.13365 | 6.10793 | 0.28686 | 10.02232 | 1.19446 | 2.16021 | 0.00000 | 1.34438 |
| **Htra3** | 0.00624 | 0.00419 | 0.00354 | 0.29925 | 0.21118 | 0.00986 | 0.00428 | 0.01060 | 0.01089 | 0.00699 | 0.00640 | 0.00684 | 0.00728 | 0.00814 |
| **Htt** | 0.82303 | 0.03071 | 2.02387 | 0.79333 | 0.97464 | 0.81455 | 0.02012 | 2.15987 | 0.00828 | 0.27919 | 1.18904 | 1.91896 | 1.67837 | 1.47678 |
| **Hunk** | 0.00000 | 0.00375 | 0.48258 | 0.15494 | 0.32126 | 0.00000 | 0.00366 | 0.00392 | 0.02226 | 0.00449 | 0.00501 | 1.06009 | 0.20486 | 0.01393 |
| **Hus1** | 1.57228 | 0.00244 | 2.59238 | 2.42746 | 0.93120 | 0.70159 | 0.00901 | 0.34834 | 6.29733 | 0.00717 | 0.38784 | 1.63887 | 0.22007 | 4.98349 |
| **Hus1b** | 0.00264 | 0.00427 | 0.07791 | 0.56365 | 0.00233 | 0.03116 | 0.00413 | 0.00438 | 0.02039 | 0.00465 | 0.44526 | 0.00446 | 0.00731 | 0.00000 |
| **Huwe1** | 22.36570 | 0.40998 | 28.38869 | 15.74440 | 17.19712 | 19.00376 | 4.81025 | 4.51058 | 29.49774 | 0.28622 | 11.86818 | 7.56272 | 6.33395 | 15.94644 |
| **Hvcn1** | 0.56050 | 0.00364 | 1.71410 | 0.28603 | 3.76964 | 0.49161 | 1.23380 | 1.11320 | 0.00710 | 0.02872 | 0.17840 | 0.31462 | 1.94041 | 0.60683 |
| **Hyal1** | 0.01787 | 0.00349 | 0.01616 | 1.40293 | 0.45753 | 0.07318 | 0.00372 | 0.00287 | 0.05204 | 0.00476 | 0.16297 | 0.72658 | 0.11580 | 0.18996 |
| **Hyal2** | 0.00000 | 0.01131 | 0.84152 | 1.49906 | 0.00000 | 1.00421 | 0.00411 | 0.67880 | 0.00702 | 0.01182 | 0.10061 | 1.57202 | 0.04553 | 0.03134 |
| **Hyal3** | 0.18477 | 0.05391 | 0.32070 | 0.60424 | 0.50140 | 0.55328 | 0.13311 | 1.14606 | 0.86131 | 0.31663 | 0.03105 | 0.08957 | 0.94316 | 0.97375 |
| **Hydin** | 0.00000 | 0.00420 | 0.14812 | 0.41650 | 0.18016 | 0.17074 | 0.00400 | 0.00325 | 0.00906 | 0.00500 | 0.00499 | 0.01402 | 0.15766 | 0.00552 |
| **Hyi** | 0.00509 | 0.01528 | 0.00116 | 0.57037 | 0.01844 | 0.11186 | 0.01114 | 0.40556 | 0.00914 | 0.00889 | 0.00362 | 0.01138 | 0.03163 | 0.05557 |
| **Hykk** | 0.34160 | 0.04050 | 0.06444 | 0.00000 | 0.12755 | 1.36877 | 0.00371 | 0.29380 | 0.18082 | 0.08489 | 0.05302 | 0.81490 | 0.17656 | 0.12129 |
| **Hyls1** | 0.00000 | 0.00395 | 0.08128 | 0.19656 | 0.01403 | 0.45056 | 0.00467 | 0.00846 | 0.07713 | 0.00651 | 0.05702 | 0.34370 | 0.47387 | 0.73509 |
| **Hyou1** | 0.00000 | 0.83194 | 3.26673 | 4.00357 | 1.12035 | 2.33125 | 1.49444 | 2.17872 | 1.17648 | 0.04909 | 1.96288 | 0.55544 | 1.07519 | 4.17100 |

| | | | | | | | | | | | | | | |
|---|---|---|---|---|---|---|---|---|---|---|---|---|---|---|
| **Hypk** | 5.93464 | 3.21167 | 0.00000 | 5.31325 | 13.17823 | 11.17661 | 1.42860 | 11.33441 | 3.72141 | 0.40285 | 0.47073 | 9.74763 | 13.81066 | 7.54981 |
| **Iah1** | 1.19176 | 0.04012 | 0.41272 | 1.30758 | 0.53958 | 1.12242 | 0.00853 | 0.40370 | 0.01302 | 0.03907 | 2.50126 | 2.85404 | 0.01418 | 2.17430 |
| **Iars** | 13.12480 | 7.85199 | 9.40839 | 4.83535 | 11.25072 | 14.73634 | 2.93205 | 9.49150 | 8.01748 | 0.08591 | 15.76178 | 10.26197 | 10.67100 | 6.33939 |
| **Iars2** | 2.02511 | 0.02973 | 1.41492 | 0.60838 | 0.19006 | 1.25432 | 0.01314 | 1.66000 | 1.67838 | 0.02052 | 5.42857 | 4.37038 | 0.00587 | 0.67551 |
| **Iba57** | 0.04260 | 0.03809 | 0.26535 | 0.29018 | 0.04619 | 0.10218 | 0.01004 | 0.00693 | 0.03569 | 0.00641 | 0.00613 | 0.02910 | 0.03969 | 0.43761 |
| **Ibtk** | 3.12875 | 0.09319 | 11.21488 | 2.41614 | 8.83869 | 5.02205 | 5.22565 | 2.90836 | 9.81554 | 0.12467 | 7.76318 | 6.09345 | 1.83051 | 11.44317 |
| **Ica1** | 0.31909 | 0.00368 | 0.10172 | 0.00026 | 0.31324 | 0.01229 | 0.00386 | 0.00607 | 0.00055 | 0.00752 | 0.00554 | 0.00538 | 3.77542 | 2.75388 |
| **Ica1l** | 1.02404 | 0.00380 | 0.01076 | 0.31694 | 1.12666 | 0.00000 | 0.00367 | 0.00470 | 1.07705 | 0.00560 | 0.01018 | 0.01048 | 1.03365 | 0.63563 |
| **Icam1** | 5.48364 | 0.00887 | 5.69833 | 1.09930 | 1.48858 | 0.66277 | 0.00777 | 0.00000 | 1.18143 | 0.00560 | 0.00613 | 0.83277 | 8.23814 | 1.24978 |
| **Icam2** | 0.00371 | 0.00382 | 0.00000 | 1.47193 | 0.24015 | 0.00000 | 0.00360 | 0.00269 | 0.00769 | 0.00604 | 0.34782 | 0.15761 | 0.04202 | 0.02244 |
| **Icam4** | 0.12350 | 0.00419 | 4.27776 | 3.42174 | 0.60103 | 1.47956 | 0.01888 | 1.06750 | 0.00672 | 0.01384 | 1.97196 | 0.00000 | 0.00000 | 0.00000 |
| **Ice1** | 4.70079 | 0.09900 | 6.81295 | 2.20447 | 7.20079 | 4.55987 | 1.69282 | 2.24351 | 6.95528 | 0.03411 | 2.12328 | 5.64327 | 11.17076 | 9.58129 |
| **Ice2** | 1.01365 | 0.10248 | 1.23004 | 0.33950 | 1.96304 | 0.37438 | 0.00911 | 0.69967 | 0.00594 | 0.01006 | 0.95104 | 2.08067 | 1.73228 | 1.80693 |
| **Ick** | 0.12908 | 0.00324 | 0.05844 | 0.08056 | 0.89875 | 0.76038 | 0.01679 | 0.10598 | 0.00699 | 0.01209 | 0.00000 | 1.39464 | 0.10472 | 0.61982 |
| **Icmt** | 1.09313 | 0.01688 | 3.55183 | 1.17384 | 3.15500 | 4.54292 | 0.97189 | 1.61038 | 0.02704 | 0.03493 | 2.90724 | 2.06133 | 1.77640 | 4.83815 |
| **Icosl** | 0.39985 | 2.66333 | 0.13823 | 0.16597 | 0.23991 | 1.00569 | 0.66458 | 0.01058 | 0.13556 | 0.01836 | 0.01557 | 0.94539 | 0.41296 | 0.01110 |
| **Ict1** | 4.56802 | 0.24668 | 7.29949 | 16.07827 | 13.04523 | 4.11767 | 4.94712 | 5.55302 | 0.28713 | 2.66639 | 0.43800 | 8.49786 | 1.37541 | 0.53985 |
| **Id1** | 0.00000 | 3.61177 | 3.58215 | 0.00000 | 4.60608 | 0.06216 | 0.00145 | 0.00000 | 0.00179 | 0.01648 | 0.03050 | 3.83445 | 1.84140 | 0.00100 |
| **Id2** | 0.00265 | 4.54975 | 0.00000 | 0.00000 | 0.00377 | 3.15541 | 2.12758 | 5.60235 | 0.01060 | 0.93648 | 0.83402 | 0.02002 | 0.84667 | 0.02472 |
| **Id3** | 0.45108 | 0.07882 | 26.10718 | 5.90976 | 10.99063 | 0.12327 | 2.57354 | 0.87246 | 0.00340 | 0.15294 | 1.33474 | 3.51581 | 0.07715 | 0.63620 |
| **Id4** | 0.00572 | 0.00426 | 0.01151 | 0.01435 | 0.08422 | 0.01303 | 0.00460 | 0.00884 | 0.01318 | 0.00730 | 0.00889 | 0.00831 | 0.69715 | 0.00151 |
| **Ide** | 4.15557 | 6.91292 | 10.65709 | 3.57402 | 3.37551 | 11.27579 | 1.33858 | 9.37127 | 8.61705 | 0.14637 | 17.09716 | 13.87319 | 1.09124 | 4.66656 |
| **Idh1** | 3.14048 | 0.12157 | 9.27978 | 7.59009 | 3.41719 | 7.74041 | 2.77072 | 7.18813 | 14.75172 | 0.05446 | 9.38888 | 11.98924 | 0.00000 | 6.48210 |
| **Idh2** | 2.43815 | 5.44812 | 12.73116 | 22.93892 | 8.25380 | 9.95921 | 3.85558 | 1.14330 | 1.08500 | 0.22362 | 1.56610 | 4.07874 | 0.34466 | 3.78110 |
| **Idh3a** | 4.00949 | 7.56912 | 10.21357 | 17.15976 | 4.80515 | 31.24274 | 10.34919 | 18.34742 | 3.51028 | 0.78607 | 20.98409 | 7.91213 | 0.83589 | 2.05255 |
| **Idh3b** | 10.18984 | 6.85849 | 23.67508 | 11.38003 | 14.65501 | 16.72560 | 7.66526 | 21.18230 | 3.38926 | 11.47889 | 16.79727 | 11.25555 | 4.65555 | 8.51710 |
| **Idh3g** | 20.00134 | 9.22714 | 31.80435 | 20.74420 | 18.94952 | 19.45983 | 5.48467 | 20.85990 | 15.18252 | 1.56081 | 14.40492 | 17.40897 | 3.13297 | 16.45334 |
| **Idi1** | 1.54694 | 0.01773 | 2.48531 | 1.45797 | 0.00000 | 2.88191 | 0.00199 | 0.00744 | 6.77897 | 0.01243 | 2.30875 | 1.44460 | 0.22909 | 0.20293 |
| **Idi2** | 0.03836 | 0.00337 | 2.35714 | 0.65717 | 1.36627 | 0.61317 | 0.00308 | 0.00000 | 0.90986 | 2.31001 | 0.00000 | 0.00000 | 0.06406 | 2.77816 |
| **Idnk** | 0.90655 | 6.49823 | 0.00000 | 2.65194 | 1.74368 | 0.64030 | 0.01352 | 0.63695 | 0.17344 | 0.02366 | 0.01432 | 0.30719 | 0.08035 | 2.00709 |
| **Ids** | 1.64470 | 0.02275 | 5.97396 | 1.84262 | 3.68145 | 2.15556 | 2.44568 | 0.00000 | 2.72210 | 0.05081 | 1.59681 | 3.47048 | 2.78425 | 1.81938 |
| **Idua** | 5.10411 | 0.00287 | 3.12126 | 0.25732 | 0.26373 | 0.00000 | 1.58237 | 0.04811 | 0.27224 | 0.01620 | 0.00000 | 0.47429 | 0.94429 | 4.08767 |
| **Ier2** | 0.00296 | 0.53645 | 0.08777 | 4.60110 | 0.62486 | 0.61155 | 1.39209 | 4.08984 | 0.00531 | 2.41539 | 0.08196 | 0.88170 | 0.06007 | 0.00000 |
| **Ier3** | 0.00856 | 0.00408 | 0.00933 | 2.44071 | 0.36531 | 0.88606 | 0.00466 | 0.12698 | 0.00588 | 0.01240 | 0.00000 | 1.01717 | 0.01751 | 0.03118 |
| **Ier3ip1** | 2.14455 | 0.22827 | 0.64461 | 1.73790 | 3.75485 | 8.56756 | 0.03147 | 6.29956 | 0.06443 | 3.23886 | 2.79724 | 7.47747 | 3.12044 | 0.22572 |
| **Ier5** | 0.00092 | 0.00395 | 0.13353 | 0.00000 | 0.18498 | 0.00819 | 0.00420 | 0.00874 | 0.00965 | 0.00635 | 0.00731 | 0.03185 | 3.23194 | 0.00376 |
| **Ier5l** | 0.01336 | 0.00472 | 0.02099 | 0.43395 | 0.16107 | 0.00335 | 0.00443 | 0.01602 | 0.01101 | 0.00633 | 0.01443 | 0.01886 | 0.03043 | 0.00211 |
| **Iffo1** | 0.03521 | 0.03610 | 0.04763 | 0.00000 | 0.01071 | 0.03831 | 0.01455 | 0.01965 | 0.01003 | 0.01763 | 0.00279 | 1.46230 | 0.50761 | 0.00782 |
| **Iffo2** | 0.30644 | 3.00906 | 0.68213 | 0.44220 | 1.80247 | 2.44057 | 0.82404 | 9.20691 | 3.90380 | 0.08103 | 0.05664 | 0.03299 | 6.80160 | 8.37717 |
| **Ifi203** | 1.05957 | 0.02802 | 0.26616 | 0.01412 | 0.77812 | 0.07149 | 0.05828 | 0.00785 | 0.92883 | 0.00652 | 0.03218 | 0.01064 | 0.77617 | 0.53829 |
| **Ifi27** | 0.34661 | 0.00387 | 0.26053 | 0.00000 | 1.63629 | 0.00000 | 0.00392 | 0.00658 | 0.52223 | 0.00600 | 0.00377 | 0.00287 | 2.81168 | 2.48517 |
| **Ifi30** | 3.04119 | 0.24714 | 7.25448 | 13.14460 | 1.33360 | 10.91987 | 3.63038 | 14.59255 | 0.47300 | 0.31470 | 6.66604 | 4.31651 | 1.53142 | 1.71989 |
| **Ifi35** | 3.39564 | 0.00953 | 7.32484 | 1.09235 | 12.39269 | 0.77609 | 0.00725 | 1.82217 | 2.79914 | 0.00000 | 0.06060 | 0.00000 | 5.05934 | 8.31944 |
| **Ifit1bl2** | 0.44161 | 0.01262 | 0.11806 | 0.17045 | 0.10838 | 0.27490 | 0.40897 | 0.05898 | 0.22204 | 0.03754 | 0.01359 | 0.04652 | 0.15458 | 0.37594 |
| **Ifitm1** | 0.33371 | 0.00424 | 0.06891 | 0.00963 | 0.08308 | 0.00546 | 0.00441 | 0.00959 | 0.02267 | 0.00646 | 0.00733 | 0.02064 | 0.27982 | 0.62673 |
| **Ifitm10** | 0.13480 | 0.00403 | 0.06335 | 0.00468 | 0.45994 | 0.05317 | 0.00424 | 0.00547 | 0.32785 | 0.00599 | 0.00339 | 0.18397 | 0.38346 | 0.00000 |
| **Ifitm2** | 0.24982 | 0.02808 | 0.03335 | 2.73222 | 3.43021 | 0.21873 | 0.00371 | 0.74246 | 0.13102 | 0.01392 | 0.32203 | 0.78387 | 0.00620 | 0.77734 |
| **Ifitm3** | 0.05329 | 0.00427 | 0.00789 | 0.00000 | 2.01979 | 0.01108 | 0.00467 | 0.01735 | 0.03154 | 0.00742 | 0.00421 | 0.61365 | 0.00548 | 0.31611 |
| **Ifitm6** | 0.01325 | 0.00442 | 0.01308 | 0.01562 | 0.01487 | 0.01220 | 0.00439 | 0.01014 | 0.01266 | 0.00754 | 0.00866 | 0.00962 | 0.03102 | 0.06935 |
| **Ifitm7** | 0.02607 | 0.00444 | 0.02273 | 0.03643 | 0.01187 | 0.02309 | 0.00438 | 0.00753 | 0.01427 | 0.00674 | 0.00722 | 0.01914 | 0.08329 | 0.01611 |
| **Ifnar1** | 4.25612 | 0.75989 | 0.72989 | 5.46806 | 3.90331 | 8.15522 | 1.69839 | 2.11060 | 2.91405 | 2.07874 | 2.69531 | 4.10598 | 6.54394 | 5.09779 |
| **Ifnar2** | 0.02004 | 0.00383 | 0.15680 | 0.00000 | 0.02898 | 0.00000 | 0.00364 | 0.69619 | 0.01097 | 0.00488 | 0.00526 | 0.01405 | 0.00638 | 0.02300 |
| **Ifngr1** | 0.00685 | 0.01120 | 0.00000 | 0.00000 | 2.20642 | 0.03607 | 1.08137 | 1.18396 | 0.15370 | 0.01509 | 0.00000 | 0.43097 | 0.00844 | 1.14132 |
| **Ifngr2** | 0.20144 | 0.02089 | 0.02166 | 0.05140 | 0.04414 | 0.22184 | 0.00416 | 1.12547 | 0.17359 | 0.00742 | 0.00180 | 0.10548 | 0.04446 | 0.68297 |
| **Ifnlr1** | 0.00823 | 0.00444 | 0.01341 | 0.01356 | 0.00922 | 0.01257 | 0.00436 | 0.01021 | 0.01020 | 0.00695 | 0.00872 | 0.00995 | 0.00818 | 0.11433 |

| | | | | | | | | | | | | | | |
|---|---|---|---|---|---|---|---|---|---|---|---|---|---|---|
| **Ifrd1** | 21.98459 | 0.06595 | 8.24283 | 4.65600 | 15.27769 | 7.46994 | 0.18838 | 3.99401 | 8.46770 | 0.51956 | 4.03120 | 4.86971 | 15.76946 | 9.50391 |
| **Ifrd2** | 0.79365 | 0.14493 | 2.63350 | 1.83150 | 1.29691 | 1.55479 | 0.00894 | 1.09066 | 0.04641 | 0.27298 | 0.84135 | 1.10564 | 0.43714 | 0.89300 |
| **Ift122** | 0.42217 | 0.00431 | 0.15386 | 1.07974 | 1.02317 | 1.65685 | 0.00369 | 0.01029 | 0.02148 | 0.00834 | 1.37085 | 0.10978 | 0.24690 | 1.30071 |
| **Ift140** | 0.05469 | 4.35739 | 0.05882 | 0.00000 | 0.57616 | 2.76567 | 3.09478 | 3.88214 | 0.00000 | 0.13897 | 0.00000 | 2.46427 | 0.20408 | 1.61098 |
| **Ift172** | 0.00000 | 0.02196 | 1.45388 | 1.99315 | 0.93964 | 2.24819 | 3.38719 | 0.75819 | 0.04425 | 0.09064 | 0.10976 | 1.38392 | 0.39360 | 0.63866 |
| **Ift20** | 5.00617 | 9.92488 | 5.08718 | 3.95760 | 6.76009 | 5.88177 | 4.98547 | 11.71843 | 0.07700 | 0.27118 | 3.09186 | 6.07724 | 8.33716 | 2.84377 |
| **Ift22** | 0.01340 | 0.00459 | 0.00000 | 0.35917 | 0.19144 | 1.65685 | 0.00402 | 0.08647 | 0.20390 | 0.05896 | 1.97707 | 0.42010 | 0.00801 | 0.00000 |
| **Ift27** | 0.00000 | 0.01018 | 0.61464 | 0.68601 | 0.54627 | 0.05928 | 0.00408 | 0.44040 | 0.00806 | 0.00484 | 0.05383 | 1.00955 | 0.00108 | 0.06190 |
| **Ift43** | 0.45905 | 7.33259 | 0.76862 | 0.68891 | 1.48182 | 1.34909 | 1.65954 | 2.06332 | 0.78288 | 0.04206 | 0.50037 | 1.94478 | 2.70356 | 0.81308 |
| **Ift46** | 2.78215 | 2.42409 | 7.00822 | 3.20300 | 5.15549 | 5.15905 | 3.03841 | 9.32689 | 3.87563 | 3.10545 | 7.68813 | 6.59231 | 1.16200 | 7.57096 |
| **Ift52** | 0.61038 | 0.14732 | 0.37384 | 0.00000 | 1.35088 | 3.77982 | 0.00230 | 0.64647 | 0.00000 | 0.01510 | 5.70197 | 4.96519 | 2.66115 | 2.22857 |
| **Ift74** | 0.44982 | 0.00508 | 0.27646 | 0.00000 | 0.46247 | 0.43629 | 0.00431 | 0.17579 | 0.00581 | 0.00508 | 0.59685 | 0.00338 | 0.61791 | 2.74039 |
| **Ift80** | 0.78521 | 0.00466 | 0.00000 | 0.47951 | 0.29535 | 0.11172 | 0.00379 | 0.00946 | 3.59734 | 0.00833 | 0.00929 | 0.10966 | 0.06751 | 0.00000 |
| **Ift81** | 0.11892 | 0.00367 | 0.03791 | 0.00284 | 0.86947 | 0.56417 | 2.23832 | 0.00709 | 0.01164 | 0.06330 | 0.00217 | 0.00000 | 1.57792 | 3.30804 |
| **Ift88** | 1.80504 | 0.00298 | 0.24651 | 0.57797 | 0.15762 | 0.84372 | 0.00328 | 0.01158 | 2.54535 | 0.00095 | 1.65591 | 0.57777 | 0.73214 | 1.59942 |
| **Igbp1** | 11.63882 | 1.35995 | 18.86363 | 7.49328 | 11.52654 | 13.23505 | 3.22854 | 12.84028 | 14.31358 | 0.13281 | 14.29671 | 13.50319 | 0.00000 | 6.22248 |
| **Igdcc3** | 0.48203 | 0.01316 | 0.00000 | 0.12672 | 0.44277 | 0.00000 | 1.47811 | 0.92751 | 0.00490 | 0.08959 | 0.00342 | 0.02490 | 0.56698 | 0.24450 |
| **Igf1** | 0.01468 | 0.00446 | 0.00962 | 0.01097 | 0.15880 | 0.01344 | 0.00494 | 0.01022 | 0.02028 | 0.00691 | 0.00878 | 0.00917 | 0.00485 | 0.01039 |
| **Igf1r** | 0.33822 | 0.00400 | 0.00000 | 0.01063 | 0.09530 | 0.00000 | 0.00425 | 0.01095 | 0.06583 | 0.00341 | 0.74896 | 0.00562 | 0.20805 | 0.48550 |
| **Igf2** | 0.00383 | 0.02084 | 0.16938 | 0.00000 | 0.14837 | 0.37153 | 0.00378 | 0.86897 | 0.01454 | 0.01506 | 0.01547 | 0.21825 | 0.01045 | 0.00694 |
| **Igf2bp1** | 1.26330 | 0.16568 | 14.59746 | 12.77503 | 6.37391 | 13.34028 | 0.03044 | 5.21351 | 0.73038 | 7.43920 | 7.24486 | 10.34137 | 6.26982 | 3.82339 |
| **Igf2bp2** | 4.94542 | 4.75515 | 16.71951 | 8.22101 | 19.27761 | 12.63892 | 2.05200 | 9.25069 | 7.45937 | 12.84322 | 12.04467 | 12.18915 | 0.09528 | 14.72099 |
| **Igf2bp3** | 0.57376 | 5.36184 | 1.08936 | 8.92997 | 1.87494 | 14.27029 | 3.09938 | 9.03963 | 4.49706 | 2.51123 | 6.47598 | 5.99312 | 6.43241 | 4.73805 |
| **Igf2r** | 3.44205 | 4.74291 | 5.49493 | 1.34398 | 3.16107 | 2.44563 | 0.25304 | 0.57763 | 7.76322 | 0.27276 | 4.78175 | 1.50627 | 7.51872 | 17.55112 |
| **Igfbp2** | 1.05180 | 0.00439 | 0.05006 | 0.00564 | 0.11480 | 0.01705 | 0.00417 | 0.00903 | 0.01168 | 0.01383 | 0.00638 | 0.00571 | 0.47344 | 0.32865 |
| **Igfbp3** | 0.00343 | 0.00430 | 0.00845 | 0.01398 | 0.00000 | 0.01056 | 0.00424 | 0.00880 | 0.02183 | 0.00695 | 0.00722 | 0.00861 | 0.25137 | 0.94905 |
| **Igfbp4** | 0.02551 | 0.00447 | 0.00000 | 0.00827 | 1.11708 | 0.01290 | 0.00471 | 0.01042 | 0.00959 | 0.00630 | 0.00901 | 0.00749 | 0.00512 | 0.16135 |
| **Igfbp5** | 0.22344 | 0.00443 | 0.00919 | 0.00983 | 0.14117 | 0.01095 | 0.00448 | 0.01219 | 0.00966 | 0.00645 | 0.00736 | 0.00837 | 0.00470 | 0.01240 |
| **Igfbp6** | 0.60190 | 0.00418 | 0.02766 | 0.16931 | 0.34674 | 0.00541 | 0.00432 | 0.00666 | 0.01066 | 0.00558 | 0.01518 | 0.01309 | 0.01664 | 0.08242 |
| **Igfbp7** | 0.20613 | 0.00442 | 0.00731 | 0.00973 | 0.18861 | 0.01057 | 0.00433 | 0.01206 | 0.03014 | 0.00649 | 0.00715 | 0.00809 | 0.00360 | 0.02320 |
| **Igflr1** | 0.04162 | 0.02047 | 0.15661 | 0.36900 | 0.00000 | 0.20396 | 0.06408 | 1.71249 | 0.00650 | 0.02248 | 2.97354 | 0.21469 | 0.00464 | 0.16776 |
| **Ighmbp2** | 0.32510 | 1.17425 | 4.00478 | 4.75248 | 2.34846 | 0.23501 | 0.00762 | 0.00000 | 0.00000 | 0.03469 | 0.52342 | 3.37633 | 1.70094 | 0.14639 |
| **Igip** | 0.73269 | 0.00433 | 0.12175 | 0.00000 | 0.75233 | 0.00000 | 0.00403 | 0.01001 | 2.20257 | 0.00597 | 0.14748 | 0.00548 | 0.00000 | 0.00000 |
| **Igsf1** | 0.00673 | 0.00431 | 0.07619 | 0.17520 | 0.00738 | 0.00897 | 0.00428 | 0.00749 | 0.01272 | 0.00593 | 0.00717 | 0.00784 | 0.00670 | 0.01004 |
| **Igsf23** | 0.00680 | 0.00462 | 0.01696 | 0.14401 | 0.00887 | 0.01652 | 0.00472 | 0.05045 | 0.01363 | 0.01891 | 0.01052 | 0.00636 | 0.00676 | 0.02435 |
| **Igsf3** | 0.00000 | 0.01184 | 1.68039 | 0.62504 | 0.71417 | 3.54838 | 1.54070 | 1.88868 | 2.37362 | 0.00196 | 0.03121 | 0.28236 | 0.91117 | 2.42040 |
| **Igsf5** | 0.00664 | 0.01133 | 0.00776 | 0.00000 | 0.00509 | 0.00264 | 0.00406 | 1.07352 | 0.14013 | 0.00578 | 0.00139 | 0.20746 | 0.00382 | 0.04382 |
| **Igsf8** | 0.01380 | 0.00484 | 0.00458 | 0.70555 | 0.07982 | 0.06362 | 0.00478 | 0.05436 | 0.00546 | 0.00734 | 0.00435 | 0.79384 | 0.21350 | 0.18072 |
| **Igsf9** | 0.92403 | 0.15663 | 0.51792 | 2.34541 | 0.87134 | 0.36449 | 3.09381 | 1.96324 | 0.07381 | 1.21368 | 0.00000 | 0.58302 | 0.01168 | 0.00005 |
| **Igsf9b** | 0.00768 | 0.00432 | 0.00000 | 1.06423 | 0.09975 | 0.08605 | 0.68508 | 0.00467 | 0.01179 | 0.02345 | 0.00170 | 0.31465 | 0.00275 | 0.14805 |
| **Ihh** | 0.01009 | 0.00440 | 0.11767 | 0.00182 | 0.07469 | 0.01391 | 0.00432 | 0.01134 | 0.00926 | 0.00618 | 0.00755 | 0.00982 | 0.02143 | 0.12910 |
| **Ik** | 5.76787 | 7.31654 | 9.65678 | 7.89545 | 6.08052 | 12.51784 | 3.20237 | 5.43589 | 14.08031 | 5.23572 | 10.80213 | 8.92088 | 9.79331 | 8.44715 |
| **Ikbip** | 1.65175 | 0.29219 | 0.42119 | 0.68863 | 0.98882 | 1.79206 | 2.72645 | 3.30630 | 2.90073 | 0.06592 | 0.01054 | 0.75372 | 0.95482 | 1.66684 |
| **Ikbkap** | 6.27745 | 7.85060 | 8.85221 | 4.23882 | 6.76039 | 6.99102 | 0.00787 | 4.31263 | 2.53067 | 0.01102 | 7.87707 | 8.16235 | 0.82274 | 6.75127 |
| **Ikbkb** | 0.18927 | 0.01902 | 0.00000 | 0.14654 | 0.18408 | 0.00000 | 0.00366 | 2.91380 | 0.01244 | 0.00554 | 0.05167 | 0.84055 | 0.00325 | 0.15219 |
| **Ikbkg** | 5.18814 | 0.06607 | 2.79871 | 0.57967 | 5.03470 | 4.02906 | 0.34188 | 0.56655 | 10.46927 | 0.98974 | 3.92292 | 1.38536 | 8.69412 | 6.56802 |
| **Ikzf2** | 0.01216 | 0.00504 | 0.00943 | 0.00877 | 0.25012 | 0.01095 | 0.00439 | 0.02946 | 0.01445 | 0.01533 | 0.00913 | 0.00855 | 0.00537 | 0.00746 |
| **Ikzf4** | 0.00366 | 0.00426 | 0.00930 | 0.06603 | 0.09469 | 0.14316 | 0.00440 | 0.00518 | 0.00810 | 0.00550 | 0.00609 | 0.00607 | 0.31916 | 0.00831 |
| **Ikzf5** | 1.44340 | 0.01093 | 0.56641 | 0.76335 | 0.76850 | 0.64179 | 0.07072 | 0.00243 | 2.69987 | 0.03527 | 0.68711 | 1.17186 | 0.64079 | 4.62427 |
| **Il10rb** | 0.01978 | 0.00421 | 0.00000 | 0.08454 | 0.17641 | 0.52970 | 0.00409 | 0.00388 | 0.00896 | 0.00636 | 0.03924 | 0.02013 | 0.13205 | 0.00002 |
| **Il11** | 0.17987 | 0.05161 | 0.00000 | 0.43349 | 0.29328 | 0.53491 | 0.07182 | 0.26745 | 0.34300 | 0.23450 | 0.02511 | 0.07333 | 1.99447 | 0.68621 |
| **Il11ra1** | 0.00225 | 0.01172 | 0.09431 | 0.00000 | 0.13827 | 0.26027 | 2.22439 | 0.50705 | 0.01215 | 0.09668 | 0.00407 | 0.01225 | 0.23781 | 0.05160 |
| **Il12rb1** | 0.00559 | 0.00415 | 0.01828 | 0.27695 | 0.15304 | 0.00655 | 0.00389 | 0.00950 | 0.02399 | 0.00838 | 0.00659 | 0.01154 | 0.07869 | 0.00000 |
| **Il12rb2** | 0.00208 | 0.00456 | 0.15207 | 0.00023 | 0.22295 | 0.09528 | 0.00424 | 0.00731 | 0.85656 | 0.00642 | 0.00553 | 0.02331 | 0.00133 | 0.08018 |
| **Il13ra1** | 0.33152 | 0.00418 | 0.00154 | 0.16060 | 0.16190 | 0.00484 | 0.00455 | 0.00701 | 0.02596 | 0.00657 | 0.00621 | 0.01391 | 0.00416 | 0.22274 |

| | | | | | | | | | | | | | | |
|---|---|---|---|---|---|---|---|---|---|---|---|---|---|---|
| **Il17b** | 0.00730 | 0.00548 | 0.00000 | 0.14442 | 0.96490 | 0.02485 | 0.45648 | 0.48459 | 0.01211 | 0.01006 | 0.01934 | 0.00703 | 0.00415 | 0.00580 |
| **Il17d** | 0.00855 | 0.00441 | 0.01586 | 0.02489 | 0.01120 | 0.02720 | 0.03435 | 0.00671 | 0.02822 | 0.00605 | 0.01449 | 0.00851 | 0.04430 | 0.00790 |
| **Il17ra** | 0.23555 | 2.03698 | 0.75678 | 0.77517 | 0.58592 | 1.54259 | 0.00296 | 0.22876 | 0.50079 | 0.00530 | 0.48873 | 1.34605 | 6.75609 | 1.96058 |
| **Il17rb** | 0.00134 | 0.00386 | 0.03209 | 0.00000 | 0.25966 | 0.00000 | 0.00426 | 0.42328 | 0.03419 | 1.46385 | 0.00109 | 0.02088 | 0.27492 | 0.00000 |
| **Il17rc** | 0.02241 | 0.00408 | 0.00776 | 0.43692 | 0.01219 | 0.02264 | 0.00412 | 0.00832 | 0.01204 | 0.00641 | 0.00521 | 0.02084 | 0.00722 | 0.00762 |
| **Il17rd** | 0.86927 | 0.00243 | 0.14952 | 1.65250 | 1.17842 | 0.45568 | 0.00909 | 0.01195 | 4.29253 | 0.08589 | 0.35195 | 1.93344 | 2.76794 | 1.78665 |
| **Il17re** | 0.00338 | 0.00418 | 0.00000 | 0.98079 | 0.06171 | 0.18051 | 0.00455 | 0.10955 | 0.00957 | 0.01465 | 0.00266 | 0.00344 | 0.05647 | 0.00000 |
| **Il18** | 0.01630 | 0.00428 | 0.32345 | 0.00000 | 0.06694 | 0.04675 | 0.00405 | 0.00544 | 0.00670 | 0.00624 | 0.00631 | 0.00866 | 0.34060 | 0.45140 |
| **Il18bp** | 0.06832 | 0.02780 | 0.00844 | 0.48747 | 0.14233 | 0.24831 | 0.00462 | 0.73856 | 0.00463 | 0.02278 | 0.02484 | 0.67317 | 0.00359 | 0.92023 |
| **Il1bos** | 0.00377 | 0.00443 | 0.04552 | 0.03443 | 0.04664 | 0.01505 | 0.00429 | 0.00876 | 0.15116 | 0.00698 | 0.00706 | 0.00947 | 0.00892 | 0.26141 |
| **Il1f9** | 0.56145 | 0.00381 | 0.54694 | 0.00000 | 0.13844 | 1.41470 | 0.20714 | 0.00159 | 1.15129 | 0.04404 | 0.00611 | 0.03469 | 3.87418 | 0.52214 |
| **Il1r1** | 0.01652 | 0.00494 | 0.00000 | 0.01116 | 0.19486 | 0.00622 | 0.00406 | 0.02423 | 1.66299 | 0.00781 | 0.00536 | 0.00804 | 0.00121 | 0.00000 |
| **Il1rap** | 0.06468 | 0.00421 | 0.16092 | 0.08243 | 0.01691 | 0.00366 | 0.00399 | 0.00841 | 0.00743 | 0.01478 | 0.00733 | 0.00884 | 0.21703 | 0.02954 |
| **Il1rapl2** | 0.02092 | 0.02693 | 0.01792 | 0.05252 | 0.00974 | 0.02517 | 0.01620 | 0.05331 | 0.22219 | 0.00660 | 0.01662 | 0.02852 | 0.01913 | 0.10721 |
| **Il1rl1** | 0.09681 | 0.04416 | 0.00000 | 0.35507 | 0.00574 | 0.24281 | 0.01127 | 0.05698 | 0.04759 | 0.11175 | 0.01147 | 0.10410 | 0.23374 | 0.17529 |
| **Il20rb** | 0.00692 | 0.00435 | 0.08133 | 0.00931 | 0.36032 | 0.01490 | 0.00410 | 0.00795 | 0.14820 | 0.01099 | 0.35525 | 0.00549 | 0.14171 | 0.00000 |
| **Il23a** | 0.00437 | 0.00425 | 0.46362 | 0.16913 | 0.00219 | 0.00634 | 0.00405 | 0.00622 | 0.01430 | 0.00683 | 0.00548 | 0.01183 | 0.00657 | 0.01795 |
| **Il2ra** | 0.00446 | 0.00477 | 0.01677 | 0.01478 | 0.01677 | 0.01664 | 0.00425 | 0.01657 | 0.20131 | 0.00731 | 0.00922 | 0.03063 | 0.07718 | 0.11316 |
| **Il2rg** | 0.00000 | 0.01518 | 0.28317 | 1.51057 | 0.20254 | 0.32302 | 0.00349 | 0.99995 | 1.90780 | 0.00856 | 0.00000 | 0.44782 | 0.62284 | 1.70671 |
| **Il31** | 0.00513 | 0.00436 | 0.01395 | 0.01240 | 0.00915 | 0.01302 | 0.00426 | 0.00938 | 0.00586 | 0.00637 | 0.00913 | 0.00940 | 0.25619 | 0.08575 |
| **Il33** | 0.68982 | 0.01071 | 0.77113 | 0.00000 | 0.00000 | 0.54998 | 0.00325 | 1.79002 | 0.63871 | 0.01433 | 0.00342 | 0.00070 | 0.11480 | 0.00000 |
| **Il34** | 0.00000 | 0.00369 | 0.45565 | 0.56327 | 0.45599 | 1.86860 | 0.00911 | 0.00210 | 0.10358 | 0.00724 | 0.00000 | 0.61135 | 0.00127 | 0.69808 |
| **Il3ra** | 0.03421 | 0.21455 | 0.29674 | 0.53484 | 0.05135 | 0.00000 | 0.00366 | 0.06239 | 0.00898 | 0.00978 | 0.00469 | 0.00614 | 0.02084 | 0.00203 |
| **Il4** | 0.01419 | 0.00444 | 0.01444 | 0.01277 | 0.01402 | 0.01301 | 0.00427 | 0.01017 | 0.00904 | 0.00714 | 0.00875 | 0.00994 | 0.03912 | 0.02212 |
| **Il4ra** | 0.04785 | 0.00398 | 0.60609 | 1.44084 | 0.18137 | 0.00000 | 0.00409 | 0.00640 | 0.00477 | 0.00468 | 0.00457 | 0.01407 | 0.00280 | 0.10133 |
| **Il5ra** | 0.04495 | 0.02079 | 0.00976 | 0.11543 | 0.02939 | 0.09333 | 0.03984 | 0.02533 | 0.18357 | 0.01770 | 0.02631 | 0.01676 | 0.08576 | 0.08303 |
| **Il6** | 0.00968 | 0.00396 | 0.01305 | 0.00151 | 0.01376 | 0.12941 | 0.00401 | 0.09006 | 0.01293 | 0.00646 | 0.01229 | 0.01023 | 0.00738 | 0.00999 |
| **Il6ra** | 0.10442 | 0.00836 | 1.76049 | 0.00000 | 0.87834 | 1.05443 | 0.00245 | 0.00000 | 0.17010 | 0.00230 | 0.00005 | 4.39480 | 0.58624 | 4.57099 |
| **Il6st** | 0.00094 | 0.13161 | 0.00791 | 0.00000 | 0.52283 | 0.29362 | 0.00299 | 0.10911 | 0.00534 | 0.01584 | 0.00000 | 5.07685 | 0.16257 | 2.24338 |
| **Il7** | 0.00856 | 0.00408 | 0.00379 | 0.00458 | 0.39122 | 0.01466 | 0.00471 | 0.00662 | 0.01766 | 0.00627 | 0.00458 | 0.41960 | 0.02314 | 0.71286 |
| **Ildr1** | 0.00226 | 0.05490 | 0.00753 | 0.00000 | 0.00904 | 0.03274 | 2.92022 | 3.94833 | 0.01245 | 0.03879 | 0.00000 | 6.68712 | 0.00578 | 0.01043 |
| **Ilf2** | 6.07144 | 0.12024 | 12.90395 | 7.73513 | 11.80422 | 6.87033 | 3.12783 | 5.62715 | 23.58595 | 0.18217 | 13.83059 | 12.86729 | 0.87994 | 24.72074 |
| **Ilf3** | 4.86123 | 0.15415 | 11.16283 | 4.76929 | 10.88628 | 8.35709 | 0.69388 | 4.61857 | 3.86458 | 0.14762 | 5.71958 | 9.06879 | 2.45504 | 17.20244 |
| **Ilk** | 1.90533 | 0.02915 | 3.38158 | 0.08116 | 2.90025 | 3.78729 | 4.03357 | 2.77687 | 1.66160 | 12.18252 | 6.81599 | 2.48336 | 0.62467 | 2.02446 |
| **Ilkap** | 3.89215 | 0.05714 | 1.92880 | 2.39452 | 1.89257 | 2.29558 | 2.52833 | 0.93904 | 0.05008 | 0.04859 | 0.00000 | 3.45272 | 0.40858 | 1.29247 |
| **Ilvbl** | 0.01403 | 0.00423 | 0.26076 | 1.29856 | 0.00444 | 0.29791 | 0.33079 | 0.00840 | 0.00576 | 0.02167 | 0.00336 | 0.01099 | 0.00477 | 0.04632 |
| **Immp1l** | 2.11249 | 0.13450 | 0.17873 | 2.80725 | 1.79842 | 2.41524 | 2.11287 | 5.50467 | 2.38305 | 0.84909 | 1.63454 | 1.90564 | 0.79843 | 0.85768 |
| **Immp2l** | 0.05114 | 0.00437 | 0.01747 | 0.00805 | 0.04919 | 0.01273 | 0.00426 | 0.00963 | 0.00688 | 0.00711 | 0.00800 | 0.02733 | 0.04229 | 0.04728 |
| **Immt** | 8.17360 | 0.28875 | 9.83640 | 7.88818 | 5.69582 | 7.83277 | 4.83944 | 9.31832 | 9.02184 | 7.43808 | 11.68158 | 13.78660 | 2.92492 | 10.12591 |
| **Imp3** | 3.67238 | 0.02566 | 0.00000 | 2.72423 | 14.13129 | 2.98614 | 0.02096 | 0.81824 | 0.50300 | 0.10858 | 0.32579 | 2.91509 | 12.82153 | 0.15029 |
| **Imp4** | 13.90799 | 0.28605 | 26.32069 | 15.51976 | 24.43323 | 19.50041 | 3.19489 | 4.09552 | 4.05586 | 11.91423 | 7.82635 | 14.31698 | 11.28389 | 17.82122 |
| **Impa1** | 0.00000 | 0.01625 | 0.33499 | 2.55029 | 4.24747 | 3.93727 | 0.02058 | 0.00502 | 2.46342 | 0.01497 | 2.33070 | 4.71085 | 1.96913 | 3.26980 |
| **Impa2** | 1.44159 | 0.09118 | 3.29659 | 12.56594 | 4.97451 | 3.72531 | 2.03714 | 3.79102 | 0.00000 | 0.17112 | 2.75455 | 3.31319 | 3.89022 | 1.37895 |
| **Impact** | 1.39232 | 0.02420 | 5.70031 | 8.43582 | 7.00042 | 8.11286 | 0.00000 | 0.18669 | 17.29337 | 0.00000 | 3.18457 | 3.02119 | 0.00000 | 9.42189 |
| **Impad1** | 1.09818 | 0.00385 | 0.31225 | 0.00000 | 0.20429 | 1.76846 | 0.01554 | 0.22341 | 0.96474 | 0.02821 | 0.00000 | 0.45699 | 0.00266 | 0.00000 |
| **Impdh1** | 8.79149 | 0.87362 | 18.66133 | 22.55146 | 16.38317 | 7.35454 | 0.07608 | 2.08965 | 0.00000 | 0.15225 | 7.54921 | 6.77591 | 4.84020 | 2.52156 |
| **Impdh2** | 48.87690 | 11.38266 | 63.55171 | 44.97640 | 56.80062 | 24.63786 | 9.54483 | 19.82049 | 32.88830 | 19.78159 | 26.99900 | 23.72783 | 25.67367 | 44.26765 |
| **Impg2** | 0.02828 | 0.00404 | 0.15647 | 0.43111 | 0.00121 | 0.00000 | 0.00389 | 0.00627 | 0.08936 | 0.00592 | 0.01518 | 0.02013 | 0.11743 | 0.00007 |
| **Ina** | 0.60322 | 0.00408 | 1.10625 | 0.72764 | 0.26938 | 1.17821 | 0.00369 | 0.00000 | 0.02339 | 0.00320 | 0.08511 | 0.01068 | 0.00064 | 3.53824 |
| **Inadl** | 0.58846 | 0.00298 | 1.93089 | 0.14912 | 0.54601 | 0.61459 | 3.37367 | 0.55453 | 0.00081 | 0.10834 | 2.82080 | 0.24956 | 0.18304 | 1.25544 |
| **Inafm1** | 0.27191 | 0.00321 | 0.18605 | 0.49708 | 0.76250 | 0.10559 | 0.00919 | 0.00224 | 0.00396 | 0.00704 | 0.00814 | 0.91152 | 0.29586 | 0.23075 |
| **Incenp** | 4.97350 | 1.72387 | 9.66530 | 9.19177 | 7.10830 | 4.77653 | 2.29496 | 7.76256 | 3.05757 | 13.84027 | 5.13130 | 7.49430 | 14.68629 | 6.82239 |
| **Inf2** | 0.20969 | 0.03867 | 0.01107 | 1.20966 | 0.00214 | 0.00000 | 0.01042 | 1.22652 | 0.01492 | 0.05781 | 0.16257 | 0.05490 | 0.00641 | 0.12306 |
| **Ing1** | 9.56583 | 0.17788 | 10.35044 | 6.52991 | 12.75503 | 8.97391 | 6.33080 | 3.20029 | 9.01036 | 0.25237 | 3.05074 | 5.55440 | 11.59568 | 14.81340 |
| **Ing2** | 4.43638 | 0.07886 | 3.66630 | 2.19365 | 2.21961 | 3.80454 | 2.23040 | 4.14355 | 0.96180 | 0.03101 | 2.12432 | 3.84102 | 3.98252 | 1.56056 |

| | | | | | | | | | | | | | | |
|---|---|---|---|---|---|---|---|---|---|---|---|---|---|---|
| **Ing3** | 0.00000 | 0.00987 | 0.87459 | 0.07121 | 1.91569 | 1.76923 | 1.89516 | 0.44357 | 0.33253 | 0.02702 | 0.32795 | 1.77675 | 0.25923 | 3.44219 |
| **Ing4** | 6.90610 | 5.96167 | 9.41483 | 9.15044 | 13.29901 | 9.81488 | 2.98168 | 3.49035 | 0.19454 | 0.09985 | 8.41022 | 5.49492 | 4.23114 | 10.21968 |
| **Ing5** | 6.75931 | 0.13146 | 4.53578 | 4.05723 | 4.00170 | 5.28785 | 3.45323 | 2.65143 | 3.06869 | 15.18020 | 6.06793 | 4.50841 | 0.00000 | 6.87599 |
| **Inha** | 0.01801 | 0.00450 | 0.00333 | 0.00869 | 1.13447 | 0.01346 | 0.00451 | 0.01042 | 0.01329 | 0.00703 | 0.00848 | 0.00751 | 0.00212 | 0.03274 |
| **Inhba** | 0.59328 | 0.00443 | 0.04325 | 0.02737 | 0.01751 | 0.04148 | 0.00435 | 0.04181 | 0.02059 | 0.00677 | 0.01905 | 0.04861 | 0.00514 | 0.07286 |
| **Inhbb** | 0.00000 | 0.00949 | 2.02989 | 0.00000 | 0.64028 | 0.00000 | 0.00291 | 0.00000 | 0.66021 | 0.00363 | 0.00057 | 2.12583 | 0.08695 | 0.00000 |
| **Inip** | 2.89113 | 3.16697 | 4.36428 | 6.46103 | 3.24941 | 4.03939 | 1.05702 | 1.63070 | 0.59956 | 0.09508 | 0.00732 | 2.75740 | 2.95250 | 1.35050 |
| **Ino80** | 1.21153 | 0.01009 | 6.94951 | 3.77514 | 7.65822 | 2.06061 | 0.37126 | 0.75938 | 2.01962 | 0.00624 | 5.04145 | 0.79935 | 2.22233 | 7.74837 |
| **Ino80b** | 0.00893 | 0.00460 | 0.01675 | 0.01905 | 0.07773 | 0.01091 | 0.00433 | 0.01362 | 0.01188 | 0.00703 | 0.01366 | 0.01040 | 0.01948 | 0.01495 |
| **Ino80c** | 9.93709 | 0.40683 | 10.19884 | 7.83515 | 7.12766 | 10.84847 | 0.97890 | 8.05825 | 8.60207 | 0.03646 | 10.88764 | 5.17323 | 6.67436 | 12.77525 |
| **Ino80d** | 5.87580 | 0.03843 | 4.04665 | 4.40692 | 10.74674 | 6.50507 | 0.03247 | 1.83194 | 8.45930 | 0.54656 | 0.18880 | 2.79568 | 6.35118 | 12.91310 |
| **Ino80dos** | 0.01573 | 0.00444 | 0.08174 | 0.10891 | 0.02366 | 0.00745 | 0.00414 | 0.01092 | 0.10475 | 0.00646 | 0.00925 | 0.00931 | 0.49080 | 0.04545 |
| **Ino80e** | 0.00000 | 3.06999 | 4.54043 | 10.41523 | 4.16129 | 0.84961 | 0.00690 | 6.12503 | 0.00000 | 0.04031 | 3.85853 | 3.61509 | 0.85516 | 0.08289 |
| **Inpp1** | 0.00000 | 0.00370 | 0.54295 | 1.18516 | 0.00000 | 0.28991 | 0.05965 | 0.39045 | 0.01022 | 0.01782 | 0.00000 | 0.55338 | 0.10158 | 0.00000 |
| **Inpp4a** | 0.34617 | 0.01837 | 0.27974 | 0.00000 | 0.36043 | 0.96543 | 0.01255 | 0.00359 | 0.00570 | 9.94813 | 0.00000 | 0.56871 | 0.20548 | 1.09148 |
| **Inpp5a** | 0.25519 | 0.00420 | 0.01715 | 0.00000 | 0.79105 | 0.56993 | 0.00449 | 0.00902 | 0.01492 | 0.00714 | 0.01514 | 0.61566 | 0.42295 | 0.52440 |
| **Inpp5b** | 0.33355 | 0.01107 | 1.70834 | 0.00000 | 0.78867 | 1.86729 | 2.67238 | 0.70643 | 2.70926 | 0.04078 | 2.42048 | 0.67359 | 0.00771 | 3.13716 |
| **Inpp5d** | 0.58648 | 2.12794 | 2.39040 | 1.81021 | 0.53898 | 0.45233 | 0.01349 | 0.00000 | 0.61068 | 0.00000 | 5.55624 | 0.95773 | 0.47204 | 1.67840 |
| **Inpp5e** | 0.00000 | 0.00425 | 0.02059 | 0.60279 | 0.00000 | 0.00000 | 0.94959 | 1.02838 | 1.04524 | 0.03943 | 0.01774 | 1.06195 | 0.40962 | 0.71986 |
| **Inpp5f** | 0.83945 | 0.01021 | 1.35620 | 1.58057 | 0.40928 | 2.69296 | 0.25959 | 1.56085 | 0.00746 | 0.02049 | 1.51175 | 0.63649 | 0.41062 | 2.38766 |
| **Inpp5k** | 5.66325 | 0.01596 | 7.36733 | 2.04753 | 6.05016 | 3.51567 | 2.15767 | 0.42609 | 0.78820 | 0.04944 | 3.67909 | 1.95994 | 3.40536 | 2.26315 |
| **Inppl1** | 0.07655 | 0.00978 | 0.40903 | 0.50251 | 0.80329 | 0.00000 | 0.00348 | 0.45865 | 0.01204 | 0.01674 | 0.00390 | 1.26771 | 0.24227 | 0.08493 |
| **Insig1** | 0.00000 | 1.01048 | 5.08120 | 6.97206 | 0.79650 | 3.75537 | 0.00632 | 6.94765 | 5.83133 | 0.09817 | 5.16335 | 1.38446 | 1.33906 | 3.52883 |
| **Insig2** | 1.00555 | 0.02351 | 0.09201 | 0.00000 | 0.24051 | 0.98212 | 0.00358 | 1.39604 | 0.04510 | 0.01806 | 0.54970 | 1.10954 | 0.00617 | 0.83930 |
| **Insm1** | 0.00840 | 0.00423 | 0.06493 | 0.01424 | 0.01202 | 0.03079 | 0.00422 | 0.05625 | 0.02628 | 0.00747 | 0.01633 | 0.03201 | 0.63311 | 0.00746 |
| **Insr** | 0.01487 | 0.03722 | 0.00690 | 0.00000 | 0.01916 | 0.09699 | 0.00288 | 0.19912 | 0.04228 | 0.00537 | 0.00384 | 3.47749 | 0.01861 | 0.02659 |
| **Ints1** | 6.85191 | 0.15085 | 14.53814 | 12.38773 | 14.19646 | 2.86281 | 0.02807 | 6.71187 | 0.04057 | 1.66295 | 1.52812 | 10.67846 | 4.13942 | 2.74917 |
| **Ints10** | 2.75398 | 0.03381 | 0.91983 | 2.44536 | 4.76578 | 4.45686 | 3.92265 | 1.30457 | 0.03194 | 0.04138 | 8.10524 | 8.11117 | 3.60285 | 6.64192 |
| **Ints12** | 9.16309 | 0.02890 | 6.26221 | 3.04090 | 5.28365 | 6.90448 | 1.27795 | 0.60187 | 9.42568 | 0.02886 | 4.04653 | 4.77129 | 2.26942 | 8.42275 |
| **Ints2** | 5.47456 | 0.00218 | 2.25689 | 0.92101 | 5.32525 | 1.62887 | 3.08211 | 0.29672 | 10.63866 | 0.02601 | 0.00000 | 1.21475 | 5.44867 | 5.44666 |
| **Ints3** | 6.15736 | 0.10421 | 14.58436 | 5.35506 | 16.71792 | 12.68377 | 1.75287 | 4.72697 | 4.24923 | 0.09903 | 6.68392 | 8.47410 | 6.08639 | 17.63938 |
| **Ints4** | 8.41719 | 0.05622 | 10.84148 | 3.90056 | 10.59457 | 6.94121 | 0.51034 | 0.00128 | 13.41899 | 0.00000 | 10.78069 | 7.92534 | 5.90167 | 12.84325 |
| **Ints5** | 2.27468 | 0.39249 | 3.42096 | 5.42074 | 5.28292 | 5.35855 | 2.31327 | 5.82648 | 0.05378 | 0.13826 | 2.60493 | 2.52989 | 5.78006 | 0.32483 |
| **Ints6** | 1.53059 | 0.00000 | 3.84421 | 0.00000 | 5.15550 | 0.86167 | 2.46566 | 0.00000 | 5.04490 | 0.02267 | 2.82493 | 3.27976 | 3.85066 | 4.18083 |
| **Ints7** | 6.93137 | 0.08116 | 9.56235 | 4.36390 | 9.94161 | 4.67182 | 0.02286 | 4.25635 | 1.20097 | 0.05019 | 8.79257 | 5.02588 | 2.61429 | 10.95295 |
| **Ints8** | 2.45680 | 0.00171 | 2.45775 | 0.00000 | 1.45588 | 1.96081 | 0.91594 | 0.44753 | 10.55581 | 0.01215 | 0.60634 | 1.99656 | 1.64210 | 1.96820 |
| **Ints9** | 5.00911 | 0.04839 | 6.43761 | 1.50164 | 2.64177 | 2.99388 | 1.51901 | 1.54729 | 1.40075 | 0.02410 | 6.53573 | 2.34369 | 1.89133 | 7.02098 |
| **Intu** | 0.47095 | 0.02658 | 0.00000 | 0.36251 | 6.48740 | 0.23662 | 0.02134 | 0.22254 | 9.05758 | 0.02597 | 0.22649 | 1.04417 | 16.02834 | 20.26649 |
| **Invs** | 1.15999 | 0.49091 | 3.42815 | 1.79182 | 2.02563 | 2.38229 | 0.81629 | 0.82769 | 0.75268 | 0.03823 | 1.84747 | 1.91258 | 0.77884 | 3.54678 |
| **Ip6k1** | 1.24840 | 0.15001 | 2.56375 | 0.00000 | 0.69983 | 3.47233 | 3.79872 | 1.18818 | 0.00000 | 0.07350 | 2.02333 | 2.08332 | 3.78486 | 2.02528 |
| **Ip6k2** | 0.01153 | 0.00411 | 0.23275 | 1.08564 | 0.04023 | 0.79069 | 0.38542 | 0.77800 | 0.01028 | 0.02759 | 0.67187 | 0.55879 | 0.01277 | 0.00973 |
| **Ipcef1** | 0.01273 | 0.01227 | 0.01016 | 0.01176 | 0.08853 | 0.01070 | 0.00483 | 0.00957 | 0.05647 | 0.00681 | 0.00756 | 0.00885 | 0.01463 | 0.42315 |
| **Ipmk** | 0.61221 | 0.01828 | 0.64412 | 1.13045 | 2.36433 | 1.28113 | 0.00190 | 2.19190 | 8.85340 | 0.00751 | 2.18332 | 0.59239 | 6.42383 | 7.50743 |
| **Ipo11** | 10.42014 | 0.04493 | 8.85463 | 2.03506 | 13.07834 | 4.10661 | 0.70313 | 0.34341 | 17.28984 | 0.02283 | 9.23103 | 4.48769 | 7.33135 | 8.22595 |
| **Ipo13** | 1.66085 | 0.01829 | 1.03880 | 1.32913 | 0.71512 | 3.27032 | 0.22045 | 0.27238 | 0.08212 | 0.03569 | 0.50063 | 2.46090 | 0.58635 | 3.71712 |
| **Ipo4** | 17.97871 | 0.15920 | 32.26865 | 12.91923 | 28.96656 | 8.26743 | 1.15546 | 5.94465 | 0.18888 | 0.03100 | 7.98651 | 11.39404 | 6.85588 | 7.19836 |
| **Ipo5** | 31.33918 | 3.55676 | 41.03999 | 24.00754 | 36.38001 | 19.32956 | 3.65964 | 12.40358 | 36.95649 | 7.60636 | 19.33083 | 9.74553 | 16.21672 | 35.33496 |
| **Ipo7** | 2.27283 | 0.01072 | 2.85825 | 0.84429 | 1.76617 | 0.41517 | 0.00131 | 0.43931 | 13.10748 | 0.00452 | 4.71294 | 0.08203 | 0.91526 | 0.66725 |
| **Ipo8** | 5.12932 | 0.03525 | 5.58653 | 1.46168 | 5.68365 | 3.00870 | 2.07348 | 0.63209 | 11.39424 | 0.00000 | 6.64816 | 5.30251 | 5.53346 | 1.27555 |
| **Ipo9** | 0.00000 | 0.02357 | 4.90213 | 4.15334 | 3.37683 | 3.95039 | 0.00028 | 4.27205 | 6.73280 | 0.06724 | 6.88439 | 1.25148 | 2.39154 | 3.34370 |
| **Ipp** | 1.43213 | 0.00422 | 0.18164 | 0.51602 | 0.00000 | 0.73202 | 1.30710 | 0.02117 | 2.93936 | 0.00569 | 0.00245 | 0.04237 | 0.54526 | 1.43859 |
| **Ippk** | 1.36706 | 0.00807 | 4.91054 | 2.13439 | 2.32118 | 2.26715 | 0.03039 | 2.63714 | 1.36286 | 0.00003 | 4.23368 | 2.92658 | 1.55248 | 10.00477 |
| **Iqca** | 0.00003 | 0.00371 | 0.24107 | 0.00758 | 0.00000 | 1.66276 | 0.00375 | 0.00123 | 0.01265 | 0.01076 | 0.05735 | 0.00246 | 1.48659 | 3.68183 |
| **Iqcb1** | 0.00578 | 0.01842 | 0.01084 | 0.00000 | 0.01237 | 0.14383 | 0.01419 | 1.82822 | 0.03017 | 0.00706 | 0.00404 | 0.02162 | 0.30785 | 0.01891 |
| **Iqcc** | 1.53504 | 0.01197 | 1.10935 | 0.85589 | 0.00000 | 0.00000 | 0.01039 | 2.71892 | 0.01331 | 0.02942 | 0.42757 | 0.00738 | 0.36237 | 0.02308 |

| | | | | | | | | | | | | | | |
|---|---|---|---|---|---|---|---|---|---|---|---|---|---|---|
| **Iqcd** | 0.01441 | 0.00453 | 0.17863 | 0.01220 | 0.10766 | 0.03470 | 0.00496 | 0.07445 | 0.00832 | 0.00611 | 0.00946 | 0.00815 | 0.16115 | 0.04995 |
| **Iqce** | 0.00000 | 0.00229 | 0.30036 | 1.61142 | 0.00000 | 0.43531 | 0.00204 | 0.00000 | 0.00379 | 9.16333 | 0.82967 | 0.00000 | 0.00657 | 0.39768 |
| **Iqcf1** | 0.95284 | 0.00408 | 0.09318 | 0.45398 | 0.03414 | 0.01186 | 0.00399 | 0.00828 | 0.00384 | 0.00852 | 0.00914 | 0.00470 | 0.81135 | 0.00000 |
| **Iqcf5** | 0.00903 | 0.00425 | 0.01051 | 0.06610 | 0.20353 | 0.00938 | 0.00418 | 0.00725 | 0.01480 | 0.00785 | 0.00734 | 0.00956 | 0.43319 | 0.02414 |
| **Iqcg** | 3.38064 | 0.00835 | 6.38405 | 2.52659 | 9.91037 | 0.56675 | 0.00118 | 0.00000 | 5.72952 | 0.00000 | 4.27518 | 0.79510 | 4.49070 | 10.91708 |
| **Iqgap1** | 1.51276 | 8.80145 | 5.17213 | 5.35373 | 2.98888 | 10.34435 | 4.06276 | 13.80065 | 11.74657 | 0.30323 | 12.74147 | 3.68327 | 2.45180 | 3.95872 |
| **Iqgap2** | 0.00000 | 0.00398 | 0.56737 | 0.15555 | 0.53319 | 0.00000 | 0.00393 | 0.20119 | 1.56252 | 0.00414 | 0.00302 | 0.00713 | 0.63337 | 0.06628 |
| **Iqgap3** | 4.75375 | 0.08675 | 15.13966 | 4.66400 | 7.07905 | 5.14604 | 3.48561 | 7.59932 | 0.31534 | 0.43892 | 5.66114 | 1.09298 | 0.00000 | 5.46957 |
| **Iqsec1** | 1.55063 | 0.11429 | 2.35008 | 2.25545 | 0.97486 | 2.00852 | 0.01314 | 5.54874 | 0.00490 | 8.83472 | 0.00000 | 2.42614 | 1.20401 | 0.00000 |
| **Iqsec2** | 0.01491 | 0.00486 | 0.01856 | 0.16248 | 0.03114 | 0.10539 | 0.00477 | 0.01323 | 0.01434 | 0.00530 | 0.01581 | 0.01284 | 0.01046 | 0.02385 |
| **Iqub** | 2.73952 | 0.00304 | 3.43219 | 0.00000 | 7.92426 | 0.35922 | 0.00310 | 0.43001 | 8.39252 | 0.00000 | 0.10553 | 0.00000 | 6.20802 | 10.21963 |
| **Irak1** | 4.72391 | 0.32567 | 0.34566 | 2.40349 | 1.13848 | 9.37449 | 1.22278 | 1.14904 | 5.03203 | 10.18171 | 2.18207 | 4.96720 | 21.63802 | 5.44176 |
| **Irak1bp1** | 0.51009 | 0.00385 | 0.36956 | 0.17756 | 0.02524 | 0.00331 | 0.00381 | 0.00457 | 0.45070 | 0.00609 | 0.00700 | 0.00871 | 2.56115 | 0.12385 |
| **Irak2** | 0.88883 | 0.12826 | 2.76782 | 3.65716 | 4.61554 | 6.54291 | 0.24325 | 1.93641 | 3.36623 | 0.34179 | 0.80131 | 0.11785 | 6.88179 | 2.26109 |
| **Irak3** | 0.80714 | 0.00375 | 0.20454 | 1.59067 | 0.13076 | 0.00000 | 0.00389 | 0.00601 | 0.01321 | 0.00554 | 0.00740 | 0.01269 | 1.34498 | 0.43426 |
| **Irak4** | 0.02784 | 0.00391 | 0.00289 | 1.02330 | 0.19607 | 0.00000 | 0.00373 | 0.00963 | 0.03500 | 0.00574 | 0.00968 | 0.01478 | 0.55469 | 0.00557 |
| **Ireb2** | 9.68486 | 0.03838 | 4.77103 | 0.00000 | 3.95383 | 5.19454 | 0.01065 | 0.54584 | 20.98379 | 0.02336 | 7.16501 | 4.70842 | 6.55241 | 4.71816 |
| **Irf1** | 0.00000 | 0.00372 | 0.00000 | 0.00574 | 0.71446 | 0.01107 | 0.00402 | 0.00625 | 0.22335 | 0.00613 | 0.00385 | 0.38119 | 2.49638 | 0.42307 |
| **Irf2bp1** | 0.26421 | 0.05831 | 2.08508 | 5.49751 | 0.00217 | 1.14186 | 2.25062 | 4.78269 | 0.00000 | 1.04133 | 0.25195 | 3.30950 | 1.96074 | 0.00000 |
| **Irf2bp2** | 0.01048 | 0.01157 | 0.00751 | 0.00654 | 0.01018 | 0.46112 | 0.00377 | 0.73809 | 0.00568 | 0.00605 | 0.00506 | 0.01329 | 0.81195 | 0.07651 |
| **Irf2bpl** | 1.55103 | 3.12702 | 9.89043 | 1.20058 | 10.03594 | 1.99483 | 3.77825 | 1.95419 | 4.33911 | 0.08943 | 0.09899 | 0.43381 | 1.91842 | 8.10415 |
| **Irf3** | 0.00000 | 2.84223 | 3.67731 | 7.13508 | 0.78581 | 0.20598 | 0.01289 | 6.16252 | 0.00000 | 0.17557 | 3.42409 | 2.99935 | 0.85345 | 0.00000 |
| **Irf5** | 0.00214 | 0.00206 | 0.09846 | 0.00460 | 0.15144 | 0.00000 | 0.00196 | 0.00202 | 0.00370 | 10.15825 | 0.00375 | 0.00000 | 0.00642 | 0.30492 |
| **Irf6** | 0.00000 | 0.00546 | 0.40588 | 0.28253 | 1.47832 | 0.12037 | 0.00487 | 0.11259 | 0.00689 | 0.00627 | 0.00354 | 0.00296 | 0.00212 | 0.09222 |
| **Irf7** | 0.03243 | 0.00431 | 0.00000 | 0.13712 | 0.09647 | 0.18180 | 0.00398 | 0.00456 | 0.28273 | 0.00884 | 0.00486 | 0.00709 | 0.29864 | 0.01963 |
| **Irf8** | 0.00251 | 0.00421 | 0.00496 | 0.01116 | 0.33222 | 0.00987 | 0.00403 | 0.02152 | 0.00782 | 0.00587 | 0.00756 | 0.00733 | 0.00405 | 0.90411 |
| **Irf9** | 5.24161 | 0.00249 | 2.73661 | 1.15533 | 6.89424 | 0.43451 | 0.04778 | 0.07216 | 0.50361 | 0.00982 | 1.04439 | 0.98335 | 1.99852 | 7.28027 |
| **Irgm1** | 0.07228 | 0.00425 | 0.01305 | 0.02446 | 0.00873 | 0.03053 | 0.01617 | 0.02475 | 0.08226 | 0.01863 | 0.00782 | 0.01918 | 0.45965 | 0.00936 |
| **Irgm2** | 0.01384 | 0.01314 | 0.02494 | 0.13203 | 0.08593 | 0.06314 | 0.01611 | 0.02881 | 0.15718 | 0.00708 | 0.01963 | 0.02495 | 0.09131 | 0.14767 |
| **Irgq** | 4.81814 | 0.04667 | 7.09959 | 0.00000 | 5.07472 | 5.10046 | 2.35060 | 2.40934 | 0.11066 | 0.20643 | 0.19915 | 0.68083 | 8.88377 | 3.42788 |
| **Irs3** | 0.00876 | 0.00447 | 0.01381 | 0.00000 | 0.08222 | 0.14137 | 0.00486 | 0.00598 | 0.03072 | 0.00603 | 0.00794 | 0.01169 | 0.00587 | 0.00281 |
| **Irx2** | 0.09218 | 0.00408 | 0.00500 | 0.19173 | 0.00504 | 0.00367 | 0.00409 | 1.29205 | 0.01087 | 0.00606 | 0.00459 | 0.03932 | 0.13885 | 0.00000 |
| **Isca1** | 4.27587 | 2.24962 | 8.79763 | 6.26439 | 8.64204 | 7.98294 | 0.69036 | 9.50783 | 0.64671 | 11.18462 | 1.87895 | 8.50209 | 3.09980 | 4.98399 |
| **Isca2** | 0.94898 | 0.02678 | 1.24492 | 2.45604 | 2.18170 | 3.05362 | 6.06825 | 0.80725 | 0.40705 | 0.14255 | 1.27660 | 2.80716 | 0.00000 | 1.70013 |
| **Iscu** | 7.16608 | 0.05409 | 1.35090 | 0.00000 | 5.12115 | 4.00681 | 0.09984 | 2.04466 | 0.72219 | 10.04565 | 0.00000 | 4.16879 | 4.36669 | 0.74168 |
| **Isg15** | 0.00299 | 0.00411 | 0.00634 | 0.01307 | 0.31846 | 0.01383 | 0.00466 | 0.00894 | 0.00768 | 0.00663 | 0.00957 | 0.00682 | 2.55091 | 0.00586 |
| **Isg20** | 0.20755 | 0.00384 | 0.21178 | 0.00000 | 1.89061 | 0.19076 | 0.00386 | 0.35031 | 0.00622 | 0.00694 | 0.00268 | 0.00392 | 0.13585 | 0.65421 |
| **Isg20l2** | 0.31146 | 0.06821 | 2.52053 | 1.75047 | 3.80771 | 1.21872 | 0.20324 | 0.24490 | 12.31475 | 0.05312 | 3.18335 | 2.28061 | 1.60908 | 11.61070 |
| **Isl2** | 0.00472 | 0.00386 | 0.54495 | 0.09486 | 2.77484 | 0.02531 | 0.01014 | 0.34170 | 0.02860 | 0.00495 | 0.00186 | 0.00668 | 0.00000 | 0.00065 |
| **Islr2** | 0.01182 | 0.00483 | 0.01439 | 0.00558 | 0.01345 | 0.01342 | 0.00416 | 0.01583 | 0.01131 | 0.00666 | 0.00678 | 0.07342 | 0.04220 | 0.00702 |
| **Ism2** | 0.00921 | 0.00529 | 0.00801 | 0.08890 | 0.14616 | 0.00587 | 0.00419 | 0.04168 | 0.00688 | 0.00690 | 0.01132 | 0.04471 | 0.14203 | 0.21900 |
| **Isoc1** | 0.99675 | 0.03420 | 0.81789 | 3.45541 | 1.13275 | 3.16068 | 0.00241 | 2.54564 | 1.50937 | 0.00860 | 4.35120 | 1.58519 | 0.00311 | 0.93121 |
| **Isoc2b** | 0.69780 | 0.00405 | 2.44432 | 0.24716 | 0.90810 | 0.00000 | 0.00371 | 0.00094 | 0.10543 | 0.02798 | 0.18020 | 0.00794 | 0.08173 | 0.73444 |
| **Israa** | 0.00163 | 0.00436 | 0.00000 | 0.29430 | 0.16231 | 0.00000 | 0.00425 | 0.00972 | 1.46901 | 0.00654 | 0.00283 | 0.00685 | 0.00170 | 0.33869 |
| **Ist1** | 10.19763 | 0.13822 | 22.04909 | 13.78982 | 17.01185 | 16.01005 | 6.50351 | 7.97297 | 15.50344 | 0.21704 | 7.50538 | 8.33868 | 9.88117 | 17.63950 |
| **Isy1** | 6.05892 | 7.61822 | 14.83790 | 8.71310 | 5.19174 | 9.38015 | 0.03314 | 4.36236 | 0.95650 | 0.05713 | 6.17258 | 9.32046 | 2.19211 | 5.97695 |
| **Isyna1** | 33.59156 | 8.11004 | 58.44924 | 65.14251 | 47.24806 | 37.25911 | 7.23253 | 13.53114 | 0.00000 | 19.49634 | 21.81088 | 18.73361 | 0.81012 | 8.05839 |
| **Itch** | 4.14581 | 0.02028 | 10.20374 | 3.46224 | 13.35695 | 10.23220 | 1.42425 | 3.00332 | 19.24136 | 0.01690 | 6.33657 | 2.85861 | 2.31406 | 11.53154 |
| **Itfg1** | 3.54079 | 0.04148 | 0.67127 | 0.40656 | 0.16481 | 0.80601 | 0.04425 | 0.62266 | 0.35697 | 0.00313 | 3.43179 | 2.09591 | 0.01814 | 0.48970 |
| **Itfg2** | 3.19500 | 0.02729 | 3.27602 | 1.29058 | 3.89658 | 1.48129 | 0.00775 | 2.84171 | 0.38592 | 0.02383 | 4.45500 | 1.12347 | 0.82868 | 6.34915 |
| **Itga1** | 0.24153 | 0.00416 | 0.80438 | 0.10980 | 0.12247 | 0.01097 | 0.00397 | 0.00265 | 0.25527 | 0.00455 | 0.00336 | 0.41966 | 0.00087 | 0.01590 |
| **Itga10** | 0.00354 | 0.00401 | 0.02436 | 0.00088 | 0.00975 | 0.02650 | 0.00429 | 0.02261 | 0.01074 | 0.00630 | 0.00718 | 0.03828 | 0.71136 | 0.00542 |
| **Itga11** | 0.31098 | 0.00460 | 0.56328 | 0.00000 | 1.23242 | 0.00002 | 0.00408 | 0.01877 | 0.05754 | 0.00658 | 0.00489 | 0.00720 | 0.00214 | 0.62466 |
| **Itga2b** | 0.00000 | 0.00468 | 0.47900 | 0.53334 | 0.00418 | 0.00000 | 0.00428 | 0.04430 | 0.00911 | 0.00737 | 0.00628 | 0.55872 | 0.00416 | 0.02193 |
| **Itga3** | 0.92198 | 2.99171 | 4.27176 | 3.78682 | 0.60543 | 5.33459 | 2.87233 | 3.94934 | 0.06127 | 0.14778 | 3.30613 | 5.47758 | 0.11117 | 0.48331 |

| | | | | | | | | | | | | | | |
|---|---|---|---|---|---|---|---|---|---|---|---|---|---|---|
| **Itga4** | 0.07253 | 0.04697 | 0.09247 | 0.03563 | 0.05174 | 0.09476 | 0.00430 | 0.01212 | 0.48051 | 0.00775 | 0.02535 | 0.13048 | 0.13699 | 0.28951 |
| **Itga5** | 1.19009 | 0.04969 | 3.35162 | 4.35438 | 0.00000 | 7.29889 | 7.30531 | 0.96535 | 1.40468 | 3.55465 | 0.00000 | 7.17151 | 0.00001 | 0.00971 |
| **Itga6** | 0.53883 | 0.22387 | 2.94808 | 0.87027 | 2.33200 | 5.47976 | 3.26061 | 0.15882 | 2.04280 | 0.05041 | 9.38159 | 11.49000 | 3.63505 | 6.19614 |
| **Itga7** | 0.01412 | 0.00451 | 0.05100 | 0.00076 | 0.08729 | 0.39208 | 0.00416 | 0.00541 | 0.00514 | 0.00700 | 0.00514 | 0.00942 | 0.00373 | 0.17063 |
| **Itga9** | 0.02531 | 0.00423 | 0.00868 | 0.12947 | 0.00801 | 0.01830 | 0.00403 | 0.00580 | 0.01036 | 0.00674 | 0.00637 | 0.01367 | 0.13596 | 0.07161 |
| **Itgae** | 0.78869 | 0.00338 | 1.87049 | 0.32283 | 0.91144 | 0.77428 | 0.00304 | 0.00000 | 2.92592 | 0.00730 | 1.15499 | 0.00588 | 1.66210 | 3.86427 |
| **Itgal** | 0.00304 | 0.00383 | 0.00000 | 0.00134 | 0.27406 | 0.00000 | 0.00451 | 0.01111 | 0.02354 | 0.00324 | 0.72093 | 0.26632 | 0.54605 | 0.41659 |
| **Itgam** | 0.01467 | 0.00469 | 0.00939 | 0.02576 | 0.01550 | 0.04788 | 0.00427 | 0.00762 | 0.00922 | 0.01082 | 0.01130 | 0.00787 | 0.06543 | 0.21857 |
| **Itgav** | 2.88783 | 0.00218 | 0.24116 | 0.33812 | 4.52022 | 0.35976 | 0.00882 | 1.03581 | 10.57315 | 0.00615 | 0.00000 | 1.26195 | 5.32508 | 3.01957 |
| **Itgb1** | 4.92431 | 0.20973 | 6.36307 | 6.06550 | 0.52792 | 9.10127 | 0.00000 | 2.11905 | 4.96594 | 0.41041 | 7.39855 | 5.62663 | 0.01096 | 6.58142 |
| **Itgb1bp1** | 6.19752 | 0.09574 | 11.73976 | 14.21919 | 5.63957 | 4.23775 | 0.00471 | 0.22253 | 1.98838 | 2.79834 | 4.34216 | 8.70168 | 5.09729 | 1.19685 |
| **Itgb3** | 0.02847 | 0.00443 | 0.22153 | 0.07164 | 0.00228 | 0.09021 | 0.00999 | 0.00414 | 0.00615 | 0.00681 | 0.01487 | 0.01782 | 0.30919 | 0.10798 |
| **Itgb3bp** | 0.82592 | 0.00412 | 0.07891 | 0.32798 | 0.35316 | 0.66285 | 0.00446 | 0.00349 | 0.10036 | 0.00701 | 0.00570 | 0.05976 | 0.00937 | 0.00000 |
| **Itgb4** | 0.00000 | 0.00385 | 0.43879 | 1.54293 | 2.61477 | 0.00000 | 0.00384 | 0.00449 | 0.03197 | 0.00450 | 0.46293 | 0.00493 | 0.31506 | 1.57154 |
| **Itgb5** | 5.28537 | 0.14374 | 17.52801 | 9.45235 | 14.72864 | 6.86979 | 4.99176 | 6.52643 | 2.45827 | 0.24497 | 2.56350 | 4.01475 | 2.60766 | 18.18977 |
| **Itgb7** | 0.90676 | 0.00369 | 3.64937 | 1.99259 | 6.72193 | 0.89449 | 0.00313 | 0.00000 | 0.06366 | 0.00994 | 0.00275 | 0.00554 | 1.68274 | 1.24489 |
| **Itih5** | 0.03112 | 0.00445 | 0.04613 | 0.00903 | 0.01094 | 0.01148 | 0.01627 | 0.01025 | 0.01067 | 0.00699 | 0.00827 | 0.00967 | 0.00481 | 0.02589 |
| **Itln1** | 0.23423 | 0.00435 | 0.01099 | 0.16927 | 0.04305 | 0.10099 | 0.00423 | 0.00515 | 0.01200 | 0.00591 | 0.00489 | 0.00648 | 0.00555 | 0.00267 |
| **Itm2a** | 0.00000 | 0.00421 | 0.47208 | 0.06150 | 0.36317 | 0.01218 | 0.00497 | 0.79475 | 0.18834 | 0.00817 | 1.09471 | 0.01508 | 0.01900 | 1.32290 |
| **Itm2b** | 0.66228 | 0.01489 | 1.28725 | 6.06798 | 3.71909 | 1.23386 | 0.63461 | 3.38727 | 0.00142 | 4.35881 | 0.58524 | 3.06184 | 0.20984 | 1.86140 |
| **Itm2c** | 0.02282 | 0.00293 | 0.99852 | 1.15933 | 2.42385 | 0.55346 | 0.00355 | 0.00000 | 0.04820 | 0.00296 | 0.00000 | 1.56692 | 0.00000 | 0.46510 |
| **Itpa** | 11.05846 | 10.43933 | 22.47440 | 16.29157 | 16.07535 | 16.12103 | 7.28014 | 9.21800 | 10.17918 | 11.53043 | 8.89377 | 9.08269 | 3.46816 | 14.48095 |
| **Itpk1** | 1.31971 | 0.02812 | 0.96540 | 0.00000 | 1.96251 | 0.83971 | 0.00843 | 0.01798 | 0.01783 | 0.00605 | 1.57041 | 5.53073 | 3.54068 | 0.00000 |
| **Itpka** | 1.90020 | 0.00351 | 0.31486 | 0.42970 | 0.88812 | 0.02734 | 0.00398 | 0.06410 | 0.00823 | 0.01333 | 0.00636 | 0.70390 | 0.00613 | 0.00000 |
| **Itpkb** | 0.00824 | 0.00441 | 0.02588 | 0.00000 | 0.47301 | 0.01807 | 0.00412 | 0.00851 | 0.02226 | 0.00482 | 0.39200 | 0.00601 | 0.00334 | 0.01290 |
| **Itpkc** | 2.99008 | 0.00974 | 5.27730 | 0.00000 | 5.16780 | 0.05387 | 0.00855 | 1.29179 | 1.97808 | 0.00501 | 2.23720 | 0.45096 | 7.75884 | 7.29434 |
| **Itpr1** | 0.00000 | 0.00395 | 0.33147 | 0.01322 | 0.98454 | 0.00000 | 0.00421 | 0.00719 | 1.29466 | 0.00565 | 0.00336 | 0.00375 | 2.54812 | 1.72683 |
| **Itpr2** | 0.03290 | 0.00401 | 0.53414 | 0.22639 | 0.58201 | 0.00000 | 0.00364 | 0.00469 | 0.54959 | 0.00593 | 0.00372 | 0.01218 | 1.38126 | 0.70741 |
| **Itpr3** | 1.03731 | 0.02793 | 2.28929 | 0.76723 | 3.10218 | 2.32873 | 0.01075 | 1.94502 | 0.05048 | 0.00000 | 1.18995 | 0.31953 | 0.71747 | 0.16990 |
| **Itpripl1** | 0.00000 | 0.00261 | 0.28643 | 0.21333 | 0.00000 | 0.47618 | 0.00335 | 0.00000 | 0.34257 | 0.00219 | 2.91713 | 0.66779 | 0.70478 | 0.13166 |
| **Itpripl2** | 0.15611 | 0.00429 | 0.00767 | 0.00792 | 0.00419 | 0.53503 | 0.70708 | 1.43416 | 0.00758 | 0.04151 | 0.00320 | 0.00000 | 0.02546 | 0.04741 |
| **Itsn1** | 5.50112 | 0.13634 | 5.34294 | 1.50255 | 8.33848 | 2.31280 | 0.13009 | 2.42176 | 3.99643 | 1.30671 | 1.39413 | 7.27607 | 6.65357 | 9.49210 |
| **Itsn2** | 0.00000 | 0.01756 | 1.82310 | 0.34609 | 0.73476 | 2.37556 | 0.00321 | 0.56843 | 1.29325 | 0.00594 | 1.25740 | 0.80961 | 1.05654 | 2.16093 |
| **Ivd** | 0.19609 | 0.06102 | 3.13956 | 2.17987 | 1.15655 | 2.34259 | 0.01644 | 1.97360 | 0.16897 | 0.02442 | 1.62356 | 3.03340 | 0.76989 | 3.79424 |
| **Ivns1abp** | 24.01170 | 3.34745 | 22.69790 | 17.52076 | 23.48444 | 19.79266 | 3.94257 | 6.87413 | 30.34681 | 0.20691 | 14.97729 | 4.85932 | 12.11534 | 15.71469 |
| **Iws1** | 10.42300 | 0.06433 | 7.65317 | 2.63763 | 12.22143 | 5.29830 | 2.50780 | 4.33047 | 12.88086 | 1.46645 | 7.79369 | 3.95070 | 11.51279 | 13.10972 |
| **Izumo1** | 0.00791 | 0.00402 | 0.00686 | 0.16608 | 0.13408 | 0.02293 | 0.01065 | 0.12328 | 0.01314 | 0.00553 | 0.01294 | 0.01701 | 0.00712 | 0.01563 |
| **Izumo1r** | 0.16624 | 0.00423 | 0.01286 | 0.00999 | 0.36247 | 0.01412 | 0.00408 | 0.00813 | 0.03189 | 0.00988 | 0.00702 | 0.00661 | 1.16396 | 0.00000 |
| **Izumo4** | 0.00282 | 0.00535 | 0.24220 | 0.14324 | 0.24020 | 0.02819 | 0.00535 | 0.03518 | 0.11911 | 0.00700 | 0.00561 | 0.00492 | 0.25190 | 0.00000 |
| **Jade1** | 0.00000 | 1.72185 | 3.89326 | 0.47482 | 1.39761 | 7.29376 | 5.10087 | 4.68835 | 8.97490 | 0.27869 | 5.06949 | 3.55547 | 7.69395 | 9.24400 |
| **Jade2** | 0.27041 | 0.00395 | 0.00540 | 0.21288 | 0.29665 | 0.00273 | 0.00389 | 0.00593 | 0.03246 | 0.00669 | 0.00555 | 0.01220 | 0.84447 | 0.77968 |
| **Jade3** | 0.83476 | 0.01195 | 0.12918 | 1.46692 | 1.13357 | 2.09362 | 2.48276 | 1.19336 | 6.26102 | 0.02678 | 1.46918 | 0.99246 | 0.14494 | 1.02892 |
| **Jag1** | 0.18185 | 0.00411 | 0.20010 | 0.04996 | 0.00918 | 0.00552 | 0.00394 | 0.00709 | 0.02157 | 0.00853 | 0.00590 | 0.01025 | 0.72189 | 0.00756 |
| **Jag2** | 0.18679 | 0.00408 | 0.70376 | 2.18551 | 0.51164 | 0.00000 | 0.00380 | 0.23777 | 0.00699 | 0.00357 | 0.00198 | 0.01110 | 0.03081 | 0.00000 |
| **Jagn1** | 5.63844 | 5.56367 | 11.90566 | 8.46139 | 7.28476 | 10.34715 | 3.29040 | 6.88476 | 2.00838 | 0.10505 | 3.99390 | 6.35462 | 4.00722 | 2.81973 |
| **Jak1** | 3.71974 | 0.04486 | 1.32297 | 1.66650 | 5.15398 | 6.25094 | 3.08598 | 4.99090 | 2.28786 | 0.16941 | 4.91562 | 3.60301 | 1.77309 | 2.18591 |
| **Jak2** | 2.39038 | 0.00974 | 0.66035 | 0.00000 | 3.86912 | 0.15146 | 0.00287 | 0.43781 | 9.06792 | 0.00375 | 0.03926 | 1.31077 | 11.12112 | 12.81361 |
| **Jak3** | 0.00428 | 0.00397 | 0.03216 | 1.99011 | 0.40867 | 0.00000 | 0.00386 | 0.00781 | 0.00927 | 0.00588 | 0.00505 | 0.77002 | 0.03634 | 0.00000 |
| **Jakmip1** | 0.69189 | 0.00393 | 2.57634 | 0.00000 | 3.33632 | 0.42791 | 0.00338 | 0.00000 | 0.02814 | 0.00984 | 1.68297 | 0.19963 | 0.00000 | 1.72791 |
| **Jakmip3** | 0.00426 | 0.00429 | 0.01445 | 0.03220 | 0.19551 | 0.01109 | 0.00439 | 0.00750 | 0.01543 | 0.00630 | 0.00797 | 0.01020 | 0.55547 | 0.00392 |
| **Jam2** | 0.22536 | 0.06889 | 1.04633 | 2.71854 | 0.94476 | 0.00000 | 0.00084 | 0.00000 | 3.26141 | 0.01152 | 1.26658 | 8.17992 | 3.82184 | 0.86329 |
| **Jam3** | 0.00000 | 0.00359 | 0.05751 | 0.35742 | 0.02509 | 0.05469 | 0.00356 | 0.00353 | 0.00327 | 0.00507 | 0.01151 | 0.14701 | 6.01924 | 0.49023 |
| **Jarid2** | 11.34030 | 8.87841 | 33.50760 | 10.90952 | 21.06923 | 23.14063 | 3.57660 | 12.15626 | 19.67177 | 0.57830 | 17.25745 | 17.32090 | 3.79691 | 16.83960 |
| **Jazf1** | 0.00000 | 0.00416 | 0.12919 | 0.02886 | 0.10886 | 0.30559 | 0.00404 | 0.00396 | 0.12835 | 0.00445 | 0.07268 | 0.00282 | 2.68887 | 0.06609 |
| **Jchain** | 0.00742 | 0.00437 | 0.01383 | 0.01923 | 0.02790 | 0.33064 | 0.01575 | 0.00552 | 0.04033 | 0.00765 | 0.00574 | 0.00868 | 0.01033 | 0.02073 |

| | | | | | | | | | | | | | | |
|---|---|---|---|---|---|---|---|---|---|---|---|---|---|---|
| **Jdp2** | 0.02054 | 0.00474 | 0.01141 | 0.00801 | 0.01854 | 0.08204 | 0.00435 | 0.00724 | 0.07289 | 0.02257 | 0.00742 | 0.00941 | 0.01377 | 0.02371 |
| **Jkamp** | 3.22336 | 0.65327 | 1.40464 | 5.63222 | 3.37727 | 3.15744 | 2.93643 | 3.16853 | 1.14011 | 0.13447 | 3.23537 | 3.89213 | 1.51994 | 1.57386 |
| **Jmjd1c** | 1.60017 | 0.05245 | 1.38896 | 1.07669 | 1.20561 | 1.41253 | 0.01031 | 0.88057 | 0.92924 | 0.01051 | 2.05874 | 2.82520 | 2.00768 | 3.23653 |
| **Jmjd4** | 0.29069 | 0.02423 | 2.53478 | 2.72479 | 0.00000 | 1.99980 | 0.01475 | 0.86928 | 0.61096 | 0.00169 | 0.21955 | 1.86112 | 0.58863 | 1.43054 |
| **Jmjd6** | 10.66103 | 6.48035 | 16.09858 | 9.95522 | 17.84528 | 8.10151 | 3.83662 | 0.72004 | 2.06305 | 1.34015 | 4.92343 | 6.95985 | 16.25897 | 11.42866 |
| **Jmjd7** | 0.05209 | 0.00425 | 0.10283 | 0.16509 | 0.13864 | 0.09772 | 0.03954 | 0.03604 | 0.00825 | 0.00643 | 0.10710 | 0.26444 | 0.13132 | 0.04447 |
| **Jmjd8** | 0.21918 | 0.01102 | 0.60633 | 0.77657 | 0.43645 | 1.43806 | 0.00362 | 1.03052 | 0.03003 | 0.00558 | 0.00000 | 0.65252 | 0.14485 | 0.39218 |
| **Jmy** | 8.81979 | 0.00915 | 6.45331 | 1.92249 | 7.12390 | 1.44280 | 2.00149 | 0.26425 | 19.64913 | 0.02800 | 0.12065 | 0.69586 | 10.67336 | 14.79272 |
| **Josd1** | 2.57157 | 0.04708 | 5.36122 | 1.49213 | 1.45739 | 6.34012 | 0.41535 | 2.64080 | 0.11199 | 0.00000 | 3.09814 | 5.03027 | 2.13227 | 4.22375 |
| **Josd2** | 0.00000 | 0.03457 | 3.48913 | 7.37272 | 1.66967 | 2.08837 | 0.02122 | 5.02231 | 0.01046 | 3.06033 | 2.63242 | 2.33110 | 0.00000 | 0.00000 |
| **Jph1** | 0.43780 | 0.01418 | 0.03607 | 0.14303 | 0.10896 | 0.00000 | 0.00410 | 0.33382 | 1.33544 | 0.01773 | 0.00062 | 0.18203 | 0.00372 | 0.00000 |
| **Jpx** | 0.11793 | 0.04758 | 0.03577 | 0.25535 | 0.07829 | 0.18514 | 0.05428 | 0.16815 | 0.22224 | 0.14538 | 0.03150 | 0.05278 | 0.74630 | 0.30760 |
| **Jrk** | 0.00000 | 0.00412 | 1.75773 | 0.36699 | 0.00000 | 0.70868 | 0.00932 | 0.19485 | 0.00262 | 0.01031 | 0.04316 | 0.00000 | 1.02555 | 0.16421 |
| **Jsrp1** | 0.00521 | 0.00431 | 0.02787 | 0.16416 | 0.56789 | 0.00862 | 0.00407 | 0.00691 | 0.01151 | 0.00606 | 0.00817 | 0.00747 | 0.00460 | 0.00439 |
| **Jtb** | 4.01528 | 0.03518 | 13.17745 | 13.61959 | 17.92340 | 0.00000 | 3.90686 | 4.78507 | 2.59519 | 8.68039 | 0.21369 | 5.67046 | 4.32097 | 0.38304 |
| **Jun** | 0.01631 | 0.12459 | 0.52758 | 0.23567 | 0.39354 | 0.07982 | 0.05034 | 3.99735 | 0.04913 | 0.01586 | 0.11667 | 0.04673 | 0.15327 | 0.00159 |
| **Junb** | 0.02770 | 0.03044 | 0.00770 | 0.36399 | 0.48931 | 0.00019 | 0.00467 | 0.98186 | 0.02011 | 0.00746 | 0.00805 | 0.06366 | 0.07772 | 0.00000 |
| **Jund** | 0.14798 | 0.28470 | 0.00911 | 1.26031 | 1.58996 | 0.07106 | 0.22190 | 1.71623 | 0.06345 | 0.10230 | 0.00000 | 0.11865 | 1.37868 | 0.00000 |
| **Jup** | 1.51217 | 3.03673 | 3.73196 | 1.62263 | 3.58525 | 3.79582 | 0.01239 | 6.80311 | 0.32224 | 0.01420 | 1.47888 | 2.78973 | 0.46127 | 1.39675 |
| **Kalrn** | 0.02334 | 0.03910 | 0.74860 | 0.00000 | 1.36969 | 0.78837 | 0.01079 | 1.95320 | 0.02045 | 0.03256 | 0.00000 | 0.49193 | 0.31815 | 0.51532 |
| **Kank1** | 0.17220 | 6.25954 | 2.06406 | 0.00000 | 1.34785 | 3.19316 | 2.20517 | 5.31684 | 0.00000 | 0.06687 | 0.00000 | 0.29271 | 0.28318 | 0.98731 |
| **Kank2** | 0.47094 | 0.04072 | 0.20186 | 0.33493 | 1.77650 | 0.67622 | 0.01534 | 0.87779 | 0.65756 | 0.03679 | 0.00000 | 1.66095 | 2.13563 | 2.26270 |
| **Kank3** | 1.29413 | 0.01129 | 0.76110 | 4.42817 | 0.39289 | 1.52450 | 0.00345 | 0.00000 | 0.00756 | 0.01681 | 1.29417 | 0.01040 | 0.65431 | 0.09611 |
| **Kank4** | 1.17321 | 0.00404 | 0.04691 | 0.00000 | 2.30634 | 0.06640 | 0.00405 | 0.00593 | 1.46248 | 0.00634 | 0.00174 | 0.00299 | 2.00417 | 0.26793 |
| **Kank4os** | 0.06345 | 0.01260 | 0.08345 | 0.00000 | 0.10149 | 0.16578 | 0.00388 | 0.64620 | 3.23066 | 0.00654 | 0.11507 | 0.15893 | 0.01000 | 0.40898 |
| **Kansl1** | 4.52216 | 0.03835 | 6.31791 | 2.30714 | 10.30296 | 3.78828 | 3.61662 | 1.57546 | 3.91703 | 0.12144 | 4.46237 | 2.86601 | 6.19034 | 7.88884 |
| **Kansl1l** | 0.62442 | 0.00431 | 0.14844 | 0.12946 | 0.00000 | 0.09680 | 0.01130 | 0.09550 | 1.27969 | 0.00590 | 0.93977 | 0.00197 | 0.02018 | 0.00000 |
| **Kansl2** | 7.40732 | 0.11359 | 15.32497 | 7.35690 | 17.74031 | 9.86862 | 2.34340 | 5.71200 | 8.67880 | 0.28491 | 13.38670 | 9.44325 | 13.06787 | 23.59219 |
| **Kansl3** | 6.90373 | 2.64416 | 7.46794 | 4.95345 | 7.43708 | 4.67208 | 3.39013 | 0.00000 | 8.25294 | 0.09362 | 2.59603 | 2.95204 | 9.21891 | 11.54998 |
| **Kantr** | 0.02434 | 0.00439 | 0.06801 | 0.09626 | 0.00918 | 0.01588 | 0.00431 | 0.02877 | 0.02472 | 0.00834 | 0.00880 | 0.00592 | 0.28953 | 0.07131 |
| **Kars** | 19.36447 | 0.32162 | 33.18487 | 19.25473 | 29.82452 | 23.87279 | 7.81927 | 11.45580 | 11.76618 | 0.50856 | 14.74773 | 17.69947 | 8.73054 | 28.07175 |
| **Kat2a** | 2.03543 | 0.10029 | 11.67916 | 5.93289 | 6.68091 | 8.28525 | 0.99896 | 5.53402 | 1.19419 | 0.03331 | 8.52451 | 8.93187 | 4.00177 | 9.75700 |
| **Kat2b** | 2.85069 | 0.01749 | 1.36440 | 0.75006 | 0.59759 | 0.00000 | 0.00204 | 0.14343 | 9.30826 | 0.00847 | 0.71895 | 1.71522 | 3.55328 | 0.00000 |
| **Kat5** | 4.97526 | 0.06588 | 9.61744 | 6.20556 | 7.74985 | 6.14470 | 3.46020 | 0.92694 | 0.90038 | 0.10579 | 4.40064 | 4.24544 | 1.01230 | 8.01833 |
| **Kat6a** | 3.49816 | 0.01102 | 0.00000 | 1.30326 | 1.88431 | 2.43257 | 0.71833 | 0.70458 | 0.89895 | 0.03215 | 1.06653 | 1.28289 | 5.87987 | 4.56803 |
| **Kat6b** | 7.01330 | 3.10593 | 9.30543 | 0.94563 | 7.87242 | 9.83672 | 4.84850 | 2.53464 | 7.91049 | 0.10629 | 4.66711 | 5.58069 | 4.35237 | 9.02759 |
| **Kat7** | 10.49252 | 0.74614 | 15.51366 | 1.80860 | 19.05402 | 17.53756 | 4.93277 | 6.68227 | 11.60441 | 0.17490 | 10.35643 | 7.46067 | 8.61089 | 13.44673 |
| **Kat8** | 6.91274 | 0.02299 | 13.14764 | 3.71050 | 13.08691 | 3.09717 | 3.45681 | 0.14826 | 6.48645 | 0.12286 | 4.42503 | 2.54403 | 3.43255 | 4.69068 |
| **Katna1** | 7.24646 | 6.56509 | 9.44855 | 5.07921 | 10.35629 | 7.92496 | 6.44231 | 1.57988 | 9.31516 | 0.33457 | 7.38403 | 4.49849 | 20.06949 | 19.52071 |
| **Katnal1** | 0.01866 | 0.00452 | 0.01181 | 0.01737 | 0.01229 | 0.02522 | 0.00421 | 0.01089 | 0.01119 | 0.00667 | 0.00703 | 0.02888 | 0.82257 | 0.19415 |
| **Katnal2** | 0.01294 | 0.00423 | 0.00695 | 0.29393 | 0.15177 | 0.23319 | 0.00398 | 0.00342 | 0.01038 | 0.00486 | 0.01786 | 0.01176 | 0.00643 | 0.00781 |
| **Katnb1** | 1.10252 | 0.05155 | 4.13862 | 5.16907 | 1.61081 | 4.74146 | 0.00302 | 1.42076 | 0.00000 | 0.02368 | 3.36918 | 0.84182 | 0.24328 | 1.16961 |
| **Katnbl1** | 4.13086 | 1.46134 | 2.08639 | 0.74527 | 3.44492 | 1.20245 | 2.17252 | 2.91660 | 8.64679 | 3.50718 | 0.61844 | 0.97216 | 4.12098 | 1.27189 |
| **Kazald1** | 0.00953 | 0.00437 | 0.00862 | 0.04916 | 0.18265 | 0.01005 | 0.00405 | 0.00814 | 0.01230 | 0.00663 | 0.01381 | 0.01207 | 0.00552 | 0.00799 |
| **Kazn** | 1.29208 | 4.63572 | 5.54897 | 0.84691 | 1.29885 | 2.98184 | 0.00778 | 1.59341 | 0.77770 | 0.00000 | 2.07404 | 2.38904 | 0.01286 | 2.08788 |
| **Kbtbd11** | 0.01108 | 0.00435 | 0.01765 | 0.01963 | 0.03475 | 0.01239 | 0.01635 | 0.00916 | 0.02452 | 0.00657 | 0.00819 | 0.00845 | 0.32999 | 0.01509 |
| **Kbtbd13** | 0.01783 | 0.04069 | 0.98746 | 0.59008 | 0.30901 | 2.21053 | 0.00245 | 5.88892 | 0.25839 | 0.08121 | 2.07053 | 0.00000 | 0.00000 | 0.34619 |
| **Kbtbd2** | 4.64477 | 0.01877 | 0.70332 | 0.20110 | 0.95762 | 1.37491 | 0.01080 | 0.07348 | 0.86681 | 0.03713 | 1.93960 | 0.29504 | 4.34952 | 5.41795 |
| **Kbtbd3** | 0.01363 | 0.01326 | 0.00459 | 0.26075 | 0.15012 | 0.02565 | 0.00396 | 0.01488 | 0.00870 | 0.01642 | 0.00674 | 0.01989 | 0.00745 | 0.18839 |
| **Kbtbd4** | 3.79775 | 5.97438 | 3.07239 | 2.41426 | 3.20300 | 3.55348 | 2.83873 | 0.28200 | 3.21844 | 0.02380 | 3.92415 | 1.65689 | 1.56528 | 3.65737 |
| **Kbtbd6** | 0.02301 | 0.00406 | 0.01789 | 0.00000 | 0.00381 | 0.03081 | 0.01034 | 0.00570 | 0.00759 | 0.00628 | 0.04056 | 0.10792 | 0.24924 | 0.37439 |
| **Kbtbd7** | 0.16878 | 0.13324 | 0.25164 | 0.38244 | 0.00000 | 1.78210 | 0.00344 | 0.01968 | 2.14100 | 0.01995 | 0.07423 | 0.21998 | 4.58697 | 0.22952 |
| **Kbtbd8** | 1.19767 | 0.00245 | 1.71768 | 0.47691 | 2.84551 | 1.57068 | 0.00262 | 0.00000 | 5.81996 | 0.00283 | 0.00000 | 0.89887 | 7.67479 | 1.74102 |
| **Kcmf1** | 19.59280 | 5.10477 | 16.62860 | 11.30804 | 16.93751 | 15.97774 | 5.87504 | 7.64927 | 2.50662 | 11.81704 | 12.78720 | 11.99671 | 13.73505 | 11.43945 |
| **Kcna2** | 0.02523 | 0.01860 | 0.01463 | 0.01998 | 0.08794 | 0.02508 | 0.00413 | 0.03735 | 0.11140 | 0.00690 | 0.01023 | 0.02582 | 0.02171 | 0.05588 |

| | | | | | | | | | | | | | | |
|---|---|---|---|---|---|---|---|---|---|---|---|---|---|---|
| **Kcna7** | 0.00219 | 0.00430 | 0.04747 | 0.01015 | 0.00768 | 0.01133 | 0.00421 | 0.00913 | 0.01447 | 0.00659 | 0.00853 | 0.00865 | 0.59220 | 0.00899 |
| **Kcnab1** | 0.03620 | 0.00445 | 0.18170 | 0.00000 | 0.00417 | 0.06670 | 0.00426 | 0.00557 | 0.00895 | 0.00553 | 0.00862 | 0.01133 | 0.20575 | 0.06373 |
| **Kcnab2** | 0.01640 | 0.00413 | 0.00314 | 0.30960 | 0.22066 | 0.00672 | 0.00408 | 0.01359 | 0.02181 | 0.00524 | 0.00636 | 0.00628 | 0.02872 | 0.16995 |
| **Kcnb1** | 0.00713 | 0.00390 | 0.36991 | 0.02401 | 0.00399 | 0.05659 | 0.00408 | 1.06659 | 0.01214 | 0.00627 | 0.00417 | 0.00495 | 0.00596 | 0.01651 |
| **Kcnc3** | 0.00888 | 0.00446 | 0.00977 | 0.01150 | 0.16138 | 0.02356 | 0.00416 | 0.00918 | 0.01198 | 0.00711 | 0.00818 | 0.00910 | 0.00520 | 0.01366 |
| **Kcnd1** | 0.00245 | 0.00412 | 0.02939 | 0.52474 | 0.27256 | 0.00456 | 0.00404 | 0.00725 | 0.01091 | 0.00614 | 0.00600 | 0.00870 | 0.00607 | 0.02089 |
| **Kcnd3** | 0.04487 | 0.00444 | 0.05257 | 0.01209 | 0.00018 | 0.00765 | 0.00423 | 0.01070 | 0.08223 | 0.00768 | 0.00691 | 0.00907 | 0.00990 | 0.26812 |
| **Kcne1** | 0.00585 | 0.00438 | 0.01498 | 0.01179 | 0.01126 | 0.01626 | 0.00428 | 0.00900 | 0.01119 | 0.00652 | 0.00894 | 0.00962 | 0.22320 | 0.00370 |
| **Kcne3** | 0.00000 | 0.00417 | 0.08278 | 0.24218 | 0.61070 | 0.00000 | 0.00392 | 0.00674 | 0.30227 | 0.00811 | 0.00426 | 0.01391 | 0.00130 | 0.10430 |
| **Kcnf1** | 0.00000 | 0.00451 | 2.02653 | 0.06596 | 4.07781 | 0.74017 | 2.61690 | 0.32405 | 3.15781 | 0.10621 | 0.00475 | 0.00000 | 0.00000 | 1.55762 |
| **Kcng3** | 0.00538 | 0.00438 | 0.01383 | 0.01355 | 0.07105 | 0.01320 | 0.00419 | 0.00933 | 0.00967 | 0.00697 | 0.00890 | 0.00906 | 0.23042 | 0.00308 |
| **Kcng4** | 0.01171 | 0.00421 | 0.36012 | 0.41558 | 0.11613 | 0.00917 | 0.00418 | 0.00573 | 0.01560 | 0.00470 | 0.17418 | 0.00489 | 0.02345 | 0.01461 |
| **Kcnh1** | 0.00000 | 0.00408 | 0.06154 | 0.24298 | 0.34681 | 0.00000 | 0.00393 | 0.00603 | 1.01858 | 0.00904 | 0.00325 | 0.01096 | 0.46136 | 0.00000 |
| **Kcnh2** | 0.01828 | 0.00462 | 0.07487 | 0.09888 | 0.00781 | 0.04229 | 0.00477 | 0.03815 | 0.01165 | 0.01873 | 0.00948 | 0.00580 | 0.00651 | 0.01216 |
| **Kcnh3** | 0.04806 | 0.00449 | 0.88548 | 0.12062 | 0.73541 | 0.12968 | 0.00398 | 0.00282 | 0.01400 | 0.00502 | 0.11724 | 0.00519 | 0.00230 | 0.00235 |
| **Kcnh4** | 0.00607 | 0.00437 | 0.33773 | 0.00088 | 0.06779 | 0.00554 | 0.00406 | 0.01092 | 0.00947 | 0.00788 | 0.00641 | 0.00942 | 0.02740 | 0.03412 |
| **Kcnip2** | 0.35925 | 0.00446 | 0.26207 | 0.00000 | 0.78267 | 0.00102 | 0.00429 | 0.01197 | 0.00777 | 0.00602 | 0.00569 | 0.00720 | 0.00306 | 0.11826 |
| **Kcnip3** | 0.02029 | 0.00447 | 0.02789 | 0.04903 | 0.13861 | 0.94479 | 0.00411 | 0.00492 | 0.03381 | 0.01156 | 0.01761 | 0.00266 | 0.04933 | 0.17233 |
| **Kcnip4** | 0.00918 | 0.00435 | 0.01058 | 0.05883 | 0.04477 | 0.00991 | 0.00409 | 0.00823 | 0.00945 | 0.00820 | 0.00749 | 0.01067 | 0.02570 | 0.09906 |
| **Kcnj10** | 0.01397 | 0.01987 | 0.14070 | 0.00337 | 0.01531 | 0.03000 | 0.01016 | 0.01975 | 0.09731 | 0.00693 | 0.01749 | 0.03168 | 0.01667 | 0.01019 |
| **Kcnj13** | 0.00341 | 0.00432 | 0.05077 | 0.05443 | 0.00077 | 0.04979 | 0.00400 | 0.00521 | 0.66964 | 0.00636 | 0.00648 | 0.01167 | 0.13595 | 0.00000 |
| **Kcnj14** | 0.00318 | 0.00407 | 0.00158 | 0.00000 | 0.28520 | 0.39745 | 0.00399 | 0.00364 | 0.00356 | 0.00604 | 0.00209 | 0.27652 | 0.20109 | 0.55242 |
| **Kcnj15** | 0.02360 | 0.06787 | 0.07695 | 0.07696 | 0.02102 | 0.19608 | 0.04712 | 0.09974 | 0.09939 | 0.03437 | 0.03955 | 0.12750 | 0.03835 | 0.10295 |
| **Kcnj16** | 0.02377 | 0.00413 | 0.02530 | 0.01805 | 0.01815 | 0.02397 | 0.00414 | 0.03036 | 0.06159 | 0.02944 | 0.01816 | 0.04237 | 0.01594 | 0.06016 |
| **Kcnj2** | 0.05154 | 0.00456 | 0.01153 | 0.02501 | 0.01020 | 0.01697 | 0.00425 | 0.01453 | 0.10185 | 0.00846 | 0.01006 | 0.01284 | 0.20693 | 0.01555 |
| **Kcnk4** | 0.19440 | 0.00437 | 0.11108 | 0.01386 | 0.08930 | 0.00782 | 0.00441 | 0.01229 | 0.00895 | 0.00662 | 0.00719 | 0.00793 | 0.01228 | 0.02421 |
| **Kcnk5** | 12.11572 | 8.04286 | 9.14134 | 2.46971 | 16.59416 | 2.25135 | 2.56988 | 4.72063 | 4.12090 | 0.10477 | 0.44839 | 1.63750 | 4.30411 | 19.60642 |
| **Kcnk6** | 9.00328 | 0.05800 | 4.25994 | 5.78124 | 9.56034 | 4.75869 | 1.36142 | 3.34399 | 17.32731 | 0.14784 | 0.54191 | 1.05053 | 12.09550 | 13.05444 |
| **Kcnmb3** | 0.00780 | 0.00518 | 0.03386 | 0.00342 | 0.09260 | 0.00811 | 0.08966 | 0.04470 | 0.01218 | 0.02063 | 0.00821 | 0.01029 | 0.06818 | 0.02940 |
| **Kcnn1** | 0.17844 | 0.33621 | 0.25521 | 0.61056 | 0.10575 | 0.40472 | 0.05390 | 0.20305 | 0.30575 | 0.15631 | 0.01485 | 0.07291 | 0.35339 | 0.41734 |
| **Kcnn2** | 0.02782 | 0.00433 | 0.01000 | 0.02843 | 0.18113 | 0.07100 | 0.00417 | 0.00539 | 0.01514 | 0.00572 | 0.00991 | 0.00984 | 0.33602 | 0.00384 |
| **Kcnn4** | 0.00000 | 0.03905 | 1.21959 | 0.00000 | 0.02478 | 0.00000 | 3.55815 | 0.00000 | 0.00513 | 0.11259 | 0.00000 | 2.08730 | 0.00444 | 0.01967 |
| **Kcnq1** | 0.00309 | 0.00415 | 0.00917 | 0.10577 | 0.01283 | 0.00834 | 0.00400 | 0.00638 | 0.01825 | 0.00615 | 0.00695 | 0.01397 | 0.33344 | 0.16290 |
| **Kcnq1ot1** | 6.75865 | 1.37146 | 3.46955 | 2.51034 | 3.06518 | 2.29349 | 0.76725 | 1.78758 | 14.22415 | 0.07012 | 0.22509 | 0.86169 | 3.76567 | 1.89182 |
| **Kcnq5** | 0.00992 | 0.00444 | 0.01598 | 0.01341 | 0.01272 | 0.01286 | 0.00433 | 0.01030 | 0.01125 | 0.00701 | 0.00882 | 0.01000 | 0.00813 | 0.02963 |
| **Kcnrg** | 0.00889 | 0.00453 | 0.01030 | 0.02482 | 0.10624 | 0.01612 | 0.00433 | 0.01547 | 0.01240 | 0.00801 | 0.00977 | 0.00795 | 0.00607 | 0.00943 |
| **Kcns3** | 4.79593 | 0.00929 | 5.81771 | 2.52013 | 2.71284 | 6.61928 | 0.01985 | 2.43389 | 1.89861 | 0.06298 | 0.22034 | 2.03219 | 1.98809 | 0.51521 |
| **Kcnu1** | 0.00000 | 0.00485 | 0.33919 | 0.07024 | 0.00000 | 0.53253 | 0.00390 | 0.13550 | 0.03423 | 0.01454 | 1.68541 | 0.12321 | 0.39161 | 1.07338 |
| **Kcnv2** | 0.00305 | 0.00402 | 0.00829 | 0.00001 | 0.06260 | 0.32428 | 0.00415 | 0.27382 | 0.01877 | 0.00517 | 0.00583 | 0.00940 | 0.40609 | 0.09383 |
| **Kcp** | 0.00530 | 0.00435 | 0.10456 | 0.34650 | 0.00786 | 0.01163 | 0.00420 | 0.00714 | 0.00866 | 0.00676 | 0.00636 | 0.00997 | 0.03499 | 0.01995 |
| **Kctd1** | 0.32588 | 0.01260 | 0.26594 | 0.00154 | 0.10197 | 0.42502 | 0.00460 | 0.10003 | 0.00526 | 0.00545 | 0.06885 | 0.00461 | 0.00308 | 0.19881 |
| **Kctd10** | 0.49618 | 0.07561 | 8.60316 | 4.72130 | 4.24088 | 3.75494 | 0.91323 | 1.35775 | 0.21488 | 0.01875 | 2.77460 | 2.96655 | 0.22450 | 3.18241 |
| **Kctd12** | 0.05619 | 0.01165 | 0.01184 | 0.08888 | 0.06997 | 0.01198 | 0.00411 | 0.01519 | 0.00560 | 0.00613 | 0.01078 | 0.00522 | 1.37741 | 0.00380 |
| **Kctd12b** | 0.42754 | 0.00445 | 0.01053 | 0.03586 | 0.00435 | 0.02257 | 0.00437 | 0.01145 | 0.01500 | 0.01117 | 0.00611 | 0.01144 | 0.04709 | 0.00213 |
| **Kctd13** | 2.01631 | 0.00341 | 2.78093 | 0.30536 | 0.86849 | 0.48462 | 1.50114 | 0.00000 | 0.14820 | 0.03704 | 0.01441 | 0.65113 | 3.86298 | 1.68752 |
| **Kctd14** | 0.03437 | 0.00456 | 0.01532 | 0.04946 | 0.88912 | 0.08046 | 0.01050 | 0.03730 | 0.07738 | 0.02019 | 0.01812 | 0.02702 | 0.45052 | 0.17761 |
| **Kctd15** | 0.07737 | 0.00876 | 1.15236 | 0.14615 | 3.86594 | 0.00000 | 1.20983 | 1.42074 | 0.37730 | 5.41977 | 0.00000 | 1.60210 | 0.55524 | 1.53355 |
| **Kctd17** | 0.09861 | 0.00426 | 0.00973 | 0.06312 | 0.01663 | 0.01207 | 0.00414 | 0.00987 | 0.00913 | 0.00719 | 0.00702 | 0.00730 | 0.26986 | 0.00263 |
| **Kctd18** | 1.68213 | 0.02169 | 1.44680 | 3.75774 | 2.96548 | 1.35056 | 0.53872 | 0.47012 | 0.42901 | 0.03177 | 0.04043 | 1.09961 | 4.35733 | 3.34102 |
| **Kctd2** | 0.56554 | 0.01122 | 0.85939 | 2.67043 | 1.95477 | 1.49261 | 1.87188 | 0.29740 | 0.01194 | 0.03687 | 0.08349 | 1.32338 | 0.35426 | 0.74172 |
| **Kctd20** | 6.42740 | 0.08679 | 5.63905 | 5.53414 | 5.54021 | 7.68465 | 0.06156 | 0.59804 | 4.36083 | 0.03448 | 1.97603 | 5.74173 | 6.75581 | 6.28325 |
| **Kctd21** | 0.14424 | 0.00432 | 0.32325 | 0.00000 | 0.00000 | 0.18720 | 0.00421 | 0.00497 | 0.00859 | 0.01241 | 0.00656 | 0.03212 | 0.06935 | 0.00141 |
| **Kctd3** | 4.82232 | 2.02974 | 4.47022 | 0.00000 | 3.83781 | 5.50190 | 3.48088 | 0.00000 | 0.18607 | 0.12045 | 0.00000 | 6.70217 | 7.43902 | 7.64938 |
| **Kctd4** | 0.00249 | 0.00435 | 0.00406 | 0.06574 | 0.00000 | 0.00882 | 0.00413 | 0.00899 | 0.06402 | 0.00625 | 0.00621 | 0.00917 | 0.00875 | 0.81676 |
| **Kctd5** | 7.32487 | 0.04625 | 6.84024 | 2.15655 | 7.35831 | 7.15514 | 3.14889 | 3.81300 | 0.00000 | 0.09449 | 4.99134 | 3.78766 | 1.96942 | 5.53333 |

| | | | | | | | | | | | | | | |
|---|---|---|---|---|---|---|---|---|---|---|---|---|---|---|
| **Kctd6** | 0.03815 | 0.08421 | 0.12989 | 1.89184 | 0.21892 | 1.69000 | 0.04302 | 2.50063 | 0.14440 | 0.02401 | 0.60813 | 0.13042 | 2.03017 | 1.22693 |
| **Kctd7** | 0.00000 | 0.00399 | 0.39295 | 0.08842 | 0.10309 | 0.00000 | 0.00386 | 0.00603 | 0.40463 | 0.00760 | 0.00507 | 0.01016 | 1.34671 | 0.21830 |
| **Kctd9** | 3.13525 | 0.02469 | 6.40914 | 2.35852 | 1.49563 | 8.35996 | 3.92372 | 2.83353 | 6.72206 | 0.09864 | 0.93175 | 1.39252 | 1.71614 | 0.00000 |
| **Kdelc1** | 0.00000 | 0.01025 | 0.29399 | 0.00150 | 0.00105 | 0.00000 | 0.00351 | 0.00398 | 0.63812 | 0.00743 | 0.14941 | 1.87985 | 0.00739 | 0.00158 |
| **Kdelc2** | 0.34688 | 0.15070 | 0.00241 | 0.37305 | 0.17073 | 1.26050 | 0.07246 | 0.24859 | 0.30867 | 0.10183 | 0.20209 | 0.12964 | 2.32992 | 0.62193 |
| **Kdelr1** | 0.00000 | 0.08963 | 2.42307 | 6.54513 | 1.34446 | 1.39190 | 1.42122 | 6.47641 | 0.12688 | 10.32958 | 6.37898 | 4.73621 | 1.03999 | 0.00000 |
| **Kdelr2** | 1.36753 | 0.03328 | 1.18430 | 6.78814 | 1.31903 | 4.47275 | 2.44529 | 1.52254 | 0.07237 | 0.07620 | 0.24075 | 2.73056 | 0.21259 | 0.81569 |
| **Kdelr3** | 1.10836 | 0.13319 | 1.25966 | 3.86425 | 0.65450 | 2.12163 | 0.80322 | 0.00494 | 0.00001 | 0.06664 | 0.00000 | 3.38344 | 0.00000 | 2.01550 |
| **Kdf1** | 0.00000 | 0.11164 | 1.11456 | 3.15362 | 1.81723 | 0.00000 | 0.26541 | 4.13874 | 0.01034 | 0.02713 | 0.00000 | 2.53477 | 0.51502 | 0.13869 |
| **Kdm1a** | 1.98299 | 0.00000 | 4.95192 | 0.00000 | 1.60807 | 3.18095 | 1.07804 | 0.59551 | 14.87180 | 0.02967 | 4.91326 | 3.40227 | 1.60652 | 16.09909 |
| **Kdm1b** | 0.94632 | 0.01718 | 0.00000 | 0.49582 | 1.37468 | 0.32702 | 0.00355 | 1.02995 | 1.63180 | 0.00875 | 3.17190 | 0.04954 | 2.00693 | 1.15700 |
| **Kdm2a** | 0.19543 | 0.01145 | 0.82419 | 0.00000 | 1.09153 | 2.32627 | 0.00360 | 0.95967 | 0.14258 | 0.01761 | 0.00000 | 1.46948 | 2.96518 | 1.15563 |
| **Kdm2b** | 18.40196 | 3.47169 | 23.95850 | 15.80780 | 23.32075 | 12.24072 | 1.47098 | 4.31293 | 19.15243 | 0.07142 | 11.45762 | 6.60356 | 10.41663 | 15.40682 |
| **Kdm3a** | 7.47434 | 0.00000 | 7.64479 | 0.85519 | 7.54086 | 1.50564 | 3.22460 | 0.01898 | 11.88168 | 0.03256 | 4.11316 | 3.64774 | 7.76155 | 7.70759 |
| **Kdm3b** | 3.45976 | 0.75658 | 12.64472 | 3.57164 | 8.59300 | 6.01998 | 0.78864 | 3.27958 | 11.52376 | 0.08144 | 10.06885 | 1.96943 | 0.07562 | 12.66167 |
| **Kdm4a** | 0.12196 | 0.00439 | 0.01126 | 0.02651 | 0.11084 | 0.20485 | 0.01782 | 0.07677 | 0.07262 | 0.00811 | 0.06542 | 0.07304 | 3.47617 | 0.16041 |
| **Kdm4b** | 0.76279 | 0.02795 | 2.08462 | 1.58047 | 0.83965 | 1.24890 | 0.00397 | 1.57031 | 0.00426 | 0.03362 | 0.21356 | 0.98414 | 1.33721 | 0.06347 |
| **Kdm4c** | 5.77757 | 0.01945 | 1.94056 | 0.00000 | 2.48539 | 1.32761 | 0.19717 | 0.00000 | 16.77520 | 0.00754 | 0.84937 | 0.83924 | 4.76671 | 11.50817 |
| **Kdm4d** | 0.39691 | 0.00429 | 0.00867 | 0.01113 | 0.07236 | 0.00987 | 0.00431 | 0.00975 | 0.00808 | 0.00668 | 0.00690 | 0.00658 | 0.80150 | 0.09567 |
| **Kdm5a** | 4.93387 | 3.95586 | 2.12958 | 2.40087 | 4.70046 | 2.56024 | 1.04082 | 2.34727 | 15.39139 | 0.01050 | 1.44954 | 1.07811 | 4.48871 | 8.02610 |
| **Kdm5b** | 8.09299 | 10.74826 | 10.27484 | 5.54863 | 16.95658 | 2.75472 | 2.79456 | 0.35620 | 20.04074 | 0.11456 | 4.53040 | 3.93730 | 14.18571 | 19.70105 |
| **Kdm5c** | 1.13548 | 1.50870 | 3.31021 | 4.42933 | 5.38569 | 8.07759 | 0.01246 | 1.58685 | 9.07824 | 0.01269 | 0.53526 | 3.28265 | 1.39112 | 6.84987 |
| **Kdm5d** | 0.01257 | 0.00400 | 0.00821 | 0.31502 | 0.01298 | 0.00653 | 0.01577 | 0.00473 | 0.01073 | 0.01540 | 0.00808 | 0.08088 | 0.01075 | 0.02418 |
| **Kdm6a** | 2.74019 | 0.00017 | 5.69066 | 0.71883 | 2.04691 | 3.36676 | 0.20801 | 0.47120 | 17.76719 | 0.00000 | 5.78441 | 2.56261 | 0.11515 | 4.50991 |
| **Kdm6b** | 0.06022 | 0.61499 | 0.29454 | 2.14866 | 1.43270 | 1.08702 | 0.10453 | 3.38245 | 0.01536 | 0.08396 | 0.15343 | 2.83265 | 0.19524 | 0.03034 |
| **Kdm7a** | 4.95058 | 0.02155 | 1.50310 | 0.58867 | 6.44469 | 1.06404 | 2.35853 | 0.03707 | 16.21786 | 0.20830 | 0.62899 | 1.43486 | 7.14878 | 11.48826 |
| **Kdm8** | 0.08931 | 0.00421 | 0.19249 | 0.00000 | 0.02033 | 0.07289 | 0.02736 | 0.00612 | 0.23722 | 0.00883 | 0.08815 | 0.03229 | 0.35568 | 1.07770 |
| **Kdsr** | 6.43875 | 0.08678 | 1.14725 | 0.52896 | 0.29911 | 4.37688 | 2.41572 | 1.03417 | 2.38333 | 0.14065 | 3.13212 | 0.50359 | 0.93710 | 2.71952 |
| **Keap1** | 5.40287 | 1.84425 | 5.53604 | 5.71059 | 1.95319 | 4.28439 | 6.10124 | 8.93279 | 2.00509 | 0.85911 | 1.70397 | 3.12854 | 0.26009 | 1.74836 |
| **Kel** | 0.00469 | 0.01276 | 0.00082 | 0.36013 | 0.00549 | 0.54926 | 0.00519 | 0.35546 | 0.01769 | 0.00475 | 0.13712 | 0.00987 | 0.00766 | 0.00296 |
| **Khdc1a** | 25.54620 | 2.99857 | 13.80157 | 7.62932 | 26.11626 | 7.63081 | 0.87601 | 13.59423 | 28.59921 | 12.14233 | 0.66208 | 0.00000 | 11.96211 | 16.03273 |
| **Khdc1b** | 0.54633 | 0.00244 | 0.28107 | 1.20347 | 5.05106 | 0.00000 | 0.00304 | 0.22153 | 10.04962 | 0.01371 | 0.00000 | 0.00000 | 9.53907 | 21.34396 |
| **Khdc1c** | 3.62458 | 0.00351 | 0.29196 | 1.81913 | 3.24908 | 0.13294 | 0.00323 | 0.44615 | 5.87147 | 0.01684 | 0.05654 | 0.00000 | 5.42878 | 6.26481 |
| **Khdc3** | 3.63450 | 5.70541 | 0.67169 | 20.95926 | 4.89620 | 6.08661 | 2.97254 | 5.57111 | 0.35006 | 7.55895 | 1.95933 | 14.04436 | 7.71291 | 0.00000 |
| **Khdrbs1** | 5.90357 | 0.06556 | 9.79889 | 5.22375 | 9.44506 | 7.11884 | 2.24714 | 2.80051 | 9.18804 | 0.05533 | 5.64419 | 5.30741 | 14.14527 | 8.97061 |
| **Khdrbs3** | 0.35188 | 0.00445 | 0.00434 | 0.04585 | 0.07561 | 0.00836 | 0.00435 | 0.01257 | 0.00734 | 0.00617 | 0.00646 | 0.00657 | 0.00396 | 0.50678 |
| **Khk** | 0.02408 | 0.01917 | 0.56635 | 0.06135 | 0.32626 | 0.35620 | 0.00380 | 0.55227 | 0.00719 | 0.00655 | 0.01527 | 0.00728 | 0.37848 | 0.03381 |
| **Khnyn** | 0.92428 | 0.03744 | 2.49170 | 5.24154 | 2.30452 | 2.85445 | 0.02112 | 1.69710 | 0.44924 | 0.03458 | 2.40609 | 0.53021 | 0.07070 | 2.21204 |
| **Khsrp** | 1.36458 | 0.52394 | 2.13011 | 1.31312 | 2.17958 | 2.69651 | 0.04936 | 0.97685 | 1.06741 | 0.03794 | 0.04385 | 1.54904 | 9.58102 | 1.37892 |
| **Kidins220** | 7.16978 | 0.00098 | 7.32521 | 3.02919 | 5.32092 | 2.65308 | 0.98230 | 0.44096 | 13.06039 | 0.00300 | 1.67271 | 2.13688 | 1.82054 | 6.29599 |
| **Kif11** | 6.97748 | 0.05800 | 5.93441 | 0.90967 | 2.87964 | 4.07435 | 0.07055 | 0.08504 | 14.50929 | 0.85581 | 1.57691 | 3.87522 | 7.97432 | 9.02315 |
| **Kif13a** | 0.00162 | 0.00393 | 0.25802 | 0.00000 | 0.00000 | 1.80126 | 0.04278 | 1.37403 | 0.20592 | 0.00000 | 2.91005 | 0.38217 | 0.00402 | 1.35964 |
| **Kif13b** | 1.22703 | 0.00373 | 0.72751 | 1.13215 | 0.03771 | 0.00000 | 0.00391 | 0.40801 | 0.46672 | 0.00463 | 0.00374 | 0.00836 | 0.58000 | 0.55762 |
| **Kif14** | 0.76555 | 0.00275 | 0.77861 | 1.25312 | 1.69620 | 0.76362 | 0.01005 | 1.12431 | 2.07549 | 0.01983 | 2.27077 | 0.65871 | 3.01601 | 0.00000 |
| **Kif15** | 0.00000 | 0.03900 | 1.21444 | 1.42031 | 2.60545 | 0.57458 | 0.00968 | 0.16089 | 2.00696 | 0.02112 | 0.11043 | 1.28988 | 0.08878 | 1.29397 |
| **Kif16b** | 4.67565 | 0.01284 | 8.40837 | 2.71513 | 8.80254 | 2.29374 | 0.15792 | 0.00000 | 8.58272 | 0.00652 | 3.55437 | 5.04470 | 0.74122 | 8.11777 |
| **Kif17** | 0.00000 | 0.00378 | 0.22100 | 0.24612 | 0.01245 | 0.00000 | 0.00373 | 0.00523 | 0.40305 | 0.00551 | 0.00512 | 0.01320 | 2.33001 | 0.13357 |
| **Kif18a** | 7.73352 | 0.00130 | 1.80761 | 1.85502 | 4.56595 | 0.55537 | 0.00662 | 1.31693 | 17.26928 | 0.01119 | 1.44708 | 0.65394 | 11.11213 | 2.13184 |
| **Kif18b** | 2.23603 | 0.00343 | 1.81787 | 2.02669 | 3.88105 | 0.85283 | 0.00335 | 0.01310 | 0.58476 | 0.01671 | 1.13985 | 0.23295 | 6.94161 | 1.05894 |
| **Kif1a** | 0.06155 | 0.00463 | 0.02975 | 0.00625 | 0.35076 | 0.00994 | 0.00418 | 0.01423 | 0.00857 | 0.00673 | 0.00790 | 0.00820 | 0.03482 | 0.11354 |
| **Kif1b** | 6.27403 | 0.05181 | 7.33264 | 1.54652 | 6.23202 | 6.27810 | 1.47605 | 0.44210 | 11.32265 | 0.05160 | 4.11366 | 2.14270 | 6.36501 | 10.34636 |
| **Kif1bp** | 5.18191 | 0.00000 | 3.46246 | 0.63671 | 5.63970 | 6.81807 | 3.21431 | 1.80353 | 2.49711 | 0.09716 | 5.83127 | 2.10858 | 6.96115 | 5.83175 |
| **Kif1c** | 0.13243 | 2.99833 | 0.05134 | 0.00000 | 1.15550 | 0.88267 | 2.11680 | 2.26302 | 0.14442 | 0.07207 | 0.00000 | 0.87532 | 0.95653 | 0.35594 |
| **Kif20a** | 4.50723 | 0.19024 | 19.50504 | 1.29564 | 15.62454 | 7.29087 | 5.95680 | 5.88504 | 0.52833 | 0.16453 | 12.45862 | 3.67767 | 0.07453 | 19.19187 |
| **Kif20b** | 1.52294 | 0.01763 | 0.15723 | 0.00000 | 1.08196 | 0.50910 | 0.00754 | 0.00000 | 11.34275 | 0.00483 | 0.10038 | 0.47383 | 8.20361 | 8.84044 |

| | | | | | | | | | | | | | | |
|---|---|---|---|---|---|---|---|---|---|---|---|---|---|---|
| **Kif21a** | 0.01431 | 0.00506 | 0.07235 | 0.00000 | 0.01242 | 0.00253 | 0.00437 | 0.04727 | 1.11255 | 0.00811 | 0.00305 | 0.21276 | 0.00437 | 0.00000 |
| **Kif21b** | 0.40724 | 0.00870 | 0.56738 | 0.00000 | 0.19914 | 1.03277 | 0.00316 | 0.09498 | 0.00843 | 0.00182 | 1.12117 | 3.26112 | 0.00550 | 0.00420 |
| **Kif22** | 12.31357 | 0.22573 | 21.27377 | 8.40749 | 24.49167 | 10.06436 | 3.37463 | 7.33633 | 2.02885 | 0.16051 | 7.24215 | 8.24276 | 12.57870 | 26.46525 |
| **Kif23** | 16.08268 | 0.06579 | 24.42191 | 3.69468 | 16.55550 | 8.46761 | 7.05922 | 7.64019 | 21.82697 | 6.25480 | 8.40224 | 3.55012 | 10.76818 | 23.46866 |
| **Kif24** | 0.80400 | 0.05981 | 0.13208 | 0.00000 | 0.96878 | 1.50621 | 0.04883 | 0.04946 | 1.08398 | 0.01798 | 1.57756 | 0.82785 | 0.75728 | 1.86946 |
| **Kif26a** | 0.00731 | 0.00475 | 0.18308 | 0.03107 | 0.00726 | 0.00599 | 0.00419 | 0.01343 | 0.01332 | 0.00588 | 0.00490 | 0.18281 | 0.11675 | 0.01079 |
| **Kif27** | 0.00170 | 0.00484 | 0.00000 | 0.02448 | 0.28185 | 0.35803 | 0.03226 | 0.00831 | 0.07236 | 0.00816 | 0.00724 | 0.00316 | 1.24568 | 0.34774 |
| **Kif2a** | 5.40629 | 0.13014 | 5.78354 | 1.02969 | 5.92833 | 5.45240 | 0.92939 | 1.29303 | 17.09421 | 0.06138 | 2.40655 | 4.41881 | 7.78567 | 7.06494 |
| **Kif2c** | 1.43350 | 0.03482 | 6.74046 | 2.93796 | 7.19854 | 1.76111 | 4.06004 | 3.60539 | 1.05673 | 13.32192 | 8.84962 | 3.17574 | 7.87150 | 9.01679 |
| **Kif3a** | 1.60093 | 0.02698 | 1.04251 | 5.11684 | 1.28783 | 1.91004 | 0.38825 | 0.63604 | 0.48919 | 0.07123 | 0.14026 | 1.42664 | 1.21559 | 3.42380 |
| **Kif3b** | 0.00000 | 0.00326 | 0.65197 | 0.64713 | 0.25550 | 1.74654 | 0.02612 | 0.00000 | 0.00169 | 0.00437 | 0.00000 | 0.78379 | 1.78751 | 3.13812 |
| **Kif3c** | 0.90433 | 0.00249 | 4.21792 | 0.62758 | 3.17229 | 0.11881 | 0.00313 | 0.59480 | 4.65736 | 0.00000 | 2.91571 | 0.00053 | 5.18192 | 12.32398 |
| **Kif4** | 12.64889 | 0.04803 | 10.43846 | 2.63083 | 12.30930 | 3.95010 | 0.00525 | 1.80012 | 13.55488 | 0.02425 | 2.75313 | 5.03396 | 7.94746 | 13.85103 |
| **Kif5a** | 0.04713 | 0.00386 | 0.19036 | 0.00899 | 0.11776 | 0.07568 | 0.00982 | 0.82924 | 0.00967 | 0.01847 | 0.01623 | 0.00474 | 0.17886 | 0.00268 |
| **Kif5b** | 9.95580 | 3.41736 | 12.49521 | 3.83223 | 13.95585 | 12.09342 | 0.38196 | 2.94144 | 22.35891 | 0.11881 | 9.75960 | 6.06736 | 3.39647 | 17.72742 |
| **Kif5c** | 0.00000 | 0.00400 | 0.00000 | 0.01725 | 0.00000 | 0.00000 | 0.00402 | 0.00835 | 2.29886 | 0.00811 | 0.00403 | 0.00381 | 2.71794 | 3.60610 |
| **Kif7** | 0.06114 | 0.00481 | 0.01750 | 0.13959 | 0.01767 | 0.05002 | 0.00441 | 0.04340 | 0.01212 | 0.01331 | 0.00474 | 0.06510 | 0.13370 | 0.01725 |
| **Kif9** | 1.31439 | 0.02540 | 1.77165 | 0.00000 | 1.06077 | 1.00261 | 0.00320 | 0.00390 | 1.96259 | 0.00739 | 0.00000 | 1.11223 | 2.50874 | 2.07516 |
| **Kifap3** | 1.04363 | 0.00296 | 0.50300 | 1.27502 | 2.36572 | 1.48275 | 0.00362 | 0.37524 | 0.00298 | 0.00000 | 4.82493 | 0.00217 | 0.00000 | 2.66237 |
| **Kifc1** | 2.86427 | 0.05592 | 4.31322 | 3.41586 | 1.48226 | 2.25181 | 0.03435 | 1.68543 | 0.08363 | 0.02171 | 0.05811 | 2.30806 | 6.37844 | 1.96891 |
| **Kifc2** | 1.64727 | 0.50746 | 0.25262 | 0.65526 | 0.53551 | 0.24083 | 0.00872 | 0.92917 | 0.00171 | 0.02233 | 0.00000 | 5.18302 | 2.48101 | 0.77623 |
| **Kifc3** | 6.58832 | 0.07403 | 18.06733 | 4.78677 | 13.08654 | 0.42180 | 0.41020 | 1.45062 | 0.22606 | 0.00000 | 0.44265 | 5.33402 | 5.35192 | 1.51033 |
| **Kifc5b** | 1.39494 | 0.03754 | 1.96453 | 2.77271 | 0.77059 | 1.22217 | 0.01699 | 1.17731 | 0.11335 | 0.01534 | 0.39518 | 1.54567 | 3.05690 | 1.55935 |
| **Kin** | 0.80458 | 0.01795 | 7.00923 | 2.55093 | 4.07671 | 1.28748 | 0.64853 | 2.43717 | 6.08548 | 3.96154 | 5.44393 | 1.44451 | 3.76466 | 5.23556 |
| **Kirrel** | 0.07073 | 0.02213 | 0.00959 | 0.00000 | 0.01163 | 0.13940 | 0.00403 | 0.00938 | 0.01235 | 0.00561 | 0.00331 | 0.59356 | 0.00520 | 0.01497 |
| **Kirrel2** | 0.00526 | 0.00431 | 0.11046 | 0.00932 | 0.05410 | 0.01066 | 0.00412 | 0.01298 | 0.00913 | 0.00713 | 0.00903 | 0.01009 | 0.09219 | 0.02556 |
| **Kiss1r** | 0.00377 | 0.00403 | 0.00520 | 0.68716 | 0.03641 | 0.00372 | 0.00399 | 0.00623 | 0.01193 | 0.00528 | 0.00635 | 0.01939 | 0.00821 | 0.00853 |
| **Kit** | 0.31447 | 0.00338 | 0.54015 | 0.02471 | 0.24679 | 0.00000 | 0.00393 | 0.00529 | 0.24954 | 0.00603 | 0.00388 | 0.49694 | 1.35225 | 4.12069 |
| **Kitl** | 0.00000 | 0.00430 | 0.00151 | 0.01440 | 0.00076 | 0.01007 | 0.00402 | 0.00837 | 1.22084 | 0.00684 | 0.00618 | 0.01116 | 0.49591 | 0.00000 |
| **Kiz** | 4.72417 | 0.01554 | 2.83404 | 0.32273 | 3.55669 | 1.29406 | 0.00153 | 2.36186 | 5.25313 | 0.01582 | 3.08076 | 1.10627 | 3.95252 | 1.17730 |
| **Kl** | 0.01745 | 0.13321 | 0.10640 | 0.01362 | 0.05822 | 0.13059 | 0.03627 | 0.02110 | 0.31096 | 0.03134 | 0.00877 | 0.03724 | 0.20623 | 0.76006 |
| **Klb** | 0.00000 | 0.06393 | 3.16095 | 4.19400 | 2.00146 | 0.83068 | 0.01541 | 0.78763 | 1.12847 | 0.00000 | 5.55752 | 3.92624 | 4.02772 | 4.30766 |
| **Klc1** | 0.02859 | 0.04546 | 1.15393 | 2.38131 | 1.32020 | 0.00000 | 0.46549 | 0.79072 | 0.01733 | 0.02902 | 0.01586 | 2.53136 | 0.14489 | 0.00000 |
| **Klc2** | 0.01008 | 0.00439 | 0.07691 | 0.00000 | 0.24952 | 0.00777 | 0.00433 | 0.01486 | 0.01365 | 0.00790 | 0.00684 | 0.01281 | 0.14657 | 0.45760 |
| **Klc3** | 0.00334 | 0.00385 | 0.29843 | 0.00000 | 0.34190 | 0.00273 | 0.00412 | 0.00518 | 0.00863 | 0.00568 | 0.00748 | 0.65946 | 0.01090 | 0.02200 |
| **Klc4** | 3.95908 | 0.03050 | 5.75052 | 4.80135 | 4.50638 | 3.98890 | 0.02955 | 1.00979 | 0.00172 | 0.07474 | 1.04251 | 0.53702 | 2.13865 | 1.44104 |
| **Klf1** | 0.00964 | 0.00530 | 0.01023 | 0.09861 | 0.03628 | 0.01120 | 0.00446 | 0.08475 | 0.01017 | 0.00757 | 0.00712 | 0.00784 | 0.01841 | 0.02574 |
| **Klf10** | 0.43307 | 0.65964 | 7.91840 | 0.90085 | 8.24454 | 2.08827 | 0.00655 | 6.24386 | 7.93680 | 0.03923 | 1.16806 | 0.26478 | 1.39917 | 1.09736 |
| **Klf11** | 7.27635 | 0.00100 | 3.64101 | 0.57921 | 6.16078 | 1.06480 | 1.83971 | 1.54335 | 0.29751 | 0.06418 | 0.00000 | 1.95380 | 9.71676 | 5.58757 |
| **Klf13** | 0.62516 | 5.31178 | 5.19959 | 1.24864 | 4.32062 | 3.35724 | 2.37328 | 1.81115 | 5.97262 | 2.48118 | 1.41714 | 4.89640 | 2.67984 | 6.36942 |
| **Klf15** | 0.00143 | 0.01534 | 0.08912 | 0.00000 | 0.04365 | 0.85875 | 0.01066 | 0.62886 | 0.00777 | 0.00626 | 0.33274 | 0.00822 | 0.21617 | 0.00000 |
| **Klf16** | 0.45320 | 0.01075 | 0.74883 | 3.10752 | 5.11406 | 1.38605 | 0.78307 | 1.31375 | 1.82296 | 0.03249 | 0.00000 | 0.66357 | 6.91044 | 5.75988 |
| **Klf17** | 11.64506 | 0.00000 | 3.09931 | 0.00000 | 24.04957 | 0.00000 | 2.30006 | 0.00000 | 24.86651 | 0.03557 | 0.00000 | 0.00000 | 18.38371 | 37.14209 |
| **Klf2** | 9.46627 | 0.00479 | 1.82225 | 0.78899 | 17.41988 | 0.00000 | 0.06514 | 0.90555 | 0.26407 | 0.01093 | 0.07488 | 6.76091 | 10.47115 | 1.67258 |
| **Klf3** | 0.62249 | 0.02626 | 1.58772 | 0.00000 | 1.32653 | 1.31158 | 1.69166 | 0.29435 | 0.01415 | 0.03251 | 2.06756 | 1.35034 | 1.26156 | 0.01950 |
| **Klf4** | 10.51259 | 8.04022 | 13.46617 | 6.50779 | 13.61282 | 2.07388 | 2.67200 | 2.84935 | 5.46170 | 0.05287 | 5.91832 | 10.94412 | 6.25634 | 8.40316 |
| **Klf5** | 0.00000 | 6.17293 | 13.62380 | 8.45608 | 25.35794 | 0.00000 | 4.59682 | 10.86162 | 0.48347 | 7.50850 | 1.33115 | 9.23578 | 0.00000 | 0.00000 |
| **Klf6** | 0.03619 | 5.17228 | 0.47843 | 0.48652 | 0.00982 | 3.02263 | 6.70636 | 6.68943 | 0.00822 | 0.31789 | 1.81402 | 0.56426 | 0.00396 | 0.00000 |
| **Klf7** | 0.01023 | 0.00403 | 0.01384 | 0.00000 | 0.02474 | 0.00356 | 0.00388 | 0.67786 | 0.02558 | 0.00531 | 0.00826 | 0.01359 | 0.26206 | 0.04228 |
| **Klf8** | 1.40249 | 0.00828 | 2.49700 | 0.66668 | 3.90279 | 1.64492 | 0.00132 | 1.29104 | 6.40911 | 0.00814 | 0.19365 | 4.34474 | 0.01292 | 0.50462 |
| **Klf9** | 0.34557 | 4.25252 | 5.48605 | 3.79847 | 5.81797 | 5.75169 | 0.89286 | 2.79639 | 6.97153 | 0.67547 | 0.00000 | 6.54642 | 5.18582 | 9.09133 |
| **Klhdc1** | 0.01472 | 0.00431 | 0.25986 | 0.00000 | 0.02043 | 0.39762 | 2.07370 | 0.00343 | 1.85735 | 0.00375 | 0.00472 | 0.00284 | 0.00000 | 0.99449 |
| **Klhdc10** | 2.86235 | 0.11557 | 7.19360 | 4.44677 | 5.57881 | 6.35570 | 2.55088 | 2.14125 | 5.38436 | 0.13906 | 0.29409 | 3.31222 | 2.34620 | 6.80880 |
| **Klhdc2** | 4.91428 | 0.02087 | 3.32250 | 3.78502 | 3.19468 | 0.91847 | 0.01318 | 4.50700 | 0.25441 | 0.01786 | 4.16635 | 3.80047 | 8.09378 | 7.23102 |
| **Klhdc3** | 0.34468 | 0.64151 | 7.57378 | 5.61643 | 1.24582 | 7.59996 | 0.02931 | 8.27933 | 0.00000 | 0.04792 | 1.15408 | 6.58146 | 0.43454 | 0.01255 |

| | | | | | | | | | | | | | | |
|---|---|---|---|---|---|---|---|---|---|---|---|---|---|---|
| **Klhdc4** | 11.16095 | 0.06478 | 11.43913 | 16.82893 | 9.99696 | 14.67855 | 0.74327 | 2.45063 | 0.83869 | 0.10534 | 6.45381 | 5.41430 | 4.16111 | 10.58358 |
| **Klhdc9** | 2.73345 | 0.02053 | 0.56764 | 1.84421 | 1.44610 | 1.14369 | 0.01791 | 1.21574 | 0.50555 | 0.02171 | 0.08188 | 1.03244 | 0.12466 | 2.01029 |
| **Klhl10** | 1.05556 | 0.00367 | 2.79425 | 0.52827 | 5.72110 | 0.00834 | 0.00305 | 0.00000 | 3.30794 | 0.01208 | 0.00000 | 0.00223 | 0.37429 | 7.07199 |
| **Klhl11** | 1.25022 | 0.00240 | 0.98238 | 0.08060 | 0.78715 | 0.39238 | 0.00356 | 0.00000 | 0.72973 | 0.00285 | 0.40657 | 0.14135 | 12.00283 | 4.13019 |
| **Klhl12** | 5.17453 | 7.70270 | 5.16192 | 1.91036 | 8.10784 | 5.07454 | 2.40563 | 0.06696 | 13.14866 | 0.05876 | 0.60137 | 3.16877 | 5.37338 | 10.85258 |
| **Klhl13** | 0.01167 | 0.00470 | 0.03273 | 0.00398 | 0.01206 | 0.01121 | 0.00393 | 0.01961 | 0.01899 | 0.00806 | 0.00436 | 0.41317 | 0.04449 | 0.02030 |
| **Klhl15** | 10.09242 | 0.00898 | 0.66098 | 0.00000 | 1.59066 | 0.46379 | 0.02053 | 0.37444 | 0.03737 | 0.00558 | 0.02454 | 1.22205 | 2.30569 | 3.62854 |
| **Klhl17** | 0.00460 | 0.09016 | 0.07970 | 0.00000 | 0.48273 | 0.00000 | 0.00387 | 0.83485 | 0.00874 | 0.00261 | 1.24123 | 0.09086 | 0.00387 | 0.40332 |
| **Klhl18** | 0.16023 | 0.00364 | 0.68267 | 0.00000 | 2.13744 | 0.02181 | 0.06889 | 1.45051 | 0.08199 | 0.00114 | 1.44634 | 0.00168 | 1.78427 | 1.13490 |
| **Klhl2** | 1.53813 | 0.01521 | 3.15458 | 1.50552 | 1.90224 | 2.09120 | 0.00156 | 1.79918 | 2.25559 | 0.02066 | 2.57705 | 3.31690 | 0.88003 | 4.24255 |
| **Klhl20** | 0.19976 | 0.00361 | 1.85192 | 0.00000 | 0.93156 | 2.24666 | 0.00352 | 0.27759 | 0.12724 | 0.02044 | 0.00000 | 2.13904 | 0.09819 | 3.10856 |
| **Klhl21** | 5.91257 | 0.52589 | 8.94255 | 5.38376 | 9.00268 | 8.26235 | 2.75277 | 1.84125 | 7.64552 | 0.02204 | 0.00000 | 5.45602 | 6.98342 | 10.88288 |
| **Klhl22** | 0.51549 | 0.02888 | 4.77618 | 7.45053 | 0.00000 | 6.62350 | 0.00257 | 0.54516 | 0.00000 | 0.01747 | 2.50471 | 2.42285 | 0.12196 | 3.02362 |
| **Klhl23** | 0.00000 | 0.01149 | 0.00000 | 0.15905 | 0.00000 | 0.00000 | 1.25130 | 0.01040 | 1.36658 | 2.77892 | 0.00000 | 0.00682 | 0.85744 | 0.00000 |
| **Klhl24** | 2.31312 | 0.00228 | 0.89069 | 0.00000 | 1.89673 | 2.77343 | 0.00307 | 1.83143 | 0.09708 | 0.00615 | 1.49569 | 0.75192 | 4.70179 | 2.97852 |
| **Klhl25** | 0.21269 | 0.01171 | 0.43702 | 0.97503 | 0.89186 | 0.00000 | 0.02608 | 0.40314 | 0.00886 | 0.01069 | 0.99741 | 0.23972 | 0.00955 | 0.57271 |
| **Klhl26** | 0.01123 | 0.00431 | 0.00000 | 1.49213 | 0.00399 | 1.64201 | 0.37746 | 0.60706 | 0.02998 | 0.01963 | 0.00490 | 0.05845 | 0.00707 | 0.00679 |
| **Klhl28** | 0.00000 | 0.00444 | 0.38642 | 0.03485 | 0.11631 | 0.00000 | 0.01080 | 0.04866 | 0.00760 | 0.01691 | 0.00829 | 0.00423 | 0.56168 | 1.48126 |
| **Klhl3** | 0.00875 | 0.00410 | 0.00469 | 0.10704 | 0.00685 | 0.00944 | 0.00432 | 0.00608 | 0.14227 | 0.00603 | 0.01007 | 0.17205 | 0.00611 | 0.00128 |
| **Klhl30** | 0.02621 | 0.00450 | 0.01190 | 0.00189 | 0.01360 | 0.30825 | 0.00418 | 0.00573 | 0.01070 | 0.00647 | 0.00616 | 0.01075 | 0.00737 | 0.00812 |
| **Klhl32** | 0.00582 | 0.00427 | 0.01943 | 0.21106 | 0.00444 | 0.00739 | 0.00419 | 0.00813 | 0.01349 | 0.00680 | 0.00698 | 0.01351 | 0.02064 | 0.16295 |
| **Klhl35** | 0.01342 | 0.00444 | 0.01161 | 0.01430 | 0.05222 | 0.01248 | 0.00418 | 0.01011 | 0.00819 | 0.00734 | 0.00857 | 0.00948 | 0.01512 | 0.08214 |
| **Klhl36** | 0.00650 | 0.00363 | 0.30409 | 0.00000 | 0.18212 | 0.01576 | 0.00929 | 0.48241 | 0.00956 | 0.00657 | 0.00513 | 0.48091 | 0.01751 | 0.00975 |
| **Klhl41** | 1.47670 | 0.27752 | 0.00000 | 3.80927 | 0.81336 | 4.14429 | 0.58189 | 1.88821 | 0.74595 | 1.65846 | 0.05210 | 0.83370 | 3.78773 | 1.85590 |
| **Klhl42** | 0.06565 | 0.02545 | 0.31004 | 0.43570 | 0.00000 | 0.11033 | 0.00332 | 0.01099 | 0.01067 | 0.00614 | 0.00313 | 2.31531 | 0.00753 | 0.08970 |
| **Klhl5** | 0.00553 | 0.00412 | 0.02686 | 0.57733 | 0.29899 | 0.00709 | 0.00404 | 0.00800 | 0.12722 | 0.00663 | 0.00475 | 0.00812 | 0.00226 | 0.05061 |
| **Klhl6** | 0.00426 | 0.02076 | 0.01550 | 0.00704 | 0.00852 | 0.02285 | 0.00504 | 0.08162 | 0.01400 | 0.00585 | 0.00906 | 0.00726 | 0.77156 | 0.00992 |
| **Klhl7** | 1.04488 | 0.11617 | 1.94580 | 1.72671 | 0.56188 | 1.39102 | 0.17306 | 0.36416 | 1.50499 | 0.26296 | 0.13998 | 0.45084 | 7.08190 | 3.67056 |
| **Klhl8** | 0.00000 | 0.00298 | 0.73181 | 0.00000 | 1.49694 | 0.38051 | 0.00356 | 0.00034 | 0.06768 | 0.00459 | 0.00142 | 0.30060 | 10.74555 | 3.87026 |
| **Klhl9** | 0.89597 | 0.00341 | 0.45892 | 0.17407 | 0.72950 | 0.96629 | 0.01939 | 0.09821 | 3.05416 | 0.02682 | 0.78417 | 0.85883 | 0.51151 | 1.14209 |
| **Klk1b24** | 0.00749 | 0.00443 | 0.01322 | 0.01132 | 0.07335 | 0.01350 | 0.00421 | 0.01000 | 0.01305 | 0.00653 | 0.00909 | 0.00955 | 0.07646 | 0.00388 |
| **Klrb1a** | 0.03037 | 0.00422 | 0.00000 | 0.35874 | 0.31891 | 0.66249 | 0.00403 | 0.00063 | 0.03005 | 0.00421 | 0.68324 | 0.00500 | 0.09213 | 0.00000 |
| **Klrg2** | 1.55488 | 0.00057 | 3.52526 | 0.00302 | 5.59766 | 0.17388 | 0.00193 | 0.00000 | 1.20857 | 0.00570 | 0.00401 | 3.18999 | 7.05317 | 6.21276 |
| **Klri2** | 0.00854 | 0.00460 | 0.01572 | 0.02613 | 0.01464 | 0.01400 | 0.02218 | 0.01788 | 0.03441 | 0.00772 | 0.00995 | 0.00912 | 0.06141 | 0.02904 |
| **Kmt2a** | 0.05031 | 0.00369 | 0.00000 | 0.01878 | 0.00734 | 1.87117 | 0.04498 | 0.65381 | 0.01013 | 0.01327 | 0.00000 | 0.88756 | 0.13963 | 0.00652 |
| **Kmt2b** | 4.58560 | 6.44598 | 2.71050 | 1.27343 | 1.93683 | 5.40488 | 0.93547 | 7.00417 | 0.00631 | 0.02953 | 3.92286 | 4.44534 | 1.82698 | 2.01304 |
| **Kmt2c** | 5.79660 | 0.03718 | 10.76459 | 2.97599 | 13.77105 | 5.24749 | 0.03727 | 3.81207 | 24.23568 | 0.00858 | 8.29556 | 1.91797 | 7.81677 | 17.75600 |
| **Kmt2d** | 0.00000 | 0.03424 | 0.55937 | 2.69925 | 0.46963 | 1.56457 | 1.31068 | 1.22152 | 1.81625 | 0.04794 | 0.00000 | 2.14468 | 2.39415 | 0.51258 |
| **Kmt2e** | 0.32202 | 0.01182 | 0.39089 | 0.20792 | 0.09652 | 0.31782 | 0.04609 | 0.17393 | 0.25920 | 0.01148 | 0.01944 | 0.14754 | 1.24536 | 0.06851 |
| **Kmt5a** | 8.91820 | 0.48975 | 16.76918 | 5.55636 | 19.57505 | 12.96153 | 3.39755 | 8.98531 | 3.84683 | 0.27965 | 6.44532 | 7.47081 | 6.82723 | 13.82282 |
| **Kncn** | 0.01376 | 0.00421 | 0.06143 | 0.16478 | 0.01877 | 0.00599 | 0.00403 | 0.00741 | 0.01257 | 0.00832 | 0.00652 | 0.01280 | 0.01098 | 0.01098 |
| **Kndc1** | 0.00884 | 0.02479 | 0.00492 | 0.19299 | 0.21026 | 0.00202 | 0.00436 | 0.65320 | 0.01358 | 0.00994 | 0.00512 | 0.07582 | 0.10094 | 0.03081 |
| **Kng1** | 5.92041 | 0.01738 | 7.86076 | 5.03750 | 0.00000 | 3.73698 | 0.00156 | 0.00000 | 4.27911 | 0.01433 | 3.14706 | 0.80442 | 0.17503 | 1.80384 |
| **Kng2** | 1.36901 | 0.01082 | 1.34478 | 0.99941 | 0.00000 | 1.09458 | 0.00342 | 0.00000 | 0.81313 | 0.03032 | 0.99278 | 0.09600 | 0.01561 | 0.21235 |
| **Knop1** | 12.86208 | 0.23723 | 15.22833 | 10.97616 | 14.31806 | 10.51199 | 5.12627 | 6.52892 | 13.70443 | 0.36107 | 4.01788 | 10.57129 | 9.17118 | 19.78773 |
| **Knstrn** | 0.39298 | 5.34640 | 3.99791 | 1.35444 | 1.65835 | 3.53572 | 5.33089 | 0.12168 | 3.39070 | 0.16548 | 1.89468 | 5.16574 | 3.77304 | 2.49238 |
| **Kntc1** | 4.69922 | 0.01535 | 4.84988 | 0.00000 | 1.38320 | 4.42357 | 0.00663 | 2.30360 | 3.58835 | 0.02451 | 2.21781 | 4.02496 | 4.76267 | 5.44015 |
| **Kpna1** | 7.22012 | 0.02170 | 9.28266 | 3.52517 | 17.78908 | 9.21549 | 1.39707 | 1.44150 | 29.35086 | 0.04981 | 9.84701 | 4.28831 | 8.30958 | 14.29953 |
| **Kpna2** | 33.45124 | 10.16408 | 41.07348 | 20.60710 | 39.05844 | 29.26003 | 6.17590 | 19.21378 | 40.32792 | 0.68570 | 25.84082 | 9.84703 | 22.07106 | 41.81613 |
| **Kpna3** | 5.02965 | 0.02594 | 5.26409 | 4.25776 | 4.62672 | 6.54052 | 0.39514 | 0.39667 | 14.31756 | 0.02803 | 7.31598 | 5.05114 | 2.47581 | 4.20012 |
| **Kpna4** | 6.45453 | 0.15833 | 1.83904 | 1.87939 | 4.30903 | 2.82507 | 1.21566 | 0.83570 | 6.90459 | 0.03671 | 1.37016 | 1.91890 | 6.87364 | 0.91968 |
| **Kpna6** | 5.33353 | 0.12674 | 11.81951 | 5.99287 | 6.05266 | 9.90468 | 2.64714 | 6.45781 | 9.25626 | 0.09782 | 5.55798 | 6.76899 | 8.22056 | 2.55639 |
| **Kpna7** | 0.27527 | 0.00444 | 0.00000 | 0.00000 | 0.46578 | 0.09401 | 0.00415 | 0.00619 | 0.00597 | 0.00659 | 0.00623 | 0.00765 | 0.06998 | 0.21790 |
| **Kpnb1** | 8.47416 | 5.24491 | 27.40228 | 19.27393 | 22.97452 | 29.48099 | 5.64466 | 12.56685 | 22.05653 | 7.10338 | 17.97280 | 19.86230 | 10.77113 | 27.52945 |
| **Kptn** | 0.00000 | 0.02646 | 4.20155 | 6.43157 | 1.31892 | 0.50507 | 0.00268 | 1.97094 | 0.35398 | 0.01525 | 0.00668 | 3.47053 | 0.32873 | 0.00000 |

| | | | | | | | | | | | | | | |
|---|---|---|---|---|---|---|---|---|---|---|---|---|---|---|
| **Kras** | 3.97963 | 0.21332 | 0.92223 | 0.00000 | 3.12665 | 4.43202 | 0.83553 | 0.81682 | 7.89314 | 0.02397 | 0.67167 | 2.13790 | 1.12208 | 5.41865 |
| **Krba1** | 0.00000 | 0.00361 | 0.56690 | 1.94546 | 1.16379 | 1.03348 | 0.00381 | 0.00000 | 0.01048 | 0.00998 | 0.06684 | 0.54336 | 0.30922 | 0.73014 |
| **Krcc1** | 1.78316 | 0.02992 | 0.93546 | 0.26160 | 1.25161 | 2.96083 | 3.33099 | 0.05298 | 3.68987 | 9.74837 | 0.00000 | 4.04001 | 1.50636 | 0.00000 |
| **Kremen1** | 0.08598 | 0.02920 | 0.28041 | 0.09920 | 0.55425 | 0.40802 | 0.57065 | 3.52196 | 0.00488 | 0.01190 | 0.87975 | 0.01228 | 1.27494 | 0.02921 |
| **Kremen2** | 0.09204 | 0.05794 | 0.29571 | 2.48443 | 2.45461 | 1.20065 | 0.80369 | 3.63431 | 0.67292 | 0.26283 | 0.00665 | 0.10402 | 1.02352 | 0.35600 |
| **Kri1** | 0.26434 | 2.86013 | 3.85418 | 0.83128 | 3.40305 | 2.69117 | 0.02481 | 2.57487 | 1.73475 | 4.28488 | 2.18075 | 0.63797 | 0.00000 | 5.60725 |
| **Krit1** | 2.88970 | 0.00153 | 1.91247 | 0.50957 | 3.84255 | 1.44971 | 3.10043 | 0.06895 | 8.08301 | 0.03490 | 0.73849 | 2.30268 | 0.54293 | 0.45052 |
| **Krr1** | 11.35971 | 0.14853 | 12.57793 | 3.18845 | 10.53147 | 12.05637 | 2.61299 | 2.97516 | 22.51336 | 0.10065 | 7.02805 | 6.14945 | 6.06960 | 14.02347 |
| **Krt10** | 0.00701 | 0.00362 | 0.02279 | 0.30818 | 0.00502 | 0.35630 | 0.40027 | 0.00903 | 0.01192 | 0.00514 | 0.00906 | 0.57993 | 0.22612 | 0.00000 |
| **Krt12** | 0.00446 | 0.00434 | 0.01052 | 0.01075 | 0.16885 | 0.01317 | 0.00414 | 0.00911 | 0.00891 | 0.00634 | 0.00935 | 0.00859 | 0.40802 | 0.02602 |
| **Krt18** | 0.00000 | 20.11865 | 12.02255 | 8.77994 | 1.58814 | 26.58541 | 16.87169 | 55.03985 | 0.42049 | 32.15987 | 4.26362 | 3.37937 | 0.23662 | 0.00000 |
| **Krt19** | 0.01282 | 0.00426 | 0.00000 | 0.00267 | 1.37569 | 0.00001 | 0.00433 | 0.01193 | 0.01186 | 0.00614 | 0.54279 | 0.00446 | 0.00282 | 0.00866 |
| **Krt222** | 0.00911 | 0.01235 | 0.00000 | 0.02395 | 0.39271 | 0.03209 | 0.01032 | 0.01743 | 0.44816 | 0.00696 | 0.00649 | 0.01143 | 0.37198 | 0.34537 |
| **Krt25** | 0.03518 | 0.00456 | 0.22099 | 0.03519 | 0.56286 | 0.03189 | 0.00412 | 0.00800 | 0.01014 | 0.00667 | 0.04186 | 0.00574 | 0.02819 | 0.17287 |
| **Krt27** | 0.09044 | 0.00501 | 1.32305 | 0.52906 | 0.00757 | 0.08130 | 0.00370 | 0.00505 | 0.10661 | 0.01030 | 0.35051 | 0.00378 | 0.00763 | 0.10104 |
| **Krt28** | 1.04953 | 0.00365 | 5.22811 | 1.49497 | 6.24109 | 0.00000 | 0.00303 | 0.00164 | 0.00000 | 0.01008 | 0.00346 | 0.00287 | 3.75037 | 0.60272 |
| **Krt7** | 0.01586 | 0.02221 | 0.31321 | 0.05935 | 0.52278 | 0.31236 | 0.01005 | 2.41317 | 0.00870 | 0.02155 | 0.00346 | 0.00342 | 0.00758 | 0.00225 |
| **Krt8** | 0.00000 | 18.97226 | 4.57406 | 14.99602 | 0.57999 | 26.13711 | 16.45195 | 51.91989 | 0.00000 | 26.34157 | 9.20945 | 4.94512 | 0.03043 | 0.00000 |
| **Krt84** | 0.00352 | 0.00419 | 0.01470 | 0.01144 | 0.00819 | 0.01182 | 0.00407 | 0.01574 | 0.02548 | 0.00615 | 0.00916 | 0.00854 | 0.59783 | 0.00234 |
| **Krtcap2** | 0.98246 | 2.60780 | 0.25323 | 8.18973 | 5.64501 | 6.63951 | 2.04813 | 5.02812 | 0.01290 | 4.44828 | 0.27713 | 7.79371 | 0.01273 | 0.32803 |
| **Krtcap3** | 0.00000 | 0.03914 | 0.13529 | 0.95440 | 0.53657 | 0.00000 | 0.00464 | 0.77235 | 0.00774 | 0.00412 | 0.78510 | 0.46623 | 0.04054 | 0.29680 |
| **Krtdap** | 0.00458 | 0.01234 | 0.00000 | 0.27601 | 0.10256 | 0.49781 | 0.08915 | 0.23753 | 0.02231 | 0.00729 | 0.00370 | 0.00333 | 0.00643 | 0.00000 |
| **Ksr1** | 0.01180 | 0.00372 | 0.06052 | 0.00000 | 0.00741 | 0.10832 | 0.00474 | 0.18841 | 0.01158 | 0.00658 | 0.71687 | 0.74954 | 0.25838 | 0.07997 |
| **Kti12** | 9.02443 | 0.02943 | 8.05778 | 4.47170 | 14.70247 | 2.84517 | 4.48035 | 1.16942 | 2.25357 | 0.10993 | 0.12195 | 5.79282 | 12.45401 | 2.18558 |
| **Ktn1** | 2.86197 | 0.02955 | 1.25566 | 5.34621 | 0.84424 | 7.85126 | 0.03651 | 1.17974 | 4.00981 | 0.04006 | 2.36410 | 4.40636 | 3.24486 | 8.11932 |
| **Kxd1** | 9.14190 | 8.41044 | 20.20353 | 16.80686 | 15.21397 | 18.22439 | 5.29312 | 9.48335 | 4.19511 | 13.36700 | 10.30566 | 6.97412 | 1.66063 | 9.42725 |
| **Kyat1** | 0.55751 | 0.02822 | 1.23020 | 1.22687 | 4.02318 | 0.53908 | 0.00356 | 1.11801 | 0.35585 | 0.00237 | 1.45403 | 1.90948 | 0.00000 | 3.55578 |
| **Kyat3** | 0.00000 | 0.08211 | 1.27753 | 7.10509 | 2.35221 | 1.64583 | 0.00508 | 0.09083 | 0.00000 | 0.04213 | 3.66875 | 12.55620 | 4.88668 | 6.12418 |
| **L1cam** | 0.00000 | 0.00468 | 0.31287 | 0.54842 | 0.49174 | 0.17443 | 0.00386 | 0.00509 | 0.47975 | 0.00636 | 0.00204 | 0.10498 | 0.29099 | 2.46827 |
| **L1td1** | 0.99502 | 11.45634 | 5.01586 | 6.02928 | 0.00000 | 9.96773 | 6.21125 | 12.00781 | 2.63737 | 1.07611 | 9.25247 | 15.71614 | 9.15004 | 0.00000 |
| **L2hgdh** | 18.35714 | 0.04694 | 11.74475 | 6.06813 | 12.03138 | 0.95732 | 0.00000 | 0.81097 | 19.05081 | 0.21105 | 2.63449 | 3.18943 | 10.99795 | 20.23463 |
| **L3hypdh** | 0.00410 | 0.00416 | 0.09822 | 0.38819 | 0.48322 | 0.02145 | 0.00403 | 0.00675 | 0.01113 | 0.00647 | 0.00525 | 0.01065 | 0.00738 | 0.00205 |
| **L3mbtl2** | 1.82291 | 0.04085 | 9.03107 | 3.03743 | 11.10864 | 7.74996 | 0.31949 | 4.34855 | 0.00509 | 12.40984 | 3.80658 | 3.56843 | 9.69954 | 3.74155 |
| **L3mbtl3** | 0.66394 | 0.19259 | 0.17997 | 0.09199 | 0.54256 | 0.50869 | 0.00387 | 0.00825 | 2.38929 | 0.00715 | 0.00213 | 0.08439 | 0.03398 | 0.28466 |
| **l7Rn6** | 9.61979 | 4.33366 | 11.61373 | 14.10900 | 15.11956 | 10.72001 | 4.82579 | 8.52356 | 1.97848 | 17.28358 | 2.95522 | 11.56255 | 3.42933 | 0.04606 |
| **Lacc1** | 0.07289 | 0.00417 | 0.03555 | 0.02154 | 0.03883 | 0.11027 | 0.13845 | 0.00333 | 0.01154 | 0.00780 | 0.00806 | 0.00786 | 0.02204 | 0.00787 |
| **Lace1** | 0.00000 | 0.00402 | 0.38338 | 0.20228 | 0.34345 | 0.01345 | 0.00423 | 0.24357 | 0.54880 | 0.01420 | 0.00205 | 1.08277 | 0.00477 | 0.00000 |
| **Lactb** | 1.99847 | 0.00964 | 1.30538 | 0.60892 | 3.94433 | 4.39283 | 3.41300 | 6.04428 | 3.76838 | 0.05685 | 0.11154 | 1.68643 | 2.78666 | 3.42360 |
| **Lactb2** | 9.05916 | 0.07901 | 7.62852 | 3.85930 | 6.52876 | 3.40024 | 0.35892 | 1.27676 | 5.74394 | 0.07234 | 5.48515 | 6.78922 | 3.73185 | 4.65808 |
| **Lad1** | 0.00000 | 0.03395 | 0.89555 | 3.59079 | 0.03349 | 2.33104 | 1.37164 | 1.95779 | 0.01869 | 0.01973 | 0.00233 | 0.19831 | 0.00917 | 0.00000 |
| **Lag3** | 0.02632 | 0.00537 | 0.04309 | 0.00755 | 0.01992 | 0.01105 | 0.01706 | 0.20321 | 0.03631 | 0.00984 | 0.03204 | 0.00992 | 0.01980 | 0.01765 |
| **Lage3** | 4.61134 | 0.18510 | 0.53443 | 3.72109 | 5.27685 | 7.12466 | 0.00718 | 5.75373 | 0.52456 | 0.09132 | 2.56384 | 6.23913 | 1.09618 | 4.91287 |
| **Lama1** | 0.05517 | 0.44505 | 0.20840 | 0.00000 | 0.06754 | 0.00000 | 0.01341 | 2.67891 | 0.06488 | 0.05771 | 0.00000 | 11.27417 | 0.00450 | 0.12393 |
| **Lama2** | 0.16774 | 0.03451 | 0.01296 | 0.01136 | 0.01201 | 0.01148 | 0.00424 | 0.00873 | 0.00889 | 0.00638 | 0.00743 | 0.00734 | 0.68523 | 0.04483 |
| **Lama3** | 0.00000 | 0.02919 | 0.46059 | 1.70627 | 0.03140 | 0.58910 | 1.02937 | 0.04240 | 0.00518 | 0.00000 | 4.93112 | 1.87766 | 0.10331 | 0.00000 |
| **Lama4** | 0.02516 | 0.01079 | 0.14620 | 0.04062 | 0.03460 | 0.01342 | 0.00409 | 0.00744 | 0.01258 | 0.00568 | 0.00333 | 1.25028 | 0.00535 | 1.73241 |
| **Lama5** | 0.73453 | 6.01330 | 4.82147 | 10.70462 | 0.00000 | 2.82310 | 2.07124 | 6.93870 | 0.00000 | 0.15694 | 2.54406 | 8.37783 | 0.30788 | 0.00000 |
| **Lamb1** | 1.65977 | 8.20234 | 5.01012 | 5.57542 | 2.73976 | 7.48913 | 3.42607 | 3.76251 | 1.62708 | 0.12939 | 0.00000 | 20.24633 | 3.18972 | 4.67903 |
| **Lamb2** | 0.10628 | 0.01201 | 1.07430 | 0.96803 | 0.53314 | 1.73467 | 0.00366 | 1.81712 | 0.00488 | 1.58069 | 1.03397 | 0.00000 | 0.13663 | 0.00951 |
| **Lamc1** | 0.58042 | 4.19912 | 0.61738 | 0.00000 | 0.63082 | 5.55278 | 1.11227 | 2.44474 | 0.02415 | 0.08176 | 4.41106 | 11.86364 | 0.44368 | 0.09005 |
| **Lamc2** | 0.54436 | 5.58929 | 0.00000 | 0.08459 | 0.58970 | 0.36192 | 0.04056 | 0.76827 | 0.41143 | 0.00626 | 0.02871 | 0.00000 | 0.00522 | 0.16014 |
| **Lamc3** | 0.25728 | 0.00432 | 0.21644 | 0.00000 | 0.63998 | 0.00064 | 0.00393 | 0.01016 | 0.37423 | 0.00695 | 0.00473 | 0.00658 | 0.36578 | 0.18728 |
| **Lamp1** | 24.36875 | 8.96632 | 29.90354 | 29.76296 | 17.89224 | 32.09191 | 7.90978 | 20.03474 | 4.06840 | 6.39452 | 17.33968 | 20.13054 | 1.44933 | 25.00733 |
| **Lamp2** | 14.79838 | 0.02503 | 12.46675 | 8.21993 | 15.68265 | 8.86978 | 0.17183 | 0.11228 | 28.14714 | 0.03177 | 5.63361 | 4.95434 | 10.01033 | 16.54124 |
| **Lamp3** | 0.77007 | 0.14995 | 1.40023 | 0.93568 | 0.67087 | 1.63235 | 0.31088 | 0.23663 | 0.75111 | 0.00100 | 0.90333 | 1.29838 | 0.67697 | 0.32611 |

| | | | | | | | | | | | | | | |
|---|---|---|---|---|---|---|---|---|---|---|---|---|---|---|
| **Lamtor1** | 7.52410 | 0.16708 | 15.61470 | 20.71566 | 15.84857 | 14.50545 | 3.64388 | 9.74927 | 3.01561 | 0.39272 | 6.83156 | 11.55614 | 4.54283 | 3.45276 |
| **Lamtor2** | 0.79423 | 0.02511 | 0.00062 | 0.12104 | 0.00307 | 1.89063 | 0.94343 | 2.22352 | 0.00000 | 5.65400 | 0.24034 | 1.26015 | 1.05016 | 1.01534 |
| **Lamtor3** | 9.17839 | 5.76817 | 6.68262 | 3.65541 | 6.93518 | 12.44997 | 0.14649 | 10.13309 | 4.18255 | 0.04420 | 6.48971 | 4.19813 | 1.55149 | 5.84207 |
| **Lamtor4** | 1.30393 | 0.09812 | 0.00000 | 4.17057 | 2.68811 | 2.85168 | 0.97188 | 6.35575 | 0.12730 | 6.01203 | 0.33944 | 3.47178 | 0.19840 | 1.42262 |
| **Lamtor5** | 5.25866 | 0.15525 | 1.38517 | 13.14124 | 3.92597 | 6.48666 | 2.33243 | 4.71995 | 1.54955 | 1.00516 | 3.85835 | 13.44964 | 0.00000 | 5.21598 |
| **Lancl1** | 5.69976 | 0.24528 | 1.16537 | 4.91036 | 2.53369 | 6.99699 | 2.57261 | 7.07481 | 0.08937 | 0.11910 | 4.61830 | 7.24834 | 0.82328 | 1.39353 |
| **Lancl2** | 0.00000 | 3.92461 | 0.18579 | 0.00000 | 0.36634 | 0.20538 | 0.00271 | 0.00459 | 0.00008 | 0.04131 | 1.21076 | 0.71569 | 0.62096 | 0.48618 |
| **Lap3** | 8.74721 | 2.74860 | 12.97506 | 16.67850 | 7.00320 | 18.23814 | 9.52828 | 15.23941 | 5.74132 | 13.78090 | 12.85010 | 21.28718 | 3.75136 | 9.53873 |
| **Laptm4a** | 2.23806 | 0.07752 | 3.84747 | 7.11226 | 4.34113 | 6.54496 | 3.43381 | 1.23299 | 2.07069 | 0.17070 | 0.82650 | 6.80129 | 4.20467 | 1.66325 |
| **Laptm4b** | 2.72875 | 0.05344 | 0.00000 | 6.82131 | 3.80897 | 5.33044 | 0.00000 | 6.89796 | 1.61394 | 0.10616 | 1.29858 | 6.47048 | 10.30443 | 0.21667 |
| **Laptm5** | 0.02160 | 0.01894 | 0.01612 | 0.45551 | 0.00994 | 0.01084 | 0.00384 | 0.00466 | 0.01136 | 0.00704 | 0.00408 | 0.67997 | 0.02430 | 0.00390 |
| **Large** | 0.04171 | 0.01851 | 0.33641 | 0.15204 | 0.66797 | 0.00000 | 0.00938 | 1.86795 | 0.00885 | 0.00798 | 0.77485 | 0.21653 | 0.36210 | 0.50158 |
| **Larp1** | 11.58153 | 3.65368 | 19.92173 | 19.22529 | 20.64220 | 13.91603 | 4.10673 | 9.32695 | 10.36194 | 0.30099 | 13.53462 | 12.40768 | 11.17281 | 24.06943 |
| **Larp1b** | 25.61477 | 0.04712 | 13.48129 | 9.57305 | 30.25138 | 4.72028 | 4.00774 | 4.87783 | 21.34320 | 0.20005 | 5.70454 | 1.76627 | 19.81685 | 30.16151 |
| **Larp4** | 27.84359 | 0.01123 | 19.44807 | 6.65960 | 20.21300 | 2.36646 | 1.14446 | 1.00314 | 46.98492 | 0.00000 | 9.00679 | 1.18523 | 15.71627 | 23.23743 |
| **Larp4b** | 14.43718 | 4.78691 | 11.88785 | 2.52539 | 15.16994 | 18.31916 | 5.11209 | 2.06765 | 14.88887 | 0.25384 | 13.30923 | 6.84472 | 7.96232 | 15.01703 |
| **Larp7** | 9.13073 | 0.25516 | 14.51292 | 9.52055 | 13.07181 | 13.53010 | 7.92896 | 9.61872 | 6.80479 | 0.33872 | 16.19942 | 14.05552 | 6.91591 | 19.14282 |
| **Lars** | 6.37169 | 0.87632 | 8.41280 | 2.74455 | 8.63245 | 7.48395 | 0.33025 | 2.82883 | 5.85847 | 0.01894 | 11.54189 | 8.53001 | 0.00000 | 8.64076 |
| **Lars2** | 0.32292 | 0.03523 | 0.89108 | 1.03386 | 0.15104 | 1.30056 | 0.00874 | 3.11217 | 1.23084 | 0.22076 | 3.29467 | 2.69556 | 1.25124 | 2.11325 |
| **Las1l** | 6.52104 | 0.11431 | 5.70950 | 9.68949 | 11.50198 | 11.00665 | 1.70349 | 3.95338 | 9.13465 | 8.82761 | 4.92552 | 5.94559 | 7.51451 | 10.56391 |
| **Lasp1** | 1.50281 | 10.98895 | 20.96538 | 11.82721 | 11.39669 | 18.16684 | 1.77343 | 10.04600 | 3.30025 | 1.40534 | 8.79983 | 6.90490 | 0.00000 | 4.28650 |
| **Lat** | 1.52008 | 0.00362 | 0.64218 | 0.56141 | 0.48509 | 0.00000 | 0.00392 | 0.00180 | 0.50376 | 0.00534 | 0.00857 | 0.14464 | 0.64460 | 0.00000 |
| **Lat2** | 0.43437 | 0.00332 | 3.23116 | 0.92439 | 1.14647 | 0.51426 | 0.00829 | 0.00000 | 0.23653 | 0.00000 | 3.03818 | 0.00172 | 1.57621 | 1.82403 |
| **Lats1** | 3.04184 | 0.02851 | 1.75062 | 0.25077 | 4.33802 | 0.58102 | 0.60856 | 0.20326 | 8.82337 | 0.03652 | 0.41866 | 2.21199 | 0.34985 | 6.07888 |
| **Lats2** | 4.39755 | 0.00260 | 4.35113 | 0.29720 | 2.47021 | 2.58556 | 2.89641 | 0.15286 | 3.23428 | 6.79061 | 0.15840 | 0.00000 | 0.00000 | 3.54556 |
| **Lbh** | 0.09608 | 0.00415 | 0.01464 | 0.01087 | 0.08254 | 0.02095 | 0.00440 | 0.01999 | 0.01367 | 0.00622 | 0.00669 | 0.01169 | 0.97961 | 0.00262 |
| **Lbhd1** | 0.35056 | 0.00432 | 0.78938 | 0.00000 | 1.74027 | 0.00000 | 0.00381 | 0.00737 | 0.44106 | 0.00487 | 0.00373 | 0.00451 | 0.00001 | 1.78350 |
| **Lbp** | 0.00167 | 0.00463 | 0.11009 | 0.00000 | 1.25374 | 0.33525 | 0.00429 | 0.00548 | 0.01891 | 0.01081 | 0.01554 | 0.00899 | 0.11186 | 0.06257 |
| **Lbr** | 10.96790 | 0.04485 | 15.25536 | 5.22305 | 10.02328 | 10.45318 | 2.88793 | 2.02391 | 20.23923 | 0.15685 | 5.32242 | 5.79676 | 7.48350 | 12.28208 |
| **Lbx2** | 0.01837 | 0.00414 | 0.04675 | 0.03930 | 0.61272 | 0.01692 | 0.00403 | 0.00681 | 0.01055 | 0.00659 | 0.00870 | 0.00983 | 0.82854 | 0.02729 |
| **Lca5** | 0.01951 | 0.00402 | 0.01039 | 0.47209 | 0.00103 | 0.00000 | 0.84773 | 0.00903 | 0.00937 | 0.00438 | 0.03053 | 0.01626 | 0.00308 | 0.29550 |
| **Lca5l** | 0.00374 | 0.00392 | 0.00766 | 0.09996 | 0.07464 | 0.00977 | 0.00381 | 0.07025 | 0.00682 | 0.00806 | 0.00668 | 0.00983 | 0.51256 | 0.38088 |
| **Lcat** | 0.05400 | 0.00444 | 0.79642 | 0.22679 | 0.11480 | 0.23909 | 0.00390 | 0.00128 | 0.02332 | 0.00545 | 0.00707 | 0.00756 | 0.00701 | 0.05881 |
| **Lck** | 2.97776 | 0.00308 | 6.01020 | 0.13297 | 1.06520 | 0.00000 | 0.00288 | 0.00000 | 0.45942 | 0.00000 | 1.63255 | 0.07410 | 0.91627 | 1.63419 |
| **Lclat1** | 13.99418 | 0.01237 | 9.70795 | 0.66646 | 16.50931 | 2.66008 | 0.03004 | 2.51051 | 13.12556 | 0.06906 | 2.86653 | 2.87661 | 9.59528 | 22.35579 |
| **Lcmt1** | 0.94377 | 0.00775 | 2.56096 | 3.47551 | 1.51826 | 4.41042 | 3.16917 | 6.04563 | 0.48612 | 0.17439 | 0.70678 | 5.79238 | 0.07421 | 1.58009 |
| **Lcmt2** | 2.87844 | 0.15541 | 2.90764 | 2.04559 | 2.28173 | 1.34051 | 2.44489 | 2.34039 | 1.56060 | 1.09677 | 0.86862 | 2.56346 | 11.87183 | 1.27377 |
| **Lcn10** | 0.00000 | 0.00421 | 0.37829 | 0.57558 | 0.00000 | 0.00000 | 0.00422 | 1.00858 | 0.01376 | 0.08708 | 0.00124 | 0.00432 | 0.00611 | 0.00570 |
| **Lcn2** | 0.21132 | 0.00543 | 0.00000 | 0.00000 | 0.85576 | 0.00238 | 0.00437 | 0.39327 | 0.31003 | 0.01475 | 0.00392 | 0.00564 | 0.20669 | 0.00000 |
| **Lcn9** | 0.00453 | 0.03709 | 0.16305 | 0.28649 | 0.00000 | 0.09671 | 0.00395 | 1.45215 | 0.21188 | 0.00838 | 0.66897 | 0.00661 | 0.00415 | 0.01630 |
| **Lcor** | 0.14487 | 0.01314 | 0.63037 | 0.27958 | 0.04370 | 0.05061 | 1.41997 | 0.02903 | 0.48769 | 0.01074 | 0.10439 | 0.21677 | 0.00081 | 0.93294 |
| **Lcorl** | 0.31370 | 0.00497 | 0.16175 | 0.35958 | 0.14545 | 0.07986 | 0.14751 | 0.05863 | 0.04825 | 0.00639 | 0.05176 | 0.07359 | 0.03491 | 0.00000 |
| **Lcp1** | 1.66068 | 7.72608 | 4.89718 | 0.86649 | 0.00000 | 4.89004 | 6.20504 | 21.60103 | 0.00000 | 0.44428 | 0.46851 | 1.08068 | 0.48095 | 0.10579 |
| **Lct** | 0.03302 | 0.00413 | 0.17340 | 0.29767 | 0.00000 | 1.66223 | 0.44445 | 0.47002 | 0.00432 | 0.03320 | 0.00134 | 0.29140 | 0.19584 | 0.29994 |
| **Lctl** | 5.00034 | 0.00299 | 4.26999 | 0.00000 | 3.85184 | 2.95193 | 2.12942 | 0.00000 | 4.11507 | 0.05461 | 1.48530 | 0.00000 | 0.99045 | 3.32103 |
| **Ldah** | 3.66724 | 2.23247 | 12.69389 | 7.05787 | 8.26876 | 10.18904 | 0.87330 | 0.83249 | 11.73083 | 0.03758 | 6.40414 | 5.35987 | 0.88101 | 4.43437 |
| **Ldb1** | 4.88270 | 0.55002 | 2.60107 | 0.60034 | 1.39518 | 1.32235 | 0.01601 | 0.73147 | 0.10727 | 0.02606 | 0.53183 | 1.78542 | 1.27427 | 2.28778 |
| **Ldha** | 1.64006 | 0.14735 | 3.27356 | 5.32867 | 3.25773 | 2.83242 | 0.03230 | 0.78555 | 0.60317 | 0.11554 | 1.40061 | 9.35112 | 1.88361 | 1.99318 |
| **Ldhb** | 3.90400 | 0.00171 | 6.02991 | 1.68292 | 6.37665 | 0.13829 | 0.00232 | 0.00000 | 2.74064 | 0.00526 | 1.84252 | 0.24358 | 8.34301 | 14.10518 |
| **Ldhc** | 0.16865 | 0.00398 | 0.13450 | 0.42131 | 0.19730 | 0.00000 | 0.00429 | 0.01583 | 0.00000 | 0.00751 | 0.00895 | 0.00356 | 3.31529 | 2.59729 |
| **Ldhd** | 0.03131 | 0.00435 | 0.03611 | 0.00752 | 0.07249 | 0.01134 | 0.00463 | 0.01032 | 0.02361 | 0.00646 | 0.00799 | 0.00952 | 0.27047 | 0.02460 |
| **Ldlr** | 0.66285 | 6.22960 | 1.79409 | 1.91031 | 2.45915 | 2.25480 | 3.13835 | 2.86165 | 0.40974 | 0.11151 | 0.00000 | 2.98140 | 0.55998 | 1.70360 |
| **Ldlrad2** | 0.20150 | 0.07738 | 0.05260 | 0.68904 | 0.17154 | 0.56198 | 0.09864 | 0.46607 | 0.34375 | 0.52203 | 0.03520 | 0.04392 | 0.39409 | 0.30930 |
| **Ldlrad3** | 0.01979 | 0.00434 | 0.57035 | 0.00690 | 0.20590 | 0.00377 | 0.00970 | 0.00923 | 0.00863 | 0.00654 | 0.00875 | 0.00971 | 0.01830 | 0.02403 |
| **Ldlrad4** | 0.01943 | 0.00438 | 0.08803 | 0.00990 | 0.00811 | 0.00957 | 0.00421 | 0.01015 | 0.00941 | 0.00667 | 0.00788 | 0.00876 | 0.17194 | 0.01771 |

| | | | | | | | | | | | | | | |
|---|---|---|---|---|---|---|---|---|---|---|---|---|---|---|
| **Ldlrap1** | 4.00901 | 0.87568 | 10.74121 | 2.81245 | 6.71099 | 4.31191 | 0.01889 | 1.53867 | 0.46433 | 5.57343 | 0.71369 | 3.33423 | 0.88781 | 4.19543 |
| **Ldoc1** | 0.00651 | 0.01143 | 0.00553 | 0.00716 | 0.00830 | 3.05051 | 2.75695 | 0.82543 | 0.01016 | 0.12255 | 0.00371 | 0.00000 | 0.00639 | 0.00722 |
| **Ldoc1l** | 0.00000 | 0.00388 | 0.96599 | 0.87193 | 0.63960 | 0.26389 | 0.00351 | 0.00000 | 0.16944 | 0.00000 | 2.15176 | 0.00641 | 0.31747 | 1.84726 |
| **Lect1** | 0.08932 | 0.00407 | 0.49415 | 0.59908 | 0.10699 | 0.00000 | 0.00395 | 0.00673 | 0.00616 | 0.00571 | 0.00695 | 0.01153 | 0.00136 | 0.22446 |
| **Lef1** | 0.08502 | 0.00442 | 0.01259 | 0.01336 | 0.01236 | 0.01096 | 0.00423 | 0.01068 | 0.01102 | 0.00694 | 0.00761 | 0.00923 | 0.02110 | 0.14368 |
| **Lekr1** | 0.00783 | 0.01200 | 0.00649 | 0.03279 | 0.01327 | 0.31396 | 0.00408 | 0.00516 | 0.00960 | 0.00767 | 0.00571 | 0.01138 | 0.00854 | 0.04836 |
| **Lemd1** | 0.00000 | 0.00467 | 0.24074 | 0.00000 | 0.38098 | 0.00006 | 0.00422 | 0.01468 | 1.10685 | 0.00826 | 0.00434 | 0.00653 | 0.33619 | 0.28211 |
| **Lemd2** | 0.15996 | 5.68440 | 0.57356 | 1.55856 | 0.08885 | 2.18065 | 1.13845 | 0.36956 | 0.00787 | 0.03980 | 0.32841 | 2.73605 | 0.17574 | 0.01379 |
| **Lemd3** | 2.99214 | 0.00298 | 0.46916 | 0.00000 | 0.22641 | 0.00000 | 0.06519 | 0.07559 | 8.31646 | 0.00548 | 0.07234 | 0.36476 | 5.54267 | 2.61178 |
| **Lenep** | 0.33525 | 0.00410 | 0.00000 | 0.74786 | 0.56047 | 0.08541 | 0.00409 | 0.34227 | 0.10421 | 0.00713 | 0.00508 | 0.00701 | 0.01056 | 0.19858 |
| **Leng1** | 1.31902 | 0.02529 | 3.27021 | 3.49039 | 6.82882 | 0.84679 | 3.23441 | 3.77568 | 0.86541 | 0.12958 | 0.00000 | 1.03973 | 10.18340 | 0.09880 |
| **Leng8** | 0.65167 | 0.42875 | 0.09368 | 2.63997 | 2.99468 | 0.17107 | 2.68318 | 1.17230 | 0.08080 | 0.01976 | 0.25416 | 0.01426 | 2.72247 | 0.00000 |
| **Leng9** | 0.01427 | 0.00497 | 0.05531 | 0.33059 | 0.02692 | 0.00036 | 0.00528 | 0.53175 | 0.01589 | 0.00531 | 0.46922 | 0.00409 | 0.16964 | 0.00000 |
| **Leo1** | 12.62791 | 7.11005 | 17.62694 | 5.98008 | 16.77167 | 9.91510 | 2.17603 | 4.06296 | 0.28648 | 0.01510 | 12.62956 | 5.21124 | 5.76859 | 15.48932 |
| **Lepr** | 0.61890 | 0.44238 | 0.28766 | 0.10724 | 0.25669 | 0.97672 | 0.13201 | 0.07415 | 1.25482 | 0.73784 | 0.09998 | 0.19526 | 5.99282 | 1.74446 |
| **Leprot** | 1.93388 | 0.07217 | 2.80166 | 1.91901 | 0.77648 | 5.04956 | 4.09227 | 2.82870 | 0.00651 | 1.54493 | 4.15468 | 6.70650 | 0.03332 | 5.12706 |
| **Leprotl1** | 0.37286 | 0.02882 | 1.15890 | 1.45807 | 1.64660 | 3.83639 | 0.00366 | 0.53948 | 0.00233 | 0.03611 | 0.16712 | 1.32636 | 1.61331 | 1.12064 |
| **Letm1** | 5.45334 | 3.93429 | 11.23244 | 4.25715 | 2.88664 | 9.77554 | 2.11420 | 8.02294 | 1.59886 | 0.07931 | 6.96342 | 6.28516 | 2.98585 | 3.80794 |
| **Letm2** | 1.90904 | 0.00397 | 0.45847 | 0.01414 | 0.34312 | 0.39220 | 0.00966 | 0.00123 | 0.00822 | 0.00444 | 0.24475 | 0.00698 | 0.94753 | 0.27597 |
| **Letmd1** | 0.90714 | 0.01636 | 5.92819 | 2.69626 | 3.22785 | 0.35406 | 0.00242 | 0.00000 | 0.48619 | 0.00697 | 0.21901 | 1.44997 | 0.00000 | 3.35629 |
| **Lexm** | 0.00000 | 0.00450 | 0.22375 | 0.22651 | 0.41288 | 0.10247 | 0.00399 | 0.00266 | 0.94856 | 0.00571 | 0.01497 | 0.02528 | 0.04035 | 0.75797 |
| **Lgals1** | 0.32416 | 5.26849 | 0.00000 | 0.00000 | 0.16386 | 0.23083 | 0.87491 | 16.34005 | 0.00000 | 11.83585 | 0.00000 | 0.54388 | 0.00940 | 0.40885 |
| **Lgals12** | 0.01295 | 0.00440 | 0.20322 | 0.00000 | 0.01983 | 0.01425 | 0.01175 | 0.10493 | 0.01248 | 0.00687 | 0.00650 | 0.00892 | 0.00720 | 0.01456 |
| **Lgals2** | 0.01364 | 0.00410 | 0.02823 | 0.04371 | 0.05070 | 0.08665 | 0.00394 | 0.00447 | 0.00732 | 0.00606 | 0.00599 | 0.01020 | 0.94210 | 0.00749 |
| **Lgals3** | 0.83336 | 2.66648 | 1.86804 | 3.88930 | 1.15683 | 0.53704 | 4.74672 | 15.99563 | 0.00000 | 5.97184 | 2.13010 | 0.04496 | 1.33296 | 0.54248 |
| **Lgals3bp** | 0.04200 | 0.00477 | 0.01057 | 0.00917 | 0.17401 | 0.03542 | 0.00444 | 0.00980 | 0.01028 | 0.00600 | 0.00789 | 0.00862 | 0.00555 | 0.01082 |
| **Lgals4** | 0.00118 | 0.00387 | 0.04308 | 0.00627 | 0.00388 | 1.09100 | 0.69097 | 1.52473 | 0.01343 | 1.72519 | 0.03009 | 0.00000 | 0.00681 | 0.00000 |
| **Lgals7** | 0.00895 | 0.00451 | 0.02982 | 0.15071 | 0.01533 | 0.20953 | 0.00426 | 0.00732 | 0.01000 | 0.00945 | 0.05882 | 0.00499 | 0.01027 | 0.02023 |
| **Lgals8** | 9.25718 | 0.02980 | 13.97205 | 16.44927 | 11.18723 | 9.52623 | 0.04694 | 2.11366 | 12.80494 | 0.12185 | 9.59919 | 1.84842 | 0.53261 | 12.28560 |
| **Lgals9** | 0.00000 | 0.00807 | 5.98675 | 5.30000 | 1.96110 | 10.80848 | 0.00146 | 0.00000 | 1.02314 | 0.01689 | 7.41476 | 1.08922 | 0.00000 | 2.86664 |
| **Lgalsl** | 7.58131 | 0.01746 | 2.39069 | 1.22008 | 3.99030 | 1.64339 | 2.11482 | 1.91207 | 8.96631 | 0.05879 | 2.01130 | 0.70584 | 2.30556 | 3.91882 |
| **Lgi2** | 0.00728 | 0.00462 | 0.01970 | 0.01013 | 0.01161 | 0.03650 | 0.00435 | 0.00815 | 0.01079 | 0.00632 | 0.00841 | 0.00949 | 0.18079 | 0.00609 |
| **Lgi3** | 0.00415 | 0.00415 | 0.27493 | 0.17239 | 0.00534 | 0.00001 | 0.00401 | 0.00840 | 0.00782 | 0.00780 | 0.00589 | 0.01528 | 0.05866 | 0.03003 |
| **Lgmn** | 5.02429 | 5.96732 | 15.30714 | 9.83521 | 11.29441 | 9.67154 | 3.55528 | 9.55249 | 0.04755 | 4.88837 | 8.57966 | 7.21400 | 0.89981 | 8.96336 |
| **Lgr4** | 0.29544 | 0.01106 | 0.00000 | 0.00000 | 0.61121 | 0.84868 | 0.00305 | 1.09309 | 6.05126 | 0.00507 | 0.22934 | 1.08754 | 1.37706 | 0.00000 |
| **Lgr5** | 0.03985 | 0.02991 | 0.03009 | 0.05300 | 0.30420 | 0.13142 | 0.02482 | 0.05528 | 0.05927 | 0.01573 | 0.00968 | 0.02276 | 0.64784 | 0.05539 |
| **Lhb** | 0.00721 | 0.00439 | 0.01275 | 0.01208 | 0.10920 | 0.01312 | 0.00438 | 0.01137 | 0.00991 | 0.00648 | 0.00891 | 0.00969 | 0.05524 | 0.00741 |
| **Lhfp** | 0.07125 | 0.01244 | 0.01459 | 0.00240 | 0.00982 | 0.00395 | 0.01275 | 0.29963 | 0.01150 | 0.00591 | 0.00799 | 0.00948 | 0.00506 | 0.01389 |
| **Lhfpl2** | 0.67521 | 0.00000 | 0.60994 | 0.00000 | 0.82921 | 2.52778 | 2.75642 | 3.60740 | 0.01717 | 0.04375 | 0.00000 | 7.87080 | 0.63966 | 0.91246 |
| **Lhfpl5** | 0.58503 | 0.00397 | 1.19314 | 0.96506 | 0.81964 | 0.00000 | 0.00378 | 0.01173 | 0.00929 | 0.00505 | 0.00366 | 0.00747 | 0.29919 | 0.00000 |
| **Lhpp** | 0.00000 | 0.00510 | 0.19402 | 0.50681 | 0.11421 | 0.10990 | 0.00416 | 0.73675 | 0.01860 | 0.00817 | 0.01167 | 0.01928 | 0.00485 | 0.41184 |
| **Lhx2** | 0.50718 | 0.00406 | 1.37893 | 0.28516 | 0.10699 | 0.00000 | 0.00368 | 0.00576 | 0.01040 | 0.00749 | 0.01011 | 0.01284 | 0.04843 | 0.00000 |
| **Lhx8** | 0.00359 | 0.00407 | 0.01401 | 0.00473 | 0.00564 | 0.07328 | 0.00409 | 0.00497 | 0.00970 | 0.00527 | 0.00986 | 0.00854 | 1.73455 | 0.05467 |
| **Lias** | 10.86915 | 0.07258 | 16.08776 | 7.69087 | 17.39651 | 10.09098 | 2.15164 | 8.99394 | 18.17538 | 0.08485 | 13.48991 | 6.21020 | 3.03459 | 24.55174 |
| **Lif** | 0.38235 | 1.68443 | 1.15380 | 0.00000 | 1.21517 | 0.00000 | 0.36893 | 4.25640 | 0.00576 | 0.00418 | 0.00151 | 0.00410 | 0.17544 | 0.62367 |
| **Lifr** | 0.10157 | 0.15478 | 0.35828 | 0.00000 | 0.14069 | 0.65125 | 0.20300 | 0.34397 | 0.62468 | 0.39861 | 0.31229 | 3.79999 | 0.87416 | 0.41129 |
| **Lig1** | 2.36621 | 4.61427 | 9.58739 | 11.34969 | 3.40973 | 11.00388 | 5.21472 | 3.68628 | 0.75414 | 6.57065 | 6.33718 | 9.36960 | 0.00000 | 4.19085 |
| **Lig3** | 2.36606 | 0.02788 | 12.02590 | 8.47340 | 7.80185 | 5.41354 | 3.04356 | 4.87175 | 6.51516 | 0.04583 | 12.97735 | 5.04095 | 1.06089 | 11.94956 |
| **Lig4** | 0.12669 | 0.00393 | 0.16728 | 0.94393 | 0.55859 | 0.00000 | 0.00404 | 0.02195 | 0.00711 | 0.00604 | 0.01305 | 0.00742 | 0.50890 | 1.96985 |
| **Lilra6** | 0.17253 | 0.00427 | 0.08843 | 0.00206 | 0.00654 | 0.00871 | 0.00422 | 0.00998 | 0.05518 | 0.00775 | 0.00579 | 0.07369 | 0.06755 | 0.05540 |
| **Lim2** | 0.27599 | 0.00416 | 0.00860 | 0.24330 | 0.00982 | 0.00421 | 0.00401 | 0.00710 | 0.01119 | 0.00791 | 0.00969 | 0.01267 | 0.00749 | 0.00207 |
| **Lima1** | 1.81897 | 0.12774 | 8.39824 | 2.31048 | 1.95636 | 0.95693 | 1.77485 | 4.42321 | 1.10755 | 0.04021 | 4.85220 | 3.49867 | 2.37134 | 5.73450 |
| **Limch1** | 0.00000 | 0.00354 | 0.39756 | 0.00000 | 0.42849 | 0.00000 | 0.00418 | 1.40070 | 0.01147 | 0.00098 | 1.49754 | 0.00289 | 1.49925 | 0.41052 |
| **Limd1** | 0.62510 | 0.06864 | 0.51729 | 2.98796 | 1.08207 | 1.79444 | 0.00320 | 3.64700 | 0.00000 | 0.01403 | 0.00000 | 1.42785 | 1.47934 | 2.01630 |
| **Limd2** | 1.21958 | 3.41143 | 0.91149 | 1.45271 | 2.15317 | 3.17214 | 0.23462 | 0.69543 | 0.63352 | 0.06521 | 0.17370 | 2.81032 | 7.59142 | 1.48531 |

| | | | | | | | | | | | | | | |
|---|---|---|---|---|---|---|---|---|---|---|---|---|---|---|
| **Lime1** | 0.88096 | 0.00453 | 1.81686 | 0.39000 | 1.14690 | 0.87823 | 0.00407 | 0.33642 | 0.02614 | 0.04765 | 0.00000 | 0.71343 | 1.85916 | 0.55674 |
| **Limk1** | 0.04389 | 0.01265 | 0.22549 | 0.00000 | 0.81045 | 0.09213 | 0.00388 | 0.74148 | 0.01559 | 0.01853 | 0.00109 | 0.09877 | 0.72378 | 0.07998 |
| **Limk2** | 4.29621 | 0.61241 | 12.73686 | 12.06214 | 4.71044 | 12.06430 | 1.83305 | 12.69856 | 2.10890 | 1.05751 | 2.45066 | 3.96589 | 3.09134 | 4.29009 |
| **Lims1** | 0.63611 | 1.23104 | 1.36835 | 0.84104 | 4.39073 | 4.73254 | 0.26027 | 2.18430 | 5.49259 | 0.00431 | 1.84731 | 2.25292 | 1.74130 | 4.34048 |
| **Lims2** | 1.13052 | 0.02762 | 1.80456 | 1.84259 | 0.73893 | 4.26459 | 0.01174 | 0.80337 | 0.00458 | 0.04023 | 1.55173 | 0.00000 | 0.10014 | 0.29022 |
| **Lin28a** | 0.00000 | 0.04037 | 1.00470 | 1.40620 | 2.28913 | 0.86889 | 0.02094 | 0.18812 | 1.82543 | 0.01263 | 0.00000 | 0.50139 | 0.81096 | 1.06917 |
| **Lin28b** | 0.77197 | 0.00809 | 2.27843 | 1.06111 | 0.51730 | 3.70974 | 0.00265 | 0.00000 | 0.00678 | 0.00466 | 2.63360 | 2.96039 | 0.28477 | 3.69334 |
| **Lin37** | 0.00000 | 7.53793 | 8.15986 | 9.23537 | 4.98292 | 1.24999 | 4.01890 | 6.61447 | 0.07963 | 10.80443 | 3.77192 | 4.97225 | 0.00000 | 0.00000 |
| **Lin52** | 2.70190 | 0.01043 | 0.49545 | 0.00000 | 0.00000 | 0.50533 | 0.00914 | 0.00000 | 2.98894 | 0.00579 | 0.00000 | 0.45972 | 3.71665 | 9.07831 |
| **Lin54** | 2.83212 | 0.01010 | 1.35829 | 0.00000 | 2.05115 | 2.43807 | 0.91725 | 0.03972 | 9.08774 | 0.00484 | 1.73718 | 0.04919 | 6.28993 | 4.04340 |
| **Lin7b** | 0.00672 | 0.00424 | 0.00926 | 0.08029 | 0.03363 | 0.03621 | 0.01615 | 0.00709 | 0.00953 | 0.00712 | 0.00704 | 0.00800 | 0.19235 | 0.00329 |
| **Lin7c** | 10.04570 | 0.52568 | 8.30562 | 2.11019 | 12.51103 | 10.22888 | 3.34204 | 6.51196 | 24.16098 | 13.06189 | 9.38100 | 2.55536 | 5.22305 | 14.19422 |
| **Lin9** | 0.30297 | 0.00449 | 0.10354 | 0.00000 | 0.00000 | 0.05701 | 0.00312 | 0.19349 | 7.77893 | 0.00742 | 0.78171 | 0.01510 | 1.85904 | 5.03000 |
| **Lincred1** | 0.00409 | 0.00459 | 0.00343 | 0.01665 | 0.18927 | 0.00943 | 0.00445 | 0.01498 | 0.15544 | 0.00735 | 0.00713 | 0.00845 | 0.00385 | 0.28253 |
| **Lingo1** | 0.02107 | 0.00409 | 0.31480 | 0.23302 | 0.00427 | 0.01811 | 0.10250 | 0.00298 | 0.00787 | 0.02020 | 0.00936 | 0.01148 | 0.06214 | 0.02336 |
| **Lingo2** | 0.01055 | 0.00444 | 0.01279 | 0.01402 | 0.00842 | 0.01331 | 0.00430 | 0.00994 | 0.00997 | 0.00770 | 0.00827 | 0.00989 | 0.00480 | 0.13835 |
| **Lins1** | 0.40139 | 0.01935 | 0.21011 | 0.87087 | 0.22024 | 0.15084 | 2.20297 | 0.00206 | 0.33711 | 0.03752 | 0.01626 | 0.25584 | 0.01026 | 0.28834 |
| **Lipe** | 0.01505 | 0.00461 | 0.06981 | 0.01982 | 0.27504 | 0.01641 | 0.00483 | 0.01086 | 0.01903 | 0.00858 | 0.00757 | 0.01002 | 0.07577 | 0.00227 |
| **Lipg** | 0.02449 | 0.00473 | 0.01112 | 0.00000 | 0.05932 | 0.14403 | 0.00421 | 0.01208 | 0.02392 | 0.00664 | 0.01700 | 0.11375 | 0.00940 | 0.02238 |
| **Liph** | 0.08680 | 0.02764 | 0.14300 | 0.44548 | 0.22709 | 0.01989 | 0.00947 | 0.00722 | 1.15772 | 0.00965 | 0.00353 | 0.02710 | 0.70618 | 0.79487 |
| **Lipo2** | 0.03687 | 0.00425 | 0.04014 | 0.09270 | 0.00888 | 0.00578 | 0.01033 | 0.02949 | 0.01165 | 0.00659 | 0.09679 | 0.02489 | 0.03793 | 0.00948 |
| **Lipo3** | 2.24238 | 0.00316 | 0.18304 | 0.35511 | 0.00000 | 0.05517 | 0.00371 | 0.44137 | 0.01605 | 0.00835 | 1.77634 | 0.00125 | 1.82207 | 0.00000 |
| **Lipt1** | 0.68183 | 0.01992 | 1.89336 | 1.23789 | 1.47650 | 1.01250 | 1.13427 | 0.62906 | 2.26317 | 0.03506 | 0.02400 | 0.00000 | 0.51496 | 3.28613 |
| **Lipt2** | 1.03744 | 0.00405 | 0.00000 | 0.18830 | 0.62023 | 0.00258 | 0.00397 | 0.00641 | 0.00556 | 0.00676 | 0.01473 | 0.02452 | 0.01117 | 0.10149 |
| **Litaf** | 6.19489 | 10.85058 | 14.69166 | 15.05334 | 15.35961 | 17.52982 | 9.50384 | 17.03199 | 8.06666 | 4.06755 | 5.36909 | 4.61497 | 3.56417 | 23.15239 |
| **Lix1** | 0.12972 | 0.08913 | 0.13222 | 0.37787 | 0.16489 | 0.39795 | 0.07395 | 0.23705 | 0.39682 | 0.04407 | 0.04908 | 0.17656 | 0.70169 | 0.70678 |
| **Lix1l** | 0.02301 | 0.00430 | 0.01210 | 0.00909 | 0.00791 | 0.02123 | 0.00421 | 0.00777 | 0.02268 | 0.00575 | 0.08564 | 0.00773 | 0.32843 | 0.04134 |
| **Llgl1** | 3.73877 | 0.15389 | 9.95393 | 5.11909 | 3.31522 | 4.01782 | 3.33353 | 1.95692 | 0.57450 | 0.19279 | 5.93391 | 4.22525 | 3.90782 | 2.62177 |
| **Llgl2** | 0.00000 | 5.03516 | 6.62387 | 2.09316 | 7.01901 | 0.00000 | 0.75066 | 9.37592 | 0.00000 | 0.07341 | 0.41257 | 0.73292 | 0.52346 | 0.00000 |
| **Llph** | 22.16530 | 0.29406 | 13.16303 | 24.92198 | 24.55939 | 17.00427 | 6.94046 | 9.28761 | 14.58812 | 1.00756 | 11.08295 | 10.77427 | 9.11288 | 19.89565 |
| **Lman1** | 1.33324 | 0.03183 | 1.23515 | 3.69292 | 0.92236 | 4.69302 | 0.25010 | 1.76170 | 0.77947 | 0.02999 | 4.45685 | 3.66252 | 0.00000 | 3.35286 |
| **Lman1l** | 0.01059 | 0.00430 | 0.01394 | 0.02335 | 0.01042 | 0.01240 | 0.00421 | 0.00867 | 0.01254 | 0.00712 | 0.00847 | 0.00819 | 0.41577 | 0.03635 |
| **Lman2** | 10.87608 | 5.92205 | 10.27348 | 14.27204 | 12.07825 | 19.79810 | 5.62926 | 11.08266 | 0.23298 | 0.42574 | 6.49747 | 14.90301 | 4.36221 | 15.38491 |
| **Lman2l** | 0.00001 | 0.11841 | 1.25033 | 0.00000 | 0.10104 | 2.23109 | 0.00294 | 2.34375 | 0.06325 | 0.00904 | 0.00000 | 1.47676 | 0.13053 | 1.48226 |
| **Lmbr1** | 0.73864 | 0.00273 | 1.85584 | 0.31080 | 1.19301 | 1.37123 | 0.75006 | 0.00000 | 4.64601 | 0.02595 | 1.35496 | 0.83880 | 1.76538 | 0.52360 |
| **Lmbr1l** | 0.01820 | 0.01160 | 0.42399 | 0.30233 | 0.35318 | 0.15851 | 0.02936 | 0.11222 | 0.04270 | 0.02007 | 0.00255 | 0.24446 | 2.70094 | 0.41164 |
| **Lmbrd1** | 7.27912 | 0.07068 | 5.35973 | 0.42949 | 5.72462 | 7.30958 | 2.23013 | 0.53514 | 11.19151 | 0.11204 | 5.07498 | 2.36679 | 0.69833 | 14.74157 |
| **Lmbrd2** | 0.00000 | 0.01251 | 0.08118 | 0.14368 | 0.00000 | 0.00000 | 0.58816 | 0.01216 | 2.18739 | 0.01029 | 1.10206 | 0.46771 | 0.05677 | 0.40287 |
| **Lmf1** | 0.00968 | 0.01880 | 0.00000 | 0.98734 | 0.00986 | 1.46145 | 3.37411 | 0.18255 | 0.00776 | 0.06441 | 0.04055 | 0.84261 | 0.00593 | 0.06208 |
| **Lmf2** | 0.51226 | 0.09781 | 4.78205 | 2.68248 | 2.54603 | 2.06065 | 2.88525 | 3.23687 | 0.01370 | 0.06345 | 4.08280 | 5.26229 | 2.07088 | 0.00000 |
| **Lmln** | 0.00000 | 0.00414 | 0.12930 | 0.27035 | 0.00664 | 0.00000 | 0.00399 | 0.00792 | 0.02599 | 0.00579 | 0.00567 | 0.01619 | 0.02403 | 0.26524 |
| **Lmna** | 1.92611 | 9.04185 | 5.42741 | 10.97045 | 8.96034 | 2.25995 | 7.35755 | 15.96420 | 0.02024 | 12.78900 | 2.51328 | 8.24752 | 2.68170 | 0.00000 |
| **Lmnb1** | 0.82438 | 0.10239 | 2.62149 | 1.82834 | 0.26254 | 1.34370 | 0.27126 | 2.16038 | 2.37908 | 0.67044 | 0.15702 | 1.89404 | 2.43988 | 4.08435 |
| **Lmnb2** | 0.85025 | 0.03293 | 2.36331 | 6.22885 | 2.53322 | 2.67169 | 2.47050 | 4.38583 | 0.12747 | 0.14820 | 0.00000 | 2.43942 | 0.00265 | 4.56354 |
| **Lmntd2** | 0.00137 | 0.00325 | 0.00000 | 0.22112 | 0.00000 | 0.34812 | 0.00354 | 0.00250 | 0.07236 | 0.00460 | 0.01792 | 0.94748 | 0.00297 | 0.42220 |
| **Lmo1** | 0.00899 | 0.00408 | 0.08459 | 0.27832 | 1.09812 | 0.00000 | 0.00398 | 0.00606 | 0.00802 | 0.00560 | 0.02012 | 0.02035 | 0.14753 | 0.00000 |
| **Lmo2** | 0.43169 | 0.00161 | 0.64486 | 0.00000 | 0.00000 | 0.00410 | 0.00293 | 0.01004 | 0.00765 | 0.01430 | 0.00289 | 3.53504 | 0.00277 | 0.00438 |
| **Lmo4** | 0.07973 | 0.00353 | 0.02075 | 0.36371 | 0.12470 | 0.00606 | 0.01054 | 0.00800 | 0.01822 | 0.01107 | 0.01537 | 1.08607 | 0.08179 | 0.21069 |
| **Lmo7** | 1.64016 | 0.02798 | 9.58649 | 3.21707 | 8.23253 | 2.41283 | 1.63697 | 0.38655 | 7.64539 | 0.04009 | 2.33832 | 0.55304 | 2.29564 | 4.49749 |
| **Lmod1** | 0.12523 | 0.00436 | 0.00634 | 0.01005 | 0.38858 | 0.01280 | 0.00428 | 0.01064 | 0.01890 | 0.00699 | 0.00742 | 0.00759 | 0.21587 | 0.01028 |
| **Lmod3** | 0.25416 | 0.00437 | 0.00000 | 0.00576 | 0.79506 | 0.02249 | 0.00413 | 0.00853 | 0.00832 | 0.00758 | 0.00557 | 0.00500 | 0.00262 | 3.02005 |
| **Lmtk2** | 1.07913 | 0.01836 | 4.25570 | 1.49887 | 2.43659 | 0.00000 | 0.01939 | 2.13594 | 1.70320 | 0.01413 | 0.69052 | 0.33978 | 1.04993 | 2.59519 |
| **Lmx1a** | 0.00297 | 0.00417 | 0.00327 | 0.00960 | 0.58846 | 0.01387 | 0.00408 | 0.00798 | 0.01979 | 0.00717 | 0.00932 | 0.00737 | 1.59495 | 0.00000 |
| **Lmx1b** | 0.00939 | 0.00450 | 0.01743 | 0.09088 | 0.01395 | 0.00208 | 0.00524 | 0.18755 | 0.01228 | 0.00563 | 0.01627 | 0.01114 | 0.00669 | 0.01084 |
| **Lncenc1** | 1.90588 | 0.00204 | 0.00000 | 0.00000 | 0.00000 | 5.43320 | 0.00294 | 0.00000 | 0.00756 | 0.00000 | 4.18742 | 0.72844 | 1.00995 | 0.00000 |

| | | | | | | | | | | | | | | |
|---|---|---|---|---|---|---|---|---|---|---|---|---|---|---|
| **Lnpep** | 0.00000 | 0.00402 | 0.00000 | 0.00680 | 0.29638 | 0.03001 | 0.00397 | 0.00546 | 0.95070 | 0.00567 | 0.00642 | 0.00608 | 1.83290 | 0.82649 |
| **Lnpk1** | 1.96293 | 0.01119 | 1.95424 | 0.45123 | 1.00009 | 3.95804 | 0.00305 | 0.71080 | 2.96312 | 0.01894 | 1.68750 | 0.59665 | 0.62508 | 1.66759 |
| **Lnx1** | 0.03728 | 0.02149 | 0.10500 | 0.03859 | 0.19924 | 0.58246 | 0.11842 | 0.00986 | 0.00797 | 0.00403 | 1.06749 | 0.00910 | 0.01463 | 0.10314 |
| **Lnx2** | 0.07681 | 0.00288 | 0.30536 | 0.46004 | 0.65322 | 1.33216 | 0.00888 | 1.08923 | 0.00739 | 0.02149 | 1.19039 | 1.04512 | 1.50188 | 0.00000 |
| **LOC10050470** | 0.00897 | 0.00438 | 0.01509 | 0.01669 | 0.09358 | 0.01208 | 0.00423 | 0.01124 | 0.01022 | 0.00893 | 0.04560 | 0.00968 | 0.23224 | 0.02133 |
| **LOC10050502** | 0.01526 | 0.00450 | 0.00802 | 0.02392 | 0.01928 | 0.06540 | 0.00439 | 0.03310 | 0.07669 | 0.01503 | 0.01045 | 0.01795 | 0.62215 | 0.04736 |
| **LOC10086161** | 0.02236 | 0.00459 | 0.01270 | 0.05161 | 0.05581 | 0.01012 | 0.00397 | 0.00923 | 0.01905 | 0.00708 | 0.00800 | 0.08931 | 0.01282 | 0.01197 |
| **LOC10105607** | 0.07906 | 0.01118 | 1.36399 | 0.73899 | 0.76252 | 0.61451 | 0.16998 | 0.17685 | 5.51523 | 0.30957 | 0.84530 | 1.07357 | 0.03889 | 0.15493 |
| **LOC10263481** | 2.06417 | 0.16031 | 0.00000 | 0.47647 | 0.52541 | 1.44431 | 0.28746 | 0.85559 | 1.47164 | 1.14543 | 0.00092 | 0.94421 | 3.00109 | 0.25526 |
| **LOC10524239** | 0.07480 | 0.00408 | 0.18734 | 0.05217 | 0.22702 | 0.06468 | 0.00400 | 0.00375 | 0.33037 | 0.00517 | 0.01057 | 0.01356 | 0.87349 | 0.00000 |
| **LOC10524257** | 0.28097 | 0.07871 | 0.10126 | 0.38190 | 0.27979 | 0.63337 | 0.05525 | 0.09765 | 0.37765 | 0.21230 | 0.07136 | 0.14449 | 0.97744 | 0.77526 |
| **LOC10524586** | 0.80443 | 0.00339 | 0.48251 | 0.00000 | 0.83413 | 0.04563 | 0.00393 | 0.00000 | 0.39809 | 0.00219 | 0.80466 | 0.39460 | 0.57214 | 1.32416 |
| **LOC10524605** | 0.00579 | 0.00442 | 0.00861 | 0.01458 | 0.00000 | 0.01048 | 0.00420 | 0.00996 | 0.06989 | 0.00742 | 0.00749 | 0.00959 | 0.01313 | 0.42481 |
| **LOC10524650** | 0.00655 | 0.00994 | 0.04373 | 0.00000 | 0.02237 | 0.14763 | 0.00935 | 0.00721 | 0.01366 | 0.00670 | 0.00411 | 1.82811 | 0.05106 | 0.03778 |
| **LOC10602923** | 0.02192 | 0.00408 | 0.04286 | 0.03249 | 0.00126 | 0.08252 | 0.45654 | 0.00769 | 0.00912 | 0.01383 | 0.00613 | 0.00472 | 4.14133 | 0.13997 |
| **LOC106740** | 0.01259 | 0.00432 | 0.00757 | 0.02609 | 0.28077 | 0.02827 | 0.00414 | 0.00698 | 0.02210 | 0.00923 | 0.00741 | 0.00690 | 0.44983 | 0.12209 |
| **LOC215458** | 0.14483 | 0.01253 | 0.21586 | 0.14790 | 0.47369 | 0.00000 | 0.00468 | 0.04525 | 0.09161 | 0.00687 | 0.12258 | 0.00674 | 0.02931 | 0.84882 |
| **LOC666331** | 0.00457 | 0.00442 | 0.02251 | 0.01129 | 0.01920 | 0.01045 | 0.00437 | 0.00979 | 0.10431 | 0.00629 | 0.00790 | 0.00918 | 0.22575 | 0.11786 |
| **Lockd** | 0.00901 | 0.00435 | 0.05266 | 0.00952 | 0.00976 | 0.01113 | 0.00432 | 0.00918 | 0.00667 | 0.00723 | 0.00766 | 0.03979 | 0.00639 | 0.09673 |
| **Lonp1** | 8.90466 | 1.58850 | 15.59960 | 14.92025 | 13.29777 | 10.18833 | 4.85829 | 6.83610 | 1.39171 | 0.18065 | 10.25399 | 7.81684 | 4.27002 | 15.59142 |
| **Lonp2** | 4.60029 | 0.08131 | 9.07827 | 4.15205 | 8.81215 | 6.15857 | 0.09214 | 3.37306 | 4.31585 | 0.02982 | 4.07580 | 3.30463 | 6.39337 | 8.93273 |
| **Lonrf1** | 0.01116 | 0.04584 | 0.29523 | 0.19888 | 0.00000 | 1.22373 | 1.13243 | 0.50544 | 0.00837 | 0.03267 | 0.96800 | 0.88325 | 0.45354 | 0.95347 |
| **Lonrf2** | 0.00817 | 0.00441 | 0.05766 | 0.00606 | 0.24295 | 0.01261 | 0.00422 | 0.01000 | 0.12685 | 0.00639 | 0.01813 | 0.01184 | 0.00624 | 0.02047 |
| **Lonrf3** | 2.79798 | 0.01088 | 3.24616 | 1.15835 | 2.53468 | 0.66191 | 0.00308 | 0.15400 | 2.39490 | 0.00000 | 0.00000 | 0.05190 | 6.05262 | 6.37814 |
| **Loxhd1** | 0.00359 | 0.00427 | 0.30323 | 0.01532 | 0.36391 | 0.00074 | 0.00453 | 0.00995 | 0.01052 | 0.00664 | 0.01014 | 0.01163 | 0.05479 | 0.50344 |
| **Loxl1** | 0.13344 | 0.00435 | 1.47987 | 0.00000 | 3.58902 | 0.00745 | 0.00385 | 0.00425 | 0.00646 | 0.00350 | 0.00391 | 0.00556 | 0.32505 | 0.01119 |
| **Loxl2** | 2.35252 | 0.03482 | 0.00000 | 0.44186 | 0.22338 | 2.03496 | 0.24950 | 0.29014 | 1.73371 | 1.08749 | 0.03770 | 0.70182 | 8.10029 | 1.43536 |
| **Lpar1** | 2.27387 | 0.02028 | 1.97528 | 1.82596 | 1.78286 | 2.46595 | 0.13875 | 0.34135 | 4.13215 | 0.01209 | 0.02463 | 0.46911 | 0.17630 | 4.01426 |
| **Lpar2** | 0.00000 | 0.02481 | 0.75161 | 0.27733 | 0.00000 | 0.34576 | 0.01057 | 1.15938 | 0.00965 | 0.00770 | 0.00681 | 0.14930 | 0.00491 | 0.01547 |
| **Lpar5** | 0.00381 | 0.00405 | 0.47312 | 0.09449 | 0.02456 | 0.00000 | 0.00406 | 0.38403 | 0.00777 | 0.00692 | 0.00458 | 0.00774 | 0.02158 | 0.08009 |
| **Lpar6** | 0.00275 | 0.00425 | 0.07749 | 0.00553 | 0.00601 | 0.01156 | 0.00419 | 0.00913 | 0.02842 | 0.00634 | 0.00978 | 0.00866 | 0.77745 | 0.00140 |
| **Lpcat1** | 0.00000 | 0.01892 | 1.59983 | 1.47372 | 0.30853 | 0.89458 | 0.16438 | 1.54896 | 0.00770 | 0.00483 | 1.43724 | 0.88445 | 0.00000 | 0.46768 |
| **Lpcat2** | 0.01225 | 0.00433 | 0.36906 | 0.00536 | 0.28605 | 0.00819 | 0.00395 | 0.00833 | 0.09648 | 0.00850 | 0.00639 | 0.00718 | 0.24539 | 0.05755 |
| **Lpcat3** | 6.48018 | 1.93395 | 17.37529 | 17.36297 | 7.30546 | 14.78850 | 0.56174 | 11.26495 | 2.03948 | 6.58846 | 11.27166 | 16.49557 | 0.64711 | 10.11533 |
| **Lpcat4** | 0.03150 | 0.00415 | 0.05702 | 2.14252 | 0.40277 | 0.21157 | 0.00374 | 2.27658 | 0.06068 | 0.05377 | 0.00000 | 0.02726 | 0.15437 | 0.07333 |
| **Lpgat1** | 5.63935 | 0.14845 | 7.56843 | 2.53033 | 6.60753 | 7.24939 | 4.37648 | 3.24636 | 8.20932 | 0.17455 | 4.34894 | 2.29358 | 3.59421 | 7.72764 |
| **Lpin1** | 0.01389 | 0.00299 | 2.47858 | 0.00000 | 1.00047 | 0.79075 | 0.06577 | 0.00000 | 0.00586 | 0.00116 | 0.00544 | 1.53660 | 0.00000 | 0.06869 |
| **Lpin2** | 0.00000 | 0.00341 | 0.00000 | 0.00816 | 0.09406 | 0.01049 | 0.00389 | 0.00472 | 0.33929 | 0.01194 | 0.00293 | 0.66959 | 2.51248 | 3.18095 |
| **Lpin3** | 0.24922 | 0.00268 | 1.15895 | 1.87698 | 0.51648 | 0.29551 | 0.00864 | 0.26314 | 0.40820 | 0.02321 | 2.79069 | 1.16358 | 0.87394 | 1.20179 |
| **Lpl** | 0.00227 | 0.00446 | 0.01787 | 0.03218 | 0.01073 | 0.00726 | 0.01920 | 0.38489 | 0.04506 | 0.00585 | 0.01275 | 0.02174 | 0.30643 | 0.58194 |
| **Lpo** | 0.00632 | 0.00443 | 0.00892 | 0.01456 | 0.00000 | 0.01109 | 0.00420 | 0.01002 | 0.08764 | 0.00702 | 0.00788 | 0.00975 | 0.00500 | 0.28391 |
| **Lpp** | 0.27734 | 0.07118 | 2.13578 | 0.80771 | 1.18336 | 4.37039 | 1.05930 | 0.31627 | 1.25212 | 0.08236 | 2.81960 | 4.69496 | 0.43873 | 0.68848 |
| **Lppos** | 0.33721 | 0.01754 | 0.01133 | 0.12264 | 0.38479 | 0.35317 | 0.59611 | 0.24524 | 0.00672 | 0.03659 | 0.03025 | 0.00222 | 0.45258 | 0.00000 |
| **Lrba** | 1.00294 | 5.34766 | 3.09555 | 3.76263 | 1.66193 | 4.50034 | 0.82712 | 2.08175 | 0.86145 | 0.05276 | 2.05735 | 3.54301 | 1.58252 | 0.99923 |
| **Lrch1** | 0.02515 | 0.00563 | 0.00441 | 0.00000 | 0.58291 | 0.02538 | 0.00481 | 0.13019 | 0.18825 | 0.00720 | 0.00314 | 0.21082 | 0.00510 | 0.19935 |
| **Lrch3** | 0.01088 | 0.00406 | 0.02642 | 0.05837 | 0.09980 | 0.02102 | 0.00419 | 0.00598 | 0.34679 | 0.00568 | 0.01355 | 0.06143 | 0.55233 | 0.12159 |
| **Lrcol1** | 0.17416 | 0.00440 | 0.06148 | 0.95912 | 0.55840 | 0.00000 | 0.00394 | 0.00872 | 0.05634 | 0.01156 | 0.00508 | 0.00579 | 0.05178 | 0.00000 |
| **Lrfn1** | 0.05742 | 1.93627 | 0.12881 | 0.00087 | 0.00929 | 0.12814 | 0.00364 | 0.20911 | 0.01817 | 0.00771 | 0.00000 | 0.81889 | 0.00913 | 0.00348 |
| **Lrfn4** | 0.03154 | 0.86766 | 0.01686 | 0.76438 | 0.27904 | 0.00000 | 0.00491 | 0.11899 | 0.01700 | 0.00660 | 0.00601 | 0.17469 | 0.00590 | 0.00000 |
| **Lrguk** | 0.08146 | 0.01251 | 0.01791 | 0.00567 | 0.11742 | 0.04380 | 0.00432 | 0.00788 | 0.01019 | 0.00576 | 0.00727 | 0.00900 | 0.00650 | 0.01467 |
| **Lrif1** | 2.10819 | 0.00330 | 0.83514 | 0.00000 | 0.00000 | 0.29793 | 0.56194 | 0.52636 | 5.73286 | 0.01306 | 0.03004 | 0.06950 | 3.67714 | 0.00000 |
| **Lrig1** | 0.09898 | 0.01208 | 0.67608 | 0.06410 | 0.00000 | 0.97640 | 0.00435 | 0.05672 | 0.05309 | 0.00668 | 0.98550 | 0.20240 | 1.12682 | 1.17002 |
| **Lrig2** | 0.03278 | 0.01972 | 0.75692 | 0.37294 | 0.30127 | 0.50040 | 0.00562 | 0.53092 | 0.03801 | 0.01227 | 0.31309 | 0.68245 | 1.95618 | 1.44059 |
| **Lrig3** | 0.02054 | 0.00450 | 0.00976 | 0.00596 | 0.01276 | 0.00330 | 0.00406 | 0.01209 | 0.01141 | 0.00691 | 0.12039 | 0.28394 | 0.00682 | 0.01090 |
| **Lrmp** | 0.00687 | 0.00440 | 1.11046 | 0.00000 | 0.18784 | 0.00379 | 0.00396 | 0.00981 | 0.00897 | 0.00680 | 0.00558 | 0.00835 | 0.00602 | 0.01015 |

| | | | | | | | | | | | | | | |
|---|---|---|---|---|---|---|---|---|---|---|---|---|---|---|
| **Lrp1** | 0.48990 | 0.00377 | 0.00000 | 0.45963 | 0.36124 | 0.00000 | 0.00406 | 0.06569 | 0.09170 | 0.01666 | 1.86930 | 0.14621 | 0.28997 | 0.38682 |
| **Lrp10** | 1.77732 | 0.10176 | 2.72217 | 3.75373 | 0.17107 | 2.90007 | 4.08997 | 10.20521 | 0.15057 | 0.70387 | 1.30959 | 2.89326 | 0.14818 | 0.44527 |
| **Lrp11** | 2.19134 | 0.00987 | 4.18344 | 1.11334 | 1.84101 | 5.89256 | 0.02242 | 0.63134 | 2.44669 | 0.01569 | 3.18634 | 3.35462 | 0.00000 | 0.72597 |
| **Lrp12** | 0.00061 | 0.00407 | 0.11563 | 0.00000 | 0.20264 | 0.00013 | 0.01189 | 0.26029 | 0.16917 | 0.00613 | 0.00229 | 0.23739 | 3.23691 | 0.00000 |
| **Lrp2** | 0.00000 | 1.02189 | 5.13492 | 6.22962 | 0.92162 | 39.49702 | 7.50289 | 12.17933 | 0.58965 | 16.89801 | 9.24849 | 0.00003 | 0.00000 | 0.00000 |
| **Lrp3** | 0.04217 | 0.00443 | 0.03021 | 0.00000 | 0.02021 | 0.24823 | 0.00421 | 0.00491 | 0.01047 | 0.00910 | 0.00342 | 0.18768 | 0.01327 | 0.00587 |
| **Lrp4** | 0.01821 | 0.00388 | 0.00000 | 0.17076 | 0.01635 | 0.60060 | 0.01590 | 0.07081 | 0.73523 | 0.00425 | 0.26411 | 0.02004 | 0.28701 | 0.00000 |
| **Lrp5** | 0.06272 | 0.00395 | 0.00988 | 0.00735 | 0.04698 | 0.02478 | 0.00396 | 0.15141 | 0.00856 | 0.00670 | 0.00544 | 0.03337 | 0.08093 | 0.18906 |
| **Lrp6** | 1.63939 | 0.00948 | 3.61884 | 0.43571 | 2.48928 | 1.55354 | 0.00199 | 0.05115 | 8.35071 | 0.00557 | 0.00000 | 2.56959 | 2.69835 | 6.78320 |
| **Lrp8** | 0.19459 | 0.03650 | 0.41890 | 0.00000 | 0.00000 | 0.28296 | 0.02585 | 0.00644 | 0.05752 | 0.02372 | 0.00072 | 3.16955 | 0.00356 | 0.75008 |
| **Lrpap1** | 19.72497 | 9.15028 | 43.76451 | 45.01577 | 20.91486 | 36.87981 | 12.56312 | 26.58757 | 17.09891 | 2.64632 | 18.90915 | 17.84205 | 3.17793 | 15.63242 |
| **Lrpprc** | 7.65397 | 0.09584 | 18.30177 | 9.87657 | 8.51849 | 13.46302 | 2.76523 | 3.49527 | 10.57805 | 16.27765 | 9.99744 | 8.29303 | 0.55591 | 7.21691 |
| **Lrr1** | 19.79633 | 2.65086 | 22.83371 | 15.12025 | 35.85259 | 7.15499 | 2.66635 | 9.68408 | 17.25200 | 12.01557 | 4.56026 | 2.44399 | 11.05710 | 5.98222 |
| **Lrrc1** | 0.09694 | 0.00459 | 0.00828 | 0.00734 | 0.09870 | 0.08851 | 0.00484 | 0.00614 | 0.02185 | 0.00845 | 0.00663 | 0.01029 | 0.33934 | 0.01855 |
| **Lrrc10b** | 0.01668 | 0.00426 | 0.01287 | 0.01445 | 0.09888 | 0.01752 | 0.00417 | 0.02196 | 0.01408 | 0.00709 | 0.01163 | 0.01144 | 0.23502 | 0.03227 |
| **Lrrc14** | 3.27867 | 0.02576 | 1.14189 | 2.78326 | 1.88915 | 5.28128 | 0.03144 | 0.01419 | 0.00000 | 0.04115 | 0.00000 | 3.16604 | 0.08627 | 0.95476 |
| **Lrrc14b** | 0.09887 | 0.01286 | 0.03756 | 0.09482 | 0.10127 | 0.31991 | 0.03685 | 0.09018 | 0.21599 | 0.02795 | 0.01389 | 0.17805 | 0.32717 | 0.33980 |
| **Lrrc15** | 0.67034 | 0.00372 | 0.65990 | 0.13071 | 0.26274 | 0.00000 | 0.00991 | 0.00875 | 0.00702 | 0.00734 | 0.00528 | 0.00592 | 1.61418 | 7.34303 |
| **Lrrc16a** | 0.00000 | 6.63551 | 2.39554 | 0.39240 | 0.29078 | 1.59403 | 3.36369 | 1.63096 | 0.01060 | 0.05205 | 3.47019 | 1.02918 | 0.78626 | 0.40697 |
| **Lrrc17** | 0.17405 | 0.02051 | 0.12101 | 0.03709 | 0.08224 | 0.12838 | 0.00446 | 0.09602 | 0.03247 | 0.02407 | 0.04441 | 0.01945 | 0.81555 | 1.15053 |
| **Lrrc2** | 0.30703 | 0.03240 | 0.75560 | 0.84588 | 0.05956 | 0.68135 | 0.09219 | 0.03753 | 0.91775 | 0.42418 | 0.00858 | 1.84654 | 5.34757 | 0.97529 |
| **Lrrc20** | 0.02162 | 0.00385 | 0.13159 | 0.12357 | 0.46722 | 1.20734 | 0.01731 | 0.01023 | 0.12900 | 0.02203 | 0.07038 | 0.16562 | 0.23148 | 0.10515 |
| **Lrrc23** | 0.00333 | 0.01259 | 0.07397 | 0.00000 | 0.42925 | 0.11109 | 0.00390 | 0.00797 | 0.00828 | 0.00531 | 0.00320 | 0.83419 | 0.12190 | 0.04939 |
| **Lrrc25** | 0.00920 | 0.00447 | 0.08751 | 0.00000 | 0.95589 | 0.01002 | 0.00425 | 0.01091 | 0.01131 | 0.00769 | 0.00706 | 0.00901 | 0.00561 | 0.01221 |
| **Lrrc27** | 0.00833 | 0.00499 | 0.01121 | 0.00000 | 0.09194 | 0.01768 | 0.00446 | 0.02477 | 0.00772 | 0.00613 | 0.00490 | 0.32017 | 0.00551 | 0.03801 |
| **Lrrc28** | 1.08909 | 0.01020 | 1.55280 | 0.00000 | 0.92885 | 1.59700 | 0.00373 | 0.33813 | 0.55296 | 0.01051 | 0.07017 | 1.31048 | 0.95072 | 0.00000 |
| **Lrrc3** | 0.11757 | 0.12061 | 0.04176 | 0.19881 | 0.06196 | 0.15226 | 0.09215 | 0.02973 | 0.29864 | 0.02581 | 0.01194 | 0.08788 | 0.30795 | 0.24819 |
| **Lrrc31** | 0.03321 | 0.02117 | 0.02503 | 0.19334 | 0.06008 | 0.09387 | 0.03619 | 0.06575 | 0.09154 | 0.03631 | 0.02549 | 0.03847 | 0.68551 | 0.17156 |
| **Lrrc32** | 0.03056 | 0.00441 | 0.01288 | 0.01924 | 0.01695 | 0.01180 | 0.00421 | 0.00990 | 0.02199 | 0.00862 | 0.00821 | 0.00911 | 0.06859 | 0.00573 |
| **Lrrc34** | 0.00521 | 0.00419 | 0.11475 | 0.24558 | 0.00541 | 0.00320 | 0.00402 | 0.00667 | 0.01374 | 0.01188 | 0.00632 | 0.01630 | 0.00650 | 0.00916 |
| **Lrrc39** | 0.53776 | 0.02741 | 0.42677 | 0.35312 | 0.44781 | 0.97294 | 0.36426 | 0.23137 | 1.28465 | 0.50780 | 0.30169 | 0.39849 | 0.73121 | 0.65514 |
| **Lrrc4** | 0.04762 | 0.00410 | 0.00000 | 0.60199 | 0.16037 | 0.20335 | 0.00394 | 0.00326 | 0.34874 | 0.00549 | 0.00484 | 0.25911 | 0.65471 | 0.13651 |
| **Lrrc40** | 2.51050 | 0.02792 | 5.80408 | 4.85845 | 5.57925 | 4.89036 | 1.12604 | 2.49496 | 4.70248 | 0.04772 | 4.33878 | 6.01887 | 3.98100 | 8.85435 |
| **Lrrc41** | 2.93759 | 0.05798 | 3.57183 | 1.11866 | 3.56442 | 2.52465 | 1.39773 | 2.06921 | 1.31663 | 0.00550 | 5.05789 | 1.47108 | 1.13541 | 2.87053 |
| **Lrrc42** | 18.76698 | 0.02353 | 20.69799 | 6.45825 | 17.61849 | 1.34854 | 3.38514 | 2.27729 | 8.10543 | 0.06112 | 6.34181 | 5.55324 | 15.43373 | 13.12522 |
| **Lrrc43** | 0.00349 | 0.00436 | 0.06568 | 0.00590 | 0.02692 | 0.01014 | 0.00432 | 0.01152 | 0.00736 | 0.00654 | 0.00790 | 0.00979 | 0.14650 | 0.12170 |
| **Lrrc45** | 0.64849 | 0.01924 | 3.11729 | 1.81929 | 0.41960 | 1.13561 | 1.98302 | 0.02989 | 0.00000 | 0.11130 | 1.89910 | 0.52681 | 0.63415 | 0.00000 |
| **Lrrc46** | 0.04914 | 0.01077 | 0.83881 | 0.00678 | 0.91420 | 0.10786 | 0.00322 | 1.92021 | 0.00386 | 0.00960 | 0.00504 | 0.01467 | 1.14564 | 0.17986 |
| **Lrrc47** | 0.33318 | 0.00425 | 1.31454 | 1.16571 | 1.04393 | 1.98891 | 0.05222 | 0.07504 | 0.84608 | 0.01381 | 1.45390 | 0.39291 | 0.00403 | 2.41655 |
| **Lrrc48** | 0.60064 | 0.00378 | 0.46524 | 0.00000 | 0.42284 | 0.06934 | 0.00349 | 0.22546 | 0.06887 | 0.00586 | 0.00380 | 0.00381 | 0.03245 | 0.34913 |
| **Lrrc49** | 0.01126 | 0.00437 | 0.01199 | 0.01289 | 0.00906 | 0.01039 | 0.00427 | 0.00950 | 0.00927 | 0.01296 | 0.02167 | 0.00939 | 0.12645 | 0.20438 |
| **Lrrc51** | 1.43850 | 0.00333 | 1.71443 | 0.91618 | 0.77036 | 0.43163 | 0.01396 | 0.00000 | 0.01889 | 0.01994 | 0.00000 | 0.66033 | 0.42432 | 0.94281 |
| **Lrrc56** | 0.03160 | 0.01156 | 0.07184 | 0.68659 | 0.00699 | 0.84048 | 0.01224 | 0.21967 | 0.00619 | 0.01835 | 0.00000 | 1.13011 | 1.06350 | 0.00000 |
| **Lrrc57** | 0.12293 | 0.00200 | 3.14233 | 0.46940 | 1.71158 | 3.71130 | 0.62095 | 0.23512 | 6.76909 | 0.06430 | 5.89480 | 0.53148 | 3.54063 | 7.92289 |
| **Lrrc58** | 6.29465 | 4.64980 | 2.31267 | 7.46405 | 11.82098 | 13.97898 | 2.41764 | 12.85598 | 24.19952 | 5.32709 | 5.33559 | 6.63629 | 4.54672 | 14.10826 |
| **Lrrc59** | 15.22333 | 7.54462 | 26.20824 | 21.28413 | 15.42195 | 31.19559 | 7.64356 | 20.53812 | 6.25686 | 5.78970 | 15.51819 | 18.49412 | 5.48060 | 10.95374 |
| **Lrrc61** | 0.01610 | 0.02742 | 0.03392 | 0.02792 | 0.06731 | 0.46045 | 0.03418 | 0.35926 | 0.02229 | 0.05192 | 0.05548 | 0.03216 | 0.05750 | 0.10982 |
| **Lrrc73** | 0.02713 | 0.00414 | 0.01581 | 0.05069 | 0.01930 | 0.13248 | 0.00404 | 0.00385 | 0.00539 | 0.00656 | 0.00675 | 0.01430 | 0.46599 | 0.04293 |
| **Lrrc74a** | 0.00906 | 0.00491 | 0.02648 | 0.00055 | 0.01338 | 0.23756 | 0.00435 | 0.00664 | 0.00781 | 0.00630 | 0.00719 | 0.01150 | 0.00613 | 0.02869 |
| **Lrrc74b** | 0.01419 | 0.00440 | 0.00946 | 0.02550 | 0.33895 | 0.03704 | 0.01111 | 0.00809 | 0.05872 | 0.00659 | 0.01007 | 0.01955 | 0.16869 | 0.25380 |
| **Lrrc75a** | 0.01234 | 0.00475 | 0.00286 | 0.01084 | 0.14909 | 0.79495 | 2.62242 | 0.04754 | 0.00896 | 0.07845 | 0.00504 | 0.00000 | 0.00684 | 0.00580 |
| **Lrrc75b** | 1.17861 | 0.01053 | 0.52812 | 0.11035 | 0.12239 | 0.40577 | 0.00997 | 1.83903 | 1.86958 | 0.01611 | 0.03061 | 0.79131 | 0.01217 | 0.00000 |
| **Lrrc8a** | 0.63610 | 4.04620 | 0.15507 | 0.68436 | 1.59730 | 1.96391 | 1.97042 | 0.09292 | 1.40157 | 0.02973 | 0.00000 | 2.00578 | 1.31312 | 1.58993 |
| **Lrrc8b** | 0.01657 | 0.00412 | 0.21595 | 0.00000 | 0.12852 | 0.08457 | 0.00401 | 0.00385 | 0.07234 | 0.00501 | 0.00471 | 0.21660 | 0.02310 | 0.47476 |
| **Lrrc8d** | 0.02984 | 0.01908 | 0.76292 | 0.22012 | 0.42474 | 0.52064 | 0.00376 | 1.07472 | 0.93671 | 0.00780 | 1.75402 | 0.99812 | 1.00800 | 1.86633 |

| | | | | | | | | | | | | | | |
|---|---|---|---|---|---|---|---|---|---|---|---|---|---|---|
| **Lrrc8e** | 0.00000 | 0.04210 | 0.95023 | 0.71788 | 0.14529 | 0.05261 | 0.02101 | 0.05881 | 0.97458 | 0.02511 | 0.02632 | 0.00966 | 1.03443 | 3.17268 |
| **Lrrcc1** | 0.00000 | 0.00415 | 0.07636 | 0.00000 | 0.55371 | 1.14650 | 0.00446 | 0.18121 | 2.94944 | 0.00633 | 0.00000 | 0.21082 | 1.04443 | 0.10160 |
| **Lrrfip1** | 0.00000 | 0.06237 | 0.58438 | 0.63331 | 1.02298 | 7.24300 | 3.91100 | 10.97699 | 0.49454 | 5.77944 | 4.01384 | 0.00000 | 1.57676 | 5.71964 |
| **Lrrfip2** | 8.02625 | 0.04765 | 8.20599 | 2.30439 | 7.17978 | 6.29421 | 0.02469 | 2.89716 | 4.43801 | 0.07037 | 5.80953 | 8.10965 | 0.56113 | 5.14243 |
| **Lrriq3** | 0.00888 | 0.00449 | 0.01523 | 0.00953 | 0.27398 | 0.01335 | 0.00493 | 0.01029 | 0.02039 | 0.00680 | 0.00878 | 0.00866 | 0.00646 | 0.02038 |
| **Lrrn4** | 0.00000 | 0.00383 | 0.52210 | 0.00364 | 0.33055 | 0.50621 | 0.01447 | 1.90065 | 0.01545 | 0.01269 | 0.00243 | 0.00378 | 0.95218 | 0.64951 |
| **Lrrn4cl** | 0.17040 | 0.03721 | 0.01348 | 0.50432 | 0.10982 | 0.32436 | 0.05197 | 0.10824 | 0.25367 | 0.10472 | 0.01081 | 0.05698 | 0.75596 | 0.36107 |
| **Lrrtm2** | 0.00599 | 0.00407 | 0.13753 | 0.16571 | 0.03844 | 0.04395 | 0.00417 | 0.00209 | 0.01033 | 0.00398 | 0.29554 | 0.08573 | 0.00955 | 0.18269 |
| **Lrsam1** | 1.25162 | 0.00977 | 1.33331 | 1.34642 | 0.84452 | 1.38592 | 0.71015 | 1.03495 | 0.00573 | 0.00401 | 0.00997 | 3.22412 | 0.00284 | 0.38650 |
| **Lrwd1** | 1.55944 | 0.06795 | 5.97730 | 7.00558 | 2.79652 | 2.38694 | 0.00068 | 5.26112 | 0.08309 | 0.04244 | 2.32827 | 3.56777 | 1.46822 | 0.37160 |
| **Lsg1** | 5.44467 | 0.06934 | 16.23092 | 1.50552 | 11.12277 | 7.30043 | 1.59558 | 1.44377 | 0.18544 | 0.07607 | 7.62521 | 5.58325 | 1.08318 | 6.41342 |
| **Lsm1** | 2.37278 | 0.02662 | 0.96543 | 6.80189 | 5.17046 | 2.80424 | 0.02271 | 1.17961 | 0.09631 | 0.11625 | 2.07809 | 2.64491 | 3.63224 | 0.10570 |
| **Lsm10** | 0.00000 | 0.06926 | 1.01492 | 0.55574 | 1.90501 | 0.66994 | 0.00732 | 1.55799 | 0.73065 | 12.28859 | 0.33975 | 1.18765 | 2.10673 | 2.06216 |
| **Lsm11** | 1.34586 | 0.02545 | 0.61876 | 1.74161 | 1.27620 | 0.63229 | 0.43123 | 0.20551 | 0.23680 | 0.05115 | 0.00846 | 0.03313 | 0.56280 | 0.52199 |
| **Lsm12** | 6.03609 | 0.05524 | 5.87739 | 14.09798 | 16.15331 | 3.27371 | 2.59077 | 3.03045 | 3.31178 | 0.09558 | 2.39165 | 1.75645 | 3.43385 | 4.35975 |
| **Lsm14a** | 5.53096 | 0.08965 | 11.23282 | 5.91468 | 5.56290 | 10.69348 | 6.35627 | 6.59720 | 4.43159 | 0.26619 | 7.55146 | 8.74781 | 7.26091 | 5.04163 |
| **Lsm14b** | 0.29831 | 0.01308 | 0.26718 | 0.72403 | 0.49723 | 0.48164 | 0.00393 | 0.70187 | 0.52116 | 0.00799 | 0.00702 | 0.47641 | 5.85301 | 0.49767 |
| **Lsm2** | 1.90005 | 0.10665 | 0.65609 | 2.50582 | 13.11575 | 3.86540 | 0.82945 | 3.54066 | 1.12680 | 0.04474 | 2.87353 | 4.29448 | 6.81007 | 5.01819 |
| **Lsm3** | 1.84192 | 0.09876 | 1.02422 | 5.02630 | 6.82217 | 11.82845 | 5.33554 | 2.12234 | 4.22693 | 9.45753 | 1.81152 | 11.94209 | 0.00000 | 4.73964 |
| **Lsm4** | 0.00891 | 0.06101 | 0.00000 | 18.12199 | 11.46568 | 0.35712 | 0.05197 | 0.71975 | 0.00943 | 0.08452 | 0.01781 | 1.14375 | 0.38245 | 0.00000 |
| **Lsm5** | 3.54837 | 4.03373 | 0.00000 | 6.07863 | 8.39576 | 5.60178 | 1.60989 | 5.08084 | 7.67023 | 1.28875 | 0.40497 | 5.38186 | 0.13448 | 6.24996 |
| **Lsm6** | 2.58531 | 0.16954 | 0.69554 | 1.69405 | 4.49944 | 6.19113 | 0.85307 | 6.86752 | 3.72665 | 2.52521 | 2.05207 | 5.26259 | 2.17188 | 2.85071 |
| **Lsm7** | 2.98694 | 0.11682 | 0.00000 | 3.13142 | 5.33508 | 6.08927 | 0.92875 | 3.75469 | 0.00000 | 4.32487 | 0.17157 | 5.79422 | 2.54627 | 1.70969 |
| **Lsm8** | 5.95181 | 0.23942 | 3.05058 | 6.62311 | 9.12788 | 9.34266 | 2.96900 | 3.87786 | 5.67253 | 0.98586 | 3.46983 | 6.08269 | 0.63696 | 9.77809 |
| **Lsp1** | 0.01757 | 0.00437 | 0.09086 | 0.00372 | 0.09225 | 0.01097 | 0.00445 | 0.00987 | 0.01127 | 0.00620 | 0.00770 | 0.00900 | 0.35271 | 0.02281 |
| **Lsr** | 0.00000 | 0.13378 | 0.24638 | 9.87317 | 1.28417 | 11.76156 | 1.97886 | 10.78460 | 0.02278 | 0.22060 | 5.31019 | 8.06501 | 0.45651 | 0.63391 |
| **Lss** | 3.04154 | 0.00935 | 7.77909 | 2.00265 | 4.13661 | 6.58507 | 0.01635 | 0.42985 | 5.05016 | 0.00418 | 0.00000 | 3.05297 | 0.22234 | 7.58067 |
| **Lst1** | 0.39611 | 0.00398 | 0.00000 | 0.11749 | 0.98819 | 0.10105 | 0.00399 | 0.01940 | 0.20240 | 0.01023 | 0.00509 | 0.00104 | 3.98054 | 0.10499 |
| **Lta** | 0.00591 | 0.00431 | 0.01209 | 0.01216 | 0.00395 | 0.01108 | 0.00447 | 0.01321 | 0.00662 | 0.00608 | 0.00858 | 0.00804 | 0.79738 | 0.28037 |
| **Lta4h** | 0.66730 | 9.37735 | 0.94858 | 4.16933 | 0.94852 | 8.97747 | 4.27806 | 3.13777 | 0.09946 | 11.76888 | 4.07502 | 3.23345 | 0.08231 | 0.33335 |
| **Ltb** | 0.01107 | 0.00407 | 0.00737 | 0.40151 | 1.22080 | 0.00000 | 0.00393 | 0.01210 | 0.01336 | 0.00523 | 0.02067 | 0.01505 | 0.00274 | 0.00462 |
| **Ltbp1** | 0.11714 | 0.00440 | 0.16238 | 0.00197 | 0.00414 | 0.00850 | 0.00422 | 0.01186 | 0.00948 | 0.00681 | 0.00804 | 0.00936 | 0.00371 | 0.02809 |
| **Ltbp2** | 0.55623 | 0.01228 | 1.25828 | 0.16471 | 0.46527 | 0.46007 | 0.00467 | 1.44307 | 0.00866 | 0.01146 | 0.00385 | 0.00202 | 0.09513 | 0.08714 |
| **Ltbp3** | 0.00777 | 0.00329 | 0.67982 | 0.00000 | 1.36932 | 0.00213 | 0.00346 | 0.12646 | 0.00444 | 0.01060 | 0.00601 | 0.73234 | 1.09124 | 0.00431 |
| **Ltbp4** | 1.53523 | 0.12292 | 3.86389 | 1.17041 | 1.96624 | 0.00000 | 0.00290 | 0.29191 | 0.01608 | 0.03363 | 0.00000 | 2.63860 | 0.91283 | 0.57705 |
| **Ltf** | 0.00666 | 0.00450 | 0.22964 | 0.00000 | 4.71193 | 0.00450 | 0.00434 | 0.01039 | 0.01132 | 0.00714 | 0.00472 | 0.00596 | 0.00000 | 0.00508 |
| **Ltk** | 0.02674 | 0.00435 | 0.00654 | 0.00434 | 0.42049 | 0.00495 | 0.00524 | 0.26893 | 0.00830 | 0.00619 | 0.00911 | 0.00700 | 0.53827 | 0.02993 |
| **Ltn1** | 6.23201 | 0.04244 | 2.34028 | 0.00000 | 2.66634 | 3.10867 | 1.36793 | 1.34966 | 8.13290 | 0.02324 | 4.46006 | 1.65960 | 4.99225 | 3.38381 |
| **Ltv1** | 23.85053 | 3.33690 | 25.28217 | 15.70842 | 27.27409 | 17.29531 | 7.09204 | 8.67691 | 17.95691 | 12.93253 | 12.83819 | 9.00865 | 13.96313 | 21.14296 |
| **Luc7l** | 8.21626 | 0.11950 | 23.22800 | 9.49651 | 19.00243 | 13.15144 | 0.66984 | 3.44028 | 9.09334 | 12.56580 | 10.49496 | 5.76234 | 5.39435 | 12.92821 |
| **Luc7l2** | 6.67833 | 0.09697 | 4.21190 | 1.29640 | 5.91700 | 4.45108 | 0.98418 | 1.17776 | 11.05825 | 1.07406 | 0.15786 | 3.84707 | 14.62120 | 8.61274 |
| **Luc7l3** | 12.59396 | 0.07703 | 12.68285 | 2.11697 | 11.94877 | 10.21334 | 2.10810 | 2.65315 | 15.56764 | 0.07109 | 9.92913 | 5.74729 | 8.78201 | 11.05424 |
| **Lurap1** | 0.41855 | 0.00477 | 0.09318 | 0.00340 | 0.16734 | 0.69429 | 0.00428 | 0.59912 | 0.00554 | 0.03202 | 0.00276 | 0.00499 | 0.05926 | 0.00000 |
| **Lurap1l** | 0.05622 | 0.06711 | 0.00229 | 0.00000 | 0.00664 | 0.31623 | 2.65904 | 2.16444 | 0.01024 | 0.09604 | 0.00000 | 0.59206 | 0.18861 | 0.00000 |
| **Luzp1** | 0.07032 | 4.73213 | 3.66219 | 2.63214 | 8.20400 | 5.48434 | 2.87707 | 2.78003 | 1.80329 | 0.15142 | 5.02108 | 1.19206 | 7.99142 | 7.63918 |
| **Luzp2** | 0.20645 | 0.00456 | 0.02968 | 0.01589 | 0.00356 | 0.00784 | 0.00419 | 0.01438 | 0.04030 | 0.01057 | 0.00666 | 0.00835 | 0.00442 | 0.13093 |
| **Luzp4** | 0.00893 | 0.00443 | 0.02896 | 0.01113 | 0.00808 | 0.01228 | 0.00428 | 0.01028 | 0.00792 | 0.00702 | 0.00894 | 0.00981 | 0.00980 | 0.15433 |
| **Lxn** | 0.03147 | 0.00451 | 0.29500 | 0.03458 | 0.39095 | 0.80144 | 0.00423 | 0.62382 | 0.00730 | 0.00740 | 0.51269 | 0.02754 | 0.09625 | 1.20024 |
| **Ly6a** | 6.20676 | 6.56458 | 1.44887 | 2.51505 | 1.19973 | 15.06795 | 1.49324 | 2.71467 | 0.42068 | 0.05225 | 3.98032 | 1.65588 | 2.31739 | 0.36332 |
| **Ly6c1** | 0.10150 | 0.13912 | 0.28595 | 1.39145 | 0.08724 | 0.47624 | 0.00949 | 0.03109 | 0.01039 | 0.01102 | 0.19563 | 0.44397 | 0.27100 | 0.15841 |
| **Ly6c2** | 0.00389 | 0.01129 | 0.08682 | 0.31846 | 0.01546 | 0.00743 | 0.00401 | 0.00479 | 0.01123 | 0.00592 | 0.01418 | 0.10676 | 0.00451 | 0.03555 |
| **Ly6e** | 8.25836 | 0.04353 | 0.95967 | 6.55582 | 0.61967 | 3.78353 | 0.00976 | 0.01893 | 0.00746 | 0.05599 | 0.00000 | 3.07043 | 1.47425 | 0.00000 |
| **Ly6f** | 0.00000 | 0.83552 | 0.92754 | 4.76497 | 0.00000 | 0.75612 | 3.51867 | 5.01306 | 0.00794 | 0.16397 | 0.00000 | 0.00000 | 0.00609 | 0.00000 |
| **Ly6g** | 0.00833 | 0.02696 | 0.05750 | 0.08257 | 0.01169 | 0.01241 | 0.01104 | 0.04912 | 0.01250 | 0.00592 | 0.01546 | 0.00674 | 0.00670 | 0.01092 |
| **Ly6g5b** | 0.05680 | 0.02400 | 0.34042 | 0.01747 | 0.46535 | 0.00000 | 0.00400 | 0.84502 | 0.00983 | 0.01130 | 1.15913 | 0.00242 | 0.00517 | 0.61505 |

| | | | | | | | | | | | | | | |
|---|---|---|---|---|---|---|---|---|---|---|---|---|---|---|
| **Ly6g5c** | 0.01578 | 0.00446 | 0.00755 | 0.01166 | 0.13020 | 0.00963 | 0.00478 | 0.01041 | 0.25446 | 0.00743 | 0.00699 | 0.00879 | 0.00547 | 0.13361 |
| **Ly6g6c** | 0.02261 | 0.00444 | 0.00000 | 0.68399 | 0.02441 | 0.00000 | 0.00407 | 0.02778 | 0.01792 | 0.00737 | 0.52676 | 0.88279 | 0.31976 | 0.00000 |
| **Ly6g6e** | 4.32661 | 0.02322 | 0.43790 | 0.00000 | 0.24561 | 0.00000 | 0.00297 | 0.00004 | 0.01587 | 0.02444 | 0.66143 | 3.99858 | 0.57715 | 0.62570 |
| **Ly6i** | 0.06362 | 0.06676 | 0.08782 | 0.01093 | 0.18951 | 0.13710 | 0.00928 | 1.11825 | 0.10275 | 0.00588 | 0.05034 | 0.01462 | 0.14620 | 0.00000 |
| **Ly75** | 0.02213 | 0.00445 | 0.02694 | 0.01177 | 0.03022 | 0.01123 | 0.00434 | 0.01051 | 0.01056 | 0.00690 | 0.00768 | 0.00980 | 0.03823 | 0.21352 |
| **Ly96** | 1.07686 | 0.00383 | 0.18722 | 0.59907 | 0.58320 | 0.01501 | 0.00380 | 0.00367 | 1.39178 | 0.02267 | 0.06203 | 0.00934 | 1.17054 | 0.00000 |
| **Lyar** | 19.48638 | 0.84193 | 23.23818 | 17.48185 | 29.10851 | 14.65695 | 3.68514 | 5.99837 | 1.09503 | 5.26631 | 9.49613 | 12.91744 | 16.60744 | 12.00939 |
| **Lyn** | 0.00000 | 0.02989 | 0.92001 | 0.22055 | 0.94081 | 3.04407 | 0.00390 | 2.14331 | 0.37522 | 0.01827 | 0.00687 | 0.00191 | 0.00000 | 0.00000 |
| **Lypd1** | 0.19454 | 0.03576 | 0.02677 | 0.14749 | 0.10748 | 0.45026 | 0.07743 | 0.06988 | 0.44891 | 0.12368 | 0.04851 | 0.13603 | 3.34212 | 0.34890 |
| **Lypd3** | 0.14910 | 0.00409 | 0.83907 | 0.94229 | 0.00000 | 0.00000 | 0.00379 | 0.00677 | 0.08127 | 0.00922 | 0.00446 | 0.00664 | 0.00893 | 0.08941 |
| **Lypd5** | 0.01507 | 0.00455 | 0.47099 | 0.00000 | 0.00827 | 0.10197 | 0.00418 | 0.00510 | 0.08269 | 0.00535 | 0.00649 | 0.00818 | 0.00697 | 0.01117 |
| **Lypd6** | 0.00660 | 0.00440 | 0.02263 | 0.01081 | 0.01104 | 0.01337 | 0.00427 | 0.00980 | 0.02001 | 0.00673 | 0.00924 | 0.00966 | 0.15115 | 0.00146 |
| **Lypd8** | 0.02111 | 0.00438 | 0.43248 | 0.01245 | 0.19993 | 0.00289 | 0.00400 | 0.01075 | 0.00962 | 0.01184 | 0.00616 | 0.00668 | 0.15353 | 0.03235 |
| **Lypla1** | 10.57094 | 0.06808 | 9.14022 | 2.72855 | 10.12299 | 3.79474 | 1.03569 | 1.68024 | 9.21512 | 0.06779 | 10.26917 | 3.36342 | 2.55993 | 9.93605 |
| **Lypla2** | 0.00000 | 0.04885 | 4.46639 | 6.28906 | 4.97623 | 0.00000 | 0.50318 | 4.70419 | 0.01236 | 0.03076 | 0.00000 | 1.57079 | 0.11431 | 0.00000 |
| **Lyplal1** | 0.00438 | 0.00399 | 0.24776 | 0.02710 | 0.06825 | 0.00000 | 0.00410 | 0.00885 | 0.00372 | 0.00511 | 0.41206 | 0.00631 | 1.29705 | 0.02941 |
| **Lyrm1** | 0.45332 | 0.00830 | 0.60596 | 3.19681 | 0.07915 | 4.58081 | 0.01189 | 0.00713 | 1.49636 | 11.97386 | 2.49577 | 0.53762 | 1.42305 | 0.57374 |
| **Lyrm2** | 0.74263 | 0.02892 | 1.26244 | 2.55484 | 2.42809 | 1.72655 | 3.35671 | 1.59822 | 2.98191 | 0.02380 | 2.36706 | 1.00389 | 1.04002 | 2.45033 |
| **Lyrm4** | 2.12949 | 0.01827 | 0.32482 | 2.15488 | 0.63543 | 3.55040 | 1.63804 | 2.09001 | 2.26689 | 0.06490 | 1.80082 | 0.99441 | 0.48773 | 0.00000 |
| **Lyrm5** | 2.25440 | 0.02317 | 0.76102 | 2.58754 | 1.39439 | 2.52923 | 0.82783 | 0.54528 | 1.77942 | 0.02709 | 5.45422 | 2.94048 | 1.56453 | 0.95359 |
| **Lyrm7** | 0.32224 | 0.00473 | 0.08808 | 0.06452 | 0.17963 | 0.65833 | 0.00413 | 0.59801 | 0.02447 | 0.01052 | 0.00000 | 0.61186 | 0.16172 | 0.03131 |
| **Lyrm9** | 0.18861 | 0.01141 | 0.01151 | 0.45930 | 0.51940 | 0.40494 | 0.00976 | 1.50507 | 0.15148 | 0.04829 | 0.00237 | 0.41674 | 0.27717 | 0.57384 |
| **Lysmd1** | 0.00000 | 0.00241 | 0.62078 | 0.82109 | 0.74180 | 1.42683 | 0.00954 | 0.16845 | 0.03584 | 0.01138 | 1.03132 | 1.05727 | 6.29860 | 1.84222 |
| **Lysmd2** | 0.69485 | 0.04999 | 0.27139 | 0.00000 | 0.15912 | 0.93693 | 1.45457 | 0.64178 | 0.00600 | 0.01789 | 0.00000 | 2.48448 | 0.01143 | 0.00680 |
| **Lysmd3** | 0.00781 | 0.00442 | 0.23758 | 0.30707 | 0.25365 | 0.71037 | 0.01412 | 0.10354 | 0.30938 | 0.01027 | 0.15560 | 0.00131 | 0.00263 | 0.17940 |
| **Lysmd4** | 0.46208 | 0.01129 | 0.14442 | 0.14505 | 0.34388 | 0.84612 | 0.01478 | 1.81532 | 0.02913 | 0.02825 | 1.45607 | 0.12015 | 1.35484 | 0.70185 |
| **Lyst** | 0.01210 | 0.00436 | 0.00000 | 0.06889 | 0.01435 | 0.08744 | 0.00414 | 0.00391 | 0.91554 | 0.00522 | 0.47549 | 0.04150 | 0.00198 | 0.66143 |
| **Lyz1** | 0.00000 | 0.00416 | 0.23134 | 0.01147 | 0.16149 | 0.02182 | 0.00427 | 0.01561 | 0.00978 | 0.00490 | 0.03567 | 0.01035 | 0.37189 | 0.00388 |
| **Lyz2** | 0.00469 | 0.00424 | 0.56674 | 0.01846 | 0.52441 | 0.00760 | 0.00414 | 0.00891 | 0.01981 | 0.00414 | 0.11156 | 0.00894 | 0.27258 | 0.00705 |
| **Lzic** | 2.86302 | 1.94314 | 3.85935 | 1.48456 | 0.36568 | 2.15480 | 0.02938 | 0.05366 | 0.49631 | 0.00000 | 4.49040 | 2.19899 | 3.71625 | 2.49873 |
| **Lztfl1** | 7.73462 | 0.00960 | 0.34831 | 0.94253 | 0.63895 | 1.63722 | 0.00192 | 0.10457 | 11.84709 | 0.00466 | 2.49328 | 0.47056 | 3.18546 | 4.55342 |
| **Lztr1** | 3.94461 | 0.89746 | 4.20709 | 4.76888 | 4.08188 | 2.39827 | 0.00767 | 2.76223 | 0.00522 | 0.02108 | 0.37373 | 3.79925 | 3.99653 | 1.10640 |
| **Lzts2** | 0.27026 | 0.21679 | 1.24931 | 0.37835 | 3.53158 | 0.63721 | 0.98945 | 2.36236 | 0.02079 | 0.06706 | 0.00000 | 0.00000 | 0.18481 | 0.16075 |
| **Lzts3** | 0.00447 | 0.01252 | 0.00804 | 0.03252 | 0.21206 | 0.04661 | 0.01078 | 0.08183 | 0.05131 | 0.02144 | 0.02049 | 0.00969 | 0.20161 | 0.20950 |
| **M6pr** | 10.34209 | 7.13636 | 16.23752 | 11.79922 | 9.26047 | 16.26182 | 3.15063 | 5.52085 | 12.46884 | 0.25469 | 10.09031 | 9.99741 | 6.34243 | 16.61307 |
| **Maats1** | 0.02310 | 0.00421 | 0.00365 | 0.12762 | 0.40955 | 0.00995 | 0.00393 | 0.00602 | 0.00802 | 0.00657 | 0.00700 | 0.02563 | 0.00796 | 0.02298 |
| **Mab21l3** | 0.00221 | 0.03711 | 0.04731 | 0.20335 | 0.06028 | 0.14604 | 0.35421 | 0.00687 | 0.00933 | 0.00764 | 0.04379 | 0.00317 | 0.00297 | 0.16692 |
| **Macf1** | 0.00000 | 1.21624 | 13.39780 | 7.23919 | 14.57164 | 5.78357 | 1.50476 | 4.77487 | 7.47581 | 0.07608 | 3.61382 | 5.79006 | 6.65500 | 8.08591 |
| **Macrod1** | 0.05515 | 0.00501 | 0.04697 | 0.03403 | 0.03472 | 0.01494 | 0.00440 | 0.06198 | 0.00974 | 0.00735 | 0.00702 | 0.02502 | 0.00673 | 0.03142 |
| **Macrod2** | 0.04533 | 0.00309 | 0.00000 | 0.19363 | 0.26436 | 0.00000 | 0.26602 | 0.00075 | 0.00838 | 0.01814 | 0.40815 | 1.05634 | 0.15012 | 0.01022 |
| **Mad1l1** | 1.30653 | 0.07566 | 5.89349 | 3.81626 | 2.37594 | 4.35388 | 4.25649 | 5.12308 | 0.39510 | 0.21886 | 4.54237 | 5.71992 | 1.01822 | 1.96742 |
| **Mad2l1** | 11.54705 | 0.08987 | 18.04932 | 11.11646 | 11.46681 | 9.63838 | 0.49141 | 0.08577 | 13.20115 | 6.76926 | 5.86514 | 6.99114 | 7.59703 | 13.80486 |
| **Mad2l1bp** | 15.59243 | 1.41702 | 20.03648 | 12.14761 | 27.82342 | 12.85891 | 9.55829 | 16.49673 | 15.56826 | 1.48358 | 4.30235 | 1.91255 | 19.22107 | 29.68333 |
| **Mad2l2** | 7.03744 | 0.17960 | 11.74525 | 8.01520 | 26.01906 | 2.77874 | 1.31030 | 6.69120 | 0.00000 | 8.40419 | 6.39126 | 8.59336 | 1.82079 | 10.18637 |
| **Madd** | 1.26298 | 0.00432 | 3.59714 | 1.98834 | 0.70756 | 0.95739 | 0.00877 | 0.00000 | 0.00362 | 0.00731 | 0.00000 | 0.04085 | 0.14489 | 1.38419 |
| **Maea** | 4.15187 | 1.36778 | 16.51972 | 11.56522 | 9.00188 | 14.04512 | 7.54123 | 7.75491 | 0.00000 | 0.45824 | 7.24912 | 8.96888 | 1.43645 | 0.62733 |
| **Mael** | 0.10471 | 0.00385 | 0.52413 | 0.00000 | 1.45310 | 0.06927 | 0.00402 | 0.01373 | 0.00116 | 0.00532 | 0.00613 | 0.00510 | 2.42862 | 0.35592 |
| **Maf1** | 2.61322 | 0.10968 | 4.08983 | 4.77385 | 4.68314 | 6.50385 | 2.28327 | 4.56481 | 1.82097 | 0.10838 | 4.29782 | 9.42548 | 1.75162 | 1.64153 |
| **Mafb** | 0.38453 | 0.00467 | 0.01054 | 0.03244 | 0.32447 | 0.01235 | 0.00428 | 0.01712 | 0.02879 | 0.01125 | 0.00605 | 0.00644 | 0.08108 | 0.00000 |
| **Maff** | 0.02239 | 0.01359 | 0.00000 | 0.37456 | 1.34651 | 1.15456 | 0.00432 | 0.72370 | 0.00433 | 0.01426 | 0.03202 | 0.00264 | 0.38084 | 0.12963 |
| **Mafg** | 0.86937 | 5.94231 | 1.37228 | 3.34496 | 3.14560 | 2.40478 | 1.78497 | 0.20656 | 0.89173 | 0.05564 | 1.52326 | 2.12883 | 3.69322 | 5.05243 |
| **Mafk** | 1.38187 | 0.01256 | 0.32714 | 1.89656 | 0.64821 | 0.15673 | 0.63013 | 0.61316 | 0.14005 | 0.00640 | 0.00000 | 0.47322 | 2.94219 | 1.15186 |
| **Magea1** | 0.00000 | 0.00467 | 0.00000 | 0.22891 | 0.12266 | 0.00000 | 0.00459 | 0.06814 | 0.26535 | 0.00610 | 0.03104 | 0.05797 | 0.00754 | 0.69458 |
| **Magea10** | 0.00539 | 0.00446 | 0.00693 | 0.00985 | 0.37960 | 0.01235 | 0.00417 | 0.01000 | 0.00661 | 0.00624 | 0.00866 | 0.00827 | 0.00926 | 0.31218 |
| **Magea2** | 0.00802 | 0.00460 | 0.00000 | 0.57022 | 0.23677 | 0.00000 | 0.00467 | 0.15944 | 0.59513 | 0.01167 | 0.02331 | 0.10494 | 0.00686 | 1.12689 |

| | | | | | | | | | | | | | | |
|---|---|---|---|---|---|---|---|---|---|---|---|---|---|---|
| **Magea3** | 0.00000 | 0.00446 | 0.00000 | 0.41544 | 0.04954 | 0.00000 | 0.00427 | 0.01559 | 0.38990 | 0.00557 | 0.01553 | 0.03915 | 0.00393 | 0.32547 |
| **Magea4** | 0.00892 | 0.00448 | 0.01259 | 0.04345 | 0.01209 | 0.00812 | 0.00472 | 0.02533 | 0.06911 | 0.00628 | 0.01030 | 0.01040 | 0.00472 | 0.05956 |
| **Magea5** | 0.04193 | 0.02151 | 0.00000 | 2.57692 | 0.50226 | 0.00000 | 0.01150 | 0.92257 | 2.61225 | 0.01579 | 0.00000 | 0.49087 | 0.00000 | 2.65125 |
| **Magea6** | 0.00323 | 0.00455 | 0.00378 | 0.17871 | 0.03267 | 0.00663 | 0.00462 | 0.02522 | 0.11397 | 0.00625 | 0.00931 | 0.01053 | 0.00438 | 0.18239 |
| **Magea8** | 0.00000 | 0.00456 | 0.00000 | 0.77487 | 0.03712 | 0.00000 | 0.00444 | 0.05963 | 1.27597 | 0.00552 | 0.00551 | 0.09914 | 0.00189 | 1.57887 |
| **Mageb16** | 5.47567 | 0.03589 | 6.15663 | 0.96480 | 4.55744 | 8.73182 | 5.18492 | 1.54200 | 8.05240 | 0.24068 | 4.54215 | 6.37013 | 1.10031 | 0.93486 |
| **Mageb16.ps1** | 0.46568 | 0.03374 | 0.70042 | 0.00000 | 0.40031 | 0.98266 | 0.77455 | 0.08173 | 1.07040 | 0.02718 | 0.51469 | 0.81223 | 0.03068 | 0.13305 |
| **Mageb3** | 0.00000 | 0.00425 | 0.08176 | 0.09396 | 0.19688 | 0.07365 | 0.00513 | 0.22308 | 0.00771 | 0.00805 | 0.00498 | 0.00462 | 0.27960 | 1.13558 |
| **Mageb4** | 0.00000 | 0.00372 | 0.00902 | 0.00249 | 0.00000 | 0.22083 | 0.00388 | 0.00378 | 0.16690 | 0.00499 | 0.00868 | 0.00761 | 4.61029 | 0.00384 |
| **Maged1** | 0.00000 | 0.06828 | 1.37534 | 0.77029 | 2.08333 | 0.44343 | 0.04225 | 0.08743 | 0.00000 | 0.03209 | 2.43880 | 6.68761 | 0.00000 | 1.73341 |
| **Maged2** | 0.05257 | 0.00417 | 0.00728 | 0.14741 | 0.10283 | 0.00554 | 0.00417 | 0.00507 | 0.01911 | 0.01108 | 0.01584 | 0.34364 | 0.00541 | 0.02799 |
| **Magee2** | 0.00509 | 0.00353 | 0.01687 | 0.08358 | 0.01141 | 0.00739 | 0.00392 | 0.00430 | 0.01160 | 0.00559 | 0.01142 | 0.78185 | 0.00659 | 0.01180 |
| **Magef1** | 0.00230 | 0.01880 | 0.06408 | 0.36019 | 0.00291 | 0.00494 | 0.51529 | 0.00374 | 0.05808 | 0.01124 | 0.00502 | 0.00673 | 0.00517 | 0.01050 |
| **Mageh1** | 0.39355 | 0.00428 | 0.00000 | 0.52568 | 0.19424 | 0.24458 | 0.00439 | 0.00397 | 0.00967 | 0.00547 | 0.00338 | 0.00458 | 0.00711 | 0.00000 |
| **Magi1** | 0.06368 | 0.01137 | 0.02154 | 0.25742 | 0.00000 | 0.24625 | 2.40146 | 2.08406 | 0.23122 | 0.05271 | 0.00000 | 1.24938 | 0.41987 | 1.10664 |
| **Magi2** | 0.04023 | 0.01943 | 0.00413 | 0.43591 | 0.00262 | 0.02985 | 0.02728 | 0.09912 | 0.13935 | 0.04319 | 0.38327 | 0.02648 | 0.06318 | 0.00000 |
| **Magi3** | 0.01212 | 0.00451 | 0.00622 | 0.17622 | 0.00468 | 0.05918 | 0.00458 | 0.02839 | 0.03263 | 0.02695 | 0.00894 | 0.00667 | 0.56751 | 0.00000 |
| **Magoh** | 12.35905 | 0.40356 | 2.38923 | 16.41818 | 23.40829 | 12.19417 | 2.40297 | 8.80694 | 20.06806 | 13.70343 | 3.70316 | 9.57452 | 22.86481 | 20.41469 |
| **Magohb** | 1.99786 | 0.01191 | 0.00000 | 2.09789 | 3.23230 | 1.20518 | 0.00881 | 1.71822 | 3.71093 | 0.04725 | 0.16578 | 1.94774 | 1.47073 | 2.60502 |
| **Magt1** | 6.94937 | 0.06112 | 5.22658 | 3.79939 | 3.91734 | 5.85522 | 0.74990 | 5.36239 | 13.53490 | 0.14152 | 0.25369 | 4.65371 | 2.65719 | 6.12515 |
| **Majin** | 0.09784 | 0.00422 | 5.29121 | 1.68896 | 3.52250 | 2.13488 | 0.00285 | 0.00000 | 2.32369 | 0.01545 | 0.03588 | 0.00000 | 0.00000 | 4.35722 |
| **Mak** | 0.00909 | 0.00446 | 0.00445 | 0.00967 | 0.69458 | 0.01279 | 0.00425 | 0.01013 | 0.01616 | 0.00642 | 0.00862 | 0.00781 | 0.02089 | 0.15201 |
| **Mak16** | 4.77310 | 7.77789 | 10.90375 | 4.82118 | 12.36565 | 13.47119 | 3.48169 | 0.43142 | 16.99140 | 0.12050 | 6.89157 | 10.38360 | 9.76118 | 11.46809 |
| **Mal** | 0.02461 | 0.00441 | 0.01203 | 0.01249 | 0.01817 | 0.01189 | 0.00434 | 0.01066 | 0.00580 | 0.00697 | 0.00823 | 0.00981 | 0.03463 | 0.20946 |
| **Malat1** | 1.27216 | 6.00321 | 2.65361 | 2.75975 | 0.34726 | 4.59484 | 1.09898 | 2.11998 | 8.38618 | 11.10762 | 3.62124 | 5.69343 | 1.92786 | 8.03442 |
| **Malsu1** | 1.48161 | 0.01023 | 0.00000 | 0.71035 | 3.13119 | 1.26255 | 0.93468 | 0.05941 | 0.22687 | 0.02993 | 0.00000 | 1.24346 | 0.89856 | 0.24354 |
| **Malt1** | 3.08595 | 0.06518 | 0.59606 | 0.59094 | 0.80539 | 0.77924 | 0.10295 | 0.01242 | 0.40617 | 0.12197 | 0.07190 | 0.16637 | 1.58677 | 3.08120 |
| **Mamdc2** | 0.00137 | 0.00881 | 0.12771 | 0.14374 | 0.00086 | 0.00347 | 0.00295 | 0.00230 | 0.01135 | 0.00494 | 0.00730 | 2.94266 | 0.82173 | 0.00849 |
| **Mamdc4** | 0.00000 | 0.01355 | 0.05439 | 0.34764 | 0.14844 | 0.47074 | 0.00398 | 0.01674 | 0.26542 | 0.00831 | 0.00594 | 0.00327 | 0.02336 | 0.00000 |
| **Maml1** | 0.28122 | 0.02714 | 1.12414 | 0.00000 | 0.34888 | 1.67464 | 2.11707 | 2.02739 | 2.27942 | 0.01964 | 1.79887 | 1.00745 | 2.21644 | 0.24430 |
| **Maml2** | 0.05109 | 0.02090 | 0.17630 | 0.00000 | 0.14586 | 0.01126 | 0.00447 | 0.01198 | 0.00924 | 0.00779 | 0.00810 | 0.00931 | 0.02519 | 0.02058 |
| **Man1a2** | 5.23433 | 0.02294 | 6.14082 | 0.00000 | 1.56959 | 6.27660 | 0.00087 | 0.85047 | 9.17589 | 0.00000 | 5.46179 | 3.65047 | 2.23158 | 6.17074 |
| **Man1b1** | 4.56697 | 0.63915 | 3.49137 | 6.38810 | 2.23543 | 2.93016 | 0.00204 | 1.05383 | 3.70724 | 0.03549 | 1.92050 | 2.35333 | 4.10003 | 2.95950 |
| **Man1c1** | 0.02928 | 0.00444 | 0.43947 | 0.18508 | 0.06369 | 0.20178 | 0.01005 | 0.00338 | 0.02908 | 0.02788 | 0.00696 | 0.00539 | 0.33894 | 0.06386 |
| **Man2a1** | 0.01879 | 0.08158 | 0.07913 | 0.00000 | 0.37516 | 4.80989 | 0.13756 | 3.32180 | 0.00837 | 0.00104 | 1.95581 | 6.04250 | 0.42830 | 0.05038 |
| **Man2a2** | 0.00000 | 0.03038 | 0.00000 | 0.82179 | 1.23617 | 0.37105 | 0.01400 | 0.04641 | 0.43060 | 1.75870 | 0.28061 | 0.93514 | 3.75969 | 0.00000 |
| **Man2b1** | 1.23972 | 0.01507 | 7.08508 | 5.45688 | 3.13929 | 5.26740 | 0.00677 | 0.48275 | 0.33622 | 0.03791 | 4.99044 | 3.12684 | 1.35768 | 5.42191 |
| **Man2b2** | 0.03020 | 0.00426 | 0.10168 | 0.78286 | 0.12692 | 0.00000 | 0.00425 | 0.06362 | 0.02483 | 0.00552 | 0.01220 | 0.01098 | 0.00300 | 0.14917 |
| **Man2c1** | 0.07721 | 0.09880 | 0.74770 | 3.75711 | 1.62977 | 0.60466 | 0.01267 | 4.78180 | 0.00064 | 12.04034 | 2.66509 | 0.70229 | 1.12320 | 0.00000 |
| **Man2c1os** | 0.06406 | 0.00468 | 0.10586 | 0.00481 | 0.65015 | 0.02695 | 0.00431 | 0.01254 | 0.00903 | 0.00881 | 0.00602 | 0.01563 | 0.00937 | 0.04408 |
| **Manbal** | 2.00495 | 0.08467 | 6.60997 | 3.71694 | 4.31386 | 7.65899 | 2.27814 | 3.18696 | 0.01207 | 0.08910 | 3.90306 | 0.00000 | 0.38319 | 3.34437 |
| **Manea** | 0.24569 | 0.01834 | 0.12961 | 0.10587 | 0.00000 | 0.02331 | 0.00349 | 0.07631 | 1.88450 | 0.00625 | 0.07135 | 0.01219 | 0.79654 | 0.00000 |
| **Maneal** | 1.11832 | 0.02493 | 0.50115 | 0.30974 | 0.56519 | 0.11955 | 0.00323 | 0.08029 | 3.22641 | 0.02934 | 0.00000 | 1.07652 | 0.10396 | 3.25782 |
| **Manf** | 2.94677 | 0.10196 | 5.73474 | 24.81095 | 9.61104 | 9.98166 | 5.24053 | 1.92213 | 3.04425 | 0.40653 | 0.12611 | 7.38143 | 8.03770 | 2.36684 |
| **Mansc1** | 0.00664 | 0.00446 | 0.00400 | 0.02067 | 0.06145 | 0.00995 | 0.00416 | 0.00972 | 0.43272 | 0.00677 | 0.00705 | 0.00894 | 0.00506 | 0.00999 |
| **Mansc4** | 0.00677 | 0.00451 | 0.00469 | 0.04618 | 0.00984 | 0.37891 | 0.00412 | 0.00404 | 0.01298 | 0.00585 | 0.28869 | 0.01208 | 0.00388 | 0.06366 |
| **Maoa** | 0.61798 | 1.06516 | 1.01389 | 0.00000 | 0.00003 | 0.00000 | 0.02419 | 0.84387 | 0.68865 | 0.01283 | 0.01026 | 0.26062 | 0.48425 | 0.19516 |
| **Map10** | 0.00000 | 0.00399 | 0.34038 | 0.00108 | 0.01104 | 0.01532 | 0.00980 | 0.00622 | 0.02149 | 0.00551 | 0.01293 | 0.00712 | 2.79884 | 0.01508 |
| **Map1a** | 0.09350 | 0.00508 | 0.00517 | 0.19369 | 0.00515 | 0.21419 | 0.00425 | 0.00725 | 0.01589 | 0.00669 | 0.00508 | 0.01239 | 0.12390 | 0.00000 |
| **Map1b** | 0.17711 | 0.00352 | 0.39048 | 0.00000 | 0.69059 | 0.10303 | 0.00393 | 0.00224 | 0.00783 | 0.00439 | 0.00748 | 0.79250 | 0.03844 | 0.41043 |
| **Map1lc3a** | 12.17581 | 0.05809 | 5.22070 | 9.87513 | 22.72292 | 2.87902 | 0.58823 | 3.24186 | 0.51955 | 0.11525 | 0.01327 | 1.47037 | 9.35720 | 12.58065 |
| **Map1lc3b** | 42.09644 | 5.55774 | 48.10727 | 55.86509 | 57.32844 | 48.53955 | 13.65134 | 29.40689 | 27.78762 | 22.18311 | 29.86918 | 34.81373 | 27.20472 | 52.03142 |
| **Map1s** | 0.20776 | 3.73557 | 0.54654 | 3.38156 | 7.15483 | 0.13495 | 0.82355 | 0.00686 | 0.00877 | 0.03726 | 0.00000 | 0.36201 | 0.26197 | 0.00000 |
| **Map2** | 0.02630 | 0.00434 | 0.06629 | 0.00453 | 0.00428 | 0.00988 | 0.00423 | 0.00952 | 0.12501 | 0.00753 | 0.00605 | 0.05708 | 0.00593 | 0.08653 |
| **Map2k1** | 8.22798 | 0.02815 | 7.28198 | 0.79777 | 9.72170 | 7.25551 | 2.33885 | 1.59071 | 9.64965 | 0.06860 | 8.26875 | 6.45684 | 3.30951 | 12.29363 |

| | | | | | | | | | | | | | | |
|---|---|---|---|---|---|---|---|---|---|---|---|---|---|---|
| **Map2k2** | 3.57189 | 4.20217 | 25.57376 | 17.33550 | 21.61903 | 2.17329 | 5.90686 | 9.10041 | 0.00000 | 0.87733 | 5.98429 | 9.58644 | 2.42891 | 0.85866 |
| **Map2k3** | 2.91596 | 0.03540 | 5.99139 | 8.43653 | 5.47690 | 10.74895 | 4.91360 | 4.14449 | 0.08494 | 0.16717 | 3.17523 | 7.73069 | 2.07659 | 2.13968 |
| **Map2k3os** | 0.44075 | 0.00437 | 0.08123 | 0.00652 | 0.00323 | 0.02350 | 0.00444 | 0.01150 | 0.01018 | 0.00632 | 0.06188 | 0.04037 | 0.00421 | 0.00440 |
| **Map2k4** | 6.42736 | 0.02299 | 3.12577 | 2.17025 | 2.44836 | 8.86073 | 2.33279 | 1.84122 | 2.07753 | 0.07188 | 10.43091 | 4.05762 | 2.37562 | 1.79027 |
| **Map2k5** | 0.01616 | 0.00411 | 0.00630 | 0.00000 | 0.16464 | 0.42694 | 0.00416 | 0.01270 | 0.00623 | 0.01763 | 0.00675 | 0.02436 | 0.01510 | 0.13879 |
| **Map2k6** | 0.39509 | 0.05971 | 4.83628 | 1.88327 | 2.13790 | 1.67136 | 2.54956 | 3.99513 | 0.34028 | 0.07114 | 3.17746 | 7.42633 | 0.00000 | 0.96349 |
| **Map2k7** | 1.04362 | 0.10183 | 5.53594 | 3.29671 | 9.06736 | 1.59451 | 3.98141 | 6.03427 | 0.28436 | 0.10781 | 1.74626 | 6.11712 | 0.54815 | 7.59896 |
| **Map3k1** | 0.00000 | 0.00346 | 0.11542 | 0.00000 | 0.18958 | 0.19094 | 0.00380 | 0.41010 | 0.29465 | 0.00656 | 0.00000 | 1.03628 | 0.16523 | 0.72564 |
| **Map3k10** | 0.01814 | 0.01510 | 0.06769 | 0.23213 | 0.03762 | 0.00000 | 0.00483 | 0.18094 | 0.01043 | 0.00705 | 0.04849 | 0.02782 | 0.15127 | 0.05230 |
| **Map3k11** | 0.38200 | 0.01780 | 1.38440 | 1.67013 | 1.06799 | 0.32202 | 2.43813 | 2.30696 | 0.00997 | 0.04644 | 0.12210 | 1.90910 | 0.00000 | 0.26651 |
| **Map3k12** | 0.57925 | 0.37618 | 0.11863 | 0.43818 | 0.10541 | 0.00000 | 0.00381 | 0.00796 | 0.21739 | 0.00461 | 0.39533 | 0.58032 | 0.00909 | 0.33150 |
| **Map3k13** | 0.00513 | 0.00435 | 0.02003 | 0.01123 | 0.00904 | 0.01224 | 0.00432 | 0.00952 | 0.00989 | 0.00643 | 0.01113 | 0.00998 | 0.25527 | 0.05549 |
| **Map3k14** | 0.00168 | 0.00428 | 0.10712 | 0.00381 | 0.34359 | 0.00469 | 0.00443 | 0.00990 | 0.00493 | 0.00667 | 0.00624 | 0.00809 | 0.31677 | 1.30857 |
| **Map3k15** | 0.00869 | 0.00388 | 0.32869 | 0.00000 | 0.22407 | 0.80825 | 0.00390 | 0.00462 | 0.01673 | 0.00495 | 0.03257 | 0.52390 | 0.00324 | 0.01119 |
| **Map3k2** | 2.07136 | 0.00351 | 0.13165 | 0.00000 | 1.12093 | 1.32043 | 0.01023 | 0.85699 | 2.20454 | 0.00595 | 0.02555 | 0.81281 | 1.17993 | 2.61809 |
| **Map3k3** | 0.18762 | 0.00992 | 1.74171 | 1.77600 | 3.52703 | 4.03167 | 0.00344 | 0.00715 | 0.95670 | 0.00634 | 1.27324 | 1.25626 | 1.20199 | 4.85497 |
| **Map3k4** | 0.00000 | 0.08345 | 0.60441 | 1.54147 | 0.62342 | 6.36709 | 0.00773 | 1.51418 | 1.35839 | 0.03969 | 2.77155 | 3.42575 | 0.62494 | 0.21719 |
| **Map3k5** | 0.08504 | 0.00906 | 2.20834 | 0.14310 | 2.78172 | 1.10110 | 0.00792 | 0.20072 | 2.45726 | 0.01285 | 2.43078 | 1.53370 | 0.93009 | 3.69093 |
| **Map3k6** | 0.00198 | 0.01320 | 0.30594 | 2.40402 | 0.49299 | 0.00000 | 0.00377 | 0.15889 | 0.02787 | 0.01442 | 0.01368 | 1.63950 | 0.91145 | 0.00000 |
| **Map3k7** | 2.29726 | 0.00000 | 3.22872 | 1.87017 | 5.09104 | 4.67818 | 0.05129 | 0.98353 | 6.29427 | 0.04213 | 2.25049 | 4.99793 | 1.83798 | 8.41710 |
| **Map3k9** | 1.35018 | 2.12542 | 0.00000 | 0.00000 | 0.90326 | 0.97975 | 0.00203 | 0.00000 | 1.38862 | 8.16126 | 1.13252 | 0.00000 | 0.00658 | 0.21303 |
| **Map4** | 3.05296 | 5.27568 | 16.94660 | 7.11460 | 6.41037 | 15.35621 | 2.44226 | 4.80161 | 5.75871 | 0.19255 | 7.01432 | 9.90465 | 9.02525 | 4.53055 |
| **Map4k1** | 0.39604 | 0.00444 | 0.12452 | 2.64717 | 2.46669 | 0.72930 | 0.00392 | 0.08433 | 0.00138 | 0.02202 | 0.04210 | 0.66639 | 0.92384 | 0.00000 |
| **Map4k2** | 0.12611 | 0.02931 | 0.39656 | 2.71880 | 0.90781 | 0.00000 | 0.01047 | 0.69327 | 0.82747 | 1.02997 | 2.49669 | 0.87147 | 0.08232 | 0.00000 |
| **Map4k3** | 3.78155 | 0.00706 | 1.43978 | 0.34629 | 2.53695 | 0.75123 | 0.00083 | 2.42259 | 7.38344 | 0.00113 | 7.38462 | 1.80291 | 1.88229 | 4.33628 |
| **Map4k4** | 2.66882 | 0.35017 | 1.88326 | 2.05033 | 1.56853 | 4.00913 | 0.01799 | 4.11922 | 1.95794 | 0.05454 | 0.00000 | 8.70807 | 0.54785 | 0.17237 |
| **Map4k5** | 7.68486 | 0.02181 | 2.76309 | 0.57727 | 11.38658 | 0.82104 | 1.75370 | 1.08346 | 18.51292 | 0.03089 | 2.18530 | 2.01399 | 8.16643 | 13.31577 |
| **Map6** | 0.12596 | 0.00432 | 0.00000 | 0.00443 | 0.25889 | 0.02802 | 0.00408 | 0.00656 | 0.00773 | 0.00572 | 0.62829 | 0.00507 | 0.07210 | 0.09342 |
| **Map6d1** | 0.88998 | 0.00422 | 0.13050 | 0.01418 | 0.76483 | 0.00128 | 0.00445 | 0.01143 | 0.00727 | 0.00705 | 0.00537 | 0.00408 | 0.54969 | 1.30684 |
| **Map7** | 16.34032 | 0.03697 | 21.49629 | 10.43609 | 18.77894 | 10.37531 | 3.17370 | 1.25627 | 8.90464 | 11.75825 | 10.02879 | 6.73585 | 1.50575 | 16.26112 |
| **Map7d1** | 0.14646 | 0.03262 | 0.45730 | 0.00000 | 0.67084 | 0.50254 | 0.37317 | 2.56513 | 0.02088 | 0.02256 | 0.00000 | 0.85404 | 0.24796 | 0.00653 |
| **Map7d2** | 4.39470 | 0.00277 | 6.01303 | 2.42950 | 7.67090 | 0.74071 | 1.89273 | 0.00000 | 19.08369 | 0.01046 | 0.00000 | 0.00000 | 0.00000 | 8.44249 |
| **Map9** | 0.05786 | 0.00396 | 0.00000 | 0.26943 | 0.00000 | 0.38123 | 0.00388 | 0.00270 | 0.00810 | 0.01307 | 0.01038 | 0.03530 | 0.11145 | 1.37786 |
| **Mapk1** | 7.45088 | 0.04221 | 4.71642 | 2.50216 | 5.73341 | 14.11552 | 0.89398 | 6.60143 | 10.67422 | 0.08522 | 5.21916 | 7.83025 | 4.89825 | 6.02023 |
| **Mapk10** | 0.00458 | 0.00359 | 0.44297 | 0.50925 | 0.12502 | 0.00000 | 0.00363 | 0.00062 | 0.01658 | 0.00424 | 0.18221 | 0.51782 | 0.01172 | 0.01481 |
| **Mapk11** | 0.00727 | 0.00408 | 0.01347 | 0.14712 | 0.02408 | 0.04101 | 0.00400 | 0.00379 | 0.00451 | 0.00479 | 0.02085 | 0.01221 | 0.59553 | 0.49453 |
| **Mapk12** | 0.23279 | 0.00382 | 0.01580 | 0.00000 | 0.01833 | 0.00000 | 0.85777 | 0.00000 | 0.01611 | 0.02403 | 0.79014 | 0.00771 | 0.02869 | 0.16065 |
| **Mapk13** | 1.22591 | 0.02086 | 1.88086 | 2.57879 | 2.61136 | 0.00000 | 0.00346 | 0.08683 | 0.00539 | 0.01746 | 2.82845 | 1.11261 | 0.07614 | 1.03306 |
| **Mapk14** | 3.21192 | 6.25212 | 8.52086 | 0.06112 | 10.23854 | 5.83467 | 1.54749 | 2.95512 | 4.25660 | 0.04893 | 5.40657 | 6.48358 | 5.61247 | 12.60544 |
| **Mapk1ip1** | 2.32596 | 0.04396 | 0.44046 | 2.67883 | 0.65143 | 2.30201 | 3.49097 | 1.05715 | 0.01889 | 0.03914 | 0.14444 | 2.66670 | 0.24269 | 0.00000 |
| **Mapk1ip1l** | 8.40451 | 0.04162 | 9.16838 | 11.02107 | 9.99194 | 11.59849 | 1.65449 | 5.03486 | 13.32796 | 0.18767 | 8.42885 | 7.88807 | 6.99608 | 10.13329 |
| **Mapk3** | 0.40600 | 0.05074 | 4.06194 | 3.11948 | 2.64244 | 1.19626 | 0.02120 | 3.30502 | 0.21997 | 0.03003 | 1.73117 | 0.93809 | 2.55781 | 0.00000 |
| **Mapk6** | 18.84529 | 0.08086 | 18.36377 | 6.91437 | 18.53059 | 13.06833 | 3.40802 | 6.59553 | 11.77529 | 0.86859 | 11.54593 | 4.24158 | 10.27506 | 15.70947 |
| **Mapk7** | 0.48176 | 0.01824 | 1.79032 | 3.19462 | 2.45523 | 1.28416 | 0.00337 | 1.53757 | 0.02740 | 0.01136 | 0.00000 | 3.74175 | 2.68357 | 0.00000 |
| **Mapk8** | 0.44476 | 0.00190 | 0.15376 | 0.50030 | 0.00000 | 2.69084 | 0.00320 | 0.00061 | 0.00150 | 0.01763 | 2.69598 | 1.47445 | 1.08010 | 4.95897 |
| **Mapk8ip1** | 0.32022 | 0.00501 | 0.07843 | 0.22637 | 0.37619 | 0.06014 | 0.00426 | 0.09353 | 0.00366 | 0.01824 | 0.00379 | 0.05315 | 0.78726 | 0.37600 |
| **Mapk8ip3** | 0.04788 | 0.01423 | 0.60868 | 0.77765 | 0.28825 | 2.00494 | 0.00147 | 0.01449 | 0.00155 | 4.39222 | 0.67454 | 4.89557 | 1.63582 | 0.00000 |
| **Mapk9** | 2.05155 | 1.74749 | 0.40455 | 1.04848 | 0.46182 | 1.37615 | 0.04487 | 1.17961 | 0.19768 | 0.00640 | 0.38498 | 2.06439 | 3.49907 | 3.41720 |
| **Mapkap1** | 3.20056 | 0.03085 | 4.48341 | 0.00000 | 7.45022 | 1.90855 | 1.03855 | 2.13984 | 0.00000 | 7.46855 | 2.92805 | 2.37155 | 2.09228 | 2.95573 |
| **Mapkapk2** | 0.96942 | 0.06656 | 1.02861 | 0.00000 | 0.56055 | 4.38563 | 3.10239 | 3.65056 | 0.38761 | 0.07034 | 0.75518 | 2.95959 | 5.62609 | 0.00000 |
| **Mapkapk3** | 1.11079 | 0.00878 | 1.88038 | 0.64062 | 2.60575 | 2.16397 | 0.00215 | 0.84885 | 4.14869 | 0.00864 | 0.00000 | 2.75523 | 0.24272 | 0.54469 |
| **Mapkapk5** | 7.72118 | 0.02181 | 7.70910 | 2.03647 | 6.32883 | 5.10533 | 1.43327 | 0.65392 | 2.54883 | 0.02392 | 4.38772 | 1.45114 | 10.14058 | 12.11789 |
| **Mapkbp1** | 1.11954 | 0.01401 | 0.24920 | 0.23309 | 1.25375 | 0.00000 | 0.00994 | 0.28794 | 2.36140 | 0.02718 | 1.68015 | 0.39837 | 2.37202 | 3.62588 |
| **Mapre1** | 16.60834 | 9.35775 | 23.42541 | 24.18715 | 26.06345 | 23.63740 | 5.62446 | 8.60232 | 17.92105 | 5.29522 | 15.26898 | 13.14241 | 10.97772 | 23.78758 |
| **Mapre2** | 1.11171 | 0.00977 | 1.30270 | 1.23130 | 0.71456 | 0.84738 | 0.00300 | 0.00000 | 1.60469 | 0.00839 | 0.03737 | 2.09679 | 0.01080 | 0.41562 |

| | | | | | | | | | | | | | | |
|---|---|---|---|---|---|---|---|---|---|---|---|---|---|---|
| **Mapre3** | 0.00000 | 0.02033 | 0.33643 | 0.58411 | 0.02678 | 0.42064 | 0.01048 | 2.96206 | 0.00952 | 0.00954 | 1.16168 | 0.00231 | 0.00843 | 0.00000 |
| **Mapt** | 1.05992 | 0.05177 | 3.48661 | 4.10082 | 2.28300 | 1.81287 | 0.00855 | 0.83152 | 2.32661 | 0.00808 | 0.93783 | 3.54823 | 0.00000 | 3.01800 |
| **X2.Mar** | 6.61680 | 0.00952 | 2.17556 | 1.04308 | 5.23949 | 1.71282 | 3.46499 | 1.06498 | 0.78008 | 0.09540 | 0.13342 | 0.00000 | 5.29337 | 5.55627 |
| **X2.Mar.1** | 0.65878 | 4.24707 | 0.37946 | 0.00000 | 0.45593 | 0.46660 | 0.02495 | 0.80594 | 0.44701 | 0.01163 | 0.18119 | 2.50926 | 2.48448 | 0.12358 |
| **X3.Mar** | 0.00536 | 0.00419 | 0.01557 | 0.09281 | 0.06097 | 0.00825 | 0.00461 | 0.00658 | 0.01597 | 0.00594 | 0.01638 | 0.01470 | 0.25691 | 0.00000 |
| **X5.Mar** | 12.53520 | 0.40267 | 5.64426 | 4.46606 | 12.50617 | 9.09738 | 3.99967 | 4.08715 | 5.71084 | 0.59175 | 1.94418 | 5.97012 | 10.79825 | 11.72516 |
| **X6.Mar** | 3.55292 | 0.03214 | 9.63462 | 6.74114 | 5.86949 | 8.44115 | 2.66392 | 3.89406 | 12.03238 | 9.68104 | 6.06287 | 5.54220 | 2.86339 | 7.60704 |
| **X7.Mar** | 1.23329 | 0.00330 | 0.64955 | 0.12423 | 0.92726 | 0.13575 | 0.00326 | 0.00000 | 4.05816 | 0.01118 | 0.69915 | 0.21615 | 0.06287 | 7.10005 |
| **X8.Mar** | 0.15666 | 0.00480 | 0.29772 | 0.99852 | 0.68776 | 0.00000 | 0.00426 | 0.87480 | 0.12626 | 0.00900 | 0.00000 | 0.45451 | 0.31876 | 0.01251 |
| **X9.Mar** | 0.14189 | 0.00413 | 0.00365 | 0.01187 | 0.70856 | 0.01096 | 0.00393 | 0.01502 | 0.00647 | 0.00903 | 0.02567 | 0.00549 | 0.91240 | 0.00322 |
| **Marcks** | 1.24990 | 0.85365 | 5.71498 | 8.99065 | 0.30410 | 11.36523 | 3.30819 | 4.83578 | 0.84987 | 0.24290 | 9.28748 | 13.37658 | 0.02451 | 0.00000 |
| **Marcksl1** | 4.61237 | 7.18759 | 18.06210 | 22.25416 | 20.96081 | 22.94567 | 7.50493 | 27.74452 | 3.61395 | 16.43335 | 10.91826 | 10.19312 | 2.95099 | 10.28044 |
| **Marf1** | 2.20217 | 0.01504 | 2.29320 | 1.35661 | 3.97411 | 2.27233 | 0.00759 | 0.18502 | 3.51402 | 0.00918 | 1.27074 | 3.24495 | 3.94500 | 2.90906 |
| **Mark1** | 0.00000 | 0.01253 | 8.65185 | 2.55159 | 5.59208 | 1.53422 | 0.00282 | 0.27041 | 1.49922 | 0.00000 | 0.02117 | 0.42193 | 0.33663 | 3.35812 |
| **Mark2** | 0.49331 | 0.18955 | 7.17929 | 3.60353 | 10.10573 | 1.79879 | 1.08783 | 3.43731 | 0.08728 | 0.39396 | 2.38120 | 3.55679 | 3.85656 | 5.59642 |
| **Mark3** | 6.31419 | 7.00412 | 14.12728 | 3.34211 | 11.28890 | 10.01602 | 3.77404 | 4.70299 | 2.99999 | 0.09617 | 4.99708 | 5.42241 | 7.46524 | 14.22971 |
| **Mark4** | 0.02439 | 0.00445 | 0.04091 | 0.06725 | 0.05090 | 0.00568 | 0.00485 | 0.04484 | 0.01202 | 0.00631 | 0.00720 | 0.02034 | 0.00795 | 0.01238 |
| **Mars** | 11.51546 | 0.35040 | 20.08320 | 16.02192 | 10.91152 | 18.58946 | 5.63594 | 7.13683 | 1.68159 | 0.32539 | 12.72777 | 13.65425 | 2.71159 | 10.45745 |
| **Mars2** | 0.00000 | 0.01172 | 2.00624 | 2.38207 | 2.88024 | 1.03429 | 2.48784 | 0.02683 | 1.51367 | 0.11165 | 0.00382 | 0.09106 | 4.04493 | 3.23941 |
| **Marveld1** | 0.04077 | 0.00431 | 0.37109 | 0.00056 | 0.75953 | 0.00001 | 0.00423 | 0.01305 | 0.00701 | 0.00693 | 0.00564 | 0.00950 | 0.05688 | 0.74103 |
| **Marveld2** | 0.82407 | 0.04634 | 1.93852 | 0.00000 | 2.47942 | 0.00000 | 0.00277 | 3.65994 | 2.52345 | 0.00936 | 0.00060 | 0.05595 | 2.36093 | 3.20861 |
| **Marveld3** | 0.00000 | 0.00433 | 0.09465 | 0.00000 | 0.32778 | 0.37120 | 0.00410 | 0.00448 | 0.03720 | 0.00574 | 0.00447 | 0.17496 | 0.00543 | 0.79642 |
| **Masp2** | 0.00368 | 0.00420 | 0.01831 | 0.00000 | 0.00708 | 0.13012 | 0.00428 | 0.00476 | 0.00806 | 0.01253 | 0.00377 | 0.35717 | 0.00528 | 0.34683 |
| **Mast1** | 0.08411 | 0.00380 | 1.74070 | 0.21481 | 5.50884 | 0.00000 | 0.00376 | 0.53975 | 0.24169 | 0.01009 | 0.00000 | 0.16352 | 1.70391 | 0.04503 |
| **Mast2** | 3.34646 | 0.03164 | 2.29560 | 1.83941 | 3.24011 | 6.08769 | 2.23728 | 2.50373 | 0.45403 | 0.07145 | 1.36658 | 1.98083 | 4.52707 | 1.46835 |
| **Mast3** | 5.57021 | 0.02773 | 7.14929 | 3.38527 | 8.93523 | 1.95120 | 0.02008 | 2.51067 | 1.02761 | 0.00440 | 1.91818 | 0.00000 | 1.03350 | 6.79313 |
| **Mast4** | 0.01482 | 0.00319 | 0.62682 | 0.00000 | 0.87955 | 0.51843 | 0.00320 | 0.00000 | 0.04369 | 2.02413 | 0.00319 | 0.39248 | 0.01446 | 0.94270 |
| **Mastl** | 5.94067 | 0.00257 | 3.42721 | 0.98076 | 4.79860 | 0.25583 | 0.00835 | 0.50453 | 8.15747 | 0.00000 | 2.52995 | 0.82081 | 0.87448 | 4.74394 |
| **Mat1a** | 0.01067 | 0.00445 | 0.01453 | 0.01367 | 0.01683 | 0.01313 | 0.00430 | 0.01036 | 0.01160 | 0.00699 | 0.00881 | 0.00997 | 0.00749 | 0.01230 |
| **Mat2a** | 17.96352 | 7.46373 | 32.65209 | 26.54675 | 33.93595 | 24.04169 | 7.84754 | 13.84503 | 35.63018 | 1.55108 | 19.43173 | 10.47672 | 16.63191 | 39.53555 |
| **Mat2b** | 10.72809 | 0.05154 | 8.59337 | 4.09274 | 15.26254 | 3.05063 | 0.65263 | 3.22194 | 7.45353 | 0.04063 | 7.67716 | 6.84205 | 14.91343 | 20.91915 |
| **Matk** | 0.46625 | 0.00431 | 0.03774 | 0.00715 | 0.14234 | 0.00770 | 0.04047 | 0.01086 | 0.01538 | 0.00643 | 0.00846 | 0.00861 | 0.00471 | 0.00411 |
| **Matn2** | 0.01676 | 0.00446 | 0.04273 | 0.00089 | 0.02939 | 0.14232 | 0.00425 | 0.00730 | 0.00650 | 0.00641 | 0.00627 | 0.01094 | 0.00885 | 0.04950 |
| **Matr3** | 7.26877 | 1.60918 | 13.40376 | 4.17406 | 12.24561 | 10.78561 | 1.16874 | 5.08789 | 25.65338 | 0.04120 | 13.53619 | 6.43904 | 4.60757 | 4.73442 |
| **Mau2** | 0.78036 | 0.06595 | 4.94886 | 2.14948 | 5.81968 | 4.21939 | 0.20435 | 2.65174 | 6.72439 | 0.18241 | 4.67178 | 3.02111 | 1.79562 | 8.99432 |
| **Mavs** | 0.89047 | 0.04456 | 1.56120 | 3.28944 | 3.20416 | 3.85401 | 3.31594 | 1.13168 | 0.00000 | 0.09030 | 0.26848 | 1.55063 | 1.78944 | 6.15197 |
| **Max** | 3.26918 | 0.82188 | 8.04099 | 3.44609 | 3.21507 | 7.25848 | 2.34404 | 1.48467 | 1.86893 | 5.98948 | 3.94936 | 3.48761 | 4.98845 | 4.75124 |
| **Maz** | 0.11144 | 0.03598 | 0.00000 | 0.42369 | 0.26391 | 0.21922 | 0.35603 | 1.47623 | 0.32411 | 0.01470 | 0.00531 | 0.75085 | 1.03410 | 0.40311 |
| **Mb21d1** | 0.06747 | 0.21274 | 0.02846 | 0.21341 | 0.44863 | 0.33296 | 0.06417 | 0.09175 | 1.78827 | 0.09383 | 0.02034 | 0.04793 | 0.76457 | 0.48687 |
| **Mb21d2** | 0.25655 | 0.00291 | 0.24137 | 0.00000 | 0.15057 | 0.29108 | 0.00348 | 0.01497 | 0.00902 | 0.01372 | 0.96189 | 1.14596 | 0.01456 | 0.61893 |
| **Mbd1** | 0.16300 | 0.01164 | 0.43350 | 1.65712 | 0.95873 | 1.07185 | 0.88137 | 0.41733 | 0.01369 | 0.01025 | 0.00000 | 0.12528 | 0.44043 | 0.98331 |
| **Mbd2** | 1.39460 | 0.02077 | 0.06962 | 1.49440 | 0.62819 | 0.75612 | 0.00386 | 2.98237 | 0.01509 | 0.00246 | 1.27637 | 0.54590 | 0.55203 | 0.05211 |
| **Mbd3** | 8.12648 | 0.17058 | 20.95471 | 9.99626 | 21.07717 | 12.97594 | 3.46182 | 6.50228 | 0.00000 | 0.25562 | 2.42450 | 11.98687 | 2.64915 | 3.13460 |
| **Mbd3l2** | 2.37388 | 0.00283 | 0.89801 | 0.00000 | 3.96816 | 0.00000 | 0.00247 | 0.00161 | 8.31079 | 0.00476 | 0.00000 | 0.00000 | 4.63950 | 8.21033 |
| **Mbd4** | 0.60110 | 0.05741 | 1.02575 | 0.58180 | 0.68229 | 0.34537 | 0.02004 | 0.07677 | 3.25769 | 0.03880 | 0.05794 | 0.13022 | 0.14936 | 0.93419 |
| **Mbd5** | 0.95202 | 0.00392 | 0.97546 | 0.24434 | 0.47529 | 0.19969 | 0.00365 | 0.00000 | 0.15198 | 0.00322 | 0.00882 | 0.01453 | 0.36400 | 1.28931 |
| **Mbd6** | 0.70527 | 0.29845 | 1.20990 | 2.26555 | 0.51087 | 1.66900 | 0.24767 | 2.11475 | 0.00157 | 2.87055 | 0.00000 | 0.00000 | 1.41022 | 0.11349 |
| **Mbip** | 4.31894 | 0.03152 | 1.39093 | 0.26962 | 4.76714 | 2.32162 | 1.02930 | 2.35018 | 3.87587 | 0.03663 | 2.54808 | 3.25407 | 2.77608 | 4.22487 |
| **Mbl2** | 0.00000 | 0.27520 | 1.22172 | 0.11156 | 1.77310 | 0.00000 | 0.00336 | 0.00761 | 2.28906 | 0.00467 | 0.00178 | 0.00278 | 0.18570 | 2.80358 |
| **Mblac1** | 0.01417 | 0.00398 | 0.16165 | 0.84811 | 0.11538 | 0.15986 | 0.04984 | 0.35238 | 0.05717 | 0.00824 | 0.00223 | 0.00388 | 0.05075 | 0.05651 |
| **Mblac2** | 0.00937 | 0.00332 | 0.06999 | 0.01294 | 0.21962 | 0.00223 | 0.02741 | 0.00370 | 0.35125 | 0.01345 | 0.00277 | 1.10135 | 0.22724 | 0.55814 |
| **Mbnl1** | 0.15771 | 0.00384 | 0.32459 | 0.07186 | 0.21238 | 0.00000 | 0.00374 | 0.18965 | 0.00811 | 0.00414 | 0.00819 | 0.00684 | 0.00241 | 0.27099 |
| **Mbnl2** | 0.06098 | 0.00430 | 0.58785 | 0.32120 | 0.00000 | 0.93266 | 0.31200 | 1.00555 | 0.89618 | 0.81606 | 0.00000 | 0.00000 | 0.00017 | 0.00000 |
| **Mbnl3** | 0.00000 | 10.98439 | 0.87337 | 3.16324 | 2.10250 | 0.00000 | 6.86239 | 10.54607 | 2.61608 | 7.30823 | 0.00000 | 0.08785 | 1.16633 | 3.53831 |
| **Mboat1** | 1.30613 | 0.00262 | 2.72975 | 0.39127 | 0.00000 | 0.86025 | 0.00246 | 0.00000 | 4.66941 | 0.02214 | 0.45252 | 1.51929 | 0.00702 | 1.93384 |

| | | | | | | | | | | | | | | |
|---|---|---|---|---|---|---|---|---|---|---|---|---|---|---|
| **Mboat2** | 4.92340 | 0.03707 | 7.84461 | 9.05681 | 4.52354 | 4.19037 | 0.00681 | 0.31815 | 7.37187 | 0.00781 | 3.74885 | 5.94815 | 1.10891 | 7.61040 |
| **Mboat4** | 0.06819 | 0.00384 | 0.00000 | 0.26787 | 0.28398 | 0.97081 | 0.00385 | 0.00000 | 0.03036 | 0.01864 | 0.67807 | 0.05926 | 0.02076 | 0.17783 |
| **Mboat7** | 4.39267 | 0.03707 | 3.85836 | 4.26889 | 4.82992 | 4.29360 | 0.01829 | 0.00000 | 1.06239 | 9.77788 | 0.00000 | 1.70670 | 1.23265 | 1.93624 |
| **Mbp** | 0.00639 | 0.12914 | 0.00592 | 0.00000 | 0.00849 | 4.69023 | 0.01954 | 7.93074 | 0.04824 | 0.05247 | 0.02307 | 0.07641 | 0.24079 | 0.25063 |
| **Mbtd1** | 11.04151 | 0.07644 | 9.71784 | 1.09264 | 14.38763 | 7.24952 | 0.00154 | 3.35557 | 20.16485 | 0.02810 | 6.23868 | 8.05511 | 11.03841 | 13.66370 |
| **Mbtps1** | 5.31165 | 0.22455 | 9.99963 | 4.98350 | 6.93961 | 4.58078 | 3.75510 | 2.92862 | 1.36937 | 0.09778 | 5.21380 | 8.40908 | 5.47685 | 5.46937 |
| **Mbtps2** | 4.16877 | 0.02345 | 3.59646 | 2.22015 | 0.36063 | 2.04958 | 0.00833 | 0.25718 | 6.76592 | 0.01179 | 2.26295 | 3.88225 | 2.27211 | 0.00000 |
| **Mc1r** | 0.07264 | 0.03994 | 0.21332 | 0.17225 | 0.03156 | 0.07559 | 0.02194 | 0.04794 | 0.19625 | 0.01910 | 0.01708 | 0.06345 | 0.30302 | 0.11733 |
| **Mcam** | 0.08296 | 0.00445 | 0.03184 | 0.01423 | 0.02421 | 0.01106 | 0.00430 | 0.01093 | 0.02093 | 0.00691 | 0.00738 | 0.00921 | 0.00494 | 0.02004 |
| **Mcat** | 0.69662 | 0.08107 | 1.04997 | 3.48211 | 0.05424 | 2.50400 | 1.62944 | 3.15763 | 0.00000 | 0.03459 | 5.14120 | 3.99395 | 0.34133 | 0.38101 |
| **Mcc** | 0.32618 | 0.10751 | 0.05441 | 0.59890 | 0.18806 | 1.57295 | 0.13201 | 0.17541 | 0.57665 | 0.17423 | 0.02851 | 0.26354 | 1.84382 | 1.54618 |
| **Mccc1** | 3.84368 | 0.53853 | 3.21682 | 2.54340 | 1.14050 | 3.58712 | 1.12423 | 2.05142 | 2.09416 | 0.05203 | 2.51740 | 2.02524 | 0.13375 | 2.01152 |
| **Mccc2** | 0.84760 | 10.34061 | 4.57073 | 3.49067 | 0.70770 | 8.09513 | 0.53105 | 5.19481 | 1.72734 | 0.08460 | 3.63147 | 6.27720 | 0.03372 | 2.27193 |
| **Mcee** | 1.75895 | 3.49863 | 0.35651 | 1.21865 | 0.96107 | 2.08423 | 1.85291 | 4.33935 | 1.76933 | 0.10169 | 0.36542 | 2.35677 | 1.12137 | 0.55161 |
| **Mcf2** | 0.50619 | 0.00404 | 0.08882 | 0.00000 | 1.36357 | 0.00059 | 0.00410 | 0.00624 | 0.00644 | 0.00593 | 0.00629 | 0.27352 | 0.10899 | 1.59337 |
| **Mcfd2** | 4.09750 | 6.12169 | 3.74091 | 11.40067 | 7.34714 | 6.50332 | 4.53255 | 2.70602 | 7.79394 | 0.11564 | 2.82674 | 7.10524 | 3.71864 | 16.83312 |
| **Mcl1** | 22.46811 | 0.17644 | 19.52834 | 16.08300 | 31.35913 | 19.36601 | 5.21044 | 6.65664 | 24.33997 | 0.24168 | 9.74328 | 11.19260 | 13.14385 | 25.29777 |
| **Mcm10** | 11.94462 | 0.10747 | 15.47969 | 12.77520 | 20.58381 | 12.50129 | 1.54722 | 2.12046 | 11.18989 | 0.08873 | 14.93626 | 11.03383 | 9.92236 | 20.47099 |
| **Mcm2** | 10.86100 | 2.67233 | 21.12311 | 14.60971 | 18.30498 | 20.21669 | 0.92393 | 2.90903 | 3.77024 | 0.18207 | 9.65127 | 14.93282 | 9.82869 | 15.86785 |
| **Mcm3** | 12.53154 | 0.12853 | 23.48746 | 11.80503 | 10.24712 | 15.19241 | 2.86076 | 0.98967 | 4.39008 | 0.09078 | 9.82736 | 12.31100 | 6.34517 | 19.45466 |
| **Mcm3ap** | 1.78245 | 0.05793 | 4.02270 | 3.76237 | 3.81182 | 1.95147 | 1.66061 | 1.37345 | 0.34952 | 0.04919 | 1.52626 | 5.68126 | 7.38125 | 2.17974 |
| **Mcm4** | 10.03583 | 0.38523 | 14.03099 | 8.26755 | 7.93461 | 11.44129 | 5.81456 | 0.98099 | 11.23948 | 6.92324 | 8.90514 | 10.23462 | 9.04092 | 14.14965 |
| **Mcm5** | 15.10258 | 0.42683 | 23.17458 | 9.99508 | 17.45963 | 23.55641 | 4.29843 | 10.60265 | 3.56283 | 0.32705 | 14.10414 | 17.89684 | 1.86474 | 15.42468 |
| **Mcm6** | 24.43529 | 2.28549 | 21.28352 | 12.95785 | 24.52857 | 12.89566 | 0.48713 | 6.86969 | 15.48586 | 0.10446 | 10.28190 | 11.76170 | 15.31406 | 24.73237 |
| **Mcm7** | 13.22095 | 7.88298 | 25.77088 | 22.82531 | 16.43276 | 16.70602 | 4.54397 | 6.88364 | 4.58856 | 0.31122 | 16.79335 | 13.84613 | 4.77999 | 16.88228 |
| **Mcm8** | 4.54283 | 0.05997 | 1.40543 | 4.17042 | 3.98678 | 4.10076 | 0.61866 | 2.49967 | 1.14582 | 0.02306 | 3.02337 | 4.69040 | 0.56623 | 3.91705 |
| **Mcm9** | 0.02383 | 0.00400 | 0.53587 | 0.10878 | 0.02350 | 0.42039 | 0.01392 | 0.65935 | 0.00648 | 0.00567 | 0.00404 | 0.00842 | 0.00269 | 0.46629 |
| **Mcmbp** | 7.55714 | 1.20293 | 10.79442 | 5.77586 | 8.99252 | 9.41757 | 1.32407 | 3.33674 | 6.08999 | 0.08604 | 10.87395 | 8.34252 | 10.15640 | 7.60609 |
| **Mcmdc2** | 0.03823 | 0.16621 | 0.23433 | 0.10317 | 0.15021 | 0.07037 | 0.06230 | 0.12004 | 0.99677 | 0.02993 | 0.02000 | 0.14213 | 0.20580 | 0.37980 |
| **Mcoln1** | 0.00000 | 0.01180 | 0.39477 | 2.24085 | 0.00000 | 1.07949 | 0.00354 | 0.81476 | 0.33178 | 0.00490 | 0.37236 | 0.15289 | 0.66821 | 0.09598 |
| **Mcph1** | 2.80404 | 0.00990 | 5.11861 | 1.53833 | 3.14830 | 3.33508 | 0.00994 | 0.34536 | 2.33584 | 0.02212 | 1.90226 | 1.38828 | 2.32532 | 8.04431 |
| **Mcrs1** | 9.23218 | 5.68557 | 30.85181 | 16.54282 | 22.41786 | 19.10666 | 2.09380 | 6.38112 | 0.21354 | 9.97094 | 9.79453 | 13.60251 | 3.47062 | 16.69000 |
| **Mctp1** | 0.00415 | 0.00429 | 0.01060 | 0.01043 | 0.18081 | 0.01423 | 0.00413 | 0.00855 | 0.01294 | 0.00744 | 0.00909 | 0.00848 | 0.66044 | 0.00266 |
| **Mctp2** | 0.01333 | 0.00444 | 0.02000 | 0.01340 | 0.01268 | 0.01228 | 0.00429 | 0.01030 | 0.00901 | 0.00737 | 0.00872 | 0.00968 | 0.03799 | 0.05337 |
| **Mcts1** | 6.70252 | 1.49235 | 2.81476 | 8.41419 | 6.00076 | 10.36770 | 4.54035 | 5.63791 | 6.62413 | 10.46454 | 4.15543 | 3.23610 | 1.43813 | 2.62781 |
| **Mcts2** | 4.61807 | 0.04019 | 1.51216 | 3.92395 | 3.56340 | 1.76509 | 2.77742 | 1.66723 | 1.86561 | 0.20798 | 2.70518 | 1.36191 | 1.56079 | 3.05939 |
| **Mcu** | 0.00369 | 0.00317 | 0.00000 | 0.00000 | 0.31943 | 1.73334 | 0.00367 | 0.03581 | 0.00368 | 0.01222 | 0.01177 | 1.13060 | 0.33596 | 0.14201 |
| **Mcur1** | 2.13420 | 4.26469 | 2.49452 | 1.81057 | 1.51791 | 3.03116 | 2.45497 | 5.65483 | 1.42658 | 0.20527 | 2.87824 | 0.69818 | 1.77140 | 2.85475 |
| **Mdc1** | 2.13787 | 0.18829 | 0.61400 | 1.01411 | 2.74887 | 5.98346 | 0.10280 | 2.38772 | 4.06552 | 0.11234 | 2.86488 | 3.60631 | 8.86659 | 6.38062 |
| **Mdfic** | 0.04315 | 0.00402 | 0.01304 | 0.00000 | 0.01010 | 0.00638 | 0.00402 | 0.99050 | 0.01177 | 0.00653 | 0.00658 | 0.01104 | 0.00535 | 0.01435 |
| **Mdga1** | 0.01485 | 0.01234 | 0.01390 | 0.01627 | 0.12106 | 0.01249 | 0.00442 | 0.03463 | 0.06075 | 0.00788 | 0.00766 | 0.01834 | 0.01692 | 0.01339 |
| **Mdh1** | 2.12777 | 8.00044 | 8.96554 | 4.54063 | 2.15524 | 19.16028 | 7.54059 | 12.43609 | 1.27888 | 13.92631 | 5.49740 | 9.82999 | 0.58893 | 1.93159 |
| **Mdh2** | 14.90918 | 10.92417 | 36.10087 | 32.70806 | 23.51225 | 21.91490 | 12.30011 | 28.72979 | 3.06709 | 13.18130 | 17.35745 | 26.43652 | 4.78369 | 14.92823 |
| **Mdk** | 0.06531 | 0.07385 | 0.15671 | 0.79377 | 0.09018 | 0.00000 | 0.01850 | 1.81661 | 0.16950 | 0.03040 | 0.59780 | 6.16028 | 0.19747 | 0.00000 |
| **Mdm1** | 0.37006 | 0.00417 | 0.00000 | 0.45394 | 0.20500 | 1.04538 | 0.01012 | 0.00441 | 0.00581 | 0.14638 | 0.01173 | 0.02532 | 0.00754 | 0.84502 |
| **Mdm2** | 3.53405 | 0.06458 | 7.00565 | 2.78691 | 6.58472 | 9.68133 | 3.89888 | 1.68565 | 5.57657 | 0.13630 | 2.11227 | 5.12507 | 15.24433 | 11.99405 |
| **Mdm4** | 5.19469 | 0.02668 | 6.23337 | 4.22737 | 11.27211 | 5.94082 | 3.51408 | 0.23188 | 18.99529 | 7.29505 | 6.37249 | 8.43758 | 7.45014 | 21.96357 |
| **Mdn1** | 12.07493 | 0.11071 | 17.61246 | 7.74397 | 18.74759 | 8.56012 | 0.06908 | 0.98725 | 24.95575 | 0.00000 | 5.29928 | 7.95822 | 8.60571 | 18.77437 |
| **Mdp1** | 0.40971 | 0.07006 | 1.39010 | 6.17562 | 6.74144 | 5.54863 | 4.69042 | 1.07268 | 1.59552 | 0.26631 | 2.39735 | 1.52773 | 1.48804 | 5.28586 |
| **Mdrl** | 0.39695 | 0.01094 | 1.22214 | 0.95006 | 1.29897 | 0.44324 | 0.00891 | 0.01203 | 0.57061 | 0.00416 | 0.68446 | 0.31947 | 0.00000 | 0.37800 |
| **Me1** | 4.57428 | 0.12410 | 3.65408 | 2.43517 | 8.88052 | 7.51615 | 1.74418 | 6.58869 | 1.01352 | 0.20346 | 7.58719 | 14.21456 | 2.69751 | 7.53899 |
| **Me2** | 0.18920 | 0.01024 | 0.33217 | 3.93291 | 2.14526 | 2.86784 | 0.00328 | 0.05694 | 1.26436 | 0.20954 | 2.44699 | 0.00000 | 1.43616 | 1.86332 |
| **Me3** | 0.00644 | 0.01312 | 0.00621 | 0.03019 | 0.14089 | 0.01664 | 0.01099 | 0.01421 | 0.01410 | 0.00655 | 0.00980 | 0.01080 | 0.15571 | 0.50074 |
| **Mea1** | 2.26067 | 0.51708 | 2.75371 | 9.90157 | 4.61937 | 9.82141 | 0.95686 | 6.48401 | 0.40255 | 11.75146 | 3.76690 | 7.47143 | 2.22057 | 1.32495 |
| **Meaf6** | 5.50674 | 0.10574 | 11.56229 | 8.21484 | 5.76926 | 6.92462 | 1.32060 | 5.37504 | 2.11624 | 7.22686 | 2.55298 | 4.23741 | 2.71340 | 2.84001 |

| | | | | | | | | | | | | | | |
|---|---|---|---|---|---|---|---|---|---|---|---|---|---|---|
| **Mecom** | 0.01421 | 0.00451 | 0.01256 | 0.01383 | 0.09688 | 0.01286 | 0.00436 | 0.01082 | 0.01049 | 0.00643 | 0.00877 | 0.00918 | 0.00639 | 0.01420 |
| **Mecp2** | 0.01070 | 0.00439 | 0.01242 | 0.49862 | 0.45660 | 0.52527 | 0.00992 | 0.18227 | 0.45613 | 0.00649 | 0.05500 | 0.05051 | 0.67349 | 0.71266 |
| **Mecr** | 4.32467 | 0.51173 | 2.99031 | 0.87501 | 2.92046 | 5.15911 | 2.67069 | 3.90841 | 0.00000 | 0.12105 | 2.81372 | 5.36879 | 1.22197 | 0.83685 |
| **Med1** | 1.80335 | 2.05462 | 4.66982 | 6.03500 | 7.37005 | 4.37918 | 0.06384 | 1.30267 | 10.76211 | 0.05645 | 5.59154 | 4.43209 | 7.05587 | 7.73833 |
| **Med10** | 14.41912 | 4.83396 | 15.43415 | 14.70659 | 15.54459 | 13.90708 | 6.57555 | 9.94880 | 1.56831 | 6.68790 | 6.29626 | 8.83395 | 10.25500 | 9.91775 |
| **Med11** | 3.31094 | 0.77544 | 3.91770 | 5.87263 | 7.08655 | 4.45437 | 2.50825 | 2.70094 | 0.42125 | 0.07953 | 2.97932 | 1.36432 | 2.40565 | 4.07140 |
| **Med12** | 0.21513 | 0.08074 | 1.21929 | 2.30072 | 2.48778 | 0.22774 | 0.59710 | 1.57860 | 0.04331 | 0.01647 | 0.35258 | 1.49288 | 0.78853 | 0.39778 |
| **Med12l** | 0.14285 | 0.00455 | 1.00605 | 1.34832 | 0.46971 | 0.75827 | 0.30035 | 0.83863 | 1.10567 | 0.01202 | 0.14985 | 0.11380 | 0.02199 | 0.20548 |
| **Med13** | 0.10681 | 0.00201 | 1.20388 | 0.95302 | 3.05591 | 2.99657 | 2.13096 | 0.00487 | 9.08043 | 0.84609 | 2.90131 | 0.30019 | 8.92598 | 4.15736 |
| **Med13l** | 0.09882 | 0.08718 | 1.63439 | 0.23902 | 1.34711 | 2.20859 | 0.00354 | 0.66687 | 0.27104 | 0.03747 | 0.01305 | 1.52464 | 1.66832 | 5.65414 |
| **Med14** | 14.81024 | 0.00717 | 6.99285 | 0.29480 | 12.75900 | 1.93531 | 0.04626 | 0.68288 | 27.14269 | 0.00000 | 2.46079 | 1.64819 | 8.31535 | 10.98962 |
| **Med15** | 2.66828 | 0.05057 | 6.69109 | 4.42664 | 4.17872 | 2.31312 | 2.63679 | 3.11391 | 0.88167 | 0.03598 | 3.93977 | 5.45545 | 1.86937 | 0.86868 |
| **Med16** | 0.33607 | 0.01738 | 0.81638 | 5.88014 | 5.28329 | 0.35491 | 0.04538 | 0.37416 | 0.08964 | 0.05706 | 0.04244 | 1.29147 | 1.72280 | 0.26832 |
| **Med17** | 14.32316 | 0.04852 | 15.25579 | 4.20089 | 12.71814 | 6.79649 | 0.81544 | 4.18742 | 8.29035 | 10.72615 | 10.54015 | 3.95122 | 3.67466 | 10.69667 |
| **Med18** | 0.00000 | 0.00458 | 0.05758 | 0.56181 | 0.02144 | 0.00000 | 0.25526 | 0.16798 | 1.27335 | 0.01357 | 0.04629 | 0.05121 | 0.95390 | 0.00000 |
| **Med19** | 3.49658 | 1.87752 | 7.86324 | 6.47124 | 10.91997 | 3.67264 | 0.00672 | 1.46916 | 5.89127 | 0.02310 | 2.23874 | 2.56658 | 3.60557 | 2.72635 |
| **Med20** | 14.82586 | 0.03674 | 13.23947 | 4.87907 | 9.93567 | 8.41514 | 3.18965 | 1.18974 | 4.53240 | 0.07132 | 8.83470 | 5.81417 | 5.69688 | 7.59890 |
| **Med21** | 7.32847 | 0.09957 | 2.51617 | 7.18544 | 13.75017 | 7.93508 | 3.48358 | 3.75762 | 17.52468 | 0.19932 | 2.95589 | 7.46353 | 10.69137 | 10.39472 |
| **Med22** | 4.91003 | 0.07303 | 7.28296 | 3.21754 | 8.82068 | 10.14887 | 2.34649 | 2.04845 | 1.13586 | 0.06582 | 3.02526 | 2.78462 | 3.29902 | 4.09507 |
| **Med23** | 9.67041 | 0.00525 | 5.80301 | 1.14376 | 11.74874 | 3.90396 | 0.17852 | 0.38866 | 6.01507 | 0.04277 | 0.14966 | 6.14056 | 7.69141 | 10.65214 |
| **Med24** | 2.06140 | 6.01713 | 10.32474 | 6.78970 | 10.54479 | 5.89566 | 1.72692 | 0.22951 | 0.00735 | 0.03045 | 7.54994 | 3.17602 | 1.47207 | 2.94800 |
| **Med25** | 0.00000 | 0.08557 | 0.99122 | 12.06825 | 6.27479 | 0.00000 | 0.03513 | 0.49377 | 0.00212 | 0.02104 | 0.00724 | 1.99417 | 0.11687 | 0.00000 |
| **Med26** | 5.16898 | 0.05215 | 6.66140 | 1.15346 | 7.53626 | 2.11955 | 0.00781 | 1.84030 | 0.78803 | 0.02331 | 6.34577 | 2.42383 | 6.06183 | 4.26447 |
| **Med27** | 15.65559 | 7.19459 | 19.52815 | 9.12608 | 23.25614 | 13.24213 | 3.96382 | 5.15814 | 1.47644 | 0.20214 | 10.04283 | 7.59833 | 5.26308 | 19.36316 |
| **Med28** | 3.92863 | 0.07518 | 2.88769 | 19.57715 | 21.17999 | 5.82269 | 0.57563 | 2.08601 | 1.59057 | 5.32465 | 0.79569 | 5.93119 | 7.69133 | 2.60011 |
| **Med29** | 4.04049 | 5.62741 | 6.18565 | 9.12860 | 19.09777 | 3.84561 | 0.74075 | 4.12237 | 1.53796 | 0.05126 | 1.07278 | 2.15978 | 13.35454 | 3.07719 |
| **Med30** | 1.27666 | 0.00362 | 0.00000 | 0.11975 | 0.46527 | 0.42032 | 0.22856 | 0.01902 | 0.12463 | 0.02165 | 0.07386 | 0.57040 | 0.09563 | 0.00000 |
| **Med31** | 3.44086 | 0.00997 | 0.23483 | 0.63291 | 7.59314 | 1.79580 | 0.14411 | 0.29550 | 4.94001 | 0.03376 | 0.39744 | 0.71414 | 11.87940 | 9.96131 |
| **Med4** | 11.21510 | 0.14293 | 16.63685 | 7.12263 | 9.02427 | 11.81184 | 4.15303 | 6.76704 | 6.21214 | 0.12221 | 9.93157 | 7.55647 | 9.81232 | 17.73614 |
| **Med6** | 14.34331 | 0.00997 | 6.98750 | 0.00000 | 7.20867 | 3.64116 | 0.52523 | 0.00000 | 16.36900 | 12.45181 | 8.04323 | 2.50963 | 9.49626 | 14.07929 |
| **Med7** | 6.14550 | 0.00684 | 4.33723 | 3.84003 | 4.43282 | 5.69778 | 5.34447 | 4.60784 | 2.19541 | 0.15660 | 9.02655 | 1.33856 | 8.51855 | 3.48764 |
| **Med8** | 12.58143 | 9.62071 | 9.04662 | 5.32316 | 7.06182 | 13.20129 | 2.99551 | 11.30826 | 6.33759 | 7.26309 | 10.17894 | 5.60250 | 1.92680 | 7.29597 |
| **Med9** | 0.98819 | 0.04632 | 0.00000 | 1.99173 | 6.89154 | 2.26132 | 0.41024 | 2.47894 | 0.37442 | 0.01310 | 2.41815 | 2.77835 | 4.94578 | 0.00000 |
| **Medag** | 0.02055 | 0.01208 | 0.01053 | 0.02051 | 0.00775 | 0.03599 | 0.00424 | 0.01017 | 0.03545 | 0.00787 | 0.00780 | 0.01791 | 0.03500 | 0.22302 |
| **Mef2a** | 1.43710 | 0.01528 | 0.11168 | 0.00000 | 0.00000 | 0.22473 | 0.00381 | 0.25548 | 0.91440 | 0.00714 | 0.00147 | 0.39683 | 1.44020 | 0.16892 |
| **Mef2d** | 0.63877 | 0.03474 | 5.36117 | 3.52439 | 4.37035 | 4.35702 | 1.13433 | 3.22411 | 3.70582 | 0.10639 | 0.90547 | 0.16039 | 1.57097 | 0.69300 |
| **Meg3** | 0.02230 | 0.01146 | 0.16123 | 1.35509 | 0.01988 | 0.08746 | 0.00372 | 0.00228 | 0.02172 | 0.00474 | 0.00303 | 0.56738 | 0.00486 | 0.03129 |
| **Megf11** | 2.74326 | 0.00491 | 1.17245 | 0.36974 | 0.12113 | 1.95319 | 0.00399 | 0.04293 | 0.00359 | 0.01497 | 0.72516 | 0.00000 | 0.39109 | 0.61931 |
| **Megf6** | 0.00000 | 0.00494 | 1.30449 | 0.68829 | 0.23017 | 0.10498 | 0.00391 | 0.02160 | 0.03243 | 0.00418 | 0.00396 | 0.11150 | 0.00320 | 0.20559 |
| **Megf8** | 0.30561 | 9.26453 | 0.55781 | 1.30291 | 0.38994 | 0.00000 | 0.00790 | 0.88174 | 0.11072 | 0.02217 | 0.65049 | 0.68596 | 0.00000 | 1.47418 |
| **Megf9** | 0.00920 | 0.00443 | 0.01080 | 0.01540 | 0.03574 | 0.01202 | 0.00483 | 0.00983 | 0.01404 | 0.00671 | 0.00834 | 0.01026 | 0.00766 | 0.19642 |
| **Mei1** | 2.27379 | 0.00360 | 1.60362 | 0.00000 | 2.96129 | 0.76575 | 0.00337 | 0.00000 | 1.07222 | 0.03336 | 0.00081 | 0.01000 | 1.78148 | 1.55917 |
| **Mei4** | 0.06028 | 0.00432 | 0.00996 | 0.05660 | 0.03186 | 0.02697 | 0.00410 | 0.01759 | 0.08451 | 0.02468 | 0.01565 | 0.03945 | 0.05186 | 0.08428 |
| **Meikin** | 0.97888 | 0.00426 | 0.66414 | 0.31395 | 0.70993 | 0.38446 | 0.00411 | 0.00283 | 0.03375 | 0.00661 | 0.00309 | 0.00339 | 0.20818 | 0.57624 |
| **Meioc** | 0.00524 | 0.00414 | 0.07971 | 0.11634 | 0.00000 | 0.00663 | 0.00396 | 0.00572 | 0.03962 | 0.00415 | 0.49059 | 0.00994 | 0.00319 | 0.50559 |
| **Meis1** | 0.02689 | 0.00467 | 0.00335 | 0.00000 | 0.01239 | 0.37730 | 0.00394 | 0.37089 | 0.00944 | 0.00807 | 0.00000 | 1.17969 | 0.00594 | 0.02720 |
| **Meis2** | 0.04810 | 0.00374 | 0.00000 | 0.00000 | 0.70695 | 0.00158 | 0.01473 | 0.19684 | 0.00375 | 0.00490 | 0.01006 | 0.00610 | 0.86966 | 2.62798 |
| **Meis3** | 0.01234 | 0.00403 | 0.00425 | 0.75767 | 0.00584 | 0.04129 | 0.02042 | 0.00396 | 0.01540 | 0.01956 | 0.01765 | 0.01159 | 0.00528 | 0.01122 |
| **Melk** | 11.57653 | 3.34270 | 12.26981 | 5.44984 | 8.78921 | 9.11495 | 3.15373 | 4.16258 | 12.04056 | 0.13289 | 4.71692 | 5.34775 | 3.37042 | 12.06927 |
| **Memo1** | 15.89690 | 0.03437 | 13.85770 | 3.58747 | 20.41076 | 3.81220 | 5.05854 | 0.86404 | 8.61335 | 0.09990 | 3.62489 | 6.03002 | 10.67008 | 13.24803 |
| **Men1** | 2.92565 | 0.05782 | 8.64565 | 1.39130 | 5.08742 | 4.75197 | 4.58893 | 3.11048 | 0.43578 | 0.13261 | 2.74886 | 6.24181 | 1.90669 | 1.69260 |
| **Mep1b** | 6.21909 | 0.01335 | 4.88248 | 1.71263 | 3.80690 | 0.00000 | 0.00829 | 0.07640 | 9.00761 | 0.00712 | 0.09496 | 0.68314 | 0.00000 | 7.75823 |
| **Mepce** | 3.05747 | 0.25548 | 11.12987 | 4.19160 | 6.69712 | 6.18271 | 3.39235 | 4.41923 | 0.79791 | 0.08316 | 3.83874 | 2.20508 | 10.86788 | 6.17764 |
| **Mertk** | 0.65168 | 0.52614 | 0.66147 | 0.00000 | 2.99029 | 0.77804 | 0.00696 | 1.00385 | 9.19578 | 0.00118 | 0.96876 | 1.47619 | 1.12292 | 2.45550 |
| **Mesdc1** | 2.22428 | 0.17848 | 0.51319 | 0.77423 | 1.43978 | 0.64223 | 0.00304 | 0.01267 | 0.04100 | 0.12740 | 0.03454 | 3.60019 | 4.30442 | 0.00000 |

| | | | | | | | | | | | | | | |
|---|---|---|---|---|---|---|---|---|---|---|---|---|---|---|
| **Mesdc2** | 9.36040 | 4.92881 | 16.45880 | 16.94498 | 9.87222 | 17.41977 | 6.30432 | 5.73495 | 7.04322 | 1.84995 | 13.70154 | 18.99053 | 3.88258 | 13.06143 |
| **Mesp2** | 0.08212 | 0.00369 | 0.16101 | 1.91047 | 0.77785 | 0.00000 | 0.00402 | 0.00623 | 0.00902 | 0.00560 | 0.00543 | 0.00632 | 3.27723 | 0.07294 |
| **Mest** | 0.00656 | 0.00309 | 0.00000 | 0.25503 | 0.00000 | 0.49663 | 0.00365 | 0.00000 | 1.29407 | 0.00188 | 1.16313 | 0.70251 | 0.01028 | 0.40494 |
| **Metap1** | 8.96395 | 5.21179 | 11.32420 | 7.85941 | 12.49107 | 13.96616 | 2.06318 | 3.15081 | 9.88496 | 0.15248 | 9.54775 | 6.25275 | 2.47884 | 13.71926 |
| **Metap1d** | 1.39540 | 0.03512 | 2.62139 | 0.81211 | 0.78387 | 0.30533 | 0.00812 | 2.32704 | 4.00005 | 0.04138 | 3.77266 | 1.60034 | 0.00000 | 1.07556 |
| **Metap2** | 27.78904 | 0.53691 | 39.81937 | 24.79955 | 39.54554 | 24.90948 | 6.88696 | 13.15516 | 33.66719 | 13.90091 | 18.25236 | 16.69663 | 17.43580 | 34.80807 |
| **Metrn** | 0.01359 | 0.00468 | 0.00497 | 0.10937 | 0.11087 | 0.14256 | 0.00447 | 0.00905 | 0.00723 | 0.00588 | 0.00581 | 0.00566 | 0.02035 | 0.02994 |
| **Mettl1** | 0.60566 | 0.00258 | 1.55517 | 1.29678 | 3.43415 | 0.91593 | 0.00904 | 0.96256 | 0.11678 | 0.00982 | 2.12935 | 1.38969 | 0.21237 | 0.58039 |
| **Mettl10** | 1.11178 | 0.01809 | 0.79653 | 0.58667 | 0.77643 | 0.78092 | 0.00285 | 2.73733 | 0.28333 | 0.00871 | 1.29256 | 1.70768 | 0.00000 | 1.07239 |
| **Mettl13** | 1.97276 | 0.01570 | 5.66082 | 5.83187 | 4.50511 | 5.26754 | 2.13388 | 0.07142 | 4.92088 | 0.08693 | 5.08217 | 2.85288 | 2.26191 | 5.95953 |
| **Mettl14** | 0.41690 | 5.65545 | 1.21931 | 4.97086 | 4.49975 | 3.03088 | 0.01456 | 2.29517 | 3.51090 | 0.00881 | 0.56697 | 0.30894 | 0.47172 | 6.89830 |
| **Mettl15** | 0.38686 | 0.00411 | 0.32136 | 0.59757 | 0.86175 | 0.38515 | 0.00373 | 0.00226 | 0.31942 | 0.00560 | 0.00426 | 0.03366 | 0.73863 | 0.54825 |
| **Mettl16** | 8.46568 | 2.47752 | 9.22981 | 2.53389 | 8.89080 | 7.38232 | 0.01039 | 0.24437 | 3.55949 | 12.52686 | 7.30568 | 4.61795 | 4.76822 | 9.80743 |
| **Mettl17** | 0.00000 | 0.04944 | 6.06567 | 6.53838 | 2.55007 | 5.68648 | 4.70135 | 3.33615 | 0.00000 | 0.14267 | 3.84502 | 0.96674 | 0.32566 | 3.09761 |
| **Mettl18** | 0.24542 | 8.97754 | 1.66651 | 2.36205 | 0.54432 | 0.22370 | 0.00179 | 1.34452 | 0.38244 | 0.01026 | 0.00519 | 0.24627 | 0.01021 | 2.24104 |
| **Mettl2** | 1.58661 | 3.85481 | 6.34363 | 6.68342 | 6.35737 | 7.82407 | 1.46464 | 4.27438 | 6.36023 | 0.17472 | 1.75113 | 1.24350 | 4.21849 | 7.17801 |
| **Mettl20** | 1.62970 | 0.01115 | 0.88520 | 0.00000 | 0.14054 | 1.36757 | 0.89766 | 0.16590 | 0.01183 | 0.01159 | 0.00874 | 0.55444 | 0.02077 | 0.29316 |
| **Mettl21a** | 0.24772 | 0.07561 | 0.86552 | 1.03905 | 0.12541 | 0.35607 | 1.13767 | 0.00348 | 0.06664 | 0.02048 | 2.09883 | 0.35142 | 0.00434 | 3.11998 |
| **Mettl21c** | 0.00774 | 0.00444 | 0.02639 | 0.01044 | 0.01129 | 0.02798 | 0.00429 | 0.00851 | 0.01109 | 0.00667 | 0.03955 | 0.00834 | 0.04126 | 0.03509 |
| **Mettl22** | 0.34598 | 0.05859 | 2.40580 | 4.69614 | 0.30789 | 4.97736 | 3.70908 | 1.09378 | 0.01249 | 4.05631 | 0.37928 | 0.45957 | 0.13878 | 0.93198 |
| **Mettl23** | 0.00000 | 0.01051 | 0.38580 | 1.84293 | 0.88447 | 0.95568 | 0.00982 | 0.77044 | 0.46194 | 0.02390 | 0.00000 | 2.05548 | 0.20805 | 0.19253 |
| **Mettl25** | 0.48785 | 0.00309 | 1.49482 | 0.72760 | 0.87926 | 0.52362 | 0.01018 | 0.39830 | 0.01248 | 0.00000 | 3.72951 | 0.00191 | 0.50171 | 0.33794 |
| **Mettl26** | 0.31827 | 4.59224 | 3.54014 | 3.93564 | 1.47656 | 0.24592 | 0.03472 | 3.63639 | 0.06058 | 11.49225 | 0.03490 | 5.15043 | 0.68982 | 0.10049 |
| **Mettl3** | 1.68444 | 0.00067 | 2.02598 | 1.26170 | 2.60883 | 2.58717 | 0.25915 | 0.59164 | 4.98206 | 0.01639 | 4.90644 | 2.76840 | 1.16342 | 3.73228 |
| **Mettl4** | 0.27654 | 0.00325 | 0.75129 | 0.30329 | 0.71188 | 0.32870 | 0.01568 | 0.07894 | 2.67038 | 0.00000 | 3.00977 | 1.14397 | 1.35871 | 1.49052 |
| **Mettl5** | 2.04846 | 0.03693 | 2.51712 | 4.54706 | 3.87625 | 1.93062 | 3.31607 | 4.96060 | 2.89721 | 0.14202 | 0.82546 | 4.95911 | 0.53656 | 2.76859 |
| **Mettl6** | 0.16344 | 0.05654 | 2.58466 | 3.15132 | 1.77246 | 1.45224 | 0.01846 | 2.84947 | 3.51572 | 0.05359 | 1.68969 | 2.02009 | 0.93587 | 3.72669 |
| **Mettl7a1** | 0.00556 | 0.02113 | 0.00000 | 0.00000 | 0.07577 | 0.21439 | 0.02340 | 0.87662 | 0.00902 | 0.00603 | 0.79008 | 0.56829 | 0.10737 | 0.03765 |
| **Mettl7a2** | 0.00381 | 0.00473 | 0.00972 | 0.00000 | 0.05334 | 0.01938 | 0.01234 | 0.55595 | 0.00640 | 0.00539 | 0.05551 | 0.04099 | 0.61491 | 0.12484 |
| **Mettl7a3** | 0.00558 | 0.00446 | 0.00971 | 0.57645 | 0.00680 | 0.00000 | 0.00483 | 0.08701 | 0.01281 | 0.00680 | 0.06794 | 0.01312 | 0.01199 | 0.01639 |
| **Mettl8** | 0.00000 | 0.04171 | 1.46693 | 0.00000 | 0.29413 | 1.65386 | 0.00323 | 0.29554 | 0.87113 | 0.02232 | 0.00000 | 1.59144 | 0.00000 | 0.00000 |
| **Mettl9** | 2.20655 | 0.02118 | 5.56869 | 0.92263 | 7.37396 | 2.12121 | 0.00591 | 1.87739 | 5.59624 | 0.00000 | 4.02416 | 4.04491 | 8.07304 | 3.60738 |
| **Mex3a** | 0.00299 | 0.03471 | 0.04818 | 1.01274 | 0.00000 | 0.03526 | 0.00398 | 0.21656 | 0.02656 | 0.03076 | 1.68797 | 0.00712 | 0.01102 | 1.41839 |
| **Mex3b** | 0.84965 | 0.00356 | 1.49500 | 0.14663 | 0.49528 | 0.37149 | 0.00391 | 0.00000 | 0.00600 | 0.00180 | 1.48162 | 0.09306 | 0.77530 | 0.50210 |
| **Mex3c** | 3.09920 | 0.22630 | 7.14912 | 0.15834 | 7.08646 | 3.12895 | 3.65665 | 0.74204 | 16.89552 | 0.09461 | 3.63413 | 1.46173 | 9.99154 | 6.77304 |
| **Mex3d** | 0.43768 | 0.02055 | 0.35079 | 0.51284 | 0.97394 | 0.49201 | 0.00466 | 0.20128 | 0.07227 | 0.00695 | 0.02122 | 0.00465 | 0.02528 | 0.60341 |
| **Mfap1a** | 3.74607 | 0.73105 | 4.19216 | 2.50119 | 5.70304 | 5.50321 | 0.13557 | 0.87330 | 7.01477 | 0.02390 | 3.61153 | 3.01518 | 4.47687 | 7.37929 |
| **Mfap1b** | 0.20579 | 0.41344 | 1.07429 | 0.45043 | 1.41533 | 1.13129 | 0.41202 | 0.06896 | 2.52052 | 0.00642 | 1.44997 | 0.49380 | 0.00000 | 2.28031 |
| **Mfap2** | 0.00534 | 0.00442 | 0.00588 | 0.00468 | 0.25755 | 0.07627 | 0.00413 | 0.00584 | 0.00881 | 0.00699 | 0.00857 | 0.00818 | 1.51595 | 0.00294 |
| **Mfap3** | 0.16797 | 0.08752 | 1.76471 | 0.55026 | 0.00000 | 2.16756 | 0.00148 | 0.66055 | 5.47405 | 0.01290 | 0.63818 | 5.11581 | 1.21016 | 5.42056 |
| **Mfap3l** | 0.01091 | 0.00445 | 0.01450 | 0.01340 | 0.01413 | 0.01318 | 0.00429 | 0.01028 | 0.01145 | 0.00711 | 0.00881 | 0.01003 | 0.00831 | 0.01285 |
| **Mfap4** | 0.00000 | 0.01512 | 0.50134 | 0.64076 | 0.56802 | 0.45531 | 0.00400 | 0.09671 | 0.78078 | 0.00611 | 1.11181 | 0.01014 | 0.36709 | 0.02357 |
| **Mfap5** | 0.00000 | 0.00360 | 0.66236 | 2.08403 | 1.03210 | 0.82053 | 0.00352 | 0.00000 | 1.70398 | 0.01058 | 0.00000 | 0.01286 | 4.44967 | 0.79741 |
| **Mff** | 5.31105 | 0.03393 | 1.88733 | 0.00000 | 2.21364 | 7.39191 | 2.31630 | 5.09767 | 2.37205 | 0.03435 | 3.80055 | 6.66120 | 2.87746 | 0.96416 |
| **Mfge8** | 1.35821 | 0.31445 | 14.29026 | 13.50314 | 7.93236 | 3.92484 | 2.49629 | 8.86377 | 0.02571 | 0.15660 | 6.24229 | 12.77175 | 0.40973 | 3.17217 |
| **Mfhas1** | 0.01347 | 0.00415 | 0.00000 | 0.43239 | 0.13296 | 0.58215 | 0.00404 | 0.00907 | 0.01298 | 0.00489 | 0.00307 | 0.01186 | 0.00795 | 0.00645 |
| **Mfi2** | 0.01317 | 0.00425 | 0.36462 | 0.24902 | 0.02434 | 0.02141 | 0.00959 | 0.00348 | 0.01071 | 0.00567 | 0.61719 | 0.00662 | 0.01610 | 0.00000 |
| **Mfn1** | 12.59102 | 6.84898 | 23.00203 | 13.24377 | 14.76082 | 13.75075 | 2.67808 | 9.11701 | 6.44091 | 0.50949 | 13.38479 | 10.98941 | 3.73805 | 13.52490 |
| **Mfn2** | 0.81222 | 0.04085 | 3.66788 | 4.53167 | 4.62119 | 6.08027 | 0.07658 | 3.22960 | 0.61877 | 0.04828 | 2.00086 | 1.97800 | 1.49539 | 7.04476 |
| **Mfng** | 0.01048 | 0.00449 | 0.12888 | 0.00000 | 0.00941 | 0.18983 | 0.00421 | 0.00533 | 0.01104 | 0.00593 | 0.01203 | 0.00979 | 0.00768 | 0.01486 |
| **Mfsd1** | 7.95860 | 0.14355 | 15.77721 | 21.04071 | 14.32580 | 20.25270 | 5.29309 | 6.41613 | 17.96796 | 1.61367 | 16.87851 | 17.49580 | 2.14456 | 9.89654 |
| **Mfsd10** | 1.00435 | 1.05158 | 8.60692 | 6.30603 | 2.34756 | 1.05253 | 2.10174 | 7.62720 | 0.00000 | 0.07054 | 1.63386 | 3.86707 | 0.22735 | 0.00000 |
| **Mfsd11** | 2.42301 | 0.16746 | 0.97112 | 1.64068 | 1.32773 | 5.47955 | 3.33492 | 1.09103 | 3.93731 | 0.92654 | 1.85817 | 1.94366 | 7.36754 | 6.18888 |
| **Mfsd12** | 0.01179 | 0.00418 | 0.10285 | 0.08544 | 0.24001 | 0.00488 | 0.00392 | 0.00715 | 0.01568 | 0.00778 | 0.01227 | 0.01265 | 0.32147 | 0.01797 |
| **Mfsd13a** | 0.00265 | 4.60209 | 0.00000 | 3.66322 | 0.09706 | 0.26432 | 0.05658 | 5.48059 | 0.00723 | 0.00645 | 0.00000 | 2.90322 | 0.25166 | 0.00000 |

| | | | | | | | | | | | | | | |
|---|---|---|---|---|---|---|---|---|---|---|---|---|---|---|
| **Mfsd14a** | 10.57741 | 0.01388 | 9.32906 | 3.08975 | 4.02552 | 4.29668 | 2.33795 | 2.17473 | 22.23917 | 0.01875 | 6.95813 | 2.13926 | 14.82669 | 20.98235 |
| **Mfsd14b** | 4.04763 | 2.13470 | 8.92590 | 1.07818 | 9.53320 | 2.69841 | 0.00000 | 1.22212 | 23.29449 | 0.00000 | 4.92735 | 0.68539 | 2.56186 | 14.19266 |
| **Mfsd2a** | 0.01443 | 0.00409 | 0.16769 | 0.00388 | 0.00000 | 0.00556 | 0.00402 | 0.00933 | 0.02145 | 0.00609 | 0.00654 | 0.00709 | 1.77293 | 0.64296 |
| **Mfsd2b** | 0.00000 | 0.01932 | 0.76889 | 0.69740 | 0.42922 | 0.40196 | 0.01177 | 1.38198 | 0.03436 | 0.00531 | 0.00634 | 0.00000 | 1.24167 | 0.25946 |
| **Mfsd3** | 0.00808 | 0.01116 | 0.00113 | 0.35697 | 0.03772 | 0.91103 | 0.00422 | 0.03646 | 0.00980 | 0.00680 | 0.00243 | 0.00259 | 0.02379 | 0.01555 |
| **Mfsd4a** | 0.03739 | 0.00462 | 0.02360 | 0.13429 | 0.18725 | 0.13975 | 0.04259 | 0.07577 | 0.08961 | 0.04013 | 0.01095 | 0.05517 | 0.35542 | 0.31479 |
| **Mfsd4b1** | 0.00386 | 0.00396 | 0.00300 | 0.53481 | 0.00538 | 0.04217 | 1.05909 | 0.02173 | 0.01295 | 0.01111 | 0.06927 | 0.00792 | 0.00698 | 0.00633 |
| **Mfsd4b2** | 0.01062 | 0.00385 | 0.01214 | 0.05557 | 0.01062 | 0.00000 | 0.04620 | 0.16610 | 0.01280 | 0.00477 | 0.12423 | 0.00878 | 0.00683 | 0.01040 |
| **Mfsd4b3** | 0.00771 | 0.00399 | 0.01078 | 0.25700 | 0.00823 | 0.00000 | 0.00949 | 0.02834 | 0.01266 | 0.00577 | 0.10875 | 0.01258 | 0.00673 | 0.00881 |
| **Mfsd4b5** | 0.02915 | 0.00401 | 0.03960 | 0.04355 | 0.03046 | 0.06083 | 0.02060 | 0.14056 | 0.09655 | 0.01200 | 0.17457 | 0.03501 | 0.04038 | 0.01753 |
| **Mfsd5** | 18.09826 | 0.07961 | 19.09438 | 7.53862 | 21.51444 | 1.59051 | 4.12136 | 4.86333 | 0.96287 | 0.14066 | 6.89992 | 5.73768 | 11.37818 | 9.69370 |
| **Mfsd6** | 0.00341 | 0.00410 | 0.01457 | 0.01117 | 0.00696 | 0.01661 | 0.00412 | 0.00765 | 0.00648 | 0.00612 | 0.01109 | 0.00766 | 1.92062 | 0.02534 |
| **Mfsd7a** | 0.17762 | 0.00393 | 1.00917 | 1.25993 | 0.15712 | 0.00000 | 0.00370 | 0.02159 | 0.10261 | 0.00376 | 0.00866 | 0.01087 | 0.00362 | 0.45600 |
| **Mfsd7b** | 0.45793 | 0.00366 | 0.41123 | 0.00000 | 0.20381 | 0.84444 | 0.05792 | 0.29655 | 3.36946 | 0.01347 | 0.00000 | 0.94467 | 2.80207 | 0.64222 |
| **Mfsd7c** | 11.95558 | 0.06527 | 23.69804 | 11.77640 | 12.89538 | 19.33031 | 6.97299 | 4.17566 | 19.41512 | 7.59822 | 7.52924 | 6.14112 | 3.08174 | 18.36231 |
| **Mfsd8** | 1.48674 | 0.00315 | 1.22222 | 1.08446 | 0.00000 | 0.53131 | 0.00307 | 0.23860 | 2.52025 | 0.00563 | 0.00000 | 0.79270 | 0.00477 | 1.19768 |
| **Mfsd9** | 0.00099 | 0.00329 | 0.09501 | 0.00000 | 0.20181 | 0.00295 | 1.41322 | 0.00227 | 0.04631 | 0.04894 | 0.00000 | 0.88590 | 0.23394 | 0.42083 |
| **Mga** | 0.62602 | 0.24179 | 3.67093 | 4.27865 | 4.59883 | 2.50524 | 0.57888 | 0.73237 | 20.18470 | 0.86904 | 3.02253 | 1.93272 | 10.69701 | 7.53718 |
| **Mgarp** | 0.62166 | 0.11070 | 4.41117 | 7.20186 | 2.50289 | 2.77698 | 4.27146 | 0.00021 | 0.00000 | 0.18828 | 8.54792 | 11.36695 | 0.23003 | 1.85186 |
| **Mgat1** | 0.03005 | 0.05136 | 0.00816 | 0.00000 | 0.20794 | 0.74097 | 4.94383 | 1.95699 | 0.01496 | 0.18531 | 1.21005 | 1.51770 | 0.33186 | 0.08821 |
| **Mgat2** | 2.91896 | 7.33296 | 7.44405 | 4.10765 | 6.80140 | 5.25905 | 0.44239 | 2.32131 | 6.01057 | 0.07050 | 4.45646 | 3.18776 | 13.77699 | 7.49879 |
| **Mgat3** | 0.99304 | 0.00382 | 2.37044 | 0.50732 | 0.03168 | 0.93308 | 0.00253 | 0.17125 | 6.21833 | 0.01556 | 1.50568 | 0.02727 | 1.46899 | 1.04740 |
| **Mgat4a** | 1.74379 | 0.01172 | 4.23224 | 0.19749 | 1.25375 | 1.22443 | 0.00276 | 0.00000 | 1.50255 | 0.01166 | 1.55059 | 0.50661 | 0.38296 | 2.44243 |
| **Mgat4b** | 0.02317 | 0.00411 | 0.02269 | 0.15704 | 0.07497 | 0.02361 | 0.00423 | 0.05303 | 0.01910 | 0.03130 | 0.00869 | 0.06834 | 0.17463 | 0.00000 |
| **Mgat4e** | 0.02741 | 0.00443 | 0.01414 | 0.01171 | 0.02024 | 0.01303 | 0.00430 | 0.01009 | 0.01455 | 0.00662 | 0.00865 | 0.00970 | 0.08413 | 0.00637 |
| **Mgat5** | 0.01237 | 0.00494 | 0.01118 | 0.01456 | 0.08171 | 0.01517 | 0.00427 | 0.03539 | 0.01287 | 0.00814 | 0.00804 | 0.05149 | 0.00663 | 0.01028 |
| **Mgea5** | 10.23937 | 0.02850 | 6.41328 | 2.27923 | 17.71246 | 1.40596 | 4.52467 | 4.95629 | 20.52275 | 0.23087 | 2.00580 | 6.25464 | 5.57866 | 15.26102 |
| **Mgl2** | 0.21871 | 0.02071 | 3.22993 | 3.98032 | 0.31281 | 2.42389 | 1.23174 | 0.00481 | 0.78929 | 0.04810 | 0.20875 | 0.09458 | 1.25214 | 2.35334 |
| **Mgll** | 0.57327 | 0.02061 | 0.00000 | 0.00000 | 0.00430 | 2.84769 | 0.00362 | 0.06297 | 0.02139 | 0.04320 | 0.25620 | 1.77810 | 0.00453 | 0.00682 |
| **Mgme1** | 2.61589 | 0.03025 | 10.54635 | 0.00008 | 8.57689 | 6.84648 | 0.00000 | 0.57929 | 3.15212 | 0.01021 | 6.46719 | 2.56864 | 0.00000 | 8.99742 |
| **Mgmt** | 0.00647 | 0.03974 | 0.00178 | 0.00000 | 0.00974 | 0.76069 | 0.01410 | 0.21828 | 0.00934 | 0.00674 | 0.03009 | 3.03926 | 0.02434 | 0.03312 |
| **Mgp** | 0.01389 | 0.00450 | 0.00457 | 0.01002 | 0.60290 | 0.01427 | 0.00435 | 0.01007 | 0.01270 | 0.00736 | 0.00879 | 0.00804 | 0.00369 | 0.01257 |
| **Mgrn1** | 1.92861 | 0.01585 | 6.92288 | 4.94826 | 9.70206 | 2.75450 | 2.70751 | 2.31514 | 0.15182 | 0.11981 | 0.00000 | 5.34850 | 2.12985 | 4.78975 |
| **Mgst1** | 0.01637 | 0.00450 | 0.00443 | 0.00990 | 0.60876 | 0.01357 | 0.00474 | 0.01023 | 0.01573 | 0.00679 | 0.00883 | 0.00798 | 0.00358 | 0.01061 |
| **Mgst2** | 0.00239 | 0.02264 | 0.01263 | 1.12814 | 0.00433 | 0.00000 | 0.82897 | 1.63844 | 0.01436 | 0.08344 | 0.00171 | 0.14848 | 0.00658 | 0.00539 |
| **Mgst3** | 0.00986 | 2.25996 | 0.00834 | 0.00000 | 0.01035 | 0.37312 | 0.08496 | 9.17939 | 0.08233 | 3.55963 | 0.00000 | 0.30681 | 1.26397 | 0.00000 |
| **Mia** | 0.13464 | 0.00521 | 0.11619 | 0.01063 | 0.10211 | 0.02837 | 0.00434 | 0.02945 | 0.01061 | 0.01247 | 0.00647 | 0.00704 | 0.11752 | 0.00966 |
| **Mia2** | 0.24614 | 0.00443 | 0.00000 | 0.01142 | 0.21488 | 0.00425 | 0.00403 | 0.01029 | 1.62732 | 0.00760 | 0.00417 | 0.00691 | 0.10696 | 0.25493 |
| **Mia3** | 3.43056 | 0.06951 | 5.10453 | 3.30157 | 2.38127 | 6.76252 | 1.09392 | 3.52210 | 2.69833 | 0.13774 | 7.22567 | 3.75070 | 1.45125 | 5.51378 |
| **Mib1** | 0.18625 | 0.00325 | 0.69295 | 0.00000 | 1.47862 | 0.00000 | 0.00324 | 0.00460 | 4.57003 | 0.00000 | 1.82266 | 0.00593 | 1.83637 | 2.88255 |
| **Mib2** | 0.00000 | 0.00269 | 0.36832 | 3.46047 | 1.98374 | 0.00000 | 0.01039 | 1.04934 | 0.01266 | 0.01588 | 0.00000 | 1.87338 | 0.00369 | 0.00186 |
| **Mical1** | 0.00000 | 0.00982 | 1.68468 | 2.44840 | 0.54709 | 2.16252 | 4.82004 | 1.36350 | 0.01246 | 0.16122 | 0.00000 | 3.81003 | 0.00000 | 0.00000 |
| **Mical2** | 0.20262 | 0.00439 | 0.31726 | 0.00492 | 0.00287 | 0.00747 | 0.00411 | 0.01077 | 0.01034 | 0.00683 | 0.00601 | 0.00814 | 0.00362 | 0.00879 |
| **Mical3** | 3.96345 | 0.00189 | 9.49759 | 3.04150 | 6.39662 | 1.48213 | 1.66868 | 1.37091 | 0.95055 | 0.00775 | 1.51384 | 1.17850 | 3.44238 | 6.70998 |
| **Micall1** | 0.17754 | 0.01173 | 0.94846 | 0.00000 | 0.83277 | 0.00000 | 1.52233 | 0.02723 | 0.10468 | 0.02952 | 0.00000 | 0.66867 | 4.57483 | 1.69517 |
| **Micall2** | 0.04271 | 0.01055 | 0.00930 | 0.37633 | 0.00419 | 0.00000 | 0.00346 | 1.53464 | 0.00619 | 0.01478 | 0.01102 | 0.03093 | 0.00442 | 0.13220 |
| **Micu1** | 1.72224 | 0.01880 | 1.42959 | 0.74944 | 0.76479 | 3.61162 | 0.51177 | 1.12971 | 0.06118 | 0.51246 | 0.53339 | 1.28052 | 1.49341 | 0.58090 |
| **Micu2** | 0.00159 | 0.01749 | 0.00000 | 0.33003 | 0.12599 | 0.53968 | 1.10568 | 1.46060 | 0.83414 | 0.39408 | 0.00000 | 2.27360 | 1.30112 | 1.36653 |
| **Mid1** | 0.35357 | 0.01223 | 0.07719 | 0.00000 | 0.00000 | 0.13380 | 1.16162 | 0.84185 | 0.15533 | 0.02439 | 0.00000 | 0.61966 | 0.03154 | 0.11407 |
| **Mid1ip1** | 0.00427 | 0.02822 | 0.00000 | 1.95178 | 1.03609 | 0.89922 | 2.62385 | 0.34779 | 0.01397 | 0.11447 | 1.02153 | 1.96294 | 0.91885 | 0.00000 |
| **Midn** | 0.05333 | 0.09586 | 0.09565 | 0.72861 | 0.19185 | 0.40460 | 0.08211 | 1.98935 | 0.19566 | 0.00506 | 0.62517 | 0.16625 | 2.15142 | 0.03896 |
| **Mief1** | 5.49844 | 0.02040 | 4.38254 | 4.09021 | 5.30765 | 5.61410 | 2.26685 | 1.34893 | 9.56963 | 0.03240 | 3.31341 | 0.74464 | 2.78529 | 4.75137 |
| **Mief2** | 0.20010 | 0.05045 | 2.25429 | 0.20473 | 1.58573 | 1.40290 | 0.39383 | 5.62170 | 1.45757 | 0.05438 | 1.59837 | 0.80131 | 1.40147 | 1.23545 |
| **Mien1** | 1.53545 | 5.10854 | 2.46062 | 19.27181 | 8.40639 | 15.37495 | 5.58923 | 13.95494 | 0.26287 | 12.57393 | 8.46593 | 4.99278 | 0.00000 | 1.76636 |
| **Mier1** | 4.01899 | 0.00279 | 1.86366 | 1.01537 | 1.04240 | 1.72816 | 1.00778 | 0.32480 | 1.76746 | 0.01980 | 0.00000 | 1.74065 | 0.33261 | 1.34732 |

| | | | | | | | | | | | | | | |
|---|---|---|---|---|---|---|---|---|---|---|---|---|---|---|
| **Mier2** | 0.00202 | 0.01801 | 0.04098 | 3.85044 | 3.34582 | 0.21855 | 0.00377 | 1.07984 | 0.00566 | 0.02483 | 0.00106 | 1.63006 | 0.07237 | 0.00000 |
| **Mier3** | 25.61889 | 0.04723 | 19.34117 | 10.67400 | 29.45167 | 14.29113 | 3.40309 | 4.25903 | 35.87043 | 0.10735 | 8.87132 | 5.13617 | 13.50532 | 39.27685 |
| **Mif** | 0.58313 | 0.62710 | 1.06024 | 12.35724 | 11.30278 | 4.11225 | 3.53880 | 6.39033 | 0.00508 | 7.32503 | 0.42159 | 9.54129 | 1.23480 | 0.00000 |
| **Mif4gd** | 2.29235 | 0.13202 | 5.52351 | 2.98627 | 3.48225 | 2.80070 | 0.39823 | 5.48915 | 0.00337 | 0.01251 | 2.39727 | 5.22933 | 2.78532 | 1.11306 |
| **Miip** | 4.00596 | 2.44088 | 11.18522 | 17.09354 | 2.56666 | 3.54352 | 1.76552 | 2.87477 | 0.00000 | 0.15313 | 4.85441 | 4.75462 | 0.29062 | 3.13555 |
| **Milr1** | 0.32928 | 0.01266 | 0.31492 | 0.51392 | 0.66541 | 0.00000 | 0.02552 | 0.01389 | 0.04930 | 0.01627 | 0.68445 | 0.23739 | 0.70401 | 1.53448 |
| **Mina** | 0.64544 | 0.02489 | 5.71136 | 1.96274 | 2.43306 | 2.43834 | 0.00144 | 1.11206 | 3.73027 | 0.00445 | 1.70065 | 2.87190 | 0.16476 | 8.18612 |
| **Mink1** | 0.93370 | 0.00406 | 0.60427 | 1.73184 | 1.52235 | 0.18787 | 0.04432 | 0.00035 | 0.02770 | 0.00584 | 0.33658 | 0.20346 | 0.27468 | 0.18871 |
| **Minos1** | 4.20287 | 0.22097 | 0.00000 | 3.19049 | 8.48970 | 11.83752 | 2.11338 | 8.70402 | 1.07704 | 10.93563 | 2.77866 | 8.49321 | 7.28073 | 2.96458 |
| **Minpp1** | 0.60947 | 0.07164 | 3.56897 | 0.71964 | 2.58187 | 4.82800 | 0.00160 | 4.72347 | 1.57923 | 0.00000 | 2.11174 | 3.57407 | 0.05060 | 6.06580 |
| **Mios** | 3.06176 | 0.07122 | 1.43549 | 0.71629 | 6.46841 | 1.31442 | 0.00192 | 1.01728 | 4.72989 | 0.00054 | 3.75244 | 1.36175 | 7.20021 | 10.36393 |
| **Miox** | 0.08129 | 0.00342 | 0.58242 | 0.78599 | 1.64409 | 0.01417 | 0.00373 | 0.01656 | 0.08258 | 0.00579 | 0.00646 | 0.00282 | 5.30591 | 3.48016 |
| **Mip** | 0.14829 | 0.02040 | 0.00924 | 0.01067 | 0.25936 | 0.00909 | 0.00435 | 0.01016 | 0.15845 | 0.00636 | 0.00643 | 0.00705 | 0.16480 | 0.02344 |
| **Mipep** | 1.77572 | 0.03217 | 1.15069 | 0.59252 | 1.80090 | 2.88622 | 0.02108 | 4.02973 | 0.15731 | 0.00475 | 4.66813 | 2.10239 | 0.63289 | 4.17803 |
| **Mipol1** | 0.00325 | 0.00423 | 1.46449 | 0.00000 | 0.00133 | 0.00987 | 0.00393 | 0.00480 | 0.05277 | 0.00442 | 0.01332 | 0.01641 | 0.25588 | 0.00000 |
| **Mir1291** | 0.00966 | 0.00445 | 0.04293 | 0.00831 | 0.04612 | 0.03890 | 0.00421 | 0.00826 | 0.02876 | 0.00668 | 0.00895 | 0.00982 | 0.00585 | 0.04220 |
| **Mir17hg** | 0.01913 | 0.00544 | 0.50934 | 1.09771 | 0.65270 | 0.00000 | 0.00389 | 0.49117 | 2.63679 | 0.01437 | 0.00000 | 0.20812 | 0.44143 | 0.00000 |
| **Mir1949** | 0.02653 | 0.00460 | 0.00718 | 0.04153 | 0.00282 | 0.01297 | 0.00418 | 0.01098 | 0.10647 | 0.00964 | 0.00831 | 0.01072 | 0.00897 | 0.08027 |
| **Mir1982** | 0.00594 | 0.00512 | 0.02645 | 0.14249 | 0.01746 | 0.02527 | 0.10810 | 0.03832 | 0.01043 | 0.00783 | 0.05463 | 0.00868 | 0.04375 | 0.01562 |
| **Mir1a.1hg** | 0.01172 | 0.00456 | 0.01989 | 0.02011 | 0.02862 | 0.01365 | 0.01022 | 0.01046 | 0.04050 | 0.00660 | 0.00944 | 0.01261 | 0.03232 | 0.04811 |
| **Mir22hg** | 0.00000 | 0.02247 | 0.45945 | 0.92101 | 0.19971 | 0.32557 | 1.14181 | 0.17620 | 0.58814 | 0.05020 | 1.18006 | 0.00000 | 0.29257 | 0.00000 |
| **Mir25** | 0.01578 | 0.00409 | 0.05372 | 0.52038 | 0.06409 | 0.00970 | 0.00393 | 0.00733 | 0.02541 | 0.00639 | 0.07300 | 0.02713 | 0.00514 | 0.06269 |
| **Mir3064** | 0.01054 | 0.00406 | 0.14051 | 0.17811 | 0.02069 | 0.04000 | 0.07633 | 0.02103 | 0.20246 | 0.00492 | 0.01888 | 0.03189 | 0.00581 | 0.29219 |
| **Mir3068** | 0.01010 | 0.00445 | 0.01466 | 0.01333 | 0.01059 | 0.01204 | 0.00425 | 0.01039 | 0.06752 | 0.00708 | 0.00825 | 0.01009 | 0.00824 | 0.01748 |
| **Mir3091** | 0.01895 | 0.00410 | 0.00144 | 0.17320 | 0.13759 | 0.12218 | 0.00415 | 0.06690 | 0.01355 | 0.00827 | 0.01990 | 0.12562 | 0.00671 | 0.01438 |
| **Mir3471.1** | 0.03061 | 0.00455 | 0.02299 | 0.04821 | 0.03401 | 0.03770 | 0.03732 | 0.06243 | 0.10320 | 0.00659 | 0.04348 | 0.02335 | 0.23906 | 0.18386 |
| **Mir3471.2** | 0.00750 | 0.00442 | 0.01365 | 0.01208 | 0.02500 | 0.01570 | 0.00427 | 0.00966 | 0.01198 | 0.00670 | 0.00863 | 0.00985 | 0.07472 | 0.04050 |
| **Mir3473f** | 0.01980 | 0.04334 | 0.01460 | 0.02579 | 0.02173 | 0.03115 | 0.01032 | 0.01796 | 0.06059 | 0.00780 | 0.00735 | 0.02084 | 0.31179 | 0.07323 |
| **Mir425** | 0.01453 | 0.00428 | 0.04284 | 0.05573 | 0.01054 | 0.00279 | 0.00427 | 0.01005 | 0.01184 | 0.00663 | 0.12820 | 0.01091 | 0.00671 | 0.01067 |
| **Mir6236** | 0.01560 | 0.00467 | 0.01001 | 0.00000 | 0.07008 | 0.02978 | 0.00444 | 0.03893 | 0.01544 | 0.00703 | 0.07365 | 0.03954 | 0.09008 | 0.01808 |
| **Mir6385** | 0.05278 | 0.00391 | 0.05630 | 0.10671 | 0.10696 | 0.04100 | 0.28325 | 0.10891 | 0.00809 | 0.01061 | 0.00411 | 0.02216 | 0.00338 | 0.12594 |
| **Mir6516** | 0.02429 | 0.07078 | 0.01122 | 0.00000 | 0.01160 | 0.00474 | 0.00445 | 0.20963 | 0.07498 | 0.00780 | 0.00524 | 0.10110 | 0.00933 | 0.03580 |
| **Mir677** | 0.00909 | 0.00449 | 0.01318 | 0.05516 | 0.01850 | 0.01381 | 0.00455 | 0.01284 | 0.01100 | 0.00702 | 0.00875 | 0.00862 | 0.00724 | 0.02647 |
| **Mir680.2** | 0.02515 | 0.00464 | 0.02943 | 0.01216 | 0.02430 | 0.01451 | 0.00441 | 0.01294 | 0.02311 | 0.00789 | 0.00795 | 0.00919 | 0.07581 | 0.02385 |
| **Mir682** | 0.02849 | 0.00444 | 0.01726 | 0.03569 | 0.03298 | 0.05137 | 0.00434 | 0.05021 | 0.00975 | 0.04861 | 0.00618 | 0.02766 | 0.19711 | 0.02492 |
| **Mir684.1** | 0.94585 | 0.01162 | 0.89031 | 0.56638 | 1.36295 | 0.58969 | 0.17302 | 0.13505 | 0.60189 | 0.06083 | 0.27620 | 0.18157 | 0.56054 | 0.92716 |
| **Mir684.2** | 0.22224 | 0.00416 | 0.19616 | 0.10879 | 0.41903 | 0.14930 | 0.05048 | 0.05744 | 0.17134 | 0.00716 | 0.03941 | 0.06267 | 0.04210 | 0.10486 |
| **Mir6910** | 0.00986 | 0.00440 | 0.05038 | 0.03872 | 0.01012 | 0.01439 | 0.00420 | 0.00795 | 0.01205 | 0.00604 | 0.01170 | 0.01211 | 0.00450 | 0.06538 |
| **Mir692.1** | 0.09146 | 0.58239 | 0.65713 | 2.48624 | 0.56497 | 1.35837 | 0.30029 | 1.52305 | 0.00736 | 0.63035 | 0.68533 | 0.66718 | 0.03889 | 0.08146 |
| **Mir6935** | 0.00868 | 0.60144 | 0.04020 | 0.03170 | 0.01166 | 0.02017 | 0.00392 | 0.00360 | 0.01195 | 0.00622 | 0.00631 | 0.08915 | 0.00701 | 0.01132 |
| **Mir6951** | 0.00973 | 0.00432 | 0.02618 | 0.00494 | 0.01268 | 0.02624 | 0.00431 | 0.00787 | 0.01640 | 0.00668 | 0.04558 | 0.02974 | 0.00962 | 0.01869 |
| **Mir697** | 0.00814 | 0.00433 | 0.04234 | 0.22325 | 0.02001 | 0.00678 | 0.00432 | 0.00969 | 0.03316 | 0.00739 | 0.00682 | 0.00896 | 0.01042 | 0.00592 |
| **Mir7013** | 0.00948 | 0.00437 | 0.01676 | 0.00934 | 0.01270 | 0.01350 | 0.00438 | 0.00955 | 0.01424 | 0.00692 | 0.00818 | 0.03201 | 0.01229 | 0.03028 |
| **Mir7014** | 0.00627 | 0.00449 | 0.04158 | 0.02882 | 0.00860 | 0.01255 | 0.00440 | 0.01466 | 0.01341 | 0.00715 | 0.00951 | 0.00820 | 0.10965 | 0.04554 |
| **Mir7024** | 0.01151 | 0.00442 | 0.05022 | 0.01599 | 0.06667 | 0.01080 | 0.00413 | 0.01015 | 0.01366 | 0.00921 | 0.00773 | 0.00866 | 0.00644 | 0.04381 |
| **Mir703** | 10.09867 | 4.18298 | 6.40601 | 13.27342 | 9.29225 | 16.28027 | 3.22254 | 12.49632 | 10.06468 | 6.41744 | 7.26930 | 11.68482 | 2.40151 | 6.65363 |
| **Mir7036** | 0.00495 | 0.00502 | 0.04446 | 0.31631 | 0.01691 | 0.00883 | 0.00418 | 0.15168 | 0.01310 | 0.00592 | 0.03097 | 0.01744 | 0.00587 | 0.01261 |
| **Mir7036b** | 0.00856 | 0.00443 | 0.01250 | 0.01578 | 0.09101 | 0.01262 | 0.00431 | 0.01082 | 0.00914 | 0.00723 | 0.00876 | 0.00919 | 0.05553 | 0.01029 |
| **Mir7051** | 0.02920 | 0.00461 | 0.03555 | 0.12676 | 0.03780 | 0.10645 | 0.00445 | 0.01878 | 0.02554 | 0.00821 | 0.00540 | 0.06897 | 0.01340 | 0.05247 |
| **Mir7067** | 0.06945 | 0.00431 | 0.05689 | 0.03172 | 0.04181 | 0.02865 | 0.00432 | 0.00739 | 0.01192 | 0.01132 | 0.07426 | 0.04366 | 0.07256 | 0.01952 |
| **Mir7069** | 0.01174 | 0.00437 | 0.05696 | 0.04629 | 0.02927 | 0.01091 | 0.00425 | 0.00882 | 0.01366 | 0.00674 | 0.00821 | 0.00955 | 0.00881 | 0.02145 |
| **Mir7079** | 0.47578 | 0.00523 | 0.17985 | 0.12622 | 0.02292 | 0.17151 | 0.00429 | 0.09982 | 0.23210 | 0.01215 | 0.00497 | 0.13594 | 0.01072 | 0.05084 |
| **Mir763** | 0.00587 | 0.00440 | 0.03212 | 0.01792 | 0.00499 | 0.01119 | 0.00429 | 0.01041 | 0.01130 | 0.00787 | 0.00734 | 0.00895 | 0.00485 | 0.38588 |
| **Mir8091** | 0.09547 | 0.00432 | 0.06293 | 0.00912 | 0.08553 | 0.01059 | 0.00413 | 0.00960 | 0.01098 | 0.00757 | 0.00719 | 0.00781 | 0.39340 | 0.00871 |
| **Mir8112** | 0.00480 | 0.00404 | 0.02068 | 0.06780 | 0.05295 | 0.02783 | 0.07613 | 0.02685 | 0.01727 | 0.61526 | 0.00905 | 0.03739 | 0.00564 | 0.01210 |

| | | | | | | | | | | | | | | |
|---|---|---|---|---|---|---|---|---|---|---|---|---|---|---|
| **Mira** | 0.00532 | 0.00469 | 0.00418 | 0.01059 | 0.03387 | 0.00901 | 0.00405 | 0.01332 | 0.44775 | 0.00621 | 0.00624 | 0.02715 | 0.40743 | 0.00000 |
| **Mirc35hg** | 0.34528 | 0.12044 | 0.03896 | 1.08026 | 0.23925 | 0.72191 | 0.08596 | 0.17485 | 0.43260 | 0.22631 | 0.02844 | 0.07430 | 0.82470 | 0.34839 |
| **Mirt1** | 0.13881 | 0.01982 | 0.04907 | 0.16049 | 0.12118 | 0.14206 | 0.02240 | 0.09622 | 0.11684 | 0.04186 | 0.01332 | 0.03604 | 0.34611 | 0.26243 |
| **Mis12** | 3.45347 | 2.17702 | 2.59039 | 0.58898 | 0.46862 | 2.98754 | 0.30787 | 0.00000 | 7.62092 | 0.03809 | 0.34420 | 1.62283 | 2.37545 | 2.69807 |
| **Mis18a** | 8.42740 | 0.04266 | 11.55311 | 8.49478 | 15.08875 | 2.63858 | 0.02929 | 5.26631 | 3.68830 | 0.29155 | 3.60562 | 3.35300 | 11.40896 | 2.38072 |
| **Mis18bp1** | 1.97951 | 0.00110 | 2.91996 | 1.11292 | 2.57739 | 2.27643 | 0.00185 | 0.18709 | 10.91964 | 0.00000 | 3.80945 | 1.51316 | 8.05347 | 0.00000 |
| **Misp** | 0.00855 | 0.00350 | 0.00570 | 0.15726 | 0.01450 | 0.00643 | 0.00372 | 0.00332 | 0.01367 | 0.00739 | 0.00506 | 0.71963 | 0.01258 | 0.00767 |
| **Mitd1** | 2.89778 | 6.21557 | 3.83566 | 0.95856 | 3.42230 | 3.65685 | 2.17271 | 0.43039 | 1.66369 | 0.11835 | 4.39394 | 1.89095 | 0.00000 | 2.59997 |
| **Mitf** | 0.00570 | 0.00446 | 0.01262 | 0.01231 | 0.10350 | 0.01547 | 0.00440 | 0.00975 | 0.00848 | 0.00603 | 0.00891 | 0.01197 | 0.10871 | 0.04004 |
| **Mixl1** | 0.00000 | 0.00386 | 0.29964 | 0.00000 | 2.01382 | 0.00000 | 0.00358 | 0.00372 | 2.77091 | 0.00595 | 0.00000 | 0.16637 | 0.82751 | 2.46125 |
| **Mki67** | 8.73668 | 0.21722 | 4.17317 | 6.04949 | 4.44134 | 3.30879 | 2.33345 | 0.76452 | 19.50289 | 11.38470 | 3.04124 | 6.15036 | 14.30981 | 6.86962 |
| **Mkks** | 6.61453 | 0.07128 | 4.00235 | 3.74592 | 5.20299 | 7.09650 | 1.34227 | 3.90782 | 5.82483 | 0.06135 | 1.22121 | 2.42799 | 5.40506 | 7.42226 |
| **Mkl1** | 0.12372 | 0.02055 | 1.82755 | 1.62806 | 0.44036 | 2.65101 | 1.83324 | 2.12413 | 0.00279 | 0.04195 | 0.23074 | 0.65183 | 0.00681 | 0.28989 |
| **Mkl2** | 0.13294 | 0.04773 | 0.68000 | 1.06488 | 0.70490 | 1.50673 | 0.02209 | 0.03835 | 0.10896 | 0.04091 | 0.01168 | 0.09274 | 0.38118 | 0.60094 |
| **Mkln1** | 5.95832 | 0.01667 | 3.21471 | 0.62142 | 2.07374 | 5.41521 | 0.01373 | 4.64694 | 5.57931 | 0.02515 | 2.00527 | 2.35333 | 0.98420 | 0.96538 |
| **Mknk1** | 4.00365 | 0.00095 | 2.46309 | 1.13582 | 3.33561 | 0.11149 | 1.39585 | 1.21830 | 0.35390 | 0.02744 | 3.23992 | 1.44044 | 6.11002 | 5.93667 |
| **Mknk2** | 2.73436 | 0.05228 | 6.77160 | 3.31189 | 8.45541 | 1.59406 | 0.93518 | 5.22620 | 0.40969 | 0.03789 | 0.06664 | 1.58312 | 10.49200 | 2.26805 |
| **Mkrn1** | 23.14323 | 7.63213 | 26.80563 | 19.91015 | 27.71984 | 21.77597 | 5.84928 | 15.78859 | 11.02254 | 11.20766 | 14.92749 | 18.83665 | 16.25177 | 16.91130 |
| **Mkrn2** | 4.25404 | 0.05203 | 7.09225 | 5.17752 | 14.29848 | 5.92175 | 0.01924 | 2.36739 | 10.77171 | 0.04123 | 4.68474 | 0.88709 | 9.89853 | 16.12222 |
| **Mkrn2os** | 0.54258 | 0.00392 | 1.24384 | 0.00000 | 4.36204 | 0.00575 | 0.00379 | 0.00539 | 1.79499 | 0.00640 | 0.00753 | 0.00265 | 0.61366 | 5.37440 |
| **Mks1** | 0.00102 | 1.49885 | 0.04175 | 0.69576 | 0.00000 | 1.62148 | 1.33621 | 0.00000 | 0.00619 | 0.02854 | 5.17519 | 2.63229 | 0.00779 | 0.00525 |
| **Mlec** | 2.28368 | 0.27070 | 4.22052 | 0.00000 | 1.23235 | 12.05767 | 5.37556 | 5.32148 | 1.65865 | 0.21865 | 1.85371 | 12.60251 | 2.31641 | 3.26643 |
| **Mlf1** | 0.00527 | 0.01168 | 0.00000 | 0.06367 | 0.98245 | 0.02933 | 0.00437 | 0.00081 | 0.01193 | 0.01674 | 1.01035 | 0.00730 | 0.39003 | 0.00000 |
| **Mlf2** | 19.18256 | 8.37462 | 38.52520 | 34.82101 | 41.99771 | 32.73028 | 5.66848 | 18.66276 | 1.81595 | 0.87313 | 14.59563 | 19.23072 | 10.07045 | 9.84754 |
| **Mlh1** | 1.94081 | 0.02241 | 1.42965 | 3.89163 | 0.83240 | 1.42882 | 0.03670 | 2.05371 | 0.00919 | 0.01782 | 0.82223 | 1.33177 | 0.00796 | 1.02566 |
| **Mlh3** | 0.05521 | 0.12965 | 0.21000 | 0.24862 | 0.98047 | 0.11467 | 0.01505 | 0.00276 | 5.51759 | 0.09205 | 0.00000 | 0.53260 | 0.21434 | 0.63264 |
| **Mlkl** | 0.05288 | 0.01187 | 0.45588 | 0.26611 | 0.01020 | 0.00000 | 0.00406 | 0.01139 | 0.01198 | 0.00536 | 0.00593 | 0.00612 | 0.00446 | 0.00712 |
| **Mllt1** | 0.41276 | 0.00409 | 1.73121 | 0.10202 | 1.37980 | 1.23418 | 0.00344 | 0.94939 | 0.25191 | 0.03316 | 0.15483 | 0.30594 | 1.60116 | 0.72988 |
| **Mllt10** | 4.62586 | 0.00590 | 6.00335 | 0.06971 | 4.76165 | 5.93432 | 1.14206 | 0.78036 | 10.72380 | 8.35338 | 1.83043 | 2.80127 | 3.36029 | 13.22873 |
| **Mllt11** | 0.17090 | 0.00385 | 0.14523 | 0.10067 | 0.05639 | 0.00635 | 0.00953 | 0.13035 | 0.08505 | 0.00731 | 0.00540 | 0.02044 | 1.87714 | 0.52457 |
| **Mllt3** | 0.00000 | 0.02604 | 0.00000 | 0.00000 | 0.05262 | 0.85196 | 0.00336 | 0.07683 | 0.99057 | 0.01003 | 0.00000 | 1.55518 | 1.94157 | 1.87343 |
| **Mllt4** | 7.90164 | 4.46447 | 3.30077 | 1.08413 | 5.81517 | 7.85613 | 0.17093 | 5.03685 | 10.39889 | 0.02971 | 2.24599 | 10.34694 | 2.48291 | 5.27324 |
| **Mllt6** | 2.54030 | 7.62940 | 7.40759 | 4.41747 | 6.54565 | 7.45074 | 3.71968 | 4.17768 | 1.14484 | 0.16240 | 0.00000 | 4.14163 | 0.00000 | 6.34759 |
| **Mlph** | 0.03594 | 0.00477 | 0.03145 | 0.04576 | 0.04119 | 0.03811 | 0.01044 | 0.03502 | 0.06071 | 0.01568 | 0.02947 | 0.03684 | 0.00591 | 0.05432 |
| **Mlst8** | 1.07190 | 0.03742 | 6.91489 | 5.23546 | 5.84596 | 3.60584 | 0.61490 | 1.42689 | 5.30338 | 0.04380 | 3.44287 | 4.49508 | 1.80908 | 4.97886 |
| **Mlx** | 2.48365 | 0.02844 | 4.94910 | 1.58121 | 0.30144 | 4.97994 | 0.03995 | 0.31639 | 4.77574 | 0.01781 | 0.15854 | 1.27940 | 0.34952 | 3.19829 |
| **Mlxip** | 0.65307 | 0.62881 | 0.97539 | 0.00000 | 2.80314 | 1.41846 | 0.00862 | 0.00000 | 2.25118 | 0.00998 | 0.01872 | 1.47190 | 2.50034 | 10.10684 |
| **Mlxipl** | 1.92973 | 0.00373 | 2.49879 | 3.20663 | 1.51798 | 0.08195 | 0.00339 | 0.00000 | 0.50033 | 0.01750 | 0.00258 | 0.00661 | 0.84430 | 0.49017 |
| **Mlycd** | 0.06665 | 0.00382 | 0.61761 | 0.00000 | 0.02297 | 0.48309 | 0.00361 | 0.71438 | 0.00996 | 0.00489 | 0.36576 | 0.01138 | 0.11548 | 0.00216 |
| **Mmaa** | 2.41134 | 0.02518 | 3.65479 | 2.61462 | 2.49852 | 4.19125 | 3.22948 | 1.13927 | 1.80740 | 0.08266 | 2.92273 | 3.01140 | 0.36451 | 4.58342 |
| **Mmab** | 0.46697 | 0.01970 | 0.06558 | 0.16950 | 0.00000 | 1.34703 | 0.00482 | 0.24429 | 0.17937 | 0.01329 | 0.17955 | 0.79824 | 0.33837 | 0.00000 |
| **Mmachc** | 1.93197 | 0.11602 | 2.57207 | 0.69302 | 3.30828 | 4.40397 | 0.06144 | 1.44904 | 5.76979 | 0.08893 | 2.70184 | 3.97181 | 0.02868 | 0.76879 |
| **Mmadhc** | 12.05059 | 7.16150 | 9.38283 | 2.60018 | 6.27586 | 10.76091 | 2.56775 | 7.37778 | 4.33802 | 0.15208 | 8.38119 | 10.79127 | 0.06086 | 2.74570 |
| **Mmd** | 0.03772 | 0.00446 | 0.00981 | 0.01009 | 0.14586 | 0.01391 | 0.00419 | 0.01029 | 0.01486 | 0.00771 | 0.00843 | 0.00917 | 0.00520 | 0.00813 |
| **Mmd2** | 0.00380 | 0.01247 | 0.06792 | 0.28377 | 0.70618 | 0.00696 | 0.00450 | 0.01998 | 0.01665 | 0.00778 | 0.00806 | 0.01021 | 0.00300 | 0.11414 |
| **Mme** | 0.44163 | 0.02460 | 0.40912 | 0.12083 | 1.47177 | 0.76648 | 0.00397 | 0.02835 | 0.20904 | 0.00936 | 0.38539 | 1.01667 | 0.00406 | 0.00000 |
| **Mmel1** | 0.00808 | 0.01930 | 0.01029 | 0.00000 | 0.00865 | 0.02571 | 0.93022 | 2.81362 | 0.01293 | 0.04087 | 0.01565 | 0.01338 | 0.00703 | 0.00981 |
| **Mmgt1** | 9.02997 | 0.00089 | 6.71837 | 1.13433 | 5.47190 | 5.18453 | 0.38141 | 0.66402 | 10.69755 | 0.01858 | 4.73321 | 0.88446 | 2.76066 | 8.30818 |
| **Mmgt2** | 2.12045 | 0.01106 | 1.37341 | 1.08435 | 1.10971 | 3.42655 | 2.91242 | 0.00000 | 0.00214 | 0.09732 | 0.06449 | 0.84004 | 0.77500 | 0.33153 |
| **Mmp11** | 0.01944 | 0.00459 | 0.26642 | 0.06410 | 0.00381 | 0.01227 | 0.00474 | 0.04285 | 0.01041 | 0.00983 | 0.01765 | 0.01908 | 0.00433 | 0.01988 |
| **Mmp12** | 0.00433 | 0.01435 | 0.26223 | 0.00000 | 0.00239 | 0.13086 | 0.00411 | 0.56357 | 0.00962 | 0.02420 | 0.57503 | 0.68036 | 0.01709 | 0.13857 |
| **Mmp14** | 0.26443 | 0.00451 | 0.01713 | 0.02053 | 0.00817 | 0.01797 | 0.00458 | 0.03223 | 0.03587 | 0.02776 | 0.01876 | 0.01402 | 0.01753 | 0.03231 |
| **Mmp15** | 0.00167 | 0.05488 | 0.07605 | 0.00000 | 0.05906 | 0.00100 | 0.00974 | 1.33633 | 0.00741 | 0.00805 | 0.00218 | 3.09157 | 0.20258 | 0.01941 |
| **Mmp16** | 0.02213 | 0.00450 | 0.01210 | 0.00766 | 0.29429 | 0.01028 | 0.01080 | 0.01891 | 0.24448 | 0.00661 | 0.00802 | 0.00769 | 0.06124 | 0.00094 |
| **Mmp17** | 0.01043 | 0.00425 | 0.47185 | 0.00000 | 0.00000 | 0.51206 | 0.00391 | 0.18078 | 0.03340 | 0.00442 | 0.10054 | 0.05414 | 0.68133 | 0.01412 |

| | | | | | | | | | | | | | | |
|---|---|---|---|---|---|---|---|---|---|---|---|---|---|---|
| **Mmp19** | 0.00000 | 0.00413 | 0.00000 | 0.01344 | 0.00000 | 0.00517 | 0.00395 | 0.00786 | 0.89593 | 0.00655 | 0.00548 | 0.00641 | 1.64631 | 0.83721 |
| **Mmp1b** | 0.00302 | 0.00387 | 0.01134 | 0.02246 | 0.00578 | 0.00170 | 0.00405 | 0.00599 | 0.03615 | 0.00651 | 0.11899 | 0.17203 | 1.05124 | 0.00666 |
| **Mmp2** | 0.09172 | 0.00429 | 0.00893 | 0.07923 | 0.04358 | 0.00756 | 0.00408 | 0.00784 | 0.01390 | 0.00641 | 0.00677 | 0.01399 | 0.01008 | 0.16769 |
| **Mmp21** | 0.00484 | 0.00437 | 0.06064 | 0.34774 | 0.00652 | 0.00674 | 0.00433 | 0.01180 | 0.01310 | 0.00723 | 0.00730 | 0.00976 | 0.00665 | 0.00908 |
| **Mmp23** | 0.02869 | 0.00428 | 0.10845 | 0.24027 | 0.01733 | 0.05261 | 0.01533 | 0.29430 | 0.01101 | 0.01090 | 0.00486 | 0.03198 | 0.00830 | 0.00926 |
| **Mmp24** | 0.01635 | 0.00434 | 0.10999 | 0.00000 | 0.59565 | 0.00870 | 0.00434 | 0.01010 | 0.00609 | 0.00603 | 0.00858 | 0.00835 | 0.52954 | 0.08684 |
| **Mmp25** | 0.01896 | 0.03624 | 0.77351 | 0.00000 | 0.61566 | 0.00600 | 0.00382 | 0.01743 | 0.02501 | 0.00647 | 0.00596 | 0.02688 | 0.10876 | 0.22643 |
| **Mmp9** | 0.13278 | 0.00417 | 0.09199 | 0.23963 | 0.17262 | 0.00285 | 0.00394 | 0.00671 | 0.00820 | 0.00671 | 0.00685 | 0.01851 | 0.23170 | 0.01055 |
| **Mmrn2** | 0.34168 | 0.00415 | 0.75782 | 0.42908 | 0.00000 | 0.00000 | 0.00412 | 0.00757 | 0.16400 | 0.00480 | 0.00439 | 0.00579 | 0.16623 | 0.88397 |
| **Mms19** | 4.78708 | 0.49661 | 9.42313 | 2.11251 | 5.82935 | 5.13445 | 0.61153 | 5.02934 | 2.00169 | 0.00005 | 3.64316 | 6.46955 | 3.20989 | 4.09714 |
| **Mms22l** | 7.58602 | 0.00605 | 5.19414 | 2.49573 | 8.35761 | 2.37355 | 0.70190 | 0.50764 | 5.46287 | 0.00724 | 0.06896 | 5.02443 | 4.17214 | 4.94011 |
| **Mn1** | 0.95436 | 0.00408 | 0.07634 | 0.00418 | 0.07489 | 0.00465 | 0.00469 | 0.01094 | 0.05390 | 0.01389 | 0.00625 | 0.00471 | 2.44334 | 0.00000 |
| **Mnat1** | 2.86722 | 4.47496 | 9.19663 | 7.11706 | 7.17253 | 5.93549 | 0.35725 | 2.32778 | 5.22427 | 0.04284 | 4.29522 | 6.75038 | 3.90157 | 3.23096 |
| **Mnd1** | 1.72509 | 2.88378 | 0.01408 | 3.06230 | 0.32623 | 1.11993 | 0.99300 | 2.86027 | 0.32847 | 0.06872 | 2.44923 | 0.00000 | 0.02690 | 0.76667 |
| **Mnd1.ps** | 0.53605 | 0.93422 | 0.01260 | 0.67645 | 0.13881 | 0.30171 | 0.25400 | 0.76606 | 0.08801 | 0.03327 | 0.72957 | 0.00477 | 0.00835 | 0.19038 |
| **Mns1** | 0.02919 | 0.01123 | 0.52579 | 0.64354 | 0.49556 | 0.01648 | 0.00349 | 0.00173 | 1.36976 | 0.00609 | 0.00490 | 0.00576 | 1.36165 | 0.84989 |
| **Mnt** | 2.02379 | 0.00795 | 2.25818 | 3.47015 | 2.00843 | 3.17591 | 0.20823 | 0.51926 | 0.04788 | 10.06486 | 1.64054 | 3.17178 | 1.39677 | 2.13938 |
| **Mob1a** | 0.33732 | 0.00402 | 0.47461 | 0.00000 | 0.80934 | 0.04619 | 0.00356 | 0.02362 | 0.97666 | 0.00436 | 0.01427 | 0.02920 | 1.49748 | 0.38536 |
| **Mob1b** | 0.97125 | 0.00366 | 0.73768 | 0.19180 | 2.45785 | 0.51379 | 0.00344 | 0.00000 | 3.14843 | 0.00521 | 0.19764 | 0.02173 | 1.75693 | 0.00000 |
| **Mob2** | 3.40060 | 2.18015 | 7.57537 | 5.78659 | 7.12214 | 6.67729 | 4.07900 | 1.23921 | 0.00000 | 0.15208 | 3.03456 | 3.45288 | 0.34845 | 2.16933 |
| **Mob3a** | 0.16621 | 0.01461 | 0.07417 | 0.60451 | 0.00000 | 0.70868 | 0.01613 | 0.98585 | 0.00336 | 0.02367 | 0.72800 | 0.36425 | 2.25286 | 0.02971 |
| **Mob3b** | 0.10764 | 0.09840 | 0.00000 | 1.57555 | 0.05985 | 1.25193 | 0.08890 | 0.82155 | 0.22810 | 0.14009 | 0.15449 | 0.54796 | 1.20752 | 0.07539 |
| **Mob3c** | 0.98508 | 0.10954 | 0.20043 | 0.93794 | 0.00000 | 0.51609 | 0.05609 | 0.22508 | 0.10877 | 0.02157 | 0.06222 | 0.53228 | 0.04762 | 0.31194 |
| **Mob4** | 10.16955 | 0.17687 | 10.11042 | 8.31731 | 12.60239 | 11.56995 | 1.96372 | 2.24908 | 8.67720 | 12.29541 | 6.26490 | 4.05129 | 4.64265 | 7.14911 |
| **Mocos** | 0.06271 | 0.00393 | 0.95475 | 1.44599 | 0.00000 | 1.43898 | 0.00380 | 0.02207 | 0.00461 | 0.02598 | 0.60913 | 0.00349 | 0.09666 | 0.07477 |
| **Mocs1** | 0.00000 | 0.00229 | 0.08944 | 0.53940 | 0.21136 | 0.45366 | 0.08825 | 2.52332 | 0.05860 | 0.00614 | 2.09246 | 1.52640 | 0.00855 | 0.00000 |
| **Mocs2** | 0.78263 | 0.02017 | 0.24369 | 0.10054 | 0.00000 | 0.96556 | 1.81933 | 1.38907 | 1.65009 | 0.03446 | 0.04668 | 1.00131 | 0.04795 | 0.00000 |
| **Mocs3** | 1.49607 | 0.00328 | 1.07657 | 1.25357 | 2.16269 | 0.66892 | 0.31427 | 0.34163 | 0.39440 | 0.04019 | 0.80415 | 1.07015 | 5.58451 | 0.00000 |
| **Mog** | 0.10386 | 0.00429 | 0.65183 | 0.00000 | 0.44509 | 0.12349 | 0.00390 | 0.00245 | 0.26579 | 0.00496 | 0.00444 | 0.00540 | 0.81378 | 0.00000 |
| **Mogat1** | 0.01074 | 0.00446 | 0.00916 | 0.00792 | 0.43214 | 0.01190 | 0.00493 | 0.01017 | 0.10075 | 0.00668 | 0.00833 | 0.00825 | 0.07530 | 0.02321 |
| **Mogat2** | 0.02370 | 0.02064 | 2.05907 | 0.04770 | 0.23603 | 0.05993 | 0.01045 | 0.00017 | 0.01561 | 0.02181 | 0.00273 | 2.83352 | 0.00000 | 0.00853 |
| **Mogs** | 4.07624 | 0.05953 | 5.65611 | 2.76159 | 1.97832 | 3.98317 | 3.18565 | 3.69619 | 0.08434 | 7.76365 | 0.00000 | 4.58563 | 5.30731 | 1.81509 |
| **Mok** | 0.00000 | 0.00345 | 0.93185 | 0.54732 | 0.99975 | 1.24622 | 0.04023 | 0.89065 | 0.00215 | 0.03086 | 0.00000 | 1.18931 | 0.06652 | 0.03761 |
| **Mon1a** | 8.98104 | 0.00846 | 5.56225 | 12.46041 | 21.16745 | 2.73095 | 3.22622 | 0.19704 | 0.00000 | 0.21850 | 0.00000 | 2.11122 | 6.65970 | 1.29706 |
| **Mon1b** | 0.99551 | 0.07241 | 2.87334 | 1.48484 | 2.09598 | 2.06740 | 1.32232 | 3.56211 | 0.97295 | 0.07371 | 1.63365 | 2.72211 | 2.61566 | 1.21469 |
| **Mon2** | 7.96309 | 0.07549 | 8.42527 | 2.85559 | 7.87610 | 6.65237 | 4.09166 | 1.09809 | 2.78909 | 0.14377 | 2.05047 | 7.47198 | 3.85947 | 5.11786 |
| **Morc1** | 0.25372 | 0.03389 | 0.31674 | 0.28181 | 0.58787 | 0.66470 | 0.00368 | 0.00534 | 0.41876 | 0.00583 | 0.00000 | 1.90318 | 1.08787 | 0.00000 |
| **Morc2a** | 7.28857 | 4.17311 | 9.13400 | 2.67268 | 8.33026 | 3.12308 | 4.54605 | 2.47963 | 13.05361 | 0.19725 | 5.50750 | 6.03096 | 1.94230 | 11.15430 |
| **Morc3** | 4.32711 | 0.00788 | 12.44477 | 3.31538 | 9.31706 | 5.25231 | 0.01756 | 1.03041 | 8.82967 | 0.00884 | 5.10539 | 2.29205 | 6.77458 | 12.02806 |
| **Morc4** | 0.02169 | 0.00442 | 0.00887 | 0.00680 | 0.18344 | 0.04347 | 0.00417 | 0.00793 | 0.00939 | 0.00635 | 0.00732 | 0.01521 | 0.01799 | 0.01880 |
| **Morf4l1** | 4.86361 | 6.15778 | 17.30630 | 15.16713 | 12.24559 | 19.23063 | 4.47305 | 16.94590 | 20.36984 | 0.42861 | 17.21065 | 16.53342 | 4.45622 | 17.37882 |
| **Morf4l1.ps1** | 2.12377 | 3.94007 | 10.02617 | 8.23645 | 6.07309 | 11.14487 | 2.59449 | 10.31241 | 12.91822 | 0.17866 | 10.82361 | 10.05120 | 2.01718 | 10.42622 |
| **Morf4l1b** | 1.87301 | 3.51553 | 9.03341 | 7.60434 | 6.18210 | 10.64939 | 2.29265 | 9.59608 | 12.61781 | 0.15289 | 10.17120 | 9.39588 | 2.23952 | 9.99055 |
| **Morf4l2** | 4.92412 | 0.84438 | 15.65750 | 17.63666 | 7.54963 | 29.98485 | 4.62200 | 15.26601 | 8.87504 | 0.33157 | 17.29845 | 15.15510 | 5.27987 | 12.35995 |
| **Morn1** | 0.33848 | 0.00434 | 0.45360 | 0.10225 | 0.42662 | 0.03034 | 0.00465 | 0.08571 | 0.00495 | 0.02000 | 0.00712 | 0.09552 | 0.44911 | 0.08642 |
| **Morn2** | 0.37980 | 0.01073 | 0.01907 | 0.05592 | 0.35320 | 0.62712 | 0.02628 | 0.00877 | 0.00788 | 0.01360 | 0.24316 | 0.81830 | 0.00410 | 0.08523 |
| **Morn4** | 0.00000 | 0.00355 | 0.43819 | 1.32578 | 0.42691 | 0.00000 | 0.00410 | 0.98729 | 0.00183 | 0.00229 | 1.17880 | 0.96880 | 1.15191 | 0.19042 |
| **Morn5** | 0.00648 | 0.00462 | 0.04293 | 0.14731 | 0.16330 | 0.00911 | 0.00414 | 0.01135 | 0.01071 | 0.00676 | 0.00738 | 0.01239 | 0.10400 | 0.11480 |
| **Mos** | 0.02802 | 0.00441 | 0.01370 | 0.01690 | 0.01225 | 0.01464 | 0.00424 | 0.00922 | 0.01174 | 0.00874 | 0.00815 | 0.00934 | 0.08494 | 0.00640 |
| **Mospd1** | 6.59073 | 0.00778 | 9.79266 | 2.91262 | 9.96350 | 6.07555 | 1.21453 | 3.33837 | 26.62709 | 0.60755 | 2.49046 | 1.28930 | 5.85118 | 19.12009 |
| **Mospd2** | 2.40933 | 0.00386 | 1.07824 | 0.00000 | 0.00000 | 1.94884 | 0.00292 | 0.03119 | 4.42486 | 0.00346 | 0.00000 | 0.59780 | 0.00000 | 0.00000 |
| **Mospd3** | 0.88405 | 0.03429 | 1.74540 | 6.40583 | 6.41635 | 0.96236 | 0.16886 | 1.49957 | 0.01901 | 0.02635 | 0.00010 | 1.79263 | 0.46542 | 0.26831 |
| **Mospd4** | 0.04047 | 0.01190 | 0.00660 | 0.28302 | 0.01256 | 0.03251 | 0.00425 | 0.00584 | 0.01139 | 0.00559 | 0.00611 | 0.00645 | 0.25321 | 0.00000 |
| **Mov10** | 0.00000 | 1.23278 | 8.26148 | 2.58536 | 5.96827 | 0.33988 | 0.00690 | 2.98955 | 0.05649 | 0.00000 | 0.99667 | 4.83828 | 0.00000 | 5.54713 |
| **Mov10l1** | 0.00432 | 0.01095 | 0.06197 | 0.45460 | 0.87408 | 0.28593 | 0.00373 | 0.00258 | 0.20827 | 0.01854 | 0.02020 | 0.00656 | 2.64611 | 1.07353 |

| | | | | | | | | | | | | | | |
|---|---|---|---|---|---|---|---|---|---|---|---|---|---|---|
| **Mpc1** | 17.43356 | 6.15917 | 15.17231 | 20.27396 | 25.77056 | 9.93471 | 3.25740 | 6.15565 | 4.59069 | 0.19648 | 11.02076 | 4.64585 | 13.27665 | 13.23700 |
| **Mpc2** | 3.36778 | 0.01602 | 0.00000 | 20.23518 | 27.68214 | 0.90552 | 2.05064 | 0.44388 | 1.41480 | 0.13602 | 0.01536 | 0.81145 | 12.01436 | 1.15488 |
| **Mpdu1** | 3.76111 | 0.24398 | 18.91307 | 17.17178 | 11.55177 | 13.96895 | 3.40419 | 10.62269 | 2.14612 | 2.85765 | 8.47850 | 12.45424 | 0.58134 | 10.10315 |
| **Mpdz** | 0.00000 | 0.00327 | 0.40318 | 0.00000 | 0.30476 | 0.00000 | 0.00357 | 0.00250 | 1.01092 | 0.00565 | 0.00093 | 0.38720 | 4.40263 | 2.89474 |
| **Mpeg1** | 0.00790 | 0.00429 | 0.04764 | 0.00000 | 0.01216 | 0.02036 | 0.00444 | 0.01557 | 0.01157 | 0.00703 | 0.00477 | 0.33445 | 0.00770 | 0.01133 |
| **Mpg** | 0.81236 | 0.01103 | 0.68991 | 4.45389 | 0.91387 | 0.84977 | 0.00400 | 0.93396 | 0.04753 | 0.02649 | 0.01143 | 1.85369 | 0.03814 | 0.00000 |
| **Mphosph10** | 11.76054 | 0.01219 | 9.74373 | 1.42803 | 9.44854 | 7.11079 | 3.77625 | 0.71308 | 16.93070 | 0.03403 | 7.00825 | 6.20460 | 7.64173 | 14.27431 |
| **Mphosph6** | 5.31533 | 0.08843 | 10.00402 | 4.99382 | 10.71963 | 4.98918 | 4.69333 | 5.59674 | 12.91214 | 0.16464 | 8.11145 | 5.35568 | 5.50666 | 13.69766 |
| **Mphosph8** | 1.85503 | 0.02182 | 4.42385 | 1.43089 | 2.96846 | 6.60518 | 2.02801 | 1.23842 | 1.71608 | 0.02990 | 9.06990 | 4.39343 | 0.06539 | 4.31240 |
| **Mphosph9** | 0.57254 | 0.00299 | 1.44642 | 0.37942 | 3.90650 | 0.94417 | 0.00385 | 0.73516 | 5.73788 | 0.00824 | 0.00000 | 1.32665 | 3.53472 | 4.98650 |
| **Mpi** | 0.48939 | 0.03870 | 2.96936 | 0.45316 | 1.44346 | 1.51112 | 3.16385 | 2.06653 | 2.08241 | 0.11771 | 2.92054 | 2.30235 | 0.00000 | 3.23768 |
| **Mpl** | 0.38757 | 0.00444 | 0.30915 | 0.00000 | 1.46333 | 0.00000 | 0.00417 | 0.00947 | 0.78719 | 0.00625 | 0.00391 | 0.00486 | 0.01184 | 0.96906 |
| **Mplkip** | 1.33553 | 0.14033 | 0.12675 | 0.96945 | 2.60971 | 0.62488 | 0.00427 | 0.06929 | 0.99303 | 0.01601 | 0.04626 | 0.23890 | 1.92858 | 1.02359 |
| **Mpnd** | 0.05044 | 0.02674 | 0.45535 | 1.17082 | 0.29663 | 1.09569 | 1.73993 | 0.11335 | 0.00179 | 0.05801 | 1.21347 | 5.27452 | 0.00055 | 0.45910 |
| **Mpp1** | 9.07980 | 5.65779 | 10.38226 | 7.59801 | 12.30437 | 23.09192 | 6.82229 | 15.17328 | 10.67739 | 7.28763 | 5.11967 | 6.49650 | 3.36358 | 11.95791 |
| **Mpp3** | 0.02293 | 0.00420 | 1.37089 | 0.23910 | 0.24344 | 0.00000 | 0.00370 | 0.00552 | 0.20583 | 0.00700 | 0.00503 | 0.00725 | 0.11041 | 0.04143 |
| **Mpp4** | 0.13534 | 0.00449 | 0.76446 | 0.09290 | 1.28292 | 0.05257 | 0.00405 | 0.00559 | 0.29090 | 0.01378 | 0.00438 | 0.00650 | 0.00000 | 0.23414 |
| **Mpp5** | 0.32364 | 0.02907 | 0.07139 | 0.44100 | 0.46623 | 1.74349 | 0.99615 | 0.03303 | 3.74002 | 0.03640 | 0.00000 | 1.69126 | 2.22129 | 0.84656 |
| **Mpp6** | 4.77071 | 0.00769 | 0.70075 | 3.65465 | 3.24227 | 1.62313 | 1.12679 | 0.00326 | 3.24882 | 0.02699 | 0.15372 | 3.99697 | 1.77451 | 3.05357 |
| **Mpp7** | 0.24478 | 0.08796 | 0.23283 | 0.53978 | 0.37141 | 2.12267 | 0.15326 | 0.58341 | 0.58149 | 0.10117 | 0.15097 | 0.55313 | 1.10460 | 0.39675 |
| **Mppe1** | 0.38600 | 0.00262 | 4.36653 | 1.28215 | 1.41272 | 1.02343 | 0.98872 | 1.07832 | 0.08099 | 0.01100 | 2.04641 | 1.92003 | 0.00000 | 2.15197 |
| **Mpped1** | 0.01040 | 0.00439 | 0.03270 | 0.02842 | 0.01178 | 0.00880 | 0.00416 | 0.00979 | 0.01239 | 0.01112 | 0.00705 | 0.00827 | 0.01411 | 0.59319 |
| **Mpped2** | 0.23603 | 0.00285 | 0.36690 | 0.00000 | 0.00000 | 0.23163 | 0.26199 | 1.62914 | 0.00839 | 0.02197 | 0.00000 | 1.53169 | 0.02232 | 0.00000 |
| **Mprip** | 5.20143 | 5.46544 | 13.25256 | 10.40754 | 11.23076 | 14.45332 | 6.44968 | 12.93913 | 1.29020 | 6.34957 | 9.57865 | 5.67769 | 6.15931 | 8.69898 |
| **Mpst** | 0.01053 | 0.02051 | 0.00663 | 0.49774 | 0.02234 | 0.00000 | 0.00534 | 0.72247 | 0.01225 | 0.00524 | 0.00538 | 0.10332 | 0.00649 | 0.00989 |
| **Mpv17** | 1.14858 | 0.20220 | 1.69985 | 0.32894 | 1.01597 | 2.05811 | 0.28140 | 1.07734 | 2.01814 | 1.00946 | 1.69969 | 1.51345 | 5.86978 | 3.09557 |
| **Mpv17l** | 3.50569 | 0.13650 | 0.25581 | 2.07481 | 0.90043 | 4.83004 | 0.68976 | 1.08741 | 2.87698 | 1.72370 | 0.07233 | 1.13487 | 15.40763 | 3.10577 |
| **Mpv17l2** | 0.86313 | 0.01878 | 0.42315 | 3.53969 | 1.80270 | 1.76212 | 1.53459 | 3.20506 | 0.00338 | 0.08464 | 0.62318 | 0.62224 | 0.01025 | 0.57158 |
| **Mpzl1** | 0.00000 | 0.02348 | 0.81997 | 1.81000 | 0.07674 | 0.84535 | 0.01053 | 0.40750 | 0.02132 | 0.00940 | 1.23551 | 0.56799 | 1.07267 | 2.15438 |
| **Mpzl2** | 4.18704 | 0.09567 | 1.34458 | 3.35987 | 0.00000 | 6.08473 | 2.92895 | 6.25327 | 5.75418 | 0.09323 | 6.21641 | 2.84911 | 0.10551 | 0.17882 |
| **Mpzl3** | 0.00503 | 0.01309 | 0.01249 | 0.01722 | 0.03223 | 0.33765 | 0.00433 | 0.31717 | 0.00970 | 0.01799 | 0.00963 | 0.00341 | 0.01299 | 0.37125 |
| **Mrap** | 0.34516 | 0.00446 | 0.41084 | 0.01121 | 0.39965 | 0.42211 | 0.00417 | 0.00720 | 0.00510 | 0.00569 | 0.00681 | 0.00353 | 0.00530 | 0.43709 |
| **Mras** | 0.53703 | 1.55632 | 1.51832 | 4.94595 | 0.34049 | 14.19427 | 4.21086 | 5.01754 | 0.14366 | 0.20349 | 5.10686 | 6.23022 | 0.25034 | 0.09664 |
| **Mrc1** | 0.33037 | 0.00444 | 0.00329 | 0.00835 | 0.23954 | 0.00957 | 0.00425 | 0.01110 | 0.00897 | 0.00639 | 0.00726 | 0.00725 | 0.00918 | 0.28543 |
| **Mre11a** | 2.76469 | 0.05641 | 2.98486 | 3.40431 | 2.63153 | 2.13079 | 0.05769 | 0.25439 | 0.12513 | 0.05096 | 1.56219 | 5.03959 | 2.90186 | 0.59933 |
| **Mreg** | 0.00000 | 0.30571 | 8.14859 | 11.18167 | 8.94408 | 7.13716 | 1.81109 | 7.40290 | 15.61883 | 0.09144 | 8.12156 | 4.33671 | 1.32392 | 11.29918 |
| **Mrfap1** | 9.27348 | 4.56658 | 15.28393 | 20.37630 | 19.63200 | 19.75434 | 0.65387 | 16.60899 | 14.30796 | 10.95155 | 4.90710 | 15.65452 | 5.97280 | 8.28260 |
| **Mrgbp** | 0.72327 | 0.01338 | 0.02432 | 0.68197 | 0.03510 | 0.50143 | 0.00429 | 0.86773 | 0.00291 | 0.00277 | 1.33026 | 0.52268 | 0.12536 | 0.02538 |
| **Mrgpra6** | 1.74424 | 3.40940 | 1.87095 | 0.96496 | 1.25086 | 1.15454 | 0.33938 | 0.15905 | 14.47992 | 0.02481 | 0.19704 | 1.02390 | 1.25283 | 3.18757 |
| **Mrgprb1** | 0.04448 | 0.00479 | 0.01234 | 0.05001 | 0.02753 | 0.07621 | 0.01702 | 0.01126 | 0.06959 | 0.07094 | 0.00823 | 0.02412 | 0.03918 | 0.18503 |
| **Mrgpre** | 0.01570 | 0.00492 | 0.04105 | 0.02379 | 0.01691 | 0.03027 | 0.02450 | 0.04780 | 0.06584 | 0.01447 | 0.01511 | 0.02980 | 0.05643 | 0.05979 |
| **Mri1** | 1.22596 | 2.10380 | 0.00000 | 1.82391 | 1.00916 | 1.94409 | 0.00329 | 0.08930 | 0.00000 | 0.02346 | 0.00000 | 3.30146 | 1.65470 | 0.47583 |
| **Mrln** | 0.10963 | 0.00415 | 0.01407 | 0.00505 | 0.00828 | 0.00726 | 0.12912 | 0.00434 | 0.01114 | 0.14181 | 0.00704 | 0.00892 | 0.01285 | 0.00549 |
| **Mrm1** | 0.05083 | 0.01047 | 0.57260 | 1.20342 | 0.77543 | 3.63973 | 0.00321 | 2.72354 | 2.14339 | 0.01401 | 0.40756 | 1.32467 | 3.07677 | 0.26184 |
| **Mrm2** | 2.15794 | 0.06355 | 6.61671 | 4.25497 | 3.44700 | 1.94650 | 2.87022 | 1.23964 | 0.05013 | 0.09174 | 2.21544 | 3.61670 | 0.04170 | 0.95839 |
| **Mro** | 0.01948 | 0.00437 | 0.01429 | 0.01920 | 0.02271 | 0.01839 | 0.00421 | 0.01158 | 0.01061 | 0.00692 | 0.02527 | 0.01153 | 0.16837 | 0.14758 |
| **Mroh1** | 0.02569 | 0.10384 | 0.56129 | 4.18539 | 0.05684 | 2.41120 | 0.69748 | 9.79016 | 0.38638 | 0.10774 | 1.66186 | 0.97639 | 0.06877 | 1.03391 |
| **Mroh5** | 0.00617 | 0.00462 | 0.00424 | 0.17312 | 0.22971 | 0.01417 | 0.00462 | 0.03245 | 0.01017 | 0.00927 | 0.00967 | 0.00550 | 0.00764 | 0.02201 |
| **Mroh8** | 0.00390 | 0.00404 | 0.78850 | 0.29559 | 0.83187 | 0.00000 | 0.00358 | 0.00444 | 3.35976 | 0.00519 | 0.00141 | 0.00449 | 0.46600 | 1.68292 |
| **Mrpl1** | 4.45933 | 0.06477 | 6.52773 | 4.54622 | 1.63452 | 2.43597 | 3.30883 | 2.09501 | 7.62070 | 13.16583 | 5.39808 | 8.32401 | 0.05725 | 1.94806 |
| **Mrpl10** | 0.73705 | 1.61208 | 7.58063 | 9.25801 | 3.27063 | 5.30831 | 4.53142 | 3.82613 | 1.34628 | 4.34678 | 5.96110 | 6.12440 | 2.06739 | 3.98962 |
| **Mrpl11** | 5.40196 | 0.12296 | 5.80800 | 11.39898 | 9.52915 | 11.29862 | 3.42477 | 9.00414 | 0.82011 | 0.41427 | 6.00676 | 7.77807 | 1.69263 | 2.42514 |
| **Mrpl12** | 8.82344 | 2.04679 | 15.08433 | 22.75975 | 22.81269 | 8.77337 | 2.87965 | 14.98030 | 0.53803 | 6.00256 | 1.04033 | 11.66780 | 8.80490 | 0.79238 |
| **Mrpl13** | 9.67904 | 0.18929 | 4.83303 | 8.21943 | 8.87443 | 10.75594 | 3.57645 | 9.12179 | 5.02171 | 3.09791 | 7.61988 | 11.82623 | 0.30094 | 6.36639 |
| **Mrpl14** | 1.43558 | 0.06941 | 1.28335 | 2.71092 | 4.95761 | 4.33287 | 1.96157 | 4.67054 | 0.00000 | 1.81481 | 0.39625 | 6.52322 | 0.20707 | 0.33341 |

| | | | | | | | | | | | | | | |
|---|---|---|---|---|---|---|---|---|---|---|---|---|---|---|
| **Mrpl15** | 15.69255 | 0.31070 | 20.49992 | 18.33424 | 22.52660 | 6.34730 | 6.26394 | 11.49821 | 4.82665 | 0.56048 | 5.42631 | 11.55948 | 3.65572 | 4.24879 |
| **Mrpl16** | 8.72426 | 0.10669 | 8.20841 | 8.03937 | 7.14096 | 8.56183 | 0.77204 | 5.65902 | 1.50569 | 0.09858 | 4.76158 | 7.12764 | 0.45045 | 3.94545 |
| **Mrpl17** | 5.74840 | 4.77083 | 12.31438 | 28.53549 | 11.87775 | 9.06310 | 9.21938 | 12.69298 | 2.64812 | 11.95198 | 6.62246 | 15.34229 | 0.81446 | 1.76969 |
| **Mrpl18** | 12.37142 | 5.30481 | 22.46052 | 18.57539 | 16.61900 | 20.81718 | 7.68678 | 18.96914 | 1.27264 | 15.69966 | 11.98171 | 15.56500 | 5.60511 | 13.95211 |
| **Mrpl19** | 7.53734 | 3.31363 | 9.76041 | 11.88508 | 9.59292 | 8.50724 | 0.30701 | 9.53711 | 2.06188 | 0.46714 | 5.07766 | 8.95438 | 0.27044 | 8.12803 |
| **Mrpl2** | 2.92069 | 4.11350 | 13.11266 | 18.22937 | 6.59802 | 6.74944 | 3.06757 | 10.38338 | 0.00000 | 14.51960 | 6.37938 | 7.07190 | 0.18959 | 2.47114 |
| **Mrpl20** | 10.64240 | 8.26980 | 6.49258 | 17.63298 | 22.50301 | 12.22530 | 5.99136 | 14.09533 | 1.18842 | 4.59951 | 4.42323 | 9.84063 | 8.87984 | 8.30108 |
| **Mrpl21** | 5.59406 | 1.73457 | 5.21817 | 7.59466 | 10.63099 | 5.16141 | 1.31881 | 4.44338 | 0.60972 | 2.60219 | 5.77182 | 8.67781 | 0.80384 | 8.18830 |
| **Mrpl22** | 10.36210 | 6.34231 | 12.20030 | 20.38287 | 23.89563 | 13.62428 | 3.56080 | 8.37268 | 13.47410 | 1.36068 | 8.40294 | 8.71922 | 0.00000 | 20.16425 |
| **Mrpl23** | 3.61129 | 2.29490 | 0.00000 | 3.65230 | 6.28692 | 5.00792 | 0.96240 | 6.38275 | 1.32095 | 9.00753 | 0.56034 | 8.70752 | 1.35185 | 1.48334 |
| **Mrpl24** | 1.97674 | 6.08531 | 8.39677 | 11.46052 | 4.01689 | 11.37174 | 1.79342 | 4.81007 | 2.02800 | 15.90370 | 8.63447 | 10.07703 | 0.00000 | 4.89343 |
| **Mrpl27** | 4.29722 | 2.66288 | 3.24431 | 4.56708 | 8.01328 | 4.29149 | 0.01001 | 5.15128 | 0.54285 | 14.12683 | 5.06811 | 5.04148 | 2.01670 | 2.63104 |
| **Mrpl28** | 4.08988 | 0.33346 | 20.97927 | 17.02113 | 11.71772 | 10.87472 | 5.28325 | 13.34637 | 0.00000 | 2.53581 | 5.99781 | 9.78182 | 0.00000 | 0.70220 |
| **Mrpl3** | 7.15462 | 0.18443 | 6.22981 | 1.36183 | 8.08127 | 7.40068 | 4.43463 | 4.41197 | 4.20016 | 0.21162 | 3.13993 | 8.17004 | 2.40340 | 3.26994 |
| **Mrpl30** | 6.12552 | 0.20480 | 3.18347 | 14.87052 | 9.80777 | 11.27013 | 1.99488 | 5.16348 | 1.96224 | 11.05077 | 4.90580 | 8.15869 | 0.41854 | 4.34735 |
| **Mrpl32** | 7.73798 | 0.05947 | 3.77057 | 5.67001 | 10.05969 | 6.08859 | 1.83994 | 3.72260 | 1.72079 | 8.30565 | 5.07968 | 6.85195 | 3.37837 | 6.56905 |
| **Mrpl33** | 3.05373 | 0.15140 | 0.00000 | 1.08621 | 2.61990 | 4.36804 | 1.78230 | 5.35635 | 2.43970 | 8.47301 | 0.25044 | 4.34285 | 0.75963 | 2.44633 |
| **Mrpl34** | 0.00000 | 1.67321 | 0.81631 | 6.35761 | 8.59025 | 0.00322 | 3.49021 | 6.72914 | 0.00000 | 0.26670 | 0.01879 | 5.70493 | 3.16515 | 0.00000 |
| **Mrpl35** | 8.98520 | 0.43140 | 6.13870 | 12.11289 | 9.89542 | 10.23011 | 6.65844 | 2.41183 | 7.63012 | 0.55373 | 4.64964 | 9.40659 | 2.06371 | 8.07986 |
| **Mrpl36** | 4.82200 | 0.17928 | 6.33534 | 14.13119 | 5.02122 | 7.19337 | 4.79374 | 9.48517 | 0.00000 | 2.32998 | 3.00663 | 8.12460 | 0.48504 | 0.13318 |
| **Mrpl37** | 5.76816 | 0.10357 | 10.71054 | 12.83824 | 8.50020 | 3.61363 | 2.05360 | 8.71652 | 0.00000 | 0.12094 | 1.85213 | 9.13298 | 1.20241 | 2.24961 |
| **Mrpl38** | 2.15870 | 7.07253 | 2.08142 | 14.89886 | 4.93412 | 3.68377 | 0.02948 | 6.31671 | 0.00453 | 0.04360 | 0.44629 | 5.18252 | 1.98543 | 1.57578 |
| **Mrpl39** | 13.30854 | 0.04350 | 10.29557 | 9.81555 | 7.47079 | 2.70101 | 6.05333 | 0.84566 | 4.52834 | 0.36789 | 4.68884 | 7.56872 | 1.80714 | 1.26950 |
| **Mrpl4** | 5.46383 | 0.90262 | 21.13542 | 24.70814 | 19.35463 | 6.58207 | 0.04810 | 15.53762 | 0.25357 | 0.17490 | 3.64501 | 11.51157 | 6.62542 | 3.84553 |
| **Mrpl40** | 13.72264 | 6.15448 | 11.82531 | 18.26420 | 24.10111 | 11.59000 | 3.86692 | 10.78342 | 4.53239 | 2.50528 | 11.14261 | 9.32117 | 13.02482 | 10.60140 |
| **Mrpl41** | 4.74110 | 3.51995 | 0.98847 | 9.93171 | 8.96736 | 5.37527 | 0.01659 | 6.19757 | 1.26045 | 7.18446 | 2.88575 | 7.31956 | 5.97073 | 6.35393 |
| **Mrpl42** | 8.19335 | 0.60389 | 1.66685 | 5.89052 | 11.41125 | 10.80364 | 3.85077 | 10.24171 | 3.21498 | 7.84718 | 3.44839 | 14.14091 | 3.86890 | 7.15868 |
| **Mrpl43** | 2.51597 | 4.17128 | 8.78225 | 11.50146 | 15.35115 | 2.71078 | 3.06750 | 10.13833 | 0.84171 | 0.62014 | 2.09168 | 8.55365 | 3.56004 | 1.48064 |
| **Mrpl44** | 8.17463 | 1.38874 | 19.29337 | 11.63700 | 15.11473 | 10.26228 | 5.07295 | 8.51830 | 3.08499 | 8.65741 | 9.90389 | 9.75424 | 1.08333 | 11.03677 |
| **Mrpl45** | 8.97559 | 0.14131 | 15.23858 | 9.41896 | 15.45877 | 14.93073 | 4.99921 | 11.74600 | 2.90413 | 0.42941 | 7.85607 | 8.52571 | 1.75955 | 14.53582 |
| **Mrpl46** | 9.06777 | 6.90631 | 14.47774 | 8.41797 | 15.25062 | 10.99815 | 2.94889 | 9.22931 | 2.23159 | 12.38585 | 7.73967 | 5.84555 | 0.02207 | 5.94062 |
| **Mrpl47** | 4.81546 | 0.13838 | 7.93982 | 15.79729 | 8.54714 | 9.57146 | 3.66392 | 6.29324 | 12.35117 | 13.09348 | 9.16327 | 7.56903 | 0.72773 | 5.45357 |
| **Mrpl48** | 11.89474 | 0.07083 | 10.13843 | 11.20067 | 6.15346 | 7.03182 | 0.00763 | 7.16195 | 8.08221 | 12.70357 | 3.57857 | 6.48586 | 5.14486 | 6.95414 |
| **Mrpl49** | 5.39110 | 0.13947 | 14.06117 | 10.23281 | 13.25873 | 12.64985 | 3.17449 | 6.30552 | 1.45658 | 12.75312 | 3.40615 | 9.69256 | 3.17778 | 6.44666 |
| **Mrpl50** | 3.51169 | 1.13116 | 3.62289 | 6.50333 | 4.75568 | 5.70687 | 0.01446 | 1.07832 | 5.18959 | 0.01070 | 5.94117 | 7.16173 | 1.81426 | 7.42856 |
| **Mrpl51** | 5.25418 | 0.22827 | 4.49779 | 11.79136 | 13.50361 | 9.17517 | 4.23316 | 9.06481 | 3.44349 | 14.89449 | 2.97527 | 12.76588 | 9.52939 | 7.84327 |
| **Mrpl52** | 0.04744 | 1.74013 | 0.04096 | 1.91131 | 1.86556 | 0.37164 | 0.74107 | 3.49186 | 0.10876 | 0.09190 | 0.00000 | 2.74187 | 0.24121 | 0.00000 |
| **Mrpl53** | 0.83365 | 0.17138 | 0.41885 | 9.36588 | 8.54699 | 5.79896 | 2.14347 | 6.79729 | 0.00000 | 4.53545 | 2.74448 | 9.00955 | 1.60175 | 0.14776 |
| **Mrpl54** | 0.21005 | 0.26312 | 0.00000 | 6.35110 | 7.20659 | 1.26074 | 0.77525 | 1.18853 | 0.08492 | 1.95001 | 0.19650 | 2.20247 | 2.34454 | 0.00000 |
| **Mrpl55** | 3.73202 | 3.26280 | 1.65070 | 8.27993 | 5.30033 | 8.58628 | 3.29203 | 7.87690 | 0.02518 | 0.81092 | 3.10771 | 7.64224 | 0.55000 | 4.42641 |
| **Mrpl57** | 0.05115 | 0.43617 | 0.00000 | 6.32788 | 7.97840 | 1.91511 | 1.20428 | 4.05174 | 1.43930 | 3.17363 | 0.09634 | 5.48180 | 1.30480 | 0.92292 |
| **Mrpl9** | 6.67310 | 0.03853 | 10.89485 | 0.65997 | 4.80371 | 4.80301 | 0.75884 | 2.24547 | 2.44364 | 0.07328 | 2.54230 | 4.36616 | 4.20826 | 6.03115 |
| **Mrps10** | 0.75430 | 5.73205 | 9.59962 | 7.54453 | 12.98363 | 2.49252 | 4.06454 | 8.04168 | 0.07565 | 0.13019 | 1.41098 | 6.09059 | 0.54120 | 0.00000 |
| **Mrps11** | 2.34740 | 1.36792 | 7.71757 | 7.72791 | 6.18956 | 8.05051 | 5.64666 | 6.56749 | 0.00000 | 2.35568 | 7.48333 | 7.09731 | 0.22398 | 1.60605 |
| **Mrps12** | 0.10895 | 5.57887 | 3.17444 | 13.29952 | 8.00657 | 2.12650 | 4.76833 | 10.20398 | 0.00000 | 0.27962 | 1.12168 | 10.54992 | 1.48706 | 0.00000 |
| **Mrps14** | 6.59364 | 0.09976 | 1.59524 | 5.02243 | 10.04637 | 9.84536 | 1.91649 | 8.54286 | 1.48926 | 4.62806 | 6.11309 | 7.77622 | 1.84193 | 5.59423 |
| **Mrps15** | 4.43713 | 2.27955 | 0.16928 | 15.61055 | 6.86523 | 2.53667 | 0.03050 | 3.65406 | 0.52135 | 0.11719 | 0.00000 | 4.09976 | 2.90459 | 0.00000 |
| **Mrps16** | 0.99108 | 4.69913 | 0.00000 | 6.42947 | 3.93366 | 4.77172 | 1.62509 | 5.67541 | 0.30737 | 0.16025 | 1.13004 | 6.21875 | 1.51916 | 0.48582 |
| **Mrps17** | 5.23390 | 9.34761 | 1.65703 | 8.52277 | 12.26807 | 10.35576 | 2.17402 | 0.93588 | 1.65629 | 3.72150 | 3.58632 | 6.77064 | 2.05197 | 7.07410 |
| **Mrps18a** | 6.76653 | 5.60517 | 9.47604 | 14.67014 | 9.56636 | 5.82474 | 4.07771 | 7.72377 | 0.11951 | 0.18703 | 3.05910 | 7.80900 | 0.03316 | 1.14363 |
| **Mrps18b** | 21.68332 | 4.27238 | 31.68518 | 29.87995 | 23.70191 | 25.13086 | 7.76568 | 14.34499 | 18.03713 | 13.42867 | 19.81407 | 12.52152 | 4.92713 | 19.25490 |
| **Mrps18c** | 3.07617 | 1.21240 | 0.00000 | 2.02217 | 5.23278 | 2.31080 | 2.02324 | 2.62073 | 2.69999 | 8.07658 | 0.61205 | 4.13833 | 2.26816 | 3.13314 |
| **Mrps2** | 5.32537 | 0.05047 | 7.94920 | 10.66488 | 9.41881 | 9.01921 | 4.08097 | 2.72707 | 7.13821 | 0.13351 | 2.52932 | 9.27819 | 2.32222 | 6.73607 |
| **Mrps21** | 1.83134 | 0.22899 | 0.00000 | 1.34277 | 3.33156 | 4.12289 | 1.86325 | 6.22622 | 1.50923 | 4.38577 | 0.07023 | 4.38300 | 1.80504 | 2.34091 |
| **Mrps22** | 8.09175 | 0.43559 | 3.25547 | 14.42248 | 14.20745 | 5.64747 | 0.04628 | 0.68750 | 2.74897 | 0.03413 | 0.18357 | 3.50738 | 2.06711 | 2.44860 |

| | | | | | | | | | | | | | | |
|---|---|---|---|---|---|---|---|---|---|---|---|---|---|---|
| **Mrps23** | 4.22686 | 1.27691 | 16.67003 | 16.47907 | 15.01424 | 15.12322 | 6.33469 | 11.98081 | 0.61096 | 0.41153 | 7.17286 | 8.45566 | 0.57973 | 2.17315 |
| **Mrps24** | 2.66616 | 0.04845 | 0.00000 | 1.08664 | 11.39683 | 2.88118 | 0.08933 | 1.97596 | 0.16003 | 3.05097 | 0.04162 | 3.95241 | 6.67617 | 0.17751 |
| **Mrps25** | 3.30983 | 0.07027 | 2.59548 | 5.19369 | 1.25101 | 6.49189 | 0.02961 | 5.17679 | 0.82527 | 9.97161 | 4.21511 | 4.27413 | 0.70142 | 2.71849 |
| **Mrps26** | 3.86193 | 0.04625 | 2.81274 | 6.81209 | 13.48581 | 1.85579 | 1.92463 | 5.31499 | 0.13007 | 0.11379 | 0.04261 | 2.83983 | 6.13403 | 0.89302 |
| **Mrps27** | 2.95857 | 4.68965 | 7.17481 | 10.31150 | 6.14995 | 7.32608 | 1.59844 | 2.89629 | 0.92861 | 0.07178 | 8.10804 | 3.60162 | 0.46056 | 6.57235 |
| **Mrps28** | 0.51430 | 4.98958 | 1.09735 | 2.02283 | 1.89295 | 0.39742 | 0.24518 | 0.99081 | 0.00416 | 3.07803 | 0.82762 | 3.89797 | 1.50823 | 0.13473 |
| **Mrps30** | 9.44129 | 0.28477 | 3.29368 | 0.93934 | 9.35821 | 4.24283 | 0.22533 | 1.44101 | 6.11521 | 0.02533 | 2.22648 | 2.86209 | 8.25442 | 2.88444 |
| **Mrps31** | 3.61695 | 3.22599 | 12.16509 | 4.24938 | 3.86844 | 8.31181 | 2.56450 | 4.98767 | 7.86219 | 9.44049 | 7.95705 | 2.76362 | 0.00000 | 8.86163 |
| **Mrps33** | 3.69266 | 0.76282 | 0.11109 | 3.43339 | 7.65419 | 6.26525 | 2.46259 | 3.97063 | 2.98585 | 11.06444 | 1.32131 | 6.71031 | 1.65421 | 2.95198 |
| **Mrps34** | 1.09963 | 0.03944 | 0.00000 | 7.37839 | 6.72977 | 1.33927 | 0.00360 | 1.19415 | 0.00185 | 0.02174 | 0.02325 | 1.70684 | 0.00000 | 1.09836 |
| **Mrps35** | 5.90134 | 0.58873 | 1.29872 | 4.97170 | 5.01535 | 8.37366 | 1.80377 | 0.49693 | 0.05494 | 0.08700 | 1.65418 | 5.63794 | 0.00000 | 1.29127 |
| **Mrps36** | 2.00500 | 0.15335 | 1.73731 | 0.95829 | 3.54906 | 5.30033 | 2.61421 | 7.35575 | 1.36910 | 11.42211 | 0.95701 | 5.14409 | 0.73897 | 0.92606 |
| **Mrps5** | 3.10914 | 0.08298 | 8.48030 | 4.74335 | 1.89967 | 6.12500 | 0.13619 | 6.97171 | 0.32697 | 0.02271 | 6.80271 | 6.58785 | 0.00000 | 4.52655 |
| **Mrps6** | 4.04147 | 1.87044 | 3.37898 | 13.45184 | 9.68762 | 2.71352 | 1.46209 | 1.33260 | 1.92478 | 8.05997 | 0.00030 | 2.18107 | 8.33338 | 0.44315 |
| **Mrps7** | 12.93507 | 5.09325 | 19.25507 | 17.12065 | 23.77692 | 8.57600 | 4.54394 | 9.52983 | 3.02764 | 6.83933 | 5.76346 | 9.15230 | 7.38415 | 11.22457 |
| **Mrps9** | 8.76003 | 0.27035 | 11.45482 | 10.21012 | 8.62681 | 4.89020 | 3.78686 | 9.07006 | 0.06594 | 12.34596 | 8.98457 | 7.20994 | 0.81292 | 3.78637 |
| **Mrrf** | 3.79853 | 0.09011 | 6.92511 | 3.69394 | 4.40227 | 6.11661 | 1.03185 | 5.34026 | 1.40127 | 0.01494 | 4.55342 | 5.17789 | 1.61978 | 4.44432 |
| **Mrs2** | 3.05167 | 0.00314 | 1.98310 | 1.73889 | 1.48033 | 1.58091 | 0.04724 | 1.73844 | 0.01020 | 0.01659 | 0.00000 | 1.12692 | 0.01217 | 1.85805 |
| **Mrto4** | 22.39587 | 1.80002 | 29.10867 | 22.64673 | 22.18939 | 21.42507 | 4.93455 | 7.88987 | 7.90910 | 13.46754 | 12.97531 | 13.59151 | 8.60276 | 15.44223 |
| **Ms4a1** | 0.02241 | 0.00443 | 0.01430 | 0.01240 | 0.01200 | 0.01263 | 0.00426 | 0.01040 | 0.02077 | 0.00670 | 0.00851 | 0.00994 | 0.03839 | 0.01560 |
| **Ms4a6c** | 0.01175 | 0.32200 | 0.01392 | 0.06120 | 0.18545 | 0.01205 | 0.00414 | 0.01187 | 0.15762 | 0.00642 | 0.01530 | 0.05439 | 0.02217 | 0.09907 |
| **Msantd1** | 0.37849 | 0.00440 | 0.01712 | 0.00614 | 0.21751 | 0.01096 | 0.00490 | 0.01113 | 0.01491 | 0.00694 | 0.00731 | 0.00748 | 0.07832 | 0.00093 |
| **Msantd2** | 6.62375 | 0.02439 | 10.41542 | 1.52333 | 10.01931 | 2.75703 | 3.81171 | 0.00000 | 12.15993 | 0.00898 | 2.12373 | 1.45515 | 2.36752 | 17.88029 |
| **Msantd3** | 0.67699 | 0.02660 | 0.12868 | 0.00000 | 0.00855 | 0.00301 | 0.00361 | 0.02329 | 0.00829 | 0.00691 | 0.00329 | 1.87612 | 0.06944 | 0.00459 |
| **Msantd4** | 1.60789 | 0.01175 | 1.65903 | 1.47004 | 1.07079 | 4.06080 | 0.00347 | 0.46276 | 0.54971 | 0.01817 | 0.80046 | 0.50819 | 0.53077 | 2.61914 |
| **Msc** | 0.00000 | 0.01002 | 0.59164 | 1.80633 | 1.07467 | 2.84121 | 1.90295 | 2.76183 | 0.00354 | 0.13180 | 0.00000 | 2.12570 | 0.00236 | 0.09293 |
| **Msh2** | 11.01593 | 0.51363 | 11.99427 | 5.48116 | 5.59168 | 6.00395 | 2.99089 | 6.31601 | 4.73754 | 0.09825 | 9.25276 | 14.37987 | 1.77620 | 5.77233 |
| **Msh3** | 2.58872 | 0.01052 | 2.06457 | 0.74759 | 2.45509 | 0.98489 | 0.00332 | 1.03500 | 3.48235 | 0.01583 | 0.00000 | 1.79758 | 2.26709 | 5.05075 |
| **Msh4** | 3.35869 | 0.00219 | 3.83330 | 3.45634 | 1.86617 | 0.98469 | 3.04267 | 0.74943 | 0.82073 | 0.01089 | 4.89715 | 0.43322 | 0.07557 | 5.15374 |
| **Msh5** | 0.01529 | 0.01253 | 0.26424 | 0.21139 | 0.00000 | 0.32973 | 0.00382 | 0.00365 | 0.00705 | 0.01721 | 0.00194 | 0.33715 | 0.69573 | 0.00845 |
| **Msh6** | 13.70556 | 4.49124 | 15.42105 | 4.86677 | 12.47831 | 9.18231 | 0.51217 | 3.22307 | 14.35244 | 0.04325 | 9.14337 | 14.18165 | 3.19404 | 20.34587 |
| **Msi2** | 0.04110 | 0.00449 | 0.07811 | 0.00265 | 0.23663 | 0.00825 | 0.00410 | 0.01122 | 0.30222 | 0.00903 | 0.00647 | 0.00791 | 0.00492 | 0.03203 |
| **Msl1** | 5.41715 | 0.04910 | 3.35849 | 2.50318 | 6.43134 | 5.87168 | 3.61508 | 3.15183 | 2.39034 | 0.08391 | 0.00000 | 3.61495 | 2.35724 | 6.79612 |
| **Msl2** | 1.60549 | 0.01015 | 0.99659 | 0.47222 | 2.76392 | 0.83873 | 2.58699 | 1.29391 | 7.31061 | 0.07003 | 3.40063 | 1.48415 | 9.00988 | 2.70204 |
| **Msl3** | 2.26259 | 0.98147 | 8.17260 | 3.61018 | 5.28720 | 7.52913 | 0.19209 | 1.07278 | 2.45039 | 0.00517 | 2.54023 | 2.11695 | 2.96602 | 8.35147 |
| **Msln** | 0.00571 | 0.01448 | 0.01233 | 0.06691 | 0.01204 | 0.00591 | 0.00468 | 0.26178 | 0.00644 | 0.00706 | 0.00747 | 0.01011 | 0.09464 | 0.07465 |
| **Mslnl** | 0.00000 | 0.00436 | 0.44942 | 0.98886 | 0.86886 | 0.14002 | 0.00489 | 0.10880 | 0.28539 | 0.00589 | 0.00201 | 0.00251 | 0.08455 | 0.17340 |
| **Msmo1** | 10.74616 | 0.01609 | 7.17898 | 8.02871 | 0.68694 | 5.60957 | 2.81229 | 0.08656 | 13.51090 | 0.14016 | 1.22397 | 0.83075 | 0.14557 | 9.07025 |
| **Msn** | 0.37051 | 5.33172 | 0.02655 | 0.69378 | 0.44945 | 2.33675 | 1.24815 | 14.06414 | 0.94851 | 1.10077 | 0.97201 | 0.00000 | 0.36126 | 0.00000 |
| **Msra** | 0.00000 | 0.03407 | 1.12898 | 0.38721 | 0.09985 | 0.45958 | 0.00311 | 2.97371 | 0.00267 | 0.02305 | 0.00000 | 0.86772 | 0.00002 | 0.90299 |
| **Msrb1** | 0.29199 | 0.00364 | 0.10413 | 0.15900 | 0.32445 | 0.16118 | 0.03402 | 0.02729 | 0.05423 | 0.00672 | 0.23038 | 0.15366 | 6.73267 | 0.14966 |
| **Msrb2** | 0.22208 | 0.01693 | 0.22220 | 0.00000 | 0.01524 | 0.46239 | 0.00350 | 0.02777 | 0.00971 | 0.01190 | 0.23080 | 2.49752 | 0.00883 | 0.01081 |
| **Msrb3** | 0.15499 | 0.00407 | 0.00000 | 0.34453 | 0.00000 | 0.17827 | 0.00435 | 0.27028 | 1.19662 | 0.11714 | 0.48345 | 0.04542 | 0.03279 | 0.00000 |
| **Mss51** | 0.01093 | 0.00513 | 0.01392 | 0.00215 | 0.01396 | 0.13357 | 0.00441 | 0.02065 | 0.01199 | 0.00732 | 0.00923 | 0.01127 | 0.01690 | 0.00789 |
| **Mst1r** | 0.17245 | 0.00396 | 0.41544 | 0.16100 | 0.00000 | 0.00257 | 0.00384 | 0.00393 | 0.00902 | 0.01281 | 0.00848 | 0.09915 | 0.03397 | 0.00346 |
| **Msto1** | 1.34151 | 0.04447 | 2.73204 | 2.23204 | 1.47371 | 1.68327 | 0.00960 | 1.61128 | 0.08022 | 0.04982 | 0.34428 | 2.48610 | 2.18764 | 2.94108 |
| **Msx2** | 0.00694 | 0.01005 | 0.03653 | 0.00000 | 0.00545 | 0.00600 | 0.90208 | 0.59402 | 0.03145 | 2.25958 | 0.00000 | 0.77962 | 0.00550 | 0.00356 |
| **Mt1** | 8.05695 | 7.27647 | 3.19811 | 17.70015 | 33.15564 | 5.13525 | 8.65332 | 21.84417 | 0.00727 | 13.37423 | 1.39183 | 1.13588 | 6.29738 | 4.24505 |
| **Mt2** | 12.92982 | 9.16739 | 0.91411 | 18.68026 | 22.73462 | 6.24779 | 8.96374 | 29.40305 | 0.37844 | 13.29644 | 2.05168 | 1.25789 | 19.87753 | 18.15685 |
| **Mt4** | 0.00851 | 0.01245 | 0.00503 | 0.05832 | 0.30014 | 0.01878 | 0.00480 | 0.01833 | 0.01485 | 0.00995 | 0.00861 | 0.10195 | 0.01477 | 0.01926 |
| **Mta1** | 0.12092 | 0.05571 | 0.15181 | 0.60172 | 0.92183 | 0.08914 | 0.01082 | 1.21785 | 0.00698 | 0.02302 | 0.03325 | 1.07225 | 0.00284 | 0.05614 |
| **Mta2** | 0.00000 | 5.19629 | 8.32894 | 7.12826 | 8.85104 | 3.49945 | 1.39052 | 3.47428 | 4.96035 | 0.06562 | 1.13265 | 4.00831 | 13.13619 | 8.20435 |
| **Mta3** | 16.00043 | 8.40886 | 43.21270 | 24.54410 | 40.02796 | 13.30348 | 9.20660 | 18.77096 | 0.00000 | 0.62564 | 17.29506 | 12.59349 | 1.31873 | 3.60127 |
| **Mtap** | 0.50631 | 0.05528 | 0.13252 | 0.83726 | 0.82271 | 1.80514 | 0.00970 | 2.00456 | 2.00694 | 0.09811 | 2.44665 | 3.61746 | 1.43889 | 0.00000 |
| **Mtap7d3** | 0.01176 | 0.00450 | 0.00790 | 0.01140 | 0.30164 | 0.01379 | 0.00424 | 0.01039 | 0.01248 | 0.00714 | 0.00885 | 0.00854 | 0.00668 | 0.01095 |

| | | | | | | | | | | | | | | |
|---|---|---|---|---|---|---|---|---|---|---|---|---|---|---|
| **Mtbp** | 0.95533 | 0.00322 | 0.90188 | 0.68548 | 0.00000 | 0.19197 | 0.14231 | 0.01544 | 2.78486 | 0.01396 | 1.51582 | 0.24685 | 1.02228 | 2.36439 |
| **Mtch1** | 5.82133 | 4.67725 | 1.39398 | 0.00000 | 1.35507 | 3.43122 | 0.38871 | 3.78626 | 0.00168 | 11.46616 | 3.80762 | 2.66137 | 0.56565 | 2.19038 |
| **Mtch2** | 15.12812 | 1.49063 | 28.26027 | 27.63609 | 21.21436 | 27.88895 | 6.49729 | 18.77314 | 11.34429 | 3.99917 | 18.95496 | 19.22775 | 3.57741 | 10.74648 |
| **Mtcl1** | 0.54016 | 2.20180 | 1.52617 | 0.54612 | 0.00000 | 0.49032 | 0.00880 | 2.85075 | 0.00426 | 0.05649 | 3.55378 | 1.73435 | 0.00185 | 0.26642 |
| **Mtcp1** | 0.24259 | 0.00387 | 0.06266 | 0.00000 | 0.00000 | 0.16992 | 0.00411 | 0.00365 | 0.00621 | 0.00574 | 0.00368 | 0.31454 | 0.00324 | 0.13239 |
| **Mtdh** | 3.53734 | 0.08033 | 3.87230 | 5.76681 | 1.51632 | 8.31500 | 1.79811 | 6.59453 | 8.46032 | 0.12652 | 5.66744 | 6.85311 | 2.42996 | 6.67528 |
| **Mterf1a** | 0.00000 | 0.00437 | 0.01334 | 0.02705 | 0.00000 | 0.00934 | 0.00407 | 0.00948 | 0.28839 | 0.00653 | 0.10367 | 0.00708 | 0.30991 | 0.41380 |
| **Mterf1b** | 0.00039 | 0.00430 | 0.01713 | 0.01189 | 0.00351 | 0.01672 | 0.00413 | 0.00802 | 0.16228 | 0.00650 | 0.04239 | 0.00724 | 0.33687 | 0.36523 |
| **Mterf2** | 0.00000 | 0.01122 | 1.18667 | 0.55305 | 0.18431 | 0.00000 | 0.00940 | 0.20697 | 0.01715 | 0.01189 | 0.01339 | 0.00611 | 0.00306 | 0.39529 |
| **Mterf3** | 1.65369 | 0.01173 | 2.29107 | 0.81388 | 0.00000 | 0.92330 | 0.03671 | 0.25027 | 3.15591 | 0.00521 | 2.27822 | 0.42226 | 0.78230 | 3.58073 |
| **Mterf4** | 5.48737 | 5.92703 | 4.06862 | 3.72259 | 2.63431 | 3.23858 | 0.07284 | 5.23077 | 7.53630 | 0.02614 | 1.41194 | 2.95129 | 0.04383 | 3.42951 |
| **Mtf1** | 4.06049 | 0.25334 | 2.65075 | 4.12863 | 9.47284 | 3.18307 | 1.55199 | 5.35152 | 7.91466 | 0.11827 | 0.16179 | 3.06556 | 6.30986 | 10.54177 |
| **Mtf2** | 16.89771 | 0.22666 | 20.87724 | 7.98034 | 23.33109 | 34.11041 | 8.71319 | 9.44494 | 43.04019 | 0.51510 | 16.56090 | 6.15087 | 16.39013 | 19.75421 |
| **Mtfmt** | 0.61889 | 0.02530 | 1.25357 | 0.51901 | 0.90387 | 1.05591 | 0.04530 | 3.64117 | 0.48080 | 0.01787 | 1.20765 | 1.25503 | 1.23334 | 1.22731 |
| **Mtfp1** | 0.00000 | 0.03250 | 0.45902 | 0.78923 | 0.00000 | 2.64572 | 0.02685 | 5.57596 | 0.00726 | 0.02670 | 0.83438 | 3.10171 | 0.00887 | 0.00000 |
| **Mtfr1** | 4.14789 | 0.10954 | 4.89649 | 4.44765 | 1.85994 | 10.86667 | 4.13596 | 4.29213 | 3.42874 | 0.21437 | 5.22080 | 10.07453 | 2.14829 | 4.58393 |
| **Mtfr1l** | 4.37752 | 0.05636 | 6.11666 | 4.95288 | 7.32748 | 9.88246 | 2.98514 | 5.92947 | 1.17603 | 0.14639 | 8.56589 | 7.55945 | 1.05148 | 1.49780 |
| **Mtfr2** | 0.06438 | 0.02552 | 0.00741 | 0.12424 | 0.00177 | 0.06532 | 0.00401 | 0.00346 | 0.00746 | 0.00635 | 0.00555 | 0.03782 | 0.56406 | 0.47680 |
| **Mtg1** | 1.33509 | 1.24986 | 10.14268 | 16.23092 | 4.57558 | 7.96493 | 1.64010 | 1.34347 | 1.91671 | 0.16969 | 5.67367 | 5.65740 | 0.33259 | 0.93656 |
| **Mtg2** | 0.00000 | 0.04471 | 4.39939 | 2.07177 | 1.95538 | 0.00000 | 0.00275 | 2.40472 | 0.01850 | 0.03375 | 1.43939 | 2.80918 | 0.20130 | 0.03072 |
| **Mthfd1** | 9.99170 | 4.65278 | 21.77980 | 16.03221 | 20.83283 | 18.94161 | 3.41113 | 13.20939 | 1.16384 | 0.52184 | 16.82134 | 16.38558 | 1.29975 | 7.47806 |
| **Mthfd1l** | 0.50242 | 0.01558 | 1.37943 | 0.92538 | 0.49218 | 1.42862 | 0.00826 | 0.00000 | 0.66527 | 0.00000 | 3.76567 | 1.95451 | 0.66519 | 0.82430 |
| **Mthfd2** | 1.25889 | 0.04243 | 5.54765 | 6.48823 | 3.90607 | 1.41649 | 3.83530 | 0.28936 | 1.29024 | 0.10680 | 9.84418 | 2.53389 | 0.79763 | 2.45904 |
| **Mthfr** | 5.53150 | 0.06331 | 10.56647 | 6.23717 | 9.11506 | 4.76625 | 3.11501 | 3.41418 | 2.15414 | 0.11765 | 3.01114 | 4.66317 | 5.76179 | 9.70218 |
| **Mthfs** | 0.79555 | 0.00992 | 0.23211 | 0.00000 | 0.37615 | 1.15385 | 0.00341 | 0.02612 | 0.03702 | 0.01230 | 0.23737 | 1.90422 | 0.80073 | 0.00799 |
| **Mthfsd** | 0.00000 | 0.00994 | 3.89614 | 9.20404 | 3.81301 | 5.83315 | 2.82581 | 0.00000 | 0.00150 | 0.07559 | 3.20195 | 0.38504 | 0.00000 | 0.93937 |
| **Mthfsl** | 0.91971 | 0.01030 | 0.17382 | 0.00000 | 0.42604 | 1.07801 | 0.00358 | 0.00641 | 0.04224 | 0.01982 | 0.35299 | 1.51866 | 0.63097 | 0.04389 |
| **Mtif2** | 1.93056 | 0.15340 | 0.81054 | 3.08880 | 2.76900 | 1.88216 | 0.16537 | 1.34343 | 3.63538 | 0.33192 | 1.56043 | 4.23429 | 2.00826 | 2.68310 |
| **Mtif3** | 0.38650 | 6.13063 | 3.16393 | 2.82611 | 2.91784 | 1.95842 | 0.04368 | 1.84098 | 0.00000 | 0.05031 | 3.52810 | 1.82767 | 0.85523 | 0.33807 |
| **Mtl5** | 0.00705 | 0.00434 | 0.39810 | 0.00000 | 0.15896 | 0.00663 | 0.00404 | 0.00852 | 0.08813 | 0.00615 | 0.00719 | 0.03629 | 0.00372 | 0.00025 |
| **Mtm1** | 1.36706 | 0.00374 | 0.63720 | 0.06769 | 0.91208 | 1.81109 | 2.40592 | 0.42251 | 2.33898 | 0.06721 | 0.90152 | 0.00000 | 0.00812 | 0.42867 |
| **Mtmr1** | 1.22943 | 0.01883 | 0.33966 | 0.00000 | 1.26755 | 1.47947 | 0.01554 | 1.08900 | 1.47455 | 0.00162 | 1.52053 | 0.43949 | 1.78279 | 0.40817 |
| **Mtmr10** | 2.66125 | 0.02015 | 2.65270 | 1.52019 | 5.69660 | 1.66513 | 4.00430 | 2.74068 | 0.80403 | 0.08977 | 3.16498 | 1.03898 | 0.00000 | 4.11923 |
| **Mtmr11** | 0.01082 | 0.00465 | 0.01982 | 0.00618 | 0.01082 | 0.05231 | 0.00436 | 0.00751 | 0.01061 | 0.00621 | 0.00838 | 0.00986 | 0.24538 | 0.00211 |
| **Mtmr12** | 3.02528 | 0.03276 | 0.82015 | 1.20808 | 2.25849 | 1.35528 | 0.01599 | 2.16067 | 0.57816 | 0.01509 | 2.66165 | 4.00105 | 3.05426 | 4.70532 |
| **Mtmr14** | 1.51605 | 0.00663 | 6.04108 | 0.00000 | 8.69265 | 3.17878 | 0.01797 | 3.93452 | 0.00000 | 0.02465 | 6.26958 | 3.38605 | 4.60248 | 2.77968 |
| **Mtmr2** | 0.33693 | 0.08365 | 0.51785 | 1.43791 | 0.39991 | 1.19492 | 0.23932 | 1.12663 | 3.87820 | 0.05660 | 0.06283 | 1.64213 | 2.19857 | 1.07051 |
| **Mtmr3** | 5.22576 | 0.01604 | 6.57927 | 0.61482 | 3.94801 | 4.82679 | 0.01551 | 1.60688 | 1.86759 | 0.00394 | 2.82622 | 2.79468 | 4.82213 | 7.63520 |
| **Mtmr4** | 1.83220 | 0.03274 | 2.19437 | 5.11681 | 0.96190 | 4.43352 | 4.76794 | 2.65486 | 5.99799 | 0.12432 | 2.26185 | 4.58295 | 0.95917 | 0.62486 |
| **Mtmr6** | 0.89683 | 0.01891 | 1.57132 | 0.88555 | 1.53962 | 1.49048 | 0.00168 | 2.74352 | 8.58676 | 0.00414 | 1.98071 | 1.85041 | 0.52074 | 3.58906 |
| **Mtmr7** | 1.62806 | 0.03496 | 3.16102 | 0.24438 | 1.02557 | 1.15819 | 0.00268 | 2.02572 | 0.60625 | 0.00572 | 0.29139 | 2.67227 | 0.17531 | 0.56104 |
| **Mtmr9** | 0.42207 | 0.01062 | 1.10142 | 0.96833 | 0.71114 | 0.41003 | 0.00340 | 0.86951 | 0.25581 | 0.02403 | 0.00000 | 0.00711 | 0.08907 | 0.13269 |
| **Mto1** | 4.05269 | 0.01548 | 1.82563 | 1.62124 | 0.44232 | 1.74966 | 3.46077 | 1.88342 | 2.97860 | 0.03779 | 0.00000 | 3.68530 | 1.14670 | 0.78794 |
| **Mtor** | 4.25321 | 8.12113 | 3.60344 | 4.27609 | 5.25780 | 4.53493 | 1.60626 | 0.62363 | 1.32000 | 0.07606 | 3.42217 | 2.66944 | 5.79876 | 6.54513 |
| **Mtpap** | 5.76908 | 0.04890 | 6.31978 | 1.17140 | 7.31233 | 4.05960 | 4.57995 | 1.05103 | 5.54265 | 0.09220 | 5.17220 | 4.64389 | 0.18613 | 7.57466 |
| **Mtpn** | 9.10884 | 2.31880 | 4.28244 | 4.95961 | 5.29632 | 15.32504 | 0.58890 | 4.80226 | 14.93646 | 0.60241 | 7.90672 | 8.72611 | 7.64306 | 3.15671 |
| **Mtr** | 1.85130 | 0.03246 | 1.34423 | 3.02218 | 0.89385 | 1.34106 | 0.00256 | 0.00000 | 0.00336 | 0.01499 | 5.04624 | 1.80422 | 0.51418 | 1.36880 |
| **Mtrf1** | 0.07749 | 0.01902 | 0.20124 | 0.00000 | 0.18262 | 0.33704 | 2.13268 | 0.00000 | 0.01847 | 0.05783 | 3.71131 | 1.03470 | 1.04873 | 0.33821 |
| **Mtrf1l** | 2.68194 | 0.00229 | 0.63104 | 0.58453 | 1.95930 | 0.34362 | 0.00328 | 0.00000 | 0.00343 | 0.01101 | 0.43443 | 1.72086 | 0.40191 | 2.70453 |
| **Mtrr** | 3.01745 | 0.03115 | 7.77967 | 5.30275 | 4.35626 | 6.48024 | 1.51567 | 2.79060 | 2.18679 | 0.03821 | 5.81895 | 1.75315 | 1.48936 | 11.22703 |
| **Mtss1** | 0.03699 | 0.00919 | 2.23327 | 2.70447 | 1.88020 | 3.04894 | 1.80439 | 1.23539 | 0.00000 | 0.08203 | 1.23140 | 3.16308 | 1.33944 | 0.21205 |
| **Mtss1l** | 0.03373 | 0.00446 | 0.25888 | 0.13769 | 0.13254 | 0.02153 | 0.00405 | 0.00639 | 0.00710 | 0.00527 | 0.00578 | 0.01061 | 0.01481 | 0.09954 |
| **Mttp** | 0.00000 | 0.00376 | 0.43093 | 0.14269 | 0.74456 | 0.00000 | 0.99436 | 0.00000 | 1.67711 | 0.01135 | 0.00198 | 0.00351 | 0.24348 | 4.88734 |
| **Mturn** | 0.08570 | 0.00403 | 0.90010 | 0.15560 | 0.00000 | 0.18324 | 0.60452 | 1.62348 | 0.00637 | 0.01980 | 0.00648 | 0.00141 | 0.11306 | 0.03756 |
| **Mtus1** | 9.96297 | 0.01397 | 9.02050 | 1.04138 | 17.18317 | 8.17428 | 5.60235 | 2.89873 | 17.64821 | 0.28639 | 9.59119 | 0.18579 | 12.86019 | 20.37896 |

| | | | | | | | | | | | | | | |
|---|---|---|---|---|---|---|---|---|---|---|---|---|---|---|
| **Mtus2** | 1.01167 | 0.00329 | 0.09261 | 1.09828 | 1.24037 | 1.06816 | 0.00360 | 0.00798 | 0.40129 | 0.00522 | 0.01608 | 0.11894 | 5.33861 | 1.27384 |
| **Mtx1** | 2.95612 | 0.12347 | 15.66871 | 14.14521 | 10.27067 | 2.97285 | 7.32872 | 10.47040 | 0.00000 | 0.32501 | 4.76730 | 6.75127 | 0.36325 | 0.00000 |
| **Mtx2** | 8.67412 | 1.73330 | 0.43026 | 0.99556 | 2.18847 | 2.43804 | 0.00605 | 1.29839 | 0.11908 | 11.06274 | 1.22080 | 2.71756 | 1.86968 | 0.00000 |
| **Mtx3** | 0.00000 | 0.00291 | 0.42306 | 0.88801 | 1.29164 | 1.55964 | 0.04928 | 0.00000 | 4.68531 | 0.01259 | 0.06104 | 1.23193 | 0.81263 | 1.80596 |
| **Muc1** | 0.00169 | 0.02633 | 0.02422 | 1.80634 | 0.00260 | 2.42304 | 0.00327 | 0.00347 | 0.01455 | 0.01626 | 0.00000 | 1.65389 | 0.00849 | 0.00001 |
| **Muc13** | 0.00583 | 0.01072 | 0.00000 | 0.05702 | 0.00000 | 0.00575 | 0.00361 | 0.00437 | 0.21629 | 0.01319 | 0.00291 | 0.97119 | 0.17607 | 0.38089 |
| **Muc4** | 0.00949 | 0.00444 | 0.01420 | 0.01337 | 0.01191 | 0.01287 | 0.00434 | 0.01028 | 0.01184 | 0.00694 | 0.00885 | 0.01001 | 0.00952 | 0.04320 |
| **Mul1** | 1.48228 | 0.09182 | 5.93415 | 10.98248 | 4.53562 | 4.01373 | 2.89936 | 4.44851 | 0.04409 | 0.12966 | 2.59790 | 3.45354 | 4.96602 | 1.39544 |
| **Mum1** | 1.30194 | 11.23929 | 12.93006 | 16.63337 | 7.13373 | 14.05537 | 1.49815 | 5.99521 | 0.62604 | 0.25925 | 9.89167 | 5.84987 | 2.67416 | 0.93973 |
| **Mum1l1** | 0.01210 | 0.00443 | 0.10622 | 0.00195 | 0.13357 | 0.01366 | 0.00414 | 0.01007 | 0.00623 | 0.00672 | 0.00764 | 0.01034 | 0.00499 | 0.08485 |
| **Murc** | 0.02091 | 0.00458 | 0.14163 | 0.00000 | 0.01542 | 0.50428 | 0.00397 | 0.00577 | 0.01849 | 0.01048 | 0.00765 | 0.02624 | 0.01218 | 0.00320 |
| **Mus81** | 0.41510 | 2.96361 | 0.70652 | 0.00000 | 2.26176 | 1.31398 | 3.30609 | 3.82071 | 0.00774 | 0.12899 | 1.97881 | 4.42186 | 1.69064 | 0.54246 |
| **Musk** | 0.00366 | 0.00445 | 0.00736 | 0.01364 | 0.19121 | 0.01175 | 0.00415 | 0.00989 | 0.02515 | 0.00649 | 0.00800 | 0.00848 | 0.00502 | 0.31102 |
| **Mustn1** | 0.01641 | 0.00436 | 0.00886 | 0.00541 | 0.22564 | 0.01459 | 0.00502 | 0.00915 | 0.01902 | 0.00664 | 0.00780 | 0.05212 | 0.00526 | 0.00586 |
| **Mut** | 1.22073 | 0.01725 | 0.44784 | 0.00000 | 0.00000 | 1.34186 | 0.00366 | 0.37583 | 1.75649 | 0.00000 | 2.03652 | 0.73049 | 0.70227 | 0.11794 |
| **Mutyh** | 2.01914 | 0.09907 | 6.98520 | 2.50458 | 2.50169 | 1.80605 | 2.17667 | 4.29591 | 0.00000 | 10.24248 | 4.24263 | 0.90355 | 0.61839 | 0.27469 |
| **Mvb12a** | 0.10307 | 0.08654 | 0.75967 | 5.25129 | 2.40712 | 0.94361 | 2.40427 | 0.77900 | 0.01494 | 0.11032 | 0.01598 | 0.34580 | 0.48319 | 0.00000 |
| **Mvb12b** | 0.00962 | 0.12692 | 0.16414 | 0.00000 | 0.49055 | 0.01505 | 0.00383 | 0.00409 | 0.05824 | 0.00544 | 0.02423 | 0.00467 | 2.04147 | 0.80713 |
| **Mvd** | 0.32754 | 6.23501 | 9.48881 | 0.53617 | 2.29855 | 5.31574 | 0.00612 | 1.50904 | 0.04535 | 0.02144 | 1.00336 | 2.17803 | 0.10311 | 0.20515 |
| **Mvk** | 0.00000 | 0.04606 | 0.68407 | 0.58914 | 0.51694 | 1.43811 | 0.02474 | 4.15820 | 0.00900 | 0.03676 | 0.00000 | 4.86978 | 1.43371 | 0.00000 |
| **Mvp** | 1.54551 | 5.97787 | 3.68426 | 4.14623 | 3.21584 | 6.87433 | 2.72178 | 5.70273 | 0.01009 | 9.15384 | 5.06483 | 3.10120 | 1.79295 | 1.08636 |
| **Mxd1** | 0.62709 | 0.01522 | 0.42240 | 0.11998 | 0.04337 | 0.27670 | 0.22853 | 0.37788 | 0.10123 | 0.03258 | 0.26468 | 0.00239 | 0.52296 | 0.82726 |
| **Mxd3** | 0.79039 | 0.02928 | 3.70530 | 3.89822 | 0.55337 | 0.67444 | 0.00866 | 0.24267 | 0.00741 | 0.00739 | 2.41815 | 1.24243 | 0.43711 | 0.00000 |
| **Mxd4** | 0.01114 | 0.00439 | 0.16799 | 0.00000 | 0.14486 | 0.02125 | 0.00417 | 0.00791 | 0.00685 | 0.00577 | 0.00783 | 0.01983 | 0.02809 | 0.10229 |
| **Mxi1** | 2.14438 | 0.01961 | 1.60309 | 0.51533 | 1.40257 | 0.00000 | 0.00469 | 0.70422 | 0.00623 | 0.00317 | 0.00000 | 1.75979 | 2.10315 | 0.61441 |
| **Mxra7** | 0.13982 | 0.00422 | 0.33507 | 0.26206 | 0.75857 | 0.00000 | 0.00387 | 0.00823 | 0.77401 | 0.00664 | 0.00406 | 0.01202 | 0.66927 | 0.35201 |
| **Mxra8** | 0.01851 | 0.01158 | 0.00000 | 1.08053 | 0.49450 | 0.00000 | 0.00424 | 0.09313 | 1.42049 | 0.00716 | 0.00184 | 0.00743 | 0.64018 | 1.16152 |
| **Myadm** | 0.83599 | 0.01823 | 3.07136 | 4.21871 | 0.77388 | 3.03577 | 0.00169 | 0.02480 | 10.58326 | 0.02318 | 0.56365 | 1.95689 | 3.92086 | 7.11163 |
| **Myadml2** | 0.08339 | 0.00444 | 0.02215 | 0.01590 | 0.00992 | 0.01083 | 0.00420 | 0.01065 | 0.01439 | 0.00795 | 0.00774 | 0.00904 | 0.01148 | 0.08188 |
| **Mybbp1a** | 27.74718 | 2.46195 | 34.35188 | 20.63340 | 28.89888 | 17.23671 | 5.34969 | 5.56687 | 11.31713 | 1.11680 | 14.07718 | 17.37845 | 16.67688 | 17.95880 |
| **Mybl1** | 0.00000 | 0.00415 | 0.10453 | 0.00203 | 0.00000 | 0.04946 | 0.00402 | 0.00543 | 0.02633 | 0.00652 | 0.00575 | 0.00738 | 1.41980 | 0.79575 |
| **Mybl2** | 2.82341 | 8.21282 | 19.80386 | 16.44310 | 12.18594 | 19.80617 | 4.63103 | 15.26175 | 2.06715 | 0.38461 | 9.11680 | 15.75312 | 8.02051 | 10.01455 |
| **Mybpc1** | 0.28028 | 0.00351 | 0.39157 | 0.01992 | 0.01526 | 0.00000 | 0.00360 | 0.00825 | 0.42186 | 0.00532 | 0.42046 | 0.71421 | 0.13859 | 0.82704 |
| **Mybpc2** | 3.65083 | 0.02530 | 9.21125 | 5.69980 | 2.38638 | 3.06420 | 0.01646 | 0.51923 | 0.16077 | 0.01455 | 5.66387 | 0.81907 | 0.40134 | 3.32082 |
| **Myc** | 6.61039 | 3.30546 | 9.21727 | 0.00000 | 3.82655 | 0.48868 | 3.61697 | 0.48414 | 1.49987 | 0.05533 | 0.43830 | 3.92545 | 2.85528 | 7.33555 |
| **Mycbp** | 4.67352 | 7.66437 | 8.15069 | 2.11567 | 8.98013 | 10.15876 | 4.48855 | 3.65900 | 4.23667 | 0.37697 | 8.50802 | 5.17385 | 6.50761 | 2.95374 |
| **Mycbp2** | 2.78479 | 0.00615 | 8.69242 | 0.00000 | 4.34667 | 5.93910 | 1.06566 | 1.38993 | 7.07344 | 0.04690 | 6.07033 | 4.26362 | 2.36798 | 9.90562 |
| **Mycbpap** | 0.10847 | 0.08287 | 0.30212 | 0.43190 | 0.00000 | 3.93336 | 2.75256 | 1.63311 | 0.01052 | 0.07710 | 0.09035 | 1.06956 | 0.00365 | 0.00500 |
| **Mycl** | 0.17867 | 0.00323 | 1.36729 | 0.62167 | 0.04583 | 0.00000 | 0.00338 | 0.00000 | 0.04259 | 0.01958 | 0.61889 | 0.61048 | 0.00228 | 1.92919 |
| **Mycn** | 0.00457 | 7.22583 | 0.00479 | 0.00000 | 0.14392 | 0.37499 | 0.00259 | 2.47630 | 0.00259 | 0.00626 | 0.00000 | 0.40113 | 0.44463 | 0.03573 |
| **Mycs** | 0.00591 | 0.00435 | 0.01503 | 0.01200 | 0.00992 | 0.01303 | 0.00426 | 0.00931 | 0.01471 | 0.00634 | 0.00900 | 0.00932 | 0.33543 | 0.01113 |
| **Myd88** | 2.05679 | 0.06347 | 0.99507 | 3.44892 | 0.39556 | 2.19994 | 0.00306 | 4.60939 | 1.24298 | 0.01143 | 0.00000 | 0.11527 | 2.33187 | 0.77094 |
| **Mydgf** | 0.75282 | 6.99753 | 4.57448 | 7.96673 | 5.09191 | 4.50658 | 2.36957 | 8.26825 | 0.01154 | 1.43382 | 3.60468 | 3.78411 | 0.00000 | 0.79529 |
| **Myef2** | 1.50108 | 0.00863 | 3.36540 | 3.13628 | 1.91926 | 2.92247 | 0.28023 | 0.64804 | 5.03937 | 0.50571 | 2.86232 | 3.41999 | 2.86505 | 5.52528 |
| **Myeov2** | 0.35988 | 0.80556 | 0.00000 | 0.00000 | 0.48560 | 1.88621 | 0.73589 | 1.96200 | 0.42171 | 5.30589 | 0.00000 | 1.32701 | 5.42564 | 0.91375 |
| **Myg1** | 4.95485 | 0.05700 | 9.66328 | 2.09126 | 7.18676 | 3.66277 | 2.87758 | 2.60034 | 0.01482 | 0.06929 | 0.32698 | 1.49732 | 2.72922 | 5.00475 |
| **Myh10** | 6.44405 | 7.26368 | 14.83290 | 6.83388 | 7.66413 | 19.02761 | 8.49974 | 21.67231 | 3.98068 | 15.54374 | 13.58564 | 1.65378 | 0.73584 | 5.36607 |
| **Myh11** | 0.74781 | 0.00407 | 0.16954 | 0.37221 | 0.13715 | 0.00000 | 0.00400 | 0.00698 | 0.05680 | 0.00540 | 0.01108 | 0.01251 | 0.00551 | 1.22286 |
| **Myh13** | 0.00651 | 0.00393 | 0.00596 | 0.00000 | 0.01381 | 0.11440 | 0.01056 | 0.00501 | 0.01476 | 0.00606 | 0.23951 | 0.46638 | 0.00739 | 0.00817 |
| **Myh14** | 0.00000 | 0.01180 | 1.03174 | 0.69326 | 0.66871 | 0.40636 | 1.23639 | 0.00000 | 0.01904 | 0.06219 | 0.01064 | 0.00313 | 0.01342 | 0.08186 |
| **Myh15** | 0.00281 | 0.00433 | 0.34333 | 0.00462 | 0.00000 | 0.00163 | 0.00404 | 0.01136 | 0.07071 | 0.00677 | 0.00603 | 0.00766 | 0.02877 | 0.63077 |
| **Myh3** | 0.01566 | 0.00445 | 0.00971 | 0.01055 | 0.17697 | 0.01619 | 0.00453 | 0.00929 | 0.01172 | 0.00660 | 0.01187 | 0.01509 | 0.00572 | 0.01094 |
| **Myh6** | 0.00503 | 0.00437 | 0.82865 | 0.05411 | 0.78923 | 0.00073 | 0.00420 | 0.01397 | 0.01150 | 0.00691 | 0.00557 | 0.00618 | 0.00079 | 0.02187 |
| **Myh7** | 0.00000 | 0.00435 | 0.90476 | 0.16847 | 1.38466 | 0.00000 | 0.00413 | 0.01024 | 0.06222 | 0.00580 | 0.00842 | 0.00589 | 0.02603 | 0.03536 |
| **Myh7b** | 0.00889 | 0.02589 | 0.02027 | 0.00000 | 0.02528 | 0.04797 | 0.00371 | 1.07840 | 0.01191 | 0.03031 | 0.00000 | 1.30091 | 0.01805 | 0.00690 |

| | | | | | | | | | | | | | | |
|---|---|---|---|---|---|---|---|---|---|---|---|---|---|---|
| **Myh9** | 1.41467 | 0.82219 | 0.11766 | 1.40518 | 0.01422 | 0.81353 | 0.05277 | 3.65030 | 0.02825 | 0.10653 | 0.00738 | 0.29602 | 1.04553 | 0.32180 |
| **Myl12a** | 11.93471 | 13.84237 | 17.57160 | 15.67222 | 12.74840 | 27.42398 | 11.34022 | 23.70520 | 6.19339 | 22.14195 | 16.24241 | 16.81002 | 1.51216 | 11.08407 |
| **Myl12b** | 6.75954 | 7.77712 | 12.05947 | 11.36820 | 7.12817 | 23.45804 | 9.61668 | 23.18135 | 5.33203 | 20.67555 | 13.91054 | 9.22370 | 1.61589 | 5.31691 |
| **Myl2** | 0.01217 | 0.00433 | 0.04270 | 0.02477 | 0.02660 | 0.00907 | 0.00415 | 0.00908 | 0.00663 | 0.01325 | 0.00718 | 0.00865 | 0.15359 | 0.26415 |
| **Myl4** | 0.00481 | 0.00441 | 0.17244 | 0.06689 | 0.00459 | 0.14135 | 0.00421 | 0.00546 | 0.01305 | 0.00634 | 0.00589 | 0.00595 | 0.07517 | 0.04251 |
| **Myl6** | 9.53281 | 10.34996 | 6.38789 | 26.33122 | 21.71974 | 30.54132 | 10.60160 | 32.84075 | 9.93761 | 24.18943 | 7.11688 | 15.64454 | 4.36049 | 10.23761 |
| **Myl6b** | 0.03290 | 0.02303 | 0.03897 | 0.11294 | 0.07200 | 0.05061 | 0.06395 | 1.02246 | 0.04008 | 0.14739 | 0.01534 | 0.07642 | 0.01310 | 0.06189 |
| **Myl9** | 0.23893 | 8.88615 | 0.00000 | 0.00000 | 1.21637 | 0.75951 | 0.00333 | 0.32761 | 0.00641 | 0.00593 | 0.00000 | 0.49766 | 0.00000 | 0.00687 |
| **Mylip** | 0.00000 | 0.00318 | 0.35212 | 0.37497 | 0.00000 | 0.87104 | 0.00434 | 0.61789 | 0.03502 | 0.00776 | 1.01382 | 1.17402 | 1.24684 | 0.00086 |
| **Mylk** | 0.02717 | 0.00458 | 0.00437 | 0.00980 | 0.66899 | 0.01403 | 0.00450 | 0.02015 | 0.01732 | 0.00631 | 0.00844 | 0.01589 | 0.29444 | 0.01131 |
| **Mylk2** | 0.01069 | 0.00436 | 0.01070 | 0.04240 | 0.05899 | 0.01115 | 0.00408 | 0.00850 | 0.01154 | 0.00954 | 0.00790 | 0.00964 | 0.08933 | 0.00362 |
| **Mylk3** | 0.00733 | 0.00435 | 0.07256 | 0.00748 | 0.00802 | 0.01307 | 0.00422 | 0.00918 | 0.01103 | 0.00765 | 0.00815 | 0.00882 | 0.36603 | 0.00236 |
| **Mylk4** | 0.05118 | 0.02767 | 0.02501 | 0.00975 | 0.02658 | 0.03221 | 0.01555 | 0.05641 | 0.18890 | 0.00650 | 0.01895 | 0.02904 | 0.07223 | 0.04831 |
| **Mylpf** | 11.04462 | 0.24451 | 2.11811 | 8.44040 | 5.31064 | 17.75539 | 5.03877 | 12.35363 | 2.98584 | 15.75843 | 2.85034 | 4.06032 | 2.91712 | 2.31634 |
| **Mynn** | 1.38227 | 2.00901 | 2.90972 | 0.95142 | 4.00592 | 1.05237 | 2.16586 | 0.65436 | 6.60072 | 0.02152 | 4.01292 | 2.05987 | 3.03082 | 5.82664 |
| **Myo10** | 0.60045 | 0.00317 | 0.75656 | 0.50260 | 1.83106 | 1.50147 | 0.00332 | 0.20813 | 0.30578 | 0.01228 | 0.80059 | 0.82938 | 0.00000 | 4.91328 |
| **Myo15** | 0.00404 | 0.00428 | 0.17426 | 0.00000 | 0.29878 | 0.14545 | 0.00424 | 0.00380 | 0.00759 | 0.00530 | 0.00756 | 0.00998 | 0.36492 | 0.32266 |
| **Myo16** | 0.01620 | 0.00444 | 0.01421 | 0.01377 | 0.01326 | 0.01335 | 0.00428 | 0.01018 | 0.01491 | 0.00725 | 0.00860 | 0.00995 | 0.01447 | 0.01216 |
| **Myo18a** | 0.00000 | 0.03970 | 1.69827 | 0.35756 | 0.18630 | 0.81271 | 0.00933 | 2.08904 | 0.02912 | 0.00364 | 0.03891 | 1.19281 | 2.34366 | 1.17705 |
| **Myo19** | 1.83645 | 2.69741 | 1.27471 | 0.00000 | 0.00000 | 3.50752 | 0.00176 | 2.61782 | 0.02842 | 0.02595 | 2.16042 | 2.61713 | 2.06631 | 1.67093 |
| **Myo1a** | 0.00083 | 0.00315 | 0.00346 | 0.00000 | 0.34967 | 0.01272 | 0.00873 | 3.79453 | 0.08756 | 3.88864 | 0.00000 | 0.29588 | 0.11526 | 0.00000 |
| **Myo1b** | 10.81981 | 0.03035 | 15.97822 | 7.26181 | 16.91446 | 8.30468 | 0.82469 | 0.00000 | 17.76605 | 0.04661 | 6.13185 | 9.71411 | 7.11108 | 13.47368 |
| **Myo1c** | 5.33043 | 1.25595 | 12.91198 | 5.07162 | 5.39424 | 14.46874 | 2.50194 | 12.28362 | 0.61898 | 5.61666 | 2.07425 | 2.60989 | 2.96564 | 3.95065 |
| **Myo1d** | 0.01207 | 0.03465 | 1.08578 | 0.00000 | 0.00000 | 0.48226 | 0.01769 | 3.08413 | 0.01537 | 9.15481 | 0.00000 | 0.54029 | 0.42985 | 0.45257 |
| **Myo1e** | 0.01001 | 0.02517 | 0.00958 | 0.03681 | 0.21585 | 0.27096 | 0.01975 | 7.28791 | 0.09274 | 10.75352 | 0.00399 | 0.02935 | 0.21438 | 0.07198 |
| **Myo1g** | 0.00000 | 0.00407 | 0.47964 | 0.65116 | 0.59324 | 0.00000 | 0.00390 | 0.01149 | 0.00705 | 0.00723 | 0.01028 | 0.00808 | 0.20814 | 0.22075 |
| **Myo1h** | 0.00586 | 0.00533 | 0.08272 | 0.00000 | 0.02095 | 0.07852 | 0.00421 | 0.03154 | 0.01117 | 0.00580 | 0.00643 | 0.32273 | 0.00707 | 0.01023 |
| **Myo3b** | 0.00000 | 0.00432 | 0.00000 | 0.01346 | 0.14923 | 0.00669 | 0.00417 | 0.01014 | 1.24251 | 0.00676 | 0.00561 | 0.00799 | 0.47439 | 0.17108 |
| **Myo5a** | 0.40819 | 0.09926 | 0.84756 | 1.20876 | 1.87141 | 1.56915 | 0.00336 | 0.00000 | 0.84916 | 0.00737 | 1.59826 | 0.00746 | 0.79531 | 2.49887 |
| **Myo5b** | 0.27122 | 0.04018 | 1.17098 | 0.00000 | 0.00000 | 3.80208 | 0.84379 | 6.87606 | 0.16558 | 6.91057 | 0.36690 | 0.28130 | 1.36174 | 1.76198 |
| **Myo5c** | 0.00676 | 0.00457 | 0.00843 | 0.17713 | 0.01078 | 0.00945 | 0.00401 | 0.00874 | 0.02198 | 0.00686 | 0.00630 | 0.05763 | 0.01424 | 0.01557 |
| **Myo6** | 0.17017 | 0.01006 | 1.13546 | 3.13680 | 2.68855 | 0.68743 | 3.20095 | 0.21629 | 0.04258 | 0.03825 | 0.10234 | 1.45106 | 1.76899 | 1.26274 |
| **Myo7a** | 0.00000 | 0.00386 | 0.00004 | 0.76421 | 0.00000 | 0.48289 | 0.00388 | 0.00240 | 0.90528 | 0.01069 | 0.00073 | 0.24953 | 0.00393 | 0.43523 |
| **Myo7b** | 0.00424 | 0.00386 | 0.05725 | 0.24300 | 0.00000 | 0.00000 | 0.00387 | 0.48320 | 0.02261 | 0.01421 | 0.01875 | 0.02049 | 0.00490 | 0.28885 |
| **Myo9a** | 4.20192 | 0.03036 | 4.12275 | 0.00000 | 7.46121 | 0.65136 | 2.28522 | 1.26395 | 26.16825 | 0.03851 | 1.38594 | 1.54187 | 1.52364 | 7.05844 |
| **Myo9b** | 1.37174 | 1.77215 | 0.99237 | 1.99383 | 0.00001 | 2.07415 | 2.13614 | 1.90101 | 0.00000 | 0.06353 | 0.00000 | 1.67388 | 1.43737 | 1.49332 |
| **Myod1** | 0.00390 | 0.01818 | 0.01426 | 0.00390 | 0.01158 | 0.01496 | 0.00393 | 0.00556 | 0.01937 | 0.00662 | 0.00396 | 1.50518 | 0.00937 | 0.01526 |
| **Myof** | 0.08377 | 0.01159 | 0.00924 | 0.00000 | 0.01063 | 0.00404 | 0.00896 | 3.04081 | 0.01338 | 0.01674 | 0.00502 | 0.02211 | 0.00492 | 0.01182 |
| **Myom1** | 0.03918 | 0.00447 | 0.10592 | 0.00000 | 0.84872 | 0.00815 | 0.00412 | 0.01072 | 0.01338 | 0.00719 | 0.00667 | 0.00768 | 0.00806 | 0.16547 |
| **Myom2** | 0.00855 | 0.00421 | 0.00000 | 0.19096 | 0.00000 | 0.03671 | 0.00373 | 0.04456 | 1.01132 | 0.00676 | 0.00376 | 0.01044 | 0.20405 | 1.68322 |
| **Myom3** | 0.01290 | 0.00395 | 0.01624 | 0.50684 | 0.00410 | 0.01956 | 0.00386 | 0.00349 | 0.01121 | 0.00501 | 0.03735 | 0.01531 | 0.47065 | 0.02683 |
| **Myoz1** | 0.00637 | 0.00436 | 0.41628 | 0.00527 | 0.01757 | 0.05716 | 0.00419 | 0.00446 | 0.00924 | 0.00355 | 0.70101 | 0.00795 | 0.04714 | 0.02534 |
| **Myoz3** | 0.00941 | 0.01287 | 0.02562 | 0.09114 | 0.41408 | 0.01412 | 0.00450 | 0.14542 | 0.23630 | 0.02387 | 0.06865 | 0.04831 | 0.07238 | 0.22121 |
| **Mypn** | 4.62779 | 0.02624 | 4.83514 | 3.30559 | 2.25526 | 3.00321 | 0.49206 | 2.15114 | 1.92283 | 0.03652 | 4.96043 | 0.11662 | 0.49550 | 4.44412 |
| **Mypop** | 0.27597 | 0.02427 | 0.04744 | 0.28550 | 0.40349 | 0.28633 | 0.02040 | 1.09338 | 0.05496 | 0.01106 | 0.00226 | 0.00186 | 0.71685 | 0.08424 |
| **Myrf** | 0.11903 | 0.00445 | 0.03761 | 0.01353 | 0.03540 | 0.00970 | 0.00427 | 0.01105 | 0.03472 | 0.00625 | 0.00724 | 0.00921 | 0.06920 | 0.02525 |
| **Myrfl** | 0.75470 | 0.01063 | 0.46979 | 0.80005 | 0.88519 | 0.00000 | 0.00384 | 0.00234 | 0.08713 | 0.00613 | 0.02494 | 0.11393 | 1.99484 | 0.07243 |
| **Myrip** | 0.04421 | 0.00444 | 0.01336 | 0.01245 | 0.01205 | 0.01263 | 0.00427 | 0.01055 | 0.01196 | 0.00666 | 0.00822 | 0.00985 | 0.00498 | 0.05824 |
| **Mysm1** | 7.05893 | 0.05274 | 8.11247 | 2.39305 | 10.89149 | 1.52915 | 1.02025 | 0.65115 | 17.53252 | 0.30438 | 2.44005 | 2.75621 | 5.11232 | 17.75464 |
| **Myt1** | 0.01067 | 0.00458 | 0.01343 | 0.01227 | 0.00430 | 0.01105 | 0.00435 | 0.01321 | 0.01089 | 0.00707 | 0.00807 | 0.00967 | 0.05750 | 0.36392 |
| **Myt1l** | 0.00000 | 0.00365 | 0.24798 | 0.00000 | 0.15408 | 0.55038 | 0.00369 | 0.00076 | 0.01197 | 0.00452 | 0.28131 | 0.31911 | 0.45420 | 0.72608 |
| **Myzap** | 0.00656 | 0.00444 | 0.00534 | 0.29414 | 0.12934 | 0.01027 | 0.00435 | 0.01655 | 0.01077 | 0.00746 | 0.00832 | 0.01168 | 0.01009 | 0.02807 |
| **Mzb1** | 0.00970 | 0.00442 | 0.35214 | 0.00341 | 0.01971 | 0.00730 | 0.00429 | 0.01062 | 0.01021 | 0.00674 | 0.00819 | 0.00853 | 0.01555 | 0.01031 |
| **Mzf1** | 0.00629 | 0.00442 | 0.00580 | 0.11546 | 0.00984 | 0.02007 | 0.00429 | 0.01042 | 0.06503 | 0.00740 | 0.01293 | 0.00859 | 0.00666 | 0.04483 |
| **Mzt1** | 5.02832 | 2.50430 | 2.76291 | 4.10491 | 3.18351 | 3.81254 | 1.19457 | 2.43074 | 5.95562 | 0.03490 | 1.91388 | 2.46925 | 3.47760 | 3.13670 |

| | | | | | | | | | | | | | | |
|---|---|---|---|---|---|---|---|---|---|---|---|---|---|---|
| **Mzt2** | 0.82536 | 0.23651 | 1.59238 | 2.64181 | 3.29607 | 2.78509 | 1.95885 | 2.36954 | 0.05401 | 0.07341 | 0.93361 | 3.53408 | 0.45824 | 0.00000 |
| **N4bp1** | 4.01432 | 0.06194 | 7.58071 | 1.36433 | 8.62516 | 3.91032 | 0.59510 | 0.87481 | 2.36184 | 0.02615 | 5.24616 | 5.77146 | 4.82100 | 9.00413 |
| **N4bp2** | 0.00000 | 0.00511 | 0.10973 | 0.00000 | 0.00000 | 0.33468 | 0.00378 | 0.05100 | 1.52318 | 0.00413 | 0.57059 | 0.06844 | 2.45913 | 4.03150 |
| **N4bp2l1** | 0.01423 | 0.00446 | 0.18236 | 0.00000 | 0.31433 | 0.00656 | 0.00412 | 0.01113 | 0.12033 | 0.00695 | 0.00641 | 0.00874 | 0.00567 | 0.01139 |
| **N4bp2l2** | 7.30441 | 0.00366 | 10.19812 | 3.25301 | 12.36392 | 4.91147 | 1.57705 | 1.56777 | 24.23763 | 4.81558 | 8.29850 | 3.92451 | 9.30356 | 16.44011 |
| **N4bp3** | 0.05139 | 0.04156 | 1.38776 | 2.05249 | 0.93017 | 0.00000 | 0.95963 | 3.96176 | 0.01359 | 0.02074 | 0.00000 | 0.72483 | 0.02384 | 0.03076 |
| **N6amt1** | 3.10341 | 0.01800 | 1.14318 | 1.72208 | 1.53880 | 0.92441 | 0.01374 | 1.26053 | 1.66585 | 0.05471 | 0.34600 | 0.51943 | 1.88778 | 1.85851 |
| **Naa10** | 7.80205 | 2.15336 | 13.56597 | 9.60157 | 13.73199 | 12.95181 | 3.56597 | 9.31607 | 0.14598 | 9.52393 | 8.69143 | 8.08686 | 4.39621 | 1.57131 |
| **Naa11** | 1.56521 | 0.03364 | 2.99826 | 1.32272 | 0.10100 | 7.98364 | 3.02197 | 2.72836 | 0.00000 | 0.10331 | 1.84079 | 0.00000 | 0.79984 | 1.86456 |
| **Naa15** | 10.25780 | 0.11621 | 13.88158 | 6.55068 | 10.25728 | 12.01645 | 3.46552 | 4.11796 | 27.39267 | 0.12966 | 12.13885 | 6.58838 | 5.83215 | 9.50712 |
| **Naa16** | 11.78438 | 0.02222 | 8.25221 | 5.21304 | 7.13203 | 4.30777 | 0.00090 | 0.29762 | 5.13807 | 0.03763 | 4.49816 | 4.45649 | 2.18257 | 10.57302 |
| **Naa20** | 6.11518 | 0.00249 | 2.02250 | 0.00000 | 0.26074 | 1.68908 | 0.04768 | 0.38464 | 4.10545 | 0.03551 | 1.95355 | 0.72638 | 0.33814 | 2.40319 |
| **Naa25** | 3.62407 | 0.03684 | 9.75994 | 0.58731 | 7.68966 | 6.77499 | 0.82172 | 4.95870 | 9.65019 | 0.02993 | 10.23724 | 2.89024 | 1.57874 | 9.21762 |
| **Naa30** | 2.50714 | 0.00167 | 3.58497 | 1.06473 | 1.99303 | 2.33803 | 0.27335 | 1.43003 | 10.38448 | 0.00912 | 4.34565 | 0.76104 | 3.66733 | 1.51490 |
| **Naa35** | 14.45216 | 0.07781 | 18.62329 | 6.13173 | 13.62949 | 8.99944 | 1.93966 | 1.31243 | 15.64992 | 0.06677 | 4.72653 | 7.34206 | 3.97238 | 13.03846 |
| **Naa38** | 1.10261 | 0.08944 | 0.00000 | 3.02958 | 4.38506 | 1.93059 | 0.77711 | 6.73494 | 0.03653 | 9.85333 | 0.54349 | 4.06729 | 1.08358 | 0.23599 |
| **Naa40** | 4.52662 | 3.55855 | 6.59145 | 5.17388 | 4.65914 | 4.44072 | 0.05069 | 2.85517 | 0.35486 | 0.01372 | 7.51304 | 3.81826 | 3.13980 | 5.25007 |
| **Naa50** | 13.29119 | 0.55013 | 12.61970 | 11.91613 | 16.01397 | 15.29829 | 0.45160 | 4.64640 | 18.44136 | 1.96087 | 11.46007 | 7.95813 | 11.03982 | 5.66167 |
| **Naa60** | 4.32462 | 0.05613 | 4.49121 | 3.82177 | 5.07910 | 4.23027 | 0.03286 | 4.31047 | 0.07325 | 0.04779 | 0.12023 | 3.95633 | 1.60757 | 0.48724 |
| **Naaa** | 0.00000 | 0.04176 | 4.52293 | 7.10899 | 1.90104 | 4.11096 | 2.31301 | 4.06800 | 2.33956 | 0.09320 | 0.66989 | 0.01651 | 0.29794 | 3.25332 |
| **Naalad2** | 16.25400 | 0.00000 | 17.45767 | 9.48509 | 10.29915 | 8.48230 | 0.07928 | 0.00000 | 29.37984 | 0.00000 | 8.38811 | 0.00000 | 4.55300 | 14.42132 |
| **Naaladl1** | 0.00000 | 0.01200 | 0.13792 | 0.16940 | 0.00000 | 0.08430 | 0.01012 | 0.13639 | 0.09486 | 0.01875 | 0.01135 | 1.22772 | 0.00977 | 0.61726 |
| **Nab1** | 2.82127 | 0.27781 | 0.00000 | 0.00000 | 0.18050 | 0.47565 | 0.00374 | 0.58611 | 0.13021 | 0.00000 | 2.71305 | 0.51293 | 1.41831 | 0.00000 |
| **Nab2** | 0.03258 | 0.00441 | 0.03678 | 0.17595 | 0.00341 | 0.00900 | 1.09985 | 0.00439 | 0.04199 | 0.01835 | 0.00972 | 0.09187 | 0.11966 | 0.01662 |
| **Nabp1** | 3.26047 | 0.04630 | 7.12955 | 0.00000 | 9.38006 | 4.62306 | 0.95424 | 5.33137 | 17.67460 | 0.03610 | 3.86990 | 0.76408 | 13.10903 | 7.68071 |
| **Nabp2** | 0.66165 | 0.02666 | 1.78250 | 2.44840 | 0.37228 | 1.75365 | 3.55687 | 3.68508 | 0.00000 | 0.05783 | 2.02868 | 1.27173 | 0.42534 | 0.00000 |
| **Naca** | 32.90448 | 13.89070 | 38.95715 | 50.08497 | 40.85578 | 43.18534 | 13.97805 | 36.29512 | 25.21339 | 27.01442 | 28.04555 | 31.56074 | 12.85645 | 26.84624 |
| **Nacad** | 0.01149 | 0.00551 | 0.26021 | 0.55285 | 0.00494 | 0.00000 | 0.00447 | 0.32330 | 0.01124 | 0.00927 | 0.00542 | 0.02247 | 0.00964 | 0.03759 |
| **Nacc1** | 0.53004 | 0.05019 | 2.44683 | 3.09205 | 2.19979 | 1.58348 | 0.47419 | 2.91011 | 0.00570 | 0.04138 | 0.89611 | 3.16608 | 2.71328 | 0.38899 |
| **Nacc2** | 3.39298 | 0.01836 | 6.58665 | 4.22277 | 8.66801 | 2.82111 | 0.01557 | 1.41541 | 3.51270 | 0.05795 | 0.05657 | 0.34841 | 1.71125 | 8.15211 |
| **Nadk** | 12.06395 | 0.11168 | 15.49598 | 12.30542 | 16.54577 | 11.85794 | 5.30938 | 6.70004 | 6.31617 | 3.83189 | 8.72421 | 7.21455 | 5.99373 | 13.50538 |
| **Nadk2** | 0.27894 | 0.01027 | 0.79643 | 0.00000 | 1.13506 | 0.40126 | 0.00908 | 0.22510 | 0.00093 | 0.02109 | 0.00000 | 0.79313 | 2.88382 | 2.02326 |
| **Nadsyn1** | 0.01426 | 0.01221 | 0.33030 | 0.00000 | 0.19133 | 0.02113 | 0.00512 | 0.03249 | 0.68215 | 0.00635 | 0.00179 | 0.44263 | 0.00208 | 0.01135 |
| **Nae1** | 7.99000 | 0.08096 | 5.22208 | 1.72476 | 7.57271 | 8.13391 | 4.51175 | 2.79663 | 2.96362 | 0.12418 | 4.56608 | 6.47993 | 0.44316 | 7.03086 |
| **Naf1** | 8.88599 | 1.09090 | 0.37786 | 0.56786 | 3.02944 | 0.72249 | 0.01766 | 0.00000 | 5.35495 | 6.53482 | 0.00000 | 1.01512 | 3.78122 | 0.57470 |
| **Naga** | 0.00000 | 0.44439 | 0.89458 | 1.52065 | 0.00000 | 2.19203 | 0.84905 | 7.52175 | 0.01616 | 0.03691 | 3.23896 | 0.72518 | 0.00830 | 0.00000 |
| **Nagk** | 0.53616 | 7.42533 | 3.52291 | 9.49034 | 1.13793 | 5.30837 | 4.60513 | 12.86545 | 0.02299 | 12.98962 | 6.55533 | 1.75091 | 0.00000 | 1.22314 |
| **Naglu** | 0.04206 | 0.00431 | 0.47830 | 0.01668 | 0.01108 | 0.00423 | 0.00407 | 0.00998 | 0.01055 | 0.01554 | 0.00797 | 0.00681 | 0.06058 | 0.03341 |
| **Nagpa** | 1.20266 | 0.05399 | 3.00072 | 7.48753 | 0.07225 | 6.41045 | 0.03062 | 3.78888 | 0.00000 | 0.07645 | 2.11897 | 2.98696 | 1.35965 | 0.97183 |
| **Nags** | 0.22750 | 0.00410 | 0.61617 | 0.06167 | 0.61987 | 0.00000 | 0.00995 | 0.00685 | 0.00503 | 0.00691 | 0.00502 | 0.00757 | 0.46550 | 0.73050 |
| **Naif1** | 0.01134 | 0.00450 | 0.16055 | 0.15339 | 0.09067 | 0.06764 | 0.00430 | 0.00885 | 0.00894 | 0.00559 | 0.13612 | 0.01305 | 0.11236 | 0.07623 |
| **Naip2** | 0.02806 | 0.01178 | 0.02171 | 0.04067 | 0.03988 | 0.03912 | 0.01541 | 0.04503 | 0.13314 | 0.02134 | 0.00979 | 0.02641 | 0.16132 | 0.07191 |
| **Naip5** | 0.20241 | 0.00416 | 0.06322 | 0.07320 | 0.05303 | 0.32033 | 0.12249 | 0.02301 | 0.70446 | 0.07342 | 0.02510 | 0.12122 | 2.57216 | 0.44764 |
| **Nampt** | 4.35598 | 0.00775 | 2.72066 | 1.22544 | 5.73339 | 4.92935 | 0.01365 | 0.15135 | 8.42528 | 0.00919 | 0.30188 | 4.32206 | 8.41623 | 4.54624 |
| **Nanog** | 2.13965 | 0.35473 | 8.25547 | 15.06788 | 5.10184 | 20.53988 | 1.85798 | 0.59645 | 8.38551 | 1.94081 | 17.51973 | 13.92897 | 12.87755 | 5.21988 |
| **Nanos1** | 0.00000 | 0.00422 | 0.17225 | 1.87779 | 0.52565 | 0.00000 | 0.01026 | 0.11166 | 1.14505 | 0.03369 | 0.89893 | 0.39039 | 4.86845 | 0.00000 |
| **Nanos2** | 0.00388 | 0.00407 | 0.00607 | 0.58566 | 0.60663 | 0.00812 | 0.00401 | 0.00548 | 0.00836 | 0.00581 | 0.00631 | 0.00730 | 0.57944 | 0.00000 |
| **Nanos3** | 0.77223 | 0.00424 | 0.23857 | 0.00000 | 0.39435 | 0.00135 | 0.00400 | 0.01094 | 0.00643 | 0.00668 | 0.00582 | 0.00674 | 0.32315 | 0.27266 |
| **Nanp** | 0.32744 | 0.00428 | 0.42005 | 0.83829 | 0.00000 | 0.14497 | 0.00414 | 1.03315 | 0.08833 | 0.00180 | 1.88631 | 0.54702 | 0.14719 | 0.35052 |
| **Nans** | 0.00000 | 0.14388 | 1.67915 | 1.34866 | 3.24052 | 0.00000 | 0.00276 | 4.57505 | 0.01914 | 0.00545 | 1.47774 | 0.65106 | 1.10502 | 0.00000 |
| **Nap1l1** | 33.90638 | 1.85581 | 29.74269 | 23.09525 | 38.81290 | 28.05083 | 2.90090 | 15.03896 | 28.94405 | 7.85360 | 22.15893 | 22.43941 | 17.82215 | 28.93110 |
| **Nap1l2** | 0.01129 | 0.00480 | 0.00631 | 0.00511 | 0.32310 | 0.08329 | 0.00430 | 0.00875 | 0.01084 | 0.00597 | 0.00918 | 0.00835 | 0.09839 | 0.05894 |
| **Nap1l3** | 0.33575 | 0.00423 | 0.10844 | 0.09112 | 0.00000 | 0.06679 | 0.00398 | 0.00429 | 0.00903 | 0.00596 | 0.00633 | 0.02034 | 0.01608 | 0.02628 |
| **Nap1l4** | 2.21137 | 5.22936 | 11.83454 | 2.99671 | 11.45805 | 7.77717 | 0.02254 | 4.85780 | 4.29836 | 0.09768 | 5.82494 | 4.42670 | 11.23439 | 6.89114 |
| **Napa** | 5.57255 | 7.32468 | 14.17317 | 18.80321 | 4.56657 | 14.14760 | 4.07443 | 10.47938 | 0.73320 | 0.18955 | 10.40774 | 10.76425 | 2.83832 | 2.84361 |

| | | | | | | | | | | | | | | |
|---|---|---|---|---|---|---|---|---|---|---|---|---|---|---|
| **Napb** | 0.66176 | 0.01150 | 0.35672 | 0.52103 | 0.95794 | 0.00000 | 0.00371 | 0.00617 | 1.97651 | 0.00940 | 0.00312 | 0.00554 | 0.00553 | 0.17650 |
| **Napepld** | 0.22814 | 0.02370 | 0.67678 | 1.38647 | 2.78488 | 1.35865 | 0.18456 | 2.34898 | 0.01971 | 0.06739 | 0.15840 | 1.54958 | 0.60776 | 1.74448 |
| **Napg** | 8.34135 | 0.03995 | 4.88136 | 3.03196 | 3.75841 | 4.56785 | 5.05743 | 2.44579 | 6.77150 | 0.10301 | 2.64811 | 3.93914 | 2.51936 | 7.57301 |
| **Naprt** | 0.00000 | 0.02086 | 0.57950 | 0.82689 | 0.42868 | 0.09474 | 0.00416 | 0.01749 | 0.00877 | 0.01374 | 0.06688 | 1.24533 | 0.02745 | 0.00000 |
| **Napsa** | 0.00000 | 0.03106 | 1.71189 | 0.73152 | 0.27659 | 0.57832 | 0.00320 | 0.00000 | 0.13485 | 0.00876 | 0.43196 | 0.72482 | 0.00000 | 0.79615 |
| **Narf** | 0.00000 | 0.00396 | 0.88330 | 0.44162 | 0.00000 | 0.89373 | 0.00459 | 0.61160 | 0.00262 | 0.00550 | 0.00000 | 0.44939 | 0.98339 | 0.68327 |
| **Narfl** | 3.51484 | 5.83992 | 9.06073 | 7.77917 | 10.60113 | 11.93451 | 4.04822 | 5.67062 | 0.00000 | 0.22156 | 11.29883 | 5.90229 | 3.23812 | 5.01255 |
| **Nars** | 13.02934 | 5.18185 | 18.26565 | 16.47303 | 12.29656 | 21.11628 | 2.76389 | 12.09592 | 15.62574 | 0.24085 | 23.62372 | 16.48529 | 4.49975 | 19.53033 |
| **Nars2** | 0.27559 | 0.00307 | 0.88927 | 0.00000 | 0.14612 | 1.40230 | 0.01559 | 0.00372 | 0.02131 | 0.00185 | 0.77832 | 0.97303 | 0.45094 | 0.84653 |
| **Nasp** | 33.16424 | 6.07461 | 34.53603 | 25.68057 | 38.20811 | 26.26629 | 6.56547 | 7.98566 | 34.28573 | 15.91286 | 19.49875 | 17.31055 | 22.85722 | 37.66566 |
| **Nat1** | 0.00551 | 0.00414 | 0.00625 | 0.00991 | 0.00298 | 0.00887 | 0.00423 | 0.00735 | 0.62991 | 0.00665 | 0.00354 | 0.24564 | 0.01022 | 0.02049 |
| **Nat10** | 3.03237 | 0.04367 | 12.03899 | 12.14246 | 11.13127 | 5.97732 | 0.68069 | 2.43811 | 0.52160 | 0.04199 | 11.15816 | 3.61087 | 0.86780 | 9.40870 |
| **Nat14** | 0.02780 | 0.00431 | 0.02978 | 0.01369 | 0.37009 | 0.02347 | 0.01129 | 0.01077 | 0.05765 | 0.00883 | 0.00766 | 0.00878 | 0.19621 | 0.00000 |
| **Nat6** | 0.05406 | 0.01983 | 0.22997 | 0.91389 | 0.25124 | 1.25458 | 0.00391 | 1.47730 | 0.00958 | 0.02083 | 0.13765 | 0.12755 | 0.02012 | 0.00000 |
| **Nat8** | 0.01566 | 0.00400 | 0.01245 | 0.00000 | 0.01266 | 0.03542 | 0.00426 | 0.00602 | 0.01141 | 0.00616 | 0.00513 | 0.35779 | 0.05947 | 0.00960 |
| **Nat8f1** | 0.02719 | 0.00378 | 0.01364 | 0.00000 | 0.00795 | 0.35434 | 0.01946 | 0.03316 | 0.01197 | 0.01077 | 0.00376 | 0.90178 | 0.81375 | 0.00687 |
| **Nat8f2** | 0.11583 | 0.01158 | 0.50385 | 1.27045 | 0.42688 | 0.41465 | 0.05599 | 0.53321 | 1.18055 | 0.00524 | 0.00000 | 0.19994 | 3.21681 | 2.00252 |
| **Nat8f3** | 0.00655 | 0.00479 | 0.00880 | 0.01638 | 0.00962 | 0.00962 | 0.00424 | 0.01615 | 0.15025 | 0.00704 | 0.00717 | 0.01478 | 0.00583 | 0.12647 |
| **Nat8f4** | 0.01692 | 0.00442 | 0.02916 | 0.61003 | 0.24236 | 0.05490 | 0.24291 | 0.00821 | 0.09590 | 0.00612 | 0.00806 | 0.03487 | 0.33166 | 0.05079 |
| **Nat8f7** | 0.00902 | 0.00455 | 0.01319 | 0.01353 | 0.01593 | 0.01212 | 0.00424 | 0.01141 | 0.02839 | 0.00685 | 0.00818 | 0.01274 | 0.00396 | 0.07982 |
| **Nat8l** | 0.00992 | 0.00426 | 0.00711 | 0.35291 | 0.00763 | 0.14249 | 0.00420 | 0.00393 | 0.01159 | 0.00536 | 0.00497 | 0.05624 | 0.04757 | 0.00000 |
| **Nat9** | 0.29404 | 0.06330 | 1.43114 | 3.33668 | 2.00036 | 4.29634 | 0.02100 | 2.09414 | 0.00400 | 8.95015 | 6.64350 | 4.96074 | 0.15685 | 1.68161 |
| **Natd1** | 4.44865 | 0.42285 | 6.33644 | 2.20333 | 6.87178 | 9.64275 | 3.46473 | 6.35431 | 8.12314 | 0.15861 | 2.87116 | 2.39257 | 8.51964 | 7.83494 |
| **Nav1** | 0.55591 | 0.11267 | 3.48954 | 0.86693 | 7.60916 | 0.21988 | 0.08327 | 1.78239 | 10.23424 | 0.06138 | 1.27393 | 0.30215 | 3.99460 | 2.73833 |
| **Nav2** | 0.10362 | 0.00361 | 1.37271 | 0.23395 | 0.68924 | 1.02163 | 0.01422 | 0.78150 | 0.00736 | 0.01995 | 0.36095 | 0.00000 | 1.43723 | 1.57457 |
| **Nav3** | 0.00000 | 0.00414 | 0.35142 | 0.07380 | 0.26440 | 0.00892 | 0.00406 | 0.00719 | 1.22639 | 0.00615 | 0.01581 | 0.16637 | 0.31010 | 0.97699 |
| **Nbas** | 0.09541 | 0.01290 | 0.00000 | 1.50894 | 0.44857 | 0.84995 | 2.40313 | 0.09889 | 0.43143 | 0.03011 | 1.77920 | 0.92270 | 0.01641 | 1.55727 |
| **Nbea** | 0.03425 | 0.00437 | 0.00709 | 0.01262 | 0.07370 | 0.00938 | 0.00435 | 0.01182 | 0.05594 | 0.00629 | 0.00710 | 0.00922 | 0.00494 | 0.62079 |
| **Nbeal1** | 0.00714 | 0.00360 | 0.03203 | 0.00000 | 0.29078 | 0.21496 | 0.00357 | 0.00212 | 3.73834 | 0.01364 | 0.00000 | 0.29278 | 1.83064 | 0.95394 |
| **Nbeal2** | 0.00000 | 0.14921 | 2.51720 | 1.54480 | 1.76866 | 0.00000 | 1.48995 | 6.15485 | 0.00865 | 8.45406 | 0.05863 | 0.27656 | 0.00078 | 0.00000 |
| **Nbl1** | 0.02791 | 0.01220 | 0.00000 | 0.52454 | 0.61274 | 0.79013 | 0.00504 | 0.42207 | 0.01140 | 0.00555 | 0.08997 | 0.00566 | 0.00604 | 0.00000 |
| **Nbn** | 4.95555 | 0.00110 | 3.02644 | 1.20752 | 5.55144 | 0.18709 | 0.00108 | 0.00142 | 9.65583 | 0.00000 | 3.59809 | 1.60533 | 1.00852 | 3.75869 |
| **Nbr1** | 6.51869 | 0.89942 | 7.75102 | 4.57119 | 9.80302 | 10.10262 | 0.08612 | 3.22661 | 3.58898 | 0.03570 | 8.27875 | 5.11843 | 7.03729 | 7.53647 |
| **Ncald** | 0.00965 | 0.00436 | 0.01979 | 0.01180 | 0.03155 | 0.01338 | 0.00433 | 0.01061 | 0.00997 | 0.00674 | 0.00867 | 0.00978 | 0.15549 | 0.00434 |
| **Ncam1** | 0.33446 | 0.00433 | 0.01903 | 0.01477 | 0.02634 | 0.01021 | 0.00438 | 0.01057 | 0.03893 | 0.00649 | 0.00993 | 0.02857 | 0.01870 | 0.03084 |
| **Ncan** | 0.01392 | 0.02022 | 0.08570 | 0.39432 | 0.01525 | 0.33477 | 0.01631 | 0.03980 | 0.07877 | 0.02473 | 0.01531 | 0.00261 | 0.08185 | 0.09843 |
| **Ncapd2** | 2.97799 | 0.27719 | 11.70819 | 7.26037 | 7.50323 | 10.84306 | 3.67019 | 8.86342 | 4.84437 | 0.09989 | 9.38890 | 7.93106 | 6.47617 | 15.77224 |
| **Ncapd3** | 1.94704 | 0.00160 | 3.18827 | 0.00000 | 0.63852 | 0.43166 | 0.00863 | 0.00000 | 2.24528 | 0.00000 | 2.54041 | 2.43182 | 2.54593 | 4.92569 |
| **Ncapg** | 1.27696 | 0.08634 | 1.28935 | 1.05786 | 0.51793 | 3.23519 | 0.00653 | 0.69921 | 7.63922 | 0.01999 | 4.37205 | 3.71309 | 0.93071 | 0.00000 |
| **Ncapg2** | 1.64709 | 0.00206 | 2.52781 | 1.39985 | 1.38110 | 1.61045 | 0.00223 | 0.00000 | 5.93012 | 0.00686 | 2.26152 | 0.70065 | 3.30816 | 6.69112 |
| **Ncaph** | 1.47122 | 0.06195 | 7.33230 | 4.29159 | 3.22491 | 8.00330 | 0.00581 | 5.17259 | 0.49830 | 0.06216 | 8.68280 | 2.07515 | 1.39994 | 4.79848 |
| **Ncaph2** | 3.21770 | 0.22768 | 17.55818 | 17.82668 | 11.47975 | 14.30331 | 7.10665 | 12.58828 | 0.68898 | 0.80239 | 13.72272 | 13.77858 | 2.47574 | 10.86731 |
| **Ncbp1** | 10.12794 | 0.11930 | 19.42392 | 8.99788 | 19.24788 | 13.03741 | 3.72145 | 9.85630 | 21.88044 | 0.14423 | 12.31873 | 6.31507 | 11.59935 | 21.17473 |
| **Ncbp2** | 17.79238 | 0.22599 | 13.65347 | 10.89865 | 18.47045 | 16.86592 | 2.30761 | 9.34227 | 8.93201 | 9.80237 | 10.54216 | 10.19843 | 12.56183 | 21.51656 |
| **Ncbp3** | 5.17619 | 0.16653 | 2.97842 | 2.76812 | 6.56646 | 8.78311 | 2.53807 | 1.26197 | 9.78680 | 0.48206 | 4.05916 | 5.31825 | 10.18321 | 13.71750 |
| **Ncdn** | 0.59692 | 0.09270 | 2.16980 | 3.89223 | 2.10849 | 0.27793 | 1.91301 | 2.87410 | 0.00156 | 0.11495 | 0.00000 | 1.78617 | 0.21162 | 0.00000 |
| **Nceh1** | 0.07166 | 0.01030 | 0.38215 | 0.71820 | 0.35750 | 0.02115 | 0.00353 | 0.00000 | 0.00844 | 0.00247 | 0.99983 | 0.77394 | 0.95430 | 1.43604 |
| **Ncf1** | 0.00000 | 0.00427 | 0.48828 | 0.01486 | 0.57336 | 0.00000 | 0.01027 | 0.01434 | 0.04684 | 0.00664 | 0.02372 | 0.00940 | 0.86190 | 1.14963 |
| **Ncf2** | 0.12817 | 0.01886 | 0.27303 | 1.08837 | 0.00000 | 0.38244 | 0.00312 | 4.50623 | 0.00568 | 0.00617 | 0.00000 | 0.26936 | 0.02746 | 0.11580 |
| **Nck1** | 1.33864 | 0.01714 | 1.13625 | 1.53009 | 2.89900 | 2.93305 | 0.18976 | 1.20363 | 1.92516 | 0.01894 | 2.23289 | 0.04401 | 0.66083 | 1.61008 |
| **Nck2** | 0.07914 | 0.01211 | 0.00138 | 0.00000 | 0.00747 | 0.04428 | 0.00353 | 1.05104 | 0.62466 | 0.02035 | 0.05523 | 0.03962 | 0.12458 | 0.00000 |
| **Nckap1** | 13.75089 | 0.07362 | 8.46534 | 4.93460 | 10.78770 | 12.70389 | 2.33036 | 4.00266 | 16.64724 | 0.07451 | 8.86051 | 9.29605 | 8.87996 | 16.23405 |
| **Nckap1l** | 0.01669 | 0.00487 | 0.00802 | 0.19649 | 0.04011 | 0.01797 | 0.00414 | 0.00976 | 0.01013 | 0.00517 | 0.00483 | 0.23561 | 0.12906 | 0.32793 |
| **Nckap5** | 0.00312 | 0.00447 | 0.00000 | 0.01686 | 0.00000 | 0.00777 | 0.00408 | 0.00973 | 0.84714 | 0.00713 | 0.00621 | 0.00950 | 0.00380 | 0.19517 |
| **Nckap5l** | 0.09218 | 0.00439 | 0.14333 | 0.01711 | 0.00533 | 0.01081 | 0.00421 | 0.01038 | 0.01018 | 0.01268 | 0.00711 | 0.01017 | 0.17404 | 0.00229 |

| | | | | | | | | | | | | | | |
|---|---|---|---|---|---|---|---|---|---|---|---|---|---|---|
| **Nckipsd** | 0.00000 | 0.03922 | 1.03528 | 0.00000 | 0.72532 | 1.13407 | 1.82466 | 3.30640 | 0.00542 | 0.02400 | 0.00000 | 2.12780 | 0.00209 | 0.05298 |
| **Ncl** | 32.24519 | 6.23439 | 42.57798 | 34.54661 | 31.80863 | 33.59442 | 12.03222 | 18.29113 | 29.65325 | 11.58371 | 26.24138 | 26.65038 | 16.28117 | 39.74305 |
| **Ncln** | 0.00000 | 0.15726 | 10.17366 | 8.07715 | 3.41015 | 0.00000 | 3.14774 | 8.43146 | 0.00000 | 0.09696 | 1.26077 | 7.07602 | 0.24757 | 0.00000 |
| **Ncmap** | 0.11914 | 0.00384 | 2.13992 | 0.39425 | 0.00000 | 0.00000 | 0.00361 | 0.01402 | 0.00000 | 0.00358 | 0.06876 | 0.00366 | 1.95888 | 0.24311 |
| **Ncoa1** | 1.62171 | 0.01892 | 1.79020 | 0.54499 | 0.08806 | 4.08821 | 2.99545 | 2.47070 | 1.19777 | 0.06170 | 0.00000 | 1.37133 | 2.71977 | 0.46123 |
| **Ncoa2** | 0.99275 | 0.00280 | 0.15903 | 0.93982 | 0.67194 | 1.04703 | 0.61350 | 1.59166 | 0.92455 | 0.02678 | 0.00000 | 1.27332 | 1.55137 | 1.55987 |
| **Ncoa3** | 20.12077 | 4.15533 | 19.94249 | 10.01313 | 19.63359 | 11.93727 | 3.21785 | 1.85078 | 9.88082 | 0.37484 | 5.74341 | 8.07272 | 13.47847 | 18.54407 |
| **Ncoa4** | 9.09379 | 0.04067 | 12.70514 | 3.28085 | 8.10787 | 13.78072 | 3.25421 | 6.03480 | 9.59485 | 0.10523 | 11.86187 | 3.74417 | 5.55697 | 6.42187 |
| **Ncoa5** | 1.05557 | 0.01074 | 0.18497 | 1.25049 | 0.99213 | 1.09785 | 0.06487 | 0.55096 | 0.43671 | 1.02687 | 0.51561 | 1.68734 | 0.38994 | 1.51835 |
| **Ncoa6** | 5.39795 | 0.08285 | 2.56075 | 0.00000 | 8.00067 | 6.11763 | 0.08857 | 1.06891 | 8.90588 | 0.00000 | 3.95434 | 5.43617 | 7.43449 | 12.54283 |
| **Ncoa7** | 0.00949 | 0.08374 | 0.01674 | 0.00942 | 0.02360 | 0.04364 | 0.00420 | 0.00415 | 0.02160 | 0.00543 | 0.00612 | 0.11401 | 0.26997 | 0.02927 |
| **Ncor1** | 2.54707 | 4.14430 | 6.85920 | 0.00000 | 4.05967 | 10.11769 | 2.20058 | 7.18547 | 8.76504 | 0.09562 | 9.38787 | 6.45199 | 2.53101 | 8.46169 |
| **Ncor2** | 0.14194 | 0.03178 | 0.72084 | 1.77842 | 0.32784 | 0.21767 | 0.00288 | 0.68015 | 0.00800 | 0.00954 | 0.06361 | 2.99624 | 0.00658 | 0.00764 |
| **Ncs1** | 0.09687 | 0.00405 | 0.25720 | 0.38459 | 0.61960 | 0.00000 | 0.00376 | 0.00597 | 0.02061 | 0.00666 | 0.00463 | 0.01339 | 0.01220 | 0.68468 |
| **Ncstn** | 0.91322 | 0.00144 | 2.45053 | 2.61329 | 0.59797 | 1.06108 | 0.00730 | 3.07522 | 7.22477 | 0.00000 | 3.02028 | 2.53394 | 0.49702 | 3.37620 |
| **Ndc1** | 4.17104 | 0.04698 | 6.22200 | 6.09663 | 2.21088 | 5.25160 | 0.99165 | 2.19749 | 9.02828 | 0.04224 | 5.97611 | 5.91903 | 2.11101 | 12.98615 |
| **Ndc80** | 10.22031 | 0.05296 | 11.84453 | 7.05726 | 13.31132 | 5.16552 | 2.60430 | 1.78176 | 14.15257 | 16.12877 | 7.29776 | 3.36579 | 5.24133 | 17.13347 |
| **Nde1** | 0.45587 | 0.01962 | 2.37429 | 3.47958 | 1.22899 | 4.22981 | 0.01021 | 2.26667 | 0.24523 | 0.01427 | 0.08827 | 0.66940 | 1.22970 | 3.40068 |
| **Ndel1** | 5.81654 | 0.03742 | 1.96809 | 7.05336 | 7.49127 | 0.87470 | 1.39905 | 1.55365 | 0.09023 | 0.04437 | 0.81513 | 4.91216 | 6.80686 | 0.64348 |
| **Ndfip1** | 1.69158 | 0.02206 | 1.27571 | 1.31548 | 1.08984 | 2.71443 | 2.66038 | 0.67392 | 0.79568 | 0.16399 | 2.40017 | 3.87170 | 3.40018 | 0.78958 |
| **Ndfip2** | 2.45437 | 0.01654 | 1.17948 | 0.22658 | 0.41928 | 3.75133 | 2.15153 | 0.03136 | 0.02493 | 0.05847 | 1.56453 | 1.85608 | 2.98562 | 0.00712 |
| **Ndnl2** | 3.90528 | 0.01757 | 2.96979 | 2.54515 | 7.60035 | 1.91574 | 0.20188 | 1.79189 | 0.08597 | 1.49961 | 2.38109 | 1.34206 | 4.68426 | 6.87290 |
| **Ndor1** | 9.29191 | 8.59182 | 9.32511 | 4.35749 | 8.10547 | 2.77909 | 0.93078 | 3.00082 | 3.74471 | 0.05280 | 2.02760 | 2.24766 | 4.51800 | 3.60157 |
| **Ndrg1** | 1.31430 | 10.46879 | 0.00000 | 0.00000 | 0.00474 | 2.73764 | 4.28621 | 18.39794 | 0.00649 | 4.98336 | 0.00000 | 0.22029 | 0.70055 | 0.00000 |
| **Ndrg2** | 0.00000 | 0.00498 | 0.18178 | 1.09300 | 0.01807 | 0.58765 | 0.00427 | 0.04893 | 0.01283 | 0.00482 | 0.45009 | 0.00158 | 0.01047 | 0.00000 |
| **Ndrg3** | 6.11084 | 3.82042 | 8.97858 | 7.07270 | 6.35220 | 12.07285 | 4.37929 | 2.23136 | 10.78778 | 0.18029 | 7.80286 | 8.17458 | 0.98790 | 6.76445 |
| **Ndrg4** | 0.13144 | 0.00443 | 0.12315 | 0.00524 | 0.00537 | 0.00611 | 0.00432 | 0.01153 | 0.25293 | 0.00689 | 0.00611 | 0.01153 | 0.00465 | 0.12127 |
| **Ndst1** | 0.18890 | 0.00358 | 1.94349 | 0.00000 | 2.96991 | 1.70230 | 0.00306 | 2.57354 | 1.75137 | 0.00340 | 0.00000 | 0.66190 | 0.15122 | 5.42247 |
| **Ndst2** | 0.03628 | 0.00392 | 0.29630 | 0.00564 | 0.01700 | 0.04297 | 0.00435 | 0.00518 | 0.03785 | 0.00921 | 0.91649 | 0.32715 | 0.02634 | 0.18354 |
| **Ndufa1** | 0.40368 | 1.61627 | 0.01039 | 0.00000 | 0.67030 | 2.51819 | 0.36403 | 3.37650 | 0.30822 | 0.86264 | 0.00000 | 2.66622 | 3.20393 | 0.53601 |
| **Ndufa10** | 8.20368 | 1.71331 | 5.55044 | 12.74248 | 5.23019 | 8.96520 | 1.12018 | 6.42577 | 2.09999 | 6.94476 | 2.56651 | 9.19527 | 0.25369 | 4.50962 |
| **Ndufa11** | 3.19668 | 3.74143 | 0.94679 | 5.12262 | 5.79357 | 5.17368 | 4.04640 | 14.11071 | 0.00000 | 12.99023 | 0.89989 | 7.45621 | 2.21003 | 4.67639 |
| **Ndufa12** | 1.19176 | 2.72347 | 0.40449 | 4.22320 | 7.94353 | 6.69363 | 3.88589 | 6.37377 | 0.20468 | 4.32676 | 0.87917 | 7.74036 | 0.78473 | 0.32467 |
| **Ndufa13** | 1.51726 | 3.86221 | 0.07673 | 2.09175 | 4.08983 | 6.02392 | 4.39596 | 11.51421 | 0.00827 | 15.87229 | 0.23959 | 6.66856 | 0.25449 | 1.82867 |
| **Ndufa2** | 0.57532 | 3.38073 | 0.00000 | 0.51370 | 2.09141 | 2.85473 | 0.68488 | 5.80936 | 0.08879 | 12.21658 | 0.18602 | 3.26730 | 5.63408 | 0.16940 |
| **Ndufa3** | 1.38246 | 4.60303 | 0.27284 | 2.34491 | 1.71898 | 2.80960 | 0.15531 | 1.97831 | 0.25757 | 9.08006 | 0.47318 | 1.21627 | 1.45436 | 1.33714 |
| **Ndufa4** | 2.13547 | 4.96716 | 0.00000 | 0.95834 | 4.69989 | 12.24817 | 2.39937 | 11.48843 | 4.74993 | 14.82797 | 0.74624 | 10.03712 | 1.51570 | 2.94302 |
| **Ndufa5** | 2.62167 | 2.97570 | 0.00000 | 0.32920 | 1.23878 | 7.43960 | 0.68822 | 8.04343 | 0.62817 | 2.65493 | 0.01246 | 6.39820 | 0.21902 | 0.95361 |
| **Ndufa6** | 0.07245 | 0.10786 | 0.00000 | 0.81373 | 3.83317 | 2.25458 | 3.21193 | 8.63596 | 0.37207 | 4.49098 | 0.00540 | 7.50868 | 1.64390 | 0.00000 |
| **Ndufa7** | 2.68707 | 2.94141 | 0.14516 | 0.54369 | 4.41572 | 6.74963 | 3.08934 | 6.77764 | 0.42709 | 10.23336 | 0.51630 | 8.40460 | 3.80983 | 1.91825 |
| **Ndufa8** | 1.72500 | 0.29129 | 4.15642 | 10.86803 | 6.69671 | 10.24970 | 4.35356 | 14.70150 | 0.15960 | 14.30611 | 6.43292 | 7.62031 | 0.00000 | 1.73972 |
| **Ndufa9** | 10.98386 | 0.32314 | 15.19605 | 22.35201 | 9.36625 | 16.61289 | 7.69667 | 20.66538 | 10.17213 | 0.55552 | 14.61593 | 11.10795 | 3.26755 | 11.64036 |
| **Ndufab1** | 0.64283 | 0.04370 | 0.08563 | 1.69629 | 4.17447 | 1.65243 | 2.19970 | 1.80952 | 0.40159 | 0.17294 | 0.04639 | 3.52778 | 4.64926 | 0.00725 |
| **Ndufaf1** | 2.96080 | 0.01292 | 3.27529 | 0.54651 | 8.02400 | 5.40423 | 1.87861 | 2.74391 | 2.48986 | 0.05973 | 8.45394 | 3.39233 | 1.40435 | 5.72178 |
| **Ndufaf2** | 1.96298 | 0.02334 | 0.29126 | 3.80608 | 2.58437 | 2.10462 | 0.66460 | 2.59179 | 1.30054 | 9.91333 | 0.61526 | 3.63384 | 3.83988 | 3.63716 |
| **Ndufaf3** | 0.00000 | 0.32245 | 0.35362 | 1.65259 | 0.08324 | 1.49162 | 0.00234 | 1.92136 | 0.01403 | 7.48545 | 0.64471 | 2.06618 | 0.00525 | 0.00000 |
| **Ndufaf4** | 9.19574 | 2.41798 | 4.89554 | 4.08047 | 3.71369 | 4.37018 | 2.24144 | 2.33142 | 8.35167 | 10.48757 | 2.92296 | 6.86705 | 2.42918 | 7.50673 |
| **Ndufaf5** | 3.37585 | 0.01743 | 4.05696 | 0.23652 | 5.42502 | 1.83716 | 0.02809 | 1.38893 | 2.42475 | 0.01055 | 1.27393 | 2.82916 | 2.88960 | 1.27974 |
| **Ndufaf6** | 1.41363 | 0.00272 | 0.40854 | 0.54515 | 1.36311 | 0.06296 | 0.00346 | 0.00320 | 0.04678 | 0.01378 | 0.11515 | 1.40444 | 0.86679 | 0.03122 |
| **Ndufaf7** | 8.22085 | 6.43235 | 9.53066 | 10.56470 | 7.48589 | 7.79217 | 0.13727 | 2.78807 | 6.60757 | 0.23436 | 11.47747 | 3.28199 | 5.82340 | 11.49438 |
| **Ndufb10** | 4.78564 | 1.46517 | 2.78812 | 8.68914 | 11.91006 | 10.03650 | 4.75206 | 14.38890 | 0.40343 | 11.54841 | 4.73791 | 12.81376 | 3.94217 | 4.13691 |
| **Ndufb11** | 7.44137 | 5.47234 | 7.57361 | 11.51612 | 17.01817 | 11.63927 | 5.22264 | 13.66768 | 0.49939 | 16.31416 | 1.10437 | 9.65432 | 5.62025 | 5.18456 |
| **Ndufb2** | 0.02745 | 0.03505 | 0.00000 | 1.05108 | 3.59486 | 1.04393 | 0.16316 | 1.09829 | 0.01077 | 1.75889 | 0.00000 | 1.23113 | 2.83481 | 0.00000 |
| **Ndufb3** | 3.53437 | 0.21034 | 0.04655 | 1.22492 | 3.24224 | 4.88005 | 2.01810 | 7.26378 | 1.08589 | 10.53406 | 0.58529 | 7.06761 | 2.38882 | 2.24488 |
| **Ndufb4** | 2.07246 | 1.96542 | 0.00000 | 0.13928 | 2.51267 | 3.47693 | 2.53813 | 6.07068 | 0.52534 | 9.43856 | 0.00000 | 5.20704 | 1.10508 | 0.37340 |

| | | | | | | | | | | | | | | |
|---|---|---|---|---|---|---|---|---|---|---|---|---|---|---|
| **Ndufb5** | 4.25556 | 5.70685 | 1.94147 | 10.30495 | 6.41361 | 8.46104 | 6.26767 | 15.83605 | 0.38616 | 11.35934 | 3.20647 | 13.26413 | 0.35593 | 0.78265 |
| **Ndufb6** | 2.07040 | 4.58075 | 0.13528 | 4.29158 | 8.13789 | 6.92234 | 4.90992 | 9.76599 | 0.05217 | 12.40034 | 0.82523 | 10.07138 | 4.92407 | 0.09109 |
| **Ndufb7** | 2.68744 | 4.15847 | 0.13530 | 2.09865 | 5.95042 | 3.79851 | 0.37351 | 7.11442 | 0.00805 | 9.61895 | 0.30106 | 4.98529 | 1.07290 | 0.78781 |
| **Ndufb8** | 2.22166 | 11.96969 | 1.07577 | 11.30787 | 7.21807 | 8.71214 | 4.22722 | 16.39935 | 0.38717 | 17.68992 | 2.75393 | 11.36979 | 0.40725 | 1.70825 |
| **Ndufb9** | 1.96259 | 6.26256 | 1.56732 | 11.32337 | 8.62070 | 9.11423 | 7.48662 | 21.31336 | 0.59459 | 12.08021 | 2.98544 | 17.87833 | 3.11159 | 0.05594 |
| **Ndufc1** | 0.96194 | 0.22728 | 0.00000 | 2.04377 | 3.61993 | 3.14757 | 1.41430 | 8.56190 | 0.04070 | 3.72020 | 0.00490 | 5.94991 | 2.24340 | 0.00000 |
| **Ndufc2** | 1.00762 | 0.14099 | 0.18200 | 5.69306 | 3.89764 | 6.15141 | 1.60080 | 10.06330 | 0.01715 | 3.99443 | 1.21595 | 7.95355 | 0.05969 | 0.02015 |
| **Ndufs1** | 12.39204 | 2.49370 | 13.90671 | 8.48344 | 11.81979 | 12.72099 | 5.53817 | 2.80933 | 18.36952 | 0.71383 | 11.25363 | 8.75367 | 3.31628 | 8.61265 |
| **Ndufs2** | 8.38163 | 3.28301 | 9.70802 | 14.63594 | 4.63971 | 13.46532 | 4.57284 | 13.18522 | 2.54274 | 5.04720 | 7.73763 | 12.69087 | 2.54517 | 2.23164 |
| **Ndufs3** | 8.80159 | 6.49853 | 13.43435 | 23.95567 | 14.40158 | 7.91009 | 6.56701 | 17.93528 | 2.19947 | 0.60459 | 5.13021 | 12.68974 | 1.08709 | 1.42203 |
| **Ndufs4** | 1.79567 | 0.18843 | 0.00000 | 2.39883 | 0.57186 | 2.80192 | 3.83164 | 6.45773 | 1.01945 | 9.08303 | 1.18015 | 8.98202 | 0.48348 | 1.58750 |
| **Ndufs6** | 1.89822 | 3.73099 | 0.09665 | 8.97359 | 6.07397 | 7.61551 | 0.74452 | 15.55595 | 0.30979 | 11.34363 | 0.78163 | 8.45911 | 0.11360 | 1.34029 |
| **Ndufs7** | 1.30060 | 8.66275 | 8.62000 | 15.35106 | 8.68627 | 9.98290 | 2.11232 | 10.54739 | 0.00000 | 11.72181 | 4.45291 | 10.26021 | 0.54835 | 0.00000 |
| **Ndufs8** | 11.35217 | 0.22982 | 6.13527 | 15.82376 | 12.33050 | 11.40703 | 5.79574 | 9.86618 | 0.00000 | 17.06398 | 6.15914 | 9.19908 | 2.56377 | 5.07093 |
| **Ndufv1** | 4.50248 | 10.87174 | 13.56138 | 18.70010 | 9.04861 | 14.00437 | 8.19276 | 15.31830 | 0.26044 | 0.70638 | 14.04159 | 11.60536 | 0.00000 | 3.43299 |
| **Ndufv2** | 3.77542 | 1.19296 | 3.87943 | 10.33165 | 8.30139 | 6.95982 | 5.56531 | 14.15605 | 0.36718 | 0.95625 | 4.14974 | 14.43269 | 4.38948 | 2.69940 |
| **Ndufv3** | 1.20846 | 3.03309 | 0.15666 | 0.78435 | 2.88850 | 3.21909 | 0.37122 | 7.24487 | 0.40182 | 4.18344 | 0.13142 | 2.90034 | 2.20626 | 1.24832 |
| **Neat1** | 8.74471 | 12.07614 | 3.20553 | 9.29290 | 0.00000 | 4.16333 | 6.42619 | 11.62078 | 1.25248 | 0.26762 | 0.88626 | 2.32180 | 0.57559 | 2.56782 |
| **Neb** | 0.02593 | 0.01181 | 0.05938 | 0.00440 | 0.01968 | 0.00523 | 0.00408 | 0.00602 | 0.00675 | 0.08338 | 0.00538 | 0.14591 | 0.02750 | 0.72235 |
| **Necab2** | 0.33560 | 0.00426 | 0.07635 | 0.62367 | 0.00000 | 0.79745 | 0.05153 | 0.00344 | 0.01878 | 0.02254 | 0.00289 | 0.00586 | 1.08561 | 0.00000 |
| **Necab3** | 0.98519 | 0.00411 | 0.38826 | 1.77376 | 0.34313 | 0.04275 | 0.00376 | 0.71305 | 0.00694 | 0.00942 | 0.33267 | 0.03221 | 0.00235 | 0.00000 |
| **Necap1** | 9.61486 | 3.08200 | 15.04344 | 5.27315 | 11.44260 | 10.21430 | 5.66433 | 1.91728 | 8.82061 | 0.21046 | 8.79493 | 7.67072 | 4.34231 | 13.74611 |
| **Necap2** | 3.75571 | 4.02740 | 6.01313 | 5.07752 | 11.38697 | 4.94082 | 3.23848 | 15.66870 | 0.00000 | 0.91860 | 1.42531 | 2.89323 | 2.95783 | 0.00000 |
| **Nectin1** | 0.18532 | 0.02696 | 0.39471 | 0.19680 | 0.00000 | 0.20216 | 0.56675 | 1.44452 | 0.00681 | 0.10426 | 1.59992 | 2.22489 | 0.19972 | 0.23133 |
| **Nectin2** | 0.58461 | 0.09342 | 7.20361 | 9.62314 | 0.93419 | 6.36429 | 0.12264 | 10.82923 | 0.73901 | 0.09670 | 1.06559 | 4.01827 | 2.01394 | 0.00000 |
| **Nectin3** | 0.00005 | 0.00176 | 0.00000 | 0.36637 | 0.84462 | 0.29289 | 0.13578 | 0.00212 | 5.03679 | 0.00551 | 0.00000 | 2.45131 | 4.44217 | 0.37189 |
| **Nedd1** | 8.56783 | 0.01658 | 3.60623 | 0.38765 | 5.59822 | 1.33756 | 1.07709 | 0.25601 | 16.49220 | 0.02684 | 1.91392 | 1.15853 | 1.55835 | 11.71095 |
| **Nedd4** | 2.01733 | 0.04459 | 0.00000 | 1.03839 | 2.20314 | 1.56585 | 0.10228 | 0.56730 | 0.83244 | 0.02911 | 3.81082 | 3.45156 | 5.13656 | 0.75103 |
| **Nedd4l** | 0.65681 | 0.13455 | 18.22347 | 5.55086 | 10.34618 | 0.44751 | 5.47230 | 10.15901 | 0.13544 | 13.77120 | 6.04937 | 4.57690 | 0.01424 | 0.39339 |
| **Nedd8** | 8.41380 | 0.92763 | 3.15040 | 13.57952 | 12.41021 | 13.76509 | 2.42977 | 12.41701 | 1.88510 | 15.25446 | 4.79239 | 12.57632 | 8.51964 | 5.62101 |
| **Nedd9** | 0.00386 | 0.00407 | 0.11954 | 0.03709 | 0.00316 | 0.00591 | 0.00410 | 0.00782 | 0.00240 | 0.00603 | 0.00751 | 0.01092 | 1.11564 | 0.36653 |
| **Nefh** | 0.00000 | 0.00367 | 0.30955 | 0.80837 | 0.00000 | 0.20646 | 0.00366 | 0.00000 | 0.01532 | 0.00000 | 1.86024 | 0.01149 | 0.00299 | 1.60054 |
| **Nefl** | 0.04376 | 0.00428 | 0.15416 | 0.18349 | 0.00275 | 0.11917 | 0.00421 | 0.00639 | 0.01191 | 0.00605 | 0.00480 | 0.00909 | 0.00551 | 0.00904 |
| **Nefm** | 0.00368 | 0.00456 | 0.00429 | 0.24130 | 0.32501 | 0.01000 | 0.00402 | 0.01870 | 0.01159 | 0.00522 | 0.00480 | 0.34957 | 0.00500 | 0.00646 |
| **Neil1** | 0.02961 | 0.00446 | 1.48498 | 0.46308 | 0.00000 | 0.53595 | 0.00388 | 0.26915 | 0.18158 | 0.01070 | 0.14227 | 1.12432 | 0.00000 | 0.06158 |
| **Neil2** | 0.00837 | 0.00431 | 0.12706 | 0.01196 | 0.01425 | 0.01069 | 0.00410 | 0.00916 | 0.02029 | 0.01004 | 0.00769 | 0.00769 | 0.50312 | 0.00660 |
| **Neil3** | 0.01274 | 0.00345 | 0.19499 | 0.00000 | 0.50180 | 0.24997 | 0.00366 | 0.74749 | 0.01480 | 0.01989 | 0.00000 | 0.63990 | 0.26478 | 0.49280 |
| **Nek1** | 0.70724 | 0.00357 | 1.39280 | 0.14796 | 0.49391 | 0.61712 | 0.00398 | 0.28193 | 3.94872 | 0.01763 | 0.00439 | 0.72098 | 2.71376 | 1.45683 |
| **Nek11** | 0.00283 | 0.00424 | 0.24293 | 0.17452 | 0.00000 | 0.60781 | 0.00398 | 0.00281 | 0.00977 | 0.00542 | 0.04749 | 0.00917 | 0.00513 | 0.47161 |
| **Nek2** | 12.10013 | 1.43809 | 20.21077 | 5.72920 | 14.17950 | 8.87151 | 0.19925 | 2.25862 | 6.90086 | 0.04849 | 6.45268 | 4.92126 | 16.84422 | 13.85419 |
| **Nek3** | 0.59820 | 3.52544 | 0.00112 | 1.11432 | 0.86107 | 0.04541 | 0.00287 | 0.00246 | 0.07570 | 0.00675 | 0.00677 | 0.92204 | 0.08247 | 1.50905 |
| **Nek4** | 0.36066 | 0.06221 | 0.00000 | 0.27469 | 0.13880 | 0.33266 | 0.02054 | 0.22538 | 1.83711 | 0.02481 | 1.57291 | 0.78854 | 1.64220 | 0.58445 |
| **Nek5** | 0.00462 | 0.00419 | 0.00000 | 0.14748 | 0.36695 | 0.00702 | 0.00417 | 0.00609 | 0.01964 | 0.00553 | 0.00640 | 0.01329 | 0.23755 | 0.35279 |
| **Nek6** | 0.07280 | 4.20154 | 0.40723 | 4.24983 | 0.00000 | 11.82179 | 4.70570 | 4.43936 | 0.15700 | 0.10782 | 0.98765 | 0.00000 | 0.00527 | 0.02458 |
| **Nek7** | 0.39222 | 0.19712 | 0.05423 | 0.69851 | 0.38991 | 1.59517 | 0.63584 | 0.32857 | 0.39310 | 0.23806 | 0.66045 | 0.37990 | 0.99534 | 1.02219 |
| **Nek8** | 1.08766 | 4.43038 | 2.63844 | 1.94770 | 1.65156 | 0.74774 | 0.00292 | 0.93676 | 0.00000 | 0.00188 | 0.00000 | 0.23292 | 1.39040 | 2.12943 |
| **Nek9** | 3.31122 | 2.62834 | 3.61601 | 2.29571 | 2.02051 | 2.66480 | 0.69097 | 2.58247 | 0.25976 | 0.04564 | 5.06867 | 4.77646 | 1.49468 | 2.66371 |
| **Nelfa** | 1.66189 | 0.04600 | 10.51552 | 5.73452 | 8.88590 | 5.34101 | 1.70897 | 2.75579 | 6.00653 | 0.01656 | 1.81437 | 6.04127 | 6.80781 | 10.56850 |
| **Nelfb** | 1.24294 | 0.36853 | 8.03262 | 6.40226 | 6.67334 | 11.67162 | 4.07143 | 7.83752 | 2.69035 | 0.19758 | 6.97219 | 6.61599 | 5.27573 | 7.81830 |
| **Nelfcd** | 5.54867 | 0.07689 | 13.28014 | 12.32443 | 14.77526 | 3.92121 | 4.15983 | 3.41173 | 1.29548 | 0.17368 | 5.97793 | 5.87967 | 5.51842 | 8.52406 |
| **Nelfe** | 8.13275 | 0.07674 | 13.00973 | 14.13108 | 15.50872 | 5.46113 | 3.13493 | 5.04369 | 0.99534 | 0.13437 | 5.52956 | 5.27706 | 8.07862 | 8.23339 |
| **Nell1** | 0.00311 | 0.00378 | 0.01423 | 0.00001 | 0.00670 | 0.04107 | 0.00406 | 0.15865 | 0.03140 | 0.00529 | 0.00726 | 0.01142 | 1.17324 | 0.00265 |
| **Nemf** | 4.62060 | 0.02446 | 5.92356 | 0.00000 | 4.99113 | 2.83650 | 0.84607 | 2.35676 | 13.14253 | 0.02631 | 4.39060 | 2.96605 | 3.60105 | 5.43731 |
| **Nemp1** | 7.78007 | 0.00036 | 0.91877 | 1.01545 | 9.40336 | 2.87005 | 0.00148 | 0.87326 | 12.23354 | 0.00971 | 0.00000 | 1.74097 | 8.45476 | 17.69760 |
| **Nemp2** | 0.00642 | 0.00488 | 0.01590 | 0.00487 | 0.01281 | 0.00936 | 0.01027 | 0.01605 | 0.24977 | 0.01390 | 0.00433 | 0.16977 | 0.00677 | 0.02900 |

| | | | | | | | | | | | | | | |
|---|---|---|---|---|---|---|---|---|---|---|---|---|---|---|
| **Nenf** | 0.05665 | 0.00556 | 0.00917 | 0.00000 | 0.06230 | 0.00434 | 0.00442 | 0.26545 | 0.00562 | 0.00730 | 0.00430 | 0.18841 | 0.63828 | 0.44310 |
| **Neo1** | 0.28462 | 0.02301 | 0.57874 | 0.12665 | 0.00000 | 3.00566 | 0.00382 | 2.55392 | 0.59107 | 0.04698 | 0.00548 | 0.08275 | 0.80435 | 0.00000 |
| **Nepro** | 2.39021 | 0.02440 | 4.57404 | 1.96754 | 7.13233 | 1.04832 | 3.16780 | 1.15280 | 4.99321 | 0.01898 | 0.69484 | 1.60915 | 7.93077 | 8.21292 |
| **Nes** | 0.17259 | 0.00370 | 0.00086 | 0.00000 | 0.00000 | 0.14947 | 0.00971 | 0.00374 | 0.25224 | 0.00517 | 0.00309 | 0.43010 | 0.43501 | 0.00000 |
| **Nespas** | 0.71448 | 0.00377 | 0.01152 | 0.55131 | 0.15441 | 0.06231 | 0.00367 | 0.00474 | 0.00415 | 0.00618 | 0.00321 | 0.62455 | 0.00347 | 0.25728 |
| **Net1** | 0.70446 | 0.02173 | 5.32096 | 0.72885 | 2.47864 | 2.25780 | 0.43706 | 3.18957 | 3.00883 | 14.23007 | 0.00000 | 3.80537 | 0.63779 | 1.70728 |
| **Neto2** | 1.05215 | 0.00298 | 0.79161 | 0.01992 | 2.34965 | 2.25857 | 0.00790 | 0.00000 | 11.14696 | 0.00724 | 1.44235 | 0.00000 | 2.83931 | 3.93670 |
| **Neu1** | 4.01311 | 0.08609 | 2.30218 | 9.38628 | 3.71543 | 14.51965 | 9.22374 | 16.70245 | 1.48531 | 0.47968 | 4.01896 | 0.00000 | 0.00000 | 2.78999 |
| **Neu3** | 0.28427 | 0.15536 | 0.04298 | 0.36935 | 0.20874 | 0.96061 | 0.10806 | 0.17259 | 0.73433 | 0.14620 | 0.16936 | 0.34827 | 3.08133 | 0.66717 |
| **Neurl1a** | 0.01031 | 0.00432 | 0.35885 | 0.23585 | 0.03378 | 0.20356 | 0.00428 | 0.00270 | 0.01247 | 0.00456 | 0.07742 | 0.00841 | 0.00601 | 0.00933 |
| **Neurl2** | 0.17545 | 0.02412 | 1.19788 | 0.32760 | 1.22541 | 0.04587 | 0.00376 | 0.00627 | 0.00743 | 0.01959 | 0.01589 | 0.09310 | 0.16227 | 0.09012 |
| **Neurl3** | 0.04558 | 0.00436 | 0.10562 | 0.03733 | 0.05740 | 0.01912 | 0.01154 | 0.02717 | 0.22124 | 0.00899 | 0.00669 | 0.00831 | 0.43524 | 0.12182 |
| **Neurl4** | 0.38328 | 2.03913 | 7.44345 | 6.74242 | 1.59869 | 7.41621 | 0.00730 | 3.58356 | 0.00000 | 0.05919 | 8.58588 | 2.21639 | 0.60069 | 1.53762 |
| **Nexn** | 2.34224 | 0.14546 | 1.26207 | 0.00000 | 3.51248 | 0.30016 | 0.00344 | 0.47458 | 9.35051 | 0.00443 | 0.07480 | 0.00000 | 1.98215 | 7.19985 |
| **Nf1** | 0.66930 | 0.01099 | 0.36002 | 0.29469 | 1.15345 | 0.39364 | 0.00369 | 0.00148 | 1.07908 | 0.00578 | 0.00326 | 0.08137 | 2.07438 | 0.63791 |
| **Nf2** | 6.98348 | 0.27244 | 4.48386 | 4.92088 | 15.58552 | 8.22232 | 0.00660 | 4.38916 | 9.40066 | 0.18663 | 3.68830 | 0.81220 | 9.51349 | 15.82005 |
| **Nfat5** | 1.16322 | 0.00262 | 0.94439 | 0.00000 | 0.93802 | 0.44106 | 0.36784 | 0.20474 | 0.28508 | 0.00478 | 0.00000 | 1.80258 | 0.93065 | 0.24221 |
| **Nfatc1** | 1.15870 | 0.00393 | 0.05215 | 0.18415 | 0.00000 | 0.46912 | 0.00379 | 0.00409 | 0.00027 | 0.01037 | 0.00354 | 0.00272 | 2.18241 | 0.00000 |
| **Nfatc2** | 0.00000 | 0.00411 | 0.06937 | 0.81263 | 0.64696 | 0.00000 | 0.00405 | 0.00682 | 0.01129 | 0.00602 | 0.00693 | 0.00690 | 0.08755 | 0.15499 |
| **Nfatc2ip** | 4.05192 | 0.05782 | 2.95099 | 6.06023 | 5.99487 | 5.01089 | 0.68350 | 1.99232 | 7.27510 | 0.13256 | 1.93974 | 3.29508 | 4.66889 | 1.88528 |
| **Nfatc3** | 7.78218 | 2.22736 | 5.21659 | 2.33651 | 5.75599 | 6.33548 | 0.50868 | 3.19384 | 4.90272 | 0.02641 | 4.90392 | 2.47697 | 7.74259 | 10.80919 |
| **Nfatc4** | 0.01235 | 0.04118 | 0.00866 | 0.03965 | 0.11750 | 0.01166 | 0.00974 | 0.01493 | 0.01802 | 0.01788 | 0.00325 | 3.57469 | 0.00465 | 0.01268 |
| **Nfe2** | 0.00000 | 0.01213 | 0.80446 | 0.00000 | 0.84806 | 0.27579 | 0.00399 | 0.23024 | 0.00765 | 0.00466 | 0.00000 | 0.82256 | 1.75593 | 0.19906 |
| **Nfe2l1** | 2.46115 | 0.29465 | 12.22333 | 9.31980 | 5.13523 | 9.24912 | 0.47264 | 8.18506 | 0.36340 | 0.08993 | 5.83029 | 5.73417 | 3.48548 | 10.97170 |
| **Nfe2l2** | 0.30500 | 0.02248 | 1.75205 | 2.89298 | 0.50696 | 6.69833 | 2.31667 | 4.77306 | 0.00000 | 0.04180 | 10.95553 | 2.27184 | 2.02283 | 2.47245 |
| **Nfe2l3** | 0.00000 | 0.00508 | 0.19407 | 0.20370 | 0.00000 | 0.03094 | 0.00400 | 0.01663 | 0.04087 | 0.00868 | 0.00729 | 0.00506 | 0.58177 | 0.00000 |
| **Nfib** | 1.04242 | 0.00416 | 0.00872 | 0.18317 | 0.48832 | 0.00383 | 0.00451 | 0.00677 | 0.01584 | 0.00681 | 0.00638 | 0.01082 | 0.00430 | 0.00000 |
| **Nfic** | 0.09614 | 0.00416 | 0.77937 | 0.41945 | 0.13057 | 0.05704 | 0.35123 | 0.00431 | 0.02933 | 0.03678 | 0.00432 | 0.04526 | 0.01988 | 0.00000 |
| **Nfil3** | 0.79474 | 0.00386 | 1.45308 | 0.16583 | 0.33896 | 0.56527 | 0.00360 | 0.00000 | 0.08259 | 0.00531 | 0.00313 | 0.00461 | 1.96419 | 1.07817 |
| **Nfix** | 0.07804 | 0.00420 | 0.31200 | 0.00000 | 0.92562 | 0.00632 | 0.00434 | 0.02174 | 0.02327 | 0.00564 | 0.00556 | 0.00751 | 0.00094 | 0.01138 |
| **Nfkb1** | 0.03373 | 0.00434 | 0.01031 | 0.00000 | 0.21650 | 0.02661 | 0.00522 | 0.48802 | 0.00982 | 0.00547 | 0.00805 | 0.01033 | 0.61205 | 0.01724 |
| **Nfkb2** | 0.00727 | 0.00395 | 0.01355 | 1.42328 | 0.29881 | 0.00000 | 0.00383 | 0.03552 | 0.00660 | 0.00588 | 0.00702 | 0.21223 | 0.23177 | 0.02414 |
| **Nfkbia** | 0.01860 | 0.01089 | 0.65501 | 0.00000 | 2.74610 | 0.07397 | 0.01957 | 0.06768 | 0.38968 | 0.00446 | 0.02973 | 0.65448 | 1.08502 | 0.29352 |
| **Nfkbib** | 0.00000 | 2.12838 | 6.86538 | 7.07262 | 7.14195 | 0.00000 | 0.73505 | 5.85046 | 0.03738 | 0.06581 | 0.00000 | 3.38040 | 0.52911 | 0.00000 |
| **Nfkbie** | 0.00345 | 0.01208 | 0.00519 | 0.64491 | 0.00150 | 0.00000 | 0.00428 | 0.01388 | 0.08145 | 0.00680 | 0.02168 | 0.01016 | 0.00553 | 0.02088 |
| **Nfkbil1** | 0.09745 | 0.00431 | 0.04023 | 0.24140 | 1.40844 | 0.06697 | 0.00466 | 0.88234 | 0.00634 | 0.01355 | 0.01681 | 0.00725 | 0.19941 | 0.00000 |
| **Nfkbiz** | 0.33823 | 0.01458 | 0.00000 | 0.74991 | 0.41734 | 0.00000 | 0.00397 | 1.20557 | 0.85532 | 0.00382 | 0.92271 | 0.00744 | 0.06245 | 0.00000 |
| **Nfrkb** | 1.92183 | 7.12651 | 1.60615 | 1.66131 | 3.08606 | 1.15737 | 0.30646 | 0.54160 | 0.59357 | 0.00977 | 0.53101 | 0.46984 | 3.21580 | 3.31613 |
| **Nfs1** | 2.57091 | 0.03896 | 7.18719 | 5.76133 | 4.53898 | 5.47581 | 0.00103 | 1.56911 | 2.35080 | 0.02360 | 4.04378 | 4.15875 | 4.84589 | 3.64365 |
| **Nfu1** | 7.44982 | 10.94514 | 5.66189 | 8.55464 | 6.97638 | 9.60739 | 2.98235 | 5.05514 | 0.76503 | 0.17447 | 8.32092 | 15.22832 | 3.31487 | 3.58545 |
| **Nfx1** | 8.08636 | 0.07954 | 12.57427 | 6.73518 | 7.98275 | 10.69249 | 3.40356 | 6.24633 | 5.14716 | 0.06003 | 8.86055 | 8.72092 | 4.92021 | 9.24133 |
| **Nfxl1** | 0.22947 | 0.02648 | 0.12132 | 2.08484 | 1.49771 | 1.44094 | 0.46737 | 0.36085 | 1.52289 | 0.01563 | 1.25195 | 2.21265 | 1.10159 | 7.34877 |
| **Nfya** | 2.53816 | 0.00410 | 1.47558 | 2.16455 | 0.40067 | 0.70493 | 0.00387 | 0.02567 | 0.12024 | 0.01189 | 0.62348 | 0.03045 | 3.63939 | 1.03035 |
| **Nfyb** | 13.24410 | 0.05229 | 10.06403 | 8.18294 | 13.19911 | 6.40042 | 0.00000 | 1.54471 | 10.48063 | 0.05591 | 3.38591 | 6.88429 | 11.66235 | 19.02532 |
| **Nfyc** | 9.25644 | 1.96740 | 26.10794 | 10.88713 | 22.22916 | 12.90729 | 5.26155 | 9.87855 | 5.40032 | 11.68981 | 14.45084 | 9.38120 | 4.91005 | 17.54843 |
| **Ngdn** | 8.59584 | 7.87579 | 26.35108 | 17.18008 | 15.20891 | 16.12397 | 6.88014 | 11.17371 | 3.94351 | 13.61365 | 14.46982 | 12.53857 | 4.86874 | 15.46430 |
| **Ngef** | 0.15336 | 0.00395 | 0.01504 | 0.55051 | 0.43411 | 0.00000 | 0.00382 | 0.00628 | 0.01054 | 0.00540 | 0.01878 | 0.01475 | 0.52824 | 0.11301 |
| **Ngfr** | 0.02878 | 0.00433 | 0.02129 | 0.03297 | 0.02838 | 0.02167 | 0.00420 | 0.01275 | 0.02669 | 0.00715 | 0.01053 | 0.01850 | 0.03648 | 0.02527 |
| **Ngfrap1** | 0.19854 | 0.93154 | 0.44280 | 1.96149 | 0.76341 | 1.93036 | 0.01629 | 2.42360 | 0.00616 | 0.02718 | 0.00000 | 0.06216 | 0.00192 | 0.05766 |
| **Ngly1** | 0.43972 | 0.02550 | 0.33602 | 0.19340 | 0.63431 | 0.74600 | 0.07428 | 1.33614 | 0.01074 | 0.00413 | 0.00000 | 2.38852 | 0.63952 | 1.09745 |
| **Ngrn** | 3.03240 | 0.03320 | 5.44284 | 10.31569 | 6.12970 | 6.56233 | 0.00828 | 3.04244 | 0.26179 | 0.08251 | 3.31585 | 4.00128 | 4.97758 | 0.00000 |
| **Nhej1** | 3.36834 | 0.02139 | 3.54755 | 1.23768 | 2.61117 | 2.06946 | 0.00929 | 1.78904 | 0.86237 | 0.02257 | 0.07241 | 0.06007 | 0.26605 | 1.47251 |
| **Nhlrc2** | 0.00000 | 0.02404 | 1.43030 | 1.18443 | 2.09844 | 0.00000 | 3.66862 | 1.42892 | 0.02204 | 0.05602 | 0.64131 | 2.91425 | 0.96336 | 1.02828 |
| **Nhlrc3** | 0.01786 | 0.00360 | 0.00873 | 0.00000 | 0.01276 | 0.00000 | 0.00354 | 0.73192 | 0.01344 | 0.01408 | 0.22361 | 0.36212 | 0.01151 | 0.01190 |
| **Nhlrc4** | 0.01251 | 0.00407 | 0.01007 | 0.00432 | 0.15935 | 0.00727 | 0.00413 | 0.60286 | 0.02033 | 0.00651 | 0.00730 | 0.01045 | 0.00529 | 0.01532 |

| | | | | | | | | | | | | | | |
|---|---|---|---|---|---|---|---|---|---|---|---|---|---|---|
| **Nhp2** | 14.72098 | 8.13029 | 21.23224 | 39.00794 | 35.07230 | 19.44537 | 6.74986 | 18.75611 | 1.14019 | 16.56046 | 9.78537 | 15.67079 | 16.74026 | 5.79422 |
| **Nhp2l1** | 27.88168 | 9.51520 | 33.10859 | 28.45272 | 39.11818 | 28.51915 | 9.94703 | 15.17360 | 25.71833 | 3.09649 | 16.91771 | 19.93416 | 22.39915 | 39.25283 |
| **Nhsl1** | 0.00404 | 0.01146 | 0.08426 | 0.31962 | 0.70040 | 0.32712 | 1.04169 | 0.38491 | 0.01440 | 0.06088 | 0.51835 | 0.01047 | 1.31843 | 0.77967 |
| **Nhsl2** | 0.01433 | 0.01416 | 0.01077 | 0.03165 | 0.02486 | 0.05976 | 0.01704 | 0.02703 | 0.06522 | 0.00811 | 0.01218 | 0.02379 | 0.01667 | 0.01029 |
| **Nicn1** | 0.21801 | 0.01822 | 2.99755 | 3.06266 | 4.83026 | 1.40961 | 1.78485 | 2.16802 | 3.13644 | 0.03093 | 0.01885 | 2.37867 | 1.84160 | 4.28386 |
| **Nid1** | 6.29554 | 0.02329 | 9.77298 | 2.68154 | 1.56112 | 1.47908 | 0.00542 | 3.79624 | 6.54663 | 0.03796 | 5.07347 | 4.15092 | 0.29780 | 1.75583 |
| **Nid2** | 0.00000 | 0.03533 | 2.00778 | 2.18342 | 0.73830 | 2.97617 | 0.00297 | 2.60742 | 2.17197 | 0.01600 | 0.00000 | 0.00000 | 0.49940 | 0.99670 |
| **Nif3l1** | 8.38316 | 1.07644 | 9.59549 | 2.49353 | 5.95997 | 10.83656 | 0.00000 | 1.88341 | 6.37940 | 0.02562 | 4.46593 | 5.60365 | 0.92213 | 9.72539 |
| **Nifk** | 17.97352 | 2.54238 | 25.11334 | 11.71783 | 21.49672 | 22.73393 | 6.89831 | 8.59175 | 20.38795 | 0.36531 | 14.51506 | 12.93031 | 12.57130 | 23.00733 |
| **Nim1k** | 0.01378 | 0.00430 | 0.07478 | 0.00360 | 0.00735 | 0.02118 | 0.00421 | 0.00799 | 0.01049 | 0.00634 | 0.00824 | 0.00937 | 0.47150 | 0.00452 |
| **Nin** | 17.36548 | 0.00000 | 13.86686 | 2.57748 | 22.92571 | 0.15492 | 0.44628 | 0.03514 | 14.69716 | 0.00000 | 4.00700 | 0.00777 | 12.27996 | 30.31815 |
| **Ninj1** | 0.00000 | 6.54651 | 0.50619 | 0.14578 | 1.40410 | 0.21325 | 0.00881 | 3.16128 | 0.00453 | 0.02699 | 0.00007 | 0.07069 | 0.94065 | 0.11151 |
| **Ninj2** | 0.00323 | 0.00518 | 0.01219 | 0.00000 | 0.02588 | 0.17790 | 0.00438 | 0.02294 | 0.01051 | 0.00677 | 0.00794 | 0.01428 | 0.70258 | 0.01945 |
| **Ninl** | 0.63828 | 0.00307 | 2.62770 | 0.51732 | 1.56716 | 1.79102 | 0.00337 | 0.00000 | 0.07722 | 0.00000 | 1.32343 | 0.20368 | 2.81458 | 3.60709 |
| **Nip7** | 13.53866 | 2.08630 | 22.28815 | 15.63490 | 20.08240 | 17.30073 | 6.72156 | 6.59918 | 17.52458 | 2.55521 | 10.69640 | 12.19758 | 18.94847 | 23.89948 |
| **Nipa2** | 1.66215 | 0.03436 | 2.46045 | 1.48643 | 4.40706 | 3.39763 | 0.38883 | 0.83292 | 11.52776 | 1.00024 | 1.79902 | 1.68798 | 6.98600 | 4.47854 |
| **Nipal1** | 6.73793 | 0.00761 | 12.82911 | 3.98995 | 14.38246 | 0.00000 | 0.47795 | 2.38989 | 11.42756 | 0.00000 | 2.11164 | 1.45474 | 3.91714 | 24.54413 |
| **Nipal3** | 0.06210 | 0.00370 | 1.53676 | 0.28522 | 1.23678 | 0.00000 | 0.00344 | 0.09690 | 0.40231 | 0.00371 | 0.99944 | 0.00565 | 0.46162 | 0.00000 |
| **Nipbl** | 2.75785 | 0.03453 | 6.88018 | 0.43725 | 10.83895 | 5.72530 | 0.00000 | 1.40884 | 13.77015 | 0.00955 | 6.97469 | 3.01559 | 3.32160 | 12.09087 |
| **Nipsnap1** | 0.16827 | 0.06652 | 1.14350 | 0.41640 | 1.90823 | 0.00000 | 4.82338 | 3.38338 | 0.81364 | 0.11537 | 0.00000 | 0.19175 | 1.69140 | 5.04602 |
| **Nipsnap3b** | 1.96818 | 0.06025 | 1.76174 | 5.62597 | 3.83760 | 5.98809 | 0.33808 | 5.30909 | 1.52409 | 0.04292 | 4.69221 | 4.41849 | 0.00000 | 2.32710 |
| **Nisch** | 0.26596 | 0.00809 | 0.63065 | 1.57156 | 2.22975 | 3.57786 | 1.21624 | 1.19930 | 0.00105 | 0.03887 | 0.00000 | 4.56935 | 1.61045 | 0.33309 |
| **Nit1** | 0.58273 | 4.56755 | 3.01733 | 0.64880 | 1.40216 | 3.55562 | 0.10925 | 0.70982 | 0.15167 | 0.02806 | 3.82719 | 3.66209 | 0.06297 | 2.39893 |
| **Nit2** | 2.14657 | 0.53189 | 1.91856 | 2.48158 | 0.95136 | 4.08962 | 3.01270 | 1.44162 | 1.40226 | 0.11202 | 1.82414 | 3.31529 | 0.24130 | 0.63479 |
| **Nkain1** | 0.00000 | 0.02487 | 1.43229 | 0.89603 | 0.00000 | 0.83263 | 0.01313 | 0.00317 | 0.00117 | 0.04349 | 0.00000 | 3.12890 | 2.58463 | 0.25983 |
| **Nkain3** | 0.01593 | 0.00443 | 0.02319 | 0.02408 | 0.02886 | 0.02685 | 0.01076 | 0.02116 | 0.04177 | 0.00757 | 0.02789 | 0.04379 | 0.07396 | 0.04085 |
| **Nkap** | 6.19877 | 0.02287 | 5.31784 | 2.25065 | 7.34906 | 4.04287 | 1.40772 | 1.27924 | 12.53904 | 0.02218 | 2.13266 | 1.96802 | 0.00000 | 1.93817 |
| **Nkapl** | 0.00453 | 0.00489 | 0.01256 | 0.01226 | 0.08583 | 0.01287 | 0.00444 | 0.02235 | 0.01935 | 0.00665 | 0.00866 | 0.00873 | 0.21350 | 0.04490 |
| **Nkd1** | 0.00622 | 0.01225 | 0.04480 | 0.05972 | 0.20038 | 0.01883 | 0.00430 | 0.00805 | 0.00581 | 0.00551 | 0.01274 | 0.00908 | 0.38618 | 0.09732 |
| **Nkd2** | 0.02905 | 0.00447 | 0.08364 | 0.00631 | 0.02204 | 0.00649 | 0.00412 | 0.01124 | 0.45391 | 0.00752 | 0.00618 | 0.00898 | 0.01066 | 0.01303 |
| **Nkiras1** | 4.96407 | 0.14418 | 5.67588 | 5.37701 | 11.49071 | 4.08428 | 0.10594 | 1.83646 | 4.35174 | 0.05220 | 0.01841 | 1.08992 | 6.37243 | 7.36973 |
| **Nkiras2** | 0.00000 | 0.01934 | 3.19242 | 0.00000 | 1.70567 | 2.51473 | 0.00345 | 1.05858 | 0.00472 | 0.01723 | 1.07609 | 0.61990 | 0.00000 | 0.46054 |
| **Nkpd1** | 0.00711 | 0.00441 | 0.02543 | 0.11999 | 0.02051 | 0.01569 | 0.00405 | 0.00597 | 0.02561 | 0.01181 | 0.00952 | 0.01712 | 0.04667 | 0.00786 |
| **Nkrf** | 1.58997 | 0.00306 | 1.85379 | 0.00000 | 4.97385 | 0.21097 | 0.00216 | 1.10058 | 6.84185 | 0.00601 | 0.65445 | 0.46682 | 0.20411 | 5.81250 |
| **Nktr** | 5.81708 | 0.03426 | 5.12304 | 3.73806 | 5.92560 | 2.55592 | 1.99189 | 1.25654 | 8.06316 | 0.03550 | 0.83442 | 2.78536 | 7.75445 | 8.63316 |
| **Nkx2.3** | 0.00739 | 0.00439 | 0.14861 | 0.00847 | 0.33150 | 0.00679 | 0.00412 | 0.01052 | 0.00601 | 0.00837 | 0.00727 | 0.00828 | 0.15319 | 0.02455 |
| **Nkx6.2** | 0.00234 | 0.00232 | 0.01410 | 0.87046 | 0.33966 | 0.00000 | 0.00333 | 0.00262 | 0.01590 | 0.00498 | 0.00727 | 2.22114 | 0.00565 | 0.00042 |
| **Nle1** | 2.71146 | 0.97366 | 5.83024 | 6.05016 | 6.17262 | 1.28868 | 1.78794 | 0.00000 | 0.10204 | 0.05696 | 3.65105 | 1.46918 | 3.22034 | 2.88029 |
| **Nlgn1** | 0.00909 | 0.00439 | 0.01336 | 0.01128 | 0.01116 | 0.01320 | 0.00433 | 0.00959 | 0.00997 | 0.00686 | 0.00815 | 0.02170 | 0.01229 | 0.12160 |
| **Nlgn2** | 0.02971 | 0.00410 | 0.00565 | 0.34535 | 0.00867 | 0.00550 | 0.00399 | 0.00592 | 0.01442 | 0.00559 | 0.00672 | 0.01879 | 0.00639 | 0.00704 |
| **Nlk** | 0.65681 | 0.00328 | 1.67396 | 1.34248 | 2.31901 | 4.51341 | 0.00353 | 1.45701 | 3.37825 | 0.00000 | 0.98315 | 0.90637 | 2.15691 | 2.05276 |
| **Nln** | 1.74818 | 0.04102 | 3.95432 | 3.04092 | 1.55585 | 2.03090 | 0.59810 | 3.61244 | 0.67723 | 0.05972 | 4.04323 | 3.12627 | 0.87045 | 1.84086 |
| **Nlrc3** | 0.52726 | 0.00432 | 0.29571 | 0.00000 | 0.08003 | 0.52145 | 0.00397 | 0.00380 | 0.28901 | 0.01748 | 0.26362 | 0.00896 | 0.66381 | 0.02666 |
| **Nlrc4** | 0.00000 | 0.00404 | 0.21855 | 0.11382 | 0.31836 | 0.25719 | 0.00402 | 0.00402 | 0.02348 | 0.00588 | 0.01093 | 0.00943 | 0.60102 | 0.59850 |
| **Nlrp12** | 0.00000 | 0.00345 | 0.00000 | 1.08576 | 2.31092 | 0.65583 | 0.00346 | 0.00000 | 0.34097 | 0.00160 | 1.27343 | 0.00295 | 2.07273 | 2.64449 |
| **Nlrp14** | 7.76282 | 0.00951 | 0.73066 | 0.46288 | 3.75956 | 0.57127 | 0.00274 | 0.00000 | 5.58161 | 0.00466 | 0.00220 | 0.00000 | 6.97684 | 10.54240 |
| **Nlrp1a** | 2.91494 | 0.55316 | 1.02609 | 1.60832 | 2.06328 | 1.27117 | 0.39251 | 0.30661 | 1.72761 | 0.93039 | 0.69540 | 1.78086 | 3.20386 | 1.17024 |
| **Nlrp2** | 0.00262 | 0.00416 | 0.01344 | 0.01257 | 0.00450 | 0.01233 | 0.00412 | 0.00833 | 0.02466 | 0.00615 | 0.00893 | 0.00789 | 1.46799 | 0.06760 |
| **Nlrp3** | 0.00682 | 0.00438 | 0.01462 | 0.01715 | 0.02323 | 0.01330 | 0.00435 | 0.01027 | 0.01037 | 0.00668 | 0.00865 | 0.00952 | 0.09214 | 0.02709 |
| **Nlrp4a** | 0.04167 | 0.00437 | 0.00000 | 0.02112 | 0.02871 | 0.14819 | 0.00400 | 0.00451 | 1.22052 | 0.00603 | 0.00480 | 0.03913 | 0.59136 | 0.15671 |
| **Nlrp4b** | 0.04980 | 0.00429 | 0.00000 | 0.01104 | 0.43781 | 0.01081 | 0.00417 | 0.00880 | 0.02014 | 0.00639 | 0.00759 | 0.00704 | 0.69132 | 0.54190 |
| **Nlrp4c** | 0.46506 | 0.00000 | 1.15296 | 2.81436 | 2.13899 | 4.72444 | 0.00627 | 0.00000 | 8.21300 | 0.03683 | 0.00000 | 5.25744 | 1.13519 | 4.64803 |
| **Nlrp4e** | 0.00000 | 0.00399 | 0.19246 | 0.00248 | 0.31499 | 0.11042 | 0.00396 | 0.00439 | 0.70277 | 0.00599 | 0.00654 | 0.16146 | 0.33999 | 0.62285 |
| **Nlrp4f** | 0.00000 | 0.01028 | 0.11108 | 0.23419 | 0.17692 | 0.02653 | 0.00353 | 0.00279 | 1.08393 | 0.00530 | 0.00076 | 0.81996 | 4.04238 | 3.30557 |
| **Nlrp4g** | 0.00631 | 0.00445 | 0.00858 | 0.00404 | 0.08319 | 0.09875 | 0.00446 | 0.00578 | 0.01586 | 0.00564 | 0.00887 | 0.01010 | 0.06355 | 0.30445 |

| | | | | | | | | | | | | | | |
|---|---|---|---|---|---|---|---|---|---|---|---|---|---|---|
| **Nlrp5** | 3.32132 | 0.00256 | 0.34007 | 0.04486 | 2.47325 | 0.17709 | 0.00813 | 0.00000 | 5.49987 | 0.00482 | 0.00000 | 0.16175 | 5.84726 | 6.97205 |
| **Nlrp5.ps** | 1.58846 | 0.00367 | 0.00000 | 0.10108 | 0.65653 | 0.12837 | 0.00375 | 0.00286 | 1.17775 | 0.01259 | 0.00040 | 0.43653 | 0.77948 | 0.70148 |
| **Nlrp6** | 0.00537 | 0.00435 | 0.30345 | 0.00298 | 0.00474 | 0.00626 | 0.00417 | 0.01051 | 0.01458 | 0.00677 | 0.00690 | 0.00863 | 0.25206 | 0.00833 |
| **Nlrp9a** | 0.00000 | 0.00405 | 0.00000 | 0.05364 | 0.00000 | 0.02735 | 0.00378 | 0.00423 | 1.53638 | 0.00684 | 0.00661 | 0.00729 | 1.54044 | 0.22544 |
| **Nlrp9b** | 0.00000 | 0.00426 | 0.00000 | 0.38225 | 0.00000 | 0.16727 | 0.00389 | 0.00187 | 0.36701 | 0.00400 | 0.29688 | 0.00616 | 0.17973 | 1.08000 |
| **Nlrp9c** | 0.02501 | 0.00406 | 0.13639 | 0.00000 | 0.09727 | 0.18564 | 0.00388 | 0.00319 | 0.09110 | 0.00499 | 0.00670 | 0.00619 | 1.21983 | 0.72127 |
| **Nlrx1** | 0.03231 | 0.81614 | 0.05759 | 2.85185 | 0.12739 | 1.82000 | 0.01473 | 3.49010 | 0.00692 | 0.03081 | 0.00000 | 0.03505 | 0.55872 | 0.00000 |
| **Nmb** | 0.51266 | 0.02434 | 0.13294 | 0.64372 | 0.30404 | 0.25136 | 0.00412 | 0.34732 | 0.57660 | 0.01738 | 0.07999 | 0.00105 | 4.45998 | 0.14250 |
| **Nmd3** | 15.45487 | 2.58228 | 20.24570 | 5.98102 | 12.83481 | 9.33865 | 2.96326 | 2.63974 | 13.81900 | 0.14012 | 12.33757 | 7.82672 | 10.56470 | 14.56625 |
| **Nme1** | 19.52876 | 1.89195 | 19.95051 | 30.94410 | 29.12067 | 32.94206 | 10.03231 | 19.97906 | 11.19052 | 21.82554 | 14.27778 | 26.52156 | 9.04888 | 18.81881 |
| **Nme2** | 27.55065 | 13.49143 | 22.13333 | 46.47842 | 43.41698 | 43.27818 | 12.33614 | 29.78683 | 9.09247 | 26.68280 | 15.02860 | 30.38236 | 23.51893 | 29.92852 |
| **Nme3** | 0.06521 | 0.04088 | 0.00000 | 2.37179 | 0.39327 | 0.77897 | 1.37573 | 5.21237 | 0.03233 | 0.23001 | 1.37318 | 0.80853 | 0.00648 | 0.06070 |
| **Nme4** | 0.42868 | 0.00700 | 0.42565 | 2.59569 | 1.45380 | 1.03799 | 0.22302 | 0.00000 | 0.00916 | 0.02664 | 1.85565 | 4.93313 | 0.64820 | 0.00000 |
| **Nme5** | 0.23127 | 0.00464 | 0.00000 | 0.14647 | 2.13052 | 0.00325 | 0.00405 | 0.01385 | 0.74361 | 0.00921 | 0.00425 | 0.00429 | 1.45093 | 0.48462 |
| **Nme6** | 3.15004 | 0.07852 | 9.50745 | 8.80080 | 5.44740 | 5.13225 | 1.22583 | 5.22075 | 2.35738 | 0.11247 | 6.54204 | 7.43796 | 1.20432 | 3.64472 |
| **Nme7** | 1.10169 | 0.03823 | 3.75516 | 0.42562 | 0.67082 | 0.86319 | 0.01861 | 2.21602 | 0.08690 | 0.04872 | 3.17353 | 3.05737 | 0.65874 | 1.74508 |
| **Nmi** | 0.00098 | 0.00413 | 0.09810 | 0.23989 | 0.05064 | 0.00677 | 0.00415 | 0.00925 | 0.00499 | 0.00710 | 0.00576 | 0.00609 | 0.49234 | 0.29326 |
| **Nmnat1** | 0.20894 | 0.00367 | 0.72218 | 1.34719 | 0.17427 | 0.92208 | 0.00358 | 0.27880 | 0.46235 | 0.00333 | 0.31247 | 0.00000 | 0.94485 | 2.21287 |
| **Nmnat2** | 0.00392 | 0.01512 | 0.01739 | 0.01477 | 0.00835 | 0.01497 | 0.00450 | 0.92952 | 0.03329 | 0.00839 | 0.00761 | 0.00699 | 0.36186 | 0.00548 |
| **Nmnat3** | 0.01069 | 0.00445 | 0.01446 | 0.01336 | 0.01368 | 0.01323 | 0.00429 | 0.01028 | 0.01009 | 0.00701 | 0.00889 | 0.01004 | 0.00694 | 0.01946 |
| **Nmral1** | 0.00000 | 0.04504 | 1.99022 | 1.19337 | 1.48736 | 1.30807 | 0.17194 | 1.74710 | 0.54592 | 0.01559 | 0.00000 | 3.21743 | 0.87714 | 1.74869 |
| **Nmrk1** | 1.14709 | 0.06324 | 0.26981 | 0.28838 | 0.13580 | 2.85431 | 0.34255 | 2.80126 | 1.69623 | 0.36353 | 0.17418 | 2.03510 | 7.33907 | 0.95794 |
| **Nmrk2** | 0.17636 | 0.06583 | 0.04306 | 0.60166 | 0.11220 | 0.45140 | 0.11874 | 0.14672 | 0.36560 | 0.24291 | 0.01360 | 0.18783 | 0.18377 | 0.28500 |
| **Nmt1** | 1.11667 | 0.00239 | 0.54936 | 1.99088 | 0.71823 | 1.89777 | 0.02089 | 0.34033 | 4.72154 | 3.15924 | 0.99089 | 1.13580 | 0.44560 | 4.02237 |
| **Nmt2** | 10.02749 | 0.09536 | 6.11083 | 2.87660 | 4.56798 | 6.89009 | 0.03704 | 3.51042 | 8.62891 | 0.01322 | 3.27698 | 4.28748 | 2.94345 | 5.10968 |
| **Nmu** | 0.00543 | 0.00440 | 0.01245 | 0.01270 | 0.00614 | 0.01194 | 0.00425 | 0.00994 | 0.04575 | 0.00649 | 0.00834 | 0.00975 | 0.11148 | 0.09331 |
| **Nnat** | 0.00845 | 0.00456 | 0.00975 | 0.23863 | 0.00935 | 0.00920 | 0.00518 | 0.07745 | 0.01154 | 0.01285 | 0.00731 | 0.00833 | 0.21475 | 0.00000 |
| **Nnt** | 0.11271 | 0.02061 | 0.99938 | 0.11820 | 0.00000 | 4.11282 | 0.00401 | 1.34576 | 0.45627 | 0.04216 | 0.01162 | 0.01740 | 0.00088 | 0.37798 |
| **Noa1** | 7.51575 | 0.00184 | 5.17648 | 1.56850 | 8.13511 | 0.16951 | 0.00241 | 0.00000 | 1.57177 | 0.00000 | 1.18791 | 0.39740 | 10.01478 | 10.88508 |
| **Nob1** | 11.51894 | 5.03336 | 23.84421 | 12.26704 | 19.18729 | 9.15697 | 6.05206 | 5.05808 | 3.48773 | 0.35153 | 10.56528 | 8.33851 | 9.53488 | 29.49121 |
| **Nobox** | 0.00392 | 0.00402 | 0.00000 | 0.00432 | 0.32687 | 0.00000 | 0.00436 | 0.00993 | 0.01578 | 0.00317 | 0.84669 | 0.00438 | 0.75338 | 0.06637 |
| **Noc2l** | 6.48171 | 3.07315 | 18.85492 | 13.98016 | 15.80777 | 10.15800 | 5.52967 | 8.55513 | 6.54753 | 0.22378 | 9.86501 | 10.25467 | 0.28947 | 18.24528 |
| **Noc3l** | 1.31110 | 0.04513 | 3.94797 | 0.91655 | 5.61636 | 2.71151 | 0.01978 | 0.31006 | 2.89577 | 0.00000 | 3.41508 | 2.57486 | 2.28349 | 7.90599 |
| **Noc4l** | 11.49321 | 0.18083 | 15.98328 | 18.35357 | 13.17040 | 8.92491 | 0.29649 | 6.98235 | 0.39468 | 0.08818 | 1.18667 | 9.40000 | 6.70820 | 8.62278 |
| **Noct** | 6.88736 | 0.08432 | 11.14646 | 2.38467 | 2.99067 | 6.86905 | 6.70308 | 5.22990 | 3.15149 | 0.14789 | 4.01113 | 1.82534 | 12.67415 | 8.35375 |
| **Nod1** | 0.00000 | 0.00323 | 0.49145 | 0.39226 | 0.04257 | 0.00000 | 0.00362 | 0.00458 | 0.01199 | 0.00000 | 2.66659 | 0.02396 | 0.51747 | 0.11932 |
| **Nol10** | 2.34307 | 0.00951 | 0.56373 | 1.63979 | 3.82642 | 2.18732 | 0.02010 | 0.00000 | 4.13120 | 0.01890 | 2.98467 | 1.14607 | 3.12019 | 6.07553 |
| **Nol11** | 7.19878 | 0.01140 | 16.50223 | 4.07854 | 15.27496 | 8.96527 | 1.18883 | 3.54218 | 22.64026 | 0.00000 | 14.21127 | 1.81094 | 7.85382 | 19.22961 |
| **Nol12** | 13.08119 | 0.18961 | 17.86954 | 7.90994 | 17.70411 | 9.25344 | 2.73777 | 6.12006 | 1.92764 | 14.62488 | 10.33474 | 8.34251 | 5.61059 | 14.52550 |
| **Nol3** | 0.00574 | 0.00386 | 0.00500 | 0.41053 | 0.11515 | 0.00000 | 0.00363 | 0.00374 | 0.01397 | 0.00554 | 0.01904 | 0.39325 | 0.00582 | 0.00521 |
| **Nol4l** | 4.16995 | 0.02045 | 8.43826 | 3.86580 | 7.51479 | 0.99623 | 0.00000 | 0.07196 | 7.80745 | 11.99500 | 2.21194 | 2.52180 | 4.63987 | 9.66531 |
| **Nol6** | 1.01440 | 5.05946 | 6.90573 | 5.51628 | 8.17080 | 4.01440 | 0.35831 | 1.39796 | 1.36579 | 0.03158 | 4.09553 | 2.74710 | 1.16256 | 5.27267 |
| **Nol7** | 12.41099 | 0.09623 | 1.37517 | 14.32701 | 26.80056 | 5.93263 | 1.00092 | 5.27426 | 7.89614 | 2.22069 | 1.24636 | 6.55680 | 11.96575 | 2.18543 |
| **Nol8** | 7.93755 | 0.00000 | 7.31636 | 2.73700 | 7.89241 | 3.16105 | 3.44906 | 0.00000 | 10.30099 | 0.06664 | 2.62210 | 1.99090 | 9.69693 | 12.83938 |
| **Nol9** | 8.90826 | 0.06936 | 11.72239 | 1.61973 | 6.74360 | 3.52382 | 1.99449 | 0.68889 | 2.64022 | 4.89598 | 6.61334 | 4.71213 | 8.90411 | 12.12213 |
| **Nolc1** | 24.88092 | 7.65564 | 39.91896 | 23.53650 | 35.67929 | 16.86569 | 5.16440 | 8.02978 | 23.92531 | 0.31220 | 15.17032 | 13.80214 | 13.01020 | 41.40871 |
| **Nom1** | 8.61430 | 2.02932 | 4.31466 | 4.72217 | 6.00956 | 2.63693 | 0.98722 | 0.00000 | 11.09832 | 0.02726 | 0.10460 | 1.30544 | 11.51069 | 8.82995 |
| **Nomo1** | 3.48096 | 0.08879 | 7.13427 | 5.69688 | 3.35600 | 12.10521 | 2.60808 | 4.77076 | 0.20502 | 0.10947 | 2.37885 | 7.72225 | 1.65216 | 5.41029 |
| **Nono** | 19.71257 | 4.09046 | 31.91920 | 14.83555 | 28.19096 | 17.34991 | 7.35366 | 13.52770 | 30.04835 | 3.96652 | 13.93610 | 10.52029 | 8.58869 | 27.28978 |
| **Nop10** | 4.97542 | 0.18117 | 0.00000 | 3.66359 | 6.44333 | 8.68534 | 1.40631 | 9.20270 | 3.01708 | 6.19055 | 0.28329 | 6.80908 | 7.74306 | 6.70930 |
| **Nop14** | 15.25303 | 0.19991 | 18.75510 | 6.46440 | 10.22162 | 10.02591 | 0.02439 | 6.32116 | 5.92648 | 0.88449 | 13.77183 | 8.97067 | 4.66538 | 13.15835 |
| **Nop16** | 11.47624 | 0.21215 | 17.49912 | 15.23617 | 15.28845 | 17.20667 | 0.07894 | 6.81542 | 8.72768 | 0.13887 | 10.87806 | 10.19622 | 9.71838 | 14.38294 |
| **Nop2** | 17.67805 | 2.64850 | 27.29564 | 10.56441 | 20.02817 | 13.26747 | 4.98449 | 2.08960 | 5.35050 | 0.10740 | 8.16802 | 11.02484 | 12.93616 | 20.97452 |
| **Nop56** | 21.33164 | 11.28761 | 33.34974 | 29.31310 | 27.56803 | 25.54879 | 11.51919 | 20.47126 | 20.38167 | 9.06375 | 22.06352 | 20.27079 | 15.53619 | 31.72745 |
| **Nop58** | 11.08998 | 2.58002 | 11.74437 | 3.31322 | 13.99106 | 9.92543 | 0.62488 | 1.71838 | 22.78445 | 0.13420 | 9.97160 | 9.70032 | 9.54348 | 8.73563 |

| | | | | | | | | | | | | | | |
|---|---|---|---|---|---|---|---|---|---|---|---|---|---|---|
| **Nop9** | 7.80333 | 0.09762 | 16.63911 | 5.05422 | 19.27325 | 6.37034 | 0.01418 | 5.75381 | 1.60445 | 0.12978 | 5.90687 | 6.24920 | 11.59769 | 8.86477 |
| **Nos1** | 2.28272 | 0.03730 | 4.64725 | 0.68377 | 2.13062 | 1.31145 | 0.35427 | 2.56737 | 0.04748 | 10.85429 | 1.60972 | 3.03219 | 3.10635 | 2.20865 |
| **Nos3** | 0.01469 | 0.00432 | 0.00000 | 0.20216 | 0.13338 | 0.19849 | 0.00412 | 0.00576 | 0.01356 | 0.01310 | 0.21224 | 0.00878 | 0.00613 | 0.00325 |
| **Nosip** | 3.30304 | 0.16781 | 11.27746 | 10.21321 | 7.93631 | 3.48585 | 4.27218 | 9.81352 | 1.50939 | 2.34860 | 7.41493 | 4.40614 | 4.19077 | 0.23338 |
| **Nostrin** | 0.07733 | 0.00512 | 0.06418 | 0.36602 | 0.08693 | 0.00000 | 0.00417 | 0.05047 | 0.01962 | 0.00760 | 0.21511 | 0.05053 | 0.45078 | 0.02027 |
| **Notch1** | 0.56488 | 0.01091 | 0.87938 | 0.84501 | 1.20258 | 0.00000 | 0.08036 | 0.01307 | 0.09971 | 0.01481 | 0.00626 | 2.06496 | 0.00857 | 0.36243 |
| **Notch2** | 0.05417 | 0.00458 | 0.00484 | 0.11824 | 0.00753 | 0.20024 | 0.00452 | 0.01538 | 0.00767 | 0.00793 | 0.00708 | 0.00812 | 0.53574 | 0.04986 |
| **Notch3** | 0.01525 | 0.00425 | 0.01308 | 0.00859 | 0.00693 | 0.01279 | 0.00434 | 0.00900 | 0.01791 | 0.00653 | 0.00758 | 0.03331 | 0.22398 | 0.11479 |
| **Notch4** | 0.08261 | 0.00441 | 0.09645 | 0.08882 | 0.64964 | 0.07360 | 0.00419 | 0.00550 | 0.02360 | 0.00603 | 0.00558 | 0.06477 | 0.00601 | 0.09781 |
| **Noto** | 0.00000 | 0.00444 | 0.91757 | 0.02926 | 1.13133 | 0.01052 | 0.00383 | 0.00623 | 1.95144 | 0.00596 | 0.00285 | 0.00753 | 0.00000 | 0.61494 |
| **Notum** | 0.52638 | 0.00401 | 0.85589 | 0.13392 | 0.00000 | 0.00000 | 0.00383 | 0.00769 | 0.00828 | 0.00847 | 0.01698 | 0.00810 | 0.59146 | 0.00628 |
| **Noxo1** | 0.01554 | 0.00370 | 0.50407 | 0.03947 | 0.40338 | 0.10123 | 0.00410 | 0.01471 | 0.01233 | 0.00565 | 0.01617 | 0.10798 | 2.08273 | 0.00210 |
| **Noxred1** | 0.20049 | 0.01023 | 0.00000 | 0.00000 | 0.07921 | 0.77023 | 0.64091 | 0.01368 | 0.75631 | 0.06565 | 0.14726 | 2.39387 | 0.05702 | 0.73595 |
| **Npas1** | 0.01921 | 0.00410 | 0.22047 | 0.49315 | 0.10671 | 0.05615 | 0.00378 | 0.00367 | 0.00749 | 0.00696 | 0.00610 | 0.00818 | 0.28028 | 0.00000 |
| **Npas2** | 0.00000 | 0.00380 | 0.96640 | 0.00000 | 0.18613 | 0.02511 | 0.00371 | 0.11707 | 0.25694 | 0.00532 | 0.00371 | 0.18036 | 0.00000 | 0.98696 |
| **Npat** | 5.99397 | 4.61347 | 4.40452 | 0.00000 | 2.91231 | 2.67886 | 1.58165 | 0.57309 | 13.18033 | 0.05544 | 2.81981 | 1.76384 | 8.99084 | 2.56225 |
| **Npc1** | 2.97034 | 0.03829 | 8.29397 | 2.00451 | 1.39633 | 5.21842 | 1.63605 | 2.73016 | 2.34619 | 0.00849 | 6.18398 | 4.04253 | 2.05333 | 2.55055 |
| **Npc1l1** | 3.17655 | 0.00321 | 1.18126 | 0.00000 | 4.50426 | 0.00000 | 0.00872 | 0.09463 | 1.37831 | 0.00278 | 0.00000 | 0.00066 | 0.56694 | 7.97104 |
| **Npc2** | 25.97535 | 7.99935 | 39.87412 | 39.51932 | 25.89358 | 32.24910 | 11.26430 | 27.72415 | 11.05427 | 3.65301 | 19.97123 | 8.39012 | 6.99500 | 22.95709 |
| **Npdc1** | 0.14101 | 0.01954 | 2.75395 | 4.14515 | 4.11627 | 0.98725 | 0.03285 | 1.59502 | 0.01781 | 0.01505 | 0.00000 | 0.00000 | 0.00924 | 1.03136 |
| **Npepl1** | 0.18062 | 7.75829 | 0.26260 | 0.95374 | 0.49106 | 0.92523 | 0.00754 | 4.31084 | 0.03885 | 0.03067 | 0.00000 | 0.95339 | 0.00000 | 0.11254 |
| **Npepps** | 3.42144 | 0.65072 | 0.82379 | 0.00000 | 1.16958 | 6.02764 | 0.01327 | 1.72915 | 3.29861 | 0.05203 | 5.10517 | 5.42551 | 5.18496 | 2.84715 |
| **Npff** | 0.00610 | 0.01390 | 0.02802 | 0.00000 | 0.02907 | 0.00967 | 0.00459 | 0.27137 | 0.01288 | 0.00613 | 0.01399 | 0.06259 | 0.17100 | 0.00337 |
| **Nphp1** | 0.00000 | 0.00423 | 0.00000 | 0.21495 | 0.53830 | 0.21262 | 0.00397 | 0.01668 | 0.71642 | 0.01626 | 0.00027 | 0.45409 | 0.35382 | 0.09611 |
| **Nphp3** | 0.02786 | 0.01108 | 0.01459 | 0.05987 | 0.07281 | 0.11409 | 0.00396 | 0.00432 | 0.35503 | 0.00582 | 0.00403 | 0.19449 | 0.12489 | 0.56106 |
| **Nphp4** | 0.53005 | 0.00458 | 0.00523 | 0.08659 | 0.05128 | 0.01217 | 0.00509 | 0.04510 | 0.00707 | 0.01934 | 0.00940 | 0.00503 | 0.03054 | 0.11282 |
| **Nphs1** | 0.08874 | 0.07610 | 0.05137 | 0.11112 | 0.04019 | 0.06615 | 0.02175 | 0.00918 | 0.34760 | 0.08368 | 0.00505 | 0.00877 | 0.36457 | 0.34351 |
| **Nphs2** | 0.00000 | 0.00448 | 0.12912 | 0.00590 | 0.18554 | 0.00140 | 0.00402 | 0.01120 | 0.95470 | 0.00894 | 0.00491 | 0.00691 | 0.03486 | 0.49483 |
| **Npl** | 19.37747 | 0.10609 | 8.49505 | 0.51959 | 3.80054 | 2.58276 | 0.15837 | 4.99177 | 23.43559 | 3.21010 | 4.74287 | 3.42915 | 12.06085 | 23.41268 |
| **Nploc4** | 9.29244 | 2.74539 | 8.52123 | 5.76798 | 7.18191 | 9.68080 | 3.07560 | 2.74485 | 0.92372 | 0.09993 | 10.21917 | 4.99507 | 2.13882 | 6.57777 |
| **Npm1** | 49.16928 | 13.63272 | 64.11270 | 62.78443 | 57.16324 | 54.94369 | 17.45307 | 35.97011 | 60.21096 | 27.00541 | 38.17841 | 40.89682 | 21.12872 | 53.66185 |
| **Npm2** | 2.30629 | 0.00369 | 0.63668 | 0.32648 | 3.65831 | 0.05178 | 0.00357 | 0.00346 | 0.81229 | 0.00687 | 0.00261 | 0.00000 | 6.55147 | 3.06655 |
| **Npm3** | 0.18060 | 0.00419 | 0.21647 | 0.68238 | 0.42024 | 0.37120 | 0.07017 | 0.36773 | 0.04918 | 0.00814 | 0.00339 | 0.09726 | 1.53598 | 0.00000 |
| **Npm3.ps1** | 0.08498 | 0.01190 | 0.14707 | 0.30356 | 0.21162 | 0.23253 | 0.04995 | 0.21769 | 0.02675 | 0.00757 | 0.00273 | 0.06229 | 0.55429 | 0.00000 |
| **Npnt** | 0.01249 | 0.00499 | 0.09294 | 0.34164 | 0.02599 | 0.01460 | 0.00432 | 0.03860 | 0.00567 | 0.02051 | 0.00878 | 0.00574 | 0.00405 | 0.12068 |
| **Nppa** | 0.00246 | 0.02898 | 0.06289 | 0.00000 | 0.45776 | 0.17744 | 0.00366 | 0.00590 | 0.02059 | 0.00658 | 0.00247 | 0.81058 | 0.00298 | 0.06672 |
| **Nppb** | 0.00168 | 0.00445 | 0.22319 | 0.00483 | 0.02209 | 0.04238 | 0.00411 | 0.01121 | 0.02594 | 0.00663 | 0.55938 | 0.00539 | 0.00750 | 0.48896 |
| **Npr1** | 1.04532 | 0.02700 | 0.80666 | 4.33879 | 2.97698 | 2.45554 | 1.01640 | 1.25068 | 0.08489 | 1.09376 | 2.44241 | 0.57918 | 1.47854 | 2.63796 |
| **Npr2** | 0.02277 | 0.68645 | 1.70515 | 0.00000 | 0.00187 | 0.28410 | 0.00311 | 1.89271 | 0.00744 | 0.00248 | 0.00000 | 0.54763 | 0.00156 | 0.00785 |
| **Npr3** | 0.08995 | 0.00426 | 0.11522 | 0.30630 | 0.00000 | 0.03460 | 0.00417 | 0.00556 | 0.01364 | 0.00653 | 0.01881 | 0.00840 | 0.00552 | 0.01390 |
| **Nprl2** | 0.89802 | 0.02605 | 4.69031 | 1.90716 | 0.85515 | 2.45484 | 3.89482 | 3.00362 | 0.41371 | 0.14818 | 3.08519 | 1.79723 | 0.36059 | 0.73837 |
| **Nprl3** | 0.03685 | 0.00426 | 0.53177 | 0.00000 | 0.18232 | 0.05397 | 0.53598 | 0.02934 | 0.11082 | 0.01718 | 0.83299 | 0.10828 | 0.43582 | 0.55489 |
| **Npsr1** | 0.00177 | 0.00443 | 0.01627 | 0.01423 | 0.12630 | 0.00981 | 0.00427 | 0.01001 | 0.10175 | 0.00592 | 0.00778 | 0.00813 | 0.28575 | 0.18181 |
| **Nptn** | 1.37170 | 0.00281 | 0.87127 | 0.03857 | 0.07867 | 0.08463 | 0.00306 | 0.57477 | 3.40866 | 0.01602 | 0.17158 | 0.00243 | 8.71794 | 0.00000 |
| **Nptx1** | 0.00633 | 0.01317 | 0.01243 | 0.00490 | 0.02116 | 0.01146 | 0.00389 | 0.02055 | 0.01057 | 0.00670 | 0.00402 | 0.86433 | 1.05591 | 0.09279 |
| **Nptx2** | 0.06879 | 0.01092 | 0.44883 | 0.00000 | 0.15566 | 0.00088 | 0.00373 | 0.00423 | 0.15156 | 0.00707 | 0.00344 | 0.55810 | 4.19278 | 0.25058 |
| **Npw** | 0.01534 | 0.00445 | 0.00790 | 0.03000 | 0.49640 | 0.01303 | 0.00425 | 0.00939 | 0.01237 | 0.00653 | 0.00835 | 0.00706 | 0.00790 | 0.00979 |
| **Npy** | 0.01028 | 0.00378 | 0.03677 | 0.75675 | 3.21766 | 0.00000 | 0.00390 | 0.00382 | 0.01070 | 0.02069 | 0.01842 | 0.37029 | 0.38512 | 0.12949 |
| **Npy1r** | 0.13363 | 0.39139 | 0.04004 | 0.36324 | 0.16415 | 0.23374 | 0.06252 | 0.16986 | 0.68900 | 0.05918 | 0.02345 | 0.15034 | 0.22571 | 0.52598 |
| **Nqo1** | 0.00000 | 0.00373 | 0.07000 | 1.02015 | 0.11417 | 0.68326 | 0.15589 | 0.00856 | 0.21415 | 0.10159 | 0.00884 | 0.52045 | 0.27281 | 0.58992 |
| **Nqo2** | 1.36543 | 0.02039 | 2.10736 | 4.39106 | 1.31472 | 0.95177 | 3.81174 | 1.27091 | 0.02859 | 0.26883 | 0.05631 | 1.39596 | 0.07817 | 1.33962 |
| **Nr0b2** | 0.02840 | 0.00434 | 0.35199 | 0.00000 | 0.21324 | 0.00444 | 0.00398 | 0.01054 | 0.00544 | 0.00566 | 0.00639 | 0.00894 | 0.13473 | 0.06033 |
| **Nr1d1** | 0.00000 | 0.00525 | 2.62760 | 0.69499 | 2.19104 | 0.04431 | 1.07400 | 0.00742 | 0.00681 | 0.03243 | 0.00232 | 0.01854 | 0.01017 | 0.01962 |
| **Nr1d2** | 5.66283 | 1.17888 | 2.20904 | 0.00000 | 5.47110 | 0.29228 | 0.00739 | 0.76685 | 8.71040 | 0.00218 | 0.00000 | 0.00000 | 4.12337 | 2.93872 |
| **Nr1h2** | 3.46459 | 0.06776 | 19.10488 | 10.49333 | 13.62918 | 2.31960 | 0.05439 | 4.90435 | 0.00678 | 0.11978 | 0.44935 | 6.69234 | 1.92309 | 0.36924 |

| | | | | | | | | | | | | | | |
|---|---|---|---|---|---|---|---|---|---|---|---|---|---|---|
| **Nr1h3** | 2.93932 | 0.00998 | 7.13951 | 0.41108 | 10.66468 | 0.00000 | 0.00247 | 0.00075 | 0.13115 | 0.00000 | 1.13274 | 0.00575 | 4.32003 | 2.77177 |
| **Nr1h4** | 0.00506 | 0.00434 | 0.00000 | 0.01553 | 0.35869 | 0.26433 | 0.00429 | 0.00941 | 0.03371 | 0.00813 | 0.12469 | 0.00441 | 0.00491 | 0.00537 |
| **Nr2c1** | 0.05518 | 0.00937 | 1.29962 | 0.00000 | 0.38400 | 2.43951 | 0.00314 | 1.53560 | 0.05327 | 0.01578 | 1.77862 | 1.41253 | 0.31518 | 0.36918 |
| **Nr2c2** | 1.03823 | 0.00234 | 0.00014 | 0.72084 | 3.10326 | 0.18186 | 0.01312 | 0.00000 | 10.02432 | 0.00367 | 0.75828 | 0.44774 | 5.35129 | 9.24504 |
| **Nr2c2ap** | 0.33394 | 1.23674 | 1.97591 | 2.83089 | 3.35046 | 4.11405 | 0.00890 | 0.54208 | 0.74485 | 6.45259 | 1.13120 | 1.80313 | 3.48595 | 1.61834 |
| **Nr2f2** | 0.19072 | 0.00413 | 0.00172 | 0.00000 | 0.03298 | 0.47991 | 0.01576 | 0.23491 | 0.11541 | 0.00888 | 0.03082 | 0.36583 | 0.50800 | 0.01014 |
| **Nr2f6** | 0.14995 | 0.28450 | 0.01509 | 0.00000 | 0.01176 | 0.06098 | 0.86608 | 1.20194 | 0.00868 | 0.01170 | 0.00000 | 1.11904 | 0.00922 | 0.01580 |
| **Nr3c1** | 0.02790 | 0.00397 | 0.00000 | 0.00000 | 0.10428 | 0.59006 | 0.00369 | 0.29780 | 0.02085 | 0.01107 | 0.00000 | 1.86372 | 0.92192 | 0.05906 |
| **Nr4a1** | 0.04984 | 0.00346 | 0.20038 | 0.61623 | 0.05939 | 0.33770 | 0.02308 | 0.71683 | 0.00636 | 0.01939 | 1.82185 | 0.50234 | 0.24796 | 0.00000 |
| **Nr4a2** | 0.00784 | 0.00428 | 0.10990 | 0.00006 | 0.35342 | 0.01074 | 0.00408 | 0.00890 | 0.01699 | 0.00761 | 0.01116 | 0.00913 | 0.68212 | 0.00176 |
| **Nr5a2** | 13.71903 | 0.07283 | 12.05715 | 0.00000 | 13.59422 | 2.95713 | 0.14668 | 0.55557 | 8.73056 | 0.00547 | 2.00734 | 0.54004 | 13.68547 | 12.56488 |
| **Nr6a1** | 0.62378 | 0.00148 | 5.88604 | 6.13593 | 2.53998 | 10.82254 | 0.00761 | 0.00000 | 2.43062 | 0.04577 | 4.40950 | 1.30639 | 1.94439 | 5.61992 |
| **Nradd** | 0.05956 | 0.01318 | 1.59394 | 0.00000 | 0.29452 | 0.10463 | 0.00363 | 0.56657 | 0.01472 | 0.00247 | 0.23068 | 0.29389 | 0.00000 | 0.00758 |
| **Nrap** | 0.00825 | 0.01082 | 0.01249 | 0.00000 | 0.00700 | 0.10307 | 0.00390 | 0.00513 | 0.01007 | 0.00624 | 0.00271 | 0.86791 | 0.01471 | 0.18565 |
| **Nrarp** | 0.00311 | 0.00422 | 0.01564 | 0.16238 | 0.06965 | 0.01117 | 0.00480 | 0.01176 | 0.01989 | 0.00796 | 0.00816 | 0.00753 | 1.59744 | 0.00000 |
| **Nras** | 7.49687 | 0.08053 | 3.93559 | 5.24395 | 3.69229 | 10.00180 | 0.08528 | 5.68634 | 6.64195 | 0.13854 | 7.59250 | 7.43843 | 3.94195 | 6.98145 |
| **Nrbf2** | 4.85974 | 0.02138 | 4.86592 | 2.36392 | 4.34445 | 5.88764 | 4.06286 | 0.95627 | 1.04475 | 0.09730 | 1.14533 | 5.35040 | 5.94938 | 3.13748 |
| **Nrbp1** | 10.32178 | 7.57966 | 15.91274 | 7.62532 | 25.12018 | 9.36762 | 4.36794 | 7.28116 | 0.00000 | 0.15670 | 8.24637 | 4.20523 | 1.67895 | 12.99633 |
| **Nrbp2** | 0.29287 | 0.00342 | 0.00231 | 0.00197 | 0.18805 | 0.09753 | 0.00405 | 0.00533 | 0.00761 | 0.00637 | 0.00230 | 0.90661 | 0.00696 | 0.00842 |
| **Nrcam** | 0.00066 | 0.00473 | 0.03426 | 0.34350 | 0.00061 | 0.03541 | 0.00409 | 0.01663 | 0.71611 | 0.00579 | 0.00930 | 0.02290 | 0.09123 | 0.00000 |
| **Nrd1** | 12.65287 | 0.87540 | 9.72763 | 5.37081 | 4.66632 | 7.87504 | 3.29696 | 1.62734 | 2.06587 | 0.15174 | 9.11355 | 6.21867 | 0.87234 | 4.31229 |
| **Nrde2** | 0.19586 | 0.05244 | 5.76555 | 3.82626 | 3.50012 | 1.69962 | 3.33419 | 0.67213 | 0.00106 | 0.06118 | 0.39601 | 3.10758 | 0.04040 | 0.03369 |
| **Nrep** | 0.05386 | 0.00435 | 0.01403 | 0.01177 | 0.00851 | 0.01278 | 0.00422 | 0.00960 | 0.02034 | 0.00673 | 0.00810 | 0.00905 | 0.30756 | 0.00374 |
| **Nrf1** | 2.77960 | 0.08248 | 13.56852 | 3.73192 | 12.34574 | 7.06827 | 2.18365 | 2.64635 | 0.41453 | 0.07700 | 1.73720 | 3.26345 | 5.79434 | 9.47620 |
| **Nrg1** | 0.01273 | 0.01100 | 0.00876 | 0.05400 | 0.43254 | 0.00667 | 0.00355 | 0.00470 | 0.00822 | 0.00595 | 0.00361 | 1.46520 | 0.01329 | 0.40082 |
| **Nrg2** | 0.00672 | 0.00428 | 0.09023 | 0.20175 | 0.07383 | 0.00000 | 0.00413 | 0.00908 | 0.15265 | 0.00632 | 0.00530 | 0.03801 | 0.01160 | 0.06263 |
| **Nrg4** | 0.00512 | 0.00403 | 0.36031 | 0.07136 | 0.02641 | 0.04874 | 0.00408 | 0.04311 | 1.46666 | 0.00510 | 0.07736 | 0.00984 | 0.00071 | 0.00265 |
| **Nrip1** | 0.00000 | 0.00541 | 0.44569 | 0.81626 | 0.00000 | 0.00000 | 0.00445 | 0.30912 | 0.02543 | 0.00680 | 0.00471 | 0.01736 | 0.13028 | 1.60038 |
| **Nrip2** | 0.00829 | 0.00453 | 0.01199 | 0.33404 | 0.01295 | 0.13703 | 0.00436 | 0.00511 | 0.01680 | 0.00626 | 0.00633 | 0.00608 | 0.01171 | 0.05297 |
| **Nrip3** | 0.39123 | 0.00424 | 0.29921 | 0.00737 | 0.60681 | 0.01065 | 0.00411 | 0.00734 | 0.00601 | 0.00650 | 0.00753 | 0.00568 | 1.11072 | 0.10224 |
| **Nrl** | 0.01121 | 0.00403 | 0.00876 | 0.09967 | 0.00859 | 0.00000 | 0.00400 | 0.00698 | 0.01861 | 0.01388 | 0.36934 | 0.02682 | 0.11799 | 0.01082 |
| **Nrm** | 0.20656 | 0.01270 | 0.01800 | 3.66304 | 0.00000 | 1.72178 | 0.52090 | 2.88169 | 0.01326 | 0.23813 | 0.52072 | 0.05952 | 0.00970 | 0.56505 |
| **Nrn1l** | 0.36731 | 0.00435 | 0.10316 | 0.13318 | 0.51625 | 0.05413 | 0.00402 | 0.00488 | 0.01812 | 0.00623 | 0.00603 | 0.00606 | 0.03076 | 0.00223 |
| **Nrp** | 0.00000 | 0.00474 | 0.00000 | 0.08732 | 0.26042 | 0.00000 | 0.51722 | 0.14664 | 0.78523 | 0.04437 | 0.00000 | 0.77310 | 0.08378 | 0.22109 |
| **Nrp1** | 0.09133 | 0.00437 | 0.00975 | 0.01240 | 0.00315 | 0.01013 | 0.00430 | 0.01016 | 0.01008 | 0.00686 | 0.01092 | 0.01346 | 0.01909 | 0.49564 |
| **Nrp2** | 0.05835 | 0.00410 | 0.00876 | 0.74567 | 0.00522 | 0.01166 | 0.00397 | 0.02134 | 0.08224 | 0.00575 | 0.01387 | 0.02838 | 0.01700 | 0.01439 |
| **Nrros** | 0.04458 | 0.00432 | 1.85862 | 0.68342 | 0.65054 | 0.00000 | 0.00364 | 0.00615 | 0.03555 | 0.00284 | 0.00782 | 0.01521 | 0.28395 | 0.07295 |
| **Nrsn2** | 0.41843 | 0.00421 | 0.24088 | 0.31594 | 0.07182 | 0.02958 | 0.00405 | 0.00382 | 0.00760 | 0.00655 | 0.02206 | 0.00562 | 0.13508 | 0.02119 |
| **Nrxn2** | 0.00653 | 0.00436 | 0.06698 | 0.00395 | 0.25131 | 0.01155 | 0.00413 | 0.01411 | 0.01973 | 0.00691 | 0.00768 | 0.00950 | 0.02450 | 0.01355 |
| **Nsa2** | 18.97422 | 1.50176 | 29.00464 | 24.89896 | 26.25010 | 18.20059 | 7.46481 | 20.25569 | 27.74156 | 9.22648 | 16.10966 | 19.66502 | 11.85108 | 29.45643 |
| **Nsd1** | 4.14835 | 0.05743 | 9.55059 | 4.27607 | 7.24646 | 10.20280 | 6.32137 | 3.26507 | 10.79606 | 0.25731 | 2.42310 | 6.69355 | 4.91566 | 10.78171 |
| **Nsdhl** | 1.23941 | 0.01413 | 0.73369 | 0.42430 | 0.67887 | 7.89640 | 0.00164 | 1.54378 | 0.23089 | 0.02949 | 3.17006 | 4.65531 | 0.00000 | 1.17544 |
| **Nsf** | 2.41121 | 0.09726 | 12.48199 | 7.14588 | 3.96909 | 8.27896 | 2.48464 | 6.68971 | 6.72131 | 0.03515 | 9.13755 | 5.33196 | 0.76031 | 6.84047 |
| **Nsfl1c** | 6.39516 | 5.88473 | 16.06042 | 17.23422 | 10.54029 | 23.93492 | 5.48821 | 9.67871 | 4.14612 | 1.52803 | 15.41039 | 16.63303 | 1.65383 | 12.68730 |
| **Nsl1** | 1.15127 | 0.04790 | 3.43971 | 0.00000 | 3.21060 | 1.52611 | 1.52416 | 2.39514 | 0.96709 | 0.04761 | 2.84550 | 1.44653 | 0.77434 | 2.75607 |
| **Nsmaf** | 0.00000 | 0.06080 | 0.24202 | 0.00000 | 0.00244 | 0.92871 | 0.00975 | 2.15002 | 0.03709 | 0.00046 | 1.87060 | 4.96723 | 0.04690 | 0.00000 |
| **Nsmce1** | 7.74448 | 4.86776 | 12.94465 | 18.71913 | 11.22460 | 18.78222 | 5.07389 | 7.61712 | 0.91756 | 13.21893 | 14.23283 | 8.18832 | 2.91451 | 8.66533 |
| **Nsmce2** | 11.55524 | 0.02475 | 11.11036 | 10.70593 | 10.73548 | 11.68542 | 6.07462 | 5.56725 | 4.27479 | 0.28341 | 7.83370 | 7.28966 | 8.38366 | 10.91083 |
| **Nsmce4a** | 28.97058 | 0.13377 | 12.76752 | 9.71409 | 28.32561 | 8.98586 | 4.46456 | 1.76267 | 4.45541 | 0.16859 | 9.47820 | 10.15000 | 8.75615 | 16.34228 |
| **Nsmf** | 0.05953 | 0.00453 | 0.28747 | 0.00083 | 0.10899 | 0.20987 | 0.00409 | 0.00627 | 0.01715 | 0.00696 | 0.00471 | 0.00717 | 0.00774 | 0.10300 |
| **Nsrp1** | 6.66736 | 0.09281 | 7.77078 | 5.43387 | 6.69683 | 5.99870 | 0.50003 | 1.11265 | 7.95754 | 0.02873 | 3.58448 | 4.51024 | 4.51653 | 9.09301 |
| **Nsun2** | 23.30646 | 0.13969 | 25.40908 | 15.05484 | 24.22647 | 15.78970 | 5.71279 | 6.86059 | 9.62920 | 10.72688 | 11.47076 | 11.91441 | 8.21996 | 24.86800 |
| **Nsun3** | 0.17268 | 0.00357 | 0.00000 | 0.00000 | 0.18399 | 0.26181 | 0.00389 | 0.00245 | 0.00921 | 0.00518 | 0.48990 | 0.52260 | 0.00619 | 0.01356 |
| **Nsun4** | 1.65704 | 0.02841 | 0.27020 | 1.50820 | 3.24173 | 0.57491 | 0.00993 | 0.59036 | 1.12344 | 0.00000 | 2.65475 | 1.05701 | 3.16328 | 5.06734 |
| **Nsun5** | 7.68425 | 0.21799 | 16.99036 | 10.52008 | 14.40886 | 3.57854 | 2.14700 | 2.15398 | 0.00000 | 0.07728 | 6.36823 | 4.18813 | 8.96315 | 4.70459 |

| | | | | | | | | | | | | | | |
|---|---|---|---|---|---|---|---|---|---|---|---|---|---|---|
| **Nsun6** | 0.92444 | 0.01946 | 0.59535 | 0.45002 | 1.70420 | 1.54441 | 0.41706 | 0.48617 | 0.29899 | 0.02754 | 1.04582 | 0.73792 | 0.00491 | 2.68923 |
| **Nsun7** | 0.29726 | 0.00367 | 0.05555 | 0.60670 | 0.85135 | 0.00000 | 0.00326 | 0.00350 | 3.64935 | 0.00616 | 0.00295 | 0.02003 | 1.11691 | 1.99517 |
| **Nt5c** | 3.00349 | 0.04932 | 3.65000 | 12.81473 | 5.40512 | 0.53722 | 0.10866 | 5.41330 | 0.30941 | 8.12458 | 0.69336 | 2.19654 | 0.01194 | 1.76632 |
| **Nt5c2** | 1.82811 | 0.04718 | 3.74601 | 4.37767 | 2.44701 | 2.97895 | 1.13792 | 1.01403 | 1.56919 | 0.03815 | 0.60337 | 1.56979 | 3.25475 | 3.45360 |
| **Nt5c3** | 3.95849 | 0.00231 | 1.32725 | 0.00000 | 2.10271 | 1.03990 | 0.00321 | 0.00000 | 0.02205 | 0.00265 | 0.04228 | 1.62303 | 0.74915 | 1.41169 |
| **Nt5c3b** | 8.65801 | 0.14768 | 18.06641 | 11.53227 | 11.42905 | 2.07504 | 0.01157 | 1.75302 | 0.14717 | 0.10588 | 6.81855 | 5.14171 | 0.28957 | 7.45800 |
| **Nt5dc1** | 1.38474 | 0.00332 | 0.63024 | 1.06751 | 1.09782 | 2.22050 | 0.01551 | 0.01259 | 0.06866 | 0.08723 | 0.80369 | 0.19907 | 1.16087 | 0.76124 |
| **Nt5dc2** | 0.21004 | 0.00325 | 0.90930 | 0.10593 | 0.00000 | 0.00000 | 0.00347 | 0.57069 | 0.00588 | 0.01300 | 0.00000 | 1.12742 | 0.00491 | 0.05493 |
| **Nt5dc3** | 0.10395 | 7.14412 | 0.00000 | 0.92393 | 0.51424 | 3.01076 | 2.81825 | 2.24893 | 0.23896 | 0.57283 | 0.00000 | 1.26119 | 0.55590 | 1.26008 |
| **Nt5e** | 0.00612 | 0.03056 | 0.01146 | 0.00555 | 0.00876 | 0.01244 | 0.00384 | 0.01195 | 0.00833 | 0.00732 | 0.00369 | 1.04635 | 0.00374 | 0.11033 |
| **Nt5m** | 0.23754 | 0.00347 | 0.42832 | 0.25922 | 0.00000 | 1.05718 | 0.00364 | 0.85586 | 0.00812 | 0.01384 | 0.02171 | 0.74288 | 0.00437 | 0.00413 |
| **Ntan1** | 0.93281 | 0.10210 | 1.96382 | 0.84459 | 0.87102 | 2.81741 | 0.34842 | 5.54440 | 0.00180 | 11.86834 | 0.00000 | 3.30172 | 3.22406 | 0.00000 |
| **Ntf5** | 0.31169 | 0.01300 | 0.01864 | 0.03022 | 0.35135 | 0.04833 | 0.00416 | 0.79434 | 0.12836 | 0.00759 | 0.06979 | 0.00713 | 0.05328 | 0.07047 |
| **Nthl1** | 1.00116 | 0.02998 | 1.84979 | 3.45520 | 0.51250 | 2.61522 | 0.00260 | 0.06018 | 0.00660 | 0.01229 | 5.55779 | 2.53140 | 0.00402 | 0.61075 |
| **Ntmt1** | 3.60831 | 5.95052 | 6.97549 | 6.15609 | 12.20410 | 3.12336 | 0.00000 | 4.66364 | 0.00000 | 0.00543 | 2.31192 | 1.58332 | 6.03837 | 0.44360 |
| **Ntn1** | 0.03836 | 0.00439 | 0.01828 | 0.03147 | 0.01166 | 0.01101 | 0.00425 | 0.00899 | 0.00952 | 0.00703 | 0.01107 | 0.01168 | 0.01149 | 0.02098 |
| **Ntn3** | 0.04823 | 0.00434 | 0.02009 | 0.05105 | 0.00487 | 0.00976 | 0.00418 | 0.00827 | 0.12715 | 0.00616 | 0.01217 | 0.01332 | 0.00618 | 0.00770 |
| **Ntpcr** | 0.00231 | 0.01016 | 0.04309 | 0.06678 | 0.15171 | 0.20221 | 0.00379 | 0.18082 | 0.01010 | 0.00555 | 0.00000 | 1.87572 | 0.00722 | 0.00968 |
| **Ntrk2** | 0.02741 | 0.00450 | 0.02155 | 0.01436 | 0.01441 | 0.04290 | 0.02175 | 0.02289 | 0.11070 | 0.01979 | 0.01428 | 0.01388 | 0.32591 | 0.01091 |
| **Nuak1** | 0.05808 | 0.06223 | 0.08393 | 0.00000 | 0.00294 | 0.99442 | 0.00242 | 2.39723 | 0.02144 | 0.02304 | 0.27653 | 2.94938 | 0.00734 | 0.00000 |
| **Nuak2** | 0.01638 | 0.02559 | 0.01455 | 0.00000 | 0.01168 | 0.00377 | 0.00436 | 0.71140 | 0.01258 | 0.00862 | 0.00841 | 0.01226 | 0.00741 | 0.01108 |
| **Nub1** | 23.56277 | 3.97381 | 33.84437 | 19.17194 | 34.53668 | 12.70192 | 3.29233 | 8.70137 | 24.88507 | 13.30447 | 17.94744 | 6.73823 | 15.19971 | 35.75010 |
| **Nubp1** | 2.25329 | 0.17813 | 2.94280 | 2.50002 | 5.58152 | 3.01503 | 0.00774 | 3.60551 | 0.89452 | 0.52668 | 1.98500 | 1.80432 | 1.53151 | 4.10321 |
| **Nubp2** | 10.57439 | 0.47701 | 21.27148 | 21.28205 | 17.67291 | 19.16468 | 3.54884 | 11.02829 | 2.59607 | 0.35562 | 14.04638 | 13.45566 | 5.52479 | 16.23814 |
| **Nubpl** | 1.18373 | 0.16000 | 0.92034 | 0.00000 | 0.94386 | 0.83771 | 0.04655 | 1.86223 | 0.06054 | 0.03110 | 0.00000 | 2.61257 | 0.35875 | 0.00000 |
| **Nucb1** | 0.28936 | 6.16607 | 5.10069 | 9.84242 | 2.22593 | 4.98648 | 1.51575 | 3.84820 | 0.00000 | 0.06858 | 0.00000 | 5.70434 | 1.59564 | 0.17238 |
| **Nucb2** | 2.84336 | 0.01860 | 6.63398 | 4.14412 | 2.58814 | 1.67212 | 0.00466 | 2.53581 | 5.18214 | 0.00000 | 3.08892 | 6.87928 | 1.55112 | 5.38925 |
| **Nucks1** | 13.45235 | 1.39940 | 12.90590 | 14.73967 | 12.53528 | 21.53037 | 4.58637 | 10.64425 | 21.19797 | 10.02158 | 12.73428 | 17.83224 | 10.15918 | 9.34289 |
| **Nudc** | 16.42823 | 4.40482 | 18.92014 | 24.31478 | 18.94649 | 16.28201 | 2.49171 | 10.70544 | 2.21070 | 15.72335 | 9.53205 | 12.87462 | 11.90829 | 5.99939 |
| **Nudcd1** | 6.59848 | 0.00000 | 5.35635 | 3.65692 | 3.37990 | 3.39432 | 1.22167 | 0.51079 | 4.79557 | 0.00000 | 9.31578 | 2.68832 | 4.72269 | 7.60386 |
| **Nudcd2** | 7.32681 | 0.11130 | 11.80654 | 11.26997 | 14.03574 | 2.60810 | 5.66813 | 5.29146 | 8.65575 | 0.28787 | 4.35421 | 6.21731 | 2.94346 | 0.00000 |
| **Nudcd3** | 3.56865 | 6.77507 | 7.41537 | 6.56579 | 1.94532 | 8.81239 | 2.99086 | 3.11619 | 3.87110 | 1.10485 | 5.57314 | 3.85112 | 2.18791 | 2.59668 |
| **Nudt1** | 1.41141 | 0.03303 | 0.13089 | 4.18209 | 0.92974 | 3.92321 | 0.02488 | 2.17991 | 0.50918 | 7.24496 | 4.96411 | 0.81500 | 0.00000 | 1.72566 |
| **Nudt10** | 0.06958 | 0.00396 | 0.02857 | 0.31257 | 0.00369 | 0.08091 | 2.07781 | 0.00876 | 0.00940 | 0.00961 | 0.00428 | 0.01730 | 0.01050 | 0.00769 |
| **Nudt11** | 0.82713 | 0.00360 | 0.00000 | 0.22402 | 0.00000 | 0.31844 | 0.10531 | 0.01651 | 0.01655 | 0.01177 | 0.14509 | 0.98022 | 0.00626 | 0.05933 |
| **Nudt12** | 0.00000 | 0.00359 | 0.00000 | 0.75263 | 0.08907 | 0.63221 | 0.00356 | 0.00425 | 2.51064 | 0.00573 | 0.00000 | 0.76039 | 0.00000 | 0.00000 |
| **Nudt13** | 0.41033 | 0.00386 | 0.28585 | 0.00000 | 1.45752 | 0.74427 | 0.00385 | 1.69904 | 0.75081 | 0.00908 | 1.43860 | 0.20656 | 0.00000 | 0.88412 |
| **Nudt14** | 0.37252 | 0.07088 | 0.10118 | 0.81985 | 0.08540 | 1.14470 | 1.15249 | 1.64426 | 0.92140 | 0.05342 | 0.00000 | 1.02749 | 1.14785 | 0.80127 |
| **Nudt15** | 1.52684 | 0.00322 | 3.35070 | 3.02090 | 1.16589 | 1.02345 | 0.00838 | 0.00000 | 0.21030 | 0.00673 | 0.00000 | 0.70890 | 0.08209 | 3.28621 |
| **Nudt16l1** | 0.75708 | 7.65798 | 4.10442 | 6.48592 | 1.88258 | 6.31415 | 5.27667 | 6.57322 | 0.57418 | 0.26471 | 5.09904 | 4.69101 | 0.00000 | 0.04253 |
| **Nudt17** | 0.45990 | 0.02080 | 0.35253 | 0.73972 | 0.00000 | 0.62798 | 0.00363 | 0.30874 | 0.00648 | 0.01820 | 0.00000 | 2.07873 | 0.00633 | 0.15782 |
| **Nudt18** | 0.00372 | 0.00447 | 0.03157 | 0.29791 | 0.32457 | 0.00000 | 0.00517 | 0.21830 | 0.00987 | 0.04392 | 0.00713 | 0.01024 | 0.52667 | 0.03487 |
| **Nudt19** | 11.02794 | 0.00742 | 3.36493 | 7.34223 | 4.28119 | 4.53822 | 3.90864 | 0.91552 | 0.90802 | 0.10827 | 3.63492 | 2.40511 | 1.08228 | 1.55650 |
| **Nudt2** | 5.89526 | 0.04877 | 4.28820 | 8.05719 | 5.45220 | 5.95146 | 3.22562 | 4.01770 | 1.35333 | 0.14329 | 4.31240 | 3.32636 | 0.00000 | 4.53450 |
| **Nudt21** | 10.19646 | 2.37623 | 8.26953 | 18.44898 | 18.90714 | 4.08668 | 0.93134 | 4.62405 | 7.24519 | 7.22129 | 2.29259 | 11.18648 | 6.96756 | 7.29145 |
| **Nudt22** | 0.95652 | 5.86115 | 2.35153 | 2.39697 | 1.58107 | 1.43481 | 0.00693 | 7.06958 | 0.28516 | 0.05420 | 0.11214 | 0.41833 | 0.09148 | 0.70749 |
| **Nudt3** | 5.18244 | 0.04597 | 11.02684 | 5.14610 | 8.49103 | 8.33816 | 2.08690 | 3.66658 | 6.35479 | 0.10603 | 2.08830 | 3.59809 | 5.13999 | 7.90397 |
| **Nudt4** | 30.50126 | 0.82111 | 28.62619 | 13.20577 | 36.22853 | 26.13337 | 7.05004 | 9.94601 | 36.53811 | 10.10445 | 16.62628 | 14.61470 | 23.10674 | 31.08735 |
| **Nudt5** | 0.05087 | 0.20256 | 0.32917 | 2.68390 | 0.59975 | 8.14698 | 0.02346 | 5.47691 | 0.93892 | 0.05178 | 7.12436 | 9.34217 | 3.04613 | 0.00000 |
| **Nudt6** | 0.01103 | 0.00380 | 0.43767 | 0.05400 | 0.68422 | 0.62572 | 0.00401 | 0.03416 | 0.01496 | 0.01343 | 0.37929 | 0.03495 | 0.37663 | 0.00484 |
| **Nudt7** | 0.02439 | 0.00372 | 0.05343 | 0.18136 | 0.13207 | 0.19431 | 0.01371 | 0.84203 | 0.02396 | 0.00648 | 0.27091 | 0.00278 | 0.53342 | 0.07514 |
| **Nudt8** | 0.00240 | 0.01101 | 0.03657 | 0.26215 | 0.27637 | 0.57139 | 0.00413 | 0.05786 | 0.02368 | 0.00487 | 0.00349 | 0.00586 | 0.01196 | 0.04909 |
| **Nudt9** | 3.03617 | 0.04160 | 16.32758 | 9.06750 | 5.72385 | 3.34475 | 0.49303 | 2.83252 | 0.00000 | 9.91423 | 4.62551 | 4.45179 | 0.00000 | 1.71266 |
| **Nuf2** | 5.39738 | 0.04430 | 2.32243 | 2.64961 | 4.37889 | 4.87412 | 1.34933 | 0.00000 | 4.86755 | 0.04593 | 2.72969 | 3.91979 | 1.16824 | 6.21377 |
| **Nufip1** | 12.27430 | 0.03869 | 12.11714 | 10.01219 | 25.10547 | 3.32806 | 2.57589 | 4.74045 | 9.81980 | 0.04016 | 6.83402 | 4.96039 | 14.90052 | 17.60636 |

| | | | | | | | | | | | | | | |
|---|---|---|---|---|---|---|---|---|---|---|---|---|---|---|
| **Nufip2** | 2.67242 | 0.03201 | 7.15050 | 2.18322 | 3.07409 | 2.01686 | 0.35654 | 2.92005 | 20.84607 | 0.05406 | 1.98579 | 1.44041 | 10.08774 | 6.46806 |
| **Numa1** | 1.38313 | 0.25686 | 7.85989 | 4.86218 | 3.44405 | 11.23981 | 0.03695 | 7.86164 | 0.31636 | 0.41058 | 7.52569 | 4.91131 | 2.59412 | 4.48910 |
| **Numb** | 1.01625 | 0.01902 | 0.01121 | 0.00000 | 0.15380 | 0.01434 | 0.00396 | 1.88402 | 0.22952 | 0.01088 | 0.00000 | 1.00682 | 1.51931 | 0.37282 |
| **Numbl** | 0.07429 | 0.00389 | 0.27799 | 0.93619 | 2.02131 | 0.04823 | 0.00504 | 0.06745 | 0.01024 | 0.00493 | 0.07333 | 0.82847 | 0.05880 | 0.00000 |
| **Nup107** | 2.92302 | 0.02329 | 4.09053 | 0.00000 | 4.17393 | 2.26128 | 0.34574 | 0.06700 | 11.44583 | 0.01432 | 4.90239 | 1.91267 | 7.66570 | 16.32090 |
| **Nup133** | 15.02797 | 0.93430 | 17.13917 | 7.41115 | 14.06285 | 10.92720 | 2.90388 | 2.18764 | 10.66597 | 0.11861 | 6.47400 | 5.85210 | 9.01485 | 14.51036 |
| **Nup153** | 10.39310 | 0.71242 | 14.45007 | 4.07188 | 12.81020 | 7.40663 | 3.26897 | 2.61447 | 19.36383 | 0.04461 | 8.59745 | 5.30103 | 5.16002 | 13.65915 |
| **Nup155** | 7.09238 | 0.35129 | 9.24202 | 4.82724 | 8.40049 | 3.48974 | 0.00373 | 0.00000 | 17.75440 | 0.00000 | 7.81360 | 2.87455 | 8.32696 | 13.57970 |
| **Nup160** | 5.83083 | 0.48149 | 9.62984 | 0.67431 | 9.67862 | 6.37326 | 0.03807 | 0.00000 | 16.33945 | 0.00000 | 7.52755 | 3.41680 | 3.33680 | 13.94001 |
| **Nup188** | 0.89672 | 0.03799 | 2.64806 | 4.48083 | 6.16670 | 2.34894 | 0.01190 | 1.90346 | 0.02658 | 4.35239 | 4.17104 | 4.46518 | 2.37043 | 1.25883 |
| **Nup205** | 8.11170 | 0.04132 | 13.64153 | 1.60757 | 14.37838 | 10.08730 | 0.49144 | 0.41022 | 15.66911 | 0.02753 | 4.23712 | 9.08891 | 4.46702 | 12.17864 |
| **Nup210** | 0.00000 | 0.03396 | 1.81162 | 0.81629 | 0.00000 | 0.58206 | 0.00395 | 0.29830 | 0.02225 | 0.00659 | 0.00164 | 0.25234 | 0.32460 | 0.18567 |
| **Nup210l** | 0.46874 | 0.00422 | 0.03359 | 0.00273 | 0.50301 | 0.00885 | 0.00413 | 0.00975 | 0.00584 | 0.00604 | 0.00769 | 0.00621 | 1.06096 | 0.11711 |
| **Nup214** | 3.73443 | 0.09467 | 6.53901 | 5.44365 | 7.38603 | 6.30404 | 0.00678 | 0.80975 | 1.02253 | 4.14559 | 4.04139 | 6.18986 | 7.81916 | 3.63786 |
| **Nup35** | 17.59089 | 0.12819 | 22.40131 | 11.40767 | 26.88293 | 14.10252 | 2.95909 | 6.16313 | 24.48684 | 0.03293 | 14.23865 | 7.06776 | 14.16207 | 24.20128 |
| **Nup37** | 9.30548 | 0.06888 | 12.35153 | 4.83227 | 6.56210 | 5.01605 | 0.09130 | 6.27145 | 10.45637 | 9.71258 | 9.34829 | 5.75549 | 1.17222 | 6.75918 |
| **Nup43** | 12.67035 | 2.56381 | 13.53494 | 10.93945 | 20.97885 | 9.99700 | 3.04833 | 5.18529 | 11.95605 | 0.14374 | 11.18974 | 6.61517 | 8.30394 | 16.13825 |
| **Nup50** | 10.68056 | 0.11651 | 12.43766 | 9.61051 | 10.89128 | 11.83148 | 6.13499 | 6.12360 | 8.17762 | 0.22206 | 6.47391 | 7.39783 | 8.70366 | 11.62013 |
| **Nup54** | 4.47191 | 1.65179 | 6.12044 | 0.00000 | 3.34559 | 2.79387 | 0.52822 | 2.49320 | 6.44321 | 0.00000 | 9.00315 | 4.56008 | 6.42262 | 9.19564 |
| **Nup62** | 4.47726 | 8.98733 | 17.69366 | 11.35309 | 8.11344 | 12.26351 | 4.00816 | 7.22633 | 0.25146 | 0.21944 | 9.23703 | 10.27683 | 9.89617 | 5.94348 |
| **Nup62cl** | 0.03356 | 0.00416 | 0.55190 | 0.00000 | 0.01357 | 0.27107 | 0.00416 | 0.00132 | 0.04006 | 0.01254 | 0.00421 | 0.14581 | 0.00673 | 1.90976 |
| **Nup85** | 17.38097 | 3.49620 | 23.74626 | 10.00100 | 20.89862 | 10.66571 | 0.89411 | 4.62233 | 8.56792 | 0.03710 | 7.36902 | 10.61261 | 0.00000 | 20.01875 |
| **Nup88** | 7.57246 | 0.05149 | 8.40975 | 2.58352 | 3.84520 | 3.80258 | 1.92917 | 2.50359 | 8.63890 | 1.37818 | 9.11409 | 4.34424 | 3.14626 | 9.30347 |
| **Nup93** | 4.39510 | 2.48433 | 9.01008 | 8.79449 | 8.46677 | 8.32743 | 4.81039 | 3.51423 | 4.55176 | 0.38622 | 6.54085 | 3.54360 | 2.00080 | 8.48680 |
| **Nup98** | 5.28702 | 0.12575 | 1.62100 | 0.10340 | 3.13204 | 4.04162 | 2.65182 | 0.91248 | 11.25185 | 1.70768 | 0.43978 | 2.50121 | 13.69895 | 6.73154 |
| **Nupl1** | 0.42055 | 0.38781 | 1.90780 | 1.12001 | 0.93336 | 2.01092 | 0.00983 | 0.16547 | 2.21575 | 0.00403 | 2.36549 | 1.27601 | 0.54653 | 2.05298 |
| **Nupl2** | 3.20983 | 4.42421 | 1.10095 | 0.35702 | 1.40162 | 3.33941 | 0.18325 | 0.66632 | 3.06137 | 0.01356 | 0.15876 | 0.14967 | 2.56837 | 0.48013 |
| **Nupr1** | 0.39740 | 0.02002 | 0.10075 | 1.58554 | 1.04561 | 0.00000 | 0.00364 | 1.11413 | 0.85167 | 0.00799 | 0.01686 | 0.01066 | 5.86777 | 1.24838 |
| **Nupr1l** | 0.00817 | 0.00444 | 0.01528 | 0.01312 | 0.03280 | 0.01376 | 0.00422 | 0.00986 | 0.01533 | 0.00659 | 0.00843 | 0.00965 | 0.05321 | 0.05631 |
| **Nus1** | 8.22563 | 0.01026 | 3.51524 | 0.53185 | 2.48730 | 9.44861 | 3.55478 | 2.82476 | 14.31977 | 0.11385 | 4.79853 | 8.14939 | 10.33016 | 4.89529 |
| **Nusap1** | 5.72860 | 0.02270 | 2.14010 | 0.37049 | 6.18551 | 4.50824 | 3.96074 | 6.32326 | 8.49542 | 0.29143 | 6.78865 | 0.73609 | 6.19236 | 9.40773 |
| **Nutf2** | 0.08930 | 0.01049 | 0.67289 | 0.80262 | 0.34974 | 1.49639 | 0.00330 | 2.17545 | 0.00718 | 0.05443 | 2.87386 | 0.94319 | 0.02087 | 0.12495 |
| **Nutm1** | 0.54536 | 0.00410 | 0.27427 | 0.09426 | 0.13113 | 0.00489 | 0.36666 | 0.00300 | 0.67637 | 0.02222 | 0.00385 | 0.01649 | 2.27465 | 0.10472 |
| **Nvl** | 6.80747 | 0.41591 | 10.08403 | 4.47601 | 9.84720 | 4.85851 | 0.39139 | 0.62890 | 14.71353 | 0.05540 | 4.92972 | 4.53368 | 5.70006 | 13.64481 |
| **Nwd1** | 0.01725 | 0.02346 | 0.31072 | 0.00000 | 0.10806 | 0.00000 | 0.00468 | 0.74664 | 0.04738 | 0.08625 | 0.00576 | 0.01282 | 0.39923 | 0.04277 |
| **Nxf1** | 11.59758 | 4.98510 | 17.51184 | 7.81929 | 24.38209 | 9.78254 | 4.43540 | 7.41150 | 22.43420 | 0.22667 | 7.53041 | 7.17635 | 11.98543 | 25.86216 |
| **Nxf2** | 1.71110 | 0.00413 | 0.42589 | 0.00000 | 0.11624 | 0.00000 | 0.00420 | 0.01222 | 0.12624 | 0.00604 | 0.00505 | 0.00607 | 0.00767 | 1.23154 |
| **Nxf7** | 0.00689 | 1.63540 | 0.00314 | 0.00308 | 0.14075 | 0.50128 | 0.00343 | 3.38007 | 0.01056 | 0.00584 | 0.00477 | 0.01088 | 0.00629 | 0.00401 |
| **Nxn** | 0.85278 | 0.00361 | 1.83629 | 0.00000 | 1.22328 | 0.52798 | 0.00328 | 0.50029 | 0.02271 | 0.00198 | 0.30940 | 0.71261 | 0.15190 | 0.95529 |
| **Nxnl1** | 0.00503 | 0.00435 | 0.01491 | 0.01250 | 0.01084 | 0.01312 | 0.00427 | 0.00926 | 0.01003 | 0.00658 | 0.00911 | 0.00929 | 0.33235 | 0.00987 |
| **Nxnl2** | 0.00717 | 0.01160 | 0.01232 | 0.00367 | 0.00794 | 0.02352 | 0.00434 | 0.01657 | 0.00795 | 0.00646 | 0.00645 | 0.06550 | 0.00695 | 0.19262 |
| **Nxpe3** | 0.16398 | 0.00259 | 0.00000 | 0.39038 | 0.03115 | 0.26354 | 0.00358 | 0.20367 | 0.66239 | 0.02033 | 0.02532 | 1.84387 | 0.04489 | 0.15252 |
| **Nxpe4** | 0.01316 | 0.00419 | 0.00755 | 0.15700 | 0.00887 | 0.01192 | 0.00424 | 0.00853 | 0.01630 | 0.00745 | 0.00682 | 0.19495 | 0.06026 | 0.00470 |
| **Nxpe5** | 0.00704 | 0.00450 | 0.05676 | 0.00302 | 0.40962 | 0.01000 | 0.00413 | 0.01081 | 0.19503 | 0.00762 | 0.00750 | 0.00802 | 0.00246 | 0.00000 |
| **Nxt1** | 3.02300 | 0.11807 | 8.33524 | 4.79416 | 11.75761 | 1.80865 | 3.09367 | 10.26687 | 0.24869 | 0.07195 | 0.95329 | 4.87344 | 8.39423 | 0.22889 |
| **Nxt2** | 0.26588 | 0.00420 | 0.22897 | 0.00000 | 0.00000 | 0.02280 | 0.04418 | 0.00691 | 3.66507 | 0.06239 | 0.00439 | 0.01045 | 1.58582 | 2.73821 |
| **Nyap1** | 0.01701 | 0.01239 | 0.00626 | 0.36112 | 0.77185 | 0.00467 | 0.00526 | 0.25483 | 0.00629 | 0.00663 | 0.00590 | 0.00528 | 0.16579 | 0.14007 |
| **Nynrin** | 0.32434 | 0.19471 | 0.00000 | 2.47235 | 0.00000 | 6.29288 | 0.64973 | 5.28917 | 0.00281 | 0.09262 | 4.57978 | 4.32243 | 0.00979 | 0.00000 |
| **Nyx** | 0.02288 | 0.00436 | 0.02964 | 0.01782 | 0.00719 | 0.01127 | 0.00423 | 0.00921 | 0.01065 | 0.00639 | 0.01203 | 0.01079 | 0.11833 | 0.15753 |
| **Oacyl** | 0.01722 | 0.00441 | 0.01504 | 0.01431 | 0.00843 | 0.03958 | 0.00438 | 0.00726 | 0.01041 | 0.00633 | 0.00770 | 0.01349 | 0.00811 | 0.12387 |
| **Oaf** | 0.05012 | 0.00425 | 0.01582 | 0.00804 | 0.01037 | 0.01508 | 0.00434 | 0.01038 | 0.00973 | 0.00689 | 0.00763 | 0.03682 | 0.22087 | 0.00915 |
| **Oard1** | 5.00809 | 4.94140 | 5.22094 | 14.28247 | 5.47379 | 12.22018 | 2.22388 | 4.50154 | 2.08901 | 0.72041 | 9.30386 | 8.81090 | 0.73371 | 3.34245 |
| **Oas1a** | 0.00000 | 0.02632 | 0.86867 | 1.07756 | 0.00000 | 1.60234 | 1.90225 | 3.07929 | 0.21019 | 0.06399 | 1.05592 | 3.86826 | 0.00148 | 0.00000 |
| **Oas1b** | 0.04397 | 0.00453 | 0.03132 | 0.03962 | 0.03233 | 0.04030 | 0.05058 | 0.06522 | 0.08684 | 0.01396 | 0.02752 | 0.02609 | 0.27527 | 0.08556 |
| **Oas1c** | 0.00000 | 0.00389 | 0.23964 | 0.00000 | 0.94084 | 0.35839 | 0.00376 | 0.00259 | 0.42480 | 0.00658 | 0.27289 | 0.57703 | 1.22608 | 1.77802 |

| | | | | | | | | | | | | | | |
|---|---|---|---|---|---|---|---|---|---|---|---|---|---|---|
| **Oas1d** | 0.00000 | 0.00871 | 1.53845 | 0.00000 | 0.80881 | 1.03011 | 0.00813 | 0.17787 | 1.21965 | 0.00834 | 1.85821 | 1.22242 | 5.63374 | 6.60159 |
| **Oas1e** | 0.00000 | 0.00277 | 0.22707 | 0.00000 | 0.05820 | 0.17338 | 0.00352 | 0.00830 | 0.04119 | 0.01816 | 0.04589 | 1.53816 | 1.49678 | 0.42214 |
| **Oas1f** | 0.00359 | 0.01234 | 0.01125 | 0.00000 | 0.01657 | 0.03215 | 0.00369 | 0.11061 | 0.03239 | 0.00520 | 0.00360 | 0.81658 | 0.43606 | 0.10392 |
| **Oas1g** | 0.00000 | 0.03466 | 0.63739 | 0.99055 | 0.00000 | 1.40258 | 1.52986 | 2.60747 | 0.17709 | 0.07220 | 1.30957 | 3.21818 | 0.00282 | 0.00000 |
| **Oas1h** | 0.33628 | 0.00421 | 0.01249 | 0.01431 | 0.00499 | 0.01011 | 0.00418 | 0.00970 | 0.00791 | 0.00653 | 0.01071 | 0.00650 | 1.02234 | 0.01352 |
| **Oas3** | 0.12312 | 0.22574 | 0.09064 | 0.14649 | 0.30673 | 0.08938 | 0.09205 | 0.07703 | 0.41514 | 0.00735 | 0.14262 | 0.09039 | 0.69182 | 0.27842 |
| **Oasl1** | 0.00528 | 0.00435 | 0.01500 | 0.01245 | 0.01061 | 0.01351 | 0.00425 | 0.00923 | 0.01106 | 0.00664 | 0.00899 | 0.00936 | 0.31074 | 0.00282 |
| **Oasl2** | 2.39982 | 0.00340 | 0.95175 | 0.00000 | 2.68203 | 0.00000 | 0.00374 | 0.00000 | 0.02988 | 0.00438 | 0.00334 | 0.49060 | 0.94009 | 0.82925 |
| **Oat** | 0.01310 | 7.06732 | 0.00000 | 0.00000 | 3.36666 | 3.89935 | 1.32169 | 10.70954 | 0.26137 | 0.07693 | 4.46685 | 2.71133 | 8.57579 | 4.89323 |
| **Oaz1** | 0.25892 | 0.42235 | 0.57059 | 1.36040 | 0.89330 | 0.69352 | 0.14284 | 1.23805 | 0.02941 | 0.13818 | 0.53159 | 1.10667 | 1.04248 | 0.12760 |
| **Oaz2** | 2.53242 | 1.91815 | 4.64068 | 7.78761 | 7.59238 | 5.67847 | 4.25227 | 1.99363 | 1.34070 | 0.18219 | 2.25610 | 5.12880 | 8.10932 | 4.94156 |
| **Oaz3** | 0.00779 | 0.00436 | 0.04329 | 0.00607 | 0.09705 | 0.01219 | 0.00418 | 0.00983 | 0.00928 | 0.00632 | 0.00846 | 0.00919 | 0.30601 | 0.00626 |
| **Obfc1** | 2.06826 | 0.01788 | 2.52149 | 1.69110 | 2.27657 | 1.71373 | 1.33943 | 3.65425 | 1.04213 | 0.02997 | 3.42990 | 0.55197 | 1.09512 | 2.45074 |
| **Obox1** | 0.00000 | 0.00384 | 0.00000 | 0.06140 | 0.33136 | 0.00356 | 0.00389 | 0.00643 | 0.48290 | 0.00760 | 0.00775 | 0.00360 | 2.71116 | 2.45688 |
| **Obox2** | 0.00270 | 0.00385 | 0.00000 | 0.11102 | 0.53714 | 0.00419 | 0.00389 | 0.00618 | 0.25002 | 0.00774 | 0.00474 | 0.00342 | 1.97238 | 3.17405 |
| **Obox3** | 0.00000 | 0.00293 | 0.00000 | 0.01908 | 0.08973 | 0.00770 | 0.00943 | 0.00678 | 0.26157 | 0.00573 | 0.01123 | 0.00642 | 12.54889 | 0.64157 |
| **Obox5** | 0.00000 | 0.00354 | 0.06736 | 0.00000 | 1.17327 | 0.00235 | 0.00397 | 0.00663 | 0.17980 | 0.00679 | 0.00372 | 0.35712 | 4.75647 | 2.76341 |
| **Obox6** | 32.69771 | 0.00973 | 28.93057 | 3.10862 | 39.86292 | 0.00000 | 0.00000 | 0.00000 | 35.14535 | 0.00000 | 2.38310 | 0.12751 | 18.54449 | 38.30097 |
| **Obox7** | 0.00123 | 0.00404 | 0.02025 | 0.01003 | 0.07279 | 0.01121 | 0.00407 | 0.00717 | 0.01715 | 0.00668 | 0.00721 | 0.01516 | 1.62246 | 0.90490 |
| **Obp2a** | 0.00855 | 1.32071 | 0.01031 | 0.00474 | 0.01312 | 0.39693 | 0.00372 | 0.00702 | 0.01079 | 0.00674 | 0.01008 | 0.00324 | 0.00783 | 0.00812 |
| **Obsl1** | 0.01749 | 0.04610 | 0.00944 | 0.00003 | 0.02908 | 0.01219 | 0.00407 | 0.30147 | 0.01122 | 0.00666 | 0.00079 | 1.01302 | 0.25930 | 0.00788 |
| **Oc90** | 0.00000 | 0.00487 | 0.56150 | 0.00000 | 0.24533 | 2.86445 | 0.00382 | 0.00536 | 0.34102 | 0.00596 | 0.00307 | 0.00157 | 0.00025 | 0.00000 |
| **Ocel1** | 0.21491 | 0.00508 | 0.02663 | 0.42962 | 0.14061 | 0.01225 | 0.03556 | 0.21231 | 0.00650 | 0.02493 | 0.00559 | 0.09739 | 0.21445 | 0.17961 |
| **Ociad1** | 9.66860 | 0.16506 | 16.53869 | 11.29459 | 9.58951 | 15.32084 | 4.09177 | 13.20133 | 8.64634 | 0.21918 | 14.06144 | 12.05456 | 2.65524 | 11.70294 |
| **Ociad2** | 0.15786 | 0.03046 | 0.08046 | 0.16658 | 0.09975 | 0.46369 | 0.05432 | 0.21983 | 0.99275 | 0.06190 | 0.01593 | 0.77426 | 2.77197 | 0.46678 |
| **Ocln** | 0.35432 | 0.01025 | 2.16501 | 0.62342 | 1.86225 | 0.55583 | 2.76363 | 3.60455 | 8.35673 | 0.05611 | 2.59769 | 0.00805 | 0.37738 | 1.96491 |
| **Ocm** | 5.47322 | 0.00370 | 3.47645 | 0.76081 | 1.99691 | 0.65498 | 0.00311 | 1.00944 | 2.55761 | 0.03007 | 0.24271 | 0.00000 | 2.32209 | 5.27175 |
| **Ocrl** | 2.97813 | 0.02495 | 4.40627 | 0.91961 | 1.50490 | 4.95584 | 2.55545 | 1.45689 | 5.33469 | 0.03908 | 7.38182 | 3.95562 | 0.00000 | 0.56937 |
| **Odc1** | 49.13189 | 13.00473 | 45.31242 | 34.67909 | 51.31289 | 38.85909 | 9.83710 | 27.07404 | 17.21143 | 6.28093 | 21.40600 | 20.57031 | 26.08704 | 46.78954 |
| **Odf2** | 0.07593 | 0.43978 | 1.15287 | 2.06897 | 0.72000 | 1.67521 | 3.63275 | 2.95736 | 0.23872 | 3.59927 | 2.50919 | 4.26121 | 3.86965 | 1.59333 |
| **Odf2l** | 1.30146 | 0.00394 | 0.08162 | 0.12563 | 0.15879 | 0.29068 | 0.03068 | 0.16641 | 0.13771 | 0.01288 | 0.02233 | 0.08736 | 0.00488 | 3.25631 |
| **Odf3b** | 0.00543 | 0.00433 | 0.00499 | 0.19780 | 0.15004 | 0.01079 | 0.00421 | 0.00995 | 0.01089 | 0.00671 | 0.00666 | 0.00688 | 0.00591 | 0.01148 |
| **Ofd1** | 3.24229 | 0.26834 | 3.44416 | 0.41163 | 2.85756 | 0.10864 | 0.01197 | 2.49212 | 4.76462 | 0.01601 | 0.00000 | 2.94902 | 0.62093 | 3.65573 |
| **Ogdh** | 8.48883 | 0.48599 | 13.78551 | 10.84589 | 10.25018 | 17.31159 | 1.34029 | 7.54607 | 7.87370 | 3.12708 | 6.28815 | 11.12260 | 20.00086 | 15.14338 |
| **Ogdhl** | 0.00452 | 0.00993 | 0.00000 | 2.00521 | 0.00409 | 0.23332 | 0.00332 | 0.16362 | 0.01149 | 0.00582 | 0.00367 | 2.04253 | 0.01056 | 0.03865 |
| **Ogfod1** | 3.60244 | 0.02556 | 0.00000 | 1.84564 | 6.76159 | 5.74479 | 0.00912 | 1.34057 | 2.08853 | 0.06700 | 4.92580 | 0.36295 | 1.28167 | 11.97545 |
| **Ogfod2** | 0.10523 | 0.01233 | 0.00004 | 2.83971 | 1.49114 | 0.67127 | 0.00443 | 1.32628 | 0.00674 | 0.01190 | 2.02942 | 0.59590 | 0.50838 | 0.00000 |
| **Ogfod3** | 0.99648 | 0.03597 | 2.37120 | 1.80050 | 0.70514 | 0.99393 | 0.00878 | 1.67881 | 0.22177 | 0.02418 | 0.00000 | 2.31413 | 0.59234 | 0.00000 |
| **Ogfr** | 0.31483 | 0.03335 | 0.62873 | 2.37817 | 1.96229 | 1.47125 | 0.01986 | 5.20735 | 0.00114 | 0.08217 | 1.96950 | 1.27694 | 0.07865 | 0.00000 |
| **Ogfrl1** | 0.77157 | 0.00360 | 0.03234 | 1.40726 | 0.00000 | 1.84048 | 0.00334 | 0.13193 | 2.25551 | 0.01141 | 0.22794 | 0.63210 | 1.16908 | 0.00000 |
| **Ogg1** | 0.20935 | 0.47877 | 0.42815 | 1.09955 | 0.57447 | 0.60333 | 0.66298 | 0.36026 | 0.02337 | 0.01185 | 0.64656 | 1.10383 | 0.27171 | 0.00000 |
| **Ogn** | 0.02826 | 0.00449 | 0.00485 | 0.01389 | 0.12370 | 0.01019 | 0.00424 | 0.01059 | 0.19658 | 0.00644 | 0.00723 | 0.00868 | 0.00472 | 0.10698 |
| **Ogt** | 3.87362 | 0.05076 | 10.94552 | 7.89373 | 2.99554 | 9.30534 | 3.75133 | 1.52192 | 19.61526 | 0.06860 | 7.69464 | 5.56169 | 1.14506 | 3.07496 |
| **Oip5** | 0.17915 | 0.01848 | 1.01588 | 0.41864 | 1.53104 | 0.46283 | 0.06296 | 1.17989 | 0.30516 | 0.04000 | 0.00000 | 1.28227 | 2.26267 | 0.07687 |
| **Ola1** | 10.06300 | 0.14788 | 15.57249 | 13.12058 | 14.39143 | 7.70697 | 2.86891 | 5.42422 | 16.76897 | 0.51145 | 4.36884 | 8.09598 | 1.92006 | 13.93348 |
| **Olah** | 0.00443 | 0.00436 | 0.01294 | 0.01221 | 0.00775 | 0.01229 | 0.00430 | 0.00936 | 0.00671 | 0.00662 | 0.00865 | 0.00931 | 0.25752 | 0.18369 |
| **Olfm1** | 1.25901 | 0.00562 | 0.01539 | 0.78414 | 0.85415 | 0.03013 | 0.00455 | 0.29832 | 0.00873 | 0.00844 | 0.01533 | 0.00480 | 0.36341 | 1.65989 |
| **Olfml1** | 0.01282 | 0.00445 | 0.00041 | 0.01049 | 0.00000 | 0.02004 | 0.00403 | 0.00820 | 1.20130 | 0.00645 | 0.04878 | 0.00771 | 0.00692 | 0.18770 |
| **Olfml2a** | 0.04948 | 0.00429 | 0.01060 | 0.02274 | 0.12592 | 0.01050 | 0.00425 | 0.00889 | 0.00859 | 0.00859 | 0.00775 | 0.00736 | 0.55572 | 0.10205 |
| **Olfr1033** | 0.02573 | 0.01203 | 0.02831 | 0.04555 | 0.06285 | 0.00000 | 0.00419 | 0.00811 | 0.07786 | 0.00509 | 0.22017 | 0.14382 | 0.01030 | 0.14924 |
| **Olfr1055** | 4.89316 | 0.11506 | 0.65011 | 0.12794 | 1.92618 | 1.18752 | 0.18572 | 0.26550 | 1.38846 | 0.06086 | 0.28236 | 0.17904 | 2.29042 | 0.53498 |
| **Olfr118** | 0.01445 | 0.00445 | 0.01192 | 0.01068 | 0.11195 | 0.01287 | 0.00418 | 0.01035 | 0.00974 | 0.00630 | 0.00903 | 0.00931 | 0.01178 | 0.06464 |
| **Olfr1300.ps1** | 0.01079 | 0.00462 | 0.01453 | 0.01302 | 0.01377 | 0.01256 | 0.00433 | 0.01380 | 0.01324 | 0.00742 | 0.00883 | 0.01003 | 0.00950 | 0.01203 |
| **Olfr1372.ps1** | 2.16686 | 0.00464 | 0.76075 | 0.00000 | 0.60700 | 0.15490 | 0.00462 | 0.05230 | 0.82387 | 0.00478 | 0.01549 | 0.00380 | 0.02826 | 0.32605 |
| **Olfr287** | 0.01494 | 0.00459 | 0.01294 | 0.01084 | 0.01320 | 0.01955 | 0.00434 | 0.01191 | 0.03757 | 0.00674 | 0.00871 | 0.00979 | 0.09904 | 0.07774 |

| | | | | | | | | | | | | | | |
|---|---|---|---|---|---|---|---|---|---|---|---|---|---|---|
| **Olfr376** | 1.29029 | 0.00426 | 0.01212 | 0.00995 | 0.00183 | 0.00783 | 0.00426 | 0.01093 | 0.00609 | 0.00662 | 0.00686 | 0.00625 | 0.35344 | 0.00189 |
| **Olfr456** | 0.00599 | 0.00441 | 0.00464 | 0.01987 | 0.01506 | 0.00895 | 0.00424 | 0.01054 | 0.23296 | 0.00829 | 0.00709 | 0.00938 | 0.02257 | 0.18816 |
| **Olfr613** | 0.29841 | 0.89380 | 0.27351 | 0.10976 | 0.20879 | 0.08693 | 0.03554 | 0.19762 | 1.19921 | 0.00666 | 0.10866 | 0.09418 | 0.45464 | 0.36964 |
| **Olfr787** | 0.00790 | 0.00442 | 0.01486 | 0.01265 | 0.01267 | 0.01336 | 0.00428 | 0.00992 | 0.01134 | 0.00677 | 0.00892 | 0.00988 | 0.09453 | 0.00455 |
| **Olfr815** | 0.00938 | 0.00443 | 0.01425 | 0.01298 | 0.01241 | 0.01289 | 0.00428 | 0.01022 | 0.02348 | 0.00683 | 0.00872 | 0.00999 | 0.03953 | 0.00960 |
| **Olfr856.ps1** | 0.29244 | 0.12465 | 0.08461 | 0.05424 | 0.26524 | 0.03853 | 0.00420 | 0.09378 | 0.98548 | 0.01544 | 0.04857 | 0.29394 | 0.33729 | 0.31201 |
| **Olfr976** | 0.00723 | 0.00447 | 0.00230 | 0.01622 | 0.00470 | 0.00828 | 0.00412 | 0.01015 | 0.95479 | 0.00744 | 0.00630 | 0.00976 | 0.00695 | 0.00000 |
| **Oma1** | 0.64440 | 0.00261 | 0.75471 | 2.22164 | 0.15613 | 3.66669 | 0.00416 | 0.54579 | 0.07516 | 0.03772 | 0.60829 | 2.08117 | 0.51893 | 3.31807 |
| **Omd** | 0.86277 | 0.04332 | 0.00000 | 0.31537 | 0.10530 | 1.26205 | 0.16555 | 0.16529 | 1.03342 | 0.37333 | 0.01570 | 0.30826 | 5.96287 | 1.08811 |
| **Omt2a** | 0.22046 | 0.00170 | 0.00000 | 0.29654 | 2.31858 | 0.01008 | 0.00326 | 0.00355 | 2.17104 | 0.00788 | 0.00031 | 0.00000 | 18.06254 | 10.62796 |
| **Omt2b** | 0.13690 | 0.00187 | 0.00000 | 0.43742 | 2.26869 | 0.00000 | 0.00320 | 0.00501 | 2.91972 | 0.00760 | 0.00000 | 0.00000 | 16.47610 | 10.65230 |
| **Onecut2** | 0.03175 | 0.00435 | 0.02055 | 0.00979 | 0.01375 | 0.01753 | 0.00433 | 0.01350 | 0.02068 | 0.00635 | 0.01042 | 0.01011 | 0.01347 | 0.08008 |
| **Onecut3** | 0.01794 | 0.00405 | 0.01478 | 0.40349 | 0.00705 | 0.00974 | 0.00388 | 0.00482 | 0.02005 | 0.00564 | 0.02143 | 0.01810 | 0.15860 | 0.04030 |
| **Ooep** | 0.13630 | 0.42175 | 0.00000 | 1.11319 | 0.39214 | 2.87206 | 2.28287 | 8.25856 | 0.47938 | 3.63640 | 2.38924 | 0.75095 | 2.53625 | 0.38451 |
| **Oog2** | 0.00000 | 0.00423 | 0.03850 | 0.04759 | 0.82409 | 0.00389 | 0.00406 | 0.00807 | 0.14430 | 0.00743 | 0.00884 | 0.00787 | 0.30756 | 1.86945 |
| **Oog3** | 0.00000 | 0.00339 | 0.07427 | 0.15367 | 3.27803 | 0.00000 | 0.00339 | 0.00756 | 5.91974 | 0.02486 | 0.00000 | 0.00000 | 5.38925 | 6.06190 |
| **Oog4** | 0.57137 | 0.01879 | 0.10637 | 0.06263 | 0.65664 | 0.12061 | 0.01574 | 0.36152 | 0.77118 | 0.43562 | 0.00008 | 0.00000 | 1.15685 | 0.79976 |
| **Oosp1** | 0.00000 | 0.01057 | 1.48163 | 0.11892 | 0.70697 | 0.12921 | 0.00316 | 0.00000 | 3.02562 | 0.00612 | 0.00528 | 0.00000 | 9.92391 | 4.28510 |
| **Oosp2** | 0.00394 | 0.00423 | 0.02095 | 0.00000 | 0.51616 | 0.13959 | 0.00401 | 0.00524 | 0.01861 | 0.00735 | 0.13836 | 0.00682 | 0.51696 | 0.00000 |
| **Oosp3** | 0.03800 | 0.00411 | 0.00000 | 0.02409 | 0.64520 | 0.01030 | 0.00474 | 0.00714 | 0.13978 | 0.00699 | 0.00974 | 0.00812 | 1.54274 | 0.00000 |
| **Opa1** | 4.13817 | 3.27957 | 2.98071 | 4.29741 | 2.45037 | 2.53792 | 0.00000 | 5.82768 | 8.93132 | 0.66013 | 1.43299 | 6.93646 | 1.31858 | 2.08046 |
| **Opa3** | 0.00000 | 0.24412 | 2.35514 | 0.62658 | 2.06205 | 2.16379 | 0.01458 | 1.22834 | 2.17064 | 0.00704 | 0.34639 | 2.91122 | 1.36379 | 3.81866 |
| **Ophn1** | 0.17365 | 0.00417 | 0.01307 | 0.01487 | 0.00770 | 0.01138 | 0.00412 | 0.00924 | 0.01005 | 0.00615 | 0.00753 | 0.00683 | 1.34487 | 0.01126 |
| **Oplah** | 0.03403 | 4.28263 | 1.22190 | 0.42427 | 0.04823 | 0.62465 | 0.00309 | 3.99453 | 0.00811 | 0.01708 | 0.00093 | 0.05693 | 0.00637 | 0.03555 |
| **Opn3** | 0.02116 | 0.00474 | 0.14127 | 0.00197 | 0.01200 | 0.03912 | 0.00434 | 0.00865 | 0.01420 | 0.00740 | 0.00696 | 0.00947 | 0.00779 | 0.01632 |
| **Oprd1** | 0.00554 | 0.00441 | 0.01899 | 0.01902 | 0.01513 | 0.02022 | 0.00432 | 0.01025 | 0.01078 | 0.00678 | 0.00933 | 0.00856 | 0.25620 | 0.00312 |
| **Optn** | 1.70080 | 0.06959 | 3.71600 | 0.63341 | 2.06035 | 0.17263 | 0.00179 | 3.55123 | 0.42235 | 0.00000 | 5.76076 | 2.13393 | 0.02838 | 3.32035 |
| **Orai1** | 0.11560 | 0.00335 | 0.00000 | 0.00000 | 0.45417 | 0.72525 | 1.11329 | 0.11301 | 0.00584 | 0.01610 | 0.03075 | 0.31831 | 3.44048 | 0.00250 |
| **Orai2** | 0.01112 | 0.00366 | 0.12876 | 0.60835 | 0.00071 | 0.15332 | 0.00368 | 0.00208 | 0.01087 | 0.00472 | 0.01544 | 0.39525 | 0.17507 | 0.00000 |
| **Orai3** | 0.00504 | 0.04331 | 0.00803 | 0.00000 | 0.02917 | 0.00446 | 0.00557 | 0.52338 | 0.04176 | 0.00584 | 0.00051 | 0.50299 | 0.01056 | 0.17388 |
| **Oraov1** | 1.87912 | 0.01682 | 2.82877 | 3.56324 | 4.27452 | 2.19960 | 3.03254 | 1.24701 | 3.99173 | 1.37390 | 2.01790 | 0.89101 | 5.97415 | 6.60020 |
| **Orc1** | 3.33002 | 0.02977 | 4.19406 | 1.29559 | 4.80338 | 4.06748 | 2.02744 | 0.00000 | 0.37735 | 0.01743 | 1.10548 | 3.01443 | 5.85226 | 4.55279 |
| **Orc2** | 12.36043 | 0.61001 | 7.17351 | 3.27183 | 8.04778 | 4.64229 | 0.09081 | 4.72217 | 10.64912 | 0.00000 | 6.08071 | 6.09729 | 6.18581 | 6.51897 |
| **Orc3** | 5.18710 | 0.06260 | 7.72597 | 3.61610 | 5.73448 | 7.88094 | 2.62929 | 6.06272 | 8.08275 | 0.07606 | 11.40438 | 3.76739 | 0.76451 | 5.71019 |
| **Orc4** | 5.56710 | 0.04868 | 5.45987 | 4.90103 | 10.97369 | 5.37646 | 4.27878 | 2.03105 | 12.23748 | 5.33665 | 7.19669 | 3.12593 | 6.55053 | 6.16364 |
| **Orc5** | 3.94701 | 0.02884 | 6.06872 | 7.93411 | 2.46688 | 7.43098 | 2.39129 | 5.09907 | 7.47992 | 0.07393 | 6.93018 | 4.97539 | 1.05706 | 5.75908 |
| **Orc6** | 5.79056 | 0.15775 | 14.70078 | 1.53562 | 10.27109 | 11.23915 | 4.17585 | 4.45191 | 15.94763 | 16.90059 | 8.25496 | 9.14230 | 5.12930 | 25.71663 |
| **Ormdl1** | 2.38225 | 0.00751 | 2.51836 | 1.96660 | 2.13752 | 6.07979 | 1.27310 | 2.17207 | 8.17873 | 0.03092 | 3.66188 | 3.39229 | 0.00000 | 0.00000 |
| **Ormdl2** | 2.46882 | 0.07468 | 5.48874 | 4.54034 | 2.25760 | 5.38188 | 2.94062 | 4.81884 | 2.65812 | 0.02387 | 3.70959 | 2.11529 | 0.00000 | 2.32230 |
| **Ormdl3** | 3.12666 | 0.04823 | 5.11347 | 4.19576 | 7.19737 | 3.65964 | 4.79840 | 2.34540 | 0.36195 | 4.95683 | 1.66405 | 3.43543 | 8.07353 | 4.66130 |
| **Os9** | 0.51774 | 0.00330 | 0.28840 | 1.04093 | 0.00000 | 2.18078 | 2.49796 | 0.13008 | 0.08321 | 0.03439 | 0.82556 | 0.43506 | 1.28237 | 3.09932 |
| **Osbp** | 2.90768 | 0.00937 | 2.18794 | 0.00000 | 2.03806 | 3.61851 | 1.76542 | 0.39286 | 2.29941 | 1.58294 | 3.90522 | 1.33466 | 2.66755 | 3.97648 |
| **Osbp2** | 0.01757 | 0.00473 | 0.01389 | 0.07239 | 0.05070 | 0.01367 | 0.00428 | 0.01340 | 0.01297 | 0.00728 | 0.01379 | 0.01227 | 0.00577 | 0.01457 |
| **Osbpl10** | 0.20168 | 0.00443 | 5.23520 | 2.48756 | 1.44771 | 0.49812 | 0.00378 | 0.23735 | 0.05732 | 0.00383 | 0.37658 | 0.00370 | 0.00000 | 0.34003 |
| **Osbpl11** | 3.56870 | 0.00794 | 8.96073 | 1.33017 | 9.21233 | 4.28476 | 1.85579 | 1.41238 | 0.35865 | 0.04065 | 3.72515 | 2.43082 | 5.10879 | 8.81601 |
| **Osbpl1a** | 0.43828 | 0.02979 | 0.00000 | 1.55778 | 0.39275 | 0.44460 | 0.00303 | 0.04182 | 0.58731 | 0.01525 | 0.41977 | 3.86703 | 3.46686 | 0.27336 |
| **Osbpl2** | 11.96454 | 4.13665 | 14.02450 | 2.18051 | 6.75928 | 6.43466 | 3.82551 | 0.26220 | 3.12058 | 13.85028 | 2.27056 | 4.17101 | 5.89005 | 6.81968 |
| **Osbpl3** | 0.39767 | 0.01207 | 0.66581 | 0.04664 | 0.23675 | 0.68720 | 0.11280 | 0.19224 | 0.14200 | 0.00458 | 0.00329 | 0.00233 | 0.54780 | 0.39434 |
| **Osbpl5** | 0.07920 | 0.00516 | 0.43985 | 0.98355 | 0.00000 | 0.00000 | 0.02280 | 0.04858 | 0.00975 | 0.03622 | 0.00380 | 0.01113 | 0.50566 | 0.00536 |
| **Osbpl6** | 4.44583 | 1.18099 | 13.89949 | 3.99594 | 12.59859 | 0.83040 | 0.00000 | 1.10269 | 14.12493 | 0.01618 | 2.72577 | 0.00000 | 3.71873 | 10.56912 |
| **Osbpl7** | 0.98743 | 0.10905 | 6.64116 | 5.42280 | 5.01659 | 10.79383 | 3.05539 | 5.48350 | 0.64195 | 5.79687 | 0.14663 | 1.18662 | 1.18930 | 5.89691 |
| **Osbpl8** | 7.75749 | 1.50325 | 2.93906 | 1.24545 | 1.76322 | 3.64424 | 0.07248 | 2.67176 | 3.48292 | 0.12557 | 6.56215 | 1.47334 | 12.10852 | 2.55633 |
| **Osbpl9** | 2.17561 | 0.02277 | 11.17598 | 5.45184 | 6.08199 | 3.05945 | 0.30165 | 1.26509 | 9.41721 | 0.00000 | 4.82279 | 2.95487 | 1.49477 | 12.42319 |
| **Oscar** | 0.84899 | 0.00498 | 1.00235 | 0.01602 | 0.16558 | 0.00000 | 0.00411 | 0.04139 | 0.00470 | 0.01515 | 0.01305 | 0.01528 | 0.81289 | 0.63518 |
| **Oscp1** | 0.34457 | 0.02376 | 0.35215 | 0.52005 | 0.00000 | 0.14256 | 0.00428 | 0.38409 | 0.09008 | 0.02056 | 0.90033 | 0.06794 | 0.87725 | 0.09138 |

| | | | | | | | | | | | | | | |
|---|---|---|---|---|---|---|---|---|---|---|---|---|---|---|
| **Oser1** | 12.47829 | 1.93819 | 18.16942 | 10.06847 | 15.57734 | 10.07693 | 7.91307 | 4.63537 | 7.77230 | 0.27927 | 11.36694 | 6.32626 | 15.53657 | 11.49670 |
| **Osgep** | 7.48778 | 0.04259 | 2.57152 | 10.96451 | 5.06703 | 3.03300 | 0.81539 | 4.80295 | 1.61160 | 0.11700 | 0.36672 | 6.83547 | 0.05868 | 1.78540 |
| **Osgepl1** | 4.98800 | 0.00157 | 5.07994 | 4.20169 | 3.81240 | 2.82373 | 0.05934 | 0.82163 | 4.14924 | 0.00000 | 2.24053 | 2.44904 | 0.01617 | 3.82073 |
| **Osgin1** | 0.01358 | 0.00438 | 0.32567 | 0.00188 | 0.00471 | 0.00713 | 0.00418 | 0.01103 | 0.01935 | 0.00697 | 0.00801 | 0.00888 | 0.01949 | 0.03125 |
| **Osgin2** | 2.82067 | 0.01805 | 0.70491 | 0.29432 | 0.00000 | 1.06434 | 1.03072 | 3.06288 | 1.53201 | 0.02543 | 2.82179 | 0.27161 | 0.92738 | 0.00000 |
| **Osm** | 0.55423 | 0.01107 | 0.81740 | 0.05876 | 0.53552 | 0.26084 | 0.00369 | 0.52944 | 0.31125 | 0.00839 | 0.00566 | 0.00230 | 0.89405 | 0.00000 |
| **Osr2** | 0.00506 | 0.00367 | 0.32094 | 0.86617 | 0.64216 | 0.00000 | 0.00411 | 0.00216 | 0.02212 | 0.00574 | 0.00396 | 0.34447 | 0.00370 | 0.00000 |
| **Ost4** | 1.71048 | 0.15878 | 0.00000 | 0.81260 | 0.83765 | 0.47980 | 1.50630 | 1.41060 | 1.31530 | 0.28982 | 0.14205 | 1.19725 | 3.25527 | 1.88418 |
| **Ostc** | 4.01713 | 8.28417 | 7.22221 | 9.81819 | 9.30621 | 18.83173 | 5.27188 | 8.10660 | 4.65444 | 0.28025 | 13.25739 | 13.98877 | 1.59669 | 9.40483 |
| **Ostf1** | 3.48371 | 3.76026 | 1.74234 | 13.80205 | 3.49325 | 12.19082 | 4.06780 | 6.32449 | 0.68712 | 9.42518 | 6.26566 | 9.32949 | 1.00799 | 1.29486 |
| **Ostm1** | 7.83135 | 0.12062 | 2.65147 | 5.53924 | 2.87340 | 5.46293 | 0.91599 | 0.00076 | 9.59512 | 0.03506 | 5.60992 | 6.36648 | 2.82127 | 5.64317 |
| **Otoa** | 0.00000 | 0.00414 | 1.39645 | 0.30724 | 0.66225 | 0.00000 | 0.00371 | 0.00555 | 0.00651 | 0.00308 | 0.00970 | 0.00996 | 0.00000 | 0.39055 |
| **Otog** | 0.72784 | 0.00401 | 0.54101 | 0.01468 | 0.58632 | 0.00618 | 0.00432 | 0.00408 | 0.05132 | 0.01076 | 0.00404 | 0.27818 | 0.01127 | 0.05743 |
| **Otop1** | 0.00750 | 0.00450 | 0.00610 | 0.09126 | 0.01202 | 0.34048 | 0.00404 | 0.00420 | 0.01297 | 0.00659 | 0.00541 | 0.01099 | 0.08113 | 0.02321 |
| **Otub1** | 2.89126 | 0.05475 | 5.70785 | 3.67202 | 8.16195 | 3.34206 | 0.01339 | 1.73603 | 0.15716 | 0.12962 | 4.39761 | 1.76275 | 2.40703 | 8.79681 |
| **Otub2** | 0.00000 | 0.00403 | 0.38404 | 0.44931 | 0.07364 | 0.14560 | 0.09394 | 0.00187 | 0.00685 | 0.01097 | 0.00438 | 0.00666 | 0.49718 | 0.36824 |
| **Otud1** | 0.44328 | 1.19178 | 7.77197 | 3.39808 | 2.63755 | 1.36339 | 1.09886 | 3.05985 | 4.94116 | 0.07211 | 0.55513 | 2.71747 | 2.31107 | 5.76062 |
| **Otud3** | 1.74319 | 0.01036 | 0.36720 | 0.49333 | 1.06877 | 1.89370 | 0.00982 | 0.59298 | 0.00842 | 0.01236 | 0.00000 | 0.69077 | 1.63688 | 1.32328 |
| **Otud4** | 5.06223 | 0.07097 | 9.43750 | 2.31949 | 7.63388 | 10.41290 | 3.39115 | 0.97710 | 20.53229 | 0.05505 | 4.10915 | 4.62991 | 11.32778 | 5.64208 |
| **Otud5** | 10.93237 | 8.70534 | 15.77326 | 8.11500 | 18.39238 | 11.81991 | 1.62174 | 3.39022 | 3.63672 | 0.07227 | 7.36958 | 5.49104 | 6.86487 | 11.19066 |
| **Otud6b** | 18.22751 | 0.04108 | 18.87832 | 8.93463 | 16.24393 | 17.17469 | 3.80527 | 1.70169 | 26.40108 | 0.15390 | 13.12629 | 6.09272 | 15.37737 | 20.46278 |
| **Otud7a** | 0.01190 | 0.00445 | 0.01304 | 0.01326 | 0.01106 | 0.01230 | 0.00431 | 0.01035 | 0.04251 | 0.00687 | 0.00851 | 0.00998 | 0.00662 | 0.05194 |
| **Otud7b** | 0.82891 | 0.04368 | 0.57078 | 0.00000 | 0.10064 | 3.23942 | 2.61664 | 0.39142 | 1.46585 | 0.06276 | 2.25871 | 1.37444 | 0.08013 | 0.01646 |
| **Otulin** | 9.30666 | 0.00000 | 4.67394 | 1.51356 | 6.00309 | 1.57749 | 3.53099 | 1.84593 | 0.33032 | 0.10672 | 1.97886 | 3.36537 | 1.39450 | 4.71949 |
| **Otx1** | 0.00440 | 0.00396 | 0.00583 | 0.38041 | 0.00744 | 0.01231 | 0.00417 | 0.00605 | 0.02560 | 0.00523 | 0.00727 | 0.08082 | 0.08171 | 0.00462 |
| **Otx2** | 0.00000 | 0.01454 | 0.86116 | 1.06196 | 0.00000 | 0.00000 | 0.15141 | 0.52031 | 0.24428 | 0.04360 | 1.30459 | 0.25875 | 0.13166 | 0.28029 |
| **Ovca2** | 0.09781 | 0.01128 | 0.64605 | 1.92842 | 0.17403 | 1.88108 | 0.21391 | 0.01800 | 0.58411 | 0.01323 | 0.00332 | 0.56599 | 0.00823 | 1.97916 |
| **Ovgp1** | 0.85789 | 0.00435 | 0.00000 | 0.00000 | 13.71659 | 0.05742 | 0.00425 | 0.00807 | 0.34456 | 0.00936 | 0.00137 | 0.00383 | 0.00000 | 0.00000 |
| **Ovol1** | 0.24267 | 6.95628 | 3.49996 | 0.00000 | 3.59991 | 0.00000 | 0.01897 | 4.76459 | 1.52517 | 0.02197 | 0.00000 | 0.00000 | 0.47257 | 4.16650 |
| **Ovol2** | 7.22979 | 0.00923 | 3.56740 | 1.91833 | 13.63059 | 0.67167 | 0.00861 | 1.60893 | 4.84233 | 5.63723 | 0.00000 | 0.00000 | 6.59440 | 10.43128 |
| **Ovol3** | 0.01206 | 0.00443 | 0.01472 | 0.01272 | 0.01320 | 0.01325 | 0.00428 | 0.01013 | 0.01132 | 0.00683 | 0.00884 | 0.00995 | 0.05895 | 0.00580 |
| **Oxa1l** | 3.03398 | 3.74116 | 13.47384 | 7.18869 | 7.35720 | 7.68101 | 4.94504 | 6.87728 | 0.56848 | 0.21242 | 6.55638 | 8.02938 | 1.15374 | 2.28649 |
| **Oxct1** | 2.63432 | 0.07135 | 0.00000 | 0.58142 | 3.57891 | 9.83928 | 3.21771 | 9.24950 | 0.05513 | 0.11310 | 0.81142 | 3.09648 | 0.26566 | 2.20938 |
| **Oxgr1** | 0.02715 | 0.00412 | 0.03087 | 0.04325 | 0.02978 | 0.06571 | 0.00998 | 0.01602 | 0.03349 | 0.06780 | 0.02803 | 0.04129 | 0.02551 | 0.02152 |
| **Oxld1** | 0.18687 | 0.01495 | 0.00000 | 0.54402 | 0.62897 | 0.17413 | 0.00418 | 0.38740 | 0.03379 | 0.01289 | 0.32520 | 0.57131 | 0.00238 | 0.11898 |
| **Oxnad1** | 0.91528 | 0.78750 | 1.53593 | 0.26449 | 0.77843 | 1.82868 | 0.37976 | 1.61459 | 0.04336 | 0.08320 | 1.16725 | 0.80628 | 0.93066 | 2.25249 |
| **Oxr1** | 0.00000 | 0.01760 | 0.65339 | 0.00000 | 0.20531 | 2.15518 | 2.20612 | 0.37228 | 0.00514 | 0.02344 | 0.47639 | 0.81229 | 2.00985 | 0.02190 |
| **Oxsm** | 0.02508 | 0.00375 | 0.00991 | 0.00000 | 0.00586 | 0.30250 | 0.00386 | 0.12494 | 1.54305 | 0.00590 | 0.02271 | 0.17674 | 0.05212 | 0.00000 |
| **Oxsr1** | 3.51216 | 6.54701 | 6.34223 | 4.24306 | 6.84185 | 7.13790 | 3.16598 | 5.83051 | 6.47162 | 0.18398 | 5.21691 | 6.90394 | 2.72334 | 8.05313 |
| **P2rx2** | 0.00931 | 0.00444 | 0.01150 | 0.08004 | 0.01483 | 0.00880 | 0.00419 | 0.00998 | 0.00949 | 0.00686 | 0.00898 | 0.01378 | 0.13527 | 0.03254 |
| **P2rx3** | 0.11551 | 0.04343 | 0.00058 | 0.27217 | 0.00326 | 2.61888 | 0.01509 | 5.30231 | 0.01122 | 0.04354 | 0.03572 | 0.03282 | 0.00946 | 0.00229 |
| **P2rx4** | 0.28995 | 0.00311 | 5.64103 | 2.19985 | 0.80094 | 3.04837 | 0.04933 | 0.00000 | 1.45412 | 0.04476 | 1.78362 | 0.32106 | 1.75715 | 1.40060 |
| **P2rx5** | 0.00000 | 0.00454 | 0.60840 | 0.07511 | 0.32865 | 0.00000 | 0.00475 | 0.04394 | 0.00493 | 0.00881 | 0.00754 | 0.00461 | 0.10604 | 0.60954 |
| **P2rx7** | 0.00000 | 0.00339 | 0.71257 | 0.09655 | 0.42119 | 0.00000 | 0.00370 | 0.01546 | 0.32676 | 0.00609 | 0.00412 | 1.19499 | 0.00203 | 0.77864 |
| **P2ry14** | 0.00774 | 0.01539 | 0.32159 | 0.00000 | 0.00000 | 0.00000 | 0.00391 | 0.83623 | 1.84343 | 0.00797 | 0.52157 | 0.05773 | 0.00175 | 0.03896 |
| **P2ry4** | 0.03195 | 0.01298 | 0.01768 | 0.04909 | 0.03966 | 0.05486 | 0.00424 | 0.03624 | 0.07953 | 0.00936 | 0.02132 | 0.01355 | 0.25227 | 0.13798 |
| **P3h1** | 0.03559 | 0.09214 | 0.02557 | 0.00000 | 1.00853 | 0.20547 | 0.00357 | 0.03288 | 0.00125 | 0.00536 | 0.00066 | 3.64921 | 0.82900 | 1.46998 |
| **P3h3** | 0.56346 | 0.06827 | 0.00000 | 0.15199 | 0.10468 | 0.87079 | 0.13738 | 0.09909 | 0.68540 | 0.12419 | 0.02941 | 0.15384 | 3.99635 | 0.35847 |
| **P3h4** | 2.59985 | 0.00451 | 2.08013 | 0.07003 | 1.32498 | 0.32681 | 0.00391 | 0.00016 | 0.01124 | 0.01085 | 0.00277 | 0.04074 | 0.00000 | 1.71877 |
| **P4ha1** | 0.23354 | 0.04648 | 7.57704 | 1.89773 | 3.55041 | 3.00841 | 1.22879 | 0.49015 | 0.13499 | 0.00000 | 2.82747 | 6.54693 | 0.69910 | 0.48926 |
| **P4ha2** | 2.33888 | 0.12778 | 0.47244 | 6.28386 | 1.03801 | 4.62907 | 0.71884 | 0.61144 | 0.73696 | 0.06108 | 0.24161 | 8.16848 | 0.90818 | 0.00004 |
| **P4ha3** | 3.76771 | 0.00322 | 1.34831 | 0.08567 | 0.17072 | 1.75242 | 0.00344 | 0.00000 | 0.00000 | 0.01718 | 0.00481 | 0.00000 | 8.41177 | 1.44087 |
| **P4hb** | 13.96395 | 7.56785 | 24.84841 | 18.47105 | 17.30683 | 20.89507 | 9.04190 | 12.52031 | 1.69489 | 5.08001 | 9.28261 | 18.22485 | 12.43731 | 20.54673 |
| **P4htm** | 1.66298 | 0.00346 | 2.63803 | 2.36279 | 1.26735 | 0.46150 | 0.00353 | 0.00000 | 0.01030 | 0.00000 | 0.00454 | 0.45177 | 0.00000 | 0.15974 |
| **Pa2g4** | 29.00595 | 6.21330 | 32.59735 | 29.61653 | 42.97589 | 29.31400 | 6.65389 | 12.86913 | 21.00615 | 5.17411 | 23.20389 | 18.10048 | 12.92954 | 30.22804 |

| | | | | | | | | | | | | | | |
|---|---|---|---|---|---|---|---|---|---|---|---|---|---|---|
| **Pabpc1** | 35.68008 | 9.76253 | 39.55562 | 19.83774 | 32.73534 | 34.03960 | 8.96821 | 18.20995 | 30.22135 | 17.31779 | 24.52482 | 26.97272 | 17.31609 | 35.75343 |
| **Pabpc1l** | 0.00786 | 0.00445 | 0.05041 | 0.02375 | 0.46284 | 0.01527 | 0.00438 | 0.01767 | 0.04516 | 0.00969 | 0.00829 | 0.00679 | 0.37121 | 0.00462 |
| **Pabpc4** | 8.42498 | 8.89130 | 28.22791 | 43.37163 | 41.85673 | 4.45798 | 4.24567 | 12.73988 | 3.71691 | 2.00598 | 1.09865 | 13.58403 | 8.12591 | 5.51072 |
| **Pabpn1** | 2.48324 | 9.18452 | 15.33416 | 13.81293 | 23.62312 | 13.88485 | 3.31207 | 18.62787 | 10.31788 | 0.64364 | 10.29947 | 12.46099 | 9.72172 | 26.63362 |
| **Pabpn1l** | 0.00822 | 0.00436 | 0.10233 | 0.00455 | 0.02269 | 0.00988 | 0.00428 | 0.01127 | 0.01032 | 0.00692 | 0.00769 | 0.01056 | 0.13817 | 0.00523 |
| **Pacrgl** | 3.12005 | 0.04440 | 1.04411 | 1.44107 | 3.90520 | 1.84054 | 0.23424 | 2.99625 | 0.91759 | 0.01013 | 0.00000 | 2.82295 | 2.90638 | 2.46057 |
| **Pacs1** | 0.37052 | 0.01586 | 1.29564 | 0.05161 | 0.38256 | 0.69271 | 0.00283 | 0.13936 | 0.00266 | 0.01315 | 0.00321 | 3.27975 | 1.45718 | 0.21419 |
| **Pacs2** | 0.08143 | 0.04555 | 1.05018 | 0.45601 | 1.21828 | 1.54353 | 0.18974 | 1.67566 | 0.14214 | 0.09625 | 0.02249 | 2.49322 | 4.76398 | 3.04739 |
| **Pacsin1** | 0.00552 | 0.00427 | 0.33664 | 0.21418 | 0.00000 | 0.96276 | 0.00395 | 0.01176 | 0.00644 | 0.01000 | 0.01074 | 0.00274 | 0.43047 | 0.00000 |
| **Pacsin2** | 1.26877 | 2.27126 | 2.81425 | 3.00935 | 3.40766 | 13.49893 | 0.87574 | 5.05613 | 0.77078 | 0.06248 | 5.37041 | 7.78158 | 7.15150 | 13.62407 |
| **Pacsin3** | 0.11940 | 5.42590 | 1.32166 | 1.93375 | 2.01346 | 1.25245 | 1.00043 | 1.26996 | 0.15789 | 0.04736 | 1.29934 | 1.04798 | 1.19818 | 0.09865 |
| **Padi1** | 0.04259 | 0.00463 | 0.05370 | 0.12382 | 0.15306 | 0.04405 | 0.00412 | 0.00583 | 0.00922 | 0.00800 | 0.00595 | 0.03068 | 0.03353 | 0.00440 |
| **Padi2** | 0.14643 | 0.22560 | 0.58361 | 0.09760 | 0.54710 | 0.24341 | 0.05237 | 0.07215 | 1.36714 | 0.09007 | 0.03418 | 0.06641 | 1.37033 | 1.14741 |
| **Padi3** | 0.21031 | 0.00385 | 0.59114 | 0.00107 | 0.00000 | 0.00618 | 0.00389 | 0.00357 | 0.00771 | 0.01308 | 0.00360 | 0.55224 | 0.02817 | 0.00725 |
| **Padi4** | 0.09445 | 0.00395 | 0.24299 | 0.02119 | 0.01184 | 0.06116 | 0.00997 | 0.00581 | 0.11532 | 0.04531 | 0.00386 | 0.21198 | 0.56717 | 0.07114 |
| **Padi6** | 8.55504 | 0.00197 | 7.90405 | 2.11822 | 6.96734 | 0.39101 | 0.00198 | 0.75046 | 3.33397 | 0.00000 | 0.04374 | 0.00000 | 7.49675 | 11.18423 |
| **Paf1** | 4.28702 | 0.10569 | 8.87746 | 10.23524 | 6.96589 | 4.95187 | 5.66412 | 7.04518 | 0.00000 | 0.35894 | 8.27429 | 4.64064 | 4.30665 | 2.90552 |
| **Pafah1b1** | 15.53551 | 0.79428 | 14.52213 | 7.68788 | 9.91068 | 14.39301 | 5.39234 | 6.04767 | 15.49416 | 0.26219 | 9.08292 | 9.20573 | 9.15770 | 11.64941 |
| **Pafah1b2** | 13.20823 | 0.02167 | 8.47247 | 1.86905 | 15.76991 | 9.58220 | 1.06279 | 0.00000 | 14.52248 | 0.04831 | 7.05832 | 2.00801 | 10.21284 | 12.32176 |
| **Pafah1b3** | 4.12744 | 0.09168 | 5.46746 | 10.74291 | 6.05031 | 2.89935 | 0.09521 | 5.87838 | 0.00000 | 8.96395 | 0.21623 | 4.76143 | 2.45817 | 2.82243 |
| **Pafah2** | 0.00000 | 0.01131 | 3.12247 | 3.16305 | 0.43989 | 7.13096 | 3.28877 | 0.57553 | 0.00537 | 0.11013 | 7.67670 | 0.09214 | 0.45342 | 0.15595 |
| **Pag1** | 0.97110 | 0.01334 | 0.00024 | 0.00000 | 0.00000 | 0.12668 | 0.01517 | 0.05144 | 1.07822 | 0.00220 | 1.09839 | 0.49312 | 0.21228 | 0.67554 |
| **Pagr1a** | 2.05071 | 0.09497 | 5.98053 | 7.40747 | 4.20471 | 2.62017 | 0.03548 | 7.37212 | 0.00000 | 14.39485 | 2.70623 | 8.27463 | 0.74188 | 0.58645 |
| **Paics** | 23.37250 | 0.53060 | 31.46974 | 16.27118 | 30.31302 | 22.59640 | 7.64191 | 13.45351 | 32.90419 | 0.67136 | 18.93094 | 18.08556 | 10.03223 | 28.51425 |
| **Paip1** | 0.11468 | 0.00446 | 0.14248 | 0.23183 | 0.11649 | 1.63332 | 0.00377 | 0.54378 | 2.54673 | 0.02126 | 0.02909 | 0.48063 | 4.65613 | 0.00000 |
| **Paip2** | 24.34230 | 0.07028 | 17.06054 | 7.49869 | 16.64892 | 13.44746 | 4.54897 | 7.04995 | 16.95086 | 0.18176 | 8.16095 | 7.22273 | 5.74717 | 14.37508 |
| **Paip2b** | 3.35464 | 0.46610 | 7.67944 | 4.64557 | 11.64574 | 7.90888 | 4.92990 | 10.21109 | 19.07230 | 6.10142 | 0.03558 | 2.86273 | 8.12051 | 15.44473 |
| **Pak1** | 2.29203 | 0.01849 | 5.85340 | 1.29908 | 3.81120 | 1.36559 | 0.00164 | 0.00000 | 7.76691 | 0.01001 | 0.15467 | 1.50496 | 4.75209 | 3.45944 |
| **Pak1ip1** | 14.47022 | 8.32020 | 25.51348 | 13.74494 | 22.02699 | 13.08208 | 2.11413 | 8.99261 | 9.21515 | 0.28349 | 15.21036 | 9.81636 | 14.90126 | 14.93241 |
| **Pak2** | 2.60724 | 0.94225 | 2.48613 | 0.00000 | 5.97890 | 7.90613 | 3.22862 | 4.67567 | 9.93969 | 0.10015 | 4.11811 | 4.29197 | 5.51923 | 8.74940 |
| **Pak4** | 0.37108 | 0.24784 | 9.69656 | 7.08266 | 9.56576 | 0.40285 | 2.55563 | 6.49430 | 0.23468 | 9.25139 | 2.34310 | 8.29290 | 2.19602 | 1.16499 |
| **Pak6** | 0.00241 | 0.57475 | 0.63910 | 0.00333 | 0.13679 | 2.59061 | 2.87878 | 3.09125 | 0.10744 | 0.11587 | 0.12293 | 0.00000 | 0.01213 | 0.36123 |
| **Pak7** | 0.00403 | 0.00448 | 0.00000 | 0.01619 | 0.12452 | 0.00686 | 0.00421 | 0.01000 | 0.87661 | 0.00705 | 0.00585 | 0.00807 | 0.16921 | 0.19009 |
| **Palb2** | 0.07469 | 0.00975 | 1.59613 | 0.70225 | 1.70886 | 0.00000 | 0.00280 | 0.79503 | 1.93513 | 0.00000 | 1.17773 | 0.60387 | 2.41827 | 2.53051 |
| **Pald1** | 0.00000 | 0.00211 | 2.26499 | 1.00955 | 0.80805 | 0.70057 | 0.00825 | 0.00000 | 0.00216 | 0.00000 | 1.80354 | 1.95684 | 0.01523 | 1.00462 |
| **Palld** | 0.37470 | 6.77666 | 2.68351 | 1.20941 | 0.48760 | 2.00366 | 0.00144 | 0.00000 | 0.46458 | 0.00444 | 3.60770 | 0.21089 | 2.71722 | 1.10986 |
| **Palm** | 0.18709 | 0.00409 | 1.81181 | 1.83997 | 3.14753 | 0.18338 | 0.10293 | 0.29899 | 0.00398 | 0.02394 | 0.01539 | 0.16207 | 0.28549 | 0.00000 |
| **Palm3** | 0.60195 | 1.50557 | 7.01754 | 8.26191 | 1.89239 | 9.93783 | 0.00760 | 1.45708 | 0.00000 | 0.13020 | 0.21169 | 6.12594 | 0.24956 | 1.19565 |
| **Pam16** | 0.31825 | 3.07816 | 0.62354 | 9.19804 | 12.83133 | 3.05747 | 3.57974 | 6.81427 | 0.00218 | 0.31248 | 0.40831 | 6.11554 | 1.49971 | 0.00000 |
| **Pan2** | 4.04958 | 0.00853 | 7.98719 | 1.38859 | 2.68166 | 4.42011 | 3.25997 | 0.12392 | 1.61631 | 0.05898 | 5.18793 | 1.99870 | 1.14672 | 6.37671 |
| **Pan3** | 0.83459 | 0.70412 | 0.67189 | 1.70700 | 1.45501 | 1.03605 | 0.19231 | 0.85787 | 9.96035 | 0.01552 | 0.00000 | 0.56991 | 5.89238 | 1.26049 |
| **Panct2** | 1.57568 | 0.02628 | 0.35985 | 0.00000 | 0.07197 | 0.72379 | 0.06027 | 0.39311 | 2.15540 | 0.01648 | 0.00675 | 0.53724 | 0.64361 | 0.00000 |
| **Pank1** | 1.66296 | 0.01882 | 0.02907 | 0.02654 | 0.08878 | 0.05924 | 0.00400 | 0.02538 | 0.04587 | 0.05761 | 0.54006 | 0.00948 | 0.70334 | 0.03441 |
| **Pank2** | 0.66925 | 0.00285 | 0.47235 | 0.00000 | 1.48572 | 0.75670 | 0.00409 | 0.27863 | 0.32941 | 0.00472 | 0.00000 | 1.18756 | 2.55792 | 2.22430 |
| **Pank3** | 24.98829 | 3.24949 | 15.12741 | 4.11359 | 22.61067 | 7.40154 | 1.62003 | 4.51223 | 37.20872 | 6.40147 | 8.69045 | 1.23364 | 19.42849 | 26.94106 |
| **Pank4** | 0.00041 | 0.00422 | 0.51827 | 0.21610 | 0.45921 | 0.00446 | 0.01595 | 0.00812 | 0.01129 | 0.00893 | 0.01721 | 0.53602 | 0.34236 | 0.12631 |
| **Panx1** | 0.90122 | 0.00359 | 0.42662 | 0.32401 | 0.96228 | 0.07383 | 0.00929 | 1.02474 | 0.22853 | 0.01251 | 0.00000 | 0.31925 | 0.57574 | 0.00000 |
| **Paox** | 0.00410 | 0.00888 | 0.00000 | 1.34672 | 0.70388 | 1.43679 | 0.00278 | 0.95318 | 0.00225 | 0.01022 | 0.88175 | 2.27546 | 0.05233 | 1.18850 |
| **Papd4** | 8.76702 | 0.00002 | 5.94173 | 2.08328 | 5.52031 | 1.88594 | 1.54447 | 1.46059 | 11.10566 | 0.04063 | 5.17296 | 2.10035 | 3.19841 | 0.89710 |
| **Papd5** | 13.74695 | 0.00410 | 12.94430 | 4.69147 | 21.45913 | 9.26812 | 0.00830 | 4.22388 | 22.18185 | 0.04203 | 6.70835 | 3.81235 | 8.97934 | 23.45281 |
| **Papd7** | 2.99512 | 3.17672 | 3.90349 | 0.00000 | 1.69359 | 3.88566 | 2.86567 | 1.25603 | 0.87481 | 0.02474 | 2.55444 | 4.16009 | 11.01232 | 8.59333 |
| **Papln** | 0.01263 | 0.00443 | 0.01641 | 0.00979 | 0.58263 | 0.01324 | 0.00437 | 0.01325 | 0.00885 | 0.00629 | 0.00825 | 0.00813 | 0.00671 | 0.02707 |
| **Papola** | 20.60325 | 7.53511 | 23.73572 | 18.94920 | 29.25319 | 19.45386 | 5.87928 | 7.81898 | 27.26877 | 0.55280 | 14.21242 | 14.25900 | 13.27392 | 17.78705 |
| **Papolb** | 0.47196 | 0.00393 | 0.11852 | 0.01397 | 0.00000 | 0.00671 | 0.00423 | 0.01070 | 0.85942 | 0.00624 | 0.02722 | 0.00701 | 1.89298 | 1.21844 |
| **Papolg** | 1.05202 | 0.00215 | 0.62550 | 0.79899 | 0.13324 | 1.98658 | 2.03864 | 0.38950 | 1.16942 | 0.10093 | 2.77828 | 1.14394 | 0.26687 | 0.18403 |

| | | | | | | | | | | | | | | |
|---|---|---|---|---|---|---|---|---|---|---|---|---|---|---|
| **Pappa** | 0.17626 | 0.02130 | 0.01418 | 0.01469 | 0.00677 | 0.00984 | 0.00430 | 0.01680 | 0.04103 | 0.00713 | 0.00699 | 0.00858 | 0.01533 | 0.01165 |
| **Papss1** | 0.28798 | 0.00366 | 0.85001 | 0.84989 | 1.74355 | 0.08427 | 0.00409 | 0.30307 | 4.17635 | 0.01716 | 0.00406 | 0.01474 | 2.45147 | 11.83876 |
| **Papss2** | 0.01463 | 0.00380 | 0.01676 | 0.04423 | 0.01607 | 0.08525 | 0.00925 | 0.47658 | 0.03630 | 0.00577 | 0.26180 | 0.00880 | 0.00744 | 0.03388 |
| **Paqr4** | 0.08292 | 0.00352 | 1.65810 | 2.15119 | 2.71885 | 2.47474 | 0.01536 | 1.46305 | 0.00152 | 0.75265 | 1.20117 | 0.00000 | 0.00000 | 0.52962 |
| **Paqr6** | 0.18699 | 0.00440 | 0.01212 | 0.02011 | 0.00904 | 0.01447 | 0.00419 | 0.00914 | 0.01104 | 0.01139 | 0.00692 | 0.00850 | 0.04558 | 0.00440 |
| **Paqr7** | 0.06768 | 0.01421 | 0.02249 | 0.03612 | 0.22652 | 0.16335 | 0.06433 | 0.03160 | 0.25434 | 0.00747 | 0.01943 | 0.02266 | 0.37032 | 0.01803 |
| **Pard3** | 3.60530 | 0.06764 | 3.10496 | 0.56812 | 2.33334 | 3.31495 | 0.03588 | 1.24182 | 2.06209 | 12.36537 | 3.66601 | 4.43892 | 3.07964 | 5.63113 |
| **Pard3b** | 0.02522 | 0.01271 | 0.02384 | 0.02761 | 0.00342 | 0.01145 | 0.02259 | 0.01516 | 0.03472 | 0.00657 | 0.00900 | 0.01683 | 0.43979 | 0.24294 |
| **Pard6a** | 0.73197 | 0.01809 | 1.48434 | 1.18312 | 1.53634 | 1.85679 | 1.65522 | 2.72749 | 0.00939 | 0.06781 | 3.59867 | 1.51173 | 0.00160 | 0.30750 |
| **Pard6b** | 2.89466 | 2.63803 | 8.50588 | 5.39408 | 2.59041 | 8.71634 | 3.59444 | 4.27322 | 0.35082 | 9.47460 | 0.36053 | 0.82823 | 0.50978 | 0.97960 |
| **Pard6g** | 0.01512 | 0.00443 | 0.17019 | 0.13605 | 0.00296 | 0.01007 | 0.00465 | 0.02753 | 0.01797 | 0.00852 | 0.00814 | 0.13977 | 0.00589 | 0.06286 |
| **Parg** | 2.16550 | 0.03503 | 4.17465 | 2.29629 | 6.12294 | 5.84518 | 0.00077 | 0.30105 | 12.54858 | 0.00356 | 7.36778 | 0.33423 | 2.13754 | 13.80960 |
| **Park2** | 0.73303 | 0.02784 | 0.00000 | 0.23602 | 1.31572 | 1.63718 | 0.14347 | 0.48478 | 0.56159 | 0.37106 | 0.01785 | 0.10322 | 4.47906 | 0.45196 |
| **Park7** | 9.12187 | 5.52866 | 4.76498 | 10.57543 | 7.92246 | 14.81122 | 1.87660 | 13.80276 | 0.09908 | 4.88917 | 8.68215 | 11.88417 | 1.60575 | 3.86946 |
| **Parl** | 10.69913 | 2.09192 | 6.83745 | 12.80229 | 7.32353 | 7.06122 | 0.05907 | 4.19674 | 3.22719 | 0.06372 | 4.72158 | 8.15682 | 2.09290 | 3.47598 |
| **Parn** | 4.01146 | 0.02522 | 9.29433 | 0.36484 | 4.72822 | 4.05483 | 1.65762 | 1.41766 | 5.19960 | 0.10041 | 2.95413 | 3.67423 | 2.36091 | 4.77809 |
| **Parp1** | 18.33162 | 7.13980 | 30.40132 | 15.87073 | 24.74010 | 15.01573 | 7.36438 | 10.31469 | 2.31764 | 9.60461 | 19.68496 | 20.22568 | 8.74223 | 17.34198 |
| **Parp10** | 0.00497 | 0.00434 | 0.01270 | 0.01202 | 0.11698 | 0.01311 | 0.00438 | 0.01095 | 0.00887 | 0.00635 | 0.00897 | 0.00941 | 0.22446 | 0.00447 |
| **Parp11** | 0.00798 | 0.00411 | 0.00806 | 0.11498 | 0.11740 | 0.63306 | 0.00939 | 0.00302 | 0.13506 | 0.01245 | 0.01775 | 0.02744 | 0.01995 | 0.31871 |
| **Parp12** | 0.00000 | 0.00231 | 0.20446 | 0.32854 | 1.97481 | 0.00000 | 0.00347 | 0.00311 | 0.88881 | 0.00317 | 1.05203 | 0.00344 | 10.39223 | 8.48939 |
| **Parp14** | 0.00516 | 0.00420 | 0.00944 | 0.01271 | 0.12493 | 0.01255 | 0.00471 | 0.00839 | 0.02938 | 0.00692 | 0.00887 | 0.00783 | 1.22891 | 0.00684 |
| **Parp16** | 0.55275 | 0.00451 | 0.83607 | 0.44924 | 1.77814 | 1.37129 | 0.01031 | 0.01232 | 0.07373 | 0.00466 | 0.21057 | 0.09706 | 0.34008 | 0.45147 |
| **Parp2** | 9.80030 | 0.06715 | 11.44245 | 3.10849 | 7.80796 | 6.85932 | 1.78285 | 1.39093 | 10.03370 | 0.06273 | 13.54056 | 2.79771 | 2.98096 | 10.30856 |
| **Parp4** | 1.13012 | 0.00275 | 1.71114 | 0.11522 | 0.89921 | 0.62445 | 3.59463 | 1.08545 | 0.57471 | 5.67016 | 0.15643 | 0.00000 | 2.81107 | 0.00000 |
| **Parp6** | 0.96058 | 0.01063 | 0.14147 | 0.17749 | 1.78276 | 0.36935 | 0.00439 | 1.02133 | 0.67939 | 0.02354 | 5.16708 | 0.83347 | 0.00383 | 1.72871 |
| **Parp8** | 18.33151 | 0.01458 | 23.47618 | 5.28986 | 20.16458 | 5.45350 | 2.60827 | 0.00000 | 13.59537 | 0.00000 | 14.24383 | 1.90423 | 4.14238 | 10.00465 |
| **Parp9** | 3.50040 | 0.00231 | 0.00000 | 0.00000 | 1.81344 | 0.08237 | 0.00267 | 0.00000 | 6.05499 | 0.01091 | 0.00000 | 1.06139 | 3.83853 | 11.81188 |
| **Parpbp** | 0.96342 | 0.00327 | 1.53466 | 0.89518 | 1.80705 | 1.84451 | 0.00377 | 0.59906 | 0.13613 | 0.00163 | 3.38060 | 0.07664 | 0.00000 | 1.72501 |
| **Pars2** | 0.67103 | 0.03112 | 1.98216 | 3.29153 | 0.30245 | 1.73621 | 0.00394 | 1.30178 | 0.00244 | 0.01321 | 0.00000 | 0.00000 | 0.00000 | 0.92679 |
| **Particl** | 0.07269 | 0.00451 | 0.13067 | 0.00000 | 0.00100 | 0.16561 | 0.00429 | 0.00549 | 0.00690 | 0.00588 | 0.00544 | 0.00895 | 0.17401 | 0.21296 |
| **Parva** | 0.11306 | 0.00330 | 0.00000 | 0.00005 | 0.45492 | 2.41904 | 0.68670 | 2.48800 | 0.01159 | 0.01689 | 2.01648 | 0.00000 | 0.00390 | 0.00000 |
| **Parvb** | 2.42410 | 0.42647 | 7.29221 | 5.13552 | 3.66716 | 4.58333 | 2.33122 | 6.89935 | 0.03334 | 0.16638 | 4.11521 | 3.96469 | 0.00000 | 1.78734 |
| **Pask** | 2.68366 | 0.01809 | 1.94792 | 2.58825 | 1.34529 | 1.50258 | 0.00879 | 2.08296 | 1.31629 | 0.02036 | 3.84182 | 0.24530 | 1.24257 | 1.66151 |
| **Pate2** | 0.00801 | 0.00441 | 0.04423 | 0.00820 | 0.03615 | 0.00835 | 0.00430 | 0.01205 | 0.23576 | 0.00708 | 0.00703 | 0.01019 | 0.01719 | 0.09866 |
| **Pate3** | 0.00880 | 0.00444 | 0.01485 | 0.00880 | 0.00796 | 0.04317 | 0.00422 | 0.00852 | 0.07949 | 0.00764 | 0.00745 | 0.01008 | 0.00397 | 0.05003 |
| **Pate4** | 0.01557 | 0.00441 | 0.03244 | 0.00441 | 0.00936 | 0.13338 | 0.00410 | 0.00711 | 0.16385 | 0.01120 | 0.00625 | 0.00933 | 0.00509 | 0.04281 |
| **Patl1** | 5.61016 | 0.66349 | 5.14393 | 0.15151 | 1.91246 | 4.06785 | 1.24110 | 0.00000 | 3.20649 | 0.05694 | 0.01541 | 3.02432 | 4.79192 | 4.40090 |
| **Patl2** | 0.00000 | 0.01986 | 0.06892 | 0.11216 | 0.00000 | 3.91590 | 0.01061 | 2.24082 | 0.01867 | 0.00628 | 1.57090 | 0.49409 | 0.09882 | 0.03376 |
| **Patz1** | 5.32375 | 0.01534 | 9.69276 | 2.93444 | 8.44628 | 1.35874 | 1.20131 | 0.50476 | 1.39280 | 0.02574 | 2.03826 | 2.09406 | 2.39394 | 11.09910 |
| **Pawr** | 0.36417 | 2.10241 | 3.48875 | 0.69916 | 0.24175 | 1.95183 | 1.55671 | 2.07174 | 1.58345 | 10.60745 | 1.40437 | 2.61497 | 0.00000 | 0.78965 |
| **Pax3** | 0.02228 | 0.01189 | 0.05164 | 0.00948 | 0.24847 | 0.02516 | 0.01607 | 0.04463 | 0.02966 | 0.00644 | 0.01512 | 0.00876 | 0.30910 | 0.04061 |
| **Pax6** | 0.03547 | 0.00393 | 2.13296 | 0.00000 | 7.89833 | 0.00000 | 0.00242 | 0.00397 | 8.17029 | 0.00392 | 0.00000 | 0.00469 | 0.08075 | 3.15723 |
| **Pax8** | 0.26405 | 0.00444 | 0.00959 | 0.00954 | 0.07986 | 0.01148 | 0.00486 | 0.01094 | 0.01711 | 0.00712 | 0.00714 | 0.00847 | 0.00443 | 0.00373 |
| **Pax9** | 0.01603 | 0.07455 | 0.05840 | 0.08198 | 0.50876 | 0.04288 | 0.01026 | 0.03929 | 0.04773 | 0.01783 | 0.00600 | 0.01425 | 0.28674 | 0.11971 |
| **Paxbp1** | 4.03628 | 0.00212 | 3.20806 | 1.74077 | 4.15012 | 2.82485 | 1.52949 | 0.66242 | 4.47123 | 0.02757 | 2.73086 | 1.17555 | 5.41168 | 7.69571 |
| **Paxip1** | 5.58703 | 0.10196 | 6.68439 | 5.66864 | 1.10918 | 6.55896 | 3.33778 | 1.62737 | 0.94446 | 0.11155 | 3.98054 | 4.69330 | 2.12122 | 8.54303 |
| **Pbdc1** | 11.99292 | 0.11768 | 19.25332 | 14.58848 | 18.13893 | 20.31572 | 6.55885 | 0.71226 | 15.73781 | 1.77990 | 11.35565 | 13.94925 | 1.26889 | 20.26469 |
| **Pbk** | 12.80057 | 0.03912 | 13.18094 | 8.64216 | 16.89344 | 6.70765 | 0.03675 | 1.68774 | 11.98768 | 0.00000 | 9.03483 | 9.84690 | 9.09988 | 11.92095 |
| **Pbld1** | 1.93112 | 0.06719 | 0.00000 | 1.28575 | 0.57557 | 2.56446 | 0.42075 | 0.22978 | 2.43845 | 0.83133 | 0.06436 | 0.68417 | 13.21795 | 2.39401 |
| **Pbld2** | 0.04033 | 0.00445 | 0.12074 | 0.16818 | 0.29973 | 0.25656 | 0.00431 | 0.00687 | 0.00562 | 0.00581 | 0.39991 | 0.00487 | 0.21539 | 0.72813 |
| **Pbrm1** | 3.43578 | 0.05555 | 0.46371 | 0.46823 | 4.76942 | 4.38659 | 0.02986 | 2.31848 | 7.14678 | 0.02417 | 3.47821 | 4.24012 | 2.24208 | 6.12295 |
| **Pbx1** | 0.02209 | 0.00442 | 0.01426 | 0.01654 | 0.09745 | 0.01142 | 0.00430 | 0.01009 | 0.04170 | 0.00613 | 0.00802 | 0.00922 | 0.21317 | 0.00381 |
| **Pbx2** | 0.00000 | 0.02941 | 0.50857 | 2.02157 | 0.63900 | 0.71316 | 0.00424 | 0.03334 | 0.01151 | 0.00545 | 0.02816 | 0.87234 | 0.00678 | 0.00512 |
| **Pbx3** | 2.35803 | 0.01817 | 0.01183 | 0.00000 | 1.07259 | 0.00006 | 2.52493 | 0.00271 | 1.86030 | 0.06226 | 0.00000 | 0.95518 | 0.76139 | 1.23904 |
| **Pbx4** | 0.00000 | 0.01286 | 0.93836 | 1.18874 | 1.50752 | 0.73149 | 0.10350 | 0.05459 | 0.00735 | 0.60691 | 0.28514 | 0.00000 | 0.16137 | 0.16146 |

| | | | | | | | | | | | | | | |
|---|---|---|---|---|---|---|---|---|---|---|---|---|---|---|
| **Pbxip1** | 0.06642 | 0.06455 | 0.20213 | 0.70429 | 0.43018 | 0.00000 | 0.01015 | 2.45231 | 0.65251 | 0.00381 | 0.67119 | 0.91988 | 0.15920 | 0.15530 |
| **Pcbd1** | 0.00802 | 0.01158 | 0.05363 | 0.61274 | 0.03670 | 0.93607 | 1.13722 | 0.39081 | 0.00740 | 0.04059 | 0.67848 | 0.25111 | 0.70335 | 0.00000 |
| **Pcbd2** | 0.34914 | 1.21794 | 0.00000 | 0.00000 | 0.20397 | 0.33235 | 0.00316 | 0.78921 | 0.02775 | 1.32288 | 0.00000 | 0.47621 | 0.11461 | 0.13121 |
| **Pcbp1** | 33.71856 | 2.45122 | 35.70940 | 44.77333 | 46.11515 | 22.61907 | 11.68130 | 20.35411 | 9.55218 | 12.58638 | 12.60954 | 26.73351 | 21.69467 | 16.29335 |
| **Pcbp2** | 12.56398 | 0.17052 | 10.18217 | 0.73277 | 15.33639 | 11.78482 | 1.73306 | 8.96849 | 7.03740 | 2.52049 | 11.34299 | 9.34609 | 3.46506 | 9.62673 |
| **Pcbp3** | 0.00000 | 0.00393 | 0.17097 | 0.41576 | 0.59854 | 0.16733 | 0.00368 | 0.00009 | 0.00778 | 0.00463 | 0.64756 | 0.00637 | 1.09815 | 0.00000 |
| **Pcbp4** | 0.03529 | 2.88278 | 0.00342 | 0.00000 | 0.27108 | 0.00304 | 0.79921 | 6.57449 | 0.01103 | 0.15866 | 0.00347 | 0.02986 | 0.00488 | 0.00309 |
| **Pcca** | 0.77122 | 0.01942 | 0.00000 | 0.00000 | 0.47588 | 0.25733 | 0.00359 | 0.00445 | 0.18933 | 0.00593 | 0.00213 | 1.05810 | 0.04806 | 0.32855 |
| **Pccb** | 2.99153 | 0.07023 | 8.96138 | 4.67814 | 8.57575 | 3.96171 | 0.00104 | 3.79860 | 2.58280 | 1.42020 | 0.00000 | 1.82619 | 1.79928 | 11.79087 |
| **Pcdh1** | 0.04445 | 0.01541 | 0.01250 | 0.00000 | 0.01117 | 0.00558 | 0.00439 | 0.24516 | 0.01019 | 0.00811 | 0.00521 | 0.17006 | 0.01934 | 0.01572 |
| **Pcdh10** | 0.00976 | 0.00445 | 0.00538 | 0.01661 | 0.00000 | 0.00996 | 0.00416 | 0.00994 | 0.28242 | 0.00758 | 0.00702 | 0.00969 | 0.00461 | 0.11367 |
| **Pcdh15** | 0.00910 | 0.00443 | 0.01387 | 0.01252 | 0.03424 | 0.01292 | 0.00431 | 0.01058 | 0.01113 | 0.00668 | 0.00896 | 0.00996 | 0.00893 | 0.04401 |
| **Pcdh19** | 0.03439 | 0.00534 | 0.01198 | 0.00000 | 0.01174 | 0.00638 | 0.01040 | 0.20459 | 0.01098 | 0.00763 | 0.00329 | 0.48512 | 0.00526 | 0.01996 |
| **Pcdh8** | 0.00274 | 0.00436 | 0.13830 | 0.00371 | 0.00092 | 0.00671 | 0.00425 | 0.01113 | 0.02408 | 0.00717 | 0.00687 | 0.01022 | 0.06940 | 0.29556 |
| **Pcdh9** | 0.00621 | 0.00450 | 0.00223 | 0.01104 | 0.35046 | 0.01071 | 0.00416 | 0.01027 | 0.34580 | 0.00694 | 0.00718 | 0.00812 | 0.00349 | 0.00000 |
| **Pced1a** | 0.11632 | 0.01140 | 0.55923 | 1.50933 | 0.00000 | 0.58384 | 0.00385 | 0.00563 | 0.00808 | 0.00527 | 0.27615 | 0.13392 | 0.01324 | 0.49713 |
| **Pced1b** | 0.11577 | 3.25945 | 0.03033 | 0.38529 | 0.82985 | 0.15201 | 0.03299 | 0.05464 | 1.08132 | 0.02045 | 0.09826 | 0.29575 | 1.80317 | 2.23480 |
| **Pcf11** | 4.25833 | 0.01784 | 3.19745 | 0.00000 | 2.41842 | 1.48489 | 0.93188 | 0.10377 | 4.63730 | 6.76315 | 4.40968 | 1.96903 | 1.34706 | 1.15087 |
| **Pcgf1** | 5.03226 | 0.01649 | 3.41076 | 0.68825 | 13.20392 | 1.56247 | 0.58132 | 1.11035 | 6.63417 | 0.03455 | 2.31196 | 0.53981 | 11.13258 | 7.67384 |
| **Pcgf2** | 0.02053 | 0.00455 | 0.01274 | 0.01745 | 0.01520 | 0.01025 | 0.00433 | 0.07295 | 0.01177 | 0.01128 | 0.00723 | 0.62182 | 0.02044 | 0.00803 |
| **Pcgf3** | 0.82367 | 0.02589 | 1.12012 | 0.00000 | 0.58630 | 0.38319 | 0.00222 | 1.99081 | 3.40530 | 0.01633 | 3.85902 | 1.23762 | 1.98626 | 0.87050 |
| **Pcgf5** | 4.19849 | 0.01775 | 9.48632 | 1.93953 | 12.17045 | 1.60126 | 1.05755 | 1.37068 | 8.06334 | 0.03754 | 3.64743 | 0.00000 | 3.80278 | 10.36218 |
| **Pcgf6** | 13.22113 | 0.10057 | 7.29721 | 1.40148 | 14.98482 | 5.04549 | 0.00000 | 2.87666 | 8.12935 | 0.01586 | 2.93151 | 3.36012 | 13.09984 | 13.10455 |
| **Pcid2** | 0.13782 | 0.01816 | 4.10684 | 1.91369 | 1.34485 | 2.21267 | 0.00963 | 1.08185 | 0.33979 | 0.01098 | 1.35342 | 1.81006 | 0.13224 | 3.18740 |
| **Pcif1** | 0.40245 | 0.04690 | 1.20731 | 1.33934 | 0.44947 | 0.37285 | 0.00901 | 0.95905 | 0.08343 | 2.41583 | 1.04029 | 2.56466 | 0.73065 | 0.00000 |
| **Pck2** | 0.00000 | 0.03641 | 2.62490 | 9.02786 | 1.61143 | 2.01388 | 0.02166 | 0.89696 | 0.00000 | 0.00257 | 8.90509 | 2.84586 | 0.87598 | 2.32824 |
| **Pclo** | 0.06536 | 0.01982 | 0.01807 | 0.09099 | 0.49317 | 0.10457 | 0.05772 | 0.07340 | 0.17129 | 0.00773 | 0.28298 | 0.03456 | 0.46547 | 0.27175 |
| **Pcm1** | 1.78507 | 0.01304 | 2.85349 | 0.58501 | 2.38469 | 2.78113 | 0.00000 | 0.13331 | 14.96484 | 6.98458 | 2.50850 | 4.23581 | 1.92232 | 1.41498 |
| **Pcmt1** | 9.66361 | 3.31516 | 13.76038 | 6.71133 | 11.96136 | 7.10957 | 4.25636 | 5.42406 | 5.38751 | 4.69623 | 3.29677 | 6.64443 | 6.40514 | 7.79980 |
| **Pcmtd1** | 1.89828 | 0.99238 | 3.41506 | 1.47247 | 2.68700 | 6.94066 | 2.48604 | 1.33866 | 6.10527 | 5.74719 | 3.77423 | 3.93074 | 1.01144 | 0.00000 |
| **Pcmtd2** | 0.03031 | 0.01089 | 0.22598 | 0.08786 | 0.56703 | 0.14263 | 0.01085 | 1.09240 | 0.23004 | 0.01873 | 0.01718 | 1.49399 | 0.05346 | 0.00417 |
| **Pcna** | 22.30782 | 0.45131 | 23.18201 | 15.27728 | 16.04176 | 25.61525 | 9.33539 | 10.89053 | 10.78663 | 3.33638 | 16.01070 | 21.14049 | 15.08305 | 21.88037 |
| **Pcnp** | 6.07074 | 0.04980 | 5.74592 | 6.72645 | 3.95287 | 10.95837 | 5.24196 | 5.78666 | 9.06740 | 0.18326 | 8.51263 | 6.92187 | 1.77072 | 5.25048 |
| **Pcnt** | 4.82307 | 0.00895 | 11.06299 | 7.07179 | 13.70550 | 3.90681 | 4.68561 | 1.33478 | 0.00000 | 0.03813 | 1.89801 | 2.11917 | 3.15453 | 2.75348 |
| **Pcnx** | 0.67498 | 0.01705 | 1.31950 | 0.31928 | 1.73820 | 0.59234 | 0.00863 | 2.35155 | 0.41353 | 0.00811 | 0.44883 | 2.09053 | 1.37167 | 0.25990 |
| **Pcnx2** | 0.00837 | 0.00440 | 0.00259 | 0.54855 | 0.04941 | 0.00750 | 0.00380 | 0.00629 | 0.00671 | 0.00576 | 0.00507 | 0.29675 | 0.03191 | 0.26984 |
| **Pcnx3** | 0.00000 | 5.74915 | 2.20869 | 2.57876 | 2.37323 | 0.78759 | 0.11397 | 3.24831 | 0.00000 | 0.03274 | 0.92461 | 3.29189 | 0.00000 | 0.41497 |
| **Pcnx4** | 0.00000 | 0.00389 | 0.10827 | 0.00000 | 0.37554 | 0.00000 | 0.73177 | 0.00287 | 0.98265 | 0.00645 | 0.03327 | 0.00525 | 1.91167 | 0.87425 |
| **Pcolce** | 0.06996 | 0.00441 | 0.22078 | 0.94570 | 0.13571 | 0.00000 | 0.00436 | 0.01703 | 0.00506 | 0.00726 | 0.00606 | 0.00572 | 0.06429 | 0.11647 |
| **Pcolce2** | 0.01904 | 0.00457 | 0.00324 | 0.01941 | 0.00000 | 0.00707 | 0.00419 | 0.01675 | 0.25224 | 0.00774 | 0.00332 | 0.47720 | 0.00879 | 0.10700 |
| **Pcp2** | 0.08172 | 0.00433 | 0.01460 | 0.02358 | 0.01010 | 0.00600 | 0.00430 | 0.00997 | 0.01102 | 0.00656 | 0.06697 | 0.01437 | 0.00558 | 0.00887 |
| **Pcp4l1** | 0.01295 | 0.00404 | 0.19411 | 0.41724 | 0.01927 | 0.00000 | 0.00410 | 0.00713 | 0.01321 | 0.00566 | 0.01256 | 0.01938 | 0.05025 | 0.00735 |
| **Pcsk1n** | 0.00771 | 0.00452 | 0.00940 | 0.09106 | 0.09929 | 0.01403 | 0.00451 | 0.01462 | 0.01077 | 0.00626 | 0.00768 | 0.00889 | 0.00737 | 0.01282 |
| **Pcsk6** | 4.65476 | 2.63549 | 12.16067 | 5.46119 | 1.20375 | 6.95579 | 2.37090 | 8.93573 | 1.79579 | 0.08654 | 0.47278 | 9.02657 | 0.12690 | 0.93410 |
| **Pcsk7** | 3.99528 | 2.04898 | 4.84855 | 3.42143 | 4.56340 | 3.47668 | 0.38020 | 1.95069 | 2.48108 | 0.03053 | 6.29258 | 3.56493 | 0.53901 | 5.99858 |
| **Pctp** | 0.02957 | 0.00441 | 0.18166 | 0.03079 | 0.02085 | 0.03085 | 0.00413 | 0.00897 | 0.02684 | 0.03371 | 0.00729 | 0.02654 | 0.06105 | 0.09015 |
| **Pcx** | 0.00000 | 0.01085 | 0.26306 | 0.24032 | 0.02051 | 0.93812 | 0.00355 | 1.30542 | 0.00266 | 0.01475 | 0.00940 | 0.59630 | 0.38490 | 0.69231 |
| **Pcyox1** | 0.41352 | 0.08705 | 0.03048 | 0.29730 | 0.00000 | 0.00000 | 0.00307 | 1.68153 | 0.95335 | 0.00753 | 3.27912 | 0.58706 | 0.00972 | 1.08824 |
| **Pcyox1l** | 0.00000 | 0.00316 | 0.52548 | 0.09814 | 0.62845 | 0.16695 | 0.01082 | 0.23308 | 0.03843 | 0.00365 | 0.65208 | 1.07518 | 0.00982 | 0.00000 |
| **Pcyt1a** | 7.40018 | 0.01546 | 4.23577 | 2.55212 | 8.14446 | 10.01197 | 1.46241 | 1.70197 | 13.22263 | 0.03147 | 0.00000 | 1.92398 | 13.74164 | 14.64628 |
| **Pcyt1b** | 0.00000 | 0.01076 | 2.69603 | 1.02092 | 0.09238 | 1.80628 | 0.00247 | 2.80888 | 1.89035 | 0.03820 | 1.15310 | 1.23754 | 0.36716 | 2.60187 |
| **Pcyt2** | 7.76801 | 5.64448 | 21.98870 | 21.78771 | 15.41451 | 9.93092 | 1.51309 | 1.73714 | 0.17565 | 0.09247 | 8.15257 | 6.13835 | 1.58487 | 9.94708 |
| **Pdap1** | 1.16530 | 0.19032 | 0.46243 | 4.86627 | 5.35935 | 5.64791 | 0.30562 | 4.86463 | 0.82815 | 0.28770 | 0.22569 | 4.77489 | 8.13215 | 1.74958 |
| **Pdcd10** | 11.75866 | 3.12273 | 7.82504 | 3.05234 | 4.78328 | 14.16220 | 3.26783 | 7.41882 | 9.34246 | 0.08787 | 8.44609 | 5.77352 | 5.29111 | 3.66833 |
| **Pdcd11** | 0.00000 | 0.15383 | 11.48914 | 9.49317 | 12.61646 | 1.93620 | 4.05401 | 3.99554 | 0.58810 | 0.10184 | 1.82210 | 5.95468 | 1.28314 | 0.37487 |

| | | | | | | | | | | | | | | |
|---|---|---|---|---|---|---|---|---|---|---|---|---|---|---|
| **Pdcd2** | 3.96740 | 0.22946 | 3.07070 | 1.94300 | 3.30122 | 0.40415 | 0.00229 | 0.91993 | 2.81396 | 0.61207 | 2.83157 | 3.51659 | 5.32421 | 2.48235 |
| **Pdcd2l** | 6.32752 | 0.03683 | 7.33800 | 9.73962 | 11.65099 | 3.15825 | 0.41174 | 1.38228 | 0.96260 | 0.02762 | 2.16667 | 5.17349 | 6.33887 | 5.20637 |
| **Pdcd4** | 0.11501 | 0.00471 | 0.00438 | 0.00000 | 0.20745 | 0.19704 | 0.06178 | 0.35515 | 0.18849 | 0.01167 | 0.02378 | 0.61243 | 0.29780 | 0.32444 |
| **Pdcd5** | 2.01476 | 0.04312 | 0.00000 | 2.20354 | 2.13752 | 5.11318 | 1.70228 | 2.13642 | 0.71618 | 0.18276 | 0.19375 | 2.80412 | 2.45843 | 0.00000 |
| **Pdcd6** | 12.33228 | 4.62546 | 18.71733 | 15.75026 | 14.77235 | 18.29491 | 3.39523 | 10.70747 | 4.60675 | 15.71342 | 9.63234 | 11.18448 | 0.00000 | 1.28258 |
| **Pdcd6ip** | 5.64472 | 0.26502 | 5.65148 | 3.29112 | 6.81794 | 1.78979 | 2.00253 | 5.67573 | 1.24675 | 0.06933 | 2.47389 | 7.62479 | 5.64229 | 1.54189 |
| **Pdcd7** | 4.34275 | 0.02350 | 13.14724 | 1.39120 | 10.75827 | 3.89949 | 3.60154 | 5.84176 | 10.66264 | 0.09762 | 0.75249 | 2.77760 | 5.65526 | 10.12725 |
| **Pdcl** | 0.07623 | 0.03455 | 5.31177 | 3.12552 | 7.67896 | 5.28441 | 0.00836 | 1.62039 | 8.16752 | 0.01647 | 3.25710 | 0.89056 | 6.79051 | 10.30492 |
| **Pdcl3** | 6.15771 | 0.16361 | 7.24735 | 3.59180 | 4.29468 | 9.99453 | 7.04527 | 12.06680 | 2.40965 | 4.48676 | 10.22147 | 4.41476 | 0.30917 | 5.57688 |
| **Pddc1** | 0.03491 | 0.01797 | 1.63265 | 1.07464 | 0.22425 | 0.42881 | 1.84409 | 1.50463 | 0.82838 | 0.04113 | 2.35325 | 2.38852 | 0.40257 | 1.17568 |
| **Pde12** | 5.38687 | 0.03126 | 7.65907 | 3.01729 | 5.45890 | 3.49081 | 4.66037 | 1.07650 | 1.18591 | 0.15472 | 2.01686 | 3.11105 | 7.93490 | 1.69714 |
| **Pde1b** | 0.17062 | 0.00402 | 0.00500 | 0.00000 | 0.00663 | 0.34848 | 0.00950 | 0.25645 | 0.00911 | 0.01373 | 0.00856 | 0.23434 | 0.00430 | 0.01536 |
| **Pde2a** | 0.01266 | 0.00429 | 0.23065 | 0.00000 | 0.00831 | 0.00911 | 0.00430 | 0.00999 | 0.00959 | 0.00672 | 0.00844 | 0.02348 | 0.18527 | 0.00396 |
| **Pde3a** | 0.01140 | 0.00442 | 0.04663 | 0.01397 | 0.00392 | 0.01038 | 0.00425 | 0.01014 | 0.02613 | 0.00750 | 0.00759 | 0.00922 | 0.00603 | 0.19338 |
| **Pde3b** | 0.00000 | 0.00388 | 0.01808 | 0.01019 | 0.00000 | 0.00341 | 0.00365 | 0.00739 | 2.60386 | 0.00783 | 0.00410 | 0.00530 | 3.61978 | 0.89735 |
| **Pde4a** | 0.97865 | 0.00405 | 0.00000 | 0.10608 | 0.31090 | 0.00000 | 0.00374 | 0.00519 | 0.85589 | 0.01526 | 0.57097 | 0.00669 | 0.02163 | 0.56026 |
| **Pde4c** | 0.10417 | 0.13217 | 0.04992 | 0.30338 | 0.16552 | 0.19921 | 0.02889 | 0.16013 | 1.07090 | 0.07220 | 0.01083 | 0.07365 | 0.45516 | 0.26860 |
| **Pde4d** | 0.04784 | 0.00442 | 0.00932 | 0.00495 | 0.00658 | 0.00348 | 0.00399 | 0.01244 | 0.00910 | 0.00704 | 0.12884 | 0.25042 | 1.35636 | 0.00500 |
| **Pde4dip** | 1.59265 | 0.01042 | 0.00000 | 1.16144 | 0.44409 | 5.04898 | 0.00982 | 2.04677 | 0.00824 | 0.01981 | 0.07283 | 1.50171 | 1.33426 | 0.30342 |
| **Pde5a** | 0.24578 | 0.00444 | 0.13676 | 0.16593 | 0.23427 | 0.20574 | 0.04694 | 0.04698 | 0.20744 | 0.04589 | 0.02942 | 0.11011 | 0.26721 | 0.30434 |
| **Pde6a** | 0.38689 | 0.00347 | 0.09812 | 0.41380 | 0.34546 | 0.00000 | 0.00372 | 0.00467 | 0.00000 | 0.00502 | 0.29467 | 0.02539 | 3.25332 | 1.88802 |
| **Pde6b** | 0.00425 | 0.00432 | 0.29676 | 0.15667 | 0.00225 | 0.11056 | 0.00405 | 0.00279 | 0.01042 | 0.00641 | 0.00528 | 0.01152 | 0.00553 | 0.03060 |
| **Pde6d** | 1.15050 | 0.03972 | 0.42789 | 1.08518 | 1.06588 | 2.42677 | 0.00318 | 2.23849 | 0.72741 | 0.02809 | 1.88629 | 1.38285 | 1.99992 | 1.05538 |
| **Pde6g** | 0.00000 | 3.71703 | 0.57501 | 0.00000 | 0.76355 | 0.03434 | 0.00306 | 0.00000 | 0.35287 | 0.00452 | 0.00385 | 0.00000 | 0.22999 | 0.57344 |
| **Pde7a** | 1.42290 | 0.06323 | 0.22240 | 1.15772 | 0.68204 | 1.54630 | 0.00318 | 0.43928 | 2.37364 | 0.02492 | 0.20654 | 2.02316 | 0.02688 | 1.09849 |
| **Pde8a** | 0.02099 | 0.00410 | 0.36886 | 0.09723 | 0.32193 | 0.00000 | 0.00405 | 0.02095 | 0.06111 | 0.00632 | 0.00339 | 0.53238 | 0.00540 | 2.11815 |
| **Pdf** | 6.16820 | 0.04498 | 3.61891 | 0.00000 | 5.48408 | 1.06785 | 0.00744 | 3.53639 | 0.41900 | 0.00009 | 1.66577 | 2.75639 | 1.64569 | 4.40633 |
| **Pdgfa** | 0.00000 | 1.45129 | 0.69931 | 5.79413 | 1.18206 | 6.72156 | 0.59413 | 4.80414 | 0.07895 | 0.07309 | 1.86289 | 0.70218 | 0.20628 | 0.30636 |
| **Pdgfb** | 0.34117 | 0.00366 | 0.39565 | 0.06287 | 1.10163 | 0.01077 | 0.00378 | 0.00243 | 0.01128 | 0.00539 | 0.00470 | 0.40015 | 0.55976 | 0.31705 |
| **Pdgfra** | 1.05394 | 0.08204 | 1.98967 | 0.00000 | 0.47292 | 0.00173 | 0.00000 | 0.00000 | 1.09398 | 0.02622 | 0.00020 | 11.87579 | 0.00047 | 0.12316 |
| **Pdgfrb** | 0.17032 | 0.00444 | 0.01376 | 0.01070 | 0.00895 | 0.01151 | 0.00431 | 0.01123 | 0.01089 | 0.00659 | 0.00750 | 0.00934 | 0.00520 | 0.00709 |
| **Pdha1** | 16.34134 | 6.01295 | 28.85570 | 18.89946 | 21.35120 | 24.65665 | 4.60784 | 11.75984 | 18.87079 | 8.92022 | 12.81326 | 8.18886 | 3.29775 | 15.09031 |
| **Pdha2** | 0.01714 | 0.00395 | 0.00789 | 0.00055 | 0.00371 | 0.39860 | 0.00387 | 0.00303 | 0.02413 | 0.00528 | 0.18236 | 0.00690 | 1.92352 | 0.07452 |
| **Pdhb** | 19.49002 | 5.40900 | 25.66885 | 28.27255 | 18.42324 | 26.60616 | 5.16479 | 15.21478 | 5.49479 | 14.16148 | 19.64502 | 17.88438 | 8.02949 | 12.28087 |
| **Pdhx** | 5.33243 | 0.05498 | 9.61676 | 3.78036 | 7.96513 | 8.78634 | 2.42951 | 1.72076 | 7.26193 | 0.15342 | 5.99185 | 5.30403 | 1.90239 | 8.30621 |
| **Pdia2** | 0.33221 | 0.00454 | 0.11174 | 0.00000 | 0.00208 | 0.38880 | 0.00424 | 0.00496 | 0.01033 | 0.00558 | 0.00484 | 0.00819 | 0.00445 | 0.00417 |
| **Pdia3** | 12.15878 | 0.02932 | 1.53431 | 3.31935 | 3.96607 | 4.64890 | 3.90202 | 1.53653 | 12.90268 | 0.11496 | 4.03583 | 2.27452 | 9.20597 | 7.38877 |
| **Pdia4** | 0.00000 | 0.03487 | 1.08361 | 0.83923 | 0.00000 | 0.00000 | 0.00202 | 1.94654 | 6.82934 | 0.02092 | 1.20274 | 2.25809 | 0.70240 | 6.37042 |
| **Pdia5** | 0.12636 | 0.11702 | 14.94674 | 20.12161 | 5.32973 | 11.47593 | 1.90713 | 0.12871 | 4.22354 | 0.05535 | 9.24256 | 7.46670 | 0.00000 | 7.11787 |
| **Pdia6** | 6.08726 | 7.21152 | 7.83096 | 14.97837 | 2.90738 | 25.99411 | 11.14616 | 5.81147 | 0.76701 | 13.58222 | 11.18520 | 14.55020 | 1.93079 | 2.74139 |
| **Pdik1l** | 0.00000 | 0.00306 | 0.18506 | 0.26501 | 0.00000 | 1.04738 | 0.00877 | 1.28140 | 1.17178 | 0.01866 | 0.13932 | 1.30506 | 0.00538 | 0.71692 |
| **Pdk1** | 0.00245 | 0.00357 | 0.07862 | 0.00000 | 0.18450 | 1.15125 | 0.00374 | 0.11863 | 0.14621 | 0.00730 | 0.00456 | 0.10131 | 1.57048 | 2.87943 |
| **Pdk2** | 0.00173 | 0.01888 | 0.33775 | 0.03555 | 0.00489 | 0.00096 | 0.01530 | 0.02761 | 0.00955 | 0.01239 | 0.00383 | 1.25627 | 0.00986 | 0.01532 |
| **Pdk3** | 1.73069 | 0.00369 | 0.95249 | 0.00000 | 3.10039 | 0.01837 | 0.00356 | 0.00000 | 0.03928 | 0.00298 | 0.22561 | 0.00602 | 2.89235 | 2.14840 |
| **Pdlim1** | 4.85467 | 11.64263 | 10.57671 | 10.99691 | 3.93154 | 14.61538 | 2.28005 | 12.79713 | 0.00000 | 0.20914 | 7.44074 | 6.09603 | 3.31769 | 3.43883 |
| **Pdlim2** | 0.22252 | 0.00395 | 0.54476 | 0.20253 | 0.50781 | 0.00000 | 0.00552 | 0.30898 | 0.00631 | 3.10574 | 0.00332 | 0.00000 | 0.08392 | 0.05909 |
| **Pdlim3** | 0.12066 | 0.00457 | 0.75516 | 0.00000 | 1.97587 | 0.14068 | 0.00409 | 0.00357 | 0.08597 | 0.00459 | 0.00391 | 0.01074 | 0.00000 | 0.42342 |
| **Pdlim4** | 0.00668 | 0.08424 | 0.03096 | 0.00000 | 0.43977 | 0.00764 | 0.00368 | 0.46324 | 0.00762 | 0.00638 | 0.00000 | 1.82049 | 0.01071 | 0.02485 |
| **Pdlim5** | 3.41428 | 0.01686 | 3.12010 | 0.00000 | 3.64993 | 8.40494 | 1.89387 | 2.67516 | 2.11803 | 0.05587 | 0.84892 | 2.00848 | 1.29201 | 3.15250 |
| **Pdlim7** | 1.23950 | 0.04001 | 1.84318 | 0.67868 | 1.78250 | 1.61139 | 0.91672 | 4.07217 | 0.00646 | 8.17524 | 2.34820 | 0.69567 | 0.12538 | 0.02261 |
| **Pdp1** | 0.49240 | 0.00456 | 0.98273 | 0.00000 | 0.19687 | 0.24791 | 0.00399 | 0.00259 | 0.00924 | 0.02295 | 0.00393 | 0.01463 | 0.30170 | 0.00933 |
| **Pdp2** | 1.05095 | 0.05775 | 7.07012 | 4.89754 | 8.48050 | 3.17274 | 0.00588 | 0.50118 | 13.60525 | 0.00000 | 2.67644 | 1.27550 | 6.51233 | 11.64085 |
| **Pdpk1** | 5.67385 | 0.02876 | 5.27585 | 1.32373 | 7.29357 | 6.45025 | 4.19030 | 0.52586 | 14.97600 | 8.47645 | 0.00000 | 4.55434 | 2.57637 | 7.68857 |
| **Pdpn** | 0.25350 | 0.69528 | 4.48971 | 6.09048 | 1.67205 | 3.46894 | 0.24757 | 5.14544 | 3.55372 | 0.10589 | 0.50984 | 9.34711 | 3.45430 | 3.95952 |
| **Pdpr** | 0.00000 | 0.00432 | 0.15076 | 0.01523 | 0.02614 | 0.00000 | 0.00531 | 0.73334 | 0.35250 | 0.00507 | 0.00598 | 0.02066 | 1.34354 | 1.95179 |

| | | | | | | | | | | | | | | |
|---|---|---|---|---|---|---|---|---|---|---|---|---|---|---|
| **Pdrg1** | 8.64006 | 0.06034 | 7.13146 | 3.67897 | 9.98990 | 10.40869 | 2.58172 | 2.27703 | 1.62696 | 0.07027 | 2.01028 | 3.77411 | 12.87752 | 9.69373 |
| **Pds5a** | 1.43396 | 0.03133 | 3.58506 | 0.00000 | 2.75183 | 1.91200 | 0.08543 | 0.64931 | 12.25295 | 0.00620 | 1.76440 | 1.78109 | 2.27811 | 4.97164 |
| **Pds5b** | 1.56123 | 3.08886 | 0.71965 | 0.00000 | 4.92434 | 2.34501 | 0.01051 | 0.04379 | 9.79020 | 6.50609 | 2.96963 | 2.05059 | 5.88335 | 5.33016 |
| **Pdss1** | 3.79571 | 0.00076 | 3.11310 | 2.80076 | 3.16291 | 2.48411 | 0.24741 | 0.99718 | 0.81649 | 0.00832 | 2.04632 | 3.40411 | 0.06016 | 2.66876 |
| **Pdss2** | 0.00000 | 0.00416 | 3.68868 | 1.50310 | 2.42629 | 0.00000 | 0.00997 | 0.57969 | 0.71031 | 0.01723 | 0.00000 | 0.27991 | 0.06345 | 0.79559 |
| **Pdxdc1** | 3.52189 | 0.13890 | 2.31271 | 3.15889 | 2.43825 | 3.38063 | 0.16897 | 1.26127 | 1.28788 | 0.08470 | 2.32868 | 2.96696 | 2.59190 | 2.77613 |
| **Pdxk** | 17.56563 | 0.16487 | 44.39616 | 30.01963 | 45.61681 | 20.86602 | 0.05641 | 2.73249 | 1.45760 | 0.24341 | 12.43672 | 5.69809 | 8.88242 | 31.06259 |
| **Pdxk.ps** | 0.01939 | 0.01268 | 0.04282 | 0.02955 | 0.07397 | 0.03235 | 0.00432 | 0.01209 | 0.05836 | 0.00788 | 0.01437 | 0.02267 | 0.01249 | 0.06064 |
| **Pdxp** | 2.43159 | 0.06168 | 3.30001 | 2.98187 | 2.02095 | 5.09463 | 0.59485 | 7.40431 | 0.84851 | 0.06398 | 3.04107 | 1.11275 | 8.26843 | 0.00000 |
| **Pdzd11** | 7.28593 | 0.07852 | 9.32235 | 5.38558 | 11.21402 | 10.77967 | 0.02075 | 4.11704 | 9.59406 | 8.84045 | 8.12362 | 5.50046 | 2.51773 | 8.82206 |
| **Pdzd2** | 0.04302 | 0.01978 | 0.12123 | 0.46529 | 0.12916 | 1.38140 | 0.02816 | 0.41233 | 0.12604 | 0.05585 | 0.00226 | 0.00351 | 2.55848 | 1.53746 |
| **Pdzd3** | 3.64851 | 5.81840 | 11.23224 | 7.14423 | 0.37785 | 15.64966 | 3.77330 | 4.88224 | 0.47406 | 0.18138 | 3.53418 | 0.93050 | 0.00000 | 0.00000 |
| **Pdzd4** | 0.78050 | 0.00342 | 0.00686 | 0.47118 | 0.51702 | 0.00000 | 0.00358 | 0.00272 | 1.26525 | 0.00636 | 0.00000 | 0.55007 | 0.02465 | 0.47421 |
| **Pdzd8** | 4.80160 | 0.10484 | 3.60482 | 1.20675 | 3.83594 | 3.05031 | 0.00000 | 1.80452 | 12.08561 | 0.03224 | 2.78701 | 4.56403 | 1.64380 | 2.71143 |
| **Pdzk1** | 0.58913 | 7.40349 | 0.00000 | 0.00000 | 0.14182 | 6.05234 | 7.08136 | 10.41974 | 0.00000 | 0.33112 | 0.89012 | 0.53538 | 0.32924 | 0.50074 |
| **Pdzk1ip1** | 0.00474 | 0.15089 | 0.00000 | 0.57600 | 0.11356 | 0.86203 | 0.02695 | 8.41740 | 0.00876 | 0.09177 | 0.00660 | 0.01518 | 0.00750 | 0.00384 |
| **Pdzph1** | 0.00987 | 0.00444 | 0.01399 | 0.01337 | 0.01197 | 0.01319 | 0.00429 | 0.01022 | 0.00747 | 0.00694 | 0.00893 | 0.01002 | 0.00572 | 0.05759 |
| **Pdzrn3** | 0.02833 | 0.00435 | 0.01149 | 0.08324 | 0.00702 | 0.01153 | 0.00440 | 0.00840 | 0.01134 | 0.00579 | 0.00721 | 0.00819 | 0.00976 | 0.11117 |
| **Pea15a** | 0.21168 | 5.14723 | 0.51963 | 0.23575 | 1.11461 | 1.05202 | 0.00229 | 1.62631 | 0.05889 | 0.01308 | 0.00000 | 0.95310 | 0.00000 | 1.91848 |
| **Peak1** | 1.85460 | 0.02054 | 1.92088 | 2.10896 | 0.00000 | 2.72690 | 0.12620 | 3.44988 | 3.45679 | 0.00000 | 1.18740 | 0.25052 | 2.22569 | 1.53106 |
| **Pebp1** | 5.96825 | 4.33465 | 15.15107 | 24.62513 | 20.17800 | 13.25478 | 2.88015 | 9.32665 | 1.67679 | 9.22258 | 5.23887 | 15.71135 | 11.03168 | 8.39605 |
| **Pecam1** | 1.71643 | 0.00000 | 4.35249 | 0.19218 | 9.27538 | 0.99086 | 0.00533 | 0.00000 | 10.81447 | 0.00000 | 0.89977 | 6.45393 | 2.38069 | 12.23792 |
| **Pecr** | 1.40226 | 0.04081 | 1.08235 | 1.16492 | 0.00000 | 1.09277 | 0.00276 | 0.00000 | 0.20602 | 0.01609 | 1.91861 | 2.30680 | 4.04921 | 0.22310 |
| **Pef1** | 4.28784 | 6.91002 | 14.24358 | 15.08673 | 8.36022 | 11.34306 | 3.67227 | 6.23124 | 4.85815 | 17.17772 | 12.24793 | 8.47910 | 0.24774 | 6.19557 |
| **Peg13** | 0.04284 | 0.00429 | 0.00901 | 0.00911 | 0.01235 | 0.00062 | 0.00423 | 0.01285 | 0.01992 | 0.00606 | 0.28483 | 0.00567 | 0.00958 | 0.02220 |
| **Peg3** | 0.00324 | 0.00422 | 0.00468 | 0.12777 | 0.00000 | 0.02699 | 0.00393 | 0.00442 | 0.51991 | 0.00523 | 0.01849 | 0.02149 | 0.00742 | 0.13495 |
| **Peli1** | 1.35897 | 2.90946 | 3.06388 | 0.00000 | 1.27648 | 6.04275 | 0.01417 | 0.43803 | 1.63032 | 0.01717 | 4.48124 | 2.22262 | 3.46500 | 2.96258 |
| **Peli2** | 0.00489 | 0.00521 | 0.00860 | 0.01707 | 0.00997 | 0.80994 | 0.00440 | 0.85143 | 0.00950 | 0.01773 | 0.00474 | 0.00434 | 0.09294 | 0.00000 |
| **Pelo** | 6.96077 | 0.06483 | 8.31268 | 4.76784 | 2.72537 | 4.22554 | 2.46346 | 3.21176 | 1.36253 | 0.07277 | 2.12984 | 4.53458 | 0.00000 | 1.76608 |
| **Pelp1** | 5.79302 | 0.02262 | 1.90961 | 6.88763 | 17.16426 | 3.05658 | 0.02628 | 0.29851 | 0.01062 | 0.09395 | 0.57298 | 3.78612 | 7.44939 | 4.66782 |
| **Pemt** | 33.89907 | 0.13293 | 41.73038 | 46.32559 | 29.27647 | 26.32450 | 6.59636 | 1.21936 | 22.39463 | 2.19886 | 10.18668 | 1.91172 | 13.72379 | 22.74500 |
| **Peo1** | 3.02134 | 0.03484 | 4.14864 | 1.66081 | 7.32845 | 3.58191 | 3.85050 | 0.78873 | 1.69608 | 0.13307 | 4.91648 | 2.43322 | 3.43660 | 7.56668 |
| **Pepd** | 0.11956 | 4.37701 | 2.54914 | 5.36518 | 2.78298 | 7.37344 | 3.40008 | 11.95642 | 0.00000 | 0.09985 | 7.80095 | 5.76296 | 0.00000 | 1.10132 |
| **Per1** | 0.02413 | 0.00340 | 0.49080 | 1.59860 | 0.75494 | 0.00000 | 0.00352 | 0.00002 | 0.34046 | 0.00536 | 0.24997 | 0.75556 | 0.24156 | 0.40855 |
| **Per2** | 0.20791 | 0.00419 | 0.16481 | 0.17068 | 0.40993 | 0.00273 | 0.00469 | 0.00906 | 0.01969 | 0.00600 | 0.00616 | 0.00574 | 0.00405 | 0.73598 |
| **Per3** | 0.69066 | 0.00393 | 0.00000 | 0.14640 | 0.97398 | 0.00767 | 0.00424 | 0.00846 | 0.00521 | 0.00701 | 0.00668 | 0.00381 | 1.86217 | 0.10271 |
| **Perm1** | 0.00000 | 0.00417 | 0.00461 | 0.01583 | 0.00000 | 0.00960 | 0.00403 | 0.00798 | 0.37014 | 0.00684 | 0.00964 | 0.00817 | 1.48036 | 0.00000 |
| **Perp** | 3.81008 | 0.25215 | 5.23958 | 4.68128 | 2.06061 | 3.00894 | 0.15837 | 10.37227 | 1.18538 | 6.58894 | 4.04831 | 1.71779 | 0.00000 | 0.00000 |
| **Pes1** | 6.06145 | 7.79185 | 11.06191 | 14.02231 | 11.68356 | 7.25661 | 0.04271 | 7.63872 | 10.56043 | 0.10340 | 8.42946 | 6.58721 | 7.02204 | 12.33152 |
| **Pet100** | 0.84265 | 0.00406 | 0.00000 | 0.53557 | 0.07075 | 1.33046 | 0.04299 | 0.60384 | 0.19119 | 0.12875 | 0.02591 | 0.49445 | 0.44881 | 1.13358 |
| **Pet2** | 0.15903 | 0.00365 | 0.00486 | 0.00000 | 0.00264 | 1.12724 | 0.00380 | 0.00193 | 0.00963 | 0.00263 | 1.04924 | 0.02852 | 2.48480 | 0.00000 |
| **Pex1** | 2.90863 | 0.02465 | 1.51472 | 1.79061 | 3.89308 | 1.26502 | 1.59301 | 1.69398 | 1.41096 | 0.04446 | 2.33908 | 1.42560 | 0.51668 | 5.43044 |
| **Pex10** | 0.24274 | 0.00340 | 0.11320 | 0.22965 | 0.05089 | 0.32486 | 0.00399 | 0.00890 | 0.00781 | 0.00617 | 0.68950 | 0.42657 | 0.30000 | 0.12844 |
| **Pex11a** | 0.00000 | 0.02343 | 0.58443 | 0.00000 | 0.19254 | 0.56551 | 0.00402 | 0.98987 | 0.00572 | 0.00701 | 0.06173 | 0.03226 | 0.58319 | 1.35429 |
| **Pex11b** | 0.19728 | 0.08014 | 0.38566 | 0.36429 | 1.63679 | 2.94750 | 2.15276 | 1.12042 | 0.00000 | 10.05401 | 0.72737 | 2.04648 | 1.95501 | 0.00000 |
| **Pex11g** | 0.02053 | 0.00486 | 0.31320 | 0.00087 | 0.00268 | 0.23897 | 0.00425 | 0.00493 | 0.00859 | 0.00679 | 0.00569 | 0.01004 | 0.09257 | 0.16118 |
| **Pex12** | 1.12187 | 0.01899 | 2.68023 | 2.76749 | 3.44836 | 0.83383 | 2.16808 | 2.29293 | 2.07259 | 0.00700 | 3.98415 | 0.93145 | 2.55758 | 2.02314 |
| **Pex13** | 0.78692 | 0.01154 | 0.00039 | 1.18765 | 0.57902 | 1.45715 | 0.01992 | 0.27414 | 0.06060 | 0.05700 | 2.84365 | 0.86929 | 2.54820 | 0.70875 |
| **Pex14** | 0.00000 | 0.12681 | 2.75002 | 4.64394 | 2.83698 | 1.63045 | 0.00255 | 1.35297 | 0.01451 | 0.03260 | 0.44105 | 3.05700 | 1.16542 | 1.26111 |
| **Pex16** | 5.00742 | 0.14418 | 19.50003 | 19.17970 | 12.99827 | 3.26689 | 4.45797 | 12.37786 | 0.08036 | 9.33615 | 5.40499 | 2.70329 | 0.88149 | 7.59067 |
| **Pex19** | 8.18427 | 0.05797 | 9.64680 | 7.09463 | 14.28717 | 5.15581 | 4.38228 | 2.97714 | 14.02650 | 0.20361 | 6.13450 | 6.14594 | 11.23603 | 18.11845 |
| **Pex2** | 3.06737 | 0.04535 | 1.89796 | 4.32158 | 2.22103 | 5.25958 | 0.61106 | 5.00782 | 0.04361 | 0.01460 | 2.85457 | 5.36543 | 0.07941 | 1.67894 |
| **Pex26** | 3.14174 | 6.08808 | 1.04566 | 0.37533 | 0.90198 | 4.75944 | 0.68319 | 0.35646 | 0.00000 | 0.03175 | 0.46553 | 2.08892 | 2.15353 | 0.00000 |
| **Pex3** | 8.81144 | 0.03993 | 3.26316 | 4.90658 | 6.78706 | 3.93760 | 4.72657 | 0.42052 | 5.18437 | 0.05563 | 4.85564 | 6.82699 | 2.31897 | 6.06383 |
| **Pex5** | 0.55784 | 0.07306 | 9.97504 | 4.38366 | 3.58055 | 0.86422 | 0.09439 | 2.98089 | 0.19797 | 0.04592 | 0.42707 | 2.41982 | 0.00000 | 0.93732 |

| | | | | | | | | | | | | | | |
|---|---|---|---|---|---|---|---|---|---|---|---|---|---|---|
| **Pex6** | 0.69202 | 7.69459 | 3.35251 | 6.61344 | 4.56019 | 2.51776 | 0.24763 | 1.35616 | 0.00000 | 0.01245 | 0.68507 | 3.54266 | 0.05421 | 0.00000 |
| **Pex7** | 0.46656 | 0.01019 | 0.30694 | 0.22359 | 0.16067 | 1.39594 | 0.00334 | 0.00323 | 1.88655 | 0.00510 | 0.00000 | 1.09614 | 0.00169 | 0.30377 |
| **Pfas** | 0.43944 | 0.11718 | 0.77613 | 3.78104 | 1.38784 | 1.69894 | 0.13626 | 0.84417 | 1.76178 | 0.06952 | 0.31703 | 2.52523 | 1.17462 | 1.62768 |
| **Pfdn1** | 4.70856 | 2.09824 | 1.47202 | 12.32990 | 6.76855 | 9.57556 | 4.97222 | 12.74028 | 0.51950 | 9.78000 | 4.34548 | 11.54001 | 0.34033 | 2.92648 |
| **Pfdn2** | 0.77120 | 5.49489 | 0.31253 | 9.22583 | 12.90059 | 7.09047 | 1.29371 | 4.91959 | 8.93028 | 6.92194 | 1.79103 | 7.13365 | 0.32998 | 11.32667 |
| **Pfdn4** | 4.49952 | 0.08608 | 0.82463 | 7.38719 | 5.63576 | 4.70075 | 2.92349 | 1.13304 | 5.46508 | 13.42169 | 2.64694 | 5.50028 | 2.24912 | 3.87381 |
| **Pfdn5** | 10.01416 | 6.69749 | 1.50911 | 9.21101 | 14.16771 | 7.67356 | 4.01671 | 7.96070 | 5.25041 | 15.76811 | 0.76133 | 9.60162 | 8.41490 | 11.22892 |
| **Pfdn6** | 8.64883 | 5.26863 | 5.39759 | 16.17654 | 14.72400 | 6.19057 | 4.02993 | 8.15235 | 3.70948 | 0.32243 | 2.87814 | 7.50962 | 2.66434 | 14.21132 |
| **Pfkfb1** | 0.91011 | 0.01142 | 0.59609 | 1.53770 | 0.62368 | 0.00000 | 0.01826 | 2.31259 | 0.03972 | 0.00303 | 0.73315 | 0.00613 | 0.53937 | 0.59679 |
| **Pfkfb2** | 0.17915 | 0.01053 | 4.02927 | 2.78986 | 3.61829 | 0.97944 | 0.00794 | 2.55798 | 5.14125 | 0.01827 | 0.00000 | 2.42234 | 0.03024 | 0.00000 |
| **Pfkfb3** | 6.76194 | 0.00105 | 10.03498 | 1.37398 | 18.03633 | 0.00000 | 0.00459 | 0.00000 | 20.79358 | 0.00000 | 0.00000 | 0.67205 | 6.54504 | 16.38727 |
| **Pfkfb4** | 0.09464 | 0.01156 | 2.70553 | 1.10005 | 0.00000 | 3.95093 | 1.12174 | 0.92575 | 0.04884 | 0.07587 | 2.48647 | 0.00000 | 0.00000 | 0.17157 |
| **Pfkl** | 0.44961 | 7.02446 | 2.95682 | 3.99269 | 1.42738 | 7.14540 | 0.29222 | 4.74651 | 0.00000 | 0.07899 | 2.18238 | 3.75318 | 0.20219 | 0.49784 |
| **Pfkm** | 0.10743 | 0.00396 | 0.35926 | 0.32702 | 0.16856 | 0.00000 | 0.08742 | 0.00314 | 0.00776 | 0.01944 | 0.00458 | 0.08505 | 0.02475 | 0.52165 |
| **Pfkp** | 0.06249 | 2.35273 | 0.11208 | 1.00565 | 1.04688 | 0.00000 | 0.00104 | 0.00180 | 0.01777 | 7.78503 | 0.00024 | 3.39028 | 1.10253 | 0.03546 |
| **Pfn1** | 0.00000 | 1.51372 | 2.96919 | 14.12218 | 19.84834 | 0.69475 | 1.98454 | 11.18265 | 5.50489 | 4.59721 | 1.67842 | 2.83375 | 0.00000 | 7.93791 |
| **Pfn2** | 0.03774 | 0.01919 | 0.03266 | 2.51612 | 0.04269 | 0.85855 | 3.20143 | 0.01542 | 0.32781 | 0.04899 | 0.83570 | 1.26186 | 0.60839 | 0.27591 |
| **Pfn4** | 0.35653 | 0.00394 | 0.59379 | 0.00000 | 0.57826 | 0.02729 | 0.01795 | 0.02764 | 0.02627 | 0.00000 | 1.97575 | 0.00466 | 0.00761 | 0.13905 |
| **Pga5** | 0.36942 | 0.01349 | 0.04001 | 0.00000 | 0.45667 | 0.02249 | 0.01557 | 0.01187 | 0.00510 | 0.00672 | 0.00468 | 0.20931 | 0.67381 | 0.02249 |
| **Pgam1** | 19.98542 | 3.86482 | 32.91937 | 31.44386 | 28.69598 | 17.32700 | 1.48027 | 11.24266 | 12.41759 | 0.25984 | 11.13909 | 21.30700 | 10.45313 | 14.92531 |
| **Pgam2** | 0.22424 | 4.58702 | 0.84874 | 3.72364 | 0.57929 | 5.88863 | 0.01858 | 2.43774 | 0.00560 | 0.02460 | 1.74904 | 2.72014 | 0.41657 | 2.12206 |
| **Pgam5** | 10.10795 | 0.07978 | 10.20403 | 5.51940 | 4.11333 | 12.40472 | 5.84980 | 5.10760 | 3.01795 | 0.19298 | 8.17788 | 9.05436 | 0.45079 | 8.84254 |
| **Pgap1** | 0.00329 | 0.00388 | 0.00000 | 0.29171 | 0.00447 | 0.84975 | 0.00990 | 0.00437 | 0.00404 | 0.00513 | 0.00271 | 0.39007 | 0.00629 | 0.35938 |
| **Pgap2** | 5.73464 | 0.02984 | 4.94130 | 2.44738 | 7.87565 | 1.50227 | 5.34659 | 4.70586 | 8.64850 | 0.44267 | 1.57103 | 2.10066 | 2.87563 | 13.63959 |
| **Pgap3** | 0.00435 | 0.01277 | 0.13329 | 0.00000 | 0.04416 | 0.00000 | 0.00554 | 0.71363 | 0.00743 | 0.00542 | 0.00686 | 0.01850 | 0.04291 | 0.14224 |
| **Pgbd5** | 0.00468 | 0.00241 | 0.67749 | 0.00000 | 1.20293 | 2.77031 | 0.00879 | 0.00407 | 0.00269 | 0.00363 | 0.18887 | 1.78978 | 2.59153 | 0.01764 |
| **Pgc** | 0.02976 | 0.01141 | 0.03293 | 0.08453 | 0.01226 | 0.00000 | 0.00410 | 0.40914 | 0.01265 | 0.00780 | 0.60478 | 0.11945 | 0.01085 | 0.00480 |
| **Pgd** | 8.82692 | 5.38535 | 26.34835 | 26.80489 | 19.70576 | 24.82668 | 3.89979 | 12.86938 | 0.51243 | 8.20225 | 16.50092 | 20.27059 | 6.12971 | 16.99372 |
| **Pggt1b** | 0.00000 | 0.00396 | 0.36646 | 1.34319 | 0.17475 | 1.65615 | 0.00349 | 2.02387 | 3.16718 | 0.02825 | 0.00420 | 0.00000 | 0.00241 | 1.38357 |
| **Pgk1** | 3.56940 | 9.68400 | 6.27922 | 13.08079 | 2.40654 | 16.49287 | 5.80042 | 14.48415 | 8.17587 | 0.34351 | 13.98716 | 3.31568 | 0.09519 | 7.15524 |
| **Pgls** | 0.00442 | 0.01074 | 0.08380 | 0.38550 | 0.69802 | 0.05738 | 0.00395 | 0.13131 | 0.01335 | 0.01727 | 0.00233 | 1.34576 | 0.00634 | 0.01133 |
| **Pglyrp1** | 1.15699 | 0.00366 | 0.91183 | 2.21873 | 1.17211 | 0.62326 | 0.02060 | 0.00444 | 0.01488 | 0.00456 | 0.00000 | 0.09746 | 1.21733 | 0.07105 |
| **Pgm1** | 0.15674 | 0.59938 | 1.20573 | 1.94938 | 0.08057 | 1.26010 | 0.00184 | 0.00000 | 0.54901 | 0.00000 | 1.81027 | 5.44232 | 1.02551 | 2.99686 |
| **Pgm2** | 0.09279 | 0.14220 | 0.00000 | 0.00000 | 0.41803 | 6.84007 | 4.08595 | 12.28163 | 0.00275 | 0.08690 | 2.69431 | 2.24999 | 1.20541 | 0.90808 |
| **Pgm2l1** | 3.21684 | 0.00318 | 1.07467 | 0.92103 | 0.76732 | 0.45117 | 0.00282 | 0.00000 | 7.19723 | 0.00375 | 0.00000 | 0.14842 | 2.34248 | 3.74606 |
| **Pgm3** | 2.37910 | 0.04887 | 2.19852 | 2.76476 | 5.06593 | 3.60703 | 0.16071 | 3.34658 | 6.80463 | 0.00428 | 4.21110 | 2.69580 | 3.39282 | 7.00666 |
| **Pgm5** | 0.75571 | 0.00451 | 0.00000 | 0.00529 | 0.51231 | 0.01093 | 0.00421 | 0.01585 | 0.11529 | 0.00663 | 0.00598 | 0.00885 | 0.08662 | 0.16090 |
| **Pgp** | 0.21497 | 0.00952 | 0.38115 | 1.59484 | 6.96067 | 0.74615 | 0.44834 | 1.08394 | 0.00460 | 0.05951 | 0.05780 | 2.82755 | 2.23401 | 0.00000 |
| **Pgpep1** | 0.18122 | 0.01197 | 0.38323 | 0.00000 | 0.00000 | 0.00000 | 0.01699 | 0.80966 | 0.01119 | 6.66770 | 0.12450 | 0.84257 | 0.09588 | 0.24391 |
| **Pgpep1l** | 0.03051 | 0.00446 | 0.00893 | 0.04201 | 0.01219 | 0.05718 | 0.00441 | 0.01006 | 0.04658 | 0.00778 | 0.00674 | 0.00749 | 0.08047 | 0.07409 |
| **Pgrmc1** | 7.24548 | 2.15414 | 5.06689 | 11.21468 | 11.89925 | 11.04122 | 4.20676 | 4.79554 | 6.46088 | 0.14750 | 3.06218 | 3.03212 | 3.04821 | 1.86798 |
| **Pgrmc2** | 5.69656 | 0.00920 | 2.91604 | 3.40033 | 5.13043 | 4.32600 | 1.29647 | 1.44673 | 5.57338 | 0.06288 | 1.04013 | 2.75319 | 8.90261 | 6.24206 |
| **Pgs1** | 8.97312 | 2.54325 | 8.72431 | 3.29304 | 13.44002 | 4.40287 | 2.98167 | 3.48806 | 0.00830 | 0.06748 | 4.28904 | 5.61823 | 6.91365 | 3.99750 |
| **Phactr1** | 0.00939 | 0.00476 | 0.00957 | 0.02810 | 0.11042 | 0.01834 | 0.01727 | 0.02283 | 0.05169 | 0.00688 | 0.00989 | 0.01392 | 0.00553 | 0.05650 |
| **Phactr2** | 0.01712 | 0.00410 | 0.00000 | 0.23461 | 0.03318 | 0.25942 | 0.00385 | 0.00311 | 0.01122 | 0.00613 | 0.00433 | 0.02153 | 0.01871 | 0.52138 |
| **Phactr4** | 0.02527 | 0.00903 | 0.68793 | 0.00000 | 1.54306 | 0.11258 | 2.34899 | 0.38600 | 0.02130 | 0.02485 | 0.77997 | 2.56417 | 1.16299 | 0.00889 |
| **Phax** | 8.91486 | 0.05860 | 13.04198 | 10.55037 | 17.01468 | 6.86693 | 5.73693 | 3.71076 | 4.01586 | 0.29171 | 6.89166 | 5.42961 | 3.59439 | 16.77100 |
| **Phb** | 3.22141 | 6.01614 | 19.96592 | 19.93132 | 21.96489 | 2.58122 | 6.08111 | 16.74595 | 0.00000 | 14.73432 | 5.52816 | 18.08511 | 0.43809 | 1.32630 |
| **Phb2** | 8.72761 | 6.91403 | 20.90195 | 23.38665 | 18.03056 | 16.31357 | 3.83213 | 11.97327 | 5.85726 | 9.69182 | 9.69236 | 14.72649 | 3.19772 | 13.24177 |
| **Phc1** | 3.47049 | 0.01664 | 3.21969 | 3.32455 | 4.30647 | 0.78267 | 0.00217 | 2.02135 | 1.40463 | 0.00805 | 1.97597 | 2.09740 | 1.33828 | 6.01123 |
| **Phc2** | 0.00000 | 3.49485 | 3.24692 | 0.55184 | 4.59988 | 2.00810 | 0.06456 | 3.38400 | 0.04478 | 0.00000 | 3.28479 | 2.14166 | 9.84421 | 9.08219 |
| **Phc3** | 0.00000 | 0.00331 | 0.15313 | 0.59437 | 0.00000 | 0.41115 | 0.00345 | 0.14738 | 0.68555 | 0.00546 | 0.00000 | 0.93990 | 0.00356 | 0.30116 |
| **Phex** | 0.01161 | 0.01307 | 0.03001 | 0.02695 | 0.05059 | 0.01485 | 0.00411 | 0.02034 | 0.45178 | 0.00685 | 0.04352 | 0.01730 | 0.32031 | 0.00000 |
| **Phf1** | 0.04979 | 0.00359 | 0.07096 | 1.54161 | 0.78623 | 0.17159 | 0.25282 | 0.71997 | 0.08306 | 0.01376 | 0.00000 | 0.38446 | 3.92976 | 1.99756 |
| **Phf10** | 2.31523 | 0.00261 | 0.00000 | 0.49552 | 0.97534 | 1.21597 | 0.01445 | 0.00000 | 4.04543 | 0.00155 | 3.78219 | 0.55383 | 0.39206 | 3.20303 |

| | | | | | | | | | | | | | | |
|---|---|---|---|---|---|---|---|---|---|---|---|---|---|---|
| **Phf11a** | 0.36396 | 1.82233 | 0.94920 | 0.10300 | 0.40967 | 3.33383 | 0.75952 | 1.03635 | 0.36369 | 0.01831 | 0.38543 | 1.06760 | 0.20897 | 0.00000 |
| **Phf11b** | 0.04952 | 0.21365 | 0.28727 | 0.02049 | 0.03777 | 1.55083 | 0.11425 | 0.09652 | 0.07435 | 0.00651 | 0.31423 | 0.36384 | 0.01426 | 0.06725 |
| **Phf11c** | 0.00767 | 0.00417 | 0.01511 | 0.05439 | 0.00972 | 0.04364 | 0.40559 | 0.00507 | 0.01396 | 0.01115 | 0.03351 | 0.00636 | 0.00685 | 0.01042 |
| **Phf11d** | 0.83208 | 0.64232 | 3.22004 | 2.17295 | 0.00000 | 12.75900 | 2.46401 | 1.92050 | 0.54101 | 0.12046 | 5.04068 | 5.11675 | 0.02719 | 0.71389 |
| **Phf12** | 2.00134 | 5.36741 | 6.41990 | 2.84095 | 7.02649 | 4.91847 | 0.85605 | 4.58189 | 0.45067 | 0.04198 | 4.25497 | 7.02550 | 3.62035 | 3.19356 |
| **Phf13** | 8.19473 | 0.14790 | 11.24443 | 7.52566 | 15.54147 | 5.89647 | 5.26424 | 6.90161 | 10.91090 | 0.23498 | 4.28795 | 3.99947 | 9.36417 | 17.09154 |
| **Phf14** | 4.24780 | 0.00791 | 2.40100 | 0.00000 | 1.67468 | 3.10852 | 0.00133 | 0.40836 | 7.40138 | 0.00000 | 4.55510 | 1.88618 | 1.44374 | 5.87351 |
| **Phf19** | 0.00936 | 5.30692 | 0.28955 | 0.00000 | 0.00609 | 0.00000 | 0.00393 | 0.47149 | 0.01054 | 0.00582 | 0.00425 | 0.00882 | 0.01457 | 0.00872 |
| **Phf2** | 0.19462 | 0.00343 | 0.00000 | 0.67900 | 0.00212 | 0.34241 | 0.04922 | 0.20276 | 0.00746 | 0.01594 | 0.00000 | 0.82611 | 0.16694 | 0.00000 |
| **Phf20** | 1.96510 | 0.07660 | 4.25624 | 3.20994 | 6.31859 | 3.97322 | 0.01789 | 5.31649 | 4.80462 | 0.03548 | 2.83932 | 3.49977 | 8.92295 | 10.40445 |
| **Phf20l1** | 0.28643 | 0.00390 | 0.48794 | 0.50658 | 0.10395 | 0.97736 | 0.00415 | 0.43841 | 0.91815 | 0.00561 | 0.00000 | 1.26819 | 0.86826 | 1.51051 |
| **Phf21a** | 0.06846 | 0.00448 | 0.37803 | 0.39588 | 2.84099 | 0.08490 | 0.00949 | 0.00391 | 0.70343 | 0.00685 | 0.00477 | 0.00700 | 0.06571 | 1.29756 |
| **Phf21b** | 0.00000 | 0.02919 | 1.61769 | 1.22672 | 0.93723 | 0.51672 | 1.27816 | 0.56997 | 0.00796 | 0.03275 | 0.00000 | 0.20712 | 0.09935 | 0.00000 |
| **Phf23** | 5.13006 | 10.69858 | 11.16234 | 16.81077 | 8.50412 | 13.20284 | 7.78002 | 10.45064 | 4.63950 | 0.60924 | 8.46699 | 6.99433 | 12.36992 | 9.08260 |
| **Phf3** | 9.88077 | 0.20613 | 13.06513 | 6.01653 | 11.52833 | 8.29948 | 6.37074 | 0.00000 | 21.00017 | 0.28387 | 6.41176 | 6.51915 | 7.49023 | 15.09001 |
| **Phf5a** | 14.83089 | 6.44978 | 17.20661 | 22.83867 | 20.60476 | 20.44556 | 2.89703 | 11.49015 | 9.61486 | 21.11584 | 13.51184 | 12.73015 | 8.81967 | 19.81145 |
| **Phf6** | 0.00000 | 0.00894 | 1.66183 | 6.05130 | 0.48024 | 1.82010 | 0.03581 | 0.00000 | 0.00403 | 0.01158 | 6.64909 | 1.28970 | 0.79438 | 0.00000 |
| **Phf7** | 0.01819 | 0.02071 | 0.00000 | 1.16878 | 0.00427 | 0.61325 | 0.00384 | 0.00535 | 0.02658 | 0.01880 | 0.00000 | 0.61014 | 1.22838 | 0.00000 |
| **Phf8** | 2.76242 | 0.02937 | 4.37483 | 0.00000 | 2.59983 | 1.82054 | 0.00284 | 1.02089 | 3.60073 | 0.00288 | 1.08192 | 0.86404 | 1.40642 | 2.87002 |
| **Phgdh** | 18.96914 | 11.72005 | 46.83296 | 43.52296 | 34.30229 | 30.17242 | 7.80216 | 18.69104 | 0.45071 | 11.05063 | 24.11276 | 8.75926 | 3.34306 | 17.69310 |
| **Phip** | 0.94206 | 0.01901 | 0.48341 | 0.87261 | 0.49984 | 2.51774 | 0.00375 | 0.11832 | 2.55627 | 0.01237 | 0.34317 | 1.38809 | 1.62678 | 0.24932 |
| **Phka1** | 0.07130 | 0.00451 | 0.01059 | 0.01315 | 0.11176 | 0.01154 | 0.00436 | 0.01115 | 0.00787 | 0.00633 | 0.00797 | 0.00861 | 0.00642 | 0.10846 |
| **Phka2** | 0.01492 | 5.72266 | 0.16449 | 0.74017 | 0.00000 | 0.62506 | 0.00243 | 0.01018 | 0.03552 | 0.00456 | 0.00705 | 0.03295 | 0.63351 | 0.17404 |
| **Phkb** | 0.02056 | 0.01029 | 0.27059 | 0.00000 | 0.39263 | 1.10697 | 0.00312 | 1.08934 | 0.00906 | 0.00420 | 0.07432 | 1.09128 | 0.58253 | 0.02257 |
| **Phkg1** | 0.49493 | 0.00379 | 2.14273 | 0.00000 | 1.37962 | 0.02962 | 0.00332 | 0.00000 | 1.22265 | 0.00804 | 0.01286 | 0.00563 | 3.45538 | 0.11637 |
| **Phkg2** | 0.26297 | 0.00370 | 1.33823 | 0.58561 | 0.99087 | 0.27843 | 0.00937 | 1.26965 | 0.02231 | 0.01773 | 0.06163 | 0.70470 | 0.00000 | 1.39171 |
| **Phlda1** | 1.52841 | 0.01084 | 1.96649 | 1.48643 | 1.27867 | 0.77946 | 1.81263 | 0.00415 | 0.00543 | 0.07369 | 1.22351 | 2.01711 | 0.18695 | 0.03682 |
| **Phlda2** | 0.01860 | 0.00414 | 0.13481 | 0.59803 | 1.27252 | 0.00000 | 0.00396 | 0.00930 | 0.02496 | 0.00705 | 0.00669 | 0.00583 | 0.03384 | 0.00426 |
| **Phlda3** | 0.00000 | 0.00325 | 1.19049 | 1.21951 | 0.06259 | 0.08903 | 0.00403 | 0.03558 | 0.01298 | 0.01153 | 0.00407 | 1.33251 | 0.02806 | 0.15982 |
| **Phldb1** | 0.06959 | 0.00418 | 0.00969 | 0.21045 | 0.11448 | 0.00586 | 0.00422 | 0.00716 | 0.01352 | 0.00637 | 0.00692 | 0.01524 | 0.00572 | 0.00807 |
| **Phldb2** | 0.14947 | 0.02209 | 0.00774 | 0.04519 | 0.00293 | 0.00018 | 1.28468 | 0.09060 | 0.01704 | 0.02009 | 0.58656 | 0.07472 | 0.34353 | 0.00000 |
| **Phldb3** | 0.01088 | 0.48421 | 0.13085 | 0.54485 | 5.86537 | 0.00000 | 0.08810 | 0.88300 | 0.02012 | 0.01152 | 0.00000 | 0.00256 | 0.01078 | 0.00000 |
| **Phlpp1** | 0.72994 | 0.01124 | 1.15064 | 0.07637 | 0.00000 | 0.00000 | 0.00375 | 1.46607 | 0.00807 | 0.01134 | 0.00479 | 0.00436 | 0.00359 | 0.00590 |
| **Phlpp2** | 1.16244 | 0.00379 | 0.01895 | 0.58177 | 0.15916 | 1.58394 | 0.00399 | 0.20563 | 0.37640 | 0.02160 | 0.00358 | 0.26779 | 3.46533 | 0.57127 |
| **Phospho2** | 2.64088 | 0.01643 | 1.21829 | 1.58744 | 5.94779 | 0.17081 | 1.79374 | 1.32441 | 6.29798 | 0.02356 | 0.00000 | 1.23366 | 6.12899 | 5.10871 |
| **Phox2a** | 0.02697 | 0.00443 | 0.01045 | 0.02072 | 0.10064 | 0.01195 | 0.00412 | 0.00999 | 0.01422 | 0.00932 | 0.00841 | 0.00855 | 0.00956 | 0.00718 |
| **Phpt1** | 1.74148 | 3.45581 | 0.00691 | 0.50682 | 0.36130 | 3.61158 | 1.55451 | 4.35597 | 0.13139 | 10.51667 | 0.13032 | 5.30185 | 0.01371 | 0.66919 |
| **Phrf1** | 7.69311 | 7.01080 | 14.48527 | 8.01550 | 11.21411 | 6.73181 | 2.66531 | 3.90893 | 0.68535 | 0.08692 | 4.51503 | 2.11321 | 10.11157 | 11.65113 |
| **Phtf1** | 0.03516 | 0.00205 | 0.50729 | 0.00000 | 0.57755 | 1.66438 | 0.00349 | 0.71606 | 0.01454 | 0.00386 | 0.17524 | 2.10279 | 3.49832 | 1.60776 |
| **Phtf2** | 3.35997 | 0.00689 | 4.01770 | 0.00000 | 0.66926 | 1.91573 | 0.00169 | 0.28555 | 8.42329 | 0.00927 | 2.87589 | 4.23181 | 10.25740 | 6.13383 |
| **Phyh** | 10.16726 | 1.31148 | 10.42115 | 8.68182 | 6.35092 | 5.67393 | 4.01516 | 3.92629 | 2.58385 | 0.10398 | 3.95113 | 5.73814 | 2.51773 | 1.93556 |
| **Phyhd1** | 0.43915 | 0.00389 | 0.12017 | 0.00000 | 0.00604 | 0.05557 | 0.00977 | 0.00386 | 0.08969 | 0.01340 | 0.00389 | 0.18549 | 0.37212 | 0.40524 |
| **Phyhipl** | 2.85482 | 0.00392 | 0.00000 | 0.00684 | 0.56266 | 0.00000 | 0.00390 | 0.01011 | 1.41550 | 0.00695 | 0.00338 | 0.00182 | 1.76357 | 0.76771 |
| **Phykpl** | 0.75471 | 0.00491 | 0.52357 | 0.00000 | 1.23154 | 0.67912 | 0.00423 | 0.07548 | 0.19778 | 0.00314 | 1.35118 | 0.06578 | 0.03574 | 0.08240 |
| **Pi15** | 0.01775 | 0.04344 | 0.02929 | 0.04128 | 0.03875 | 0.06986 | 0.01015 | 0.05433 | 0.21203 | 0.00635 | 0.00864 | 0.04638 | 0.46642 | 0.08342 |
| **Pi16** | 0.32550 | 0.00436 | 0.59104 | 0.00000 | 0.70785 | 0.00144 | 0.00393 | 0.00988 | 0.00789 | 0.00794 | 0.00580 | 0.00681 | 0.01264 | 0.10826 |
| **Pi4k2a** | 4.89236 | 0.37813 | 4.30399 | 4.70564 | 5.11705 | 7.62264 | 0.80408 | 6.72267 | 0.34083 | 0.11998 | 3.62164 | 4.56279 | 6.37307 | 4.82313 |
| **Pi4k2b** | 5.87063 | 0.00621 | 3.87430 | 1.29602 | 5.58812 | 5.50753 | 2.75411 | 3.12359 | 6.43234 | 11.59254 | 4.60346 | 1.23873 | 6.37168 | 7.31197 |
| **Pi4ka** | 4.52002 | 9.12784 | 4.68788 | 1.00354 | 2.71327 | 4.93958 | 0.00478 | 5.91654 | 1.58200 | 0.03953 | 1.75179 | 4.85945 | 1.71414 | 1.49605 |
| **Pi4kb** | 0.00236 | 0.00429 | 0.02304 | 0.04714 | 0.00000 | 0.02250 | 0.00484 | 0.08136 | 0.05212 | 0.00659 | 0.00731 | 0.17402 | 1.99412 | 1.19014 |
| **Pianp** | 0.00000 | 0.00389 | 0.90703 | 0.40451 | 0.00000 | 2.37447 | 0.00375 | 0.00000 | 0.00394 | 0.01001 | 0.00000 | 0.28404 | 0.07503 | 0.00000 |
| **Pias1** | 3.19380 | 0.00943 | 1.24021 | 0.00000 | 5.35261 | 2.52599 | 1.21709 | 1.51771 | 3.02419 | 0.03669 | 0.97731 | 2.65254 | 3.35424 | 7.23287 |
| **Pias2** | 5.76977 | 0.10132 | 8.75652 | 6.27867 | 3.56887 | 13.00255 | 5.06954 | 3.13873 | 9.21634 | 0.14111 | 6.28344 | 5.38441 | 1.91989 | 3.50910 |
| **Pias3** | 0.00000 | 0.02489 | 0.71183 | 1.72704 | 0.85162 | 0.86181 | 0.00317 | 1.52622 | 0.01348 | 0.00450 | 0.27060 | 2.07956 | 1.23887 | 0.47649 |
| **Pias4** | 0.00191 | 0.00335 | 0.00278 | 4.80659 | 8.25297 | 0.05884 | 0.00920 | 0.03160 | 0.02110 | 0.00483 | 0.00935 | 0.72790 | 0.04025 | 0.00000 |

| | | | | | | | | | | | | | | |
|---|---|---|---|---|---|---|---|---|---|---|---|---|---|---|
| **Pibf1** | 2.26298 | 0.70895 | 4.13076 | 0.27344 | 3.92405 | 2.05241 | 0.47599 | 0.00000 | 1.88918 | 0.03958 | 4.75941 | 1.00303 | 1.18506 | 5.66929 |
| **Picalm** | 4.58250 | 0.04798 | 6.81137 | 4.10384 | 6.71432 | 4.28423 | 1.23595 | 2.06061 | 12.74276 | 0.03510 | 6.10377 | 3.37720 | 2.89357 | 6.57145 |
| **Pick1** | 0.00000 | 0.03485 | 1.92992 | 1.73952 | 1.01252 | 3.93317 | 2.20409 | 1.28747 | 0.00370 | 0.11379 | 2.61006 | 1.77933 | 0.00373 | 0.25407 |
| **Pid1** | 0.00477 | 0.00433 | 0.00000 | 0.18127 | 0.10608 | 0.03852 | 0.00415 | 0.00421 | 0.01701 | 0.00465 | 0.33566 | 0.00443 | 0.00414 | 0.08341 |
| **Pidd1** | 1.17559 | 0.02373 | 4.47649 | 0.92415 | 1.43650 | 1.26694 | 0.00266 | 0.00000 | 0.00445 | 0.00221 | 0.18743 | 2.50835 | 0.00000 | 2.10706 |
| **Piezo1** | 0.23192 | 2.98756 | 1.55820 | 0.94622 | 0.48337 | 0.94902 | 0.00873 | 0.58188 | 0.03200 | 0.01185 | 0.89452 | 0.61289 | 0.00000 | 0.02790 |
| **Pif1** | 0.10060 | 0.04171 | 2.82140 | 2.31592 | 1.22860 | 0.33302 | 0.02109 | 2.11413 | 0.00000 | 0.01937 | 0.71057 | 2.74493 | 2.32233 | 0.69943 |
| **Piga** | 1.00444 | 0.28928 | 1.82594 | 1.34558 | 0.83011 | 1.53217 | 0.00334 | 0.89073 | 3.58559 | 0.00909 | 3.26334 | 0.07825 | 2.20226 | 4.50785 |
| **Pigb** | 0.00000 | 0.00395 | 1.33514 | 0.56991 | 0.55461 | 1.39106 | 0.00464 | 0.00574 | 0.07264 | 0.00288 | 0.00000 | 0.57450 | 0.00000 | 0.62938 |
| **Pigc** | 0.00000 | 0.02364 | 0.99343 | 1.51890 | 0.48609 | 1.32145 | 0.01222 | 1.34316 | 0.01517 | 0.01324 | 0.69027 | 0.32617 | 0.00000 | 1.32060 |
| **Pigf** | 2.23136 | 0.02728 | 1.75908 | 2.57660 | 1.22970 | 6.34781 | 1.40407 | 3.77500 | 4.76208 | 0.05207 | 1.30013 | 0.86705 | 0.44838 | 2.11933 |
| **Pigg** | 0.22484 | 0.01048 | 0.52502 | 0.00000 | 0.02731 | 0.50433 | 0.00343 | 0.00000 | 1.03208 | 0.00201 | 1.08614 | 0.49114 | 0.45401 | 0.36137 |
| **Pigh** | 0.18888 | 0.00351 | 0.34191 | 0.00000 | 0.58571 | 0.26315 | 0.00965 | 0.35508 | 0.00503 | 0.00462 | 0.00000 | 0.46421 | 0.19932 | 0.62502 |
| **Pigk** | 3.71423 | 0.00832 | 3.47741 | 6.62820 | 1.31578 | 5.91332 | 0.00181 | 2.81269 | 0.38009 | 0.02955 | 0.89829 | 3.15086 | 0.00000 | 3.66625 |
| **Pigl** | 1.36817 | 0.02752 | 0.29642 | 0.86688 | 1.44548 | 2.94667 | 0.58210 | 0.02315 | 1.25109 | 0.04676 | 2.20349 | 4.14215 | 0.05080 | 1.23753 |
| **Pigm** | 1.15409 | 0.03087 | 2.58721 | 1.79588 | 3.20608 | 1.42092 | 0.00066 | 2.75142 | 9.13551 | 0.01620 | 2.21447 | 2.98364 | 3.19974 | 6.95672 |
| **Pign** | 0.41341 | 0.00330 | 0.53828 | 0.75146 | 0.45463 | 0.90555 | 0.00349 | 0.00332 | 0.00341 | 0.00594 | 0.00999 | 1.55026 | 1.05020 | 0.10194 |
| **Pigo** | 2.58707 | 5.59513 | 6.15394 | 3.26087 | 3.67904 | 0.00000 | 0.76831 | 3.65064 | 0.68877 | 6.32210 | 3.53412 | 1.11914 | 1.98497 | 2.41247 |
| **Pigp** | 6.46006 | 4.04306 | 0.10603 | 3.01519 | 5.27547 | 12.09000 | 2.57203 | 10.70578 | 2.47183 | 12.92644 | 1.37234 | 11.37749 | 9.84758 | 2.51171 |
| **Pigq** | 2.75091 | 1.66385 | 10.78176 | 7.73894 | 3.05805 | 7.53084 | 2.94032 | 5.45059 | 2.85844 | 0.08465 | 1.79534 | 8.77436 | 5.91524 | 9.23838 |
| **Pigs** | 6.08686 | 0.13303 | 13.54910 | 9.51425 | 17.40640 | 5.81612 | 2.50443 | 3.97547 | 1.01950 | 0.06146 | 0.11868 | 8.05920 | 4.11861 | 5.26006 |
| **Pigt** | 0.08197 | 0.00399 | 0.24845 | 0.85399 | 0.62360 | 0.02943 | 0.01030 | 0.09350 | 0.00635 | 0.03399 | 0.00281 | 0.34678 | 0.00090 | 0.27515 |
| **Pigu** | 0.05927 | 1.45581 | 4.37311 | 0.00009 | 3.02736 | 2.15705 | 0.02052 | 3.53896 | 0.01899 | 0.02723 | 1.26632 | 2.32522 | 0.00000 | 1.33268 |
| **Pigv** | 0.44869 | 0.24071 | 0.95628 | 0.89007 | 0.97551 | 1.87029 | 0.62418 | 0.18614 | 0.48959 | 0.01931 | 0.40968 | 0.33935 | 0.43003 | 0.61240 |
| **Pigw** | 0.35875 | 0.00446 | 0.00000 | 0.43203 | 0.00000 | 0.10807 | 0.00433 | 0.00789 | 0.60370 | 0.00608 | 0.00368 | 0.00997 | 0.31116 | 0.09469 |
| **Pigx** | 9.90997 | 0.02644 | 0.62048 | 3.78224 | 11.02140 | 2.03705 | 2.20455 | 1.08948 | 4.91835 | 0.00388 | 0.05860 | 0.86769 | 6.07249 | 1.29647 |
| **Pigyl** | 0.70538 | 0.18017 | 0.94102 | 11.99909 | 4.92983 | 4.19537 | 3.79096 | 7.68177 | 0.00000 | 8.46965 | 1.89510 | 5.09128 | 1.03819 | 0.00000 |
| **Pigz** | 0.02418 | 0.00421 | 1.34385 | 0.09548 | 0.00002 | 0.57367 | 0.99359 | 0.04133 | 0.01694 | 0.00648 | 0.05741 | 0.00000 | 0.02227 | 0.00000 |
| **Pih1d1** | 6.03093 | 0.24743 | 7.17428 | 6.57704 | 5.47963 | 4.18170 | 0.49871 | 8.88507 | 0.00000 | 1.99004 | 3.71442 | 7.03830 | 2.26870 | 2.15121 |
| **Pih1d2** | 0.00715 | 0.00408 | 0.00274 | 0.29325 | 0.14399 | 0.00782 | 0.00418 | 0.00722 | 0.00798 | 0.00630 | 0.00563 | 0.04562 | 0.01074 | 0.46646 |
| **Pik3ap1** | 0.00999 | 0.00472 | 0.15485 | 0.01193 | 0.01008 | 0.03932 | 0.00435 | 0.00769 | 0.01092 | 0.00651 | 0.00980 | 0.01223 | 0.00697 | 0.01150 |
| **Pik3c2a** | 6.21939 | 0.01799 | 1.84724 | 0.00000 | 6.63995 | 1.81117 | 0.01905 | 1.88383 | 17.80434 | 0.16967 | 1.46956 | 1.24920 | 6.18585 | 5.24832 |
| **Pik3c2b** | 0.11441 | 0.00416 | 0.02179 | 0.91832 | 2.27422 | 0.13665 | 0.01629 | 0.74263 | 0.86792 | 0.02273 | 0.00000 | 1.03410 | 1.23554 | 1.20321 |
| **Pik3c2g** | 0.04305 | 0.00439 | 0.01037 | 0.07885 | 0.00995 | 0.03016 | 0.00425 | 0.00625 | 0.01294 | 0.00782 | 0.01882 | 0.01768 | 0.02643 | 0.00545 |
| **Pik3c3** | 8.93028 | 0.01464 | 8.62592 | 3.66648 | 6.96860 | 5.29430 | 0.00000 | 1.07333 | 6.80214 | 0.01383 | 3.15710 | 3.03257 | 3.60260 | 9.71960 |
| **Pik3ca** | 0.24015 | 0.02579 | 2.61039 | 0.00000 | 1.60452 | 3.24611 | 0.01676 | 0.86262 | 5.63401 | 0.01576 | 1.69842 | 1.37600 | 3.93891 | 2.76742 |
| **Pik3cb** | 4.32803 | 0.00318 | 0.97552 | 0.00000 | 4.11188 | 0.58483 | 0.00300 | 0.85841 | 5.49029 | 0.00521 | 0.00200 | 0.01446 | 5.53607 | 3.77642 |
| **Pik3cd** | 0.00000 | 0.26996 | 0.16430 | 1.12079 | 2.40944 | 2.24145 | 0.00355 | 3.63756 | 0.12966 | 0.03229 | 2.68089 | 0.00000 | 3.27182 | 3.15876 |
| **Pik3ip1** | 0.00000 | 0.00396 | 0.89313 | 0.02350 | 0.82764 | 0.77284 | 0.00425 | 0.02671 | 0.01791 | 0.01101 | 0.00602 | 0.00373 | 0.40345 | 0.42424 |
| **Pik3r1** | 0.00000 | 0.04572 | 0.51288 | 0.00000 | 0.00000 | 1.00887 | 2.02139 | 2.73088 | 0.01509 | 0.05660 | 0.97648 | 0.30709 | 1.01196 | 2.23084 |
| **Pik3r2** | 0.00000 | 0.03554 | 2.95272 | 6.84260 | 1.93133 | 0.00000 | 4.15450 | 4.00106 | 0.00117 | 0.04155 | 0.12714 | 1.10721 | 0.06145 | 0.00000 |
| **Pik3r3** | 0.61902 | 0.00393 | 0.58007 | 0.00000 | 1.17701 | 0.29076 | 0.00380 | 0.00000 | 0.83727 | 0.00424 | 0.00367 | 0.00000 | 3.43701 | 2.65352 |
| **Pik3r4** | 4.33322 | 0.00704 | 3.16863 | 3.17764 | 2.06943 | 1.49491 | 3.02597 | 0.09688 | 1.87739 | 0.08542 | 2.77168 | 3.44562 | 2.62350 | 4.28551 |
| **Pik3r6** | 0.16327 | 0.00302 | 1.48346 | 1.63243 | 0.00000 | 1.62318 | 0.40705 | 0.00000 | 0.01949 | 1.34519 | 3.35867 | 0.00000 | 0.05749 | 0.00000 |
| **Pikfyve** | 4.51968 | 0.03514 | 0.95950 | 1.37383 | 0.78043 | 1.78577 | 0.33919 | 1.41136 | 3.24791 | 0.01714 | 0.21832 | 2.56830 | 4.07515 | 4.75173 |
| **Pilra** | 0.12549 | 0.00428 | 0.06276 | 0.12152 | 0.01063 | 0.32189 | 0.02157 | 0.05136 | 0.25293 | 0.14138 | 0.00575 | 0.18259 | 1.98717 | 0.18947 |
| **Pilrb2** | 0.00800 | 0.00452 | 0.05330 | 0.00796 | 0.00835 | 0.01159 | 0.00424 | 0.01147 | 0.00981 | 0.00805 | 0.00757 | 0.01299 | 0.00468 | 0.11606 |
| **Pim1** | 0.43596 | 0.01027 | 2.56539 | 1.85589 | 3.50679 | 2.71789 | 0.00819 | 1.65526 | 1.28251 | 0.03588 | 0.80708 | 0.54352 | 3.00193 | 4.33559 |
| **Pim2** | 2.48267 | 0.01286 | 3.70843 | 0.15981 | 4.47800 | 1.78271 | 0.00288 | 2.18593 | 0.98248 | 0.00660 | 0.00000 | 0.42578 | 1.60700 | 3.52398 |
| **Pim3** | 0.98135 | 0.60788 | 10.32230 | 5.02531 | 12.32713 | 2.22654 | 3.17908 | 9.07451 | 0.73995 | 0.04030 | 2.15265 | 2.16446 | 7.58893 | 0.89314 |
| **Pin1** | 4.59764 | 6.83589 | 13.79026 | 13.78972 | 15.99558 | 14.00030 | 5.38213 | 13.53576 | 0.08035 | 0.31749 | 8.36468 | 17.46508 | 4.36018 | 1.82658 |
| **Pin1rt1** | 0.80608 | 0.17205 | 4.81828 | 0.54812 | 9.59030 | 0.85569 | 0.11039 | 0.10327 | 0.04029 | 0.01296 | 0.14573 | 0.37362 | 3.42218 | 0.51829 |
| **Pin4** | 2.95872 | 0.06003 | 0.00000 | 2.02972 | 5.30104 | 5.04530 | 0.76298 | 2.95130 | 2.44339 | 0.07721 | 0.00000 | 3.13716 | 5.74978 | 1.97980 |
| **Pink1** | 4.54542 | 2.26821 | 13.52141 | 8.81424 | 13.25183 | 3.29969 | 3.02977 | 5.15550 | 1.33431 | 10.93239 | 1.15042 | 1.58235 | 1.42823 | 6.87543 |
| **Pinlyp** | 1.02589 | 0.00374 | 1.11759 | 3.83174 | 1.78492 | 0.48087 | 0.00370 | 0.00123 | 0.13141 | 0.01793 | 0.65101 | 0.00585 | 0.36368 | 2.37143 |

| | | | | | | | | | | | | | | |
|---|---|---|---|---|---|---|---|---|---|---|---|---|---|---|
| **Pinx1** | 5.99702 | 0.02580 | 7.17164 | 1.50991 | 1.41623 | 6.56578 | 0.24208 | 1.43514 | 5.10487 | 0.04523 | 7.63359 | 2.26090 | 0.09353 | 4.80745 |
| **Pip4k2a** | 0.30945 | 4.67847 | 0.00000 | 0.00000 | 0.46870 | 1.83892 | 0.00199 | 1.38194 | 1.68229 | 0.00230 | 1.34820 | 1.11333 | 1.07176 | 0.67025 |
| **Pip4k2b** | 0.05897 | 0.01227 | 0.24076 | 0.00000 | 1.14984 | 0.32266 | 0.00429 | 0.77336 | 0.00463 | 0.00798 | 0.54733 | 0.38245 | 1.68779 | 0.13338 |
| **Pip4k2c** | 7.15979 | 0.07002 | 14.69654 | 7.57270 | 7.21473 | 13.34645 | 1.22986 | 2.62383 | 2.91332 | 0.05703 | 6.82035 | 1.70757 | 1.41365 | 14.48284 |
| **Pip5k1a** | 0.84730 | 0.01931 | 1.50767 | 0.00000 | 4.91176 | 1.91218 | 0.00286 | 4.03909 | 1.46994 | 0.00315 | 0.04019 | 0.78430 | 3.91484 | 6.78403 |
| **Pip5k1b** | 0.00000 | 0.01204 | 0.20188 | 0.00000 | 0.00079 | 0.00237 | 0.00415 | 0.35242 | 0.00888 | 0.00566 | 0.00000 | 0.83977 | 1.71058 | 0.01878 |
| **Pip5k1c** | 0.98923 | 7.80321 | 2.36121 | 1.94643 | 0.45493 | 2.81269 | 2.99676 | 2.59692 | 0.00000 | 0.11735 | 0.00000 | 0.49126 | 2.91560 | 2.01058 |
| **Pipox** | 0.03058 | 0.00394 | 0.00289 | 0.25489 | 0.00000 | 0.45765 | 0.00419 | 0.15320 | 0.05073 | 0.02279 | 0.24481 | 0.00099 | 0.47107 | 0.05320 |
| **Pir** | 0.00000 | 0.01070 | 0.31345 | 0.63231 | 0.00000 | 1.45545 | 0.00364 | 0.01086 | 0.01240 | 0.02100 | 0.02658 | 1.60172 | 0.00802 | 0.00659 |
| **Pira6** | 0.70388 | 0.00420 | 0.10685 | 0.00244 | 0.00000 | 0.00488 | 0.00416 | 0.00997 | 0.09407 | 0.00799 | 0.00508 | 0.10413 | 0.01605 | 0.17608 |
| **Pirb** | 0.00000 | 0.00425 | 0.50154 | 0.00000 | 0.12466 | 0.00000 | 0.00409 | 0.00943 | 0.34034 | 0.00701 | 0.00470 | 0.03576 | 0.01576 | 1.09146 |
| **Pisd** | 4.00662 | 0.08510 | 5.53426 | 4.53690 | 6.65117 | 6.14821 | 0.31707 | 3.24140 | 1.43034 | 0.07778 | 4.01671 | 2.54620 | 2.96066 | 5.76141 |
| **Pisd.ps1** | 1.87193 | 0.14279 | 2.44033 | 7.26313 | 1.25641 | 5.60587 | 1.23958 | 3.26105 | 0.41487 | 0.03298 | 2.74427 | 3.78043 | 5.54588 | 5.26944 |
| **Pisd.ps2** | 0.19100 | 0.17123 | 0.61233 | 0.96902 | 0.18627 | 0.43540 | 0.01604 | 0.55628 | 0.09885 | 0.00142 | 1.15982 | 0.27764 | 0.06914 | 1.27499 |
| **Pithd1** | 1.88744 | 0.09262 | 2.12695 | 3.00647 | 3.19348 | 3.84898 | 0.19844 | 1.30117 | 0.00000 | 0.03964 | 6.68463 | 2.73823 | 4.40451 | 1.89269 |
| **Pitpna** | 20.04435 | 4.55021 | 16.19129 | 7.74964 | 21.89669 | 10.54748 | 2.67641 | 4.59121 | 7.40772 | 0.62379 | 3.04745 | 8.26839 | 10.45442 | 13.25850 |
| **Pitpnb** | 15.78052 | 0.08762 | 20.50747 | 5.33631 | 21.92456 | 12.95863 | 2.93892 | 2.80479 | 8.52589 | 0.08839 | 8.26011 | 6.53228 | 8.02660 | 20.87870 |
| **Pitpnc1** | 0.00000 | 0.01965 | 6.31467 | 3.23803 | 1.61707 | 6.17747 | 4.04960 | 2.00886 | 7.54167 | 0.12407 | 0.83777 | 0.38372 | 2.71174 | 0.73435 |
| **Pitpnm1** | 0.01626 | 0.00417 | 0.00410 | 0.43494 | 0.16843 | 0.01553 | 0.00403 | 0.00502 | 0.01467 | 0.00524 | 0.00602 | 0.00832 | 0.00939 | 0.00498 |
| **Pitpnm2** | 0.22585 | 0.00297 | 0.47108 | 0.21976 | 0.20668 | 0.18215 | 0.00342 | 0.01283 | 0.00396 | 0.00482 | 0.00243 | 1.35054 | 0.38022 | 0.00565 |
| **Pitrm1** | 0.53235 | 0.05434 | 0.59763 | 4.94289 | 0.20106 | 8.85819 | 2.77792 | 3.51285 | 1.65911 | 0.07700 | 5.72394 | 3.56977 | 0.06165 | 4.29102 |
| **Pitx1** | 0.00346 | 0.65579 | 0.01505 | 0.41659 | 1.21062 | 0.05114 | 0.00399 | 0.00747 | 0.01297 | 0.00648 | 0.00432 | 0.92320 | 0.00231 | 0.00669 |
| **Piwil1** | 0.00818 | 0.01287 | 0.00983 | 0.39480 | 0.05956 | 0.02336 | 0.00469 | 0.04582 | 0.03788 | 0.00826 | 0.01767 | 0.00738 | 0.26725 | 0.00000 |
| **Piwil2** | 2.27528 | 0.01076 | 5.74940 | 4.57543 | 11.39432 | 6.28222 | 1.44420 | 0.00000 | 2.33807 | 0.00901 | 1.60619 | 0.89139 | 0.82268 | 9.38282 |
| **Pja1** | 0.11687 | 0.00897 | 0.05320 | 1.03105 | 1.04481 | 0.93476 | 1.10548 | 1.17080 | 0.00189 | 0.00844 | 3.29390 | 1.18392 | 1.27072 | 0.15643 |
| **Pja2** | 1.06441 | 0.00333 | 0.24540 | 0.80304 | 0.23398 | 1.04877 | 0.54092 | 0.97639 | 0.55046 | 0.01780 | 3.03870 | 0.33247 | 0.04954 | 0.32348 |
| **Pkd1** | 0.33401 | 1.55351 | 0.05432 | 0.11970 | 0.25341 | 0.02580 | 0.18018 | 0.17047 | 0.00284 | 0.01046 | 0.52717 | 0.38800 | 1.16999 | 0.38329 |
| **Pkd1l2** | 0.55741 | 0.01303 | 2.25259 | 0.45995 | 2.18311 | 0.00000 | 0.00371 | 1.26243 | 0.06067 | 0.00313 | 0.00021 | 0.06450 | 0.69680 | 2.40778 |
| **Pkd1l3** | 0.01615 | 0.00416 | 0.05765 | 0.00421 | 0.16846 | 0.08286 | 0.00389 | 0.00413 | 0.73967 | 0.01525 | 0.00919 | 0.00555 | 1.00416 | 0.69803 |
| **Pkd2** | 0.09278 | 0.03282 | 0.00000 | 0.05949 | 0.07441 | 0.01339 | 0.01730 | 0.14370 | 0.59014 | 0.02505 | 0.00549 | 0.05158 | 0.01538 | 2.69836 |
| **Pkd2l1** | 0.00641 | 0.00400 | 0.03535 | 0.13607 | 0.00504 | 0.01832 | 0.00411 | 0.00521 | 0.01455 | 0.00626 | 0.00627 | 0.17050 | 0.12274 | 0.07151 |
| **Pkd2l2** | 0.02543 | 0.00412 | 0.03107 | 0.00107 | 0.03171 | 0.03931 | 0.00400 | 0.00587 | 0.72427 | 0.00567 | 0.00791 | 0.00660 | 2.29670 | 0.44900 |
| **Pkig** | 4.88820 | 0.19146 | 6.86040 | 10.46049 | 14.95261 | 9.55551 | 1.52128 | 10.58957 | 0.00221 | 4.16229 | 3.48595 | 6.43463 | 2.13459 | 1.76348 |
| **Pklr** | 0.01799 | 0.00426 | 0.36220 | 0.22926 | 0.30210 | 0.26023 | 0.00377 | 0.00201 | 0.00522 | 0.00723 | 0.00464 | 0.00801 | 0.01528 | 0.16121 |
| **Pkm** | 1.56448 | 10.60082 | 10.40017 | 17.12240 | 4.46361 | 16.50168 | 6.82921 | 15.31727 | 0.18952 | 0.57897 | 14.65125 | 22.74107 | 7.58603 | 0.58437 |
| **Pkmyt1** | 0.00000 | 0.01820 | 1.08830 | 1.76680 | 2.43307 | 0.00000 | 0.00228 | 0.57732 | 0.00317 | 8.29613 | 0.08551 | 0.22832 | 0.14931 | 0.00000 |
| **Pkn1** | 0.01670 | 0.03218 | 0.23285 | 2.34188 | 0.22149 | 0.00000 | 0.01029 | 3.11252 | 0.01798 | 0.01247 | 2.32346 | 1.47953 | 0.00284 | 0.03726 |
| **Pkn2** | 8.02527 | 0.02209 | 2.70741 | 2.60783 | 5.31685 | 3.01545 | 1.47719 | 0.07900 | 16.13377 | 4.69506 | 3.03723 | 3.49388 | 8.30082 | 2.72444 |
| **Pkn3** | 1.08825 | 0.10094 | 0.37831 | 0.69365 | 1.47356 | 0.28891 | 0.00401 | 0.08916 | 0.00200 | 0.01427 | 0.00055 | 0.80914 | 1.77960 | 1.02314 |
| **Pknox1** | 3.41882 | 5.87329 | 17.38932 | 0.75807 | 16.30734 | 6.26536 | 4.37134 | 1.58389 | 5.80957 | 0.07299 | 3.00164 | 2.77270 | 2.75476 | 7.46196 |
| **Pknox2** | 0.00798 | 0.00437 | 0.00290 | 0.00000 | 0.39265 | 0.41297 | 0.00439 | 0.00668 | 0.01160 | 0.00734 | 0.00635 | 0.00943 | 0.00634 | 0.00499 |
| **Pkp2** | 1.98743 | 0.14305 | 6.10527 | 0.46553 | 0.49373 | 5.70889 | 2.65022 | 4.96297 | 0.18871 | 0.03404 | 4.91646 | 0.00000 | 2.24034 | 0.00000 |
| **Pkp3** | 0.02612 | 0.01284 | 0.01017 | 2.66393 | 0.15591 | 0.00000 | 0.34990 | 1.23122 | 0.00900 | 0.03430 | 0.00259 | 0.11134 | 0.17286 | 0.00578 |
| **Pkp4** | 0.88552 | 0.03312 | 4.94947 | 3.60555 | 1.81378 | 5.52183 | 0.00120 | 4.64247 | 2.02471 | 0.02516 | 0.00000 | 4.47924 | 4.49236 | 6.11080 |
| **Pla1a** | 1.18767 | 0.00307 | 2.98622 | 0.66450 | 0.72757 | 0.00000 | 0.00824 | 0.26556 | 2.20260 | 2.32813 | 0.00000 | 0.24664 | 0.34578 | 1.12524 |
| **Pla2g10** | 0.32174 | 5.50753 | 2.96472 | 1.41490 | 0.59104 | 11.37742 | 0.00202 | 1.33197 | 0.00343 | 0.04257 | 2.62605 | 0.47395 | 1.21859 | 0.04530 |
| **Pla2g10os** | 0.02294 | 0.00445 | 0.01313 | 0.02074 | 0.02440 | 0.01788 | 0.00453 | 0.02498 | 0.01196 | 0.00723 | 0.00886 | 0.00824 | 0.17519 | 0.00935 |
| **Pla2g12a** | 5.86260 | 0.00115 | 2.60775 | 4.26171 | 4.61365 | 2.00844 | 0.00778 | 2.75928 | 0.16050 | 0.03247 | 2.52964 | 1.74468 | 1.55971 | 4.39644 |
| **Pla2g12b** | 0.00300 | 0.02152 | 0.00621 | 0.00000 | 0.00753 | 0.24615 | 0.00380 | 0.00578 | 0.01785 | 0.00504 | 0.00203 | 0.50283 | 0.95432 | 0.00281 |
| **Pla2g15** | 0.00000 | 7.83239 | 11.02060 | 7.79228 | 14.06809 | 11.34265 | 4.93412 | 10.03120 | 3.75351 | 0.23676 | 4.26805 | 4.82618 | 9.91167 | 15.64456 |
| **Pla2g16** | 12.53994 | 3.06307 | 6.27041 | 8.58396 | 14.16729 | 1.48862 | 3.12069 | 0.33948 | 6.60851 | 0.11194 | 2.66255 | 0.76358 | 6.45097 | 14.56721 |
| **Pla2g1b** | 0.00614 | 0.00456 | 0.01647 | 0.01128 | 0.23650 | 0.01145 | 0.00403 | 0.01150 | 0.00808 | 0.00739 | 0.00798 | 0.02026 | 0.66679 | 0.06322 |
| **Pla2g2d** | 0.04435 | 0.02861 | 0.01904 | 0.14103 | 0.08110 | 0.09257 | 0.04056 | 0.07599 | 0.15431 | 0.03164 | 0.01499 | 0.09527 | 0.12212 | 0.23182 |
| **Pla2g2f** | 0.21174 | 0.01827 | 0.00794 | 0.00870 | 0.00604 | 0.01657 | 0.00387 | 0.00535 | 0.00967 | 0.00970 | 0.00384 | 1.18309 | 0.03451 | 0.00370 |
| **Pla2g4a** | 0.05045 | 0.00448 | 0.00000 | 0.01147 | 0.15187 | 0.00741 | 0.00398 | 0.01056 | 1.57394 | 0.00667 | 0.00497 | 0.00824 | 0.00159 | 0.00000 |

| | | | | | | | | | | | | | | |
|---|---|---|---|---|---|---|---|---|---|---|---|---|---|---|
| **Pla2g4c** | 0.00000 | 0.00540 | 0.00000 | 0.09026 | 0.33275 | 0.31571 | 0.00423 | 0.09526 | 1.19914 | 0.00895 | 0.00407 | 0.00423 | 1.03813 | 1.62194 |
| **Pla2g4d** | 0.00227 | 0.00438 | 0.00905 | 0.01349 | 0.07023 | 0.01058 | 0.00434 | 0.01071 | 0.00645 | 0.00612 | 0.00772 | 0.00902 | 0.06542 | 0.73778 |
| **Pla2g4e** | 0.11754 | 0.00241 | 1.24942 | 0.00000 | 9.52586 | 0.08997 | 0.00153 | 0.00000 | 12.00767 | 0.00271 | 0.00000 | 1.18284 | 0.48428 | 6.38235 |
| **Pla2g4f** | 0.03886 | 0.00327 | 0.22640 | 0.99332 | 0.51665 | 0.00000 | 0.00340 | 1.03379 | 0.01002 | 0.00669 | 1.27599 | 0.46291 | 0.22024 | 0.00000 |
| **Pla2g5** | 0.02460 | 0.00393 | 1.24583 | 0.43404 | 0.19294 | 1.87664 | 0.02739 | 0.00628 | 0.54576 | 0.00372 | 0.21246 | 0.07385 | 0.00393 | 0.63176 |
| **Pla2g6** | 0.46681 | 0.03434 | 1.29040 | 2.71594 | 0.32199 | 3.57749 | 0.10771 | 0.70023 | 0.00333 | 0.00984 | 3.23090 | 1.49389 | 0.00638 | 0.19726 |
| **Pla2g7** | 1.03708 | 0.00372 | 0.16601 | 0.74287 | 0.00000 | 1.40870 | 0.00355 | 0.00000 | 0.00941 | 0.00000 | 3.09814 | 0.00293 | 0.37653 | 0.00000 |
| **Plaa** | 20.86924 | 0.37648 | 15.87999 | 5.60326 | 19.23204 | 14.00253 | 1.89300 | 8.23003 | 27.55961 | 2.38980 | 11.27106 | 13.31524 | 6.68070 | 15.86189 |
| **Plac1** | 0.00000 | 0.00509 | 0.24161 | 0.00000 | 0.00000 | 0.30536 | 0.00407 | 0.70103 | 0.00997 | 0.45391 | 0.00015 | 0.00757 | 0.14780 | 0.82214 |
| **Plac8** | 0.24513 | 0.00445 | 0.00171 | 0.08331 | 0.53988 | 0.19060 | 0.00543 | 0.42996 | 0.00952 | 0.01887 | 0.00638 | 0.00606 | 0.02390 | 1.13763 |
| **Plac9b** | 2.54499 | 0.03070 | 1.99216 | 6.18827 | 5.14199 | 3.26341 | 0.59390 | 2.45993 | 0.90900 | 0.14685 | 4.40830 | 2.88799 | 0.59704 | 2.07422 |
| **Plag1** | 0.00335 | 0.00484 | 0.00000 | 0.21424 | 0.46348 | 0.26163 | 0.00406 | 0.01325 | 0.00774 | 0.01006 | 0.23908 | 0.00403 | 1.04708 | 0.00609 |
| **Plagl1** | 0.00643 | 0.00411 | 0.15323 | 0.39356 | 0.01642 | 0.48376 | 0.00395 | 0.01286 | 0.00831 | 0.00585 | 0.01210 | 0.01283 | 0.28800 | 0.00000 |
| **Plagl2** | 0.43203 | 0.02000 | 1.53555 | 0.00000 | 1.56514 | 0.76089 | 3.19156 | 0.00001 | 2.26214 | 0.05109 | 2.44013 | 1.11032 | 0.04413 | 1.66811 |
| **Plat** | 0.09019 | 0.00408 | 0.33841 | 0.43338 | 0.28854 | 0.32472 | 0.01057 | 0.31112 | 0.38943 | 0.03790 | 0.00175 | 0.00000 | 1.22051 | 0.35011 |
| **Platr10** | 0.12111 | 0.00409 | 0.07875 | 0.23905 | 0.38774 | 0.01914 | 0.06996 | 0.06475 | 0.17268 | 0.00546 | 0.01011 | 0.05197 | 0.21347 | 0.30876 |
| **Platr14** | 0.78327 | 0.00853 | 3.17476 | 0.00000 | 1.23487 | 4.04479 | 1.16448 | 1.07634 | 2.04658 | 0.03647 | 2.72221 | 3.62253 | 0.00000 | 3.89265 |
| **Platr17** | 8.05247 | 0.00228 | 10.48549 | 2.26817 | 12.76456 | 3.45781 | 0.95376 | 3.58228 | 1.01154 | 0.14701 | 0.01478 | 0.00000 | 9.38072 | 9.16537 |
| **Platr20** | 0.01380 | 0.00442 | 0.01642 | 0.01445 | 0.01290 | 0.00731 | 0.00476 | 0.03074 | 0.01164 | 0.02058 | 0.12910 | 0.00947 | 0.00756 | 0.01240 |
| **Platr21** | 0.00485 | 0.00476 | 0.09229 | 0.00000 | 0.14106 | 0.67634 | 0.00402 | 0.00597 | 0.01597 | 0.00583 | 0.00529 | 0.01062 | 0.00565 | 0.00271 |
| **Platr22** | 0.00750 | 0.00444 | 0.01288 | 0.00780 | 0.00868 | 0.06310 | 0.00433 | 0.00714 | 0.00891 | 0.00711 | 0.00777 | 0.01030 | 0.00745 | 0.14262 |
| **Platr26** | 0.00948 | 0.00448 | 0.03567 | 0.00000 | 0.01401 | 0.04044 | 0.00446 | 0.00728 | 0.01087 | 0.00678 | 0.00916 | 0.11170 | 0.00785 | 0.01168 |
| **Platr4** | 0.30236 | 0.00343 | 0.00000 | 0.40383 | 0.15188 | 0.66272 | 0.00387 | 0.69172 | 0.00790 | 0.00702 | 0.38005 | 1.06662 | 0.07482 | 0.56288 |
| **Platr5** | 0.03199 | 0.00423 | 0.01054 | 0.00000 | 0.28528 | 0.07324 | 0.00549 | 0.01051 | 0.04328 | 0.00625 | 0.00462 | 0.20273 | 0.04650 | 0.05618 |
| **Platr9** | 0.01375 | 0.01226 | 0.01323 | 0.02247 | 0.02614 | 0.08226 | 0.01108 | 0.01843 | 0.14617 | 0.00732 | 0.00723 | 0.45837 | 0.35573 | 0.08620 |
| **Plaur** | 0.12689 | 0.00466 | 0.37091 | 0.00273 | 0.07148 | 0.35025 | 0.04480 | 0.02709 | 0.06155 | 0.01648 | 0.01787 | 0.35511 | 0.02817 | 0.00000 |
| **Plb1** | 0.01948 | 0.02091 | 0.00000 | 0.00000 | 0.46231 | 1.61990 | 0.00377 | 0.00415 | 0.00529 | 0.00417 | 0.49114 | 0.80932 | 0.01663 | 1.15004 |
| **Plbd1** | 16.92608 | 4.71958 | 12.00034 | 3.78746 | 6.27530 | 2.88471 | 0.00000 | 0.00000 | 8.49920 | 0.00542 | 0.00000 | 3.18427 | 0.61840 | 10.53379 |
| **Plbd2** | 0.42681 | 0.01761 | 3.90313 | 7.65550 | 0.08329 | 5.74626 | 0.04039 | 0.19145 | 0.98002 | 0.04361 | 1.55094 | 2.84699 | 0.36141 | 2.58743 |
| **Plcb1** | 0.00862 | 0.00442 | 0.01183 | 0.01148 | 0.00812 | 0.01965 | 0.00424 | 0.00888 | 0.02096 | 0.00661 | 0.00900 | 0.00955 | 0.13745 | 0.19462 |
| **Plcb3** | 1.04605 | 0.12918 | 1.98320 | 6.56069 | 0.35866 | 1.34655 | 0.00923 | 3.36714 | 0.08686 | 0.02299 | 0.10725 | 2.44842 | 0.13725 | 0.00000 |
| **Plcd1** | 1.98372 | 0.09497 | 14.61380 | 6.67752 | 4.03750 | 5.73285 | 0.02410 | 6.90734 | 0.62319 | 0.06825 | 5.03545 | 2.64548 | 0.00000 | 3.34531 |
| **Plcd3** | 1.88653 | 0.03249 | 8.76647 | 0.24572 | 12.41580 | 5.34838 | 0.01270 | 0.70757 | 0.01111 | 10.40448 | 0.23980 | 0.00000 | 1.26068 | 5.23262 |
| **Plcd4** | 2.45023 | 0.00288 | 1.42717 | 0.38788 | 4.84005 | 0.08549 | 0.00256 | 0.00000 | 5.63673 | 0.01358 | 0.00000 | 0.00189 | 5.68683 | 8.11225 |
| **Plce1** | 0.00415 | 0.00425 | 0.00000 | 0.01218 | 0.00000 | 0.00523 | 0.00404 | 0.00828 | 1.51075 | 0.00673 | 0.00395 | 0.05145 | 0.19227 | 0.84633 |
| **Plcg1** | 0.00000 | 0.00382 | 1.80431 | 1.13232 | 0.14026 | 0.63415 | 0.00883 | 0.00000 | 0.62840 | 0.00679 | 0.12370 | 0.19616 | 0.00000 | 2.56083 |
| **Plcg2** | 0.00000 | 0.04724 | 6.85817 | 5.90210 | 5.45191 | 0.27777 | 0.00267 | 2.12016 | 2.94295 | 0.01608 | 0.00000 | 0.63747 | 0.00000 | 2.57001 |
| **Plch1** | 0.00802 | 0.00495 | 0.33661 | 0.02408 | 0.00259 | 0.01162 | 0.00406 | 0.01974 | 0.07841 | 0.01039 | 0.00795 | 0.00704 | 0.00965 | 0.00277 |
| **Plcxd1** | 0.00000 | 0.00432 | 0.38761 | 0.21928 | 0.02549 | 0.02018 | 0.00417 | 0.01391 | 0.00846 | 0.00467 | 0.00783 | 0.01794 | 0.40118 | 0.00398 |
| **Pld1** | 0.28010 | 0.01193 | 0.00000 | 0.00000 | 0.10136 | 0.00000 | 1.34465 | 0.00267 | 3.21420 | 0.01550 | 0.00000 | 0.22717 | 4.77354 | 5.26634 |
| **Pld2** | 0.11638 | 0.00383 | 1.39303 | 0.00005 | 1.56946 | 0.00000 | 0.00390 | 1.62464 | 0.03881 | 0.01717 | 0.00774 | 0.02357 | 1.71251 | 2.08827 |
| **Pld3** | 1.70456 | 2.32147 | 13.10853 | 15.63675 | 3.46702 | 10.16890 | 2.56231 | 5.87852 | 0.17601 | 0.22147 | 6.59693 | 4.49216 | 0.01455 | 7.53957 |
| **Pld4** | 0.00659 | 0.00439 | 0.01716 | 0.01207 | 0.01349 | 0.01320 | 0.00428 | 0.00972 | 0.01080 | 0.00657 | 0.00895 | 0.00963 | 0.18956 | 0.00468 |
| **Pld5** | 0.38540 | 0.00456 | 0.01678 | 0.00708 | 0.13258 | 0.01010 | 0.00451 | 0.01601 | 0.04808 | 0.00640 | 0.00647 | 0.01168 | 0.00463 | 0.07620 |
| **Pld6** | 0.00000 | 0.00402 | 0.00626 | 0.03021 | 0.13788 | 0.00962 | 0.00427 | 0.00726 | 0.18500 | 0.00587 | 0.00681 | 0.02352 | 2.03530 | 0.00000 |
| **Plec** | 0.38872 | 2.73203 | 4.66111 | 0.90294 | 0.00760 | 4.29607 | 0.98747 | 8.30332 | 0.00000 | 0.06826 | 0.22825 | 0.09391 | 0.52174 | 0.67839 |
| **Plek** | 0.01031 | 0.07469 | 0.01825 | 0.05847 | 0.22064 | 0.16642 | 0.00405 | 0.00692 | 0.04080 | 0.01454 | 0.00689 | 0.00809 | 0.62047 | 0.16165 |
| **Plek2** | 0.68647 | 0.02976 | 1.54266 | 0.87650 | 3.83214 | 0.35145 | 0.00373 | 1.23537 | 4.81632 | 0.00387 | 0.01649 | 0.29692 | 0.00000 | 2.85215 |
| **Plekha1** | 7.99859 | 0.00831 | 7.34428 | 2.01285 | 5.12735 | 9.59644 | 1.14797 | 1.06420 | 8.11885 | 0.03454 | 8.99790 | 0.52462 | 3.25424 | 5.83004 |
| **Plekha2** | 0.00633 | 0.00420 | 0.22732 | 0.00000 | 0.22641 | 0.01041 | 0.00398 | 0.01014 | 0.02329 | 0.00729 | 0.01229 | 0.08021 | 0.99786 | 0.79876 |
| **Plekha3** | 0.92335 | 0.00281 | 1.23194 | 1.42574 | 2.64341 | 1.53065 | 0.00282 | 0.24796 | 3.47560 | 0.00768 | 0.70805 | 0.29544 | 8.62706 | 1.61035 |
| **Plekha4** | 0.61218 | 0.00483 | 0.13593 | 2.09023 | 3.34423 | 0.92840 | 0.00425 | 0.16208 | 0.00309 | 0.03965 | 0.88317 | 0.00003 | 1.10180 | 0.58409 |
| **Plekha5** | 0.77875 | 0.04861 | 1.76748 | 0.09019 | 1.61648 | 1.17367 | 1.41323 | 0.16089 | 3.10336 | 0.01735 | 1.71095 | 1.70233 | 1.77982 | 6.76866 |
| **Plekha6** | 1.01635 | 0.00246 | 0.96905 | 0.00000 | 1.81683 | 0.00000 | 0.00312 | 0.27187 | 0.74950 | 0.00670 | 0.00157 | 1.72889 | 0.81219 | 2.12445 |
| **Plekha7** | 0.06708 | 0.03509 | 1.02919 | 0.03917 | 0.92624 | 3.06329 | 3.16988 | 3.38567 | 0.01011 | 0.07715 | 2.95917 | 1.12680 | 0.73276 | 1.00177 |

| | | | | | | | | | | | | | | |
|---|---|---|---|---|---|---|---|---|---|---|---|---|---|---|
| **Plekha8** | 0.00000 | 0.07587 | 0.97622 | 0.00000 | 1.30607 | 2.86904 | 0.67882 | 1.75490 | 5.31910 | 0.04534 | 3.38282 | 2.64366 | 3.60800 | 8.59636 |
| **Plekhb1** | 0.69761 | 0.00420 | 1.75299 | 0.00000 | 1.29202 | 0.00000 | 0.00397 | 0.00542 | 0.10921 | 0.00322 | 0.00445 | 0.00430 | 0.17616 | 1.65713 |
| **Plekhb2** | 0.07909 | 0.16425 | 0.45916 | 0.60925 | 1.17526 | 3.02403 | 2.18905 | 2.92440 | 0.51538 | 10.22626 | 0.17990 | 0.00000 | 1.54992 | 0.44572 |
| **Plekhd1** | 0.01570 | 0.00520 | 0.00000 | 0.16018 | 0.30982 | 0.09565 | 1.33365 | 0.26195 | 0.01655 | 0.05633 | 0.01790 | 0.12707 | 0.35469 | 0.18062 |
| **Plekhf1** | 0.52015 | 5.49280 | 6.75708 | 7.96208 | 0.00055 | 6.87389 | 7.26676 | 7.98523 | 0.00000 | 13.71394 | 2.55668 | 6.30710 | 0.10163 | 0.40593 |
| **Plekhf2** | 9.10452 | 0.10533 | 9.61407 | 7.22756 | 13.58346 | 11.77215 | 3.31720 | 7.12973 | 16.31431 | 0.09022 | 6.88543 | 5.56867 | 9.11764 | 15.09867 |
| **Plekhg1** | 0.00885 | 0.00435 | 0.05234 | 0.00786 | 0.00000 | 0.01166 | 0.00436 | 0.00911 | 0.00795 | 0.00645 | 0.00763 | 0.00857 | 0.27517 | 0.69575 |
| **Plekhg2** | 0.41174 | 5.45114 | 3.36801 | 1.59872 | 2.78209 | 0.22077 | 0.02237 | 0.03478 | 0.03021 | 0.00000 | 0.21561 | 0.42629 | 0.10603 | 0.00000 |
| **Plekhg3** | 2.71865 | 0.30759 | 7.73908 | 5.01264 | 6.63665 | 6.69271 | 2.02649 | 3.63364 | 0.48261 | 0.06810 | 4.88704 | 2.80144 | 2.61941 | 6.19787 |
| **Plekhg4** | 0.01011 | 0.00443 | 0.01385 | 0.01686 | 0.01310 | 0.01311 | 0.00428 | 0.01038 | 0.01097 | 0.00725 | 0.00860 | 0.00972 | 0.00545 | 0.02477 |
| **Plekhg5** | 4.73093 | 0.03974 | 3.03106 | 0.42445 | 8.09015 | 0.00000 | 0.01549 | 0.40232 | 0.10709 | 0.00368 | 0.00000 | 3.91515 | 5.84468 | 2.35621 |
| **Plekhg6** | 0.48062 | 0.00416 | 0.47494 | 0.00000 | 0.87281 | 0.00000 | 0.00402 | 0.00810 | 0.02005 | 0.00949 | 0.00530 | 0.00557 | 0.90247 | 0.99900 |
| **Plekhh1** | 1.02669 | 0.02550 | 1.55786 | 1.59822 | 1.10970 | 2.20635 | 0.00923 | 1.88048 | 0.10050 | 0.00545 | 3.25989 | 1.70936 | 0.80648 | 0.79294 |
| **Plekhh2** | 0.01990 | 0.00497 | 0.01787 | 0.00728 | 0.17886 | 0.03543 | 0.00406 | 0.02841 | 0.05814 | 0.01414 | 0.01375 | 0.13764 | 0.04746 | 0.06227 |
| **Plekhh3** | 0.00056 | 3.94918 | 0.10951 | 0.25363 | 0.30037 | 0.04316 | 0.05469 | 0.70622 | 0.00634 | 0.00656 | 0.00242 | 0.00000 | 0.00171 | 0.07226 |
| **Plekhj1** | 2.25735 | 0.07046 | 0.76777 | 7.93601 | 5.60367 | 1.14407 | 0.00855 | 2.12984 | 0.40015 | 0.03002 | 0.15482 | 2.69422 | 4.52388 | 0.57632 |
| **Plekhm1** | 6.97444 | 0.14918 | 7.46929 | 3.51071 | 12.96926 | 3.60429 | 1.21766 | 0.17407 | 2.13222 | 4.96261 | 3.81596 | 0.00000 | 7.21187 | 10.68040 |
| **Plekhm2** | 0.48861 | 0.01201 | 2.20170 | 1.88841 | 4.79529 | 0.23002 | 0.03022 | 0.34375 | 0.00537 | 0.00159 | 0.01034 | 0.98406 | 0.57088 | 0.37226 |
| **Plekhm3** | 6.28061 | 0.00179 | 6.14549 | 0.00000 | 8.56515 | 1.27774 | 0.34794 | 0.00000 | 8.30289 | 0.00000 | 1.42579 | 0.86326 | 2.69929 | 10.84806 |
| **Plekhn1** | 0.02044 | 0.00406 | 0.02509 | 0.08314 | 0.00484 | 0.00718 | 0.00373 | 0.00531 | 0.02979 | 0.00665 | 0.00901 | 0.63967 | 0.33627 | 0.00901 |
| **Plekho1** | 0.00000 | 0.00361 | 0.22869 | 0.00000 | 0.50713 | 1.16183 | 0.21083 | 0.00616 | 0.00438 | 0.01707 | 0.00000 | 0.93382 | 0.00357 | 0.21902 |
| **Plekho2** | 0.70848 | 0.02590 | 3.64453 | 2.02637 | 1.57336 | 2.66628 | 4.33412 | 2.70419 | 0.00000 | 0.07968 | 2.29103 | 0.17780 | 2.41930 | 1.04684 |
| **Plekhs1** | 0.00529 | 0.00450 | 0.00001 | 0.00742 | 1.24976 | 0.01124 | 0.00525 | 0.01959 | 0.08839 | 0.00618 | 0.00847 | 0.00697 | 0.11034 | 0.00000 |
| **Plet1** | 0.00000 | 0.18760 | 0.38734 | 4.69448 | 1.18018 | 12.01272 | 4.80058 | 8.88771 | 0.02947 | 0.10068 | 2.21385 | 1.90873 | 0.00000 | 0.00000 |
| **Plgrkt** | 0.00298 | 0.01235 | 0.00000 | 0.79463 | 0.33371 | 0.80057 | 0.69325 | 0.79191 | 0.01069 | 0.02218 | 0.00000 | 0.26287 | 0.00893 | 0.00563 |
| **Plin1** | 0.00417 | 0.00436 | 0.09847 | 0.24179 | 0.20679 | 0.00608 | 0.00417 | 0.00884 | 0.00831 | 0.00623 | 0.00633 | 0.00732 | 0.00432 | 0.07742 |
| **Plin2** | 1.24194 | 0.27225 | 5.06483 | 13.34354 | 4.60179 | 11.22828 | 1.41199 | 15.27049 | 0.51562 | 0.84773 | 10.48445 | 11.23678 | 4.33436 | 2.48496 |
| **Plin3** | 0.00000 | 8.81909 | 7.34466 | 9.31087 | 4.73615 | 1.52271 | 2.74543 | 8.49698 | 0.30267 | 0.21655 | 7.08808 | 9.29688 | 2.60832 | 1.13631 |
| **Plin5** | 0.02885 | 0.00373 | 0.00000 | 0.00000 | 0.25470 | 0.00000 | 0.01113 | 1.52100 | 0.01552 | 0.01185 | 0.52592 | 0.15709 | 0.00477 | 0.00651 |
| **Plk1** | 1.68386 | 3.26207 | 19.79358 | 9.36330 | 22.06573 | 0.36560 | 2.27404 | 9.16886 | 0.00000 | 0.08387 | 6.24741 | 5.66162 | 1.83565 | 0.00000 |
| **Plk2** | 0.30397 | 0.00373 | 0.42505 | 0.00000 | 0.00000 | 0.03958 | 0.00414 | 0.00364 | 0.01200 | 0.02435 | 0.22181 | 0.57594 | 0.02412 | 0.91406 |
| **Plk3** | 0.14099 | 0.00342 | 0.00159 | 0.49924 | 0.00195 | 0.76284 | 1.59567 | 0.01683 | 0.02155 | 0.00748 | 1.27513 | 0.10177 | 1.30153 | 0.01556 |
| **Plk4** | 5.40679 | 0.00192 | 2.84427 | 0.20190 | 2.72267 | 0.67679 | 0.00780 | 0.06222 | 8.60800 | 0.00180 | 0.00000 | 2.40145 | 1.15122 | 6.00851 |
| **Plk5** | 0.16688 | 0.00410 | 4.62452 | 4.96115 | 0.05980 | 0.24661 | 0.00311 | 0.00000 | 0.00455 | 0.01189 | 1.03963 | 0.00407 | 0.00760 | 0.00000 |
| **Pllp** | 0.00512 | 0.00492 | 0.05574 | 0.00906 | 0.28104 | 0.00850 | 0.00443 | 0.02213 | 0.00771 | 0.00732 | 0.00793 | 0.00795 | 0.00733 | 0.10892 |
| **Plod1** | 1.68144 | 0.08726 | 8.34876 | 4.02621 | 0.00000 | 2.90363 | 0.74229 | 3.66700 | 0.59139 | 0.06234 | 0.52516 | 3.74155 | 0.20600 | 1.80298 |
| **Plod2** | 0.02805 | 0.00191 | 0.00000 | 0.52802 | 0.00302 | 0.00000 | 0.02091 | 0.00003 | 0.00941 | 0.00917 | 0.78226 | 2.75454 | 0.01573 | 0.26218 |
| **Plod3** | 1.26734 | 0.01632 | 5.97145 | 4.75797 | 2.17741 | 4.70438 | 0.00197 | 0.00000 | 0.04012 | 0.00851 | 0.43820 | 3.36878 | 0.94341 | 2.82052 |
| **Plp1** | 0.02776 | 0.02753 | 0.00000 | 0.23637 | 0.00000 | 0.00000 | 0.01004 | 0.04576 | 0.30112 | 0.00377 | 1.06987 | 0.37530 | 0.05560 | 0.08658 |
| **Plp2** | 3.29126 | 10.18481 | 19.78179 | 14.03080 | 9.23034 | 18.91447 | 3.46434 | 16.72870 | 2.53162 | 12.24717 | 3.92547 | 11.76981 | 0.00000 | 3.44645 |
| **Plpp1** | 4.11907 | 5.64752 | 8.94918 | 6.69155 | 5.44788 | 3.62143 | 3.47875 | 4.04794 | 1.88749 | 0.14274 | 2.22144 | 13.65741 | 0.44195 | 2.27556 |
| **Plpp2** | 0.77793 | 0.02463 | 1.30373 | 7.05758 | 1.88266 | 2.46107 | 2.66518 | 0.77111 | 0.04383 | 7.64948 | 0.00000 | 1.80677 | 0.00000 | 0.08702 |
| **Plpp3** | 0.42579 | 0.00436 | 0.15272 | 0.62802 | 1.28134 | 2.89862 | 0.00343 | 0.94837 | 5.12595 | 0.01513 | 0.19711 | 0.26029 | 2.06621 | 1.23322 |
| **Plpp5** | 10.05106 | 0.02963 | 8.87964 | 0.25739 | 11.43552 | 1.87669 | 3.02804 | 4.69482 | 6.07955 | 1.61481 | 2.00259 | 3.19974 | 7.47411 | 13.86530 |
| **Plpp6** | 0.99838 | 0.01373 | 0.00000 | 0.81209 | 0.08517 | 0.10523 | 0.00394 | 1.08517 | 0.00726 | 0.01367 | 0.00000 | 0.71069 | 0.00620 | 0.05692 |
| **Plppr3** | 0.06896 | 0.00420 | 0.01030 | 0.20867 | 0.31316 | 0.01101 | 0.09681 | 0.00957 | 0.01094 | 0.01803 | 0.01082 | 0.03135 | 0.02661 | 0.00378 |
| **Plppr4** | 0.01211 | 0.00454 | 0.01184 | 0.01867 | 0.01019 | 0.02053 | 0.01104 | 0.02058 | 0.10018 | 0.00710 | 0.00896 | 0.02285 | 0.01975 | 0.01752 |
| **Plrg1** | 8.98366 | 3.22019 | 9.53280 | 7.03085 | 10.09716 | 12.47639 | 3.80570 | 8.07771 | 13.38439 | 0.18704 | 8.66362 | 9.30158 | 10.53403 | 14.31465 |
| **Pls1** | 0.00000 | 0.00394 | 0.46195 | 0.01320 | 0.56709 | 1.57154 | 0.01758 | 0.09564 | 0.00570 | 0.00344 | 1.04351 | 0.00000 | 0.56832 | 0.42491 |
| **Pls3** | 0.07793 | 0.01087 | 1.29105 | 1.25776 | 0.45348 | 4.62832 | 0.00822 | 3.65130 | 4.11467 | 0.02782 | 0.32760 | 1.49649 | 0.69988 | 0.05347 |
| **Plscr1** | 1.20334 | 0.06294 | 0.00000 | 0.59159 | 0.00255 | 0.94272 | 0.82052 | 0.00000 | 0.01178 | 0.02018 | 1.18971 | 4.42844 | 0.30408 | 0.00000 |
| **Plscr3** | 0.01647 | 0.00457 | 0.22428 | 0.06469 | 0.18700 | 0.00000 | 0.00469 | 0.06278 | 0.01437 | 0.00753 | 0.00614 | 0.51847 | 0.00520 | 0.00632 |
| **Pltp** | 0.00000 | 0.08161 | 0.30354 | 0.00035 | 0.06353 | 0.01349 | 0.11089 | 2.26394 | 0.05008 | 0.01597 | 0.00000 | 3.17606 | 0.05996 | 0.16765 |
| **Plvap** | 0.18651 | 0.00436 | 0.03424 | 0.01120 | 0.01644 | 0.01031 | 0.00425 | 0.00995 | 0.01689 | 0.00634 | 0.00747 | 0.00897 | 0.27543 | 0.01140 |
| **Plxdc1** | 0.02164 | 0.00438 | 0.64277 | 0.00126 | 0.01187 | 0.00796 | 0.00404 | 0.01033 | 0.00959 | 0.00677 | 0.00918 | 0.00843 | 0.04174 | 0.00664 |

| | | | | | | | | | | | | | | |
|---|---|---|---|---|---|---|---|---|---|---|---|---|---|---|
| **Plxdc2** | 0.01960 | 0.00433 | 0.01028 | 0.00000 | 0.00536 | 0.00386 | 0.00520 | 0.87290 | 0.00435 | 0.00530 | 0.00184 | 0.29413 | 0.40564 | 0.27857 |
| **Plxna1** | 0.55562 | 0.03882 | 1.34011 | 2.38289 | 0.10952 | 3.39337 | 3.65589 | 1.49362 | 0.16496 | 0.88233 | 0.00000 | 4.62448 | 1.04429 | 0.00000 |
| **Plxna2** | 0.29504 | 0.01180 | 0.02139 | 0.00000 | 0.00000 | 0.01302 | 0.00351 | 1.27312 | 0.01477 | 0.01050 | 0.00363 | 0.03393 | 0.01752 | 0.52593 |
| **Plxna3** | 0.12925 | 0.00422 | 0.35564 | 0.16826 | 0.31207 | 0.00000 | 0.00389 | 0.00470 | 0.00984 | 0.00606 | 0.00388 | 0.28220 | 0.01920 | 0.28465 |
| **Plxna4** | 0.01584 | 0.00443 | 0.01562 | 0.01050 | 0.07451 | 0.01307 | 0.00418 | 0.01021 | 0.00870 | 0.00647 | 0.00933 | 0.00948 | 0.06890 | 0.02055 |
| **Plxnb1** | 0.04689 | 0.02476 | 0.48633 | 1.34755 | 0.01374 | 0.43537 | 0.00342 | 1.37357 | 0.00533 | 0.00480 | 0.00009 | 0.04249 | 0.41758 | 0.04388 |
| **Plxnb2** | 0.00000 | 0.14816 | 12.72108 | 13.81651 | 4.83410 | 6.42854 | 3.74736 | 10.54500 | 0.00000 | 1.44337 | 1.31683 | 7.47955 | 0.38244 | 0.00000 |
| **Plxnb3** | 0.00000 | 0.01396 | 0.00720 | 0.00450 | 0.00000 | 0.00610 | 0.00389 | 0.03282 | 0.31355 | 0.01337 | 0.00288 | 0.41486 | 0.05291 | 0.67370 |
| **Plxnc1** | 0.00000 | 0.00404 | 0.00323 | 0.26395 | 0.00000 | 0.00680 | 0.00425 | 0.00739 | 0.01734 | 0.00539 | 0.00567 | 0.00471 | 1.35478 | 1.91423 |
| **Plxnd1** | 0.41749 | 0.00398 | 0.70302 | 0.79840 | 0.00000 | 0.37718 | 0.00437 | 0.12286 | 0.00561 | 0.00797 | 0.00191 | 0.00264 | 0.07444 | 0.00000 |
| **Pm20d2** | 0.00364 | 0.00435 | 0.27309 | 0.00065 | 0.00579 | 0.01460 | 0.00421 | 0.00863 | 0.00990 | 0.00667 | 0.00775 | 0.00940 | 0.19436 | 0.00843 |
| **Pmaip1** | 1.03333 | 0.04149 | 0.81501 | 0.00000 | 0.00000 | 1.18842 | 0.00300 | 0.00060 | 0.49691 | 0.00675 | 0.00000 | 2.53829 | 0.02621 | 0.00000 |
| **Pmch** | 0.37280 | 0.00436 | 0.03988 | 0.05336 | 0.24193 | 0.00801 | 0.00426 | 0.00962 | 0.01017 | 0.00602 | 0.00689 | 0.00616 | 0.01095 | 0.02603 |
| **Pmel** | 0.30884 | 0.06671 | 4.65574 | 0.81655 | 2.23120 | 0.00000 | 0.00493 | 5.06319 | 0.52611 | 15.21015 | 0.00000 | 1.59914 | 1.29761 | 3.46181 |
| **Pmepa1** | 0.16524 | 0.02512 | 0.00000 | 0.31041 | 0.00343 | 0.15344 | 0.00394 | 0.07496 | 0.01235 | 0.02041 | 0.01335 | 0.34976 | 0.00616 | 0.00036 |
| **Pmf1** | 0.05058 | 4.62472 | 10.52373 | 6.06595 | 7.37991 | 8.34887 | 3.66377 | 10.05914 | 0.00000 | 12.61534 | 2.54322 | 8.29928 | 1.00070 | 0.00000 |
| **Pmfbp1** | 0.56326 | 0.00425 | 0.80232 | 0.00000 | 0.42566 | 0.00000 | 0.00387 | 0.00748 | 3.22381 | 0.00548 | 0.00306 | 0.00447 | 0.11035 | 1.06466 |
| **Pmis2** | 0.00961 | 0.00452 | 0.07797 | 0.00000 | 0.01104 | 0.33753 | 0.00420 | 0.00522 | 0.08727 | 0.00737 | 0.00650 | 0.00844 | 0.01509 | 0.01277 |
| **Pml** | 0.70820 | 0.04929 | 6.17048 | 1.28796 | 4.68885 | 0.64139 | 1.85763 | 3.89050 | 1.24309 | 0.01783 | 4.40372 | 1.84286 | 1.55538 | 5.86375 |
| **Pmm1** | 19.76466 | 2.90557 | 46.57357 | 39.48503 | 28.06409 | 25.55346 | 0.25686 | 7.48445 | 0.64142 | 0.48167 | 16.32552 | 25.02842 | 8.95631 | 13.10231 |
| **Pmm2** | 3.84304 | 0.10154 | 14.72563 | 5.96739 | 9.22956 | 11.53219 | 0.03744 | 6.94150 | 7.03435 | 0.17013 | 5.10128 | 9.88809 | 0.27795 | 9.67427 |
| **Pmp22** | 0.06663 | 0.00445 | 0.01220 | 0.01164 | 0.08532 | 0.01224 | 0.00443 | 0.01152 | 0.01078 | 0.00643 | 0.00811 | 0.00925 | 0.00603 | 0.01002 |
| **Pmpca** | 1.90767 | 0.02223 | 4.11140 | 7.79515 | 1.90356 | 8.53305 | 3.12212 | 0.46352 | 2.84779 | 0.03090 | 4.07446 | 5.82750 | 0.84027 | 5.13024 |
| **Pmpcb** | 11.08307 | 5.44768 | 11.97481 | 6.07094 | 9.65394 | 9.87488 | 3.17120 | 5.01221 | 9.93648 | 0.12052 | 7.08187 | 10.38563 | 4.61289 | 4.36687 |
| **Pms1** | 0.73331 | 0.00390 | 0.36615 | 0.00000 | 0.00000 | 0.21266 | 1.11139 | 0.02961 | 2.01713 | 0.00424 | 0.00000 | 0.45495 | 0.00612 | 0.28183 |
| **Pms2** | 1.63613 | 0.00972 | 2.71510 | 0.78700 | 3.46828 | 5.45476 | 2.66970 | 0.19659 | 2.06916 | 0.10046 | 3.96153 | 0.45247 | 6.04104 | 7.77397 |
| **Pmvk** | 0.33924 | 7.23083 | 0.00000 | 0.00000 | 0.59552 | 11.59900 | 3.06059 | 6.93263 | 0.00000 | 0.18897 | 5.04286 | 11.46559 | 1.27216 | 0.09360 |
| **Pnck** | 0.00923 | 0.06872 | 0.01119 | 0.00000 | 0.01107 | 0.00507 | 0.00384 | 2.25902 | 0.01228 | 0.00849 | 0.00593 | 0.03380 | 0.00787 | 0.01347 |
| **Pnisr** | 2.06265 | 2.58334 | 4.25278 | 2.26877 | 3.45597 | 3.37213 | 0.61318 | 2.45699 | 8.10638 | 0.00000 | 5.17842 | 2.99098 | 1.52358 | 15.63893 |
| **Pnkd** | 0.95137 | 0.20727 | 0.39249 | 6.13754 | 3.01699 | 3.42321 | 2.99666 | 8.52496 | 0.69490 | 5.14038 | 0.80173 | 7.37806 | 0.06307 | 0.32979 |
| **Pnkp** | 1.31559 | 0.01625 | 10.58475 | 5.59721 | 3.23484 | 1.66128 | 0.00092 | 1.77993 | 0.04557 | 0.02991 | 3.41391 | 3.59862 | 0.48560 | 0.84879 |
| **Pnldc1** | 1.35721 | 0.03141 | 1.84317 | 0.51561 | 2.31225 | 3.27977 | 0.02737 | 0.92955 | 0.46457 | 0.04001 | 2.32838 | 2.14517 | 3.19372 | 5.65584 |
| **Pnlip** | 0.00437 | 0.00437 | 0.01126 | 0.01276 | 0.00587 | 0.01439 | 0.00425 | 0.00919 | 0.09560 | 0.00718 | 0.01012 | 0.01008 | 0.19815 | 0.04231 |
| **Pnliprp2** | 0.00000 | 0.01769 | 3.05439 | 5.68853 | 1.00610 | 0.30689 | 0.01423 | 0.00000 | 1.29947 | 0.00000 | 0.00052 | 2.32697 | 0.00000 | 0.00000 |
| **Pnma1** | 0.00789 | 0.00388 | 0.01583 | 0.02698 | 0.04780 | 0.04189 | 0.00403 | 0.25930 | 0.01195 | 0.01070 | 0.01112 | 0.02011 | 0.00645 | 0.01000 |
| **Pnma2** | 0.00485 | 0.00441 | 0.00272 | 0.00580 | 0.09580 | 0.10384 | 0.00426 | 0.00817 | 0.78746 | 0.00644 | 0.00562 | 0.00939 | 0.00350 | 0.00000 |
| **Pnma3** | 0.00000 | 0.04886 | 0.29311 | 0.97851 | 0.08077 | 1.44207 | 1.03656 | 1.67601 | 0.00863 | 0.00463 | 0.35691 | 0.08259 | 0.81556 | 0.15174 |
| **Pnma5** | 0.00000 | 0.05091 | 1.69145 | 0.00000 | 0.38409 | 6.61290 | 2.45388 | 6.46697 | 0.02620 | 0.08985 | 6.48351 | 0.86575 | 8.27372 | 0.59214 |
| **Pnmal2** | 0.00384 | 0.00438 | 0.21298 | 0.00583 | 0.00655 | 0.00696 | 0.00423 | 0.01076 | 0.00639 | 0.00686 | 0.00735 | 0.00884 | 0.15162 | 0.05829 |
| **Pnn** | 9.21756 | 0.05830 | 7.56727 | 5.98017 | 12.12228 | 7.64960 | 1.53242 | 2.98204 | 3.52754 | 0.08594 | 4.97445 | 3.92233 | 14.27630 | 10.62754 |
| **Pno1** | 14.48311 | 7.35887 | 20.55734 | 13.18711 | 23.16288 | 8.97272 | 5.26333 | 9.65577 | 10.89648 | 0.27512 | 3.61828 | 10.67829 | 8.89349 | 5.94039 |
| **Pnoc** | 0.00824 | 0.00443 | 0.01534 | 0.01343 | 0.01493 | 0.01811 | 0.00437 | 0.01035 | 0.03242 | 0.00734 | 0.00865 | 0.00944 | 0.11734 | 0.00865 |
| **Pnp** | 11.01904 | 5.70206 | 17.30703 | 10.80657 | 12.21711 | 19.78004 | 2.87043 | 2.85223 | 10.23934 | 3.37713 | 8.18300 | 6.01216 | 12.61784 | 11.42901 |
| **Pnp2** | 4.28881 | 3.96486 | 9.82137 | 2.65190 | 5.53987 | 4.80413 | 2.25335 | 0.39375 | 6.09870 | 11.31045 | 1.99031 | 0.63473 | 3.03990 | 7.48241 |
| **Pnpla1** | 0.37090 | 0.00404 | 0.12304 | 0.12832 | 0.27628 | 0.20023 | 0.00952 | 0.54699 | 0.21801 | 0.00617 | 0.02886 | 0.00216 | 0.63880 | 0.19696 |
| **Pnpla2** | 0.52422 | 4.94965 | 4.51715 | 4.30943 | 3.76156 | 0.17439 | 2.39398 | 3.97241 | 0.00000 | 0.07144 | 1.47716 | 0.72362 | 0.17850 | 0.00000 |
| **Pnpla3** | 2.26530 | 0.01976 | 7.59384 | 3.65020 | 3.50098 | 0.39639 | 0.02417 | 0.39091 | 5.10928 | 0.00000 | 0.00000 | 0.45036 | 0.88920 | 4.34363 |
| **Pnpla6** | 2.09289 | 0.15249 | 5.26370 | 7.22947 | 1.53449 | 6.20776 | 0.00000 | 0.59560 | 1.14245 | 0.08012 | 4.89519 | 12.20613 | 2.94940 | 1.12218 |
| **Pnpla7** | 0.51383 | 0.00380 | 1.56051 | 1.13978 | 0.15782 | 0.18223 | 0.00381 | 0.73060 | 0.00000 | 0.01469 | 0.00130 | 0.00166 | 2.03871 | 2.31962 |
| **Pnpla8** | 17.64357 | 0.01246 | 8.13356 | 3.75195 | 7.28663 | 5.56878 | 0.79393 | 3.08293 | 23.09886 | 0.00790 | 3.44254 | 4.51932 | 4.09540 | 10.16490 |
| **Pnpo** | 0.16612 | 0.04229 | 1.66264 | 3.94220 | 0.00000 | 2.28518 | 0.00782 | 4.93533 | 0.00000 | 1.84102 | 3.14758 | 0.00000 | 0.07448 | 1.58257 |
| **Pnpt1** | 2.67352 | 0.00324 | 0.56823 | 0.41439 | 0.02393 | 1.25842 | 1.44539 | 0.00948 | 3.65998 | 0.02388 | 0.16340 | 0.82108 | 0.07429 | 0.71030 |
| **Pnrc1** | 0.14485 | 0.01170 | 0.15802 | 0.36371 | 0.45881 | 0.00000 | 0.04187 | 0.74562 | 0.33575 | 0.00465 | 0.31101 | 0.05426 | 5.48796 | 0.25465 |
| **Pnrc2** | 0.00000 | 6.71723 | 1.10979 | 0.64120 | 1.54298 | 4.33541 | 1.61594 | 0.73358 | 1.98472 | 6.15483 | 2.49347 | 2.31434 | 0.00000 | 5.44636 |
| **Poc1a** | 1.43074 | 0.11820 | 5.03557 | 5.42721 | 4.80604 | 3.33811 | 2.78074 | 4.85467 | 2.68796 | 0.09345 | 4.39769 | 4.36066 | 3.00352 | 5.08365 |

| | | | | | | | | | | | | | | |
|---|---|---|---|---|---|---|---|---|---|---|---|---|---|---|
| **Poc1b** | 0.00000 | 0.00323 | 0.24237 | 0.00000 | 0.15715 | 0.18448 | 0.00367 | 0.00122 | 0.00483 | 0.00000 | 1.73536 | 0.67061 | 0.15936 | 1.19376 |
| **Poc5** | 9.02667 | 3.08092 | 3.34946 | 0.00000 | 3.89630 | 5.74724 | 0.00147 | 0.00000 | 0.12400 | 0.05382 | 0.57990 | 4.05091 | 2.41104 | 3.43955 |
| **Podnl1** | 0.00892 | 0.00445 | 0.01436 | 0.00757 | 0.47596 | 0.01312 | 0.00420 | 0.01006 | 0.01400 | 0.00629 | 0.00915 | 0.00821 | 0.11376 | 0.03024 |
| **Podxl** | 0.21130 | 0.00447 | 0.00589 | 0.01067 | 0.10716 | 0.00852 | 0.00446 | 0.01115 | 0.13788 | 0.00640 | 0.00633 | 0.00823 | 0.01060 | 0.17953 |
| **Pofut1** | 0.77680 | 0.01956 | 0.32350 | 0.00000 | 0.69338 | 1.27952 | 0.00348 | 0.14600 | 0.00230 | 0.00000 | 2.27869 | 1.00820 | 0.56150 | 1.41302 |
| **Pofut2** | 1.35073 | 0.05699 | 2.80208 | 2.12158 | 1.07802 | 5.41267 | 0.09499 | 0.60680 | 0.00000 | 0.01280 | 1.29015 | 3.87731 | 0.64881 | 1.93038 |
| **Pogk** | 0.09968 | 0.01129 | 0.23921 | 0.42382 | 0.89848 | 0.05871 | 0.00403 | 0.00242 | 0.09438 | 0.08476 | 0.00928 | 0.00615 | 0.02732 | 1.16958 |
| **Poglut1** | 0.40898 | 0.03672 | 4.92038 | 3.39705 | 1.20010 | 6.35985 | 0.00121 | 0.00000 | 3.20292 | 0.01937 | 3.67754 | 6.43506 | 2.74814 | 1.39053 |
| **Pogz** | 1.94868 | 0.03937 | 4.31594 | 1.88464 | 4.76885 | 2.71153 | 2.61416 | 3.65814 | 7.11778 | 0.11165 | 2.69141 | 3.19311 | 4.27508 | 4.94867 |
| **Pola1** | 4.16241 | 0.02602 | 4.62845 | 2.71359 | 2.97023 | 2.70199 | 0.00087 | 0.00000 | 10.22667 | 0.00000 | 1.23802 | 2.94954 | 1.24473 | 4.03455 |
| **Pola2** | 0.65022 | 5.38207 | 4.25574 | 4.36240 | 3.58967 | 1.31508 | 2.26303 | 1.37056 | 0.76030 | 0.26523 | 0.49771 | 2.67010 | 4.55604 | 0.28445 |
| **Polb** | 9.94867 | 0.09305 | 11.85173 | 5.03675 | 8.38276 | 9.99531 | 3.48883 | 4.08728 | 4.28616 | 0.11311 | 3.04370 | 3.46412 | 2.87318 | 6.79206 |
| **Pold1** | 0.70768 | 0.14003 | 14.24760 | 10.15363 | 8.16847 | 4.52530 | 0.80827 | 1.36129 | 0.00000 | 0.09676 | 6.13631 | 8.21724 | 0.05081 | 0.31466 |
| **Pold2** | 3.29075 | 0.08834 | 7.47965 | 8.51262 | 9.09997 | 7.10706 | 1.66475 | 0.20744 | 0.00128 | 0.11004 | 8.95854 | 8.66130 | 0.12980 | 0.59187 |
| **Pold3** | 14.64113 | 0.15431 | 21.55505 | 11.23094 | 19.92833 | 11.21306 | 6.57365 | 6.32596 | 12.16679 | 0.25224 | 9.59873 | 6.88974 | 4.14841 | 20.45685 |
| **Pold4** | 0.00574 | 0.00536 | 0.00463 | 0.06389 | 0.11290 | 0.33618 | 0.00478 | 0.04918 | 0.01109 | 0.00858 | 0.10784 | 0.18368 | 0.00447 | 0.02071 |
| **Poldip2** | 2.40346 | 9.63306 | 4.56804 | 4.61465 | 2.66824 | 5.68039 | 2.55849 | 3.35939 | 0.88475 | 0.04961 | 4.76532 | 7.43394 | 1.33191 | 0.50826 |
| **Poldip3** | 5.86388 | 7.16398 | 16.70800 | 14.08188 | 13.30649 | 24.25556 | 4.78167 | 10.79198 | 13.37114 | 1.24846 | 11.94809 | 10.56600 | 10.56615 | 16.20432 |
| **Pole** | 6.19951 | 0.09216 | 5.57396 | 3.51305 | 5.11157 | 4.32446 | 3.90101 | 0.61703 | 4.60486 | 6.10378 | 2.12702 | 7.01999 | 1.48300 | 6.09094 |
| **Pole2** | 4.05724 | 0.01567 | 7.62967 | 0.00344 | 1.48643 | 5.76722 | 0.88452 | 0.14754 | 0.44201 | 0.00669 | 4.40464 | 4.06612 | 2.54213 | 5.09230 |
| **Pole3** | 5.17772 | 0.77598 | 4.08784 | 9.55566 | 8.17743 | 8.15274 | 4.33186 | 5.20437 | 2.06048 | 0.32103 | 2.25082 | 6.79196 | 6.54643 | 5.56164 |
| **Pole4** | 5.31264 | 0.12021 | 2.95951 | 11.47528 | 19.97299 | 4.11940 | 0.18505 | 3.23014 | 8.04514 | 3.51126 | 0.88152 | 4.29000 | 9.96502 | 4.98990 |
| **Polg** | 0.27795 | 0.02636 | 7.25714 | 3.15922 | 0.72694 | 6.05609 | 1.64487 | 1.70935 | 0.57324 | 2.00424 | 2.15255 | 2.14269 | 0.41473 | 1.45800 |
| **Polg2** | 0.42920 | 0.01255 | 1.31211 | 0.00000 | 1.52813 | 1.45249 | 0.01160 | 1.10310 | 0.23559 | 0.01885 | 0.00000 | 0.98700 | 0.95004 | 0.23645 |
| **Polh** | 0.00000 | 0.00441 | 0.22862 | 0.10335 | 0.23099 | 0.01974 | 0.00467 | 0.03892 | 0.90777 | 0.00613 | 0.00427 | 0.00820 | 0.36391 | 0.25678 |
| **Poli** | 0.01130 | 0.03899 | 0.00000 | 0.00000 | 0.01448 | 0.97578 | 2.55167 | 0.45940 | 0.04070 | 0.03049 | 0.00000 | 0.60881 | 0.74911 | 0.20187 |
| **Polk** | 0.86323 | 0.00391 | 0.75690 | 0.00000 | 0.68165 | 0.76746 | 0.01095 | 0.04212 | 0.53855 | 0.00497 | 0.34176 | 0.28289 | 0.00000 | 0.86275 |
| **Poll** | 0.31951 | 0.00173 | 3.59378 | 1.24137 | 2.82450 | 0.97582 | 0.09029 | 3.45536 | 0.00000 | 0.02381 | 3.28569 | 2.13909 | 1.36033 | 1.25996 |
| **Polm** | 0.00000 | 0.02602 | 4.19579 | 2.59962 | 1.84046 | 3.52938 | 0.02773 | 1.84406 | 0.39157 | 0.03473 | 2.01322 | 2.77737 | 4.57977 | 0.56162 |
| **Poln** | 0.00987 | 0.00442 | 0.01073 | 0.01232 | 0.12304 | 0.01359 | 0.00486 | 0.00970 | 0.02009 | 0.00686 | 0.00914 | 0.01420 | 0.00594 | 0.00665 |
| **Polq** | 6.42397 | 0.01617 | 4.62454 | 1.16273 | 9.22756 | 3.78392 | 0.63999 | 0.00000 | 16.61892 | 0.00704 | 0.12175 | 1.01485 | 5.77029 | 13.50471 |
| **Polr1a** | 6.05251 | 0.08774 | 22.40113 | 14.55011 | 15.76473 | 9.91645 | 1.14162 | 4.44518 | 5.41901 | 0.07428 | 10.02170 | 7.62825 | 8.28523 | 16.76872 |
| **Polr1b** | 11.75264 | 0.61335 | 23.67760 | 14.95564 | 17.03104 | 14.94018 | 6.53904 | 3.99234 | 10.86681 | 0.21912 | 13.53369 | 10.87715 | 4.97023 | 16.16687 |
| **Polr1c** | 15.43049 | 7.19515 | 30.72654 | 19.19952 | 27.82833 | 16.51409 | 5.80177 | 17.42914 | 18.95363 | 0.39112 | 12.43501 | 11.11008 | 11.13009 | 26.76137 |
| **Polr1d** | 12.78551 | 0.95480 | 8.11377 | 19.80177 | 23.17030 | 13.39615 | 6.35121 | 14.43166 | 16.05244 | 17.89143 | 4.40694 | 14.57163 | 13.77251 | 19.10131 |
| **Polr1e** | 4.24739 | 0.03069 | 15.13314 | 3.44315 | 12.92630 | 2.78036 | 3.38466 | 1.29702 | 10.13832 | 0.03247 | 6.75697 | 2.85396 | 9.91102 | 23.42389 |
| **Polr2a** | 11.68440 | 0.35123 | 16.40976 | 6.50205 | 15.20793 | 9.46201 | 0.03676 | 10.28857 | 1.11902 | 0.07343 | 6.07012 | 13.78947 | 6.04001 | 12.55779 |
| **Polr2b** | 14.44727 | 0.01557 | 17.97364 | 5.81364 | 13.99634 | 12.41464 | 0.23890 | 1.36595 | 20.72013 | 0.03484 | 12.57087 | 5.99303 | 9.41794 | 16.09734 |
| **Polr2c** | 9.68213 | 8.60155 | 25.46316 | 21.67313 | 25.90149 | 15.22852 | 5.91988 | 10.13907 | 4.37557 | 7.98377 | 11.04827 | 8.84219 | 6.47548 | 19.63247 |
| **Polr2d** | 1.22387 | 0.06106 | 12.82857 | 6.23861 | 12.00605 | 9.68599 | 1.59638 | 6.97331 | 7.01752 | 0.16126 | 4.64927 | 3.93891 | 2.99232 | 15.37325 |
| **Polr2e** | 19.56333 | 6.04192 | 34.78242 | 26.23084 | 29.47702 | 25.54676 | 8.53999 | 16.05610 | 1.13704 | 9.81762 | 13.85297 | 18.11473 | 10.01823 | 25.78341 |
| **Polr2f** | 5.39589 | 4.32939 | 2.37092 | 12.05083 | 30.91890 | 9.04908 | 2.83883 | 8.38012 | 0.00000 | 9.83558 | 1.06974 | 4.43622 | 2.29530 | 4.50924 |
| **Polr2g** | 5.17640 | 0.67414 | 7.11767 | 8.58542 | 8.21361 | 7.30535 | 2.23019 | 0.60819 | 5.96330 | 9.41804 | 5.35623 | 6.88331 | 9.03366 | 8.18892 |
| **Polr2h** | 7.44005 | 0.13752 | 10.39540 | 12.56751 | 21.06396 | 13.68715 | 0.02222 | 3.35248 | 7.02952 | 0.19921 | 8.53664 | 8.07007 | 7.97276 | 17.80128 |
| **Polr2i** | 0.97300 | 0.30734 | 1.44293 | 5.18081 | 4.88041 | 2.81564 | 1.34107 | 3.14367 | 0.00000 | 0.20227 | 0.56218 | 3.94251 | 2.76955 | 0.79610 |
| **Polr2j** | 9.28807 | 5.31947 | 1.56919 | 6.86033 | 15.66393 | 8.40230 | 0.18528 | 8.56592 | 6.20962 | 5.59137 | 0.38365 | 6.70074 | 14.05792 | 11.25891 |
| **Polr2k** | 4.05945 | 0.00089 | 0.00000 | 1.52994 | 5.64181 | 3.52782 | 0.59644 | 0.48339 | 4.39481 | 0.09300 | 0.09351 | 2.94503 | 8.99254 | 2.50463 |
| **Polr2l** | 6.54234 | 2.28043 | 2.01681 | 5.83382 | 13.49135 | 7.42492 | 2.07532 | 5.96057 | 2.25122 | 8.84427 | 0.12651 | 3.54468 | 17.35921 | 7.27845 |
| **Polr2m** | 6.87110 | 0.18432 | 9.80704 | 10.54266 | 12.29048 | 13.05009 | 1.43747 | 11.88729 | 7.30469 | 0.20458 | 8.08911 | 8.49603 | 4.84517 | 18.49242 |
| **Polr3a** | 3.00882 | 0.02185 | 6.78883 | 2.05013 | 9.14564 | 2.81192 | 0.76702 | 3.47420 | 1.41014 | 0.03640 | 7.13745 | 2.47657 | 9.33645 | 15.14053 |
| **Polr3b** | 4.22764 | 0.03693 | 3.43096 | 1.63815 | 7.68814 | 2.48590 | 0.01242 | 0.00000 | 4.00131 | 0.00211 | 1.70251 | 5.02341 | 3.86659 | 11.21111 |
| **Polr3c** | 3.43342 | 2.39307 | 6.60243 | 2.28992 | 6.41409 | 3.30621 | 2.73320 | 0.74648 | 6.24669 | 0.08352 | 4.91075 | 2.16601 | 0.00000 | 7.24008 |
| **Polr3d** | 3.55554 | 0.01655 | 8.62165 | 4.84258 | 10.79306 | 5.97923 | 0.16758 | 1.00625 | 1.70688 | 0.03210 | 3.07708 | 3.03321 | 9.80518 | 9.31219 |
| **Polr3e** | 4.86713 | 0.09351 | 7.25830 | 1.83409 | 4.63153 | 6.70642 | 0.01387 | 2.32721 | 0.13830 | 0.00802 | 2.91762 | 4.42810 | 2.92031 | 3.23272 |
| **Polr3f** | 4.51023 | 0.01521 | 4.95651 | 2.43703 | 4.35686 | 3.10394 | 2.52687 | 0.00000 | 11.77572 | 0.16036 | 5.67552 | 2.27309 | 3.78112 | 5.34064 |

| | | | | | | | | | | | | | | |
|---|---|---|---|---|---|---|---|---|---|---|---|---|---|---|
| **Polr3g** | 9.08004 | 0.01150 | 12.76358 | 8.53176 | 14.69830 | 11.50871 | 1.46996 | 5.55979 | 12.78759 | 0.09558 | 13.97144 | 2.98384 | 5.55261 | 16.89202 |
| **Polr3gl** | 0.06323 | 0.02141 | 0.01572 | 0.12816 | 0.27366 | 0.53272 | 0.01636 | 0.04694 | 0.05365 | 0.00888 | 0.06118 | 0.03803 | 0.32962 | 0.18851 |
| **Polr3h** | 4.74034 | 0.04454 | 3.29495 | 11.19720 | 5.21679 | 2.84319 | 0.00163 | 1.96105 | 0.97084 | 0.16193 | 0.38773 | 3.64871 | 7.35871 | 2.91001 |
| **Polr3k** | 3.27575 | 0.04340 | 0.94822 | 0.60020 | 4.13959 | 2.90436 | 0.34722 | 0.03792 | 4.28592 | 0.04283 | 1.33687 | 1.97896 | 8.53193 | 10.34819 |
| **Polrmt** | 9.03213 | 0.04862 | 13.56649 | 6.35547 | 15.68396 | 1.42657 | 0.00000 | 2.45926 | 0.46407 | 0.01272 | 1.67725 | 4.21907 | 6.07002 | 6.72956 |
| **Pom121** | 3.25666 | 2.31557 | 9.68040 | 1.76077 | 1.74969 | 5.24035 | 1.54689 | 1.36194 | 1.75442 | 0.02225 | 2.53421 | 2.65387 | 9.21754 | 12.91027 |
| **Pomc** | 0.03619 | 0.00473 | 0.02467 | 0.22152 | 0.57561 | 0.07552 | 0.00397 | 0.00719 | 0.01041 | 0.01561 | 0.03898 | 0.00669 | 0.02721 | 0.02204 |
| **Pomgnt1** | 0.94268 | 0.03163 | 2.61086 | 3.31028 | 2.34275 | 5.89664 | 2.47522 | 3.72598 | 3.61177 | 0.17061 | 0.00000 | 4.19868 | 0.46636 | 0.53421 |
| **Pomgnt2** | 0.51798 | 0.00286 | 0.65478 | 0.10623 | 0.00000 | 0.00557 | 0.00339 | 0.00002 | 0.00926 | 0.00666 | 0.12237 | 1.42620 | 0.11063 | 0.00115 |
| **Pomk** | 0.47109 | 0.01995 | 0.00000 | 0.97093 | 0.06435 | 0.33985 | 0.00332 | 0.99086 | 0.01783 | 0.00518 | 2.14645 | 0.79980 | 0.71619 | 0.00000 |
| **Pomp** | 15.52289 | 8.10856 | 10.41503 | 24.98379 | 19.49348 | 26.57735 | 5.35819 | 15.84694 | 11.15478 | 23.33022 | 11.60900 | 15.96742 | 6.75711 | 13.36256 |
| **Pomt1** | 0.00467 | 0.02777 | 0.00000 | 0.44992 | 0.10825 | 1.30979 | 0.00378 | 1.49243 | 0.00927 | 0.13205 | 0.30254 | 2.12159 | 0.00907 | 0.03188 |
| **Pomt2** | 1.12296 | 0.01004 | 1.05898 | 1.57285 | 0.31400 | 3.93928 | 0.00313 | 0.00000 | 2.70655 | 0.00866 | 0.82046 | 2.03354 | 0.00000 | 1.12222 |
| **Pon2** | 0.00000 | 2.61025 | 0.13293 | 0.13563 | 0.00000 | 1.34821 | 4.38086 | 3.07589 | 0.07819 | 0.11763 | 1.51874 | 0.99466 | 0.43178 | 0.00000 |
| **Pop1** | 1.16871 | 0.03989 | 6.53214 | 3.61997 | 2.80946 | 2.78123 | 0.00849 | 0.85558 | 0.05179 | 0.00687 | 2.88187 | 4.36342 | 3.82355 | 0.71756 |
| **Pop4** | 11.17157 | 0.26229 | 17.59908 | 18.55157 | 15.87140 | 19.62705 | 6.80547 | 7.69448 | 10.90073 | 0.47516 | 13.96812 | 10.05092 | 4.05359 | 20.18713 |
| **Pop5** | 7.05298 | 1.54892 | 0.07085 | 8.31986 | 17.02288 | 3.50899 | 1.53306 | 0.07267 | 2.73708 | 0.11552 | 0.30132 | 2.30929 | 8.32715 | 2.98379 |
| **Pop7** | 4.68987 | 0.09395 | 6.61551 | 11.18379 | 9.48538 | 6.29277 | 2.35242 | 1.79942 | 0.00000 | 0.19144 | 5.13132 | 6.61472 | 0.40213 | 1.84729 |
| **Popdc3** | 0.00528 | 0.00439 | 0.00000 | 0.91504 | 0.14535 | 1.35514 | 0.00400 | 0.02169 | 0.00485 | 0.00400 | 1.10206 | 0.18143 | 0.19442 | 0.00781 |
| **Por** | 0.56007 | 0.08014 | 8.98755 | 8.93448 | 3.65020 | 5.51039 | 2.76326 | 4.77318 | 0.02478 | 0.09392 | 2.87592 | 3.22855 | 1.09263 | 1.48883 |
| **Porcn** | 5.66748 | 0.03185 | 6.21076 | 8.40316 | 2.63987 | 0.59854 | 0.01329 | 0.00000 | 0.63838 | 0.00000 | 0.00000 | 2.41023 | 0.06357 | 2.83078 |
| **Postn** | 0.53655 | 0.00437 | 0.14624 | 0.00199 | 0.12666 | 0.00501 | 0.00411 | 0.01214 | 0.00867 | 0.00753 | 0.00614 | 0.00878 | 0.00352 | 0.00477 |
| **Pot1a** | 1.01836 | 0.00392 | 1.77282 | 1.18710 | 2.39267 | 0.66897 | 0.03292 | 0.25012 | 5.72341 | 0.00000 | 1.34549 | 0.31393 | 0.00000 | 6.15793 |
| **Pot1b** | 0.13544 | 0.00378 | 0.02925 | 0.36257 | 0.16660 | 0.05792 | 0.04135 | 0.12257 | 0.97568 | 0.00683 | 0.12628 | 0.15179 | 0.36398 | 0.47769 |
| **Pou2f1** | 0.61787 | 0.03229 | 3.25914 | 0.15893 | 3.42490 | 2.63555 | 0.01823 | 3.00947 | 0.75341 | 0.02213 | 0.00000 | 3.31011 | 0.77568 | 0.32541 |
| **Pou2f2** | 0.41865 | 0.00408 | 0.03849 | 0.01056 | 0.13568 | 0.00904 | 0.00424 | 0.01606 | 0.00339 | 0.00667 | 0.00657 | 0.00547 | 1.86805 | 0.06219 |
| **Pou2f3** | 0.00000 | 0.03209 | 0.15315 | 0.00000 | 2.27998 | 0.00000 | 0.00921 | 0.05661 | 2.50391 | 0.01440 | 0.00000 | 3.42664 | 0.46004 | 6.49968 |
| **Pou3f1** | 0.00323 | 0.00409 | 0.22189 | 0.00303 | 0.09958 | 0.00553 | 0.00421 | 0.00814 | 0.00830 | 0.00903 | 0.00518 | 0.31164 | 0.11835 | 0.06672 |
| **Pou3f3** | 0.03869 | 0.00437 | 0.01081 | 0.03259 | 0.07206 | 0.04507 | 0.00432 | 0.00720 | 0.01099 | 0.00558 | 0.01144 | 0.01135 | 0.00392 | 0.15601 |
| **Pou4f1** | 0.00000 | 0.00400 | 0.21388 | 0.01226 | 0.77320 | 0.00051 | 0.00977 | 0.00777 | 0.00803 | 0.00534 | 0.00962 | 0.00838 | 1.98820 | 0.87856 |
| **Pou4f2** | 0.41072 | 0.00428 | 0.04581 | 0.00453 | 0.28142 | 0.01500 | 0.00413 | 0.00802 | 0.00928 | 0.01171 | 0.00581 | 0.05143 | 0.03060 | 0.00267 |
| **Pou5f1** | 12.02683 | 0.48018 | 26.28010 | 28.57731 | 16.73238 | 18.69229 | 5.61453 | 11.50877 | 1.62474 | 0.51721 | 13.72448 | 18.57159 | 2.99701 | 11.54663 |
| **Pou6f1** | 2.08069 | 0.00975 | 2.03922 | 3.61617 | 7.07307 | 2.05521 | 0.00227 | 0.00000 | 5.62009 | 0.00241 | 0.01153 | 1.72551 | 0.99686 | 8.28242 |
| **Pp2d1** | 0.02116 | 0.00437 | 0.03255 | 0.00888 | 0.01722 | 0.00743 | 0.00426 | 0.01078 | 0.78146 | 0.00729 | 0.00546 | 0.03150 | 0.00702 | 0.00000 |
| **Ppa1** | 10.71523 | 0.09278 | 8.89689 | 6.19143 | 12.43799 | 12.79481 | 0.37261 | 5.19076 | 1.14869 | 0.06374 | 6.85870 | 9.31045 | 7.70165 | 9.84111 |
| **Ppa2** | 5.76438 | 0.05138 | 1.78291 | 0.00000 | 3.86893 | 2.74831 | 1.63646 | 1.09139 | 1.41065 | 0.07075 | 3.89413 | 1.96407 | 1.55886 | 1.75570 |
| **Ppan** | 8.19655 | 4.72912 | 25.56197 | 24.77507 | 27.18688 | 7.39334 | 5.16282 | 6.50649 | 0.00000 | 0.16611 | 4.74199 | 12.29970 | 5.86185 | 1.15373 |
| **Ppara** | 0.03412 | 0.00469 | 0.01378 | 0.00297 | 0.09502 | 0.01524 | 0.00446 | 0.01520 | 0.01437 | 0.00814 | 0.00443 | 0.38451 | 0.94801 | 0.02713 |
| **Ppard** | 0.95858 | 2.48585 | 2.22362 | 0.00000 | 1.16335 | 0.00459 | 1.14896 | 1.77299 | 0.00120 | 0.01230 | 1.84405 | 1.72536 | 0.28377 | 0.43170 |
| **Ppargc1b** | 0.00000 | 7.56866 | 0.89988 | 2.12932 | 0.05174 | 4.81445 | 0.02051 | 0.07814 | 0.01196 | 8.81811 | 0.83190 | 0.00000 | 0.11409 | 0.00000 |
| **Ppat** | 17.95198 | 0.17647 | 24.95120 | 9.43428 | 16.96555 | 9.20821 | 5.71379 | 9.11458 | 25.21418 | 0.53692 | 12.65164 | 10.80999 | 23.45451 | 19.15017 |
| **Ppcdc** | 1.42068 | 0.01001 | 1.93638 | 3.57300 | 0.60131 | 7.62954 | 0.00274 | 0.19411 | 0.00000 | 0.00052 | 5.95231 | 1.02626 | 0.40790 | 1.73830 |
| **Ppcs** | 2.04555 | 6.52245 | 3.29417 | 8.53593 | 4.23092 | 5.18738 | 0.84888 | 2.33975 | 0.00000 | 0.05901 | 1.83224 | 6.67379 | 5.93271 | 0.00000 |
| **Ppdpf** | 0.21411 | 0.07777 | 0.01073 | 4.52667 | 3.06760 | 0.16721 | 0.03619 | 4.69808 | 0.00261 | 0.05050 | 0.00000 | 1.97113 | 0.45013 | 0.06064 |
| **Ppef1** | 0.00390 | 0.00442 | 0.00469 | 0.00290 | 0.04204 | 0.41094 | 0.00408 | 0.00455 | 0.00755 | 0.01115 | 0.00648 | 0.00904 | 0.00572 | 0.75518 |
| **Ppef2** | 0.00839 | 0.00470 | 0.03335 | 0.02252 | 0.03550 | 0.23892 | 0.00427 | 0.00794 | 0.01045 | 0.00610 | 0.07816 | 0.04788 | 0.00699 | 0.00933 |
| **Ppfia1** | 10.13002 | 4.33972 | 16.35690 | 7.27354 | 10.98226 | 8.89092 | 3.11585 | 7.10904 | 7.73374 | 1.36821 | 6.77604 | 2.95823 | 9.90658 | 15.51144 |
| **Ppfia3** | 0.00632 | 0.00449 | 0.00954 | 0.26182 | 0.01063 | 0.00641 | 0.00543 | 0.24115 | 0.01253 | 0.00585 | 0.00689 | 0.00615 | 0.00665 | 0.00999 |
| **Ppfia4** | 1.14970 | 0.00407 | 1.08460 | 1.14934 | 0.96381 | 0.00001 | 0.00369 | 0.00365 | 0.30406 | 0.01805 | 0.00345 | 0.00564 | 0.00000 | 0.00000 |
| **Ppfibp1** | 0.52235 | 0.92911 | 0.34150 | 0.00000 | 0.13635 | 0.74826 | 0.03552 | 0.89246 | 3.46123 | 0.00794 | 1.33347 | 0.78588 | 2.01306 | 3.23853 |
| **Ppfibp2** | 13.46313 | 0.49219 | 20.80308 | 2.69620 | 17.35348 | 10.60652 | 3.53057 | 2.77260 | 14.41679 | 0.09161 | 7.97493 | 0.98350 | 0.00000 | 13.90750 |
| **Pphln1** | 10.00750 | 0.11408 | 13.24181 | 9.16066 | 20.02027 | 11.59877 | 3.40519 | 4.72630 | 15.39505 | 5.45446 | 4.51803 | 5.61673 | 10.27875 | 12.83913 |
| **Ppia** | 37.00847 | 15.88016 | 31.71787 | 45.15234 | 40.54829 | 48.82574 | 14.66206 | 37.81455 | 20.20601 | 30.10505 | 26.58934 | 37.40012 | 18.83134 | 32.74928 |
| **Ppib** | 0.45696 | 4.71404 | 1.31786 | 5.67851 | 1.34261 | 5.11178 | 2.89072 | 5.20007 | 0.00999 | 3.07536 | 6.78553 | 4.29300 | 0.00000 | 1.51337 |
| **Ppic** | 0.50532 | 0.02735 | 1.51343 | 1.77300 | 0.23859 | 1.07346 | 0.00346 | 0.49534 | 0.19359 | 0.00000 | 1.52611 | 2.70239 | 0.69765 | 0.00128 |

| | | | | | | | | | | | | | | |
|---|---|---|---|---|---|---|---|---|---|---|---|---|---|---|
| **Ppid** | 10.52620 | 3.03418 | 21.05131 | 13.38912 | 12.40859 | 20.17247 | 4.77519 | 6.03425 | 12.67534 | 0.19887 | 12.88901 | 14.48843 | 8.54230 | 8.23184 |
| **Ppie** | 0.00000 | 5.71588 | 12.29567 | 9.12423 | 12.21751 | 12.74763 | 0.17804 | 8.03399 | 1.33535 | 0.16257 | 12.41983 | 3.32308 | 0.61055 | 9.88617 |
| **Ppif** | 3.29271 | 0.07373 | 2.27329 | 4.09349 | 4.49490 | 6.20334 | 2.45373 | 4.34130 | 0.34025 | 0.15135 | 2.47879 | 5.62119 | 4.32938 | 1.00134 |
| **Ppig** | 21.42733 | 4.43325 | 20.45522 | 8.42217 | 21.16023 | 13.22053 | 0.02479 | 5.09457 | 34.90404 | 0.07641 | 10.38847 | 10.64697 | 5.25299 | 28.28142 |
| **Ppih** | 4.90615 | 0.12624 | 10.33139 | 13.92939 | 7.03803 | 10.69916 | 3.96485 | 8.29565 | 9.82907 | 3.01337 | 6.43095 | 8.22126 | 2.28034 | 11.19268 |
| **Ppil1** | 4.42270 | 4.00753 | 18.23043 | 18.41992 | 18.22887 | 16.73488 | 5.51051 | 8.22021 | 8.03248 | 3.32200 | 7.17397 | 11.73350 | 2.39400 | 20.94038 |
| **Ppil2** | 3.40755 | 5.34999 | 9.65867 | 2.83657 | 16.17701 | 5.03406 | 3.35036 | 7.52405 | 0.02117 | 0.06973 | 9.30057 | 6.44137 | 0.52392 | 9.20683 |
| **Ppil3** | 0.64738 | 0.02652 | 2.12796 | 1.12726 | 3.12977 | 3.73186 | 0.01449 | 1.92437 | 0.00248 | 0.11208 | 2.64249 | 4.05177 | 0.41865 | 1.36247 |
| **Ppil4** | 6.20485 | 0.04150 | 8.54714 | 2.94350 | 8.54110 | 8.22299 | 1.40637 | 0.43137 | 5.61299 | 0.01496 | 1.59868 | 3.77608 | 3.64729 | 11.57116 |
| **Ppil6** | 0.00161 | 0.01279 | 0.00565 | 0.19601 | 0.21198 | 0.01440 | 0.00477 | 0.05619 | 0.00951 | 0.00774 | 0.00932 | 0.00979 | 0.27611 | 0.50275 |
| **Ppip5k1** | 0.42040 | 0.01364 | 0.64328 | 1.48017 | 0.28372 | 0.04777 | 0.00421 | 2.55007 | 1.46366 | 0.00553 | 0.28591 | 0.02894 | 0.01117 | 1.05211 |
| **Ppip5k2** | 1.99783 | 0.02526 | 3.86517 | 1.88586 | 1.60918 | 2.99327 | 0.25166 | 0.38907 | 10.43490 | 0.01310 | 2.32056 | 2.48671 | 1.94500 | 5.62125 |
| **Ppl** | 0.00000 | 0.01987 | 0.99540 | 0.89652 | 0.00000 | 0.15412 | 3.93137 | 0.67406 | 1.26955 | 0.20799 | 0.04948 | 0.00000 | 0.00000 | 0.00000 |
| **Ppm1a** | 29.07023 | 0.15057 | 40.21760 | 17.36925 | 24.23905 | 8.06243 | 0.50443 | 6.13606 | 28.54692 | 0.03049 | 16.77155 | 18.15655 | 13.77452 | 24.48553 |
| **Ppm1b** | 12.21480 | 2.81601 | 10.86427 | 5.55559 | 7.83697 | 9.17354 | 2.81487 | 2.35760 | 15.33468 | 0.03739 | 9.46841 | 7.82806 | 6.53058 | 17.11335 |
| **Ppm1d** | 1.22855 | 0.01006 | 1.20922 | 0.44306 | 0.87916 | 2.14874 | 3.07975 | 1.86459 | 0.02349 | 0.12294 | 0.04287 | 0.19860 | 6.66283 | 1.08949 |
| **Ppm1e** | 0.57036 | 0.00399 | 0.20101 | 0.09204 | 0.00000 | 0.00372 | 0.00410 | 0.00817 | 0.01853 | 0.00509 | 0.00607 | 0.00430 | 1.75484 | 0.17659 |
| **Ppm1f** | 2.41208 | 0.03193 | 2.73999 | 1.68146 | 3.90309 | 2.79733 | 1.05909 | 1.81729 | 4.21903 | 0.07503 | 1.26312 | 1.82657 | 0.45370 | 1.77678 |
| **Ppm1g** | 17.04964 | 1.91021 | 23.26036 | 10.35562 | 26.24905 | 17.48893 | 4.92216 | 11.81509 | 3.81148 | 12.84732 | 11.61070 | 11.06266 | 8.44731 | 30.11897 |
| **Ppm1h** | 0.00000 | 0.00402 | 0.22812 | 0.00831 | 0.11398 | 0.00824 | 0.00416 | 0.01758 | 0.00822 | 0.00633 | 0.01751 | 0.02560 | 1.61967 | 1.03218 |
| **Ppm1j** | 0.27437 | 0.02646 | 0.40226 | 0.00000 | 0.04391 | 0.00000 | 0.00332 | 1.21828 | 0.00726 | 0.00602 | 0.00000 | 0.65291 | 0.00282 | 0.01430 |
| **Ppm1k** | 4.06242 | 0.23324 | 0.47557 | 0.85797 | 0.27859 | 4.29457 | 0.49375 | 0.90072 | 5.68766 | 1.15850 | 0.04203 | 2.11940 | 15.48654 | 2.33499 |
| **Ppm1l** | 1.13152 | 0.00339 | 3.02549 | 2.45178 | 0.85185 | 1.85301 | 0.00258 | 0.00000 | 5.29480 | 0.01584 | 0.00000 | 0.45410 | 0.00000 | 4.23161 |
| **Ppm1m** | 0.00158 | 0.01840 | 0.09867 | 0.76441 | 0.10525 | 0.44362 | 0.00362 | 0.00301 | 0.02933 | 0.00293 | 1.22828 | 1.22375 | 0.01758 | 0.02322 |
| **Ppm1n** | 0.40856 | 0.01052 | 0.31460 | 1.13717 | 0.03841 | 0.00023 | 0.00371 | 0.67288 | 0.09005 | 0.01464 | 0.94153 | 0.00583 | 0.03698 | 0.00000 |
| **Ppme1** | 3.35705 | 0.08011 | 8.03720 | 3.34510 | 1.99761 | 5.20783 | 0.39731 | 0.43457 | 0.00000 | 0.00546 | 4.74434 | 3.95797 | 0.29476 | 0.32436 |
| **Ppox** | 0.00000 | 0.02429 | 1.40886 | 0.10971 | 0.00000 | 1.07219 | 1.14460 | 1.83501 | 0.00509 | 0.00903 | 0.54786 | 3.77415 | 0.00000 | 0.57517 |
| **Ppp1ca** | 15.73405 | 9.13068 | 25.29884 | 26.01821 | 20.25361 | 25.26837 | 7.24735 | 27.74894 | 0.76815 | 1.06213 | 11.87635 | 15.98303 | 2.23597 | 11.83066 |
| **Ppp1cb** | 0.03104 | 0.20660 | 7.82327 | 3.81078 | 4.11568 | 11.55630 | 1.55814 | 9.56718 | 10.64133 | 14.68298 | 12.03451 | 4.77154 | 7.33395 | 1.78703 |
| **Ppp1cc** | 26.07722 | 2.83548 | 36.96578 | 19.39259 | 41.14079 | 27.28895 | 11.56842 | 22.39548 | 15.99819 | 9.94066 | 8.22993 | 19.69076 | 20.03160 | 23.94273 |
| **Ppp1r10** | 5.99112 | 5.32402 | 7.88979 | 0.00000 | 7.01294 | 3.23091 | 3.49619 | 3.55871 | 0.76061 | 0.13685 | 2.98384 | 2.78434 | 3.64092 | 4.11130 |
| **Ppp1r11** | 1.06008 | 3.44456 | 3.06673 | 8.68836 | 3.40581 | 10.98133 | 2.57719 | 6.39357 | 1.65618 | 0.40368 | 5.27581 | 4.87241 | 0.26423 | 3.11227 |
| **Ppp1r12a** | 3.04433 | 0.01576 | 3.24531 | 0.00000 | 1.91828 | 2.69552 | 0.81041 | 0.00000 | 1.80885 | 0.01233 | 4.52277 | 2.05902 | 2.24659 | 9.07442 |
| **Ppp1r12b** | 0.12734 | 0.21671 | 0.08701 | 0.45915 | 0.32460 | 0.19157 | 0.16533 | 0.14526 | 1.06408 | 0.01475 | 0.08227 | 0.12368 | 1.47260 | 1.46888 |
| **Ppp1r12c** | 3.46581 | 5.85687 | 4.02466 | 8.89142 | 5.13755 | 3.78496 | 1.43752 | 1.05763 | 0.71110 | 0.13081 | 0.51757 | 5.89103 | 4.08646 | 0.70868 |
| **Ppp1r13b** | 0.89831 | 0.01834 | 2.55795 | 0.00000 | 1.27452 | 0.46565 | 0.00844 | 3.64939 | 0.00293 | 0.02151 | 1.45004 | 1.83166 | 1.81762 | 1.40978 |
| **Ppp1r13l** | 0.16703 | 0.00397 | 0.23600 | 0.98505 | 0.30192 | 0.80431 | 0.00379 | 0.75769 | 0.01118 | 0.01628 | 0.00000 | 0.07861 | 1.09567 | 0.00000 |
| **Ppp1r14a** | 0.00840 | 0.00434 | 0.00749 | 0.18302 | 0.12674 | 0.01120 | 0.00410 | 0.00789 | 0.01137 | 0.00622 | 0.00707 | 0.01012 | 0.00652 | 0.00916 |
| **Ppp1r14b** | 3.74760 | 0.69731 | 1.28423 | 7.51094 | 13.39756 | 5.45007 | 4.75571 | 8.41552 | 0.00000 | 5.87721 | 0.04757 | 5.59672 | 4.48775 | 1.05793 |
| **Ppp1r14c** | 0.06307 | 0.00514 | 0.13770 | 0.07834 | 0.10453 | 0.24990 | 0.22667 | 0.02788 | 0.00980 | 0.04734 | 0.00922 | 0.00223 | 0.03960 | 0.00744 |
| **Ppp1r14d** | 0.26317 | 0.05104 | 0.46187 | 0.00000 | 1.00612 | 0.34167 | 0.01424 | 0.31402 | 0.01204 | 0.02689 | 0.00000 | 6.11996 | 0.53126 | 0.01755 |
| **Ppp1r15a** | 4.24693 | 0.03971 | 4.27422 | 2.40247 | 4.01539 | 0.79909 | 1.64927 | 1.81045 | 0.00000 | 0.03222 | 0.02911 | 0.30103 | 10.27282 | 6.18410 |
| **Ppp1r15b** | 6.99001 | 0.67892 | 7.37295 | 0.46388 | 1.59826 | 2.28666 | 3.09195 | 4.62581 | 22.32411 | 0.32069 | 5.73901 | 3.40626 | 13.77454 | 18.76583 |
| **Ppp1r16a** | 0.24988 | 0.02804 | 1.22173 | 2.12499 | 3.01675 | 0.23108 | 0.00323 | 2.43511 | 0.00000 | 0.00545 | 1.71760 | 0.78399 | 1.75783 | 0.00000 |
| **Ppp1r16b** | 1.84887 | 0.07573 | 0.08524 | 0.58946 | 3.24246 | 1.86682 | 0.49526 | 0.41730 | 6.25480 | 0.88232 | 0.03096 | 0.57496 | 10.68844 | 10.71917 |
| **Ppp1r18** | 0.79249 | 0.40946 | 0.79944 | 2.80805 | 0.79521 | 4.02553 | 0.17017 | 7.72247 | 0.27991 | 1.40127 | 0.00000 | 1.17914 | 0.00000 | 0.00251 |
| **Ppp1r1b** | 0.01066 | 0.00445 | 0.01454 | 0.01353 | 0.01701 | 0.01315 | 0.00430 | 0.01044 | 0.01154 | 0.00700 | 0.00881 | 0.01002 | 0.00763 | 0.01222 |
| **Ppp1r2** | 10.03711 | 6.04334 | 5.76894 | 4.54100 | 12.31335 | 4.89499 | 3.17994 | 2.96012 | 6.57517 | 0.11155 | 5.38301 | 7.03405 | 11.26242 | 2.02990 |
| **Ppp1r2.ps3** | 1.18184 | 0.96112 | 0.61587 | 0.41998 | 2.13940 | 0.27028 | 0.43817 | 0.01713 | 0.30967 | 0.00425 | 0.28930 | 1.01817 | 1.40472 | 0.54345 |
| **Ppp1r2.ps7** | 0.00185 | 0.00429 | 0.05099 | 0.00811 | 0.00632 | 0.01127 | 0.00420 | 0.00930 | 0.01938 | 0.00632 | 0.01273 | 0.00809 | 0.82503 | 0.00173 |
| **Ppp1r21** | 1.23506 | 0.01662 | 3.75179 | 0.22710 | 1.46341 | 1.73950 | 0.01387 | 2.77055 | 0.13339 | 0.00000 | 3.84349 | 0.57832 | 0.93350 | 0.49258 |
| **Ppp1r26** | 1.54424 | 0.02108 | 2.12186 | 1.87895 | 2.02611 | 2.61136 | 0.12611 | 2.07629 | 0.53180 | 0.04705 | 0.02525 | 0.00000 | 0.00498 | 1.22987 |
| **Ppp1r35** | 0.00266 | 0.00338 | 0.00000 | 0.54161 | 1.03373 | 0.10946 | 0.02211 | 0.31571 | 0.01831 | 1.22434 | 0.00090 | 0.46944 | 0.01149 | 0.04603 |
| **Ppp1r36** | 0.00463 | 0.00444 | 0.18797 | 0.00172 | 0.02653 | 0.07549 | 0.00438 | 0.00528 | 0.02454 | 0.00617 | 0.00674 | 0.01184 | 0.00397 | 0.68516 |
| **Ppp1r37** | 0.00297 | 0.00364 | 1.02699 | 0.59535 | 1.04620 | 0.02748 | 0.00394 | 0.01526 | 0.01438 | 0.00246 | 0.49487 | 0.66087 | 0.00275 | 0.01171 |

| | | | | | | | | | | | | | | |
|---|---|---|---|---|---|---|---|---|---|---|---|---|---|---|
| **Ppp1r3b** | 0.00000 | 0.13605 | 2.08590 | 1.09433 | 0.12598 | 11.33699 | 6.32152 | 8.34186 | 0.00000 | 2.97775 | 4.90861 | 0.00000 | 0.80694 | 1.16463 |
| **Ppp1r3c** | 0.00397 | 0.00441 | 0.09621 | 0.00485 | 0.00698 | 0.00960 | 0.00427 | 0.01055 | 0.06616 | 0.00656 | 0.00798 | 0.00903 | 0.24515 | 0.00000 |
| **Ppp1r3d** | 0.01430 | 0.00426 | 0.00854 | 0.00983 | 0.26849 | 0.01266 | 0.00412 | 0.00853 | 0.00555 | 0.00677 | 0.00814 | 0.00787 | 0.69142 | 0.38863 |
| **Ppp1r3e** | 0.01062 | 0.00541 | 0.01478 | 0.00333 | 0.01202 | 0.00461 | 0.00436 | 0.59304 | 0.01248 | 0.00872 | 0.00815 | 0.01276 | 0.00782 | 0.01148 |
| **Ppp1r3f** | 0.01085 | 0.00440 | 0.00757 | 0.13918 | 0.08366 | 0.05397 | 0.00505 | 0.07859 | 0.01102 | 0.00557 | 0.00591 | 0.02791 | 0.00604 | 0.00828 |
| **Ppp1r3fos** | 0.00618 | 0.00557 | 0.00873 | 0.20412 | 0.00861 | 0.26343 | 0.00420 | 0.02545 | 0.01106 | 0.01016 | 0.00628 | 0.05242 | 0.00825 | 0.00896 |
| **Ppp1r3g** | 0.00479 | 0.00417 | 0.01158 | 0.08908 | 0.00959 | 0.01071 | 0.00403 | 0.00953 | 0.01170 | 0.00698 | 0.00711 | 0.01354 | 0.25566 | 0.00108 |
| **Ppp1r7** | 7.89612 | 4.85688 | 7.06324 | 6.17807 | 5.72281 | 12.49918 | 2.70632 | 4.91688 | 3.65369 | 0.17749 | 8.97155 | 5.42683 | 5.96028 | 8.29739 |
| **Ppp1r8** | 4.97863 | 0.07037 | 7.96408 | 7.41019 | 11.89227 | 8.48360 | 4.17209 | 5.41719 | 4.26607 | 0.23291 | 2.53251 | 5.49981 | 12.49043 | 8.86266 |
| **Ppp1r9a** | 0.93277 | 0.01185 | 0.43106 | 0.01289 | 1.76541 | 1.05308 | 0.00464 | 0.77753 | 0.01705 | 0.01131 | 0.00559 | 0.00678 | 0.00000 | 1.18933 |
| **Ppp1r9b** | 0.02593 | 0.02277 | 0.05651 | 0.15714 | 0.14955 | 0.83558 | 0.01765 | 1.12770 | 0.01331 | 1.38002 | 0.00000 | 0.00000 | 0.00613 | 0.00345 |
| **Ppp2ca** | 20.57704 | 0.33137 | 17.01253 | 11.13407 | 17.85260 | 18.64253 | 4.73293 | 7.52333 | 13.49153 | 0.21846 | 9.34052 | 13.37273 | 7.41930 | 13.75284 |
| **Ppp2cb** | 4.59313 | 0.06799 | 0.27349 | 0.04616 | 0.17003 | 0.19117 | 0.01957 | 0.03094 | 6.11740 | 0.02219 | 0.73676 | 0.20184 | 9.22949 | 4.60549 |
| **Ppp2r1a** | 1.96751 | 3.11858 | 20.77176 | 18.40443 | 11.84494 | 25.41261 | 5.28943 | 11.85907 | 10.15709 | 11.22376 | 17.64555 | 17.76696 | 1.13425 | 17.43272 |
| **Ppp2r1b** | 3.02276 | 0.02544 | 1.24611 | 2.36969 | 0.86562 | 3.30667 | 0.02242 | 0.38499 | 1.12309 | 0.02227 | 1.37746 | 2.16296 | 1.72024 | 1.64975 |
| **Ppp2r2a** | 13.04174 | 0.09195 | 12.42336 | 4.19241 | 19.08932 | 8.56502 | 4.32466 | 5.34477 | 15.36087 | 0.13220 | 9.05243 | 6.04869 | 13.06862 | 17.11314 |
| **Ppp2r2b** | 0.58656 | 0.00419 | 0.00000 | 0.45999 | 0.72547 | 0.32392 | 1.95741 | 0.00418 | 2.35734 | 0.01298 | 0.00067 | 0.00098 | 1.57093 | 0.23934 |
| **Ppp2r2c** | 1.00614 | 3.92485 | 3.59320 | 0.00000 | 3.18069 | 3.59545 | 0.00760 | 1.36469 | 0.34821 | 0.01098 | 5.75219 | 3.30495 | 0.00000 | 1.63651 |
| **Ppp2r2d** | 7.24158 | 0.02992 | 6.93576 | 2.10262 | 9.57898 | 5.22449 | 0.96729 | 6.70694 | 4.58432 | 0.03824 | 5.27458 | 4.36243 | 6.39749 | 3.22942 |
| **Ppp2r3a** | 0.96281 | 0.00412 | 0.25893 | 0.05546 | 0.00000 | 0.03221 | 0.00422 | 0.00407 | 0.01393 | 0.00558 | 0.00991 | 0.05024 | 0.69637 | 0.00406 |
| **Ppp2r3c** | 2.04942 | 0.02079 | 3.30203 | 1.47787 | 4.44197 | 4.04901 | 0.00644 | 0.00000 | 7.29355 | 0.01734 | 3.97056 | 4.95922 | 1.50500 | 2.33385 |
| **Ppp2r3d** | 1.01212 | 0.11806 | 8.55196 | 3.12980 | 2.74133 | 4.89814 | 4.27893 | 0.19682 | 1.07053 | 0.47735 | 1.54828 | 2.84488 | 0.50491 | 1.01677 |
| **Ppp2r5a** | 0.31590 | 0.01786 | 1.51165 | 0.76493 | 0.69226 | 1.31984 | 0.00336 | 1.94510 | 0.00343 | 0.01407 | 0.82467 | 0.34967 | 0.00657 | 0.06479 |
| **Ppp2r5b** | 0.00000 | 0.02249 | 1.48593 | 1.07175 | 0.00000 | 0.60780 | 0.85807 | 3.20512 | 0.01174 | 10.42093 | 2.75039 | 2.22686 | 0.01940 | 0.00000 |
| **Ppp2r5c** | 2.58766 | 0.10450 | 11.05744 | 3.13180 | 5.47823 | 12.41988 | 3.47905 | 10.43650 | 6.62390 | 0.47709 | 9.15022 | 8.40347 | 5.67955 | 8.24335 |
| **Ppp2r5d** | 0.57541 | 6.44624 | 2.30773 | 2.73992 | 0.75062 | 3.94658 | 0.83368 | 5.00385 | 0.13882 | 0.10913 | 1.82750 | 5.23733 | 4.34102 | 0.40819 |
| **Ppp2r5e** | 10.12201 | 0.00581 | 5.67098 | 0.64888 | 5.06964 | 5.10247 | 0.00109 | 0.00000 | 1.31374 | 0.00000 | 8.85474 | 2.54239 | 4.04235 | 3.21194 |
| **Ppp3ca** | 0.62850 | 0.01663 | 2.43216 | 0.54057 | 0.46578 | 2.16237 | 0.00248 | 1.30050 | 1.75279 | 0.01394 | 0.01670 | 1.80109 | 0.44198 | 0.10338 |
| **Ppp3cb** | 0.00000 | 0.02735 | 0.49914 | 0.00000 | 0.51592 | 0.63952 | 1.74614 | 0.06912 | 1.88740 | 0.05590 | 2.83397 | 2.92004 | 1.21068 | 2.27361 |
| **Ppp3cc** | 0.05198 | 0.00451 | 0.00000 | 0.01554 | 0.04542 | 0.00733 | 0.00420 | 0.01092 | 0.83789 | 0.00670 | 0.00590 | 0.00886 | 0.00565 | 0.09199 |
| **Ppp3r1** | 6.24821 | 8.05284 | 2.47793 | 5.52375 | 0.26477 | 12.48321 | 1.80511 | 4.37186 | 3.32620 | 0.07637 | 2.51799 | 6.45230 | 3.47472 | 1.65629 |
| **Ppp4c** | 0.00000 | 0.92945 | 11.37091 | 7.21932 | 13.23416 | 5.54773 | 2.00101 | 4.56848 | 0.01627 | 0.06739 | 5.36021 | 5.05051 | 0.00000 | 2.89185 |
| **Ppp4r1** | 3.19823 | 2.15547 | 8.53207 | 4.50765 | 0.21065 | 10.87085 | 5.12084 | 9.48158 | 0.06181 | 0.15709 | 7.03766 | 8.25855 | 2.48435 | 2.19766 |
| **Ppp4r1l.ps** | 0.06084 | 0.00436 | 0.02476 | 0.06575 | 0.01196 | 0.04819 | 0.40541 | 0.00758 | 0.07340 | 0.01289 | 0.02976 | 0.02044 | 0.01967 | 0.05431 |
| **Ppp4r2** | 18.75664 | 0.93663 | 12.63047 | 0.00000 | 21.93825 | 8.15818 | 5.83742 | 1.86724 | 29.29181 | 0.17879 | 7.41125 | 6.10883 | 15.98524 | 24.03354 |
| **Ppp4r4** | 0.15134 | 0.00438 | 0.01396 | 0.01095 | 0.00638 | 0.01206 | 0.00433 | 0.00995 | 0.00915 | 0.00658 | 0.00755 | 0.00860 | 0.25511 | 0.00255 |
| **Ppp5c** | 0.95784 | 5.21309 | 14.77021 | 11.95342 | 13.75673 | 3.71131 | 0.35374 | 5.58083 | 0.00000 | 0.02928 | 4.50912 | 8.95772 | 1.95355 | 0.16617 |
| **Ppp6c** | 9.25942 | 0.07047 | 14.74330 | 7.47408 | 8.96804 | 3.93666 | 0.37197 | 2.34114 | 12.40069 | 0.01936 | 7.38584 | 2.14377 | 6.74744 | 9.58786 |
| **Ppp6r1** | 0.12473 | 0.00344 | 0.12020 | 0.24704 | 0.42728 | 0.15352 | 0.26578 | 1.14498 | 0.01355 | 0.04651 | 0.94110 | 0.24224 | 0.54500 | 0.09863 |
| **Ppp6r2** | 0.43463 | 0.00421 | 0.00000 | 0.03883 | 0.00000 | 0.00325 | 0.00389 | 0.00778 | 0.98028 | 0.00731 | 0.00434 | 0.00984 | 0.42606 | 0.00000 |
| **Ppp6r3** | 2.30447 | 1.04381 | 22.89019 | 13.21706 | 19.34933 | 2.88863 | 7.37485 | 10.36336 | 1.91667 | 0.88520 | 11.38842 | 9.38236 | 3.58067 | 3.16780 |
| **Pprc1** | 12.61740 | 8.58990 | 17.12453 | 15.17328 | 23.26502 | 4.83715 | 3.43041 | 3.76766 | 3.55967 | 0.11336 | 5.98830 | 10.08902 | 15.33065 | 12.30017 |
| **Ppt1** | 1.40839 | 0.18720 | 0.89187 | 0.57018 | 0.00000 | 0.00004 | 1.63509 | 3.17994 | 2.37063 | 0.08254 | 1.54922 | 5.96838 | 0.41897 | 0.00000 |
| **Ppt2** | 9.13983 | 0.38550 | 1.28618 | 1.27114 | 16.57515 | 0.79159 | 0.00343 | 0.15601 | 3.28580 | 0.03038 | 0.06194 | 0.00000 | 5.93283 | 6.51640 |
| **Pptc7** | 1.39790 | 0.00311 | 1.44022 | 0.00000 | 0.57372 | 3.09440 | 0.80020 | 0.35993 | 1.18161 | 0.07828 | 0.00000 | 1.83935 | 5.32351 | 4.39158 |
| **Ppwd1** | 0.15313 | 0.03318 | 0.79074 | 0.20377 | 1.43385 | 0.10210 | 0.00397 | 0.98273 | 3.07070 | 0.00578 | 0.01672 | 0.18199 | 0.16027 | 2.30435 |
| **Pqbp1** | 8.31613 | 0.45661 | 21.56129 | 10.40786 | 17.26826 | 7.99227 | 3.93887 | 3.49474 | 0.00000 | 0.11874 | 4.94466 | 10.43973 | 0.28025 | 4.96518 |
| **Pqlc1** | 10.94205 | 0.03004 | 20.58566 | 6.27505 | 15.93335 | 6.49852 | 2.70713 | 3.42907 | 1.04762 | 15.39620 | 3.03977 | 0.00000 | 0.56037 | 13.51359 |
| **Pqlc2** | 0.00000 | 0.01683 | 1.88735 | 4.36163 | 0.27578 | 1.59777 | 0.65014 | 5.66930 | 0.00300 | 0.03118 | 1.09499 | 2.69757 | 0.40631 | 0.00000 |
| **Pqlc3** | 0.00406 | 0.00936 | 0.00000 | 1.65436 | 0.02460 | 2.56754 | 3.60155 | 3.29974 | 0.01332 | 0.08797 | 2.44075 | 2.45421 | 0.19992 | 0.23108 |
| **Pradc1** | 2.48898 | 0.05850 | 2.78153 | 0.08014 | 4.13306 | 6.19205 | 2.18847 | 1.68975 | 2.79498 | 0.09439 | 0.43355 | 4.49846 | 3.19409 | 1.93306 |
| **Praf2** | 3.01291 | 0.20954 | 1.77055 | 0.00000 | 1.38820 | 2.53473 | 0.00083 | 5.56219 | 0.01589 | 7.18890 | 0.01585 | 3.04552 | 2.00563 | 0.09544 |
| **Pram1** | 0.00923 | 0.00449 | 0.05233 | 0.00000 | 0.01621 | 0.11539 | 0.00430 | 0.00616 | 0.00917 | 0.00635 | 0.00853 | 0.01430 | 0.00454 | 0.05253 |
| **Pramef12** | 7.50838 | 0.01900 | 1.87772 | 2.39440 | 8.04519 | 6.68696 | 2.17020 | 0.04088 | 12.19557 | 0.09473 | 0.93108 | 1.70726 | 6.16971 | 9.24402 |
| **Pramef17** | 0.38621 | 0.57260 | 0.21165 | 0.00000 | 0.38438 | 2.28541 | 3.24265 | 0.00153 | 0.72132 | 0.12136 | 0.21702 | 0.20241 | 0.78306 | 1.18373 |

| | | | | | | | | | | | | | | |
|---|---|---|---|---|---|---|---|---|---|---|---|---|---|---|
| **Pramef25** | 0.00000 | 0.00406 | 0.02509 | 0.00990 | 0.01758 | 0.01279 | 0.00404 | 0.00819 | 0.11372 | 0.00638 | 0.00807 | 0.00731 | 2.45813 | 0.00000 |
| **Pramef6** | 0.01680 | 0.02755 | 1.43040 | 0.28918 | 0.16385 | 1.27556 | 0.09331 | 0.06441 | 1.23945 | 0.09874 | 0.53946 | 0.12574 | 0.48992 | 0.16857 |
| **Pramef8** | 2.52623 | 0.22386 | 2.29264 | 2.31124 | 2.52492 | 1.34218 | 1.14512 | 0.24373 | 2.88538 | 0.07705 | 0.00000 | 1.62581 | 1.56504 | 6.42170 |
| **Pramel1** | 0.00000 | 0.00457 | 0.00000 | 0.19333 | 0.91114 | 0.03457 | 0.00455 | 0.03263 | 0.30782 | 0.01637 | 0.00776 | 0.00582 | 0.16078 | 0.00000 |
| **Pramel3** | 0.00993 | 0.00412 | 0.01376 | 0.00650 | 0.01520 | 0.01547 | 0.35225 | 0.00415 | 0.01098 | 0.00570 | 0.00848 | 0.01029 | 0.01658 | 0.14371 |
| **Pramel4** | 22.69208 | 7.22899 | 27.14752 | 18.67849 | 17.40054 | 28.69449 | 6.93875 | 3.55777 | 29.90947 | 6.73529 | 12.71665 | 11.11808 | 9.47474 | 16.52252 |
| **Pramel5** | 27.34525 | 1.90022 | 39.17545 | 26.95682 | 22.68157 | 38.68773 | 10.23005 | 12.88071 | 23.87489 | 15.61176 | 17.81597 | 11.70112 | 7.91943 | 11.68376 |
| **Pramel6** | 25.14739 | 11.72881 | 24.81707 | 31.10785 | 31.02028 | 27.57210 | 10.08909 | 23.57600 | 33.76146 | 2.49958 | 13.87290 | 18.21201 | 14.01630 | 27.58687 |
| **Pramel7** | 14.09847 | 0.02190 | 12.44873 | 0.00000 | 2.57815 | 4.09951 | 1.88319 | 3.69750 | 34.87969 | 1.51042 | 3.77297 | 2.20573 | 2.93569 | 35.31901 |
| **Prc1** | 9.86546 | 0.24730 | 11.24610 | 1.22304 | 16.44246 | 6.90487 | 2.21116 | 1.18954 | 28.82079 | 0.12768 | 9.88321 | 2.59544 | 15.28916 | 21.89523 |
| **Prcc** | 0.00000 | 0.02842 | 1.20842 | 0.28737 | 1.93550 | 0.00000 | 0.00358 | 0.50527 | 2.85028 | 0.00147 | 0.37089 | 0.31304 | 0.00000 | 8.12227 |
| **Prcp** | 1.98494 | 0.00203 | 4.25915 | 0.75408 | 3.57022 | 2.03233 | 0.00962 | 0.72100 | 5.51929 | 0.01167 | 0.00000 | 2.33696 | 0.59747 | 6.34106 |
| **Prdm10** | 0.22867 | 0.00384 | 0.97989 | 0.03155 | 1.18756 | 0.53312 | 0.00369 | 1.04938 | 0.32062 | 0.01685 | 0.29985 | 0.00221 | 0.27986 | 2.94612 |
| **Prdm14** | 0.15564 | 0.00247 | 0.98963 | 1.11344 | 7.91804 | 0.00000 | 0.00266 | 0.00000 | 3.85942 | 0.00341 | 0.00000 | 1.25444 | 2.18586 | 3.66082 |
| **Prdm15** | 0.66773 | 2.47126 | 1.90770 | 1.45093 | 1.85391 | 0.19168 | 0.00304 | 0.00638 | 0.00166 | 0.00204 | 0.00455 | 0.00230 | 2.01789 | 1.70497 |
| **Prdm16** | 0.20812 | 0.00475 | 0.29599 | 0.70792 | 0.50392 | 0.00000 | 0.00420 | 0.14189 | 0.11662 | 0.00754 | 0.00426 | 0.62423 | 0.00326 | 0.35853 |
| **Prdm2** | 4.19424 | 0.07508 | 8.48435 | 2.26733 | 3.29217 | 7.13145 | 3.24186 | 3.14438 | 5.53375 | 0.10692 | 2.13890 | 2.44903 | 8.03532 | 9.53640 |
| **Prdm4** | 3.70645 | 0.01580 | 6.62707 | 2.64684 | 9.13727 | 5.50466 | 0.00080 | 0.50931 | 11.09125 | 0.00000 | 3.78229 | 2.30622 | 7.59096 | 14.00253 |
| **Prdm6** | 0.01932 | 0.00513 | 0.12156 | 0.05796 | 0.08830 | 0.01122 | 0.00426 | 0.04222 | 0.00952 | 0.02155 | 0.00773 | 0.00649 | 0.00864 | 0.01587 |
| **Prdm9** | 0.07091 | 0.05383 | 0.25656 | 0.45711 | 0.43823 | 0.61576 | 0.06958 | 0.06233 | 2.30634 | 0.11465 | 0.03914 | 0.02635 | 1.80749 | 0.79897 |
| **Prdx1** | 15.73506 | 9.51356 | 17.14058 | 27.75894 | 18.49602 | 33.39960 | 9.40703 | 21.85953 | 21.55484 | 2.86423 | 21.19483 | 36.20907 | 5.54743 | 16.47774 |
| **Prdx2** | 3.90921 | 8.08931 | 7.89198 | 9.77888 | 7.49804 | 7.04968 | 6.22497 | 12.03930 | 2.54314 | 14.70889 | 6.62198 | 13.41134 | 5.51573 | 6.22768 |
| **Prdx3** | 0.71459 | 3.40869 | 13.21539 | 2.32066 | 6.20566 | 6.48160 | 4.53483 | 2.54595 | 13.91849 | 0.17908 | 3.63734 | 2.64154 | 0.04662 | 12.50326 |
| **Prdx4** | 6.28719 | 0.12682 | 8.34098 | 6.74828 | 10.01584 | 9.60743 | 4.16184 | 7.69670 | 0.85837 | 0.31012 | 4.75388 | 12.01661 | 3.35328 | 3.56699 |
| **Prdx5** | 7.37317 | 6.45970 | 6.01519 | 15.59083 | 11.48111 | 9.16913 | 4.96057 | 10.72825 | 0.00000 | 17.06465 | 6.27688 | 6.26175 | 2.20221 | 2.38153 |
| **Prdx6** | 8.45671 | 0.85798 | 29.77717 | 33.47208 | 17.09598 | 28.59705 | 10.22007 | 19.76837 | 8.47417 | 21.78970 | 13.51794 | 22.59793 | 0.50589 | 10.57867 |
| **Prdx6b** | 0.00975 | 0.00421 | 0.23830 | 0.39546 | 0.03965 | 0.23062 | 0.14170 | 0.10358 | 0.03427 | 0.28012 | 0.08422 | 0.14619 | 0.00576 | 0.05537 |
| **Preb** | 2.99158 | 0.14284 | 9.69336 | 9.58490 | 4.09301 | 9.79748 | 2.88406 | 5.36007 | 1.37541 | 0.13716 | 2.09577 | 7.93987 | 0.00000 | 7.17728 |
| **Prelid1** | 10.61952 | 1.42812 | 18.22387 | 23.98104 | 22.47384 | 17.73810 | 3.46650 | 15.63855 | 0.02832 | 9.93303 | 5.09097 | 17.13084 | 1.87907 | 4.59739 |
| **Prelid2** | 8.80391 | 0.07417 | 10.27374 | 19.36553 | 6.40828 | 18.22913 | 6.13894 | 4.67038 | 5.51139 | 10.87061 | 11.61102 | 2.31060 | 2.05342 | 6.78630 |
| **Prelp** | 0.05948 | 0.00429 | 0.00797 | 0.01216 | 0.01312 | 0.01148 | 0.00454 | 0.00885 | 0.02429 | 0.00615 | 0.00721 | 0.00822 | 0.45044 | 0.51072 |
| **Prep** | 0.32760 | 8.12003 | 6.00918 | 10.23276 | 1.68801 | 13.48832 | 5.61394 | 7.52733 | 0.00000 | 9.12540 | 8.07389 | 9.07176 | 1.88343 | 0.00000 |
| **Prepl** | 0.70157 | 0.01166 | 1.93852 | 0.00000 | 0.00000 | 2.00689 | 0.07945 | 0.07301 | 3.86079 | 0.00770 | 3.49554 | 0.44708 | 0.23131 | 3.07923 |
| **Prex1** | 0.23709 | 0.02021 | 0.35761 | 1.57308 | 0.06017 | 0.11520 | 0.01564 | 4.53540 | 0.03072 | 0.01851 | 0.59841 | 0.06539 | 0.42206 | 0.48676 |
| **Prex2** | 1.60002 | 0.26257 | 0.50141 | 1.09038 | 1.38423 | 0.81879 | 0.17206 | 0.98434 | 4.45628 | 0.35702 | 1.10017 | 0.97596 | 13.72048 | 6.16799 |
| **Prg4** | 0.23213 | 0.00512 | 0.10714 | 0.09120 | 0.14598 | 0.10492 | 0.00457 | 0.04711 | 0.01981 | 0.00837 | 0.00775 | 0.11461 | 0.47655 | 0.02002 |
| **Prickle1** | 0.01630 | 0.00306 | 0.98054 | 0.67055 | 3.51685 | 0.00000 | 0.00327 | 0.02814 | 1.72378 | 0.00904 | 0.00000 | 0.15962 | 3.48497 | 13.00382 |
| **Prickle2** | 0.02039 | 0.00517 | 0.01314 | 0.00428 | 0.01351 | 0.07607 | 0.00432 | 0.03982 | 0.01256 | 0.00825 | 0.00657 | 0.01293 | 0.00916 | 0.01092 |
| **Prickle3** | 0.00739 | 0.01583 | 0.00616 | 0.00000 | 0.47608 | 0.00669 | 0.00429 | 0.27967 | 0.01303 | 0.00743 | 0.00717 | 0.03722 | 0.28237 | 0.00386 |
| **Prim1** | 4.69613 | 3.05174 | 10.06437 | 3.27140 | 8.36229 | 7.18513 | 1.15556 | 5.67692 | 10.28266 | 0.08436 | 10.05236 | 8.26792 | 2.27689 | 10.17140 |
| **Prim2** | 5.96963 | 0.02762 | 8.57350 | 2.88852 | 7.03975 | 5.19577 | 4.88755 | 3.77246 | 7.41520 | 2.28701 | 7.42003 | 7.41974 | 2.90277 | 11.07252 |
| **Primpol** | 1.38168 | 0.00292 | 1.31776 | 0.00000 | 1.00924 | 1.21082 | 0.00325 | 0.15931 | 0.03025 | 0.01305 | 0.40858 | 3.22855 | 0.21134 | 2.01577 |
| **Prkaa1** | 7.01863 | 0.02252 | 14.00477 | 7.30990 | 7.12450 | 5.30857 | 0.24816 | 3.38574 | 15.79017 | 0.07295 | 4.64834 | 1.04130 | 4.12477 | 9.36116 |
| **Prkaa2** | 0.00000 | 0.00343 | 0.29784 | 0.99424 | 0.91626 | 1.95976 | 0.01442 | 0.00433 | 0.70034 | 0.00513 | 0.48948 | 0.42522 | 0.00406 | 2.07806 |
| **Prkab1** | 17.66915 | 7.44953 | 21.47343 | 10.10705 | 24.12349 | 17.47310 | 6.26086 | 11.20013 | 4.31574 | 1.57938 | 13.48047 | 11.06363 | 7.02725 | 14.88369 |
| **Prkab2** | 0.01436 | 0.01346 | 0.00000 | 0.35610 | 0.00532 | 1.77599 | 1.75430 | 0.07686 | 0.00595 | 0.05549 | 0.00000 | 0.77238 | 0.33485 | 0.06200 |
| **Prkaca** | 0.00787 | 0.03917 | 0.24465 | 0.44772 | 0.00000 | 2.25434 | 0.01452 | 3.69514 | 0.00000 | 0.04426 | 1.64697 | 3.73398 | 0.06347 | 1.41532 |
| **Prkacb** | 3.72582 | 0.03970 | 2.94033 | 0.00000 | 5.23142 | 2.51590 | 0.00107 | 2.31319 | 5.06934 | 0.00000 | 3.72640 | 5.22400 | 1.44743 | 4.70178 |
| **Prkag1** | 1.67694 | 0.06683 | 9.84493 | 12.59370 | 9.34052 | 7.45401 | 0.05694 | 6.06815 | 0.06155 | 1.28463 | 7.86398 | 4.96146 | 1.99021 | 1.77299 |
| **Prkag2** | 0.01284 | 0.00407 | 0.00791 | 0.77011 | 0.02768 | 0.48719 | 0.00414 | 1.54657 | 0.00692 | 0.02750 | 0.00000 | 0.53072 | 0.02997 | 0.01400 |
| **Prkag2os1** | 0.01261 | 0.00435 | 0.01344 | 0.02474 | 0.00884 | 0.05027 | 0.67265 | 0.01066 | 0.01046 | 0.02951 | 0.00634 | 0.00602 | 0.10322 | 0.00366 |
| **Prkar1a** | 3.19981 | 1.37798 | 5.65655 | 10.93648 | 6.14325 | 8.88855 | 1.26764 | 1.72044 | 0.04328 | 0.07008 | 7.32338 | 9.71574 | 0.57958 | 5.37148 |
| **Prkar1b** | 0.01227 | 0.01482 | 0.00175 | 0.00000 | 0.01639 | 0.00000 | 0.00412 | 0.15130 | 0.01236 | 0.00536 | 1.02315 | 0.47332 | 0.00737 | 0.00692 |
| **Prkar2a** | 1.63428 | 0.39720 | 2.94138 | 3.60631 | 2.90141 | 5.73322 | 0.01459 | 2.19151 | 2.82760 | 0.01982 | 4.34383 | 3.05853 | 0.62959 | 11.57884 |
| **Prkar2b** | 0.02799 | 0.00413 | 0.00000 | 0.00788 | 0.00000 | 0.01262 | 0.00411 | 0.00666 | 0.18447 | 0.00605 | 0.00533 | 0.11472 | 0.58649 | 1.00502 |

| | | | | | | | | | | | | | | |
|---|---|---|---|---|---|---|---|---|---|---|---|---|---|---|
| **Prkca** | 0.10247 | 0.00425 | 0.32459 | 0.09781 | 0.17063 | 0.00150 | 0.00429 | 0.00812 | 0.00866 | 0.00562 | 0.01332 | 0.01445 | 0.00752 | 0.01398 |
| **Prkcb** | 0.00681 | 0.01313 | 0.01173 | 0.03394 | 0.08314 | 0.01355 | 0.01096 | 0.02068 | 0.03673 | 0.00686 | 0.02369 | 0.01173 | 0.00547 | 0.06523 |
| **Prkcd** | 4.60238 | 0.01018 | 1.81177 | 0.00000 | 3.44186 | 0.26031 | 0.00323 | 0.50017 | 2.45445 | 0.00326 | 0.03559 | 0.42671 | 8.18340 | 15.60111 |
| **Prkce** | 0.00000 | 0.00470 | 1.03750 | 0.15353 | 2.08556 | 0.08730 | 0.00430 | 0.01927 | 0.03510 | 0.01592 | 0.05011 | 0.01447 | 0.01119 | 0.46493 |
| **Prkcg** | 1.88342 | 0.01520 | 8.18270 | 0.68869 | 7.34531 | 0.00000 | 5.52598 | 7.25714 | 0.97858 | 14.79096 | 2.72228 | 0.75749 | 1.55850 | 4.61787 |
| **Prkch** | 0.00168 | 0.00354 | 0.00896 | 0.00609 | 0.00677 | 0.01134 | 0.00404 | 0.00585 | 0.04143 | 0.00663 | 0.00363 | 0.93896 | 0.04729 | 0.22317 |
| **Prkci** | 1.44400 | 0.06071 | 2.08080 | 1.09525 | 0.04368 | 3.18865 | 1.46079 | 0.98910 | 0.08731 | 0.04087 | 2.80923 | 5.47647 | 1.44561 | 1.16754 |
| **Prkcq** | 0.00508 | 0.00437 | 0.01368 | 0.01219 | 0.00782 | 0.01244 | 0.00426 | 0.00955 | 0.02651 | 0.00632 | 0.00878 | 0.00950 | 0.24053 | 0.05795 |
| **Prkcsh** | 1.79511 | 1.08342 | 13.20472 | 17.36456 | 6.11951 | 12.01416 | 5.27945 | 9.82317 | 0.00000 | 0.31999 | 6.48353 | 10.42681 | 1.15564 | 1.25029 |
| **Prkcz** | 2.53369 | 0.15273 | 6.30952 | 4.58618 | 2.92749 | 8.21768 | 2.43110 | 8.81008 | 0.00000 | 3.75125 | 2.63188 | 4.72648 | 2.74555 | 1.00939 |
| **Prkd1** | 0.00464 | 0.00432 | 0.01363 | 0.01349 | 0.05243 | 0.01150 | 0.00430 | 0.00929 | 0.00505 | 0.00583 | 0.00849 | 0.00809 | 0.69042 | 0.25341 |
| **Prkd2** | 0.00000 | 0.00376 | 0.58099 | 0.51331 | 0.22972 | 0.18950 | 0.00352 | 0.00005 | 0.00763 | 0.00947 | 0.00506 | 0.24308 | 0.24467 | 0.10757 |
| **Prkd3** | 0.42769 | 0.00427 | 0.22527 | 0.02271 | 0.06732 | 0.00109 | 0.00401 | 0.00765 | 0.00677 | 0.00589 | 0.00676 | 0.52103 | 0.23577 | 0.20140 |
| **Prkdc** | 2.86925 | 0.00120 | 2.87666 | 0.44386 | 2.83671 | 2.42953 | 0.00807 | 0.00000 | 1.98789 | 0.00694 | 0.08381 | 3.10771 | 2.55618 | 5.22537 |
| **Prkg1** | 0.00510 | 0.00445 | 0.00000 | 0.03161 | 0.24117 | 0.00113 | 0.00604 | 0.10739 | 0.40640 | 0.01277 | 0.00977 | 0.00867 | 0.00815 | 0.40634 |
| **Prkra** | 0.23912 | 0.00286 | 0.20900 | 1.32174 | 0.00000 | 0.00000 | 0.00280 | 0.00000 | 0.00970 | 3.96061 | 1.08026 | 0.00000 | 0.00613 | 0.00000 |
| **Prkrip1** | 5.82937 | 7.41719 | 13.38309 | 12.37969 | 11.60289 | 9.83806 | 2.01131 | 1.22818 | 2.81289 | 0.14981 | 2.38445 | 6.80459 | 5.68022 | 10.16300 |
| **Prkrir** | 1.23297 | 0.05290 | 2.14604 | 0.00000 | 3.09997 | 4.64461 | 0.03067 | 0.41304 | 3.70423 | 0.00000 | 4.92882 | 3.94408 | 8.95770 | 3.91177 |
| **Prkx** | 0.04174 | 0.00414 | 0.79784 | 0.86294 | 0.45976 | 1.51510 | 1.07412 | 0.15395 | 1.31234 | 0.03184 | 0.14900 | 0.18972 | 0.79921 | 1.01144 |
| **Prl3b1** | 4.63890 | 0.00401 | 1.53999 | 0.67996 | 3.80054 | 0.30297 | 0.00362 | 0.00000 | 2.27102 | 0.00711 | 0.00062 | 0.00085 | 0.00000 | 2.61674 |
| **Prlr** | 0.00000 | 0.16282 | 0.01044 | 0.05406 | 0.00000 | 0.00000 | 0.00373 | 0.00673 | 1.85626 | 0.00667 | 0.00331 | 0.02858 | 0.43675 | 2.46149 |
| **Prm2** | 0.01718 | 0.00435 | 0.01872 | 0.13663 | 0.02116 | 0.05957 | 0.00413 | 0.00558 | 0.01192 | 0.00701 | 0.00603 | 0.01094 | 0.00712 | 0.00920 |
| **Prm3** | 0.00803 | 0.00393 | 0.01682 | 0.03165 | 0.02613 | 0.05061 | 0.00385 | 0.11498 | 0.01327 | 0.02753 | 0.05629 | 0.00576 | 0.00677 | 0.01165 |
| **Prmt1** | 0.52620 | 0.00381 | 0.20249 | 0.46162 | 1.36598 | 0.26643 | 0.02721 | 0.00338 | 0.06507 | 0.01113 | 0.18592 | 0.37076 | 0.47848 | 0.37670 |
| **Prmt2** | 0.12249 | 0.00422 | 0.16188 | 0.00115 | 0.17980 | 0.00434 | 0.00415 | 0.01098 | 0.00390 | 0.00676 | 0.00646 | 0.00914 | 0.55998 | 0.38436 |
| **Prmt3** | 7.85684 | 0.01270 | 3.71746 | 3.45374 | 8.54838 | 5.96132 | 1.39482 | 1.60305 | 6.18634 | 0.00128 | 11.25487 | 3.66769 | 0.58125 | 9.93352 |
| **Prmt5** | 11.72244 | 7.69258 | 29.41084 | 13.41729 | 17.53608 | 18.75131 | 3.81648 | 8.66327 | 10.35643 | 0.24101 | 16.98655 | 12.71758 | 4.30842 | 21.39306 |
| **Prmt6** | 4.46325 | 0.04296 | 2.37173 | 2.28088 | 10.57591 | 0.89045 | 0.66112 | 2.27319 | 6.36293 | 0.04276 | 1.00161 | 0.51740 | 17.51628 | 4.80176 |
| **Prmt7** | 1.58691 | 0.07443 | 8.37048 | 10.80160 | 2.35961 | 8.77531 | 3.38344 | 1.59229 | 0.47523 | 0.18135 | 12.05383 | 6.21978 | 0.91259 | 3.37428 |
| **Prmt9** | 11.35018 | 0.01723 | 6.56176 | 1.48313 | 7.09784 | 3.38865 | 0.00851 | 1.06569 | 3.93137 | 0.00525 | 0.48863 | 0.00000 | 0.00000 | 7.56947 |
| **Prnp** | 41.87396 | 12.17085 | 35.63411 | 18.84864 | 41.90779 | 41.71677 | 9.82971 | 13.11270 | 45.16064 | 8.40309 | 22.92131 | 13.16144 | 16.42820 | 36.27211 |
| **Prob1** | 0.03046 | 0.00438 | 0.01164 | 0.01512 | 0.00626 | 0.01406 | 0.00427 | 0.00895 | 0.01044 | 0.00795 | 0.00767 | 0.01293 | 0.07314 | 0.22930 |
| **Proc** | 0.00673 | 0.05690 | 0.01195 | 0.00000 | 0.00914 | 0.03454 | 0.00448 | 1.32426 | 0.00854 | 0.00820 | 0.00615 | 0.01150 | 0.03148 | 0.13426 |
| **Proca1** | 0.50343 | 0.01147 | 0.13010 | 0.98332 | 1.59018 | 0.03669 | 0.00876 | 2.25950 | 1.54139 | 0.00591 | 0.00000 | 0.00349 | 1.56063 | 0.00000 |
| **Procr** | 0.01295 | 0.03158 | 0.36401 | 0.00000 | 0.00000 | 0.33945 | 0.00285 | 0.00185 | 0.14353 | 0.00509 | 0.00652 | 3.49392 | 1.41643 | 0.00000 |
| **Prodh** | 0.50422 | 0.20855 | 3.49901 | 5.22404 | 0.00000 | 1.50790 | 0.00270 | 0.02244 | 0.01724 | 0.01594 | 2.35739 | 1.51122 | 0.00968 | 0.00000 |
| **Prokr2** | 0.17758 | 0.00438 | 0.29946 | 0.24564 | 0.20612 | 0.00000 | 2.12529 | 0.02254 | 0.03132 | 0.15465 | 0.00210 | 0.00274 | 0.37668 | 0.00000 |
| **Prom1** | 0.00000 | 0.01069 | 0.23588 | 0.14930 | 0.03424 | 0.00000 | 0.01484 | 2.73644 | 0.41701 | 0.01282 | 0.00239 | 0.00644 | 0.01389 | 0.29723 |
| **Prom2** | 0.56301 | 0.00435 | 0.48368 | 2.18172 | 0.86057 | 0.06239 | 0.12741 | 0.03199 | 1.19783 | 0.04980 | 0.00000 | 0.00364 | 0.00727 | 0.91703 |
| **Prorsd1** | 3.12058 | 0.00093 | 2.20424 | 2.04688 | 1.78384 | 2.23975 | 0.00237 | 0.00001 | 2.04591 | 0.00000 | 6.35945 | 0.90954 | 4.13783 | 5.65049 |
| **Pros1** | 0.07419 | 2.10753 | 0.02095 | 1.65863 | 0.01983 | 2.71574 | 3.16809 | 0.02290 | 0.02974 | 0.04384 | 0.60694 | 8.29114 | 0.00441 | 0.00000 |
| **Prosc** | 7.33398 | 0.07383 | 14.61363 | 3.52207 | 18.19837 | 5.57238 | 1.52652 | 0.58017 | 12.39576 | 0.02196 | 2.13272 | 4.75275 | 8.75289 | 19.33495 |
| **Proser1** | 0.00000 | 0.00980 | 2.31126 | 0.62331 | 0.66734 | 0.82796 | 1.73736 | 0.10387 | 0.00232 | 0.03561 | 2.31873 | 2.69624 | 0.90883 | 0.74224 |
| **Proser2** | 0.52142 | 0.01406 | 0.59482 | 1.55165 | 2.17264 | 0.83440 | 1.26528 | 0.86964 | 0.73940 | 0.02016 | 2.04879 | 0.00304 | 0.10768 | 0.67283 |
| **Proser3** | 0.11528 | 0.00339 | 0.02193 | 0.58916 | 0.24563 | 0.23122 | 0.00365 | 0.00221 | 0.00784 | 0.00594 | 0.15472 | 0.55961 | 0.81003 | 0.00000 |
| **Prox2** | 0.03912 | 0.00448 | 0.01148 | 0.04496 | 0.02108 | 0.03137 | 0.00417 | 0.01235 | 0.01602 | 0.01318 | 0.00762 | 0.01756 | 0.10667 | 0.02767 |
| **Prpf18** | 3.65799 | 0.02290 | 10.26989 | 4.34778 | 10.98765 | 5.27010 | 0.05159 | 0.51100 | 8.71088 | 0.00000 | 8.09864 | 3.00393 | 4.11794 | 15.76171 |
| **Prpf19** | 0.77768 | 0.48407 | 0.22921 | 1.07435 | 1.09204 | 2.02123 | 0.17292 | 1.02030 | 0.57957 | 1.37439 | 0.05024 | 1.94929 | 5.46385 | 0.24620 |
| **Prpf3** | 2.61297 | 0.02292 | 5.74646 | 1.16824 | 6.69403 | 6.71983 | 0.00658 | 1.49011 | 8.50076 | 0.02363 | 3.50760 | 4.02364 | 1.89225 | 13.67275 |
| **Prpf31** | 3.82400 | 2.94858 | 8.74227 | 14.60535 | 6.71420 | 7.91771 | 0.03912 | 0.85017 | 1.65884 | 0.07467 | 5.95986 | 6.12464 | 2.68418 | 9.27849 |
| **Prpf38a** | 9.71750 | 6.51118 | 14.09721 | 12.00402 | 11.75172 | 11.27435 | 0.02547 | 7.21826 | 14.55006 | 0.09712 | 8.66103 | 6.73254 | 10.63771 | 13.00353 |
| **Prpf38b** | 9.83028 | 0.23412 | 6.08490 | 2.78644 | 8.46709 | 8.72337 | 0.02411 | 0.22037 | 1.33807 | 0.00000 | 5.72052 | 4.08177 | 4.50872 | 8.61597 |
| **Prpf39** | 1.34055 | 0.00291 | 0.72780 | 1.97800 | 0.80923 | 2.03411 | 1.30695 | 0.35645 | 5.33089 | 1.45706 | 0.42872 | 0.38836 | 3.65367 | 1.61851 |
| **Prpf4** | 5.44937 | 0.11527 | 7.71818 | 3.24636 | 8.49460 | 9.97995 | 2.58750 | 5.45229 | 6.44180 | 0.04446 | 6.54064 | 7.39465 | 5.84987 | 11.36648 |
| **Prpf40a** | 11.51517 | 0.01477 | 9.35656 | 4.67424 | 12.44658 | 9.94749 | 1.67408 | 2.27470 | 21.18363 | 0.08979 | 3.50472 | 3.05218 | 10.99761 | 16.17553 |

| | | | | | | | | | | | | | | |
|---|---|---|---|---|---|---|---|---|---|---|---|---|---|---|
| **Prpf40b** | 0.00000 | 0.00293 | 0.44751 | 1.13257 | 0.24562 | 0.00000 | 0.00345 | 0.19687 | 0.00848 | 0.00493 | 1.22224 | 1.15194 | 0.12991 | 0.00000 |
| **Prpf4b** | 7.15341 | 0.02454 | 5.70060 | 0.85952 | 5.57349 | 3.21274 | 0.35901 | 0.15807 | 14.92141 | 0.05328 | 4.67601 | 3.40538 | 8.38159 | 9.17872 |
| **Prpf6** | 12.25992 | 0.11708 | 9.95357 | 5.03201 | 8.26018 | 4.17850 | 1.63631 | 2.39238 | 1.36874 | 8.88290 | 7.91637 | 3.69122 | 4.21465 | 9.32889 |
| **Prpf8** | 14.42066 | 7.37403 | 22.62906 | 9.50448 | 20.54425 | 20.98436 | 6.09383 | 2.84220 | 11.46733 | 5.84698 | 15.05132 | 13.04110 | 6.08769 | 16.36400 |
| **Prph** | 0.00536 | 0.00386 | 0.01413 | 0.00298 | 0.00982 | 0.02293 | 0.00375 | 0.00302 | 0.01116 | 0.76068 | 0.00530 | 0.00741 | 0.01517 | 0.02178 |
| **Prph2** | 0.00315 | 0.00428 | 0.01510 | 0.01225 | 0.00887 | 0.01337 | 0.00420 | 0.00873 | 0.01074 | 0.00646 | 0.00911 | 0.00877 | 0.67012 | 0.00231 |
| **Prps1** | 35.84859 | 4.10538 | 39.67051 | 24.44612 | 41.28979 | 25.77527 | 7.47499 | 12.17486 | 40.98340 | 3.32915 | 15.69748 | 17.26718 | 16.91413 | 40.09788 |
| **Prps1l3** | 5.95321 | 0.12184 | 13.50666 | 7.62091 | 12.12621 | 5.63540 | 2.54688 | 2.38501 | 14.74926 | 0.06875 | 6.32651 | 5.11878 | 12.52028 | 16.51051 |
| **Prps2** | 0.09898 | 0.01434 | 0.01572 | 0.40360 | 0.31278 | 0.58549 | 0.00381 | 0.04158 | 0.01391 | 0.00687 | 1.35320 | 1.21725 | 0.19387 | 0.12080 |
| **Prpsap1** | 17.09185 | 0.43346 | 14.89433 | 5.76567 | 13.40289 | 6.24109 | 3.66794 | 10.26057 | 0.47215 | 10.17419 | 1.06424 | 12.19547 | 10.46070 | 10.26264 |
| **Prpsap2** | 14.79343 | 7.44797 | 23.70699 | 14.05907 | 21.05062 | 20.17274 | 8.42107 | 5.89116 | 11.72701 | 0.55264 | 18.52079 | 9.76127 | 5.39216 | 21.69137 |
| **Prr11** | 3.02385 | 0.33057 | 0.21987 | 1.51401 | 6.37016 | 0.73189 | 2.15005 | 1.14393 | 11.33966 | 0.13064 | 0.57131 | 0.73105 | 8.33314 | 7.08583 |
| **Prr12** | 0.53064 | 0.01264 | 1.04592 | 0.00000 | 0.97986 | 0.61760 | 0.00391 | 0.96348 | 0.24541 | 0.02817 | 0.02530 | 0.67487 | 2.10532 | 0.48933 |
| **Prr13** | 13.23758 | 0.33463 | 27.23796 | 21.63309 | 23.45575 | 13.92591 | 3.64194 | 13.37464 | 10.28227 | 5.23730 | 14.82375 | 9.06213 | 9.46654 | 25.08682 |
| **Prr14** | 3.29464 | 0.59577 | 7.47883 | 1.15782 | 1.95442 | 2.28500 | 7.33264 | 0.87501 | 0.04760 | 0.38174 | 4.09916 | 3.30419 | 0.51197 | 3.71719 |
| **Prr14l** | 1.43228 | 0.04317 | 2.63720 | 1.91803 | 4.62323 | 3.01389 | 2.79638 | 3.07787 | 4.68544 | 0.11418 | 0.07862 | 2.03706 | 6.53322 | 3.49444 |
| **Prr15l** | 0.00204 | 0.00384 | 0.00000 | 1.14287 | 1.75802 | 0.00000 | 2.47834 | 0.00053 | 0.02107 | 0.04501 | 0.55539 | 0.00459 | 0.54146 | 0.25380 |
| **Prr18** | 0.32114 | 0.00393 | 0.13454 | 0.24884 | 0.00000 | 0.05684 | 0.50091 | 0.00222 | 0.02048 | 0.00653 | 0.00749 | 0.00953 | 0.41703 | 0.00000 |
| **Prr19** | 1.08287 | 0.00413 | 0.28085 | 0.34718 | 1.14981 | 0.02034 | 0.00403 | 0.00476 | 0.01175 | 0.00677 | 0.00637 | 0.00735 | 0.00472 | 0.00000 |
| **Prr22** | 0.02941 | 0.00370 | 0.46016 | 0.12087 | 0.00000 | 0.03838 | 0.00394 | 0.35269 | 0.01158 | 0.00629 | 0.00131 | 0.50969 | 0.00563 | 0.00664 |
| **Prr23a1** | 4.36351 | 0.00343 | 0.31712 | 0.00000 | 2.37437 | 0.12145 | 0.00384 | 0.00317 | 0.00756 | 0.00497 | 0.01293 | 0.00469 | 4.56066 | 0.20031 |
| **Prr23a2** | 0.47280 | 0.00433 | 0.06672 | 0.00877 | 0.27421 | 0.03109 | 0.00418 | 0.00688 | 0.00782 | 0.01275 | 0.01466 | 0.00997 | 0.20138 | 0.01899 |
| **Prr23a3** | 1.09764 | 0.00216 | 1.10803 | 0.39313 | 1.76344 | 0.00000 | 0.00732 | 0.79311 | 0.11886 | 9.49908 | 0.00000 | 0.00000 | 0.75123 | 0.60871 |
| **Prr3** | 0.18032 | 0.05220 | 2.79353 | 5.98154 | 2.30429 | 3.67267 | 1.21164 | 2.17445 | 0.96149 | 0.04727 | 1.46023 | 1.39391 | 0.03738 | 5.27262 |
| **Prr36** | 0.02161 | 0.00439 | 0.23255 | 0.00336 | 0.00330 | 0.01046 | 0.00423 | 0.00921 | 0.01281 | 0.00665 | 0.00696 | 0.00844 | 0.47891 | 0.00627 |
| **Prr5** | 0.30101 | 0.02819 | 0.49803 | 2.34319 | 0.94023 | 0.89189 | 2.42878 | 3.48310 | 0.01073 | 0.09038 | 0.02850 | 0.29179 | 0.02513 | 0.19227 |
| **Prr5l** | 0.00518 | 0.00421 | 0.00443 | 0.28771 | 0.36792 | 0.02711 | 0.02160 | 0.01390 | 0.26990 | 0.00568 | 0.00965 | 0.01094 | 0.17414 | 0.18269 |
| **Prrc1** | 0.16918 | 0.06458 | 3.67142 | 2.52058 | 2.86879 | 2.75220 | 0.22779 | 1.44798 | 0.00000 | 0.06170 | 6.69022 | 5.75431 | 0.00000 | 2.51295 |
| **Prrc2a** | 0.39367 | 1.94371 | 6.98044 | 4.52101 | 6.66534 | 0.56636 | 0.09627 | 2.81873 | 0.08452 | 0.04512 | 2.40433 | 3.01561 | 1.32737 | 0.36545 |
| **Prrc2b** | 0.93404 | 0.15686 | 3.63825 | 3.67288 | 4.67658 | 4.03744 | 0.13678 | 4.26318 | 1.38997 | 3.44624 | 3.63382 | 7.04113 | 0.66571 | 1.54106 |
| **Prrc2c** | 0.00000 | 0.02512 | 9.11902 | 4.79262 | 9.67824 | 6.40582 | 0.00497 | 3.08368 | 17.04096 | 0.02164 | 3.76017 | 1.58973 | 12.46143 | 8.75859 |
| **Prrg1** | 0.01855 | 0.00445 | 0.01413 | 0.01485 | 0.01611 | 0.01276 | 0.00428 | 0.01035 | 0.01176 | 0.00739 | 0.00864 | 0.00979 | 0.00782 | 0.01132 |
| **Prrg2** | 0.07266 | 0.03470 | 1.98720 | 0.81625 | 1.53665 | 2.14087 | 1.58384 | 1.34196 | 0.00397 | 0.10091 | 0.84066 | 1.46066 | 0.30130 | 0.18034 |
| **Prrg4** | 0.00444 | 0.00396 | 0.00551 | 0.28161 | 0.09412 | 0.04390 | 0.00411 | 0.04748 | 0.00489 | 0.01645 | 0.00926 | 0.01025 | 0.00386 | 0.21053 |
| **Prrt3** | 0.00474 | 0.00478 | 0.00988 | 0.19420 | 0.00910 | 0.00912 | 0.00449 | 0.01699 | 0.00603 | 0.01348 | 0.00734 | 0.00734 | 0.08702 | 0.12563 |
| **Prrt4** | 0.57516 | 0.00415 | 0.26283 | 0.06652 | 0.80043 | 0.00000 | 0.00398 | 0.00731 | 0.01365 | 0.00461 | 0.29983 | 0.01453 | 0.00684 | 0.15099 |
| **Prrx1** | 0.15379 | 0.00438 | 0.01953 | 0.02076 | 0.01135 | 0.01398 | 0.00425 | 0.02384 | 0.05065 | 0.00678 | 0.00833 | 0.00819 | 0.01869 | 0.02902 |
| **Prrx2** | 0.00000 | 0.00417 | 0.22758 | 0.00369 | 0.00018 | 0.00643 | 0.00407 | 0.00941 | 0.01965 | 0.00658 | 0.00725 | 0.00773 | 1.33122 | 0.00595 |
| **Prss16** | 0.21626 | 0.00366 | 0.00000 | 0.00000 | 0.00543 | 0.66409 | 0.00368 | 0.00852 | 0.02799 | 0.00508 | 0.00000 | 0.62143 | 0.18705 | 0.00000 |
| **Prss23** | 0.17451 | 0.00448 | 0.01043 | 0.01083 | 0.08928 | 0.01149 | 0.00453 | 0.01093 | 0.01048 | 0.00655 | 0.00749 | 0.00876 | 0.00492 | 0.00797 |
| **Prss3** | 0.00876 | 0.00435 | 0.01583 | 0.06436 | 0.01051 | 0.01049 | 0.00426 | 0.00822 | 0.01592 | 0.00616 | 0.01091 | 0.01104 | 0.01054 | 0.02618 |
| **Prss32** | 0.00781 | 6.74674 | 0.01497 | 0.03362 | 0.00375 | 0.00000 | 2.61479 | 0.27226 | 0.00487 | 0.09848 | 0.01128 | 0.01166 | 0.00193 | 0.15545 |
| **Prss35** | 0.14039 | 0.01128 | 0.03186 | 0.00000 | 0.09092 | 0.60448 | 0.14342 | 0.04312 | 0.16179 | 0.03378 | 0.07139 | 1.98522 | 0.09533 | 0.08589 |
| **Prss36** | 1.34890 | 0.00993 | 1.12099 | 1.77618 | 0.97423 | 2.79644 | 0.00336 | 0.00000 | 0.00326 | 0.00587 | 1.61544 | 1.23807 | 0.46759 | 0.33772 |
| **Prss38** | 0.03714 | 0.00387 | 0.08178 | 1.16677 | 0.00000 | 0.66975 | 0.00359 | 0.00420 | 0.00664 | 0.01638 | 0.00000 | 0.87334 | 0.03283 | 0.14121 |
| **Prss41** | 0.00863 | 0.00395 | 0.00892 | 0.03805 | 0.01112 | 0.07998 | 0.00371 | 0.68373 | 0.01214 | 0.00593 | 0.00495 | 0.01251 | 0.00693 | 0.01045 |
| **Prss45** | 0.01393 | 0.00441 | 0.00426 | 0.00142 | 0.00000 | 0.37638 | 0.00412 | 0.00466 | 0.06165 | 0.00596 | 0.00614 | 0.00990 | 0.01373 | 0.82743 |
| **Prss48** | 0.00703 | 0.00443 | 0.02699 | 0.00532 | 0.06967 | 0.03715 | 0.00417 | 0.00862 | 0.01710 | 0.00794 | 0.00844 | 0.00989 | 0.00352 | 0.07813 |
| **Prss53** | 0.00641 | 0.00408 | 0.00566 | 0.18581 | 0.00896 | 0.10831 | 0.00383 | 0.50405 | 0.01198 | 0.00775 | 0.00349 | 0.01082 | 0.00702 | 0.00912 |
| **Prss57** | 0.01071 | 0.00432 | 0.04754 | 0.00917 | 0.00759 | 0.01233 | 0.00421 | 0.00917 | 0.01092 | 0.00666 | 0.00860 | 0.00848 | 0.61305 | 0.00532 |
| **Prss8** | 0.00284 | 5.91822 | 0.00000 | 0.93776 | 0.00263 | 3.80766 | 4.15968 | 13.03820 | 0.00000 | 1.53554 | 1.48569 | 2.12447 | 0.06281 | 0.28500 |
| **Prtg** | 0.00000 | 0.01468 | 0.00000 | 0.04124 | 0.00030 | 0.75876 | 0.00417 | 0.48049 | 0.55138 | 0.02091 | 0.28961 | 0.88210 | 0.00529 | 0.00000 |
| **Prtn3** | 0.00000 | 0.00467 | 0.38206 | 0.88150 | 0.73524 | 0.00000 | 0.00408 | 0.08387 | 0.02589 | 0.01582 | 0.00558 | 0.00428 | 0.00792 | 0.00751 |
| **Prune** | 6.26453 | 0.23367 | 7.24551 | 6.20732 | 7.23774 | 7.19436 | 4.28722 | 7.86701 | 8.29995 | 0.13664 | 8.23058 | 2.42058 | 3.51105 | 12.11246 |
| **Prune2** | 0.03697 | 0.00434 | 0.01470 | 0.01118 | 0.00984 | 0.01310 | 0.00427 | 0.00933 | 0.01050 | 0.00655 | 0.00845 | 0.01229 | 0.30181 | 0.00350 |

| | | | | | | | | | | | | | | |
|---|---|---|---|---|---|---|---|---|---|---|---|---|---|---|
| **Prx** | 0.35501 | 0.01471 | 1.06546 | 0.98687 | 0.65919 | 0.00000 | 0.00426 | 0.51368 | 0.02534 | 0.01143 | 0.11424 | 0.64891 | 0.48699 | 0.62573 |
| **Psap** | 5.50489 | 12.55293 | 23.11572 | 30.34558 | 10.80631 | 30.32065 | 6.90073 | 29.89034 | 3.15066 | 7.65033 | 17.40424 | 13.36050 | 7.03127 | 13.87660 |
| **Psat1** | 31.91520 | 6.93847 | 48.17662 | 45.20843 | 39.27471 | 31.97950 | 7.78082 | 4.17656 | 36.95028 | 1.74829 | 25.90234 | 16.18485 | 13.36834 | 36.85584 |
| **Psd** | 0.03604 | 0.00443 | 0.13913 | 0.00393 | 0.00807 | 0.00688 | 0.00451 | 0.02013 | 0.01010 | 0.00652 | 0.00831 | 0.01467 | 0.00543 | 0.02036 |
| **Psd2** | 0.30832 | 0.00422 | 0.26962 | 0.00000 | 0.16120 | 0.00026 | 0.00392 | 0.01007 | 0.51368 | 0.00660 | 0.00471 | 0.00568 | 0.56117 | 0.80710 |
| **Psd3** | 0.04927 | 0.01970 | 0.02391 | 0.03075 | 0.01274 | 0.02097 | 0.01630 | 0.03658 | 0.06921 | 0.00654 | 0.01876 | 0.00928 | 0.41867 | 0.40152 |
| **Psd4** | 0.18367 | 0.03326 | 1.46029 | 5.08943 | 3.26555 | 2.39624 | 3.03594 | 3.94512 | 0.23025 | 0.16167 | 1.86469 | 0.29737 | 0.08553 | 1.54750 |
| **Psen1** | 3.17108 | 5.26776 | 8.57978 | 4.39807 | 3.55105 | 10.13655 | 3.39900 | 6.34695 | 6.38360 | 0.13794 | 7.01390 | 4.06588 | 0.64422 | 5.31459 |
| **Psen2** | 0.02337 | 0.00391 | 0.00210 | 0.00000 | 0.31762 | 0.12633 | 0.00330 | 2.78317 | 0.00649 | 0.01767 | 0.00342 | 0.01636 | 0.23505 | 0.50438 |
| **Psenen** | 1.33948 | 5.86297 | 0.48144 | 5.26139 | 2.38625 | 5.62665 | 3.25719 | 9.93569 | 2.79169 | 13.60634 | 0.98647 | 5.16297 | 0.40532 | 0.40014 |
| **Psg16** | 10.42591 | 1.44440 | 14.20891 | 4.34756 | 24.46936 | 3.57382 | 0.62510 | 1.14862 | 41.79447 | 2.44796 | 2.87598 | 2.55964 | 21.18157 | 51.75554 |
| **Psg17** | 0.00745 | 0.00439 | 0.01438 | 0.01154 | 0.02721 | 0.01312 | 0.00424 | 0.00974 | 0.00956 | 0.00633 | 0.00936 | 0.00964 | 0.17001 | 0.01555 |
| **Psg23** | 8.80222 | 0.53496 | 2.36878 | 0.49903 | 4.90621 | 4.00819 | 0.50183 | 0.73681 | 3.89685 | 0.26968 | 0.93586 | 0.82693 | 10.16100 | 1.51232 |
| **Psg28** | 0.01780 | 0.03332 | 0.05235 | 0.02554 | 0.01420 | 0.01528 | 0.00450 | 0.05752 | 0.21745 | 0.00788 | 0.01332 | 0.01980 | 0.03301 | 0.00000 |
| **Psg29** | 2.09095 | 0.12732 | 0.41738 | 0.11635 | 0.69968 | 0.84598 | 0.16369 | 0.14642 | 0.71746 | 0.04785 | 0.15939 | 0.10985 | 1.78292 | 0.31359 |
| **Psip1** | 0.24522 | 0.02046 | 1.82664 | 0.00000 | 1.67384 | 1.30933 | 0.03112 | 2.97949 | 12.26463 | 3.15602 | 5.05244 | 6.59109 | 5.23772 | 9.30674 |
| **Pskh1** | 0.06843 | 0.04345 | 1.06003 | 0.00000 | 0.48272 | 2.55901 | 1.51205 | 2.19098 | 0.89716 | 0.05309 | 0.00000 | 1.97427 | 1.49956 | 2.84436 |
| **Psma1** | 20.43737 | 9.58819 | 23.86750 | 24.26641 | 21.62934 | 35.36566 | 9.91047 | 17.58098 | 16.11800 | 15.98640 | 19.41376 | 23.43257 | 1.61768 | 15.14357 |
| **Psma2** | 20.07966 | 6.94062 | 18.61355 | 29.53136 | 19.83246 | 29.22006 | 7.01502 | 19.41635 | 22.48540 | 4.39631 | 21.02328 | 22.35464 | 7.53063 | 16.14403 |
| **Psma3** | 20.07045 | 0.45795 | 24.22547 | 22.64975 | 20.43470 | 32.78793 | 9.07421 | 11.46170 | 17.39959 | 11.12940 | 19.55951 | 21.48377 | 8.95271 | 14.74476 |
| **Psma4** | 21.84594 | 7.28208 | 29.98804 | 26.94548 | 20.06887 | 30.93064 | 10.26763 | 20.58772 | 32.55713 | 18.66880 | 25.87256 | 24.34634 | 10.28397 | 17.96602 |
| **Psma5** | 11.71622 | 0.52409 | 22.38420 | 16.54905 | 15.54989 | 30.62470 | 8.06674 | 12.34283 | 14.67928 | 14.98214 | 22.87531 | 20.22799 | 3.07275 | 16.03392 |
| **Psma6** | 31.62250 | 9.68345 | 35.04112 | 33.85299 | 34.38641 | 38.12991 | 9.71991 | 18.90639 | 30.62848 | 18.34882 | 24.55145 | 21.22149 | 8.40040 | 24.90742 |
| **Psma7** | 13.95999 | 7.81771 | 28.87657 | 30.96302 | 19.29962 | 33.92777 | 10.56042 | 22.46748 | 7.62419 | 21.31459 | 21.59332 | 22.94567 | 0.51463 | 13.48141 |
| **Psma8** | 0.12389 | 0.00427 | 0.00784 | 0.01347 | 0.00551 | 0.01106 | 0.00410 | 0.00850 | 0.10424 | 0.00473 | 0.61166 | 0.00594 | 0.00411 | 0.00000 |
| **Psmb1** | 19.64156 | 7.53435 | 20.98954 | 31.30376 | 22.69136 | 24.36313 | 9.31616 | 20.65325 | 12.06490 | 17.47277 | 17.94711 | 21.98975 | 8.14176 | 21.28449 |
| **Psmb10** | 0.98594 | 0.03461 | 0.43790 | 1.63285 | 7.36832 | 1.17260 | 0.00354 | 0.45094 | 0.05022 | 0.97153 | 0.29482 | 1.38644 | 1.27949 | 0.36226 |
| **Psmb11** | 0.05967 | 0.02943 | 0.41464 | 1.47875 | 0.23091 | 0.34014 | 0.06128 | 0.08072 | 0.46318 | 0.02614 | 0.01436 | 0.10912 | 0.32606 | 0.40770 |
| **Psmb2** | 6.89085 | 2.98226 | 19.71408 | 27.50496 | 18.60158 | 24.15082 | 8.93893 | 18.51278 | 7.03742 | 9.54296 | 17.93687 | 19.81545 | 0.43865 | 19.87321 |
| **Psmb3** | 10.73886 | 7.91313 | 7.73113 | 25.16476 | 16.54279 | 25.17251 | 6.09713 | 15.57904 | 6.31030 | 16.07469 | 10.58446 | 20.64785 | 2.70442 | 8.35887 |
| **Psmb4** | 14.40969 | 3.96462 | 19.12904 | 39.20392 | 30.01062 | 19.19470 | 8.42353 | 19.35944 | 4.78631 | 21.04858 | 7.75914 | 24.21677 | 10.73195 | 5.76073 |
| **Psmb5** | 16.71570 | 8.04104 | 25.73273 | 28.53047 | 20.43867 | 20.83296 | 9.67837 | 19.49828 | 1.92236 | 15.83698 | 16.05764 | 17.44039 | 12.49944 | 3.16915 |
| **Psmb6** | 14.95913 | 8.30751 | 17.69045 | 31.48504 | 19.95700 | 25.38886 | 11.38266 | 21.86435 | 2.41342 | 15.86495 | 20.09866 | 20.26154 | 3.39638 | 14.79051 |
| **Psmb7** | 22.46087 | 7.84278 | 26.31630 | 21.78942 | 18.50226 | 20.70971 | 6.10955 | 14.27481 | 24.43252 | 1.54299 | 19.81788 | 17.82438 | 5.25180 | 18.95065 |
| **Psmb8** | 0.01614 | 0.00450 | 0.00956 | 0.01271 | 0.21173 | 0.01321 | 0.00448 | 0.01057 | 0.01051 | 0.00659 | 0.00872 | 0.00885 | 0.00548 | 0.01450 |
| **Psmb9** | 0.00000 | 0.01307 | 0.12789 | 0.66939 | 0.00293 | 0.00000 | 0.00400 | 0.08252 | 0.01384 | 0.00756 | 0.29675 | 1.36392 | 0.00577 | 0.01181 |
| **Psmc1** | 17.00161 | 3.94743 | 29.93079 | 27.37252 | 21.54047 | 27.00829 | 9.53740 | 13.26723 | 23.29201 | 17.25659 | 21.28381 | 20.50078 | 6.88103 | 22.36352 |
| **Psmc2** | 21.61854 | 6.71188 | 35.45882 | 24.78785 | 24.13551 | 30.62108 | 10.27358 | 13.92861 | 19.18490 | 2.03847 | 23.50088 | 22.48321 | 3.01819 | 20.67499 |
| **Psmc3** | 9.68257 | 6.18062 | 30.59615 | 30.06132 | 13.44884 | 30.91973 | 11.65026 | 18.94729 | 6.66253 | 16.19128 | 21.01493 | 20.39385 | 1.76960 | 12.29782 |
| **Psmc3ip** | 4.16131 | 0.01113 | 5.02919 | 2.39301 | 7.38617 | 0.71590 | 0.00295 | 0.40900 | 1.06892 | 0.02708 | 1.67662 | 1.69906 | 5.51117 | 6.44556 |
| **Psmc4** | 17.90524 | 3.40399 | 36.38115 | 29.51500 | 25.17273 | 33.02590 | 8.38708 | 14.73236 | 5.59500 | 16.39337 | 20.04332 | 19.53086 | 4.64343 | 14.37807 |
| **Psmc5** | 17.23854 | 3.01717 | 31.24047 | 29.51882 | 14.01208 | 27.06204 | 10.00015 | 19.78231 | 11.22337 | 17.95081 | 19.76409 | 18.81273 | 1.72151 | 18.95565 |
| **Psmc6** | 15.34666 | 4.10746 | 24.00303 | 21.99251 | 15.63857 | 29.37035 | 6.93925 | 11.33462 | 26.90615 | 0.62744 | 22.40357 | 18.45968 | 4.06100 | 12.44453 |
| **Psmd1** | 11.27067 | 6.33896 | 23.33654 | 17.54795 | 15.23302 | 26.64476 | 5.07417 | 11.90086 | 16.28984 | 0.49520 | 21.23263 | 18.20305 | 2.67724 | 18.81100 |
| **Psmd10** | 1.45676 | 0.00577 | 0.62710 | 3.41790 | 1.45851 | 5.59844 | 0.59913 | 3.95697 | 2.43747 | 0.06644 | 7.98447 | 4.59242 | 0.00000 | 3.44327 |
| **Psmd11** | 13.59879 | 6.13825 | 29.19059 | 27.84511 | 22.32054 | 29.03472 | 9.12862 | 5.78517 | 21.52817 | 14.73209 | 18.68111 | 20.26799 | 7.98769 | 21.41780 |
| **Psmd12** | 24.86414 | 7.45688 | 30.57326 | 27.56084 | 24.75151 | 29.11071 | 7.47200 | 13.58765 | 13.12020 | 1.56399 | 19.31365 | 18.82710 | 8.42559 | 17.68198 |
| **Psmd13** | 5.82462 | 6.12165 | 22.34066 | 23.46292 | 15.04351 | 20.27448 | 7.44534 | 13.22484 | 8.90187 | 12.38767 | 15.23787 | 15.93872 | 0.81092 | 18.46310 |
| **Psmd14** | 18.93892 | 8.76503 | 24.74825 | 20.03621 | 22.39063 | 29.68625 | 7.66316 | 11.95183 | 16.46962 | 4.88417 | 19.75916 | 17.09158 | 11.78958 | 18.13417 |
| **Psmd2** | 18.09062 | 10.77075 | 25.08618 | 24.12789 | 12.91621 | 29.28761 | 7.73263 | 11.96879 | 12.55286 | 7.98560 | 21.46313 | 17.55921 | 3.02512 | 11.46600 |
| **Psmd3** | 15.87812 | 5.62729 | 34.70853 | 28.53741 | 25.46652 | 26.03182 | 6.61311 | 14.03464 | 0.00000 | 0.61963 | 14.93560 | 16.57490 | 5.32477 | 14.47340 |
| **Psmd4** | 14.31679 | 5.45537 | 36.23877 | 35.09030 | 29.54734 | 32.47478 | 9.07072 | 22.58153 | 1.26110 | 9.32278 | 21.84141 | 21.46894 | 12.51279 | 11.12011 |
| **Psmd5** | 5.71725 | 0.03552 | 5.14169 | 8.66710 | 9.04396 | 7.60978 | 2.76822 | 2.59099 | 5.81672 | 0.10970 | 6.40698 | 6.88752 | 3.81246 | 4.81566 |
| **Psmd6** | 21.47528 | 0.32325 | 25.99319 | 24.44924 | 18.13945 | 17.36748 | 7.02194 | 16.53922 | 11.73742 | 0.44986 | 13.28603 | 15.85108 | 9.11245 | 5.67050 |
| **Psmd7** | 16.47291 | 6.90547 | 31.33863 | 25.94563 | 31.45292 | 25.29410 | 8.53175 | 14.10582 | 16.48272 | 0.78296 | 18.44442 | 17.81621 | 11.51512 | 23.57760 |

| | | | | | | | | | | | | | | |
|---|---|---|---|---|---|---|---|---|---|---|---|---|---|---|
| **Psmd8** | 17.30213 | 5.10072 | 26.51599 | 34.46818 | 28.46506 | 19.79233 | 2.96342 | 14.43446 | 7.28455 | 6.61079 | 8.45919 | 13.21328 | 5.92404 | 12.16697 |
| **Psmd9** | 6.54715 | 7.93631 | 11.25259 | 9.32610 | 6.65551 | 5.10806 | 2.80790 | 0.65231 | 0.12196 | 15.28406 | 9.53794 | 3.89632 | 2.00222 | 2.13244 |
| **Psme1** | 6.28408 | 7.65882 | 12.94730 | 13.40724 | 14.83676 | 7.12998 | 0.00876 | 8.68254 | 0.01326 | 0.09808 | 6.28303 | 3.75784 | 0.69286 | 2.42029 |
| **Psme2** | 0.67676 | 0.04551 | 0.75792 | 0.00000 | 1.65311 | 3.04524 | 1.08902 | 3.58486 | 0.95441 | 6.36824 | 3.66093 | 4.59752 | 2.04827 | 3.03833 |
| **Psme2b** | 0.18245 | 0.01833 | 0.12301 | 0.00000 | 0.29400 | 0.65117 | 0.20008 | 0.70133 | 0.15759 | 1.58785 | 0.74782 | 0.97737 | 0.65940 | 0.59404 |
| **Psme3** | 11.11498 | 0.11469 | 15.42621 | 8.15279 | 15.50376 | 15.69473 | 3.16440 | 5.05589 | 1.77116 | 0.18513 | 14.12111 | 11.15786 | 11.36942 | 6.18277 |
| **Psme4** | 8.04620 | 2.54530 | 4.96468 | 1.67766 | 4.34394 | 5.07556 | 0.82374 | 2.73213 | 11.73443 | 0.04329 | 5.58364 | 6.84423 | 7.07574 | 5.67718 |
| **Psmf1** | 5.11993 | 0.07738 | 7.97057 | 5.79427 | 1.42823 | 6.07564 | 6.15141 | 5.01656 | 1.36339 | 0.23584 | 8.72026 | 5.18053 | 2.22314 | 2.35197 |
| **Psmg1** | 1.50912 | 0.27590 | 8.24039 | 2.91498 | 5.35754 | 5.96459 | 2.83174 | 5.07407 | 2.02765 | 10.05139 | 1.74843 | 7.52478 | 0.58222 | 1.44432 |
| **Psmg2** | 10.66561 | 0.11026 | 15.81443 | 9.82682 | 9.39674 | 16.05813 | 3.41171 | 6.21295 | 1.91398 | 15.87896 | 8.96340 | 14.97949 | 1.47591 | 7.62012 |
| **Psmg3** | 5.09766 | 0.09878 | 8.50230 | 6.43025 | 5.69633 | 6.54491 | 4.59874 | 3.25799 | 0.82282 | 0.12890 | 9.81978 | 5.19577 | 1.53647 | 10.97908 |
| **Psmg4** | 0.10987 | 0.05075 | 0.00000 | 1.30611 | 1.53079 | 0.87514 | 0.84198 | 1.82272 | 0.00692 | 0.07942 | 0.00000 | 1.44875 | 0.02753 | 0.03121 |
| **Psors1c2** | 0.04683 | 0.00417 | 0.27026 | 0.00891 | 0.49226 | 0.00303 | 0.00394 | 0.00848 | 0.14880 | 0.01304 | 0.00538 | 0.00620 | 0.96122 | 0.19811 |
| **Pspc1** | 2.06085 | 0.04183 | 2.04907 | 2.02279 | 3.14324 | 2.72405 | 1.11758 | 2.02515 | 1.00558 | 0.03314 | 3.62858 | 1.48567 | 1.95123 | 2.89227 |
| **Psph** | 4.83477 | 0.08878 | 9.62177 | 5.70593 | 7.37238 | 2.79473 | 2.89053 | 1.70341 | 1.74503 | 0.01161 | 8.00026 | 2.08567 | 0.49004 | 6.73812 |
| **Psrc1** | 0.79279 | 0.06030 | 8.49579 | 5.51261 | 1.95221 | 2.55577 | 3.06495 | 2.52526 | 0.00000 | 0.06803 | 4.67156 | 12.65583 | 0.00000 | 1.13377 |
| **Pstk** | 2.08149 | 0.02484 | 0.68061 | 2.95784 | 0.45023 | 1.36797 | 3.75408 | 1.99671 | 0.00298 | 0.10753 | 0.18760 | 2.49885 | 0.34204 | 0.12852 |
| **Pstpip1** | 0.00472 | 0.63259 | 0.49888 | 1.00169 | 0.00000 | 0.38249 | 0.00408 | 0.00176 | 0.00407 | 0.00554 | 0.00284 | 0.01966 | 0.00128 | 0.45846 |
| **Pstpip2** | 0.00873 | 0.00417 | 0.23714 | 0.03266 | 0.08726 | 0.03920 | 0.00407 | 0.02505 | 0.01994 | 0.00523 | 0.03374 | 0.00679 | 0.14764 | 0.53227 |
| **Ptafr** | 0.07114 | 0.02087 | 0.03503 | 0.15488 | 0.07801 | 0.49461 | 0.04765 | 0.11696 | 0.18035 | 0.07395 | 0.03168 | 0.10040 | 0.93816 | 0.23813 |
| **Ptar1** | 0.05610 | 0.03518 | 0.58706 | 0.00000 | 0.00000 | 0.91473 | 0.00304 | 0.16458 | 2.01894 | 0.00509 | 0.00000 | 1.32610 | 0.01235 | 1.03050 |
| **Ptbp1** | 22.37017 | 7.17620 | 41.98143 | 23.25962 | 35.96905 | 28.27503 | 10.38157 | 19.94946 | 8.35665 | 12.75859 | 16.92902 | 21.82633 | 12.40051 | 23.69568 |
| **Ptbp2** | 0.33570 | 0.00344 | 0.10694 | 0.00177 | 0.32733 | 0.24483 | 0.56709 | 0.00358 | 0.21277 | 0.02073 | 0.29401 | 0.38001 | 2.10320 | 0.55485 |
| **Ptbp3** | 7.62499 | 1.35315 | 5.26723 | 1.40292 | 2.49403 | 16.33695 | 1.80897 | 4.74657 | 7.48560 | 0.09782 | 6.64361 | 11.07371 | 3.53751 | 0.00000 |
| **Ptcd1** | 0.28597 | 0.03572 | 1.38166 | 1.31322 | 0.80748 | 0.00000 | 2.70345 | 1.82436 | 0.06202 | 0.12827 | 0.48324 | 1.55048 | 0.00000 | 1.25131 |
| **Ptcd2** | 2.76932 | 1.38349 | 5.68510 | 7.24959 | 4.98591 | 5.66467 | 5.12581 | 8.54673 | 2.98102 | 0.16115 | 2.77780 | 5.19176 | 0.00000 | 4.21909 |
| **Ptcd3** | 7.58153 | 1.56323 | 15.27551 | 6.12637 | 12.62244 | 11.57270 | 1.34725 | 2.19677 | 15.42031 | 7.51795 | 14.04150 | 9.86520 | 4.13864 | 14.06746 |
| **Ptch1** | 0.21789 | 0.01112 | 0.24349 | 0.22194 | 0.00000 | 0.70054 | 0.02656 | 0.18055 | 0.00797 | 0.03485 | 1.76826 | 1.10875 | 0.01489 | 0.04443 |
| **Ptch2** | 0.00000 | 0.00436 | 0.39750 | 0.59531 | 0.32814 | 0.07072 | 0.00362 | 0.22356 | 0.00602 | 0.01090 | 0.81207 | 0.05813 | 0.00247 | 0.09050 |
| **Ptcra** | 0.14110 | 0.01963 | 0.02323 | 0.06844 | 0.10498 | 0.27886 | 0.02800 | 0.02085 | 0.27351 | 0.06603 | 0.01792 | 0.06583 | 2.29475 | 0.16415 |
| **Ptdss1** | 7.65418 | 7.89035 | 14.89590 | 9.13333 | 8.97136 | 16.44352 | 4.41567 | 5.77732 | 11.10924 | 1.15258 | 8.28701 | 12.75826 | 0.95260 | 9.79157 |
| **Ptdss2** | 16.69882 | 0.14515 | 17.47343 | 16.62416 | 18.47580 | 5.27606 | 0.02411 | 6.88998 | 5.54358 | 8.08060 | 3.82056 | 9.17417 | 9.58968 | 32.52875 |
| **Pten** | 10.22021 | 0.28440 | 6.01162 | 3.76291 | 10.86531 | 4.64913 | 0.39729 | 0.96190 | 18.42922 | 0.01922 | 4.34323 | 3.32655 | 6.54618 | 5.35643 |
| **Pter** | 0.00270 | 0.00268 | 0.02834 | 0.00000 | 0.00456 | 0.41952 | 0.00353 | 0.00345 | 0.00529 | 0.00758 | 0.00000 | 1.83203 | 0.00751 | 0.35676 |
| **Ptger1** | 0.22235 | 0.00409 | 0.01044 | 0.39498 | 0.16455 | 0.28098 | 0.00376 | 0.00857 | 0.01082 | 0.00507 | 0.00609 | 0.01034 | 0.00536 | 0.00000 |
| **Ptger3** | 0.00280 | 0.00418 | 0.00764 | 0.17816 | 0.00000 | 0.00310 | 0.00398 | 0.00670 | 0.17347 | 0.00562 | 0.01809 | 0.01700 | 0.01084 | 0.25402 |
| **Ptger4** | 0.00434 | 0.00379 | 0.01556 | 0.17828 | 0.00000 | 0.00480 | 0.00392 | 0.00597 | 0.00708 | 0.00516 | 0.01732 | 0.01233 | 2.24758 | 0.59383 |
| **Ptges** | 7.22500 | 13.37974 | 20.31495 | 37.59686 | 0.00000 | 43.41385 | 13.04903 | 29.11591 | 0.60253 | 2.19248 | 23.52712 | 20.55257 | 2.81413 | 0.26689 |
| **Ptges2** | 0.60051 | 0.12193 | 3.39339 | 2.90151 | 4.86916 | 6.01857 | 1.25642 | 2.44725 | 0.13676 | 0.89334 | 2.60777 | 2.18814 | 1.18484 | 2.07612 |
| **Ptges3** | 18.17103 | 2.54811 | 13.69969 | 21.46719 | 16.82842 | 22.83222 | 4.37977 | 13.14985 | 36.09184 | 4.55052 | 12.93527 | 11.41943 | 8.73518 | 10.11805 |
| **Ptges3l** | 1.76942 | 1.12516 | 0.57956 | 1.21515 | 1.27786 | 0.00000 | 0.00442 | 0.46522 | 0.01664 | 0.00717 | 0.00027 | 0.00000 | 0.17929 | 0.37053 |
| **Ptgfrn** | 0.00000 | 0.00885 | 0.94693 | 0.00000 | 0.00000 | 5.27615 | 0.00250 | 0.85533 | 0.00000 | 0.37691 | 0.17671 | 4.42375 | 0.00111 | 0.20302 |
| **Ptgis** | 8.03878 | 0.05021 | 16.48721 | 12.69179 | 10.77044 | 1.66065 | 2.62532 | 2.17432 | 1.03169 | 0.20248 | 9.60765 | 2.65337 | 0.17974 | 12.82085 |
| **Ptgr1** | 0.20124 | 15.18536 | 8.64134 | 12.98330 | 1.38392 | 24.02510 | 12.29724 | 33.23957 | 2.34763 | 23.50362 | 17.52547 | 11.39148 | 0.00000 | 1.85818 |
| **Ptgr2** | 0.02986 | 0.03915 | 1.07642 | 2.95647 | 0.90332 | 2.97083 | 3.10061 | 1.72721 | 1.06494 | 0.12534 | 3.92834 | 1.99564 | 0.07300 | 2.42674 |
| **Ptgs1** | 0.01175 | 0.00412 | 0.00955 | 0.00284 | 0.23353 | 0.01561 | 0.00445 | 0.00884 | 0.00960 | 0.00623 | 0.00559 | 0.24212 | 0.00780 | 0.01119 |
| **Pth1r** | 0.00599 | 0.21778 | 0.01409 | 0.00000 | 0.00992 | 0.00885 | 0.00273 | 0.12782 | 0.01033 | 0.01366 | 0.00211 | 5.44565 | 0.00411 | 0.04369 |
| **Ptk2** | 2.39698 | 0.02984 | 3.99309 | 0.00000 | 1.53136 | 5.07494 | 0.01368 | 1.29759 | 3.58815 | 0.00972 | 7.10689 | 3.21715 | 4.10573 | 5.63573 |
| **Ptk2b** | 0.00741 | 0.01070 | 0.01030 | 0.00000 | 0.12828 | 0.00417 | 0.00352 | 0.97133 | 0.00982 | 0.00550 | 0.00879 | 0.01270 | 0.06430 | 0.00778 |
| **Ptk6** | 0.32636 | 6.57858 | 0.01624 | 0.00000 | 0.02990 | 0.02801 | 0.03479 | 2.97554 | 0.07098 | 0.05420 | 0.00000 | 1.12723 | 0.14387 | 0.14979 |
| **Ptk7** | 0.14880 | 0.01245 | 0.02017 | 0.15558 | 0.00067 | 0.00000 | 0.00529 | 0.08947 | 0.03839 | 0.00567 | 0.03589 | 0.00659 | 0.00762 | 0.25065 |
| **Ptma** | 37.20171 | 17.84741 | 61.14528 | 62.38833 | 56.83321 | 52.81498 | 16.94724 | 43.22914 | 31.39550 | 28.85656 | 33.36971 | 41.31887 | 13.90223 | 35.29694 |
| **Ptms** | 0.07489 | 0.16299 | 1.91315 | 0.75506 | 0.14595 | 1.38085 | 0.49754 | 7.87694 | 0.00868 | 12.50098 | 1.82741 | 3.05198 | 0.03038 | 0.00309 |
| **Ptov1** | 0.00000 | 0.02270 | 5.17051 | 7.46916 | 1.93350 | 0.00000 | 2.10751 | 5.31172 | 0.00000 | 1.99752 | 0.00000 | 6.42779 | 0.51476 | 0.00000 |
| **Ptp4a1** | 3.31248 | 0.02335 | 0.63455 | 0.49654 | 0.76358 | 0.19103 | 0.00310 | 0.80947 | 8.69356 | 0.01498 | 0.36816 | 0.51750 | 0.78182 | 0.00000 |

| | | | | | | | | | | | | | | |
|---|---|---|---|---|---|---|---|---|---|---|---|---|---|---|
| **Ptp4a2** | 14.11978 | 8.82797 | 23.86089 | 13.64010 | 25.67727 | 25.06496 | 3.46336 | 14.29729 | 31.53322 | 0.41090 | 17.39932 | 13.39968 | 11.46341 | 21.24233 |
| **Ptp4a3** | 0.15479 | 0.03268 | 0.19972 | 1.49934 | 0.74378 | 0.00000 | 0.03723 | 0.00000 | 0.17391 | 0.03232 | 0.89593 | 1.65391 | 0.58417 | 0.26494 |
| **Ptpa** | 4.09675 | 3.79823 | 9.39900 | 1.69942 | 7.18021 | 8.83377 | 3.81104 | 9.47593 | 0.60624 | 0.53807 | 1.98214 | 5.20131 | 0.71951 | 5.44146 |
| **Ptpdc1** | 0.00000 | 0.00391 | 0.09722 | 0.00000 | 0.17228 | 0.00000 | 0.00388 | 0.00786 | 0.24456 | 0.00428 | 0.95424 | 0.00418 | 0.73514 | 0.54464 |
| **Ptpmt1** | 1.45788 | 0.16430 | 0.77915 | 5.61128 | 2.14157 | 1.94109 | 2.40235 | 2.91431 | 1.95127 | 4.40582 | 2.03015 | 3.15232 | 0.98312 | 0.00000 |
| **Ptpn1** | 2.89675 | 2.81053 | 4.49957 | 4.53186 | 4.85711 | 5.88664 | 0.18151 | 1.63878 | 0.00203 | 0.04442 | 1.66834 | 6.21020 | 7.32066 | 2.20397 |
| **Ptpn11** | 5.54746 | 0.04040 | 6.84977 | 11.24225 | 3.98355 | 8.25378 | 2.73863 | 0.61379 | 8.40034 | 0.22643 | 5.33150 | 11.34474 | 12.89413 | 13.86222 |
| **Ptpn12** | 4.93876 | 0.06437 | 5.71464 | 2.00181 | 4.67172 | 8.81154 | 4.62012 | 1.93264 | 7.39089 | 0.18242 | 7.01689 | 3.69530 | 2.00917 | 5.43392 |
| **Ptpn13** | 10.50879 | 0.00963 | 10.41786 | 1.40930 | 13.29682 | 1.60696 | 0.21599 | 0.40405 | 1.51773 | 0.00000 | 6.99100 | 2.68231 | 2.70233 | 10.11108 |
| **Ptpn14** | 0.05856 | 0.01996 | 0.01975 | 0.00000 | 0.10480 | 0.00924 | 0.00518 | 0.82085 | 0.00973 | 0.00593 | 0.00630 | 0.01361 | 0.00671 | 0.01037 |
| **Ptpn18** | 0.00371 | 0.01172 | 0.00118 | 1.06548 | 0.62825 | 0.02337 | 0.02132 | 0.03173 | 0.09556 | 0.01197 | 0.00431 | 0.54998 | 0.20200 | 0.00000 |
| **Ptpn2** | 19.53310 | 0.05295 | 15.57202 | 7.92062 | 21.48650 | 3.14726 | 0.09057 | 2.06283 | 21.14541 | 0.02686 | 13.20145 | 5.57785 | 11.02527 | 21.06716 |
| **Ptpn21** | 7.02322 | 0.02602 | 5.00585 | 2.52166 | 13.44913 | 0.90759 | 0.02644 | 2.15452 | 3.94604 | 0.00208 | 4.46899 | 0.56886 | 4.99479 | 13.58856 |
| **Ptpn22** | 2.93304 | 0.01155 | 7.59005 | 0.70269 | 6.22241 | 0.42442 | 0.00206 | 0.00000 | 4.90531 | 0.00000 | 0.00469 | 0.00067 | 0.42466 | 3.26653 |
| **Ptpn23** | 3.78597 | 0.08732 | 6.44565 | 2.94213 | 3.35635 | 1.82162 | 0.00792 | 3.42211 | 0.01085 | 0.03767 | 0.77825 | 1.57083 | 0.36059 | 4.72416 |
| **Ptpn3** | 0.00000 | 0.05435 | 1.14457 | 1.43785 | 2.79871 | 2.97796 | 2.64896 | 0.99248 | 2.88033 | 0.06537 | 3.88290 | 2.14638 | 0.09251 | 1.99776 |
| **Ptpn4** | 0.01772 | 0.00377 | 0.06875 | 0.00000 | 0.23139 | 0.50416 | 0.00952 | 0.00295 | 0.02397 | 0.00627 | 0.01218 | 0.29946 | 0.99148 | 0.09895 |
| **Ptpn6** | 0.00000 | 0.02265 | 11.89677 | 5.75001 | 0.79488 | 0.85550 | 0.00080 | 2.30758 | 0.00000 | 0.01539 | 3.02004 | 2.26305 | 0.00000 | 0.57846 |
| **Ptpn7** | 0.07557 | 0.09366 | 0.02560 | 0.25814 | 0.08269 | 0.26301 | 0.04015 | 0.13345 | 0.24270 | 0.09937 | 0.02316 | 0.04262 | 0.26640 | 0.28744 |
| **Ptpn9** | 1.66547 | 0.00981 | 0.11022 | 0.55575 | 1.92499 | 0.27578 | 0.00312 | 0.00113 | 2.49371 | 0.00497 | 0.00000 | 1.06538 | 3.74530 | 7.58165 |
| **Ptpra** | 2.55417 | 0.04791 | 2.80861 | 2.25483 | 2.04056 | 4.86880 | 3.49815 | 4.55391 | 0.19252 | 0.05857 | 5.87277 | 3.38717 | 2.20576 | 1.43502 |
| **Ptprb** | 0.02362 | 0.00441 | 0.40208 | 0.00000 | 0.30082 | 0.17137 | 0.00458 | 0.01024 | 0.01283 | 0.00495 | 0.02387 | 0.04545 | 0.01344 | 0.00942 |
| **Ptprcap** | 0.31441 | 0.00421 | 0.43996 | 0.49606 | 0.07323 | 0.00000 | 0.00479 | 0.40507 | 0.00834 | 0.00322 | 0.75853 | 0.13957 | 0.01841 | 0.00000 |
| **Ptpre** | 0.01937 | 0.00430 | 0.00371 | 0.00759 | 0.31426 | 0.18083 | 0.00972 | 0.00516 | 0.02744 | 0.04296 | 0.00887 | 0.00879 | 0.02289 | 0.06036 |
| **Ptprf** | 0.06523 | 6.98944 | 0.23321 | 2.36666 | 0.71649 | 2.59832 | 2.89527 | 8.91968 | 0.03656 | 0.10335 | 0.00000 | 0.04326 | 0.00224 | 0.23475 |
| **Ptprg** | 0.08796 | 0.01096 | 0.81682 | 1.90994 | 0.40602 | 0.96674 | 0.08686 | 0.00000 | 0.01550 | 0.00830 | 0.16850 | 0.96710 | 0.12083 | 2.87156 |
| **Ptprh** | 0.00228 | 0.00404 | 0.01161 | 0.02795 | 0.05843 | 0.01531 | 0.00408 | 0.03277 | 0.00747 | 0.00780 | 0.00814 | 0.00696 | 1.58835 | 0.12162 |
| **Ptprj** | 0.45928 | 0.00489 | 0.64749 | 0.23437 | 0.74755 | 0.10565 | 0.00398 | 0.00891 | 1.26340 | 0.00583 | 0.28255 | 0.02778 | 0.48150 | 1.75703 |
| **Ptprk** | 0.03845 | 0.00379 | 0.46326 | 0.00000 | 0.00928 | 0.62496 | 0.29046 | 3.49171 | 0.01169 | 0.03190 | 0.00000 | 0.30078 | 0.10210 | 0.00000 |
| **Ptprm** | 0.03926 | 0.00441 | 0.02242 | 0.01080 | 0.00954 | 0.01217 | 0.00425 | 0.01024 | 0.01907 | 0.00646 | 0.00829 | 0.00941 | 0.15487 | 0.01145 |
| **Ptprn** | 0.00000 | 0.01132 | 1.56748 | 3.66600 | 0.24755 | 0.89218 | 0.00942 | 0.03374 | 0.00597 | 0.00272 | 0.00000 | 0.00302 | 0.00176 | 0.01135 |
| **Ptpro** | 0.00548 | 0.00432 | 0.01372 | 0.01277 | 0.00996 | 0.01307 | 0.00424 | 0.00890 | 0.00610 | 0.00702 | 0.00882 | 0.00898 | 0.43170 | 0.08045 |
| **Ptprr** | 0.00280 | 0.00422 | 0.01460 | 0.01260 | 0.00819 | 0.01276 | 0.00420 | 0.00869 | 0.02289 | 0.00635 | 0.00904 | 0.00838 | 0.99148 | 0.00347 |
| **Ptprs** | 0.10514 | 0.03139 | 0.05301 | 0.16317 | 0.94636 | 0.12858 | 0.00492 | 0.77232 | 0.01012 | 0.00569 | 0.04820 | 0.02649 | 0.00142 | 0.41688 |
| **Ptprt** | 0.00424 | 0.00434 | 0.08229 | 0.01377 | 0.14239 | 0.01236 | 0.00430 | 0.01030 | 0.01066 | 0.00658 | 0.01117 | 0.02094 | 0.69685 | 0.08030 |
| **Ptpru** | 0.00000 | 0.00419 | 0.36586 | 1.23106 | 0.05062 | 0.00000 | 0.00410 | 0.00992 | 0.00905 | 0.00537 | 0.00475 | 0.00684 | 0.00518 | 0.02864 |
| **Ptprz1** | 0.08163 | 0.00425 | 0.00968 | 0.07299 | 0.00456 | 0.01502 | 0.00994 | 0.00652 | 0.02361 | 0.01354 | 0.00644 | 0.01187 | 0.14648 | 0.13468 |
| **Ptrf** | 0.05211 | 0.00375 | 1.58094 | 2.39521 | 2.13649 | 0.03374 | 0.00362 | 0.00000 | 0.12660 | 0.02118 | 0.64150 | 0.00905 | 0.00000 | 1.76263 |
| **Ptrh1** | 1.54457 | 0.01024 | 2.27437 | 8.93054 | 7.30948 | 0.00000 | 0.53781 | 0.06784 | 0.47395 | 0.01804 | 0.00000 | 0.60176 | 3.61647 | 0.34794 |
| **Ptrh2** | 7.57490 | 0.01221 | 2.88405 | 5.06608 | 11.63094 | 2.86887 | 1.78549 | 2.94615 | 5.78577 | 10.50701 | 3.18844 | 3.81377 | 5.20900 | 8.98329 |
| **Ptrhd1** | 0.62828 | 0.01796 | 0.75257 | 1.62921 | 1.98790 | 1.91734 | 0.28368 | 1.04242 | 0.21778 | 0.06090 | 0.31530 | 1.67585 | 0.79395 | 1.00298 |
| **Pts** | 0.00495 | 0.05984 | 0.05488 | 0.33347 | 0.33515 | 0.49904 | 2.74651 | 2.03221 | 0.06940 | 0.11060 | 0.01137 | 0.97839 | 0.35738 | 0.01995 |
| **Pttg1** | 12.12311 | 7.53740 | 4.34590 | 17.33993 | 17.74821 | 6.42731 | 6.34695 | 3.36529 | 8.28070 | 14.73837 | 4.95264 | 17.92088 | 20.87455 | 15.21692 |
| **Pttg1ip** | 26.73211 | 0.24172 | 40.23476 | 25.77021 | 35.27107 | 14.99953 | 3.22165 | 7.69298 | 23.27169 | 0.18238 | 10.09118 | 9.73653 | 19.04982 | 36.10649 |
| **Puf60** | 10.31268 | 9.57535 | 24.02182 | 23.31948 | 21.56982 | 5.89106 | 3.04201 | 16.04390 | 2.70150 | 13.66858 | 7.88644 | 12.88861 | 5.68923 | 6.95893 |
| **Pum1** | 10.12554 | 0.05630 | 4.35992 | 0.00000 | 11.80937 | 6.82174 | 1.30431 | 3.09765 | 5.68749 | 0.22962 | 2.28166 | 3.52762 | 7.82473 | 12.69967 |
| **Pum2** | 23.25798 | 0.02324 | 11.20137 | 3.09750 | 17.76483 | 10.85165 | 4.36579 | 3.19859 | 29.51443 | 0.07351 | 5.03192 | 7.45082 | 9.67604 | 21.50273 |
| **Pum3** | 19.64558 | 0.11099 | 21.19912 | 6.14325 | 18.71849 | 12.02273 | 1.48203 | 3.64639 | 8.74796 | 0.04254 | 14.68288 | 12.96832 | 10.90352 | 22.54126 |
| **Pura** | 0.11682 | 0.02685 | 1.38369 | 1.09970 | 0.58920 | 0.58974 | 2.94457 | 1.51138 | 0.20014 | 0.13614 | 0.30374 | 1.57636 | 0.00003 | 0.24151 |
| **Purb** | 10.23737 | 0.03929 | 7.79424 | 4.89386 | 13.50484 | 12.85403 | 0.17862 | 1.93074 | 28.41035 | 0.01602 | 6.88651 | 2.63986 | 8.32405 | 10.09891 |
| **Purg** | 0.25131 | 0.00413 | 0.05911 | 0.00000 | 0.00100 | 0.41112 | 0.00396 | 0.00322 | 0.01172 | 0.00506 | 0.16114 | 0.01132 | 1.14979 | 0.00213 |
| **Pus1** | 2.37873 | 0.99705 | 9.19960 | 12.62245 | 5.23148 | 8.20618 | 3.99159 | 8.81757 | 0.36011 | 0.15448 | 11.62544 | 7.69889 | 5.32295 | 6.60094 |
| **Pus10** | 1.43590 | 0.60723 | 1.71720 | 0.41455 | 1.31987 | 1.55068 | 4.42713 | 0.45921 | 6.31049 | 0.03654 | 7.85454 | 3.05165 | 0.00000 | 0.13012 |
| **Pus3** | 5.78635 | 0.03074 | 3.56278 | 0.00000 | 5.78420 | 4.36757 | 0.91500 | 0.00000 | 7.42048 | 0.03166 | 4.63230 | 3.60678 | 0.00000 | 7.42045 |
| **Pus7** | 3.62786 | 0.02231 | 6.14152 | 1.27856 | 3.34187 | 5.22200 | 0.02485 | 0.00000 | 6.33263 | 0.13993 | 3.08165 | 2.25145 | 6.34518 | 6.96672 |

| | | | | | | | | | | | | | | |
|---|---|---|---|---|---|---|---|---|---|---|---|---|---|---|
| **Pus7l** | 0.00000 | 5.25732 | 0.29417 | 1.08915 | 1.34141 | 0.04476 | 0.00278 | 0.00615 | 0.04148 | 0.00534 | 0.02626 | 0.01022 | 0.20075 | 1.70445 |
| **Pusl1** | 0.44881 | 4.32404 | 4.21356 | 6.53740 | 2.83430 | 0.71304 | 0.36547 | 3.64818 | 0.09405 | 3.31655 | 0.00000 | 1.66696 | 0.84699 | 0.27763 |
| **Pvr** | 0.07526 | 0.00306 | 0.08585 | 0.00000 | 0.18884 | 0.26004 | 0.00296 | 1.57013 | 1.47267 | 0.00592 | 0.20617 | 0.96898 | 0.14430 | 1.30562 |
| **Pvt1** | 0.86384 | 0.00257 | 2.88907 | 0.00000 | 3.54183 | 0.31929 | 0.44724 | 0.00000 | 0.71063 | 0.00000 | 0.37080 | 1.57918 | 0.00000 | 1.45803 |
| **Pwp1** | 13.54266 | 6.29804 | 18.28101 | 14.02148 | 21.51938 | 9.66767 | 3.20211 | 6.74940 | 10.97281 | 0.11786 | 10.44771 | 7.21553 | 13.88745 | 20.86584 |
| **Pwp2** | 4.38677 | 0.05863 | 14.62620 | 1.84696 | 9.39230 | 10.46126 | 0.63929 | 1.55565 | 1.55201 | 0.06954 | 3.01531 | 7.33755 | 6.10702 | 12.15081 |
| **Pwwp2a** | 2.21630 | 0.21963 | 4.77775 | 1.22271 | 6.65398 | 3.54577 | 2.44994 | 0.26899 | 4.44892 | 0.23562 | 1.26842 | 2.40340 | 4.34406 | 5.48094 |
| **Pwwp2b** | 0.00000 | 0.06768 | 1.84224 | 0.01146 | 0.00000 | 1.36569 | 3.12254 | 3.20716 | 0.00677 | 0.03785 | 1.57457 | 0.00282 | 0.09839 | 0.00000 |
| **Pxdc1** | 0.46539 | 0.00416 | 0.01012 | 0.17506 | 0.00000 | 0.00802 | 0.00398 | 0.00634 | 0.00985 | 0.01037 | 0.00975 | 0.01072 | 0.07370 | 0.07833 |
| **Pxdn** | 0.11167 | 0.01209 | 0.01122 | 0.00000 | 0.02640 | 0.06921 | 0.00447 | 0.00547 | 0.01344 | 0.00566 | 0.00338 | 0.67228 | 0.00534 | 0.00571 |
| **Pxk** | 3.53881 | 0.07358 | 7.13205 | 4.59896 | 6.22283 | 7.51659 | 2.05784 | 6.89732 | 3.26211 | 0.03905 | 4.27524 | 2.95226 | 6.59817 | 4.99966 |
| **Pxmp2** | 3.12937 | 0.06326 | 0.66168 | 4.27149 | 6.90365 | 1.12205 | 3.24293 | 1.32504 | 0.62650 | 0.14723 | 0.05550 | 2.16950 | 3.78961 | 4.55626 |
| **Pxmp4** | 1.55723 | 0.10024 | 2.74871 | 1.25809 | 0.64937 | 0.70713 | 2.02667 | 6.75381 | 1.06199 | 0.08204 | 2.18883 | 1.14409 | 0.25458 | 0.92972 |
| **Pxn** | 0.00000 | 0.02715 | 0.39391 | 3.58402 | 2.47514 | 1.18790 | 0.05386 | 2.34381 | 0.83067 | 2.55735 | 0.00000 | 1.09370 | 1.23025 | 0.66305 |
| **Pxt1** | 0.00000 | 0.00446 | 0.38135 | 0.05293 | 0.80338 | 0.35070 | 0.55349 | 0.00809 | 0.02276 | 0.04800 | 0.00447 | 0.00401 | 3.68911 | 0.00135 |
| **Pxylp1** | 0.00285 | 0.00427 | 0.00980 | 0.18910 | 0.24937 | 0.00659 | 0.00410 | 0.00694 | 0.00576 | 0.00615 | 0.01346 | 0.01232 | 0.00345 | 0.47175 |
| **Pycard** | 0.26687 | 0.04283 | 0.00000 | 4.92357 | 1.33343 | 4.51834 | 0.05462 | 1.93783 | 0.03323 | 0.03148 | 1.29565 | 1.78255 | 0.00392 | 0.68948 |
| **Pycr1** | 0.03084 | 0.04805 | 0.15670 | 0.47711 | 0.05261 | 0.44198 | 0.03254 | 0.08916 | 0.10261 | 0.00736 | 0.03258 | 0.02776 | 0.08702 | 0.19545 |
| **Pycr2** | 0.27949 | 7.47379 | 4.03140 | 7.11697 | 0.95283 | 1.61924 | 0.00000 | 3.50767 | 0.10809 | 0.03659 | 2.76860 | 6.42128 | 0.83438 | 0.62599 |
| **Pycrl** | 2.35286 | 0.04845 | 2.32743 | 4.95606 | 0.41847 | 5.36509 | 3.77803 | 3.50169 | 0.00000 | 0.06921 | 5.81949 | 2.26909 | 0.29808 | 0.03486 |
| **Pydc3** | 0.00504 | 0.00434 | 0.01385 | 0.01691 | 0.01084 | 0.01226 | 0.00420 | 0.01093 | 0.12105 | 0.00680 | 0.00791 | 0.00997 | 0.20048 | 0.00000 |
| **Pygb** | 1.29835 | 0.03720 | 3.68704 | 4.85317 | 2.40244 | 1.28067 | 2.29714 | 3.72295 | 0.10283 | 0.04699 | 3.75944 | 3.88342 | 0.16932 | 0.25497 |
| **Pygl** | 0.83542 | 0.00528 | 0.54084 | 0.19685 | 1.29312 | 0.53983 | 0.00397 | 0.02058 | 0.36373 | 0.01957 | 0.00620 | 0.00106 | 0.16417 | 0.00000 |
| **Pygm** | 0.81544 | 0.01358 | 0.76092 | 0.20384 | 0.33092 | 0.00000 | 0.00464 | 0.99584 | 0.03297 | 0.00478 | 0.01680 | 0.00852 | 0.55114 | 0.22941 |
| **Pygo1** | 0.00129 | 0.00423 | 0.47199 | 0.00345 | 0.44647 | 0.00352 | 0.00424 | 0.00826 | 0.00864 | 0.00570 | 0.00591 | 0.01118 | 0.59334 | 0.00996 |
| **Pygo2** | 3.48755 | 4.62031 | 6.59496 | 3.92650 | 5.59696 | 4.92368 | 3.97970 | 6.16888 | 0.02580 | 0.07083 | 2.86271 | 5.04754 | 1.94034 | 3.81544 |
| **Pym1** | 0.00000 | 0.00374 | 0.13995 | 0.34450 | 0.15197 | 0.34919 | 0.00967 | 0.00181 | 0.01892 | 0.00463 | 0.11914 | 0.28404 | 0.15174 | 0.46412 |
| **Pyroxd1** | 0.61808 | 0.01247 | 0.71524 | 1.61272 | 1.08565 | 0.33534 | 0.00916 | 0.59184 | 0.33512 | 0.00486 | 0.00000 | 0.31751 | 0.00441 | 1.24019 |
| **Pyurf** | 0.36176 | 0.00394 | 0.01441 | 0.67594 | 0.00000 | 0.39245 | 0.68997 | 0.15695 | 0.01370 | 0.03156 | 0.18255 | 0.29697 | 0.02293 | 0.07933 |
| **Pyy** | 0.00247 | 0.12769 | 0.01634 | 0.00743 | 0.01022 | 0.02299 | 0.00282 | 0.01968 | 0.01178 | 0.00792 | 0.00318 | 5.28051 | 0.00767 | 0.01320 |
| **Pzp** | 1.16353 | 0.00337 | 0.25167 | 2.06010 | 2.54276 | 0.93420 | 0.00857 | 0.02215 | 3.96375 | 0.02515 | 0.09829 | 0.02940 | 2.40291 | 0.36350 |
| **Qars** | 10.51976 | 8.58898 | 30.44244 | 22.95180 | 15.63890 | 22.19692 | 2.70492 | 9.28542 | 10.81993 | 11.35808 | 12.69875 | 13.34504 | 8.57011 | 13.63624 |
| **Qdpr** | 0.17853 | 0.08579 | 0.00000 | 5.99357 | 2.55588 | 5.34354 | 0.00878 | 1.51438 | 0.00280 | 0.04246 | 0.10721 | 5.99963 | 0.63538 | 0.60920 |
| **Qk** | 4.30986 | 0.05311 | 6.66833 | 3.00371 | 6.78592 | 10.98500 | 5.25029 | 0.70211 | 17.98619 | 0.12750 | 6.67066 | 5.80984 | 8.51234 | 5.32112 |
| **Qpctl** | 0.33872 | 0.00447 | 0.15340 | 1.31600 | 0.61889 | 0.09752 | 0.00427 | 0.41677 | 0.01111 | 0.00769 | 2.11638 | 0.00476 | 0.88141 | 0.00000 |
| **Qrich1** | 0.00000 | 0.00047 | 5.13646 | 4.44675 | 3.73252 | 2.33092 | 1.06667 | 3.64269 | 5.82112 | 0.02334 | 3.93761 | 3.31289 | 1.59263 | 5.93200 |
| **Qrich2** | 0.00183 | 0.00456 | 0.46980 | 0.00000 | 0.49287 | 0.00318 | 0.00412 | 0.01336 | 0.00728 | 0.00559 | 0.00667 | 0.01019 | 0.07957 | 0.04730 |
| **Qrsl1** | 1.42841 | 0.04160 | 2.13650 | 3.71152 | 1.81394 | 2.70480 | 1.17109 | 2.52806 | 0.36763 | 0.02017 | 1.39531 | 1.77902 | 0.19572 | 1.06931 |
| **Qser1** | 0.35720 | 0.00357 | 0.26649 | 0.00000 | 0.68587 | 0.15991 | 0.00410 | 0.01908 | 2.02351 | 0.00547 | 0.00494 | 1.04330 | 1.78177 | 0.97241 |
| **Qsox1** | 0.09555 | 0.98598 | 0.14079 | 0.09411 | 0.27680 | 0.69011 | 0.00923 | 2.34386 | 0.00269 | 0.02424 | 0.00000 | 1.12594 | 0.35966 | 0.34672 |
| **Qsox2** | 6.03005 | 0.02566 | 3.95605 | 0.73934 | 2.58469 | 2.12474 | 4.28967 | 0.00000 | 4.36265 | 0.03074 | 0.00000 | 1.25105 | 3.36223 | 8.91411 |
| **Qtrt1** | 1.07213 | 0.35631 | 11.92220 | 9.66569 | 7.83716 | 2.46250 | 0.31576 | 7.92926 | 0.18133 | 0.03375 | 7.15028 | 1.67777 | 1.76138 | 3.45433 |
| **Qtrtd1** | 0.00000 | 0.01085 | 2.19202 | 0.94419 | 0.73312 | 1.23325 | 0.00312 | 0.60377 | 2.17872 | 0.02345 | 0.00000 | 1.72150 | 1.36033 | 8.44386 |
| **R3hcc1** | 10.72386 | 6.14881 | 16.16382 | 14.46872 | 11.99402 | 8.40490 | 7.41911 | 1.32453 | 0.35115 | 0.25769 | 9.03548 | 4.87267 | 0.29340 | 3.47119 |
| **R3hcc1l** | 1.86453 | 0.00289 | 1.01320 | 0.29038 | 2.90913 | 1.85525 | 2.39501 | 0.00373 | 0.02118 | 0.02497 | 0.26219 | 0.96070 | 0.92975 | 2.91338 |
| **R3hdm1** | 9.55431 | 0.01445 | 11.22802 | 1.87140 | 12.45852 | 4.92523 | 2.60818 | 0.00000 | 10.29606 | 0.05530 | 2.29010 | 2.51285 | 8.62384 | 17.50776 |
| **R3hdm2** | 2.60831 | 5.25653 | 9.24791 | 3.50135 | 6.16094 | 1.02030 | 0.00722 | 0.61340 | 0.33914 | 0.02433 | 1.11843 | 3.71717 | 5.23061 | 1.75429 |
| **R3hdm4** | 0.00000 | 0.32435 | 4.48390 | 4.85879 | 0.00000 | 4.03143 | 4.27960 | 8.53351 | 0.19300 | 6.66365 | 2.81821 | 7.07988 | 0.00000 | 1.84154 |
| **Rab10** | 12.86199 | 0.04695 | 9.53324 | 6.97978 | 15.05376 | 7.88270 | 1.41308 | 2.14275 | 17.13883 | 0.13177 | 5.81176 | 2.93028 | 12.09091 | 11.76515 |
| **Rab10os** | 0.58584 | 0.00390 | 0.96753 | 2.29391 | 4.31611 | 0.61029 | 0.00360 | 0.00010 | 0.26244 | 0.09989 | 0.00000 | 0.00443 | 3.04120 | 0.00000 |
| **Rab11a** | 12.07019 | 0.40551 | 2.63290 | 13.65749 | 10.85557 | 15.57364 | 5.83435 | 8.99063 | 11.16770 | 1.09921 | 2.70774 | 6.87774 | 5.49940 | 0.38599 |
| **Rab11b** | 7.11506 | 0.22084 | 16.09167 | 11.85507 | 18.40173 | 8.66491 | 4.54627 | 6.61716 | 1.48571 | 2.67318 | 4.39363 | 9.42034 | 5.30618 | 3.43043 |
| **Rab11fip1** | 6.52653 | 3.01596 | 11.24811 | 2.64171 | 7.94691 | 3.92291 | 0.06699 | 3.88103 | 7.88876 | 6.75787 | 3.43129 | 0.40529 | 3.85188 | 16.43619 |
| **Rab11fip2** | 0.04197 | 0.00426 | 0.01265 | 0.00090 | 0.00907 | 0.16251 | 0.00414 | 0.00468 | 0.00640 | 0.00640 | 0.00700 | 0.01525 | 1.24365 | 0.00830 |
| **Rab11fip3** | 0.42921 | 0.00382 | 0.04401 | 1.07712 | 0.14253 | 0.01493 | 0.00955 | 0.01155 | 0.03107 | 0.00588 | 0.06290 | 0.03485 | 0.41750 | 0.24524 |

| | | | | | | | | | | | | | | |
|---|---|---|---|---|---|---|---|---|---|---|---|---|---|---|
| **Rab11fip4** | 0.00000 | 0.01161 | 0.23228 | 2.08108 | 0.00000 | 1.62895 | 0.98256 | 1.82165 | 0.29608 | 0.03641 | 1.41723 | 0.01972 | 0.01154 | 0.11845 |
| **Rab11fip5** | 7.52265 | 9.46151 | 4.76429 | 7.89937 | 3.51462 | 4.64153 | 2.76556 | 9.90925 | 2.75463 | 0.06973 | 0.00000 | 1.62782 | 8.76233 | 5.74862 |
| **Rab12** | 0.71344 | 0.05103 | 1.32117 | 0.88819 | 3.43970 | 2.44939 | 3.04447 | 2.65617 | 1.88385 | 0.03107 | 0.00000 | 2.81113 | 5.00533 | 2.77442 |
| **Rab13** | 0.80508 | 0.00230 | 0.16609 | 0.18915 | 0.98485 | 0.36373 | 0.01705 | 1.10800 | 0.00000 | 0.01243 | 0.00000 | 2.67966 | 3.20526 | 0.07779 |
| **Rab14** | 5.25933 | 5.55788 | 6.41608 | 6.10212 | 6.57326 | 12.30004 | 0.07287 | 7.10669 | 9.46723 | 0.08152 | 8.96199 | 7.94643 | 7.76064 | 6.43618 |
| **Rab15** | 0.53647 | 7.88987 | 0.00000 | 0.77581 | 0.00000 | 0.00000 | 0.01208 | 2.81792 | 0.95804 | 0.02164 | 0.00000 | 0.85931 | 0.01339 | 0.24003 |
| **Rab17** | 0.00000 | 7.38128 | 3.28412 | 0.22292 | 0.00665 | 11.58924 | 2.78684 | 10.10717 | 0.00000 | 0.15264 | 3.08234 | 0.70179 | 0.08469 | 0.00000 |
| **Rab18** | 11.22902 | 0.08631 | 13.36649 | 10.61850 | 6.33282 | 13.11370 | 1.86150 | 9.48095 | 7.77571 | 0.14198 | 15.57390 | 11.95649 | 3.01507 | 8.85539 |
| **Rab19** | 0.00981 | 0.00462 | 0.00701 | 0.01188 | 0.00000 | 0.02592 | 0.00430 | 0.00870 | 0.07244 | 0.00658 | 0.00714 | 0.00931 | 0.02503 | 0.79090 |
| **Rab1a** | 12.50583 | 0.36779 | 20.98292 | 19.17347 | 15.65348 | 25.23593 | 8.70559 | 12.19360 | 13.41535 | 7.59826 | 12.47693 | 16.24948 | 9.27713 | 8.82722 |
| **Rab1b** | 0.06975 | 0.20517 | 2.12014 | 12.40379 | 4.22584 | 1.46398 | 0.01897 | 7.79591 | 0.41717 | 0.10819 | 0.02245 | 4.57027 | 0.00000 | 0.00000 |
| **Rab20** | 11.37354 | 6.88092 | 19.03757 | 2.29614 | 17.08561 | 4.09711 | 2.34183 | 7.78617 | 0.61325 | 11.10853 | 2.96554 | 1.91228 | 2.38610 | 10.22053 |
| **Rab21** | 2.16365 | 0.10986 | 0.85688 | 0.00000 | 0.53727 | 2.68991 | 0.00916 | 2.95489 | 0.40473 | 0.05001 | 1.23765 | 2.33649 | 0.75075 | 0.50229 |
| **Rab22a** | 1.93033 | 0.00250 | 0.42315 | 1.77653 | 2.24004 | 3.08630 | 0.84671 | 0.48399 | 0.00000 | 0.04039 | 1.47787 | 1.17802 | 1.68458 | 2.16050 |
| **Rab23** | 0.11900 | 0.01771 | 1.17837 | 1.35436 | 0.38764 | 2.46620 | 0.00291 | 0.01087 | 1.40713 | 0.01342 | 2.04808 | 1.94554 | 2.27322 | 2.54303 |
| **Rab24** | 6.42491 | 0.16149 | 3.18007 | 5.71491 | 5.09316 | 3.03860 | 3.13557 | 2.86366 | 1.95013 | 0.16842 | 4.52077 | 3.83295 | 1.53382 | 3.94597 |
| **Rab25** | 0.00381 | 0.03488 | 0.21331 | 0.12669 | 0.23004 | 0.18398 | 0.00470 | 1.19090 | 0.01815 | 0.00841 | 0.00401 | 0.00439 | 0.00498 | 0.00136 |
| **Rab26** | 0.29547 | 0.00364 | 0.05785 | 2.17638 | 0.18761 | 0.00000 | 0.00362 | 0.00416 | 0.00549 | 0.00598 | 0.01837 | 0.32845 | 0.02360 | 0.30655 |
| **Rab26os** | 0.09276 | 0.02449 | 0.02732 | 0.00956 | 0.03370 | 0.19373 | 0.00462 | 0.18480 | 0.10333 | 0.01135 | 0.00591 | 0.00494 | 0.74261 | 0.16032 |
| **Rab27a** | 0.14448 | 0.04552 | 0.27790 | 0.27996 | 0.39898 | 0.18176 | 0.03197 | 0.31149 | 0.05161 | 0.01469 | 0.01353 | 1.00105 | 1.10261 | 0.48215 |
| **Rab27b** | 0.01932 | 0.00434 | 0.02608 | 0.06956 | 0.02136 | 0.05252 | 0.00988 | 0.02062 | 0.03967 | 0.02434 | 0.01442 | 0.00738 | 0.47486 | 0.01466 |
| **Rab28** | 1.67325 | 0.01122 | 0.55031 | 0.00000 | 0.51659 | 0.54443 | 0.83075 | 0.69797 | 2.25612 | 0.01780 | 0.17344 | 0.66647 | 0.08702 | 0.00000 |
| **Rab29** | 2.89342 | 0.00000 | 8.42928 | 3.64481 | 2.33349 | 10.15851 | 0.01085 | 2.21419 | 1.70463 | 11.79169 | 3.05214 | 0.89372 | 0.00000 | 0.73107 |
| **Rab2a** | 16.31050 | 0.17809 | 6.41458 | 5.38715 | 9.61239 | 14.80487 | 7.24788 | 9.93128 | 2.30992 | 0.53450 | 6.25811 | 8.30310 | 4.12602 | 5.00546 |
| **Rab2b** | 3.28846 | 0.00253 | 1.30587 | 0.98306 | 2.17510 | 0.92151 | 0.00291 | 0.00132 | 3.70003 | 0.00364 | 0.00000 | 0.63372 | 5.07723 | 1.37790 |
| **Rab30** | 0.58786 | 0.03818 | 0.65456 | 1.12954 | 0.32878 | 1.39072 | 0.00933 | 2.10779 | 0.73902 | 0.00116 | 3.10853 | 1.49666 | 0.48226 | 0.00000 |
| **Rab31** | 0.40567 | 0.15530 | 1.43580 | 0.61654 | 0.28619 | 1.73372 | 0.00324 | 0.00000 | 0.81901 | 0.02205 | 3.10678 | 0.32634 | 0.27411 | 0.10094 |
| **Rab32** | 0.00840 | 0.00441 | 0.00919 | 0.01193 | 0.19789 | 0.01321 | 0.00476 | 0.01012 | 0.01323 | 0.01417 | 0.00871 | 0.00897 | 0.01253 | 0.07055 |
| **Rab33b** | 1.92972 | 0.13581 | 1.56702 | 1.86733 | 1.61408 | 2.37380 | 0.72252 | 3.77164 | 2.16651 | 0.65476 | 0.00663 | 0.92058 | 12.51358 | 2.11161 |
| **Rab34** | 0.99640 | 0.00371 | 0.87157 | 1.53155 | 3.87013 | 0.00000 | 0.00359 | 0.00341 | 0.77206 | 0.00410 | 0.01171 | 0.01629 | 2.18199 | 2.88806 |
| **Rab35** | 6.59945 | 0.10589 | 7.88550 | 5.27532 | 6.48971 | 8.44398 | 4.39047 | 5.50940 | 0.97066 | 0.15247 | 0.24403 | 4.87848 | 7.15620 | 1.29506 |
| **Rab38** | 0.00000 | 0.00367 | 2.94383 | 1.57424 | 0.55563 | 0.00000 | 0.00328 | 0.00000 | 1.14310 | 0.00000 | 0.00509 | 0.29529 | 0.00000 | 0.00000 |
| **Rab39** | 5.13293 | 0.00222 | 3.98602 | 2.18871 | 2.23622 | 0.00000 | 0.00171 | 0.00000 | 9.28228 | 0.00000 | 0.94366 | 0.02717 | 6.12767 | 2.85461 |
| **Rab39b** | 0.00637 | 0.00425 | 0.06976 | 0.00001 | 0.00655 | 0.08369 | 0.00417 | 0.00513 | 0.01201 | 0.00577 | 0.00724 | 0.01037 | 0.49895 | 0.04115 |
| **Rab3a** | 0.00000 | 0.01123 | 0.44767 | 0.84134 | 1.14922 | 0.01332 | 0.00412 | 0.01328 | 0.00903 | 0.02161 | 0.00388 | 0.12803 | 0.78941 | 0.00000 |
| **Rab3b** | 0.26204 | 0.00308 | 3.09306 | 1.20229 | 0.00000 | 1.20227 | 0.00310 | 0.00000 | 0.00000 | 0.00000 | 1.32623 | 0.65625 | 0.56338 | 0.76318 |
| **Rab3c** | 0.02241 | 0.04220 | 0.01460 | 0.00255 | 0.01786 | 0.04605 | 0.01022 | 0.01764 | 0.10520 | 0.16526 | 0.01284 | 0.01409 | 0.02386 | 0.05234 |
| **Rab3d** | 0.00802 | 0.00364 | 0.46297 | 0.81738 | 0.40763 | 1.14885 | 0.01150 | 0.16197 | 0.28772 | 0.02486 | 0.00727 | 0.40751 | 4.55330 | 4.19936 |
| **Rab3gap1** | 3.51914 | 0.02530 | 3.06973 | 1.45597 | 2.56506 | 2.13803 | 0.03140 | 0.00000 | 1.59835 | 0.00000 | 4.35975 | 1.96771 | 2.33969 | 5.40021 |
| **Rab3gap2** | 0.88104 | 0.01917 | 1.22094 | 0.00000 | 0.07748 | 1.07262 | 0.00999 | 0.16735 | 1.27467 | 0.01012 | 1.58433 | 1.29475 | 0.69400 | 2.52588 |
| **Rab3il1** | 4.87403 | 5.75440 | 18.68741 | 10.05358 | 17.45312 | 7.95935 | 0.71484 | 2.72046 | 1.84843 | 0.35891 | 9.12311 | 1.98975 | 0.00000 | 15.67979 |
| **Rab3ip** | 5.78665 | 2.23686 | 5.41508 | 5.15224 | 3.81082 | 6.97272 | 3.67773 | 5.20086 | 0.22800 | 0.11979 | 3.95296 | 3.22949 | 3.82561 | 5.25406 |
| **Rab40b** | 0.48726 | 0.00374 | 0.50916 | 0.33888 | 0.17810 | 0.04455 | 0.00386 | 0.06001 | 0.00862 | 0.00578 | 0.00416 | 0.00674 | 1.28497 | 0.49743 |
| **Rab40c** | 3.13331 | 0.03894 | 7.19108 | 3.56305 | 7.81948 | 9.47367 | 2.69884 | 4.35530 | 2.62791 | 0.02720 | 1.97614 | 3.53513 | 5.82444 | 8.62975 |
| **Rab42** | 0.00240 | 0.00436 | 0.08384 | 0.00000 | 0.25888 | 0.00000 | 0.00526 | 0.53634 | 0.06176 | 0.00533 | 0.00608 | 0.00986 | 0.39256 | 0.00000 |
| **Rab43** | 0.13514 | 0.02647 | 0.64987 | 0.00000 | 1.70277 | 0.46530 | 0.00932 | 1.22955 | 0.04221 | 0.02034 | 0.00000 | 0.67529 | 3.89264 | 3.33514 |
| **Rab4a** | 0.00000 | 0.02014 | 0.46614 | 1.87738 | 0.61606 | 2.02016 | 2.20283 | 0.00266 | 0.00332 | 0.06252 | 0.00000 | 0.20118 | 0.72434 | 0.00000 |
| **Rab4b** | 0.00000 | 0.01605 | 1.19425 | 1.53237 | 0.00084 | 0.15447 | 0.00846 | 1.08250 | 0.00554 | 0.01332 | 0.00000 | 3.56180 | 0.28049 | 0.00000 |
| **Rab5a** | 24.71374 | 0.28454 | 16.87063 | 6.76818 | 20.81557 | 10.73050 | 2.12850 | 3.99615 | 20.83446 | 0.10905 | 9.45076 | 7.21810 | 15.99526 | 12.94376 |
| **Rab5b** | 2.17606 | 1.21858 | 2.61018 | 4.17982 | 3.95106 | 4.02345 | 0.00162 | 0.85078 | 9.93808 | 0.32327 | 2.54488 | 0.50781 | 1.15201 | 5.91367 |
| **Rab5c** | 1.18523 | 0.42282 | 7.12821 | 5.81940 | 10.09472 | 3.19411 | 0.06617 | 6.63426 | 0.13892 | 0.03738 | 6.11226 | 3.04470 | 1.27476 | 0.54203 |
| **Rab6a** | 5.87497 | 0.05071 | 6.01557 | 0.00000 | 2.33840 | 9.82370 | 0.19654 | 3.35989 | 13.07452 | 9.31423 | 6.36895 | 5.67969 | 9.17745 | 2.41342 |
| **Rab6b** | 0.00747 | 0.00429 | 0.08794 | 0.05552 | 0.13760 | 1.21930 | 0.00408 | 0.27319 | 0.00833 | 0.01450 | 0.00326 | 0.00000 | 0.00548 | 0.00961 |
| **Rab7** | 21.53694 | 9.50921 | 33.32260 | 21.52973 | 35.18625 | 26.03420 | 8.25691 | 17.50716 | 5.70462 | 10.78387 | 17.72573 | 14.02535 | 15.46694 | 26.77120 |
| **Rab8a** | 4.88611 | 7.67484 | 15.78049 | 10.09524 | 13.97855 | 14.66472 | 2.10873 | 6.70758 | 1.30572 | 6.36548 | 8.40332 | 6.04067 | 3.61921 | 6.74251 |

| | | | | | | | | | | | | | | |
|---|---|---|---|---|---|---|---|---|---|---|---|---|---|---|
| **Rab8b** | 0.67928 | 0.01149 | 0.39891 | 0.48732 | 0.77850 | 0.68230 | 0.00992 | 0.06071 | 1.50118 | 0.00480 | 0.00000 | 0.68071 | 0.55413 | 0.14453 |
| **Rab9** | 3.69301 | 6.40847 | 8.11434 | 6.65507 | 8.37685 | 11.86773 | 2.77734 | 6.71482 | 6.08718 | 12.47435 | 8.71515 | 5.33764 | 4.77447 | 10.63964 |
| **Rabac1** | 1.43522 | 0.05222 | 4.52695 | 16.27328 | 7.15678 | 1.50939 | 0.74068 | 6.83823 | 0.00000 | 0.14344 | 0.35393 | 9.41309 | 0.51684 | 0.00000 |
| **Rabep1** | 5.17971 | 0.03364 | 2.29550 | 1.27007 | 9.42103 | 4.80840 | 0.01370 | 1.99596 | 16.64325 | 0.01003 | 0.00000 | 3.26302 | 1.92551 | 7.51296 |
| **Rabep2** | 3.53254 | 0.03306 | 7.34295 | 3.01973 | 10.07230 | 1.89665 | 0.00717 | 4.38593 | 0.00584 | 0.32597 | 2.85591 | 0.54367 | 2.98791 | 1.93283 |
| **Rabepk** | 0.66863 | 6.13766 | 1.99172 | 1.85625 | 2.43145 | 3.74861 | 0.00688 | 1.41451 | 0.44478 | 0.02054 | 2.24813 | 5.41615 | 1.20824 | 1.86388 |
| **Rabgap1** | 2.63646 | 0.00313 | 0.21379 | 0.23655 | 1.44948 | 1.10617 | 0.00328 | 0.00000 | 3.34922 | 0.01153 | 0.00471 | 0.42037 | 1.63572 | 3.06211 |
| **Rabgap1l** | 0.00760 | 0.00551 | 0.00875 | 0.02737 | 0.00376 | 0.00266 | 0.00437 | 0.30078 | 0.01949 | 0.01065 | 0.01858 | 0.01287 | 0.43750 | 0.43248 |
| **Rabgef1** | 1.18468 | 0.00904 | 4.45683 | 3.19819 | 6.99229 | 4.05583 | 1.18001 | 0.00000 | 6.75078 | 0.04615 | 2.12100 | 0.83760 | 6.86269 | 15.06401 |
| **Rabggta** | 1.43876 | 0.05984 | 8.79626 | 9.11766 | 5.31637 | 0.48149 | 0.05375 | 2.64305 | 0.55588 | 0.06357 | 4.09935 | 3.99412 | 1.20650 | 4.62030 |
| **Rabggtb** | 23.44878 | 2.38655 | 15.33422 | 0.95825 | 6.46195 | 11.66841 | 8.22066 | 9.50832 | 17.72182 | 13.26197 | 13.83284 | 8.58320 | 6.12665 | 19.63086 |
| **Rabif** | 7.02312 | 0.05152 | 9.32717 | 1.12769 | 9.10279 | 4.74766 | 3.45207 | 4.93012 | 1.22918 | 0.08894 | 6.57756 | 3.16039 | 5.91604 | 7.93545 |
| **Rabl2** | 1.80407 | 0.01781 | 1.22472 | 2.31537 | 1.64960 | 2.76957 | 0.21675 | 0.17258 | 0.74553 | 0.02073 | 0.04218 | 2.25601 | 0.58127 | 3.05456 |
| **Rabl3** | 0.00000 | 0.04563 | 2.18028 | 1.56921 | 0.67091 | 4.68447 | 0.00000 | 2.89387 | 2.18225 | 9.73515 | 2.09656 | 3.62794 | 0.19716 | 2.63170 |
| **Rabl6** | 0.00000 | 0.04991 | 4.42084 | 3.29684 | 0.29779 | 4.15578 | 2.48160 | 1.12827 | 0.13234 | 0.10162 | 1.98219 | 5.83102 | 0.00000 | 0.28168 |
| **Rac1** | 8.32862 | 0.65758 | 5.57211 | 7.50604 | 10.33083 | 9.57829 | 1.42812 | 8.20395 | 8.28202 | 2.88592 | 7.64549 | 9.25331 | 12.73446 | 7.10121 |
| **Rac2** | 0.04675 | 0.01996 | 0.03779 | 0.16023 | 0.04573 | 0.12240 | 0.05700 | 0.06164 | 0.11285 | 0.01676 | 0.01079 | 0.03567 | 0.26528 | 0.27942 |
| **Rac3** | 0.00000 | 0.00392 | 2.35060 | 2.81535 | 1.99968 | 0.59872 | 0.00341 | 0.09845 | 0.00301 | 0.08026 | 0.02877 | 0.04339 | 0.13609 | 0.11282 |
| **Racgap1** | 18.00465 | 2.26506 | 18.25067 | 9.50237 | 30.91923 | 15.16860 | 2.25910 | 13.87852 | 14.70574 | 0.15739 | 7.35656 | 6.20311 | 21.78616 | 25.00244 |
| **Rack1** | 41.46555 | 20.43354 | 63.90000 | 61.00376 | 54.31114 | 50.70260 | 15.74727 | 45.50232 | 24.39693 | 30.11379 | 32.59842 | 42.66557 | 16.25879 | 43.97876 |
| **Rad1** | 7.47648 | 0.09909 | 4.85751 | 5.20983 | 6.61575 | 4.74161 | 1.95582 | 2.78134 | 5.55250 | 0.33734 | 2.29467 | 3.21655 | 8.23505 | 3.10096 |
| **Rad17** | 2.97826 | 0.01292 | 7.72382 | 2.49396 | 11.47154 | 2.35061 | 3.43170 | 0.12458 | 0.61952 | 0.12784 | 4.21568 | 5.03011 | 5.18401 | 0.00000 |
| **Rad18** | 0.32167 | 0.00316 | 0.76909 | 0.58417 | 0.94586 | 0.45538 | 0.00365 | 0.01254 | 0.16059 | 0.00535 | 2.36282 | 0.63275 | 0.42628 | 1.02321 |
| **Rad21** | 24.77028 | 0.33619 | 26.03071 | 16.35275 | 31.76525 | 14.24639 | 5.66610 | 10.34783 | 31.80644 | 0.31967 | 16.75880 | 16.34654 | 16.69531 | 30.24422 |
| **Rad23a** | 0.07108 | 0.01455 | 0.30175 | 0.00000 | 0.02123 | 0.02852 | 0.00418 | 0.55233 | 0.01934 | 0.00734 | 0.75422 | 0.18359 | 0.00337 | 0.39833 |
| **Rad23b** | 13.60906 | 1.43414 | 7.96794 | 12.45900 | 10.10650 | 12.91616 | 2.60281 | 6.50294 | 14.48439 | 1.23823 | 9.01529 | 10.37724 | 8.88478 | 5.57688 |
| **Rad50** | 0.28751 | 2.59468 | 1.53270 | 1.30842 | 2.91877 | 2.34678 | 0.00173 | 0.00000 | 3.26492 | 0.00000 | 2.70356 | 1.44130 | 3.38062 | 2.32869 |
| **Rad51** | 5.19665 | 0.35251 | 8.98407 | 4.63698 | 10.55910 | 9.62303 | 2.89159 | 4.36063 | 7.47651 | 0.07940 | 4.90340 | 9.82865 | 2.61048 | 9.86696 |
| **Rad51ap1** | 5.35866 | 2.78468 | 8.36702 | 2.54040 | 8.12227 | 5.66282 | 0.02694 | 2.83110 | 1.92187 | 11.35290 | 8.81717 | 4.10284 | 8.63285 | 13.77532 |
| **Rad51b** | 0.61478 | 0.02462 | 0.06412 | 0.00000 | 0.00000 | 0.47521 | 0.00298 | 1.33638 | 0.66214 | 0.01479 | 1.79701 | 1.32308 | 0.00371 | 0.41257 |
| **Rad51c** | 0.02066 | 0.01106 | 1.19343 | 0.44257 | 1.34659 | 1.03288 | 0.00344 | 0.00330 | 0.94585 | 0.00346 | 0.10115 | 0.26633 | 2.17844 | 2.25845 |
| **Rad51d** | 2.58408 | 0.29465 | 2.30274 | 6.65143 | 1.82965 | 4.58300 | 0.74769 | 6.10438 | 1.14323 | 0.40659 | 1.49633 | 3.20388 | 0.86841 | 2.53339 |
| **Rad52** | 4.80885 | 0.02443 | 4.71755 | 3.96489 | 4.23008 | 3.39113 | 0.00925 | 0.32683 | 0.00073 | 0.02236 | 1.44066 | 0.74570 | 1.10896 | 0.40194 |
| **Rad54b** | 3.66028 | 0.00000 | 8.09746 | 2.34999 | 5.14760 | 5.94837 | 2.89945 | 0.23005 | 4.25391 | 0.06968 | 3.64696 | 3.50898 | 3.51387 | 3.09039 |
| **Rad54l** | 2.52890 | 0.00023 | 2.87887 | 0.00000 | 3.26260 | 4.83980 | 1.37062 | 1.67803 | 2.55443 | 0.01254 | 4.60628 | 3.05980 | 0.98634 | 6.92040 |
| **Rad54l2** | 0.38457 | 0.03728 | 3.63225 | 2.58135 | 7.80527 | 5.37941 | 0.02002 | 1.98174 | 4.18204 | 0.02725 | 3.89515 | 2.70036 | 3.96387 | 8.49567 |
| **Rad9a** | 1.07503 | 0.03386 | 6.35978 | 2.99145 | 2.91549 | 1.78112 | 5.37308 | 5.84336 | 0.44016 | 0.15265 | 5.41826 | 1.17009 | 1.83341 | 6.11327 |
| **Rad9b** | 0.00000 | 0.00406 | 0.07415 | 0.02924 | 0.32349 | 0.57734 | 0.00388 | 0.00857 | 0.21774 | 0.01035 | 0.00748 | 0.00618 | 1.79775 | 0.00000 |
| **Radil** | 0.25544 | 0.00411 | 0.52442 | 0.71992 | 0.82176 | 0.00000 | 0.00389 | 0.00626 | 0.08502 | 0.00582 | 0.00410 | 0.00706 | 0.00755 | 0.12512 |
| **Rae1** | 9.13646 | 8.65731 | 12.42522 | 7.24830 | 7.91141 | 7.82785 | 3.01416 | 2.43348 | 2.06021 | 0.11002 | 10.04749 | 9.07182 | 6.29315 | 7.89979 |
| **Raet1a** | 0.00685 | 0.00500 | 0.13833 | 0.00000 | 0.01927 | 0.01050 | 0.00431 | 0.05134 | 0.01094 | 0.00760 | 0.06095 | 0.41091 | 0.00737 | 0.01123 |
| **Raet1d** | 0.01058 | 0.01271 | 0.03344 | 0.00000 | 0.01464 | 0.01405 | 0.00425 | 0.04530 | 0.01160 | 0.00782 | 0.02748 | 0.06597 | 0.00773 | 0.01162 |
| **Raet1e** | 0.21553 | 0.00470 | 0.09220 | 0.00305 | 0.00334 | 0.06386 | 0.00405 | 0.09408 | 0.01973 | 0.02343 | 0.01645 | 0.67280 | 0.13484 | 0.00582 |
| **Raf1** | 3.66994 | 0.02546 | 4.50992 | 1.16194 | 1.30898 | 3.35331 | 0.53581 | 1.65668 | 3.78218 | 0.00000 | 3.34987 | 3.21645 | 6.12216 | 15.56969 |
| **Rai1** | 0.04394 | 0.00412 | 0.03624 | 0.39605 | 0.00407 | 0.00000 | 0.00396 | 0.00503 | 0.01300 | 0.00533 | 0.15848 | 0.21281 | 0.00616 | 0.00474 |
| **Rai14** | 0.00000 | 0.01477 | 1.83326 | 1.49662 | 0.67708 | 2.06796 | 3.35959 | 2.36241 | 0.01050 | 0.06678 | 0.61317 | 5.80926 | 0.73015 | 1.38995 |
| **Rala** | 1.50067 | 0.01712 | 0.47000 | 1.18114 | 1.85615 | 3.99689 | 1.54788 | 0.41634 | 1.64961 | 0.97763 | 0.39292 | 2.91769 | 3.16572 | 2.40326 |
| **Ralb** | 0.00000 | 0.05890 | 0.94170 | 3.43500 | 0.00000 | 0.37544 | 0.01967 | 0.82055 | 0.15960 | 0.06244 | 2.95139 | 6.02217 | 0.29722 | 0.00000 |
| **Ralbp1** | 5.92081 | 0.09173 | 11.03334 | 1.50076 | 10.19321 | 6.90011 | 3.88151 | 7.18613 | 1.23278 | 0.11084 | 2.15695 | 4.27542 | 1.66289 | 5.87174 |
| **Ralgapa1** | 2.70301 | 0.00225 | 1.72618 | 1.12202 | 0.15886 | 1.42766 | 0.00292 | 0.00000 | 1.07685 | 0.01321 | 2.16415 | 1.24651 | 0.71781 | 2.49303 |
| **Ralgapa2** | 0.23909 | 0.01882 | 1.04276 | 0.00000 | 0.32091 | 4.43400 | 0.00907 | 0.38901 | 1.17099 | 0.04124 | 0.03630 | 0.52921 | 0.29077 | 1.49886 |
| **Ralgapb** | 3.87321 | 0.00181 | 3.52468 | 0.01292 | 2.26763 | 4.10922 | 0.00237 | 0.75063 | 4.94517 | 0.00239 | 1.71932 | 2.05894 | 2.90221 | 2.48621 |
| **Ralgds** | 1.79349 | 5.61619 | 6.71609 | 2.57331 | 6.65069 | 6.46569 | 4.55270 | 4.67032 | 0.00101 | 0.21998 | 0.00000 | 0.00000 | 1.42399 | 2.59456 |
| **Ralgps1** | 0.00000 | 0.00395 | 0.19089 | 0.54696 | 0.17536 | 0.00000 | 0.01002 | 0.00755 | 0.01507 | 0.00516 | 0.00866 | 0.01015 | 1.14599 | 0.00000 |
| **Ralgps2** | 3.14631 | 0.05754 | 6.27193 | 2.24485 | 4.41009 | 2.10673 | 2.92703 | 2.60743 | 3.56426 | 0.09154 | 2.68643 | 4.75889 | 4.97441 | 4.10680 |

| | | | | | | | | | | | | | | |
|---|---|---|---|---|---|---|---|---|---|---|---|---|---|---|
| **Raly** | 0.75469 | 0.08216 | 4.33067 | 0.00000 | 4.69640 | 1.54826 | 0.00686 | 5.17661 | 0.00000 | 0.03553 | 0.59969 | 7.35304 | 1.11002 | 0.27433 |
| **Ramp1** | 0.00762 | 0.00435 | 0.22693 | 0.15968 | 0.93121 | 0.00164 | 0.00459 | 0.00861 | 0.01814 | 0.00585 | 0.00606 | 0.00603 | 0.00650 | 0.05019 |
| **Ramp2** | 3.15391 | 0.00337 | 0.09987 | 0.53924 | 4.43164 | 0.00000 | 0.00355 | 0.00851 | 1.27299 | 0.01175 | 0.00000 | 0.68560 | 3.44536 | 1.09117 |
| **Ramp3** | 0.00747 | 0.00383 | 0.01152 | 0.00425 | 0.00957 | 0.01329 | 0.00429 | 0.00652 | 0.00921 | 0.00608 | 0.00530 | 0.61289 | 0.19963 | 0.06921 |
| **Ran** | 31.30803 | 8.91091 | 45.89159 | 38.71476 | 39.90489 | 41.01721 | 13.79870 | 26.31420 | 16.78609 | 15.43174 | 23.85278 | 33.01245 | 19.02759 | 19.34838 |
| **Ranbp1** | 16.61358 | 5.87134 | 6.28331 | 26.91025 | 26.44903 | 15.09140 | 5.36586 | 8.60845 | 2.06035 | 2.86254 | 5.29393 | 15.18436 | 3.98145 | 6.15093 |
| **Ranbp10** | 5.77852 | 5.81712 | 13.41061 | 4.16536 | 14.84980 | 5.82839 | 2.77558 | 3.73459 | 5.02528 | 0.09105 | 2.16303 | 1.51706 | 11.99626 | 15.59706 |
| **Ranbp17** | 1.37433 | 0.03747 | 2.50637 | 0.00000 | 2.67695 | 1.26031 | 0.00707 | 0.23413 | 2.54517 | 0.00830 | 0.65462 | 6.21880 | 0.24611 | 3.25799 |
| **Ranbp2** | 3.82520 | 0.04778 | 5.75888 | 6.15522 | 4.67913 | 5.06096 | 2.13561 | 1.33241 | 25.52314 | 0.05087 | 5.49779 | 2.22984 | 14.45271 | 3.36101 |
| **Ranbp3** | 5.69757 | 0.12288 | 13.03440 | 6.23852 | 18.14786 | 4.97283 | 3.22940 | 3.70449 | 0.13679 | 10.07147 | 3.43675 | 8.00034 | 7.54861 | 5.25089 |
| **Ranbp3l** | 0.07887 | 0.19944 | 0.02228 | 0.33023 | 0.18060 | 0.53928 | 0.16224 | 0.18789 | 0.28410 | 0.07026 | 0.01437 | 0.07714 | 0.26024 | 0.42139 |
| **Ranbp6** | 0.11697 | 0.00414 | 0.49351 | 0.38764 | 0.17769 | 0.69848 | 0.00371 | 0.00115 | 0.04465 | 0.01301 | 0.01013 | 0.00696 | 0.02501 | 0.96838 |
| **Ranbp9** | 6.87099 | 0.01275 | 4.58440 | 1.24759 | 5.95788 | 3.93732 | 2.92101 | 2.48680 | 2.04348 | 0.03495 | 5.44772 | 3.67644 | 8.81654 | 9.39792 |
| **Rangap1** | 6.06346 | 8.01198 | 28.38794 | 19.58762 | 18.40753 | 24.67116 | 6.19085 | 11.38550 | 11.86894 | 12.09796 | 15.66855 | 11.78507 | 4.18286 | 19.57532 |
| **Rangrf** | 1.31028 | 0.12634 | 1.57811 | 7.46523 | 4.15722 | 1.40520 | 3.09831 | 3.24885 | 0.00000 | 0.09013 | 2.72230 | 7.70090 | 1.13047 | 1.36388 |
| **Rap1a** | 14.08557 | 0.10261 | 14.69553 | 6.62674 | 13.67917 | 11.45719 | 0.02229 | 0.55579 | 18.32086 | 7.97881 | 7.51686 | 9.45642 | 0.62498 | 6.53973 |
| **Rap1b** | 6.70118 | 0.04396 | 6.08001 | 6.89547 | 7.65370 | 13.53091 | 3.86973 | 3.91449 | 12.32751 | 0.20050 | 5.84116 | 5.60856 | 10.70265 | 2.80159 |
| **Rap1gap** | 0.01803 | 0.00432 | 0.13650 | 0.06834 | 0.55929 | 0.00444 | 0.00403 | 0.00833 | 0.01033 | 0.00650 | 0.00671 | 0.01123 | 0.00824 | 0.02951 |
| **Rap1gap2** | 0.30651 | 0.00322 | 0.99878 | 0.27572 | 0.61325 | 1.70299 | 1.68443 | 0.00000 | 0.65551 | 0.01517 | 0.00000 | 0.97068 | 0.81212 | 0.03807 |
| **Rap1gds1** | 0.97260 | 0.00269 | 1.42049 | 0.00000 | 0.00000 | 2.28556 | 0.00438 | 0.29335 | 2.87388 | 0.00254 | 0.00000 | 1.85161 | 0.95108 | 2.94561 |
| **Rap2a** | 1.01634 | 0.01154 | 0.24229 | 0.00000 | 0.96560 | 1.84329 | 0.01589 | 0.81097 | 0.32999 | 0.00828 | 0.00199 | 0.13895 | 3.66574 | 0.81871 |
| **Rap2b** | 0.01382 | 0.01220 | 0.07548 | 2.04441 | 0.06231 | 0.23942 | 1.88944 | 0.00403 | 0.01118 | 0.11827 | 0.00060 | 0.33857 | 0.01058 | 0.03488 |
| **Rap2c** | 2.95468 | 0.01782 | 1.67772 | 0.44793 | 4.39503 | 2.41013 | 1.33665 | 2.64165 | 10.57712 | 0.03732 | 0.00000 | 3.26269 | 0.00000 | 0.91284 |
| **Rapgef1** | 0.56309 | 0.05683 | 1.62379 | 2.69875 | 0.79612 | 4.05930 | 0.49090 | 2.33409 | 0.82837 | 0.00000 | 4.55963 | 4.47808 | 0.77982 | 0.49632 |
| **Rapgef2** | 0.47206 | 3.14922 | 0.94143 | 0.00000 | 0.20629 | 2.83540 | 0.00220 | 0.05204 | 3.50998 | 0.00436 | 0.00000 | 1.53489 | 1.34207 | 0.25152 |
| **Rapgef4** | 0.15947 | 0.06853 | 0.01660 | 0.65870 | 0.13494 | 0.46737 | 0.14077 | 0.09968 | 0.21352 | 0.35791 | 0.02109 | 0.11398 | 0.46072 | 0.35611 |
| **Rapgef5** | 0.00400 | 0.00284 | 0.00000 | 0.47636 | 0.47378 | 0.00000 | 0.00301 | 1.45450 | 0.00727 | 0.01028 | 0.00000 | 1.20731 | 0.73963 | 1.78992 |
| **Rapgef6** | 1.79987 | 4.83492 | 2.74640 | 1.05672 | 3.40611 | 3.27180 | 0.00665 | 2.55257 | 4.99186 | 0.00600 | 2.27138 | 1.97934 | 3.09375 | 7.47037 |
| **Rapgefl1** | 0.13949 | 0.01903 | 0.14247 | 1.46019 | 0.57915 | 0.42334 | 0.00410 | 1.79252 | 0.27847 | 0.01542 | 0.17040 | 0.00262 | 0.92318 | 0.31604 |
| **Raph1** | 2.13502 | 0.27104 | 0.57175 | 0.29079 | 0.56330 | 3.78851 | 0.08969 | 2.86287 | 1.06927 | 0.05264 | 1.90098 | 2.13319 | 1.00181 | 1.59568 |
| **Rapsn** | 0.03016 | 0.00399 | 0.01401 | 0.00101 | 0.00942 | 0.00552 | 1.77490 | 0.00285 | 0.01186 | 0.03754 | 0.00685 | 0.01002 | 0.00748 | 0.01012 |
| **Rarb** | 0.68897 | 0.00390 | 1.57448 | 0.00000 | 0.00005 | 0.40688 | 0.42723 | 0.11125 | 0.01541 | 0.02139 | 0.01261 | 1.09044 | 0.00000 | 0.18035 |
| **Rarg** | 2.40832 | 0.01054 | 5.68058 | 1.84800 | 9.53856 | 0.00000 | 0.54758 | 0.53021 | 0.01096 | 0.01009 | 0.00356 | 0.00000 | 4.43761 | 0.94060 |
| **Rarres2** | 0.00854 | 0.00460 | 0.01075 | 0.06035 | 0.02518 | 0.01736 | 0.00458 | 0.02844 | 0.01137 | 0.00878 | 0.01034 | 0.00720 | 0.00591 | 0.01685 |
| **Rars** | 20.71184 | 9.92626 | 24.56536 | 16.94727 | 17.08477 | 19.35446 | 7.12930 | 11.21956 | 23.70356 | 10.41371 | 18.30052 | 12.30435 | 10.09158 | 27.14691 |
| **Rars2** | 3.09613 | 0.05416 | 5.71540 | 1.77320 | 2.35340 | 7.94134 | 1.21842 | 1.98040 | 2.07078 | 0.03887 | 4.64602 | 7.13264 | 0.00000 | 2.94732 |
| **Rasa1** | 6.40231 | 0.00000 | 3.24987 | 0.00000 | 1.91505 | 5.38086 | 3.05221 | 1.30484 | 6.22449 | 0.06771 | 2.97405 | 4.57281 | 0.78850 | 4.24955 |
| **Rasa2** | 0.34235 | 0.00258 | 0.00000 | 1.30230 | 0.54669 | 1.08990 | 0.00311 | 0.12387 | 3.10871 | 0.03473 | 0.27739 | 1.00043 | 4.99283 | 2.68237 |
| **Rasa3** | 0.01011 | 0.00453 | 0.01127 | 0.00212 | 0.00777 | 0.20520 | 0.00423 | 0.00519 | 0.01336 | 0.00581 | 0.00803 | 0.01031 | 0.29005 | 0.11275 |
| **Rasa4** | 0.19943 | 0.00420 | 0.30247 | 0.46904 | 0.44450 | 0.01034 | 0.00388 | 0.00477 | 0.01916 | 0.00563 | 0.00972 | 0.01098 | 0.16630 | 0.00000 |
| **Rasal1** | 0.02802 | 0.00413 | 0.01449 | 0.37422 | 0.00702 | 0.00865 | 0.00397 | 0.00545 | 0.01253 | 0.00571 | 0.01303 | 0.01750 | 0.00657 | 0.00727 |
| **Rasal2** | 1.07705 | 0.02236 | 1.74601 | 0.00000 | 1.32360 | 0.81193 | 0.00364 | 0.01959 | 0.35617 | 0.01299 | 0.86400 | 0.22032 | 1.19050 | 2.73506 |
| **Rasal3** | 0.01074 | 0.00454 | 0.01757 | 0.01216 | 0.01394 | 0.01790 | 0.00424 | 0.01070 | 0.01157 | 0.00710 | 0.00806 | 0.01311 | 0.00746 | 0.01204 |
| **Rasd1** | 0.16953 | 1.23531 | 0.16969 | 0.00629 | 0.00000 | 0.00000 | 0.00400 | 0.01970 | 0.01549 | 0.03541 | 0.38769 | 0.00096 | 0.12187 | 0.00928 |
| **Rasd2** | 0.00470 | 0.01117 | 0.00000 | 0.02702 | 0.16007 | 0.23400 | 0.02182 | 1.35658 | 0.07966 | 0.02371 | 0.33262 | 0.00217 | 0.74124 | 0.14477 |
| **Rasef** | 0.00686 | 0.00461 | 0.01411 | 0.01159 | 0.00833 | 0.01170 | 0.00433 | 0.01441 | 0.02674 | 0.00676 | 0.00892 | 0.00932 | 0.32855 | 0.02974 |
| **Rasgef1a** | 0.00301 | 0.00396 | 0.87724 | 0.00000 | 0.08713 | 0.00348 | 0.00387 | 0.00409 | 0.00852 | 0.00902 | 0.00427 | 0.20718 | 0.09118 | 0.00420 |
| **Rasgef1b** | 0.00757 | 0.00429 | 0.01166 | 0.01327 | 0.08623 | 0.01455 | 0.00412 | 0.00835 | 0.01243 | 0.00703 | 0.00899 | 0.00841 | 0.59041 | 0.01555 |
| **Rasgrf1** | 0.00325 | 0.00426 | 0.12176 | 0.00365 | 0.00492 | 0.00877 | 0.00416 | 0.00978 | 0.02097 | 0.00646 | 0.00753 | 0.00946 | 0.63612 | 0.03800 |
| **Rasgrp2** | 0.30387 | 0.00417 | 0.04530 | 0.00515 | 0.38147 | 0.00298 | 0.00425 | 0.01094 | 0.01196 | 0.00549 | 0.39235 | 0.00993 | 0.03337 | 0.01988 |
| **Rasgrp3** | 0.03603 | 0.10670 | 0.01903 | 0.01521 | 0.01886 | 0.02592 | 0.03887 | 0.01060 | 0.03115 | 0.00640 | 0.01075 | 0.02263 | 0.05255 | 0.03011 |
| **Rasgrp4** | 1.01726 | 1.11236 | 5.33183 | 3.94205 | 3.07767 | 2.27755 | 0.01380 | 5.65550 | 0.21112 | 0.04726 | 3.04756 | 0.18395 | 0.95003 | 3.49454 |
| **Rasip1** | 0.05364 | 0.00391 | 0.46965 | 0.00000 | 0.66950 | 0.16369 | 0.00361 | 1.35357 | 0.00592 | 0.01106 | 0.00712 | 0.00661 | 0.00000 | 0.16652 |
| **Rasl10a** | 0.00638 | 0.01951 | 0.51749 | 1.07290 | 1.08372 | 0.00000 | 2.79214 | 0.72911 | 0.00425 | 0.05049 | 0.00000 | 0.00226 | 0.26410 | 0.00000 |
| **Rasl11a** | 1.62580 | 0.00428 | 0.03744 | 0.00000 | 1.37850 | 0.00239 | 0.00432 | 0.01177 | 0.03491 | 0.00652 | 0.00655 | 0.00412 | 0.01588 | 1.36846 |

| | | | | | | | | | | | | | | |
|---|---|---|---|---|---|---|---|---|---|---|---|---|---|---|
| **Rasl11b** | 0.00762 | 0.00410 | 0.01760 | 0.17357 | 0.00522 | 0.00454 | 0.00406 | 0.00649 | 0.00699 | 0.00569 | 0.02822 | 0.01594 | 0.27738 | 0.19354 |
| **Rasl12** | 0.01117 | 0.00440 | 0.01360 | 0.04203 | 0.02071 | 0.04624 | 0.00414 | 0.00644 | 0.05449 | 0.00706 | 0.00964 | 0.01724 | 0.06674 | 0.07170 |
| **Rasl2.9** | 3.84270 | 0.43215 | 5.01665 | 3.96174 | 4.34015 | 4.93496 | 1.53926 | 2.10047 | 0.60946 | 1.31865 | 2.16811 | 4.35974 | 1.93968 | 0.91766 |
| **Rassf1** | 0.37212 | 0.04167 | 2.11629 | 0.00000 | 1.68157 | 2.63880 | 6.96023 | 2.91033 | 0.14077 | 0.23281 | 0.00000 | 2.44226 | 0.87208 | 0.41290 |
| **Rassf3** | 0.00000 | 0.00994 | 5.53563 | 0.00000 | 6.17942 | 1.29329 | 0.00165 | 1.03987 | 5.62169 | 0.00451 | 1.60203 | 0.95877 | 4.47901 | 3.98940 |
| **Rassf4** | 0.00904 | 0.00446 | 0.01078 | 0.00476 | 0.01745 | 0.30539 | 0.00421 | 0.00670 | 0.01039 | 0.00678 | 0.00609 | 0.00986 | 0.00805 | 0.00980 |
| **Rassf5** | 0.00445 | 0.00414 | 0.29828 | 0.05716 | 0.02166 | 0.02681 | 0.00414 | 0.00366 | 0.01628 | 0.00746 | 0.01006 | 0.00830 | 0.57352 | 1.26765 |
| **Rassf7** | 0.63421 | 0.11759 | 12.47277 | 11.48115 | 4.46955 | 0.78998 | 4.40825 | 11.08355 | 0.00206 | 0.20054 | 0.94308 | 1.20109 | 0.78918 | 0.04708 |
| **Rassf8** | 0.00000 | 0.00384 | 0.35589 | 0.56130 | 0.00000 | 0.16291 | 2.44839 | 0.54769 | 0.00566 | 0.01588 | 0.00119 | 0.00561 | 0.69799 | 0.06799 |
| **Raver1** | 2.71405 | 0.02347 | 7.01039 | 3.90669 | 6.07338 | 3.62857 | 0.23077 | 1.08457 | 1.35946 | 0.02991 | 2.09242 | 3.54214 | 4.61315 | 1.84249 |
| **Raver2** | 0.00212 | 0.00388 | 0.06765 | 0.12853 | 0.00000 | 0.00359 | 0.00390 | 0.03745 | 0.02290 | 0.00612 | 0.00523 | 0.17312 | 0.00750 | 0.45456 |
| **Rb1** | 0.17563 | 0.00375 | 0.00000 | 0.00359 | 0.36193 | 0.01704 | 0.00382 | 0.00332 | 3.38913 | 0.00603 | 0.00845 | 0.10431 | 1.54771 | 2.66799 |
| **Rb1cc1** | 26.75985 | 0.00503 | 14.69037 | 8.28055 | 33.73902 | 4.25037 | 1.67448 | 0.06315 | 46.82795 | 0.00000 | 2.27940 | 0.39837 | 12.58082 | 31.31553 |
| **Rbak** | 0.16304 | 0.05896 | 0.70905 | 0.36737 | 1.18020 | 0.48792 | 0.07225 | 0.26093 | 0.35851 | 0.12279 | 0.03035 | 0.73376 | 0.54340 | 1.05225 |
| **Rbakdn** | 0.00926 | 0.00446 | 0.06900 | 0.00000 | 0.01033 | 0.13378 | 0.00420 | 0.33819 | 0.01135 | 0.00648 | 0.00534 | 0.00954 | 0.00750 | 0.00983 |
| **Rbbp4** | 0.85908 | 0.05560 | 2.34564 | 1.79890 | 4.66866 | 4.84671 | 1.66506 | 2.85952 | 8.79582 | 0.07663 | 3.95543 | 2.03579 | 9.18069 | 3.43394 |
| **Rbbp5** | 5.83240 | 0.01996 | 7.70747 | 0.00000 | 6.97871 | 4.57408 | 4.14184 | 2.52292 | 7.24666 | 0.11247 | 6.49810 | 4.31785 | 1.31246 | 11.19719 |
| **Rbbp6** | 9.93619 | 3.45067 | 6.97839 | 1.24557 | 6.31716 | 8.78178 | 5.10005 | 1.73655 | 11.15785 | 0.09634 | 7.48536 | 3.77867 | 12.56429 | 8.05996 |
| **Rbbp7** | 25.70147 | 8.39501 | 37.78212 | 19.67338 | 26.31460 | 35.21958 | 4.56454 | 11.75840 | 27.53802 | 0.46009 | 23.78403 | 21.28376 | 7.89280 | 18.66119 |
| **Rbbp8** | 12.40518 | 0.23164 | 5.76356 | 1.79036 | 11.79401 | 4.57325 | 2.84322 | 0.00000 | 19.20666 | 0.08813 | 1.46047 | 0.02579 | 4.80110 | 9.20748 |
| **Rbbp9** | 2.02581 | 0.04293 | 1.18342 | 0.34226 | 0.67848 | 0.71343 | 1.21834 | 0.15908 | 1.42470 | 0.02286 | 3.49204 | 1.64021 | 0.01539 | 2.22425 |
| **Rbck1** | 0.81943 | 0.09394 | 5.47098 | 3.85972 | 2.82996 | 0.22581 | 0.00169 | 3.51273 | 0.00000 | 0.00479 | 2.09717 | 3.27074 | 0.88393 | 0.00000 |
| **Rbfa** | 0.05709 | 0.01260 | 0.60127 | 0.64138 | 1.53312 | 0.49629 | 0.00404 | 0.06447 | 0.01055 | 0.01258 | 0.15440 | 0.48102 | 0.00345 | 0.00000 |
| **Rbfox2** | 1.38232 | 0.02295 | 2.12410 | 1.81933 | 2.38977 | 3.87843 | 0.66846 | 3.68081 | 0.06218 | 0.02828 | 2.92553 | 2.61537 | 2.41291 | 2.62287 |
| **Rbks** | 0.32594 | 0.02956 | 1.51660 | 1.42226 | 0.32888 | 0.63856 | 1.94581 | 0.82793 | 0.29680 | 0.06932 | 0.06140 | 1.11412 | 0.00000 | 0.41941 |
| **Rbl1** | 0.19691 | 0.17303 | 0.32275 | 0.00000 | 0.00000 | 0.00757 | 0.00393 | 1.41281 | 0.88062 | 0.01139 | 0.15834 | 0.03714 | 1.86811 | 1.16649 |
| **Rbl2** | 0.11087 | 0.00454 | 0.37796 | 0.00000 | 0.62793 | 0.29434 | 0.00374 | 0.08920 | 0.01472 | 0.00537 | 0.00472 | 0.01601 | 0.00355 | 0.30738 |
| **Rbm10** | 7.53291 | 0.04753 | 10.25331 | 4.52247 | 5.13151 | 7.71291 | 0.25227 | 5.47772 | 1.71401 | 0.04095 | 5.12603 | 2.51128 | 5.07357 | 8.41447 |
| **Rbm11** | 0.04688 | 0.02094 | 0.00000 | 0.13884 | 0.09320 | 0.15898 | 0.02991 | 0.07249 | 0.19086 | 0.02871 | 0.01221 | 0.04364 | 0.32727 | 0.21121 |
| **Rbm12** | 7.70816 | 0.04398 | 3.31436 | 2.58946 | 1.79095 | 2.03062 | 4.00044 | 1.06904 | 1.83164 | 0.14545 | 0.37797 | 2.21318 | 5.33615 | 2.89242 |
| **Rbm12b1** | 0.03424 | 0.02978 | 0.47342 | 0.06380 | 0.32832 | 0.00621 | 0.00375 | 0.70990 | 1.95127 | 0.00630 | 0.00000 | 0.78245 | 0.00066 | 1.37230 |
| **Rbm12b2** | 0.00000 | 0.01357 | 0.19921 | 0.22863 | 1.07048 | 0.59891 | 0.00393 | 0.02625 | 1.86410 | 0.01733 | 0.00000 | 0.56340 | 1.63149 | 1.94108 |
| **Rbm14** | 3.34785 | 7.45404 | 9.21885 | 10.52510 | 9.97937 | 2.31487 | 2.23909 | 5.47235 | 2.23630 | 8.23268 | 3.13193 | 6.78206 | 4.31282 | 1.58006 |
| **Rbm15** | 1.04703 | 0.06993 | 1.77605 | 2.58721 | 0.49119 | 2.92533 | 2.48788 | 0.55724 | 1.44523 | 0.01954 | 3.23863 | 0.36139 | 5.40447 | 0.59178 |
| **Rbm15b** | 0.00000 | 0.01763 | 2.40098 | 0.12018 | 1.17645 | 0.50213 | 0.00811 | 3.69833 | 0.03463 | 0.01532 | 0.28766 | 1.41016 | 1.07058 | 0.20950 |
| **Rbm17** | 13.77312 | 3.71593 | 15.87782 | 3.95705 | 17.22256 | 14.80700 | 5.04253 | 0.35848 | 14.69385 | 7.27773 | 7.57521 | 9.76665 | 11.60270 | 20.62998 |
| **Rbm18** | 5.47575 | 5.45087 | 4.27777 | 4.34419 | 6.85600 | 6.60516 | 0.80836 | 0.00000 | 8.09653 | 0.07186 | 1.42318 | 4.19378 | 4.32690 | 9.27518 |
| **Rbm19** | 1.30071 | 0.01826 | 7.38079 | 6.05177 | 7.03232 | 6.42417 | 0.80665 | 0.69118 | 0.07336 | 0.11248 | 5.10445 | 3.63065 | 4.53572 | 4.13871 |
| **Rbm20** | 0.00877 | 0.00437 | 0.00288 | 0.10054 | 0.00890 | 0.26167 | 0.15317 | 0.00682 | 0.01350 | 0.01295 | 0.08269 | 0.29295 | 0.00969 | 0.01054 |
| **Rbm22** | 9.54345 | 1.59951 | 3.52194 | 3.85137 | 9.63722 | 2.45749 | 0.31394 | 1.86465 | 3.71625 | 0.04135 | 4.01381 | 0.90698 | 9.99079 | 6.98560 |
| **Rbm25** | 11.88270 | 6.89629 | 14.09313 | 8.89145 | 11.53064 | 12.40249 | 4.61574 | 3.01269 | 22.73689 | 4.94860 | 7.17617 | 8.49120 | 10.89029 | 13.60144 |
| **Rbm26** | 1.84173 | 0.07759 | 4.51648 | 2.92491 | 5.33959 | 2.52710 | 0.00519 | 3.19569 | 3.81528 | 0.00590 | 7.54770 | 5.94069 | 2.12934 | 5.47142 |
| **Rbm27** | 2.30968 | 0.03336 | 1.29956 | 2.87963 | 3.53109 | 2.72655 | 0.00792 | 2.13367 | 11.46139 | 0.00564 | 2.25946 | 1.12639 | 9.70473 | 3.69658 |
| **Rbm28** | 11.25795 | 0.05218 | 13.54498 | 4.85801 | 15.64745 | 11.17191 | 2.00625 | 1.86448 | 6.26684 | 0.09721 | 6.11560 | 4.54842 | 11.48313 | 20.55167 |
| **Rbm3** | 27.51454 | 8.44579 | 29.64621 | 24.20862 | 25.36298 | 20.70887 | 4.04356 | 18.59182 | 32.85363 | 4.93619 | 13.93739 | 15.98573 | 8.66579 | 28.42204 |
| **Rbm33** | 9.28241 | 0.03972 | 6.58050 | 1.15115 | 8.14988 | 4.96455 | 2.89586 | 3.32558 | 3.82796 | 7.62342 | 3.91033 | 6.35682 | 9.74970 | 12.36510 |
| **Rbm34** | 4.84920 | 0.08518 | 5.70419 | 4.30828 | 7.13174 | 2.90761 | 3.25508 | 2.93135 | 12.64779 | 0.12116 | 5.16862 | 1.64573 | 2.88242 | 4.95979 |
| **Rbm38** | 0.16396 | 1.20075 | 2.79287 | 0.15124 | 0.21003 | 0.10434 | 0.88970 | 0.00000 | 0.01128 | 0.07018 | 0.58587 | 1.15317 | 2.66095 | 0.04186 |
| **Rbm39** | 21.73791 | 0.21099 | 16.60252 | 9.55268 | 23.93729 | 16.93431 | 4.52860 | 7.03756 | 23.17980 | 0.09051 | 11.96116 | 13.12588 | 13.60796 | 16.80738 |
| **Rbm3os** | 0.01138 | 0.00448 | 0.05152 | 0.00493 | 0.10456 | 0.01232 | 0.00419 | 0.01072 | 0.18681 | 0.00760 | 0.00738 | 0.00902 | 0.03439 | 0.02011 |
| **Rbm4** | 0.00689 | 0.02033 | 0.19016 | 0.06845 | 0.03049 | 0.07600 | 0.00439 | 0.16320 | 0.00773 | 0.00572 | 0.12416 | 0.05157 | 0.00537 | 0.20529 |
| **Rbm41** | 4.39815 | 0.00229 | 1.63600 | 0.00000 | 8.47636 | 0.57502 | 1.02536 | 0.00000 | 11.13858 | 0.08476 | 1.94874 | 0.35174 | 0.11767 | 10.72143 |
| **Rbm42** | 9.37239 | 4.17561 | 7.46104 | 33.85848 | 28.96928 | 2.77883 | 0.43913 | 5.63473 | 0.74651 | 7.30178 | 1.44985 | 9.67458 | 16.58645 | 3.78443 |
| **Rbm44** | 0.79878 | 0.00400 | 0.89879 | 0.29668 | 0.51465 | 0.44661 | 0.21869 | 0.32320 | 1.92726 | 0.00303 | 1.01868 | 0.48371 | 1.67053 | 0.54450 |
| **Rbm45** | 0.04978 | 0.00400 | 1.07565 | 0.46095 | 0.46979 | 0.00000 | 0.00352 | 0.18101 | 1.80933 | 0.02429 | 0.01250 | 0.00557 | 0.37532 | 1.02504 |

| | | | | | | | | | | | | | | |
|---|---|---|---|---|---|---|---|---|---|---|---|---|---|---|
| **Rbm46os** | 0.00394 | 0.00435 | 0.01688 | 0.01108 | 0.01068 | 0.01157 | 0.01620 | 0.00890 | 0.06938 | 0.00648 | 0.00793 | 0.01288 | 0.16316 | 0.15008 |
| **Rbm47** | 0.88228 | 0.31431 | 9.35143 | 3.16605 | 3.65453 | 1.72756 | 0.64738 | 4.34100 | 7.55371 | 0.05558 | 5.22137 | 1.02318 | 0.62481 | 7.50613 |
| **Rbm48** | 3.97788 | 0.00864 | 2.78437 | 1.37139 | 5.40868 | 4.74969 | 4.06782 | 0.31629 | 1.53891 | 0.07134 | 0.00000 | 1.92569 | 1.58529 | 9.97010 |
| **Rbm4b** | 0.15281 | 2.04770 | 0.80442 | 0.27855 | 0.39494 | 0.64786 | 0.83906 | 1.33813 | 1.35510 | 0.01029 | 0.22907 | 0.21663 | 0.37850 | 0.40181 |
| **Rbm5** | 9.21467 | 0.05825 | 7.72337 | 5.82374 | 6.57707 | 4.72719 | 5.32699 | 4.10022 | 13.16274 | 0.12100 | 7.84239 | 4.44435 | 8.64416 | 10.07677 |
| **Rbm6** | 7.44863 | 0.11290 | 11.79370 | 5.03789 | 15.30000 | 8.99723 | 1.12019 | 5.57497 | 15.85847 | 0.09088 | 9.41975 | 4.94942 | 8.98062 | 16.26115 |
| **Rbm7** | 4.34633 | 0.03543 | 9.02916 | 2.90539 | 10.63314 | 8.09924 | 0.07694 | 1.41034 | 6.52952 | 0.00000 | 7.39717 | 4.63193 | 2.89505 | 6.81655 |
| **Rbm8a** | 11.47825 | 7.61524 | 9.45502 | 22.19670 | 22.92868 | 18.92876 | 8.69584 | 13.66700 | 5.50832 | 19.28984 | 11.21704 | 15.94117 | 16.45237 | 20.30249 |
| **Rbms1** | 0.54326 | 0.03682 | 0.36962 | 0.29545 | 0.08780 | 1.76062 | 0.00485 | 1.14281 | 0.00607 | 0.01268 | 0.00475 | 0.00000 | 0.00429 | 0.00000 |
| **Rbms2** | 1.16488 | 5.71834 | 7.60374 | 2.11813 | 9.69922 | 6.46180 | 2.86231 | 3.52790 | 10.32670 | 0.62716 | 1.52631 | 5.45006 | 1.30903 | 8.74121 |
| **Rbmx** | 2.71225 | 0.03099 | 5.44468 | 8.87099 | 8.13373 | 1.97608 | 0.64286 | 2.54498 | 10.29967 | 0.04131 | 3.47780 | 0.97363 | 4.49948 | 8.14122 |
| **Rbmx2** | 2.44604 | 0.04110 | 4.07244 | 4.42874 | 5.88483 | 7.33786 | 2.36636 | 3.57540 | 7.10760 | 0.08553 | 0.00000 | 3.41533 | 5.61393 | 6.40781 |
| **Rbmxl1** | 14.29604 | 0.35678 | 27.49807 | 22.09985 | 21.96189 | 21.17828 | 7.89328 | 12.44917 | 9.64763 | 0.54242 | 17.06125 | 16.88258 | 10.17402 | 13.43480 |
| **Rbmxl2** | 0.01261 | 0.00419 | 0.01127 | 0.02006 | 0.02781 | 0.01180 | 0.00419 | 0.00747 | 0.00995 | 0.00805 | 0.01433 | 0.15040 | 0.11613 | 0.03468 |
| **Rbp1** | 0.00778 | 0.00439 | 0.00657 | 0.00992 | 0.66153 | 0.01421 | 0.00430 | 0.01448 | 0.01008 | 0.00665 | 0.00871 | 0.00795 | 0.00629 | 0.01032 |
| **Rbp7** | 0.00000 | 0.00426 | 0.19620 | 0.00000 | 0.30027 | 0.00716 | 0.00404 | 0.00905 | 0.02123 | 0.00784 | 0.00700 | 0.00885 | 0.79739 | 0.00361 |
| **Rbpj** | 12.13166 | 4.12635 | 11.94007 | 7.31239 | 12.04020 | 13.76236 | 2.32945 | 2.95512 | 9.84853 | 0.20549 | 14.18962 | 12.66573 | 5.75412 | 13.17192 |
| **Rbpms** | 18.58913 | 0.18673 | 7.79778 | 4.55278 | 14.73607 | 8.40498 | 2.89141 | 3.03639 | 5.32922 | 0.17617 | 6.75823 | 5.52646 | 11.80449 | 8.56162 |
| **Rbpms2** | 19.28719 | 0.38026 | 20.16332 | 19.58784 | 20.06841 | 8.80414 | 4.52012 | 8.53137 | 4.67170 | 0.33361 | 13.67157 | 16.07740 | 14.31764 | 18.15808 |
| **Rbsn** | 6.31102 | 0.02110 | 5.75770 | 2.15334 | 17.37770 | 1.17430 | 0.01692 | 0.80754 | 12.23191 | 0.00002 | 2.14335 | 1.64212 | 11.89355 | 20.15963 |
| **Rbx1** | 10.42700 | 7.02440 | 3.42686 | 8.01296 | 11.11639 | 12.43668 | 2.95590 | 8.02588 | 6.32047 | 10.50761 | 2.81659 | 12.35503 | 13.24706 | 9.24319 |
| **Rc3h1** | 4.37107 | 0.00296 | 5.86942 | 0.39941 | 4.54120 | 3.80138 | 1.32442 | 0.54570 | 14.20360 | 0.06293 | 0.65982 | 0.42096 | 5.41576 | 4.25079 |
| **Rc3h2** | 5.62507 | 0.01052 | 1.54352 | 0.00000 | 3.15478 | 0.90751 | 0.00082 | 2.21907 | 17.52897 | 0.01087 | 0.45605 | 1.98915 | 1.21573 | 1.07243 |
| **Rcan1** | 2.45085 | 0.05308 | 16.24389 | 10.46463 | 5.15588 | 6.45382 | 1.26796 | 3.56536 | 5.30802 | 0.04892 | 5.38033 | 2.19696 | 2.26379 | 3.33449 |
| **Rcan3** | 0.31071 | 0.00335 | 0.16910 | 0.06463 | 2.07046 | 0.09549 | 0.00332 | 0.00000 | 4.18830 | 0.00705 | 0.00092 | 0.00058 | 2.74661 | 7.84407 |
| **Rcbtb1** | 0.34280 | 0.07694 | 0.02937 | 0.00000 | 0.10241 | 1.13227 | 0.00439 | 0.08680 | 0.00218 | 0.00508 | 0.00545 | 2.48351 | 1.32843 | 0.59974 |
| **Rcbtb2** | 0.00601 | 0.00447 | 0.00502 | 0.01267 | 0.42148 | 0.01225 | 0.00423 | 0.01020 | 0.01022 | 0.00642 | 0.00796 | 0.00786 | 0.00514 | 0.40163 |
| **Rcc1** | 3.88674 | 6.97425 | 18.65598 | 16.22015 | 21.18702 | 13.79108 | 1.95881 | 8.31486 | 9.38697 | 0.18888 | 10.75100 | 7.85506 | 5.24379 | 18.42122 |
| **Rcc2** | 9.00557 | 7.33552 | 27.75790 | 25.00897 | 20.75812 | 15.57741 | 2.43231 | 8.10849 | 0.13117 | 5.45655 | 12.59283 | 16.37746 | 2.84282 | 14.94650 |
| **Rccd1** | 0.63502 | 0.08552 | 0.00294 | 0.02160 | 0.52581 | 0.04486 | 0.00434 | 0.00485 | 0.00639 | 0.00600 | 0.67491 | 0.00761 | 0.23677 | 0.41198 |
| **Rce1** | 4.90278 | 0.00289 | 4.78409 | 8.89102 | 5.35625 | 4.19299 | 3.40938 | 1.31334 | 0.98085 | 0.19687 | 0.60641 | 1.34117 | 0.08072 | 3.16022 |
| **Rchy1** | 7.37015 | 0.17276 | 11.26445 | 8.00622 | 11.35287 | 8.89555 | 2.92416 | 3.10388 | 7.69314 | 0.11194 | 6.06252 | 8.89998 | 7.62704 | 7.57764 |
| **Rcl1** | 7.22799 | 0.05186 | 14.42727 | 13.20313 | 16.31031 | 9.93355 | 2.70347 | 5.88952 | 6.81754 | 0.13930 | 5.74696 | 4.97312 | 6.73082 | 11.46278 |
| **Rcn1** | 0.01292 | 0.08480 | 0.01715 | 0.00590 | 0.01595 | 0.00408 | 0.00944 | 0.03455 | 0.01939 | 0.02648 | 0.04146 | 6.37893 | 0.48938 | 0.02564 |
| **Rcn2** | 1.45700 | 0.01884 | 1.52234 | 0.00000 | 0.41890 | 2.75477 | 0.78608 | 3.75205 | 2.65539 | 0.06208 | 0.00000 | 2.01804 | 1.40038 | 0.73343 |
| **Rcn3** | 0.14358 | 0.02863 | 0.81510 | 0.77910 | 1.15776 | 0.52145 | 0.00349 | 0.00209 | 0.04134 | 0.00637 | 0.10599 | 1.46744 | 1.04266 | 0.38253 |
| **Rcor1** | 0.00000 | 0.01087 | 0.28923 | 0.47198 | 2.90549 | 0.21646 | 0.00413 | 0.41345 | 0.12190 | 0.00425 | 0.29298 | 0.08000 | 1.41772 | 5.36338 |
| **Rcor2** | 0.29719 | 0.46024 | 0.15749 | 1.77311 | 3.50487 | 0.00000 | 0.11000 | 0.35170 | 0.02535 | 0.04188 | 0.00000 | 1.68788 | 4.98735 | 1.35979 |
| **Rcor3** | 0.25529 | 0.00389 | 0.65411 | 0.75215 | 0.00125 | 0.07657 | 0.80334 | 0.73043 | 1.15696 | 0.01255 | 0.00053 | 0.00285 | 0.00272 | 0.14806 |
| **Rcsd1** | 0.03054 | 0.00433 | 0.01265 | 0.01171 | 0.00569 | 0.01173 | 0.00423 | 0.00944 | 0.00795 | 0.00627 | 0.00984 | 0.00955 | 0.29905 | 0.19938 |
| **Rd3** | 0.01541 | 0.00444 | 0.07650 | 0.02516 | 0.06605 | 0.01046 | 0.00426 | 0.01090 | 0.01086 | 0.00708 | 0.00740 | 0.00837 | 0.00667 | 0.01299 |
| **Rdh1** | 0.03620 | 0.02010 | 0.33447 | 0.16005 | 0.03005 | 0.01873 | 0.03798 | 0.10731 | 0.63924 | 0.07106 | 0.00416 | 0.14066 | 0.06864 | 0.33448 |
| **Rdh10** | 11.61453 | 0.92846 | 4.25247 | 4.56042 | 4.49299 | 1.71804 | 0.00119 | 0.00000 | 6.59623 | 0.00000 | 3.11428 | 1.26016 | 8.35882 | 6.61486 |
| **Rdh11** | 0.01532 | 0.00099 | 0.00000 | 0.00004 | 0.00000 | 8.66239 | 0.00201 | 2.39400 | 0.04279 | 0.00572 | 3.62822 | 2.38086 | 2.66343 | 0.55191 |
| **Rdh12** | 0.00000 | 0.00393 | 0.08461 | 0.00000 | 0.14341 | 0.00000 | 0.00394 | 0.00730 | 0.50774 | 0.00496 | 0.85632 | 0.00365 | 0.67821 | 1.25427 |
| **Rdh13** | 7.67099 | 0.03287 | 1.13320 | 4.29653 | 4.29011 | 3.05059 | 2.01903 | 3.26390 | 5.14830 | 0.57230 | 1.30949 | 2.69853 | 2.21649 | 3.71768 |
| **Rdh14** | 0.00702 | 0.00396 | 0.16532 | 0.00000 | 0.22001 | 0.54466 | 1.13025 | 0.00339 | 0.95311 | 0.00973 | 0.00637 | 0.07378 | 0.00080 | 0.00000 |
| **Rdh16** | 0.51153 | 0.00433 | 0.02304 | 0.01249 | 0.00501 | 0.00655 | 0.05830 | 0.04228 | 0.01586 | 0.13957 | 0.00760 | 0.01457 | 0.07143 | 0.01977 |
| **Rdh18.ps** | 0.02358 | 0.02081 | 0.07234 | 0.00000 | 0.04005 | 0.08172 | 0.04038 | 0.31941 | 0.77437 | 0.86234 | 0.01065 | 0.00511 | 0.16175 | 0.05840 |
| **Rdh19** | 0.00888 | 0.00434 | 0.02261 | 0.00517 | 0.01911 | 0.01126 | 0.04471 | 0.14991 | 0.02081 | 0.45347 | 0.03102 | 0.01457 | 0.00612 | 0.07536 |
| **Rdh5** | 0.15961 | 0.02618 | 0.00924 | 0.00256 | 0.05815 | 0.15229 | 1.52427 | 0.17827 | 0.00906 | 0.00674 | 0.00432 | 0.00425 | 0.00543 | 0.00631 |
| **Rdh7** | 0.01968 | 0.00404 | 0.01169 | 0.00072 | 0.01046 | 0.00000 | 0.41814 | 0.16230 | 0.01923 | 1.56298 | 0.02267 | 0.00676 | 0.11041 | 0.01702 |
| **Rdh9** | 0.01840 | 0.01961 | 0.55186 | 0.18896 | 0.06754 | 0.02005 | 0.00407 | 0.02883 | 0.00872 | 0.00578 | 0.00572 | 0.00448 | 0.57996 | 0.08297 |
| **Rdm1** | 2.07992 | 0.04758 | 9.71864 | 6.67236 | 4.40954 | 7.58461 | 1.19093 | 4.75019 | 0.00000 | 0.04269 | 0.16341 | 0.00000 | 6.26493 | 4.53097 |
| **Rdx** | 3.99970 | 0.04174 | 4.15354 | 2.12823 | 2.17894 | 6.36211 | 0.14211 | 6.12517 | 5.20578 | 0.14531 | 0.48508 | 10.12158 | 3.27971 | 5.31044 |

| | | | | | | | | | | | | | | |
|---|---|---|---|---|---|---|---|---|---|---|---|---|---|---|
| **Rec114** | 0.00000 | 0.00423 | 0.40204 | 1.23864 | 1.04449 | 0.87842 | 0.00402 | 0.00200 | 0.00547 | 0.00467 | 0.00158 | 0.00825 | 0.15236 | 0.00000 |
| **Rec8** | 0.00000 | 0.19371 | 4.28566 | 5.51742 | 0.00000 | 0.00000 | 1.10289 | 0.00000 | 0.00849 | 0.00000 | 5.56763 | 2.99380 | 0.00000 | 0.00000 |
| **Reck** | 0.00355 | 0.00395 | 0.15529 | 0.06809 | 0.01536 | 0.00000 | 0.00392 | 0.00617 | 0.00780 | 0.00705 | 0.09704 | 0.07280 | 0.43578 | 0.42740 |
| **Recql** | 0.49247 | 0.00344 | 1.17132 | 1.15938 | 0.00000 | 1.80929 | 0.00376 | 0.10036 | 0.00471 | 0.00310 | 2.69814 | 0.37816 | 0.00612 | 2.30109 |
| **Recql4** | 0.01697 | 0.01837 | 0.45912 | 2.79094 | 0.37994 | 1.35196 | 0.00330 | 1.64895 | 0.19570 | 0.01513 | 0.00000 | 0.97753 | 0.01625 | 0.48274 |
| **Recql5** | 1.54874 | 0.06472 | 5.32413 | 3.86165 | 2.62511 | 4.40576 | 2.84260 | 1.21759 | 0.69246 | 0.06370 | 1.95050 | 2.06401 | 1.31143 | 1.81358 |
| **Reep1** | 3.77240 | 0.03762 | 20.19036 | 7.10233 | 8.00229 | 17.86248 | 2.90378 | 1.63153 | 11.66453 | 0.11283 | 11.76977 | 4.25738 | 0.27630 | 9.10298 |
| **Reep2** | 0.02051 | 0.00549 | 0.39809 | 0.00000 | 0.46617 | 0.30813 | 0.00409 | 0.24642 | 0.00362 | 0.00576 | 0.00481 | 0.02280 | 2.34468 | 0.16510 |
| **Reep3** | 2.68113 | 0.12588 | 2.36876 | 3.22000 | 2.56247 | 5.26570 | 2.62393 | 1.15448 | 4.04057 | 0.11058 | 3.79153 | 3.21364 | 0.97143 | 1.19690 |
| **Reep4** | 0.00000 | 6.88786 | 2.50564 | 6.58203 | 0.00000 | 3.61288 | 0.00064 | 4.62401 | 0.00000 | 0.04721 | 1.27772 | 2.16391 | 0.56897 | 0.73611 |
| **Reep5** | 0.22089 | 0.05776 | 0.49221 | 0.00000 | 0.80650 | 0.35461 | 1.18535 | 4.31625 | 0.00000 | 5.89539 | 0.36614 | 1.78431 | 1.74462 | 0.17022 |
| **Reep6** | 0.15828 | 0.00405 | 0.01889 | 1.93648 | 0.17958 | 0.17067 | 0.00376 | 0.01250 | 0.01329 | 0.00602 | 0.00295 | 0.00841 | 0.82323 | 0.00000 |
| **Reg1** | 1.92462 | 0.00368 | 0.05810 | 0.00461 | 0.00000 | 0.00479 | 0.00391 | 0.00782 | 0.01594 | 0.00316 | 0.76534 | 0.00319 | 2.48531 | 0.00593 |
| **Reg2** | 0.02713 | 0.00419 | 0.02266 | 0.16179 | 0.01781 | 0.15690 | 0.00390 | 0.00360 | 0.16656 | 0.00572 | 0.02276 | 0.02497 | 0.00313 | 0.02796 |
| **Rel** | 0.00000 | 0.00354 | 0.00000 | 0.00000 | 0.57609 | 0.36983 | 0.00428 | 0.03328 | 0.57453 | 0.00616 | 0.00000 | 0.58918 | 0.13347 | 0.00000 |
| **Rela** | 1.34742 | 0.04122 | 5.58401 | 4.61381 | 2.54527 | 2.21537 | 0.00185 | 4.23263 | 0.00000 | 0.02849 | 3.09522 | 0.82964 | 1.67561 | 0.74160 |
| **Relb** | 0.66503 | 3.63463 | 5.64258 | 0.00000 | 8.85483 | 0.00000 | 0.00195 | 0.46242 | 0.18821 | 0.00304 | 0.60843 | 0.14038 | 4.57764 | 3.77043 |
| **Rell1** | 0.08928 | 0.39822 | 1.25209 | 2.97301 | 0.00000 | 1.81187 | 0.93735 | 2.53936 | 1.94094 | 0.02519 | 1.61443 | 5.53311 | 2.28671 | 1.58282 |
| **Reln** | 0.00000 | 0.00383 | 0.00000 | 0.17456 | 0.70181 | 0.00000 | 0.00414 | 0.00556 | 0.19877 | 0.00392 | 0.89758 | 0.00282 | 0.55534 | 1.69045 |
| **Relt** | 0.00271 | 0.00373 | 0.01397 | 0.65100 | 0.50516 | 0.00000 | 0.10357 | 0.01356 | 0.00912 | 0.01283 | 0.00521 | 0.49632 | 0.32793 | 0.93136 |
| **Rem1** | 0.00529 | 0.00441 | 0.00581 | 0.50088 | 0.01049 | 0.01554 | 0.00494 | 0.04000 | 0.01288 | 0.00679 | 0.00569 | 0.00656 | 0.00724 | 0.00851 |
| **Renbp** | 15.62988 | 10.02333 | 20.03387 | 17.56313 | 9.20166 | 25.80579 | 6.90756 | 13.13230 | 0.00000 | 0.58824 | 14.81844 | 11.72765 | 1.86114 | 3.08415 |
| **Repin1** | 0.04462 | 0.00445 | 0.01285 | 0.00594 | 0.04000 | 0.04218 | 0.00444 | 0.01348 | 0.01209 | 0.00831 | 0.00658 | 0.10884 | 0.01420 | 0.01631 |
| **Reps1** | 6.61734 | 0.07223 | 6.39175 | 3.08028 | 8.05748 | 4.87386 | 0.00429 | 6.84964 | 4.05723 | 0.01162 | 7.38110 | 7.05414 | 5.90598 | 7.71769 |
| **Reps2** | 1.34122 | 0.00345 | 1.35823 | 0.77098 | 2.66552 | 3.14882 | 1.11822 | 0.00000 | 3.46731 | 0.00000 | 3.87971 | 0.01012 | 0.44580 | 0.00000 |
| **Rer1** | 13.94730 | 0.25377 | 6.16083 | 10.58049 | 9.33568 | 6.04839 | 3.77419 | 2.67556 | 0.00000 | 1.13630 | 0.66150 | 7.28300 | 5.84384 | 2.93036 |
| **Rere** | 2.03564 | 0.09077 | 7.14988 | 1.48726 | 8.46412 | 4.22696 | 0.14529 | 6.14280 | 0.82905 | 0.00000 | 6.12229 | 6.57551 | 4.07843 | 5.12349 |
| **Resp18** | 0.00724 | 0.00417 | 0.12213 | 0.04605 | 0.01258 | 0.02235 | 0.01075 | 0.35674 | 0.01113 | 0.00742 | 0.00641 | 0.00784 | 0.00581 | 0.01472 |
| **Rest** | 0.00000 | 0.01924 | 5.46559 | 0.71663 | 4.25726 | 1.80605 | 0.00175 | 2.31672 | 7.11570 | 0.00000 | 0.14748 | 0.18854 | 0.81948 | 7.34702 |
| **Ret** | 0.00810 | 0.00406 | 0.00675 | 0.31655 | 0.00837 | 0.00890 | 0.00383 | 0.29976 | 0.01314 | 0.00825 | 0.00523 | 0.01088 | 0.00675 | 0.00900 |
| **Retsat** | 0.16939 | 0.08897 | 2.69106 | 2.33581 | 0.39041 | 6.26972 | 0.06194 | 5.32412 | 1.67718 | 0.25098 | 3.02636 | 7.97177 | 0.05433 | 1.44404 |
| **Rev1** | 0.00000 | 0.00346 | 0.43275 | 0.35940 | 1.27121 | 0.24662 | 0.00340 | 0.20545 | 2.95584 | 0.00391 | 0.20966 | 0.49256 | 1.11826 | 1.43601 |
| **Rev3l** | 1.71314 | 0.00235 | 1.43467 | 0.63197 | 2.03671 | 0.51584 | 0.01339 | 0.00000 | 9.85889 | 0.00879 | 0.00000 | 1.15542 | 3.22829 | 1.29407 |
| **Rex2** | 1.51937 | 0.00189 | 2.83184 | 2.15662 | 1.69025 | 0.75352 | 0.18681 | 0.39937 | 15.19514 | 0.05366 | 1.21678 | 1.52815 | 5.91102 | 0.00000 |
| **Rexo1** | 1.52310 | 0.04783 | 6.53019 | 6.36991 | 6.40981 | 1.73172 | 1.57975 | 1.47612 | 0.00000 | 0.05523 | 1.79248 | 3.23819 | 2.87962 | 0.06513 |
| **Rexo2** | 8.44597 | 0.81360 | 2.41830 | 1.83718 | 3.25862 | 9.71104 | 0.10334 | 2.29234 | 0.42689 | 0.12547 | 1.24956 | 4.61858 | 0.05989 | 0.42807 |
| **Rexo4** | 8.54249 | 6.17001 | 10.93997 | 6.79035 | 13.77831 | 8.48595 | 3.47019 | 1.67263 | 6.05581 | 0.07310 | 2.54804 | 3.15923 | 8.07344 | 9.86979 |
| **Rfc1** | 9.65194 | 0.08893 | 13.87965 | 9.38687 | 19.11732 | 8.96493 | 0.00177 | 0.13323 | 21.02771 | 0.05248 | 9.24847 | 6.60286 | 6.66676 | 21.95752 |
| **Rfc2** | 4.16433 | 0.22376 | 13.22003 | 7.72159 | 7.49923 | 13.65466 | 4.50859 | 7.20149 | 2.45902 | 7.29739 | 3.88458 | 12.80624 | 1.05450 | 4.40844 |
| **Rfc3** | 11.79685 | 0.13639 | 9.98054 | 6.99321 | 6.51685 | 10.69721 | 0.00929 | 2.23983 | 2.75260 | 0.06327 | 7.88339 | 11.03724 | 1.46299 | 4.75030 |
| **Rfc4** | 3.24541 | 0.14307 | 12.04324 | 8.49742 | 9.81823 | 9.33735 | 3.07226 | 7.14110 | 8.17410 | 0.11671 | 8.52316 | 7.13732 | 1.11588 | 3.60989 |
| **Rfc5** | 6.90658 | 0.02839 | 4.42700 | 6.46438 | 5.82445 | 8.71658 | 2.43376 | 1.15286 | 1.59492 | 0.10010 | 3.46102 | 7.76319 | 1.14178 | 8.50549 |
| **Rfesd** | 0.00980 | 0.01692 | 0.87610 | 0.00000 | 0.30299 | 2.53058 | 2.41374 | 1.36078 | 0.00149 | 0.03700 | 1.44932 | 2.15366 | 0.07262 | 0.25832 |
| **Rffl** | 8.03514 | 1.05199 | 14.01279 | 3.44605 | 6.54590 | 6.12131 | 2.63473 | 3.55486 | 5.99413 | 6.77208 | 1.61802 | 1.44665 | 5.45639 | 7.05044 |
| **Rfk** | 7.59525 | 3.81739 | 11.17590 | 10.87655 | 11.28632 | 19.22523 | 5.00410 | 12.59769 | 9.29091 | 1.53404 | 10.65813 | 6.75930 | 8.52637 | 7.83514 |
| **Rfng** | 6.42582 | 4.09532 | 13.47459 | 8.62401 | 7.93447 | 5.80740 | 3.16789 | 2.08306 | 0.08251 | 0.09854 | 4.83845 | 4.26825 | 9.07257 | 6.36603 |
| **Rfpl4** | 0.00000 | 0.00402 | 0.19146 | 0.00000 | 0.14398 | 0.00376 | 0.00394 | 0.00899 | 0.03533 | 0.00652 | 0.00630 | 0.00744 | 1.95157 | 0.96033 |
| **Rfpl4b** | 0.00173 | 0.00415 | 0.01438 | 0.01626 | 0.00523 | 0.01375 | 0.00415 | 0.00992 | 0.01792 | 0.00662 | 0.00953 | 0.00694 | 2.17405 | 0.00061 |
| **Rft1** | 0.07603 | 0.08096 | 0.81142 | 0.17829 | 0.36766 | 2.35592 | 0.34215 | 0.18711 | 0.34755 | 0.08525 | 0.53854 | 2.29524 | 0.51645 | 2.81879 |
| **Rftn1** | 0.02454 | 0.00418 | 0.02096 | 0.14169 | 0.14483 | 0.00466 | 0.00423 | 0.00629 | 0.06055 | 0.00550 | 0.01651 | 0.02510 | 0.17756 | 0.06752 |
| **Rftn2** | 2.99857 | 0.00422 | 2.58536 | 0.00000 | 4.27539 | 0.35141 | 0.00482 | 0.18683 | 0.04469 | 0.00704 | 0.00756 | 0.00149 | 0.21705 | 2.58600 |
| **Rfwd2** | 13.38622 | 0.08029 | 9.02082 | 3.85412 | 12.53216 | 9.51124 | 1.21885 | 3.92739 | 17.79043 | 1.07969 | 6.09413 | 3.58435 | 12.90210 | 20.09180 |
| **Rfwd3** | 9.28675 | 0.09210 | 8.36737 | 4.28844 | 12.65668 | 7.54741 | 2.51106 | 5.98078 | 21.57050 | 0.09308 | 7.48239 | 8.75997 | 12.11633 | 10.91713 |
| **Rfx1** | 0.91184 | 6.08249 | 2.26231 | 2.96805 | 2.22864 | 0.00000 | 0.91055 | 3.07116 | 0.04243 | 0.03989 | 0.12636 | 1.74414 | 1.00911 | 0.00000 |
| **Rfx2** | 0.00000 | 3.16424 | 2.37656 | 7.02996 | 0.14866 | 0.48942 | 0.01395 | 0.56962 | 0.00000 | 0.02744 | 3.65274 | 7.72854 | 2.97774 | 3.58600 |

| | | | | | | | | | | | | | | |
|---|---|---|---|---|---|---|---|---|---|---|---|---|---|---|
| **Rfx3** | 0.00995 | 0.01352 | 0.01240 | 0.03877 | 0.01789 | 0.01134 | 0.00428 | 0.01845 | 0.14034 | 0.00822 | 0.00720 | 0.01829 | 0.02677 | 0.03121 |
| **Rfx4** | 0.00500 | 0.00414 | 0.01452 | 0.08422 | 0.00426 | 0.90304 | 0.00451 | 0.87319 | 0.01166 | 0.01300 | 1.63806 | 0.00000 | 0.00916 | 0.00000 |
| **Rfx5** | 0.00164 | 0.00422 | 0.09313 | 0.03294 | 0.37523 | 0.32947 | 0.00411 | 0.00386 | 0.03946 | 0.00626 | 0.00452 | 0.00799 | 0.60712 | 0.00000 |
| **Rfx7** | 1.03094 | 0.00911 | 0.76982 | 0.00000 | 0.75411 | 0.55777 | 0.00287 | 0.00000 | 1.09906 | 0.00000 | 1.49786 | 1.88185 | 0.80852 | 1.81847 |
| **Rfxank** | 1.16513 | 0.20644 | 2.02133 | 3.57395 | 2.43478 | 3.36539 | 0.39269 | 1.02081 | 0.99338 | 1.23308 | 0.02311 | 2.05654 | 1.15138 | 3.89272 |
| **Rfxap** | 2.44693 | 0.01223 | 0.13811 | 1.96389 | 3.02008 | 1.31231 | 1.06746 | 0.03799 | 0.67983 | 0.03164 | 0.00000 | 0.04048 | 3.92258 | 1.39318 |
| **Rgcc** | 0.01150 | 0.00384 | 0.01899 | 0.90939 | 0.40167 | 0.00000 | 0.00383 | 0.00377 | 0.01169 | 0.01137 | 0.15725 | 0.08607 | 0.00627 | 0.00284 |
| **Rgl1** | 0.00529 | 0.01483 | 0.01155 | 0.15028 | 0.02283 | 1.05834 | 0.00429 | 0.18129 | 0.00873 | 0.02394 | 0.02443 | 0.54252 | 0.25539 | 0.00000 |
| **Rgl2** | 0.00000 | 0.01160 | 1.17196 | 2.12276 | 0.00000 | 1.29469 | 0.00983 | 1.03017 | 0.00669 | 0.03313 | 0.00927 | 0.81754 | 0.76946 | 0.00000 |
| **Rgmb** | 0.02028 | 0.00408 | 0.04323 | 0.00000 | 0.30365 | 0.13693 | 0.00435 | 0.00224 | 0.08542 | 0.00397 | 0.56725 | 0.00451 | 1.46256 | 0.69752 |
| **Rgn** | 0.95581 | 0.01825 | 2.62658 | 0.00000 | 5.76321 | 6.59312 | 0.01355 | 2.99494 | 9.42612 | 0.04205 | 2.96385 | 0.00000 | 1.03185 | 3.24382 |
| **Rgp1** | 0.54842 | 0.07028 | 0.94417 | 0.75549 | 0.72455 | 2.67059 | 0.00317 | 1.47245 | 0.00102 | 0.00450 | 0.44045 | 2.44765 | 0.96083 | 0.97591 |
| **Rgs10** | 0.03819 | 0.00333 | 0.02039 | 0.00000 | 0.21545 | 0.21875 | 0.00401 | 0.00335 | 0.00911 | 0.00542 | 0.00226 | 0.95655 | 0.34121 | 0.34673 |
| **Rgs11** | 5.12783 | 0.00154 | 4.84352 | 5.37567 | 3.39763 | 3.18726 | 0.00298 | 0.43066 | 0.31188 | 0.00000 | 2.76755 | 1.32221 | 5.78552 | 2.03938 |
| **Rgs12** | 0.01365 | 0.00437 | 0.01154 | 0.01242 | 0.13822 | 0.01284 | 0.00439 | 0.01144 | 0.01259 | 0.00680 | 0.00848 | 0.01268 | 0.01820 | 0.01194 |
| **Rgs14** | 0.00448 | 0.00443 | 0.57611 | 0.35985 | 0.00000 | 0.04046 | 0.00432 | 0.00322 | 0.01340 | 0.00511 | 0.11103 | 0.02253 | 0.00301 | 0.05754 |
| **Rgs16** | 0.05142 | 0.00445 | 0.00568 | 0.01370 | 0.01254 | 0.00932 | 0.00433 | 0.01021 | 0.33841 | 0.00671 | 0.00668 | 0.00935 | 0.09684 | 0.00000 |
| **Rgs17** | 0.01039 | 0.02567 | 0.00938 | 0.00000 | 0.23210 | 0.17110 | 0.00428 | 0.00428 | 0.05935 | 0.00591 | 0.00734 | 0.28110 | 1.89559 | 0.12411 |
| **Rgs19** | 1.32966 | 0.00111 | 0.01473 | 0.06824 | 1.24760 | 0.16130 | 0.00275 | 0.00368 | 0.00295 | 0.00571 | 0.00280 | 4.34911 | 1.13648 | 0.07065 |
| **Rgs2** | 0.33670 | 0.00427 | 0.06892 | 0.00363 | 0.01841 | 0.00794 | 0.00416 | 0.01107 | 0.87902 | 0.00665 | 0.00508 | 0.00651 | 0.77091 | 0.00000 |
| **Rgs20** | 0.02578 | 0.02819 | 0.02652 | 0.02817 | 0.03117 | 0.05803 | 0.02318 | 0.06648 | 0.04831 | 0.00625 | 0.03199 | 0.02239 | 0.01637 | 0.07120 |
| **Rgs3** | 0.00000 | 0.00377 | 0.51598 | 1.30549 | 0.45367 | 0.18413 | 0.00343 | 1.33862 | 0.03190 | 0.02888 | 0.06272 | 0.24531 | 0.55876 | 0.56513 |
| **Rgs7bp** | 0.02647 | 0.04506 | 0.01777 | 0.03836 | 0.02784 | 0.02782 | 0.02195 | 0.01502 | 0.21270 | 0.02541 | 0.01750 | 0.04175 | 0.01361 | 0.08607 |
| **Rhag** | 0.01342 | 0.00449 | 0.01180 | 0.04661 | 0.01359 | 0.02936 | 0.01050 | 0.01471 | 0.03230 | 0.00801 | 0.01061 | 0.01563 | 0.09003 | 0.09238 |
| **Rhbdd1** | 0.15952 | 0.01160 | 1.80754 | 0.82796 | 1.06190 | 2.13967 | 0.00332 | 0.48608 | 2.18807 | 0.01814 | 0.00802 | 0.00000 | 0.00309 | 0.00000 |
| **Rhbdd2** | 0.62259 | 0.04933 | 0.48590 | 1.61304 | 0.15897 | 2.47932 | 1.47985 | 0.87242 | 1.29143 | 0.05163 | 0.01139 | 3.42015 | 5.21444 | 1.94385 |
| **Rhbdd3** | 0.25858 | 0.03134 | 1.32981 | 1.71848 | 0.75901 | 0.97664 | 0.00815 | 3.49025 | 0.00454 | 0.00175 | 2.68847 | 4.01568 | 0.58528 | 0.26339 |
| **Rhbdf1** | 0.14830 | 0.01148 | 0.57165 | 1.32132 | 0.33741 | 0.53781 | 0.01104 | 0.51463 | 0.00405 | 0.00725 | 0.01347 | 0.00135 | 0.06884 | 0.18693 |
| **Rhbdf2** | 0.02904 | 0.04996 | 0.46006 | 0.00000 | 0.48284 | 0.76517 | 0.00352 | 0.32265 | 0.01014 | 0.00779 | 0.00000 | 1.91296 | 0.02801 | 0.00025 |
| **Rhbdl1** | 0.06876 | 0.00422 | 0.08018 | 0.27970 | 0.03445 | 0.00980 | 0.00404 | 0.00615 | 0.01051 | 0.00984 | 0.00694 | 0.00719 | 0.03483 | 0.00000 |
| **Rhbdl2** | 0.59891 | 0.03062 | 5.07146 | 0.06794 | 1.96894 | 3.12765 | 3.73847 | 5.71686 | 0.07173 | 10.94518 | 0.00000 | 0.00000 | 0.06824 | 2.47733 |
| **Rhbdl3** | 0.01729 | 0.00382 | 0.07602 | 0.00000 | 0.82024 | 0.20291 | 0.00427 | 0.24943 | 0.11034 | 0.00621 | 0.00355 | 0.00368 | 0.01609 | 1.87678 |
| **Rhbg** | 0.00000 | 0.00404 | 0.17503 | 0.53058 | 0.60345 | 0.00000 | 0.00386 | 0.00634 | 0.00980 | 0.00596 | 0.00511 | 0.01160 | 0.07601 | 0.94300 |
| **Rhcg** | 1.63239 | 0.00386 | 0.59491 | 0.45057 | 0.89815 | 0.00000 | 0.00398 | 0.00427 | 0.14176 | 0.00531 | 0.00609 | 0.01018 | 0.00000 | 2.70377 |
| **Rhd** | 0.00095 | 0.00421 | 0.49034 | 0.07638 | 0.03544 | 0.00000 | 0.00396 | 0.00918 | 0.00776 | 0.00697 | 0.00562 | 0.01201 | 0.04771 | 0.56964 |
| **Rheb** | 9.45187 | 5.89967 | 5.45516 | 6.45476 | 10.88628 | 13.42163 | 1.92494 | 1.57642 | 3.12380 | 0.09223 | 4.36954 | 6.06533 | 8.13193 | 4.31196 |
| **Rhebl1** | 1.65432 | 0.02493 | 8.28512 | 13.15302 | 4.99073 | 2.44675 | 0.03288 | 0.02144 | 2.92477 | 0.02485 | 1.91539 | 1.03024 | 0.11542 | 1.31254 |
| **Rhno1** | 1.73703 | 11.85245 | 2.92235 | 4.49074 | 4.52222 | 4.51899 | 0.00283 | 5.29749 | 3.06263 | 5.32076 | 1.66026 | 5.72862 | 5.09097 | 2.35688 |
| **Rhoa** | 15.88469 | 2.30700 | 17.75252 | 19.43564 | 16.68225 | 10.81352 | 1.02111 | 12.70366 | 11.00747 | 0.24985 | 11.20959 | 11.17883 | 11.84502 | 13.21679 |
| **Rhob** | 1.45355 | 0.01743 | 1.49349 | 1.84245 | 0.46586 | 0.00000 | 1.04599 | 2.26788 | 0.08230 | 0.15709 | 0.06243 | 3.60518 | 0.00412 | 0.33011 |
| **Rhobtb1** | 0.00000 | 0.00423 | 0.42469 | 0.00000 | 0.36389 | 0.00000 | 0.00405 | 0.01038 | 0.66378 | 0.00745 | 0.00432 | 0.00529 | 0.72359 | 1.29533 |
| **Rhobtb2** | 0.32993 | 0.01793 | 2.31134 | 0.72546 | 3.12167 | 1.75348 | 0.01329 | 4.52580 | 0.14637 | 1.51117 | 0.07982 | 0.49042 | 3.83936 | 1.44308 |
| **Rhoc** | 0.20871 | 3.43022 | 0.38452 | 1.00793 | 0.00000 | 2.72101 | 0.08452 | 8.87908 | 0.00097 | 2.62697 | 0.00000 | 2.94882 | 0.00497 | 0.10618 |
| **Rhod** | 0.00000 | 0.00394 | 0.38890 | 0.34163 | 0.38945 | 1.03763 | 0.00393 | 0.14222 | 0.02806 | 0.01598 | 0.00707 | 0.47435 | 0.00631 | 0.29161 |
| **Rhof** | 0.82008 | 0.03492 | 0.74693 | 0.00000 | 0.00000 | 0.26461 | 0.00301 | 0.04342 | 0.24198 | 3.45308 | 0.03909 | 0.39637 | 0.65080 | 1.18219 |
| **Rhog** | 0.00000 | 0.03332 | 5.15665 | 2.94926 | 1.43241 | 0.42072 | 0.00814 | 7.04044 | 0.05660 | 0.02581 | 0.00000 | 1.26707 | 0.00000 | 0.27258 |
| **Rhoh** | 0.01186 | 0.07654 | 0.03634 | 0.09147 | 0.00936 | 0.07791 | 0.01664 | 0.00954 | 0.06783 | 0.00792 | 0.01240 | 0.03269 | 1.05034 | 0.71093 |
| **Rhoq** | 0.04867 | 0.00397 | 0.00000 | 0.26310 | 0.57774 | 0.04600 | 0.00444 | 0.00484 | 0.02162 | 0.00548 | 0.00522 | 0.64074 | 0.00327 | 0.00296 |
| **Rhot1** | 5.43705 | 0.03473 | 4.40610 | 3.29576 | 7.01129 | 5.95442 | 1.62885 | 4.67341 | 7.34234 | 0.11625 | 3.52447 | 5.56798 | 7.71792 | 11.19108 |
| **Rhot2** | 1.29820 | 0.03135 | 5.77448 | 7.09478 | 2.48035 | 2.28841 | 0.02492 | 2.87901 | 0.06175 | 0.04841 | 1.97403 | 2.95923 | 0.81447 | 2.57448 |
| **Rhou** | 0.00000 | 6.68247 | 0.00000 | 0.08289 | 0.42793 | 0.93129 | 0.00235 | 0.64124 | 0.80658 | 0.01879 | 0.00000 | 2.34214 | 0.32362 | 0.00081 |
| **Rhov** | 0.04918 | 0.00401 | 0.00000 | 0.55557 | 0.34018 | 0.04031 | 0.00381 | 0.00366 | 0.01218 | 0.00899 | 0.00564 | 0.02424 | 0.01614 | 0.12691 |
| **Rhox13** | 0.00998 | 0.00430 | 0.03720 | 0.04690 | 0.00955 | 0.00970 | 0.00413 | 0.00803 | 0.03569 | 0.00607 | 0.00763 | 0.01398 | 0.09735 | 0.00370 |
| **Rhox3c** | 0.01789 | 0.00442 | 0.01182 | 0.01313 | 0.03118 | 0.01294 | 0.00419 | 0.00996 | 0.01878 | 0.00682 | 0.00837 | 0.00964 | 0.03925 | 0.11977 |
| **Rhox3e** | 0.03653 | 0.00444 | 0.01808 | 0.01132 | 0.03821 | 0.01206 | 0.00419 | 0.01047 | 0.01529 | 0.00653 | 0.00826 | 0.00950 | 0.02486 | 0.12163 |

| | | | | | | | | | | | | | | |
|---|---|---|---|---|---|---|---|---|---|---|---|---|---|---|
| **Rhox3g** | 0.03037 | 0.00445 | 0.01833 | 0.01158 | 0.03550 | 0.01245 | 0.00420 | 0.01046 | 0.01270 | 0.00666 | 0.00830 | 0.00960 | 0.00796 | 0.11412 |
| **Rhox3h** | 0.09890 | 0.00444 | 0.01509 | 0.01069 | 0.03888 | 0.01174 | 0.00419 | 0.01075 | 0.01420 | 0.00658 | 0.00764 | 0.00924 | 0.00731 | 0.13830 |
| **Rhox5** | 0.00013 | 0.02507 | 2.26160 | 0.00000 | 0.00000 | 8.80314 | 0.00351 | 0.15340 | 0.01123 | 0.03904 | 0.00000 | 0.43496 | 0.60719 | 0.80796 |
| **Rhox6** | 0.09280 | 0.00448 | 0.17602 | 0.00000 | 0.17815 | 0.04974 | 0.00426 | 0.00725 | 0.06872 | 0.00606 | 0.00582 | 0.00931 | 0.02040 | 0.05410 |
| **Rhox7a** | 0.19388 | 0.00440 | 0.00607 | 0.01070 | 0.09979 | 0.00919 | 0.00420 | 0.01105 | 0.00756 | 0.00605 | 0.00705 | 0.00810 | 0.01500 | 0.53126 |
| **Rhox8** | 0.03745 | 0.10060 | 0.00762 | 0.04212 | 0.04439 | 0.04057 | 0.00410 | 0.03816 | 0.21466 | 0.00723 | 0.00976 | 0.01196 | 0.06661 | 0.03933 |
| **Rhox9** | 1.52025 | 0.00446 | 0.00000 | 0.00000 | 1.47293 | 0.37765 | 0.00405 | 0.00577 | 0.77353 | 0.00576 | 0.00299 | 0.00298 | 0.04012 | 0.22496 |
| **Rhpn2** | 0.44341 | 0.00979 | 2.25775 | 0.00000 | 1.89971 | 1.44864 | 1.93011 | 4.15122 | 3.80570 | 0.04123 | 0.00000 | 1.76914 | 7.14596 | 9.56789 |
| **Ribc1** | 1.36245 | 0.00104 | 2.49660 | 0.32182 | 5.22952 | 0.15738 | 0.00222 | 0.00000 | 0.06748 | 0.00183 | 2.41219 | 3.60916 | 1.11771 | 1.25626 |
| **Ric1** | 0.07688 | 0.00396 | 0.56223 | 0.16656 | 0.00000 | 0.67807 | 0.00420 | 0.02008 | 0.34375 | 0.00567 | 0.00635 | 0.00383 | 1.20576 | 2.42092 |
| **Ric3** | 0.46882 | 0.00395 | 0.14698 | 0.67661 | 0.06189 | 0.00000 | 0.00394 | 0.00753 | 0.01625 | 0.00553 | 0.00550 | 0.00696 | 0.71739 | 0.43315 |
| **Ric8a** | 1.31361 | 0.05217 | 4.68322 | 2.82481 | 2.42267 | 4.34621 | 4.37425 | 5.13813 | 0.00000 | 0.07634 | 3.49870 | 6.27759 | 0.77440 | 1.07223 |
| **Ric8b** | 0.81074 | 0.00404 | 0.27130 | 0.06573 | 0.77416 | 0.40458 | 0.00936 | 0.00249 | 0.00654 | 0.00778 | 0.00301 | 0.10614 | 0.01068 | 4.00524 |
| **Rictor** | 2.34931 | 0.00290 | 1.24779 | 1.19221 | 1.15923 | 1.37805 | 0.02759 | 0.28455 | 1.66566 | 0.01199 | 0.31528 | 1.79725 | 1.19389 | 5.87991 |
| **Rida** | 0.78914 | 0.00190 | 0.70850 | 0.00000 | 0.72456 | 1.27553 | 0.00833 | 1.34986 | 0.63421 | 0.00416 | 3.02469 | 2.10981 | 0.00690 | 1.38016 |
| **Rif1** | 5.88711 | 0.00402 | 10.25774 | 1.27465 | 15.88141 | 3.80178 | 1.95384 | 0.28202 | 43.07696 | 0.00485 | 5.31902 | 4.16000 | 16.93840 | 13.96148 |
| **Rilp** | 0.33211 | 0.00447 | 0.23949 | 0.00000 | 0.22263 | 0.42462 | 0.00440 | 0.00563 | 0.00576 | 0.00668 | 0.00431 | 0.00639 | 0.01134 | 0.19187 |
| **Rilpl1** | 0.24465 | 0.01411 | 0.33667 | 0.00000 | 0.53773 | 0.11239 | 0.00399 | 0.17139 | 0.00573 | 0.00653 | 0.00196 | 0.40171 | 1.09494 | 0.45871 |
| **Rilpl2** | 0.00000 | 0.00987 | 0.44412 | 0.00000 | 1.81148 | 0.00000 | 0.00303 | 0.80706 | 0.37250 | 0.03106 | 0.00000 | 1.66699 | 4.14501 | 0.06960 |
| **Rimbp2** | 0.00734 | 0.01233 | 0.02306 | 0.05596 | 0.00751 | 0.03695 | 0.02309 | 0.12153 | 0.07571 | 0.01376 | 0.03428 | 0.03684 | 0.26192 | 0.28338 |
| **Rimbp3** | 0.00839 | 0.00384 | 0.07151 | 0.00000 | 0.07251 | 0.63144 | 0.00388 | 0.00272 | 0.00576 | 0.01723 | 0.01818 | 0.22054 | 0.49022 | 0.14904 |
| **Rimkla** | 0.24496 | 0.01012 | 0.86350 | 0.56910 | 0.10029 | 3.01217 | 0.00344 | 0.00425 | 0.13823 | 0.00415 | 0.11873 | 1.61898 | 3.16945 | 0.79852 |
| **Rimklb** | 1.09393 | 0.01221 | 2.07427 | 0.00037 | 2.45084 | 0.13529 | 0.00343 | 0.26464 | 3.96625 | 0.01682 | 0.00119 | 0.00000 | 8.18378 | 5.97284 |
| **Rims2** | 0.00841 | 0.00415 | 0.01076 | 0.00068 | 0.00879 | 0.28178 | 0.00402 | 0.00488 | 0.01187 | 0.00702 | 0.00701 | 0.01092 | 0.58564 | 0.02212 |
| **Rims3** | 0.00000 | 0.00506 | 1.28290 | 1.29134 | 1.46586 | 0.20005 | 0.00440 | 0.05559 | 0.01791 | 0.02928 | 0.00299 | 0.00195 | 0.02709 | 1.17848 |
| **Rin1** | 0.69315 | 0.06349 | 0.00000 | 0.42488 | 0.69494 | 2.38852 | 0.11690 | 1.85755 | 1.32085 | 0.27570 | 0.00000 | 0.21243 | 5.21210 | 0.17342 |
| **Rin2** | 0.78627 | 0.00396 | 0.46297 | 0.54839 | 1.32492 | 0.06042 | 0.00458 | 0.50046 | 4.61176 | 0.00561 | 0.88811 | 0.00394 | 0.00000 | 0.25920 |
| **Rin3** | 0.00633 | 4.44872 | 0.00000 | 0.00407 | 0.02110 | 0.42535 | 2.03025 | 5.24367 | 0.01377 | 1.84444 | 0.81526 | 0.00000 | 0.00538 | 0.00000 |
| **Ring1** | 0.20870 | 0.02948 | 0.58153 | 2.36540 | 0.73913 | 0.40403 | 0.03056 | 1.88619 | 0.23610 | 0.02432 | 0.86154 | 0.90066 | 0.37632 | 0.94928 |
| **Rinl** | 0.08364 | 0.02076 | 3.53546 | 2.32415 | 3.85250 | 2.75033 | 0.00914 | 3.17337 | 0.49271 | 0.00000 | 0.80551 | 0.00000 | 2.74912 | 1.24482 |
| **Rint1** | 1.53247 | 0.04522 | 1.38368 | 2.56018 | 1.45581 | 0.81317 | 0.05901 | 1.06907 | 0.87057 | 0.13201 | 0.38622 | 2.31369 | 0.60483 | 2.08566 |
| **Riok1** | 3.45372 | 3.86574 | 5.41963 | 2.43654 | 5.88713 | 5.63182 | 1.14374 | 0.91931 | 6.64595 | 0.01832 | 5.29696 | 5.05496 | 5.10020 | 5.06780 |
| **Riok2** | 12.57013 | 0.05338 | 10.40454 | 2.03996 | 11.77291 | 5.18515 | 2.86338 | 3.34850 | 12.59345 | 13.09447 | 12.65013 | 7.76072 | 3.42780 | 15.86256 |
| **Riok3** | 13.38583 | 0.12269 | 16.99731 | 10.02175 | 18.77289 | 9.94080 | 1.49818 | 5.75538 | 9.73591 | 0.31269 | 11.04292 | 5.80064 | 8.66703 | 14.25319 |
| **Ripk1** | 2.15518 | 0.00322 | 1.09278 | 0.40339 | 0.27402 | 1.34964 | 1.35956 | 0.45157 | 0.00162 | 0.03277 | 0.00000 | 1.04672 | 2.10366 | 2.27878 |
| **Ripk2** | 0.01958 | 0.00419 | 0.00958 | 0.01200 | 0.07698 | 0.00870 | 0.00408 | 0.02541 | 0.01115 | 0.00734 | 0.06031 | 0.01113 | 0.02754 | 0.13603 |
| **Ripk3** | 0.10042 | 0.00400 | 0.87661 | 0.01136 | 0.57216 | 0.00000 | 0.00406 | 0.00686 | 0.00497 | 0.00531 | 0.00865 | 0.00880 | 0.68741 | 1.99668 |
| **Ripk4** | 0.22651 | 0.00306 | 0.15085 | 1.37218 | 1.02449 | 0.89737 | 0.00337 | 0.00763 | 0.04689 | 0.01282 | 2.37346 | 0.44120 | 0.06889 | 0.00000 |
| **Ripply3** | 0.36837 | 0.06077 | 0.05500 | 0.05821 | 0.10995 | 0.77458 | 0.07533 | 0.12172 | 0.70174 | 0.54162 | 0.01866 | 0.41560 | 2.76013 | 0.49023 |
| **Rit1** | 0.39451 | 6.43340 | 2.11607 | 0.31322 | 3.31622 | 1.85601 | 1.93928 | 7.28885 | 0.19636 | 0.02239 | 2.08080 | 0.93754 | 7.35691 | 0.44606 |
| **Rita1** | 0.24232 | 0.00412 | 0.24772 | 0.03165 | 1.60038 | 0.98420 | 1.53562 | 0.24312 | 0.01685 | 0.05777 | 0.03574 | 0.49391 | 2.13090 | 0.44836 |
| **Rlbp1** | 0.01725 | 0.00371 | 0.78181 | 1.20706 | 0.07520 | 0.18536 | 0.00368 | 0.16557 | 0.00857 | 0.00922 | 0.35223 | 0.00665 | 0.68375 | 0.00000 |
| **Rlf** | 6.27458 | 0.01944 | 3.76369 | 1.37005 | 5.67998 | 1.27288 | 0.00143 | 0.30707 | 12.60181 | 0.00000 | 0.00000 | 0.28541 | 6.34278 | 4.52398 |
| **Rlim** | 5.45341 | 0.17930 | 11.09141 | 5.78983 | 9.50736 | 13.53380 | 1.00000 | 7.81236 | 22.45417 | 6.42043 | 11.20025 | 9.43736 | 9.33970 | 11.12918 |
| **Rln3** | 0.00745 | 0.00445 | 0.05194 | 0.00835 | 0.01422 | 0.01201 | 0.00432 | 0.01040 | 0.01104 | 0.00677 | 0.00859 | 0.00955 | 0.10387 | 0.00491 |
| **Rltpr** | 0.13154 | 0.02867 | 0.35446 | 0.16038 | 0.29476 | 0.01144 | 0.00367 | 0.24114 | 0.01043 | 0.01596 | 0.00341 | 0.26221 | 0.00636 | 0.00485 |
| **Rmdn1** | 0.06928 | 0.00444 | 0.00000 | 0.46072 | 0.00000 | 0.31273 | 0.03486 | 0.01355 | 0.19730 | 0.00657 | 0.00382 | 0.65882 | 0.18830 | 0.00650 |
| **Rmdn2** | 0.00804 | 0.00430 | 0.00555 | 0.00907 | 0.48058 | 0.01325 | 0.00413 | 0.00871 | 0.00786 | 0.00635 | 0.00942 | 0.00787 | 0.69556 | 0.02478 |
| **Rmdn3** | 4.38150 | 0.40359 | 8.46688 | 9.22369 | 3.97227 | 7.91334 | 5.76397 | 6.84921 | 0.68646 | 0.18455 | 2.87139 | 5.42752 | 2.57088 | 5.49899 |
| **Rmi1** | 1.14894 | 0.00062 | 3.20248 | 0.60961 | 1.53225 | 2.14445 | 0.02529 | 0.00000 | 2.92234 | 10.56144 | 3.16934 | 0.22802 | 2.61273 | 4.66766 |
| **Rmi2** | 2.88384 | 0.28795 | 0.00000 | 2.14597 | 1.63828 | 3.90071 | 0.61728 | 0.35982 | 2.90771 | 1.05436 | 0.25185 | 1.31819 | 12.82380 | 3.28438 |
| **Rmnd5a** | 12.36184 | 0.04975 | 7.98743 | 5.82255 | 12.72383 | 9.72494 | 4.73202 | 4.74879 | 13.64741 | 0.21436 | 1.66522 | 6.64810 | 14.09773 | 13.62878 |
| **Rmnd5b** | 16.14955 | 0.02808 | 21.84339 | 5.92179 | 25.62192 | 2.24392 | 2.43368 | 3.07827 | 2.66942 | 0.08604 | 1.65602 | 2.67228 | 11.24994 | 13.77963 |
| **Rn4.5s** | 0.06021 | 0.06117 | 0.01027 | 0.00803 | 0.01381 | 0.17395 | 0.02277 | 0.43757 | 0.10627 | 0.10009 | 0.02516 | 0.36517 | 0.76319 | 0.03575 |
| **Rn45s** | 0.52121 | 7.86285 | 0.60256 | 19.93841 | 19.31118 | 3.43660 | 0.77613 | 14.38061 | 2.37560 | 1.92026 | 6.91921 | 8.52254 | 2.06112 | 4.03071 |

| | | | | | | | | | | | | | | |
|---|---|---|---|---|---|---|---|---|---|---|---|---|---|---|
| **Rnase4** | 0.04247 | 0.00528 | 0.00749 | 0.04177 | 0.01117 | 0.56428 | 0.00440 | 0.37323 | 0.01041 | 0.00905 | 0.00493 | 0.00432 | 0.00871 | 0.00916 |
| **Rnaseh1** | 8.77505 | 0.00788 | 8.72902 | 2.18731 | 2.80310 | 2.72634 | 0.01896 | 0.03473 | 5.56055 | 0.03156 | 1.71199 | 4.25986 | 2.41679 | 7.25516 |
| **Rnaseh2a** | 5.16311 | 6.88249 | 10.06564 | 10.06327 | 4.81524 | 5.46904 | 1.96534 | 3.95446 | 0.00000 | 10.72341 | 5.24302 | 6.52786 | 0.06105 | 3.65690 |
| **Rnaseh2b** | 8.03972 | 1.00882 | 15.54046 | 17.15087 | 6.20648 | 9.09413 | 2.48357 | 6.35203 | 2.96904 | 11.28726 | 6.15049 | 10.26922 | 0.00000 | 4.85966 |
| **Rnaseh2c** | 0.07830 | 0.09150 | 0.00000 | 4.24770 | 1.20237 | 0.66834 | 0.25667 | 0.85190 | 0.00521 | 0.15887 | 0.00000 | 1.64414 | 0.22981 | 0.00000 |
| **Rnasek** | 3.06271 | 0.93421 | 1.92646 | 14.65741 | 11.08344 | 14.91254 | 5.90006 | 9.42833 | 0.00000 | 0.91291 | 1.98462 | 12.37533 | 6.07718 | 3.83310 |
| **Rnasel** | 0.02594 | 0.00353 | 0.00000 | 0.57870 | 0.03655 | 0.84738 | 0.00894 | 0.73715 | 0.54341 | 0.05929 | 2.83249 | 0.00442 | 0.10319 | 0.03336 |
| **Rnaset2b** | 3.37862 | 0.04052 | 2.16366 | 2.47535 | 3.68595 | 4.53633 | 0.68656 | 1.51779 | 1.10698 | 0.07941 | 1.40086 | 1.60682 | 1.14170 | 0.90969 |
| **Rnd1** | 0.00000 | 0.02131 | 1.65513 | 4.29493 | 0.32330 | 1.36053 | 0.06451 | 0.00000 | 0.57486 | 0.00906 | 9.15513 | 1.81182 | 0.53835 | 1.77296 |
| **Rnd2** | 0.84962 | 0.78773 | 7.80449 | 4.46900 | 2.98754 | 2.09552 | 0.98600 | 6.72199 | 0.01417 | 0.01769 | 1.24624 | 0.46933 | 0.00000 | 2.82650 |
| **Rnd3** | 0.00000 | 0.09101 | 2.58185 | 2.62675 | 0.16664 | 1.37443 | 0.00076 | 0.12878 | 0.18973 | 0.06495 | 5.19218 | 9.68295 | 0.90347 | 0.78232 |
| **Rnf10** | 5.80788 | 2.77465 | 18.11392 | 12.97315 | 13.23972 | 17.22278 | 0.10246 | 16.38983 | 12.43675 | 2.14329 | 13.92019 | 11.59613 | 10.54971 | 16.97364 |
| **Rnf103** | 1.47379 | 0.16042 | 0.00000 | 0.30331 | 0.50858 | 1.32967 | 1.68908 | 0.52363 | 4.58317 | 0.02407 | 1.39269 | 0.00000 | 0.48322 | 0.60322 |
| **Rnf11** | 13.43422 | 0.00295 | 6.72105 | 2.51023 | 12.99078 | 6.62861 | 3.91784 | 4.03247 | 20.49976 | 10.53143 | 7.07484 | 1.99612 | 13.53752 | 11.30967 |
| **Rnf111** | 6.38959 | 0.02341 | 6.42143 | 3.70414 | 7.14905 | 3.49231 | 2.97779 | 1.89739 | 10.28627 | 0.02611 | 5.30681 | 3.08180 | 5.17274 | 12.18931 |
| **Rnf113a1** | 0.06373 | 0.01290 | 0.64737 | 1.01095 | 0.39399 | 0.66437 | 0.01122 | 0.82872 | 0.00741 | 0.00392 | 0.00000 | 0.50747 | 0.06120 | 0.07908 |
| **Rnf113a2** | 6.09498 | 0.03541 | 7.41503 | 6.43822 | 6.38073 | 8.70190 | 0.00220 | 0.18664 | 0.97802 | 0.00482 | 3.45511 | 0.91724 | 3.13127 | 4.44401 |
| **Rnf114** | 8.83334 | 0.15438 | 13.12012 | 10.56619 | 11.76557 | 9.02064 | 4.62696 | 7.53386 | 2.52720 | 0.24595 | 6.77102 | 6.53378 | 3.96793 | 11.40221 |
| **Rnf115** | 9.49718 | 0.78251 | 12.17886 | 10.02312 | 9.81859 | 7.77796 | 2.70537 | 1.65457 | 6.64512 | 0.02475 | 11.30915 | 3.74278 | 4.44750 | 9.98119 |
| **Rnf121** | 0.38265 | 0.01490 | 4.62920 | 2.45098 | 1.72112 | 3.97684 | 0.01167 | 2.34255 | 5.28197 | 0.02444 | 4.99918 | 1.94830 | 0.28264 | 5.25685 |
| **Rnf122** | 0.00000 | 0.00303 | 0.00000 | 0.00000 | 0.28314 | 0.67533 | 0.00296 | 0.03272 | 5.20460 | 0.00748 | 0.00260 | 0.02830 | 5.01546 | 7.96921 |
| **Rnf123** | 0.00000 | 0.00998 | 0.58395 | 0.96856 | 0.28182 | 0.81891 | 0.00305 | 2.04451 | 0.00414 | 0.01223 | 1.37309 | 1.37905 | 0.00523 | 0.08866 |
| **Rnf125** | 2.88630 | 0.01954 | 1.67196 | 0.63246 | 7.12824 | 1.92942 | 0.00229 | 0.91592 | 3.77088 | 0.00626 | 0.00000 | 2.97564 | 11.94128 | 5.30134 |
| **Rnf126** | 0.00823 | 0.00512 | 0.00105 | 0.74532 | 3.72878 | 0.00774 | 0.04863 | 0.16608 | 0.00660 | 0.00700 | 0.00407 | 0.36942 | 0.26978 | 0.00000 |
| **Rnf128** | 0.00000 | 5.08753 | 0.09617 | 0.00000 | 0.59475 | 1.49847 | 0.92425 | 3.20883 | 2.15439 | 0.02677 | 0.00000 | 2.40310 | 0.23407 | 0.00000 |
| **Rnf13** | 2.75795 | 0.00000 | 6.57400 | 1.37010 | 1.77091 | 9.29837 | 1.88782 | 2.90077 | 7.33556 | 0.10903 | 7.15285 | 3.35356 | 0.00000 | 3.99739 |
| **Rnf130** | 1.39829 | 0.01299 | 1.50718 | 1.27378 | 0.00000 | 1.27101 | 1.62282 | 0.00000 | 1.59531 | 0.01994 | 2.06486 | 6.37030 | 0.40185 | 0.33653 |
| **Rnf135** | 0.48285 | 0.02673 | 1.81031 | 0.53592 | 0.39848 | 2.54693 | 0.00316 | 2.19147 | 0.17844 | 0.02963 | 0.03383 | 1.11801 | 2.08612 | 0.40121 |
| **Rnf138** | 3.40598 | 0.48148 | 3.06493 | 5.83015 | 2.33196 | 2.09609 | 2.07967 | 1.27560 | 4.30674 | 9.47058 | 3.05726 | 4.23224 | 0.82084 | 0.00000 |
| **Rnf138rt1** | 0.00593 | 0.00441 | 0.00563 | 0.01279 | 0.16920 | 0.01062 | 0.00438 | 0.01185 | 0.21428 | 0.00667 | 0.00754 | 0.00922 | 0.02661 | 0.00455 |
| **Rnf139** | 2.14838 | 0.23287 | 1.79735 | 1.34756 | 4.21117 | 3.23209 | 0.05019 | 2.21170 | 2.43287 | 0.04857 | 2.69127 | 0.44567 | 3.26164 | 3.08864 |
| **Rnf14** | 8.59002 | 0.03422 | 5.73984 | 2.24087 | 8.28711 | 8.49645 | 1.46810 | 1.58305 | 7.84284 | 0.06466 | 5.78662 | 5.13514 | 8.84213 | 14.11541 |
| **Rnf141** | 0.00000 | 0.00333 | 0.38056 | 0.04589 | 0.00000 | 0.38714 | 0.00366 | 0.08118 | 0.00238 | 0.02185 | 1.30635 | 0.00239 | 2.25491 | 0.38609 |
| **Rnf145** | 0.83276 | 0.00445 | 1.02089 | 2.05821 | 0.00000 | 0.00000 | 0.01137 | 0.34473 | 1.14015 | 0.00368 | 0.00000 | 0.12816 | 0.50970 | 3.74522 |
| **Rnf146** | 0.13656 | 0.01219 | 0.45068 | 0.16074 | 0.19701 | 0.00000 | 0.07889 | 0.00387 | 0.63180 | 0.01789 | 0.01076 | 0.20048 | 0.52184 | 0.13578 |
| **Rnf149** | 8.88220 | 7.14993 | 12.99226 | 6.26419 | 5.66093 | 8.86150 | 1.96153 | 2.91832 | 8.88343 | 0.02598 | 7.67649 | 11.25115 | 0.00000 | 9.72439 |
| **Rnf150** | 0.02981 | 0.01904 | 0.01075 | 0.01267 | 0.02236 | 0.03576 | 0.00411 | 0.00647 | 0.11280 | 0.00633 | 0.00505 | 0.29680 | 0.01343 | 0.05046 |
| **Rnf152** | 0.00906 | 0.00490 | 0.03158 | 0.02211 | 0.01423 | 0.01525 | 0.00996 | 0.08662 | 0.02893 | 0.00699 | 0.00903 | 0.35566 | 0.02942 | 0.01442 |
| **Rnf157** | 0.01412 | 0.00447 | 0.10236 | 0.00421 | 0.21177 | 0.00713 | 0.00422 | 0.01258 | 0.01168 | 0.00681 | 0.00724 | 0.00785 | 0.45612 | 0.19872 |
| **Rnf165** | 0.00783 | 0.01339 | 0.00572 | 0.00000 | 0.36664 | 0.05805 | 0.00470 | 0.02477 | 0.04977 | 0.00650 | 0.00432 | 0.48415 | 0.17661 | 0.00177 |
| **Rnf166** | 0.33271 | 0.01236 | 1.47892 | 0.44428 | 0.72243 | 2.00416 | 0.00355 | 0.74280 | 0.00317 | 0.02409 | 2.49981 | 0.77148 | 1.15088 | 0.15498 |
| **Rnf167** | 9.12419 | 9.80130 | 20.95480 | 14.77241 | 13.51147 | 18.62771 | 4.58755 | 9.30413 | 1.65036 | 0.85473 | 11.73254 | 10.48072 | 8.35133 | 9.94774 |
| **Rnf168** | 2.75852 | 0.27852 | 0.46579 | 1.91670 | 3.13439 | 1.91082 | 1.66058 | 1.39666 | 6.94224 | 0.35345 | 0.16657 | 1.74596 | 7.99469 | 3.44903 |
| **Rnf169** | 0.40028 | 0.01128 | 0.06376 | 0.14674 | 0.23335 | 3.10999 | 0.00350 | 1.18184 | 0.00268 | 0.01474 | 1.42761 | 0.25129 | 0.51939 | 0.36083 |
| **Rnf17** | 0.02632 | 0.65199 | 0.00000 | 0.09619 | 0.63721 | 0.58453 | 0.00853 | 0.00000 | 1.85591 | 0.01248 | 1.83387 | 0.00000 | 0.59797 | 0.00000 |
| **Rnf170** | 0.22407 | 0.00160 | 0.00000 | 0.26410 | 0.24692 | 1.56454 | 0.00245 | 0.00000 | 5.35514 | 0.00660 | 1.84642 | 1.94271 | 2.08199 | 3.74011 |
| **Rnf181** | 7.96829 | 5.39140 | 17.79244 | 7.05404 | 5.78813 | 17.90327 | 6.24352 | 12.87481 | 2.06556 | 0.37042 | 11.10436 | 7.62887 | 5.54419 | 16.02617 |
| **Rnf182** | 0.00658 | 0.00439 | 0.10882 | 0.00369 | 0.00806 | 0.01169 | 0.00422 | 0.01026 | 0.05223 | 0.00696 | 0.00742 | 0.01039 | 0.10270 | 0.00195 |
| **Rnf185** | 2.26948 | 0.00050 | 6.14782 | 3.89607 | 3.96044 | 5.89005 | 3.20805 | 0.37601 | 2.73143 | 0.07134 | 5.42512 | 1.53527 | 5.49591 | 6.28457 |
| **Rnf187** | 2.24010 | 0.30454 | 3.46594 | 3.32594 | 2.13672 | 5.87603 | 0.04777 | 8.09219 | 0.22860 | 0.06921 | 3.57206 | 7.48970 | 4.86671 | 1.31129 |
| **Rnf19a** | 1.54403 | 0.02736 | 1.26210 | 0.91569 | 0.38669 | 2.44656 | 0.00362 | 0.92946 | 3.10913 | 0.02154 | 1.13587 | 1.23903 | 3.85914 | 0.28806 |
| **Rnf19b** | 9.52870 | 0.12489 | 12.34011 | 7.89941 | 16.44775 | 7.41488 | 4.73236 | 4.86720 | 1.24786 | 0.21399 | 8.12949 | 9.64500 | 8.93185 | 0.14167 |
| **Rnf2** | 9.62657 | 0.03087 | 8.51684 | 3.33960 | 11.65188 | 2.76811 | 3.31207 | 2.88723 | 15.98736 | 0.09793 | 3.54214 | 3.88844 | 11.81709 | 8.36098 |
| **Rnf20** | 2.58317 | 0.04204 | 7.66322 | 2.14103 | 6.48428 | 3.85312 | 2.46566 | 1.41552 | 15.97857 | 0.00597 | 6.84049 | 0.80136 | 9.52807 | 15.48056 |
| **Rnf207** | 0.05932 | 0.00424 | 0.57996 | 0.18784 | 0.15737 | 0.16519 | 0.00438 | 0.70099 | 0.00398 | 0.00428 | 0.00551 | 0.02791 | 0.00171 | 0.29022 |

| | | | | | | | | | | | | | | |
|---|---|---|---|---|---|---|---|---|---|---|---|---|---|---|
| **Rnf208** | 0.97986 | 0.03721 | 1.26984 | 2.30493 | 0.17587 | 1.11965 | 0.58379 | 2.35214 | 0.00387 | 0.03882 | 0.00000 | 0.00000 | 0.03178 | 0.15540 |
| **Rnf214** | 5.95612 | 0.01764 | 3.95645 | 0.26099 | 4.80029 | 4.43221 | 1.87516 | 2.79915 | 3.54957 | 0.03667 | 2.19902 | 2.32267 | 5.55908 | 5.68734 |
| **Rnf215** | 1.12884 | 0.00342 | 0.20006 | 0.94080 | 0.01007 | 0.39974 | 2.50684 | 0.93600 | 0.00293 | 0.02528 | 0.40454 | 0.52445 | 1.19347 | 0.12225 |
| **Rnf216** | 7.48496 | 0.00693 | 13.43514 | 5.14615 | 25.27421 | 2.18281 | 0.00000 | 0.00000 | 2.49329 | 0.00000 | 4.13899 | 1.94662 | 13.07825 | 17.50939 |
| **Rnf219** | 0.01423 | 0.01967 | 0.00000 | 0.10917 | 0.00006 | 0.00687 | 0.00432 | 0.00636 | 0.42413 | 0.00677 | 0.00496 | 0.01677 | 0.00427 | 0.63228 |
| **Rnf220** | 0.58076 | 2.77370 | 3.91530 | 2.31466 | 8.25837 | 3.91545 | 2.52698 | 3.23087 | 0.06971 | 0.06042 | 4.66841 | 4.74334 | 2.36242 | 1.52768 |
| **Rnf223** | 0.46197 | 0.00465 | 0.14412 | 0.02432 | 0.44541 | 0.15395 | 0.00415 | 0.01406 | 0.01409 | 0.00765 | 0.70349 | 0.00249 | 0.67124 | 0.04115 |
| **Rnf225** | 0.34751 | 0.00380 | 0.00000 | 0.02950 | 0.04284 | 0.22499 | 0.01574 | 0.16190 | 1.10323 | 0.00535 | 0.00719 | 0.03922 | 0.34348 | 0.77164 |
| **Rnf24** | 1.02869 | 0.01279 | 1.92139 | 0.03615 | 2.86373 | 1.67603 | 2.29296 | 0.34839 | 9.07879 | 0.00987 | 1.28460 | 0.16120 | 4.95967 | 5.77131 |
| **Rnf25** | 1.21980 | 0.01156 | 1.04819 | 0.49776 | 3.41226 | 1.96393 | 0.00924 | 1.79305 | 0.84315 | 0.00898 | 1.22568 | 0.23147 | 0.23684 | 3.86392 |
| **Rnf26** | 1.14747 | 0.03868 | 4.48796 | 3.97608 | 2.50715 | 6.49761 | 2.32326 | 4.10415 | 0.28987 | 0.13158 | 1.26122 | 3.04208 | 3.28669 | 2.28916 |
| **Rnf31** | 1.36442 | 0.06123 | 4.19285 | 6.36406 | 2.95626 | 1.60364 | 1.77110 | 2.40511 | 0.14095 | 0.04419 | 3.18532 | 3.16470 | 3.69718 | 2.02440 |
| **Rnf32** | 0.00346 | 0.00432 | 0.00717 | 0.02143 | 0.00740 | 1.03160 | 0.00509 | 0.75296 | 0.01295 | 0.01310 | 0.05474 | 0.00000 | 0.68811 | 0.00152 |
| **Rnf34** | 2.07387 | 0.04733 | 6.00422 | 1.53176 | 10.14102 | 4.05066 | 2.60890 | 2.57114 | 7.07905 | 0.11415 | 6.53674 | 2.33272 | 4.08569 | 11.08023 |
| **Rnf38** | 2.62725 | 1.25177 | 6.02695 | 2.41450 | 7.78883 | 4.10032 | 4.51736 | 2.61431 | 0.24196 | 0.09809 | 5.22875 | 1.54584 | 7.50060 | 5.74353 |
| **Rnf39** | 0.01214 | 0.00422 | 0.00616 | 0.28848 | 0.01093 | 0.00969 | 0.00414 | 0.00857 | 0.01310 | 0.00870 | 0.00680 | 0.01579 | 0.00941 | 0.00813 |
| **Rnf4** | 19.92596 | 5.21865 | 23.79381 | 12.26675 | 21.77872 | 18.60500 | 6.02404 | 7.22392 | 24.80879 | 0.30579 | 13.00269 | 9.02837 | 18.38253 | 25.98254 |
| **Rnf40** | 1.31593 | 5.63316 | 6.70733 | 2.94139 | 6.79425 | 2.80761 | 0.07394 | 3.31663 | 0.36998 | 0.01106 | 3.78007 | 0.57285 | 7.02059 | 7.51307 |
| **Rnf41** | 2.08082 | 0.04689 | 7.66913 | 1.07742 | 6.14311 | 4.95194 | 0.00956 | 4.31094 | 5.93062 | 14.27386 | 2.32422 | 1.19538 | 1.44600 | 3.03783 |
| **Rnf43** | 0.00637 | 0.00437 | 0.01293 | 0.01101 | 0.06717 | 0.01287 | 0.00419 | 0.00964 | 0.01583 | 0.00619 | 0.00928 | 0.00927 | 0.24145 | 0.02714 |
| **Rnf44** | 5.56847 | 1.34489 | 4.27926 | 1.44201 | 3.24157 | 2.35065 | 0.90983 | 2.36469 | 3.49116 | 5.82247 | 2.44981 | 2.33859 | 9.88468 | 4.04295 |
| **Rnf5** | 0.78430 | 0.04834 | 2.04134 | 2.04686 | 0.65833 | 1.14499 | 0.12762 | 3.04393 | 0.14750 | 1.18098 | 1.50604 | 2.62619 | 1.00496 | 0.52027 |
| **Rnf6** | 1.65220 | 0.94016 | 2.52778 | 1.05040 | 8.61283 | 3.93675 | 2.72359 | 0.72748 | 1.89522 | 0.07369 | 3.31308 | 2.12684 | 5.74298 | 15.52222 |
| **Rnf7** | 21.87015 | 0.13449 | 36.76150 | 33.10054 | 30.19444 | 19.93631 | 7.79865 | 8.94694 | 14.24886 | 13.10051 | 11.72258 | 8.02503 | 11.87728 | 21.76453 |
| **Rnf8** | 8.86505 | 0.03393 | 3.49610 | 1.77669 | 7.37455 | 2.74956 | 0.00950 | 1.12974 | 1.18421 | 0.02607 | 2.13466 | 1.17285 | 6.45654 | 4.10394 |
| **Rnft1** | 5.44502 | 0.01096 | 4.05858 | 4.63767 | 5.25113 | 1.06445 | 0.00186 | 0.72279 | 7.05323 | 0.00624 | 3.65349 | 1.70577 | 1.06621 | 5.05189 |
| **Rnft2** | 0.03263 | 0.00435 | 0.01660 | 0.00887 | 0.01269 | 0.00418 | 0.00422 | 0.01211 | 0.01111 | 0.00639 | 0.14665 | 0.00674 | 0.03481 | 0.00644 |
| **Rngtt** | 6.07328 | 0.01520 | 3.37829 | 0.51542 | 6.08637 | 2.73220 | 0.00189 | 0.44915 | 0.99335 | 0.00000 | 3.28656 | 2.32565 | 8.88405 | 5.92090 |
| **Rnh1** | 0.59309 | 0.08382 | 5.40657 | 6.22513 | 0.68927 | 3.20477 | 0.01466 | 8.66680 | 0.08234 | 0.02684 | 2.75247 | 0.33902 | 0.00000 | 0.80478 |
| **Rnmt** | 17.64388 | 2.48360 | 14.05523 | 7.99512 | 14.56402 | 14.61449 | 1.13062 | 4.67542 | 18.73824 | 0.12195 | 11.36910 | 7.96057 | 6.39745 | 17.64615 |
| **Rnmtl1** | 5.08523 | 0.13148 | 10.43868 | 3.37668 | 9.57083 | 1.72827 | 5.10713 | 0.73328 | 0.15100 | 0.20679 | 1.24311 | 3.13605 | 1.77250 | 1.82667 |
| **Rnpc3** | 0.78594 | 0.00291 | 1.02290 | 0.51507 | 2.97907 | 0.72253 | 1.68640 | 0.00000 | 3.89890 | 0.01600 | 0.23948 | 0.35129 | 3.57549 | 5.74756 |
| **Rnpep** | 8.63045 | 5.12373 | 10.58718 | 5.41775 | 10.65049 | 15.40894 | 4.46619 | 9.20323 | 0.24843 | 1.93291 | 2.11121 | 2.73668 | 0.60506 | 7.81221 |
| **Rnpepl1** | 1.76201 | 0.01833 | 3.06456 | 3.75691 | 1.13022 | 1.49383 | 1.80555 | 3.50926 | 0.00000 | 0.09495 | 0.77221 | 2.65512 | 0.56942 | 0.00000 |
| **Rnps1** | 17.14385 | 4.45508 | 30.88728 | 21.92543 | 29.81290 | 19.06104 | 7.46466 | 10.54037 | 3.71797 | 12.77487 | 13.68857 | 18.75916 | 11.24425 | 7.76078 |
| **Rnu73b** | 0.00934 | 0.00441 | 0.01367 | 0.02792 | 0.00820 | 0.01020 | 0.00420 | 0.00928 | 0.11573 | 0.00685 | 0.00729 | 0.01174 | 0.00643 | 0.01252 |
| **Robo3** | 0.00965 | 0.00432 | 0.01354 | 0.03035 | 0.01333 | 0.01495 | 0.00416 | 0.01155 | 0.01166 | 0.00739 | 0.00782 | 0.01189 | 0.01592 | 0.01000 |
| **Robo4** | 0.00642 | 0.00413 | 0.15401 | 0.34324 | 0.00000 | 0.00000 | 0.00395 | 0.00756 | 0.04872 | 0.00714 | 0.00883 | 0.01189 | 0.00430 | 0.04915 |
| **Rock1** | 1.78047 | 0.10351 | 4.37809 | 0.23861 | 3.33098 | 3.35922 | 0.00597 | 0.22675 | 19.72978 | 0.00000 | 3.49120 | 1.92266 | 11.73866 | 7.53482 |
| **Rock2** | 1.88012 | 1.21322 | 2.04158 | 0.29912 | 1.22658 | 4.04688 | 2.85904 | 0.00000 | 12.52608 | 0.08544 | 0.19866 | 0.47702 | 1.75252 | 4.47374 |
| **Rogdi** | 2.92745 | 13.52996 | 19.31398 | 23.02445 | 6.87411 | 18.11097 | 9.17314 | 23.22647 | 0.00000 | 4.03857 | 12.55076 | 16.16530 | 1.59708 | 1.83003 |
| **Rom1** | 0.06052 | 0.00409 | 0.32700 | 0.09634 | 1.91453 | 0.20335 | 0.00388 | 0.00202 | 0.00421 | 0.00521 | 0.00509 | 0.06349 | 0.45305 | 0.24924 |
| **Romo1** | 0.24347 | 0.04882 | 0.07893 | 0.50761 | 1.85457 | 1.65082 | 0.11609 | 1.70900 | 0.01331 | 3.25396 | 0.00000 | 2.08631 | 1.17316 | 0.00000 |
| **Ropn1l** | 0.00700 | 0.00409 | 0.01525 | 0.00000 | 0.01386 | 0.03323 | 0.00439 | 0.00649 | 0.01156 | 0.00647 | 0.00459 | 0.47617 | 0.00789 | 0.01113 |
| **Ror1** | 0.00908 | 0.00388 | 0.01074 | 0.02357 | 0.00888 | 0.00422 | 0.00383 | 0.00364 | 0.01475 | 0.98682 | 0.00613 | 0.00520 | 0.00791 | 0.00730 |
| **Rora** | 0.52660 | 0.00543 | 0.00000 | 0.00350 | 0.23400 | 0.00000 | 0.00446 | 0.22255 | 0.64529 | 0.00870 | 0.00417 | 0.00633 | 0.00372 | 0.00000 |
| **Rp2** | 0.02898 | 0.01652 | 0.41335 | 0.00000 | 0.34407 | 3.15994 | 2.10682 | 0.29343 | 0.06623 | 0.05858 | 1.64233 | 1.75275 | 0.17290 | 0.05061 |
| **Rp9** | 1.21121 | 1.37547 | 1.90882 | 1.72814 | 2.14012 | 1.26744 | 1.69219 | 3.75434 | 0.16907 | 0.01496 | 3.55246 | 1.34093 | 1.44476 | 0.43834 |
| **Rpa1** | 0.98965 | 5.97960 | 7.30448 | 11.75361 | 1.37675 | 15.20175 | 1.08659 | 4.05501 | 0.38887 | 0.04558 | 10.38952 | 11.18978 | 0.15038 | 7.29753 |
| **Rpa2** | 16.70566 | 9.08881 | 25.01108 | 13.54955 | 18.61036 | 19.95763 | 6.66451 | 8.37406 | 4.25251 | 0.42581 | 12.10318 | 12.89930 | 10.77280 | 17.53301 |
| **Rpa3** | 5.20253 | 0.94695 | 0.28798 | 0.00000 | 0.64802 | 2.98215 | 0.72969 | 2.11790 | 2.90459 | 0.10284 | 0.51795 | 2.86002 | 1.98295 | 2.76351 |
| **Rpain** | 2.79042 | 0.03632 | 1.62694 | 2.80077 | 1.70709 | 3.96409 | 0.00272 | 0.36322 | 0.86658 | 0.00953 | 1.19288 | 3.55168 | 0.95489 | 2.65495 |
| **Rpap1** | 1.72180 | 0.11148 | 5.24659 | 6.08705 | 3.40876 | 5.65533 | 2.89002 | 0.69293 | 0.21987 | 0.04597 | 7.18891 | 9.48897 | 1.43725 | 3.51052 |
| **Rpap2** | 2.97217 | 0.01646 | 2.34656 | 3.66257 | 2.54808 | 2.84252 | 1.31475 | 1.38812 | 2.56558 | 0.00916 | 2.25364 | 1.73663 | 0.00000 | 1.12155 |
| **Rpap3** | 8.17481 | 5.93390 | 12.26496 | 3.79490 | 21.94357 | 7.79112 | 3.42603 | 3.07435 | 0.56517 | 0.10293 | 5.72202 | 6.57118 | 10.77410 | 9.47331 |

| | | | | | | | | | | | | | | |
|---|---|---|---|---|---|---|---|---|---|---|---|---|---|---|
| **Rpe** | 2.18353 | 0.03864 | 2.04390 | 2.36705 | 2.49867 | 3.02662 | 0.00780 | 1.69936 | 4.45220 | 0.05918 | 5.30322 | 3.29311 | 1.24036 | 2.45449 |
| **Rpf1** | 7.85463 | 0.02281 | 9.46534 | 1.93868 | 10.75040 | 1.89547 | 2.16973 | 1.03449 | 5.93742 | 0.03438 | 1.54824 | 3.29361 | 8.10446 | 4.24433 |
| **Rpf2** | 13.00617 | 5.55427 | 17.38993 | 9.63021 | 19.06962 | 11.08988 | 5.39765 | 4.39369 | 25.23043 | 15.33944 | 14.10385 | 8.70706 | 7.00128 | 17.58524 |
| **Rpgr** | 0.77818 | 0.00223 | 0.53528 | 0.26424 | 0.36512 | 1.22325 | 0.00269 | 0.00000 | 3.01917 | 0.00000 | 5.68947 | 0.61898 | 0.00000 | 0.33965 |
| **Rpgrip1** | 1.70533 | 6.41854 | 5.05912 | 0.45018 | 3.13806 | 3.58324 | 0.00066 | 0.00000 | 1.12710 | 0.01217 | 2.22809 | 2.43459 | 1.90630 | 1.72761 |
| **Rpgrip1l** | 0.00495 | 0.00423 | 0.23459 | 0.02472 | 0.13867 | 1.92821 | 0.00399 | 0.00443 | 0.00373 | 0.01018 | 0.00306 | 0.01038 | 0.12722 | 1.13234 |
| **Rph3a** | 0.00233 | 0.00417 | 0.12525 | 0.00501 | 0.00739 | 0.01385 | 0.00411 | 0.00825 | 0.05974 | 0.00620 | 0.00660 | 0.00936 | 1.77110 | 0.82116 |
| **Rph3al** | 0.03369 | 0.00416 | 0.10881 | 0.00000 | 0.33485 | 0.01936 | 0.00409 | 0.00707 | 0.00429 | 0.00578 | 0.02430 | 0.00780 | 1.48990 | 0.01684 |
| **Rpia** | 1.45673 | 0.03475 | 0.00000 | 0.00000 | 4.61444 | 3.26152 | 0.50316 | 1.16556 | 1.48941 | 0.06998 | 0.21058 | 0.25543 | 9.93242 | 3.43839 |
| **Rpl10** | 44.09152 | 17.08755 | 44.52696 | 45.80430 | 49.43565 | 49.89744 | 15.17087 | 38.22230 | 37.97015 | 25.30653 | 22.17973 | 35.82701 | 17.50507 | 40.37773 |
| **Rpl10a** | 28.50406 | 13.46025 | 25.22001 | 40.55972 | 43.14382 | 37.15150 | 10.72494 | 29.91747 | 5.71499 | 19.09285 | 18.34068 | 28.49160 | 17.76967 | 21.32022 |
| **Rpl10l** | 12.08368 | 6.39207 | 12.21424 | 13.47511 | 15.97953 | 13.69874 | 1.04884 | 0.46034 | 1.97621 | 0.09731 | 4.55007 | 5.70648 | 9.64954 | 3.78306 |
| **Rpl11** | 29.15576 | 14.76556 | 15.28648 | 36.54378 | 39.84453 | 36.99694 | 11.56308 | 30.64549 | 13.54685 | 26.86355 | 10.54111 | 30.46490 | 21.74481 | 23.20281 |
| **Rpl12** | 28.43680 | 15.58166 | 22.95030 | 47.70715 | 48.94626 | 41.31971 | 14.00034 | 36.22562 | 3.79955 | 31.79181 | 14.91636 | 37.44624 | 17.18430 | 13.45609 |
| **Rpl13** | 13.22736 | 4.65633 | 0.79161 | 58.86910 | 53.61533 | 20.58612 | 3.73253 | 12.46772 | 2.53254 | 16.75733 | 4.28436 | 22.44849 | 18.96793 | 4.92880 |
| **Rpl13a** | 34.08824 | 17.66220 | 38.49774 | 61.91523 | 48.21708 | 43.13525 | 14.80088 | 39.65449 | 4.65961 | 26.09792 | 26.55710 | 36.77268 | 16.79976 | 32.24737 |
| **Rpl14** | 31.02707 | 13.83870 | 29.61989 | 54.06189 | 42.16446 | 40.29669 | 14.09008 | 38.09171 | 0.98544 | 26.88104 | 21.78542 | 33.25386 | 7.32405 | 33.08329 |
| **Rpl14.ps1** | 26.30792 | 11.71935 | 23.71820 | 47.37423 | 35.06148 | 33.49599 | 12.24995 | 33.20708 | 0.71695 | 23.42477 | 17.91782 | 28.45187 | 4.21642 | 27.29616 |
| **Rpl15** | 33.67220 | 13.88455 | 34.60439 | 48.18071 | 50.33343 | 41.20050 | 12.12257 | 36.54936 | 19.93254 | 18.21585 | 24.28213 | 34.85643 | 15.76790 | 33.95863 |
| **Rpl17** | 33.29060 | 14.03678 | 21.59955 | 42.51820 | 44.54963 | 38.49901 | 12.70319 | 35.25687 | 27.45696 | 28.68015 | 14.22176 | 30.09348 | 22.19747 | 29.48493 |
| **Rpl18** | 22.75043 | 16.04670 | 25.29194 | 53.29351 | 43.60810 | 33.78422 | 13.58825 | 40.73125 | 0.19058 | 27.74687 | 17.77713 | 34.57258 | 7.33236 | 20.38527 |
| **Rpl18a** | 24.71739 | 7.15617 | 12.64355 | 32.31789 | 41.34289 | 33.83229 | 9.52788 | 27.63857 | 1.70580 | 19.41091 | 8.85374 | 29.87179 | 16.62721 | 23.60452 |
| **Rpl19** | 30.49196 | 16.76614 | 29.74525 | 48.56319 | 39.09087 | 38.98834 | 14.61926 | 33.94608 | 16.86712 | 27.52931 | 23.24731 | 32.63338 | 16.69147 | 36.58552 |
| **Rpl21** | 25.04653 | 13.64580 | 12.19410 | 39.74547 | 38.32141 | 32.83233 | 9.66060 | 31.36850 | 22.96368 | 24.74734 | 6.75303 | 28.40720 | 20.39545 | 20.96463 |
| **Rpl22** | 17.73221 | 5.86246 | 3.41315 | 22.01443 | 24.16494 | 27.09147 | 7.62562 | 22.08271 | 14.97848 | 18.42519 | 3.65044 | 19.32961 | 18.30754 | 12.00724 |
| **Rpl22l1** | 1.25719 | 0.33544 | 0.07403 | 3.88623 | 3.63152 | 2.68096 | 3.23443 | 6.46911 | 3.70072 | 3.62366 | 0.31937 | 4.93795 | 4.39878 | 0.86530 |
| **Rpl23** | 24.73735 | 14.51028 | 16.48554 | 37.11579 | 37.41693 | 34.43398 | 10.89140 | 33.31548 | 24.95829 | 23.48846 | 10.66789 | 29.36245 | 27.16140 | 23.26956 |
| **Rpl23a** | 24.32760 | 14.12991 | 13.38394 | 39.00451 | 38.58899 | 37.12814 | 14.36825 | 34.44459 | 17.21867 | 29.06396 | 8.63730 | 34.89330 | 24.94800 | 24.56157 |
| **Rpl24** | 24.59634 | 11.64639 | 14.17842 | 39.85182 | 38.31755 | 31.98447 | 12.44724 | 30.06389 | 26.00854 | 23.63086 | 8.97809 | 29.32232 | 24.06081 | 26.31593 |
| **Rpl26** | 0.04601 | 0.00471 | 0.04288 | 0.09713 | 0.02301 | 0.03378 | 0.00427 | 0.01698 | 0.03128 | 0.01015 | 0.03225 | 0.03503 | 0.01149 | 0.01636 |
| **Rpl27** | 11.63415 | 9.81855 | 7.54146 | 25.80253 | 32.89484 | 26.44128 | 7.96787 | 28.85369 | 7.26726 | 23.00544 | 4.40797 | 26.12176 | 16.50237 | 17.89264 |
| **Rpl27a** | 15.20657 | 13.20326 | 13.92877 | 46.26803 | 39.66489 | 22.29832 | 12.09034 | 39.70791 | 5.93385 | 25.74458 | 7.27032 | 30.21293 | 17.20446 | 4.95072 |
| **Rpl28** | 19.87781 | 10.33459 | 7.57856 | 24.13133 | 27.76794 | 25.26947 | 7.84830 | 27.30886 | 2.13970 | 19.11147 | 3.71186 | 23.40069 | 15.44699 | 16.01389 |
| **Rpl29** | 27.71557 | 13.39085 | 25.52408 | 40.15234 | 42.80297 | 36.77749 | 12.46454 | 32.26844 | 0.09647 | 25.87129 | 14.75048 | 33.01081 | 5.55469 | 23.12237 |
| **Rpl3** | 34.25051 | 17.94259 | 60.46151 | 53.62499 | 45.11535 | 50.67875 | 13.87694 | 44.21830 | 38.93385 | 23.60910 | 36.68825 | 38.44957 | 11.47881 | 45.26024 |
| **Rpl30** | 0.81480 | 0.01224 | 0.09186 | 1.10697 | 1.18755 | 0.42242 | 0.19036 | 0.57642 | 0.24661 | 0.04225 | 0.05054 | 0.43955 | 0.49194 | 0.19555 |
| **Rpl31** | 15.47382 | 4.77607 | 2.84906 | 14.73752 | 21.46500 | 22.02970 | 7.27816 | 21.77702 | 11.03560 | 20.13246 | 1.55840 | 19.47205 | 19.72555 | 9.38811 |
| **Rpl31.ps12** | 7.25085 | 1.67046 | 0.38585 | 4.92511 | 8.19984 | 9.03741 | 3.05084 | 10.24880 | 3.92190 | 11.76511 | 0.17859 | 8.45122 | 13.87535 | 2.67158 |
| **Rpl32** | 23.54057 | 8.27415 | 7.92786 | 24.63628 | 31.40226 | 27.58274 | 11.02622 | 28.17633 | 14.85799 | 24.57972 | 4.84781 | 24.72463 | 18.36546 | 15.77373 |
| **Rpl32l** | 4.45162 | 0.92455 | 0.23607 | 3.35410 | 4.94777 | 4.37728 | 1.97731 | 5.62066 | 1.68141 | 6.59520 | 0.25388 | 3.99692 | 4.40307 | 2.01285 |
| **Rpl34** | 0.24264 | 0.05614 | 0.34681 | 2.67681 | 3.34471 | 0.26232 | 0.52801 | 0.79741 | 0.04551 | 0.09182 | 0.12449 | 0.35809 | 0.04291 | 0.16797 |
| **Rpl35** | 17.21534 | 8.11425 | 5.46546 | 22.49182 | 21.54997 | 25.46798 | 7.56715 | 27.57526 | 9.08858 | 24.19533 | 2.75060 | 21.33208 | 22.97133 | 12.56508 |
| **Rpl35a** | 13.89065 | 8.61497 | 3.08191 | 23.87223 | 20.97163 | 27.87976 | 8.43363 | 25.47949 | 16.27100 | 21.42220 | 2.33625 | 21.24201 | 18.42266 | 13.09185 |
| **Rpl36** | 4.01954 | 3.06378 | 2.18097 | 10.25299 | 17.23745 | 14.50763 | 4.71613 | 16.13119 | 0.00000 | 17.13856 | 0.63950 | 15.17500 | 1.17592 | 0.28105 |
| **Rpl36a** | 16.73637 | 8.21261 | 3.64961 | 26.13622 | 21.30921 | 23.34714 | 7.38482 | 25.43763 | 19.98655 | 16.93376 | 1.65399 | 20.77447 | 13.83427 | 14.34726 |
| **Rpl36al** | 11.57674 | 4.38662 | 3.43070 | 17.14938 | 21.00827 | 20.56546 | 6.89824 | 19.86484 | 11.62471 | 14.61620 | 2.71457 | 15.80753 | 20.76041 | 12.98272 |
| **Rpl37** | 8.40582 | 3.18647 | 1.60569 | 16.47927 | 10.00165 | 12.30870 | 4.17991 | 17.74108 | 5.25449 | 12.95126 | 0.53910 | 10.49310 | 19.54925 | 5.89271 |
| **Rpl37a** | 12.99010 | 7.93195 | 3.58956 | 19.71454 | 17.70302 | 27.74837 | 6.80293 | 21.91824 | 7.92163 | 19.95802 | 1.87043 | 18.91470 | 23.19333 | 11.27232 |
| **Rpl37rt** | 7.30695 | 2.84894 | 0.89643 | 14.14664 | 9.47634 | 11.46393 | 4.19411 | 17.40697 | 4.64593 | 12.65662 | 0.41875 | 10.15012 | 16.34099 | 4.81784 |
| **Rpl38** | 8.14992 | 4.81350 | 1.32054 | 15.74052 | 9.87369 | 19.52168 | 3.56412 | 19.48073 | 2.88360 | 17.82429 | 0.77942 | 14.61631 | 16.99304 | 5.40174 |
| **Rpl39** | 18.97128 | 3.02050 | 0.89039 | 8.15852 | 9.70555 | 23.95925 | 4.79335 | 10.97812 | 17.45206 | 16.01136 | 0.58720 | 13.42314 | 20.02591 | 6.89527 |
| **Rpl39l** | 3.88399 | 4.57157 | 1.84137 | 9.33410 | 22.69566 | 9.48471 | 5.05902 | 8.14277 | 3.52057 | 8.43522 | 0.31043 | 11.53539 | 14.46941 | 8.71107 |
| **Rpl3l** | 0.01964 | 0.00346 | 0.65980 | 0.21972 | 2.20670 | 0.00000 | 0.00396 | 0.00148 | 0.00511 | 0.00454 | 0.00405 | 0.43775 | 2.38890 | 0.24431 |
| **Rpl4** | 42.39113 | 19.90639 | 64.12718 | 63.30962 | 51.62469 | 56.73882 | 17.58054 | 47.12124 | 47.65077 | 25.63086 | 37.35084 | 44.57795 | 19.22642 | 45.42906 |
| **Rpl41** | 27.43745 | 9.42761 | 9.51522 | 35.30664 | 30.49050 | 37.35839 | 9.48636 | 30.60114 | 16.15640 | 25.48484 | 5.93109 | 26.42147 | 28.71152 | 24.46647 |

| | | | | | | | | | | | | | | |
|---|---|---|---|---|---|---|---|---|---|---|---|---|---|---|
| **Rpl5** | 44.60779 | 18.59219 | 57.03966 | 56.94464 | 49.15180 | 50.49491 | 15.71121 | 43.79230 | 43.64491 | 29.48824 | 34.31545 | 39.59191 | 16.53165 | 39.98391 |
| **Rpl6** | 43.35945 | 18.02170 | 58.31065 | 61.83663 | 50.85370 | 47.83097 | 17.02843 | 44.67881 | 33.75469 | 29.47010 | 36.00809 | 39.06162 | 20.15723 | 41.37341 |
| **Rpl7** | 41.78990 | 16.89846 | 50.02522 | 57.81578 | 48.64693 | 48.09936 | 15.57391 | 40.36847 | 44.14579 | 27.01211 | 32.78113 | 36.76026 | 13.37456 | 34.29013 |
| **Rpl7a** | 29.10449 | 18.99787 | 48.83719 | 45.87545 | 35.90526 | 38.15010 | 13.10040 | 35.96226 | 33.02836 | 29.27298 | 28.23985 | 35.07008 | 11.35530 | 38.31258 |
| **Rpl7l1** | 10.38440 | 0.39502 | 22.38997 | 17.46870 | 24.40488 | 15.88130 | 4.21887 | 10.31170 | 21.74439 | 2.05510 | 12.30610 | 12.55264 | 7.12121 | 23.42960 |
| **Rpl8** | 35.01332 | 16.92227 | 51.80113 | 60.35074 | 48.06043 | 44.92916 | 13.46356 | 43.46504 | 2.00306 | 30.01042 | 30.08285 | 36.59326 | 15.75653 | 15.64254 |
| **Rpl9** | 32.95634 | 15.27065 | 29.41722 | 47.59398 | 44.20648 | 40.56557 | 14.19559 | 33.83988 | 36.82840 | 29.46843 | 20.06685 | 34.22613 | 20.39705 | 32.75161 |
| **Rplp0** | 35.77295 | 20.61943 | 67.89228 | 59.36638 | 62.54941 | 51.91869 | 15.55837 | 50.28120 | 0.00000 | 31.91788 | 37.80879 | 43.69015 | 4.01471 | 21.53941 |
| **Rplp1** | 10.59879 | 14.53471 | 17.45095 | 43.47380 | 53.69200 | 31.60158 | 13.97114 | 41.11573 | 0.00000 | 25.74019 | 7.74660 | 35.17375 | 3.59096 | 0.43209 |
| **Rplp2** | 2.88443 | 9.50336 | 6.35600 | 27.60006 | 27.28192 | 14.67756 | 9.20795 | 28.46426 | 0.00000 | 22.10607 | 0.42729 | 25.10885 | 4.04259 | 0.58065 |
| **Rpn1** | 5.76967 | 0.23207 | 16.19148 | 19.89820 | 5.62951 | 19.72997 | 6.98421 | 12.22125 | 5.39245 | 0.35870 | 12.47233 | 11.07809 | 0.71245 | 8.62540 |
| **Rpn2** | 14.66344 | 11.59142 | 18.36396 | 20.72379 | 8.25113 | 21.29197 | 8.13807 | 13.65815 | 14.95022 | 0.61919 | 11.33441 | 17.18704 | 2.54554 | 15.23551 |
| **Rpp14** | 6.34916 | 0.08437 | 9.58144 | 6.04567 | 5.99962 | 9.49592 | 1.12468 | 1.00560 | 11.72727 | 10.46137 | 9.62295 | 6.92963 | 6.13593 | 12.33059 |
| **Rpp21** | 1.52792 | 4.88584 | 1.19484 | 4.77731 | 5.64337 | 2.96322 | 3.63211 | 5.10600 | 0.58279 | 11.92357 | 0.34064 | 3.64938 | 1.16601 | 0.72621 |
| **Rpp25** | 0.12807 | 0.00280 | 0.23049 | 3.27082 | 0.75872 | 0.14710 | 0.55955 | 0.30287 | 0.00114 | 0.01273 | 0.82664 | 0.99245 | 1.94757 | 0.11082 |
| **Rpp25l** | 0.48451 | 4.40916 | 4.09562 | 9.23200 | 1.80222 | 4.56815 | 0.02821 | 1.87397 | 0.00000 | 12.40540 | 0.88840 | 5.21945 | 0.00000 | 1.35362 |
| **Rpp30** | 10.00523 | 0.07044 | 11.25560 | 8.49291 | 12.76612 | 13.76076 | 3.31197 | 4.61376 | 9.32081 | 0.16221 | 7.67377 | 7.44015 | 0.87003 | 11.14823 |
| **Rpp38** | 6.26278 | 0.00846 | 6.09970 | 3.20299 | 6.64524 | 3.36367 | 0.00861 | 1.81957 | 1.44762 | 0.09135 | 6.00663 | 2.96380 | 0.27656 | 7.16562 |
| **Rpp40** | 5.64420 | 0.07232 | 7.70776 | 5.69586 | 10.57005 | 8.11139 | 0.62260 | 1.38055 | 0.54677 | 0.05947 | 7.08358 | 6.57156 | 4.59038 | 7.96806 |
| **Rprd1a** | 4.42885 | 0.03826 | 7.43577 | 0.00000 | 12.65691 | 9.44825 | 4.33544 | 4.79679 | 11.31819 | 0.11463 | 1.16395 | 4.45106 | 9.52394 | 15.37773 |
| **Rprd1b** | 9.64838 | 0.07724 | 9.27113 | 2.11069 | 8.09847 | 4.93837 | 5.13016 | 2.71370 | 11.63511 | 0.16515 | 5.00808 | 2.71556 | 7.10964 | 10.76245 |
| **Rprd2** | 0.13677 | 0.01001 | 0.74246 | 1.21346 | 2.01691 | 2.46881 | 0.29593 | 1.60795 | 5.81259 | 0.02686 | 0.00000 | 2.41953 | 3.82529 | 1.09491 |
| **Rps10** | 1.68210 | 10.10275 | 14.99234 | 36.60252 | 44.09753 | 8.97012 | 11.42913 | 37.36692 | 0.00000 | 24.04375 | 4.92029 | 28.35330 | 6.28940 | 0.00000 |
| **Rps11** | 28.45289 | 11.32570 | 13.87724 | 40.27502 | 38.94240 | 41.63601 | 11.40725 | 32.05520 | 22.86716 | 27.57668 | 9.94040 | 30.37886 | 23.96388 | 25.72328 |
| **Rps12** | 23.50549 | 14.56724 | 13.89686 | 39.40485 | 40.14393 | 42.75745 | 12.58346 | 35.07838 | 6.09422 | 27.89127 | 9.12103 | 35.67931 | 23.26620 | 22.13018 |
| **Rps13** | 21.52686 | 11.43144 | 8.62805 | 27.97526 | 40.11104 | 28.23423 | 9.33437 | 29.89528 | 2.62940 | 21.80077 | 5.06597 | 28.51289 | 24.07379 | 11.34479 |
| **Rps14** | 19.37844 | 7.83982 | 7.97932 | 32.16645 | 38.69813 | 34.16575 | 7.61017 | 31.47773 | 11.81693 | 22.75805 | 8.29485 | 28.28457 | 12.65563 | 24.13868 |
| **Rps15** | 14.63318 | 6.71503 | 2.81217 | 25.95168 | 28.33583 | 26.84800 | 6.20216 | 22.13698 | 1.96668 | 18.44356 | 4.93651 | 18.80131 | 16.89716 | 12.46502 |
| **Rps15a** | 11.39397 | 7.36632 | 1.67012 | 25.56827 | 17.54281 | 16.59305 | 5.42596 | 21.86457 | 18.54583 | 16.21394 | 2.13569 | 15.85611 | 16.89542 | 11.62127 |
| **Rps15a.ps4** | 3.86915 | 2.86751 | 0.00000 | 11.64037 | 5.90675 | 5.27065 | 1.96514 | 9.62113 | 7.13409 | 7.19889 | 0.39881 | 5.86649 | 8.21169 | 3.19122 |
| **Rps15a.ps6** | 3.22680 | 2.69602 | 0.00000 | 10.05394 | 5.19293 | 4.70134 | 1.67169 | 8.21098 | 6.66419 | 6.16745 | 0.39037 | 4.99278 | 6.95434 | 3.24593 |
| **Rps16** | 22.19915 | 13.08585 | 17.32440 | 44.00180 | 45.48891 | 38.05161 | 13.13449 | 38.72324 | 3.99909 | 28.97721 | 10.08933 | 34.44518 | 17.77346 | 16.37384 |
| **Rps17** | 16.45020 | 8.65302 | 5.04170 | 29.10093 | 28.87601 | 29.31088 | 10.57399 | 29.54990 | 16.94663 | 24.74112 | 4.01359 | 25.04688 | 24.33882 | 13.52356 |
| **Rps18** | 20.85352 | 12.89706 | 12.73604 | 36.99196 | 40.96605 | 38.34560 | 12.37187 | 33.34695 | 7.79761 | 27.99269 | 8.36699 | 33.25049 | 19.98321 | 24.04311 |
| **Rps19** | 22.02977 | 9.87701 | 9.01719 | 26.82773 | 37.05696 | 30.56649 | 11.52515 | 30.30226 | 10.56451 | 25.95226 | 5.78025 | 29.37619 | 21.54381 | 19.79366 |
| **Rps19.ps3** | 7.38113 | 3.03000 | 1.25790 | 7.75436 | 13.17062 | 9.20950 | 4.15899 | 12.06934 | 2.05678 | 11.56994 | 0.77175 | 10.28856 | 6.70002 | 5.37634 |
| **Rps19bp1** | 1.21070 | 0.05402 | 0.20219 | 7.65070 | 17.68020 | 2.14368 | 0.03389 | 0.73283 | 0.65970 | 0.14885 | 0.72028 | 2.25462 | 10.52109 | 1.45882 |
| **Rps2** | 34.34568 | 9.29097 | 16.31155 | 39.68553 | 56.12259 | 38.95531 | 7.57068 | 20.60363 | 5.47473 | 27.00557 | 13.85163 | 33.31615 | 21.18902 | 16.97532 |
| **Rps20** | 21.89908 | 11.09404 | 9.12217 | 29.09717 | 38.26969 | 34.43240 | 12.68837 | 36.27503 | 2.29041 | 26.79719 | 6.33082 | 32.73421 | 21.61326 | 7.23057 |
| **Rps21** | 12.83855 | 7.03008 | 1.57050 | 15.15396 | 14.64036 | 25.50215 | 5.61682 | 23.52001 | 2.77298 | 21.42784 | 1.59768 | 20.90837 | 23.12121 | 8.19536 |
| **Rps23** | 13.95662 | 9.02855 | 4.08208 | 16.89388 | 25.35219 | 23.27314 | 8.52832 | 20.76379 | 6.28947 | 19.09464 | 2.06587 | 20.94739 | 14.98488 | 13.24654 |
| **Rps24** | 27.37014 | 12.22074 | 8.72946 | 30.65368 | 37.39067 | 35.56300 | 10.50335 | 30.55029 | 29.85378 | 25.32232 | 7.84933 | 30.00762 | 25.92023 | 20.91908 |
| **Rps25** | 21.98325 | 10.96719 | 4.65078 | 26.05578 | 33.07092 | 28.31939 | 9.79393 | 24.58579 | 23.08923 | 21.22072 | 3.51096 | 22.93723 | 25.20295 | 15.74744 |
| **Rps26** | 18.36752 | 9.12058 | 7.31926 | 24.58754 | 24.94458 | 29.86588 | 8.76823 | 28.59742 | 1.48961 | 22.83873 | 3.98186 | 22.78906 | 19.85837 | 17.26312 |
| **Rps27** | 0.08831 | 0.01185 | 0.42790 | 0.15154 | 0.65926 | 0.09046 | 0.00442 | 0.16456 | 0.00793 | 0.03960 | 0.02508 | 0.07389 | 0.10776 | 0.01353 |
| **Rps27a** | 26.87356 | 12.22203 | 14.93218 | 45.47730 | 40.36788 | 37.22821 | 12.87099 | 39.92510 | 30.51959 | 26.26315 | 11.34481 | 35.41266 | 20.52080 | 27.12303 |
| **Rps27l** | 7.64068 | 2.29213 | 0.72490 | 4.71931 | 17.58858 | 21.47401 | 6.91492 | 14.59658 | 1.72954 | 16.73157 | 1.91072 | 18.30399 | 14.79823 | 4.17749 |
| **Rps28** | 9.21127 | 4.52657 | 1.21063 | 4.01608 | 8.04549 | 19.94447 | 3.94455 | 12.86882 | 1.93607 | 19.37470 | 0.52532 | 14.37242 | 18.02699 | 4.69643 |
| **Rps29** | 14.51784 | 4.13089 | 2.21355 | 12.79315 | 13.11627 | 30.18923 | 5.65243 | 16.85475 | 0.84077 | 22.43363 | 1.95276 | 20.30789 | 21.86675 | 9.32305 |
| **Rps3** | 40.70569 | 16.34408 | 48.87415 | 59.22656 | 44.14745 | 46.34566 | 14.09537 | 42.24926 | 35.82760 | 28.28601 | 31.33940 | 36.92797 | 18.95951 | 38.08008 |
| **Rps3a1** | 44.09494 | 17.40002 | 51.85498 | 55.81756 | 51.89156 | 50.00147 | 16.52881 | 43.22792 | 38.88753 | 30.52125 | 32.99868 | 39.57960 | 21.01857 | 38.76854 |
| **Rps4l** | 8.48677 | 8.32712 | 19.09717 | 37.51929 | 27.55706 | 9.59748 | 6.07913 | 18.88875 | 0.00000 | 14.67621 | 3.89040 | 19.91133 | 4.62572 | 1.86614 |
| **Rps4x** | 46.98384 | 15.62999 | 54.94676 | 47.34376 | 50.13334 | 49.42847 | 15.69429 | 38.39120 | 47.60622 | 26.45635 | 27.17839 | 37.13460 | 15.85007 | 40.17520 |
| **Rps5** | 33.90240 | 15.02074 | 34.66926 | 51.82523 | 46.56477 | 42.81696 | 13.29002 | 39.22315 | 8.15304 | 28.64837 | 22.88842 | 36.77009 | 16.43579 | 37.62231 |
| **Rps6** | 42.47506 | 15.68911 | 49.56208 | 59.18753 | 47.24509 | 47.94408 | 15.62483 | 41.89130 | 37.18775 | 25.89653 | 31.71780 | 39.08545 | 21.68592 | 42.01392 |

| | | | | | | | | | | | | | | |
|---|---|---|---|---|---|---|---|---|---|---|---|---|---|---|
| **Rps6ka1** | 1.28402 | 0.17551 | 11.33552 | 7.48856 | 6.81307 | 13.68069 | 1.65109 | 8.93215 | 0.02518 | 0.19652 | 5.69977 | 9.28272 | 0.35842 | 3.43933 |
| **Rps6ka2** | 0.03224 | 0.00394 | 0.69167 | 0.83341 | 0.80040 | 1.32862 | 0.00382 | 0.00003 | 0.00329 | 0.00616 | 0.00143 | 0.13574 | 1.41659 | 0.22956 |
| **Rps6ka3** | 1.01096 | 0.00161 | 0.54227 | 1.24476 | 0.48394 | 1.04428 | 1.14931 | 0.00000 | 1.79639 | 0.00108 | 4.33372 | 1.18930 | 3.02959 | 0.72652 |
| **Rps6ka4** | 0.13700 | 0.00445 | 0.80993 | 1.42025 | 1.64546 | 0.20351 | 0.00496 | 0.26350 | 0.31549 | 0.00665 | 0.17619 | 0.00226 | 0.17345 | 0.21517 |
| **Rps6ka5** | 1.60206 | 0.00351 | 1.18968 | 0.15653 | 1.01959 | 0.35994 | 0.00322 | 0.11351 | 2.40415 | 0.00295 | 0.52199 | 0.01931 | 0.49140 | 1.18475 |
| **Rps6ka6** | 0.29059 | 0.00357 | 0.06811 | 0.03584 | 1.68799 | 0.00699 | 0.00271 | 0.00374 | 7.46740 | 0.00550 | 0.00633 | 0.08776 | 4.95101 | 0.44480 |
| **Rps6kb1** | 3.95265 | 0.00182 | 3.99051 | 2.78340 | 3.52161 | 4.07095 | 0.00190 | 0.51743 | 8.30526 | 0.00000 | 2.83754 | 0.59450 | 4.37677 | 5.17872 |
| **Rps6kb2** | 1.44357 | 9.18353 | 7.15397 | 8.46816 | 1.76796 | 4.21093 | 2.06555 | 3.48542 | 0.17429 | 0.17192 | 3.91787 | 1.63769 | 1.71538 | 1.36491 |
| **Rps6kc1** | 0.11608 | 0.05338 | 1.38608 | 0.00000 | 0.39380 | 0.52635 | 0.00289 | 2.82813 | 0.18355 | 0.01949 | 0.02145 | 0.73979 | 0.54093 | 2.25964 |
| **Rps6kl1** | 0.11953 | 0.01180 | 0.03987 | 0.86669 | 0.00000 | 0.08494 | 0.00425 | 0.01723 | 0.01001 | 0.01347 | 0.00525 | 0.06035 | 0.07191 | 0.08642 |
| **Rps7** | 31.93343 | 16.73452 | 28.24932 | 50.14157 | 47.66207 | 36.77805 | 13.83783 | 34.08016 | 23.00269 | 28.68071 | 17.74925 | 34.52088 | 22.37890 | 34.11180 |
| **Rps8** | 40.01146 | 16.84785 | 37.24157 | 58.16917 | 51.80612 | 46.47660 | 16.75447 | 40.50409 | 21.74653 | 31.82856 | 23.69479 | 39.71796 | 25.05436 | 39.29619 |
| **Rps9** | 31.80892 | 17.68770 | 34.46619 | 52.91566 | 43.72412 | 41.25705 | 13.84257 | 40.25098 | 21.34918 | 30.93303 | 21.60660 | 36.79333 | 21.95313 | 35.57490 |
| **Rpsa** | 39.14155 | 19.08915 | 57.92977 | 55.35922 | 51.90521 | 53.94867 | 12.88128 | 45.68924 | 2.13624 | 34.60590 | 35.81116 | 40.51713 | 7.70233 | 41.42729 |
| **Rptor** | 1.04518 | 0.02597 | 4.58459 | 1.73023 | 6.01680 | 3.23873 | 2.51780 | 2.19077 | 0.00000 | 0.06364 | 3.09365 | 3.42169 | 0.71106 | 1.18829 |
| **Rpusd1** | 0.92654 | 0.02881 | 3.41175 | 1.80552 | 0.91042 | 4.32796 | 6.04928 | 1.97353 | 0.00000 | 0.20397 | 1.97219 | 1.26653 | 1.91362 | 0.00000 |
| **Rpusd2** | 1.07332 | 0.01945 | 2.42995 | 2.92010 | 3.60447 | 2.27695 | 0.52277 | 0.51461 | 1.42441 | 0.02406 | 0.00000 | 0.72691 | 1.92506 | 2.14864 |
| **Rpusd3** | 0.00000 | 0.05258 | 5.23799 | 7.82068 | 1.79750 | 0.99631 | 1.61923 | 4.49301 | 0.00000 | 0.05499 | 0.05179 | 3.66347 | 0.80225 | 0.63254 |
| **Rpusd4** | 2.78190 | 0.06791 | 7.20131 | 4.85258 | 4.80498 | 5.99816 | 0.00098 | 2.75871 | 1.23358 | 0.07623 | 5.38264 | 2.71869 | 0.94473 | 8.76207 |
| **Rrad** | 0.00875 | 0.03223 | 0.01125 | 0.04967 | 0.01101 | 0.00699 | 0.00467 | 0.55737 | 0.01219 | 0.00926 | 0.01123 | 0.01793 | 0.00716 | 0.01142 |
| **Rraga** | 0.01253 | 0.96056 | 0.10610 | 3.03476 | 0.00000 | 3.07368 | 4.25919 | 0.76038 | 0.24574 | 0.09837 | 2.48706 | 2.45610 | 0.01236 | 0.64998 |
| **Rragc** | 4.07049 | 0.05565 | 3.96858 | 4.23995 | 4.04705 | 5.72479 | 0.05279 | 3.88933 | 2.14424 | 0.05792 | 3.43658 | 2.07176 | 8.78195 | 10.99457 |
| **Rragd** | 11.85192 | 0.46839 | 25.04063 | 12.05173 | 11.00951 | 26.85264 | 6.61206 | 7.12486 | 22.74523 | 0.42785 | 11.54175 | 0.99570 | 0.10458 | 0.86543 |
| **Rras** | 0.14739 | 0.61840 | 0.36677 | 0.00000 | 0.04529 | 1.06692 | 0.00406 | 1.23058 | 0.05512 | 0.03270 | 0.01161 | 0.65077 | 0.74096 | 0.00000 |
| **Rras2** | 10.64135 | 0.04803 | 6.15424 | 0.00129 | 9.34460 | 9.73268 | 2.54658 | 4.33146 | 14.78010 | 0.04844 | 2.44602 | 7.97254 | 5.44502 | 7.69303 |
| **Rrbp1** | 10.71175 | 0.22236 | 7.74234 | 4.98639 | 4.15031 | 15.62430 | 4.67986 | 4.91027 | 1.98786 | 0.14001 | 6.66221 | 12.50044 | 8.25923 | 10.39118 |
| **Rreb1** | 3.58927 | 0.01612 | 8.67154 | 2.11576 | 9.76487 | 1.05438 | 0.69886 | 4.48666 | 4.19659 | 0.02812 | 0.00000 | 2.15515 | 8.89836 | 5.91443 |
| **Rrm1** | 14.83320 | 0.84207 | 21.15318 | 12.24226 | 18.41866 | 20.80459 | 6.75879 | 9.19532 | 18.24110 | 12.10009 | 19.16551 | 14.76178 | 9.78417 | 16.30288 |
| **Rrm2** | 13.19904 | 6.92104 | 27.14608 | 22.10075 | 17.15964 | 15.60774 | 3.55705 | 10.93522 | 18.98945 | 0.75134 | 14.54118 | 11.72960 | 15.87821 | 18.69585 |
| **Rrm2b** | 0.84434 | 0.00358 | 0.00000 | 0.14884 | 0.22581 | 0.00000 | 0.40088 | 0.00383 | 4.39995 | 0.03165 | 0.00000 | 0.27540 | 2.64644 | 4.94757 |
| **Rrn3** | 11.68218 | 1.53787 | 12.81810 | 10.05429 | 18.05681 | 11.70368 | 3.38468 | 7.30114 | 21.10175 | 0.16304 | 4.82445 | 7.93360 | 11.22060 | 20.58864 |
| **Rrnad1** | 4.21619 | 0.04112 | 4.63322 | 1.06459 | 3.94444 | 2.86690 | 0.29461 | 0.00000 | 1.71460 | 0.88536 | 0.00000 | 2.67513 | 0.67056 | 1.99763 |
| **Rrp1** | 12.52589 | 7.60475 | 22.65466 | 20.00615 | 14.00183 | 15.20782 | 3.22579 | 11.52417 | 1.65472 | 0.28370 | 8.68660 | 10.84735 | 5.57261 | 12.40833 |
| **Rrp12** | 3.01227 | 3.42211 | 9.49998 | 8.48672 | 11.28786 | 7.07643 | 1.66058 | 5.55579 | 2.36825 | 0.22021 | 7.24717 | 5.67182 | 10.26293 | 8.02645 |
| **Rrp15** | 0.26658 | 0.09600 | 23.47828 | 11.79943 | 22.73470 | 0.00000 | 0.02674 | 4.66932 | 0.00000 | 0.02029 | 1.11845 | 10.52421 | 0.66667 | 0.10629 |
| **Rrp1b** | 3.46184 | 0.04538 | 7.52339 | 3.32573 | 5.79401 | 3.56728 | 0.01846 | 1.95121 | 0.19139 | 0.00000 | 5.40480 | 4.37730 | 2.69593 | 7.67313 |
| **Rrp36** | 4.73623 | 0.03938 | 6.40432 | 3.77688 | 8.23563 | 7.13173 | 3.56940 | 3.78452 | 0.17721 | 0.16152 | 7.20643 | 1.97981 | 1.49854 | 4.31515 |
| **Rrp7a** | 1.51522 | 2.44037 | 3.66395 | 4.41362 | 5.31806 | 3.07873 | 2.20908 | 6.10866 | 1.02378 | 0.07542 | 4.71554 | 3.21203 | 8.16131 | 0.45115 |
| **Rrp8** | 6.44131 | 2.13431 | 13.15513 | 8.96724 | 9.44775 | 4.13258 | 3.25279 | 3.31541 | 2.61093 | 0.13756 | 7.69193 | 5.25504 | 4.87411 | 8.79315 |
| **Rrp9** | 3.13742 | 6.71047 | 11.83624 | 12.86961 | 14.60221 | 5.33955 | 2.09951 | 2.15721 | 0.00000 | 10.58797 | 2.47024 | 3.31789 | 7.39601 | 0.21399 |
| **Rrs1** | 6.88778 | 0.08375 | 11.88098 | 7.21867 | 9.24133 | 8.04448 | 5.27157 | 1.03239 | 9.22616 | 0.09290 | 2.55374 | 2.03777 | 10.48679 | 8.87862 |
| **Rs1** | 0.01753 | 0.00444 | 0.02565 | 0.01675 | 0.13649 | 0.00563 | 0.00385 | 0.00955 | 1.95197 | 0.00656 | 0.00530 | 0.00863 | 0.03761 | 0.00000 |
| **Rsad1** | 1.29274 | 0.02393 | 0.28581 | 0.06773 | 0.27706 | 0.00000 | 0.01041 | 0.77512 | 0.48504 | 0.01531 | 0.10772 | 0.71603 | 0.24517 | 1.25435 |
| **Rsad2** | 0.00742 | 0.00456 | 0.00449 | 0.01200 | 0.78795 | 0.01320 | 0.00434 | 0.01073 | 0.01029 | 0.00626 | 0.00887 | 0.00752 | 0.00453 | 0.00946 |
| **Rsbn1** | 0.14547 | 0.00443 | 0.23212 | 0.03689 | 0.41645 | 0.00987 | 0.00451 | 0.01401 | 0.53845 | 0.00736 | 0.00547 | 0.00552 | 1.52818 | 0.00000 |
| **Rsbn1l** | 0.44338 | 0.01043 | 1.39891 | 0.42362 | 1.51466 | 1.35624 | 0.04130 | 0.72964 | 4.83576 | 3.96808 | 1.36421 | 0.87430 | 5.03779 | 2.20413 |
| **Rsf1** | 2.83292 | 0.01081 | 2.72754 | 0.78510 | 2.17370 | 2.20353 | 0.00184 | 1.81349 | 10.59795 | 0.00466 | 1.80319 | 0.87897 | 3.83536 | 4.34288 |
| **Rsl1** | 0.11640 | 0.01301 | 0.11166 | 0.00000 | 0.30338 | 1.16697 | 0.15076 | 0.23560 | 0.02234 | 0.01045 | 0.03613 | 0.02783 | 0.00265 | 0.43938 |
| **Rsl1d1** | 24.47510 | 9.81256 | 31.62391 | 17.36798 | 32.31898 | 25.69668 | 7.02512 | 9.44649 | 29.70715 | 12.56872 | 21.63321 | 17.76276 | 16.29304 | 32.36356 |
| **Rsl24d1** | 2.37034 | 4.54942 | 10.69862 | 10.07188 | 16.00749 | 6.79432 | 2.66632 | 5.02685 | 30.02452 | 13.14942 | 7.99296 | 6.44510 | 8.86662 | 25.04606 |
| **Rslcan18** | 0.01010 | 0.00383 | 0.02962 | 0.00000 | 0.00785 | 0.35141 | 0.00398 | 0.00395 | 0.21891 | 0.00589 | 0.00551 | 0.53978 | 0.00542 | 0.00000 |
| **Rsph1** | 0.24199 | 0.01106 | 0.87537 | 0.16991 | 1.07943 | 0.50592 | 0.08951 | 1.24152 | 0.41063 | 0.11395 | 0.77065 | 0.47340 | 1.51219 | 1.13389 |
| **Rsph3a** | 0.15286 | 0.00440 | 0.00770 | 0.00446 | 0.06011 | 0.02624 | 0.00482 | 0.01249 | 0.00748 | 0.00569 | 0.10946 | 0.16247 | 0.00349 | 0.10254 |
| **Rsph3b** | 0.14422 | 0.00448 | 0.00814 | 0.00000 | 0.05946 | 0.02920 | 0.00451 | 0.00830 | 0.00862 | 0.00557 | 0.09622 | 0.17210 | 0.00401 | 0.07800 |
| **Rsph9** | 1.09601 | 0.00421 | 0.03759 | 0.38826 | 0.32894 | 0.01114 | 0.00401 | 0.00718 | 0.00875 | 0.00738 | 0.00585 | 0.00467 | 0.01319 | 0.16872 |

| | | | | | | | | | | | | | | |
|---|---|---|---|---|---|---|---|---|---|---|---|---|---|---|
| **Rspo1** | 0.00611 | 0.00430 | 0.01250 | 0.07345 | 0.29561 | 0.01002 | 0.00472 | 0.00713 | 0.02021 | 0.00697 | 0.00752 | 0.01265 | 0.05260 | 0.00000 |
| **Rspo2** | 0.00000 | 0.00381 | 0.00000 | 0.00000 | 0.00000 | 0.30724 | 0.00371 | 0.00227 | 2.78067 | 0.00793 | 0.00774 | 0.00782 | 0.92285 | 3.38248 |
| **Rspry1** | 0.00000 | 0.02462 | 0.59244 | 0.31236 | 1.47389 | 1.27869 | 0.67269 | 2.81290 | 4.46328 | 0.02765 | 0.00000 | 1.04270 | 6.27234 | 0.10624 |
| **Rsrc1** | 5.26253 | 0.06506 | 2.14033 | 3.02490 | 1.78004 | 5.71229 | 1.35699 | 2.73935 | 3.51071 | 0.05389 | 4.41607 | 4.98684 | 0.00000 | 0.24647 |
| **Rsrc2** | 11.17001 | 0.08231 | 15.26711 | 5.03468 | 15.61938 | 13.53293 | 3.24208 | 4.37987 | 14.52136 | 13.26077 | 9.63799 | 8.62712 | 14.21185 | 17.07847 |
| **Rsrp1** | 4.79804 | 0.72747 | 3.16317 | 4.12361 | 4.94971 | 4.52466 | 0.51853 | 2.85688 | 2.04782 | 7.66961 | 2.57493 | 3.48029 | 18.86926 | 5.58764 |
| **Rsu1** | 0.44630 | 0.06240 | 1.94686 | 0.00000 | 0.55273 | 2.66981 | 0.00734 | 4.02663 | 0.02844 | 0.01897 | 2.01714 | 6.42949 | 0.00000 | 1.59861 |
| **Rtbdn** | 0.03625 | 0.00485 | 0.26893 | 0.02185 | 1.82156 | 0.00285 | 0.00418 | 0.01906 | 0.00563 | 0.00895 | 0.00613 | 0.00532 | 0.00665 | 0.70193 |
| **Rtca** | 7.33585 | 0.02455 | 3.09385 | 5.18844 | 4.00865 | 2.18455 | 0.00841 | 2.69316 | 0.00000 | 0.04376 | 2.59143 | 3.39821 | 3.71807 | 1.88051 |
| **Rtcb** | 5.00680 | 6.18548 | 8.33710 | 10.00954 | 5.71814 | 17.26597 | 4.55263 | 5.52861 | 7.11069 | 0.32998 | 4.15054 | 11.59544 | 1.75813 | 14.21121 |
| **Rtel1** | 8.26588 | 0.21528 | 12.36815 | 12.72610 | 6.54592 | 6.55542 | 3.59263 | 5.63004 | 3.31802 | 0.20282 | 5.77022 | 6.20074 | 3.32529 | 12.08998 |
| **Rtf1** | 5.51512 | 3.31578 | 3.88057 | 1.55763 | 6.90068 | 6.24109 | 1.52365 | 2.08380 | 3.45817 | 11.06023 | 2.98870 | 2.15411 | 4.59514 | 8.92992 |
| **Rtfdc1** | 8.67141 | 0.09711 | 14.95430 | 9.16145 | 20.40944 | 14.71942 | 3.58387 | 4.08385 | 6.99914 | 0.20888 | 8.38775 | 4.69277 | 8.25071 | 19.86285 |
| **Rtkn** | 0.00255 | 0.00888 | 0.00900 | 2.14628 | 0.01973 | 0.37767 | 0.00299 | 0.00284 | 0.00739 | 0.01777 | 0.00043 | 2.97557 | 0.00368 | 0.07057 |
| **Rtkn2** | 0.07796 | 0.01162 | 0.16731 | 0.43968 | 1.03122 | 0.01369 | 0.00416 | 0.00477 | 0.64637 | 0.05430 | 0.11893 | 0.05675 | 0.65033 | 0.34654 |
| **Rtn1** | 0.00844 | 0.02152 | 0.00200 | 0.00000 | 0.36386 | 0.23926 | 0.00443 | 0.77883 | 0.02191 | 0.00527 | 0.00058 | 0.16894 | 0.00466 | 0.00087 |
| **Rtn2** | 2.56956 | 0.00359 | 1.69140 | 0.00000 | 10.41356 | 0.29910 | 0.00366 | 0.00000 | 0.00000 | 0.00291 | 0.00114 | 0.00170 | 1.75906 | 2.16139 |
| **Rtn3** | 0.71035 | 0.10522 | 2.41229 | 2.86956 | 3.20912 | 3.14286 | 0.02965 | 9.35420 | 2.95108 | 0.27799 | 3.22421 | 4.51303 | 0.75006 | 0.95510 |
| **Rtn4** | 27.19518 | 0.25746 | 24.45577 | 26.63422 | 21.25623 | 27.46165 | 7.79017 | 13.67882 | 30.61347 | 4.44288 | 12.26230 | 13.24551 | 19.81258 | 9.77264 |
| **Rtn4ip1** | 0.08810 | 0.00369 | 1.16768 | 1.06253 | 0.00000 | 2.42752 | 1.48656 | 0.00000 | 1.19270 | 0.04970 | 0.00000 | 0.31651 | 0.26283 | 0.37381 |
| **Rtn4r** | 0.00000 | 0.00431 | 3.40410 | 0.88245 | 5.23797 | 0.00000 | 0.00369 | 0.00558 | 0.00803 | 0.00000 | 0.00659 | 0.01349 | 0.16876 | 0.11034 |
| **Rtn4rl1** | 0.01080 | 0.00434 | 0.04953 | 0.00444 | 0.33919 | 0.01376 | 0.00422 | 0.00934 | 0.01047 | 0.00687 | 0.00827 | 0.00828 | 0.49340 | 0.10057 |
| **Rtp3** | 3.99234 | 0.16479 | 2.09658 | 0.12172 | 1.58978 | 2.15693 | 0.13086 | 0.16957 | 2.12780 | 0.15273 | 0.76380 | 2.31092 | 0.74736 | 1.15677 |
| **Rttn** | 0.00000 | 0.00441 | 0.50061 | 0.03211 | 0.47477 | 0.00000 | 0.00516 | 0.28662 | 0.50509 | 0.00460 | 0.13647 | 0.02193 | 0.03677 | 1.45866 |
| **Rubcn** | 0.11062 | 0.00372 | 1.36979 | 0.00000 | 0.47467 | 1.77027 | 0.01159 | 0.13672 | 2.15116 | 0.00521 | 0.06048 | 0.89231 | 0.69408 | 2.11759 |
| **Rufy1** | 0.52638 | 0.01397 | 1.38196 | 2.80253 | 2.09349 | 1.21403 | 3.02529 | 0.06968 | 0.00000 | 0.11167 | 1.73524 | 0.25280 | 1.07228 | 0.37946 |
| **Rufy2** | 0.01401 | 0.02138 | 0.00000 | 0.46412 | 0.38200 | 0.42086 | 0.01060 | 0.41859 | 0.07010 | 0.00614 | 0.00232 | 0.00190 | 2.15499 | 0.07326 |
| **Rufy3** | 1.16447 | 2.57787 | 1.59846 | 3.06917 | 1.79324 | 1.53435 | 1.34502 | 2.24784 | 2.38999 | 0.10977 | 0.39645 | 2.65295 | 0.85750 | 3.35795 |
| **Rufy4** | 0.00845 | 0.00450 | 0.01055 | 0.00139 | 0.01025 | 0.42206 | 0.00423 | 0.00513 | 0.00962 | 0.00589 | 0.00832 | 0.01051 | 0.00761 | 0.07603 |
| **Rundc1** | 2.17964 | 0.04695 | 1.52870 | 0.83566 | 2.46089 | 2.23736 | 0.69293 | 0.00000 | 0.06205 | 0.03322 | 1.53754 | 2.34251 | 2.31661 | 6.56383 |
| **Rundc3a** | 0.05284 | 0.04363 | 5.25116 | 3.36247 | 1.84419 | 0.00000 | 0.00334 | 0.49097 | 0.00138 | 0.02350 | 0.00866 | 0.94899 | 0.13484 | 0.42389 |
| **Rundc3b** | 0.00164 | 0.00414 | 0.00692 | 0.00435 | 0.00301 | 0.17427 | 0.15983 | 0.00339 | 0.00945 | 0.00523 | 0.00601 | 0.00865 | 0.56747 | 0.62996 |
| **Runx1** | 0.03922 | 0.06690 | 0.18243 | 0.00000 | 0.01568 | 1.09830 | 0.00418 | 0.45232 | 0.00876 | 0.00530 | 0.00000 | 0.43380 | 0.15843 | 0.00010 |
| **Runx1t1** | 0.21134 | 0.24963 | 0.68406 | 0.00000 | 1.41372 | 0.49731 | 0.05403 | 0.00000 | 1.30266 | 0.00923 | 1.09206 | 1.85814 | 1.52537 | 2.12010 |
| **Runx2** | 0.01405 | 0.00430 | 0.01605 | 0.02898 | 0.05068 | 0.01118 | 0.00424 | 0.00957 | 0.03401 | 0.00613 | 0.00856 | 0.00844 | 0.68425 | 0.08968 |
| **Rusc1** | 9.71592 | 0.00916 | 13.58112 | 0.94089 | 19.06334 | 0.00000 | 2.36706 | 0.00000 | 0.79588 | 0.03838 | 1.17012 | 0.01916 | 1.62334 | 7.13948 |
| **Rusc2** | 0.15971 | 0.00401 | 1.53467 | 0.62798 | 0.27647 | 0.12067 | 0.00894 | 0.00000 | 0.00771 | 0.00258 | 0.31661 | 0.00785 | 0.46777 | 0.21538 |
| **Ruvbl1** | 14.88457 | 0.30506 | 20.99346 | 11.82878 | 18.67550 | 18.56915 | 4.51176 | 8.18694 | 4.71060 | 0.24815 | 12.77496 | 14.96711 | 12.27175 | 13.88286 |
| **Ruvbl2** | 13.77138 | 11.72826 | 30.73598 | 25.15016 | 20.67248 | 18.89543 | 6.85044 | 15.86289 | 0.09323 | 3.21240 | 16.51695 | 18.04093 | 2.50817 | 6.65222 |
| **Rwdd1** | 25.67176 | 0.08782 | 7.36604 | 10.20556 | 29.45061 | 11.12288 | 5.41013 | 2.51281 | 14.43947 | 1.65173 | 2.49742 | 9.25976 | 12.81225 | 11.62866 |
| **Rwdd2a** | 0.81840 | 0.02035 | 0.01116 | 0.70022 | 1.45029 | 0.00000 | 0.28560 | 0.00964 | 0.00822 | 0.00746 | 0.01102 | 0.38703 | 1.24880 | 0.03512 |
| **Rwdd2b** | 0.58077 | 0.01912 | 2.43532 | 0.54982 | 2.60606 | 1.18092 | 0.00371 | 0.37176 | 0.00475 | 0.01964 | 0.80728 | 1.31610 | 1.10683 | 1.73060 |
| **Rwdd3** | 0.10559 | 0.01330 | 0.30728 | 0.54808 | 0.04867 | 0.65717 | 0.00413 | 0.32076 | 0.01154 | 0.00642 | 0.00636 | 0.13638 | 1.08326 | 0.02977 |
| **Rwdd4a** | 3.62510 | 3.32468 | 1.55330 | 9.56659 | 13.79489 | 5.25392 | 1.80461 | 0.86864 | 7.27261 | 0.07411 | 1.99359 | 1.62050 | 11.84728 | 6.06873 |
| **Rxra** | 0.56670 | 7.44106 | 12.32912 | 7.10560 | 8.47097 | 4.91189 | 4.54006 | 0.21555 | 0.18480 | 0.16059 | 0.98638 | 0.13931 | 0.63792 | 1.78190 |
| **Rxrb** | 1.05939 | 0.10106 | 7.73379 | 8.53926 | 5.50970 | 5.53478 | 5.37828 | 1.73662 | 0.00820 | 0.19668 | 2.10179 | 4.57874 | 0.10094 | 1.65057 |
| **Rxrg** | 0.01022 | 0.02732 | 0.01720 | 0.00000 | 0.02472 | 0.15686 | 0.03573 | 0.15139 | 0.02063 | 0.02216 | 0.04030 | 0.31725 | 0.00815 | 0.01204 |
| **Rybp** | 13.33949 | 5.57991 | 15.22993 | 0.00000 | 17.31255 | 8.69370 | 3.12486 | 2.15125 | 24.35543 | 0.14164 | 6.01636 | 6.30138 | 12.88271 | 17.63716 |
| **Ryk** | 1.90724 | 6.18472 | 4.07934 | 2.50754 | 4.09401 | 9.67357 | 2.70072 | 7.30973 | 0.64215 | 2.99794 | 6.89100 | 8.51554 | 2.31910 | 10.87631 |
| **Ryr1** | 0.40475 | 0.01605 | 1.25345 | 0.87546 | 0.24792 | 1.52450 | 0.00364 | 0.81135 | 0.02994 | 0.00687 | 0.09190 | 0.70619 | 0.05846 | 0.79069 |
| **Ryr2** | 0.00000 | 0.00419 | 1.46742 | 0.16823 | 0.01483 | 0.00261 | 1.77535 | 0.00000 | 0.11380 | 0.05770 | 0.00165 | 0.22665 | 0.02308 | 0.30302 |
| **Ryr3** | 0.01552 | 0.00487 | 0.30613 | 0.00000 | 0.00673 | 0.06562 | 0.00425 | 0.00736 | 0.59801 | 0.00572 | 0.00676 | 0.00705 | 0.01723 | 0.00000 |
| **S100a1** | 0.01304 | 0.00446 | 0.01077 | 0.01126 | 0.11373 | 0.01355 | 0.00492 | 0.01016 | 0.02032 | 0.00694 | 0.00880 | 0.00941 | 0.00579 | 0.00654 |
| **S100a10** | 0.11502 | 7.58699 | 0.00000 | 0.00000 | 1.31170 | 4.78435 | 2.71701 | 7.48143 | 0.00000 | 11.95016 | 0.00000 | 1.58559 | 0.00000 | 0.09475 |
| **S100a11** | 0.00541 | 2.58572 | 0.06122 | 0.34431 | 0.66222 | 10.03184 | 6.12736 | 22.00007 | 1.84670 | 15.88812 | 0.85127 | 1.06968 | 0.22780 | 0.46024 |

| | | | | | | | | | | | | | | |
|---|---|---|---|---|---|---|---|---|---|---|---|---|---|---|
| **S100a13** | 0.00371 | 0.01095 | 0.00155 | 0.34783 | 0.13093 | 0.03250 | 0.00411 | 0.20296 | 0.06039 | 0.00828 | 0.00381 | 0.00744 | 0.82284 | 0.01758 |
| **S100a14** | 0.00363 | 0.06357 | 0.00000 | 0.58240 | 0.43232 | 0.28026 | 0.01585 | 4.54148 | 0.00682 | 0.02592 | 0.00633 | 0.00507 | 0.31705 | 0.06237 |
| **S100a16** | 0.00418 | 0.14132 | 0.00000 | 0.00000 | 1.21826 | 0.00302 | 0.00383 | 4.79103 | 0.01866 | 0.91890 | 0.00000 | 0.07801 | 0.00282 | 0.00000 |
| **S100a3** | 0.00000 | 0.00449 | 0.31530 | 0.06027 | 1.35237 | 0.00000 | 0.00408 | 0.01176 | 0.11385 | 0.00540 | 0.00473 | 0.00778 | 0.08466 | 0.50738 |
| **S100a4** | 0.16283 | 0.00445 | 0.00856 | 0.00873 | 0.17711 | 0.01192 | 0.00420 | 0.01111 | 0.00970 | 0.00635 | 0.00789 | 0.00827 | 0.00442 | 0.00716 |
| **S100a6** | 0.21200 | 0.00416 | 0.00000 | 0.00000 | 0.44961 | 0.31369 | 0.00450 | 0.07577 | 0.00772 | 0.01080 | 0.00359 | 0.19229 | 0.00396 | 0.04067 |
| **S100g** | 0.00860 | 0.00446 | 0.01049 | 0.01189 | 0.15911 | 0.01347 | 0.00465 | 0.01060 | 0.01006 | 0.00660 | 0.00902 | 0.00919 | 0.00583 | 0.02856 |
| **S100pbp** | 0.45442 | 0.42741 | 4.67520 | 0.00000 | 3.50265 | 1.51811 | 0.02290 | 0.96366 | 1.50414 | 0.00880 | 0.00000 | 0.81138 | 0.77459 | 0.17490 |
| **S1pr1** | 0.01714 | 0.00530 | 0.15161 | 0.00000 | 0.00682 | 0.41169 | 0.04281 | 0.22256 | 0.01237 | 0.00726 | 0.01061 | 0.00634 | 0.00594 | 0.01212 |
| **S1pr2** | 1.85690 | 0.00327 | 6.68224 | 4.29427 | 2.63775 | 0.30322 | 0.00236 | 0.00000 | 1.83825 | 0.00078 | 1.49152 | 0.03609 | 0.00000 | 2.88938 |
| **S1pr3** | 0.03401 | 0.00439 | 0.01396 | 0.01217 | 0.00955 | 0.01280 | 0.00425 | 0.00984 | 0.02532 | 0.00671 | 0.00827 | 0.00947 | 0.17272 | 0.00721 |
| **S1pr4** | 0.04031 | 0.01248 | 0.00435 | 0.27068 | 0.06788 | 0.25107 | 0.10465 | 0.00779 | 0.04334 | 0.05182 | 0.00563 | 0.05591 | 0.36709 | 0.13233 |
| **S1pr5** | 0.59519 | 0.03496 | 1.17246 | 1.67568 | 0.25798 | 1.24186 | 1.01338 | 2.94011 | 1.16256 | 0.03190 | 0.00000 | 0.00000 | 0.01840 | 0.16445 |
| **Saa1** | 0.20726 | 0.00449 | 0.01675 | 0.01536 | 0.20130 | 0.01123 | 0.00425 | 0.01176 | 0.06423 | 0.00855 | 0.00666 | 0.00737 | 0.02000 | 0.00221 |
| **Saa2** | 0.49448 | 0.00442 | 0.04821 | 0.05946 | 0.03130 | 0.00967 | 0.00408 | 0.00823 | 0.10852 | 0.00802 | 0.00560 | 0.01096 | 0.00491 | 0.00000 |
| **Saa3** | 0.19114 | 0.00459 | 0.00269 | 0.17508 | 0.05531 | 0.09718 | 0.00464 | 0.05645 | 0.00683 | 0.01251 | 0.00762 | 0.00416 | 0.06983 | 0.03185 |
| **Saal1** | 0.12125 | 0.05438 | 5.64322 | 8.28975 | 3.26687 | 5.74698 | 0.00907 | 2.17660 | 0.18474 | 13.23275 | 4.74651 | 6.32097 | 0.00000 | 3.57058 |
| **Sac3d1** | 0.69074 | 0.03466 | 0.52116 | 0.00014 | 3.58204 | 0.63134 | 1.17786 | 1.94180 | 0.09609 | 0.13178 | 0.00000 | 2.50487 | 10.72523 | 0.22610 |
| **Sacm1l** | 3.53363 | 0.01550 | 1.95455 | 1.29127 | 0.61086 | 4.65877 | 0.00165 | 1.51797 | 6.03501 | 0.94285 | 4.02619 | 3.22976 | 0.19334 | 1.20555 |
| **Sae1** | 14.25365 | 4.12885 | 23.82793 | 18.76036 | 13.83072 | 12.44190 | 1.58254 | 5.08473 | 12.68684 | 9.53165 | 10.29913 | 15.40789 | 5.74635 | 17.04666 |
| **Safb** | 2.31457 | 0.15349 | 13.03353 | 11.92460 | 11.26526 | 1.32602 | 1.71541 | 5.43167 | 0.00990 | 3.35007 | 1.14725 | 6.96350 | 1.30245 | 1.47141 |
| **Safb2** | 0.80795 | 0.03247 | 3.90322 | 7.62735 | 4.89247 | 0.82695 | 0.56215 | 4.07414 | 0.21200 | 0.05377 | 0.08746 | 1.98439 | 0.77787 | 0.22124 |
| **Sag** | 0.07883 | 0.00452 | 0.13869 | 0.00500 | 0.00903 | 0.01359 | 0.00423 | 0.01086 | 0.00840 | 0.00827 | 0.00689 | 0.00896 | 0.00639 | 0.06575 |
| **Sall1** | 1.04086 | 0.02175 | 3.24042 | 1.64942 | 2.87658 | 2.02624 | 0.06095 | 0.09125 | 2.19276 | 0.01979 | 1.23647 | 4.90220 | 0.00000 | 2.75745 |
| **Sall2** | 0.03164 | 0.00444 | 0.01553 | 0.01274 | 0.01020 | 0.01264 | 0.00433 | 0.01036 | 0.01096 | 0.00703 | 0.00838 | 0.00996 | 0.00908 | 0.06004 |
| **Sall4** | 21.63061 | 5.98282 | 27.17774 | 13.40863 | 24.73898 | 14.41399 | 8.35363 | 15.39249 | 16.57129 | 1.61779 | 13.95180 | 15.86430 | 5.53146 | 21.23339 |
| **Samd1** | 1.54677 | 0.03361 | 0.66374 | 0.59067 | 6.04054 | 0.97192 | 0.15412 | 0.84426 | 0.65515 | 0.20306 | 0.76499 | 0.72640 | 0.13672 | 2.77581 |
| **Samd10** | 0.00639 | 0.02974 | 0.00976 | 0.00000 | 0.01079 | 0.00743 | 0.00354 | 2.55257 | 0.01403 | 0.06410 | 0.02085 | 0.03674 | 0.00838 | 0.04390 |
| **Samd4b** | 3.11054 | 0.15158 | 12.54567 | 4.28189 | 8.99665 | 13.72525 | 3.46760 | 8.23030 | 0.16851 | 0.20089 | 2.15114 | 7.56895 | 2.75311 | 2.77391 |
| **Samd7** | 0.00681 | 0.00442 | 0.06287 | 0.00577 | 0.00000 | 0.00939 | 0.00419 | 0.01072 | 0.02460 | 0.00687 | 0.00711 | 0.01179 | 0.31232 | 0.45367 |
| **Samd8** | 0.59377 | 0.03342 | 0.33475 | 0.74742 | 0.21420 | 2.49683 | 0.07195 | 2.21086 | 0.50187 | 0.11382 | 2.70648 | 0.54010 | 3.17052 | 1.27638 |
| **Samhd1** | 1.27450 | 0.01858 | 3.86278 | 2.25269 | 5.04028 | 0.17932 | 1.39058 | 0.00000 | 0.87809 | 0.03140 | 0.26072 | 0.91389 | 3.69823 | 6.05949 |
| **Samm50** | 1.25499 | 1.12686 | 2.14140 | 0.00000 | 3.60894 | 4.06127 | 1.74419 | 8.28298 | 0.01147 | 0.06955 | 1.24539 | 5.13399 | 0.85550 | 0.41053 |
| **Sap130** | 2.14242 | 0.02080 | 2.82438 | 0.59963 | 5.06328 | 4.11483 | 1.88521 | 0.79424 | 2.29902 | 12.59595 | 3.85540 | 2.77030 | 5.71400 | 4.72341 |
| **Sap18** | 3.07272 | 0.76408 | 0.75590 | 0.31240 | 0.53307 | 4.05775 | 1.33788 | 1.09401 | 5.32445 | 0.67121 | 0.12164 | 1.19958 | 6.46540 | 2.44366 |
| **Sap30** | 0.16631 | 0.00336 | 0.04265 | 0.26553 | 1.34001 | 0.47679 | 0.00393 | 0.01335 | 1.52418 | 0.01923 | 0.51439 | 0.69292 | 0.87707 | 1.38345 |
| **Sap30bp** | 10.39432 | 0.06698 | 12.06408 | 5.66397 | 15.61040 | 10.54754 | 4.03530 | 4.31950 | 0.16810 | 0.17943 | 4.55870 | 4.48226 | 6.37719 | 12.73688 |
| **Sap30l** | 3.32756 | 0.00848 | 0.29580 | 2.33871 | 5.87792 | 0.83521 | 0.06792 | 1.02761 | 0.31116 | 0.04445 | 0.01398 | 3.47446 | 14.15814 | 1.26007 |
| **Sapcd1** | 0.01245 | 0.04499 | 0.01153 | 0.00000 | 0.01705 | 0.04066 | 0.00419 | 0.18384 | 0.00905 | 0.01059 | 0.00234 | 0.48918 | 0.03101 | 0.01870 |
| **Sapcd2** | 6.98261 | 0.00879 | 15.23049 | 11.40906 | 20.93342 | 2.83752 | 5.04678 | 1.98497 | 2.31807 | 0.15811 | 3.89230 | 0.00000 | 11.94691 | 6.25101 |
| **Sar1a** | 11.83802 | 0.21690 | 13.23998 | 13.26486 | 15.44495 | 13.79492 | 6.67995 | 8.27758 | 8.14657 | 0.43222 | 7.02403 | 10.69293 | 10.14148 | 9.93231 |
| **Sar1b** | 12.27270 | 1.66205 | 18.78138 | 15.23476 | 13.77979 | 18.29873 | 1.83996 | 2.48246 | 9.64550 | 11.83616 | 8.82640 | 11.61158 | 0.95550 | 14.05984 |
| **Saraf** | 7.87333 | 0.04959 | 8.74536 | 9.49158 | 7.29248 | 9.79528 | 4.57801 | 3.06962 | 2.89146 | 1.46407 | 9.99671 | 5.98349 | 2.18793 | 10.00851 |
| **Sardh** | 0.08868 | 0.01040 | 2.36220 | 0.57853 | 3.95418 | 0.00000 | 0.00324 | 0.35176 | 0.07233 | 0.01161 | 0.83934 | 0.00371 | 2.39944 | 1.71864 |
| **Sarm1** | 0.53525 | 0.00910 | 0.68233 | 0.00000 | 2.42925 | 3.95226 | 0.05685 | 7.63182 | 2.53268 | 0.05254 | 1.26579 | 2.18931 | 0.47636 | 1.92904 |
| **Sarnp** | 8.14700 | 0.85688 | 10.85452 | 10.39435 | 7.33148 | 13.28832 | 8.17529 | 8.24764 | 12.34460 | 12.09739 | 8.03184 | 12.24759 | 5.43567 | 5.00237 |
| **Sars** | 7.85501 | 4.32085 | 12.41316 | 15.59970 | 4.08740 | 8.92372 | 4.22055 | 10.69554 | 0.00000 | 0.19532 | 12.34694 | 10.02457 | 0.30114 | 4.90393 |
| **Sars2** | 1.65478 | 0.14024 | 8.09341 | 8.50820 | 3.86809 | 5.68097 | 0.03910 | 1.62715 | 0.00000 | 0.05140 | 3.01953 | 6.01410 | 1.24990 | 1.06917 |
| **Sart1** | 5.59812 | 8.75920 | 14.06882 | 11.71134 | 11.11343 | 6.34717 | 1.47198 | 6.00734 | 1.47974 | 1.82188 | 5.50511 | 5.19790 | 5.98243 | 4.05737 |
| **Sart3** | 2.85415 | 0.11523 | 7.89472 | 14.85849 | 9.53083 | 4.17728 | 3.56063 | 5.28743 | 0.39447 | 0.20353 | 2.50886 | 6.15626 | 3.98340 | 1.93912 |
| **Sash1** | 0.38421 | 0.00384 | 0.05851 | 0.20710 | 0.01259 | 0.70782 | 0.67455 | 0.01226 | 0.00775 | 0.04957 | 0.00000 | 0.34142 | 0.68950 | 0.00415 |
| **Sass6** | 1.44549 | 0.19086 | 1.62992 | 1.05303 | 1.13292 | 2.82588 | 0.21732 | 3.39661 | 4.06158 | 0.02919 | 5.16903 | 0.14420 | 1.59728 | 1.80546 |
| **Sat1** | 20.23662 | 8.96700 | 35.85269 | 24.47464 | 30.72402 | 19.89259 | 4.39167 | 0.00000 | 36.90440 | 11.47760 | 18.55588 | 22.44630 | 1.16583 | 25.07275 |
| **Sat2** | 2.00924 | 0.00360 | 0.72818 | 0.18532 | 0.96193 | 0.00000 | 0.00396 | 0.00504 | 0.13238 | 0.01444 | 0.00448 | 0.00202 | 0.48913 | 6.49374 |
| **Satb1** | 7.83925 | 0.07180 | 8.29659 | 3.38773 | 14.30891 | 3.92035 | 0.82813 | 8.24989 | 13.03105 | 0.05391 | 4.94114 | 6.60905 | 1.38623 | 3.41736 |

| | | | | | | | | | | | | | | |
|---|---|---|---|---|---|---|---|---|---|---|---|---|---|---|
| **Satb2** | 2.07526 | 0.06080 | 6.54946 | 1.26678 | 3.00462 | 11.06886 | 0.74881 | 3.43670 | 0.36295 | 0.01406 | 0.09213 | 2.69805 | 1.06504 | 2.13736 |
| **Sav1** | 4.28722 | 0.00923 | 0.81698 | 0.40727 | 2.09112 | 0.77366 | 0.83878 | 0.00000 | 3.22962 | 0.00944 | 0.00000 | 2.38575 | 2.25602 | 2.86353 |
| **Saxo1** | 0.01482 | 0.00212 | 0.39245 | 0.00000 | 0.55709 | 1.89897 | 0.00934 | 0.13940 | 0.00144 | 0.03092 | 4.95285 | 0.79218 | 0.92461 | 1.57214 |
| **Saxo1os** | 0.00563 | 0.00441 | 0.01767 | 0.05955 | 0.01070 | 0.02279 | 0.00434 | 0.01057 | 0.01116 | 0.00784 | 0.00840 | 0.00725 | 0.26210 | 0.00168 |
| **Saxo2** | 0.02449 | 0.00451 | 0.01367 | 0.00397 | 0.01857 | 0.08160 | 0.00428 | 0.00698 | 0.01190 | 0.00631 | 0.00862 | 0.01081 | 0.01461 | 0.02323 |
| **Saysd1** | 0.09623 | 0.02225 | 0.21376 | 0.00000 | 1.29656 | 0.51447 | 0.02271 | 0.70411 | 0.28056 | 0.02932 | 0.00000 | 0.46268 | 0.69503 | 0.05026 |
| **Sbds** | 3.70739 | 0.07290 | 14.62926 | 3.89463 | 8.25722 | 7.90325 | 4.18054 | 6.56364 | 3.92364 | 0.81044 | 3.31200 | 4.91051 | 3.12871 | 18.99690 |
| **Sbf1** | 0.00000 | 0.02889 | 0.97360 | 1.54018 | 1.32010 | 0.00000 | 0.83755 | 1.42047 | 0.00794 | 0.02693 | 0.18203 | 0.80701 | 0.03567 | 0.00000 |
| **Sbf2** | 0.04894 | 0.01236 | 0.26436 | 0.00000 | 0.16165 | 1.58417 | 0.00473 | 0.56802 | 0.30922 | 0.01958 | 0.00000 | 0.59296 | 0.38357 | 2.07859 |
| **Sbk1** | 0.02319 | 0.00363 | 1.69478 | 1.13040 | 0.09643 | 2.18864 | 0.00418 | 0.56609 | 0.03585 | 0.01386 | 0.00000 | 0.83271 | 2.78724 | 0.40930 |
| **Sbk3** | 0.01214 | 0.00440 | 0.17971 | 0.00376 | 0.10488 | 0.00690 | 0.00419 | 0.01186 | 0.00864 | 0.00705 | 0.00697 | 0.00909 | 0.02509 | 0.08350 |
| **Sbno1** | 6.74178 | 0.02747 | 9.25981 | 4.18684 | 12.76671 | 13.69940 | 0.00000 | 4.71433 | 24.24827 | 0.06878 | 4.08283 | 4.84034 | 11.55062 | 17.84183 |
| **Sbno2** | 1.42630 | 0.43089 | 4.30296 | 3.24183 | 5.69824 | 0.61793 | 2.04443 | 5.13592 | 0.00000 | 0.06788 | 1.17626 | 2.17325 | 0.43486 | 0.08735 |
| **Sbp** | 0.89985 | 0.02438 | 1.21253 | 1.28540 | 0.25280 | 0.19971 | 0.00404 | 0.38163 | 0.36516 | 0.00946 | 0.00094 | 0.00385 | 0.00000 | 0.90837 |
| **Sbpl** | 0.41600 | 0.00532 | 0.68291 | 0.62361 | 0.06090 | 0.12987 | 0.01046 | 0.19467 | 0.10668 | 0.01018 | 0.00368 | 0.00373 | 0.00032 | 0.36995 |
| **Sbsn** | 0.19937 | 0.00466 | 0.32882 | 0.19963 | 0.06433 | 0.45949 | 1.16484 | 0.00502 | 0.01220 | 0.03609 | 0.00331 | 0.00194 | 0.03552 | 0.07303 |
| **Sbspon** | 0.00370 | 0.01225 | 0.00262 | 0.00867 | 0.18152 | 0.01216 | 0.00429 | 0.01013 | 0.51951 | 0.00640 | 0.00339 | 0.21158 | 0.00681 | 0.00000 |
| **Sc5d** | 0.94335 | 0.00294 | 0.00395 | 2.64484 | 0.30279 | 3.23842 | 1.62367 | 0.00286 | 1.75885 | 0.01945 | 0.00000 | 1.11235 | 0.97175 | 0.00000 |
| **Scaf1** | 0.07652 | 0.02835 | 0.63302 | 1.23270 | 1.00129 | 0.00000 | 0.14035 | 1.47514 | 0.00853 | 0.01105 | 0.32863 | 0.29866 | 0.15681 | 0.00000 |
| **Scaf11** | 5.13398 | 0.01246 | 5.89199 | 2.05745 | 8.89435 | 5.84198 | 1.43115 | 3.07282 | 5.14879 | 0.03145 | 5.41260 | 4.31687 | 6.93250 | 11.96055 |
| **Scaf4** | 0.19560 | 0.03867 | 2.93133 | 0.91937 | 4.12140 | 1.72498 | 0.00165 | 3.84455 | 0.63647 | 0.01857 | 0.48014 | 4.38820 | 3.29736 | 0.11868 |
| **Scaf8** | 3.37678 | 0.02050 | 6.87059 | 4.61845 | 5.69063 | 7.07358 | 2.36862 | 4.32389 | 6.87077 | 0.04042 | 2.76987 | 4.67083 | 7.51584 | 3.80289 |
| **Scai** | 1.37315 | 0.00352 | 0.56785 | 0.00000 | 0.80960 | 0.00000 | 0.00321 | 0.00436 | 6.10440 | 0.00595 | 0.00192 | 0.02861 | 3.56820 | 0.00000 |
| **Scamp1** | 10.66745 | 0.15294 | 3.44976 | 1.95488 | 3.99693 | 7.32382 | 0.46086 | 5.53859 | 5.98635 | 0.10621 | 6.44579 | 2.54373 | 3.94945 | 4.36338 |
| **Scamp2** | 7.95629 | 0.06276 | 12.96722 | 3.08660 | 5.73056 | 11.39317 | 3.70937 | 6.94276 | 1.99873 | 0.13486 | 3.56043 | 2.46560 | 0.01889 | 5.67125 |
| **Scamp3** | 0.00000 | 0.22117 | 1.43490 | 1.05090 | 0.20659 | 0.27816 | 0.00383 | 0.43165 | 0.01245 | 0.00149 | 0.82605 | 0.54224 | 0.00097 | 0.00776 |
| **Scamp4** | 6.52594 | 8.69751 | 21.88419 | 12.47137 | 10.99753 | 13.20083 | 6.35420 | 12.04923 | 0.43858 | 7.16159 | 8.70548 | 8.04510 | 0.71770 | 8.23195 |
| **Scamp5** | 1.69358 | 0.01840 | 3.18452 | 3.98710 | 3.41888 | 1.06130 | 0.33568 | 2.90770 | 0.29882 | 0.02735 | 0.12459 | 0.05929 | 1.39820 | 1.35221 |
| **Scand1** | 0.03694 | 0.00473 | 0.01261 | 0.01613 | 0.11085 | 0.01457 | 0.00455 | 0.07246 | 0.00991 | 0.00752 | 0.00778 | 0.01797 | 0.08376 | 0.00831 |
| **Scap** | 2.88922 | 0.01614 | 9.22354 | 2.66043 | 5.27960 | 2.60576 | 0.00097 | 4.01687 | 1.55776 | 0.01110 | 1.56887 | 2.28538 | 5.88683 | 6.16729 |
| **Scaper** | 0.00000 | 0.00443 | 0.49385 | 0.00000 | 0.44902 | 0.39650 | 0.00331 | 0.01306 | 2.25515 | 0.00434 | 0.00207 | 0.05576 | 1.58649 | 0.00000 |
| **Scarb1** | 0.08536 | 0.01760 | 1.31765 | 0.72256 | 0.10948 | 3.52416 | 0.00924 | 0.08797 | 0.00391 | 0.00549 | 1.88400 | 1.45219 | 0.60117 | 1.46939 |
| **Scarb2** | 2.95964 | 0.02761 | 9.36077 | 6.12191 | 0.24294 | 10.54809 | 2.28751 | 3.40144 | 0.71866 | 0.04884 | 5.25685 | 6.79632 | 0.00000 | 4.54330 |
| **Scarf1** | 0.01027 | 0.00432 | 0.19486 | 0.00701 | 0.01515 | 0.00401 | 0.00418 | 0.01030 | 0.06463 | 0.00810 | 0.00658 | 0.00794 | 0.32470 | 0.35419 |
| **Sccpdh** | 0.00234 | 0.03980 | 0.88261 | 0.46313 | 0.00000 | 0.00000 | 0.37570 | 4.39806 | 0.14934 | 0.00659 | 1.44956 | 3.52223 | 0.86079 | 0.00000 |
| **Scd1** | 3.09967 | 0.06760 | 10.33473 | 12.35340 | 3.27529 | 14.03803 | 0.02675 | 1.84400 | 11.85876 | 0.22941 | 8.03024 | 12.08054 | 7.55304 | 9.97675 |
| **Scd2** | 0.00000 | 0.21124 | 7.34402 | 12.60936 | 1.24165 | 24.05293 | 1.60089 | 6.06656 | 0.00000 | 0.33510 | 13.34967 | 21.51809 | 0.18106 | 0.67836 |
| **Scd3** | 0.47056 | 0.01015 | 1.26086 | 1.78072 | 0.01755 | 6.71952 | 0.09077 | 0.07876 | 0.34985 | 0.06938 | 0.27844 | 1.26511 | 4.34157 | 0.16141 |
| **Scd4** | 0.28520 | 0.08655 | 0.25364 | 0.14592 | 0.16068 | 0.44451 | 0.15813 | 0.01309 | 0.20814 | 0.04252 | 0.07990 | 0.57923 | 0.17689 | 0.11112 |
| **Scel** | 0.00920 | 0.00434 | 0.01364 | 0.01184 | 0.00832 | 0.01257 | 0.00425 | 0.00934 | 0.00963 | 0.00631 | 0.00918 | 0.00939 | 0.31573 | 0.08898 |
| **Scfd1** | 2.54648 | 1.22015 | 3.49708 | 2.19269 | 3.43279 | 5.97366 | 0.98737 | 3.43492 | 5.75394 | 0.04442 | 8.73768 | 4.77140 | 0.75689 | 5.73442 |
| **Scfd2** | 4.33866 | 0.02513 | 3.67889 | 0.76977 | 2.56950 | 0.73599 | 0.00208 | 0.71377 | 2.33761 | 0.02160 | 2.34844 | 1.44697 | 2.58841 | 1.99791 |
| **Scg3** | 0.00000 | 0.00403 | 0.05433 | 0.50627 | 0.12502 | 0.00000 | 0.00406 | 0.00765 | 0.02138 | 0.00650 | 0.00501 | 0.00861 | 0.37070 | 1.23220 |
| **Scgb2b23.ps** | 0.02653 | 0.00449 | 0.01606 | 0.03394 | 0.01155 | 0.02256 | 0.00502 | 0.03578 | 0.01236 | 0.00598 | 0.00821 | 0.00827 | 0.00685 | 0.01543 |
| **Scimp** | 0.00275 | 0.00451 | 0.00000 | 0.01074 | 0.78720 | 0.01021 | 0.00418 | 0.01009 | 0.27525 | 0.00797 | 0.00674 | 0.00724 | 0.00218 | 0.21719 |
| **Scin** | 0.01735 | 0.01167 | 0.01257 | 0.01486 | 0.01551 | 0.01345 | 0.00404 | 0.03668 | 0.05501 | 0.04320 | 0.00757 | 0.01213 | 0.01917 | 0.01605 |
| **Sclt1** | 0.00774 | 0.00413 | 0.00000 | 0.24645 | 0.33676 | 0.36300 | 0.00377 | 0.00316 | 0.00710 | 0.00561 | 0.00327 | 0.11914 | 0.08965 | 0.77255 |
| **Scly** | 0.02838 | 0.01268 | 0.01287 | 0.02027 | 0.02322 | 0.00355 | 0.10886 | 0.08459 | 0.06070 | 0.01752 | 0.28308 | 0.00707 | 0.10418 | 0.00000 |
| **Scmh1** | 2.89589 | 6.69528 | 4.17620 | 0.46801 | 1.30339 | 1.93337 | 0.00051 | 0.90108 | 0.20472 | 0.01957 | 2.98164 | 3.72012 | 1.11277 | 0.64567 |
| **Scml2** | 0.09466 | 0.00441 | 0.00000 | 0.01294 | 0.00000 | 0.00662 | 0.00411 | 0.00959 | 0.79541 | 0.00694 | 0.00532 | 0.00847 | 0.09735 | 0.51543 |
| **Scml4** | 0.03296 | 0.00355 | 0.01565 | 0.00000 | 0.03544 | 0.25257 | 0.02078 | 0.02946 | 0.14607 | 0.03479 | 0.54344 | 0.47987 | 0.46440 | 0.07668 |
| **Scn10a** | 0.00160 | 0.00447 | 0.20314 | 0.00000 | 0.25017 | 0.00521 | 0.00424 | 0.01216 | 0.00995 | 0.00615 | 0.00693 | 0.00893 | 0.04500 | 0.21775 |
| **Scn11a** | 0.00517 | 0.00431 | 0.04346 | 0.14279 | 0.00703 | 0.06767 | 0.00412 | 0.00408 | 0.01031 | 0.00587 | 0.00614 | 0.01388 | 0.00708 | 0.07929 |
| **Scn2b** | 0.11297 | 0.09033 | 0.00995 | 1.32506 | 0.49114 | 0.85053 | 0.08969 | 0.29465 | 0.09603 | 0.35635 | 0.00761 | 0.09162 | 0.55844 | 0.26007 |
| **Scn3a** | 0.01024 | 0.00441 | 0.08953 | 0.00415 | 0.20394 | 0.01254 | 0.00416 | 0.01346 | 0.00660 | 0.00759 | 0.00875 | 0.00865 | 0.06528 | 0.03174 |

| | | | | | | | | | | | | | | |
|---|---|---|---|---|---|---|---|---|---|---|---|---|---|---|
| **Scn8a** | 0.10812 | 0.00543 | 0.00738 | 0.15655 | 0.00359 | 0.00341 | 0.00447 | 0.09130 | 0.00750 | 0.00691 | 0.00596 | 0.02525 | 0.00450 | 0.63620 |
| **Scnm1** | 6.99774 | 0.02835 | 5.83464 | 7.11155 | 8.00529 | 4.45173 | 0.13666 | 3.33682 | 4.45938 | 0.02729 | 5.25283 | 4.38537 | 7.42290 | 4.08615 |
| **Sco1** | 0.00000 | 0.10785 | 1.07152 | 1.57935 | 0.94465 | 1.07767 | 0.05023 | 0.84887 | 2.98997 | 0.00239 | 0.65891 | 0.01944 | 1.53553 | 1.29943 |
| **Scoc** | 1.55034 | 0.00113 | 0.31636 | 0.42943 | 3.52941 | 2.45076 | 1.21938 | 0.26969 | 6.59845 | 0.00475 | 2.73106 | 2.34725 | 5.75613 | 3.50736 |
| **Scp2** | 2.71462 | 7.27117 | 10.22071 | 12.15453 | 5.90677 | 16.84901 | 4.18334 | 7.08169 | 4.11781 | 8.98855 | 10.83079 | 11.20176 | 2.50654 | 6.18444 |
| **Scpep1** | 10.11154 | 0.12249 | 20.87285 | 11.63436 | 14.55730 | 20.74299 | 5.33370 | 5.97455 | 20.38745 | 16.63576 | 10.60787 | 7.67710 | 5.64445 | 25.54859 |
| **Scpep1os** | 0.22098 | 0.03700 | 0.06807 | 0.09439 | 0.03611 | 0.32032 | 0.04123 | 0.14370 | 0.65001 | 0.07397 | 0.05849 | 0.09625 | 1.63434 | 0.46528 |
| **Scrib** | 0.00000 | 0.06144 | 1.07449 | 8.43928 | 1.84093 | 0.00000 | 0.28091 | 2.71739 | 0.00296 | 0.01415 | 0.00000 | 4.84671 | 0.21534 | 0.00000 |
| **Scrn2** | 0.00000 | 0.03222 | 2.24020 | 2.32851 | 1.69412 | 0.94430 | 0.00355 | 0.77528 | 0.00521 | 0.02000 | 1.20253 | 0.72716 | 0.01061 | 0.00000 |
| **Scrn3** | 0.00000 | 0.00538 | 0.42451 | 1.45016 | 0.64624 | 0.32247 | 0.00393 | 0.88914 | 0.20546 | 0.00771 | 0.08681 | 0.03906 | 1.03669 | 0.55086 |
| **Sct** | 0.01284 | 2.03301 | 0.03119 | 0.22886 | 0.13914 | 0.28545 | 0.01130 | 0.30146 | 0.01151 | 2.24564 | 0.12158 | 0.03948 | 0.00951 | 0.00000 |
| **Scube1** | 0.05781 | 0.00433 | 0.00449 | 0.03790 | 0.00546 | 0.09575 | 0.01033 | 0.01763 | 0.11563 | 0.02479 | 0.00779 | 0.02747 | 0.43901 | 0.08952 |
| **Scyl1** | 2.57452 | 6.13379 | 12.92797 | 8.77288 | 9.41594 | 6.75621 | 0.02999 | 3.55061 | 0.01596 | 0.03988 | 5.26826 | 2.54082 | 0.57416 | 4.34961 |
| **Scyl2** | 0.18238 | 0.01771 | 2.93462 | 0.36735 | 2.56035 | 1.50250 | 0.00287 | 1.09095 | 1.40485 | 0.00000 | 2.34398 | 2.21179 | 1.00961 | 4.04119 |
| **Scyl3** | 3.01635 | 0.02472 | 2.00356 | 1.86476 | 1.67924 | 2.69059 | 3.64335 | 1.37610 | 0.00452 | 0.12460 | 1.13043 | 1.84546 | 0.55388 | 0.56757 |
| **Sdad1** | 4.97720 | 0.06040 | 5.66667 | 7.99207 | 4.34304 | 3.35272 | 0.00068 | 0.61316 | 7.05702 | 0.00000 | 2.11950 | 5.22893 | 3.66840 | 7.69515 |
| **Sdc1** | 0.81720 | 0.00305 | 1.94456 | 2.12497 | 0.23460 | 1.00871 | 0.00326 | 0.00000 | 0.00376 | 0.00270 | 0.00000 | 0.34391 | 4.56545 | 2.39105 |
| **Sdc2** | 0.06931 | 0.00441 | 0.00772 | 0.00979 | 0.31817 | 0.01235 | 0.00442 | 0.01241 | 0.00959 | 0.00632 | 0.00814 | 0.00855 | 0.00613 | 0.00884 |
| **Sdc3** | 0.41339 | 0.00477 | 0.05932 | 0.00000 | 0.58967 | 0.00785 | 0.00397 | 0.01601 | 0.01913 | 0.00982 | 0.00492 | 0.19520 | 0.17668 | 0.00812 |
| **Sdc4** | 0.00000 | 6.05760 | 5.09477 | 5.76402 | 0.00000 | 20.99099 | 3.28301 | 21.82600 | 0.00000 | 0.54326 | 10.59900 | 12.79207 | 0.16084 | 0.00000 |
| **Sdcbp** | 0.55260 | 0.04648 | 1.56544 | 4.40647 | 1.56910 | 7.64621 | 2.54402 | 4.86514 | 7.12186 | 0.03529 | 5.81491 | 5.03492 | 0.35493 | 3.26339 |
| **Sdcbp2** | 0.00000 | 0.00938 | 0.13611 | 0.00000 | 0.35181 | 2.75542 | 0.00866 | 0.89490 | 0.00053 | 0.00373 | 2.19623 | 0.97527 | 1.73285 | 0.00000 |
| **Sdccag3** | 0.00000 | 4.71456 | 3.91158 | 2.01230 | 1.07469 | 4.30970 | 1.69306 | 4.00152 | 1.39037 | 0.03190 | 4.02722 | 3.79100 | 0.66707 | 7.32480 |
| **Sdccag8** | 0.01130 | 0.00486 | 0.90276 | 0.16151 | 0.04763 | 0.14217 | 0.00420 | 0.78604 | 0.03012 | 0.00454 | 0.00460 | 0.01134 | 0.78431 | 0.53248 |
| **Sde2** | 6.37996 | 3.15291 | 6.59039 | 1.08512 | 4.28347 | 3.96247 | 0.42297 | 2.40115 | 8.33508 | 0.08393 | 4.11171 | 3.39404 | 7.29081 | 12.13849 |
| **Sdf2** | 5.14798 | 0.08781 | 6.87289 | 1.61525 | 2.76450 | 5.27666 | 2.53784 | 5.85602 | 5.80003 | 0.14716 | 5.27237 | 6.68374 | 2.38312 | 8.52892 |
| **Sdf2l1** | 0.01922 | 0.01243 | 0.03644 | 0.33003 | 0.08324 | 0.05472 | 0.00382 | 0.00486 | 0.06968 | 0.00537 | 0.04531 | 0.36287 | 0.30452 | 0.28572 |
| **Sdf4** | 11.13974 | 0.21055 | 13.61230 | 16.79123 | 11.50699 | 15.27568 | 1.44259 | 10.33817 | 7.73963 | 0.30012 | 10.36972 | 11.59296 | 11.32273 | 12.58810 |
| **Sdha** | 8.62144 | 0.37436 | 15.21911 | 8.37513 | 8.88686 | 18.72966 | 4.43292 | 11.72940 | 10.30473 | 0.22652 | 12.46002 | 14.24748 | 0.00000 | 10.43820 |
| **Sdhaf1** | 0.17840 | 0.00406 | 0.00873 | 1.57409 | 0.10659 | 0.02749 | 0.00386 | 0.00367 | 0.01576 | 0.01247 | 0.00554 | 0.01410 | 0.16754 | 0.03191 |
| **Sdhaf2** | 3.61772 | 0.03045 | 4.61627 | 3.87645 | 3.33224 | 8.76919 | 2.19217 | 3.23246 | 4.00769 | 0.03924 | 4.32428 | 5.16683 | 0.67065 | 6.83792 |
| **Sdhaf3** | 0.67452 | 0.01056 | 0.27942 | 0.37501 | 1.04594 | 2.13863 | 0.00335 | 0.01539 | 0.63335 | 0.01162 | 0.05578 | 1.69058 | 0.00178 | 0.55792 |
| **Sdhaf4** | 0.16738 | 0.03832 | 0.09072 | 0.36485 | 0.13579 | 2.61822 | 0.02013 | 3.00099 | 0.32175 | 0.03003 | 1.23929 | 4.12757 | 0.00494 | 0.37155 |
| **Sdhb** | 8.31051 | 8.11827 | 11.35640 | 6.84143 | 5.70958 | 16.10196 | 8.82294 | 13.29977 | 0.01582 | 11.85142 | 10.77253 | 14.17091 | 0.59819 | 3.29254 |
| **Sdhc** | 2.53872 | 7.35532 | 8.27182 | 9.50919 | 2.30641 | 14.26411 | 0.52067 | 8.89346 | 3.22260 | 12.49349 | 13.00590 | 12.13648 | 0.37388 | 5.28397 |
| **Sdhd** | 4.21096 | 0.02960 | 1.85773 | 0.00000 | 0.00002 | 2.78631 | 1.41514 | 1.12763 | 2.41838 | 0.05796 | 6.46133 | 3.10605 | 1.29197 | 3.44753 |
| **Sdk1** | 0.00000 | 0.00409 | 0.79165 | 0.96469 | 0.05557 | 0.34673 | 0.00388 | 0.10194 | 0.00427 | 0.00711 | 0.00392 | 0.00223 | 0.35688 | 0.00000 |
| **Sdk2** | 0.00876 | 0.00440 | 0.25133 | 0.00318 | 0.09339 | 0.00640 | 0.00409 | 0.01098 | 0.01151 | 0.00880 | 0.00684 | 0.00905 | 0.00933 | 0.02116 |
| **Sdr39u1** | 0.00000 | 0.58009 | 1.44424 | 1.85524 | 0.93828 | 0.59739 | 0.12025 | 0.38589 | 0.03196 | 0.00937 | 0.00000 | 0.07144 | 0.00000 | 0.49802 |
| **Sdsl** | 0.00909 | 0.00427 | 0.01146 | 0.19717 | 0.23137 | 0.00702 | 0.00398 | 0.00671 | 0.01041 | 0.00665 | 0.00674 | 0.01233 | 0.00505 | 0.01136 |
| **Sebox** | 0.07380 | 0.00452 | 0.06549 | 0.01326 | 0.38839 | 0.00912 | 0.00405 | 0.01284 | 0.11197 | 0.00769 | 0.00695 | 0.00539 | 1.65693 | 0.51308 |
| **Sec1** | 0.00000 | 0.00298 | 0.55057 | 3.20354 | 1.96665 | 1.82349 | 0.00334 | 0.00000 | 0.01819 | 0.00987 | 0.16240 | 1.09422 | 2.03552 | 0.00000 |
| **Sec11a** | 16.51107 | 1.91703 | 12.77675 | 9.16247 | 12.26816 | 10.70150 | 0.47752 | 10.54938 | 9.24031 | 0.11664 | 2.46114 | 6.17236 | 3.61469 | 12.60990 |
| **Sec11c** | 3.48374 | 0.07200 | 0.87143 | 5.07399 | 3.25134 | 3.74664 | 3.13468 | 6.09986 | 0.00000 | 13.18282 | 0.27684 | 1.10723 | 0.01205 | 1.05239 |
| **Sec13** | 12.81906 | 0.35047 | 21.58109 | 15.85960 | 14.45116 | 18.58687 | 3.02720 | 12.77824 | 5.67718 | 14.19106 | 11.08945 | 13.09298 | 3.51738 | 14.50157 |
| **Sec14l1** | 0.21711 | 0.41069 | 0.23366 | 0.22309 | 1.15337 | 2.27631 | 0.00270 | 2.64580 | 0.00441 | 0.03455 | 1.19260 | 2.01668 | 0.07122 | 0.00000 |
| **Sec14l2** | 0.12064 | 0.02549 | 0.57523 | 0.03310 | 0.25566 | 0.42106 | 0.00395 | 0.42673 | 0.00816 | 0.01716 | 0.08577 | 0.54930 | 0.45652 | 0.16298 |
| **Sec14l3** | 0.71735 | 0.01277 | 0.30636 | 0.18820 | 0.00000 | 0.00000 | 0.01108 | 0.09890 | 0.01033 | 0.00940 | 0.00813 | 0.00416 | 0.56073 | 0.02983 |
| **Sec14l4** | 0.47409 | 0.01116 | 0.94220 | 0.00000 | 0.74648 | 0.07531 | 0.00340 | 0.40396 | 0.46304 | 0.01203 | 0.00194 | 0.82757 | 0.00040 | 1.07846 |
| **Sec16a** | 4.54691 | 0.13782 | 3.68633 | 4.64198 | 5.88465 | 8.68099 | 0.03246 | 0.09725 | 4.94495 | 0.00986 | 3.10803 | 6.08123 | 6.86964 | 8.02336 |
| **Sec16b** | 0.44926 | 0.00396 | 0.28622 | 0.00000 | 3.09206 | 0.00000 | 0.00400 | 0.00588 | 0.64823 | 0.00544 | 0.00274 | 0.02663 | 1.40519 | 1.49887 |
| **Sec22a** | 0.68349 | 0.11155 | 2.44625 | 1.66270 | 2.27924 | 4.76953 | 3.38850 | 3.52870 | 0.52237 | 0.07838 | 0.00000 | 4.09980 | 0.42610 | 3.84443 |
| **Sec22b** | 5.50463 | 0.20634 | 6.80696 | 4.94800 | 4.43979 | 4.92560 | 0.01824 | 7.48152 | 1.07742 | 0.03914 | 2.37330 | 6.75062 | 0.49999 | 1.41709 |
| **Sec22c** | 0.02362 | 0.01209 | 0.50896 | 1.28210 | 0.36719 | 0.64037 | 0.00474 | 0.74273 | 0.08232 | 0.00632 | 0.00000 | 0.86188 | 0.00396 | 1.89742 |
| **Sec23a** | 4.08063 | 0.04485 | 3.30350 | 1.17266 | 7.30055 | 5.57372 | 1.50639 | 2.10361 | 12.81799 | 0.05115 | 2.82749 | 4.56732 | 1.58722 | 9.67687 |

| | | | | | | | | | | | | | | |
|---|---|---|---|---|---|---|---|---|---|---|---|---|---|---|
| **Sec23b** | 2.43624 | 7.99457 | 6.49415 | 4.60073 | 7.32851 | 8.16459 | 2.93475 | 3.10240 | 3.30743 | 0.13356 | 6.78473 | 5.63970 | 0.96511 | 7.74885 |
| **Sec23ip** | 4.42407 | 0.02154 | 2.87505 | 1.16566 | 5.31024 | 4.73233 | 4.32016 | 0.00016 | 9.88793 | 0.24539 | 9.79199 | 1.12562 | 6.84214 | 3.55710 |
| **Sec24a** | 2.82172 | 0.00361 | 2.29106 | 0.89174 | 1.48855 | 0.92915 | 2.90990 | 0.73460 | 10.40257 | 0.06442 | 0.82059 | 0.18887 | 5.84464 | 2.58975 |
| **Sec24b** | 6.31231 | 0.03343 | 5.15064 | 1.60037 | 4.60269 | 7.62476 | 1.59707 | 0.00000 | 0.73790 | 10.96801 | 0.16295 | 5.56486 | 1.32959 | 2.85511 |
| **Sec24c** | 2.75128 | 6.28384 | 9.84483 | 9.54974 | 9.33616 | 15.00516 | 0.23091 | 3.67271 | 4.46317 | 0.10408 | 1.90729 | 9.10883 | 3.34932 | 9.73467 |
| **Sec24d** | 0.04354 | 0.01985 | 0.80024 | 0.16876 | 0.02981 | 0.89624 | 0.00362 | 0.01750 | 0.00370 | 0.00370 | 0.00000 | 0.79327 | 1.03039 | 0.00000 |
| **Sec31a** | 6.92605 | 1.19699 | 14.40845 | 7.72745 | 9.33458 | 8.92875 | 1.33140 | 6.96146 | 0.64742 | 0.15068 | 5.45626 | 12.49064 | 5.51804 | 3.22200 |
| **Sec31b** | 0.00000 | 0.00429 | 0.40762 | 0.73256 | 0.00395 | 0.00000 | 0.00392 | 0.01175 | 0.00822 | 0.00751 | 0.00436 | 0.06024 | 0.05244 | 0.01843 |
| **Sec61a1** | 3.80846 | 0.24492 | 13.86929 | 10.34887 | 9.27830 | 20.07644 | 5.31118 | 8.22577 | 0.00731 | 0.45573 | 9.16741 | 12.97889 | 1.30740 | 9.44328 |
| **Sec61a2** | 1.94771 | 4.72762 | 0.30495 | 0.40966 | 1.14807 | 0.00000 | 0.03649 | 1.86465 | 2.68163 | 0.00405 | 0.50661 | 3.67177 | 1.24194 | 1.52767 |
| **Sec61b** | 0.28884 | 3.48491 | 0.47051 | 1.88534 | 5.48390 | 3.46650 | 1.90235 | 9.26062 | 0.06681 | 10.95990 | 0.12968 | 8.34140 | 1.18782 | 0.00000 |
| **Sec61g** | 1.68405 | 1.82225 | 0.00000 | 0.43154 | 1.04598 | 6.26721 | 2.09399 | 5.97219 | 1.45793 | 5.96254 | 0.14907 | 5.40799 | 4.16432 | 0.60471 |
| **Sec62** | 13.76526 | 6.83926 | 15.50681 | 3.72991 | 6.85259 | 14.99560 | 4.70068 | 3.85796 | 8.51877 | 0.26434 | 10.52817 | 4.70926 | 4.25698 | 16.28258 |
| **Sec63** | 5.86483 | 0.36563 | 1.33025 | 3.20691 | 3.44337 | 6.36840 | 2.63550 | 3.61463 | 5.08971 | 0.07105 | 2.61918 | 3.32446 | 7.17563 | 8.79095 |
| **Secisbp2** | 1.60157 | 0.01597 | 3.89900 | 3.20148 | 2.66503 | 2.33395 | 0.00176 | 0.43284 | 1.25923 | 0.02318 | 2.88037 | 3.98659 | 1.19086 | 5.29078 |
| **Secisbp2l** | 0.04295 | 0.01326 | 0.21178 | 0.61399 | 0.00000 | 0.17172 | 0.00507 | 0.15441 | 1.19295 | 0.00544 | 0.00539 | 0.02214 | 1.26649 | 0.73725 |
| **Sectm1a** | 0.00318 | 0.00430 | 0.00845 | 0.00915 | 0.34735 | 0.01401 | 0.00415 | 0.00857 | 0.00921 | 0.00614 | 0.00899 | 0.01250 | 0.50724 | 0.00197 |
| **Seh1l** | 12.56817 | 0.26990 | 15.71419 | 5.24235 | 14.55884 | 10.30282 | 1.03488 | 1.78274 | 22.28948 | 9.16706 | 11.20109 | 7.29508 | 6.74598 | 13.56607 |
| **Sel1l** | 2.88429 | 0.04767 | 1.55739 | 0.00000 | 0.75513 | 4.73450 | 3.20778 | 1.28734 | 1.17638 | 0.10449 | 2.45594 | 4.70337 | 0.80690 | 2.85113 |
| **Selenbp1** | 0.00000 | 0.00447 | 1.22179 | 0.06695 | 0.70770 | 0.29007 | 0.00399 | 0.00149 | 0.00878 | 0.00480 | 0.00535 | 0.01020 | 0.07342 | 1.15549 |
| **Selenbp2** | 0.01267 | 0.00429 | 1.49138 | 0.22026 | 0.69667 | 0.08571 | 0.00939 | 0.00000 | 0.01248 | 0.00372 | 0.00547 | 0.00733 | 0.21368 | 0.53864 |
| **Selk** | 5.02539 | 1.46930 | 1.94532 | 8.85156 | 8.27747 | 9.57067 | 3.54811 | 3.91850 | 6.23554 | 9.99123 | 0.40985 | 6.09989 | 9.41241 | 6.10739 |
| **Selm** | 0.02743 | 0.00313 | 0.03224 | 0.00000 | 0.65382 | 0.01966 | 0.00401 | 0.71000 | 0.01261 | 0.01455 | 0.15324 | 1.29783 | 0.09568 | 0.00146 |
| **Selo** | 0.00231 | 0.00454 | 0.24644 | 0.00000 | 0.19123 | 0.27878 | 0.01149 | 0.22212 | 0.00653 | 0.00664 | 0.00000 | 0.52662 | 0.10377 | 0.15161 |
| **Selplg** | 0.17123 | 0.00443 | 0.04801 | 0.20760 | 0.16773 | 0.16499 | 0.05409 | 0.04548 | 0.16093 | 0.00706 | 0.10751 | 0.04769 | 0.09557 | 0.20693 |
| **Selt** | 29.16761 | 3.05622 | 27.17109 | 20.20417 | 29.36549 | 20.03216 | 1.73224 | 10.29588 | 38.94967 | 0.16265 | 14.59390 | 16.55554 | 16.04805 | 27.54607 |
| **Sema3b** | 0.00296 | 0.03414 | 0.00185 | 0.72694 | 0.00567 | 0.04195 | 0.00994 | 3.45530 | 0.01094 | 0.02431 | 0.00000 | 0.26373 | 0.00426 | 0.03609 |
| **Sema3c** | 0.01608 | 0.00441 | 0.01077 | 0.01258 | 0.14341 | 0.01304 | 0.00443 | 0.01006 | 0.01877 | 0.00648 | 0.00864 | 0.00872 | 0.22602 | 0.00442 |
| **Sema3e** | 0.18907 | 0.06355 | 0.00000 | 0.49297 | 0.12179 | 0.43539 | 0.08190 | 1.52982 | 0.21305 | 0.44569 | 0.00968 | 0.14123 | 0.70500 | 0.49675 |
| **Sema4a** | 0.41070 | 0.00432 | 0.92880 | 0.28715 | 0.31356 | 0.02381 | 0.00421 | 0.01699 | 0.00653 | 0.00511 | 0.21827 | 0.00378 | 0.22200 | 0.35169 |
| **Sema4b** | 0.06170 | 3.75679 | 0.00000 | 2.43830 | 0.04387 | 1.66068 | 0.11342 | 0.51791 | 0.08242 | 0.03575 | 0.00000 | 1.30170 | 0.24861 | 0.03664 |
| **Sema4c** | 0.03103 | 0.00438 | 0.01309 | 0.02076 | 0.01492 | 0.01244 | 0.00419 | 0.00947 | 0.01223 | 0.00932 | 0.00819 | 0.00892 | 0.12068 | 0.00334 |
| **Sema4d** | 0.16627 | 0.00329 | 0.87185 | 0.23134 | 0.00000 | 0.45853 | 2.93305 | 0.83043 | 0.00874 | 0.13714 | 0.98594 | 0.41770 | 0.54998 | 1.03825 |
| **Sema4g** | 1.90742 | 0.00351 | 1.16954 | 2.21692 | 1.12632 | 1.53677 | 0.00342 | 3.60646 | 0.00792 | 0.01745 | 0.60370 | 0.53838 | 0.00266 | 0.64711 |
| **Sema5a** | 0.00264 | 0.02091 | 0.00000 | 0.90067 | 0.57868 | 0.00000 | 0.00502 | 0.24030 | 0.00939 | 0.00064 | 1.86252 | 0.88388 | 0.58658 | 0.53566 |
| **Sema5b** | 0.00000 | 0.00397 | 0.08407 | 1.12866 | 0.00084 | 0.00000 | 0.01005 | 0.02764 | 0.03944 | 0.00509 | 0.00784 | 0.02083 | 0.14621 | 0.00000 |
| **Sema6a** | 0.24230 | 0.00405 | 0.00584 | 0.34624 | 0.06282 | 0.00315 | 0.00414 | 0.00600 | 0.01553 | 0.00546 | 0.01732 | 0.01689 | 0.28321 | 0.01207 |
| **Sema6b** | 0.00589 | 0.00443 | 0.09168 | 0.00320 | 0.42957 | 0.01021 | 0.00423 | 0.01105 | 0.02222 | 0.00696 | 0.00864 | 0.00884 | 0.03349 | 0.01904 |
| **Sema6c** | 0.01064 | 0.00411 | 0.00797 | 0.25228 | 0.13824 | 0.00469 | 0.00387 | 0.00613 | 0.02434 | 0.00561 | 0.01810 | 0.01642 | 0.12616 | 0.00000 |
| **Sema6d** | 0.00727 | 0.05759 | 0.01925 | 0.00000 | 0.01883 | 0.05211 | 0.01532 | 0.06367 | 0.05185 | 0.01341 | 0.01451 | 0.56980 | 0.03966 | 0.02362 |
| **Sema7a** | 0.22561 | 0.00435 | 0.01240 | 0.01626 | 0.13030 | 0.01389 | 0.00461 | 0.00923 | 0.01505 | 0.01079 | 0.00646 | 0.06334 | 0.01304 | 0.00520 |
| **Senp1** | 1.51848 | 0.01701 | 4.78032 | 0.00000 | 4.42504 | 4.81180 | 0.01486 | 2.31133 | 2.07045 | 0.02452 | 0.77548 | 1.71588 | 2.64344 | 3.86155 |
| **Senp2** | 4.16738 | 0.12425 | 2.83043 | 0.74400 | 2.17874 | 2.00726 | 0.00738 | 0.00000 | 9.43804 | 0.02988 | 2.05024 | 2.64673 | 7.04905 | 8.20096 |
| **Senp3** | 16.50662 | 0.00000 | 9.11528 | 2.35184 | 5.23816 | 0.40472 | 1.80829 | 0.29687 | 12.48890 | 2.26167 | 1.84748 | 2.23531 | 11.66901 | 17.37137 |
| **Senp5** | 5.63911 | 0.00108 | 4.41779 | 0.00004 | 4.33446 | 5.70212 | 1.62426 | 0.26699 | 5.74387 | 0.08754 | 0.00000 | 2.80737 | 4.64587 | 9.68951 |
| **Senp6** | 14.00152 | 0.06360 | 9.81957 | 0.56543 | 17.06482 | 7.27879 | 2.60546 | 3.22975 | 6.53015 | 0.03287 | 0.87310 | 4.23710 | 8.27030 | 19.36959 |
| **Senp7** | 0.16107 | 0.05184 | 0.57018 | 0.29631 | 1.29375 | 0.30823 | 0.00967 | 0.03652 | 0.16099 | 0.00448 | 0.03985 | 1.12301 | 2.09843 | 2.51896 |
| **Senp8** | 1.03001 | 0.01466 | 1.53201 | 0.53513 | 0.95383 | 2.66611 | 0.00355 | 0.20297 | 0.71530 | 0.00864 | 1.70795 | 0.28271 | 0.00538 | 0.00000 |
| **X15.Sep** | 13.03258 | 7.75373 | 18.60415 | 18.23893 | 8.06488 | 18.23384 | 9.33367 | 18.32922 | 2.62047 | 13.72424 | 9.07622 | 20.20726 | 0.86148 | 6.12260 |
| **Sephs1** | 9.27966 | 0.09752 | 14.07063 | 5.08102 | 6.60788 | 8.24704 | 0.76738 | 0.50771 | 3.08425 | 0.12884 | 11.49871 | 6.68512 | 2.68331 | 9.76459 |
| **Sephs2** | 0.00000 | 0.17853 | 4.10083 | 4.50971 | 2.91590 | 3.48609 | 2.68960 | 8.38406 | 0.68898 | 5.60699 | 4.50993 | 2.98655 | 6.57634 | 3.60908 |
| **Sepn1** | 0.21357 | 0.00386 | 0.74849 | 0.00000 | 0.21088 | 0.51927 | 0.00374 | 0.00324 | 0.00594 | 0.01944 | 0.02507 | 0.78100 | 0.01709 | 0.00177 |
| **Sepp1** | 0.01826 | 0.00450 | 0.00000 | 0.01300 | 0.36303 | 0.00767 | 0.00433 | 0.01120 | 0.69725 | 0.00660 | 0.00616 | 0.00775 | 0.01842 | 0.10326 |
| **Sepsecs** | 2.19673 | 0.00371 | 0.34646 | 1.30252 | 0.13152 | 1.05239 | 0.00386 | 0.11154 | 0.46242 | 0.01156 | 0.00000 | 0.23121 | 0.58359 | 0.33896 |
| **X1.Sep** | 0.16217 | 0.01308 | 0.21871 | 0.13402 | 0.38734 | 0.31854 | 0.00966 | 1.56241 | 0.00812 | 0.33244 | 0.00000 | 0.00254 | 0.14142 | 0.00000 |

| | | | | | | | | | | | | | | |
|---|---|---|---|---|---|---|---|---|---|---|---|---|---|---|
| **X10.Sep** | 0.41514 | 0.00398 | 0.07977 | 0.09751 | 0.19266 | 0.02366 | 0.00419 | 0.27230 | 4.38569 | 0.00596 | 0.00000 | 0.32072 | 0.60333 | 1.52659 |
| **X11.Sep** | 0.71925 | 0.16275 | 0.03712 | 0.97352 | 1.66725 | 6.05533 | 0.07742 | 0.78283 | 0.21762 | 0.17633 | 2.62935 | 14.94758 | 1.81044 | 3.21336 |
| **X12.Sep** | 0.00000 | 0.01241 | 2.80224 | 3.07197 | 0.22811 | 1.39836 | 0.00254 | 0.16056 | 0.01545 | 4.79007 | 4.27791 | 0.00000 | 0.01414 | 0.09561 |
| **X2.Sep** | 6.87953 | 4.97529 | 9.35636 | 2.41819 | 14.73173 | 16.01962 | 4.40351 | 3.32726 | 17.57304 | 0.14526 | 12.80237 | 9.02055 | 6.99770 | 8.91721 |
| **X3.Sep** | 2.76997 | 0.01697 | 2.82197 | 1.31189 | 4.18384 | 0.90867 | 0.00249 | 0.00000 | 5.18820 | 0.00000 | 0.00000 | 0.00524 | 4.64976 | 3.77490 |
| **X4.Sep** | 0.00000 | 0.00429 | 0.07376 | 0.49526 | 0.00000 | 0.06593 | 0.00407 | 0.00252 | 0.01111 | 0.00436 | 0.40450 | 0.00389 | 0.83875 | 0.01632 |
| **X6.Sep** | 0.00205 | 0.62980 | 0.00000 | 0.33451 | 0.00102 | 3.99524 | 0.48783 | 0.30693 | 0.02928 | 0.04336 | 0.00157 | 0.00000 | 0.03175 | 0.09521 |
| **X7.Sep** | 1.50842 | 0.03033 | 6.97350 | 1.65124 | 1.71579 | 4.42745 | 0.85175 | 0.73509 | 9.51821 | 0.00583 | 10.27906 | 2.31309 | 3.90300 | 0.00000 |
| **X8.Sep** | 0.13162 | 0.03692 | 6.28891 | 5.57546 | 1.47442 | 6.42064 | 0.62138 | 1.42568 | 2.37187 | 0.15459 | 0.96545 | 0.03299 | 0.00000 | 1.79252 |
| **X9.Sep** | 0.09238 | 0.01118 | 0.14441 | 0.53498 | 0.01840 | 0.00000 | 0.01477 | 0.00674 | 0.01085 | 0.01459 | 0.00367 | 0.88600 | 0.00488 | 0.04078 |
| **Sepw1** | 5.23176 | 2.00510 | 2.44544 | 9.79061 | 12.15960 | 4.06089 | 2.07064 | 9.21824 | 0.31163 | 12.26022 | 1.09082 | 3.65331 | 2.26389 | 4.16139 |
| **Serac1** | 0.99704 | 0.00332 | 1.66719 | 0.29907 | 2.60769 | 1.35001 | 0.00347 | 0.96800 | 0.36291 | 0.00116 | 2.62978 | 0.00000 | 0.00938 | 2.09305 |
| **Serbp1** | 34.31412 | 8.20928 | 22.99604 | 28.07210 | 33.94295 | 21.23324 | 6.99174 | 9.85573 | 15.94051 | 16.32849 | 13.91395 | 18.78947 | 16.50528 | 16.29490 |
| **Serf1** | 0.07143 | 0.00375 | 0.00327 | 0.48671 | 0.65278 | 0.04924 | 0.00913 | 0.00319 | 0.01601 | 0.01287 | 0.02201 | 0.35266 | 0.80055 | 0.00000 |
| **Serf2** | 8.73763 | 9.42793 | 3.93484 | 27.37284 | 27.00235 | 19.44569 | 5.24118 | 23.55950 | 6.96251 | 17.03784 | 4.64209 | 15.04420 | 14.95438 | 11.22777 |
| **Sergef** | 0.30579 | 0.02849 | 0.92430 | 0.27461 | 0.49240 | 0.00000 | 0.00341 | 0.00041 | 0.00866 | 0.00157 | 0.93889 | 1.27457 | 0.00000 | 0.07856 |
| **Serhl** | 0.16405 | 2.56264 | 1.65881 | 2.01526 | 1.23665 | 1.83944 | 0.03863 | 4.37782 | 0.75600 | 0.69984 | 0.43346 | 0.74920 | 0.00000 | 3.33712 |
| **Serinc1** | 14.90137 | 0.05778 | 12.67835 | 5.63817 | 10.13140 | 7.46228 | 0.46554 | 1.40514 | 23.17172 | 0.00000 | 1.40972 | 4.82147 | 1.60382 | 6.31289 |
| **Serinc2** | 0.15999 | 0.03384 | 1.51357 | 0.87168 | 0.86148 | 3.27159 | 0.01445 | 4.26675 | 0.30356 | 0.03384 | 0.08269 | 0.61525 | 0.01922 | 1.04218 |
| **Serinc3** | 16.72832 | 2.71056 | 25.34890 | 17.69378 | 12.06689 | 20.85808 | 5.41499 | 10.87455 | 20.15301 | 3.60292 | 13.71557 | 10.12529 | 13.90615 | 18.86068 |
| **Serinc5** | 0.27618 | 0.00373 | 0.41969 | 0.00000 | 0.78422 | 0.55451 | 0.00380 | 0.07076 | 2.18405 | 0.00239 | 0.63554 | 0.00309 | 0.00000 | 0.03991 |
| **Serp1** | 5.80944 | 0.10088 | 7.00591 | 3.85043 | 8.95212 | 9.52041 | 5.92835 | 4.97556 | 11.11941 | 11.01844 | 4.32775 | 1.65041 | 14.73237 | 9.01199 |
| **Serpina1a** | 0.01091 | 0.00438 | 0.01561 | 0.00412 | 0.11742 | 0.03240 | 0.01618 | 0.03216 | 0.02588 | 0.05919 | 0.01578 | 0.02550 | 0.01514 | 0.02500 |
| **Serpina1b** | 0.01120 | 0.00446 | 0.01358 | 0.01297 | 0.05765 | 0.01311 | 0.00429 | 0.01071 | 0.01161 | 0.00682 | 0.00872 | 0.00968 | 0.00920 | 0.02083 |
| **Serpina1c** | 0.00969 | 0.00446 | 0.01141 | 0.01211 | 0.11495 | 0.01363 | 0.00447 | 0.01042 | 0.01403 | 0.00705 | 0.00884 | 0.00943 | 0.00645 | 0.00973 |
| **Serpina1d** | 0.01003 | 0.00443 | 0.01017 | 0.02376 | 0.10656 | 0.01237 | 0.00443 | 0.00942 | 0.01484 | 0.00670 | 0.00821 | 0.01113 | 0.00715 | 0.01232 |
| **Serpina1e** | 0.00828 | 0.00451 | 0.00157 | 0.00742 | 1.72661 | 0.01397 | 0.00459 | 0.01016 | 0.01531 | 0.00689 | 0.00884 | 0.00734 | 0.00186 | 0.00437 |
| **Serpina3m** | 0.00528 | 0.00326 | 0.01082 | 0.01022 | 0.01158 | 0.01107 | 0.00396 | 0.00533 | 0.01259 | 0.00637 | 0.00518 | 1.38590 | 0.00723 | 0.01293 |
| **Serpinb1a** | 0.00268 | 0.01012 | 0.00000 | 0.00000 | 0.00600 | 0.49114 | 0.00912 | 0.00802 | 0.01204 | 0.00304 | 0.89069 | 2.68439 | 0.51824 | 0.00000 |
| **Serpinb1b** | 0.01062 | 0.00438 | 0.01396 | 0.01550 | 0.01402 | 0.01443 | 0.00436 | 0.00974 | 0.01204 | 0.00724 | 0.01390 | 0.03103 | 0.00753 | 0.01174 |
| **Serpinb1c** | 0.00525 | 0.00379 | 0.00000 | 0.53461 | 0.32296 | 0.26159 | 0.00365 | 0.00503 | 0.02209 | 0.00726 | 0.00412 | 0.19989 | 0.19126 | 0.20473 |
| **Serpinb6a** | 3.74549 | 0.41912 | 14.48060 | 16.88554 | 1.56512 | 23.65301 | 10.19591 | 20.36082 | 1.90452 | 3.33353 | 9.92965 | 18.19851 | 0.00000 | 3.99981 |
| **Serpinb6b** | 0.01748 | 0.03585 | 0.06523 | 0.19599 | 0.00531 | 0.19923 | 0.15916 | 5.04298 | 0.01300 | 0.02869 | 0.05151 | 0.11154 | 0.48937 | 0.05275 |
| **Serpinb6c** | 0.50008 | 0.06800 | 0.89138 | 6.20526 | 0.00000 | 7.38548 | 3.55219 | 12.69205 | 0.01734 | 0.16192 | 1.49456 | 0.21350 | 0.00000 | 0.00000 |
| **Serpinb6d** | 0.02234 | 0.00420 | 0.04193 | 0.13122 | 0.00817 | 0.13678 | 0.03301 | 0.26909 | 0.01309 | 0.00639 | 0.04636 | 0.00603 | 0.00671 | 0.00734 |
| **Serpinb6e** | 0.00796 | 0.00444 | 0.03948 | 0.08110 | 0.01063 | 0.11349 | 0.04621 | 0.14609 | 0.01268 | 0.00719 | 0.01481 | 0.00676 | 0.00698 | 0.00945 |
| **Serpinb9** | 0.00798 | 0.00477 | 0.03098 | 0.11639 | 0.01135 | 0.12167 | 0.00464 | 0.04167 | 0.02414 | 0.00658 | 0.02521 | 0.00995 | 0.00649 | 0.01602 |
| **Serpinb9b** | 0.00000 | 0.02703 | 3.10314 | 2.97477 | 0.00000 | 7.90217 | 2.06724 | 6.09938 | 0.00447 | 0.07673 | 3.02483 | 0.34455 | 0.00310 | 0.00000 |
| **Serpinc1** | 0.77539 | 0.03228 | 0.56339 | 0.56052 | 3.00485 | 0.06269 | 0.00382 | 0.00070 | 0.18973 | 0.01013 | 0.00338 | 0.00521 | 1.22368 | 1.67734 |
| **Serpind1** | 0.00000 | 0.00429 | 0.00000 | 0.01683 | 0.01887 | 0.00744 | 0.00396 | 0.00837 | 1.06784 | 0.00677 | 0.00623 | 0.00804 | 0.78425 | 0.00000 |
| **Serpine1** | 1.08073 | 0.00434 | 0.01210 | 0.01005 | 0.00341 | 0.00773 | 0.00424 | 0.01233 | 0.00889 | 0.00677 | 0.00688 | 0.00750 | 0.00434 | 0.00374 |
| **Serpine2** | 0.35420 | 0.01935 | 2.85959 | 2.76256 | 0.39731 | 3.05399 | 2.49940 | 2.00682 | 0.00000 | 4.20116 | 0.00000 | 8.20476 | 3.80792 | 0.47087 |
| **Serpine3** | 0.06200 | 0.00433 | 0.00891 | 0.02141 | 0.00887 | 0.00759 | 0.00416 | 0.01491 | 0.10796 | 0.00735 | 0.05682 | 0.00737 | 0.00540 | 0.08817 |
| **Serpinf1** | 0.40348 | 0.00434 | 0.01083 | 0.01834 | 0.01648 | 0.01217 | 0.00415 | 0.00912 | 0.01495 | 0.00621 | 0.09797 | 0.00951 | 0.00422 | 0.00843 |
| **Serpinf2** | 0.02760 | 0.00436 | 0.12947 | 0.00915 | 0.53538 | 0.00470 | 0.00406 | 0.01071 | 0.00998 | 0.00736 | 0.00681 | 0.00643 | 0.35677 | 0.42205 |
| **Serping1** | 9.37341 | 0.00000 | 2.60110 | 1.48866 | 2.92845 | 0.00000 | 0.00176 | 0.00000 | 3.72537 | 0.00000 | 3.46877 | 2.55269 | 4.61248 | 5.08744 |
| **Serpinh1** | 0.52356 | 1.52429 | 0.01677 | 0.00000 | 0.23043 | 0.00264 | 0.00000 | 0.42233 | 0.00623 | 0.02223 | 0.00178 | 14.20156 | 0.00353 | 0.11012 |
| **Serpini1** | 0.00000 | 0.00376 | 0.47735 | 0.00000 | 0.31738 | 0.00484 | 0.00380 | 0.00619 | 0.00000 | 0.00466 | 0.03504 | 0.00628 | 3.64124 | 1.02032 |
| **Sertad1** | 1.78399 | 5.22976 | 7.76582 | 4.75355 | 10.52184 | 0.00000 | 3.33746 | 8.23471 | 0.00000 | 6.88671 | 2.77844 | 1.37571 | 9.25461 | 0.18639 |
| **Sertad2** | 0.58561 | 0.01937 | 0.84077 | 0.00000 | 1.54032 | 1.40571 | 0.00307 | 1.40647 | 3.07614 | 0.00000 | 2.35040 | 1.42672 | 1.23633 | 6.48061 |
| **Sertad3** | 0.00939 | 0.00565 | 0.17404 | 0.40616 | 0.75278 | 1.76568 | 0.00411 | 0.72137 | 0.03262 | 0.00909 | 0.02154 | 0.00000 | 0.00590 | 0.00831 |
| **Sesn1** | 2.28097 | 0.01396 | 5.85239 | 2.67661 | 3.44719 | 1.50205 | 0.01261 | 1.73804 | 2.15106 | 0.03211 | 3.72730 | 3.21333 | 1.55467 | 7.02917 |
| **Sesn2** | 0.26729 | 1.76952 | 2.72015 | 1.81250 | 3.53068 | 1.49016 | 0.15092 | 0.60577 | 0.07324 | 0.00970 | 1.24335 | 1.07414 | 0.32384 | 4.54052 |
| **Sesn3** | 0.37458 | 0.00311 | 0.25121 | 1.62080 | 1.38369 | 1.59794 | 0.00874 | 0.01542 | 1.27449 | 0.00118 | 1.51805 | 0.24649 | 0.76169 | 3.46908 |
| **Sestd1** | 0.00143 | 0.01765 | 0.05682 | 0.42704 | 0.00000 | 0.00000 | 0.00344 | 0.24272 | 0.00000 | 0.00625 | 0.00455 | 1.72664 | 2.19588 | 0.96942 |

| | | | | | | | | | | | | | | |
|---|---|---|---|---|---|---|---|---|---|---|---|---|---|---|
| **Set** | 29.04290 | 9.56135 | 46.51367 | 39.63524 | 47.23079 | 34.96233 | 7.91824 | 22.78019 | 38.97590 | 10.68461 | 21.24146 | 27.74972 | 10.63385 | 40.97867 |
| **Setd1a** | 3.61387 | 0.03406 | 9.15521 | 5.14301 | 8.52878 | 1.25036 | 0.09049 | 2.67411 | 0.00000 | 0.01008 | 4.44178 | 2.35980 | 3.34000 | 3.48322 |
| **Setd1b** | 0.70549 | 0.24831 | 0.25760 | 0.91224 | 2.52467 | 1.18548 | 0.00370 | 0.35532 | 0.90551 | 0.03894 | 0.88857 | 0.90742 | 1.76740 | 0.27028 |
| **Setd2** | 19.68725 | 6.34112 | 23.81146 | 5.06706 | 20.91790 | 10.18111 | 6.52622 | 1.94290 | 27.38450 | 0.15759 | 5.78851 | 4.66244 | 16.21261 | 22.15583 |
| **Setd3** | 5.35048 | 0.05736 | 9.27819 | 8.07006 | 8.11507 | 11.28468 | 5.80205 | 9.44366 | 6.02014 | 0.80654 | 10.97826 | 6.91524 | 4.67929 | 3.57360 |
| **Setd4** | 1.14532 | 0.01803 | 4.87144 | 3.42561 | 1.48645 | 1.59341 | 0.01499 | 2.08847 | 0.26219 | 0.06395 | 0.58544 | 0.00000 | 1.04462 | 1.02398 |
| **Setd5** | 0.24155 | 0.01648 | 0.55080 | 0.00202 | 0.63258 | 0.66947 | 0.03572 | 0.39908 | 1.06634 | 0.00296 | 0.53122 | 2.03954 | 0.24458 | 2.42352 |
| **Setd6** | 2.04891 | 0.01242 | 2.13716 | 3.71295 | 2.07168 | 2.20388 | 0.36288 | 1.02811 | 0.00370 | 0.05539 | 0.00000 | 1.52332 | 0.71908 | 1.10651 |
| **Setd7** | 0.03557 | 0.00419 | 0.03905 | 1.58703 | 0.00000 | 0.00000 | 1.29332 | 0.00434 | 0.03198 | 0.02826 | 0.00957 | 0.02730 | 0.01851 | 0.00000 |
| **Setdb1** | 7.44341 | 0.02083 | 8.55386 | 0.69113 | 3.04378 | 3.41430 | 1.68175 | 1.84635 | 7.91521 | 10.08878 | 3.48219 | 3.33560 | 7.72467 | 8.08567 |
| **Setdb2** | 0.12312 | 0.00408 | 0.00738 | 0.01073 | 0.00022 | 0.00487 | 0.00407 | 0.00896 | 0.26784 | 0.00675 | 0.17118 | 0.00483 | 1.84190 | 0.01689 |
| **Setmar** | 1.30441 | 0.00462 | 0.33197 | 0.16998 | 0.06890 | 0.18409 | 0.00399 | 0.00310 | 0.11393 | 0.00513 | 0.00467 | 0.01224 | 0.01811 | 0.02811 |
| **Setx** | 12.14658 | 8.00089 | 11.55737 | 5.44897 | 18.50655 | 14.70215 | 2.10061 | 4.75928 | 24.32716 | 0.00000 | 14.54470 | 10.78705 | 10.86681 | 17.80036 |
| **Sez6l** | 0.00260 | 0.00428 | 0.02262 | 0.00972 | 0.19004 | 0.01242 | 0.00428 | 0.01006 | 0.01616 | 0.00648 | 0.00864 | 0.00866 | 0.55463 | 0.00170 |
| **Sez6l2** | 1.63832 | 0.00326 | 1.77536 | 0.91698 | 1.75290 | 0.46314 | 0.00370 | 0.01779 | 0.16479 | 0.01053 | 0.21062 | 0.83329 | 0.93872 | 0.86659 |
| **Sf1** | 8.57823 | 9.84224 | 11.77676 | 8.13295 | 16.44877 | 8.29961 | 3.98430 | 7.08799 | 2.29188 | 11.24645 | 4.85432 | 9.89265 | 11.13923 | 3.33330 |
| **Sf3a1** | 5.59251 | 5.71009 | 15.24285 | 15.98032 | 10.24339 | 9.42396 | 5.81067 | 7.91315 | 3.18702 | 0.32367 | 10.59327 | 9.95460 | 9.69047 | 4.81144 |
| **Sf3a2** | 0.07903 | 0.11899 | 0.57720 | 1.74302 | 1.61919 | 0.00000 | 0.07646 | 0.83026 | 0.00595 | 0.03701 | 0.20343 | 0.91556 | 0.23420 | 0.00000 |
| **Sf3a3** | 12.35485 | 5.20895 | 18.62564 | 9.97749 | 17.57774 | 12.39096 | 0.06897 | 6.08458 | 14.37538 | 10.17896 | 12.11200 | 10.41090 | 4.91824 | 14.11899 |
| **Sf3b1** | 23.73633 | 6.54711 | 21.40266 | 7.43358 | 27.46144 | 15.83129 | 5.85034 | 4.20591 | 33.26453 | 0.19070 | 14.20468 | 11.85422 | 18.63534 | 22.58146 |
| **Sf3b2** | 16.00416 | 0.34822 | 28.14157 | 17.47135 | 23.81951 | 16.04905 | 2.67536 | 10.92609 | 9.13900 | 14.63207 | 18.04656 | 15.10887 | 9.21439 | 16.29969 |
| **Sf3b3** | 13.43536 | 2.58740 | 26.79592 | 4.82875 | 23.79752 | 20.06795 | 7.40976 | 7.89249 | 19.32239 | 0.47278 | 15.73219 | 13.44405 | 4.84333 | 15.38607 |
| **Sf3b4** | 1.25096 | 2.35666 | 18.49952 | 15.36504 | 20.06315 | 0.00000 | 4.79506 | 10.49242 | 0.06069 | 1.33845 | 2.11914 | 8.90272 | 2.91356 | 0.34427 |
| **Sf3b5** | 0.00000 | 8.55734 | 4.88814 | 17.39521 | 16.31136 | 0.95910 | 6.47271 | 14.50504 | 0.00000 | 7.48643 | 0.84436 | 15.82483 | 1.94064 | 0.00000 |
| **Sf3b6** | 12.23766 | 4.37222 | 8.61491 | 13.48452 | 16.14391 | 14.22760 | 5.65572 | 5.36175 | 11.78757 | 9.99401 | 8.65914 | 10.59104 | 9.73253 | 13.77983 |
| **Sfi1** | 0.99483 | 0.06208 | 2.57891 | 8.35408 | 3.32847 | 5.98661 | 2.13399 | 5.21772 | 2.30348 | 7.63312 | 1.31848 | 2.13476 | 14.62094 | 0.75843 |
| **Sfmbt1** | 0.00000 | 0.00008 | 0.32231 | 0.28933 | 0.95786 | 2.22708 | 0.00236 | 0.00008 | 1.66435 | 0.01560 | 1.56293 | 3.19445 | 6.44030 | 3.19238 |
| **Sfmbt2** | 0.21779 | 4.13460 | 1.19005 | 1.95382 | 0.21547 | 2.01944 | 0.87483 | 2.18007 | 1.61528 | 0.01845 | 2.49474 | 3.11440 | 0.00000 | 1.30511 |
| **Sfn** | 13.07825 | 6.69342 | 29.42233 | 25.41760 | 32.24470 | 8.29831 | 2.43184 | 10.10610 | 0.33639 | 8.54359 | 14.89746 | 6.60221 | 2.34941 | 3.09707 |
| **Sfpq** | 10.84191 | 5.45347 | 19.88393 | 14.90376 | 18.93794 | 18.17681 | 6.64463 | 9.93387 | 22.19630 | 0.93790 | 10.30536 | 14.11422 | 5.74056 | 16.76873 |
| **Sfr1** | 11.84412 | 11.23625 | 19.48339 | 14.58500 | 13.52490 | 15.56849 | 6.98067 | 7.88034 | 7.99535 | 0.40259 | 13.31559 | 13.77421 | 3.96885 | 15.99859 |
| **Sfrp1** | 0.43444 | 0.11682 | 0.61849 | 1.64676 | 0.00000 | 4.20801 | 0.02932 | 0.00352 | 0.00153 | 0.01405 | 0.00000 | 3.97691 | 0.51210 | 0.12772 |
| **Sfrp5** | 0.00749 | 0.00407 | 0.01748 | 0.45044 | 0.00183 | 0.00636 | 0.00400 | 0.01408 | 0.05198 | 0.00507 | 0.01540 | 0.02706 | 0.01584 | 0.04625 |
| **Sfswap** | 0.00000 | 0.01612 | 7.17331 | 4.02849 | 9.11912 | 1.39303 | 4.04504 | 3.03764 | 0.15080 | 0.17551 | 2.40828 | 2.53883 | 1.07003 | 1.41129 |
| **Sft2d1** | 5.02851 | 3.43538 | 6.63767 | 9.26111 | 5.23272 | 4.93095 | 2.38174 | 1.54936 | 0.00393 | 10.12564 | 1.38596 | 4.85335 | 0.86232 | 0.00000 |
| **Sft2d2** | 0.01081 | 0.03385 | 0.00000 | 0.15826 | 0.20523 | 2.06090 | 1.73397 | 1.59031 | 0.01403 | 0.06065 | 0.00080 | 0.05482 | 0.00542 | 0.00000 |
| **Sft2d3** | 0.00327 | 0.00428 | 0.00000 | 0.14126 | 0.14399 | 0.03326 | 0.00441 | 0.00938 | 0.25579 | 0.00751 | 0.00539 | 0.08826 | 0.28098 | 0.00000 |
| **Sfta2** | 0.03054 | 0.00500 | 0.00813 | 0.09870 | 0.01143 | 0.01988 | 0.00410 | 0.01816 | 0.00781 | 0.01230 | 0.00844 | 0.00730 | 0.00797 | 0.05143 |
| **Sfxn1** | 3.67645 | 0.07238 | 1.95466 | 0.92806 | 5.38184 | 4.52152 | 0.00223 | 1.12391 | 0.04420 | 0.01238 | 3.40889 | 3.95651 | 0.00000 | 2.95897 |
| **Sfxn2** | 2.26522 | 0.05805 | 2.05814 | 2.51715 | 1.05656 | 1.70612 | 0.03112 | 3.54660 | 0.38416 | 0.03354 | 2.99615 | 2.01877 | 1.92649 | 2.10677 |
| **Sfxn3** | 0.07954 | 0.01366 | 0.00507 | 0.02778 | 0.21855 | 0.03840 | 0.00431 | 0.09654 | 0.35025 | 0.00697 | 0.03494 | 0.03796 | 0.03409 | 0.03468 |
| **Sfxn4** | 0.55098 | 0.01007 | 1.27166 | 0.39035 | 2.03259 | 2.07998 | 0.13515 | 0.27292 | 1.61231 | 0.24138 | 2.46702 | 0.09435 | 7.08692 | 1.19728 |
| **Sfxn5** | 0.00234 | 0.01083 | 0.25089 | 0.01796 | 0.00542 | 0.34783 | 0.00390 | 0.17215 | 0.01014 | 0.00862 | 0.02520 | 0.36134 | 0.00713 | 0.00984 |
| **Sgcb** | 0.04410 | 0.00406 | 0.00363 | 0.02687 | 0.05035 | 0.00000 | 0.00396 | 0.01091 | 0.01239 | 0.01161 | 1.13012 | 0.00360 | 0.05607 | 0.00412 |
| **Sgce** | 0.06026 | 0.00445 | 0.26294 | 0.00000 | 0.74653 | 0.00000 | 0.00445 | 0.01025 | 0.84473 | 0.00698 | 0.00406 | 0.00587 | 0.00000 | 0.70164 |
| **Sgf29** | 8.22450 | 0.04622 | 14.58278 | 6.49212 | 12.20129 | 7.87618 | 0.19047 | 3.60593 | 0.88802 | 0.07064 | 2.24049 | 1.57873 | 1.89880 | 10.42553 |
| **Sgk1** | 17.06027 | 2.80222 | 20.24765 | 11.35806 | 22.77443 | 16.39444 | 10.40042 | 9.32998 | 6.92979 | 5.26824 | 9.12012 | 8.76566 | 12.23398 | 3.79592 |
| **Sgk2** | 0.00638 | 0.00522 | 0.12802 | 0.00000 | 0.00905 | 0.12491 | 0.00423 | 0.01733 | 0.01062 | 0.00612 | 0.00412 | 0.19416 | 0.00691 | 0.03145 |
| **Sgk3** | 15.03813 | 0.00670 | 11.19402 | 0.87631 | 13.02712 | 4.71769 | 0.87596 | 3.52287 | 24.77299 | 0.05293 | 3.36596 | 1.06707 | 4.75680 | 13.72543 |
| **Sgms1** | 0.00000 | 0.00263 | 0.44733 | 0.56737 | 0.00000 | 0.00000 | 0.00766 | 0.01090 | 0.00624 | 5.42240 | 0.58077 | 0.00000 | 1.06881 | 0.68773 |
| **Sgms2** | 0.01628 | 0.03531 | 0.02672 | 0.34546 | 0.99970 | 0.14727 | 0.00476 | 0.08598 | 0.09015 | 0.01455 | 0.03149 | 0.04798 | 0.42416 | 0.04692 |
| **Sgol1** | 6.38353 | 3.92461 | 5.79796 | 3.82345 | 1.81016 | 5.10534 | 2.76353 | 0.47777 | 4.20032 | 0.15115 | 5.01067 | 4.88742 | 4.46041 | 6.11690 |
| **Sgol2a** | 1.50237 | 0.03426 | 0.84230 | 2.33306 | 2.66520 | 0.70534 | 1.98206 | 0.41908 | 12.03097 | 0.16292 | 4.04169 | 1.02725 | 6.46912 | 5.11454 |
| **Sgol2b** | 0.07218 | 0.00508 | 0.01663 | 0.07001 | 0.02745 | 0.02893 | 0.03451 | 0.02174 | 0.18095 | 0.00756 | 0.04509 | 0.04363 | 0.03306 | 0.33294 |
| **Sgpl1** | 3.58585 | 0.08421 | 3.42743 | 7.98272 | 0.39377 | 4.77127 | 2.32312 | 4.43120 | 3.98406 | 0.08247 | 0.58666 | 13.50194 | 0.46028 | 1.78616 |

| | | | | | | | | | | | | | | |
|---|---|---|---|---|---|---|---|---|---|---|---|---|---|---|
| **Sgpp1** | 19.11834 | 4.60466 | 9.52745 | 0.63774 | 13.74301 | 6.16146 | 3.12731 | 5.65889 | 27.04850 | 0.12535 | 3.35690 | 2.54058 | 9.85117 | 18.85924 |
| **Sgpp2** | 0.07961 | 0.01991 | 0.00804 | 0.23400 | 0.18824 | 0.26446 | 0.11588 | 0.07302 | 0.41402 | 0.10651 | 0.01838 | 0.05087 | 0.31208 | 0.30266 |
| **Sgsh** | 0.41384 | 0.00547 | 0.00779 | 0.00294 | 0.25942 | 0.01029 | 0.00448 | 0.11019 | 0.00793 | 0.00826 | 0.00633 | 0.00602 | 0.71221 | 0.15322 |
| **Sgsm1** | 0.08332 | 0.00398 | 1.18668 | 0.00000 | 1.16016 | 0.25778 | 0.01516 | 0.08651 | 1.63444 | 0.04072 | 0.00452 | 0.54107 | 0.47583 | 0.23891 |
| **Sgsm2** | 0.20897 | 0.01188 | 0.79608 | 0.12560 | 0.46612 | 0.37058 | 0.00393 | 0.41644 | 0.35981 | 0.00483 | 0.19716 | 0.37797 | 0.06418 | 0.45719 |
| **Sgsm3** | 1.58062 | 0.04839 | 5.24453 | 3.68211 | 4.90672 | 6.03283 | 0.25250 | 3.17069 | 0.34087 | 0.05958 | 2.26183 | 4.03686 | 6.04103 | 4.93678 |
| **Sgta** | 2.88569 | 0.32928 | 22.49942 | 18.94109 | 25.60197 | 9.31543 | 0.79634 | 13.96148 | 0.07770 | 0.13839 | 7.81202 | 11.38642 | 4.78417 | 0.94747 |
| **Sgtb** | 0.00627 | 0.00403 | 0.12830 | 0.18864 | 0.04802 | 0.05085 | 0.15192 | 0.00322 | 0.08250 | 0.00510 | 0.00433 | 0.07549 | 0.22876 | 0.00000 |
| **Sh2b1** | 1.31167 | 0.03314 | 0.75589 | 2.32342 | 4.59262 | 1.29487 | 0.02023 | 2.32277 | 0.04955 | 0.04623 | 2.80402 | 3.28167 | 2.62557 | 0.42634 |
| **Sh2b2** | 0.59058 | 0.00411 | 0.06910 | 0.36101 | 0.33401 | 0.00000 | 0.00476 | 0.03730 | 0.00267 | 0.00539 | 0.00556 | 0.02996 | 1.04899 | 0.11827 |
| **Sh2b3** | 4.11740 | 3.75500 | 8.74970 | 1.70868 | 6.30098 | 4.67987 | 2.05677 | 2.14054 | 0.54248 | 0.05667 | 3.29646 | 2.46856 | 5.90583 | 3.82796 |
| **Sh2d3c** | 0.43853 | 0.00360 | 1.03743 | 0.22190 | 2.70494 | 0.00000 | 0.00413 | 0.01396 | 0.00643 | 0.02342 | 0.01298 | 0.00719 | 7.59831 | 0.04831 |
| **Sh2d4a** | 0.01933 | 0.01152 | 0.00701 | 0.00482 | 0.01784 | 0.50547 | 0.01067 | 1.02952 | 0.02088 | 0.01423 | 0.01436 | 0.00591 | 0.00965 | 0.01804 |
| **Sh2d4b** | 0.00000 | 0.01920 | 0.50448 | 0.07109 | 0.00000 | 0.22500 | 0.01229 | 0.64859 | 0.01083 | 0.00555 | 0.50501 | 0.65226 | 0.00467 | 0.00990 |
| **Sh2d5** | 0.07600 | 0.00443 | 0.00000 | 0.02403 | 0.20536 | 0.00555 | 0.00395 | 0.00979 | 1.53048 | 0.00883 | 0.00479 | 0.00858 | 0.00751 | 0.00000 |
| **Sh2d7** | 0.00841 | 0.00412 | 0.00915 | 0.00512 | 0.01339 | 0.28025 | 0.00417 | 0.08208 | 0.01172 | 0.01273 | 0.11738 | 0.00472 | 0.00772 | 0.00807 |
| **Sh3bgr** | 0.00127 | 0.00377 | 0.07494 | 0.03735 | 0.40714 | 0.15107 | 0.00396 | 0.00326 | 0.00932 | 0.00510 | 0.00353 | 0.35642 | 0.28565 | 0.00053 |
| **Sh3bgrl** | 0.06526 | 0.00475 | 0.00855 | 0.00868 | 0.21268 | 0.01795 | 0.00457 | 0.01462 | 0.02742 | 0.00714 | 0.01472 | 0.01167 | 0.00472 | 0.01109 |
| **Sh3bgrl2** | 0.00041 | 6.30707 | 0.00000 | 0.24400 | 0.00000 | 2.61536 | 2.76794 | 9.11074 | 0.00885 | 0.12830 | 2.03772 | 2.41726 | 0.06927 | 1.25051 |
| **Sh3bgrl3** | 2.26546 | 0.20764 | 1.04882 | 10.17241 | 4.16314 | 10.68141 | 2.71847 | 5.04939 | 0.23868 | 7.48250 | 1.93004 | 14.73484 | 1.68743 | 1.09202 |
| **Sh3bp1** | 0.03804 | 3.36225 | 0.21263 | 4.71008 | 0.15568 | 0.80058 | 3.49602 | 4.69905 | 0.00895 | 0.10185 | 0.15890 | 4.71442 | 0.00883 | 0.00000 |
| **Sh3bp2** | 0.00012 | 0.02021 | 0.40682 | 0.00000 | 1.49565 | 0.00000 | 0.01243 | 0.66239 | 0.06353 | 0.00513 | 0.00000 | 0.38249 | 0.50394 | 0.33835 |
| **Sh3bp4** | 0.30294 | 0.01021 | 0.10956 | 1.55826 | 0.34323 | 0.11127 | 2.34361 | 0.00342 | 0.01044 | 0.10107 | 0.00000 | 2.56555 | 0.00918 | 0.00082 |
| **Sh3bp5** | 0.00311 | 0.37731 | 0.00000 | 0.63679 | 0.00298 | 0.26436 | 0.00346 | 2.35477 | 0.00944 | 0.01998 | 0.84879 | 0.00105 | 0.14868 | 0.00000 |
| **Sh3bp5l** | 2.22893 | 0.04370 | 9.22788 | 1.94112 | 6.10096 | 2.90218 | 0.40548 | 0.15039 | 1.15260 | 0.00000 | 1.27377 | 1.19151 | 3.67727 | 5.05797 |
| **Sh3d19** | 0.32356 | 0.05286 | 2.78099 | 0.00000 | 0.78667 | 2.38803 | 0.01691 | 0.72988 | 1.89527 | 0.03874 | 0.09002 | 1.49703 | 0.37101 | 2.31532 |
| **Sh3d21** | 2.15193 | 0.19777 | 4.43943 | 3.32088 | 2.55273 | 6.20262 | 1.65363 | 8.06358 | 0.50827 | 0.07901 | 0.28746 | 0.00000 | 0.08219 | 1.38063 |
| **Sh3gl1** | 0.99515 | 4.70491 | 3.04120 | 0.45946 | 1.69899 | 2.23339 | 1.78162 | 0.38911 | 0.24522 | 3.47902 | 3.30111 | 2.89759 | 2.33936 | 4.37891 |
| **Sh3gl2** | 0.01545 | 0.00449 | 0.01428 | 0.03532 | 0.01816 | 0.02233 | 0.01615 | 0.02579 | 0.04745 | 0.01637 | 0.00813 | 0.01475 | 0.00916 | 0.05222 |
| **Sh3gl3** | 0.14873 | 0.01206 | 0.00415 | 0.06372 | 0.00004 | 0.00525 | 0.00374 | 0.00544 | 0.02180 | 0.01244 | 0.00377 | 0.68411 | 0.01203 | 0.45567 |
| **Sh3glb1** | 7.51832 | 0.03984 | 4.49416 | 0.99825 | 6.28401 | 10.10960 | 4.43114 | 1.90345 | 2.60826 | 0.09485 | 5.83057 | 3.68719 | 0.41353 | 2.11156 |
| **Sh3glb2** | 0.00000 | 0.02554 | 2.25641 | 3.41975 | 2.27788 | 0.36544 | 0.61058 | 2.49644 | 0.06022 | 0.02418 | 1.82092 | 2.89641 | 0.13334 | 0.00000 |
| **Sh3kbp1** | 0.17791 | 0.18539 | 0.08170 | 0.04320 | 0.00000 | 0.76780 | 0.00385 | 0.40446 | 0.40169 | 0.00638 | 0.00425 | 0.00936 | 0.00667 | 0.21254 |
| **Sh3pxd2a** | 0.92240 | 0.00373 | 0.39028 | 0.19935 | 1.28863 | 0.45385 | 0.00391 | 0.00000 | 0.02661 | 0.01774 | 0.47226 | 0.01301 | 1.75432 | 0.48512 |
| **Sh3pxd2b** | 0.07708 | 0.00447 | 0.00559 | 0.00812 | 0.33514 | 0.01601 | 0.00461 | 0.01028 | 0.01865 | 0.00712 | 0.00782 | 0.00820 | 0.00408 | 0.00391 |
| **Sh3rf1** | 0.00019 | 0.00398 | 0.01012 | 0.00000 | 0.01379 | 0.03409 | 3.38683 | 0.71688 | 0.12491 | 0.08003 | 0.00000 | 0.47155 | 0.86452 | 0.69851 |
| **Sh3rf3** | 0.00222 | 0.00416 | 0.11140 | 0.03526 | 0.29380 | 0.00822 | 0.00402 | 0.01386 | 0.00496 | 0.00823 | 0.00660 | 0.00569 | 0.76905 | 0.42593 |
| **Sh3tc1** | 0.10619 | 0.05485 | 3.55499 | 1.88523 | 0.43116 | 2.54353 | 3.27754 | 3.31775 | 0.00630 | 9.87012 | 1.17749 | 1.22567 | 0.00786 | 0.00000 |
| **Sh3tc2** | 0.00000 | 0.02711 | 0.99752 | 0.05255 | 0.00000 | 4.15829 | 1.05205 | 0.52550 | 0.00704 | 0.03832 | 1.66080 | 0.72504 | 0.00408 | 0.01170 |
| **Sh3yl1** | 0.88527 | 0.01822 | 1.00994 | 0.00000 | 0.45013 | 1.72660 | 1.44139 | 0.01968 | 0.00195 | 0.03043 | 0.00000 | 1.80708 | 1.47240 | 0.01690 |
| **Sharpin** | 2.71683 | 0.01174 | 3.08812 | 4.23497 | 1.68343 | 1.41079 | 0.01524 | 1.18282 | 0.00035 | 0.10906 | 0.88364 | 0.87538 | 0.00026 | 0.02594 |
| **Shb** | 0.00000 | 0.00346 | 0.50223 | 0.09709 | 0.19142 | 0.30854 | 0.22953 | 0.00107 | 0.00375 | 0.00530 | 0.00375 | 0.55115 | 1.65851 | 0.00000 |
| **Shbg** | 0.31480 | 0.00373 | 1.66438 | 0.00000 | 2.87267 | 0.00000 | 0.00346 | 0.00415 | 0.52560 | 0.00265 | 0.00293 | 0.00302 | 4.05140 | 0.71755 |
| **Shc1** | 0.11123 | 0.12046 | 5.75095 | 2.45880 | 6.01677 | 7.09264 | 0.09136 | 5.61045 | 2.56376 | 0.09999 | 4.90426 | 3.42206 | 1.11331 | 5.39954 |
| **Shc3** | 1.10339 | 0.04313 | 5.68256 | 0.39506 | 2.00051 | 0.67001 | 0.00297 | 1.41958 | 0.02557 | 0.04185 | 4.09605 | 1.31587 | 0.00000 | 0.43525 |
| **Shcbp1** | 6.33445 | 0.01778 | 5.35807 | 2.32590 | 4.90351 | 5.90355 | 3.25098 | 1.98447 | 8.32476 | 8.21751 | 11.15673 | 4.69896 | 0.27468 | 3.43775 |
| **Shcbp1l** | 0.92282 | 0.00398 | 0.17310 | 0.09643 | 0.59395 | 0.00000 | 0.00989 | 0.00541 | 1.50082 | 0.00655 | 0.00582 | 0.01012 | 0.77499 | 0.00000 |
| **Shf** | 0.00220 | 0.02848 | 0.02011 | 0.05806 | 0.00841 | 0.01590 | 0.00352 | 0.02205 | 0.01502 | 0.00744 | 0.00927 | 1.96956 | 0.12479 | 0.00293 |
| **Shfm1** | 4.41264 | 0.70848 | 0.19654 | 0.00000 | 4.86557 | 10.91268 | 4.03505 | 4.62236 | 0.97813 | 8.93681 | 0.14427 | 7.26627 | 8.16274 | 2.89022 |
| **Shisa5** | 0.00000 | 0.07898 | 0.56464 | 0.94413 | 0.14213 | 2.23426 | 0.00792 | 0.78648 | 0.00714 | 0.01791 | 0.00000 | 7.15398 | 0.00296 | 0.06275 |
| **Shisa7** | 0.31567 | 0.01717 | 1.25139 | 2.51274 | 1.07649 | 0.97648 | 0.00878 | 0.00000 | 0.38153 | 0.00438 | 0.00000 | 1.38539 | 0.66830 | 0.09038 |
| **Shisa8** | 0.01309 | 0.00422 | 0.00658 | 0.02608 | 0.37569 | 0.01320 | 0.00426 | 0.00896 | 0.00903 | 0.00627 | 0.00890 | 0.00913 | 0.82461 | 0.00311 |
| **Shkbp1** | 0.00000 | 0.07945 | 5.20343 | 3.26882 | 0.26222 | 0.09032 | 0.02329 | 1.02101 | 0.00000 | 1.01547 | 0.92005 | 8.42527 | 0.10960 | 0.00000 |
| **Shmt1** | 4.26580 | 3.22480 | 12.07856 | 9.12483 | 7.25440 | 8.32494 | 1.75156 | 1.76304 | 6.03283 | 0.54022 | 10.68180 | 14.59914 | 12.37440 | 11.72972 |
| **Shmt2** | 0.54782 | 3.05106 | 8.31860 | 11.57178 | 0.77206 | 7.25224 | 0.52877 | 3.16527 | 0.00000 | 0.08709 | 6.65317 | 8.42393 | 2.63490 | 0.57043 |

| | | | | | | | | | | | | | | |
|---|---|---|---|---|---|---|---|---|---|---|---|---|---|---|
| **Shoc2** | 3.18196 | 0.54882 | 1.96018 | 0.00000 | 0.86284 | 3.63630 | 1.65927 | 2.38844 | 2.39682 | 0.08120 | 0.72277 | 2.79083 | 1.40870 | 0.24266 |
| **Shox2** | 0.69455 | 0.00441 | 0.00298 | 0.01151 | 0.00000 | 0.00512 | 0.00429 | 0.01205 | 0.44623 | 0.00684 | 0.00531 | 0.00790 | 0.00307 | 0.00000 |
| **Shprh** | 3.05916 | 0.00336 | 2.25505 | 0.40350 | 1.29803 | 0.54419 | 0.00331 | 0.00000 | 4.28111 | 0.01374 | 0.08626 | 0.28476 | 0.00000 | 2.89090 |
| **Shq1** | 3.54388 | 1.94884 | 1.20232 | 3.13119 | 2.77141 | 2.93900 | 0.83222 | 0.30269 | 0.00004 | 0.02481 | 0.08544 | 3.88054 | 0.98056 | 2.45976 |
| **Shroom1** | 0.21884 | 0.00441 | 0.10051 | 0.00337 | 0.17806 | 0.00690 | 0.00444 | 0.01367 | 0.00530 | 0.00610 | 0.00691 | 0.00858 | 0.00460 | 0.10737 |
| **Shroom2** | 3.38674 | 0.00392 | 0.00022 | 0.03355 | 0.63889 | 2.17657 | 0.00369 | 1.80551 | 3.53080 | 0.01182 | 0.00921 | 0.00000 | 1.30313 | 0.37588 |
| **Shroom3** | 0.53871 | 1.72303 | 7.32860 | 4.60928 | 0.50743 | 10.31392 | 1.90116 | 8.72249 | 0.02472 | 0.14519 | 9.50664 | 6.62040 | 0.38603 | 0.24430 |
| **Shroom4** | 0.00000 | 0.01125 | 0.33150 | 0.37135 | 0.24515 | 0.24553 | 0.00370 | 0.51844 | 1.12235 | 0.00594 | 0.00339 | 0.00177 | 1.17033 | 0.41461 |
| **Shtn1** | 0.00000 | 0.03204 | 0.00000 | 0.10124 | 0.34428 | 0.00000 | 0.02162 | 0.05043 | 1.32242 | 0.00722 | 0.04544 | 0.43764 | 0.00070 | 1.25851 |
| **Siae** | 0.00424 | 0.00463 | 0.16703 | 0.00000 | 0.32247 | 0.66813 | 0.00406 | 0.00524 | 0.29032 | 0.01712 | 0.02802 | 0.01337 | 0.00609 | 0.00000 |
| **Siah1a** | 4.60397 | 0.00239 | 1.52432 | 0.44841 | 7.92529 | 1.51336 | 0.00216 | 1.33001 | 12.41430 | 0.00274 | 0.16161 | 0.21389 | 6.47759 | 12.40919 |
| **Siah1b** | 6.16286 | 0.00903 | 1.03972 | 0.99453 | 11.26209 | 3.16112 | 0.00277 | 0.30619 | 7.88137 | 0.00481 | 1.95009 | 1.03263 | 8.01204 | 6.45231 |
| **Siah2** | 0.00000 | 4.30917 | 0.30963 | 0.00000 | 1.46537 | 0.89763 | 2.68298 | 1.55100 | 0.92314 | 0.05663 | 1.14265 | 0.31912 | 13.42038 | 6.11910 |
| **Sidt2** | 5.33260 | 6.34204 | 19.53654 | 18.06559 | 7.71364 | 15.92713 | 7.49756 | 10.69194 | 1.76231 | 0.94672 | 8.94321 | 2.53314 | 1.91064 | 13.03807 |
| **Sigirr** | 0.97219 | 7.18368 | 3.38627 | 2.87565 | 1.03585 | 1.19296 | 0.01417 | 1.32727 | 0.00007 | 0.01005 | 2.05792 | 0.00012 | 0.97599 | 0.54695 |
| **Siglecg** | 1.70648 | 0.00348 | 1.77353 | 0.89720 | 1.50909 | 0.00000 | 0.00345 | 0.00000 | 0.20594 | 0.00000 | 1.10936 | 0.07447 | 0.00000 | 0.29415 |
| **Sigmar1** | 0.83411 | 0.02037 | 0.61131 | 3.30139 | 1.28638 | 1.76148 | 0.00337 | 0.00000 | 0.11789 | 0.00228 | 1.24088 | 1.56822 | 1.86844 | 0.72166 |
| **Sik1** | 0.02108 | 0.02027 | 0.12323 | 1.48991 | 1.00133 | 0.38083 | 0.02179 | 0.00835 | 0.01822 | 0.00847 | 0.01001 | 2.16626 | 0.21479 | 0.41781 |
| **Sik2** | 0.00628 | 0.00356 | 0.00000 | 0.01645 | 0.23115 | 0.00000 | 0.00399 | 0.09344 | 0.01777 | 0.00486 | 0.93492 | 0.18227 | 0.00659 | 0.00676 |
| **Sik3** | 1.95817 | 0.19304 | 4.57790 | 0.00000 | 4.37389 | 0.93646 | 0.00262 | 2.13062 | 0.86502 | 0.00086 | 1.89300 | 2.33599 | 0.53079 | 2.99839 |
| **Sike1** | 4.46759 | 0.05262 | 2.53037 | 1.01613 | 4.85037 | 2.39593 | 2.83626 | 2.87793 | 2.49458 | 0.08926 | 2.11652 | 1.94969 | 2.70069 | 4.40310 |
| **Sil1** | 1.02049 | 0.00336 | 2.25056 | 3.77258 | 2.73184 | 3.15091 | 0.90289 | 0.00000 | 0.02375 | 0.02584 | 0.74891 | 0.67690 | 0.89944 | 0.56032 |
| **Simc1** | 2.89399 | 0.00296 | 2.66350 | 0.59779 | 3.06231 | 1.08602 | 0.21132 | 0.43981 | 6.42885 | 0.02805 | 2.32794 | 0.79480 | 1.98622 | 4.26975 |
| **Sin3a** | 5.30437 | 0.11235 | 3.73821 | 1.34232 | 3.99278 | 2.66695 | 2.77228 | 4.05799 | 5.19879 | 0.12041 | 1.15452 | 2.14260 | 5.90123 | 6.80513 |
| **Sin3b** | 4.52130 | 11.53919 | 12.04360 | 16.13920 | 6.62271 | 10.70479 | 4.77369 | 15.90197 | 0.90178 | 0.43830 | 4.67769 | 14.53310 | 4.02580 | 0.78350 |
| **Sipa1** | 0.22411 | 0.18154 | 2.27489 | 1.11243 | 3.41801 | 0.92996 | 0.02007 | 0.00000 | 0.00043 | 0.00768 | 0.01962 | 3.13349 | 0.00000 | 0.63185 |
| **Sipa1l1** | 0.01000 | 0.11588 | 1.07537 | 0.83878 | 0.14330 | 1.13865 | 0.06675 | 1.24248 | 0.00143 | 0.01788 | 0.09441 | 0.97817 | 0.20941 | 1.89975 |
| **Sipa1l2** | 0.00000 | 0.11289 | 0.23135 | 0.00000 | 0.00000 | 4.30661 | 2.78874 | 6.25066 | 0.10924 | 0.07403 | 5.28214 | 5.34296 | 1.29821 | 1.28226 |
| **Sipa1l3** | 0.05573 | 0.01245 | 0.12438 | 0.35748 | 0.13263 | 0.04951 | 0.00424 | 0.06669 | 0.15315 | 0.00509 | 0.00994 | 1.01239 | 0.63860 | 0.00000 |
| **Sirpa** | 0.01907 | 0.00463 | 0.00779 | 0.00841 | 0.27422 | 0.01232 | 0.00432 | 0.01283 | 0.05019 | 0.00632 | 0.00662 | 0.06289 | 0.00443 | 0.00619 |
| **Sirt1** | 11.97019 | 2.35602 | 6.71584 | 2.00418 | 10.27186 | 4.74845 | 1.44724 | 0.66146 | 24.85202 | 0.05289 | 1.94811 | 2.42140 | 15.71308 | 10.14486 |
| **Sirt2** | 3.49675 | 0.02947 | 4.79939 | 1.21566 | 1.96894 | 4.82337 | 4.79280 | 4.23760 | 0.01583 | 0.13508 | 1.77067 | 3.62241 | 0.84731 | 3.76445 |
| **Sirt3** | 0.00344 | 0.02768 | 0.00000 | 0.59870 | 0.17525 | 0.66582 | 0.01642 | 1.69217 | 0.07800 | 0.09187 | 0.81988 | 0.70643 | 0.01521 | 0.10645 |
| **Sirt4** | 0.00000 | 0.00408 | 0.59192 | 0.35978 | 0.60541 | 0.00000 | 0.00407 | 0.01353 | 0.13930 | 0.01221 | 0.01266 | 0.09586 | 0.48574 | 0.10585 |
| **Sirt5** | 1.47758 | 0.01951 | 0.53611 | 1.87392 | 0.42441 | 0.48345 | 1.94094 | 0.03319 | 0.03274 | 0.04204 | 0.04148 | 0.61325 | 0.95650 | 0.10172 |
| **Sirt6** | 3.87886 | 0.04870 | 9.93739 | 4.99617 | 7.22171 | 2.16856 | 0.01368 | 1.57670 | 0.00000 | 0.01941 | 2.82223 | 0.06558 | 0.14536 | 0.87191 |
| **Sirt7** | 2.07461 | 0.05124 | 7.28543 | 3.04574 | 7.72012 | 1.96399 | 3.95789 | 2.95680 | 0.00000 | 0.09386 | 3.20131 | 1.24067 | 4.30904 | 0.00000 |
| **Siva1** | 2.91045 | 0.13942 | 1.16280 | 7.53191 | 12.11968 | 5.22455 | 0.04158 | 6.48680 | 0.00000 | 6.29483 | 0.68961 | 5.94164 | 4.06489 | 1.48106 |
| **Six1** | 2.14481 | 0.00354 | 0.00474 | 0.21061 | 0.09432 | 0.01134 | 1.70936 | 0.00759 | 0.00771 | 0.01695 | 0.00319 | 0.21547 | 0.00762 | 0.00186 |
| **Six2** | 0.02228 | 0.02145 | 0.04240 | 0.04896 | 0.14103 | 0.02931 | 0.00429 | 0.04763 | 0.04313 | 0.00626 | 0.00952 | 0.04146 | 0.01298 | 0.01231 |
| **Six4** | 0.00000 | 0.00385 | 0.00000 | 0.00539 | 0.01289 | 0.01859 | 0.00999 | 0.00562 | 0.63322 | 0.00668 | 0.00225 | 0.51098 | 0.51743 | 0.18332 |
| **Six5** | 0.00613 | 0.00380 | 0.00182 | 0.05859 | 0.00702 | 0.53369 | 2.05533 | 0.01000 | 0.03599 | 0.00594 | 0.00194 | 0.10460 | 0.08756 | 0.00660 |
| **Ska1** | 10.14752 | 4.86959 | 7.41009 | 7.22055 | 12.08078 | 7.49363 | 3.97244 | 5.45300 | 5.68798 | 0.19113 | 4.76664 | 4.86355 | 11.29002 | 1.38538 |
| **Ska2** | 0.26599 | 0.01615 | 0.94331 | 1.51134 | 0.78659 | 2.67182 | 0.00240 | 0.14671 | 5.58713 | 0.01608 | 0.42205 | 1.38732 | 5.91624 | 2.91371 |
| **Ska3** | 4.29340 | 0.12217 | 5.01312 | 0.79698 | 5.21970 | 5.93445 | 0.32121 | 4.38177 | 5.35480 | 0.02268 | 0.34938 | 4.44328 | 5.99710 | 6.49911 |
| **Skap1** | 0.30872 | 0.00461 | 0.06291 | 0.00000 | 0.16932 | 0.17556 | 0.00413 | 0.00583 | 0.19675 | 0.00642 | 0.00586 | 0.00634 | 0.01883 | 0.50246 |
| **Skap2** | 1.80750 | 0.00609 | 0.96626 | 0.81996 | 0.32118 | 4.15287 | 0.33589 | 0.89837 | 3.08923 | 0.02624 | 2.22584 | 5.38476 | 6.01709 | 4.82276 |
| **Ski** | 9.35384 | 8.08293 | 13.41694 | 6.23807 | 7.81553 | 3.09414 | 2.19854 | 4.98173 | 0.18740 | 6.13436 | 3.69275 | 3.86686 | 4.22402 | 1.40307 |
| **Skida1** | 0.06863 | 0.02713 | 0.49465 | 0.31019 | 0.13040 | 0.55637 | 1.17915 | 1.35892 | 0.00806 | 0.01678 | 0.00000 | 0.16276 | 0.00768 | 0.03989 |
| **Skil** | 0.84721 | 0.00307 | 0.59005 | 0.00000 | 0.00000 | 0.06288 | 0.02171 | 0.91869 | 1.71354 | 0.01460 | 0.93795 | 1.13839 | 0.29910 | 0.00000 |
| **Skint2** | 0.01587 | 0.00449 | 0.01585 | 0.02831 | 0.01200 | 0.01055 | 0.00488 | 0.03227 | 0.01673 | 0.00642 | 0.00828 | 0.00804 | 0.01716 | 0.03132 |
| **Skint6** | 0.00723 | 0.00440 | 0.01484 | 0.01253 | 0.01207 | 0.01333 | 0.00428 | 0.00979 | 0.01280 | 0.00671 | 0.00892 | 0.00979 | 0.13079 | 0.00401 |
| **Skiv2l** | 0.00000 | 0.05903 | 2.83066 | 7.09682 | 1.41728 | 2.47199 | 1.09560 | 2.06869 | 0.01896 | 0.03140 | 0.35608 | 2.78519 | 0.52508 | 0.38374 |
| **Skiv2l2** | 20.37705 | 6.92376 | 17.42074 | 5.99652 | 23.49548 | 11.17027 | 3.84939 | 6.47893 | 28.24037 | 0.12242 | 6.78568 | 6.03717 | 9.96542 | 23.11454 |
| **Skp1a** | 9.89807 | 1.62968 | 4.85820 | 10.66531 | 6.14791 | 19.31606 | 3.99595 | 7.27202 | 8.01358 | 15.56344 | 8.01704 | 10.78383 | 7.41783 | 7.57197 |

| | | | | | | | | | | | | | | |
|---|---|---|---|---|---|---|---|---|---|---|---|---|---|---|
| **Skp2** | 7.22499 | 0.00899 | 7.89672 | 1.17510 | 8.45019 | 1.96849 | 0.04910 | 0.00000 | 0.50228 | 0.00000 | 1.13236 | 0.51111 | 9.09973 | 10.45907 |
| **Sla2** | 0.02291 | 0.00446 | 0.03823 | 0.11753 | 0.10245 | 0.29432 | 0.00456 | 0.00542 | 0.01288 | 0.00586 | 0.00459 | 0.00519 | 0.00467 | 0.04530 |
| **Slain1** | 0.41221 | 0.00273 | 2.03939 | 0.00000 | 5.14564 | 0.00000 | 0.00342 | 0.00980 | 0.00000 | 0.00284 | 0.00375 | 0.00000 | 12.77843 | 2.16990 |
| **Slain2** | 2.72568 | 0.06577 | 4.55069 | 1.32960 | 6.77818 | 4.14722 | 0.06645 | 0.49549 | 3.64184 | 0.00119 | 4.71534 | 4.44140 | 6.90251 | 5.04700 |
| **Slamf7** | 0.00000 | 0.00422 | 0.31126 | 0.00000 | 0.12217 | 0.00187 | 0.00450 | 0.00865 | 1.03750 | 0.00713 | 0.00542 | 0.00634 | 0.79583 | 0.28812 |
| **Slamf8** | 0.01707 | 0.01183 | 0.02304 | 0.02466 | 0.01795 | 0.04924 | 0.01021 | 0.00911 | 0.08898 | 0.00799 | 0.01858 | 0.03775 | 0.29099 | 0.02927 |
| **Slamf9** | 0.00000 | 0.01246 | 1.14746 | 0.05066 | 0.24825 | 0.20407 | 0.00324 | 0.68132 | 0.00682 | 6.68031 | 0.01037 | 0.00772 | 0.04476 | 0.00000 |
| **Slbp** | 18.47147 | 0.28609 | 21.90885 | 15.85213 | 20.96519 | 13.24740 | 5.48234 | 14.40063 | 14.46563 | 0.25254 | 6.04967 | 8.32708 | 15.17073 | 12.43086 |
| **Slc10a1** | 0.02885 | 0.00539 | 0.00000 | 0.21199 | 0.06353 | 0.21834 | 0.00434 | 0.22301 | 0.82780 | 0.01687 | 0.11754 | 0.01235 | 0.04012 | 0.18349 |
| **Slc10a2** | 0.02951 | 0.00467 | 0.02330 | 0.02178 | 0.02144 | 0.03190 | 0.01046 | 0.01755 | 0.20039 | 0.00697 | 0.01239 | 0.00992 | 0.03677 | 0.00402 |
| **Slc10a3** | 1.73699 | 0.00272 | 0.87232 | 0.64465 | 1.81138 | 0.96644 | 0.00850 | 0.70708 | 1.87390 | 0.01744 | 0.00000 | 1.00432 | 2.46723 | 5.99766 |
| **Slc10a4** | 0.00700 | 0.00435 | 0.01808 | 0.17580 | 0.00933 | 0.01066 | 0.00421 | 0.00893 | 0.00995 | 0.00640 | 0.00748 | 0.00950 | 0.00587 | 0.02168 |
| **Slc10a6** | 1.14658 | 0.00417 | 3.32631 | 1.37237 | 0.60242 | 1.06346 | 1.57453 | 0.00000 | 0.01061 | 0.00000 | 0.00000 | 0.00000 | 1.94882 | 1.44156 |
| **Slc10a7** | 0.50791 | 0.00991 | 4.29450 | 1.96889 | 1.21881 | 3.10889 | 2.24489 | 0.00000 | 3.01963 | 0.05602 | 2.11893 | 2.57117 | 0.58225 | 0.39554 |
| **Slc11a1** | 0.06242 | 0.00423 | 0.58828 | 0.00000 | 0.10337 | 0.14443 | 0.00412 | 0.00560 | 0.19390 | 0.02482 | 0.01954 | 0.02785 | 1.10685 | 0.09178 |
| **Slc11a2** | 6.28579 | 0.07585 | 16.47557 | 2.57667 | 10.12057 | 8.84533 | 1.45139 | 4.02901 | 3.83889 | 8.01055 | 4.26903 | 3.08403 | 5.58469 | 13.76821 |
| **Slc12a2** | 24.50041 | 0.00000 | 16.99795 | 2.20001 | 19.07707 | 0.28322 | 0.00000 | 1.26740 | 35.12103 | 0.01090 | 0.66699 | 2.01187 | 13.17137 | 31.12194 |
| **Slc12a4** | 0.39903 | 0.00295 | 7.91068 | 6.04184 | 1.34943 | 2.95130 | 0.01322 | 0.37766 | 0.01192 | 0.00000 | 0.26677 | 1.37970 | 0.00000 | 1.84093 |
| **Slc12a5** | 0.00828 | 0.00415 | 0.00601 | 0.25424 | 0.01287 | 0.02062 | 0.00396 | 0.00499 | 0.01507 | 0.00557 | 0.00549 | 0.01649 | 0.05875 | 0.07908 |
| **Slc12a6** | 0.02949 | 0.01911 | 1.26744 | 0.85783 | 1.50841 | 1.40351 | 0.00313 | 1.30400 | 4.72892 | 0.01207 | 0.00000 | 1.99716 | 0.29084 | 0.31286 |
| **Slc12a7** | 8.37211 | 0.08319 | 24.80441 | 6.69712 | 3.98326 | 4.79409 | 2.92212 | 3.16995 | 6.64866 | 0.00000 | 4.57572 | 8.85359 | 0.20052 | 8.08958 |
| **Slc12a8** | 0.09410 | 0.07438 | 0.41781 | 0.00000 | 0.48587 | 0.05017 | 0.00428 | 0.02336 | 0.55149 | 0.01265 | 0.00929 | 0.01959 | 0.49700 | 0.19896 |
| **Slc12a9** | 0.17371 | 0.01853 | 1.49872 | 0.81170 | 0.84911 | 0.35400 | 0.00336 | 0.00000 | 0.13515 | 0.00583 | 0.00106 | 0.88147 | 1.06136 | 0.00000 |
| **Slc13a2** | 6.35773 | 4.66670 | 9.79021 | 0.00000 | 5.63500 | 1.54575 | 0.33707 | 4.26456 | 2.93296 | 0.01139 | 0.07610 | 0.00000 | 0.00000 | 11.50276 |
| **Slc13a2os** | 0.20877 | 0.03374 | 0.24347 | 0.01459 | 0.36443 | 0.06367 | 0.00398 | 0.05627 | 0.11572 | 0.00813 | 0.00550 | 0.00432 | 0.55568 | 0.58732 |
| **Slc13a3** | 0.01145 | 0.00408 | 0.14475 | 0.00431 | 0.03367 | 0.00722 | 0.00415 | 0.01027 | 0.01065 | 0.00691 | 0.00742 | 0.00868 | 1.67475 | 0.03988 |
| **Slc13a5** | 0.00343 | 0.11059 | 0.01413 | 0.00000 | 0.00920 | 0.00479 | 0.00284 | 0.27189 | 0.01052 | 0.00690 | 0.00541 | 5.31843 | 0.00638 | 0.02518 |
| **Slc14a1** | 0.00735 | 0.00445 | 0.00327 | 0.03334 | 0.83269 | 0.01363 | 0.00455 | 0.00917 | 0.01863 | 0.00612 | 0.00790 | 0.00672 | 0.00465 | 0.00592 |
| **Slc15a1** | 0.07010 | 0.01301 | 0.05129 | 0.09932 | 0.04359 | 0.07157 | 0.05126 | 0.09852 | 0.15474 | 0.01582 | 0.01795 | 0.03265 | 0.08869 | 0.15519 |
| **Slc15a2** | 0.00266 | 2.77793 | 0.00000 | 0.00000 | 0.00471 | 0.00000 | 4.38192 | 15.14679 | 0.00510 | 1.71204 | 0.20862 | 0.12215 | 0.26918 | 0.02244 |
| **Slc15a4** | 1.19477 | 0.04279 | 3.45118 | 0.00000 | 1.15464 | 6.33902 | 1.54275 | 0.94323 | 0.32336 | 0.06486 | 3.46149 | 1.61671 | 0.09578 | 3.56692 |
| **Slc16a1** | 0.00000 | 4.21155 | 2.66974 | 4.24737 | 0.00000 | 5.35380 | 2.36113 | 2.48045 | 5.67790 | 0.03298 | 0.31821 | 0.08650 | 2.35908 | 8.58790 |
| **Slc16a10** | 0.13301 | 0.00873 | 1.75035 | 0.00000 | 1.36773 | 1.70787 | 0.00827 | 0.00000 | 0.42317 | 0.00727 | 0.16889 | 2.44645 | 4.62499 | 4.10419 |
| **Slc16a11** | 0.01685 | 0.02469 | 0.01334 | 0.00000 | 0.04636 | 0.00459 | 0.01746 | 0.38932 | 0.01220 | 0.01614 | 0.00777 | 0.01660 | 0.00475 | 0.07627 |
| **Slc16a13** | 0.01533 | 0.05218 | 0.56853 | 1.78836 | 0.07163 | 2.72864 | 0.54322 | 1.81272 | 0.03987 | 0.01131 | 0.00000 | 1.10361 | 0.13195 | 0.49037 |
| **Slc16a14** | 0.04098 | 0.00437 | 0.01070 | 0.01227 | 0.07010 | 0.01241 | 0.00422 | 0.00937 | 0.02953 | 0.00768 | 0.00771 | 0.00867 | 0.21063 | 0.18341 |
| **Slc16a3** | 0.15313 | 0.16369 | 0.04090 | 0.25310 | 0.14085 | 0.03292 | 0.00390 | 0.61380 | 0.01362 | 0.01751 | 0.02592 | 0.00314 | 0.00739 | 0.07350 |
| **Slc16a4** | 0.00784 | 0.00407 | 0.01724 | 0.08152 | 0.06940 | 0.01076 | 0.00409 | 0.00566 | 0.01879 | 0.00681 | 0.00564 | 0.19499 | 0.00573 | 0.09723 |
| **Slc16a5** | 0.66627 | 0.01228 | 1.87774 | 0.00000 | 2.19972 | 0.54193 | 0.00465 | 1.12749 | 0.10463 | 0.00331 | 0.00179 | 0.32531 | 0.73349 | 2.93048 |
| **Slc16a6** | 0.00000 | 0.02045 | 5.55223 | 0.79344 | 3.63475 | 0.48106 | 0.00979 | 0.43129 | 1.75547 | 0.00547 | 0.00000 | 0.29929 | 1.07638 | 16.35431 |
| **Slc17a5** | 0.16429 | 0.01241 | 2.23141 | 1.02563 | 2.48034 | 0.85678 | 0.04013 | 0.18004 | 1.77761 | 0.03494 | 0.60500 | 1.32504 | 0.21729 | 3.33884 |
| **Slc17a7** | 0.00798 | 0.00439 | 0.01295 | 0.08671 | 0.03662 | 0.01362 | 0.00428 | 0.00830 | 0.01074 | 0.00614 | 0.00855 | 0.00897 | 0.00673 | 0.01323 |
| **Slc17a8** | 0.05564 | 0.11710 | 0.01234 | 0.32354 | 0.06281 | 0.31146 | 0.04529 | 0.06295 | 0.20652 | 0.03845 | 0.01557 | 0.06656 | 0.05072 | 0.23430 |
| **Slc17a9** | 0.00485 | 0.00429 | 0.01081 | 0.12804 | 0.05415 | 0.01273 | 0.00416 | 0.00701 | 0.00836 | 0.00616 | 0.00732 | 0.00747 | 0.10415 | 0.02505 |
| **Slc18a2** | 0.02689 | 0.00421 | 0.01215 | 0.02410 | 0.06490 | 0.03908 | 0.00377 | 0.00553 | 3.89329 | 0.00665 | 0.00293 | 0.10332 | 0.28070 | 0.19364 |
| **Slc18b1** | 0.56679 | 0.24208 | 0.00017 | 0.37187 | 0.47413 | 0.00000 | 0.03774 | 0.04494 | 0.99202 | 0.00660 | 0.00074 | 0.32819 | 0.80562 | 0.28625 |
| **Slc19a1** | 0.05707 | 0.01011 | 0.89218 | 0.01216 | 0.66267 | 0.50516 | 1.43538 | 0.00417 | 0.01388 | 0.02022 | 0.00000 | 1.37741 | 0.00818 | 1.12301 |
| **Slc19a2** | 1.34462 | 0.00389 | 0.55266 | 0.17583 | 0.00000 | 0.47886 | 0.00417 | 0.07355 | 1.63810 | 0.00712 | 0.02039 | 0.37617 | 0.02183 | 0.50800 |
| **Slc19a3** | 3.85440 | 0.00223 | 0.07052 | 0.00002 | 1.37631 | 0.00000 | 0.00270 | 0.06041 | 9.41507 | 0.00620 | 0.00197 | 0.00000 | 10.00561 | 10.08641 |
| **Slc1a1** | 10.85944 | 0.01860 | 17.17308 | 12.30599 | 9.05100 | 5.17519 | 2.14399 | 4.35062 | 10.75334 | 0.12066 | 5.97251 | 8.02623 | 4.48761 | 5.61468 |
| **Slc1a3** | 0.00000 | 0.05549 | 0.26252 | 0.00000 | 0.54449 | 0.17691 | 0.00310 | 0.00987 | 0.84775 | 0.00537 | 0.00000 | 3.69545 | 0.00075 | 3.65628 |
| **Slc1a4** | 0.00532 | 0.02371 | 0.00148 | 0.38584 | 0.00307 | 0.00000 | 0.00311 | 0.00234 | 0.01500 | 0.00450 | 0.11816 | 2.36417 | 1.70662 | 0.00000 |
| **Slc1a5** | 2.79939 | 2.30784 | 2.71299 | 2.99781 | 3.21655 | 1.02855 | 2.33073 | 1.48058 | 0.52794 | 0.07774 | 0.00000 | 0.97810 | 6.09472 | 4.47319 |
| **Slc1a7** | 0.04033 | 0.00450 | 0.02179 | 0.06229 | 0.05550 | 0.12839 | 0.01692 | 0.03696 | 0.15904 | 0.02119 | 0.02450 | 0.03410 | 0.44644 | 0.15163 |
| **Slc20a1** | 2.53583 | 0.13004 | 14.49738 | 25.06660 | 9.29440 | 29.81328 | 10.13919 | 12.00706 | 15.31178 | 1.85610 | 15.94036 | 5.77494 | 0.89179 | 12.93201 |

| | | | | | | | | | | | | | | |
|---|---|---|---|---|---|---|---|---|---|---|---|---|---|---|
| **Slc20a2** | 0.00000 | 0.04085 | 0.37237 | 1.26753 | 0.00000 | 1.28370 | 0.00302 | 0.43566 | 0.00000 | 0.02585 | 2.64995 | 3.15105 | 1.02604 | 1.08077 |
| **Slc22a13** | 0.99351 | 0.00422 | 5.11823 | 0.69297 | 1.82285 | 0.40297 | 0.00308 | 0.00000 | 0.96012 | 0.00000 | 0.00000 | 0.12803 | 0.00000 | 0.57269 |
| **Slc22a13b.ps** | 2.15580 | 0.01146 | 6.29217 | 0.77488 | 2.55093 | 0.65128 | 0.00264 | 0.00000 | 1.81580 | 0.00000 | 0.00181 | 0.48207 | 0.00000 | 1.09805 |
| **Slc22a15** | 0.07449 | 0.01237 | 0.03951 | 0.08004 | 0.05179 | 0.13096 | 0.02167 | 0.01214 | 0.12488 | 0.01801 | 0.00874 | 0.05607 | 0.43218 | 0.24571 |
| **Slc22a16** | 0.00918 | 0.00390 | 0.07478 | 0.00130 | 0.01153 | 0.00019 | 0.00425 | 0.00859 | 0.01383 | 0.00661 | 0.60327 | 0.16781 | 0.00766 | 0.00821 |
| **Slc22a17** | 0.10879 | 0.01095 | 2.78936 | 0.00627 | 0.08563 | 0.78643 | 0.01407 | 2.31817 | 0.00251 | 0.01640 | 0.03549 | 0.00959 | 0.12627 | 0.00000 |
| **Slc22a20** | 0.29371 | 0.01237 | 0.05807 | 0.94268 | 1.27207 | 0.20500 | 0.00353 | 0.00233 | 1.87315 | 0.00562 | 0.31671 | 0.24916 | 1.49326 | 1.43029 |
| **Slc22a21** | 0.00000 | 0.01301 | 0.08241 | 0.00000 | 0.00392 | 0.13275 | 0.08070 | 0.09475 | 0.01020 | 0.00558 | 0.27269 | 0.39316 | 0.62090 | 0.04170 |
| **Slc22a23** | 0.00242 | 0.06804 | 0.00000 | 0.07159 | 0.14531 | 3.69127 | 3.63043 | 2.54869 | 0.00079 | 0.08404 | 0.00000 | 0.32633 | 0.58250 | 0.31166 |
| **Slc22a28** | 0.00628 | 0.00439 | 0.01495 | 0.01244 | 0.01157 | 0.01343 | 0.00427 | 0.00957 | 0.01122 | 0.00668 | 0.00897 | 0.00964 | 0.19184 | 0.00318 |
| **Slc22a5** | 0.00000 | 0.00419 | 0.01912 | 0.00000 | 0.00000 | 1.11378 | 5.00203 | 2.09755 | 0.00371 | 0.13785 | 0.00000 | 0.30090 | 0.37631 | 1.33685 |
| **Slc22a7** | 0.00481 | 0.00439 | 0.00592 | 0.29615 | 0.09018 | 0.00656 | 0.00404 | 0.01093 | 0.01213 | 0.00647 | 0.00677 | 0.03449 | 0.00431 | 0.06434 |
| **Slc23a1** | 0.64889 | 0.02826 | 0.00000 | 0.39383 | 0.11531 | 0.79613 | 0.09584 | 0.15748 | 1.10235 | 0.20611 | 0.04356 | 0.21421 | 5.29535 | 0.88724 |
| **Slc23a2** | 2.43959 | 0.00124 | 3.27774 | 3.55156 | 2.00279 | 3.36147 | 0.01453 | 0.87657 | 4.37528 | 0.02591 | 0.24459 | 2.62983 | 4.82929 | 3.75273 |
| **Slc23a3** | 0.71635 | 0.00458 | 0.33296 | 0.00000 | 0.23383 | 0.04523 | 0.00422 | 0.00666 | 0.00615 | 0.00571 | 0.00568 | 0.00734 | 0.52089 | 0.00890 |
| **Slc24a3** | 0.03146 | 0.00437 | 0.00000 | 0.08276 | 0.00221 | 0.00448 | 0.00401 | 0.00838 | 0.74583 | 0.00772 | 0.00514 | 0.01258 | 0.00430 | 0.00000 |
| **Slc24a5** | 0.00000 | 0.03892 | 1.30506 | 2.18854 | 0.00000 | 2.01219 | 0.00821 | 1.34881 | 0.28590 | 0.06699 | 5.70598 | 5.01545 | 0.00000 | 0.96029 |
| **Slc25a1** | 0.04091 | 0.05707 | 0.00000 | 0.11661 | 0.02674 | 0.86392 | 0.00266 | 0.09340 | 0.00751 | 0.02518 | 0.00000 | 5.81971 | 0.01675 | 0.01050 |
| **Slc25a10** | 1.28822 | 0.10262 | 4.19859 | 7.55158 | 3.56056 | 1.88213 | 2.89118 | 6.62659 | 0.00069 | 5.67395 | 2.61077 | 3.96126 | 0.67259 | 0.19161 |
| **Slc25a11** | 2.16022 | 2.47257 | 4.94506 | 10.99292 | 5.82114 | 10.01191 | 1.65626 | 10.66405 | 0.58181 | 0.12864 | 0.24515 | 8.96301 | 2.57327 | 3.06919 |
| **Slc25a12** | 5.33273 | 0.01677 | 8.90162 | 0.32258 | 13.57343 | 0.00000 | 0.00171 | 1.06437 | 3.22101 | 0.00353 | 0.00000 | 0.02120 | 1.37063 | 11.48514 |
| **Slc25a13** | 1.00949 | 0.01833 | 3.48497 | 3.88978 | 5.69137 | 7.03687 | 3.47377 | 0.41131 | 6.84286 | 0.06540 | 4.73065 | 10.33127 | 1.46391 | 7.82463 |
| **Slc25a14** | 4.64309 | 0.02656 | 1.01632 | 1.97820 | 1.74792 | 5.45185 | 1.38183 | 1.35929 | 2.98798 | 0.02770 | 2.91470 | 1.02551 | 0.80277 | 3.50330 |
| **Slc25a15** | 3.54752 | 0.04207 | 1.62408 | 2.85407 | 2.35009 | 2.23309 | 0.00253 | 1.15076 | 5.98345 | 0.00737 | 3.27452 | 2.87869 | 1.91582 | 7.65588 |
| **Slc25a16** | 3.32639 | 0.00798 | 2.99511 | 0.98169 | 2.56576 | 3.02522 | 0.22889 | 0.32648 | 6.08535 | 0.00598 | 0.12572 | 3.77821 | 3.83018 | 7.92172 |
| **Slc25a17** | 6.74875 | 0.05026 | 8.43389 | 6.39717 | 8.21258 | 8.43769 | 3.18624 | 0.52437 | 9.24726 | 0.07843 | 5.87370 | 9.03985 | 3.18840 | 11.46657 |
| **Slc25a18** | 0.25951 | 0.00392 | 0.48565 | 0.00000 | 0.40802 | 0.00000 | 0.00497 | 0.12221 | 0.04143 | 0.00544 | 0.00208 | 0.72016 | 0.37926 | 0.01251 |
| **Slc25a19** | 1.70170 | 0.04202 | 5.32966 | 5.31387 | 6.53113 | 4.17095 | 0.05926 | 2.77269 | 0.00000 | 0.04190 | 1.60433 | 1.44262 | 0.99428 | 2.83329 |
| **Slc25a2** | 0.06120 | 1.74235 | 0.32807 | 0.00194 | 0.00284 | 0.13617 | 0.00333 | 0.00019 | 0.01044 | 0.00531 | 0.00617 | 0.00425 | 0.00956 | 0.00997 |
| **Slc25a20** | 4.54871 | 0.01689 | 11.14972 | 5.81874 | 11.39637 | 8.34446 | 0.91741 | 1.83652 | 10.42438 | 0.07518 | 4.80493 | 3.09243 | 1.94038 | 14.71004 |
| **Slc25a21** | 0.00000 | 0.00353 | 1.52428 | 2.72445 | 0.29679 | 0.22654 | 0.00343 | 0.00000 | 0.11866 | 0.00000 | 1.87407 | 0.02615 | 0.03780 | 0.00000 |
| **Slc25a22** | 2.00832 | 5.91894 | 4.29811 | 3.21293 | 2.52439 | 0.28280 | 0.86458 | 0.09937 | 0.16766 | 0.03350 | 1.02334 | 0.92220 | 2.33599 | 1.99598 |
| **Slc25a23** | 1.84981 | 0.01117 | 0.52667 | 0.00000 | 0.95850 | 0.08103 | 0.06699 | 0.90159 | 0.10230 | 0.01217 | 0.00156 | 0.07591 | 1.02998 | 1.01708 |
| **Slc25a24** | 0.04332 | 0.00487 | 0.07988 | 0.00000 | 0.01174 | 0.00000 | 0.00457 | 0.42336 | 0.00593 | 0.00549 | 0.00135 | 0.42969 | 0.00500 | 0.23732 |
| **Slc25a25** | 5.74202 | 0.06136 | 4.48697 | 1.35538 | 13.44524 | 2.10916 | 3.11477 | 2.69624 | 0.29223 | 0.08394 | 1.11314 | 0.00000 | 10.30027 | 16.16699 |
| **Slc25a26** | 0.61945 | 0.04285 | 0.93367 | 2.16548 | 1.18752 | 2.01225 | 0.65457 | 0.93951 | 0.10229 | 9.48184 | 1.69282 | 1.89206 | 0.42312 | 3.09690 |
| **Slc25a27** | 0.02899 | 0.01196 | 0.02454 | 0.05039 | 0.05049 | 0.01175 | 0.00423 | 0.00742 | 0.03351 | 0.00592 | 0.01893 | 0.01153 | 0.20816 | 0.00797 |
| **Slc25a28** | 6.56273 | 2.19570 | 6.57184 | 3.89642 | 8.13377 | 1.44134 | 1.31242 | 3.80208 | 1.11255 | 0.05291 | 2.14832 | 0.50670 | 10.99401 | 7.14434 |
| **Slc25a3** | 32.66197 | 10.37015 | 25.86998 | 41.79990 | 35.22488 | 22.14818 | 7.27701 | 20.55466 | 10.76064 | 13.03332 | 13.26340 | 18.05832 | 12.59719 | 16.55852 |
| **Slc25a30** | 0.00000 | 0.00414 | 0.05240 | 0.02894 | 0.00000 | 0.00000 | 0.00394 | 0.00826 | 0.23306 | 0.00511 | 0.33974 | 0.00680 | 0.27480 | 0.48662 |
| **Slc25a31** | 3.96686 | 0.00388 | 0.86641 | 0.00000 | 1.51563 | 0.00000 | 0.00393 | 0.00609 | 0.19290 | 0.00170 | 0.97534 | 0.00558 | 0.00473 | 0.31797 |
| **Slc25a32** | 1.72254 | 0.00171 | 0.36926 | 0.00000 | 1.18075 | 0.54219 | 0.00303 | 0.00000 | 3.48606 | 0.00000 | 4.24330 | 0.83316 | 5.11975 | 0.69724 |
| **Slc25a33** | 0.16353 | 0.00229 | 0.93925 | 0.64840 | 0.76493 | 1.33689 | 0.00297 | 0.12231 | 1.08496 | 0.00143 | 1.18351 | 1.75997 | 1.25257 | 3.03742 |
| **Slc25a35** | 0.01592 | 0.01076 | 0.49662 | 0.10794 | 0.16281 | 0.03503 | 0.00350 | 0.26801 | 0.01329 | 0.00652 | 0.51546 | 1.41876 | 0.00649 | 0.00349 |
| **Slc25a36** | 0.68640 | 0.00193 | 2.04523 | 0.59102 | 3.87315 | 0.04978 | 0.00175 | 0.03037 | 9.77035 | 0.00275 | 0.00000 | 1.54714 | 5.92181 | 5.99714 |
| **Slc25a37** | 0.58331 | 0.00437 | 0.00000 | 0.00294 | 1.21077 | 0.00000 | 0.00437 | 0.01136 | 0.00733 | 0.01721 | 0.18385 | 0.00847 | 0.00275 | 0.07269 |
| **Slc25a38** | 2.11864 | 0.01746 | 4.89685 | 0.32245 | 7.37656 | 3.24976 | 0.61042 | 0.99235 | 0.31409 | 0.03526 | 0.56203 | 2.86357 | 4.87514 | 5.29431 |
| **Slc25a39** | 2.61396 | 8.85053 | 11.26533 | 17.12654 | 6.31054 | 11.21557 | 6.00186 | 11.94258 | 0.00000 | 2.86917 | 3.51538 | 12.55643 | 2.31032 | 0.00000 |
| **Slc25a4** | 16.37602 | 12.20124 | 21.14372 | 28.79465 | 18.82450 | 29.83060 | 9.71294 | 25.65203 | 1.72350 | 15.01306 | 15.86708 | 17.00698 | 0.32859 | 7.39680 |
| **Slc25a40** | 0.19524 | 0.00303 | 0.13589 | 0.00000 | 0.00000 | 1.12729 | 0.84053 | 0.00000 | 0.85036 | 0.02490 | 0.89528 | 0.82764 | 0.06157 | 0.00000 |
| **Slc25a41** | 1.40997 | 0.00290 | 6.51751 | 0.14373 | 4.77559 | 0.31126 | 0.00250 | 0.00000 | 0.29705 | 0.00000 | 2.66303 | 0.00539 | 1.11468 | 3.83592 |
| **Slc25a42** | 0.85404 | 0.03223 | 0.48924 | 0.11104 | 0.31125 | 0.13917 | 0.04151 | 0.06006 | 0.45586 | 0.02615 | 0.01868 | 0.06058 | 0.43767 | 0.74833 |
| **Slc25a43** | 0.04416 | 0.00481 | 0.05745 | 1.03108 | 0.00381 | 0.07902 | 0.00394 | 0.00474 | 0.00961 | 0.00720 | 0.00789 | 0.01720 | 0.39452 | 0.00000 |
| **Slc25a44** | 16.17909 | 6.51574 | 16.11490 | 12.55799 | 16.64515 | 15.25079 | 5.14014 | 5.16623 | 14.55883 | 0.23036 | 3.42720 | 3.06853 | 15.95408 | 22.35290 |
| **Slc25a45** | 0.06696 | 0.00436 | 0.49776 | 0.03536 | 0.23568 | 0.17975 | 0.00408 | 0.00298 | 0.22826 | 0.00517 | 0.00600 | 0.00973 | 0.25321 | 0.00000 |

| | | | | | | | | | | | | | | |
|---|---|---|---|---|---|---|---|---|---|---|---|---|---|---|
| **Slc25a46** | 3.16062 | 0.00816 | 1.61566 | 1.23504 | 1.94360 | 3.54722 | 0.01358 | 0.88079 | 5.59659 | 0.01789 | 3.60108 | 3.34876 | 2.78779 | 0.33332 |
| **Slc25a47** | 0.00653 | 0.00439 | 0.16374 | 0.00349 | 0.00765 | 0.01767 | 0.00422 | 0.00837 | 0.00980 | 0.00684 | 0.00742 | 0.00926 | 0.22341 | 0.00338 |
| **Slc25a48** | 0.02194 | 0.00399 | 0.01188 | 0.00308 | 0.01038 | 0.01800 | 0.00421 | 0.00870 | 0.01203 | 0.00664 | 0.00457 | 0.44490 | 0.01169 | 0.05331 |
| **Slc25a5** | 28.11376 | 9.60779 | 40.21586 | 45.24037 | 25.81249 | 56.72651 | 10.93299 | 36.00624 | 15.41374 | 22.76286 | 34.32821 | 36.91271 | 2.29010 | 24.64168 |
| **Slc25a51** | 6.44780 | 0.15727 | 4.10921 | 4.18471 | 9.12177 | 1.07864 | 0.09839 | 3.37057 | 4.70217 | 0.02385 | 1.50176 | 2.02333 | 5.91287 | 3.93598 |
| **Slc25a53** | 0.15661 | 0.00425 | 0.25583 | 0.00000 | 0.53480 | 0.05027 | 0.00414 | 0.00399 | 1.12434 | 0.00549 | 0.00463 | 0.04097 | 0.33603 | 0.30867 |
| **Slc26a1** | 0.00726 | 0.00443 | 0.01394 | 0.01316 | 0.03083 | 0.01263 | 0.00428 | 0.01008 | 0.00670 | 0.00681 | 0.00882 | 0.00945 | 0.13368 | 0.04974 |
| **Slc26a10** | 0.01493 | 0.00448 | 0.12111 | 0.00413 | 0.01073 | 0.03204 | 0.00426 | 0.00803 | 0.01134 | 0.00685 | 0.00804 | 0.00953 | 0.00708 | 0.01117 |
| **Slc26a11** | 0.00000 | 2.73106 | 3.03362 | 1.37220 | 2.91003 | 0.00000 | 0.14682 | 0.00000 | 0.70400 | 0.00000 | 0.00554 | 0.18660 | 0.11624 | 2.81086 |
| **Slc26a2** | 0.00000 | 0.02515 | 0.20582 | 0.98309 | 0.83728 | 0.79766 | 0.00447 | 0.53667 | 0.03326 | 0.00986 | 0.00969 | 0.00840 | 1.62900 | 0.42055 |
| **Slc26a4** | 0.23299 | 0.00407 | 0.09817 | 0.00232 | 0.00000 | 0.00149 | 0.00376 | 0.00915 | 2.24337 | 0.00801 | 0.00368 | 0.00482 | 2.06009 | 1.32780 |
| **Slc26a5** | 0.71012 | 0.00450 | 0.83316 | 0.55682 | 0.00000 | 0.09336 | 0.00416 | 0.00446 | 0.03879 | 0.01356 | 0.00507 | 0.00438 | 0.00069 | 0.72609 |
| **Slc26a6** | 0.00963 | 0.00382 | 0.05860 | 0.01155 | 0.00795 | 0.06846 | 0.00411 | 0.00453 | 0.00690 | 0.01365 | 0.00399 | 0.40524 | 0.00922 | 0.44196 |
| **Slc26a8** | 0.00308 | 0.00402 | 0.00335 | 0.00926 | 0.53656 | 0.03595 | 0.40304 | 0.00922 | 0.00653 | 0.01075 | 0.00858 | 0.00656 | 0.29646 | 0.30231 |
| **Slc26a9** | 0.00713 | 0.00450 | 0.00799 | 0.01299 | 0.22483 | 0.01239 | 0.00462 | 0.00991 | 0.00817 | 0.00645 | 0.00858 | 0.00876 | 0.00657 | 0.18139 |
| **Slc27a1** | 0.12203 | 0.05235 | 0.84439 | 0.09916 | 0.21840 | 0.38158 | 0.06556 | 0.10027 | 0.06062 | 0.04567 | 0.67461 | 0.84649 | 1.15124 | 0.75124 |
| **Slc27a2** | 0.00410 | 0.00556 | 0.04381 | 0.00000 | 0.18721 | 0.00192 | 0.00438 | 0.36288 | 0.01063 | 0.00808 | 0.00625 | 0.01266 | 0.00647 | 0.49470 |
| **Slc27a3** | 0.11614 | 0.00228 | 1.40107 | 0.20813 | 0.38038 | 0.54420 | 0.00329 | 0.29637 | 0.00474 | 0.00386 | 0.19497 | 2.31369 | 0.15672 | 0.01206 |
| **Slc27a4** | 3.05963 | 0.08436 | 6.29529 | 5.57771 | 1.28662 | 4.84336 | 0.02084 | 7.38022 | 0.18324 | 12.87936 | 2.27072 | 2.61908 | 6.39648 | 4.70659 |
| **Slc27a5** | 1.63678 | 0.00325 | 2.18422 | 0.13316 | 2.99299 | 0.00000 | 0.00310 | 0.00190 | 2.91161 | 0.00267 | 0.28084 | 0.00346 | 3.53725 | 3.46386 |
| **Slc28a1** | 0.00992 | 0.00423 | 0.01235 | 0.25798 | 0.00833 | 0.00906 | 0.00425 | 0.00688 | 0.01088 | 0.00581 | 0.00670 | 0.00878 | 0.00653 | 0.03539 |
| **Slc28a2** | 0.10228 | 0.00441 | 0.13171 | 0.00560 | 0.26195 | 0.01129 | 0.00407 | 0.00925 | 0.01176 | 0.01450 | 0.00627 | 0.00859 | 0.02101 | 0.00452 |
| **Slc28a3** | 2.61628 | 0.21813 | 3.34424 | 5.51173 | 1.73619 | 6.45289 | 3.40886 | 5.16301 | 1.12669 | 2.15364 | 0.16064 | 1.00244 | 1.37206 | 1.62749 |
| **Slc29a1** | 0.33998 | 6.16584 | 2.68768 | 2.70094 | 0.94499 | 2.56889 | 0.03653 | 6.08530 | 0.00000 | 0.03880 | 0.10251 | 2.32260 | 0.00000 | 1.30819 |
| **Slc29a2** | 0.28605 | 0.01009 | 2.71046 | 5.59993 | 2.03732 | 2.67451 | 0.00291 | 1.80014 | 0.54231 | 0.00594 | 0.14090 | 1.28279 | 8.48318 | 1.01335 |
| **Slc29a3** | 3.94115 | 0.01159 | 5.43700 | 0.16754 | 6.55135 | 1.28076 | 0.03249 | 0.76408 | 0.16335 | 0.00000 | 2.78149 | 0.00440 | 0.09880 | 2.83280 |
| **Slc29a4** | 0.07515 | 0.00388 | 0.36304 | 0.00000 | 0.00051 | 0.04516 | 0.00401 | 0.04064 | 0.00815 | 0.01694 | 0.65510 | 0.05598 | 0.29296 | 0.00000 |
| **Slc2a1** | 0.00000 | 6.60841 | 3.36736 | 6.35583 | 0.00000 | 3.23102 | 3.15277 | 1.11429 | 0.00000 | 0.07728 | 0.02598 | 7.73019 | 0.00968 | 0.02778 |
| **Slc2a12** | 0.00007 | 0.01107 | 0.64914 | 0.00155 | 0.16620 | 0.44678 | 0.00379 | 0.67216 | 0.01741 | 0.00269 | 0.40847 | 0.00313 | 0.11081 | 0.01165 |
| **Slc2a3** | 4.95044 | 0.34197 | 10.34536 | 22.73672 | 4.70708 | 17.69789 | 7.45956 | 20.90699 | 2.93796 | 0.65873 | 9.79942 | 0.15460 | 2.42249 | 4.37915 |
| **Slc2a4** | 0.01049 | 0.01361 | 0.54227 | 0.24382 | 0.36427 | 0.36580 | 3.39560 | 1.06019 | 0.11597 | 0.14837 | 0.08213 | 0.00028 | 1.34058 | 1.13664 |
| **Slc2a4rg.ps** | 0.10273 | 0.00452 | 0.23846 | 0.47111 | 0.00000 | 1.27102 | 0.00403 | 0.00329 | 0.00544 | 0.00711 | 0.00642 | 0.05666 | 0.19039 | 0.02910 |
| **Slc2a7** | 0.00695 | 0.00432 | 0.01008 | 0.22775 | 0.00941 | 0.01117 | 0.00438 | 0.00811 | 0.01271 | 0.00592 | 0.00713 | 0.00790 | 0.00682 | 0.01000 |
| **Slc2a8** | 3.54068 | 0.13337 | 18.53158 | 17.50692 | 3.36308 | 2.43966 | 3.66498 | 2.81941 | 0.15633 | 0.17739 | 1.86233 | 5.78469 | 0.00000 | 1.19048 |
| **Slc2a9** | 0.00629 | 0.00417 | 0.16333 | 0.00000 | 0.00446 | 0.29453 | 0.00393 | 0.02069 | 0.20669 | 0.00680 | 0.15125 | 0.00545 | 0.00578 | 0.00000 |
| **Slc30a1** | 0.22449 | 6.06873 | 0.07564 | 8.83480 | 0.60736 | 6.11274 | 5.09810 | 10.61639 | 1.56301 | 0.29454 | 1.38953 | 1.90295 | 7.49688 | 0.00000 |
| **Slc30a2** | 0.70114 | 4.48439 | 0.67697 | 3.41403 | 0.00000 | 0.87721 | 2.01068 | 2.55941 | 0.00000 | 0.12077 | 2.88050 | 0.82096 | 1.53752 | 1.18453 |
| **Slc30a3** | 0.11529 | 0.00333 | 0.20459 | 1.13584 | 0.03620 | 0.00000 | 0.00341 | 0.00182 | 0.07807 | 0.00506 | 0.00393 | 0.75210 | 0.02575 | 0.00000 |
| **Slc30a4** | 2.50045 | 0.01312 | 3.90115 | 7.41899 | 0.50615 | 3.67932 | 1.32804 | 4.37184 | 8.79536 | 3.18287 | 4.57841 | 5.87367 | 0.21272 | 1.05188 |
| **Slc30a5** | 4.75744 | 0.11921 | 5.29726 | 0.00000 | 4.32379 | 4.96404 | 2.98214 | 0.04179 | 10.24235 | 0.06931 | 3.15921 | 4.01894 | 3.77929 | 9.97061 |
| **Slc30a6** | 2.53335 | 0.00583 | 3.34082 | 1.46839 | 4.20359 | 6.54196 | 1.26587 | 1.51484 | 3.22205 | 0.00667 | 6.80821 | 4.27183 | 2.68051 | 5.97235 |
| **Slc30a7** | 1.65350 | 0.02412 | 2.51929 | 1.97002 | 2.61399 | 1.32481 | 0.07794 | 1.34463 | 2.41851 | 0.04001 | 0.12975 | 1.92157 | 0.00000 | 3.24406 |
| **Slc30a9** | 0.75746 | 0.02222 | 1.21596 | 0.00000 | 0.28160 | 3.89924 | 0.05164 | 2.13055 | 0.00033 | 0.01320 | 1.21648 | 4.63349 | 0.51324 | 0.00595 |
| **Slc31a1** | 3.34220 | 0.68197 | 3.51535 | 0.90066 | 2.31792 | 7.88046 | 0.07993 | 6.34905 | 4.94910 | 0.03110 | 1.68237 | 6.01082 | 2.42451 | 4.77401 |
| **Slc31a2** | 0.91829 | 0.02624 | 0.00000 | 0.44105 | 0.00000 | 0.23989 | 0.00402 | 1.32280 | 1.65554 | 0.01860 | 0.00000 | 1.89813 | 0.00290 | 0.00000 |
| **Slc33a1** | 0.59577 | 0.00390 | 1.62355 | 0.76613 | 0.00000 | 1.86902 | 2.62779 | 0.07849 | 0.96655 | 0.06236 | 0.92615 | 0.13528 | 0.77822 | 1.53596 |
| **Slc34a2** | 10.06027 | 6.10836 | 14.72116 | 5.27701 | 5.01220 | 17.94312 | 5.93540 | 7.08896 | 20.98992 | 0.31374 | 7.54183 | 0.29839 | 18.75973 | 19.86307 |
| **Slc35a1** | 0.46972 | 0.00929 | 0.50116 | 0.00000 | 0.36568 | 0.15827 | 0.00282 | 0.29991 | 0.52495 | 0.00000 | 4.05482 | 2.28919 | 0.01138 | 0.00000 |
| **Slc35a2** | 1.41240 | 0.00309 | 2.93340 | 2.14120 | 0.69923 | 2.56769 | 1.63322 | 0.00000 | 4.76195 | 0.05581 | 0.49323 | 0.89284 | 1.04689 | 2.08885 |
| **Slc35a3** | 1.49650 | 0.48209 | 1.17395 | 0.48781 | 1.96642 | 1.22271 | 0.01469 | 0.69460 | 12.17803 | 0.00726 | 2.71016 | 0.80036 | 1.07318 | 2.90910 |
| **Slc35a4** | 2.11775 | 5.90936 | 9.18117 | 7.95333 | 5.40944 | 7.31759 | 5.08441 | 7.59489 | 0.39395 | 4.45726 | 9.95554 | 3.61277 | 1.58819 | 2.87624 |
| **Slc35a5** | 1.08151 | 0.00389 | 0.21625 | 0.05617 | 0.04293 | 0.15082 | 0.00443 | 0.10526 | 0.64243 | 0.00613 | 0.03884 | 0.00582 | 1.81468 | 0.49248 |
| **Slc35b1** | 6.37807 | 1.96759 | 12.25870 | 13.76578 | 4.88880 | 15.97370 | 5.91870 | 6.37954 | 1.83425 | 0.32042 | 7.57018 | 7.65145 | 3.81411 | 5.39520 |
| **Slc35b2** | 2.33329 | 0.05318 | 2.15690 | 6.98534 | 2.93274 | 3.68990 | 0.00708 | 1.85723 | 0.10252 | 0.03126 | 1.36307 | 7.03885 | 0.75006 | 3.78411 |
| **Slc35b3** | 0.00000 | 0.05295 | 0.34746 | 0.57464 | 0.07570 | 0.03449 | 0.00921 | 0.01561 | 0.00347 | 0.00438 | 0.41463 | 1.79909 | 0.37286 | 0.85809 |

| | | | | | | | | | | | | | | |
|---|---|---|---|---|---|---|---|---|---|---|---|---|---|---|
| **Slc35b4** | 2.12395 | 0.00882 | 3.98039 | 1.84682 | 1.83725 | 1.01902 | 0.61283 | 1.93408 | 2.40474 | 0.01121 | 3.24070 | 1.29588 | 0.85287 | 4.82613 |
| **Slc35c1** | 0.04218 | 0.02699 | 0.01437 | 0.54156 | 0.00000 | 0.17065 | 0.14120 | 2.13747 | 0.00500 | 0.02403 | 1.20158 | 0.02835 | 0.00417 | 0.96440 |
| **Slc35c2** | 1.04937 | 0.02813 | 5.45873 | 8.35649 | 6.37940 | 8.20462 | 0.00247 | 0.05109 | 0.00000 | 0.02023 | 2.43657 | 2.35209 | 0.50547 | 4.09061 |
| **Slc35d1** | 0.34389 | 0.01819 | 0.07696 | 0.00000 | 0.02522 | 1.47528 | 0.00325 | 1.26820 | 1.17233 | 0.00709 | 0.00000 | 1.09217 | 0.00635 | 0.00000 |
| **Slc35d2** | 0.05033 | 0.01150 | 0.10688 | 0.54493 | 0.00839 | 0.06716 | 0.00959 | 0.07718 | 0.24236 | 0.00695 | 0.03093 | 0.07010 | 0.64046 | 0.17109 |
| **Slc35e1** | 2.01278 | 0.11637 | 5.02256 | 4.33324 | 1.68597 | 9.82186 | 1.80285 | 4.47204 | 5.28382 | 0.09795 | 4.59272 | 5.12349 | 3.98423 | 6.39518 |
| **Slc35e2** | 3.64513 | 0.23421 | 0.44879 | 3.24274 | 1.30669 | 4.82086 | 1.13288 | 1.23592 | 2.15049 | 1.69879 | 0.53145 | 3.12116 | 5.69850 | 6.50159 |
| **Slc35e3** | 2.44915 | 0.02011 | 3.81416 | 1.10756 | 0.15128 | 2.05999 | 0.85369 | 0.57781 | 0.30604 | 0.00653 | 0.11761 | 0.67063 | 5.03451 | 2.78345 |
| **Slc35e4** | 4.58770 | 0.02714 | 11.04781 | 6.18426 | 9.09058 | 2.68051 | 0.77205 | 4.68486 | 0.64503 | 0.03089 | 0.04904 | 0.05766 | 2.65310 | 4.44757 |
| **Slc35f1** | 0.03226 | 0.00447 | 0.01552 | 0.02371 | 0.03038 | 0.01613 | 0.00417 | 0.01162 | 0.15160 | 0.00755 | 0.00629 | 0.02068 | 0.01943 | 0.07624 |
| **Slc35f2** | 15.77603 | 0.08852 | 13.88798 | 10.04749 | 11.80347 | 10.18353 | 3.85905 | 1.69283 | 15.21891 | 4.79880 | 6.40191 | 4.59339 | 13.22298 | 10.76761 |
| **Slc35f4** | 0.01004 | 0.03524 | 0.01438 | 0.00000 | 0.01585 | 0.00547 | 0.00436 | 0.32830 | 0.01244 | 0.00902 | 0.00567 | 0.09919 | 0.00844 | 0.01271 |
| **Slc35f5** | 0.78419 | 0.00296 | 0.39123 | 0.00000 | 0.00000 | 1.07577 | 2.04692 | 0.74152 | 2.20636 | 0.05078 | 0.46598 | 1.27781 | 1.36532 | 0.00000 |
| **Slc35f6** | 2.07509 | 0.16590 | 8.85344 | 7.67582 | 4.07070 | 5.43162 | 1.99731 | 4.19760 | 2.99739 | 0.15707 | 0.14044 | 5.21458 | 0.00000 | 2.27723 |
| **Slc35g1** | 9.33598 | 0.24191 | 8.38270 | 6.67716 | 10.62208 | 12.68738 | 3.34720 | 10.04781 | 22.84669 | 0.17389 | 6.88751 | 3.85926 | 4.57132 | 19.57516 |
| **Slc36a1** | 0.00000 | 0.01784 | 0.18155 | 0.49860 | 0.40751 | 0.49689 | 1.67922 | 0.81833 | 0.00862 | 0.05924 | 0.00056 | 0.00000 | 0.38245 | 1.05514 |
| **Slc36a1os** | 6.29213 | 0.49954 | 1.81091 | 0.18942 | 2.64540 | 3.17592 | 0.42000 | 0.49677 | 3.09362 | 0.26010 | 0.65619 | 0.49340 | 6.11720 | 1.29831 |
| **Slc36a3** | 0.00909 | 0.00459 | 0.01674 | 0.02136 | 0.01745 | 0.11917 | 0.00493 | 0.03030 | 0.02704 | 0.00646 | 0.00745 | 0.00933 | 0.00712 | 0.01730 |
| **Slc37a1** | 0.81932 | 5.44571 | 9.47056 | 2.24098 | 4.11174 | 5.06521 | 1.82476 | 3.48619 | 1.05555 | 0.05353 | 0.35856 | 2.68978 | 2.01053 | 7.72782 |
| **Slc37a2** | 0.00230 | 5.72533 | 0.00000 | 0.96995 | 0.00216 | 4.22068 | 2.80669 | 9.05164 | 0.00000 | 0.06742 | 0.00000 | 6.51228 | 0.00222 | 0.21224 |
| **Slc37a3** | 0.43999 | 0.06323 | 1.67658 | 1.51615 | 0.82663 | 5.86679 | 0.73707 | 0.00748 | 5.19755 | 0.03375 | 3.21782 | 5.84400 | 3.83915 | 6.61581 |
| **Slc37a4** | 0.16791 | 2.63029 | 0.42262 | 2.04228 | 0.71786 | 0.00000 | 0.00967 | 1.76392 | 1.64771 | 0.01863 | 0.00000 | 0.00615 | 0.00312 | 1.33972 |
| **Slc38a1** | 0.00436 | 0.00382 | 0.00000 | 0.38665 | 0.00000 | 1.52765 | 1.87674 | 2.96361 | 0.02077 | 0.02848 | 1.39981 | 0.00000 | 0.00776 | 0.00000 |
| **Slc38a10** | 1.55963 | 2.41521 | 3.58306 | 3.22991 | 3.45779 | 3.43199 | 0.00097 | 3.32080 | 0.22655 | 0.02855 | 1.91791 | 4.13080 | 0.00000 | 2.87363 |
| **Slc38a2** | 0.06240 | 0.00364 | 0.47821 | 0.24271 | 0.12025 | 0.61959 | 0.00362 | 0.43351 | 0.00761 | 0.00561 | 0.60331 | 0.36419 | 0.00495 | 0.01709 |
| **Slc38a3** | 0.00155 | 0.00457 | 0.00358 | 0.13249 | 0.00101 | 0.01512 | 0.00483 | 0.05541 | 0.01596 | 0.00940 | 0.00940 | 0.00629 | 0.00341 | 0.91982 |
| **Slc38a4** | 0.00000 | 0.06634 | 7.75831 | 5.22050 | 1.40210 | 5.96873 | 3.47867 | 7.20578 | 1.55957 | 0.12373 | 4.87362 | 7.55509 | 0.32264 | 2.62122 |
| **Slc38a6** | 1.31573 | 0.03371 | 0.44672 | 0.69312 | 1.21260 | 1.22735 | 0.48661 | 1.74340 | 0.28409 | 0.08552 | 0.00613 | 2.15312 | 0.57798 | 1.11872 |
| **Slc38a7** | 2.42008 | 0.04465 | 3.03197 | 6.21397 | 0.78667 | 4.14156 | 0.31449 | 4.90378 | 1.10209 | 0.04571 | 2.49528 | 6.04807 | 1.80969 | 1.07430 |
| **Slc38a8** | 0.34668 | 0.00432 | 0.22587 | 0.00595 | 0.56791 | 0.92798 | 0.00399 | 0.00291 | 0.03046 | 0.00502 | 0.01841 | 0.00254 | 0.37564 | 0.99827 |
| **Slc38a9** | 0.73519 | 0.00370 | 1.00011 | 0.00000 | 0.53416 | 1.34950 | 0.00345 | 0.01214 | 0.86680 | 0.00586 | 1.55265 | 0.00743 | 0.00000 | 0.67556 |
| **Slc39a1** | 3.22355 | 0.67600 | 0.00000 | 1.74956 | 5.05328 | 2.83516 | 0.26261 | 3.37631 | 7.07807 | 0.05002 | 0.00000 | 1.18758 | 6.05654 | 3.18428 |
| **Slc39a10** | 2.38188 | 0.02948 | 3.46517 | 1.34688 | 2.62594 | 2.83994 | 0.00062 | 0.00000 | 9.58322 | 0.01427 | 3.57297 | 3.13056 | 4.53407 | 4.66625 |
| **Slc39a11** | 0.00000 | 0.00407 | 2.55586 | 1.04911 | 0.26142 | 0.00000 | 0.00315 | 1.26855 | 0.05451 | 0.00000 | 0.00427 | 0.01569 | 0.97104 | 1.69871 |
| **Slc39a13** | 0.18110 | 0.01083 | 0.79552 | 0.00000 | 1.90264 | 0.46234 | 0.00330 | 1.10645 | 0.01334 | 0.00737 | 2.16484 | 0.00630 | 0.12873 | 0.15009 |
| **Slc39a14** | 2.05971 | 0.02210 | 2.92481 | 5.51612 | 1.23591 | 7.36259 | 4.31818 | 2.96267 | 3.83853 | 0.31374 | 5.32434 | 5.93759 | 0.49445 | 4.50029 |
| **Slc39a2** | 1.58749 | 0.01113 | 4.02519 | 1.82603 | 3.63969 | 2.54459 | 1.11892 | 5.09290 | 0.03262 | 0.13138 | 0.00000 | 0.00000 | 0.00444 | 1.35556 |
| **Slc39a3** | 0.20118 | 0.01119 | 0.78363 | 1.12667 | 0.27127 | 1.13057 | 0.01107 | 0.11229 | 0.01140 | 0.01050 | 0.01383 | 0.52638 | 0.38947 | 0.29288 |
| **Slc39a4** | 10.58907 | 0.26604 | 20.51713 | 14.68882 | 14.07779 | 16.84054 | 3.03644 | 7.37617 | 0.00000 | 0.14498 | 6.00338 | 2.92133 | 7.61425 | 4.64572 |
| **Slc39a5** | 0.01636 | 0.00473 | 0.01653 | 0.03106 | 0.04689 | 0.02146 | 0.01036 | 0.02618 | 0.01936 | 0.00903 | 0.00796 | 0.02361 | 0.09475 | 0.01079 |
| **Slc39a6** | 1.68048 | 0.00849 | 4.68836 | 1.12647 | 1.00684 | 1.85233 | 0.00128 | 0.89500 | 5.20441 | 0.01148 | 3.09821 | 1.93450 | 8.34398 | 3.53902 |
| **Slc39a7** | 2.15755 | 0.56549 | 12.37083 | 15.49876 | 2.44393 | 10.18379 | 2.52789 | 6.76982 | 0.16430 | 0.13242 | 5.90260 | 8.92930 | 1.46303 | 3.84691 |
| **Slc39a9** | 1.80462 | 0.03936 | 3.23511 | 1.60933 | 2.76352 | 6.80833 | 0.28515 | 1.24567 | 6.30077 | 1.21043 | 4.47313 | 3.00653 | 3.65785 | 9.47607 |
| **Slc3a1** | 0.00778 | 0.00388 | 0.01815 | 0.00262 | 0.02600 | 0.00512 | 0.00376 | 0.20942 | 0.00916 | 0.00553 | 0.00900 | 0.00996 | 0.01164 | 0.11794 |
| **Slc3a2** | 4.91948 | 0.54854 | 21.59476 | 20.58062 | 5.87008 | 8.91735 | 7.05946 | 5.70831 | 0.81144 | 0.54231 | 11.68561 | 6.72794 | 5.95101 | 5.86498 |
| **Slc41a1** | 0.09291 | 0.00960 | 0.09288 | 0.75948 | 0.21619 | 1.26452 | 0.00951 | 0.00265 | 0.01455 | 0.01154 | 0.00000 | 2.28597 | 0.43837 | 0.62735 |
| **Slc41a2** | 0.26112 | 0.00408 | 0.33668 | 0.08769 | 0.09374 | 0.69424 | 0.00435 | 0.01062 | 0.05958 | 0.00787 | 0.00396 | 0.01357 | 0.00336 | 0.93000 |
| **Slc41a3** | 0.01484 | 0.02733 | 0.01460 | 0.01255 | 0.01566 | 0.27742 | 0.00409 | 0.00418 | 0.03241 | 0.00552 | 0.01591 | 0.01083 | 0.00784 | 0.00702 |
| **Slc43a1** | 0.00280 | 0.01738 | 0.00747 | 0.06158 | 0.11111 | 0.00715 | 0.00350 | 0.00351 | 0.00947 | 0.00499 | 0.00674 | 1.91962 | 0.49479 | 0.00492 |
| **Slc43a2** | 0.00325 | 0.02640 | 0.02183 | 0.06423 | 0.00811 | 0.04175 | 0.00387 | 0.12020 | 0.43656 | 0.01801 | 0.02128 | 0.01447 | 0.39736 | 0.00000 |
| **Slc43a3** | 11.80697 | 4.53875 | 19.82874 | 15.40908 | 6.56596 | 10.96275 | 0.00139 | 0.98994 | 7.75245 | 0.05622 | 13.84381 | 7.31025 | 0.71810 | 12.75139 |
| **Slc44a1** | 0.00000 | 0.04500 | 0.25544 | 0.52245 | 0.02378 | 0.07053 | 0.02895 | 1.82089 | 1.19173 | 0.00651 | 0.02149 | 0.03785 | 1.00331 | 0.01408 |
| **Slc44a2** | 0.00000 | 0.03383 | 0.83482 | 2.39514 | 2.24719 | 1.36697 | 0.76127 | 2.26980 | 0.26683 | 0.05030 | 0.97804 | 0.00000 | 4.51891 | 5.26862 |
| **Slc44a3** | 0.02490 | 0.00327 | 0.55372 | 0.22489 | 0.05561 | 0.01446 | 0.00349 | 0.56245 | 0.00718 | 0.03243 | 0.00000 | 0.82371 | 0.45415 | 0.00203 |
| **Slc44a4** | 0.74721 | 3.46064 | 1.45211 | 0.01014 | 0.26798 | 3.67902 | 1.75913 | 2.65529 | 0.07020 | 0.04864 | 0.00241 | 0.00000 | 0.87057 | 1.15927 |

| | | | | | | | | | | | | | | |
|---|---|---|---|---|---|---|---|---|---|---|---|---|---|---|
| **Slc45a3** | 0.51111 | 0.00194 | 2.78373 | 0.27106 | 3.28713 | 0.00000 | 0.00217 | 0.11046 | 5.91461 | 0.02133 | 0.50588 | 1.10286 | 11.68393 | 8.12182 |
| **Slc45a4** | 4.92454 | 4.14599 | 17.51207 | 4.51272 | 3.06245 | 2.95776 | 1.11775 | 7.06382 | 0.10305 | 2.03809 | 0.65171 | 13.50003 | 0.24221 | 3.20916 |
| **Slc46a1** | 7.33335 | 3.31433 | 15.01967 | 10.80540 | 5.43205 | 17.44163 | 0.40669 | 15.36000 | 1.45171 | 2.22736 | 12.04205 | 2.92274 | 0.39039 | 1.13859 |
| **Slc46a3** | 0.02077 | 0.04465 | 0.01876 | 0.00135 | 0.01830 | 0.00005 | 1.38574 | 0.00298 | 0.01385 | 0.10139 | 0.00481 | 2.24619 | 0.03458 | 0.00574 |
| **Slc48a1** | 0.03269 | 0.00331 | 0.01696 | 0.00000 | 0.73613 | 0.54130 | 0.00376 | 0.01503 | 1.05876 | 0.00534 | 0.00000 | 0.90224 | 0.12224 | 0.00000 |
| **Slc4a1** | 0.00700 | 0.00455 | 0.00544 | 0.01417 | 0.30185 | 0.01144 | 0.00431 | 0.01094 | 0.12653 | 0.00643 | 0.00792 | 0.00816 | 0.00384 | 0.00000 |
| **Slc4a11** | 2.81732 | 0.07114 | 3.40929 | 5.17596 | 6.36246 | 7.40184 | 0.34379 | 1.79608 | 1.35605 | 0.07650 | 0.55706 | 0.00000 | 9.89867 | 6.13159 |
| **Slc4a1ap** | 2.67139 | 0.00587 | 3.41158 | 3.27684 | 9.43798 | 3.45972 | 0.01841 | 0.00000 | 3.87999 | 11.60266 | 5.03976 | 2.06072 | 3.01804 | 11.38946 |
| **Slc4a2** | 1.24770 | 0.13082 | 3.67029 | 2.25172 | 0.00000 | 8.66775 | 2.25926 | 7.73612 | 0.00000 | 0.10772 | 1.09788 | 2.41033 | 0.03335 | 0.00000 |
| **Slc4a3** | 0.00000 | 0.00307 | 3.46497 | 0.72691 | 0.29230 | 0.00000 | 0.00309 | 0.00000 | 0.00884 | 0.00000 | 0.00777 | 1.50985 | 0.65513 | 0.00000 |
| **Slc4a5** | 0.00417 | 0.00389 | 0.03766 | 1.94711 | 0.00151 | 0.75835 | 0.00369 | 0.77418 | 0.02843 | 0.00644 | 0.00000 | 0.00571 | 0.00759 | 0.00613 |
| **Slc4a7** | 0.31430 | 0.00325 | 0.28282 | 0.42667 | 0.11686 | 0.01373 | 0.01420 | 0.00000 | 1.91500 | 0.00331 | 2.22194 | 0.26436 | 1.37038 | 0.00002 |
| **Slc4a8** | 0.06948 | 0.00368 | 0.00000 | 0.54333 | 0.00354 | 0.00000 | 0.00376 | 0.61833 | 0.01030 | 0.00648 | 0.68636 | 0.03178 | 0.40282 | 0.07419 |
| **Slc50a1** | 0.00979 | 0.00458 | 0.00000 | 0.04353 | 0.17310 | 1.41598 | 2.04936 | 0.00838 | 0.00594 | 0.07529 | 0.00617 | 0.00000 | 0.00503 | 0.02613 |
| **Slc51a** | 0.00000 | 0.01910 | 3.74969 | 3.37355 | 0.00000 | 0.55872 | 0.00336 | 0.00000 | 0.00935 | 0.00000 | 2.17003 | 0.00454 | 0.00000 | 0.49784 |
| **Slc51b** | 0.21233 | 0.01153 | 0.00000 | 0.00000 | 0.11940 | 0.09609 | 0.00344 | 1.91281 | 0.06546 | 0.00385 | 0.66253 | 0.34351 | 0.14819 | 0.00000 |
| **Slc52a2** | 0.00694 | 0.01663 | 0.24705 | 0.25505 | 0.33134 | 0.93337 | 0.01582 | 1.68270 | 0.28614 | 0.01798 | 0.00000 | 3.52153 | 3.17038 | 0.32780 |
| **Slc52a3** | 0.04214 | 0.01453 | 0.88289 | 0.76117 | 0.00000 | 1.06872 | 0.65771 | 0.09475 | 0.79129 | 0.06393 | 0.00351 | 0.00000 | 0.00959 | 0.00000 |
| **Slc5a11** | 4.06080 | 0.07018 | 6.41373 | 11.23435 | 0.00000 | 14.32312 | 0.78252 | 12.08936 | 2.59791 | 0.28765 | 1.78472 | 0.00000 | 1.27037 | 5.58052 |
| **Slc5a12** | 0.01129 | 0.01245 | 0.00567 | 0.00971 | 0.28684 | 0.01969 | 0.00416 | 0.00875 | 0.04446 | 0.00734 | 0.00803 | 0.00842 | 0.00370 | 0.13274 |
| **Slc5a2** | 0.00767 | 0.00412 | 0.01240 | 0.45962 | 0.00629 | 0.01078 | 0.00392 | 0.00484 | 0.01749 | 0.00540 | 0.13456 | 0.01195 | 0.00665 | 0.00592 |
| **Slc5a3** | 2.40799 | 0.00264 | 3.25897 | 0.59000 | 2.96900 | 2.14217 | 0.07374 | 0.00000 | 12.06761 | 0.01107 | 0.63090 | 0.53398 | 1.72626 | 4.65996 |
| **Slc5a4b** | 0.01926 | 0.00420 | 1.21013 | 0.33955 | 0.15191 | 0.93641 | 0.00382 | 0.00336 | 1.27043 | 0.00960 | 0.00294 | 0.00146 | 0.92673 | 0.71459 |
| **Slc5a5** | 0.80571 | 0.00391 | 0.76095 | 1.04624 | 0.68174 | 0.02537 | 0.00891 | 0.00005 | 0.02731 | 0.00938 | 0.01199 | 0.01368 | 0.01856 | 0.41396 |
| **Slc5a6** | 29.51897 | 8.33261 | 42.06161 | 22.84403 | 34.47946 | 24.65588 | 7.66041 | 13.26232 | 24.40437 | 16.16547 | 7.52997 | 3.09182 | 14.10954 | 41.86634 |
| **Slc5a8** | 0.00517 | 0.00445 | 0.09025 | 0.01404 | 0.08762 | 0.01025 | 0.00410 | 0.00987 | 0.09131 | 0.01261 | 0.00638 | 0.00799 | 0.00378 | 0.32480 |
| **Slc6a12** | 0.00000 | 0.00426 | 0.51283 | 0.01499 | 0.00061 | 0.51714 | 1.52047 | 1.03798 | 0.01033 | 0.03080 | 0.76645 | 0.00000 | 0.00810 | 0.00122 |
| **Slc6a13** | 0.00000 | 0.02785 | 0.31320 | 0.00850 | 0.00000 | 3.39573 | 0.01087 | 4.14890 | 0.00190 | 0.00573 | 0.02211 | 0.00513 | 0.00190 | 0.62978 |
| **Slc6a14** | 0.48137 | 1.35525 | 0.39993 | 0.00000 | 0.00000 | 2.69807 | 0.01461 | 2.04021 | 0.00167 | 0.02081 | 1.21420 | 0.02989 | 0.69963 | 0.00000 |
| **Slc6a15** | 0.00000 | 0.00290 | 0.00000 | 0.01180 | 0.48385 | 0.00657 | 0.00351 | 0.00404 | 0.15403 | 0.00697 | 0.00284 | 1.31973 | 3.34697 | 3.53105 |
| **Slc6a16** | 0.00698 | 0.00433 | 0.01463 | 0.04366 | 0.00335 | 0.00824 | 0.00415 | 0.00827 | 0.10527 | 0.00607 | 0.01232 | 0.01316 | 0.07468 | 0.06621 |
| **Slc6a18** | 0.10847 | 0.01379 | 0.02501 | 0.07982 | 0.03892 | 0.07362 | 0.01735 | 0.04783 | 0.68737 | 0.00915 | 0.03614 | 0.02503 | 0.03477 | 0.07292 |
| **Slc6a2** | 0.04080 | 0.01262 | 0.03105 | 0.11612 | 0.06792 | 0.10421 | 0.02786 | 0.03162 | 0.11153 | 0.03904 | 0.01196 | 0.03469 | 0.09553 | 0.18642 |
| **Slc6a20b** | 0.02090 | 0.02181 | 0.03823 | 0.07851 | 0.03885 | 0.09513 | 0.01671 | 0.02388 | 0.12363 | 0.04387 | 0.00922 | 0.02740 | 0.02263 | 0.10504 |
| **Slc6a4** | 0.32461 | 0.00403 | 0.87260 | 0.00000 | 0.00000 | 0.27072 | 0.00365 | 0.00000 | 1.96719 | 0.00390 | 0.00279 | 0.09423 | 0.16137 | 0.00000 |
| **Slc6a6** | 10.16257 | 0.04507 | 16.28512 | 3.90659 | 11.87285 | 2.46188 | 2.62942 | 2.64310 | 16.12644 | 0.08499 | 3.26522 | 4.40086 | 1.95347 | 20.18965 |
| **Slc6a7** | 0.00232 | 0.00409 | 0.01592 | 0.01199 | 0.01211 | 0.01161 | 0.00408 | 0.00820 | 0.00686 | 0.00600 | 0.00861 | 0.00733 | 1.99500 | 0.25968 |
| **Slc6a8** | 21.60865 | 9.68907 | 31.99048 | 12.98495 | 17.16053 | 29.21050 | 6.24410 | 27.06327 | 17.57405 | 12.95178 | 8.93781 | 15.00170 | 1.92875 | 30.88454 |
| **Slc6a9** | 0.16274 | 0.00405 | 0.24747 | 0.00000 | 0.86049 | 0.00000 | 0.00399 | 0.00790 | 0.00927 | 0.00646 | 0.02401 | 0.00942 | 1.12801 | 1.31103 |
| **Slc7a1** | 0.10032 | 1.08291 | 0.41593 | 0.00000 | 0.19555 | 0.13348 | 0.00303 | 0.00326 | 0.00817 | 0.01351 | 0.00358 | 1.56706 | 0.25951 | 0.05355 |
| **Slc7a11** | 0.53782 | 1.14103 | 3.91162 | 2.15956 | 1.10547 | 4.18889 | 1.11350 | 4.41682 | 0.95811 | 1.46287 | 2.26337 | 4.29553 | 0.46171 | 1.59887 |
| **Slc7a14** | 1.70345 | 0.16297 | 0.00000 | 0.90041 | 0.94859 | 2.49254 | 0.49360 | 0.51069 | 5.01640 | 1.72490 | 0.12690 | 0.86405 | 12.26424 | 8.66912 |
| **Slc7a2** | 1.99361 | 0.23283 | 0.12548 | 4.33302 | 1.26975 | 2.71613 | 0.38709 | 1.02215 | 2.02694 | 1.03552 | 0.05871 | 1.68674 | 1.47196 | 1.66486 |
| **Slc7a3** | 0.01613 | 0.00495 | 0.12760 | 0.00000 | 0.01370 | 0.01145 | 0.00414 | 0.02095 | 0.00835 | 0.00718 | 0.00662 | 0.06189 | 0.00734 | 0.08889 |
| **Slc7a4** | 0.43909 | 0.00428 | 0.44419 | 0.01528 | 0.00000 | 0.00312 | 0.00417 | 0.01036 | 0.00509 | 0.00583 | 0.00604 | 0.00586 | 0.27739 | 0.08545 |
| **Slc7a5** | 2.33599 | 0.05778 | 2.48432 | 4.03674 | 2.72995 | 1.81811 | 0.00192 | 1.66591 | 3.24777 | 0.00759 | 1.72931 | 3.68345 | 4.87250 | 4.67406 |
| **Slc7a6** | 3.73404 | 2.95690 | 9.24185 | 10.11375 | 4.88357 | 15.65232 | 8.36591 | 13.34658 | 16.01742 | 15.14521 | 11.81434 | 11.15311 | 0.77891 | 8.62724 |
| **Slc7a6os** | 4.12052 | 4.86470 | 2.34283 | 4.47731 | 9.07046 | 1.66525 | 0.17433 | 2.21240 | 2.23254 | 0.01356 | 0.32479 | 1.95279 | 7.07548 | 0.68093 |
| **Slc7a7** | 0.79410 | 0.02765 | 2.86523 | 0.35512 | 1.67788 | 1.89765 | 4.51249 | 1.07628 | 0.92716 | 0.11004 | 0.00000 | 0.58773 | 0.00000 | 0.94858 |
| **Slc7a8** | 0.01814 | 0.01094 | 0.00184 | 0.78608 | 0.00462 | 0.00000 | 0.00386 | 0.00548 | 0.01459 | 0.00361 | 1.18038 | 0.00904 | 0.02679 | 0.00000 |
| **Slc7a9** | 0.01878 | 0.00438 | 0.01171 | 0.03194 | 0.02413 | 0.01118 | 0.00412 | 0.00922 | 0.00992 | 0.01289 | 0.00813 | 0.00855 | 0.07128 | 0.01020 |
| **Slc8b1** | 2.47268 | 0.87326 | 7.11633 | 5.48244 | 9.15995 | 6.91850 | 0.02860 | 5.20054 | 1.81909 | 10.33635 | 1.20177 | 2.35399 | 1.34297 | 12.02800 |
| **Slc9a1** | 0.60905 | 4.82752 | 2.26609 | 1.08074 | 2.24639 | 3.30130 | 1.01111 | 6.33655 | 0.07900 | 0.05003 | 2.56562 | 2.21591 | 2.67904 | 0.40911 |
| **Slc9a3r1** | 0.00000 | 2.59031 | 0.30598 | 0.92638 | 0.29093 | 3.26442 | 6.05414 | 16.89738 | 0.00133 | 10.54298 | 1.23767 | 0.58576 | 0.22413 | 0.05192 |
| **Slc9a3r2** | 0.01833 | 0.00433 | 0.00744 | 0.01263 | 0.20647 | 0.01455 | 0.00457 | 0.01496 | 0.01303 | 0.00631 | 0.00813 | 0.01090 | 0.00527 | 0.25855 |

| | | | | | | | | | | | | | | |
|---|---|---|---|---|---|---|---|---|---|---|---|---|---|---|
| **Slc9a5** | 0.01443 | 0.00562 | 0.15408 | 0.00000 | 0.00694 | 0.00000 | 0.60546 | 0.34438 | 0.04520 | 0.00863 | 0.00078 | 0.28783 | 0.01118 | 0.00828 |
| **Slc9a6** | 1.13757 | 0.05722 | 0.21876 | 1.62695 | 0.00000 | 5.66737 | 3.42706 | 1.74528 | 1.94647 | 0.09158 | 0.54780 | 1.94961 | 0.10582 | 0.00000 |
| **Slc9a8** | 0.13762 | 0.01028 | 2.68464 | 2.69332 | 0.58083 | 3.16510 | 0.27697 | 1.37107 | 0.29854 | 0.01462 | 0.00000 | 2.06216 | 0.00000 | 1.10301 |
| **Slc9b1** | 2.14473 | 0.01120 | 0.28368 | 1.79402 | 0.82498 | 1.54772 | 0.00415 | 0.19905 | 0.00457 | 0.00009 | 1.65444 | 1.54784 | 0.00271 | 2.66034 |
| **Slc9b2** | 0.00577 | 0.01072 | 0.00692 | 0.00000 | 0.00804 | 0.31035 | 0.00370 | 0.42102 | 0.02529 | 0.00666 | 0.00202 | 0.15819 | 0.43499 | 0.02199 |
| **Slco1a5** | 0.00000 | 0.00425 | 0.06698 | 0.00552 | 0.00000 | 0.01063 | 0.00424 | 0.00899 | 0.01394 | 0.00670 | 0.00716 | 0.00831 | 0.58006 | 0.49376 |
| **Slco2a1** | 0.06948 | 0.05640 | 1.02991 | 2.55721 | 0.24265 | 1.20533 | 1.25690 | 1.26044 | 0.00894 | 0.02306 | 3.42281 | 0.00000 | 0.00099 | 0.00000 |
| **Slco2b1** | 0.00893 | 0.01180 | 0.01441 | 0.06962 | 0.01492 | 0.00885 | 0.00409 | 0.00617 | 0.01195 | 0.04096 | 0.00722 | 0.01353 | 0.00722 | 0.01546 |
| **Slco3a1** | 0.01987 | 0.00435 | 0.00000 | 0.01166 | 1.01012 | 0.00837 | 0.00408 | 0.00841 | 0.03375 | 0.00627 | 0.00668 | 0.00619 | 0.15258 | 1.57040 |
| **Slco4a1** | 0.00000 | 0.00418 | 0.00150 | 0.11852 | 0.00000 | 0.00912 | 0.00410 | 0.00653 | 0.18135 | 0.00575 | 0.00634 | 0.00682 | 0.48404 | 0.31660 |
| **Slco4c1** | 2.54285 | 0.01927 | 2.90106 | 0.08081 | 2.22848 | 0.00000 | 0.00309 | 0.46881 | 12.66838 | 0.00121 | 0.00054 | 0.00155 | 0.87199 | 2.50764 |
| **Slco5a1** | 0.00132 | 0.00436 | 0.16380 | 0.11138 | 0.29949 | 0.00000 | 0.00401 | 0.00834 | 0.43633 | 0.00726 | 0.00490 | 0.01068 | 0.00237 | 0.06067 |
| **Slf1** | 0.14032 | 0.00397 | 0.17447 | 0.15624 | 0.00000 | 0.20863 | 0.08876 | 0.21358 | 0.33013 | 0.00567 | 0.00682 | 0.01377 | 0.18086 | 0.18731 |
| **Slfn2** | 0.00000 | 0.00946 | 0.23460 | 1.17605 | 0.00000 | 0.00000 | 0.00325 | 0.00000 | 0.01962 | 0.00211 | 1.30889 | 1.29663 | 0.11328 | 0.06739 |
| **Slfn4** | 0.01230 | 0.00477 | 0.03707 | 0.06939 | 0.01232 | 0.05411 | 0.00475 | 0.02145 | 0.01584 | 0.00738 | 0.05329 | 0.02177 | 0.01185 | 0.16268 |
| **Slfn5** | 1.66596 | 0.07940 | 0.00000 | 1.16366 | 0.34865 | 2.84910 | 0.53659 | 0.47622 | 2.44565 | 1.08474 | 0.09319 | 1.11130 | 13.94835 | 2.06939 |
| **Slfn9** | 0.01289 | 0.00414 | 0.02775 | 0.05874 | 0.03090 | 0.01485 | 0.00408 | 0.02689 | 0.14402 | 0.00606 | 0.01074 | 0.00728 | 0.01448 | 0.02854 |
| **Slirp** | 0.18326 | 0.55012 | 0.02644 | 0.00016 | 0.98766 | 2.67390 | 1.26563 | 2.46737 | 1.01390 | 6.75497 | 0.01076 | 2.53833 | 2.02199 | 0.62958 |
| **Slit3** | 0.10388 | 0.00446 | 0.01603 | 0.01239 | 0.03209 | 0.01201 | 0.00444 | 0.01057 | 0.00818 | 0.00701 | 0.00756 | 0.00930 | 0.00383 | 0.05310 |
| **Slk** | 0.77629 | 1.64755 | 1.72265 | 0.87506 | 0.89836 | 4.01161 | 1.86593 | 2.65985 | 0.50177 | 0.05641 | 0.42938 | 2.42201 | 2.29694 | 0.00000 |
| **Slmap** | 12.00870 | 0.18933 | 6.08298 | 3.08733 | 8.23437 | 7.33367 | 1.40963 | 0.83870 | 18.85608 | 0.04451 | 8.86111 | 3.34948 | 9.36873 | 9.94286 |
| **Slmo1** | 0.01181 | 0.00394 | 1.75780 | 0.70573 | 0.00000 | 0.00000 | 0.00359 | 0.01250 | 0.02141 | 0.00294 | 0.01113 | 0.01634 | 0.00000 | 0.35957 |
| **Slmo2** | 3.51837 | 8.34560 | 5.29374 | 6.66438 | 3.54798 | 9.85314 | 6.39788 | 0.57814 | 0.44104 | 0.16891 | 4.85892 | 6.49113 | 1.19294 | 3.70260 |
| **Sln** | 0.00753 | 0.00441 | 0.02653 | 0.01103 | 0.01128 | 0.01210 | 0.00427 | 0.00999 | 0.00699 | 0.00714 | 0.00848 | 0.00966 | 0.10781 | 0.08744 |
| **Sltm** | 15.29402 | 0.06768 | 6.69394 | 2.65441 | 16.88917 | 3.07719 | 5.26236 | 3.78559 | 18.34107 | 0.05494 | 3.47576 | 3.70246 | 7.16711 | 21.49850 |
| **Slu7** | 11.72183 | 0.08108 | 14.32380 | 3.78244 | 11.01205 | 4.99121 | 6.31901 | 0.69732 | 16.40681 | 0.18614 | 6.04253 | 4.99913 | 11.18583 | 18.55619 |
| **Slx1b** | 4.00461 | 0.64653 | 3.09389 | 11.83876 | 3.42041 | 8.21920 | 1.24882 | 4.45733 | 1.56339 | 3.73799 | 0.47117 | 3.60718 | 6.02709 | 4.20996 |
| **Slx4** | 0.57933 | 0.00319 | 2.29858 | 2.32384 | 0.10657 | 3.46887 | 0.01715 | 0.98121 | 0.14497 | 0.03718 | 3.00456 | 0.38238 | 1.00536 | 2.22351 |
| **Slx4ip** | 0.00000 | 0.00285 | 0.23494 | 0.00000 | 0.65754 | 0.60398 | 0.15607 | 0.00570 | 2.07015 | 0.00831 | 0.72669 | 0.88229 | 2.57178 | 4.20067 |
| **Smad1** | 0.02620 | 0.00462 | 0.00901 | 0.00000 | 0.08436 | 0.25676 | 0.00422 | 0.00619 | 0.01159 | 0.00567 | 0.02828 | 0.01755 | 0.00818 | 0.01770 |
| **Smad2** | 6.83966 | 0.05124 | 3.04968 | 1.68088 | 3.26934 | 3.08662 | 0.03756 | 1.61703 | 8.72772 | 0.03503 | 5.88643 | 2.23809 | 1.52898 | 3.71175 |
| **Smad3** | 0.30110 | 0.01025 | 3.17939 | 0.36683 | 1.85498 | 0.91959 | 0.00393 | 1.06979 | 0.10699 | 0.00000 | 1.56201 | 2.08010 | 1.11210 | 1.58540 |
| **Smad4** | 2.98454 | 0.12523 | 6.56640 | 2.30299 | 3.05344 | 3.98400 | 0.00565 | 4.25963 | 1.56661 | 5.56714 | 1.04059 | 4.68157 | 0.57727 | 4.60367 |
| **Smad5** | 0.48526 | 0.00894 | 0.77629 | 0.00000 | 0.27079 | 0.97764 | 0.82875 | 3.45739 | 0.73656 | 0.05537 | 0.06318 | 3.32588 | 0.71391 | 0.40870 |
| **Smad7** | 0.00394 | 0.02147 | 0.08295 | 0.03055 | 0.18313 | 0.07942 | 0.01670 | 0.37091 | 0.41620 | 0.00974 | 0.24090 | 0.00066 | 3.68440 | 0.74326 |
| **Smad9** | 0.07564 | 0.03359 | 0.00004 | 0.45902 | 0.02660 | 0.37657 | 0.00385 | 0.01108 | 0.01274 | 0.00863 | 0.01308 | 0.01258 | 0.10053 | 0.26120 |
| **Smagp** | 2.43591 | 0.02476 | 0.00000 | 1.15181 | 0.97007 | 4.98700 | 0.12394 | 1.82938 | 0.56628 | 0.05360 | 0.44046 | 2.59178 | 2.28814 | 1.88746 |
| **Smap1** | 0.79445 | 0.08245 | 8.55037 | 2.77651 | 1.70494 | 5.57650 | 0.33489 | 0.60160 | 1.83901 | 0.05104 | 4.39717 | 4.99568 | 0.00000 | 2.81038 |
| **Smap2** | 11.22178 | 0.09063 | 12.36386 | 3.62919 | 9.78134 | 4.35964 | 0.03560 | 3.61110 | 3.59848 | 0.02264 | 0.11397 | 6.48562 | 4.36950 | 7.08338 |
| **Smarca2** | 0.04234 | 0.03028 | 2.51975 | 0.13729 | 0.39901 | 0.50755 | 0.01551 | 0.36523 | 0.60552 | 0.00455 | 0.00000 | 4.22809 | 3.27593 | 2.04061 |
| **Smarca4** | 1.94917 | 0.17474 | 29.06746 | 19.50548 | 27.44268 | 2.23300 | 3.38691 | 10.15261 | 0.99096 | 4.18041 | 7.04481 | 10.47565 | 5.66398 | 7.77534 |
| **Smarca5** | 14.75086 | 0.37249 | 15.39772 | 4.41528 | 12.50630 | 8.28106 | 0.37966 | 1.87144 | 33.44185 | 0.04481 | 10.05137 | 9.47516 | 13.83680 | 10.49443 |
| **Smarca5.ps** | 5.77044 | 0.06034 | 5.69093 | 0.68652 | 4.20851 | 2.63090 | 0.16852 | 0.49629 | 17.73355 | 0.00000 | 2.92804 | 3.58474 | 7.14564 | 3.98526 |
| **Smarcad1** | 12.41834 | 0.07050 | 2.02190 | 3.11444 | 4.14331 | 2.22908 | 0.76339 | 0.26792 | 12.40578 | 5.71529 | 1.22838 | 3.13415 | 8.16654 | 4.92970 |
| **Smarcal1** | 0.85714 | 0.01781 | 3.10375 | 2.40558 | 2.43882 | 4.17491 | 0.30626 | 4.65697 | 0.00000 | 0.01670 | 2.57770 | 1.32753 | 0.17569 | 1.84191 |
| **Smarcb1** | 1.48385 | 0.65591 | 6.35485 | 13.19727 | 9.93046 | 7.42146 | 2.20773 | 6.41218 | 2.60334 | 0.14249 | 4.15928 | 10.91325 | 4.40929 | 10.39790 |
| **Smarcc1** | 6.25884 | 0.18773 | 8.98749 | 8.35485 | 11.15513 | 7.78096 | 0.75859 | 5.47281 | 8.06343 | 0.07503 | 5.16624 | 7.37773 | 5.91939 | 7.70119 |
| **Smarcc2** | 0.00640 | 0.04903 | 1.20892 | 0.00017 | 3.32114 | 1.19627 | 0.45412 | 0.91292 | 0.77231 | 0.03056 | 0.24942 | 1.28108 | 3.22245 | 0.16819 |
| **Smarcd1** | 1.29099 | 0.01133 | 1.63805 | 1.31068 | 3.15501 | 0.18450 | 0.00328 | 2.90465 | 0.00913 | 0.00167 | 1.24873 | 0.42951 | 1.46079 | 0.70849 |
| **Smarcd2** | 8.37941 | 0.04456 | 24.22288 | 15.88284 | 13.36622 | 2.06882 | 4.68975 | 5.49542 | 3.88861 | 0.21183 | 3.28240 | 3.52055 | 10.32631 | 16.88811 |
| **Smarcd3** | 1.42596 | 0.01356 | 2.88202 | 2.48456 | 0.79798 | 0.39233 | 0.00391 | 0.31915 | 0.00609 | 0.00165 | 0.00316 | 0.00197 | 0.00000 | 0.00000 |
| **Smarce1** | 4.90610 | 0.09456 | 16.55847 | 6.15244 | 11.98944 | 6.56256 | 4.97189 | 7.52602 | 8.11861 | 13.74895 | 9.51194 | 7.26566 | 3.20775 | 17.28920 |
| **Smc1a** | 14.77284 | 0.07311 | 10.76766 | 3.05566 | 14.07899 | 8.73073 | 2.45137 | 5.75029 | 18.00506 | 0.10998 | 5.79158 | 8.63196 | 1.20853 | 11.73551 |
| **Smc1b** | 0.28637 | 0.00376 | 0.25966 | 0.38400 | 0.62202 | 0.44608 | 0.00366 | 0.00000 | 0.13264 | 0.00471 | 0.81658 | 0.00855 | 2.14124 | 1.14032 |
| **Smc2** | 6.44298 | 0.03763 | 3.71523 | 1.78789 | 5.13833 | 6.31129 | 2.37309 | 0.61183 | 11.07508 | 0.03234 | 4.70181 | 7.20885 | 2.63650 | 3.69000 |

| | | | | | | | | | | | | | | |
|---|---|---|---|---|---|---|---|---|---|---|---|---|---|---|
| **Smc3** | 11.45613 | 0.62777 | 10.41932 | 2.16171 | 14.65510 | 7.84659 | 2.00871 | 2.11343 | 20.53390 | 0.08077 | 11.73886 | 7.56341 | 8.27189 | 14.12387 |
| **Smc4** | 11.29297 | 0.05794 | 11.20138 | 5.03850 | 12.96668 | 8.59498 | 2.76051 | 5.41326 | 31.58481 | 0.17559 | 9.14358 | 7.00515 | 9.39628 | 6.21743 |
| **Smc5** | 7.10876 | 0.01335 | 9.69483 | 3.67929 | 9.60702 | 4.09036 | 0.00366 | 0.71421 | 14.34424 | 0.00557 | 8.16156 | 8.26001 | 1.01490 | 7.57560 |
| **Smc6** | 11.87156 | 0.04814 | 7.72459 | 2.38934 | 10.17021 | 4.54858 | 0.02803 | 4.08351 | 18.83984 | 0.03097 | 5.89847 | 7.31542 | 10.18703 | 14.14158 |
| **Smchd1** | 6.74426 | 0.04720 | 2.45998 | 3.19007 | 13.23810 | 3.23690 | 3.21111 | 2.19643 | 16.07651 | 0.07397 | 2.45652 | 2.07677 | 7.10586 | 11.14921 |
| **Smco1** | 0.35869 | 0.11986 | 0.07099 | 0.70872 | 0.39008 | 0.59567 | 0.15875 | 0.25081 | 0.62595 | 0.15675 | 0.04553 | 0.22029 | 1.69109 | 1.07560 |
| **Smco2** | 0.00701 | 0.00417 | 0.00977 | 0.27770 | 0.00371 | 0.44198 | 0.00397 | 0.00304 | 0.00987 | 0.00518 | 0.01442 | 0.01202 | 0.00657 | 0.18629 |
| **Smco4** | 0.03005 | 0.00443 | 0.00613 | 0.00000 | 0.00809 | 0.58403 | 0.00427 | 0.16854 | 0.00729 | 0.00707 | 0.00510 | 0.23513 | 0.81904 | 0.00522 |
| **Smcr8** | 2.48530 | 0.00838 | 2.89297 | 1.23459 | 4.01158 | 3.86792 | 0.70950 | 0.66769 | 2.48802 | 0.02926 | 4.75847 | 2.01592 | 5.93340 | 6.36027 |
| **Smdt1** | 0.25173 | 4.36970 | 0.11506 | 7.71827 | 8.88201 | 2.45215 | 0.84202 | 8.13854 | 0.30080 | 1.62509 | 0.02002 | 5.90411 | 1.21813 | 0.00000 |
| **Smek1** | 0.16304 | 0.09579 | 3.49160 | 1.00797 | 7.01167 | 3.78857 | 0.23725 | 5.34620 | 13.26578 | 0.01449 | 2.17508 | 1.90051 | 10.44143 | 10.31603 |
| **Smek2** | 6.26461 | 0.03067 | 4.92729 | 0.00000 | 4.13046 | 2.84327 | 0.64229 | 2.28683 | 15.34917 | 0.03941 | 1.00842 | 3.07620 | 3.22356 | 4.20315 |
| **Smg1** | 3.97219 | 4.87965 | 4.79221 | 3.72962 | 10.21924 | 6.93694 | 2.31378 | 5.43792 | 20.32482 | 0.05929 | 4.54520 | 3.06029 | 6.55343 | 6.08304 |
| **Smg5** | 4.23467 | 0.05696 | 9.47452 | 10.79982 | 17.24539 | 3.80194 | 0.60477 | 4.64225 | 0.63984 | 0.07837 | 6.56142 | 5.45961 | 7.25672 | 7.38026 |
| **Smg6** | 0.93476 | 0.00300 | 2.70211 | 2.02390 | 2.70070 | 0.91918 | 0.00336 | 0.39834 | 1.24136 | 0.01774 | 2.22180 | 0.63303 | 2.28329 | 4.29410 |
| **Smg7** | 3.75972 | 1.07000 | 2.98723 | 3.26966 | 7.61490 | 3.96985 | 2.61455 | 5.83214 | 11.27318 | 0.18868 | 4.45792 | 2.96439 | 9.10242 | 5.43287 |
| **Smg8** | 6.64479 | 0.01856 | 6.02239 | 2.67874 | 12.50339 | 5.42245 | 0.00092 | 0.22468 | 6.67118 | 0.00000 | 6.71604 | 2.15820 | 9.06435 | 12.85249 |
| **Smg9** | 0.00000 | 0.03017 | 6.32070 | 0.73211 | 3.29571 | 1.36588 | 0.00308 | 0.44274 | 0.00428 | 0.00000 | 1.61010 | 0.53029 | 0.00000 | 0.14557 |
| **Smim1** | 0.50132 | 1.39172 | 0.33359 | 1.01726 | 0.12279 | 0.37586 | 0.00340 | 3.33862 | 0.00727 | 0.04150 | 0.00000 | 0.82414 | 0.00527 | 0.00430 |
| **Smim10l1** | 0.01416 | 0.00425 | 0.00515 | 0.06430 | 0.20291 | 0.00851 | 0.00421 | 0.00796 | 0.06508 | 0.00633 | 0.00633 | 0.02995 | 0.00729 | 0.18205 |
| **Smim10l2a** | 0.01266 | 0.00413 | 0.00965 | 0.01889 | 0.09742 | 0.01487 | 0.00402 | 0.02054 | 0.00993 | 0.01131 | 0.00698 | 0.04828 | 0.02934 | 0.05730 |
| **Smim11** | 0.57284 | 0.51969 | 0.00000 | 0.00001 | 0.30785 | 1.51769 | 0.43340 | 1.89048 | 0.01697 | 0.07276 | 0.01470 | 0.65608 | 1.12565 | 0.00503 |
| **Smim12** | 0.18045 | 0.09329 | 1.80430 | 0.85430 | 0.21779 | 3.70770 | 1.90317 | 3.19129 | 0.00000 | 0.15337 | 2.61834 | 2.59993 | 1.35708 | 0.05895 |
| **Smim13** | 7.72839 | 0.00000 | 6.77081 | 3.81979 | 12.92673 | 3.70916 | 1.69632 | 0.11873 | 21.53113 | 0.08535 | 2.70567 | 2.18921 | 7.00633 | 8.68278 |
| **Smim14** | 0.04416 | 0.02428 | 0.12185 | 0.00000 | 0.21746 | 0.02656 | 0.79571 | 1.21211 | 1.12739 | 9.76211 | 0.00000 | 0.07688 | 3.22286 | 1.42381 |
| **Smim15** | 0.71119 | 0.01666 | 2.99603 | 0.30046 | 0.38069 | 3.45378 | 1.99961 | 1.26839 | 8.84120 | 4.03561 | 3.57086 | 2.28097 | 2.18805 | 4.41528 |
| **Smim19** | 0.40089 | 0.00391 | 0.07564 | 1.44829 | 0.83916 | 0.09025 | 0.96974 | 0.94989 | 0.00702 | 0.02229 | 0.33006 | 0.06443 | 0.70147 | 0.00000 |
| **Smim20** | 2.35224 | 2.30483 | 4.54997 | 6.25748 | 3.89653 | 4.37914 | 1.55194 | 7.42639 | 1.07662 | 0.09281 | 2.13390 | 2.86933 | 0.62527 | 0.00000 |
| **Smim22** | 0.00476 | 0.03067 | 0.00492 | 0.00000 | 0.20762 | 0.86356 | 0.00414 | 0.24108 | 0.01592 | 0.32685 | 0.02084 | 0.22125 | 0.00775 | 0.00283 |
| **Smim24** | 0.00788 | 0.14024 | 0.35618 | 0.01491 | 0.32885 | 0.45864 | 0.00383 | 0.00256 | 0.01582 | 0.00848 | 0.00426 | 0.00185 | 0.02359 | 0.00731 |
| **Smim4** | 0.32480 | 0.01162 | 0.04786 | 0.01215 | 0.67857 | 0.18708 | 0.12206 | 0.61519 | 0.00392 | 1.65611 | 0.00409 | 0.18431 | 0.75704 | 0.63273 |
| **Smim5** | 0.00974 | 0.00444 | 0.01412 | 0.01351 | 0.03477 | 0.01275 | 0.00434 | 0.01102 | 0.01737 | 0.00673 | 0.00865 | 0.00997 | 0.00589 | 0.02273 |
| **Smim6** | 0.01077 | 0.01252 | 0.14151 | 0.03178 | 0.07769 | 0.00000 | 0.00521 | 0.39202 | 0.01670 | 0.00554 | 0.06675 | 0.02271 | 0.00628 | 0.32837 |
| **Smim7** | 4.39623 | 0.12911 | 9.84450 | 4.95956 | 13.78752 | 14.47643 | 3.47172 | 10.24193 | 11.62289 | 13.07960 | 7.98663 | 3.54016 | 7.06405 | 17.27759 |
| **Smim8** | 0.52465 | 0.02519 | 0.00073 | 0.71362 | 0.59579 | 1.66619 | 0.31844 | 1.40155 | 0.03224 | 0.01738 | 0.85478 | 2.54699 | 0.40555 | 0.18051 |
| **Smim9** | 0.13835 | 0.00424 | 0.00414 | 0.01790 | 0.13788 | 0.16194 | 0.00429 | 0.01911 | 0.14348 | 0.00813 | 0.00592 | 0.00428 | 0.60103 | 0.14549 |
| **Smkr.ps** | 0.00788 | 0.01372 | 0.00729 | 0.25398 | 0.00501 | 0.00000 | 0.00472 | 0.45120 | 0.06947 | 0.00598 | 0.02080 | 0.01394 | 0.05161 | 0.00075 |
| **Smn1** | 8.00109 | 0.11772 | 10.27943 | 12.27997 | 16.19230 | 5.88964 | 2.08182 | 3.45773 | 1.76902 | 0.07086 | 12.20165 | 6.22318 | 5.03863 | 7.70340 |
| **Smndc1** | 10.04559 | 2.56442 | 8.48640 | 5.49111 | 9.90906 | 5.37002 | 3.95627 | 0.00000 | 13.96899 | 0.14480 | 8.03074 | 3.99074 | 9.50471 | 8.71558 |
| **Smo** | 0.03281 | 0.00397 | 0.31469 | 1.20297 | 0.07542 | 0.00000 | 0.00396 | 0.00623 | 0.00843 | 0.00441 | 0.01384 | 0.01963 | 0.00527 | 0.17055 |
| **Smoc1** | 0.52570 | 0.01153 | 1.26180 | 0.00000 | 0.65993 | 3.86774 | 0.01078 | 0.19605 | 0.01410 | 0.00971 | 0.38723 | 0.80002 | 0.00000 | 0.28785 |
| **Smoc2** | 0.00828 | 0.00435 | 0.15456 | 0.03133 | 0.09103 | 0.00400 | 0.00428 | 0.01212 | 0.29529 | 0.00690 | 0.00552 | 0.00820 | 0.05114 | 0.17592 |
| **Smox** | 0.01310 | 0.00578 | 0.25542 | 1.05536 | 0.27869 | 0.03883 | 0.00457 | 0.19827 | 0.00553 | 0.00853 | 0.00591 | 0.02624 | 0.58250 | 0.00000 |
| **Smpd1** | 4.41680 | 0.74058 | 25.30024 | 25.33776 | 5.13096 | 17.59791 | 10.54763 | 26.63610 | 0.00000 | 13.81223 | 17.56238 | 0.14347 | 4.76468 | 2.45735 |
| **Smpd2** | 3.22678 | 5.00509 | 1.14944 | 6.18021 | 0.53990 | 0.00000 | 5.37930 | 1.78898 | 0.00000 | 0.09297 | 2.50615 | 3.12238 | 0.84606 | 0.00000 |
| **Smpd3** | 0.07073 | 0.00392 | 0.11408 | 0.00000 | 0.01129 | 0.36387 | 0.00376 | 0.05103 | 0.28710 | 0.00603 | 0.05933 | 0.47284 | 0.42831 | 0.11941 |
| **Smpd4** | 7.92663 | 0.43745 | 13.08114 | 4.74789 | 13.51654 | 9.85977 | 0.02776 | 4.35412 | 4.08287 | 0.18569 | 8.69102 | 6.95201 | 11.18494 | 13.56750 |
| **Smpdl3a** | 4.50526 | 8.28827 | 7.20152 | 10.76354 | 9.33819 | 8.12458 | 2.54568 | 4.08158 | 11.64356 | 0.04301 | 0.05644 | 0.19998 | 0.00000 | 1.58964 |
| **Smpdl3b** | 0.19482 | 0.07244 | 3.73473 | 4.07788 | 1.05341 | 2.43380 | 0.02941 | 2.87391 | 0.07325 | 0.02374 | 3.13361 | 4.23072 | 0.79960 | 0.36261 |
| **Sms** | 33.52077 | 4.05036 | 35.04151 | 23.79495 | 35.92441 | 18.03055 | 4.84340 | 10.49508 | 34.57096 | 3.98231 | 9.68005 | 18.63543 | 5.09214 | 23.31568 |
| **Smtn** | 0.00000 | 2.72059 | 2.49263 | 1.47030 | 2.11290 | 0.00000 | 1.60976 | 10.24895 | 0.00877 | 0.07860 | 0.00975 | 0.07253 | 0.00000 | 0.00000 |
| **Smtnl1** | 0.01009 | 0.01220 | 0.26568 | 0.00101 | 0.00820 | 0.01454 | 0.00422 | 0.00873 | 0.01051 | 0.00790 | 0.00652 | 0.02190 | 0.00873 | 0.01372 |
| **Smtnl2** | 0.68587 | 0.08151 | 0.04905 | 0.79426 | 0.23153 | 0.95253 | 1.00474 | 0.28137 | 0.41784 | 0.25361 | 0.03677 | 0.27389 | 0.85808 | 0.64400 |
| **Smu1** | 15.45136 | 8.62937 | 19.16236 | 9.31687 | 14.68939 | 18.82467 | 1.90168 | 10.96229 | 13.66168 | 5.38765 | 16.70019 | 12.79809 | 10.81800 | 19.29571 |
| **Smug1** | 2.52296 | 0.02234 | 1.67516 | 2.10039 | 2.62066 | 4.12453 | 0.00924 | 1.28482 | 3.75906 | 0.03047 | 0.13322 | 0.81499 | 0.00900 | 2.13054 |

| | | | | | | | | | | | | | | |
|---|---|---|---|---|---|---|---|---|---|---|---|---|---|---|
| **Smurf1** | 2.20101 | 3.72588 | 1.03085 | 1.64392 | 3.10606 | 2.19893 | 0.00000 | 4.40035 | 0.08218 | 9.52261 | 2.39467 | 3.19366 | 0.91811 | 0.69498 |
| **Smurf2** | 9.57673 | 0.03188 | 3.41476 | 0.00000 | 9.94031 | 6.40803 | 1.61457 | 2.48773 | 5.46017 | 0.08080 | 0.81137 | 4.54991 | 7.93918 | 13.48358 |
| **Smyd1** | 1.01679 | 0.03399 | 0.14919 | 0.56686 | 0.07840 | 0.00000 | 0.01524 | 3.69914 | 0.01341 | 0.01782 | 0.00000 | 1.22563 | 0.15048 | 0.28080 |
| **Smyd2** | 5.17621 | 0.11551 | 2.87684 | 1.38949 | 1.55452 | 1.20576 | 0.00670 | 2.55194 | 0.70625 | 11.08257 | 0.00000 | 0.26647 | 0.00000 | 0.00000 |
| **Smyd3** | 1.65199 | 0.05845 | 3.29792 | 0.00000 | 0.97859 | 2.83875 | 3.25962 | 2.72078 | 0.00492 | 0.03576 | 1.82572 | 1.85932 | 0.32817 | 1.04674 |
| **Smyd4** | 0.87438 | 0.04001 | 0.98325 | 0.12202 | 0.99391 | 2.07107 | 0.01623 | 0.27972 | 0.31110 | 0.00948 | 2.08912 | 0.91831 | 0.02821 | 2.27854 |
| **Smyd5** | 11.32074 | 8.35705 | 28.91474 | 24.56874 | 19.96773 | 12.37208 | 3.29500 | 6.19370 | 6.34401 | 0.13371 | 16.22211 | 12.47159 | 3.37724 | 20.07990 |
| **Snai1** | 1.40922 | 0.00795 | 4.01295 | 1.76507 | 3.35477 | 0.00000 | 0.00215 | 0.00000 | 0.25809 | 0.00000 | 0.00266 | 2.83389 | 8.26705 | 1.09142 |
| **Snai2** | 0.06142 | 0.01212 | 0.02342 | 0.01517 | 0.03222 | 0.02699 | 0.01616 | 0.00869 | 0.06089 | 0.00668 | 0.01079 | 0.01912 | 0.01494 | 0.03897 |
| **Snai3** | 0.00445 | 0.00443 | 0.00000 | 0.30672 | 0.71661 | 0.45225 | 0.00410 | 0.00458 | 0.00565 | 0.00569 | 0.00323 | 0.00317 | 0.13338 | 0.04106 |
| **Snap23** | 6.27682 | 0.00000 | 5.69775 | 2.18461 | 11.65243 | 6.33491 | 0.01512 | 0.88158 | 17.86342 | 0.02932 | 7.67673 | 3.11253 | 10.03400 | 10.26962 |
| **Snap29** | 4.98827 | 0.00783 | 5.20609 | 3.11636 | 1.74909 | 0.54614 | 2.96690 | 4.02286 | 0.27533 | 0.65561 | 1.05400 | 3.35425 | 4.61223 | 1.19466 |
| **Snap47** | 0.17502 | 0.68459 | 3.85417 | 4.85539 | 0.51902 | 5.53838 | 0.35607 | 0.43102 | 0.00000 | 0.00751 | 7.27405 | 4.57248 | 2.22380 | 2.10743 |
| **Snap91** | 1.11134 | 0.00393 | 0.50745 | 0.25512 | 1.67521 | 0.43625 | 0.00363 | 0.00000 | 1.48305 | 0.00725 | 0.00532 | 0.00244 | 0.08907 | 4.54008 |
| **Snapc1** | 2.31760 | 0.00199 | 1.65661 | 1.33413 | 2.45844 | 2.53126 | 0.00299 | 0.00000 | 0.16752 | 0.02654 | 0.74474 | 1.08933 | 7.72802 | 1.36914 |
| **Snapc2** | 2.69490 | 0.37189 | 4.52102 | 5.41841 | 7.06463 | 2.10278 | 0.80841 | 3.53368 | 0.03760 | 0.06053 | 4.18607 | 4.43076 | 8.52707 | 2.15682 |
| **Snapc3** | 3.44355 | 0.00137 | 5.10016 | 3.72064 | 15.33687 | 0.00000 | 0.13648 | 1.64390 | 2.49799 | 0.01168 | 0.25853 | 2.45140 | 8.35547 | 4.01283 |
| **Snapc4** | 3.07025 | 0.05296 | 6.83000 | 3.04397 | 3.64315 | 3.75352 | 5.22760 | 3.50913 | 0.34456 | 0.14612 | 4.60192 | 2.18155 | 8.19009 | 4.19902 |
| **Snapc5** | 3.07054 | 0.08423 | 1.20258 | 3.41392 | 4.73884 | 5.00070 | 0.00794 | 5.85094 | 0.70801 | 0.24048 | 1.69135 | 3.74734 | 6.59462 | 3.69895 |
| **Snapin** | 0.87534 | 0.01869 | 2.13034 | 1.75684 | 3.88676 | 3.46776 | 1.12990 | 0.18985 | 0.86282 | 0.02864 | 0.88119 | 2.30876 | 0.13405 | 4.09256 |
| **Snca** | 0.00774 | 0.00454 | 0.00284 | 0.00977 | 1.14633 | 0.01351 | 0.00457 | 0.01042 | 0.01318 | 0.00646 | 0.00889 | 0.00740 | 0.00339 | 0.00628 |
| **Sncb** | 0.00756 | 0.00505 | 0.03500 | 0.06402 | 0.14377 | 0.00783 | 0.00438 | 0.03426 | 0.01388 | 0.00729 | 0.00961 | 0.01073 | 0.00593 | 0.00740 |
| **Sncg** | 0.00355 | 0.00418 | 0.01095 | 0.35979 | 0.00438 | 0.00670 | 0.00406 | 0.00828 | 0.03910 | 0.00677 | 0.00627 | 0.00900 | 0.01967 | 0.08304 |
| **Snd1** | 0.03941 | 5.92360 | 1.78403 | 1.10208 | 1.65671 | 0.57689 | 2.39563 | 1.03706 | 0.23116 | 0.07662 | 0.22454 | 2.43043 | 0.92987 | 3.17372 |
| **Snf8** | 10.65080 | 3.73463 | 13.30946 | 13.75153 | 14.08897 | 12.23006 | 1.11626 | 9.56013 | 6.16066 | 2.11295 | 5.94577 | 7.25983 | 8.30889 | 14.14454 |
| **Snhg1** | 13.30775 | 8.07778 | 3.12070 | 7.24514 | 7.72297 | 7.36204 | 3.68235 | 5.89652 | 10.78454 | 0.56299 | 3.48527 | 2.93784 | 12.54018 | 2.20044 |
| **Snhg10** | 0.01644 | 0.00424 | 0.06012 | 0.01524 | 0.00457 | 0.01532 | 0.00405 | 0.00747 | 0.20706 | 0.00767 | 0.00706 | 0.00744 | 0.82148 | 0.00000 |
| **Snhg11** | 0.16598 | 0.03320 | 0.47250 | 3.51649 | 0.00000 | 5.02996 | 0.04585 | 0.03170 | 4.20512 | 0.16328 | 1.17527 | 1.45318 | 2.43848 | 0.66813 |
| **Snhg12** | 5.26926 | 0.10075 | 1.16586 | 4.77305 | 9.65354 | 3.27471 | 3.10036 | 7.02956 | 2.50079 | 0.20654 | 3.26790 | 5.12797 | 5.23050 | 2.35716 |
| **Snhg14** | 1.20288 | 0.01250 | 3.60302 | 1.24076 | 4.56956 | 0.00000 | 0.00077 | 0.58495 | 19.72473 | 0.00003 | 0.00000 | 0.04693 | 0.00000 | 0.20885 |
| **Snhg15** | 1.97349 | 0.01294 | 0.59333 | 2.24280 | 4.86274 | 1.65133 | 0.19571 | 0.63923 | 0.25524 | 0.41829 | 0.21333 | 1.54928 | 4.21436 | 0.77579 |
| **Snhg17** | 2.72434 | 0.13791 | 4.59525 | 7.47693 | 7.25930 | 0.00000 | 1.55078 | 3.78777 | 5.39523 | 0.45643 | 3.33903 | 1.31784 | 2.02230 | 3.87257 |
| **Snhg18** | 0.25511 | 0.00436 | 0.23079 | 0.00000 | 0.79362 | 0.06782 | 0.00397 | 0.00435 | 0.00318 | 0.00532 | 0.00574 | 0.01063 | 0.50297 | 0.83337 |
| **Snhg20** | 0.05614 | 0.32160 | 0.00771 | 0.78905 | 0.40889 | 0.14499 | 0.00392 | 0.48308 | 0.00484 | 0.00836 | 0.00000 | 0.67435 | 1.35324 | 0.02458 |
| **Snhg3** | 1.91055 | 0.03667 | 0.38278 | 2.01414 | 2.13501 | 2.43598 | 1.10543 | 2.78526 | 1.48919 | 1.29840 | 0.04781 | 0.79742 | 5.11954 | 0.50118 |
| **Snhg4** | 2.26074 | 0.02143 | 0.65111 | 0.79949 | 1.19362 | 0.14738 | 0.00459 | 0.36212 | 2.72777 | 0.02788 | 0.00232 | 0.00000 | 1.96264 | 0.63385 |
| **Snhg5** | 11.67225 | 2.15818 | 14.73716 | 11.39291 | 15.10168 | 9.13143 | 3.39926 | 11.94198 | 13.24701 | 0.21190 | 6.03032 | 8.99160 | 6.11429 | 15.19990 |
| **Snhg6** | 0.50009 | 0.00400 | 0.06278 | 0.00051 | 0.24577 | 0.10706 | 0.09494 | 0.09139 | 0.09720 | 0.52371 | 0.00515 | 0.21878 | 1.43665 | 0.10284 |
| **Snhg7** | 0.93226 | 0.01870 | 0.87513 | 1.55866 | 1.12894 | 0.68020 | 1.74776 | 1.34900 | 0.48663 | 0.06906 | 0.15941 | 0.62620 | 0.71717 | 2.26439 |
| **Snhg8** | 0.88451 | 0.02136 | 0.16522 | 0.00417 | 1.50594 | 1.00125 | 0.15896 | 0.67936 | 0.00724 | 0.04681 | 0.08135 | 0.67140 | 1.13378 | 1.17869 |
| **Snhg9** | 0.06575 | 0.00468 | 0.02097 | 0.06303 | 0.05150 | 0.08178 | 0.01051 | 0.05114 | 0.01822 | 0.02674 | 0.00597 | 0.02526 | 0.40590 | 0.02278 |
| **Snip1** | 6.05393 | 0.01106 | 5.10002 | 1.49749 | 3.69479 | 1.95792 | 4.07653 | 0.24655 | 3.01412 | 0.06647 | 0.41289 | 0.69840 | 12.11023 | 5.45705 |
| **Snn** | 0.12909 | 0.00371 | 0.06487 | 0.81037 | 0.00000 | 0.00000 | 0.00367 | 0.00286 | 0.70305 | 0.03103 | 0.10601 | 0.06253 | 1.39694 | 0.00000 |
| **Snora15** | 0.01324 | 0.00446 | 0.01532 | 0.03228 | 0.01217 | 0.01313 | 0.00432 | 0.01001 | 0.02126 | 0.00668 | 0.00808 | 0.00896 | 0.01711 | 0.01755 |
| **Snora16a** | 0.02063 | 0.00461 | 0.01954 | 0.00000 | 0.01316 | 0.03764 | 0.00517 | 0.06410 | 0.01445 | 0.00615 | 0.00611 | 0.08929 | 0.00625 | 0.01397 |
| **Snora21** | 0.16429 | 0.00455 | 0.09331 | 0.05186 | 0.37293 | 0.00659 | 0.00497 | 0.18762 | 0.13476 | 0.00746 | 0.04544 | 0.00518 | 0.00752 | 0.22042 |
| **Snora23** | 0.01181 | 0.00467 | 0.03572 | 0.00272 | 0.07191 | 0.03213 | 0.00423 | 0.00938 | 0.20000 | 0.00683 | 0.00843 | 0.00956 | 0.00567 | 0.00000 |
| **Snora28** | 0.00247 | 0.00436 | 0.09064 | 0.00000 | 0.03026 | 0.12116 | 0.00464 | 0.06556 | 0.32783 | 0.00518 | 0.24328 | 0.02051 | 0.00427 | 0.27556 |
| **Snora2b** | 0.06384 | 0.00420 | 0.02086 | 0.02812 | 0.00566 | 0.01241 | 0.00405 | 0.00727 | 0.04066 | 0.00643 | 0.00705 | 0.01013 | 0.71435 | 0.19205 |
| **Snora3** | 0.07253 | 0.00418 | 0.22701 | 0.00000 | 0.15205 | 0.00000 | 0.00402 | 0.47576 | 0.02116 | 0.00663 | 0.20333 | 0.12007 | 0.01365 | 0.30411 |
| **Snora31** | 0.31291 | 0.01194 | 0.26402 | 0.26541 | 0.56992 | 0.03346 | 0.00418 | 0.00510 | 0.01910 | 0.00636 | 0.00396 | 0.27174 | 0.25948 | 0.16987 |
| **Snora33** | 0.51741 | 0.14686 | 0.48136 | 0.19192 | 0.33554 | 0.34675 | 0.00422 | 0.22148 | 0.13197 | 0.01972 | 0.06019 | 0.32465 | 0.48841 | 0.32402 |
| **Snora34** | 0.04892 | 0.00458 | 0.08041 | 0.03217 | 0.06214 | 0.05299 | 0.00425 | 0.00971 | 0.05762 | 0.00690 | 0.00960 | 0.01226 | 0.00527 | 0.10770 |
| **Snora41** | 0.72549 | 0.00398 | 0.28061 | 0.04383 | 0.26436 | 0.32046 | 0.65809 | 0.00226 | 0.04724 | 0.02876 | 0.01519 | 0.31475 | 0.13304 | 0.20670 |
| **Snora43** | 0.01138 | 0.00474 | 0.24298 | 0.06821 | 0.02306 | 0.01610 | 0.00490 | 0.15742 | 0.09997 | 0.00559 | 0.00854 | 0.00754 | 0.00480 | 0.11499 |

| | | | | | | | | | | | | | | |
|---|---|---|---|---|---|---|---|---|---|---|---|---|---|---|
| **Snora44** | 0.00758 | 0.00520 | 0.34890 | 0.06657 | 0.10307 | 0.17135 | 0.00415 | 0.12861 | 0.54596 | 0.00709 | 0.38042 | 0.01220 | 0.00185 | 0.00000 |
| **Snora47** | 0.00475 | 0.00436 | 0.01463 | 0.01245 | 0.00979 | 0.01296 | 0.00425 | 0.00936 | 0.01662 | 0.00656 | 0.00896 | 0.00938 | 0.29806 | 0.01382 |
| **Snora52** | 0.00731 | 0.16126 | 0.06587 | 0.06477 | 0.04138 | 0.00651 | 0.00397 | 0.00679 | 0.00963 | 0.00746 | 0.04469 | 0.00506 | 0.00558 | 0.01672 |
| **Snora5c** | 0.00871 | 0.00457 | 0.04012 | 0.04524 | 0.01865 | 0.02989 | 0.00455 | 0.01486 | 0.00989 | 0.00696 | 0.00688 | 0.01863 | 0.00756 | 0.01703 |
| **Snora61** | 0.16979 | 0.01110 | 0.37062 | 0.58973 | 0.33031 | 0.15620 | 0.00348 | 0.39781 | 0.60865 | 0.00293 | 1.09804 | 0.32278 | 0.00292 | 0.16373 |
| **Snora62** | 0.01544 | 0.00473 | 0.07847 | 0.01144 | 0.07850 | 0.03059 | 0.00441 | 0.00886 | 0.01198 | 0.00901 | 0.01339 | 0.00820 | 0.00766 | 0.07094 |
| **Snora64** | 1.04441 | 0.00366 | 0.71342 | 0.18077 | 1.29175 | 0.38513 | 0.04234 | 0.13085 | 0.02382 | 0.01012 | 0.15185 | 0.11664 | 0.00244 | 0.53683 |
| **Snora65** | 1.07336 | 1.54151 | 1.48080 | 0.99436 | 0.90530 | 0.80664 | 1.23993 | 0.71309 | 0.16019 | 0.04059 | 0.02681 | 0.89706 | 0.00000 | 0.68898 |
| **Snora68** | 0.90394 | 0.44113 | 0.61459 | 0.41065 | 0.66837 | 0.28292 | 0.02649 | 0.41557 | 0.07174 | 0.02557 | 0.62048 | 0.02983 | 0.03851 | 0.24768 |
| **Snora69** | 0.30369 | 0.00418 | 0.14668 | 0.00000 | 0.04058 | 0.11891 | 0.00429 | 0.00454 | 0.01926 | 0.00590 | 0.00520 | 0.11647 | 0.00532 | 0.32944 |
| **Snora70** | 0.30924 | 0.01297 | 0.93272 | 1.19558 | 0.22681 | 0.75547 | 0.00965 | 0.20396 | 1.42805 | 0.03405 | 0.25952 | 0.21628 | 0.05522 | 0.91304 |
| **Snora74a** | 0.08705 | 0.00445 | 0.04544 | 0.00780 | 0.02385 | 0.00901 | 0.00419 | 0.01107 | 0.11110 | 0.00713 | 0.00692 | 0.00912 | 0.00412 | 0.13130 |
| **Snora75** | 0.00659 | 0.00432 | 0.08211 | 0.05971 | 0.03904 | 0.01766 | 0.00422 | 0.00745 | 0.00741 | 0.00616 | 0.01612 | 0.01613 | 0.02746 | 0.09123 |
| **Snora78** | 0.01581 | 0.00436 | 0.01827 | 0.05490 | 0.04257 | 0.02793 | 0.00424 | 0.05956 | 0.01354 | 0.00722 | 0.00730 | 0.01197 | 0.00852 | 0.01121 |
| **Snora7a** | 0.11697 | 0.00426 | 0.05501 | 0.00000 | 0.25209 | 0.06432 | 0.00463 | 0.14560 | 0.01167 | 0.00685 | 0.30913 | 0.34808 | 0.02294 | 0.24173 |
| **Snora81** | 0.03219 | 0.00447 | 0.27326 | 0.05391 | 0.01345 | 0.04667 | 0.00410 | 0.00526 | 0.58915 | 0.00550 | 0.02608 | 0.00750 | 0.01259 | 0.12909 |
| **Snord100** | 0.48502 | 0.07610 | 0.79250 | 0.56176 | 0.81716 | 0.23717 | 0.00401 | 0.02427 | 0.29150 | 0.00738 | 0.50977 | 0.53202 | 0.28512 | 0.35036 |
| **Snord104** | 0.03005 | 0.00434 | 0.01003 | 0.03160 | 0.05051 | 0.03169 | 0.00420 | 0.00641 | 0.01008 | 0.01065 | 0.01213 | 0.05109 | 0.14568 | 0.00842 |
| **Snord12** | 0.70083 | 0.00515 | 0.70387 | 0.00001 | 0.68457 | 0.26044 | 0.00428 | 0.24598 | 0.46171 | 0.00980 | 0.41074 | 0.13206 | 0.00110 | 0.21594 |
| **Snord15a** | 0.01367 | 0.00518 | 0.08174 | 0.25609 | 0.05066 | 0.00581 | 0.00443 | 0.10862 | 0.00918 | 0.00820 | 0.00886 | 0.00729 | 0.00671 | 0.09196 |
| **Snord15b** | 0.24460 | 0.03416 | 0.30418 | 0.00000 | 0.09724 | 0.00000 | 0.00433 | 0.30663 | 0.18505 | 0.00597 | 0.09940 | 0.07869 | 0.00397 | 0.01352 |
| **Snord16a** | 0.00000 | 0.01209 | 0.67375 | 0.40161 | 0.29204 | 0.22884 | 0.06764 | 0.16295 | 0.78036 | 0.00979 | 0.49365 | 0.35384 | 0.01425 | 0.59332 |
| **Snord17** | 0.00000 | 0.00423 | 0.11684 | 0.34750 | 0.18281 | 0.00000 | 0.00398 | 0.31265 | 0.12169 | 0.00365 | 0.44048 | 0.03855 | 0.00360 | 0.47031 |
| **Snord1b** | 0.01525 | 0.00445 | 0.08504 | 0.01022 | 0.01196 | 0.01115 | 0.00429 | 0.01171 | 0.01213 | 0.00794 | 0.00839 | 0.00899 | 0.00819 | 0.01073 |
| **Snord2** | 0.01015 | 0.00449 | 0.06040 | 0.03514 | 0.01355 | 0.02333 | 0.00414 | 0.00807 | 0.23129 | 0.00645 | 0.02862 | 0.00947 | 0.00470 | 0.04070 |
| **Snord22** | 1.95827 | 1.28720 | 0.98996 | 0.64917 | 0.45734 | 0.84801 | 0.00352 | 0.07247 | 1.44487 | 0.00339 | 0.67529 | 0.92804 | 0.04559 | 0.08168 |
| **Snord32a** | 0.05974 | 0.00396 | 0.30591 | 0.33843 | 0.05665 | 0.07738 | 0.00401 | 0.03539 | 0.02483 | 0.00534 | 0.40563 | 0.19813 | 0.00406 | 0.12909 |
| **Snord33** | 0.38069 | 0.01191 | 0.71963 | 0.29455 | 0.40105 | 0.12946 | 0.43031 | 0.20548 | 0.00815 | 0.01068 | 0.50078 | 0.37283 | 0.00162 | 0.29260 |
| **Snord34** | 0.51917 | 0.00402 | 0.82666 | 0.39153 | 0.18061 | 0.17478 | 0.00466 | 0.34484 | 0.00589 | 0.00230 | 0.78551 | 0.33432 | 0.16002 | 0.43504 |
| **Snord35a** | 0.79183 | 0.01375 | 1.44753 | 0.30972 | 1.07473 | 0.28971 | 0.00397 | 0.32927 | 0.00340 | 0.01064 | 0.49508 | 0.46325 | 0.18683 | 0.73405 |
| **Snord35b** | 0.41074 | 0.00547 | 0.22243 | 0.19548 | 0.01725 | 0.10371 | 0.00430 | 0.17410 | 0.01115 | 0.01056 | 0.00482 | 0.00718 | 0.01826 | 0.09903 |
| **Snord38a** | 0.03762 | 0.00425 | 0.02075 | 0.06828 | 0.02304 | 0.01463 | 0.00412 | 0.00637 | 0.01637 | 0.00896 | 0.00937 | 0.04990 | 0.00735 | 0.04510 |
| **Snord42a** | 0.00813 | 1.10728 | 0.02275 | 0.05291 | 0.01072 | 0.09612 | 0.00411 | 0.00724 | 0.01489 | 0.00705 | 0.11501 | 0.17645 | 0.00474 | 0.03695 |
| **Snord42b** | 0.11170 | 1.27009 | 0.13336 | 0.22951 | 0.15143 | 0.00000 | 0.00360 | 0.00258 | 0.01188 | 0.37061 | 0.16035 | 0.00548 | 0.01052 | 0.14978 |
| **Snord45b** | 0.00621 | 0.00445 | 0.04056 | 0.00604 | 0.02002 | 0.01212 | 0.00496 | 0.03654 | 0.09976 | 0.00620 | 0.01410 | 0.03633 | 0.00722 | 0.11296 |
| **Snord47** | 1.30589 | 0.00375 | 0.48713 | 0.26114 | 0.25979 | 0.29305 | 0.32822 | 0.06280 | 0.46020 | 0.01928 | 0.95343 | 0.14949 | 0.09332 | 0.18289 |
| **Snord49a** | 0.02507 | 0.01326 | 0.08402 | 0.05068 | 0.00670 | 0.00227 | 0.00451 | 0.46356 | 0.12106 | 0.00614 | 0.07402 | 0.07932 | 0.00396 | 0.01516 |
| **Snord49b** | 0.03527 | 0.00477 | 0.02189 | 0.17738 | 0.06980 | 0.02158 | 0.00446 | 0.02388 | 0.01300 | 0.01044 | 0.01625 | 0.01774 | 0.03426 | 0.00784 |
| **Snord4a** | 0.21752 | 0.73886 | 0.11297 | 0.07322 | 0.16363 | 0.04896 | 0.00422 | 0.03491 | 0.01088 | 0.00834 | 0.08836 | 0.18904 | 0.05299 | 0.09909 |
| **Snord52** | 0.01862 | 1.32255 | 0.55821 | 0.17049 | 0.02590 | 0.30887 | 0.04192 | 0.10877 | 0.45424 | 0.82315 | 0.09172 | 0.04157 | 0.00268 | 0.20221 |
| **Snord53** | 0.00940 | 0.00444 | 0.05707 | 0.01984 | 0.03815 | 0.01003 | 0.00418 | 0.01045 | 0.07259 | 0.00761 | 0.00951 | 0.00997 | 0.00457 | 0.04320 |
| **Snord55** | 0.01639 | 0.00437 | 0.01120 | 0.10476 | 0.17062 | 0.01154 | 0.00414 | 0.00749 | 0.01301 | 0.00578 | 0.01029 | 0.01004 | 0.00574 | 0.00850 |
| **Snord57** | 0.01105 | 0.00538 | 0.13236 | 0.02233 | 0.08888 | 0.12051 | 0.00436 | 0.15585 | 0.02321 | 0.01018 | 0.00636 | 0.02347 | 0.00448 | 0.21640 |
| **Snord58b** | 0.29775 | 0.00441 | 0.18027 | 0.10131 | 0.10011 | 0.19954 | 0.00982 | 0.00361 | 0.04359 | 0.00577 | 0.21231 | 0.00598 | 0.00995 | 0.22640 |
| **Snord65** | 0.14510 | 0.00411 | 0.10092 | 0.23866 | 0.14411 | 0.09127 | 0.00392 | 0.00331 | 0.22923 | 0.00670 | 0.00548 | 0.06289 | 0.12326 | 0.08335 |
| **Snord68** | 0.00845 | 0.10172 | 0.01573 | 0.10142 | 0.18696 | 0.02065 | 0.00415 | 0.00789 | 0.01331 | 0.00835 | 0.00818 | 0.04456 | 0.00669 | 0.00952 |
| **Snord69** | 0.01083 | 0.01498 | 0.07708 | 0.09038 | 0.03720 | 0.05344 | 0.00412 | 0.03834 | 0.32994 | 0.00720 | 0.00632 | 0.05600 | 0.00463 | 0.14785 |
| **Snord72** | 0.37677 | 0.00397 | 0.20695 | 0.41137 | 0.39964 | 0.17165 | 0.00409 | 0.00628 | 0.00711 | 0.00617 | 0.04729 | 0.00924 | 0.49827 | 0.32134 |
| **Snord73a** | 0.00505 | 0.00439 | 0.05964 | 0.06019 | 0.05946 | 0.00003 | 0.01001 | 0.00952 | 0.22797 | 0.00561 | 0.33470 | 0.00897 | 0.00356 | 0.01357 |
| **Snord8** | 0.03252 | 0.00457 | 0.06175 | 0.00685 | 0.01663 | 0.01830 | 0.00431 | 0.01035 | 0.03892 | 0.00719 | 0.00712 | 0.00927 | 0.01167 | 0.19464 |
| **Snord83b** | 0.21668 | 0.00537 | 0.21112 | 0.31554 | 0.02936 | 0.05388 | 0.00414 | 0.07695 | 0.27724 | 0.00854 | 0.01423 | 0.04775 | 0.00717 | 0.10678 |
| **Snord87** | 0.02685 | 0.00439 | 0.02219 | 0.01280 | 0.00733 | 0.01071 | 0.00424 | 0.01017 | 0.01747 | 0.00708 | 0.01562 | 0.01151 | 0.02707 | 0.11844 |
| **Snord89** | 0.00566 | 0.00421 | 0.08233 | 0.00000 | 0.02525 | 0.05339 | 0.09897 | 0.00575 | 0.03174 | 0.00705 | 0.00661 | 0.01056 | 0.01161 | 0.15612 |
| **Snord92** | 0.01030 | 0.00449 | 0.02684 | 0.00592 | 0.06347 | 0.07210 | 0.00422 | 0.00806 | 0.12601 | 0.00665 | 0.00664 | 0.00972 | 0.00595 | 0.00687 |
| **Snord95** | 0.00816 | 0.00413 | 0.25203 | 0.00000 | 0.12711 | 0.06439 | 0.05591 | 0.00573 | 0.01509 | 0.00524 | 0.02232 | 0.04694 | 0.00303 | 0.17705 |

| | | | | | | | | | | | | | | |
|---|---|---|---|---|---|---|---|---|---|---|---|---|---|---|
| **Snord96a** | 0.04559 | 0.00414 | 0.56846 | 0.80072 | 0.17461 | 0.05336 | 0.32204 | 0.26959 | 0.37279 | 0.01119 | 0.09799 | 0.01635 | 0.00785 | 0.30068 |
| **Snord98** | 0.00872 | 0.00489 | 0.02560 | 0.01433 | 0.01095 | 0.01105 | 0.00439 | 0.02697 | 0.13540 | 0.00892 | 0.00836 | 0.04515 | 0.02031 | 0.04069 |
| **Snord99** | 0.07296 | 0.00450 | 0.17474 | 0.23240 | 0.07682 | 0.24886 | 0.00399 | 0.49058 | 0.47179 | 0.00600 | 0.77476 | 0.00596 | 0.00371 | 0.00000 |
| **Snph** | 0.00713 | 0.00432 | 0.01351 | 0.01466 | 0.00987 | 0.01269 | 0.00421 | 0.00905 | 0.00560 | 0.00690 | 0.00904 | 0.00885 | 0.39686 | 0.07583 |
| **Snrk** | 0.14326 | 0.01105 | 1.21599 | 3.30758 | 0.21673 | 0.73577 | 2.96970 | 1.01380 | 0.01724 | 0.01875 | 1.33001 | 2.35301 | 0.75974 | 0.00000 |
| **Snrnp200** | 14.36703 | 0.12571 | 17.40197 | 4.92028 | 13.92322 | 13.22069 | 3.84358 | 0.41666 | 15.80351 | 0.10714 | 12.11783 | 11.52730 | 5.33318 | 14.70282 |
| **Snrnp25** | 5.15338 | 1.08618 | 2.43751 | 7.13810 | 9.70734 | 4.63828 | 0.01160 | 7.06476 | 0.75912 | 0.05510 | 3.48363 | 7.35285 | 8.55788 | 9.71146 |
| **Snrnp27** | 4.97933 | 4.81192 | 1.92150 | 7.14215 | 10.14725 | 5.13791 | 4.29124 | 6.03422 | 1.34386 | 0.53950 | 2.26325 | 6.03440 | 3.29956 | 7.97702 |
| **Snrnp35** | 0.53816 | 0.02949 | 0.18161 | 0.68832 | 0.68923 | 0.42688 | 0.00377 | 1.99126 | 1.12779 | 0.02356 | 2.91680 | 0.26137 | 2.73545 | 1.42539 |
| **Snrnp40** | 7.74401 | 0.07162 | 11.80391 | 16.54288 | 14.02946 | 13.15847 | 0.90848 | 4.06501 | 2.90529 | 4.53017 | 2.81967 | 8.39740 | 2.93597 | 6.85351 |
| **Snrnp48** | 2.89607 | 0.00991 | 5.82734 | 3.28097 | 5.11964 | 5.39067 | 1.45938 | 0.35495 | 2.27617 | 0.02323 | 0.17980 | 2.50765 | 4.23163 | 5.01432 |
| **Snrnp70** | 0.20326 | 1.32320 | 0.70287 | 3.13757 | 2.33596 | 1.14273 | 0.12849 | 3.72325 | 0.31216 | 4.61246 | 0.88611 | 3.86157 | 2.90776 | 2.06958 |
| **Snrpa** | 0.00000 | 3.44761 | 10.33562 | 18.22869 | 18.10857 | 0.64743 | 3.59509 | 9.22739 | 0.00000 | 13.39381 | 0.00941 | 10.85251 | 6.87999 | 0.00000 |
| **Snrpa1** | 11.40663 | 0.93748 | 16.81760 | 21.38944 | 20.40930 | 11.93751 | 3.64433 | 10.89916 | 1.59280 | 1.68976 | 11.22089 | 8.71445 | 7.03213 | 8.69574 |
| **Snrpb** | 6.20642 | 10.23011 | 40.27456 | 37.08658 | 30.40459 | 17.01722 | 10.14610 | 20.39038 | 0.08872 | 1.23169 | 14.63235 | 22.67277 | 5.05641 | 0.00000 |
| **Snrpb2** | 12.55124 | 8.65974 | 15.66276 | 13.73799 | 15.79551 | 17.74212 | 6.32397 | 9.69168 | 25.46700 | 3.22423 | 15.47530 | 12.15394 | 2.04402 | 16.38471 |
| **Snrpc** | 2.22838 | 8.96675 | 18.15749 | 25.76757 | 28.02299 | 1.34072 | 6.54239 | 16.73384 | 0.27699 | 6.59045 | 5.96357 | 15.63881 | 9.35412 | 0.52052 |
| **Snrpd1** | 10.80076 | 4.32436 | 1.50813 | 17.26766 | 14.34056 | 14.21321 | 6.09519 | 8.31248 | 7.22056 | 16.88714 | 4.12273 | 9.77187 | 15.24031 | 11.43372 |
| **Snrpd2** | 8.17668 | 4.29465 | 0.50587 | 7.85781 | 14.17101 | 15.41572 | 5.06123 | 13.52844 | 0.15204 | 6.00501 | 1.37053 | 13.28194 | 12.10063 | 6.38949 |
| **Snrpd3** | 12.53043 | 0.16263 | 1.04719 | 10.32140 | 6.82949 | 5.47626 | 3.88958 | 4.14325 | 5.21672 | 13.61740 | 1.51469 | 8.60699 | 4.63810 | 6.79008 |
| **Snrpe** | 7.46930 | 3.21002 | 0.00000 | 8.64002 | 13.32569 | 12.66231 | 4.05277 | 10.53432 | 9.62674 | 11.29939 | 1.23715 | 9.84930 | 17.06392 | 7.88253 |
| **Snrpf** | 0.00000 | 0.34663 | 0.00000 | 0.98864 | 1.52080 | 0.64234 | 0.25367 | 0.24833 | 0.16199 | 0.07230 | 0.00625 | 0.84579 | 0.36154 | 0.18715 |
| **Snrpg** | 6.15181 | 0.33194 | 0.00000 | 3.84753 | 9.98250 | 16.16034 | 3.02345 | 9.08521 | 2.35640 | 8.77862 | 0.66568 | 11.04234 | 16.96721 | 3.64403 |
| **Snta1** | 0.56744 | 0.01050 | 1.20072 | 1.11833 | 1.62301 | 0.00000 | 0.01413 | 0.00656 | 0.00717 | 0.00984 | 0.00336 | 0.32092 | 1.05826 | 0.06409 |
| **Sntb2** | 0.00000 | 0.00247 | 0.65920 | 0.53131 | 0.21971 | 0.01406 | 0.00390 | 1.09948 | 1.05502 | 0.01176 | 0.00000 | 1.90474 | 6.17525 | 3.42041 |
| **Sntg2** | 0.00848 | 0.00441 | 0.11958 | 0.00671 | 0.00824 | 0.00803 | 0.00422 | 0.01123 | 0.01059 | 0.00879 | 0.00725 | 0.00926 | 0.00558 | 0.14356 |
| **Snupn** | 10.08443 | 0.01399 | 6.10048 | 0.95773 | 13.50309 | 3.55271 | 3.10685 | 5.67295 | 15.52647 | 0.11775 | 1.87158 | 1.42491 | 9.45845 | 19.52122 |
| **Snw1** | 0.45680 | 0.04638 | 0.82972 | 0.41532 | 2.89882 | 7.81135 | 0.20943 | 2.21469 | 8.69929 | 0.06003 | 5.42433 | 3.59111 | 3.07346 | 4.54098 |
| **Snx1** | 14.68658 | 10.93658 | 21.71382 | 13.39705 | 13.68003 | 28.42011 | 9.97160 | 15.56203 | 1.02112 | 11.48842 | 22.85968 | 9.01879 | 4.33938 | 9.74328 |
| **Snx10** | 1.07214 | 0.01554 | 2.09027 | 3.07613 | 3.56613 | 2.92878 | 0.00188 | 0.35596 | 4.78380 | 0.02188 | 2.77762 | 4.95203 | 0.44085 | 2.56373 |
| **Snx11** | 0.00000 | 0.00481 | 2.50239 | 1.08269 | 3.78280 | 0.02697 | 0.01011 | 0.50119 | 0.00390 | 0.01154 | 0.01975 | 0.00353 | 0.33596 | 0.28389 |
| **Snx12** | 2.55063 | 4.61890 | 1.27864 | 1.21637 | 5.08054 | 2.86287 | 3.79079 | 4.57738 | 4.93629 | 0.08803 | 1.45022 | 0.89084 | 2.22356 | 4.02721 |
| **Snx13** | 0.59641 | 0.01884 | 0.23817 | 1.14476 | 0.72691 | 2.73750 | 0.00305 | 0.46078 | 3.93282 | 0.00725 | 2.70114 | 0.71278 | 0.48054 | 0.45288 |
| **Snx14** | 5.08306 | 0.01232 | 2.72106 | 0.00000 | 3.93745 | 1.46789 | 0.00221 | 0.18873 | 8.87216 | 0.00235 | 0.48403 | 1.27677 | 0.32536 | 3.03541 |
| **Snx15** | 0.29449 | 2.12036 | 1.85060 | 2.96637 | 1.04104 | 1.20398 | 4.75453 | 4.00240 | 0.27053 | 0.19467 | 1.50412 | 4.49541 | 1.43219 | 0.75201 |
| **Snx16** | 14.91846 | 0.00849 | 7.12836 | 3.05713 | 13.67568 | 3.74026 | 2.14754 | 0.00000 | 14.15942 | 0.00000 | 0.08198 | 1.51118 | 1.22315 | 3.17930 |
| **Snx17** | 0.00000 | 0.23686 | 3.93846 | 0.10378 | 3.43251 | 5.52157 | 5.50655 | 7.92782 | 0.00841 | 0.19598 | 8.32516 | 6.50236 | 2.10248 | 2.00306 |
| **Snx18** | 0.53375 | 0.02252 | 1.28665 | 0.08251 | 4.79258 | 2.00239 | 0.98537 | 1.91179 | 1.96425 | 0.01604 | 5.87112 | 4.88341 | 5.25806 | 6.06337 |
| **Snx19** | 0.11182 | 0.00394 | 0.00000 | 0.00012 | 0.00000 | 0.47142 | 1.40030 | 0.00199 | 2.14075 | 0.01269 | 0.00348 | 0.00731 | 0.03641 | 0.18645 |
| **Snx2** | 14.39710 | 11.06087 | 21.04054 | 26.12890 | 12.32659 | 36.42144 | 11.44534 | 24.60173 | 17.62343 | 5.13382 | 25.19985 | 22.81353 | 1.93725 | 10.29468 |
| **Snx21** | 0.01309 | 0.00441 | 0.01054 | 0.10202 | 0.12957 | 0.05401 | 0.00409 | 0.13047 | 0.01253 | 0.00556 | 0.00647 | 0.13399 | 0.00665 | 0.01044 |
| **Snx22** | 0.12994 | 0.03287 | 1.06687 | 2.32043 | 0.73503 | 1.17589 | 0.00377 | 1.70029 | 0.00222 | 0.02227 | 0.00000 | 0.06563 | 0.00000 | 1.41744 |
| **Snx24** | 0.03499 | 0.01357 | 0.44779 | 0.05073 | 0.00194 | 0.83797 | 0.00421 | 0.04937 | 0.00982 | 0.01362 | 0.00000 | 0.39226 | 0.00590 | 0.00335 |
| **Snx25** | 0.01142 | 0.00296 | 0.00000 | 0.00000 | 0.00764 | 1.71496 | 3.52407 | 0.45084 | 0.01267 | 0.23957 | 1.01563 | 0.78142 | 0.55928 | 0.00000 |
| **Snx27** | 7.45914 | 8.92851 | 14.33237 | 5.86996 | 10.49924 | 11.27436 | 0.06838 | 3.42675 | 9.65350 | 0.02904 | 8.78826 | 10.70693 | 2.52106 | 13.36407 |
| **Snx29** | 0.00000 | 0.00435 | 0.22987 | 0.00822 | 0.11252 | 0.00443 | 0.00414 | 0.01191 | 0.05220 | 0.00651 | 0.00682 | 0.00859 | 0.35953 | 0.25872 |
| **Snx3** | 25.57050 | 9.04317 | 16.56108 | 23.96569 | 31.06592 | 16.02077 | 4.82558 | 11.05149 | 3.46451 | 0.25214 | 6.51503 | 10.73662 | 11.63100 | 9.31121 |
| **Snx30** | 6.35523 | 7.87849 | 10.78982 | 1.20947 | 6.53705 | 6.38487 | 2.73093 | 0.85202 | 7.01093 | 0.17743 | 0.09511 | 2.07626 | 1.05489 | 6.94850 |
| **Snx32** | 0.01423 | 0.00432 | 0.01665 | 0.01160 | 0.00876 | 0.01267 | 0.00428 | 0.00928 | 0.00972 | 0.00624 | 0.00901 | 0.00904 | 0.45841 | 0.03108 |
| **Snx33** | 0.01931 | 0.00490 | 0.00750 | 0.00187 | 0.22697 | 0.01406 | 0.00404 | 0.01690 | 0.04898 | 0.00736 | 0.00570 | 0.12311 | 0.03685 | 0.00152 |
| **Snx4** | 6.62650 | 4.73791 | 10.59227 | 3.16433 | 10.31634 | 5.35118 | 3.70438 | 2.12015 | 4.16676 | 0.08183 | 6.75771 | 6.64962 | 3.88583 | 12.96675 |
| **Snx5** | 26.76439 | 0.57674 | 24.73379 | 24.17985 | 25.33675 | 15.89110 | 7.54464 | 11.20733 | 25.49564 | 0.51783 | 19.52207 | 14.00570 | 9.35958 | 19.83364 |
| **Snx6** | 14.23623 | 0.45159 | 22.80926 | 13.61364 | 24.18159 | 16.15277 | 4.22390 | 7.73828 | 7.57578 | 0.24604 | 12.73662 | 12.60349 | 6.39979 | 22.25239 |
| **Snx7** | 0.00766 | 0.00415 | 0.00111 | 0.11588 | 0.00569 | 0.33337 | 0.79004 | 0.00431 | 0.01769 | 0.02985 | 0.00085 | 0.30444 | 0.40753 | 0.00000 |
| **Snx8** | 2.10084 | 0.15881 | 4.69406 | 9.71071 | 7.64487 | 2.78765 | 2.95256 | 2.15497 | 0.06897 | 0.05456 | 3.47833 | 0.03578 | 2.09443 | 3.71125 |

| | | | | | | | | | | | | | | |
|---|---|---|---|---|---|---|---|---|---|---|---|---|---|---|
| **Snx9** | 0.84294 | 0.01850 | 0.00000 | 0.00000 | 1.12144 | 3.10866 | 2.30131 | 0.59000 | 0.00672 | 0.04078 | 1.14007 | 1.68158 | 0.27679 | 1.88403 |
| **Soat1** | 0.00000 | 4.08828 | 3.37243 | 2.00216 | 0.90452 | 1.41904 | 1.31980 | 2.24977 | 8.23619 | 0.06459 | 0.51015 | 2.31294 | 1.80972 | 5.19118 |
| **Soat2** | 0.75077 | 0.02716 | 0.93625 | 0.30770 | 1.06793 | 0.01623 | 0.01493 | 0.01051 | 0.87929 | 0.03255 | 0.00609 | 0.05699 | 1.47573 | 0.53891 |
| **Socs1** | 0.02406 | 0.58821 | 0.01485 | 0.05149 | 0.03419 | 0.22670 | 0.17897 | 0.37560 | 0.01141 | 0.06634 | 0.02185 | 0.01020 | 1.14211 | 0.00000 |
| **Socs2** | 0.55200 | 0.02994 | 1.51533 | 2.90026 | 0.33113 | 0.45831 | 0.00242 | 0.04447 | 0.46924 | 0.00000 | 1.13882 | 4.36288 | 0.59491 | 1.89591 |
| **Socs3** | 11.91593 | 0.12261 | 5.02311 | 2.30181 | 13.84035 | 0.00000 | 0.01204 | 1.51073 | 2.12297 | 0.00000 | 0.49000 | 7.86393 | 13.85193 | 11.77962 |
| **Socs4** | 0.54162 | 0.09662 | 0.77617 | 0.32657 | 1.49207 | 0.77545 | 0.00375 | 0.00000 | 1.12411 | 0.00498 | 0.00579 | 0.11849 | 4.92746 | 2.82028 |
| **Socs5** | 0.06573 | 0.00418 | 0.03109 | 0.13231 | 0.00112 | 0.21922 | 1.77104 | 0.00641 | 0.00137 | 0.11292 | 0.02135 | 0.05933 | 2.70783 | 0.04464 |
| **Socs6** | 4.21729 | 0.00877 | 1.71509 | 1.65748 | 0.00000 | 1.76306 | 1.46504 | 2.30162 | 3.66662 | 0.03548 | 0.00000 | 2.05809 | 1.16161 | 2.22196 |
| **Socs7** | 0.00000 | 0.00178 | 1.10494 | 0.43257 | 4.21888 | 1.75270 | 2.28283 | 2.26163 | 4.23006 | 0.03449 | 0.47326 | 1.28557 | 7.80271 | 11.60276 |
| **Sod1** | 3.25758 | 6.14273 | 0.84727 | 14.13958 | 7.71210 | 23.62349 | 8.84285 | 15.20652 | 1.10913 | 15.37729 | 4.58176 | 23.54451 | 6.63515 | 1.08523 |
| **Sod2** | 12.04093 | 5.95245 | 7.21089 | 28.14060 | 15.92316 | 19.24426 | 4.42747 | 14.60582 | 13.48254 | 10.34064 | 8.64393 | 11.25427 | 13.40976 | 5.16614 |
| **Soga1** | 0.10034 | 0.12888 | 0.46391 | 0.38157 | 0.48325 | 0.62804 | 0.00958 | 0.00593 | 0.37532 | 0.01231 | 0.01040 | 0.62738 | 0.00156 | 1.56674 |
| **Sohlh1** | 0.19854 | 0.00457 | 0.36067 | 0.00139 | 0.00195 | 0.01895 | 0.00416 | 0.00920 | 0.00934 | 0.00776 | 0.00765 | 0.00939 | 0.00791 | 0.00984 |
| **Son** | 15.33653 | 1.86145 | 12.94021 | 7.18701 | 13.30675 | 7.78394 | 0.74460 | 3.44312 | 17.37239 | 0.03504 | 6.57215 | 6.34412 | 11.85515 | 11.97787 |
| **Sorbs1** | 3.42751 | 0.00963 | 8.85133 | 3.01572 | 5.55638 | 7.01666 | 2.99430 | 1.80439 | 11.20595 | 0.03688 | 1.13094 | 1.75776 | 2.61453 | 8.13576 |
| **Sorbs2** | 0.04978 | 0.00447 | 0.01306 | 0.00868 | 0.02342 | 0.00765 | 0.00543 | 0.06794 | 0.03728 | 0.00655 | 0.00805 | 0.00807 | 0.19176 | 0.00000 |
| **Sorbs3** | 0.01940 | 0.02332 | 0.34707 | 0.00000 | 0.00797 | 0.16829 | 0.00390 | 1.80397 | 0.00379 | 0.05342 | 0.00410 | 0.03306 | 0.50583 | 0.16687 |
| **Sorcs2** | 0.07716 | 0.00454 | 0.25114 | 0.01647 | 0.07989 | 0.00975 | 0.00438 | 0.02727 | 0.01088 | 0.00863 | 0.00842 | 0.00623 | 0.00557 | 0.01084 |
| **Sorcs3** | 0.00897 | 0.00509 | 0.01136 | 0.04773 | 0.01321 | 0.00839 | 0.00439 | 0.03432 | 0.01514 | 0.00757 | 0.00775 | 0.01272 | 0.01003 | 0.01687 |
| **Sord** | 27.97090 | 15.49325 | 50.75749 | 47.90697 | 39.66232 | 37.61551 | 14.68624 | 36.23477 | 40.19663 | 21.90166 | 26.20518 | 15.97699 | 12.19356 | 42.98981 |
| **Sorl1** | 0.34857 | 0.00425 | 0.71232 | 0.02818 | 0.08192 | 0.00000 | 0.00400 | 0.00983 | 0.00564 | 0.00512 | 0.01017 | 0.01189 | 0.00119 | 0.85746 |
| **Sort1** | 0.01069 | 0.01215 | 0.01855 | 0.38349 | 0.06253 | 0.19310 | 0.00485 | 0.57587 | 0.00864 | 0.00725 | 0.00562 | 0.00940 | 0.03814 | 0.03793 |
| **Sos1** | 13.05769 | 0.39812 | 9.93048 | 1.65591 | 13.16405 | 1.52616 | 1.39680 | 0.00000 | 17.35438 | 2.35894 | 1.01472 | 1.01202 | 10.48168 | 17.56748 |
| **Sos2** | 3.25181 | 0.00065 | 2.48859 | 0.00000 | 1.82287 | 2.14747 | 0.00193 | 0.00000 | 3.06679 | 0.00000 | 5.92422 | 1.80171 | 0.41949 | 1.75820 |
| **Sowaha** | 0.00000 | 0.00404 | 0.37508 | 0.07064 | 0.07899 | 0.00232 | 0.00392 | 0.00647 | 0.50583 | 0.00735 | 0.00508 | 0.00619 | 1.51093 | 0.00000 |
| **Sowahb** | 0.00281 | 0.01199 | 0.07636 | 0.00000 | 0.01115 | 1.30220 | 0.00388 | 0.01462 | 0.01058 | 0.00729 | 0.00686 | 0.53327 | 0.01237 | 0.00768 |
| **Sowahc** | 0.29241 | 0.00465 | 0.11995 | 0.50663 | 0.38078 | 0.54348 | 0.00442 | 0.72470 | 1.37342 | 0.01419 | 0.12780 | 0.10230 | 0.00157 | 2.39703 |
| **Sox11** | 0.06277 | 0.00397 | 0.00537 | 0.62267 | 0.11752 | 0.00000 | 0.00375 | 0.01187 | 0.01941 | 0.00509 | 0.02723 | 0.01844 | 0.00563 | 0.00000 |
| **Sox12** | 0.90564 | 3.42513 | 1.91077 | 0.93885 | 2.34909 | 0.19131 | 0.01321 | 0.00000 | 1.98223 | 0.01016 | 1.02588 | 1.22353 | 0.44352 | 3.02910 |
| **Sox13** | 3.61251 | 0.10444 | 10.03723 | 7.40953 | 3.63682 | 10.94875 | 0.23746 | 7.09655 | 0.00864 | 0.04941 | 3.40893 | 1.42297 | 4.81777 | 1.75131 |
| **Sox15** | 15.45012 | 0.08078 | 26.74961 | 6.34053 | 27.40740 | 1.56908 | 0.03949 | 1.09553 | 0.22645 | 13.33370 | 3.73137 | 2.44636 | 6.76183 | 6.36260 |
| **Sox17** | 0.00379 | 0.08516 | 0.00824 | 0.00000 | 0.44774 | 0.00488 | 0.00429 | 0.42839 | 0.02402 | 0.01376 | 0.00000 | 2.10625 | 0.00486 | 0.00994 |
| **Sox2** | 0.00000 | 0.47687 | 1.77313 | 1.15010 | 0.16205 | 0.00000 | 0.00000 | 0.00000 | 0.00384 | 0.02803 | 3.65417 | 9.00850 | 0.09514 | 0.00000 |
| **Sox21** | 7.11981 | 0.00841 | 7.24414 | 0.00000 | 13.10276 | 0.00000 | 1.05680 | 0.88226 | 7.09208 | 0.02560 | 0.00000 | 2.14367 | 0.00000 | 7.21695 |
| **Sox2ot** | 0.01183 | 0.01266 | 0.02379 | 0.03404 | 0.02994 | 0.03186 | 0.01727 | 0.02532 | 0.06558 | 0.01512 | 0.01654 | 0.04096 | 0.09013 | 0.14448 |
| **Sox30** | 0.00839 | 0.00431 | 0.00910 | 0.00965 | 0.17814 | 0.01483 | 0.00484 | 0.01069 | 0.01736 | 0.00632 | 0.00919 | 0.00890 | 0.35072 | 0.01486 |
| **Sox4** | 0.02339 | 0.00433 | 0.00951 | 0.01097 | 0.21037 | 0.01246 | 0.00439 | 0.01106 | 0.00531 | 0.00606 | 0.00876 | 0.00887 | 0.27110 | 0.07857 |
| **Sox6** | 0.00374 | 0.01247 | 0.26287 | 0.32331 | 0.00261 | 0.00000 | 0.00511 | 0.06533 | 0.01264 | 0.00636 | 0.00793 | 0.00551 | 0.00628 | 0.00834 |
| **Sox7** | 0.00676 | 0.02209 | 0.00889 | 0.00000 | 0.01618 | 0.02240 | 0.00393 | 0.03024 | 0.04605 | 0.00550 | 0.26528 | 0.40700 | 0.01074 | 0.01968 |
| **Sp1** | 7.05128 | 0.10164 | 11.56629 | 2.28955 | 13.59016 | 8.14201 | 0.03403 | 0.26976 | 12.74576 | 0.02288 | 3.10503 | 3.54255 | 6.93088 | 10.46743 |
| **Sp100** | 0.06526 | 0.04167 | 0.03848 | 0.22632 | 0.69551 | 0.17522 | 0.02694 | 0.07295 | 0.52197 | 0.04686 | 0.01356 | 0.07105 | 0.24378 | 1.19443 |
| **Sp110** | 41.00436 | 3.52840 | 43.96411 | 17.25135 | 46.26645 | 16.85308 | 7.94657 | 1.46508 | 39.60868 | 2.39080 | 6.53027 | 1.88440 | 23.19895 | 44.65971 |
| **Sp140** | 13.63964 | 1.05881 | 7.89216 | 1.10526 | 15.51164 | 1.65530 | 3.40228 | 1.11693 | 16.15517 | 4.00998 | 3.21820 | 0.61141 | 10.33127 | 13.41450 |
| **Sp2** | 0.17593 | 3.49067 | 2.67946 | 0.78762 | 6.75187 | 0.54467 | 0.00248 | 1.29385 | 0.34404 | 0.00000 | 1.03155 | 0.90152 | 2.52980 | 0.00000 |
| **Sp3** | 1.22176 | 0.00219 | 0.41441 | 0.17108 | 0.45566 | 0.09399 | 0.14791 | 0.53164 | 21.11298 | 0.00867 | 2.04038 | 0.94630 | 4.52073 | 2.72350 |
| **Sp3os** | 1.27284 | 0.01145 | 0.00000 | 0.48814 | 2.38578 | 0.39701 | 0.00390 | 0.17804 | 2.89945 | 0.01083 | 0.11725 | 0.25815 | 4.92989 | 2.14285 |
| **Sp4** | 0.20061 | 0.00309 | 0.30968 | 0.22612 | 1.30900 | 0.00000 | 0.22058 | 0.76866 | 1.03366 | 0.00284 | 2.56336 | 0.00451 | 0.54262 | 0.00000 |
| **Sp5** | 0.01119 | 0.00443 | 0.01410 | 0.01094 | 0.01402 | 0.02629 | 0.00427 | 0.00933 | 0.01150 | 0.00759 | 0.00829 | 0.01019 | 0.00878 | 0.01363 |
| **Sp9** | 0.01048 | 0.00441 | 0.01480 | 0.01250 | 0.01244 | 0.01333 | 0.00427 | 0.00989 | 0.01103 | 0.00673 | 0.00889 | 0.00983 | 0.11495 | 0.00448 |
| **Spa17** | 0.11723 | 0.00480 | 0.00000 | 0.38949 | 0.07158 | 0.24465 | 0.00519 | 0.34764 | 0.35481 | 0.00751 | 0.01322 | 0.00781 | 0.04344 | 0.03529 |
| **Spaca6** | 0.10340 | 0.00413 | 0.33206 | 0.00000 | 0.00000 | 0.00569 | 0.00437 | 0.35880 | 0.08151 | 0.00571 | 0.00507 | 0.03476 | 0.18592 | 0.00721 |
| **Spag1** | 1.44256 | 0.00478 | 0.27005 | 0.59261 | 1.28365 | 0.48206 | 0.01068 | 0.02286 | 1.59726 | 0.00608 | 0.00574 | 0.00000 | 2.78557 | 4.73306 |
| **Spag16** | 0.00546 | 0.00430 | 0.00878 | 0.01416 | 0.14344 | 0.01252 | 0.00414 | 0.00847 | 0.06946 | 0.00739 | 0.00920 | 0.01646 | 0.27512 | 0.00000 |
| **Spag17** | 0.00932 | 0.00442 | 0.06325 | 0.04680 | 0.07474 | 0.03053 | 0.00435 | 0.00613 | 0.01098 | 0.00552 | 0.01376 | 0.01372 | 0.00653 | 0.03785 |

| | | | | | | | | | | | | | | |
|---|---|---|---|---|---|---|---|---|---|---|---|---|---|---|
| **Spag4** | 1.11996 | 0.02127 | 0.42002 | 1.50207 | 0.12053 | 0.51534 | 0.00362 | 0.00354 | 0.58081 | 0.00537 | 0.00000 | 0.45307 | 0.68978 | 0.35964 |
| **Spag5** | 2.53692 | 9.08241 | 2.48312 | 2.95720 | 3.87456 | 0.46929 | 1.59441 | 3.18008 | 3.07083 | 0.13771 | 3.81716 | 2.47901 | 11.04954 | 8.27208 |
| **Spag6l** | 0.15492 | 0.00428 | 0.34648 | 0.14183 | 0.18611 | 0.02026 | 0.00409 | 0.00476 | 0.00780 | 0.00749 | 0.00564 | 0.00563 | 0.62550 | 0.01085 |
| **Spag7** | 5.08450 | 0.37475 | 19.98873 | 19.14875 | 15.46052 | 17.57010 | 3.44785 | 18.54495 | 4.35751 | 1.99828 | 15.24421 | 12.85397 | 2.16908 | 13.61301 |
| **Spag9** | 9.55293 | 0.00583 | 8.48441 | 3.86283 | 2.72759 | 4.05429 | 2.51773 | 0.65578 | 11.22660 | 0.05081 | 4.92391 | 4.12984 | 3.88999 | 3.81194 |
| **Spam1** | 0.00405 | 0.00441 | 0.01037 | 0.01342 | 0.00023 | 0.01105 | 0.00425 | 0.00995 | 0.04176 | 0.00664 | 0.00779 | 0.00967 | 0.04597 | 0.37106 |
| **Sparc** | 2.03197 | 0.98745 | 0.41277 | 0.00000 | 0.05365 | 1.92119 | 0.00000 | 0.57236 | 0.29403 | 0.04200 | 1.20196 | 15.75889 | 0.00525 | 0.51350 |
| **Sparcl1** | 0.01808 | 0.00451 | 0.01008 | 0.01378 | 0.20059 | 0.01298 | 0.00432 | 0.01087 | 0.01043 | 0.00655 | 0.00856 | 0.00860 | 0.00452 | 0.03012 |
| **Spast** | 5.87502 | 0.02747 | 2.15153 | 2.02523 | 6.59058 | 1.99074 | 2.95208 | 3.50635 | 7.02040 | 0.05984 | 5.94086 | 5.10539 | 9.73536 | 9.70727 |
| **Spata1** | 0.04971 | 0.00515 | 0.03232 | 0.00000 | 0.00493 | 0.00936 | 0.00399 | 0.04863 | 0.20584 | 0.00688 | 0.00349 | 0.29475 | 0.00470 | 0.00065 |
| **Spata13** | 0.00380 | 0.01215 | 0.03582 | 0.00295 | 0.31314 | 2.72034 | 0.00430 | 1.47554 | 0.02833 | 0.01100 | 0.24934 | 0.00000 | 0.00591 | 0.00620 |
| **Spata17** | 0.05469 | 0.00401 | 0.17611 | 0.07307 | 0.02035 | 0.00352 | 0.00402 | 0.00537 | 0.02592 | 0.00609 | 0.04632 | 0.16576 | 0.02043 | 0.12520 |
| **Spata2** | 5.91012 | 0.11269 | 6.43422 | 4.48208 | 7.17494 | 6.48918 | 2.39375 | 4.27725 | 4.90857 | 0.13834 | 0.78641 | 3.46112 | 14.83415 | 10.11993 |
| **Spata21** | 0.01286 | 0.00406 | 0.29242 | 0.01747 | 0.41765 | 0.34622 | 0.00419 | 0.00465 | 0.00000 | 0.01215 | 0.00674 | 0.00202 | 2.58356 | 1.99783 |
| **Spata22** | 0.01452 | 0.00447 | 0.07408 | 0.00000 | 0.00966 | 0.19536 | 0.00425 | 0.00505 | 0.01146 | 0.00576 | 0.11999 | 0.01253 | 0.01207 | 0.03201 |
| **Spata24** | 0.36892 | 0.00308 | 0.07353 | 0.80483 | 0.38535 | 0.66839 | 0.00920 | 0.00327 | 0.00392 | 0.02417 | 0.00560 | 1.62562 | 0.33416 | 1.20722 |
| **Spata25** | 0.03435 | 0.00428 | 0.10922 | 0.01388 | 0.00617 | 0.01089 | 0.00413 | 0.00829 | 0.01156 | 0.00918 | 0.00693 | 0.00984 | 0.39082 | 0.01521 |
| **Spata2l** | 0.10866 | 0.00415 | 0.01045 | 0.01149 | 0.00713 | 0.00000 | 0.00407 | 0.01118 | 0.00946 | 0.00910 | 0.55154 | 0.00465 | 0.15810 | 0.00261 |
| **Spata33** | 0.02573 | 0.00443 | 0.01303 | 0.01192 | 0.02314 | 0.01255 | 0.00454 | 0.01045 | 0.01957 | 0.00676 | 0.00855 | 0.00994 | 0.01760 | 0.03896 |
| **Spata5** | 7.94639 | 0.00954 | 3.54886 | 3.27099 | 6.47720 | 3.47678 | 0.02768 | 1.77169 | 4.16810 | 0.01734 | 0.06089 | 1.57177 | 11.35264 | 8.23599 |
| **Spata5l1** | 0.00000 | 0.00367 | 0.31185 | 0.11742 | 0.16143 | 0.00697 | 0.00375 | 0.00271 | 0.01014 | 0.00540 | 0.00640 | 0.49694 | 0.00412 | 0.01180 |
| **Spata6** | 0.00853 | 0.00442 | 0.01105 | 0.01300 | 0.03536 | 0.01189 | 0.00421 | 0.00998 | 0.01863 | 0.00669 | 0.00794 | 0.00947 | 0.02784 | 0.34342 |
| **Spata7** | 4.26145 | 0.00235 | 3.92862 | 2.80649 | 4.33653 | 3.04931 | 0.10297 | 0.35070 | 0.82808 | 0.00000 | 1.41409 | 1.62336 | 0.75836 | 8.45971 |
| **Spats1** | 0.00551 | 0.00376 | 0.00000 | 0.08211 | 1.06203 | 0.12280 | 0.14243 | 0.09736 | 0.35065 | 0.00557 | 0.04920 | 0.06685 | 0.80056 | 0.00000 |
| **Spats2** | 0.00000 | 0.04766 | 1.36254 | 0.88057 | 1.21967 | 1.11029 | 1.66230 | 1.03183 | 0.00968 | 4.27744 | 2.65156 | 3.91499 | 0.30666 | 0.53373 |
| **Spats2l** | 0.02812 | 0.00444 | 0.05659 | 0.00852 | 0.01737 | 0.00930 | 0.00427 | 0.01111 | 0.01021 | 0.00676 | 0.01903 | 0.00904 | 0.00689 | 0.05610 |
| **Spc24** | 1.21267 | 4.73025 | 5.03767 | 9.70731 | 7.70531 | 4.23192 | 1.59745 | 4.95476 | 0.28323 | 8.85717 | 0.79834 | 7.48106 | 1.12893 | 1.30404 |
| **Spc25** | 6.91234 | 1.63985 | 2.58203 | 2.88988 | 3.91643 | 8.48234 | 0.09844 | 2.53825 | 0.20287 | 2.32659 | 8.22823 | 4.56312 | 13.60120 | 2.33247 |
| **Spcs1** | 6.64273 | 5.41985 | 3.04550 | 13.40002 | 11.64049 | 12.59378 | 4.77174 | 10.60134 | 8.45385 | 0.41837 | 3.94860 | 10.50472 | 2.59399 | 6.91010 |
| **Spcs2** | 3.22022 | 0.23253 | 6.44504 | 9.56150 | 5.11110 | 13.26716 | 4.52107 | 3.97428 | 1.24482 | 0.29838 | 3.35671 | 9.40842 | 1.52109 | 3.78353 |
| **Spcs3** | 2.43221 | 0.15763 | 2.61838 | 3.34259 | 1.34499 | 7.99671 | 1.69299 | 1.63390 | 5.59950 | 0.02383 | 4.64915 | 6.05475 | 0.00000 | 2.29241 |
| **Spdef** | 0.00352 | 0.00431 | 0.01316 | 0.01244 | 0.09908 | 0.01305 | 0.00434 | 0.01027 | 0.00849 | 0.00624 | 0.00901 | 0.00898 | 0.45326 | 0.00371 |
| **Spdl1** | 8.38973 | 0.04799 | 5.95073 | 2.73058 | 7.94734 | 3.52539 | 2.73067 | 2.84126 | 4.15600 | 0.06160 | 3.55279 | 3.12045 | 10.64926 | 18.25747 |
| **Spdya** | 0.00000 | 0.00433 | 0.04110 | 0.00000 | 0.49959 | 0.07471 | 0.15538 | 0.00325 | 1.98964 | 0.00689 | 0.00449 | 0.00640 | 0.12918 | 0.42300 |
| **Spdye4a** | 0.05704 | 0.00440 | 0.07503 | 0.00612 | 0.00433 | 0.00884 | 0.00425 | 0.01065 | 0.09556 | 0.00679 | 0.00688 | 0.00949 | 0.17245 | 0.00000 |
| **Spdye4b** | 0.72395 | 0.00424 | 0.28516 | 0.00000 | 0.12804 | 0.00190 | 0.00388 | 0.00927 | 1.10888 | 0.00840 | 0.00388 | 0.00545 | 0.64611 | 0.00000 |
| **Specc1** | 0.14967 | 0.01159 | 0.66753 | 0.82212 | 0.22279 | 1.07223 | 0.00383 | 1.13574 | 0.65861 | 0.02600 | 0.00000 | 0.86366 | 0.46001 | 0.00000 |
| **Specc1l** | 0.08770 | 0.00182 | 1.83772 | 0.65424 | 1.75765 | 2.79135 | 0.00279 | 0.83374 | 1.09637 | 0.00000 | 5.20251 | 0.56258 | 3.19105 | 3.33210 |
| **Speer1** | 0.17900 | 0.00437 | 0.01729 | 0.00733 | 0.05280 | 0.02250 | 0.00440 | 0.01100 | 0.01211 | 0.00674 | 0.00681 | 0.00916 | 0.06034 | 0.02209 |
| **Speer2** | 0.01588 | 0.00440 | 0.03723 | 0.01982 | 0.01895 | 0.07014 | 0.00421 | 0.00723 | 0.01202 | 0.00710 | 0.00647 | 0.01268 | 0.00954 | 0.01135 |
| **Speer3** | 0.00662 | 0.00437 | 0.09945 | 0.05191 | 0.02173 | 0.11733 | 0.00415 | 0.00563 | 0.01335 | 0.00690 | 0.00555 | 0.01394 | 0.01563 | 0.01138 |
| **Speer4b** | 0.02258 | 0.00432 | 0.01350 | 0.03710 | 0.01153 | 0.01423 | 0.00435 | 0.01083 | 0.01171 | 0.00586 | 0.00771 | 0.00847 | 0.07167 | 0.07382 |
| **Speer4c** | 0.07030 | 0.00431 | 0.06141 | 0.03636 | 0.14995 | 0.03035 | 0.00414 | 0.00648 | 0.03017 | 0.01216 | 0.01008 | 0.01046 | 0.09817 | 0.06437 |
| **Speer4cos** | 0.00290 | 0.00465 | 0.00684 | 0.03063 | 0.00430 | 0.03320 | 0.00426 | 0.00866 | 0.23273 | 0.00657 | 0.00749 | 0.00961 | 0.23528 | 0.00000 |
| **Speer4d** | 0.06250 | 0.00425 | 0.07711 | 0.07245 | 0.18010 | 0.02354 | 0.00414 | 0.00732 | 0.04860 | 0.00558 | 0.00915 | 0.01135 | 0.13434 | 0.05815 |
| **Speer4e** | 0.11910 | 0.00422 | 0.09516 | 0.05289 | 0.17016 | 0.05282 | 0.00403 | 0.00501 | 0.13124 | 0.00560 | 0.01141 | 0.00961 | 0.29191 | 0.11362 |
| **Speer5.ps1** | 0.14873 | 0.00412 | 0.11788 | 0.00000 | 0.33286 | 0.08229 | 0.00407 | 0.01615 | 0.33565 | 0.00602 | 0.00745 | 0.02132 | 0.34569 | 0.46516 |
| **Speer6.ps1** | 0.02204 | 0.00439 | 0.09113 | 0.03006 | 0.01807 | 0.12730 | 0.00420 | 0.00603 | 0.01177 | 0.00723 | 0.00683 | 0.01293 | 0.00924 | 0.01013 |
| **Speer8.ps1** | 0.00889 | 0.00438 | 0.01303 | 0.01294 | 0.00980 | 0.01105 | 0.00425 | 0.00983 | 0.04563 | 0.00682 | 0.02117 | 0.01168 | 0.11181 | 0.01127 |
| **Spef1** | 0.00376 | 0.00397 | 0.00578 | 0.09767 | 0.21576 | 0.02375 | 0.00400 | 0.00519 | 0.00759 | 0.00605 | 0.00523 | 0.21328 | 0.34926 | 0.00467 |
| **Speg** | 0.01363 | 0.00439 | 0.20151 | 0.00000 | 2.05659 | 0.00399 | 0.00413 | 0.00975 | 0.01877 | 0.00619 | 0.00648 | 0.00701 | 0.27116 | 0.06729 |
| **Spen** | 1.95935 | 1.05289 | 3.85052 | 2.56398 | 2.98993 | 0.62534 | 0.00354 | 1.01783 | 0.08722 | 0.01295 | 0.58274 | 0.77131 | 0.21009 | 1.20510 |
| **Spesp1** | 0.00000 | 0.00344 | 0.74934 | 0.00988 | 0.04697 | 0.00265 | 0.00378 | 0.00180 | 0.08183 | 0.00478 | 0.00355 | 0.64230 | 1.44112 | 0.00404 |
| **Spg11** | 2.16180 | 0.02666 | 2.76758 | 1.33769 | 3.09072 | 3.24394 | 0.00203 | 2.63252 | 0.57707 | 0.00449 | 4.21975 | 2.56423 | 0.43514 | 1.95725 |
| **Spg20** | 1.09738 | 0.09419 | 1.03821 | 1.92262 | 0.43232 | 3.56255 | 2.37597 | 2.78703 | 0.92888 | 0.09527 | 0.12298 | 6.34272 | 1.38725 | 2.26769 |

| | | | | | | | | | | | | | | |
|---|---|---|---|---|---|---|---|---|---|---|---|---|---|---|
| **Spg21** | 6.17000 | 2.07162 | 5.88621 | 0.06461 | 6.42203 | 6.27656 | 3.24921 | 1.72314 | 0.00855 | 0.10497 | 1.36646 | 7.19180 | 0.51992 | 0.84040 |
| **Spg7** | 2.73632 | 0.05289 | 2.75808 | 3.11062 | 2.36384 | 0.90802 | 2.67274 | 4.31585 | 0.23098 | 0.05859 | 6.25602 | 3.48638 | 3.53713 | 1.71186 |
| **Sphk1** | 0.02351 | 0.07768 | 0.33269 | 0.25658 | 1.00872 | 0.00000 | 0.56933 | 0.52025 | 0.00570 | 0.00990 | 0.00916 | 1.00885 | 0.00367 | 0.44877 |
| **Sphk2** | 0.69531 | 0.03382 | 2.12424 | 3.32883 | 3.64268 | 2.21928 | 2.77535 | 6.33203 | 0.00000 | 0.20391 | 1.02067 | 1.89049 | 2.73599 | 0.25648 |
| **Spi1** | 0.00493 | 0.00449 | 0.01474 | 0.22515 | 0.97580 | 0.10525 | 0.00410 | 0.06390 | 0.01137 | 0.00619 | 0.01109 | 0.00729 | 0.00293 | 0.00781 |
| **Spib** | 0.04225 | 0.00454 | 0.00000 | 0.06027 | 0.06193 | 0.02917 | 0.01627 | 0.01558 | 0.62584 | 0.01792 | 0.00509 | 0.01080 | 0.04360 | 0.00000 |
| **Spic** | 25.21106 | 0.03591 | 16.19875 | 5.51401 | 41.56970 | 0.00000 | 0.00000 | 0.00000 | 31.20284 | 0.05623 | 7.04451 | 16.18843 | 15.05010 | 31.17093 |
| **Spice1** | 3.79798 | 0.01113 | 0.32086 | 1.42628 | 2.28320 | 0.46485 | 0.00342 | 0.00006 | 0.23870 | 0.00199 | 1.59122 | 1.77845 | 0.68467 | 3.52489 |
| **Spidr** | 1.28821 | 0.01730 | 1.18641 | 0.18268 | 1.33556 | 0.65993 | 0.00314 | 0.13047 | 0.98380 | 0.00293 | 0.00000 | 1.06643 | 0.77796 | 2.30405 |
| **Spin1** | 9.55539 | 3.51019 | 10.02027 | 4.71701 | 10.47400 | 13.60189 | 3.48367 | 2.61607 | 16.48553 | 0.18895 | 8.25631 | 7.28282 | 12.54696 | 16.88497 |
| **Spin2c** | 0.00742 | 0.00426 | 0.00310 | 0.55543 | 0.00570 | 0.84960 | 2.17579 | 0.00253 | 0.03470 | 0.03773 | 0.15521 | 0.00351 | 0.12899 | 0.12047 |
| **Spin2d** | 0.49666 | 0.00380 | 0.23786 | 0.00000 | 0.37648 | 0.03743 | 0.00375 | 0.16092 | 0.47853 | 0.00631 | 0.00413 | 0.00321 | 0.75273 | 0.23719 |
| **Spink2** | 0.62856 | 0.01679 | 0.05287 | 2.18940 | 4.05830 | 0.10160 | 0.66861 | 0.34665 | 2.49314 | 0.04162 | 0.11595 | 2.23706 | 0.77028 | 2.35529 |
| **Spink4** | 0.08005 | 0.00425 | 0.04261 | 0.03852 | 0.01654 | 0.02995 | 0.00408 | 0.00663 | 0.00960 | 0.00987 | 0.00701 | 0.01427 | 0.16896 | 0.00424 |
| **Spint1** | 8.53160 | 4.82907 | 14.55797 | 5.63483 | 9.37319 | 8.62556 | 7.23087 | 8.79828 | 3.36991 | 0.18330 | 5.66640 | 11.15608 | 3.08339 | 7.15375 |
| **Spint2** | 7.97486 | 4.62293 | 6.61559 | 7.33649 | 7.75439 | 4.37665 | 2.31937 | 4.04955 | 2.05135 | 0.11847 | 4.92006 | 3.91538 | 1.03693 | 9.59495 |
| **Spint5** | 0.01619 | 0.01158 | 0.00487 | 0.00000 | 0.16868 | 0.18091 | 0.00409 | 0.00524 | 0.01582 | 0.00685 | 0.00361 | 0.37208 | 0.00529 | 0.00503 |
| **Spire1** | 0.31132 | 0.00317 | 0.21442 | 0.14927 | 1.65932 | 0.00000 | 0.00321 | 0.01056 | 2.78438 | 0.00803 | 1.66977 | 0.13455 | 5.88985 | 5.22880 |
| **Spire2** | 0.74482 | 0.00426 | 0.22289 | 0.01026 | 0.05157 | 0.02823 | 0.00429 | 0.00780 | 0.19689 | 0.00677 | 0.00556 | 0.00772 | 0.33154 | 0.00000 |
| **Spn** | 0.00319 | 0.00430 | 0.01337 | 0.01191 | 0.00634 | 0.01276 | 0.00423 | 0.00890 | 0.00948 | 0.00608 | 0.00902 | 0.00890 | 0.50402 | 0.14095 |
| **Spns1** | 4.99734 | 6.87068 | 37.17353 | 30.29648 | 32.16680 | 3.51872 | 1.87750 | 9.85224 | 0.00000 | 0.16917 | 8.08529 | 15.20078 | 4.11183 | 1.08098 |
| **Spns2** | 1.28165 | 0.83770 | 0.93114 | 0.62720 | 1.58313 | 0.28298 | 0.00350 | 0.31138 | 0.48223 | 0.43925 | 0.11147 | 0.04033 | 2.77003 | 3.22448 |
| **Spns3** | 0.00000 | 0.00386 | 0.23865 | 0.85177 | 0.00162 | 0.00000 | 0.00402 | 0.01170 | 0.01448 | 0.01308 | 0.92293 | 0.00551 | 0.00713 | 0.00000 |
| **Spock1** | 0.01059 | 0.00444 | 0.07119 | 0.00606 | 0.01147 | 0.01226 | 0.00427 | 0.01076 | 0.01836 | 0.00726 | 0.00940 | 0.00972 | 0.00746 | 0.00983 |
| **Spock2** | 0.06536 | 0.00421 | 0.56829 | 0.19704 | 0.18725 | 0.00002 | 0.00406 | 0.02501 | 0.02526 | 0.01959 | 0.00771 | 0.00517 | 0.02137 | 0.03436 |
| **Spock3** | 0.01057 | 0.00444 | 0.01465 | 0.02008 | 0.01368 | 0.01331 | 0.00431 | 0.00990 | 0.01185 | 0.00690 | 0.00865 | 0.00949 | 0.00742 | 0.01182 |
| **Spon1** | 0.00000 | 0.00419 | 0.57681 | 0.00000 | 0.60152 | 0.01464 | 0.00381 | 0.00360 | 0.18733 | 0.00490 | 0.21934 | 0.00490 | 0.60074 | 0.40546 |
| **Spon2** | 0.00495 | 0.00385 | 0.42094 | 0.13200 | 0.00597 | 0.00437 | 0.00404 | 0.90469 | 0.01118 | 0.01523 | 0.00957 | 0.01478 | 0.00664 | 0.01305 |
| **Spop** | 1.96818 | 0.02215 | 8.97626 | 1.41131 | 6.43529 | 4.15030 | 2.32503 | 1.20795 | 11.97956 | 0.06721 | 4.87549 | 2.85191 | 3.12290 | 12.33483 |
| **Spopl** | 0.78463 | 0.00352 | 0.06946 | 0.26318 | 0.60138 | 0.16969 | 0.00342 | 0.21330 | 3.16381 | 0.02442 | 0.00000 | 0.45118 | 0.00000 | 0.02031 |
| **Spp1** | 0.10084 | 8.03908 | 0.00000 | 0.38087 | 0.00000 | 3.49062 | 0.00000 | 2.91482 | 0.00452 | 0.11646 | 10.22494 | 24.63153 | 0.22866 | 0.01429 |
| **Sppl2a** | 1.25277 | 0.05404 | 1.28006 | 0.69181 | 0.24869 | 2.07313 | 0.03366 | 0.86705 | 2.85999 | 1.52048 | 1.46079 | 4.96503 | 0.92712 | 0.78173 |
| **Sppl2b** | 0.06968 | 0.01141 | 0.67805 | 1.23711 | 0.47162 | 0.15688 | 0.02572 | 0.95072 | 0.00753 | 0.01236 | 0.00000 | 0.73302 | 0.19993 | 0.04774 |
| **Sppl3** | 4.55733 | 2.04439 | 15.27219 | 3.46351 | 17.84863 | 6.13560 | 2.41117 | 0.91860 | 2.37751 | 2.31951 | 3.32611 | 4.22242 | 2.99241 | 16.36674 |
| **Spr** | 0.37499 | 1.83378 | 0.03620 | 4.69205 | 6.56032 | 1.02535 | 0.00840 | 0.65468 | 0.27877 | 0.02098 | 1.20580 | 2.10625 | 0.22757 | 1.53781 |
| **Spred1** | 0.11928 | 0.00378 | 0.80749 | 0.83370 | 2.61949 | 0.49666 | 0.00255 | 0.00000 | 7.52974 | 0.00490 | 0.00000 | 0.26647 | 1.15548 | 0.86694 |
| **Spred2** | 0.03840 | 0.00416 | 0.03998 | 0.21011 | 0.04070 | 0.18534 | 0.00398 | 0.00255 | 0.02059 | 0.00451 | 0.24414 | 0.10394 | 0.02671 | 0.00296 |
| **Spred3** | 0.06838 | 0.00413 | 0.09838 | 0.00000 | 0.12607 | 0.00749 | 0.00421 | 0.00793 | 0.16064 | 0.00729 | 0.00476 | 0.27026 | 0.02296 | 0.08342 |
| **Sprtn** | 0.00430 | 0.00393 | 0.18451 | 0.00000 | 0.45254 | 1.16144 | 0.00387 | 0.00000 | 0.21046 | 0.00364 | 0.80161 | 0.01150 | 0.60696 | 0.24864 |
| **Spry1** | 0.13574 | 0.02071 | 0.23578 | 1.06737 | 0.01776 | 0.53068 | 0.00398 | 1.46193 | 0.00788 | 0.00546 | 0.37462 | 0.00265 | 0.01460 | 0.13426 |
| **Spry2** | 4.67046 | 0.04223 | 0.04086 | 0.47660 | 0.00000 | 1.09761 | 1.14925 | 0.02711 | 4.66309 | 0.02613 | 0.00000 | 2.45497 | 2.72569 | 0.00000 |
| **Spry4** | 0.27579 | 0.00364 | 0.02879 | 0.23322 | 0.73222 | 0.00000 | 0.00367 | 0.01257 | 0.59333 | 0.01366 | 0.00355 | 0.75982 | 6.46447 | 1.76671 |
| **Spryd3** | 0.07104 | 0.05574 | 1.29604 | 2.44279 | 1.63856 | 1.20138 | 0.02746 | 1.89656 | 1.25584 | 0.01787 | 1.58673 | 0.23879 | 1.05039 | 2.79507 |
| **Spryd4** | 1.66644 | 0.11929 | 4.35518 | 0.00000 | 3.25418 | 3.93520 | 2.26934 | 3.67437 | 0.70336 | 5.58505 | 4.29255 | 4.87505 | 0.00000 | 3.11587 |
| **Spryd7** | 7.29439 | 5.40780 | 2.77210 | 4.25401 | 5.11007 | 3.01684 | 1.14571 | 0.01835 | 6.19367 | 0.10968 | 3.59727 | 0.87302 | 2.32351 | 0.36895 |
| **Spsb1** | 0.16243 | 0.00410 | 0.25173 | 0.02614 | 0.09398 | 0.01854 | 0.78375 | 0.01615 | 0.00842 | 0.01105 | 0.00608 | 0.00511 | 0.00538 | 0.01701 |
| **Spsb2** | 0.00293 | 0.06680 | 0.76699 | 0.00000 | 0.26080 | 0.02792 | 2.07172 | 4.93489 | 0.00168 | 0.05882 | 0.06009 | 1.08408 | 0.25914 | 0.06791 |
| **Spsb3** | 3.39212 | 0.15915 | 7.62482 | 3.53864 | 9.19581 | 3.63533 | 0.41795 | 5.00082 | 0.00000 | 0.04490 | 0.40599 | 2.30174 | 2.25851 | 3.38221 |
| **Spsb4** | 1.44150 | 0.10068 | 2.06424 | 0.00000 | 4.12835 | 5.64354 | 5.54854 | 8.57720 | 1.63866 | 10.99499 | 2.04156 | 0.98620 | 6.82786 | 2.95006 |
| **Sptan1** | 19.26773 | 2.32189 | 14.27500 | 12.44637 | 12.41101 | 14.72864 | 3.43164 | 9.59586 | 4.28385 | 0.31625 | 7.88822 | 5.19274 | 5.27652 | 17.62173 |
| **Sptb** | 0.21255 | 0.00419 | 0.24499 | 0.51126 | 0.86506 | 0.16537 | 0.00397 | 0.00386 | 0.03270 | 0.00590 | 0.01607 | 0.00916 | 0.00167 | 0.07962 |
| **Sptbn1** | 1.74447 | 2.80420 | 1.38026 | 0.38730 | 0.70726 | 4.14539 | 2.31347 | 1.29671 | 0.32483 | 0.06305 | 1.18615 | 4.35179 | 2.15601 | 0.00000 |
| **Sptbn2** | 3.90910 | 0.00290 | 14.32902 | 6.42879 | 11.31553 | 2.46670 | 0.00670 | 0.61290 | 0.72312 | 0.00000 | 0.06601 | 0.14891 | 1.28254 | 9.97298 |
| **Sptbn4** | 0.00254 | 0.00423 | 0.01737 | 1.30161 | 0.33890 | 0.00000 | 0.00394 | 0.00617 | 0.00884 | 0.00513 | 0.00512 | 0.19343 | 0.01004 | 0.05568 |
| **Sptlc1** | 3.95830 | 0.05744 | 8.78672 | 3.82939 | 3.56085 | 5.42547 | 2.55891 | 5.53478 | 5.21296 | 0.03196 | 7.34626 | 4.90607 | 0.35268 | 9.23301 |

| | | | | | | | | | | | | | | |
|---|---|---|---|---|---|---|---|---|---|---|---|---|---|---|
| **Sptlc2** | 6.07567 | 4.26465 | 5.60516 | 3.06297 | 4.92614 | 11.09698 | 1.10830 | 6.18370 | 9.03295 | 0.02495 | 3.59342 | 6.48353 | 8.41829 | 10.97969 |
| **Sptssa** | 4.10443 | 0.16596 | 15.60466 | 12.16609 | 10.02631 | 10.79885 | 6.72624 | 9.86477 | 6.05560 | 3.84052 | 7.18594 | 8.07878 | 0.74184 | 11.07784 |
| **Spty2d1** | 12.17716 | 3.97908 | 8.42075 | 3.96658 | 8.60121 | 1.52348 | 1.27961 | 2.34569 | 20.67043 | 0.02318 | 6.48608 | 0.02643 | 7.41868 | 7.65990 |
| **Spz1** | 0.84584 | 0.00360 | 0.23824 | 0.00391 | 0.02108 | 0.00000 | 0.00392 | 0.01012 | 0.00000 | 0.00625 | 0.09904 | 0.00881 | 4.38517 | 0.28338 |
| **Sqle** | 7.66304 | 0.05213 | 7.15825 | 0.00000 | 12.95745 | 11.02163 | 0.00016 | 0.00000 | 5.72095 | 0.00609 | 0.00000 | 4.27004 | 11.75546 | 14.07991 |
| **Sqstm1** | 6.75999 | 0.21880 | 4.30045 | 15.01438 | 5.86886 | 25.50892 | 6.42571 | 8.82801 | 1.40890 | 0.46501 | 12.63133 | 12.91973 | 0.43700 | 5.09923 |
| **Sra1** | 2.43231 | 2.36247 | 4.22189 | 6.55260 | 4.75274 | 6.18130 | 4.62802 | 10.20011 | 0.45516 | 12.66386 | 5.14481 | 7.69945 | 0.97517 | 2.79481 |
| **Srbd1** | 2.30483 | 0.02505 | 1.80726 | 2.78263 | 3.19107 | 2.67376 | 1.68444 | 0.73328 | 1.36483 | 0.01725 | 1.26704 | 1.87133 | 1.33335 | 4.56414 |
| **Src** | 0.09691 | 0.06295 | 0.06568 | 0.00000 | 0.35718 | 0.15672 | 0.00313 | 3.67792 | 0.00947 | 0.03206 | 0.00000 | 1.21074 | 0.27880 | 0.96599 |
| **Srcap** | 0.02097 | 0.00325 | 0.71880 | 0.75992 | 1.70095 | 0.30460 | 0.00340 | 0.25556 | 1.21242 | 0.02427 | 0.08737 | 0.71437 | 3.02495 | 1.25822 |
| **Srcin1** | 0.05169 | 0.00428 | 0.03734 | 0.09860 | 0.05964 | 0.00664 | 0.00395 | 0.00808 | 0.00924 | 0.01282 | 0.00929 | 0.00991 | 0.01525 | 0.03332 |
| **Srd5a1** | 0.57244 | 0.00407 | 0.31149 | 0.00000 | 0.34836 | 0.01481 | 0.00409 | 0.00492 | 0.00660 | 0.02182 | 0.00493 | 0.12545 | 0.12900 | 0.01533 |
| **Srd5a3** | 0.50449 | 0.01028 | 0.19416 | 0.44700 | 0.93250 | 0.60096 | 1.95805 | 0.52290 | 1.50996 | 0.07378 | 0.00000 | 1.04375 | 1.21364 | 0.00000 |
| **Srebf1** | 0.01319 | 0.58333 | 1.31012 | 2.51108 | 0.80079 | 0.20316 | 2.70876 | 2.28300 | 0.12703 | 0.08053 | 1.18561 | 2.87766 | 0.11545 | 0.04639 |
| **Srebf2** | 0.16935 | 4.35428 | 1.43088 | 0.00038 | 0.00000 | 2.29448 | 0.00693 | 1.41925 | 0.00618 | 0.02142 | 2.63490 | 3.00684 | 0.01249 | 0.02005 |
| **Srek1** | 3.55910 | 6.77447 | 6.00019 | 2.55235 | 7.05162 | 6.78304 | 0.00000 | 0.00000 | 5.26942 | 0.01316 | 2.93064 | 2.29844 | 4.68985 | 12.71662 |
| **Srek1ip1** | 0.00000 | 0.03294 | 1.26435 | 1.63934 | 0.84952 | 1.36994 | 1.17581 | 4.60178 | 0.00689 | 0.02309 | 0.45934 | 2.07516 | 0.00000 | 1.14224 |
| **Srf** | 0.25953 | 0.08195 | 2.93715 | 0.00000 | 2.88133 | 2.39681 | 2.49980 | 5.02590 | 2.34077 | 0.10437 | 1.37584 | 0.09758 | 1.64341 | 0.54721 |
| **Srfbp1** | 9.41601 | 4.99369 | 12.63791 | 5.95466 | 11.40878 | 7.08827 | 0.00000 | 0.24585 | 3.57668 | 0.00829 | 5.16926 | 2.91058 | 3.36057 | 16.84766 |
| **Srgap1** | 0.00000 | 0.00350 | 0.88165 | 0.00000 | 0.34225 | 0.00702 | 0.00320 | 1.08187 | 0.57487 | 0.01576 | 0.00000 | 1.08143 | 0.89606 | 0.03982 |
| **Srgap2** | 0.84758 | 0.06246 | 0.37536 | 1.82303 | 0.00000 | 3.12201 | 4.56281 | 1.37668 | 0.14314 | 0.14173 | 0.69426 | 2.61759 | 0.72923 | 0.30370 |
| **Srgap3** | 0.03313 | 0.00398 | 0.02496 | 0.00000 | 0.09244 | 0.00221 | 0.00354 | 0.67900 | 0.01953 | 0.01067 | 0.07349 | 0.08757 | 0.01506 | 0.01898 |
| **Srgn** | 0.00000 | 0.09058 | 1.04867 | 0.26474 | 0.09452 | 0.26742 | 0.00924 | 0.00534 | 0.09285 | 0.00444 | 0.00114 | 2.02442 | 0.90713 | 0.49703 |
| **Sri** | 0.00000 | 10.28423 | 0.23113 | 0.32184 | 0.39242 | 0.31928 | 0.01657 | 4.03937 | 0.00000 | 9.09295 | 0.29365 | 3.80632 | 0.00521 | 0.32657 |
| **Srl** | 1.19213 | 0.00343 | 0.57145 | 0.00000 | 2.82726 | 0.10084 | 0.01030 | 0.00107 | 3.42715 | 0.00563 | 0.19495 | 0.55369 | 1.86850 | 0.00000 |
| **Srm** | 16.44713 | 0.18653 | 12.71887 | 12.18996 | 13.10677 | 12.69940 | 3.19554 | 4.21573 | 0.12471 | 0.13706 | 4.93155 | 6.96850 | 2.31941 | 6.80057 |
| **Srp14** | 7.58064 | 7.76374 | 7.66648 | 16.84220 | 12.74030 | 20.41394 | 5.58054 | 7.06337 | 3.41574 | 0.74144 | 10.91999 | 13.49405 | 12.05285 | 9.16495 |
| **Srp19** | 5.78247 | 3.96238 | 2.62854 | 12.72046 | 9.96173 | 4.75076 | 0.25379 | 7.36692 | 1.73218 | 3.67227 | 1.60727 | 8.45932 | 3.91723 | 2.03662 |
| **Srp54a** | 8.79432 | 0.08313 | 7.63462 | 3.52243 | 6.70024 | 7.33450 | 1.96874 | 3.06278 | 7.20951 | 0.06669 | 6.48862 | 5.31322 | 3.21962 | 2.83110 |
| **Srp54b** | 7.93487 | 0.06252 | 7.47223 | 3.35708 | 5.95069 | 5.67801 | 2.10414 | 1.84368 | 5.53375 | 0.07259 | 5.29007 | 4.79858 | 2.70411 | 2.48443 |
| **Srp68** | 5.36612 | 11.57233 | 9.47921 | 9.11334 | 5.97387 | 11.96960 | 4.57350 | 9.55428 | 2.16047 | 4.85437 | 10.56412 | 10.75638 | 0.57999 | 5.02377 |
| **Srp72** | 4.90101 | 0.05402 | 5.87759 | 6.08500 | 7.39178 | 5.84459 | 1.50582 | 6.04322 | 10.69239 | 0.10828 | 6.63528 | 9.83233 | 4.11772 | 7.59200 |
| **Srp9** | 13.33919 | 0.08455 | 6.70742 | 21.78048 | 29.60093 | 10.68922 | 8.00973 | 2.68200 | 6.78583 | 0.39576 | 4.32288 | 7.54489 | 13.85467 | 6.23273 |
| **Srpk1** | 9.18802 | 2.82685 | 20.89124 | 10.03898 | 18.83476 | 9.16829 | 2.38520 | 5.00750 | 13.29414 | 0.14375 | 7.29769 | 3.80510 | 9.95033 | 22.23025 |
| **Srpk2** | 2.67139 | 0.03783 | 1.13779 | 0.00000 | 3.11880 | 2.61291 | 0.93415 | 1.04665 | 5.61352 | 0.05103 | 0.06692 | 4.61561 | 3.70609 | 2.82504 |
| **Srpk3** | 0.01027 | 0.48402 | 0.01207 | 0.02919 | 0.00995 | 0.02150 | 0.00397 | 0.00676 | 0.01040 | 0.00769 | 0.00939 | 0.00419 | 0.20827 | 0.02185 |
| **Srpr** | 10.69354 | 0.07513 | 16.45896 | 4.65135 | 7.89471 | 9.38560 | 2.32608 | 2.00259 | 14.48951 | 11.69100 | 10.66861 | 7.70666 | 11.09524 | 19.28437 |
| **Srprb** | 9.42760 | 4.70206 | 17.04393 | 14.82007 | 9.28298 | 16.39488 | 2.40428 | 3.93825 | 1.04657 | 0.14982 | 8.05085 | 7.75575 | 4.07596 | 14.46242 |
| **Srr** | 1.59611 | 0.02224 | 1.14803 | 0.99786 | 1.23139 | 1.82526 | 0.00366 | 0.50146 | 0.15139 | 0.00000 | 3.08697 | 0.23751 | 2.12892 | 1.65094 |
| **Srrd** | 0.72242 | 0.01042 | 0.54001 | 0.33434 | 0.77234 | 0.59810 | 0.00889 | 0.19744 | 0.05661 | 0.00984 | 0.01932 | 0.41750 | 0.12988 | 1.05162 |
| **Srrm1** | 4.79708 | 1.86894 | 14.20065 | 5.97692 | 12.83252 | 13.95056 | 2.02995 | 6.69642 | 7.55030 | 0.08752 | 8.37963 | 11.54287 | 8.16886 | 14.26693 |
| **Srrm2** | 0.08815 | 0.09295 | 1.68482 | 2.49254 | 7.58937 | 4.96720 | 0.38011 | 1.77716 | 6.83168 | 0.07454 | 2.26275 | 3.82651 | 7.77158 | 5.48180 |
| **Srrt** | 4.82116 | 0.31049 | 18.74311 | 14.42930 | 15.15791 | 15.22750 | 0.47831 | 11.18718 | 0.09578 | 0.03866 | 13.08208 | 16.30277 | 2.36986 | 9.08253 |
| **Srsf1** | 16.18950 | 5.52099 | 24.40749 | 11.66910 | 24.42801 | 16.68196 | 2.61111 | 8.25972 | 41.11841 | 11.59356 | 9.22525 | 11.30100 | 13.81016 | 19.58291 |
| **Srsf10** | 8.24741 | 0.02742 | 6.92898 | 11.45270 | 8.72299 | 10.60644 | 2.78418 | 0.41530 | 13.17587 | 2.84695 | 7.59529 | 6.05812 | 11.48107 | 1.31244 |
| **Srsf11** | 14.45522 | 0.08619 | 13.44108 | 2.98808 | 12.70107 | 16.46569 | 2.63254 | 7.60939 | 12.46982 | 0.39545 | 9.17603 | 8.65093 | 7.93733 | 15.04934 |
| **Srsf2** | 28.32814 | 8.10331 | 24.99416 | 16.63761 | 27.04244 | 20.62336 | 7.04731 | 10.54118 | 21.22457 | 12.32226 | 11.20179 | 15.92453 | 19.21192 | 22.80172 |
| **Srsf3** | 22.53088 | 5.68982 | 38.10575 | 22.92283 | 39.41237 | 35.16003 | 9.38620 | 22.22965 | 33.30690 | 2.84827 | 23.53441 | 23.97610 | 14.81439 | 30.63623 |
| **Srsf4** | 0.01148 | 0.88063 | 0.44038 | 0.12148 | 0.38995 | 0.00000 | 0.00440 | 0.17982 | 0.01347 | 0.00502 | 0.09966 | 0.26032 | 0.00343 | 0.00636 |
| **Srsf5** | 13.84059 | 4.64652 | 15.43223 | 9.17287 | 6.96829 | 7.72938 | 1.53550 | 4.96852 | 19.12313 | 9.70030 | 10.88527 | 5.46790 | 16.24384 | 10.53883 |
| **Srsf6** | 21.15844 | 4.68822 | 17.00667 | 9.91766 | 29.85331 | 12.84248 | 6.53787 | 8.49602 | 19.02963 | 9.93792 | 13.19122 | 14.91801 | 15.59392 | 14.74096 |
| **Srsf7** | 24.21731 | 11.81493 | 29.86876 | 24.99074 | 34.04658 | 29.88494 | 6.42874 | 17.28218 | 18.76679 | 13.86864 | 19.79924 | 17.59700 | 13.49760 | 27.26627 |
| **Srsf9** | 1.82069 | 0.01097 | 3.60568 | 0.00000 | 5.33185 | 0.68506 | 0.02245 | 0.29899 | 0.83644 | 3.35335 | 0.56690 | 1.07268 | 1.74756 | 4.89073 |
| **Srxn1** | 0.70049 | 0.02454 | 0.18394 | 1.42229 | 0.00000 | 2.19653 | 3.79420 | 2.20230 | 0.00312 | 0.11644 | 0.00000 | 2.48558 | 0.00502 | 0.07650 |
| **Ss18** | 8.16858 | 4.81341 | 19.24246 | 10.79431 | 11.10068 | 19.58302 | 6.67297 | 9.42535 | 12.16746 | 0.48153 | 11.85383 | 7.09036 | 2.46188 | 14.46600 |

| | | | | | | | | | | | | | | |
|---|---|---|---|---|---|---|---|---|---|---|---|---|---|---|
| **Ss18l1** | 0.66974 | 0.01004 | 0.65347 | 1.73080 | 0.09053 | 0.53209 | 0.00343 | 0.00000 | 0.08714 | 0.02049 | 0.03273 | 0.76141 | 1.40283 | 1.35831 |
| **Ssb** | 11.62410 | 4.31737 | 15.92576 | 6.97462 | 5.68316 | 18.61619 | 6.61656 | 4.40965 | 18.88975 | 0.23525 | 17.03664 | 12.41716 | 1.87438 | 8.03252 |
| **Ssbp1** | 1.28717 | 0.05135 | 1.49486 | 2.56245 | 2.73957 | 5.88982 | 4.32492 | 1.11185 | 4.01553 | 0.63156 | 1.00075 | 8.47237 | 4.47911 | 4.92252 |
| **Ssbp2** | 0.88792 | 0.00361 | 0.24147 | 0.03182 | 2.66919 | 0.77602 | 0.00380 | 0.00105 | 1.37080 | 0.00515 | 0.00143 | 0.00000 | 1.19824 | 5.02643 |
| **Ssbp3** | 0.80698 | 0.09354 | 2.68835 | 2.30123 | 1.67468 | 6.24677 | 0.43934 | 4.95802 | 0.02191 | 0.06627 | 0.89008 | 4.01599 | 3.28725 | 0.37211 |
| **Ssbp4** | 0.04155 | 0.00410 | 0.00515 | 0.48263 | 0.01710 | 0.01188 | 0.00400 | 0.00436 | 0.02461 | 0.00761 | 0.00738 | 0.17350 | 0.02757 | 0.00000 |
| **Ssc4d** | 0.93520 | 0.00418 | 1.53538 | 0.00000 | 3.40380 | 0.54337 | 0.00481 | 1.04274 | 0.09033 | 0.00313 | 0.00370 | 0.00000 | 2.13772 | 0.92374 |
| **Ssfa2** | 1.29338 | 0.00451 | 0.47580 | 0.28047 | 0.39267 | 0.80083 | 0.00394 | 0.00208 | 0.00498 | 0.00331 | 0.65128 | 0.04328 | 0.01394 | 2.01339 |
| **Ssh1** | 0.00812 | 0.01246 | 0.07111 | 0.00000 | 0.00665 | 0.02644 | 0.30619 | 0.81551 | 0.00741 | 0.02303 | 0.04939 | 0.01116 | 0.10656 | 0.03133 |
| **Ssh2** | 0.41485 | 0.01285 | 1.07383 | 0.08350 | 1.55826 | 0.78661 | 0.00350 | 0.00202 | 1.33564 | 0.00613 | 0.00000 | 0.49092 | 0.23984 | 1.55788 |
| **Ssh3** | 0.40453 | 0.02962 | 0.08841 | 0.15352 | 0.17489 | 0.03315 | 0.30616 | 0.22148 | 0.00475 | 0.00658 | 0.00023 | 0.53079 | 0.35552 | 0.03361 |
| **Ssmem1** | 0.00939 | 0.00413 | 0.09341 | 0.00790 | 0.04943 | 0.00000 | 0.11969 | 0.00637 | 0.00921 | 0.00549 | 0.35378 | 0.01180 | 0.18198 | 0.20371 |
| **Ssna1** | 10.23311 | 0.15388 | 10.16438 | 11.20116 | 18.40800 | 8.87635 | 4.40153 | 7.65036 | 0.48202 | 3.54803 | 6.21577 | 8.78254 | 2.70461 | 5.80925 |
| **Sspn** | 0.00000 | 0.00401 | 0.00000 | 0.14823 | 0.00000 | 0.00000 | 0.00380 | 0.00634 | 0.32004 | 0.00671 | 0.00484 | 0.01215 | 0.48925 | 2.29703 |
| **Sspo** | 0.00251 | 0.00380 | 0.40012 | 0.05202 | 0.00440 | 0.00403 | 0.00410 | 0.00313 | 0.01113 | 0.00593 | 0.00424 | 0.42314 | 0.00567 | 0.01119 |
| **Ssr1** | 2.67120 | 0.29050 | 10.70273 | 11.55300 | 6.47117 | 15.29261 | 1.87163 | 6.65217 | 7.64778 | 0.30119 | 8.45457 | 9.83092 | 1.44327 | 8.37894 |
| **Ssr2** | 10.33594 | 0.42298 | 17.86645 | 17.47003 | 12.46610 | 26.06587 | 4.96208 | 9.31105 | 12.48846 | 3.41119 | 11.15849 | 19.90735 | 6.09319 | 19.57996 |
| **Ssr3** | 5.53416 | 2.27627 | 10.05018 | 6.12012 | 8.45286 | 18.78747 | 6.27428 | 8.06160 | 22.91469 | 0.88588 | 10.01068 | 12.26612 | 3.47559 | 11.73543 |
| **Ssr4** | 12.43243 | 7.06833 | 6.41648 | 16.52866 | 17.56515 | 14.44214 | 5.19642 | 12.52154 | 5.62113 | 8.75425 | 3.90212 | 14.28002 | 1.58428 | 8.73716 |
| **Ssrp1** | 13.18138 | 9.08792 | 18.76404 | 15.54120 | 11.46251 | 22.97836 | 8.71183 | 19.09938 | 13.56360 | 0.63163 | 17.53927 | 14.69593 | 4.75915 | 14.96809 |
| **Sssca1** | 0.06640 | 0.12699 | 5.74894 | 17.94801 | 17.31798 | 0.48312 | 1.96799 | 4.52749 | 0.00285 | 8.70369 | 0.73351 | 7.17742 | 0.02608 | 0.00000 |
| **Ssu72** | 19.05736 | 0.24343 | 11.06047 | 15.46230 | 18.12673 | 16.68162 | 4.12924 | 8.38677 | 4.05328 | 0.27410 | 8.31532 | 10.07596 | 6.49847 | 7.98655 |
| **Ssx2ip** | 1.09431 | 0.04084 | 3.33888 | 1.08312 | 4.55799 | 0.22359 | 0.00814 | 3.68061 | 1.82724 | 0.00000 | 0.01646 | 2.47367 | 9.14184 | 5.27152 |
| **Ssxa1** | 0.00710 | 0.00440 | 0.01416 | 0.00529 | 0.01736 | 0.07595 | 0.00423 | 0.00663 | 0.01148 | 0.00622 | 0.00784 | 0.01061 | 0.12866 | 0.00352 |
| **St13** | 8.20166 | 0.18278 | 16.29651 | 22.47566 | 11.91912 | 23.27468 | 4.52828 | 9.80876 | 7.40131 | 8.75765 | 15.60866 | 15.98752 | 8.07833 | 7.79498 |
| **St14** | 0.00000 | 0.06658 | 3.55063 | 2.53929 | 0.45250 | 4.73008 | 2.01054 | 4.82436 | 0.25096 | 0.08493 | 2.79556 | 2.07704 | 0.00000 | 0.46033 |
| **St3gal1** | 0.03221 | 0.02111 | 0.01118 | 0.01935 | 0.16602 | 0.01255 | 0.00451 | 0.03378 | 0.06070 | 0.00678 | 0.00728 | 0.01709 | 0.06260 | 0.03687 |
| **St3gal2** | 0.92336 | 0.01137 | 0.00000 | 0.02156 | 1.04547 | 0.04276 | 0.00426 | 0.18083 | 0.47625 | 0.01802 | 0.00000 | 1.92795 | 0.01179 | 2.38847 |
| **St3gal3** | 0.00020 | 0.05781 | 0.31599 | 0.00000 | 0.00535 | 0.35803 | 0.00405 | 0.94403 | 0.00546 | 0.00878 | 0.00000 | 0.63808 | 0.03007 | 0.11695 |
| **St3gal4** | 0.00536 | 0.00274 | 0.01099 | 0.00000 | 0.01550 | 0.02117 | 0.00386 | 0.00457 | 0.01306 | 0.00645 | 0.18317 | 1.98811 | 0.00925 | 0.08498 |
| **St3gal5** | 0.00000 | 0.00437 | 0.30132 | 0.69683 | 0.17008 | 0.32564 | 0.00393 | 0.00289 | 0.00536 | 0.02441 | 0.01321 | 0.00439 | 0.11612 | 0.13668 |
| **St3gal6** | 0.08260 | 0.00452 | 1.98485 | 1.61245 | 0.00308 | 1.98044 | 0.49102 | 1.95992 | 1.14009 | 0.02310 | 0.02029 | 0.01967 | 1.88512 | 0.43384 |
| **St5** | 0.27499 | 0.00347 | 0.54249 | 2.19342 | 2.03745 | 1.74919 | 0.00373 | 0.00162 | 0.78582 | 0.01201 | 0.01445 | 0.82267 | 0.04629 | 1.28998 |
| **St6gal1** | 0.00625 | 0.00428 | 0.00860 | 0.01389 | 0.00000 | 0.01095 | 0.00427 | 0.00844 | 0.01177 | 0.00627 | 0.00718 | 0.00835 | 0.29685 | 1.10410 |
| **St6galnac2** | 1.14534 | 0.00649 | 3.67854 | 10.42081 | 1.79061 | 5.38340 | 2.54514 | 0.17141 | 1.84202 | 0.06928 | 6.85816 | 5.52846 | 0.01839 | 1.78323 |
| **St6galnac4** | 0.01960 | 0.00434 | 0.00948 | 0.07808 | 0.00405 | 0.01123 | 0.00440 | 0.00841 | 0.01609 | 0.00586 | 0.00704 | 0.00820 | 0.01242 | 0.25837 |
| **St6galnac5** | 0.00487 | 0.00468 | 0.01198 | 0.00000 | 0.01068 | 0.02721 | 0.00410 | 0.02448 | 0.01086 | 0.00579 | 0.00431 | 0.48138 | 0.21661 | 0.01116 |
| **St6galnac6** | 0.04803 | 0.00445 | 0.00987 | 0.01376 | 0.13475 | 0.01255 | 0.00415 | 0.01018 | 0.01058 | 0.00744 | 0.00831 | 0.00861 | 0.02016 | 0.00608 |
| **St7** | 0.68053 | 0.00435 | 0.00000 | 0.07928 | 0.00000 | 0.00000 | 0.00484 | 0.22724 | 0.83332 | 0.00652 | 0.00347 | 0.01273 | 0.12519 | 0.22666 |
| **St7l** | 1.79634 | 0.01155 | 1.99510 | 0.14520 | 2.21914 | 1.18175 | 0.01119 | 0.02523 | 1.38602 | 0.04747 | 0.00000 | 0.80628 | 0.88424 | 2.75545 |
| **St8sia1** | 0.00000 | 0.02384 | 0.08527 | 0.09098 | 0.35947 | 0.07081 | 0.01596 | 0.03889 | 0.11874 | 0.01924 | 0.01542 | 0.24473 | 0.27655 | 0.23810 |
| **Stab1** | 0.09088 | 0.00426 | 0.49101 | 0.00355 | 0.08723 | 0.00589 | 0.00404 | 0.00872 | 0.01236 | 0.00926 | 0.00643 | 0.00709 | 0.65499 | 0.06131 |
| **Stac2** | 4.38876 | 0.05054 | 10.91396 | 6.17782 | 12.35266 | 4.96468 | 1.13761 | 0.98311 | 0.00000 | 0.04951 | 0.49948 | 1.17742 | 0.00000 | 4.90007 |
| **Stag1** | 2.59735 | 0.00596 | 3.92278 | 0.00000 | 2.07412 | 4.20726 | 1.29241 | 0.61183 | 2.56246 | 0.01676 | 4.79137 | 5.60133 | 0.98587 | 3.39398 |
| **Stag2** | 19.14286 | 0.04508 | 12.61569 | 0.83841 | 12.46338 | 6.10361 | 0.00345 | 0.13780 | 31.75691 | 0.70008 | 3.69106 | 0.22695 | 6.48220 | 7.44020 |
| **Stag3** | 2.29015 | 0.01728 | 2.61140 | 4.26319 | 2.36635 | 2.80488 | 0.00285 | 0.00000 | 1.86082 | 0.01827 | 1.36204 | 0.50915 | 0.98212 | 0.32197 |
| **Stam** | 4.79793 | 0.02321 | 4.57343 | 0.00021 | 3.54538 | 3.10594 | 2.56618 | 1.97677 | 1.08228 | 0.02044 | 2.49897 | 3.76772 | 1.62811 | 3.87676 |
| **Stam2** | 13.04130 | 0.01934 | 12.14043 | 8.91621 | 10.10466 | 6.72201 | 0.33014 | 0.92527 | 7.29184 | 0.03655 | 5.65622 | 7.18957 | 4.52452 | 13.18366 |
| **Stambp** | 0.04114 | 0.00293 | 0.00000 | 1.09672 | 1.33798 | 2.28034 | 0.01492 | 2.77591 | 2.16812 | 0.05404 | 3.10483 | 0.00000 | 2.24578 | 0.32875 |
| **Stambpl1** | 0.05006 | 0.00312 | 0.01259 | 0.00461 | 0.00923 | 0.01212 | 0.00398 | 0.00619 | 0.00959 | 0.00627 | 0.00416 | 1.67393 | 0.00513 | 0.02009 |
| **Stamos** | 0.00707 | 0.00376 | 0.01826 | 0.05294 | 0.00800 | 0.43749 | 0.00418 | 0.10659 | 0.01241 | 0.02672 | 0.89269 | 0.00134 | 0.00802 | 0.00261 |
| **Stap2** | 0.00635 | 0.00436 | 0.03093 | 0.24274 | 0.20973 | 0.00076 | 0.00468 | 0.00936 | 0.01646 | 0.00602 | 0.03272 | 0.01569 | 0.00643 | 0.00826 |
| **Star** | 1.02393 | 0.00396 | 0.35123 | 0.00000 | 1.33140 | 0.43814 | 0.00432 | 0.00288 | 0.01701 | 0.00499 | 0.00227 | 0.24501 | 0.73564 | 0.05540 |
| **Stard10** | 0.00000 | 5.32018 | 14.95110 | 18.27265 | 0.20232 | 13.44127 | 5.52556 | 25.61071 | 0.08010 | 8.42211 | 4.84290 | 7.92603 | 0.00000 | 0.00000 |
| **Stard13** | 0.01318 | 0.00432 | 0.02650 | 0.01046 | 0.00697 | 0.01250 | 0.00424 | 0.00920 | 0.00514 | 0.00620 | 0.00890 | 0.00876 | 0.49469 | 0.09614 |

| | | | | | | | | | | | | | | |
|---|---|---|---|---|---|---|---|---|---|---|---|---|---|---|
| **Stard3** | 0.71617 | 0.02389 | 2.58967 | 2.38341 | 1.95210 | 3.48177 | 0.00255 | 2.88771 | 0.02142 | 0.03999 | 0.43020 | 2.60180 | 0.25476 | 0.28559 |
| **Stard3nl** | 2.52488 | 1.32935 | 3.63270 | 4.28148 | 0.97712 | 7.58369 | 0.32061 | 2.35036 | 0.00673 | 0.06909 | 4.29411 | 3.56793 | 0.28059 | 2.63785 |
| **Stard4** | 0.16804 | 0.00418 | 1.92972 | 1.40061 | 0.00000 | 3.47030 | 3.71847 | 0.54271 | 0.01176 | 0.11503 | 0.00000 | 0.00000 | 0.00504 | 0.00000 |
| **Stard5** | 0.00385 | 0.00395 | 0.00000 | 0.25664 | 0.00000 | 0.72037 | 0.01063 | 0.00773 | 0.21404 | 0.01256 | 0.00321 | 0.00527 | 1.02248 | 0.56810 |
| **Stard7** | 0.93375 | 0.07488 | 2.49930 | 3.81207 | 2.08932 | 4.27857 | 2.40704 | 9.37174 | 1.85087 | 2.93504 | 4.98663 | 4.17323 | 5.67145 | 2.03856 |
| **Stat1** | 0.15483 | 0.08039 | 0.20806 | 0.14590 | 0.08613 | 0.26642 | 0.02738 | 0.13234 | 0.13900 | 0.23638 | 0.00077 | 1.18633 | 0.59666 | 0.22342 |
| **Stat2** | 0.47441 | 0.01007 | 1.32632 | 2.20870 | 3.09601 | 0.37957 | 0.00370 | 0.67277 | 0.00000 | 0.00000 | 0.98704 | 2.52934 | 4.89753 | 7.18782 |
| **Stat3** | 3.47364 | 0.15762 | 6.36206 | 5.68394 | 2.87422 | 13.42989 | 6.18216 | 5.16531 | 2.12520 | 1.23766 | 9.69953 | 9.86773 | 3.97140 | 2.36247 |
| **Stat4** | 0.00314 | 0.00427 | 0.01091 | 0.01189 | 0.12370 | 0.01209 | 0.00433 | 0.01016 | 0.00418 | 0.00638 | 0.00869 | 0.00873 | 0.53994 | 0.15872 |
| **Stat5a** | 0.24589 | 0.00474 | 1.39388 | 1.77953 | 1.74387 | 1.72187 | 0.00394 | 0.31002 | 0.35486 | 0.02179 | 0.55512 | 0.11549 | 0.01511 | 0.49789 |
| **Stat5b** | 0.01859 | 0.00395 | 0.55984 | 0.15505 | 0.73577 | 0.13755 | 0.00396 | 0.00083 | 0.42052 | 0.00463 | 0.00186 | 0.43725 | 0.00000 | 0.01478 |
| **Stat6** | 0.19162 | 0.04313 | 0.14454 | 7.50440 | 0.04674 | 3.68215 | 0.03194 | 4.15547 | 0.00726 | 1.07262 | 0.00000 | 0.28269 | 0.00186 | 1.00308 |
| **Stau1** | 2.32484 | 0.77822 | 3.96155 | 1.96394 | 5.71072 | 2.35928 | 0.10212 | 3.34598 | 4.40517 | 0.04632 | 6.37026 | 3.95540 | 8.49775 | 12.01659 |
| **Stau2** | 0.13162 | 0.00360 | 0.45129 | 0.95231 | 0.32848 | 0.05797 | 2.68600 | 0.00745 | 0.02064 | 0.04555 | 0.01009 | 0.43340 | 0.00318 | 0.03228 |
| **Stbd1** | 0.24285 | 0.00403 | 0.42791 | 0.05498 | 0.00000 | 0.00000 | 0.00377 | 0.00739 | 0.00426 | 0.00690 | 0.00477 | 0.05063 | 1.37991 | 2.89245 |
| **Stc2** | 0.00560 | 0.00437 | 0.01426 | 0.01136 | 0.03664 | 0.01320 | 0.00424 | 0.00953 | 0.00947 | 0.00623 | 0.00941 | 0.00942 | 0.23913 | 0.01183 |
| **Steap1** | 0.10903 | 0.00230 | 0.00000 | 0.25291 | 0.13799 | 0.62316 | 0.00369 | 0.00248 | 0.01806 | 0.00642 | 0.00000 | 2.18635 | 0.54188 | 0.00000 |
| **Steap2** | 0.08807 | 0.00387 | 0.24610 | 0.10111 | 0.00000 | 0.00730 | 0.03897 | 0.03803 | 0.03467 | 0.00556 | 0.01339 | 0.49480 | 0.01475 | 0.72377 |
| **Steap3** | 0.51213 | 0.94701 | 12.68241 | 6.96104 | 4.85521 | 4.16496 | 0.00729 | 4.50308 | 2.57802 | 0.01746 | 4.01647 | 0.00000 | 0.00000 | 1.67598 |
| **Stil** | 1.46875 | 0.03526 | 1.90359 | 0.00000 | 0.00000 | 1.26032 | 0.00262 | 0.92154 | 1.30524 | 0.00498 | 1.94260 | 2.64418 | 4.71344 | 0.00000 |
| **Stim1** | 0.89103 | 0.01283 | 1.95365 | 0.00000 | 1.83364 | 1.01902 | 0.00385 | 0.09644 | 2.86591 | 0.01671 | 0.88190 | 0.06368 | 2.04063 | 1.28520 |
| **Stim2** | 0.16078 | 0.00476 | 0.07220 | 0.00000 | 0.00000 | 0.47649 | 0.00405 | 0.06913 | 0.01141 | 0.00704 | 0.10810 | 1.24616 | 0.47767 | 0.57766 |
| **Stip1** | 27.89162 | 7.86626 | 52.91679 | 46.88898 | 41.15111 | 44.21087 | 8.95630 | 16.98128 | 25.95899 | 2.91240 | 32.54379 | 23.09272 | 8.44758 | 42.39304 |
| **Stk10** | 1.17734 | 0.00324 | 0.12332 | 0.00000 | 3.03261 | 0.67087 | 0.00375 | 0.02027 | 0.01262 | 0.00379 | 1.27797 | 0.05322 | 1.83623 | 3.06455 |
| **Stk11** | 1.18675 | 0.32851 | 17.42050 | 12.81900 | 15.85602 | 1.03221 | 6.56190 | 9.91202 | 0.00000 | 0.89690 | 6.89608 | 8.43210 | 1.17604 | 1.98268 |
| **Stk11ip** | 0.25224 | 0.00321 | 1.31858 | 4.91370 | 3.48666 | 1.14993 | 0.01921 | 1.48501 | 0.01043 | 0.00430 | 1.67171 | 0.00000 | 1.74058 | 1.47011 |
| **Stk16** | 9.20751 | 0.08526 | 8.81494 | 13.05595 | 12.28594 | 6.71228 | 1.04901 | 2.37940 | 2.57862 | 0.18409 | 3.60301 | 7.32150 | 6.51240 | 3.55776 |
| **Stk17b** | 2.35635 | 0.00296 | 1.50973 | 1.30257 | 1.83262 | 0.85202 | 0.00357 | 0.00453 | 0.62168 | 0.01535 | 2.30564 | 0.54097 | 1.59383 | 3.37965 |
| **Stk19** | 3.01870 | 0.10042 | 6.15734 | 3.55209 | 7.38182 | 2.00892 | 3.07957 | 3.16690 | 0.02661 | 0.14882 | 0.11236 | 3.78021 | 2.74507 | 1.05093 |
| **Stk24** | 10.13734 | 3.71143 | 21.60181 | 10.87215 | 12.95162 | 13.29878 | 4.09751 | 4.06885 | 2.33586 | 0.11897 | 11.81609 | 6.30317 | 4.30188 | 5.59686 |
| **Stk25** | 0.98011 | 8.25332 | 5.97785 | 4.31837 | 4.33801 | 2.97366 | 1.28718 | 4.75581 | 3.23740 | 0.04811 | 4.11365 | 1.51900 | 0.71795 | 10.20939 |
| **Stk3** | 0.20084 | 0.00239 | 0.11776 | 0.00000 | 0.14455 | 3.13256 | 0.92234 | 0.00050 | 1.42860 | 0.01503 | 0.30905 | 1.92168 | 1.37266 | 0.00054 |
| **Stk31** | 2.97813 | 0.02568 | 1.44424 | 0.05205 | 2.91141 | 2.59324 | 0.00304 | 0.67378 | 1.76247 | 0.00185 | 5.88057 | 2.50280 | 1.45913 | 3.13938 |
| **Stk35** | 5.80709 | 0.00131 | 4.46598 | 0.00000 | 4.00318 | 0.80431 | 0.00817 | 0.04327 | 2.43308 | 0.00000 | 0.00000 | 0.39665 | 17.70981 | 1.93426 |
| **Stk36** | 0.00000 | 0.00444 | 0.31925 | 0.25907 | 0.76185 | 0.20628 | 0.00421 | 0.01211 | 0.00349 | 0.01074 | 0.00409 | 0.00310 | 0.54356 | 0.31184 |
| **Stk38** | 8.97895 | 0.06941 | 9.56733 | 6.60948 | 6.34318 | 3.44612 | 2.21747 | 2.19748 | 9.28908 | 0.03390 | 8.63677 | 7.41013 | 5.01405 | 11.63066 |
| **Stk38l** | 1.89618 | 0.01608 | 1.63827 | 2.24819 | 3.27607 | 1.58407 | 1.46879 | 1.46968 | 10.43908 | 0.04656 | 1.84117 | 2.56688 | 4.77706 | 3.60232 |
| **Stk39** | 0.31996 | 0.00457 | 0.01252 | 0.01368 | 0.00626 | 0.01173 | 0.00429 | 0.01317 | 0.01743 | 0.00708 | 0.00728 | 0.00844 | 0.00462 | 0.00368 |
| **Stk4** | 8.99131 | 0.26701 | 9.07953 | 2.25983 | 7.99357 | 5.21338 | 4.51982 | 0.25915 | 4.63923 | 0.44110 | 1.87473 | 0.66271 | 1.56774 | 4.86336 |
| **Stk40** | 0.99898 | 2.56315 | 5.85975 | 6.95768 | 2.15205 | 8.19956 | 0.00555 | 6.92327 | 0.00000 | 0.05329 | 2.64897 | 5.41538 | 4.89024 | 3.04254 |
| **Stkld1** | 0.00773 | 0.00450 | 0.00933 | 0.01330 | 0.00729 | 0.00891 | 0.00461 | 0.02403 | 0.20903 | 0.00668 | 0.00772 | 0.00927 | 0.03706 | 0.00000 |
| **Stmn1** | 7.30020 | 0.41249 | 8.08381 | 8.99305 | 3.52054 | 7.20983 | 3.59465 | 9.19730 | 0.07449 | 0.14599 | 10.47308 | 7.54658 | 6.37326 | 4.99607 |
| **Stmn1.rs1** | 0.03674 | 0.00463 | 0.03076 | 0.06042 | 0.03250 | 0.04102 | 0.02771 | 0.05995 | 0.01464 | 0.00612 | 0.04531 | 0.08306 | 0.06073 | 0.01791 |
| **Stmn3** | 0.93157 | 1.17449 | 22.72921 | 26.30760 | 27.42497 | 0.00000 | 0.22561 | 0.19321 | 0.00000 | 0.13182 | 3.58879 | 1.17400 | 3.13077 | 0.00000 |
| **Stom** | 0.00535 | 0.00495 | 0.00487 | 0.00000 | 0.32939 | 0.15527 | 0.00427 | 0.02125 | 0.00736 | 0.00543 | 0.00366 | 0.20499 | 0.00588 | 0.07466 |
| **Stoml1** | 0.04255 | 0.02087 | 0.15912 | 1.55744 | 0.28812 | 0.45866 | 4.03232 | 0.37557 | 0.14839 | 0.17018 | 1.29071 | 1.17513 | 0.66638 | 0.06652 |
| **Stoml2** | 11.35146 | 7.49354 | 13.97120 | 15.18533 | 16.74846 | 4.30660 | 3.76621 | 7.69141 | 3.67989 | 2.91910 | 6.10596 | 10.09380 | 1.94824 | 5.06264 |
| **Stoml3** | 0.02111 | 0.00439 | 0.01148 | 0.01675 | 0.19307 | 0.01408 | 0.00422 | 0.00885 | 0.01944 | 0.00593 | 0.01124 | 0.00999 | 0.11896 | 0.01620 |
| **Ston2** | 0.70856 | 0.38687 | 0.58079 | 0.76669 | 0.50333 | 0.08580 | 0.01564 | 0.08913 | 0.96835 | 0.02774 | 0.02581 | 0.02162 | 1.92944 | 0.79036 |
| **Stox1** | 0.05243 | 0.00448 | 0.19649 | 0.38853 | 0.00000 | 0.00000 | 0.00435 | 0.03260 | 0.28704 | 0.00839 | 0.00676 | 0.00535 | 0.39341 | 0.00000 |
| **Stox2** | 0.00627 | 0.00442 | 0.00857 | 0.01194 | 0.25863 | 0.01202 | 0.00436 | 0.01098 | 0.00711 | 0.00603 | 0.00890 | 0.00866 | 0.11061 | 0.16273 |
| **Stra13** | 3.90542 | 5.53964 | 0.08621 | 2.77285 | 3.43448 | 9.09967 | 2.81179 | 4.50782 | 0.32711 | 0.22818 | 1.87106 | 5.38292 | 1.39691 | 1.20702 |
| **Stra6** | 0.01222 | 0.00491 | 0.01182 | 0.01785 | 0.02125 | 0.21555 | 0.00452 | 0.00754 | 0.01107 | 0.00623 | 0.00719 | 0.00740 | 0.00739 | 0.01620 |
| **Stra8** | 0.00256 | 0.00439 | 0.04186 | 0.87635 | 0.14572 | 0.63302 | 0.00471 | 1.07854 | 0.01695 | 0.02587 | 0.00434 | 0.00000 | 0.00751 | 0.00617 |
| **Strada** | 3.98052 | 0.08550 | 11.44917 | 10.90545 | 7.47287 | 9.16840 | 0.27642 | 5.31594 | 0.54714 | 0.24464 | 3.54859 | 2.37478 | 0.00000 | 1.65932 |

| | | | | | | | | | | | | | | |
|---|---|---|---|---|---|---|---|---|---|---|---|---|---|---|
| **Stradb** | 0.26243 | 5.00438 | 0.15614 | 0.42256 | 0.05731 | 2.48547 | 0.00915 | 0.83433 | 0.06218 | 0.01212 | 0.25117 | 1.50659 | 7.42951 | 1.16972 |
| **Strap** | 24.14775 | 3.56683 | 21.85403 | 10.32277 | 28.99221 | 15.31259 | 6.77982 | 8.64880 | 17.07415 | 0.27863 | 12.41374 | 9.78569 | 13.68730 | 23.41995 |
| **Strbp** | 4.60341 | 0.09668 | 5.63779 | 4.28833 | 5.18072 | 7.28557 | 0.09297 | 1.24478 | 2.66304 | 0.07646 | 4.55071 | 10.08651 | 6.99053 | 9.18761 |
| **Strc** | 0.00656 | 0.00436 | 0.52040 | 0.00000 | 0.12385 | 0.00539 | 0.00422 | 0.01253 | 0.14469 | 0.00635 | 0.00678 | 0.00817 | 0.00535 | 0.00229 |
| **Strip1** | 2.88326 | 0.05849 | 5.82257 | 3.38318 | 5.26490 | 8.35043 | 0.01255 | 4.24745 | 0.00000 | 0.13024 | 2.59391 | 3.41375 | 9.09932 | 6.15268 |
| **Strip2** | 0.00293 | 0.00409 | 0.00770 | 0.00997 | 0.07399 | 0.17033 | 0.00440 | 0.00798 | 0.01868 | 0.00647 | 0.00625 | 0.03981 | 1.55106 | 0.06455 |
| **Strn** | 0.29590 | 0.00376 | 0.81084 | 0.45895 | 1.05426 | 0.01921 | 0.00365 | 0.00000 | 0.63713 | 0.00345 | 0.19555 | 0.00570 | 1.02581 | 2.95639 |
| **Strn3** | 7.76395 | 0.15110 | 6.65887 | 0.80884 | 10.72054 | 5.16677 | 1.64282 | 0.00000 | 31.91660 | 0.03227 | 6.23609 | 4.57218 | 2.64248 | 9.99731 |
| **Strn4** | 0.86263 | 4.77407 | 6.71230 | 5.10466 | 4.32181 | 3.27000 | 1.71979 | 1.70513 | 0.00000 | 8.19505 | 1.82042 | 7.11961 | 0.83735 | 0.00000 |
| **Stt3a** | 3.44330 | 3.88334 | 7.70345 | 7.92764 | 2.04228 | 9.95798 | 0.15816 | 2.26930 | 20.45387 | 0.06529 | 6.82714 | 7.10137 | 2.27374 | 11.46784 |
| **Stt3b** | 10.05394 | 1.25588 | 9.45007 | 6.23397 | 10.68408 | 14.29895 | 2.79550 | 0.77929 | 14.91763 | 0.10166 | 6.54698 | 7.08068 | 11.49046 | 12.77575 |
| **Stub1** | 0.41132 | 0.00999 | 2.74702 | 5.52062 | 2.31098 | 0.80105 | 0.01391 | 1.31976 | 0.00109 | 4.25819 | 0.00000 | 1.38493 | 0.00000 | 0.18203 |
| **Stx11** | 0.00868 | 0.00438 | 0.01403 | 0.01212 | 0.00923 | 0.01279 | 0.00427 | 0.00963 | 0.02256 | 0.00635 | 0.00886 | 0.00957 | 0.20736 | 0.03208 |
| **Stx12** | 5.49206 | 1.93844 | 7.53451 | 0.85343 | 7.51003 | 8.42707 | 2.76227 | 3.70444 | 9.36344 | 0.04270 | 4.71055 | 4.00566 | 1.27359 | 10.16331 |
| **Stx16** | 0.76525 | 0.00914 | 2.45260 | 3.80003 | 0.14868 | 2.76211 | 0.31615 | 2.56436 | 0.12129 | 0.05717 | 0.00000 | 2.82855 | 0.67261 | 2.79040 |
| **Stx17** | 7.69147 | 1.76257 | 5.82149 | 3.35323 | 5.97686 | 3.89108 | 0.44913 | 0.00000 | 5.00542 | 0.02821 | 5.68659 | 2.07705 | 0.00000 | 11.74340 |
| **Stx18** | 1.62275 | 3.43878 | 2.55728 | 2.77993 | 2.21128 | 3.69087 | 2.32843 | 6.50763 | 0.03438 | 0.05906 | 3.74258 | 2.55537 | 0.00000 | 0.23261 |
| **Stx1a** | 1.51155 | 0.00386 | 6.60848 | 3.66616 | 3.63725 | 0.52250 | 3.49137 | 0.30193 | 0.00249 | 0.09178 | 1.08875 | 0.00000 | 1.56248 | 2.05923 |
| **Stx1b** | 0.00877 | 0.00413 | 0.00448 | 0.23983 | 0.03436 | 0.00512 | 0.00393 | 0.00669 | 0.14517 | 0.00586 | 0.00593 | 0.01193 | 0.24533 | 0.00000 |
| **Stx2** | 0.15450 | 0.02252 | 0.25559 | 0.00000 | 0.90452 | 0.00034 | 0.01579 | 0.01612 | 0.01763 | 0.00602 | 0.00483 | 0.20582 | 0.27884 | 0.36834 |
| **Stx3** | 1.99259 | 3.48225 | 3.55995 | 3.46343 | 1.63959 | 8.82216 | 1.40610 | 6.34679 | 3.42235 | 0.09955 | 0.28453 | 2.62247 | 0.66846 | 2.79017 |
| **Stx4a** | 0.13182 | 0.05007 | 10.38998 | 4.57968 | 4.18961 | 7.37032 | 0.01332 | 3.05743 | 0.07177 | 0.18387 | 3.97911 | 4.19625 | 0.00000 | 2.07349 |
| **Stx5a** | 0.94746 | 0.06401 | 6.01216 | 2.18216 | 3.06299 | 4.75658 | 0.03685 | 5.01943 | 2.61125 | 0.06324 | 5.11649 | 3.52504 | 7.25980 | 2.35341 |
| **Stx6** | 0.04560 | 0.00370 | 0.31283 | 1.23837 | 0.53637 | 0.11855 | 0.81045 | 0.00000 | 0.01179 | 0.01188 | 1.27876 | 0.42934 | 0.32198 | 0.94229 |
| **Stx7** | 1.77764 | 0.05037 | 10.09528 | 3.44384 | 5.83316 | 17.65713 | 3.99263 | 4.77275 | 3.06013 | 0.11028 | 5.40993 | 3.22732 | 0.00000 | 2.05508 |
| **Stx8** | 0.47049 | 0.10174 | 2.24513 | 5.78712 | 0.08270 | 2.89590 | 0.28864 | 6.57725 | 0.00000 | 0.41501 | 4.50737 | 5.58298 | 0.00000 | 1.62610 |
| **Stxbp1** | 0.22421 | 0.01220 | 0.53551 | 0.43600 | 1.29819 | 0.00700 | 0.01111 | 0.66533 | 0.00540 | 0.00598 | 0.01414 | 2.00065 | 0.39775 | 0.57450 |
| **Stxbp2** | 0.66562 | 5.21431 | 4.04483 | 7.42077 | 1.44635 | 4.40139 | 2.60809 | 12.01618 | 0.76541 | 12.37902 | 4.38344 | 4.78204 | 2.24413 | 0.40974 |
| **Stxbp3** | 0.01363 | 0.04550 | 0.90283 | 0.77410 | 0.00000 | 3.49339 | 0.00338 | 0.68239 | 1.89347 | 0.00000 | 2.67271 | 0.86149 | 0.36936 | 1.15965 |
| **Stxbp3.ps** | 0.00947 | 0.00441 | 0.05315 | 0.02127 | 0.02648 | 0.09758 | 0.00464 | 0.03798 | 0.24323 | 0.14476 | 0.17223 | 0.06018 | 0.22247 | 0.10646 |
| **Stxbp4** | 0.22200 | 0.08860 | 0.13581 | 0.29738 | 0.19018 | 0.65637 | 0.07880 | 0.21686 | 0.35304 | 0.40262 | 0.07288 | 0.59683 | 0.62747 | 0.87517 |
| **Stxbp5** | 0.56335 | 0.00202 | 0.43701 | 0.00000 | 1.02209 | 3.02343 | 0.00292 | 1.57308 | 1.24849 | 0.04277 | 1.55093 | 1.39934 | 4.99376 | 5.38710 |
| **Stxbp5l** | 0.00000 | 0.00442 | 0.40503 | 0.00000 | 1.06052 | 0.00741 | 0.03822 | 0.02839 | 1.96947 | 0.00613 | 0.00460 | 0.01157 | 0.41905 | 0.28610 |
| **Stxbp6** | 1.07934 | 0.01920 | 0.29274 | 1.00689 | 0.00000 | 1.05419 | 0.00414 | 0.00175 | 0.09372 | 0.01535 | 0.23288 | 0.00511 | 0.36567 | 0.00000 |
| **Styk1** | 0.02217 | 0.00439 | 0.01452 | 0.01861 | 0.00943 | 0.01256 | 0.00431 | 0.00970 | 0.01211 | 0.00621 | 0.00824 | 0.00908 | 0.10462 | 0.06335 |
| **Styx** | 0.19781 | 0.00316 | 1.14002 | 1.14177 | 1.61628 | 1.57566 | 0.37058 | 0.86038 | 5.08012 | 0.03028 | 1.61047 | 0.21417 | 0.80393 | 2.38136 |
| **Styxl1** | 0.01611 | 0.00455 | 0.01363 | 0.00837 | 0.01757 | 0.01556 | 0.00424 | 0.00961 | 0.01096 | 0.00687 | 0.27464 | 0.00587 | 0.03053 | 0.05663 |
| **Sub1** | 8.72792 | 7.21454 | 0.28058 | 13.05302 | 13.49164 | 19.58845 | 4.88583 | 13.21660 | 13.56951 | 12.78404 | 4.15684 | 16.42193 | 20.50159 | 8.49159 |
| **Sucla2** | 9.97228 | 0.22221 | 15.06993 | 13.16734 | 9.89057 | 14.59083 | 3.89362 | 2.84413 | 7.05487 | 4.63498 | 8.89611 | 12.43700 | 5.83185 | 5.51839 |
| **Suclg1** | 9.88411 | 9.51123 | 21.29124 | 19.10063 | 10.56993 | 21.49725 | 9.59401 | 23.84391 | 2.30749 | 19.65567 | 18.35957 | 23.91527 | 1.29564 | 4.69661 |
| **Suclg2** | 2.79457 | 3.54362 | 14.15136 | 10.12972 | 6.46541 | 14.20523 | 3.49676 | 10.94393 | 1.93761 | 0.19380 | 11.90942 | 11.22892 | 1.82224 | 5.36466 |
| **Suco** | 3.89295 | 0.11131 | 1.04437 | 0.07104 | 0.82777 | 4.46864 | 2.69143 | 1.05722 | 1.05400 | 0.03296 | 1.06720 | 2.06798 | 2.41022 | 1.77024 |
| **Suds3** | 11.50574 | 7.05386 | 10.40914 | 4.05792 | 17.08901 | 6.92766 | 4.06032 | 3.38759 | 0.70687 | 0.24052 | 3.96549 | 3.54379 | 8.58167 | 12.43120 |
| **Sufu** | 0.05869 | 0.01024 | 0.67933 | 0.73219 | 0.00000 | 0.14851 | 2.08649 | 0.00000 | 0.69783 | 0.03670 | 0.00000 | 2.53952 | 1.42408 | 0.99220 |
| **Sugct** | 0.00705 | 0.00426 | 0.01405 | 0.01530 | 0.01152 | 0.01460 | 0.00419 | 0.00809 | 0.00971 | 0.00757 | 0.00875 | 0.00839 | 0.71811 | 0.00549 |
| **Sugp1** | 1.38728 | 0.07822 | 8.62052 | 6.86391 | 3.28686 | 2.89144 | 3.06527 | 1.26615 | 0.15348 | 0.11076 | 1.12782 | 3.44519 | 5.00768 | 1.44381 |
| **Sugp2** | 0.46474 | 0.00405 | 0.79485 | 0.00000 | 0.71496 | 1.05801 | 0.00448 | 0.83177 | 0.01560 | 0.01979 | 0.00000 | 0.70257 | 0.04941 | 0.55938 |
| **Sugt1** | 38.80684 | 10.57025 | 48.07576 | 36.94887 | 44.76090 | 33.17138 | 9.49370 | 21.92378 | 35.58147 | 21.11566 | 21.58246 | 19.57571 | 16.47276 | 36.66787 |
| **Sulf2** | 0.12875 | 0.13119 | 2.84641 | 4.42182 | 0.45187 | 5.48856 | 0.34483 | 0.06871 | 0.00430 | 0.05719 | 0.00000 | 11.14314 | 0.00000 | 0.26825 |
| **Sult1a1** | 0.06658 | 0.00443 | 0.00895 | 0.01417 | 0.23952 | 0.02444 | 0.00412 | 0.00856 | 0.28915 | 0.00679 | 0.00644 | 0.00804 | 0.12334 | 0.00000 |
| **Sult2a6** | 0.30530 | 0.00440 | 0.00940 | 0.01121 | 0.00002 | 0.00871 | 0.00420 | 0.01131 | 0.11258 | 0.00660 | 0.00657 | 0.00880 | 0.03945 | 0.04472 |
| **Sult2b1** | 1.23117 | 0.03186 | 3.68564 | 1.18110 | 3.47490 | 0.00000 | 0.00279 | 0.00000 | 0.01136 | 0.00000 | 0.19690 | 1.61714 | 0.58588 | 0.97813 |
| **Sult4a1** | 0.04995 | 0.00424 | 0.98715 | 0.71707 | 0.96798 | 0.33085 | 0.00383 | 0.00609 | 0.27977 | 0.00633 | 0.01611 | 0.00453 | 0.18163 | 0.34434 |
| **Sult5a1** | 0.42660 | 0.03298 | 1.06309 | 1.04712 | 1.31628 | 3.94116 | 0.01351 | 1.54677 | 1.85664 | 0.02221 | 1.06077 | 0.64349 | 0.09925 | 2.03238 |
| **Sult6b1** | 1.18960 | 0.00344 | 0.45969 | 0.49866 | 0.28907 | 0.77627 | 2.12864 | 0.44475 | 0.35940 | 0.11928 | 0.00000 | 0.29175 | 1.15359 | 0.00000 |

| | | | | | | | | | | | | | | |
|---|---|---|---|---|---|---|---|---|---|---|---|---|---|---|
| **Sumf1** | 1.54549 | 0.02850 | 0.20261 | 2.26868 | 0.99837 | 1.63525 | 0.01981 | 0.79837 | 1.50877 | 0.00346 | 0.77458 | 0.77719 | 0.96288 | 2.29076 |
| **Sumf2** | 0.00000 | 0.01885 | 1.34799 | 0.00000 | 0.29174 | 2.76538 | 0.00331 | 1.40196 | 0.32233 | 0.00000 | 2.84424 | 1.52064 | 0.30201 | 0.32916 |
| **Sumo1** | 16.58615 | 6.73549 | 34.59158 | 34.24124 | 31.47945 | 27.05449 | 8.53961 | 6.66896 | 31.16374 | 13.83670 | 17.52010 | 14.29542 | 18.09363 | 27.58261 |
| **Sumo2** | 33.97651 | 0.47546 | 43.78984 | 35.38662 | 47.24343 | 28.77789 | 10.48766 | 19.92423 | 26.45724 | 2.28738 | 15.75908 | 18.11432 | 10.34063 | 31.82281 |
| **Sumo3** | 6.74054 | 0.10338 | 7.87496 | 5.83480 | 13.49832 | 7.54699 | 1.25101 | 8.37399 | 8.82233 | 0.12201 | 1.16970 | 7.14293 | 3.77434 | 8.02522 |
| **Sun1** | 14.77027 | 3.23710 | 24.29683 | 12.72034 | 16.72532 | 13.17439 | 1.73551 | 5.17745 | 25.28759 | 6.01084 | 8.64513 | 9.83828 | 9.97285 | 27.76741 |
| **Sun2** | 0.03995 | 0.01251 | 0.00767 | 0.00000 | 0.01508 | 0.08691 | 0.22398 | 0.71941 | 0.02413 | 0.02767 | 0.00267 | 0.07203 | 0.14131 | 0.06176 |
| **Suox** | 0.28183 | 6.86465 | 4.18021 | 2.20292 | 0.00000 | 6.72841 | 2.76213 | 0.08683 | 0.00000 | 0.04582 | 3.53903 | 1.24863 | 0.00000 | 0.65369 |
| **Supt16** | 0.52614 | 0.02253 | 1.66642 | 0.68819 | 0.21341 | 0.56173 | 0.00367 | 0.81985 | 0.71023 | 0.00347 | 1.67194 | 0.11934 | 1.43653 | 3.05667 |
| **Supt20** | 0.10291 | 0.03481 | 2.33072 | 0.90650 | 2.59478 | 3.42062 | 1.05329 | 1.98221 | 2.08287 | 0.00554 | 3.01373 | 2.41350 | 0.79402 | 4.70081 |
| **Supt3** | 5.48273 | 0.00197 | 5.16404 | 0.88318 | 1.24534 | 0.00000 | 0.00251 | 0.01655 | 0.00835 | 0.00000 | 1.60059 | 2.48554 | 0.43735 | 1.91671 |
| **Supt4a** | 10.99496 | 5.34186 | 4.88065 | 14.53371 | 18.79189 | 11.15419 | 2.18098 | 1.06560 | 10.92083 | 0.62828 | 5.59434 | 7.16721 | 15.89505 | 12.68453 |
| **Supt4b** | 4.68831 | 2.69614 | 1.52309 | 6.13668 | 8.56313 | 3.99024 | 0.78959 | 0.30708 | 4.53407 | 0.04617 | 1.89298 | 2.59671 | 7.60837 | 5.25241 |
| **Supt5** | 14.77329 | 0.29286 | 24.24420 | 18.22125 | 24.05328 | 9.26203 | 4.12294 | 11.74351 | 5.68425 | 1.62860 | 8.87381 | 8.97732 | 12.22658 | 17.08211 |
| **Supt6** | 2.76809 | 8.30012 | 12.65567 | 7.32021 | 11.62169 | 10.15966 | 0.36652 | 6.98328 | 0.64094 | 0.10092 | 4.90819 | 7.46366 | 5.74420 | 8.46564 |
| **Supt7l** | 3.69883 | 0.27648 | 3.36609 | 0.00000 | 3.38770 | 5.19896 | 2.25507 | 0.68432 | 2.57806 | 0.09256 | 4.53180 | 3.87780 | 3.57184 | 7.84606 |
| **Supv3l1** | 3.41081 | 0.04304 | 12.69640 | 6.49180 | 10.25728 | 7.86811 | 0.08570 | 3.09616 | 3.39488 | 0.07469 | 5.34945 | 5.57921 | 3.99533 | 14.96314 |
| **Surf1** | 7.48144 | 0.05050 | 10.36461 | 11.52601 | 1.44000 | 7.35745 | 0.03982 | 4.85998 | 1.21103 | 0.01683 | 9.15480 | 2.38718 | 0.73270 | 2.76627 |
| **Surf2** | 9.76404 | 0.16947 | 22.17415 | 8.03476 | 14.76586 | 11.84761 | 3.31971 | 3.33201 | 6.52528 | 0.17643 | 10.50366 | 10.73779 | 2.49749 | 13.96233 |
| **Surf4** | 5.37502 | 5.48410 | 8.20540 | 8.51547 | 3.44774 | 15.25863 | 4.14944 | 5.81843 | 2.55324 | 0.31036 | 3.55488 | 8.89760 | 4.57648 | 8.46254 |
| **Surf6** | 1.88855 | 0.00992 | 4.34729 | 2.11795 | 5.73208 | 1.80479 | 0.00291 | 1.25542 | 1.24858 | 0.00363 | 0.09700 | 0.19270 | 10.40268 | 4.45151 |
| **Susd1** | 0.00838 | 0.00436 | 0.01333 | 0.04080 | 0.01267 | 0.01122 | 0.00418 | 0.00838 | 0.01135 | 0.00692 | 0.00787 | 0.01256 | 0.04930 | 0.00651 |
| **Susd2** | 0.11708 | 0.03691 | 0.62077 | 0.00000 | 0.00000 | 1.84491 | 0.00362 | 2.57746 | 0.30143 | 0.02556 | 0.00000 | 1.16672 | 0.31907 | 0.00000 |
| **Susd3** | 0.00059 | 0.00433 | 1.33898 | 1.66556 | 2.87994 | 0.00000 | 0.01536 | 0.01322 | 0.01606 | 0.00512 | 0.00374 | 0.00381 | 1.91943 | 0.07148 |
| **Susd4** | 0.00628 | 0.00490 | 0.16367 | 0.00000 | 0.18390 | 0.10872 | 0.00498 | 0.00686 | 0.02473 | 0.00594 | 0.00640 | 0.00900 | 0.00576 | 0.00557 |
| **Susd6** | 1.19244 | 0.00300 | 1.50237 | 0.00000 | 0.62835 | 2.97544 | 0.91984 | 0.04305 | 2.22902 | 0.01722 | 0.98154 | 0.32241 | 3.56966 | 4.58547 |
| **Suv39h1** | 3.45067 | 3.38083 | 6.92346 | 2.69252 | 6.93363 | 2.96374 | 2.67793 | 0.40459 | 4.24279 | 0.05759 | 4.32440 | 4.10359 | 5.28016 | 8.21714 |
| **Suv39h2** | 0.81785 | 0.00380 | 0.51186 | 0.14379 | 0.74665 | 0.70910 | 2.26279 | 0.67253 | 2.84428 | 0.01722 | 1.69562 | 0.09894 | 4.94752 | 1.63652 |
| **Suv420h1** | 6.61485 | 0.12506 | 6.10271 | 2.55528 | 9.52972 | 6.65057 | 2.39637 | 1.71858 | 6.82063 | 0.05589 | 0.45375 | 4.81887 | 10.27906 | 6.75443 |
| **Suv420h2** | 0.00000 | 0.10512 | 0.29135 | 2.02835 | 0.51348 | 0.35335 | 0.05264 | 2.54029 | 0.00102 | 0.03492 | 1.87259 | 3.13391 | 0.29361 | 0.08558 |
| **Suz12** | 22.97508 | 1.54326 | 19.40321 | 7.98694 | 21.77974 | 13.08547 | 3.55817 | 5.01403 | 29.76580 | 0.25848 | 16.54391 | 11.11843 | 15.96493 | 25.06807 |
| **Sva** | 0.00460 | 0.00434 | 0.01504 | 0.01231 | 0.01032 | 0.01346 | 0.00424 | 0.00915 | 0.01092 | 0.00658 | 0.00903 | 0.00928 | 0.35854 | 0.00251 |
| **Svbp** | 0.05203 | 0.00223 | 0.00351 | 0.00000 | 0.59386 | 0.28118 | 1.20118 | 0.81596 | 0.00990 | 6.58779 | 0.00000 | 0.98308 | 0.00601 | 0.00000 |
| **Svil** | 0.32753 | 0.00439 | 0.06979 | 0.07138 | 0.01363 | 0.00804 | 0.00425 | 0.01162 | 0.01454 | 0.00918 | 0.00896 | 0.01203 | 0.00420 | 0.25161 |
| **Svip** | 0.15711 | 0.00360 | 0.00000 | 0.05105 | 0.08353 | 0.61494 | 0.00371 | 1.10209 | 0.00464 | 0.03356 | 0.00000 | 0.40823 | 0.90616 | 2.46066 |
| **Svop** | 0.00957 | 0.00432 | 0.03574 | 0.00897 | 0.00931 | 0.01229 | 0.00422 | 0.00942 | 0.00916 | 0.00648 | 0.00875 | 0.00905 | 0.44647 | 0.01339 |
| **Svopl** | 0.02290 | 0.00456 | 0.01872 | 0.06187 | 0.03710 | 0.02534 | 0.00458 | 0.02359 | 0.08964 | 0.00862 | 0.01077 | 0.02180 | 0.44751 | 0.00000 |
| **Swap70** | 1.15485 | 3.49234 | 6.04480 | 3.65371 | 2.41255 | 5.97661 | 4.86757 | 3.36986 | 0.05420 | 0.17597 | 6.26028 | 1.70782 | 0.57111 | 2.74657 |
| **Swi5** | 3.95357 | 0.80466 | 0.77282 | 5.13104 | 2.92697 | 9.73429 | 1.56791 | 8.19567 | 1.71941 | 13.69522 | 4.07530 | 9.49067 | 1.63510 | 3.94801 |
| **Swsap1** | 0.00000 | 0.00426 | 1.44324 | 1.78312 | 1.11982 | 2.51021 | 0.00475 | 0.29397 | 0.00945 | 0.01287 | 0.00000 | 0.00000 | 0.73591 | 0.71201 |
| **Swt1** | 10.81421 | 0.04353 | 7.95452 | 0.51681 | 11.10403 | 4.86180 | 1.06311 | 0.00017 | 17.21905 | 0.01453 | 7.26774 | 2.89942 | 10.44985 | 16.38058 |
| **Syap1** | 15.51247 | 0.07641 | 14.23736 | 3.41323 | 20.00552 | 9.43158 | 2.17427 | 0.18922 | 22.00932 | 0.02564 | 2.34096 | 2.14931 | 5.74626 | 12.70460 |
| **Sybu** | 0.00358 | 0.00432 | 0.00863 | 0.01024 | 0.31831 | 0.01298 | 0.00434 | 0.01013 | 0.00882 | 0.00597 | 0.00934 | 0.00855 | 0.38623 | 0.02630 |
| **Syce1** | 0.69211 | 0.00448 | 0.00000 | 0.00000 | 0.18706 | 0.44724 | 0.00423 | 0.00626 | 0.00538 | 0.00625 | 0.00367 | 0.00758 | 0.00611 | 0.15765 |
| **Syce1l** | 0.20151 | 0.00364 | 0.15035 | 0.01971 | 0.00000 | 0.50561 | 2.29499 | 0.00205 | 0.00573 | 0.02847 | 0.00249 | 0.30676 | 0.00472 | 0.27861 |
| **Syce2** | 3.43214 | 7.19935 | 3.72708 | 21.20632 | 12.88869 | 9.57829 | 2.01344 | 4.54986 | 1.97921 | 0.23024 | 5.57538 | 1.30971 | 9.45476 | 0.00000 |
| **Syce3** | 0.01969 | 0.00418 | 0.00000 | 0.09282 | 0.23052 | 0.07331 | 0.00384 | 0.00450 | 0.33639 | 0.00998 | 0.01138 | 0.00710 | 0.52815 | 0.00000 |
| **Sycn** | 2.30051 | 0.00379 | 0.00000 | 1.40102 | 5.36945 | 0.49232 | 0.00441 | 0.99985 | 0.09882 | 0.01716 | 0.10962 | 0.60166 | 1.47838 | 0.82665 |
| **Sycp1** | 0.04977 | 0.00365 | 1.84002 | 0.84338 | 0.31670 | 0.00000 | 0.00230 | 0.00450 | 12.37355 | 0.00306 | 0.03508 | 0.00794 | 0.33924 | 3.84277 |
| **Sycp1.ps1** | 0.00769 | 0.00433 | 0.03934 | 0.06388 | 0.00000 | 0.00893 | 0.00416 | 0.00791 | 0.48693 | 0.00664 | 0.01278 | 0.01076 | 0.00528 | 0.25526 |
| **Sycp2** | 0.52275 | 0.00409 | 0.00000 | 0.01195 | 0.19761 | 0.00978 | 0.00422 | 0.00864 | 0.00565 | 0.00671 | 0.00638 | 0.00512 | 1.33648 | 1.32852 |
| **Sycp2l** | 0.00767 | 0.00447 | 0.00911 | 0.01258 | 0.26139 | 0.01219 | 0.00435 | 0.01114 | 0.00891 | 0.00615 | 0.00851 | 0.00852 | 0.01377 | 0.14234 |
| **Sycp3** | 3.92745 | 0.01134 | 1.60799 | 2.31502 | 0.30955 | 2.85845 | 0.01025 | 1.12317 | 0.95149 | 0.02941 | 2.36803 | 0.44536 | 3.37697 | 2.84363 |
| **Syde1** | 0.07657 | 0.00437 | 0.01341 | 0.00487 | 0.00821 | 0.06851 | 0.00421 | 0.00666 | 0.03495 | 0.00653 | 0.00682 | 0.00982 | 0.16813 | 0.00076 |
| **Syde2** | 2.80747 | 0.04761 | 0.47216 | 0.52367 | 0.00000 | 0.27510 | 0.80637 | 0.04505 | 0.79836 | 0.01826 | 0.82777 | 0.73049 | 1.03145 | 0.32202 |

| | | | | | | | | | | | | | | |
|---|---|---|---|---|---|---|---|---|---|---|---|---|---|---|
| **Syf2** | 13.03356 | 8.75394 | 19.87903 | 17.48457 | 23.73557 | 14.79010 | 7.10938 | 8.58442 | 3.30806 | 2.40361 | 8.19405 | 7.23126 | 10.53328 | 9.07722 |
| **Sympk** | 0.20835 | 0.01230 | 0.00442 | 1.06174 | 0.33788 | 0.18540 | 0.00419 | 0.01241 | 0.00719 | 0.01813 | 0.01543 | 0.80561 | 0.00682 | 0.08770 |
| **Syn1** | 0.01870 | 0.00438 | 0.01317 | 0.26300 | 0.01010 | 0.07407 | 0.00423 | 0.00577 | 0.01411 | 0.00719 | 0.01962 | 0.00784 | 0.00868 | 0.00719 |
| **Syn2** | 0.01743 | 0.00446 | 0.00813 | 0.01042 | 0.38188 | 0.01277 | 0.00425 | 0.01053 | 0.00963 | 0.00622 | 0.00887 | 0.00835 | 0.00641 | 0.05029 |
| **Syna** | 0.02410 | 0.00455 | 0.24708 | 0.03805 | 0.41617 | 0.01032 | 0.00478 | 0.03324 | 0.00744 | 0.00865 | 0.00879 | 0.00501 | 0.35351 | 0.05628 |
| **Sync** | 0.00000 | 0.01476 | 1.28430 | 0.26745 | 0.60522 | 0.29980 | 0.00445 | 0.29029 | 0.44317 | 0.00550 | 0.00261 | 0.00187 | 0.00000 | 0.47079 |
| **Syncrip** | 17.39914 | 0.09652 | 24.24313 | 16.43130 | 22.60812 | 15.53247 | 3.18502 | 4.20376 | 26.74310 | 5.28879 | 8.82895 | 6.23278 | 11.71248 | 18.86246 |
| **Syne1** | 0.04795 | 0.02592 | 0.01705 | 0.10774 | 0.07221 | 0.79137 | 0.66690 | 0.81676 | 0.22479 | 0.01328 | 0.03711 | 0.01666 | 0.03894 | 0.05161 |
| **Syne2** | 4.22900 | 0.00221 | 8.22884 | 2.48333 | 11.85619 | 1.10099 | 1.40305 | 1.72605 | 4.24625 | 0.02481 | 0.85491 | 0.20498 | 6.23558 | 7.35375 |
| **Syne3** | 0.01205 | 0.00428 | 0.71776 | 0.14102 | 4.43940 | 0.04934 | 0.00404 | 0.00234 | 0.09003 | 0.00624 | 0.27873 | 0.01860 | 0.00000 | 0.03033 |
| **Syngr1** | 8.92985 | 5.67857 | 6.29363 | 22.06482 | 12.40246 | 6.02932 | 4.85638 | 7.14383 | 0.58202 | 2.21932 | 1.52252 | 0.89462 | 2.27767 | 5.82701 |
| **Syngr2** | 2.59356 | 3.96262 | 10.97525 | 14.19669 | 4.07291 | 12.75977 | 4.12525 | 14.80501 | 0.36621 | 1.81807 | 2.10320 | 4.60270 | 5.06139 | 3.87540 |
| **Syngr3** | 1.17546 | 0.00382 | 0.08424 | 0.97442 | 0.61741 | 0.35919 | 0.94679 | 0.00431 | 0.00853 | 0.04853 | 0.00188 | 0.00110 | 4.80097 | 0.29072 |
| **Syngr4** | 0.00926 | 0.00508 | 0.01067 | 0.00321 | 0.01389 | 0.16524 | 0.00433 | 0.02180 | 0.01107 | 0.00798 | 0.17166 | 0.00725 | 0.00795 | 0.00908 |
| **Synj1** | 4.32002 | 2.23055 | 2.84947 | 0.89342 | 3.46088 | 6.04744 | 5.41218 | 0.32552 | 5.94577 | 4.79049 | 4.95976 | 1.13757 | 4.54656 | 5.01579 |
| **Synj2** | 0.03509 | 0.00435 | 0.01124 | 0.06439 | 0.01297 | 0.01347 | 0.00426 | 0.00850 | 0.01103 | 0.00720 | 0.00861 | 0.00887 | 0.02687 | 0.02142 |
| **Synj2bp** | 2.44074 | 0.63579 | 1.61145 | 0.00000 | 1.48428 | 2.04100 | 0.05228 | 0.91957 | 5.66354 | 0.05145 | 1.42036 | 2.09298 | 0.24778 | 1.71436 |
| **Synm** | 0.00000 | 0.00444 | 0.00125 | 0.01529 | 0.00000 | 0.00609 | 0.00404 | 0.00940 | 0.88911 | 0.00746 | 0.00545 | 0.00897 | 0.01009 | 0.79184 |
| **Synpo2l** | 0.08728 | 0.00431 | 1.39949 | 0.08912 | 0.76556 | 1.72826 | 0.00460 | 0.59694 | 0.00534 | 0.00906 | 0.00266 | 0.00000 | 1.24848 | 0.06656 |
| **Synpr** | 0.20419 | 0.00365 | 0.04448 | 0.13041 | 0.04291 | 0.00000 | 0.00379 | 0.00370 | 0.57276 | 0.00651 | 0.17127 | 0.39456 | 0.45613 | 0.08394 |
| **Synrg** | 2.45778 | 5.61407 | 2.38942 | 1.04062 | 1.63262 | 4.01665 | 0.33102 | 1.48477 | 0.63192 | 0.15244 | 3.29343 | 3.10068 | 4.61581 | 6.09653 |
| **Syp** | 1.41292 | 0.00382 | 0.66011 | 0.12003 | 0.00000 | 0.50687 | 0.00381 | 0.04272 | 0.05587 | 0.00532 | 0.01819 | 0.00593 | 0.30042 | 0.13046 |
| **Sypl** | 4.04221 | 0.03623 | 3.82064 | 5.96473 | 0.00000 | 12.49066 | 0.01191 | 4.58578 | 7.82476 | 0.02879 | 6.58995 | 4.55123 | 1.36104 | 2.03364 |
| **Sys1** | 7.01150 | 0.45172 | 1.36561 | 5.23594 | 10.29134 | 6.65853 | 0.02024 | 2.48545 | 0.51702 | 0.01647 | 1.39362 | 3.69926 | 8.41475 | 2.40176 |
| **Syt11** | 0.02815 | 0.00384 | 0.44161 | 0.23388 | 0.17258 | 0.50378 | 0.06292 | 0.04166 | 0.08732 | 0.02182 | 0.02543 | 0.01435 | 4.40019 | 0.30370 |
| **Syt12** | 0.05718 | 0.06993 | 0.03287 | 0.06017 | 0.04726 | 0.12801 | 0.01652 | 0.03470 | 0.10819 | 0.03657 | 0.01795 | 0.04373 | 0.12872 | 0.19793 |
| **Syt14** | 0.01466 | 0.00439 | 0.02798 | 0.02420 | 0.00949 | 0.01149 | 0.00429 | 0.00966 | 0.04389 | 0.00641 | 0.00769 | 0.00924 | 0.07420 | 0.01916 |
| **Syt15** | 0.45216 | 0.20148 | 0.00000 | 0.39407 | 0.26551 | 0.66463 | 0.22761 | 0.22965 | 0.48081 | 0.07846 | 0.01143 | 0.32748 | 0.91316 | 0.73450 |
| **Syt16** | 0.00871 | 0.00444 | 0.01376 | 0.01349 | 0.01027 | 0.01270 | 0.00435 | 0.01024 | 0.01046 | 0.00695 | 0.00879 | 0.00998 | 0.00819 | 0.08308 |
| **Syt17** | 0.02712 | 0.00444 | 0.02701 | 0.00127 | 0.00862 | 0.17520 | 0.00422 | 0.00520 | 0.01103 | 0.00617 | 0.00714 | 0.01020 | 0.17673 | 0.04629 |
| **Syt2** | 0.00680 | 0.01165 | 0.60150 | 0.14850 | 0.24691 | 0.00000 | 0.00388 | 0.02072 | 0.01509 | 0.00497 | 0.44303 | 0.00819 | 0.02771 | 0.00399 |
| **Syt5** | 0.02802 | 0.00406 | 0.93675 | 0.16689 | 0.41254 | 0.00000 | 0.00386 | 0.01532 | 0.01376 | 0.00451 | 0.01828 | 0.01272 | 0.28684 | 0.02510 |
| **Syt7** | 0.03856 | 0.00433 | 0.04570 | 0.01913 | 0.28950 | 0.00741 | 0.00409 | 0.00900 | 0.01148 | 0.00738 | 0.00658 | 0.00954 | 0.14252 | 0.51342 |
| **Syt9** | 0.00000 | 0.00892 | 1.03079 | 0.00000 | 0.00134 | 0.79483 | 0.00283 | 0.00000 | 0.00501 | 0.01550 | 0.00000 | 2.99525 | 0.10276 | 0.00126 |
| **Sytl1** | 0.01072 | 0.00445 | 0.01397 | 0.01212 | 0.02586 | 0.01906 | 0.00455 | 0.00932 | 0.01448 | 0.00705 | 0.00857 | 0.01003 | 0.00717 | 0.01021 |
| **Sytl2** | 0.02835 | 0.00443 | 0.08889 | 0.00511 | 0.07324 | 0.00706 | 0.00428 | 0.01196 | 0.00809 | 0.00651 | 0.00684 | 0.00873 | 0.03872 | 0.51050 |
| **Sytl3** | 0.00000 | 0.02537 | 0.56409 | 5.45506 | 0.44122 | 0.00000 | 0.01414 | 2.71688 | 0.00880 | 0.02390 | 0.00000 | 1.01848 | 0.00000 | 0.31225 |
| **Sytl4** | 0.00400 | 0.00426 | 0.01631 | 0.01502 | 0.01186 | 0.01349 | 0.00415 | 0.00828 | 0.05565 | 0.00643 | 0.01730 | 0.00865 | 0.71513 | 0.00000 |
| **Syvn1** | 0.61758 | 2.84924 | 2.18897 | 4.13532 | 0.44975 | 1.65680 | 1.10294 | 3.29154 | 0.00000 | 0.06857 | 0.06058 | 3.64178 | 0.30296 | 0.09751 |
| **Szrd1** | 1.96954 | 1.70476 | 10.75328 | 8.57730 | 11.52118 | 13.22611 | 4.77069 | 9.31010 | 0.22016 | 0.45615 | 6.72824 | 10.29948 | 0.65646 | 2.05896 |
| **Szt2** | 0.00000 | 0.07383 | 2.55968 | 2.99592 | 0.55235 | 1.60197 | 5.44883 | 0.50527 | 0.00000 | 0.15868 | 0.00000 | 4.69379 | 1.40948 | 0.00000 |
| **T2** | 0.05964 | 0.01184 | 0.03081 | 0.03833 | 0.00775 | 0.06031 | 0.02781 | 0.01414 | 0.08392 | 0.01559 | 0.02251 | 0.01351 | 1.38482 | 0.09599 |
| **Tab1** | 1.69322 | 0.05432 | 2.77287 | 1.10437 | 1.35988 | 3.65494 | 0.54497 | 0.23073 | 0.00333 | 0.07578 | 1.80201 | 1.18404 | 0.00000 | 0.33804 |
| **Tab2** | 2.16309 | 0.04758 | 6.46709 | 0.00000 | 4.55697 | 5.64838 | 0.60974 | 3.58013 | 11.35778 | 0.00970 | 1.33930 | 2.83725 | 4.28331 | 9.05253 |
| **Tab3** | 2.62454 | 0.01063 | 1.07265 | 0.00649 | 0.42813 | 2.87742 | 0.31785 | 0.20370 | 1.81741 | 0.03182 | 0.00000 | 0.57853 | 0.09928 | 1.97888 |
| **Tac2** | 0.00507 | 0.01947 | 0.00885 | 0.01637 | 0.04520 | 0.14009 | 0.41712 | 1.23534 | 0.01302 | 0.03116 | 0.00322 | 0.02277 | 0.07775 | 0.02311 |
| **Tacc1** | 0.77869 | 0.03875 | 1.32033 | 0.82582 | 4.59626 | 3.35053 | 1.16012 | 3.19499 | 0.33511 | 0.04383 | 1.66598 | 3.66017 | 2.49389 | 4.50203 |
| **Tacc2** | 1.54263 | 0.05315 | 1.36166 | 1.39445 | 1.82710 | 7.26563 | 0.01545 | 4.07147 | 0.02385 | 0.00355 | 5.96552 | 1.97982 | 1.07998 | 2.37906 |
| **Tacc3** | 6.60976 | 0.23354 | 15.76899 | 9.04409 | 10.71319 | 11.40657 | 0.06385 | 11.01965 | 6.15252 | 0.08488 | 9.40197 | 7.00194 | 8.97535 | 13.94894 |
| **Taco1** | 0.34640 | 0.01129 | 1.74422 | 2.23790 | 1.50549 | 1.34371 | 0.00364 | 0.31623 | 0.00075 | 0.03576 | 0.93430 | 1.22096 | 0.46983 | 0.18921 |
| **Tacr3** | 0.00964 | 0.00434 | 0.01331 | 0.01190 | 0.00710 | 0.01230 | 0.00423 | 0.00938 | 0.01922 | 0.00630 | 0.00891 | 0.00935 | 0.30584 | 0.10401 |
| **Tacstd2** | 0.00389 | 9.59904 | 0.00000 | 0.01583 | 0.00635 | 3.37618 | 1.27392 | 10.26920 | 0.00933 | 0.07869 | 0.62071 | 0.07015 | 0.00867 | 0.00000 |
| **Tada1** | 9.09039 | 2.88317 | 5.77629 | 6.83116 | 15.01847 | 7.83244 | 7.10367 | 6.86154 | 7.37594 | 0.27482 | 5.59863 | 3.63118 | 4.66554 | 6.14497 |
| **Tada2a** | 7.11082 | 0.10811 | 10.75956 | 6.40470 | 10.37643 | 6.07438 | 2.78355 | 2.57380 | 8.53169 | 0.03617 | 5.33743 | 7.11957 | 6.84010 | 14.47951 |
| **Tada2b** | 1.33371 | 0.01278 | 0.67034 | 2.31276 | 0.52249 | 0.16417 | 1.22901 | 1.15923 | 1.75817 | 0.04587 | 1.31486 | 0.37024 | 1.14214 | 0.00000 |

| | | | | | | | | | | | | | | |
|---|---|---|---|---|---|---|---|---|---|---|---|---|---|---|
| **Tada3** | 1.03159 | 0.17026 | 8.32091 | 0.67024 | 6.23237 | 11.08073 | 6.74667 | 7.22141 | 0.00000 | 15.05538 | 7.17965 | 3.37157 | 0.84942 | 1.26782 |
| **Taf1** | 7.88430 | 0.06764 | 8.07705 | 1.54797 | 7.45193 | 5.54098 | 2.53205 | 1.91481 | 21.60560 | 0.13013 | 3.95119 | 2.57484 | 7.48007 | 8.48052 |
| **Taf10** | 1.54195 | 0.02518 | 0.04061 | 0.00000 | 2.86439 | 0.88106 | 1.14349 | 0.41579 | 0.04372 | 4.25058 | 0.00356 | 1.84501 | 7.56131 | 0.08231 |
| **Taf11** | 6.24519 | 0.21994 | 11.70842 | 11.28369 | 11.38258 | 9.74743 | 5.67116 | 5.67819 | 1.73385 | 0.35896 | 5.87339 | 10.10837 | 3.75589 | 9.90468 |
| **Taf12** | 7.08868 | 0.04414 | 10.13683 | 4.46318 | 8.79696 | 7.62827 | 2.73017 | 5.41938 | 6.50014 | 0.09648 | 1.82513 | 4.49971 | 6.43853 | 8.75713 |
| **Taf13** | 7.42215 | 6.63887 | 13.72618 | 12.39064 | 13.02594 | 5.01696 | 1.62459 | 3.53011 | 20.73087 | 0.00000 | 5.71490 | 2.69951 | 5.28610 | 12.78369 |
| **Taf15** | 0.02325 | 0.00460 | 0.00000 | 0.15921 | 0.00786 | 0.21726 | 0.01712 | 0.11835 | 0.09056 | 0.01146 | 0.00131 | 0.53439 | 0.02949 | 0.53722 |
| **Taf1a** | 3.62982 | 0.05675 | 5.67791 | 3.81622 | 7.45269 | 1.50815 | 0.00655 | 0.00000 | 5.36187 | 0.00002 | 4.29444 | 2.66444 | 8.85331 | 7.80591 |
| **Taf1b** | 3.52709 | 4.10500 | 5.72279 | 1.34973 | 6.84663 | 5.06514 | 4.96262 | 0.21353 | 10.67125 | 1.29221 | 5.55900 | 0.97334 | 6.38958 | 7.40014 |
| **Taf1c** | 2.03229 | 0.09724 | 16.94070 | 4.47047 | 11.08768 | 1.87188 | 0.08231 | 1.03146 | 0.03745 | 0.00000 | 2.87034 | 1.24662 | 0.00000 | 1.69145 |
| **Taf1d** | 15.26975 | 3.42355 | 16.63599 | 12.71255 | 14.41244 | 1.89334 | 7.93130 | 7.94260 | 28.80311 | 10.89271 | 6.54954 | 7.75583 | 14.55655 | 14.68425 |
| **Taf2** | 5.95439 | 0.01584 | 5.30868 | 2.49869 | 6.56611 | 1.75579 | 1.75722 | 0.02159 | 9.53258 | 0.00479 | 6.57901 | 3.78443 | 3.94663 | 12.78340 |
| **Taf3** | 0.10803 | 0.03714 | 2.00488 | 0.92566 | 2.94728 | 1.37832 | 0.00893 | 0.17636 | 0.01443 | 0.02276 | 1.81712 | 1.89363 | 6.99807 | 3.71500 |
| **Taf4** | 3.58478 | 1.08079 | 4.50689 | 1.64239 | 2.83088 | 3.28701 | 4.89784 | 1.10923 | 0.00000 | 0.13833 | 2.43589 | 2.31527 | 5.05559 | 3.65253 |
| **Taf4b** | 3.59583 | 0.01651 | 5.12737 | 0.65013 | 8.21680 | 0.08841 | 0.20239 | 0.15545 | 22.06660 | 0.00000 | 3.19556 | 1.12252 | 8.93037 | 6.65730 |
| **Taf5** | 3.60268 | 0.00078 | 0.94460 | 0.00000 | 1.61519 | 2.32203 | 0.00258 | 0.91269 | 8.19655 | 0.01049 | 2.07018 | 2.47286 | 9.47969 | 1.50368 |
| **Taf5l** | 6.68977 | 0.00000 | 5.63165 | 0.00000 | 3.95950 | 0.28041 | 1.58247 | 0.16518 | 8.51696 | 0.02405 | 3.27303 | 1.58083 | 13.04398 | 11.02234 |
| **Taf6** | 3.08473 | 0.18381 | 10.89803 | 2.60966 | 8.92459 | 4.85687 | 2.09103 | 3.39430 | 1.15966 | 0.02660 | 4.88510 | 3.03954 | 5.31292 | 11.07456 |
| **Taf6l** | 0.00206 | 0.00428 | 0.00267 | 0.29770 | 0.62851 | 0.01036 | 0.00414 | 0.00806 | 0.13243 | 0.00590 | 0.02168 | 0.01103 | 0.05487 | 0.00000 |
| **Taf7** | 0.00000 | 6.19485 | 13.06658 | 5.20688 | 9.82814 | 4.53105 | 0.01967 | 4.74243 | 7.26678 | 0.03137 | 8.15869 | 5.43331 | 0.00000 | 14.16063 |
| **Taf8** | 4.39671 | 0.06073 | 5.15354 | 1.62564 | 3.82174 | 1.44256 | 0.01607 | 1.14033 | 4.08460 | 0.00000 | 2.34111 | 1.53857 | 2.89865 | 8.75317 |
| **Taf9** | 24.60616 | 0.18800 | 19.87641 | 11.28422 | 31.75462 | 10.28872 | 5.42792 | 11.85570 | 25.38897 | 3.06491 | 9.57818 | 10.74185 | 20.01986 | 23.90395 |
| **Taf9b** | 0.00000 | 0.00423 | 0.00000 | 0.01690 | 0.00000 | 0.00358 | 0.00386 | 0.00814 | 1.85419 | 0.00698 | 0.05188 | 0.01049 | 0.47328 | 0.16642 |
| **Tagap** | 0.97776 | 2.81125 | 0.36175 | 1.29230 | 1.00433 | 1.40238 | 0.00977 | 0.04102 | 1.71261 | 0.01509 | 0.88442 | 0.02779 | 0.55866 | 1.64946 |
| **Tagap1** | 1.67039 | 4.10398 | 0.51722 | 2.08985 | 1.77432 | 1.69216 | 0.00290 | 0.03054 | 2.87455 | 0.02291 | 1.57538 | 0.00000 | 1.06917 | 1.99424 |
| **Tagln** | 1.24157 | 0.00397 | 1.19052 | 0.28450 | 0.38663 | 0.24642 | 0.00360 | 0.00559 | 0.39967 | 0.00394 | 0.00180 | 0.50593 | 0.10372 | 0.06388 |
| **Tagln2** | 1.34117 | 11.53282 | 15.48751 | 18.52124 | 10.75794 | 35.89031 | 10.24196 | 36.27398 | 2.42354 | 19.19028 | 19.55122 | 8.22635 | 0.00000 | 5.98251 |
| **Taldo1** | 13.92142 | 10.41657 | 36.73059 | 30.29642 | 23.45514 | 32.00192 | 6.73426 | 20.91415 | 1.41224 | 17.38283 | 21.34565 | 19.69627 | 8.27141 | 21.17432 |
| **Tamm41** | 0.26323 | 0.03893 | 1.02765 | 0.82510 | 1.60831 | 1.68822 | 0.01780 | 0.87488 | 0.00303 | 0.00630 | 0.00000 | 0.76414 | 0.37471 | 0.41355 |
| **Tanc1** | 1.85095 | 1.66034 | 4.33982 | 2.13850 | 1.62612 | 4.35189 | 0.00753 | 2.62172 | 3.85828 | 0.00000 | 5.30904 | 0.16870 | 0.92825 | 0.00000 |
| **Tanc2** | 1.06086 | 0.00358 | 0.70185 | 0.00000 | 2.45703 | 0.23984 | 0.00318 | 0.76006 | 4.22784 | 0.00659 | 0.00819 | 0.31443 | 0.83822 | 1.82752 |
| **Tango2** | 3.71348 | 0.01350 | 5.49568 | 1.67610 | 5.03593 | 4.17209 | 2.30514 | 0.00000 | 2.94651 | 0.04659 | 3.42108 | 4.24767 | 2.65992 | 7.91998 |
| **Tango6** | 1.38546 | 0.03279 | 1.82657 | 2.13950 | 3.29212 | 1.42380 | 0.25941 | 0.44924 | 0.22152 | 0.00000 | 3.70297 | 2.37803 | 1.13141 | 3.31085 |
| **Tank** | 0.68858 | 0.01136 | 1.25376 | 0.00000 | 1.26432 | 1.30847 | 0.00384 | 0.63812 | 1.37116 | 0.00030 | 1.11496 | 1.12922 | 0.74219 | 0.56902 |
| **Taok1** | 16.89436 | 0.02505 | 10.07586 | 0.75179 | 17.33849 | 8.36191 | 0.27479 | 3.20721 | 29.54105 | 0.09811 | 3.76229 | 4.92584 | 10.87796 | 12.11051 |
| **Taok2** | 3.08024 | 1.45055 | 8.66289 | 5.20099 | 4.40167 | 5.08560 | 4.54554 | 3.93832 | 0.00000 | 0.28109 | 0.54704 | 3.97848 | 3.01781 | 3.04737 |
| **Taok3** | 0.37552 | 0.03442 | 0.51258 | 0.37468 | 1.42115 | 0.65895 | 0.00376 | 0.00123 | 0.36892 | 0.00627 | 0.08801 | 0.00985 | 0.01388 | 0.63531 |
| **Tap1** | 0.00000 | 0.04243 | 0.69139 | 0.52639 | 0.14313 | 0.10329 | 0.00394 | 0.72511 | 0.00633 | 0.00058 | 1.66067 | 1.38515 | 0.00113 | 0.07615 |
| **Tapbp** | 0.61255 | 0.01908 | 0.40179 | 0.68255 | 0.62074 | 1.78480 | 0.03724 | 0.54734 | 0.24653 | 0.04589 | 0.01215 | 0.02688 | 0.85190 | 0.20241 |
| **Tapbpl** | 1.15974 | 0.04261 | 0.04018 | 0.57548 | 0.17023 | 2.01559 | 0.29432 | 0.31668 | 1.38591 | 0.89052 | 0.03300 | 0.44022 | 9.35970 | 0.95823 |
| **Tapt1** | 1.02654 | 0.00299 | 1.34095 | 0.13774 | 1.46231 | 1.05060 | 0.03230 | 0.00000 | 3.53539 | 0.00915 | 0.57395 | 1.05115 | 0.00000 | 3.05020 |
| **Tarbp1** | 0.01452 | 0.01075 | 0.82592 | 0.44721 | 0.01199 | 0.03772 | 0.27126 | 0.00376 | 0.11844 | 0.01161 | 0.03246 | 0.34035 | 0.46309 | 0.59918 |
| **Tarbp2** | 12.18149 | 4.95364 | 21.91096 | 17.01278 | 15.39552 | 15.65663 | 6.83172 | 8.17451 | 1.65908 | 0.35881 | 7.00044 | 11.45470 | 4.49193 | 8.46371 |
| **Tardbp** | 18.07284 | 2.85787 | 31.59504 | 17.67804 | 23.29368 | 25.50782 | 6.05487 | 12.60236 | 26.17402 | 15.95770 | 18.41899 | 17.55627 | 7.88865 | 24.22800 |
| **Tarm1** | 0.64699 | 0.00441 | 0.01765 | 0.00000 | 0.16042 | 0.23494 | 0.00400 | 0.01271 | 0.01375 | 0.00614 | 0.00427 | 0.00744 | 0.09011 | 0.29469 |
| **Tars** | 3.56224 | 0.22402 | 9.92436 | 16.19308 | 11.43976 | 8.86813 | 6.65493 | 10.04998 | 0.06292 | 0.39218 | 14.94270 | 13.45980 | 0.44282 | 0.00036 |
| **Tars2** | 2.00584 | 0.06098 | 8.53476 | 9.24947 | 2.94913 | 5.32030 | 0.72342 | 4.51407 | 0.01953 | 0.05296 | 8.49782 | 3.07497 | 1.60252 | 3.72761 |
| **Tarsl2** | 0.23642 | 0.00324 | 1.58826 | 0.97726 | 1.61114 | 2.91660 | 0.00320 | 0.10079 | 0.00129 | 0.00000 | 2.75648 | 0.36563 | 0.00000 | 0.61948 |
| **Tas1r1** | 0.00000 | 4.51866 | 0.10942 | 1.88984 | 0.32282 | 0.81275 | 0.00288 | 0.18916 | 0.00817 | 0.00485 | 0.22235 | 1.18889 | 0.79316 | 0.00000 |
| **Tas1r3** | 0.01027 | 0.00431 | 0.15893 | 0.00730 | 0.00648 | 0.01178 | 0.00415 | 0.01271 | 0.01066 | 0.00668 | 0.01111 | 0.00949 | 0.25562 | 0.00507 |
| **Tasp1** | 1.12851 | 0.01070 | 3.46508 | 0.13170 | 2.04223 | 0.00000 | 0.00202 | 0.49829 | 10.94518 | 0.00246 | 3.30117 | 1.23295 | 1.42889 | 10.56649 |
| **Tat** | 1.35791 | 0.00073 | 0.47371 | 1.72281 | 0.31478 | 1.03693 | 0.00215 | 0.00000 | 2.30712 | 0.00000 | 5.58976 | 2.22501 | 0.81669 | 2.90951 |
| **Tatdn1** | 2.12871 | 0.00907 | 1.93287 | 1.69267 | 4.65284 | 3.78403 | 0.00781 | 0.26103 | 7.68705 | 0.03667 | 0.00000 | 2.93622 | 0.48568 | 2.42299 |
| **Tatdn2** | 1.66355 | 0.01918 | 0.00000 | 0.12778 | 0.68854 | 0.11645 | 0.00913 | 1.16550 | 2.67603 | 0.02106 | 0.00244 | 0.06887 | 1.81190 | 1.32482 |
| **Tatdn3** | 1.07118 | 0.00325 | 1.52922 | 0.16351 | 1.82066 | 0.00000 | 0.00382 | 1.88830 | 0.11822 | 0.01515 | 0.93902 | 0.39623 | 1.91705 | 5.15900 |

| | | | | | | | | | | | | | | |
|---|---|---|---|---|---|---|---|---|---|---|---|---|---|---|
| **Tax1bp1** | 21.38292 | 1.91229 | 21.01758 | 5.28802 | 19.94473 | 13.16355 | 1.04241 | 6.19927 | 23.88151 | 0.07553 | 14.56057 | 12.31469 | 8.38777 | 20.41908 |
| **Tax1bp3** | 3.19155 | 7.77210 | 17.76069 | 9.73407 | 2.57149 | 22.25009 | 6.08855 | 14.60513 | 1.56447 | 2.17947 | 8.44258 | 8.85811 | 1.54111 | 5.24054 |
| **Taz** | 3.59099 | 0.12334 | 2.21643 | 4.82477 | 1.31561 | 3.19770 | 0.00752 | 4.76066 | 3.31826 | 0.04157 | 1.90552 | 3.88812 | 0.49986 | 0.88142 |
| **Tbc1d1** | 1.16194 | 0.03115 | 4.47426 | 1.14115 | 6.31024 | 1.22628 | 0.92272 | 2.46430 | 2.68322 | 0.03819 | 1.50422 | 4.37710 | 4.14069 | 3.66996 |
| **Tbc1d10a** | 0.32496 | 4.26882 | 1.30273 | 0.06976 | 1.52802 | 1.01497 | 0.00302 | 2.11794 | 0.02621 | 0.00868 | 0.95706 | 0.01506 | 0.07640 | 0.22264 |
| **Tbc1d10b** | 5.55709 | 0.04776 | 7.27641 | 7.14565 | 8.75497 | 7.85065 | 0.41584 | 6.90454 | 1.95567 | 0.04903 | 1.56437 | 4.37804 | 4.44790 | 4.40642 |
| **Tbc1d10c** | 0.03314 | 0.00396 | 1.15252 | 0.23132 | 0.62615 | 0.40644 | 0.00388 | 0.90799 | 0.00534 | 0.00312 | 0.00746 | 0.00074 | 0.54071 | 0.48631 |
| **Tbc1d12** | 0.51545 | 0.01285 | 0.11134 | 0.26796 | 0.28666 | 0.04264 | 0.00476 | 0.11574 | 0.87228 | 0.00535 | 0.05645 | 0.69341 | 0.65940 | 0.00000 |
| **Tbc1d13** | 1.78485 | 1.72351 | 2.36488 | 3.09710 | 7.42084 | 8.38382 | 0.06435 | 0.28981 | 2.09919 | 0.04208 | 2.87082 | 3.32971 | 6.70626 | 3.04409 |
| **Tbc1d14** | 0.50158 | 0.03414 | 3.55015 | 0.00000 | 2.55812 | 3.75731 | 1.02874 | 1.14353 | 0.00538 | 0.05427 | 0.14709 | 2.44902 | 2.48510 | 2.86616 |
| **Tbc1d15** | 9.30701 | 4.49991 | 14.17444 | 4.11098 | 9.14115 | 8.04427 | 1.70933 | 0.60183 | 18.21991 | 0.00874 | 10.08506 | 4.17386 | 5.85722 | 13.26339 |
| **Tbc1d16** | 0.01310 | 0.00438 | 0.00015 | 0.02371 | 0.35462 | 0.01023 | 0.00407 | 0.00893 | 0.00820 | 0.00939 | 0.00708 | 0.00860 | 0.00700 | 0.84054 |
| **Tbc1d17** | 2.94705 | 0.05581 | 7.65906 | 5.79770 | 7.23550 | 1.68632 | 2.89158 | 4.03695 | 0.00316 | 0.04649 | 3.06532 | 2.70002 | 0.00000 | 1.95342 |
| **Tbc1d19** | 0.18880 | 0.00357 | 0.07839 | 0.05172 | 0.48380 | 0.31629 | 0.00348 | 0.37376 | 0.01324 | 0.06667 | 1.40534 | 0.29329 | 0.04225 | 0.00000 |
| **Tbc1d2** | 0.00291 | 0.04544 | 0.19660 | 0.70404 | 0.00000 | 2.63350 | 0.25530 | 3.11709 | 0.01299 | 0.03600 | 0.12212 | 0.00000 | 0.04251 | 0.24920 |
| **Tbc1d20** | 0.00000 | 0.01167 | 2.18710 | 0.06348 | 2.51577 | 1.11237 | 0.00307 | 0.01215 | 2.24367 | 0.08349 | 0.45727 | 0.53298 | 0.11031 | 5.05427 |
| **Tbc1d22a** | 2.16869 | 0.03419 | 5.80132 | 4.88746 | 8.52088 | 2.53794 | 0.00235 | 0.62548 | 0.29487 | 0.00000 | 2.84007 | 2.23057 | 0.63485 | 3.29622 |
| **Tbc1d22b** | 0.29661 | 0.00139 | 1.56041 | 0.24437 | 2.27874 | 1.78543 | 0.00239 | 0.95525 | 0.04441 | 0.02428 | 0.02643 | 3.65098 | 0.68718 | 0.63800 |
| **Tbc1d23** | 5.09349 | 3.03105 | 6.09692 | 4.06418 | 5.52631 | 6.50258 | 1.75761 | 0.00000 | 9.28723 | 0.04069 | 2.64675 | 2.50911 | 7.73414 | 4.31230 |
| **Tbc1d24** | 0.41355 | 0.05766 | 0.40918 | 0.75314 | 0.03398 | 0.70360 | 0.04002 | 0.23021 | 0.70286 | 0.02340 | 0.01178 | 0.53615 | 1.85233 | 0.66742 |
| **Tbc1d25** | 2.78588 | 0.04646 | 4.62161 | 3.72398 | 8.39474 | 2.61737 | 0.00095 | 2.39786 | 0.36750 | 0.02656 | 5.50328 | 3.70056 | 2.99915 | 4.72636 |
| **Tbc1d2b** | 0.00000 | 0.01909 | 0.60408 | 0.33281 | 0.00000 | 0.97585 | 2.86785 | 1.88137 | 0.02229 | 0.09224 | 0.00000 | 0.19650 | 1.00979 | 0.37590 |
| **Tbc1d31** | 1.52381 | 0.04559 | 2.69426 | 1.59254 | 0.95486 | 1.98056 | 3.26022 | 0.64186 | 3.09753 | 0.03517 | 0.18531 | 2.22847 | 0.00000 | 1.79887 |
| **Tbc1d32** | 0.02587 | 0.00435 | 0.00712 | 0.02205 | 0.04001 | 0.02361 | 0.00472 | 0.00733 | 0.02398 | 0.00475 | 0.55827 | 0.00491 | 0.00581 | 0.00325 |
| **Tbc1d5** | 0.00000 | 0.00985 | 0.50226 | 0.00000 | 0.42908 | 0.94901 | 0.75868 | 0.01207 | 0.00496 | 0.02341 | 0.21951 | 1.52281 | 0.00158 | 1.08554 |
| **Tbc1d7** | 0.80178 | 0.04315 | 0.64683 | 0.77056 | 0.12300 | 0.11697 | 0.00869 | 1.74190 | 0.00373 | 0.02664 | 3.01550 | 3.71338 | 0.02086 | 0.04577 |
| **Tbc1d8** | 0.00000 | 0.01226 | 0.57304 | 0.00000 | 0.00173 | 0.00000 | 1.62402 | 1.14378 | 0.00001 | 0.06267 | 0.26059 | 0.40047 | 1.65220 | 1.05561 |
| **Tbc1d8b** | 0.03183 | 0.06177 | 0.18107 | 0.00000 | 0.02106 | 0.01062 | 0.00451 | 0.03094 | 0.02976 | 0.00682 | 0.02427 | 0.02317 | 0.01381 | 0.07868 |
| **Tbc1d9** | 8.77563 | 0.00124 | 3.27652 | 3.13256 | 11.97690 | 1.54932 | 1.81786 | 0.00000 | 7.61400 | 0.06308 | 0.31290 | 0.23261 | 10.07756 | 11.81631 |
| **Tbc1d9b** | 0.19973 | 0.05007 | 0.00000 | 0.00000 | 1.34282 | 1.80578 | 1.00923 | 1.34044 | 0.01078 | 0.03175 | 4.19941 | 3.51885 | 1.67350 | 0.68002 |
| **Tbca** | 1.64913 | 3.22154 | 0.00000 | 0.00000 | 2.44345 | 5.17493 | 2.23294 | 2.72056 | 1.81270 | 12.17979 | 0.30086 | 6.90276 | 3.98536 | 0.00000 |
| **Tbcb** | 2.94981 | 0.11660 | 7.24329 | 9.88608 | 11.52630 | 3.79811 | 4.93842 | 8.52563 | 0.00000 | 11.56596 | 4.43391 | 5.60859 | 3.04501 | 3.82747 |
| **Tbcc** | 0.53839 | 0.21686 | 0.54980 | 3.42138 | 5.17190 | 0.42861 | 0.76431 | 0.13104 | 0.04973 | 8.20205 | 0.31442 | 1.13829 | 6.28647 | 0.00000 |
| **Tbccd1** | 3.31488 | 0.01791 | 7.23475 | 1.47705 | 4.08335 | 3.32181 | 0.72062 | 0.14162 | 4.08017 | 0.01320 | 2.51342 | 2.36025 | 0.82224 | 4.24278 |
| **Tbcd** | 0.00000 | 0.01580 | 1.42719 | 1.07852 | 0.28330 | 0.60949 | 2.07274 | 1.19989 | 0.01226 | 0.01458 | 1.09661 | 2.45079 | 0.23026 | 0.76893 |
| **Tbce** | 5.38063 | 0.02189 | 2.95306 | 0.50822 | 2.74666 | 0.96158 | 0.00202 | 0.49859 | 7.22001 | 0.00141 | 4.55020 | 0.76292 | 0.03769 | 3.36431 |
| **Tbcel** | 0.02040 | 0.00442 | 0.00984 | 0.04370 | 0.32851 | 0.01179 | 0.00437 | 0.01338 | 0.08848 | 0.00599 | 0.00700 | 0.00684 | 0.00412 | 0.01022 |
| **Tbck** | 0.24362 | 0.02477 | 1.16688 | 1.28066 | 0.94508 | 3.04561 | 0.03395 | 0.97095 | 0.34893 | 0.06886 | 2.34226 | 1.09491 | 0.08692 | 1.76985 |
| **Tbk1** | 1.68773 | 0.01231 | 0.28095 | 0.65071 | 0.38844 | 1.03314 | 0.00422 | 0.02870 | 0.41998 | 0.00041 | 2.38057 | 0.58515 | 2.59124 | 1.93742 |
| **Tbkbp1** | 0.00863 | 0.00440 | 0.05821 | 0.00833 | 0.68953 | 0.01127 | 0.00439 | 0.00981 | 0.01045 | 0.00662 | 0.00712 | 0.02296 | 0.00535 | 0.01025 |
| **Tbl1x** | 0.20048 | 0.00388 | 0.43836 | 0.84057 | 0.00000 | 0.12672 | 0.00947 | 1.65514 | 0.02847 | 0.00480 | 0.00138 | 0.00264 | 0.22910 | 0.39528 |
| **Tbl1xr1** | 1.30445 | 4.69948 | 6.68637 | 1.41298 | 1.61351 | 7.58390 | 4.14820 | 6.90584 | 4.63658 | 10.51965 | 5.69676 | 6.05267 | 0.62395 | 1.26568 |
| **Tbl2** | 4.38800 | 2.24244 | 5.76141 | 6.28613 | 5.97060 | 6.97033 | 0.00835 | 0.00000 | 3.74656 | 0.00988 | 0.00000 | 1.97577 | 2.99190 | 10.20388 |
| **Tbl3** | 5.72987 | 0.08134 | 21.58928 | 11.17825 | 27.67330 | 6.32312 | 1.61602 | 4.38513 | 0.12162 | 0.09057 | 8.60532 | 8.91903 | 1.26906 | 3.80428 |
| **Tbp** | 3.17390 | 0.02181 | 5.46130 | 3.68040 | 6.81855 | 5.03405 | 0.00000 | 0.99070 | 6.94797 | 0.02312 | 4.65301 | 3.28276 | 10.25618 | 15.36056 |
| **Tbpl1** | 2.12933 | 0.08689 | 4.33640 | 3.26459 | 4.39431 | 5.24441 | 2.99434 | 0.76449 | 1.03462 | 0.06999 | 8.76201 | 7.60483 | 1.20393 | 4.27228 |
| **Tbrg1** | 19.70876 | 9.29596 | 20.07896 | 21.62379 | 19.46274 | 15.13929 | 6.75780 | 14.75881 | 8.29866 | 15.12325 | 10.73013 | 11.61160 | 3.87478 | 13.21493 |
| **Tbrg3** | 0.07975 | 0.03035 | 0.09178 | 0.43741 | 0.03431 | 0.35969 | 0.10785 | 0.51342 | 0.57456 | 0.34822 | 0.02521 | 0.06600 | 0.24532 | 0.53607 |
| **Tbrg4** | 4.87106 | 0.19497 | 14.42785 | 13.08981 | 11.51569 | 4.08689 | 0.03496 | 4.04749 | 0.48685 | 0.03850 | 6.10774 | 7.03990 | 6.35228 | 4.26251 |
| **Tbx15** | 0.01463 | 0.04797 | 0.01313 | 0.00242 | 0.15581 | 0.01755 | 0.00280 | 0.04137 | 0.00989 | 0.00554 | 0.00316 | 6.03909 | 0.00571 | 0.01591 |
| **Tbx18** | 0.03003 | 0.01925 | 0.00796 | 0.01427 | 0.04243 | 0.02662 | 0.00984 | 0.02901 | 0.26575 | 0.01988 | 0.00716 | 1.18670 | 0.03076 | 0.07627 |
| **Tbx19** | 0.00580 | 0.00438 | 0.01218 | 0.01272 | 0.11360 | 0.01271 | 0.00434 | 0.01066 | 0.01745 | 0.00621 | 0.00888 | 0.00915 | 0.20283 | 0.01354 |
| **Tbx2** | 0.01667 | 0.00443 | 0.01661 | 0.02328 | 0.45975 | 0.01102 | 0.00433 | 0.00915 | 0.01571 | 0.00642 | 0.01075 | 0.01040 | 0.00433 | 0.00512 |
| **Tbx20** | 2.64341 | 0.04534 | 2.36945 | 2.70693 | 0.00000 | 3.44088 | 1.43651 | 0.92990 | 2.07412 | 0.09180 | 7.10147 | 2.11323 | 0.07230 | 0.87280 |
| **Tbx21** | 0.00813 | 0.00436 | 0.01218 | 0.00666 | 0.81718 | 0.01291 | 0.00418 | 0.00921 | 0.01069 | 0.00635 | 0.00880 | 0.00814 | 0.38041 | 0.01146 |

| | | | | | | | | | | | | | | |
|---|---|---|---|---|---|---|---|---|---|---|---|---|---|---|
| **Tbx3** | 8.63450 | 0.29604 | 12.77382 | 14.05086 | 16.01590 | 8.32739 | 1.17804 | 3.78413 | 6.25273 | 0.09232 | 5.52877 | 7.99073 | 5.28101 | 3.40133 |
| **Tbx4** | 0.00811 | 0.00437 | 0.01312 | 0.01339 | 0.02648 | 0.01277 | 0.00419 | 0.00931 | 0.02953 | 0.00687 | 0.00866 | 0.00922 | 0.26783 | 0.00598 |
| **Tbx6** | 0.00689 | 0.00448 | 0.03186 | 0.00738 | 0.05394 | 0.02194 | 0.00432 | 0.00910 | 0.01520 | 0.00635 | 0.00894 | 0.00952 | 0.00351 | 0.20541 |
| **Tbxa2r** | 0.00328 | 0.00538 | 0.00405 | 0.48392 | 0.49859 | 0.00000 | 0.00543 | 0.64934 | 0.01416 | 0.00703 | 0.00702 | 0.00497 | 0.92199 | 0.00000 |
| **Tc2n** | 0.72077 | 0.22977 | 0.00000 | 0.58120 | 0.31595 | 0.72107 | 0.08349 | 0.10040 | 3.36051 | 0.13175 | 0.02091 | 0.20085 | 4.30832 | 0.19471 |
| **Tcaim** | 0.31444 | 0.00298 | 0.11662 | 0.96434 | 0.43639 | 0.53101 | 0.00330 | 0.03688 | 0.68357 | 0.00623 | 5.35722 | 0.98285 | 0.52640 | 1.19893 |
| **Tcam1** | 0.90551 | 0.00366 | 2.81174 | 1.54681 | 1.55543 | 0.06437 | 0.00310 | 0.00000 | 1.35766 | 0.00000 | 0.82325 | 0.01500 | 0.00570 | 3.09584 |
| **Tcea1** | 18.60108 | 2.88174 | 14.47174 | 7.87028 | 24.45524 | 7.37662 | 0.12514 | 5.38961 | 23.23177 | 0.42911 | 10.94070 | 10.62915 | 15.70339 | 23.47013 |
| **Tcea3** | 0.00000 | 0.03093 | 1.51082 | 1.11075 | 0.23309 | 3.08799 | 0.00374 | 0.42092 | 0.01779 | 0.01880 | 1.53539 | 0.72806 | 0.49026 | 0.00000 |
| **Tceal8** | 1.32515 | 0.01169 | 0.00127 | 0.00000 | 0.00000 | 0.92077 | 0.00376 | 0.00314 | 1.43558 | 0.00580 | 0.00315 | 0.01093 | 0.17347 | 0.42354 |
| **Tceanc** | 0.00902 | 0.01359 | 0.14749 | 0.21611 | 1.09761 | 0.57996 | 0.01839 | 0.25907 | 0.01049 | 0.00760 | 0.13682 | 0.02866 | 0.82951 | 0.00000 |
| **Tceanc2** | 2.87253 | 0.03460 | 5.07060 | 1.43514 | 3.80799 | 4.02728 | 0.00880 | 5.01642 | 0.02975 | 0.01765 | 0.02236 | 0.76629 | 2.08107 | 0.13807 |
| **Tceb1** | 30.99310 | 4.14726 | 22.22471 | 19.60009 | 26.25537 | 25.03588 | 9.00697 | 9.09967 | 37.13903 | 15.37955 | 19.44803 | 10.52648 | 22.35814 | 18.79884 |
| **Tceb2** | 3.02433 | 3.33842 | 0.00000 | 3.69962 | 11.67438 | 10.16807 | 2.82584 | 15.01075 | 0.03746 | 11.74333 | 0.34422 | 9.96928 | 5.25849 | 2.15490 |
| **Tceb3** | 11.27670 | 0.02851 | 10.89531 | 3.64415 | 18.75038 | 8.66908 | 2.12017 | 1.99533 | 16.39814 | 0.10587 | 3.08630 | 4.18292 | 19.90728 | 20.71442 |
| **Tcerg1** | 9.86977 | 0.05611 | 9.63669 | 3.34046 | 14.95030 | 6.10265 | 3.96845 | 3.97391 | 14.85511 | 0.04008 | 10.82299 | 5.01343 | 10.21040 | 11.26596 |
| **Tcerg1l** | 0.00198 | 0.00430 | 0.01052 | 0.01311 | 0.19821 | 0.01179 | 0.00426 | 0.00915 | 0.00969 | 0.00567 | 0.00877 | 0.00763 | 1.00583 | 0.11969 |
| **Tcf12** | 1.00829 | 0.00294 | 0.19842 | 0.61661 | 0.22992 | 0.26824 | 0.04940 | 0.00017 | 2.85949 | 0.00467 | 0.00000 | 1.12794 | 0.65857 | 3.75500 |
| **Tcf15** | 0.00594 | 0.00410 | 0.02169 | 0.00000 | 0.11181 | 0.05014 | 0.03971 | 0.11986 | 0.02399 | 1.22007 | 0.02532 | 0.17027 | 0.74561 | 0.00481 |
| **Tcf19** | 0.17923 | 0.01074 | 0.77195 | 0.00000 | 0.74508 | 1.57132 | 0.00311 | 2.20091 | 0.00000 | 0.01957 | 0.20422 | 0.93845 | 3.24708 | 0.79123 |
| **Tcf20** | 0.59401 | 0.01238 | 0.13365 | 0.00042 | 0.57435 | 2.09125 | 0.12447 | 1.42890 | 0.39001 | 0.06145 | 0.04761 | 0.24351 | 1.10079 | 1.59921 |
| **Tcf23** | 1.25542 | 0.03263 | 2.45692 | 0.90300 | 0.55923 | 0.00000 | 0.00870 | 0.00000 | 0.17347 | 0.02352 | 0.00269 | 2.49855 | 0.00000 | 2.19070 |
| **Tcf24** | 1.56326 | 0.00374 | 1.09646 | 1.66083 | 0.13171 | 0.00000 | 0.00939 | 1.55440 | 0.00762 | 0.01175 | 0.00000 | 0.00488 | 0.02022 | 0.00000 |
| **Tcf25** | 9.55424 | 0.14018 | 23.71073 | 13.74117 | 21.59342 | 13.81917 | 3.76556 | 6.98551 | 7.61689 | 4.44271 | 6.05485 | 5.84823 | 4.56472 | 18.42127 |
| **Tcf3** | 0.06138 | 0.01818 | 1.83184 | 5.47343 | 1.64791 | 0.00000 | 0.00262 | 2.86857 | 0.04149 | 0.00858 | 0.00000 | 4.12770 | 1.77133 | 0.08244 |
| **Tcf4** | 0.46058 | 0.00433 | 0.29147 | 0.02118 | 0.37494 | 0.17478 | 0.00397 | 0.00388 | 1.12638 | 0.00586 | 0.00795 | 0.02192 | 0.00825 | 0.08541 |
| **Tcf7** | 0.00133 | 0.00275 | 0.04108 | 0.00000 | 0.68643 | 0.26722 | 0.00350 | 0.00000 | 0.00638 | 0.00071 | 2.13293 | 0.77184 | 1.16899 | 1.14663 |
| **Tcf7l1** | 0.00803 | 0.01991 | 0.03286 | 0.00000 | 0.40586 | 0.13902 | 0.00385 | 0.01051 | 0.01382 | 0.00607 | 0.00236 | 2.05337 | 0.15799 | 0.00786 |
| **Tcf7l2** | 0.02237 | 0.03907 | 0.38059 | 0.04559 | 1.86493 | 0.00000 | 0.00348 | 0.00317 | 0.04918 | 0.00491 | 0.01549 | 3.10348 | 0.00205 | 0.08997 |
| **Tcfl5** | 1.81068 | 0.05613 | 0.98004 | 4.41995 | 4.94124 | 3.17317 | 2.32988 | 3.36378 | 0.03251 | 0.07873 | 3.05890 | 0.03222 | 1.75087 | 0.70598 |
| **Tchh** | 0.00000 | 0.00370 | 0.01198 | 0.01253 | 0.00000 | 0.00819 | 0.00398 | 0.00878 | 0.28379 | 0.00605 | 0.00839 | 0.00625 | 5.95829 | 0.00000 |
| **Tchp** | 0.24653 | 0.02838 | 0.79317 | 0.00000 | 0.74824 | 1.80540 | 0.00324 | 0.22335 | 0.00263 | 0.00374 | 1.22202 | 2.70062 | 2.70305 | 0.69637 |
| **Tcirg1** | 0.67314 | 0.02063 | 4.05198 | 1.74050 | 4.67162 | 0.06478 | 0.00415 | 0.37427 | 0.00003 | 0.00711 | 0.19435 | 0.00993 | 2.28107 | 0.83728 |
| **Tcl1** | 0.48324 | 1.33846 | 0.00000 | 0.00000 | 1.23078 | 0.00000 | 0.00227 | 0.02780 | 0.34639 | 0.02042 | 0.79173 | 4.58641 | 7.87151 | 1.74193 |
| **Tcl1b1** | 0.00000 | 0.00402 | 0.06550 | 0.03759 | 0.32929 | 0.00262 | 0.00377 | 0.00857 | 0.69007 | 0.00860 | 0.00494 | 0.00634 | 3.84243 | 0.71406 |
| **Tcl1b2** | 0.00495 | 0.00515 | 0.02445 | 0.26168 | 0.00000 | 0.02898 | 0.00396 | 0.02645 | 0.11857 | 0.01058 | 0.00830 | 0.00396 | 1.40072 | 0.27786 |
| **Tcl1b3** | 0.00000 | 0.00448 | 0.01766 | 0.01901 | 0.02689 | 0.00837 | 0.00399 | 0.01057 | 0.60816 | 0.00738 | 0.00682 | 0.00675 | 0.99792 | 0.29717 |
| **Tcl1b4** | 0.01270 | 0.00426 | 0.01244 | 0.01795 | 0.00633 | 0.01327 | 0.00400 | 0.01746 | 0.00945 | 0.00690 | 0.00864 | 0.00665 | 1.29279 | 0.22827 |
| **Tcl1b5** | 0.00438 | 0.00442 | 0.01592 | 0.01812 | 0.01387 | 0.01326 | 0.00408 | 0.01028 | 0.06987 | 0.00671 | 0.00866 | 0.00706 | 1.20972 | 0.12610 |
| **Tcn2** | 1.07610 | 8.65548 | 3.40144 | 4.19273 | 0.84178 | 3.09532 | 3.80740 | 2.62801 | 0.00740 | 0.13295 | 1.10589 | 3.18700 | 0.02019 | 1.93599 |
| **Tcof1** | 3.33825 | 6.56365 | 13.00354 | 10.33082 | 11.39282 | 6.58952 | 1.86825 | 3.65120 | 0.21275 | 0.05094 | 8.66043 | 6.22146 | 8.90116 | 4.19625 |
| **Tcp1** | 24.85867 | 8.72494 | 34.89563 | 26.68545 | 26.90880 | 30.37073 | 9.69428 | 16.52535 | 22.27259 | 4.68544 | 20.64089 | 26.79261 | 7.03099 | 30.06075 |
| **Tcp10a** | 0.01022 | 0.00442 | 0.14494 | 0.00404 | 0.01092 | 0.00899 | 0.00423 | 0.01114 | 0.01162 | 0.00785 | 0.00739 | 0.00987 | 0.00747 | 0.01156 |
| **Tcp10b** | 0.01039 | 0.00443 | 0.09838 | 0.00571 | 0.01204 | 0.01022 | 0.00425 | 0.01099 | 0.01168 | 0.00765 | 0.00775 | 0.00997 | 0.00749 | 0.01173 |
| **Tcp10c** | 0.01038 | 0.00443 | 0.09719 | 0.00575 | 0.01213 | 0.01024 | 0.00425 | 0.01098 | 0.01165 | 0.00765 | 0.00775 | 0.01006 | 0.00747 | 0.01183 |
| **Tcp11** | 0.37047 | 0.00400 | 0.53555 | 0.10666 | 0.49181 | 0.00275 | 0.00384 | 0.00552 | 0.00623 | 0.00709 | 0.00517 | 0.01407 | 0.77484 | 0.00883 |
| **Tcp11l1** | 0.14079 | 0.01983 | 0.00951 | 0.01265 | 0.23683 | 0.15381 | 0.00447 | 0.01928 | 0.10705 | 0.01419 | 0.00844 | 0.00664 | 0.58565 | 0.00000 |
| **Tcp11l2** | 1.51543 | 0.00312 | 1.68254 | 0.00000 | 1.27538 | 0.25108 | 0.00857 | 0.00000 | 1.03440 | 0.00126 | 0.80782 | 0.29491 | 1.87177 | 0.31034 |
| **Tcstv1** | 0.13221 | 0.02033 | 0.01506 | 0.10038 | 0.00532 | 0.35383 | 0.02867 | 0.59853 | 0.03779 | 0.00640 | 0.00390 | 0.00000 | 1.64158 | 0.04362 |
| **Tcstv3** | 2.12215 | 0.00429 | 0.31041 | 0.56919 | 0.37475 | 0.66768 | 0.01723 | 0.02767 | 0.53641 | 0.02372 | 0.06291 | 0.00000 | 3.84050 | 0.64870 |
| **Tcta** | 0.64683 | 0.00318 | 0.50339 | 0.00000 | 0.00000 | 1.08923 | 0.00389 | 0.59814 | 0.00778 | 0.00535 | 0.00000 | 1.32453 | 0.39145 | 0.00000 |
| **Tcte2** | 0.00538 | 0.00529 | 0.00526 | 0.00000 | 0.01192 | 0.58057 | 0.00435 | 0.09834 | 0.02404 | 0.00778 | 0.01442 | 0.37319 | 0.00834 | 0.00372 |
| **Tctex1d2** | 0.66384 | 0.00499 | 0.23821 | 0.06704 | 0.15243 | 0.27636 | 0.00414 | 0.49647 | 0.65824 | 0.03113 | 0.34834 | 0.00120 | 0.00794 | 0.31811 |
| **Tctn1** | 0.03510 | 0.01227 | 0.20444 | 0.80901 | 0.04019 | 0.54898 | 0.11535 | 0.00325 | 0.01162 | 0.04197 | 0.30284 | 0.32976 | 0.01266 | 0.00246 |
| **Tctn2** | 0.00000 | 0.01960 | 1.49133 | 1.78593 | 0.94455 | 0.49587 | 0.00346 | 1.38561 | 0.01240 | 0.01194 | 0.34442 | 0.68290 | 0.00000 | 1.74962 |

| | | | | | | | | | | | | | | |
|---|---|---|---|---|---|---|---|---|---|---|---|---|---|---|
| **Tctn3** | 1.28609 | 0.02951 | 2.41165 | 1.30052 | 0.78729 | 0.53319 | 1.08841 | 1.85865 | 2.38036 | 0.02443 | 2.78628 | 0.01085 | 0.00978 | 2.75590 |
| **Tdgf1** | 7.98354 | 0.08913 | 10.37736 | 5.66097 | 10.09959 | 0.00000 | 0.00442 | 0.00000 | 16.28787 | 0.01896 | 3.22382 | 22.03858 | 0.26029 | 16.65106 |
| **Tdh** | 9.55455 | 0.05026 | 19.00375 | 5.68950 | 8.53469 | 2.08793 | 0.00000 | 0.00000 | 0.00000 | 0.01452 | 5.24635 | 10.04755 | 3.69393 | 5.31279 |
| **Tdp1** | 0.11747 | 0.02170 | 0.92800 | 0.00000 | 0.36408 | 4.25729 | 1.08835 | 0.25833 | 0.27052 | 0.02855 | 1.85778 | 1.24358 | 0.00173 | 0.00000 |
| **Tdp2** | 3.82196 | 0.02197 | 3.04304 | 3.60376 | 8.60873 | 3.21950 | 1.23631 | 0.28677 | 6.98608 | 0.03913 | 0.82456 | 3.37289 | 7.91959 | 5.13469 |
| **Tdpoz1** | 0.51069 | 0.00304 | 0.36367 | 0.13567 | 0.14971 | 0.00000 | 0.00315 | 0.25302 | 2.12766 | 0.00551 | 0.00272 | 0.00301 | 6.82792 | 1.94528 |
| **Tdpoz2** | 0.47629 | 0.00440 | 2.57430 | 0.25276 | 2.84521 | 0.98814 | 0.00300 | 0.38234 | 4.11430 | 0.00298 | 0.00000 | 0.00000 | 0.41176 | 1.94730 |
| **Tdpoz3** | 0.01016 | 0.00444 | 0.01230 | 0.01261 | 0.03425 | 0.01167 | 0.00424 | 0.01032 | 0.04163 | 0.00693 | 0.00801 | 0.00965 | 0.01355 | 0.21986 |
| **Tdpoz4** | 0.00000 | 0.00507 | 0.04781 | 0.01743 | 0.00000 | 0.00169 | 0.01043 | 0.03469 | 0.77417 | 0.00808 | 0.00498 | 0.10769 | 0.04696 | 2.31615 |
| **Tdrd1** | 0.01229 | 0.00422 | 0.17060 | 0.00537 | 0.15166 | 0.01266 | 0.00417 | 0.00783 | 0.00471 | 0.00796 | 0.01098 | 0.01034 | 1.10102 | 0.69439 |
| **Tdrd12** | 0.00000 | 0.01144 | 0.09805 | 0.18313 | 0.37020 | 2.20526 | 0.00414 | 0.00000 | 0.01274 | 0.00538 | 2.68567 | 0.00151 | 0.27201 | 0.79889 |
| **Tdrd3** | 0.86577 | 0.00829 | 3.90750 | 0.87848 | 1.90692 | 3.32729 | 1.58707 | 1.92568 | 2.11821 | 0.10340 | 3.92764 | 2.79342 | 0.85407 | 0.40639 |
| **Tdrd5** | 0.54134 | 0.00339 | 0.67961 | 0.00000 | 1.82260 | 0.60139 | 2.59886 | 0.00000 | 0.33309 | 0.05637 | 0.00000 | 0.61656 | 0.03873 | 0.81419 |
| **Tdrd7** | 0.00090 | 0.00381 | 0.15188 | 0.01878 | 0.25322 | 0.00000 | 0.00355 | 0.52057 | 0.30165 | 0.01187 | 0.00658 | 0.01021 | 0.00471 | 0.38353 |
| **Tdrd9** | 1.11021 | 0.01075 | 4.44470 | 1.37884 | 2.84994 | 0.00000 | 0.00298 | 0.00000 | 0.37826 | 0.00000 | 1.78974 | 0.37612 | 0.97603 | 3.48617 |
| **Tdrkh** | 0.00843 | 0.00367 | 0.00935 | 0.39249 | 0.17039 | 0.26729 | 0.00347 | 0.00200 | 1.16837 | 0.00490 | 0.04751 | 0.26199 | 1.69942 | 0.00000 |
| **Tdrp** | 4.69011 | 6.95871 | 18.97225 | 11.31311 | 1.24460 | 30.88006 | 10.57062 | 22.81111 | 4.35484 | 18.63612 | 7.39060 | 6.22913 | 0.28504 | 0.00000 |
| **Tead1** | 3.59999 | 0.05209 | 6.02119 | 0.48650 | 2.11077 | 8.50925 | 2.38742 | 2.14746 | 3.52838 | 0.02674 | 5.48551 | 6.28227 | 3.25890 | 4.40230 |
| **Tead2** | 0.01430 | 0.08747 | 0.00008 | 1.84202 | 0.96138 | 1.34487 | 3.67398 | 9.31256 | 0.00000 | 12.02985 | 0.96022 | 1.23795 | 0.27757 | 1.29996 |
| **Tead3** | 0.63401 | 0.00428 | 0.15304 | 0.03766 | 0.47385 | 0.08599 | 0.00413 | 0.10576 | 0.05910 | 0.00704 | 0.08412 | 0.03262 | 0.37089 | 0.11955 |
| **Tead4** | 10.01006 | 0.08399 | 16.98151 | 8.17025 | 23.46564 | 11.94834 | 0.01760 | 4.06183 | 4.51391 | 0.09070 | 6.63380 | 4.88225 | 6.84876 | 12.11697 |
| **Tec** | 0.00933 | 0.00440 | 0.02602 | 0.01194 | 0.01944 | 0.01204 | 0.00422 | 0.01001 | 0.02128 | 0.00704 | 0.00836 | 0.00995 | 0.10349 | 0.01474 |
| **Tecpr1** | 0.00000 | 0.00399 | 1.47551 | 1.14319 | 0.57674 | 0.62937 | 0.00409 | 0.13617 | 0.18981 | 0.00995 | 0.40040 | 1.09432 | 0.00000 | 0.00000 |
| **Tecpr2** | 0.57006 | 0.00383 | 1.47595 | 0.00000 | 0.99470 | 1.38837 | 0.08130 | 0.13188 | 2.97970 | 0.03566 | 0.64366 | 0.35004 | 0.14838 | 2.44845 |
| **Tecr** | 5.93843 | 0.30361 | 12.01288 | 12.21872 | 5.25121 | 14.41940 | 1.81551 | 11.87321 | 1.06263 | 0.25201 | 7.68333 | 13.86085 | 0.47307 | 4.82333 |
| **Teddm1b** | 0.51586 | 0.01117 | 0.02968 | 0.36435 | 0.12387 | 0.79448 | 0.31530 | 0.07015 | 1.08425 | 0.22818 | 0.08170 | 0.41368 | 6.35549 | 0.52070 |
| **Teddm3** | 0.01037 | 0.01391 | 0.01221 | 0.05316 | 0.01521 | 0.06213 | 0.00415 | 0.00834 | 0.01230 | 0.00617 | 0.00975 | 0.04542 | 0.00694 | 0.01031 |
| **Tef** | 0.66579 | 0.01016 | 0.19677 | 0.18236 | 1.43923 | 0.64189 | 0.00356 | 0.00175 | 0.00000 | 0.00565 | 0.00000 | 0.31635 | 3.57080 | 1.21516 |
| **Tefm** | 3.94783 | 0.84871 | 6.75805 | 3.27393 | 9.90505 | 3.00174 | 0.00968 | 3.51036 | 5.90597 | 13.16895 | 5.29426 | 1.88482 | 1.68408 | 8.50171 |
| **Tek** | 0.00626 | 0.00437 | 0.01168 | 0.01239 | 0.00355 | 0.01161 | 0.00431 | 0.00964 | 0.02361 | 0.00631 | 0.00830 | 0.00945 | 0.17284 | 0.27093 |
| **Tekt1** | 0.00446 | 0.00459 | 0.00243 | 0.19789 | 0.38420 | 0.01634 | 0.00482 | 0.06252 | 0.01284 | 0.00858 | 0.01016 | 0.00479 | 0.09295 | 0.00315 |
| **Tekt2** | 5.68567 | 0.02413 | 1.04878 | 0.00000 | 0.92037 | 0.00000 | 0.00404 | 0.49280 | 0.11059 | 0.00535 | 0.00244 | 0.00000 | 3.30656 | 2.09173 |
| **Telo2** | 0.04876 | 5.35545 | 3.28269 | 3.85576 | 0.98166 | 1.05933 | 0.00783 | 0.04770 | 0.05073 | 0.01196 | 1.32646 | 1.53708 | 0.52464 | 0.77414 |
| **Ten1** | 1.21356 | 5.99500 | 2.94451 | 6.14123 | 4.26345 | 4.79370 | 0.61435 | 5.99119 | 0.01169 | 0.06013 | 1.00829 | 2.91227 | 0.95596 | 1.19803 |
| **Tenm3** | 0.01093 | 0.00365 | 0.00974 | 0.00674 | 0.01000 | 0.00382 | 0.00415 | 0.00566 | 0.01283 | 0.00670 | 0.09213 | 0.72222 | 0.25612 | 0.00330 |
| **Tenm4** | 0.03025 | 0.00391 | 0.02096 | 0.02136 | 0.11427 | 0.01214 | 0.00404 | 0.56664 | 0.00884 | 0.00677 | 0.00625 | 0.00661 | 0.00475 | 0.03151 |
| **Tep1** | 1.43316 | 0.00400 | 5.56712 | 4.79727 | 6.19221 | 1.26231 | 0.00912 | 0.06206 | 1.56773 | 0.00801 | 1.67205 | 0.89454 | 0.37900 | 4.29997 |
| **Terb1** | 0.02216 | 0.03503 | 0.15485 | 0.00230 | 0.06491 | 0.04289 | 0.00450 | 0.00561 | 0.00837 | 0.00667 | 0.00607 | 0.00798 | 0.02410 | 0.62213 |
| **Terb2** | 1.90102 | 3.69580 | 0.05858 | 0.00000 | 2.31513 | 3.14139 | 0.00304 | 0.48293 | 3.15128 | 0.00000 | 3.00068 | 0.44747 | 1.28085 | 0.89350 |
| **Terf1** | 14.36064 | 0.06139 | 12.72914 | 4.98523 | 9.41589 | 2.95814 | 2.21388 | 3.83621 | 7.50212 | 0.02451 | 1.38786 | 3.84995 | 6.04905 | 1.95735 |
| **Terf2** | 2.73601 | 0.05329 | 3.13760 | 3.79115 | 3.13284 | 3.31874 | 0.00203 | 2.22115 | 1.50713 | 0.00866 | 3.24976 | 3.23343 | 0.92149 | 2.06750 |
| **Terf2ip** | 2.96234 | 0.21543 | 4.28809 | 2.03592 | 2.90339 | 1.62828 | 0.02653 | 2.56512 | 1.55296 | 0.01679 | 2.87106 | 0.86238 | 4.48091 | 1.23658 |
| **Tert** | 0.37345 | 0.00390 | 0.62231 | 0.29103 | 0.01315 | 0.55727 | 0.00457 | 0.17996 | 0.00331 | 0.00417 | 0.00071 | 0.55291 | 0.00127 | 0.43159 |
| **Tes** | 0.74146 | 0.31412 | 6.70669 | 8.48725 | 2.53953 | 11.46323 | 4.38960 | 11.26910 | 0.00000 | 0.20890 | 4.02917 | 15.80547 | 1.49848 | 1.41558 |
| **Tesc** | 0.00000 | 0.02573 | 0.93210 | 3.54921 | 2.00262 | 1.76498 | 0.00373 | 0.02304 | 0.00000 | 0.01287 | 0.55489 | 0.00623 | 2.54173 | 1.97859 |
| **Tescl** | 0.01054 | 0.00435 | 0.06593 | 0.09643 | 0.04794 | 0.02505 | 0.00404 | 0.00615 | 0.00990 | 0.00577 | 0.01264 | 0.01393 | 0.00765 | 0.02593 |
| **Tesk1** | 0.07936 | 0.02450 | 0.21106 | 3.43777 | 0.96589 | 0.00000 | 0.33186 | 2.62420 | 0.01550 | 0.02556 | 0.00000 | 1.09290 | 0.14415 | 0.00000 |
| **Tesk2** | 0.70732 | 0.01104 | 3.00186 | 0.00000 | 3.97473 | 2.64250 | 0.00342 | 0.21344 | 3.26523 | 0.00000 | 0.00000 | 1.62195 | 2.06063 | 6.04878 |
| **Tet1** | 6.75965 | 0.37709 | 1.95216 | 0.00000 | 10.02239 | 4.96145 | 1.19376 | 0.62226 | 25.76518 | 0.01032 | 10.32354 | 8.64035 | 1.03200 | 15.81081 |
| **Tet2** | 0.00000 | 0.08091 | 0.38167 | 3.86769 | 0.00000 | 6.48638 | 3.64717 | 3.35744 | 0.70316 | 0.06472 | 3.92213 | 4.59822 | 0.01136 | 0.91091 |
| **Tet3** | 0.01508 | 0.00358 | 0.01947 | 0.08223 | 0.42524 | 0.91715 | 0.00381 | 0.02031 | 0.31492 | 0.01204 | 0.00359 | 0.24839 | 2.10373 | 0.00000 |
| **Tex10** | 8.34539 | 0.01210 | 8.37219 | 6.32508 | 7.76775 | 4.22649 | 0.15684 | 2.77268 | 10.47045 | 0.00000 | 8.85679 | 5.02224 | 3.13598 | 10.10259 |
| **Tex101** | 0.00848 | 0.00434 | 0.46423 | 0.00000 | 0.00607 | 0.00487 | 0.00424 | 0.01070 | 0.01954 | 0.00891 | 0.00544 | 0.09598 | 0.00721 | 0.00828 |
| **Tex11** | 0.56982 | 0.00365 | 0.17518 | 0.00000 | 0.05949 | 0.07258 | 0.00386 | 0.00352 | 1.57016 | 0.00623 | 0.00482 | 0.00220 | 4.66469 | 1.16562 |
| **Tex12** | 0.29188 | 0.00403 | 0.75844 | 0.10797 | 1.11211 | 0.34921 | 0.00364 | 0.00020 | 0.95342 | 0.00620 | 0.10404 | 0.00692 | 0.97588 | 0.40840 |

| | | | | | | | | | | | | | | |
|---|---|---|---|---|---|---|---|---|---|---|---|---|---|---|
| **Tex13** | 0.00400 | 0.00440 | 0.00812 | 0.01364 | 0.02351 | 0.01117 | 0.00470 | 0.00955 | 0.00885 | 0.00689 | 0.00764 | 0.00928 | 0.00511 | 0.89499 |
| **Tex14** | 0.59036 | 0.00337 | 0.66893 | 0.57456 | 0.20113 | 0.85985 | 0.00354 | 0.00000 | 0.01134 | 0.00466 | 0.00000 | 0.67961 | 0.26154 | 1.63451 |
| **Tex15** | 2.32691 | 0.00346 | 0.00000 | 0.17361 | 3.51194 | 0.14458 | 0.00336 | 0.00174 | 3.54286 | 0.00566 | 0.00000 | 0.00532 | 2.00428 | 2.24911 |
| **Tex19.1** | 3.24309 | 5.53682 | 12.51440 | 19.49893 | 10.76876 | 17.17833 | 3.59723 | 4.51113 | 9.94670 | 0.17417 | 16.76281 | 10.21160 | 6.31068 | 8.55898 |
| **Tex19.2** | 0.00000 | 0.45663 | 6.98378 | 14.67714 | 0.96796 | 6.51131 | 1.50127 | 2.25998 | 0.16023 | 0.44212 | 13.05579 | 3.16350 | 0.09392 | 0.32221 |
| **Tex2** | 3.71622 | 0.14083 | 5.06280 | 3.60274 | 5.86685 | 7.65341 | 0.02448 | 2.66711 | 11.00617 | 0.04193 | 0.61505 | 3.45430 | 1.07271 | 11.34454 |
| **Tex21** | 0.00195 | 0.01303 | 0.20679 | 0.00000 | 0.00686 | 0.00000 | 0.00370 | 0.01444 | 0.01357 | 0.00372 | 1.24711 | 0.79660 | 0.01030 | 0.00639 |
| **Tex26** | 0.00764 | 0.00439 | 0.01425 | 0.01233 | 0.01054 | 0.01281 | 0.00426 | 0.00977 | 0.02658 | 0.00654 | 0.00886 | 0.00973 | 0.16351 | 0.00863 |
| **Tex261** | 3.31183 | 0.07816 | 3.58650 | 4.46691 | 2.85360 | 6.19921 | 1.48213 | 3.75006 | 0.10396 | 0.09237 | 0.07797 | 4.13447 | 0.49021 | 2.75412 |
| **Tex264** | 3.53288 | 1.53964 | 10.62409 | 12.67475 | 5.17845 | 9.94673 | 2.72113 | 4.31814 | 0.03217 | 0.17560 | 1.67043 | 9.00225 | 3.64399 | 0.49645 |
| **Tex30** | 6.93283 | 0.03044 | 3.22610 | 0.90013 | 2.39114 | 3.78010 | 2.15126 | 3.07735 | 6.48776 | 0.02106 | 4.26362 | 4.74422 | 5.75694 | 1.05129 |
| **Tex37** | 0.21630 | 0.00447 | 0.00442 | 0.00934 | 0.27639 | 0.01046 | 0.00455 | 0.01066 | 0.00993 | 0.00619 | 0.00744 | 0.00759 | 0.06272 | 0.16258 |
| **Tex38** | 0.00630 | 0.00420 | 0.17162 | 0.04698 | 0.08110 | 0.00530 | 0.00426 | 0.00907 | 0.00941 | 0.00568 | 0.01290 | 0.01416 | 0.22769 | 0.00882 |
| **Tex40** | 0.22963 | 6.33385 | 0.15486 | 1.33047 | 0.27475 | 0.39988 | 0.47015 | 0.07800 | 0.26119 | 0.03872 | 0.01579 | 0.05191 | 0.00173 | 0.01212 |
| **Tex43** | 0.00466 | 0.00424 | 0.01375 | 0.01390 | 0.00731 | 0.01218 | 0.00420 | 0.00861 | 0.01740 | 0.00669 | 0.00887 | 0.00821 | 0.89017 | 0.05545 |
| **Tex9** | 0.94399 | 0.03839 | 2.57306 | 0.96195 | 2.17017 | 0.41602 | 0.04905 | 0.18282 | 1.15140 | 0.07835 | 2.55880 | 0.28724 | 1.56971 | 2.61495 |
| **Tfam** | 4.85190 | 0.05679 | 11.50614 | 6.18129 | 8.78994 | 7.81821 | 2.69750 | 5.75815 | 5.28422 | 4.54963 | 6.43585 | 7.40893 | 1.81116 | 5.95952 |
| **Tfap2a** | 0.00000 | 0.01186 | 0.01071 | 0.00000 | 0.00962 | 0.00000 | 1.05134 | 1.40206 | 0.17437 | 1.50383 | 0.00000 | 0.16882 | 0.02079 | 0.00803 |
| **Tfap2c** | 5.13066 | 0.18571 | 18.48638 | 9.32444 | 21.22012 | 10.49309 | 4.44953 | 7.79950 | 5.41114 | 0.25933 | 7.62617 | 5.00901 | 1.93638 | 13.12441 |
| **Tfap2e** | 0.00241 | 0.00450 | 0.28278 | 0.00339 | 0.04179 | 0.00000 | 0.00411 | 0.03728 | 0.00000 | 0.00723 | 0.00597 | 0.00503 | 4.92289 | 1.05409 |
| **Tfap4** | 0.00000 | 0.00356 | 1.39132 | 0.46964 | 2.21248 | 0.00821 | 0.37429 | 0.50382 | 0.09933 | 0.00365 | 0.40482 | 0.38706 | 0.38126 | 0.55842 |
| **Tfb1m** | 1.66834 | 0.01685 | 1.90618 | 0.23381 | 0.82376 | 3.82190 | 5.09524 | 0.07036 | 1.56082 | 0.10590 | 0.04122 | 2.98976 | 0.00000 | 1.38425 |
| **Tfb2m** | 11.44683 | 4.41191 | 9.69602 | 1.47021 | 16.00291 | 5.06918 | 2.79315 | 1.15875 | 16.32518 | 0.01185 | 1.57496 | 0.08965 | 13.97061 | 10.37015 |
| **Tfcp2** | 1.31291 | 0.00298 | 2.89492 | 0.88268 | 4.32097 | 0.25564 | 0.00171 | 0.00000 | 11.15157 | 0.00000 | 0.00000 | 0.16819 | 3.04157 | 8.97340 |
| **Tfcp2l1** | 4.93835 | 8.08635 | 23.49079 | 22.31346 | 9.25172 | 31.12831 | 4.55322 | 14.67251 | 10.29429 | 11.01065 | 19.81786 | 17.66405 | 0.52449 | 1.14263 |
| **Tfdp1** | 7.32880 | 0.14579 | 13.51532 | 6.22048 | 11.87156 | 12.85695 | 2.12126 | 4.81870 | 14.16656 | 0.10542 | 8.29082 | 10.16417 | 4.08860 | 14.97077 |
| **Tfdp2** | 2.58299 | 0.00132 | 1.36588 | 0.38911 | 4.00005 | 0.47270 | 0.00273 | 0.32940 | 1.19194 | 0.00000 | 1.83323 | 1.93685 | 3.94400 | 4.23249 |
| **Tfe3** | 4.03626 | 0.01385 | 3.72531 | 5.18135 | 3.43865 | 6.66912 | 2.13289 | 3.16341 | 0.00000 | 0.12491 | 3.65732 | 5.03478 | 1.90182 | 4.37813 |
| **Tfeb** | 2.81834 | 0.17593 | 9.68892 | 7.94044 | 12.22660 | 1.32654 | 4.60855 | 15.18386 | 0.00000 | 1.43762 | 1.19128 | 1.04446 | 0.40218 | 0.00000 |
| **Tfg** | 12.54098 | 0.19794 | 9.99303 | 8.27546 | 20.50805 | 11.98203 | 0.02714 | 5.96758 | 4.20210 | 0.59170 | 5.44527 | 10.62681 | 13.58844 | 10.94916 |
| **Tfip11** | 2.63575 | 0.06667 | 6.70271 | 4.63646 | 6.88672 | 7.86627 | 1.36778 | 0.85194 | 0.08558 | 1.21554 | 1.08869 | 2.02032 | 7.39113 | 1.93034 |
| **Tfpi** | 0.02624 | 0.22192 | 0.00810 | 0.00000 | 0.01895 | 0.19679 | 0.00249 | 0.00824 | 0.20337 | 0.00558 | 0.00000 | 5.65222 | 2.23834 | 0.00000 |
| **Tfpt** | 0.00000 | 0.02633 | 6.25259 | 9.03716 | 9.04190 | 0.00000 | 0.02644 | 3.62606 | 0.00000 | 0.01692 | 0.01083 | 0.82517 | 2.39517 | 0.00000 |
| **Tfr2** | 0.00443 | 0.00371 | 0.00687 | 0.47538 | 0.03556 | 0.00655 | 0.00400 | 0.00381 | 0.01109 | 0.00594 | 0.00565 | 0.38961 | 0.00906 | 0.01024 |
| **Tfrc** | 22.85503 | 0.00000 | 18.81506 | 5.18961 | 20.75306 | 6.92547 | 0.00000 | 0.29446 | 46.80608 | 0.00000 | 4.85067 | 2.64011 | 14.80776 | 21.61100 |
| **Tgds** | 1.21103 | 0.00282 | 0.00000 | 0.74192 | 0.48151 | 0.00000 | 0.00428 | 0.59942 | 0.90279 | 0.00000 | 2.26408 | 0.87509 | 1.95208 | 2.57418 |
| **Tgfa** | 0.00605 | 0.00443 | 0.00264 | 0.00000 | 0.62435 | 0.07803 | 0.00508 | 0.00565 | 0.01524 | 0.00612 | 0.00475 | 0.17495 | 0.00262 | 0.03879 |
| **Tgfb1i1** | 0.11720 | 0.00444 | 0.01332 | 0.01039 | 0.02748 | 0.01205 | 0.00425 | 0.01098 | 0.01253 | 0.00655 | 0.00781 | 0.00941 | 0.00523 | 0.00710 |
| **Tgfb2** | 0.17330 | 0.00391 | 0.02274 | 0.00828 | 0.00331 | 0.00000 | 0.00410 | 0.00953 | 0.00708 | 0.00540 | 0.28244 | 0.00937 | 1.37909 | 0.74539 |
| **Tgfb3** | 0.32948 | 0.00426 | 0.01170 | 0.01004 | 0.00000 | 0.00885 | 0.00417 | 0.00993 | 0.00632 | 0.00609 | 0.00670 | 0.00741 | 0.50304 | 0.39864 |
| **Tgfbr1** | 1.47440 | 0.03776 | 1.24286 | 0.61110 | 1.23070 | 3.81307 | 0.10544 | 0.00000 | 5.83593 | 0.01492 | 0.42943 | 3.05466 | 4.05907 | 0.00000 |
| **Tgfbr2** | 0.03301 | 0.00379 | 0.00120 | 0.00000 | 0.11415 | 0.84870 | 0.00461 | 0.00466 | 0.01885 | 0.00674 | 0.00000 | 0.73214 | 0.00608 | 0.00161 |
| **Tgfbr3** | 0.01268 | 0.00481 | 0.02003 | 0.00028 | 0.00682 | 0.09723 | 0.00430 | 0.00787 | 0.65909 | 0.00711 | 0.00667 | 0.00988 | 0.02086 | 0.00000 |
| **Tgfbr3l** | 0.02090 | 0.00438 | 0.01253 | 0.08367 | 0.01236 | 0.01244 | 0.00440 | 0.00913 | 0.01627 | 0.00625 | 0.00776 | 0.00821 | 0.01096 | 0.02210 |
| **Tgfbrap1** | 3.10652 | 0.00098 | 2.44391 | 0.39708 | 3.19567 | 0.87277 | 2.13643 | 0.64095 | 1.01439 | 0.09557 | 3.30362 | 1.93007 | 2.14071 | 5.43094 |
| **Tgif1** | 0.07731 | 0.03109 | 4.14458 | 0.00000 | 1.64068 | 2.78667 | 2.21062 | 1.47157 | 1.07662 | 0.06501 | 1.38077 | 4.11263 | 1.24432 | 2.89357 |
| **Tgif2** | 2.26582 | 0.02691 | 4.88801 | 3.37664 | 6.96115 | 3.30723 | 2.07369 | 0.87331 | 2.25855 | 0.06428 | 6.20442 | 4.74522 | 2.35148 | 5.19447 |
| **Tgm2** | 0.04984 | 0.00431 | 0.45415 | 0.11408 | 0.00641 | 0.11366 | 0.00389 | 0.00226 | 0.01574 | 0.00586 | 0.01215 | 0.00917 | 0.10331 | 0.00510 |
| **Tgm4** | 2.25665 | 0.01224 | 1.71915 | 0.00000 | 1.53862 | 1.81382 | 0.00375 | 1.74423 | 0.00000 | 0.00346 | 0.00280 | 0.07318 | 3.20383 | 0.86251 |
| **Tgoln1** | 5.55915 | 0.02654 | 5.52659 | 1.09481 | 2.47964 | 5.96500 | 1.32203 | 2.75649 | 14.85998 | 0.03084 | 5.77343 | 4.68079 | 8.58863 | 9.04601 |
| **Tgs1** | 2.89989 | 0.03069 | 2.51975 | 0.28955 | 2.64830 | 3.30638 | 0.00885 | 0.42688 | 4.29486 | 0.01471 | 2.26747 | 2.01461 | 8.34985 | 4.35036 |
| **Th** | 0.26101 | 0.00398 | 0.00635 | 0.01062 | 0.26656 | 0.01081 | 0.00400 | 0.00861 | 0.00927 | 0.00705 | 0.00729 | 0.00557 | 3.21127 | 0.00365 |
| **Thada** | 0.18027 | 0.00256 | 1.60822 | 2.53449 | 0.67178 | 2.22345 | 0.00259 | 0.94071 | 0.75880 | 0.00000 | 4.47747 | 0.49358 | 0.00000 | 3.47114 |
| **Thap1** | 6.15499 | 0.00199 | 0.89029 | 0.00000 | 1.44429 | 0.78778 | 0.00335 | 0.00804 | 1.75823 | 0.02218 | 0.03554 | 0.96088 | 7.20731 | 1.65560 |
| **Thap11** | 3.76283 | 6.39505 | 5.17965 | 6.74954 | 8.75703 | 2.53270 | 4.12156 | 4.09859 | 0.00000 | 1.41409 | 3.49451 | 4.00741 | 16.43741 | 5.52191 |

| | | | | | | | | | | | | | | |
|---|---|---|---|---|---|---|---|---|---|---|---|---|---|---|
| **Thap2** | 0.01324 | 0.01299 | 0.78860 | 0.20100 | 0.34575 | 0.64245 | 0.01873 | 0.10536 | 0.08651 | 0.01790 | 0.01130 | 0.13665 | 0.36982 | 0.01969 |
| **Thap3** | 0.52855 | 0.00272 | 0.68334 | 1.88991 | 2.21506 | 1.85043 | 0.01537 | 0.61617 | 0.03855 | 0.00321 | 2.93637 | 0.70005 | 0.35701 | 0.07041 |
| **Thap4** | 4.25999 | 3.48701 | 3.72469 | 4.12718 | 10.82556 | 4.65977 | 3.90738 | 9.43238 | 1.76601 | 10.31971 | 7.30690 | 5.53106 | 4.14503 | 8.30644 |
| **Thap6** | 0.03149 | 0.07470 | 0.18893 | 0.14840 | 0.03713 | 0.19139 | 0.16823 | 0.46369 | 0.26056 | 0.31358 | 1.21153 | 0.70567 | 0.80718 | 1.64912 |
| **Thap7** | 0.00000 | 3.87208 | 3.99478 | 4.84513 | 5.46033 | 0.00000 | 2.07719 | 3.00201 | 0.00003 | 0.07265 | 0.00000 | 1.59401 | 0.35887 | 0.00000 |
| **Thbd** | 0.26579 | 0.00345 | 0.38824 | 0.00000 | 0.17393 | 0.25517 | 0.00370 | 0.00205 | 0.00579 | 0.00441 | 0.00159 | 0.89160 | 0.03138 | 0.25328 |
| **Thbs1** | 3.34591 | 0.00532 | 0.00671 | 0.00000 | 0.01190 | 0.38683 | 0.00442 | 0.18262 | 0.00508 | 0.00693 | 0.00303 | 0.10380 | 0.00392 | 0.01193 |
| **Thbs3** | 0.00535 | 0.00446 | 0.03033 | 0.70125 | 0.14364 | 0.00000 | 0.00503 | 0.62629 | 0.00761 | 0.00524 | 0.01358 | 0.01449 | 0.05878 | 0.49260 |
| **Theg** | 0.02697 | 0.00445 | 0.01566 | 0.01392 | 0.00392 | 0.00920 | 0.00469 | 0.02286 | 0.03202 | 0.00698 | 0.00782 | 0.00820 | 0.21274 | 0.26487 |
| **Thegl** | 0.01197 | 0.00444 | 0.01330 | 0.00832 | 0.01149 | 0.03166 | 0.01046 | 0.00795 | 0.01277 | 0.00638 | 0.03680 | 0.00899 | 0.00836 | 0.04865 |
| **Them4** | 4.24318 | 0.01069 | 8.19908 | 2.50852 | 6.52513 | 0.51443 | 0.00279 | 0.41472 | 1.03323 | 0.00913 | 0.29326 | 1.53559 | 3.19951 | 5.11814 |
| **Them5** | 0.35712 | 0.00387 | 0.23650 | 0.42459 | 0.00000 | 0.00000 | 0.00376 | 0.00422 | 0.00956 | 0.00687 | 0.00459 | 0.10721 | 0.00517 | 0.00289 |
| **Themis** | 0.08541 | 0.01276 | 0.02888 | 0.04849 | 0.04355 | 0.02987 | 0.01070 | 0.02849 | 0.16588 | 0.00868 | 0.01211 | 0.02190 | 0.02985 | 0.13512 |
| **Themis2** | 0.00431 | 0.00446 | 0.00000 | 0.01101 | 1.33197 | 0.01082 | 0.00420 | 0.00921 | 0.00619 | 0.00702 | 0.00736 | 0.00683 | 0.01051 | 0.79783 |
| **Themis3** | 0.00000 | 0.00432 | 0.06529 | 0.27633 | 0.30494 | 0.00607 | 0.00397 | 0.00780 | 0.16100 | 0.00652 | 0.00686 | 0.00595 | 0.00269 | 0.66034 |
| **Thg1l** | 0.01220 | 0.01033 | 0.39178 | 0.00000 | 2.24948 | 0.24720 | 0.00837 | 1.80432 | 3.15209 | 0.00173 | 1.43058 | 0.89705 | 0.43651 | 1.47219 |
| **Thnsl1** | 0.00363 | 0.01252 | 0.00032 | 0.24146 | 0.00337 | 0.00690 | 0.00507 | 0.97902 | 0.00693 | 0.00648 | 0.00286 | 0.02479 | 0.01035 | 1.07101 |
| **Thoc1** | 14.42061 | 0.00000 | 9.90134 | 3.50669 | 12.94099 | 6.27341 | 3.71744 | 2.52887 | 21.79689 | 0.10872 | 4.95588 | 6.39508 | 3.68436 | 7.16703 |
| **Thoc2** | 3.63323 | 0.01704 | 3.48621 | 3.14408 | 5.57491 | 5.45267 | 0.00000 | 0.00000 | 30.21418 | 0.03459 | 1.46973 | 2.99009 | 2.14735 | 2.44898 |
| **Thoc3** | 7.78386 | 0.01940 | 7.80403 | 4.41188 | 16.31764 | 5.44006 | 2.49591 | 4.15364 | 5.29368 | 0.04961 | 4.15628 | 4.15572 | 6.39538 | 11.17032 |
| **Thoc5** | 4.77648 | 7.56015 | 10.37242 | 6.27893 | 10.08041 | 5.27900 | 0.03054 | 5.67327 | 6.48675 | 3.38314 | 10.73928 | 6.01981 | 3.21277 | 13.67073 |
| **Thoc6** | 3.19864 | 2.89618 | 13.52833 | 8.17326 | 8.56390 | 10.59596 | 3.79432 | 2.01762 | 0.01741 | 0.14709 | 9.67419 | 7.91231 | 2.15906 | 1.94709 |
| **Thoc7** | 16.83994 | 6.40311 | 10.70456 | 13.31855 | 15.34516 | 13.43362 | 2.74499 | 4.27770 | 8.28615 | 11.67978 | 6.18463 | 8.99550 | 6.29813 | 7.22907 |
| **Thop1** | 0.00000 | 0.07468 | 5.82337 | 12.75998 | 6.47174 | 0.00000 | 0.01404 | 1.59611 | 0.00000 | 0.03048 | 0.02928 | 2.99605 | 0.33250 | 0.00000 |
| **Thra** | 0.02277 | 0.01231 | 1.10053 | 0.06222 | 3.18046 | 0.52172 | 0.00372 | 1.00076 | 0.16121 | 0.02960 | 0.00000 | 0.00400 | 0.00000 | 1.07206 |
| **Thrap3** | 0.81869 | 4.59123 | 8.94262 | 12.45464 | 4.24556 | 7.44212 | 0.26131 | 2.93688 | 1.79982 | 0.08443 | 4.44895 | 6.38285 | 3.99583 | 1.83314 |
| **Thrsp** | 0.00000 | 0.00482 | 1.99207 | 0.00606 | 0.62025 | 1.29467 | 0.01116 | 0.00799 | 0.01439 | 0.00192 | 0.00337 | 0.00173 | 0.00000 | 0.00000 |
| **Thsd1** | 0.00386 | 0.00429 | 0.00405 | 0.24528 | 0.00594 | 0.10185 | 0.00410 | 0.00557 | 0.00640 | 0.00632 | 0.00504 | 0.00599 | 0.19845 | 0.10469 |
| **Thsd4** | 0.01134 | 0.00487 | 0.01179 | 0.00009 | 0.00910 | 0.20992 | 0.00443 | 0.00659 | 0.01786 | 0.00616 | 0.01111 | 0.01520 | 0.00848 | 0.10189 |
| **Thsd7a** | 0.00835 | 0.00426 | 0.01268 | 0.06401 | 0.00628 | 0.01163 | 0.00421 | 0.00742 | 0.04753 | 0.00596 | 0.00818 | 0.00806 | 0.38135 | 0.00000 |
| **Thsd7b** | 0.00356 | 0.00431 | 0.01418 | 0.01194 | 0.00873 | 0.01306 | 0.00424 | 0.00896 | 0.00512 | 0.00609 | 0.00923 | 0.00895 | 0.50970 | 0.08300 |
| **Thtpa** | 0.00571 | 0.00432 | 0.80288 | 0.27258 | 0.00000 | 0.02680 | 0.00493 | 0.06242 | 0.05328 | 0.00510 | 0.00773 | 0.01333 | 0.00253 | 0.77372 |
| **Thumpd1** | 9.87353 | 2.98399 | 10.09705 | 7.78863 | 13.77915 | 6.21037 | 2.83190 | 2.99992 | 10.86591 | 0.06045 | 1.42739 | 6.77761 | 3.61984 | 13.97485 |
| **Thumpd2** | 0.00757 | 0.00352 | 0.00697 | 0.11519 | 0.34624 | 0.00622 | 0.00448 | 0.01988 | 0.00963 | 0.00580 | 0.00662 | 1.25434 | 0.00633 | 0.01020 |
| **Thumpd3** | 4.19129 | 0.02542 | 3.64010 | 4.82865 | 2.56580 | 5.21512 | 1.69569 | 3.24085 | 5.01190 | 0.03287 | 3.70077 | 0.99788 | 1.52723 | 0.92546 |
| **Thy1** | 0.20714 | 0.00441 | 0.01129 | 0.01407 | 0.00419 | 0.01012 | 0.00424 | 0.01067 | 0.00569 | 0.00659 | 0.00731 | 0.00982 | 0.00614 | 0.16370 |
| **Thyn1** | 1.41764 | 0.00049 | 4.09279 | 3.30332 | 0.53303 | 2.67793 | 1.80097 | 1.98920 | 1.40751 | 5.77698 | 3.43494 | 2.17804 | 0.61397 | 2.99479 |
| **Tia1** | 0.01611 | 0.66818 | 0.00000 | 0.29127 | 0.51134 | 0.65639 | 0.04957 | 0.91760 | 0.20135 | 0.02118 | 0.00000 | 0.78762 | 0.00539 | 0.09600 |
| **Tial1** | 12.33693 | 0.69516 | 8.37842 | 2.96912 | 11.28649 | 6.48276 | 1.24200 | 4.52739 | 13.68635 | 0.09906 | 4.69584 | 4.58939 | 7.24262 | 9.24583 |
| **Tiam1** | 1.90678 | 0.00348 | 0.06154 | 0.81140 | 2.31590 | 0.33863 | 1.35429 | 0.40975 | 3.04477 | 0.01322 | 0.00097 | 0.01245 | 4.18846 | 2.25862 |
| **Tiam2** | 0.03258 | 0.00380 | 0.01221 | 0.00000 | 1.14187 | 0.41433 | 0.43395 | 0.00350 | 0.00000 | 0.00530 | 0.00334 | 0.21256 | 3.16129 | 1.85000 |
| **Ticam1** | 0.34972 | 0.02856 | 0.96567 | 1.88404 | 0.88463 | 0.41198 | 1.09323 | 3.30951 | 1.14611 | 0.08062 | 0.00000 | 0.00000 | 2.40993 | 0.88270 |
| **Ticam2** | 0.00000 | 0.00431 | 0.26448 | 0.06903 | 0.93723 | 0.00000 | 0.00392 | 0.00889 | 1.04271 | 0.00575 | 0.00375 | 0.00561 | 0.11227 | 0.64082 |
| **Ticrr** | 11.31598 | 0.06851 | 14.26872 | 8.97428 | 13.17695 | 6.71577 | 3.67273 | 0.19344 | 13.80460 | 0.21902 | 5.29574 | 7.63935 | 10.38847 | 15.02579 |
| **Tie1** | 0.17774 | 0.02608 | 1.22353 | 1.70684 | 0.00000 | 1.55028 | 0.00366 | 0.45057 | 0.00234 | 0.02959 | 1.81388 | 0.00000 | 0.73979 | 0.06643 |
| **Tifab** | 0.28147 | 0.02307 | 0.11081 | 0.00000 | 0.17444 | 0.02728 | 0.00434 | 0.88589 | 0.00896 | 0.00561 | 0.00000 | 0.29685 | 0.00300 | 0.32002 |
| **Tigar** | 9.39843 | 0.17091 | 33.43544 | 23.95042 | 24.00424 | 17.49563 | 7.06628 | 7.10027 | 10.67222 | 14.30771 | 18.02565 | 12.26172 | 13.51124 | 24.00604 |
| **Tigd2** | 0.01586 | 0.00315 | 0.00169 | 0.00000 | 0.00271 | 0.00000 | 1.77395 | 0.00000 | 0.01006 | 0.01085 | 1.56598 | 0.30392 | 1.31390 | 0.08726 |
| **Tigd3** | 0.08502 | 0.23015 | 0.03716 | 0.27525 | 0.01799 | 0.11410 | 0.01550 | 0.00548 | 0.52241 | 0.08362 | 0.01434 | 0.04562 | 2.30422 | 0.99390 |
| **Tigd5** | 0.03099 | 0.00455 | 0.33412 | 0.00000 | 0.00217 | 0.09021 | 0.00521 | 0.19445 | 0.00961 | 0.00535 | 0.00580 | 0.07156 | 0.00699 | 0.07672 |
| **Tigit** | 0.84795 | 0.06410 | 0.24150 | 0.01919 | 0.40971 | 0.05786 | 0.00389 | 0.03708 | 1.29084 | 0.00711 | 0.01956 | 0.00382 | 0.57786 | 0.38485 |
| **Timd2** | 44.39293 | 8.07886 | 53.92595 | 48.36607 | 40.01422 | 51.10483 | 11.73160 | 13.46491 | 52.30030 | 8.06106 | 27.01286 | 5.95834 | 18.58320 | 39.49967 |
| **Timd4** | 0.00642 | 0.00449 | 0.00416 | 0.00777 | 0.06693 | 0.04145 | 0.00413 | 0.00757 | 0.77204 | 0.00617 | 0.00785 | 0.00919 | 0.00371 | 0.00000 |
| **Timeless** | 2.73428 | 0.02074 | 3.29669 | 0.23522 | 1.75612 | 4.16851 | 0.55104 | 3.37634 | 1.33144 | 0.00765 | 6.95426 | 4.60917 | 1.66589 | 6.64454 |
| **Timm10** | 7.46855 | 1.36790 | 2.12136 | 9.77050 | 11.02280 | 12.22161 | 1.21711 | 3.60743 | 2.63466 | 1.14387 | 3.19419 | 5.49362 | 10.89813 | 6.61862 |

| | | | | | | | | | | | | | | |
|---|---|---|---|---|---|---|---|---|---|---|---|---|---|---|
| **Timm10b** | 0.00456 | 0.03382 | 0.04479 | 0.15958 | 1.03646 | 0.29839 | 1.95435 | 0.94036 | 0.00890 | 0.07306 | 0.00000 | 0.77544 | 0.46175 | 0.00000 |
| **Timm13** | 2.27248 | 0.37482 | 2.59855 | 16.56987 | 18.02099 | 7.59662 | 3.05207 | 15.63706 | 0.00000 | 5.87344 | 2.54957 | 11.97615 | 2.21298 | 1.09856 |
| **Timm17a** | 28.99225 | 11.23565 | 27.24756 | 24.91933 | 36.45529 | 18.71993 | 6.89888 | 8.15478 | 26.37959 | 0.61370 | 14.71279 | 12.72141 | 16.64459 | 30.12698 |
| **Timm17b** | 4.64839 | 10.13931 | 13.16692 | 13.01678 | 13.42684 | 15.21153 | 2.23945 | 7.52953 | 0.57288 | 4.40128 | 4.47627 | 9.42344 | 1.74601 | 5.46537 |
| **Timm21** | 0.00000 | 0.00338 | 0.00000 | 0.00000 | 0.00000 | 0.12128 | 0.00293 | 2.40113 | 1.66760 | 0.01821 | 1.56275 | 0.48587 | 0.00910 | 0.96813 |
| **Timm22** | 9.24362 | 10.80712 | 24.02574 | 18.13263 | 22.39475 | 15.98273 | 5.08188 | 2.81558 | 8.00060 | 0.54815 | 7.76508 | 7.25256 | 6.21978 | 12.87845 |
| **Timm23** | 29.51285 | 9.99182 | 45.41977 | 39.01216 | 44.83737 | 26.18794 | 10.70304 | 16.77269 | 20.13190 | 14.61891 | 17.22488 | 23.25113 | 13.05493 | 19.78586 |
| **Timm44** | 7.78741 | 7.93278 | 14.86578 | 11.01399 | 15.49433 | 12.13488 | 1.02187 | 9.25645 | 0.47608 | 1.16006 | 9.14657 | 10.15948 | 2.02587 | 8.24099 |
| **Timm50** | 5.43257 | 0.17693 | 10.84944 | 16.01696 | 12.30038 | 5.24470 | 0.45747 | 7.06196 | 0.00000 | 1.70897 | 6.33818 | 7.28787 | 0.00000 | 0.05922 |
| **Timm8a1** | 24.53695 | 0.25911 | 29.55157 | 17.37170 | 40.78753 | 8.69119 | 3.76817 | 11.37656 | 13.85746 | 0.09685 | 7.33123 | 7.89448 | 1.73844 | 15.36441 |
| **Timm8a2** | 4.72413 | 0.00106 | 6.94045 | 5.77062 | 8.01818 | 6.15843 | 0.00842 | 0.20240 | 0.00000 | 0.03021 | 5.25046 | 2.17010 | 0.02526 | 3.51309 |
| **Timm8b** | 0.49828 | 0.05671 | 0.00000 | 0.80728 | 3.59304 | 3.38867 | 2.28436 | 5.70567 | 0.36517 | 0.18385 | 0.18195 | 2.61838 | 4.54480 | 0.26419 |
| **Timm9** | 9.27668 | 0.05519 | 6.86383 | 4.57271 | 6.72846 | 6.35594 | 4.14530 | 8.26803 | 0.29744 | 11.17438 | 5.63185 | 6.08236 | 4.60652 | 5.22415 |
| **Timmdc1** | 2.11456 | 0.04551 | 0.51375 | 2.06441 | 0.69110 | 2.05844 | 0.06740 | 0.35936 | 0.59498 | 0.29549 | 0.29697 | 0.48385 | 0.80373 | 1.01929 |
| **Timp1** | 0.84053 | 0.07524 | 0.31401 | 0.00000 | 0.02580 | 0.04809 | 0.00805 | 0.95268 | 0.00265 | 0.01833 | 0.00000 | 6.06138 | 0.35650 | 0.94062 |
| **Timp2** | 0.22364 | 0.00450 | 0.01081 | 0.01045 | 0.08825 | 0.01268 | 0.00437 | 0.01184 | 0.00725 | 0.00616 | 0.00740 | 0.00843 | 0.00499 | 0.02790 |
| **Timp3** | 0.16213 | 0.00441 | 0.00416 | 0.00754 | 0.22435 | 0.00573 | 0.00510 | 0.22555 | 0.16199 | 0.00549 | 0.01080 | 0.01247 | 0.00324 | 0.00000 |
| **Tinagl1** | 0.00000 | 0.10961 | 3.58335 | 3.73221 | 0.00000 | 4.26602 | 4.45623 | 7.52147 | 0.00274 | 0.17671 | 3.97258 | 1.07268 | 0.00543 | 0.00000 |
| **Tinf2** | 5.01529 | 0.98158 | 9.42269 | 5.06638 | 11.58686 | 4.15199 | 2.59136 | 4.58273 | 0.45311 | 5.92013 | 1.33092 | 5.02591 | 9.19880 | 3.14151 |
| **Tiparp** | 5.37985 | 4.35083 | 0.05736 | 1.86171 | 5.82231 | 1.57790 | 0.00000 | 0.00408 | 19.88308 | 2.51281 | 2.20141 | 0.67413 | 8.63225 | 5.36223 |
| **Tipin** | 21.21244 | 0.24860 | 18.05085 | 10.73611 | 17.83298 | 13.34292 | 1.80152 | 7.45307 | 18.06747 | 15.83264 | 8.69249 | 12.92326 | 13.93336 | 22.46405 |
| **Tiprl** | 4.89170 | 0.04179 | 9.64771 | 6.00724 | 10.59488 | 5.58264 | 4.35341 | 0.45828 | 2.90636 | 0.12729 | 8.23794 | 5.87911 | 13.70158 | 12.72094 |
| **Tirap** | 0.24169 | 0.01878 | 0.12142 | 0.23164 | 0.09629 | 1.01798 | 0.05623 | 2.10456 | 0.75351 | 0.50612 | 0.04758 | 0.71174 | 1.78353 | 0.71094 |
| **Tjap1** | 1.19519 | 0.04076 | 3.15779 | 3.24613 | 5.90851 | 1.86803 | 0.00896 | 0.52921 | 0.01708 | 0.01447 | 0.00000 | 2.68071 | 2.88556 | 0.00000 |
| **Tjp1** | 4.10116 | 0.07765 | 3.08964 | 2.30909 | 7.07735 | 7.39078 | 2.35767 | 3.57312 | 14.21718 | 0.10797 | 4.71677 | 1.36470 | 2.41079 | 2.51965 |
| **Tjp2** | 0.00000 | 9.82286 | 6.01841 | 3.29037 | 0.09597 | 12.47022 | 5.37456 | 10.72599 | 1.42400 | 0.19615 | 4.19988 | 3.43628 | 2.78977 | 4.29346 |
| **Tjp3** | 0.00000 | 0.01135 | 0.23081 | 0.13911 | 0.70994 | 0.09907 | 1.47840 | 1.96530 | 0.00804 | 0.61817 | 0.00000 | 0.00000 | 0.48938 | 0.03085 |
| **Tk1** | 17.58352 | 4.15083 | 26.34299 | 13.89123 | 17.70691 | 22.56926 | 7.07583 | 20.44908 | 4.22339 | 9.13941 | 15.33799 | 8.20288 | 3.39405 | 18.91361 |
| **Tk2** | 0.70848 | 0.01932 | 0.00000 | 2.10851 | 0.02382 | 2.78513 | 0.00441 | 1.69451 | 0.03021 | 0.01217 | 0.38043 | 0.00000 | 1.11185 | 1.30294 |
| **Tkfc** | 0.01342 | 0.01229 | 0.04747 | 0.02281 | 0.36143 | 0.02614 | 0.00413 | 0.00753 | 0.02477 | 0.00747 | 0.00742 | 0.00754 | 0.03028 | 0.02557 |
| **Tkt** | 6.50963 | 0.05522 | 15.76060 | 2.78502 | 17.15815 | 5.96690 | 0.00000 | 1.87407 | 1.67666 | 2.34668 | 4.23578 | 5.10114 | 5.13473 | 9.51742 |
| **Tktl1** | 7.84372 | 0.00167 | 6.36113 | 1.97477 | 4.86440 | 2.81053 | 0.00148 | 0.00000 | 5.78933 | 0.02375 | 1.10685 | 0.67872 | 3.26490 | 6.12286 |
| **Tktl2** | 0.00290 | 0.00475 | 0.22733 | 0.00000 | 0.12699 | 0.00486 | 0.00395 | 0.01508 | 0.00387 | 0.00545 | 0.00793 | 0.04880 | 0.37214 | 0.21453 |
| **Tlcd1** | 2.92495 | 0.01525 | 1.64962 | 7.20448 | 3.50147 | 1.35596 | 0.36612 | 0.00000 | 0.72655 | 0.05074 | 1.25292 | 2.25900 | 1.43038 | 0.58157 |
| **Tlcd2** | 0.04738 | 0.00305 | 0.04603 | 0.00178 | 3.05156 | 0.42980 | 0.03560 | 0.00495 | 5.45112 | 0.00681 | 0.01093 | 0.04014 | 4.52217 | 16.23345 |
| **Tldc1** | 0.00000 | 0.00317 | 0.37795 | 0.35280 | 0.42558 | 0.67296 | 0.00918 | 0.00000 | 0.00550 | 0.00200 | 0.84071 | 0.95009 | 0.00292 | 0.31696 |
| **Tle1** | 1.26954 | 0.02963 | 0.38340 | 2.94997 | 0.47053 | 3.09977 | 2.53027 | 0.00313 | 0.05304 | 0.04425 | 0.00000 | 4.21380 | 2.09350 | 0.00194 |
| **Tle2** | 0.01184 | 0.00475 | 0.01736 | 0.01681 | 0.15477 | 0.02075 | 0.01032 | 0.01954 | 0.00861 | 0.00686 | 0.00622 | 0.05614 | 0.03209 | 0.06975 |
| **Tle3** | 0.48195 | 0.02582 | 1.09484 | 1.18621 | 2.69223 | 1.68619 | 0.00258 | 1.40980 | 1.53274 | 0.03784 | 3.07963 | 1.89063 | 0.06644 | 1.87799 |
| **Tle4** | 13.21052 | 1.00411 | 14.38121 | 0.00000 | 4.52874 | 10.96453 | 2.72495 | 1.74523 | 15.24344 | 5.48358 | 5.35821 | 1.43919 | 9.94792 | 6.84558 |
| **Tle6** | 2.41353 | 0.01245 | 6.05630 | 2.92616 | 4.68040 | 1.22248 | 0.00266 | 0.61994 | 0.02764 | 0.02677 | 0.55492 | 0.10216 | 2.53617 | 5.74424 |
| **Tlk1** | 4.67231 | 0.00597 | 4.59516 | 0.59089 | 4.54849 | 2.71412 | 3.07622 | 0.00000 | 12.12117 | 0.11586 | 5.60062 | 4.69983 | 1.62003 | 4.13432 |
| **Tlk2** | 6.01872 | 0.03910 | 5.11670 | 1.14568 | 10.25522 | 6.43790 | 3.08738 | 2.27822 | 13.96180 | 0.05003 | 3.49083 | 2.96221 | 9.22482 | 10.02592 |
| **Tll1** | 0.06110 | 0.00450 | 0.01368 | 0.01208 | 0.00705 | 0.01944 | 0.01729 | 0.02747 | 0.02156 | 0.00669 | 0.01113 | 0.00881 | 0.36600 | 0.15650 |
| **Tln1** | 1.39801 | 0.08746 | 6.14919 | 7.40030 | 1.92970 | 5.54029 | 2.50528 | 7.20275 | 0.00000 | 1.32076 | 3.94593 | 3.76976 | 0.17509 | 0.00000 |
| **Tln2** | 0.05267 | 0.17376 | 1.86953 | 1.57205 | 0.64360 | 2.01795 | 0.01963 | 3.54475 | 0.09871 | 0.03816 | 4.11354 | 4.08754 | 1.06365 | 0.15678 |
| **Tlr3** | 0.01048 | 0.00437 | 0.01467 | 0.01172 | 0.01153 | 0.01557 | 0.00425 | 0.00941 | 0.01087 | 0.00671 | 0.00889 | 0.00968 | 0.20260 | 0.00626 |
| **Tm2d1** | 0.76391 | 0.01130 | 0.00000 | 1.18136 | 1.17566 | 2.08469 | 0.08735 | 0.07034 | 0.55454 | 0.72380 | 0.03193 | 0.95753 | 0.00903 | 0.00000 |
| **Tm2d2** | 1.82529 | 0.03699 | 4.15083 | 2.27488 | 1.55136 | 2.61333 | 3.22430 | 8.87738 | 0.00162 | 0.06908 | 2.10323 | 2.78789 | 1.17244 | 0.00000 |
| **Tm2d3** | 3.32095 | 0.02413 | 5.41476 | 6.81429 | 4.73914 | 5.91847 | 2.69339 | 0.99885 | 0.00507 | 0.19057 | 5.20068 | 2.77341 | 2.07353 | 2.00983 |
| **Tm4sf1** | 0.01101 | 0.00443 | 0.00769 | 0.01085 | 0.35168 | 0.01341 | 0.00460 | 0.01123 | 0.01241 | 0.00656 | 0.00889 | 0.00873 | 0.00627 | 0.00828 |
| **Tm4sf5** | 0.01322 | 0.01945 | 0.17611 | 0.73142 | 0.50721 | 0.00000 | 0.00348 | 0.00414 | 0.00462 | 0.00705 | 0.00382 | 0.81794 | 0.14251 | 0.35632 |
| **Tm6sf1** | 0.00000 | 0.00350 | 1.07390 | 0.00000 | 0.32214 | 0.36834 | 0.00376 | 0.00000 | 0.00585 | 0.00342 | 0.37209 | 0.58840 | 0.09530 | 0.07457 |
| **Tm6sf2** | 0.00425 | 0.00435 | 0.42325 | 0.00000 | 0.02459 | 0.35556 | 0.13276 | 0.00157 | 0.13626 | 0.02295 | 0.00575 | 0.00664 | 0.00341 | 0.00000 |
| **Tm7sf2** | 0.00327 | 0.00400 | 0.00000 | 0.02316 | 0.36248 | 0.50505 | 0.00464 | 1.71051 | 0.00250 | 0.00566 | 0.00000 | 0.90712 | 0.08166 | 0.37666 |

| | | | | | | | | | | | | | | |
|---|---|---|---|---|---|---|---|---|---|---|---|---|---|---|
| **Tm7sf3** | 1.72937 | 0.05246 | 1.07911 | 1.18126 | 0.21926 | 3.19461 | 1.47154 | 0.69636 | 0.85106 | 0.09280 | 3.98300 | 4.32509 | 2.97711 | 1.01390 |
| **Tm9sf1** | 4.23108 | 0.00000 | 3.26600 | 0.54638 | 1.75331 | 3.47046 | 1.92945 | 0.97280 | 1.25835 | 8.34482 | 2.15611 | 1.65433 | 1.41282 | 3.28287 |
| **Tm9sf2** | 11.76489 | 0.04771 | 9.09337 | 0.93811 | 3.72929 | 12.97926 | 0.21013 | 2.16634 | 18.77440 | 5.11548 | 6.83091 | 7.48946 | 4.09420 | 3.57653 |
| **Tm9sf3** | 4.64359 | 4.52976 | 4.89380 | 2.25099 | 5.43229 | 13.21865 | 2.81376 | 1.65431 | 11.00539 | 0.01588 | 9.68612 | 8.85660 | 1.10669 | 6.40583 |
| **Tm9sf4** | 1.75536 | 0.03648 | 7.96625 | 3.33898 | 1.51499 | 7.23514 | 2.28706 | 8.00677 | 2.95356 | 0.28470 | 6.62251 | 4.67877 | 1.50355 | 3.79373 |
| **Tma16** | 12.13111 | 0.00000 | 12.89939 | 3.74292 | 14.01261 | 1.94729 | 2.55990 | 0.13196 | 7.19229 | 7.79127 | 8.29196 | 2.69823 | 7.27618 | 14.27142 |
| **Tma7** | 14.88350 | 2.40879 | 3.78380 | 22.32019 | 21.84575 | 19.84485 | 8.07881 | 16.16889 | 11.56393 | 15.37136 | 2.91225 | 11.45016 | 13.93594 | 13.03680 |
| **Tmbim1** | 10.27205 | 5.12031 | 9.85845 | 13.68497 | 3.13573 | 14.08703 | 3.95827 | 10.54616 | 2.09670 | 0.30788 | 7.58659 | 9.48323 | 5.41484 | 1.47099 |
| **Tmbim4** | 2.33184 | 0.02910 | 4.42671 | 6.43992 | 1.18813 | 2.65375 | 0.16635 | 3.19812 | 0.55150 | 0.04361 | 2.66468 | 6.93930 | 0.06350 | 0.00069 |
| **Tmbim6** | 3.64040 | 5.60611 | 13.71070 | 10.86535 | 5.81782 | 13.74446 | 7.01886 | 15.44517 | 0.00000 | 10.75795 | 11.71901 | 12.02108 | 6.27395 | 0.81343 |
| **Tmc2** | 0.08512 | 0.02636 | 0.05744 | 0.07980 | 0.04024 | 0.20050 | 1.73814 | 0.03989 | 0.08503 | 0.17948 | 0.00782 | 0.06959 | 0.19785 | 0.28443 |
| **Tmc4** | 0.73263 | 0.00364 | 0.69934 | 0.00000 | 1.46470 | 0.08082 | 0.00395 | 0.00102 | 0.08929 | 0.00489 | 0.00474 | 0.27361 | 1.28131 | 0.28772 |
| **Tmc5** | 0.34432 | 0.00447 | 0.00000 | 0.00754 | 0.78002 | 0.00515 | 0.00449 | 0.01217 | 0.87329 | 0.00765 | 0.00424 | 0.00585 | 0.00265 | 0.00000 |
| **Tmc6** | 0.17916 | 0.23170 | 6.80277 | 11.31060 | 2.01038 | 4.75798 | 3.36709 | 0.49832 | 0.19840 | 0.09405 | 6.19021 | 2.09653 | 0.58034 | 0.41866 |
| **Tmc7** | 0.01038 | 0.00443 | 0.12978 | 0.00688 | 0.00443 | 0.00674 | 0.00414 | 0.01061 | 0.24574 | 0.00800 | 0.00689 | 0.01293 | 0.00550 | 0.00235 |
| **Tmc8** | 0.00305 | 0.00426 | 0.00600 | 0.07115 | 0.20121 | 0.05764 | 0.01223 | 0.29101 | 0.05041 | 0.00635 | 0.01007 | 0.01681 | 0.35992 | 0.01309 |
| **Tmcc1** | 0.94458 | 0.00272 | 0.19872 | 0.71983 | 3.13175 | 1.14196 | 1.79597 | 0.01022 | 3.36231 | 0.06573 | 0.00000 | 0.94304 | 1.66879 | 5.80580 |
| **Tmcc2** | 0.91887 | 0.00409 | 0.01169 | 0.00757 | 0.02411 | 0.02188 | 0.00461 | 0.00758 | 0.00594 | 0.00644 | 0.00642 | 0.00573 | 1.53839 | 0.06628 |
| **Tmcc3** | 0.00000 | 0.00380 | 0.00000 | 0.29115 | 0.00000 | 0.00000 | 0.00371 | 0.00564 | 3.18717 | 0.01323 | 0.00253 | 0.00404 | 0.47852 | 6.82555 |
| **Tmco1** | 9.69025 | 8.75699 | 16.55235 | 15.70646 | 11.33218 | 18.08358 | 6.32617 | 13.77618 | 6.75908 | 10.28966 | 12.53834 | 12.25451 | 8.64414 | 11.82048 |
| **Tmco2** | 0.38226 | 0.00419 | 0.10713 | 0.00000 | 0.18140 | 0.09996 | 0.00431 | 0.00490 | 0.01272 | 0.00575 | 0.00518 | 0.01785 | 0.80032 | 0.18528 |
| **Tmco3** | 3.03071 | 0.05902 | 1.35369 | 0.72328 | 0.47818 | 2.31402 | 0.04847 | 0.98699 | 0.58308 | 0.05449 | 0.00000 | 0.60932 | 2.52631 | 1.03854 |
| **Tmco4** | 0.32854 | 0.03273 | 0.38295 | 0.80257 | 0.00000 | 3.92897 | 1.96583 | 1.79958 | 0.00071 | 0.10140 | 0.04553 | 0.61298 | 0.79609 | 0.62835 |
| **Tmco6** | 0.00492 | 0.00438 | 0.04812 | 0.08967 | 0.09852 | 0.03390 | 0.00421 | 0.00815 | 0.01784 | 0.00578 | 0.17528 | 0.00488 | 0.02766 | 0.03640 |
| **Tmed1** | 2.89708 | 4.22065 | 13.86487 | 5.81629 | 6.00925 | 6.79101 | 2.40970 | 1.47308 | 0.12999 | 0.07166 | 2.85199 | 4.87582 | 0.00000 | 3.74312 |
| **Tmed10** | 13.02085 | 7.32712 | 21.68955 | 18.09257 | 17.19920 | 20.64875 | 5.86086 | 13.50077 | 5.95290 | 13.64190 | 11.65229 | 15.79389 | 6.36507 | 15.74824 |
| **Tmed2** | 10.91564 | 7.20485 | 14.27959 | 9.39741 | 12.30786 | 16.47099 | 5.66957 | 6.02960 | 15.14525 | 0.15989 | 3.95276 | 9.83414 | 6.19289 | 18.57758 |
| **Tmed3** | 0.91626 | 0.00419 | 0.29143 | 0.00000 | 0.94352 | 0.36945 | 0.00344 | 1.91594 | 0.02466 | 0.00694 | 0.09628 | 0.40579 | 0.14194 | 0.15627 |
| **Tmed4** | 3.82000 | 0.16886 | 5.80654 | 4.27900 | 3.73461 | 2.09065 | 1.91713 | 5.46134 | 5.79724 | 0.11134 | 4.02989 | 3.31093 | 2.84090 | 7.99644 |
| **Tmed5** | 8.74655 | 2.90209 | 11.61173 | 5.25719 | 12.09826 | 8.75696 | 4.55409 | 1.13951 | 28.03376 | 0.13005 | 3.31413 | 2.12864 | 10.72883 | 3.01367 |
| **Tmed6** | 0.00000 | 0.00303 | 0.22424 | 0.59415 | 2.21909 | 0.00000 | 0.00847 | 0.00118 | 1.15983 | 2.92399 | 0.00000 | 0.00000 | 1.66969 | 4.45606 |
| **Tmed7** | 4.69622 | 0.00070 | 4.55232 | 0.88929 | 2.14260 | 5.63702 | 1.46880 | 3.53513 | 7.50994 | 0.02148 | 5.39560 | 1.83139 | 0.49956 | 1.68214 |
| **Tmed8** | 1.49483 | 0.00325 | 0.82409 | 1.55888 | 0.40183 | 2.13165 | 0.00315 | 0.00115 | 3.62547 | 0.00432 | 0.00000 | 0.59710 | 1.74483 | 0.23852 |
| **Tmed9** | 1.39459 | 0.17760 | 3.97583 | 0.64484 | 4.28295 | 5.17671 | 1.11910 | 0.79202 | 1.15185 | 0.04138 | 0.04760 | 2.80776 | 9.45762 | 2.57898 |
| **Tmeff1** | 0.50751 | 0.00000 | 1.03792 | 0.00000 | 0.00000 | 1.73729 | 1.92575 | 0.00000 | 0.10946 | 10.04375 | 2.42288 | 3.62075 | 1.11186 | 0.00000 |
| **Tmem101** | 0.49135 | 0.06082 | 5.46662 | 6.19891 | 1.97164 | 3.13503 | 1.61830 | 0.45531 | 0.01300 | 0.01582 | 2.13454 | 4.44931 | 0.00000 | 5.62057 |
| **Tmem104** | 0.19692 | 0.04165 | 0.00000 | 2.05528 | 0.40212 | 0.67733 | 0.01053 | 0.68654 | 0.00681 | 0.00747 | 0.10109 | 0.80158 | 0.04851 | 0.03192 |
| **Tmem106a** | 0.00000 | 0.03880 | 1.31744 | 1.71090 | 1.19131 | 0.00000 | 0.00278 | 0.08195 | 0.01471 | 0.04521 | 0.68106 | 2.83176 | 1.37802 | 1.54855 |
| **Tmem106b** | 0.58390 | 0.00458 | 0.30071 | 0.60829 | 0.34888 | 0.87487 | 0.48625 | 0.29515 | 1.91991 | 0.02796 | 1.32212 | 0.38452 | 0.00000 | 0.50140 |
| **Tmem106c** | 0.00343 | 0.03211 | 0.00000 | 0.74840 | 0.62140 | 0.94823 | 0.01110 | 0.29468 | 0.02588 | 0.01448 | 0.01072 | 1.33657 | 1.05741 | 0.01721 |
| **Tmem107** | 0.24421 | 0.01812 | 0.41359 | 0.70987 | 0.06129 | 0.52869 | 0.00946 | 0.00000 | 0.00674 | 0.00385 | 1.08556 | 1.48843 | 0.98563 | 0.00000 |
| **Tmem108** | 0.02857 | 0.00482 | 0.01457 | 0.01217 | 0.01374 | 0.01132 | 0.00444 | 0.02039 | 0.01098 | 0.00771 | 0.00865 | 0.00964 | 0.05913 | 0.00794 |
| **Tmem109** | 1.29648 | 0.02453 | 1.31615 | 2.36894 | 0.74193 | 3.58999 | 0.03147 | 5.64005 | 0.00329 | 5.47811 | 0.00000 | 1.36162 | 0.99069 | 0.08339 |
| **Tmem11** | 8.02198 | 2.47284 | 16.70719 | 13.12007 | 11.43571 | 13.94266 | 3.75468 | 9.78457 | 2.71605 | 0.20646 | 11.16612 | 5.10857 | 7.82661 | 10.09598 |
| **Tmem110** | 0.37689 | 0.00323 | 0.36386 | 0.84321 | 0.37756 | 2.13673 | 0.00383 | 0.24061 | 0.00389 | 0.01542 | 2.63576 | 0.02351 | 0.36075 | 0.53917 |
| **Tmem115** | 0.67126 | 0.07334 | 2.71005 | 1.44451 | 0.60308 | 3.57525 | 4.31715 | 4.53506 | 0.20214 | 0.21065 | 0.09430 | 2.70295 | 0.31778 | 1.04764 |
| **Tmem116** | 0.58468 | 0.01860 | 0.35271 | 0.12927 | 0.00000 | 1.40489 | 2.15339 | 0.79943 | 0.00190 | 0.05664 | 0.61389 | 2.10940 | 0.45629 | 0.30512 |
| **Tmem117** | 0.00000 | 0.00431 | 0.00111 | 0.01565 | 0.00000 | 0.00745 | 0.00396 | 0.00848 | 1.29587 | 0.00669 | 0.00622 | 0.00822 | 0.66265 | 0.00000 |
| **Tmem120a** | 1.74291 | 4.26135 | 9.45425 | 17.26997 | 15.81316 | 5.16219 | 2.58342 | 6.89453 | 0.09301 | 0.06895 | 0.27283 | 0.38575 | 4.00900 | 10.21743 |
| **Tmem120b** | 0.03757 | 0.02882 | 1.59406 | 1.51845 | 1.62359 | 0.04764 | 0.00341 | 0.00000 | 0.00838 | 0.00525 | 1.00356 | 0.29465 | 0.86937 | 0.00000 |
| **Tmem123** | 2.40883 | 0.01040 | 0.32686 | 1.23979 | 3.13371 | 0.78429 | 0.00252 | 1.18530 | 6.07446 | 0.00118 | 1.87789 | 0.59737 | 1.00605 | 1.48892 |
| **Tmem125** | 0.00032 | 0.95484 | 0.06874 | 0.25863 | 0.00101 | 0.06588 | 3.56572 | 4.22657 | 0.01477 | 0.04715 | 0.37462 | 0.34676 | 0.00608 | 0.00532 |
| **Tmem126a** | 0.00000 | 0.03196 | 0.29529 | 0.00000 | 1.42326 | 0.19232 | 1.00921 | 3.82230 | 0.26450 | 5.67605 | 0.86800 | 4.82231 | 0.02445 | 0.55340 |
| **Tmem126b** | 0.89587 | 0.00225 | 0.15841 | 1.94753 | 0.98891 | 0.44185 | 0.43620 | 1.60844 | 0.14853 | 0.01761 | 1.35198 | 1.90112 | 0.01181 | 0.35607 |
| **Tmem127** | 0.17377 | 0.01178 | 0.03998 | 0.39615 | 0.63971 | 2.96734 | 0.01016 | 1.34403 | 1.43875 | 1.35585 | 0.34052 | 0.96112 | 0.00787 | 1.35596 |

| | | | | | | | | | | | | | | |
|---|---|---|---|---|---|---|---|---|---|---|---|---|---|---|
| **Tmem128** | 2.68226 | 0.08972 | 2.07410 | 2.72493 | 1.51011 | 2.99592 | 0.41902 | 2.99712 | 2.13430 | 0.03476 | 1.75159 | 2.31500 | 0.62716 | 7.28663 |
| **Tmem129** | 1.93241 | 0.01734 | 2.83667 | 0.00000 | 1.83765 | 1.58847 | 0.00253 | 2.59789 | 1.29475 | 0.01071 | 0.58273 | 1.24972 | 2.91122 | 0.28936 |
| **Tmem130** | 0.01007 | 0.01184 | 0.35170 | 0.03305 | 0.00000 | 0.00000 | 0.00413 | 0.01034 | 0.01104 | 0.01081 | 0.00582 | 0.00718 | 0.20075 | 1.42005 |
| **Tmem131** | 2.38017 | 0.38681 | 4.87985 | 4.22607 | 0.30167 | 4.22240 | 0.01073 | 3.34603 | 0.06308 | 0.03267 | 2.75042 | 9.91686 | 0.00000 | 1.17721 |
| **Tmem132b** | 0.00000 | 0.00394 | 0.23388 | 0.14219 | 0.18439 | 0.00124 | 0.00388 | 0.00658 | 0.00532 | 0.00533 | 0.01680 | 0.01328 | 1.33599 | 0.02819 |
| **Tmem132c** | 0.00197 | 0.00446 | 0.00000 | 0.49020 | 0.06406 | 0.56314 | 0.00408 | 0.00448 | 0.08651 | 0.00519 | 0.00276 | 0.00693 | 0.20945 | 0.00000 |
| **Tmem134** | 0.08892 | 0.05404 | 0.00000 | 2.79630 | 0.85610 | 0.27424 | 0.01060 | 2.84245 | 0.01859 | 0.04946 | 0.06239 | 0.76690 | 2.45338 | 0.22047 |
| **Tmem135** | 0.50714 | 0.00484 | 0.92080 | 2.73046 | 0.00100 | 1.50513 | 0.01795 | 0.39234 | 2.12548 | 0.00569 | 0.00000 | 0.41113 | 0.80082 | 0.00000 |
| **Tmem136** | 0.00521 | 0.06094 | 0.00599 | 0.21478 | 0.02271 | 0.11649 | 0.00457 | 0.05965 | 0.02323 | 0.00686 | 0.02314 | 0.16965 | 0.07088 | 0.19602 |
| **Tmem138** | 0.37829 | 8.03517 | 2.19018 | 2.31638 | 2.28785 | 2.46657 | 0.00644 | 2.56913 | 1.23233 | 0.02912 | 1.87351 | 2.01909 | 3.74850 | 1.69132 |
| **Tmem139** | 0.00000 | 0.01936 | 0.28707 | 0.47931 | 0.01124 | 2.71324 | 0.90601 | 3.45888 | 0.00681 | 0.05993 | 0.19452 | 0.00000 | 0.02602 | 0.00000 |
| **Tmem141** | 0.02514 | 0.02498 | 0.61212 | 3.24458 | 1.44110 | 2.20528 | 0.67417 | 0.02924 | 0.27497 | 0.03459 | 0.26049 | 2.62127 | 0.00000 | 1.55244 |
| **Tmem143** | 0.00533 | 0.00405 | 0.23384 | 0.00000 | 0.17335 | 0.09160 | 0.13936 | 0.00229 | 0.00771 | 0.01060 | 0.01718 | 0.14655 | 0.16281 | 0.00445 |
| **Tmem144** | 0.29604 | 0.00291 | 1.23205 | 1.69144 | 0.03910 | 3.33129 | 0.00254 | 3.04121 | 1.28161 | 0.01612 | 5.69827 | 0.27311 | 0.01728 | 0.58428 |
| **Tmem145** | 0.04852 | 0.00456 | 0.03179 | 0.14885 | 0.37122 | 0.01481 | 0.00449 | 0.02486 | 0.04396 | 0.01058 | 0.00827 | 0.00539 | 0.03892 | 0.04035 |
| **Tmem147** | 3.02785 | 0.13520 | 2.85820 | 14.07186 | 0.89285 | 7.70259 | 2.62383 | 7.40888 | 0.00000 | 0.76546 | 3.42818 | 11.07826 | 0.06399 | 0.35495 |
| **Tmem14a** | 0.01202 | 0.00396 | 0.00697 | 0.00059 | 0.00627 | 0.38719 | 2.15280 | 0.00175 | 0.01150 | 0.01095 | 0.37955 | 0.00619 | 0.36167 | 0.00123 |
| **Tmem14c** | 2.58127 | 6.10238 | 4.18315 | 9.33867 | 6.90006 | 4.41261 | 5.25100 | 12.86163 | 0.18680 | 0.16652 | 4.03379 | 8.90694 | 1.66114 | 4.22232 |
| **Tmem150a** | 0.42059 | 0.06851 | 0.69355 | 1.73422 | 0.00000 | 3.92292 | 1.87157 | 3.15096 | 0.00000 | 0.05296 | 0.21938 | 3.47005 | 0.35046 | 0.00000 |
| **Tmem150c** | 0.00000 | 0.00416 | 0.00000 | 0.01407 | 0.18067 | 0.00649 | 0.00411 | 0.00793 | 0.39790 | 0.00637 | 0.00623 | 0.00649 | 1.41277 | 0.84310 |
| **Tmem158** | 0.02526 | 0.00447 | 0.05250 | 0.02322 | 0.10439 | 0.01437 | 0.01085 | 0.01548 | 0.01047 | 0.00800 | 0.00781 | 0.04915 | 0.00689 | 0.01221 |
| **Tmem159** | 11.26575 | 7.87407 | 21.79265 | 16.42129 | 11.93747 | 25.42785 | 2.16048 | 6.51817 | 1.43707 | 0.50176 | 13.69324 | 8.30667 | 7.18688 | 16.47513 |
| **Tmem160** | 0.04123 | 0.00452 | 0.02129 | 0.01083 | 0.24809 | 0.03505 | 0.00462 | 0.03052 | 0.00694 | 0.01669 | 0.00674 | 0.03920 | 0.23440 | 0.01056 |
| **Tmem161a** | 0.85482 | 0.01869 | 0.09058 | 4.00795 | 0.11137 | 0.43956 | 0.00981 | 0.74341 | 0.00452 | 0.01923 | 0.00000 | 3.52893 | 0.11241 | 0.17972 |
| **Tmem161b** | 0.06183 | 0.00502 | 0.00000 | 0.22368 | 0.00000 | 1.61214 | 0.00360 | 0.03500 | 6.85607 | 0.02931 | 0.00465 | 0.00000 | 0.71289 | 1.16505 |
| **Tmem164** | 0.00000 | 0.00389 | 1.25650 | 1.13791 | 1.03277 | 0.10922 | 1.93768 | 0.00000 | 1.34699 | 0.01862 | 0.00000 | 0.51410 | 4.78730 | 2.27161 |
| **Tmem165** | 1.06037 | 0.02739 | 0.00000 | 2.08983 | 0.58324 | 3.56358 | 0.00903 | 1.55316 | 1.13329 | 0.02298 | 0.61495 | 3.47224 | 1.31672 | 0.68014 |
| **Tmem167** | 10.15887 | 0.05809 | 5.03232 | 8.52200 | 11.25571 | 11.93672 | 2.13683 | 5.51905 | 12.94215 | 10.20271 | 2.85145 | 5.18778 | 3.67438 | 4.53196 |
| **Tmem167b** | 4.15518 | 0.02802 | 2.70060 | 3.47339 | 0.89569 | 1.42356 | 0.00272 | 1.40143 | 4.18096 | 0.00000 | 3.99680 | 0.48845 | 1.22031 | 7.46037 |
| **Tmem168** | 0.71261 | 0.01045 | 1.37425 | 1.16972 | 0.06221 | 3.15770 | 2.27659 | 1.37950 | 6.14633 | 0.06055 | 0.46080 | 1.32231 | 0.43551 | 1.60834 |
| **Tmem169** | 0.00954 | 0.00443 | 0.01152 | 0.01543 | 0.00595 | 0.01368 | 0.00426 | 0.00961 | 0.01776 | 0.00721 | 0.00790 | 0.00969 | 0.01076 | 0.21594 |
| **Tmem17** | 0.00000 | 0.00463 | 0.69169 | 2.55854 | 0.00000 | 0.32887 | 0.00361 | 0.12511 | 0.55019 | 0.00591 | 0.00000 | 1.13426 | 0.00247 | 0.28597 |
| **Tmem170** | 0.21661 | 0.00334 | 0.34675 | 0.63675 | 1.42090 | 0.56658 | 0.00356 | 0.60211 | 0.00000 | 0.01722 | 1.49063 | 0.00459 | 3.75078 | 0.95029 |
| **Tmem170b** | 0.40299 | 0.02070 | 0.00000 | 0.06564 | 0.00000 | 0.45622 | 0.92460 | 0.62479 | 0.97614 | 0.02418 | 0.00298 | 0.18229 | 0.01610 | 0.01546 |
| **Tmem171** | 0.00409 | 0.00404 | 0.00000 | 0.00765 | 0.00651 | 1.18341 | 0.03284 | 1.19792 | 0.00938 | 0.02654 | 1.35526 | 0.05838 | 0.00720 | 0.02073 |
| **Tmem173** | 0.00176 | 0.00393 | 0.01358 | 0.01314 | 0.02383 | 0.01178 | 0.00405 | 0.00838 | 0.02332 | 0.00614 | 0.00910 | 0.00701 | 3.53972 | 0.00000 |
| **Tmem175** | 0.06729 | 0.01025 | 0.80560 | 3.34701 | 0.00000 | 4.13543 | 0.14919 | 3.77444 | 0.01165 | 0.02145 | 0.11444 | 1.41570 | 0.26231 | 2.52088 |
| **Tmem176b** | 0.02274 | 0.00450 | 0.00377 | 0.00777 | 1.55749 | 0.01292 | 0.00466 | 0.01062 | 0.01347 | 0.00649 | 0.00833 | 0.00730 | 0.00220 | 0.00515 |
| **Tmem177** | 0.00000 | 0.01238 | 0.41564 | 2.21828 | 0.15711 | 2.26989 | 0.00472 | 0.85292 | 0.00590 | 0.01787 | 0.00000 | 0.00000 | 0.15935 | 2.41636 |
| **Tmem179b** | 0.90921 | 0.03259 | 0.86319 | 2.93849 | 0.16814 | 2.72033 | 0.03402 | 5.64650 | 0.00688 | 0.05927 | 1.66134 | 4.55764 | 0.92081 | 0.33158 |
| **Tmem18** | 1.26126 | 0.04831 | 0.35077 | 0.00000 | 0.17184 | 1.98969 | 0.00360 | 0.52616 | 0.30687 | 0.01117 | 2.49094 | 1.82493 | 0.21441 | 0.90201 |
| **Tmem181a** | 0.25241 | 0.03635 | 0.67236 | 1.30880 | 0.25503 | 0.52682 | 0.00441 | 0.04044 | 0.03473 | 0.00483 | 0.05108 | 0.13091 | 0.00184 | 0.23821 |
| **Tmem181b.p** | 2.14378 | 1.21037 | 1.46734 | 2.26615 | 0.27438 | 1.86402 | 2.23593 | 0.45215 | 2.41494 | 0.05026 | 0.52139 | 0.39976 | 0.13584 | 0.59695 |
| **Tmem181c.p** | 0.03443 | 0.00431 | 0.39975 | 0.52666 | 0.06449 | 0.41767 | 1.31580 | 0.13580 | 0.24083 | 0.00530 | 0.05437 | 0.24023 | 0.42101 | 0.10453 |
| **Tmem183a** | 6.59617 | 7.03728 | 9.73527 | 8.25469 | 10.07878 | 10.47156 | 0.02793 | 4.87128 | 5.44559 | 0.56852 | 0.55524 | 8.76087 | 3.05393 | 9.17065 |
| **Tmem184a** | 0.54624 | 0.00413 | 2.14122 | 0.03928 | 1.20519 | 0.00000 | 0.00372 | 0.01148 | 0.00904 | 0.00490 | 0.00482 | 0.00501 | 0.02360 | 0.91894 |
| **Tmem184b** | 0.00000 | 0.01701 | 2.17852 | 0.61264 | 0.00000 | 0.09023 | 0.00957 | 1.26168 | 0.00728 | 0.00304 | 0.01880 | 3.23436 | 0.01184 | 0.25787 |
| **Tmem184c** | 2.26324 | 0.00193 | 8.98248 | 3.01526 | 1.92937 | 1.94809 | 0.00715 | 0.50114 | 14.84628 | 0.02711 | 0.44509 | 0.99145 | 10.61971 | 7.28454 |
| **Tmem185a** | 0.00000 | 0.02266 | 0.70070 | 1.95669 | 1.58456 | 2.58719 | 0.00371 | 0.66102 | 2.83116 | 0.02685 | 0.21327 | 0.76396 | 0.86481 | 2.81752 |
| **Tmem185b** | 2.48411 | 0.01774 | 3.91181 | 4.05654 | 1.83450 | 2.19933 | 2.41008 | 0.75502 | 7.85319 | 0.52487 | 0.05311 | 2.04395 | 0.00000 | 2.12800 |
| **Tmem186** | 0.61734 | 0.02326 | 1.17028 | 1.69876 | 1.75335 | 1.13621 | 3.54811 | 1.28240 | 0.04988 | 0.04699 | 0.65111 | 0.04508 | 0.96445 | 3.46793 |
| **Tmem189** | 0.03795 | 0.04114 | 2.88883 | 3.69440 | 2.56824 | 1.27824 | 0.00283 | 4.57802 | 0.63281 | 0.01569 | 0.33840 | 0.01828 | 2.63651 | 2.71160 |
| **Tmem19** | 0.76803 | 0.01570 | 1.73504 | 2.94533 | 1.41666 | 4.45409 | 0.23092 | 1.04582 | 0.00783 | 0.02087 | 1.61891 | 4.68985 | 0.57504 | 1.87605 |
| **Tmem190** | 0.00716 | 0.00441 | 0.01481 | 0.01265 | 0.01306 | 0.01325 | 0.00428 | 0.00983 | 0.01366 | 0.00666 | 0.00891 | 0.00976 | 0.13195 | 0.00427 |
| **Tmem191c** | 0.33231 | 0.02377 | 5.93921 | 13.79045 | 1.54807 | 0.00001 | 1.13086 | 0.50683 | 0.00000 | 0.05164 | 0.01486 | 1.13930 | 0.50307 | 0.00000 |

| | | | | | | | | | | | | | | |
|---|---|---|---|---|---|---|---|---|---|---|---|---|---|---|
| **Tmem192** | 4.78599 | 0.04083 | 3.78112 | 5.91534 | 7.99202 | 4.43751 | 0.80367 | 0.25613 | 0.01247 | 10.08359 | 3.31087 | 3.30195 | 0.69278 | 2.09980 |
| **Tmem198** | 0.00000 | 0.00362 | 0.69505 | 1.05787 | 0.00000 | 0.00016 | 0.00376 | 0.00084 | 0.01507 | 0.00503 | 0.01043 | 0.42853 | 0.00635 | 0.00425 |
| **Tmem198b** | 0.00476 | 0.00377 | 0.25808 | 0.17282 | 0.00000 | 0.17917 | 0.00374 | 0.45447 | 0.43162 | 0.00435 | 0.00000 | 0.26406 | 0.01905 | 0.00000 |
| **Tmem199** | 8.83281 | 0.01450 | 9.46464 | 4.84516 | 4.14480 | 11.65396 | 0.97101 | 0.05580 | 3.36361 | 0.06268 | 7.94743 | 5.14623 | 2.36575 | 4.54952 |
| **Tmem2** | 0.35705 | 0.00387 | 0.70517 | 0.30551 | 0.95325 | 0.00000 | 0.01140 | 0.73552 | 2.83228 | 0.00276 | 0.93568 | 0.25002 | 0.96958 | 0.54895 |
| **Tmem201** | 0.66806 | 0.05739 | 6.94770 | 4.87079 | 13.14285 | 2.83666 | 0.10597 | 3.46589 | 2.87021 | 0.01677 | 1.44789 | 0.15186 | 8.22668 | 4.00765 |
| **Tmem203** | 0.23885 | 0.01045 | 0.14870 | 0.45067 | 0.46031 | 0.10858 | 0.00381 | 0.00698 | 0.00642 | 0.00658 | 0.00385 | 0.73056 | 0.96165 | 0.08347 |
| **Tmem205** | 0.01303 | 0.02851 | 0.00000 | 0.95393 | 0.09286 | 0.38224 | 0.00412 | 0.22845 | 0.02174 | 0.04528 | 1.13589 | 1.92556 | 0.00597 | 0.00000 |
| **Tmem206** | 0.91102 | 0.05084 | 0.00000 | 0.73107 | 0.83759 | 2.14769 | 0.25974 | 0.20144 | 1.01622 | 0.90116 | 0.01118 | 0.34172 | 7.73390 | 1.70366 |
| **Tmem208** | 6.43147 | 4.99717 | 4.26219 | 9.94861 | 6.64406 | 9.67477 | 5.07624 | 2.99503 | 1.97846 | 9.82482 | 1.17156 | 9.93924 | 4.28935 | 4.19276 |
| **Tmem209** | 11.45038 | 0.03771 | 9.68738 | 3.48783 | 3.34350 | 7.75649 | 0.79159 | 2.16506 | 15.75229 | 7.65120 | 6.74987 | 7.09939 | 2.04732 | 13.03939 |
| **Tmem213** | 0.00587 | 0.00430 | 0.00754 | 0.00000 | 0.00650 | 0.37035 | 0.00402 | 0.72478 | 0.00745 | 0.01351 | 0.00471 | 0.01217 | 0.00459 | 0.18404 |
| **Tmem214** | 0.95497 | 0.08102 | 5.70563 | 4.99986 | 0.68383 | 10.20594 | 3.65691 | 0.60520 | 0.00560 | 15.47263 | 3.45818 | 3.75533 | 0.74895 | 2.90722 |
| **Tmem215** | 0.00571 | 0.00437 | 0.01498 | 0.01234 | 0.01111 | 0.01344 | 0.00426 | 0.00941 | 0.01089 | 0.00661 | 0.00902 | 0.00952 | 0.24704 | 0.00341 |
| **Tmem216** | 1.37564 | 0.03260 | 1.41556 | 0.73907 | 4.45910 | 5.41899 | 2.24671 | 3.87051 | 4.53792 | 0.06713 | 1.66415 | 1.76439 | 0.73724 | 5.14017 |
| **Tmem218** | 0.00310 | 0.00380 | 0.02082 | 0.00000 | 1.05218 | 0.63306 | 0.00452 | 0.00413 | 0.01372 | 0.01156 | 0.00000 | 0.58530 | 0.00476 | 0.05209 |
| **Tmem219** | 0.35774 | 0.04093 | 1.27804 | 2.20630 | 0.00000 | 1.72664 | 3.08503 | 2.58508 | 0.00131 | 0.06622 | 0.01203 | 0.81182 | 0.02889 | 0.61862 |
| **Tmem222** | 1.21682 | 8.08430 | 7.67722 | 15.38929 | 3.22838 | 11.68175 | 6.11889 | 5.71114 | 3.08326 | 0.37591 | 6.69980 | 6.63830 | 2.65763 | 4.34592 |
| **Tmem223** | 1.40699 | 0.03350 | 2.42652 | 1.61956 | 7.78557 | 1.61777 | 0.33361 | 3.89991 | 0.06473 | 0.04601 | 1.25017 | 3.71070 | 4.70928 | 0.02271 |
| **Tmem229b** | 11.38112 | 0.00783 | 6.81694 | 3.02555 | 17.37669 | 2.65088 | 0.04875 | 0.00000 | 11.06581 | 0.00585 | 1.52867 | 1.11479 | 3.99077 | 26.72253 |
| **Tmem230** | 0.02192 | 0.00854 | 1.08925 | 5.31093 | 0.40041 | 4.62120 | 2.55397 | 0.52898 | 0.00283 | 0.05250 | 5.98186 | 2.07216 | 0.00000 | 2.97841 |
| **Tmem231** | 2.55379 | 0.22426 | 7.14532 | 13.11517 | 5.52264 | 6.90463 | 2.26960 | 9.22512 | 4.15716 | 0.19185 | 11.03015 | 7.20149 | 6.91543 | 6.29461 |
| **Tmem232** | 0.00494 | 0.00437 | 0.00115 | 0.06113 | 0.00610 | 0.00828 | 0.00438 | 0.00926 | 0.33343 | 0.00624 | 0.00628 | 0.00875 | 0.00980 | 0.10650 |
| **Tmem234** | 0.89961 | 0.82792 | 3.70875 | 4.09769 | 2.06710 | 1.21713 | 0.09847 | 2.63957 | 0.22271 | 0.04367 | 0.58912 | 2.28760 | 0.46120 | 0.83104 |
| **Tmem235** | 0.17339 | 0.00418 | 0.06456 | 0.18220 | 0.00000 | 0.00686 | 0.00409 | 0.00622 | 0.00620 | 0.01054 | 0.01132 | 0.01175 | 0.00812 | 0.46637 |
| **Tmem237** | 0.02607 | 0.00477 | 0.14292 | 0.33500 | 0.37195 | 0.59885 | 0.00399 | 0.41659 | 0.00498 | 0.01682 | 0.00257 | 0.03621 | 0.02411 | 0.29199 |
| **Tmem241** | 0.00000 | 0.00444 | 0.54989 | 0.00000 | 0.09347 | 0.60001 | 1.51408 | 0.00352 | 0.49370 | 0.00972 | 0.00469 | 0.00257 | 0.00092 | 0.00000 |
| **Tmem242** | 2.68749 | 0.11816 | 4.95094 | 2.07095 | 5.43708 | 6.43014 | 0.01996 | 4.51688 | 0.74881 | 0.05464 | 5.22728 | 5.19105 | 0.08275 | 3.63862 |
| **Tmem243** | 0.00603 | 0.00367 | 0.01770 | 0.03631 | 0.03563 | 0.00929 | 0.00392 | 0.00737 | 0.01537 | 0.00734 | 0.25774 | 0.75279 | 0.01575 | 0.01549 |
| **Tmem245** | 0.14076 | 0.04271 | 1.50609 | 2.28701 | 0.28668 | 0.33195 | 2.88826 | 0.27706 | 2.05171 | 0.08374 | 0.89973 | 1.71405 | 0.54003 | 1.07050 |
| **Tmem246** | 0.00982 | 0.00447 | 0.03978 | 0.00799 | 0.07222 | 0.01199 | 0.00422 | 0.01074 | 0.01103 | 0.00651 | 0.00882 | 0.00949 | 0.01549 | 0.03837 |
| **Tmem248** | 2.41511 | 0.41269 | 2.13873 | 1.28388 | 3.34815 | 5.74249 | 4.86518 | 6.43782 | 7.40770 | 0.30819 | 1.89963 | 6.51685 | 3.46678 | 8.32432 |
| **Tmem25** | 0.00652 | 0.00427 | 0.00889 | 0.00910 | 0.04388 | 0.00108 | 0.00430 | 0.01376 | 0.01630 | 0.00585 | 0.21432 | 0.00614 | 0.03838 | 0.06532 |
| **Tmem251** | 1.51087 | 0.00265 | 1.30725 | 1.20797 | 1.00454 | 1.93799 | 0.01020 | 2.19844 | 2.60793 | 0.03010 | 0.08374 | 1.86183 | 4.09128 | 3.07738 |
| **Tmem253** | 0.07479 | 0.00442 | 0.34521 | 0.22245 | 0.00000 | 0.23763 | 0.00386 | 0.00241 | 0.08381 | 0.00532 | 0.00558 | 0.04185 | 0.00314 | 0.00000 |
| **Tmem254a** | 0.00557 | 2.17260 | 0.01411 | 0.11085 | 0.04993 | 0.05205 | 0.00386 | 0.00395 | 0.00516 | 0.00547 | 0.01216 | 0.00642 | 0.00281 | 0.21222 |
| **Tmem255b** | 0.00541 | 0.00442 | 0.00760 | 0.01135 | 0.37625 | 0.01268 | 0.00434 | 0.01070 | 0.00588 | 0.00649 | 0.00896 | 0.00838 | 0.15473 | 0.05002 |
| **Tmem256** | 0.00359 | 0.05353 | 0.00000 | 0.00000 | 1.00919 | 1.08597 | 2.47919 | 1.88637 | 0.01833 | 1.21373 | 0.04584 | 3.30302 | 0.01964 | 0.00000 |
| **Tmem258** | 1.28255 | 2.01658 | 0.03053 | 0.33365 | 0.91555 | 3.28105 | 0.31008 | 3.59446 | 0.23176 | 8.00344 | 0.00000 | 3.33253 | 3.68814 | 0.90822 |
| **Tmem259** | 0.03825 | 0.02838 | 0.03053 | 0.51132 | 1.39443 | 0.07787 | 0.01092 | 0.04267 | 0.00798 | 0.03219 | 0.01298 | 0.06330 | 0.13865 | 0.06656 |
| **Tmem260** | 0.00334 | 0.00474 | 0.00435 | 1.22350 | 0.09903 | 0.10567 | 0.00424 | 0.00734 | 0.00832 | 0.00588 | 0.00686 | 0.00884 | 0.18274 | 0.00000 |
| **Tmem261** | 0.79734 | 0.04651 | 0.62781 | 2.68511 | 1.72545 | 1.32841 | 2.45306 | 3.05268 | 0.00000 | 0.17760 | 1.17476 | 4.37554 | 0.00000 | 0.18434 |
| **Tmem263** | 1.16760 | 0.06693 | 1.19908 | 1.06434 | 1.18562 | 7.90520 | 3.13042 | 1.57371 | 5.70902 | 0.13302 | 2.79253 | 3.16144 | 1.76210 | 0.58412 |
| **Tmem265** | 1.59032 | 0.01827 | 2.76130 | 1.60715 | 2.00875 | 2.74521 | 0.87094 | 0.00000 | 0.37695 | 0.05027 | 1.75313 | 0.87530 | 0.13714 | 1.64993 |
| **Tmem266** | 0.01065 | 0.00371 | 0.03415 | 0.00000 | 0.01024 | 0.01383 | 0.00364 | 0.64008 | 0.01217 | 0.00705 | 0.01565 | 0.30658 | 0.00633 | 0.01367 |
| **Tmem267** | 9.07014 | 0.00267 | 1.65605 | 0.28561 | 6.41379 | 0.96289 | 0.00244 | 0.86757 | 7.31701 | 0.00804 | 0.00000 | 0.00000 | 1.42896 | 5.94693 |
| **Tmem268** | 0.30169 | 0.02602 | 0.67833 | 0.49131 | 0.69439 | 0.05064 | 0.00400 | 0.00000 | 0.26950 | 0.00714 | 0.17684 | 0.00573 | 0.03272 | 1.27450 |
| **Tmem29** | 2.37629 | 0.01128 | 0.10128 | 1.10363 | 4.85543 | 2.25611 | 0.00290 | 1.42525 | 3.61914 | 0.02547 | 0.07120 | 2.10594 | 2.83348 | 1.92919 |
| **Tmem30a** | 6.36406 | 0.03187 | 2.16382 | 1.16908 | 2.48309 | 6.36589 | 0.00057 | 1.58161 | 16.57027 | 0.01306 | 3.34570 | 1.99706 | 2.06270 | 0.58764 |
| **Tmem30b** | 0.00000 | 0.00421 | 0.47507 | 2.64200 | 0.07078 | 1.05019 | 2.57368 | 0.00068 | 0.00519 | 0.07115 | 0.20399 | 0.05215 | 0.08130 | 1.06388 |
| **Tmem33** | 7.23499 | 1.04390 | 10.54409 | 7.67482 | 4.94678 | 7.20948 | 2.06712 | 5.55508 | 6.42691 | 0.12687 | 4.76523 | 3.78265 | 3.84384 | 3.57358 |
| **Tmem35b** | 0.02059 | 0.02846 | 0.00000 | 3.54722 | 1.00378 | 1.98707 | 3.62400 | 0.00763 | 0.00321 | 0.17150 | 2.10036 | 0.66837 | 1.13833 | 1.07040 |
| **Tmem37** | 5.26817 | 0.01729 | 7.44474 | 4.84680 | 16.24260 | 0.92209 | 0.00856 | 0.63333 | 0.32768 | 0.00000 | 0.00000 | 0.38034 | 1.73381 | 9.39299 |
| **Tmem38a** | 1.15424 | 0.00482 | 0.27342 | 0.01171 | 0.32392 | 0.81521 | 0.00431 | 0.02163 | 0.00875 | 0.00986 | 0.00578 | 0.00517 | 0.57028 | 0.95248 |
| **Tmem38b** | 2.97323 | 1.00314 | 3.56214 | 1.96699 | 1.57128 | 6.49466 | 2.04651 | 2.93750 | 5.40035 | 0.09289 | 0.03767 | 0.77518 | 0.61977 | 0.84972 |

| | | | | | | | | | | | | | | |
|---|---|---|---|---|---|---|---|---|---|---|---|---|---|---|
| **Tmem39a** | 4.71622 | 0.06771 | 9.09020 | 2.08951 | 4.08298 | 8.79798 | 2.24638 | 0.55386 | 6.92557 | 0.05160 | 10.65403 | 7.86160 | 3.85270 | 9.17826 |
| **Tmem39b** | 0.07021 | 0.01147 | 4.18096 | 0.00000 | 3.56119 | 0.67160 | 0.00297 | 2.33276 | 0.00000 | 0.01992 | 0.00913 | 0.19947 | 1.10718 | 2.93090 |
| **Tmem40** | 0.00000 | 0.01208 | 0.12633 | 1.35664 | 0.17876 | 1.77198 | 0.01237 | 0.29129 | 0.01132 | 0.01898 | 0.00000 | 0.59071 | 0.00620 | 0.00251 |
| **Tmem41a** | 1.33986 | 0.00490 | 0.63177 | 3.78873 | 0.05795 | 0.54238 | 0.00421 | 0.08133 | 0.38374 | 0.02450 | 0.01268 | 0.28570 | 2.16194 | 0.12020 |
| **Tmem41b** | 35.84962 | 4.66853 | 46.75730 | 38.90449 | 38.45780 | 31.43629 | 7.01074 | 11.17044 | 35.73577 | 9.69210 | 21.83025 | 26.22036 | 19.42783 | 26.49032 |
| **Tmem42** | 0.40113 | 0.00388 | 0.06495 | 0.30455 | 0.00000 | 0.04355 | 0.04818 | 0.32459 | 0.00484 | 0.02306 | 0.01621 | 0.00270 | 0.90254 | 0.00000 |
| **Tmem43** | 3.07845 | 0.03106 | 3.40863 | 2.79619 | 0.14227 | 5.60143 | 2.36766 | 0.26956 | 0.11209 | 0.04377 | 0.00000 | 2.98532 | 2.27495 | 1.52614 |
| **Tmem44** | 0.13609 | 0.00430 | 0.35902 | 0.01286 | 0.08983 | 0.00254 | 0.00460 | 0.00899 | 0.04763 | 0.01378 | 0.00706 | 0.00971 | 0.00934 | 0.59288 |
| **Tmem45a** | 0.02496 | 0.02020 | 0.01143 | 0.04118 | 0.01102 | 0.02024 | 0.00446 | 0.02057 | 0.01173 | 0.00859 | 0.00636 | 0.56327 | 0.00649 | 0.01004 |
| **Tmem45b** | 0.00000 | 0.06730 | 1.38011 | 2.98358 | 0.03697 | 5.96719 | 2.54406 | 4.36536 | 0.46949 | 0.12819 | 0.07985 | 0.00000 | 0.04229 | 0.00000 |
| **Tmem47** | 0.09543 | 0.01152 | 0.01108 | 0.00630 | 0.03274 | 0.01518 | 0.00431 | 0.00684 | 0.01137 | 0.00709 | 0.00627 | 0.11036 | 1.87747 | 0.01442 |
| **Tmem5** | 4.52300 | 0.00246 | 0.33671 | 1.39295 | 0.51438 | 0.85250 | 0.26294 | 0.00111 | 1.10381 | 0.00511 | 0.00769 | 1.40918 | 1.30008 | 0.65795 |
| **Tmem50a** | 3.75591 | 0.08481 | 4.45336 | 9.89036 | 3.03451 | 2.70399 | 0.06558 | 3.36859 | 2.43102 | 0.17378 | 5.42403 | 4.89568 | 0.49490 | 3.08323 |
| **Tmem50b** | 22.15353 | 0.09737 | 21.02596 | 7.62314 | 25.97733 | 9.82609 | 3.06774 | 9.55759 | 12.58172 | 0.17365 | 2.33923 | 0.17983 | 14.19427 | 28.41240 |
| **Tmem51** | 0.00000 | 0.00449 | 0.37061 | 0.68690 | 0.26492 | 1.44529 | 0.00503 | 0.10482 | 0.01977 | 0.00544 | 0.01743 | 0.00000 | 0.00449 | 0.00000 |
| **Tmem52** | 0.58156 | 0.00499 | 0.82887 | 0.63996 | 0.28251 | 0.91330 | 0.00954 | 0.00681 | 0.01127 | 0.00825 | 0.00251 | 0.00498 | 0.00572 | 0.07881 |
| **Tmem53** | 0.00953 | 0.00384 | 0.02070 | 0.00000 | 0.52933 | 0.00567 | 0.00367 | 0.46523 | 0.02195 | 0.00652 | 0.00725 | 0.01192 | 0.00764 | 0.00502 |
| **Tmem54** | 0.01111 | 0.00393 | 0.01279 | 0.00000 | 0.01169 | 0.08752 | 0.01486 | 0.49459 | 0.01127 | 0.00745 | 0.00191 | 0.15341 | 0.00704 | 0.01062 |
| **Tmem55a** | 3.89847 | 4.14720 | 6.87632 | 8.55961 | 2.03497 | 5.67733 | 0.48572 | 5.62782 | 5.93561 | 8.30548 | 6.05913 | 1.04111 | 0.53467 | 3.80092 |
| **Tmem55b** | 1.32676 | 0.06533 | 5.99342 | 3.76025 | 5.40496 | 6.54787 | 0.00588 | 2.24596 | 0.96941 | 1.72510 | 1.89277 | 5.46589 | 7.28071 | 3.74430 |
| **Tmem56** | 11.86227 | 0.00154 | 7.01352 | 0.25507 | 17.64151 | 0.00000 | 0.00000 | 0.00000 | 27.75019 | 0.00000 | 0.00000 | 0.00000 | 2.05812 | 13.09947 |
| **Tmem57** | 1.75127 | 5.21757 | 7.89461 | 1.57697 | 0.31371 | 7.31616 | 0.76868 | 0.55425 | 4.65391 | 0.02222 | 0.00000 | 3.96343 | 4.98789 | 7.34596 |
| **Tmem59** | 20.44670 | 3.93652 | 31.53421 | 25.15780 | 19.70083 | 24.94617 | 6.88114 | 19.49486 | 16.84638 | 2.04053 | 15.41057 | 13.69477 | 7.55387 | 14.05370 |
| **Tmem60** | 1.64363 | 0.02930 | 1.09720 | 4.03360 | 1.28431 | 3.42834 | 0.01504 | 0.83198 | 2.35802 | 1.64332 | 2.92553 | 3.40008 | 5.05069 | 0.72668 |
| **Tmem62** | 5.08090 | 6.30528 | 3.46515 | 0.48919 | 0.77476 | 18.43219 | 6.44492 | 10.38429 | 11.23483 | 0.31111 | 10.81520 | 0.00000 | 0.93445 | 4.10490 |
| **Tmem63a** | 0.09940 | 0.02146 | 0.53783 | 0.30722 | 1.34932 | 0.00000 | 0.00373 | 0.00870 | 0.08296 | 0.00572 | 0.00477 | 0.61535 | 0.01010 | 1.42176 |
| **Tmem63b** | 0.00601 | 0.01097 | 0.00000 | 0.81640 | 0.65057 | 2.28434 | 0.00393 | 0.90188 | 0.00369 | 0.01345 | 0.00635 | 0.38807 | 0.17110 | 1.20030 |
| **Tmem64** | 15.72890 | 0.00283 | 14.43369 | 0.62669 | 2.88804 | 5.42416 | 5.39439 | 1.37277 | 5.07457 | 8.66289 | 9.99596 | 3.74456 | 0.00798 | 6.92271 |
| **Tmem65** | 5.76045 | 0.01520 | 4.89568 | 3.06062 | 3.26592 | 10.66407 | 3.74095 | 0.09269 | 2.07880 | 0.42559 | 4.38473 | 4.44544 | 3.46157 | 4.24933 |
| **Tmem67** | 0.06833 | 0.00412 | 0.08532 | 0.00000 | 0.55526 | 0.07509 | 0.00411 | 0.00458 | 0.05246 | 0.00551 | 0.00392 | 0.42618 | 0.48319 | 0.38900 |
| **Tmem68** | 5.16485 | 0.00256 | 2.34958 | 0.00000 | 1.46935 | 1.08602 | 0.16899 | 1.22744 | 6.08065 | 0.01360 | 3.18733 | 0.76481 | 3.63166 | 1.98034 |
| **Tmem69** | 2.76050 | 0.16830 | 0.26262 | 1.27275 | 0.97518 | 2.59322 | 0.55280 | 0.77227 | 3.77658 | 1.22475 | 0.22888 | 1.47338 | 10.82826 | 2.15268 |
| **Tmem70** | 12.31008 | 3.47616 | 3.41762 | 2.92739 | 9.94287 | 5.14419 | 4.35339 | 0.44783 | 4.68376 | 0.14880 | 2.30285 | 9.15509 | 4.28365 | 8.41648 |
| **Tmem72** | 0.00000 | 0.00467 | 0.00000 | 0.01688 | 0.00000 | 0.01234 | 0.00415 | 0.01479 | 1.21754 | 0.00737 | 0.00641 | 0.00765 | 0.57390 | 0.09966 |
| **Tmem79** | 0.00000 | 0.08727 | 2.40273 | 2.11995 | 1.08701 | 1.24693 | 5.25781 | 12.87659 | 0.18258 | 0.17618 | 1.92040 | 0.00000 | 2.24124 | 1.78023 |
| **Tmem8** | 3.45480 | 6.56258 | 8.05189 | 1.61107 | 2.95420 | 3.16586 | 0.00000 | 0.60703 | 1.73044 | 0.04463 | 3.85062 | 7.80927 | 0.82871 | 3.47533 |
| **Tmem80** | 0.00298 | 0.01179 | 0.02385 | 0.00000 | 0.15616 | 0.99932 | 0.00438 | 0.05362 | 0.00535 | 0.01752 | 0.22514 | 0.75457 | 0.01391 | 0.32556 |
| **Tmem81** | 0.00000 | 0.01180 | 0.91958 | 2.07129 | 0.40182 | 0.23722 | 0.00362 | 0.00000 | 0.51093 | 0.00914 | 0.01386 | 0.01586 | 0.00015 | 1.07960 |
| **Tmem82** | 0.07018 | 0.01127 | 0.06245 | 1.67261 | 0.30127 | 0.30914 | 0.09221 | 0.13637 | 0.22017 | 0.07375 | 0.35410 | 1.56129 | 0.34532 | 0.53527 |
| **Tmem86a** | 0.23041 | 0.02752 | 0.45331 | 2.34855 | 0.26196 | 0.00000 | 0.00480 | 1.10552 | 0.00233 | 0.00587 | 0.00000 | 0.87471 | 0.75802 | 0.17434 |
| **Tmem86b** | 1.07998 | 0.01279 | 0.30642 | 0.00000 | 0.67059 | 0.00000 | 2.68737 | 0.26253 | 0.05289 | 0.06297 | 0.25329 | 0.39434 | 0.00611 | 0.07095 |
| **Tmem87a** | 0.00000 | 0.00431 | 1.94744 | 1.49390 | 0.83960 | 0.00000 | 0.00250 | 0.28350 | 9.30615 | 0.01269 | 0.20916 | 0.68748 | 0.97314 | 3.76053 |
| **Tmem87b** | 5.73662 | 0.01894 | 4.43230 | 2.97899 | 6.20293 | 1.75301 | 2.95861 | 0.00000 | 4.62912 | 0.06440 | 2.10824 | 0.00000 | 1.96276 | 11.74706 |
| **Tmem88** | 0.00913 | 0.01231 | 0.10660 | 0.00000 | 0.10892 | 0.00077 | 0.00486 | 0.29699 | 0.00950 | 0.01362 | 0.00769 | 0.32589 | 0.00462 | 0.01913 |
| **Tmem88b** | 0.04120 | 0.00441 | 0.01243 | 0.02654 | 0.02468 | 0.02670 | 0.01016 | 0.00772 | 0.14727 | 0.04166 | 0.00729 | 0.03402 | 0.29698 | 0.04515 |
| **Tmem8b** | 0.01714 | 0.04318 | 0.01838 | 0.00000 | 0.02018 | 0.68302 | 0.00419 | 0.01840 | 0.02638 | 0.00755 | 0.01315 | 0.02926 | 0.17579 | 0.00472 |
| **Tmem9** | 0.00000 | 0.37402 | 2.34873 | 10.98904 | 0.15057 | 5.51116 | 5.16854 | 16.81043 | 0.00413 | 0.42235 | 6.02803 | 8.47935 | 0.00935 | 0.00000 |
| **Tmem91** | 0.00746 | 0.00446 | 0.00936 | 0.01273 | 0.12867 | 0.01180 | 0.00429 | 0.01100 | 0.10205 | 0.00659 | 0.00809 | 0.00922 | 0.00951 | 0.00458 |
| **Tmem92** | 0.91969 | 0.28766 | 3.94764 | 1.51239 | 0.85634 | 1.35886 | 0.34959 | 0.09112 | 3.31121 | 0.00227 | 0.91794 | 0.06746 | 5.69228 | 3.10428 |
| **Tmem94** | 0.01816 | 0.13620 | 0.02227 | 0.00842 | 0.54020 | 0.02346 | 0.00410 | 0.00441 | 0.02669 | 0.00519 | 0.00580 | 0.25686 | 0.23104 | 0.27538 |
| **Tmem97** | 11.56236 | 10.15764 | 24.58522 | 20.73346 | 10.05676 | 17.16646 | 2.33771 | 10.94198 | 3.88054 | 0.15015 | 12.08554 | 14.69421 | 5.19584 | 15.66763 |
| **Tmem98** | 0.00798 | 0.04080 | 0.00000 | 0.00000 | 0.05405 | 0.00000 | 0.00414 | 0.99251 | 0.00936 | 0.00219 | 1.17011 | 0.55538 | 0.00929 | 0.01248 |
| **Tmem9b** | 5.05004 | 0.02995 | 1.57505 | 1.82833 | 4.80085 | 7.14045 | 3.90372 | 3.27543 | 1.89086 | 1.61476 | 5.00501 | 1.36677 | 0.01681 | 2.59782 |
| **Tmf1** | 1.94976 | 0.00284 | 0.62168 | 0.00000 | 0.10104 | 1.83909 | 0.00302 | 0.63378 | 3.31142 | 0.00893 | 0.43422 | 0.64344 | 2.04831 | 2.10801 |
| **Tmigd1** | 1.60385 | 0.00368 | 1.34216 | 0.42294 | 0.35896 | 0.01999 | 0.00322 | 0.00000 | 2.91031 | 0.00444 | 0.00286 | 0.00398 | 2.49352 | 0.63319 |

| | | | | | | | | | | | | | | |
|---|---|---|---|---|---|---|---|---|---|---|---|---|---|---|
| **Tmod1** | 0.00953 | 1.50322 | 0.04028 | 0.00773 | 0.01337 | 0.07540 | 0.00367 | 0.00267 | 0.01144 | 0.00570 | 0.00804 | 0.00380 | 0.00710 | 0.01042 |
| **Tmod2** | 0.61141 | 0.00396 | 0.53999 | 0.46933 | 0.08911 | 0.00000 | 0.00420 | 0.00547 | 0.04464 | 0.01190 | 0.00824 | 0.01406 | 0.08095 | 0.70346 |
| **Tmod3** | 6.17299 | 7.57695 | 5.95108 | 2.50446 | 5.00335 | 7.70498 | 2.92024 | 2.11718 | 5.53004 | 0.19932 | 4.86165 | 3.50973 | 2.19740 | 8.52847 |
| **Tmod4** | 0.01427 | 0.00425 | 0.01399 | 0.14431 | 0.07841 | 0.01073 | 0.00412 | 0.01008 | 0.01800 | 0.00723 | 0.00662 | 0.00769 | 0.03241 | 0.01293 |
| **Tmpo** | 14.22188 | 0.08083 | 12.14831 | 5.82346 | 9.12102 | 12.33606 | 3.15264 | 2.85907 | 9.07979 | 0.12974 | 4.93597 | 10.58301 | 12.52301 | 12.02882 |
| **Tmppe** | 4.76621 | 0.41767 | 7.72563 | 13.95112 | 6.98167 | 9.11042 | 0.98181 | 4.16098 | 16.34745 | 2.36117 | 0.12061 | 3.27647 | 7.45232 | 23.80808 |
| **Tmprss11a** | 0.00196 | 0.00425 | 0.00550 | 0.01305 | 0.20264 | 0.00985 | 0.00442 | 0.00790 | 0.02039 | 0.00588 | 0.00731 | 0.00733 | 0.72776 | 1.02687 |
| **Tmprss12** | 0.02001 | 0.00494 | 0.06944 | 0.08504 | 0.00744 | 0.23120 | 0.00468 | 0.17664 | 0.00990 | 0.01027 | 0.03957 | 0.00218 | 0.50033 | 0.00000 |
| **Tmprss2** | 0.00227 | 2.62773 | 0.00000 | 0.00000 | 0.00249 | 3.49498 | 4.80671 | 8.66789 | 0.01013 | 5.91996 | 0.86841 | 0.01326 | 0.00583 | 0.00000 |
| **Tmprss9** | 0.00617 | 0.00443 | 0.09059 | 0.00505 | 0.09734 | 0.01767 | 0.00430 | 0.01131 | 0.00804 | 0.00673 | 0.00755 | 0.00984 | 0.13739 | 0.04821 |
| **Tmsb10** | 2.48480 | 5.70705 | 0.00000 | 4.11754 | 4.65884 | 12.74555 | 6.39680 | 30.54949 | 0.00274 | 23.80058 | 0.60834 | 4.75803 | 2.56175 | 0.38473 |
| **Tmsb15a** | 0.01856 | 0.00444 | 0.01617 | 0.01245 | 0.01188 | 0.01262 | 0.00427 | 0.01033 | 0.01980 | 0.00672 | 0.00859 | 0.00989 | 0.03492 | 0.01574 |
| **Tmsb15b1** | 0.00999 | 0.00482 | 0.01138 | 0.03809 | 0.01314 | 0.02370 | 0.00419 | 0.01761 | 0.01326 | 0.00906 | 0.01006 | 0.00804 | 0.00650 | 0.01948 |
| **Tmsb4x** | 0.04575 | 11.04220 | 0.00000 | 0.00000 | 5.30818 | 1.23315 | 7.00085 | 20.53842 | 0.60282 | 12.67971 | 0.28905 | 0.94491 | 0.05748 | 0.00000 |
| **Tmtc1** | 0.00000 | 0.00412 | 0.11879 | 0.90726 | 0.61079 | 0.34795 | 0.00374 | 0.00247 | 0.11241 | 0.00524 | 0.08724 | 0.01318 | 0.11388 | 1.58504 |
| **Tmtc2** | 0.02143 | 0.00395 | 0.01135 | 0.06671 | 0.00743 | 0.16426 | 0.00427 | 1.18064 | 0.01057 | 0.01028 | 0.01075 | 0.00396 | 0.13510 | 0.05279 |
| **Tmtc3** | 1.42996 | 0.00950 | 0.88194 | 0.06291 | 1.89316 | 1.63986 | 0.00309 | 0.00000 | 2.64031 | 0.00455 | 0.00000 | 1.80195 | 2.56715 | 5.02939 |
| **Tmtc4** | 0.02682 | 0.00330 | 0.35793 | 0.02419 | 0.28275 | 1.21206 | 0.00349 | 0.00000 | 0.02089 | 0.00578 | 2.75896 | 0.47106 | 0.00264 | 1.10462 |
| **Tmub1** | 0.02937 | 0.01305 | 1.28698 | 0.10139 | 0.00000 | 0.42565 | 0.55009 | 1.51288 | 0.00951 | 0.01522 | 0.22065 | 0.00109 | 0.04394 | 0.00000 |
| **Tmub2** | 4.84379 | 0.05254 | 7.93943 | 3.78101 | 2.88004 | 4.18406 | 2.69530 | 3.19535 | 0.64755 | 0.06725 | 1.56326 | 3.81688 | 0.74451 | 6.60619 |
| **Tmx1** | 0.10790 | 0.01159 | 0.52270 | 2.13316 | 0.53484 | 1.70468 | 0.25549 | 1.01918 | 1.66335 | 0.00101 | 3.51231 | 0.69684 | 1.21643 | 0.18909 |
| **Tmx2** | 3.01263 | 0.12751 | 9.74087 | 9.18565 | 5.42485 | 11.34440 | 1.76432 | 5.25326 | 13.68273 | 0.11753 | 7.22648 | 8.93445 | 1.14024 | 9.54432 |
| **Tmx3** | 1.27524 | 0.00302 | 1.34470 | 0.00000 | 1.15486 | 1.15519 | 0.00875 | 1.63114 | 9.45184 | 0.00137 | 0.87436 | 0.97602 | 1.40951 | 0.00000 |
| **Tmx4** | 0.06669 | 0.02243 | 0.12619 | 0.87291 | 0.00000 | 3.08460 | 0.94595 | 5.98852 | 0.81458 | 0.11066 | 7.03475 | 4.21418 | 0.27169 | 0.22123 |
| **Tnc** | 0.25425 | 0.00442 | 0.01255 | 0.01008 | 0.00539 | 0.01078 | 0.00427 | 0.01137 | 0.00641 | 0.00635 | 0.00743 | 0.00912 | 0.00460 | 0.06444 |
| **Tnfaip1** | 4.23961 | 0.03246 | 7.93603 | 5.31160 | 2.59682 | 11.68609 | 2.66145 | 6.40828 | 4.73942 | 0.05631 | 6.11198 | 0.87064 | 1.81496 | 6.76413 |
| **Tnfaip2** | 0.05450 | 0.00440 | 0.01083 | 0.01156 | 0.11796 | 0.01263 | 0.00439 | 0.01195 | 0.00689 | 0.00691 | 0.00791 | 0.00934 | 0.00406 | 0.09033 |
| **Tnfaip3** | 0.01614 | 0.00447 | 0.01369 | 0.01376 | 0.02961 | 0.01224 | 0.00429 | 0.01077 | 0.02680 | 0.00655 | 0.00843 | 0.00971 | 0.01171 | 0.03202 |
| **Tnfaip6** | 0.02475 | 0.00428 | 0.01986 | 0.08775 | 0.01912 | 0.07852 | 0.01584 | 0.05461 | 0.05612 | 0.04619 | 0.01678 | 0.01988 | 0.51918 | 0.16500 |
| **Tnfaip8** | 0.21325 | 0.03037 | 0.26286 | 0.24589 | 0.41176 | 0.08913 | 0.00941 | 0.73530 | 1.14031 | 0.00516 | 0.00000 | 0.88777 | 0.88825 | 1.63106 |
| **Tnfaip8l2** | 0.00053 | 0.00350 | 0.00000 | 0.10599 | 2.02901 | 0.00000 | 0.00371 | 0.00504 | 0.24538 | 0.01406 | 0.00410 | 0.00624 | 3.76688 | 4.50790 |
| **Tnfrsf10b** | 0.28104 | 0.18453 | 0.60005 | 3.48894 | 0.42433 | 2.69265 | 0.14396 | 0.32959 | 1.45185 | 0.28979 | 0.05059 | 1.47750 | 2.24278 | 0.88625 |
| **Tnfrsf11a** | 0.47466 | 0.00437 | 0.04117 | 0.03738 | 0.14891 | 0.01548 | 0.05347 | 0.16949 | 1.88901 | 0.00690 | 0.00650 | 0.00819 | 0.35587 | 0.01412 |
| **Tnfrsf12a** | 0.19604 | 0.01945 | 0.95102 | 2.06026 | 1.32687 | 1.31633 | 1.10682 | 0.76368 | 0.69931 | 0.07450 | 0.56628 | 0.00000 | 1.39092 | 0.21054 |
| **Tnfrsf13c** | 0.00600 | 0.00446 | 0.00902 | 0.01068 | 0.26165 | 0.02476 | 0.00421 | 0.00872 | 0.03095 | 0.00708 | 0.00860 | 0.00795 | 0.49142 | 0.04060 |
| **Tnfrsf17** | 1.54750 | 0.00377 | 1.27183 | 0.01222 | 0.05052 | 0.00000 | 0.00362 | 0.00381 | 0.76038 | 0.00523 | 0.00746 | 0.00475 | 0.92682 | 3.08334 |
| **Tnfrsf19** | 0.21778 | 0.02313 | 0.06196 | 0.66030 | 0.43826 | 0.00000 | 0.00379 | 0.02836 | 0.41490 | 0.01016 | 0.00305 | 0.13802 | 0.51675 | 0.53115 |
| **Tnfrsf1a** | 0.09914 | 0.00416 | 0.22127 | 0.21859 | 1.36368 | 0.00519 | 0.00406 | 0.00526 | 0.01636 | 0.00602 | 0.01349 | 0.01508 | 0.00161 | 0.00376 |
| **Tnfrsf1b** | 0.01763 | 0.00467 | 0.09033 | 0.00000 | 0.00933 | 0.00866 | 0.00428 | 0.01435 | 0.00638 | 0.00710 | 0.00572 | 0.14473 | 0.00820 | 0.35388 |
| **Tnfrsf21** | 0.12619 | 0.00426 | 0.05490 | 0.00000 | 0.25282 | 0.15987 | 0.00393 | 0.00347 | 0.01029 | 0.01113 | 2.20392 | 0.71472 | 0.00395 | 0.73992 |
| **Tnfrsf22** | 4.49189 | 0.00078 | 1.39009 | 0.11033 | 5.42890 | 0.12431 | 0.36833 | 0.03776 | 29.30158 | 0.01656 | 0.15078 | 0.32996 | 5.27056 | 47.76964 |
| **Tnfrsf23** | 0.03422 | 0.00410 | 0.13732 | 0.00001 | 0.03113 | 0.02929 | 0.03747 | 0.01439 | 0.02936 | 0.00738 | 0.00751 | 0.02230 | 0.17463 | 0.23736 |
| **Tnfrsf25** | 0.02040 | 0.00392 | 0.10462 | 0.00000 | 0.51250 | 0.00782 | 0.00442 | 0.00621 | 0.01086 | 0.00635 | 0.00433 | 0.55788 | 0.00477 | 0.01066 |
| **Tnfrsf26** | 0.00336 | 0.00429 | 0.11456 | 0.32102 | 0.02204 | 0.02457 | 0.00414 | 0.00950 | 0.01144 | 0.00614 | 0.00755 | 0.00692 | 0.01394 | 0.01402 |
| **Tnfrsf4** | 0.01713 | 0.01476 | 0.00483 | 0.00365 | 0.27121 | 0.01546 | 0.00503 | 0.59423 | 0.02030 | 0.01123 | 0.33222 | 0.01124 | 0.00587 | 0.00293 |
| **Tnfrsf8** | 0.00496 | 0.00419 | 0.01766 | 0.33356 | 0.00356 | 0.01894 | 0.00410 | 0.00487 | 0.01756 | 0.00567 | 0.01072 | 0.01106 | 0.00357 | 0.17430 |
| **Tnfrsf9** | 0.43760 | 0.02637 | 0.12092 | 0.00000 | 0.20849 | 0.00000 | 1.27183 | 3.55438 | 1.27704 | 0.01451 | 0.00579 | 0.00628 | 0.42314 | 0.00000 |
| **Tnfsf10** | 0.84895 | 0.03542 | 0.08730 | 0.36450 | 0.32727 | 0.98935 | 0.13883 | 0.09182 | 1.27659 | 0.16273 | 0.04262 | 0.28547 | 5.47964 | 0.75697 |
| **Tnfsf12** | 0.19865 | 0.00397 | 0.08967 | 0.00000 | 1.00528 | 0.00000 | 0.00407 | 0.00668 | 0.34830 | 0.20975 | 0.00395 | 0.00413 | 1.66221 | 0.12015 |
| **Tnfsf13b** | 10.93701 | 0.00700 | 18.35056 | 5.64841 | 21.25192 | 0.91623 | 0.00281 | 0.00000 | 22.06483 | 0.00000 | 1.61596 | 0.00000 | 9.73838 | 20.19557 |
| **Tnik** | 0.00647 | 0.00557 | 0.00682 | 0.00000 | 0.00239 | 0.00063 | 0.00407 | 0.50423 | 1.90805 | 0.00970 | 0.00508 | 0.01505 | 0.01191 | 0.00070 |
| **Tnip1** | 10.21747 | 0.04045 | 4.55105 | 6.07513 | 16.86791 | 1.90707 | 0.56706 | 3.41119 | 2.19331 | 1.31739 | 3.68627 | 2.48642 | 8.15175 | 10.92700 |
| **Tnip2** | 3.45217 | 0.01108 | 0.27659 | 0.79461 | 2.57688 | 0.08228 | 0.00963 | 0.01467 | 0.70049 | 0.00552 | 0.00151 | 0.54023 | 0.76446 | 0.27697 |
| **Tnk1** | 0.00432 | 0.00420 | 0.00063 | 0.29580 | 0.00353 | 0.00510 | 0.00401 | 0.00813 | 0.16518 | 0.00705 | 0.00576 | 0.01003 | 0.00532 | 0.00163 |
| **Tnk2** | 0.72674 | 0.00453 | 0.35628 | 2.32706 | 1.17900 | 0.31303 | 0.01160 | 0.13136 | 0.36157 | 0.04178 | 0.36530 | 0.49731 | 3.02670 | 0.00000 |

| | | | | | | | | | | | | | | |
|---|---|---|---|---|---|---|---|---|---|---|---|---|---|---|
| **Tnks** | 0.00000 | 0.01065 | 0.79772 | 0.95243 | 0.04437 | 0.69504 | 0.00341 | 0.00000 | 1.41688 | 0.00304 | 0.35898 | 0.49263 | 1.71340 | 0.79021 |
| **Tnks1bp1** | 0.08771 | 2.86397 | 0.07781 | 0.96871 | 0.13230 | 0.37989 | 0.78878 | 4.00866 | 0.01029 | 0.02523 | 0.88106 | 0.00000 | 0.52857 | 0.00000 |
| **Tnks2** | 14.79209 | 0.03999 | 11.89068 | 4.08928 | 18.76712 | 7.49246 | 0.02438 | 5.90110 | 27.03233 | 0.02971 | 12.22746 | 7.24386 | 7.00264 | 19.34155 |
| **Tnnc2** | 1.08082 | 0.00399 | 0.29714 | 0.58549 | 0.45260 | 0.00000 | 0.00402 | 0.00783 | 0.02950 | 0.00612 | 0.00815 | 0.00570 | 0.62308 | 1.27710 |
| **Tnni3** | 0.00001 | 0.02209 | 0.43037 | 0.05681 | 0.10525 | 0.10974 | 0.00456 | 0.01088 | 0.02339 | 0.00666 | 0.06440 | 0.29543 | 0.06332 | 0.02604 |
| **Tnnt1** | 0.01953 | 0.00455 | 0.01092 | 0.03743 | 0.02141 | 0.01156 | 0.00405 | 0.01063 | 0.01074 | 0.01214 | 0.00983 | 0.01435 | 0.03292 | 0.04372 |
| **Tnpo1** | 6.27644 | 0.03024 | 1.14041 | 2.18732 | 7.57340 | 5.34300 | 0.34934 | 0.02963 | 10.21229 | 0.80950 | 0.02387 | 2.16859 | 4.92628 | 6.49759 |
| **Tnpo2** | 0.73920 | 0.04958 | 0.66859 | 0.76545 | 0.47191 | 0.55939 | 0.06418 | 0.62536 | 6.09514 | 0.02879 | 0.39451 | 0.39245 | 5.34301 | 0.95166 |
| **Tnpo3** | 3.40218 | 6.47429 | 5.57184 | 1.09516 | 7.24030 | 4.26063 | 0.90838 | 0.59762 | 10.32336 | 0.03076 | 2.30625 | 1.96317 | 11.05549 | 9.27362 |
| **Tnrc18** | 2.53138 | 5.05547 | 13.11707 | 7.27605 | 12.03589 | 4.94678 | 1.00995 | 0.69006 | 0.69472 | 0.07070 | 2.17328 | 4.94861 | 0.00000 | 3.87133 |
| **Tnrc6a** | 2.70764 | 0.04004 | 6.74930 | 3.15671 | 4.72585 | 6.98664 | 3.38241 | 4.83102 | 3.00571 | 0.13354 | 0.00000 | 2.07035 | 0.24958 | 3.67195 |
| **Tnrc6b** | 0.00000 | 0.00305 | 0.71370 | 0.00000 | 1.12750 | 0.42302 | 0.01399 | 0.64101 | 0.56751 | 0.00499 | 0.00000 | 1.31295 | 1.93501 | 1.72456 |
| **Tnrc6c** | 0.11277 | 0.00358 | 0.14341 | 0.97100 | 0.78052 | 0.49218 | 0.04171 | 2.91559 | 0.09169 | 0.01079 | 0.03193 | 0.81851 | 0.15421 | 1.22434 |
| **Tns1** | 0.48982 | 0.02009 | 0.29064 | 0.89147 | 0.29425 | 0.78479 | 0.00448 | 0.29875 | 0.02020 | 0.01288 | 0.59071 | 2.07569 | 0.00585 | 0.02820 |
| **Tns2** | 0.04416 | 0.00433 | 0.01157 | 0.01435 | 0.04788 | 0.02039 | 0.00443 | 0.00894 | 0.02946 | 0.00852 | 0.00758 | 0.00872 | 0.25824 | 0.00000 |
| **Tns3** | 2.00313 | 0.05910 | 1.05602 | 0.26188 | 0.70889 | 2.86451 | 1.84602 | 2.35931 | 0.45497 | 2.85173 | 0.69433 | 1.83383 | 0.76862 | 0.00000 |
| **Tns4** | 0.03185 | 0.04451 | 0.24153 | 0.66766 | 0.68721 | 1.22528 | 0.09259 | 0.07835 | 1.16447 | 0.06977 | 0.02480 | 0.06075 | 1.64597 | 0.62319 |
| **Tob1** | 0.02697 | 0.00482 | 0.03282 | 0.52643 | 0.00000 | 0.00000 | 0.00451 | 0.03400 | 0.02713 | 0.00673 | 0.00564 | 0.04351 | 0.13825 | 0.29872 |
| **Tob2** | 0.00777 | 0.00389 | 0.39999 | 0.19281 | 1.07015 | 0.00000 | 0.00441 | 0.12751 | 0.01692 | 0.00513 | 0.01176 | 0.07860 | 8.03560 | 0.09840 |
| **Toe1** | 9.45850 | 0.13835 | 14.79503 | 5.74101 | 6.55345 | 7.03151 | 2.73775 | 10.00037 | 6.94511 | 0.06785 | 6.95314 | 5.84426 | 1.80774 | 13.52944 |
| **Tollip** | 8.74284 | 1.64583 | 13.78888 | 7.73574 | 7.47027 | 11.99125 | 1.50661 | 6.57599 | 5.52599 | 0.07567 | 9.14573 | 10.01317 | 1.32644 | 10.22873 |
| **Tom1** | 4.39641 | 4.03603 | 8.31597 | 5.97448 | 6.59237 | 10.72786 | 0.01585 | 5.93549 | 0.31585 | 0.08310 | 4.08453 | 7.42588 | 0.12935 | 3.75135 |
| **Tom1l1** | 0.47842 | 0.01895 | 1.48683 | 2.08397 | 1.13949 | 1.50158 | 1.98995 | 0.77864 | 1.17798 | 0.02463 | 2.43355 | 0.52372 | 0.00000 | 1.35318 |
| **Tom1l2** | 1.18827 | 0.01766 | 5.04546 | 2.86886 | 7.41093 | 4.78579 | 0.00927 | 0.97366 | 0.00000 | 0.00301 | 3.01646 | 0.57049 | 4.45948 | 2.30980 |
| **Tomm20** | 34.15711 | 0.46894 | 12.56382 | 25.08310 | 41.41365 | 22.85109 | 6.40113 | 4.79497 | 18.13793 | 0.53098 | 8.14207 | 14.84938 | 20.03991 | 18.50653 |
| **Tomm20l** | 0.32444 | 0.00490 | 0.00984 | 0.12596 | 0.44317 | 0.01124 | 0.00438 | 0.01847 | 0.10443 | 0.00893 | 0.00566 | 0.00543 | 0.01403 | 0.02630 |
| **Tomm22** | 20.78784 | 8.69679 | 34.48194 | 29.20879 | 36.07483 | 23.90189 | 9.99799 | 16.74935 | 8.47111 | 15.39298 | 16.59638 | 16.60737 | 13.24235 | 22.59505 |
| **Tomm34** | 1.08019 | 0.66273 | 1.61643 | 3.50622 | 4.42897 | 1.59119 | 0.73868 | 2.31275 | 0.18367 | 0.01510 | 1.89454 | 3.62366 | 5.17743 | 3.90575 |
| **Tomm40** | 1.70205 | 5.37441 | 18.64683 | 15.72617 | 21.23686 | 6.37397 | 1.85366 | 10.15038 | 0.00000 | 10.08550 | 2.97269 | 9.28547 | 2.23369 | 0.00000 |
| **Tomm40l** | 0.00418 | 0.01966 | 0.00000 | 0.63439 | 0.18003 | 0.00000 | 0.00378 | 2.46987 | 0.00846 | 0.00742 | 1.83285 | 2.29810 | 0.00664 | 0.01685 |
| **Tomm5** | 20.88768 | 3.92861 | 9.01104 | 25.84122 | 33.53800 | 21.68228 | 5.48827 | 11.01882 | 9.94047 | 4.58417 | 5.79974 | 12.83048 | 18.61972 | 22.63337 |
| **Tomm6** | 2.80527 | 2.54934 | 0.00000 | 1.98762 | 7.84854 | 7.51995 | 1.42022 | 3.45072 | 0.53626 | 3.91704 | 0.50353 | 5.99264 | 5.66154 | 0.50404 |
| **Tomm7** | 4.88450 | 3.62890 | 0.00000 | 11.08050 | 5.93910 | 10.22812 | 2.47149 | 16.58419 | 3.31591 | 11.49196 | 0.76669 | 8.48443 | 9.43725 | 7.08253 |
| **Tomm70a** | 17.98757 | 0.05714 | 15.77750 | 9.25360 | 17.71315 | 10.34564 | 6.03046 | 5.63551 | 21.22889 | 0.47704 | 9.80200 | 11.27112 | 4.77693 | 12.85099 |
| **Tomt** | 0.12378 | 0.01280 | 0.04774 | 0.28098 | 0.05730 | 0.09910 | 0.00383 | 0.60319 | 1.36119 | 0.00570 | 0.01314 | 0.01358 | 0.00550 | 0.50042 |
| **Tonsl** | 2.19743 | 0.06142 | 5.88784 | 1.03370 | 4.88934 | 2.34248 | 0.01390 | 5.42769 | 0.00000 | 0.01986 | 2.58074 | 4.61061 | 2.35098 | 0.00000 |
| **Top1** | 3.90937 | 0.93139 | 15.78777 | 5.09022 | 11.23995 | 12.23084 | 1.31215 | 0.81060 | 33.06139 | 0.04445 | 6.06416 | 6.52101 | 13.38108 | 14.57275 |
| **Top1mt** | 1.14461 | 0.01200 | 1.47708 | 3.07455 | 4.18954 | 0.12503 | 0.20352 | 0.00000 | 0.04103 | 0.00988 | 0.00159 | 0.91995 | 0.00000 | 0.58118 |
| **Top2a** | 14.16695 | 0.19069 | 14.43606 | 4.07971 | 15.93387 | 10.54680 | 0.00000 | 8.20243 | 25.82074 | 5.45791 | 15.99856 | 8.71379 | 12.12992 | 18.07753 |
| **Top2b** | 0.12657 | 0.02903 | 0.00000 | 0.00000 | 0.04661 | 1.54319 | 1.83356 | 2.28030 | 0.00795 | 0.05438 | 0.72296 | 4.73995 | 0.00566 | 0.00385 |
| **Top3a** | 2.85339 | 0.01976 | 4.66889 | 2.68813 | 4.03033 | 2.48569 | 2.26936 | 1.99856 | 0.14825 | 0.08530 | 3.80702 | 0.34882 | 2.07277 | 3.14803 |
| **Top3b** | 0.00000 | 0.75783 | 2.62001 | 3.44957 | 0.67885 | 1.23367 | 4.50075 | 3.04359 | 0.00628 | 0.10634 | 3.52334 | 2.30081 | 0.00725 | 0.00000 |
| **Topaz1** | 0.18423 | 0.00382 | 0.11840 | 0.00008 | 0.00000 | 0.00638 | 0.00403 | 0.00746 | 0.28140 | 0.00861 | 0.00365 | 0.55557 | 0.00482 | 0.00000 |
| **Topbp1** | 13.67396 | 0.24317 | 15.82154 | 3.43234 | 16.60884 | 8.38307 | 2.22139 | 3.42749 | 12.22335 | 0.06688 | 9.69433 | 8.71158 | 9.06801 | 17.45739 |
| **Topors** | 9.32221 | 0.07193 | 12.77546 | 2.82936 | 18.01425 | 7.13927 | 0.00000 | 5.02902 | 10.72402 | 0.02478 | 9.65971 | 3.65310 | 11.26948 | 21.09611 |
| **Toporsos** | 0.03050 | 0.00484 | 0.01612 | 0.00000 | 0.04734 | 0.03555 | 0.06474 | 0.08902 | 0.01128 | 0.01900 | 0.00706 | 0.02287 | 0.00832 | 0.00938 |
| **Tor1a** | 5.31796 | 2.81642 | 4.10549 | 1.94588 | 5.34884 | 3.62569 | 4.92100 | 1.09843 | 2.47559 | 0.23340 | 0.48605 | 4.86107 | 9.82762 | 6.18771 |
| **Tor1aip1** | 3.86663 | 0.04976 | 7.73888 | 3.22055 | 9.19586 | 2.97787 | 0.00778 | 1.72600 | 4.07731 | 0.00530 | 3.12092 | 2.09600 | 9.28123 | 11.20651 |
| **Tor1aip2** | 4.91528 | 3.95689 | 15.07137 | 13.10247 | 7.98036 | 11.16510 | 7.21891 | 7.43209 | 16.27070 | 2.44361 | 11.25972 | 10.30349 | 10.38706 | 10.46100 |
| **Tor1b** | 15.07340 | 0.05555 | 12.60272 | 2.63699 | 12.49956 | 6.53200 | 0.49695 | 2.66933 | 17.20418 | 0.02302 | 2.88569 | 2.48464 | 12.33310 | 24.14891 |
| **Tor2a** | 7.51573 | 0.06565 | 15.83303 | 5.39050 | 18.04825 | 3.58489 | 0.00000 | 6.29854 | 0.31599 | 0.02180 | 2.10913 | 3.78011 | 5.35092 | 3.59584 |
| **Tor3a** | 0.00000 | 0.00342 | 0.40229 | 0.23418 | 1.25327 | 0.00000 | 0.00353 | 0.00000 | 0.94672 | 0.00373 | 0.41573 | 0.33554 | 0.49026 | 1.98863 |
| **Tor4a** | 1.42110 | 0.15357 | 0.00000 | 1.11761 | 0.93479 | 2.95495 | 0.37480 | 2.21852 | 2.22561 | 0.97128 | 0.07445 | 1.49279 | 12.77130 | 1.04485 |
| **Tox3** | 0.00878 | 0.00438 | 0.28674 | 0.00172 | 0.00663 | 0.00628 | 0.00424 | 0.01151 | 0.01140 | 0.00766 | 0.00671 | 0.00944 | 0.00843 | 0.03516 |
| **Tox4** | 5.23550 | 0.19714 | 6.40321 | 9.18867 | 9.67144 | 8.96111 | 0.04816 | 4.01271 | 10.77443 | 0.15877 | 4.46065 | 6.41806 | 8.08713 | 11.60893 |

| | | | | | | | | | | | | | | |
|---|---|---|---|---|---|---|---|---|---|---|---|---|---|---|
| **Tpbg** | 0.06464 | 0.00439 | 0.01165 | 0.01043 | 0.00541 | 0.01186 | 0.00431 | 0.01020 | 0.00559 | 0.00654 | 0.00789 | 0.01641 | 0.00447 | 0.24366 |
| **Tpbgl** | 0.06926 | 0.00404 | 0.00982 | 0.00000 | 0.06007 | 0.02499 | 0.00409 | 0.79496 | 0.01128 | 0.00576 | 0.00411 | 0.02810 | 0.00615 | 0.00998 |
| **Tpcn1** | 0.09169 | 0.00396 | 0.02231 | 0.00000 | 0.03176 | 0.37875 | 0.00410 | 0.30117 | 0.00743 | 0.00548 | 0.00443 | 0.01314 | 0.21617 | 0.03245 |
| **Tpcn2** | 0.15216 | 0.00254 | 0.04002 | 1.48911 | 0.00000 | 0.63901 | 0.00327 | 0.00000 | 0.01040 | 0.01112 | 2.12998 | 1.31752 | 0.00539 | 0.00000 |
| **Tpd52** | 21.14973 | 0.32836 | 28.08954 | 23.78824 | 25.32687 | 21.39878 | 6.33081 | 13.08184 | 10.23522 | 0.48598 | 17.04114 | 12.15868 | 0.87593 | 17.96088 |
| **Tpd52l1** | 0.00509 | 0.00449 | 0.00604 | 0.01208 | 0.48043 | 0.01250 | 0.00436 | 0.01093 | 0.00654 | 0.00619 | 0.00873 | 0.00800 | 0.00571 | 0.16727 |
| **Tpd52l2** | 18.58309 | 5.84539 | 23.39109 | 17.48738 | 22.52512 | 18.27448 | 5.81312 | 6.81806 | 9.37441 | 19.09562 | 13.84131 | 9.91087 | 9.64163 | 19.77956 |
| **Tpgs1** | 0.00167 | 0.01920 | 0.02807 | 5.89851 | 1.97435 | 0.00000 | 0.00403 | 0.05809 | 0.00525 | 0.15137 | 0.03260 | 0.14441 | 0.46561 | 0.00000 |
| **Tpgs2** | 2.65555 | 0.00176 | 0.00000 | 0.53402 | 1.05217 | 0.19792 | 0.00292 | 0.82438 | 1.21601 | 0.93243 | 1.93726 | 1.58132 | 0.86879 | 1.04117 |
| **Tph2** | 0.00866 | 0.00444 | 0.01372 | 0.01348 | 0.01021 | 0.01268 | 0.00434 | 0.01024 | 0.01135 | 0.00694 | 0.00876 | 0.00998 | 0.00897 | 0.08217 |
| **Tpi1** | 2.31457 | 12.31582 | 3.93634 | 7.50576 | 1.12091 | 7.85825 | 6.04629 | 28.89953 | 0.47283 | 0.51045 | 10.07341 | 6.04780 | 3.62868 | 0.31290 |
| **Tpk1** | 0.00055 | 0.71445 | 0.75113 | 0.00000 | 0.38925 | 0.40274 | 1.77914 | 1.40818 | 1.01156 | 0.01960 | 2.38949 | 0.84379 | 0.00000 | 0.00000 |
| **Tpm1** | 4.47708 | 8.70377 | 12.14437 | 18.32254 | 4.18261 | 16.97976 | 8.55566 | 20.36060 | 2.85023 | 11.80978 | 6.39775 | 6.53332 | 0.00000 | 0.00000 |
| **Tpm2** | 0.57553 | 0.05371 | 0.00000 | 0.62292 | 1.46957 | 3.59899 | 0.06148 | 1.74178 | 0.00242 | 0.03385 | 0.02947 | 0.00000 | 0.33051 | 0.00000 |
| **Tpm3** | 17.64124 | 8.61955 | 34.51894 | 22.41805 | 29.87840 | 31.67803 | 10.01600 | 28.01548 | 15.89358 | 1.17592 | 21.85752 | 17.05457 | 11.33388 | 27.33000 |
| **Tpm4** | 8.91384 | 9.46581 | 21.01063 | 12.74500 | 7.09561 | 21.18099 | 10.17322 | 22.08050 | 10.71463 | 8.33602 | 15.42112 | 15.16195 | 0.15583 | 1.91297 |
| **Tpmt** | 1.00261 | 0.01140 | 0.67083 | 0.46009 | 1.28431 | 3.19482 | 0.07368 | 2.06866 | 2.26130 | 0.09365 | 0.65799 | 0.02397 | 3.06522 | 1.90563 |
| **Tpo** | 0.11346 | 0.00442 | 0.01396 | 0.01177 | 0.00906 | 0.01250 | 0.00426 | 0.01042 | 0.01005 | 0.00701 | 0.00770 | 0.00934 | 0.05607 | 0.00476 |
| **Tpp1** | 0.00654 | 0.07526 | 0.00000 | 2.66758 | 0.00000 | 4.57294 | 0.55529 | 6.56673 | 0.00982 | 9.95902 | 1.74975 | 3.73930 | 0.00560 | 0.00000 |
| **Tpp2** | 8.90727 | 0.01384 | 6.87575 | 1.84009 | 4.84003 | 3.15419 | 1.14435 | 0.22379 | 10.13329 | 0.00564 | 8.83423 | 4.65721 | 3.80902 | 6.10948 |
| **Tppp** | 0.81193 | 0.00434 | 0.10097 | 0.00183 | 0.13608 | 0.00159 | 0.00405 | 0.01179 | 0.42447 | 0.00749 | 0.00586 | 0.00567 | 0.00331 | 0.84912 |
| **Tppp3** | 0.59021 | 0.03976 | 4.70004 | 3.64282 | 1.24234 | 0.41920 | 0.00348 | 0.03075 | 0.00972 | 0.01010 | 0.00038 | 1.15216 | 0.00000 | 0.00000 |
| **Tpr** | 16.49369 | 0.10526 | 13.49155 | 5.12517 | 23.96996 | 13.06903 | 3.17829 | 6.24605 | 19.70344 | 0.14881 | 8.06913 | 8.77814 | 17.58290 | 20.14941 |
| **Tpra1** | 0.02690 | 0.01154 | 1.60958 | 1.06446 | 0.20869 | 1.24636 | 1.10651 | 1.12276 | 0.00212 | 0.02871 | 0.00000 | 1.98721 | 0.39744 | 0.00000 |
| **Tprgl** | 3.29955 | 7.69905 | 9.38387 | 5.80685 | 4.52861 | 4.44996 | 1.77409 | 5.12608 | 0.69767 | 0.63175 | 2.78012 | 4.33163 | 2.61723 | 3.05549 |
| **Tprkb** | 3.27595 | 0.08424 | 4.72910 | 4.46031 | 4.13472 | 5.95531 | 4.14659 | 3.62360 | 5.21146 | 0.12995 | 6.14168 | 7.51445 | 2.65776 | 8.03575 |
| **Tprn** | 6.47372 | 9.55105 | 12.43239 | 2.85638 | 15.45960 | 3.49964 | 6.87838 | 4.37359 | 5.74004 | 0.32040 | 0.68843 | 3.86572 | 8.48955 | 16.57619 |
| **Tpst1** | 1.06674 | 0.01528 | 2.50199 | 0.00000 | 1.02341 | 0.00000 | 0.00257 | 0.00179 | 0.39122 | 0.00836 | 0.13620 | 3.34597 | 0.73014 | 0.73215 |
| **Tpst2** | 0.58072 | 0.14149 | 2.09374 | 4.69372 | 1.68666 | 3.21779 | 4.74596 | 5.60248 | 0.00052 | 0.14273 | 0.04496 | 1.07721 | 0.20497 | 0.15940 |
| **Tpt1** | 27.59336 | 9.60520 | 10.13110 | 59.69577 | 53.01046 | 25.59638 | 7.54439 | 16.77759 | 14.57210 | 14.60811 | 13.38948 | 27.23798 | 23.73237 | 13.67344 |
| **Tpx2** | 19.12203 | 5.92883 | 20.32069 | 4.57620 | 19.51726 | 10.54453 | 4.21919 | 4.71778 | 22.17156 | 0.13963 | 14.96501 | 10.58069 | 17.62510 | 20.41249 |
| **Tra2a** | 21.25801 | 4.22848 | 16.82118 | 14.71496 | 18.67286 | 21.54397 | 4.49854 | 16.35782 | 27.28133 | 10.69507 | 17.88375 | 12.08353 | 12.37591 | 9.36800 |
| **Tra2b** | 14.19191 | 0.29958 | 21.47735 | 20.44054 | 20.84742 | 20.33894 | 4.73634 | 9.15198 | 24.63780 | 2.79045 | 17.65461 | 11.59671 | 16.41101 | 11.20548 |
| **Trabd** | 0.00000 | 4.73404 | 2.26691 | 2.42776 | 0.28644 | 2.14027 | 2.46585 | 2.70064 | 0.00919 | 0.07031 | 0.28040 | 1.39133 | 0.01509 | 0.00000 |
| **Trabd2b** | 0.02682 | 0.00434 | 0.01167 | 0.02189 | 0.10899 | 0.01612 | 0.00437 | 0.00919 | 0.04018 | 0.00613 | 0.01143 | 0.00956 | 0.38326 | 0.00290 |
| **Tradd** | 0.00000 | 0.00455 | 0.37108 | 2.67603 | 0.80434 | 0.29622 | 0.00419 | 0.66231 | 0.03443 | 0.00589 | 0.00339 | 0.25934 | 0.03259 | 0.07170 |
| **Traf1** | 0.00801 | 0.01762 | 0.01997 | 0.00000 | 0.41126 | 0.00607 | 0.01405 | 0.79872 | 0.01860 | 0.01059 | 0.01117 | 0.01761 | 0.11164 | 0.02636 |
| **Traf2** | 3.85610 | 0.05116 | 3.94060 | 4.35759 | 3.53419 | 4.97600 | 3.16549 | 5.71912 | 1.14421 | 0.06924 | 3.78784 | 2.07312 | 2.63024 | 5.27921 |
| **Traf3** | 0.23501 | 0.00835 | 0.20477 | 0.00000 | 1.35083 | 1.85840 | 0.00888 | 0.00000 | 1.04175 | 0.01427 | 1.07578 | 2.70981 | 3.39473 | 1.85250 |
| **Traf3ip1** | 0.32558 | 0.01006 | 1.51822 | 2.52501 | 0.35279 | 0.78373 | 0.00805 | 2.31729 | 0.21821 | 0.02698 | 2.52839 | 0.36182 | 0.12138 | 2.06995 |
| **Traf3ip2** | 0.00000 | 0.00410 | 0.40774 | 0.91404 | 0.04210 | 0.02709 | 0.00397 | 0.00288 | 0.00568 | 0.02026 | 0.00494 | 0.00971 | 0.22702 | 0.02609 |
| **Traf4** | 0.58038 | 5.99645 | 8.48284 | 3.51263 | 5.73125 | 8.13888 | 2.65969 | 5.18406 | 0.72698 | 0.08596 | 1.21962 | 2.03587 | 0.98265 | 3.75138 |
| **Traf5** | 0.01800 | 0.00440 | 0.12666 | 0.00552 | 0.01087 | 0.00905 | 0.00422 | 0.01101 | 0.01289 | 0.00735 | 0.00812 | 0.01076 | 0.00854 | 0.01195 |
| **Traf6** | 0.00727 | 0.00402 | 0.00000 | 0.24940 | 0.00087 | 0.45182 | 0.00380 | 0.00335 | 0.01630 | 0.00639 | 0.22199 | 0.42849 | 0.09777 | 0.64293 |
| **Traf7** | 2.67660 | 0.10523 | 12.33001 | 10.82993 | 7.47024 | 12.50573 | 2.63961 | 5.80984 | 0.35150 | 0.15351 | 9.27254 | 8.94751 | 4.04409 | 7.69318 |
| **Trafd1** | 2.15343 | 0.03139 | 3.63790 | 0.00000 | 5.37986 | 1.68677 | 0.27371 | 1.95980 | 0.01027 | 0.02124 | 1.58233 | 1.32393 | 0.80362 | 0.86871 |
| **Traip** | 4.77783 | 0.05810 | 6.24176 | 2.61793 | 4.82606 | 3.93576 | 4.19327 | 3.16512 | 5.49009 | 0.08309 | 3.01825 | 1.39896 | 8.57773 | 6.83666 |
| **Trak1** | 0.64607 | 6.67002 | 5.94041 | 2.39985 | 4.25547 | 1.15333 | 0.00742 | 4.11270 | 2.66057 | 0.04227 | 0.14563 | 0.13150 | 1.59316 | 3.92017 |
| **Trak2** | 3.28420 | 0.01630 | 4.79859 | 2.65947 | 4.56354 | 5.88677 | 0.00136 | 1.36078 | 5.70572 | 0.02115 | 2.34438 | 3.00489 | 5.10271 | 5.06323 |
| **Tram1** | 3.22264 | 0.13763 | 3.27880 | 2.87866 | 1.32531 | 5.63175 | 0.01551 | 0.68458 | 1.62871 | 0.03251 | 2.82239 | 2.10088 | 0.75539 | 1.80009 |
| **Tram2** | 0.14695 | 0.00388 | 0.00189 | 0.19979 | 0.00506 | 0.62691 | 0.00382 | 0.00062 | 0.00878 | 0.00730 | 0.46696 | 0.06676 | 0.02395 | 0.00000 |
| **Trank1** | 0.01370 | 0.00420 | 0.00962 | 0.11695 | 0.01209 | 0.00932 | 0.00399 | 0.00666 | 0.01274 | 0.00845 | 0.00698 | 0.01518 | 0.20792 | 0.00774 |
| **Trap1** | 3.04203 | 8.17995 | 7.96658 | 13.41071 | 2.90579 | 6.35774 | 2.52978 | 7.01372 | 0.12974 | 12.17220 | 6.93087 | 10.52996 | 2.69077 | 5.67896 |
| **Trap1a** | 0.00000 | 3.24377 | 1.96518 | 11.27485 | 0.24553 | 7.75111 | 7.98434 | 19.88019 | 2.14524 | 12.89464 | 4.74044 | 2.17237 | 0.34646 | 4.49997 |
| **Trappc1** | 2.30782 | 0.07434 | 0.00000 | 2.53361 | 2.02759 | 5.43389 | 0.09496 | 7.75658 | 0.43159 | 0.06408 | 3.00356 | 6.14814 | 0.00000 | 0.81846 |

| | | | | | | | | | | | | | | |
|---|---|---|---|---|---|---|---|---|---|---|---|---|---|---|
| **Trappc10** | 1.03545 | 0.02257 | 0.57660 | 0.24962 | 3.61186 | 2.23767 | 1.38633 | 0.42255 | 0.89788 | 0.07105 | 1.79431 | 0.28282 | 8.29941 | 4.30088 |
| **Trappc11** | 1.67097 | 0.08914 | 5.56095 | 1.63771 | 3.24825 | 8.70963 | 3.99335 | 3.17976 | 1.36579 | 0.05194 | 5.53379 | 2.79613 | 1.24364 | 4.55142 |
| **Trappc12** | 1.13733 | 0.03229 | 3.99673 | 1.06986 | 3.51961 | 3.01562 | 0.00155 | 4.52145 | 0.00000 | 0.03916 | 4.25673 | 2.57863 | 0.56198 | 3.08735 |
| **Trappc13** | 9.79882 | 0.33990 | 9.42059 | 3.51381 | 7.93266 | 9.48032 | 4.69637 | 3.36198 | 7.65742 | 1.56634 | 7.25179 | 9.25935 | 1.42916 | 6.60861 |
| **Trappc2** | 0.68260 | 0.01065 | 0.96412 | 0.87404 | 0.16329 | 1.32971 | 1.58463 | 0.45809 | 0.04874 | 0.02588 | 2.12947 | 1.55614 | 0.01957 | 0.34902 |
| **Trappc2l** | 2.99226 | 0.13481 | 0.27157 | 6.39713 | 3.81519 | 3.13749 | 2.88585 | 6.85977 | 1.11262 | 0.30106 | 0.98072 | 3.17922 | 1.17957 | 4.16418 |
| **Trappc3** | 0.74443 | 0.01990 | 1.67648 | 0.34728 | 3.79731 | 3.03710 | 0.02815 | 2.55188 | 1.12151 | 0.04306 | 4.64031 | 1.00028 | 0.00000 | 0.84533 |
| **Trappc4** | 7.76908 | 0.10394 | 14.26747 | 11.31299 | 13.17949 | 12.35726 | 4.78340 | 5.57645 | 3.74474 | 13.18774 | 7.92713 | 5.52383 | 6.63304 | 5.83264 |
| **Trappc5** | 0.98095 | 3.89874 | 0.93122 | 4.26927 | 1.27677 | 3.45642 | 1.92888 | 3.65120 | 0.01501 | 7.50715 | 0.93273 | 3.73563 | 0.01231 | 0.01335 |
| **Trappc6a** | 0.07628 | 0.02594 | 0.47455 | 2.42373 | 1.25208 | 0.49487 | 0.00277 | 2.86272 | 0.00323 | 0.04026 | 0.00000 | 2.56045 | 0.22860 | 0.00000 |
| **Trappc6b** | 2.49740 | 0.08527 | 2.18758 | 1.27975 | 3.54437 | 3.69355 | 3.68161 | 2.21136 | 2.06515 | 0.05935 | 1.76017 | 1.92420 | 4.15352 | 3.28018 |
| **Trappc8** | 8.37563 | 0.00000 | 7.47061 | 0.00000 | 1.62335 | 7.05175 | 4.71653 | 0.00500 | 7.52083 | 0.11530 | 10.07360 | 2.79809 | 6.07981 | 10.30610 |
| **Trappc9** | 2.23930 | 0.03282 | 3.06391 | 1.51130 | 6.21025 | 1.06486 | 2.88993 | 1.31233 | 7.30297 | 0.10012 | 4.30780 | 1.81662 | 10.61449 | 13.85986 |
| **Trdmt1** | 1.30763 | 0.00371 | 0.63648 | 0.00000 | 0.19072 | 0.32255 | 0.35247 | 0.01405 | 4.57511 | 0.00341 | 0.00299 | 0.16685 | 0.00120 | 0.00000 |
| **Trerf1** | 0.01679 | 0.00442 | 0.06384 | 0.00512 | 0.10289 | 0.01181 | 0.00415 | 0.01056 | 0.01135 | 0.00647 | 0.00805 | 0.00921 | 0.07568 | 0.00855 |
| **Trex1** | 0.00788 | 0.07097 | 0.11742 | 1.21212 | 0.43760 | 1.66348 | 0.01491 | 3.56488 | 0.00175 | 0.04151 | 0.96221 | 0.72992 | 0.00301 | 0.59480 |
| **Trf** | 0.00768 | 0.00438 | 0.25555 | 0.00000 | 0.19076 | 0.00671 | 0.00452 | 0.01189 | 0.01201 | 0.00645 | 0.00680 | 0.00941 | 0.00870 | 0.04202 |
| **Trhr2** | 0.00967 | 0.00475 | 0.01419 | 0.00165 | 0.01415 | 0.30580 | 0.00429 | 0.00600 | 0.01110 | 0.00629 | 0.00728 | 0.01075 | 0.00804 | 0.00880 |
| **Triap1** | 6.53025 | 0.72783 | 14.93944 | 12.68523 | 8.94102 | 12.48217 | 5.18024 | 10.79665 | 6.09210 | 13.40316 | 9.96393 | 5.15331 | 0.69345 | 16.07729 |
| **Trib1** | 6.18324 | 0.05268 | 4.89901 | 3.15305 | 5.47760 | 4.57903 | 2.49378 | 5.86685 | 7.25437 | 0.06854 | 0.00000 | 0.31205 | 2.53664 | 1.50213 |
| **Trib2** | 0.00626 | 0.01200 | 0.01497 | 0.01207 | 0.00709 | 0.01235 | 0.00421 | 0.00863 | 0.01460 | 0.00632 | 0.00870 | 0.00866 | 0.56671 | 0.08997 |
| **Trib3** | 0.53237 | 0.00368 | 2.01755 | 1.36508 | 1.47431 | 0.00000 | 0.00360 | 0.01949 | 0.51514 | 0.00925 | 0.90120 | 0.01090 | 2.56105 | 0.73127 |
| **Trim11** | 1.57825 | 0.01708 | 3.14571 | 2.32130 | 0.19200 | 2.38552 | 0.00270 | 2.46467 | 0.00000 | 0.04005 | 0.04282 | 1.39853 | 2.61311 | 0.00000 |
| **Trim12c** | 0.03405 | 0.02672 | 0.10212 | 0.38077 | 0.34062 | 0.32396 | 0.00432 | 0.04576 | 0.17804 | 0.01402 | 0.03095 | 0.07179 | 0.04542 | 0.12724 |
| **Trim13** | 2.84188 | 0.16932 | 5.49279 | 3.93497 | 6.51024 | 1.92974 | 5.61852 | 0.95839 | 8.05603 | 0.12806 | 2.14862 | 3.56848 | 11.34995 | 8.51264 |
| **Trim14** | 0.53787 | 0.00415 | 0.37945 | 0.87258 | 0.00000 | 0.19607 | 0.06235 | 0.03365 | 0.14151 | 0.03344 | 0.65038 | 0.18637 | 0.52306 | 0.46414 |
| **Trim16** | 0.03417 | 0.00418 | 0.15757 | 0.34814 | 0.00479 | 0.00421 | 0.00422 | 0.00740 | 0.01173 | 0.00583 | 0.00615 | 0.00792 | 0.00573 | 0.00761 |
| **Trim17** | 0.03409 | 0.00471 | 0.01041 | 0.16059 | 0.00474 | 0.13018 | 0.00420 | 0.00975 | 0.07339 | 0.01660 | 0.00768 | 0.00766 | 0.01088 | 0.00000 |
| **Trim2** | 0.02708 | 0.02022 | 0.01529 | 0.00000 | 0.00552 | 0.03214 | 0.03021 | 2.51846 | 0.13283 | 0.02955 | 0.01471 | 0.26249 | 0.01469 | 0.06022 |
| **Trim23** | 0.06400 | 0.00447 | 0.69025 | 0.06613 | 0.92051 | 0.32003 | 0.00376 | 0.00191 | 0.66302 | 0.00302 | 0.30834 | 0.14406 | 0.04455 | 0.93833 |
| **Trim24** | 16.94316 | 7.27697 | 17.71779 | 10.03090 | 17.49383 | 9.07858 | 1.70715 | 0.00000 | 8.45539 | 10.89255 | 9.75949 | 9.72739 | 6.97551 | 14.56898 |
| **Trim25** | 2.93042 | 0.11966 | 5.74726 | 1.45455 | 1.60632 | 5.71387 | 1.48322 | 1.50824 | 3.66974 | 7.59351 | 0.40331 | 8.81865 | 1.71875 | 4.69799 |
| **Trim26** | 0.00184 | 0.06296 | 0.25582 | 0.40706 | 0.47604 | 0.81440 | 0.00998 | 3.56957 | 0.00186 | 0.01424 | 1.98219 | 1.10029 | 0.99507 | 1.10784 |
| **Trim27** | 12.01382 | 0.89624 | 8.73739 | 7.56304 | 9.26529 | 10.87005 | 1.94757 | 5.61279 | 5.44987 | 0.33903 | 4.57101 | 9.05852 | 8.12946 | 11.23325 |
| **Trim28** | 27.90126 | 10.64077 | 42.91373 | 33.65500 | 32.67793 | 29.79582 | 4.82288 | 19.90335 | 8.34755 | 16.98210 | 22.54867 | 29.02735 | 20.76855 | 24.80469 |
| **Trim3** | 0.66113 | 0.02786 | 0.00000 | 1.90584 | 0.51379 | 0.29483 | 0.00423 | 1.20169 | 0.00766 | 0.00245 | 1.36234 | 0.33386 | 0.06403 | 0.00000 |
| **Trim31** | 0.57888 | 0.00442 | 0.01365 | 0.01769 | 0.00392 | 0.01740 | 0.00424 | 0.01014 | 0.00875 | 0.01030 | 0.00657 | 0.00765 | 0.01819 | 0.00322 |
| **Trim32** | 0.36598 | 0.01976 | 1.00108 | 1.72624 | 1.03776 | 1.05997 | 1.49546 | 0.67036 | 0.24457 | 0.04183 | 0.00000 | 0.61923 | 1.18672 | 0.03826 |
| **Trim33** | 9.36462 | 0.15631 | 11.09818 | 0.07382 | 14.30930 | 8.16975 | 2.21022 | 1.62690 | 19.75125 | 0.01781 | 4.51009 | 6.76236 | 12.19663 | 20.58595 |
| **Trim34a** | 0.00559 | 0.00400 | 0.01217 | 0.00435 | 0.01081 | 0.01573 | 0.00431 | 0.00733 | 0.01063 | 0.00640 | 0.00499 | 0.39990 | 0.02945 | 0.05009 |
| **Trim35** | 0.07737 | 0.00444 | 0.01417 | 0.01149 | 0.01151 | 0.01232 | 0.00430 | 0.01086 | 0.01144 | 0.00669 | 0.00803 | 0.00972 | 0.00582 | 0.00903 |
| **Trim36** | 0.23178 | 0.07372 | 0.33088 | 0.65670 | 0.78717 | 0.68144 | 0.06685 | 0.03329 | 0.23631 | 0.10416 | 0.02035 | 0.21113 | 1.29049 | 0.45311 |
| **Trim37** | 6.23368 | 0.03821 | 2.37248 | 1.21952 | 4.90903 | 2.51001 | 0.00107 | 1.14364 | 13.38417 | 0.00703 | 2.91518 | 2.59268 | 2.25095 | 4.14698 |
| **Trim38** | 0.11286 | 0.01376 | 7.76885 | 2.87765 | 0.00000 | 3.58151 | 0.05597 | 0.05725 | 0.00000 | 0.00000 | 3.33296 | 4.92232 | 0.13730 | 0.16678 |
| **Trim39** | 3.66938 | 0.07292 | 5.95216 | 6.91059 | 6.13223 | 3.67354 | 3.98303 | 3.88045 | 11.88709 | 0.22251 | 1.37016 | 1.99262 | 2.71409 | 7.43209 |
| **Trim40** | 0.36480 | 0.00438 | 0.00000 | 0.01029 | 0.14697 | 0.00000 | 0.00404 | 0.01088 | 2.04459 | 0.00703 | 0.00369 | 0.00630 | 0.00058 | 1.30791 |
| **Trim41** | 1.38669 | 0.07878 | 3.67064 | 5.55778 | 1.73456 | 6.11016 | 0.46951 | 6.61199 | 0.36209 | 0.03613 | 0.55650 | 4.79808 | 4.96129 | 0.55957 |
| **Trim43a** | 34.41452 | 6.43617 | 41.53102 | 30.34168 | 46.31420 | 28.01087 | 4.10310 | 9.06239 | 49.61761 | 3.81831 | 15.68954 | 14.87387 | 14.52749 | 46.00751 |
| **Trim43b** | 33.49824 | 4.37771 | 36.84250 | 23.99775 | 46.06091 | 17.95046 | 2.33752 | 4.06993 | 49.48475 | 2.38223 | 11.97084 | 9.45260 | 18.25767 | 47.18707 |
| **Trim43c** | 29.96568 | 2.57820 | 23.93271 | 14.48769 | 35.60537 | 10.58652 | 1.42030 | 2.61454 | 39.45981 | 2.55732 | 7.44157 | 6.28519 | 14.04031 | 36.90789 |
| **Trim44** | 2.71817 | 0.06831 | 0.44641 | 0.00000 | 2.63350 | 0.00000 | 0.00737 | 0.07095 | 7.60260 | 0.05576 | 1.98449 | 3.65024 | 3.72684 | 4.53078 |
| **Trim45** | 0.01343 | 0.00433 | 0.00420 | 0.17431 | 0.00792 | 0.06634 | 0.00399 | 0.00242 | 0.01716 | 0.00454 | 0.57467 | 0.00937 | 0.00687 | 0.00417 |
| **Trim47** | 0.09598 | 0.00407 | 0.73275 | 1.19974 | 0.23844 | 1.89082 | 0.00387 | 0.48871 | 0.00329 | 0.00637 | 0.00000 | 1.19394 | 0.00734 | 0.08386 |
| **Trim5** | 0.01427 | 0.00457 | 0.03080 | 0.07025 | 0.01748 | 0.01718 | 0.01021 | 0.02909 | 0.07214 | 0.00681 | 0.02314 | 0.01486 | 0.01905 | 0.02624 |
| **Trim52** | 0.31893 | 0.00518 | 0.06614 | 0.00000 | 0.06595 | 0.10032 | 0.00417 | 0.02267 | 0.24058 | 0.00653 | 0.00507 | 0.14944 | 0.16972 | 0.06026 |

| | | | | | | | | | | | | | | |
|---|---|---|---|---|---|---|---|---|---|---|---|---|---|---|
| **Trim54** | 0.10419 | 0.00439 | 0.70077 | 0.49172 | 1.33059 | 0.00000 | 0.00437 | 0.01617 | 0.01534 | 0.00716 | 0.00538 | 0.00591 | 0.06277 | 0.00000 |
| **Trim56** | 0.02477 | 0.00469 | 0.02582 | 0.00770 | 0.14730 | 0.01348 | 0.00426 | 0.03062 | 0.00638 | 0.00676 | 0.00863 | 0.03930 | 0.10049 | 0.19865 |
| **Trim59** | 0.45282 | 1.12416 | 0.80119 | 0.36227 | 0.00000 | 0.91912 | 0.11591 | 1.67513 | 6.18726 | 0.01767 | 3.02119 | 0.13989 | 5.50992 | 0.00000 |
| **Trim6** | 5.95687 | 0.02992 | 7.02332 | 2.17910 | 5.66847 | 3.57680 | 0.01726 | 0.89466 | 14.17944 | 0.03268 | 2.05818 | 5.66850 | 8.91473 | 7.14155 |
| **Trim60** | 0.00000 | 0.00418 | 0.00527 | 0.01340 | 0.00000 | 0.00867 | 0.00414 | 0.00805 | 0.04350 | 0.00611 | 0.00676 | 0.00757 | 0.43150 | 2.17943 |
| **Trim61** | 0.00000 | 0.00271 | 0.00000 | 0.00815 | 1.50696 | 0.00000 | 0.00303 | 0.00459 | 4.78649 | 0.00757 | 0.00198 | 0.00000 | 9.19098 | 11.79834 |
| **Trim62** | 0.01017 | 0.00409 | 0.00496 | 0.01209 | 0.09950 | 1.28500 | 0.17484 | 0.07597 | 0.00483 | 0.00692 | 0.00440 | 0.00000 | 0.08924 | 0.02228 |
| **Trim63** | 0.27590 | 0.00413 | 0.03405 | 0.01227 | 0.00153 | 0.01235 | 0.00405 | 0.00828 | 0.01249 | 0.00817 | 0.00698 | 0.00609 | 1.84403 | 0.00102 |
| **Trim65** | 0.00761 | 0.00455 | 0.01402 | 0.00000 | 0.00576 | 0.04840 | 0.00474 | 0.01139 | 0.03841 | 0.00603 | 0.00614 | 0.10228 | 0.24030 | 0.10584 |
| **Trim66** | 0.00000 | 0.00402 | 0.44818 | 0.33402 | 0.77787 | 0.00000 | 0.00365 | 0.00469 | 1.37738 | 0.00489 | 0.01058 | 0.01389 | 0.00000 | 1.52971 |
| **Trim67** | 0.00650 | 0.02190 | 0.11423 | 0.00000 | 0.82991 | 0.00817 | 0.00422 | 0.01976 | 0.03408 | 0.00622 | 0.00664 | 0.02223 | 0.00616 | 0.09415 |
| **Trim68** | 0.00635 | 0.00376 | 0.00491 | 0.07593 | 0.42412 | 0.01184 | 0.00421 | 0.00445 | 0.01026 | 0.00509 | 0.00480 | 0.57333 | 0.00524 | 0.01014 |
| **Trim69** | 0.00641 | 0.00430 | 0.00935 | 0.00969 | 0.00404 | 0.02674 | 0.00418 | 0.00754 | 0.20737 | 0.00628 | 0.00790 | 0.00896 | 0.49449 | 0.00000 |
| **Trim7** | 1.01964 | 0.00370 | 10.78339 | 1.97607 | 4.36193 | 0.42413 | 0.00797 | 0.62737 | 5.31438 | 0.00000 | 0.24840 | 0.13867 | 0.00000 | 8.39853 |
| **Trim71** | 0.00000 | 0.00360 | 0.00000 | 0.00000 | 0.15361 | 0.00082 | 0.00352 | 0.94768 | 0.54572 | 0.00480 | 0.04007 | 0.01522 | 2.13330 | 0.09897 |
| **Trim75** | 0.02123 | 0.00409 | 0.10941 | 0.00323 | 0.03401 | 0.00989 | 0.00410 | 0.00911 | 0.00596 | 0.00615 | 0.00683 | 0.00760 | 1.54982 | 0.49502 |
| **Trim8** | 0.20488 | 1.10759 | 0.68733 | 0.47320 | 0.20681 | 0.88318 | 0.00409 | 0.34297 | 0.00476 | 0.01426 | 0.23845 | 0.29921 | 0.32003 | 0.00000 |
| **Triml1** | 0.01815 | 0.01891 | 0.24086 | 0.27414 | 0.31438 | 0.01994 | 0.00403 | 0.00475 | 0.41482 | 0.00545 | 0.03098 | 0.59280 | 0.01035 | 0.24089 |
| **Triml2** | 0.12564 | 0.17615 | 11.95147 | 11.39582 | 5.53896 | 0.00000 | 3.22250 | 6.42993 | 7.11939 | 0.19726 | 0.26342 | 2.99186 | 0.71639 | 4.88820 |
| **Trio** | 0.05191 | 1.15238 | 0.21419 | 0.32464 | 0.03498 | 0.79846 | 0.00388 | 0.83275 | 0.06931 | 0.00495 | 0.33504 | 0.24149 | 0.01252 | 0.19935 |
| **Triobp** | 0.18363 | 0.03395 | 1.83965 | 3.81255 | 1.59933 | 0.00000 | 0.05850 | 3.47414 | 0.00194 | 0.04442 | 0.00000 | 1.72878 | 0.28180 | 0.30774 |
| **Trip10** | 0.18990 | 0.26862 | 0.15877 | 0.17103 | 0.12633 | 0.00000 | 0.09383 | 0.13570 | 0.05378 | 0.02089 | 0.69801 | 0.03100 | 0.00600 | 0.21112 |
| **Trip11** | 0.09023 | 0.00389 | 0.11793 | 0.05524 | 0.00000 | 0.03096 | 0.00408 | 0.00474 | 1.08115 | 0.00623 | 0.00524 | 0.17692 | 0.85616 | 0.45486 |
| **Trip12** | 17.35111 | 0.08080 | 22.11003 | 13.04239 | 22.79670 | 22.23312 | 6.55977 | 10.39132 | 32.50248 | 7.83757 | 18.18880 | 17.18508 | 16.10464 | 22.33113 |
| **Trip13** | 12.42313 | 5.63030 | 16.28282 | 13.04459 | 23.49541 | 10.43665 | 4.83637 | 4.69470 | 13.08398 | 0.15289 | 11.71958 | 4.50659 | 6.86686 | 21.06105 |
| **Trip4** | 2.44326 | 5.28602 | 5.17224 | 2.16962 | 6.88826 | 1.56104 | 0.12164 | 5.97562 | 4.38171 | 0.59292 | 5.03534 | 2.87120 | 4.22947 | 7.64139 |
| **Trip6** | 0.26715 | 0.04292 | 0.01436 | 0.10359 | 0.27555 | 0.86004 | 0.00397 | 0.60988 | 0.00767 | 0.01610 | 0.02369 | 2.13634 | 0.01189 | 0.22380 |
| **Trit1** | 0.65970 | 0.00788 | 2.84186 | 5.11303 | 1.84102 | 2.29672 | 0.00799 | 0.03325 | 1.61541 | 0.00294 | 4.37229 | 4.51816 | 0.01307 | 2.41489 |
| **Trmo** | 0.83295 | 0.00448 | 2.48750 | 0.14730 | 3.14486 | 0.26215 | 0.00368 | 0.23638 | 0.47315 | 0.02154 | 0.81241 | 0.21757 | 0.16427 | 2.29608 |
| **Trmt1** | 1.50015 | 5.95974 | 13.25253 | 18.63603 | 5.30587 | 5.36603 | 0.10406 | 8.33570 | 0.00000 | 0.11270 | 9.23905 | 9.20571 | 0.04434 | 0.10552 |
| **Trmt10a** | 0.64920 | 0.03740 | 2.41351 | 6.42273 | 5.13719 | 4.24751 | 0.44658 | 0.12487 | 1.21990 | 0.09367 | 5.67406 | 4.37641 | 2.88211 | 10.26182 |
| **Trmt10b** | 3.57922 | 0.01008 | 3.87851 | 2.81248 | 0.69959 | 3.37411 | 0.01373 | 2.69448 | 0.82358 | 0.03168 | 0.10011 | 2.65271 | 0.00000 | 2.25744 |
| **Trmt10c** | 1.10869 | 0.17695 | 5.32796 | 1.19197 | 4.81396 | 1.28762 | 1.65536 | 0.59291 | 3.22250 | 0.04839 | 0.62595 | 1.82570 | 3.80590 | 7.47155 |
| **Trmt11** | 3.54851 | 0.00813 | 4.26479 | 0.00000 | 1.71718 | 2.38593 | 3.80833 | 2.79176 | 10.74743 | 0.04127 | 3.02610 | 2.22427 | 1.78184 | 1.50351 |
| **Trmt112** | 0.63627 | 0.13530 | 7.57010 | 10.79442 | 12.33119 | 1.96778 | 0.64297 | 5.62986 | 0.08241 | 6.58672 | 0.84551 | 10.81915 | 4.38256 | 0.00000 |
| **Trmt12** | 0.09872 | 0.01835 | 2.66620 | 1.71272 | 1.95543 | 0.34084 | 0.00115 | 0.02530 | 0.03655 | 12.55245 | 1.81043 | 0.09180 | 4.42772 | 0.61901 |
| **Trmt13** | 0.00000 | 0.00381 | 1.75706 | 0.09745 | 0.66301 | 1.31565 | 1.23624 | 0.34619 | 1.52545 | 0.03316 | 0.13530 | 0.14927 | 0.00000 | 1.34048 |
| **Trmt1l** | 10.71127 | 0.00846 | 7.52826 | 0.49255 | 6.03322 | 5.03698 | 1.65104 | 1.99422 | 5.04038 | 0.01129 | 5.49653 | 2.26724 | 1.10183 | 5.71828 |
| **Trmt2a** | 9.25394 | 0.11481 | 11.88241 | 5.73424 | 4.74481 | 7.08242 | 3.15513 | 3.99892 | 1.10463 | 0.10179 | 3.95406 | 6.14727 | 0.63723 | 4.82396 |
| **Trmt2b** | 1.53573 | 0.01341 | 0.00000 | 1.03073 | 4.13303 | 0.21205 | 0.00413 | 0.85046 | 1.85058 | 0.01546 | 0.00000 | 0.00160 | 0.05265 | 4.78001 |
| **Trmt44** | 0.50326 | 0.01079 | 1.31463 | 0.78510 | 0.50492 | 0.36343 | 0.00335 | 0.43049 | 0.49284 | 0.02580 | 0.35620 | 0.74642 | 0.05877 | 0.27012 |
| **Trmt5** | 2.73882 | 0.05072 | 7.60051 | 1.92368 | 3.86301 | 5.08665 | 2.48803 | 2.67290 | 3.18395 | 9.79395 | 7.16441 | 1.76871 | 0.99363 | 10.64128 |
| **Trmt6** | 9.46660 | 0.03854 | 13.31194 | 10.98550 | 10.73138 | 7.86864 | 0.96966 | 0.62238 | 8.40849 | 0.10422 | 10.30709 | 5.36245 | 2.25142 | 15.01396 |
| **Trmt61a** | 3.62471 | 0.00176 | 3.56416 | 0.92760 | 6.99332 | 1.95030 | 0.16728 | 0.25825 | 0.63485 | 0.20636 | 1.19523 | 2.20822 | 6.26362 | 2.80219 |
| **Trmt61b** | 2.51891 | 0.03008 | 4.64228 | 0.00000 | 8.49023 | 0.51486 | 0.00296 | 1.48108 | 1.17556 | 0.47658 | 0.00000 | 0.41109 | 1.03639 | 2.06510 |
| **Trmu** | 0.53506 | 0.00344 | 0.88461 | 2.28419 | 0.25470 | 0.46718 | 2.59137 | 0.53991 | 0.08173 | 0.05902 | 0.00000 | 0.62641 | 0.00612 | 0.11299 |
| **Trnau1ap** | 1.91715 | 0.00186 | 0.72747 | 3.25163 | 6.37830 | 1.59185 | 0.00295 | 0.21488 | 1.18518 | 0.00731 | 0.00000 | 2.97267 | 6.66905 | 0.89478 |
| **Trnp1** | 0.91723 | 0.00368 | 0.40104 | 0.88078 | 0.45153 | 0.00000 | 0.00391 | 0.00523 | 0.01278 | 0.00618 | 0.00703 | 0.00827 | 2.77400 | 0.00000 |
| **Trnt1** | 11.48592 | 0.03780 | 4.71120 | 1.51666 | 4.72544 | 3.26986 | 2.25670 | 1.15280 | 10.92406 | 0.07677 | 3.96853 | 2.69561 | 3.72875 | 7.74715 |
| **Tro** | 0.00765 | 0.00434 | 0.01641 | 0.04761 | 0.02641 | 0.02379 | 0.00432 | 0.00700 | 0.03125 | 0.00582 | 0.00808 | 0.00766 | 0.24368 | 0.00274 |
| **Troap** | 0.18627 | 0.05856 | 0.99015 | 2.45161 | 2.03406 | 0.12294 | 0.00944 | 2.74978 | 0.23561 | 0.01964 | 0.32597 | 2.24035 | 0.69493 | 0.00000 |
| **Trove2** | 1.32918 | 0.04975 | 0.26594 | 0.63657 | 0.78045 | 0.70039 | 0.08055 | 0.01088 | 4.11471 | 0.04179 | 0.21958 | 0.62993 | 0.28860 | 1.82112 |
| **Trp53** | 1.08332 | 5.80046 | 6.16564 | 0.88213 | 2.41248 | 2.34367 | 0.01581 | 2.91758 | 0.11211 | 0.00053 | 3.92379 | 7.14477 | 0.00000 | 4.69668 |
| **Trp53bp1** | 7.29225 | 0.03688 | 5.80239 | 1.98751 | 8.77772 | 2.46307 | 0.03905 | 3.03642 | 4.95529 | 4.39237 | 0.30527 | 5.11658 | 5.03880 | 15.73112 |
| **Trp53bp2** | 0.00413 | 0.01677 | 0.02225 | 3.11270 | 0.00000 | 18.46929 | 1.66131 | 1.94917 | 0.00750 | 0.07053 | 4.71671 | 0.74475 | 0.41029 | 0.00000 |

| | | | | | | | | | | | | | | |
|---|---|---|---|---|---|---|---|---|---|---|---|---|---|---|
| **Trp53i11** | 0.00000 | 0.01140 | 3.02543 | 0.86234 | 0.92109 | 1.94905 | 0.02612 | 0.78653 | 0.04566 | 0.00370 | 0.00000 | 1.25551 | 0.00000 | 0.30132 |
| **Trp53i13** | 0.00406 | 0.00408 | 0.00591 | 1.08230 | 0.17172 | 0.07192 | 0.00403 | 0.00452 | 0.01878 | 0.00495 | 0.00730 | 0.00841 | 0.01188 | 0.00104 |
| **Trp53inp1** | 1.00980 | 0.01642 | 1.56968 | 1.77539 | 0.64067 | 1.99047 | 1.52887 | 0.01625 | 3.07416 | 0.02683 | 0.08262 | 1.63312 | 4.83992 | 0.68157 |
| **Trp53inp2** | 0.91378 | 0.01967 | 4.75305 | 2.08825 | 1.46519 | 7.44010 | 0.42281 | 1.78531 | 0.65648 | 0.08486 | 0.29717 | 0.94446 | 4.86336 | 7.51754 |
| **Trp53rka** | 0.71878 | 0.01831 | 0.14953 | 0.67903 | 0.43941 | 1.45395 | 1.13998 | 1.07139 | 0.16479 | 0.04272 | 0.00000 | 1.65829 | 1.32235 | 1.52728 |
| **Trp53rkb** | 1.39235 | 0.01144 | 0.18853 | 0.89270 | 0.32137 | 0.74565 | 2.39273 | 0.25790 | 0.21414 | 0.02725 | 0.00629 | 0.44749 | 1.34775 | 1.14398 |
| **Trp73** | 0.00000 | 0.00414 | 0.88712 | 1.01128 | 0.00000 | 1.08745 | 0.00397 | 0.00000 | 0.00614 | 0.00917 | 0.01598 | 0.00616 | 0.35247 | 0.00000 |
| **Trpa1** | 0.00000 | 0.00395 | 0.17280 | 1.01342 | 0.00000 | 0.00000 | 0.00399 | 0.00653 | 0.01497 | 0.00527 | 0.00475 | 0.01303 | 0.42563 | 0.00982 |
| **Trpc1** | 0.04092 | 0.00464 | 0.03365 | 0.37895 | 0.07087 | 0.03792 | 0.01685 | 0.03635 | 0.08312 | 0.01804 | 0.03858 | 0.02715 | 0.80110 | 0.16263 |
| **Trpc4ap** | 1.10831 | 8.73120 | 5.59308 | 5.81734 | 9.14890 | 4.59657 | 3.08148 | 2.39661 | 0.06377 | 0.12961 | 2.80134 | 0.68045 | 1.30377 | 2.37805 |
| **Trpc5** | 0.04862 | 0.00410 | 0.00684 | 0.03569 | 0.01424 | 0.12666 | 0.02807 | 0.03551 | 0.52322 | 0.00760 | 0.00364 | 0.17467 | 0.61268 | 0.38484 |
| **Trpc5os** | 0.00883 | 0.00467 | 0.22529 | 0.16918 | 0.00005 | 0.00626 | 0.00423 | 0.01387 | 0.01230 | 0.00947 | 0.00707 | 0.00657 | 0.01619 | 0.29336 |
| **Trpc6** | 21.45694 | 2.49737 | 6.41475 | 1.69436 | 11.94331 | 11.43045 | 1.52869 | 1.97489 | 11.55574 | 1.14299 | 3.82984 | 1.83053 | 14.88506 | 5.84037 |
| **Trpd52l3** | 0.00533 | 0.00444 | 0.00584 | 0.01341 | 0.22656 | 0.01130 | 0.00445 | 0.01004 | 0.00554 | 0.00660 | 0.00777 | 0.00847 | 0.00789 | 0.67687 |
| **Trpm3** | 0.01092 | 0.01378 | 0.01769 | 0.04776 | 0.01659 | 0.02813 | 0.03540 | 0.05256 | 0.06093 | 0.00961 | 0.01059 | 0.01858 | 0.02734 | 0.05004 |
| **Trpm4** | 0.00000 | 0.05508 | 0.16554 | 1.46605 | 0.00000 | 0.10555 | 0.01071 | 3.40154 | 0.01370 | 0.02284 | 0.89820 | 0.87494 | 0.00571 | 0.00000 |
| **Trpm6** | 0.00074 | 0.00379 | 0.00000 | 0.00321 | 0.00000 | 1.14316 | 0.00337 | 2.51587 | 0.04084 | 0.00900 | 1.27347 | 0.00185 | 0.03509 | 0.49729 |
| **Trpm7** | 3.44083 | 0.03596 | 2.00878 | 0.00000 | 2.06677 | 4.58327 | 0.26423 | 0.63097 | 10.68582 | 0.03899 | 5.97614 | 6.66104 | 6.49072 | 7.08727 |
| **Trps1** | 0.22202 | 0.01222 | 0.00000 | 0.00000 | 0.00000 | 0.02162 | 0.00384 | 0.02028 | 0.79347 | 0.00662 | 0.00001 | 1.26271 | 0.00808 | 0.03468 |
| **Trpt1** | 0.75045 | 0.11308 | 2.22533 | 0.64703 | 0.53084 | 1.21481 | 0.22913 | 2.98015 | 0.00000 | 0.52935 | 0.00000 | 1.00299 | 1.60013 | 0.69436 |
| **Trpv1** | 0.00542 | 0.00435 | 0.01027 | 0.55109 | 0.00683 | 0.00000 | 0.00421 | 0.01017 | 0.01506 | 0.00686 | 0.01442 | 0.11666 | 0.00565 | 0.01195 |
| **Trpv2** | 0.02278 | 0.00483 | 0.00856 | 0.01750 | 0.00352 | 0.00823 | 0.00429 | 0.01967 | 0.21405 | 0.00761 | 0.00714 | 0.01048 | 0.05906 | 0.05670 |
| **Trpv4** | 0.01639 | 0.00443 | 0.01817 | 0.00764 | 0.01329 | 0.16493 | 0.01014 | 0.00533 | 0.01373 | 0.00571 | 0.02246 | 0.01147 | 0.00734 | 0.00860 |
| **Trpv5** | 0.21299 | 0.00355 | 0.74838 | 0.00000 | 0.70068 | 0.22863 | 0.38738 | 1.55890 | 0.00850 | 0.02039 | 0.03439 | 0.51884 | 0.00280 | 0.16128 |
| **Trpv6** | 0.01024 | 0.01068 | 0.01251 | 0.00000 | 0.01254 | 0.00466 | 0.00359 | 0.60779 | 0.01252 | 0.00561 | 0.00499 | 0.10202 | 0.00794 | 0.01139 |
| **Trrap** | 0.93720 | 0.34567 | 2.45988 | 1.74604 | 0.94678 | 1.69273 | 0.00330 | 0.18759 | 0.29890 | 0.00117 | 2.29683 | 2.18041 | 2.97277 | 8.47870 |
| **Trub1** | 3.40501 | 6.35090 | 3.91494 | 2.72439 | 0.93238 | 4.75163 | 1.51625 | 5.16689 | 0.20068 | 0.16840 | 4.15483 | 6.80325 | 0.41226 | 0.94849 |
| **Trub2** | 2.56330 | 0.05598 | 6.70844 | 5.00272 | 2.62051 | 6.64558 | 3.45389 | 3.01873 | 3.67096 | 0.11719 | 4.83214 | 6.75084 | 10.32115 | 5.01934 |
| **Tsacc** | 0.04389 | 0.00417 | 0.00000 | 0.39245 | 0.39518 | 0.26112 | 0.00425 | 0.37831 | 0.62587 | 0.01595 | 0.56728 | 0.06051 | 0.00961 | 0.23382 |
| **Tsc1** | 0.87109 | 0.00144 | 1.88977 | 1.20732 | 2.98880 | 2.98425 | 0.00257 | 0.00000 | 0.80650 | 0.01617 | 2.86064 | 1.55100 | 4.26541 | 4.07651 |
| **Tsc2** | 5.28159 | 0.89905 | 10.66976 | 5.51972 | 6.21925 | 4.80457 | 0.04340 | 4.54025 | 1.17784 | 0.07467 | 3.83168 | 7.22326 | 1.98560 | 3.24845 |
| **Tsc22d1** | 0.79681 | 9.40471 | 6.52221 | 2.23135 | 6.33457 | 2.17093 | 1.85601 | 11.71341 | 0.80446 | 0.09631 | 2.75124 | 9.53838 | 5.52455 | 0.66145 |
| **Tsc22d2** | 0.15756 | 0.00102 | 1.07325 | 0.31212 | 2.77270 | 0.46751 | 0.16334 | 0.65995 | 13.33948 | 0.04344 | 0.52557 | 1.31123 | 14.97173 | 6.46944 |
| **Tsc22d3** | 0.01802 | 0.00471 | 0.00661 | 0.00000 | 0.21016 | 0.25205 | 0.00434 | 0.00865 | 0.02255 | 0.00581 | 0.00904 | 0.01031 | 0.00654 | 0.01160 |
| **Tsc22d4** | 1.67456 | 2.20281 | 4.66669 | 3.95646 | 9.80027 | 0.73991 | 1.26338 | 2.13674 | 0.08264 | 4.69085 | 0.00000 | 1.00621 | 7.69044 | 1.04601 |
| **Tsen15** | 1.28766 | 0.05175 | 0.64296 | 3.55080 | 3.99661 | 0.50472 | 1.82021 | 2.30974 | 0.23190 | 0.04645 | 1.88847 | 6.93285 | 0.89902 | 1.28090 |
| **Tsen2** | 5.61904 | 0.04324 | 6.77432 | 3.54247 | 9.11504 | 3.18279 | 0.00807 | 2.21144 | 0.32310 | 0.02804 | 4.69744 | 2.57220 | 5.77271 | 6.52549 |
| **Tsen34** | 8.37161 | 0.14648 | 6.36407 | 5.49997 | 9.17713 | 5.45949 | 4.93373 | 2.30902 | 0.14205 | 10.44878 | 3.33710 | 4.50453 | 3.56261 | 1.36630 |
| **Tsen54** | 0.57603 | 4.75258 | 1.35134 | 0.72151 | 1.90974 | 2.18938 | 0.00251 | 1.14047 | 0.00484 | 0.03565 | 0.13930 | 0.72253 | 0.87146 | 1.26026 |
| **Tsfm** | 0.00000 | 0.06297 | 3.39621 | 8.67542 | 3.80122 | 0.81544 | 0.04735 | 8.05922 | 0.01122 | 0.00788 | 3.76636 | 2.72516 | 0.00000 | 0.08382 |
| **Tsg101** | 12.65577 | 0.20871 | 6.98225 | 6.46844 | 6.19771 | 10.19289 | 3.53348 | 8.74282 | 8.82611 | 0.10008 | 6.97093 | 7.71231 | 4.45544 | 5.86542 |
| **Tsga10** | 0.00380 | 0.00416 | 0.01485 | 0.00619 | 0.11195 | 0.03282 | 0.00429 | 0.00715 | 0.00824 | 0.00543 | 0.32934 | 0.01067 | 0.43921 | 0.00281 |
| **Tsga8** | 0.01000 | 0.00447 | 0.00969 | 0.01035 | 0.28493 | 0.01353 | 0.00421 | 0.01043 | 0.01132 | 0.00647 | 0.00916 | 0.00876 | 0.00543 | 0.00891 |
| **Tshz1** | 0.00065 | 0.00507 | 0.39432 | 0.00000 | 0.23529 | 0.09243 | 0.00411 | 0.06004 | 0.02530 | 0.02067 | 0.00547 | 0.00771 | 0.74311 | 0.03638 |
| **Tshz2** | 0.00537 | 0.00432 | 0.01658 | 0.01252 | 0.00911 | 0.01314 | 0.00422 | 0.00901 | 0.01876 | 0.00682 | 0.00878 | 0.00917 | 0.43014 | 0.00981 |
| **Tsix** | 0.01675 | 0.00539 | 0.02555 | 0.00000 | 0.01337 | 0.00616 | 0.00455 | 0.13890 | 0.01357 | 0.00903 | 0.00646 | 0.10791 | 0.01366 | 0.01533 |
| **Tsks** | 0.12542 | 0.00424 | 0.09185 | 0.00000 | 0.70281 | 0.00796 | 0.00409 | 0.00921 | 0.09366 | 0.00667 | 0.00631 | 0.00680 | 1.32159 | 0.00000 |
| **Tsku** | 0.01433 | 0.00430 | 0.07912 | 0.29903 | 0.00755 | 0.00619 | 0.00423 | 0.01124 | 0.01205 | 0.00770 | 0.00656 | 0.00937 | 0.00627 | 0.00957 |
| **Tslp** | 0.00127 | 0.00444 | 0.00189 | 0.01183 | 0.30561 | 0.00807 | 0.00424 | 0.01000 | 0.32879 | 0.00754 | 0.00657 | 0.00766 | 0.01241 | 0.74553 |
| **Tsn** | 8.53598 | 9.75880 | 13.98401 | 16.75783 | 14.46675 | 15.43564 | 4.08493 | 6.83841 | 9.60296 | 0.22068 | 11.58338 | 12.40071 | 1.62680 | 13.76792 |
| **Tsnax** | 6.77674 | 0.40081 | 6.45684 | 1.50199 | 5.98622 | 8.66297 | 0.51078 | 1.64103 | 3.42368 | 2.79048 | 3.21404 | 5.76851 | 1.00017 | 9.03519 |
| **Tsnaxip1** | 0.31886 | 0.00394 | 1.31213 | 0.00000 | 0.48684 | 0.17380 | 0.00378 | 0.06051 | 2.44164 | 0.00424 | 0.00281 | 0.00476 | 0.38196 | 0.71642 |
| **Tspan1** | 0.05903 | 0.11587 | 6.51265 | 8.62894 | 2.20271 | 2.96302 | 2.66745 | 6.69192 | 2.82646 | 1.66636 | 2.76067 | 1.85743 | 0.00000 | 0.60440 |
| **Tspan10** | 0.57979 | 0.00405 | 0.64367 | 0.00331 | 0.00000 | 1.11635 | 0.01186 | 0.32420 | 0.00728 | 0.00066 | 1.35312 | 0.00000 | 0.09428 | 0.00326 |
| **Tspan13** | 0.03302 | 0.00406 | 0.04076 | 0.09795 | 0.52572 | 0.01289 | 0.00436 | 0.00487 | 0.01198 | 0.01675 | 0.00633 | 0.01185 | 0.97534 | 0.00000 |

| | | | | | | | | | | | | | | |
|---|---|---|---|---|---|---|---|---|---|---|---|---|---|---|
| **Tspan14** | 2.30061 | 0.03050 | 1.94665 | 0.00003 | 0.74008 | 1.65858 | 0.13764 | 1.41811 | 0.15142 | 0.01321 | 3.95669 | 3.11957 | 1.03946 | 2.69263 |
| **Tspan15** | 0.00910 | 0.04571 | 0.03676 | 0.13303 | 0.04684 | 0.58344 | 0.01509 | 2.86025 | 0.08536 | 0.03428 | 0.00418 | 0.00744 | 0.17040 | 0.76210 |
| **Tspan17** | 0.16964 | 0.02841 | 7.41881 | 4.30364 | 1.23605 | 0.55933 | 0.00286 | 1.69746 | 0.09872 | 0.00000 | 0.17502 | 0.87079 | 0.15274 | 0.00000 |
| **Tspan3** | 0.75274 | 0.02330 | 1.08820 | 1.54020 | 0.17459 | 6.00588 | 1.39989 | 0.15138 | 0.00001 | 0.04523 | 7.79274 | 2.42461 | 0.00042 | 0.02972 |
| **Tspan31** | 1.73881 | 7.63354 | 7.38562 | 1.53526 | 4.60424 | 12.13211 | 4.97023 | 5.51811 | 3.06206 | 0.26926 | 4.30866 | 9.25782 | 0.66220 | 5.96226 |
| **Tspan33** | 0.02566 | 0.55700 | 0.01112 | 0.00000 | 0.03208 | 0.00992 | 0.00416 | 1.64645 | 0.01329 | 0.01146 | 0.00492 | 0.02370 | 0.00783 | 0.01191 |
| **Tspan4** | 3.24765 | 0.02027 | 7.28991 | 1.56491 | 12.94231 | 0.42373 | 0.00309 | 0.96849 | 0.92448 | 0.00000 | 0.02122 | 0.58934 | 0.11116 | 6.69859 |
| **Tspan5** | 0.00590 | 0.00431 | 0.01120 | 0.12763 | 0.01596 | 0.08599 | 0.00405 | 0.00415 | 0.00932 | 0.00637 | 0.00604 | 0.01016 | 1.40616 | 0.02683 |
| **Tspan6** | 0.01693 | 0.00416 | 0.06053 | 0.00732 | 0.00249 | 0.00000 | 0.00400 | 0.00974 | 0.39685 | 0.00500 | 0.76080 | 0.00971 | 0.03159 | 0.00000 |
| **Tspan7** | 0.00838 | 0.00492 | 0.16091 | 0.00000 | 0.00923 | 0.23643 | 0.00431 | 0.00629 | 0.01135 | 0.00608 | 0.00729 | 0.00885 | 0.00793 | 0.00836 |
| **Tspan8** | 1.32599 | 12.96530 | 0.47720 | 0.00000 | 0.19677 | 7.14261 | 11.18139 | 32.73406 | 0.13332 | 11.49160 | 0.30596 | 0.40155 | 1.35660 | 0.55274 |
| **Tspan9** | 0.01061 | 0.02482 | 3.03685 | 0.00002 | 0.76350 | 0.94761 | 0.00926 | 0.33349 | 0.00029 | 0.02380 | 0.63447 | 1.46596 | 0.29834 | 0.05503 |
| **Tspo** | 0.00334 | 0.02502 | 0.01623 | 0.18758 | 0.00945 | 0.00592 | 0.00350 | 2.05260 | 0.01147 | 0.00644 | 0.12674 | 2.26256 | 0.00666 | 0.00804 |
| **Tspyl1** | 1.45333 | 0.02565 | 1.02947 | 2.54977 | 3.15381 | 4.19625 | 2.74434 | 0.60046 | 1.81645 | 0.13241 | 1.66160 | 2.42235 | 2.02431 | 3.16049 |
| **Tspyl2** | 0.54300 | 0.00430 | 0.00000 | 0.95447 | 0.12764 | 0.07186 | 0.00494 | 0.40404 | 1.08620 | 0.00629 | 0.00299 | 0.00667 | 0.69417 | 1.68386 |
| **Tspyl3** | 0.01250 | 0.00448 | 0.01618 | 0.00878 | 0.01149 | 0.05502 | 0.00427 | 0.00734 | 0.00969 | 0.00677 | 0.00813 | 0.01022 | 0.03079 | 0.05459 |
| **Tspyl4** | 0.00810 | 0.00433 | 0.01178 | 0.01200 | 0.00431 | 0.01491 | 0.00420 | 0.00895 | 0.07183 | 0.00645 | 0.00914 | 0.00905 | 0.35362 | 0.08148 |
| **Tsr1** | 6.99880 | 0.00767 | 11.90700 | 2.12474 | 10.26966 | 4.55027 | 3.15102 | 1.93315 | 11.41766 | 0.09150 | 5.85346 | 2.34405 | 3.38825 | 10.48166 |
| **Tsr2** | 8.76587 | 0.04275 | 24.11604 | 8.63560 | 20.75811 | 11.78535 | 1.36075 | 3.25026 | 14.49540 | 6.91412 | 4.21160 | 4.50310 | 0.00000 | 23.22783 |
| **Tsr3** | 5.52688 | 0.00217 | 2.14606 | 7.83136 | 7.53969 | 1.30123 | 2.70001 | 0.14148 | 0.31603 | 3.37320 | 0.26193 | 0.70961 | 3.47456 | 2.54110 |
| **Tssc1** | 6.97683 | 0.01549 | 9.53899 | 3.21490 | 4.92873 | 5.99133 | 2.16737 | 1.29024 | 0.66708 | 0.02135 | 7.11042 | 2.62594 | 0.16748 | 4.23456 |
| **Tssc4** | 2.15146 | 8.86479 | 3.97990 | 3.38507 | 3.26393 | 2.89829 | 0.00113 | 0.39609 | 0.00000 | 0.00000 | 4.32601 | 1.60596 | 2.98591 | 2.33403 |
| **Tst** | 0.00000 | 0.00386 | 0.17615 | 0.90850 | 0.32899 | 0.00012 | 0.00441 | 0.00893 | 0.01299 | 0.00818 | 0.79041 | 0.75689 | 0.00148 | 0.13421 |
| **Tsta3** | 0.65355 | 2.23932 | 5.05356 | 7.14225 | 0.69947 | 3.52625 | 3.17564 | 6.88887 | 0.05549 | 0.10145 | 4.30933 | 4.18920 | 0.87016 | 0.97257 |
| **Tstd1** | 0.54455 | 0.02318 | 0.07443 | 0.76530 | 0.51408 | 0.58163 | 0.56320 | 0.76642 | 0.97359 | 0.05087 | 0.00000 | 0.00000 | 2.52046 | 0.22391 |
| **Tstd2** | 0.53875 | 0.00363 | 1.61407 | 0.59990 | 0.46965 | 0.15246 | 0.00328 | 0.70499 | 0.50810 | 0.00795 | 0.39359 | 0.36001 | 0.08920 | 0.34341 |
| **Tstd3** | 0.10122 | 0.03511 | 0.94922 | 0.14024 | 1.16267 | 2.45020 | 2.02091 | 0.53387 | 0.77481 | 0.13461 | 0.00000 | 1.48291 | 1.16728 | 0.86718 |
| **Tsx** | 1.16952 | 0.00291 | 0.15749 | 0.55393 | 0.42433 | 0.16716 | 0.01003 | 0.31308 | 0.00641 | 0.02756 | 0.02843 | 1.05753 | 1.80731 | 0.97214 |
| **Ttbk1** | 0.00872 | 0.00418 | 0.00868 | 0.16804 | 0.00574 | 0.00791 | 0.00402 | 0.00628 | 0.01744 | 0.00686 | 0.00699 | 0.01436 | 0.11441 | 0.11391 |
| **Ttbk2** | 1.69914 | 0.02877 | 1.66244 | 0.17681 | 0.90643 | 2.76377 | 0.08698 | 0.05563 | 2.88713 | 0.05706 | 0.00000 | 1.23738 | 1.71389 | 4.22631 |
| **Ttc1** | 19.51884 | 0.47593 | 25.53685 | 16.32749 | 23.21699 | 17.69340 | 2.78543 | 4.70249 | 18.63224 | 0.19916 | 16.42258 | 16.51353 | 13.32191 | 21.47945 |
| **Ttc12** | 0.00935 | 0.00434 | 0.00395 | 0.05640 | 0.57218 | 0.01028 | 0.00411 | 0.00774 | 0.01374 | 0.01334 | 0.00759 | 0.01141 | 0.00454 | 0.00547 |
| **Ttc13** | 6.01517 | 0.01519 | 10.40954 | 3.86200 | 3.49223 | 6.09664 | 1.88219 | 2.89511 | 0.61019 | 0.00000 | 4.56717 | 4.74570 | 0.20990 | 3.25210 |
| **Ttc14** | 3.33439 | 0.00822 | 1.10016 | 0.61209 | 1.55411 | 2.84751 | 1.63245 | 0.42748 | 8.38159 | 0.01338 | 2.12919 | 1.44378 | 5.10030 | 1.30817 |
| **Ttc16** | 0.06797 | 0.00422 | 0.00831 | 0.01220 | 0.31675 | 0.01056 | 0.00431 | 0.00951 | 0.19638 | 0.00734 | 0.00676 | 0.00710 | 0.70279 | 0.44557 |
| **Ttc17** | 0.86559 | 0.01821 | 3.43667 | 0.75782 | 0.00000 | 1.74645 | 0.63233 | 0.00000 | 2.15477 | 0.04618 | 1.10429 | 0.49184 | 0.00003 | 2.03350 |
| **Ttc19** | 2.20901 | 0.01016 | 1.97020 | 1.70019 | 2.63900 | 3.20539 | 1.23300 | 1.72922 | 3.91769 | 0.08068 | 3.54245 | 1.45445 | 0.70919 | 3.37348 |
| **Ttc21a** | 0.00967 | 0.00443 | 0.08357 | 0.00474 | 0.04436 | 0.01098 | 0.00418 | 0.01085 | 0.01132 | 0.00712 | 0.00814 | 0.01031 | 0.00839 | 0.01799 |
| **Ttc23** | 0.01522 | 0.00466 | 0.00857 | 0.01086 | 0.12680 | 0.01789 | 0.00452 | 0.01360 | 0.13180 | 0.00678 | 0.00738 | 0.00879 | 0.32449 | 0.00000 |
| **Ttc23l** | 0.04389 | 0.00444 | 0.02666 | 0.01080 | 0.01206 | 0.01226 | 0.00427 | 0.01059 | 0.01017 | 0.00707 | 0.00807 | 0.01022 | 0.00603 | 0.01566 |
| **Ttc25** | 0.00859 | 0.02075 | 1.73776 | 0.32898 | 1.73938 | 0.00843 | 0.00372 | 0.01100 | 0.52972 | 0.00353 | 0.03751 | 0.01059 | 0.04033 | 0.00023 |
| **Ttc26** | 0.08576 | 0.02883 | 0.02963 | 0.60456 | 0.34481 | 0.33914 | 0.04588 | 0.05994 | 0.29713 | 0.05715 | 0.01834 | 0.08858 | 1.09625 | 1.17073 |
| **Ttc27** | 10.34206 | 0.03537 | 16.09479 | 6.26490 | 10.97225 | 5.48047 | 0.01594 | 1.03601 | 7.36669 | 0.00000 | 5.38521 | 3.11041 | 2.56952 | 9.47552 |
| **Ttc28** | 0.10292 | 0.00443 | 0.11498 | 0.00543 | 0.41840 | 0.00731 | 0.00423 | 0.01094 | 0.01216 | 0.00597 | 0.00681 | 0.00871 | 0.06418 | 0.11575 |
| **Ttc3** | 0.00000 | 0.02525 | 0.55469 | 1.13487 | 2.42377 | 1.81465 | 0.00229 | 0.25921 | 5.91142 | 0.00000 | 1.57581 | 1.70647 | 0.04713 | 6.10740 |
| **Ttc30a1** | 0.10987 | 0.00504 | 0.07650 | 0.00000 | 0.75745 | 0.16067 | 0.00421 | 0.06602 | 0.15753 | 0.00874 | 0.00662 | 0.10072 | 2.76096 | 0.05131 |
| **Ttc30a2** | 0.01408 | 0.00425 | 0.01580 | 0.00936 | 0.03535 | 0.04544 | 0.00420 | 0.00652 | 0.07508 | 0.00633 | 0.00971 | 0.04611 | 0.09198 | 0.03888 |
| **Ttc30b** | 0.91427 | 0.00365 | 0.03121 | 0.45294 | 0.17290 | 0.42010 | 0.00382 | 0.00300 | 0.32702 | 0.00579 | 0.00837 | 0.10109 | 1.81176 | 0.00000 |
| **Ttc32** | 0.12476 | 0.00394 | 0.01783 | 0.45886 | 0.67911 | 0.09147 | 0.01586 | 0.76296 | 0.01531 | 0.01436 | 0.14542 | 0.30551 | 0.11050 | 0.00437 |
| **Ttc33** | 0.72875 | 0.02305 | 6.84238 | 2.21471 | 3.80741 | 7.21349 | 0.00623 | 1.46747 | 5.59778 | 0.01562 | 2.94268 | 2.82911 | 0.63117 | 8.82842 |
| **Ttc34** | 0.00889 | 0.00448 | 0.03622 | 0.00954 | 0.00660 | 0.00914 | 0.00419 | 0.01085 | 0.29089 | 0.00738 | 0.00727 | 0.00957 | 0.03992 | 0.00365 |
| **Ttc37** | 0.03094 | 0.00353 | 0.88844 | 0.79085 | 0.07271 | 0.37087 | 0.01012 | 0.28765 | 2.16304 | 0.00276 | 0.70649 | 0.62689 | 0.03370 | 1.15563 |
| **Ttc38** | 0.08032 | 0.03588 | 0.01051 | 0.94343 | 0.41232 | 0.15489 | 0.02866 | 0.14176 | 0.46003 | 0.02335 | 0.10868 | 0.01653 | 0.24703 | 0.64931 |
| **Ttc39a** | 0.01178 | 0.01201 | 0.01140 | 0.20078 | 0.16659 | 0.00000 | 0.00503 | 1.08574 | 0.05681 | 0.00375 | 0.65017 | 0.00241 | 0.20186 | 0.00000 |
| **Ttc39b** | 2.59106 | 0.02799 | 4.01162 | 1.72709 | 0.81119 | 1.48122 | 0.03107 | 0.57826 | 3.77228 | 0.01073 | 0.87283 | 0.70651 | 0.25524 | 1.22979 |

| | | | | | | | | | | | | | | |
|---|---|---|---|---|---|---|---|---|---|---|---|---|---|---|
| **Ttc39c** | 0.00000 | 0.00442 | 0.48360 | 0.45967 | 0.08131 | 0.00000 | 0.00438 | 0.08497 | 0.01516 | 0.00093 | 1.75032 | 0.05263 | 0.00409 | 0.20573 |
| **Ttc4** | 3.32564 | 0.07071 | 9.56772 | 7.33013 | 5.14216 | 10.38451 | 5.72348 | 3.16324 | 5.23971 | 0.18274 | 7.98708 | 7.29023 | 1.06730 | 9.05772 |
| **Ttc41** | 0.30735 | 0.00470 | 0.18326 | 0.57748 | 0.00000 | 0.65420 | 0.00400 | 0.00538 | 0.32824 | 0.00419 | 0.69851 | 0.05942 | 0.00510 | 0.00000 |
| **Ttc5** | 0.00000 | 0.44727 | 2.32717 | 0.74414 | 2.70723 | 0.54610 | 0.02171 | 2.09678 | 0.55686 | 0.10570 | 1.21876 | 0.97036 | 0.13895 | 1.45659 |
| **Ttc6** | 0.00432 | 0.00439 | 0.02185 | 0.08533 | 0.04445 | 0.00331 | 0.00418 | 0.00744 | 0.81902 | 0.00711 | 0.00489 | 0.01333 | 0.00341 | 0.05329 |
| **Ttc7** | 0.55273 | 3.42844 | 0.00000 | 0.19939 | 0.06796 | 0.07516 | 0.00310 | 0.11002 | 0.01591 | 0.00490 | 0.42992 | 0.00000 | 0.12890 | 0.07338 |
| **Ttc7b** | 1.26094 | 0.09484 | 11.74808 | 7.15680 | 6.82597 | 1.97569 | 2.84094 | 10.30257 | 1.64642 | 0.10713 | 4.51496 | 8.78263 | 0.68028 | 1.45501 |
| **Ttc8** | 0.00822 | 0.00372 | 0.00000 | 0.00561 | 0.12937 | 0.00000 | 0.00413 | 0.00367 | 0.00562 | 0.00634 | 0.47286 | 0.41939 | 0.00325 | 0.64738 |
| **Ttc9** | 0.19033 | 0.00446 | 0.00450 | 0.27779 | 0.05333 | 0.00926 | 0.00392 | 0.05985 | 0.01011 | 0.00568 | 0.00495 | 0.31975 | 0.00490 | 0.00416 |
| **Ttc9c** | 0.24294 | 8.06570 | 1.85690 | 0.73782 | 0.85049 | 3.23275 | 0.07792 | 0.63680 | 5.09870 | 0.03726 | 1.37744 | 1.62950 | 1.83985 | 1.83683 |
| **Ttf1** | 2.10192 | 0.16974 | 2.42778 | 1.69491 | 1.52646 | 3.63358 | 0.45895 | 1.56902 | 9.60117 | 0.79552 | 0.57285 | 1.39172 | 14.40829 | 5.32535 |
| **Ttf2** | 2.33939 | 0.00208 | 3.41768 | 3.30789 | 0.33063 | 3.79891 | 0.01493 | 1.47298 | 0.00137 | 0.01635 | 3.55069 | 1.57329 | 0.00000 | 0.96247 |
| **Tti1** | 3.99003 | 0.02265 | 6.38936 | 0.92370 | 5.25773 | 6.39338 | 0.00135 | 0.00000 | 0.71133 | 0.00000 | 5.71891 | 2.62636 | 1.34530 | 5.17534 |
| **Tti2** | 2.19432 | 0.00272 | 3.97027 | 2.32444 | 1.82210 | 2.27257 | 2.19905 | 0.02088 | 1.13927 | 0.06899 | 3.84043 | 0.76622 | 1.38176 | 4.14227 |
| **Ttk** | 6.09489 | 0.01223 | 4.19658 | 1.04872 | 4.16877 | 3.65988 | 2.69852 | 0.03657 | 8.32835 | 0.03907 | 4.91553 | 5.26517 | 1.00338 | 4.44670 |
| **Ttl** | 4.75275 | 0.03124 | 9.47018 | 7.67877 | 6.40378 | 5.04729 | 0.73886 | 2.36653 | 0.82927 | 0.04982 | 4.09752 | 2.68269 | 0.00000 | 6.45741 |
| **Ttll1** | 1.32448 | 0.01161 | 1.53880 | 1.16084 | 1.92325 | 2.62249 | 0.00376 | 0.74329 | 0.04642 | 0.01671 | 0.04647 | 0.35068 | 1.28325 | 2.35446 |
| **Ttll10** | 0.34378 | 0.00494 | 2.00547 | 0.28263 | 0.17649 | 0.03349 | 0.00395 | 0.00349 | 0.18996 | 0.00330 | 0.00823 | 0.00552 | 0.00000 | 0.00000 |
| **Ttll11** | 0.02798 | 0.00438 | 0.00963 | 0.01578 | 0.00557 | 0.01020 | 0.00425 | 0.00988 | 0.00967 | 0.00755 | 0.00749 | 0.00905 | 0.04999 | 0.57249 |
| **Ttll12** | 0.34456 | 0.09591 | 3.38892 | 7.91812 | 2.69024 | 6.34156 | 0.01701 | 5.88550 | 0.19388 | 0.04546 | 1.10134 | 6.13638 | 0.95897 | 2.20885 |
| **Ttll13** | 0.00413 | 0.00420 | 0.48776 | 0.09126 | 0.04481 | 0.12295 | 0.00469 | 0.00276 | 0.01277 | 0.00463 | 0.00476 | 0.20702 | 1.07939 | 0.00207 |
| **Ttll2** | 0.00473 | 0.01078 | 0.01469 | 0.00000 | 0.00786 | 0.03245 | 0.47566 | 0.00330 | 0.00596 | 0.01797 | 0.00261 | 0.44814 | 0.01313 | 0.10027 |
| **Ttll3** | 0.02641 | 0.01145 | 0.97398 | 0.26237 | 0.44535 | 0.46910 | 1.17861 | 0.00000 | 0.81179 | 0.05512 | 2.26404 | 0.00940 | 0.90813 | 0.00000 |
| **Ttll4** | 1.45614 | 0.01162 | 3.41741 | 0.47092 | 1.11164 | 1.94583 | 0.00313 | 1.51191 | 3.08934 | 0.00000 | 4.14791 | 0.66689 | 1.33718 | 4.78470 |
| **Ttll5** | 0.77113 | 0.05192 | 1.34275 | 0.39730 | 1.40865 | 3.68104 | 0.00337 | 0.58953 | 0.30390 | 0.03474 | 0.00000 | 3.38217 | 0.51891 | 2.18986 |
| **Ttll6** | 1.43152 | 0.02186 | 3.80567 | 0.62456 | 1.90702 | 2.98271 | 0.00302 | 2.29665 | 1.39521 | 0.00609 | 0.80501 | 0.87592 | 0.38230 | 0.78783 |
| **Ttll9** | 0.00890 | 0.00445 | 0.00000 | 0.00000 | 0.89031 | 0.09479 | 0.00417 | 0.00578 | 0.00714 | 0.00590 | 0.01096 | 0.00892 | 1.44264 | 0.04032 |
| **Ttn** | 0.00000 | 0.00406 | 0.14973 | 0.18676 | 1.09761 | 0.00000 | 0.00405 | 0.06318 | 7.94767 | 0.00571 | 0.00395 | 0.00376 | 0.91081 | 4.53645 |
| **Ttpal** | 2.15696 | 0.01901 | 4.61577 | 0.00000 | 3.03152 | 4.53328 | 0.01452 | 1.95344 | 1.56748 | 0.00000 | 2.79552 | 0.44984 | 1.31194 | 4.16868 |
| **Ttyh1** | 0.14322 | 0.00448 | 0.00702 | 0.00912 | 0.34447 | 0.01165 | 0.00431 | 0.01063 | 0.00771 | 0.00666 | 0.00759 | 0.00759 | 0.02679 | 0.03311 |
| **Ttyh2** | 0.78578 | 3.24770 | 1.25286 | 5.56504 | 2.92048 | 3.98821 | 0.40953 | 6.79820 | 3.13457 | 0.11324 | 1.15978 | 2.40156 | 4.04415 | 7.83164 |
| **Ttyh3** | 0.05155 | 0.01781 | 0.01188 | 0.05524 | 0.06291 | 0.08330 | 0.01447 | 0.60090 | 0.04277 | 0.00566 | 0.00470 | 0.05542 | 1.56723 | 0.05120 |
| **Tub** | 0.00000 | 0.00361 | 1.07308 | 0.00000 | 0.61819 | 0.00000 | 0.00389 | 1.37885 | 0.02259 | 0.00516 | 0.00000 | 0.55514 | 0.50924 | 3.11810 |
| **Tuba1a** | 6.75263 | 6.16868 | 23.04853 | 16.33640 | 10.57358 | 25.19384 | 7.50713 | 22.25395 | 6.54003 | 9.45632 | 14.32745 | 22.02263 | 7.61255 | 9.56451 |
| **Tuba1b** | 8.91321 | 9.27273 | 28.26063 | 22.04899 | 14.74281 | 29.41570 | 8.96319 | 25.70482 | 10.22630 | 14.31608 | 17.02416 | 25.65358 | 2.99416 | 14.11388 |
| **Tuba1c** | 10.48810 | 8.38162 | 30.52473 | 22.26373 | 14.14081 | 32.84745 | 9.85174 | 28.28203 | 10.04824 | 13.40172 | 19.81326 | 27.02247 | 5.83817 | 14.68014 |
| **Tuba3a** | 2.55013 | 0.00336 | 4.89192 | 0.00000 | 1.57189 | 1.69898 | 0.01950 | 0.00000 | 0.00247 | 0.00257 | 0.00165 | 0.16178 | 0.92704 | 2.79366 |
| **Tuba3b** | 3.00527 | 0.00291 | 7.01472 | 0.25954 | 3.53936 | 1.19077 | 0.00811 | 0.00000 | 0.00000 | 0.00000 | 0.00445 | 0.37788 | 2.07434 | 5.07126 |
| **Tuba4a** | 3.99761 | 3.24062 | 17.24973 | 26.12176 | 1.62182 | 10.51016 | 5.21883 | 8.67044 | 0.51992 | 14.55982 | 9.40354 | 8.76264 | 5.17553 | 5.44752 |
| **Tuba8** | 0.00000 | 0.00457 | 0.33085 | 0.00531 | 0.00000 | 0.72523 | 3.20584 | 0.80338 | 0.00652 | 0.13010 | 0.00225 | 0.00000 | 0.20489 | 0.01871 |
| **Tubb2a** | 0.02736 | 0.01226 | 0.34573 | 2.85725 | 0.36404 | 0.15005 | 0.01121 | 0.65268 | 0.05995 | 0.00420 | 0.71073 | 0.43557 | 0.53849 | 0.00000 |
| **Tubb2b** | 0.02374 | 0.00470 | 0.03027 | 0.42358 | 0.02191 | 0.01497 | 0.00455 | 0.04915 | 0.00843 | 0.00805 | 0.05014 | 0.01450 | 0.03384 | 0.02752 |
| **Tubb3** | 0.65017 | 0.02660 | 2.17826 | 0.25434 | 3.64979 | 0.00000 | 0.00923 | 0.01176 | 0.08744 | 0.00227 | 0.00825 | 0.01344 | 1.69563 | 0.01380 |
| **Tubb4b** | 25.97067 | 16.30235 | 41.54307 | 32.91709 | 38.15816 | 30.52221 | 10.64405 | 24.56764 | 6.91444 | 14.58056 | 23.19673 | 18.59194 | 21.15413 | 35.86084 |
| **Tubb5** | 3.31127 | 10.05608 | 15.47379 | 19.60015 | 12.97210 | 25.81225 | 9.20550 | 22.26751 | 1.02436 | 1.09977 | 15.73791 | 25.24018 | 9.67391 | 9.03997 |
| **Tubb6** | 2.45660 | 0.08037 | 5.91914 | 7.72518 | 3.89875 | 2.80509 | 4.14393 | 3.66891 | 0.02406 | 0.14610 | 2.03390 | 3.03667 | 1.88560 | 2.04972 |
| **Tubd1** | 1.79612 | 0.01899 | 1.32009 | 0.52111 | 1.02030 | 2.29515 | 0.01555 | 0.01103 | 1.46042 | 0.01922 | 0.00000 | 1.58487 | 0.00000 | 0.58934 |
| **Tube1** | 0.54847 | 0.03834 | 0.00000 | 2.20158 | 0.00000 | 1.56256 | 0.01865 | 0.74615 | 1.86425 | 7.70352 | 3.94609 | 1.49690 | 0.01797 | 1.44583 |
| **Tubg1** | 4.53202 | 0.08665 | 10.55076 | 7.87721 | 8.61417 | 7.09927 | 0.01893 | 1.12625 | 0.02295 | 0.04871 | 5.48236 | 2.93057 | 1.37636 | 8.73934 |
| **Tubg2** | 0.31306 | 0.00401 | 2.28844 | 0.85261 | 1.85082 | 1.10383 | 0.00353 | 0.20471 | 0.34224 | 0.01040 | 0.51692 | 0.21404 | 2.39447 | 2.44618 |
| **Tubgcp2** | 0.98585 | 0.05016 | 4.05446 | 3.63004 | 3.60466 | 4.05749 | 2.70108 | 4.10373 | 0.22013 | 0.10353 | 3.64998 | 8.13460 | 6.01651 | 1.34368 |
| **Tubgcp3** | 5.39245 | 2.23875 | 7.78863 | 0.00000 | 8.05767 | 6.04149 | 2.16336 | 4.19212 | 0.49060 | 0.05478 | 5.44689 | 4.52803 | 1.55829 | 7.03873 |
| **Tubgcp4** | 0.01218 | 0.01894 | 0.81582 | 1.03971 | 0.27640 | 0.38974 | 0.00364 | 1.23705 | 0.05249 | 0.01829 | 0.00002 | 0.44367 | 0.36121 | 1.34677 |
| **Tubgcp5** | 2.18632 | 0.01104 | 2.36458 | 1.58801 | 0.30170 | 0.71568 | 0.04265 | 0.00000 | 0.30198 | 0.00000 | 4.11105 | 0.08693 | 0.61163 | 6.14641 |
| **Tubgcp6** | 0.20773 | 0.00430 | 1.27330 | 0.47147 | 1.31489 | 1.15639 | 0.01050 | 0.12942 | 0.82505 | 0.00313 | 0.61505 | 0.33546 | 0.93290 | 1.21181 |

| | | | | | | | | | | | | | | |
|---|---|---|---|---|---|---|---|---|---|---|---|---|---|---|
| **Tufm** | 16.76487 | 10.98428 | 29.70128 | 29.05183 | 21.42613 | 17.50506 | 5.71633 | 13.23228 | 3.42204 | 13.36619 | 13.24816 | 14.92772 | 2.80648 | 16.00914 |
| **Tuft1** | 2.47770 | 0.08502 | 4.08174 | 7.85482 | 4.34152 | 6.79455 | 5.66203 | 6.35926 | 0.45911 | 0.29366 | 1.49830 | 2.13495 | 4.36869 | 3.90907 |
| **Tug1** | 7.89144 | 0.00872 | 3.45377 | 3.33112 | 3.67681 | 4.38777 | 1.58214 | 1.03380 | 18.04283 | 0.04759 | 2.08112 | 2.54580 | 1.39427 | 3.42131 |
| **Tulp1** | 0.31815 | 0.00427 | 2.58974 | 0.00000 | 3.81205 | 0.00000 | 0.00366 | 0.00507 | 0.05329 | 0.00323 | 0.01533 | 0.00672 | 0.09492 | 0.61582 |
| **Tulp2** | 0.01677 | 0.00437 | 0.12947 | 0.00468 | 0.01572 | 0.00121 | 0.00515 | 0.09893 | 0.00872 | 0.00542 | 0.00882 | 0.00708 | 0.45067 | 0.21571 |
| **Tulp3** | 0.00000 | 0.03760 | 0.56262 | 0.13870 | 0.00000 | 0.06725 | 0.00399 | 0.12605 | 1.35976 | 0.01052 | 0.04390 | 0.84637 | 3.47385 | 1.73930 |
| **Tulp4** | 1.34930 | 0.00800 | 1.98180 | 0.21002 | 4.65789 | 1.41261 | 0.01829 | 0.44977 | 3.42913 | 1.53286 | 0.00000 | 3.41062 | 1.25428 | 3.55642 |
| **Tusc2** | 0.56414 | 0.02476 | 2.88471 | 0.00000 | 1.15864 | 3.97908 | 2.15603 | 3.25377 | 0.89311 | 0.06882 | 1.18095 | 3.11103 | 0.90541 | 1.66188 |
| **Tusc3** | 2.16257 | 0.00497 | 0.18160 | 0.92255 | 0.85965 | 0.15726 | 0.00414 | 0.52511 | 0.99289 | 0.00868 | 0.00256 | 0.15325 | 0.13642 | 1.95982 |
| **Tut1** | 0.71951 | 1.81925 | 7.82035 | 5.43852 | 5.84874 | 3.51639 | 1.50797 | 4.32253 | 0.00727 | 1.18994 | 5.03651 | 1.76898 | 2.79414 | 8.25720 |
| **Tvp23a** | 0.01192 | 0.00444 | 0.17256 | 0.00000 | 0.01396 | 0.39792 | 0.00408 | 0.00451 | 0.01035 | 0.01288 | 0.00581 | 0.01038 | 0.00725 | 0.00807 |
| **Tvp23b** | 1.30379 | 5.12910 | 2.70614 | 0.71231 | 1.30196 | 6.73826 | 0.00686 | 3.47830 | 1.24104 | 0.05889 | 2.03496 | 5.18639 | 0.05151 | 5.82058 |
| **Twf1** | 6.20852 | 0.04466 | 4.27343 | 5.28759 | 5.84277 | 7.29413 | 2.34042 | 8.05537 | 3.72686 | 0.10110 | 5.30562 | 5.89328 | 1.64152 | 9.58948 |
| **Twf2** | 0.00000 | 5.49588 | 2.42445 | 2.47138 | 1.32253 | 0.00000 | 0.61818 | 1.80952 | 0.00239 | 9.90738 | 1.10050 | 1.88702 | 0.29528 | 0.06251 |
| **Twistnb** | 8.64422 | 0.02133 | 5.40192 | 1.59457 | 8.23713 | 4.99637 | 1.98577 | 0.00000 | 12.62581 | 0.44946 | 5.27203 | 5.04181 | 5.18786 | 1.50979 |
| **Twsg1** | 0.00000 | 0.03361 | 0.49577 | 0.45939 | 0.37732 | 0.91688 | 1.02890 | 1.35851 | 0.58555 | 0.03338 | 0.00000 | 0.47634 | 1.04949 | 0.00000 |
| **Txk** | 0.56172 | 0.00467 | 0.43365 | 0.00000 | 0.77244 | 0.15197 | 0.00423 | 0.00473 | 1.88045 | 0.00688 | 0.00326 | 0.00418 | 0.00000 | 0.39420 |
| **Txlna** | 8.61728 | 0.35818 | 12.02283 | 11.07916 | 14.74847 | 11.28621 | 6.19762 | 8.47949 | 1.38876 | 0.27274 | 9.25627 | 6.63501 | 10.41082 | 11.66828 |
| **Txlng** | 11.02667 | 3.69401 | 8.86997 | 0.00000 | 14.96432 | 5.00288 | 0.98293 | 1.94389 | 24.15227 | 0.00000 | 0.90710 | 4.32423 | 5.65835 | 13.74874 |
| **Txn1** | 1.25929 | 5.77109 | 0.00000 | 2.77842 | 1.57195 | 15.07837 | 6.62601 | 18.48513 | 0.79463 | 17.77343 | 2.49377 | 20.21738 | 0.39964 | 2.19954 |
| **Txn2** | 11.79124 | 10.39834 | 27.74838 | 18.18539 | 14.41483 | 23.53944 | 3.42946 | 20.10859 | 0.00000 | 10.58615 | 10.46082 | 15.55078 | 1.03384 | 13.08150 |
| **Txndc11** | 0.00000 | 1.28620 | 1.83383 | 1.54936 | 1.62435 | 0.15526 | 2.09298 | 0.49317 | 0.00063 | 0.07504 | 0.00214 | 3.21318 | 0.75145 | 0.06783 |
| **Txndc12** | 0.85329 | 4.32317 | 9.72397 | 7.42009 | 1.08005 | 13.57397 | 5.43085 | 7.56195 | 1.90306 | 8.78786 | 10.06604 | 18.55682 | 0.00000 | 1.21432 |
| **Txndc15** | 5.56720 | 0.02859 | 5.48353 | 0.45499 | 7.98080 | 1.99350 | 0.23005 | 3.51940 | 3.92016 | 0.00000 | 1.91902 | 5.11003 | 2.73571 | 3.60090 |
| **Txndc16** | 0.00000 | 0.03948 | 0.08404 | 0.00003 | 0.00000 | 7.26481 | 1.87076 | 8.00765 | 2.01538 | 0.07534 | 0.14013 | 0.00000 | 1.31316 | 0.56808 |
| **Txndc17** | 4.20240 | 0.25663 | 0.73232 | 6.82271 | 4.56993 | 9.35763 | 1.75922 | 12.01451 | 0.09935 | 6.67045 | 3.10231 | 10.37990 | 1.27212 | 2.80114 |
| **Txndc2** | 0.00000 | 0.00431 | 0.20418 | 0.00000 | 0.29845 | 0.00440 | 0.00429 | 0.00947 | 0.00851 | 0.00676 | 0.00626 | 0.00833 | 0.36850 | 0.75047 |
| **Txndc5** | 0.02483 | 0.17338 | 0.00000 | 1.92512 | 0.29771 | 3.45620 | 6.26628 | 2.51119 | 0.68547 | 0.07565 | 0.27942 | 1.75675 | 0.31322 | 1.06069 |
| **Txndc9** | 16.78783 | 10.37996 | 16.29428 | 10.23427 | 14.05059 | 17.76336 | 4.47851 | 12.11464 | 15.63044 | 2.71720 | 13.22651 | 13.75218 | 5.21718 | 13.09080 |
| **Txnip** | 0.41699 | 7.21774 | 0.00000 | 0.00000 | 1.98143 | 0.00000 | 0.00225 | 3.03266 | 0.00000 | 0.01517 | 0.00142 | 0.00000 | 2.10252 | 0.50630 |
| **Txnl1** | 22.25654 | 6.10685 | 18.29764 | 26.62002 | 19.50951 | 13.34728 | 5.30286 | 6.25354 | 6.79119 | 0.19603 | 11.00850 | 13.17655 | 8.11124 | 2.05302 |
| **Txnl4a** | 11.57059 | 3.94750 | 16.85600 | 19.81948 | 24.21660 | 16.00582 | 5.66768 | 13.07191 | 2.63276 | 13.32241 | 9.35973 | 13.90035 | 10.52759 | 12.68913 |
| **Txnl4b** | 0.00000 | 0.33639 | 0.51787 | 2.63309 | 0.42853 | 0.28902 | 0.04969 | 1.44381 | 0.00977 | 0.00334 | 1.04860 | 0.45565 | 0.10806 | 0.10260 |
| **Txnrd1** | 1.51416 | 0.00388 | 3.59150 | 0.20074 | 4.05435 | 8.67608 | 0.05330 | 0.90638 | 0.26050 | 7.29835 | 6.00453 | 7.31564 | 0.42365 | 2.76087 |
| **Txnrd2** | 2.26838 | 0.04100 | 4.66157 | 3.38761 | 0.97345 | 5.48151 | 0.06034 | 0.55236 | 0.73403 | 0.01751 | 5.47864 | 2.10180 | 0.00000 | 1.58760 |
| **Txnrd3** | 0.00841 | 0.00378 | 0.01644 | 0.08353 | 0.32145 | 0.00626 | 0.00408 | 0.00503 | 0.01086 | 0.00648 | 0.00848 | 0.29171 | 0.51372 | 0.35889 |
| **Tyk2** | 4.21024 | 0.02824 | 15.13453 | 7.91221 | 9.73139 | 2.65957 | 0.01467 | 0.96871 | 1.52753 | 0.00000 | 1.27903 | 0.01206 | 3.41546 | 9.91771 |
| **Tymp** | 0.00837 | 0.00432 | 0.01903 | 0.08386 | 0.09783 | 0.00987 | 0.00407 | 0.00837 | 0.01144 | 0.00633 | 0.01138 | 0.01061 | 0.00584 | 0.00956 |
| **Tyms** | 12.94728 | 0.40322 | 22.16160 | 17.55937 | 17.05009 | 6.16520 | 2.59883 | 6.34223 | 8.82916 | 10.91437 | 6.65837 | 5.45778 | 0.82131 | 7.94825 |
| **Tyms.ps** | 1.43384 | 0.01177 | 2.57517 | 2.02020 | 1.88451 | 0.53119 | 0.13198 | 0.56911 | 0.33758 | 1.36485 | 0.70777 | 0.50585 | 0.08824 | 0.58454 |
| **Tyr** | 0.63378 | 0.02072 | 0.05940 | 0.48601 | 0.22298 | 0.87127 | 0.13937 | 0.13698 | 1.44809 | 0.21220 | 0.03872 | 0.21885 | 5.25771 | 0.71530 |
| **Tyro3** | 0.70347 | 8.92652 | 3.20379 | 0.91041 | 2.22143 | 4.73684 | 2.55194 | 2.96570 | 0.34137 | 11.53957 | 0.43332 | 8.59994 | 1.13521 | 0.60439 |
| **Tysnd1** | 1.57098 | 0.03923 | 0.30238 | 2.32122 | 0.65211 | 1.01938 | 2.18155 | 1.37915 | 0.00068 | 2.99659 | 0.47654 | 1.73172 | 2.37973 | 0.09787 |
| **Tyw1** | 1.59762 | 0.05658 | 8.10072 | 5.53439 | 5.70368 | 7.93883 | 1.83748 | 0.26134 | 0.60201 | 0.07974 | 1.03356 | 4.46321 | 0.71799 | 6.65310 |
| **Tyw3** | 1.61263 | 0.02639 | 3.56478 | 1.06769 | 3.31845 | 1.83705 | 2.05712 | 0.66481 | 0.30141 | 3.28148 | 0.69797 | 0.89494 | 0.61039 | 0.75935 |
| **Tyw5** | 3.47243 | 0.01487 | 4.24900 | 3.31972 | 3.03898 | 1.78007 | 0.00110 | 1.30912 | 5.00116 | 0.00678 | 4.15606 | 4.13329 | 1.62159 | 1.04553 |
| **U2af1** | 0.21755 | 3.09378 | 7.23242 | 10.21077 | 7.46999 | 6.00646 | 3.73170 | 8.61510 | 0.62473 | 12.96265 | 6.37767 | 6.08582 | 4.68976 | 6.65894 |
| **U2af1l4** | 1.88270 | 0.07726 | 3.84828 | 5.46292 | 2.95930 | 5.63135 | 1.04543 | 5.32672 | 0.00000 | 7.88301 | 0.08882 | 3.57480 | 0.39101 | 1.75241 |
| **U2af2** | 0.98475 | 0.36355 | 0.00000 | 0.70688 | 6.73968 | 5.51485 | 0.14515 | 2.65838 | 1.80872 | 0.07922 | 0.48348 | 2.93942 | 10.08976 | 1.81317 |
| **U2surp** | 7.36767 | 3.49034 | 8.03347 | 1.62751 | 11.45954 | 8.24691 | 4.08697 | 2.83829 | 20.80455 | 0.12517 | 5.41645 | 6.67210 | 7.65741 | 11.04876 |
| **Uaca** | 0.91464 | 0.00394 | 0.49966 | 0.01280 | 1.98217 | 1.37854 | 0.00366 | 0.00217 | 0.78316 | 0.00926 | 0.14232 | 0.78718 | 0.00000 | 0.83554 |
| **Uap1** | 10.94337 | 4.55993 | 25.30292 | 10.39316 | 24.43709 | 8.55454 | 2.93184 | 0.75347 | 26.66991 | 0.20443 | 8.30012 | 14.46364 | 2.90081 | 24.11056 |
| **Uap1l1** | 3.16791 | 4.44263 | 10.28867 | 11.95743 | 6.77212 | 4.74797 | 0.05959 | 1.09663 | 0.02254 | 0.05005 | 6.36065 | 0.33709 | 3.08790 | 3.58615 |
| **Uba1** | 24.38143 | 9.11272 | 40.54723 | 23.75054 | 28.86585 | 31.41669 | 4.68826 | 15.47770 | 9.71418 | 5.40219 | 16.90247 | 20.31818 | 9.20786 | 22.34641 |
| **Uba1y** | 0.08325 | 0.08202 | 0.00000 | 0.10224 | 2.05697 | 0.00213 | 0.06775 | 0.00000 | 0.35479 | 0.00306 | 1.40437 | 0.54244 | 4.35098 | 0.85791 |

| | | | | | | | | | | | | | | |
|---|---|---|---|---|---|---|---|---|---|---|---|---|---|---|
| **Uba2** | 16.05300 | 3.37104 | 20.30135 | 10.33209 | 19.46975 | 16.34942 | 4.63806 | 7.36010 | 13.25984 | 0.28478 | 13.94157 | 18.94777 | 10.19876 | 24.70178 |
| **Uba3** | 2.42552 | 1.29783 | 2.48883 | 1.66856 | 1.18749 | 3.21983 | 0.74273 | 0.33574 | 10.12802 | 0.02134 | 6.75564 | 1.89992 | 3.21901 | 1.95792 |
| **Uba5** | 16.80610 | 0.04780 | 8.35143 | 4.48483 | 12.89418 | 6.71130 | 2.83908 | 2.65163 | 6.80240 | 0.04227 | 1.37771 | 5.11017 | 4.97771 | 5.75554 |
| **Uba52** | 24.94015 | 9.99856 | 7.84523 | 24.59379 | 34.88035 | 34.33520 | 9.35531 | 30.02972 | 11.10227 | 25.93056 | 5.22012 | 28.23833 | 22.70856 | 20.12069 |
| **Uba6** | 2.36215 | 0.00295 | 1.02948 | 0.00000 | 1.58108 | 1.68253 | 0.00272 | 0.00000 | 6.97134 | 0.00375 | 0.00000 | 0.78013 | 1.03305 | 0.78143 |
| **Ubac1** | 4.10276 | 7.69720 | 4.23233 | 5.76864 | 3.30534 | 9.64059 | 1.50409 | 3.23510 | 3.02209 | 7.06427 | 4.77965 | 7.67796 | 2.34705 | 6.05401 |
| **Ubac2** | 1.13684 | 0.12289 | 5.90044 | 4.92858 | 5.49443 | 3.00588 | 0.19845 | 6.42827 | 0.00000 | 0.02477 | 2.41531 | 6.90314 | 0.94287 | 1.37892 |
| **Ubald1** | 1.20627 | 5.09115 | 1.42428 | 3.43894 | 4.17192 | 1.23959 | 2.90433 | 5.19637 | 0.11674 | 12.26414 | 0.02015 | 2.67875 | 2.38904 | 0.73237 |
| **Ubap1** | 6.59103 | 0.06079 | 8.77063 | 6.50535 | 9.68267 | 3.67796 | 0.43425 | 2.77448 | 1.74502 | 0.05224 | 6.86702 | 5.38788 | 8.24856 | 6.15646 |
| **Ubap2** | 9.85386 | 0.27888 | 7.60012 | 14.60071 | 16.21404 | 9.10768 | 3.49594 | 3.74332 | 0.92041 | 2.17151 | 1.48539 | 7.99731 | 11.29514 | 2.71416 |
| **Ubap2l** | 6.93909 | 0.09154 | 9.04206 | 7.88415 | 9.19666 | 7.23364 | 4.59022 | 3.70767 | 1.23122 | 1.43827 | 3.39108 | 6.65427 | 7.48667 | 6.67350 |
| **Ubash3a** | 0.00729 | 0.00467 | 0.10885 | 0.34958 | 0.00561 | 0.00387 | 0.00423 | 0.01126 | 0.01289 | 0.00536 | 0.00606 | 0.03231 | 0.00664 | 0.00849 |
| **Ubash3b** | 0.00000 | 0.01772 | 0.39823 | 0.00000 | 0.58732 | 1.48159 | 0.00299 | 2.67038 | 0.23445 | 0.02446 | 0.86612 | 0.32978 | 0.55240 | 1.91528 |
| **Ubb** | 22.81897 | 13.02697 | 34.59376 | 35.74980 | 21.14492 | 41.09490 | 11.45640 | 33.93815 | 10.57564 | 12.00237 | 28.10799 | 30.20203 | 10.44339 | 26.41422 |
| **Ubc** | 15.49641 | 11.76458 | 24.71732 | 27.18332 | 23.10781 | 26.26989 | 5.24196 | 22.71896 | 15.34001 | 16.33766 | 15.69971 | 15.38606 | 16.29706 | 21.55909 |
| **Ube2a** | 5.97903 | 6.85254 | 4.83722 | 9.76025 | 12.92965 | 4.05503 | 4.33531 | 10.98435 | 3.64629 | 9.80952 | 1.44698 | 17.31975 | 9.95403 | 4.27016 |
| **Ube2b** | 15.33561 | 6.25910 | 12.55055 | 8.32276 | 14.82383 | 8.38692 | 5.43144 | 4.82367 | 4.63191 | 1.32584 | 9.07057 | 4.81236 | 8.66945 | 12.89182 |
| **Ube2c** | 20.54145 | 7.87326 | 32.99489 | 33.52170 | 43.64710 | 12.49471 | 8.77048 | 19.32719 | 5.35401 | 15.82706 | 3.57353 | 13.25083 | 14.00988 | 13.74321 |
| **Ube2cbp** | 0.98299 | 0.00277 | 3.58614 | 2.31599 | 1.69460 | 1.07020 | 0.00254 | 1.43056 | 0.02237 | 0.00352 | 2.93615 | 0.48534 | 0.00000 | 0.23765 |
| **Ube2d.ps** | 0.00000 | 0.00238 | 0.82467 | 1.45217 | 0.96204 | 0.00000 | 1.65534 | 0.04759 | 0.39779 | 0.01869 | 3.23219 | 0.88713 | 0.15162 | 0.75668 |
| **Ube2d1** | 0.82963 | 0.01056 | 0.14692 | 0.00000 | 1.03423 | 0.27020 | 0.05364 | 0.60167 | 0.01677 | 0.00520 | 0.02801 | 1.23523 | 0.15956 | 0.07602 |
| **Ube2d2a** | 12.72383 | 0.07897 | 5.77907 | 1.93378 | 10.08993 | 8.16375 | 1.50698 | 7.17694 | 10.31139 | 4.49209 | 6.27001 | 6.11502 | 20.92413 | 6.43792 |
| **Ube2d3** | 12.66393 | 3.29255 | 20.47957 | 20.46273 | 20.83623 | 21.05217 | 2.80852 | 8.57627 | 15.94277 | 0.32071 | 14.25319 | 14.99067 | 16.11581 | 12.33061 |
| **Ube2dnl1** | 0.00510 | 0.00388 | 0.07495 | 0.00000 | 0.03925 | 0.17944 | 0.00499 | 0.06867 | 0.07064 | 0.00478 | 0.01444 | 0.70880 | 0.37713 | 0.23446 |
| **Ube2dnl2** | 0.00568 | 0.00407 | 0.01332 | 0.00000 | 0.01314 | 0.05935 | 0.00475 | 0.01287 | 0.02253 | 0.00574 | 0.00580 | 0.41406 | 0.05052 | 0.14894 |
| **Ube2e1** | 4.48172 | 0.04074 | 11.33741 | 6.91714 | 8.29503 | 8.77212 | 2.73324 | 4.31066 | 9.60417 | 0.08059 | 4.09420 | 2.78258 | 0.00000 | 8.89266 |
| **Ube2e2** | 0.00422 | 0.00471 | 0.00003 | 0.48561 | 0.25402 | 0.01386 | 0.00400 | 0.67131 | 0.00546 | 0.00873 | 0.00903 | 0.00811 | 0.18362 | 0.25326 |
| **Ube2e3** | 17.80903 | 2.93001 | 11.29673 | 9.04230 | 21.39668 | 8.82579 | 6.09482 | 2.37154 | 22.90633 | 0.16771 | 8.45516 | 5.14340 | 6.88855 | 17.67046 |
| **Ube2f** | 4.29791 | 0.06742 | 7.88483 | 7.11972 | 6.55833 | 8.17271 | 3.10798 | 2.90534 | 0.90629 | 0.07034 | 1.76928 | 3.57718 | 0.33097 | 3.36733 |
| **Ube2g1** | 2.46259 | 0.27595 | 4.83905 | 0.59273 | 5.78079 | 1.99079 | 0.01437 | 1.56706 | 7.42761 | 0.00000 | 2.18649 | 4.40081 | 3.16658 | 0.55619 |
| **Ube2g2** | 3.04510 | 1.01178 | 4.92334 | 2.70269 | 6.06893 | 4.58973 | 0.03040 | 3.46130 | 0.06814 | 0.05395 | 0.00000 | 1.07990 | 2.15829 | 3.26861 |
| **Ube2h** | 4.83828 | 5.00423 | 11.68604 | 6.40042 | 8.20001 | 25.44480 | 0.47974 | 6.86229 | 8.79524 | 8.62812 | 8.07724 | 5.37815 | 3.56261 | 5.48214 |
| **Ube2i** | 18.44004 | 0.33954 | 19.41798 | 17.13520 | 18.45916 | 17.67314 | 3.87525 | 10.27957 | 3.24126 | 1.99595 | 6.54043 | 17.32337 | 8.60130 | 9.66984 |
| **Ube2j1** | 4.82252 | 0.44344 | 3.31977 | 2.50963 | 2.01493 | 3.39923 | 0.31678 | 2.15015 | 0.72910 | 0.01664 | 0.03021 | 0.72370 | 1.47381 | 2.25098 |
| **Ube2j2** | 0.20637 | 0.08626 | 0.28255 | 0.61055 | 1.08147 | 0.30270 | 0.13377 | 0.31021 | 0.57119 | 0.05840 | 0.10200 | 0.42239 | 1.52871 | 0.28458 |
| **Ube2k** | 15.19125 | 0.31568 | 12.40253 | 10.80499 | 14.36165 | 15.89992 | 4.76300 | 14.42667 | 13.10283 | 0.32284 | 10.35071 | 15.09590 | 10.76926 | 14.87012 |
| **Ube2l3** | 16.34834 | 5.26995 | 14.73496 | 20.06152 | 28.84594 | 25.38473 | 5.48651 | 15.28807 | 16.05592 | 14.56976 | 10.11325 | 17.36406 | 10.73724 | 19.39972 |
| **Ube2l6** | 1.39023 | 6.11111 | 1.57820 | 1.97261 | 1.30556 | 7.23955 | 3.04528 | 2.82904 | 1.20599 | 2.78160 | 1.62269 | 10.85993 | 1.34221 | 0.41457 |
| **Ube2m** | 2.29639 | 5.01171 | 3.22089 | 5.03328 | 0.96739 | 5.08122 | 1.65306 | 6.96021 | 0.04121 | 0.15651 | 4.47405 | 5.77717 | 0.00428 | 0.90860 |
| **Ube2n** | 6.15299 | 0.17504 | 14.34425 | 9.87648 | 8.44113 | 15.74904 | 5.98776 | 8.26468 | 13.58372 | 13.09422 | 11.99100 | 11.42214 | 8.76277 | 13.73677 |
| **Ube2o** | 4.23073 | 1.39692 | 15.86279 | 6.35056 | 12.18531 | 10.49594 | 0.79550 | 1.24352 | 0.00000 | 1.23130 | 7.72868 | 6.69630 | 7.72657 | 7.67347 |
| **Ube2q1** | 3.32668 | 5.03149 | 4.24673 | 4.85336 | 2.29489 | 7.27088 | 5.77592 | 3.45482 | 4.54174 | 0.22667 | 3.62676 | 6.48965 | 0.69953 | 2.40141 |
| **Ube2q2** | 1.88846 | 0.02731 | 6.39254 | 6.22162 | 1.80710 | 6.68532 | 0.10812 | 1.48496 | 7.08614 | 0.00000 | 5.22043 | 4.34121 | 1.46183 | 4.67712 |
| **Ube2r2** | 1.47469 | 0.01287 | 3.62175 | 2.77782 | 1.87863 | 2.59915 | 4.68371 | 6.68295 | 5.90969 | 1.22489 | 4.32057 | 6.08791 | 2.04375 | 3.33710 |
| **Ube2s** | 26.02063 | 2.23712 | 11.65445 | 13.47786 | 33.82663 | 16.33525 | 2.70650 | 6.21202 | 0.22372 | 5.91583 | 6.90827 | 12.29256 | 12.07557 | 9.55449 |
| **Ube2t** | 5.11069 | 0.01662 | 3.69532 | 1.30037 | 3.13556 | 2.16561 | 0.58628 | 0.37717 | 1.68907 | 0.06398 | 2.77647 | 3.86765 | 2.51712 | 2.70508 |
| **Ube2u** | 0.87065 | 0.00392 | 0.14595 | 0.10465 | 0.00000 | 0.14637 | 0.00387 | 0.00184 | 0.70756 | 0.00612 | 0.28877 | 0.06896 | 0.00459 | 0.49491 |
| **Ube2v1** | 6.19630 | 7.34322 | 26.37950 | 20.87415 | 25.43993 | 17.47771 | 4.99738 | 13.53089 | 0.00000 | 14.96891 | 13.27408 | 14.08606 | 3.56116 | 4.85713 |
| **Ube2v2** | 10.70415 | 3.88209 | 9.30478 | 7.80486 | 9.56659 | 12.47157 | 3.42363 | 5.11664 | 19.08450 | 9.53057 | 7.99380 | 7.56368 | 6.12072 | 7.32470 |
| **Ube2w** | 0.00000 | 0.01789 | 1.92508 | 0.70860 | 2.34706 | 2.84704 | 2.16245 | 0.45602 | 2.81792 | 0.07005 | 1.43517 | 2.24734 | 2.23030 | 1.29111 |
| **Ube2z** | 9.24293 | 6.91905 | 16.53940 | 8.62554 | 17.66983 | 14.17341 | 3.96974 | 10.08223 | 13.02220 | 5.16571 | 6.83061 | 9.97892 | 8.08207 | 14.13983 |
| **Ube3a** | 0.92009 | 0.01484 | 4.48279 | 0.85858 | 0.89962 | 7.24802 | 0.03346 | 1.88169 | 12.94562 | 0.00720 | 0.72099 | 3.88251 | 3.42947 | 0.00000 |
| **Ube3b** | 2.30583 | 0.08214 | 3.49463 | 5.31182 | 2.89620 | 7.84717 | 4.85864 | 4.22772 | 0.04271 | 13.20590 | 9.07257 | 6.36133 | 0.98375 | 5.12279 |
| **Ube3c** | 6.59645 | 0.05835 | 10.34978 | 2.46714 | 7.93732 | 8.28488 | 0.13410 | 2.98118 | 8.67968 | 0.97373 | 1.54992 | 6.55528 | 4.14857 | 10.98007 |
| **Ube4a** | 2.30084 | 0.02205 | 2.20358 | 0.00000 | 2.10740 | 3.64795 | 1.22074 | 0.19840 | 3.05881 | 0.02043 | 4.17577 | 0.78861 | 1.25007 | 2.01374 |

| | | | | | | | | | | | | | | |
|---|---|---|---|---|---|---|---|---|---|---|---|---|---|---|
| **Ube4b** | 2.52232 | 0.06698 | 9.25643 | 3.44459 | 9.11032 | 3.59716 | 0.30435 | 5.20107 | 2.03133 | 0.04386 | 2.62773 | 7.08390 | 8.88882 | 3.12344 |
| **Ubfd1** | 13.51423 | 0.06824 | 8.04399 | 6.65813 | 15.65632 | 13.96814 | 0.06526 | 2.93842 | 9.08558 | 0.04969 | 10.17292 | 8.85373 | 12.44789 | 13.83737 |
| **Ubiad1** | 3.81762 | 0.00127 | 2.85081 | 0.00000 | 3.62539 | 4.47815 | 3.35288 | 0.56584 | 3.88554 | 0.36122 | 2.38522 | 1.83404 | 1.86170 | 6.97144 |
| **Ubl3** | 4.28075 | 4.56065 | 8.96722 | 1.14813 | 4.13020 | 4.61639 | 1.62094 | 4.53026 | 1.14788 | 0.04868 | 7.04425 | 6.06613 | 3.08088 | 0.64655 |
| **Ubl4a** | 3.31081 | 0.04366 | 2.59605 | 0.88749 | 2.62562 | 6.23636 | 2.85001 | 0.36237 | 2.41546 | 0.00985 | 1.06053 | 1.99587 | 3.65960 | 2.56016 |
| **Ubl5** | 3.62969 | 4.65208 | 0.00000 | 7.42584 | 10.00332 | 5.65231 | 1.37162 | 9.57902 | 6.81353 | 6.90185 | 0.57970 | 5.96745 | 17.68612 | 6.96996 |
| **Ubl7** | 13.01548 | 6.64662 | 18.88341 | 9.65817 | 25.99445 | 8.05939 | 1.52521 | 2.27386 | 0.52449 | 0.07502 | 5.35673 | 2.04345 | 4.63378 | 4.80248 |
| **Ublcp1** | 6.10190 | 0.02024 | 7.11039 | 4.81733 | 4.40559 | 4.89212 | 0.00575 | 0.00000 | 9.37332 | 0.00000 | 6.27487 | 4.67060 | 1.92188 | 10.38619 |
| **Ubn1** | 6.24448 | 0.02275 | 2.22740 | 6.10279 | 6.58080 | 4.94965 | 1.82319 | 2.58780 | 1.51337 | 0.11154 | 1.17571 | 2.42137 | 6.14136 | 10.05908 |
| **Ubn2** | 4.30830 | 0.17204 | 1.22540 | 0.54560 | 4.97182 | 3.53631 | 0.03046 | 0.15552 | 9.38192 | 0.01919 | 1.43703 | 2.61813 | 5.72496 | 8.32744 |
| **Ubox5** | 1.86413 | 0.02847 | 2.80873 | 1.43140 | 3.93685 | 2.16757 | 0.00276 | 0.86598 | 0.10833 | 0.01144 | 3.95428 | 2.28532 | 1.12528 | 2.51389 |
| **Ubp1** | 2.28407 | 0.12465 | 3.43385 | 5.59672 | 2.75820 | 9.27268 | 1.49802 | 3.43851 | 3.82239 | 8.53336 | 4.09152 | 5.49142 | 9.53163 | 8.21239 |
| **Ubqln1** | 13.17956 | 2.63927 | 18.42781 | 13.95524 | 13.55217 | 18.48243 | 5.00384 | 6.05248 | 8.69577 | 4.56125 | 14.63818 | 11.86350 | 8.41821 | 13.40486 |
| **Ubqln2** | 0.78511 | 0.01979 | 0.76105 | 0.76930 | 0.08854 | 1.37363 | 2.58497 | 1.54734 | 0.13581 | 0.66521 | 0.04418 | 0.63816 | 3.65804 | 0.00000 |
| **Ubqln4** | 2.01506 | 0.59506 | 3.06506 | 4.61203 | 1.75082 | 8.18590 | 5.03592 | 6.54014 | 1.35145 | 0.17795 | 3.69808 | 4.49232 | 0.90874 | 4.34454 |
| **Ubr1** | 1.20973 | 0.02100 | 1.40141 | 0.00000 | 0.67906 | 2.44900 | 0.10086 | 1.50128 | 1.43458 | 0.01571 | 0.92090 | 0.60488 | 0.10851 | 0.66116 |
| **Ubr2** | 3.04829 | 0.00195 | 3.43216 | 0.00000 | 1.83281 | 2.08120 | 0.00296 | 0.78564 | 2.87177 | 0.01094 | 4.28983 | 1.24151 | 1.35825 | 3.69761 |
| **Ubr3** | 8.17439 | 0.05054 | 10.34091 | 5.18558 | 6.26492 | 10.94875 | 0.01743 | 2.55935 | 10.97088 | 0.05653 | 4.24639 | 7.28991 | 7.93550 | 4.80741 |
| **Ubr4** | 5.51712 | 0.18238 | 16.91727 | 15.47206 | 9.16357 | 18.23787 | 6.19985 | 7.25264 | 2.60070 | 2.56760 | 7.97555 | 15.74478 | 4.16661 | 4.76913 |
| **Ubr5** | 13.19713 | 0.03134 | 13.81596 | 3.26218 | 24.18538 | 6.19859 | 1.76456 | 5.27527 | 27.44112 | 0.00000 | 10.58473 | 6.37555 | 14.34711 | 22.29935 |
| **Ubr7** | 4.50246 | 0.03937 | 5.30765 | 2.25399 | 3.38383 | 5.97322 | 1.89838 | 1.48498 | 0.13419 | 0.07232 | 4.37386 | 4.56209 | 1.96962 | 3.64945 |
| **Ubtd1** | 1.57313 | 0.07381 | 6.90738 | 3.67323 | 9.68122 | 0.57934 | 0.01834 | 0.06398 | 0.00000 | 0.00000 | 0.45598 | 1.31196 | 0.35343 | 0.04064 |
| **Ubtd2** | 4.20824 | 0.00171 | 1.16096 | 0.40245 | 2.24796 | 0.26403 | 2.27928 | 0.00602 | 1.34835 | 0.02933 | 1.42263 | 0.98360 | 5.83200 | 7.43032 |
| **Ubtf** | 6.55920 | 4.53725 | 28.71094 | 11.62183 | 32.07236 | 12.90186 | 2.76660 | 7.24988 | 6.76398 | 0.16005 | 12.27808 | 14.28606 | 5.43415 | 28.00899 |
| **Ubtfl1** | 1.16272 | 0.01311 | 0.84481 | 0.51386 | 1.53854 | 0.19624 | 0.00113 | 0.14914 | 20.75937 | 0.00722 | 0.23705 | 0.00251 | 0.01943 | 12.49337 |
| **Ubxn1** | 24.09297 | 3.21665 | 36.28918 | 47.37308 | 38.35421 | 21.54475 | 8.96713 | 15.01279 | 1.03562 | 12.41742 | 15.13233 | 19.30668 | 6.88666 | 20.90629 |
| **Ubxn11** | 0.82365 | 0.00365 | 0.54109 | 0.18991 | 0.12390 | 0.00000 | 0.00369 | 0.00226 | 0.00393 | 0.00551 | 0.00430 | 0.18499 | 0.71836 | 1.31738 |
| **Ubxn2a** | 9.58728 | 0.09040 | 5.53801 | 2.59090 | 8.90364 | 6.65154 | 0.02434 | 2.73033 | 15.52341 | 0.03136 | 6.84196 | 3.00498 | 3.88773 | 10.45889 |
| **Ubxn2b** | 0.93501 | 0.00261 | 1.29308 | 0.55090 | 1.29923 | 0.98206 | 0.00410 | 0.27234 | 1.42293 | 0.01037 | 0.00000 | 2.45031 | 0.00000 | 0.21796 |
| **Ubxn4** | 9.42149 | 0.16651 | 7.47083 | 7.22943 | 10.56023 | 15.31002 | 1.38128 | 8.85656 | 5.21052 | 0.14669 | 8.29551 | 4.85178 | 11.04864 | 9.65246 |
| **Ubxn6** | 4.03498 | 4.66622 | 11.31482 | 15.48041 | 11.99507 | 0.00000 | 0.03939 | 6.11377 | 0.02939 | 0.05122 | 0.75166 | 6.07358 | 3.65016 | 0.92404 |
| **Ubxn7** | 2.46634 | 0.00902 | 2.66570 | 0.73385 | 1.30531 | 1.83309 | 2.82100 | 0.31346 | 2.28114 | 0.05572 | 0.00000 | 2.41321 | 5.01788 | 2.21714 |
| **Ubxn8** | 13.98928 | 0.05396 | 16.02600 | 10.47534 | 11.63622 | 12.36199 | 1.86217 | 2.29665 | 6.59534 | 0.02594 | 9.19215 | 4.34342 | 2.47783 | 13.11407 |
| **Uchl1** | 0.68427 | 0.02780 | 19.81950 | 28.37740 | 5.56355 | 6.71299 | 1.12738 | 0.00000 | 0.00000 | 0.15363 | 13.17755 | 5.23140 | 2.39157 | 2.07903 |
| **Uchl3** | 8.34972 | 0.03613 | 4.22498 | 7.83788 | 9.89214 | 6.54174 | 0.03453 | 3.11976 | 3.63284 | 11.43670 | 5.45378 | 5.31565 | 6.10023 | 3.22120 |
| **Uchl4** | 1.81268 | 0.00266 | 0.74085 | 1.52366 | 1.95939 | 1.34356 | 0.00911 | 0.39315 | 0.69102 | 2.86935 | 1.13424 | 1.07143 | 0.75396 | 0.75872 |
| **Uchl5** | 12.74219 | 0.09867 | 10.17723 | 11.59687 | 3.77799 | 8.69734 | 2.38489 | 9.49344 | 7.58911 | 6.25211 | 7.08614 | 7.46689 | 0.05568 | 2.70893 |
| **Uck1** | 0.96988 | 0.01554 | 1.66365 | 1.50719 | 2.62327 | 3.20131 | 0.59837 | 0.10388 | 0.14864 | 0.05843 | 1.33414 | 3.65475 | 0.36423 | 1.24759 |
| **Uck2** | 7.87068 | 0.08326 | 7.13888 | 8.59007 | 9.30570 | 7.15333 | 0.33302 | 2.52848 | 0.20973 | 7.60729 | 7.39049 | 8.89737 | 12.45144 | 4.41835 |
| **Uckl1** | 0.59381 | 0.00311 | 0.42364 | 0.00000 | 0.00000 | 0.92036 | 2.01704 | 0.56232 | 0.00683 | 0.05553 | 1.46287 | 1.02916 | 0.01560 | 0.07133 |
| **Ucn2** | 0.00402 | 0.00456 | 0.00549 | 0.25541 | 0.00629 | 0.19725 | 0.00529 | 0.17549 | 0.03218 | 0.00577 | 0.00449 | 0.00389 | 0.19181 | 0.00000 |
| **Ucp2** | 0.02123 | 0.02084 | 0.12354 | 0.00000 | 2.02105 | 0.00000 | 0.00400 | 1.69063 | 0.00739 | 0.00696 | 0.05033 | 0.01201 | 0.29454 | 0.00757 |
| **Uevld** | 0.44075 | 0.02674 | 1.39176 | 0.32432 | 0.22571 | 1.22913 | 0.03570 | 2.34999 | 0.00674 | 0.03045 | 0.90974 | 0.24385 | 0.37935 | 2.51145 |
| **Ufc1** | 12.18987 | 5.54480 | 17.28739 | 19.41727 | 11.27312 | 11.56420 | 2.62770 | 10.96863 | 6.08826 | 0.25515 | 10.02610 | 9.64429 | 2.65775 | 15.72438 |
| **Ufd1l** | 6.83554 | 9.11319 | 13.64046 | 12.37082 | 11.68849 | 15.00798 | 2.93505 | 2.41494 | 13.30731 | 0.09117 | 11.29333 | 10.22424 | 4.58936 | 14.68034 |
| **Ufl1** | 1.05195 | 0.00922 | 0.00000 | 1.56847 | 0.21079 | 0.87800 | 0.00290 | 0.27465 | 1.45832 | 0.00472 | 0.59575 | 3.53698 | 0.02057 | 2.10300 |
| **Ufm1** | 8.35847 | 0.17410 | 5.42924 | 6.96520 | 11.57475 | 9.04551 | 2.31382 | 5.12311 | 7.14668 | 7.45103 | 7.07053 | 5.30403 | 3.41593 | 10.77587 |
| **Ufsp1** | 0.30726 | 0.00377 | 0.46468 | 0.59174 | 0.98281 | 0.00000 | 0.00360 | 0.90786 | 0.00737 | 0.00759 | 0.00305 | 0.33662 | 0.03382 | 0.00000 |
| **Ufsp2** | 3.78968 | 0.01535 | 4.23354 | 4.09566 | 4.47564 | 7.64337 | 1.46030 | 1.60252 | 4.46184 | 0.02607 | 8.73972 | 2.34658 | 0.26877 | 1.91459 |
| **Ugcg** | 22.28057 | 2.95477 | 18.61984 | 5.07435 | 21.21368 | 10.55013 | 0.62905 | 2.57911 | 32.80620 | 0.01054 | 7.90211 | 4.69323 | 7.99189 | 18.33897 |
| **Ugdh** | 0.44233 | 0.06561 | 0.65369 | 4.54980 | 4.24459 | 3.90028 | 4.31406 | 5.43344 | 5.61148 | 0.14299 | 5.97209 | 4.96213 | 1.63537 | 9.51961 |
| **Uggt1** | 7.61996 | 3.47611 | 18.09312 | 2.20886 | 14.10606 | 8.81839 | 5.30006 | 4.15679 | 15.43330 | 0.06997 | 4.38677 | 7.38001 | 1.28386 | 18.45620 |
| **Uggt2** | 0.02510 | 0.01152 | 0.00641 | 0.00000 | 0.00935 | 0.34885 | 0.00454 | 0.02002 | 0.01498 | 0.00497 | 0.00226 | 0.62104 | 0.15040 | 0.03310 |
| **Ugp2** | 4.72284 | 4.15364 | 1.24401 | 3.50865 | 3.71781 | 8.15145 | 5.37126 | 5.99244 | 4.59493 | 0.33536 | 2.94772 | 1.49260 | 0.35854 | 1.93486 |
| **Ugt2b36** | 0.35746 | 0.00384 | 0.00000 | 0.00742 | 0.61250 | 0.00180 | 0.00376 | 0.00937 | 2.96853 | 0.00706 | 0.00303 | 0.00373 | 4.29981 | 0.54126 |

| | | | | | | | | | | | | | | |
|---|---|---|---|---|---|---|---|---|---|---|---|---|---|---|
| **Ugt8a** | 0.00251 | 0.00429 | 0.01384 | 0.01216 | 0.00649 | 0.01239 | 0.00421 | 0.00894 | 0.03475 | 0.00616 | 0.00881 | 0.00883 | 0.60224 | 0.03121 |
| **Uhmk1** | 3.82923 | 0.00327 | 1.08271 | 0.32766 | 2.63270 | 0.00000 | 0.00332 | 0.00358 | 3.85326 | 0.00440 | 0.00070 | 0.00567 | 5.20002 | 1.60800 |
| **Uhrf1** | 0.00000 | 1.39507 | 1.44186 | 5.17232 | 0.42716 | 1.93202 | 0.81185 | 1.83064 | 0.01018 | 0.06863 | 1.98883 | 15.08097 | 0.93386 | 1.32704 |
| **Uhrf1bp1** | 3.23504 | 2.87521 | 3.03537 | 1.81971 | 3.53839 | 3.34582 | 0.65248 | 2.95929 | 0.39860 | 0.04138 | 7.48642 | 3.49317 | 5.65030 | 4.00175 |
| **Uhrf1bp1l** | 5.47888 | 0.00000 | 1.07767 | 0.25828 | 1.61147 | 3.17427 | 0.75401 | 1.50786 | 9.68632 | 0.02279 | 2.39898 | 2.93158 | 4.69214 | 5.13321 |
| **Uhrf2** | 0.08086 | 0.03074 | 0.00000 | 0.00000 | 0.18655 | 0.30092 | 0.00388 | 0.03022 | 0.92062 | 0.00841 | 0.00000 | 1.15479 | 0.51077 | 0.00000 |
| **Uimc1** | 3.82734 | 3.00509 | 0.98935 | 0.13840 | 1.42215 | 1.66619 | 0.00146 | 0.50015 | 7.12590 | 0.01023 | 1.05038 | 1.62153 | 2.16087 | 4.33272 |
| **Ulbp1** | 0.00736 | 0.00424 | 0.30126 | 0.04001 | 0.00180 | 0.00320 | 0.00411 | 0.00923 | 0.01804 | 0.00854 | 0.00639 | 0.00700 | 0.30917 | 0.13558 |
| **Ulk1** | 18.07650 | 4.89025 | 33.15663 | 13.89552 | 23.33795 | 9.13921 | 1.78024 | 4.81303 | 12.34140 | 10.65171 | 12.73595 | 18.53757 | 11.52016 | 28.04736 |
| **Ulk2** | 0.37657 | 0.02886 | 0.45387 | 0.00000 | 4.15464 | 0.91968 | 0.00345 | 0.26949 | 1.71379 | 0.00668 | 0.00722 | 1.25626 | 2.61466 | 4.67852 |
| **Ulk3** | 0.08169 | 0.03182 | 0.29459 | 0.36350 | 0.00074 | 0.28681 | 0.00932 | 0.94969 | 0.17769 | 0.00477 | 0.01147 | 0.00510 | 0.00623 | 0.08689 |
| **Ulk4** | 0.56285 | 0.00385 | 0.22149 | 0.02598 | 0.28778 | 0.00206 | 0.00368 | 0.11645 | 0.00643 | 0.00928 | 0.00454 | 0.07665 | 0.18062 | 0.00287 |
| **Umad1** | 0.14299 | 0.00329 | 0.58534 | 0.00000 | 0.92222 | 0.03237 | 0.02447 | 0.87382 | 1.09985 | 0.00354 | 0.92325 | 0.71669 | 0.79623 | 0.00000 |
| **Umodl1** | 0.66925 | 0.00427 | 0.31869 | 0.00000 | 3.02170 | 0.00000 | 0.00504 | 0.26952 | 0.27639 | 0.00456 | 0.00478 | 0.00520 | 1.16021 | 0.45298 |
| **Umps** | 10.76129 | 5.06181 | 19.53646 | 14.61515 | 13.15265 | 15.16118 | 5.91500 | 8.48302 | 7.34146 | 0.27029 | 12.93863 | 9.38462 | 4.66371 | 18.32715 |
| **Unc119** | 0.27914 | 2.59723 | 0.17345 | 1.12630 | 1.47335 | 0.75870 | 0.15597 | 0.86269 | 0.00051 | 0.00604 | 0.03453 | 2.23794 | 0.72246 | 0.33696 |
| **Unc119b** | 3.99049 | 0.01488 | 2.17986 | 3.92440 | 8.17248 | 3.49638 | 0.02413 | 2.06697 | 1.89887 | 0.03182 | 1.47429 | 2.72973 | 8.40717 | 4.19742 |
| **Unc13a** | 0.01212 | 0.00447 | 0.03276 | 0.01272 | 0.28643 | 0.02975 | 0.00448 | 0.00877 | 0.01276 | 0.00707 | 0.00711 | 0.01254 | 0.00975 | 0.08513 |
| **Unc13b** | 1.63701 | 0.00367 | 1.52284 | 0.73404 | 1.77447 | 1.18422 | 0.00390 | 0.01148 | 3.75116 | 0.00356 | 0.00193 | 0.02857 | 3.19685 | 4.83926 |
| **Unc13c** | 0.84623 | 0.00391 | 0.28642 | 0.18507 | 0.94191 | 0.18299 | 0.00372 | 0.00000 | 3.22414 | 0.00564 | 0.00174 | 0.00256 | 0.01344 | 7.40652 |
| **Unc13d** | 0.00758 | 0.00503 | 0.09400 | 0.00000 | 0.25960 | 0.13890 | 0.00444 | 0.16885 | 0.00870 | 0.00746 | 0.00556 | 0.01027 | 0.01073 | 0.11628 |
| **Unc45a** | 0.73902 | 3.42926 | 7.29500 | 9.21662 | 2.19987 | 2.45640 | 3.15756 | 7.05588 | 0.01480 | 5.04327 | 2.64024 | 0.38305 | 1.15951 | 0.00000 |
| **Unc45b** | 0.18154 | 0.00383 | 2.83352 | 0.00000 | 2.45235 | 0.00000 | 0.00346 | 0.00335 | 0.04724 | 0.00094 | 0.00719 | 0.01140 | 1.68853 | 0.56929 |
| **Unc50** | 6.60034 | 0.68414 | 3.03901 | 0.53457 | 0.29441 | 3.36740 | 0.02611 | 1.46667 | 2.93052 | 0.02849 | 0.72247 | 2.36937 | 4.52718 | 4.35483 |
| **Unc5b** | 0.12761 | 0.00402 | 0.23002 | 0.40909 | 0.04288 | 0.35015 | 2.06864 | 0.00221 | 0.04338 | 0.03776 | 0.00213 | 0.01577 | 0.00483 | 0.25449 |
| **Unc5c** | 0.01807 | 0.00417 | 0.00000 | 0.01548 | 0.33585 | 0.00738 | 0.00397 | 0.00694 | 1.20539 | 0.00948 | 0.00456 | 0.00648 | 1.30307 | 0.53812 |
| **Unc5cl** | 0.01242 | 0.00443 | 0.12284 | 0.01465 | 0.07628 | 0.00774 | 0.00416 | 0.01096 | 0.01026 | 0.00988 | 0.00728 | 0.00862 | 0.01714 | 0.01260 |
| **Unc80** | 0.01250 | 0.00437 | 0.01062 | 0.00000 | 0.01425 | 0.36556 | 0.00419 | 0.00464 | 0.01117 | 0.00575 | 0.00604 | 0.03016 | 0.00804 | 0.00821 |
| **Unc93a** | 0.15170 | 0.04866 | 0.39514 | 0.00000 | 0.27880 | 0.80263 | 2.20533 | 5.93413 | 0.02192 | 0.04060 | 0.00000 | 0.91597 | 0.26280 | 0.30120 |
| **Unc93b1** | 0.02342 | 0.00442 | 0.00726 | 0.00993 | 0.55974 | 0.02139 | 0.00438 | 0.00994 | 0.00821 | 0.00699 | 0.01125 | 0.00786 | 0.06804 | 0.03518 |
| **Uncx** | 0.06231 | 0.00427 | 0.07854 | 0.00653 | 0.32969 | 0.01346 | 0.00443 | 0.01528 | 0.02009 | 0.00851 | 0.00749 | 0.01052 | 0.67442 | 0.02095 |
| **Ung** | 3.34126 | 1.25011 | 4.35459 | 7.81621 | 10.19122 | 4.02827 | 0.07304 | 1.64397 | 2.34398 | 0.07950 | 2.11126 | 5.99575 | 6.14187 | 10.18082 |
| **Unk** | 1.45259 | 0.01938 | 4.54428 | 2.12854 | 2.67075 | 1.68706 | 0.01741 | 0.76237 | 0.00170 | 0.00517 | 0.00000 | 0.42570 | 2.78528 | 0.67574 |
| **Unkl** | 2.33343 | 0.05889 | 7.14079 | 6.48230 | 7.15464 | 7.26433 | 0.00765 | 1.07184 | 1.47860 | 0.04301 | 5.05252 | 3.25273 | 3.89783 | 6.07455 |
| **Upb1** | 0.00526 | 0.00417 | 0.00297 | 0.20941 | 0.01174 | 0.01377 | 0.00397 | 0.00515 | 0.06760 | 0.00668 | 0.00731 | 0.01406 | 0.05607 | 0.35725 |
| **Upf1** | 0.00000 | 0.01084 | 0.75161 | 1.12926 | 1.49370 | 0.04244 | 0.06228 | 0.87982 | 0.00241 | 0.02202 | 0.00000 | 1.58711 | 1.48730 | 0.00000 |
| **Upf2** | 9.18597 | 0.06038 | 7.08514 | 4.54680 | 9.26116 | 8.14308 | 1.26140 | 3.06185 | 7.40910 | 0.07488 | 4.08154 | 5.33343 | 4.81265 | 4.63793 |
| **Upf3a** | 1.90172 | 0.06517 | 3.95717 | 4.29144 | 5.40042 | 1.93431 | 0.81693 | 1.21831 | 0.39317 | 0.03728 | 1.25840 | 5.91776 | 1.35025 | 3.66451 |
| **Upf3b** | 12.58444 | 0.03957 | 10.81931 | 2.19449 | 13.92303 | 7.39331 | 0.00471 | 0.00000 | 12.68345 | 1.87739 | 4.57803 | 3.67641 | 0.87297 | 11.08327 |
| **Upk1a** | 0.00000 | 0.00427 | 0.48345 | 0.27322 | 0.25624 | 0.09654 | 0.00448 | 0.00617 | 0.00973 | 0.01321 | 0.00574 | 0.00701 | 0.92154 | 0.00000 |
| **Upk3a** | 0.32462 | 0.00381 | 0.36142 | 0.77246 | 0.66228 | 0.98019 | 0.00358 | 0.00000 | 0.00390 | 0.01005 | 0.61541 | 0.00568 | 0.49856 | 0.95706 |
| **Upk3b** | 0.19024 | 0.00423 | 0.09765 | 0.11209 | 0.57782 | 0.41643 | 0.00392 | 0.00000 | 0.06776 | 0.01491 | 1.06647 | 0.00897 | 0.00522 | 0.00000 |
| **Upk3bl** | 0.06436 | 0.00424 | 0.02722 | 0.04266 | 0.17879 | 0.01519 | 0.00400 | 0.00674 | 0.00851 | 0.01171 | 0.00677 | 0.00829 | 0.47248 | 0.00339 |
| **Upp1** | 0.00167 | 0.29948 | 0.17979 | 0.32778 | 0.29862 | 0.08241 | 0.05780 | 1.11087 | 0.00414 | 0.10012 | 1.92153 | 21.45758 | 3.02360 | 0.48976 |
| **Uqcc1** | 6.68896 | 0.07064 | 13.17592 | 9.14987 | 10.24121 | 8.59388 | 0.20770 | 2.26111 | 14.63216 | 1.59385 | 9.73407 | 4.58859 | 2.02948 | 12.70511 |
| **Uqcc2** | 0.38461 | 0.01237 | 0.00000 | 1.20010 | 1.88103 | 2.42965 | 1.73357 | 1.44016 | 0.04637 | 0.14920 | 0.01404 | 0.86563 | 0.56459 | 0.00150 |
| **Uqcc3** | 7.03373 | 0.07854 | 1.89284 | 9.15494 | 10.48522 | 1.45167 | 0.82204 | 2.97743 | 0.15799 | 11.04578 | 4.52464 | 3.36107 | 10.14225 | 8.41658 |
| **Uqcr10** | 6.07543 | 4.96922 | 0.13175 | 2.60118 | 7.03137 | 9.87497 | 5.61169 | 12.71105 | 0.66887 | 16.28547 | 0.21236 | 10.12240 | 6.73301 | 3.58491 |
| **Uqcr11** | 0.68937 | 4.15963 | 0.00000 | 2.07908 | 2.08570 | 2.78966 | 2.41714 | 10.48334 | 0.01284 | 10.98223 | 0.00000 | 5.79395 | 0.28526 | 0.30075 |
| **Uqcrb** | 8.35926 | 0.13769 | 0.01462 | 7.11717 | 4.58929 | 11.14969 | 2.43075 | 9.74704 | 6.34638 | 9.02875 | 1.28295 | 6.97632 | 1.70689 | 1.75396 |
| **Uqcrc1** | 14.21791 | 12.46704 | 26.11918 | 29.76033 | 23.36213 | 15.07519 | 9.58774 | 23.43996 | 0.00000 | 0.84488 | 11.40201 | 20.18156 | 0.04730 | 2.78722 |
| **Uqcrc2** | 14.57682 | 4.77659 | 18.74310 | 19.43930 | 9.50546 | 25.90609 | 7.91070 | 15.84252 | 19.55435 | 0.47731 | 15.40385 | 20.11696 | 2.45945 | 15.07071 |
| **Uqcrfs1** | 13.39319 | 0.15696 | 4.44039 | 6.47748 | 12.72615 | 10.35007 | 2.52574 | 6.75450 | 2.01376 | 8.39960 | 2.85241 | 8.29545 | 7.88614 | 3.29045 |
| **Uqcrh** | 7.17164 | 5.95627 | 0.00000 | 2.43497 | 8.74073 | 10.74955 | 5.65483 | 9.80344 | 4.34201 | 14.20005 | 0.38369 | 8.80293 | 6.24853 | 4.79941 |
| **Uqcrq** | 0.35726 | 0.80280 | 0.25795 | 4.25123 | 9.25000 | 3.50267 | 3.12176 | 11.55804 | 0.00000 | 6.57997 | 0.02543 | 8.28594 | 0.61670 | 0.00000 |

| | | | | | | | | | | | | | | |
|---|---|---|---|---|---|---|---|---|---|---|---|---|---|---|
| **Urah** | 0.00383 | 0.01916 | 0.02276 | 0.00000 | 0.01107 | 0.06173 | 0.00402 | 0.00828 | 0.01156 | 0.00694 | 0.13373 | 1.25129 | 0.00725 | 0.01379 |
| **Urb1** | 4.70710 | 0.04537 | 7.56326 | 3.94488 | 8.91839 | 4.65635 | 0.38484 | 1.20182 | 0.31081 | 0.00577 | 2.69174 | 1.98626 | 5.53591 | 6.87587 |
| **Urb2** | 0.92777 | 0.00324 | 2.41082 | 2.16649 | 2.20574 | 0.70324 | 0.00294 | 0.00000 | 2.36214 | 0.00627 | 0.55173 | 0.02550 | 1.17521 | 6.17230 |
| **Urgcp** | 0.11015 | 8.66748 | 6.11468 | 0.00000 | 7.98684 | 0.00000 | 3.56867 | 2.04157 | 0.34009 | 0.14334 | 0.28442 | 1.67002 | 1.41795 | 1.84732 |
| **Uri1** | 3.69661 | 0.04930 | 3.12356 | 0.00000 | 4.46354 | 2.78846 | 2.94126 | 2.06991 | 1.01243 | 0.03354 | 7.20973 | 8.39131 | 1.08174 | 1.48957 |
| **Urm1** | 3.59741 | 0.33081 | 2.05871 | 1.56113 | 7.29151 | 1.06683 | 0.04985 | 0.90599 | 0.68063 | 0.02207 | 2.04244 | 0.64357 | 3.65468 | 1.36019 |
| **Urod** | 2.62761 | 0.20472 | 13.61381 | 11.90668 | 10.62000 | 6.08627 | 0.00996 | 5.30818 | 4.32994 | 0.02928 | 7.39677 | 8.45989 | 3.87663 | 13.20318 |
| **Uros** | 0.81457 | 0.02232 | 1.04326 | 0.38084 | 0.00000 | 1.32108 | 0.00950 | 0.56659 | 0.07205 | 0.00447 | 0.13569 | 0.58605 | 0.62454 | 1.26349 |
| **Usb1** | 0.34207 | 0.10367 | 2.47788 | 2.09510 | 2.49123 | 2.47265 | 0.00323 | 1.71058 | 2.24060 | 0.00122 | 0.00000 | 2.23207 | 0.67222 | 0.14475 |
| **Use1** | 5.34814 | 0.07417 | 5.25563 | 10.22904 | 5.26353 | 3.34947 | 4.09765 | 6.38096 | 0.08731 | 11.07931 | 3.73363 | 3.39845 | 2.20076 | 1.81968 |
| **Usf1** | 0.00000 | 0.01021 | 1.82359 | 0.53350 | 0.85909 | 0.00000 | 0.00948 | 1.96147 | 0.24186 | 0.01310 | 3.92655 | 0.58886 | 0.89472 | 7.96937 |
| **Usf2** | 0.01338 | 0.04624 | 0.60697 | 5.93603 | 0.28675 | 2.32651 | 2.03168 | 6.40813 | 0.00000 | 0.07553 | 4.25364 | 3.74208 | 0.26154 | 0.45343 |
| **Usf3** | 0.84498 | 0.00180 | 0.38622 | 0.11634 | 2.21613 | 0.18463 | 0.00834 | 0.39846 | 5.52539 | 0.01310 | 2.91944 | 1.60305 | 1.24849 | 4.19788 |
| **Ush2a** | 0.00418 | 0.00454 | 0.00774 | 0.01183 | 0.19103 | 0.01038 | 0.00442 | 0.01559 | 0.04752 | 0.00651 | 0.00787 | 0.00896 | 0.02574 | 0.31597 |
| **Usmg5** | 2.28902 | 0.42365 | 0.00000 | 0.64889 | 3.07762 | 6.88484 | 1.55208 | 5.95372 | 1.46067 | 8.74014 | 0.00000 | 5.20761 | 9.36247 | 1.35101 |
| **Uso1** | 5.56142 | 0.05046 | 7.85312 | 8.15507 | 6.91208 | 8.32748 | 2.64827 | 5.76839 | 2.27043 | 0.09443 | 5.41412 | 5.26235 | 1.28780 | 11.43033 |
| **Usp1** | 21.36097 | 0.48301 | 19.10940 | 9.53899 | 27.41485 | 6.43435 | 2.20815 | 2.10234 | 35.19526 | 0.04174 | 10.87536 | 4.05637 | 16.90025 | 22.99715 |
| **Usp10** | 8.52594 | 5.22510 | 25.83168 | 14.87466 | 31.87466 | 3.19083 | 3.69441 | 8.43525 | 0.94253 | 0.14029 | 4.14271 | 8.10323 | 11.33230 | 2.70301 |
| **Usp11** | 0.61929 | 0.01435 | 0.04030 | 0.00000 | 0.12762 | 0.49522 | 0.00371 | 0.31389 | 0.00692 | 0.00849 | 1.46956 | 1.25820 | 2.01920 | 0.00000 |
| **Usp12** | 1.40649 | 0.00965 | 1.17175 | 4.64033 | 4.05806 | 5.13783 | 0.78538 | 0.15284 | 2.17094 | 0.11370 | 3.47070 | 2.64318 | 5.42678 | 6.47628 |
| **Usp13** | 0.06916 | 0.00403 | 0.31623 | 0.67544 | 1.00129 | 0.00000 | 0.00386 | 0.00541 | 0.03674 | 0.00792 | 0.00502 | 0.00999 | 0.14212 | 0.00000 |
| **Usp14** | 19.93989 | 0.08420 | 23.16858 | 14.66819 | 14.90988 | 20.14366 | 4.37678 | 6.61840 | 23.48807 | 3.08239 | 18.71875 | 13.21707 | 3.44585 | 10.27613 |
| **Usp15** | 10.82566 | 0.00000 | 7.94012 | 3.18054 | 10.41610 | 6.86216 | 2.62585 | 6.38209 | 10.88651 | 0.09041 | 11.77111 | 3.91858 | 4.65958 | 5.31177 |
| **Usp16** | 16.74840 | 0.03926 | 16.79151 | 6.15264 | 23.32830 | 9.89613 | 1.69769 | 0.00671 | 25.59636 | 0.04796 | 6.74488 | 6.46323 | 11.63625 | 25.23906 |
| **Usp17la** | 0.00000 | 0.05341 | 0.04979 | 0.05438 | 0.02611 | 0.01503 | 0.16090 | 0.01092 | 0.22071 | 0.00658 | 0.03043 | 0.04455 | 1.10396 | 0.00000 |
| **Usp17lb** | 0.00000 | 0.00402 | 0.17409 | 0.07360 | 0.00000 | 0.00226 | 0.00373 | 0.00580 | 0.90789 | 0.00636 | 0.00542 | 0.01230 | 1.10585 | 0.00000 |
| **Usp17lc** | 0.00577 | 0.00441 | 0.02651 | 0.01066 | 0.00903 | 0.01235 | 0.00426 | 0.00994 | 0.04940 | 0.00658 | 0.00873 | 0.00950 | 0.17542 | 0.00000 |
| **Usp17ld** | 0.00648 | 0.00439 | 0.02436 | 0.01754 | 0.00937 | 0.01189 | 0.00423 | 0.00930 | 0.04946 | 0.00672 | 0.00865 | 0.01072 | 0.12505 | 0.00000 |
| **Usp17le** | 0.00846 | 0.00437 | 0.01425 | 0.01615 | 0.00995 | 0.01262 | 0.00422 | 0.00916 | 0.02907 | 0.00668 | 0.00850 | 0.01008 | 0.22019 | 0.00000 |
| **Usp19** | 0.33139 | 0.02647 | 6.42359 | 9.52142 | 7.54690 | 0.00000 | 0.02223 | 2.46726 | 0.37896 | 0.01385 | 0.76698 | 2.32628 | 7.63268 | 0.29297 |
| **Usp2** | 0.00000 | 2.10395 | 1.55785 | 2.87689 | 0.60567 | 0.66347 | 0.02150 | 0.00952 | 0.03317 | 0.00421 | 0.00896 | 0.46310 | 0.51603 | 0.51948 |
| **Usp20** | 0.23823 | 0.00367 | 1.01175 | 1.26531 | 0.90589 | 1.53413 | 0.00376 | 0.00000 | 0.00278 | 0.00992 | 0.01806 | 0.42034 | 0.65940 | 0.44513 |
| **Usp21** | 1.85071 | 3.65132 | 2.19645 | 0.47836 | 3.43182 | 0.74628 | 0.00169 | 3.40497 | 0.38240 | 0.01627 | 2.39482 | 2.89388 | 1.59395 | 2.19074 |
| **Usp22** | 11.18676 | 0.10397 | 16.68488 | 5.71375 | 15.30961 | 8.63528 | 3.03544 | 2.92505 | 17.18250 | 0.06478 | 4.05145 | 4.63072 | 9.75510 | 17.57600 |
| **Usp24** | 3.65066 | 0.11008 | 5.79550 | 3.05720 | 2.26086 | 10.06463 | 0.67360 | 8.53391 | 8.17458 | 1.72820 | 6.56400 | 6.82676 | 0.28954 | 4.80691 |
| **Usp25** | 7.70468 | 5.36951 | 2.21800 | 3.60402 | 2.37819 | 0.00000 | 0.00161 | 0.21372 | 2.79813 | 0.00395 | 2.47904 | 1.27481 | 0.00000 | 2.32760 |
| **Usp26** | 0.00735 | 0.00443 | 0.01169 | 0.01391 | 0.00535 | 0.01211 | 0.00428 | 0.01008 | 0.02298 | 0.00697 | 0.00837 | 0.00983 | 0.00583 | 0.23391 |
| **Usp27x** | 0.00689 | 0.00401 | 0.00510 | 0.00000 | 0.19281 | 0.38671 | 0.00422 | 0.01955 | 0.01547 | 0.00643 | 0.01009 | 0.06361 | 0.00570 | 0.00407 |
| **Usp28** | 0.00000 | 0.00769 | 4.47877 | 1.69226 | 0.61605 | 2.39881 | 0.01663 | 0.46327 | 3.06571 | 0.03197 | 0.33326 | 5.52155 | 4.38748 | 7.65014 |
| **Usp29** | 2.50104 | 0.00209 | 0.29426 | 0.85047 | 8.90703 | 2.77404 | 0.00216 | 0.00000 | 11.11578 | 0.02489 | 0.00000 | 0.00000 | 11.68880 | 5.86972 |
| **Usp3** | 0.93625 | 0.18366 | 0.25804 | 3.41684 | 1.48845 | 2.73097 | 0.10138 | 1.79003 | 0.43108 | 0.24414 | 0.00000 | 2.13452 | 2.07931 | 3.13991 |
| **Usp30** | 2.11578 | 0.01935 | 3.71249 | 2.33246 | 0.67943 | 3.72946 | 1.39520 | 0.69699 | 0.57523 | 0.01186 | 0.88088 | 1.09501 | 0.01232 | 1.19433 |
| **Usp31** | 0.38795 | 0.00227 | 1.23447 | 0.00000 | 0.24626 | 3.94375 | 0.00301 | 0.00000 | 0.65504 | 0.00210 | 2.49521 | 1.26322 | 0.38425 | 1.29369 |
| **Usp32** | 2.29095 | 0.00653 | 1.83749 | 0.27328 | 2.28050 | 1.46979 | 0.01364 | 0.20927 | 3.28220 | 0.00000 | 2.55019 | 5.25303 | 2.77842 | 3.04863 |
| **Usp33** | 16.51961 | 0.00000 | 10.52708 | 2.66114 | 17.86182 | 3.63358 | 1.86706 | 2.29054 | 28.74286 | 0.01034 | 2.34636 | 2.15587 | 11.79618 | 12.28341 |
| **Usp34** | 1.07057 | 0.01091 | 2.08679 | 0.00000 | 3.97481 | 2.40837 | 0.16935 | 0.53522 | 19.41290 | 0.00000 | 2.22629 | 1.08089 | 4.39396 | 8.54091 |
| **Usp35** | 0.78655 | 0.00508 | 0.00406 | 0.35402 | 0.00000 | 0.02059 | 0.00408 | 0.02058 | 0.01691 | 0.01100 | 0.00715 | 0.00516 | 0.10600 | 0.00000 |
| **Usp36** | 1.57939 | 1.00288 | 3.75212 | 5.41879 | 6.06009 | 6.35259 | 0.00850 | 1.38119 | 0.11007 | 0.03137 | 0.44101 | 1.83610 | 4.28718 | 9.15683 |
| **Usp37** | 11.58789 | 0.04747 | 8.51614 | 3.35331 | 19.25938 | 5.76922 | 0.00690 | 2.30742 | 17.50163 | 9.28493 | 7.47328 | 3.08591 | 10.58306 | 14.67106 |
| **Usp38** | 14.43054 | 0.02421 | 8.63193 | 1.28100 | 13.89247 | 3.95409 | 1.29233 | 0.80555 | 8.19157 | 0.01939 | 3.03661 | 3.09809 | 14.93809 | 18.24583 |
| **Usp39** | 3.37736 | 0.06897 | 8.70299 | 5.47121 | 11.83776 | 7.63690 | 4.85678 | 1.80893 | 6.38954 | 0.09730 | 3.90758 | 4.73164 | 6.58431 | 11.38171 |
| **Usp4** | 4.56499 | 4.95314 | 7.41085 | 3.21994 | 5.79557 | 5.60545 | 3.85191 | 4.84159 | 5.06871 | 0.06384 | 6.26911 | 4.59939 | 4.57133 | 8.77941 |
| **Usp40** | 0.86162 | 0.02798 | 0.00000 | 0.61489 | 0.30491 | 1.85632 | 0.00403 | 0.64032 | 3.45613 | 0.01904 | 1.07904 | 1.83106 | 0.00000 | 0.92852 |
| **Usp42** | 3.15990 | 0.01030 | 3.04293 | 2.05373 | 3.67456 | 3.05004 | 0.00304 | 0.96470 | 2.82959 | 0.02553 | 0.71528 | 1.66488 | 3.38421 | 2.62128 |
| **Usp43** | 1.30133 | 5.59418 | 6.63243 | 4.42280 | 1.56655 | 6.43619 | 4.76874 | 0.00000 | 0.02200 | 0.11672 | 2.22826 | 1.28888 | 0.00291 | 0.63377 |

| | | | | | | | | | | | | | | |
|---|---|---|---|---|---|---|---|---|---|---|---|---|---|---|
| **Usp44** | 0.00032 | 0.00446 | 0.02305 | 0.01125 | 0.00000 | 0.00876 | 0.00415 | 0.01006 | 0.31868 | 0.00753 | 0.00670 | 0.00919 | 0.06361 | 0.33130 |
| **Usp45** | 2.87897 | 0.01860 | 1.79923 | 1.18416 | 2.89769 | 3.38486 | 0.02136 | 0.06935 | 7.22268 | 0.01053 | 1.91645 | 1.01881 | 1.32310 | 1.47692 |
| **Usp46** | 0.51979 | 0.01935 | 0.79579 | 0.56919 | 0.00000 | 1.25836 | 0.00325 | 1.93481 | 0.09789 | 0.02700 | 1.66515 | 0.02729 | 2.74747 | 0.02319 |
| **Usp47** | 1.68235 | 0.02386 | 1.19117 | 0.00000 | 1.57950 | 2.43195 | 0.95157 | 2.43428 | 6.46075 | 0.02242 | 0.52652 | 4.18490 | 1.72053 | 4.44226 |
| **Usp48** | 21.73016 | 2.77419 | 32.47252 | 23.19053 | 30.36281 | 22.82028 | 7.46646 | 13.39520 | 21.14648 | 5.80761 | 18.06840 | 17.01294 | 6.42624 | 23.56618 |
| **Usp49** | 0.42959 | 0.02687 | 0.82919 | 0.00000 | 2.45047 | 1.65843 | 0.04427 | 0.00258 | 0.59072 | 0.00526 | 0.00000 | 1.97246 | 3.47676 | 0.92380 |
| **Usp5** | 6.24502 | 3.36664 | 14.84420 | 9.54200 | 10.53135 | 12.17353 | 2.76316 | 7.24482 | 1.72477 | 0.15354 | 12.64498 | 11.44193 | 2.31590 | 7.25855 |
| **Usp50** | 0.00409 | 0.00442 | 0.00917 | 0.01290 | 0.05208 | 0.01118 | 0.00417 | 0.00987 | 0.01116 | 0.00678 | 0.00784 | 0.00920 | 0.01503 | 0.59009 |
| **Usp53** | 0.49632 | 0.00463 | 1.20415 | 0.00000 | 0.00000 | 0.17523 | 0.00360 | 0.37504 | 4.75213 | 0.00270 | 0.48353 | 0.00217 | 4.18912 | 1.62787 |
| **Usp54** | 3.93712 | 0.01943 | 2.58948 | 2.30280 | 4.68442 | 1.21229 | 0.00233 | 2.54275 | 7.44360 | 0.00164 | 0.30817 | 0.40955 | 6.07542 | 7.90428 |
| **Usp6nl** | 0.35357 | 0.02006 | 0.13987 | 0.07670 | 0.00000 | 0.05127 | 0.00504 | 0.82358 | 0.00763 | 0.00683 | 0.01082 | 0.00635 | 0.06244 | 0.13871 |
| **Usp7** | 15.15534 | 0.10171 | 19.58548 | 6.39276 | 17.91330 | 9.64075 | 2.98316 | 6.79176 | 11.54227 | 0.26594 | 10.75396 | 12.81770 | 11.90943 | 14.87195 |
| **Usp8** | 7.77539 | 0.05045 | 6.38636 | 1.69367 | 5.79899 | 7.97760 | 2.90587 | 1.81632 | 3.92352 | 0.05503 | 6.70136 | 6.37171 | 2.25784 | 4.47590 |
| **Usp9x** | 9.91852 | 0.01017 | 6.24105 | 3.44180 | 8.26605 | 8.66347 | 4.04680 | 0.02596 | 22.37007 | 0.05630 | 6.98431 | 7.65637 | 6.78199 | 8.25465 |
| **Uspl1** | 4.77783 | 0.02307 | 6.68726 | 2.55643 | 5.03548 | 6.15049 | 2.04073 | 0.69177 | 4.28117 | 3.22619 | 1.33497 | 3.31067 | 12.21501 | 8.58318 |
| **Utf1** | 0.00116 | 0.06136 | 0.08885 | 12.69659 | 4.18522 | 0.00000 | 0.02168 | 0.26512 | 0.00535 | 0.03213 | 0.04945 | 4.03895 | 0.02497 | 0.01277 |
| **Utp11l** | 9.72734 | 0.15752 | 10.84258 | 10.01972 | 16.36838 | 12.25730 | 3.76314 | 5.84744 | 7.17211 | 0.09299 | 10.77437 | 9.10284 | 10.56721 | 12.02008 |
| **Utp14a** | 3.62442 | 0.09425 | 11.53913 | 1.81953 | 8.02202 | 3.82276 | 2.46456 | 2.76080 | 17.10977 | 0.03055 | 4.67170 | 4.54702 | 0.09450 | 14.31474 |
| **Utp14b** | 0.31112 | 0.00901 | 1.61398 | 1.90608 | 2.04803 | 0.22493 | 0.00293 | 1.24254 | 1.08877 | 0.01891 | 1.88010 | 1.67713 | 7.80295 | 7.07337 |
| **Utp15** | 6.33741 | 0.06082 | 9.97942 | 4.73325 | 8.88143 | 7.11710 | 1.21818 | 0.14348 | 10.68458 | 0.05563 | 5.13082 | 6.31093 | 11.00667 | 13.65023 |
| **Utp18** | 8.95486 | 0.01038 | 10.50178 | 6.43318 | 17.54021 | 5.32698 | 3.19253 | 2.58542 | 13.22845 | 0.14849 | 5.21753 | 7.62231 | 4.24879 | 10.78472 |
| **Utp20** | 3.92269 | 3.28133 | 10.12750 | 2.25352 | 4.71186 | 2.78301 | 0.00463 | 1.87453 | 10.42535 | 0.02414 | 3.87202 | 3.90836 | 5.18523 | 10.10247 |
| **Utp23** | 7.54594 | 0.15128 | 5.46476 | 3.55106 | 13.69516 | 5.30313 | 2.80225 | 2.00585 | 8.01545 | 0.19254 | 3.56251 | 1.38940 | 14.18167 | 16.60125 |
| **Utp3** | 18.17340 | 0.17104 | 21.95313 | 11.97625 | 24.96051 | 9.00057 | 4.28423 | 6.88842 | 15.13761 | 0.07015 | 4.71161 | 8.24243 | 17.03291 | 17.53081 |
| **Utp6** | 6.97379 | 0.06153 | 6.64932 | 1.49497 | 5.89130 | 6.85643 | 0.01203 | 0.29893 | 12.20878 | 18.29419 | 7.85177 | 6.20576 | 6.08878 | 11.01789 |
| **Utrn** | 0.41766 | 0.01113 | 0.68268 | 0.75443 | 0.11516 | 1.75520 | 3.05513 | 0.67673 | 2.86070 | 0.03049 | 0.06015 | 0.72501 | 0.69650 | 0.42243 |
| **Uts2r** | 0.05104 | 0.00423 | 0.01046 | 0.00000 | 0.35312 | 0.10330 | 0.00417 | 0.00501 | 0.01129 | 0.00528 | 0.00725 | 0.04829 | 0.24456 | 0.00303 |
| **Uty** | 0.00385 | 0.00420 | 0.01159 | 0.19981 | 0.00614 | 0.00000 | 0.00474 | 0.02071 | 0.01844 | 0.00572 | 0.14094 | 0.06949 | 0.57055 | 0.00000 |
| **Uvrag** | 0.43487 | 0.01774 | 0.56517 | 0.90672 | 2.63738 | 0.98884 | 0.00884 | 2.43458 | 0.14601 | 0.15192 | 0.00000 | 1.96193 | 0.44326 | 0.79357 |
| **Uvssa** | 1.19751 | 0.04049 | 0.04747 | 0.15946 | 0.00000 | 0.19804 | 0.00364 | 0.00571 | 2.93486 | 0.00744 | 0.00993 | 0.18407 | 0.20289 | 0.05757 |
| **Uxs1** | 0.05437 | 0.00398 | 0.51397 | 0.39660 | 0.19881 | 0.00640 | 0.00347 | 0.00209 | 1.65946 | 0.00699 | 0.01353 | 0.07891 | 0.23399 | 0.58450 |
| **Uxt** | 3.94532 | 3.27833 | 0.19592 | 5.06224 | 7.51184 | 3.53112 | 2.22648 | 4.01444 | 1.54210 | 0.14184 | 0.44241 | 2.68397 | 2.14870 | 3.29121 |
| **Vac14** | 1.99838 | 7.24117 | 6.52390 | 1.21883 | 5.93995 | 8.77094 | 4.21961 | 4.40955 | 0.79576 | 0.08220 | 5.70122 | 6.07651 | 4.03779 | 7.02390 |
| **Vamp1** | 0.00362 | 0.00432 | 0.00954 | 0.15034 | 0.17440 | 0.01110 | 0.00440 | 0.00712 | 0.00680 | 0.01264 | 0.00641 | 0.00684 | 0.00339 | 0.30750 |
| **Vamp2** | 0.00000 | 0.00468 | 0.05746 | 1.31569 | 0.91563 | 0.08681 | 0.00524 | 0.03702 | 0.12851 | 0.02251 | 0.00367 | 0.00412 | 0.00065 | 0.53317 |
| **Vamp3** | 7.52621 | 0.06083 | 11.60801 | 11.61842 | 12.44263 | 13.99063 | 3.81236 | 4.02560 | 10.66502 | 1.57633 | 9.71299 | 3.48909 | 0.45359 | 7.04489 |
| **Vamp4** | 0.38614 | 0.03535 | 0.58067 | 0.00000 | 0.69908 | 0.71969 | 0.00368 | 1.90620 | 0.06312 | 0.00279 | 0.64137 | 0.00724 | 0.26146 | 1.19590 |
| **Vamp5** | 0.01870 | 0.01165 | 0.00246 | 0.13294 | 0.77552 | 0.01404 | 0.00410 | 0.01632 | 0.02001 | 0.00657 | 0.00686 | 0.00572 | 0.02799 | 0.00498 |
| **Vamp8** | 8.61430 | 0.58280 | 3.41643 | 10.56310 | 16.97766 | 5.52784 | 2.57812 | 12.20189 | 8.55455 | 13.57598 | 1.19121 | 2.92748 | 4.94443 | 8.65948 |
| **Vangl1** | 1.12736 | 0.11290 | 1.32467 | 0.83754 | 0.87680 | 2.15386 | 0.22553 | 0.67866 | 0.14655 | 0.02327 | 2.58885 | 9.19436 | 0.00000 | 0.65777 |
| **Vangl2** | 0.11370 | 0.01312 | 0.56717 | 0.00000 | 0.39800 | 0.20967 | 0.01145 | 0.02265 | 0.03096 | 0.02124 | 0.00743 | 0.05908 | 0.00250 | 0.00427 |
| **Vapa** | 7.05165 | 0.42302 | 14.35136 | 14.08289 | 17.58089 | 16.12531 | 3.34646 | 7.55804 | 19.27745 | 0.34281 | 11.73982 | 10.74607 | 11.29366 | 14.18782 |
| **Vapb** | 0.16670 | 0.00385 | 0.42449 | 0.43793 | 1.15070 | 0.54397 | 0.00364 | 0.79707 | 0.00505 | 0.05038 | 0.01314 | 0.03267 | 0.35106 | 0.06814 |
| **Vars** | 0.00000 | 0.04004 | 1.71166 | 18.47122 | 4.45303 | 0.43087 | 0.12827 | 1.27022 | 0.00594 | 0.06705 | 0.00000 | 3.89223 | 0.02063 | 0.00000 |
| **Vars2** | 0.48191 | 0.01114 | 2.73876 | 2.64653 | 0.69583 | 0.52175 | 2.11677 | 0.00617 | 0.11639 | 0.02622 | 1.01911 | 1.35465 | 0.24141 | 0.00000 |
| **Vash1** | 0.00394 | 0.00441 | 0.01178 | 0.01670 | 0.00789 | 0.01160 | 0.00432 | 0.01129 | 0.00788 | 0.00653 | 0.00838 | 0.00875 | 0.58721 | 0.27066 |
| **Vasn** | 0.10171 | 0.00384 | 0.42010 | 1.18271 | 0.13342 | 0.00000 | 0.00390 | 0.01071 | 0.00605 | 0.00564 | 0.00578 | 0.00600 | 1.98683 | 0.00000 |
| **Vasp** | 0.00000 | 0.54908 | 2.80494 | 3.07750 | 0.23572 | 4.70993 | 4.43445 | 12.29208 | 0.49780 | 2.06921 | 6.45609 | 2.37166 | 2.34676 | 1.30313 |
| **Vat1** | 0.56459 | 0.03835 | 0.41424 | 0.43580 | 3.71358 | 0.35301 | 1.45757 | 0.04682 | 0.53315 | 0.11264 | 0.15784 | 0.28702 | 1.68440 | 0.82137 |
| **Vat1l** | 0.00784 | 0.00505 | 0.00777 | 0.03511 | 0.01053 | 0.03785 | 0.00428 | 0.37334 | 0.01423 | 0.00738 | 0.19408 | 0.10989 | 0.00676 | 0.00953 |
| **Vav1** | 0.41561 | 0.01330 | 1.08329 | 0.61072 | 0.43079 | 0.21466 | 0.01708 | 0.01708 | 0.29475 | 0.00726 | 0.00308 | 0.04720 | 0.07966 | 0.35317 |
| **Vav2** | 0.52658 | 0.01579 | 8.33638 | 2.63646 | 3.37244 | 5.32010 | 3.34786 | 2.76659 | 0.99959 | 0.08032 | 3.69240 | 2.24865 | 0.00060 | 3.99660 |
| **Vax2os** | 0.00349 | 0.01108 | 0.00974 | 0.02255 | 0.01115 | 0.01019 | 0.00363 | 0.01632 | 0.01241 | 0.00644 | 0.00417 | 1.56123 | 0.00680 | 0.01077 |
| **Vbp1** | 3.80227 | 6.32574 | 12.86232 | 9.84820 | 10.64605 | 17.68230 | 3.48380 | 8.27252 | 20.58109 | 0.16508 | 3.40837 | 7.42913 | 3.55154 | 11.82061 |
| **Vcl** | 4.66370 | 4.83593 | 0.95126 | 1.48708 | 3.27315 | 7.97115 | 6.53538 | 4.74888 | 3.49037 | 10.84604 | 3.25990 | 1.56070 | 2.27617 | 2.71888 |

| | | | | | | | | | | | | | | |
|---|---|---|---|---|---|---|---|---|---|---|---|---|---|---|
| **Vcp** | 18.34692 | 6.86782 | 34.01283 | 29.20090 | 25.48055 | 30.95063 | 8.13459 | 17.59027 | 1.60613 | 14.84071 | 20.86459 | 23.90100 | 10.64985 | 12.29766 |
| **Vcpip1** | 2.16845 | 0.00253 | 1.83246 | 0.00000 | 2.66987 | 1.31325 | 0.03595 | 0.28682 | 7.74072 | 0.00753 | 0.38169 | 0.95439 | 5.52662 | 0.00000 |
| **Vcpkmt** | 0.38776 | 0.00377 | 0.07892 | 1.29017 | 0.84402 | 0.46357 | 0.00389 | 0.30755 | 0.15462 | 0.01369 | 0.04794 | 0.13806 | 0.01789 | 0.01264 |
| **Vdac1** | 3.08999 | 4.67219 | 16.21162 | 17.45002 | 8.03642 | 6.34992 | 3.69440 | 11.84430 | 13.74161 | 13.88900 | 11.79415 | 11.84851 | 3.78129 | 18.93275 |
| **Vdac2** | 9.66960 | 6.50194 | 19.01356 | 23.22890 | 11.99144 | 18.85251 | 7.56717 | 22.12423 | 6.14682 | 16.63516 | 14.90665 | 18.95855 | 1.38559 | 6.01747 |
| **Vdac3** | 13.59361 | 10.39704 | 16.53623 | 16.26896 | 12.59115 | 21.77083 | 5.21197 | 23.18676 | 17.30624 | 9.34113 | 18.66730 | 12.82668 | 1.63671 | 12.05513 |
| **Vdr** | 0.09591 | 0.00432 | 0.46925 | 0.00000 | 0.02860 | 0.00148 | 0.00430 | 0.01244 | 0.00572 | 0.00652 | 0.00578 | 0.00934 | 0.00476 | 0.51256 |
| **Vegfa** | 0.10731 | 0.00422 | 0.00859 | 0.20178 | 0.00609 | 0.01001 | 0.00413 | 0.01244 | 0.01060 | 0.00658 | 0.00922 | 0.01270 | 0.02902 | 0.01868 |
| **Vegfb** | 0.00077 | 0.76993 | 0.06202 | 0.58340 | 0.18999 | 0.00000 | 0.01676 | 0.48036 | 0.00873 | 1.91827 | 0.00346 | 0.15410 | 0.12416 | 0.00000 |
| **Veph1** | 0.01603 | 0.00480 | 0.01756 | 0.07781 | 0.04552 | 0.25198 | 0.02376 | 0.05761 | 0.34599 | 0.05536 | 0.01148 | 0.08411 | 0.22865 | 0.09013 |
| **Vezf1** | 0.62304 | 0.00258 | 1.21175 | 0.67649 | 0.46965 | 2.43678 | 0.07195 | 0.27146 | 0.00000 | 0.00822 | 3.82872 | 0.57037 | 3.22625 | 3.95190 |
| **Vezt** | 0.68047 | 0.01797 | 1.30649 | 1.10520 | 1.35909 | 2.48650 | 0.00956 | 0.51452 | 0.01443 | 0.02013 | 2.70544 | 1.86344 | 0.25341 | 1.12714 |
| **Vgf** | 0.02937 | 0.00445 | 0.01371 | 0.01216 | 0.01678 | 0.01260 | 0.00441 | 0.01033 | 0.02583 | 0.00666 | 0.00841 | 0.00982 | 0.03819 | 0.00891 |
| **Vgll1** | 0.01180 | 0.01382 | 0.01511 | 0.00234 | 0.01087 | 0.01492 | 0.00405 | 0.01576 | 0.01220 | 0.00636 | 0.00451 | 0.39524 | 0.01441 | 0.05389 |
| **Vgll3** | 0.10220 | 0.01123 | 0.00274 | 0.01333 | 0.00589 | 0.90124 | 2.51975 | 2.43306 | 0.00992 | 0.02816 | 0.00252 | 0.01258 | 0.00608 | 0.00000 |
| **Vgll4** | 0.01612 | 0.00470 | 0.06840 | 0.05162 | 0.56975 | 0.75342 | 0.00417 | 0.00528 | 0.00833 | 0.00562 | 0.00728 | 0.00474 | 0.00781 | 0.00626 |
| **Vhl** | 8.28599 | 0.03190 | 11.36305 | 10.81354 | 12.52494 | 8.91706 | 2.54386 | 4.65656 | 12.83964 | 0.06251 | 7.48749 | 0.72006 | 13.73255 | 17.45290 |
| **Vil1** | 0.43420 | 0.00380 | 0.00000 | 0.02469 | 0.32742 | 0.00000 | 0.00435 | 0.01590 | 1.53984 | 0.00817 | 0.00484 | 0.00060 | 5.35882 | 3.80068 |
| **Vill** | 0.00000 | 4.84008 | 2.09087 | 3.30942 | 0.85376 | 3.28262 | 0.01296 | 4.48979 | 0.01098 | 0.03496 | 1.38507 | 0.00000 | 0.00000 | 0.04448 |
| **Vim** | 0.65932 | 0.02635 | 0.00067 | 0.00000 | 2.15160 | 0.00967 | 0.00377 | 0.09124 | 0.00861 | 0.00636 | 0.00321 | 1.41262 | 0.00000 | 0.02659 |
| **Vimp** | 17.92268 | 0.20783 | 20.72010 | 15.76824 | 17.49957 | 15.47994 | 4.85707 | 10.52197 | 8.85705 | 0.21093 | 5.40483 | 11.59525 | 17.77156 | 18.89017 |
| **Vipas39** | 0.14101 | 0.05973 | 4.54469 | 4.73641 | 2.02633 | 7.06818 | 1.34446 | 4.61523 | 4.50663 | 0.06950 | 4.79081 | 4.04094 | 0.94145 | 1.78524 |
| **Vkorc1** | 1.04143 | 0.01145 | 2.14354 | 9.63016 | 5.47637 | 3.62700 | 0.06579 | 3.25447 | 0.00000 | 9.35594 | 0.93367 | 7.65906 | 4.78633 | 0.00000 |
| **Vkorc1l1** | 5.56109 | 0.04596 | 3.47984 | 4.51609 | 5.75263 | 5.16300 | 4.12532 | 2.83690 | 13.40132 | 0.14803 | 2.99558 | 3.29322 | 8.64058 | 9.40669 |
| **Vldlr** | 0.07416 | 0.00480 | 0.01133 | 0.00620 | 0.00914 | 0.14642 | 0.00510 | 0.24029 | 0.00670 | 0.00686 | 0.00591 | 0.00481 | 0.63744 | 0.04859 |
| **Vma21** | 3.33918 | 0.04185 | 0.38834 | 0.81323 | 2.13591 | 1.98396 | 0.00868 | 2.76132 | 2.23214 | 0.06602 | 1.51786 | 1.95699 | 0.37804 | 0.79035 |
| **Vmac** | 0.78842 | 5.93740 | 0.20598 | 0.05327 | 0.00000 | 0.03009 | 0.00332 | 0.11794 | 0.01109 | 0.02451 | 0.01410 | 0.31615 | 0.07007 | 0.61770 |
| **Vmn1r132** | 0.00850 | 0.00444 | 0.01371 | 0.01165 | 0.05254 | 0.01344 | 0.00423 | 0.01010 | 0.01319 | 0.00662 | 0.00905 | 0.00973 | 0.04587 | 0.00538 |
| **Vmn1r148** | 0.00802 | 0.00441 | 0.01457 | 0.01324 | 0.01813 | 0.01325 | 0.00425 | 0.00985 | 0.01184 | 0.00695 | 0.00892 | 0.00972 | 0.10161 | 0.00406 |
| **Vmn1r32** | 0.00000 | 0.00443 | 0.00242 | 0.01604 | 0.00000 | 0.00847 | 0.00410 | 0.00939 | 0.32804 | 0.00708 | 0.00627 | 0.00913 | 0.00487 | 0.80723 |
| **Vmn1r51** | 0.01180 | 0.00443 | 0.01203 | 0.01307 | 0.00797 | 0.01179 | 0.00426 | 0.01035 | 0.05484 | 0.00677 | 0.00849 | 0.01000 | 0.03102 | 0.08651 |
| **Vmn1r58** | 0.17946 | 0.27893 | 0.20982 | 0.23707 | 0.27046 | 0.23644 | 0.12130 | 0.23539 | 1.77460 | 0.01396 | 0.06772 | 0.26186 | 0.27896 | 0.47324 |
| **Vmn1r65** | 0.02193 | 0.98753 | 0.16044 | 0.10799 | 0.17649 | 0.09285 | 0.00379 | 0.01264 | 0.33941 | 0.01375 | 0.03773 | 0.11658 | 0.86644 | 0.26616 |
| **Vmn2r.ps129** | 0.56769 | 0.04470 | 0.00023 | 0.34739 | 0.15337 | 1.39535 | 0.09632 | 0.09050 | 1.07312 | 0.80643 | 0.01268 | 0.24935 | 5.15179 | 0.45038 |
| **Vmn2r100** | 0.00939 | 0.00448 | 0.01382 | 0.01464 | 0.02464 | 0.01331 | 0.00430 | 0.01039 | 0.01208 | 0.00667 | 0.00858 | 0.00975 | 0.00485 | 0.04483 |
| **Vmn2r12** | 0.01080 | 0.00431 | 0.03654 | 0.02617 | 0.02445 | 0.01766 | 0.00418 | 0.01508 | 0.01101 | 0.00684 | 0.01064 | 0.03552 | 0.00830 | 0.03821 |
| **Vmn2r121** | 0.00964 | 0.00438 | 0.04782 | 0.03429 | 0.03229 | 0.01447 | 0.00420 | 0.01321 | 0.01999 | 0.00706 | 0.00979 | 0.05619 | 0.00764 | 0.04727 |
| **Vmn2r28** | 0.01078 | 0.00438 | 0.01478 | 0.01106 | 0.01566 | 0.01348 | 0.00432 | 0.00971 | 0.01201 | 0.00669 | 0.00871 | 0.01720 | 0.09267 | 0.00471 |
| **Vmn2r29** | 0.00815 | 0.00403 | 0.05283 | 0.02972 | 0.09808 | 0.01495 | 0.75017 | 0.01061 | 0.01770 | 0.02144 | 0.01245 | 0.02031 | 0.00553 | 0.00668 |
| **Vmn2r55** | 0.01786 | 0.00437 | 0.02866 | 0.01796 | 0.02000 | 0.01104 | 0.01013 | 0.00844 | 0.22041 | 0.00767 | 0.02154 | 0.01693 | 0.00971 | 0.02294 |
| **Vmn2r62** | 0.01156 | 0.00446 | 0.00970 | 0.08445 | 0.01172 | 0.01235 | 0.00432 | 0.01240 | 0.03730 | 0.00894 | 0.00817 | 0.00889 | 0.00751 | 0.00902 |
| **Vmn2r94** | 0.00411 | 0.00445 | 0.00319 | 0.01342 | 0.19915 | 0.01032 | 0.00450 | 0.00940 | 0.25311 | 0.00653 | 0.00758 | 0.00852 | 0.22629 | 0.00000 |
| **Vmn2r98** | 0.00831 | 0.00448 | 0.01195 | 0.01517 | 0.02949 | 0.01266 | 0.00430 | 0.01037 | 0.04302 | 0.00665 | 0.00811 | 0.00961 | 0.00446 | 0.08540 |
| **Vmp1** | 19.74277 | 0.21774 | 29.90251 | 20.47683 | 22.25462 | 18.52930 | 1.13993 | 4.24085 | 24.61088 | 11.80422 | 14.33650 | 10.04784 | 8.03928 | 28.41817 |
| **Vnn1** | 1.05294 | 0.04816 | 29.22032 | 9.07303 | 11.33976 | 11.08834 | 3.00101 | 2.95404 | 27.07329 | 0.12617 | 6.74071 | 3.17060 | 0.00000 | 15.86937 |
| **Vprbp** | 4.31073 | 0.01666 | 1.88821 | 0.00000 | 0.97278 | 0.10966 | 0.00231 | 0.07936 | 6.11147 | 0.00000 | 3.63088 | 0.63595 | 5.28013 | 8.88793 |
| **Vpreb3** | 1.37466 | 0.00377 | 1.33339 | 0.13503 | 5.85648 | 0.44414 | 0.00366 | 0.24993 | 1.23985 | 0.01312 | 0.00046 | 0.00071 | 0.41012 | 1.85230 |
| **Vps11** | 2.80775 | 0.05942 | 12.23252 | 6.46577 | 7.16545 | 6.07900 | 1.53541 | 2.29817 | 1.49858 | 0.01541 | 6.66736 | 3.88941 | 1.89880 | 7.37562 |
| **Vps13a** | 0.05633 | 0.00389 | 0.39774 | 1.34918 | 0.66259 | 0.06017 | 0.00379 | 0.00084 | 0.36566 | 0.00504 | 0.31982 | 0.27593 | 0.14547 | 4.34194 |
| **Vps13b** | 2.89336 | 0.00978 | 2.51268 | 1.26509 | 1.24389 | 1.99486 | 0.00254 | 1.05621 | 1.56861 | 0.00666 | 0.28173 | 2.88338 | 0.13666 | 1.24957 |
| **Vps13c** | 0.56416 | 0.01121 | 1.70772 | 1.44961 | 0.00000 | 1.10087 | 0.00416 | 0.01714 | 0.00200 | 0.00181 | 2.10253 | 1.02745 | 0.48198 | 2.44677 |
| **Vps13d** | 2.18196 | 7.67286 | 6.53721 | 4.58124 | 4.77530 | 5.74531 | 0.63133 | 0.12030 | 5.01221 | 0.03498 | 1.48252 | 0.73854 | 1.71209 | 8.54777 |
| **Vps16** | 6.73210 | 6.54443 | 8.36101 | 9.72294 | 8.12584 | 7.42575 | 3.30726 | 0.60098 | 6.40707 | 0.09333 | 8.79452 | 5.73495 | 3.49823 | 9.21043 |
| **Vps18** | 0.24660 | 0.02583 | 3.98621 | 3.12454 | 2.68062 | 0.00000 | 2.55151 | 1.17461 | 0.06978 | 0.04050 | 0.84884 | 2.79171 | 2.07348 | 0.42107 |
| **Vps25** | 4.80197 | 0.31463 | 8.29717 | 8.87375 | 3.10989 | 14.85615 | 5.03688 | 18.06350 | 0.47760 | 4.31297 | 9.74020 | 15.55412 | 0.81224 | 3.04071 |

| | | | | | | | | | | | | | | |
|---|---|---|---|---|---|---|---|---|---|---|---|---|---|---|
| **Vps26a** | 9.84593 | 0.06026 | 16.20046 | 10.64679 | 8.03243 | 12.46373 | 0.06233 | 3.26033 | 1.44390 | 0.02256 | 8.49934 | 6.18780 | 3.92876 | 11.13072 |
| **Vps26b** | 1.33248 | 0.01673 | 4.36305 | 2.05620 | 3.26682 | 2.36286 | 1.76697 | 1.94136 | 3.18494 | 0.00865 | 2.45168 | 2.27722 | 2.70527 | 1.90735 |
| **Vps28** | 4.33758 | 7.64844 | 4.25149 | 4.51007 | 3.67634 | 12.57862 | 1.36712 | 10.48955 | 0.14652 | 11.21692 | 5.03824 | 4.95108 | 0.50536 | 1.74636 |
| **Vps29** | 1.81292 | 2.52752 | 3.55506 | 10.01463 | 0.93046 | 16.08108 | 4.96310 | 11.01720 | 1.60844 | 6.34252 | 7.99462 | 10.74360 | 2.05402 | 0.82087 |
| **Vps33a** | 1.01831 | 0.08948 | 8.63120 | 7.93811 | 6.69500 | 10.45008 | 3.88738 | 0.55853 | 2.53620 | 0.09522 | 7.49737 | 6.87608 | 4.06329 | 5.09434 |
| **Vps33b** | 1.05401 | 0.01959 | 11.50880 | 1.50550 | 1.86659 | 4.10220 | 2.62300 | 0.60231 | 2.84441 | 0.10211 | 3.67597 | 0.45134 | 0.49231 | 2.04156 |
| **Vps35** | 16.58934 | 2.21295 | 13.72370 | 9.46898 | 11.08307 | 17.15446 | 0.53291 | 4.98049 | 18.31256 | 0.03709 | 14.05056 | 11.70142 | 4.18353 | 10.54104 |
| **Vps36** | 1.56066 | 0.04586 | 1.51200 | 4.23921 | 1.25337 | 1.71922 | 1.16103 | 4.35444 | 0.00462 | 2.57292 | 0.99882 | 3.03722 | 0.00000 | 0.67426 |
| **Vps37a** | 7.11747 | 0.03267 | 3.14984 | 2.48364 | 3.47260 | 3.56848 | 1.04452 | 0.16850 | 1.99852 | 0.04271 | 1.33547 | 3.30490 | 2.58044 | 5.18796 |
| **Vps37b** | 0.72692 | 0.07250 | 11.93513 | 7.16231 | 14.90549 | 0.32664 | 4.07863 | 6.62171 | 0.00000 | 0.10921 | 4.72238 | 2.07454 | 1.83929 | 0.01432 |
| **Vps37c** | 0.98812 | 0.01864 | 0.97120 | 0.30892 | 0.55452 | 1.65942 | 0.10957 | 1.69671 | 0.00000 | 0.00000 | 3.33136 | 1.41682 | 0.00017 | 1.18211 |
| **Vps37d** | 0.04621 | 0.00434 | 0.05383 | 0.47911 | 1.58969 | 0.00881 | 0.00401 | 0.00568 | 0.01003 | 0.00660 | 0.00949 | 0.07894 | 0.03207 | 0.00022 |
| **Vps39** | 1.61619 | 0.00232 | 1.48026 | 1.59506 | 0.60950 | 3.77247 | 0.00281 | 0.00000 | 1.89747 | 0.00300 | 3.00063 | 0.99322 | 0.36604 | 1.90294 |
| **Vps41** | 9.86051 | 0.06915 | 9.36784 | 5.23804 | 5.47124 | 8.17394 | 1.91368 | 0.07829 | 6.87032 | 7.40613 | 4.99857 | 7.27141 | 0.00000 | 5.02206 |
| **Vps45** | 0.17871 | 0.02611 | 0.54288 | 0.22862 | 0.16598 | 3.71197 | 0.01551 | 1.18272 | 0.00799 | 0.01289 | 2.10010 | 2.96279 | 0.55037 | 0.00000 |
| **Vps4a** | 4.87942 | 0.05799 | 9.30129 | 2.86260 | 6.15314 | 8.52566 | 0.15700 | 5.62533 | 1.23563 | 5.54844 | 0.56542 | 7.27976 | 7.49246 | 7.06083 |
| **Vps4b** | 8.48111 | 0.05761 | 5.40166 | 3.30605 | 8.15070 | 6.40783 | 2.18374 | 4.14771 | 5.10019 | 0.07150 | 3.13237 | 4.72146 | 5.08675 | 7.35535 |
| **Vps50** | 3.91054 | 0.02917 | 4.23521 | 0.00000 | 3.29309 | 4.24035 | 0.04870 | 0.09883 | 7.61793 | 0.02314 | 3.75233 | 2.95164 | 0.94044 | 3.26897 |
| **Vps51** | 1.97496 | 0.27470 | 3.51843 | 9.79736 | 1.58244 | 1.52626 | 0.00826 | 5.11760 | 0.00614 | 0.03110 | 2.98665 | 3.12821 | 0.17476 | 0.41388 |
| **Vps52** | 1.84536 | 0.09999 | 8.00692 | 7.68254 | 5.37221 | 5.96815 | 0.00584 | 3.52094 | 0.09516 | 1.37304 | 5.47814 | 6.04644 | 0.23341 | 1.00763 |
| **Vps53** | 1.06033 | 0.07889 | 4.41781 | 1.77168 | 2.38368 | 4.25110 | 1.28051 | 3.68488 | 1.03184 | 0.00000 | 4.27206 | 6.74212 | 0.19323 | 4.17160 |
| **Vps54** | 1.98933 | 0.02960 | 6.04951 | 3.93905 | 4.39363 | 9.65111 | 2.37492 | 3.68536 | 6.54548 | 0.06059 | 6.24187 | 4.57736 | 1.48087 | 0.14818 |
| **Vps72** | 0.00000 | 3.05861 | 6.25420 | 8.34751 | 10.81550 | 0.00000 | 0.24921 | 3.56999 | 0.24034 | 0.03269 | 0.50814 | 4.45930 | 0.16435 | 0.00000 |
| **Vps8** | 1.31000 | 0.00983 | 9.74413 | 2.61817 | 7.76996 | 0.76851 | 0.00788 | 0.12692 | 3.22199 | 0.00660 | 1.93403 | 1.64217 | 0.19905 | 0.61631 |
| **Vps9d1** | 0.37474 | 0.02172 | 1.79751 | 0.69809 | 0.78388 | 0.08650 | 2.47372 | 1.17014 | 0.00000 | 0.03304 | 0.06427 | 0.21306 | 1.90941 | 0.00000 |
| **Vrk1** | 10.91360 | 0.20313 | 12.62185 | 8.19551 | 14.50073 | 6.33311 | 2.48153 | 5.23675 | 8.98624 | 0.13506 | 12.75843 | 10.60337 | 5.93029 | 17.21232 |
| **Vrk2** | 0.51482 | 0.00346 | 2.25087 | 1.13559 | 0.73421 | 0.92654 | 1.31987 | 0.39064 | 0.00434 | 0.01835 | 0.18392 | 0.73493 | 0.00000 | 1.27288 |
| **Vrk3** | 4.22176 | 0.01064 | 9.67947 | 4.44184 | 2.81244 | 0.00000 | 0.16589 | 4.25885 | 1.00926 | 0.01841 | 2.73364 | 0.95621 | 1.03099 | 2.64003 |
| **Vrtn** | 0.00925 | 0.00436 | 0.01018 | 0.01241 | 0.16481 | 0.01270 | 0.00414 | 0.00928 | 0.01375 | 0.00612 | 0.00902 | 0.00856 | 0.30650 | 0.02639 |
| **Vsig10** | 0.36327 | 0.03440 | 0.07700 | 0.07885 | 1.30740 | 0.55056 | 0.03088 | 0.85633 | 0.13238 | 0.04190 | 0.00123 | 0.00829 | 0.03307 | 0.37128 |
| **Vsig10l** | 0.84005 | 0.00390 | 1.25235 | 0.87737 | 0.20743 | 1.18910 | 0.00385 | 0.00000 | 0.00058 | 0.00402 | 0.00010 | 0.04570 | 0.68043 | 0.95766 |
| **Vsir** | 0.03415 | 0.00438 | 0.00000 | 0.00000 | 0.51020 | 0.41041 | 0.00485 | 0.00538 | 0.00975 | 0.00680 | 0.00372 | 0.07511 | 0.00266 | 0.29000 |
| **Vstm5** | 0.00000 | 0.00397 | 0.40808 | 0.00000 | 0.00058 | 0.00000 | 0.00369 | 0.39848 | 0.01137 | 0.00690 | 0.00000 | 2.28029 | 0.01672 | 0.13909 |
| **Vsx2** | 0.00000 | 0.00422 | 0.59344 | 0.44723 | 0.02649 | 0.02393 | 0.00399 | 0.00166 | 0.00655 | 0.00444 | 0.00904 | 0.01351 | 1.31160 | 0.00000 |
| **Vta1** | 4.35401 | 0.13646 | 4.67263 | 2.83177 | 4.87014 | 7.74971 | 3.22849 | 4.71698 | 0.62746 | 0.09432 | 6.05011 | 6.78998 | 2.75117 | 0.54928 |
| **Vtcn1** | 0.29940 | 0.02564 | 0.95893 | 1.79181 | 0.16089 | 14.39456 | 0.00313 | 1.58670 | 0.00211 | 0.05418 | 4.98146 | 0.00000 | 0.00000 | 0.20618 |
| **Vti1a** | 0.00392 | 0.03034 | 0.10406 | 1.00452 | 0.33239 | 0.51442 | 0.43891 | 0.65003 | 0.00917 | 0.01580 | 0.02079 | 0.44096 | 0.01398 | 0.45668 |
| **Vti1b** | 8.43046 | 0.54078 | 4.69600 | 8.22419 | 4.61798 | 4.97231 | 1.61834 | 4.96766 | 1.62466 | 0.12094 | 1.33793 | 4.67799 | 4.17235 | 0.62374 |
| **Vwa2** | 0.01709 | 0.00434 | 0.01380 | 0.01228 | 0.00919 | 0.01240 | 0.00431 | 0.00959 | 0.00922 | 0.00637 | 0.00880 | 0.00927 | 0.30704 | 0.07944 |
| **Vwa3a** | 0.00728 | 0.00454 | 0.15710 | 0.00000 | 0.25568 | 0.00000 | 0.00402 | 0.01068 | 1.26780 | 0.00746 | 0.00397 | 0.00737 | 0.01783 | 2.63853 |
| **Vwa5b1** | 0.01135 | 0.00495 | 3.13071 | 0.10510 | 0.50263 | 0.06130 | 0.00371 | 0.00006 | 0.01774 | 0.00000 | 0.01928 | 0.00623 | 0.00000 | 0.35519 |
| **Vwa5b2** | 0.37281 | 0.00403 | 0.14746 | 0.23998 | 0.25668 | 0.00215 | 0.00418 | 0.00837 | 0.00329 | 0.00623 | 0.00611 | 0.00550 | 0.98417 | 0.01266 |
| **Vwa7** | 0.05593 | 0.01547 | 0.09156 | 0.00000 | 0.00708 | 0.02252 | 0.01603 | 0.28632 | 0.10290 | 0.01908 | 0.00374 | 0.15021 | 0.29368 | 0.02592 |
| **Vwa8** | 0.30069 | 9.00913 | 0.00000 | 1.06082 | 0.75366 | 1.18659 | 0.00136 | 0.23341 | 0.00206 | 0.02899 | 0.00000 | 4.81068 | 0.12885 | 1.31893 |
| **Vwa9** | 4.64716 | 0.01986 | 4.60769 | 1.47852 | 4.51522 | 4.65377 | 2.71796 | 1.31313 | 0.72727 | 0.07523 | 5.31417 | 4.84057 | 2.98361 | 0.83704 |
| **Vwc2** | 0.00853 | 0.00443 | 0.01453 | 0.01286 | 0.01259 | 0.01314 | 0.00428 | 0.01007 | 0.02024 | 0.00685 | 0.00881 | 0.00994 | 0.06423 | 0.00349 |
| **Vwce** | 0.02375 | 0.00441 | 0.08211 | 0.17047 | 0.01934 | 0.00000 | 0.00477 | 0.08979 | 0.01413 | 0.01671 | 0.00706 | 0.00599 | 0.02397 | 0.00443 |
| **Vwf** | 0.00930 | 0.00444 | 0.01350 | 0.01172 | 0.01215 | 0.02701 | 0.00426 | 0.00904 | 0.01857 | 0.00689 | 0.00778 | 0.01019 | 0.01491 | 0.02876 |
| **Wac** | 9.16988 | 0.10818 | 13.31827 | 4.99438 | 13.31583 | 4.98293 | 0.42014 | 4.09359 | 27.01759 | 0.05010 | 9.46087 | 1.04016 | 11.38560 | 12.14236 |
| **Wapl** | 12.55799 | 0.92361 | 9.83868 | 2.22734 | 8.89496 | 8.59969 | 4.74752 | 2.01657 | 23.71825 | 8.19748 | 8.91274 | 8.16782 | 12.93937 | 11.33526 |
| **Wars** | 2.29834 | 1.80454 | 11.13782 | 12.44919 | 7.29478 | 9.87746 | 4.13126 | 6.01956 | 0.16115 | 0.19557 | 9.71333 | 14.43883 | 0.08601 | 8.75933 |
| **Wars2** | 9.72803 | 0.02252 | 6.46921 | 4.10073 | 9.59651 | 2.00729 | 0.86259 | 1.57912 | 3.82629 | 0.02490 | 4.37786 | 2.39363 | 1.80985 | 3.81395 |
| **Was** | 0.08578 | 0.04651 | 0.14768 | 0.13280 | 0.94865 | 0.17337 | 0.04722 | 0.07749 | 0.37320 | 0.00681 | 0.02286 | 0.04553 | 0.29283 | 0.71344 |
| **Wasf1** | 0.02473 | 0.10434 | 0.00000 | 1.13913 | 0.13171 | 0.67219 | 0.00393 | 0.25127 | 0.19690 | 0.00343 | 0.81425 | 0.17955 | 1.13977 | 0.00000 |
| **Wasf2** | 0.00000 | 0.76178 | 2.40101 | 4.99789 | 3.68673 | 3.37842 | 0.52637 | 1.59356 | 0.02698 | 0.04420 | 0.65534 | 0.42017 | 1.81201 | 1.89878 |

| | | | | | | | | | | | | | | |
|---|---|---|---|---|---|---|---|---|---|---|---|---|---|---|
| **Wash1** | 1.62351 | 0.77963 | 0.14059 | 1.12057 | 3.24887 | 1.99646 | 0.02184 | 1.95268 | 0.13003 | 0.01035 | 1.27706 | 4.01843 | 1.07948 | 0.24089 |
| **Wasl** | 10.94490 | 0.04070 | 4.87629 | 0.69617 | 3.79484 | 8.88963 | 4.35831 | 4.31827 | 13.24359 | 0.19354 | 4.96032 | 1.86514 | 7.80717 | 3.63517 |
| **Wbp1** | 0.01738 | 0.10072 | 1.01426 | 3.33166 | 1.39370 | 4.59513 | 0.00903 | 2.27954 | 0.00000 | 0.04798 | 1.97215 | 4.59031 | 0.00783 | 0.33394 |
| **Wbp11** | 9.62322 | 5.81127 | 11.42525 | 5.89985 | 14.34777 | 4.42730 | 2.90538 | 9.84703 | 0.79389 | 9.42271 | 9.52422 | 9.48292 | 11.86248 | 2.47385 |
| **Wbp1l** | 3.43741 | 4.26807 | 7.53870 | 2.48978 | 5.86306 | 8.08193 | 5.52780 | 5.49648 | 0.24030 | 0.13958 | 3.29929 | 10.25463 | 3.74640 | 8.96462 |
| **Wbp2** | 3.14648 | 0.06761 | 7.80477 | 8.94358 | 4.82548 | 6.70377 | 4.27795 | 10.52207 | 0.38082 | 0.35316 | 1.74731 | 5.02426 | 2.50322 | 1.77698 |
| **Wbp2nl** | 0.00000 | 0.00441 | 0.73408 | 0.00000 | 0.66193 | 0.61806 | 0.00405 | 0.00211 | 0.02900 | 0.01247 | 0.00388 | 0.00466 | 0.25880 | 0.19606 |
| **Wbp4** | 7.33630 | 0.06897 | 3.09158 | 2.62403 | 6.19883 | 2.83231 | 4.06452 | 1.81298 | 4.05496 | 0.07695 | 3.19387 | 4.40936 | 4.36571 | 7.16254 |
| **Wbp5** | 0.01524 | 0.03265 | 0.00000 | 0.11453 | 1.25526 | 2.26658 | 0.00309 | 0.29533 | 0.05927 | 9.99076 | 0.00000 | 0.00000 | 0.00612 | 0.00000 |
| **Wbscr16** | 2.93112 | 0.00914 | 5.19118 | 5.44552 | 4.05237 | 2.53008 | 0.00266 | 0.72995 | 0.00448 | 0.00951 | 5.47103 | 2.00881 | 1.60351 | 1.37964 |
| **Wbscr17** | 0.01512 | 0.00448 | 0.00944 | 0.01301 | 0.19889 | 0.01182 | 0.00437 | 0.01046 | 0.00746 | 0.00591 | 0.00865 | 0.00834 | 0.13612 | 0.15038 |
| **Wbscr22** | 7.45985 | 5.08758 | 17.23201 | 10.08741 | 17.48573 | 14.91758 | 3.16114 | 7.06173 | 0.03870 | 0.89435 | 10.81032 | 9.48172 | 1.37330 | 9.72420 |
| **Wbscr25** | 0.00532 | 0.00440 | 0.11624 | 0.00652 | 0.00259 | 0.00750 | 0.00431 | 0.01093 | 0.00839 | 0.00862 | 0.00692 | 0.00901 | 0.00358 | 0.38181 |
| **Wbscr27** | 0.74646 | 0.01613 | 2.44525 | 5.72627 | 0.83894 | 0.30948 | 1.09922 | 4.66729 | 0.81060 | 0.42084 | 1.41486 | 2.17273 | 1.20470 | 0.55375 |
| **Wdfy1** | 2.09894 | 0.05791 | 10.43121 | 7.73747 | 7.28432 | 12.32538 | 2.17402 | 2.88520 | 13.36324 | 0.14761 | 6.67998 | 4.95382 | 0.51488 | 7.47859 |
| **Wdfy2** | 0.57253 | 0.00368 | 0.69679 | 0.93918 | 0.00000 | 0.14480 | 0.00499 | 0.39140 | 0.01296 | 0.00240 | 0.86823 | 0.72990 | 0.53238 | 0.37747 |
| **Wdfy3** | 1.29346 | 0.03691 | 2.11705 | 0.02572 | 4.86863 | 0.13466 | 0.00280 | 1.60411 | 3.02704 | 0.00308 | 0.15606 | 1.36528 | 1.54523 | 3.61240 |
| **Wdfy4** | 0.00669 | 0.00434 | 0.00000 | 0.00904 | 0.38287 | 0.22388 | 0.00413 | 0.00386 | 0.01670 | 0.01356 | 0.63142 | 0.00839 | 0.00550 | 0.02888 |
| **Wdhd1** | 2.11856 | 0.05036 | 3.42603 | 2.03420 | 4.67771 | 5.69833 | 0.00547 | 0.00000 | 4.19669 | 9.60158 | 2.46282 | 3.60583 | 0.87850 | 9.50361 |
| **Wdpcp** | 0.01323 | 0.00415 | 0.01430 | 0.01724 | 0.01266 | 0.01502 | 0.00407 | 0.02908 | 0.01102 | 0.00738 | 0.00815 | 0.01056 | 0.05810 | 0.01071 |
| **Wdr1** | 0.08921 | 7.28073 | 8.01235 | 9.41109 | 5.32518 | 15.55767 | 4.59464 | 18.61571 | 0.52451 | 0.34262 | 12.07072 | 6.80664 | 1.69689 | 2.48626 |
| **Wdr11** | 1.70305 | 0.02506 | 3.95871 | 1.96174 | 3.26722 | 4.32066 | 2.76657 | 0.46777 | 1.33669 | 0.07808 | 6.56846 | 5.50932 | 0.00871 | 3.41008 |
| **Wdr12** | 8.51722 | 0.05730 | 17.42581 | 16.20168 | 17.03970 | 8.89208 | 3.35207 | 3.50602 | 16.57160 | 8.70627 | 14.80161 | 7.88795 | 2.65576 | 16.88779 |
| **Wdr13** | 2.63916 | 0.03185 | 2.24395 | 0.00000 | 1.30520 | 0.06649 | 0.28735 | 3.01984 | 1.06270 | 0.00818 | 0.02027 | 0.36184 | 0.00000 | 2.96429 |
| **Wdr17** | 0.00863 | 0.00450 | 0.00907 | 0.01310 | 0.22442 | 0.01205 | 0.00431 | 0.01069 | 0.00752 | 0.00607 | 0.00895 | 0.00849 | 0.05271 | 0.13600 |
| **Wdr18** | 8.46508 | 3.31273 | 23.31697 | 21.50302 | 23.15600 | 7.99076 | 2.02084 | 10.35798 | 0.23985 | 0.14368 | 6.19960 | 11.08741 | 2.21304 | 5.96125 |
| **Wdr19** | 0.07567 | 0.01181 | 0.03170 | 0.06548 | 0.00530 | 0.02932 | 0.00415 | 0.00738 | 0.11607 | 0.00617 | 0.00607 | 0.01098 | 0.18652 | 0.00000 |
| **Wdr20** | 0.11938 | 0.01749 | 2.19375 | 0.16304 | 2.15270 | 1.49398 | 2.61698 | 0.34411 | 4.76485 | 0.20145 | 2.14927 | 1.11718 | 4.10257 | 7.71685 |
| **Wdr20rt** | 0.02896 | 0.00365 | 0.60333 | 0.00000 | 0.26082 | 0.20191 | 0.42677 | 0.02200 | 0.79900 | 0.01554 | 0.24148 | 0.55747 | 0.08005 | 1.24620 |
| **Wdr24** | 2.17343 | 0.02246 | 4.52482 | 0.73662 | 6.46386 | 2.53248 | 0.00284 | 1.29127 | 0.00000 | 0.00514 | 1.18551 | 1.18108 | 0.00000 | 3.01282 |
| **Wdr25** | 0.01260 | 0.01836 | 0.00000 | 0.44502 | 0.06086 | 0.18150 | 0.73983 | 0.00351 | 0.01103 | 0.01042 | 0.05825 | 0.82254 | 0.34437 | 0.60295 |
| **Wdr26** | 11.57271 | 0.10872 | 12.31159 | 4.08360 | 13.20638 | 15.25855 | 6.01510 | 6.61550 | 19.18628 | 2.12820 | 8.68752 | 5.24618 | 13.58052 | 20.80388 |
| **Wdr27** | 0.19144 | 0.00438 | 0.00990 | 0.01488 | 0.22765 | 0.01010 | 0.00412 | 0.00906 | 0.24055 | 0.01001 | 0.00577 | 0.00697 | 0.20119 | 0.11173 |
| **Wdr3** | 14.37451 | 0.03848 | 15.82222 | 6.10523 | 13.83568 | 10.05208 | 3.57167 | 0.86860 | 16.03068 | 0.09854 | 11.86993 | 7.46010 | 7.26819 | 13.19126 |
| **Wdr31** | 0.03515 | 0.00409 | 0.10003 | 2.10892 | 0.13622 | 0.00000 | 0.00405 | 0.02501 | 0.00583 | 0.00600 | 0.00717 | 0.00722 | 0.61754 | 0.04560 |
| **Wdr33** | 10.17806 | 0.12023 | 12.16813 | 6.67814 | 9.99819 | 9.53067 | 4.26626 | 3.25715 | 4.88956 | 0.23959 | 6.19959 | 3.64724 | 8.29865 | 8.80420 |
| **Wdr34** | 4.73713 | 0.01078 | 8.40645 | 0.55240 | 3.64106 | 0.87136 | 0.03336 | 0.36036 | 0.68666 | 0.01391 | 1.23488 | 0.37756 | 4.60066 | 2.85937 |
| **Wdr35** | 0.06066 | 0.01075 | 0.00000 | 0.00000 | 0.55659 | 0.76092 | 0.00403 | 0.00768 | 0.00887 | 0.00799 | 0.00345 | 0.99743 | 0.49301 | 0.01519 |
| **Wdr36** | 17.09053 | 0.03178 | 15.11133 | 7.99934 | 19.99459 | 4.41508 | 2.50412 | 2.65947 | 15.15218 | 0.00000 | 6.94183 | 5.34171 | 12.39868 | 20.48738 |
| **Wdr37** | 3.30225 | 0.01642 | 1.13768 | 0.00000 | 1.71596 | 1.12638 | 0.34146 | 1.24672 | 1.35735 | 0.00336 | 2.64309 | 1.73138 | 6.90136 | 3.71915 |
| **Wdr4** | 7.24686 | 9.91727 | 10.06683 | 10.61393 | 12.77133 | 6.05862 | 2.24471 | 4.65733 | 1.32047 | 0.14650 | 1.07901 | 4.80763 | 2.12437 | 3.44840 |
| **Wdr41** | 0.05664 | 0.00430 | 0.00776 | 0.07282 | 0.01324 | 0.06534 | 0.00402 | 0.00583 | 0.02768 | 0.00812 | 0.00596 | 0.01347 | 0.00507 | 0.07819 |
| **Wdr43** | 32.59363 | 0.61978 | 25.52885 | 9.93074 | 37.63501 | 10.00754 | 5.48263 | 2.72777 | 25.13674 | 0.15206 | 8.16941 | 7.98414 | 19.36920 | 35.50524 |
| **Wdr44** | 0.02672 | 0.00364 | 0.49943 | 0.00000 | 0.30822 | 0.06553 | 0.00341 | 0.02462 | 4.88518 | 0.00530 | 0.08530 | 0.09242 | 3.60103 | 0.00000 |
| **Wdr45** | 23.59638 | 8.96504 | 33.46818 | 15.05572 | 25.44467 | 25.33629 | 2.91485 | 6.58803 | 6.87295 | 9.30827 | 12.76434 | 7.81667 | 7.20058 | 27.68255 |
| **Wdr45b** | 18.91818 | 7.11533 | 28.73706 | 17.60824 | 33.08349 | 3.34687 | 5.75052 | 11.62774 | 7.29838 | 6.07027 | 7.88357 | 5.85738 | 13.77997 | 10.89344 |
| **Wdr46** | 11.32577 | 4.05349 | 18.96635 | 12.49380 | 21.66066 | 3.50472 | 0.01451 | 4.75550 | 2.96331 | 0.28876 | 5.81345 | 7.74511 | 7.03758 | 3.71431 |
| **Wdr47** | 2.15349 | 0.01349 | 2.60247 | 0.30134 | 2.86610 | 1.41421 | 0.00966 | 0.57708 | 4.30827 | 0.00416 | 0.00000 | 0.52763 | 3.98371 | 6.21265 |
| **Wdr48** | 0.32996 | 0.00994 | 4.49517 | 1.98173 | 5.30437 | 2.00002 | 0.74537 | 1.78697 | 2.58146 | 0.05701 | 0.38140 | 2.02724 | 1.45971 | 3.19562 |
| **Wdr5** | 14.72228 | 0.09650 | 23.36513 | 16.64388 | 21.41572 | 21.60263 | 4.60336 | 5.13388 | 14.64189 | 5.41436 | 11.88714 | 8.39765 | 11.72187 | 17.39778 |
| **Wdr53** | 5.51787 | 0.06358 | 5.72900 | 1.98607 | 7.29595 | 5.17052 | 0.02260 | 4.43885 | 4.86657 | 0.03933 | 4.71168 | 2.01763 | 2.57915 | 7.17957 |
| **Wdr54** | 0.00870 | 0.00408 | 0.27417 | 0.15731 | 0.19152 | 0.48588 | 0.00416 | 0.00000 | 0.01468 | 0.00306 | 1.22481 | 0.01685 | 0.01001 | 0.03767 |
| **Wdr55** | 1.97293 | 6.03362 | 7.36835 | 4.19625 | 9.50987 | 3.34949 | 2.09723 | 3.22795 | 0.53439 | 0.09591 | 2.60615 | 2.71464 | 4.72959 | 9.94611 |
| **Wdr59** | 1.84619 | 0.00195 | 2.61839 | 0.00000 | 4.83655 | 2.53478 | 0.01316 | 0.04401 | 3.53251 | 0.00097 | 3.86889 | 1.16466 | 0.01252 | 2.23547 |
| **Wdr5b** | 0.01374 | 0.00421 | 0.00000 | 0.45886 | 0.00145 | 0.10660 | 0.00409 | 0.00265 | 0.01523 | 0.00386 | 0.46499 | 0.00463 | 0.01563 | 0.51013 |

| | | | | | | | | | | | | | | |
|---|---|---|---|---|---|---|---|---|---|---|---|---|---|---|
| **Wdr6** | 0.00000 | 0.07770 | 7.88084 | 3.56322 | 1.45389 | 1.90590 | 4.16726 | 2.20320 | 0.20563 | 0.16863 | 2.09426 | 4.12377 | 0.07243 | 2.30628 |
| **Wdr60** | 0.30211 | 0.00406 | 0.05340 | 0.06930 | 0.00000 | 0.51213 | 0.00467 | 0.14768 | 0.01241 | 0.00515 | 0.02162 | 0.00000 | 1.57386 | 0.04566 |
| **Wdr61** | 14.52913 | 0.13900 | 12.79037 | 5.48290 | 13.61491 | 16.33542 | 3.84835 | 4.51335 | 12.64587 | 14.12195 | 9.81680 | 11.60448 | 0.89965 | 9.98870 |
| **Wdr62** | 0.85543 | 4.71339 | 4.45481 | 1.76946 | 9.15020 | 1.78269 | 0.00705 | 0.00000 | 0.04768 | 0.03106 | 1.35609 | 2.10658 | 1.88561 | 4.93667 |
| **Wdr63** | 0.04058 | 0.00441 | 0.01444 | 0.01160 | 0.01066 | 0.01270 | 0.00430 | 0.01004 | 0.01056 | 0.00650 | 0.00861 | 0.00959 | 0.12956 | 0.01341 |
| **Wdr7** | 2.01407 | 0.00362 | 0.88610 | 0.00000 | 1.62093 | 1.72563 | 0.00362 | 0.19335 | 3.19526 | 0.01828 | 1.73929 | 0.59238 | 0.00000 | 4.09888 |
| **Wdr70** | 0.12352 | 0.00938 | 6.73133 | 1.30237 | 4.35739 | 1.95994 | 1.02677 | 0.00000 | 0.76057 | 0.00000 | 1.82487 | 2.03900 | 0.85947 | 4.01365 |
| **Wdr72** | 1.01725 | 0.03065 | 1.02525 | 0.35302 | 1.52768 | 0.00000 | 1.98781 | 0.98958 | 5.26523 | 0.01030 | 1.39459 | 0.00000 | 0.00000 | 0.65365 |
| **Wdr73** | 10.73633 | 10.41512 | 16.66988 | 5.92492 | 16.33855 | 9.65236 | 2.26183 | 6.42741 | 1.35782 | 0.09247 | 5.60823 | 5.63190 | 3.46590 | 11.05578 |
| **Wdr74** | 17.17521 | 8.72403 | 30.27932 | 20.39980 | 22.82095 | 12.39675 | 5.07254 | 12.20814 | 7.34833 | 16.61834 | 12.31176 | 11.55717 | 10.08570 | 14.22808 |
| **Wdr75** | 18.01104 | 0.07166 | 17.70809 | 3.59578 | 12.24489 | 7.63554 | 3.09524 | 0.35043 | 21.78051 | 0.03023 | 10.34874 | 5.30287 | 9.30809 | 17.70034 |
| **Wdr76** | 0.00000 | 0.00409 | 0.00000 | 1.44273 | 0.15021 | 1.20327 | 0.21821 | 0.00924 | 0.54943 | 0.02889 | 1.37030 | 0.59250 | 0.00577 | 1.00125 |
| **Wdr77** | 20.51455 | 7.22296 | 27.01682 | 12.27847 | 23.94903 | 12.90837 | 5.66730 | 11.63486 | 14.28870 | 6.10222 | 13.17171 | 14.64057 | 11.47001 | 21.80046 |
| **Wdr78** | 0.03666 | 0.00463 | 0.03887 | 0.00000 | 0.00735 | 0.20154 | 0.00428 | 0.00580 | 0.01927 | 0.00586 | 0.00671 | 0.01043 | 0.08271 | 0.06188 |
| **Wdr8** | 0.00000 | 0.02417 | 2.73643 | 2.75415 | 0.00000 | 1.64832 | 0.00249 | 0.08486 | 0.00038 | 0.02191 | 1.95093 | 2.86817 | 0.00423 | 0.36629 |
| **Wdr81** | 3.43740 | 0.02562 | 10.20245 | 7.83575 | 8.76142 | 1.53996 | 0.00923 | 0.88088 | 1.03904 | 0.00080 | 4.60469 | 0.04923 | 7.46351 | 2.79621 |
| **Wdr82** | 6.02132 | 0.09544 | 8.57512 | 6.54369 | 8.48252 | 12.37648 | 1.10333 | 8.39159 | 10.11268 | 0.08571 | 10.28017 | 8.76207 | 4.47623 | 6.09672 |
| **Wdr83** | 0.19609 | 0.03872 | 6.63107 | 5.25154 | 1.48817 | 2.05608 | 2.85737 | 1.91906 | 0.35215 | 1.22221 | 4.18926 | 2.47973 | 0.00000 | 0.44935 |
| **Wdr83os** | 3.77683 | 6.79630 | 1.76164 | 7.03877 | 5.53860 | 5.92506 | 3.89117 | 6.37141 | 1.46441 | 12.67564 | 2.57249 | 7.63968 | 1.10056 | 2.02191 |
| **Wdr89** | 0.42812 | 0.02243 | 2.16198 | 0.00000 | 1.38879 | 2.09267 | 0.02849 | 0.29211 | 3.62557 | 0.00345 | 0.21866 | 0.00000 | 0.48996 | 1.83455 |
| **Wdr90** | 1.74247 | 0.00247 | 4.29692 | 1.95285 | 1.28312 | 0.22753 | 0.00234 | 1.26341 | 0.83544 | 0.00000 | 2.73697 | 0.78832 | 0.02722 | 1.38982 |
| **Wdr91** | 2.96751 | 0.01794 | 9.70541 | 5.72923 | 6.47177 | 4.56119 | 2.23594 | 2.82080 | 0.15996 | 0.05829 | 5.02562 | 0.00000 | 0.97297 | 3.01854 |
| **Wdr92** | 6.84884 | 0.02495 | 4.74895 | 0.40190 | 2.44198 | 1.97638 | 0.53305 | 1.52569 | 4.60395 | 0.93376 | 3.42444 | 2.76104 | 6.21570 | 5.89877 |
| **Wdr93** | 0.00000 | 0.00488 | 0.76171 | 0.45303 | 0.43608 | 0.02265 | 0.00397 | 0.01657 | 0.03474 | 0.00842 | 0.00597 | 0.00496 | 0.08353 | 0.97730 |
| **Wdr95** | 0.07719 | 0.02326 | 0.00000 | 0.87779 | 0.54010 | 1.47323 | 0.00863 | 0.00000 | 0.00622 | 0.02523 | 2.60888 | 1.66032 | 0.17409 | 0.36523 |
| **Wdsub1** | 0.10119 | 0.01016 | 2.24046 | 0.00000 | 0.35325 | 1.14288 | 0.00861 | 0.29185 | 0.03446 | 0.02754 | 1.63331 | 1.46500 | 0.02405 | 1.46609 |
| **Wdtc1** | 6.74145 | 7.59901 | 17.37273 | 5.56273 | 22.90452 | 6.06760 | 1.79413 | 5.92545 | 1.23141 | 0.05551 | 3.07594 | 2.51853 | 10.42712 | 5.70771 |
| **Wdyhv1** | 0.29882 | 0.00288 | 1.17984 | 0.00000 | 1.54743 | 0.99556 | 0.04265 | 0.25835 | 0.52304 | 0.00939 | 0.49230 | 0.86956 | 3.45709 | 1.39440 |
| **Wee1** | 8.90066 | 0.01999 | 5.85624 | 0.00000 | 7.99546 | 3.05902 | 0.00518 | 3.66923 | 9.93104 | 0.00000 | 7.95092 | 6.75870 | 1.78954 | 10.33880 |
| **Wee2** | 0.07563 | 0.00425 | 0.01519 | 0.01155 | 0.00000 | 0.00988 | 0.00411 | 0.01097 | 0.05094 | 0.00656 | 0.00751 | 0.00678 | 2.08474 | 0.14190 |
| **Wfdc15a** | 0.35863 | 0.00422 | 0.00333 | 0.00621 | 0.01865 | 0.45325 | 0.00420 | 0.48597 | 0.00639 | 0.00913 | 0.12111 | 0.13594 | 0.14498 | 0.27923 |
| **Wfdc15b** | 0.00967 | 0.00447 | 0.01575 | 0.01213 | 0.06842 | 0.01334 | 0.00457 | 0.01053 | 0.01371 | 0.00679 | 0.00900 | 0.00959 | 0.00683 | 0.01072 |
| **Wfdc18** | 0.00937 | 0.00445 | 0.01149 | 0.01206 | 0.11385 | 0.01343 | 0.00467 | 0.01058 | 0.01500 | 0.00681 | 0.00885 | 0.00946 | 0.00662 | 0.00917 |
| **Wfdc2** | 0.02534 | 4.66328 | 0.03149 | 0.00000 | 1.83210 | 0.24472 | 0.00769 | 4.70004 | 0.02633 | 2.27935 | 0.00000 | 0.46789 | 0.01405 | 0.00000 |
| **Wfdc3** | 0.01825 | 0.00416 | 0.13763 | 0.39386 | 0.00000 | 0.01270 | 0.00414 | 0.00499 | 0.03150 | 0.00516 | 0.01676 | 0.00828 | 0.02227 | 0.20631 |
| **Wfikkn1** | 0.00000 | 0.00425 | 0.06040 | 0.38647 | 0.00000 | 0.17307 | 0.00378 | 0.00377 | 0.39114 | 0.00624 | 0.01428 | 0.02926 | 0.19908 | 0.05192 |
| **Wfikkn2** | 0.01706 | 0.00418 | 0.11025 | 0.01108 | 0.01543 | 0.02223 | 0.00952 | 0.04656 | 0.08046 | 0.00636 | 0.00740 | 0.01840 | 0.01948 | 0.06070 |
| **Wfs1** | 0.01253 | 0.00381 | 0.36316 | 1.62859 | 0.00782 | 0.64254 | 0.00373 | 0.60742 | 0.00646 | 0.00408 | 0.81094 | 0.00220 | 0.40026 | 0.00000 |
| **Whamm** | 1.56006 | 0.04512 | 3.02840 | 6.50688 | 0.00000 | 6.27154 | 1.65183 | 3.51528 | 0.00000 | 0.12720 | 7.06184 | 4.31238 | 1.78597 | 0.49885 |
| **Whrn** | 0.00000 | 0.01242 | 3.99206 | 3.03351 | 0.17191 | 0.90350 | 0.01126 | 0.40725 | 0.01875 | 0.00726 | 0.02444 | 0.89506 | 0.07898 | 0.00000 |
| **Whsc1** | 0.36237 | 0.22162 | 2.25322 | 0.71443 | 1.11198 | 3.29415 | 0.05387 | 0.55562 | 2.34533 | 0.00995 | 0.00000 | 2.29679 | 1.20241 | 4.41381 |
| **Whsc1l1** | 0.34841 | 0.01380 | 0.66333 | 1.14847 | 1.70829 | 1.27412 | 2.50590 | 0.13665 | 3.39934 | 0.03023 | 0.00000 | 0.55425 | 1.32623 | 4.35294 |
| **Wif1** | 0.04224 | 0.00420 | 0.01183 | 0.21420 | 0.01062 | 0.00506 | 0.00400 | 0.00713 | 0.01469 | 0.00804 | 0.01839 | 0.01219 | 0.00985 | 0.05661 |
| **Wipf1** | 0.96048 | 0.02798 | 0.97779 | 0.00000 | 4.88695 | 3.56902 | 4.97162 | 5.55949 | 6.41601 | 0.25925 | 0.00013 | 0.00000 | 2.12122 | 6.51822 |
| **Wipf2** | 3.64210 | 0.01923 | 3.14144 | 1.13136 | 5.67486 | 4.04368 | 1.14217 | 2.49338 | 3.00175 | 0.05842 | 0.06351 | 0.04356 | 5.53684 | 9.95043 |
| **Wipi1** | 0.05511 | 0.00356 | 1.71455 | 0.11484 | 2.91564 | 0.53580 | 0.13490 | 0.08418 | 0.70293 | 0.00299 | 0.00000 | 0.61724 | 0.02080 | 0.13957 |
| **Wipi2** | 16.49186 | 0.96519 | 19.82021 | 8.45692 | 24.27016 | 13.42117 | 5.27210 | 5.94964 | 10.88049 | 0.26229 | 8.92686 | 2.10058 | 13.26657 | 23.13651 |
| **Wisp1** | 0.41233 | 0.00441 | 0.01301 | 0.01008 | 0.00589 | 0.01000 | 0.00429 | 0.01177 | 0.00984 | 0.00658 | 0.00706 | 0.00871 | 0.00470 | 0.00509 |
| **Wisp3** | 0.00472 | 0.00423 | 0.14841 | 0.17004 | 0.10834 | 0.00229 | 0.00416 | 0.00763 | 0.01852 | 0.00553 | 0.01436 | 0.01542 | 0.03819 | 0.01316 |
| **Wiz** | 6.89235 | 0.02291 | 7.21342 | 5.25755 | 5.79614 | 3.71610 | 4.24939 | 3.88414 | 0.60268 | 0.09109 | 3.50329 | 3.46022 | 2.68046 | 1.49461 |
| **Wls** | 0.08643 | 0.02119 | 0.02905 | 0.00000 | 0.61664 | 0.01001 | 0.04692 | 0.34820 | 0.01183 | 0.00606 | 0.00602 | 0.00965 | 0.07503 | 0.08234 |
| **Wnk1** | 14.13274 | 5.17805 | 16.84787 | 8.61843 | 26.25546 | 14.55158 | 1.21120 | 3.10666 | 33.35745 | 15.74565 | 5.96296 | 9.67781 | 15.55701 | 18.26021 |
| **Wnk2** | 0.44955 | 0.09693 | 1.44591 | 0.00000 | 0.91792 | 0.87845 | 1.58208 | 11.47195 | 0.51896 | 2.39947 | 0.26463 | 0.20493 | 0.26159 | 0.00000 |
| **Wnk3** | 1.89156 | 0.01116 | 3.09548 | 1.46606 | 4.51891 | 0.16801 | 0.00204 | 0.37780 | 15.06972 | 0.00000 | 0.05556 | 1.03320 | 0.62501 | 3.26701 |
| **Wnk4** | 0.03006 | 0.00400 | 0.00736 | 0.41257 | 0.03980 | 0.00540 | 0.00420 | 0.00464 | 0.01389 | 0.00558 | 0.01015 | 0.20167 | 0.00908 | 0.02057 |

| | | | | | | | | | | | | | | |
|---|---|---|---|---|---|---|---|---|---|---|---|---|---|---|
| **Wnt10b** | 0.49728 | 0.00499 | 0.52443 | 0.22945 | 0.99936 | 0.00000 | 0.00434 | 0.09992 | 0.00647 | 0.00866 | 0.00397 | 0.00399 | 0.00504 | 0.21827 |
| **Wnt2b** | 0.06969 | 0.00445 | 0.03255 | 0.01066 | 0.03363 | 0.01294 | 0.00473 | 0.01026 | 0.07661 | 0.00674 | 0.00689 | 0.01003 | 0.00380 | 0.00930 |
| **Wnt3a** | 0.00000 | 0.01302 | 1.14049 | 0.12030 | 0.13308 | 1.88638 | 0.01074 | 3.71750 | 0.00741 | 0.01086 | 0.16324 | 0.00000 | 0.00538 | 0.00000 |
| **Wnt4** | 0.13491 | 0.09698 | 0.02692 | 0.43440 | 0.55706 | 0.37341 | 0.06039 | 0.27768 | 0.49264 | 0.20483 | 0.04301 | 0.11242 | 0.52064 | 0.52346 |
| **Wnt6** | 0.00509 | 3.89401 | 0.00617 | 0.00267 | 0.01006 | 0.29285 | 0.00282 | 0.62262 | 0.01182 | 0.02117 | 0.00000 | 0.49452 | 0.00627 | 0.00771 |
| **Wnt7a** | 0.00734 | 0.00440 | 0.01009 | 0.01176 | 0.23494 | 0.01301 | 0.00444 | 0.01185 | 0.00915 | 0.00643 | 0.00893 | 0.00922 | 0.00927 | 0.02686 |
| **Wnt7b** | 0.00000 | 5.53061 | 1.64798 | 1.43387 | 0.00000 | 2.54662 | 5.39445 | 14.02494 | 0.00406 | 0.24342 | 0.04393 | 0.00000 | 0.00594 | 0.00000 |
| **Wrap53** | 3.03393 | 0.03759 | 11.77390 | 5.10170 | 5.30397 | 2.53706 | 0.00979 | 2.10170 | 0.26920 | 5.92937 | 2.41917 | 6.90452 | 1.25356 | 8.76104 |
| **Wrb** | 0.99903 | 0.01564 | 1.43747 | 3.06565 | 3.09761 | 3.75908 | 0.04238 | 0.00000 | 2.95743 | 0.01548 | 0.00000 | 3.44635 | 0.50074 | 3.60656 |
| **Wrn** | 0.08294 | 0.00424 | 0.02037 | 0.13779 | 0.09612 | 0.03527 | 0.00401 | 0.00493 | 0.01560 | 0.00627 | 0.01402 | 0.01158 | 0.42918 | 0.19856 |
| **Wrnip1** | 2.65904 | 0.02812 | 2.11566 | 4.68780 | 2.18320 | 5.82746 | 1.91379 | 0.00000 | 0.00607 | 0.01769 | 2.59252 | 5.33283 | 1.41876 | 3.25705 |
| **Wsb1** | 7.32598 | 9.29743 | 25.05713 | 5.95945 | 26.53903 | 12.37437 | 7.59631 | 9.58242 | 29.83384 | 6.00799 | 5.35190 | 5.71432 | 20.65794 | 29.44296 |
| **Wsb2** | 1.93261 | 0.05777 | 4.19555 | 1.59736 | 7.91041 | 6.96011 | 1.28424 | 2.03711 | 1.22148 | 0.05822 | 2.19754 | 3.35668 | 0.26539 | 10.56521 |
| **Wscd2** | 0.00000 | 0.00403 | 0.25734 | 0.49242 | 0.16943 | 2.55664 | 0.00377 | 0.60172 | 0.00957 | 0.00490 | 0.00272 | 0.00000 | 0.03258 | 0.00000 |
| **Wt1** | 0.01108 | 0.00452 | 0.00625 | 0.01242 | 0.51602 | 0.01314 | 0.00434 | 0.01098 | 0.01012 | 0.00633 | 0.00883 | 0.00794 | 0.00534 | 0.01048 |
| **Wtap** | 15.91097 | 10.00078 | 32.05322 | 30.74122 | 35.74113 | 28.92119 | 12.11734 | 15.92644 | 39.14139 | 20.59328 | 22.73580 | 9.67379 | 16.17153 | 31.80432 |
| **Wtip** | 0.01701 | 0.01937 | 0.73094 | 1.41891 | 0.42334 | 0.15389 | 1.66684 | 0.52645 | 0.03998 | 0.02220 | 0.00000 | 2.91616 | 0.08056 | 0.12263 |
| **Wwc1** | 0.00000 | 0.04663 | 3.18998 | 0.44024 | 0.28342 | 1.58753 | 0.01076 | 1.17272 | 0.00029 | 0.00899 | 1.57660 | 0.50872 | 0.03076 | 0.16549 |
| **Wwc2** | 0.49810 | 0.06376 | 11.63875 | 7.36348 | 2.80953 | 3.89854 | 2.68203 | 4.72000 | 0.62745 | 0.12789 | 6.53789 | 1.30224 | 4.23028 | 3.38243 |
| **Wwox** | 0.00918 | 0.00392 | 0.00000 | 0.11322 | 0.35691 | 0.00000 | 0.00412 | 0.01203 | 0.00871 | 0.00459 | 0.96052 | 0.42572 | 0.26791 | 0.00392 |
| **Wwp1** | 0.29492 | 0.00950 | 0.06613 | 1.43866 | 0.06557 | 2.86318 | 0.00399 | 0.48826 | 0.00398 | 0.01217 | 0.00000 | 3.48957 | 0.42960 | 1.96551 |
| **Wwp2** | 13.67248 | 0.05856 | 9.76788 | 4.78593 | 10.35089 | 6.34905 | 2.31737 | 4.10366 | 2.09832 | 0.08070 | 9.55733 | 4.02238 | 6.54123 | 11.78165 |
| **Wwtr1** | 9.86247 | 0.02727 | 6.71042 | 3.20736 | 8.01625 | 3.11517 | 1.09635 | 0.45058 | 5.00205 | 0.03477 | 7.50784 | 6.53870 | 6.67963 | 6.04703 |
| **Xab2** | 6.81715 | 6.53305 | 14.39839 | 11.37171 | 11.15939 | 10.62306 | 2.90274 | 5.24962 | 0.00000 | 9.04285 | 8.36874 | 7.45119 | 0.84346 | 11.26724 |
| **Xaf1** | 0.56062 | 0.04374 | 0.00000 | 0.35241 | 0.10494 | 1.12393 | 0.15481 | 0.05843 | 0.77689 | 0.31830 | 0.02041 | 0.35657 | 5.23826 | 0.46860 |
| **Xbp1** | 1.16271 | 3.21316 | 2.43830 | 5.59448 | 0.22009 | 4.52479 | 7.29341 | 6.31544 | 0.12211 | 0.32708 | 3.26464 | 1.30737 | 0.00000 | 0.00000 |
| **Xcr1** | 0.06662 | 0.05373 | 0.02879 | 0.35014 | 0.02330 | 0.06785 | 0.03311 | 0.00959 | 0.19269 | 0.02901 | 0.01139 | 0.07519 | 0.21136 | 1.50844 |
| **Xdh** | 0.00804 | 0.00429 | 0.01585 | 0.35042 | 0.16126 | 0.00612 | 0.00421 | 0.00723 | 0.08711 | 0.00558 | 0.00613 | 0.00711 | 0.00433 | 0.00000 |
| **Xiap** | 7.16422 | 0.01655 | 3.62368 | 1.56152 | 3.56109 | 6.01530 | 0.11681 | 0.29458 | 21.07385 | 0.01609 | 1.99271 | 3.82690 | 1.60721 | 4.85192 |
| **Xist** | 2.07801 | 14.02646 | 5.37863 | 7.19317 | 12.72000 | 6.70737 | 0.02211 | 17.49523 | 14.79928 | 0.09693 | 0.00000 | 4.82366 | 0.08167 | 1.56973 |
| **Xk** | 0.99869 | 0.00404 | 0.01577 | 0.00000 | 0.00295 | 0.78520 | 0.00397 | 0.01480 | 0.01493 | 0.00605 | 0.00343 | 0.03839 | 0.02950 | 0.00152 |
| **Xkr8** | 0.53618 | 0.02523 | 1.89562 | 3.34409 | 0.13270 | 3.43916 | 0.04355 | 0.13332 | 0.62597 | 0.05877 | 1.66787 | 4.78532 | 0.34172 | 1.91367 |
| **Xkr9** | 1.36756 | 0.00461 | 1.10322 | 0.01386 | 0.09662 | 0.02337 | 0.00294 | 0.00443 | 8.88670 | 0.00549 | 0.00276 | 0.00362 | 0.02387 | 1.07829 |
| **Xkrx** | 0.13786 | 0.02604 | 0.00292 | 0.02567 | 0.00000 | 0.06902 | 0.01510 | 0.29132 | 0.89530 | 0.02624 | 0.00409 | 0.01924 | 0.01456 | 0.40011 |
| **Xlr** | 0.00403 | 0.00466 | 0.58624 | 0.00653 | 0.65517 | 0.07873 | 0.00394 | 0.00783 | 0.55434 | 0.01263 | 0.00445 | 0.01091 | 0.03185 | 0.50255 |
| **Xlr3a** | 0.00000 | 0.02615 | 0.01385 | 3.04020 | 0.38370 | 8.35812 | 0.00317 | 0.00000 | 1.26313 | 0.00222 | 3.87473 | 0.69883 | 0.98486 | 0.26415 |
| **Xlr3b** | 0.00000 | 0.09959 | 0.12831 | 1.45110 | 0.48252 | 7.26549 | 0.00298 | 0.00134 | 1.86600 | 0.01538 | 5.21440 | 0.98930 | 0.82087 | 0.77872 |
| **Xlr3c** | 0.00000 | 0.02589 | 0.00000 | 0.66722 | 0.55807 | 7.74407 | 0.00299 | 0.01824 | 2.06284 | 0.01513 | 4.73710 | 0.90961 | 0.62190 | 0.49848 |
| **Xlr4a** | 0.58470 | 0.19566 | 0.46713 | 0.00000 | 0.26986 | 2.57801 | 0.00384 | 0.01464 | 0.70391 | 0.00142 | 1.47323 | 0.53016 | 0.42652 | 0.39875 |
| **Xlr4b** | 1.61645 | 0.77510 | 1.53975 | 0.99184 | 0.53828 | 8.00715 | 0.00266 | 0.38249 | 2.37351 | 0.03950 | 3.17342 | 1.71626 | 1.06352 | 1.28569 |
| **Xlr4c** | 0.62220 | 0.56750 | 0.44229 | 0.24323 | 0.28283 | 2.95014 | 0.00379 | 0.21586 | 0.58750 | 0.00852 | 1.53769 | 0.57858 | 0.47377 | 0.60745 |
| **Xlr5a** | 0.25649 | 0.01012 | 0.70139 | 0.51780 | 1.81090 | 0.12049 | 0.00325 | 0.24964 | 1.95783 | 0.00000 | 2.93532 | 0.40668 | 0.16098 | 3.24862 |
| **Xlr5b** | 0.11799 | 0.00302 | 0.88433 | 0.64371 | 1.42109 | 0.03388 | 0.00325 | 0.21396 | 2.30278 | 0.00000 | 2.61378 | 0.23747 | 0.12865 | 3.54447 |
| **Xlr5c** | 0.04182 | 0.00374 | 0.36412 | 0.15881 | 0.45013 | 0.00835 | 0.00383 | 0.06672 | 0.98154 | 0.00385 | 0.86851 | 0.11921 | 0.00057 | 1.22796 |
| **Xndc1** | 0.17551 | 0.01063 | 0.02313 | 0.64134 | 0.81150 | 0.73650 | 0.06609 | 1.12524 | 0.06880 | 0.00627 | 0.21229 | 1.05405 | 2.12499 | 0.15098 |
| **Xpa** | 0.88555 | 0.01146 | 0.29430 | 0.00000 | 0.44421 | 0.15107 | 0.00402 | 0.18200 | 0.29168 | 0.00600 | 0.00000 | 0.53838 | 0.89530 | 1.21115 |
| **Xpc** | 0.00000 | 0.01928 | 0.85747 | 1.88200 | 0.46502 | 0.98536 | 4.31719 | 0.03290 | 0.02675 | 0.03809 | 0.00000 | 1.79166 | 0.54686 | 0.37588 |
| **Xpnpep1** | 3.92237 | 0.41602 | 22.68617 | 14.15572 | 14.89318 | 17.97677 | 9.35507 | 12.85285 | 0.32122 | 17.16353 | 12.19758 | 13.09414 | 3.99412 | 14.00195 |
| **Xpnpep2** | 0.04348 | 0.01771 | 0.40559 | 0.00000 | 0.06729 | 0.00000 | 0.00370 | 0.52740 | 0.08738 | 0.01178 | 0.00521 | 0.01039 | 0.00742 | 0.69564 |
| **Xpnpep3** | 0.38918 | 0.07695 | 0.59122 | 1.37674 | 0.88844 | 1.50735 | 0.06312 | 0.61678 | 2.53518 | 0.15057 | 0.47401 | 1.20584 | 1.50640 | 3.02925 |
| **Xpo1** | 17.17200 | 3.23352 | 23.50710 | 10.71799 | 23.66563 | 15.71782 | 4.53985 | 1.94104 | 46.78817 | 0.17534 | 9.74399 | 8.36055 | 16.34084 | 8.31489 |
| **Xpo4** | 3.38597 | 0.39735 | 2.68881 | 0.00000 | 4.01883 | 2.37134 | 0.00144 | 0.00000 | 9.16540 | 0.00135 | 0.00000 | 1.84577 | 1.93119 | 4.92069 |
| **Xpo5** | 1.82850 | 0.05879 | 10.49332 | 9.16024 | 6.10457 | 4.12142 | 1.84592 | 4.05340 | 0.76176 | 0.09103 | 3.71247 | 5.32162 | 0.46156 | 5.97705 |
| **Xpo6** | 4.60120 | 5.71551 | 16.01583 | 3.19007 | 12.90722 | 11.58783 | 1.44990 | 3.59319 | 0.43518 | 0.05177 | 7.34288 | 6.63288 | 3.45666 | 6.39595 |
| **Xpo7** | 6.29010 | 0.13627 | 7.86124 | 5.86882 | 7.70827 | 7.66021 | 0.10341 | 3.43441 | 10.03346 | 0.28872 | 5.26383 | 5.03365 | 2.69177 | 3.15128 |

| | | | | | | | | | | | | | | |
|---|---|---|---|---|---|---|---|---|---|---|---|---|---|---|
| **Xpot** | 23.28065 | 0.13501 | 21.45147 | 4.68657 | 21.88467 | 14.19962 | 4.79853 | 4.89433 | 20.44327 | 0.13661 | 19.35310 | 11.90274 | 8.22871 | 18.06906 |
| **Xpr1** | 13.83466 | 0.01906 | 11.96981 | 1.17197 | 15.17374 | 4.87105 | 0.00000 | 0.44915 | 21.02082 | 0.00000 | 5.73039 | 0.23283 | 4.78403 | 13.12258 |
| **Xrcc1** | 6.10207 | 0.16182 | 15.96171 | 19.90202 | 12.00905 | 5.73706 | 2.38720 | 2.67366 | 0.23141 | 11.50850 | 6.47228 | 8.75878 | 0.11790 | 2.83114 |
| **Xrcc2** | 0.00000 | 0.04823 | 0.14096 | 1.75838 | 0.66021 | 0.27227 | 0.10639 | 1.06292 | 1.67151 | 0.10981 | 0.09506 | 0.16073 | 0.81531 | 4.96467 |
| **Xrcc3** | 0.19086 | 0.04497 | 1.45430 | 1.56207 | 1.48267 | 2.10002 | 0.00373 | 2.29747 | 0.02259 | 0.00479 | 0.77559 | 1.02998 | 0.58424 | 0.10773 |
| **Xrcc4** | 2.04356 | 0.00398 | 0.00013 | 4.65498 | 0.93109 | 3.07440 | 0.01104 | 0.28410 | 3.68860 | 0.00559 | 0.39895 | 0.00000 | 0.20754 | 0.00000 |
| **Xrcc5** | 0.00000 | 0.03088 | 0.87726 | 1.49391 | 1.34964 | 1.77911 | 0.00199 | 0.13434 | 0.00616 | 0.03053 | 5.98260 | 3.36013 | 1.71275 | 3.05114 |
| **Xrcc6** | 1.64376 | 6.55883 | 4.34042 | 1.50614 | 5.42933 | 5.29367 | 4.20804 | 0.24739 | 1.41552 | 0.16338 | 6.68141 | 8.62020 | 0.50474 | 6.33393 |
| **Xrcc6bp1** | 0.20221 | 0.00392 | 0.00000 | 0.00000 | 2.34059 | 0.06611 | 0.00420 | 0.00506 | 0.00027 | 0.00640 | 0.00916 | 0.07972 | 2.30427 | 0.12718 |
| **Xrn1** | 4.91668 | 0.06314 | 1.72798 | 1.30262 | 2.96426 | 1.12630 | 0.04725 | 0.23174 | 4.37901 | 0.06371 | 0.00000 | 1.67466 | 1.30209 | 2.47784 |
| **Xrn2** | 11.11871 | 0.02432 | 13.40303 | 3.65052 | 13.15418 | 7.82938 | 2.34699 | 1.74731 | 9.26641 | 0.02665 | 9.70136 | 7.94235 | 4.59345 | 13.80504 |
| **Xxylt1** | 0.00331 | 0.00323 | 0.00000 | 0.63982 | 0.00000 | 0.58953 | 0.00351 | 0.00477 | 0.34560 | 0.00676 | 0.00000 | 1.63375 | 0.00306 | 0.07949 |
| **Xylb** | 2.69669 | 0.04730 | 13.78316 | 4.91084 | 13.67269 | 3.86083 | 0.51404 | 6.87965 | 4.82805 | 0.17874 | 0.36376 | 1.60270 | 4.67370 | 13.06003 |
| **Xylt2** | 0.00000 | 0.04135 | 0.74623 | 1.50067 | 0.30891 | 0.84414 | 0.03281 | 3.60037 | 0.00158 | 0.01755 | 0.00000 | 2.98542 | 0.56098 | 0.56184 |
| **Yae1d1** | 9.38709 | 0.01189 | 7.33464 | 3.74332 | 7.88642 | 5.16589 | 1.93463 | 3.84929 | 10.18231 | 0.07954 | 8.12210 | 4.28881 | 1.54985 | 12.71097 |
| **Yaf2** | 0.57694 | 0.00340 | 1.41968 | 0.55220 | 0.79806 | 0.51964 | 0.32459 | 0.48597 | 0.51454 | 0.01226 | 0.08334 | 0.97374 | 0.78983 | 0.33859 |
| **Yap1** | 5.40169 | 0.06577 | 8.17842 | 5.34476 | 10.03344 | 4.22508 | 2.36380 | 4.64254 | 11.32707 | 0.06978 | 3.68908 | 5.31910 | 15.23109 | 14.07114 |
| **Yars** | 12.52239 | 4.05901 | 22.63899 | 13.62646 | 17.14026 | 15.17030 | 2.54729 | 7.72664 | 5.31321 | 0.20539 | 17.29132 | 11.32725 | 3.47862 | 14.94469 |
| **Yars2** | 6.26202 | 0.04037 | 1.46907 | 2.15239 | 2.36421 | 2.85757 | 1.62782 | 0.44628 | 1.44979 | 0.03204 | 2.97782 | 1.93249 | 1.98397 | 1.46307 |
| **Ybey** | 2.21760 | 0.01111 | 0.62950 | 1.07655 | 2.43400 | 1.53185 | 0.06136 | 0.81977 | 3.30532 | 0.37004 | 1.21077 | 0.48046 | 1.83065 | 1.00988 |
| **Ybx1** | 36.54627 | 7.44783 | 34.46440 | 21.68018 | 37.55115 | 28.24327 | 9.56648 | 17.94144 | 23.88835 | 23.23176 | 22.19017 | 20.99158 | 21.19036 | 33.59737 |
| **Ybx2** | 0.00000 | 5.83361 | 0.73076 | 3.81447 | 0.80329 | 0.00000 | 0.00000 | 3.52385 | 0.04604 | 8.98232 | 0.12031 | 2.26929 | 1.79791 | 0.02659 |
| **Ybx3** | 25.00503 | 2.63752 | 30.73261 | 27.91836 | 26.84503 | 24.70272 | 10.57830 | 21.10894 | 5.14654 | 18.36693 | 15.61913 | 17.50071 | 3.95196 | 7.17089 |
| **Ydjc** | 0.24309 | 0.02091 | 0.17844 | 1.83675 | 0.07879 | 0.12193 | 0.00392 | 0.00388 | 0.00340 | 0.01087 | 0.00549 | 0.13866 | 2.28268 | 0.00000 |
| **Yeats2** | 0.83624 | 0.07009 | 4.45582 | 3.15094 | 3.92464 | 4.23089 | 3.44881 | 2.25304 | 3.97464 | 0.08913 | 7.58456 | 3.39197 | 3.42954 | 4.83405 |
| **Yeats4** | 0.68075 | 0.03002 | 4.38633 | 7.73992 | 7.69623 | 2.76602 | 2.04502 | 2.56826 | 0.11865 | 0.07458 | 1.72348 | 5.41244 | 1.43527 | 0.00000 |
| **Yes1** | 2.17442 | 0.00264 | 0.12819 | 0.02918 | 0.00006 | 0.00000 | 0.00290 | 0.00640 | 5.58835 | 0.00549 | 0.36097 | 1.76268 | 0.96632 | 0.00000 |
| **Yif1a** | 6.50801 | 0.21879 | 10.60961 | 8.11888 | 4.40920 | 12.84239 | 3.06696 | 11.03710 | 0.06964 | 0.17049 | 8.80645 | 8.49896 | 0.04081 | 3.55163 |
| **Yif1b** | 0.00000 | 0.21991 | 18.26328 | 14.76265 | 10.67935 | 1.78544 | 2.74281 | 12.02415 | 0.00000 | 12.71810 | 5.22679 | 10.36451 | 0.01420 | 0.00000 |
| **Yipf1** | 2.49000 | 0.01331 | 7.23658 | 2.31015 | 6.74208 | 6.32490 | 0.00576 | 2.52218 | 4.02169 | 0.05883 | 4.43318 | 3.44857 | 7.96108 | 13.18364 |
| **Yipf2** | 0.00000 | 0.01018 | 1.26842 | 2.35940 | 0.07951 | 1.43068 | 0.01423 | 1.97669 | 0.00000 | 0.04564 | 2.00695 | 2.84647 | 0.00274 | 0.55796 |
| **Yipf3** | 3.57390 | 0.04801 | 8.39785 | 12.37212 | 7.30416 | 3.40949 | 3.43437 | 4.04476 | 0.00000 | 0.08513 | 0.43690 | 3.70733 | 3.15745 | 0.89513 |
| **Yipf4** | 3.18723 | 0.00252 | 1.09624 | 2.77003 | 1.99710 | 1.09777 | 0.03106 | 0.00000 | 12.65167 | 0.00000 | 1.93104 | 0.07340 | 7.43101 | 0.40989 |
| **Yipf5** | 10.22632 | 0.12678 | 7.22239 | 8.69833 | 9.75982 | 10.87954 | 3.57315 | 5.01832 | 6.72216 | 0.17150 | 4.19371 | 5.82540 | 2.17914 | 10.56290 |
| **Yipf6** | 5.93495 | 4.16036 | 1.83592 | 5.26593 | 2.41284 | 2.74454 | 0.85328 | 0.88275 | 12.38452 | 0.02665 | 3.05781 | 1.30516 | 0.60102 | 3.71080 |
| **Ykt6** | 5.90803 | 6.57899 | 8.14018 | 5.38935 | 6.61666 | 14.31190 | 5.37638 | 3.67418 | 4.09929 | 0.40112 | 2.42526 | 6.71108 | 0.16885 | 2.74782 |
| **Ylpm1** | 2.47956 | 0.02685 | 3.40777 | 2.86836 | 12.04065 | 2.66441 | 1.96102 | 0.25774 | 4.58767 | 0.04722 | 1.55459 | 3.25281 | 6.57643 | 7.03444 |
| **Yme1l1** | 14.26711 | 3.74746 | 13.62780 | 7.55737 | 14.51607 | 6.94027 | 1.59868 | 5.13946 | 32.99570 | 0.04045 | 12.17842 | 3.84133 | 9.09666 | 11.92466 |
| **Yod1** | 0.00000 | 0.00456 | 0.00000 | 0.10583 | 0.42085 | 0.01331 | 0.00444 | 0.03749 | 0.12984 | 0.00905 | 0.00810 | 0.00475 | 0.00407 | 0.95392 |
| **Ypel1** | 0.00443 | 0.00415 | 0.01631 | 0.63763 | 0.21328 | 0.59924 | 0.00376 | 0.00321 | 0.01533 | 0.00554 | 0.00639 | 0.00770 | 0.03771 | 0.09206 |
| **Ypel2** | 15.68799 | 0.10122 | 12.90392 | 4.11817 | 22.78952 | 8.21066 | 0.25178 | 4.21360 | 19.29720 | 0.50441 | 4.19901 | 5.63643 | 10.20606 | 24.81329 |
| **Ypel3** | 3.65825 | 0.00264 | 1.33938 | 3.31166 | 5.32153 | 0.55072 | 0.00324 | 0.04839 | 0.00010 | 0.01067 | 0.14757 | 0.95624 | 3.24670 | 0.21164 |
| **Ypel5** | 11.45746 | 2.83211 | 11.93097 | 7.10066 | 16.52648 | 2.68130 | 3.83836 | 0.53573 | 20.86230 | 0.12247 | 6.42803 | 4.10946 | 9.33862 | 12.89232 |
| **Yrdc** | 12.86085 | 6.90992 | 7.70309 | 6.51633 | 11.99221 | 3.41772 | 0.68312 | 0.00000 | 7.99156 | 0.01472 | 2.93190 | 4.67569 | 13.84461 | 7.37991 |
| **Ythdc1** | 9.31326 | 0.06484 | 6.31465 | 5.06988 | 12.20497 | 7.34214 | 0.24094 | 2.27320 | 10.95043 | 0.00293 | 5.19001 | 6.49303 | 7.57968 | 9.31125 |
| **Ythdc2** | 0.08302 | 0.21311 | 0.12743 | 0.10093 | 0.00000 | 0.00000 | 0.08421 | 0.00000 | 3.97257 | 0.00233 | 1.48696 | 0.50141 | 0.19436 | 0.32046 |
| **Ythdf1** | 9.40830 | 0.11723 | 15.40234 | 5.08050 | 13.15269 | 11.50634 | 2.87224 | 4.91879 | 7.52739 | 0.07675 | 7.59404 | 9.32017 | 6.88407 | 11.49256 |
| **Ythdf2** | 3.39761 | 0.89841 | 4.03537 | 5.07682 | 12.78938 | 5.24046 | 3.39085 | 1.62619 | 14.38314 | 0.16289 | 1.22176 | 2.82921 | 21.19309 | 12.80557 |
| **Ythdf3** | 3.04264 | 1.52541 | 3.49783 | 0.72615 | 1.03046 | 2.59588 | 0.14400 | 3.29468 | 6.25199 | 0.01494 | 3.40833 | 1.88853 | 3.29605 | 0.92083 |
| **Ywhab** | 10.38976 | 4.00649 | 17.03577 | 16.85690 | 20.63738 | 21.13107 | 3.02158 | 5.88974 | 20.45591 | 9.43805 | 10.07826 | 13.79728 | 7.65299 | 16.79075 |
| **Ywhae** | 18.06254 | 8.14074 | 32.81818 | 32.10587 | 27.11740 | 31.89041 | 7.31190 | 21.37156 | 23.57086 | 4.20573 | 21.54811 | 23.78022 | 9.65230 | 29.71333 |
| **Ywhag** | 1.65826 | 8.33027 | 24.67871 | 18.08342 | 20.30731 | 15.02101 | 2.83865 | 15.24127 | 5.56203 | 12.17417 | 11.89989 | 13.78681 | 10.40873 | 3.66201 |
| **Ywhah** | 19.60313 | 0.43988 | 16.92369 | 10.74688 | 14.83714 | 16.66347 | 7.43333 | 9.21511 | 11.31301 | 5.75673 | 4.27390 | 11.53981 | 7.32479 | 14.46717 |
| **Ywhaq** | 33.89893 | 10.29994 | 47.38569 | 41.02844 | 36.73281 | 31.45350 | 8.39865 | 21.51312 | 23.06538 | 10.34937 | 17.36415 | 19.96498 | 19.53863 | 17.57287 |
| **Ywhaz** | 8.28407 | 11.14703 | 20.06766 | 18.86251 | 6.02500 | 27.28606 | 2.35060 | 14.16494 | 12.25225 | 10.87067 | 16.28143 | 16.57806 | 2.82824 | 9.40810 |

| | | | | | | | | | | | | | | |
|---|---|---|---|---|---|---|---|---|---|---|---|---|---|---|
| **Yy1** | 16.06414 | 0.04834 | 13.37992 | 4.54567 | 17.49863 | 11.10358 | 1.39361 | 3.22639 | 14.94566 | 0.04466 | 3.92078 | 4.29546 | 12.03245 | 15.91614 |
| **Yy2** | 0.20394 | 0.00273 | 1.84833 | 0.00000 | 1.28569 | 0.41763 | 0.00131 | 0.00000 | 15.63808 | 0.00000 | 0.86257 | 0.62231 | 0.36566 | 5.39967 |
| **Zadh2** | 0.01435 | 0.00384 | 0.00000 | 0.44502 | 0.12769 | 0.36968 | 0.00384 | 0.01662 | 0.00768 | 0.01298 | 0.00209 | 0.14907 | 0.00618 | 0.07546 |
| **Zak** | 0.10999 | 0.01165 | 0.08534 | 0.10871 | 0.04513 | 2.24483 | 0.00983 | 0.03740 | 1.13407 | 0.05001 | 1.61572 | 0.06222 | 0.03511 | 0.41587 |
| **Zan** | 0.00000 | 0.00395 | 2.09956 | 0.40277 | 0.79823 | 0.02945 | 0.00351 | 0.00000 | 0.00875 | 0.00155 | 0.26048 | 0.02036 | 0.99681 | 0.09866 |
| **Zap70** | 0.66747 | 0.00335 | 0.62822 | 1.00307 | 3.31933 | 0.76309 | 0.00335 | 0.00000 | 0.02522 | 0.00421 | 2.29635 | 0.01014 | 0.00000 | 1.42809 |
| **Zar1** | 0.53178 | 0.00419 | 0.01331 | 0.06050 | 0.01251 | 0.01246 | 0.00410 | 0.00778 | 0.00760 | 0.00837 | 0.00626 | 0.00871 | 0.53761 | 0.02353 |
| **Zar1l** | 0.00000 | 0.00396 | 0.20636 | 0.00000 | 0.15357 | 0.00254 | 0.81293 | 0.00649 | 0.02748 | 0.02114 | 0.00655 | 0.00837 | 3.07761 | 0.08631 |
| **Zbed3** | 0.11719 | 0.00379 | 0.00000 | 0.07135 | 0.50137 | 0.00401 | 0.00384 | 0.00606 | 0.39191 | 0.00705 | 0.00521 | 0.00727 | 3.66376 | 0.82955 |
| **Zbed4** | 4.72939 | 1.28429 | 6.25738 | 1.30460 | 6.48785 | 7.58068 | 1.35793 | 2.26828 | 11.46643 | 0.02638 | 6.57883 | 3.90588 | 3.84536 | 15.52461 |
| **Zbed5** | 0.99936 | 0.10330 | 0.15782 | 17.98202 | 16.37354 | 1.70039 | 0.27174 | 3.43569 | 0.06928 | 1.16770 | 0.00000 | 5.21937 | 8.04476 | 0.00000 |
| **Zbed6** | 0.01647 | 0.00401 | 0.00000 | 0.04056 | 0.02994 | 0.01035 | 0.00368 | 0.00639 | 1.30252 | 0.00713 | 0.00000 | 0.79706 | 0.71187 | 0.00000 |
| **Zbp1** | 0.00719 | 0.00442 | 0.04737 | 0.00526 | 0.21810 | 0.01223 | 0.00416 | 0.00996 | 0.00933 | 0.00837 | 0.00804 | 0.00963 | 0.09940 | 0.05062 |
| **Zbtb1** | 0.12523 | 0.00325 | 0.27095 | 0.26636 | 0.69420 | 0.33040 | 1.34963 | 0.00707 | 1.91903 | 0.03314 | 0.00000 | 0.37260 | 2.57178 | 2.65222 |
| **Zbtb10** | 19.13134 | 2.65191 | 9.58287 | 0.43998 | 13.45559 | 6.29195 | 2.41805 | 5.61261 | 26.53034 | 0.25140 | 5.81239 | 4.62258 | 14.65607 | 24.31112 |
| **Zbtb11** | 2.19438 | 0.02180 | 1.28705 | 0.16927 | 1.75862 | 0.26204 | 0.00791 | 0.08262 | 11.76637 | 0.00221 | 1.29954 | 0.19300 | 12.34877 | 7.64561 |
| **Zbtb12** | 0.02432 | 0.00441 | 0.02094 | 0.05008 | 0.00728 | 0.02473 | 0.00421 | 0.00726 | 0.02208 | 0.00615 | 0.00966 | 0.01078 | 0.01888 | 0.06281 |
| **Zbtb14** | 0.12711 | 0.94633 | 3.09600 | 2.80215 | 6.11486 | 2.63260 | 0.00323 | 0.20748 | 0.00000 | 0.00149 | 1.74222 | 0.85858 | 3.36517 | 2.85031 |
| **Zbtb16** | 0.00502 | 0.01994 | 0.01200 | 0.00252 | 0.00986 | 0.44532 | 0.00432 | 0.00595 | 0.01525 | 0.00588 | 0.00707 | 0.00878 | 0.26438 | 0.00633 |
| **Zbtb17** | 6.38495 | 3.57526 | 16.52954 | 6.61748 | 18.58194 | 5.19505 | 3.24792 | 8.50659 | 0.00000 | 2.10339 | 9.04857 | 1.37153 | 3.25025 | 9.62603 |
| **Zbtb18** | 0.08755 | 0.01847 | 0.23959 | 0.00000 | 1.69286 | 0.59824 | 0.00384 | 0.25123 | 0.17395 | 0.00153 | 1.38091 | 1.03298 | 0.31081 | 0.09993 |
| **Zbtb2** | 2.21639 | 3.09890 | 1.49272 | 3.15705 | 3.05472 | 1.62158 | 1.94639 | 0.67387 | 0.28967 | 0.01439 | 1.60055 | 2.71841 | 5.62944 | 0.02334 |
| **Zbtb20** | 0.00822 | 0.00421 | 0.00202 | 0.26323 | 0.36148 | 0.01744 | 0.00453 | 0.00676 | 0.02072 | 0.00739 | 0.00598 | 0.00876 | 0.00402 | 0.00783 |
| **Zbtb21** | 0.23247 | 0.01189 | 2.24082 | 0.17802 | 1.45176 | 1.25122 | 0.01801 | 0.09375 | 1.74381 | 0.00108 | 0.46115 | 0.93182 | 2.10134 | 1.08766 |
| **Zbtb22** | 0.57808 | 0.04723 | 2.03234 | 3.22720 | 2.78633 | 0.33457 | 0.02312 | 1.13531 | 0.01419 | 0.03345 | 1.19648 | 1.20285 | 0.24011 | 0.09655 |
| **Zbtb24** | 1.56711 | 0.14038 | 2.72198 | 1.65543 | 2.74018 | 1.78808 | 2.51022 | 0.28182 | 0.83902 | 0.09184 | 4.27791 | 2.01030 | 2.62779 | 0.27692 |
| **Zbtb25** | 0.01342 | 0.00472 | 0.54168 | 0.00000 | 0.28673 | 1.20797 | 0.00439 | 1.26896 | 0.00737 | 0.00672 | 0.00000 | 0.39932 | 0.01011 | 0.00387 |
| **Zbtb26** | 0.73295 | 0.00401 | 0.00000 | 0.00000 | 0.25601 | 0.33116 | 0.00399 | 0.00969 | 2.57295 | 0.00943 | 0.00185 | 0.22084 | 0.00864 | 1.31433 |
| **Zbtb3** | 0.03396 | 0.02385 | 0.16777 | 0.74491 | 0.71532 | 0.41008 | 0.96822 | 0.10094 | 0.00729 | 0.05422 | 1.16349 | 0.00001 | 0.00943 | 0.89608 |
| **Zbtb32** | 0.00341 | 0.00400 | 0.04175 | 1.07166 | 0.02858 | 0.01374 | 0.00403 | 0.00622 | 0.00766 | 0.00626 | 0.00550 | 0.01040 | 0.40775 | 0.03473 |
| **Zbtb33** | 0.00634 | 0.00387 | 0.08230 | 0.19228 | 0.03234 | 0.10157 | 0.00403 | 0.00390 | 0.00733 | 0.00600 | 0.00453 | 0.15082 | 0.72438 | 0.06072 |
| **Zbtb34** | 7.76347 | 0.01824 | 5.53095 | 1.82914 | 8.15043 | 4.04457 | 0.03777 | 0.45570 | 1.03955 | 0.08126 | 0.45218 | 0.51002 | 1.55864 | 6.70935 |
| **Zbtb37** | 1.68506 | 0.00368 | 0.39979 | 0.24232 | 1.87317 | 0.03186 | 0.00333 | 0.00182 | 2.39692 | 0.00359 | 0.88505 | 0.02481 | 3.58311 | 3.98170 |
| **Zbtb39** | 0.46333 | 0.01125 | 1.03377 | 0.12117 | 0.36635 | 2.00154 | 0.90500 | 0.00908 | 0.00668 | 0.05291 | 0.00000 | 1.19275 | 0.47839 | 0.53081 |
| **Zbtb4** | 0.04173 | 0.00403 | 0.01189 | 0.01056 | 0.00883 | 0.01158 | 0.00435 | 0.00654 | 0.01040 | 0.00628 | 0.00572 | 0.32712 | 0.03070 | 0.07783 |
| **Zbtb40** | 1.40464 | 0.03960 | 5.97482 | 4.41960 | 2.44212 | 3.48316 | 0.32351 | 0.01174 | 0.11058 | 0.01449 | 0.00000 | 3.15265 | 0.27936 | 3.10883 |
| **Zbtb41** | 2.94757 | 0.00396 | 0.90725 | 0.00000 | 0.94164 | 1.31035 | 0.00306 | 0.00000 | 6.37032 | 0.00576 | 0.07185 | 0.02185 | 2.62139 | 2.07725 |
| **Zbtb42** | 0.02128 | 0.00489 | 0.31190 | 0.15065 | 0.02702 | 0.25515 | 0.00413 | 0.01327 | 0.00909 | 0.02668 | 0.00512 | 0.00417 | 0.01125 | 0.04350 |
| **Zbtb43** | 5.83789 | 0.00129 | 6.52443 | 2.84818 | 6.06580 | 3.69760 | 1.46337 | 1.38986 | 11.30045 | 0.07627 | 2.42202 | 0.84032 | 8.09331 | 5.25093 |
| **Zbtb44** | 5.83783 | 0.05893 | 4.87739 | 2.41633 | 3.86864 | 0.99411 | 1.50826 | 3.29444 | 8.92145 | 0.09753 | 7.43965 | 2.11895 | 5.88521 | 7.23793 |
| **Zbtb45** | 1.23994 | 0.01611 | 0.74406 | 1.31492 | 0.31396 | 1.74964 | 0.00888 | 0.03589 | 0.00332 | 0.03854 | 0.07336 | 3.85996 | 2.38204 | 0.00000 |
| **Zbtb46** | 0.43581 | 0.00383 | 1.20927 | 0.47051 | 1.21479 | 0.00066 | 0.00379 | 0.06224 | 0.58396 | 0.00329 | 0.30678 | 0.00518 | 0.17338 | 0.16785 |
| **Zbtb48** | 0.90418 | 0.02791 | 5.21966 | 2.29658 | 2.48958 | 0.71005 | 5.44038 | 3.13840 | 0.02534 | 0.30798 | 1.06966 | 2.06904 | 0.49033 | 1.15132 |
| **Zbtb49** | 0.03663 | 0.00505 | 0.11962 | 0.01052 | 0.05209 | 0.00865 | 0.00433 | 0.07306 | 0.07188 | 0.00837 | 0.00521 | 0.24058 | 0.46554 | 0.00000 |
| **Zbtb5** | 2.67220 | 0.01228 | 2.74588 | 0.08281 | 2.68390 | 0.87989 | 0.00322 | 1.59814 | 4.48505 | 0.03398 | 1.02778 | 0.01366 | 3.72969 | 1.10753 |
| **Zbtb6** | 0.23001 | 0.01964 | 0.23379 | 2.06891 | 1.16710 | 0.30522 | 0.23601 | 0.45255 | 4.00781 | 0.00543 | 0.29290 | 0.09796 | 2.87569 | 0.91746 |
| **Zbtb7a** | 1.05173 | 0.02731 | 0.67161 | 0.43707 | 1.96395 | 0.03865 | 0.94474 | 0.66627 | 0.02781 | 0.00715 | 0.00317 | 0.45191 | 2.82326 | 0.49418 |
| **Zbtb7b** | 2.65036 | 0.11030 | 7.15988 | 8.80473 | 5.96518 | 4.89488 | 3.69637 | 11.72701 | 0.09338 | 3.97519 | 1.67385 | 3.56173 | 0.56471 | 0.61187 |
| **Zbtb7c** | 0.00000 | 0.00428 | 1.21153 | 0.00000 | 0.25887 | 0.24168 | 0.00372 | 0.00000 | 0.00421 | 0.00351 | 0.00539 | 0.00771 | 0.93299 | 0.07305 |
| **Zbtb8a** | 0.14026 | 0.01850 | 3.29747 | 0.09671 | 4.03387 | 1.76212 | 0.04488 | 2.54056 | 0.62376 | 0.02990 | 0.00822 | 0.00222 | 1.77451 | 6.00197 |
| **Zbtb8b** | 0.00000 | 0.03397 | 3.92583 | 0.00000 | 1.54551 | 0.68511 | 0.00928 | 1.98187 | 1.73652 | 0.00248 | 0.66017 | 0.04738 | 1.03376 | 1.68981 |
| **Zbtb8os** | 1.07859 | 0.09282 | 0.44391 | 2.49268 | 1.11877 | 3.13395 | 0.02385 | 2.75993 | 1.92687 | 10.43608 | 2.31514 | 2.72801 | 0.10487 | 2.15049 |
| **Zbtb9** | 0.61585 | 0.00361 | 1.58835 | 1.81703 | 0.76906 | 1.76761 | 0.00900 | 0.00000 | 0.13370 | 0.00306 | 0.01329 | 0.34397 | 2.81742 | 2.70960 |
| **Zc2hc1a** | 0.01923 | 0.00440 | 0.05809 | 0.02188 | 0.17959 | 0.14693 | 0.00412 | 0.00475 | 0.00886 | 0.00600 | 0.00828 | 0.03069 | 0.01163 | 0.10198 |
| **Zc2hc1c** | 0.28547 | 0.00438 | 0.14809 | 0.00301 | 0.02429 | 0.00594 | 0.00433 | 0.01120 | 0.01904 | 0.01486 | 0.00624 | 0.00971 | 0.02568 | 0.02145 |

| | | | | | | | | | | | | | | |
|---|---|---|---|---|---|---|---|---|---|---|---|---|---|---|
| **Zc3h10** | 0.52774 | 2.80063 | 1.60584 | 3.59924 | 1.97544 | 1.46324 | 3.17173 | 1.39902 | 0.00008 | 0.05058 | 2.23592 | 0.00000 | 1.21029 | 0.28614 |
| **Zc3h11a** | 6.10264 | 0.36557 | 7.59828 | 6.29324 | 10.53902 | 15.99053 | 4.31536 | 6.18231 | 11.24671 | 0.15383 | 9.26215 | 7.41305 | 12.24698 | 14.03319 |
| **Zc3h12a** | 1.82157 | 0.10835 | 1.02006 | 0.65160 | 2.12602 | 0.18975 | 0.02230 | 0.01591 | 1.10548 | 0.08407 | 0.00474 | 0.05081 | 3.64825 | 0.39821 |
| **Zc3h12c** | 3.51213 | 0.00333 | 1.29548 | 0.00000 | 0.65904 | 0.00000 | 0.00249 | 0.15098 | 11.28378 | 0.00886 | 0.00255 | 0.00000 | 1.09041 | 4.65908 |
| **Zc3h13** | 2.86163 | 0.04652 | 5.91774 | 2.54605 | 6.30786 | 4.08885 | 0.00103 | 1.09144 | 4.58059 | 0.00000 | 3.78836 | 3.84486 | 6.87099 | 8.99299 |
| **Zc3h14** | 10.28153 | 0.01082 | 10.24269 | 5.76025 | 10.92375 | 4.44701 | 1.89331 | 0.29544 | 9.97861 | 0.07791 | 7.13052 | 6.39455 | 2.49974 | 10.87695 |
| **Zc3h15** | 23.12135 | 3.49884 | 14.31978 | 13.89653 | 22.00543 | 14.26881 | 4.82254 | 3.54881 | 17.87331 | 0.24166 | 8.06423 | 10.52725 | 13.80384 | 17.71004 |
| **Zc3h18** | 5.25506 | 4.00455 | 6.72129 | 3.97225 | 5.21823 | 2.77856 | 0.09932 | 1.40259 | 0.63749 | 0.01364 | 0.47304 | 2.95150 | 6.56157 | 3.83211 |
| **Zc3h3** | 1.43724 | 0.09430 | 1.71752 | 1.55353 | 3.22254 | 1.06116 | 0.02861 | 1.09549 | 0.05368 | 0.00124 | 0.46608 | 1.10031 | 1.68136 | 0.72760 |
| **Zc3h4** | 0.06669 | 0.01020 | 0.63736 | 0.68592 | 3.80676 | 2.59890 | 1.33122 | 2.61394 | 2.79204 | 0.01526 | 0.00000 | 2.18554 | 1.52744 | 3.72254 |
| **Zc3h6** | 0.02296 | 0.00449 | 0.02690 | 0.00621 | 0.05011 | 0.06873 | 0.00416 | 0.00661 | 0.01034 | 0.00697 | 0.00840 | 0.01244 | 0.20664 | 0.02177 |
| **Zc3h7a** | 2.69482 | 0.01337 | 1.12743 | 1.64017 | 1.41195 | 1.50011 | 0.05965 | 0.50175 | 2.83542 | 0.00000 | 3.11006 | 0.77133 | 0.00000 | 3.51340 |
| **Zc3h7b** | 0.26920 | 0.00373 | 0.11642 | 0.00000 | 0.43072 | 0.19288 | 0.00421 | 0.00314 | 0.09318 | 0.00528 | 0.00243 | 0.41885 | 0.76238 | 0.09675 |
| **Zc3h8** | 5.54444 | 0.00256 | 5.33547 | 1.17666 | 7.06545 | 3.80390 | 0.00257 | 0.52311 | 6.63482 | 0.03172 | 0.10080 | 1.52962 | 0.88880 | 10.04497 |
| **Zc3hav1** | 0.98353 | 7.92786 | 0.00000 | 0.32784 | 0.96717 | 0.56678 | 0.00576 | 2.48273 | 0.00000 | 0.00110 | 1.58120 | 4.14751 | 1.07369 | 0.00000 |
| **Zc3hav1l** | 0.00924 | 0.02505 | 0.01561 | 0.00000 | 0.00643 | 0.42874 | 0.01004 | 0.03277 | 0.03009 | 0.01241 | 0.00608 | 0.05024 | 1.40484 | 0.00000 |
| **Zc3hc1** | 15.32220 | 6.27846 | 30.95360 | 12.27090 | 17.75128 | 6.91916 | 2.76811 | 2.98782 | 6.95669 | 0.14148 | 14.40143 | 7.89750 | 11.03592 | 18.51280 |
| **Zcchc10** | 3.93934 | 0.04367 | 6.27400 | 5.60089 | 6.69508 | 7.69506 | 1.12357 | 0.00000 | 6.84941 | 0.01622 | 3.65780 | 4.27075 | 7.91836 | 8.21460 |
| **Zcchc11** | 1.44866 | 0.00000 | 1.39180 | 0.05062 | 0.47363 | 1.78222 | 0.06476 | 0.12678 | 2.09042 | 6.48279 | 6.95162 | 1.51610 | 0.78846 | 0.23192 |
| **Zcchc13** | 0.00993 | 0.00444 | 0.01369 | 0.00278 | 0.01315 | 0.11501 | 0.00430 | 0.00620 | 0.01751 | 0.00594 | 0.00938 | 0.01120 | 0.02875 | 0.03110 |
| **Zcchc14** | 0.12740 | 0.01173 | 0.90150 | 0.00000 | 0.40606 | 0.60884 | 0.00434 | 0.47578 | 0.03419 | 0.02825 | 0.93608 | 1.08547 | 0.92148 | 3.06208 |
| **Zcchc16** | 0.00291 | 0.00412 | 0.01343 | 0.00563 | 0.00942 | 0.01537 | 0.00441 | 0.00841 | 0.01267 | 0.00655 | 0.00658 | 0.20264 | 0.57101 | 0.00299 |
| **Zcchc17** | 2.58290 | 0.06099 | 10.12380 | 5.68154 | 3.61132 | 3.46564 | 2.00021 | 5.28920 | 0.57108 | 2.82748 | 0.93707 | 6.30809 | 1.67439 | 1.62669 |
| **Zcchc2** | 0.95433 | 0.01087 | 0.71165 | 1.14382 | 0.40092 | 0.50932 | 1.14688 | 1.18201 | 0.59095 | 0.01017 | 0.54050 | 0.85327 | 4.19733 | 0.00000 |
| **Zcchc3** | 3.78523 | 0.41047 | 5.71151 | 2.97894 | 4.98717 | 5.97792 | 1.37356 | 2.13694 | 0.64717 | 0.06026 | 2.44247 | 2.48765 | 1.32365 | 1.81377 |
| **Zcchc4** | 1.13854 | 0.09851 | 2.98841 | 1.15841 | 3.31305 | 2.36720 | 0.00254 | 1.04109 | 1.76486 | 0.04814 | 1.96349 | 2.14618 | 1.05614 | 3.74739 |
| **Zcchc6** | 0.07137 | 0.00213 | 3.00784 | 0.00000 | 0.75868 | 2.07995 | 1.17115 | 0.56108 | 2.51243 | 4.28614 | 1.09957 | 0.83008 | 0.56866 | 5.05624 |
| **Zcchc7** | 4.17317 | 0.02410 | 4.73483 | 1.87064 | 3.33773 | 3.11722 | 0.01870 | 0.00000 | 5.42727 | 0.00000 | 4.55251 | 2.99657 | 3.57030 | 6.25906 |
| **Zcchc8** | 2.80391 | 0.24256 | 7.08293 | 3.79510 | 9.31453 | 4.59003 | 2.28735 | 4.14130 | 0.89439 | 0.02790 | 7.79148 | 7.96642 | 1.20421 | 6.88754 |
| **Zcchc9** | 5.34893 | 0.12635 | 5.19880 | 0.91948 | 3.80134 | 2.39549 | 2.27825 | 3.25982 | 4.63185 | 0.05065 | 0.59921 | 3.85157 | 2.36615 | 7.25127 |
| **Zcrb1** | 6.30447 | 0.03311 | 0.62855 | 4.37866 | 11.10992 | 4.56186 | 1.24945 | 0.00007 | 0.31773 | 0.05714 | 1.92387 | 1.57826 | 8.19866 | 2.70566 |
| **Zcwpw1** | 1.46801 | 0.00307 | 3.25188 | 0.35774 | 1.61195 | 2.58924 | 0.00248 | 0.00000 | 4.85264 | 0.00300 | 3.24002 | 0.13391 | 0.00000 | 3.36311 |
| **Zdbf2** | 0.01768 | 0.02670 | 0.13546 | 0.27619 | 0.15633 | 0.30208 | 0.00351 | 0.13294 | 0.00649 | 0.01642 | 0.55196 | 0.15869 | 0.12405 | 0.39474 |
| **Zdhhc11** | 0.01294 | 0.00469 | 0.13475 | 0.00750 | 0.00615 | 0.00501 | 0.00413 | 0.01812 | 0.30794 | 0.01225 | 0.00615 | 0.00858 | 0.01658 | 0.00433 |
| **Zdhhc12** | 0.30272 | 5.37160 | 3.25652 | 6.35068 | 0.53219 | 3.23063 | 0.97787 | 3.74572 | 0.00263 | 11.75161 | 1.20958 | 7.29277 | 0.25200 | 0.00000 |
| **Zdhhc13** | 0.20740 | 0.00358 | 0.55026 | 0.00000 | 1.07535 | 1.52480 | 0.00387 | 0.29633 | 0.81377 | 0.00451 | 0.27568 | 1.03723 | 0.00260 | 1.65128 |
| **Zdhhc14** | 0.00895 | 0.00429 | 0.01272 | 0.05686 | 0.01858 | 0.01053 | 0.00419 | 0.00799 | 0.01063 | 0.00670 | 0.00755 | 0.01322 | 0.12433 | 0.00361 |
| **Zdhhc15** | 0.00394 | 0.01019 | 0.00000 | 0.34402 | 0.00425 | 0.00000 | 0.00379 | 0.00000 | 0.01389 | 0.00198 | 1.00244 | 0.72967 | 0.00745 | 0.00679 |
| **Zdhhc16** | 1.13659 | 0.03863 | 2.90626 | 1.47368 | 0.69637 | 3.19366 | 0.53962 | 3.53681 | 0.32252 | 0.04583 | 3.06667 | 6.23940 | 1.04723 | 0.12244 |
| **Zdhhc17** | 0.00000 | 0.01894 | 0.95030 | 0.71396 | 1.60431 | 2.61137 | 2.11165 | 3.10455 | 5.62809 | 0.01901 | 1.33578 | 0.93716 | 0.47608 | 2.07151 |
| **Zdhhc18** | 1.71009 | 1.34172 | 5.27984 | 0.92036 | 8.55076 | 3.23353 | 0.06096 | 3.25184 | 0.00000 | 0.04698 | 0.20316 | 2.12177 | 1.44114 | 0.53421 |
| **Zdhhc19** | 0.00775 | 0.00419 | 0.00547 | 0.02197 | 0.12703 | 0.00000 | 0.00422 | 0.01194 | 0.01077 | 0.00414 | 0.80245 | 0.00391 | 0.00708 | 0.02218 |
| **Zdhhc2** | 0.01077 | 0.00257 | 0.28980 | 0.00000 | 0.20394 | 1.86834 | 0.00302 | 0.00317 | 1.94249 | 0.01755 | 0.00000 | 2.33337 | 0.02737 | 0.01549 |
| **Zdhhc20** | 13.74610 | 0.93012 | 8.74793 | 1.46843 | 7.04455 | 9.16527 | 1.58500 | 1.82160 | 22.75342 | 0.19312 | 2.70932 | 4.74457 | 4.13951 | 15.29828 |
| **Zdhhc21** | 0.02934 | 0.00471 | 0.00238 | 0.00000 | 0.01395 | 0.77154 | 1.65027 | 0.02466 | 0.38222 | 0.05255 | 0.00124 | 0.14266 | 0.11771 | 0.00000 |
| **Zdhhc24** | 0.19569 | 0.02268 | 0.00000 | 1.01160 | 0.00000 | 0.35526 | 0.09402 | 0.00234 | 1.35612 | 0.01044 | 0.00000 | 4.05577 | 0.20657 | 0.10284 |
| **Zdhhc3** | 2.79469 | 0.28847 | 4.71778 | 7.49548 | 4.87214 | 7.85247 | 0.92204 | 5.62207 | 5.93753 | 0.15955 | 3.77715 | 2.43934 | 1.15675 | 4.82007 |
| **Zdhhc4** | 5.47169 | 0.13693 | 3.43585 | 4.11503 | 1.85044 | 5.21969 | 0.13964 | 8.36730 | 0.00689 | 13.52013 | 4.82135 | 3.64915 | 0.42781 | 0.77979 |
| **Zdhhc5** | 1.38651 | 0.02316 | 1.05917 | 0.84883 | 4.98650 | 3.21413 | 0.02516 | 0.97484 | 2.62960 | 0.03167 | 3.23504 | 3.24852 | 8.63624 | 4.34746 |
| **Zdhhc6** | 4.30035 | 0.04318 | 3.16433 | 2.39410 | 1.37657 | 7.02460 | 3.09799 | 3.86162 | 1.36491 | 2.91863 | 4.82747 | 8.35866 | 3.74386 | 2.33992 |
| **Zdhhc7** | 7.30460 | 0.05247 | 12.03962 | 4.75289 | 18.79201 | 3.69802 | 3.01608 | 2.84786 | 0.01002 | 0.09480 | 0.87493 | 2.18623 | 10.34067 | 9.16158 |
| **Zdhhc8** | 0.11404 | 0.00461 | 0.11626 | 0.25945 | 0.07232 | 0.00000 | 0.00547 | 0.22269 | 0.01117 | 0.00841 | 0.00802 | 0.00988 | 0.17265 | 0.01085 |
| **Zdhhc9** | 0.04367 | 0.00397 | 0.19051 | 0.00000 | 1.06105 | 0.34035 | 0.00406 | 0.12916 | 0.03932 | 0.00641 | 0.01279 | 0.00390 | 0.00331 | 0.58836 |
| **Zeb1** | 0.01505 | 0.00294 | 0.00729 | 0.02147 | 0.09238 | 0.00000 | 0.00893 | 0.00257 | 0.00763 | 3.56092 | 0.00484 | 0.46744 | 0.63210 | 0.06145 |
| **Zeb2** | 0.07439 | 0.00446 | 0.00548 | 0.01165 | 0.39094 | 0.01257 | 0.00473 | 0.01097 | 0.02616 | 0.00648 | 0.00796 | 0.00777 | 0.00403 | 0.00261 |

| | | | | | | | | | | | | | | |
|---|---|---|---|---|---|---|---|---|---|---|---|---|---|---|
| **Zer1** | 0.55122 | 0.02563 | 0.92115 | 0.00000 | 0.85257 | 2.19922 | 0.01443 | 2.77853 | 0.00154 | 0.00381 | 0.00000 | 2.13231 | 0.72948 | 0.00442 |
| **Zfa.ps** | 0.07726 | 0.00417 | 0.00932 | 0.04937 | 0.16572 | 0.01279 | 0.09101 | 0.01942 | 0.22147 | 0.00753 | 0.03234 | 0.00741 | 0.14578 | 0.15395 |
| **Zfand1** | 1.18964 | 0.04052 | 1.42714 | 2.60875 | 2.98978 | 3.33276 | 0.36833 | 1.91005 | 0.31259 | 0.01583 | 4.14457 | 6.77651 | 0.38897 | 1.86071 |
| **Zfand2a** | 9.87011 | 0.06054 | 16.78766 | 15.28238 | 18.53252 | 10.92643 | 7.42721 | 1.50106 | 15.86732 | 14.38544 | 10.99506 | 4.53050 | 17.15785 | 17.55227 |
| **Zfand2b** | 3.57456 | 0.10247 | 11.30155 | 8.90726 | 6.09896 | 4.42984 | 1.73466 | 5.58789 | 0.25853 | 7.32631 | 2.20904 | 3.34937 | 1.15239 | 0.71567 |
| **Zfand3** | 2.03081 | 5.06524 | 5.17130 | 5.94186 | 8.31594 | 2.86116 | 0.66604 | 3.58836 | 0.02682 | 0.10401 | 3.15841 | 3.47370 | 1.21073 | 1.32019 |
| **Zfand4** | 0.01062 | 0.00429 | 0.13074 | 0.01076 | 0.08619 | 0.00424 | 0.00419 | 0.00955 | 0.01417 | 0.00774 | 0.17403 | 0.00935 | 0.35276 | 0.25336 |
| **Zfand5** | 20.03462 | 0.12387 | 9.36487 | 6.43221 | 18.66670 | 5.27470 | 1.28863 | 1.79211 | 24.77325 | 0.02857 | 4.90359 | 2.84006 | 27.28541 | 10.81591 |
| **Zfand6** | 1.05185 | 0.03195 | 1.01661 | 3.16605 | 6.53262 | 3.39948 | 1.96228 | 5.78530 | 9.29614 | 0.06848 | 2.71640 | 2.27895 | 5.84244 | 1.71805 |
| **Zfat** | 0.09001 | 0.00455 | 0.12723 | 0.13315 | 0.56975 | 0.88626 | 0.00393 | 0.37166 | 0.00699 | 0.00479 | 0.42937 | 0.07023 | 0.00837 | 1.04867 |
| **Zfc3h1** | 2.21434 | 0.00182 | 0.41896 | 0.00000 | 0.15821 | 1.52894 | 2.27420 | 0.00000 | 8.73552 | 0.02718 | 1.40883 | 1.18204 | 5.16428 | 1.54337 |
| **Zfhx2** | 0.01066 | 0.00371 | 0.16065 | 0.01025 | 0.02896 | 0.00480 | 0.76443 | 0.00256 | 0.00885 | 0.02121 | 0.00275 | 0.38034 | 0.02005 | 0.01108 |
| **Zfhx3** | 0.09906 | 0.03700 | 0.13750 | 0.08086 | 0.71571 | 0.53901 | 0.85597 | 1.42676 | 0.78556 | 4.50937 | 0.46188 | 0.13812 | 0.61480 | 0.00000 |
| **Zfp1** | 1.46710 | 0.03787 | 3.51631 | 2.27727 | 4.91144 | 2.38875 | 1.57049 | 0.00000 | 0.79745 | 0.00985 | 3.45520 | 6.09842 | 3.63696 | 4.13189 |
| **Zfp101** | 0.95934 | 0.00374 | 0.25421 | 1.21140 | 0.81136 | 0.00003 | 0.00288 | 0.01765 | 6.99028 | 0.00000 | 2.98066 | 0.08483 | 1.79597 | 0.66296 |
| **Zfp105** | 3.91634 | 0.02340 | 0.74671 | 2.62592 | 0.42772 | 0.99005 | 0.00251 | 1.15188 | 0.00516 | 0.01375 | 0.62792 | 3.24177 | 0.00000 | 1.04877 |
| **Zfp106** | 8.77311 | 0.97623 | 13.65208 | 8.54408 | 19.85605 | 10.37736 | 2.61871 | 7.98565 | 23.03287 | 0.15007 | 15.81753 | 10.16393 | 4.40700 | 25.28302 |
| **Zfp108** | 0.00000 | 0.00407 | 0.51260 | 0.00000 | 0.00066 | 0.17566 | 0.00400 | 0.00189 | 0.00675 | 0.00530 | 0.04868 | 0.01665 | 0.81755 | 0.01469 |
| **Zfp109** | 0.00758 | 0.00500 | 0.20410 | 0.37665 | 0.05530 | 0.02750 | 0.10857 | 0.03574 | 0.01866 | 0.00640 | 0.00443 | 0.17537 | 0.01064 | 0.04989 |
| **Zfp11** | 0.06855 | 0.00406 | 0.11923 | 0.01195 | 0.01685 | 0.24887 | 0.02866 | 0.00680 | 0.65031 | 0.31672 | 0.02615 | 0.07196 | 1.16411 | 0.12366 |
| **Zfp110** | 2.33438 | 2.26577 | 6.20075 | 5.31348 | 5.84480 | 5.46756 | 1.03145 | 4.04329 | 8.61359 | 0.05450 | 7.20652 | 4.52104 | 1.55461 | 3.82228 |
| **Zfp111** | 0.01681 | 0.20224 | 0.75244 | 0.35263 | 0.33739 | 0.70281 | 0.04594 | 0.06983 | 0.08939 | 0.01810 | 0.00000 | 1.02223 | 0.17355 | 0.04330 |
| **Zfp112** | 0.00502 | 0.00449 | 0.26025 | 0.17112 | 0.33216 | 0.39493 | 0.00489 | 0.00341 | 0.02017 | 0.00497 | 0.00461 | 0.00390 | 0.00377 | 0.00000 |
| **Zfp113** | 1.40053 | 0.10654 | 0.75361 | 0.82939 | 1.41166 | 2.36003 | 0.23955 | 0.12273 | 5.50118 | 0.59286 | 1.25738 | 0.81537 | 8.74472 | 3.40036 |
| **Zfp114** | 0.01198 | 0.00440 | 0.03906 | 0.00000 | 0.01279 | 0.03327 | 0.00426 | 0.01314 | 0.01184 | 0.01162 | 0.66232 | 0.21767 | 0.02506 | 0.00887 |
| **Zfp119a** | 0.00890 | 0.00404 | 0.03925 | 0.21708 | 0.02726 | 0.02749 | 0.01487 | 0.00502 | 0.31850 | 0.02949 | 0.03128 | 0.06317 | 0.00343 | 0.02884 |
| **Zfp119b** | 0.04862 | 0.01178 | 0.37174 | 0.00000 | 0.30058 | 0.07538 | 0.02856 | 0.00816 | 0.55641 | 0.18265 | 0.10950 | 0.20348 | 0.04727 | 0.26350 |
| **Zfp12** | 1.78352 | 0.09583 | 2.36212 | 0.00000 | 1.33093 | 0.85906 | 0.08796 | 0.18049 | 1.58849 | 0.06770 | 0.15866 | 3.18083 | 3.33259 | 2.14917 |
| **Zfp120** | 0.15581 | 0.09721 | 0.71988 | 1.09241 | 0.78254 | 1.07908 | 0.04549 | 0.01328 | 2.31293 | 0.00497 | 0.16092 | 1.24807 | 2.96732 | 2.13530 |
| **Zfp13** | 0.00515 | 0.00321 | 0.38598 | 0.39806 | 0.14591 | 0.04299 | 0.00376 | 0.82324 | 0.00781 | 0.01254 | 0.00000 | 1.14372 | 0.01244 | 0.02824 |
| **Zfp131** | 11.89952 | 0.02709 | 6.41185 | 1.60258 | 12.93312 | 4.81682 | 0.76104 | 1.67135 | 20.52641 | 0.00483 | 1.88993 | 0.76110 | 14.67568 | 15.26162 |
| **Zfp133.ps** | 0.03348 | 0.00450 | 0.09213 | 0.00000 | 0.01042 | 0.18087 | 0.00419 | 0.00671 | 0.01066 | 0.00767 | 0.00618 | 0.01073 | 0.00769 | 0.00887 |
| **Zfp14** | 0.00807 | 0.00491 | 0.19014 | 0.00000 | 0.00000 | 0.50700 | 0.00380 | 0.00695 | 0.77048 | 0.00687 | 0.00721 | 0.08815 | 0.25953 | 0.00000 |
| **Zfp141** | 1.89097 | 0.05913 | 2.17942 | 2.92473 | 2.66307 | 2.94368 | 0.52760 | 0.55814 | 5.70829 | 0.03981 | 4.08213 | 0.93921 | 3.23065 | 2.82224 |
| **Zfp142** | 0.95851 | 8.64548 | 3.75360 | 2.25522 | 4.65758 | 1.83328 | 0.00054 | 1.21094 | 0.62209 | 0.01991 | 4.68744 | 0.50811 | 2.88502 | 1.84360 |
| **Zfp143** | 14.88460 | 0.00518 | 12.03441 | 2.03131 | 13.63137 | 4.15633 | 0.13730 | 0.96679 | 21.46797 | 6.61464 | 4.83802 | 1.42030 | 7.42826 | 20.07007 |
| **Zfp146** | 0.04590 | 0.00460 | 0.04016 | 0.01889 | 0.00267 | 0.00765 | 0.00417 | 0.02412 | 0.01371 | 0.00867 | 0.16919 | 0.00501 | 0.00402 | 0.27940 |
| **Zfp148** | 0.83558 | 0.00000 | 6.12993 | 0.96859 | 7.48001 | 4.84925 | 0.11038 | 0.50124 | 15.87982 | 0.00000 | 3.84331 | 3.36454 | 5.07799 | 6.17874 |
| **Zfp157** | 0.00000 | 0.00272 | 1.01568 | 0.02767 | 0.00000 | 1.56358 | 0.00338 | 0.01715 | 0.00431 | 0.00161 | 1.84643 | 1.18782 | 0.00085 | 0.15200 |
| **Zfp160** | 0.78395 | 0.00311 | 0.50195 | 1.22315 | 0.54135 | 0.47259 | 0.55442 | 0.00000 | 5.49783 | 0.02517 | 0.00581 | 1.00621 | 0.23243 | 0.00000 |
| **Zfp174** | 0.00697 | 0.02108 | 0.00000 | 0.10194 | 0.00981 | 0.01164 | 0.00430 | 0.03055 | 1.16059 | 0.00937 | 0.02179 | 0.02558 | 0.04395 | 0.01437 |
| **Zfp180** | 8.09170 | 0.00974 | 4.90838 | 1.05437 | 6.82629 | 4.20366 | 1.11092 | 0.70025 | 7.58480 | 0.02399 | 0.56569 | 1.35246 | 0.05958 | 11.02774 |
| **Zfp182** | 0.00246 | 0.00382 | 0.07517 | 0.00000 | 0.60750 | 0.00000 | 0.00337 | 0.07326 | 1.52331 | 0.00000 | 3.13722 | 0.25104 | 0.19460 | 0.00000 |
| **Zfp184** | 0.00000 | 0.01425 | 0.21974 | 0.99432 | 0.90465 | 0.30553 | 0.00377 | 0.21697 | 3.32591 | 0.00433 | 1.20891 | 0.34867 | 0.53500 | 0.00000 |
| **Zfp185** | 0.00708 | 0.00445 | 0.00853 | 0.01966 | 0.01145 | 0.40384 | 0.00414 | 0.00636 | 0.02001 | 0.00633 | 0.00515 | 0.00651 | 0.07052 | 0.01030 |
| **Zfp189** | 0.01339 | 0.00436 | 0.01488 | 0.01222 | 0.01079 | 0.01333 | 0.00424 | 0.00945 | 0.01108 | 0.00663 | 0.00883 | 0.00942 | 0.26614 | 0.00271 |
| **Zfp2** | 0.32604 | 0.00344 | 0.13865 | 0.19957 | 0.01886 | 0.45581 | 0.00395 | 0.00224 | 1.13104 | 0.00481 | 0.00000 | 0.75111 | 0.00207 | 0.00000 |
| **Zfp202** | 0.05887 | 0.01983 | 0.01480 | 0.00900 | 0.02675 | 0.68169 | 0.01124 | 0.00899 | 0.17898 | 0.04000 | 0.01467 | 0.02286 | 1.09451 | 0.06057 |
| **Zfp207** | 11.56373 | 4.44718 | 28.16696 | 6.59679 | 29.34005 | 20.89724 | 4.55323 | 9.50356 | 24.02221 | 0.13575 | 14.10704 | 13.76213 | 13.07762 | 32.76292 |
| **Zfp212** | 1.37031 | 0.02116 | 3.44659 | 2.67140 | 2.13923 | 2.86560 | 3.75804 | 2.42219 | 1.91467 | 0.04159 | 0.97281 | 0.21482 | 1.72497 | 0.56677 |
| **Zfp213** | 0.91356 | 0.00307 | 0.14878 | 1.32438 | 0.36979 | 0.09678 | 0.01033 | 1.12316 | 0.63246 | 0.00655 | 0.14141 | 1.31206 | 0.00546 | 0.00000 |
| **Zfp217** | 2.81899 | 0.04589 | 3.69098 | 3.44501 | 10.99146 | 1.82621 | 2.00691 | 2.15724 | 9.44890 | 0.05395 | 3.25716 | 3.45300 | 7.35935 | 7.72554 |
| **Zfp219** | 0.00000 | 2.63009 | 1.18310 | 0.78532 | 1.26944 | 0.01557 | 0.34852 | 1.57082 | 0.00987 | 0.02315 | 0.00000 | 1.00611 | 0.00419 | 0.09390 |
| **Zfp229** | 0.33030 | 0.34486 | 0.47781 | 0.75075 | 0.28487 | 1.20887 | 0.97308 | 0.24272 | 4.84610 | 0.06551 | 0.00000 | 0.38196 | 2.25193 | 1.13269 |
| **Zfp235** | 0.58853 | 0.00379 | 0.11693 | 0.20465 | 0.45130 | 0.57622 | 0.00413 | 0.78888 | 0.01974 | 0.03547 | 0.00000 | 0.36377 | 0.32745 | 0.27511 |

| | | | | | | | | | | | | | | |
|---|---|---|---|---|---|---|---|---|---|---|---|---|---|---|
| **Zfp236** | 3.08696 | 0.02472 | 0.94627 | 0.95592 | 0.63305 | 0.99147 | 0.34590 | 0.45158 | 3.85285 | 0.03568 | 2.41633 | 2.18901 | 1.50538 | 1.60145 |
| **Zfp239** | 9.44694 | 0.05676 | 3.57948 | 2.87910 | 10.68484 | 4.23178 | 0.00184 | 0.04477 | 4.00635 | 0.00000 | 5.97629 | 1.08017 | 4.98141 | 13.63869 |
| **Zfp24** | 1.25195 | 0.02996 | 1.50559 | 0.19655 | 0.47752 | 1.54282 | 0.00277 | 1.09799 | 2.59958 | 0.00000 | 3.11084 | 1.21379 | 2.99252 | 0.00000 |
| **Zfp248** | 0.00951 | 0.01911 | 0.16468 | 0.11891 | 0.00589 | 0.15285 | 0.00399 | 0.49274 | 0.06855 | 0.00400 | 0.91646 | 0.16533 | 0.13841 | 0.13154 |
| **Zfp251** | 0.55510 | 0.02978 | 2.07831 | 0.00000 | 2.43111 | 2.75157 | 0.00322 | 0.46622 | 0.95676 | 0.01987 | 1.60150 | 0.35136 | 0.00000 | 3.13931 |
| **Zfp26** | 0.00000 | 0.02489 | 0.00000 | 2.21283 | 0.06003 | 1.07336 | 1.07752 | 0.01029 | 4.64518 | 0.01179 | 0.19915 | 1.15969 | 0.97391 | 0.00000 |
| **Zfp260** | 1.94683 | 0.07658 | 0.82248 | 0.12341 | 0.00000 | 1.70290 | 0.00333 | 0.01602 | 2.84271 | 0.00457 | 0.00000 | 0.68224 | 0.00000 | 1.60880 |
| **Zfp263** | 0.00000 | 0.01121 | 1.50644 | 1.57466 | 0.80568 | 1.30098 | 2.71246 | 0.49447 | 2.73301 | 0.02776 | 0.56492 | 0.65484 | 1.68915 | 2.23193 |
| **Zfp266** | 0.04200 | 0.00436 | 0.00976 | 0.06189 | 0.03568 | 0.01036 | 0.00434 | 0.00781 | 0.00965 | 0.00647 | 0.00876 | 0.01331 | 0.00609 | 0.10643 |
| **Zfp27** | 0.00000 | 0.00515 | 0.64761 | 0.29413 | 0.47724 | 0.33209 | 0.00422 | 0.15441 | 0.01593 | 0.00599 | 0.00101 | 0.36509 | 0.53712 | 0.00000 |
| **Zfp273** | 0.08222 | 0.00401 | 0.23568 | 0.48817 | 0.17957 | 0.08094 | 0.01491 | 0.00000 | 1.55857 | 0.00343 | 0.73314 | 0.01302 | 0.43117 | 0.22822 |
| **Zfp275** | 0.11329 | 0.01153 | 0.52160 | 0.00000 | 0.02283 | 1.05730 | 0.01347 | 0.72281 | 0.01047 | 0.00426 | 0.00000 | 0.68024 | 0.07844 | 0.00000 |
| **Zfp276** | 0.00000 | 0.00389 | 0.78619 | 1.93132 | 0.52388 | 1.16813 | 0.00397 | 1.26408 | 0.41293 | 0.01196 | 0.00000 | 1.21227 | 0.99056 | 0.17324 |
| **Zfp277** | 4.63614 | 0.38070 | 4.58894 | 1.51031 | 4.71789 | 4.60163 | 0.07101 | 4.47778 | 2.35913 | 0.01330 | 6.42390 | 1.42175 | 0.61158 | 1.14039 |
| **Zfp28** | 0.00000 | 0.00431 | 1.51549 | 0.01137 | 0.51550 | 0.43675 | 0.00396 | 0.00000 | 0.00184 | 0.01146 | 0.00395 | 0.00319 | 0.42197 | 1.06387 |
| **Zfp280b** | 11.60272 | 0.99973 | 14.53714 | 6.95618 | 18.80862 | 14.14504 | 4.18108 | 8.28130 | 3.60134 | 10.82945 | 9.07797 | 7.14267 | 13.60001 | 19.45369 |
| **Zfp280c** | 0.00204 | 0.55506 | 0.00516 | 0.00523 | 0.00000 | 0.21587 | 0.00388 | 0.14632 | 0.06697 | 0.00629 | 0.00619 | 0.00577 | 0.49280 | 0.37068 |
| **Zfp280d** | 5.06672 | 0.00227 | 4.26331 | 0.41006 | 5.98086 | 1.77424 | 0.00190 | 0.43279 | 11.48848 | 0.00791 | 3.04789 | 0.46930 | 0.84406 | 3.70176 |
| **Zfp281** | 6.28690 | 3.04658 | 6.58320 | 4.85037 | 6.62100 | 8.54423 | 1.58067 | 0.00000 | 15.88937 | 0.08928 | 3.83626 | 1.71717 | 9.36496 | 6.52873 |
| **Zfp282** | 0.38617 | 0.05317 | 0.69669 | 1.48998 | 0.15669 | 0.58274 | 0.03826 | 2.98671 | 0.07900 | 0.02247 | 0.11474 | 0.99014 | 0.68754 | 1.44786 |
| **Zfp287** | 0.03717 | 0.01198 | 0.01712 | 0.05663 | 0.01638 | 0.04990 | 0.01012 | 0.01670 | 0.10335 | 0.02747 | 0.02828 | 0.02258 | 0.15283 | 0.04577 |
| **Zfp292** | 2.41651 | 0.00212 | 1.23118 | 0.90375 | 3.29178 | 1.04276 | 0.00137 | 0.05153 | 14.47237 | 0.02435 | 0.05914 | 2.51978 | 1.97914 | 5.45514 |
| **Zfp296** | 2.07559 | 3.83005 | 26.90912 | 6.28337 | 30.47920 | 0.59674 | 9.41574 | 12.88331 | 0.00000 | 12.69809 | 2.83714 | 8.56382 | 9.46730 | 2.36854 |
| **Zfp3** | 0.02145 | 0.00486 | 0.35402 | 0.09039 | 0.00000 | 0.03641 | 1.09926 | 0.00470 | 0.14424 | 0.06514 | 0.01253 | 0.00349 | 0.00304 | 0.66227 |
| **Zfp30** | 0.00149 | 0.00450 | 0.17602 | 0.00293 | 0.17586 | 0.07461 | 0.00476 | 0.00743 | 0.01139 | 0.00883 | 0.00685 | 0.03475 | 0.39067 | 0.00928 |
| **Zfp300** | 0.07488 | 0.01255 | 0.09077 | 0.14386 | 0.05639 | 0.07321 | 0.02805 | 0.01999 | 0.42556 | 0.00720 | 0.02165 | 0.07896 | 0.07742 | 0.26141 |
| **Zfp316** | 1.46685 | 0.42737 | 0.21566 | 1.23539 | 1.79172 | 1.41078 | 0.28557 | 0.06867 | 1.49857 | 0.54637 | 0.29899 | 0.30685 | 3.45228 | 2.03152 |
| **Zfp317** | 0.30231 | 3.41800 | 1.15416 | 1.70444 | 1.18194 | 0.35443 | 0.01416 | 0.03516 | 3.85061 | 0.02935 | 0.36946 | 2.24972 | 0.62912 | 2.34302 |
| **Zfp318** | 3.44593 | 0.01761 | 0.29037 | 0.00000 | 0.66055 | 0.70834 | 0.00850 | 0.00131 | 1.22377 | 1.81740 | 0.20640 | 1.86441 | 3.37977 | 0.58715 |
| **Zfp319** | 0.00000 | 0.00397 | 0.34936 | 0.56436 | 0.03871 | 0.37711 | 0.00398 | 0.00193 | 0.00338 | 0.00436 | 0.00574 | 0.05229 | 1.39276 | 0.02702 |
| **Zfp322a** | 1.39789 | 0.00134 | 0.28366 | 1.87740 | 0.00000 | 3.09513 | 0.03488 | 0.00000 | 3.17747 | 0.03701 | 2.02032 | 2.48107 | 1.30701 | 1.49399 |
| **Zfp324** | 0.00469 | 0.00399 | 0.06806 | 0.01427 | 0.30483 | 0.02748 | 0.00430 | 0.00203 | 0.00554 | 0.01321 | 1.54100 | 0.00640 | 0.99691 | 0.03433 |
| **Zfp326** | 0.12706 | 0.02748 | 0.00000 | 0.54294 | 0.03989 | 2.82367 | 0.12683 | 0.01166 | 0.34823 | 0.03898 | 0.00000 | 1.91589 | 0.08092 | 0.00007 |
| **Zfp329** | 1.17076 | 0.02794 | 0.62403 | 0.70819 | 0.00000 | 0.04570 | 0.26910 | 0.01806 | 2.54334 | 0.01420 | 0.60051 | 0.01193 | 1.61940 | 0.00000 |
| **Zfp330** | 7.43873 | 0.07368 | 4.00834 | 7.35099 | 3.06988 | 7.79256 | 3.86301 | 1.26449 | 4.88363 | 0.19492 | 6.89142 | 6.18319 | 7.18074 | 10.27151 |
| **Zfp334** | 0.04552 | 0.00426 | 0.01242 | 0.01438 | 0.00941 | 0.05519 | 0.00415 | 0.00571 | 0.01934 | 0.00830 | 0.00820 | 0.00773 | 0.55715 | 0.01209 |
| **Zfp335** | 2.80385 | 3.01712 | 6.96531 | 2.75107 | 10.25020 | 1.83899 | 0.61131 | 0.07328 | 0.23734 | 0.00000 | 1.47811 | 1.58022 | 4.38352 | 0.22550 |
| **Zfp341** | 0.05120 | 0.00343 | 0.32343 | 2.14768 | 2.22267 | 0.35119 | 0.00336 | 1.65490 | 0.00387 | 0.00120 | 1.16032 | 0.07349 | 0.45844 | 0.40182 |
| **Zfp345** | 1.71290 | 0.15580 | 1.56037 | 2.47157 | 1.27233 | 2.40429 | 2.20826 | 0.14081 | 6.16222 | 0.45837 | 0.69201 | 1.26372 | 0.41063 | 1.41010 |
| **Zfp346** | 0.16349 | 0.01126 | 0.79722 | 2.06048 | 1.05981 | 2.39575 | 0.00316 | 0.00000 | 1.03315 | 0.03864 | 1.55081 | 1.65395 | 1.00324 | 0.58263 |
| **Zfp35** | 0.81823 | 0.00242 | 4.09626 | 0.89182 | 3.93114 | 3.22331 | 0.00781 | 0.44947 | 1.44111 | 0.16556 | 2.26986 | 1.08597 | 0.53585 | 5.48198 |
| **Zfp352** | 0.00242 | 0.00405 | 0.01433 | 0.01440 | 0.00518 | 0.01484 | 0.00411 | 0.00753 | 0.02265 | 0.00615 | 0.01402 | 0.00818 | 2.30846 | 0.00180 |
| **Zfp354a** | 0.00764 | 0.00390 | 0.00497 | 0.51040 | 0.00491 | 0.06289 | 0.00383 | 0.00000 | 0.02719 | 0.00313 | 0.91934 | 0.01947 | 0.00497 | 0.00641 |
| **Zfp354b** | 0.00497 | 0.00416 | 0.00242 | 0.27687 | 0.00229 | 0.00008 | 0.00414 | 0.00833 | 0.14665 | 0.00622 | 0.03984 | 0.01127 | 0.00345 | 0.05977 |
| **Zfp354c** | 0.01596 | 0.00479 | 0.01588 | 0.03636 | 0.01105 | 0.01813 | 0.00405 | 0.01409 | 0.01178 | 0.00593 | 0.00450 | 0.32216 | 0.00728 | 0.01719 |
| **Zfp358** | 0.02027 | 0.00428 | 0.00629 | 0.07779 | 0.18235 | 0.01212 | 0.00433 | 0.01157 | 0.01007 | 0.01434 | 0.00712 | 0.00751 | 0.00724 | 0.01003 |
| **Zfp36** | 3.77322 | 5.22056 | 3.11082 | 0.48537 | 7.75188 | 0.00000 | 0.69216 | 4.78740 | 2.72606 | 0.13174 | 2.90202 | 0.00000 | 6.58816 | 1.95419 |
| **Zfp362** | 0.08100 | 0.00406 | 0.11954 | 0.00000 | 0.06326 | 0.90208 | 0.00448 | 0.01227 | 0.01897 | 0.01046 | 0.00000 | 0.38381 | 0.00491 | 0.00082 |
| **Zfp367** | 0.00000 | 0.00404 | 0.49539 | 0.24979 | 0.23027 | 0.56872 | 0.67262 | 0.31916 | 8.81785 | 0.02967 | 0.50233 | 0.11213 | 1.32433 | 1.30969 |
| **Zfp369** | 0.54404 | 0.00442 | 0.10065 | 0.00000 | 0.05961 | 0.28593 | 0.02995 | 0.03319 | 0.10363 | 0.00611 | 0.06643 | 0.05717 | 0.00635 | 0.09127 |
| **Zfp36l1** | 0.01786 | 0.02145 | 0.00000 | 0.22928 | 0.00420 | 0.14713 | 0.00285 | 0.65253 | 0.01564 | 0.04156 | 0.28561 | 5.17518 | 0.00550 | 0.01474 |
| **Zfp36l2** | 1.08839 | 1.62537 | 1.00448 | 3.08838 | 3.32528 | 2.17715 | 4.02329 | 10.29615 | 3.77403 | 0.32888 | 2.26354 | 1.27618 | 0.86584 | 2.13329 |
| **Zfp37** | 0.01603 | 0.01292 | 0.21144 | 0.06369 | 0.00491 | 0.36284 | 0.11371 | 0.01636 | 0.04589 | 0.00857 | 0.00818 | 0.00319 | 0.00636 | 0.01849 |
| **Zfp382** | 0.02046 | 0.00440 | 0.25590 | 0.00143 | 0.00697 | 0.00827 | 0.00421 | 0.01115 | 0.01101 | 0.00737 | 0.00827 | 0.00937 | 0.00588 | 0.01330 |
| **Zfp383** | 0.00000 | 0.00351 | 0.09030 | 0.13606 | 0.28272 | 0.00000 | 0.00367 | 0.38694 | 1.01035 | 0.01252 | 1.09196 | 0.09352 | 0.11713 | 0.42513 |

| | | | | | | | | | | | | | | |
|---|---|---|---|---|---|---|---|---|---|---|---|---|---|---|
| **Zfp384** | 0.03850 | 0.00535 | 0.12047 | 0.56531 | 0.21656 | 0.00000 | 0.00456 | 0.18066 | 0.15281 | 0.00863 | 0.00786 | 0.02985 | 0.66137 | 0.05933 |
| **Zfp385a** | 0.07077 | 0.00378 | 0.03950 | 0.17973 | 0.38724 | 0.09143 | 0.03216 | 0.05560 | 0.00831 | 0.01059 | 0.01823 | 0.84766 | 0.43203 | 0.00000 |
| **Zfp385c** | 0.13670 | 0.01192 | 0.01247 | 0.14683 | 0.14562 | 0.18799 | 0.02743 | 0.03192 | 0.15149 | 0.04208 | 0.01693 | 0.03963 | 1.07331 | 0.28626 |
| **Zfp386** | 0.33010 | 0.01151 | 2.83252 | 1.71333 | 3.30711 | 1.04394 | 0.00907 | 0.00000 | 2.99581 | 0.00000 | 0.04706 | 0.75987 | 0.00000 | 1.67674 |
| **Zfp39** | 0.01031 | 0.00384 | 0.01678 | 0.07926 | 0.08589 | 0.00000 | 0.00369 | 0.04167 | 0.01818 | 0.00181 | 1.53549 | 0.00726 | 0.36440 | 0.00000 |
| **Zfp395** | 0.01809 | 0.03396 | 1.08186 | 1.40976 | 0.69575 | 0.36400 | 0.02572 | 0.82033 | 0.04868 | 0.00997 | 2.00512 | 0.22985 | 0.01199 | 0.12365 |
| **Zfp397** | 0.60670 | 0.02569 | 0.36732 | 0.90075 | 0.11125 | 0.03779 | 0.01558 | 0.00000 | 1.54899 | 0.00104 | 1.65585 | 0.00970 | 0.97655 | 0.00000 |
| **Zfp398** | 0.00593 | 0.00373 | 0.00000 | 1.07360 | 0.23271 | 0.78187 | 2.74540 | 0.37767 | 0.00383 | 0.13527 | 0.08591 | 0.00000 | 1.73671 | 0.00000 |
| **Zfp40** | 0.00177 | 0.00439 | 0.01998 | 0.01386 | 0.07198 | 0.01433 | 0.01031 | 0.00870 | 0.02894 | 0.00572 | 0.01909 | 0.00816 | 0.23080 | 0.36672 |
| **Zfp407** | 0.00000 | 0.00000 | 3.10745 | 1.81119 | 3.31749 | 0.00000 | 1.06396 | 0.00000 | 1.45383 | 9.94533 | 2.98197 | 1.41394 | 4.97880 | 2.11057 |
| **Zfp408** | 0.91807 | 0.03841 | 0.33051 | 0.26446 | 0.34636 | 0.60310 | 0.44151 | 0.56303 | 1.72631 | 0.01451 | 1.71178 | 0.04611 | 0.54016 | 0.53494 |
| **Zfp41** | 0.00724 | 0.01971 | 0.01302 | 0.00002 | 0.01172 | 0.01830 | 0.00429 | 0.01706 | 0.01211 | 0.00672 | 0.00359 | 1.51464 | 0.00703 | 0.01070 |
| **Zfp410** | 4.54686 | 0.05977 | 8.25369 | 3.37888 | 6.87761 | 6.40291 | 4.45889 | 5.30368 | 6.08640 | 0.07264 | 6.54186 | 5.16659 | 8.05569 | 12.58393 |
| **Zfp414** | 0.71866 | 0.03581 | 3.92959 | 3.65331 | 2.88157 | 3.02113 | 0.02915 | 0.91901 | 0.00000 | 6.73765 | 3.49391 | 3.61232 | 0.00474 | 1.62418 |
| **Zfp418** | 0.00000 | 0.00398 | 0.64585 | 0.02249 | 0.17312 | 0.02977 | 0.00403 | 0.00262 | 0.01809 | 0.00392 | 0.50404 | 0.23491 | 0.28036 | 0.07031 |
| **Zfp42** | 1.10446 | 0.89409 | 12.77490 | 8.08303 | 4.36334 | 19.45212 | 6.09103 | 12.61902 | 3.62190 | 0.49832 | 17.87141 | 8.11370 | 7.19966 | 3.46913 |
| **Zfp420** | 0.32139 | 0.00407 | 1.25980 | 0.03790 | 0.89829 | 0.43226 | 0.00357 | 0.00000 | 1.90996 | 0.00381 | 0.00409 | 0.10356 | 0.62565 | 1.29293 |
| **Zfp422** | 0.24398 | 0.02084 | 0.05994 | 0.23743 | 0.30846 | 0.54736 | 0.03311 | 0.05404 | 0.83422 | 0.02543 | 0.01706 | 0.07487 | 0.15078 | 1.25104 |
| **Zfp423** | 0.00859 | 0.00440 | 0.00192 | 0.06761 | 0.49378 | 0.01200 | 0.00421 | 0.01053 | 0.05625 | 0.00681 | 0.00696 | 0.00662 | 0.00324 | 0.00000 |
| **Zfp426** | 0.97545 | 0.01122 | 1.43756 | 0.18223 | 0.13095 | 0.61094 | 0.96031 | 0.35132 | 3.15758 | 0.00955 | 1.09973 | 0.00804 | 0.89037 | 0.84884 |
| **Zfp428** | 0.08558 | 0.00403 | 0.01083 | 1.41816 | 1.09781 | 0.00922 | 0.00384 | 0.00520 | 0.02236 | 0.00571 | 0.00729 | 0.02657 | 0.03054 | 0.00653 |
| **Zfp429** | 0.00000 | 0.00409 | 0.17096 | 0.00000 | 0.34213 | 0.87074 | 1.06466 | 0.00304 | 0.00424 | 0.02299 | 0.11360 | 0.24664 | 0.34684 | 0.35741 |
| **Zfp433** | 2.57077 | 0.00240 | 3.79361 | 1.81050 | 6.93819 | 1.60235 | 0.01199 | 1.26851 | 13.03836 | 0.43137 | 2.30000 | 0.03901 | 2.01559 | 0.00000 |
| **Zfp438** | 0.21538 | 0.01000 | 0.53417 | 0.90438 | 0.39444 | 0.81425 | 0.00328 | 0.51973 | 0.75597 | 0.00252 | 2.87373 | 0.15335 | 1.33856 | 1.00989 |
| **Zfp442** | 0.22738 | 0.32431 | 0.44633 | 0.51996 | 2.44118 | 0.37273 | 0.22508 | 0.10392 | 5.38928 | 0.03679 | 0.20271 | 0.25408 | 1.71416 | 13.50645 |
| **Zfp444** | 0.08390 | 0.04131 | 0.17845 | 1.02768 | 0.40892 | 0.51409 | 0.06143 | 0.01741 | 0.84688 | 0.01046 | 0.10771 | 0.34857 | 1.22662 | 0.48895 |
| **Zfp445** | 3.05449 | 0.01000 | 2.70248 | 3.48486 | 0.82448 | 1.39840 | 0.50275 | 0.19690 | 13.74237 | 0.02495 | 1.77630 | 1.75461 | 4.88320 | 2.41134 |
| **Zfp446** | 2.21840 | 0.04336 | 1.22438 | 3.70446 | 1.02401 | 2.35849 | 0.01770 | 1.02364 | 0.02192 | 0.02268 | 0.00000 | 1.83042 | 1.99929 | 0.18704 |
| **Zfp449** | 0.00706 | 0.00450 | 0.00000 | 0.01400 | 0.19174 | 0.00853 | 0.00419 | 0.00985 | 1.00942 | 0.00811 | 0.00604 | 0.00844 | 0.00737 | 0.00943 |
| **Zfp451** | 6.48980 | 0.03203 | 3.43785 | 0.27587 | 3.78543 | 4.52431 | 0.00281 | 0.07485 | 0.03459 | 0.01358 | 0.00000 | 2.99252 | 2.82639 | 1.93542 |
| **Zfp454** | 0.58189 | 0.00406 | 0.00000 | 0.26844 | 0.00154 | 0.57593 | 0.00406 | 0.00327 | 0.00323 | 0.01096 | 0.00635 | 0.00516 | 0.61359 | 0.22367 |
| **Zfp455** | 0.14361 | 0.00469 | 0.02713 | 0.02269 | 0.09895 | 0.11892 | 0.00482 | 0.02965 | 0.13021 | 0.00762 | 0.00931 | 0.04569 | 0.01701 | 0.01972 |
| **Zfp456** | 0.17308 | 0.00456 | 0.01252 | 0.00000 | 0.12358 | 0.35793 | 0.06417 | 0.05451 | 0.00762 | 0.00690 | 0.05432 | 0.44485 | 0.04568 | 0.09772 |
| **Zfp457** | 0.01556 | 0.00463 | 0.18062 | 0.00390 | 0.01138 | 0.01393 | 0.00433 | 0.01793 | 0.03263 | 0.00711 | 0.01559 | 0.02012 | 0.06854 | 0.06127 |
| **Zfp458** | 0.00477 | 0.00357 | 0.32625 | 0.21354 | 0.52916 | 0.17210 | 0.00400 | 0.02384 | 0.01505 | 0.00432 | 0.51085 | 0.40848 | 0.00599 | 0.00000 |
| **Zfp459** | 0.49382 | 0.07307 | 0.27155 | 1.40312 | 0.14245 | 0.29075 | 0.02724 | 0.06715 | 1.45623 | 0.03460 | 0.43734 | 2.20331 | 0.53518 | 1.14406 |
| **Zfp46** | 0.00313 | 0.00390 | 0.21464 | 0.00000 | 0.04954 | 0.08771 | 0.00380 | 0.10241 | 0.02682 | 0.00618 | 0.22889 | 0.00510 | 0.59395 | 0.10071 |
| **Zfp462** | 0.00463 | 0.01699 | 0.18877 | 0.00000 | 0.47164 | 0.22907 | 0.00398 | 0.44498 | 0.00676 | 0.00601 | 0.00000 | 0.56245 | 0.26665 | 0.23674 |
| **Zfp467** | 0.00662 | 0.00458 | 0.00607 | 0.01577 | 0.49084 | 0.01380 | 0.01027 | 0.01072 | 0.01285 | 0.00674 | 0.01199 | 0.00838 | 0.00588 | 0.07166 |
| **Zfp472** | 3.36394 | 0.00967 | 3.87033 | 4.12650 | 6.42693 | 2.10300 | 4.40921 | 0.26008 | 7.03294 | 0.15441 | 2.37840 | 1.83986 | 0.00000 | 2.81671 |
| **Zfp473** | 0.00000 | 0.00447 | 0.30104 | 0.96384 | 0.24187 | 0.25157 | 0.01187 | 0.13779 | 0.00620 | 0.01384 | 0.03320 | 0.00420 | 0.00433 | 0.76297 |
| **Zfp488** | 0.81169 | 1.91994 | 0.43053 | 0.20176 | 0.52407 | 0.27290 | 0.14092 | 0.18372 | 3.27966 | 0.02840 | 0.15648 | 0.36234 | 0.38051 | 1.11727 |
| **Zfp493** | 0.00000 | 0.00352 | 0.15272 | 0.53140 | 0.08478 | 0.68903 | 1.98784 | 0.00074 | 0.05818 | 0.05650 | 1.95191 | 0.70079 | 0.00584 | 0.93089 |
| **Zfp503** | 0.05529 | 0.00448 | 0.00560 | 0.01218 | 0.39204 | 0.01251 | 0.00424 | 0.01052 | 0.00870 | 0.00765 | 0.00819 | 0.00772 | 0.04064 | 0.04844 |
| **Zfp507** | 0.00000 | 0.00456 | 0.19190 | 1.33233 | 0.66349 | 2.70962 | 0.19124 | 0.02387 | 1.30286 | 0.01411 | 0.21587 | 0.66659 | 0.74396 | 1.07570 |
| **Zfp51** | 1.78930 | 0.00322 | 2.84117 | 1.38452 | 2.13412 | 2.62517 | 2.42212 | 0.00723 | 10.98796 | 0.03825 | 0.84784 | 0.00087 | 5.42683 | 0.42767 |
| **Zfp511** | 1.42684 | 0.04356 | 0.84792 | 5.89649 | 4.54921 | 1.52578 | 0.15108 | 1.32351 | 0.36613 | 0.02023 | 0.05678 | 2.71078 | 0.41410 | 1.58223 |
| **Zfp512** | 0.00000 | 0.02921 | 1.20816 | 0.00000 | 0.03794 | 1.51480 | 0.00324 | 1.22698 | 0.00127 | 0.03374 | 2.29461 | 0.65877 | 0.00000 | 0.98689 |
| **Zfp512b** | 1.30624 | 0.01208 | 1.16328 | 0.52274 | 0.00000 | 0.00000 | 0.89335 | 1.01834 | 2.33368 | 0.00839 | 0.14107 | 0.90744 | 1.11250 | 1.58845 |
| **Zfp513** | 3.81642 | 0.08250 | 4.01791 | 4.48101 | 6.55669 | 3.11091 | 3.30629 | 3.96754 | 0.00025 | 0.11014 | 0.68034 | 1.65217 | 8.00456 | 0.24092 |
| **Zfp516** | 0.04230 | 0.00446 | 0.00000 | 0.42479 | 0.11867 | 0.79522 | 0.00508 | 0.56431 | 0.00936 | 0.00555 | 0.00162 | 0.00819 | 0.00804 | 0.00368 |
| **Zfp518a** | 0.44531 | 0.08532 | 0.00000 | 2.45282 | 1.73356 | 0.95121 | 0.08790 | 0.30818 | 13.97493 | 0.00229 | 1.59470 | 0.74834 | 2.51913 | 1.55712 |
| **Zfp518b** | 0.15624 | 2.63899 | 0.65403 | 1.72498 | 2.24782 | 0.74455 | 0.00268 | 1.22788 | 2.56975 | 0.00901 | 1.36379 | 0.54388 | 2.05142 | 3.97419 |
| **Zfp52** | 0.00000 | 0.00463 | 0.91970 | 0.72604 | 0.41532 | 0.02372 | 0.00392 | 0.07435 | 1.91420 | 0.00604 | 0.00099 | 0.18515 | 1.07100 | 0.11618 |
| **Zfp523** | 0.01296 | 0.03314 | 0.31751 | 0.00000 | 0.00870 | 0.00403 | 0.00382 | 0.02517 | 0.00984 | 0.00819 | 0.00567 | 0.49823 | 0.01120 | 0.01801 |

| | | | | | | | | | | | | | | |
|---|---|---|---|---|---|---|---|---|---|---|---|---|---|---|
| **Zfp524** | 0.00000 | 0.01739 | 0.73048 | 0.81958 | 0.90930 | 1.06471 | 0.01451 | 2.29849 | 0.00463 | 11.77211 | 0.00467 | 0.74597 | 0.00144 | 0.92118 |
| **Zfp526** | 0.00000 | 0.02010 | 0.28288 | 0.83412 | 0.46594 | 0.00000 | 0.00382 | 0.08933 | 0.00460 | 0.00517 | 0.00789 | 0.52504 | 0.05306 | 0.18674 |
| **Zfp53** | 0.00000 | 0.00530 | 0.43703 | 0.23094 | 0.63417 | 0.15507 | 0.00408 | 0.56931 | 0.02756 | 0.03076 | 0.09671 | 0.02484 | 0.64812 | 0.13396 |
| **Zfp532** | 1.70532 | 0.01206 | 3.19537 | 0.40075 | 2.71334 | 0.93772 | 0.55424 | 1.77879 | 0.02832 | 0.00000 | 1.76409 | 0.50403 | 0.51263 | 0.87039 |
| **Zfp534** | 0.02697 | 0.00302 | 2.14824 | 1.42853 | 1.58640 | 0.79259 | 0.13267 | 0.22233 | 11.18315 | 0.70711 | 1.04654 | 1.22849 | 0.19723 | 0.16675 |
| **Zfp54** | 0.00000 | 0.02169 | 0.00000 | 1.69596 | 1.09758 | 0.81112 | 0.00411 | 0.31983 | 1.76642 | 0.00074 | 2.43009 | 0.00000 | 2.13586 | 0.17881 |
| **Zfp541** | 0.03278 | 0.00409 | 1.64201 | 0.03402 | 0.00000 | 0.02188 | 0.00377 | 0.00000 | 0.00933 | 0.00243 | 0.01914 | 0.02330 | 0.42033 | 0.27024 |
| **Zfp553** | 0.15114 | 0.06645 | 4.39173 | 2.54746 | 2.95785 | 1.19356 | 4.69733 | 0.88853 | 1.65686 | 0.15740 | 2.68503 | 3.57889 | 4.39368 | 1.61988 |
| **Zfp558** | 1.29423 | 0.01189 | 0.40948 | 0.01488 | 0.53893 | 0.02170 | 0.00419 | 0.01099 | 1.00908 | 0.00749 | 0.00158 | 0.66026 | 0.86010 | 0.00000 |
| **Zfp560** | 0.00000 | 0.00430 | 0.10359 | 0.21578 | 0.53684 | 0.05592 | 0.80578 | 0.12897 | 0.00536 | 0.01342 | 0.00260 | 0.19299 | 0.39817 | 0.07292 |
| **Zfp563** | 0.00568 | 0.00433 | 0.10945 | 0.00000 | 0.00000 | 0.35939 | 0.00364 | 0.23396 | 0.02948 | 0.00528 | 1.16455 | 0.03041 | 0.30144 | 0.46253 |
| **Zfp566** | 0.71537 | 0.00936 | 3.21594 | 2.80392 | 3.30867 | 3.70716 | 2.71573 | 0.00125 | 0.00000 | 0.04672 | 1.62108 | 2.68541 | 2.92934 | 4.92432 |
| **Zfp568** | 10.41957 | 0.20740 | 13.88843 | 4.75286 | 14.54676 | 12.72775 | 0.63752 | 5.42315 | 19.17295 | 8.06182 | 10.24852 | 4.03315 | 5.50069 | 19.69011 |
| **Zfp57** | 0.16649 | 0.01630 | 1.27100 | 0.56370 | 1.09000 | 0.47420 | 0.35985 | 0.60061 | 0.35027 | 0.01719 | 4.15480 | 2.47359 | 3.28500 | 1.01708 |
| **Zfp572** | 0.01297 | 0.00445 | 0.00762 | 0.01226 | 0.22967 | 0.01168 | 0.00446 | 0.01059 | 0.00746 | 0.00637 | 0.00829 | 0.00878 | 0.01525 | 0.28636 |
| **Zfp574** | 0.38429 | 2.93953 | 1.94782 | 8.82721 | 8.04237 | 0.95229 | 0.04523 | 0.54335 | 0.00000 | 0.03465 | 0.16218 | 3.52977 | 3.54708 | 0.00000 |
| **Zfp575** | 14.66957 | 0.87174 | 3.30901 | 0.79158 | 7.78573 | 6.18997 | 0.59321 | 0.99917 | 6.24860 | 0.30546 | 2.99293 | 0.71093 | 7.69239 | 2.88862 |
| **Zfp579** | 0.29923 | 0.00319 | 0.15403 | 0.78976 | 0.94321 | 0.00000 | 0.00362 | 0.00258 | 0.01027 | 0.00599 | 0.00388 | 1.18028 | 0.01441 | 0.02937 |
| **Zfp58** | 1.03678 | 2.37912 | 0.00000 | 0.29988 | 0.42306 | 0.08255 | 0.00258 | 0.49859 | 0.40372 | 0.00629 | 0.00000 | 1.79985 | 0.00363 | 1.52299 |
| **Zfp580** | 0.00811 | 0.00419 | 0.01128 | 0.12990 | 0.05895 | 0.01406 | 0.00418 | 0.00553 | 0.01162 | 0.00576 | 0.01519 | 0.01509 | 0.24916 | 0.00600 |
| **Zfp583** | 0.06639 | 0.00431 | 0.00791 | 0.29810 | 0.82912 | 0.30948 | 2.84486 | 0.00517 | 0.00889 | 0.11995 | 0.53299 | 0.51709 | 0.84687 | 0.04188 |
| **Zfp59** | 0.00000 | 0.00455 | 0.87267 | 0.03851 | 0.46520 | 0.27381 | 0.00415 | 0.20927 | 0.01385 | 0.00481 | 0.00000 | 0.45053 | 0.81748 | 2.56441 |
| **Zfp592** | 2.78521 | 7.26991 | 3.65782 | 5.49877 | 4.62302 | 1.93364 | 0.39247 | 2.49509 | 3.55890 | 9.92254 | 2.59506 | 1.54932 | 3.73755 | 2.68800 |
| **Zfp593** | 1.32143 | 0.02700 | 1.34659 | 2.24031 | 5.39584 | 2.32558 | 0.02165 | 0.67547 | 0.08162 | 0.05767 | 0.20689 | 1.61105 | 2.18052 | 0.16092 |
| **Zfp595** | 0.00000 | 0.00443 | 0.00000 | 0.56212 | 0.00153 | 0.13636 | 0.00420 | 0.00575 | 0.98839 | 0.00520 | 0.03132 | 0.01735 | 0.00327 | 0.00000 |
| **Zfp597** | 1.23806 | 0.00305 | 0.41954 | 0.00000 | 1.31742 | 0.96317 | 2.76281 | 0.00000 | 3.37675 | 0.00000 | 2.47727 | 0.35074 | 1.22892 | 0.71690 |
| **Zfp598** | 9.12732 | 3.32636 | 9.68520 | 7.33815 | 14.31389 | 4.12165 | 1.85507 | 8.03618 | 0.00000 | 0.14324 | 3.25863 | 7.62550 | 10.35165 | 2.52880 |
| **Zfp599** | 1.10942 | 0.01136 | 5.67513 | 0.00000 | 0.51326 | 0.84957 | 0.00240 | 0.44431 | 10.69677 | 0.00000 | 1.11308 | 0.02603 | 5.98599 | 2.26069 |
| **Zfp60** | 0.76154 | 0.00360 | 0.31978 | 0.81132 | 0.39080 | 1.20100 | 0.00366 | 0.00000 | 0.03445 | 0.00423 | 0.35133 | 0.20273 | 0.47881 | 1.06574 |
| **Zfp600** | 3.13064 | 0.04008 | 4.93705 | 3.98307 | 3.36631 | 1.54465 | 0.27400 | 0.86780 | 21.02601 | 0.06819 | 2.07468 | 3.34924 | 8.88404 | 0.00000 |
| **Zfp605** | 0.46180 | 0.01283 | 0.09093 | 0.26484 | 0.59067 | 1.38306 | 0.47002 | 0.10526 | 2.35596 | 0.16231 | 0.00216 | 0.16778 | 2.26790 | 0.03438 |
| **Zfp606** | 0.00000 | 0.00429 | 0.30481 | 0.98146 | 0.01552 | 0.06097 | 0.00410 | 0.00210 | 0.00967 | 0.01022 | 0.34132 | 0.01394 | 0.00346 | 0.08228 |
| **Zfp607a** | 0.41966 | 0.01149 | 0.07031 | 0.19208 | 0.12009 | 0.87449 | 0.07579 | 0.12457 | 1.44212 | 0.10596 | 0.13095 | 0.27801 | 4.42994 | 0.60879 |
| **Zfp607b** | 0.31041 | 0.02091 | 0.11138 | 0.55295 | 0.89023 | 0.31265 | 0.03356 | 0.11784 | 2.12371 | 0.09468 | 0.00155 | 0.07536 | 0.45569 | 0.00000 |
| **Zfp609** | 0.24120 | 0.00386 | 0.70479 | 0.00000 | 0.37277 | 0.00000 | 0.00364 | 0.57318 | 0.07893 | 0.00677 | 0.00572 | 0.02223 | 0.00112 | 0.41313 |
| **Zfp61** | 0.11093 | 2.79843 | 2.46278 | 2.83901 | 0.96099 | 1.61112 | 0.00923 | 0.80537 | 0.00000 | 0.01667 | 0.00000 | 1.29453 | 2.54775 | 1.16636 |
| **Zfp612** | 0.93428 | 0.00375 | 0.13844 | 0.00000 | 0.44484 | 0.29861 | 0.00371 | 0.47804 | 0.43629 | 0.00694 | 0.00440 | 0.00801 | 0.00563 | 1.47152 |
| **Zfp616** | 0.00987 | 0.00442 | 0.01328 | 0.01539 | 0.01495 | 0.01227 | 0.00425 | 0.01005 | 0.02044 | 0.00763 | 0.00864 | 0.00962 | 0.04751 | 0.04227 |
| **Zfp617** | 0.60943 | 0.02948 | 1.41075 | 0.68515 | 1.66505 | 2.87284 | 1.97859 | 0.00000 | 2.54624 | 0.39425 | 8.21000 | 1.64837 | 6.08794 | 1.43870 |
| **Zfp619** | 0.00497 | 0.00418 | 0.10735 | 0.14160 | 0.00553 | 0.00475 | 0.00415 | 0.01086 | 0.01178 | 0.00743 | 0.08796 | 0.00564 | 0.23203 | 0.00000 |
| **Zfp62** | 0.29705 | 0.00293 | 0.07655 | 0.92780 | 0.20440 | 0.52655 | 0.01042 | 0.10390 | 0.23879 | 0.00296 | 2.58832 | 0.25442 | 2.66295 | 0.75238 |
| **Zfp622** | 19.26250 | 2.26344 | 11.17833 | 13.85613 | 21.62896 | 10.87461 | 2.44817 | 0.27120 | 12.92672 | 0.09904 | 3.02930 | 5.46133 | 22.24054 | 12.08773 |
| **Zfp623** | 1.27118 | 0.01045 | 1.78360 | 0.70174 | 1.85292 | 2.90766 | 0.01431 | 0.78938 | 0.00000 | 0.00229 | 0.00000 | 1.05159 | 4.78733 | 0.77473 |
| **Zfp626** | 0.71051 | 0.06950 | 0.78030 | 1.83030 | 0.70710 | 1.64880 | 0.13067 | 0.63767 | 4.29245 | 1.90576 | 0.22448 | 0.20472 | 4.45343 | 1.81769 |
| **Zfp628** | 0.00808 | 0.01756 | 0.15355 | 0.54991 | 0.67155 | 0.34518 | 0.71445 | 0.01010 | 0.00724 | 0.00508 | 0.00000 | 1.82541 | 0.39362 | 0.00000 |
| **Zfp629** | 0.01753 | 0.00451 | 0.02609 | 0.00590 | 0.22226 | 0.01948 | 0.00413 | 0.00999 | 0.56108 | 0.00700 | 0.00624 | 0.00812 | 0.00226 | 0.00313 |
| **Zfp637** | 3.14657 | 0.00203 | 4.22032 | 2.72781 | 4.46932 | 4.51736 | 1.93103 | 0.00000 | 0.68743 | 0.01537 | 0.03649 | 2.04533 | 0.60259 | 2.55732 |
| **Zfp638** | 11.04947 | 0.00000 | 5.66905 | 1.24644 | 9.14018 | 3.85193 | 1.97926 | 1.56568 | 24.92560 | 0.03654 | 3.82011 | 3.37074 | 2.26729 | 4.34677 |
| **Zfp639** | 0.00000 | 0.03877 | 2.65359 | 2.37775 | 2.35445 | 1.50888 | 4.45490 | 1.10572 | 0.62866 | 0.01686 | 6.00096 | 1.09958 | 6.81659 | 11.55519 |
| **Zfp64** | 0.00000 | 8.31382 | 4.76030 | 4.03581 | 2.55009 | 1.99522 | 0.53370 | 4.96080 | 0.00657 | 0.03440 | 0.41580 | 3.06374 | 0.03274 | 0.00000 |
| **Zfp644** | 0.63369 | 0.00248 | 0.00000 | 0.60175 | 3.96763 | 0.00000 | 0.00123 | 0.00000 | 16.85568 | 0.01240 | 0.00000 | 0.08069 | 6.34902 | 7.70824 |
| **Zfp646** | 0.45917 | 0.02672 | 2.18267 | 0.26586 | 0.36518 | 1.41837 | 0.00312 | 1.12154 | 0.01218 | 0.00553 | 0.79896 | 0.96125 | 0.05694 | 2.10958 |
| **Zfp647** | 0.01502 | 0.00420 | 0.00295 | 0.10261 | 0.18295 | 0.13504 | 0.00414 | 0.00366 | 0.00862 | 0.00504 | 0.00576 | 0.01201 | 0.37008 | 0.02507 |
| **Zfp65** | 0.38789 | 0.13212 | 0.64056 | 1.32589 | 0.21026 | 0.26373 | 0.08409 | 0.07655 | 2.99255 | 0.00469 | 0.52918 | 0.24021 | 0.44607 | 0.08811 |
| **Zfp651** | 0.01540 | 0.00456 | 0.01075 | 0.00000 | 0.02430 | 0.00818 | 0.00467 | 0.09327 | 0.01535 | 0.00847 | 0.00331 | 0.99178 | 0.00684 | 0.00901 |

| | | | | | | | | | | | | | | |
|---|---|---|---|---|---|---|---|---|---|---|---|---|---|---|
| **Zfp652** | 0.23067 | 0.01734 | 0.32610 | 0.19490 | 0.34484 | 0.46143 | 0.00360 | 0.72584 | 0.00591 | 0.00655 | 0.00000 | 0.98541 | 0.18257 | 0.36655 |
| **Zfp653** | 2.00113 | 5.71680 | 1.11087 | 0.00870 | 2.40034 | 1.20018 | 0.00262 | 0.52909 | 0.01950 | 0.00507 | 0.81144 | 0.23968 | 0.64001 | 0.16142 |
| **Zfp654** | 1.17048 | 0.00287 | 1.71844 | 0.65864 | 1.25459 | 1.03432 | 0.01069 | 0.16760 | 5.57416 | 0.01071 | 1.56949 | 0.80612 | 7.24508 | 0.00000 |
| **Zfp655** | 9.03952 | 3.43600 | 8.98528 | 9.11453 | 13.75646 | 8.75711 | 6.00028 | 1.67548 | 17.32554 | 0.12989 | 8.66873 | 3.78830 | 11.28541 | 12.52119 |
| **Zfp658** | 0.50363 | 0.00375 | 0.62692 | 0.48344 | 0.13888 | 0.73755 | 0.00940 | 1.78122 | 0.01787 | 0.02438 | 0.58864 | 0.00000 | 0.01376 | 0.74282 |
| **Zfp661** | 0.35747 | 0.01335 | 0.54890 | 0.19250 | 0.08390 | 0.00000 | 0.00399 | 0.02305 | 0.25141 | 0.01372 | 0.00406 | 0.20075 | 0.05880 | 0.00000 |
| **Zfp664** | 2.11760 | 0.04361 | 2.55384 | 1.39402 | 2.42507 | 9.77573 | 1.40863 | 2.20811 | 2.52480 | 0.09021 | 4.89361 | 4.90442 | 12.25723 | 6.44297 |
| **Zfp667** | 0.02016 | 0.00470 | 0.33764 | 0.00000 | 0.00000 | 0.40841 | 1.61649 | 0.42132 | 0.00848 | 0.01313 | 0.05605 | 0.01680 | 0.49375 | 0.00000 |
| **Zfp668** | 1.10356 | 0.02583 | 4.41482 | 2.76783 | 3.09970 | 1.58679 | 0.00311 | 1.52836 | 0.01922 | 0.74671 | 1.80080 | 0.35944 | 2.56753 | 0.00000 |
| **Zfp672** | 1.18888 | 0.00338 | 0.94165 | 0.00000 | 0.96810 | 0.65654 | 1.97069 | 0.00000 | 1.32908 | 0.05072 | 0.44383 | 0.29580 | 2.22288 | 1.45002 |
| **Zfp677** | 0.00000 | 0.00455 | 0.20159 | 0.00556 | 0.01275 | 0.00230 | 0.00379 | 0.01299 | 1.63510 | 0.00637 | 0.00307 | 0.06418 | 1.25364 | 0.00000 |
| **Zfp68** | 2.32924 | 0.00365 | 1.09055 | 0.94873 | 0.83183 | 0.44427 | 0.01406 | 1.37416 | 1.55283 | 0.01038 | 0.00078 | 0.00082 | 1.70080 | 2.78775 |
| **Zfp687** | 0.04302 | 0.00345 | 0.13529 | 0.92702 | 0.07833 | 1.11432 | 0.05612 | 0.00000 | 0.37709 | 0.00166 | 1.30119 | 0.37467 | 0.20610 | 0.00000 |
| **Zfp688** | 0.00000 | 0.00302 | 0.37654 | 1.33219 | 0.69136 | 0.67014 | 2.14163 | 0.03790 | 0.00849 | 0.07759 | 0.01604 | 2.43324 | 0.00563 | 0.00000 |
| **Zfp689** | 0.00552 | 0.01553 | 0.11297 | 0.00000 | 0.01315 | 0.00000 | 0.01605 | 0.68825 | 0.02869 | 0.00951 | 0.47358 | 0.51818 | 0.79617 | 0.01039 |
| **Zfp69** | 0.00000 | 0.00395 | 0.20577 | 0.00000 | 0.35037 | 0.00940 | 0.00388 | 0.15480 | 0.00471 | 0.00700 | 0.00920 | 0.00703 | 0.13777 | 0.32040 |
| **Zfp691** | 0.04203 | 0.02014 | 1.52544 | 1.39872 | 0.18319 | 0.26760 | 0.00325 | 2.21066 | 0.10215 | 0.00770 | 1.81048 | 0.11942 | 0.00196 | 0.00000 |
| **Zfp692** | 1.26387 | 0.03056 | 1.82226 | 4.48386 | 0.93993 | 4.20652 | 0.13035 | 3.05054 | 0.13999 | 0.02250 | 5.60855 | 0.11860 | 0.14561 | 1.07032 |
| **Zfp7** | 1.41202 | 0.09282 | 5.90802 | 2.49214 | 1.74098 | 4.75642 | 4.44529 | 3.39919 | 0.27841 | 0.09579 | 0.34343 | 0.70520 | 8.31327 | 5.57097 |
| **Zfp703** | 0.05317 | 0.01927 | 0.16824 | 1.69682 | 0.16498 | 0.00000 | 0.01642 | 3.05157 | 0.00635 | 0.05617 | 0.00000 | 0.10589 | 0.24928 | 0.09548 |
| **Zfp704** | 0.19038 | 0.21026 | 4.44932 | 0.43289 | 0.67079 | 5.43292 | 3.20926 | 6.46839 | 0.32086 | 0.15314 | 1.35497 | 11.02331 | 0.00000 | 2.24781 |
| **Zfp706** | 25.13697 | 6.96069 | 21.56742 | 16.62348 | 32.68761 | 26.16787 | 7.36565 | 11.15301 | 19.06832 | 10.93175 | 9.09444 | 15.81229 | 20.34073 | 17.47213 |
| **Zfp707** | 1.79121 | 1.12456 | 6.69238 | 7.04015 | 5.88858 | 1.75033 | 1.86773 | 4.93269 | 0.07533 | 0.08284 | 0.22383 | 1.15372 | 10.84284 | 0.35057 |
| **Zfp708** | 0.02509 | 0.00388 | 1.12081 | 0.62186 | 0.90704 | 0.02836 | 1.51212 | 0.00000 | 3.06949 | 0.07759 | 0.14302 | 0.09342 | 1.19860 | 1.59077 |
| **Zfp709** | 0.00073 | 0.00371 | 0.05027 | 0.00000 | 0.00288 | 0.16621 | 0.00390 | 0.00362 | 0.38804 | 0.00525 | 0.00210 | 0.52880 | 0.28923 | 0.00000 |
| **Zfp710** | 3.80209 | 6.51152 | 2.80869 | 2.00808 | 8.37496 | 1.83268 | 0.00078 | 6.23335 | 0.05012 | 0.06504 | 0.00000 | 1.01906 | 3.02360 | 6.11644 |
| **Zfp712** | 0.00829 | 0.00443 | 0.06889 | 0.00000 | 0.10896 | 0.00792 | 0.00506 | 0.03230 | 0.01392 | 0.00636 | 0.00684 | 0.05787 | 0.09353 | 0.05300 |
| **Zfp715** | 0.56701 | 1.51703 | 0.80050 | 0.17755 | 0.56718 | 0.12505 | 0.00290 | 0.48814 | 3.88054 | 0.00407 | 1.13419 | 0.00205 | 1.05567 | 0.00000 |
| **Zfp719** | 0.01823 | 0.00388 | 0.49284 | 0.00000 | 0.92700 | 1.38422 | 0.00407 | 0.00438 | 0.01709 | 0.00588 | 0.00191 | 0.04185 | 1.11397 | 0.01238 |
| **Zfp72** | 0.01914 | 0.00368 | 0.44378 | 0.00000 | 0.54091 | 0.11515 | 0.01412 | 0.09204 | 1.68800 | 0.01853 | 0.67649 | 0.72129 | 0.00000 | 0.00000 |
| **Zfp729a** | 0.00000 | 0.01776 | 0.86863 | 1.03899 | 1.73300 | 1.44990 | 0.62311 | 0.00000 | 6.35758 | 0.01698 | 2.39605 | 0.27356 | 0.67587 | 0.78169 |
| **Zfp729b** | 2.24009 | 0.00156 | 2.08353 | 1.09808 | 5.20492 | 1.20083 | 0.89191 | 0.00000 | 11.32460 | 0.01357 | 3.33609 | 1.35093 | 1.41270 | 0.29240 |
| **Zfp738** | 0.16670 | 0.63101 | 0.22830 | 0.00000 | 0.64573 | 0.19821 | 0.01471 | 0.02504 | 0.54160 | 0.00396 | 1.24072 | 0.03058 | 1.12288 | 0.48156 |
| **Zfp74** | 0.44490 | 0.00320 | 0.03204 | 0.44382 | 0.00000 | 0.24567 | 0.00349 | 0.00000 | 0.02898 | 0.00000 | 2.06521 | 0.12125 | 0.80244 | 1.47093 |
| **Zfp740** | 2.55078 | 0.07764 | 8.85682 | 3.34182 | 11.32945 | 6.15214 | 0.17669 | 2.27498 | 5.25917 | 0.10021 | 2.42748 | 4.54279 | 1.12179 | 12.45976 |
| **Zfp746** | 1.64037 | 0.00311 | 1.43458 | 3.24132 | 1.42115 | 0.75164 | 0.02639 | 4.23502 | 1.79972 | 0.01824 | 1.84745 | 0.36112 | 0.75834 | 0.06951 |
| **Zfp747** | 0.00000 | 0.03471 | 1.18675 | 0.00000 | 0.13905 | 0.59082 | 1.18260 | 0.41165 | 0.25743 | 7.01389 | 0.20578 | 0.89222 | 0.00000 | 0.25595 |
| **Zfp748** | 1.51727 | 0.01014 | 0.87138 | 1.05891 | 0.76969 | 1.26526 | 0.10857 | 0.53040 | 0.74956 | 0.00000 | 3.59542 | 1.75147 | 2.41862 | 0.62884 |
| **Zfp750** | 0.00460 | 0.00432 | 0.01082 | 0.01032 | 0.14640 | 0.01295 | 0.00415 | 0.00905 | 0.01894 | 0.00598 | 0.00959 | 0.00862 | 0.50876 | 0.01625 |
| **Zfp758** | 0.00000 | 0.00393 | 0.16694 | 0.02701 | 0.40277 | 0.27428 | 0.00948 | 0.06680 | 1.29861 | 0.00714 | 0.01841 | 0.03758 | 0.68398 | 0.48590 |
| **Zfp759** | 0.06356 | 0.00368 | 0.57909 | 0.68164 | 0.22689 | 0.00000 | 0.47450 | 0.00000 | 0.93373 | 0.00000 | 2.03791 | 0.27944 | 0.93576 | 0.00000 |
| **Zfp760** | 0.45109 | 0.00488 | 0.13463 | 0.23817 | 0.00000 | 0.03033 | 0.00427 | 0.26848 | 0.00887 | 0.00640 | 0.46225 | 0.00498 | 0.05497 | 0.36280 |
| **Zfp763** | 0.01875 | 0.03213 | 0.84800 | 0.00000 | 0.58809 | 0.01508 | 0.00346 | 0.01593 | 0.26127 | 0.01009 | 1.79519 | 0.79659 | 0.15635 | 0.41532 |
| **Zfp764** | 0.00000 | 0.00422 | 0.22996 | 0.09625 | 0.11075 | 0.11849 | 0.25519 | 0.06504 | 0.04944 | 1.32814 | 0.05823 | 0.14760 | 0.00376 | 0.16089 |
| **Zfp768** | 0.03373 | 0.00396 | 0.08005 | 0.00000 | 0.00815 | 0.15745 | 0.00416 | 0.30778 | 0.01054 | 0.01210 | 0.55601 | 0.02000 | 0.00645 | 0.00578 |
| **Zfp770** | 0.18473 | 0.00450 | 0.63154 | 0.00000 | 0.20194 | 0.62152 | 0.00460 | 0.06739 | 0.04855 | 0.00613 | 0.02266 | 0.13628 | 2.62473 | 0.25038 |
| **Zfp771** | 0.02289 | 0.00442 | 0.01083 | 0.02131 | 0.06365 | 0.04574 | 0.00417 | 0.00804 | 0.01443 | 0.00692 | 0.00762 | 0.00942 | 0.01875 | 0.01185 |
| **Zfp772** | 1.57802 | 0.01237 | 1.30499 | 1.47050 | 2.10680 | 1.11751 | 0.00377 | 0.51239 | 0.32496 | 0.02605 | 0.31802 | 0.00547 | 0.03393 | 0.74370 |
| **Zfp773** | 0.15323 | 0.00428 | 0.06882 | 0.01533 | 0.01767 | 0.07470 | 0.00423 | 0.00562 | 0.05788 | 0.00616 | 0.01123 | 0.04114 | 0.22869 | 0.13928 |
| **Zfp775** | 3.06627 | 0.01633 | 1.41771 | 1.43170 | 14.63095 | 0.56875 | 0.00240 | 0.08241 | 2.18675 | 0.02923 | 1.79718 | 1.19717 | 10.88278 | 6.01889 |
| **Zfp777** | 0.00000 | 0.54895 | 2.41338 | 4.57190 | 6.58699 | 0.00000 | 0.15836 | 2.14313 | 0.31107 | 0.09176 | 0.00485 | 1.69649 | 1.39857 | 0.08908 |
| **Zfp78** | 0.00000 | 0.01716 | 0.70083 | 0.36155 | 0.55995 | 0.37155 | 0.00341 | 0.00000 | 0.49641 | 0.00000 | 1.87090 | 0.60910 | 2.39881 | 1.93328 |
| **Zfp780b** | 0.07965 | 0.12766 | 0.13660 | 0.70923 | 0.41400 | 0.37047 | 0.08883 | 0.20741 | 1.39320 | 0.09999 | 0.07506 | 0.16848 | 1.86954 | 2.02926 |
| **Zfp781** | 0.36741 | 0.03663 | 2.15806 | 1.07461 | 1.57906 | 1.86563 | 0.84424 | 0.07345 | 6.14791 | 0.03947 | 0.58319 | 0.21942 | 2.01098 | 0.83453 |
| **Zfp783** | 0.02216 | 0.00444 | 0.01190 | 0.00000 | 0.00360 | 0.01266 | 0.00522 | 0.21547 | 0.00830 | 0.00571 | 0.00561 | 0.14757 | 0.00784 | 0.48593 |

| | | | | | | | | | | | | | | |
|---|---|---|---|---|---|---|---|---|---|---|---|---|---|---|
| **Zfp784** | 0.75162 | 0.02402 | 0.39386 | 0.20529 | 0.00000 | 0.27610 | 0.00422 | 0.73142 | 0.00260 | 0.00863 | 0.00108 | 0.29616 | 0.54279 | 0.00629 |
| **Zfp786** | 0.00716 | 0.00407 | 0.43133 | 0.04364 | 0.00303 | 0.08818 | 0.00393 | 0.04127 | 0.01033 | 0.00525 | 0.00521 | 0.00449 | 0.00633 | 0.01744 |
| **Zfp787** | 0.00548 | 0.00365 | 0.00258 | 0.00000 | 0.29970 | 0.40578 | 0.32105 | 0.50913 | 0.00774 | 0.03686 | 0.02027 | 0.64434 | 0.49726 | 0.04287 |
| **Zfp788** | 0.01950 | 0.00428 | 2.01304 | 2.26292 | 0.24820 | 1.05291 | 0.42401 | 1.20208 | 4.02162 | 0.01155 | 0.24554 | 0.00000 | 0.00000 | 0.37847 |
| **Zfp790** | 0.45116 | 0.00308 | 1.92438 | 0.60114 | 1.36187 | 0.84555 | 0.00329 | 0.28243 | 0.49050 | 0.00620 | 1.15532 | 0.31104 | 2.47551 | 0.47125 |
| **Zfp791** | 0.00000 | 0.01912 | 2.07044 | 0.66212 | 0.95003 | 0.26233 | 0.00347 | 0.21322 | 1.87372 | 0.00356 | 0.00147 | 0.00000 | 5.38252 | 0.00000 |
| **Zfp799** | 0.37131 | 0.10436 | 0.15891 | 0.31114 | 0.33745 | 0.67068 | 0.10649 | 0.11900 | 0.64457 | 0.08138 | 0.05609 | 0.11429 | 0.75710 | 0.75848 |
| **Zfp800** | 0.13787 | 0.00368 | 1.24374 | 0.00000 | 0.24554 | 0.32775 | 0.00363 | 0.43983 | 0.93345 | 0.02516 | 0.09517 | 0.07926 | 2.56683 | 0.00000 |
| **Zfp808** | 1.06336 | 0.10389 | 1.56308 | 2.69725 | 2.66749 | 3.12280 | 1.68506 | 0.08753 | 6.37640 | 0.48363 | 2.27851 | 1.51329 | 2.00065 | 0.00000 |
| **Zfp809** | 0.95949 | 0.02101 | 3.83637 | 2.40389 | 2.42019 | 3.15966 | 0.27425 | 0.54872 | 2.07014 | 0.01132 | 5.65993 | 0.45777 | 2.21331 | 3.79983 |
| **Zfp81** | 0.73519 | 0.00375 | 0.72967 | 1.20393 | 0.60156 | 0.00000 | 0.00408 | 0.00308 | 1.13078 | 0.00504 | 0.01098 | 0.02460 | 0.95064 | 0.16444 |
| **Zfp810** | 0.23033 | 3.77955 | 0.49074 | 1.62706 | 0.20116 | 1.18038 | 0.00267 | 0.00000 | 1.66403 | 0.02006 | 0.53252 | 0.00697 | 3.94773 | 0.37672 |
| **Zfp811** | 0.00823 | 0.00410 | 0.00751 | 0.35583 | 0.00598 | 0.01714 | 0.08437 | 0.00418 | 0.09768 | 0.01715 | 0.00998 | 0.01098 | 0.04471 | 0.00944 |
| **Zfp819** | 0.68841 | 0.06102 | 1.60168 | 0.20618 | 0.17721 | 0.00000 | 0.00318 | 1.01163 | 0.00433 | 0.01719 | 0.03235 | 2.82969 | 1.53794 | 0.03104 |
| **Zfp82** | 0.06750 | 0.00425 | 2.12475 | 0.00000 | 0.92172 | 0.14683 | 0.48042 | 0.90059 | 0.60049 | 0.01223 | 0.93735 | 0.36272 | 1.92278 | 5.34135 |
| **Zfp820** | 3.99988 | 0.00212 | 5.57392 | 1.62695 | 4.02307 | 0.14913 | 0.00000 | 0.00000 | 21.67663 | 0.00000 | 0.00174 | 0.54804 | 5.76118 | 0.87722 |
| **Zfp821** | 5.02152 | 0.00301 | 8.00796 | 4.33396 | 6.34757 | 1.38290 | 1.79425 | 1.73904 | 0.00000 | 0.00853 | 0.04294 | 0.29370 | 6.57635 | 2.18462 |
| **Zfp825** | 2.55836 | 0.04585 | 5.59589 | 3.91311 | 1.67660 | 3.69015 | 2.32865 | 0.27822 | 10.95468 | 0.01134 | 1.32907 | 3.39376 | 0.00000 | 4.66222 |
| **Zfp827** | 0.04572 | 0.00443 | 0.12032 | 0.00905 | 0.04559 | 0.00838 | 0.00429 | 0.01070 | 0.01303 | 0.00751 | 0.00715 | 0.01001 | 0.00557 | 0.00992 |
| **Zfp830** | 1.59109 | 0.18784 | 4.99408 | 1.61519 | 1.98855 | 2.63373 | 1.95351 | 2.24378 | 0.44247 | 0.06330 | 3.01079 | 1.36613 | 6.54921 | 6.50614 |
| **Zfp839** | 2.20723 | 0.00965 | 0.81339 | 0.00000 | 1.37190 | 0.42857 | 1.14442 | 0.00000 | 0.34402 | 0.01585 | 1.06577 | 0.88932 | 0.64558 | 1.71102 |
| **Zfp84** | 1.23540 | 0.00236 | 0.48265 | 0.11197 | 0.92569 | 1.21663 | 0.01411 | 0.13922 | 2.92274 | 0.02374 | 1.93114 | 1.10604 | 1.21765 | 2.29035 |
| **Zfp846** | 0.48729 | 0.00329 | 0.91463 | 0.76317 | 1.41680 | 0.49831 | 0.00344 | 0.00000 | 0.00695 | 0.02031 | 0.00893 | 0.60220 | 0.53002 | 1.79468 |
| **Zfp85** | 0.00805 | 0.00417 | 0.20395 | 0.11547 | 0.10917 | 0.28538 | 0.00396 | 0.00000 | 0.23422 | 0.00212 | 1.92043 | 0.00751 | 0.03639 | 0.04277 |
| **Zfp850** | 0.07701 | 0.27608 | 1.05862 | 1.67709 | 0.80545 | 1.43406 | 0.04044 | 0.30700 | 3.75684 | 0.13056 | 0.04631 | 1.33328 | 0.58743 | 0.92718 |
| **Zfp862.ps** | 0.24522 | 0.00400 | 0.00130 | 0.86156 | 0.32384 | 0.41810 | 0.00396 | 0.00887 | 0.01158 | 0.00494 | 0.04918 | 0.02077 | 0.00934 | 0.05648 |
| **Zfp865** | 0.30276 | 0.00421 | 0.55346 | 1.39262 | 0.40450 | 0.31794 | 0.00413 | 0.00142 | 0.32772 | 0.01302 | 0.20891 | 0.00768 | 0.21815 | 0.00000 |
| **Zfp866** | 0.82341 | 0.06395 | 2.04624 | 0.00000 | 0.33862 | 0.53882 | 0.03529 | 0.74997 | 0.16089 | 0.01289 | 0.00000 | 0.51914 | 1.12091 | 0.97315 |
| **Zfp867** | 0.00000 | 0.01465 | 0.17061 | 0.37892 | 0.00000 | 0.00000 | 0.01080 | 0.52634 | 0.03353 | 0.01101 | 0.00300 | 0.00550 | 0.45018 | 0.69328 |
| **Zfp868** | 2.58584 | 0.00839 | 1.95485 | 0.84075 | 2.04496 | 2.49021 | 1.44267 | 0.42354 | 2.62828 | 0.03001 | 2.62902 | 1.39693 | 4.10977 | 3.44524 |
| **Zfp869** | 3.24568 | 0.01539 | 3.64780 | 1.75255 | 2.49687 | 3.19395 | 4.68984 | 2.58849 | 2.17939 | 0.04315 | 6.43427 | 1.31673 | 0.97412 | 8.13080 |
| **Zfp87** | 0.00283 | 0.01000 | 0.75778 | 0.49846 | 0.02814 | 2.16308 | 0.00350 | 0.00000 | 0.00717 | 0.00410 | 0.00000 | 2.12958 | 0.36089 | 2.29018 |
| **Zfp870** | 0.28122 | 0.06256 | 1.08836 | 0.00000 | 0.67622 | 1.97381 | 0.04054 | 1.15499 | 0.61155 | 0.17614 | 0.00000 | 0.54419 | 4.85690 | 0.59534 |
| **Zfp871** | 1.02844 | 0.01141 | 1.12162 | 0.63843 | 2.25222 | 0.76679 | 1.72321 | 0.01183 | 3.92070 | 0.04060 | 0.01545 | 0.74561 | 0.01714 | 0.72658 |
| **Zfp872** | 0.02205 | 0.00435 | 0.16435 | 0.01542 | 0.05720 | 0.01448 | 0.00404 | 0.00709 | 0.80671 | 0.00952 | 0.01405 | 0.01753 | 0.15466 | 0.00000 |
| **Zfp873** | 0.00000 | 0.00338 | 1.17576 | 0.00000 | 1.13924 | 0.10361 | 0.00336 | 0.00793 | 1.65879 | 0.00284 | 0.01422 | 0.52835 | 0.90600 | 2.76924 |
| **Zfp874a** | 3.00467 | 0.00213 | 0.49342 | 1.04589 | 1.84429 | 1.52739 | 0.00735 | 0.00000 | 10.45427 | 0.00000 | 3.83848 | 0.21582 | 0.74494 | 1.92694 |
| **Zfp874b** | 0.50828 | 0.16817 | 0.51090 | 0.76641 | 0.38369 | 1.12373 | 0.11722 | 0.24203 | 2.43039 | 0.09447 | 1.30652 | 1.45621 | 3.49597 | 2.08497 |
| **Zfp882** | 0.28548 | 0.12822 | 0.24323 | 0.63699 | 0.37582 | 0.59403 | 0.04718 | 0.04394 | 0.41964 | 0.08993 | 0.01190 | 0.73938 | 1.90693 | 0.75448 |
| **Zfp9** | 0.26191 | 0.04573 | 0.27067 | 0.59728 | 0.08262 | 0.15616 | 0.01688 | 0.06280 | 0.40550 | 0.35418 | 0.03813 | 0.12583 | 0.92330 | 0.23031 |
| **Zfp90** | 0.01149 | 0.03049 | 0.00603 | 0.00000 | 0.00707 | 0.16937 | 0.00376 | 0.00846 | 0.00417 | 0.00546 | 0.00177 | 0.80107 | 0.26883 | 0.14251 |
| **Zfp91** | 7.42774 | 0.32660 | 1.94520 | 4.08770 | 5.31019 | 5.22777 | 0.02422 | 3.44155 | 13.92312 | 0.02357 | 6.52949 | 3.23024 | 4.36508 | 3.93661 |
| **Zfp93** | 0.27420 | 0.00316 | 0.00708 | 0.00000 | 0.00974 | 0.38575 | 0.00357 | 0.00890 | 0.29465 | 0.00511 | 0.00000 | 1.17191 | 0.25839 | 0.00000 |
| **Zfp930** | 0.00000 | 0.05306 | 2.88370 | 1.71529 | 1.29084 | 2.37815 | 1.63934 | 3.16851 | 5.65754 | 0.19142 | 1.09249 | 2.47186 | 0.39819 | 1.42323 |
| **Zfp931** | 0.14067 | 0.17006 | 0.07427 | 0.18517 | 0.07535 | 0.32221 | 0.27660 | 0.12427 | 0.26485 | 0.54464 | 0.16375 | 0.26411 | 0.08300 | 0.00000 |
| **Zfp932** | 13.74682 | 0.04813 | 12.35962 | 2.36272 | 15.56882 | 8.77520 | 1.94159 | 3.64071 | 31.70434 | 2.22372 | 2.47489 | 1.43173 | 0.00000 | 9.45125 |
| **Zfp933** | 0.15781 | 0.05601 | 0.57372 | 0.55157 | 0.54312 | 1.02263 | 0.04066 | 0.07026 | 2.07270 | 0.03475 | 0.14421 | 0.29393 | 0.42994 | 0.78062 |
| **Zfp934** | 0.46444 | 0.11320 | 1.73424 | 1.40764 | 1.63420 | 4.01249 | 0.53513 | 0.06619 | 5.68142 | 0.48436 | 1.93313 | 1.42668 | 0.66940 | 0.00000 |
| **Zfp935** | 1.52012 | 0.05355 | 1.36934 | 2.32722 | 2.56055 | 0.64632 | 0.18336 | 0.18517 | 5.03695 | 0.39881 | 0.26660 | 0.85869 | 4.93574 | 2.29100 |
| **Zfp936** | 6.69042 | 0.00841 | 4.30177 | 2.82449 | 6.12036 | 3.30539 | 0.00000 | 0.46117 | 23.08682 | 1.67611 | 0.18871 | 1.90612 | 9.15807 | 2.33895 |
| **Zfp937** | 0.51075 | 0.17445 | 0.72029 | 0.33930 | 0.30874 | 0.40258 | 0.15681 | 1.19061 | 1.03859 | 0.02694 | 0.23623 | 0.70106 | 0.84036 | 0.95480 |
| **Zfp938** | 0.08928 | 0.01339 | 0.01771 | 0.34943 | 0.08100 | 0.02802 | 0.08328 | 0.05675 | 0.11725 | 0.00818 | 0.00939 | 0.01955 | 0.00762 | 0.01886 |
| **Zfp939** | 0.22949 | 0.01037 | 4.10838 | 0.58512 | 2.45763 | 0.00000 | 0.00132 | 0.00000 | 7.29509 | 4.66264 | 1.86319 | 0.00000 | 0.98454 | 2.55139 |
| **Zfp94** | 0.03757 | 0.00457 | 0.93003 | 0.03385 | 0.00000 | 1.38309 | 0.00417 | 0.00184 | 0.00526 | 0.00498 | 0.00496 | 0.01492 | 0.00263 | 0.13549 |
| **Zfp940** | 0.00404 | 0.00402 | 0.07307 | 0.62100 | 0.00000 | 0.44237 | 0.00522 | 0.28898 | 0.00409 | 0.00091 | 1.42062 | 0.24294 | 0.04403 | 0.15195 |

| | | | | | | | | | | | | | | |
|---|---|---|---|---|---|---|---|---|---|---|---|---|---|---|
| **Zfp941** | 0.08875 | 0.03964 | 0.01336 | 0.28965 | 0.05164 | 0.28114 | 0.02963 | 0.08902 | 0.16197 | 0.08809 | 0.00309 | 0.07580 | 0.70429 | 0.28828 |
| **Zfp942** | 3.10637 | 0.00329 | 0.64631 | 1.53963 | 0.70057 | 0.25807 | 1.08600 | 0.45838 | 5.66928 | 0.02894 | 1.37734 | 0.23879 | 0.76896 | 0.00000 |
| **Zfp943** | 0.53186 | 0.00373 | 0.49501 | 0.02093 | 0.52243 | 0.19986 | 0.00414 | 0.01877 | 0.16227 | 0.01268 | 0.00275 | 0.18135 | 0.83102 | 0.00000 |
| **Zfp944** | 0.91214 | 0.01184 | 1.12568 | 0.00000 | 0.27231 | 0.24447 | 0.01045 | 0.70266 | 3.55481 | 0.00435 | 0.01183 | 0.33376 | 1.39521 | 0.05618 |
| **Zfp945** | 0.00768 | 0.00432 | 0.03860 | 0.04893 | 0.03483 | 0.01931 | 0.01019 | 0.01298 | 0.21254 | 0.00665 | 0.01167 | 0.01050 | 0.00632 | 0.00827 |
| **Zfp946** | 0.00974 | 0.00428 | 0.00253 | 0.07317 | 0.04152 | 0.04681 | 0.00416 | 0.01000 | 0.26596 | 0.00697 | 0.01244 | 0.01224 | 0.00449 | 0.21912 |
| **Zfp947** | 0.75681 | 0.00378 | 1.11270 | 1.11613 | 3.10195 | 0.17040 | 0.03233 | 1.02111 | 8.55914 | 0.01662 | 0.00274 | 0.03092 | 2.16727 | 1.12965 |
| **Zfp948** | 0.87584 | 0.00424 | 0.73638 | 0.28991 | 0.16505 | 0.00000 | 0.04115 | 0.05832 | 2.52451 | 0.01037 | 0.56895 | 0.00520 | 5.78220 | 0.00000 |
| **Zfp949** | 0.19820 | 1.01070 | 1.72065 | 1.96351 | 2.80846 | 5.70630 | 0.41226 | 0.58461 | 5.38212 | 0.01132 | 0.66068 | 1.05024 | 3.78179 | 3.85552 |
| **Zfp950** | 0.60283 | 0.29547 | 1.42076 | 1.39804 | 1.13346 | 0.90740 | 2.64481 | 0.05182 | 7.49720 | 0.20610 | 0.19421 | 0.17358 | 2.36381 | 3.42573 |
| **Zfp951** | 2.92334 | 0.02310 | 5.45676 | 4.89410 | 3.11418 | 2.55320 | 0.17737 | 1.00949 | 12.95069 | 12.25921 | 0.43691 | 1.42909 | 3.99066 | 0.41974 |
| **Zfp952** | 0.14227 | 0.00963 | 0.00000 | 1.63651 | 0.34591 | 0.31777 | 1.40749 | 0.00000 | 0.01331 | 0.00826 | 0.79030 | 1.23605 | 0.03093 | 1.93705 |
| **Zfp953** | 0.03147 | 2.29163 | 0.46059 | 0.46550 | 0.17690 | 0.51556 | 0.00478 | 0.77803 | 0.03064 | 0.00540 | 0.22889 | 0.04794 | 0.44692 | 0.11863 |
| **Zfp954** | 3.51389 | 0.45292 | 5.78929 | 3.94793 | 5.26240 | 3.54448 | 3.01925 | 1.84288 | 1.38607 | 0.04080 | 2.84254 | 1.68719 | 5.06830 | 5.95227 |
| **Zfp955a** | 0.69897 | 0.00396 | 1.08609 | 0.14214 | 1.08514 | 1.43630 | 0.06909 | 0.00273 | 4.84804 | 0.01347 | 0.42619 | 0.22590 | 1.68218 | 2.05376 |
| **Zfp955b** | 0.95075 | 0.01182 | 0.00965 | 0.11374 | 1.72176 | 0.27972 | 0.38187 | 0.16352 | 7.07074 | 0.00381 | 0.63985 | 0.90729 | 0.71789 | 2.70742 |
| **Zfp956** | 0.00379 | 0.04614 | 0.04057 | 2.24156 | 0.00000 | 2.23039 | 1.30647 | 1.37295 | 0.00000 | 0.02205 | 4.52254 | 0.20994 | 1.91422 | 0.40737 |
| **Zfp957** | 0.00365 | 0.00433 | 0.00692 | 0.01315 | 0.00000 | 0.01023 | 0.00422 | 0.00911 | 0.15177 | 0.00638 | 0.00750 | 0.00879 | 0.37000 | 0.27060 |
| **Zfp958** | 2.15265 | 0.02370 | 5.30170 | 1.39799 | 4.64440 | 4.80260 | 0.02144 | 1.92902 | 5.31557 | 0.19166 | 6.22867 | 1.97449 | 5.37630 | 3.31485 |
| **Zfp959** | 0.12688 | 0.01905 | 0.82513 | 0.34726 | 0.62508 | 0.17375 | 0.04493 | 0.01073 | 0.71620 | 0.01044 | 0.03215 | 0.50504 | 0.32344 | 1.03033 |
| **Zfp960** | 9.26153 | 0.16769 | 11.12615 | 7.15237 | 6.36351 | 2.85928 | 0.79968 | 0.21961 | 22.21872 | 0.06173 | 2.99631 | 1.53566 | 2.39076 | 1.50263 |
| **Zfp961** | 2.89881 | 0.09171 | 4.50827 | 0.66478 | 1.42977 | 7.19834 | 0.19115 | 4.23985 | 13.21442 | 0.03773 | 3.59852 | 3.72102 | 2.90606 | 6.90391 |
| **Zfp963** | 0.00000 | 0.00391 | 0.25699 | 1.04797 | 0.25932 | 0.48383 | 0.00383 | 0.00229 | 0.00769 | 0.00458 | 0.00775 | 0.23128 | 0.02499 | 0.23298 |
| **Zfp97** | 7.64056 | 0.10743 | 8.43945 | 5.18935 | 4.78900 | 2.43404 | 0.32917 | 0.15604 | 17.67729 | 0.01988 | 2.10869 | 1.06318 | 1.57908 | 0.22283 |
| **Zfp970** | 4.37098 | 0.47144 | 6.93576 | 0.76930 | 10.62515 | 4.20382 | 1.01868 | 0.48844 | 18.61514 | 2.64340 | 2.02453 | 2.48365 | 21.15809 | 24.45259 |
| **Zfp971** | 0.23891 | 0.07871 | 1.00753 | 0.65446 | 0.63833 | 1.21044 | 0.31489 | 0.91569 | 2.77905 | 0.43421 | 1.30625 | 0.44140 | 0.32715 | 3.11114 |
| **Zfp972** | 0.30761 | 0.05197 | 0.35013 | 0.11348 | 0.76765 | 0.12352 | 0.08569 | 0.00000 | 1.93460 | 3.00458 | 0.11278 | 0.18455 | 4.17920 | 0.24762 |
| **Zfp973** | 0.01477 | 0.00442 | 0.05502 | 0.02690 | 0.01588 | 0.04751 | 0.05002 | 0.05991 | 0.08793 | 0.01652 | 0.03535 | 0.02565 | 0.01923 | 0.01421 |
| **Zfp974** | 0.10817 | 0.00404 | 0.00925 | 0.00000 | 0.19834 | 0.45401 | 0.00304 | 0.00320 | 6.58307 | 0.00661 | 0.00402 | 0.01892 | 2.82204 | 0.00000 |
| **Zfp975** | 0.58084 | 0.05988 | 0.36369 | 0.79550 | 0.93352 | 0.56997 | 0.02049 | 0.61757 | 3.31164 | 0.39414 | 0.44122 | 1.78445 | 1.04287 | 0.92290 |
| **Zfp976** | 0.08274 | 0.07094 | 0.43874 | 0.28158 | 0.41418 | 0.47724 | 0.04650 | 0.19325 | 2.64699 | 0.05059 | 0.41284 | 0.26183 | 0.47182 | 0.49254 |
| **Zfp979** | 1.02930 | 0.24512 | 3.76963 | 3.46113 | 2.34991 | 1.68200 | 0.03670 | 0.84380 | 16.07409 | 0.10629 | 1.50475 | 3.88615 | 3.86018 | 0.60884 |
| **Zfp980** | 2.39921 | 0.01582 | 4.53306 | 2.84831 | 2.32524 | 0.83803 | 0.37892 | 1.03903 | 19.02653 | 0.29879 | 1.59451 | 2.02792 | 8.00402 | 0.00000 |
| **Zfp981** | 4.28797 | 0.00010 | 4.58124 | 2.60120 | 2.27861 | 1.12810 | 1.81379 | 0.41409 | 17.79473 | 2.78778 | 2.02792 | 1.68141 | 8.83416 | 0.00000 |
| **Zfp982** | 0.15901 | 0.01199 | 0.66268 | 1.21510 | 1.27501 | 0.53485 | 0.15962 | 0.33645 | 9.23061 | 0.08798 | 0.78156 | 0.26779 | 0.00464 | 0.13307 |
| **Zfp983** | 0.00237 | 0.00343 | 0.25465 | 0.01261 | 0.01723 | 0.09167 | 0.00997 | 0.35680 | 0.96829 | 0.00000 | 2.04329 | 0.00738 | 0.39108 | 0.00000 |
| **Zfp984** | 2.62824 | 0.26776 | 8.87402 | 5.50448 | 7.80973 | 4.54260 | 0.78933 | 0.41417 | 23.78443 | 3.38193 | 3.71779 | 5.17287 | 6.32345 | 0.92905 |
| **Zfp985** | 1.51482 | 0.07921 | 5.36420 | 6.19625 | 3.25935 | 3.68498 | 1.16801 | 1.37058 | 11.00577 | 0.63918 | 3.39459 | 4.75224 | 4.38778 | 0.00000 |
| **Zfp987** | 0.42459 | 0.07085 | 4.34511 | 5.09040 | 4.70072 | 1.47604 | 0.44851 | 0.15194 | 17.23646 | 0.81151 | 1.93321 | 3.50131 | 6.48834 | 0.00000 |
| **Zfp991** | 0.07757 | 0.06368 | 0.50369 | 1.01185 | 0.07570 | 0.78172 | 0.53697 | 1.34224 | 1.03582 | 3.73136 | 0.58467 | 0.64470 | 0.00020 | 0.06129 |
| **Zfp992** | 1.86170 | 2.48484 | 0.49680 | 0.25176 | 0.41035 | 1.30657 | 0.03900 | 0.33096 | 1.07878 | 0.00000 | 2.81864 | 1.32132 | 0.75458 | 0.13271 |
| **Zfp994** | 1.52247 | 0.17047 | 1.08688 | 0.90111 | 0.59599 | 0.65352 | 0.31707 | 0.52001 | 2.18643 | 0.11001 | 0.57168 | 0.91225 | 2.69581 | 0.32241 |
| **Zfp995** | 0.05411 | 0.00387 | 0.26987 | 0.16668 | 0.10976 | 0.13095 | 0.00432 | 0.01970 | 0.75290 | 0.00486 | 0.19848 | 0.38891 | 0.00701 | 0.09014 |
| **Zfpl1** | 0.00000 | 0.00339 | 0.28910 | 9.04551 | 7.47574 | 0.15929 | 0.23019 | 0.29867 | 0.01011 | 0.02844 | 0.02672 | 0.72085 | 0.48393 | 0.00000 |
| **Zfpm1** | 0.02638 | 0.01132 | 0.03029 | 0.16550 | 0.00939 | 0.00501 | 0.00407 | 0.61159 | 0.01112 | 0.01944 | 0.02514 | 0.01048 | 0.08408 | 0.04509 |
| **Zfr** | 8.63390 | 0.62873 | 7.88162 | 4.82126 | 5.20119 | 5.98910 | 2.03542 | 2.53241 | 8.65434 | 0.00000 | 8.25111 | 6.72723 | 4.30228 | 1.81212 |
| **Zfr2** | 0.00305 | 0.00423 | 0.15611 | 0.31837 | 0.15855 | 0.00354 | 0.00440 | 0.00918 | 0.01104 | 0.00525 | 0.00610 | 0.00697 | 0.01014 | 0.00859 |
| **Zfx** | 6.12505 | 0.00111 | 2.62992 | 0.48421 | 5.88205 | 2.53979 | 3.46285 | 0.00000 | 10.61292 | 0.08617 | 3.78392 | 0.98436 | 3.15275 | 4.78777 |
| **Zfyve1** | 0.59806 | 0.00990 | 5.20332 | 5.30100 | 1.63855 | 0.22297 | 1.46747 | 1.48951 | 0.01359 | 0.05065 | 3.46168 | 0.96150 | 0.55827 | 0.26087 |
| **Zfyve16** | 0.00000 | 0.00287 | 0.19890 | 0.56084 | 0.05721 | 1.16313 | 0.15544 | 0.00964 | 1.79712 | 0.01511 | 0.41110 | 1.33119 | 0.38976 | 1.03461 |
| **Zfyve19** | 0.49189 | 0.07315 | 3.06691 | 6.38350 | 1.53789 | 3.82881 | 0.30060 | 4.18308 | 0.41350 | 0.01689 | 5.28511 | 2.99689 | 0.19379 | 1.89418 |
| **Zfyve21** | 0.00000 | 0.07594 | 0.99761 | 3.20432 | 0.20672 | 1.35415 | 1.06984 | 2.90065 | 0.00161 | 0.04052 | 0.06964 | 3.27067 | 1.20801 | 0.00000 |
| **Zfyve26** | 4.91229 | 0.02225 | 8.70406 | 6.19011 | 1.60932 | 9.21690 | 5.76490 | 2.68606 | 4.28289 | 0.22372 | 5.55772 | 0.80813 | 5.39856 | 10.36134 |
| **Zfyve27** | 2.88145 | 0.81153 | 2.54399 | 2.58371 | 2.87509 | 3.01146 | 0.83338 | 0.04759 | 2.34391 | 0.10047 | 0.97522 | 0.87928 | 2.91760 | 1.69166 |
| **Zfyve9** | 0.04996 | 0.00444 | 0.34383 | 0.08657 | 0.09368 | 0.00875 | 0.00422 | 0.01092 | 0.01056 | 0.00778 | 0.01208 | 0.01797 | 1.14651 | 0.36444 |

| | | | | | | | | | | | | | | |
|---|---|---|---|---|---|---|---|---|---|---|---|---|---|---|
| **Zg16** | 0.00773 | 0.00434 | 0.10977 | 0.12620 | 0.00932 | 0.00823 | 0.00435 | 0.00821 | 0.01206 | 0.00614 | 0.00707 | 0.00776 | 0.00721 | 0.01037 |
| **Zglp1** | 0.00135 | 0.00460 | 0.10272 | 0.37880 | 0.28498 | 0.00000 | 0.00495 | 0.13839 | 0.01022 | 0.00868 | 0.00660 | 0.00608 | 0.00324 | 0.14888 |
| **Zgpat** | 0.17610 | 0.01751 | 2.32177 | 3.45247 | 7.11662 | 1.14283 | 0.00311 | 0.00000 | 0.00000 | 0.01959 | 1.05076 | 1.49297 | 1.78038 | 1.91690 |
| **Zgrf1** | 0.28357 | 0.00349 | 0.98393 | 1.26261 | 0.94112 | 1.50713 | 2.19780 | 0.10471 | 0.78185 | 0.00084 | 0.53027 | 0.29339 | 0.46656 | 2.05241 |
| **Zhx1** | 0.00660 | 0.00444 | 0.05158 | 0.00493 | 0.04044 | 0.01194 | 0.00439 | 0.01016 | 0.35846 | 0.00685 | 0.00688 | 0.00906 | 0.14786 | 0.00000 |
| **Zhx3** | 0.61313 | 0.00391 | 0.68816 | 0.61569 | 0.16873 | 0.10804 | 0.00381 | 0.00042 | 0.00012 | 0.00440 | 0.00422 | 0.03038 | 2.03371 | 0.41870 |
| **Zic3** | 0.01357 | 0.01368 | 0.01780 | 0.00294 | 0.02485 | 0.01366 | 0.00420 | 0.01839 | 0.01416 | 0.00662 | 0.00535 | 0.18809 | 0.00602 | 0.01643 |
| **Zik1** | 0.01646 | 0.00439 | 0.01265 | 0.02663 | 0.01264 | 0.01091 | 0.00419 | 0.01970 | 0.01141 | 0.00724 | 0.05063 | 0.15524 | 0.00935 | 0.01425 |
| **Zkscan1** | 0.52882 | 0.00295 | 2.05162 | 0.65286 | 3.04689 | 1.83465 | 0.10383 | 0.00000 | 5.42812 | 0.00198 | 0.13726 | 0.67916 | 2.56529 | 3.23041 |
| **Zkscan14** | 1.56043 | 0.04375 | 0.69810 | 1.81193 | 4.50994 | 1.20894 | 1.69222 | 0.98992 | 0.89294 | 10.34175 | 0.09456 | 0.00000 | 2.24856 | 1.09577 |
| **Zkscan16** | 0.00747 | 0.00446 | 0.00799 | 0.01540 | 0.00396 | 0.01035 | 0.00420 | 0.01029 | 0.19940 | 0.00744 | 0.00750 | 0.00986 | 0.00356 | 0.06006 |
| **Zkscan17** | 0.00000 | 0.01746 | 1.60244 | 2.21715 | 2.10065 | 3.76728 | 0.03061 | 2.83142 | 0.01287 | 0.01919 | 2.64875 | 1.74041 | 1.22864 | 2.19566 |
| **Zkscan3** | 0.00000 | 0.01169 | 0.09552 | 0.27654 | 0.05237 | 0.00760 | 0.01047 | 0.06251 | 0.03400 | 0.01550 | 0.00626 | 1.17815 | 0.00826 | 0.99200 |
| **Zkscan4** | 0.00678 | 0.00429 | 0.01606 | 0.02526 | 0.00747 | 0.01067 | 0.00419 | 0.00835 | 0.04053 | 0.00609 | 0.01138 | 0.01119 | 0.33229 | 0.01163 |
| **Zkscan5** | 0.00000 | 0.02892 | 3.80721 | 0.00000 | 2.35749 | 0.59876 | 0.00266 | 0.35836 | 0.25775 | 0.00755 | 1.67027 | 3.49120 | 2.18833 | 2.51947 |
| **Zkscan6** | 0.98708 | 0.01946 | 1.18698 | 1.12305 | 1.19292 | 1.41975 | 2.84529 | 0.72540 | 0.53085 | 0.03591 | 0.00000 | 2.05172 | 0.70181 | 2.14650 |
| **Zkscan7** | 0.01403 | 0.17534 | 0.05099 | 0.00895 | 0.00860 | 0.00000 | 0.00377 | 0.00866 | 0.00989 | 0.00090 | 1.72224 | 0.09611 | 0.16245 | 0.06419 |
| **Zkscan8** | 0.03014 | 0.37076 | 0.01645 | 0.42986 | 0.00729 | 0.01298 | 0.00921 | 0.00339 | 0.01124 | 0.01118 | 0.01085 | 0.08003 | 0.02207 | 0.09341 |
| **Zmat2** | 0.00000 | 0.08572 | 1.37620 | 5.49539 | 1.89693 | 3.00495 | 0.37615 | 4.01862 | 3.63197 | 0.12483 | 3.80264 | 2.70245 | 3.08582 | 4.30391 |
| **Zmat3** | 1.45313 | 0.06887 | 0.49024 | 2.32272 | 0.78778 | 1.23938 | 0.04026 | 0.09605 | 0.71462 | 0.05881 | 1.55367 | 0.46880 | 0.04767 | 1.43855 |
| **Zmat4** | 0.16526 | 0.00437 | 0.01270 | 0.04679 | 0.00613 | 0.01070 | 0.00431 | 0.00949 | 0.00865 | 0.00636 | 0.00741 | 0.00811 | 0.01174 | 0.07284 |
| **Zmat5** | 5.14092 | 0.44678 | 6.57130 | 13.99692 | 11.39891 | 8.78293 | 2.62132 | 5.58140 | 0.00000 | 0.23666 | 4.56271 | 4.80492 | 3.90929 | 3.00550 |
| **Zmiz1** | 6.59518 | 0.12860 | 17.00664 | 3.34155 | 21.52105 | 16.40879 | 1.78089 | 9.04285 | 5.65929 | 9.71216 | 7.98336 | 10.06548 | 10.14583 | 21.19107 |
| **Zmiz2** | 2.57639 | 0.11150 | 6.43995 | 10.59062 | 7.01063 | 4.15561 | 3.12934 | 6.52092 | 0.11885 | 0.12789 | 3.78178 | 2.25548 | 1.26086 | 1.05937 |
| **Zmpste24** | 29.78447 | 2.13406 | 37.34096 | 24.85315 | 31.42239 | 28.50498 | 5.61440 | 6.00271 | 40.40747 | 0.35073 | 21.82579 | 4.95088 | 12.86491 | 36.27619 |
| **Zmym1** | 3.22150 | 0.01859 | 10.32691 | 4.73039 | 7.45505 | 3.20859 | 2.29769 | 1.50651 | 1.21061 | 0.02070 | 3.59297 | 8.68449 | 6.11655 | 0.90615 |
| **Zmym2** | 5.40521 | 0.04562 | 1.53488 | 1.69100 | 4.06808 | 0.84552 | 0.00658 | 2.08399 | 11.98149 | 0.00240 | 6.17039 | 2.12890 | 4.16535 | 1.74239 |
| **Zmym3** | 7.18380 | 0.12027 | 5.41287 | 3.88893 | 7.37138 | 2.22826 | 4.10543 | 0.63074 | 6.71662 | 0.16014 | 0.00000 | 1.96833 | 0.46273 | 7.70221 |
| **Zmym4** | 3.09313 | 0.03877 | 2.93265 | 0.48335 | 4.10544 | 3.65636 | 0.25225 | 0.00000 | 4.24816 | 0.01299 | 4.94218 | 3.92671 | 2.45712 | 4.01004 |
| **Zmym5** | 1.45574 | 0.02597 | 0.50360 | 0.38048 | 0.62004 | 1.41754 | 2.53851 | 0.03519 | 4.84100 | 0.11506 | 1.23037 | 0.81803 | 0.77994 | 2.34914 |
| **Zmym6** | 0.10292 | 0.11101 | 0.91194 | 0.82631 | 0.09145 | 1.83738 | 0.05181 | 0.07091 | 0.64197 | 0.03001 | 1.65429 | 0.88306 | 0.93762 | 0.23303 |
| **Zmynd11** | 0.72856 | 0.00419 | 0.03036 | 0.06681 | 1.25024 | 0.64476 | 0.00495 | 0.23332 | 0.03055 | 0.00357 | 0.80618 | 0.09197 | 0.00201 | 0.50640 |
| **Zmynd12** | 0.00876 | 0.00455 | 0.02132 | 0.01867 | 0.25083 | 0.01666 | 0.00425 | 0.00973 | 0.01292 | 0.00686 | 0.01049 | 0.00829 | 0.01908 | 0.00836 |
| **Zmynd19** | 8.87629 | 0.07103 | 1.89950 | 2.98059 | 7.36738 | 5.02285 | 0.72916 | 2.26963 | 10.41091 | 0.04201 | 4.35753 | 3.65869 | 11.64070 | 11.30593 |
| **Zmynd8** | 2.02828 | 0.06543 | 5.76655 | 0.50284 | 10.58090 | 2.24933 | 0.35231 | 1.81245 | 1.82148 | 0.26126 | 3.44904 | 2.54736 | 7.99991 | 5.81051 |
| **Znf41.ps** | 1.59356 | 1.74571 | 0.40195 | 0.47762 | 0.40877 | 1.80964 | 0.06736 | 0.33987 | 0.40820 | 0.00000 | 2.49624 | 1.28586 | 0.44734 | 0.43700 |
| **Znfx1** | 0.15575 | 0.02294 | 0.66935 | 0.78646 | 1.07883 | 1.16967 | 0.00407 | 0.23246 | 0.61040 | 0.01244 | 0.00810 | 0.32620 | 2.87524 | 0.00000 |
| **Znhit1** | 4.77481 | 0.14615 | 2.02839 | 12.27063 | 9.89238 | 4.64447 | 5.17368 | 4.09754 | 0.13842 | 0.32198 | 2.16599 | 5.13970 | 0.62905 | 2.80501 |
| **Znhit2** | 0.05331 | 0.00378 | 0.09513 | 1.23873 | 2.20513 | 0.21214 | 0.01013 | 0.00488 | 0.02557 | 0.00614 | 0.01283 | 0.34481 | 3.18007 | 0.00000 |
| **Znhit3** | 16.59490 | 0.05721 | 13.01219 | 6.38358 | 25.85436 | 14.23476 | 5.22292 | 5.26467 | 16.66192 | 0.32453 | 7.52617 | 3.72975 | 18.75319 | 24.12876 |
| **Znhit6** | 12.89151 | 8.01594 | 10.68778 | 15.25116 | 14.02542 | 7.88338 | 2.96238 | 3.12062 | 11.43000 | 11.48758 | 5.86445 | 10.25727 | 12.56538 | 13.37639 |
| **Znrd1** | 3.35624 | 0.30101 | 3.09079 | 9.24397 | 16.43977 | 10.36216 | 5.18404 | 2.71138 | 7.38568 | 11.42080 | 3.47030 | 8.09587 | 12.78494 | 16.15088 |
| **Znrd1as** | 0.16571 | 0.02179 | 0.67151 | 1.26346 | 0.79628 | 1.12423 | 0.00390 | 1.11034 | 1.54523 | 0.03668 | 0.07912 | 2.94723 | 0.00000 | 0.45640 |
| **Znrf1** | 1.84746 | 0.07405 | 6.23887 | 3.13250 | 8.29553 | 2.18350 | 0.00145 | 2.97987 | 0.25016 | 0.00000 | 1.69361 | 3.39750 | 3.50888 | 3.21619 |
| **Znrf2** | 8.42413 | 4.68818 | 8.21449 | 3.75382 | 14.65851 | 5.73610 | 6.12668 | 5.72237 | 21.66746 | 3.37943 | 8.06977 | 5.69544 | 9.28540 | 13.36405 |
| **Znrf3** | 0.14879 | 0.01773 | 1.28653 | 0.76202 | 0.88179 | 0.76948 | 0.00882 | 0.00552 | 3.52912 | 0.00438 | 0.08334 | 1.68253 | 2.25705 | 3.40812 |
| **Zp1** | 1.33432 | 0.00401 | 0.47224 | 0.14219 | 0.29573 | 0.00000 | 0.00409 | 0.00827 | 0.00491 | 0.00611 | 0.00582 | 0.00330 | 0.27383 | 1.70616 |
| **Zp2** | 2.47503 | 0.01010 | 0.34500 | 0.18127 | 3.31155 | 0.09557 | 0.00417 | 0.02955 | 0.61715 | 0.00470 | 0.00166 | 0.00000 | 6.21002 | 2.00933 |
| **Zp3** | 3.08205 | 0.00238 | 6.27449 | 1.36372 | 7.62795 | 0.00000 | 0.00238 | 0.00078 | 1.95036 | 0.00468 | 0.00400 | 0.00833 | 10.99445 | 8.06118 |
| **Zpbp2** | 0.33976 | 0.00474 | 0.17761 | 0.05899 | 0.30380 | 0.05821 | 0.00416 | 0.00557 | 0.05335 | 0.00509 | 0.00915 | 0.01143 | 0.00314 | 0.03721 |
| **Zpr1** | 11.75604 | 0.16265 | 17.45467 | 8.45278 | 19.24057 | 10.57005 | 4.21159 | 1.95602 | 7.76036 | 0.10773 | 6.55614 | 3.64531 | 9.91842 | 19.87427 |
| **Zranb1** | 1.86704 | 7.69315 | 1.39130 | 2.41819 | 2.01998 | 1.36548 | 2.48989 | 0.14098 | 9.73748 | 0.03084 | 0.81713 | 0.99287 | 1.48348 | 7.26233 |
| **Zranb2** | 7.92403 | 0.00550 | 10.06978 | 5.66533 | 9.67180 | 8.78603 | 2.39239 | 0.79003 | 11.41550 | 0.10572 | 2.96368 | 5.47833 | 4.48454 | 14.40468 |
| **Zranb3** | 0.30316 | 0.02424 | 3.27805 | 0.00000 | 1.28056 | 1.47456 | 0.00261 | 0.41421 | 1.29000 | 0.00000 | 2.09861 | 1.12288 | 0.82774 | 1.81515 |
| **Zrsr1** | 0.67929 | 0.00457 | 2.25359 | 0.45567 | 0.96273 | 1.61603 | 0.11940 | 0.10381 | 3.13220 | 0.00492 | 0.00242 | 0.00000 | 3.52477 | 4.41329 |

| | | | | | | | | | | | | | | |
|---|---|---|---|---|---|---|---|---|---|---|---|---|---|---|
| **Zrsr2** | 3.88701 | 0.02558 | 3.04188 | 3.45654 | 4.46515 | 6.54123 | 0.01000 | 0.97937 | 4.01981 | 0.42799 | 0.00000 | 3.34117 | 0.67581 | 2.87125 |
| **Zscan10** | 0.39119 | 0.08581 | 0.97222 | 0.48005 | 2.17279 | 0.00000 | 1.14767 | 0.00253 | 0.00061 | 0.01973 | 0.00000 | 7.65409 | 3.38127 | 0.03836 |
| **Zscan12** | 0.93506 | 0.01725 | 0.83224 | 2.69295 | 0.79326 | 1.81041 | 0.04585 | 0.00479 | 0.83760 | 0.01014 | 0.00000 | 2.29272 | 1.51702 | 0.86467 |
| **Zscan18** | 0.00000 | 0.00315 | 1.14648 | 1.33124 | 0.38730 | 0.42400 | 0.00333 | 0.00000 | 0.00171 | 0.00177 | 0.84809 | 0.41347 | 1.72032 | 2.02004 |
| **Zscan20** | 0.84178 | 0.00364 | 0.92015 | 0.00000 | 0.73970 | 0.00000 | 0.03999 | 0.00000 | 0.06289 | 0.01351 | 1.13044 | 0.21525 | 0.12526 | 0.72031 |
| **Zscan21** | 6.31183 | 0.56046 | 6.80949 | 6.07919 | 3.71235 | 9.48642 | 0.48042 | 1.31146 | 1.02877 | 0.04556 | 9.70707 | 3.69118 | 5.07701 | 10.45230 |
| **Zscan22** | 1.07164 | 0.00483 | 1.16281 | 0.00000 | 1.49166 | 1.32150 | 0.00389 | 0.68932 | 0.00400 | 0.00541 | 0.00000 | 0.75467 | 0.00286 | 0.54548 |
| **Zscan25** | 0.00000 | 0.00426 | 0.50177 | 0.11200 | 0.58715 | 0.08661 | 0.04535 | 0.01623 | 0.00909 | 0.00449 | 0.09335 | 0.09760 | 0.01984 | 0.13113 |
| **Zscan26** | 0.73693 | 6.82251 | 0.67369 | 1.68997 | 1.59824 | 1.85694 | 2.14203 | 0.50696 | 0.70789 | 0.09724 | 0.25879 | 0.07213 | 1.46976 | 1.35977 |
| **Zscan29** | 0.22558 | 0.02234 | 0.72112 | 0.20969 | 0.80919 | 1.65969 | 2.01030 | 0.53489 | 0.38693 | 0.05874 | 1.72696 | 0.02444 | 4.64613 | 0.23746 |
| **Zscan4a** | 0.00000 | 0.00410 | 0.02503 | 0.00714 | 0.15319 | 0.00869 | 0.00416 | 0.00895 | 0.21567 | 0.00598 | 0.00737 | 0.00701 | 2.10513 | 0.24542 |
| **Zscan4b** | 0.00207 | 0.00428 | 0.01420 | 0.01147 | 0.03882 | 0.01116 | 0.00424 | 0.00906 | 0.10666 | 0.00629 | 0.00827 | 0.00851 | 0.70074 | 0.03259 |
| **Zscan4c** | 0.00000 | 0.00408 | 0.01844 | 0.01255 | 0.09319 | 0.00794 | 0.00428 | 0.00849 | 0.53158 | 0.00629 | 0.00682 | 0.00669 | 2.32435 | 0.04998 |
| **Zscan4d** | 0.00000 | 0.00415 | 0.04856 | 0.00483 | 0.11647 | 0.00711 | 0.00412 | 0.00938 | 0.48931 | 0.00667 | 0.00678 | 0.00713 | 1.90382 | 0.00000 |
| **Zscan4f** | 0.00000 | 0.00407 | 0.01024 | 0.01409 | 0.09196 | 0.00759 | 0.00436 | 0.00832 | 0.58439 | 0.00633 | 0.00666 | 0.00666 | 2.37904 | 0.08506 |
| **Zscan5b** | 0.96957 | 0.00113 | 3.02664 | 0.21055 | 5.54102 | 0.17049 | 0.00106 | 0.00000 | 15.23314 | 0.00000 | 0.00586 | 0.00000 | 15.50477 | 22.16914 |
| **Zswim1** | 1.63206 | 0.15569 | 8.64597 | 9.63798 | 6.13665 | 3.35163 | 1.87496 | 9.10357 | 0.14281 | 0.05672 | 4.23821 | 3.57755 | 8.46419 | 1.20514 |
| **Zswim3** | 0.00000 | 0.00956 | 5.45055 | 0.00000 | 6.98507 | 3.91287 | 4.24952 | 2.20795 | 0.39629 | 0.07756 | 2.02192 | 1.10358 | 4.85483 | 6.19072 |
| **Zswim4** | 0.33501 | 0.00281 | 0.95648 | 0.59123 | 1.21709 | 0.24663 | 0.50475 | 0.77939 | 0.14273 | 0.02361 | 0.00000 | 1.50227 | 2.36237 | 1.77384 |
| **Zswim5** | 0.00000 | 0.01237 | 0.24568 | 0.16220 | 0.00000 | 0.18913 | 0.00970 | 0.01028 | 1.75428 | 0.00595 | 0.00455 | 0.50520 | 0.11430 | 0.00000 |
| **Zswim6** | 0.44914 | 0.00417 | 0.18450 | 0.00000 | 0.51557 | 0.63006 | 0.00390 | 0.01843 | 1.01810 | 0.00618 | 0.00479 | 0.24381 | 0.00221 | 0.00000 |
| **Zswim7** | 0.00557 | 0.01039 | 0.00000 | 2.44665 | 0.46332 | 1.02489 | 0.00399 | 0.38042 | 0.01120 | 0.89641 | 1.03239 | 2.34439 | 0.02413 | 0.01398 |
| **Zswim8** | 0.97726 | 0.02817 | 0.57736 | 0.00000 | 0.59867 | 1.98822 | 1.50026 | 2.38719 | 0.00476 | 1.39197 | 0.03899 | 1.31494 | 0.30433 | 0.34920 |
| **Zufsp** | 9.09937 | 0.00885 | 4.86635 | 1.05329 | 1.93786 | 1.97277 | 0.01465 | 1.44416 | 8.20601 | 0.00000 | 3.82253 | 1.65088 | 0.00000 | 0.72579 |
| **Zw10** | 1.31074 | 0.06400 | 5.96667 | 5.59559 | 1.56666 | 4.11200 | 1.05210 | 4.61004 | 2.94380 | 0.03075 | 4.67122 | 2.86866 | 3.70299 | 2.44214 |
| **Zwilch** | 1.36791 | 0.06180 | 5.67100 | 5.68174 | 4.49438 | 5.70103 | 0.97885 | 1.08762 | 5.87505 | 0.05309 | 9.41198 | 3.44377 | 2.50638 | 6.32457 |
| **Zwint** | 7.29727 | 2.25043 | 14.20145 | 7.03079 | 4.06310 | 16.31742 | 3.05504 | 4.87051 | 9.94709 | 8.84083 | 13.27278 | 11.03357 | 2.95575 | 6.00021 |
| **Zxda** | 0.05679 | 0.06335 | 0.02200 | 0.14665 | 0.04154 | 0.32628 | 0.38932 | 0.02698 | 0.26382 | 0.04896 | 0.02354 | 0.04429 | 0.17895 | 0.14466 |
| **Zxdc** | 0.20134 | 0.00298 | 0.74889 | 1.11391 | 0.71942 | 2.98157 | 0.05225 | 1.71465 | 1.37333 | 0.00000 | 2.62004 | 0.36243 | 0.00000 | 0.87336 |
| **Zyg11a** | 3.33954 | 2.44347 | 3.88463 | 2.09088 | 5.23876 | 0.00000 | 0.00208 | 0.00000 | 2.37145 | 0.00448 | 0.75598 | 1.13240 | 0.49228 | 6.63057 |
| **Zyg11b** | 0.91686 | 0.11202 | 1.02296 | 1.69158 | 2.17844 | 2.28069 | 2.60095 | 0.36556 | 6.94332 | 0.45046 | 3.57425 | 3.30060 | 6.04761 | 1.70939 |
| **Zyx** | 1.30430 | 7.54783 | 5.40206 | 3.95472 | 0.10257 | 3.75111 | 8.32785 | 15.85535 | 0.03083 | 6.10526 | 0.00000 | 5.79778 | 0.91468 | 0.00000 |
| **Zzef1** | 3.24487 | 0.04079 | 3.23709 | 2.13267 | 4.24466 | 2.62449 | 0.01959 | 3.42437 | 2.13395 | 0.04176 | 2.42038 | 2.80238 | 3.71450 | 6.84060 |
| **Zzz3** | 12.26663 | 0.03416 | 9.47385 | 1.57572 | 15.88563 | 5.19022 | 0.01486 | 0.38280 | 19.62325 | 0.03548 | 6.45114 | 5.75114 | 9.73422 | 15.67501 |

### Supplementary Table S10

#### Human male NMF coefficient matrix

| Gene | Cluster 1 | Cluster 2 | Cluster 3 | Cluster 4 | Cluster 5 | Cluster 6 | Cluster 7 | Cluster 8 | Cluster 9 | Cluster 10 | Cluster 11 | Cluster 12 |
|---|---|---|---|---|---|---|---|---|---|---|---|---|
| A1BG | 0.01065 | 0.12519 | 0.16888 | 0.58816 | 0.00000 | 0.03306 | 0.66803 | 10.01036 | 0.00000 | 0.00000 | 0.15939 | 1.66466 |
| A1BG-AS1 | 0.10918 | 0.06425 | 0.03971 | 0.01151 | 0.03362 | 0.00000 | 0.04057 | 0.11597 | 0.07577 | 0.00000 | 0.03480 | 0.21834 |
| A1CF | 1.18030 | 0.01045 | 0.01545 | 0.00810 | 0.03140 | 0.00000 | 0.00000 | 0.78403 | 0.08200 | 0.08622 | 0.01647 | 0.13157 |
| A2M | 0.00592 | 0.00623 | 0.30270 | 0.69115 | 0.00119 | 0.00271 | 0.00000 | 0.01111 | 0.12010 | 0.07883 | 0.00447 | 0.03990 |
| A2M-AS1 | 0.00512 | 0.00823 | 0.14641 | 0.00467 | 0.00000 | 0.13577 | 0.00000 | 0.01371 | 0.00000 | 0.06345 | 0.01519 | 0.11460 |
| A2ML1 | 2.61229 | 0.00000 | 1.47952 | 0.00095 | 0.04502 | 0.39744 | 0.00000 | 0.18380 | 1.03566 | 0.00000 | 0.25789 | 0.08482 |
| A2MP1 | 0.00626 | 0.01351 | 0.02981 | 0.00966 | 0.01318 | 0.02855 | 0.01673 | 0.01044 | 0.01896 | 0.01070 | 0.01458 | 0.01581 |
| A3GALT2 | 0.01021 | 0.03415 | 0.01682 | 0.02016 | 0.02022 | 0.00010 | 0.01572 | 0.09310 | 0.02481 | 0.02052 | 0.01540 | 0.22080 |
| A4GALT | 4.35830 | 22.89782 | 3.81468 | 13.04783 | 23.56960 | 0.00000 | 1.88494 | 0.00000 | 27.01796 | 19.11778 | 11.84582 | 12.36201 |
| A4GNT | 0.00282 | 0.02861 | 0.01253 | 0.09646 | 0.03661 | 0.00000 | 0.02007 | 0.03782 | 0.02880 | 0.01044 | 0.03486 | 0.03767 |
| AAAS | 3.86798 | 20.07037 | 2.65980 | 11.32142 | 15.07429 | 5.55037 | 3.53599 | 2.09597 | 8.93952 | 0.00000 | 13.96547 | 12.48683 |
| AACS | 3.90753 | 0.44186 | 1.55554 | 4.89749 | 8.65239 | 0.06112 | 4.38063 | 0.36418 | 6.58750 | 0.00000 | 3.05308 | 0.88543 |
| AACSP1 | 0.10501 | 0.04014 | 0.01149 | 0.00000 | 0.00000 | 0.02691 | 0.03248 | 0.59499 | 0.03222 | 0.09304 | 0.01485 | 0.04279 |
| AADAC | 0.00365 | 0.00053 | 0.08088 | 0.00000 | 0.01933 | 0.09177 | 0.00000 | 0.01038 | 0.09633 | 0.13947 | 0.00002 | 0.00628 |
| AADACL2 | 0.00564 | 0.00580 | 0.14467 | 0.01270 | 0.04625 | 0.07459 | 0.00000 | 0.01063 | 0.08786 | 0.02354 | 0.00584 | 0.01283 |
| AADACL2-AS1 | 0.00282 | 0.06763 | 0.48953 | 0.08198 | 0.19951 | 0.14757 | 0.16338 | 0.04521 | 0.58045 | 0.15807 | 0.05832 | 0.16266 |
| AADACL3 | 0.00360 | 0.00000 | 0.00000 | 0.10398 | 2.78186 | 0.02053 | 0.00000 | 0.00853 | 0.00000 | 0.17143 | 1.33419 | 0.00644 |
| AADACP1 | 0.00708 | 0.01073 | 0.03279 | 0.00566 | 0.01909 | 0.00000 | 0.00182 | 0.01145 | 0.11558 | 0.12498 | 0.01276 | 0.02370 |
| AADAT | 0.00000 | 0.00045 | 1.99622 | 0.59401 | 1.20702 | 3.11431 | 0.31569 | 0.22794 | 1.42440 | 0.97490 | 2.31981 | 0.09645 |
| AAED1 | 0.23381 | 0.98978 | 2.93629 | 0.00248 | 0.00000 | 1.67737 | 1.19193 | 0.64856 | 0.31857 | 0.00000 | 0.53184 | 1.19731 |
| AAGAB | 3.41612 | 7.75736 | 12.08216 | 4.41568 | 4.67888 | 17.34156 | 9.85124 | 1.89202 | 14.27172 | 0.00000 | 9.36463 | 6.35921 |
| AAK1 | 2.65122 | 0.39171 | 14.25561 | 1.05634 | 8.13374 | 4.74878 | 13.60588 | 3.61868 | 11.54405 | 22.17256 | 3.08297 | 3.47375 |
| AAMDC | 0.09698 | 6.77052 | 0.08972 | 3.58066 | 3.30438 | 4.96844 | 0.00000 | 3.28796 | 0.00000 | 2.34753 | 0.60322 | 12.05993 |
| AAMP | 0.20433 | 33.43126 | 27.42113 | 24.19008 | 26.76964 | 11.84315 | 37.11492 | 12.80448 | 37.49436 | 0.00000 | 28.10009 | 29.52437 |
| AANAT | 0.01958 | 0.01629 | 0.03705 | 0.02563 | 0.00522 | 0.00000 | 0.02988 | 0.01001 | 0.01619 | 0.04711 | 0.07070 | 0.04139 |
| AAR2 | 0.98700 | 12.49638 | 5.97693 | 6.92383 | 7.97533 | 8.49386 | 9.42327 | 9.24849 | 16.25870 | 0.00000 | 9.77999 | 15.13857 |
| AARD | 0.03141 | 0.00615 | 0.05540 | 0.05495 | 0.00000 | 0.13892 | 0.00000 | 0.04070 | 0.03942 | 0.10697 | 0.04384 | 0.04630 |
| AARS | 2.60138 | 3.98580 | 12.99391 | 15.71051 | 15.92766 | 1.73940 | 11.53758 | 0.71157 | 23.83032 | 0.00000 | 26.04039 | 9.52825 |
| AARS2 | 0.32640 | 12.29041 | 1.70688 | 11.91209 | 4.25417 | 0.00000 | 0.05203 | 0.43441 | 5.90089 | 9.07073 | 1.13250 | 7.31074 |
| AARSD1 | 0.00834 | 0.36799 | 0.09088 | 0.26285 | 0.11730 | 0.41519 | 0.20365 | 0.17055 | 0.15608 | 0.00000 | 0.54495 | 0.22210 |
| AASDH | 1.81221 | 1.40073 | 1.77544 | 1.72609 | 0.45616 | 8.75811 | 0.00000 | 2.16438 | 2.17392 | 5.91160 | 1.87076 | 3.48480 |
| AASDHPPT | 1.08149 | 9.59760 | 5.19483 | 2.62504 | 0.00000 | 24.45669 | 10.20785 | 0.87449 | 10.19697 | 3.32029 | 10.15962 | 6.44283 |
| AASS | 0.14851 | 1.21754 | 5.99296 | 2.75146 | 0.00000 | 1.24449 | 3.16880 | 0.55874 | 6.01323 | 10.09079 | 6.12182 | 0.66395 |
| AATF | 1.33723 | 17.75527 | 14.82085 | 10.21871 | 17.79898 | 12.68044 | 21.20229 | 9.83448 | 17.35037 | 0.00000 | 13.87781 | 14.72757 |
| AATK | 0.01505 | 0.03866 | 0.01477 | 0.00539 | 0.00000 | 0.06298 | 0.01592 | 0.01001 | 0.00234 | 0.04527 | 0.01577 | 0.10975 |
| ABAT | 1.85080 | 0.47278 | 0.20078 | 0.58489 | 0.00000 | 0.00000 | 0.06309 | 0.37812 | 0.00000 | 0.13245 | 0.00247 | 1.69065 |
| ABCA1 | 4.14062 | 1.31195 | 1.25174 | 9.94120 | 0.00000 | 6.72432 | 0.00901 | 0.19817 | 5.15780 | 2.36132 | 0.00000 | 0.13217 |
| ABCA10 | 0.35176 | 0.00000 | 0.01898 | 0.00000 | 0.01062 | 0.02623 | 0.01451 | 0.13076 | 0.01168 | 0.24630 | 0.00283 | 0.68522 |
| ABCA11P | 0.47787 | 1.00689 | 1.31640 | 0.67891 | 0.70555 | 1.59706 | 0.38490 | 2.32899 | 1.10467 | 0.06209 | 0.79280 | 1.53766 |
| ABCA12 | 0.53360 | 0.01548 | 0.02712 | 0.08053 | 0.05258 | 0.00000 | 0.00065 | 0.26463 | 0.05894 | 0.42407 | 0.00307 | 0.01017 |
| ABCA13 | 0.61796 | 2.21395 | 0.26933 | 0.89781 | 1.21908 | 0.00009 | 0.00172 | 0.20393 | 0.83015 | 0.00000 | 0.57813 | 1.09651 |
| ABCA17P | 0.29737 | 0.72368 | 0.06065 | 0.08936 | 0.00000 | 0.00000 | 0.02565 | 3.22974 | 0.00000 | 0.04170 | 0.00694 | 11.32334 |
| ABCA2 | 1.16109 | 0.22446 | 0.29524 | 0.01862 | 0.00000 | 0.00000 | 0.00726 | 0.01013 | 0.00000 | 0.29875 | 0.11471 | 0.19521 |
| ABCA3 | 2.68308 | 0.86822 | 0.04188 | 1.43338 | 0.27996 | 0.00000 | 0.01314 | 0.44926 | 0.00019 | 0.08075 | 0.04027 | 1.20057 |
| ABCA4 | 0.44453 | 0.01298 | 0.18889 | 0.01012 | 1.21300 | 0.00000 | 0.00065 | 0.00000 | 1.30693 | 0.02148 | 0.76680 | 0.08388 |
| ABCA5 | 1.49335 | 0.17668 | 0.07655 | 0.59153 | 0.12292 | 1.02847 | 0.00000 | 0.66615 | 0.45152 | 0.00000 | 0.00031 | 0.56415 |

| | | | | | | | | | | | | |
|---|---|---|---|---|---|---|---|---|---|---|---|---|
| **ABCA6** | 0.21601 | 0.00939 | 0.01714 | 0.01474 | 0.00801 | 0.01239 | 0.00925 | 0.02457 | 0.01431 | 0.07106 | 0.01318 | 0.05087 |
| **ABCA7** | 0.04196 | 0.07472 | 1.15744 | 1.02879 | 0.52072 | 0.00000 | 0.00001 | 0.17015 | 1.16883 | 2.08621 | 0.08788 | 0.29731 |
| **ABCA8** | 0.01660 | 0.02204 | 0.01984 | 0.01708 | 0.01099 | 0.01295 | 0.02583 | 0.01917 | 0.00235 | 0.01223 | 0.06633 | 0.01569 |
| **ABCA9** | 0.03660 | 0.04650 | 0.02904 | 0.00935 | 0.01187 | 0.00192 | 0.01050 | 0.05385 | 0.01605 | 0.06657 | 0.01694 | 0.01704 |
| **ABCB1** | 0.14357 | 0.27630 | 0.02394 | 0.00000 | 0.00000 | 0.00314 | 0.02486 | 0.91074 | 0.00237 | 0.03223 | 0.03602 | 0.76111 |
| **ABCB10** | 0.00000 | 0.44524 | 0.00000 | 0.00000 | 3.34647 | 9.00413 | 0.00001 | 0.04470 | 1.54630 | 11.87098 | 2.02959 | 0.46443 |
| **ABCB11** | 0.03096 | 0.01554 | 0.01538 | 0.01185 | 0.01572 | 0.01868 | 0.01689 | 0.01332 | 0.01999 | 0.01413 | 0.01421 | 0.01530 |
| **ABCB4** | 8.29632 | 0.02143 | 0.01611 | 0.00704 | 0.05897 | 0.00000 | 0.00750 | 0.54361 | 0.20702 | 0.00000 | 0.00261 | 0.00000 |
| **ABCB5** | 0.09530 | 0.09093 | 0.02622 | 0.70398 | 0.00000 | 0.06260 | 0.00000 | 0.07662 | 0.00000 | 0.05835 | 0.04118 | 0.32293 |
| **ABCB6** | 0.34650 | 12.86253 | 18.24454 | 8.17934 | 11.79296 | 0.00000 | 3.59749 | 0.06348 | 28.92085 | 0.96785 | 14.77185 | 4.84850 |
| **ABCB7** | 0.22314 | 2.85818 | 0.00000 | 3.74747 | 1.87185 | 6.54290 | 1.91358 | 4.46256 | 6.54893 | 5.80271 | 7.13692 | 2.62384 |
| **ABCB8** | 0.28923 | 21.74547 | 6.69942 | 11.69978 | 16.72708 | 0.00000 | 1.35872 | 0.00000 | 7.88803 | 1.54531 | 5.21032 | 11.52137 |
| **ABCB9** | 2.78357 | 0.31927 | 0.39917 | 0.30898 | 0.00177 | 0.00000 | 0.07161 | 1.25635 | 0.95678 | 0.00000 | 0.01312 | 0.75056 |
| **ABCC1** | 3.53518 | 1.23009 | 2.34225 | 0.86087 | 0.00000 | 0.14627 | 0.17762 | 0.11556 | 0.00000 | 1.39650 | 0.07368 | 0.95878 |
| **ABCC10** | 0.11569 | 1.70502 | 1.78305 | 5.86486 | 0.88697 | 0.00000 | 0.00000 | 0.00000 | 6.12221 | 2.10546 | 0.15862 | 0.08910 |
| **ABCC11** | 0.00936 | 0.01177 | 0.03353 | 0.03587 | 0.00488 | 0.00000 | 0.00930 | 0.05552 | 0.01725 | 0.13070 | 0.02127 | 0.02112 |
| **ABCC12** | 0.25006 | 0.06938 | 0.02044 | 0.18091 | 0.02150 | 0.03346 | 0.02426 | 0.15514 | 0.01397 | 0.00000 | 0.01020 | 0.05703 |
| **ABCC13** | 0.00592 | 3.08390 | 0.13716 | 0.26834 | 0.18831 | 0.09586 | 0.00000 | 0.03843 | 0.00000 | 0.01372 | 0.01888 | 0.03047 |
| **ABCC2** | 0.10941 | 0.00000 | 0.04279 | 6.11442 | 0.05602 | 1.14063 | 0.00000 | 0.08322 | 0.00000 | 0.45305 | 0.30801 | 0.55694 |
| **ABCC3** | 2.55908 | 0.00413 | 0.04471 | 0.01655 | 0.01197 | 0.02080 | 0.05212 | 0.19379 | 0.00423 | 0.00000 | 0.01272 | 0.10629 |
| **ABCC4** | 4.00358 | 3.14430 | 0.00000 | 0.34879 | 10.70329 | 5.16066 | 0.00000 | 2.46961 | 13.04027 | 6.15603 | 10.06048 | 0.13829 |
| **ABCC5** | 1.19035 | 5.75965 | 12.99941 | 2.98767 | 0.59422 | 9.53668 | 4.63144 | 3.01297 | 22.99132 | 10.22337 | 5.12224 | 13.26201 |
| **ABCC5-AS1** | 0.00448 | 0.00629 | 0.02629 | 0.01449 | 0.00000 | 0.02574 | 0.04614 | 0.01140 | 0.00118 | 0.13471 | 0.07687 | 0.02126 |
| **ABCC6** | 7.51650 | 0.07765 | 0.03013 | 0.00000 | 0.00000 | 0.01202 | 0.00016 | 5.12547 | 0.05709 | 0.13196 | 0.02169 | 0.06684 |
| **ABCC6P1** | 10.31901 | 0.13180 | 0.00000 | 0.00000 | 0.00000 | 0.00000 | 0.00000 | 12.97624 | 0.00305 | 0.10999 | 0.01555 | 0.77493 |
| **ABCC8** | 0.00270 | 0.00352 | 0.00451 | 0.00000 | 0.06873 | 0.06511 | 0.02425 | 0.16021 | 0.00000 | 0.08609 | 0.04586 | 0.04099 |
| **ABCC9** | 0.27607 | 0.00867 | 0.05179 | 0.01382 | 0.03232 | 0.01924 | 0.00000 | 0.00000 | 0.10617 | 0.20641 | 0.00000 | 0.08071 |
| **ABCD1** | 0.47895 | 0.18093 | 3.64139 | 0.00000 | 0.47729 | 0.00000 | 2.05607 | 0.00000 | 2.99448 | 0.00000 | 0.12555 | 0.22517 |
| **ABCD2** | 0.00549 | 0.02632 | 0.00691 | 0.00391 | 0.00000 | 0.04567 | 0.03169 | 0.01010 | 0.00000 | 0.05734 | 0.02883 | 0.01020 |
| **ABCD3** | 2.56613 | 4.86295 | 5.76368 | 0.75835 | 2.55918 | 24.46203 | 9.77090 | 1.22306 | 11.36248 | 9.82735 | 6.13168 | 4.10743 |
| **ABCD4** | 8.21505 | 3.34939 | 0.15571 | 4.64097 | 3.12141 | 0.00000 | 0.00000 | 2.78760 | 4.62711 | 0.86686 | 3.95993 | 3.35167 |
| **ABCE1** | 3.72148 | 8.86120 | 14.22034 | 2.53030 | 0.33002 | 41.84155 | 18.52073 | 3.65976 | 14.45834 | 9.42474 | 16.13267 | 10.98262 |
| **ABCF1** | 3.64958 | 15.36161 | 20.33914 | 12.39908 | 21.11019 | 10.78067 | 36.95158 | 17.54001 | 19.25552 | 30.06536 | 14.10606 | 36.29594 |
| **ABCF2** | 3.54466 | 23.03704 | 16.48937 | 10.79037 | 20.21609 | 9.56986 | 26.10459 | 25.94914 | 26.50482 | 0.00000 | 20.02253 | 38.32583 |
| **ABCF3** | 0.94539 | 14.06330 | 12.59484 | 10.64124 | 10.74942 | 3.15816 | 9.32102 | 4.22818 | 15.16260 | 0.00000 | 14.81003 | 12.87561 |
| **ABCG1** | 0.00352 | 0.02686 | 0.22002 | 0.00000 | 0.00047 | 0.04381 | 0.00000 | 0.01088 | 0.00000 | 0.16056 | 0.01893 | 0.03202 |
| **ABCG2** | 0.01225 | 0.00000 | 18.86650 | 0.08134 | 20.00290 | 0.74834 | 29.47965 | 0.02150 | 54.46542 | 0.00574 | 22.37912 | 0.00000 |
| **ABCG4** | 0.38529 | 0.00501 | 0.04753 | 0.01971 | 0.09082 | 0.01273 | 0.02703 | 0.10459 | 0.01976 | 0.00000 | 0.01472 | 0.50617 |
| **ABCG5** | 0.38894 | 0.00550 | 0.00257 | 0.01503 | 0.00215 | 0.04256 | 0.05109 | 0.00953 | 0.00441 | 0.15232 | 0.00412 | 0.00767 |
| **ABCG8** | 1.93028 | 0.18291 | 0.01271 | 0.00000 | 0.08528 | 0.00000 | 0.01517 | 0.15991 | 0.02063 | 0.01308 | 0.00689 | 0.03904 |
| **ABHD1** | 0.32758 | 0.00571 | 0.02236 | 0.00328 | 0.05582 | 0.03850 | 0.00310 | 0.00506 | 0.14264 | 0.26412 | 0.00482 | 0.04752 |
| **ABHD10** | 0.19460 | 3.50851 | 3.95087 | 2.02257 | 1.06292 | 9.46734 | 6.36234 | 0.00000 | 1.36586 | 0.00000 | 11.79569 | 2.74382 |
| **ABHD11** | 0.90012 | 8.33903 | 12.14205 | 5.94624 | 11.87800 | 4.73451 | 6.40217 | 2.09617 | 11.45111 | 0.00000 | 15.27991 | 3.63260 |
| **ABHD12** | 1.65821 | 8.63527 | 10.62291 | 7.56067 | 19.14028 | 0.00542 | 2.29291 | 1.70146 | 15.62922 | 13.57957 | 3.02789 | 1.48302 |
| **ABHD12B** | 1.51933 | 3.32834 | 1.09756 | 5.76893 | 1.43640 | 4.32052 | 0.73570 | 0.00000 | 0.00000 | 1.45464 | 3.33438 | 1.54052 |
| **ABHD13** | 0.30049 | 2.69835 | 3.74210 | 0.85182 | 0.02439 | 9.27410 | 0.00000 | 1.56239 | 4.59821 | 1.90205 | 3.09840 | 5.26513 |
| **ABHD14A** | 0.32315 | 0.09082 | 1.00155 | 0.67669 | 0.00000 | 0.00000 | 0.54178 | 0.02034 | 0.90909 | 0.00000 | 0.87659 | 0.32868 |
| **ABHD14B** | 1.28341 | 9.01728 | 1.42622 | 0.79053 | 0.00000 | 0.00000 | 3.21675 | 0.00000 | 2.98129 | 1.14823 | 6.18609 | 12.37794 |

| | | | | | | | | | | | | |
|---|---|---|---|---|---|---|---|---|---|---|---|---|
| **ABHD15** | 0.70770 | 2.09221 | 0.00000 | 2.05509 | 0.96463 | 3.71871 | 0.00000 | 0.05888 | 0.70491 | 2.23207 | 1.55846 | 2.30244 |
| **ABHD16A** | 4.57934 | 5.65273 | 3.93418 | 9.74635 | 3.01814 | 0.00000 | 0.00000 | 0.13926 | 7.49532 | 0.60640 | 3.29155 | 3.01102 |
| **ABHD16B** | 0.15362 | 0.00000 | 0.08607 | 0.03289 | 0.01187 | 0.04112 | 0.00000 | 0.00290 | 0.10499 | 0.40894 | 0.12981 | 0.10451 |
| **ABHD17A** | 0.01439 | 0.03696 | 0.15597 | 0.00395 | 0.35271 | 0.00000 | 0.29126 | 0.00603 | 0.00000 | 1.08553 | 0.00000 | 0.07463 |
| **ABHD17B** | 1.57451 | 1.57571 | 3.09500 | 2.31079 | 1.20108 | 3.89836 | 3.28746 | 3.10592 | 2.51293 | 7.78368 | 1.02213 | 0.33160 |
| **ABHD17C** | 0.00000 | 4.26000 | 3.40719 | 0.00000 | 4.44717 | 8.59567 | 10.07446 | 0.09458 | 0.30463 | 15.67670 | 2.99536 | 1.97943 |
| **ABHD2** | 7.96257 | 4.98824 | 3.78898 | 3.92309 | 3.47676 | 9.73000 | 2.52395 | 3.77679 | 6.96177 | 0.34334 | 4.97339 | 5.09143 |
| **ABHD3** | 1.48181 | 9.66555 | 6.37637 | 1.54199 | 2.83410 | 9.58960 | 0.00000 | 12.36315 | 8.29773 | 1.41360 | 2.51108 | 11.47780 |
| **ABHD4** | 0.07676 | 0.13664 | 14.77584 | 2.57981 | 7.62409 | 0.00096 | 8.02824 | 0.06451 | 12.28723 | 0.00000 | 2.56938 | 0.00145 |
| **ABHD5** | 0.89459 | 5.26775 | 13.03731 | 3.08089 | 3.61166 | 13.30873 | 8.90257 | 2.00777 | 9.90865 | 4.93073 | 10.76699 | 2.43919 |
| **ABHD6** | 0.00000 | 3.03934 | 3.95104 | 5.33367 | 3.84857 | 3.83971 | 0.00000 | 0.03328 | 1.72654 | 4.52551 | 2.98827 | 0.85206 |
| **ABHD8** | 0.00215 | 0.03554 | 1.20383 | 0.05662 | 0.15030 | 0.00000 | 1.39140 | 0.53710 | 2.06427 | 0.00000 | 1.35138 | 0.13086 |
| **ABI1** | 2.80503 | 7.06347 | 9.33875 | 7.60474 | 0.00000 | 20.11522 | 12.33581 | 4.53669 | 9.35099 | 4.27482 | 11.54622 | 3.80269 |
| **ABI2** | 0.98144 | 1.83405 | 3.45823 | 3.37950 | 0.69732 | 4.52350 | 2.66036 | 1.67079 | 1.67290 | 2.00003 | 2.11645 | 1.48878 |
| **ABI3** | 0.02521 | 0.02161 | 0.02649 | 0.15213 | 0.35182 | 0.12483 | 0.00247 | 0.16205 | 0.00000 | 0.00035 | 0.00429 | 0.23118 |
| **ABI3BP** | 3.59645 | 0.04789 | 0.14977 | 0.07206 | 0.00030 | 0.03971 | 0.02738 | 0.57075 | 0.00258 | 0.33940 | 0.01717 | 0.02772 |
| **ABL1** | 6.04879 | 3.15585 | 2.00796 | 4.84335 | 3.04940 | 0.00000 | 4.63562 | 0.19422 | 4.87918 | 8.32807 | 1.19931 | 1.88008 |
| **ABL2** | 2.14511 | 0.73951 | 0.00000 | 4.69630 | 3.07038 | 9.14541 | 6.41779 | 2.34056 | 0.56222 | 13.40352 | 2.44351 | 2.47002 |
| **ABLIM1** | 2.15666 | 2.21590 | 0.00000 | 0.64723 | 0.00000 | 0.99554 | 0.52756 | 0.13671 | 0.00000 | 0.94164 | 0.32367 | 1.28233 |
| **ABLIM2** | 1.41820 | 0.04726 | 0.32108 | 0.00000 | 0.07854 | 0.00000 | 0.00000 | 0.11789 | 0.00000 | 0.28384 | 0.03062 | 0.07241 |
| **ABLIM3** | 0.13242 | 0.06623 | 1.83691 | 0.40873 | 0.00000 | 0.03144 | 0.88901 | 0.00320 | 0.03510 | 0.00000 | 0.00000 | 0.07206 |
| **ABO** | 0.09605 | 0.08351 | 0.01079 | 0.06695 | 0.08133 | 0.00000 | 0.02161 | 0.00000 | 0.00592 | 0.02707 | 0.06086 | 0.28981 |
| **ABR** | 4.72786 | 3.70100 | 8.05949 | 10.42300 | 7.30172 | 0.00000 | 11.97247 | 1.49212 | 7.41724 | 9.36250 | 2.71637 | 0.97087 |
| **ABRA** | 0.00540 | 0.00985 | 0.14253 | 0.02871 | 0.01340 | 0.00000 | 0.00000 | 0.00895 | 0.03423 | 0.05668 | 0.10878 | 0.04846 |
| **ABRACL** | 0.14192 | 16.90840 | 4.85413 | 3.35650 | 4.35736 | 25.05484 | 10.57506 | 0.00000 | 8.38856 | 2.78170 | 7.81298 | 1.43520 |
| **ABT1** | 0.00000 | 19.30925 | 20.84068 | 15.55134 | 14.65784 | 19.03286 | 26.88082 | 21.30738 | 20.28262 | 0.00000 | 21.76722 | 28.96443 |
| **ABTB1** | 0.09972 | 0.60758 | 0.62492 | 0.49624 | 0.32104 | 0.00000 | 0.55645 | 0.36031 | 0.64642 | 0.08979 | 0.17210 | 1.84028 |
| **ABTB2** | 5.64879 | 6.99051 | 3.42563 | 5.03194 | 1.11324 | 0.00000 | 0.74828 | 1.55666 | 3.48423 | 2.85428 | 0.01049 | 6.69255 |
| **AC093375.1** | 0.00445 | 0.26009 | 0.06439 | 0.00000 | 0.14937 | 0.16010 | 0.00000 | 0.00544 | 0.03722 | 0.12171 | 0.04458 | 0.08179 |
| **ACAA1** | 4.23159 | 8.94522 | 13.43089 | 9.87054 | 10.05237 | 0.00000 | 8.25826 | 3.00524 | 15.36039 | 0.00000 | 16.75333 | 1.60102 |
| **ACAA2** | 3.48171 | 18.36869 | 3.65531 | 16.63684 | 15.31319 | 28.10130 | 20.44280 | 2.08450 | 17.72760 | 1.17179 | 21.23384 | 5.70060 |
| **ACACA** | 1.72285 | 2.90405 | 10.60326 | 4.87728 | 12.61122 | 6.49509 | 12.91812 | 0.66974 | 20.68213 | 17.13302 | 9.06541 | 3.70663 |
| **ACACB** | 0.71218 | 0.21435 | 0.00000 | 0.54995 | 0.00000 | 0.20860 | 0.00000 | 0.40911 | 0.00000 | 0.45975 | 0.02316 | 0.51970 |
| **ACAD10** | 0.47829 | 3.91598 | 1.00626 | 2.43336 | 3.64760 | 0.00000 | 1.03143 | 1.90582 | 1.32696 | 0.19542 | 2.97609 | 2.58301 |
| **ACAD11** | 0.67768 | 0.20181 | 0.07210 | 0.21495 | 0.00000 | 0.02836 | 0.01539 | 0.41192 | 0.24525 | 0.00010 | 0.13711 | 0.03006 |
| **ACAD8** | 1.90080 | 5.95459 | 2.82663 | 5.99106 | 3.14265 | 6.29446 | 7.01817 | 0.75306 | 4.54288 | 0.00000 | 9.15536 | 3.53347 |
| **ACAD9** | 2.10525 | 5.12383 | 4.62025 | 8.79101 | 3.03930 | 5.79470 | 5.20148 | 2.35731 | 8.49969 | 0.00215 | 9.25392 | 5.27160 |
| **ACADL** | 0.00462 | 0.00131 | 0.05402 | 0.39936 | 0.04217 | 0.12882 | 0.01820 | 0.02166 | 0.00000 | 0.01099 | 0.00984 | 0.01308 |
| **ACADM** | 0.21958 | 1.82034 | 7.15786 | 5.74176 | 0.00000 | 10.51521 | 11.39562 | 0.00000 | 10.89069 | 4.77949 | 13.13210 | 0.14328 |
| **ACADS** | 0.00000 | 10.15912 | 2.98419 | 7.27648 | 6.16645 | 0.00000 | 0.08969 | 0.33932 | 0.00000 | 1.24174 | 1.61876 | 0.84588 |
| **ACADSB** | 1.23886 | 0.73098 | 2.02629 | 0.67768 | 2.02805 | 8.66043 | 3.71616 | 5.12238 | 3.14356 | 2.57767 | 10.00953 | 0.18645 |
| **ACADVL** | 0.68163 | 4.32959 | 14.29361 | 4.68069 | 4.30879 | 0.00000 | 4.38656 | 0.55115 | 7.47629 | 4.93067 | 6.28187 | 0.18391 |
| **ACAN** | 0.13361 | 0.01728 | 0.10570 | 0.04533 | 0.10328 | 0.04568 | 0.00583 | 0.01207 | 0.00950 | 0.00000 | 0.01710 | 0.05155 |
| **ACAP1** | 0.00130 | 0.68794 | 0.04881 | 0.09143 | 0.00000 | 0.00000 | 0.16223 | 0.43300 | 0.00000 | 0.43145 | 0.31836 | 0.75465 |
| **ACAP2** | 3.95157 | 1.27738 | 3.54313 | 3.02461 | 0.00000 | 15.21644 | 6.34932 | 1.38059 | 5.98861 | 13.16520 | 2.36880 | 1.95281 |
| **ACAP3** | 0.00161 | 0.07837 | 0.74649 | 0.03624 | 0.75191 | 0.00000 | 0.19806 | 0.07440 | 1.00614 | 4.80537 | 0.00011 | 0.00261 |
| **ACAT1** | 1.84069 | 8.01315 | 5.67243 | 1.76926 | 1.03640 | 17.62753 | 11.72835 | 1.49251 | 16.46552 | 0.00000 | 16.54704 | 2.84569 |
| **ACAT2** | 3.04825 | 2.59714 | 11.81748 | 22.76191 | 27.47887 | 26.99725 | 18.79109 | 6.45035 | 26.52629 | 0.00000 | 40.10344 | 0.58859 |

| | | | | | | | | | | | | |
|---|---|---|---|---|---|---|---|---|---|---|---|---|
| **ACBD3** | 2.50962 | 1.58803 | 4.41532 | 2.87057 | 1.06290 | 11.93056 | 8.73831 | 10.61700 | 0.00000 | 14.42866 | 3.54626 | 3.71680 |
| **ACBD4** | 0.05905 | 0.06069 | 0.58475 | 0.01771 | 0.00000 | 0.00203 | 0.05119 | 0.02060 | 0.09114 | 0.02895 | 0.13249 | 0.09966 |
| **ACBD5** | 0.00725 | 2.72247 | 1.73028 | 2.01821 | 0.00000 | 7.56874 | 0.00000 | 6.92491 | 4.86946 | 1.26036 | 4.32857 | 5.86628 |
| **ACBD6** | 0.13569 | 5.61357 | 4.48960 | 2.29197 | 3.82296 | 4.42393 | 3.23707 | 0.00000 | 4.07003 | 2.84831 | 3.19675 | 2.49886 |
| **ACBD7** | 0.11959 | 0.13416 | 0.03940 | 0.15319 | 0.00000 | 0.00000 | 0.17506 | 0.64444 | 0.00000 | 0.26622 | 0.04819 | 0.30872 |
| **ACCS** | 0.00255 | 0.04585 | 0.05596 | 0.01073 | 0.01198 | 0.00000 | 0.00828 | 0.04700 | 0.04019 | 0.09955 | 0.03283 | 0.00889 |
| **ACCSL** | 20.89981 | 0.00000 | 0.00746 | 0.00749 | 0.00701 | 0.00000 | 0.00671 | 3.85231 | 0.00392 | 0.00835 | 0.04559 | 0.57471 |
| **ACD** | 2.53854 | 9.02484 | 8.30713 | 6.64784 | 8.51505 | 0.00000 | 6.20765 | 1.36703 | 6.98692 | 0.00000 | 8.77630 | 5.06099 |
| **ACE** | 0.05894 | 5.93939 | 0.00000 | 4.16872 | 2.03997 | 0.13257 | 0.00000 | 0.00000 | 0.00000 | 0.50043 | 0.00000 | 7.55190 |
| **ACE2** | 4.72425 | 3.61209 | 0.00000 | 0.00000 | 30.31944 | 1.77120 | 0.00000 | 0.00000 | 0.00000 | 0.58649 | 13.84540 | 0.00000 |
| **ACER1** | 0.11741 | 0.14198 | 0.01271 | 0.00185 | 0.00760 | 0.02217 | 0.01687 | 0.00000 | 0.01791 | 0.01254 | 0.01056 | 0.23604 |
| **ACER2** | 0.65034 | 1.12318 | 1.32416 | 0.81817 | 0.31483 | 0.00000 | 0.00000 | 1.43446 | 1.31595 | 0.37205 | 0.00000 | 3.15882 |
| **ACER3** | 1.35896 | 2.07438 | 6.67443 | 0.02445 | 0.75373 | 5.03943 | 1.46318 | 0.37217 | 14.79297 | 1.67270 | 2.77382 | 5.11377 |
| **ACHE** | 0.14846 | 0.53547 | 0.10080 | 0.89268 | 0.54385 | 0.00000 | 0.00000 | 0.09548 | 0.39596 | 0.28969 | 0.02976 | 0.43633 |
| **ACIN1** | 3.41482 | 4.75666 | 10.30311 | 8.35779 | 16.69859 | 0.66757 | 24.03763 | 5.39680 | 11.19333 | 32.33155 | 7.04664 | 10.65990 |
| **ACKR2** | 0.67842 | 0.00000 | 3.44003 | 0.00000 | 5.40778 | 0.00000 | 0.45059 | 0.00000 | 14.84801 | 0.89343 | 4.92914 | 0.10669 |
| **ACKR3** | 0.00433 | 0.04838 | 0.58939 | 2.20087 | 3.79920 | 0.00000 | 0.14800 | 0.00871 | 0.00000 | 0.04032 | 0.26276 | 0.00968 |
| **ACKR4** | 0.18475 | 0.02656 | 0.00560 | 0.00276 | 0.00000 | 0.06537 | 0.02375 | 0.00394 | 0.02746 | 0.11390 | 0.01898 | 0.01704 |
| **ACLY** | 10.70157 | 12.80914 | 17.32608 | 19.05566 | 32.52326 | 18.78895 | 31.48702 | 3.43102 | 26.93463 | 12.79024 | 29.67128 | 6.81097 |
| **ACMSD** | 0.01005 | 0.00784 | 0.04686 | 0.01977 | 0.04221 | 0.04394 | 0.01180 | 0.01014 | 0.04074 | 0.02245 | 0.01775 | 0.02313 |
| **ACN9** | 1.72816 | 12.76419 | 7.58698 | 0.87816 | 0.68537 | 22.93976 | 5.78620 | 2.92887 | 9.25687 | 2.56143 | 9.73512 | 15.40432 |
| **ACO1** | 2.68901 | 4.07784 | 5.73315 | 9.53536 | 5.53393 | 13.68097 | 6.44868 | 0.52326 | 6.92446 | 0.59502 | 13.92706 | 0.16119 |
| **ACO2** | 0.59435 | 13.88456 | 11.52997 | 14.30911 | 5.46499 | 6.38091 | 2.18649 | 2.72233 | 21.14626 | 0.00000 | 17.46494 | 5.69815 |
| **ACOT1** | 0.08347 | 0.01785 | 0.18729 | 0.21862 | 0.02510 | 0.16259 | 0.00000 | 0.09666 | 0.14376 | 0.00000 | 0.23098 | 0.11512 |
| **ACOT11** | 0.00403 | 1.30813 | 0.46670 | 6.61551 | 0.08566 | 0.00000 | 0.03240 | 1.65216 | 0.00000 | 0.00000 | 0.13677 | 5.07293 |
| **ACOT12** | 0.01231 | 0.00946 | 0.00793 | 0.01054 | 0.01176 | 0.04112 | 0.00920 | 0.03672 | 0.01315 | 0.05527 | 0.02331 | 0.03421 |
| **ACOT13** | 2.42439 | 3.72068 | 9.56415 | 6.70070 | 4.43471 | 10.45275 | 3.14103 | 3.48425 | 3.04487 | 0.00000 | 5.97431 | 6.05721 |
| **ACOT2** | 0.54433 | 0.24781 | 2.03001 | 0.23750 | 0.00000 | 1.06380 | 0.00000 | 1.09428 | 0.37779 | 0.00000 | 1.35043 | 4.21680 |
| **ACOT4** | 0.93255 | 0.00000 | 1.78723 | 0.00000 | 0.86532 | 2.51857 | 1.02679 | 1.52668 | 1.22996 | 0.00000 | 3.82496 | 1.56878 |
| **ACOT6** | 0.00910 | 0.00969 | 0.04722 | 0.01289 | 0.05060 | 0.04965 | 0.01589 | 0.08884 | 0.10263 | 0.00000 | 0.06517 | 0.03671 |
| **ACOT7** | 1.53407 | 15.97607 | 10.32803 | 5.57989 | 13.93856 | 6.81537 | 21.93724 | 1.00918 | 12.77603 | 0.00000 | 8.10490 | 3.37932 |
| **ACOT8** | 0.56341 | 6.49560 | 1.85913 | 5.34536 | 3.37897 | 5.38603 | 0.00000 | 0.00000 | 2.57341 | 0.00000 | 2.37441 | 4.85943 |
| **ACOT9** | 3.85120 | 3.07008 | 1.38901 | 0.53801 | 0.47882 | 0.98452 | 1.09541 | 2.20810 | 1.84984 | 0.76807 | 1.19806 | 8.90011 |
| **ACOX1** | 0.88831 | 3.21629 | 15.39517 | 4.40235 | 11.51278 | 7.29811 | 10.77721 | 0.36460 | 10.61336 | 0.68828 | 18.10252 | 1.33485 |
| **ACOX2** | 0.01176 | 0.00235 | 0.00495 | 0.25791 | 0.01044 | 0.07340 | 0.01652 | 0.03408 | 0.00264 | 0.05513 | 0.01008 | 0.01196 |
| **ACOX3** | 0.92992 | 2.37931 | 3.85993 | 0.83193 | 0.83102 | 1.97763 | 0.75535 | 1.84949 | 1.06159 | 0.00000 | 3.48500 | 3.43203 |
| **ACOXL** | 0.07306 | 1.24475 | 0.76192 | 5.26480 | 1.96184 | 1.92650 | 4.14738 | 0.37153 | 8.94410 | 0.00000 | 3.29920 | 0.00000 |
| **ACP1** | 3.55182 | 20.77722 | 17.60307 | 13.05662 | 15.58795 | 29.53688 | 28.31984 | 12.80846 | 20.14158 | 5.00732 | 16.26458 | 29.78514 |
| **ACP2** | 0.50217 | 4.79502 | 1.21575 | 3.26909 | 4.34709 | 3.92016 | 0.00000 | 0.00000 | 3.35369 | 0.00000 | 3.23821 | 1.70667 |
| **ACP5** | 0.00692 | 13.76939 | 4.58444 | 4.96201 | 6.00577 | 2.56215 | 2.54603 | 0.13642 | 5.43358 | 0.00000 | 9.33963 | 5.04856 |
| **ACP6** | 0.13191 | 0.87642 | 5.65074 | 1.19119 | 0.02512 | 1.65773 | 1.39387 | 0.20052 | 0.00000 | 0.00000 | 2.58568 | 0.46644 |
| **ACPP** | 0.00553 | 0.00000 | 0.94890 | 4.72198 | 0.00000 | 0.63686 | 0.83790 | 0.00000 | 2.11673 | 0.00000 | 0.36206 | 0.10709 |
| **ACPT** | 0.01425 | 0.02409 | 0.09768 | 0.20870 | 0.24636 | 0.00000 | 0.00000 | 0.05235 | 0.17052 | 0.01919 | 0.00184 | 0.17252 |
| **ACR** | 0.02640 | 0.02711 | 0.01609 | 0.00506 | 0.01438 | 0.02184 | 0.00510 | 0.00778 | 0.01692 | 0.11452 | 0.01073 | 0.01086 |
| **ACRBP** | 0.16214 | 0.13575 | 0.03712 | 0.01849 | 0.08352 | 0.00000 | 0.00191 | 0.00000 | 0.06350 | 0.04977 | 0.00251 | 0.28948 |
| **ACRC** | 0.00000 | 0.12101 | 0.16592 | 0.16690 | 0.03155 | 0.48975 | 0.27987 | 0.17227 | 0.69740 | 0.57232 | 0.05906 | 0.21814 |
| **ACRV1** | 0.00140 | 0.08599 | 0.15171 | 0.43193 | 0.06639 | 0.09348 | 0.20752 | 0.09676 | 0.00177 | 0.02051 | 0.06136 | 0.20205 |
| **ACSBG1** | 0.14991 | 0.00000 | 0.00464 | 0.02136 | 0.04381 | 0.20930 | 0.02281 | 0.17053 | 0.16571 | 0.45633 | 0.05041 | 0.00169 |

| | | | | | | | | | | | | |
|---|---|---|---|---|---|---|---|---|---|---|---|---|
| **ACSBG2** | 0.01883 | 0.03414 | 0.07527 | 0.01063 | 0.02022 | 0.00000 | 0.03671 | 0.00733 | 0.05043 | 0.06739 | 0.04186 | 0.14436 |
| **ACSF2** | 0.00409 | 0.13548 | 0.62439 | 1.88105 | 0.32517 | 0.52125 | 0.05152 | 0.00705 | 0.12800 | 0.00000 | 0.12098 | 0.00143 |
| **ACSF3** | 0.47151 | 3.71713 | 0.13324 | 1.46152 | 3.97792 | 0.00000 | 0.12461 | 0.00000 | 4.72714 | 0.81774 | 4.21338 | 1.74839 |
| **ACSL1** | 0.65346 | 0.94551 | 3.81502 | 0.89926 | 0.00000 | 0.03537 | 1.36981 | 2.59715 | 0.00301 | 0.00000 | 0.00000 | 0.61905 |
| **ACSL3** | 2.48164 | 21.31898 | 8.04722 | 9.18730 | 14.04473 | 34.33203 | 0.00000 | 5.40960 | 40.96067 | 20.88027 | 24.37546 | 23.21167 |
| **ACSL4** | 0.28078 | 2.76829 | 0.00000 | 3.06487 | 3.49826 | 12.48457 | 1.37664 | 1.11272 | 0.00000 | 9.64784 | 12.47626 | 8.35453 |
| **ACSL5** | 0.14546 | 0.04900 | 0.03327 | 0.04121 | 0.00000 | 0.06717 | 0.03270 | 0.00782 | 0.02283 | 0.16783 | 0.04148 | 0.01160 |
| **ACSL6** | 0.00215 | 0.00000 | 0.00000 | 0.00000 | 0.34120 | 0.00000 | 0.01960 | 0.00630 | 5.47714 | 0.70668 | 6.70163 | 0.00697 |
| **ACSM1** | 0.00371 | 0.01213 | 0.01129 | 0.00529 | 0.04328 | 0.01032 | 0.05407 | 0.01277 | 0.06243 | 0.06612 | 0.00674 | 0.10073 |
| **ACSM2A** | 0.00774 | 0.01541 | 0.01747 | 0.01265 | 0.01719 | 0.01107 | 0.02570 | 0.01062 | 0.00790 | 0.01679 | 0.03154 | 0.01969 |
| **ACSM3** | 0.03686 | 0.00000 | 0.00000 | 0.89993 | 0.03143 | 0.26192 | 0.36845 | 0.07144 | 0.00000 | 0.00000 | 0.00453 | 0.01612 |
| **ACSM4** | 0.00591 | 0.02782 | 0.01410 | 0.01349 | 0.01962 | 0.01251 | 0.01777 | 0.00877 | 0.02096 | 0.01241 | 0.01659 | 0.06159 |
| **ACSM5** | 0.01384 | 0.04831 | 0.01298 | 0.00798 | 0.01042 | 0.01448 | 0.02328 | 0.13154 | 0.03660 | 0.00964 | 0.01214 | 0.05014 |
| **ACSM6** | 0.01250 | 0.01535 | 0.02436 | 0.00739 | 0.10297 | 0.00000 | 0.00672 | 0.00765 | 0.06238 | 0.09273 | 0.03903 | 0.04593 |
| **ACSS1** | 0.00850 | 0.00800 | 0.08832 | 0.03634 | 0.00000 | 0.04927 | 0.00485 | 0.01045 | 0.00000 | 0.04048 | 0.01737 | 0.01251 |
| **ACSS2** | 1.65091 | 1.68769 | 2.23917 | 6.37573 | 21.17104 | 0.00000 | 2.52335 | 1.50187 | 4.55311 | 3.78038 | 9.98803 | 9.25003 |
| **ACSS3** | 0.96232 | 0.00000 | 0.03185 | 0.05200 | 0.01305 | 0.10828 | 0.28413 | 0.08876 | 0.00753 | 0.00000 | 0.14124 | 0.00246 |
| **ACTA1** | 0.43193 | 0.07451 | 0.53698 | 0.04048 | 0.00000 | 0.07944 | 0.00000 | 0.06855 | 0.00000 | 1.09253 | 0.23433 | 0.10663 |
| **ACTA2** | 0.03058 | 0.02749 | 0.65025 | 0.08102 | 0.00010 | 0.12438 | 0.00000 | 0.02043 | 0.00000 | 0.16939 | 0.02926 | 0.00268 |
| **ACTB** | 12.40971 | 79.74472 | 65.41752 | 51.08747 | 86.89505 | 70.25426 | 83.67737 | 18.70669 | 90.62102 | 22.38223 | 78.05805 | 46.92440 |
| **ACTBL2** | 0.00626 | 0.01589 | 0.87208 | 0.05878 | 0.00000 | 0.12158 | 0.00000 | 0.17877 | 0.00000 | 0.10603 | 0.02740 | 0.00000 |
| **ACTC1** | 0.19370 | 0.05856 | 5.58358 | 0.00000 | 0.00000 | 0.09787 | 2.99695 | 0.04419 | 0.00000 | 0.18269 | 0.23745 | 0.00566 |
| **ACTG1** | 17.89213 | 66.91304 | 57.28683 | 50.69478 | 58.13847 | 52.31184 | 51.64357 | 22.48791 | 57.49679 | 6.67535 | 25.89481 | 50.70800 |
| **ACTG1P4** | 0.03777 | 0.07336 | 0.18555 | 0.15280 | 0.00000 | 0.32724 | 0.00000 | 0.03087 | 0.00000 | 0.06674 | 0.07697 | 0.02570 |
| **ACTG2** | 3.18341 | 4.98981 | 0.55435 | 1.04088 | 8.12379 | 1.43967 | 0.71764 | 1.23941 | 0.00000 | 0.00000 | 5.99814 | 0.00000 |
| **ACTL10** | 0.00666 | 0.01163 | 0.07012 | 0.00541 | 0.10901 | 0.00000 | 0.00049 | 0.00742 | 0.02277 | 0.10748 | 0.04456 | 0.07252 |
| **ACTL6A** | 1.18130 | 13.82892 | 13.41701 | 6.22859 | 2.45468 | 30.59532 | 16.85773 | 2.50491 | 15.78452 | 0.00000 | 14.57367 | 11.41416 |
| **ACTL6B** | 0.00542 | 0.01747 | 0.01083 | 0.01317 | 0.00802 | 0.02418 | 0.01766 | 0.01037 | 0.01139 | 0.05447 | 0.01406 | 0.01315 |
| **ACTL7A** | 0.00671 | 0.01813 | 0.01616 | 0.03331 | 0.01730 | 0.00440 | 0.01837 | 0.01055 | 0.03755 | 0.00006 | 0.01626 | 0.01859 |
| **ACTL7B** | 0.13040 | 0.01558 | 0.02263 | 0.00869 | 0.01332 | 0.00833 | 0.00549 | 0.37745 | 0.03531 | 0.03764 | 0.01487 | 0.00767 |
| **ACTL8** | 49.10202 | 0.29034 | 0.00000 | 0.00000 | 0.10102 | 0.00000 | 0.00247 | 25.34031 | 0.00000 | 0.00353 | 0.00000 | 11.37031 |
| **ACTL9** | 0.00542 | 0.01257 | 0.02445 | 0.02085 | 0.00630 | 0.02222 | 0.04502 | 0.01035 | 0.00054 | 0.05250 | 0.01326 | 0.01388 |
| **ACTN1** | 6.86794 | 16.01258 | 25.01691 | 3.77715 | 29.70670 | 0.00000 | 21.37545 | 3.02535 | 31.06943 | 9.21726 | 32.25912 | 2.59924 |
| **ACTN1-AS1** | 0.01846 | 0.03752 | 0.02527 | 0.00914 | 0.02426 | 0.00804 | 0.01195 | 0.07898 | 0.02132 | 0.02415 | 0.03808 | 0.02267 |
| **ACTN2** | 0.43427 | 0.18615 | 0.43517 | 0.00336 | 0.08265 | 0.12100 | 0.82260 | 0.51721 | 1.40355 | 0.00000 | 0.34086 | 0.37528 |
| **ACTN3** | 0.27999 | 0.00000 | 0.00002 | 0.68986 | 0.00000 | 0.06338 | 0.00842 | 0.01421 | 0.00000 | 0.20985 | 0.00363 | 0.08381 |
| **ACTN4** | 0.76040 | 13.43961 | 15.55217 | 17.47343 | 25.27455 | 0.00000 | 11.11835 | 1.24904 | 19.67040 | 25.08805 | 22.50330 | 8.28627 |
| **ACTR10** | 0.38232 | 5.55312 | 1.41192 | 3.13032 | 2.61505 | 24.29148 | 4.87207 | 0.28709 | 3.34449 | 0.00020 | 8.45283 | 10.41790 |
| **ACTR1A** | 1.45300 | 15.90590 | 18.32694 | 13.48574 | 15.89640 | 12.63636 | 21.72673 | 2.91671 | 19.87496 | 0.00000 | 18.91069 | 10.38689 |
| **ACTR1B** | 0.78281 | 2.02244 | 2.55069 | 2.53205 | 2.99632 | 1.50181 | 0.21510 | 0.00000 | 0.96804 | 4.84349 | 4.10504 | 0.51456 |
| **ACTR2** | 6.75506 | 0.98738 | 7.54276 | 4.01175 | 5.16941 | 17.19330 | 15.70320 | 3.02962 | 0.90295 | 17.20395 | 9.62211 | 6.11370 |
| **ACTR3** | 1.57354 | 12.25090 | 14.63086 | 8.43014 | 6.63422 | 34.24140 | 19.42156 | 4.32699 | 11.76565 | 9.94823 | 18.33152 | 10.06211 |
| **ACTR3B** | 1.93847 | 1.53390 | 1.94116 | 0.91818 | 1.62813 | 0.00008 | 2.35340 | 0.71921 | 0.08784 | 1.53466 | 1.13226 | 0.38174 |
| **ACTR3C** | 0.00740 | 0.01783 | 0.02456 | 0.13898 | 0.10195 | 0.00000 | 0.03080 | 0.01059 | 0.02947 | 0.07310 | 0.02279 | 0.01705 |
| **ACTR5** | 1.46548 | 4.28200 | 4.06974 | 2.65052 | 2.32205 | 1.43095 | 0.79360 | 4.34483 | 6.89498 | 0.00000 | 3.10074 | 5.11594 |
| **ACTR6** | 0.00000 | 1.54151 | 13.03981 | 0.75646 | 1.50883 | 25.23816 | 15.31793 | 3.62246 | 14.23943 | 0.47330 | 7.98976 | 4.08850 |
| **ACTR8** | 1.11870 | 6.15503 | 3.69004 | 4.45873 | 1.14195 | 15.73494 | 5.85999 | 1.28351 | 4.55060 | 0.00000 | 6.52612 | 9.63757 |
| **ACTRT3** | 0.00057 | 0.00000 | 0.89807 | 0.49467 | 0.15803 | 0.78996 | 1.00920 | 0.17534 | 0.75524 | 0.00000 | 1.71004 | 0.57945 |

| | | | | | | | | | | | | |
|---|---|---|---|---|---|---|---|---|---|---|---|---|
| **ACVR1** | 1.25875 | 0.68382 | 0.25555 | 1.24480 | 5.35307 | 6.74181 | 1.87646 | 0.17520 | 3.46494 | 3.34895 | 2.02720 | 0.00000 |
| **ACVR1B** | 8.00024 | 0.94351 | 0.17179 | 4.32066 | 1.32751 | 1.25511 | 1.19353 | 0.61531 | 0.17022 | 3.30068 | 0.25067 | 2.39279 |
| **ACVR1C** | 0.54697 | 0.24095 | 0.07979 | 0.00000 | 0.21676 | 0.00000 | 0.25074 | 0.00000 | 0.28148 | 0.17916 | 0.00000 | 0.61158 |
| **ACVR2A** | 1.15030 | 2.63022 | 1.10549 | 0.99457 | 2.55126 | 4.61906 | 0.00000 | 6.49675 | 6.91255 | 1.46877 | 0.24569 | 1.72320 |
| **ACVR2B** | 5.87453 | 0.84472 | 6.31344 | 7.24908 | 1.31600 | 6.64032 | 12.04470 | 2.99654 | 0.00158 | 23.77843 | 2.89426 | 3.37944 |
| **ACVR2B-AS1** | 0.09625 | 0.07670 | 0.10530 | 0.07832 | 0.01456 | 0.07171 | 0.04454 | 0.00741 | 0.11465 | 0.13720 | 0.00000 | 0.24469 |
| **ACVRL1** | 0.00518 | 0.00571 | 0.43076 | 1.34941 | 0.02561 | 0.00000 | 0.12847 | 0.00943 | 0.00757 | 0.07405 | 0.02240 | 0.01638 |
| **ACY1** | 0.02371 | 0.32475 | 0.80527 | 0.34584 | 0.47243 | 0.00000 | 0.01355 | 0.01120 | 1.38204 | 0.00000 | 1.27344 | 0.22187 |
| **ACY3** | 0.00668 | 0.01119 | 0.01336 | 0.00793 | 0.10039 | 0.00005 | 0.03999 | 0.01035 | 0.00000 | 0.05834 | 0.01240 | 0.01634 |
| **ACYP1** | 1.24568 | 1.26686 | 6.95980 | 0.00000 | 0.00000 | 3.33481 | 6.87754 | 0.57920 | 3.92911 | 0.16861 | 1.84379 | 0.11977 |
| **ACYP2** | 0.14659 | 0.08291 | 0.22864 | 0.30203 | 0.00000 | 0.49985 | 0.30546 | 0.13228 | 0.02797 | 0.47335 | 0.04863 | 0.12767 |
| **ADA** | 6.86979 | 10.49357 | 6.68585 | 4.37539 | 1.10061 | 5.63757 | 2.46050 | 0.39183 | 2.63966 | 6.11759 | 8.61026 | 0.56488 |
| **ADAD1** | 7.30734 | 0.02694 | 0.03111 | 0.00000 | 0.00969 | 0.04433 | 0.00000 | 3.25669 | 0.01485 | 0.00000 | 0.05547 | 3.86065 |
| **ADAD2** | 1.46038 | 0.02275 | 0.08691 | 0.00587 | 0.00588 | 0.00000 | 0.00793 | 0.15620 | 0.02882 | 0.37247 | 0.00808 | 0.24319 |
| **ADAL** | 3.05185 | 4.39157 | 1.46536 | 0.64052 | 1.06399 | 6.96419 | 1.39029 | 3.65916 | 0.00000 | 5.17024 | 4.04611 | 3.88052 |
| **ADAM10** | 3.44880 | 6.73323 | 7.70476 | 6.15384 | 4.57545 | 28.07384 | 0.00000 | 6.73781 | 21.71001 | 10.55414 | 9.18391 | 8.93217 |
| **ADAM11** | 0.26432 | 0.00000 | 0.10238 | 0.00119 | 0.27744 | 0.00000 | 0.15796 | 0.01071 | 0.14420 | 0.04131 | 0.22278 | 0.05717 |
| **ADAM12** | 1.56714 | 0.02766 | 0.28611 | 0.00000 | 0.04191 | 0.10788 | 0.03001 | 0.32498 | 0.00006 | 0.28499 | 0.00000 | 0.10994 |
| **ADAM15** | 1.80029 | 0.83025 | 8.05448 | 0.65995 | 14.80910 | 0.00000 | 3.80239 | 0.02982 | 31.80470 | 5.60880 | 13.25155 | 0.78696 |
| **ADAM17** | 4.70425 | 4.04723 | 7.80412 | 2.51248 | 2.99043 | 18.53824 | 0.00000 | 10.92892 | 17.13853 | 12.01241 | 5.79943 | 3.15139 |
| **ADAM18** | 0.00712 | 0.08283 | 1.64298 | 2.79418 | 1.27196 | 1.05108 | 0.00000 | 0.01988 | 0.54160 | 0.66201 | 0.58788 | 0.00000 |
| **ADAM19** | 9.32709 | 0.32419 | 3.39065 | 1.33864 | 2.11469 | 1.75607 | 0.18275 | 0.92937 | 0.00000 | 2.55115 | 2.06801 | 6.03535 |
| **ADAM1A** | 0.00124 | 0.35675 | 0.02177 | 0.49836 | 0.00000 | 0.00000 | 0.55971 | 0.00385 | 1.17056 | 2.18347 | 0.98955 | 0.31320 |
| **ADAM2** | 0.01036 | 0.01589 | 0.08841 | 0.01587 | 0.02291 | 0.00457 | 0.01199 | 0.01227 | 0.00943 | 0.01608 | 0.01858 | 0.01892 |
| **ADAM20** | 0.01146 | 0.00000 | 0.13880 | 0.05467 | 0.00000 | 0.11375 | 0.00000 | 0.01851 | 0.03440 | 0.28003 | 0.00627 | 0.15611 |
| **ADAM20P1** | 0.00211 | 0.01991 | 0.04047 | 0.06206 | 0.01705 | 0.00000 | 0.00000 | 0.14926 | 0.10423 | 0.65827 | 0.00000 | 0.06048 |
| **ADAM21** | 0.00599 | 0.05386 | 0.08859 | 0.03083 | 0.00192 | 0.02278 | 0.01892 | 0.14618 | 0.14922 | 0.00000 | 0.11402 | 0.43793 |
| **ADAM21P1** | 0.00519 | 0.02398 | 0.01749 | 0.01205 | 0.01547 | 0.01996 | 0.01798 | 0.02936 | 0.03678 | 0.00287 | 0.01582 | 0.05498 |
| **ADAM22** | 0.54865 | 0.02543 | 0.05278 | 0.00348 | 0.00000 | 0.00000 | 0.08734 | 0.00000 | 0.00000 | 0.31235 | 0.27628 | 0.06215 |
| **ADAM23** | 0.08941 | 1.34645 | 0.40011 | 2.75184 | 1.74065 | 0.93755 | 0.00000 | 0.23574 | 0.00000 | 1.70600 | 0.00000 | 0.64950 |
| **ADAM28** | 0.27430 | 0.01061 | 0.15701 | 0.01239 | 0.00709 | 0.01879 | 0.00526 | 0.02966 | 0.01380 | 0.05029 | 0.02745 | 0.00935 |
| **ADAM32** | 0.10322 | 0.04512 | 0.00823 | 0.09002 | 0.00000 | 0.16555 | 0.00000 | 0.17848 | 0.09996 | 0.31431 | 0.11389 | 0.06451 |
| **ADAM33** | 0.00331 | 0.01361 | 0.01735 | 0.01799 | 0.03377 | 0.00519 | 0.02119 | 0.07076 | 0.02246 | 0.00000 | 0.01691 | 0.08790 |
| **ADAM3A** | 0.00482 | 0.00940 | 0.03709 | 0.00296 | 0.00000 | 0.01950 | 0.10991 | 0.01008 | 0.00000 | 0.14059 | 0.01208 | 0.01084 |
| **ADAM5** | 0.00579 | 0.01943 | 0.01522 | 0.01253 | 0.02144 | 0.01173 | 0.02060 | 0.01142 | 0.01801 | 0.00748 | 0.02113 | 0.03832 |
| **ADAM8** | 0.82695 | 0.24303 | 1.84678 | 0.00000 | 0.04912 | 0.00000 | 0.00006 | 0.15355 | 0.40587 | 0.94118 | 0.04928 | 0.28813 |
| **ADAM9** | 0.11483 | 5.47619 | 10.37686 | 0.80888 | 7.99012 | 26.44494 | 0.00000 | 2.54711 | 31.55974 | 3.70329 | 15.63704 | 3.78423 |
| **ADAMTS1** | 0.00808 | 0.41319 | 15.15699 | 0.24581 | 2.30233 | 2.16289 | 0.53127 | 0.40576 | 8.18476 | 0.00004 | 1.23335 | 0.71142 |
| **ADAMTS10** | 0.19950 | 0.01315 | 0.03505 | 0.09322 | 0.00000 | 0.03439 | 0.02681 | 0.00945 | 0.00000 | 0.05918 | 0.01514 | 0.00903 |
| **ADAMTS12** | 0.27854 | 0.25348 | 0.03928 | 0.71555 | 0.03829 | 0.44548 | 0.00000 | 0.00000 | 0.00000 | 0.01556 | 0.57167 | 0.00943 |
| **ADAMTS13** | 0.13509 | 0.10189 | 0.03058 | 0.11811 | 0.00000 | 0.00000 | 0.01276 | 0.09865 | 0.13502 | 0.07960 | 0.16959 | 0.15745 |
| **ADAMTS14** | 1.41334 | 0.20113 | 0.02599 | 0.00535 | 0.09663 | 0.00275 | 0.02749 | 0.30293 | 0.01152 | 0.00202 | 0.02031 | 0.40551 |
| **ADAMTS15** | 0.68099 | 0.04355 | 0.02052 | 0.05798 | 0.00000 | 0.00745 | 0.00000 | 0.00047 | 0.02588 | 0.18729 | 0.01056 | 0.01875 |
| **ADAMTS16** | 0.01400 | 0.00577 | 0.03776 | 0.09662 | 0.00000 | 0.01461 | 0.01339 | 0.00964 | 0.00000 | 0.06150 | 0.06824 | 0.03919 |
| **ADAMTS17** | 3.53193 | 0.30506 | 0.06171 | 0.10687 | 0.67215 | 0.00014 | 0.00000 | 0.46625 | 0.02886 | 0.19362 | 0.05134 | 0.34628 |
| **ADAMTS18** | 0.96206 | 0.01771 | 0.03014 | 0.04220 | 0.37318 | 0.00000 | 0.01285 | 0.10155 | 0.04175 | 0.00025 | 0.14686 | 0.10082 |
| **ADAMTS19** | 0.14046 | 0.14477 | 0.06673 | 0.07766 | 0.00000 | 0.00000 | 0.00000 | 0.03574 | 0.00000 | 0.11860 | 0.11293 | 0.20186 |
| **ADAMTS2** | 0.02400 | 1.35296 | 0.00000 | 0.00000 | 0.40288 | 0.09474 | 0.00000 | 0.10842 | 0.00000 | 0.29756 | 0.08224 | 0.00000 |

| | | | | | | | | | | | | |
|---|---|---|---|---|---|---|---|---|---|---|---|---|
| **ADAMTS20** | 0.00738 | 0.00443 | 3.81405 | 0.21136 | 0.00016 | 0.00000 | 0.00000 | 0.13658 | 0.00000 | 0.35534 | 0.01524 | 0.47434 |
| **ADAMTS3** | 0.79817 | 0.45371 | 0.54331 | 0.17498 | 0.52263 | 0.82915 | 0.11688 | 0.00000 | 0.00000 | 0.28395 | 0.03683 | 0.21821 |
| **ADAMTS4** | 0.20903 | 0.00550 | 0.05987 | 2.12903 | 0.00000 | 0.00000 | 0.00317 | 0.05282 | 0.01274 | 0.23590 | 0.09413 | 0.05247 |
| **ADAMTS5** | 0.12651 | 0.08660 | 0.11451 | 0.00000 | 0.12732 | 0.13732 | 0.00000 | 0.00920 | 0.19499 | 0.13656 | 0.05303 | 0.00529 |
| **ADAMTS6** | 0.00227 | 0.03876 | 0.49507 | 0.00000 | 0.09129 | 0.30486 | 0.00000 | 0.16994 | 0.56751 | 1.00243 | 1.17570 | 0.53157 |
| **ADAMTS7** | 0.01548 | 0.01096 | 0.02856 | 0.01302 | 0.01913 | 0.01172 | 0.01891 | 0.03635 | 0.02523 | 0.01610 | 0.02359 | 0.05070 |
| **ADAMTS8** | 0.00809 | 0.00036 | 0.02736 | 0.04743 | 0.01956 | 0.13469 | 0.05618 | 0.00745 | 0.00000 | 0.00335 | 0.05186 | 0.05905 |
| **ADAMTS9** | 8.32079 | 0.26588 | 0.04972 | 1.11150 | 0.09822 | 0.88292 | 0.00000 | 3.16881 | 0.00000 | 0.72440 | 0.28698 | 0.00211 |
| **ADAMTS9-AS2** | 0.00622 | 0.06404 | 0.01050 | 0.00924 | 0.01224 | 0.01071 | 0.01463 | 0.01109 | 0.06605 | 0.00788 | 0.02643 | 0.01339 |
| **ADAMTSL1** | 3.27615 | 0.00904 | 0.04639 | 0.02022 | 0.01710 | 0.00000 | 0.02110 | 0.67641 | 0.02415 | 0.00000 | 0.02980 | 0.02579 |
| **ADAMTSL2** | 0.10221 | 0.18959 | 0.02222 | 0.05479 | 0.03922 | 0.00000 | 0.01595 | 0.00360 | 0.03273 | 0.00000 | 0.02149 | 0.10492 |
| **ADAMTSL3** | 4.24882 | 0.62204 | 0.00000 | 1.93697 | 1.30902 | 0.00000 | 0.48699 | 0.26882 | 0.10624 | 0.28397 | 0.00000 | 0.84941 |
| **ADAMTSL4** | 0.00001 | 0.00000 | 1.78272 | 0.00000 | 1.33713 | 0.00000 | 0.85892 | 0.02446 | 5.84585 | 3.02097 | 4.13886 | 0.00000 |
| **ADAMTSL4-AS1** | 0.04195 | 0.06798 | 0.08728 | 0.01928 | 0.00611 | 0.00000 | 0.00019 | 0.06200 | 0.02021 | 0.10488 | 0.03904 | 0.09906 |
| **ADAMTSL5** | 0.71790 | 5.49383 | 0.04542 | 1.62998 | 0.00000 | 0.00000 | 0.14650 | 0.00000 | 0.81629 | 1.62280 | 0.28768 | 2.74210 |
| **ADAP1** | 4.90778 | 2.13781 | 0.04103 | 0.00000 | 0.13172 | 0.00137 | 0.00593 | 0.73589 | 0.00000 | 0.19081 | 0.02471 | 0.22568 |
| **ADAP2** | 0.00256 | 0.89343 | 7.11883 | 1.43206 | 0.00000 | 0.00000 | 7.54364 | 1.58783 | 0.00000 | 0.00000 | 0.96077 | 0.96456 |
| **ADAR** | 20.17298 | 8.60969 | 13.27701 | 14.53831 | 11.58094 | 13.17820 | 19.05457 | 1.97887 | 12.61932 | 13.54708 | 8.39506 | 5.46552 |
| **ADARB1** | 1.01145 | 0.65722 | 0.20066 | 0.00000 | 1.65331 | 0.81732 | 1.46632 | 0.01883 | 2.08578 | 4.55044 | 0.69663 | 0.04680 |
| **ADARB2** | 0.07327 | 0.00852 | 0.01257 | 0.00529 | 0.00000 | 0.05477 | 0.02420 | 0.07405 | 0.00033 | 0.07220 | 0.01025 | 0.01512 |
| **ADAT1** | 4.05388 | 2.12474 | 3.67929 | 0.34062 | 2.85428 | 1.79876 | 2.89712 | 3.92363 | 0.23544 | 1.54305 | 4.09072 | 4.81913 |
| **ADAT2** | 0.00000 | 1.32782 | 0.47588 | 0.01233 | 0.00000 | 6.47099 | 1.40522 | 0.42902 | 2.78026 | 2.64103 | 3.61675 | 3.46661 |
| **ADAT3** | 0.09066 | 0.38200 | 3.01053 | 1.13678 | 1.97798 | 0.00000 | 2.59743 | 0.52237 | 1.60039 | 6.71831 | 0.88111 | 0.65447 |
| **ADCK1** | 0.00000 | 16.36074 | 0.00000 | 2.27144 | 7.23218 | 1.83441 | 0.10176 | 0.22564 | 1.71862 | 2.05692 | 3.57015 | 3.45080 |
| **ADCK2** | 0.80452 | 19.09203 | 3.74062 | 2.25115 | 4.52290 | 3.32579 | 3.06696 | 0.55033 | 1.00352 | 11.21080 | 1.53402 | 7.73356 |
| **ADCK3** | 0.54206 | 10.95103 | 3.99578 | 2.43892 | 3.80585 | 0.43532 | 6.85082 | 0.04772 | 12.22713 | 2.21903 | 7.54396 | 9.33229 |
| **ADCK4** | 0.11240 | 2.99506 | 2.72334 | 1.95024 | 2.73418 | 0.00000 | 2.60660 | 0.00000 | 2.63509 | 0.00618 | 2.12698 | 2.24709 |
| **ADCK5** | 0.08423 | 3.37657 | 0.35488 | 3.77413 | 4.35005 | 0.00000 | 0.06577 | 0.37278 | 1.52734 | 2.86916 | 2.53258 | 1.42380 |
| **ADCY1** | 0.21729 | 0.08035 | 0.57132 | 0.03830 | 0.07908 | 0.00037 | 0.00000 | 0.04250 | 0.00000 | 0.20896 | 0.02967 | 0.01209 |
| **ADCY10** | 0.00005 | 0.05081 | 1.16651 | 0.52782 | 0.41058 | 1.17472 | 0.00000 | 0.30558 | 0.50573 | 2.62546 | 0.25820 | 0.44394 |
| **ADCY10P1** | 0.40870 | 0.00000 | 0.32216 | 0.26911 | 0.00000 | 0.14587 | 0.07182 | 0.73682 | 0.84033 | 0.90737 | 0.05277 | 0.19434 |
| **ADCY2** | 1.17183 | 0.00864 | 0.03243 | 0.04657 | 0.02782 | 0.00000 | 0.05058 | 0.17444 | 0.00000 | 0.04507 | 0.04024 | 0.00343 |
| **ADCY3** | 2.36852 | 1.02999 | 0.86232 | 0.43292 | 0.51681 | 0.03932 | 0.99772 | 0.70549 | 0.88784 | 0.89551 | 0.42891 | 0.54803 |
| **ADCY4** | 0.00546 | 0.01535 | 0.01915 | 0.08389 | 0.00000 | 0.02836 | 0.03165 | 0.01117 | 0.00015 | 0.05798 | 0.02034 | 0.01217 |
| **ADCY5** | 11.42266 | 0.13970 | 1.40648 | 0.02889 | 0.00000 | 0.00000 | 0.00000 | 2.14419 | 0.29156 | 0.04831 | 0.06285 | 0.48320 |
| **ADCY6** | 0.08634 | 0.02837 | 0.06524 | 0.90739 | 0.00000 | 0.22037 | 0.00000 | 0.16198 | 0.04104 | 0.03077 | 0.02144 | 0.17297 |
| **ADCY7** | 3.24197 | 0.04244 | 0.45150 | 0.05262 | 0.00000 | 1.18422 | 0.00000 | 0.22829 | 0.08583 | 2.22111 | 0.00000 | 0.09490 |
| **ADCY8** | 0.00850 | 0.01822 | 0.04081 | 0.02381 | 0.03067 | 0.00117 | 0.02525 | 0.02039 | 0.00402 | 0.00708 | 0.05644 | 0.01699 |
| **ADCY9** | 8.02396 | 0.26405 | 0.28792 | 0.04102 | 2.53506 | 0.00000 | 0.00000 | 1.48445 | 1.14112 | 1.56996 | 0.16925 | 0.42916 |
| **ADCYAP1** | 0.00250 | 0.01577 | 0.01091 | 0.02237 | 0.01702 | 0.04799 | 0.01235 | 0.08581 | 0.00779 | 0.03171 | 0.01243 | 0.01047 |
| **ADCYAP1R1** | 0.01253 | 0.03246 | 0.03533 | 0.01234 | 0.00397 | 0.01634 | 0.01374 | 0.03847 | 0.00000 | 0.07450 | 0.02538 | 0.01656 |
| **ADD1** | 3.15168 | 4.63965 | 8.11236 | 9.41309 | 6.70501 | 3.65578 | 17.50081 | 0.59066 | 5.82026 | 19.91783 | 5.61349 | 4.35008 |
| **ADD2** | 3.46272 | 0.25229 | 0.32391 | 1.27928 | 0.06635 | 0.00000 | 0.00927 | 0.07514 | 0.01796 | 0.00000 | 0.02027 | 0.12948 |
| **ADD3** | 15.22861 | 3.44329 | 0.51697 | 11.26450 | 0.64819 | 17.92108 | 5.25017 | 1.88404 | 1.22173 | 7.14618 | 7.19481 | 1.83201 |
| **ADD3-AS1** | 0.03908 | 0.01376 | 0.04363 | 0.02074 | 0.08344 | 0.00000 | 0.03424 | 0.01385 | 0.02177 | 0.00000 | 0.03003 | 0.01770 |
| **ADGB** | 0.02560 | 0.01466 | 0.00687 | 0.01044 | 0.00977 | 0.01310 | 0.10429 | 0.03288 | 0.00000 | 0.03632 | 0.01081 | 0.01299 |
| **ADH1A** | 0.00740 | 0.01525 | 0.01408 | 0.01202 | 0.01545 | 0.01948 | 0.01834 | 0.03911 | 0.01974 | 0.01199 | 0.01516 | 0.01448 |
| **ADH1B** | 0.00554 | 0.00856 | 0.04150 | 0.00623 | 0.03170 | 0.05293 | 0.01188 | 0.01307 | 0.00000 | 0.01972 | 0.02359 | 0.01345 |

| | | | | | | | | | | | | |
|---|---|---|---|---|---|---|---|---|---|---|---|---|
| **ADH1C** | 0.00233 | 0.01434 | 0.01491 | 0.01133 | 0.01237 | 0.02118 | 0.01630 | 0.04823 | 0.01771 | 0.01665 | 0.02218 | 0.01411 |
| **ADH4** | 0.00716 | 0.02070 | 0.02003 | 0.01289 | 0.01631 | 0.01747 | 0.01537 | 0.01019 | 0.02764 | 0.01563 | 0.01830 | 0.01652 |
| **ADH5** | 4.14445 | 10.15783 | 10.95046 | 8.46702 | 2.87572 | 14.41177 | 14.65915 | 1.02728 | 10.82546 | 0.67868 | 12.17771 | 8.17934 |
| **ADH6** | 0.00871 | 0.01410 | 0.01135 | 0.01617 | 0.02049 | 0.01950 | 0.01908 | 0.03200 | 0.05213 | 0.02321 | 0.00904 | 0.01694 |
| **ADH7** | 0.01654 | 0.00753 | 0.01815 | 0.08574 | 0.00000 | 0.05255 | 0.00839 | 0.06816 | 0.00690 | 0.02611 | 0.01573 | 0.01034 |
| **ADHFE1** | 0.00806 | 0.16140 | 0.21226 | 0.16174 | 0.04994 | 0.00000 | 0.04736 | 0.01630 | 0.00935 | 0.00000 | 0.04675 | 0.84496 |
| **ADI1** | 3.09809 | 24.57587 | 10.82636 | 13.23500 | 24.77535 | 24.27498 | 22.20816 | 0.04301 | 28.19648 | 3.07551 | 33.95077 | 12.61892 |
| **ADIG** | 0.00489 | 0.00763 | 0.01449 | 0.00231 | 0.07092 | 0.01750 | 0.00597 | 0.01028 | 0.01269 | 0.09729 | 0.00646 | 0.01270 |
| **ADIPOQ** | 0.00654 | 0.01907 | 0.01531 | 0.01945 | 0.01281 | 0.00810 | 0.02019 | 0.01015 | 0.02304 | 0.02121 | 0.02449 | 0.02050 |
| **ADIPOQ-AS1** | 0.00587 | 0.01409 | 0.01962 | 0.01160 | 0.01259 | 0.01285 | 0.02492 | 0.00964 | 0.01115 | 0.02015 | 0.03306 | 0.03530 |
| **ADIPOR1** | 2.32267 | 15.74377 | 19.57062 | 10.98070 | 9.09789 | 18.21895 | 10.76002 | 8.36316 | 25.74288 | 0.00000 | 12.79159 | 14.01101 |
| **ADIPOR2** | 3.02898 | 1.38818 | 6.02687 | 3.93697 | 8.69643 | 6.74073 | 10.37125 | 7.04804 | 7.31354 | 23.08595 | 5.33226 | 6.14214 |
| **ADIRF** | 0.01828 | 0.01721 | 0.12695 | 0.01984 | 0.10338 | 0.00000 | 0.00667 | 0.04400 | 0.01865 | 0.00434 | 0.06019 | 0.02828 |
| **ADK** | 1.28024 | 0.00000 | 10.62547 | 0.12250 | 8.08231 | 7.23329 | 13.47880 | 0.16076 | 20.21478 | 0.91189 | 21.01220 | 0.26441 |
| **ADM** | 0.93264 | 1.98695 | 32.69828 | 5.89243 | 18.74959 | 13.01811 | 0.00000 | 2.35635 | 19.03671 | 0.00000 | 8.99925 | 25.87848 |
| **ADM2** | 0.53034 | 0.00000 | 0.59689 | 0.00000 | 0.00000 | 0.03118 | 0.56189 | 0.01675 | 0.15654 | 0.27414 | 0.26663 | 0.11519 |
| **ADM5** | 0.00109 | 0.19574 | 0.74318 | 1.34097 | 0.13085 | 0.00000 | 2.74547 | 0.07763 | 1.90025 | 3.70486 | 0.00000 | 0.18065 |
| **ADNP** | 1.73830 | 5.79547 | 9.97297 | 10.29316 | 3.63210 | 20.55522 | 18.14745 | 4.94356 | 8.85418 | 20.86871 | 4.86246 | 8.49993 |
| **ADNP2** | 4.19399 | 2.22333 | 6.67439 | 2.47226 | 0.00000 | 7.30840 | 8.95082 | 5.52777 | 3.74669 | 13.70186 | 0.00000 | 3.99024 |
| **ADO** | 1.93679 | 0.91665 | 0.83360 | 0.99713 | 0.22867 | 4.25139 | 2.18756 | 0.00000 | 0.00000 | 6.04882 | 0.89745 | 1.61197 |
| **ADORA1** | 0.00238 | 0.01202 | 0.04716 | 0.03857 | 0.00225 | 0.01162 | 0.01017 | 0.04631 | 0.00000 | 0.06283 | 0.06172 | 0.01222 |
| **ADORA2A-AS1** | 0.12037 | 0.07382 | 0.03103 | 0.03451 | 0.12086 | 0.00000 | 0.00050 | 0.00866 | 0.05900 | 0.14272 | 0.00000 | 0.02828 |
| **ADORA2A.uid-2** | 0.02471 | 0.03728 | 0.01456 | 0.01825 | 0.01564 | 0.00696 | 0.01600 | 0.01270 | 0.02297 | 0.01975 | 0.01457 | 0.02751 |
| **ADORA2B** | 0.00000 | 5.94014 | 0.00000 | 6.30063 | 0.57976 | 0.00000 | 0.00115 | 0.09849 | 2.59952 | 1.25865 | 1.04528 | 2.66591 |
| **ADORA3** | 0.00231 | 1.36426 | 0.00053 | 0.65472 | 0.28026 | 0.44168 | 0.00000 | 0.02525 | 0.00000 | 0.00217 | 0.02779 | 0.21991 |
| **ADPGK** | 16.83013 | 11.60825 | 13.35724 | 9.53574 | 7.57543 | 18.46717 | 0.00000 | 5.74547 | 23.80303 | 0.00000 | 11.44670 | 10.33489 |
| **ADPGK-AS1** | 0.00724 | 0.12891 | 0.41098 | 0.06716 | 0.00000 | 0.00000 | 0.00000 | 0.00666 | 0.19957 | 0.38277 | 0.27425 | 0.11502 |
| **ADPRH** | 0.08249 | 0.09615 | 1.17851 | 0.10688 | 0.13069 | 0.00000 | 0.02809 | 0.00872 | 0.00000 | 0.00000 | 0.08233 | 0.01325 |
| **ADPRHL1** | 0.79079 | 0.18622 | 0.00000 | 0.78452 | 0.34747 | 0.00000 | 0.43222 | 0.19143 | 0.52619 | 0.00000 | 0.60489 | 0.42972 |
| **ADPRHL2** | 0.24220 | 12.97967 | 6.65713 | 7.20093 | 6.77298 | 6.17291 | 12.76220 | 2.11257 | 7.49924 | 0.00000 | 13.30772 | 12.88951 |
| **ADPRM** | 1.12426 | 0.41532 | 2.90073 | 1.30765 | 0.00000 | 1.62133 | 2.82396 | 0.98977 | 0.54824 | 0.00000 | 2.53323 | 3.11720 |
| **ADRA1D** | 0.00586 | 0.02307 | 0.22245 | 0.04313 | 0.01452 | 0.01922 | 0.06252 | 0.01089 | 0.00155 | 0.00000 | 0.02927 | 0.02096 |
| **ADRA2A** | 0.00355 | 0.01828 | 0.01634 | 0.02211 | 0.02382 | 0.00000 | 0.03922 | 0.03786 | 0.02122 | 0.01234 | 0.01621 | 0.01583 |
| **ADRA2B** | 0.97944 | 0.01437 | 0.01828 | 0.06623 | 0.01748 | 0.01146 | 0.01291 | 0.03693 | 0.02937 | 0.00000 | 0.01256 | 0.00910 |
| **ADRA2C** | 0.00558 | 0.09579 | 0.01438 | 0.00000 | 0.01069 | 0.09093 | 0.00000 | 0.00977 | 0.00244 | 0.09744 | 0.00828 | 0.00493 |
| **ADRB1** | 0.02288 | 0.01540 | 0.03799 | 0.01818 | 0.01148 | 0.02205 | 0.06137 | 0.01953 | 0.00000 | 0.03988 | 0.02338 | 0.01376 |
| **ADRB2** | 0.40751 | 0.01408 | 0.01241 | 0.01403 | 0.01244 | 0.00000 | 0.02192 | 0.09332 | 0.10034 | 0.03124 | 0.00507 | 0.03767 |
| **ADRB3** | 0.00660 | 0.01509 | 0.07459 | 0.17488 | 0.01875 | 0.01605 | 0.01452 | 0.00987 | 0.00739 | 0.00069 | 0.02402 | 0.04280 |
| **ADRBK1** | 1.92791 | 2.53889 | 2.29440 | 3.36780 | 7.00990 | 0.00000 | 2.13652 | 0.00000 | 3.83051 | 13.73269 | 0.64772 | 3.56432 |
| **ADRBK2** | 1.36440 | 0.62025 | 7.99773 | 2.18293 | 2.66350 | 5.05601 | 4.19798 | 0.00000 | 0.39324 | 8.01862 | 0.70440 | 0.04500 |
| **ADRM1** | 6.43471 | 34.48883 | 29.12369 | 26.71786 | 39.97911 | 8.61665 | 36.40617 | 12.91911 | 33.51244 | 0.39061 | 25.45518 | 25.15180 |
| **ADSL** | 1.75294 | 16.58044 | 17.45403 | 13.41417 | 10.24981 | 19.79849 | 22.64071 | 4.00560 | 25.18957 | 0.00000 | 27.78998 | 17.18806 |
| **ADSS** | 1.09578 | 2.60635 | 12.30980 | 2.56666 | 0.00000 | 15.39288 | 13.64883 | 4.60033 | 8.68698 | 3.97212 | 7.46867 | 3.64911 |
| **ADSSL1** | 0.04268 | 0.00000 | 0.13100 | 0.00000 | 0.00000 | 0.01935 | 0.48116 | 0.17028 | 0.04130 | 0.25376 | 0.00000 | 0.00821 |
| **ADTRP** | 1.65757 | 5.72850 | 5.71219 | 0.00000 | 5.93103 | 13.22515 | 0.00000 | 0.13954 | 14.58242 | 0.81089 | 5.02565 | 0.03984 |
| **AEBP1** | 0.00361 | 0.08193 | 0.06316 | 0.29643 | 0.07498 | 0.00000 | 0.01022 | 0.00903 | 0.00002 | 0.18080 | 0.06291 | 0.02051 |
| **AEBP2** | 0.11303 | 4.69884 | 14.83897 | 1.53272 | 0.00000 | 28.50692 | 7.92298 | 10.97661 | 4.32859 | 12.40103 | 6.56918 | 5.71179 |
| **AEN** | 3.64229 | 26.49172 | 15.15433 | 19.86572 | 19.77186 | 5.63769 | 15.58210 | 9.88136 | 21.94168 | 10.12942 | 16.39308 | 24.77876 |

| | | | | | | | | | | | | |
|---|---|---|---|---|---|---|---|---|---|---|---|---|
| **AES** | 0.00273 | 3.59942 | 4.07572 | 0.61151 | 1.19942 | 0.00000 | 0.05683 | 3.14575 | 0.00000 | 0.54817 | 0.06362 | 19.95343 |
| **AFAP1** | 7.11801 | 1.55634 | 0.69335 | 2.98762 | 4.25584 | 0.65556 | 4.26726 | 0.23148 | 7.41244 | 3.40653 | 2.12930 | 1.14881 |
| **AFAP1-AS1** | 8.19297 | 0.32007 | 0.02345 | 0.06404 | 1.97042 | 0.69234 | 0.08198 | 0.34195 | 0.20591 | 0.00180 | 0.08512 | 0.47887 |
| **AFAP1L1** | 0.00148 | 2.09459 | 6.31288 | 0.00000 | 0.00000 | 1.24885 | 0.22042 | 0.00000 | 0.00000 | 0.70800 | 1.06314 | 1.08187 |
| **AFAP1L2** | 1.73974 | 0.10909 | 0.05004 | 0.00000 | 0.00000 | 0.04461 | 0.69361 | 0.60144 | 0.00000 | 0.20669 | 0.00000 | 0.03086 |
| **AFF1** | 9.66356 | 4.56045 | 3.71241 | 6.80611 | 5.80812 | 10.48849 | 3.21676 | 9.24088 | 3.97298 | 13.79059 | 4.95419 | 11.22722 |
| **AFF2** | 0.58942 | 0.00483 | 0.06926 | 0.10269 | 0.02902 | 0.10244 | 0.00000 | 0.00318 | 0.23200 | 0.00000 | 0.00806 | 0.02463 |
| **AFF3** | 0.73288 | 0.05136 | 0.00146 | 0.20513 | 0.13986 | 0.00000 | 0.06155 | 0.35814 | 0.00000 | 0.20239 | 0.00522 | 0.17543 |
| **AFF4** | 5.52454 | 0.83212 | 14.50096 | 4.46829 | 7.57620 | 9.77734 | 11.87870 | 2.68462 | 8.13284 | 24.98887 | 10.14529 | 4.18248 |
| **AFG3L1P** | 0.93994 | 2.62314 | 1.06281 | 2.28366 | 2.70550 | 1.25058 | 1.42123 | 0.24463 | 3.02036 | 2.85054 | 0.87436 | 3.44052 |
| **AFG3L2** | 1.11085 | 15.60828 | 10.90987 | 16.21365 | 18.54674 | 6.01992 | 14.12980 | 1.58528 | 14.87453 | 14.54857 | 16.34522 | 3.74550 |
| **AFMID** | 0.13526 | 1.31323 | 3.39465 | 2.41295 | 0.92678 | 1.47113 | 9.08864 | 16.43322 | 5.90406 | 0.00000 | 4.44179 | 2.12412 |
| **AFP** | 0.01150 | 0.02383 | 0.00566 | 0.01924 | 0.01240 | 0.00404 | 0.00586 | 0.00991 | 0.18147 | 0.00000 | 0.00468 | 0.01724 |
| **AFTPH** | 0.00000 | 2.57924 | 4.22625 | 9.81655 | 4.51651 | 3.93866 | 9.10327 | 10.31816 | 0.00000 | 13.25666 | 4.72925 | 15.23832 |
| **AGA** | 0.09175 | 0.93384 | 0.90543 | 0.81769 | 0.11645 | 2.48124 | 0.00000 | 0.07803 | 0.38893 | 1.01975 | 1.43660 | 0.21550 |
| **AGAP1** | 2.34887 | 1.88194 | 4.45412 | 1.93618 | 4.79073 | 1.90402 | 5.00158 | 0.00000 | 0.08155 | 8.40527 | 1.03015 | 0.23812 |
| **AGAP11** | 0.00947 | 0.00000 | 0.00407 | 0.00000 | 0.11930 | 0.02413 | 0.03212 | 0.03954 | 0.01865 | 0.12600 | 0.12618 | 0.01606 |
| **AGAP2** | 0.01227 | 0.01959 | 0.09257 | 0.00573 | 0.02085 | 0.02708 | 0.01677 | 0.03335 | 0.00000 | 0.14921 | 0.02177 | 0.01767 |
| **AGAP2-AS1** | 0.02219 | 0.05497 | 0.15568 | 0.01029 | 0.00000 | 0.00000 | 0.05249 | 0.00516 | 0.16024 | 0.03483 | 0.07655 | 0.08965 |
| **AGAP3** | 1.40822 | 3.56014 | 5.51135 | 1.00821 | 9.55701 | 0.00000 | 7.17423 | 0.04977 | 4.26864 | 9.55161 | 1.32473 | 0.09817 |
| **AGAP4** | 0.03351 | 0.08316 | 0.02102 | 0.00000 | 0.00000 | 0.02212 | 0.13894 | 0.09073 | 0.10068 | 0.31764 | 0.02584 | 0.11364 |
| **AGAP5** | 0.00568 | 0.05682 | 0.04318 | 0.00000 | 0.13997 | 0.00000 | 0.03559 | 0.01791 | 0.10092 | 0.16658 | 0.02822 | 0.11101 |
| **AGAP6** | 0.00023 | 0.29637 | 0.34434 | 0.07113 | 0.14110 | 0.30253 | 0.36027 | 0.35264 | 0.96767 | 2.40482 | 0.33196 | 0.59997 |
| **AGAP7P** | 0.00448 | 0.00466 | 0.02879 | 0.12418 | 0.09614 | 0.00000 | 0.05659 | 0.02131 | 0.19230 | 0.03380 | 0.18309 | 0.12656 |
| **AGBL1** | 0.20946 | 0.01135 | 0.02076 | 0.02959 | 0.00658 | 0.01596 | 0.01749 | 0.06270 | 0.00886 | 0.04320 | 0.02865 | 0.00942 |
| **AGBL2** | 0.00490 | 0.05691 | 0.02332 | 0.00867 | 0.00000 | 0.03456 | 0.02193 | 0.65157 | 0.05019 | 0.01651 | 0.02425 | 0.21791 |
| **AGBL3** | 0.34823 | 0.21667 | 0.17892 | 0.15224 | 0.11987 | 0.01921 | 0.10130 | 0.33660 | 0.18886 | 0.00989 | 0.13396 | 0.08338 |
| **AGBL4** | 0.35588 | 0.03605 | 0.05853 | 0.08269 | 0.05235 | 0.00000 | 0.02092 | 0.01190 | 0.00000 | 0.01127 | 0.11686 | 0.01351 |
| **AGBL5** | 0.10246 | 1.50529 | 1.58166 | 0.56401 | 0.43111 | 1.11578 | 1.73363 | 0.08021 | 1.40012 | 1.91938 | 0.29576 | 1.46773 |
| **AGBL5-AS1** | 0.00572 | 0.01998 | 0.01238 | 0.01265 | 0.00866 | 0.01469 | 0.01801 | 0.01050 | 0.03570 | 0.03983 | 0.04143 | 0.01418 |
| **AGER** | 0.00000 | 2.80604 | 0.00000 | 0.02319 | 10.13494 | 0.00000 | 0.29507 | 0.37168 | 4.72087 | 0.74805 | 3.88497 | 0.68111 |
| **AGFG1** | 1.26404 | 3.08157 | 8.79572 | 3.29492 | 6.58880 | 12.87098 | 12.22869 | 1.31496 | 0.00000 | 23.19525 | 0.98564 | 2.31888 |
| **AGFG2** | 3.93975 | 0.62397 | 0.00000 | 1.82338 | 0.18437 | 0.00000 | 0.62206 | 0.52214 | 0.00000 | 0.01798 | 0.23076 | 0.70842 |
| **AGGF1** | 4.23709 | 6.89548 | 2.08538 | 6.85158 | 2.69486 | 13.11646 | 2.73338 | 0.87700 | 2.43145 | 6.29704 | 2.45667 | 5.88550 |
| **AGK** | 9.16189 | 7.01441 | 0.00000 | 2.33155 | 1.84671 | 6.73870 | 0.69492 | 1.80833 | 1.64405 | 0.00000 | 5.04858 | 4.81349 |
| **AGL** | 0.00000 | 21.51595 | 4.18966 | 27.15761 | 4.90877 | 56.02084 | 22.26134 | 2.57079 | 12.64390 | 44.42677 | 27.70131 | 16.61284 |
| **AGMAT** | 0.46444 | 0.06352 | 0.08587 | 0.09050 | 0.08278 | 0.32699 | 0.01474 | 0.04395 | 0.14620 | 0.23522 | 0.12671 | 0.06356 |
| **AGMO** | 2.83071 | 0.16249 | 0.08991 | 0.00031 | 0.00822 | 0.04688 | 0.00781 | 0.18280 | 0.02432 | 0.00547 | 0.00731 | 0.03389 |
| **AGO1** | 3.08278 | 1.81085 | 3.23076 | 4.16611 | 3.53005 | 4.31337 | 8.06855 | 1.69127 | 2.40709 | 0.86645 | 2.69961 | 6.41112 |
| **AGO2** | 8.90288 | 1.81926 | 6.08566 | 1.62941 | 7.65552 | 1.69259 | 8.67018 | 1.72142 | 2.29353 | 17.29175 | 3.64345 | 3.66815 |
| **AGO3** | 0.80398 | 0.99164 | 3.39142 | 1.25136 | 0.08714 | 2.26847 | 0.00000 | 4.03116 | 4.60231 | 1.35657 | 0.53390 | 2.20997 |
| **AGO4** | 0.14330 | 0.35139 | 0.95915 | 0.65407 | 0.00000 | 1.14614 | 0.58310 | 0.12010 | 0.32735 | 0.98414 | 0.30448 | 0.05954 |
| **AGPAT1** | 1.17235 | 4.39047 | 4.25970 | 4.92048 | 2.49407 | 0.00000 | 5.49501 | 0.27487 | 9.53208 | 0.00000 | 4.93232 | 6.36982 |
| **AGPAT2** | 3.61034 | 19.69086 | 1.84669 | 19.41782 | 18.68808 | 0.00000 | 1.41579 | 1.18808 | 10.06606 | 2.82903 | 8.04046 | 1.10981 |
| **AGPAT3** | 2.08401 | 1.59009 | 0.00000 | 1.79473 | 1.29124 | 0.10310 | 0.59894 | 0.00000 | 0.00000 | 1.75714 | 0.44711 | 0.11759 |
| **AGPAT4** | 1.90116 | 0.00000 | 1.36047 | 0.94304 | 0.01209 | 0.54709 | 0.00000 | 0.61977 | 0.48308 | 1.13147 | 0.37032 | 0.15658 |
| **AGPAT4-IT1** | 0.16902 | 0.03019 | 0.05856 | 0.00021 | 0.00000 | 0.05026 | 0.00000 | 0.01749 | 0.51305 | 0.12363 | 0.01834 | 0.06166 |
| **AGPAT5** | 0.61469 | 1.14493 | 11.88207 | 0.00000 | 0.82408 | 12.30355 | 10.58586 | 0.34010 | 27.69225 | 12.00282 | 11.44739 | 0.25841 |

| | | | | | | | | | | | | |
|---|---|---|---|---|---|---|---|---|---|---|---|---|
| **AGPAT6** | 5.60313 | 15.28075 | 12.84393 | 17.25335 | 15.63499 | 1.33582 | 14.28394 | 2.17415 | 17.68523 | 6.33564 | 9.26931 | 8.94965 |
| **AGPAT9** | 2.14638 | 16.54025 | 0.00000 | 0.00000 | 14.57372 | 1.42759 | 0.00000 | 3.76605 | 0.00000 | 0.82882 | 9.39343 | 7.54797 |
| **AGPS** | 1.02540 | 2.48180 | 7.12533 | 0.62745 | 0.00000 | 14.36898 | 16.16852 | 0.87848 | 2.71154 | 11.83295 | 5.41918 | 0.17807 |
| **AGR2** | 0.11338 | 0.02070 | 0.01656 | 0.02727 | 0.04988 | 0.00000 | 0.01245 | 0.00583 | 0.03142 | 0.03249 | 0.00811 | 0.01411 |
| **AGRN** | 0.23482 | 0.26415 | 0.25914 | 0.02749 | 3.89770 | 0.00000 | 0.08040 | 0.00149 | 2.24208 | 2.45974 | 0.62139 | 0.00495 |
| **AGRP** | 0.00435 | 0.05044 | 0.01536 | 0.01292 | 0.00355 | 0.02323 | 0.02017 | 0.03260 | 0.01305 | 0.03482 | 0.01411 | 0.00960 |
| **AGT** | 0.00764 | 0.02426 | 0.01108 | 0.00837 | 0.03461 | 0.01689 | 0.02116 | 0.01029 | 0.02962 | 0.01297 | 0.01139 | 0.08322 |
| **AGTPBP1** | 2.68192 | 1.36695 | 2.37386 | 1.44242 | 0.00000 | 6.82859 | 0.00000 | 0.76184 | 3.60618 | 5.04229 | 1.65753 | 1.06650 |
| **AGTR1** | 0.00525 | 0.04114 | 0.10804 | 0.00000 | 0.05547 | 0.09697 | 0.00000 | 0.03587 | 0.00000 | 0.05499 | 0.03943 | 0.01770 |
| **AGTRAP** | 3.56169 | 6.67361 | 3.98953 | 10.18126 | 7.98103 | 0.00000 | 1.50465 | 0.74293 | 4.36960 | 0.02272 | 5.84465 | 0.17795 |
| **AGXT** | 0.01431 | 0.01314 | 0.01733 | 0.00971 | 0.02251 | 0.01830 | 0.02219 | 0.00923 | 0.00986 | 0.03009 | 0.01392 | 0.01540 |
| **AGXT2** | 0.00580 | 0.01432 | 0.02433 | 0.04222 | 0.00399 | 0.01341 | 0.03558 | 0.01101 | 0.00000 | 0.02515 | 0.05324 | 0.01789 |
| **AHCTF1** | 0.96407 | 0.31891 | 1.80292 | 0.63746 | 0.19565 | 11.20435 | 3.98188 | 8.25086 | 1.11148 | 13.26048 | 0.86242 | 2.37974 |
| **AHCTF1P1** | 0.36194 | 0.03723 | 0.47673 | 0.00000 | 0.17317 | 2.51590 | 0.36184 | 3.12146 | 0.59654 | 2.91866 | 0.09130 | 0.66121 |
| **AHCY** | 6.37126 | 25.20217 | 24.51140 | 21.28578 | 19.04856 | 14.28691 | 30.94579 | 4.64801 | 38.44125 | 0.00000 | 23.53376 | 20.84678 |
| **AHCYL1** | 0.29380 | 0.43866 | 19.10082 | 4.74959 | 6.08214 | 10.16463 | 24.37744 | 0.34268 | 0.22452 | 22.95145 | 7.22486 | 0.53700 |
| **AHCYL2** | 0.02923 | 0.22837 | 0.81022 | 5.54721 | 0.27344 | 2.17177 | 1.26099 | 1.39448 | 0.05308 | 3.41401 | 0.08494 | 0.59632 |
| **AHDC1** | 0.63193 | 0.08481 | 1.77008 | 0.19355 | 1.42908 | 0.00000 | 1.62576 | 0.00000 | 0.21049 | 4.24799 | 0.05491 | 0.05753 |
| **AHI1** | 1.66285 | 0.18010 | 0.22241 | 0.00000 | 0.00000 | 3.89833 | 2.49881 | 0.83482 | 1.07007 | 5.05354 | 2.51806 | 1.94122 |
| **AHNAK** | 0.00000 | 13.70249 | 2.13664 | 4.88983 | 32.96304 | 14.05869 | 5.32448 | 1.07967 | 23.70485 | 39.06655 | 28.88014 | 0.92885 |
| **AHNAK2** | 0.00763 | 0.00492 | 0.08832 | 0.00318 | 0.00000 | 0.11282 | 0.04338 | 0.01698 | 0.05550 | 0.03987 | 0.12168 | 0.05900 |
| **AHR** | 0.45561 | 0.74155 | 1.18174 | 0.48205 | 0.00000 | 0.88637 | 0.81258 | 0.06237 | 0.15551 | 1.83146 | 0.00000 | 0.56027 |
| **AHRR** | 0.91782 | 0.24374 | 0.02676 | 0.05671 | 0.26067 | 0.00000 | 0.00079 | 0.11709 | 0.46006 | 1.01913 | 0.00001 | 0.03563 |
| **AHSA1** | 13.81472 | 29.36874 | 24.25907 | 20.79500 | 19.33242 | 26.75502 | 43.12160 | 14.72609 | 36.97348 | 0.85896 | 28.57341 | 28.39828 |
| **AHSA2** | 0.31319 | 0.23150 | 1.79548 | 0.99037 | 0.12270 | 0.11502 | 0.87408 | 0.00000 | 0.65988 | 2.78208 | 0.03072 | 0.28823 |
| **AHSP** | 0.00466 | 0.41496 | 0.00218 | 0.00000 | 0.02422 | 0.11755 | 0.00000 | 0.01200 | 0.00000 | 0.02138 | 0.00323 | 0.44082 |
| **AICDA** | 0.00909 | 0.33070 | 0.01636 | 0.99811 | 0.00000 | 0.08131 | 0.00000 | 0.02597 | 0.00000 | 0.03111 | 0.00657 | 0.07218 |
| **AIDA** | 0.07622 | 0.11586 | 0.55812 | 0.00000 | 0.30166 | 1.43779 | 1.31211 | 0.21786 | 0.00000 | 1.49856 | 0.24191 | 0.67962 |
| **AIF1** | 0.03270 | 0.03311 | 0.01989 | 0.00958 | 0.08463 | 0.00904 | 0.01644 | 0.04476 | 0.02792 | 0.00000 | 0.02922 | 0.02103 |
| **AIF1L** | 0.00000 | 28.38097 | 37.85020 | 16.79657 | 42.59835 | 16.03917 | 29.48461 | 3.09008 | 24.04541 | 2.02146 | 33.44262 | 15.10997 |
| **AIFM1** | 6.85941 | 12.78064 | 7.31182 | 10.63022 | 9.13268 | 13.31841 | 12.14341 | 3.44457 | 15.28870 | 0.59051 | 17.23775 | 6.62440 |
| **AIFM2** | 0.00051 | 1.58497 | 2.31526 | 6.06067 | 2.57073 | 0.00000 | 2.76304 | 0.70356 | 0.00000 | 0.00000 | 1.34868 | 0.05877 |
| **AIFM3** | 0.00247 | 0.02139 | 0.02210 | 0.35573 | 0.02851 | 0.00000 | 0.20494 | 0.03216 | 0.00000 | 0.00000 | 0.06014 | 0.09195 |
| **AIG1** | 0.51103 | 0.08882 | 0.41357 | 0.00000 | 0.00000 | 0.22595 | 0.34221 | 0.60658 | 0.03108 | 0.92112 | 0.03261 | 0.03822 |
| **AIM1** | 11.36045 | 0.00000 | 8.29866 | 1.98721 | 0.00000 | 0.15798 | 1.74889 | 0.90194 | 0.08253 | 1.23405 | 0.00000 | 0.24477 |
| **AIM1L** | 0.07475 | 0.00000 | 7.63307 | 4.54388 | 18.61156 | 0.00000 | 2.42235 | 0.29899 | 11.17808 | 3.10146 | 9.37328 | 0.20477 |
| **AIM2** | 0.00534 | 0.18146 | 0.00534 | 0.00000 | 0.02386 | 0.09677 | 0.00000 | 0.01000 | 0.00000 | 0.03683 | 0.02684 | 0.00068 |
| **AIMP1** | 0.62017 | 13.60438 | 7.53402 | 7.43481 | 11.11887 | 32.60840 | 7.98533 | 0.00000 | 10.16123 | 19.07166 | 12.82863 | 6.64796 |
| **AIMP2** | 5.71569 | 13.25002 | 6.97324 | 6.57410 | 8.53092 | 10.38670 | 3.87347 | 2.57479 | 10.51616 | 0.00000 | 14.50387 | 4.80920 |
| **AIP** | 4.07090 | 3.45522 | 3.24994 | 4.80451 | 1.23577 | 0.00000 | 5.26149 | 6.72762 | 1.75093 | 0.00000 | 2.41102 | 9.90657 |
| **AIPL1** | 0.08511 | 0.00968 | 0.04573 | 0.00666 | 0.00552 | 0.03242 | 0.01086 | 0.08900 | 0.01122 | 0.04610 | 0.01340 | 0.02813 |
| **AIRE** | 0.00565 | 0.42438 | 0.53796 | 1.37687 | 2.51078 | 0.00000 | 0.00000 | 0.03483 | 0.00000 | 0.07997 | 0.00000 | 0.08517 |
| **AIRN** | 0.00264 | 0.04262 | 0.21028 | 0.17271 | 0.01409 | 0.00000 | 0.01327 | 0.07365 | 0.00541 | 0.46288 | 0.01829 | 0.00470 |
| **AJAP1** | 0.00688 | 0.00353 | 0.02821 | 0.00679 | 0.50532 | 0.04850 | 0.04690 | 0.01291 | 0.00000 | 0.00001 | 0.06182 | 0.02340 |
| **AJUBA** | 1.29668 | 1.27865 | 5.41849 | 0.00000 | 4.12799 | 0.08067 | 10.92569 | 5.57347 | 4.82375 | 0.03565 | 4.56157 | 3.14513 |
| **AK1** | 2.35847 | 4.34571 | 1.23873 | 0.73788 | 2.39575 | 3.18521 | 1.01036 | 0.57820 | 1.54072 | 1.44752 | 2.37226 | 1.26599 |
| **AK2** | 1.90937 | 15.87034 | 8.18981 | 14.05678 | 18.00746 | 17.43283 | 14.70435 | 6.51473 | 14.11708 | 3.57316 | 18.18465 | 13.30493 |
| **AK3** | 0.08563 | 0.00000 | 1.28431 | 0.65448 | 0.00000 | 0.61630 | 1.99227 | 0.08878 | 1.39173 | 2.29817 | 1.32944 | 0.00000 |

| | | | | | | | | | | | | |
|---|---|---|---|---|---|---|---|---|---|---|---|---|
| **AK4** | 0.19795 | 19.30846 | 0.00000 | 22.65928 | 22.82865 | 25.11527 | 6.42185 | 1.16438 | 9.67057 | 28.03753 | 25.29469 | 15.79777 |
| **AK5** | 3.47539 | 0.82436 | 0.35017 | 1.47126 | 0.37996 | 0.64414 | 0.50451 | 0.61291 | 0.28152 | 0.18625 | 0.21926 | 0.42746 |
| **AK6** | 0.00000 | 8.42082 | 7.34628 | 1.43807 | 0.00000 | 31.26380 | 12.88119 | 4.10440 | 12.17839 | 3.90253 | 11.45184 | 16.54607 |
| **AK7** | 0.09237 | 0.21340 | 2.73573 | 0.00001 | 0.30793 | 0.85251 | 0.00025 | 1.03630 | 0.00000 | 0.00000 | 0.21449 | 2.50361 |
| **AK8** | 0.37809 | 0.44620 | 0.07878 | 1.69918 | 0.20179 | 0.00000 | 0.05680 | 0.16741 | 0.00000 | 0.04015 | 0.01870 | 0.53320 |
| **AK9** | 1.86201 | 0.41010 | 0.57859 | 0.00000 | 0.00619 | 0.63399 | 0.00000 | 0.34863 | 0.55831 | 0.00000 | 0.00137 | 0.41486 |
| **AKAP1** | 9.28458 | 10.00196 | 7.62868 | 13.86218 | 9.71439 | 7.51272 | 8.36998 | 2.59390 | 11.93221 | 28.20336 | 8.34173 | 9.13858 |
| **AKAP10** | 0.54528 | 3.39321 | 2.63531 | 3.05239 | 0.26485 | 9.49746 | 0.00139 | 3.98780 | 6.38056 | 5.93066 | 1.55484 | 6.51938 |
| **AKAP11** | 8.50977 | 0.09822 | 1.67126 | 0.67694 | 0.32536 | 4.27068 | 1.75955 | 0.81877 | 0.65464 | 6.34994 | 0.43332 | 1.66608 |
| **AKAP12** | 0.00000 | 17.19473 | 3.31335 | 20.20316 | 15.60389 | 35.12698 | 3.78007 | 1.31846 | 17.39756 | 62.73109 | 4.08720 | 14.66003 |
| **AKAP13** | 5.90877 | 0.53149 | 15.17855 | 8.70135 | 5.40366 | 0.32687 | 11.89665 | 7.22265 | 17.70915 | 16.32759 | 3.43296 | 13.13399 |
| **AKAP14** | 0.20295 | 0.00355 | 0.01468 | 0.01243 | 0.04921 | 0.00000 | 0.03630 | 0.02231 | 0.01755 | 0.03163 | 0.01393 | 0.41828 |
| **AKAP2.uid-1139** | 0.00199 | 0.46120 | 0.04729 | 0.00000 | 0.11889 | 0.00000 | 0.00156 | 0.03130 | 0.43691 | 0.29353 | 0.46691 | 0.25338 |
| **AKAP2.uid-1192** | 0.00486 | 0.03016 | 0.05279 | 0.01538 | 0.10311 | 0.08112 | 0.00000 | 0.00982 | 0.12440 | 0.03955 | 0.26766 | 0.01109 |
| **AKAP3** | 0.00205 | 1.08213 | 0.02852 | 0.60174 | 0.01708 | 0.00000 | 0.00755 | 0.35666 | 0.35981 | 0.00000 | 0.03812 | 1.45734 |
| **AKAP5** | 0.00071 | 0.02835 | 0.53673 | 0.00000 | 0.06035 | 0.38474 | 0.00000 | 0.02146 | 0.03146 | 0.31142 | 0.23195 | 0.91298 |
| **AKAP6** | 0.76075 | 0.13910 | 0.13818 | 0.05332 | 0.22825 | 0.13943 | 0.00000 | 0.01570 | 0.01848 | 0.00000 | 0.08686 | 0.15804 |
| **AKAP7** | 0.56863 | 0.58030 | 0.00000 | 1.44681 | 0.00000 | 0.55310 | 0.00000 | 0.00000 | 0.00000 | 0.75606 | 0.01165 | 0.38128 |
| **AKAP8** | 3.91033 | 16.08663 | 12.53957 | 9.68362 | 12.74238 | 2.84732 | 13.96304 | 19.22930 | 16.60474 | 7.80718 | 8.35275 | 20.06881 |
| **AKAP8L** | 0.65561 | 0.90305 | 1.90588 | 2.17860 | 2.93884 | 0.00000 | 0.30963 | 0.74765 | 1.82792 | 4.06281 | 1.05705 | 1.66376 |
| **AKAP9** | 0.79106 | 3.71406 | 6.64059 | 3.95859 | 5.00120 | 14.43289 | 2.42733 | 0.00000 | 11.46761 | 31.53790 | 3.40335 | 4.40174 |
| **AKIP1** | 6.20662 | 14.33733 | 4.78599 | 5.10904 | 5.82088 | 7.70058 | 6.80134 | 2.26756 | 6.71660 | 1.43677 | 9.55633 | 17.62266 |
| **AKIRIN1** | 2.27909 | 19.63049 | 13.29080 | 11.99738 | 9.89556 | 16.52283 | 6.93352 | 3.42543 | 14.70260 | 17.22922 | 9.34224 | 14.39117 |
| **AKIRIN2** | 12.70145 | 13.39770 | 11.42201 | 11.23590 | 6.96177 | 8.38284 | 8.96991 | 14.95837 | 12.29938 | 1.40324 | 6.02312 | 17.22215 |
| **AKNA** | 0.28437 | 0.00154 | 0.08371 | 3.81020 | 0.86978 | 0.00000 | 1.12086 | 0.00602 | 0.00000 | 0.10568 | 0.00000 | 0.00915 |
| **AKNAD1** | 0.07096 | 0.02995 | 0.07774 | 0.00630 | 0.00401 | 0.01204 | 0.02284 | 0.01472 | 0.03126 | 0.01123 | 0.08460 | 0.01371 |
| **AKR1A1** | 3.35155 | 22.97931 | 14.77894 | 12.47023 | 26.02345 | 17.17290 | 25.94147 | 2.56990 | 21.50071 | 0.00002 | 25.32102 | 11.70165 |
| **AKR1B1** | 3.80549 | 2.69993 | 2.56766 | 3.41725 | 2.29973 | 3.85884 | 0.11381 | 0.18351 | 0.00058 | 0.00000 | 0.02912 | 0.16887 |
| **AKR1B10** | 0.66088 | 0.01117 | 0.02562 | 0.02014 | 0.00863 | 0.01416 | 0.01424 | 0.42261 | 0.00930 | 0.01929 | 0.03597 | 0.00000 |
| **AKR1B15** | 0.06501 | 0.01465 | 0.01450 | 0.01407 | 0.01700 | 0.00868 | 0.02670 | 0.12362 | 0.01594 | 0.02018 | 0.01466 | 0.01702 |
| **AKR1C1** | 0.00519 | 0.01788 | 0.01570 | 0.01558 | 0.02085 | 0.00104 | 0.01916 | 0.01263 | 0.03458 | 0.00000 | 0.01625 | 0.06077 |
| **AKR1C2** | 0.00983 | 0.01991 | 0.01306 | 0.00773 | 0.02235 | 0.02888 | 0.01976 | 0.00993 | 0.01040 | 0.02939 | 0.01337 | 0.01379 |
| **AKR1C3** | 4.10686 | 0.02366 | 0.00814 | 0.00000 | 0.15611 | 0.00715 | 0.01921 | 0.76118 | 0.00000 | 0.03191 | 0.00767 | 0.52024 |
| **AKR1C6P** | 0.00640 | 0.01440 | 0.01539 | 0.01360 | 0.01196 | 0.01238 | 0.02344 | 0.01163 | 0.00672 | 0.01794 | 0.04291 | 0.02584 |
| **AKR1CL1** | 0.00563 | 0.01842 | 0.01505 | 0.00949 | 0.02364 | 0.01989 | 0.01685 | 0.01147 | 0.01560 | 0.01878 | 0.01417 | 0.02388 |
| **AKR1D1** | 0.29993 | 0.03076 | 0.03738 | 0.00294 | 0.00583 | 0.10434 | 0.00000 | 0.07455 | 0.00000 | 0.05373 | 0.00776 | 0.29532 |
| **AKR1E2** | 0.05396 | 0.35665 | 0.00834 | 0.00000 | 0.07096 | 0.07686 | 0.06349 | 0.39974 | 0.00000 | 0.06343 | 0.00007 | 0.22772 |
| **AKR7A2** | 1.43653 | 3.51811 | 8.79905 | 3.63336 | 2.09608 | 4.96523 | 6.26183 | 0.11294 | 11.81882 | 0.00000 | 8.56282 | 1.32736 |
| **AKR7A2P1** | 0.04742 | 0.17288 | 0.43017 | 0.17823 | 0.06593 | 0.15378 | 0.06679 | 0.02193 | 0.63428 | 0.00000 | 0.51547 | 0.13457 |
| **AKR7A3** | 0.00891 | 0.22589 | 0.24186 | 0.00000 | 0.04935 | 0.37720 | 0.04550 | 0.00134 | 0.14678 | 0.00483 | 0.01074 | 0.32765 |
| **AKR7L** | 0.00757 | 0.08548 | 0.20805 | 0.00000 | 0.00000 | 0.16030 | 0.00000 | 0.00279 | 0.33944 | 0.37722 | 0.20225 | 0.14169 |
| **AKT1** | 5.24826 | 1.72725 | 7.56096 | 1.24234 | 9.86707 | 1.71597 | 7.92837 | 0.57837 | 1.25442 | 3.41747 | 0.72354 | 0.00000 |
| **AKT1S1** | 1.64162 | 4.70702 | 11.84401 | 0.99521 | 8.60218 | 0.00000 | 8.09077 | 6.84292 | 5.57615 | 18.84363 | 0.67701 | 9.89126 |
| **AKT2** | 6.62808 | 3.32007 | 7.01688 | 4.23978 | 8.68253 | 0.00000 | 6.25987 | 0.43371 | 10.79270 | 13.79017 | 3.01831 | 1.64613 |
| **AKT3** | 4.00019 | 0.64648 | 0.80342 | 0.74135 | 1.18624 | 1.82117 | 0.50911 | 0.12140 | 0.00000 | 2.92582 | 0.15080 | 1.18376 |
| **AKTIP** | 6.69484 | 5.25411 | 1.13152 | 0.00000 | 1.94736 | 4.40495 | 2.67083 | 0.52016 | 3.65365 | 2.38793 | 6.21206 | 0.98208 |
| **ALAD** | 0.00000 | 14.27611 | 6.50112 | 2.36154 | 29.45280 | 5.83916 | 9.50769 | 3.89138 | 12.12145 | 0.00000 | 25.81960 | 20.22338 |
| **ALAS1** | 6.36593 | 22.19616 | 29.52376 | 1.16511 | 15.82589 | 23.62306 | 13.39402 | 7.13632 | 10.97844 | 0.00000 | 20.16351 | 15.90035 |

| | | | | | | | | | | | | |
|---|---|---|---|---|---|---|---|---|---|---|---|---|
| **ALAS2** | 0.00943 | 0.02356 | 0.06439 | 0.02412 | 0.04821 | 0.00000 | 0.02140 | 0.01650 | 0.03652 | 0.00000 | 0.01947 | 0.02004 |
| **ALB** | 0.00440 | 0.00641 | 0.00764 | 0.00858 | 0.00000 | 0.04368 | 0.09285 | 0.00955 | 0.02187 | 0.06569 | 0.01521 | 0.06357 |
| **ALCAM** | 0.84256 | 0.02972 | 0.28324 | 1.12643 | 0.21052 | 0.00000 | 0.00000 | 0.19337 | 0.23112 | 0.63025 | 0.00000 | 0.18553 |
| **ALDH16A1** | 0.00000 | 1.96229 | 7.34097 | 2.47220 | 0.00000 | 0.00000 | 2.54844 | 0.50137 | 1.45926 | 5.18821 | 3.92520 | 1.11114 |
| **ALDH18A1** | 3.98217 | 3.81768 | 7.69485 | 0.64061 | 1.92173 | 5.87175 | 3.00762 | 1.15308 | 13.41432 | 0.00004 | 6.93878 | 0.28016 |
| **ALDH1A1** | 0.22203 | 0.00000 | 1.52676 | 7.46518 | 0.00000 | 0.00000 | 0.02014 | 0.03831 | 0.21334 | 0.15120 | 1.50725 | 0.00229 |
| **ALDH1A2** | 7.14710 | 0.00575 | 0.03758 | 0.01990 | 0.00528 | 0.21714 | 0.11215 | 0.54193 | 0.05711 | 0.00000 | 0.02953 | 0.00000 |
| **ALDH1A3** | 0.00402 | 0.37915 | 0.60951 | 1.41758 | 0.00043 | 0.01257 | 0.00000 | 0.09285 | 0.00000 | 0.02689 | 0.00995 | 0.06981 |
| **ALDH1B1** | 0.00147 | 1.26324 | 3.47943 | 2.22584 | 0.03770 | 1.01679 | 0.57063 | 0.07412 | 4.15659 | 0.00000 | 3.64488 | 0.00000 |
| **ALDH1L1** | 0.01340 | 0.03107 | 0.20778 | 0.04471 | 0.00000 | 0.03350 | 0.00000 | 0.03632 | 0.00000 | 0.05107 | 0.03667 | 0.00913 |
| **ALDH1L2** | 0.00486 | 0.00000 | 0.28709 | 0.00000 | 0.05344 | 0.24035 | 0.69714 | 0.13280 | 0.01412 | 0.24734 | 0.24755 | 0.12486 |
| **ALDH2** | 0.11751 | 2.88764 | 10.20073 | 36.48247 | 18.22402 | 2.66415 | 7.97139 | 0.00960 | 3.74744 | 0.17294 | 6.98790 | 0.00000 |
| **ALDH3A1** | 0.00568 | 0.26876 | 0.00676 | 0.00000 | 0.14648 | 0.00000 | 0.03822 | 0.00893 | 0.00000 | 0.06803 | 0.02186 | 0.03390 |
| **ALDH3A2** | 1.03636 | 12.54991 | 0.00000 | 14.18238 | 5.42797 | 19.03333 | 5.36824 | 0.79134 | 0.09117 | 5.69295 | 2.59202 | 1.63886 |
| **ALDH3B1** | 0.00679 | 0.01673 | 0.11479 | 0.01145 | 0.00900 | 0.02004 | 0.07743 | 0.01461 | 0.00008 | 0.00276 | 0.08122 | 0.02567 |
| **ALDH3B2** | 0.04672 | 0.00975 | 1.04949 | 0.00000 | 0.00127 | 0.01730 | 0.68271 | 0.02704 | 0.00000 | 0.06553 | 0.05269 | 0.00861 |
| **ALDH4A1** | 0.40983 | 2.39133 | 8.37923 | 1.50507 | 6.01451 | 0.00000 | 4.30707 | 0.08858 | 14.45924 | 1.22706 | 8.06942 | 0.00000 |
| **ALDH5A1** | 3.45853 | 0.27885 | 1.27539 | 0.27069 | 0.92436 | 0.67964 | 0.76647 | 0.06975 | 0.00000 | 0.60971 | 0.60723 | 0.00000 |
| **ALDH6A1** | 0.32246 | 7.07421 | 2.68234 | 2.48409 | 2.30030 | 10.67522 | 2.88767 | 0.39402 | 0.53875 | 4.12596 | 10.69615 | 2.20241 |
| **ALDH7A1** | 6.43334 | 6.26814 | 5.65280 | 5.82778 | 1.89871 | 7.66260 | 5.31311 | 4.92146 | 6.27361 | 0.00000 | 6.36211 | 0.87540 |
| **ALDH8A1** | 0.03766 | 0.01320 | 0.04621 | 0.01300 | 0.02158 | 0.01395 | 0.00813 | 0.56368 | 0.02712 | 0.02366 | 0.01196 | 0.02878 |
| **ALDH9A1** | 4.34139 | 8.22482 | 3.33296 | 3.92369 | 1.50420 | 10.40984 | 8.17349 | 2.50445 | 10.15522 | 2.26337 | 13.29464 | 10.93560 |
| **ALDOA** | 10.22454 | 55.79713 | 49.54375 | 45.62061 | 57.33976 | 27.28322 | 69.99712 | 26.56908 | 67.82144 | 15.81690 | 47.86486 | 51.29718 |
| **ALDOB** | 0.00597 | 0.02305 | 0.01518 | 0.01691 | 0.00973 | 0.01725 | 0.00960 | 0.01048 | 0.03560 | 0.02739 | 0.01448 | 0.01531 |
| **ALDOC** | 0.00226 | 1.06416 | 2.55908 | 6.77244 | 0.88297 | 0.00000 | 0.86901 | 3.23540 | 4.19866 | 0.00000 | 0.61841 | 1.15111 |
| **ALG1** | 0.04054 | 7.10645 | 6.51728 | 3.74738 | 3.54666 | 0.00000 | 0.00000 | 0.29174 | 14.89931 | 0.00000 | 8.12635 | 2.78212 |
| **ALG10** | 0.24047 | 0.38475 | 0.18692 | 0.24401 | 0.31159 | 0.95668 | 0.00000 | 0.06398 | 0.00000 | 0.23582 | 0.12795 | 0.32910 |
| **ALG10B** | 0.00192 | 0.46373 | 0.23040 | 0.07082 | 0.49719 | 0.74351 | 0.00000 | 0.10814 | 0.47910 | 0.37888 | 0.26040 | 0.82684 |
| **ALG11** | 0.39892 | 3.18080 | 4.13312 | 0.29367 | 0.35091 | 5.49197 | 0.00000 | 0.69810 | 4.11483 | 1.32929 | 2.18293 | 5.06861 |
| **ALG12** | 1.10127 | 6.79381 | 4.89960 | 3.37910 | 3.57523 | 0.00000 | 1.06177 | 0.36142 | 10.20017 | 1.77861 | 3.53934 | 4.72193 |
| **ALG13** | 4.07474 | 2.09173 | 5.89348 | 0.69184 | 0.00000 | 16.08801 | 7.04678 | 3.95964 | 4.59262 | 7.38431 | 6.93942 | 5.60267 |
| **ALG14** | 0.00365 | 5.60090 | 6.57149 | 0.04886 | 5.01401 | 12.45777 | 0.00000 | 0.00000 | 11.03698 | 0.23806 | 4.28876 | 2.09075 |
| **ALG1L** | 0.17310 | 1.20653 | 0.03802 | 0.05384 | 0.29808 | 0.45897 | 0.00000 | 0.00000 | 0.00000 | 0.00000 | 0.09264 | 0.27964 |
| **ALG1L2** | 0.00388 | 0.01325 | 0.04073 | 0.01156 | 0.01017 | 0.01867 | 0.00587 | 0.01692 | 0.02130 | 0.07001 | 0.01560 | 0.01924 |
| **ALG1L9P** | 0.07195 | 0.07288 | 0.03150 | 0.06037 | 0.00247 | 0.00000 | 0.01499 | 0.10997 | 0.06479 | 0.03257 | 0.03415 | 0.03488 |
| **ALG2** | 2.90273 | 15.75347 | 15.94611 | 11.13186 | 5.05982 | 16.97743 | 4.14131 | 8.08126 | 23.59318 | 0.00000 | 9.48608 | 17.04555 |
| **ALG3** | 1.64092 | 17.94190 | 14.24776 | 8.84803 | 14.95992 | 2.68824 | 5.16302 | 1.86766 | 27.58442 | 0.00000 | 10.80464 | 18.87157 |
| **ALG5** | 0.56478 | 9.68917 | 10.94636 | 3.10362 | 6.07370 | 23.24613 | 0.00000 | 3.84764 | 21.40586 | 4.20127 | 8.69500 | 12.19846 |
| **ALG6** | 0.44352 | 6.52335 | 2.73531 | 0.78859 | 2.91660 | 15.77658 | 0.00000 | 0.35895 | 5.37525 | 2.47490 | 4.60009 | 8.59950 |
| **ALG8** | 0.12687 | 10.86099 | 12.26256 | 5.38070 | 4.19556 | 14.53090 | 0.00000 | 0.19364 | 20.68394 | 0.00000 | 7.03108 | 8.50307 |
| **ALG9** | 0.28127 | 3.76427 | 2.63667 | 1.57893 | 2.17343 | 8.11935 | 0.00000 | 0.64772 | 7.29164 | 1.64349 | 2.45821 | 0.17866 |
| **ALK** | 0.11815 | 0.01643 | 0.21275 | 0.01537 | 0.02350 | 0.00000 | 0.03933 | 0.16377 | 0.00000 | 0.02559 | 0.01435 | 0.00832 |
| **ALKBH1** | 3.70221 | 11.71252 | 4.24225 | 5.74162 | 2.94636 | 14.56670 | 5.32636 | 8.79078 | 4.56210 | 0.30399 | 4.89531 | 14.52572 |
| **ALKBH2** | 7.09311 | 8.12928 | 0.92303 | 2.25173 | 0.87507 | 4.03734 | 4.18152 | 2.26080 | 4.93474 | 0.00000 | 2.57583 | 10.12983 |
| **ALKBH3** | 1.58627 | 1.06106 | 2.52033 | 0.00000 | 0.48626 | 1.56993 | 2.62295 | 4.06211 | 0.98861 | 0.00000 | 1.75218 | 1.01099 |
| **ALKBH3-AS1** | 0.06241 | 0.01702 | 0.08804 | 0.00000 | 0.02819 | 0.03223 | 0.06441 | 0.00998 | 0.01263 | 0.06461 | 0.00707 | 0.01070 |
| **ALKBH4** | 0.15623 | 7.35881 | 2.75828 | 3.24608 | 2.26176 | 1.44521 | 3.66465 | 0.55365 | 5.85605 | 0.00000 | 3.74323 | 5.11445 |
| **ALKBH5** | 12.81864 | 8.94416 | 11.00754 | 9.84283 | 12.25077 | 0.13687 | 12.88894 | 5.10541 | 8.27868 | 11.77757 | 5.33440 | 9.54949 |

| | | | | | | | | | | | | |
|---|---|---|---|---|---|---|---|---|---|---|---|---|
| **ALKBH6** | 0.00107 | 0.69783 | 4.52351 | 2.03725 | 1.95652 | 0.00000 | 2.85012 | 0.00000 | 2.09690 | 1.55201 | 3.88741 | 1.02078 |
| **ALKBH7** | 3.74945 | 17.86739 | 6.37243 | 20.29087 | 12.79053 | 0.00000 | 9.44898 | 7.08687 | 9.18080 | 1.63419 | 7.38782 | 15.74127 |
| **ALKBH8** | 0.00000 | 2.02955 | 1.11080 | 0.73384 | 1.85576 | 3.27712 | 5.69438 | 5.77190 | 1.34740 | 0.72405 | 1.47650 | 4.42950 |
| **ALLC** | 0.19398 | 0.01861 | 0.01217 | 0.01156 | 0.01437 | 0.00828 | 0.02191 | 0.03468 | 0.02240 | 0.02451 | 0.01537 | 0.01145 |
| **ALMS1** | 10.49491 | 0.68227 | 1.15230 | 0.44102 | 1.59826 | 5.80026 | 3.07378 | 4.24335 | 0.02734 | 6.73054 | 0.61738 | 2.41103 |
| **ALMS1-IT1** | 0.93283 | 0.14320 | 0.05122 | 0.02839 | 0.04250 | 0.05833 | 0.25194 | 0.08191 | 0.00000 | 0.44533 | 0.08401 | 0.01544 |
| **ALMS1P** | 0.02831 | 0.01171 | 0.01385 | 0.01721 | 0.00466 | 0.01326 | 0.04898 | 0.01054 | 0.01274 | 0.10808 | 0.01614 | 0.01403 |
| **ALOX12** | 0.00103 | 0.06762 | 3.47234 | 0.76352 | 0.00000 | 0.00731 | 2.70910 | 0.17168 | 5.04689 | 0.00000 | 0.02613 | 0.48780 |
| **ALOX12B** | 0.02067 | 0.01429 | 0.06883 | 0.00819 | 0.00889 | 0.01804 | 0.00000 | 0.00968 | 0.07791 | 0.03430 | 0.02795 | 0.01440 |
| **ALOX12P2** | 0.02307 | 0.05616 | 0.13308 | 0.00404 | 0.00353 | 0.01867 | 0.01025 | 0.11368 | 0.02451 | 0.02829 | 0.01196 | 0.05607 |
| **ALOX15** | 11.58028 | 0.14166 | 0.02444 | 0.00000 | 0.08187 | 0.00000 | 0.04022 | 3.00653 | 0.00000 | 0.00000 | 0.02457 | 0.36686 |
| **ALOX15B** | 0.46897 | 0.00567 | 0.06467 | 0.00580 | 0.03214 | 0.02930 | 0.00827 | 0.00000 | 0.01967 | 0.00000 | 0.01116 | 0.15377 |
| **ALOX5** | 0.00000 | 1.94093 | 0.00000 | 6.77550 | 0.79925 | 1.09531 | 0.00000 | 0.02106 | 0.19130 | 0.09419 | 0.98029 | 0.00000 |
| **ALOX5AP** | 0.01690 | 0.09704 | 0.00894 | 0.11994 | 0.01216 | 0.01853 | 0.01751 | 0.00845 | 0.02368 | 0.00076 | 0.01236 | 0.01049 |
| **ALOXE3** | 3.66965 | 0.00881 | 0.21561 | 0.00897 | 0.03413 | 0.00000 | 0.00110 | 0.04761 | 0.00627 | 0.01316 | 0.03551 | 0.10448 |
| **ALPI** | 0.10885 | 0.06408 | 0.03021 | 0.24001 | 0.19818 | 0.00000 | 0.02691 | 0.03975 | 0.02244 | 0.00000 | 0.04708 | 0.06403 |
| **ALPK1** | 0.00488 | 0.01036 | 0.00850 | 0.01089 | 0.03676 | 0.01529 | 0.06655 | 0.01012 | 0.00000 | 0.06965 | 0.01510 | 0.02020 |
| **ALPK2** | 0.14319 | 0.04398 | 0.03922 | 0.01153 | 0.00000 | 0.01608 | 0.05044 | 0.08635 | 0.00000 | 0.05155 | 0.03285 | 0.01631 |
| **ALPK3** | 0.10478 | 0.00000 | 0.50986 | 0.00000 | 0.40118 | 0.00000 | 0.46237 | 0.00979 | 0.96199 | 1.42006 | 1.04695 | 0.50037 |
| **ALPL** | 0.00054 | 51.85434 | 0.79508 | 34.44756 | 50.53916 | 0.00000 | 0.00000 | 0.00000 | 4.56345 | 9.42360 | 9.27388 | 27.29858 |
| **ALPP** | 1.99175 | 17.36565 | 0.00000 | 0.27630 | 43.91255 | 0.00000 | 0.00009 | 1.56390 | 40.49615 | 4.41137 | 29.83160 | 4.68497 |
| **ALPPL2** | 8.76756 | 56.50602 | 0.00000 | 10.59841 | 80.12932 | 8.10496 | 0.11540 | 22.28954 | 46.61001 | 33.01831 | 22.14551 | 43.87640 |
| **ALS2** | 2.13657 | 3.18312 | 2.38335 | 4.93240 | 1.67265 | 5.20688 | 4.19144 | 8.48118 | 1.74504 | 2.72655 | 1.37744 | 6.35178 |
| **ALS2CL** | 0.09334 | 0.25772 | 0.76165 | 0.03770 | 0.09632 | 0.00000 | 0.05503 | 0.00000 | 0.67065 | 0.00000 | 0.00000 | 0.28332 |
| **ALS2CR11** | 2.51860 | 1.49957 | 0.11192 | 0.00297 | 0.02672 | 0.29691 | 0.00000 | 0.92423 | 0.00000 | 0.00372 | 0.09089 | 1.81742 |
| **ALS2CR12** | 0.05531 | 0.00000 | 0.05055 | 0.11489 | 0.00000 | 0.09172 | 0.00000 | 0.10374 | 0.01083 | 0.24923 | 0.20948 | 0.49827 |
| **ALX1** | 0.00527 | 2.57439 | 0.28425 | 0.38898 | 0.00000 | 1.33482 | 0.82065 | 0.01711 | 0.06252 | 0.00000 | 0.20613 | 2.76422 |
| **ALX3** | 0.00430 | 0.35512 | 0.14035 | 0.27808 | 0.11237 | 0.00000 | 0.00000 | 0.08725 | 0.50289 | 0.16604 | 0.16931 | 0.07259 |
| **ALX4** | 0.00516 | 0.01803 | 0.07574 | 0.00648 | 0.01042 | 0.03683 | 0.00000 | 0.07113 | 0.00000 | 0.07913 | 0.01272 | 0.02006 |
| **ALYREF** | 3.50023 | 2.64287 | 7.42881 | 4.41790 | 7.72876 | 1.42558 | 17.83277 | 10.76414 | 0.00000 | 32.58820 | 2.41306 | 3.03441 |
| **AMACR** | 0.42401 | 0.51554 | 0.08403 | 0.27924 | 0.00000 | 0.07715 | 0.00000 | 0.31152 | 0.13719 | 0.00000 | 0.48691 | 0.18217 |
| **AMBN** | 0.01016 | 0.00068 | 0.00000 | 2.47035 | 0.07087 | 0.04173 | 0.00389 | 0.01595 | 0.28937 | 0.00000 | 0.04749 | 0.01166 |
| **AMBP** | 0.60923 | 0.01523 | 0.01933 | 0.01033 | 0.01604 | 0.00000 | 0.02218 | 1.43802 | 0.03844 | 0.00738 | 0.00990 | 0.00978 |
| **AMBRA1** | 11.65224 | 9.68534 | 1.56022 | 4.65227 | 6.40211 | 0.04986 | 4.64671 | 2.25362 | 5.41875 | 4.25956 | 1.89612 | 9.45537 |
| **1.00000** | 10.69565 | 20.94659 | 13.25111 | 10.83744 | 13.44874 | 47.45410 | 28.14872 | 27.76213 | 25.46312 | 19.41929 | 25.58396 | 32.50168 |
| **AMDHD1** | 0.00365 | 0.00000 | 0.30136 | 4.73011 | 0.07433 | 0.36091 | 0.12802 | 1.20287 | 0.31089 | 2.73192 | 0.00000 | 2.30831 |
| **AMDHD2** | 0.00000 | 3.86730 | 8.12446 | 3.95709 | 5.85393 | 0.00000 | 2.25420 | 3.71871 | 4.83044 | 1.32058 | 8.64480 | 7.38358 |
| **AMER1** | 0.00555 | 0.11486 | 0.27212 | 0.19363 | 0.00000 | 0.00000 | 0.02307 | 0.00430 | 0.39158 | 1.10130 | 0.07681 | 0.11704 |
| **AMER2** | 0.00561 | 0.00000 | 1.10820 | 0.00778 | 0.62586 | 0.11340 | 0.30102 | 0.00976 | 0.33293 | 0.00000 | 0.19602 | 0.06393 |
| **AMER3** | 0.10934 | 0.05528 | 0.01653 | 0.01056 | 0.01584 | 0.00001 | 0.02349 | 0.00392 | 0.02338 | 0.01251 | 0.01656 | 0.01341 |
| **AMFR** | 5.07879 | 4.38921 | 0.33561 | 2.23643 | 7.09265 | 12.70605 | 0.67380 | 0.48896 | 2.78131 | 20.22647 | 3.74015 | 7.17708 |
| **AMH** | 0.00860 | 0.05663 | 0.24127 | 0.18450 | 0.00763 | 0.00000 | 0.06821 | 0.02642 | 0.05965 | 0.94761 | 0.00000 | 0.00298 |
| **AMHR2** | 0.31506 | 0.42618 | 1.25145 | 2.58126 | 0.15429 | 0.26331 | 0.22384 | 0.17169 | 0.07481 | 0.00000 | 0.00000 | 0.31944 |
| **AMICA1** | 0.01065 | 0.00000 | 0.80499 | 0.93505 | 0.06839 | 0.00000 | 0.00000 | 0.01763 | 0.00000 | 0.02144 | 0.01408 | 0.59360 |
| **AMIGO1** | 1.58085 | 0.00000 | 0.06885 | 0.00000 | 0.06405 | 0.00000 | 0.07957 | 0.00100 | 0.13307 | 0.21472 | 0.00915 | 0.03982 |
| **AMIGO2** | 0.32698 | 0.00000 | 0.12368 | 0.14151 | 0.00578 | 0.00000 | 0.06595 | 0.64761 | 0.00000 | 0.03119 | 0.09592 | 1.20942 |
| **AMIGO3** | 0.05217 | 0.19034 | 1.57970 | 1.11309 | 0.86054 | 0.00000 | 0.44057 | 0.00467 | 1.25954 | 1.64368 | 0.42868 | 0.65757 |
| **AMMECR1** | 0.02995 | 0.70081 | 5.26010 | 0.39717 | 3.59056 | 4.97950 | 10.62757 | 0.00000 | 3.29397 | 2.77528 | 6.12757 | 0.16382 |

| | | | | | | | | | | | | |
|---|---|---|---|---|---|---|---|---|---|---|---|---|
| **AMMECR1L** | 5.42871 | 3.59151 | 3.75826 | 3.27999 | 5.29065 | 1.31590 | 4.59560 | 9.23428 | 4.01841 | 5.59836 | 1.18365 | 5.09638 |
| **AMN** | 0.13424 | 0.00000 | 0.00000 | 0.06307 | 0.53826 | 0.00000 | 0.05547 | 0.00734 | 0.13240 | 0.48402 | 0.14086 | 0.01497 |
| **AMN1** | 1.65700 | 0.34161 | 3.94906 | 0.33145 | 0.00000 | 1.30910 | 0.68039 | 0.34568 | 3.17660 | 0.00000 | 1.83923 | 2.40535 |
| **AMOT** | 10.14756 | 4.96978 | 0.00000 | 3.82992 | 3.49843 | 10.84660 | 12.62619 | 0.00000 | 4.31863 | 22.56814 | 3.16956 | 7.86625 |
| **AMOTL1** | 6.14715 | 0.32738 | 3.43334 | 8.68982 | 1.59508 | 0.00000 | 1.90608 | 0.32256 | 0.00000 | 0.00000 | 1.49333 | 0.67221 |
| **AMOTL2** | 0.00185 | 13.61167 | 17.04897 | 0.00000 | 18.47892 | 1.97070 | 20.86722 | 0.00000 | 12.14128 | 0.00000 | 14.12850 | 3.51699 |
| **AMPD1** | 0.17904 | 0.02772 | 0.05778 | 0.03381 | 0.01530 | 0.00000 | 0.00574 | 0.10805 | 0.04390 | 0.00124 | 0.01494 | 0.08733 |
| **AMPD2** | 5.63154 | 3.44464 | 5.51702 | 2.68371 | 3.04356 | 0.00003 | 4.28145 | 0.65358 | 5.91267 | 1.01158 | 2.00745 | 4.69902 |
| **AMPD3** | 2.92075 | 0.01008 | 0.46712 | 0.60849 | 0.00000 | 0.10983 | 0.10362 | 0.10527 | 0.01079 | 0.00000 | 0.12624 | 0.47992 |
| **AMPH** | 2.26831 | 1.18223 | 0.00000 | 0.02421 | 0.00000 | 0.15395 | 0.00000 | 0.00402 | 0.00000 | 0.01391 | 0.00000 | 1.72325 |
| **AMT** | 0.06322 | 5.56863 | 0.86767 | 9.41702 | 4.25002 | 0.00000 | 1.44313 | 0.10626 | 2.65138 | 0.85148 | 8.69017 | 0.03048 |
| **AMY2B** | 0.00139 | 0.00000 | 0.08156 | 0.02487 | 0.00000 | 0.52263 | 0.21683 | 0.15744 | 0.08186 | 0.41956 | 0.17358 | 0.05489 |
| **AMZ1** | 0.15786 | 0.05049 | 0.55019 | 0.03468 | 0.00896 | 0.00000 | 0.00958 | 0.06993 | 0.66010 | 0.00000 | 0.00000 | 0.21364 |
| **AMZ2** | 3.27322 | 7.19894 | 3.02747 | 1.10239 | 2.65741 | 8.45970 | 1.47516 | 2.51902 | 0.98797 | 1.90919 | 5.08831 | 5.98004 |
| **AMZ2P1** | 1.01174 | 1.26211 | 0.23526 | 0.93823 | 0.01961 | 1.22104 | 0.43068 | 0.92987 | 0.00000 | 0.00000 | 0.33805 | 0.95103 |
| **ANAPC1** | 1.09351 | 2.41112 | 1.09570 | 2.14048 | 1.21131 | 2.24839 | 1.48401 | 1.41267 | 3.48057 | 1.72577 | 3.21788 | 2.57073 |
| **ANAPC10** | 1.16396 | 2.48699 | 6.30437 | 1.49518 | 0.00000 | 14.60319 | 4.10922 | 2.72777 | 4.42719 | 0.00000 | 4.32188 | 7.34964 |
| **ANAPC11** | 8.60142 | 13.24989 | 21.95549 | 23.67685 | 30.14506 | 14.96350 | 28.41248 | 13.92986 | 28.36663 | 9.06880 | 25.80501 | 12.41715 |
| **ANAPC13** | 2.46563 | 5.89769 | 12.29809 | 8.70771 | 2.96881 | 21.05991 | 21.69514 | 6.09075 | 10.50464 | 2.55918 | 13.94899 | 14.71266 |
| **ANAPC15** | 1.49922 | 4.56675 | 2.51818 | 1.38077 | 2.03433 | 1.70151 | 3.75465 | 4.59548 | 2.49666 | 0.00000 | 2.76945 | 4.49276 |
| **ANAPC16** | 1.08007 | 7.20845 | 5.29453 | 7.14244 | 7.76563 | 14.90132 | 9.25513 | 0.27116 | 7.90666 | 6.02670 | 9.93684 | 11.63842 |
| **ANAPC1P1** | 0.05986 | 0.01691 | 0.01561 | 0.03713 | 0.01883 | 0.00785 | 0.02245 | 0.33699 | 0.02200 | 0.00000 | 0.01518 | 0.05351 |
| **ANAPC2** | 0.45765 | 0.33210 | 1.59188 | 0.38207 | 1.38226 | 0.00000 | 0.00000 | 0.69194 | 1.46160 | 2.35981 | 1.00187 | 2.09997 |
| **ANAPC4** | 2.78185 | 3.14903 | 8.69255 | 4.66567 | 0.00000 | 13.13182 | 4.41373 | 1.35330 | 8.59831 | 7.65947 | 8.14689 | 1.12395 |
| **ANAPC5** | 0.97465 | 21.14616 | 11.53597 | 13.89659 | 16.95140 | 21.10423 | 16.43391 | 4.12916 | 25.74969 | 8.90650 | 23.10163 | 17.61761 |
| **ANAPC7** | 5.01918 | 13.08597 | 10.51150 | 4.82198 | 1.41565 | 13.01410 | 10.70849 | 1.38791 | 16.11045 | 7.45401 | 10.30068 | 13.41558 |
| **ANG** | 0.05772 | 0.19488 | 0.41155 | 0.01943 | 0.11739 | 0.00000 | 0.06910 | 0.00031 | 0.57080 | 0.00000 | 0.00000 | 1.66261 |
| **ANGEL1** | 0.17214 | 5.52993 | 1.27997 | 5.05386 | 2.40615 | 0.00000 | 0.00000 | 0.27947 | 5.69408 | 1.12419 | 1.40811 | 2.88298 |
| **ANGEL2** | 0.05333 | 1.77909 | 4.72085 | 4.89672 | 0.00000 | 9.28744 | 5.24611 | 3.18563 | 4.02712 | 0.09706 | 4.37205 | 5.70724 |
| **ANGPT1** | 0.00000 | 0.81521 | 1.50096 | 0.00000 | 0.00000 | 6.82547 | 5.26934 | 0.25549 | 4.15669 | 2.23546 | 1.53309 | 2.10978 |
| **ANGPT2** | 0.06525 | 0.00000 | 0.04156 | 0.38340 | 0.09177 | 0.42957 | 0.00000 | 1.01322 | 0.57260 | 2.33536 | 0.50359 | 0.82758 |
| **ANGPT4** | 0.00563 | 0.01286 | 0.28118 | 0.02278 | 0.01960 | 0.02219 | 0.00154 | 0.01173 | 0.00000 | 0.01777 | 0.02200 | 0.01906 |
| **ANGPTL1** | 0.03390 | 0.08610 | 0.01791 | 0.02666 | 0.16675 | 0.00000 | 0.00441 | 0.02439 | 0.05020 | 0.35195 | 0.00979 | 0.09388 |
| **ANGPTL2** | 0.00579 | 0.00447 | 0.04903 | 0.02294 | 0.02703 | 0.04664 | 0.00000 | 0.01026 | 0.00201 | 0.07556 | 0.01914 | 0.01225 |
| **ANGPTL3** | 0.00630 | 0.00573 | 0.05179 | 0.05120 | 0.05019 | 0.03614 | 0.00000 | 0.00906 | 0.10820 | 0.01081 | 0.04354 | 0.03538 |
| **ANGPTL4** | 0.11132 | 0.07724 | 0.03462 | 0.12598 | 0.00000 | 0.00000 | 0.01106 | 0.18929 | 0.04909 | 0.09140 | 0.01308 | 0.17003 |
| **ANGPTL5** | 0.20241 | 0.00172 | 0.30884 | 0.00338 | 0.00056 | 0.04844 | 0.00000 | 0.26116 | 0.24503 | 0.03136 | 0.01499 | 0.10105 |
| **ANGPTL6** | 0.07354 | 0.01295 | 0.02275 | 0.00916 | 0.00972 | 0.03454 | 0.03030 | 0.09171 | 0.07048 | 0.00000 | 0.00453 | 0.08368 |
| **ANGPTL7** | 0.00273 | 0.02736 | 0.04276 | 0.02277 | 0.12060 | 0.00000 | 0.00231 | 0.02335 | 0.01795 | 0.10207 | 0.06710 | 0.02077 |
| **ANHX** | 0.04128 | 0.01548 | 0.01329 | 0.00957 | 0.01050 | 0.03174 | 0.01404 | 0.00605 | 0.01638 | 0.00941 | 0.03037 | 0.04763 |
| **ANK1** | 0.00584 | 2.03630 | 1.43969 | 0.14920 | 0.14015 | 0.00000 | 1.01321 | 0.24802 | 0.46338 | 0.38594 | 0.00000 | 0.08106 |
| **ANK2** | 0.74182 | 0.00495 | 0.54412 | 0.56677 | 0.09490 | 0.00000 | 0.04492 | 0.02603 | 0.09009 | 0.23140 | 0.12734 | 0.08619 |
| **ANK3** | 5.56432 | 0.14114 | 0.31017 | 1.32033 | 0.00000 | 0.00000 | 0.97942 | 1.26143 | 2.01014 | 1.92387 | 2.15307 | 0.22582 |
| **ANKAR** | 0.10938 | 0.05694 | 0.40934 | 0.15517 | 0.00000 | 0.20500 | 0.08207 | 0.19510 | 0.11402 | 1.29902 | 0.08048 | 0.25003 |
| **ANKDD1A** | 0.10506 | 0.04226 | 0.41150 | 0.02158 | 0.17111 | 0.17417 | 0.17056 | 0.00000 | 0.00000 | 0.57740 | 0.66069 | 0.12099 |
| **ANKDD1B** | 0.00000 | 1.39307 | 0.06049 | 0.78368 | 0.10495 | 0.71173 | 0.00000 | 0.31525 | 0.00000 | 0.38023 | 0.19123 | 0.54828 |
| **ANKEF1** | 0.64032 | 0.88641 | 1.51737 | 0.00000 | 0.00000 | 2.23929 | 0.64343 | 0.19381 | 0.47708 | 0.03287 | 0.53188 | 0.67580 |
| **ANKFN1** | 0.01143 | 0.06160 | 0.61663 | 0.00846 | 0.04300 | 0.00009 | 0.10459 | 0.02251 | 0.00000 | 0.00000 | 0.01388 | 0.04793 |

| | | | | | | | | | | | | |
|---|---|---|---|---|---|---|---|---|---|---|---|---|
| **ANKFY1** | 0.13203 | 4.46634 | 4.82166 | 3.68406 | 8.43566 | 5.98930 | 7.77046 | 2.18216 | 10.32173 | 6.41301 | 5.88406 | 4.17982 |
| **ANKH** | 4.45434 | 0.41144 | 1.21751 | 0.18456 | 0.63722 | 1.30690 | 0.70213 | 2.34233 | 1.14350 | 3.42418 | 0.22258 | 0.05824 |
| **ANKHD1-EIF4E** | 0.00315 | 0.06065 | 0.14108 | 0.01240 | 0.08903 | 0.00000 | 0.13961 | 0.02893 | 0.04644 | 0.20427 | 0.04899 | 0.08903 |
| **ANKHD1.uid-68** | 0.00122 | 0.00000 | 1.11969 | 0.00000 | 0.00000 | 0.60800 | 2.33458 | 0.05105 | 0.29991 | 3.40304 | 0.24712 | 0.11171 |
| **ANKHD1.uid-74** | 0.00508 | 0.36639 | 0.75533 | 0.69226 | 1.40177 | 0.67286 | 1.09608 | 0.23151 | 0.00000 | 3.95975 | 0.00000 | 0.90622 |
| **ANKIB1** | 2.24172 | 2.37314 | 4.09051 | 0.63013 | 1.43191 | 10.50052 | 5.45420 | 1.20752 | 0.73678 | 19.96296 | 2.82466 | 2.82968 |
| **ANKK1** | 0.00760 | 0.03089 | 0.00000 | 0.74841 | 0.01317 | 0.04155 | 0.03941 | 0.00333 | 0.00871 | 0.01925 | 0.01769 | 0.45333 |
| **ANKLE1** | 0.00617 | 0.14176 | 3.58243 | 0.35546 | 0.00000 | 0.00000 | 0.53006 | 0.10106 | 4.27095 | 0.00000 | 2.05877 | 0.30302 |
| **ANKLE2** | 3.86198 | 2.98511 | 4.97589 | 3.13127 | 4.96266 | 7.38553 | 1.77900 | 0.00001 | 9.40232 | 14.21465 | 2.78691 | 2.26746 |
| **ANKMY1** | 0.29207 | 0.46225 | 0.22780 | 0.17746 | 0.02625 | 0.00000 | 0.10772 | 0.02281 | 1.49347 | 0.00000 | 0.50350 | 0.23934 |
| **ANKMY2** | 0.99240 | 0.69354 | 10.32335 | 3.66138 | 0.00000 | 1.10450 | 11.00212 | 3.79688 | 8.66948 | 0.00000 | 2.58600 | 0.78323 |
| **ANKRA2** | 0.33770 | 3.65383 | 3.32551 | 1.77134 | 1.01922 | 5.75014 | 0.93334 | 0.61399 | 0.82367 | 0.00000 | 0.93610 | 2.19677 |
| **ANKRD1** | 0.03180 | 0.09538 | 0.54849 | 1.93206 | 0.90717 | 0.00000 | 0.04499 | 0.06197 | 0.00000 | 0.67049 | 0.04843 | 0.02514 |
| **ANKRD10** | 4.74093 | 3.83060 | 16.92710 | 14.63274 | 3.30637 | 15.68072 | 24.86063 | 10.98814 | 24.26060 | 2.97943 | 16.09557 | 7.15303 |
| **ANKRD11** | 3.60356 | 1.70385 | 6.46479 | 5.58428 | 14.28788 | 1.92010 | 21.67897 | 2.74459 | 6.66209 | 56.00730 | 8.42754 | 3.24397 |
| **ANKRD12** | 1.66126 | 1.42647 | 6.10991 | 1.37625 | 0.00000 | 12.00011 | 0.00002 | 9.00544 | 11.81350 | 20.57121 | 1.10102 | 3.51474 |
| **ANKRD13A** | 7.31476 | 0.80365 | 2.03910 | 2.72693 | 4.80269 | 6.09483 | 6.29323 | 1.57139 | 1.23563 | 7.22852 | 3.82012 | 2.13619 |
| **ANKRD13B** | 1.13480 | 0.37179 | 0.09220 | 0.28312 | 0.18545 | 0.00000 | 0.31371 | 0.15984 | 0.03431 | 1.86361 | 0.00000 | 0.29995 |
| **ANKRD13C** | 0.68715 | 4.73121 | 5.06124 | 4.10770 | 0.72416 | 15.51122 | 10.25294 | 10.21748 | 9.88255 | 4.36712 | 6.61305 | 3.95712 |
| **ANKRD13D** | 0.05706 | 0.45984 | 1.56422 | 0.82884 | 1.78713 | 0.00000 | 1.77895 | 0.01800 | 2.30847 | 5.22268 | 0.30653 | 0.12750 |
| **ANKRD16** | 0.04217 | 0.20698 | 0.53027 | 0.20906 | 1.26672 | 0.00002 | 1.25054 | 0.10000 | 0.63972 | 1.86400 | 0.92831 | 0.12582 |
| **ANKRD17** | 6.11932 | 5.25284 | 11.59437 | 7.96020 | 5.28523 | 25.72256 | 20.63385 | 4.70173 | 7.24072 | 29.11542 | 6.11384 | 7.35516 |
| **ANKRD18A** | 0.00564 | 0.01059 | 0.02087 | 0.01054 | 0.01107 | 0.02139 | 0.01296 | 0.00881 | 0.01303 | 0.03576 | 0.01707 | 0.05016 |
| **ANKRD18B** | 0.01381 | 0.03933 | 0.19022 | 0.34563 | 0.01306 | 0.02155 | 0.01934 | 0.01874 | 0.02925 | 0.00605 | 0.02410 | 0.04575 |
| **ANKRD18DP** | 0.03548 | 0.02016 | 0.05100 | 0.04353 | 0.02577 | 0.00000 | 0.01465 | 0.00842 | 0.02086 | 0.00000 | 0.01546 | 0.04921 |
| **ANKRD19P** | 0.20969 | 2.19270 | 2.13091 | 2.31465 | 4.07141 | 0.16675 | 7.98731 | 1.47297 | 2.66150 | 9.45818 | 2.02540 | 2.14193 |
| **ANKRD2** | 0.00715 | 0.11313 | 0.28464 | 0.41449 | 0.01912 | 0.00005 | 0.34848 | 0.00857 | 0.00000 | 0.00000 | 1.04066 | 0.03344 |
| **ANKRD20A11P** | 0.00747 | 0.02731 | 0.00000 | 0.00000 | 0.01117 | 0.00000 | 0.00132 | 0.01203 | 0.75793 | 0.44142 | 0.00000 | 0.04258 |
| **ANKRD20A12P** | 0.02529 | 0.08914 | 0.01772 | 0.00000 | 0.01474 | 0.10913 | 0.00000 | 0.01104 | 0.08170 | 0.04917 | 0.01327 | 0.02282 |
| **ANKRD20A19P** | 0.01175 | 0.02030 | 2.75154 | 0.00086 | 0.00000 | 0.02923 | 0.72887 | 0.00959 | 1.80073 | 0.00000 | 0.00000 | 0.03787 |
| **ANKRD20A4** | 0.01628 | 0.01452 | 0.01228 | 0.00907 | 0.01084 | 0.02760 | 0.01580 | 0.03166 | 0.01293 | 0.03325 | 0.01490 | 0.04578 |
| **ANKRD20A5P** | 0.04945 | 0.20656 | 0.00000 | 0.13590 | 0.03331 | 0.10818 | 0.00000 | 0.03503 | 0.00000 | 0.20992 | 0.00197 | 0.20668 |
| **ANKRD20A8P** | 0.00785 | 0.01750 | 0.02822 | 0.02322 | 0.01941 | 0.00000 | 0.01398 | 0.03857 | 0.03206 | 0.02794 | 0.01503 | 0.06646 |
| **ANKRD20A9P** | 0.05430 | 0.05286 | 0.01174 | 0.18272 | 0.18636 | 0.00468 | 0.16111 | 0.19007 | 0.02274 | 0.16706 | 0.02016 | 0.08409 |
| **ANKRD22** | 1.08649 | 0.01138 | 0.00431 | 0.03741 | 0.01831 | 0.00000 | 0.12699 | 0.06801 | 0.03861 | 0.02042 | 0.00292 | 0.01430 |
| **ANKRD23** | 0.00290 | 0.00000 | 1.72289 | 0.00000 | 0.18396 | 0.00000 | 0.00000 | 0.00435 | 0.35505 | 1.54741 | 0.17588 | 0.24902 |
| **ANKRD24** | 0.07275 | 0.02949 | 0.11293 | 0.00000 | 0.04409 | 0.00000 | 0.27928 | 0.12091 | 0.10759 | 0.20618 | 0.03632 | 0.04860 |
| **ANKRD26** | 2.74278 | 1.23572 | 1.58439 | 0.72128 | 2.07861 | 4.87783 | 0.00000 | 0.80385 | 0.00000 | 14.02012 | 1.65277 | 3.27581 |
| **ANKRD26P1** | 1.15542 | 0.28598 | 0.41270 | 0.55342 | 0.00000 | 0.00000 | 0.67665 | 0.59303 | 0.00019 | 0.74911 | 0.23451 | 0.53590 |
| **ANKRD26P3** | 0.05093 | 0.08593 | 0.01602 | 0.01163 | 0.01732 | 0.00000 | 0.02269 | 0.07795 | 0.04046 | 0.00000 | 0.01383 | 0.16576 |
| **ANKRD27** | 2.07757 | 3.15442 | 4.23399 | 2.30086 | 2.32418 | 3.51924 | 4.36680 | 1.09981 | 7.77578 | 1.17747 | 5.29867 | 1.54330 |
| **ANKRD28** | 0.94530 | 4.22821 | 0.00000 | 1.29376 | 1.68526 | 11.39623 | 2.72316 | 0.25740 | 0.00000 | 11.00610 | 1.77202 | 2.28964 |
| **ANKRD29** | 0.00670 | 0.00781 | 0.02354 | 0.02279 | 0.03369 | 0.00000 | 0.01800 | 0.00646 | 0.01551 | 0.00224 | 0.05288 | 0.18682 |
| **ANKRD30A** | 0.42490 | 0.00000 | 0.00421 | 0.00000 | 0.16218 | 0.01874 | 0.00162 | 0.01192 | 0.00000 | 0.27523 | 0.00000 | 0.11529 |
| **ANKRD30B** | 0.00725 | 0.03520 | 0.06264 | 0.01312 | 0.01732 | 0.00483 | 0.01625 | 0.02070 | 0.01121 | 0.01951 | 0.02097 | 0.01451 |
| **ANKRD30BL** | 0.02024 | 1.45672 | 3.41371 | 0.65996 | 3.05036 | 0.00000 | 0.00000 | 0.00000 | 4.17729 | 4.00313 | 0.07952 | 0.29795 |
| **ANKRD30BP3** | 0.00620 | 0.01411 | 0.01383 | 0.01280 | 0.01538 | 0.01864 | 0.01849 | 0.01105 | 0.01789 | 0.01500 | 0.01486 | 0.02092 |
| **ANKRD31** | 0.00988 | 0.00573 | 0.01290 | 0.00689 | 0.06353 | 0.01167 | 0.02328 | 0.07888 | 0.00000 | 0.11065 | 0.01929 | 0.04702 |

| | | | | | | | | | | | | |
|---|---|---|---|---|---|---|---|---|---|---|---|---|
| **ANKRD32** | 3.08679 | 1.53176 | 3.21633 | 0.00000 | 0.39377 | 12.59735 | 4.78488 | 6.83200 | 3.88054 | 14.84160 | 2.74142 | 4.84972 |
| **ANKRD33** | 0.00623 | 0.01553 | 0.10490 | 0.04790 | 0.00702 | 0.01816 | 0.02239 | 0.01048 | 0.08143 | 0.00965 | 0.00808 | 0.01624 |
| **ANKRD33B** | 0.00000 | 3.63805 | 0.00000 | 0.00000 | 10.21864 | 5.33865 | 6.55756 | 0.80940 | 1.65870 | 10.37222 | 3.82458 | 0.00000 |
| **ANKRD34A** | 0.03338 | 0.00268 | 0.14689 | 1.51688 | 0.04337 | 0.00000 | 0.03774 | 0.01051 | 0.00000 | 0.12759 | 0.00768 | 0.01329 |
| **ANKRD34B** | 0.00341 | 0.01414 | 0.02955 | 0.01763 | 0.02648 | 0.01362 | 0.02533 | 0.02614 | 0.03468 | 0.00000 | 0.01318 | 0.05149 |
| **ANKRD34C** | 0.00579 | 0.01340 | 0.01144 | 0.06804 | 0.00475 | 0.00080 | 0.01868 | 0.01058 | 0.00000 | 0.03536 | 0.08111 | 0.01669 |
| **ANKRD35** | 0.02983 | 0.00000 | 0.15415 | 0.01514 | 0.00725 | 0.00000 | 0.01698 | 9.17595 | 0.00588 | 0.06952 | 0.04504 | 0.49337 |
| **ANKRD36** | 0.01064 | 0.19648 | 0.21898 | 0.00000 | 0.00000 | 0.79746 | 0.00000 | 0.25616 | 1.15848 | 1.91901 | 0.03300 | 2.31018 |
| **ANKRD36B** | 0.03996 | 0.16935 | 0.47289 | 0.00068 | 0.00000 | 0.13594 | 0.00003 | 0.64678 | 2.46546 | 1.08707 | 0.00000 | 2.15601 |
| **ANKRD36BP1** | 1.82815 | 0.67764 | 0.09156 | 0.45968 | 0.30325 | 1.43925 | 0.00000 | 0.58136 | 0.00000 | 0.00000 | 1.75345 | 0.11623 |
| **ANKRD36BP2** | 0.59436 | 0.09049 | 0.00000 | 0.00000 | 0.09764 | 0.10446 | 0.00000 | 0.53344 | 0.00000 | 0.63637 | 0.01921 | 0.35865 |
| **ANKRD37** | 2.40121 | 0.16352 | 1.29192 | 0.36790 | 0.43930 | 0.00024 | 0.09937 | 5.78853 | 0.60389 | 0.00000 | 0.03007 | 4.15997 |
| **ANKRD39** | 0.27318 | 4.15880 | 12.42140 | 0.59345 | 3.43496 | 1.46179 | 5.96583 | 1.19567 | 5.88512 | 1.13019 | 6.94326 | 3.55986 |
| **ANKRD40** | 1.29978 | 9.21256 | 3.00579 | 6.38575 | 3.07137 | 15.75222 | 15.91700 | 9.42896 | 5.54697 | 0.00000 | 11.29919 | 13.75045 |
| **ANKRD42** | 1.45982 | 0.99674 | 0.99882 | 0.48328 | 0.16823 | 0.98265 | 0.50485 | 2.53383 | 0.76240 | 0.00000 | 0.61875 | 1.98868 |
| **ANKRD44** | 0.48792 | 0.00536 | 0.06519 | 0.09458 | 0.04234 | 0.06793 | 0.05225 | 0.12276 | 0.16888 | 0.00000 | 0.02731 | 0.01054 |
| **ANKRD44-IT1** | 0.01326 | 0.01284 | 0.01589 | 0.02160 | 0.00915 | 0.02751 | 0.03074 | 0.01136 | 0.01111 | 0.01574 | 0.01591 | 0.02170 |
| **ANKRD45** | 0.83691 | 0.11433 | 0.00000 | 4.52888 | 0.18051 | 0.00588 | 0.08059 | 0.24986 | 0.00000 | 0.00000 | 0.04670 | 0.02155 |
| **ANKRD46** | 2.18541 | 3.81372 | 0.50551 | 0.57656 | 0.97238 | 5.61109 | 0.00000 | 0.00000 | 0.50790 | 0.01001 | 3.74196 | 1.11230 |
| **ANKRD49** | 0.10719 | 3.39780 | 3.13762 | 0.00002 | 0.00000 | 10.85274 | 1.34676 | 2.86120 | 4.25724 | 5.26709 | 3.32904 | 2.99447 |
| **ANKRD50** | 0.01920 | 0.85647 | 4.55406 | 0.48306 | 1.26026 | 4.56740 | 1.99797 | 3.32480 | 1.00847 | 6.77201 | 0.06256 | 1.79678 |
| **ANKRD52** | 2.09441 | 0.38532 | 0.86011 | 0.68943 | 1.08797 | 0.00000 | 0.12187 | 0.27662 | 3.39049 | 5.77863 | 0.00000 | 1.14112 |
| **ANKRD53** | 0.17885 | 0.00859 | 0.06125 | 0.00620 | 0.11950 | 0.00000 | 0.01889 | 0.02026 | 0.00000 | 0.00000 | 0.36669 | 0.02550 |
| **ANKRD54** | 1.29197 | 4.14110 | 5.38067 | 2.35601 | 2.85942 | 0.00000 | 3.33649 | 2.39700 | 6.18434 | 4.07605 | 2.90742 | 0.84818 |
| **ANKRD55** | 0.02256 | 2.68902 | 0.20032 | 6.38877 | 0.00000 | 0.02269 | 0.00000 | 0.00000 | 0.04562 | 0.04090 | 0.00000 | 3.11445 |
| **ANKRD6** | 4.29274 | 0.24576 | 0.55772 | 0.00000 | 3.08328 | 1.01712 | 8.80455 | 0.26282 | 12.88626 | 0.31172 | 2.19111 | 0.11863 |
| **ANKRD61** | 0.00606 | 0.01567 | 0.05411 | 0.06329 | 0.24571 | 0.00000 | 0.04328 | 0.00668 | 0.07447 | 0.00000 | 0.09739 | 0.19639 |
| **ANKRD62** | 0.00545 | 0.07978 | 0.01329 | 0.01658 | 0.01652 | 0.10705 | 0.00000 | 0.19176 | 0.00000 | 0.09598 | 0.01210 | 0.65240 |
| **ANKRD62P1-PA** | 0.16794 | 0.00504 | 0.03053 | 0.09151 | 0.06897 | 0.00000 | 0.00707 | 0.02119 | 0.11702 | 0.50510 | 0.00000 | 0.07385 |
| **ANKRD65** | 0.00518 | 0.02293 | 0.96897 | 0.01529 | 0.00000 | 0.07656 | 0.09286 | 0.01174 | 0.04741 | 0.00654 | 0.14360 | 0.02424 |
| **ANKRD7** | 0.21416 | 0.27841 | 0.06779 | 0.06430 | 0.02019 | 0.08088 | 0.00000 | 0.44926 | 0.00000 | 0.09413 | 0.09028 | 0.10216 |
| **ANKRD9** | 0.00180 | 1.43838 | 0.66388 | 0.22198 | 1.42252 | 0.00000 | 0.08332 | 0.01275 | 0.56076 | 1.40071 | 0.00000 | 0.03786 |
| **ANKS1A** | 4.37720 | 7.41749 | 10.88565 | 2.15581 | 12.10299 | 0.45556 | 13.85591 | 0.35238 | 10.10283 | 3.82784 | 5.70661 | 2.34855 |
| **ANKS1B** | 0.74611 | 0.00365 | 0.03769 | 0.01684 | 0.02025 | 0.03006 | 0.00980 | 0.18815 | 0.03857 | 0.01642 | 0.00676 | 0.28560 |
| **ANKS3** | 0.11296 | 2.03786 | 3.59373 | 0.56965 | 1.66991 | 0.00000 | 0.00003 | 0.09590 | 4.75895 | 2.36876 | 0.98106 | 1.19488 |
| **ANKS4B** | 3.38719 | 0.00176 | 0.00783 | 1.42815 | 0.03052 | 0.03974 | 0.05342 | 0.15713 | 0.00000 | 0.00000 | 0.01850 | 0.00162 |
| **ANKS6** | 0.72365 | 1.16580 | 0.32923 | 0.07167 | 0.77083 | 0.00000 | 0.25146 | 0.36371 | 0.00000 | 2.97716 | 0.70718 | 0.61386 |
| **ANKUB1** | 0.00621 | 0.01008 | 0.01816 | 0.03709 | 0.03668 | 0.03505 | 0.01478 | 0.01020 | 0.01421 | 0.01548 | 0.01555 | 0.01508 |
| **ANKZF1** | 2.72285 | 11.13322 | 8.35178 | 2.98535 | 7.72537 | 0.00000 | 8.62737 | 4.51870 | 9.31549 | 3.70337 | 8.39524 | 16.77150 |
| **ANLN** | 5.00280 | 5.76240 | 8.88290 | 0.93355 | 6.78637 | 38.69561 | 0.00014 | 4.32963 | 21.20412 | 10.37177 | 19.66478 | 3.54565 |
| **ANO1** | 0.00096 | 0.50628 | 1.47596 | 0.00000 | 3.86656 | 0.00000 | 0.00000 | 0.00712 | 0.64328 | 0.94741 | 2.10520 | 0.00042 |
| **ANO1-AS2** | 0.00654 | 0.01704 | 0.10301 | 0.01513 | 0.02246 | 0.00650 | 0.01258 | 0.01061 | 0.01691 | 0.00590 | 0.01720 | 0.01765 |
| **ANO10** | 4.58583 | 3.61533 | 6.14684 | 4.45100 | 3.27130 | 0.22460 | 0.47324 | 2.76497 | 4.47113 | 1.80018 | 1.84770 | 4.29299 |
| **ANO2** | 0.00597 | 0.00981 | 0.01252 | 0.04990 | 0.18317 | 0.05411 | 0.00673 | 0.00990 | 0.03166 | 0.00000 | 0.04648 | 0.02024 |
| **ANO3** | 0.51611 | 0.00998 | 0.25565 | 0.04077 | 0.00479 | 0.02271 | 0.00000 | 0.03131 | 0.10275 | 0.01517 | 0.00824 | 0.05923 |
| **ANO4** | 0.35202 | 0.49363 | 0.00407 | 0.30429 | 0.00000 | 0.00503 | 0.00000 | 0.09770 | 0.02705 | 0.29278 | 0.00000 | 0.04297 |
| **ANO5** | 0.55889 | 0.10123 | 0.16010 | 0.29501 | 0.00000 | 0.18102 | 0.00000 | 0.47164 | 0.03342 | 0.76028 | 0.00000 | 0.00000 |
| **ANO6** | 16.53090 | 0.27301 | 7.58995 | 6.36563 | 5.28595 | 7.27735 | 3.89540 | 4.88675 | 32.10253 | 5.54959 | 5.06086 | 2.15341 |

| | | | | | | | | | | | | |
|---|---|---|---|---|---|---|---|---|---|---|---|---|
| **ANO7** | 0.72179 | 0.30707 | 1.03257 | 0.00000 | 0.60111 | 0.00000 | 0.00000 | 0.00000 | 4.03884 | 1.32939 | 1.86294 | 0.07876 |
| **ANO8** | 0.36992 | 0.00467 | 0.00000 | 0.03593 | 0.00000 | 0.00000 | 0.10927 | 0.00302 | 0.17437 | 0.17292 | 0.03477 | 0.04026 |
| **ANO9** | 0.00407 | 0.03534 | 0.22723 | 0.00000 | 0.00000 | 0.00000 | 0.00000 | 0.01054 | 0.66606 | 0.19439 | 0.33631 | 0.00718 |
| **ANP32A** | 0.97811 | 5.64582 | 12.31252 | 7.97571 | 10.56936 | 7.71888 | 17.69604 | 0.13817 | 4.47611 | 13.83297 | 7.28420 | 5.28761 |
| **ANP32A-IT1** | 0.00177 | 0.06367 | 0.14114 | 0.22881 | 0.08506 | 0.00001 | 0.50551 | 0.00317 | 0.22129 | 2.53771 | 0.09385 | 0.16342 |
| **ANP32AP1** | 0.13927 | 0.48505 | 1.81855 | 1.02218 | 1.80003 | 1.20242 | 1.24294 | 0.02350 | 0.96780 | 0.45333 | 0.99289 | 0.82136 |
| **ANP32B** | 2.93254 | 22.80182 | 16.43330 | 15.46214 | 21.62399 | 28.17875 | 41.85503 | 9.88214 | 18.63089 | 34.11967 | 20.82947 | 24.76167 |
| **ANP32C** | 0.01171 | 0.04356 | 0.27660 | 0.06848 | 0.17770 | 0.34324 | 0.27263 | 0.01255 | 0.00477 | 0.21876 | 0.08801 | 0.08677 |
| **ANP32D** | 0.00567 | 0.01502 | 0.01803 | 0.02289 | 0.01805 | 0.01243 | 0.05176 | 0.00951 | 0.01335 | 0.03300 | 0.02329 | 0.03241 |
| **ANP32E** | 0.00000 | 1.84192 | 29.63300 | 17.04111 | 7.24839 | 25.09516 | 44.58882 | 1.11842 | 15.92367 | 13.19474 | 24.54732 | 0.50155 |
| **ANPEP** | 0.00000 | 35.01262 | 2.60842 | 33.70072 | 46.10134 | 16.28373 | 0.00000 | 0.00000 | 23.68150 | 14.80662 | 21.80155 | 0.52409 |
| **ANTXR1** | 0.00886 | 0.06499 | 0.29251 | 0.16437 | 0.00476 | 0.05235 | 0.10760 | 0.01151 | 0.00000 | 0.00482 | 0.01194 | 0.00962 |
| **ANTXR2** | 0.08712 | 0.49115 | 2.92857 | 1.87276 | 0.40653 | 0.64615 | 0.00000 | 0.10225 | 0.00000 | 0.16134 | 0.00000 | 0.00049 |
| **ANTXRL** | 0.00545 | 0.01481 | 0.04551 | 0.00583 | 0.04800 | 0.00043 | 0.00164 | 0.01045 | 0.09098 | 0.06289 | 0.03538 | 0.01449 |
| **ANXA1** | 0.00787 | 0.00000 | 35.64129 | 0.00000 | 0.00000 | 0.09066 | 5.81244 | 0.00652 | 1.61879 | 0.00000 | 0.00000 | 0.03284 |
| **ANXA10** | 0.00802 | 0.01496 | 0.01481 | 0.01397 | 0.01274 | 0.01340 | 0.02153 | 0.08285 | 0.00946 | 0.02035 | 0.03287 | 0.01238 |
| **ANXA11** | 1.06185 | 14.16847 | 12.76748 | 5.03475 | 24.09532 | 0.00003 | 13.98387 | 0.11100 | 14.69794 | 10.06991 | 16.53069 | 2.07649 |
| **ANXA13** | 0.01969 | 0.01359 | 0.01683 | 0.01147 | 0.01408 | 0.02502 | 0.01607 | 0.01483 | 0.01788 | 0.01560 | 0.01627 | 0.01482 |
| **ANXA2** | 0.72518 | 26.09228 | 31.75782 | 29.73901 | 44.89473 | 38.68489 | 44.28159 | 1.52130 | 38.37844 | 15.56380 | 44.14829 | 5.73960 |
| **ANXA2P1** | 0.04869 | 1.31591 | 3.45151 | 2.75178 | 3.24249 | 5.14385 | 2.52446 | 0.01995 | 3.56291 | 0.00000 | 7.52920 | 0.15797 |
| **ANXA2P2** | 0.62199 | 17.13577 | 23.34891 | 22.46592 | 28.30964 | 28.95083 | 24.26090 | 0.22976 | 31.04714 | 0.00000 | 36.41624 | 2.53463 |
| **ANXA2P3** | 0.02466 | 1.27718 | 3.07298 | 2.43752 | 3.93496 | 4.50478 | 3.98483 | 0.01368 | 2.75628 | 0.47191 | 6.33034 | 0.13365 |
| **ANXA2R** | 0.01088 | 0.03659 | 0.06831 | 0.09057 | 0.01533 | 0.00000 | 0.17027 | 0.16165 | 0.00000 | 0.08027 | 0.00874 | 0.89653 |
| **ANXA3** | 0.10500 | 21.78357 | 7.61233 | 11.62420 | 12.38801 | 45.85344 | 23.06820 | 0.00000 | 24.93816 | 0.00000 | 26.75452 | 4.98666 |
| **ANXA4** | 0.10115 | 3.89930 | 1.23810 | 2.78378 | 0.52877 | 0.70463 | 0.04829 | 0.05586 | 0.00000 | 0.00000 | 0.47183 | 1.31828 |
| **ANXA5** | 4.79437 | 14.57772 | 36.40277 | 6.76033 | 18.84060 | 27.59419 | 41.63666 | 7.31572 | 31.48488 | 0.00000 | 34.27295 | 27.97479 |
| **ANXA6** | 0.18300 | 14.35978 | 6.93119 | 3.20106 | 18.31551 | 0.00000 | 33.93467 | 0.21398 | 29.83651 | 3.65995 | 36.99117 | 1.11694 |
| **ANXA7** | 8.83352 | 16.52411 | 16.06104 | 7.59266 | 7.06940 | 34.00322 | 18.69214 | 3.20865 | 18.07543 | 0.00000 | 22.42338 | 9.20102 |
| **ANXA8.uid-1418** | 0.00606 | 0.09087 | 0.11365 | 0.00000 | 0.23291 | 0.04186 | 0.18347 | 0.00979 | 0.00000 | 0.00806 | 0.77118 | 0.00711 |
| **ANXA8L1** | 0.00553 | 0.07944 | 0.00000 | 0.00000 | 0.38686 | 0.20497 | 0.55146 | 0.00750 | 0.56526 | 0.00000 | 1.24103 | 0.00159 |
| **ANXA9** | 0.01033 | 0.02097 | 2.11844 | 0.02673 | 0.04372 | 0.12263 | 0.36103 | 0.00764 | 1.54445 | 0.00000 | 0.35511 | 0.01163 |
| **AOAH** | 3.45004 | 0.00976 | 0.00000 | 0.02312 | 0.00000 | 0.13585 | 0.06619 | 0.31204 | 0.00857 | 0.00989 | 0.00580 | 0.04044 |
| **AOC1** | 0.00572 | 0.02274 | 0.02093 | 0.00919 | 0.00297 | 0.02962 | 0.00714 | 0.01031 | 0.00000 | 0.04681 | 0.07492 | 0.01290 |
| **AOC2** | 0.28714 | 0.57603 | 2.01218 | 0.90727 | 0.85102 | 0.00000 | 1.36462 | 0.40621 | 3.12981 | 3.23319 | 0.00000 | 1.22983 |
| **AOC3** | 0.00000 | 0.40506 | 0.85088 | 0.41892 | 0.00000 | 0.36325 | 1.95527 | 0.01848 | 1.82878 | 5.56533 | 0.28265 | 0.58753 |
| **AOC4P** | 0.05753 | 0.02144 | 0.02307 | 0.03892 | 0.07030 | 0.00000 | 0.02867 | 0.00649 | 0.05136 | 0.00000 | 0.01226 | 0.02433 |
| **AOX1** | 2.96495 | 2.59397 | 0.00000 | 0.70280 | 0.02481 | 0.00000 | 0.01547 | 0.01821 | 0.00386 | 0.01343 | 0.00445 | 7.93772 |
| **AOX2P** | 0.27168 | 0.01301 | 0.01728 | 0.04234 | 0.01350 | 0.00000 | 0.03980 | 0.01089 | 0.01285 | 0.02844 | 0.01660 | 0.03573 |
| **AP1AR** | 0.10623 | 3.01499 | 4.00686 | 3.52408 | 0.26773 | 10.45477 | 8.31085 | 1.29396 | 3.54546 | 1.04299 | 4.57177 | 2.89118 |
| **AP1B1** | 1.25993 | 9.21863 | 8.78532 | 12.31843 | 20.28892 | 0.00000 | 12.37345 | 0.61445 | 13.61786 | 5.05810 | 12.73715 | 6.41797 |
| **AP1B1P1** | 0.22120 | 0.01471 | 0.01652 | 0.01232 | 0.01470 | 0.01660 | 0.01657 | 0.01951 | 0.02280 | 0.01299 | 0.01520 | 0.01265 |
| **AP1G1** | 2.71579 | 8.96609 | 10.34893 | 4.56800 | 2.46143 | 23.63442 | 14.42288 | 13.30953 | 5.52692 | 11.93736 | 10.14757 | 11.65532 |
| **AP1G2** | 0.10048 | 0.39353 | 4.97371 | 6.55539 | 2.66475 | 0.00000 | 3.35277 | 0.03290 | 8.41483 | 0.69418 | 4.06178 | 0.30384 |
| **AP1M1** | 9.76327 | 5.18585 | 8.17322 | 4.61155 | 3.90305 | 0.32379 | 10.36488 | 2.50394 | 5.85406 | 0.00000 | 6.22846 | 8.61979 |
| **AP1M2** | 7.50278 | 12.19896 | 8.72875 | 18.17797 | 16.73094 | 4.53090 | 17.73889 | 3.45371 | 15.48968 | 0.00000 | 15.21505 | 10.13009 |
| **AP1S1** | 1.19128 | 7.89565 | 15.08641 | 9.64471 | 4.48178 | 4.14851 | 12.44275 | 0.44521 | 9.22605 | 0.00000 | 7.31768 | 1.92040 |
| **AP1S2** | 0.07032 | 0.52827 | 0.51861 | 1.92874 | 0.00000 | 0.57236 | 1.60838 | 1.49796 | 0.91752 | 0.00000 | 0.12960 | 1.06369 |
| **AP1S3** | 0.14758 | 1.31317 | 0.67871 | 0.38852 | 0.64698 | 2.11323 | 0.94188 | 0.05889 | 0.00000 | 0.33213 | 1.19381 | 4.16409 |

| | | | | | | | | | | | | |
|---|---|---|---|---|---|---|---|---|---|---|---|---|
| **AP2A1** | 1.97392 | 4.09575 | 8.74693 | 3.45965 | 19.77995 | 0.00000 | 1.98397 | 0.00000 | 12.65255 | 4.14539 | 6.11496 | 2.75206 |
| **AP2A2** | 2.04657 | 0.27081 | 1.77021 | 0.97687 | 4.39953 | 0.00000 | 2.14026 | 0.49785 | 2.55660 | 11.24520 | 0.58595 | 0.95117 |
| **AP2B1** | 7.48464 | 13.56313 | 12.61599 | 14.02592 | 10.87481 | 20.75846 | 13.04321 | 2.65111 | 21.68017 | 5.06158 | 15.36195 | 20.56860 |
| **AP2M1** | 5.54624 | 32.91270 | 28.39410 | 36.04983 | 50.55303 | 21.69214 | 33.55739 | 6.79215 | 36.79602 | 1.39667 | 33.59715 | 29.36746 |
| **AP2S1** | 2.27338 | 26.19455 | 21.23313 | 22.14713 | 38.39863 | 15.55706 | 37.22775 | 8.22363 | 30.14210 | 14.43118 | 22.26517 | 32.40596 |
| **AP3B1** | 1.98195 | 3.74069 | 7.28565 | 4.34928 | 5.63659 | 21.78669 | 12.75086 | 2.01784 | 10.28006 | 10.72198 | 8.06577 | 7.02719 |
| **AP3B2** | 2.81449 | 0.40670 | 0.64675 | 0.00365 | 0.00000 | 0.00000 | 0.35356 | 0.14065 | 0.00000 | 0.03767 | 0.05626 | 0.24305 |
| **AP3D1** | 1.75082 | 5.70660 | 12.86961 | 9.10248 | 26.31063 | 0.00000 | 26.43092 | 0.12530 | 5.71649 | 41.20549 | 8.99507 | 5.42503 |
| **AP3M1** | 0.00028 | 6.52721 | 6.66365 | 1.08731 | 0.66191 | 15.47414 | 4.21978 | 2.04889 | 11.04769 | 0.00000 | 5.43211 | 5.60251 |
| **AP3M2** | 1.72757 | 2.16415 | 1.74450 | 0.88763 | 0.39613 | 3.03214 | 1.85724 | 5.51148 | 7.06855 | 2.27134 | 2.01322 | 5.44419 |
| **AP3S1** | 1.75547 | 9.91157 | 5.68717 | 3.78600 | 1.09019 | 28.72057 | 8.89482 | 5.65283 | 1.74038 | 9.86188 | 8.79321 | 10.69307 |
| **AP3S2** | 0.07666 | 0.84916 | 0.29831 | 0.24426 | 0.33454 | 0.63724 | 0.00000 | 0.33218 | 0.70802 | 0.00000 | 0.30676 | 0.64935 |
| **AP4B1** | 0.02892 | 2.05391 | 1.81214 | 0.32310 | 1.09960 | 2.30008 | 2.03386 | 6.52073 | 2.30925 | 1.54677 | 2.93848 | 4.25873 |
| **AP4B1-AS1** | 0.00330 | 0.04697 | 0.06963 | 0.00853 | 0.02807 | 0.03294 | 0.00000 | 0.12471 | 0.11840 | 0.01112 | 0.02752 | 0.08880 |
| **AP4E1** | 0.00000 | 1.30500 | 2.61388 | 0.00000 | 3.66608 | 4.39025 | 4.74283 | 2.45766 | 0.22869 | 10.00907 | 5.21084 | 2.46451 |
| **AP4M1** | 0.05191 | 8.29943 | 4.63916 | 3.01017 | 2.78166 | 1.79770 | 2.48748 | 8.19096 | 3.40122 | 0.00000 | 4.56848 | 7.96317 |
| **AP4S1** | 0.03753 | 0.10753 | 0.84832 | 0.75302 | 0.00000 | 1.94949 | 1.54657 | 0.00976 | 0.47077 | 2.18574 | 0.40554 | 0.16828 |
| **AP5B1** | 0.01294 | 1.27454 | 6.99675 | 2.88087 | 3.23378 | 0.00000 | 2.07111 | 0.21078 | 0.59393 | 4.22615 | 0.31905 | 0.72663 |
| **AP5M1** | 0.75101 | 2.97274 | 3.20287 | 5.94664 | 2.80590 | 25.30063 | 6.68884 | 5.36584 | 4.33710 | 0.00000 | 6.67466 | 8.19942 |
| **AP5S1** | 0.85330 | 3.26963 | 2.67322 | 2.09670 | 3.73928 | 0.06069 | 2.95219 | 1.11717 | 2.15755 | 0.00000 | 2.27428 | 7.85825 |
| **AP5Z1** | 0.51310 | 3.15903 | 5.54177 | 3.46613 | 3.35368 | 0.00000 | 4.41143 | 0.81286 | 7.61422 | 3.25143 | 2.00792 | 2.57439 |
| **APAF1** | 1.16783 | 0.85483 | 0.45787 | 1.80870 | 0.27408 | 0.00000 | 0.62190 | 1.00607 | 0.60448 | 0.44867 | 0.09611 | 1.09469 |
| **APBA1** | 4.67408 | 0.32291 | 0.01679 | 0.00000 | 0.00000 | 0.00000 | 0.00000 | 0.34363 | 0.07738 | 0.09692 | 0.13517 | 0.26270 |
| **APBA2** | 1.26136 | 0.58875 | 0.00311 | 0.14289 | 0.01184 | 0.00000 | 0.05174 | 0.00616 | 0.01061 | 0.00000 | 0.04268 | 0.30268 |
| **APBA3** | 0.00242 | 8.59729 | 3.29844 | 0.51190 | 1.52947 | 0.00000 | 0.12265 | 0.00000 | 4.71590 | 1.17842 | 2.94745 | 3.54111 |
| **APBB1** | 0.39169 | 1.34667 | 1.91103 | 0.57455 | 3.71386 | 0.00000 | 2.65395 | 0.03865 | 2.27820 | 0.00000 | 2.38186 | 0.91988 |
| **APBB1IP** | 0.00176 | 0.00000 | 0.80467 | 0.00000 | 0.83325 | 0.00000 | 0.44317 | 0.02281 | 0.36703 | 0.22773 | 0.29124 | 0.14016 |
| **APBB2** | 5.02791 | 5.05520 | 4.05648 | 5.58302 | 1.31215 | 5.44057 | 4.70975 | 6.06386 | 6.73173 | 0.64138 | 4.61663 | 11.61920 |
| **APBB3** | 0.29782 | 0.43880 | 1.07246 | 1.16774 | 0.67130 | 0.00000 | 1.09914 | 0.04102 | 2.28394 | 0.66530 | 1.54461 | 0.96834 |
| **APC** | 7.15404 | 0.42326 | 3.52368 | 1.12022 | 0.00000 | 7.58343 | 1.23171 | 2.25938 | 4.91893 | 12.57044 | 2.06236 | 4.75244 |
| **APC2** | 0.90087 | 0.20887 | 0.12614 | 0.27147 | 0.10263 | 0.00000 | 0.06013 | 0.12137 | 0.16838 | 0.30436 | 0.00005 | 0.12401 |
| **APCDD1** | 0.00782 | 0.01810 | 0.10641 | 0.00523 | 0.01018 | 0.03241 | 0.00000 | 0.00684 | 0.18950 | 0.00000 | 0.08746 | 0.10494 |
| **APCDD1L** | 0.00636 | 0.01914 | 0.03195 | 0.01459 | 0.01466 | 0.00800 | 0.01323 | 0.01058 | 0.04110 | 0.01826 | 0.01194 | 0.01629 |
| **APCDD1L-AS1** | 0.54809 | 0.02799 | 0.04368 | 0.01623 | 0.04846 | 0.00000 | 0.01092 | 0.02167 | 0.04021 | 0.00000 | 0.01160 | 0.02196 |
| **APEH** | 0.01534 | 17.75027 | 16.52219 | 19.32336 | 22.85896 | 1.56898 | 10.38742 | 0.62273 | 11.36982 | 0.00000 | 24.09492 | 10.20913 |
| **APELA** | 0.00524 | 0.00981 | 0.07780 | 6.83988 | 13.33616 | 0.61704 | 0.00000 | 0.00258 | 0.00000 | 0.00000 | 3.23959 | 0.00472 |
| **APEX1** | 3.47615 | 18.74461 | 19.44546 | 23.78094 | 13.32581 | 41.13156 | 29.18707 | 13.81224 | 31.00081 | 0.00046 | 23.89355 | 34.42348 |
| **APEX2** | 3.09380 | 9.03240 | 6.90238 | 6.93696 | 7.65795 | 4.38819 | 5.14465 | 6.81893 | 13.18570 | 0.00000 | 5.10125 | 12.89452 |
| **APH1A** | 8.21353 | 7.29782 | 8.74694 | 6.50682 | 4.19983 | 4.16375 | 4.02630 | 0.41305 | 13.83920 | 0.00000 | 13.38354 | 9.57369 |
| **APH1B** | 0.14630 | 0.23513 | 0.29525 | 0.13940 | 0.17088 | 0.09405 | 0.00000 | 0.10231 | 0.30210 | 0.00000 | 1.01701 | 0.40728 |
| **API5** | 3.73683 | 8.09193 | 12.24261 | 4.14037 | 0.00000 | 25.51202 | 14.78368 | 13.30300 | 14.63042 | 0.00000 | 12.17071 | 11.16239 |
| **APIP** | 0.17629 | 1.68810 | 1.54895 | 3.22481 | 1.26522 | 7.72294 | 2.59370 | 0.00000 | 2.43962 | 0.00000 | 3.61777 | 1.41784 |
| **APITD1** | 1.70654 | 2.81751 | 0.00000 | 1.27363 | 0.24338 | 4.04514 | 0.47058 | 1.00744 | 0.21710 | 0.63991 | 0.57222 | 1.63052 |
| **APLF** | 0.02791 | 0.32607 | 0.32209 | 0.19675 | 0.27780 | 0.00000 | 0.00000 | 0.35062 | 0.00000 | 0.41951 | 0.52155 | 0.47931 |
| **APLN** | 0.14101 | 0.01571 | 0.03617 | 0.01498 | 0.00000 | 0.04774 | 0.00379 | 0.00253 | 0.00030 | 0.06464 | 0.01422 | 0.00920 |
| **APLNR** | 0.00411 | 0.00774 | 0.00525 | 0.61757 | 0.00000 | 0.00515 | 0.04856 | 0.00963 | 0.00000 | 0.15337 | 0.02562 | 0.01199 |
| **APLP1** | 0.02901 | 0.09373 | 0.04195 | 0.42155 | 0.02288 | 0.07933 | 0.00000 | 0.12147 | 0.00000 | 0.43009 | 0.00000 | 0.09468 |
| **APLP2** | 5.57307 | 31.49473 | 28.29977 | 30.43516 | 41.44133 | 48.53202 | 10.12796 | 14.86045 | 57.23557 | 41.86658 | 28.54435 | 25.94607 |

| | | | | | | | | | | | | |
|---|---|---|---|---|---|---|---|---|---|---|---|---|
| **APMAP** | 1.09715 | 14.11492 | 14.23602 | 3.79794 | 5.25565 | 15.26709 | 0.00000 | 3.13780 | 25.46917 | 0.00000 | 8.41820 | 9.00579 |
| **APOA1** | 0.00095 | 0.05657 | 26.98662 | 21.99603 | 0.85563 | 0.00000 | 0.00000 | 0.04812 | 7.57919 | 0.00000 | 0.14915 | 0.16352 |
| **APOA1BP** | 0.25337 | 9.88606 | 11.03569 | 6.37327 | 13.52590 | 2.82832 | 16.85169 | 0.59316 | 19.07723 | 0.00922 | 16.96436 | 3.29404 |
| **APOA2** | 0.03557 | 0.00000 | 3.31705 | 9.21760 | 0.00000 | 0.16780 | 1.89182 | 0.82533 | 0.00000 | 0.00000 | 0.05998 | 0.01938 |
| **APOA4** | 0.01015 | 0.20230 | 0.00000 | 0.44041 | 0.93463 | 0.00000 | 0.09905 | 0.01157 | 0.27873 | 0.00000 | 0.36051 | 0.03061 |
| **APOA5** | 0.00653 | 0.09449 | 0.34088 | 0.07332 | 0.01154 | 0.00000 | 0.00000 | 0.00665 | 0.02549 | 0.00822 | 0.07436 | 0.13246 |
| **APOB** | 0.08133 | 0.02517 | 0.07170 | 0.23646 | 0.03076 | 0.00000 | 0.09044 | 0.00588 | 0.00000 | 0.04782 | 0.01394 | 0.01207 |
| **APOBEC1** | 0.00728 | 0.01739 | 0.01407 | 0.00893 | 0.01716 | 0.02457 | 0.01804 | 0.03151 | 0.00583 | 0.01953 | 0.03824 | 0.02751 |
| **APOBEC2** | 0.18632 | 0.30146 | 2.58525 | 0.22722 | 0.00000 | 0.00000 | 0.00000 | 0.00772 | 1.29558 | 0.64423 | 0.88022 | 0.66267 |
| **APOBEC3A** | 0.00796 | 0.10074 | 0.00000 | 0.00000 | 0.00000 | 0.03666 | 0.03498 | 0.00920 | 0.00000 | 0.19938 | 0.00257 | 0.00193 |
| **APOBEC3B** | 0.00801 | 0.01780 | 0.08034 | 0.21010 | 0.00000 | 0.03166 | 0.00540 | 0.01008 | 0.01846 | 0.01655 | 0.05838 | 0.01734 |
| **APOBEC3B-AS1** | 0.01221 | 0.00361 | 0.07435 | 0.11100 | 0.00888 | 0.15317 | 0.00000 | 0.00925 | 0.01779 | 0.01384 | 0.00766 | 0.06200 |
| **APOBEC3C** | 0.00557 | 0.00772 | 0.33586 | 4.34853 | 0.03883 | 0.00210 | 0.01961 | 0.00905 | 0.00000 | 0.00151 | 0.02425 | 0.01535 |
| **APOBEC3D** | 0.02690 | 0.01408 | 0.16742 | 2.62337 | 0.09786 | 0.00190 | 0.09983 | 0.00799 | 0.00000 | 0.00000 | 0.03489 | 0.01501 |
| **APOBEC3F** | 0.00553 | 0.32349 | 0.27065 | 3.66717 | 0.00000 | 0.03002 | 0.00000 | 0.20088 | 0.05459 | 0.03657 | 0.00258 | 1.99720 |
| **APOBEC3G** | 0.12946 | 0.00330 | 0.38291 | 2.47372 | 0.01067 | 0.00000 | 0.02287 | 0.04444 | 0.00000 | 0.08095 | 0.01723 | 0.02956 |
| **APOBEC3H** | 0.00595 | 0.01211 | 0.01224 | 0.01889 | 0.01793 | 0.02149 | 0.01861 | 0.00945 | 0.01531 | 0.02095 | 0.01570 | 0.03190 |
| **APOBEC4** | 0.00629 | 0.01110 | 0.03382 | 0.00871 | 0.00582 | 0.03978 | 0.02307 | 0.01032 | 0.05347 | 0.00689 | 0.03354 | 0.01530 |
| **APOBR** | 0.00133 | 0.37437 | 0.00281 | 0.39415 | 0.00000 | 0.10380 | 0.00000 | 0.21057 | 0.00000 | 0.04374 | 0.00006 | 2.64910 |
| **APOC1** | 0.54283 | 31.69843 | 14.79746 | 43.79998 | 39.76755 | 26.89098 | 7.44487 | 0.49666 | 19.56979 | 22.53088 | 17.56475 | 25.00218 |
| **APOC1P1** | 0.00012 | 2.78816 | 0.44097 | 6.16292 | 4.82933 | 0.21879 | 0.58527 | 0.17787 | 0.00000 | 0.12054 | 0.47342 | 6.07773 |
| **APOC3** | 0.00589 | 0.04101 | 2.64551 | 0.00013 | 1.39929 | 0.18382 | 0.00000 | 0.00371 | 2.21993 | 0.00000 | 0.00077 | 0.17829 |
| **APOC4-APOC2** | 0.00639 | 0.02147 | 0.02512 | 0.02393 | 0.01413 | 0.00861 | 0.01357 | 0.01755 | 0.02004 | 0.06923 | 0.01206 | 0.03378 |
| **APOD** | 0.72838 | 0.00000 | 0.20653 | 0.29857 | 0.04554 | 0.04415 | 0.04605 | 0.24015 | 0.77683 | 0.01545 | 3.01708 | 0.00000 |
| **APOE** | 0.00000 | 0.98609 | 4.05605 | 20.79597 | 9.02868 | 0.00000 | 0.02447 | 0.04044 | 8.43516 | 7.52022 | 0.34750 | 0.00000 |
| **APOF** | 0.02212 | 0.03927 | 0.05139 | 0.00652 | 0.01058 | 0.03066 | 0.00895 | 0.03984 | 0.01627 | 0.01658 | 0.02811 | 0.04268 |
| **APOH** | 0.00363 | 0.42596 | 0.02074 | 0.00954 | 0.00000 | 0.00866 | 0.03056 | 0.02466 | 0.02228 | 0.01557 | 0.03183 | 0.19834 |
| **APOL1** | 0.00477 | 0.00454 | 0.00000 | 0.26966 | 0.04458 | 0.04064 | 0.00255 | 0.00978 | 0.06140 | 0.07517 | 0.00784 | 0.01101 |
| **APOL2** | 0.00508 | 0.00000 | 0.32401 | 0.33558 | 0.00000 | 0.02167 | 0.74311 | 0.01120 | 0.65023 | 0.00000 | 0.40531 | 0.01343 |
| **APOL3** | 0.05003 | 0.01455 | 0.01681 | 0.01710 | 0.01329 | 0.01273 | 0.01559 | 0.01237 | 0.01825 | 0.02865 | 0.01651 | 0.01442 |
| **APOL4** | 0.06509 | 0.00203 | 0.24043 | 0.11931 | 0.00000 | 0.13194 | 0.00000 | 0.00562 | 0.93702 | 0.00000 | 0.00000 | 0.01040 |
| **APOL5** | 0.00879 | 0.00452 | 0.01553 | 0.01196 | 0.07787 | 0.01529 | 0.04736 | 0.04492 | 0.00000 | 0.00000 | 0.01617 | 0.26537 |
| **APOL6** | 0.00714 | 0.00719 | 0.03579 | 0.02301 | 0.03226 | 0.00000 | 0.02142 | 0.02686 | 0.00000 | 0.07601 | 0.09677 | 0.09772 |
| **APOLD1** | 0.00832 | 0.00000 | 0.07194 | 0.05408 | 0.16833 | 0.03888 | 0.03031 | 0.00687 | 0.05389 | 0.07214 | 0.15119 | 0.09565 |
| **APOM** | 0.00318 | 0.06374 | 0.16327 | 0.76235 | 0.02474 | 0.07831 | 0.00003 | 0.01988 | 0.00004 | 0.02077 | 0.11211 | 0.04957 |
| **APOO** | 2.80108 | 4.86515 | 0.18787 | 3.92862 | 0.77432 | 6.42691 | 1.11284 | 1.26345 | 1.77627 | 0.44680 | 3.31099 | 5.09887 |
| **APOOL** | 0.00000 | 1.36832 | 1.66738 | 1.74598 | 0.00000 | 12.06516 | 8.05614 | 0.05329 | 0.03169 | 7.49658 | 11.11863 | 1.21729 |
| **APOOP5** | 0.18029 | 0.35524 | 0.09197 | 0.33539 | 0.09718 | 0.56156 | 0.13935 | 0.21466 | 0.11834 | 0.00002 | 0.32044 | 0.34299 |
| **APOPT1** | 1.18332 | 4.89724 | 5.50419 | 5.51532 | 6.95317 | 7.61331 | 11.26772 | 2.88596 | 10.51016 | 2.86321 | 7.50811 | 7.25403 |
| **APP** | 0.86671 | 3.24815 | 6.30465 | 0.00000 | 5.92126 | 0.00000 | 0.00000 | 1.52780 | 4.26683 | 11.02461 | 16.15113 | 0.76562 |
| **APPBP2** | 1.33216 | 1.76862 | 3.39269 | 1.30822 | 0.27304 | 3.63781 | 0.00000 | 1.84412 | 1.92770 | 1.47815 | 1.09401 | 1.40348 |
| **APPL1** | 1.32397 | 4.12881 | 4.46828 | 4.95792 | 4.21371 | 15.35555 | 7.58936 | 0.30087 | 0.90606 | 17.48787 | 5.66922 | 0.38984 |
| **APPL2** | 2.70985 | 1.37131 | 1.99510 | 0.55897 | 0.25616 | 2.23863 | 1.03316 | 0.15120 | 0.55280 | 0.00000 | 1.33846 | 0.57065 |
| **APRT** | 0.00000 | 17.87180 | 20.38848 | 19.21828 | 26.21390 | 1.34805 | 31.39030 | 0.00425 | 27.76678 | 1.93660 | 29.47698 | 6.38256 |
| **APTR** | 0.10911 | 2.03635 | 7.00629 | 1.74062 | 2.98903 | 1.20409 | 4.09206 | 9.52748 | 1.86301 | 1.31417 | 2.38597 | 3.34592 |
| **APTX** | 1.22457 | 16.98240 | 14.14986 | 6.05924 | 9.80807 | 22.05303 | 13.73341 | 11.10005 | 10.81395 | 0.00000 | 10.96228 | 20.01364 |
| **AQP1** | 0.00336 | 0.00570 | 0.03629 | 0.03157 | 0.01887 | 0.07082 | 0.00002 | 0.04357 | 0.06698 | 0.02301 | 0.00587 | 0.02770 |
| **AQP10** | 0.00659 | 0.01483 | 0.03380 | 0.01869 | 0.01132 | 0.01912 | 0.01847 | 0.01205 | 0.01646 | 0.00651 | 0.03455 | 0.01698 |

| | | | | | | | | | | | | |
|---|---|---|---|---|---|---|---|---|---|---|---|---|
| **AQP11** | 0.00232 | 0.66667 | 1.21214 | 0.73553 | 0.00000 | 1.39034 | 0.00000 | 0.77232 | 0.81990 | 0.47955 | 0.33574 | 1.02985 |
| **AQP2** | 0.00610 | 0.01546 | 0.01562 | 0.01642 | 0.01422 | 0.01591 | 0.01834 | 0.01338 | 0.01731 | 0.01617 | 0.02191 | 0.01666 |
| **AQP3** | 0.00000 | 62.47159 | 10.53940 | 6.06828 | 23.81202 | 12.29011 | 0.00000 | 1.09302 | 32.62429 | 13.00727 | 11.54481 | 15.01295 |
| **AQP4** | 0.00573 | 0.01165 | 0.03966 | 0.13311 | 0.00758 | 0.02906 | 0.03011 | 0.01011 | 0.00439 | 0.02411 | 0.01663 | 0.01764 |
| **AQP4-AS1** | 0.00968 | 0.02887 | 0.03160 | 0.15783 | 0.00624 | 0.10962 | 0.00000 | 0.01628 | 0.00000 | 0.05521 | 0.01139 | 0.02995 |
| **AQP5** | 0.02548 | 0.01524 | 0.01796 | 0.01152 | 0.02556 | 0.00737 | 0.05131 | 0.01258 | 0.00228 | 0.02691 | 0.01549 | 0.01724 |
| **AQP6** | 0.30688 | 0.11195 | 0.04181 | 0.25749 | 0.35700 | 0.57553 | 0.44418 | 1.30502 | 0.05750 | 0.74831 | 0.02865 | 0.19100 |
| **AQP7** | 0.02115 | 0.36455 | 0.02581 | 0.00000 | 0.07171 | 0.00000 | 0.03833 | 0.03775 | 0.00000 | 0.03547 | 0.10661 | 0.12492 |
| **AQP7P1** | 0.00601 | 0.01456 | 0.01372 | 0.01008 | 0.01358 | 0.02694 | 0.01643 | 0.01016 | 0.02058 | 0.01562 | 0.01488 | 0.02168 |
| **AQP8** | 0.06703 | 0.00445 | 0.00888 | 0.05005 | 0.00917 | 0.00000 | 0.00047 | 0.00320 | 0.03602 | 0.29041 | 0.02938 | 0.04267 |
| **AQP9** | 0.03345 | 0.00000 | 0.00451 | 0.00183 | 0.00273 | 0.12773 | 0.00966 | 0.00371 | 0.00000 | 0.03151 | 0.04338 | 0.04287 |
| **AQPEP** | 0.00000 | 0.25531 | 0.91144 | 0.61375 | 0.98192 | 1.50328 | 0.00000 | 0.00135 | 0.00000 | 1.15739 | 0.38819 | 0.00000 |
| **AQR** | 2.42846 | 12.11392 | 9.11134 | 5.55785 | 1.12545 | 22.77938 | 6.29782 | 7.40058 | 10.50474 | 4.55542 | 10.73511 | 12.03294 |
| **AR** | 0.00367 | 0.14672 | 0.00893 | 0.01440 | 0.21649 | 0.00000 | 0.02728 | 0.07494 | 0.00000 | 0.15531 | 0.00000 | 0.00060 |
| **ARAF** | 0.83767 | 6.22946 | 5.41718 | 5.31767 | 7.17822 | 0.00000 | 7.61133 | 0.39392 | 6.32219 | 0.60760 | 8.99831 | 3.45806 |
| **ARAP1** | 0.45219 | 8.41754 | 1.78405 | 1.74327 | 10.68291 | 0.00000 | 1.94470 | 0.00000 | 4.64047 | 10.16378 | 5.63784 | 2.57621 |
| **ARAP2** | 5.36783 | 0.02049 | 0.18812 | 0.03115 | 0.01114 | 0.52050 | 0.08530 | 0.56143 | 0.00000 | 0.13429 | 0.00000 | 0.23522 |
| **ARAP3** | 0.02511 | 0.01310 | 0.09575 | 0.07667 | 0.00304 | 0.00121 | 0.00409 | 0.03297 | 0.01097 | 0.05803 | 0.03795 | 0.03823 |
| **ARC** | 0.82357 | 0.01405 | 0.04531 | 0.00928 | 0.01694 | 0.01264 | 0.03084 | 0.00181 | 0.01514 | 0.00398 | 0.01314 | 0.01058 |
| **ARCN1** | 6.41867 | 16.00111 | 10.84039 | 9.76334 | 6.84643 | 25.93081 | 16.47573 | 9.52502 | 15.06797 | 3.04326 | 16.95216 | 16.80801 |
| **AREL1** | 2.53900 | 2.84195 | 3.88076 | 2.99885 | 3.53679 | 6.81632 | 1.67897 | 5.24250 | 7.78670 | 4.11593 | 1.78519 | 4.64133 |
| **ARF1** | 15.33890 | 40.34440 | 38.15303 | 33.72722 | 42.78860 | 30.62442 | 49.29678 | 23.76497 | 47.64813 | 3.42152 | 33.15421 | 34.95522 |
| **ARF3** | 9.04574 | 7.43495 | 12.10195 | 12.18362 | 15.84063 | 2.63856 | 14.54587 | 11.34951 | 5.06915 | 19.00275 | 8.11610 | 7.07191 |
| **ARF4** | 8.54861 | 14.09345 | 25.08338 | 11.27133 | 15.17275 | 46.03785 | 28.63214 | 10.08466 | 29.52164 | 4.99586 | 29.09631 | 18.11385 |
| **ARF5** | 6.19842 | 22.96711 | 14.62148 | 16.21895 | 23.60756 | 1.08026 | 20.85683 | 2.30300 | 23.41478 | 2.21420 | 16.34654 | 12.71409 |
| **ARF6** | 1.36078 | 3.89836 | 10.79728 | 6.66067 | 8.76007 | 28.49634 | 17.69898 | 16.60452 | 4.10852 | 21.40656 | 13.25410 | 9.19042 |
| **ARFGAP1** | 0.98642 | 11.14909 | 9.68557 | 7.32613 | 6.59923 | 0.00000 | 9.85517 | 0.44581 | 15.14471 | 16.32437 | 5.38102 | 9.42750 |
| **ARFGAP2** | 3.61359 | 18.96029 | 13.39006 | 12.18418 | 14.44777 | 0.00709 | 12.70853 | 6.64691 | 13.48535 | 0.00000 | 12.94748 | 16.02006 |
| **ARFGAP3** | 2.87449 | 4.79258 | 0.69395 | 6.00938 | 0.49778 | 6.72074 | 0.00000 | 4.48150 | 1.08932 | 1.82985 | 0.08103 | 4.88284 |
| **ARFGEF1** | 3.08837 | 9.40008 | 1.30220 | 11.01326 | 7.37743 | 27.61057 | 0.00000 | 1.91753 | 0.00000 | 22.83340 | 0.80945 | 6.20170 |
| **ARFGEF2** | 6.14709 | 6.23303 | 4.43061 | 9.47350 | 2.68199 | 18.16273 | 2.86998 | 2.35115 | 2.67609 | 14.69812 | 4.35615 | 3.07571 |
| **ARFIP1** | 3.10641 | 6.71437 | 5.09693 | 6.64183 | 4.90235 | 18.75533 | 5.02417 | 4.50396 | 3.47165 | 1.32736 | 14.98258 | 5.88620 |
| **ARFIP2** | 0.55526 | 9.24471 | 14.71977 | 10.14672 | 13.59047 | 0.30977 | 13.81310 | 3.62185 | 28.25898 | 0.00000 | 19.14836 | 6.88478 |
| **ARFRP1** | 0.85832 | 9.65866 | 3.80410 | 5.70884 | 5.98799 | 0.00192 | 4.27201 | 1.07587 | 6.25577 | 1.84743 | 9.09589 | 4.65345 |
| **ARG1** | 0.00625 | 0.00010 | 0.02960 | 0.06332 | 0.00000 | 0.06891 | 0.06365 | 0.01622 | 0.04182 | 0.03036 | 0.13895 | 0.07975 |
| **ARG2** | 9.17736 | 1.28968 | 7.06814 | 1.12527 | 0.00000 | 6.37157 | 5.12049 | 1.60780 | 1.30519 | 0.00000 | 1.02444 | 5.38673 |
| **ARGFX** | 0.00875 | 12.76706 | 0.23939 | 12.65470 | 0.00000 | 0.00000 | 0.00000 | 25.07568 | 0.00000 | 0.23308 | 0.00000 | 41.93174 |
| **ARGFXP2** | 0.00158 | 0.41525 | 0.00754 | 0.56597 | 0.10670 | 0.09937 | 0.00003 | 2.01041 | 0.01078 | 0.02312 | 0.09820 | 3.65991 |
| **ARGLU1** | 0.00000 | 7.74858 | 13.27262 | 6.76401 | 5.59848 | 21.98271 | 16.33932 | 18.11801 | 10.35587 | 26.64086 | 12.24121 | 14.35297 |
| **ARHGAP1** | 2.80119 | 14.72989 | 8.00088 | 10.59016 | 12.64443 | 0.00000 | 9.70567 | 6.86960 | 4.39733 | 6.70158 | 13.33276 | 11.65244 |
| **ARHGAP10** | 11.72519 | 0.00000 | 6.12200 | 0.84329 | 8.43712 | 0.00000 | 11.36925 | 2.60603 | 9.48132 | 7.33346 | 3.89538 | 1.09976 |
| **ARHGAP11A** | 4.17517 | 5.73612 | 3.68837 | 0.00000 | 0.00000 | 20.34551 | 8.03780 | 5.89793 | 6.35977 | 15.62405 | 3.72566 | 7.59137 |
| **ARHGAP11B** | 0.00123 | 0.12007 | 0.72644 | 0.10624 | 0.38831 | 0.00000 | 0.00000 | 0.17592 | 0.94095 | 1.91450 | 0.29145 | 0.57386 |
| **ARHGAP12** | 5.94037 | 2.44853 | 2.51167 | 6.39239 | 2.27523 | 15.81556 | 3.66988 | 0.95705 | 2.29327 | 7.07660 | 5.89485 | 2.10973 |
| **ARHGAP15** | 2.04585 | 0.02043 | 0.03312 | 0.00756 | 0.03066 | 0.02986 | 0.00919 | 0.23179 | 0.19628 | 0.00000 | 0.06004 | 0.00565 |
| **ARHGAP17** | 6.15549 | 3.34254 | 6.44848 | 1.79555 | 3.04336 | 0.00000 | 9.64508 | 2.09256 | 6.73594 | 7.52691 | 2.49714 | 1.42904 |
| **ARHGAP18** | 16.68644 | 8.46301 | 11.68566 | 1.68676 | 17.52948 | 24.48349 | 23.08631 | 7.80313 | 16.20657 | 15.06466 | 25.11305 | 0.00000 |
| **ARHGAP19** | 2.35867 | 3.68316 | 3.16730 | 2.66546 | 2.32448 | 6.79628 | 3.35393 | 0.49734 | 2.45875 | 2.55560 | 1.48452 | 2.90331 |

| | | | | | | | | | | | | |
|---|---|---|---|---|---|---|---|---|---|---|---|---|
| **ARHGAP20** | 0.45907 | 0.04538 | 0.08305 | 0.00727 | 0.00587 | 0.00000 | 0.00160 | 0.30860 | 0.03590 | 0.09543 | 0.01080 | 0.06579 |
| **ARHGAP21** | 1.30463 | 0.85389 | 1.70966 | 2.11845 | 4.98823 | 8.98048 | 1.93377 | 1.39132 | 0.00000 | 14.27186 | 1.33451 | 4.74177 |
| **ARHGAP22** | 0.07523 | 1.21629 | 0.00910 | 0.00000 | 0.00000 | 0.00000 | 0.05602 | 0.01318 | 0.00000 | 0.97421 | 0.02791 | 0.00339 |
| **ARHGAP23** | 0.00113 | 0.06244 | 0.67324 | 0.00000 | 6.67751 | 0.00000 | 4.57221 | 0.04845 | 0.90139 | 1.61413 | 1.87424 | 0.00761 |
| **ARHGAP24** | 0.51942 | 0.00000 | 8.00964 | 0.01253 | 0.86821 | 0.79832 | 2.11527 | 0.17196 | 2.90328 | 0.00000 | 1.11706 | 0.04375 |
| **ARHGAP25** | 0.00722 | 0.12622 | 0.02289 | 0.08232 | 0.01027 | 0.00000 | 0.00339 | 0.01063 | 0.19636 | 0.00323 | 0.01656 | 0.01151 |
| **ARHGAP26** | 6.33548 | 0.00000 | 0.14758 | 0.25466 | 0.95812 | 1.91402 | 0.54253 | 0.18434 | 0.00000 | 1.33917 | 0.07540 | 0.31200 |
| **ARHGAP26-AS1** | 0.60228 | 0.01334 | 0.04146 | 0.00060 | 0.07789 | 0.03395 | 0.00011 | 0.27593 | 0.02840 | 0.03135 | 0.00439 | 0.12561 |
| **ARHGAP27** | 1.05044 | 1.72939 | 1.75510 | 0.00000 | 0.90255 | 1.21616 | 2.08487 | 0.00000 | 2.73034 | 0.08934 | 0.69808 | 1.61496 |
| **ARHGAP28** | 0.03921 | 0.29608 | 1.19540 | 0.00000 | 0.00000 | 1.48207 | 3.00903 | 0.00330 | 0.00000 | 0.03961 | 0.18556 | 0.05546 |
| **ARHGAP29** | 3.07031 | 0.00000 | 3.16945 | 0.00000 | 3.23001 | 7.31167 | 11.81132 | 1.89256 | 6.63936 | 15.81749 | 7.89495 | 0.95133 |
| **ARHGAP30** | 0.20828 | 0.06231 | 0.65517 | 0.21565 | 0.27518 | 0.00000 | 0.00000 | 0.17993 | 0.17670 | 2.34515 | 0.00000 | 0.16746 |
| **ARHGAP31** | 0.00453 | 0.14364 | 0.37638 | 0.22722 | 0.04675 | 0.00000 | 0.20395 | 0.02872 | 0.28318 | 0.18773 | 0.08282 | 0.23800 |
| **ARHGAP31-AS1** | 0.39742 | 0.01414 | 0.02983 | 0.03621 | 0.01702 | 0.00623 | 0.01659 | 0.44481 | 0.02203 | 0.00009 | 0.01502 | 0.02125 |
| **ARHGAP32** | 6.46056 | 1.64388 | 7.50325 | 2.51629 | 2.02273 | 2.89597 | 3.75299 | 0.47990 | 0.55410 | 7.31394 | 1.08253 | 0.00000 |
| **ARHGAP33** | 1.56532 | 0.00000 | 0.31650 | 0.42152 | 0.03170 | 0.04354 | 0.07686 | 0.02046 | 0.00000 | 0.24982 | 0.05229 | 0.10212 |
| **ARHGAP35** | 2.77894 | 1.71015 | 1.02592 | 1.48392 | 5.48676 | 0.00000 | 4.24143 | 0.32461 | 2.98555 | 7.62632 | 2.80548 | 1.90581 |
| **ARHGAP36** | 0.00630 | 0.06178 | 0.01457 | 0.00000 | 0.00000 | 0.07282 | 0.03278 | 0.01411 | 0.01499 | 0.03178 | 0.04307 | 0.01173 |
| **ARHGAP39** | 2.52231 | 0.52042 | 0.09136 | 0.04946 | 0.29115 | 0.00000 | 0.47477 | 0.00000 | 0.18346 | 3.76242 | 0.00000 | 0.37958 |
| **ARHGAP4** | 0.05460 | 0.12881 | 0.12737 | 0.05156 | 0.04400 | 0.00000 | 0.08317 | 0.00645 | 0.00005 | 0.00000 | 0.03952 | 0.03579 |
| **ARHGAP40** | 0.42346 | 0.01082 | 0.00670 | 0.10494 | 0.00383 | 0.01073 | 0.01151 | 0.00179 | 0.05057 | 0.02918 | 0.01036 | 0.01203 |
| **ARHGAP42** | 0.06510 | 0.93719 | 0.34067 | 4.73790 | 0.61037 | 4.26934 | 0.97391 | 0.00000 | 0.00000 | 2.67437 | 1.23224 | 0.00017 |
| **ARHGAP44** | 1.92093 | 0.37610 | 0.28528 | 0.02595 | 1.97898 | 1.40194 | 0.23728 | 0.09906 | 0.00000 | 2.53836 | 0.89313 | 0.00000 |
| **ARHGAP5** | 3.83880 | 0.00000 | 3.04856 | 2.46745 | 2.74009 | 22.99264 | 6.50914 | 3.38073 | 4.03654 | 26.41078 | 1.82769 | 9.09146 |
| **ARHGAP5-AS1** | 0.32224 | 0.20736 | 0.24336 | 0.18652 | 0.00709 | 1.72340 | 0.00000 | 0.02804 | 0.26324 | 0.00000 | 0.00000 | 1.21127 |
| **ARHGAP6** | 0.60089 | 0.01353 | 0.03011 | 0.01123 | 0.01894 | 0.02507 | 0.04645 | 0.00675 | 0.00000 | 0.02898 | 0.06336 | 0.02028 |
| **ARHGAP8** | 0.07549 | 0.55642 | 0.37058 | 0.12859 | 0.23974 | 0.31851 | 0.00000 | 0.00761 | 0.41813 | 0.00000 | 0.58592 | 0.02722 |
| **ARHGAP9** | 0.14953 | 1.84135 | 0.19440 | 0.00000 | 0.00000 | 0.01808 | 0.12825 | 0.10136 | 0.00024 | 0.05472 | 0.03897 | 0.63373 |
| **ARHGDIA** | 7.37419 | 1.45238 | 22.24900 | 9.79333 | 14.95959 | 0.00000 | 27.70994 | 2.25665 | 16.98224 | 19.16059 | 11.84937 | 2.25507 |
| **ARHGDIB** | 0.07849 | 0.14089 | 2.62889 | 0.00000 | 1.34640 | 0.17411 | 1.20520 | 0.02619 | 1.27525 | 0.00000 | 0.29687 | 0.00269 |
| **ARHGDIG** | 0.03348 | 0.11019 | 0.01230 | 0.04152 | 0.25123 | 0.00000 | 0.01046 | 0.01762 | 0.00000 | 0.26971 | 0.00000 | 0.02597 |
| **ARHGEF1** | 0.03924 | 5.61168 | 3.71804 | 3.68452 | 3.48745 | 0.00000 | 1.14420 | 0.08133 | 6.25376 | 3.09226 | 2.84977 | 3.21612 |
| **ARHGEF10** | 0.21198 | 0.37112 | 0.56016 | 0.37708 | 0.42083 | 0.00000 | 0.00000 | 0.02977 | 0.09816 | 1.30666 | 0.22544 | 0.53748 |
| **ARHGEF10L** | 0.01547 | 0.12769 | 0.54080 | 0.01189 | 0.00000 | 0.01882 | 0.04661 | 0.01044 | 0.48858 | 0.00785 | 0.00000 | 0.01558 |
| **ARHGEF11** | 2.19091 | 0.39358 | 0.92701 | 1.05480 | 5.23277 | 0.00000 | 3.16912 | 0.00000 | 1.86440 | 8.12056 | 0.00000 | 3.99851 |
| **ARHGEF12** | 8.65345 | 0.00000 | 2.69203 | 0.87871 | 3.62680 | 6.82652 | 4.99184 | 1.07962 | 0.73303 | 6.43392 | 0.90725 | 0.27732 |
| **ARHGEF15** | 0.00504 | 0.01887 | 0.25004 | 2.30479 | 0.18748 | 0.00000 | 0.00028 | 0.01024 | 0.00000 | 0.02753 | 0.08506 | 0.03524 |
| **ARHGEF16** | 2.18644 | 2.37804 | 1.63236 | 5.32549 | 5.24449 | 0.00000 | 1.42911 | 0.04407 | 1.71126 | 3.49000 | 3.77021 | 0.00000 |
| **ARHGEF17** | 0.04064 | 0.15112 | 1.10215 | 0.00000 | 3.67018 | 0.00000 | 0.46276 | 0.04145 | 2.39607 | 0.89908 | 0.79077 | 0.00000 |
| **ARHGEF18** | 0.00000 | 6.77845 | 2.24118 | 2.30210 | 8.57266 | 0.00000 | 5.73695 | 2.31675 | 2.07071 | 10.02992 | 1.73672 | 4.57334 |
| **ARHGEF19** | 0.00225 | 0.06575 | 0.00000 | 0.46783 | 0.17517 | 0.00000 | 0.36166 | 0.32642 | 0.61762 | 0.58864 | 0.11283 | 0.05962 |
| **ARHGEF2** | 0.00193 | 0.00000 | 0.93859 | 1.28951 | 2.69402 | 0.00000 | 3.50188 | 0.00000 | 1.55187 | 2.35718 | 1.98214 | 0.11222 |
| **ARHGEF25** | 0.00204 | 0.00684 | 0.05086 | 0.04777 | 0.00371 | 0.02782 | 0.00000 | 0.04482 | 0.12340 | 0.06050 | 0.00229 | 0.05533 |
| **ARHGEF26** | 1.58391 | 2.01154 | 0.00000 | 0.67166 | 10.34955 | 3.70895 | 1.61094 | 2.02624 | 4.31343 | 2.22903 | 7.17135 | 3.38191 |
| **ARHGEF26-AS1** | 0.33675 | 0.00000 | 0.00000 | 0.00074 | 0.04566 | 0.36571 | 0.00000 | 0.00000 | 0.00000 | 0.00525 | 0.00838 | 0.03874 |
| **ARHGEF28** | 1.60800 | 0.01051 | 0.00105 | 0.06168 | 0.00404 | 0.00000 | 0.00424 | 0.06329 | 0.04180 | 0.30565 | 0.00036 | 0.15073 |
| **ARHGEF3** | 0.83543 | 0.64473 | 0.68286 | 0.98612 | 0.87711 | 2.32654 | 0.00000 | 1.02341 | 1.15527 | 0.06855 | 0.08793 | 1.15450 |
| **ARHGEF33** | 2.09356 | 0.81086 | 0.11605 | 0.00000 | 0.21430 | 0.06835 | 0.04362 | 0.33125 | 0.00000 | 0.02314 | 0.12377 | 5.80116 |

| | | | | | | | | | | | | |
|---|---|---|---|---|---|---|---|---|---|---|---|---|
| **ARHGEF34P** | 0.00903 | 0.01583 | 0.01583 | 0.01053 | 0.02413 | 0.02396 | 0.06190 | 0.01173 | 0.01967 | 0.00229 | 0.01415 | 0.02174 |
| **ARHGEF35** | 0.00531 | 0.06731 | 0.11770 | 0.07299 | 0.00000 | 0.01687 | 0.20301 | 0.00996 | 0.00000 | 0.04624 | 0.06924 | 0.01527 |
| **ARHGEF37** | 0.00086 | 0.00000 | 0.75369 | 0.00000 | 0.04778 | 0.27893 | 0.18627 | 0.14592 | 0.00000 | 0.09463 | 0.00000 | 0.39862 |
| **ARHGEF38** | 0.19703 | 0.16014 | 0.00000 | 0.16676 | 0.35794 | 0.95256 | 0.07916 | 0.08212 | 0.04619 | 0.03976 | 0.25527 | 0.02510 |
| **ARHGEF38-IT1** | 0.01238 | 0.11978 | 0.00000 | 0.08831 | 0.05746 | 0.19344 | 0.00000 | 0.02265 | 0.02304 | 0.00000 | 0.03811 | 0.00505 |
| **ARHGEF39** | 1.47621 | 10.13354 | 3.78554 | 3.73892 | 5.84348 | 0.00032 | 1.30973 | 0.00000 | 6.20600 | 0.00000 | 3.18322 | 4.49745 |
| **ARHGEF4** | 0.00617 | 0.07811 | 2.02907 | 0.01909 | 0.22904 | 0.00000 | 0.00000 | 0.00925 | 0.00000 | 0.00000 | 0.28679 | 0.03558 |
| **ARHGEF40** | 0.06124 | 0.01339 | 0.03919 | 0.43727 | 0.00000 | 0.10776 | 0.00000 | 0.00734 | 0.00000 | 0.08968 | 0.00526 | 0.00609 |
| **ARHGEF5** | 0.02169 | 0.01311 | 0.44077 | 0.22183 | 0.09258 | 0.01986 | 1.18603 | 0.01274 | 0.03642 | 0.45764 | 0.03936 | 0.14658 |
| **ARHGEF6** | 0.37913 | 0.05836 | 0.01937 | 0.01251 | 0.01187 | 0.00000 | 0.01895 | 0.08223 | 0.04212 | 0.00000 | 0.03124 | 0.20098 |
| **ARHGEF7** | 5.08775 | 7.00878 | 7.45427 | 3.12704 | 5.59085 | 5.02697 | 7.98495 | 7.54733 | 12.33368 | 3.07323 | 5.21369 | 9.48669 |
| **ARHGEF9** | 1.34689 | 0.08462 | 0.41942 | 0.36951 | 0.61130 | 0.63241 | 0.00000 | 0.24682 | 0.40237 | 0.30357 | 0.82088 | 0.36214 |
| **ARID1A** | 6.48827 | 5.89275 | 7.33432 | 11.38947 | 16.99691 | 11.25315 | 24.04816 | 4.53292 | 5.34903 | 37.64118 | 10.55808 | 9.77938 |
| **ARID1B** | 1.87964 | 2.47777 | 3.53896 | 3.34029 | 0.85072 | 10.15835 | 6.32698 | 0.41013 | 1.00145 | 9.75647 | 0.39778 | 0.91913 |
| **ARID2** | 5.84755 | 0.95008 | 3.07029 | 0.74761 | 0.00000 | 14.93212 | 12.84597 | 2.10799 | 2.99380 | 17.53189 | 4.10295 | 5.40408 |
| **ARID3A** | 0.00072 | 0.00000 | 22.17093 | 1.64077 | 4.28805 | 0.00000 | 27.42119 | 0.46313 | 6.13256 | 22.27192 | 1.35029 | 1.45840 |
| **ARID3B** | 3.49797 | 2.44804 | 9.34764 | 4.03773 | 2.65228 | 0.00000 | 7.07517 | 2.67060 | 4.31194 | 7.20843 | 2.32261 | 5.85983 |
| **ARID3C** | 0.02086 | 0.08649 | 0.01744 | 0.01666 | 0.00000 | 0.00572 | 0.01289 | 0.01787 | 0.01219 | 0.04702 | 0.03810 | 0.04279 |
| **ARID4A** | 8.49219 | 0.01163 | 3.34503 | 2.45808 | 3.26593 | 7.77635 | 2.60825 | 11.99308 | 2.64074 | 14.03253 | 1.06889 | 3.80304 |
| **ARID4B** | 3.08743 | 0.35790 | 9.10268 | 3.88634 | 3.27897 | 9.11715 | 8.53304 | 11.13778 | 4.51746 | 19.86919 | 1.91452 | 2.78768 |
| **ARID5A** | 0.88745 | 1.50628 | 0.23067 | 1.70782 | 0.96412 | 0.00000 | 0.09470 | 0.00000 | 1.75785 | 0.00000 | 1.62687 | 0.76618 |
| **ARID5B** | 0.36733 | 0.16986 | 6.30876 | 1.71605 | 0.00000 | 0.46443 | 3.06270 | 0.06883 | 0.72488 | 0.00000 | 0.41122 | 0.01575 |
| **ARIH1** | 8.16250 | 2.58663 | 5.99623 | 2.89372 | 1.07063 | 11.90634 | 12.37998 | 3.09593 | 0.09033 | 18.94528 | 2.88158 | 1.33215 |
| **ARIH2** | 3.98030 | 12.84748 | 15.77125 | 10.55766 | 14.01275 | 19.61367 | 24.46006 | 8.90580 | 22.23839 | 8.11529 | 17.34596 | 19.08635 |
| **ARIH2OS** | 0.40366 | 0.15385 | 0.22256 | 0.27460 | 0.47626 | 0.10269 | 0.19468 | 0.41939 | 0.43011 | 0.00000 | 0.61613 | 0.65289 |
| **1.00000** | 2.05817 | 9.66136 | 4.23197 | 3.64122 | 2.91174 | 29.72019 | 4.52919 | 0.33453 | 7.35978 | 4.58475 | 8.69682 | 6.60479 |
| **10.00000** | 0.01250 | 0.00758 | 0.00000 | 0.24672 | 0.00000 | 0.07582 | 0.08712 | 0.01468 | 0.00000 | 0.20000 | 0.16688 | 0.02879 |
| **11.00000** | 0.00962 | 0.12515 | 1.18732 | 0.70868 | 0.37842 | 0.31980 | 1.70235 | 0.12786 | 1.03028 | 0.00000 | 0.66882 | 0.09808 |
| **ARL13A** | 0.07661 | 0.01002 | 0.04845 | 0.01976 | 0.03808 | 0.02660 | 0.03480 | 0.00473 | 0.00830 | 0.04498 | 0.02771 | 0.09729 |
| **ARL13B** | 7.80811 | 0.35339 | 1.38927 | 1.63891 | 0.00000 | 2.43455 | 4.19263 | 1.43745 | 1.75525 | 1.82551 | 0.37280 | 0.80773 |
| **14.00000** | 0.00550 | 0.01316 | 0.01244 | 0.01172 | 0.01998 | 0.00983 | 0.07256 | 0.01153 | 0.00000 | 0.02100 | 0.01431 | 0.04684 |
| **ARL14EP** | 1.78256 | 4.65005 | 0.42702 | 1.08188 | 1.37316 | 8.14291 | 1.55337 | 5.81029 | 0.45908 | 2.30007 | 3.10565 | 7.64402 |
| **ARL14EPL** | 0.00627 | 0.01466 | 0.01828 | 0.01688 | 0.02431 | 0.01980 | 0.02390 | 0.01049 | 0.01312 | 0.01001 | 0.01577 | 0.01638 |
| **15.00000** | 0.66400 | 4.48527 | 9.06518 | 0.64663 | 3.34593 | 20.47822 | 4.57799 | 1.18885 | 3.70919 | 0.00013 | 1.78348 | 15.50529 |
| **16.00000** | 0.00000 | 8.41587 | 6.10043 | 4.69071 | 5.67268 | 4.85265 | 7.18398 | 3.52173 | 7.89983 | 0.06585 | 3.68488 | 5.95481 |
| **ARL17A.uid-210** | 0.00879 | 0.07678 | 0.02701 | 0.00936 | 0.10498 | 0.00000 | 0.00071 | 0.08200 | 0.08218 | 0.07685 | 0.00033 | 0.05952 |
| **ARL17A.uid-219** | 0.00261 | 0.05940 | 0.01757 | 0.01042 | 0.04484 | 0.07098 | 0.02214 | 0.11811 | 0.02852 | 0.28515 | 0.06224 | 0.09242 |
| **ARL17A.uid-219** | 0.00152 | 0.02401 | 0.01446 | 0.10525 | 0.06598 | 0.00000 | 0.00424 | 0.13607 | 0.09448 | 0.08164 | 0.05959 | 0.06725 |
| **2.00000** | 0.78756 | 14.10833 | 6.26604 | 12.13797 | 16.83152 | 1.26917 | 16.19865 | 0.07298 | 10.50643 | 4.06963 | 11.78010 | 5.88264 |
| **ARL2BP** | 2.58270 | 11.73350 | 1.14616 | 2.94585 | 7.80614 | 12.02849 | 2.86002 | 0.96740 | 2.68068 | 0.00000 | 9.40594 | 5.33188 |
| **3.00000** | 2.28880 | 1.80527 | 6.59080 | 2.88919 | 1.15932 | 6.23235 | 9.10514 | 2.16669 | 3.59481 | 2.67370 | 3.72540 | 7.01187 |
| **ARL4A** | 1.33205 | 1.65104 | 0.64246 | 1.52517 | 3.30110 | 5.08854 | 0.00000 | 0.83718 | 0.00000 | 0.41301 | 0.98979 | 2.46326 |
| **ARL4C** | 3.74208 | 0.22872 | 6.74806 | 0.07069 | 1.58062 | 0.06617 | 1.90781 | 1.12313 | 0.00000 | 5.10749 | 0.02886 | 0.25276 |
| **ARL4D** | 7.44945 | 22.17066 | 1.75638 | 14.78977 | 17.59258 | 7.16428 | 9.52010 | 34.44620 | 7.23681 | 0.00000 | 14.13420 | 27.66661 |
| **ARL5A** | 1.83434 | 1.46867 | 13.92307 | 3.24272 | 1.77689 | 19.13403 | 15.78010 | 0.91998 | 2.60827 | 22.84294 | 5.16693 | 1.77171 |
| **ARL5B** | 2.53241 | 1.36865 | 11.05417 | 0.47044 | 0.10810 | 11.47042 | 3.14750 | 8.10647 | 3.85850 | 0.00839 | 1.95506 | 7.09367 |
| **ARL5C** | 0.00412 | 0.02181 | 0.01729 | 0.01259 | 0.01351 | 0.01484 | 0.01624 | 0.02247 | 0.03308 | 0.01848 | 0.01247 | 0.01610 |
| **6.00000** | 0.51502 | 0.22265 | 0.57028 | 0.08428 | 0.00000 | 1.53443 | 0.75568 | 0.21148 | 0.72934 | 0.00000 | 0.50585 | 0.80107 |

| | | | | | | | | | | | | |
|---|---|---|---|---|---|---|---|---|---|---|---|---|
| **ARL6IP1** | 9.96018 | 28.69527 | 21.91058 | 9.87996 | 11.96934 | 58.02781 | 0.00000 | 25.47297 | 47.89750 | 29.20781 | 14.10889 | 30.03195 |
| **ARL6IP4** | 0.40114 | 9.66457 | 5.69392 | 10.37967 | 11.19740 | 4.45251 | 8.01476 | 0.13161 | 8.20001 | 3.72867 | 6.00468 | 3.21861 |
| **ARL6IP5** | 0.84438 | 2.63942 | 6.00174 | 0.15409 | 8.50632 | 16.28230 | 0.00000 | 0.00000 | 11.91707 | 8.25067 | 10.22724 | 15.94857 |
| **ARL6IP6** | 1.11970 | 7.83226 | 10.94979 | 4.18512 | 0.91091 | 19.64792 | 0.00000 | 4.20646 | 11.75468 | 1.55344 | 5.70320 | 10.14845 |
| **ARL8A** | 0.01974 | 0.81039 | 0.00000 | 0.96714 | 1.47409 | 0.00000 | 0.49636 | 0.01111 | 0.00000 | 4.63770 | 0.25874 | 1.17481 |
| **ARL8B** | 4.46802 | 17.62469 | 16.00696 | 9.66896 | 4.27955 | 33.33059 | 10.92528 | 20.65097 | 9.98401 | 11.24643 | 5.83936 | 30.12527 |
| **9.00000** | 0.42881 | 0.04822 | 0.07624 | 0.00622 | 0.00000 | 0.00000 | 0.05388 | 0.73801 | 0.08594 | 0.16111 | 0.00000 | 0.03818 |
| **ARMC1** | 8.66383 | 10.24042 | 1.84196 | 8.07984 | 2.51775 | 33.35333 | 6.18393 | 11.00807 | 7.35426 | 4.47382 | 8.99129 | 12.39501 |
| **ARMC10** | 0.30632 | 1.04728 | 1.07332 | 0.74891 | 1.17390 | 2.45467 | 0.90318 | 0.00916 | 1.95320 | 1.58300 | 0.48218 | 0.51663 |
| **ARMC12** | 0.18685 | 0.01161 | 0.06243 | 0.04112 | 0.09633 | 0.00000 | 0.02911 | 0.07226 | 0.00000 | 0.03257 | 0.01506 | 0.01062 |
| **ARMC2** | 13.76300 | 0.14246 | 0.24328 | 0.28579 | 0.00000 | 0.33588 | 0.11063 | 4.05390 | 0.00000 | 0.28416 | 0.63828 | 0.78246 |
| **ARMC3** | 0.00681 | 0.02603 | 0.03609 | 0.06276 | 0.04509 | 0.00000 | 0.04536 | 0.01045 | 0.02631 | 0.06298 | 0.02087 | 0.01769 |
| **ARMC4** | 0.00932 | 0.01351 | 0.00816 | 0.01006 | 0.01157 | 0.01438 | 0.09063 | 0.00975 | 0.00000 | 0.03619 | 0.01125 | 0.02163 |
| **ARMC5** | 0.99069 | 0.58464 | 0.41307 | 0.37725 | 0.60828 | 0.00000 | 0.85470 | 0.56912 | 0.55199 | 0.80975 | 0.24880 | 0.91700 |
| **ARMC6** | 0.75724 | 12.11460 | 8.66326 | 5.18014 | 9.98327 | 0.00000 | 11.88562 | 2.14345 | 18.13345 | 0.00000 | 6.82185 | 15.26099 |
| **ARMC7** | 1.62730 | 9.54885 | 3.19891 | 3.13180 | 8.52017 | 0.00000 | 6.00862 | 1.45001 | 8.75864 | 2.14554 | 7.34638 | 9.11612 |
| **ARMC8** | 6.46581 | 11.25099 | 6.48644 | 5.28059 | 1.45301 | 20.43695 | 6.13272 | 7.24394 | 7.85419 | 5.53394 | 8.73195 | 10.60862 |
| **ARMC9** | 4.03564 | 1.12044 | 0.99886 | 2.76789 | 0.00000 | 0.30997 | 1.72296 | 3.21690 | 0.00000 | 0.00000 | 3.69109 | 1.39912 |
| **ARMCX1** | 0.00624 | 0.00286 | 0.15690 | 0.11258 | 0.00705 | 0.15215 | 0.21345 | 0.01070 | 1.04657 | 0.00000 | 0.00000 | 0.02636 |
| **ARMCX2** | 0.04232 | 0.08394 | 0.13231 | 1.19308 | 0.00000 | 0.43804 | 0.06984 | 0.00000 | 0.10949 | 0.00000 | 0.00000 | 0.23228 |
| **ARMCX3** | 0.02867 | 0.00000 | 1.43819 | 0.82750 | 0.11475 | 0.32349 | 0.95598 | 0.14749 | 0.75516 | 0.00000 | 0.00000 | 0.10867 |
| **ARMCX4** | 0.08852 | 0.00000 | 0.10161 | 0.00000 | 0.69068 | 0.35381 | 0.31970 | 0.00766 | 0.00000 | 0.44443 | 0.61482 | 0.02208 |
| **ARMCX5** | 0.00701 | 0.46143 | 0.06717 | 0.69879 | 0.93111 | 0.42393 | 0.00000 | 0.00000 | 0.84810 | 0.00000 | 0.00000 | 0.44203 |
| **ARMCX5-GPRA** | 0.00931 | 0.02522 | 0.01488 | 0.09371 | 0.02262 | 0.05833 | 0.12773 | 0.00979 | 0.03549 | 0.00000 | 0.00804 | 0.04532 |
| **ARMCX6** | 0.11391 | 0.03863 | 0.00847 | 0.29324 | 0.03925 | 0.04424 | 0.09238 | 0.00808 | 0.46089 | 0.00000 | 0.00000 | 0.02143 |
| **ARMS2** | 0.00626 | 0.06647 | 0.03150 | 0.00593 | 0.01015 | 0.00922 | 0.05726 | 0.01074 | 0.00140 | 0.00000 | 0.14721 | 0.01497 |
| **ARNT** | 0.64988 | 8.41477 | 6.42050 | 5.17407 | 1.49924 | 6.00519 | 3.94185 | 1.27003 | 2.83708 | 1.54511 | 5.49984 | 4.80635 |
| **ARNT2** | 0.42778 | 0.06183 | 0.18567 | 0.04577 | 0.01478 | 0.00000 | 0.02942 | 0.00775 | 0.00000 | 0.06114 | 0.01619 | 0.02682 |
| **ARNTL** | 2.49488 | 0.44737 | 1.00555 | 0.69723 | 0.00000 | 1.29299 | 1.03568 | 1.21598 | 0.11391 | 0.00000 | 0.62669 | 0.86896 |
| **ARNTL2** | 0.51761 | 0.50008 | 0.15869 | 0.29126 | 1.88045 | 2.88198 | 0.29926 | 0.00000 | 0.00000 | 1.09245 | 1.40598 | 0.04956 |
| **ARNTL2-AS1** | 0.01081 | 0.01398 | 0.01687 | 0.00944 | 0.04769 | 0.01799 | 0.01543 | 0.05636 | 0.02077 | 0.01725 | 0.01487 | 0.01477 |
| **ARPC1A** | 1.67913 | 19.08739 | 14.71988 | 11.27224 | 14.67285 | 24.28191 | 18.01873 | 1.37426 | 21.70172 | 0.00000 | 16.78470 | 10.58061 |
| **ARPC1B** | 4.57684 | 32.30370 | 27.46481 | 17.81511 | 28.33136 | 5.81328 | 36.49394 | 2.45472 | 39.84012 | 0.00000 | 31.03420 | 19.21529 |
| **ARPC2** | 4.12494 | 30.60364 | 32.75826 | 26.18385 | 24.21661 | 48.71205 | 38.69362 | 15.08735 | 36.72695 | 17.31809 | 32.25247 | 29.24695 |
| **ARPC3** | 0.75364 | 11.40460 | 13.92418 | 8.75979 | 4.11925 | 34.52375 | 12.66611 | 2.67304 | 14.83782 | 3.00068 | 12.49742 | 16.46772 |
| **ARPC4** | 3.56192 | 15.86072 | 22.02735 | 17.96623 | 22.40407 | 10.30459 | 35.87302 | 10.54327 | 10.77113 | 8.97705 | 15.47796 | 14.78416 |
| **ARPC5** | 0.20065 | 15.56222 | 23.36542 | 10.43792 | 15.89963 | 40.89290 | 23.80751 | 2.29400 | 23.71198 | 0.00000 | 23.81242 | 13.79163 |
| **ARPC5L** | 0.52527 | 17.46335 | 16.19829 | 9.20493 | 14.03839 | 5.16146 | 15.78599 | 11.65650 | 16.75268 | 20.26499 | 11.61636 | 11.61454 |
| **ARPIN** | 0.16004 | 0.23782 | 0.16290 | 0.49153 | 0.32783 | 0.11287 | 0.00000 | 0.18821 | 0.06184 | 0.39833 | 0.34227 | 0.12637 |
| **ARPP19** | 1.44915 | 11.34901 | 16.42858 | 4.89663 | 2.80983 | 35.79814 | 15.74029 | 15.90161 | 5.37405 | 20.33937 | 11.91505 | 16.95226 |
| **ARPP21** | 0.22614 | 0.01459 | 0.01770 | 0.01293 | 0.01574 | 0.01247 | 0.01809 | 0.00552 | 0.03129 | 0.00887 | 0.01156 | 0.01613 |
| **ARR3** | 0.00671 | 0.00000 | 0.01036 | 0.14976 | 0.04647 | 0.09385 | 0.00000 | 0.24273 | 0.06334 | 0.04466 | 0.01309 | 0.30880 |
| **ARRB1** | 0.00000 | 5.28896 | 9.60475 | 11.31635 | 5.64323 | 0.00000 | 9.04971 | 3.30603 | 4.52427 | 0.00000 | 4.46608 | 5.47156 |
| **ARRB2** | 0.27087 | 0.89852 | 1.32265 | 1.17690 | 0.00000 | 0.00000 | 3.92710 | 0.00000 | 1.33309 | 2.23307 | 0.94707 | 0.33412 |
| **ARRDC1** | 0.36014 | 2.52102 | 6.04273 | 2.71750 | 4.89666 | 0.00000 | 1.46862 | 0.19309 | 7.57032 | 1.60403 | 6.08005 | 0.54709 |
| **ARRDC2** | 1.91046 | 3.92994 | 0.60224 | 5.68759 | 0.27349 | 0.18773 | 0.23627 | 7.01504 | 0.00000 | 2.07167 | 0.39890 | 5.89483 |
| **ARRDC3** | 0.47920 | 4.34725 | 2.29947 | 0.56841 | 3.47020 | 29.08194 | 0.00000 | 2.74480 | 0.00000 | 6.01129 | 8.61279 | 0.82813 |
| **ARRDC3-AS1** | 0.08525 | 0.04161 | 0.18318 | 0.02186 | 0.00524 | 0.02482 | 0.00000 | 0.00803 | 0.00937 | 0.02380 | 0.02295 | 0.06453 |

| | | | | | | | | | | | | |
|---|---|---|---|---|---|---|---|---|---|---|---|---|
| **ARRDC4** | 0.00000 | 1.46258 | 0.00000 | 0.11647 | 0.95555 | 5.17381 | 2.50648 | 0.00968 | 0.65721 | 1.61477 | 0.00000 | 0.73627 |
| **ARRDC5** | 0.00542 | 0.01167 | 0.00488 | 0.32718 | 0.00000 | 0.00009 | 0.01433 | 0.01030 | 0.00000 | 0.04555 | 0.08966 | 0.01594 |
| **ARSA** | 0.45484 | 0.95430 | 0.99991 | 1.11550 | 0.00000 | 0.00000 | 0.04364 | 0.09357 | 1.14073 | 0.24573 | 0.76218 | 1.21170 |
| **ARSB** | 0.00172 | 0.48592 | 3.76706 | 0.00000 | 0.06473 | 1.25321 | 0.89852 | 0.06809 | 2.00551 | 1.25088 | 0.25236 | 0.00000 |
| **ARSD** | 0.47818 | 0.04395 | 0.01377 | 0.00000 | 0.22789 | 0.00000 | 0.07130 | 0.00000 | 0.24397 | 0.24627 | 0.00000 | 0.20651 |
| **ARSE** | 0.00572 | 0.00860 | 0.51313 | 2.19471 | 0.03144 | 0.00000 | 0.00000 | 0.00950 | 0.01568 | 0.00758 | 0.02972 | 0.01498 |
| **ARSG** | 0.33101 | 0.52941 | 0.13922 | 0.00000 | 0.25689 | 0.00000 | 0.00000 | 0.55096 | 0.30860 | 0.04462 | 0.47713 | 0.24821 |
| **ARSH** | 0.04243 | 0.01545 | 0.01381 | 0.01101 | 0.01519 | 0.02061 | 0.01786 | 0.01248 | 0.01810 | 0.01607 | 0.01425 | 0.01450 |
| **ARSI** | 0.18622 | 0.26581 | 0.00000 | 0.18434 | 0.33266 | 0.09095 | 0.00000 | 0.00000 | 0.00000 | 0.08350 | 0.00000 | 2.67836 |
| **ARSJ** | 0.45121 | 0.00690 | 0.03604 | 0.01838 | 0.05874 | 0.00000 | 0.02756 | 0.00204 | 0.00000 | 0.10388 | 0.00782 | 0.02214 |
| **ARSK** | 0.04187 | 0.48221 | 1.81612 | 0.06277 | 1.15081 | 2.60690 | 0.41941 | 2.49550 | 7.01441 | 1.62752 | 2.39183 | 1.48284 |
| **ART1** | 0.00590 | 0.01550 | 0.06159 | 0.01930 | 0.02102 | 0.00000 | 0.03752 | 0.01014 | 0.01850 | 0.01710 | 0.01833 | 0.03172 |
| **ART3** | 0.79412 | 0.21291 | 0.00635 | 0.00000 | 0.00151 | 0.04489 | 0.01895 | 0.94441 | 0.01891 | 0.00797 | 0.00374 | 0.71870 |
| **ART4** | 0.11649 | 0.03142 | 0.19420 | 0.06002 | 0.11448 | 0.03358 | 0.02518 | 0.03439 | 0.00000 | 0.01965 | 0.01222 | 0.01119 |
| **ART5** | 0.56343 | 0.33476 | 0.01240 | 0.01352 | 0.00000 | 0.00851 | 0.00866 | 1.51809 | 0.03790 | 0.01167 | 0.00905 | 1.03306 |
| **ARTN** | 0.00852 | 0.01551 | 0.03322 | 0.01172 | 0.01504 | 0.01625 | 0.01361 | 0.01021 | 0.02421 | 0.01098 | 0.01795 | 0.01661 |
| **ARV1** | 1.40750 | 3.08697 | 4.21104 | 1.33672 | 1.10522 | 3.41009 | 0.87799 | 0.01631 | 5.72806 | 1.27561 | 5.73402 | 1.89421 |
| **ARVCF** | 0.00268 | 0.05504 | 0.16064 | 0.13125 | 0.26303 | 0.00000 | 0.18277 | 0.02393 | 0.00000 | 0.14202 | 0.16399 | 0.01205 |
| **ARX** | 0.00141 | 0.00831 | 0.14723 | 0.87862 | 0.00000 | 0.00000 | 0.07927 | 0.11777 | 0.00000 | 0.24281 | 0.10883 | 0.22067 |
| **AS3MT** | 0.00646 | 0.01558 | 0.02148 | 0.01135 | 0.01566 | 0.02318 | 0.01501 | 0.01003 | 0.02073 | 0.01043 | 0.01391 | 0.02378 |
| **ASAH1** | 3.12401 | 0.77285 | 11.43511 | 1.08315 | 9.03394 | 24.69771 | 0.00050 | 2.40322 | 51.76134 | 4.67780 | 15.38977 | 0.29938 |
| **ASAH2** | 1.39607 | 0.01523 | 0.05247 | 0.04727 | 0.07202 | 0.01565 | 0.00981 | 0.28674 | 0.01436 | 0.00000 | 0.01430 | 0.01367 |
| **ASAH2B** | 0.03632 | 0.24625 | 0.07521 | 0.01196 | 0.04463 | 0.42177 | 0.00000 | 0.12855 | 0.10338 | 0.00000 | 0.25377 | 0.36326 |
| **ASAP1** | 3.94245 | 4.07558 | 14.83929 | 1.12719 | 10.14195 | 12.39833 | 20.42706 | 0.95922 | 4.94756 | 22.87918 | 4.49593 | 0.00176 |
| **ASAP1-IT1** | 0.00686 | 0.00000 | 0.18624 | 0.00052 | 0.07788 | 0.00017 | 0.02852 | 0.00929 | 0.38241 | 1.95022 | 0.00000 | 0.03267 |
| **ASAP1-IT2** | 0.00106 | 0.00000 | 0.35492 | 0.01397 | 0.50590 | 0.08965 | 0.00000 | 0.02922 | 0.27384 | 1.99407 | 0.13504 | 0.00754 |
| **ASAP2** | 5.27926 | 0.44038 | 0.00000 | 2.16080 | 5.86556 | 5.73899 | 2.77178 | 0.28439 | 1.38044 | 7.70078 | 4.09135 | 0.00000 |
| **ASAP3** | 0.25117 | 4.17955 | 3.50843 | 1.24697 | 0.21469 | 1.43490 | 0.00000 | 1.93254 | 0.57357 | 0.00000 | 0.06647 | 1.66170 |
| **ASB1** | 0.61538 | 2.49123 | 2.61797 | 4.00703 | 1.56662 | 1.79042 | 1.99803 | 1.20008 | 1.57837 | 2.28054 | 0.28218 | 3.65814 |
| **ASB11** | 0.19973 | 0.02737 | 0.01419 | 0.01274 | 0.02397 | 0.00053 | 0.01963 | 0.01155 | 0.04625 | 0.00000 | 0.01210 | 0.01369 |
| **ASB12** | 0.00638 | 0.03923 | 0.00000 | 0.15454 | 0.00625 | 0.01335 | 0.05962 | 0.01031 | 0.17553 | 0.00000 | 0.00009 | 0.01570 |
| **ASB13** | 0.00372 | 0.04973 | 0.23493 | 0.17292 | 0.00000 | 0.14274 | 0.00000 | 0.01911 | 0.06442 | 0.00000 | 0.21312 | 0.00600 |
| **ASB14** | 0.01278 | 0.08960 | 0.01836 | 0.67664 | 0.24606 | 0.04451 | 0.13142 | 0.01482 | 0.07878 | 0.95159 | 0.01720 | 0.07172 |
| **ASB15** | 0.00842 | 0.00000 | 0.00000 | 0.23169 | 0.00973 | 0.07823 | 0.00000 | 0.00927 | 0.46433 | 0.06491 | 0.27774 | 0.07597 |
| **ASB16** | 0.02872 | 0.11692 | 0.12049 | 0.02432 | 0.09682 | 0.00000 | 0.01049 | 0.01136 | 0.03619 | 0.00000 | 0.14302 | 0.10082 |
| **ASB16-AS1** | 0.51054 | 1.27360 | 1.60503 | 0.16257 | 0.00456 | 0.22065 | 0.67632 | 1.07953 | 0.70396 | 0.47257 | 1.32605 | 1.46042 |
| **ASB18** | 0.00879 | 0.04011 | 0.01781 | 0.01533 | 0.00944 | 0.02651 | 0.01512 | 0.01248 | 0.01355 | 0.02300 | 0.01474 | 0.01907 |
| **ASB2** | 1.26470 | 0.55635 | 0.00115 | 0.08942 | 0.00000 | 0.00000 | 0.06931 | 0.17716 | 0.00000 | 0.10618 | 0.00179 | 0.03152 |
| **ASB3** | 0.00351 | 1.25069 | 1.30810 | 0.00000 | 0.00000 | 1.70289 | 2.05834 | 1.10576 | 0.14824 | 2.95786 | 0.36481 | 1.21823 |
| **ASB5** | 0.02686 | 0.01859 | 0.03009 | 0.01665 | 0.02623 | 0.00084 | 0.03137 | 0.02571 | 0.02249 | 0.00000 | 0.03151 | 0.01798 |
| **ASB6** | 4.55817 | 11.85768 | 7.31045 | 4.46140 | 6.40623 | 0.00000 | 6.79200 | 5.12876 | 12.13205 | 4.00413 | 7.54452 | 9.37850 |
| **ASB7** | 0.38436 | 2.41543 | 2.29920 | 1.86256 | 1.28555 | 3.51273 | 1.31630 | 7.18289 | 2.56414 | 4.40472 | 0.50761 | 3.88123 |
| **ASB8** | 4.79057 | 4.33821 | 2.64034 | 1.11470 | 1.96953 | 7.56766 | 4.65671 | 1.39455 | 2.60518 | 1.84451 | 4.19560 | 4.54497 |
| **ASB9** | 0.00568 | 0.01321 | 0.04227 | 0.02907 | 0.00000 | 0.02386 | 0.05626 | 0.01038 | 0.02151 | 0.03424 | 0.01396 | 0.01479 |
| **ASB9P1** | 0.01635 | 0.00816 | 0.00884 | 0.11657 | 0.02808 | 0.00818 | 0.01049 | 0.02700 | 0.04972 | 0.03458 | 0.03562 | 0.05530 |
| **ASCC1** | 0.00047 | 5.45131 | 5.19688 | 4.01295 | 2.75644 | 13.55990 | 4.83171 | 2.67781 | 9.81255 | 0.00002 | 5.22502 | 12.32096 |
| **ASCC2** | 4.00056 | 11.31359 | 8.59823 | 9.20571 | 13.19685 | 0.00000 | 8.20292 | 3.59421 | 14.17546 | 0.00000 | 9.57726 | 12.68764 |
| **ASCC3** | 4.37163 | 4.05189 | 6.29732 | 6.24521 | 2.35023 | 16.97316 | 5.61432 | 4.19862 | 7.33268 | 10.58560 | 1.83441 | 3.33082 |

| | | | | | | | | | | | | |
|---|---|---|---|---|---|---|---|---|---|---|---|---|
| **ASCL1** | 0.11274 | 0.00553 | 0.00775 | 0.00822 | 0.03376 | 0.04140 | 0.00976 | 0.15728 | 0.01508 | 0.04507 | 0.01033 | 0.05654 |
| **ASCL2** | 0.27624 | 0.04319 | 0.00312 | 0.06353 | 0.02637 | 0.00000 | 0.00097 | 0.05485 | 0.04232 | 0.13141 | 0.00450 | 0.16240 |
| **ASCL3** | 0.00629 | 0.02500 | 0.01486 | 0.01143 | 0.01286 | 0.01695 | 0.01693 | 0.01048 | 0.02015 | 0.01673 | 0.01527 | 0.01614 |
| **ASF1A** | 8.03701 | 9.88863 | 8.22681 | 0.00000 | 0.34715 | 28.29089 | 7.22497 | 3.87888 | 7.92550 | 0.37832 | 4.61633 | 14.40168 |
| **ASF1B** | 5.37534 | 6.44372 | 10.01324 | 5.72013 | 6.61915 | 6.16848 | 7.53941 | 2.74012 | 10.56311 | 0.00000 | 6.69087 | 5.73838 |
| **ASGR1** | 0.56779 | 0.04891 | 0.54700 | 0.00000 | 0.00000 | 0.01155 | 0.00000 | 0.00207 | 0.80842 | 0.08976 | 0.00031 | 0.01044 |
| **ASGR2** | 0.04666 | 1.49623 | 0.03325 | 0.00737 | 0.22413 | 0.00000 | 0.02481 | 0.00028 | 0.02932 | 0.00000 | 0.08907 | 0.38014 |
| **ASH1L** | 3.34420 | 0.00000 | 5.61588 | 3.50193 | 5.66378 | 8.75995 | 0.85379 | 6.46102 | 7.64014 | 25.77328 | 5.10231 | 4.94271 |
| **ASH1L-AS1** | 0.21765 | 1.06091 | 0.74149 | 0.07331 | 0.89563 | 0.91487 | 0.00000 | 5.44940 | 0.40241 | 1.30760 | 0.79818 | 1.56611 |
| **ASH2L** | 1.33713 | 18.85496 | 10.08706 | 26.13315 | 0.00064 | 18.21277 | 5.06616 | 1.76880 | 4.64463 | 1.96562 | 7.81996 | 14.95669 |
| **ASIC1** | 0.47204 | 0.10033 | 0.49415 | 0.02457 | 0.00000 | 0.00000 | 0.03802 | 0.03624 | 0.00000 | 0.62334 | 0.02763 | 0.02594 |
| **ASIC2** | 0.02079 | 0.07152 | 0.09384 | 0.25003 | 0.06488 | 0.04979 | 0.00000 | 0.09994 | 0.01658 | 0.03369 | 0.03702 | 0.03738 |
| **ASIC3** | 0.03057 | 0.52462 | 0.51441 | 0.17797 | 0.04577 | 0.00000 | 0.01124 | 0.00000 | 0.38170 | 1.74419 | 0.00000 | 1.57457 |
| **ASIC4** | 0.14619 | 0.00876 | 0.00965 | 0.00511 | 0.00891 | 0.04533 | 0.01440 | 0.00291 | 0.00665 | 0.04475 | 0.01763 | 0.01105 |
| **ASIP** | 0.00193 | 0.04300 | 0.03626 | 0.04990 | 0.05931 | 0.00000 | 0.02755 | 0.02531 | 0.00000 | 0.15739 | 0.06730 | 0.05485 |
| **ASL** | 0.01521 | 4.51679 | 3.94787 | 3.09741 | 5.27752 | 0.00000 | 2.53626 | 1.12209 | 4.52785 | 1.50248 | 4.10705 | 4.71733 |
| **ASNA1** | 0.02414 | 16.54358 | 14.42032 | 12.86028 | 18.64538 | 0.00000 | 19.89953 | 17.31434 | 16.33353 | 0.00000 | 18.89498 | 28.79591 |
| **ASNS** | 0.56832 | 0.00000 | 3.76820 | 6.54013 | 11.57073 | 10.75644 | 19.76792 | 0.00000 | 11.47721 | 1.33562 | 12.34992 | 0.00000 |
| **ASNSD1** | 2.20057 | 12.25143 | 16.27312 | 6.96975 | 4.77878 | 25.04744 | 19.13794 | 5.48933 | 10.03431 | 6.88823 | 12.93103 | 14.46524 |
| **ASPA** | 0.00241 | 0.28184 | 0.00000 | 1.56551 | 0.00000 | 0.12023 | 0.00000 | 0.00714 | 0.00000 | 0.23081 | 0.00961 | 0.00451 |
| **ASPDH** | 0.05579 | 0.01591 | 0.03001 | 0.01103 | 0.03680 | 0.00483 | 0.04406 | 0.01886 | 0.00933 | 0.00828 | 0.02665 | 0.01665 |
| **ASPG** | 0.00546 | 0.02548 | 0.01991 | 0.00978 | 0.01046 | 0.01661 | 0.03156 | 0.01214 | 0.01187 | 0.02717 | 0.01614 | 0.01423 |
| **ASPH** | 3.76158 | 6.33808 | 3.28056 | 9.33792 | 0.00000 | 9.88807 | 0.00000 | 2.66109 | 6.78329 | 1.76358 | 2.70169 | 3.66346 |
| **ASPHD1** | 0.00548 | 0.38696 | 0.03868 | 0.10286 | 0.12511 | 0.00000 | 0.10636 | 0.00619 | 0.06717 | 0.06366 | 0.02537 | 0.79293 |
| **ASPHD2** | 2.10708 | 0.13925 | 0.00000 | 1.21757 | 0.00002 | 0.59574 | 0.16645 | 0.71703 | 0.07114 | 0.92121 | 0.00000 | 0.49531 |
| **ASPM** | 6.76532 | 1.29230 | 3.40339 | 0.76920 | 2.91461 | 21.71830 | 0.00000 | 1.84636 | 12.16877 | 35.33825 | 0.73669 | 13.32025 |
| **ASPN** | 0.00662 | 0.00684 | 0.06472 | 0.17092 | 0.23744 | 0.00000 | 0.01554 | 0.06418 | 0.15439 | 1.39337 | 0.00402 | 0.03211 |
| **ASPRV1** | 0.01449 | 0.33707 | 3.40135 | 0.00548 | 0.09730 | 0.00000 | 0.00000 | 0.01878 | 1.44924 | 0.00000 | 0.00121 | 0.05244 |
| **ASPSCR1** | 0.82072 | 1.46558 | 2.47192 | 0.48708 | 1.28180 | 0.00000 | 0.67430 | 3.73183 | 1.51506 | 3.11891 | 0.33882 | 0.54303 |
| **ASRGL1** | 0.58364 | 27.63310 | 5.03998 | 37.32088 | 33.00200 | 21.04805 | 14.81346 | 2.48032 | 3.15208 | 9.15627 | 19.20370 | 21.00180 |
| **ASS1** | 0.02882 | 0.00000 | 5.38518 | 3.49661 | 3.25888 | 0.00540 | 11.49997 | 0.01919 | 10.73726 | 0.02430 | 7.07565 | 0.00296 |
| **ASTE1** | 0.71294 | 3.40205 | 0.31513 | 0.48333 | 0.81421 | 3.11970 | 1.05859 | 1.60092 | 1.48836 | 1.47961 | 3.49782 | 2.75996 |
| **ASTL** | 4.28798 | 0.30787 | 0.00000 | 0.00000 | 0.05656 | 0.05385 | 0.00419 | 1.11075 | 0.00000 | 0.06370 | 0.00000 | 0.51872 |
| **ASTN1** | 0.04371 | 0.01811 | 0.02386 | 0.02013 | 0.05011 | 0.00054 | 0.02308 | 0.02857 | 0.00592 | 0.01282 | 0.04799 | 0.01486 |
| **ASTN2** | 6.24105 | 5.99288 | 0.21527 | 3.08199 | 6.03386 | 2.62829 | 0.00000 | 0.65853 | 1.32139 | 0.04993 | 2.33878 | 1.78956 |
| **ASUN** | 1.24610 | 7.28717 | 4.74009 | 0.39433 | 3.26921 | 21.87652 | 8.73382 | 0.28398 | 5.29741 | 7.49379 | 7.61659 | 6.61478 |
| **ASXL1** | 7.93341 | 2.92362 | 2.63482 | 1.46347 | 9.00399 | 3.08212 | 8.88126 | 1.78238 | 4.59031 | 19.22077 | 3.74012 | 4.77964 |
| **ASXL2** | 2.39034 | 1.41834 | 3.72351 | 1.64778 | 4.02215 | 8.18002 | 6.05425 | 3.52094 | 4.06607 | 13.76390 | 1.88927 | 5.72569 |
| **ASXL3** | 1.66244 | 0.00000 | 0.29484 | 0.05375 | 0.00000 | 0.22728 | 0.00000 | 0.31730 | 0.17447 | 1.46561 | 0.00000 | 0.01005 |
| **ASZ1** | 0.12000 | 0.03838 | 0.02191 | 0.00560 | 0.02172 | 0.03836 | 0.01446 | 0.07329 | 0.03403 | 0.00000 | 0.00696 | 0.89653 |
| **ATAD1** | 5.26898 | 9.91694 | 8.70781 | 1.10802 | 1.20163 | 19.20028 | 0.00000 | 4.39394 | 21.03875 | 4.29584 | 3.52648 | 13.03402 |
| **ATAD2** | 4.09702 | 2.15854 | 6.90756 | 3.83066 | 5.20251 | 21.12601 | 6.56405 | 15.50580 | 7.05447 | 15.35194 | 5.24688 | 12.54471 |
| **ATAD2B** | 3.00038 | 0.54132 | 2.28661 | 0.12944 | 1.64653 | 9.19159 | 4.07330 | 2.81812 | 3.08486 | 12.65834 | 2.67480 | 1.55097 |
| **ATAD3A** | 0.69156 | 3.22056 | 5.05067 | 3.63356 | 11.04256 | 0.00000 | 10.52234 | 0.10270 | 0.72236 | 14.14093 | 5.65006 | 1.33463 |
| **ATAD3B** | 0.00000 | 18.92627 | 11.73346 | 14.05880 | 27.50180 | 0.00000 | 24.94074 | 1.75754 | 10.01639 | 37.76529 | 9.11703 | 20.97131 |
| **ATAD3C** | 6.93706 | 0.43163 | 0.26912 | 0.05899 | 0.00000 | 0.00000 | 0.35393 | 1.69764 | 0.37914 | 0.16186 | 0.09763 | 0.09887 |
| **ATAD5** | 1.43835 | 0.88116 | 3.31156 | 2.31362 | 4.41934 | 16.76023 | 0.00005 | 0.76447 | 6.52895 | 17.17517 | 3.71271 | 3.78228 |
| **ATAT1** | 0.43049 | 1.53774 | 2.67845 | 2.19334 | 0.36634 | 0.00000 | 2.87962 | 0.96286 | 6.08152 | 0.06760 | 3.53535 | 1.25940 |

| | | | | | | | | | | | | |
|---|---|---|---|---|---|---|---|---|---|---|---|---|
| **ATCAY** | 0.02530 | 0.00961 | 0.02982 | 0.07915 | 0.02361 | 0.01898 | 0.00281 | 0.02734 | 0.03349 | 0.09126 | 0.01242 | 0.25268 |
| **ATE1** | 2.16460 | 2.64152 | 2.00971 | 5.70082 | 0.56679 | 16.69310 | 3.82306 | 0.64156 | 8.49582 | 5.70392 | 4.61411 | 2.16532 |
| **ATE1-AS1** | 0.00912 | 0.02018 | 0.02105 | 0.04068 | 0.03045 | 0.00827 | 0.03369 | 0.01711 | 0.04374 | 0.01953 | 0.01765 | 0.02254 |
| **ATF1** | 0.31323 | 2.63237 | 8.34188 | 4.33943 | 0.00000 | 15.63624 | 17.46561 | 9.21168 | 14.20093 | 10.51393 | 7.05891 | 4.91407 |
| **ATF2** | 9.86579 | 1.98863 | 1.72325 | 1.85185 | 1.61328 | 7.39668 | 4.44939 | 6.53073 | 6.52181 | 1.69782 | 6.00125 | 3.35221 |
| **ATF3** | 2.68142 | 9.46449 | 4.17258 | 1.85243 | 6.59831 | 3.62331 | 5.84904 | 2.05914 | 3.80577 | 0.00000 | 0.68117 | 18.90965 |
| **ATF4** | 6.95331 | 17.25345 | 21.61756 | 17.02809 | 21.77080 | 17.84571 | 30.77351 | 9.91073 | 37.78900 | 23.21724 | 18.65759 | 19.22904 |
| **ATF5** | 1.01492 | 2.14573 | 1.64279 | 1.77255 | 10.73955 | 0.03698 | 11.46098 | 0.11513 | 0.00000 | 13.51364 | 2.24114 | 4.26971 |
| **ATF6** | 3.44636 | 8.95656 | 5.55171 | 0.91816 | 4.27466 | 22.96399 | 0.00000 | 0.78544 | 12.69475 | 7.76761 | 4.22754 | 2.80301 |
| **ATF6B** | 1.12619 | 2.37284 | 4.04489 | 2.41676 | 1.18648 | 0.00000 | 0.00000 | 0.14960 | 4.78150 | 0.01570 | 5.88665 | 4.02272 |
| **ATF7** | 2.97130 | 2.60360 | 2.68438 | 2.34686 | 1.47924 | 3.58565 | 4.14754 | 3.33982 | 3.72693 | 3.05443 | 3.23155 | 0.73270 |
| **ATF7IP** | 6.35542 | 14.95324 | 7.72226 | 10.31459 | 1.21949 | 23.16425 | 12.35193 | 10.38652 | 12.60922 | 10.68966 | 4.32080 | 17.51149 |
| **ATF7IP2** | 1.71869 | 0.18503 | 5.31123 | 0.00000 | 0.00000 | 1.22453 | 3.02610 | 0.28875 | 4.52283 | 3.43844 | 2.31969 | 0.43392 |
| **ATG10** | 0.28204 | 5.66272 | 0.00000 | 4.41000 | 3.43238 | 6.84729 | 5.67279 | 0.62728 | 4.40122 | 6.85063 | 8.66272 | 9.12666 |
| **ATG101** | 2.25381 | 17.27469 | 12.31500 | 8.31812 | 14.85266 | 0.00000 | 18.89507 | 24.93904 | 14.96807 | 0.55681 | 4.84172 | 24.60046 |
| **ATG12** | 0.00000 | 8.14472 | 13.22533 | 6.18700 | 4.46247 | 18.77820 | 12.78177 | 21.55017 | 6.52936 | 7.51997 | 6.98705 | 15.69236 |
| **ATG13** | 3.64095 | 15.85041 | 13.88095 | 12.20611 | 9.68660 | 0.00000 | 9.90029 | 17.24979 | 17.99004 | 5.60613 | 9.42024 | 25.31466 |
| **ATG14** | 7.74210 | 6.93364 | 1.72373 | 0.80706 | 9.27975 | 10.79407 | 5.22812 | 17.88352 | 3.00554 | 17.22298 | 4.58083 | 19.20911 |
| **ATG16L1** | 2.19193 | 14.80487 | 8.64069 | 4.52343 | 4.43026 | 4.91000 | 5.37082 | 11.43445 | 14.24075 | 4.56730 | 16.10285 | 22.23604 |
| **ATG16L2** | 0.12984 | 0.20111 | 0.23880 | 0.34082 | 0.00000 | 0.00000 | 0.00000 | 0.11825 | 0.00000 | 0.11716 | 0.19985 | 0.00000 |
| **ATG2A** | 1.10396 | 22.61343 | 4.52213 | 0.00000 | 6.47944 | 0.00000 | 1.03791 | 2.91915 | 4.69773 | 18.93683 | 0.02299 | 29.08410 |
| **ATG2B** | 3.57796 | 0.55505 | 3.96872 | 0.81099 | 0.54215 | 2.84566 | 1.28524 | 1.62648 | 3.06948 | 4.15501 | 0.08032 | 0.23782 |
| **ATG3** | 2.19337 | 39.84193 | 12.46045 | 32.37854 | 14.48256 | 47.11404 | 16.84708 | 25.34293 | 6.68096 | 6.35314 | 16.00768 | 45.87030 |
| **ATG4A** | 3.64938 | 2.48695 | 3.43527 | 0.44047 | 1.52153 | 7.19808 | 2.06992 | 2.81318 | 2.02419 | 0.00000 | 2.12397 | 2.53409 |
| **ATG4B** | 1.91962 | 10.87467 | 12.15496 | 9.80866 | 16.29240 | 0.00044 | 14.03227 | 2.63846 | 19.64655 | 3.96718 | 8.60312 | 10.88971 |
| **ATG4C** | 0.41508 | 3.72403 | 1.25274 | 0.47471 | 0.00000 | 13.13289 | 3.30163 | 0.09754 | 2.60825 | 1.21833 | 4.26959 | 3.85464 |
| **ATG4D** | 0.83043 | 1.13656 | 0.92742 | 2.15124 | 1.65048 | 0.00000 | 1.38899 | 0.65708 | 0.59208 | 0.00005 | 0.86247 | 0.88941 |
| **ATG5** | 4.05264 | 17.01163 | 14.23281 | 17.08702 | 3.98398 | 41.78115 | 15.99862 | 19.00729 | 17.73010 | 0.00000 | 12.27047 | 32.50304 |
| **ATG7** | 2.57761 | 5.87146 | 4.95773 | 3.97731 | 2.66145 | 8.46983 | 2.41961 | 3.74295 | 6.99863 | 0.00000 | 4.68974 | 9.61644 |
| **ATG9A** | 1.28690 | 15.88022 | 10.25662 | 10.05451 | 9.75057 | 0.00000 | 1.52592 | 8.18760 | 23.86228 | 9.86325 | 4.53135 | 32.82439 |
| **ATG9B** | 0.03086 | 10.62741 | 2.60477 | 4.64086 | 7.15241 | 0.00000 | 0.00000 | 3.50727 | 0.50706 | 2.51039 | 0.00000 | 18.18041 |
| **ATHL1** | 0.05394 | 0.10673 | 0.36784 | 0.62718 | 0.18437 | 0.00000 | 0.06491 | 0.00356 | 0.18325 | 0.58675 | 0.05573 | 0.04136 |
| **ATIC** | 2.35763 | 21.98782 | 25.85456 | 10.65133 | 15.40667 | 27.71656 | 30.10568 | 1.01818 | 31.33216 | 5.17764 | 26.69113 | 20.38968 |
| **ATL1** | 0.00049 | 0.00000 | 0.33079 | 0.09569 | 0.04960 | 0.31595 | 0.09928 | 0.04696 | 0.00000 | 0.22689 | 0.20463 | 0.09900 |
| **ATL2** | 6.33880 | 12.53694 | 5.64082 | 10.66167 | 1.54024 | 22.88731 | 0.00000 | 9.88346 | 18.86913 | 6.89343 | 2.81828 | 11.43909 |
| **ATL3** | 8.29052 | 6.35366 | 1.68958 | 6.45825 | 5.18982 | 14.44703 | 0.83409 | 2.17905 | 0.84044 | 1.64344 | 5.55265 | 9.53440 |
| **ATM** | 1.44663 | 0.00000 | 0.36364 | 1.93729 | 0.47276 | 3.12169 | 0.25982 | 3.11693 | 3.34202 | 8.62333 | 1.16387 | 2.04440 |
| **ATMIN** | 4.79388 | 3.69871 | 7.69242 | 0.00000 | 6.21222 | 11.52610 | 10.70196 | 0.95525 | 0.00000 | 22.35942 | 4.57529 | 4.96048 |
| **ATN1** | 1.40601 | 0.89597 | 6.30454 | 3.11693 | 5.88590 | 0.00000 | 2.93657 | 0.09329 | 6.09162 | 14.54907 | 1.72458 | 0.82591 |
| **ATOH7** | 0.00351 | 0.07565 | 0.04850 | 0.00000 | 0.01697 | 0.01974 | 0.00884 | 0.13439 | 0.01913 | 0.02344 | 0.01195 | 0.41844 |
| **ATOH8** | 0.01856 | 0.01499 | 0.08856 | 0.03306 | 0.01282 | 0.00000 | 0.02081 | 0.00580 | 0.04034 | 0.00000 | 0.04188 | 0.11843 |
| **ATOX1** | 1.51502 | 12.15802 | 15.28323 | 10.71762 | 17.52556 | 5.71688 | 18.87707 | 4.84261 | 8.09682 | 6.74030 | 11.12705 | 12.98635 |
| **ATP10A** | 0.66025 | 0.00000 | 0.01564 | 0.07812 | 0.00695 | 0.00000 | 0.00602 | 4.04505 | 0.04578 | 0.02841 | 0.02837 | 0.32902 |
| **ATP10B** | 0.37554 | 0.00421 | 0.89486 | 0.01451 | 0.02923 | 0.05626 | 0.04752 | 0.11773 | 0.49410 | 0.00000 | 0.09101 | 0.03462 |
| **ATP10D** | 4.46772 | 0.01227 | 1.98642 | 0.00522 | 0.00466 | 0.00000 | 0.00043 | 1.31384 | 0.00000 | 0.00870 | 0.10570 | 0.00813 |
| **ATP11A** | 3.74290 | 9.69753 | 2.22669 | 8.46233 | 24.48746 | 18.83898 | 4.56305 | 0.00000 | 16.19597 | 40.58566 | 12.63173 | 2.66407 |
| **ATP11A-AS1** | 0.00184 | 0.04844 | 0.00000 | 0.47857 | 2.92977 | 0.00000 | 0.00123 | 0.00000 | 0.38182 | 4.55791 | 0.50049 | 0.54918 |
| **ATP11AUN** | 0.00215 | 0.58399 | 0.09661 | 0.20191 | 0.80066 | 0.00000 | 0.00147 | 0.21573 | 0.00016 | 0.05793 | 0.82293 | 0.61697 |

| | | | | | | | | | | | | |
|---|---|---|---|---|---|---|---|---|---|---|---|---|
| **ATP11B** | 3.08647 | 1.20690 | 0.00000 | 0.00000 | 4.80913 | 13.49899 | 0.00000 | 0.91923 | 9.25377 | 11.17813 | 7.40897 | 5.25634 |
| **ATP11C** | 0.12362 | 0.00000 | 3.24418 | 7.91615 | 2.40198 | 14.36709 | 0.99328 | 0.22822 | 9.67798 | 20.10968 | 6.64892 | 0.82696 |
| **ATP12A** | 0.00132 | 0.02407 | 1.39618 | 1.73271 | 0.00000 | 0.46174 | 0.00000 | 0.00868 | 0.16481 | 0.15713 | 0.00000 | 0.00699 |
| **ATP13A1** | 1.53901 | 5.49882 | 8.02580 | 7.92100 | 8.35331 | 0.00000 | 0.02440 | 0.34175 | 13.47750 | 19.06069 | 2.75941 | 5.71166 |
| **ATP13A2** | 0.31695 | 14.21431 | 2.38085 | 7.89925 | 31.59740 | 0.00000 | 0.12293 | 0.00000 | 10.93360 | 10.97931 | 7.30926 | 4.04143 |
| **ATP13A3** | 4.39410 | 5.06237 | 5.35232 | 0.00000 | 1.08995 | 33.26656 | 5.24814 | 2.19884 | 7.90765 | 35.73808 | 2.27612 | 3.87609 |
| **ATP13A4** | 0.16373 | 0.01460 | 0.02360 | 0.01307 | 0.01542 | 0.01299 | 0.01997 | 0.00590 | 0.01793 | 0.01420 | 0.01493 | 0.01578 |
| **ATP13A5** | 0.23869 | 0.01459 | 0.01395 | 0.01273 | 0.01517 | 0.01445 | 0.01760 | 0.14103 | 0.01985 | 0.01148 | 0.01422 | 0.00803 |
| **ATP1A1** | 8.56053 | 25.12571 | 17.48156 | 10.66739 | 42.66344 | 21.79237 | 9.98157 | 6.73745 | 28.51793 | 43.82300 | 16.12323 | 22.61961 |
| **ATP1A1-AS1** | 0.45801 | 0.66729 | 0.47735 | 0.27831 | 1.83425 | 0.34043 | 0.08181 | 0.02736 | 0.66137 | 1.28392 | 0.46505 | 0.61546 |
| **ATP1A2** | 0.00556 | 0.00364 | 0.04827 | 0.04606 | 0.00000 | 0.03095 | 0.03516 | 0.01804 | 0.00000 | 0.15521 | 0.03739 | 0.02058 |
| **ATP1A3** | 0.02687 | 0.01028 | 0.14380 | 0.70007 | 0.09497 | 0.00000 | 0.16370 | 0.00963 | 0.07861 | 0.09736 | 0.00472 | 0.01619 |
| **ATP1A4** | 0.00418 | 0.01443 | 1.83147 | 0.00412 | 0.01648 | 0.03194 | 0.00000 | 0.04597 | 0.00000 | 0.01168 | 0.04301 | 0.07153 |
| **ATP1B1** | 3.93486 | 11.55660 | 0.00000 | 18.62998 | 23.09030 | 24.19923 | 0.00000 | 4.53674 | 0.00000 | 24.89243 | 4.31464 | 3.72970 |
| **ATP1B2** | 0.00760 | 0.71597 | 0.02810 | 0.00801 | 0.11283 | 0.14875 | 0.00000 | 2.67022 | 0.00000 | 0.06947 | 0.01744 | 6.81776 |
| **ATP1B3** | 1.90510 | 28.02270 | 24.06226 | 7.34272 | 21.94665 | 50.77998 | 13.49353 | 8.11798 | 47.75194 | 19.34331 | 19.36678 | 23.81161 |
| **ATP2A1** | 0.09389 | 0.98742 | 0.32267 | 0.49612 | 0.31241 | 0.00000 | 0.39074 | 0.00862 | 0.16515 | 0.74510 | 0.00000 | 1.33697 |
| **ATP2A2** | 8.59764 | 2.46042 | 6.96450 | 1.18372 | 5.02115 | 13.18284 | 3.70445 | 6.29648 | 8.82204 | 37.98049 | 1.80904 | 8.54358 |
| **ATP2A3** | 1.21283 | 0.13038 | 0.07799 | 0.00939 | 0.19107 | 0.00000 | 0.32269 | 0.58633 | 0.50012 | 0.00000 | 0.00000 | 1.17123 |
| **ATP2B1** | 9.95120 | 2.66330 | 11.51175 | 10.32238 | 10.72286 | 40.10188 | 14.03564 | 4.64419 | 45.15827 | 52.02531 | 21.92545 | 8.30790 |
| **ATP2B2** | 0.13281 | 0.00217 | 0.04494 | 0.09360 | 0.02891 | 0.06915 | 0.00000 | 0.00344 | 0.01691 | 0.10729 | 0.03005 | 0.00899 |
| **ATP2B3** | 0.01943 | 0.00499 | 0.05506 | 0.01552 | 0.00000 | 0.07026 | 0.00000 | 0.01027 | 0.00823 | 0.08589 | 0.01180 | 0.04192 |
| **ATP2B4** | 6.88734 | 0.42116 | 5.14148 | 4.62466 | 4.96431 | 3.31672 | 1.59999 | 1.29854 | 5.19755 | 3.30581 | 3.13514 | 0.00000 |
| **ATP2C1** | 3.22207 | 6.40396 | 10.12106 | 4.24561 | 2.98527 | 21.24843 | 0.00000 | 2.15652 | 12.61006 | 12.93878 | 5.63357 | 3.75321 |
| **ATP2C2** | 2.91573 | 1.19097 | 4.16988 | 0.00000 | 1.19611 | 0.00000 | 1.99420 | 1.14568 | 6.44673 | 0.85101 | 1.21819 | 0.28005 |
| **ATP4A** | 0.00505 | 0.05385 | 0.04794 | 0.00528 | 0.02632 | 0.00000 | 0.04971 | 0.01051 | 0.05390 | 0.07689 | 0.01241 | 0.01409 |
| **ATP4B** | 0.00510 | 0.00840 | 0.01712 | 0.00792 | 0.05151 | 0.02553 | 0.02029 | 0.00749 | 0.04807 | 0.04914 | 0.00868 | 0.08904 |
| **ATP5A1** | 8.80436 | 40.66449 | 34.41560 | 37.85344 | 39.08235 | 60.54919 | 53.46108 | 12.31931 | 56.55073 | 3.09356 | 51.28915 | 29.45151 |
| **ATP5B** | 8.53105 | 50.24246 | 48.10984 | 48.95433 | 50.74021 | 67.51676 | 72.04769 | 8.59630 | 74.95337 | 12.31563 | 62.34465 | 23.18054 |
| **ATP5C1** | 4.36171 | 20.46466 | 24.31567 | 23.37687 | 21.23557 | 52.92480 | 37.35107 | 7.79550 | 32.28931 | 5.99240 | 37.61522 | 12.73808 |
| **ATP5D** | 3.40143 | 23.12852 | 19.33071 | 21.17326 | 34.36817 | 0.00000 | 30.82589 | 2.12185 | 36.11514 | 16.64660 | 25.52090 | 7.77622 |
| **ATP5E** | 0.09082 | 8.51232 | 7.20836 | 6.87928 | 15.39878 | 5.48470 | 13.07182 | 1.01140 | 8.06723 | 4.77303 | 8.18454 | 8.64083 |
| **ATP5EP2** | 0.15975 | 11.72136 | 11.67881 | 10.21860 | 19.72580 | 15.16784 | 22.89920 | 1.38469 | 10.93272 | 14.04109 | 14.44310 | 12.30674 |
| **ATP5F1** | 0.76856 | 23.74654 | 20.99554 | 17.31263 | 20.30734 | 48.39991 | 28.11793 | 4.16463 | 29.35278 | 0.00000 | 29.17441 | 19.07422 |
| **ATP5G1** | 0.10814 | 13.10597 | 18.68273 | 15.93472 | 20.20329 | 7.40784 | 24.15208 | 0.81706 | 16.94897 | 6.22892 | 25.28743 | 7.62064 |
| **ATP5G2** | 3.82725 | 16.79993 | 8.07364 | 17.17781 | 26.01415 | 15.63443 | 13.09764 | 2.15612 | 14.95652 | 10.29255 | 20.88240 | 13.57566 |
| **ATP5G3** | 7.78548 | 39.69423 | 34.77255 | 38.41981 | 46.65029 | 59.65650 | 51.38527 | 10.02152 | 58.50530 | 20.04144 | 51.01848 | 29.78219 |
| **ATP5H** | 0.72423 | 10.63670 | 12.89525 | 10.19334 | 8.30526 | 23.88284 | 15.47803 | 0.79753 | 7.63020 | 10.07380 | 16.89420 | 12.79368 |
| **ATP5I** | 0.54842 | 26.65419 | 17.67558 | 25.93588 | 33.09321 | 28.13531 | 48.30887 | 1.46810 | 20.01779 | 34.50910 | 34.19350 | 22.73695 |
| **ATP5J** | 3.20907 | 23.56795 | 24.98833 | 18.52891 | 20.56342 | 48.49734 | 41.63390 | 5.95409 | 22.42819 | 15.29734 | 36.19812 | 18.02637 |
| **ATP5J2** | 2.18021 | 21.33149 | 16.11866 | 17.98973 | 23.20939 | 11.75145 | 35.10481 | 2.31836 | 21.21732 | 13.73020 | 24.63143 | 14.11458 |
| **ATP5L** | 4.81557 | 22.40363 | 21.86695 | 22.05389 | 27.64465 | 42.49788 | 37.45872 | 4.99643 | 23.28940 | 24.22751 | 30.79313 | 22.48615 |
| **ATP5L2** | 0.00265 | 0.03684 | 0.01267 | 0.04566 | 0.26387 | 0.07944 | 0.18558 | 0.01191 | 0.00000 | 0.43368 | 0.06492 | 0.02980 |
| **ATP5O** | 3.05905 | 33.56895 | 25.06744 | 30.18750 | 36.67573 | 47.80824 | 41.37373 | 1.83163 | 36.22953 | 13.94341 | 40.88413 | 19.87289 |
| **ATP5S** | 1.34698 | 1.24687 | 2.60717 | 2.41850 | 2.91684 | 4.47666 | 1.39962 | 0.36432 | 1.06079 | 0.03465 | 3.40504 | 3.45059 |
| **ATP5SL** | 3.11516 | 11.83527 | 9.62100 | 10.47727 | 12.19377 | 4.18759 | 17.13855 | 0.00000 | 13.95884 | 0.00000 | 9.10491 | 12.89650 |
| **ATP6AP1** | 1.68017 | 27.95633 | 15.44538 | 11.58540 | 28.28421 | 20.97343 | 1.15765 | 3.66302 | 42.24700 | 4.47314 | 23.03931 | 15.06588 |
| **ATP6AP1L** | 0.63570 | 0.10040 | 0.04355 | 0.14043 | 0.36005 | 0.00000 | 0.01562 | 0.27241 | 0.00000 | 0.18492 | 0.23818 | 0.12940 |

| | | | | | | | | | | | | |
|---|---|---|---|---|---|---|---|---|---|---|---|---|
| **ATP6AP2** | 0.41421 | 12.71190 | 6.22213 | 6.95205 | 9.05512 | 28.35495 | 0.00000 | 4.60284 | 15.68411 | 10.67848 | 12.87797 | 10.26682 |
| **ATP6V0A1** | 1.73820 | 5.34163 | 5.65994 | 6.33477 | 9.57522 | 0.00000 | 0.00000 | 1.11036 | 9.25453 | 6.56592 | 0.80647 | 5.12019 |
| **ATP6V0A2** | 0.01599 | 2.69176 | 0.94612 | 2.13134 | 1.00860 | 4.90306 | 0.00000 | 0.58724 | 3.86791 | 4.49238 | 1.13297 | 0.84884 |
| **ATP6V0A4** | 1.33143 | 13.47896 | 0.75171 | 0.00000 | 35.69491 | 2.00818 | 4.72125 | 0.23899 | 32.97452 | 0.00000 | 23.00425 | 0.00000 |
| **ATP6V0B** | 10.53640 | 32.95314 | 32.36006 | 21.93119 | 41.13744 | 23.20057 | 20.99974 | 10.07801 | 44.33264 | 13.77681 | 23.26272 | 21.53144 |
| **ATP6V0C** | 2.13993 | 9.53548 | 16.62308 | 3.78843 | 29.07741 | 0.00000 | 15.07729 | 1.70551 | 18.66254 | 39.27695 | 7.58850 | 6.90372 |
| **ATP6V0CP3** | 0.00524 | 0.00871 | 0.08152 | 0.01615 | 0.00257 | 0.05669 | 0.01721 | 0.01760 | 0.00000 | 0.04923 | 0.02000 | 0.01108 |
| **ATP6V0D1** | 8.08160 | 15.54641 | 20.97237 | 9.53356 | 17.02645 | 4.88182 | 21.67988 | 9.67767 | 22.27992 | 0.00000 | 17.32429 | 8.73504 |
| **ATP6V0D2** | 0.01947 | 16.33832 | 0.53479 | 0.00038 | 41.59735 | 0.16327 | 0.00000 | 0.00417 | 0.00000 | 0.00697 | 10.83575 | 0.00000 |
| **ATP6V0E1** | 3.44861 | 19.80921 | 26.84219 | 13.25827 | 22.49420 | 35.68771 | 19.80882 | 11.41544 | 31.65396 | 9.79731 | 18.84484 | 21.82716 |
| **ATP6V0E2** | 2.28018 | 1.19639 | 1.19321 | 0.29717 | 2.91259 | 0.00000 | 1.38030 | 0.09216 | 0.00000 | 1.61501 | 5.06444 | 0.14102 |
| **ATP6V0E2-AS1** | 0.00651 | 0.01398 | 0.02120 | 0.00901 | 0.01241 | 0.02611 | 0.01063 | 0.01037 | 0.11831 | 0.00503 | 0.00407 | 0.01668 |
| **ATP6V1A** | 0.65545 | 14.10968 | 20.37809 | 5.70307 | 16.99659 | 39.65265 | 15.90427 | 0.77859 | 15.53790 | 8.51357 | 20.56117 | 6.73741 |
| **ATP6V1B1** | 0.00561 | 11.90651 | 0.00000 | 0.70815 | 34.91859 | 0.10695 | 7.69649 | 0.00482 | 15.26674 | 0.00000 | 26.20907 | 0.00000 |
| **ATP6V1B2** | 10.11814 | 13.54099 | 21.35611 | 5.20473 | 6.33770 | 29.53976 | 19.28116 | 7.32612 | 14.04040 | 0.00000 | 11.50317 | 9.49961 |
| **ATP6V1C1** | 4.86506 | 10.97838 | 15.04671 | 4.02732 | 0.20593 | 29.76265 | 12.43834 | 6.62028 | 15.22057 | 0.00000 | 10.79320 | 13.59541 |
| **ATP6V1C2** | 1.50864 | 7.09950 | 15.00779 | 0.00000 | 13.37996 | 1.24041 | 11.23059 | 0.03964 | 11.82811 | 6.45262 | 10.25476 | 0.00000 |
| **ATP6V1D** | 5.84958 | 10.82989 | 13.18103 | 9.52272 | 12.02495 | 33.53222 | 17.34511 | 13.99201 | 9.71412 | 2.60293 | 14.40326 | 16.59275 |
| **ATP6V1E1** | 3.11364 | 18.22026 | 15.37115 | 9.96828 | 10.11817 | 28.04138 | 15.41218 | 6.93083 | 8.71627 | 4.52821 | 13.12922 | 20.41569 |
| **ATP6V1E2** | 0.01577 | 0.01073 | 0.09024 | 0.01203 | 0.16169 | 0.10050 | 0.00000 | 0.00874 | 0.00000 | 0.02907 | 0.00912 | 0.17934 |
| **ATP6V1F** | 5.62686 | 21.02265 | 19.26339 | 17.45043 | 21.73372 | 5.89077 | 21.33356 | 1.54986 | 21.00946 | 0.14796 | 22.84869 | 16.69121 |
| **ATP6V1G1** | 5.89227 | 16.79645 | 23.50417 | 16.36862 | 9.65892 | 50.76726 | 31.75536 | 8.21176 | 23.68140 | 15.72481 | 22.87841 | 16.39149 |
| **ATP6V1G2** | 0.00173 | 0.10966 | 0.00000 | 0.12068 | 0.01873 | 0.15574 | 0.01713 | 0.21884 | 0.04805 | 0.00000 | 0.00096 | 0.13132 |
| **ATP6V1H** | 2.87541 | 7.58506 | 11.48313 | 3.93943 | 7.55819 | 22.43023 | 12.21982 | 0.00000 | 12.77623 | 0.00000 | 12.50144 | 4.26992 |
| **ATP7A** | 1.71855 | 0.02823 | 0.75507 | 2.42946 | 0.00000 | 2.96184 | 0.00000 | 0.17954 | 0.52280 | 1.09266 | 0.00000 | 1.04164 |
| **ATP7B** | 1.60879 | 1.06870 | 12.10951 | 2.13175 | 2.34667 | 0.00000 | 1.82356 | 0.00000 | 14.32369 | 3.89307 | 3.24126 | 0.15748 |
| **ATP8A1** | 3.40784 | 0.11034 | 0.11247 | 0.63189 | 0.04534 | 0.00000 | 0.03741 | 1.48741 | 0.07388 | 0.29367 | 0.03221 | 0.03735 |
| **ATP8A2** | 0.32780 | 0.00000 | 2.23967 | 1.39925 | 0.00000 | 2.04519 | 0.71566 | 0.29849 | 1.35534 | 0.00000 | 2.39872 | 0.42278 |
| **ATP8B1** | 4.54991 | 3.09397 | 0.00000 | 0.00000 | 16.75298 | 8.51535 | 1.92568 | 5.53717 | 4.97391 | 18.77043 | 6.00834 | 6.29705 |
| **ATP8B2** | 0.03014 | 0.26241 | 0.91977 | 6.49645 | 0.00000 | 0.39254 | 0.18481 | 0.00000 | 0.00000 | 0.64036 | 0.08513 | 0.14546 |
| **ATP8B3** | 0.01065 | 0.00894 | 0.10174 | 0.44776 | 0.00000 | 0.04619 | 0.01676 | 0.19379 | 0.15166 | 0.00000 | 0.00000 | 0.92543 |
| **ATP8B4** | 0.24986 | 0.11030 | 0.27041 | 0.03844 | 0.00000 | 0.06999 | 0.09271 | 0.01228 | 0.00000 | 0.11324 | 0.02620 | 0.05069 |
| **ATP8B5P** | 0.48436 | 0.30119 | 0.54519 | 0.60001 | 0.19413 | 1.16069 | 0.40096 | 0.67744 | 0.42014 | 0.12841 | 0.44206 | 0.47365 |
| **ATP9A** | 5.56622 | 1.98705 | 0.17090 | 0.24244 | 4.88028 | 0.58447 | 1.52723 | 0.33023 | 0.00000 | 2.70065 | 0.58435 | 0.01737 |
| **ATP9B** | 0.22966 | 2.74857 | 1.79155 | 4.17727 | 2.02516 | 0.00000 | 0.58087 | 3.94381 | 2.61781 | 2.00724 | 0.63717 | 6.02015 |
| **ATPAF1** | 0.82421 | 3.09413 | 0.00000 | 3.19790 | 3.80111 | 6.03268 | 1.24061 | 0.05927 | 0.00000 | 6.33808 | 1.27455 | 1.09994 |
| **ATPAF2** | 3.63859 | 5.69843 | 6.52508 | 8.33182 | 4.98455 | 5.15587 | 8.84503 | 0.62339 | 6.55362 | 0.00000 | 9.55364 | 2.04255 |
| **ATPIF1** | 3.08811 | 12.96462 | 18.74492 | 15.18317 | 12.46352 | 20.73796 | 27.41358 | 24.66054 | 13.62372 | 13.22783 | 11.90119 | 23.32370 |
| **ATR** | 1.14259 | 0.91059 | 0.06392 | 0.47488 | 0.00000 | 4.71645 | 0.00000 | 0.47324 | 0.36540 | 3.44258 | 1.43896 | 1.22741 |
| **ATRAID** | 1.81366 | 11.97382 | 6.98410 | 14.43731 | 14.82008 | 15.68722 | 0.00000 | 0.35362 | 6.41316 | 11.82928 | 11.87239 | 6.01570 |
| **ATRIP** | 0.39422 | 2.46316 | 2.43671 | 1.68251 | 5.58444 | 1.07561 | 2.26762 | 0.80664 | 2.26541 | 4.22691 | 1.82132 | 1.70386 |
| **ATRN** | 0.18440 | 1.73624 | 1.84884 | 1.28879 | 2.03693 | 4.78161 | 0.00000 | 0.84471 | 8.67030 | 12.19417 | 1.55619 | 1.21356 |
| **ATRNL1** | 2.04836 | 0.01080 | 0.00875 | 0.02192 | 0.01976 | 0.01534 | 0.06633 | 0.27118 | 0.00381 | 0.00000 | 0.02035 | 0.02109 |
| **ATRX** | 7.02303 | 0.20396 | 3.61335 | 8.09807 | 2.57626 | 16.66453 | 7.97056 | 3.67789 | 6.87637 | 29.23615 | 5.02540 | 3.30215 |
| **ATXN1** | 1.27662 | 0.11997 | 0.10510 | 0.14818 | 0.04926 | 0.00508 | 0.03780 | 0.06656 | 0.00000 | 0.20350 | 0.01690 | 0.34396 |
| **ATXN10** | 1.66555 | 10.52623 | 8.19987 | 10.11996 | 4.42138 | 16.85714 | 7.02417 | 0.14315 | 2.94325 | 1.28870 | 10.05595 | 6.56733 |
| **ATXN1L** | 10.25420 | 3.79793 | 3.34947 | 1.91434 | 3.95944 | 2.67222 | 7.08677 | 4.38338 | 4.61650 | 6.52413 | 3.46425 | 6.95899 |
| **ATXN2** | 3.84380 | 7.58242 | 3.67682 | 8.30719 | 4.34245 | 10.82313 | 7.43436 | 3.06695 | 2.75909 | 15.06953 | 3.66509 | 6.23868 |

| | | | | | | | | | | | | |
|---|---|---|---|---|---|---|---|---|---|---|---|---|
| **ATXN2L** | 7.53356 | 4.19701 | 12.76317 | 8.29765 | 9.51276 | 0.00000 | 13.58705 | 1.25990 | 11.32366 | 30.20234 | 6.78838 | 7.77368 |
| **ATXN3** | 0.73701 | 3.98875 | 2.99127 | 3.30827 | 1.35313 | 16.70801 | 0.00000 | 2.15509 | 5.70064 | 5.06989 | 1.03674 | 4.69584 |
| **ATXN3L** | 0.01246 | 0.00007 | 0.03865 | 0.09258 | 0.02771 | 0.13832 | 0.00000 | 0.00742 | 0.00000 | 0.02959 | 0.01674 | 0.01708 |
| **ATXN7** | 2.88347 | 0.39783 | 1.54817 | 1.45441 | 1.84706 | 2.22199 | 5.08539 | 0.19388 | 0.00000 | 10.78549 | 1.73826 | 1.60757 |
| **ATXN7L1** | 1.17301 | 0.78799 | 0.16112 | 2.64032 | 0.83692 | 1.56467 | 0.57638 | 0.42009 | 1.07163 | 1.37348 | 0.77663 | 1.25617 |
| **ATXN7L2** | 3.04844 | 1.50430 | 1.93956 | 1.06223 | 0.61762 | 0.00000 | 1.21632 | 0.12987 | 2.57326 | 0.00000 | 1.49232 | 1.75552 |
| **ATXN7L3** | 0.91687 | 1.35937 | 5.11807 | 3.76020 | 5.58528 | 0.00000 | 4.62305 | 0.54719 | 2.57459 | 12.93359 | 0.42388 | 1.38488 |
| **ATXN7L3B** | 4.02760 | 7.53930 | 10.42528 | 1.27865 | 1.46784 | 14.38351 | 6.66326 | 4.13993 | 9.26974 | 1.96962 | 10.13580 | 10.90667 |
| **ATXN8OS** | 0.01392 | 0.01265 | 0.01793 | 0.01285 | 0.01133 | 0.03995 | 0.03378 | 0.00861 | 0.00986 | 0.04131 | 0.02435 | 0.02000 |
| **AUH** | 0.42598 | 3.53582 | 3.63823 | 1.90946 | 0.86624 | 4.22458 | 2.12140 | 0.38354 | 2.77641 | 3.60815 | 4.11019 | 1.71822 |
| **AUNIP** | 4.88867 | 8.31190 | 5.96361 | 0.98853 | 7.33180 | 15.90943 | 5.25219 | 4.79625 | 5.80772 | 0.00014 | 8.25200 | 8.72264 |
| **AUP1** | 3.17293 | 22.09958 | 19.63520 | 19.23775 | 24.01433 | 10.45976 | 11.84473 | 2.89152 | 45.30620 | 9.24640 | 20.75011 | 13.95544 |
| **AURKA** | 18.45954 | 25.37240 | 15.64868 | 6.63504 | 15.57728 | 31.54139 | 31.88559 | 8.74227 | 20.78743 | 0.00000 | 12.28934 | 28.58344 |
| **AURKAIP1** | 1.89602 | 27.28017 | 24.53473 | 20.82650 | 36.39398 | 7.07513 | 39.12476 | 9.73184 | 24.68945 | 18.23283 | 22.43213 | 23.28883 |
| **AURKAPS1** | 1.65984 | 1.69014 | 1.32113 | 0.25270 | 0.40979 | 2.75857 | 2.09417 | 1.02514 | 1.61016 | 1.08970 | 0.42314 | 2.15953 |
| **AURKB** | 4.61566 | 17.97405 | 18.51077 | 13.31182 | 23.26804 | 16.15002 | 18.44778 | 9.36625 | 31.09617 | 1.18235 | 15.07741 | 22.48556 |
| **AURKC** | 5.41220 | 2.78173 | 0.56223 | 0.00000 | 0.75440 | 0.37256 | 0.60910 | 10.90509 | 1.04081 | 0.00000 | 0.00000 | 11.35990 |
| **AUTS2** | 2.71325 | 0.65485 | 0.00000 | 0.68435 | 1.99472 | 1.19278 | 0.63145 | 0.32466 | 0.00000 | 3.56838 | 0.00000 | 1.43773 |
| **AVEN** | 2.83364 | 0.19729 | 2.23825 | 0.30601 | 0.76863 | 0.76300 | 1.73875 | 0.59328 | 0.14755 | 3.22679 | 0.43327 | 0.00000 |
| **AVIL** | 0.27619 | 0.16155 | 0.02128 | 0.09562 | 0.00000 | 0.12257 | 0.00000 | 0.75976 | 0.37472 | 0.43373 | 0.00000 | 0.87665 |
| **AVL9** | 2.56254 | 2.67015 | 5.64369 | 5.45012 | 2.70978 | 4.42603 | 3.60710 | 3.58959 | 1.98350 | 0.00000 | 5.98308 | 1.11037 |
| **AVP** | 0.00912 | 0.01624 | 0.01120 | 0.03377 | 0.01356 | 0.02490 | 0.01481 | 0.01450 | 0.01845 | 0.01420 | 0.02098 | 0.01951 |
| **AVPI1** | 1.03587 | 4.87708 | 16.42087 | 0.16948 | 2.38308 | 0.00000 | 7.55367 | 7.27344 | 2.36996 | 0.00000 | 5.01971 | 3.76423 |
| **AVPR1A** | 0.07793 | 0.25759 | 0.00577 | 0.06452 | 0.00000 | 0.00000 | 0.15530 | 0.00863 | 0.00000 | 0.02302 | 0.04252 | 0.00618 |
| **AXDND1** | 0.03926 | 0.14124 | 0.02375 | 0.00000 | 0.00000 | 0.42408 | 0.00000 | 0.20550 | 0.00000 | 0.12420 | 0.00111 | 0.40596 |
| **AXIN1** | 0.83002 | 2.77216 | 1.52605 | 1.30551 | 5.73962 | 0.00000 | 6.75662 | 2.54473 | 2.66174 | 13.67953 | 1.71835 | 1.02515 |
| **AXIN2** | 1.46422 | 1.96764 | 8.84750 | 5.55689 | 0.00000 | 0.01180 | 3.16504 | 0.59150 | 1.41442 | 4.02539 | 1.42448 | 0.73646 |
| **AXL** | 0.20916 | 0.09756 | 0.00000 | 0.43247 | 0.00000 | 5.87989 | 0.00000 | 0.11358 | 0.00000 | 4.18013 | 0.76741 | 0.24483 |
| **AZGP1** | 0.02586 | 0.02797 | 0.01738 | 0.01370 | 0.02877 | 0.00000 | 0.02402 | 0.18079 | 0.01679 | 0.01502 | 0.01518 | 0.03124 |
| **AZGP1P1** | 0.00592 | 0.02956 | 0.01170 | 0.01000 | 0.11839 | 0.00000 | 0.02846 | 0.03976 | 0.00000 | 0.05928 | 0.00971 | 0.01637 |
| **AZI2** | 3.53696 | 3.85856 | 6.15365 | 1.22253 | 1.12844 | 20.46471 | 7.54583 | 7.91527 | 4.67830 | 7.51619 | 3.67644 | 5.85033 |
| **AZIN1** | 5.44868 | 22.44894 | 19.72754 | 6.77532 | 4.89622 | 55.98204 | 19.89642 | 22.53004 | 23.63685 | 7.11666 | 18.74774 | 26.39018 |
| **AZIN2** | 1.60121 | 0.15427 | 0.14981 | 1.83116 | 0.19466 | 0.00000 | 0.01556 | 1.94225 | 0.06252 | 0.19440 | 0.17703 | 2.03661 |
| **AZU1** | 0.04264 | 0.01043 | 0.01499 | 0.00722 | 0.00222 | 0.02625 | 0.00118 | 0.00727 | 0.03942 | 0.08949 | 0.00722 | 0.01464 |
| **B2M** | 1.36159 | 0.00673 | 4.95146 | 13.39451 | 0.00001 | 5.80965 | 0.00000 | 1.99498 | 0.54706 | 0.00000 | 0.00317 | 1.37573 |
| **B3GALNT1** | 2.20871 | 0.56718 | 0.49066 | 0.03290 | 0.00000 | 0.34399 | 0.75991 | 0.41524 | 0.01160 | 0.00000 | 0.09007 | 0.06579 |
| **B3GALNT2** | 2.60331 | 3.40069 | 10.74040 | 0.84348 | 1.93711 | 11.02504 | 5.42143 | 2.74148 | 7.40760 | 5.37551 | 7.01042 | 3.91233 |
| **B3GALT1** | 0.07288 | 0.01724 | 0.07365 | 0.01007 | 0.01917 | 0.01868 | 0.00850 | 0.03589 | 0.04677 | 0.00000 | 0.01333 | 0.02416 |
| **B3GALT2** | 0.00162 | 0.00000 | 0.15424 | 0.00002 | 0.16359 | 0.20327 | 0.00000 | 0.08447 | 0.36669 | 0.66913 | 0.00000 | 0.22184 |
| **B3GALT4** | 0.04736 | 0.01802 | 0.43210 | 0.30674 | 0.39900 | 0.00000 | 0.01820 | 0.12978 | 0.10504 | 0.00000 | 0.17777 | 0.07632 |
| **B3GALT5** | 0.00914 | 0.01248 | 0.02936 | 0.01027 | 0.00896 | 0.02509 | 0.01538 | 0.05119 | 0.00660 | 0.01970 | 0.04994 | 0.01280 |
| **B3GALT6** | 0.02574 | 0.00000 | 2.37813 | 1.01457 | 1.09619 | 0.00000 | 4.34864 | 0.13288 | 0.74463 | 3.12853 | 0.71621 | 0.13046 |
| **B3GALTL** | 0.05726 | 0.83603 | 0.16015 | 0.95408 | 0.51713 | 2.42875 | 0.00000 | 0.13857 | 1.09489 | 0.49305 | 0.01407 | 0.00027 |
| **B3GAT1** | 0.03237 | 0.01301 | 0.02005 | 0.00907 | 0.01449 | 0.01997 | 0.03196 | 0.01039 | 0.00578 | 0.05315 | 0.01360 | 0.01367 |
| **B3GAT2** | 0.42777 | 0.07693 | 0.01484 | 0.00988 | 0.01307 | 0.01073 | 0.01961 | 0.13727 | 0.02445 | 0.00004 | 0.01223 | 0.07445 |
| **B3GAT3** | 0.66685 | 1.74752 | 5.88088 | 2.40463 | 0.93656 | 0.00000 | 2.20237 | 0.00000 | 5.12741 | 2.86498 | 1.92910 | 2.68250 |
| **B3GNT1** | 0.00000 | 5.15695 | 4.26088 | 3.30767 | 3.04103 | 0.00000 | 0.65879 | 0.15731 | 14.51549 | 3.85851 | 6.25325 | 1.94427 |
| **B3GNT2** | 22.89018 | 2.70229 | 8.24696 | 0.00000 | 5.74270 | 13.34148 | 0.00000 | 23.89912 | 30.32401 | 8.59551 | 22.46507 | 0.00000 |

| | | | | | | | | | | | | |
|---|---|---|---|---|---|---|---|---|---|---|---|---|
| **B3GNT3** | 0.06178 | 0.02726 | 0.02413 | 0.01058 | 0.00756 | 0.02202 | 0.01323 | 0.04482 | 0.01056 | 0.03029 | 0.02842 | 0.10539 |
| **B3GNT4** | 2.71494 | 0.30546 | 0.10588 | 0.43785 | 0.00000 | 0.00000 | 0.03290 | 0.61455 | 0.80080 | 0.17983 | 1.11192 | 0.21958 |
| **B3GNT5** | 1.12037 | 0.00000 | 11.68228 | 0.75942 | 0.00000 | 0.03865 | 3.11912 | 0.33305 | 5.31036 | 0.00167 | 0.29819 | 0.00000 |
| **B3GNT6** | 0.00885 | 0.02964 | 0.01545 | 0.04167 | 0.01076 | 0.01096 | 0.04929 | 0.01078 | 0.02870 | 0.05701 | 0.02063 | 0.01952 |
| **B3GNT7** | 0.00000 | 3.89794 | 0.29843 | 21.66548 | 8.60563 | 5.74126 | 0.00000 | 1.34469 | 0.00092 | 4.05645 | 20.98973 | 0.39200 |
| **B3GNT8** | 0.00604 | 0.04929 | 0.10157 | 0.05504 | 0.00000 | 0.00792 | 0.02266 | 0.00716 | 0.25404 | 0.01609 | 0.00415 | 0.09893 |
| **B3GNT9** | 0.12279 | 0.00925 | 0.05982 | 0.08468 | 0.00000 | 0.07398 | 0.00000 | 0.03461 | 0.01038 | 0.05840 | 0.01394 | 0.04687 |
| **B3GNTL1** | 0.00154 | 0.66127 | 0.71896 | 0.40777 | 0.81158 | 0.23794 | 0.28864 | 0.13273 | 0.17936 | 0.00000 | 1.68812 | 0.09936 |
| **B4GALNT1** | 0.05383 | 0.44818 | 0.41497 | 0.00346 | 0.00480 | 0.00000 | 0.00269 | 0.04855 | 0.16403 | 0.00000 | 0.05523 | 0.95452 |
| **B4GALNT2** | 0.00782 | 0.29053 | 0.00000 | 0.50096 | 0.42856 | 0.21318 | 0.00000 | 0.25482 | 0.00000 | 0.28865 | 0.00000 | 0.69820 |
| **B4GALNT3** | 0.01828 | 0.18707 | 0.72542 | 0.41700 | 1.58897 | 0.00000 | 0.01135 | 0.04750 | 2.20252 | 1.06725 | 0.00421 | 0.21681 |
| **B4GALNT4** | 0.00872 | 0.01648 | 0.21553 | 0.03269 | 0.01126 | 0.00443 | 0.00773 | 0.01091 | 0.00000 | 0.07926 | 0.03544 | 0.01457 |
| **B4GALT1** | 8.01943 | 14.69308 | 28.17394 | 14.61364 | 29.48442 | 26.14983 | 15.71440 | 15.37513 | 28.40021 | 15.09362 | 21.23594 | 7.35291 |
| **B4GALT1-AS1** | 0.00595 | 0.13271 | 0.35366 | 0.01335 | 0.25702 | 0.59993 | 0.00000 | 0.05187 | 0.00000 | 0.00000 | 0.20122 | 0.22953 |
| **B4GALT2** | 0.87041 | 1.26060 | 1.47979 | 1.25111 | 2.44459 | 0.00000 | 3.97971 | 0.01838 | 1.98209 | 2.29056 | 0.00000 | 0.23627 |
| **B4GALT3** | 4.54980 | 10.69152 | 18.62168 | 5.43297 | 12.27932 | 2.97855 | 7.98267 | 3.27843 | 19.92348 | 0.00003 | 11.78452 | 13.20009 |
| **B4GALT4** | 2.30471 | 6.31105 | 4.23219 | 9.59460 | 0.84406 | 15.82950 | 0.00000 | 1.23886 | 9.33666 | 1.44643 | 2.16877 | 2.75561 |
| **B4GALT4-AS1** | 0.00314 | 0.01227 | 0.01631 | 0.04789 | 0.06900 | 0.02092 | 0.00000 | 0.01968 | 0.06228 | 0.07676 | 0.00610 | 0.04447 |
| **B4GALT5** | 3.00966 | 7.38197 | 0.38680 | 2.17465 | 24.65376 | 15.15034 | 8.65661 | 1.12692 | 7.52663 | 29.10431 | 9.45383 | 4.50854 |
| **B4GALT6** | 1.63679 | 0.00000 | 0.00000 | 0.23578 | 0.24132 | 1.34589 | 1.15562 | 0.59294 | 0.51267 | 0.58318 | 1.90114 | 0.00000 |
| **B4GALT7** | 0.08777 | 4.73109 | 0.58745 | 1.92599 | 4.65050 | 0.00003 | 0.00000 | 0.00000 | 2.02914 | 3.59344 | 1.95606 | 2.60467 |
| **B9D1** | 1.88070 | 3.00644 | 0.80528 | 3.01127 | 1.62404 | 0.01359 | 0.10297 | 0.67633 | 0.29347 | 0.00000 | 2.32852 | 0.75760 |
| **B9D2** | 0.00283 | 2.30540 | 1.82954 | 2.09734 | 1.63771 | 0.00000 | 3.94785 | 5.02686 | 1.62373 | 0.00000 | 0.95683 | 6.29048 |
| **BAALC** | 3.12325 | 0.01234 | 0.01740 | 0.00968 | 0.01932 | 0.00201 | 0.01687 | 0.62877 | 0.02172 | 0.00000 | 0.01281 | 0.03632 |
| **BAAT** | 0.00857 | 0.01233 | 0.06830 | 0.10857 | 0.00000 | 0.35589 | 0.20666 | 0.53034 | 0.23759 | 0.05945 | 0.02849 | 0.19319 |
| **BABAM1** | 3.23086 | 14.48124 | 22.07497 | 14.66050 | 12.96695 | 6.55853 | 23.04858 | 4.88534 | 16.79783 | 0.00000 | 17.71647 | 13.40112 |
| **BACE1** | 0.10605 | 1.75305 | 0.02691 | 3.68230 | 0.49539 | 1.30540 | 0.24188 | 0.37826 | 0.00000 | 1.18398 | 0.47659 | 5.36678 |
| **BACE1-AS** | 0.03703 | 0.04723 | 0.05317 | 0.06116 | 0.00000 | 0.10490 | 0.00000 | 0.03897 | 0.11027 | 0.13900 | 0.10326 | 0.12533 |
| **BACE2** | 0.01196 | 0.04794 | 0.04849 | 0.03197 | 0.25304 | 0.00000 | 0.15948 | 0.00831 | 0.13016 | 0.02041 | 0.00000 | 0.14308 |
| **BACH1** | 12.52606 | 3.09365 | 3.71849 | 2.86953 | 0.00000 | 8.21851 | 5.27184 | 2.22325 | 3.23269 | 2.72639 | 2.25980 | 1.28118 |
| **BACH1-IT2** | 0.00368 | 0.00000 | 0.01734 | 0.00412 | 0.22499 | 0.03108 | 0.00000 | 0.00961 | 0.03099 | 0.20150 | 0.00000 | 0.01281 |
| **BACH2** | 3.04096 | 0.15042 | 0.03946 | 0.18091 | 0.00000 | 0.26509 | 0.04121 | 0.21282 | 0.00000 | 0.72617 | 0.09168 | 0.00218 |
| **BAD** | 0.92635 | 3.89553 | 6.68847 | 5.51111 | 3.33492 | 0.00000 | 6.61290 | 0.78665 | 3.05989 | 0.00000 | 3.14651 | 3.81420 |
| **BAG1** | 0.00848 | 5.27893 | 8.52404 | 4.61594 | 8.21066 | 1.36334 | 10.52577 | 2.11196 | 2.37673 | 17.78461 | 5.27415 | 0.88316 |
| **BAG2** | 0.54070 | 0.05105 | 7.13933 | 0.07650 | 0.00000 | 0.85656 | 8.61494 | 0.35005 | 2.23556 | 0.00000 | 0.67879 | 1.08857 |
| **BAG3** | 0.39527 | 2.62540 | 21.83957 | 4.65087 | 5.50877 | 0.01945 | 15.48379 | 0.84007 | 0.05535 | 7.51317 | 3.63104 | 0.93169 |
| **BAG4** | 0.71858 | 4.94297 | 6.44720 | 7.45375 | 0.93423 | 19.39309 | 9.62746 | 29.54460 | 2.73914 | 3.54828 | 9.49335 | 24.95862 |
| **BAG5** | 2.68211 | 8.28832 | 3.04850 | 3.07286 | 4.05411 | 17.77392 | 6.86087 | 10.28329 | 3.57046 | 14.85347 | 5.46847 | 12.71274 |
| **BAG6** | 3.99752 | 4.86344 | 10.31585 | 10.24731 | 15.35100 | 0.00000 | 17.48887 | 0.16706 | 3.89961 | 19.14440 | 6.56704 | 2.05483 |
| **BAGE** | 0.00252 | 0.01626 | 0.03972 | 0.02214 | 0.03238 | 0.00000 | 0.07251 | 0.22611 | 0.07911 | 0.04418 | 0.00551 | 0.52648 |
| **BAHCC1** | 0.10882 | 0.03510 | 0.00927 | 0.10608 | 0.06478 | 0.00000 | 0.03321 | 0.02223 | 0.00000 | 0.67630 | 0.01846 | 0.02996 |
| **BAHD1** | 0.40107 | 1.06624 | 3.72668 | 2.47621 | 4.32312 | 0.00000 | 5.81177 | 0.05151 | 4.71430 | 6.40820 | 0.28111 | 0.63843 |
| **BAI1** | 0.00819 | 0.00425 | 0.01456 | 0.00819 | 0.01003 | 0.03824 | 0.02364 | 0.00655 | 0.00649 | 0.02136 | 0.02932 | 0.13424 |
| **BAI2** | 0.32408 | 0.25021 | 0.03488 | 0.01728 | 0.04717 | 0.00000 | 0.06218 | 0.01959 | 0.00000 | 0.00000 | 0.03699 | 0.01364 |
| **BAI3** | 0.11619 | 0.00711 | 0.01136 | 0.00878 | 0.02458 | 0.02493 | 0.07114 | 0.00081 | 0.00000 | 0.02751 | 0.01035 | 0.07342 |
| **BAIAP2** | 12.40923 | 5.86531 | 14.66346 | 0.37562 | 10.86796 | 0.00000 | 13.26948 | 6.81929 | 23.14028 | 0.00000 | 11.66791 | 9.58535 |
| **BAIAP2-AS1** | 0.00142 | 0.00000 | 0.00000 | 0.65202 | 0.13941 | 0.00000 | 0.29797 | 0.62013 | 0.00000 | 1.03081 | 0.02379 | 0.75673 |
| **BAIAP2L1** | 8.05529 | 20.90748 | 13.15413 | 0.00000 | 22.39167 | 1.82605 | 9.79032 | 6.35384 | 15.20977 | 19.63968 | 15.15035 | 3.83668 |

| | | | | | | | | | | | | |
|---|---|---|---|---|---|---|---|---|---|---|---|---|
| **BAIAP2L2** | 0.79018 | 11.55215 | 0.00000 | 0.04934 | 0.88547 | 0.00000 | 0.00633 | 1.94836 | 0.00000 | 0.13506 | 0.00000 | 11.30939 |
| **BAIAP3** | 0.68355 | 0.55626 | 0.23455 | 0.00000 | 0.65104 | 0.00000 | 0.00000 | 0.01456 | 0.00000 | 0.93043 | 0.00000 | 0.10314 |
| **BAK1** | 0.05311 | 27.49546 | 2.44606 | 14.79189 | 2.29554 | 0.00000 | 6.00265 | 2.57793 | 4.28573 | 0.00000 | 2.32002 | 31.40838 |
| **BAMBI** | 0.00000 | 1.61662 | 2.81440 | 26.77326 | 0.87758 | 1.02980 | 0.00000 | 4.87765 | 0.87044 | 0.59840 | 0.00000 | 13.60702 |
| **BANCR** | 0.01751 | 0.05641 | 0.21249 | 1.64785 | 0.00000 | 0.00290 | 0.17419 | 0.06531 | 0.14836 | 0.01192 | 0.00000 | 0.04415 |
| **BANF1** | 6.00890 | 20.51057 | 12.90208 | 15.95027 | 19.64798 | 13.03171 | 20.75875 | 11.71790 | 16.78689 | 2.54783 | 15.82977 | 24.19441 |
| **BANF2** | 0.00627 | 0.05232 | 0.01559 | 0.01146 | 0.00918 | 0.02308 | 0.01565 | 0.01053 | 0.01838 | 0.01428 | 0.01787 | 0.01295 |
| **BANK1** | 0.75574 | 0.01323 | 0.02862 | 0.00802 | 0.00917 | 0.01154 | 0.01567 | 0.01423 | 0.01319 | 0.06497 | 0.01748 | 0.03888 |
| **BANP** | 2.64550 | 7.79830 | 3.56998 | 3.07980 | 4.46503 | 0.01070 | 4.16461 | 0.81799 | 4.45961 | 0.00010 | 3.73904 | 6.34571 |
| **BAP1** | 1.77708 | 8.09580 | 7.14968 | 8.19319 | 8.57200 | 0.00000 | 9.98580 | 4.18963 | 6.19696 | 16.92700 | 2.24499 | 6.80840 |
| **BARD1** | 7.54421 | 17.80761 | 8.65477 | 17.66023 | 8.29560 | 24.03059 | 7.13242 | 2.25434 | 0.00000 | 5.40913 | 9.32816 | 15.76183 |
| **BARHL2** | 0.03227 | 0.01243 | 0.00995 | 0.05782 | 0.01601 | 0.00971 | 0.02031 | 0.05972 | 0.01607 | 0.02521 | 0.01591 | 0.04522 |
| **BARX1** | 0.00365 | 0.23467 | 0.00000 | 0.04051 | 0.02118 | 0.03354 | 0.00000 | 0.67826 | 0.00000 | 0.11388 | 0.00283 | 1.11728 |
| **BARX2** | 2.07730 | 0.31227 | 0.00000 | 0.00000 | 0.09819 | 0.08899 | 0.00000 | 0.11195 | 0.00000 | 0.07512 | 0.04374 | 0.42139 |
| **BASP1** | 0.00000 | 10.41132 | 3.41923 | 0.00000 | 37.86231 | 21.37278 | 40.40459 | 9.13391 | 0.00058 | 74.70182 | 27.02296 | 10.67554 |
| **BASP1P1** | 0.00548 | 0.00907 | 0.02549 | 0.03314 | 0.00769 | 0.03346 | 0.00835 | 0.01016 | 0.01089 | 0.05985 | 0.02033 | 0.01343 |
| **BATF** | 0.00499 | 0.00000 | 3.51253 | 0.00872 | 1.01583 | 0.00000 | 0.96595 | 0.00779 | 0.00000 | 0.01404 | 0.67911 | 0.05009 |
| **BATF2** | 2.36392 | 0.08881 | 0.00162 | 0.00000 | 0.00826 | 0.00000 | 0.00738 | 0.40974 | 0.10586 | 0.03151 | 0.00437 | 0.87384 |
| **BATF3** | 0.15501 | 11.61464 | 0.00000 | 2.29424 | 3.31861 | 0.00000 | 2.27808 | 3.48502 | 0.00000 | 2.43099 | 2.74521 | 14.83175 |
| **BAX** | 0.38784 | 3.51703 | 7.11486 | 4.96152 | 5.31835 | 0.00000 | 3.49013 | 0.00000 | 2.10562 | 9.30956 | 6.14577 | 2.11726 |
| **BAZ1A** | 7.67766 | 0.00000 | 5.18707 | 0.33693 | 8.23198 | 14.43049 | 11.14157 | 17.70297 | 4.91199 | 33.48541 | 2.22360 | 7.22202 |
| **BAZ1B** | 11.58985 | 6.73959 | 4.66599 | 4.97723 | 8.95298 | 12.74942 | 20.68081 | 6.56134 | 5.06933 | 41.56648 | 7.92935 | 11.44541 |
| **BAZ2A** | 16.56911 | 6.19929 | 9.57546 | 8.47511 | 12.99615 | 1.80584 | 9.56120 | 4.11161 | 8.55581 | 26.07053 | 2.14015 | 17.24093 |
| **BAZ2B.uid-2643** | 0.00144 | 0.01394 | 0.80894 | 0.13672 | 0.00000 | 0.30453 | 0.07229 | 0.63377 | 0.00000 | 1.57556 | 0.00000 | 0.19977 |
| **BAZ2B.uid-2983** | 0.24412 | 0.34553 | 0.12721 | 0.00000 | 0.56342 | 0.48815 | 0.00000 | 0.78157 | 0.46189 | 0.51981 | 0.43683 | 1.30889 |
| **BBC3** | 0.02769 | 1.21429 | 0.63986 | 0.69823 | 1.87405 | 0.00000 | 0.44071 | 0.00549 | 0.25149 | 4.60688 | 0.00000 | 0.36137 |
| **BBIP1** | 0.00626 | 1.04346 | 2.15715 | 1.30564 | 0.00000 | 4.34283 | 1.38323 | 9.04317 | 1.51981 | 0.02408 | 0.28900 | 4.88252 |
| **BBOX1** | 0.02098 | 0.01459 | 0.03284 | 0.22972 | 0.00000 | 0.03187 | 0.15784 | 0.00746 | 0.00000 | 0.04867 | 0.01287 | 0.02041 |
| **BBS1** | 0.25415 | 4.08451 | 0.60441 | 1.45484 | 0.81772 | 0.05882 | 1.04538 | 0.75120 | 1.79749 | 0.15317 | 1.37214 | 2.27235 |
| **BBS10** | 0.91535 | 0.45205 | 0.27820 | 0.00000 | 0.00000 | 0.69981 | 0.67447 | 0.00000 | 0.69730 | 0.01466 | 1.52280 | 1.30765 |
| **BBS12** | 0.12264 | 0.87821 | 0.88759 | 0.00294 | 0.00000 | 0.97713 | 0.60129 | 0.07304 | 0.54478 | 0.00000 | 0.26221 | 1.30856 |
| **BBS2** | 0.10514 | 3.52456 | 3.03818 | 2.60535 | 0.00000 | 0.59332 | 2.08260 | 0.22432 | 0.99198 | 0.00000 | 3.85739 | 2.06041 |
| **BBS4** | 3.03494 | 3.50062 | 2.22836 | 1.39624 | 1.72442 | 4.56749 | 0.00000 | 2.52375 | 1.35467 | 2.26981 | 3.12155 | 4.86644 |
| **BBS5** | 1.73077 | 0.23011 | 0.13052 | 0.00000 | 0.44028 | 0.54903 | 0.00000 | 1.59084 | 0.24454 | 0.22410 | 0.13084 | 1.77387 |
| **BBS7** | 1.96108 | 2.18497 | 1.58771 | 2.45307 | 0.63957 | 3.84498 | 1.23323 | 2.46861 | 1.15933 | 0.00000 | 0.98592 | 4.35176 |
| **BBS9** | 5.72369 | 1.94686 | 1.35368 | 2.02704 | 2.46575 | 4.51925 | 0.00000 | 3.81837 | 4.26065 | 0.00000 | 0.15667 | 0.49694 |
| **BBX** | 0.80135 | 2.63950 | 3.79646 | 3.75965 | 1.51983 | 7.01859 | 6.03251 | 3.97555 | 3.54130 | 6.66680 | 0.68564 | 4.95417 |
| **BCAM** | 0.08387 | 0.70791 | 1.37525 | 0.00000 | 0.17307 | 0.00000 | 0.04891 | 0.06557 | 3.25209 | 0.61378 | 0.47349 | 0.00000 |
| **BCAN** | 0.00393 | 0.02049 | 0.00823 | 0.00340 | 0.00843 | 0.01700 | 0.05935 | 0.06084 | 0.02918 | 0.16968 | 0.02029 | 0.04037 |
| **BCAP29** | 0.10015 | 9.17607 | 0.97669 | 0.00000 | 1.66469 | 19.28431 | 0.00000 | 0.36766 | 7.25432 | 7.33196 | 5.85236 | 8.17377 |
| **BCAP31** | 4.31537 | 21.48704 | 22.39574 | 15.02181 | 21.65842 | 13.66655 | 9.50726 | 13.66719 | 39.27693 | 4.86051 | 15.48526 | 22.66229 |
| **BCAR1** | 0.94648 | 2.35832 | 5.86720 | 1.43301 | 7.90612 | 0.00000 | 11.96045 | 0.10292 | 2.00527 | 5.94923 | 1.08347 | 0.50698 |
| **BCAR3** | 0.81646 | 0.00000 | 1.20631 | 2.87793 | 0.00000 | 0.00000 | 4.68961 | 0.00000 | 3.29268 | 2.76273 | 2.44580 | 0.05313 |
| **BCAR4** | 14.48379 | 0.00000 | 14.84029 | 0.00000 | 1.98130 | 0.00000 | 2.85353 | 10.35320 | 0.00000 | 0.19578 | 0.60920 | 0.00000 |
| **BCAS1** | 0.73022 | 0.13453 | 0.19199 | 0.00048 | 0.00978 | 0.02493 | 0.10028 | 1.08279 | 0.09597 | 0.00000 | 0.00000 | 0.09976 |
| **BCAS2** | 3.06720 | 11.15168 | 19.19240 | 4.07299 | 4.39090 | 41.63563 | 15.29087 | 1.15166 | 13.50604 | 3.32062 | 14.08338 | 10.60680 |
| **BCAS3** | 0.63502 | 1.59785 | 1.63494 | 1.82312 | 2.05621 | 0.00000 | 1.27241 | 2.06604 | 0.38071 | 0.66720 | 0.63384 | 2.81616 |
| **BCAS4** | 0.35406 | 1.44613 | 0.07441 | 0.75837 | 0.00000 | 0.00000 | 0.00000 | 0.04031 | 0.13796 | 0.66931 | 0.13395 | 0.88210 |

| | | | | | | | | | | | | |
|---|---|---|---|---|---|---|---|---|---|---|---|---|
| **BCAT1** | 0.00000 | 17.06082 | 20.35091 | 21.24250 | 5.04549 | 55.84063 | 37.51730 | 6.03220 | 29.53332 | 33.49301 | 24.13957 | 29.08850 |
| **BCAT2** | 3.63090 | 7.84846 | 2.20699 | 4.04565 | 6.59141 | 0.01576 | 9.56866 | 0.42315 | 12.12367 | 0.00000 | 9.86870 | 3.78563 |
| **BCCIP** | 2.62742 | 18.72124 | 20.73819 | 5.93879 | 5.53852 | 35.95614 | 29.51775 | 15.00857 | 18.25848 | 15.78645 | 19.43582 | 22.62777 |
| **BCDIN3D** | 0.00679 | 1.17954 | 1.48348 | 0.44312 | 0.40638 | 1.52368 | 0.37643 | 0.44625 | 0.77488 | 0.00000 | 0.93821 | 3.14252 |
| **BCDIN3D-AS1** | 0.15951 | 0.01260 | 0.13229 | 0.17319 | 0.00000 | 0.00000 | 0.04179 | 0.06732 | 0.08375 | 0.08051 | 0.37793 | 0.13287 |
| **BCHE** | 0.01829 | 0.01584 | 0.01079 | 0.01197 | 0.01745 | 0.00794 | 0.07702 | 0.01654 | 0.00000 | 0.02296 | 0.01361 | 0.01562 |
| **BCKDHA** | 0.05724 | 10.41115 | 7.26891 | 5.22691 | 4.81793 | 1.08720 | 5.11269 | 0.26248 | 5.10313 | 0.00000 | 10.50272 | 10.51030 |
| **BCKDHB** | 1.77638 | 1.04339 | 0.70573 | 1.16658 | 0.00000 | 0.67523 | 0.00000 | 0.17029 | 0.83656 | 0.33395 | 1.89550 | 0.00000 |
| **BCKDK** | 1.59479 | 2.24513 | 5.80601 | 14.00015 | 4.53226 | 0.00000 | 6.00196 | 0.00000 | 13.41699 | 0.00000 | 9.57381 | 2.44392 |
| **BCL10** | 0.56675 | 6.47641 | 7.54091 | 5.40302 | 1.62681 | 20.34660 | 13.32525 | 16.05196 | 5.60207 | 1.82946 | 7.50182 | 5.95496 |
| **BCL11A** | 0.54284 | 4.71535 | 0.04683 | 2.69346 | 1.63976 | 0.57577 | 0.00000 | 0.13638 | 0.00000 | 1.95671 | 0.43899 | 5.65230 |
| **BCL11B** | 0.29258 | 0.00000 | 0.13413 | 0.17089 | 0.06202 | 0.01665 | 0.02044 | 0.08197 | 0.00000 | 0.11311 | 0.00502 | 0.09206 |
| **BCL2** | 0.01544 | 0.00786 | 0.06232 | 0.00733 | 0.01461 | 0.12400 | 0.02909 | 0.00643 | 0.06948 | 0.04572 | 0.00525 | 0.06446 |
| **BCL2A1** | 0.04998 | 0.01740 | 0.01424 | 0.01240 | 0.05225 | 0.00000 | 0.00721 | 0.01339 | 0.06679 | 0.00279 | 0.00883 | 0.04601 |
| **BCL2L1** | 3.23018 | 8.33130 | 8.73668 | 4.01350 | 12.91245 | 0.01086 | 12.36253 | 2.12335 | 7.99902 | 7.18682 | 12.28843 | 3.95362 |
| **BCL2L10** | 15.98927 | 0.00487 | 0.00920 | 0.00573 | 0.02324 | 0.00000 | 0.01290 | 1.33326 | 0.02313 | 0.00000 | 0.01040 | 0.18822 |
| **BCL2L11** | 0.75327 | 3.34325 | 1.89641 | 3.00197 | 3.72034 | 3.49073 | 4.79959 | 10.41657 | 7.20938 | 3.45041 | 1.52969 | 2.39983 |
| **BCL2L12** | 1.85889 | 14.02351 | 18.48954 | 5.83222 | 14.87512 | 0.00000 | 10.17257 | 2.57622 | 20.19924 | 4.56553 | 12.53992 | 9.81211 |
| **BCL2L13** | 6.88252 | 10.52051 | 9.74650 | 3.55849 | 4.83379 | 11.72698 | 6.77586 | 6.66885 | 6.09440 | 0.19911 | 4.77986 | 15.71218 |
| **BCL2L14** | 0.00592 | 0.10324 | 0.00010 | 0.11337 | 0.20397 | 0.12486 | 0.00000 | 0.00940 | 0.02902 | 0.00000 | 0.00663 | 0.11503 |
| **BCL2L15** | 0.00393 | 0.03273 | 0.13665 | 0.64732 | 0.00571 | 0.00000 | 0.45726 | 0.15519 | 0.00000 | 0.00000 | 0.13340 | 0.14080 |
| **BCL2L2** | 0.03908 | 0.00314 | 1.37998 | 0.81973 | 0.30858 | 0.01657 | 1.01877 | 0.11673 | 1.13505 | 0.00000 | 0.67116 | 0.08297 |
| **BCL3** | 0.00123 | 0.26246 | 0.31529 | 4.95647 | 0.33014 | 0.00000 | 0.50547 | 0.15376 | 0.00306 | 0.25018 | 0.09795 | 0.05559 |
| **BCL6** | 0.00297 | 1.18194 | 1.38532 | 0.49793 | 0.59737 | 0.00000 | 1.82825 | 0.00969 | 0.00000 | 1.90956 | 0.91727 | 0.72893 |
| **BCL6B** | 0.34945 | 0.00701 | 0.02110 | 0.00981 | 0.00353 | 0.03453 | 0.08297 | 0.00885 | 0.16028 | 0.00000 | 0.00066 | 0.09411 |
| **BCL7A** | 0.93294 | 0.89052 | 1.11251 | 2.24131 | 7.16909 | 0.00000 | 3.02668 | 0.06002 | 3.17687 | 2.63495 | 0.90995 | 0.00000 |
| **BCL7B** | 0.11652 | 5.99772 | 5.84651 | 6.07876 | 2.87126 | 3.51824 | 5.39762 | 0.41732 | 5.48847 | 4.16744 | 4.86141 | 4.63919 |
| **BCL7C** | 0.00071 | 0.36913 | 0.18421 | 0.19025 | 2.08165 | 0.00000 | 1.38198 | 0.05534 | 0.00000 | 3.24308 | 0.18615 | 0.09284 |
| **BCL9** | 3.92080 | 1.93935 | 1.60455 | 5.47989 | 4.38630 | 0.00000 | 6.04170 | 0.70474 | 1.36537 | 8.41230 | 5.00836 | 1.05806 |
| **BCL9L** | 0.60368 | 1.17697 | 9.40599 | 2.91894 | 8.27770 | 0.00000 | 14.46441 | 0.00000 | 0.03554 | 16.25157 | 0.91779 | 0.52102 |
| **BCLAF1** | 5.67952 | 10.62686 | 11.84288 | 5.13626 | 11.18997 | 35.42297 | 30.36476 | 16.59931 | 11.03645 | 30.65422 | 21.45442 | 18.02708 |
| **BCMO1** | 0.39380 | 0.12062 | 0.00867 | 0.63549 | 0.09120 | 0.00074 | 0.01454 | 0.06370 | 0.02252 | 0.00000 | 0.01224 | 0.00938 |
| **BCO2** | 0.00156 | 0.01388 | 0.17758 | 0.33837 | 0.32458 | 0.00000 | 0.00000 | 0.58323 | 0.21688 | 0.32687 | 0.20643 | 0.07601 |
| **BCOR** | 1.88200 | 3.93085 | 2.50038 | 11.58458 | 2.96156 | 5.77377 | 0.68593 | 4.58156 | 1.99339 | 7.11734 | 0.01903 | 6.55378 |
| **BCORL1** | 5.37298 | 0.88606 | 1.98552 | 0.00000 | 4.63306 | 0.00000 | 1.33870 | 0.27729 | 4.25856 | 6.94340 | 2.11230 | 2.32380 |
| **BCORP1** | 0.00577 | 0.06111 | 0.00913 | 0.02259 | 0.00000 | 0.04720 | 0.01015 | 0.01017 | 0.01223 | 0.02884 | 0.01188 | 0.01032 |
| **BCR** | 3.38619 | 2.38093 | 4.99005 | 3.85076 | 7.65009 | 0.00000 | 11.21769 | 0.11238 | 6.03398 | 15.05496 | 2.69913 | 2.64685 |
| **BCRP3** | 0.01892 | 0.01092 | 0.00550 | 0.62952 | 0.00000 | 0.00655 | 0.03011 | 0.00771 | 0.00728 | 0.03196 | 0.03328 | 0.04828 |
| **BCS1L** | 0.63801 | 15.15861 | 1.34288 | 5.78686 | 12.81526 | 8.99468 | 3.46266 | 8.63086 | 9.51302 | 5.28524 | 12.04632 | 11.83894 |
| **BCYRN1** | 0.00612 | 0.02219 | 0.01247 | 0.00987 | 0.02521 | 0.02475 | 0.02003 | 0.01055 | 0.01899 | 0.01527 | 0.01285 | 0.01519 |
| **BDH1** | 1.80957 | 0.00000 | 0.59502 | 0.12250 | 0.07023 | 0.18266 | 0.29134 | 0.76494 | 0.00000 | 0.17726 | 0.79891 | 0.00000 |
| **BDH2** | 2.16213 | 0.52254 | 0.40370 | 1.06775 | 0.10953 | 0.50854 | 0.20673 | 0.58412 | 0.79466 | 0.00000 | 0.43470 | 0.01328 |
| **BDKRB1** | 0.00490 | 0.02888 | 0.02820 | 0.00556 | 0.00823 | 0.02360 | 0.04563 | 0.00910 | 0.04773 | 0.05618 | 0.00584 | 0.04566 |
| **BDKRB2** | 0.00866 | 0.01552 | 0.02109 | 0.02036 | 0.00540 | 0.04726 | 0.01586 | 0.02081 | 0.01929 | 0.02170 | 0.01090 | 0.02327 |
| **BDNF** | 0.47363 | 0.00139 | 0.43635 | 0.88783 | 0.00000 | 0.62771 | 0.00000 | 0.08652 | 0.00000 | 0.45383 | 0.07436 | 0.18164 |
| **BDNF-AS** | 0.87507 | 0.07670 | 0.12968 | 0.14410 | 0.19336 | 0.00000 | 0.04432 | 0.13210 | 0.00000 | 0.18656 | 0.09693 | 0.25735 |
| **BDP1** | 4.37775 | 2.49824 | 5.65588 | 5.37247 | 6.27316 | 19.06975 | 7.25038 | 9.18178 | 3.56590 | 42.85124 | 4.89528 | 9.72847 |
| **BEAN1** | 1.80016 | 0.01235 | 0.35915 | 0.00386 | 0.00677 | 0.03959 | 0.01983 | 0.15943 | 0.08695 | 0.00000 | 0.00000 | 0.03732 |

| | | | | | | | | | | | | |
|---|---|---|---|---|---|---|---|---|---|---|---|---|
| **BECN1** | 2.10055 | 9.09610 | 9.09309 | 5.64118 | 3.05599 | 9.14685 | 8.90514 | 6.88167 | 8.49610 | 0.00000 | 13.72533 | 9.85540 |
| **BECN1P1** | 0.48125 | 4.60752 | 0.00000 | 0.00000 | 0.00000 | 0.00000 | 0.01160 | 20.83394 | 0.00000 | 0.02486 | 0.00000 | 19.37429 |
| **BEGAIN** | 0.00826 | 0.01346 | 0.01444 | 0.01298 | 0.01394 | 0.02129 | 0.02197 | 0.01013 | 0.01817 | 0.02117 | 0.01480 | 0.01804 |
| **BEND2** | 0.30178 | 0.00781 | 0.03174 | 0.00671 | 0.03255 | 0.02899 | 0.00613 | 0.00000 | 0.02129 | 0.02145 | 0.01107 | 0.07756 |
| **BEND3** | 1.81602 | 6.27870 | 2.22775 | 5.59004 | 3.88533 | 7.46825 | 6.16512 | 1.92689 | 2.13787 | 10.35163 | 2.79503 | 8.56793 |
| **BEND3P3** | 0.03212 | 0.02207 | 0.17719 | 0.04718 | 0.14340 | 0.00000 | 0.02104 | 0.00234 | 0.16810 | 0.23167 | 0.32569 | 0.13535 |
| **BEND4** | 0.01190 | 0.59427 | 0.36017 | 2.63737 | 0.00000 | 0.00000 | 0.04932 | 0.36442 | 0.00852 | 0.01909 | 0.06665 | 0.04799 |
| **BEND5** | 0.05230 | 0.22245 | 0.02580 | 0.05922 | 0.00000 | 0.00004 | 0.03393 | 0.01744 | 0.00000 | 0.03409 | 0.12326 | 0.00396 |
| **BEND6** | 0.41547 | 0.05735 | 0.01341 | 0.00000 | 0.00000 | 0.06385 | 0.04637 | 0.23587 | 0.01112 | 0.06920 | 0.01470 | 0.00000 |
| **BEND7** | 0.25654 | 0.01350 | 0.00000 | 0.09088 | 1.05251 | 0.40762 | 0.13149 | 0.03386 | 0.00020 | 0.63817 | 0.00000 | 0.02156 |
| **BEST1** | 0.00453 | 0.03076 | 0.02276 | 0.01136 | 0.02218 | 0.00308 | 0.08908 | 0.01049 | 0.04883 | 0.10784 | 0.01187 | 0.01603 |
| **BEST2** | 0.00320 | 0.00791 | 0.00933 | 0.00437 | 0.00000 | 0.01651 | 0.06722 | 0.02016 | 0.10628 | 0.10149 | 0.00810 | 0.06497 |
| **BEST3** | 0.02081 | 0.01581 | 0.01224 | 0.01660 | 0.00623 | 0.03779 | 0.01243 | 0.01235 | 0.01341 | 0.02851 | 0.01421 | 0.01280 |
| **BEST4** | 0.00383 | 0.03805 | 0.02836 | 0.01317 | 0.00000 | 0.01173 | 0.00662 | 0.01454 | 0.01424 | 0.19641 | 0.02823 | 0.05681 |
| **BET1** | 0.14666 | 5.60503 | 7.26288 | 0.36807 | 0.00133 | 11.90892 | 5.73937 | 2.89001 | 3.62732 | 0.00373 | 5.76254 | 7.88902 |
| **BET1L** | 1.27526 | 8.32906 | 5.80219 | 2.02980 | 11.10387 | 0.00000 | 6.22960 | 3.06510 | 9.88951 | 2.60562 | 6.86786 | 9.59350 |
| **BEX1** | 0.02937 | 0.16869 | 0.17713 | 0.41539 | 0.00000 | 0.08321 | 0.19002 | 0.01096 | 0.23651 | 0.00000 | 0.00000 | 0.15294 |
| **BEX2** | 0.07778 | 0.43451 | 5.98820 | 0.00000 | 0.84019 | 0.44593 | 11.21910 | 0.21792 | 4.16956 | 0.00104 | 0.00000 | 2.19933 |
| **BEX4** | 0.52572 | 0.00000 | 8.95194 | 3.85032 | 4.02991 | 9.14256 | 8.47903 | 0.34647 | 14.15673 | 0.00000 | 6.01464 | 5.35366 |
| **BEX5** | 0.00314 | 0.07347 | 0.08491 | 0.01301 | 0.00460 | 0.04106 | 0.01742 | 0.06587 | 0.01966 | 0.01728 | 0.02064 | 0.03145 |
| **BFAR** | 7.08902 | 17.22861 | 6.42223 | 10.10691 | 2.99987 | 13.62376 | 4.72968 | 0.63176 | 11.22060 | 5.80044 | 6.26327 | 17.88066 |
| **BFSP1** | 0.34931 | 0.06816 | 0.03527 | 0.28380 | 0.00000 | 0.02921 | 0.06047 | 0.02647 | 0.01532 | 0.12143 | 0.00463 | 0.39102 |
| **BFSP2** | 0.00723 | 0.10267 | 0.16469 | 0.15658 | 0.00000 | 0.00769 | 0.04078 | 0.04355 | 0.00006 | 0.06072 | 0.01668 | 0.34986 |
| **BGLAP** | 0.00625 | 0.01666 | 0.01957 | 0.04545 | 0.02626 | 0.00000 | 0.02238 | 0.01054 | 0.03961 | 0.01083 | 0.01692 | 0.01948 |
| **BGN** | 0.07721 | 0.01640 | 0.16241 | 0.01584 | 0.02432 | 0.00000 | 0.05136 | 0.00577 | 0.00028 | 0.01573 | 0.01549 | 0.01592 |
| **BHLHA15** | 0.00627 | 0.01347 | 0.14015 | 0.01512 | 0.01501 | 0.02223 | 0.00409 | 0.01036 | 0.00776 | 0.01725 | 0.01606 | 0.01584 |
| **BHLHB9** | 2.95738 | 0.09975 | 0.06057 | 0.16284 | 0.07184 | 0.20472 | 0.06964 | 0.34795 | 0.00000 | 0.00000 | 0.03573 | 0.05219 |
| **BHLHE22** | 0.67921 | 0.35516 | 0.00000 | 0.62054 | 0.06335 | 0.01891 | 0.00000 | 0.00000 | 0.00000 | 0.36729 | 0.03076 | 0.50519 |
| **BHLHE23** | 0.01931 | 0.01479 | 0.01612 | 0.02447 | 0.01361 | 0.01878 | 0.01784 | 0.00867 | 0.01588 | 0.01499 | 0.01530 | 0.01576 |
| **BHLHE40** | 8.36124 | 2.44151 | 16.15725 | 5.07357 | 0.00000 | 0.64006 | 2.24987 | 0.36314 | 0.00000 | 1.48813 | 0.00000 | 0.00000 |
| **BHLHE40-AS1** | 0.06342 | 0.00766 | 0.10249 | 0.00000 | 0.00463 | 0.03475 | 0.00000 | 0.00737 | 0.00000 | 0.26071 | 0.00291 | 0.00684 |
| **BHLHE41** | 0.03715 | 0.01263 | 0.10653 | 0.01693 | 0.00000 | 0.03754 | 0.01310 | 0.00698 | 0.00000 | 0.06819 | 0.02434 | 0.01399 |
| **BHMT** | 0.00209 | 3.57372 | 0.00000 | 4.57977 | 0.00000 | 0.03886 | 0.03440 | 0.77320 | 0.00066 | 0.00881 | 0.00000 | 0.94668 |
| **BHMT2** | 0.09425 | 0.02869 | 1.60631 | 0.18770 | 0.00000 | 0.00378 | 0.39848 | 0.01320 | 0.50028 | 0.03292 | 0.00000 | 0.04365 |
| **BICC1** | 2.17731 | 0.00779 | 0.01633 | 0.00909 | 0.00467 | 0.00000 | 0.01133 | 0.00051 | 0.00000 | 0.02777 | 0.20484 | 0.02945 |
| **BICD1** | 0.22776 | 1.14205 | 1.17111 | 8.88510 | 3.85623 | 3.01621 | 0.00000 | 0.21703 | 2.85182 | 3.48142 | 0.24036 | 0.14723 |
| **BICD2** | 1.41735 | 3.98071 | 4.54511 | 3.08173 | 7.78269 | 6.62717 | 6.00405 | 1.44343 | 1.71152 | 19.83177 | 2.93406 | 4.00811 |
| **BID** | 0.00224 | 0.24034 | 0.48069 | 0.26918 | 0.00000 | 0.14250 | 0.51397 | 0.21486 | 0.34956 | 0.44597 | 0.93252 | 0.09752 |
| **BIK** | 0.07702 | 19.47328 | 6.39902 | 0.00000 | 0.00000 | 0.00000 | 6.38799 | 34.66056 | 7.07248 | 5.10651 | 2.94276 | 66.35678 |
| **BIN1** | 0.00038 | 5.74631 | 4.90490 | 6.29898 | 21.50219 | 0.00000 | 11.00183 | 0.00184 | 16.43997 | 4.41358 | 10.08419 | 0.00000 |
| **BIN2** | 0.39641 | 0.76829 | 7.73565 | 0.00000 | 1.55993 | 0.09453 | 6.00302 | 0.03558 | 7.28444 | 0.76804 | 4.15676 | 0.53981 |
| **BIN3** | 3.66155 | 4.52043 | 6.21646 | 3.97112 | 5.05900 | 3.80737 | 6.34970 | 2.52706 | 7.23211 | 0.00000 | 2.16684 | 4.79936 |
| **BIN3-IT1** | 0.01779 | 0.12994 | 0.18989 | 0.13457 | 0.19363 | 0.00000 | 0.00962 | 0.06154 | 0.13877 | 1.13753 | 0.12721 | 0.05567 |
| **BIRC2** | 3.07341 | 3.37406 | 5.10173 | 3.68160 | 0.96845 | 11.71067 | 8.31617 | 11.75036 | 6.00722 | 0.00000 | 5.25806 | 11.76881 |
| **BIRC3** | 0.37150 | 0.00000 | 0.14055 | 0.07433 | 0.09606 | 0.14942 | 0.20409 | 0.10291 | 0.03663 | 0.06780 | 0.05083 | 0.09083 |
| **BIRC5** | 3.54403 | 24.04194 | 13.71565 | 16.12149 | 22.73101 | 16.45583 | 36.29236 | 19.05271 | 16.46823 | 7.03942 | 17.13419 | 21.59542 |
| **BIRC6** | 6.45378 | 2.74979 | 10.26163 | 14.97863 | 5.24508 | 18.73088 | 12.02578 | 5.36279 | 7.60546 | 33.72723 | 6.66365 | 4.67571 |
| **BIRC7** | 0.01708 | 0.01674 | 0.04739 | 0.01448 | 0.01744 | 0.01175 | 0.02508 | 0.01256 | 0.00916 | 0.01604 | 0.01653 | 0.01569 |

| | | | | | | | | | | | | |
|---|---|---|---|---|---|---|---|---|---|---|---|---|
| **BIRC8** | 0.00617 | 0.01349 | 0.02133 | 0.01421 | 0.01210 | 0.02352 | 0.01483 | 0.01042 | 0.01395 | 0.01711 | 0.03092 | 0.01585 |
| **BIVM** | 0.02446 | 0.00000 | 0.27771 | 0.18911 | 0.01260 | 0.08393 | 0.23693 | 0.00574 | 0.79138 | 0.48771 | 0.47288 | 0.01513 |
| **BLACAT1** | 0.00622 | 0.01337 | 0.05933 | 0.02148 | 0.02734 | 0.02862 | 0.01233 | 0.01040 | 0.03914 | 0.00875 | 0.01135 | 0.01572 |
| **BLACE** | 0.10080 | 1.55601 | 0.00000 | 1.28199 | 0.08254 | 0.03373 | 0.00002 | 0.03692 | 0.00000 | 0.05281 | 0.00006 | 0.70889 |
| **BLCAP** | 12.64532 | 8.80731 | 10.87527 | 2.85886 | 3.59720 | 11.60209 | 6.41998 | 9.03878 | 12.15272 | 0.00000 | 5.60766 | 14.95872 |
| **BLK** | 0.65288 | 0.01448 | 0.03143 | 0.01403 | 0.01861 | 0.00060 | 0.01714 | 0.08185 | 0.02395 | 0.00843 | 0.01457 | 0.03100 |
| **BLM** | 1.27168 | 4.56692 | 4.68092 | 1.20539 | 0.25924 | 16.00221 | 1.61930 | 3.84241 | 5.07588 | 8.25445 | 3.77545 | 4.33260 |
| **BLMH** | 1.88404 | 12.29982 | 15.04124 | 7.08645 | 16.23424 | 25.14904 | 33.08691 | 4.01003 | 18.82208 | 4.91375 | 12.41674 | 8.06415 |
| **BLNK** | 0.10625 | 0.01363 | 0.03821 | 0.01691 | 0.03002 | 0.06186 | 0.00567 | 0.07720 | 0.01804 | 0.01853 | 0.03169 | 0.33464 |
| **BLOC1S1** | 0.33474 | 4.83958 | 4.10455 | 6.20612 | 5.18096 | 0.00001 | 5.46862 | 0.56265 | 2.30827 | 2.80447 | 4.28433 | 4.05569 |
| **BLOC1S2** | 1.84404 | 14.38391 | 11.25578 | 6.39043 | 0.47775 | 30.74024 | 13.98505 | 1.02081 | 11.70644 | 8.16694 | 7.51942 | 7.72935 |
| **BLOC1S3** | 9.38016 | 3.05381 | 4.00158 | 2.50868 | 3.65614 | 3.37297 | 5.87279 | 6.28159 | 0.37024 | 0.00000 | 5.46256 | 4.68684 |
| **BLOC1S4** | 0.00000 | 0.77890 | 4.12932 | 1.47035 | 2.29169 | 0.00000 | 5.32526 | 0.47954 | 0.22867 | 4.46304 | 2.06477 | 0.85665 |
| **BLOC1S5.uid-8** | 0.00540 | 0.01688 | 0.02065 | 0.01009 | 0.01164 | 0.02135 | 0.01379 | 0.01308 | 0.02186 | 0.01911 | 0.03177 | 0.03443 |
| **BLOC1S6** | 0.63200 | 5.10362 | 8.11847 | 1.52606 | 0.00000 | 15.09924 | 9.60407 | 4.41148 | 3.44409 | 8.35815 | 3.27748 | 6.72504 |
| **BLVRA** | 0.78233 | 0.73407 | 0.03405 | 0.32298 | 0.00438 | 0.48514 | 0.07818 | 0.08671 | 0.00000 | 0.00000 | 0.00704 | 0.51491 |
| **BLVRB** | 0.74156 | 9.14686 | 10.74989 | 0.00000 | 2.81911 | 1.18724 | 7.18863 | 0.66972 | 1.49110 | 0.00000 | 1.84192 | 19.37463 |
| **BLZF1** | 0.00229 | 3.58696 | 4.92289 | 0.13014 | 0.00000 | 7.50868 | 0.00000 | 7.62942 | 4.93088 | 0.00000 | 3.67190 | 5.85653 |
| **BMF** | 0.44184 | 0.14161 | 2.08382 | 0.88347 | 0.45185 | 0.00010 | 3.08018 | 0.00184 | 0.17876 | 1.29043 | 0.00000 | 0.01826 |
| **BMI1** | 0.00366 | 0.11157 | 0.03689 | 0.01950 | 0.14705 | 0.04502 | 0.00965 | 0.04395 | 0.12360 | 0.00000 | 0.16254 | 0.02536 |
| **BMP1** | 0.08605 | 0.20091 | 0.85356 | 0.02993 | 0.44503 | 0.00000 | 0.55438 | 0.06445 | 0.79243 | 1.11491 | 0.00000 | 0.00927 |
| **BMP10** | 0.00646 | 0.05173 | 0.02924 | 0.03119 | 0.02258 | 0.00000 | 0.02460 | 0.01081 | 0.01537 | 0.00571 | 0.01937 | 0.01481 |
| **BMP15** | 15.32992 | 0.03547 | 0.00404 | 0.00209 | 0.01224 | 0.00000 | 0.00341 | 12.46936 | 0.01629 | 0.00694 | 0.00587 | 0.00000 |
| **BMP2** | 0.00321 | 0.02556 | 0.58881 | 17.96579 | 0.45936 | 0.00000 | 0.00000 | 0.00536 | 0.00000 | 0.20777 | 0.02890 | 0.04737 |
| **BMP2K** | 2.04557 | 0.54502 | 0.89574 | 2.84345 | 0.03236 | 3.33231 | 0.49494 | 0.17190 | 0.87883 | 4.31590 | 1.92508 | 1.13572 |
| **BMP3** | 0.00555 | 0.01902 | 0.01465 | 0.01107 | 0.01718 | 0.01446 | 0.03486 | 0.01213 | 0.00747 | 0.02441 | 0.01624 | 0.01513 |
| **BMP4** | 0.56337 | 0.48797 | 4.77415 | 1.98275 | 0.99261 | 1.25892 | 0.91556 | 6.29956 | 1.44505 | 1.22754 | 0.38126 | 1.36688 |
| **BMP5** | 0.00501 | 0.01616 | 0.01760 | 0.01354 | 0.01455 | 0.01009 | 0.01826 | 0.02818 | 0.01995 | 0.01199 | 0.03091 | 0.02747 |
| **BMP6** | 16.23992 | 0.00000 | 0.90178 | 6.21839 | 0.00000 | 0.00000 | 0.00008 | 2.36890 | 0.00000 | 1.04904 | 0.00000 | 0.31875 |
| **BMP7** | 0.45651 | 0.05598 | 1.38303 | 0.25310 | 0.00000 | 0.00000 | 0.38345 | 0.02529 | 0.04842 | 0.31557 | 0.00000 | 0.90234 |
| **BMP8A** | 0.07388 | 0.12578 | 0.25869 | 0.41246 | 0.70882 | 0.00000 | 0.07890 | 0.00000 | 0.42784 | 1.79958 | 0.35762 | 0.18262 |
| **BMP8B** | 0.00426 | 0.03855 | 0.07817 | 0.00000 | 0.10399 | 0.02619 | 0.01314 | 0.01250 | 0.01385 | 0.09962 | 0.02218 | 0.02517 |
| **BMPER** | 0.13959 | 0.02924 | 0.01130 | 0.02066 | 0.00000 | 0.00000 | 0.03086 | 0.26156 | 0.00000 | 0.11718 | 0.06290 | 0.03697 |
| **BMPR1A** | 0.36305 | 0.79399 | 0.00000 | 1.01392 | 2.27376 | 4.83297 | 0.00000 | 0.62157 | 4.24837 | 5.07081 | 2.02479 | 0.55051 |
| **BMPR1B** | 1.04337 | 0.23606 | 0.00000 | 0.00000 | 0.04716 | 0.15732 | 0.04953 | 0.42162 | 0.04671 | 0.27781 | 0.01684 | 0.14786 |
| **BMPR2** | 2.13718 | 0.15279 | 4.21193 | 1.89661 | 0.90447 | 7.27811 | 0.01696 | 0.05620 | 4.74430 | 7.72204 | 1.34083 | 0.65590 |
| **BMS1** | 1.72491 | 10.01457 | 9.59272 | 4.66941 | 9.97489 | 15.20263 | 11.46560 | 6.91281 | 14.55033 | 13.48881 | 6.59168 | 19.14199 |
| **BMS1P20** | 0.47465 | 2.74711 | 1.11678 | 1.32671 | 1.42278 | 2.25430 | 0.47281 | 3.13772 | 1.68379 | 1.18965 | 0.81652 | 3.35051 |
| **BMS1P4** | 0.00000 | 0.55851 | 0.00000 | 0.00000 | 0.00000 | 1.37350 | 0.00000 | 0.28310 | 0.15891 | 1.57333 | 0.39363 | 0.97956 |
| **BMS1P5.uid-14** | 0.00421 | 0.01069 | 0.00758 | 0.00477 | 0.03576 | 0.04098 | 0.03599 | 0.01715 | 0.00496 | 0.05254 | 0.01650 | 0.01867 |
| **BMS1P5.uid-14** | 0.00577 | 0.00190 | 0.01409 | 0.00754 | 0.01068 | 0.08263 | 0.01707 | 0.01472 | 0.06275 | 0.08620 | 0.00017 | 0.02942 |
| **BNC1** | 1.29812 | 0.01119 | 0.46620 | 0.00254 | 0.17896 | 0.00000 | 0.17971 | 0.04022 | 0.26840 | 0.07127 | 0.00000 | 0.06294 |
| **BNC2** | 0.66401 | 0.16926 | 0.18068 | 0.28423 | 0.00000 | 0.16785 | 0.00000 | 0.15737 | 0.51880 | 0.08889 | 0.00000 | 0.35214 |
| **BNIP1** | 2.45631 | 6.92974 | 1.21021 | 1.42980 | 3.99697 | 4.06178 | 2.51160 | 0.75289 | 0.62299 | 0.00000 | 2.15970 | 3.42301 |
| **BNIP2** | 1.77531 | 9.35635 | 13.70171 | 3.93527 | 1.69350 | 25.41146 | 6.42986 | 23.84026 | 11.18735 | 6.30100 | 6.92244 | 17.11426 |
| **BNIP3** | 4.08240 | 6.86282 | 26.77402 | 8.18845 | 5.60102 | 22.98319 | 24.31603 | 8.84961 | 17.14416 | 2.07281 | 14.15661 | 10.73000 |
| **BNIP3L** | 7.81257 | 2.49614 | 5.67269 | 1.03912 | 0.00000 | 8.23820 | 4.62204 | 12.28699 | 5.03517 | 0.05357 | 0.83832 | 7.25983 |
| **BNIPL** | 0.18494 | 0.00333 | 0.00605 | 0.00000 | 0.00000 | 0.00000 | 0.07603 | 0.00000 | 0.46277 | 0.31599 | 0.05079 | 0.09852 |

| | | | | | | | | | | | | |
|---|---|---|---|---|---|---|---|---|---|---|---|---|
| **BOC** | 0.02550 | 0.02416 | 0.00979 | 0.17274 | 0.00000 | 0.00934 | 0.05439 | 0.01246 | 0.07569 | 0.02130 | 0.00324 | 0.02586 |
| **BOD1** | 16.90349 | 0.44004 | 2.76790 | 2.19435 | 5.92292 | 0.19590 | 8.80539 | 1.36950 | 1.98064 | 8.14722 | 1.28397 | 0.09266 |
| **BOD1L1** | 5.96813 | 0.00000 | 2.97751 | 3.22746 | 2.94093 | 10.77965 | 3.44616 | 6.44745 | 1.25097 | 36.61447 | 1.90387 | 7.46226 |
| **BOD1L2** | 0.15949 | 0.00672 | 0.01707 | 0.22848 | 0.06122 | 0.02270 | 0.02465 | 0.00818 | 0.00000 | 0.06590 | 0.01920 | 0.01766 |
| **BOK** | 0.10442 | 0.37921 | 0.75646 | 0.00000 | 0.00000 | 0.28975 | 0.78154 | 0.06333 | 0.84510 | 1.30188 | 0.78229 | 0.00940 |
| **BOK-AS1** | 0.20467 | 0.18209 | 0.18566 | 0.00000 | 0.00000 | 0.00000 | 0.60383 | 0.06509 | 1.31433 | 0.05683 | 0.95339 | 0.06834 |
| **BOLA1** | 2.04128 | 4.89977 | 5.12695 | 1.20603 | 4.30053 | 1.76481 | 5.27329 | 0.67544 | 9.06459 | 0.28119 | 10.93195 | 8.46892 |
| **BOLA2+BOLA2I** | 0.00507 | 0.01905 | 0.04462 | 0.05054 | 0.01136 | 0.02442 | 0.02633 | 0.02518 | 0.02201 | 0.05404 | 0.02763 | 0.03225 |
| **BOLA2+BOLA2I** | 0.00257 | 0.04690 | 0.03358 | 0.17438 | 0.00000 | 0.09534 | 0.11403 | 0.02833 | 0.00114 | 0.36330 | 0.02594 | 0.08895 |
| **BOLA3** | 0.00000 | 14.70709 | 0.00000 | 8.61160 | 20.16600 | 20.12074 | 0.89221 | 0.00000 | 6.21612 | 6.51105 | 17.61001 | 4.59368 |
| **BOLA3-AS1** | 0.06891 | 0.35692 | 0.51925 | 0.53951 | 0.00000 | 0.00000 | 0.55499 | 0.01584 | 1.40306 | 1.47743 | 1.56697 | 0.36188 |
| **BOLL** | 0.00459 | 0.46101 | 0.06161 | 0.00000 | 0.03458 | 0.06388 | 0.00000 | 0.08092 | 0.16647 | 0.28118 | 0.00000 | 0.87270 |
| **1.00000** | 0.00706 | 0.97849 | 1.29218 | 0.02108 | 0.93000 | 0.00000 | 0.00000 | 0.07891 | 0.77548 | 0.00000 | 0.64013 | 0.20926 |
| **BORA** | 1.36155 | 5.91608 | 2.62136 | 0.93999 | 1.58367 | 14.05882 | 0.00000 | 4.45924 | 7.07333 | 0.00000 | 3.54690 | 10.33670 |
| **BPESC1** | 0.13899 | 0.06955 | 0.00000 | 0.00022 | 0.03194 | 0.35079 | 0.00840 | 0.26849 | 0.00000 | 0.28492 | 0.01289 | 0.00000 |
| **BPGM** | 12.11842 | 1.46729 | 14.49651 | 0.19006 | 0.00000 | 8.16753 | 5.41766 | 9.48405 | 3.46296 | 0.00000 | 2.22532 | 3.62898 |
| **BPHL** | 0.55483 | 2.30863 | 0.08392 | 0.59559 | 0.68724 | 2.59668 | 0.00000 | 0.12547 | 0.98682 | 0.00000 | 0.68571 | 0.97242 |
| **BPIFA2** | 0.11784 | 0.16367 | 0.11255 | 0.36640 | 0.11739 | 0.10731 | 0.00443 | 0.12815 | 0.00000 | 0.00000 | 0.18818 | 0.34112 |
| **BPIFA4P** | 0.00623 | 0.01670 | 0.01391 | 0.01092 | 0.01619 | 0.02513 | 0.02034 | 0.00923 | 0.02383 | 0.00256 | 0.01982 | 0.03903 |
| **BPIFB1** | 0.00640 | 0.01529 | 0.01315 | 0.01238 | 0.01269 | 0.02181 | 0.01618 | 0.01070 | 0.02211 | 0.01883 | 0.01433 | 0.01782 |
| **BPIFB3** | 0.01905 | 0.01531 | 0.01528 | 0.01372 | 0.01581 | 0.01684 | 0.01852 | 0.01777 | 0.01853 | 0.01150 | 0.01514 | 0.01547 |
| **BPIFB4** | 0.01274 | 0.02714 | 0.02755 | 0.02504 | 0.02217 | 0.01749 | 0.01236 | 0.06935 | 0.02092 | 0.01152 | 0.01314 | 0.24975 |
| **BPIFC** | 0.00840 | 0.00972 | 0.01516 | 0.01194 | 0.02940 | 0.00461 | 0.01752 | 0.00729 | 0.01789 | 0.02985 | 0.01607 | 0.09355 |
| **BPNT1** | 0.97059 | 2.47626 | 4.63742 | 1.40428 | 0.87158 | 5.24345 | 2.80690 | 0.30884 | 3.75613 | 0.67557 | 3.57246 | 2.97281 |
| **BPTF** | 4.47601 | 0.82650 | 4.91812 | 4.55483 | 9.77562 | 12.69306 | 16.29528 | 4.57350 | 1.89966 | 45.46522 | 5.54679 | 7.91749 |
| **BRAF** | 0.83623 | 1.19705 | 1.54935 | 1.69953 | 1.94422 | 2.17529 | 0.53315 | 0.27096 | 0.74128 | 0.68974 | 0.41963 | 1.88359 |
| **BRAP** | 2.86864 | 15.10335 | 7.59304 | 5.41482 | 6.01371 | 22.42655 | 7.29283 | 17.79214 | 8.18080 | 0.00000 | 4.99076 | 16.18061 |
| **BRAT1** | 1.36543 | 6.84619 | 3.93077 | 6.15169 | 4.73153 | 0.00000 | 3.29984 | 0.58836 | 7.19489 | 3.69286 | 4.33584 | 6.46667 |
| **BRCA1** | 4.29725 | 2.80026 | 3.18032 | 4.43303 | 5.01373 | 11.13992 | 5.37841 | 1.33838 | 0.00561 | 13.65979 | 6.09882 | 7.82797 |
| **BRCA2** | 9.68718 | 1.73444 | 3.17216 | 1.16661 | 0.26207 | 14.14049 | 0.00000 | 6.28765 | 4.20663 | 15.12108 | 5.01475 | 9.62765 |
| **BRCC3** | 0.00000 | 3.34044 | 5.69227 | 0.63934 | 2.16753 | 12.06382 | 8.19351 | 0.00000 | 8.04956 | 2.39289 | 9.86398 | 1.33151 |
| **BRD1** | 4.32869 | 1.66776 | 2.13311 | 3.71511 | 0.48878 | 2.68882 | 2.76847 | 0.19363 | 2.76760 | 10.88465 | 1.85073 | 2.06352 |
| **BRD2** | 12.01342 | 12.83364 | 19.94154 | 20.28096 | 20.38204 | 4.67903 | 29.20905 | 13.40698 | 27.44913 | 28.72876 | 15.08284 | 24.77318 |
| **BRD3** | 5.58805 | 18.67004 | 2.41457 | 24.01816 | 14.76031 | 4.24899 | 19.94861 | 0.08397 | 1.31283 | 13.20459 | 14.26144 | 10.20455 |
| **BRD4** | 10.20616 | 3.22545 | 11.14129 | 8.96008 | 17.18715 | 0.00000 | 19.36639 | 5.79706 | 3.33367 | 41.84072 | 1.55384 | 7.66194 |
| **BRD7** | 2.25996 | 1.87499 | 2.01805 | 0.35891 | 2.37744 | 4.48053 | 3.37215 | 0.92258 | 0.00000 | 11.35406 | 1.18032 | 1.30440 |
| **BRD7P3** | 0.00128 | 0.01591 | 0.09710 | 0.01158 | 0.01827 | 0.07913 | 0.00000 | 0.57119 | 0.00000 | 0.09174 | 0.05700 | 0.17141 |
| **BRD8** | 1.83865 | 13.75298 | 4.95976 | 9.28025 | 1.44133 | 13.29846 | 4.51909 | 8.56890 | 16.30745 | 9.59055 | 6.16005 | 16.99634 |
| **BRD9** | 0.83453 | 4.94981 | 5.77251 | 3.74891 | 10.08553 | 0.00000 | 3.75020 | 3.06228 | 2.20530 | 17.48245 | 5.00979 | 2.70526 |
| **BRDT** | 3.28341 | 9.83816 | 6.25737 | 19.31015 | 5.21040 | 19.75615 | 3.38450 | 30.99719 | 4.12940 | 16.96678 | 4.80450 | 29.35913 |
| **BRE** | 0.28960 | 4.85204 | 3.40085 | 2.48159 | 1.42529 | 8.04364 | 0.00000 | 0.00000 | 2.86881 | 0.01356 | 7.30831 | 1.12105 |
| **BRE-AS1** | 0.06253 | 0.19806 | 0.00000 | 0.08734 | 0.00000 | 0.12340 | 0.00000 | 0.00000 | 0.26134 | 0.18758 | 0.17674 | 0.22417 |
| **BREA2** | 0.05106 | 0.07962 | 0.01244 | 0.00101 | 0.09961 | 0.00000 | 0.06194 | 0.05993 | 0.00000 | 0.26329 | 0.00272 | 0.13405 |
| **BRF1** | 0.49418 | 1.03591 | 0.92612 | 0.41347 | 2.43139 | 0.00000 | 0.59412 | 1.03252 | 1.21952 | 5.49055 | 0.93157 | 1.25011 |
| **BRF2** | 0.13566 | 8.76704 | 7.59152 | 3.35464 | 1.66584 | 3.98399 | 5.11261 | 5.77366 | 4.23072 | 0.00000 | 4.48112 | 10.11915 |
| **BRI3** | 2.79514 | 1.35374 | 4.52810 | 0.56238 | 4.70623 | 0.00000 | 7.72717 | 0.19839 | 0.24589 | 7.74833 | 1.10844 | 0.14573 |
| **BRI3BP** | 1.32796 | 0.94064 | 2.32872 | 0.10474 | 1.10150 | 0.00000 | 1.65066 | 0.01007 | 2.06941 | 3.19870 | 1.56360 | 0.77310 |
| **BRICD5** | 0.02996 | 0.17980 | 0.85781 | 0.52222 | 0.10798 | 0.00000 | 0.05669 | 0.00094 | 0.90174 | 1.03643 | 0.00000 | 0.06206 |

| | | | | | | | | | | | | |
|---|---|---|---|---|---|---|---|---|---|---|---|---|
| **BRINP1** | 3.72567 | 0.16182 | 0.31042 | 1.17826 | 0.19555 | 0.00000 | 0.00000 | 1.11006 | 0.51076 | 0.02769 | 0.00000 | 0.00000 |
| **BRINP2** | 0.00618 | 0.05149 | 0.01515 | 0.01031 | 0.01749 | 0.00641 | 0.02346 | 0.01082 | 0.01827 | 0.01548 | 0.01914 | 0.01390 |
| **BRINP3** | 0.10772 | 0.01965 | 0.09887 | 0.01219 | 0.02391 | 0.00000 | 0.02713 | 0.06837 | 0.00828 | 0.00500 | 0.01662 | 0.01730 |
| **BRIP1** | 0.45525 | 0.40386 | 0.57040 | 0.60144 | 0.34396 | 4.76400 | 0.00000 | 0.07063 | 0.92713 | 1.73380 | 0.82828 | 1.82914 |
| **BRIX1** | 1.76747 | 11.89808 | 9.74173 | 3.36393 | 5.61105 | 49.02869 | 14.12807 | 23.66814 | 9.33693 | 12.03183 | 13.24688 | 20.30759 |
| **BRK1** | 1.77788 | 13.10261 | 16.71267 | 15.16498 | 13.15759 | 19.87450 | 20.97811 | 1.45326 | 16.88724 | 7.47307 | 15.17449 | 26.36967 |
| **BRMS1** | 1.52047 | 20.60000 | 15.98594 | 17.30413 | 26.25620 | 3.84970 | 29.13136 | 9.60438 | 17.97180 | 0.00000 | 14.12537 | 25.22969 |
| **BRMS1L** | 0.00000 | 5.46285 | 4.63767 | 5.52598 | 0.00000 | 17.93573 | 5.96385 | 3.77815 | 2.30281 | 0.90450 | 1.37717 | 5.36760 |
| **BROX** | 0.00000 | 7.02329 | 8.97830 | 2.18183 | 0.00583 | 30.08700 | 12.12899 | 8.35291 | 5.51597 | 3.50556 | 11.19677 | 11.27811 |
| **BRPF1** | 3.95730 | 3.67647 | 3.73742 | 2.03682 | 2.99716 | 1.03837 | 3.08213 | 2.37963 | 3.83416 | 12.00244 | 0.84420 | 4.42887 |
| **BRPF3** | 1.27648 | 4.84722 | 1.38864 | 3.43502 | 5.11874 | 7.83721 | 9.17657 | 0.38927 | 1.20452 | 10.49897 | 1.89383 | 7.91734 |
| **BRSK1** | 0.36945 | 0.01489 | 0.00000 | 0.16682 | 0.00000 | 0.34596 | 0.25034 | 0.15638 | 0.00000 | 1.77403 | 0.48601 | 0.06542 |
| **BRSK2** | 0.13331 | 0.01386 | 0.06593 | 0.03255 | 0.00000 | 0.02066 | 0.00000 | 0.03221 | 0.03466 | 0.25037 | 0.04991 | 0.03273 |
| **BRWD1** | 10.74520 | 5.52573 | 3.05603 | 4.44891 | 4.88820 | 14.77233 | 5.87330 | 7.85916 | 5.30091 | 22.45713 | 3.44345 | 7.51389 |
| **BRWD1-AS1** | 0.02385 | 0.01273 | 0.00806 | 0.00212 | 0.00803 | 0.05073 | 0.01134 | 0.00908 | 0.01277 | 0.06491 | 0.00923 | 0.01152 |
| **BRWD1-IT2** | 0.50993 | 0.05606 | 0.06059 | 0.04562 | 0.00000 | 0.01493 | 0.00000 | 0.00000 | 0.02827 | 0.07480 | 0.01022 | 0.05214 |
| **BRWD3** | 1.75173 | 1.56144 | 0.55226 | 4.12440 | 0.00107 | 10.42675 | 0.02002 | 1.67770 | 0.00000 | 13.60979 | 0.31442 | 10.44115 |
| **BSCL2.uid-1559** | 0.02405 | 0.10476 | 0.07452 | 0.23243 | 0.08249 | 0.16823 | 0.02173 | 0.10440 | 0.31705 | 0.00000 | 0.03916 | 0.32189 |
| **BSCL2.uid-1560** | 0.02385 | 0.10295 | 0.07728 | 0.05956 | 0.04676 | 0.00000 | 0.10634 | 0.00982 | 0.24037 | 0.04087 | 0.01064 | 0.09423 |
| **BSCL2.uid-1560** | 0.30620 | 0.36394 | 0.61196 | 0.04754 | 0.26636 | 0.04008 | 0.13455 | 0.15787 | 0.41904 | 0.00000 | 0.20929 | 0.44417 |
| **BSDC1** | 4.93461 | 7.56367 | 8.02829 | 6.94198 | 3.54107 | 4.87605 | 7.37782 | 2.49980 | 12.69447 | 2.96029 | 3.54730 | 14.34684 |
| **BSG** | 9.51123 | 47.12648 | 33.78635 | 25.27741 | 40.15391 | 23.20735 | 12.79628 | 27.29409 | 63.80360 | 28.06871 | 26.42404 | 63.34128 |
| **BSN** | 1.44906 | 0.10472 | 0.18987 | 0.00940 | 0.00000 | 0.00000 | 0.00000 | 0.00281 | 0.00000 | 0.53520 | 0.48256 | 0.14568 |
| **BSN-AS2** | 0.00644 | 0.01345 | 0.01189 | 0.03094 | 0.00801 | 0.19768 | 0.03159 | 0.00795 | 0.00000 | 0.06748 | 0.01109 | 0.03277 |
| **BSND** | 0.00944 | 0.16013 | 0.01611 | 0.04690 | 0.01698 | 0.00000 | 0.02256 | 0.00907 | 0.01836 | 0.00231 | 0.01382 | 0.05061 |
| **BSPRY** | 0.00000 | 7.35140 | 0.41687 | 0.00091 | 7.61468 | 0.69241 | 2.52908 | 0.00000 | 2.92584 | 6.61884 | 3.36331 | 2.13856 |
| **BST1** | 0.02917 | 0.14168 | 0.16373 | 0.10769 | 0.00000 | 0.27601 | 0.05576 | 0.08561 | 0.02584 | 0.00000 | 0.11393 | 0.01194 |
| **BST2** | 0.00315 | 0.02456 | 0.00090 | 0.29701 | 0.00000 | 0.23951 | 0.00000 | 0.04420 | 0.01814 | 0.02018 | 0.01285 | 0.00191 |
| **BTAF1** | 0.36166 | 1.80993 | 5.60483 | 0.80868 | 0.95421 | 15.50997 | 2.53131 | 12.41926 | 10.66922 | 17.35399 | 1.59983 | 6.25303 |
| **BTBD1** | 0.00000 | 1.49485 | 3.52888 | 2.12871 | 0.00015 | 4.81135 | 5.07605 | 0.04945 | 1.00191 | 7.89381 | 2.70290 | 0.73317 |
| **BTBD10** | 3.81022 | 3.14080 | 4.96153 | 1.62087 | 2.15002 | 10.89382 | 8.08633 | 5.96868 | 8.71707 | 0.00000 | 9.02308 | 4.57516 |
| **BTBD11** | 1.91127 | 0.03349 | 0.00000 | 0.00000 | 0.03232 | 0.32074 | 0.05465 | 0.01797 | 0.00000 | 0.00000 | 0.00687 | 0.06933 |
| **BTBD16** | 0.00528 | 0.01591 | 0.05096 | 0.00958 | 0.00587 | 0.02164 | 0.01183 | 0.01040 | 0.01564 | 0.06620 | 0.01354 | 0.01282 |
| **BTBD17** | 0.00605 | 0.01309 | 0.01637 | 0.01136 | 0.01195 | 0.01667 | 0.02095 | 0.01049 | 0.00000 | 0.02163 | 0.06375 | 0.01618 |
| **BTBD18** | 0.42803 | 0.00000 | 0.12307 | 0.21529 | 0.17752 | 0.05371 | 0.07718 | 0.57324 | 0.00000 | 0.34992 | 0.26753 | 1.22376 |
| **BTBD19** | 0.01748 | 0.05675 | 0.05693 | 0.01200 | 0.03246 | 0.01644 | 0.04327 | 0.00802 | 0.03509 | 0.00000 | 0.05604 | 0.03662 |
| **BTBD2** | 3.86677 | 0.84513 | 1.28029 | 0.95283 | 1.46753 | 0.00000 | 1.55217 | 1.54040 | 2.47633 | 3.94677 | 0.79356 | 1.05629 |
| **BTBD3** | 1.25640 | 2.46032 | 2.68039 | 2.23934 | 4.47097 | 7.39396 | 5.35338 | 0.50255 | 4.62197 | 4.17689 | 8.42171 | 0.90644 |
| **BTBD6** | 0.86227 | 2.00078 | 0.19762 | 0.00000 | 1.82234 | 0.99666 | 1.21023 | 0.10988 | 0.28469 | 6.17099 | 0.19582 | 2.00356 |
| **BTBD7** | 1.02229 | 0.72023 | 3.97597 | 3.05076 | 2.98951 | 2.84300 | 3.97335 | 2.40773 | 3.50679 | 11.20603 | 0.38344 | 2.05775 |
| **BTBD8** | 0.16012 | 0.00000 | 0.21710 | 0.35431 | 0.00569 | 0.52642 | 0.00000 | 0.00870 | 0.00000 | 0.00000 | 0.34177 | 0.50074 |
| **BTBD9** | 1.20594 | 0.12788 | 0.95206 | 0.94708 | 1.31131 | 0.00000 | 1.24100 | 0.14326 | 0.11335 | 3.83109 | 0.67330 | 1.01881 |
| **BTD** | 0.24031 | 0.75959 | 2.79121 | 2.39712 | 0.94480 | 2.04265 | 0.00000 | 0.71353 | 4.68952 | 0.00000 | 2.11956 | 1.54551 |
| **BTF3** | 12.62409 | 24.56328 | 28.56030 | 18.23743 | 23.05569 | 46.39628 | 37.67671 | 9.10694 | 42.80911 | 17.16784 | 27.61180 | 21.27342 |
| **BTF3L4** | 1.06474 | 6.38755 | 6.51461 | 4.63710 | 2.08379 | 32.81899 | 16.68167 | 1.27677 | 9.22316 | 9.42832 | 14.12631 | 8.74043 |
| **BTF3P11** | 0.00714 | 0.00311 | 0.56922 | 0.05411 | 0.00000 | 0.20202 | 0.00000 | 0.02556 | 0.00000 | 0.17954 | 0.01982 | 0.00426 |
| **BTG1** | 3.51475 | 0.00000 | 19.95860 | 2.70788 | 7.88786 | 5.21274 | 26.99989 | 5.45612 | 8.33159 | 20.46887 | 5.25623 | 1.47261 |
| **BTG2** | 0.00000 | 9.84598 | 4.45011 | 8.94424 | 2.04967 | 6.98985 | 4.72687 | 0.00000 | 4.16976 | 1.05336 | 2.19951 | 0.00000 |

| | | | | | | | | | | | | |
|---|---|---|---|---|---|---|---|---|---|---|---|---|
| **BTG3** | 0.08568 | 10.83480 | 10.41729 | 2.56012 | 3.72198 | 20.29081 | 6.19537 | 4.55897 | 7.52960 | 5.21393 | 3.07437 | 14.39257 |
| **BTG4** | 7.20689 | 0.06377 | 0.03716 | 0.00231 | 0.02342 | 0.01015 | 0.01493 | 0.13040 | 0.01884 | 0.00000 | 0.04273 | 0.04264 |
| **BTK** | 0.00053 | 0.00000 | 0.57331 | 2.29136 | 0.61520 | 0.69067 | 0.00000 | 3.24589 | 0.23075 | 0.22087 | 0.15679 | 1.91509 |
| **BTLA** | 0.15454 | 0.53380 | 0.35526 | 4.09087 | 0.00000 | 0.30651 | 0.00000 | 4.69142 | 0.00000 | 0.18966 | 0.00000 | 3.83227 |
| **BTN1A1** | 0.00296 | 3.02332 | 0.00000 | 0.00000 | 0.01505 | 0.00000 | 0.03512 | 0.05507 | 0.00000 | 0.04237 | 0.00000 | 19.47534 |
| **BTN2A1** | 0.01910 | 0.98948 | 2.44955 | 2.58833 | 0.26630 | 2.14492 | 0.19441 | 0.00006 | 3.09206 | 0.01902 | 0.54558 | 0.00000 |
| **BTN2A2** | 0.00565 | 0.00985 | 0.06957 | 0.02997 | 0.00000 | 0.02592 | 0.01285 | 0.01036 | 0.00493 | 0.02802 | 0.07985 | 0.02049 |
| **BTN2A3P** | 0.00500 | 0.01131 | 0.01088 | 0.01466 | 0.00000 | 0.01654 | 0.01920 | 0.01119 | 0.01151 | 0.09220 | 0.04674 | 0.01315 |
| **BTN3A1** | 0.00297 | 0.00363 | 0.00000 | 0.00083 | 0.00223 | 0.01926 | 0.00514 | 0.00924 | 0.01489 | 0.59102 | 0.00000 | 0.00666 |
| **BTN3A2** | 0.00611 | 0.01445 | 0.02266 | 0.02182 | 0.01440 | 0.01465 | 0.01903 | 0.01050 | 0.01442 | 0.02111 | 0.01539 | 0.01611 |
| **BTN3A3** | 0.00633 | 0.01541 | 0.01472 | 0.01311 | 0.01419 | 0.01665 | 0.01688 | 0.01057 | 0.03602 | 0.01135 | 0.01428 | 0.01658 |
| **BTNL10** | 0.00462 | 0.00828 | 0.00481 | 0.00508 | 0.00200 | 0.02529 | 0.01620 | 0.01016 | 0.01612 | 0.11974 | 0.03993 | 0.01150 |
| **BTNL2** | 0.00620 | 0.01590 | 0.01297 | 0.02386 | 0.01542 | 0.01510 | 0.01749 | 0.03917 | 0.01851 | 0.01510 | 0.01640 | 0.01585 |
| **BTNL3** | 1.20287 | 0.00752 | 0.08317 | 0.02920 | 0.05009 | 0.00000 | 0.01779 | 2.09264 | 0.00000 | 0.00000 | 0.05483 | 0.01638 |
| **BTNL8** | 0.08227 | 0.01185 | 0.07575 | 0.02283 | 0.00315 | 0.03312 | 0.00000 | 0.07528 | 0.01513 | 0.04842 | 0.01316 | 0.02292 |
| **BTNL9** | 0.13842 | 0.12943 | 0.01695 | 0.00000 | 0.01839 | 0.01347 | 0.01520 | 0.45311 | 0.01376 | 0.02418 | 0.01609 | 0.69332 |
| **BTRC** | 8.52210 | 13.49290 | 0.82841 | 1.71803 | 0.00000 | 10.18556 | 2.33514 | 11.59190 | 4.98983 | 5.28908 | 5.15719 | 21.74481 |
| **BUB1** | 5.14644 | 17.20880 | 6.81011 | 7.08086 | 12.92749 | 42.31606 | 0.01481 | 9.71794 | 17.64027 | 13.05964 | 8.23951 | 15.01183 |
| **BUB1B** | 6.26359 | 3.39258 | 3.10669 | 1.79892 | 10.85761 | 24.83758 | 6.35322 | 1.17782 | 10.80455 | 18.97584 | 6.38659 | 7.07572 |
| **BUB3** | 4.61674 | 21.49930 | 24.41420 | 17.53162 | 19.07281 | 41.57412 | 30.96031 | 21.05084 | 25.34495 | 6.48801 | 23.23792 | 27.09561 |
| **BUD13** | 3.22360 | 7.90166 | 4.01142 | 4.33950 | 3.40152 | 4.91182 | 4.53268 | 2.53005 | 3.37774 | 0.00000 | 7.30125 | 8.07632 |
| **BUD31** | 2.83986 | 9.55004 | 13.81288 | 8.40554 | 7.76825 | 25.45740 | 16.88560 | 22.33890 | 15.81681 | 0.00000 | 10.42569 | 12.23201 |
| **BVES** | 2.05804 | 0.00667 | 0.08716 | 0.12551 | 0.01112 | 0.00000 | 0.02007 | 0.31472 | 0.04575 | 0.02597 | 0.01181 | 0.12162 |
| **BVES-AS1** | 0.00859 | 0.02291 | 0.01561 | 0.01586 | 0.01877 | 0.00950 | 0.02095 | 0.01012 | 0.02125 | 0.00540 | 0.01540 | 0.02811 |
| **BYSL** | 1.81870 | 21.25862 | 19.60340 | 10.78719 | 16.35841 | 4.63771 | 26.21680 | 10.51301 | 24.90992 | 0.00000 | 17.09254 | 29.12945 |
| **BZRAP1** | 0.11884 | 0.07029 | 0.28699 | 0.00559 | 0.00642 | 0.00000 | 0.03460 | 0.00411 | 0.16628 | 0.00000 | 0.04415 | 0.06138 |
| **BZRAP1-AS1** | 0.01074 | 0.23590 | 0.21791 | 0.00000 | 0.00000 | 0.04772 | 0.30017 | 0.22781 | 0.29938 | 0.21188 | 0.01744 | 0.18206 |
| **BZW1** | 1.70809 | 11.02122 | 8.76256 | 6.60583 | 3.13931 | 29.11899 | 12.77129 | 23.79008 | 13.96490 | 0.00048 | 8.31962 | 16.37121 |
| **BZW2** | 2.33162 | 15.78415 | 32.74040 | 12.72697 | 12.80181 | 26.10315 | 28.41494 | 15.30348 | 28.16438 | 8.84901 | 19.42049 | 17.91450 |
| **C10orf10** | 0.04232 | 1.11803 | 11.59816 | 4.35688 | 27.29004 | 1.73426 | 3.09268 | 0.00000 | 0.00000 | 0.00000 | 12.21495 | 0.00000 |
| **C10orf105** | 0.00730 | 0.01822 | 0.01559 | 0.01461 | 0.01480 | 0.01801 | 0.02455 | 0.01168 | 0.01525 | 0.01062 | 0.01588 | 0.01626 |
| **C10orf11** | 0.59884 | 0.04438 | 0.05379 | 0.00432 | 0.02391 | 0.01186 | 0.00335 | 0.08008 | 0.16389 | 0.00000 | 0.00223 | 0.02547 |
| **C10orf111** | 0.00302 | 0.30232 | 0.13932 | 0.11740 | 0.07684 | 0.29560 | 0.00000 | 0.03993 | 0.44738 | 0.00000 | 0.00964 | 0.39247 |
| **C10orf113** | 0.00601 | 0.01457 | 0.05616 | 0.00903 | 0.01741 | 0.01918 | 0.01454 | 0.01043 | 0.00735 | 0.02692 | 0.01579 | 0.01556 |
| **C10orf12** | 2.00558 | 1.80358 | 1.55285 | 0.91055 | 0.00000 | 1.37827 | 2.04469 | 1.10763 | 0.19936 | 5.36107 | 1.68786 | 3.23302 |
| **C10orf120** | 0.46752 | 0.01292 | 0.01372 | 0.01142 | 0.01542 | 0.01476 | 0.01728 | 0.16254 | 0.02008 | 0.01038 | 0.01378 | 0.01318 |
| **C10orf126** | 0.00288 | 0.00000 | 0.04075 | 0.30174 | 0.00000 | 0.05762 | 0.00737 | 0.05478 | 0.00000 | 0.18179 | 0.00963 | 0.00723 |
| **C10orf131** | 0.02876 | 0.05578 | 0.10981 | 0.00431 | 0.16711 | 0.30858 | 0.07442 | 0.00415 | 0.12812 | 0.00000 | 0.40743 | 0.24957 |
| **C10orf2** | 0.57922 | 5.74460 | 2.95947 | 2.33259 | 3.48543 | 0.21093 | 8.75572 | 7.29897 | 7.65223 | 8.48716 | 3.90763 | 11.68606 |
| **C10orf25** | 0.00097 | 0.44194 | 0.34110 | 0.00000 | 0.54389 | 0.14405 | 0.18276 | 0.18403 | 0.00000 | 0.58711 | 0.53959 | 0.55949 |
| **C10orf32** | 0.00000 | 0.71566 | 3.79589 | 0.00000 | 0.00000 | 6.79266 | 6.12530 | 0.01054 | 1.15628 | 1.69613 | 1.02152 | 1.25094 |
| **C10orf35** | 1.45402 | 0.47597 | 12.06271 | 0.00000 | 0.00000 | 1.55127 | 7.49069 | 0.34260 | 3.00519 | 0.00000 | 2.05105 | 0.27449 |
| **C10orf40** | 0.00627 | 0.01544 | 0.02876 | 0.01201 | 0.01550 | 0.01601 | 0.01408 | 0.01041 | 0.01790 | 0.01680 | 0.01566 | 0.01792 |
| **C10orf54** | 0.00623 | 0.31251 | 1.97531 | 0.00154 | 0.35441 | 0.00000 | 0.00000 | 0.00740 | 0.00000 | 0.00514 | 0.00718 | 0.11556 |
| **C10orf55** | 0.00289 | 0.00742 | 0.00949 | 0.04660 | 0.00963 | 0.04347 | 0.00668 | 0.02357 | 0.01950 | 0.04387 | 0.01071 | 0.03218 |
| **C10orf67** | 0.00961 | 0.07509 | 0.02922 | 0.00000 | 0.00000 | 0.07649 | 0.00000 | 0.01232 | 0.00000 | 0.15486 | 0.02520 | 0.08709 |
| **C10orf71** | 0.01047 | 0.02308 | 0.06350 | 0.01070 | 0.03061 | 0.00000 | 0.00530 | 0.01055 | 0.13369 | 0.00000 | 0.00710 | 0.02207 |
| **C10orf76** | 0.00000 | 4.09134 | 2.48834 | 5.00709 | 0.57326 | 3.93284 | 1.87319 | 3.31402 | 2.03650 | 7.06521 | 2.51442 | 6.30907 |

| | | | | | | | | | | | | |
|---|---|---|---|---|---|---|---|---|---|---|---|---|
| **C10orf82** | 5.17922 | 0.01195 | 0.13489 | 0.00960 | 0.03664 | 0.52605 | 0.13082 | 0.59317 | 0.28671 | 0.00000 | 0.00557 | 0.00000 |
| **C10orf88** | 0.00000 | 3.34370 | 5.88425 | 0.06973 | 0.11806 | 6.40048 | 3.86090 | 2.66962 | 3.93503 | 0.00000 | 2.51104 | 4.95066 |
| **C10orf90** | 0.11269 | 0.00947 | 0.05241 | 1.27510 | 0.05236 | 0.01601 | 0.00760 | 0.00459 | 0.00000 | 0.01104 | 0.01358 | 0.01262 |
| **C10orf91** | 0.14313 | 0.02291 | 0.01799 | 0.01388 | 0.04419 | 0.00000 | 0.01950 | 0.02976 | 0.02753 | 0.00272 | 0.01316 | 0.02109 |
| **C10orf95** | 0.44274 | 0.34072 | 0.22084 | 0.29328 | 0.20373 | 0.01983 | 0.05878 | 0.00000 | 0.07193 | 0.52932 | 0.35657 | 0.66478 |
| **C10orf99** | 0.01075 | 0.00511 | 0.00333 | 0.04622 | 0.00000 | 0.09772 | 0.00505 | 0.00686 | 0.00832 | 0.01697 | 0.00961 | 0.05248 |
| **C11orf1** | 0.11795 | 1.30463 | 1.27524 | 0.71727 | 0.32798 | 1.10403 | 0.52718 | 0.00000 | 0.68898 | 0.62689 | 1.24770 | 1.35240 |
| **C11orf16** | 0.14353 | 0.10930 | 0.04989 | 0.00186 | 0.13753 | 0.00271 | 0.03724 | 0.19741 | 0.12615 | 0.00000 | 0.14758 | 0.19870 |
| **C11orf21** | 0.00994 | 0.01504 | 0.01440 | 0.01193 | 0.02124 | 0.01660 | 0.02104 | 0.02101 | 0.01639 | 0.01353 | 0.01528 | 0.01594 |
| **C11orf24** | 3.65802 | 7.36219 | 7.23761 | 2.36108 | 6.61564 | 0.00000 | 1.62273 | 0.63376 | 14.24877 | 0.00000 | 4.55075 | 8.28877 |
| **C11orf30** | 2.73188 | 1.29433 | 1.97734 | 2.66155 | 1.39633 | 3.47808 | 0.87358 | 0.72119 | 5.04492 | 6.51986 | 0.88659 | 2.66601 |
| **C11orf31** | 9.13964 | 21.82384 | 12.90418 | 11.12124 | 27.99326 | 16.03927 | 12.10900 | 16.33090 | 11.45526 | 10.83499 | 14.39053 | 22.43879 |
| **C11orf40** | 8.17210 | 0.00862 | 0.00253 | 0.06648 | 0.14856 | 0.00000 | 0.01294 | 3.24715 | 0.01207 | 0.02122 | 0.01853 | 0.64477 |
| **C11orf42** | 0.00647 | 0.02336 | 0.04603 | 0.01053 | 0.04912 | 0.00000 | 0.00744 | 0.01024 | 0.09313 | 0.00498 | 0.01182 | 0.02393 |
| **C11orf44** | 0.00850 | 0.01838 | 0.01981 | 0.01310 | 0.01423 | 0.03115 | 0.02315 | 0.01003 | 0.01176 | 0.01727 | 0.01575 | 0.01971 |
| **C11orf45** | 0.51394 | 0.42583 | 0.00204 | 0.00085 | 0.08476 | 0.27076 | 0.12201 | 0.06965 | 0.09576 | 0.00000 | 0.00000 | 0.23448 |
| **C11orf48** | 0.17364 | 0.20242 | 0.53112 | 0.19810 | 0.53784 | 0.00000 | 0.01784 | 0.82161 | 1.57258 | 0.97220 | 0.45991 | 1.23116 |
| **C11orf49** | 0.00154 | 2.53383 | 2.15287 | 1.30537 | 2.29598 | 2.62250 | 2.49122 | 0.26161 | 4.24245 | 0.00000 | 2.27162 | 1.14697 |
| **C11orf53** | 0.37372 | 0.01400 | 0.03295 | 0.01323 | 0.01484 | 0.01342 | 0.01977 | 0.00164 | 0.01593 | 0.01133 | 0.01453 | 0.01531 |
| **C11orf54** | 0.00000 | 2.38494 | 7.76532 | 1.13649 | 0.00000 | 8.90510 | 6.39637 | 2.31710 | 8.98344 | 3.65179 | 0.84711 | 2.07933 |
| **C11orf57** | 3.35971 | 3.56604 | 5.06806 | 0.67915 | 2.13495 | 6.92439 | 11.13075 | 2.13352 | 1.89763 | 11.82293 | 2.49758 | 6.86759 |
| **C11orf58** | 4.25213 | 16.52095 | 16.46151 | 10.91868 | 14.48710 | 45.54594 | 23.23900 | 2.87859 | 16.23511 | 16.47844 | 19.13652 | 21.97364 |
| **C11orf63** | 0.05550 | 0.03725 | 0.05137 | 0.03433 | 0.01279 | 0.00508 | 0.06012 | 0.05226 | 0.07718 | 0.00000 | 0.01047 | 0.02123 |
| **C11orf65** | 0.31968 | 0.84089 | 0.00945 | 0.00000 | 0.01016 | 0.29364 | 0.00000 | 0.29325 | 0.00000 | 0.01479 | 0.08266 | 1.48604 |
| **C11orf68** | 0.24386 | 0.16852 | 0.51550 | 0.87299 | 0.78726 | 0.12817 | 0.71330 | 0.09239 | 0.00000 | 3.06953 | 0.09466 | 0.58797 |
| **C11orf70** | 0.07049 | 0.00000 | 2.54512 | 0.00000 | 0.00000 | 0.01210 | 1.31424 | 0.01611 | 1.02021 | 0.00000 | 0.25168 | 0.19178 |
| **C11orf71** | 1.84510 | 3.08158 | 2.79700 | 0.00000 | 0.67185 | 5.43861 | 0.00000 | 4.67187 | 0.50025 | 2.28950 | 2.66781 | 4.20001 |
| **C11orf73** | 0.15437 | 7.08247 | 8.15736 | 5.95399 | 4.56370 | 11.26093 | 12.10610 | 0.08604 | 6.91514 | 0.00000 | 10.90197 | 2.10254 |
| **C11orf74** | 1.86029 | 2.36174 | 0.11698 | 1.14170 | 2.79136 | 5.11136 | 1.36178 | 2.00671 | 0.17027 | 0.19239 | 3.29361 | 0.69244 |
| **C11orf80** | 0.06853 | 0.78924 | 0.17502 | 0.11578 | 0.62146 | 0.00000 | 0.08979 | 0.25602 | 0.00000 | 0.93946 | 0.82446 | 0.32307 |
| **C11orf83** | 0.44033 | 16.38616 | 0.56050 | 7.34535 | 20.90131 | 0.92866 | 3.46260 | 16.39625 | 5.16173 | 4.85462 | 3.78273 | 17.60561 |
| **C11orf84** | 2.66773 | 0.59746 | 0.43225 | 0.40920 | 0.54408 | 0.00000 | 2.07009 | 0.01426 | 0.00826 | 3.81815 | 0.00111 | 0.73464 |
| **C11orf85** | 2.00538 | 0.10592 | 0.03804 | 0.01945 | 0.01317 | 0.00000 | 0.00947 | 0.22464 | 0.02436 | 0.01382 | 0.01536 | 0.50184 |
| **C11orf86** | 0.00670 | 0.01393 | 0.00739 | 0.01306 | 0.12615 | 0.05217 | 0.01523 | 0.35582 | 0.04928 | 0.00000 | 0.00369 | 0.01008 |
| **C11orf87** | 0.00224 | 0.00286 | 0.03293 | 0.05806 | 0.00623 | 0.04845 | 0.00948 | 0.03609 | 0.00698 | 0.01990 | 0.01500 | 0.16010 |
| **C11orf88** | 0.10458 | 0.00000 | 0.00000 | 0.00000 | 0.09516 | 0.03397 | 0.57839 | 0.00587 | 0.19874 | 0.11020 | 1.34502 | 0.00922 |
| **C11orf91** | 0.03353 | 0.01125 | 0.08191 | 0.00679 | 0.02812 | 0.01172 | 0.08242 | 0.06667 | 0.00000 | 0.02622 | 0.01660 | 0.04949 |
| **C11orf94** | 0.00517 | 0.01049 | 0.08705 | 0.01480 | 0.03270 | 0.00000 | 0.17443 | 0.00919 | 0.00306 | 0.08743 | 0.01119 | 0.03687 |
| **C11orf95** | 0.00000 | 0.14259 | 0.35203 | 2.08426 | 0.57204 | 2.04751 | 3.24852 | 0.44565 | 1.13969 | 7.29061 | 0.00000 | 1.42270 |
| **C11orf96** | 0.04635 | 0.01290 | 0.26144 | 0.11967 | 0.08214 | 0.00000 | 0.05348 | 0.00895 | 0.00000 | 0.01322 | 0.01361 | 0.01637 |
| **C12orf10** | 0.02526 | 12.53822 | 12.58144 | 7.73915 | 4.92319 | 0.00012 | 14.70288 | 6.97373 | 8.34778 | 0.00000 | 8.26453 | 8.33444 |
| **C12orf29** | 0.65819 | 3.35525 | 4.83532 | 0.28655 | 1.23219 | 17.07514 | 7.70023 | 0.21783 | 9.85123 | 0.00000 | 6.97946 | 7.36312 |
| **C12orf36** | 0.00862 | 0.18374 | 0.17897 | 1.01746 | 0.02899 | 0.03333 | 0.00001 | 0.00552 | 0.06456 | 0.00000 | 0.00784 | 0.52522 |
| **C12orf4** | 1.94733 | 5.13305 | 6.66798 | 1.12083 | 0.00002 | 10.63296 | 4.03692 | 2.92956 | 3.06638 | 1.17389 | 3.65020 | 6.43567 |
| **C12orf40** | 0.33146 | 0.59270 | 0.05167 | 0.00000 | 0.31046 | 0.42138 | 0.00000 | 0.70198 | 0.00000 | 0.18184 | 0.35576 | 0.59030 |
| **C12orf42** | 0.08766 | 0.08242 | 0.01049 | 0.04488 | 0.03100 | 0.02807 | 0.02831 | 0.01160 | 0.02181 | 0.00000 | 0.01249 | 0.02228 |
| **C12orf43** | 1.62768 | 8.32618 | 6.93324 | 4.55745 | 5.23578 | 5.54847 | 9.14405 | 12.88967 | 6.23142 | 5.25606 | 4.28932 | 16.80267 |
| **C12orf45** | 0.40006 | 10.56907 | 11.60753 | 4.69074 | 10.30999 | 16.95805 | 18.89532 | 11.12883 | 7.42119 | 4.01561 | 9.74056 | 16.31319 |

| | | | | | | | | | | | | |
|---|---|---|---|---|---|---|---|---|---|---|---|---|
| **C12orf49** | 1.03389 | 2.72756 | 3.26787 | 1.27722 | 2.46375 | 2.24789 | 1.16410 | 2.44148 | 3.67822 | 3.98865 | 0.00000 | 3.25557 |
| **C12orf5** | 2.21416 | 16.55652 | 22.42740 | 1.42800 | 19.57613 | 44.46556 | 32.42212 | 8.90765 | 26.26435 | 9.75565 | 24.46307 | 26.61676 |
| **C12orf50** | 0.23901 | 0.01137 | 0.01570 | 0.01481 | 0.01987 | 0.00037 | 0.04100 | 0.19625 | 0.01770 | 0.00538 | 0.01736 | 0.06049 |
| **C12orf54** | 0.03185 | 0.01103 | 0.01670 | 0.02583 | 0.00127 | 0.03665 | 0.01346 | 0.00793 | 0.02118 | 0.04455 | 0.01740 | 0.04190 |
| **C12orf56** | 0.55666 | 0.59263 | 0.00097 | 0.73808 | 0.87824 | 1.58559 | 0.11078 | 0.08194 | 0.00000 | 0.00000 | 0.13127 | 0.78611 |
| **C12orf57** | 0.01387 | 3.94610 | 11.28153 | 13.59164 | 5.84420 | 0.35957 | 8.92055 | 10.18423 | 6.28948 | 4.67502 | 2.59270 | 17.16101 |
| **C12orf60** | 1.06238 | 0.89078 | 0.81080 | 0.00000 | 0.35822 | 1.96833 | 0.73172 | 5.30290 | 0.00000 | 0.02172 | 0.21718 | 3.60415 |
| **C12orf61** | 0.08207 | 0.01139 | 0.11745 | 0.00440 | 0.01394 | 0.09033 | 0.04194 | 0.04566 | 0.01414 | 0.00000 | 0.01045 | 0.03170 |
| **C12orf65** | 5.12004 | 6.28776 | 2.16556 | 3.11681 | 2.65534 | 2.38674 | 3.64612 | 0.99219 | 3.15730 | 5.37158 | 4.19318 | 0.61738 |
| **C12orf66** | 0.59740 | 1.42480 | 0.85051 | 1.07135 | 0.00000 | 2.32013 | 0.00000 | 1.21880 | 0.00000 | 0.00000 | 1.98173 | 1.48246 |
| **C12orf71** | 0.00575 | 0.00995 | 0.01633 | 0.00891 | 0.01256 | 0.01383 | 0.01097 | 0.00966 | 0.03492 | 0.04924 | 0.03500 | 0.04181 |
| **C12orf73** | 0.00000 | 2.30320 | 3.25098 | 0.39303 | 0.00001 | 7.03866 | 3.57347 | 0.00000 | 6.04288 | 2.91029 | 3.58142 | 1.10707 |
| **C12orf74** | 0.39438 | 0.00968 | 0.01082 | 0.03178 | 0.01396 | 0.02590 | 0.01191 | 0.00145 | 0.02035 | 0.01866 | 0.01257 | 0.02946 |
| **C12orf75** | 0.98348 | 0.11933 | 4.79987 | 2.09521 | 0.00000 | 0.26045 | 2.30442 | 0.08045 | 0.00000 | 0.73137 | 0.17802 | 0.00000 |
| **C12orf76** | 0.24732 | 0.20269 | 0.76698 | 0.00000 | 0.00000 | 2.09752 | 0.81512 | 1.07527 | 2.69978 | 1.19724 | 6.40460 | 2.36423 |
| **C12orf77** | 3.86196 | 0.00755 | 0.00964 | 0.01095 | 0.01107 | 0.02456 | 0.01407 | 0.30106 | 0.01816 | 0.00611 | 0.01166 | 0.01701 |
| **C12orf79** | 0.00762 | 0.01264 | 0.12145 | 0.00127 | 0.04705 | 0.00379 | 0.04407 | 0.05352 | 0.09745 | 0.05308 | 0.00426 | 0.01272 |
| **C12orf80** | 0.00629 | 0.01525 | 0.01616 | 0.01368 | 0.01752 | 0.01638 | 0.02050 | 0.01051 | 0.01890 | 0.01286 | 0.01480 | 0.01648 |
| **C14orf1** | 5.17185 | 4.90108 | 10.65852 | 10.78350 | 26.22945 | 19.18149 | 6.01010 | 1.24365 | 26.67373 | 0.00000 | 19.65497 | 5.22102 |
| **C14orf105** | 0.02700 | 0.01580 | 0.01316 | 0.01430 | 0.01537 | 0.01565 | 0.01357 | 0.01071 | 0.02610 | 0.02354 | 0.01333 | 0.01507 |
| **C14orf119** | 3.50825 | 15.89360 | 19.69507 | 13.79092 | 13.44487 | 37.30400 | 25.44790 | 17.47879 | 20.70145 | 3.33507 | 21.36334 | 29.71272 |
| **C14orf132** | 0.00301 | 0.08152 | 0.02572 | 0.01511 | 0.01279 | 0.00000 | 0.07799 | 0.04730 | 0.00615 | 0.02202 | 0.01584 | 0.01149 |
| **C14orf142** | 0.68161 | 4.57531 | 6.54182 | 1.23016 | 0.00000 | 11.72023 | 8.89428 | 7.83353 | 3.24232 | 0.00199 | 8.07433 | 7.05814 |
| **C14orf159** | 0.00401 | 2.24384 | 0.00000 | 4.72267 | 2.09105 | 1.77906 | 0.00000 | 0.08298 | 0.92819 | 0.64620 | 0.64332 | 1.40943 |
| **C14orf166** | 14.11043 | 13.11978 | 22.21001 | 10.07402 | 16.04985 | 47.01007 | 25.88080 | 17.30542 | 22.87901 | 10.49703 | 15.26304 | 23.04892 |
| **C14orf169** | 0.19902 | 2.24703 | 2.43332 | 3.88502 | 2.47032 | 3.84190 | 6.69714 | 3.00802 | 0.27225 | 4.61438 | 1.28588 | 3.56585 |
| **C14orf178** | 0.05799 | 0.07825 | 0.02753 | 0.06216 | 0.04842 | 0.05846 | 0.06862 | 0.11129 | 0.01802 | 0.00000 | 0.05335 | 0.11028 |
| **C14orf180** | 0.00678 | 0.00862 | 0.01275 | 0.01182 | 0.00688 | 0.03359 | 0.01257 | 0.01009 | 0.01309 | 0.01485 | 0.04445 | 0.05323 |
| **C14orf182** | 0.00265 | 0.00000 | 0.26872 | 0.00000 | 0.28750 | 0.00000 | 0.00000 | 0.10227 | 0.35442 | 0.59726 | 0.26100 | 0.26376 |
| **C14orf183** | 0.05937 | 0.01701 | 0.08991 | 0.02720 | 0.04324 | 0.00000 | 0.00656 | 0.02704 | 0.03363 | 0.04714 | 0.08970 | 0.05116 |
| **C14orf2** | 7.88723 | 22.98179 | 22.76945 | 17.62504 | 24.86141 | 48.35447 | 40.71177 | 18.92310 | 25.24011 | 28.78974 | 33.27232 | 33.92479 |
| **C14orf23** | 0.01628 | 0.01244 | 0.01650 | 0.02817 | 0.01427 | 0.02431 | 0.01184 | 0.00913 | 0.03123 | 0.02108 | 0.01220 | 0.01508 |
| **C14orf28** | 0.18474 | 0.18496 | 0.27116 | 0.38404 | 0.00000 | 0.00000 | 0.00000 | 0.08550 | 0.40457 | 0.32450 | 0.28610 | 0.23688 |
| **C14orf37** | 2.42737 | 1.90824 | 0.00000 | 0.20175 | 0.68917 | 0.00000 | 0.00000 | 1.97216 | 0.00000 | 0.96111 | 0.01567 | 1.35581 |
| **C14orf39** | 0.13213 | 0.00419 | 0.00476 | 0.08037 | 0.02502 | 0.06919 | 0.00839 | 0.00574 | 0.01028 | 0.01505 | 0.01059 | 0.02706 |
| **C14orf64** | 0.00584 | 0.01774 | 0.01905 | 0.01515 | 0.02141 | 0.00466 | 0.05774 | 0.01067 | 0.00000 | 0.03095 | 0.01476 | 0.01622 |
| **C14orf79** | 0.15697 | 0.60621 | 0.13229 | 0.22353 | 0.00000 | 0.29736 | 0.06327 | 0.00181 | 0.21345 | 0.17752 | 0.02295 | 0.82217 |
| **C14orf80** | 0.00000 | 2.74665 | 0.41747 | 0.60719 | 3.47044 | 0.00000 | 1.32673 | 0.04833 | 0.00000 | 5.66699 | 0.00123 | 0.30744 |
| **C14orf93** | 0.28217 | 1.18261 | 1.44572 | 1.72411 | 2.36255 | 0.66358 | 2.23254 | 0.00011 | 3.71495 | 0.00002 | 1.75172 | 0.39425 |
| **C15orf26** | 0.18995 | 0.01527 | 0.04038 | 0.05600 | 0.01183 | 0.00787 | 0.01307 | 0.15536 | 0.01567 | 0.01492 | 0.01628 | 0.12766 |
| **C15orf27** | 0.10327 | 0.03775 | 0.06606 | 0.02178 | 0.00780 | 0.02915 | 0.01021 | 0.00928 | 0.04894 | 0.00000 | 0.10064 | 0.05109 |
| **C15orf32** | 0.00759 | 0.01539 | 0.01466 | 0.01305 | 0.01720 | 0.01374 | 0.01911 | 0.02886 | 0.01987 | 0.01073 | 0.01515 | 0.02046 |
| **C15orf37** | 0.17783 | 0.54603 | 0.26454 | 0.44974 | 0.22469 | 0.96976 | 0.00000 | 0.04485 | 0.76504 | 0.26908 | 0.08258 | 0.56202 |
| **C15orf39** | 2.13214 | 6.81355 | 2.47268 | 4.90443 | 10.16180 | 0.00000 | 7.44620 | 1.36563 | 6.53656 | 15.83710 | 4.48115 | 8.24240 |
| **C15orf40** | 0.43626 | 5.86482 | 2.30672 | 1.72495 | 3.53999 | 7.27428 | 1.13019 | 0.06853 | 2.40001 | 0.03523 | 2.73708 | 6.01105 |
| **C15orf41** | 0.00000 | 5.56708 | 2.39931 | 0.37505 | 0.73677 | 6.81459 | 2.91070 | 6.01732 | 4.95377 | 1.78922 | 1.75540 | 10.31464 |
| **C15orf43** | 0.05041 | 0.22219 | 0.03706 | 0.01171 | 0.00000 | 0.03316 | 0.00000 | 0.02127 | 0.04253 | 0.07140 | 0.01182 | 0.10216 |
| **C15orf48** | 0.02017 | 0.08107 | 0.19962 | 0.06871 | 0.22601 | 0.00000 | 0.08421 | 0.29191 | 0.00000 | 0.00000 | 0.02709 | 0.21064 |

| | | | | | | | | | | | | |
|---|---|---|---|---|---|---|---|---|---|---|---|---|
| **C15orf52** | 0.00506 | 0.01097 | 0.01128 | 0.01756 | 0.00000 | 0.01583 | 0.07032 | 0.01041 | 0.00000 | 0.08191 | 0.07154 | 0.01355 |
| **C15orf56** | 0.02962 | 0.02253 | 0.11004 | 0.02073 | 0.02143 | 0.00699 | 0.00668 | 0.00814 | 0.05487 | 0.01526 | 0.03192 | 0.01607 |
| **C15orf57** | 0.00113 | 5.09989 | 11.32045 | 0.52159 | 0.88168 | 5.64240 | 5.15300 | 1.74719 | 3.95956 | 0.00000 | 4.55920 | 1.05681 |
| **C15orf59** | 0.00861 | 0.01403 | 0.01618 | 0.01049 | 0.01218 | 0.02500 | 0.01992 | 0.01049 | 0.01928 | 0.01712 | 0.01317 | 0.01990 |
| **C15orf61** | 2.91120 | 2.93340 | 7.41945 | 1.31543 | 2.10784 | 2.29852 | 6.91131 | 0.14067 | 8.73777 | 2.17199 | 6.19446 | 1.70305 |
| **C15orf62** | 0.17562 | 0.01093 | 0.04619 | 0.00329 | 0.01279 | 0.06531 | 0.00000 | 0.03143 | 0.04679 | 0.14012 | 0.00512 | 0.02398 |
| **C15orf65** | 0.18981 | 0.43406 | 0.03968 | 0.00304 | 0.00286 | 0.03009 | 0.06170 | 1.82285 | 0.00000 | 0.00000 | 0.09629 | 5.09716 |
| **C16orf13** | 0.74388 | 7.45254 | 5.81628 | 5.99307 | 9.67944 | 0.00000 | 13.06077 | 0.29446 | 10.82118 | 0.00000 | 14.34803 | 2.35527 |
| **C16orf3** | 0.04114 | 0.01214 | 0.02072 | 0.01078 | 0.02377 | 0.01866 | 0.02874 | 0.00970 | 0.01300 | 0.00878 | 0.04457 | 0.02086 |
| **C16orf45** | 0.13397 | 0.06471 | 0.05868 | 0.00000 | 0.00000 | 0.04473 | 0.01271 | 0.24107 | 0.00756 | 0.13926 | 0.06898 | 0.08279 |
| **C16orf46** | 2.63585 | 0.73226 | 1.95039 | 0.01643 | 0.29656 | 1.46946 | 0.88985 | 2.76989 | 1.12175 | 0.00000 | 0.52777 | 2.51735 |
| **C16orf47** | 0.00877 | 0.03344 | 0.91251 | 0.00019 | 0.14823 | 0.00937 | 0.06013 | 0.03280 | 0.15526 | 0.00454 | 0.18420 | 0.01194 |
| **C16orf52** | 0.01597 | 0.00000 | 1.02672 | 0.00000 | 0.00000 | 0.99456 | 0.66575 | 0.27470 | 0.00000 | 4.37865 | 0.17027 | 0.20726 |
| **C16orf54** | 0.00384 | 1.57834 | 0.30129 | 0.00018 | 0.32309 | 0.00189 | 0.13204 | 0.48261 | 0.00000 | 0.00000 | 0.00495 | 2.03815 |
| **C16orf58** | 1.05026 | 3.70010 | 0.56259 | 0.97876 | 1.83196 | 0.00000 | 0.00000 | 0.02933 | 0.63710 | 1.87187 | 0.96386 | 1.29010 |
| **C16orf59** | 0.71311 | 2.77130 | 3.17290 | 2.64036 | 2.15589 | 0.00000 | 4.88780 | 0.16735 | 3.48495 | 0.00000 | 5.58250 | 2.14526 |
| **C16orf62** | 3.56058 | 4.69060 | 1.22243 | 4.07335 | 3.75083 | 2.66707 | 2.03206 | 1.82657 | 1.18264 | 0.00000 | 5.69018 | 3.01262 |
| **C16orf70** | 3.22131 | 8.12019 | 10.88588 | 2.57796 | 3.77902 | 4.32030 | 1.89457 | 3.72187 | 17.57803 | 0.00000 | 9.71872 | 11.34505 |
| **C16orf71** | 0.61654 | 0.03293 | 0.17428 | 0.12582 | 0.14536 | 0.00000 | 0.00000 | 0.09403 | 0.40592 | 0.34929 | 0.09510 | 0.34891 |
| **C16orf72** | 4.09905 | 1.96802 | 3.06114 | 2.45448 | 0.00000 | 15.18543 | 10.98898 | 10.24406 | 0.00000 | 17.40984 | 1.25984 | 13.50029 |
| **C16orf74** | 7.36480 | 1.26629 | 2.60894 | 0.53898 | 4.44327 | 0.00000 | 0.21981 | 0.95512 | 0.43604 | 0.22455 | 4.93446 | 0.88903 |
| **C16orf80** | 1.90232 | 14.59009 | 10.29622 | 7.81665 | 6.17204 | 15.90545 | 9.75361 | 21.25559 | 15.44238 | 0.00000 | 9.32658 | 13.17944 |
| **C16orf82** | 0.00533 | 0.01174 | 0.01025 | 0.00711 | 0.00678 | 0.02719 | 0.03824 | 0.01035 | 0.00092 | 0.05778 | 0.01442 | 0.01322 |
| **C16orf86** | 0.03395 | 0.15097 | 0.02723 | 0.28319 | 0.00000 | 0.00000 | 0.12547 | 0.01090 | 0.01747 | 0.02845 | 0.25529 | 0.00910 |
| **C16orf87** | 1.20734 | 3.60914 | 5.25438 | 0.26796 | 0.03173 | 15.06287 | 4.07356 | 1.70626 | 3.42149 | 3.30068 | 1.74359 | 7.16808 |
| **C16orf89** | 0.00769 | 0.01736 | 0.05020 | 0.01556 | 0.02547 | 0.00400 | 0.01687 | 0.01046 | 0.02368 | 0.00727 | 0.01413 | 0.01764 |
| **C16orf91** | 1.26695 | 13.18719 | 8.97376 | 2.19193 | 5.54570 | 5.94307 | 9.82117 | 18.93500 | 10.35062 | 0.00000 | 8.81434 | 20.27227 |
| **C16orf93** | 0.03948 | 0.03783 | 0.06456 | 0.02839 | 0.03432 | 0.00047 | 0.01061 | 0.03261 | 0.10206 | 0.00000 | 0.03771 | 0.05965 |
| **C16orf95** | 0.23805 | 1.01996 | 0.54807 | 0.82196 | 1.02049 | 0.00000 | 0.97454 | 0.29745 | 0.00000 | 0.56974 | 0.65617 | 0.43406 |
| **C17orf100** | 0.58351 | 0.16553 | 0.15860 | 0.07555 | 0.00000 | 0.10249 | 0.00000 | 1.49596 | 0.33948 | 0.00000 | 0.24948 | 0.49763 |
| **C17orf104** | 0.55986 | 0.21534 | 0.13745 | 0.13788 | 0.01498 | 0.66019 | 0.50896 | 0.79646 | 1.43060 | 1.42423 | 0.11892 | 0.42840 |
| **C17orf107** | 0.30904 | 0.17235 | 0.17926 | 0.40799 | 0.00002 | 0.00088 | 0.19458 | 0.18707 | 0.00264 | 0.04702 | 0.04197 | 0.28159 |
| **C17orf112** | 0.21334 | 0.01666 | 0.01351 | 0.00847 | 0.00000 | 0.00991 | 0.01127 | 0.03684 | 0.00000 | 0.03500 | 0.16197 | 0.01041 |
| **C17orf47** | 0.00566 | 0.03311 | 0.01146 | 0.00281 | 0.05501 | 0.04328 | 0.01211 | 0.00780 | 0.01641 | 0.02123 | 0.01275 | 0.07778 |
| **C17orf51** | 2.50549 | 5.08504 | 3.16129 | 3.23657 | 1.34361 | 0.00000 | 6.12781 | 8.14690 | 7.68830 | 2.99996 | 7.39237 | 9.27975 |
| **C17orf53** | 5.18462 | 7.29624 | 5.77359 | 4.16513 | 3.74157 | 2.19841 | 5.27059 | 4.86949 | 4.03010 | 0.00000 | 5.18582 | 7.73446 |
| **C17orf58** | 0.44809 | 3.77505 | 4.01055 | 0.70761 | 0.00000 | 2.71748 | 4.91754 | 0.13645 | 5.21571 | 0.00000 | 4.77330 | 1.65060 |
| **C17orf59** | 1.47635 | 1.44601 | 1.82971 | 1.38974 | 3.59890 | 0.00000 | 2.77111 | 2.00311 | 2.07671 | 5.43610 | 2.08095 | 0.60193 |
| **C17orf62** | 1.69659 | 11.26822 | 7.43685 | 5.76855 | 5.74416 | 0.00000 | 2.13700 | 0.66134 | 11.53917 | 0.00000 | 5.58121 | 7.14857 |
| **C17orf64** | 0.00632 | 0.03694 | 0.00895 | 0.00697 | 0.00868 | 0.00000 | 0.01727 | 0.00755 | 0.14700 | 0.02741 | 0.00143 | 0.08368 |
| **C17orf67** | 0.02818 | 0.00690 | 0.02313 | 0.01339 | 0.01690 | 0.00807 | 0.02093 | 0.59770 | 0.00000 | 0.03351 | 0.05514 | 0.20179 |
| **C17orf70** | 1.10930 | 7.33964 | 2.00401 | 4.90354 | 8.52780 | 0.00000 | 0.38838 | 0.00000 | 4.61317 | 1.52353 | 6.19478 | 1.56435 |
| **C17orf74** | 0.00449 | 0.01140 | 0.03485 | 0.00449 | 0.00000 | 0.01941 | 0.05655 | 0.01127 | 0.06828 | 0.07783 | 0.00000 | 0.02077 |
| **C17orf75** | 1.63922 | 4.39878 | 0.00000 | 0.10683 | 1.07853 | 11.65933 | 0.00000 | 0.63359 | 0.96521 | 2.02835 | 2.65806 | 0.54540 |
| **C17orf77** | 0.00694 | 0.01492 | 0.01595 | 0.01139 | 0.01736 | 0.01575 | 0.01787 | 0.01052 | 0.01667 | 0.01875 | 0.02192 | 0.01665 |
| **C17orf78** | 0.00624 | 0.01539 | 0.01898 | 0.01044 | 0.01249 | 0.01541 | 0.02304 | 0.01045 | 0.04771 | 0.01680 | 0.01214 | 0.01612 |
| **C17orf80** | 3.20787 | 10.53777 | 5.78385 | 3.86321 | 3.26742 | 9.33773 | 3.03067 | 5.07382 | 8.02439 | 0.00000 | 6.32815 | 14.19332 |
| **C17orf85** | 4.13194 | 7.86963 | 9.12103 | 6.49651 | 6.67246 | 3.58267 | 8.06906 | 4.53469 | 2.73332 | 18.61954 | 7.54638 | 3.87070 |

| | | | | | | | | | | | | |
|---|---|---|---|---|---|---|---|---|---|---|---|---|
| **C17orf89** | 0.00000 | 11.20167 | 6.04234 | 1.60746 | 7.84490 | 3.24133 | 4.06552 | 1.62740 | 3.49508 | 12.92323 | 11.82834 | 6.31680 |
| **C17orf96** | 1.06637 | 0.22017 | 0.82811 | 0.51325 | 2.09996 | 0.00000 | 1.65600 | 0.03513 | 0.00000 | 3.58552 | 0.00000 | 0.31384 |
| **C17orf97** | 0.00608 | 0.03098 | 0.05185 | 0.01437 | 0.15020 | 0.01082 | 0.04168 | 0.00730 | 0.02222 | 0.00000 | 0.01943 | 0.12166 |
| **C17orf98** | 0.15474 | 0.07317 | 0.13834 | 0.04548 | 0.09815 | 0.00000 | 0.00861 | 0.18700 | 0.12017 | 0.04753 | 0.10438 | 0.08860 |
| **C17orf99** | 1.48773 | 0.10623 | 0.00665 | 0.00000 | 0.01169 | 0.00000 | 0.00169 | 0.82151 | 0.00968 | 0.36175 | 0.00000 | 0.17034 |
| **C18orf21** | 0.00000 | 6.79483 | 3.70987 | 3.16529 | 2.86332 | 11.53302 | 5.74393 | 8.87756 | 4.80486 | 3.30366 | 5.59900 | 14.75692 |
| **C18orf25** | 0.74607 | 6.14448 | 4.08891 | 4.24415 | 2.80485 | 16.22522 | 7.43531 | 14.84008 | 6.13242 | 0.00000 | 4.55733 | 6.55380 |
| **C18orf32** | 0.36203 | 0.96159 | 0.74893 | 0.54081 | 0.91182 | 0.94491 | 0.00000 | 0.08767 | 0.84856 | 0.00000 | 0.86839 | 0.31703 |
| **C18orf54** | 0.62097 | 5.06070 | 2.08840 | 1.54930 | 0.00000 | 9.09832 | 2.27597 | 0.34717 | 0.38111 | 5.33815 | 2.67213 | 4.28641 |
| **C18orf61** | 0.00594 | 0.01795 | 0.00959 | 0.00113 | 0.00537 | 0.08278 | 0.02525 | 0.01802 | 0.00271 | 0.04533 | 0.01281 | 0.08421 |
| **C18orf63** | 0.00664 | 0.01367 | 0.02210 | 0.01731 | 0.01119 | 0.01354 | 0.01495 | 0.76529 | 0.01464 | 0.03011 | 0.01856 | 0.00193 |
| **C18orf64+LINC(** | 0.00631 | 0.02774 | 0.02288 | 0.01516 | 0.00723 | 0.05972 | 0.03749 | 0.00985 | 0.04620 | 0.00941 | 0.02345 | 0.02705 |
| **C18orf8** | 5.40362 | 6.57103 | 5.74298 | 2.12613 | 5.99954 | 4.48427 | 5.52860 | 2.90418 | 12.78940 | 0.00000 | 5.79665 | 4.31192 |
| **C19orf10** | 3.67840 | 18.81067 | 24.68679 | 15.50240 | 22.50132 | 15.45866 | 8.60320 | 14.72127 | 32.89652 | 6.42523 | 16.00401 | 17.32667 |
| **C19orf12** | 0.22537 | 3.22412 | 9.30241 | 0.00000 | 3.20563 | 0.25805 | 4.75493 | 0.15999 | 4.15760 | 7.41750 | 6.00172 | 1.80156 |
| **C19orf18** | 0.19504 | 0.14430 | 0.02610 | 0.00956 | 0.09111 | 0.00000 | 0.00556 | 0.20281 | 0.09071 | 0.48113 | 0.06313 | 0.35347 |
| **C19orf24** | 2.91002 | 12.92341 | 13.60745 | 6.58312 | 14.45059 | 0.00000 | 19.53294 | 2.60270 | 21.96476 | 9.01895 | 12.90952 | 13.11212 |
| **C19orf25** | 1.51276 | 6.50011 | 5.50421 | 6.38625 | 5.51865 | 0.00000 | 9.49940 | 1.13805 | 13.32968 | 0.00000 | 3.98809 | 8.49024 |
| **C19orf26** | 0.00531 | 0.02756 | 0.00598 | 0.01290 | 0.00000 | 0.03848 | 0.08661 | 0.01025 | 0.00000 | 0.05128 | 0.01377 | 0.01095 |
| **C19orf35** | 0.00468 | 0.00911 | 0.02646 | 0.02650 | 0.05865 | 0.00030 | 0.02319 | 0.01007 | 0.00751 | 0.11394 | 0.02332 | 0.01818 |
| **C19orf38** | 2.08916 | 0.13721 | 0.49671 | 0.00267 | 0.04982 | 0.00011 | 0.18823 | 0.04298 | 0.00000 | 0.00000 | 0.01808 | 1.41916 |
| **C19orf40** | 0.00177 | 1.77778 | 1.26631 | 1.27142 | 1.09506 | 3.35601 | 1.42599 | 1.08649 | 1.28051 | 0.00000 | 1.18706 | 2.45563 |
| **C19orf43** | 0.97328 | 6.46952 | 8.33590 | 4.63057 | 9.66064 | 0.00000 | 17.09167 | 8.41484 | 1.39405 | 33.66416 | 3.22071 | 12.22461 |
| **C19orf44** | 0.00142 | 0.94423 | 1.71080 | 1.38460 | 0.47211 | 0.00000 | 0.00118 | 0.71221 | 1.36410 | 2.56380 | 0.79717 | 1.30130 |
| **C19orf45** | 0.00442 | 0.01380 | 0.02735 | 0.01151 | 0.00557 | 0.01840 | 0.02804 | 0.19464 | 0.02170 | 0.05163 | 0.02305 | 0.00838 |
| **C19orf47** | 7.35547 | 4.22048 | 8.76252 | 1.32867 | 5.83275 | 0.00000 | 4.45473 | 4.56003 | 3.44689 | 2.62915 | 4.07523 | 8.23627 |
| **C19orf48** | 5.58482 | 42.31109 | 22.51400 | 27.17289 | 38.44360 | 2.20610 | 45.08593 | 20.35705 | 41.02474 | 0.00000 | 27.88463 | 54.02238 |
| **C19orf52** | 0.03960 | 1.90802 | 2.06593 | 2.31173 | 2.55580 | 0.22487 | 1.54838 | 1.26072 | 0.35114 | 4.23839 | 1.02688 | 0.70408 |
| **C19orf53** | 2.57421 | 27.25657 | 20.85082 | 19.99616 | 25.43354 | 13.43742 | 32.38244 | 25.81770 | 27.06203 | 3.94089 | 20.47184 | 40.46355 |
| **C19orf54** | 1.79440 | 7.99435 | 0.59653 | 2.77070 | 2.68090 | 0.00000 | 2.96334 | 0.67303 | 2.69262 | 1.91799 | 1.43889 | 7.32224 |
| **C19orf57** | 0.35617 | 1.18627 | 0.23847 | 0.68062 | 0.83813 | 0.00000 | 0.00225 | 0.05774 | 0.00000 | 0.21913 | 1.21791 | 0.50199 |
| **C19orf60** | 0.75952 | 7.11784 | 5.58783 | 7.13560 | 8.79863 | 0.00000 | 4.52337 | 0.19330 | 3.80468 | 1.10739 | 7.20287 | 1.43259 |
| **C19orf66** | 0.53980 | 0.01501 | 0.01543 | 0.01714 | 0.02021 | 0.01258 | 0.01697 | 0.00152 | 0.03340 | 0.00000 | 0.01296 | 0.01201 |
| **C19orf67** | 3.86000 | 0.04182 | 0.04763 | 0.01126 | 0.05489 | 0.00000 | 0.05352 | 0.51179 | 0.11875 | 0.00000 | 0.06216 | 0.04997 |
| **C19orf68** | 0.06807 | 0.00753 | 0.20671 | 0.00000 | 0.09062 | 0.00000 | 0.25901 | 0.01452 | 0.23663 | 0.64496 | 0.19165 | 0.04516 |
| **C19orf70** | 0.71870 | 11.33234 | 6.98633 | 11.76437 | 19.73151 | 0.00055 | 9.44497 | 5.07836 | 9.14897 | 1.58370 | 14.32040 | 6.94556 |
| **C19orf71** | 0.08614 | 0.02172 | 0.06499 | 0.00635 | 0.03172 | 0.00000 | 0.00000 | 0.00419 | 0.08655 | 0.15927 | 0.01410 | 0.01750 |
| **C19orf73** | 0.01051 | 0.36271 | 0.25860 | 0.00041 | 0.44583 | 0.23146 | 0.15281 | 0.46177 | 0.00000 | 0.00000 | 0.20605 | 0.54045 |
| **C19orf80** | 0.00414 | 0.01167 | 0.02651 | 0.19711 | 0.01548 | 0.00000 | 0.02040 | 0.00993 | 0.02113 | 0.14293 | 0.00644 | 0.03843 |
| **C19orf81** | 0.00399 | 0.00765 | 0.02044 | 0.02288 | 0.02925 | 0.04435 | 0.00000 | 0.18294 | 0.07006 | 0.02466 | 0.01828 | 0.09629 |
| **C19orf82** | 0.00610 | 0.35002 | 0.02300 | 0.07430 | 0.02852 | 0.07781 | 0.00000 | 0.00145 | 0.12729 | 0.07973 | 0.05477 | 1.02308 |
| **C19orf83** | 0.11046 | 0.33276 | 0.03402 | 0.00966 | 0.00000 | 0.00600 | 0.01709 | 0.71180 | 0.00963 | 0.03038 | 0.10532 | 0.50906 |
| **C19orf84** | 0.02163 | 0.02783 | 0.06302 | 0.03831 | 0.01569 | 0.00000 | 0.00622 | 0.00995 | 0.04635 | 0.01578 | 0.01671 | 0.03627 |
| **C1D** | 0.16111 | 1.78490 | 2.13683 | 0.30957 | 0.30320 | 13.95951 | 3.90590 | 3.54865 | 4.01272 | 0.92291 | 1.16406 | 7.13652 |
| **C1GALT1** | 1.91401 | 1.73154 | 7.67109 | 0.00000 | 0.48079 | 18.50986 | 0.00000 | 8.95071 | 6.68026 | 10.00756 | 5.11280 | 16.60173 |
| **C1GALT1C1** | 0.26086 | 1.53257 | 4.26823 | 2.19184 | 0.00000 | 7.82627 | 0.00000 | 0.99245 | 12.88897 | 0.74794 | 8.79128 | 0.87383 |
| **C1QA** | 0.00537 | 0.06306 | 0.00880 | 0.01266 | 0.01539 | 0.01095 | 0.00789 | 0.00731 | 0.03064 | 0.05590 | 0.01136 | 0.10464 |
| **C1QBP** | 1.50377 | 48.41660 | 34.95018 | 27.55040 | 28.44339 | 51.88071 | 46.73507 | 24.01337 | 55.70481 | 0.00287 | 40.06977 | 61.20616 |

| | | | | | | | | | | | | |
|---|---|---|---|---|---|---|---|---|---|---|---|---|
| **C1QL1** | 0.04494 | 0.18124 | 0.05136 | 0.00000 | 0.00491 | 0.01957 | 0.01418 | 0.04684 | 0.00906 | 0.07069 | 0.01218 | 0.05873 |
| **C1QL3** | 0.00566 | 0.01787 | 0.03854 | 0.00000 | 0.00309 | 0.16165 | 0.06100 | 0.00551 | 0.08389 | 0.01890 | 0.00850 | 0.15558 |
| **C1QL4** | 0.20994 | 0.00330 | 0.03977 | 0.03070 | 0.01708 | 0.02343 | 0.01215 | 0.00000 | 0.01456 | 0.04608 | 0.01178 | 0.15959 |
| **C1QTNF1** | 1.06017 | 0.01159 | 0.06357 | 0.00805 | 0.01408 | 0.02197 | 0.00493 | 0.12542 | 0.05385 | 0.00295 | 0.01069 | 0.01341 |
| **C1QTNF2** | 0.00172 | 0.15551 | 0.01636 | 0.00000 | 0.01556 | 0.01532 | 0.01781 | 0.06312 | 0.01226 | 0.02051 | 0.01108 | 0.59295 |
| **C1QTNF3** | 0.02459 | 0.00728 | 0.06844 | 0.00000 | 0.01322 | 0.07683 | 0.01088 | 0.01239 | 0.00000 | 0.13801 | 0.03144 | 0.00605 |
| **C1QTNF3-AMA(** | 0.00599 | 0.01510 | 0.01899 | 0.01573 | 0.00997 | 0.02106 | 0.01586 | 0.01137 | 0.00952 | 0.02835 | 0.03276 | 0.01856 |
| **C1QTNF4** | 0.01205 | 0.10493 | 0.48917 | 0.50952 | 0.51634 | 0.15304 | 0.23208 | 0.03193 | 0.34966 | 0.00000 | 0.22875 | 0.06795 |
| **C1QTNF5+MFR** | 0.00588 | 0.01405 | 0.03124 | 0.01406 | 0.01840 | 0.01598 | 0.03772 | 0.00952 | 0.00722 | 0.01795 | 0.01539 | 0.04893 |
| **C1QTNF6** | 6.53464 | 0.44097 | 3.71974 | 0.00000 | 7.77481 | 0.00000 | 1.95397 | 0.56820 | 10.05084 | 6.50596 | 6.33104 | 2.11448 |
| **C1QTNF7** | 0.00454 | 0.00188 | 0.00783 | 0.00570 | 0.00000 | 0.02903 | 0.01573 | 0.00924 | 0.01349 | 0.18816 | 0.08730 | 0.01538 |
| **C1QTNF8** | 0.01003 | 0.00982 | 0.04083 | 0.02865 | 0.00355 | 0.03580 | 0.00667 | 0.01000 | 0.00446 | 0.04334 | 0.01799 | 0.01384 |
| **C1QTNF9** | 0.00198 | 1.45180 | 0.15292 | 0.56577 | 0.09715 | 0.03562 | 0.00000 | 0.02660 | 0.00000 | 0.22276 | 0.00000 | 0.00000 |
| **C1QTNF9B** | 0.01017 | 0.01147 | 0.15288 | 0.02737 | 0.00000 | 0.18186 | 0.01889 | 0.01587 | 0.00000 | 0.10010 | 0.03147 | 0.01589 |
| **C1QTNF9B-AS1** | 0.02993 | 0.29759 | 0.04653 | 0.13794 | 0.15211 | 0.00002 | 0.14836 | 0.07480 | 0.03037 | 0.00000 | 0.17493 | 0.06824 |
| **C1R** | 0.00489 | 0.29108 | 0.00000 | 1.40235 | 0.06203 | 0.11632 | 0.00000 | 0.07885 | 0.00000 | 0.01020 | 0.16848 | 0.02010 |
| **C1RL** | 0.00579 | 0.01142 | 0.03135 | 0.19501 | 0.17445 | 0.05943 | 0.01546 | 0.00860 | 0.00000 | 0.00165 | 0.01965 | 0.03826 |
| **C1RL-AS1** | 0.15015 | 0.00785 | 0.00105 | 0.11018 | 0.00000 | 0.00000 | 0.07883 | 0.00298 | 0.00000 | 0.20372 | 0.00720 | 0.01366 |
| **C1S** | 0.08228 | 1.96831 | 0.04310 | 0.83657 | 0.80341 | 0.45011 | 0.00000 | 0.50209 | 1.60061 | 0.15015 | 0.02692 | 0.74454 |
| **C1orf100** | 0.10622 | 0.01089 | 0.02940 | 0.01139 | 0.01094 | 0.01849 | 0.00944 | 0.04865 | 0.02857 | 0.05401 | 0.01275 | 0.02930 |
| **C1orf101** | 0.04475 | 0.04049 | 0.36102 | 0.00722 | 0.01366 | 0.00000 | 0.00012 | 0.05188 | 0.03457 | 0.03723 | 0.01804 | 0.05063 |
| **C1orf105** | 0.00535 | 0.00000 | 1.41904 | 0.00114 | 0.07003 | 0.07744 | 1.18130 | 0.00922 | 0.06574 | 0.00000 | 0.39640 | 0.01027 |
| **C1orf106** | 0.00553 | 3.12081 | 2.54813 | 2.88547 | 5.64141 | 0.00000 | 2.53786 | 0.00333 | 5.39912 | 0.08575 | 2.82028 | 0.04856 |
| **C1orf109** | 0.00465 | 9.88225 | 8.59920 | 5.71231 | 5.83845 | 13.99656 | 11.62268 | 1.54828 | 10.46777 | 0.00000 | 9.83822 | 10.35438 |
| **C1orf110** | 0.32612 | 0.02194 | 0.01324 | 0.03801 | 0.01634 | 0.03768 | 0.01094 | 0.78749 | 0.01922 | 0.00301 | 0.01833 | 0.07105 |
| **C1orf111** | 0.00621 | 0.00647 | 0.00639 | 0.03811 | 0.01176 | 0.06459 | 0.01535 | 0.00984 | 0.01355 | 0.00910 | 0.04353 | 0.01413 |
| **C1orf112** | 0.21704 | 3.92346 | 1.70204 | 1.08247 | 1.86840 | 9.71096 | 1.96626 | 0.73114 | 7.02034 | 0.00000 | 4.33114 | 3.86712 |
| **C1orf115** | 0.21563 | 0.14935 | 4.05765 | 0.00000 | 6.09525 | 0.00000 | 2.37188 | 0.03711 | 0.00000 | 1.45643 | 0.19980 | 0.10924 |
| **C1orf116** | 0.24236 | 0.45303 | 0.05661 | 0.04978 | 1.16333 | 0.04951 | 0.08601 | 0.00679 | 0.51029 | 0.00067 | 0.09183 | 0.00512 |
| **C1orf122** | 0.11936 | 2.10501 | 2.91932 | 2.83349 | 9.05817 | 0.00000 | 8.92252 | 0.40720 | 0.00000 | 20.03103 | 0.56816 | 0.94133 |
| **C1orf123** | 0.82825 | 7.60384 | 4.79599 | 1.95028 | 3.52871 | 4.91282 | 5.58264 | 0.95711 | 6.12633 | 1.37756 | 7.99570 | 4.99552 |
| **C1orf127** | 0.49366 | 0.02869 | 0.01635 | 0.01606 | 0.14820 | 0.00000 | 0.02598 | 0.04238 | 0.01150 | 0.01965 | 0.01704 | 0.10911 |
| **C1orf131** | 3.15174 | 11.18970 | 7.96474 | 4.26849 | 6.77217 | 16.08545 | 8.80426 | 5.10680 | 8.52378 | 3.78798 | 7.41935 | 14.99362 |
| **C1orf140** | 0.04243 | 0.01603 | 0.00833 | 0.00860 | 0.01905 | 0.00908 | 0.00539 | 0.01168 | 0.12912 | 0.00676 | 0.03916 | 0.01595 |
| **C1orf141** | 0.00757 | 0.00911 | 0.01983 | 0.00935 | 0.06783 | 0.03803 | 0.01542 | 0.01050 | 0.01453 | 0.01439 | 0.02744 | 0.01676 |
| **C1orf145** | 0.00719 | 0.00622 | 0.01480 | 0.15861 | 0.00000 | 0.02682 | 0.00000 | 0.01596 | 0.01603 | 0.17214 | 0.01376 | 0.00961 |
| **C1orf146** | 0.12268 | 0.15433 | 0.16517 | 0.00000 | 0.11563 | 0.30008 | 0.20537 | 0.00079 | 0.00161 | 0.07203 | 0.06292 | 0.00314 |
| **C1orf158** | 0.00585 | 0.00995 | 0.00993 | 0.00903 | 0.01501 | 0.03358 | 0.02255 | 0.01023 | 0.00643 | 0.03231 | 0.04140 | 0.01418 |
| **C1orf159** | 0.24540 | 4.95285 | 1.05448 | 0.92896 | 1.59815 | 0.00000 | 1.23248 | 0.01062 | 4.74976 | 0.63549 | 1.36909 | 4.84987 |
| **C1orf162** | 1.07649 | 0.42210 | 0.00000 | 0.29034 | 0.07545 | 0.23325 | 0.04109 | 0.94231 | 0.00000 | 0.10420 | 0.00000 | 4.93240 |
| **C1orf167** | 0.03928 | 0.01544 | 0.04545 | 0.06540 | 0.01131 | 0.00436 | 0.01809 | 0.02961 | 0.02848 | 0.03470 | 0.01389 | 0.01384 |
| **C1orf168** | 1.34066 | 0.12999 | 0.00000 | 0.00000 | 0.12591 | 0.00000 | 0.70712 | 0.00000 | 1.62567 | 0.00000 | 2.80899 | 0.02623 |
| **C1orf174** | 4.29057 | 13.13007 | 7.42763 | 5.96355 | 5.24384 | 15.25019 | 13.02995 | 0.31041 | 13.33872 | 0.00000 | 13.72216 | 8.90235 |
| **C1orf177** | 0.01292 | 0.05719 | 0.02600 | 0.04851 | 0.06134 | 0.00000 | 0.00719 | 0.00890 | 0.07842 | 0.00000 | 0.03673 | 0.01930 |
| **C1orf186** | 0.03320 | 0.00993 | 0.05272 | 0.77204 | 0.02144 | 0.00000 | 0.02385 | 0.00762 | 0.00619 | 0.01247 | 0.02294 | 0.02042 |
| **C1orf189** | 0.00760 | 0.00484 | 0.01280 | 0.01235 | 0.01099 | 0.00933 | 0.01296 | 0.00664 | 0.01792 | 0.04540 | 0.02613 | 0.15859 |
| **C1orf192** | 0.03242 | 0.07996 | 0.05469 | 0.00000 | 0.10793 | 0.00000 | 0.00014 | 0.00163 | 0.11926 | 0.48919 | 0.00000 | 0.11821 |
| **C1orf194** | 0.03372 | 0.00000 | 0.01382 | 0.01236 | 0.02168 | 0.02047 | 0.01463 | 0.00196 | 0.02538 | 0.01081 | 0.02006 | 0.44853 |

| | | | | | | | | | | | | |
|---|---|---|---|---|---|---|---|---|---|---|---|---|
| **C1orf195** | 0.00579 | 0.01589 | 0.01161 | 0.04996 | 0.02551 | 0.03056 | 0.02611 | 0.01036 | 0.00672 | 0.01711 | 0.02363 | 0.01904 |
| **C1orf198** | 0.00286 | 3.20359 | 8.21744 | 0.00000 | 0.39742 | 0.38691 | 0.97142 | 1.34245 | 2.33581 | 0.00000 | 0.07986 | 3.44796 |
| **C1orf200** | 0.00529 | 0.01271 | 0.01254 | 0.00882 | 0.00922 | 0.01953 | 0.03496 | 0.01040 | 0.00540 | 0.06283 | 0.01638 | 0.01431 |
| **C1orf21** | 1.10795 | 0.00000 | 3.83677 | 0.00000 | 0.77980 | 0.63525 | 8.48050 | 0.44165 | 5.21055 | 0.12977 | 1.74988 | 0.03892 |
| **C1orf210** | 8.24452 | 0.75752 | 1.34330 | 0.75139 | 0.00000 | 0.46902 | 0.78891 | 0.34348 | 1.23683 | 0.00000 | 1.28739 | 0.00000 |
| **C1orf213** | 0.43724 | 0.46042 | 1.51314 | 0.37057 | 0.22899 | 0.38526 | 0.14427 | 0.53509 | 0.25577 | 0.00000 | 0.47012 | 0.39521 |
| **C1orf216** | 0.00167 | 0.93177 | 2.06248 | 2.11839 | 0.00000 | 0.21164 | 2.83023 | 3.04659 | 1.47867 | 3.22639 | 0.89400 | 4.52552 |
| **C1orf220** | 0.00358 | 0.01560 | 0.01055 | 0.02005 | 0.00000 | 0.03501 | 0.02872 | 0.03655 | 0.01360 | 0.09016 | 0.00871 | 0.05288 |
| **C1orf226** | 4.01265 | 1.93719 | 0.00000 | 2.63496 | 1.60641 | 0.00000 | 1.87320 | 0.00000 | 4.59112 | 0.48114 | 5.40414 | 1.03003 |
| **C1orf228** | 0.08381 | 0.01572 | 0.03909 | 0.03161 | 0.08165 | 0.00000 | 0.00093 | 0.00000 | 0.04916 | 0.12301 | 0.00000 | 0.24456 |
| **C1orf229** | 0.01399 | 0.01306 | 0.02277 | 0.02145 | 0.01105 | 0.01958 | 0.01525 | 0.20627 | 0.00563 | 0.03071 | 0.04049 | 0.04048 |
| **C1orf233** | 0.38639 | 0.57961 | 0.00000 | 4.62966 | 2.10517 | 0.00000 | 0.49916 | 0.00000 | 0.00000 | 1.82523 | 0.06002 | 0.38078 |
| **C1orf234** | 0.07876 | 0.01336 | 0.04841 | 0.00092 | 0.00407 | 0.08858 | 0.00000 | 0.08696 | 0.00000 | 0.03289 | 0.01107 | 0.00735 |
| **C1orf27** | 0.00086 | 1.52123 | 11.27427 | 1.15122 | 0.37096 | 14.67530 | 0.00000 | 0.00000 | 17.61076 | 7.41727 | 3.50237 | 0.45439 |
| **C1orf35** | 0.62712 | 9.24845 | 6.08698 | 1.80554 | 5.39893 | 0.00000 | 7.00651 | 1.18959 | 9.24916 | 12.70581 | 4.86501 | 3.67065 |
| **C1orf43** | 6.47545 | 17.43427 | 15.88706 | 11.09364 | 7.15066 | 23.26330 | 11.73391 | 5.33958 | 19.84000 | 0.00000 | 17.14484 | 16.79742 |
| **C1orf50** | 0.96123 | 3.78741 | 2.23817 | 2.19282 | 2.35481 | 1.55178 | 0.57046 | 0.64061 | 1.52995 | 1.55220 | 1.74706 | 2.57192 |
| **C1orf52** | 0.00000 | 12.34079 | 14.11177 | 2.59732 | 6.32504 | 24.95664 | 13.16789 | 25.70988 | 10.84176 | 6.93565 | 9.08615 | 15.51312 |
| **C1orf53** | 0.13740 | 0.18214 | 0.05984 | 0.60970 | 0.04900 | 0.29857 | 0.00000 | 0.00000 | 0.00000 | 0.17700 | 0.28651 | 0.09119 |
| **C1orf54** | 0.06424 | 0.20759 | 0.03713 | 0.06616 | 0.00000 | 0.19894 | 0.00000 | 0.00401 | 0.01998 | 0.06164 | 0.00463 | 1.14200 |
| **C1orf56** | 3.12870 | 0.11431 | 1.47149 | 0.30259 | 1.04487 | 1.72649 | 3.55662 | 0.94046 | 1.22224 | 0.00000 | 1.35931 | 0.04768 |
| **C1orf61** | 0.05296 | 0.00000 | 3.29653 | 0.00000 | 5.11008 | 0.00000 | 2.87389 | 0.15620 | 4.13962 | 0.41641 | 6.88573 | 0.79080 |
| **C1orf64** | 0.04482 | 1.51087 | 0.67171 | 0.00000 | 0.00000 | 0.00000 | 0.00000 | 0.33287 | 0.00000 | 0.25514 | 0.00000 | 1.71525 |
| **C1orf74** | 0.62341 | 2.43940 | 1.18396 | 0.28302 | 0.07061 | 1.51147 | 0.00000 | 0.05869 | 0.17644 | 0.00000 | 0.54975 | 2.49337 |
| **C1orf85** | 0.34373 | 12.40302 | 1.99715 | 7.25693 | 9.83067 | 2.06085 | 0.00000 | 0.00000 | 7.40323 | 0.00000 | 11.54230 | 3.53998 |
| **C1orf86** | 0.11080 | 0.07396 | 3.30775 | 0.80955 | 1.88806 | 0.00000 | 2.84425 | 0.49695 | 1.40776 | 5.04475 | 0.43418 | 0.24972 |
| **C1orf94** | 0.00577 | 0.05353 | 0.01848 | 0.15915 | 0.01015 | 0.00000 | 0.01761 | 0.01069 | 0.01245 | 0.02447 | 0.02768 | 0.01333 |
| **C1orf95** | 0.08388 | 0.01957 | 0.03403 | 0.01712 | 0.01610 | 0.00470 | 0.01249 | 0.06136 | 0.01882 | 0.02246 | 0.02128 | 0.01392 |
| **C2** | 0.00145 | 0.70186 | 0.19537 | 1.33667 | 0.23358 | 0.21710 | 0.00000 | 0.29194 | 0.00000 | 1.30717 | 0.00000 | 1.43219 |
| **C20orf144** | 0.01086 | 0.01729 | 0.01830 | 0.00748 | 0.01554 | 0.03815 | 0.01365 | 0.01027 | 0.02492 | 0.01032 | 0.01264 | 0.01527 |
| **C20orf166-AS1** | 0.01050 | 0.01495 | 0.01639 | 0.01257 | 0.01499 | 0.01709 | 0.02688 | 0.00958 | 0.01099 | 0.01623 | 0.01679 | 0.01610 |
| **C20orf173** | 0.05426 | 0.01476 | 0.01983 | 0.01531 | 0.01595 | 0.01560 | 0.03445 | 0.01418 | 0.02163 | 0.01210 | 0.01331 | 0.01593 |
| **C20orf194** | 3.06216 | 0.85838 | 0.00000 | 1.83039 | 0.06073 | 2.48236 | 0.00000 | 3.93113 | 0.00000 | 2.87130 | 1.08472 | 1.99892 |
| **C20orf195** | 0.00685 | 0.02064 | 0.01993 | 0.03526 | 0.04560 | 0.00000 | 0.02138 | 0.01086 | 0.02088 | 0.00000 | 0.01743 | 0.03569 |
| **C20orf196** | 0.65991 | 4.02398 | 1.14606 | 1.66052 | 1.98700 | 2.24921 | 1.25083 | 0.63928 | 0.28051 | 0.00000 | 1.60476 | 4.52902 |
| **C20orf197** | 0.07282 | 0.01288 | 0.00888 | 0.01029 | 0.00791 | 0.02943 | 0.00810 | 0.00725 | 0.05622 | 0.00473 | 0.03248 | 0.01473 |
| **C20orf202** | 0.00544 | 0.01473 | 0.05884 | 0.02737 | 0.01212 | 0.03120 | 0.00761 | 0.01178 | 0.01462 | 0.02840 | 0.01408 | 0.01401 |
| **C20orf203** | 0.00999 | 0.04757 | 0.05577 | 0.01639 | 0.05868 | 0.00000 | 0.04065 | 0.01269 | 0.06108 | 0.48273 | 0.00000 | 0.00528 |
| **C20orf24** | 0.04251 | 4.81161 | 6.00853 | 2.36059 | 2.89237 | 0.00000 | 0.30660 | 2.05714 | 4.13302 | 0.00000 | 1.49660 | 1.04784 |
| **C20orf26** | 0.32493 | 0.20883 | 0.00588 | 0.00000 | 0.00874 | 0.17408 | 0.00000 | 0.71762 | 0.00000 | 0.01566 | 0.07685 | 0.79459 |
| **C20orf27** | 1.99086 | 16.96259 | 13.59073 | 10.28555 | 15.04126 | 0.00000 | 27.67130 | 0.92576 | 30.79346 | 0.00116 | 16.87065 | 11.48328 |
| **C20orf96** | 5.26290 | 1.06046 | 0.04590 | 0.00000 | 0.62483 | 0.00000 | 0.00772 | 1.45952 | 1.42027 | 1.09943 | 0.27477 | 1.06839 |
| **C21orf119** | 0.00644 | 0.93911 | 0.98299 | 0.00000 | 0.29501 | 0.03303 | 0.38775 | 1.74232 | 0.41114 | 0.00000 | 0.55657 | 2.17312 |
| **C21orf140** | 0.00257 | 0.00883 | 0.02078 | 0.01999 | 0.00122 | 0.05382 | 0.01987 | 0.04115 | 0.00000 | 0.02908 | 0.02789 | 0.04449 |
| **C21orf15** | 0.00885 | 0.06418 | 0.00000 | 0.00552 | 0.06131 | 0.00000 | 0.03868 | 0.01494 | 0.21256 | 0.10954 | 0.00000 | 0.01494 |
| **C21orf2** | 0.11249 | 4.76444 | 1.14859 | 1.32423 | 1.76794 | 0.00000 | 0.19332 | 0.08963 | 2.82558 | 2.73557 | 1.20753 | 0.70278 |
| **C21orf33** | 1.51015 | 16.90973 | 8.91213 | 12.87042 | 11.71866 | 4.33797 | 14.37369 | 1.01329 | 9.38867 | 6.43720 | 23.93031 | 3.99546 |
| **C21orf37** | 0.00585 | 0.09211 | 0.01257 | 0.00000 | 0.03977 | 0.05268 | 0.01819 | 0.01081 | 0.00295 | 0.01958 | 0.01248 | 0.01211 |

| | | | | | | | | | | | | |
|---|---|---|---|---|---|---|---|---|---|---|---|---|
| **C21orf49** | 0.00257 | 0.04862 | 0.03583 | 0.00000 | 0.11934 | 0.02463 | 0.00155 | 0.45406 | 0.00884 | 0.04580 | 0.09289 | 0.09388 |
| **C21orf58** | 0.00067 | 0.29463 | 1.01670 | 0.49193 | 2.09914 | 0.00000 | 0.00000 | 0.19740 | 2.47014 | 3.43004 | 1.00826 | 0.90225 |
| **C21orf59** | 3.98076 | 23.36537 | 22.82868 | 7.70571 | 10.57084 | 24.05285 | 25.97608 | 3.68280 | 20.15990 | 0.00000 | 16.95064 | 21.67760 |
| **C21orf62** | 0.00800 | 0.03864 | 0.05913 | 0.10216 | 0.01533 | 0.00812 | 0.01970 | 0.01039 | 0.02189 | 0.00001 | 0.01580 | 0.02000 |
| **C21orf67** | 0.03029 | 1.14380 | 0.21476 | 0.12091 | 0.15037 | 0.00000 | 0.00000 | 0.00000 | 0.29264 | 0.05167 | 0.25025 | 1.86604 |
| **C21orf88** | 0.01594 | 0.01405 | 0.01075 | 0.01587 | 0.01378 | 0.00676 | 0.06015 | 0.01090 | 0.00571 | 0.05066 | 0.01170 | 0.01847 |
| **C21orf90** | 0.12119 | 0.03896 | 0.01803 | 0.01430 | 0.01597 | 0.00000 | 0.03212 | 0.02890 | 0.01955 | 0.00559 | 0.02413 | 0.04100 |
| **C21orf91** | 3.19854 | 2.70942 | 6.08152 | 0.91267 | 0.83948 | 8.57921 | 5.55512 | 19.61540 | 1.37758 | 0.20001 | 1.08360 | 8.54748 |
| **C21orf91-OT1** | 0.01038 | 0.01350 | 0.03593 | 0.01364 | 0.00565 | 0.02565 | 0.00954 | 0.03151 | 0.03641 | 0.02380 | 0.02902 | 0.03558 |
| **C22orf15** | 0.00428 | 0.01716 | 0.00000 | 0.00000 | 0.07427 | 0.03582 | 0.02550 | 0.00921 | 0.08464 | 0.13018 | 0.00000 | 0.01888 |
| **C22orf23** | 0.73818 | 0.06220 | 0.23729 | 0.15564 | 0.21822 | 1.02820 | 0.00000 | 0.40798 | 0.37642 | 0.00000 | 0.07104 | 0.78965 |
| **C22orf24** | 0.00600 | 0.04513 | 0.06811 | 0.01439 | 0.00000 | 0.00000 | 0.02602 | 0.00931 | 0.00000 | 0.18434 | 0.03895 | 0.02951 |
| **C22orf29** | 2.79592 | 3.47918 | 2.40834 | 1.38144 | 5.04594 | 0.02524 | 3.70056 | 0.21840 | 3.06800 | 2.25157 | 11.16789 | 5.43004 |
| **C22orf31** | 0.00521 | 0.01396 | 0.10025 | 0.01598 | 0.00004 | 0.00675 | 0.12239 | 0.01230 | 0.01714 | 0.05720 | 0.01995 | 0.02498 |
| **C22orf34** | 0.57405 | 0.24252 | 0.18062 | 0.09316 | 0.14766 | 0.15726 | 0.00000 | 0.00870 | 0.07573 | 0.00000 | 0.06100 | 1.51118 |
| **C22orf39** | 0.38969 | 1.68892 | 4.40398 | 3.90573 | 2.41401 | 1.62518 | 5.95784 | 0.52635 | 5.13572 | 1.47714 | 6.40128 | 1.72384 |
| **C22orf42** | 0.03749 | 0.01163 | 0.01458 | 0.01282 | 0.01145 | 0.03440 | 0.01962 | 0.02537 | 0.01763 | 0.00813 | 0.01298 | 0.02026 |
| **C22orf46** | 0.16768 | 1.31089 | 1.50167 | 0.76645 | 1.95280 | 0.00000 | 0.00000 | 0.27111 | 0.44652 | 1.31089 | 1.10911 | 3.21031 |
| **C2CD2** | 0.00000 | 0.43197 | 1.93318 | 0.00000 | 3.34186 | 0.43668 | 2.64477 | 0.14856 | 2.45410 | 2.25863 | 2.59753 | 0.04606 |
| **C2CD2L** | 1.20750 | 0.33799 | 1.71532 | 0.72750 | 1.26624 | 0.00000 | 0.26662 | 0.71740 | 1.46001 | 4.73124 | 0.06699 | 0.73316 |
| **C2CD3** | 2.42563 | 2.20146 | 2.55577 | 1.84072 | 1.79888 | 2.11111 | 1.86245 | 4.87820 | 4.07552 | 0.89126 | 1.50575 | 3.12550 |
| **C2CD4A** | 0.00630 | 0.01614 | 0.01494 | 0.01329 | 0.01668 | 0.01502 | 0.02479 | 0.01053 | 0.01807 | 0.01547 | 0.01483 | 0.01663 |
| **C2CD4B** | 0.04573 | 0.06541 | 0.01949 | 0.01325 | 0.00182 | 0.07151 | 0.00000 | 0.00888 | 0.00478 | 0.04289 | 0.00880 | 0.03032 |
| **C2CD4C** | 0.14559 | 0.08526 | 0.00000 | 0.75663 | 0.51062 | 0.00000 | 0.05205 | 0.00000 | 0.00000 | 0.04714 | 0.00000 | 0.15995 |
| **C2CD4D** | 0.00630 | 0.01639 | 0.03890 | 0.01533 | 0.01344 | 0.01684 | 0.01434 | 0.01046 | 0.02781 | 0.01332 | 0.01924 | 0.01621 |
| **C2CD5** | 4.59308 | 2.64009 | 0.06721 | 1.86364 | 0.70457 | 4.67236 | 1.01585 | 2.04744 | 1.58257 | 0.00000 | 1.47762 | 1.78759 |
| **C2orf15** | 3.11659 | 0.00000 | 0.28909 | 0.00001 | 0.00000 | 1.02228 | 0.00001 | 0.65367 | 1.59529 | 0.63722 | 1.68649 | 0.11574 |
| **C2orf16** | 0.51452 | 0.16269 | 0.14927 | 0.08018 | 0.00000 | 0.14068 | 0.00000 | 0.48705 | 0.18504 | 0.00002 | 0.00000 | 0.39364 |
| **C2orf27A** | 0.00369 | 0.06559 | 0.02991 | 0.01890 | 0.08107 | 0.08010 | 0.01847 | 0.01960 | 0.01407 | 0.00002 | 0.01747 | 0.03935 |
| **C2orf40** | 0.19620 | 0.08626 | 0.26509 | 0.00751 | 0.04817 | 0.00000 | 0.04645 | 0.00000 | 0.00000 | 0.00000 | 0.04748 | 0.10381 |
| **C2orf42** | 0.31294 | 2.31056 | 1.84906 | 1.13511 | 0.57676 | 1.17465 | 0.46684 | 2.51064 | 2.86032 | 0.00000 | 0.92407 | 3.12369 |
| **C2orf43** | 0.44422 | 4.04596 | 1.76162 | 1.17437 | 0.71907 | 7.18515 | 0.00000 | 1.01163 | 7.02281 | 0.48377 | 3.99384 | 3.27448 |
| **C2orf44** | 2.88808 | 3.11242 | 2.97035 | 1.34501 | 0.60554 | 2.30768 | 3.70581 | 4.34738 | 5.85759 | 0.02622 | 4.38214 | 7.34586 |
| **C2orf47** | 0.90617 | 9.14467 | 5.35748 | 3.10996 | 2.55805 | 22.30327 | 8.31923 | 0.81351 | 8.80082 | 5.23687 | 8.65796 | 5.63149 |
| **C2orf48** | 0.16228 | 0.41538 | 0.29362 | 0.86687 | 0.71217 | 0.00000 | 0.00000 | 0.24836 | 1.19445 | 0.87778 | 1.07484 | 0.43978 |
| **C2orf49** | 1.97552 | 6.49611 | 7.18164 | 3.43840 | 7.59447 | 16.33720 | 9.25559 | 13.79134 | 3.80352 | 6.77321 | 6.16984 | 15.62355 |
| **C2orf50** | 0.05492 | 0.01435 | 0.03815 | 0.03327 | 0.00530 | 0.08381 | 0.00000 | 0.04293 | 0.02396 | 0.00938 | 0.01139 | 0.00898 |
| **C2orf54** | 0.08576 | 0.01055 | 0.03045 | 0.02755 | 0.00542 | 0.03319 | 0.01258 | 0.00559 | 0.00604 | 0.03150 | 0.01779 | 0.01292 |
| **C2orf61** | 0.05360 | 0.01196 | 0.00486 | 0.06093 | 0.00914 | 0.03392 | 0.00619 | 0.08120 | 0.18868 | 0.00370 | 0.00169 | 0.01437 |
| **C2orf66** | 0.00269 | 0.00957 | 0.24955 | 0.00235 | 0.00000 | 0.03945 | 0.10337 | 0.03336 | 0.05010 | 0.19448 | 0.00203 | 0.01881 |
| **C2orf68** | 1.93055 | 10.61245 | 7.37127 | 5.77760 | 3.78297 | 8.64508 | 9.33415 | 3.75245 | 3.05606 | 9.91319 | 12.60809 | 15.13020 |
| **C2orf69** | 0.78733 | 7.27506 | 4.08245 | 0.96746 | 0.00000 | 18.11895 | 8.67527 | 16.95626 | 4.06914 | 11.31277 | 7.81589 | 13.65882 |
| **C2orf71** | 0.21449 | 0.01539 | 0.05128 | 0.02646 | 0.00618 | 0.01968 | 0.00882 | 0.01843 | 0.00842 | 0.04672 | 0.01411 | 0.01065 |
| **C2orf72** | 0.00527 | 0.05374 | 5.87111 | 0.00000 | 0.24751 | 0.00000 | 0.93588 | 0.00994 | 0.00000 | 0.02220 | 0.03424 | 0.01972 |
| **C2orf73** | 0.03923 | 0.01198 | 0.01388 | 0.01117 | 0.01265 | 0.03036 | 0.01645 | 0.01024 | 0.01709 | 0.01040 | 0.01405 | 0.02344 |
| **C2orf74** | 0.00000 | 3.78501 | 0.78547 | 0.01423 | 1.07811 | 7.63101 | 0.00000 | 0.05180 | 0.00000 | 0.54600 | 4.73758 | 4.41821 |
| **C2orf76** | 2.09087 | 2.75721 | 1.93346 | 0.00000 | 0.00000 | 3.63821 | 1.79346 | 1.65662 | 1.60398 | 0.42416 | 1.50418 | 6.26182 |
| **C2orf80** | 0.09226 | 0.30922 | 0.07049 | 0.13527 | 0.00000 | 0.04148 | 0.00000 | 0.00000 | 0.00762 | 0.04213 | 0.00781 | 0.55258 |

| | | | | | | | | | | | | |
|---|---|---|---|---|---|---|---|---|---|---|---|---|
| **C2orf81** | 0.36572 | 0.00120 | 0.01902 | 0.25890 | 0.09672 | 0.06781 | 0.00000 | 0.00000 | 0.13639 | 0.17723 | 0.00029 | 0.05900 |
| **C2orf82** | 0.00405 | 0.01092 | 0.01268 | 0.00232 | 0.00000 | 0.01948 | 0.00836 | 0.00914 | 0.01278 | 0.25808 | 0.00973 | 0.02032 |
| **C2orf83** | 0.06737 | 0.01236 | 0.02586 | 0.02741 | 0.01337 | 0.01918 | 0.02488 | 0.06542 | 0.02519 | 0.04287 | 0.00995 | 0.01189 |
| **C2orf88** | 1.33484 | 0.00000 | 0.79197 | 0.07215 | 0.00000 | 0.06348 | 0.07923 | 0.39719 | 0.28315 | 0.68179 | 0.01693 | 0.13711 |
| **C2orf91** | 0.00521 | 0.07230 | 0.01022 | 0.12657 | 0.01230 | 0.01855 | 0.01531 | 0.00828 | 0.02960 | 0.02247 | 0.01584 | 0.13668 |
| **C3** | 1.34661 | 0.00524 | 0.01194 | 0.05198 | 0.03903 | 0.00000 | 0.01632 | 4.63745 | 0.04725 | 0.05916 | 0.00851 | 0.64076 |
| **C3AR1** | 0.13742 | 0.73478 | 0.01622 | 0.00000 | 0.00000 | 0.07835 | 0.00000 | 0.00000 | 0.00364 | 0.04107 | 0.00074 | 0.91810 |
| **C3P1** | 0.00557 | 0.01401 | 0.00958 | 0.00927 | 0.01116 | 0.02554 | 0.04370 | 0.01011 | 0.02549 | 0.03995 | 0.00974 | 0.01713 |
| **C3orf14** | 1.31063 | 5.96308 | 6.32783 | 0.00000 | 0.00000 | 11.27756 | 8.38545 | 0.28891 | 1.28015 | 0.00000 | 3.40886 | 7.27481 |
| **C3orf17** | 5.61382 | 11.24866 | 3.34730 | 3.24615 | 0.00021 | 22.15441 | 2.66600 | 7.85109 | 8.33063 | 5.06689 | 2.55531 | 13.37095 |
| **C3orf18** | 0.02834 | 0.23242 | 0.42384 | 0.42779 | 0.01563 | 0.00000 | 0.01624 | 0.09342 | 0.00000 | 0.00000 | 0.14108 | 0.19827 |
| **C3orf20** | 0.01613 | 0.04114 | 0.27278 | 0.00000 | 0.00000 | 0.00000 | 0.00000 | 0.03183 | 0.23761 | 0.16242 | 0.08917 | 0.03606 |
| **C3orf27** | 0.00359 | 0.02028 | 0.00613 | 0.00190 | 0.04545 | 0.00000 | 0.00485 | 0.01018 | 0.03087 | 0.24813 | 0.00487 | 0.05024 |
| **C3orf30** | 0.00551 | 0.01351 | 0.01826 | 0.00664 | 0.01332 | 0.02086 | 0.04681 | 0.01053 | 0.01868 | 0.04327 | 0.01016 | 0.01453 |
| **C3orf33** | 1.58089 | 1.15692 | 0.26338 | 0.00000 | 0.00000 | 0.74091 | 0.20580 | 0.28601 | 0.55821 | 0.00000 | 0.00000 | 1.83108 |
| **C3orf35** | 0.00441 | 0.02217 | 0.10530 | 0.00000 | 0.00000 | 0.00760 | 0.02098 | 0.00977 | 0.36467 | 0.45586 | 0.05015 | 0.01513 |
| **C3orf36** | 0.00515 | 0.01102 | 0.00127 | 0.03381 | 0.00000 | 0.01705 | 0.29050 | 0.01025 | 0.00000 | 0.05792 | 0.00551 | 0.01644 |
| **C3orf38** | 4.69308 | 6.09825 | 6.01271 | 0.94760 | 0.00261 | 14.65520 | 4.08589 | 0.37479 | 4.88704 | 2.16857 | 3.38000 | 8.88327 |
| **C3orf43** | 0.80478 | 0.00000 | 0.00727 | 0.06994 | 0.10359 | 0.00000 | 0.01734 | 2.55233 | 0.03543 | 0.05565 | 0.01003 | 0.10881 |
| **C3orf49** | 0.67162 | 0.00886 | 0.15541 | 0.01424 | 0.00052 | 0.01379 | 0.01120 | 0.00143 | 0.00469 | 0.17662 | 0.01483 | 0.00947 |
| **C3orf52** | 0.08894 | 3.53752 | 2.44901 | 0.22071 | 6.19783 | 4.48190 | 0.00000 | 0.96455 | 10.42973 | 0.00000 | 3.05298 | 0.49020 |
| **C3orf55** | 0.00202 | 0.21815 | 0.09100 | 0.04848 | 0.00013 | 0.04439 | 0.00369 | 0.06229 | 0.01877 | 0.00615 | 0.01084 | 0.08718 |
| **C3orf56** | 22.91948 | 0.22608 | 0.00190 | 0.00000 | 0.54961 | 0.00000 | 0.00000 | 2.29075 | 0.01466 | 0.17657 | 0.10040 | 0.32790 |
| **C3orf58** | 1.84267 | 0.76496 | 2.33723 | 0.37660 | 0.00000 | 5.65109 | 0.00000 | 1.81216 | 5.79473 | 3.19155 | 0.91613 | 1.21699 |
| **C3orf62** | 0.00208 | 0.06623 | 0.00000 | 0.09732 | 0.19482 | 0.80980 | 0.76126 | 0.08204 | 0.69100 | 2.92687 | 0.44784 | 0.35019 |
| **C3orf65** | 0.13735 | 0.00000 | 0.13114 | 0.34242 | 0.04341 | 0.00000 | 0.00000 | 0.21369 | 0.00000 | 0.74055 | 0.54492 | 0.09130 |
| **C3orf67** | 1.11461 | 0.64979 | 0.15195 | 0.13533 | 0.54332 | 1.07396 | 0.00000 | 0.29753 | 0.35628 | 0.00000 | 0.37664 | 1.34633 |
| **C3orf70** | 0.29793 | 0.05298 | 0.98668 | 0.36749 | 0.00000 | 0.00000 | 0.00000 | 0.17242 | 0.00000 | 0.01696 | 0.52892 | 0.32355 |
| **C3orf80** | 0.11949 | 0.11364 | 0.49369 | 0.12474 | 1.56386 | 0.73931 | 0.00000 | 0.00168 | 0.00000 | 0.28163 | 0.00000 | 0.00000 |
| **C4A+C4B+C4B-** | 0.00608 | 0.02579 | 0.43855 | 0.05685 | 0.01923 | 0.04541 | 0.02697 | 0.00903 | 0.20908 | 0.00000 | 0.04389 | 0.03739 |
| **C4A+C4B+C4B-** | 0.00638 | 0.06499 | 0.34219 | 0.03586 | 0.09280 | 0.00000 | 0.00518 | 0.01018 | 0.12958 | 0.00000 | 0.03942 | 0.01792 |
| **C4BPA** | 0.00586 | 0.01076 | 0.00739 | 0.00710 | 0.00357 | 0.04344 | 0.03520 | 0.01034 | 0.04293 | 0.02094 | 0.01093 | 0.01407 |
| **C4BPB** | 0.00000 | 0.02061 | 0.00000 | 0.00000 | 6.69035 | 0.00000 | 1.87753 | 0.08546 | 15.01991 | 0.42702 | 14.79823 | 0.00000 |
| **C4orf17** | 0.01587 | 0.02224 | 0.01891 | 0.01265 | 0.01668 | 0.01014 | 0.01963 | 0.10017 | 0.01781 | 0.01215 | 0.01661 | 0.01192 |
| **C4orf19** | 2.02645 | 0.56909 | 8.30023 | 0.97952 | 3.32963 | 2.37400 | 5.57680 | 0.00000 | 4.60559 | 0.49056 | 9.47467 | 0.24165 |
| **C4orf22** | 0.31363 | 0.04211 | 0.01415 | 0.00623 | 0.00953 | 0.03415 | 0.01508 | 0.16186 | 0.01795 | 0.00821 | 0.01129 | 0.01157 |
| **C4orf26** | 0.00242 | 0.03292 | 0.27462 | 0.00646 | 0.03809 | 0.05855 | 0.09730 | 0.07973 | 0.00000 | 0.00977 | 0.04971 | 0.01168 |
| **C4orf27** | 10.19614 | 4.22816 | 4.81829 | 1.27012 | 2.27813 | 12.08007 | 4.88776 | 6.36426 | 0.23619 | 3.52790 | 4.26552 | 6.99023 |
| **C4orf29** | 0.71614 | 0.73431 | 2.44673 | 1.85790 | 2.01285 | 2.67846 | 0.56707 | 0.25996 | 4.64652 | 0.00000 | 3.39355 | 1.23315 |
| **C4orf3** | 0.69214 | 9.78261 | 14.09511 | 9.53538 | 13.29456 | 27.51358 | 16.54486 | 0.13517 | 8.41007 | 8.08335 | 14.77669 | 19.96974 |
| **C4orf32** | 0.00553 | 0.00000 | 0.52229 | 0.02249 | 0.53198 | 0.40760 | 0.00000 | 0.11093 | 0.11193 | 0.36985 | 0.00000 | 2.49815 |
| **C4orf33** | 0.28721 | 0.33233 | 1.62584 | 1.33048 | 0.00000 | 3.01084 | 0.00000 | 0.75752 | 1.72392 | 0.66388 | 1.96321 | 1.44552 |
| **C4orf36** | 0.00758 | 0.01131 | 0.01418 | 0.08537 | 0.00176 | 0.06582 | 0.01203 | 0.00686 | 0.00000 | 0.01974 | 0.10846 | 0.14199 |
| **C4orf45** | 0.43301 | 0.05997 | 0.05030 | 0.00496 | 0.03873 | 0.00000 | 0.08913 | 0.00174 | 0.01803 | 0.05931 | 0.12209 | 0.02220 |
| **C4orf46** | 0.00000 | 3.12504 | 6.06902 | 1.41311 | 0.49368 | 15.97920 | 7.05866 | 8.16035 | 5.56259 | 4.37495 | 5.97892 | 6.29892 |
| **C4orf47** | 2.38067 | 0.11956 | 0.31685 | 0.00000 | 0.00000 | 0.10955 | 0.15154 | 0.42138 | 0.02037 | 0.06734 | 0.04724 | 0.40467 |
| **C4orf48** | 0.18013 | 1.99584 | 4.60746 | 2.99875 | 7.67492 | 0.00000 | 3.86109 | 0.00000 | 3.44537 | 3.69880 | 3.41035 | 0.25213 |
| **C4orf51** | 0.02132 | 0.00000 | 0.15259 | 0.19646 | 0.10651 | 0.09698 | 0.22911 | 0.19209 | 0.22555 | 0.09074 | 0.05434 | 0.27153 |

| | | | | | | | | | | | | |
|---|---|---|---|---|---|---|---|---|---|---|---|---|
| **C5** | 1.82376 | 0.01014 | 0.01541 | 0.00954 | 0.01010 | 0.01411 | 0.01578 | 0.12444 | 0.01896 | 0.02205 | 0.01402 | 0.09994 |
| **C5AR1** | 0.64042 | 0.00000 | 0.00000 | 0.00000 | 0.00000 | 0.05443 | 0.77598 | 0.12504 | 0.05766 | 1.03015 | 0.00000 | 0.08742 |
| **C5AR2** | 0.00578 | 0.01333 | 0.04404 | 0.01104 | 0.01266 | 0.02084 | 0.00992 | 0.01039 | 0.00712 | 0.04205 | 0.01590 | 0.01439 |
| **C5orf15** | 0.05664 | 9.72451 | 6.31161 | 4.57818 | 0.96601 | 19.50799 | 0.00000 | 4.27952 | 17.02720 | 3.50513 | 8.32864 | 10.35644 |
| **C5orf22** | 4.93018 | 7.43010 | 5.40153 | 4.50407 | 1.03894 | 19.02684 | 7.31344 | 4.70846 | 3.73314 | 0.42657 | 4.43534 | 3.41529 |
| **C5orf24** | 0.00000 | 1.56969 | 3.87440 | 0.90766 | 0.00000 | 3.01413 | 6.67582 | 0.64602 | 5.15826 | 13.90277 | 4.18350 | 3.24815 |
| **C5orf27** | 0.14858 | 0.00735 | 0.06032 | 0.00621 | 0.01245 | 0.05984 | 0.01710 | 0.09559 | 0.00000 | 0.01447 | 0.02271 | 0.02446 |
| **C5orf28** | 0.00776 | 1.20915 | 0.14964 | 0.72942 | 0.00000 | 5.85948 | 1.14827 | 2.52833 | 2.03791 | 0.70960 | 1.01469 | 4.32436 |
| **C5orf30** | 0.00000 | 0.29141 | 14.80516 | 0.12260 | 0.00000 | 0.17015 | 4.80095 | 0.17405 | 0.37848 | 0.00000 | 0.15341 | 0.68653 |
| **C5orf34** | 1.77497 | 9.01203 | 0.47201 | 3.28473 | 0.00000 | 6.94011 | 0.84007 | 1.38551 | 0.81937 | 0.00000 | 1.17271 | 3.42156 |
| **C5orf38** | 0.00411 | 0.01033 | 0.26682 | 0.00119 | 0.00805 | 0.00000 | 0.01640 | 0.01012 | 0.05536 | 0.21467 | 0.03187 | 0.01106 |
| **C5orf42** | 1.26598 | 1.11007 | 0.45529 | 0.77835 | 0.00000 | 0.00000 | 0.00000 | 0.07605 | 0.44587 | 2.01234 | 0.13740 | 0.60132 |
| **C5orf45** | 1.28575 | 0.78428 | 0.12843 | 0.61626 | 0.29802 | 0.52621 | 0.00000 | 0.84161 | 0.40037 | 0.00000 | 0.69539 | 0.66547 |
| **C5orf47** | 1.19228 | 0.00930 | 0.10915 | 0.01087 | 0.15395 | 0.00000 | 0.00081 | 0.00995 | 0.01996 | 0.05946 | 0.00759 | 0.43212 |
| **C5orf49** | 0.08850 | 0.01127 | 0.13834 | 0.01843 | 0.04138 | 0.02240 | 0.01568 | 0.12053 | 0.00000 | 0.00876 | 0.06136 | 0.01533 |
| **C5orf51** | 1.29130 | 3.98563 | 11.15544 | 0.27762 | 0.00000 | 14.35296 | 15.68715 | 0.40754 | 8.36381 | 8.91175 | 11.56327 | 3.78321 |
| **C5orf52** | 0.00658 | 0.02244 | 0.01086 | 0.00742 | 0.00000 | 0.06924 | 0.00743 | 0.00429 | 0.00311 | 0.04812 | 0.01325 | 0.22193 |
| **C5orf55** | 0.27229 | 0.47211 | 0.11583 | 0.27554 | 0.04206 | 0.00000 | 0.00000 | 0.00000 | 0.32609 | 0.10975 | 0.06846 | 1.00774 |
| **C5orf56** | 0.02832 | 0.01886 | 0.02818 | 0.01304 | 0.01964 | 0.00563 | 0.01791 | 0.01216 | 0.06781 | 0.00351 | 0.01046 | 0.01739 |
| **C5orf58** | 0.78890 | 0.18146 | 0.00000 | 0.09478 | 0.44524 | 0.85682 | 1.18429 | 0.00000 | 0.00035 | 0.11026 | 0.62406 | 0.58430 |
| **C5orf60** | 0.04861 | 0.21602 | 0.01440 | 0.02117 | 0.21319 | 0.00000 | 0.04277 | 0.01353 | 0.01709 | 0.00000 | 0.01164 | 0.09994 |
| **C5orf63** | 0.19547 | 2.20576 | 1.46916 | 0.26529 | 0.11670 | 2.18986 | 0.51084 | 0.20487 | 0.35580 | 0.60210 | 0.64246 | 1.95216 |
| **C5orf64** | 0.01767 | 0.01313 | 0.01488 | 0.01204 | 0.01178 | 0.02021 | 0.02412 | 0.00891 | 0.00581 | 0.02117 | 0.03798 | 0.01633 |
| **C5orf66** | 0.86124 | 0.11746 | 0.29551 | 0.38497 | 0.74789 | 0.21526 | 0.35141 | 0.00000 | 0.21534 | 1.59375 | 0.15605 | 0.08560 |
| **C5orf66-AS1** | 0.00127 | 0.30583 | 0.13397 | 0.14132 | 0.00000 | 0.20639 | 0.03622 | 0.57973 | 0.00000 | 0.05073 | 0.00000 | 0.36202 |
| **C5orf66-AS2** | 0.00544 | 0.00836 | 0.02409 | 0.01013 | 0.00000 | 0.04932 | 0.01161 | 0.00987 | 0.03044 | 0.04849 | 0.00732 | 0.01726 |
| **C6orf1** | 2.03047 | 3.06283 | 4.25767 | 2.62574 | 1.12037 | 0.00429 | 0.45415 | 0.80305 | 0.66583 | 0.00000 | 0.84377 | 2.53157 |
| **C6orf10** | 0.03046 | 0.28678 | 0.03149 | 0.69740 | 0.00000 | 0.00002 | 0.05712 | 0.38112 | 0.00000 | 0.04452 | 0.00848 | 0.38718 |
| **C6orf100** | 0.01230 | 0.07171 | 0.01597 | 0.00175 | 0.05138 | 0.00000 | 0.03236 | 0.01679 | 0.00539 | 0.12062 | 0.00609 | 0.02629 |
| **C6orf106** | 4.80553 | 9.86961 | 12.06717 | 17.12857 | 20.63552 | 10.05884 | 16.81003 | 5.92903 | 2.53256 | 22.56930 | 6.14560 | 11.33896 |
| **C6orf120** | 1.30521 | 2.67803 | 3.61849 | 0.01874 | 1.03705 | 2.60745 | 0.00000 | 1.25426 | 2.45668 | 1.97000 | 1.73683 | 2.86306 |
| **C6orf123** | 0.00653 | 0.01645 | 0.01823 | 0.01290 | 0.01413 | 0.01046 | 0.01848 | 0.01050 | 0.01013 | 0.00962 | 0.06692 | 0.01731 |
| **C6orf132** | 0.00055 | 11.64961 | 2.36062 | 0.68189 | 16.91096 | 0.00000 | 7.86252 | 0.00000 | 10.43297 | 14.81416 | 9.07556 | 0.74247 |
| **C6orf136** | 3.19170 | 6.84665 | 3.70585 | 4.44639 | 4.99843 | 0.39817 | 2.88872 | 0.32416 | 5.83415 | 0.34723 | 4.30056 | 3.80694 |
| **C6orf141** | 0.00183 | 0.00394 | 0.39324 | 0.00000 | 0.00000 | 0.00760 | 0.00000 | 0.14187 | 0.41927 | 0.44407 | 0.00000 | 0.47902 |
| **C6orf147** | 0.00000 | 1.06247 | 1.45696 | 0.44572 | 0.13333 | 0.54276 | 2.30049 | 2.76480 | 6.13764 | 0.63678 | 1.55139 | 2.39467 |
| **C6orf163** | 0.10580 | 0.52601 | 0.08986 | 0.00000 | 0.36315 | 0.57363 | 0.00000 | 0.00000 | 0.30883 | 0.07617 | 0.12790 | 0.36292 |
| **C6orf164** | 0.00338 | 0.04743 | 0.00410 | 0.22001 | 0.07522 | 0.03624 | 0.00023 | 0.04917 | 0.06001 | 0.06638 | 0.04343 | 0.07290 |
| **C6orf165** | 0.04008 | 0.07919 | 0.21945 | 0.00000 | 0.00000 | 0.13968 | 0.00000 | 0.18666 | 0.00015 | 0.29269 | 0.39309 | 0.87418 |
| **C6orf201** | 0.01736 | 0.01616 | 0.02445 | 0.15898 | 0.03103 | 0.03526 | 0.13917 | 0.08069 | 0.28103 | 0.00000 | 0.13666 | 0.04993 |
| **C6orf203** | 2.66505 | 4.61388 | 0.99377 | 3.27316 | 1.95252 | 7.05851 | 1.22987 | 2.10449 | 0.00000 | 0.57074 | 3.24833 | 10.59387 |
| **C6orf211** | 1.25561 | 25.90469 | 5.94817 | 22.44019 | 6.78762 | 69.09420 | 12.78121 | 7.58113 | 13.78100 | 7.76865 | 21.11591 | 20.71465 |
| **C6orf222** | 0.02722 | 0.01596 | 0.02684 | 0.01122 | 0.00916 | 0.02220 | 0.03283 | 0.00781 | 0.00304 | 0.04203 | 0.01525 | 0.01343 |
| **C6orf223** | 0.10531 | 0.26069 | 0.32273 | 0.13384 | 0.08995 | 0.16413 | 0.98728 | 0.00499 | 0.05506 | 0.41455 | 0.30569 | 0.00917 |
| **C6orf226** | 0.39592 | 0.68298 | 1.00610 | 0.43545 | 0.00000 | 1.12056 | 0.00000 | 1.11437 | 0.54487 | 0.73674 | 0.63300 | 0.27453 |
| **C6orf229** | 0.00317 | 0.01512 | 0.06771 | 0.03192 | 0.01485 | 0.01772 | 0.03027 | 0.02988 | 0.03686 | 0.00000 | 0.00922 | 0.03743 |
| **C6orf25** | 0.39824 | 0.00708 | 0.11268 | 0.11303 | 0.00390 | 0.02220 | 0.00000 | 0.00211 | 0.02011 | 0.07715 | 0.02241 | 0.01027 |
| **C6orf47** | 0.01549 | 1.90640 | 2.05763 | 2.24071 | 2.60067 | 0.00000 | 2.12092 | 0.11452 | 2.59714 | 1.18019 | 1.32374 | 0.58998 |

| | | | | | | | | | | | | |
|---|---|---|---|---|---|---|---|---|---|---|---|---|
| **C6orf48** | 8.66830 | 13.47299 | 3.24869 | 18.29001 | 18.61512 | 19.17061 | 11.31803 | 10.95860 | 16.33687 | 14.82319 | 4.81464 | 28.01953 |
| **C6orf52** | 10.13153 | 1.00353 | 0.40223 | 0.00000 | 0.34279 | 0.00488 | 0.44189 | 10.65758 | 0.00000 | 0.00000 | 0.28777 | 2.09313 |
| **C6orf57** | 0.20656 | 0.53787 | 0.13543 | 0.26104 | 0.24429 | 1.30163 | 0.00000 | 0.19726 | 0.00000 | 0.43027 | 0.76531 | 0.92499 |
| **C6orf58** | 0.00235 | 0.04320 | 0.00957 | 0.03106 | 0.00514 | 0.03569 | 0.01072 | 0.04519 | 0.01939 | 0.02022 | 0.02062 | 0.00994 |
| **C6orf62** | 4.24080 | 19.71231 | 17.64115 | 16.66028 | 2.55248 | 68.33926 | 25.04215 | 26.60123 | 18.76105 | 3.87185 | 26.89696 | 15.45038 |
| **C6orf7** | 0.00337 | 0.01457 | 0.01306 | 0.01086 | 0.01426 | 0.02382 | 0.01673 | 0.06209 | 0.01871 | 0.01531 | 0.01609 | 0.01402 |
| **C6orf89** | 0.89038 | 8.87198 | 5.31066 | 2.09976 | 5.24101 | 5.44954 | 0.83644 | 0.81184 | 8.27663 | 0.00000 | 4.19510 | 7.51845 |
| **C6orf99** | 0.30625 | 0.13168 | 0.66479 | 0.64164 | 0.75046 | 0.33369 | 4.03778 | 0.20647 | 0.79739 | 0.00000 | 3.12634 | 0.20946 |
| **C7** | 0.02916 | 0.01423 | 0.01751 | 0.01312 | 0.02046 | 0.00961 | 0.01990 | 0.00696 | 0.02539 | 0.00609 | 0.01671 | 0.04513 |
| **C7orf13** | 1.30947 | 0.65671 | 0.17415 | 0.17154 | 0.00640 | 0.00748 | 0.22769 | 0.00000 | 0.00000 | 0.00000 | 0.75977 | 3.70267 |
| **C7orf25** | 2.27611 | 2.85330 | 1.90984 | 0.45212 | 0.87248 | 4.10227 | 0.67841 | 0.40862 | 0.96930 | 0.00000 | 1.71181 | 4.56646 |
| **C7orf26** | 2.80851 | 0.91356 | 0.27909 | 0.78574 | 3.17713 | 0.29516 | 4.43316 | 0.25753 | 2.00752 | 4.98691 | 0.44227 | 1.75931 |
| **C7orf31** | 0.11134 | 0.44282 | 0.04173 | 0.28659 | 0.29468 | 1.58514 | 0.26423 | 0.35826 | 0.00000 | 0.95675 | 0.00000 | 1.27556 |
| **C7orf33** | 0.01237 | 0.01407 | 0.27006 | 0.01468 | 0.03489 | 0.00000 | 0.00520 | 0.01460 | 0.00893 | 0.00000 | 0.01793 | 0.13456 |
| **C7orf43** | 0.32943 | 0.13203 | 1.90522 | 1.31036 | 1.25138 | 0.00000 | 2.12811 | 0.22856 | 0.78935 | 3.04757 | 0.35027 | 0.21013 |
| **C7orf49** | 1.51554 | 13.13122 | 6.18758 | 2.63329 | 1.75041 | 9.05731 | 0.00000 | 0.94203 | 7.82078 | 0.00000 | 6.76732 | 8.05936 |
| **C7orf50** | 0.35174 | 16.05295 | 9.24443 | 9.76425 | 17.17149 | 0.04202 | 13.48980 | 1.57613 | 16.77188 | 0.00000 | 10.25493 | 8.91574 |
| **C7orf55** | 0.05006 | 1.83800 | 1.14915 | 1.45806 | 1.03821 | 2.64040 | 1.42399 | 0.46453 | 0.68342 | 2.14972 | 2.22725 | 1.98998 |
| **C7orf57** | 0.00967 | 0.29798 | 0.03214 | 0.51371 | 0.00000 | 0.20360 | 0.00000 | 0.00424 | 0.05846 | 0.07861 | 0.00000 | 0.11457 |
| **C7orf60** | 1.55563 | 0.44571 | 2.35324 | 0.23625 | 0.00000 | 1.17549 | 0.00000 | 1.87977 | 0.32316 | 0.28934 | 0.15391 | 0.20239 |
| **C7orf61** | 0.00978 | 0.05779 | 0.02157 | 0.00465 | 0.02864 | 0.00000 | 0.05405 | 0.01341 | 0.01840 | 0.06863 | 0.01046 | 0.01031 |
| **C7orf63** | 0.05299 | 0.12319 | 0.00000 | 0.16422 | 0.00000 | 0.35110 | 0.24641 | 0.00000 | 0.03918 | 0.01152 | 0.07057 | 0.12400 |
| **C7orf65** | 0.00621 | 0.01466 | 0.01568 | 0.01274 | 0.01434 | 0.01995 | 0.01781 | 0.01031 | 0.01713 | 0.01888 | 0.01488 | 0.01722 |
| **C7orf69** | 0.03341 | 0.01445 | 0.02189 | 0.01353 | 0.01710 | 0.02665 | 0.01523 | 0.01767 | 0.01620 | 0.01553 | 0.01342 | 0.01465 |
| **C7orf71** | 0.08219 | 0.03161 | 0.01775 | 0.01146 | 0.01624 | 0.00000 | 0.11131 | 0.01266 | 0.00005 | 0.03981 | 0.01460 | 0.01213 |
| **C7orf73** | 0.37210 | 9.21002 | 9.50536 | 4.00891 | 4.61454 | 24.14324 | 9.43752 | 0.95071 | 4.68259 | 5.14357 | 10.29531 | 10.84036 |
| **C7orf76** | 0.00422 | 0.01594 | 0.01788 | 0.01049 | 0.02576 | 0.00488 | 0.00663 | 0.10771 | 0.04575 | 0.02608 | 0.01092 | 0.01203 |
| **C8A** | 0.06298 | 0.00448 | 0.00786 | 0.10008 | 0.03015 | 0.05976 | 0.00895 | 0.00646 | 0.00000 | 0.02295 | 0.02662 | 0.01395 |
| **C8B** | 0.01519 | 0.01207 | 0.00697 | 0.00730 | 0.00843 | 0.01623 | 0.01285 | 0.05125 | 0.02042 | 0.09434 | 0.01067 | 0.00956 |
| **C8G** | 0.01943 | 0.37928 | 0.07564 | 0.00000 | 0.00000 | 0.00000 | 0.01158 | 0.00806 | 0.00000 | 0.55820 | 0.17780 | 0.00091 |
| **C8orf12** | 5.50285 | 0.03630 | 0.03096 | 0.00000 | 0.02053 | 0.14248 | 0.24666 | 0.48360 | 0.00000 | 0.00000 | 0.00385 | 0.03130 |
| **C8orf22** | 0.08013 | 0.26547 | 0.01014 | 0.00000 | 0.00000 | 0.03420 | 0.01312 | 0.00910 | 0.01079 | 0.02983 | 0.00996 | 0.01540 |
| **C8orf31** | 0.00288 | 0.00152 | 0.06870 | 0.00000 | 0.06090 | 0.00000 | 0.15857 | 0.24638 | 0.00000 | 0.23728 | 0.55085 | 0.00400 |
| **C8orf33** | 4.42105 | 20.40026 | 5.23814 | 6.54171 | 11.73782 | 13.97483 | 13.73626 | 22.76319 | 12.08234 | 7.65167 | 8.47619 | 22.12657 |
| **C8orf34** | 0.08830 | 0.02592 | 0.01893 | 0.05227 | 0.01150 | 0.00751 | 0.01147 | 0.00298 | 0.02701 | 0.02064 | 0.01481 | 0.12391 |
| **C8orf37** | 0.48232 | 0.36036 | 0.27240 | 0.02333 | 0.03772 | 0.47955 | 0.29679 | 0.84242 | 0.74300 | 0.02197 | 0.40478 | 0.42022 |
| **C8orf37-AS1** | 1.42915 | 1.10767 | 0.00000 | 0.00000 | 0.08295 | 0.00000 | 0.05608 | 0.14201 | 0.00000 | 0.13853 | 0.01095 | 3.34991 |
| **C8orf44** | 0.06025 | 0.17611 | 0.33827 | 0.25768 | 0.00000 | 0.53314 | 0.51145 | 0.27433 | 0.16588 | 0.00013 | 0.74116 | 0.94850 |
| **C8orf44-SGK3** | 0.00603 | 0.01358 | 0.01256 | 0.01078 | 0.01201 | 0.02202 | 0.01630 | 0.01001 | 0.02006 | 0.02310 | 0.01491 | 0.02609 |
| **C8orf46** | 0.02463 | 0.01083 | 0.09559 | 0.00463 | 0.00000 | 0.02296 | 0.07185 | 0.08657 | 0.00000 | 0.10996 | 0.00734 | 0.03388 |
| **C8orf48** | 0.09552 | 0.01553 | 0.01684 | 0.12098 | 0.00652 | 0.19975 | 0.07496 | 0.00000 | 0.05664 | 0.00000 | 0.05083 | 0.17329 |
| **C8orf49** | 0.00519 | 0.01135 | 0.04281 | 0.21752 | 0.00000 | 0.01072 | 0.04423 | 0.01029 | 0.00000 | 0.05790 | 0.01353 | 0.01412 |
| **C8orf58** | 0.00390 | 0.00000 | 0.99479 | 0.00000 | 0.19259 | 0.27181 | 0.59293 | 0.87102 | 0.00000 | 0.65453 | 0.00000 | 0.08534 |
| **C8orf59** | 0.00000 | 7.11850 | 11.23541 | 1.04148 | 3.13745 | 33.52888 | 14.05640 | 1.96303 | 12.15567 | 12.90444 | 11.43898 | 12.69446 |
| **C8orf76** | 0.01460 | 2.35561 | 2.74462 | 3.24907 | 0.86462 | 7.28883 | 3.84211 | 0.71998 | 3.41221 | 1.94866 | 1.24424 | 3.55978 |
| **C8orf82** | 0.33435 | 0.00000 | 0.33435 | 0.02731 | 1.26229 | 0.00000 | 1.59601 | 0.39578 | 0.26188 | 3.95309 | 0.00000 | 0.13782 |
| **C8orf88** | 0.00315 | 0.13058 | 0.29788 | 0.06512 | 0.00000 | 0.23182 | 0.14495 | 0.32819 | 0.06572 | 0.00013 | 0.00000 | 0.44313 |
| **C9** | 0.00611 | 0.00415 | 0.15096 | 0.01514 | 0.00384 | 0.09058 | 0.00000 | 0.00865 | 0.17878 | 0.00000 | 0.01018 | 0.03569 |

| | | | | | | | | | | | | |
|---|---|---|---|---|---|---|---|---|---|---|---|---|
| **C9orf114** | 0.11618 | 10.77844 | 7.84669 | 8.18867 | 10.04892 | 1.38618 | 8.96469 | 5.33117 | 7.05966 | 5.30359 | 8.67664 | 12.12062 |
| **C9orf116** | 0.07149 | 4.40112 | 0.00000 | 0.56191 | 2.12159 | 0.19595 | 0.02003 | 0.01547 | 0.00000 | 0.00000 | 0.28203 | 0.07641 |
| **C9orf117** | 0.09758 | 0.13504 | 0.07405 | 0.09045 | 0.16237 | 0.05835 | 0.05167 | 0.05499 | 0.15828 | 0.15282 | 0.06845 | 0.17483 |
| **C9orf129** | 0.13453 | 0.01557 | 0.01608 | 0.02723 | 0.00898 | 0.03141 | 0.01711 | 0.00309 | 0.01702 | 0.03236 | 0.01682 | 0.05519 |
| **C9orf131** | 0.14178 | 0.00912 | 0.03302 | 0.16016 | 0.03411 | 0.00000 | 0.04868 | 0.06315 | 0.18422 | 0.10908 | 0.05697 | 0.20515 |
| **C9orf135** | 0.26788 | 0.09980 | 0.00000 | 3.75181 | 0.00000 | 0.41369 | 0.00016 | 0.00339 | 1.82523 | 0.00000 | 0.02682 | 0.00000 |
| **C9orf135-AS1** | 0.00404 | 0.00173 | 0.03674 | 0.00678 | 0.07103 | 0.05500 | 0.00000 | 0.01253 | 0.02025 | 0.08547 | 0.01459 | 0.01025 |
| **C9orf139** | 0.01953 | 0.01418 | 0.01176 | 0.01065 | 0.01330 | 0.01755 | 0.01680 | 0.00857 | 0.02114 | 0.03070 | 0.01330 | 0.01560 |
| **C9orf142** | 0.00696 | 8.10368 | 12.77688 | 1.70876 | 15.96411 | 0.00000 | 10.79249 | 0.00000 | 21.12699 | 5.24523 | 15.31626 | 5.15284 |
| **C9orf152** | 0.01192 | 0.00000 | 0.36493 | 0.10015 | 0.10524 | 0.05823 | 0.12291 | 0.00971 | 0.04014 | 0.00000 | 0.01013 | 0.37280 |
| **C9orf153** | 0.03125 | 0.03818 | 0.08842 | 0.00000 | 0.00006 | 0.30950 | 0.00000 | 0.27314 | 0.00000 | 0.16213 | 0.01742 | 0.13350 |
| **C9orf156** | 0.72861 | 3.81811 | 3.73915 | 1.60242 | 0.69916 | 8.16607 | 6.16561 | 2.24025 | 3.46236 | 0.00000 | 4.71234 | 3.66579 |
| **C9orf16** | 0.21555 | 6.14437 | 4.27240 | 4.50083 | 8.51691 | 0.00000 | 7.63457 | 1.87167 | 6.73102 | 0.99049 | 3.56016 | 6.28916 |
| **C9orf163** | 0.00429 | 0.01795 | 0.04495 | 0.00000 | 0.02101 | 0.06831 | 0.00000 | 0.00888 | 0.05522 | 0.11641 | 0.10851 | 0.01701 |
| **C9orf170** | 0.00616 | 0.01356 | 0.01239 | 0.01038 | 0.02011 | 0.02536 | 0.02687 | 0.01049 | 0.02009 | 0.01273 | 0.01658 | 0.01585 |
| **C9orf171** | 0.84322 | 0.01435 | 0.01919 | 0.02919 | 0.04700 | 0.00000 | 0.09121 | 0.14016 | 0.00000 | 0.01566 | 0.04023 | 0.06600 |
| **C9orf172** | 0.00567 | 0.01233 | 0.01205 | 0.02305 | 0.01778 | 0.01365 | 0.01169 | 0.00980 | 0.02685 | 0.05528 | 0.01213 | 0.02154 |
| **C9orf173** | 0.00575 | 0.01468 | 0.02403 | 0.01526 | 0.00880 | 0.02381 | 0.01253 | 0.01005 | 0.01126 | 0.04588 | 0.01428 | 0.01698 |
| **C9orf24** | 0.04104 | 0.02138 | 0.03992 | 0.05902 | 0.03590 | 0.00560 | 0.11440 | 0.01607 | 0.05178 | 0.00000 | 0.00528 | 0.01884 |
| **C9orf3** | 1.05515 | 7.81904 | 4.88168 | 2.10647 | 7.98427 | 5.38893 | 3.87560 | 2.55454 | 8.57561 | 4.41914 | 7.30638 | 3.02240 |
| **C9orf37** | 0.77622 | 3.88548 | 3.59144 | 1.23409 | 1.19966 | 0.00123 | 5.23436 | 3.01609 | 0.33536 | 0.76760 | 3.73189 | 6.17636 |
| **C9orf40** | 0.79177 | 0.39703 | 1.70048 | 0.26688 | 1.66027 | 3.97704 | 3.63115 | 0.47029 | 0.38634 | 8.02996 | 1.10332 | 2.27271 |
| **C9orf41** | 0.29045 | 0.36154 | 1.04475 | 0.86771 | 0.00000 | 2.84192 | 0.55232 | 0.67020 | 1.56621 | 1.36252 | 1.02469 | 0.21907 |
| **C9orf43** | 0.31396 | 0.67598 | 0.12308 | 0.03195 | 0.15240 | 0.08351 | 0.02430 | 0.52061 | 0.45622 | 0.00000 | 0.16858 | 0.28720 |
| **C9orf47** | 0.05862 | 0.31894 | 0.01459 | 0.42444 | 0.00000 | 0.00000 | 0.00258 | 0.49009 | 0.03111 | 0.20347 | 0.00024 | 0.31605 |
| **C9orf53** | 0.00646 | 0.01927 | 0.02267 | 0.02792 | 0.01361 | 0.00000 | 0.09049 | 0.01081 | 0.06067 | 0.00000 | 0.01310 | 0.01848 |
| **C9orf57** | 0.00591 | 0.00943 | 0.01625 | 0.01350 | 0.00912 | 0.03956 | 0.01335 | 0.00998 | 0.02207 | 0.01620 | 0.01238 | 0.03161 |
| **C9orf64** | 0.20538 | 10.60219 | 2.63567 | 3.71338 | 0.00000 | 14.81718 | 6.42685 | 5.59842 | 4.95132 | 3.19482 | 5.63687 | 16.53383 |
| **C9orf66** | 0.11263 | 0.01164 | 0.18625 | 0.01720 | 0.01188 | 0.00000 | 0.00174 | 0.02643 | 0.18318 | 0.00483 | 0.06252 | 0.13525 |
| **C9orf69** | 1.22427 | 1.17025 | 1.52172 | 1.88886 | 3.85840 | 0.00000 | 4.70757 | 0.01409 | 0.14686 | 10.51496 | 3.31377 | 0.68371 |
| **C9orf72** | 0.19951 | 0.66560 | 2.33350 | 1.46513 | 0.00000 | 4.22181 | 1.66765 | 1.88150 | 0.81733 | 0.00000 | 0.81379 | 2.33459 |
| **C9orf78** | 7.85153 | 17.52110 | 15.02871 | 9.26038 | 14.97679 | 22.34594 | 22.97318 | 8.09583 | 16.83586 | 8.52232 | 17.61201 | 18.28557 |
| **C9orf84** | 0.01099 | 0.35899 | 0.00000 | 0.12977 | 0.34918 | 0.31315 | 0.00000 | 0.31320 | 0.00000 | 0.40736 | 0.00000 | 1.06292 |
| **C9orf85** | 1.25582 | 2.28234 | 3.14556 | 2.45238 | 0.27771 | 6.55609 | 5.48176 | 11.04409 | 0.72833 | 1.74456 | 1.31836 | 3.49705 |
| **C9orf89** | 0.15063 | 2.23097 | 3.54469 | 1.93751 | 1.12219 | 0.00000 | 1.47057 | 0.26844 | 1.20131 | 1.33569 | 0.90025 | 0.51036 |
| **C9orf9** | 0.14953 | 1.12547 | 0.13256 | 0.01789 | 0.00002 | 0.10948 | 0.10322 | 0.00000 | 0.00000 | 0.08550 | 0.36202 | 0.88464 |
| **C9orf91** | 2.75987 | 0.23172 | 0.39160 | 0.17627 | 0.10876 | 1.09638 | 0.52707 | 0.19042 | 0.00000 | 0.60056 | 0.00000 | 0.04764 |
| **C9orf92** | 0.24008 | 0.05766 | 0.01051 | 0.00810 | 0.03376 | 0.04242 | 0.01253 | 0.01077 | 0.01242 | 0.01229 | 0.01240 | 0.08303 |
| **CA1** | 0.00212 | 0.01123 | 0.01521 | 0.01328 | 0.01658 | 0.01568 | 0.01811 | 0.05816 | 0.02077 | 0.00804 | 0.01528 | 0.08776 |
| **CA10** | 0.95413 | 0.00800 | 0.00439 | 0.03358 | 0.00000 | 0.03651 | 0.00978 | 0.05904 | 0.00000 | 0.11725 | 0.02710 | 0.01016 |
| **CA11** | 0.00458 | 0.01199 | 0.02921 | 0.08525 | 0.17485 | 0.00000 | 0.03708 | 0.09490 | 0.00000 | 0.08196 | 0.06999 | 0.22744 |
| **CA12** | 1.15139 | 0.00000 | 22.20642 | 0.00000 | 1.39460 | 0.00000 | 13.18402 | 1.11455 | 31.12429 | 0.97986 | 9.13772 | 0.00000 |
| **CA13** | 0.23674 | 0.00000 | 0.15900 | 0.74071 | 0.26115 | 0.08592 | 0.37209 | 0.00000 | 0.11695 | 0.03968 | 0.00000 | 0.12768 |
| **CA14** | 0.02954 | 0.00000 | 0.01023 | 0.15558 | 0.00000 | 0.00000 | 0.00919 | 0.05578 | 0.15130 | 0.49840 | 0.00135 | 0.70659 |
| **CA2** | 0.00000 | 23.40910 | 19.75561 | 12.76999 | 4.21407 | 44.39885 | 26.54580 | 1.12634 | 8.76192 | 0.47906 | 32.87146 | 11.51243 |
| **CA3** | 0.00580 | 0.01490 | 0.03443 | 0.01017 | 0.01481 | 0.01011 | 0.02708 | 0.01030 | 0.02686 | 0.03503 | 0.01325 | 0.01951 |
| **CA4** | 0.00557 | 0.00166 | 0.00000 | 5.55292 | 0.00103 | 0.08216 | 0.00000 | 0.00796 | 0.11459 | 0.00000 | 0.00486 | 0.00853 |
| **CA5A** | 0.04073 | 0.03876 | 0.02175 | 0.01920 | 0.01388 | 0.00220 | 0.01872 | 0.09532 | 0.01790 | 0.01520 | 0.02317 | 0.02438 |

| | | | | | | | | | | | | |
|---|---|---|---|---|---|---|---|---|---|---|---|---|
| **CA5B** | 0.19400 | 0.12799 | 0.50373 | 0.12842 | 0.02702 | 0.46028 | 0.00000 | 0.04088 | 0.76902 | 0.22090 | 0.45723 | 0.10960 |
| **CA5BP1** | 1.39291 | 0.99395 | 2.15181 | 0.63841 | 0.96311 | 0.13250 | 2.76216 | 1.57437 | 3.41046 | 0.00000 | 1.26432 | 2.77572 |
| **CA6** | 0.00186 | 0.04435 | 0.02066 | 0.00750 | 0.01031 | 0.01562 | 0.01434 | 1.47429 | 0.01403 | 0.01346 | 0.01653 | 0.01193 |
| **CA7** | 0.00322 | 0.11000 | 0.01591 | 0.01532 | 0.00000 | 0.00000 | 0.01900 | 0.03101 | 0.00000 | 0.02868 | 0.09845 | 0.00918 |
| **CA8** | 0.00235 | 0.42199 | 0.62337 | 1.03153 | 0.36394 | 1.64232 | 2.07054 | 0.23347 | 1.58370 | 0.00000 | 0.52112 | 0.44747 |
| **CA9** | 0.18315 | 0.21449 | 0.00130 | 0.20549 | 0.00849 | 0.00965 | 0.01571 | 0.01129 | 0.07644 | 0.01698 | 0.01692 | 0.01370 |
| **CAAP1** | 1.55476 | 4.93816 | 4.49456 | 4.56099 | 0.00000 | 18.01320 | 5.79126 | 5.12516 | 6.06795 | 5.60515 | 7.49630 | 8.47646 |
| **CAB39** | 0.29719 | 0.00000 | 5.22968 | 7.19860 | 2.55369 | 13.25561 | 10.29195 | 2.02201 | 0.00000 | 9.37700 | 0.88547 | 0.00000 |
| **CAB39L** | 0.15164 | 2.49828 | 7.07982 | 0.12983 | 0.00000 | 4.11367 | 7.31780 | 9.02619 | 5.65866 | 0.00000 | 2.27683 | 6.41698 |
| **CABIN1** | 0.99116 | 2.24142 | 7.20757 | 3.34059 | 4.58699 | 0.00000 | 6.40809 | 0.93434 | 7.04399 | 9.01978 | 1.23966 | 3.21719 |
| **CABLES1** | 0.32762 | 0.23599 | 0.77630 | 0.54908 | 0.94525 | 0.27360 | 0.25231 | 0.08750 | 0.28911 | 0.68888 | 0.00000 | 0.22958 |
| **CABLES2** | 1.24780 | 0.00000 | 0.54354 | 0.04937 | 0.45220 | 0.34413 | 0.62220 | 0.00414 | 0.00000 | 0.00000 | 0.09310 | 0.07829 |
| **CABP1** | 0.00240 | 0.05303 | 0.06826 | 0.03157 | 0.03755 | 0.00000 | 0.01973 | 0.05813 | 0.00472 | 0.01114 | 0.03053 | 0.14044 |
| **CABP2** | 0.00615 | 0.01545 | 0.01714 | 0.01160 | 0.01501 | 0.01998 | 0.01476 | 0.01063 | 0.02004 | 0.01735 | 0.01445 | 0.01725 |
| **CABP4** | 0.08806 | 0.00926 | 0.01583 | 0.02960 | 0.18399 | 0.04885 | 0.00088 | 0.04131 | 0.00000 | 0.18093 | 0.03290 | 0.26787 |
| **CABP7** | 0.02446 | 0.33599 | 0.15546 | 0.00191 | 0.27756 | 0.00000 | 0.34273 | 2.80694 | 0.00273 | 0.00000 | 0.14884 | 0.94680 |
| **CABYR** | 4.46996 | 3.71132 | 0.48670 | 1.59899 | 1.31762 | 3.48558 | 0.51896 | 4.72262 | 1.66750 | 0.00000 | 0.87534 | 8.64161 |
| **CACFD1** | 0.83828 | 2.12074 | 0.91249 | 2.43572 | 0.21553 | 0.00000 | 0.89539 | 0.29987 | 0.88064 | 0.54741 | 0.00000 | 1.28599 |
| **CACHD1** | 0.01390 | 1.85017 | 0.31729 | 3.84939 | 0.00000 | 8.43413 | 0.00000 | 0.01840 | 1.47494 | 12.58838 | 0.20610 | 3.56378 |
| **CACNA1A** | 0.33044 | 0.00000 | 0.00000 | 0.00000 | 0.00000 | 0.51843 | 0.34049 | 0.05106 | 0.00000 | 0.79955 | 0.01572 | 0.06552 |
| **CACNA1B** | 0.00093 | 0.63135 | 0.00000 | 0.14960 | 0.00000 | 0.00000 | 0.00515 | 0.03589 | 0.00000 | 1.77343 | 0.03221 | 1.76378 |
| **CACNA1C** | 2.29067 | 0.01356 | 0.02683 | 0.01023 | 0.02478 | 0.00000 | 0.02271 | 0.35632 | 0.01649 | 0.00000 | 0.03554 | 0.02966 |
| **CACNA1C-AS1** | 0.25864 | 0.01074 | 0.01579 | 0.01019 | 0.03548 | 0.04606 | 0.01362 | 0.04122 | 0.01976 | 0.00526 | 0.00894 | 0.01700 |
| **CACNA1C-AS2** | 0.01009 | 0.01384 | 0.01209 | 0.00549 | 0.02244 | 0.05879 | 0.01146 | 0.05501 | 0.00678 | 0.01391 | 0.02077 | 0.01572 |
| **CACNA1D** | 3.33717 | 0.05666 | 0.08898 | 0.18554 | 0.04059 | 0.01905 | 0.00000 | 0.37345 | 0.00000 | 0.11195 | 0.03959 | 0.52943 |
| **CACNA1E** | 1.30222 | 0.01807 | 0.01739 | 0.14931 | 0.00000 | 0.01698 | 0.00000 | 0.11849 | 0.03951 | 0.10920 | 0.00863 | 0.01923 |
| **CACNA1F** | 0.00853 | 0.01273 | 0.01365 | 0.01299 | 0.03190 | 0.02306 | 0.01238 | 0.01213 | 0.02077 | 0.03403 | 0.01503 | 0.01579 |
| **CACNA1G** | 0.06664 | 0.00530 | 0.00547 | 0.00251 | 0.00000 | 0.02841 | 0.05325 | 0.00537 | 0.00000 | 0.17194 | 0.01201 | 0.01558 |
| **CACNA1H** | 3.72203 | 0.02692 | 0.03101 | 0.01118 | 0.11497 | 0.00000 | 0.00832 | 0.00000 | 0.01155 | 0.04840 | 0.01148 | 0.24043 |
| **CACNA1I** | 0.11939 | 0.00639 | 0.01880 | 0.00876 | 0.05204 | 0.00000 | 0.01233 | 0.02521 | 0.00000 | 0.23498 | 0.01590 | 0.02407 |
| **CACNA1S** | 0.08498 | 0.09685 | 0.03926 | 0.10744 | 0.02820 | 0.00000 | 0.14481 | 0.02211 | 0.00000 | 0.00000 | 0.06797 | 0.00932 |
| **CACNA2D1** | 6.14924 | 0.68005 | 0.15904 | 0.28576 | 0.01042 | 0.00000 | 0.00000 | 0.63430 | 0.00726 | 0.01045 | 0.01641 | 0.96317 |
| **CACNA2D2** | 2.36495 | 0.95955 | 0.00000 | 5.74994 | 1.44440 | 0.34437 | 0.00000 | 0.43062 | 0.00000 | 1.83686 | 0.07573 | 0.03960 |
| **CACNA2D3** | 0.76110 | 0.57333 | 1.36382 | 0.76079 | 2.43453 | 0.94291 | 0.00000 | 0.12974 | 0.78885 | 0.42727 | 1.63943 | 0.00000 |
| **CACNA2D3-AS1** | 0.00573 | 0.01740 | 0.03344 | 0.00973 | 0.01194 | 0.01947 | 0.03110 | 0.01057 | 0.01890 | 0.03615 | 0.01364 | 0.01573 |
| **CACNA2D4** | 0.01369 | 0.01333 | 0.02494 | 0.00792 | 0.04091 | 0.01845 | 0.01614 | 0.01044 | 0.01577 | 0.01596 | 0.02643 | 0.01570 |
| **CACNB1** | 0.16207 | 0.05551 | 0.37746 | 0.29050 | 0.00000 | 0.00000 | 0.00000 | 0.71413 | 0.11369 | 0.52468 | 0.03039 | 0.22953 |
| **CACNB2** | 0.59752 | 0.86148 | 0.00000 | 0.48920 | 0.65705 | 1.81270 | 0.00000 | 0.16301 | 5.53929 | 3.04745 | 5.87532 | 0.47938 |
| **CACNB3** | 0.02056 | 7.27537 | 0.57910 | 6.32432 | 0.00000 | 0.00000 | 1.07235 | 1.29049 | 0.07428 | 0.67303 | 0.00000 | 10.53346 |
| **CACNB4** | 0.04209 | 0.09926 | 0.00323 | 0.22152 | 0.08824 | 0.08223 | 0.00000 | 0.01157 | 0.00000 | 0.20554 | 0.00000 | 0.13601 |
| **CACNG1** | 0.00274 | 0.01443 | 0.01401 | 0.01132 | 0.03486 | 0.01406 | 0.02099 | 0.04893 | 0.01719 | 0.01305 | 0.01582 | 0.01426 |
| **CACNG2** | 0.01259 | 0.01592 | 0.02202 | 0.02702 | 0.01484 | 0.01826 | 0.00934 | 0.00982 | 0.02021 | 0.03269 | 0.01548 | 0.03040 |
| **CACNG3** | 0.00500 | 0.00758 | 0.00000 | 0.26851 | 0.00000 | 0.02320 | 0.00342 | 0.00988 | 0.01521 | 0.08388 | 0.01000 | 0.01346 |
| **CACNG4** | 0.00563 | 0.01542 | 0.03401 | 0.01839 | 0.04507 | 0.04047 | 0.02075 | 0.01016 | 0.00000 | 0.05853 | 0.01943 | 0.01116 |
| **CACNG5** | 0.29182 | 0.00817 | 0.02184 | 0.01809 | 0.00482 | 0.02645 | 0.01404 | 0.00463 | 0.00931 | 0.04609 | 0.01541 | 0.02340 |
| **CACNG6** | 0.01732 | 0.02087 | 0.01634 | 0.00956 | 0.00937 | 0.03906 | 0.01460 | 0.02632 | 0.01393 | 0.02061 | 0.01259 | 0.01197 |
| **CACNG7** | 0.84906 | 0.01017 | 0.00886 | 0.00460 | 0.01201 | 0.03718 | 0.00861 | 0.03879 | 0.01644 | 0.05616 | 0.00963 | 0.05439 |
| **CACNG8** | 0.08232 | 0.00000 | 0.01438 | 0.02270 | 0.04800 | 0.00000 | 0.04798 | 0.49637 | 0.00000 | 0.15576 | 0.01043 | 0.28605 |

| | | | | | | | | | | | | |
|---|---|---|---|---|---|---|---|---|---|---|---|---|
| **CACTIN** | 1.58111 | 1.03132 | 1.19811 | 1.13370 | 2.59252 | 0.00000 | 1.58498 | 3.02413 | 1.32840 | 7.90574 | 0.27423 | 1.56764 |
| **CACTIN-AS1** | 0.02369 | 0.02513 | 0.01799 | 0.01287 | 0.03268 | 0.00000 | 0.02234 | 0.00904 | 0.03319 | 0.01086 | 0.01244 | 0.02240 |
| **CACUL1** | 7.30973 | 11.46915 | 8.42685 | 7.52413 | 3.49562 | 22.85951 | 15.29723 | 13.80039 | 8.65544 | 8.80275 | 9.84677 | 12.83005 |
| **CACYBP** | 1.15333 | 16.29317 | 26.91240 | 14.38412 | 12.79151 | 35.18938 | 37.10724 | 14.76245 | 25.68044 | 7.44814 | 26.10861 | 13.06366 |
| **CAD** | 3.17850 | 10.38537 | 11.70262 | 9.17405 | 15.11222 | 0.00000 | 12.21687 | 0.00000 | 18.13225 | 19.49871 | 12.71081 | 10.23695 |
| **CADM1** | 1.31081 | 5.78609 | 4.38271 | 22.53013 | 2.43776 | 3.45244 | 0.00000 | 5.54822 | 4.01335 | 2.40017 | 0.00000 | 2.00345 |
| **CADM2** | 0.03146 | 0.18104 | 0.13066 | 0.53237 | 0.00000 | 0.06474 | 0.01538 | 0.12668 | 0.01138 | 0.04775 | 0.00778 | 0.14388 |
| **CADM2-AS2** | 0.01006 | 0.01493 | 0.01793 | 0.01224 | 0.02148 | 0.01409 | 0.01794 | 0.04818 | 0.01579 | 0.01837 | 0.01493 | 0.01432 |
| **CADM3** | 0.00651 | 0.01552 | 0.04591 | 0.02170 | 0.01397 | 0.01230 | 0.01004 | 0.01045 | 0.05206 | 0.00641 | 0.02100 | 0.01715 |
| **CADM4** | 0.03446 | 0.02082 | 0.05811 | 0.06928 | 0.02728 | 0.00261 | 0.00039 | 0.11422 | 0.01751 | 0.24935 | 0.02504 | 0.10515 |
| **CADPS** | 0.49068 | 0.01341 | 0.09695 | 0.00339 | 0.03513 | 0.03171 | 0.04194 | 0.04449 | 0.01824 | 0.03191 | 0.02068 | 0.07523 |
| **CADPS2** | 2.39125 | 0.04582 | 0.05237 | 0.00837 | 0.18244 | 0.00000 | 0.04595 | 0.23012 | 0.14585 | 0.99030 | 0.00000 | 0.00972 |
| **CAGE1** | 0.04396 | 0.03550 | 0.06462 | 0.01338 | 0.03282 | 0.00000 | 0.05823 | 0.21028 | 0.07034 | 0.00000 | 0.04229 | 0.08116 |
| **CAHM** | 0.02507 | 0.00838 | 0.10586 | 0.00387 | 0.00641 | 0.09503 | 0.03440 | 0.03351 | 0.00000 | 0.06631 | 0.02579 | 0.05935 |
| **CALB1** | 0.70715 | 0.00000 | 0.39782 | 5.70825 | 0.72912 | 1.06382 | 0.30935 | 0.01106 | 0.55638 | 0.27085 | 3.52813 | 0.00000 |
| **CALB2** | 0.23227 | 0.00000 | 0.10111 | 0.00000 | 0.07197 | 0.05676 | 0.16469 | 0.00095 | 1.27298 | 0.37057 | 0.00000 | 0.02566 |
| **CALCA** | 0.00771 | 0.01380 | 0.01140 | 0.04733 | 0.00899 | 0.03155 | 0.02081 | 0.02930 | 0.01963 | 0.01015 | 0.01345 | 0.01633 |
| **CALCB** | 0.00263 | 0.06276 | 0.09719 | 0.01210 | 0.01268 | 0.02273 | 0.02723 | 0.35533 | 0.02202 | 0.00000 | 0.02705 | 0.02813 |
| **CALCOCO1** | 3.83907 | 2.43090 | 2.62750 | 1.42080 | 0.00000 | 0.42740 | 2.63449 | 0.04852 | 3.37315 | 0.00000 | 3.86225 | 4.41835 |
| **CALCOCO2** | 7.04750 | 7.18823 | 3.84496 | 11.78802 | 3.82943 | 11.92251 | 8.29552 | 9.37393 | 7.61852 | 0.00000 | 7.17891 | 10.62843 |
| **CALCR** | 0.00549 | 0.09102 | 0.42183 | 0.71905 | 0.07543 | 0.00000 | 0.00137 | 0.12985 | 0.00000 | 0.00462 | 0.04220 | 0.05321 |
| **CALCRL** | 0.01764 | 0.00675 | 0.02336 | 0.00135 | 0.03166 | 0.06759 | 0.00000 | 0.08250 | 0.02070 | 0.04768 | 0.02185 | 0.08313 |
| **CALD1** | 0.26878 | 0.78837 | 16.82202 | 3.59443 | 27.59267 | 0.00078 | 23.44432 | 0.17385 | 11.78031 | 3.49989 | 19.72272 | 0.00000 |
| **CALHM1** | 0.00566 | 0.00664 | 0.02046 | 0.07037 | 0.00000 | 0.05339 | 0.00001 | 0.00995 | 0.03814 | 0.04034 | 0.01125 | 0.01252 |
| **CALHM2** | 0.00489 | 0.09560 | 1.71386 | 0.96920 | 0.09192 | 0.00000 | 0.15315 | 0.00904 | 0.31650 | 0.11848 | 0.03760 | 0.02496 |
| **CALHM3** | 0.11223 | 0.01325 | 0.05243 | 0.01786 | 0.01944 | 0.01076 | 0.01843 | 0.00866 | 0.01569 | 0.01218 | 0.01711 | 0.02327 |
| **CALM1** | 21.91184 | 31.71119 | 29.73273 | 27.66860 | 32.81150 | 43.07688 | 47.21310 | 13.09209 | 32.70002 | 37.04277 | 38.77757 | 18.19585 |
| **CALM2** | 11.96690 | 34.89872 | 21.39094 | 30.16408 | 30.68709 | 85.58570 | 58.04831 | 22.33271 | 54.46551 | 22.46818 | 49.27624 | 31.27581 |
| **CALM3** | 3.50594 | 18.50213 | 12.32182 | 27.12708 | 33.46413 | 14.19142 | 16.15303 | 15.81275 | 8.09345 | 17.63594 | 12.68323 | 20.62941 |
| **CALML3** | 0.00218 | 0.02068 | 0.45643 | 0.03004 | 0.02221 | 0.05326 | 0.00000 | 0.11213 | 0.00000 | 0.03254 | 0.01383 | 0.01668 |
| **CALML3-AS1** | 0.00626 | 0.01629 | 0.17676 | 0.00687 | 0.01035 | 0.01726 | 0.00000 | 0.01036 | 0.00355 | 0.02234 | 0.04314 | 0.01546 |
| **CALML4** | 11.37288 | 0.67758 | 1.85369 | 0.35318 | 1.49115 | 4.91181 | 0.57191 | 1.30578 | 3.63077 | 5.06401 | 0.28212 | 0.14165 |
| **CALML5** | 0.11434 | 0.10718 | 0.70181 | 0.00416 | 0.08183 | 0.00000 | 0.09983 | 0.00679 | 0.00000 | 0.00000 | 0.01991 | 0.07140 |
| **CALML6** | 0.02709 | 0.01552 | 0.01995 | 0.00792 | 0.06853 | 0.00262 | 0.04278 | 0.01776 | 0.00258 | 0.04289 | 0.02005 | 0.01790 |
| **CALN1** | 5.65274 | 0.84255 | 0.00000 | 0.00000 | 0.07571 | 0.19839 | 0.00000 | 1.54548 | 0.00000 | 0.00007 | 0.03612 | 0.21344 |
| **CALR** | 2.55741 | 21.37384 | 34.73699 | 22.69927 | 33.32689 | 9.45885 | 22.80225 | 24.18238 | 52.89987 | 26.84322 | 30.51138 | 19.29732 |
| **CALR3** | 0.14114 | 0.02649 | 0.08070 | 0.15696 | 0.00000 | 0.04061 | 0.00547 | 0.00663 | 0.00658 | 0.02507 | 0.01728 | 0.31240 |
| **CALU** | 4.76874 | 14.45868 | 19.47932 | 10.30262 | 3.83340 | 34.10477 | 0.00000 | 18.36291 | 41.92305 | 17.22413 | 13.60359 | 16.26874 |
| **CALY** | 0.00547 | 0.00759 | 0.10019 | 0.00052 | 0.07853 | 0.02255 | 0.00000 | 0.01017 | 0.00000 | 0.05660 | 0.03171 | 0.01422 |
| **CAMK1** | 0.01470 | 0.02882 | 0.20554 | 0.02095 | 0.18652 | 0.00000 | 0.00000 | 0.05452 | 0.06708 | 0.29282 | 0.01781 | 0.09038 |
| **CAMK1D** | 0.91484 | 0.01730 | 0.12109 | 0.13198 | 0.03210 | 0.00480 | 0.04790 | 0.13733 | 0.00305 | 0.00000 | 0.03650 | 0.08062 |
| **CAMK2A** | 5.03883 | 0.05416 | 0.02757 | 0.00154 | 0.00000 | 0.03397 | 0.03411 | 0.41033 | 0.00000 | 0.07025 | 0.01410 | 0.26719 |
| **CAMK2B** | 0.02347 | 0.03487 | 0.00000 | 0.03853 | 0.00000 | 0.07793 | 0.12071 | 0.04283 | 0.00000 | 0.34416 | 0.02237 | 0.02234 |
| **CAMK2D** | 0.02084 | 0.00000 | 1.20500 | 5.58636 | 4.31840 | 1.42071 | 4.03142 | 0.00000 | 0.29323 | 0.03304 | 4.78379 | 0.01950 |
| **CAMK2G** | 7.17134 | 1.97652 | 5.24276 | 1.78376 | 11.38508 | 2.48508 | 12.48018 | 1.56295 | 11.36893 | 8.53822 | 6.18366 | 1.70947 |
| **CAMK2N1** | 0.00255 | 0.00000 | 0.07182 | 0.00000 | 1.27509 | 0.53501 | 0.29508 | 0.29700 | 0.00000 | 1.73370 | 0.13038 | 0.06836 |
| **CAMK2N2** | 0.12809 | 0.18662 | 0.33992 | 0.09298 | 0.00000 | 0.00000 | 0.57203 | 0.07097 | 0.11208 | 0.20373 | 0.04541 | 1.13101 |
| **CAMK4** | 0.59836 | 0.02275 | 0.02320 | 0.01081 | 0.03210 | 0.00222 | 0.04230 | 0.16382 | 0.01000 | 0.01583 | 0.02058 | 0.01566 |

| | | | | | | | | | | | | |
|---|---|---|---|---|---|---|---|---|---|---|---|---|
| **CAMKK1** | 0.08964 | 0.27843 | 0.00000 | 0.15576 | 0.24518 | 0.05808 | 0.00000 | 0.11924 | 0.06605 | 0.31051 | 0.77188 | 0.08098 |
| **CAMKK2** | 0.85702 | 6.28067 | 2.40486 | 0.00000 | 4.07633 | 0.52178 | 4.99579 | 0.00000 | 7.00705 | 3.46710 | 5.16056 | 3.27109 |
| **CAMKMT** | 0.73927 | 1.39276 | 1.61321 | 0.31754 | 1.00099 | 3.33457 | 0.00000 | 0.60691 | 1.73178 | 1.78997 | 1.64771 | 2.04300 |
| **CAMKV** | 0.00962 | 0.38465 | 0.22152 | 2.96013 | 0.00000 | 0.00357 | 0.00000 | 0.14035 | 0.43338 | 0.00205 | 0.00194 | 0.10061 |
| **CAMLG** | 3.61131 | 4.38474 | 7.93017 | 8.26513 | 2.61228 | 11.18706 | 12.04002 | 3.43216 | 9.26783 | 11.62056 | 6.84471 | 9.85688 |
| **CAMP** | 0.00120 | 0.37447 | 0.00000 | 3.20151 | 1.19947 | 0.46773 | 0.01656 | 0.20460 | 1.13577 | 3.05423 | 0.38813 | 0.29461 |
| **CAMSAP1** | 8.57329 | 2.90400 | 2.39013 | 3.34630 | 4.07717 | 0.00000 | 5.62599 | 1.34501 | 3.30621 | 7.98303 | 1.85284 | 3.15330 |
| **CAMSAP2** | 0.00000 | 1.08962 | 11.17434 | 0.00000 | 1.23419 | 11.09420 | 8.88160 | 1.04264 | 0.03767 | 16.32087 | 1.47600 | 2.79476 |
| **CAMSAP3** | 0.39381 | 0.62545 | 0.12795 | 1.88965 | 5.73846 | 0.00000 | 1.08507 | 0.01926 | 0.00000 | 4.35244 | 0.22372 | 0.00012 |
| **CAMTA1** | 1.96045 | 3.53237 | 4.60630 | 7.70990 | 4.20005 | 9.61394 | 6.59630 | 10.88636 | 3.60730 | 3.42886 | 3.74122 | 7.01233 |
| **CAMTA2** | 10.94827 | 2.11654 | 2.04324 | 3.35353 | 4.14612 | 0.00000 | 2.51847 | 3.40123 | 6.11665 | 4.32925 | 1.74792 | 1.83541 |
| **CAND1** | 0.72224 | 6.56885 | 14.03236 | 4.26717 | 4.66564 | 22.58793 | 16.48980 | 1.34587 | 9.67002 | 9.28056 | 9.29627 | 5.54474 |
| **CAND1.11** | 0.01202 | 0.02221 | 0.05079 | 0.01103 | 0.03223 | 0.05209 | 0.01061 | 0.01763 | 0.00000 | 0.01057 | 0.01967 | 0.16847 |
| **CAND2** | 0.63848 | 1.60106 | 0.12150 | 0.25325 | 0.14875 | 0.00000 | 0.00000 | 0.00000 | 0.27529 | 0.67084 | 0.00395 | 2.54539 |
| **CANT1** | 0.89833 | 2.29309 | 4.32322 | 1.29891 | 2.95475 | 1.23845 | 4.54630 | 1.60358 | 6.86890 | 7.11338 | 1.27948 | 4.71386 |
| **CANX** | 4.55482 | 22.54602 | 29.03251 | 17.05403 | 17.96100 | 58.29000 | 19.96551 | 13.15185 | 59.43329 | 40.09597 | 32.13917 | 19.70804 |
| **CAP1** | 1.13254 | 17.94181 | 24.73863 | 19.06467 | 21.51480 | 27.74141 | 36.35363 | 9.33352 | 19.69869 | 18.23166 | 25.49631 | 18.40178 |
| **CAP2** | 0.49938 | 0.32558 | 0.68064 | 0.24431 | 0.73775 | 0.00000 | 0.09861 | 0.06179 | 0.68816 | 0.03651 | 1.53091 | 0.19366 |
| **CAPG** | 0.01976 | 1.86109 | 2.30463 | 16.67905 | 6.23401 | 3.28558 | 0.90031 | 0.06730 | 7.34653 | 0.00000 | 2.73900 | 0.00000 |
| **CAPN1** | 0.85643 | 15.56255 | 15.67580 | 9.99335 | 12.90589 | 0.00000 | 11.84018 | 0.00000 | 15.80446 | 0.01025 | 10.98888 | 1.84197 |
| **CAPN10** | 1.65681 | 0.42966 | 1.94525 | 0.60545 | 1.58638 | 0.00000 | 2.82522 | 0.10010 | 2.37859 | 2.90075 | 0.43763 | 0.22105 |
| **CAPN10-AS1** | 0.14783 | 0.38107 | 0.85076 | 0.14036 | 0.18441 | 0.00000 | 0.00000 | 0.20822 | 2.42845 | 0.68867 | 1.52086 | 0.72808 |
| **CAPN11** | 0.00595 | 0.01281 | 0.01050 | 0.01123 | 0.00093 | 0.01242 | 0.03218 | 0.01040 | 0.00000 | 0.04915 | 0.08246 | 0.01491 |
| **CAPN12** | 0.37991 | 0.31569 | 0.31456 | 0.22731 | 0.77068 | 0.00000 | 0.00000 | 4.62460 | 0.49969 | 0.30021 | 0.03328 | 0.58301 |
| **CAPN13** | 5.60705 | 0.00000 | 0.04060 | 0.80261 | 0.21676 | 0.00000 | 0.00000 | 8.76254 | 0.00000 | 0.03707 | 0.02204 | 1.82976 |
| **CAPN14** | 0.01068 | 0.00659 | 0.01755 | 0.00246 | 0.00358 | 0.02494 | 0.00323 | 0.08848 | 0.02618 | 0.17781 | 0.00476 | 0.01154 |
| **CAPN15** | 0.00140 | 0.30414 | 2.65563 | 0.73652 | 3.57845 | 0.00000 | 3.39124 | 0.07747 | 3.16849 | 16.86429 | 1.41354 | 0.48010 |
| **CAPN2** | 0.60186 | 1.79494 | 2.27024 | 6.59240 | 3.97202 | 4.17818 | 11.39713 | 0.00000 | 15.12364 | 5.01877 | 17.56047 | 0.00000 |
| **CAPN3** | 2.72489 | 0.09751 | 0.00000 | 0.22925 | 0.21293 | 0.00000 | 0.05762 | 0.02611 | 0.00000 | 1.05976 | 0.08997 | 1.48133 |
| **CAPN5** | 1.79779 | 5.74173 | 0.28474 | 14.03849 | 0.00057 | 0.00000 | 0.93982 | 0.00000 | 0.66678 | 0.85593 | 0.72968 | 0.44410 |
| **CAPN6** | 0.00604 | 0.02083 | 4.28105 | 0.00426 | 0.00834 | 0.00922 | 0.01424 | 0.01088 | 0.00000 | 0.01834 | 0.01716 | 0.01688 |
| **CAPN7** | 3.30942 | 1.49934 | 2.83528 | 0.76378 | 1.22458 | 6.36949 | 3.94806 | 0.00000 | 2.03626 | 4.08982 | 2.15561 | 0.00000 |
| **CAPN8** | 0.00749 | 0.01027 | 0.01271 | 0.04773 | 0.01052 | 0.03505 | 0.02248 | 0.02834 | 0.01838 | 0.01179 | 0.01643 | 0.01982 |
| **CAPN9** | 0.00608 | 0.01372 | 0.02060 | 0.01015 | 0.01011 | 0.02020 | 0.01934 | 0.01046 | 0.01726 | 0.02041 | 0.03520 | 0.01584 |
| **CAPNS1** | 0.19529 | 5.04185 | 3.08021 | 8.65502 | 12.75347 | 0.17576 | 8.74985 | 0.44043 | 0.00475 | 18.33855 | 2.32627 | 4.13044 |
| **CAPNS2** | 0.00578 | 0.01110 | 0.02406 | 0.00558 | 0.00464 | 0.04999 | 0.05098 | 0.00983 | 0.03602 | 0.00978 | 0.00441 | 0.02671 |
| **CAPRIN1** | 6.95417 | 16.41840 | 19.80494 | 12.00663 | 8.73090 | 34.62208 | 27.69286 | 9.30706 | 30.69442 | 3.82139 | 22.13121 | 13.73523 |
| **CAPRIN2** | 5.39315 | 0.78068 | 0.47967 | 2.02354 | 0.00000 | 0.74266 | 0.26475 | 0.34285 | 0.00000 | 2.25088 | 0.08857 | 1.26350 |
| **CAPS** | 0.06303 | 0.19909 | 0.09250 | 0.03610 | 0.21336 | 0.00000 | 0.06854 | 0.03383 | 0.45590 | 0.43396 | 0.04332 | 0.10989 |
| **CAPS2** | 0.03317 | 0.00001 | 0.18568 | 0.04762 | 0.00000 | 0.21722 | 0.06138 | 0.06958 | 0.03016 | 0.00000 | 0.03354 | 0.02134 |
| **CAPSL** | 0.00615 | 0.01298 | 0.01517 | 0.01161 | 0.01063 | 0.03083 | 0.02082 | 0.01045 | 0.01942 | 0.01521 | 0.01267 | 0.01534 |
| **CAPZA1** | 4.78925 | 12.60903 | 20.99765 | 12.06846 | 0.00000 | 46.65642 | 29.27599 | 5.99960 | 9.22023 | 15.05941 | 24.88774 | 11.31369 |
| **CAPZA2** | 0.24859 | 9.96001 | 10.35482 | 5.17351 | 0.00000 | 32.30396 | 14.51987 | 0.30199 | 12.89976 | 0.01355 | 14.50626 | 3.06192 |
| **CAPZB** | 4.09165 | 15.50054 | 20.16976 | 15.06279 | 20.15293 | 13.51162 | 17.61541 | 2.76559 | 20.99493 | 2.99034 | 17.36710 | 11.49629 |
| **CARD10** | 0.09748 | 2.52286 | 1.16112 | 1.31985 | 4.95201 | 0.00000 | 2.07480 | 0.03373 | 0.86200 | 2.76796 | 0.08928 | 0.23618 |
| **CARD11** | 0.00472 | 0.12988 | 0.13774 | 0.70293 | 0.23167 | 0.00000 | 0.00751 | 0.01094 | 0.00000 | 0.04431 | 0.01150 | 0.08461 |
| **CARD14** | 0.02296 | 0.12919 | 0.19203 | 0.21779 | 0.12006 | 0.00000 | 0.01758 | 0.00654 | 0.39980 | 0.00000 | 0.13729 | 0.10457 |
| **CARD16** | 0.01814 | 0.00674 | 0.01668 | 0.01575 | 0.00933 | 0.04191 | 0.01274 | 0.00797 | 0.01265 | 0.01620 | 0.01390 | 0.07032 |

| | | | | | | | | | | | | |
|---|---|---|---|---|---|---|---|---|---|---|---|---|
| **CARD18** | 0.00369 | 0.00050 | 0.00000 | 0.00000 | 0.00213 | 0.05430 | 0.03235 | 0.00953 | 0.20857 | 0.18634 | 0.00000 | 0.00719 |
| **CARD6** | 0.05334 | 0.01198 | 0.04765 | 0.00635 | 0.00782 | 0.02565 | 0.01260 | 0.13374 | 0.00772 | 0.04279 | 0.02458 | 0.00753 |
| **CARD8** | 0.48936 | 0.52902 | 0.91699 | 0.47262 | 0.54074 | 0.60867 | 1.57699 | 0.13896 | 0.60002 | 0.00000 | 1.29519 | 0.60216 |
| **CARD9** | 0.10358 | 0.01257 | 0.19864 | 0.00000 | 0.24004 | 0.00000 | 0.01595 | 0.03250 | 0.88427 | 0.35672 | 0.05620 | 0.33451 |
| **CARF** | 0.00140 | 0.00000 | 0.21643 | 0.02498 | 0.00000 | 0.28322 | 0.00000 | 0.37160 | 0.25286 | 0.19041 | 0.00000 | 0.95952 |
| **CARHSP1** | 1.27974 | 6.15929 | 18.72268 | 18.39749 | 15.55047 | 0.99665 | 17.19689 | 9.93086 | 15.68282 | 5.67826 | 9.41740 | 2.81249 |
| **CARKD** | 4.95764 | 7.75616 | 8.68539 | 13.31521 | 1.40115 | 8.58979 | 10.80652 | 8.45441 | 8.28667 | 0.54530 | 6.40623 | 5.91147 |
| **CARM1** | 0.56809 | 1.28699 | 4.93211 | 2.53888 | 6.21439 | 0.00000 | 10.06115 | 1.12702 | 0.09695 | 21.28500 | 2.34150 | 3.00586 |
| **CARNS1** | 0.03052 | 0.01626 | 0.02963 | 0.01380 | 0.00607 | 0.02760 | 0.01200 | 0.00719 | 0.05188 | 0.03261 | 0.00759 | 0.02610 |
| **CARS** | 2.17609 | 9.59054 | 14.26528 | 6.01645 | 6.71398 | 6.15033 | 16.32745 | 1.00354 | 22.63778 | 2.22271 | 14.05542 | 8.59286 |
| **CARS2** | 0.00000 | 0.31267 | 1.39421 | 0.77822 | 1.38889 | 1.29518 | 6.17542 | 0.23112 | 2.36005 | 4.60504 | 0.72404 | 0.00031 |
| **CARTPT** | 0.00611 | 0.02063 | 0.01314 | 0.00969 | 0.01129 | 0.02507 | 0.01608 | 0.01020 | 0.01606 | 0.02131 | 0.01417 | 0.01749 |
| **CASC1** | 4.13447 | 0.06066 | 0.17852 | 0.00000 | 0.07646 | 0.07377 | 0.00000 | 0.90634 | 0.07924 | 0.02827 | 0.01702 | 0.15882 |
| **CASC10** | 0.00343 | 0.13272 | 0.54415 | 0.17960 | 0.00000 | 0.06870 | 0.14166 | 0.09186 | 0.09232 | 0.18368 | 0.16049 | 0.11441 |
| **CASC11** | 0.78525 | 0.00295 | 0.10547 | 0.03054 | 0.01009 | 0.00000 | 0.00728 | 4.69876 | 0.03519 | 0.02494 | 0.02149 | 0.40790 |
| **CASC14** | 0.00391 | 0.02113 | 0.00790 | 0.12515 | 0.00000 | 0.04150 | 0.02937 | 0.01603 | 0.01290 | 0.03866 | 0.01511 | 0.01685 |
| **CASC15** | 0.55231 | 0.12431 | 0.26248 | 0.72246 | 0.00000 | 0.41635 | 0.14580 | 1.44193 | 0.93292 | 0.11128 | 0.03169 | 0.24142 |
| **CASC17** | 0.21714 | 0.01403 | 0.01527 | 0.01156 | 0.01395 | 0.01883 | 0.01914 | 0.00290 | 0.01928 | 0.00916 | 0.01823 | 0.01367 |
| **CASC18** | 0.29301 | 0.01373 | 0.02092 | 0.01015 | 0.01153 | 0.00957 | 0.01637 | 0.00860 | 0.02361 | 0.03306 | 0.02507 | 0.01194 |
| **CASC2** | 0.19241 | 0.69108 | 0.02412 | 0.00498 | 0.01219 | 0.18306 | 0.00138 | 0.00664 | 0.00000 | 0.00803 | 0.12785 | 0.83432 |
| **CASC20** | 0.02144 | 0.04126 | 0.01059 | 0.12236 | 0.06622 | 0.00000 | 0.01180 | 0.02574 | 0.05467 | 0.01320 | 0.01584 | 0.01881 |
| **CASC21** | 0.03841 | 0.01660 | 0.01361 | 0.01254 | 0.01406 | 0.02186 | 0.01784 | 0.04940 | 0.01765 | 0.01507 | 0.01423 | 0.01258 |
| **CASC3** | 6.09253 | 7.00519 | 7.35981 | 5.28012 | 6.26714 | 11.00343 | 13.64343 | 2.97834 | 12.42303 | 7.65979 | 7.14867 | 5.09276 |
| **CASC4** | 0.43049 | 0.02728 | 6.95159 | 4.14951 | 0.00000 | 2.09912 | 1.35145 | 0.48714 | 4.81614 | 0.62362 | 1.06837 | 0.53163 |
| **CASC5** | 6.05942 | 0.00000 | 0.59530 | 0.06565 | 22.22458 | 36.83605 | 0.00000 | 7.87317 | 0.00001 | 33.76417 | 2.97731 | 11.60680 |
| **CASC8** | 1.86169 | 0.03613 | 0.08883 | 0.11724 | 0.17159 | 0.00038 | 0.02068 | 1.95624 | 0.00000 | 0.00000 | 0.03343 | 0.10169 |
| **CASC9** | 0.41734 | 0.22193 | 0.11759 | 0.66099 | 0.00000 | 0.39502 | 0.23570 | 0.00000 | 0.00000 | 0.37447 | 0.11702 | 1.04000 |
| **CASD1** | 0.69445 | 0.00000 | 7.08555 | 0.01297 | 0.00000 | 4.72759 | 3.49116 | 0.00000 | 3.71098 | 3.18217 | 1.36920 | 1.18128 |
| **CASK** | 0.00000 | 0.00000 | 0.61144 | 1.83641 | 0.40879 | 3.82717 | 2.00153 | 0.27524 | 0.00004 | 5.26182 | 0.62885 | 0.00000 |
| **CASKIN1** | 1.16366 | 0.49581 | 0.00307 | 0.08379 | 0.02828 | 0.03958 | 0.00007 | 0.00000 | 0.05262 | 0.11477 | 0.05155 | 0.11785 |
| **CASKIN2** | 0.65980 | 1.34360 | 0.06397 | 0.75719 | 2.25192 | 0.00000 | 0.89150 | 0.16090 | 1.71202 | 2.31132 | 1.02427 | 1.50765 |
| **CASP10** | 0.01684 | 1.05736 | 0.22525 | 3.12016 | 0.00000 | 1.27758 | 0.00000 | 0.00000 | 0.04948 | 0.22478 | 0.00000 | 0.39532 |
| **CASP12** | 0.05886 | 0.01455 | 0.01431 | 0.01176 | 0.01386 | 0.01722 | 0.01924 | 0.01669 | 0.01576 | 0.01631 | 0.02031 | 0.01585 |
| **CASP14** | 0.00690 | 0.02484 | 0.01226 | 0.01105 | 0.01485 | 0.01709 | 0.03912 | 0.00952 | 0.01824 | 0.04806 | 0.01320 | 0.01963 |
| **CASP2** | 2.15762 | 2.21449 | 6.37903 | 6.07672 | 1.84991 | 9.89668 | 14.62680 | 0.80172 | 8.00264 | 2.11389 | 13.49327 | 1.92183 |
| **CASP3** | 0.53463 | 8.21843 | 7.30895 | 5.55331 | 0.70209 | 23.26391 | 12.34383 | 12.62367 | 13.21901 | 0.00000 | 14.67793 | 19.53341 |
| **CASP4** | 0.00652 | 0.00733 | 0.88089 | 0.00319 | 0.05811 | 0.10739 | 0.00000 | 0.00969 | 0.00000 | 0.00405 | 0.00672 | 0.03344 |
| **CASP5** | 0.01633 | 0.01538 | 0.00804 | 0.01220 | 0.01341 | 0.01398 | 0.03551 | 0.01062 | 0.05447 | 0.00000 | 0.02359 | 0.02017 |
| **CASP6** | 0.85683 | 10.58366 | 1.18005 | 1.57285 | 3.22001 | 5.40677 | 17.96717 | 5.67975 | 9.71431 | 0.00000 | 15.75653 | 25.57932 |
| **CASP7** | 0.72355 | 4.37431 | 0.24225 | 0.61323 | 0.18271 | 0.00000 | 0.05875 | 0.02379 | 0.00000 | 0.00000 | 0.78626 | 1.55688 |
| **CASP8** | 0.72855 | 2.69811 | 0.79338 | 4.84903 | 0.00000 | 0.41440 | 0.00000 | 0.42406 | 0.00000 | 0.16961 | 0.06805 | 1.90831 |
| **CASP8AP2** | 4.36317 | 0.09757 | 3.82485 | 2.87166 | 0.00003 | 12.27122 | 4.63692 | 4.67985 | 11.73678 | 15.81783 | 4.52293 | 7.05727 |
| **CASP9** | 1.12097 | 4.06316 | 1.01132 | 3.08907 | 1.01914 | 2.54613 | 1.05053 | 0.00000 | 0.49350 | 0.00000 | 1.41668 | 5.35659 |
| **CASQ1** | 0.00681 | 0.98969 | 2.31353 | 0.39907 | 0.00000 | 0.00000 | 0.00000 | 0.22517 | 0.36226 | 1.00942 | 0.00000 | 2.12678 |
| **CASQ2** | 0.00473 | 0.10444 | 0.00000 | 0.00000 | 0.00000 | 0.05619 | 0.55002 | 0.00774 | 0.47787 | 0.05151 | 0.00000 | 0.04232 |
| **CASR** | 0.00494 | 0.03517 | 0.02067 | 0.14244 | 0.00000 | 0.01531 | 0.01439 | 0.01176 | 0.01150 | 0.03866 | 0.01916 | 0.01854 |
| **CASS4** | 0.07259 | 0.02387 | 0.05830 | 0.00000 | 0.00199 | 0.01262 | 0.04028 | 0.00078 | 0.23621 | 0.19547 | 0.00000 | 0.08965 |
| **CAST** | 4.90693 | 1.12640 | 3.88559 | 0.00000 | 12.10529 | 8.01111 | 14.65229 | 0.85727 | 11.17278 | 3.77049 | 21.24473 | 0.91386 |

| | | | | | | | | | | | | |
|---|---|---|---|---|---|---|---|---|---|---|---|---|
| **CASZ1** | 0.08725 | 0.14426 | 0.09037 | 0.00000 | 0.13900 | 0.00000 | 0.04943 | 0.34291 | 0.00000 | 0.04435 | 0.11005 | 0.48723 |
| **CAT** | 0.00631 | 0.61636 | 2.19387 | 1.76889 | 1.47990 | 4.75598 | 1.04768 | 0.00000 | 0.26516 | 0.02268 | 1.88828 | 0.30748 |
| **CATIP** | 0.03547 | 0.14770 | 0.01778 | 0.01832 | 0.03106 | 0.00000 | 0.00048 | 0.07883 | 0.08320 | 0.11423 | 0.00309 | 0.19594 |
| **CATIP-AS1** | 0.01571 | 0.10452 | 0.04658 | 0.01298 | 0.00000 | 0.03514 | 0.02599 | 0.20088 | 0.02734 | 0.11502 | 0.07535 | 0.58257 |
| **CATSPER1** | 0.29547 | 0.03150 | 0.03610 | 0.00571 | 0.00891 | 0.00206 | 0.01725 | 0.28567 | 0.00000 | 0.06112 | 0.04302 | 0.22539 |
| **CATSPER2** | 0.04444 | 0.03372 | 0.05437 | 0.01742 | 0.07806 | 0.00000 | 0.00251 | 0.20971 | 0.04067 | 0.08520 | 0.02364 | 0.16503 |
| **CATSPER2P1** | 0.03143 | 0.03089 | 0.09628 | 0.01195 | 0.05546 | 0.08269 | 0.00000 | 0.02531 | 0.06510 | 0.01632 | 0.10927 | 0.05387 |
| **CATSPER3** | 0.00433 | 0.12079 | 0.00000 | 0.47320 | 0.16104 | 0.06662 | 0.11164 | 0.00382 | 0.00000 | 0.39415 | 0.30771 | 0.13712 |
| **CATSPER4** | 0.00619 | 0.01511 | 0.01543 | 0.01246 | 0.01645 | 0.01650 | 0.02343 | 0.01050 | 0.01665 | 0.01313 | 0.01492 | 0.01841 |
| **CATSPERB** | 0.27856 | 0.09994 | 0.00953 | 0.00282 | 0.00000 | 0.00000 | 0.07616 | 0.23498 | 0.00000 | 0.24088 | 0.02391 | 0.00215 |
| **CATSPERD** | 1.11746 | 0.01125 | 0.02320 | 0.12861 | 0.02917 | 0.00000 | 0.05570 | 0.62718 | 0.03619 | 0.00000 | 0.01893 | 0.10737 |
| **CATSPERG** | 0.24113 | 0.47622 | 0.16642 | 0.25382 | 1.52589 | 0.00000 | 0.00000 | 0.34183 | 0.66642 | 1.18394 | 0.01326 | 0.36400 |
| **CAV1** | 0.00716 | 0.65047 | 15.61576 | 2.28766 | 0.00000 | 0.00185 | 0.80165 | 0.00601 | 0.00045 | 0.01615 | 0.00000 | 0.20580 |
| **CAV2** | 0.00552 | 0.00889 | 0.85138 | 0.04805 | 0.00130 | 0.04374 | 0.00000 | 0.00975 | 0.00000 | 0.02300 | 0.02326 | 0.02290 |
| **CAV3** | 0.00624 | 0.01605 | 0.01248 | 0.01084 | 0.00973 | 0.02918 | 0.01933 | 0.01040 | 0.02723 | 0.01482 | 0.01417 | 0.01528 |
| **CBFA2T2** | 2.51346 | 3.44130 | 1.46694 | 17.17217 | 5.31169 | 3.37857 | 0.00000 | 4.55608 | 2.43339 | 12.78307 | 2.63070 | 2.88236 |
| **CBFA2T3** | 2.99252 | 0.00000 | 0.00125 | 0.00545 | 0.00000 | 0.00846 | 0.08092 | 0.00260 | 0.00000 | 0.32011 | 0.00077 | 0.07388 |
| **CBFB** | 1.93353 | 2.26378 | 5.82554 | 0.43911 | 0.00000 | 16.39210 | 13.17852 | 4.73159 | 0.90653 | 12.20946 | 2.96504 | 3.06573 |
| **CBL** | 2.38815 | 5.90258 | 13.68617 | 3.54413 | 11.72284 | 9.28064 | 11.54672 | 5.70746 | 9.73481 | 18.44495 | 5.79599 | 17.05892 |
| **CBLB** | 0.57243 | 0.31217 | 30.88008 | 3.78443 | 0.00000 | 4.36419 | 17.52016 | 0.00000 | 15.74404 | 2.58549 | 0.45794 | 0.00695 |
| **CBLC** | 0.00622 | 0.76897 | 1.31625 | 0.23264 | 0.26327 | 0.00000 | 0.70821 | 0.03656 | 0.78463 | 0.00000 | 0.32159 | 0.78365 |
| **CBLL1** | 3.45952 | 11.19442 | 5.83292 | 2.36502 | 1.51236 | 22.55139 | 9.34558 | 15.62687 | 8.87205 | 0.77529 | 7.66198 | 13.67664 |
| **CBLN1** | 0.01648 | 0.21296 | 0.13320 | 0.00000 | 0.00000 | 0.00000 | 0.00000 | 6.03143 | 0.00691 | 0.33605 | 0.00000 | 2.96654 |
| **CBLN2** | 0.00507 | 0.00117 | 0.02904 | 0.00000 | 0.00000 | 0.00000 | 0.43014 | 0.00962 | 0.50179 | 0.07316 | 0.91210 | 0.01205 |
| **CBLN3** | 0.00660 | 0.01056 | 0.02023 | 0.01819 | 0.01257 | 0.03899 | 0.00185 | 0.00932 | 0.03597 | 0.03443 | 0.00889 | 0.02086 |
| **CBLN4** | 0.28128 | 0.01030 | 0.01238 | 0.00836 | 0.00960 | 0.03418 | 0.01284 | 0.26573 | 0.01597 | 0.01720 | 0.01261 | 0.02537 |
| **CBR1** | 0.01738 | 1.31458 | 14.90376 | 0.00000 | 0.00000 | 3.37019 | 20.27663 | 1.45879 | 11.67949 | 0.00000 | 5.25548 | 2.12211 |
| **CBR3** | 0.00488 | 0.82058 | 0.44311 | 2.36520 | 0.00000 | 0.02138 | 0.00000 | 0.28183 | 0.02505 | 0.10965 | 0.14263 | 1.20374 |
| **CBR3-AS1** | 1.29917 | 0.07907 | 0.54527 | 0.00000 | 0.74574 | 0.15097 | 0.43838 | 0.16832 | 0.15063 | 0.50801 | 0.59282 | 0.36569 |
| **CBR4** | 2.18368 | 3.16377 | 1.33283 | 6.54925 | 0.34707 | 15.67040 | 5.68780 | 2.26959 | 4.34563 | 3.00783 | 6.34189 | 2.83706 |
| **CBS** | 0.00007 | 0.00000 | 14.75117 | 7.97725 | 5.14712 | 0.00000 | 22.67421 | 0.01266 | 22.01184 | 6.36327 | 11.48060 | 0.00000 |
| **CBWD1** | 0.00000 | 0.06887 | 3.83280 | 0.85224 | 0.00000 | 5.89495 | 7.12514 | 1.08504 | 3.13765 | 1.14388 | 3.29766 | 1.99774 |
| **CBWD2** | 0.03662 | 0.09023 | 2.27933 | 0.46974 | 0.07349 | 3.46552 | 2.62369 | 0.88736 | 3.68027 | 0.00000 | 1.04224 | 1.04183 |
| **CBWD3** | 0.00558 | 0.01386 | 0.01481 | 0.01030 | 0.01084 | 0.02454 | 0.02703 | 0.01220 | 0.00756 | 0.02790 | 0.02427 | 0.01540 |
| **CBWD5.uid-117** | 0.14902 | 0.27676 | 1.50488 | 0.30306 | 0.70157 | 0.95592 | 1.69936 | 0.82856 | 0.62493 | 0.00003 | 0.70982 | 0.48566 |
| **CBWD6** | 0.00294 | 0.02822 | 0.40617 | 0.18001 | 0.06498 | 0.62631 | 0.53106 | 0.15647 | 0.46655 | 0.28063 | 0.09803 | 0.13154 |
| **CBX1** | 8.92083 | 8.46240 | 6.35720 | 9.27821 | 5.91060 | 14.12229 | 18.78275 | 3.94498 | 9.61088 | 10.94068 | 9.25178 | 9.79487 |
| **CBX2** | 1.62117 | 2.12609 | 9.38662 | 0.87477 | 2.11700 | 0.00000 | 10.13247 | 0.31484 | 7.90749 | 2.14513 | 0.00000 | 1.75724 |
| **CBX3** | 3.97303 | 23.65494 | 13.39315 | 13.79343 | 12.42882 | 43.12706 | 31.40551 | 15.23355 | 10.01518 | 33.56979 | 18.21748 | 19.29175 |
| **CBX3P2** | 0.04728 | 0.09425 | 0.01039 | 0.00000 | 0.01979 | 0.12432 | 0.01635 | 0.00505 | 0.00000 | 0.02191 | 0.00576 | 0.01384 |
| **CBX4** | 0.05806 | 0.04223 | 0.22023 | 0.39254 | 1.11193 | 0.00000 | 0.76997 | 0.20451 | 0.00000 | 3.30187 | 0.00000 | 0.09240 |
| **CBX5** | 8.05467 | 6.80579 | 4.26216 | 2.79677 | 3.42302 | 11.05372 | 7.04158 | 8.23560 | 2.36524 | 9.38187 | 6.39902 | 20.18243 |
| **CBX6** | 0.25375 | 0.20716 | 0.08132 | 0.98144 | 0.40068 | 0.23877 | 0.25948 | 0.33217 | 0.00000 | 1.36761 | 0.00000 | 1.21726 |
| **CBX7** | 0.01181 | 0.22447 | 0.00000 | 0.67703 | 0.00714 | 0.16252 | 0.00000 | 0.00030 | 0.05393 | 0.17009 | 0.02282 | 0.20443 |
| **CBX8** | 1.29994 | 1.02791 | 0.11165 | 0.75197 | 0.62115 | 0.16526 | 1.04961 | 1.04864 | 0.81810 | 0.00000 | 0.57472 | 2.94911 |
| **CBY1** | 4.27713 | 4.62349 | 1.30956 | 3.23885 | 4.97187 | 2.72268 | 2.94367 | 0.17427 | 0.75498 | 0.00563 | 5.44519 | 2.13846 |
| **CBY3** | 0.13045 | 0.08813 | 0.03102 | 0.09202 | 0.06656 | 0.00000 | 0.00047 | 0.00403 | 0.02136 | 0.28508 | 0.08612 | 0.02775 |
| **CC2D1A** | 0.63297 | 1.63444 | 2.81570 | 2.90847 | 2.10428 | 0.00000 | 2.71709 | 0.13535 | 3.69880 | 4.76661 | 1.75240 | 0.56466 |

| | | | | | | | | | | | | |
|---|---|---|---|---|---|---|---|---|---|---|---|---|
| **CC2D1B** | 1.04134 | 2.82918 | 7.50062 | 4.63636 | 5.93647 | 0.00000 | 4.71516 | 0.67635 | 4.15812 | 6.29636 | 4.54464 | 2.88698 |
| **CC2D2A** | 0.08844 | 0.69594 | 0.00000 | 0.05297 | 0.00000 | 0.65492 | 0.74688 | 0.19686 | 0.42871 | 0.95626 | 0.57312 | 0.00000 |
| **CC2D2B** | 0.00491 | 0.00838 | 0.01528 | 0.06347 | 0.00395 | 0.02837 | 0.00320 | 0.02191 | 0.03229 | 0.11031 | 0.03619 | 0.04355 |
| **CCAR1** | 0.68966 | 11.38079 | 14.23721 | 12.18146 | 13.15325 | 30.35638 | 25.80420 | 8.32872 | 21.88547 | 31.01649 | 13.95188 | 19.38462 |
| **CCAR2** | 2.82752 | 15.78974 | 9.31012 | 9.20142 | 10.84780 | 0.00000 | 13.62995 | 5.77123 | 20.57530 | 0.19056 | 5.17050 | 13.39214 |
| **CCAT1** | 0.00571 | 0.00640 | 0.04220 | 0.41690 | 0.00000 | 0.03722 | 0.00000 | 0.00986 | 0.00785 | 0.03175 | 0.01308 | 0.01329 |
| **CCAT2** | 0.00835 | 0.01729 | 0.01458 | 0.01170 | 0.01613 | 0.02343 | 0.01840 | 0.01023 | 0.01755 | 0.00968 | 0.01425 | 0.02348 |
| **CCBE1** | 2.29987 | 0.00000 | 2.22618 | 0.00000 | 0.77596 | 0.00000 | 1.34217 | 0.03039 | 0.00000 | 0.04180 | 3.42326 | 0.01076 |
| **CCBL1** | 0.00215 | 1.84401 | 2.38233 | 0.66218 | 0.58079 | 0.00000 | 0.00000 | 1.14432 | 8.61735 | 1.83953 | 1.72865 | 2.18938 |
| **CCBL2** | 0.05496 | 5.23015 | 8.18348 | 2.33742 | 0.24710 | 8.88291 | 1.31488 | 0.00697 | 3.26738 | 1.24225 | 5.47207 | 3.47430 |
| **CCDC101** | 1.41275 | 4.99743 | 2.18556 | 3.82649 | 5.34946 | 2.27319 | 4.90117 | 1.33458 | 2.29795 | 0.00000 | 6.19236 | 5.68719 |
| **CCDC102A** | 0.17653 | 0.08947 | 0.38277 | 0.36508 | 0.48878 | 0.30488 | 1.31748 | 0.00000 | 1.92472 | 0.00000 | 0.03956 | 0.09613 |
| **CCDC102B** | 0.42703 | 0.06560 | 0.20298 | 0.00000 | 0.00000 | 0.22750 | 0.00000 | 0.03830 | 0.04490 | 0.14609 | 0.15287 | 0.10190 |
| **CCDC103** | 0.23647 | 0.33927 | 0.10665 | 0.77891 | 0.00239 | 1.03077 | 0.00002 | 0.08741 | 0.00545 | 0.00000 | 0.89007 | 1.81960 |
| **CCDC104** | 0.63185 | 4.20483 | 1.17886 | 1.57962 | 0.00000 | 3.83782 | 2.19285 | 0.16775 | 1.96055 | 4.09993 | 1.04983 | 2.75439 |
| **CCDC105** | 0.00534 | 0.01437 | 0.00887 | 0.00993 | 0.01045 | 0.00831 | 0.02493 | 0.01065 | 0.02029 | 0.06209 | 0.02347 | 0.01476 |
| **CCDC106** | 0.27657 | 0.77929 | 0.39979 | 1.28797 | 1.19991 | 0.00000 | 0.28891 | 0.04861 | 0.80816 | 0.38582 | 1.70052 | 0.00841 |
| **CCDC107** | 0.01209 | 0.12306 | 0.84812 | 0.64228 | 0.47556 | 0.00000 | 0.01211 | 0.32001 | 1.64159 | 0.00000 | 0.25138 | 0.92002 |
| **CCDC108** | 0.18389 | 0.22287 | 0.11952 | 0.01674 | 0.00000 | 0.02385 | 0.01824 | 0.17715 | 0.00986 | 0.02622 | 0.04312 | 0.24846 |
| **CCDC109B** | 0.46840 | 8.47965 | 3.40429 | 3.97116 | 8.10054 | 6.16431 | 6.44194 | 0.46502 | 4.22038 | 1.42132 | 6.77248 | 0.12327 |
| **CCDC11** | 0.43644 | 0.23218 | 0.16641 | 0.17817 | 0.00000 | 0.11098 | 0.11266 | 0.18545 | 0.13890 | 0.22984 | 0.00344 | 0.59825 |
| **CCDC110** | 0.29440 | 0.06234 | 0.13899 | 0.00433 | 0.01299 | 0.00000 | 0.00661 | 2.66725 | 0.02648 | 0.00000 | 0.01280 | 0.15230 |
| **CCDC112** | 0.00374 | 1.66421 | 0.54284 | 0.00000 | 0.00000 | 0.68115 | 0.19868 | 0.12681 | 0.13319 | 3.07897 | 0.41031 | 0.77214 |
| **CCDC113** | 0.67253 | 1.74075 | 0.85525 | 0.00044 | 0.07948 | 0.94605 | 0.00000 | 0.23771 | 0.00000 | 0.03277 | 0.00000 | 2.53422 |
| **CCDC114** | 0.08084 | 0.18379 | 0.02370 | 0.00000 | 0.00000 | 0.02542 | 0.02503 | 0.03560 | 0.01750 | 0.05811 | 0.00796 | 0.01644 |
| **CCDC115** | 5.14232 | 6.01856 | 4.74863 | 3.52187 | 3.54932 | 5.18053 | 6.28163 | 3.71117 | 0.26999 | 0.00000 | 2.87444 | 6.12507 |
| **CCDC116** | 0.00137 | 0.02086 | 0.82374 | 0.00436 | 0.00000 | 0.00028 | 2.03276 | 0.14756 | 3.58319 | 0.00000 | 0.87084 | 0.19464 |
| **CCDC117** | 2.75889 | 11.99929 | 6.46169 | 0.74169 | 0.00000 | 14.52337 | 10.88872 | 8.36696 | 5.83569 | 11.72188 | 9.81808 | 12.34811 |
| **CCDC12** | 2.84813 | 17.53362 | 17.04192 | 14.97670 | 17.67507 | 8.39757 | 19.13014 | 26.11933 | 14.67595 | 3.45660 | 7.40513 | 19.91535 |
| **CCDC120** | 1.00364 | 0.98496 | 1.97569 | 1.20227 | 1.58401 | 0.00412 | 0.84574 | 0.09045 | 1.87703 | 1.03388 | 2.06244 | 0.41522 |
| **CCDC121** | 1.84120 | 0.70362 | 0.21421 | 0.05420 | 0.00000 | 1.15250 | 0.52202 | 0.76180 | 0.13798 | 0.00000 | 0.10184 | 0.59791 |
| **CCDC122** | 0.55790 | 0.23320 | 0.24933 | 0.00000 | 0.37462 | 0.24725 | 0.19372 | 0.32602 | 0.00243 | 0.65755 | 0.08184 | 0.87212 |
| **CCDC124** | 1.62765 | 28.68011 | 16.29073 | 17.54154 | 28.35571 | 0.00003 | 25.01304 | 3.41916 | 19.75588 | 2.91474 | 17.72589 | 30.70683 |
| **CCDC125** | 0.02947 | 0.00000 | 2.65697 | 0.00000 | 0.83890 | 2.92462 | 3.25176 | 0.00000 | 1.03853 | 1.50467 | 2.82721 | 0.32476 |
| **CCDC126** | 1.22136 | 1.16606 | 3.69151 | 1.64708 | 0.00000 | 13.94037 | 1.68063 | 4.97450 | 3.61415 | 2.34628 | 2.49026 | 4.59432 |
| **CCDC127** | 4.20071 | 4.76497 | 2.14431 | 1.29952 | 0.23964 | 6.34490 | 0.00000 | 0.72559 | 3.70165 | 2.64443 | 2.38597 | 5.71440 |
| **CCDC129** | 0.09077 | 0.01894 | 0.02500 | 0.34062 | 0.00000 | 0.05779 | 0.03190 | 0.00752 | 0.00286 | 0.02812 | 0.00681 | 0.02669 |
| **CCDC13** | 0.31041 | 0.05719 | 0.20337 | 0.04423 | 0.11800 | 0.00000 | 0.24248 | 0.30237 | 0.31167 | 0.09909 | 0.00602 | 0.62529 |
| **CCDC13-AS1** | 0.12001 | 0.01257 | 0.01494 | 0.01019 | 0.01060 | 0.02223 | 0.01297 | 0.00376 | 0.01835 | 0.03048 | 0.01403 | 0.01700 |
| **CCDC130** | 0.85593 | 5.66138 | 4.57497 | 9.47129 | 6.67945 | 0.00000 | 2.01585 | 0.01084 | 8.77435 | 1.12670 | 2.85454 | 8.38152 |
| **CCDC132** | 1.92709 | 3.90979 | 3.03735 | 0.00000 | 0.00000 | 6.27541 | 1.35019 | 0.00000 | 4.51774 | 2.54569 | 2.52624 | 0.78884 |
| **CCDC134** | 0.07107 | 1.49797 | 1.30917 | 1.32757 | 1.51096 | 0.00000 | 0.00000 | 0.15096 | 2.82509 | 0.54769 | 1.52689 | 0.42998 |
| **CCDC135** | 0.61940 | 0.23954 | 0.00000 | 0.00000 | 0.00000 | 0.00000 | 0.00588 | 0.28133 | 0.18618 | 0.22085 | 0.00568 | 0.23242 |
| **CCDC136** | 0.43163 | 0.11863 | 0.36169 | 0.55629 | 0.00000 | 0.00944 | 0.62359 | 1.78781 | 0.02822 | 1.07839 | 0.10934 | 0.70123 |
| **CCDC137** | 3.54035 | 8.85013 | 12.77444 | 7.74102 | 16.10438 | 0.87675 | 14.13802 | 9.61251 | 15.19048 | 25.53491 | 16.30350 | 9.87950 |
| **CCDC138** | 1.10393 | 1.62052 | 3.03599 | 0.30850 | 0.35957 | 6.50177 | 2.74251 | 0.06513 | 6.99970 | 0.37279 | 3.66834 | 3.75693 |
| **CCDC14** | 1.82363 | 3.62156 | 3.47130 | 1.05244 | 0.82393 | 14.89340 | 4.27532 | 2.04256 | 4.52174 | 9.94242 | 2.91097 | 5.74527 |
| **CCDC141** | 1.82247 | 0.02592 | 0.73377 | 0.39084 | 1.56805 | 0.25704 | 0.00000 | 0.55116 | 2.40810 | 0.76162 | 2.65460 | 0.44334 |

| | | | | | | | | | | | | |
|---|---|---|---|---|---|---|---|---|---|---|---|---|
| **CCDC142** | 0.00762 | 1.93653 | 6.03206 | 1.74082 | 1.53504 | 0.00000 | 1.93999 | 1.13188 | 8.59993 | 2.55566 | 1.17289 | 2.83666 |
| **CCDC144A** | 0.11131 | 0.04570 | 0.03984 | 0.01905 | 0.04382 | 0.00000 | 0.00444 | 0.14099 | 0.03731 | 0.07087 | 0.03834 | 0.72269 |
| **CCDC144B** | 0.22156 | 0.14535 | 0.21627 | 0.14860 | 0.12017 | 0.00000 | 0.00000 | 0.28202 | 0.19040 | 0.34240 | 0.20044 | 1.31074 |
| **CCDC144CP** | 0.09110 | 0.20181 | 0.48303 | 0.34736 | 0.05178 | 0.19712 | 0.12252 | 0.10488 | 1.01554 | 1.28217 | 0.17347 | 1.10041 |
| **CCDC144NL** | 0.16725 | 0.01633 | 0.14151 | 0.02431 | 0.02625 | 0.00000 | 0.00484 | 0.22409 | 0.23519 | 0.13669 | 0.01769 | 0.32158 |
| **CCDC146** | 5.88348 | 0.08225 | 0.17238 | 0.00000 | 0.09370 | 0.00001 | 0.00000 | 1.10053 | 0.06840 | 1.04493 | 0.00000 | 0.38355 |
| **CCDC147** | 16.23460 | 0.07844 | 0.01175 | 0.00400 | 0.00000 | 0.00000 | 0.00295 | 4.32435 | 0.00000 | 0.02869 | 0.07448 | 0.37226 |
| **CCDC147-AS1** | 0.00298 | 0.03242 | 0.04365 | 0.01587 | 0.01909 | 0.00383 | 0.01585 | 0.12738 | 0.01919 | 0.00000 | 0.01799 | 0.07964 |
| **CCDC148** | 2.58557 | 0.23305 | 0.01050 | 0.02244 | 0.08502 | 0.19380 | 0.00000 | 0.50068 | 0.00000 | 0.13553 | 0.00000 | 0.58621 |
| **CCDC149** | 0.90603 | 0.00000 | 0.04901 | 0.10216 | 0.05741 | 0.19996 | 0.00000 | 0.00174 | 0.00000 | 0.02127 | 0.00947 | 0.01764 |
| **CCDC15** | 5.61938 | 0.79653 | 0.00000 | 0.00000 | 0.50898 | 0.88384 | 0.00000 | 3.99302 | 0.00000 | 0.62503 | 0.07791 | 2.24012 |
| **CCDC150** | 1.39305 | 0.28074 | 0.00000 | 0.39574 | 0.00000 | 2.16581 | 2.42164 | 1.18265 | 3.27543 | 4.96243 | 0.61063 | 1.40390 |
| **CCDC151** | 0.03949 | 0.08871 | 0.02240 | 0.01253 | 0.02778 | 0.10079 | 0.00000 | 0.14663 | 0.03413 | 0.01734 | 0.00924 | 0.04553 |
| **CCDC152** | 0.00959 | 0.42247 | 0.00000 | 0.31822 | 0.83469 | 0.59450 | 0.25465 | 0.00147 | 0.41305 | 0.00000 | 0.10358 | 0.18982 |
| **CCDC153** | 0.00211 | 1.10659 | 0.12810 | 0.00000 | 0.34424 | 0.00000 | 0.00000 | 0.07206 | 0.00000 | 0.94949 | 0.28790 | 0.71143 |
| **CCDC154** | 0.00303 | 0.05870 | 0.00000 | 0.13079 | 0.00000 | 0.00000 | 0.02671 | 0.00855 | 0.14229 | 0.36293 | 0.00000 | 0.02132 |
| **CCDC155** | 0.42459 | 0.31912 | 0.10666 | 0.00000 | 0.01123 | 0.00000 | 0.00705 | 0.05793 | 0.02756 | 0.07325 | 0.02105 | 0.26533 |
| **CCDC157** | 0.35529 | 0.44478 | 0.21085 | 0.28443 | 0.36415 | 0.00000 | 0.02896 | 1.12386 | 0.50413 | 0.00000 | 0.08836 | 0.72186 |
| **CCDC158** | 0.02015 | 0.05068 | 0.01739 | 0.03256 | 0.00000 | 0.00000 | 0.00780 | 0.00216 | 0.02053 | 0.10775 | 0.01475 | 0.16947 |
| **CCDC159** | 0.04426 | 0.20234 | 0.40804 | 0.40023 | 0.00000 | 0.37516 | 0.00000 | 0.00466 | 0.00000 | 0.00874 | 0.03595 | 0.04749 |
| **CCDC160** | 0.02428 | 0.00661 | 0.10817 | 0.17821 | 0.00000 | 0.03491 | 0.00000 | 0.00957 | 0.00000 | 0.03500 | 0.02219 | 0.02732 |
| **CCDC162P** | 0.04257 | 0.01950 | 0.04940 | 0.00333 | 0.00286 | 0.10407 | 0.00000 | 0.00469 | 0.00475 | 0.05359 | 0.00675 | 0.06820 |
| **CCDC163P** | 1.35910 | 10.51466 | 0.06236 | 7.18581 | 0.00000 | 4.40660 | 0.00114 | 4.26862 | 2.02687 | 0.33099 | 0.26503 | 9.95415 |
| **CCDC166** | 0.02460 | 0.03318 | 0.01669 | 0.01540 | 0.01177 | 0.01696 | 0.01351 | 0.00850 | 0.00000 | 0.00567 | 0.14901 | 0.01691 |
| **CCDC167** | 0.59876 | 7.30466 | 13.92840 | 11.94091 | 3.73346 | 7.86626 | 18.56948 | 0.19817 | 7.05871 | 1.35600 | 8.19447 | 5.12545 |
| **CCDC168** | 0.00160 | 0.05592 | 0.21497 | 0.00000 | 0.07877 | 0.04179 | 0.12652 | 0.22335 | 0.05040 | 0.08130 | 0.03548 | 0.21792 |
| **CCDC169** | 0.11597 | 0.00000 | 0.35640 | 0.07051 | 0.00000 | 0.28373 | 0.00000 | 0.00245 | 0.20376 | 0.09939 | 0.49087 | 0.00635 |
| **CCDC17** | 0.04394 | 0.11073 | 0.34243 | 0.29086 | 0.14442 | 0.00000 | 0.00000 | 0.31204 | 0.61775 | 1.23421 | 0.04279 | 1.45434 |
| **CCDC170** | 0.00296 | 0.00000 | 0.00000 | 0.05667 | 0.00000 | 0.14924 | 0.00000 | 0.89436 | 0.02768 | 0.89025 | 0.00000 | 0.11550 |
| **CCDC171** | 1.04848 | 0.87370 | 0.37812 | 0.57565 | 0.24995 | 0.57269 | 0.00000 | 0.64490 | 0.00000 | 0.17356 | 0.26816 | 0.52704 |
| **CCDC172** | 0.15746 | 0.00370 | 0.05279 | 0.02838 | 0.01309 | 0.01558 | 0.01862 | 0.11272 | 0.01859 | 0.00000 | 0.01661 | 0.34138 |
| **CCDC173** | 0.90057 | 0.42087 | 0.00000 | 0.00000 | 0.00000 | 0.17296 | 0.43117 | 0.40356 | 0.00000 | 1.62312 | 0.95498 | 0.69751 |
| **CCDC174** | 0.70154 | 3.74961 | 2.12766 | 2.34575 | 2.72559 | 9.84292 | 4.71400 | 8.54408 | 9.05957 | 10.14124 | 1.42037 | 6.49163 |
| **CCDC175** | 1.18782 | 0.03171 | 0.02209 | 0.02250 | 0.03714 | 0.03796 | 0.00000 | 0.00000 | 0.01999 | 0.07561 | 0.01095 | 0.11022 |
| **CCDC176** | 0.91366 | 0.50887 | 0.02647 | 0.18601 | 0.22609 | 0.17199 | 0.16622 | 0.41245 | 0.73502 | 0.00181 | 0.41653 | 1.08852 |
| **CCDC177** | 0.75139 | 0.00868 | 0.02301 | 0.00941 | 0.00735 | 0.02087 | 0.02220 | 0.11529 | 0.00000 | 0.02728 | 0.06544 | 0.01954 |
| **CCDC178** | 2.34176 | 0.13176 | 0.00000 | 0.00000 | 0.62518 | 0.05316 | 0.00000 | 0.32905 | 0.00000 | 0.32636 | 0.00000 | 0.80327 |
| **CCDC18** | 1.47513 | 0.00016 | 0.33091 | 1.39702 | 0.09966 | 4.39589 | 0.57015 | 0.86567 | 0.00000 | 7.16729 | 1.29071 | 1.83367 |
| **CCDC181** | 0.06553 | 0.12298 | 0.05257 | 0.00000 | 0.00001 | 0.21275 | 0.82162 | 0.76203 | 0.00000 | 0.04672 | 0.34199 | 0.82339 |
| **CCDC183** | 0.00643 | 0.07760 | 0.20469 | 0.04165 | 0.05088 | 0.00000 | 0.10444 | 0.00914 | 0.04297 | 0.19128 | 0.12460 | 0.03539 |
| **CCDC183-AS1** | 0.10501 | 0.00208 | 0.44338 | 0.18686 | 0.00000 | 0.12294 | 0.07243 | 0.00000 | 0.00000 | 0.74855 | 0.33807 | 0.07979 |
| **CCDC184** | 0.03660 | 0.00287 | 0.11860 | 0.00144 | 0.01080 | 0.03321 | 0.00000 | 0.04227 | 0.01283 | 0.06293 | 0.01369 | 0.21670 |
| **CCDC186** | 4.86790 | 0.00000 | 8.09285 | 0.00000 | 3.57309 | 11.33833 | 2.05781 | 5.79864 | 3.81939 | 9.90934 | 2.08624 | 2.95341 |
| **CCDC19** | 0.00263 | 0.53068 | 0.40945 | 0.02445 | 1.24301 | 0.00808 | 1.07131 | 0.11475 | 0.00000 | 0.00000 | 0.20938 | 0.21448 |
| **CCDC22** | 0.42168 | 0.19587 | 0.84454 | 2.24899 | 1.73558 | 0.05837 | 0.82135 | 0.00188 | 2.46412 | 0.00000 | 3.38175 | 0.00000 |
| **CCDC23** | 0.18690 | 2.46015 | 8.56688 | 1.38538 | 1.88438 | 7.25818 | 9.35827 | 3.54411 | 5.92105 | 2.83469 | 2.82196 | 3.03944 |
| **CCDC24** | 0.49999 | 1.18928 | 0.33663 | 0.18431 | 0.14413 | 0.00000 | 0.03401 | 0.02452 | 0.64379 | 0.00000 | 0.36297 | 0.28142 |
| **CCDC25** | 19.00999 | 4.35942 | 2.98497 | 1.62041 | 0.00000 | 6.87903 | 4.20205 | 8.51779 | 0.35794 | 2.78408 | 6.61708 | 4.96377 |

| | | | | | | | | | | | | |
|---|---|---|---|---|---|---|---|---|---|---|---|---|
| **CCDC27** | 10.10480 | 0.00307 | 0.01079 | 0.00753 | 0.01731 | 0.00000 | 0.01314 | 1.81927 | 0.02400 | 0.00000 | 0.00922 | 0.26760 |
| **CCDC28A** | 2.28946 | 3.91094 | 5.86535 | 1.16916 | 0.53545 | 11.00416 | 8.65493 | 4.51283 | 4.59545 | 0.01688 | 5.55464 | 6.50690 |
| **CCDC28B** | 0.39929 | 0.10400 | 0.22188 | 1.55771 | 0.00798 | 0.01394 | 0.07582 | 0.20182 | 0.53627 | 0.00000 | 0.00000 | 0.07423 |
| **CCDC3** | 1.24956 | 0.73810 | 0.00000 | 0.36580 | 0.03615 | 0.12701 | 0.02535 | 0.16409 | 0.00000 | 0.10873 | 0.00000 | 0.05855 |
| **CCDC30** | 1.61456 | 0.00000 | 0.05122 | 0.05272 | 0.03312 | 0.00000 | 0.02971 | 0.33049 | 0.14437 | 0.29777 | 0.00000 | 0.15996 |
| **CCDC33** | 0.02954 | 0.01059 | 0.11861 | 0.01062 | 0.03661 | 0.01739 | 0.00214 | 0.01031 | 0.01360 | 0.04964 | 0.06369 | 0.01371 |
| **CCDC34** | 0.38972 | 11.32985 | 1.70060 | 0.00000 | 13.31008 | 10.27996 | 10.23110 | 5.15397 | 0.00000 | 14.52275 | 6.82251 | 18.06637 |
| **CCDC36** | 0.00121 | 0.00418 | 0.00567 | 0.00314 | 0.01229 | 0.03498 | 0.00341 | 0.48261 | 0.03002 | 0.12262 | 0.00310 | 0.05412 |
| **CCDC37** | 0.01860 | 0.01129 | 0.06358 | 0.01247 | 0.04014 | 0.00000 | 0.03401 | 0.00599 | 0.06983 | 0.07504 | 0.01694 | 0.06271 |
| **CCDC38** | 0.01908 | 0.13159 | 0.00000 | 0.31465 | 0.00000 | 0.03894 | 0.20439 | 0.09784 | 0.00000 | 0.44390 | 0.00000 | 0.08949 |
| **CCDC39** | 2.02680 | 0.01157 | 0.02703 | 0.00000 | 0.00000 | 0.37495 | 0.07160 | 0.24531 | 0.13471 | 1.21557 | 0.08898 | 0.30163 |
| **CCDC40** | 0.21851 | 0.24516 | 0.02081 | 0.06956 | 0.00000 | 0.01325 | 0.15599 | 0.02710 | 0.11090 | 0.00000 | 0.02187 | 0.40844 |
| **CCDC42** | 0.05039 | 0.08611 | 0.06374 | 0.27875 | 0.00000 | 0.00000 | 0.03167 | 0.11603 | 0.06631 | 0.00443 | 0.08622 | 0.05655 |
| **CCDC42B** | 0.00408 | 0.03738 | 0.01471 | 0.01880 | 0.00493 | 0.03586 | 0.01662 | 0.01534 | 0.03191 | 0.04618 | 0.02329 | 0.02121 |
| **CCDC43** | 2.98522 | 7.02028 | 7.84627 | 3.89798 | 4.32132 | 22.73996 | 12.72409 | 3.62629 | 9.25833 | 12.33411 | 10.62255 | 7.65222 |
| **CCDC47** | 0.69164 | 13.86574 | 25.90210 | 9.35454 | 10.61059 | 34.10805 | 0.00000 | 15.26645 | 45.78925 | 20.65339 | 24.45064 | 14.20209 |
| **CCDC50** | 4.41949 | 0.83959 | 4.61673 | 4.13043 | 3.07195 | 5.18308 | 2.56362 | 0.06667 | 5.77830 | 1.84526 | 2.56812 | 0.00000 |
| **CCDC51** | 0.83954 | 18.13993 | 9.98756 | 5.63997 | 13.42790 | 8.22510 | 18.33042 | 4.28480 | 19.31980 | 0.00000 | 26.36108 | 18.27773 |
| **CCDC53** | 1.41471 | 2.73899 | 1.33284 | 4.11423 | 2.08796 | 10.29816 | 0.84211 | 1.08631 | 0.52645 | 0.52647 | 2.77842 | 3.97085 |
| **CCDC57** | 0.00000 | 1.96810 | 0.00000 | 1.33402 | 1.27018 | 0.00000 | 0.00389 | 0.17580 | 0.90056 | 2.37847 | 0.58262 | 0.30156 |
| **CCDC58** | 0.03372 | 6.31076 | 6.70602 | 0.92383 | 1.44394 | 24.93435 | 9.78258 | 0.68919 | 11.02027 | 8.90916 | 5.83824 | 18.75061 |
| **CCDC59** | 5.89707 | 6.63163 | 11.60067 | 2.17933 | 3.19596 | 18.68116 | 13.88037 | 11.15325 | 6.87602 | 7.85792 | 8.33689 | 12.59746 |
| **CCDC6** | 10.41021 | 4.10477 | 1.76675 | 3.51783 | 6.61426 | 14.89206 | 9.29255 | 0.82472 | 7.65667 | 16.14857 | 3.69634 | 3.54766 |
| **CCDC60** | 0.51021 | 0.01124 | 0.00000 | 0.10957 | 0.16687 | 0.09700 | 0.00500 | 0.00061 | 0.00548 | 0.04274 | 0.00619 | 0.00494 |
| **CCDC61** | 0.38832 | 0.16851 | 0.34697 | 0.03596 | 0.47877 | 0.92942 | 0.06747 | 0.30775 | 0.08168 | 1.49659 | 0.00000 | 0.13220 |
| **CCDC62** | 0.00781 | 0.04398 | 0.02813 | 0.28488 | 0.04062 | 0.52178 | 0.11617 | 2.66383 | 0.00000 | 0.67190 | 0.00000 | 1.04622 |
| **CCDC63** | 0.03659 | 0.19113 | 0.01743 | 0.08230 | 0.04690 | 0.00000 | 0.05113 | 0.06392 | 0.00361 | 0.02833 | 0.03714 | 0.06832 |
| **CCDC64** | 1.16804 | 1.82781 | 1.18130 | 0.19945 | 3.94919 | 0.00000 | 0.54907 | 1.91519 | 3.22382 | 3.05426 | 0.33440 | 2.37162 |
| **CCDC64B** | 0.01433 | 0.01765 | 1.08004 | 0.02333 | 0.85571 | 0.00000 | 0.32172 | 0.05881 | 0.54938 | 1.14433 | 0.45073 | 0.01327 |
| **CCDC65** | 0.00382 | 0.62564 | 0.00000 | 0.24567 | 0.15741 | 0.26241 | 0.00430 | 0.55838 | 0.00000 | 0.68314 | 0.00000 | 1.34123 |
| **CCDC66** | 0.41031 | 1.36610 | 1.55657 | 0.17272 | 0.42834 | 6.99392 | 1.67442 | 0.06043 | 1.87216 | 6.58541 | 2.85590 | 0.17105 |
| **CCDC67** | 0.05350 | 0.01677 | 0.00645 | 0.05658 | 0.00453 | 0.05280 | 0.01136 | 0.00376 | 0.01556 | 0.01072 | 0.01281 | 0.05231 |
| **CCDC68** | 0.09083 | 0.20978 | 0.23045 | 0.13307 | 0.00000 | 0.17920 | 0.00000 | 0.00515 | 0.00000 | 0.02825 | 0.00040 | 0.01583 |
| **CCDC69** | 0.00199 | 0.19633 | 0.22063 | 0.13284 | 0.00000 | 0.00000 | 0.47768 | 0.03013 | 0.12173 | 0.06586 | 0.17502 | 1.18802 |
| **CCDC7** | 0.48869 | 0.30610 | 0.37780 | 0.16210 | 0.27328 | 1.58004 | 0.00000 | 0.80178 | 0.96484 | 0.20547 | 0.11991 | 1.15356 |
| **CCDC70** | 0.00548 | 0.01597 | 0.05310 | 0.02391 | 0.00773 | 0.00000 | 0.00189 | 0.01083 | 0.11778 | 0.04930 | 0.03575 | 0.02751 |
| **CCDC71** | 0.37901 | 0.55656 | 0.40379 | 0.28649 | 0.37981 | 0.00361 | 0.00000 | 0.00000 | 0.00000 | 0.04437 | 3.49329 | 1.06578 |
| **CCDC71L** | 0.10777 | 0.41843 | 0.75051 | 0.12538 | 0.00000 | 0.00620 | 0.47855 | 0.00000 | 0.00000 | 0.82095 | 0.00000 | 0.38508 |
| **CCDC73** | 1.67618 | 2.10856 | 0.52549 | 0.00000 | 0.94120 | 1.51387 | 0.94905 | 2.07617 | 0.00000 | 2.74972 | 0.40928 | 4.92431 |
| **CCDC74A** | 0.04234 | 0.05198 | 0.05565 | 0.03945 | 0.00908 | 0.00000 | 0.05410 | 0.04133 | 0.00000 | 0.04655 | 0.02637 | 0.03263 |
| **CCDC74B** | 0.07099 | 0.07630 | 0.02564 | 0.03104 | 0.00480 | 0.00000 | 0.03825 | 0.00188 | 0.02955 | 0.02898 | 0.01571 | 0.11015 |
| **CCDC77** | 6.54063 | 3.23646 | 5.35499 | 1.31232 | 0.93854 | 9.44541 | 7.00561 | 4.70751 | 9.84624 | 2.29108 | 3.86351 | 5.78480 |
| **CCDC78** | 0.00597 | 0.09356 | 0.04815 | 0.00708 | 0.00000 | 0.00000 | 0.12003 | 0.01078 | 0.00000 | 0.14806 | 0.01944 | 0.02464 |
| **CCDC79** | 1.37263 | 0.00000 | 0.00000 | 0.37541 | 0.00000 | 0.09236 | 0.00000 | 0.11540 | 0.05804 | 0.06891 | 0.00458 | 0.23830 |
| **CCDC8** | 1.02053 | 0.02210 | 1.27355 | 0.30081 | 0.00434 | 0.00000 | 0.17992 | 0.05408 | 0.63895 | 0.60602 | 0.90625 | 0.17795 |
| **CCDC80** | 0.03209 | 0.00000 | 0.76050 | 4.15067 | 1.70222 | 0.11951 | 0.00100 | 0.21627 | 0.10121 | 1.17352 | 0.12272 | 0.65114 |
| **CCDC81** | 0.20162 | 0.10733 | 0.01500 | 0.68163 | 0.00000 | 0.13838 | 0.18042 | 0.00000 | 0.34471 | 0.00000 | 0.29064 | 0.03264 |
| **CCDC82** | 2.02627 | 1.32869 | 3.42318 | 1.59881 | 0.00000 | 3.75207 | 2.91437 | 11.82922 | 0.67393 | 4.11362 | 2.25458 | 6.75586 |

| | | | | | | | | | | | | |
|---|---|---|---|---|---|---|---|---|---|---|---|---|
| **CCDC83** | 0.00225 | 0.02347 | 0.02585 | 0.02045 | 0.00743 | 0.02787 | 0.00000 | 0.10221 | 0.04814 | 0.15771 | 0.01175 | 0.00950 |
| **CCDC84** | 0.74661 | 5.44417 | 4.43809 | 2.50417 | 4.99974 | 3.08129 | 2.57201 | 0.11196 | 4.25249 | 5.96959 | 2.02396 | 3.08549 |
| **CCDC85A** | 0.00273 | 0.00542 | 0.03471 | 0.06786 | 0.00796 | 0.04223 | 0.01472 | 0.28826 | 0.01498 | 0.02405 | 0.01960 | 0.09423 |
| **CCDC85B** | 0.00000 | 1.39441 | 2.05244 | 0.44825 | 1.77392 | 0.00000 | 2.70555 | 0.85506 | 0.52733 | 10.53145 | 0.00000 | 1.55253 |
| **CCDC85C** | 0.60800 | 1.83721 | 2.55250 | 2.28951 | 9.46776 | 0.06894 | 10.39690 | 0.00000 | 7.57053 | 10.95434 | 2.31308 | 0.41642 |
| **CCDC86** | 1.96206 | 20.94239 | 17.69674 | 8.11436 | 16.31141 | 0.00000 | 21.07990 | 8.11555 | 31.82135 | 0.00000 | 16.43896 | 27.35016 |
| **CCDC87** | 0.39989 | 0.09246 | 0.08910 | 0.60503 | 0.26378 | 0.03763 | 0.00000 | 0.14087 | 0.00000 | 0.17208 | 0.02841 | 0.56515 |
| **CCDC88A** | 4.96998 | 0.82778 | 0.00000 | 1.86802 | 0.00000 | 2.54846 | 0.00000 | 3.20960 | 0.00000 | 11.51978 | 0.65609 | 6.43272 |
| **CCDC88B** | 0.01456 | 0.02029 | 0.51734 | 0.00000 | 0.00000 | 0.00000 | 0.20417 | 0.39730 | 0.19075 | 0.51570 | 0.03061 | 0.02742 |
| **CCDC88C** | 7.69197 | 0.14221 | 0.66408 | 0.00000 | 1.55437 | 0.00000 | 4.16431 | 0.22485 | 5.76160 | 5.40095 | 1.03188 | 0.49515 |
| **CCDC9** | 1.18910 | 0.20341 | 4.05490 | 1.93287 | 3.15990 | 0.00000 | 3.13142 | 0.67215 | 1.94493 | 6.60752 | 0.32718 | 0.86945 |
| **CCDC90B** | 1.57128 | 8.16531 | 2.27536 | 4.36103 | 4.35219 | 20.17980 | 0.00000 | 0.08463 | 1.57413 | 3.44613 | 4.14729 | 5.11977 |
| **CCDC91** | 2.93692 | 0.27341 | 2.52870 | 0.20576 | 0.77481 | 7.71739 | 3.19973 | 0.30090 | 1.05915 | 6.72656 | 0.31365 | 0.84641 |
| **CCDC92** | 1.60553 | 0.09653 | 0.14571 | 0.01303 | 0.00499 | 0.00000 | 0.00000 | 0.00000 | 0.06906 | 0.05394 | 0.03529 | 0.37438 |
| **CCDC93** | 5.20304 | 2.45733 | 5.50293 | 1.88001 | 2.62977 | 9.96469 | 2.78713 | 2.05856 | 6.05334 | 9.57209 | 4.75646 | 5.08028 |
| **CCDC94** | 0.25251 | 8.14181 | 5.79920 | 6.25022 | 9.43386 | 0.89960 | 6.27394 | 9.76489 | 9.19931 | 0.00000 | 4.44465 | 12.27531 |
| **CCDC96** | 0.76130 | 0.01080 | 0.00438 | 0.03802 | 0.07536 | 0.00000 | 0.10047 | 0.16947 | 0.00000 | 0.05095 | 0.02731 | 0.00430 |
| **CCDC97** | 5.77713 | 6.46768 | 11.10434 | 3.93364 | 8.95432 | 0.00000 | 8.24446 | 0.48940 | 10.68438 | 4.44812 | 2.87981 | 9.88946 |
| **CCER1** | 0.01774 | 0.00642 | 0.01079 | 0.00803 | 0.00637 | 0.04005 | 0.01108 | 0.00588 | 0.01297 | 0.03369 | 0.01359 | 0.08952 |
| **CCER2** | 0.01721 | 0.04950 | 0.05643 | 0.04003 | 0.00986 | 0.00000 | 0.00350 | 0.22707 | 0.08147 | 0.01845 | 0.04784 | 0.17309 |
| **CCHCR1** | 0.74887 | 6.91522 | 0.94898 | 5.14405 | 5.38845 | 0.00000 | 2.40958 | 1.35961 | 1.38027 | 3.19367 | 4.20067 | 6.78014 |
| **CCIN** | 0.12969 | 1.75674 | 0.00002 | 0.00000 | 1.79833 | 1.61278 | 0.00000 | 0.00000 | 1.31895 | 0.09837 | 1.75588 | 3.93698 |
| **CCK** | 2.30683 | 0.34895 | 0.00594 | 0.00066 | 0.04066 | 0.02394 | 0.00657 | 2.83255 | 0.01297 | 0.00495 | 0.00466 | 0.82419 |
| **CCKBR** | 0.01477 | 3.54417 | 49.72801 | 0.00000 | 0.00000 | 0.28231 | 27.47752 | 0.00000 | 62.54019 | 0.18616 | 16.68769 | 0.14495 |
| **CCL13** | 0.00608 | 0.01331 | 0.00680 | 0.00772 | 0.00639 | 0.02835 | 0.02231 | 0.01048 | 0.07358 | 0.02007 | 0.00356 | 0.01503 |
| **CCL15** | 0.00291 | 0.13153 | 0.00626 | 0.00000 | 0.01285 | 0.08179 | 0.00756 | 0.05947 | 0.00388 | 0.01385 | 0.00689 | 0.87002 |
| **CCL16** | 0.00282 | 0.00757 | 0.03019 | 0.01415 | 0.01783 | 0.00427 | 0.02232 | 0.03169 | 0.00857 | 0.00302 | 0.04778 | 0.15073 |
| **CCL19** | 0.10910 | 0.01277 | 0.01669 | 0.01617 | 0.11217 | 0.00000 | 0.03735 | 0.00529 | 0.01954 | 0.00000 | 0.01534 | 0.01825 |
| **CCL2** | 0.41604 | 0.06638 | 0.11561 | 0.04573 | 0.04822 | 0.02546 | 0.04732 | 0.18465 | 0.10700 | 0.00000 | 0.07057 | 0.04556 |
| **CCL20** | 0.91479 | 0.01397 | 0.01454 | 0.01293 | 0.01827 | 0.00463 | 0.01968 | 0.06379 | 0.02390 | 0.00296 | 0.01396 | 0.02083 |
| **CCL21** | 0.07996 | 0.01474 | 0.01495 | 0.01403 | 0.01803 | 0.00886 | 0.02212 | 0.01620 | 0.02461 | 0.01033 | 0.01355 | 0.02303 |
| **CCL22** | 0.01727 | 0.02464 | 0.02804 | 0.03499 | 0.00826 | 0.01241 | 0.01246 | 0.02084 | 0.02576 | 0.02980 | 0.01699 | 0.02339 |
| **CCL24** | 0.14221 | 0.00958 | 0.01641 | 0.00994 | 0.01551 | 0.01854 | 0.01175 | 0.00279 | 0.01784 | 0.03320 | 0.01377 | 0.04784 |
| **CCL25** | 0.01789 | 0.01435 | 0.01243 | 0.00760 | 0.02256 | 0.03366 | 0.02689 | 0.32042 | 0.01216 | 0.00000 | 0.01257 | 0.18112 |
| **CCL26** | 0.02701 | 0.08972 | 0.54969 | 0.65217 | 0.11630 | 0.00000 | 0.02595 | 0.11491 | 0.15261 | 0.14867 | 0.33786 | 0.25787 |
| **CCL27** | 0.00527 | 0.06669 | 0.12330 | 0.00658 | 0.02292 | 0.00000 | 0.00811 | 0.01024 | 0.04064 | 0.05135 | 0.06073 | 0.01980 |
| **CCL28** | 0.04041 | 0.72502 | 0.09229 | 0.00171 | 0.31302 | 0.45995 | 0.00000 | 0.03484 | 0.48768 | 0.00000 | 1.48306 | 1.40368 |
| **CCL5** | 0.00220 | 0.00000 | 0.01192 | 0.00000 | 0.17982 | 0.00000 | 0.06148 | 0.01752 | 0.04001 | 0.35790 | 0.12673 | 0.22542 |
| **CCL8** | 0.00675 | 0.01891 | 0.01074 | 0.01649 | 0.02319 | 0.00000 | 0.01322 | 0.01056 | 0.08458 | 0.00273 | 0.01070 | 0.01879 |
| **CCM2** | 0.30341 | 7.04903 | 1.56447 | 1.96353 | 8.62620 | 0.00000 | 2.51721 | 0.28316 | 0.55703 | 6.20879 | 2.12747 | 3.88623 |
| **CCM2L** | 0.91970 | 0.00000 | 0.00112 | 0.54909 | 0.03495 | 0.00000 | 0.00495 | 0.00131 | 0.00000 | 0.22247 | 0.05795 | 0.01710 |
| **CCNA1** | 10.06970 | 18.24387 | 0.00000 | 0.54695 | 3.99786 | 3.00718 | 0.00000 | 32.09411 | 0.00000 | 0.00000 | 0.26374 | 13.00994 |
| **CCNA2** | 7.57300 | 25.56392 | 15.86112 | 13.82565 | 16.06364 | 41.59189 | 20.57828 | 15.74408 | 22.94587 | 0.00000 | 17.80651 | 30.14311 |
| **CCNB1** | 17.43697 | 41.61939 | 37.22182 | 18.55311 | 26.84935 | 54.23219 | 45.52492 | 39.70669 | 35.55266 | 0.16414 | 28.76770 | 56.26398 |
| **CCNB1IP1** | 2.00787 | 10.37440 | 8.26598 | 9.63474 | 6.20332 | 15.90913 | 10.21982 | 8.36627 | 13.05603 | 7.44537 | 5.38486 | 22.98812 |
| **CCNB2** | 5.81573 | 20.27251 | 15.99673 | 9.04094 | 9.84182 | 30.09564 | 13.32151 | 5.41677 | 15.11353 | 0.00000 | 12.88113 | 16.82622 |
| **CCNB3** | 8.55552 | 0.32405 | 0.03980 | 0.13280 | 0.00000 | 0.00000 | 0.03161 | 0.83985 | 0.16320 | 0.01397 | 0.00000 | 0.30675 |
| **CCNC** | 1.17865 | 14.41965 | 17.95996 | 6.48640 | 3.66838 | 57.70797 | 24.83555 | 8.29564 | 20.94921 | 7.55554 | 16.87190 | 16.22791 |

| | | | | | | | | | | | | |
|---|---|---|---|---|---|---|---|---|---|---|---|---|
| **CCND1** | 0.69371 | 1.79185 | 3.59882 | 0.00000 | 0.00000 | 0.03624 | 2.71553 | 1.73773 | 2.35723 | 3.28314 | 1.08346 | 1.70830 |
| **CCND2** | 13.18591 | 0.00000 | 0.00000 | 0.00000 | 0.17608 | 0.04696 | 0.00000 | 2.21961 | 0.15269 | 0.00000 | 0.00000 | 3.17719 |
| **CCND3** | 2.88921 | 17.58776 | 25.13338 | 14.71557 | 19.53644 | 0.00230 | 21.30070 | 0.66961 | 21.52411 | 0.00000 | 16.30780 | 9.87768 |
| **CCNDBP1** | 1.56763 | 26.88879 | 11.65597 | 12.05624 | 12.55524 | 29.99720 | 4.05996 | 6.55351 | 3.69521 | 3.80109 | 10.32562 | 23.18140 |
| **CCNE1** | 4.58631 | 8.47062 | 31.93059 | 5.69554 | 9.98126 | 0.10039 | 19.45239 | 18.59066 | 0.88302 | 22.42291 | 1.18498 | 9.80171 |
| **CCNE2** | 0.00000 | 1.06419 | 5.21736 | 0.00000 | 0.00000 | 6.78378 | 5.36835 | 10.06046 | 4.53194 | 0.00000 | 7.80656 | 3.12330 |
| **CCNF** | 0.42710 | 23.33128 | 11.33407 | 12.19637 | 15.62574 | 3.51003 | 21.66012 | 15.35096 | 19.19776 | 11.87641 | 11.58506 | 19.07513 |
| **CCNG1** | 4.62930 | 7.09353 | 12.83421 | 6.25991 | 1.72137 | 26.75502 | 15.14356 | 2.25534 | 21.79346 | 0.97526 | 10.65294 | 6.17922 |
| **CCNG2** | 0.83749 | 0.00000 | 1.97867 | 0.00002 | 0.00000 | 2.84945 | 5.50791 | 0.00000 | 0.82926 | 2.80598 | 0.00000 | 0.00000 |
| **CCNH** | 0.83923 | 4.55994 | 7.40754 | 2.05214 | 0.07791 | 18.96494 | 0.00000 | 6.62980 | 8.63302 | 0.00000 | 5.81954 | 6.86264 |
| **CCNI** | 17.81095 | 3.36361 | 7.41645 | 6.99058 | 14.91495 | 12.43746 | 16.29050 | 5.53533 | 1.42919 | 26.94493 | 6.32068 | 3.30243 |
| **CCNI2** | 2.16803 | 0.23204 | 0.00000 | 0.00000 | 1.95126 | 0.00000 | 0.00045 | 0.14689 | 0.00000 | 0.68897 | 1.23934 | 0.35642 |
| **CCNJ** | 10.39638 | 1.25164 | 2.86401 | 1.97717 | 0.59424 | 2.21726 | 7.20742 | 2.23251 | 8.99869 | 0.13714 | 5.12077 | 0.65395 |
| **CCNJL** | 0.05093 | 0.02386 | 0.03339 | 0.01222 | 0.14084 | 0.03048 | 0.01284 | 0.00639 | 0.00722 | 0.04559 | 0.00480 | 0.01277 |
| **CCNK** | 7.70719 | 1.50037 | 6.78607 | 0.57418 | 8.19921 | 8.78775 | 8.82552 | 5.28838 | 2.37263 | 12.45734 | 5.90977 | 6.71466 |
| **CCNL1** | 2.89638 | 8.30995 | 7.20722 | 8.30191 | 4.46557 | 19.18152 | 4.79347 | 8.47144 | 11.95396 | 17.52023 | 3.65084 | 12.31834 |
| **CCNL2** | 2.77733 | 11.29617 | 9.70975 | 12.05585 | 15.09708 | 0.00000 | 2.01995 | 9.70112 | 13.24406 | 21.85682 | 7.68880 | 8.18382 |
| **CCNO** | 9.05966 | 0.51568 | 0.34418 | 1.08125 | 0.18617 | 0.00000 | 0.03740 | 0.85640 | 0.53129 | 0.07422 | 0.03638 | 1.73079 |
| **CCNT1** | 2.62099 | 0.11137 | 4.49233 | 1.95551 | 0.00000 | 10.23614 | 4.91555 | 12.24723 | 4.03785 | 15.66324 | 2.98931 | 5.58247 |
| **CCNT2** | 1.92593 | 1.56999 | 9.71890 | 2.84728 | 0.97488 | 19.53330 | 2.66319 | 8.49391 | 3.94418 | 10.41582 | 4.64276 | 6.19374 |
| **CCNT2-AS1** | 0.19988 | 0.16852 | 0.77997 | 0.00000 | 0.00000 | 0.00735 | 1.30345 | 0.07447 | 0.33053 | 0.20963 | 0.00000 | 0.03316 |
| **CCNY** | 0.66836 | 1.87588 | 2.96594 | 5.20801 | 8.41133 | 8.04718 | 11.46222 | 0.32382 | 0.00000 | 17.77451 | 0.73838 | 1.19671 |
| **CCNYL1** | 5.32249 | 0.87044 | 5.02266 | 0.73404 | 1.49068 | 6.40885 | 2.10394 | 4.16231 | 0.00000 | 6.77812 | 1.67266 | 8.32537 |
| **CCNYL2** | 0.09232 | 0.01265 | 0.01576 | 0.01713 | 0.01364 | 0.02396 | 0.01874 | 0.01121 | 0.01706 | 0.01070 | 0.01387 | 0.01934 |
| **CCP110** | 5.52018 | 3.02812 | 3.95486 | 0.71914 | 1.71942 | 11.01914 | 3.93210 | 3.32320 | 2.96014 | 6.99703 | 5.41104 | 14.81556 |
| **CCPG1** | 0.37456 | 8.99189 | 0.17359 | 2.34522 | 0.00000 | 4.49636 | 0.00000 | 3.09722 | 0.00000 | 3.09787 | 0.00000 | 15.11533 |
| **CCR1** | 0.00730 | 0.01018 | 1.13698 | 0.06639 | 0.23085 | 0.00000 | 0.00000 | 0.01413 | 0.28631 | 0.00000 | 0.03559 | 0.01820 |
| **CCR10** | 0.27634 | 0.01501 | 0.04529 | 0.03393 | 0.00057 | 0.04570 | 0.00000 | 0.05528 | 0.02581 | 0.03955 | 0.02370 | 0.00883 |
| **CCR2** | 0.00737 | 0.17516 | 0.20124 | 0.00000 | 0.00000 | 0.04918 | 0.00000 | 0.00988 | 0.00000 | 0.03643 | 0.00991 | 0.00307 |
| **CCR3** | 0.60035 | 0.00349 | 0.07493 | 0.01698 | 0.01651 | 0.02282 | 0.00002 | 0.45989 | 0.03008 | 0.19891 | 0.01007 | 0.00183 |
| **CCR4** | 0.00610 | 0.01789 | 0.01571 | 0.04224 | 0.02337 | 0.00024 | 0.04289 | 0.04414 | 0.02476 | 0.00261 | 0.01758 | 0.01649 |
| **CCR6** | 0.00367 | 0.08058 | 0.02881 | 0.00000 | 0.09379 | 0.28974 | 0.00000 | 0.01060 | 0.15418 | 0.04756 | 0.05147 | 0.00242 |
| **CCR7** | 0.00871 | 0.00000 | 36.23091 | 0.79932 | 0.77497 | 0.82845 | 0.00000 | 0.00000 | 5.18332 | 0.00000 | 0.19513 | 0.00369 |
| **CCR8** | 0.66431 | 0.03667 | 0.00000 | 8.66932 | 0.01088 | 0.02852 | 0.00545 | 0.41441 | 0.01570 | 0.00053 | 0.03942 | 0.00000 |
| **CCR9** | 0.00466 | 0.00913 | 0.00715 | 0.01868 | 0.00248 | 0.02287 | 0.01561 | 0.00976 | 0.01406 | 0.09344 | 0.01053 | 0.02858 |
| **CCRN4L** | 0.82657 | 4.41561 | 7.27520 | 4.20845 | 2.70643 | 1.59585 | 8.45513 | 20.16475 | 2.17865 | 2.67622 | 7.52990 | 12.25106 |
| **CCS** | 0.86303 | 15.79201 | 7.73482 | 12.82233 | 9.85017 | 0.00000 | 7.78719 | 1.51825 | 4.24265 | 3.02720 | 6.58412 | 10.45547 |
| **CCSAP** | 1.17717 | 0.00000 | 2.28779 | 0.65074 | 1.68444 | 4.30522 | 3.32774 | 14.09567 | 2.67601 | 6.99783 | 2.23032 | 3.05048 |
| **CCSER1** | 1.17747 | 0.02308 | 1.08895 | 0.02305 | 0.03851 | 0.00000 | 0.00000 | 0.05482 | 0.00000 | 0.01370 | 0.01899 | 0.00769 |
| **CCSER2** | 7.83265 | 4.06966 | 10.65116 | 2.60172 | 9.21183 | 15.42726 | 8.46196 | 1.16450 | 3.70391 | 9.34849 | 8.35884 | 7.03001 |
| **CCT2** | 5.68171 | 32.61591 | 31.34519 | 20.81554 | 24.53825 | 57.07590 | 49.70600 | 23.88800 | 43.28205 | 15.94980 | 38.16507 | 36.97839 |
| **CCT3** | 5.60982 | 39.97450 | 36.07273 | 35.90635 | 41.47430 | 33.44284 | 52.49892 | 17.83683 | 50.23982 | 10.43560 | 41.52814 | 38.17689 |
| **CCT4** | 0.82781 | 28.96426 | 28.32654 | 20.10922 | 22.44923 | 46.54738 | 47.66201 | 3.15721 | 38.45437 | 12.41710 | 35.19224 | 25.75406 |
| **CCT5** | 0.09223 | 34.31011 | 36.62801 | 20.57541 | 27.72109 | 53.71426 | 51.44463 | 15.52411 | 45.56873 | 13.72748 | 37.93476 | 29.37374 |
| **CCT6A** | 1.88278 | 33.92431 | 33.38342 | 21.04733 | 23.38634 | 53.24383 | 51.81729 | 7.58828 | 40.99112 | 16.88200 | 36.96829 | 29.52723 |
| **CCT6B** | 0.42256 | 0.19345 | 0.25611 | 0.09613 | 0.00000 | 0.15316 | 0.00000 | 0.58517 | 0.03122 | 0.03131 | 0.10774 | 0.22817 |
| **CCT6P1** | 0.00765 | 0.56514 | 0.66717 | 0.28729 | 0.22332 | 1.16079 | 0.00000 | 0.81458 | 0.94604 | 0.51261 | 0.33214 | 0.22807 |
| **CCT6P3** | 0.00852 | 0.20087 | 0.15398 | 0.18894 | 0.16115 | 0.62781 | 0.08129 | 0.19860 | 0.10703 | 0.90840 | 0.00530 | 0.20228 |

| | | | | | | | | | | | | |
|---|---|---|---|---|---|---|---|---|---|---|---|---|
| **CCT7** | 0.00000 | 34.51667 | 22.05190 | 28.21998 | 34.12678 | 27.70110 | 34.74917 | 8.50807 | 41.78844 | 6.08664 | 38.06223 | 36.14332 |
| **CCT8** | 11.11527 | 28.30589 | 25.79604 | 12.11212 | 12.46467 | 50.01826 | 37.34177 | 8.46670 | 39.24400 | 1.79234 | 27.80461 | 23.48034 |
| **CCZ1** | 1.05472 | 0.28935 | 0.53007 | 0.26488 | 0.21951 | 1.20165 | 1.73554 | 0.01850 | 0.00000 | 3.44687 | 0.14989 | 0.00000 |
| **CCZ1B** | 0.32830 | 0.37348 | 0.25665 | 0.00000 | 0.00009 | 0.73897 | 0.69280 | 0.00944 | 0.07133 | 1.43243 | 0.11383 | 0.09666 |
| **CD101** | 0.00329 | 0.03646 | 0.00019 | 0.32994 | 0.11432 | 0.00000 | 0.04900 | 0.00846 | 0.00000 | 0.43769 | 0.00000 | 0.02644 |
| **CD109** | 1.67763 | 0.18781 | 0.31752 | 1.09928 | 0.00186 | 0.00000 | 0.00000 | 0.62171 | 0.15833 | 0.00011 | 0.17582 | 0.04431 |
| **CD14** | 0.00255 | 0.06183 | 0.00994 | 0.37143 | 0.06723 | 0.00000 | 0.17277 | 0.28889 | 0.09459 | 0.00000 | 0.03157 | 2.30503 |
| **CD151** | 0.00000 | 3.96740 | 4.82679 | 2.10057 | 13.85237 | 0.00000 | 1.87779 | 0.06573 | 18.51759 | 13.92444 | 3.33629 | 1.47299 |
| **CD160** | 0.00677 | 0.12572 | 0.00000 | 0.11852 | 0.02027 | 0.09352 | 0.00000 | 0.00600 | 0.03263 | 0.83564 | 0.00000 | 0.04201 |
| **CD163** | 0.00513 | 0.00891 | 0.01409 | 0.20199 | 0.01133 | 0.00961 | 0.03686 | 0.00990 | 0.02757 | 0.02843 | 0.01113 | 0.06315 |
| **CD163L1** | 0.67092 | 0.00000 | 0.40512 | 0.01990 | 0.07206 | 0.69217 | 0.19249 | 0.00108 | 0.01576 | 0.00000 | 0.07102 | 0.03532 |
| **CD164** | 11.43003 | 17.57484 | 24.08710 | 5.50654 | 10.75071 | 51.45197 | 0.00000 | 13.90478 | 43.16893 | 14.42483 | 20.60322 | 19.97294 |
| **CD164L2** | 0.00289 | 0.08618 | 0.03646 | 0.00588 | 0.00735 | 0.00449 | 0.02739 | 0.11107 | 0.02751 | 0.01557 | 0.01863 | 0.08167 |
| **CD177** | 0.07009 | 0.01186 | 0.02461 | 0.01025 | 0.03100 | 0.01197 | 0.03648 | 0.03209 | 0.00028 | 0.04134 | 0.01403 | 0.01702 |
| **CD180** | 0.00204 | 0.00097 | 0.00061 | 0.00680 | 0.01147 | 0.16695 | 0.00746 | 0.05791 | 0.00000 | 0.01258 | 0.02365 | 0.01800 |
| **CD19** | 0.05838 | 0.08090 | 0.20386 | 0.67010 | 0.09806 | 0.00000 | 0.35444 | 0.06957 | 0.30088 | 0.00000 | 0.07770 | 0.04808 |
| **CD1D** | 0.00600 | 0.02308 | 0.05651 | 0.27678 | 0.00214 | 0.01480 | 0.01442 | 0.01057 | 0.04130 | 0.00981 | 0.01409 | 0.02010 |
| **CD200** | 0.00629 | 0.99335 | 0.57294 | 4.18417 | 0.00341 | 0.20896 | 0.00000 | 0.00000 | 0.00000 | 0.00000 | 0.03696 | 0.90578 |
| **CD200R1** | 0.10694 | 0.35220 | 0.00000 | 0.42392 | 0.02476 | 0.67344 | 0.00000 | 0.25204 | 0.03483 | 0.00000 | 0.37995 | 0.06209 |
| **CD200R1L** | 0.03900 | 0.01672 | 0.01847 | 0.06004 | 0.04627 | 0.00000 | 0.01260 | 0.05851 | 0.06349 | 0.02511 | 0.09903 | 0.01792 |
| **CD207** | 0.00629 | 0.02240 | 0.03450 | 0.01858 | 0.01768 | 0.00632 | 0.01732 | 0.01064 | 0.01631 | 0.01403 | 0.01636 | 0.01659 |
| **CD209** | 0.09968 | 0.01476 | 0.05567 | 0.01647 | 0.02636 | 0.04392 | 0.07747 | 0.05552 | 0.00000 | 0.06120 | 0.03366 | 0.06382 |
| **CD22** | 0.00809 | 0.61241 | 0.18351 | 0.00000 | 2.76303 | 0.00000 | 0.00000 | 0.02087 | 0.00000 | 0.10990 | 0.04155 | 0.12585 |
| **CD226** | 0.05657 | 0.01375 | 0.00625 | 0.00852 | 0.01168 | 0.03296 | 0.05606 | 0.01678 | 0.04487 | 0.00000 | 0.00400 | 0.01435 |
| **CD24** | 0.57980 | 0.00000 | 22.48073 | 5.72641 | 7.90899 | 0.22924 | 44.30662 | 0.08337 | 60.56433 | 0.00000 | 26.36921 | 0.00000 |
| **CD244** | 0.00554 | 0.03814 | 0.02480 | 0.03015 | 0.00635 | 0.08798 | 0.00000 | 0.00962 | 0.00000 | 0.03332 | 0.03291 | 0.00846 |
| **CD247** | 0.02438 | 0.04301 | 0.25029 | 0.01074 | 0.01371 | 0.00000 | 0.00000 | 0.00777 | 0.14035 | 0.00658 | 0.04726 | 0.02826 |
| **CD248** | 0.00534 | 0.00789 | 0.24136 | 0.11351 | 0.00000 | 0.02821 | 0.00000 | 0.01049 | 0.00000 | 0.05227 | 0.01592 | 0.02328 |
| **CD27** | 0.00557 | 0.01527 | 0.02605 | 0.02841 | 0.02842 | 0.01165 | 0.02819 | 0.00964 | 0.00437 | 0.03985 | 0.01550 | 0.02833 |
| **CD27-AS1** | 0.05920 | 0.20471 | 0.00000 | 0.22567 | 0.27127 | 0.12940 | 0.00000 | 0.12266 | 0.01247 | 0.56283 | 0.07571 | 0.07017 |
| **CD274** | 0.00000 | 0.00000 | 11.24997 | 0.00000 | 2.17973 | 0.01512 | 1.29531 | 0.11068 | 9.13313 | 0.07840 | 1.17941 | 0.00868 |
| **CD276** | 0.25857 | 0.17596 | 14.56760 | 0.72863 | 4.27138 | 1.74315 | 5.14759 | 0.00000 | 18.03108 | 0.00000 | 4.97479 | 1.26483 |
| **CD28** | 0.01113 | 0.00468 | 0.01414 | 0.00938 | 0.02840 | 0.03245 | 0.02903 | 0.00728 | 0.00380 | 0.01109 | 0.05109 | 0.07185 |
| **CD2AP** | 0.37285 | 5.46613 | 2.71834 | 5.09131 | 0.00000 | 26.89776 | 12.98254 | 3.70427 | 1.12400 | 25.15772 | 0.64286 | 6.40027 |
| **CD2BP2** | 0.83234 | 14.11259 | 13.55010 | 12.40953 | 11.86033 | 1.04576 | 16.63607 | 13.85827 | 14.28484 | 0.00000 | 11.53317 | 17.12608 |
| **CD300A** | 0.00620 | 0.01567 | 0.02386 | 0.01143 | 0.02749 | 0.01698 | 0.01637 | 0.01051 | 0.01921 | 0.01549 | 0.01649 | 0.01615 |
| **CD300E** | 0.04337 | 0.01855 | 0.01476 | 0.01130 | 0.06669 | 0.01020 | 0.01741 | 0.58493 | 0.01530 | 0.01716 | 0.01040 | 0.11784 |
| **CD300LB** | 0.01066 | 0.01483 | 0.01515 | 0.00655 | 0.02014 | 0.02296 | 0.01842 | 0.01170 | 0.02502 | 0.03748 | 0.01108 | 0.02399 |
| **CD300LF** | 0.00064 | 2.07405 | 0.08535 | 0.00000 | 0.79620 | 0.12450 | 0.55457 | 0.07793 | 0.00000 | 0.04426 | 0.46963 | 0.05249 |
| **CD300LG** | 0.00644 | 0.01398 | 0.03399 | 0.01085 | 0.01093 | 0.01740 | 0.01332 | 0.01040 | 0.00579 | 0.01553 | 0.05476 | 0.01643 |
| **CD302** | 0.03161 | 0.03616 | 0.01617 | 0.06798 | 0.01406 | 0.00000 | 0.01741 | 0.00890 | 0.02229 | 0.02454 | 0.02459 | 0.05028 |
| **CD320** | 0.02616 | 26.45787 | 17.91410 | 17.73997 | 14.49225 | 0.00000 | 10.68901 | 0.80190 | 33.79357 | 12.85500 | 21.22454 | 10.94725 |
| **CD33** | 0.04770 | 0.04967 | 0.00626 | 0.02340 | 0.00417 | 0.09298 | 0.00000 | 0.00665 | 0.00416 | 0.05177 | 0.00754 | 0.03440 |
| **CD34** | 0.00540 | 0.00638 | 0.01453 | 0.00277 | 0.10640 | 0.02481 | 0.01083 | 0.00975 | 0.00000 | 0.04650 | 0.04478 | 0.02112 |
| **CD36** | 6.98255 | 0.00000 | 0.00014 | 0.00000 | 0.07926 | 0.00000 | 0.11159 | 2.59778 | 0.07886 | 0.34561 | 0.22434 | 0.00000 |
| **CD37** | 1.44814 | 0.54553 | 0.00000 | 0.00000 | 0.07760 | 0.00000 | 0.09952 | 3.04260 | 0.00000 | 0.44369 | 0.01893 | 6.27768 |
| **CD38** | 1.18521 | 0.42214 | 0.18511 | 0.90468 | 0.08154 | 0.00000 | 0.01928 | 0.08240 | 0.01777 | 0.00000 | 0.00963 | 0.16084 |
| **CD3D** | 0.00560 | 0.00528 | 0.00074 | 0.10395 | 0.02540 | 0.08998 | 0.00807 | 0.01092 | 0.00643 | 0.00741 | 0.01246 | 0.01227 |

| | | | | | | | | | | | | |
|---|---|---|---|---|---|---|---|---|---|---|---|---|
| **CD3E** | 0.00465 | 0.01143 | 0.03712 | 0.01099 | 0.01387 | 0.03244 | 0.01140 | 0.01589 | 0.01749 | 0.01323 | 0.01495 | 0.04391 |
| **CD3EAP** | 1.91755 | 4.56462 | 2.46025 | 3.45427 | 6.75476 | 0.13932 | 8.92511 | 3.55611 | 4.39025 | 1.40599 | 4.95703 | 13.89603 |
| **CD3G** | 0.01367 | 0.02084 | 0.01042 | 0.06602 | 0.01080 | 0.02238 | 0.01831 | 0.04178 | 0.02172 | 0.00182 | 0.01458 | 0.03197 |
| **CD4** | 0.00606 | 0.01491 | 0.02477 | 0.01286 | 0.01129 | 0.01949 | 0.02044 | 0.01046 | 0.00726 | 0.02206 | 0.02775 | 0.01569 |
| **CD40** | 0.00229 | 0.15404 | 0.03414 | 0.01032 | 0.00597 | 0.00348 | 0.05523 | 0.24065 | 0.00567 | 0.00000 | 0.01668 | 0.94303 |
| **CD40LG** | 0.00628 | 0.02616 | 0.01450 | 0.01138 | 0.01503 | 0.01186 | 0.01879 | 0.01019 | 0.01737 | 0.01794 | 0.01625 | 0.04869 |
| **CD44** | 0.00125 | 0.58132 | 0.97605 | 3.15107 | 0.00000 | 0.00000 | 0.00000 | 0.10276 | 0.00000 | 0.22912 | 0.00000 | 0.68483 |
| **CD46** | 0.00000 | 7.08051 | 2.14419 | 5.31393 | 11.54640 | 23.75538 | 0.00000 | 1.77488 | 20.01640 | 2.22883 | 9.50425 | 7.25912 |
| **CD47** | 0.00000 | 6.51646 | 9.14779 | 0.00583 | 8.47124 | 19.89065 | 0.00000 | 0.04503 | 30.05200 | 4.57337 | 19.75152 | 4.00197 |
| **CD48** | 0.00956 | 0.17189 | 0.04309 | 0.28967 | 0.00000 | 0.02270 | 0.00660 | 0.00728 | 0.04574 | 0.00000 | 0.01215 | 0.21868 |
| **CD5** | 0.09519 | 0.19332 | 0.00368 | 0.04745 | 0.09070 | 0.00000 | 0.09404 | 0.03049 | 0.00441 | 0.10110 | 0.00413 | 0.15115 |
| **CD52** | 0.00112 | 0.21908 | 0.03908 | 0.00000 | 0.11640 | 0.00000 | 0.00000 | 0.06242 | 0.39720 | 0.70735 | 0.01620 | 0.34119 |
| **CD53** | 0.00000 | 22.50532 | 0.00000 | 5.74199 | 20.68830 | 30.24592 | 0.00000 | 0.00000 | 44.47005 | 5.56630 | 36.03692 | 2.59734 |
| **CD55** | 3.44960 | 9.58568 | 20.46444 | 0.00000 | 30.70384 | 40.29951 | 0.00000 | 0.67665 | 44.15791 | 9.83451 | 16.07743 | 0.07693 |
| **CD58** | 0.01167 | 0.78327 | 0.36069 | 0.12974 | 0.00000 | 2.06616 | 0.00000 | 0.00000 | 0.41216 | 0.00000 | 0.17899 | 0.74313 |
| **CD59** | 9.86518 | 6.65777 | 8.25133 | 2.62519 | 6.29378 | 10.47201 | 2.54110 | 7.33332 | 9.75500 | 5.47932 | 12.36989 | 2.91549 |
| **CD6** | 0.35396 | 0.03880 | 0.01381 | 0.00349 | 0.00000 | 0.04973 | 0.06738 | 0.00000 | 0.00000 | 0.02129 | 0.01151 | 0.55780 |
| **CD63** | 0.00000 | 19.88402 | 35.15062 | 27.87634 | 25.39726 | 30.80088 | 4.00028 | 8.44287 | 43.80933 | 16.78366 | 10.85741 | 15.55635 |
| **CD68** | 0.41668 | 1.02759 | 3.11382 | 0.09758 | 0.71182 | 0.00000 | 0.15030 | 0.97321 | 1.86465 | 0.61663 | 0.45015 | 1.11536 |
| **CD69** | 0.00650 | 0.01291 | 0.19188 | 0.00808 | 0.01157 | 0.02537 | 0.00000 | 0.01025 | 0.01747 | 0.00965 | 0.04386 | 0.01604 |
| **CD7** | 0.00396 | 0.02883 | 0.03962 | 0.10838 | 0.05442 | 0.00001 | 0.01551 | 0.04672 | 0.02374 | 0.00000 | 0.01306 | 0.01923 |
| **CD70** | 0.05183 | 0.15594 | 0.26299 | 0.98433 | 0.01333 | 0.00000 | 0.02463 | 0.00540 | 0.00000 | 0.06801 | 0.01444 | 0.06467 |
| **CD72** | 0.34585 | 0.00337 | 0.02577 | 0.06597 | 0.02639 | 0.00568 | 0.03586 | 0.09199 | 0.01452 | 0.00000 | 0.01571 | 1.06234 |
| **CD74** | 0.00931 | 0.02697 | 4.73459 | 0.03435 | 0.04120 | 0.00000 | 0.00000 | 0.01172 | 0.00000 | 0.08471 | 0.02334 | 0.05851 |
| **CD79A** | 0.00585 | 0.05109 | 0.00489 | 0.06700 | 0.07030 | 0.01882 | 0.04450 | 0.20738 | 0.14923 | 0.00000 | 0.02547 | 0.35839 |
| **CD79B** | 0.00412 | 0.00932 | 0.02349 | 0.00862 | 0.01100 | 0.00924 | 0.05688 | 0.01042 | 0.00124 | 0.11770 | 0.01432 | 0.04083 |
| **CD80** | 0.01176 | 0.01884 | 0.05075 | 0.02178 | 0.01983 | 0.00000 | 0.03547 | 0.02141 | 0.01567 | 0.00000 | 0.01960 | 0.06058 |
| **CD81** | 0.48303 | 7.73426 | 4.96617 | 21.00179 | 35.00113 | 15.62152 | 12.91599 | 0.00000 | 7.21780 | 37.72394 | 2.68150 | 0.44972 |
| **CD81-AS1** | 0.02165 | 0.01872 | 0.01975 | 0.01299 | 0.02486 | 0.00002 | 0.01583 | 0.05737 | 0.02213 | 0.01095 | 0.05169 | 0.01454 |
| **CD82** | 4.67570 | 3.23917 | 0.00637 | 0.00000 | 0.00649 | 0.00000 | 0.00094 | 3.46136 | 0.01367 | 0.02291 | 0.00000 | 11.46392 |
| **CD83** | 5.28304 | 1.19504 | 0.00000 | 0.73627 | 0.78448 | 0.00000 | 0.01799 | 2.97407 | 0.23532 | 0.00000 | 0.09570 | 1.92873 |
| **CD84** | 0.00617 | 0.01472 | 0.01450 | 0.02086 | 0.01049 | 0.01865 | 0.01812 | 0.01039 | 0.03097 | 0.01376 | 0.02133 | 0.01792 |
| **CD86** | 0.04629 | 0.00000 | 0.02867 | 0.01074 | 0.01469 | 0.01149 | 0.01503 | 0.16614 | 0.01326 | 0.01761 | 0.01748 | 0.76821 |
| **CD8A** | 0.00442 | 0.01417 | 0.01142 | 0.01610 | 0.02236 | 0.02563 | 0.00000 | 0.01631 | 0.12049 | 0.02742 | 0.00207 | 0.03414 |
| **CD9** | 6.91452 | 37.82716 | 9.00117 | 26.19634 | 17.16784 | 36.76800 | 0.00000 | 10.18083 | 15.17706 | 0.00000 | 1.83062 | 31.61485 |
| **CD93** | 0.00586 | 0.01188 | 0.02292 | 0.00679 | 0.00927 | 0.03473 | 0.02435 | 0.01028 | 0.02584 | 0.02630 | 0.01232 | 0.01433 |
| **CD96** | 0.00641 | 0.03718 | 0.05282 | 0.03031 | 0.00000 | 0.01681 | 0.01046 | 0.06812 | 0.00662 | 0.05873 | 0.02509 | 0.09085 |
| **CD97** | 0.00000 | 4.78683 | 0.00000 | 0.00000 | 4.35194 | 0.00000 | 0.00389 | 0.14542 | 13.19112 | 7.44443 | 7.24535 | 0.18616 |
| **CD99.uid-13523** | 0.00678 | 0.01453 | 0.01516 | 0.01166 | 0.02209 | 0.01656 | 0.01634 | 0.01064 | 0.02216 | 0.01534 | 0.01383 | 0.01665 |
| **CD99L2** | 0.14941 | 0.16439 | 0.13419 | 0.09872 | 0.13019 | 0.28667 | 0.00000 | 0.00429 | 0.00000 | 0.00000 | 0.34647 | 0.00573 |
| **CDA** | 5.35837 | 5.79715 | 0.37010 | 4.14088 | 0.74587 | 0.21130 | 0.00000 | 0.17498 | 0.00007 | 0.00253 | 0.00010 | 3.63467 |
| **CDADC1** | 0.48916 | 0.40718 | 0.29044 | 0.76954 | 1.96589 | 5.41850 | 0.00000 | 0.33272 | 2.85847 | 1.55096 | 1.56103 | 2.89954 |
| **CDAN1** | 0.00000 | 3.00677 | 1.35584 | 0.85736 | 1.88629 | 0.00000 | 0.77025 | 4.43278 | 0.57851 | 4.89310 | 0.90353 | 2.68808 |
| **CDC123** | 1.09324 | 20.21570 | 26.60226 | 13.98234 | 13.95782 | 38.94170 | 39.27554 | 6.45038 | 29.74124 | 0.00000 | 24.68069 | 23.31362 |
| **CDC14A** | 0.03600 | 1.47937 | 1.10803 | 0.30272 | 0.00000 | 1.21869 | 0.34357 | 0.06241 | 0.94109 | 0.63638 | 0.34069 | 0.33473 |
| **CDC14B** | 0.10883 | 0.95654 | 1.94967 | 0.83362 | 0.53533 | 1.18904 | 0.55281 | 0.20177 | 0.00407 | 1.10288 | 0.00000 | 0.61625 |
| **CDC14C** | 0.00592 | 0.02136 | 0.01855 | 0.02415 | 0.01497 | 0.01891 | 0.01430 | 0.01316 | 0.02881 | 0.02980 | 0.01800 | 0.02265 |
| **CDC16** | 0.69245 | 1.76383 | 2.10864 | 1.20391 | 3.97761 | 10.92179 | 8.32189 | 0.80303 | 0.00000 | 17.81040 | 4.98012 | 2.47476 |

| | | | | | | | | | | | | |
|---|---|---|---|---|---|---|---|---|---|---|---|---|
| **CDC20** | 9.77730 | 34.96450 | 22.29159 | 18.27210 | 26.91001 | 11.28370 | 37.85931 | 22.97354 | 26.20883 | 0.00525 | 20.55783 | 32.28467 |
| **CDC20B** | 2.44190 | 0.06056 | 0.01036 | 0.00000 | 0.00000 | 0.03285 | 0.04362 | 0.50626 | 0.00145 | 0.02117 | 0.02028 | 0.31565 |
| **CDC23** | 0.00000 | 7.55803 | 9.89333 | 4.02787 | 0.00000 | 12.26245 | 8.92810 | 5.00960 | 10.79617 | 2.73382 | 8.66530 | 15.46364 |
| **CDC25A** | 4.63532 | 5.24817 | 3.42094 | 3.48411 | 13.04746 | 6.89205 | 13.81158 | 7.12948 | 0.06067 | 30.00503 | 3.82167 | 14.00286 |
| **CDC25B** | 11.14103 | 10.87513 | 6.24964 | 6.62163 | 5.05043 | 0.00000 | 12.74234 | 4.13228 | 5.70029 | 15.10710 | 3.28367 | 14.71545 |
| **CDC25C** | 2.88353 | 4.13848 | 1.34005 | 3.60331 | 1.92435 | 5.42534 | 2.41785 | 2.66586 | 3.25347 | 0.00000 | 1.32876 | 7.85178 |
| **CDC26** | 0.27249 | 4.01251 | 2.41846 | 0.64481 | 1.35215 | 3.91677 | 3.10757 | 10.99776 | 1.17076 | 0.00000 | 2.00739 | 7.71992 |
| **CDC27** | 7.79587 | 4.63488 | 7.56933 | 6.23024 | 3.05100 | 20.66039 | 13.41502 | 2.59393 | 5.65941 | 9.90239 | 9.20977 | 8.44179 |
| **CDC34** | 1.27951 | 1.94519 | 4.11083 | 1.35257 | 5.14654 | 0.00000 | 2.43295 | 2.46842 | 0.00000 | 15.83691 | 0.37466 | 2.25518 |
| **CDC37** | 0.58550 | 23.21339 | 26.62649 | 18.42681 | 22.02535 | 0.00062 | 30.79566 | 22.54509 | 27.11118 | 9.92224 | 25.46129 | 23.61107 |
| **CDC37L1** | 2.32392 | 3.83637 | 4.00748 | 2.68562 | 1.24662 | 11.68958 | 3.45874 | 3.59542 | 3.30145 | 0.00000 | 3.43943 | 5.94332 |
| **CDC37L1-AS1** | 0.00729 | 0.04986 | 0.10521 | 0.05115 | 0.05903 | 0.02626 | 0.07965 | 0.00812 | 0.05190 | 0.00000 | 0.03401 | 0.05662 |
| **CDC40** | 3.69171 | 2.26186 | 5.21148 | 2.10994 | 3.18787 | 7.95791 | 6.63737 | 6.12246 | 5.31778 | 3.55137 | 3.52037 | 4.27681 |
| **CDC42** | 2.72984 | 11.23051 | 15.94565 | 12.98770 | 8.26454 | 23.67922 | 33.87287 | 14.75888 | 3.22412 | 14.88279 | 34.08446 | 14.66518 |
| **CDC42BPA** | 7.26850 | 1.56892 | 1.04244 | 2.50808 | 0.47217 | 9.95216 | 3.98193 | 2.38603 | 1.35873 | 16.01384 | 1.26954 | 2.81005 |
| **CDC42BPB** | 8.48158 | 0.00000 | 0.83562 | 2.37667 | 3.75079 | 0.00000 | 3.90144 | 1.07087 | 0.00000 | 14.43847 | 0.29823 | 1.22880 |
| **CDC42BPG** | 0.04523 | 0.73090 | 2.69932 | 0.38461 | 4.02893 | 0.00000 | 0.57544 | 0.00000 | 1.73411 | 4.54962 | 0.81310 | 1.47727 |
| **CDC42EP1** | 0.00000 | 1.69748 | 19.79534 | 1.40245 | 6.39077 | 0.00000 | 5.40659 | 0.11113 | 10.09467 | 4.34697 | 1.05980 | 7.26964 |
| **CDC42EP2** | 0.00191 | 1.90193 | 2.09148 | 0.93051 | 2.85798 | 0.00000 | 4.15316 | 1.07806 | 4.70202 | 2.03795 | 5.99744 | 3.79317 |
| **CDC42EP3** | 1.06105 | 0.74496 | 8.42685 | 1.02405 | 0.00000 | 1.69573 | 3.32204 | 0.21476 | 3.31608 | 2.27889 | 0.00000 | 0.33276 |
| **CDC42EP4** | 0.44404 | 18.38708 | 11.08866 | 33.32293 | 2.13898 | 0.00000 | 3.46535 | 0.00000 | 3.98092 | 0.00000 | 1.23949 | 5.63777 |
| **CDC42EP5** | 0.37744 | 0.01018 | 0.08778 | 0.01721 | 0.00336 | 0.03796 | 0.00913 | 0.00282 | 0.00000 | 0.02442 | 0.03168 | 0.01213 |
| **CDC42P3** | 0.09000 | 0.00213 | 0.10081 | 0.08411 | 0.00000 | 0.12972 | 0.03250 | 0.03100 | 0.00000 | 0.13977 | 0.04992 | 0.02156 |
| **CDC42SE1** | 4.40612 | 3.80824 | 15.05011 | 3.32869 | 8.12684 | 13.45562 | 7.61272 | 0.84349 | 14.06869 | 0.00000 | 4.93580 | 0.23047 |
| **CDC42SE2** | 1.66288 | 10.00418 | 5.96210 | 1.01660 | 5.08757 | 24.38343 | 21.22673 | 1.14546 | 9.73776 | 7.92916 | 18.68261 | 4.30736 |
| **CDC45** | 4.11502 | 10.53864 | 8.68389 | 5.84732 | 3.72781 | 9.86508 | 9.08434 | 1.17180 | 11.41411 | 0.00000 | 8.74694 | 11.39753 |
| **CDC5L** | 6.47651 | 11.61540 | 9.84550 | 6.74634 | 6.77372 | 30.64686 | 9.52932 | 13.61372 | 14.34092 | 9.98364 | 9.44808 | 13.56979 |
| **CDC6** | 2.98415 | 14.46378 | 12.35688 | 8.46207 | 10.50958 | 26.42177 | 13.04623 | 20.76118 | 15.55978 | 0.00000 | 16.26728 | 20.17060 |
| **CDC7** | 2.83913 | 7.22581 | 1.27395 | 3.53067 | 0.00000 | 17.70092 | 3.59788 | 1.36276 | 2.68196 | 2.37467 | 8.54477 | 7.44897 |
| **CDC73** | 0.15934 | 3.00956 | 7.40452 | 1.50501 | 0.00000 | 11.16325 | 10.64617 | 1.43619 | 4.60089 | 11.73913 | 3.17706 | 4.04745 |
| **CDCA2** | 4.28074 | 7.60556 | 3.77878 | 5.35570 | 6.34043 | 15.01606 | 9.12998 | 7.97409 | 12.83879 | 7.25165 | 5.55621 | 13.02433 |
| **CDCA3** | 3.46611 | 15.24689 | 17.59956 | 10.84612 | 16.34385 | 16.61815 | 15.55904 | 2.62586 | 15.10957 | 0.00000 | 9.18943 | 9.02906 |
| **CDCA4** | 1.70463 | 19.95682 | 6.24672 | 16.34791 | 14.65711 | 14.50526 | 18.24821 | 11.46668 | 19.82889 | 0.00000 | 19.01531 | 18.52039 |
| **CDCA5** | 2.81386 | 19.77675 | 15.15269 | 13.66298 | 15.42004 | 1.70736 | 24.22493 | 8.77699 | 32.72019 | 2.85048 | 14.75465 | 19.47876 |
| **CDCA7** | 0.00000 | 9.69030 | 2.83566 | 1.47115 | 0.12116 | 17.25713 | 5.29207 | 0.00000 | 14.10177 | 6.61076 | 11.11212 | 8.62178 |
| **CDCA7L** | 10.10402 | 0.03141 | 1.04825 | 3.65261 | 0.00000 | 0.00000 | 0.00000 | 2.20062 | 0.30186 | 0.00110 | 0.00542 | 1.60805 |
| **CDCA8** | 6.77580 | 26.33747 | 11.19823 | 8.06506 | 15.88523 | 14.69127 | 19.96194 | 18.07779 | 19.00700 | 0.00000 | 9.17850 | 23.42004 |
| **CDCP1** | 0.07276 | 4.01283 | 0.02356 | 0.40196 | 1.67982 | 0.79082 | 0.00000 | 0.00000 | 2.71063 | 1.13870 | 1.53102 | 0.57603 |
| **CDCP2** | 0.01121 | 0.01265 | 0.01969 | 0.02807 | 0.01890 | 0.04195 | 0.01439 | 0.00876 | 0.00000 | 0.04163 | 0.04127 | 0.01224 |
| **CDH1** | 5.92203 | 28.51416 | 24.05376 | 3.08827 | 17.17994 | 38.65962 | 0.00000 | 6.47099 | 64.26036 | 27.05978 | 27.68563 | 6.94923 |
| **CDH10** | 0.32564 | 0.12599 | 0.07626 | 0.20882 | 0.00000 | 0.00907 | 0.00000 | 0.44790 | 0.01542 | 0.17351 | 0.00251 | 0.07455 |
| **CDH11** | 0.00368 | 0.00373 | 1.15747 | 4.78318 | 0.00000 | 0.00077 | 0.00000 | 0.00804 | 0.01773 | 0.11883 | 0.01650 | 0.02159 |
| **CDH12** | 3.72286 | 0.01067 | 0.01442 | 0.00248 | 0.06369 | 0.00000 | 0.01370 | 0.87212 | 0.00931 | 0.05119 | 0.01064 | 0.00000 |
| **CDH13** | 0.26889 | 0.00949 | 0.02695 | 0.02397 | 0.00000 | 0.06677 | 0.02525 | 0.02918 | 0.01413 | 0.03254 | 0.01329 | 0.01252 |
| **CDH15** | 0.25162 | 0.01810 | 0.02723 | 0.02145 | 0.12749 | 0.00925 | 0.02950 | 0.04024 | 0.00000 | 0.06803 | 0.00884 | 0.06071 |
| **CDH16** | 0.03126 | 0.01553 | 0.08442 | 0.13465 | 0.01633 | 0.01983 | 0.02555 | 0.02116 | 0.03004 | 0.00000 | 0.01441 | 0.01585 |
| **CDH17** | 2.77186 | 0.09391 | 0.00291 | 0.16470 | 0.03344 | 0.00000 | 0.04511 | 2.45565 | 0.00346 | 0.00000 | 0.01479 | 0.30444 |
| **CDH18** | 1.05282 | 0.05658 | 0.57338 | 0.00060 | 0.06165 | 0.00000 | 0.12128 | 0.00843 | 0.00000 | 0.03736 | 0.00648 | 0.01043 |

| | | | | | | | | | | | | |
|---|---|---|---|---|---|---|---|---|---|---|---|---|
| **CDH19** | 0.01432 | 0.01512 | 0.01088 | 0.00784 | 0.02142 | 0.01647 | 0.00595 | 0.07581 | 0.03724 | 0.04440 | 0.01214 | 0.01057 |
| **CDH2** | 2.38536 | 0.93692 | 2.27129 | 6.50862 | 0.00000 | 0.06648 | 0.00000 | 1.56255 | 0.23651 | 0.00033 | 0.11973 | 3.23734 |
| **CDH20** | 0.02454 | 0.01215 | 0.06445 | 0.00536 | 0.10019 | 0.00000 | 0.04055 | 0.00401 | 0.00142 | 0.06642 | 0.01062 | 0.12314 |
| **CDH22** | 0.14295 | 0.01397 | 0.02556 | 0.01048 | 0.01855 | 0.02297 | 0.02049 | 0.03637 | 0.01750 | 0.01084 | 0.01226 | 0.01375 |
| **CDH23** | 0.00067 | 0.39385 | 0.06769 | 0.16145 | 0.00000 | 0.00000 | 0.00000 | 0.15717 | 0.11121 | 0.98660 | 0.12028 | 0.50586 |
| **CDH24** | 0.18231 | 0.00000 | 1.14402 | 0.00000 | 0.27594 | 0.00000 | 0.25821 | 0.40212 | 0.16093 | 0.77679 | 0.00000 | 0.55268 |
| **CDH26** | 0.03013 | 0.00999 | 0.05556 | 0.01459 | 0.02211 | 0.05649 | 0.05106 | 0.00686 | 0.07187 | 0.00923 | 0.03064 | 0.01526 |
| **CDH3** | 18.85430 | 7.59833 | 0.80170 | 1.33192 | 6.20932 | 0.00000 | 0.00000 | 8.02588 | 7.03757 | 2.56768 | 4.39154 | 15.94454 |
| **CDH4** | 0.50512 | 0.59456 | 0.00095 | 0.00697 | 0.04008 | 0.00000 | 0.01663 | 0.13858 | 0.00000 | 0.85635 | 1.05105 | 0.00001 |
| **CDH5** | 1.65623 | 0.01169 | 0.77048 | 2.12899 | 0.19736 | 0.00000 | 0.00000 | 0.00308 | 0.00000 | 0.00000 | 0.21956 | 0.00660 |
| **CDH6** | 0.01587 | 0.00396 | 0.03717 | 0.02083 | 0.03534 | 0.02045 | 0.00266 | 0.11120 | 0.02094 | 0.25059 | 0.03795 | 0.02762 |
| **CDH7** | 0.00398 | 0.01682 | 0.02842 | 0.07181 | 0.04246 | 0.00000 | 0.01220 | 0.03293 | 0.07360 | 0.00000 | 0.02255 | 0.01713 |
| **CDH8** | 0.76673 | 0.49619 | 0.30914 | 2.36672 | 0.48870 | 0.76615 | 0.00000 | 0.20421 | 0.00000 | 0.00000 | 0.18264 | 0.02307 |
| **CDH9** | 0.00485 | 0.01620 | 0.06713 | 0.03858 | 0.02801 | 0.00000 | 0.01122 | 0.07526 | 0.01309 | 0.02067 | 0.01668 | 0.01340 |
| **CDHR1** | 0.00266 | 0.41586 | 0.80941 | 14.16143 | 0.51866 | 0.00000 | 0.01489 | 0.00175 | 0.00000 | 0.89696 | 0.11076 | 0.14310 |
| **CDHR2** | 0.00428 | 0.01079 | 0.02734 | 0.09291 | 0.00000 | 0.02190 | 0.03003 | 0.05448 | 0.00045 | 0.04537 | 0.04396 | 0.02189 |
| **CDHR3** | 0.02836 | 0.02083 | 0.01410 | 0.01873 | 0.02411 | 0.02188 | 0.01694 | 0.02653 | 0.02257 | 0.02338 | 0.01292 | 0.01896 |
| **CDHR4** | 0.00626 | 0.01655 | 0.01947 | 0.02082 | 0.10069 | 0.00000 | 0.05546 | 0.01062 | 0.00014 | 0.00000 | 0.01242 | 0.02120 |
| **CDHR5** | 0.00352 | 0.00716 | 0.05479 | 0.00246 | 0.04856 | 0.00000 | 0.00291 | 0.01030 | 0.04204 | 0.30944 | 0.01126 | 0.01290 |
| **CDIP1** | 5.84815 | 2.60218 | 0.56065 | 2.65419 | 1.28186 | 0.00000 | 0.78752 | 2.13510 | 0.03079 | 0.68213 | 0.00000 | 2.08807 |
| **CDIPT** | 1.97342 | 7.33273 | 5.80754 | 4.49255 | 6.87608 | 4.26711 | 4.38442 | 2.26823 | 9.42083 | 23.78927 | 1.37410 | 8.76151 |
| **CDIPT-AS1** | 0.18284 | 0.43411 | 0.11927 | 0.33438 | 0.04612 | 0.34129 | 0.17021 | 0.92388 | 0.00000 | 0.11214 | 0.14849 | 0.61553 |
| **CDK1** | 0.99959 | 22.91081 | 13.62237 | 7.31941 | 6.43713 | 60.35046 | 16.28017 | 19.31757 | 23.65099 | 2.37495 | 21.95842 | 30.23200 |
| **CDK10** | 1.28046 | 4.26820 | 4.13788 | 5.10196 | 4.59905 | 0.00000 | 0.67635 | 1.02501 | 4.13213 | 5.24190 | 4.50891 | 3.72666 |
| **CDK11A** | 0.30773 | 0.13760 | 0.85572 | 0.46109 | 0.66089 | 0.00000 | 0.09170 | 0.47024 | 0.65361 | 2.26574 | 0.17206 | 0.47102 |
| **CDK11B** | 0.38419 | 0.40738 | 1.54631 | 0.52025 | 1.82587 | 0.00000 | 1.53155 | 1.95820 | 0.00007 | 2.92647 | 0.77515 | 1.77343 |
| **CDK12** | 13.64951 | 3.50261 | 7.99192 | 8.67183 | 11.72743 | 5.31805 | 12.34073 | 10.26171 | 7.72690 | 23.83460 | 8.03596 | 12.48188 |
| **CDK13** | 3.16717 | 0.45457 | 3.00746 | 3.69207 | 2.56295 | 7.43621 | 7.67804 | 0.73700 | 0.47983 | 15.60914 | 1.46397 | 1.17844 |
| **CDK14** | 0.01737 | 0.38291 | 0.24880 | 0.01796 | 0.20321 | 0.00000 | 0.51995 | 0.24873 | 0.00000 | 0.12334 | 0.23807 | 0.09024 |
| **CDK15** | 0.51280 | 0.00213 | 0.03061 | 0.00138 | 0.08907 | 0.02373 | 0.00054 | 0.01031 | 0.02371 | 0.06054 | 0.00589 | 0.07290 |
| **CDK16** | 2.68475 | 11.38091 | 8.20571 | 14.11257 | 15.40488 | 0.00000 | 10.70964 | 3.44889 | 9.13241 | 16.46031 | 9.39408 | 4.87887 |
| **CDK17** | 1.62094 | 2.65344 | 1.80003 | 0.76826 | 0.00000 | 3.18237 | 1.26283 | 5.67002 | 3.29465 | 0.68118 | 1.45819 | 3.43835 |
| **CDK18** | 0.14219 | 0.05103 | 0.04668 | 0.10894 | 0.00000 | 0.02031 | 0.01390 | 0.00684 | 0.00000 | 0.01885 | 0.05836 | 0.02359 |
| **CDK19** | 0.00248 | 0.00000 | 1.27442 | 0.90912 | 0.25391 | 1.09726 | 1.22243 | 0.29557 | 0.00005 | 0.25706 | 0.19806 | 0.51920 |
| **CDK2** | 0.08924 | 4.11688 | 13.07317 | 1.20508 | 5.84424 | 8.58496 | 4.81021 | 2.55995 | 6.85857 | 2.96569 | 6.04492 | 4.47629 |
| **CDK20** | 1.63323 | 0.42740 | 0.15959 | 1.12212 | 0.24403 | 0.47784 | 0.48576 | 1.75386 | 1.54962 | 0.00000 | 0.02990 | 2.76655 |
| **CDK2AP1** | 2.32026 | 8.05489 | 5.35163 | 1.39906 | 3.83560 | 12.76917 | 15.67089 | 3.07817 | 0.00000 | 15.83661 | 3.46236 | 12.25879 |
| **CDK2AP2** | 17.47786 | 11.33832 | 7.87548 | 16.74390 | 13.49775 | 4.02523 | 19.36163 | 20.60411 | 13.17139 | 0.00000 | 9.63075 | 19.10510 |
| **CDK4** | 0.43775 | 23.29497 | 17.14189 | 15.03611 | 13.13321 | 15.48739 | 28.50302 | 8.22135 | 29.27897 | 0.00000 | 21.24037 | 22.56378 |
| **CDK5** | 4.19874 | 6.81968 | 4.43422 | 6.21585 | 6.11749 | 0.79554 | 5.62167 | 0.52417 | 4.39347 | 0.00000 | 6.17432 | 2.51324 |
| **CDK5R1** | 0.63762 | 0.31621 | 0.41713 | 0.50973 | 0.00000 | 0.01814 | 3.27350 | 0.83844 | 1.51236 | 1.67807 | 0.00000 | 0.38199 |
| **CDK5R2** | 0.72978 | 0.02416 | 0.00382 | 0.00832 | 0.03403 | 0.11417 | 0.00000 | 0.00000 | 0.00000 | 0.19956 | 0.00000 | 0.08398 |
| **CDK5RAP1** | 1.03754 | 12.59259 | 6.27603 | 5.96668 | 5.94769 | 14.64744 | 10.95519 | 5.52880 | 13.16648 | 0.00000 | 14.42998 | 13.29622 |
| **CDK5RAP2** | 0.56447 | 13.88043 | 3.71287 | 2.80419 | 10.36601 | 6.40140 | 5.55851 | 2.35440 | 1.01785 | 14.05244 | 4.14966 | 16.89851 |
| **CDK5RAP3** | 0.87570 | 5.45095 | 3.30236 | 7.35505 | 7.24760 | 0.00000 | 0.63735 | 0.06101 | 4.24601 | 3.26506 | 4.72017 | 5.27262 |
| **CDK6** | 0.00000 | 2.88071 | 9.13121 | 0.00000 | 13.39893 | 4.48661 | 25.00237 | 0.30357 | 10.07354 | 17.65073 | 15.66900 | 0.00000 |
| **CDK7** | 10.73726 | 10.70493 | 19.89152 | 3.77184 | 1.78853 | 22.93919 | 19.35673 | 7.97786 | 13.73430 | 0.80311 | 7.32521 | 16.14054 |
| **CDK8** | 1.44495 | 0.64818 | 1.15979 | 1.28795 | 0.79905 | 1.36158 | 2.11303 | 0.24444 | 1.77194 | 1.33335 | 0.72596 | 1.10239 |

| | | | | | | | | | | | | |
|---|---|---|---|---|---|---|---|---|---|---|---|---|
| **CDK9** | 1.02561 | 4.69678 | 3.86333 | 6.11257 | 5.64273 | 0.00000 | 5.74156 | 11.50288 | 0.00000 | 5.76817 | 3.37643 | 3.83177 |
| **CDKAL1** | 1.35202 | 3.18570 | 4.24589 | 2.60056 | 1.83109 | 8.11834 | 3.04659 | 0.69518 | 4.32717 | 1.68705 | 3.09198 | 3.35230 |
| **CDKL1** | 0.15451 | 0.19081 | 0.10616 | 0.03537 | 0.13256 | 0.00000 | 0.00000 | 0.12492 | 0.19475 | 0.25839 | 0.03509 | 0.51321 |
| **CDKL2** | 0.32853 | 0.00000 | 0.02475 | 0.00222 | 0.15937 | 0.00000 | 0.02116 | 0.04811 | 0.07979 | 0.28244 | 0.10884 | 0.18767 |
| **CDKL3** | 0.13478 | 0.09277 | 0.24511 | 0.18534 | 0.06435 | 0.12405 | 0.07261 | 0.25514 | 0.35926 | 0.00000 | 0.04017 | 1.25781 |
| **CDKL4** | 0.06415 | 0.02085 | 0.01297 | 0.01580 | 0.03975 | 0.00000 | 0.01387 | 0.00909 | 0.08042 | 0.09746 | 0.03673 | 0.23213 |
| **CDKL5** | 2.49826 | 0.28660 | 0.35096 | 0.35335 | 0.00000 | 0.00000 | 0.10437 | 0.34246 | 0.00000 | 0.58216 | 0.61001 | 0.63204 |
| **CDKN1A** | 1.78037 | 1.68869 | 9.89658 | 0.68220 | 1.48221 | 3.65779 | 0.00000 | 1.73602 | 4.09024 | 2.01936 | 0.00000 | 0.00000 |
| **CDKN1B** | 0.25825 | 0.10147 | 0.12210 | 2.25883 | 0.00000 | 0.65754 | 3.11553 | 0.00000 | 0.00000 | 2.10833 | 2.43253 | 0.31119 |
| **CDKN1C** | 0.10395 | 0.01576 | 19.50650 | 0.99831 | 0.00000 | 0.38235 | 9.07691 | 2.60037 | 0.00000 | 0.67277 | 0.00000 | 0.16329 |
| **CDKN2A** | 0.00538 | 0.02795 | 0.13659 | 0.01164 | 0.00909 | 0.02568 | 0.09444 | 0.02226 | 0.06109 | 0.00000 | 0.00750 | 0.01359 |
| **CDKN2AIP** | 4.37724 | 2.81642 | 4.32731 | 2.12232 | 0.00000 | 8.81578 | 3.35296 | 24.85328 | 4.56666 | 0.00000 | 3.78340 | 4.91753 |
| **CDKN2AIPNL** | 0.96390 | 7.09748 | 0.00000 | 3.90225 | 5.00207 | 6.13874 | 2.45283 | 0.34581 | 2.20202 | 2.04496 | 4.40108 | 3.93488 |
| **CDKN2B** | 0.12934 | 0.18864 | 0.14899 | 0.00000 | 0.00000 | 0.02247 | 0.00000 | 0.00251 | 0.02283 | 0.12183 | 0.00249 | 0.42343 |
| **CDKN2B-AS1** | 0.02539 | 0.01194 | 0.01324 | 0.00670 | 0.00098 | 0.02605 | 0.00717 | 0.03100 | 0.01621 | 0.10138 | 0.03830 | 0.01018 |
| **CDKN2C** | 0.09934 | 0.24129 | 0.18403 | 0.58954 | 0.00000 | 0.00000 | 0.00000 | 0.26641 | 0.00000 | 0.17631 | 0.11836 | 0.39454 |
| **CDKN2D** | 0.80721 | 2.02026 | 8.26967 | 1.69419 | 3.44799 | 2.86742 | 5.50372 | 3.54028 | 3.71477 | 1.59189 | 1.39511 | 3.24461 |
| **CDKN3** | 0.22648 | 3.36945 | 2.71785 | 4.54884 | 6.67204 | 21.64416 | 1.55744 | 0.40392 | 0.52373 | 1.15528 | 4.04847 | 5.00885 |
| **CDNF** | 0.12173 | 0.44588 | 0.03770 | 0.17044 | 0.06156 | 0.51671 | 0.00000 | 0.05415 | 0.09657 | 0.24106 | 0.05431 | 0.28092 |
| **CDO1** | 0.00000 | 0.27563 | 7.94915 | 1.08978 | 0.00000 | 1.18491 | 1.18368 | 0.01571 | 2.35614 | 0.48739 | 1.66955 | 0.86995 |
| **CDON** | 2.76003 | 0.58084 | 0.49149 | 0.32229 | 0.00000 | 0.05860 | 0.16984 | 1.44915 | 0.45028 | 0.51907 | 0.10167 | 0.88842 |
| **CDPF1** | 0.97874 | 2.04688 | 1.31155 | 0.41516 | 1.38744 | 0.00175 | 2.24929 | 0.00000 | 0.78849 | 0.29102 | 3.49086 | 4.71493 |
| **CDR1** | 11.71648 | 4.94514 | 0.00000 | 0.85201 | 3.80658 | 1.77475 | 0.53244 | 1.14554 | 0.00000 | 14.33519 | 0.62006 | 14.07085 |
| **CDR2** | 1.85827 | 0.16053 | 1.75476 | 3.94492 | 0.00000 | 4.37605 | 2.41608 | 2.15013 | 0.00033 | 5.70061 | 0.00000 | 10.04632 |
| **CDR2L** | 0.81208 | 3.86675 | 2.65246 | 3.25746 | 5.43642 | 0.00000 | 11.38545 | 0.43283 | 6.90458 | 2.23642 | 0.36715 | 0.52438 |
| **CDRT1** | 0.01923 | 0.12213 | 0.04095 | 0.03009 | 0.07630 | 0.02272 | 0.05385 | 0.07734 | 0.24801 | 0.09812 | 0.06676 | 0.13561 |
| **CDRT15** | 0.00750 | 0.01668 | 0.03898 | 0.01538 | 0.01336 | 0.01710 | 0.01963 | 0.01071 | 0.04123 | 0.00784 | 0.01931 | 0.01864 |
| **CDRT15P1** | 0.00586 | 0.01236 | 0.07788 | 0.08090 | 0.00000 | 0.04987 | 0.03171 | 0.02929 | 0.14810 | 0.02279 | 0.01031 | 0.01523 |
| **CDRT4** | 0.00287 | 0.02001 | 0.00593 | 0.00503 | 0.02120 | 0.01294 | 0.00520 | 0.01510 | 0.04455 | 0.26498 | 0.03450 | 0.02678 |
| **CDS1** | 1.96914 | 0.91115 | 4.43848 | 0.00000 | 6.92501 | 7.89305 | 1.54653 | 0.54859 | 4.95914 | 8.09176 | 2.33630 | 0.83223 |
| **CDS2** | 6.65993 | 7.01378 | 8.48318 | 2.45897 | 0.00004 | 4.63843 | 3.57385 | 6.27797 | 17.43281 | 1.02771 | 10.00461 | 14.56237 |
| **CDSN** | 0.30439 | 0.05366 | 0.00501 | 0.10215 | 0.00000 | 0.03874 | 0.01011 | 0.04134 | 0.02001 | 0.01126 | 0.01066 | 0.05937 |
| **CDT1** | 4.16583 | 9.24515 | 4.74732 | 8.96385 | 22.07345 | 0.00000 | 12.77864 | 2.59864 | 2.87231 | 25.56821 | 3.97208 | 12.84462 |
| **CDV3** | 3.50137 | 11.89027 | 14.87781 | 10.63703 | 8.60896 | 39.53478 | 32.20503 | 27.28018 | 0.11584 | 29.80360 | 21.94165 | 25.69365 |
| **CDX1** | 0.00202 | 0.00000 | 1.28739 | 0.00000 | 0.00000 | 0.00000 | 10.76339 | 0.00688 | 6.64985 | 1.19109 | 1.01730 | 0.00000 |
| **CDX2** | 0.07515 | 5.48574 | 0.00000 | 6.11157 | 18.34476 | 0.00000 | 7.32469 | 0.15679 | 15.40302 | 3.12599 | 12.07283 | 0.00000 |
| **CDX4** | 0.00651 | 0.01700 | 0.03076 | 0.03427 | 0.02503 | 0.00000 | 0.02626 | 0.01054 | 0.03234 | 0.00000 | 0.01333 | 0.02881 |
| **CDYL** | 2.75872 | 3.54860 | 7.47806 | 1.09544 | 12.34472 | 9.91833 | 6.47393 | 0.00000 | 4.97003 | 13.58432 | 2.42674 | 3.86338 |
| **CDYL2** | 0.21957 | 0.13767 | 0.02344 | 0.21173 | 0.34282 | 0.09627 | 0.00000 | 0.00801 | 0.00000 | 0.31044 | 0.00000 | 0.02185 |
| **CEACAM1** | 0.02358 | 0.97514 | 0.00000 | 13.91931 | 0.75438 | 0.86642 | 0.00000 | 0.00327 | 0.49008 | 0.00000 | 0.04049 | 0.04096 |
| **CEACAM16** | 0.00730 | 0.00749 | 0.00305 | 0.08431 | 0.00001 | 0.03718 | 0.05931 | 0.00956 | 0.00024 | 0.04643 | 0.01915 | 0.01357 |
| **CEACAM19** | 0.00444 | 0.05228 | 0.51367 | 0.19524 | 0.24058 | 0.00000 | 0.00000 | 0.04944 | 0.06911 | 0.12776 | 0.00601 | 0.05659 |
| **CEACAM20** | 0.00625 | 0.00996 | 0.13631 | 0.01249 | 0.00387 | 0.05171 | 0.00827 | 0.04843 | 0.01562 | 0.01906 | 0.01351 | 0.01583 |
| **CEACAM21** | 0.00208 | 0.17692 | 0.01564 | 0.17811 | 0.02840 | 0.00000 | 0.11555 | 0.08266 | 0.00002 | 0.00000 | 0.01116 | 0.34958 |
| **CEACAM22P** | 0.28500 | 0.00000 | 0.21625 | 0.00438 | 0.20250 | 0.00000 | 0.04933 | 0.27316 | 0.06373 | 0.46245 | 0.06313 | 0.11730 |
| **CEACAM5** | 0.00739 | 0.02006 | 0.01172 | 0.07879 | 0.01072 | 0.02331 | 0.01829 | 0.00908 | 0.03372 | 0.00297 | 0.01246 | 0.03412 |
| **CEACAM6** | 0.04965 | 0.89536 | 0.20638 | 11.30282 | 0.00000 | 0.03066 | 0.00000 | 0.00000 | 0.00001 | 0.01051 | 0.02606 | 0.17279 |
| **CEACAM7** | 0.00755 | 0.03460 | 0.01064 | 0.11111 | 0.01486 | 0.00000 | 0.02852 | 0.01343 | 0.01095 | 0.00759 | 0.05767 | 0.03932 |

| | | | | | | | | | | | | |
|---|---|---|---|---|---|---|---|---|---|---|---|---|
| **CEACAM8** | 0.00619 | 0.01543 | 0.01762 | 0.01446 | 0.01350 | 0.01634 | 0.01865 | 0.01016 | 0.01646 | 0.02352 | 0.01570 | 0.01904 |
| **CEBPA** | 0.00000 | 0.91390 | 17.45265 | 0.65947 | 17.33072 | 0.31856 | 28.63640 | 0.00000 | 5.57153 | 6.63543 | 1.38930 | 0.00000 |
| **CEBPA-AS1** | 0.00549 | 0.08697 | 0.71147 | 0.00000 | 0.18397 | 0.00000 | 0.10068 | 0.00831 | 0.00000 | 0.13165 | 0.13938 | 0.05366 |
| **CEBPB** | 0.20218 | 0.08699 | 5.27739 | 0.00000 | 0.16944 | 0.00011 | 2.12396 | 0.00735 | 0.00000 | 3.54467 | 0.00000 | 0.00361 |
| **CEBPD** | 0.00120 | 0.01170 | 5.69831 | 0.23065 | 4.80937 | 0.00000 | 1.02887 | 0.02935 | 0.00000 | 0.75815 | 0.24363 | 0.00258 |
| **CEBPE** | 0.00561 | 0.03166 | 0.20202 | 0.00000 | 0.38564 | 0.00000 | 0.00114 | 0.01143 | 0.01600 | 0.06330 | 0.00236 | 0.01562 |
| **CEBPG** | 2.53609 | 9.32071 | 17.01102 | 3.16493 | 7.50274 | 15.41433 | 11.21441 | 5.44866 | 14.60421 | 6.58847 | 10.87440 | 10.18112 |
| **CEBPZ** | 1.00458 | 17.02992 | 13.94038 | 20.57166 | 11.50451 | 29.08211 | 15.71413 | 15.30106 | 13.34033 | 29.20115 | 13.89860 | 22.23633 |
| **CEBPZOS** | 0.13750 | 5.13836 | 2.11735 | 1.44659 | 0.00000 | 8.91932 | 2.97562 | 1.85798 | 1.98938 | 10.16443 | 3.50808 | 7.67823 |
| **CECR1** | 0.00268 | 1.45586 | 0.00000 | 3.95322 | 0.13783 | 0.65849 | 0.06562 | 0.05328 | 0.00000 | 0.06815 | 0.00000 | 0.23477 |
| **CECR2** | 2.82029 | 0.99772 | 0.33920 | 2.42316 | 0.62267 | 1.58975 | 0.81549 | 0.96487 | 0.00000 | 4.70440 | 0.49803 | 1.15664 |
| **CECR3** | 0.00587 | 0.02485 | 0.02922 | 0.04372 | 0.01520 | 0.00367 | 0.02033 | 0.01953 | 0.01653 | 0.00872 | 0.01714 | 0.01680 |
| **CECR5** | 0.62595 | 10.83995 | 3.81158 | 8.13743 | 5.55518 | 0.00000 | 4.86551 | 0.82653 | 2.83709 | 0.00000 | 7.72505 | 10.09152 |
| **CECR5-AS1** | 0.00581 | 0.01277 | 0.01627 | 0.08946 | 0.01232 | 0.00001 | 0.01829 | 0.00738 | 0.01770 | 0.01721 | 0.01898 | 0.11695 |
| **CECR6** | 0.06718 | 0.06829 | 0.00240 | 0.00000 | 0.02678 | 0.01460 | 0.00434 | 0.00458 | 0.02753 | 0.23635 | 0.00013 | 0.02712 |
| **CECR7** | 0.49391 | 0.01917 | 0.01408 | 0.00851 | 0.01472 | 0.01964 | 0.01657 | 0.81717 | 0.01912 | 0.00661 | 0.01694 | 0.07929 |
| **CEL** | 0.00757 | 0.01545 | 0.05626 | 0.02029 | 0.03433 | 0.00000 | 0.01150 | 0.00979 | 0.07143 | 0.04980 | 0.01272 | 0.02520 |
| **CELA1** | 0.20359 | 0.31963 | 0.32426 | 0.00000 | 0.54341 | 0.00067 | 0.81916 | 0.47902 | 1.65657 | 0.12513 | 1.24042 | 0.86035 |
| **CELA2A** | 0.19535 | 0.07891 | 0.01862 | 0.01624 | 0.04531 | 0.00331 | 0.02660 | 0.03351 | 0.01165 | 0.00000 | 0.05495 | 0.02813 |
| **CELA2B** | 0.02853 | 0.02438 | 0.01453 | 0.01257 | 0.01595 | 0.01054 | 0.02386 | 0.00701 | 0.01974 | 0.01090 | 0.01539 | 0.01884 |
| **CELA3A** | 0.40597 | 0.49386 | 0.05868 | 2.00704 | 0.06241 | 0.08523 | 0.00000 | 1.05964 | 0.01326 | 0.00000 | 0.01823 | 1.55349 |
| **CELA3B** | 0.00315 | 0.44752 | 0.08265 | 1.72795 | 0.01283 | 0.01339 | 0.00910 | 1.10714 | 0.03241 | 0.00000 | 0.00714 | 1.54502 |
| **CELF1** | 7.32232 | 11.47506 | 12.13791 | 9.46566 | 8.16883 | 3.21422 | 13.91811 | 2.66864 | 19.10491 | 29.87011 | 14.55457 | 12.70494 |
| **CELF2** | 4.59135 | 0.72488 | 0.61192 | 0.36039 | 0.00000 | 1.06270 | 0.00000 | 0.99953 | 0.00000 | 3.02145 | 0.02916 | 0.14991 |
| **CELF2-AS2** | 0.00780 | 0.03068 | 0.01451 | 0.01010 | 0.00144 | 0.04516 | 0.01230 | 0.01879 | 0.01294 | 0.04554 | 0.01063 | 0.01003 |
| **CELF3** | 0.63538 | 0.06153 | 0.13906 | 0.01678 | 0.02739 | 0.00000 | 0.12229 | 0.04298 | 0.00000 | 0.00000 | 0.14221 | 0.05913 |
| **CELF4** | 0.01666 | 0.18332 | 0.16520 | 0.33431 | 0.00000 | 0.12851 | 0.00000 | 0.05648 | 0.00000 | 0.54409 | 0.05562 | 0.04301 |
| **CELF5** | 0.35609 | 0.06399 | 0.01619 | 0.00000 | 0.05249 | 0.04577 | 0.03089 | 0.05192 | 0.00000 | 0.07953 | 0.01625 | 0.03749 |
| **CELF6** | 0.00429 | 0.21268 | 0.56534 | 0.32181 | 0.00000 | 0.00000 | 0.01609 | 0.00972 | 0.62191 | 1.29216 | 0.07226 | 0.00062 |
| **CELSR1** | 6.89214 | 0.68418 | 0.65312 | 0.00000 | 0.13800 | 0.00000 | 0.05880 | 0.12615 | 0.00000 | 2.48453 | 0.00000 | 0.54681 |
| **CELSR2** | 0.00635 | 0.85913 | 3.03933 | 0.84117 | 0.84378 | 0.00000 | 0.18821 | 0.00000 | 1.21715 | 6.08248 | 0.97872 | 1.33096 |
| **CELSR3** | 0.00851 | 0.51034 | 0.41706 | 1.10701 | 0.14432 | 0.00000 | 0.66690 | 0.10237 | 2.15663 | 1.87840 | 0.34336 | 0.56469 |
| **CELSR3-AS1** | 0.01707 | 0.03117 | 0.04930 | 0.04450 | 0.10452 | 0.00000 | 0.05639 | 0.04559 | 0.08691 | 0.00000 | 0.02353 | 0.11112 |
| **CEMIP** | 0.00433 | 0.00700 | 0.02620 | 0.00000 | 0.06363 | 0.00289 | 0.18849 | 0.00966 | 0.04043 | 0.90003 | 0.13784 | 0.04313 |
| **CEMP1** | 0.00000 | 0.64382 | 1.94094 | 0.42097 | 0.00000 | 0.00000 | 0.08946 | 0.36743 | 2.81926 | 6.64890 | 1.66280 | 0.75345 |
| **CEND1** | 0.16443 | 0.01521 | 0.01170 | 0.08923 | 0.01355 | 0.00002 | 0.02624 | 0.09515 | 0.04590 | 0.01803 | 0.01200 | 0.01914 |
| **CENPA** | 2.20339 | 9.03545 | 0.84253 | 6.98642 | 3.12702 | 4.71369 | 1.85008 | 7.99077 | 0.13725 | 9.83124 | 0.00000 | 9.69809 |
| **CENPB** | 0.11087 | 0.08800 | 1.85343 | 0.18486 | 3.27900 | 0.00000 | 3.82937 | 0.00000 | 0.00000 | 9.06352 | 0.58239 | 0.37989 |
| **CENPBD1** | 0.93738 | 0.94928 | 1.35292 | 0.25715 | 0.48242 | 2.37252 | 0.59541 | 0.48299 | 2.14662 | 0.00000 | 0.52877 | 2.32872 |
| **CENPBD1P1** | 0.03853 | 1.30844 | 1.39248 | 3.52216 | 3.15370 | 0.00000 | 5.83113 | 0.29952 | 5.09989 | 1.66869 | 2.83751 | 3.06967 |
| **CENPC** | 1.11273 | 1.04631 | 1.42849 | 0.19213 | 0.13229 | 10.46063 | 0.00000 | 3.45947 | 4.91907 | 4.35713 | 0.33031 | 1.62628 |
| **CENPE** | 10.79292 | 0.69997 | 13.13910 | 6.87162 | 4.30891 | 29.84400 | 13.18112 | 6.59969 | 7.64915 | 29.88368 | 5.53603 | 9.85093 |
| **CENPF** | 0.53666 | 9.82799 | 11.05405 | 5.57273 | 10.70366 | 28.25402 | 15.01044 | 14.79977 | 26.88941 | 46.80252 | 8.89112 | 26.15632 |
| **CENPH** | 1.24631 | 7.22970 | 1.94400 | 1.44786 | 4.09258 | 5.52714 | 0.92214 | 2.24910 | 0.00000 | 4.50609 | 3.85056 | 6.02830 |
| **CENPI** | 0.82146 | 0.24299 | 0.27576 | 0.00157 | 0.42378 | 4.89085 | 0.00000 | 0.37391 | 2.09802 | 0.00000 | 1.23902 | 0.51421 |
| **CENPJ** | 1.85614 | 3.52515 | 0.71588 | 4.85206 | 1.40818 | 13.36842 | 2.01152 | 3.26786 | 6.09723 | 12.60706 | 2.57841 | 5.70740 |
| **CENPK** | 1.13979 | 2.86761 | 4.59757 | 0.81383 | 0.00000 | 18.83294 | 5.10671 | 4.07308 | 7.11488 | 5.99271 | 5.77474 | 4.94280 |
| **CENPL** | 0.95701 | 2.66795 | 3.56358 | 1.09235 | 1.41687 | 10.68638 | 5.43577 | 7.65104 | 6.03563 | 0.00000 | 4.44245 | 6.12529 |

| | | | | | | | | | | | | |
|---|---|---|---|---|---|---|---|---|---|---|---|---|
| **CENPM** | 0.00000 | 6.47173 | 1.34846 | 6.39552 | 10.10340 | 0.09398 | 2.48589 | 1.56032 | 3.76117 | 0.93651 | 4.94668 | 4.89871 |
| **CENPN** | 1.56408 | 11.00741 | 13.34143 | 10.14653 | 12.38355 | 26.46767 | 17.17406 | 12.09110 | 9.17301 | 6.84616 | 12.28898 | 13.07173 |
| **CENPO** | 3.68136 | 4.16923 | 7.11500 | 3.03235 | 7.60373 | 2.50377 | 5.18467 | 7.38220 | 9.66657 | 1.36782 | 6.40833 | 8.09737 |
| **CENPP** | 6.72260 | 1.74588 | 1.90032 | 0.95602 | 1.04484 | 5.97757 | 0.00000 | 2.73795 | 1.87388 | 0.39586 | 3.93346 | 0.95670 |
| **CENPQ** | 1.46240 | 3.35465 | 3.33855 | 1.43038 | 0.00000 | 10.87409 | 5.60493 | 2.56574 | 7.66425 | 1.82103 | 1.16804 | 8.59851 |
| **CENPT** | 2.17643 | 8.27393 | 12.83795 | 10.19819 | 10.02224 | 0.00000 | 6.81598 | 7.15466 | 18.77092 | 3.90222 | 10.37730 | 4.92831 |
| **CENPU** | 10.58329 | 2.75288 | 7.53610 | 5.07264 | 1.90410 | 24.86792 | 11.62729 | 3.50740 | 11.47044 | 4.96522 | 5.05165 | 9.11315 |
| **CENPV** | 0.02621 | 5.32068 | 2.59877 | 6.71784 | 6.61400 | 0.00000 | 8.77152 | 0.00000 | 0.00540 | 11.44177 | 1.68943 | 2.83276 |
| **CENPW** | 1.97912 | 12.97894 | 11.19235 | 9.92233 | 12.60573 | 23.01329 | 17.79629 | 15.93053 | 8.20261 | 3.45178 | 12.27390 | 20.31810 |
| **CEP104** | 3.26566 | 1.20857 | 2.32744 | 0.82300 | 5.81750 | 3.70618 | 3.98598 | 0.46963 | 0.00000 | 10.30437 | 1.67792 | 1.09210 |
| **CEP112** | 1.17016 | 0.17161 | 0.20469 | 0.36636 | 0.54487 | 0.00000 | 0.04092 | 0.04999 | 0.08240 | 0.31697 | 0.02593 | 0.42545 |
| **CEP120** | 3.17336 | 0.70354 | 0.42663 | 0.10563 | 0.77628 | 3.97708 | 0.53159 | 0.02998 | 0.00000 | 4.06479 | 1.20019 | 0.10630 |
| **CEP128** | 3.19349 | 0.83954 | 0.89731 | 1.03413 | 1.37801 | 2.27737 | 0.00000 | 0.74803 | 0.94813 | 3.30781 | 0.55986 | 1.32392 |
| **CEP131** | 0.77013 | 2.16458 | 0.43973 | 2.08325 | 6.73874 | 0.00000 | 1.34821 | 0.29282 | 7.67355 | 6.23008 | 5.27113 | 0.56100 |
| **CEP135** | 0.61824 | 1.03138 | 0.71727 | 1.42674 | 0.00000 | 6.20921 | 1.76069 | 0.41903 | 4.68074 | 4.53569 | 2.77524 | 4.63870 |
| **CEP152** | 5.40812 | 1.72215 | 0.00477 | 2.32938 | 3.41548 | 7.21918 | 1.46347 | 1.53698 | 3.93355 | 11.85139 | 1.48440 | 3.57056 |
| **CEP162** | 2.88036 | 0.04619 | 1.94251 | 0.38061 | 0.00000 | 5.24558 | 2.05295 | 0.99224 | 1.58019 | 5.66727 | 1.00743 | 1.63118 |
| **CEP164** | 0.51092 | 5.35358 | 0.00000 | 8.75029 | 2.76027 | 0.00000 | 1.21779 | 0.52499 | 1.19962 | 7.82562 | 0.76310 | 14.23356 |
| **CEP170** | 2.14849 | 0.26729 | 0.95295 | 1.05890 | 0.00000 | 0.60267 | 1.27256 | 1.40463 | 1.31090 | 1.85059 | 0.94389 | 1.14434 |
| **CEP170B** | 1.92301 | 0.00309 | 1.95214 | 0.00000 | 2.14046 | 0.17363 | 0.94708 | 0.00000 | 1.36035 | 3.77084 | 0.00000 | 0.00000 |
| **CEP170P1** | 0.00196 | 0.01756 | 0.01170 | 0.04097 | 0.01450 | 0.03443 | 0.01575 | 0.06696 | 0.01324 | 0.00983 | 0.02947 | 0.02657 |
| **CEP19** | 0.19563 | 0.40577 | 0.16387 | 0.40415 | 0.00000 | 0.52441 | 0.20754 | 0.00000 | 0.06547 | 0.00000 | 0.10359 | 2.09640 |
| **CEP192** | 1.32769 | 0.88294 | 0.35484 | 3.26876 | 0.19384 | 8.95900 | 1.17297 | 2.45401 | 1.36190 | 9.33324 | 1.23287 | 3.11242 |
| **CEP250** | 1.69310 | 2.35053 | 3.97431 | 2.01947 | 0.00000 | 0.00000 | 3.52653 | 0.77129 | 0.94381 | 4.36507 | 3.42373 | 2.19107 |
| **CEP290** | 1.10060 | 0.00000 | 1.12514 | 0.00000 | 2.56211 | 5.62693 | 3.99525 | 0.09703 | 3.76932 | 15.57845 | 2.77647 | 0.58855 |
| **CEP350** | 2.86937 | 0.40420 | 4.50133 | 1.91816 | 3.21246 | 14.22115 | 4.21668 | 2.27746 | 2.42039 | 24.63743 | 2.28849 | 3.65550 |
| **CEP41** | 0.95400 | 3.88151 | 2.24874 | 0.18628 | 0.00000 | 4.02727 | 3.04421 | 1.34342 | 0.03236 | 1.28553 | 2.92758 | 2.40420 |
| **CEP44** | 0.04408 | 2.43475 | 1.34320 | 3.58681 | 0.10715 | 9.72377 | 2.18970 | 3.39645 | 1.30873 | 3.76246 | 1.16472 | 4.66861 |
| **CEP55** | 2.00412 | 1.56506 | 5.12947 | 1.18941 | 0.00000 | 6.16804 | 6.44248 | 2.20297 | 0.37078 | 6.62945 | 0.00000 | 3.73637 |
| **CEP57** | 5.88713 | 6.54496 | 5.76754 | 5.79270 | 4.59552 | 21.34902 | 10.46028 | 5.17756 | 7.11984 | 10.76332 | 10.81063 | 8.44477 |
| **CEP57L1** | 0.89533 | 2.12379 | 0.19964 | 0.00000 | 0.24522 | 7.09551 | 1.52039 | 0.93057 | 1.67423 | 2.56808 | 1.92573 | 2.97368 |
| **CEP63** | 2.70720 | 2.49614 | 2.55140 | 2.56592 | 3.19384 | 13.12706 | 4.00322 | 2.54905 | 2.39152 | 5.77134 | 5.78315 | 6.57849 |
| **CEP68** | 1.53923 | 5.87250 | 5.78613 | 0.00000 | 3.02896 | 1.97238 | 4.66050 | 0.18417 | 1.15651 | 0.19327 | 5.32218 | 2.46905 |
| **CEP70** | 1.90971 | 0.31381 | 0.84473 | 0.01884 | 0.23168 | 4.65924 | 1.78207 | 1.13943 | 0.47468 | 0.00000 | 0.00000 | 1.07901 |
| **CEP72** | 0.36202 | 1.72464 | 2.90336 | 1.86420 | 1.39585 | 1.08191 | 0.92549 | 0.79197 | 3.23332 | 3.09717 | 2.46210 | 0.47651 |
| **CEP76** | 2.26275 | 3.10838 | 3.84973 | 3.13292 | 1.07681 | 5.29751 | 4.57166 | 1.51999 | 6.03230 | 0.00000 | 2.43515 | 6.49863 |
| **CEP78** | 4.70819 | 6.12146 | 4.42889 | 2.38067 | 1.81473 | 3.92532 | 1.62745 | 2.68457 | 5.67683 | 2.84720 | 7.74861 | 5.71021 |
| **CEP83** | 1.00304 | 0.37181 | 1.16309 | 0.40116 | 0.22600 | 3.33922 | 0.96341 | 2.95145 | 4.08973 | 2.90720 | 1.01571 | 1.49172 |
| **CEP83-AS1** | 0.09776 | 0.00000 | 0.07373 | 0.00249 | 0.00420 | 0.09526 | 0.00000 | 0.17200 | 0.02687 | 0.01661 | 0.04115 | 0.13738 |
| **CEP85** | 3.20985 | 8.02537 | 17.27866 | 3.10931 | 4.48150 | 5.64093 | 10.15799 | 4.78768 | 20.04648 | 0.00000 | 6.59495 | 6.70156 |
| **CEP85L** | 4.24266 | 2.26867 | 3.28849 | 6.46209 | 0.00000 | 1.67207 | 0.49911 | 19.45093 | 1.02931 | 6.36259 | 0.53393 | 11.09933 |
| **CEP89** | 0.44702 | 0.10671 | 1.44548 | 1.12341 | 0.57117 | 1.00015 | 0.27191 | 0.91382 | 1.89544 | 0.13097 | 0.94361 | 0.32369 |
| **CEP95** | 3.73297 | 5.96730 | 0.50297 | 4.25510 | 1.82803 | 12.09068 | 2.08191 | 6.66769 | 5.82555 | 9.60035 | 2.20350 | 3.58702 |
| **CEP97** | 2.90443 | 0.70966 | 1.69353 | 0.00000 | 0.00000 | 6.43274 | 0.00000 | 5.48735 | 4.11174 | 2.72316 | 0.69184 | 6.65421 |
| **CEPT1** | 0.74722 | 6.70006 | 9.11970 | 2.94033 | 0.00000 | 14.77947 | 0.00000 | 0.65900 | 21.94507 | 4.32542 | 5.26907 | 7.85500 |
| **CERCAM** | 0.79979 | 0.22436 | 1.96868 | 0.00000 | 0.28216 | 0.00000 | 0.00000 | 0.62101 | 2.31806 | 0.07454 | 1.89172 | 0.79500 |
| **CERK** | 0.08267 | 0.12401 | 0.02487 | 0.11866 | 0.02310 | 0.00000 | 0.00439 | 0.00878 | 0.11583 | 0.02334 | 0.00141 | 0.00721 |
| **CERKL** | 1.04830 | 0.08771 | 0.01730 | 0.19296 | 0.02583 | 0.00000 | 0.02078 | 0.12079 | 0.01522 | 0.00000 | 0.01306 | 0.52596 |

| | | | | | | | | | | | | |
|---|---|---|---|---|---|---|---|---|---|---|---|---|
| **CERS1.uid-2288** | 0.05927 | 0.15093 | 0.16851 | 0.04828 | 0.30282 | 0.00000 | 0.00000 | 0.04628 | 0.07965 | 0.29997 | 0.00247 | 0.22657 |
| **CERS2** | 6.52435 | 8.59649 | 7.53761 | 3.92103 | 5.84381 | 12.31444 | 6.43650 | 8.29826 | 11.45451 | 15.58589 | 3.73320 | 16.38912 |
| **CERS3** | 0.16767 | 0.01261 | 0.01307 | 0.01301 | 0.01277 | 0.02093 | 0.01477 | 0.12580 | 0.01661 | 0.01908 | 0.01955 | 0.01166 |
| **CERS4** | 0.00000 | 0.16940 | 1.06229 | 0.00000 | 9.14813 | 0.00000 | 2.11854 | 0.11894 | 15.55434 | 4.36911 | 14.20342 | 2.48757 |
| **CERS5** | 0.00000 | 21.83036 | 18.92112 | 3.33252 | 19.43220 | 18.85931 | 7.38006 | 19.33634 | 30.06054 | 7.84283 | 17.64702 | 33.26501 |
| **CERS6** | 1.96795 | 0.47624 | 0.04938 | 2.36653 | 0.61308 | 0.11187 | 0.00000 | 0.06771 | 0.00000 | 0.39338 | 0.00000 | 0.24302 |
| **CES1** | 0.18646 | 0.01660 | 0.99299 | 0.00743 | 0.02320 | 0.00000 | 0.00033 | 0.02213 | 0.00000 | 0.03802 | 0.01562 | 0.01982 |
| **CES1P1** | 0.00640 | 0.01597 | 0.01368 | 0.01231 | 0.01686 | 0.01385 | 0.03506 | 0.01215 | 0.01256 | 0.01396 | 0.01567 | 0.01635 |
| **CES2** | 0.04704 | 4.01734 | 3.06947 | 4.15560 | 3.10251 | 0.00000 | 2.97828 | 0.23918 | 2.46112 | 3.98039 | 8.37890 | 1.64604 |
| **CES3** | 0.00456 | 0.73661 | 0.00000 | 0.11285 | 1.02302 | 0.05812 | 0.00000 | 0.01296 | 0.00000 | 0.17099 | 0.17822 | 0.12211 |
| **CES4A** | 0.27041 | 0.37697 | 0.07464 | 0.29465 | 0.02165 | 0.07921 | 0.05222 | 0.00000 | 0.00000 | 0.00000 | 0.15944 | 0.36867 |
| **CES5A** | 0.00542 | 0.01186 | 0.02072 | 0.01696 | 0.06714 | 0.00000 | 0.06248 | 0.01063 | 0.00000 | 0.05018 | 0.00992 | 0.01560 |
| **CES5AP1** | 0.00623 | 0.01492 | 0.03082 | 0.01202 | 0.01523 | 0.01748 | 0.01570 | 0.01047 | 0.01424 | 0.01778 | 0.01580 | 0.01607 |
| **CETN1** | 0.18165 | 0.15188 | 0.00000 | 0.00000 | 0.03570 | 0.19682 | 0.00000 | 0.00000 | 0.10197 | 0.03188 | 0.03414 | 0.38519 |
| **CETN2** | 5.26937 | 4.13697 | 3.43886 | 1.57176 | 4.98599 | 12.12665 | 6.32297 | 6.35055 | 3.16966 | 2.26438 | 7.53252 | 9.53821 |
| **CETN3** | 1.20931 | 3.84518 | 6.39443 | 1.54975 | 2.61483 | 17.40253 | 5.13522 | 0.30571 | 0.77564 | 6.23250 | 6.65578 | 6.38659 |
| **CETN4P** | 0.03845 | 0.06256 | 0.03489 | 0.01897 | 0.03863 | 0.12762 | 0.00000 | 0.01290 | 0.02008 | 0.00000 | 0.01151 | 0.15401 |
| **CETP** | 0.00891 | 0.01297 | 0.04860 | 0.02326 | 0.01138 | 0.00803 | 0.00264 | 0.01061 | 0.02999 | 0.07592 | 0.01141 | 0.01870 |
| **CFB** | 0.01628 | 0.15946 | 0.05491 | 0.14196 | 0.00000 | 0.00000 | 0.00000 | 0.24127 | 0.10500 | 0.37218 | 0.02419 | 0.34315 |
| **CFD** | 0.43395 | 2.12726 | 0.10475 | 0.73840 | 3.09535 | 0.00000 | 1.38372 | 1.06769 | 2.87300 | 0.57934 | 2.48735 | 14.98196 |
| **CFDP1** | 2.15152 | 2.99456 | 6.91335 | 6.87169 | 5.68447 | 15.23132 | 12.83993 | 8.19191 | 8.80402 | 7.21692 | 6.48972 | 10.40641 |
| **CFH** | 0.00586 | 0.01096 | 0.01719 | 0.01125 | 0.01051 | 0.02602 | 0.01987 | 0.00900 | 0.01763 | 0.01846 | 0.01771 | 0.04450 |
| **CFHR1** | 0.00798 | 0.01443 | 0.01814 | 0.03059 | 0.01183 | 0.01604 | 0.01883 | 0.01344 | 0.02867 | 0.01319 | 0.01218 | 0.01702 |
| **CFHR3** | 0.00897 | 0.02767 | 0.01517 | 0.00617 | 0.00238 | 0.06706 | 0.02088 | 0.00970 | 0.00634 | 0.01773 | 0.01174 | 0.01614 |
| **CFHR4** | 0.00575 | 0.01447 | 0.01385 | 0.00804 | 0.00702 | 0.02707 | 0.01403 | 0.01029 | 0.03895 | 0.04067 | 0.00885 | 0.01410 |
| **CFL1** | 15.41604 | 48.08951 | 47.56299 | 46.16955 | 55.88759 | 46.35580 | 64.61910 | 28.36669 | 65.91196 | 17.17402 | 45.11850 | 51.35649 |
| **CFL1P1** | 0.28832 | 0.03742 | 0.47929 | 0.25684 | 0.13270 | 0.00029 | 0.15740 | 0.09144 | 0.00000 | 0.30979 | 0.11249 | 0.09791 |
| **CFL2** | 1.18905 | 4.99551 | 2.89522 | 0.00000 | 1.32595 | 32.34461 | 6.84430 | 5.43155 | 10.95047 | 1.34748 | 10.06551 | 18.24171 |
| **CFLAR** | 0.41328 | 16.28587 | 1.28747 | 19.98588 | 0.00000 | 2.44589 | 0.00000 | 0.57227 | 0.00000 | 2.09331 | 0.45114 | 15.29268 |
| **CFLAR-AS1** | 0.00438 | 0.05731 | 0.01721 | 0.12929 | 0.00000 | 0.06729 | 0.00000 | 0.01568 | 0.00750 | 0.04576 | 0.01024 | 0.10725 |
| **CFP** | 0.01478 | 0.02361 | 0.14257 | 0.01063 | 0.03508 | 0.00000 | 0.03154 | 0.02897 | 0.07222 | 0.02476 | 0.00857 | 0.01387 |
| **CFTR** | 0.53115 | 0.04779 | 1.46715 | 0.05474 | 0.32709 | 0.00000 | 0.00000 | 0.02455 | 0.00000 | 0.06041 | 0.01136 | 0.01443 |
| **CGA** | 0.00000 | 0.00000 | 75.75761 | 0.00000 | 0.00000 | 0.10261 | 13.01717 | 0.10120 | 0.23630 | 0.32187 | 0.00000 | 1.30075 |
| **CGB** | 0.24755 | 0.01064 | 5.54520 | 0.03052 | 0.00269 | 0.00000 | 0.00000 | 0.26864 | 0.00000 | 0.08577 | 0.01400 | 0.35934 |
| **CGB1** | 0.00655 | 0.01652 | 0.04435 | 0.01399 | 0.01694 | 0.00000 | 0.07047 | 0.01094 | 0.00000 | 0.02697 | 0.02713 | 0.01853 |
| **CGB2** | 0.00586 | 0.01259 | 0.03651 | 0.01160 | 0.01274 | 0.02269 | 0.03578 | 0.01179 | 0.00306 | 0.01989 | 0.01966 | 0.02133 |
| **CGB5** | 0.47622 | 0.00997 | 5.33514 | 0.00979 | 0.01379 | 0.00000 | 0.00000 | 0.04854 | 0.00000 | 0.01970 | 0.01542 | 0.06170 |
| **CGB7** | 0.56779 | 0.01103 | 1.91848 | 0.01975 | 0.00000 | 0.01092 | 0.00570 | 0.29049 | 0.00000 | 0.07072 | 0.02481 | 0.16833 |
| **CGB8** | 0.30338 | 0.00193 | 3.88036 | 0.00745 | 0.01372 | 0.00000 | 0.00000 | 0.27784 | 0.00000 | 0.03389 | 0.01409 | 0.29741 |
| **CGGBP1** | 4.70202 | 4.66611 | 9.93722 | 5.06208 | 3.29910 | 15.76883 | 9.99940 | 4.09136 | 2.86606 | 5.11963 | 4.16412 | 6.21746 |
| **CGN** | 3.03019 | 12.27520 | 3.70442 | 10.87006 | 12.34702 | 0.00000 | 0.00000 | 0.45697 | 3.88836 | 6.39508 | 5.09771 | 0.44728 |
| **CGNL1** | 1.31998 | 1.54634 | 0.26707 | 4.73749 | 0.01130 | 0.00000 | 0.02391 | 0.25585 | 0.00000 | 0.44562 | 0.00000 | 4.77963 |
| **CGREF1** | 2.54582 | 0.23180 | 0.53037 | 0.03361 | 0.46061 | 0.00000 | 0.00000 | 0.50249 | 0.99782 | 0.00000 | 0.52451 | 0.48517 |
| **CGRRF1** | 0.00000 | 1.95926 | 0.12844 | 1.67801 | 0.00000 | 8.87003 | 0.00000 | 0.84886 | 4.56005 | 1.65023 | 1.34073 | 1.90533 |
| **CHAC1** | 5.93007 | 1.22720 | 5.15228 | 0.00000 | 0.00000 | 0.00000 | 8.57291 | 0.64022 | 18.15194 | 0.58952 | 0.71730 | 3.12125 |
| **CHAC2** | 0.04286 | 0.63585 | 0.00000 | 1.08803 | 1.59280 | 4.55854 | 0.00000 | 0.01441 | 1.56536 | 0.93402 | 4.40119 | 0.00000 |
| **CHAD** | 0.02960 | 0.08941 | 0.01496 | 0.03334 | 0.07710 | 0.00362 | 0.03661 | 0.02807 | 0.02223 | 0.00000 | 0.00888 | 0.03248 |
| **CHADL** | 0.02055 | 0.03088 | 0.14108 | 0.03865 | 0.00994 | 0.04962 | 0.05021 | 0.00828 | 0.05712 | 0.02165 | 0.05392 | 0.03930 |

| | | | | | | | | | | | | |
|---|---|---|---|---|---|---|---|---|---|---|---|---|
| **CHAF1A** | 13.25336 | 5.49013 | 4.25733 | 6.97114 | 11.10274 | 0.00749 | 12.38728 | 7.58121 | 1.36797 | 22.32953 | 4.44714 | 7.43897 |
| **CHAF1B** | 3.67527 | 10.15771 | 4.50397 | 2.64649 | 4.61398 | 7.42721 | 6.93929 | 2.57107 | 4.96352 | 0.00000 | 7.57725 | 4.02350 |
| **CHAMP1** | 7.48631 | 0.81325 | 0.96923 | 0.00000 | 1.16956 | 2.01796 | 1.51256 | 0.79945 | 0.60039 | 2.70989 | 2.68287 | 2.41361 |
| **CHAT** | 0.01257 | 0.01703 | 0.01972 | 0.01398 | 0.02272 | 0.00749 | 0.02100 | 0.04727 | 0.01990 | 0.00360 | 0.01577 | 0.05028 |
| **CHCHD1** | 0.52518 | 11.04707 | 11.31142 | 7.89284 | 8.52509 | 24.05113 | 23.94008 | 0.24665 | 20.10920 | 10.84190 | 16.36133 | 16.19321 |
| **CHCHD10** | 0.02069 | 2.46084 | 3.10910 | 5.26731 | 15.96642 | 0.00000 | 11.25466 | 0.03315 | 2.29362 | 21.96277 | 4.24150 | 0.33969 |
| **CHCHD2** | 5.85085 | 32.17338 | 20.72129 | 20.40769 | 29.48691 | 25.22189 | 32.28042 | 17.84870 | 31.32851 | 12.24739 | 30.01705 | 30.03896 |
| **CHCHD3** | 1.25457 | 13.49207 | 9.37199 | 14.71552 | 7.05963 | 20.25767 | 17.46205 | 1.14276 | 15.85517 | 0.00000 | 14.39968 | 1.20975 |
| **CHCHD4** | 0.08684 | 15.28638 | 2.90356 | 7.74605 | 7.16238 | 17.70217 | 6.93918 | 0.00000 | 8.49203 | 0.00000 | 15.21226 | 9.88881 |
| **CHCHD5** | 0.80674 | 10.46715 | 5.20774 | 7.39563 | 12.16146 | 1.12423 | 6.49638 | 1.06589 | 4.58820 | 0.00000 | 6.68580 | 9.59344 |
| **CHCHD6** | 3.30811 | 1.68593 | 1.38947 | 1.26543 | 0.95538 | 2.76757 | 2.40930 | 2.42857 | 1.75931 | 0.00000 | 4.68123 | 0.71800 |
| **CHCHD7** | 6.93095 | 10.79965 | 8.56831 | 5.58089 | 15.78159 | 27.50724 | 12.88020 | 9.90816 | 12.56869 | 6.66947 | 16.74891 | 15.02846 |
| **CHD1** | 0.28656 | 2.53207 | 7.60578 | 0.38817 | 2.52316 | 20.51971 | 12.70917 | 10.79260 | 4.31420 | 29.00242 | 7.80374 | 7.69533 |
| **CHD1L** | 0.14842 | 6.38700 | 2.98120 | 10.41595 | 1.67148 | 9.39619 | 2.35216 | 0.20844 | 5.77308 | 4.52677 | 3.26449 | 2.92202 |
| **CHD2** | 4.44486 | 0.00000 | 6.85195 | 4.47774 | 5.39182 | 4.08753 | 7.91420 | 4.18430 | 9.12619 | 23.03774 | 2.26932 | 3.57532 |
| **CHD3** | 2.94987 | 0.00000 | 2.28114 | 3.03066 | 3.73148 | 0.00000 | 2.80392 | 0.14832 | 2.62222 | 9.40822 | 0.00000 | 1.02986 |
| **CHD4** | 6.98608 | 9.06859 | 8.96533 | 15.83010 | 23.55183 | 15.32530 | 23.83257 | 2.75476 | 8.40241 | 45.14878 | 5.97743 | 11.34939 |
| **CHD5** | 0.29410 | 0.08109 | 0.05258 | 0.11116 | 0.06790 | 0.00000 | 0.04441 | 0.02381 | 0.00000 | 0.26123 | 0.05080 | 0.55594 |
| **CHD6** | 3.05080 | 4.27725 | 0.92974 | 8.07134 | 4.78597 | 5.17269 | 1.39840 | 2.71224 | 3.86155 | 10.75221 | 1.49212 | 7.41735 |
| **CHD7** | 9.71887 | 8.98728 | 12.78201 | 0.00000 | 12.88933 | 13.85558 | 18.35845 | 6.23926 | 12.86767 | 27.69549 | 9.29307 | 13.87019 |
| **CHD8** | 5.57971 | 0.00000 | 8.53999 | 3.49179 | 11.53549 | 6.46559 | 8.90370 | 7.56614 | 8.23820 | 27.22629 | 3.57389 | 9.50867 |
| **CHD9** | 0.33332 | 0.12537 | 0.33082 | 1.82200 | 0.00028 | 0.89673 | 0.16914 | 0.39744 | 0.00000 | 4.50901 | 1.10359 | 0.44474 |
| **CHDC2** | 0.02092 | 0.08349 | 0.00000 | 0.08565 | 0.00000 | 0.03915 | 0.00000 | 0.03666 | 0.11452 | 0.08159 | 0.00000 | 0.04281 |
| **CHDH** | 0.00152 | 1.02819 | 0.00000 | 4.22971 | 2.30558 | 0.00000 | 1.10342 | 0.01076 | 0.00000 | 2.27762 | 0.77253 | 0.02806 |
| **CHEK1** | 11.07967 | 8.27105 | 4.86368 | 6.51007 | 4.56591 | 15.53675 | 10.71884 | 1.65137 | 7.36577 | 5.65274 | 7.89844 | 11.35067 |
| **CHEK2** | 1.22986 | 3.99321 | 8.20142 | 4.44968 | 0.00000 | 5.77586 | 6.47023 | 0.60806 | 8.74599 | 0.00000 | 5.16171 | 2.67165 |
| **CHEK2P2** | 0.78710 | 0.01264 | 0.01152 | 0.00501 | 0.01325 | 0.05537 | 0.00845 | 0.02293 | 0.04245 | 0.00000 | 0.01236 | 0.09279 |
| **CHERP** | 4.09876 | 6.62798 | 6.90238 | 5.69137 | 6.47414 | 0.13963 | 7.17159 | 2.37011 | 11.02960 | 8.71160 | 4.86264 | 7.99476 |
| **CHFR** | 2.70908 | 4.71424 | 5.56185 | 7.41513 | 10.08520 | 3.63440 | 1.04208 | 1.94176 | 8.38186 | 1.99154 | 4.39968 | 2.63157 |
| **CHGA** | 0.44733 | 2.58827 | 0.00000 | 1.46945 | 0.95292 | 0.00000 | 0.00027 | 0.00000 | 0.16137 | 1.70610 | 0.00000 | 2.49226 |
| **CHGB** | 0.01212 | 0.01735 | 0.04128 | 0.01197 | 0.01257 | 0.01604 | 0.03154 | 0.01421 | 0.03538 | 0.00760 | 0.01140 | 0.03867 |
| **CHI3L1** | 0.01108 | 0.00000 | 0.00000 | 0.00000 | 0.04522 | 0.05166 | 0.27694 | 0.00927 | 0.56568 | 0.09063 | 0.76348 | 0.01654 |
| **CHI3L2** | 0.00076 | 2.46391 | 0.00000 | 7.16515 | 9.00442 | 14.22722 | 0.00000 | 0.84892 | 0.08151 | 0.00000 | 8.44277 | 10.45035 |
| **CHIA** | 0.00467 | 0.01772 | 0.00241 | 0.98876 | 0.02169 | 0.01859 | 0.06669 | 0.00897 | 0.00000 | 0.04958 | 0.01049 | 0.02606 |
| **CHIAP2** | 0.00406 | 0.01247 | 0.00823 | 0.00000 | 0.00000 | 0.03043 | 0.07916 | 0.00995 | 0.00000 | 0.18395 | 0.00196 | 0.00881 |
| **CHIC1** | 1.67629 | 0.00000 | 0.00000 | 0.08275 | 0.12125 | 0.20010 | 0.46122 | 0.34137 | 0.00000 | 0.11985 | 0.00000 | 1.04777 |
| **CHIC2** | 2.23214 | 3.12204 | 3.52418 | 3.64085 | 1.00777 | 4.23642 | 1.69994 | 5.75154 | 7.85410 | 2.28500 | 1.90893 | 5.01285 |
| **CHID1** | 1.44729 | 18.79471 | 12.67405 | 9.37282 | 23.65274 | 2.58265 | 0.97720 | 2.41768 | 21.53013 | 3.43113 | 19.66532 | 10.93959 |
| **CHIT1** | 0.00584 | 0.02438 | 0.02418 | 0.01740 | 0.00351 | 0.08402 | 0.02603 | 0.00983 | 0.01566 | 0.01576 | 0.01196 | 0.01039 |
| **CHKA** | 0.00679 | 0.00000 | 3.84455 | 0.16086 | 1.42263 | 0.31837 | 6.17818 | 1.07816 | 0.36238 | 3.36115 | 0.00000 | 0.06821 |
| **CHKB** | 0.00771 | 0.23236 | 0.05921 | 0.38366 | 0.61206 | 0.08555 | 0.35162 | 0.00560 | 0.00000 | 2.41353 | 0.59747 | 0.01082 |
| **CHKB-AS1** | 0.12031 | 0.10404 | 0.04958 | 0.11056 | 0.16568 | 0.00000 | 0.01947 | 0.01166 | 0.08074 | 0.23357 | 0.34475 | 0.22347 |
| **CHL1** | 0.00242 | 0.00854 | 0.04463 | 0.00077 | 0.00000 | 0.05383 | 0.04055 | 0.03759 | 0.00000 | 0.06249 | 0.01108 | 0.04077 |
| **CHM** | 0.28140 | 0.43997 | 2.63677 | 1.05386 | 0.00000 | 3.16571 | 6.63627 | 0.52048 | 0.46471 | 1.51345 | 1.95422 | 4.46972 |
| **CHML** | 0.00000 | 0.00000 | 7.11188 | 0.00000 | 0.87484 | 8.55958 | 0.92053 | 1.03736 | 0.01385 | 10.16207 | 2.76273 | 2.13206 |
| **CHMP1A** | 3.69193 | 21.82372 | 20.21901 | 11.31269 | 16.17593 | 0.00000 | 26.95103 | 13.32905 | 26.09239 | 0.00000 | 13.97535 | 23.35683 |
| **CHMP1B** | 1.80805 | 11.07997 | 21.03352 | 2.51253 | 2.55301 | 28.16784 | 14.85123 | 15.63727 | 16.71768 | 4.93823 | 16.16792 | 19.03304 |
| **CHMP1B2P** | 0.00737 | 0.27257 | 0.31587 | 0.12821 | 0.24663 | 0.00194 | 0.40186 | 0.00377 | 0.52005 | 0.51405 | 0.14127 | 0.20958 |

| | | | | | | | | | | | | |
|---|---|---|---|---|---|---|---|---|---|---|---|---|
| **CHMP2A** | 0.00000 | 8.89787 | 8.36688 | 9.82085 | 12.91564 | 6.39478 | 8.89815 | 4.72508 | 1.45338 | 3.25990 | 10.53918 | 10.70638 |
| **CHMP2B** | 5.42401 | 5.37146 | 14.50560 | 2.70505 | 5.02376 | 26.17043 | 19.28664 | 1.85084 | 11.90894 | 4.80221 | 6.60125 | 8.25261 |
| **CHMP3** | 0.73136 | 0.59233 | 0.79120 | 0.53334 | 0.84768 | 0.20370 | 0.46190 | 0.17665 | 0.66132 | 0.67385 | 0.72110 | 0.64577 |
| **CHMP4A** | 0.20395 | 4.02518 | 6.76276 | 6.71634 | 9.50627 | 12.83313 | 14.25384 | 0.00000 | 7.20823 | 1.53495 | 11.33164 | 0.42530 |
| **CHMP4B** | 0.09090 | 0.00000 | 0.00000 | 0.22824 | 11.77113 | 0.00000 | 0.45937 | 0.00000 | 0.00000 | 12.85434 | 0.00000 | 1.37113 |
| **CHMP4C** | 0.12213 | 0.18350 | 3.36057 | 2.97747 | 0.37819 | 0.60864 | 3.80673 | 0.00000 | 2.30059 | 0.00000 | 2.25300 | 0.00000 |
| **CHMP5** | 1.45602 | 12.65209 | 16.43311 | 5.97557 | 1.54443 | 26.83332 | 19.14366 | 11.44834 | 11.79506 | 0.00958 | 7.66019 | 19.76964 |
| **CHMP6** | 0.28797 | 11.06701 | 1.25851 | 3.33648 | 6.21024 | 3.63377 | 1.54341 | 1.63527 | 1.72960 | 0.00000 | 3.79242 | 7.49371 |
| **CHMP7** | 2.19105 | 7.94698 | 10.13118 | 3.31574 | 3.99877 | 0.00026 | 10.02933 | 1.42623 | 15.44637 | 0.00000 | 5.92047 | 6.09992 |
| **CHN1** | 0.15180 | 0.53066 | 0.18154 | 0.44762 | 0.43932 | 0.71226 | 0.00000 | 0.07505 | 0.58291 | 0.22037 | 0.00000 | 0.12327 |
| **CHN2** | 3.42674 | 1.10161 | 2.02089 | 0.59908 | 0.89599 | 0.00000 | 0.21843 | 0.24090 | 0.00000 | 0.22384 | 0.44881 | 1.61124 |
| **CHODL** | 0.07340 | 0.75766 | 0.01411 | 0.25962 | 0.00000 | 0.10356 | 0.08008 | 0.01191 | 0.01633 | 0.00000 | 0.04971 | 0.00415 |
| **CHORDC1** | 0.00000 | 9.42685 | 15.72731 | 3.22303 | 0.30752 | 32.40331 | 20.16270 | 9.85985 | 17.01515 | 6.22246 | 14.69329 | 10.30257 |
| **CHP1** | 4.93025 | 23.60918 | 15.18549 | 30.52422 | 20.93122 | 50.60038 | 27.07001 | 25.17168 | 10.25214 | 14.96702 | 29.37714 | 24.12619 |
| **CHP2** | 0.00391 | 0.03568 | 0.01388 | 0.00719 | 0.00359 | 0.01984 | 0.01517 | 0.14997 | 0.00978 | 0.02969 | 0.05740 | 0.01294 |
| **CHPF** | 1.21665 | 0.55473 | 5.45158 | 0.57528 | 3.40848 | 0.00000 | 0.00000 | 0.00000 | 3.52030 | 3.15373 | 0.00000 | 0.33417 |
| **CHPF2** | 0.23318 | 0.43995 | 1.92489 | 1.14215 | 1.79522 | 0.00000 | 0.00000 | 0.21852 | 1.07615 | 2.03361 | 0.13885 | 0.27892 |
| **CHPT1** | 0.36128 | 2.45217 | 4.84546 | 0.00000 | 2.31223 | 10.81169 | 2.00894 | 0.07317 | 5.92286 | 8.08314 | 1.39702 | 1.04026 |
| **CHRAC1** | 0.30688 | 11.61524 | 7.04551 | 2.73656 | 4.69713 | 6.41276 | 4.62538 | 1.50606 | 5.32599 | 5.72083 | 6.01034 | 9.77175 |
| **CHRD** | 0.01864 | 0.24013 | 0.07025 | 0.49809 | 0.54772 | 0.00000 | 0.03602 | 0.02274 | 0.00000 | 0.64759 | 0.00000 | 1.25681 |
| **CHRDL1** | 0.36928 | 0.20270 | 0.01529 | 0.00261 | 0.00204 | 0.01436 | 0.02016 | 0.04631 | 0.01904 | 0.01247 | 0.01140 | 0.00113 |
| **CHRDL2** | 0.19304 | 0.01284 | 0.00863 | 0.08939 | 0.00925 | 0.00777 | 0.01308 | 0.04463 | 0.03460 | 0.01317 | 0.01210 | 0.09107 |
| **CHRFAM7A** | 0.02901 | 0.03211 | 0.01484 | 0.00662 | 0.00645 | 0.05868 | 0.01272 | 0.02145 | 0.01779 | 0.00881 | 0.01166 | 0.08800 |
| **CHRM1** | 0.00200 | 0.01232 | 0.28091 | 0.03554 | 0.03427 | 0.00000 | 0.00769 | 0.12593 | 0.00000 | 0.01060 | 0.10399 | 0.25205 |
| **CHRM2** | 0.00236 | 0.05950 | 0.04424 | 0.00000 | 0.00000 | 0.01104 | 0.00260 | 0.03773 | 0.01621 | 0.25353 | 0.01031 | 0.00387 |
| **CHRM3** | 0.02912 | 0.14639 | 0.25373 | 0.51015 | 0.02269 | 0.00000 | 0.00798 | 0.02013 | 0.73178 | 0.00812 | 0.09671 | 0.05254 |
| **CHRM3-AS2** | 0.00633 | 0.01340 | 0.01881 | 0.00935 | 0.01053 | 0.03092 | 0.01857 | 0.01054 | 0.04601 | 0.00811 | 0.01251 | 0.01581 |
| **CHRM4** | 0.00632 | 0.06807 | 0.02552 | 0.02994 | 0.01547 | 0.01593 | 0.04569 | 0.01068 | 0.01284 | 0.00000 | 0.02022 | 0.01349 |
| **CHRM5** | 0.00770 | 0.38152 | 0.96498 | 0.00109 | 0.07380 | 0.07470 | 0.00000 | 0.01760 | 0.40747 | 0.00000 | 0.02452 | 1.56823 |
| **CHRNA1** | 0.00652 | 0.02383 | 0.01706 | 0.01366 | 0.04031 | 0.00000 | 0.03155 | 0.00942 | 0.06177 | 0.10296 | 0.05897 | 0.01902 |
| **CHRNA10** | 0.07454 | 0.00000 | 0.20296 | 0.21719 | 0.00000 | 0.00000 | 0.22475 | 0.18888 | 1.13492 | 1.77072 | 0.52822 | 5.66918 |
| **CHRNA2** | 0.00584 | 0.01289 | 0.02551 | 0.02386 | 0.01245 | 0.02451 | 0.02750 | 0.01042 | 0.01570 | 0.02953 | 0.01177 | 0.01471 |
| **CHRNA3** | 0.94440 | 0.34844 | 0.38260 | 0.07676 | 0.18563 | 0.00000 | 0.00000 | 0.06944 | 0.00000 | 0.27321 | 0.00468 | 0.02774 |
| **CHRNA4** | 0.00217 | 0.00863 | 0.00414 | 0.01296 | 0.00290 | 0.00648 | 0.01396 | 0.03633 | 0.02280 | 0.28090 | 0.00936 | 0.00868 |
| **CHRNA5** | 0.66523 | 1.79435 | 2.64260 | 0.63151 | 0.00031 | 2.51457 | 0.00000 | 0.72463 | 1.08295 | 1.00681 | 0.00000 | 1.25785 |
| **CHRNA6** | 0.09725 | 0.00793 | 0.01822 | 0.00607 | 0.04241 | 0.05056 | 0.01583 | 0.00687 | 0.00738 | 0.00420 | 0.01495 | 0.04008 |
| **CHRNA7** | 0.83273 | 0.49183 | 0.03219 | 0.10605 | 0.01229 | 0.05645 | 0.00000 | 0.60826 | 0.15559 | 0.00000 | 0.01786 | 0.48888 |
| **CHRNA9** | 0.00609 | 0.04253 | 0.01982 | 0.01248 | 0.02040 | 0.00019 | 0.04461 | 0.00942 | 0.01431 | 0.00000 | 0.01752 | 0.05166 |
| **CHRNB1** | 0.00549 | 1.07927 | 2.57382 | 0.04617 | 0.00000 | 0.00000 | 0.75711 | 6.45583 | 2.58494 | 0.43992 | 2.86249 | 4.45111 |
| **CHRNB2** | 0.01166 | 0.02719 | 0.13819 | 0.06271 | 0.00000 | 0.00000 | 0.00009 | 0.00419 | 0.03399 | 0.26717 | 0.02959 | 0.16341 |
| **CHRNB3** | 0.00590 | 0.02267 | 0.01838 | 0.02703 | 0.07409 | 0.01745 | 0.08414 | 0.00988 | 0.00000 | 0.02553 | 0.01704 | 0.22253 |
| **CHRNB4** | 1.26047 | 0.00486 | 0.04129 | 0.09734 | 0.00000 | 0.05141 | 0.00000 | 0.00000 | 0.00407 | 0.04573 | 0.01141 | 0.05939 |
| **CHRND** | 0.00405 | 0.00705 | 0.00467 | 0.01701 | 0.00853 | 0.02411 | 0.01256 | 0.00994 | 0.02759 | 0.19782 | 0.00318 | 0.00985 |
| **CHRNE** | 1.88075 | 0.17225 | 0.15479 | 0.13069 | 0.30152 | 0.00000 | 0.01173 | 0.15186 | 0.32439 | 0.08651 | 0.14265 | 0.27863 |
| **CHRNG** | 0.00594 | 0.01376 | 0.01397 | 0.01317 | 0.01226 | 0.01820 | 0.02300 | 0.01057 | 0.03317 | 0.03217 | 0.01525 | 0.01521 |
| **CHST1** | 0.00118 | 0.38219 | 0.00000 | 0.16225 | 0.03950 | 0.07099 | 0.00000 | 0.02027 | 0.00000 | 0.29014 | 0.01002 | 0.21234 |
| **CHST10** | 2.00064 | 0.43453 | 0.75635 | 0.00000 | 0.17080 | 0.00000 | 0.32361 | 0.10351 | 0.31202 | 0.00000 | 0.10652 | 0.00000 |
| **CHST11** | 0.00000 | 0.62001 | 1.19193 | 0.60748 | 0.70189 | 0.73109 | 0.00000 | 0.09119 | 0.00000 | 2.44916 | 0.00000 | 0.00000 |

| | | | | | | | | | | | | |
|---|---|---|---|---|---|---|---|---|---|---|---|---|
| **CHST12** | 3.27067 | 14.62243 | 3.12543 | 20.32220 | 6.58897 | 0.00000 | 0.00000 | 3.66951 | 7.23814 | 3.21707 | 4.90944 | 5.24915 |
| **CHST13** | 0.08513 | 0.01786 | 0.03087 | 0.01230 | 0.01777 | 0.02061 | 0.01643 | 0.03502 | 0.02205 | 0.01441 | 0.01130 | 0.01432 |
| **CHST14** | 0.02087 | 0.00000 | 2.21396 | 0.12884 | 2.63711 | 0.86933 | 0.37070 | 0.00952 | 2.83774 | 2.76480 | 0.48498 | 0.01287 |
| **CHST15** | 1.58345 | 4.46331 | 1.19339 | 1.22575 | 1.86641 | 0.00000 | 2.14380 | 1.86406 | 1.98416 | 0.00000 | 2.53045 | 7.31918 |
| **CHST2** | 0.18481 | 6.86509 | 0.00000 | 15.54901 | 11.05349 | 3.54555 | 0.00201 | 0.00000 | 0.00000 | 10.12502 | 0.90313 | 1.30725 |
| **CHST3** | 0.00563 | 0.14717 | 0.18135 | 1.44857 | 0.05956 | 0.00000 | 0.01162 | 0.00968 | 0.00000 | 0.00000 | 0.09235 | 0.01522 |
| **CHST4** | 0.88315 | 1.91848 | 0.04467 | 0.16756 | 0.00000 | 0.00000 | 0.11609 | 0.00000 | 0.00000 | 0.06768 | 0.00000 | 3.99889 |
| **CHST5** | 0.01118 | 0.40550 | 0.02038 | 0.13173 | 0.00000 | 0.00000 | 0.00441 | 0.01415 | 0.04035 | 0.03086 | 0.03548 | 0.05517 |
| **CHST6** | 0.07223 | 1.73369 | 0.00000 | 0.62826 | 0.74587 | 0.19131 | 0.00000 | 0.77418 | 0.00000 | 0.68938 | 0.09461 | 1.98421 |
| **CHST7** | 0.03651 | 0.38531 | 0.39387 | 2.57490 | 0.11672 | 0.00000 | 0.00000 | 0.08774 | 0.39179 | 0.64135 | 0.04914 | 0.49287 |
| **CHST8** | 0.00202 | 2.93381 | 0.41367 | 0.42060 | 1.48590 | 0.00000 | 0.00000 | 0.02219 | 0.00000 | 1.06197 | 0.05196 | 0.29237 |
| **CHST9** | 0.00544 | 0.03632 | 0.14119 | 1.27051 | 0.00000 | 0.05700 | 0.00000 | 0.00921 | 0.00000 | 0.01855 | 0.01418 | 0.00926 |
| **CHSY1** | 2.28001 | 1.26806 | 2.67640 | 0.00000 | 1.11348 | 6.67384 | 5.93460 | 1.65300 | 2.09831 | 6.28286 | 1.40033 | 2.15498 |
| **CHSY3** | 0.02199 | 0.01488 | 0.01735 | 0.01040 | 0.01983 | 0.01639 | 0.01802 | 0.06834 | 0.01577 | 0.02153 | 0.01391 | 0.04669 |
| **CHTF18** | 0.07168 | 4.07958 | 3.52288 | 3.69802 | 4.98895 | 0.00000 | 0.00000 | 0.26116 | 2.62280 | 7.05798 | 3.07838 | 0.79262 |
| **CHTF8** | 1.30552 | 11.47088 | 11.56989 | 7.16052 | 4.34987 | 0.00000 | 12.17994 | 3.73500 | 13.51523 | 0.00000 | 13.61198 | 14.64388 |
| **CHTOP** | 5.17709 | 28.41894 | 21.79549 | 16.50802 | 16.91882 | 18.14110 | 32.61351 | 26.46795 | 25.70057 | 6.55344 | 26.86669 | 29.52257 |
| **CHUK** | 0.58126 | 2.36889 | 5.68509 | 0.66677 | 2.36827 | 9.73442 | 8.06351 | 0.43227 | 2.46378 | 13.75261 | 3.58546 | 1.32571 |
| **CHURC1** | 0.00000 | 2.33871 | 3.93542 | 5.14359 | 5.25806 | 18.05021 | 6.26492 | 0.69717 | 1.48029 | 9.08375 | 7.33261 | 9.64660 |
| **CIAO1** | 1.25878 | 20.05116 | 20.00903 | 14.20600 | 18.01798 | 11.28205 | 18.86930 | 2.69458 | 22.84213 | 4.78732 | 19.54143 | 13.86366 |
| **CIAPIN1** | 4.65671 | 17.35570 | 15.41580 | 9.96533 | 12.45082 | 10.61638 | 15.02346 | 5.86357 | 21.37065 | 0.00000 | 19.16051 | 10.82714 |
| **CIART** | 4.69223 | 0.04136 | 0.04245 | 0.00000 | 0.00000 | 0.07859 | 0.24283 | 4.12582 | 0.00000 | 0.00000 | 0.01492 | 6.10558 |
| **CIB1** | 4.57141 | 26.01767 | 10.83949 | 13.14236 | 20.43678 | 6.96320 | 11.19959 | 2.63834 | 10.50023 | 0.31702 | 15.23064 | 18.08578 |
| **CIB2** | 0.03047 | 1.01394 | 0.13066 | 1.83437 | 3.38120 | 0.00000 | 1.19707 | 0.07316 | 0.00000 | 3.85369 | 0.00000 | 0.76028 |
| **CIB3** | 0.00963 | 0.03581 | 0.04129 | 0.01437 | 0.07267 | 0.04894 | 0.02051 | 0.05550 | 0.02223 | 0.00001 | 0.01172 | 0.04580 |
| **CIC** | 0.00149 | 0.48342 | 1.24356 | 1.58407 | 5.45077 | 0.00000 | 2.83560 | 0.00000 | 1.88844 | 14.03440 | 0.29181 | 0.62624 |
| **CIDEA** | 0.00223 | 0.02832 | 0.03879 | 0.01374 | 0.01728 | 0.00174 | 0.01773 | 0.06113 | 0.01272 | 0.02839 | 0.01719 | 0.01275 |
| **CIDEC** | 0.10761 | 0.43148 | 0.00000 | 0.28144 | 4.81433 | 0.99582 | 1.31526 | 0.07538 | 1.67726 | 0.00000 | 6.29528 | 0.00000 |
| **CIDECP** | 4.46537 | 3.80869 | 1.98725 | 1.97015 | 2.77094 | 2.12179 | 0.29030 | 7.35622 | 2.87178 | 0.00016 | 1.16080 | 6.07108 |
| **CIITA** | 0.00610 | 0.01761 | 0.01403 | 0.01157 | 0.01337 | 0.01499 | 0.01822 | 0.01009 | 0.02310 | 0.02504 | 0.01812 | 0.02062 |
| **CILP** | 0.02877 | 0.03678 | 0.04679 | 0.05007 | 0.00000 | 0.07278 | 0.00000 | 0.03526 | 0.02749 | 0.01225 | 0.01690 | 1.49656 |
| **CILP2** | 0.03397 | 0.37703 | 0.00000 | 0.26503 | 0.35842 | 0.00000 | 0.00234 | 0.04850 | 0.35803 | 0.98845 | 0.24947 | 1.08842 |
| **CINP** | 0.82999 | 14.83795 | 6.03331 | 5.83196 | 11.52373 | 13.08980 | 8.30142 | 1.92760 | 6.31445 | 0.00000 | 9.25149 | 15.28332 |
| **CIPC** | 2.42829 | 4.70363 | 3.95004 | 0.50174 | 0.49554 | 5.01009 | 3.63697 | 3.40426 | 2.61622 | 2.79792 | 4.89776 | 5.25460 |
| **CIR1** | 4.41479 | 3.55191 | 4.69851 | 2.55144 | 0.00000 | 12.18467 | 7.89060 | 6.36720 | 0.00000 | 8.39101 | 3.85532 | 10.18728 |
| **CIRBP** | 4.27213 | 12.71216 | 8.48489 | 12.68889 | 11.49106 | 4.99468 | 7.72757 | 2.22471 | 6.62553 | 8.24514 | 11.60158 | 12.25291 |
| **CIRBP-AS1** | 0.01325 | 0.15268 | 0.54171 | 0.12850 | 0.29443 | 0.00000 | 0.30137 | 0.00244 | 0.25349 | 0.00000 | 0.10510 | 0.57444 |
| **CIRH1A** | 0.41509 | 20.02653 | 21.68276 | 10.13152 | 13.47363 | 16.81299 | 21.20784 | 15.36397 | 27.08059 | 0.00000 | 21.66706 | 20.07645 |
| **CISD1** | 2.59128 | 9.90883 | 11.20995 | 11.22740 | 6.31778 | 24.01895 | 21.91910 | 7.41711 | 11.92445 | 12.24944 | 14.11542 | 12.57519 |
| **CISD2** | 0.96672 | 1.46580 | 6.44108 | 2.31777 | 0.00000 | 6.61099 | 3.05254 | 0.56510 | 9.51229 | 0.11483 | 5.00952 | 0.96080 |
| **CISD3** | 0.00071 | 5.94881 | 4.60346 | 2.17768 | 9.07912 | 0.00000 | 3.91352 | 0.00000 | 3.99797 | 3.30664 | 7.28275 | 2.27180 |
| **CISH** | 1.15751 | 4.90534 | 0.48625 | 1.35380 | 0.00000 | 1.38243 | 0.81754 | 0.27576 | 1.46325 | 0.67672 | 0.12923 | 4.53033 |
| **CIT** | 5.13226 | 7.56346 | 1.25108 | 4.92195 | 5.06107 | 6.22052 | 2.40065 | 1.43619 | 1.88464 | 5.33608 | 2.98169 | 4.33081 |
| **CITED1** | 0.14893 | 0.00923 | 0.15126 | 0.02628 | 0.01732 | 0.16589 | 0.02342 | 0.53972 | 0.00000 | 0.00112 | 0.01287 | 0.02606 |
| **CITED2** | 0.00693 | 0.00181 | 12.24409 | 0.00000 | 4.47707 | 0.00000 | 8.94399 | 0.21363 | 0.00000 | 5.07881 | 1.76646 | 0.00000 |
| **CITED4** | 0.00105 | 1.31655 | 0.00000 | 0.00000 | 9.81464 | 0.00000 | 4.12299 | 0.00000 | 0.00000 | 14.23574 | 1.81387 | 0.32727 |
| **CIZ1** | 2.83973 | 6.02843 | 2.92753 | 4.88738 | 7.40895 | 0.00000 | 3.50317 | 1.83513 | 4.77189 | 1.85364 | 2.76647 | 9.45169 |
| **CKAP2** | 5.10244 | 13.85479 | 6.55340 | 4.83851 | 5.01815 | 39.95308 | 16.79118 | 17.55607 | 10.67331 | 8.78939 | 12.74157 | 17.93085 |

| | | | | | | | | | | | | |
|---|---|---|---|---|---|---|---|---|---|---|---|---|
| **CKAP2L** | 2.13489 | 2.58580 | 1.46981 | 2.17796 | 0.97031 | 8.59800 | 0.00000 | 5.14863 | 4.22290 | 3.67339 | 3.49992 | 8.42912 |
| **CKAP4** | 0.05906 | 3.25268 | 0.38953 | 1.34165 | 3.26768 | 0.00000 | 0.00005 | 0.00318 | 0.00000 | 2.85479 | 0.00000 | 2.69033 |
| **CKAP5** | 8.90000 | 14.68473 | 15.11328 | 8.89145 | 11.08541 | 26.62514 | 22.87680 | 9.95582 | 19.81254 | 18.79665 | 11.74489 | 18.26394 |
| **CKB** | 2.71530 | 30.75727 | 27.31027 | 28.96896 | 40.81979 | 17.51417 | 31.46284 | 1.16409 | 30.41901 | 16.24308 | 15.73672 | 8.69363 |
| **CKLF** | 0.01133 | 0.27306 | 0.56808 | 0.37417 | 0.03724 | 0.40009 | 0.18568 | 0.12949 | 0.00000 | 0.49529 | 0.47772 | 0.85380 |
| **CKM** | 0.00399 | 0.07144 | 0.25617 | 0.02991 | 0.04268 | 0.00028 | 0.00583 | 0.72834 | 0.04849 | 0.00492 | 0.10472 | 0.11492 |
| **CKMT1A** | 0.02345 | 0.06013 | 0.51619 | 0.15415 | 0.00000 | 0.00000 | 0.81713 | 0.03502 | 0.00000 | 0.86203 | 0.33968 | 0.11462 |
| **CKMT1B** | 0.00481 | 0.04230 | 0.67645 | 0.25751 | 0.11865 | 0.01448 | 0.16585 | 0.01631 | 0.00000 | 0.00000 | 0.05105 | 0.19613 |
| **CKMT2** | 1.37070 | 0.06322 | 0.16434 | 0.53824 | 0.07733 | 0.00000 | 0.00554 | 0.86678 | 0.51548 | 0.39865 | 0.13699 | 0.13308 |
| **CKMT2-AS1** | 0.06532 | 1.15805 | 0.92839 | 1.02648 | 0.28403 | 0.00000 | 0.04204 | 5.22992 | 1.04229 | 0.72857 | 1.34227 | 3.30179 |
| **CKS1B** | 4.34163 | 6.31151 | 7.78331 | 8.98222 | 9.45731 | 9.89904 | 13.00637 | 4.48959 | 7.30035 | 2.41601 | 5.47814 | 9.05014 |
| **CKS2** | 8.61851 | 20.43664 | 26.36416 | 14.83194 | 17.43101 | 38.02799 | 33.87007 | 23.73628 | 24.34376 | 1.42665 | 14.90358 | 26.51079 |
| **CLASP1** | 1.32923 | 1.72527 | 1.05306 | 3.08926 | 0.90308 | 7.77051 | 2.85768 | 1.43458 | 3.24651 | 5.60393 | 2.26106 | 3.15371 |
| **CLASP2** | 1.29678 | 3.58587 | 3.91969 | 1.10564 | 0.00000 | 10.55809 | 3.49359 | 0.42122 | 5.57877 | 8.68941 | 3.99347 | 2.82095 |
| **CLASRP** | 0.90071 | 1.82112 | 2.29412 | 0.97334 | 1.97891 | 0.00000 | 0.44795 | 0.93772 | 2.96936 | 3.98769 | 0.97163 | 1.35838 |
| **CLC** | 1.89783 | 0.06888 | 0.00803 | 0.08125 | 0.04640 | 0.00000 | 0.06189 | 1.22529 | 0.00000 | 0.00730 | 0.00931 | 0.39487 |
| **CLCA1** | 0.07302 | 0.01848 | 0.04063 | 0.01740 | 0.01753 | 0.00648 | 0.02642 | 0.03667 | 0.01864 | 0.00328 | 0.01879 | 0.01437 |
| **CLCA2** | 0.00303 | 0.00000 | 0.44588 | 0.09480 | 0.00000 | 0.09151 | 0.03245 | 0.00891 | 0.00000 | 0.21720 | 0.07349 | 0.00642 |
| **CLCA3P** | 0.00889 | 0.01881 | 0.02006 | 0.00970 | 0.01660 | 0.02852 | 0.01114 | 0.00758 | 0.00930 | 0.05195 | 0.01492 | 0.06745 |
| **CLCA4** | 0.21716 | 0.01366 | 0.01451 | 0.01160 | 0.00861 | 0.00609 | 0.03818 | 0.00178 | 0.00338 | 0.07379 | 0.03001 | 0.02874 |
| **CLCC1** | 0.20627 | 6.49733 | 7.13106 | 0.15312 | 3.61991 | 6.60194 | 0.00000 | 0.60410 | 10.03514 | 1.07394 | 6.19201 | 0.33062 |
| **CLCF1** | 0.10075 | 0.42091 | 0.00346 | 0.00000 | 0.00000 | 0.05513 | 0.00240 | 0.07149 | 0.01750 | 0.06305 | 0.00944 | 0.63773 |
| **CLCN1** | 0.03076 | 0.01657 | 0.03507 | 0.00982 | 0.24910 | 0.00000 | 0.00960 | 0.01081 | 0.06681 | 0.00000 | 0.04417 | 0.07626 |
| **CLCN2** | 0.63596 | 0.26784 | 2.00735 | 0.81830 | 0.00001 | 0.00000 | 0.00000 | 0.09320 | 0.35464 | 0.33797 | 0.00002 | 0.36981 |
| **CLCN3** | 4.63461 | 5.16105 | 6.23274 | 4.63169 | 3.28074 | 14.51096 | 0.00000 | 2.88085 | 6.33820 | 9.36192 | 0.80833 | 4.11907 |
| **CLCN4** | 5.41656 | 0.10290 | 0.00000 | 0.00000 | 0.00000 | 0.00000 | 0.02037 | 1.15430 | 0.14556 | 0.36910 | 0.00000 | 1.24565 |
| **CLCN5** | 6.19753 | 2.36743 | 2.34674 | 1.66591 | 5.15633 | 5.70633 | 0.00000 | 0.60645 | 2.15622 | 5.53295 | 3.57346 | 1.04032 |
| **CLCN6** | 0.15338 | 0.00737 | 3.90920 | 1.62456 | 3.48147 | 2.16722 | 0.00000 | 0.00000 | 10.93464 | 3.44392 | 1.61068 | 0.00161 |
| **CLCN7** | 0.00341 | 2.82307 | 2.86348 | 0.00000 | 0.99213 | 0.00000 | 0.00000 | 0.01610 | 4.49573 | 10.68911 | 0.22649 | 0.71939 |
| **CLCNKA** | 0.00651 | 0.03186 | 0.05815 | 0.01049 | 0.02590 | 0.01477 | 0.04598 | 0.01054 | 0.00000 | 0.02059 | 0.01602 | 0.01510 |
| **CLCNKB** | 0.02064 | 0.05035 | 0.08891 | 0.00151 | 0.04873 | 0.03517 | 0.01353 | 0.00926 | 0.00177 | 0.05112 | 0.01706 | 0.01037 |
| **CLDN1** | 0.00451 | 0.00688 | 0.61478 | 0.00469 | 0.01065 | 0.02707 | 0.00000 | 0.01599 | 0.81324 | 0.00841 | 0.13539 | 0.01990 |
| **CLDN10** | 0.11985 | 18.46399 | 0.00000 | 0.00000 | 73.62523 | 14.12794 | 0.00000 | 1.01294 | 2.37364 | 2.00976 | 43.30398 | 0.06616 |
| **CLDN10-AS1** | 0.00545 | 0.15613 | 0.00517 | 0.00000 | 0.11542 | 0.05662 | 0.01165 | 0.01011 | 0.01402 | 0.04667 | 0.00526 | 0.00773 |
| **CLDN11** | 0.30875 | 0.00000 | 0.35746 | 0.00000 | 0.00000 | 3.05796 | 0.00029 | 0.06519 | 0.00000 | 3.85128 | 0.21229 | 5.86567 |
| **CLDN12** | 5.71689 | 7.04712 | 3.06326 | 0.01466 | 0.52757 | 9.79804 | 0.00000 | 3.30312 | 13.55333 | 0.00000 | 4.38682 | 5.98697 |
| **CLDN14** | 0.02811 | 0.02193 | 0.01858 | 0.01794 | 0.06738 | 0.01166 | 0.01487 | 0.01700 | 0.02247 | 0.00000 | 0.03286 | 0.02377 |
| **CLDN15** | 0.05206 | 0.72895 | 0.63518 | 0.84773 | 0.00000 | 0.00000 | 1.04983 | 0.06580 | 1.64388 | 3.17453 | 0.00000 | 0.62529 |
| **CLDN16** | 0.03715 | 0.00929 | 0.03221 | 0.00787 | 0.00351 | 0.04901 | 0.01380 | 0.06620 | 0.02434 | 0.03702 | 0.01078 | 0.00931 |
| **CLDN18** | 1.23733 | 0.08114 | 0.04625 | 0.14515 | 0.01519 | 0.00000 | 0.00621 | 0.37646 | 0.03157 | 0.00788 | 0.01215 | 0.23711 |
| **CLDN19** | 0.00608 | 13.13980 | 0.58102 | 14.01532 | 3.85932 | 0.00000 | 0.02415 | 0.00000 | 0.00000 | 0.22093 | 0.25327 | 1.00823 |
| **CLDN2** | 0.01637 | 0.00856 | 0.00798 | 0.00000 | 0.21928 | 0.00000 | 0.05201 | 0.01140 | 0.06590 | 0.07829 | 0.00000 | 0.03578 |
| **CLDN20** | 0.00171 | 0.06186 | 0.00000 | 0.20433 | 0.00000 | 0.25544 | 0.00000 | 0.09594 | 0.00000 | 0.02230 | 0.03543 | 0.11295 |
| **CLDN23** | 0.00289 | 0.18960 | 0.06017 | 0.00000 | 0.57303 | 0.36518 | 0.00000 | 0.04811 | 0.56705 | 0.01579 | 0.00436 | 0.30709 |
| **CLDN24** | 0.00603 | 0.01317 | 0.01312 | 0.01713 | 0.00975 | 0.02316 | 0.03812 | 0.01034 | 0.02219 | 0.01802 | 0.01371 | 0.01857 |
| **CLDN3** | 0.00084 | 0.00000 | 7.88091 | 0.00000 | 0.75463 | 0.00000 | 2.49916 | 0.00332 | 13.82440 | 0.94237 | 13.71762 | 0.00000 |
| **CLDN4** | 0.01360 | 13.37774 | 45.04445 | 2.39813 | 69.60496 | 5.13728 | 25.84970 | 0.00000 | 85.34394 | 0.00000 | 32.60165 | 0.00000 |
| **CLDN5** | 0.01187 | 0.01229 | 0.14715 | 0.01112 | 0.06452 | 0.00000 | 0.00190 | 0.00884 | 0.03130 | 0.02970 | 0.04925 | 0.05619 |

| | | | | | | | | | | | | |
|---|---|---|---|---|---|---|---|---|---|---|---|---|
| **CLDN6** | 0.01435 | 6.03350 | 11.89446 | 4.83476 | 18.45577 | 0.00000 | 2.01823 | 5.25409 | 31.20747 | 18.05866 | 8.83304 | 19.63811 |
| **CLDN7** | 0.00000 | 30.90212 | 29.69744 | 20.35287 | 34.19465 | 9.06681 | 14.50846 | 2.76409 | 56.91392 | 0.00000 | 22.98755 | 17.37239 |
| **CLDN8** | 0.00632 | 0.01412 | 0.01877 | 0.01072 | 0.03017 | 0.02749 | 0.02167 | 0.01038 | 0.01529 | 0.00555 | 0.01789 | 0.01623 |
| **CLDN9** | 0.00173 | 0.00446 | 0.04037 | 0.13577 | 0.00246 | 0.16321 | 0.00000 | 0.07114 | 0.30451 | 0.00000 | 0.00567 | 0.01863 |
| **CLDND1** | 3.13730 | 6.55817 | 19.71896 | 3.75628 | 1.19061 | 21.35571 | 0.00000 | 7.03909 | 27.97273 | 0.00000 | 8.12654 | 15.89698 |
| **CLDND2** | 0.01629 | 0.03394 | 0.05569 | 0.00000 | 0.00000 | 0.00000 | 0.13586 | 0.01494 | 0.06779 | 0.17367 | 0.13086 | 0.07501 |
| **CLEC10A** | 6.09350 | 0.02263 | 0.02152 | 0.00528 | 0.01620 | 0.00000 | 0.01389 | 2.29945 | 0.01362 | 0.00000 | 0.02304 | 0.02608 |
| **CLEC11A** | 0.31407 | 0.75618 | 0.78910 | 0.37552 | 0.45300 | 0.00000 | 0.18215 | 0.13732 | 0.18122 | 0.00000 | 0.72523 | 1.53211 |
| **CLEC12A** | 0.02009 | 0.01315 | 0.73672 | 0.32236 | 0.13351 | 0.00000 | 0.62024 | 0.01112 | 0.51191 | 0.00000 | 0.10182 | 0.09694 |
| **CLEC12B** | 0.00717 | 0.03086 | 0.17582 | 0.08488 | 0.01815 | 0.00000 | 0.01749 | 0.01969 | 0.02552 | 0.02043 | 0.01277 | 0.02280 |
| **CLEC16A** | 3.82113 | 6.57708 | 2.30111 | 5.35125 | 9.67029 | 0.00000 | 2.88795 | 1.26871 | 6.94271 | 3.74960 | 4.46027 | 4.34779 |
| **CLEC17A** | 0.78867 | 0.00151 | 0.01343 | 0.00911 | 0.00943 | 0.01113 | 0.04049 | 0.94958 | 0.01838 | 0.02155 | 0.01727 | 0.20041 |
| **CLEC18A** | 0.00622 | 0.01408 | 0.03346 | 0.01476 | 0.01301 | 0.01498 | 0.01249 | 0.01027 | 0.02454 | 0.02208 | 0.01526 | 0.02077 |
| **CLEC18B** | 0.01547 | 0.01013 | 0.00749 | 0.02242 | 0.01002 | 0.00338 | 0.03114 | 0.00841 | 0.03995 | 0.13311 | 0.01419 | 0.02197 |
| **CLEC18C** | 0.00795 | 0.00667 | 0.01727 | 0.00934 | 0.00907 | 0.05047 | 0.01974 | 0.00696 | 0.01618 | 0.00501 | 0.01318 | 0.07622 |
| **CLEC19A** | 0.00757 | 0.01435 | 0.01508 | 0.01162 | 0.01692 | 0.01672 | 0.02193 | 0.01112 | 0.01281 | 0.01854 | 0.02229 | 0.01767 |
| **CLEC1A** | 0.01358 | 0.28385 | 6.80159 | 0.25700 | 0.02203 | 0.00000 | 0.00000 | 0.00499 | 0.00000 | 0.00000 | 0.06652 | 0.22535 |
| **CLEC1B** | 0.00524 | 0.04573 | 0.62203 | 0.01990 | 0.01599 | 0.00000 | 0.00000 | 0.08528 | 0.00000 | 0.00874 | 0.03568 | 0.03736 |
| **CLEC2A** | 0.00578 | 0.00937 | 0.03927 | 0.21578 | 0.00000 | 0.03139 | 0.00857 | 0.00937 | 0.00551 | 0.04277 | 0.01564 | 0.02097 |
| **CLEC2B** | 0.06114 | 0.01349 | 0.01755 | 0.01586 | 0.01394 | 0.01438 | 0.02301 | 0.01023 | 0.01265 | 0.00742 | 0.03666 | 0.02319 |
| **CLEC2D** | 0.03005 | 0.22159 | 0.93440 | 0.39285 | 0.00000 | 1.70516 | 0.46204 | 0.25582 | 0.11041 | 0.40412 | 1.09155 | 0.59864 |
| **CLEC2L** | 0.04834 | 0.62337 | 0.59706 | 0.20769 | 0.00000 | 0.00000 | 1.36522 | 0.25252 | 0.01419 | 0.00000 | 0.41959 | 3.60162 |
| **CLEC3B** | 0.00599 | 0.01676 | 0.03399 | 0.03356 | 0.02165 | 0.00985 | 0.01417 | 0.00984 | 0.02202 | 0.01929 | 0.01569 | 0.02607 |
| **CLEC4A** | 0.71783 | 0.13563 | 0.02246 | 0.18249 | 0.00006 | 0.18019 | 0.00000 | 0.45336 | 0.00000 | 0.00742 | 0.10807 | 0.20002 |
| **CLEC4C** | 2.07314 | 0.19689 | 0.00163 | 0.00000 | 0.05270 | 0.00000 | 0.06992 | 1.48459 | 0.00000 | 0.25539 | 0.00000 | 0.00000 |
| **CLEC4D** | 0.00774 | 2.13611 | 0.00000 | 0.47638 | 0.75174 | 0.27436 | 0.00000 | 0.00000 | 0.00000 | 0.00208 | 0.00000 | 0.82731 |
| **CLEC4E** | 0.00591 | 0.00405 | 0.00000 | 2.11082 | 0.01014 | 0.05565 | 0.00000 | 0.00889 | 0.04978 | 0.00000 | 0.01393 | 0.01168 |
| **CLEC4F** | 0.00620 | 0.04358 | 0.01520 | 0.01427 | 0.02167 | 0.02315 | 0.01711 | 0.01039 | 0.01903 | 0.01149 | 0.01550 | 0.01485 |
| **CLEC4G** | 0.09544 | 0.01206 | 0.03371 | 0.04080 | 0.01302 | 0.00972 | 0.01085 | 0.01009 | 0.02412 | 0.01777 | 0.01437 | 0.03065 |
| **CLEC4GP1** | 0.00156 | 0.05085 | 1.10322 | 0.00000 | 0.00000 | 0.02470 | 0.19529 | 0.03239 | 0.96121 | 0.15123 | 0.70226 | 0.00055 |
| **CLEC4M** | 0.00470 | 0.00902 | 0.00970 | 0.00891 | 0.07667 | 0.04633 | 0.00470 | 0.03242 | 0.02685 | 0.01481 | 0.01228 | 0.01438 |
| **CLEC5A** | 0.00588 | 0.01269 | 0.01969 | 0.01079 | 0.00660 | 0.02531 | 0.01442 | 0.01032 | 0.03523 | 0.03526 | 0.01511 | 0.01669 |
| **CLEC6A** | 0.00607 | 3.18411 | 0.00000 | 0.28389 | 0.80468 | 0.52887 | 0.00000 | 0.00000 | 0.00000 | 0.00079 | 0.04505 | 1.22327 |
| **CLEC7A** | 0.05391 | 0.01240 | 1.07917 | 0.67085 | 0.00000 | 0.04111 | 0.00000 | 0.03409 | 0.05680 | 0.79751 | 0.01151 | 0.03072 |
| **CLEC9A** | 0.00542 | 0.00000 | 0.11339 | 0.34747 | 0.00000 | 0.01522 | 0.06807 | 0.00366 | 0.00000 | 0.09303 | 0.00627 | 0.67100 |
| **CLGN** | 0.66818 | 0.05261 | 0.03840 | 0.14314 | 0.09496 | 0.33935 | 0.00000 | 0.58092 | 0.00000 | 0.00000 | 0.07936 | 0.75979 |
| **CLHC1** | 0.00002 | 0.51895 | 0.11029 | 0.18288 | 0.20511 | 1.06075 | 0.24441 | 0.84540 | 0.99799 | 0.60258 | 0.32602 | 1.55756 |
| **CLIC1** | 0.17612 | 6.93890 | 29.07678 | 31.11109 | 34.76654 | 5.72197 | 42.63248 | 0.00742 | 35.84130 | 0.00000 | 25.05352 | 0.00000 |
| **CLIC2** | 0.00619 | 0.49728 | 0.00287 | 0.12379 | 0.12838 | 0.31933 | 0.00000 | 0.00273 | 0.00000 | 0.00343 | 0.02779 | 0.19122 |
| **CLIC3** | 0.00450 | 0.00000 | 4.43445 | 0.00000 | 0.00000 | 0.01766 | 5.25250 | 0.00752 | 4.75132 | 0.00000 | 4.53115 | 0.01571 |
| **CLIC4** | 6.76917 | 18.29772 | 11.58274 | 2.34393 | 9.22782 | 40.30041 | 22.32061 | 3.52189 | 20.01070 | 3.86848 | 19.28193 | 10.14626 |
| **CLIC5** | 0.00765 | 0.00000 | 0.04650 | 0.12287 | 0.07465 | 0.05960 | 0.00000 | 0.00638 | 0.00000 | 0.00000 | 0.27872 | 0.12200 |
| **CLIC6** | 0.00847 | 0.01608 | 0.02170 | 0.01845 | 0.00953 | 0.03032 | 0.01196 | 0.00997 | 0.02018 | 0.01833 | 0.01276 | 0.01471 |
| **CLINT1** | 5.47218 | 11.16480 | 13.44084 | 9.64808 | 4.33511 | 21.81493 | 18.76310 | 5.81314 | 12.64115 | 12.17901 | 12.69528 | 10.48209 |
| **CLIP1** | 8.00359 | 12.99383 | 5.96783 | 6.08673 | 14.52618 | 29.58762 | 0.00000 | 2.19238 | 10.96146 | 31.48225 | 9.66416 | 14.26050 |
| **CLIP1-AS1** | 0.00741 | 0.01972 | 0.07466 | 0.02648 | 0.12372 | 0.02305 | 0.00000 | 0.00964 | 0.12969 | 0.08495 | 0.03974 | 0.03698 |
| **CLIP2** | 0.62549 | 0.18171 | 0.03174 | 0.00979 | 0.01735 | 0.01596 | 0.00000 | 0.11327 | 0.01909 | 0.12862 | 0.01116 | 0.84663 |
| **CLIP3** | 2.29799 | 0.05415 | 0.18106 | 0.05024 | 0.11790 | 0.00000 | 0.00197 | 0.01217 | 0.18274 | 0.01154 | 0.10767 | 0.08072 |

| | | | | | | | | | | | | |
|---|---|---|---|---|---|---|---|---|---|---|---|---|
| **CLIP4** | 3.64277 | 0.07952 | 0.05681 | 0.01549 | 0.03793 | 0.00000 | 0.10715 | 0.54646 | 0.00000 | 0.17405 | 0.00019 | 0.11570 |
| **CLK1** | 6.48559 | 9.70881 | 6.63594 | 7.04794 | 2.18592 | 25.71131 | 0.00000 | 12.37364 | 11.22422 | 10.11231 | 1.40136 | 19.89114 |
| **CLK2** | 0.14950 | 0.39739 | 1.61506 | 1.56768 | 1.74027 | 4.06822 | 1.29038 | 1.21304 | 2.05571 | 7.65540 | 1.94698 | 1.70009 |
| **CLK2P** | 0.10673 | 0.04164 | 0.24624 | 0.03143 | 0.11556 | 0.53385 | 0.00000 | 0.11307 | 0.20480 | 0.44344 | 0.19646 | 0.22112 |
| **CLK3** | 4.22734 | 7.64054 | 9.57023 | 7.96765 | 6.60081 | 5.92460 | 9.96772 | 13.59819 | 12.89562 | 0.00000 | 8.64689 | 10.98706 |
| **CLK4** | 0.06556 | 1.67180 | 1.94445 | 1.68479 | 0.40482 | 4.53279 | 0.00000 | 8.31508 | 2.58270 | 0.93370 | 1.45503 | 1.95857 |
| **CLLU1** | 0.00631 | 0.01524 | 0.02793 | 0.01049 | 0.01454 | 0.01826 | 0.01836 | 0.01046 | 0.02061 | 0.01308 | 0.01531 | 0.01636 |
| **CLMN** | 0.82523 | 0.63523 | 0.30921 | 0.00000 | 0.21031 | 0.08668 | 0.26332 | 0.77189 | 0.00000 | 0.04022 | 0.56847 | 0.37939 |
| **CLMP** | 0.00787 | 0.05618 | 7.55371 | 0.00000 | 0.00000 | 0.13013 | 0.00000 | 0.00458 | 0.00000 | 0.13212 | 0.11948 | 0.07330 |
| **CLN3** | 0.20507 | 12.85675 | 15.61374 | 8.87635 | 10.54473 | 2.00699 | 13.86698 | 0.13760 | 17.81421 | 2.22532 | 9.47646 | 5.42755 |
| **CLN5** | 1.08158 | 3.01185 | 8.99492 | 0.31723 | 2.28637 | 15.08507 | 0.00000 | 1.94760 | 13.15132 | 2.67370 | 4.94136 | 0.82556 |
| **CLN6** | 1.72554 | 13.43596 | 9.39433 | 5.88039 | 7.26709 | 0.00000 | 1.55968 | 0.16494 | 5.85836 | 15.56582 | 7.75553 | 7.51652 |
| **CLN8** | 1.16315 | 5.86292 | 8.74029 | 4.49997 | 12.28179 | 4.66417 | 0.69128 | 2.77750 | 17.33798 | 5.09187 | 10.23801 | 3.83260 |
| **CLNK** | 0.00409 | 0.01366 | 0.02915 | 0.10567 | 0.01346 | 0.01322 | 0.04522 | 0.01896 | 0.00231 | 0.01021 | 0.03236 | 0.01657 |
| **CLNS1A** | 0.95614 | 16.13494 | 14.84505 | 8.00606 | 11.45443 | 30.80577 | 12.26531 | 2.19573 | 17.10284 | 0.00000 | 17.65278 | 12.95897 |
| **CLOCK** | 13.82326 | 0.08537 | 0.22229 | 0.65677 | 0.45855 | 7.26970 | 0.96568 | 6.33741 | 1.47381 | 8.77926 | 2.29280 | 3.51467 |
| **1.00000** | 0.00311 | 14.61013 | 14.52369 | 6.70495 | 4.48565 | 13.53212 | 11.01564 | 17.26134 | 15.64869 | 0.00000 | 9.36184 | 18.52053 |
| **CLPB** | 1.70500 | 8.95856 | 9.02740 | 6.49220 | 10.61152 | 0.25424 | 7.59182 | 7.24909 | 12.31418 | 0.00000 | 11.99313 | 9.48688 |
| **CLPP** | 2.22012 | 22.90428 | 12.66372 | 20.09254 | 19.28147 | 0.00000 | 23.27869 | 17.12126 | 21.03848 | 2.64800 | 16.87897 | 27.61069 |
| **CLPS** | 1.27458 | 0.08672 | 0.00000 | 0.13946 | 0.00000 | 0.00000 | 0.10524 | 0.18989 | 0.36077 | 0.25549 | 0.79771 | 0.26616 |
| **CLPSL1** | 0.00918 | 0.05342 | 0.00000 | 0.40895 | 0.76484 | 0.23629 | 0.03583 | 0.00623 | 0.00000 | 0.19492 | 0.90958 | 0.00880 |
| **CLPSL2** | 0.00631 | 0.10937 | 0.00278 | 0.09177 | 0.29198 | 0.00566 | 0.05091 | 0.00786 | 0.14225 | 0.02814 | 0.05234 | 0.08080 |
| **CLPTM1** | 2.58700 | 15.20723 | 15.96588 | 7.58774 | 11.47809 | 0.00000 | 0.00000 | 0.92782 | 17.50000 | 15.65105 | 7.05021 | 8.03064 |
| **CLPTM1L** | 7.18600 | 8.59046 | 7.99717 | 2.70056 | 7.12051 | 10.27616 | 9.12872 | 3.30584 | 8.34492 | 26.14338 | 1.78736 | 12.50922 |
| **CLPX** | 4.17766 | 13.50985 | 7.26249 | 10.40765 | 4.42098 | 26.60178 | 6.19537 | 5.39196 | 1.89943 | 7.21534 | 9.30595 | 17.58373 |
| **CLRN1** | 0.00533 | 0.01489 | 0.02230 | 0.01308 | 0.01231 | 0.02003 | 0.01373 | 0.01337 | 0.01605 | 0.02539 | 0.01514 | 0.01528 |
| **CLRN1-AS1** | 0.00774 | 0.01160 | 0.07912 | 0.00402 | 0.03061 | 0.06749 | 0.02156 | 0.01841 | 0.00000 | 0.01391 | 0.01251 | 0.02745 |
| **CLSPN** | 9.74462 | 6.75555 | 7.13788 | 7.55108 | 4.98000 | 22.23800 | 15.00623 | 17.35113 | 7.55066 | 28.21885 | 8.03043 | 13.19761 |
| **CLSTN1** | 8.31126 | 5.84477 | 0.94899 | 8.24257 | 7.53897 | 0.00005 | 2.16334 | 4.27327 | 9.24997 | 16.36605 | 0.00000 | 8.43160 |
| **CLSTN2** | 2.42194 | 0.01204 | 0.01549 | 0.00943 | 0.01220 | 0.01960 | 0.01705 | 0.67006 | 0.01959 | 0.00000 | 0.01156 | 0.03615 |
| **CLSTN2-AS1** | 0.02471 | 0.02108 | 0.02348 | 0.01126 | 0.01320 | 0.01438 | 0.01947 | 0.01024 | 0.01724 | 0.01905 | 0.01946 | 0.01810 |
| **CLSTN3** | 1.97344 | 0.17544 | 0.03922 | 0.67846 | 0.07232 | 0.00000 | 0.09542 | 0.27160 | 0.00000 | 0.43089 | 0.30668 | 0.70873 |
| **CLTA** | 7.76476 | 26.19559 | 30.95957 | 13.08594 | 28.53373 | 21.48292 | 34.41095 | 4.49420 | 34.76452 | 16.97560 | 24.87780 | 26.50373 |
| **CLTB** | 0.99174 | 9.42773 | 2.50954 | 5.72898 | 24.43086 | 0.00000 | 3.57145 | 1.42954 | 0.00000 | 14.74462 | 2.85476 | 2.69091 |
| **CLTC** | 4.46970 | 19.48971 | 18.28600 | 14.51894 | 21.76757 | 50.44939 | 38.25856 | 3.64744 | 22.44349 | 30.40103 | 24.62534 | 7.12028 |
| **CLTCL1** | 1.16674 | 1.81595 | 0.27851 | 0.54757 | 2.13195 | 0.00000 | 0.24471 | 0.10413 | 1.52275 | 0.52859 | 0.40739 | 0.98031 |
| **CLU** | 0.00153 | 0.00000 | 2.06395 | 0.02389 | 0.00000 | 0.00000 | 0.00000 | 0.22843 | 1.26880 | 1.33696 | 0.63193 | 0.04340 |
| **CLUAP1** | 2.51262 | 5.46752 | 0.07778 | 1.30544 | 1.18485 | 2.54084 | 0.46022 | 0.26000 | 0.00000 | 1.37960 | 2.24177 | 3.57961 |
| **CLUH** | 2.38208 | 12.33578 | 1.90546 | 7.43738 | 18.71184 | 0.00000 | 10.16061 | 0.69222 | 18.13239 | 33.79547 | 5.45875 | 10.63135 |
| **CLUHP3** | 0.00191 | 0.00000 | 0.52352 | 0.12371 | 0.00000 | 0.00000 | 0.00773 | 0.00521 | 2.98727 | 0.86209 | 0.27954 | 0.06757 |
| **CLUL1** | 0.80730 | 0.00311 | 0.00901 | 0.10569 | 0.05977 | 0.00000 | 0.04437 | 0.17214 | 0.00000 | 0.46024 | 0.01428 | 0.03267 |
| **CLVS1** | 0.29471 | 0.03440 | 0.15926 | 0.00171 | 0.01709 | 0.04612 | 0.30937 | 0.00000 | 0.00000 | 0.00000 | 0.01496 | 0.10800 |
| **CLVS2** | 0.01446 | 0.15724 | 0.59175 | 0.56301 | 0.00000 | 0.04377 | 0.00000 | 0.00795 | 0.50024 | 0.28930 | 0.00079 | 0.00011 |
| **CLYBL** | 0.22131 | 0.20610 | 0.09766 | 0.86567 | 0.11712 | 0.23028 | 0.00000 | 0.06890 | 0.00000 | 0.01340 | 0.62146 | 0.00000 |
| **CLYBL-AS2** | 0.00695 | 0.01635 | 0.01352 | 0.04230 | 0.02168 | 0.01666 | 0.04725 | 0.00976 | 0.00550 | 0.00288 | 0.01601 | 0.02416 |
| **CMA1** | 0.00613 | 0.01537 | 0.01616 | 0.01234 | 0.01648 | 0.01808 | 0.02356 | 0.01000 | 0.01793 | 0.00669 | 0.01529 | 0.02912 |
| **CMAHP** | 0.20110 | 0.05136 | 0.29829 | 0.01675 | 0.02355 | 0.04720 | 0.00000 | 0.00479 | 0.00000 | 0.03185 | 0.09808 | 0.01606 |
| **CMAS** | 0.59750 | 4.65185 | 15.34074 | 5.99095 | 1.95421 | 15.02329 | 11.08040 | 0.49905 | 3.76733 | 0.00000 | 6.22532 | 5.03060 |

| | | | | | | | | | | | | |
|---|---|---|---|---|---|---|---|---|---|---|---|---|
| **CMBL** | 0.42329 | 0.90923 | 1.24769 | 4.13478 | 0.00000 | 0.08494 | 0.47064 | 0.00274 | 0.73970 | 2.01859 | 0.27089 | 0.00000 |
| **CMC1** | 0.33017 | 2.89969 | 1.53399 | 2.36425 | 1.09979 | 13.72957 | 3.36270 | 0.00071 | 0.71204 | 3.59543 | 5.58672 | 3.07617 |
| **CMC2** | 7.83380 | 8.92343 | 12.36292 | 7.82357 | 7.80982 | 23.31150 | 14.05604 | 16.38194 | 10.31853 | 7.77397 | 8.86354 | 17.72212 |
| **CMC4** | 0.90426 | 0.57646 | 0.37729 | 0.44100 | 0.38234 | 0.91193 | 0.56540 | 0.17447 | 0.61458 | 0.59095 | 0.26132 | 0.23996 |
| **CMIP** | 0.67648 | 0.00000 | 0.78389 | 0.08448 | 2.33578 | 0.00000 | 3.06601 | 0.00000 | 0.00000 | 6.20313 | 0.00000 | 0.03776 |
| **CMKLR1** | 3.13677 | 0.00000 | 0.06325 | 0.00772 | 0.01986 | 0.00000 | 0.00781 | 0.36748 | 0.01200 | 0.02686 | 0.01737 | 0.30105 |
| **CMPK1** | 0.23861 | 6.50475 | 15.23648 | 1.18374 | 0.01533 | 35.11121 | 18.04372 | 9.65928 | 8.35310 | 8.25172 | 11.04345 | 5.10992 |
| **CMPK2** | 0.00386 | 0.08797 | 0.10369 | 0.00000 | 0.03104 | 0.04072 | 0.02711 | 0.00779 | 0.00000 | 0.15087 | 0.00446 | 0.04026 |
| **CMSS1** | 0.15646 | 9.59755 | 1.87948 | 1.71584 | 7.18987 | 20.22228 | 8.61433 | 0.67687 | 8.22224 | 7.79558 | 7.84939 | 7.00028 |
| **CMTM1** | 0.05598 | 0.03528 | 0.01243 | 0.04844 | 0.05113 | 0.00000 | 0.02021 | 0.03006 | 0.01631 | 0.04731 | 0.03293 | 0.01775 |
| **CMTM2** | 0.11342 | 0.01421 | 0.00798 | 0.07350 | 0.11568 | 0.03208 | 0.00000 | 0.00288 | 0.02678 | 0.05092 | 0.03938 | 0.03144 |
| **CMTM3** | 0.00171 | 0.44780 | 1.71049 | 0.74069 | 0.00000 | 0.00000 | 1.76230 | 0.00972 | 0.00015 | 0.50502 | 0.07766 | 0.00191 |
| **CMTM4** | 11.43164 | 2.09626 | 1.29585 | 3.24401 | 3.49703 | 6.25427 | 2.33309 | 0.79784 | 0.94637 | 8.21094 | 3.96151 | 0.57174 |
| **CMTM6** | 2.35024 | 13.69306 | 22.73448 | 9.70231 | 8.53239 | 41.01150 | 8.04089 | 14.27471 | 31.71489 | 23.64437 | 12.93246 | 16.04491 |
| **CMTM7** | 0.00000 | 3.61141 | 5.43559 | 12.74147 | 6.34449 | 1.12442 | 8.30788 | 0.15264 | 16.59607 | 0.93858 | 11.03296 | 0.00000 |
| **CMTM8** | 0.48535 | 0.49504 | 2.02964 | 0.15463 | 1.07639 | 0.01712 | 2.36691 | 0.00320 | 1.01064 | 1.65750 | 0.00000 | 0.01610 |
| **CMTR1** | 6.22566 | 3.98113 | 2.71857 | 2.14199 | 2.54686 | 0.00000 | 1.39349 | 0.49549 | 3.41114 | 0.54982 | 1.64499 | 2.30402 |
| **CMTR2** | 0.08614 | 1.36088 | 2.00678 | 0.00000 | 1.10879 | 6.18418 | 0.00000 | 0.88023 | 5.10464 | 2.53357 | 0.35136 | 2.62708 |
| **CMYA5** | 0.02591 | 0.21195 | 0.34160 | 0.09345 | 0.98903 | 0.85197 | 0.00000 | 0.37254 | 0.37372 | 0.43856 | 0.02082 | 0.75441 |
| **CNBD1** | 3.18578 | 0.00570 | 0.01161 | 0.00846 | 0.01446 | 0.01755 | 0.01481 | 0.00000 | 0.01641 | 0.02056 | 0.01344 | 0.10471 |
| **CNBD2** | 0.09185 | 0.27878 | 0.19595 | 0.00105 | 0.13902 | 0.00222 | 0.25859 | 0.04243 | 0.15001 | 0.00000 | 0.16855 | 0.43256 |
| **CNBP** | 16.17704 | 37.96069 | 29.74311 | 28.93381 | 22.88590 | 66.53305 | 43.92345 | 30.44701 | 40.60845 | 2.44223 | 37.57575 | 58.89765 |
| **CNDP1** | 0.56951 | 0.00060 | 0.01872 | 0.00138 | 0.25957 | 0.10116 | 0.00000 | 0.00518 | 0.09051 | 0.00000 | 0.06480 | 0.05840 |
| **CNDP2** | 1.69591 | 10.69809 | 18.91683 | 6.74143 | 11.14911 | 14.18355 | 30.61034 | 0.23098 | 30.08408 | 0.00000 | 26.53144 | 4.97457 |
| **CNEP1R1** | 0.02272 | 6.11524 | 0.99083 | 2.90044 | 0.00000 | 10.08657 | 3.64737 | 9.93402 | 3.19122 | 0.00000 | 2.79254 | 13.64821 |
| **CNFN** | 0.06596 | 1.51653 | 0.13064 | 0.07464 | 0.80552 | 0.00017 | 0.01107 | 2.16261 | 0.00000 | 0.19982 | 0.10950 | 1.94209 |
| **CNGA1** | 0.00681 | 0.17204 | 0.00000 | 1.12020 | 0.00048 | 0.05475 | 0.00000 | 0.20258 | 0.08515 | 0.24816 | 0.35460 | 0.59631 |
| **CNGA4** | 0.00356 | 0.05173 | 0.66726 | 0.00000 | 0.00000 | 0.11143 | 0.04721 | 0.08332 | 0.00000 | 0.00870 | 0.00739 | 0.00539 |
| **CNGB1** | 0.00372 | 0.01043 | 0.06442 | 0.01489 | 0.00838 | 0.00909 | 0.01232 | 0.05889 | 0.01676 | 0.12122 | 0.01318 | 0.02856 |
| **CNGB3** | 1.53771 | 0.10698 | 0.01331 | 0.00388 | 0.02615 | 0.00245 | 0.08027 | 0.11949 | 0.00027 | 0.02180 | 0.01253 | 0.00838 |
| **CNIH1** | 1.45738 | 9.12963 | 10.73592 | 6.00387 | 5.83474 | 53.73917 | 0.00000 | 11.37759 | 34.77107 | 15.94799 | 10.68866 | 20.33327 |
| **CNIH2** | 0.01433 | 0.03879 | 0.16614 | 0.02886 | 0.05532 | 0.00000 | 0.45969 | 0.06809 | 0.12692 | 0.60521 | 0.02260 | 0.01831 |
| **CNIH3** | 0.00700 | 0.00151 | 0.03521 | 4.22622 | 0.00000 | 0.08400 | 0.13222 | 0.23044 | 0.00000 | 0.00242 | 0.01853 | 0.00408 |
| **CNIH4** | 0.26323 | 18.94405 | 10.08774 | 25.40776 | 12.61665 | 47.86750 | 0.00000 | 3.04356 | 16.58245 | 17.84992 | 13.58510 | 13.94858 |
| **CNKSR1** | 0.00000 | 3.16346 | 4.38675 | 0.58062 | 0.00000 | 0.00000 | 0.72402 | 1.98313 | 3.18694 | 1.44788 | 0.00495 | 5.37592 |
| **CNKSR2** | 1.77355 | 0.00000 | 0.00000 | 0.00318 | 0.01830 | 0.20258 | 0.00344 | 0.12957 | 0.02521 | 0.38635 | 0.00421 | 0.31583 |
| **CNKSR3** | 0.57173 | 0.00000 | 0.19699 | 2.62538 | 0.00000 | 0.00000 | 0.02026 | 3.24884 | 0.02569 | 0.00750 | 0.02726 | 2.56360 |
| **CNN1** | 1.60079 | 0.00000 | 0.16424 | 0.00000 | 0.00000 | 0.04730 | 0.07090 | 0.46886 | 0.11277 | 0.38794 | 0.00000 | 0.04821 |
| **CNN2** | 0.36805 | 14.29946 | 17.80121 | 7.80239 | 5.19594 | 0.03624 | 13.07973 | 0.38354 | 14.40593 | 0.00000 | 0.00000 | 4.46466 |
| **CNN3** | 13.35481 | 24.81506 | 9.78732 | 24.61014 | 17.30770 | 44.90219 | 23.95102 | 5.88924 | 20.24801 | 1.71187 | 28.68405 | 12.72718 |
| **CNNM1** | 0.44858 | 0.17133 | 0.11570 | 0.33070 | 0.00000 | 0.00000 | 0.00000 | 0.01195 | 0.00317 | 0.47708 | 0.01215 | 0.11659 |
| **CNNM2** | 25.37393 | 0.00000 | 5.81978 | 0.30012 | 0.37454 | 0.94176 | 1.31213 | 5.06377 | 0.67583 | 6.77976 | 0.00000 | 4.11369 |
| **CNNM3** | 0.20312 | 6.90471 | 4.23990 | 2.01310 | 14.03703 | 1.85718 | 2.66806 | 0.00000 | 5.69347 | 27.76571 | 4.06132 | 4.98361 |
| **CNNM4** | 0.03950 | 5.63988 | 2.28964 | 2.83440 | 6.01549 | 6.70992 | 0.00000 | 1.19234 | 12.48913 | 10.55695 | 0.00310 | 0.73075 |
| **CNOT1** | 3.20597 | 11.93036 | 11.58697 | 10.27948 | 14.76859 | 18.35928 | 21.03884 | 7.56770 | 13.54916 | 16.71308 | 10.65578 | 8.31584 |
| **CNOT10** | 0.91065 | 12.14364 | 6.25212 | 6.52228 | 4.05944 | 17.77037 | 10.56483 | 5.70018 | 14.26676 | 3.09336 | 12.31700 | 10.41899 |
| **CNOT11** | 4.59989 | 0.22525 | 6.94182 | 3.97303 | 1.88853 | 9.80303 | 11.69784 | 0.00000 | 0.11569 | 14.98742 | 1.13333 | 1.46699 |
| **CNOT2** | 2.72651 | 11.10918 | 8.54619 | 5.39071 | 5.47287 | 22.13742 | 10.27114 | 8.11592 | 12.99236 | 6.46538 | 8.99977 | 11.69737 |

| | | | | | | | | | | | | |
|---|---|---|---|---|---|---|---|---|---|---|---|---|
| **CNOT3** | 2.58161 | 1.47681 | 5.14397 | 1.09057 | 4.78903 | 0.00000 | 0.20724 | 0.24018 | 4.04476 | 8.36623 | 1.05895 | 1.98564 |
| **CNOT4** | 6.15038 | 4.08329 | 1.57208 | 1.62237 | 2.76539 | 11.71724 | 2.47066 | 6.93326 | 3.98834 | 4.37037 | 4.05104 | 9.39482 |
| **CNOT6** | 6.98806 | 3.38222 | 5.81759 | 2.76554 | 0.00000 | 16.31423 | 13.66717 | 11.21527 | 1.57497 | 17.91433 | 4.90146 | 5.50328 |
| **CNOT6L** | 11.73875 | 1.35323 | 0.82838 | 1.09351 | 0.00000 | 4.95115 | 2.36393 | 1.02527 | 0.00000 | 5.95274 | 0.11824 | 0.00000 |
| **CNOT7** | 9.27257 | 15.03951 | 16.03540 | 10.73048 | 9.28672 | 39.95953 | 27.77300 | 8.61330 | 19.92572 | 11.46988 | 18.09991 | 18.96415 |
| **CNOT8** | 5.51287 | 7.94340 | 2.42413 | 9.07545 | 0.95661 | 20.71829 | 8.74098 | 10.09645 | 8.48809 | 0.15205 | 5.08575 | 10.74931 |
| **CNP** | 5.72114 | 12.23388 | 7.89176 | 8.38731 | 0.00000 | 0.00000 | 4.09000 | 1.66237 | 1.16562 | 2.24571 | 5.82922 | 10.27762 |
| **CNPPD1** | 0.60110 | 6.44898 | 10.24272 | 2.03599 | 5.49632 | 0.00000 | 4.46315 | 1.00578 | 9.01606 | 0.00000 | 3.50234 | 4.85516 |
| **CNPY1** | 0.01004 | 0.03947 | 0.01246 | 0.05905 | 0.00621 | 0.04457 | 0.01269 | 0.13737 | 0.01355 | 0.00648 | 0.01320 | 0.04380 |
| **CNPY2** | 1.51035 | 13.50935 | 22.84106 | 19.01504 | 17.30807 | 21.13872 | 18.62895 | 0.86015 | 33.45889 | 8.45792 | 24.21755 | 13.18528 |
| **CNPY3** | 1.43747 | 6.58427 | 6.43404 | 10.34500 | 4.73474 | 0.00000 | 6.48946 | 0.69505 | 20.11683 | 2.06366 | 7.85901 | 9.44054 |
| **CNPY4** | 2.33685 | 0.22162 | 2.94334 | 1.59864 | 0.00000 | 0.00000 | 0.41701 | 6.60016 | 0.62880 | 2.94373 | 2.92076 | 5.37824 |
| **CNR1** | 0.00573 | 0.14351 | 0.23905 | 0.20878 | 0.23726 | 0.11377 | 0.00000 | 0.01847 | 0.13055 | 0.00000 | 0.00153 | 0.00423 |
| **CNR2** | 0.07482 | 3.83865 | 0.51385 | 2.50434 | 0.19039 | 0.00000 | 0.00128 | 0.00073 | 0.15330 | 0.00000 | 0.10134 | 0.12102 |
| **CNRIP1** | 5.85790 | 0.05333 | 0.00181 | 0.00000 | 0.02563 | 0.00000 | 0.00000 | 1.79322 | 0.04257 | 0.07830 | 0.00640 | 1.11963 |
| **CNST** | 3.90822 | 3.68906 | 7.33547 | 0.66569 | 0.01994 | 5.37194 | 5.21268 | 16.47157 | 8.49452 | 3.09388 | 4.34179 | 8.75164 |
| **CNTD1** | 0.14494 | 0.07292 | 0.00052 | 0.33891 | 0.20938 | 0.44327 | 0.05756 | 0.19492 | 0.49521 | 0.94860 | 0.17342 | 0.25508 |
| **CNTD2** | 5.21242 | 0.01932 | 0.16819 | 0.00836 | 0.17054 | 0.00000 | 0.05109 | 0.53620 | 0.00000 | 0.00000 | 0.00566 | 0.16110 |
| **CNTF** | 0.00671 | 0.00883 | 0.02147 | 0.01198 | 0.01049 | 0.05974 | 0.02559 | 0.02013 | 0.00766 | 0.01158 | 0.01803 | 0.01600 |
| **CNTFR** | 0.00420 | 0.31691 | 0.07085 | 0.45028 | 0.00000 | 0.00000 | 0.03542 | 0.00418 | 0.00014 | 0.17220 | 0.00307 | 0.24631 |
| **CNTLN** | 1.50450 | 0.71050 | 0.26088 | 2.17142 | 0.17422 | 1.01457 | 0.00000 | 0.14827 | 0.00000 | 0.93273 | 0.32537 | 3.47103 |
| **CNTN1** | 0.21934 | 0.01181 | 0.00821 | 0.01414 | 0.00000 | 0.04629 | 0.04563 | 0.06921 | 0.03595 | 0.01284 | 0.02474 | 0.00992 |
| **CNTN2** | 0.12210 | 0.02744 | 0.01681 | 0.02620 | 0.10727 | 0.04695 | 0.00001 | 0.00842 | 0.03038 | 0.18705 | 0.05886 | 0.12951 |
| **CNTN3** | 0.07006 | 0.01510 | 0.02706 | 0.01260 | 0.01283 | 0.02586 | 0.02068 | 0.00595 | 0.01225 | 0.01934 | 0.01378 | 0.01386 |
| **CNTN4** | 1.40677 | 0.06435 | 0.21183 | 0.00000 | 0.00000 | 0.00000 | 0.13221 | 1.65525 | 0.70674 | 0.49349 | 0.00000 | 0.06444 |
| **CNTN4-AS1** | 0.01670 | 0.01459 | 0.01578 | 0.01129 | 0.01331 | 0.01916 | 0.01669 | 0.01901 | 0.02536 | 0.01904 | 0.01286 | 0.01494 |
| **CNTN4-AS2** | 0.00845 | 0.01424 | 0.01417 | 0.02711 | 0.01839 | 0.01764 | 0.02070 | 0.01113 | 0.03020 | 0.00659 | 0.03148 | 0.02182 |
| **CNTN5** | 1.60766 | 0.06768 | 0.13298 | 0.00217 | 0.00361 | 0.00000 | 0.00000 | 0.01737 | 0.01708 | 0.14904 | 0.02753 | 0.02453 |
| **CNTN6** | 0.74199 | 2.54557 | 0.00000 | 2.38169 | 1.35250 | 0.25201 | 0.00000 | 0.22391 | 0.00000 | 0.35654 | 2.16428 | 0.97589 |
| **CNTNAP1** | 0.30236 | 0.00000 | 0.10477 | 0.01976 | 0.09723 | 0.00000 | 0.00193 | 0.00000 | 0.09162 | 0.10973 | 0.03494 | 0.65422 |
| **CNTNAP2** | 3.73381 | 0.01109 | 6.78118 | 0.00940 | 0.00000 | 0.00000 | 1.36545 | 0.22012 | 0.00106 | 0.00000 | 0.10639 | 0.13678 |
| **CNTNAP3** | 0.00505 | 0.03581 | 0.02498 | 0.02001 | 0.02404 | 0.00767 | 0.01512 | 0.01537 | 0.01189 | 0.04063 | 0.01712 | 0.01490 |
| **CNTNAP4** | 0.45003 | 0.04451 | 0.25500 | 0.06898 | 0.01415 | 0.00000 | 0.04757 | 0.23500 | 0.30183 | 0.00000 | 0.00452 | 0.08715 |
| **CNTNAP5** | 0.00867 | 0.37360 | 0.03755 | 0.03200 | 0.00000 | 0.35742 | 0.00000 | 0.00331 | 0.01979 | 0.00508 | 0.00522 | 0.18525 |
| **CNTRL** | 5.32668 | 0.02767 | 0.11233 | 0.00000 | 0.00000 | 1.65619 | 0.69559 | 2.26535 | 0.16744 | 1.50862 | 0.94613 | 1.94520 |
| **CNTROB** | 3.24913 | 1.21806 | 2.79431 | 2.82378 | 2.37553 | 0.00000 | 2.14910 | 1.02370 | 2.33898 | 5.50372 | 0.95354 | 2.58569 |
| **COA1** | 5.96305 | 12.93003 | 4.12113 | 5.90582 | 6.93423 | 16.43706 | 6.62996 | 4.95295 | 8.84250 | 0.00000 | 12.17130 | 10.16658 |
| **COA3** | 1.20276 | 9.63578 | 9.83141 | 13.46628 | 13.47761 | 18.56506 | 17.75773 | 0.07386 | 10.72794 | 0.91005 | 19.10575 | 6.54716 |
| **COA4** | 1.39645 | 11.93566 | 19.92817 | 14.44622 | 13.69913 | 15.95157 | 19.39882 | 3.62389 | 25.16395 | 3.75611 | 18.18861 | 13.92440 |
| **COA5** | 0.49913 | 2.42269 | 4.96412 | 1.44049 | 0.48899 | 12.43434 | 4.99559 | 2.40771 | 3.00251 | 4.38417 | 5.91547 | 7.07513 |
| **COA6** | 0.00000 | 12.29738 | 15.43854 | 7.26987 | 9.90281 | 21.61998 | 27.84214 | 15.10921 | 17.20962 | 3.51161 | 12.57678 | 20.16290 |
| **COA7** | 0.96694 | 12.39734 | 12.59253 | 2.43435 | 6.12986 | 18.76049 | 11.59030 | 1.08166 | 8.76129 | 0.00000 | 16.48536 | 16.67729 |
| **COASY** | 2.57369 | 21.65131 | 19.71551 | 17.24664 | 16.81654 | 0.00000 | 19.13379 | 4.16231 | 20.06005 | 9.09390 | 18.40772 | 16.76313 |
| **COBL** | 0.00000 | 5.64719 | 1.38605 | 2.45764 | 15.86375 | 4.91207 | 10.69245 | 0.00000 | 4.40195 | 16.65257 | 5.82359 | 0.00000 |
| **COBLL1** | 4.37315 | 1.46491 | 7.78481 | 0.00000 | 14.93768 | 3.45118 | 5.04824 | 0.00000 | 1.29390 | 4.72672 | 7.91700 | 0.00000 |
| **COCH** | 1.89360 | 0.00000 | 3.47050 | 0.07507 | 0.00000 | 0.31214 | 2.64780 | 0.72998 | 10.85394 | 0.00000 | 3.23881 | 0.02866 |
| **COG1** | 0.68259 | 4.62029 | 3.22202 | 4.81159 | 2.90774 | 0.00000 | 1.58736 | 0.85030 | 4.16637 | 0.00000 | 5.67183 | 2.77475 |
| **COG2** | 0.88778 | 6.94462 | 1.66084 | 4.34568 | 5.21642 | 3.55441 | 2.77027 | 0.00000 | 4.31173 | 1.03741 | 5.07756 | 4.10472 |

| | | | | | | | | | | | | |
|---|---|---|---|---|---|---|---|---|---|---|---|---|
| **COG3** | 1.36567 | 3.00782 | 3.48746 | 2.18685 | 5.50284 | 5.63661 | 12.17956 | 3.11972 | 7.17947 | 5.78292 | 15.82534 | 2.56618 |
| **COG4** | 0.00000 | 11.44595 | 7.04913 | 8.33294 | 7.13746 | 0.18835 | 5.63222 | 0.97182 | 5.94917 | 1.07580 | 12.70743 | 5.15414 |
| **COG5** | 8.18222 | 3.83831 | 2.26288 | 1.34533 | 1.54933 | 4.64572 | 0.79255 | 1.75030 | 5.28667 | 0.67682 | 2.43856 | 2.08125 |
| **COG6** | 1.22212 | 2.01686 | 3.00807 | 0.17909 | 0.00000 | 7.59276 | 2.40947 | 0.46264 | 1.94289 | 0.27715 | 2.73085 | 1.86846 |
| **COG7** | 0.14197 | 6.08015 | 3.05080 | 4.09867 | 2.66294 | 0.00000 | 6.06767 | 2.20505 | 7.60337 | 1.32522 | 4.96141 | 3.55818 |
| **COG8** | 1.02856 | 4.41133 | 2.82964 | 1.04215 | 1.78333 | 0.00000 | 0.75283 | 1.68468 | 0.99923 | 0.60397 | 3.75140 | 1.81582 |
| **COIL** | 1.84999 | 5.31916 | 2.37427 | 1.73537 | 4.53817 | 11.81907 | 10.20973 | 4.32396 | 6.01840 | 9.35161 | 4.67763 | 6.84783 |
| **COL10A1** | 0.00538 | 0.00070 | 0.00000 | 0.14775 | 0.07603 | 0.10840 | 0.00000 | 0.01087 | 0.04344 | 0.05213 | 0.04530 | 0.07770 |
| **COL11A1** | 0.32295 | 0.01435 | 0.04664 | 0.03622 | 0.04151 | 0.03084 | 0.01130 | 0.11101 | 0.00910 | 0.06684 | 0.03413 | 0.02938 |
| **COL11A2** | 0.08906 | 0.05027 | 0.17648 | 0.79726 | 0.11283 | 0.00000 | 0.05015 | 0.07848 | 0.79259 | 0.00000 | 0.00000 | 0.90605 |
| **COL12A1** | 1.35223 | 0.02531 | 0.18607 | 0.02296 | 0.00000 | 0.00000 | 0.65701 | 0.44625 | 0.00000 | 0.12083 | 0.00920 | 0.08036 |
| **COL13A1** | 0.13272 | 0.10756 | 0.02801 | 0.01197 | 0.00021 | 0.02742 | 0.00740 | 0.00656 | 0.01513 | 0.03912 | 0.01193 | 0.08721 |
| **COL14A1** | 0.02821 | 0.02761 | 0.00325 | 0.17727 | 0.02581 | 0.00911 | 0.05682 | 0.25572 | 0.01876 | 0.10027 | 0.01640 | 0.07125 |
| **COL15A1** | 0.29821 | 0.12137 | 0.00000 | 0.00000 | 0.22814 | 0.53007 | 0.00000 | 0.01122 | 0.00000 | 0.49679 | 0.05811 | 0.01768 |
| **COL16A1** | 0.00987 | 0.00992 | 0.21844 | 0.00000 | 0.01963 | 0.03077 | 0.19081 | 0.06362 | 0.00000 | 0.03015 | 0.01380 | 0.01093 |
| **COL17A1** | 0.03289 | 0.38360 | 0.00000 | 0.58808 | 0.21561 | 0.09313 | 0.08930 | 0.03177 | 0.00000 | 0.17757 | 0.04756 | 0.07963 |
| **COL18A1** | 0.48946 | 0.05021 | 0.70311 | 1.97692 | 0.07616 | 0.05402 | 0.29664 | 0.05545 | 0.00000 | 0.00000 | 0.10301 | 0.00858 |
| **COL18A1-AS1** | 0.00417 | 0.00864 | 0.02353 | 0.00106 | 0.10972 | 0.04628 | 0.00529 | 0.02031 | 0.01065 | 0.05230 | 0.00534 | 0.01354 |
| **COL19A1** | 0.03203 | 0.00391 | 0.00000 | 0.34975 | 0.03861 | 0.78427 | 0.10928 | 0.07146 | 0.00542 | 0.52091 | 0.03565 | 0.01780 |
| **COL1A1** | 0.01117 | 0.00000 | 1.03957 | 0.01450 | 0.74721 | 0.00000 | 1.04781 | 0.00461 | 0.57001 | 0.28547 | 0.00000 | 0.24040 |
| **COL1A2** | 0.00556 | 0.04505 | 0.78072 | 0.00830 | 0.00000 | 0.06113 | 0.97384 | 0.01003 | 0.03693 | 0.00000 | 0.00000 | 0.00921 |
| **COL20A1** | 0.01150 | 0.00468 | 0.02789 | 0.00011 | 0.00430 | 0.01861 | 0.00323 | 0.00492 | 0.04005 | 0.25202 | 0.00000 | 0.09970 |
| **COL21A1** | 0.29366 | 0.00000 | 9.13649 | 0.14279 | 0.18423 | 0.38087 | 1.58256 | 0.00376 | 4.43272 | 1.69298 | 1.61006 | 0.00000 |
| **COL22A1** | 3.94108 | 0.01306 | 0.02342 | 0.01025 | 0.02581 | 0.00000 | 0.01588 | 0.30863 | 0.01683 | 0.00000 | 0.03889 | 0.21943 |
| **COL23A1** | 0.16345 | 0.00729 | 0.16980 | 1.12675 | 0.01324 | 0.00000 | 0.00000 | 0.20099 | 0.03667 | 0.01945 | 0.01366 | 0.02413 |
| **COL24A1** | 0.04055 | 0.14525 | 0.10289 | 0.02929 | 0.00000 | 0.00106 | 0.03445 | 0.45248 | 0.02381 | 0.02344 | 0.02357 | 0.10995 |
| **COL25A1** | 0.15996 | 0.02240 | 0.04812 | 0.00982 | 0.08106 | 0.00000 | 0.00942 | 0.22332 | 0.15402 | 0.00000 | 0.00359 | 0.12346 |
| **COL26A1** | 0.07439 | 1.38960 | 0.00000 | 2.21142 | 0.68358 | 0.44026 | 0.11049 | 0.21782 | 0.00000 | 0.66495 | 0.00000 | 0.15715 |
| **COL27A1** | 0.13845 | 1.60355 | 0.04575 | 1.73431 | 0.67861 | 0.00000 | 0.00000 | 0.00000 | 0.00000 | 2.38150 | 0.04152 | 2.19988 |
| **COL28A1** | 1.99783 | 0.15752 | 0.15377 | 0.00000 | 0.10201 | 0.04016 | 0.00000 | 2.24639 | 0.00000 | 0.17607 | 0.00000 | 0.08536 |
| **COL2A1** | 0.03643 | 0.21621 | 0.00265 | 0.00000 | 0.17304 | 0.00000 | 0.01004 | 0.00069 | 0.00000 | 0.60908 | 0.00000 | 0.25562 |
| **COL3A1** | 0.01885 | 0.01134 | 0.06006 | 0.00240 | 0.03059 | 0.05669 | 0.01140 | 0.05764 | 0.00000 | 0.04715 | 0.00822 | 0.00799 |
| **COL4A1** | 0.01070 | 0.00275 | 5.61845 | 12.64134 | 0.00000 | 0.00000 | 0.00000 | 0.00727 | 0.00484 | 0.12635 | 0.09909 | 0.00683 |
| **COL4A2** | 0.27099 | 0.03632 | 6.00363 | 8.53817 | 0.26827 | 0.00000 | 0.00000 | 0.05544 | 0.00000 | 0.14652 | 1.10196 | 0.00664 |
| **COL4A2-AS1** | 0.00620 | 0.01482 | 0.02882 | 0.01678 | 0.01359 | 0.01779 | 0.01509 | 0.01047 | 0.01675 | 0.01790 | 0.01500 | 0.01603 |
| **COL4A3** | 0.00154 | 0.00580 | 0.09488 | 0.02748 | 0.00000 | 0.12439 | 0.00000 | 0.04558 | 0.00623 | 0.11318 | 0.05705 | 0.01704 |
| **COL4A3BP** | 0.89854 | 3.61888 | 3.48143 | 10.62660 | 6.45155 | 12.65650 | 8.71470 | 3.75724 | 0.00000 | 14.20531 | 8.84428 | 6.03719 |
| **COL4A4** | 0.46852 | 0.12378 | 0.08648 | 0.00417 | 0.00000 | 0.01454 | 0.01723 | 0.08826 | 0.05085 | 0.04706 | 0.04164 | 0.07231 |
| **COL4A5** | 0.01300 | 0.01126 | 0.58152 | 0.30464 | 0.00000 | 0.00707 | 0.61907 | 0.01979 | 0.22662 | 0.00000 | 0.00000 | 0.01863 |
| **COL4A6** | 0.05472 | 0.01470 | 0.01875 | 0.04823 | 0.00653 | 0.02176 | 0.04830 | 0.00532 | 0.03493 | 0.00963 | 0.01595 | 0.01682 |
| **COL5A1** | 0.63125 | 0.04218 | 0.21928 | 0.09737 | 0.10659 | 0.00000 | 0.13857 | 0.00216 | 0.00000 | 0.00000 | 0.04381 | 0.05178 |
| **COL5A2** | 0.25490 | 0.60336 | 0.21032 | 0.00000 | 0.01129 | 0.00000 | 0.20080 | 0.00000 | 0.77606 | 0.30215 | 0.25631 | 1.46649 |
| **COL5A3** | 0.00761 | 0.01374 | 0.08326 | 0.01058 | 0.01997 | 0.01457 | 0.01582 | 0.00999 | 0.01698 | 0.03125 | 0.01307 | 0.01792 |
| **COL6A1** | 0.70663 | 0.30753 | 0.29693 | 1.30499 | 0.84703 | 0.00000 | 0.00000 | 0.59906 | 0.00000 | 2.69956 | 0.00000 | 3.10470 |
| **COL6A2** | 0.32914 | 0.12339 | 0.12031 | 0.32852 | 0.62978 | 0.00000 | 0.02002 | 0.11898 | 0.00000 | 1.08689 | 0.00000 | 0.11178 |
| **COL6A3** | 0.67665 | 0.02851 | 0.02951 | 0.00000 | 0.00000 | 0.04560 | 0.02765 | 0.00667 | 0.01017 | 0.07283 | 0.00649 | 0.03156 |
| **COL6A4P1** | 0.05310 | 0.07047 | 0.05385 | 0.02469 | 0.00222 | 0.00379 | 0.00846 | 0.04718 | 0.03667 | 0.02832 | 0.03800 | 0.06044 |
| **COL6A4P2** | 0.02952 | 0.00293 | 0.10166 | 0.00746 | 0.02386 | 0.02942 | 0.00496 | 0.08702 | 0.02494 | 0.07331 | 0.00677 | 0.11349 |

| | | | | | | | | | | | | |
|---|---|---|---|---|---|---|---|---|---|---|---|---|
| **COL6A5** | 0.15333 | 0.00561 | 0.13791 | 0.01278 | 0.04531 | 0.01758 | 0.05755 | 1.88573 | 0.00000 | 0.03931 | 0.01196 | 0.03781 |
| **COL6A6** | 2.47221 | 0.00000 | 0.01584 | 0.04380 | 0.11305 | 0.11983 | 0.07413 | 0.81144 | 0.00000 | 0.04380 | 0.07230 | 0.18344 |
| **COL7A1** | 0.00077 | 0.00000 | 0.85299 | 0.55864 | 0.26272 | 0.00000 | 0.04108 | 0.05725 | 2.05122 | 6.90759 | 0.01361 | 0.17850 |
| **COL8A1** | 0.00439 | 0.01924 | 0.06888 | 0.00546 | 0.08546 | 0.05268 | 0.00000 | 0.00763 | 0.00000 | 0.09777 | 0.00307 | 0.05824 |
| **COL8A2** | 0.00547 | 0.01891 | 0.06640 | 0.00642 | 0.03724 | 0.00000 | 0.00598 | 0.01078 | 0.05716 | 0.05895 | 0.00921 | 0.01495 |
| **COL9A1** | 0.01099 | 0.02973 | 0.07401 | 0.05888 | 0.00000 | 0.03373 | 0.04797 | 0.05265 | 0.00076 | 0.04943 | 0.01593 | 0.04246 |
| **COL9A2** | 0.05457 | 0.00000 | 0.21512 | 0.05594 | 0.18342 | 0.00000 | 0.05821 | 0.00843 | 0.16557 | 0.36930 | 0.03231 | 0.01097 |
| **COL9A3** | 0.00987 | 5.96992 | 2.05903 | 0.45111 | 4.77197 | 0.00000 | 0.00000 | 0.24444 | 0.83046 | 4.21529 | 1.55363 | 1.50969 |
| **COLCA1** | 0.00533 | 0.01155 | 0.01759 | 0.02227 | 0.00690 | 0.02513 | 0.04090 | 0.01051 | 0.00557 | 0.03970 | 0.01337 | 0.02088 |
| **COLEC10** | 0.00605 | 0.01902 | 0.01439 | 0.02110 | 0.02324 | 0.00048 | 0.02991 | 0.04721 | 0.02764 | 0.00705 | 0.01873 | 0.01555 |
| **COLEC11** | 0.49756 | 0.00381 | 0.15006 | 0.00000 | 0.02552 | 0.07188 | 0.06492 | 0.49886 | 0.00000 | 0.02207 | 0.01980 | 0.02640 |
| **COLEC12** | 0.00000 | 1.71115 | 5.30011 | 0.00000 | 7.74011 | 1.16111 | 0.89752 | 0.04526 | 12.61238 | 11.83491 | 4.68171 | 0.23274 |
| **COLGALT1** | 0.91744 | 16.91900 | 10.39538 | 6.66697 | 15.52887 | 8.08575 | 6.36889 | 14.54950 | 19.87527 | 6.02845 | 8.82089 | 9.79374 |
| **COLGALT2** | 0.00560 | 0.01262 | 0.02188 | 0.01260 | 0.02356 | 0.01032 | 0.01857 | 0.01066 | 0.01119 | 0.04369 | 0.03568 | 0.01538 |
| **COLQ** | 0.01265 | 0.01288 | 0.60834 | 0.15360 | 0.03802 | 0.00000 | 0.10616 | 0.02986 | 0.46054 | 0.00000 | 0.00000 | 0.14833 |
| **COMMD1** | 4.56264 | 2.84075 | 1.62931 | 3.62559 | 7.71055 | 10.27378 | 2.81932 | 4.74945 | 3.12949 | 1.01255 | 7.92923 | 0.70027 |
| **COMMD10** | 3.11483 | 2.37692 | 3.62639 | 2.70223 | 1.05823 | 14.84827 | 2.98898 | 0.49878 | 6.52066 | 1.20518 | 1.73169 | 2.07603 |
| **COMMD2** | 0.95916 | 12.71929 | 8.42162 | 4.06564 | 2.25687 | 30.50807 | 14.84731 | 1.12354 | 7.03505 | 6.36147 | 14.51206 | 10.53745 |
| **COMMD3** | 0.16550 | 3.52221 | 0.89313 | 0.00000 | 1.06621 | 8.30497 | 4.46440 | 0.00000 | 2.98457 | 3.98904 | 5.02918 | 1.41420 |
| **COMMD4** | 1.41865 | 12.71597 | 10.94218 | 11.83541 | 15.79824 | 1.83744 | 12.08235 | 0.94194 | 6.08596 | 3.60340 | 11.16032 | 6.33731 |
| **COMMD5** | 0.49883 | 3.57309 | 6.09988 | 5.87940 | 3.25606 | 1.36824 | 8.03176 | 5.76729 | 8.29337 | 0.00000 | 4.05936 | 8.85135 |
| **COMMD6** | 9.04881 | 5.29972 | 8.74907 | 3.81390 | 6.45240 | 16.93090 | 16.04074 | 8.75511 | 6.85396 | 9.34352 | 8.86083 | 6.23833 |
| **COMMD7** | 1.67370 | 8.20959 | 8.19579 | 5.05790 | 12.94288 | 7.10164 | 11.82602 | 0.28056 | 7.49042 | 7.18386 | 7.74716 | 0.44493 |
| **COMMD8** | 1.18822 | 2.02254 | 8.17284 | 1.56061 | 0.00022 | 18.75674 | 6.06801 | 1.53098 | 7.19942 | 0.29505 | 5.96386 | 5.74707 |
| **COMMD9** | 1.67017 | 8.79784 | 12.26545 | 3.49366 | 7.71107 | 11.23181 | 14.48457 | 2.96633 | 16.20686 | 0.00000 | 12.87589 | 9.98385 |
| **COMP** | 0.08721 | 0.00327 | 0.00807 | 0.32618 | 0.03036 | 0.03775 | 0.02618 | 0.93990 | 0.00000 | 0.00699 | 0.06122 | 0.10315 |
| **COMT** | 0.07109 | 3.46798 | 2.26853 | 0.00687 | 8.26283 | 0.00000 | 1.56716 | 0.02293 | 5.23761 | 0.00000 | 4.12674 | 3.40720 |
| **COMTD1** | 0.00000 | 4.26505 | 3.71413 | 1.63202 | 3.86750 | 0.00000 | 1.97272 | 0.01489 | 3.40964 | 2.70381 | 8.83332 | 0.02019 |
| **COPA** | 2.12134 | 10.21323 | 14.45091 | 6.63792 | 5.44274 | 18.12014 | 14.67781 | 0.73466 | 12.19184 | 4.48068 | 14.55778 | 8.81214 |
| **COPB1** | 0.68395 | 12.92536 | 11.48014 | 7.22883 | 2.39262 | 36.40043 | 14.48922 | 6.44950 | 14.66248 | 13.11967 | 16.70819 | 15.48239 |
| **COPB2** | 5.47340 | 11.90462 | 17.96096 | 13.32631 | 7.54644 | 28.07615 | 22.11959 | 14.31649 | 16.13027 | 0.00000 | 20.00621 | 11.21390 |
| **COPE** | 9.96241 | 23.03046 | 23.99238 | 23.29736 | 23.80733 | 4.20834 | 29.91798 | 9.80535 | 28.22372 | 0.00000 | 25.38195 | 20.56157 |
| **COPG1** | 4.18263 | 12.75203 | 11.83934 | 9.82933 | 8.71810 | 9.71226 | 13.59894 | 2.96324 | 16.75736 | 0.00000 | 21.28362 | 8.02793 |
| **COPG2** | 0.67228 | 0.77237 | 1.04136 | 0.43880 | 0.17515 | 0.91952 | 0.86267 | 0.50551 | 0.00000 | 0.49113 | 0.46652 | 0.55454 |
| **COPRS** | 5.09662 | 2.53299 | 4.43219 | 1.47361 | 5.71942 | 2.57481 | 9.76112 | 2.76697 | 2.31689 | 13.42241 | 2.05634 | 0.19454 |
| **COPS2** | 2.10967 | 7.33598 | 13.28818 | 4.50022 | 0.00000 | 30.61048 | 16.15540 | 14.64219 | 9.46498 | 9.86467 | 9.45393 | 9.80331 |
| **COPS3** | 6.79743 | 15.48416 | 18.73038 | 12.82752 | 10.60923 | 35.71579 | 21.66038 | 13.53436 | 15.91672 | 0.00000 | 17.85283 | 13.84913 |
| **COPS4** | 2.88136 | 9.16139 | 2.79391 | 6.67205 | 3.65953 | 25.47432 | 6.58491 | 9.55590 | 9.25321 | 0.00000 | 10.81019 | 11.28750 |
| **COPS5** | 1.42334 | 12.37365 | 13.82089 | 6.71947 | 3.90733 | 27.44810 | 17.06662 | 8.81384 | 18.45134 | 0.00000 | 13.90624 | 14.55170 |
| **COPS6** | 1.55080 | 17.72781 | 22.32691 | 17.01832 | 13.88716 | 9.75135 | 24.16566 | 9.36068 | 19.89485 | 0.00000 | 23.63222 | 16.30471 |
| **COPS7A** | 1.48865 | 8.97851 | 7.60217 | 5.96515 | 5.52084 | 4.17905 | 11.74800 | 7.76239 | 10.14393 | 0.00000 | 12.52580 | 16.80511 |
| **COPS7B** | 9.10066 | 13.43054 | 3.17777 | 5.57499 | 6.76627 | 0.19972 | 5.32935 | 5.79129 | 7.27821 | 0.00000 | 5.57938 | 10.50390 |
| **COPS8** | 11.26304 | 8.00756 | 5.58526 | 2.66793 | 4.41699 | 25.64118 | 8.71003 | 2.66441 | 9.63833 | 0.00077 | 8.02988 | 1.17832 |
| **COPZ1** | 1.08681 | 19.03776 | 13.52876 | 9.73044 | 9.79547 | 19.81702 | 12.14074 | 0.58375 | 11.55625 | 0.00000 | 14.95108 | 14.00629 |
| **COPZ2** | 0.01362 | 0.01410 | 0.00719 | 0.04032 | 0.00000 | 0.00630 | 0.00759 | 0.01317 | 0.01255 | 0.15031 | 0.03488 | 0.01457 |
| **COQ10A** | 0.44331 | 0.10578 | 1.25692 | 0.37344 | 0.18578 | 0.17058 | 0.60411 | 0.47883 | 1.59935 | 0.08622 | 0.00000 | 0.35808 |
| **COQ10B** | 2.38020 | 8.67476 | 11.43583 | 3.82873 | 1.57369 | 24.23640 | 6.03968 | 12.64697 | 7.42197 | 0.00000 | 6.99605 | 9.99875 |
| **COQ2** | 0.67486 | 5.29391 | 2.92046 | 3.49470 | 4.77425 | 4.45896 | 1.22664 | 0.00000 | 4.31587 | 0.53034 | 5.09976 | 3.76345 |

| | | | | | | | | | | | | |
|---|---|---|---|---|---|---|---|---|---|---|---|---|
| **COQ3** | 0.35811 | 7.45171 | 0.00000 | 1.60223 | 0.78331 | 5.79909 | 0.70675 | 0.57820 | 0.84514 | 0.42147 | 3.14724 | 3.60970 |
| **COQ4** | 0.69918 | 23.26574 | 5.83623 | 8.51620 | 13.92292 | 0.13411 | 9.84602 | 1.80125 | 7.11615 | 0.00000 | 12.97110 | 22.63977 |
| **COQ5** | 0.79116 | 10.14263 | 0.00000 | 4.83997 | 7.57028 | 13.13337 | 0.00000 | 0.36909 | 1.98941 | 0.00000 | 4.16177 | 7.30557 |
| **COQ6** | 1.96326 | 5.28735 | 3.37761 | 3.44652 | 4.08682 | 3.34810 | 3.09292 | 1.06380 | 0.00000 | 0.00000 | 6.57344 | 3.43831 |
| **COQ7** | 2.03656 | 12.64517 | 3.08827 | 2.84478 | 3.84714 | 10.32844 | 5.21809 | 6.56918 | 3.51651 | 0.00000 | 7.49929 | 11.53687 |
| **COQ9** | 1.09794 | 9.60892 | 4.71431 | 3.73919 | 7.61468 | 7.39672 | 1.58418 | 0.69513 | 4.55710 | 1.62012 | 8.58873 | 4.27579 |
| **CORIN** | 0.04173 | 0.02384 | 0.06723 | 0.16155 | 0.00459 | 0.04747 | 0.03882 | 0.04731 | 0.19092 | 0.24641 | 0.00000 | 0.04148 |
| **CORO1A** | 0.00476 | 9.31922 | 0.00000 | 8.58840 | 0.51967 | 0.43579 | 0.04989 | 1.35339 | 0.65865 | 0.00000 | 0.41579 | 16.03357 |
| **CORO1B** | 0.61887 | 7.69951 | 7.80378 | 6.45593 | 11.92358 | 0.00000 | 4.00269 | 0.65195 | 10.32169 | 0.00000 | 7.85032 | 10.24309 |
| **CORO1C** | 3.81505 | 31.68012 | 10.61626 | 15.54982 | 23.46781 | 46.12613 | 21.22578 | 12.22390 | 28.49550 | 14.65836 | 30.21918 | 24.57719 |
| **CORO2A** | 1.72092 | 5.02826 | 3.71821 | 10.51508 | 9.29368 | 3.79489 | 15.78480 | 0.00000 | 15.26759 | 1.14575 | 4.09367 | 0.00000 |
| **CORO2B** | 0.15935 | 0.01430 | 0.02716 | 0.01343 | 0.04454 | 0.01115 | 0.02316 | 0.02793 | 0.02455 | 0.00000 | 0.01343 | 0.02895 |
| **CORO6** | 0.00503 | 0.35264 | 0.07932 | 0.50115 | 0.00000 | 0.00000 | 0.02231 | 0.00437 | 0.12908 | 0.20961 | 0.00000 | 0.36942 |
| **CORO7** | 0.00855 | 0.92781 | 1.46265 | 1.76495 | 1.79286 | 0.00000 | 2.35150 | 0.00542 | 1.67958 | 0.93227 | 0.12803 | 0.00414 |
| **CORT** | 0.00864 | 0.05361 | 0.06414 | 0.07305 | 0.12274 | 0.00000 | 0.07946 | 0.32464 | 0.04341 | 0.32394 | 0.01587 | 0.06159 |
| **COTL1** | 0.42480 | 0.39262 | 11.51299 | 0.90139 | 0.30085 | 0.00000 | 3.62753 | 0.54615 | 2.69997 | 0.57332 | 1.79898 | 0.86041 |
| **COX10** | 4.47125 | 8.35452 | 7.80518 | 5.61305 | 5.10060 | 8.90261 | 9.41255 | 5.45688 | 16.47034 | 0.00000 | 10.07535 | 7.22734 |
| **COX10-AS1** | 0.05034 | 0.11610 | 1.24373 | 0.36426 | 0.21290 | 1.56340 | 0.66479 | 0.32657 | 1.61773 | 0.00000 | 0.20440 | 0.36153 |
| **COX11** | 0.03424 | 7.62143 | 5.39596 | 10.07072 | 4.25690 | 22.57186 | 10.37246 | 0.35705 | 12.79750 | 0.00000 | 10.95244 | 5.52438 |
| **COX14** | 2.21809 | 9.10769 | 13.33533 | 10.41466 | 14.68947 | 9.81419 | 19.63628 | 3.73079 | 14.83514 | 6.12150 | 15.89820 | 11.08180 |
| **COX15** | 0.81117 | 9.06774 | 4.37809 | 5.70163 | 3.95673 | 15.91201 | 8.26407 | 0.78744 | 7.28554 | 0.54145 | 9.76747 | 9.92152 |
| **COX16** | 0.30918 | 4.15505 | 4.99402 | 6.27955 | 11.53505 | 8.86991 | 9.36193 | 9.51505 | 5.09230 | 1.77113 | 6.57670 | 13.74531 |
| **COX17** | 0.27150 | 3.80788 | 3.50716 | 4.37676 | 7.03792 | 9.47875 | 8.57109 | 2.09102 | 2.14372 | 4.06922 | 4.44885 | 5.84983 |
| **COX18** | 0.23581 | 0.45837 | 0.93187 | 0.89608 | 1.82488 | 3.15493 | 0.52165 | 0.04155 | 2.71622 | 2.68464 | 2.54154 | 0.81270 |
| **COX19** | 0.93386 | 6.14867 | 5.67205 | 2.33760 | 3.46811 | 4.52983 | 10.56372 | 5.71334 | 5.01523 | 4.09560 | 2.61708 | 8.20174 |
| **COX20** | 0.00982 | 1.41380 | 1.49665 | 1.07134 | 2.62829 | 1.42355 | 1.72734 | 0.00000 | 1.31760 | 0.34662 | 1.86868 | 0.06436 |
| **COX4I1** | 5.53463 | 41.86618 | 29.75980 | 40.58725 | 52.16449 | 42.63825 | 51.52484 | 15.85433 | 49.94830 | 33.16827 | 41.99968 | 35.36385 |
| **COX4I2** | 9.07086 | 0.00000 | 0.05235 | 0.03499 | 0.00000 | 0.00000 | 0.00000 | 5.06645 | 0.01327 | 0.21759 | 0.00171 | 0.68893 |
| **COX5A** | 0.97219 | 26.71781 | 27.75247 | 26.95634 | 33.41818 | 46.34309 | 39.25287 | 4.35675 | 41.76391 | 17.86270 | 35.28990 | 22.18434 |
| **COX5B** | 7.96632 | 33.12845 | 27.11949 | 34.27245 | 41.11334 | 38.36069 | 42.95056 | 24.70705 | 36.67203 | 28.14343 | 37.76834 | 34.71218 |
| **COX6A1** | 2.26093 | 18.64632 | 15.12452 | 18.71539 | 22.99636 | 24.94402 | 16.11129 | 17.43198 | 25.18684 | 21.47076 | 19.87085 | 26.16514 |
| **COX6B1** | 8.30150 | 28.81753 | 27.19918 | 30.94216 | 42.05296 | 29.69497 | 48.08145 | 8.41352 | 35.12740 | 24.91132 | 35.95496 | 25.74039 |
| **COX6B2** | 0.00415 | 0.57114 | 0.01096 | 0.08152 | 0.00000 | 0.00289 | 0.00277 | 2.56719 | 0.01139 | 0.10561 | 0.02998 | 0.35415 |
| **COX6C** | 8.67030 | 21.29226 | 18.86488 | 23.77045 | 22.93060 | 54.34952 | 36.34195 | 16.13820 | 25.31577 | 30.27979 | 28.85314 | 26.99520 |
| **COX7A1** | 0.14139 | 0.24733 | 0.07585 | 0.47523 | 0.03861 | 0.16972 | 0.00000 | 0.01836 | 0.00000 | 0.11692 | 0.09577 | 0.00000 |
| **COX7A2** | 4.98008 | 23.80528 | 22.35261 | 25.31400 | 30.36724 | 45.98711 | 29.97955 | 12.41365 | 34.76999 | 21.33836 | 33.18785 | 26.88655 |
| **COX7A2L** | 12.00237 | 22.15341 | 16.47759 | 13.67955 | 19.53427 | 45.84207 | 25.91510 | 10.11151 | 24.31343 | 7.80997 | 21.81892 | 24.86758 |
| **COX7B** | 3.13101 | 14.68190 | 19.54262 | 23.22037 | 22.07871 | 43.22187 | 38.10064 | 17.87481 | 29.21677 | 17.74335 | 32.92110 | 23.43285 |
| **COX7B2** | 0.00000 | 7.98444 | 10.41421 | 3.09717 | 6.62778 | 25.83238 | 24.14791 | 2.96105 | 3.16798 | 6.30746 | 9.11052 | 12.44297 |
| **COX7C** | 5.23102 | 19.03188 | 23.80600 | 22.20689 | 27.01027 | 41.60274 | 37.13735 | 3.36936 | 30.62751 | 24.21479 | 27.69785 | 11.10921 |
| **COX8A** | 1.91823 | 24.81898 | 26.11110 | 31.61320 | 39.70885 | 15.01066 | 33.15479 | 10.78040 | 33.90580 | 13.87845 | 28.78033 | 22.15425 |
| **COX8C** | 0.76957 | 0.01598 | 1.82727 | 0.34033 | 0.09994 | 1.38519 | 0.00152 | 0.29072 | 2.37626 | 0.00000 | 0.57301 | 1.30154 |
| **CP** | 0.01435 | 0.12038 | 0.02881 | 0.00477 | 0.47648 | 0.22035 | 0.02194 | 0.00944 | 0.04851 | 0.00000 | 0.13411 | 0.00198 |
| **CPA1** | 1.03684 | 0.04917 | 0.64029 | 0.01491 | 0.02737 | 0.05513 | 0.00000 | 0.21956 | 0.00000 | 0.00682 | 0.01613 | 0.14690 |
| **CPA2** | 0.63902 | 0.20639 | 0.02020 | 0.00000 | 0.00000 | 0.02564 | 0.20095 | 0.03067 | 0.06126 | 0.08148 | 0.00000 | 0.00455 |
| **CPA4** | 0.07262 | 0.00707 | 0.11106 | 0.01744 | 0.02061 | 0.00000 | 0.01135 | 0.00000 | 0.01156 | 0.01926 | 0.02177 | 0.28665 |
| **CPA5** | 0.17117 | 0.01656 | 0.04196 | 0.01695 | 0.01554 | 0.01792 | 0.05617 | 0.04472 | 0.02501 | 0.00106 | 0.02033 | 0.01389 |
| **CPA6** | 0.11784 | 0.02036 | 0.02805 | 0.01833 | 0.00348 | 0.05011 | 0.00759 | 0.06093 | 0.00982 | 0.02560 | 0.01267 | 0.06579 |

| | | | | | | | | | | | | |
|---|---|---|---|---|---|---|---|---|---|---|---|---|
| **CPAMD8** | 0.00531 | 0.01147 | 0.12294 | 0.00750 | 0.01636 | 0.02320 | 0.07079 | 0.01035 | 0.00430 | 0.05146 | 0.01849 | 0.01355 |
| **CPB1** | 0.06521 | 0.00447 | 0.00874 | 0.00752 | 0.00170 | 0.06061 | 0.02609 | 0.00786 | 0.00638 | 0.03942 | 0.01231 | 0.04071 |
| **CPB2** | 0.12165 | 0.00000 | 0.01936 | 0.00000 | 0.03898 | 0.04116 | 0.01425 | 0.02974 | 0.03906 | 0.19731 | 0.02264 | 0.26683 |
| **CPB2-AS1** | 0.03655 | 0.07942 | 0.01787 | 0.00173 | 0.00000 | 0.00000 | 0.14603 | 0.08447 | 0.00000 | 0.22070 | 0.09485 | 0.22172 |
| **CPD** | 1.88577 | 2.79718 | 0.41475 | 2.70440 | 0.00000 | 7.14964 | 0.00000 | 5.29438 | 0.82547 | 8.90996 | 0.25804 | 4.03444 |
| **CPE** | 0.05316 | 0.02682 | 0.02997 | 0.04516 | 0.00866 | 0.01464 | 0.02859 | 0.05639 | 0.02120 | 0.01228 | 0.02153 | 0.05079 |
| **CPEB1** | 4.97908 | 0.56547 | 0.00000 | 0.00000 | 0.02717 | 0.08437 | 0.00792 | 1.18331 | 0.00000 | 0.16394 | 0.02042 | 1.93727 |
| **CPEB2** | 3.26605 | 0.30322 | 0.10466 | 0.75269 | 0.00080 | 2.97047 | 0.00001 | 2.84246 | 0.00000 | 3.25147 | 0.00000 | 1.00248 |
| **CPEB2-AS1** | 0.00476 | 0.01574 | 0.02082 | 0.01006 | 0.01705 | 0.01758 | 0.01520 | 0.01909 | 0.01926 | 0.02045 | 0.01509 | 0.02841 |
| **CPEB3** | 0.80292 | 0.72584 | 0.36803 | 0.00000 | 0.78466 | 1.09580 | 0.77577 | 0.77357 | 1.19188 | 2.13846 | 0.00284 | 1.43376 |
| **CPEB4** | 5.76396 | 0.25746 | 7.88714 | 0.00000 | 0.00000 | 2.64631 | 0.64270 | 8.44587 | 0.00000 | 5.05848 | 0.03084 | 3.17242 |
| **CPED1** | 5.20000 | 0.00000 | 0.09205 | 0.16360 | 0.11174 | 0.09143 | 0.04765 | 0.90742 | 0.13090 | 0.36460 | 0.04554 | 4.60865 |
| **CPLX1** | 0.17572 | 0.15235 | 0.00665 | 0.00000 | 0.00000 | 0.12876 | 0.00000 | 0.30855 | 0.00000 | 0.36716 | 0.00001 | 0.08613 |
| **CPLX2** | 0.00295 | 0.00537 | 0.25027 | 0.00000 | 0.13725 | 0.00000 | 0.14177 | 0.00701 | 0.00000 | 1.38716 | 0.19478 | 0.01839 |
| **CPLX3** | 0.00433 | 0.24020 | 0.00000 | 0.00000 | 0.02782 | 0.02222 | 0.00425 | 0.00445 | 0.01320 | 0.23762 | 0.00304 | 0.31000 |
| **CPLX4** | 2.04198 | 0.05201 | 0.00845 | 0.00334 | 0.06009 | 0.00000 | 0.00300 | 0.48192 | 0.09870 | 0.00000 | 0.00639 | 0.00000 |
| **CPM** | 0.18488 | 0.00000 | 6.07660 | 0.03881 | 0.01954 | 0.00000 | 0.00000 | 2.48920 | 0.16099 | 0.17408 | 0.00000 | 1.38590 |
| **CPN1** | 0.07378 | 0.12079 | 0.41106 | 2.29682 | 0.00000 | 0.00000 | 0.00000 | 0.00000 | 0.00000 | 0.14772 | 0.16876 | 0.21539 |
| **CPN2** | 0.00635 | 0.06339 | 0.10648 | 0.01578 | 0.04596 | 0.00000 | 0.01413 | 0.01086 | 0.01407 | 0.00783 | 0.02366 | 0.01334 |
| **CPNE1** | 1.03915 | 21.07724 | 18.89857 | 16.92264 | 16.35245 | 2.18544 | 23.01526 | 13.55239 | 21.47541 | 0.00000 | 29.05538 | 14.00392 |
| **CPNE2** | 0.17717 | 0.84903 | 0.16553 | 0.18527 | 0.48745 | 0.08043 | 0.15123 | 0.00000 | 0.09503 | 0.00000 | 0.00000 | 0.97612 |
| **CPNE3** | 0.02343 | 5.41054 | 7.50654 | 3.05043 | 1.83000 | 15.47131 | 15.47523 | 1.53508 | 3.39282 | 5.59189 | 12.24376 | 4.73472 |
| **CPNE4** | 1.44873 | 0.16183 | 0.00951 | 0.01009 | 0.00000 | 0.05874 | 0.01720 | 0.12107 | 0.00000 | 0.14950 | 0.01429 | 0.06431 |
| **CPNE5** | 0.00554 | 0.01079 | 0.19087 | 0.01048 | 0.00696 | 0.04261 | 0.00974 | 0.02832 | 0.00671 | 0.07016 | 0.01229 | 0.00990 |
| **CPNE6** | 0.03061 | 0.01020 | 0.02093 | 0.00932 | 0.00932 | 0.03207 | 0.00873 | 0.04486 | 0.01717 | 0.04009 | 0.01818 | 0.01846 |
| **CPNE7** | 0.04044 | 0.18534 | 0.05200 | 0.00000 | 0.00000 | 0.00000 | 0.00709 | 0.05440 | 0.00000 | 0.30258 | 0.02430 | 0.02675 |
| **CPNE8** | 0.14334 | 0.00000 | 2.44465 | 0.09262 | 0.00000 | 1.03553 | 0.39656 | 0.28236 | 1.53357 | 0.57350 | 3.18735 | 0.05774 |
| **CPNE9** | 4.61969 | 0.43100 | 0.10185 | 0.09158 | 0.20456 | 0.00000 | 0.09041 | 0.57277 | 0.15132 | 0.27936 | 0.00000 | 0.31674 |
| **CPO** | 0.24317 | 0.02988 | 0.00733 | 0.15933 | 0.03128 | 0.00000 | 0.05134 | 0.00213 | 0.04690 | 0.00000 | 0.01872 | 0.03117 |
| **CPOX** | 0.00000 | 1.63638 | 2.23776 | 0.00000 | 0.00000 | 5.46266 | 5.73867 | 0.55155 | 0.00000 | 10.96764 | 0.13202 | 1.86184 |
| **CPPED1** | 0.18024 | 8.21802 | 0.49987 | 7.15313 | 5.85400 | 10.75564 | 0.00000 | 0.92145 | 0.82629 | 2.26461 | 6.85550 | 9.05381 |
| **CPQ** | 0.72848 | 0.04546 | 0.00958 | 0.00427 | 0.00419 | 0.05219 | 0.01056 | 0.11651 | 0.01239 | 0.02061 | 0.01126 | 0.04678 |
| **CPS1** | 5.15622 | 0.00822 | 0.00000 | 0.09980 | 0.10591 | 0.11956 | 0.00000 | 0.37735 | 0.00000 | 0.01098 | 0.00292 | 0.57263 |
| **CPSF1** | 2.05033 | 6.23415 | 5.51520 | 11.55688 | 13.73827 | 0.00000 | 6.98488 | 0.07805 | 9.73175 | 16.31012 | 7.13066 | 5.99794 |
| **CPSF2** | 9.02977 | 4.59162 | 9.07936 | 4.55691 | 3.11626 | 22.14663 | 10.61224 | 1.47627 | 6.03069 | 7.25673 | 6.24857 | 8.13637 |
| **CPSF3** | 0.56818 | 15.93366 | 8.48501 | 5.62329 | 8.42038 | 18.43443 | 6.79002 | 0.00000 | 7.67435 | 1.93519 | 13.94550 | 7.31629 |
| **CPSF3L** | 0.09826 | 17.10139 | 13.32451 | 12.60632 | 12.78451 | 0.00000 | 14.67144 | 4.96012 | 19.18201 | 0.00000 | 14.68438 | 14.24693 |
| **CPSF4** | 2.21092 | 7.78805 | 7.27496 | 7.95429 | 6.82371 | 3.23481 | 8.17771 | 2.46891 | 6.27326 | 7.83076 | 0.00000 | 4.55101 |
| **CPSF4L** | 0.72590 | 1.09422 | 0.02103 | 0.14404 | 0.01356 | 0.04507 | 0.00000 | 0.61564 | 0.00000 | 0.13205 | 0.00044 | 0.00000 |
| **CPSF6** | 0.52152 | 15.58870 | 11.99496 | 15.55288 | 9.27709 | 29.84532 | 16.29843 | 10.97581 | 22.09777 | 6.40668 | 19.57096 | 18.98899 |
| **CPSF7** | 0.32256 | 7.43328 | 6.73003 | 4.46246 | 8.77590 | 2.39747 | 6.05209 | 3.95265 | 17.20939 | 9.58393 | 3.55336 | 9.72271 |
| **CPT1A** | 0.53272 | 2.35101 | 0.63829 | 4.19876 | 0.00000 | 0.07393 | 1.95282 | 2.66881 | 0.00000 | 0.13901 | 0.11221 | 1.46427 |
| **CPT1B** | 0.00329 | 0.01009 | 0.00363 | 0.02454 | 0.01123 | 0.06460 | 0.02100 | 0.01320 | 0.06940 | 0.13569 | 0.00111 | 0.06790 |
| **CPT1C** | 1.66070 | 0.06955 | 0.23167 | 0.13969 | 0.00000 | 0.22725 | 0.00000 | 5.71695 | 0.30698 | 1.18387 | 0.00000 | 0.80926 |
| **CPT2** | 0.39166 | 2.24898 | 3.89236 | 1.58246 | 2.12448 | 0.40326 | 1.71344 | 0.06833 | 2.84200 | 0.00000 | 3.95794 | 0.87408 |
| **CPTP** | 0.10344 | 1.84582 | 2.32301 | 0.45065 | 2.10733 | 0.00000 | 1.87677 | 0.00000 | 0.64089 | 2.56159 | 2.10335 | 1.00987 |
| **CPVL** | 1.40306 | 8.18694 | 5.29338 | 3.84863 | 13.96849 | 10.83167 | 0.02271 | 0.05541 | 19.89118 | 1.66279 | 23.57065 | 0.66406 |
| **CPXM1** | 0.03596 | 0.05684 | 0.31847 | 0.15252 | 0.00000 | 0.00000 | 0.00000 | 0.00551 | 0.01309 | 0.07507 | 0.03009 | 0.05759 |

| | | | | | | | | | | | | |
|---|---|---|---|---|---|---|---|---|---|---|---|---|
| **CPXM2** | 0.15632 | 0.00700 | 0.01402 | 0.25380 | 0.00364 | 0.05538 | 0.00000 | 0.00823 | 0.00000 | 0.11970 | 0.00888 | 0.08955 |
| **CPZ** | 0.26146 | 0.00000 | 2.94483 | 0.00000 | 0.00000 | 0.01654 | 0.19098 | 0.00734 | 0.99014 | 0.16191 | 0.14042 | 0.06042 |
| **CR1** | 0.06885 | 0.00168 | 0.00035 | 0.03002 | 0.01372 | 0.05674 | 0.01303 | 0.00915 | 0.00421 | 0.21019 | 0.00310 | 0.11639 |
| **CR1L** | 0.00238 | 0.02635 | 0.00000 | 0.78697 | 0.07069 | 0.10121 | 0.00000 | 0.10388 | 0.00698 | 0.00351 | 0.02084 | 0.01174 |
| **CR2** | 3.52786 | 0.23730 | 0.00000 | 0.00000 | 0.03535 | 0.00000 | 0.00000 | 1.76559 | 0.30399 | 0.04761 | 0.00131 | 0.07382 |
| **CRABP1** | 5.15855 | 0.00775 | 0.12209 | 0.54629 | 0.00320 | 0.01858 | 0.05823 | 0.40662 | 0.04952 | 0.00000 | 0.00440 | 0.02524 |
| **CRABP2** | 0.00268 | 0.05848 | 0.17733 | 0.63109 | 0.55582 | 0.18166 | 0.11139 | 1.19988 | 0.00001 | 0.61359 | 0.03480 | 0.31504 |
| **CRADD** | 0.06286 | 3.08792 | 2.99351 | 0.90932 | 3.42859 | 6.79045 | 2.23365 | 0.06664 | 0.88605 | 0.00000 | 3.79752 | 2.19451 |
| **CRAMP1L** | 3.18512 | 3.38196 | 1.55769 | 1.62427 | 3.09810 | 3.75039 | 2.01908 | 1.43532 | 3.52260 | 12.41359 | 0.71897 | 3.74559 |
| **CRAT** | 0.91871 | 0.27044 | 3.71682 | 0.83570 | 0.00000 | 0.00394 | 1.16120 | 0.00000 | 0.24276 | 0.00000 | 0.00000 | 0.07588 |
| **CRB1** | 0.00483 | 0.03540 | 0.03364 | 0.01413 | 0.00842 | 0.03145 | 0.01378 | 0.00926 | 0.00508 | 0.08070 | 0.02930 | 0.02930 |
| **CRB2** | 0.10236 | 0.00789 | 0.07727 | 0.00187 | 0.03634 | 0.00378 | 0.04281 | 0.00517 | 0.06495 | 0.16361 | 0.00000 | 0.01045 |
| **CRB3** | 0.35698 | 14.91222 | 8.15584 | 9.19526 | 18.08992 | 0.00000 | 6.50546 | 0.62914 | 11.00277 | 0.00000 | 9.08545 | 8.58864 |
| **CRBN** | 1.19779 | 5.84320 | 2.52175 | 2.61669 | 0.70089 | 14.49681 | 6.00971 | 0.79338 | 3.93018 | 2.07220 | 8.73569 | 4.44325 |
| **CRCP** | 2.24045 | 3.99223 | 4.38580 | 3.23699 | 6.22447 | 1.22100 | 7.05741 | 8.40920 | 5.76840 | 1.46598 | 3.19289 | 14.45686 |
| **CREB1** | 2.88950 | 4.16436 | 3.47225 | 3.67318 | 2.32470 | 10.88632 | 4.85204 | 7.37560 | 4.26070 | 6.99940 | 5.04962 | 3.69892 |
| **CREB3** | 0.00814 | 11.80788 | 2.16915 | 8.61376 | 11.89206 | 6.12195 | 3.36257 | 5.95719 | 4.78741 | 0.00000 | 4.85701 | 13.26808 |
| **CREB3L1** | 0.31835 | 0.12553 | 0.99905 | 0.60105 | 0.00000 | 0.00000 | 0.00000 | 0.16873 | 0.45056 | 0.02780 | 0.00000 | 0.00180 |
| **CREB3L2** | 7.73288 | 0.52811 | 7.85479 | 1.13282 | 0.35130 | 1.93104 | 1.42356 | 0.04697 | 0.43091 | 0.00000 | 0.46062 | 0.08233 |
| **CREB3L3** | 0.81143 | 0.98279 | 0.09848 | 1.10243 | 0.00000 | 0.00000 | 0.12777 | 0.00000 | 0.01232 | 0.05557 | 0.00000 | 0.09801 |
| **CREB3L4** | 0.20834 | 1.79297 | 1.46605 | 1.55857 | 0.62432 | 0.11504 | 1.40172 | 0.00000 | 1.97124 | 0.00000 | 1.83647 | 1.58094 |
| **CREB5** | 3.00309 | 0.00000 | 0.13754 | 0.03528 | 0.03868 | 0.00000 | 0.00005 | 0.26263 | 0.14424 | 0.49247 | 0.00000 | 0.24883 |
| **CREBBP** | 7.75914 | 0.00000 | 3.81683 | 3.86582 | 13.26434 | 8.37721 | 8.84623 | 0.97917 | 1.93578 | 34.55013 | 1.07905 | 10.56224 |
| **CREBL2** | 0.74573 | 1.85774 | 0.00000 | 9.18555 | 0.53779 | 6.84808 | 0.00000 | 0.26782 | 0.00000 | 4.54509 | 0.64976 | 0.85324 |
| **CREBRF** | 0.17666 | 0.12683 | 0.28045 | 0.32830 | 0.00000 | 2.01778 | 0.73222 | 1.56011 | 0.00000 | 3.92283 | 0.00000 | 1.62033 |
| **CREBZF** | 0.38942 | 2.60057 | 6.90583 | 1.56765 | 1.47876 | 4.50619 | 4.95070 | 2.12025 | 8.14146 | 13.67973 | 4.87390 | 0.59490 |
| **CREG1** | 0.00000 | 7.42940 | 1.92152 | 1.27414 | 0.42332 | 5.24556 | 0.00044 | 0.93726 | 2.60445 | 6.54988 | 0.22027 | 7.37294 |
| **CREG2** | 0.00545 | 0.01478 | 0.03214 | 0.01063 | 0.08293 | 0.00000 | 0.00341 | 0.01055 | 0.06890 | 0.05673 | 0.04237 | 0.01462 |
| **CRELD1** | 1.40720 | 7.24249 | 1.38486 | 3.77461 | 0.59495 | 0.00001 | 0.00000 | 0.00000 | 5.46866 | 2.08030 | 0.56821 | 12.73557 |
| **CRELD2** | 5.31415 | 5.55218 | 8.21653 | 2.47508 | 5.13953 | 0.00000 | 0.00000 | 5.17900 | 9.68620 | 1.96299 | 3.02332 | 1.87967 |
| **CREM** | 1.19667 | 7.50358 | 13.89016 | 4.30334 | 8.06110 | 20.22782 | 2.75902 | 9.93705 | 3.32684 | 0.88390 | 6.16048 | 12.84083 |
| **CRH** | 0.00413 | 0.00410 | 0.05118 | 0.00000 | 0.01025 | 0.04311 | 0.00000 | 0.00981 | 0.07311 | 0.14922 | 0.00000 | 0.01294 |
| **CRHBP** | 0.03263 | 0.00395 | 0.23154 | 2.10776 | 0.02694 | 0.03716 | 0.00000 | 1.19405 | 0.04703 | 0.08336 | 0.00588 | 0.33327 |
| **CRHR1** | 0.00598 | 0.06268 | 0.02555 | 0.01531 | 0.00687 | 0.02549 | 0.01807 | 0.03204 | 0.00848 | 0.02415 | 0.01722 | 0.00936 |
| **CRHR1-IT1** | 0.00102 | 0.12169 | 0.10502 | 0.31819 | 0.41749 | 0.09137 | 0.00000 | 0.17183 | 0.41145 | 0.65197 | 0.23625 | 0.29835 |
| **CRHR2** | 0.00440 | 0.07476 | 0.03618 | 0.00000 | 0.00000 | 0.00000 | 0.01318 | 0.00836 | 0.00000 | 0.16783 | 0.58741 | 0.05266 |
| **CRIM1** | 0.67102 | 0.00000 | 8.79272 | 0.00000 | 0.05283 | 8.59040 | 2.55062 | 0.50138 | 4.47176 | 13.44024 | 2.19653 | 0.31932 |
| **CRIP1** | 0.00000 | 17.77882 | 3.21232 | 19.85543 | 42.45854 | 8.41551 | 26.48076 | 0.00000 | 30.76783 | 9.23367 | 22.40314 | 0.00000 |
| **CRIP2** | 0.00064 | 2.15836 | 2.91878 | 0.59758 | 0.44714 | 0.00000 | 0.22793 | 0.07776 | 0.00000 | 0.06246 | 0.00000 | 0.14806 |
| **CRIP3** | 0.00232 | 0.36086 | 0.00000 | 0.85150 | 0.00000 | 0.00000 | 0.19427 | 0.01334 | 0.49563 | 0.20439 | 0.00000 | 0.41581 |
| **CRIPAK** | 0.00000 | 0.14233 | 0.28027 | 0.27074 | 0.00000 | 0.64783 | 0.66037 | 0.00196 | 0.16526 | 1.60260 | 0.47686 | 0.13664 |
| **CRIPT** | 0.10832 | 4.12595 | 6.14645 | 0.62115 | 0.00000 | 16.20663 | 8.09339 | 12.71033 | 3.23767 | 4.42814 | 4.91159 | 15.62573 |
| **CRISP2** | 0.02216 | 0.01253 | 0.01932 | 0.02873 | 0.01696 | 0.00412 | 0.03028 | 0.01001 | 0.02889 | 0.01160 | 0.01364 | 0.12201 |
| **CRISP3** | 0.00551 | 0.00919 | 0.00785 | 0.00688 | 0.00564 | 0.04644 | 0.03925 | 0.00846 | 0.01871 | 0.01686 | 0.00923 | 0.06644 |
| **CRISPLD1** | 0.46265 | 0.19648 | 0.13224 | 0.05269 | 0.00000 | 0.52508 | 0.00000 | 1.28955 | 0.00000 | 0.00000 | 0.20286 | 1.30191 |
| **CRISPLD2** | 5.91655 | 0.01164 | 1.43666 | 0.05382 | 0.00000 | 0.00000 | 0.62479 | 1.80157 | 0.00000 | 0.04255 | 0.02741 | 0.20745 |
| **CRK** | 5.21750 | 7.07163 | 7.87166 | 6.38536 | 10.71451 | 7.36874 | 21.28625 | 5.63057 | 7.51917 | 22.30510 | 8.23630 | 8.12658 |
| **CRKL** | 11.01135 | 16.32683 | 11.65747 | 13.95790 | 12.24236 | 22.24599 | 14.31974 | 25.00279 | 14.67410 | 3.51878 | 12.84240 | 19.29106 |

| | | | | | | | | | | | | |
|---|---|---|---|---|---|---|---|---|---|---|---|---|
| **CRLF1** | 0.02601 | 0.07873 | 0.11480 | 0.10315 | 0.05461 | 0.00000 | 0.11611 | 0.05537 | 0.00000 | 0.00000 | 0.02546 | 0.11203 |
| **CRLF3** | 0.46488 | 2.75943 | 7.85725 | 2.15817 | 0.00000 | 10.55125 | 3.32471 | 4.15815 | 8.15943 | 1.83522 | 7.49116 | 2.24060 |
| **CRLS1** | 6.37881 | 2.26390 | 1.89827 | 2.18638 | 3.37104 | 6.55553 | 7.68059 | 1.08412 | 0.00000 | 11.90361 | 3.23791 | 0.83757 |
| **CRMP1** | 11.09356 | 0.00367 | 0.01711 | 0.04865 | 0.00489 | 0.14600 | 0.00000 | 2.22451 | 0.00000 | 0.00189 | 0.02032 | 0.12611 |
| **CRNDE** | 0.56415 | 0.50789 | 0.00000 | 0.15597 | 0.00000 | 0.86690 | 0.00000 | 0.84775 | 0.25629 | 1.11214 | 0.30506 | 2.45611 |
| **CRNKL1** | 1.79654 | 8.02020 | 7.03524 | 2.98783 | 3.17441 | 16.71448 | 7.62778 | 12.04701 | 6.88289 | 3.59933 | 3.38113 | 11.69581 |
| **CRNN** | 0.00621 | 0.01457 | 0.01394 | 0.01308 | 0.01662 | 0.01813 | 0.02213 | 0.01014 | 0.01848 | 0.00970 | 0.01474 | 0.02265 |
| **CROCC** | 0.04711 | 0.12076 | 0.08995 | 0.46965 | 0.00000 | 0.00000 | 0.06424 | 0.14144 | 0.19164 | 0.63406 | 0.00000 | 0.03859 |
| **CROCCP2** | 0.00000 | 0.25353 | 1.53454 | 0.01999 | 0.00485 | 2.28829 | 1.60380 | 0.48055 | 3.47695 | 3.97963 | 0.00000 | 2.03859 |
| **CROCCP3** | 0.02068 | 0.01283 | 0.05612 | 0.07379 | 0.00000 | 0.11152 | 0.00000 | 0.12230 | 0.30882 | 0.10538 | 0.18506 | 0.25724 |
| **CROT** | 0.36047 | 0.00000 | 0.86972 | 0.00000 | 0.00000 | 0.64010 | 0.48453 | 0.38139 | 0.66951 | 0.92162 | 0.00000 | 0.00016 |
| **CRP** | 0.00110 | 0.00591 | 0.00006 | 0.00134 | 0.03905 | 0.03540 | 0.00000 | 0.54736 | 0.00000 | 0.20028 | 0.00136 | 0.74023 |
| **CRSP8P** | 1.30058 | 2.72883 | 1.98986 | 0.80181 | 1.00624 | 2.53379 | 1.08108 | 1.10759 | 1.92273 | 0.00000 | 1.82503 | 1.62046 |
| **CRTAC1** | 0.22323 | 0.02191 | 0.07559 | 0.00538 | 0.00000 | 0.02114 | 0.02988 | 0.73423 | 0.00816 | 0.11063 | 0.01369 | 0.05841 |
| **CRTAM** | 0.00541 | 0.07836 | 0.00839 | 0.15635 | 0.04825 | 0.00000 | 0.03620 | 0.01241 | 0.02223 | 0.00000 | 0.00950 | 0.02536 |
| **CRTAP** | 0.34504 | 3.03102 | 3.23845 | 9.06713 | 7.19093 | 5.41237 | 4.32002 | 0.12996 | 10.15020 | 4.16687 | 8.59889 | 0.31653 |
| **CRTC1** | 1.10170 | 0.98471 | 0.64452 | 1.18780 | 0.91090 | 0.00000 | 0.02995 | 0.00000 | 1.19574 | 4.52140 | 0.40266 | 0.37327 |
| **CRTC2** | 1.23310 | 1.96911 | 7.95788 | 1.45597 | 4.55736 | 0.00000 | 3.43431 | 1.25327 | 7.49302 | 15.78822 | 0.00000 | 3.98699 |
| **CRTC3** | 1.42579 | 0.79968 | 7.93615 | 0.00000 | 0.00000 | 3.51557 | 5.06896 | 0.11979 | 0.77010 | 4.04401 | 0.83456 | 0.72649 |
| **CRX** | 0.04667 | 0.13835 | 0.00975 | 0.05441 | 0.01603 | 0.00000 | 0.02259 | 0.36046 | 0.00000 | 0.15799 | 0.01385 | 0.12002 |
| **CRY1** | 4.31874 | 4.20147 | 5.51941 | 2.08380 | 1.10840 | 15.05399 | 3.35634 | 1.39333 | 4.19576 | 0.00000 | 2.41769 | 2.21787 |
| **CRY2** | 1.44610 | 9.78543 | 3.33383 | 1.18016 | 6.00169 | 0.00001 | 4.53193 | 2.65284 | 7.49123 | 0.00000 | 8.23834 | 6.98594 |
| **CRYAB** | 0.02210 | 0.02095 | 0.41675 | 0.00391 | 0.01236 | 0.11861 | 0.00000 | 0.12292 | 0.00000 | 0.00978 | 0.01587 | 0.00500 |
| **CRYBA1** | 0.05223 | 0.00351 | 0.51617 | 0.00686 | 0.25853 | 0.06654 | 0.04161 | 0.10630 | 0.00000 | 0.00000 | 0.03514 | 0.00902 |
| **CRYBA2** | 0.01418 | 0.73124 | 0.42382 | 0.00298 | 0.30133 | 0.00000 | 1.11176 | 0.01992 | 0.81873 | 0.00000 | 0.28632 | 0.19516 |
| **CRYBA4** | 0.00259 | 0.02766 | 0.14268 | 0.73487 | 0.00000 | 0.01666 | 0.00000 | 0.04498 | 0.12017 | 0.00563 | 0.04747 | 0.01363 |
| **CRYBB1** | 0.16994 | 0.01022 | 0.26101 | 1.27180 | 0.00000 | 0.00406 | 0.01736 | 0.16726 | 0.04041 | 0.01533 | 0.02287 | 0.08995 |
| **CRYBB2** | 0.00263 | 0.00236 | 0.03729 | 0.04977 | 0.02428 | 0.01878 | 0.05019 | 0.01456 | 0.00000 | 0.25606 | 0.07511 | 0.02256 |
| **CRYBB2P1** | 0.09251 | 0.72983 | 3.41942 | 1.74662 | 1.89845 | 1.00774 | 3.01741 | 0.92477 | 2.13694 | 1.21079 | 2.38117 | 1.20808 |
| **CRYBB3** | 0.00695 | 0.05166 | 0.06694 | 0.09772 | 0.00733 | 0.00498 | 0.00953 | 0.03456 | 0.03038 | 0.05877 | 0.04083 | 0.03645 |
| **CRYBG3** | 2.07547 | 0.02365 | 0.04093 | 0.36758 | 0.00000 | 0.01281 | 0.00000 | 0.76130 | 0.00000 | 0.57996 | 0.00000 | 1.23683 |
| **CRYGA** | 0.00586 | 0.01114 | 0.02021 | 0.01441 | 0.01503 | 0.01674 | 0.01617 | 0.00858 | 0.01625 | 0.02078 | 0.01564 | 0.05929 |
| **CRYGB** | 0.00173 | 0.02162 | 0.12658 | 0.32269 | 0.00000 | 0.06580 | 0.00000 | 0.05102 | 0.00000 | 0.14863 | 0.00944 | 0.05160 |
| **CRYGC** | 0.07359 | 0.12631 | 0.14174 | 0.07377 | 0.02914 | 0.34683 | 0.00000 | 0.12894 | 0.30140 | 0.15481 | 0.22820 | 0.99500 |
| **CRYGD** | 0.03304 | 0.10066 | 1.09508 | 0.59244 | 0.00000 | 0.00000 | 0.21845 | 0.03102 | 0.27609 | 0.00000 | 0.31755 | 0.19906 |
| **CRYGS** | 0.00252 | 0.06212 | 0.00139 | 0.30604 | 0.00000 | 0.00000 | 0.07387 | 0.14212 | 0.00311 | 0.62030 | 0.10952 | 0.04450 |
| **CRYL1** | 0.00622 | 0.02575 | 0.35741 | 0.03830 | 0.02318 | 0.03324 | 0.00000 | 0.01020 | 0.00000 | 0.00731 | 0.03993 | 0.01371 |
| **CRYM** | 0.50437 | 0.58258 | 0.06476 | 3.10649 | 0.00000 | 0.00000 | 0.05054 | 1.47025 | 0.01637 | 0.01032 | 0.00835 | 0.24861 |
| **CRYM-AS1** | 0.00597 | 0.01735 | 0.15459 | 0.03018 | 0.00613 | 0.00000 | 0.02034 | 0.00880 | 0.07161 | 0.04974 | 0.03162 | 0.04192 |
| **CRYZ** | 3.39015 | 3.57431 | 7.14768 | 9.99361 | 1.20503 | 8.11244 | 2.20878 | 2.43135 | 12.18018 | 0.00000 | 4.84039 | 4.15793 |
| **CRYZL1** | 0.55626 | 2.83030 | 0.66668 | 2.19082 | 0.30082 | 8.76637 | 0.00002 | 0.64169 | 2.22413 | 0.00000 | 3.76365 | 0.98605 |
| **CS** | 9.96196 | 18.48039 | 6.54726 | 19.25398 | 19.80608 | 17.70468 | 17.85020 | 11.66468 | 26.03734 | 10.96133 | 19.02294 | 18.78760 |
| **CSAD** | 0.00000 | 0.99965 | 0.36668 | 0.07391 | 2.13903 | 1.85315 | 0.00000 | 0.00000 | 1.64058 | 9.36211 | 1.91197 | 0.87362 |
| **CSAG1** | 0.03569 | 0.14645 | 0.06418 | 0.00000 | 0.00000 | 0.00823 | 0.46632 | 5.44059 | 0.03405 | 0.07045 | 0.00000 | 1.35195 |
| **CSAG3.uid-1334** | 0.00772 | 0.01535 | 0.01327 | 0.01100 | 0.01574 | 0.02322 | 0.02443 | 0.02254 | 0.01368 | 0.01322 | 0.01562 | 0.01863 |
| **CSAG4** | 0.00389 | 0.01429 | 0.01120 | 0.01213 | 0.01020 | 0.00034 | 0.03788 | 0.26882 | 0.02445 | 0.06970 | 0.01444 | 0.04423 |
| **CSDC2** | 0.06834 | 0.01407 | 0.01806 | 0.03000 | 0.02931 | 0.00247 | 0.07355 | 0.00632 | 0.01717 | 0.02624 | 0.00944 | 0.01633 |
| **CSDE1** | 16.81143 | 27.07269 | 24.31777 | 20.80416 | 15.82123 | 50.22375 | 35.16262 | 21.06959 | 32.56209 | 19.34125 | 27.37233 | 28.85772 |

| | | | | | | | | | | | | |
|---|---|---|---|---|---|---|---|---|---|---|---|---|
| **CSE1L** | 3.59959 | 27.68171 | 19.31374 | 16.36545 | 9.23531 | 53.97259 | 31.87395 | 3.43136 | 29.12469 | 13.19759 | 29.90196 | 20.60936 |
| **CSF1** | 0.00448 | 0.25190 | 0.19754 | 0.08115 | 0.00000 | 0.00099 | 0.00578 | 0.00982 | 0.00000 | 0.02443 | 0.06494 | 0.31894 |
| **CSF1R** | 11.89603 | 0.22488 | 0.21509 | 0.12240 | 0.00000 | 0.00000 | 0.00000 | 2.63231 | 0.04899 | 0.12926 | 0.38645 | 1.02073 |
| **CSF2** | 0.00620 | 0.01330 | 0.02208 | 0.02309 | 0.01358 | 0.02054 | 0.01712 | 0.01049 | 0.01222 | 0.01710 | 0.01837 | 0.01694 |
| **CSF2RB** | 0.00642 | 0.00742 | 0.06091 | 0.00365 | 0.06829 | 0.06213 | 0.00000 | 0.00966 | 0.23577 | 0.00000 | 0.00482 | 0.01445 |
| **CSF3** | 0.00538 | 0.00990 | 0.02938 | 0.01197 | 0.14983 | 0.00000 | 0.01472 | 0.01053 | 0.00720 | 0.06080 | 0.00883 | 0.01540 |
| **CSF3R** | 0.00523 | 0.32475 | 15.20802 | 0.05689 | 0.02287 | 0.00000 | 0.00000 | 0.01119 | 0.00000 | 0.19590 | 0.01083 | 0.01692 |
| **CSGALNACT1** | 2.09616 | 0.00000 | 1.40377 | 0.00000 | 0.16533 | 0.42395 | 0.97628 | 0.00000 | 1.19916 | 0.00000 | 0.94983 | 0.01181 |
| **CSGALNACT2** | 2.93609 | 0.35231 | 0.42355 | 0.08130 | 0.00000 | 1.99032 | 0.03180 | 1.95362 | 1.76096 | 0.68695 | 0.06758 | 1.92957 |
| **CSH1** | 0.00510 | 0.02898 | 0.06481 | 0.07388 | 0.00732 | 0.00000 | 0.05898 | 0.08067 | 0.00000 | 0.04941 | 0.01677 | 0.02351 |
| **CSH2** | 0.00319 | 0.01292 | 0.02613 | 0.02920 | 0.01103 | 0.01642 | 0.01649 | 0.04216 | 0.01910 | 0.02747 | 0.01517 | 0.03000 |
| **CSK** | 2.08820 | 5.08538 | 5.00797 | 6.64621 | 7.34173 | 0.00000 | 9.57586 | 1.28217 | 3.18139 | 14.70420 | 1.01024 | 4.60312 |
| **CSMD1** | 2.03461 | 0.31332 | 0.00000 | 0.18901 | 0.51288 | 0.83141 | 0.00000 | 0.24223 | 0.00000 | 0.65087 | 0.28243 | 0.00280 |
| **CSMD2** | 0.40739 | 0.00104 | 0.11504 | 0.00352 | 0.11894 | 0.00000 | 0.02896 | 0.05358 | 0.00000 | 0.32189 | 0.04271 | 0.01532 |
| **CSMD2-AS1** | 0.01741 | 0.01468 | 0.01315 | 0.01096 | 0.01177 | 0.01680 | 0.01946 | 0.05056 | 0.02742 | 0.02998 | 0.01166 | 0.01232 |
| **CSMD3** | 0.39994 | 0.01841 | 0.26205 | 0.21264 | 0.00000 | 0.16025 | 0.00000 | 0.12087 | 0.04009 | 0.47874 | 0.04818 | 0.16492 |
| **CSN3** | 0.00286 | 0.03415 | 0.14847 | 0.01464 | 0.07202 | 0.00007 | 0.10277 | 0.04521 | 0.02077 | 0.00000 | 0.00937 | 0.01635 |
| **CSNK1A1** | 2.84476 | 21.78825 | 21.02149 | 12.69155 | 13.45362 | 36.79020 | 17.70080 | 13.29026 | 24.06888 | 9.12623 | 14.00943 | 24.75129 |
| **CSNK1A1L** | 0.03476 | 0.91256 | 1.30215 | 0.37903 | 0.20110 | 2.31069 | 0.50464 | 0.65304 | 0.92108 | 0.04788 | 0.47185 | 1.09574 |
| **CSNK1A1P1** | 0.21989 | 1.14119 | 0.22132 | 0.79940 | 0.32471 | 3.61401 | 0.00000 | 0.26756 | 1.94646 | 1.55294 | 0.36584 | 1.85480 |
| **CSNK1D** | 3.32477 | 5.49560 | 8.31963 | 9.53948 | 9.40201 | 0.00046 | 10.30152 | 4.19031 | 10.43386 | 20.92748 | 3.78001 | 7.15651 |
| **CSNK1E** | 5.08357 | 9.06741 | 3.76797 | 0.80732 | 8.59034 | 0.27551 | 7.59864 | 2.79106 | 3.35707 | 15.64982 | 3.41866 | 7.82812 |
| **CSNK1G1** | 4.37188 | 3.00011 | 4.80232 | 0.00000 | 1.46278 | 6.03501 | 8.68244 | 7.99788 | 7.57666 | 7.90401 | 0.80109 | 6.45135 |
| **CSNK1G2** | 2.01678 | 0.99415 | 3.23393 | 1.49267 | 6.48928 | 0.00000 | 2.82102 | 1.78008 | 5.25185 | 11.09567 | 2.26669 | 0.75646 |
| **CSNK1G2-AS1** | 0.19970 | 0.00000 | 0.00258 | 0.00160 | 0.36553 | 0.00000 | 0.00457 | 0.00150 | 0.00000 | 0.98870 | 0.00000 | 0.04062 |
| **CSNK1G3** | 1.97405 | 3.03423 | 7.24480 | 2.02879 | 0.00000 | 11.07893 | 5.93156 | 7.25956 | 5.44054 | 4.65106 | 4.13994 | 5.02832 |
| **CSNK2A1** | 2.87066 | 13.89191 | 8.18603 | 6.91668 | 9.30363 | 20.14243 | 16.21998 | 7.93829 | 7.29189 | 10.13753 | 10.41000 | 16.59290 |
| **CSNK2A2** | 6.53797 | 1.37205 | 4.93202 | 1.14639 | 4.28984 | 1.34494 | 4.62145 | 3.25602 | 0.00000 | 9.90103 | 0.00000 | 6.06219 |
| **CSNK2A3** | 0.77408 | 5.72011 | 2.68728 | 2.05134 | 0.29639 | 8.75407 | 1.78736 | 2.69335 | 4.26848 | 0.00000 | 4.34251 | 6.29884 |
| **CSNK2B** | 9.34013 | 25.53732 | 19.99671 | 19.63987 | 29.61964 | 28.79478 | 25.21242 | 17.88279 | 26.94933 | 9.07133 | 18.33205 | 26.48682 |
| **CSPG4** | 0.07435 | 0.01513 | 0.07333 | 0.07840 | 0.00555 | 0.00641 | 0.01265 | 0.02309 | 0.28666 | 0.00267 | 0.00000 | 0.03117 |
| **CSPG5** | 0.22172 | 0.39468 | 0.00000 | 0.02916 | 0.15845 | 0.00907 | 0.01507 | 0.26377 | 0.03735 | 0.13939 | 0.00000 | 1.66870 |
| **CSPP1** | 3.94571 | 0.00000 | 2.29710 | 4.84700 | 3.99851 | 6.06263 | 3.33290 | 8.05006 | 4.88419 | 21.65562 | 3.84587 | 5.16269 |
| **CSRNP1** | 1.36862 | 6.89013 | 0.21971 | 4.92998 | 6.12106 | 0.00000 | 0.11488 | 4.28918 | 0.69116 | 5.02080 | 0.33161 | 6.07640 |
| **CSRNP2** | 4.04115 | 0.42004 | 4.01752 | 2.97994 | 1.34278 | 2.34879 | 8.10056 | 1.13018 | 3.31720 | 4.89660 | 1.15937 | 2.63050 |
| **CSRNP3** | 0.03092 | 0.04656 | 0.15100 | 0.04350 | 0.00053 | 0.37471 | 0.00000 | 0.00478 | 0.00000 | 0.13767 | 0.14852 | 0.04861 |
| **CSRP1** | 8.72330 | 2.84466 | 1.71878 | 0.15305 | 2.96699 | 0.38323 | 1.32458 | 0.89885 | 0.83216 | 0.00000 | 0.53647 | 0.60369 |
| **CSRP2** | 6.18140 | 2.67962 | 1.63318 | 2.81820 | 0.00000 | 2.74730 | 1.21961 | 1.96203 | 1.22417 | 0.00000 | 0.72935 | 2.65693 |
| **CSRP2BP** | 0.00000 | 10.09945 | 7.65847 | 0.00000 | 13.44905 | 19.06479 | 12.95911 | 2.01039 | 11.11310 | 3.13705 | 7.60416 | 6.81582 |
| **CSRP3** | 0.11099 | 0.01125 | 0.01067 | 0.02131 | 0.00673 | 0.01379 | 0.00542 | 0.11856 | 0.02262 | 0.06959 | 0.01286 | 0.01338 |
| **CST1** | 0.02810 | 0.00000 | 0.78520 | 0.10664 | 0.00000 | 0.04708 | 0.00000 | 0.09165 | 0.20986 | 1.07058 | 0.00000 | 0.00000 |
| **CST2** | 0.00391 | 0.00317 | 0.02364 | 0.07712 | 0.00000 | 0.02144 | 0.00000 | 0.01096 | 0.01338 | 0.29968 | 0.00467 | 0.01039 |
| **CST3** | 0.00000 | 1.22434 | 5.73843 | 4.15373 | 7.10039 | 0.00000 | 0.00000 | 0.07529 | 7.77425 | 3.45478 | 2.45641 | 0.19258 |
| **CST4** | 0.00647 | 0.01655 | 0.03478 | 0.09424 | 0.01816 | 0.00069 | 0.01869 | 0.01054 | 0.02489 | 0.00440 | 0.01672 | 0.01888 |
| **CST6** | 0.00870 | 0.00000 | 0.23225 | 0.01187 | 0.25064 | 0.34489 | 0.00000 | 0.01918 | 0.19402 | 0.00000 | 0.00313 | 0.00659 |
| **CST7** | 0.00654 | 0.02054 | 0.07747 | 0.00047 | 0.07310 | 0.01373 | 0.00000 | 0.00856 | 0.01432 | 0.07158 | 0.02076 | 0.06171 |
| **CST8** | 0.05625 | 0.01450 | 0.01494 | 0.01168 | 0.01481 | 0.01922 | 0.01739 | 0.04765 | 0.01813 | 0.01323 | 0.01468 | 0.01413 |
| **CSTA** | 0.05820 | 0.00000 | 0.00000 | 0.73365 | 0.62323 | 0.85003 | 0.07995 | 0.10906 | 0.00000 | 0.00000 | 0.00000 | 0.15873 |

| | | | | | | | | | | | | |
|---|---|---|---|---|---|---|---|---|---|---|---|---|
| **CSTB** | 7.47664 | 17.06749 | 17.95390 | 13.54239 | 24.50302 | 13.38498 | 31.11506 | 0.24487 | 32.81100 | 8.35211 | 16.05151 | 5.88454 |
| **CSTF1** | 5.62644 | 14.89038 | 5.70581 | 4.31528 | 5.38927 | 21.45850 | 7.15065 | 4.66664 | 11.77921 | 0.00000 | 11.09157 | 16.19626 |
| **CSTF2** | 2.26917 | 16.91582 | 10.38399 | 6.08002 | 10.35153 | 11.38104 | 15.03210 | 11.24236 | 11.12539 | 0.00000 | 19.56591 | 16.81901 |
| **CSTF2T** | 0.00116 | 6.13873 | 5.19618 | 1.10566 | 1.46643 | 9.12214 | 1.91899 | 3.76477 | 5.82389 | 0.00000 | 7.32358 | 10.88138 |
| **CSTF3** | 0.84424 | 8.61899 | 11.44075 | 3.13474 | 0.63886 | 25.86658 | 17.52652 | 12.56646 | 15.10018 | 6.25478 | 11.89908 | 13.59269 |
| **CSTF3-AS1** | 0.29755 | 0.02998 | 0.02216 | 0.02847 | 0.01681 | 0.00000 | 0.02380 | 0.41551 | 0.02613 | 0.05717 | 0.01345 | 0.01079 |
| **CSTL1** | 0.00441 | 0.08054 | 0.00604 | 0.00000 | 0.05550 | 0.03074 | 0.00156 | 0.00848 | 0.01298 | 0.09325 | 0.00440 | 0.03927 |
| **CT47A6.uid-131** | 0.00456 | 0.01186 | 0.00513 | 0.01047 | 0.00000 | 0.01608 | 0.10287 | 0.04473 | 0.00000 | 0.08432 | 0.02458 | 0.01364 |
| **CT47B1** | 0.00247 | 0.01424 | 0.00931 | 0.01609 | 0.01088 | 0.01599 | 0.01713 | 0.39210 | 0.03930 | 0.01946 | 0.01359 | 0.03203 |
| **CT55** | 0.02614 | 0.44149 | 0.00000 | 0.00149 | 0.11877 | 0.27920 | 0.14504 | 0.16733 | 0.10661 | 0.00000 | 0.04079 | 1.05340 |
| **CT62** | 0.01963 | 0.11156 | 0.02675 | 0.03686 | 0.03384 | 0.08684 | 0.00001 | 0.04334 | 0.00073 | 0.02153 | 0.00848 | 0.00651 |
| **CT83** | 0.02475 | 0.04326 | 0.03002 | 0.00739 | 0.01296 | 0.04716 | 0.16416 | 0.06950 | 0.00221 | 0.00000 | 0.01077 | 0.11752 |
| **CTAG2** | 0.00368 | 0.45795 | 0.00000 | 0.00995 | 0.00000 | 0.00000 | 0.42264 | 0.00437 | 0.21382 | 0.35346 | 0.13498 | 0.72075 |
| **CTAGE1** | 0.07320 | 0.01909 | 0.02677 | 0.03032 | 0.00488 | 0.13702 | 0.03676 | 0.01551 | 0.06228 | 0.03130 | 0.02297 | 0.01945 |
| **CTAGE10P** | 0.00956 | 0.01381 | 0.01807 | 0.01913 | 0.03746 | 0.02008 | 0.01734 | 0.01218 | 0.03555 | 0.02409 | 0.01538 | 0.02783 |
| **CTAGE11P** | 0.02739 | 0.02257 | 0.02933 | 0.01341 | 0.00817 | 0.05025 | 0.01360 | 0.01705 | 0.01950 | 0.02495 | 0.02177 | 0.01177 |
| **CTAGE4.uid-101** | 0.00622 | 0.01486 | 0.01706 | 0.01213 | 0.01464 | 0.01845 | 0.02106 | 0.01044 | 0.01741 | 0.01523 | 0.01533 | 0.01670 |
| **CTAGE5** | 3.59774 | 0.01680 | 2.34065 | 2.70767 | 0.02808 | 4.48647 | 0.00000 | 0.48649 | 4.55919 | 3.61159 | 1.46210 | 0.31801 |
| **CTAGE7P** | 0.04092 | 0.00497 | 0.01350 | 0.01447 | 0.00000 | 0.09262 | 0.04416 | 0.01230 | 0.04419 | 0.04640 | 0.01569 | 0.02875 |
| **CTAGE8** | 0.00628 | 0.01477 | 0.01585 | 0.01395 | 0.01378 | 0.02174 | 0.01781 | 0.01044 | 0.02118 | 0.01322 | 0.01398 | 0.01605 |
| **CTAGE9** | 0.00598 | 0.01365 | 0.01650 | 0.01204 | 0.01226 | 0.02560 | 0.01653 | 0.01023 | 0.02296 | 0.02683 | 0.01260 | 0.01641 |
| **CTB-113P19.1** | 0.00572 | 0.36826 | 0.10048 | 0.57859 | 0.15615 | 0.11024 | 0.00000 | 0.00803 | 0.04767 | 0.02533 | 0.08350 | 0.08457 |
| **CTB-118N6.3** | 0.00148 | 0.02756 | 0.07316 | 0.07193 | 0.00000 | 0.17862 | 0.00000 | 0.01936 | 0.32409 | 0.28247 | 0.11758 | 0.03557 |
| **CTB-12O2.1** | 0.00261 | 0.01264 | 0.01140 | 0.01443 | 0.00907 | 0.01777 | 0.05598 | 0.01881 | 0.00305 | 0.19349 | 0.03382 | 0.05593 |
| **CTB-178M22.2** | 0.21834 | 0.21475 | 0.02281 | 0.00096 | 0.00721 | 0.03762 | 0.01493 | 0.01250 | 0.01898 | 0.00962 | 0.01421 | 0.66226 |
| **CTB-7E3.1** | 0.07066 | 0.01522 | 0.02051 | 0.01148 | 0.01552 | 0.01584 | 0.01766 | 0.03326 | 0.01869 | 0.01174 | 0.01706 | 0.01395 |
| **CTBP1** | 0.55107 | 6.44428 | 1.94814 | 5.11599 | 6.16985 | 3.67619 | 9.46587 | 0.32354 | 0.00000 | 12.73057 | 8.09027 | 2.28381 |
| **CTBP1-AS** | 0.00542 | 0.09034 | 0.10045 | 0.16218 | 0.00000 | 0.00000 | 0.00089 | 0.00665 | 0.15079 | 0.36832 | 0.41410 | 0.07499 |
| **CTBP1-AS2** | 1.51605 | 0.26417 | 0.92262 | 0.32125 | 0.06206 | 0.54738 | 0.61365 | 0.00000 | 0.58297 | 1.62207 | 0.74882 | 0.20432 |
| **CTBP2** | 0.55288 | 1.56665 | 1.05380 | 5.78215 | 4.76933 | 6.31254 | 12.23276 | 0.71353 | 2.67423 | 16.77110 | 2.81532 | 0.30959 |
| **CTBS** | 0.19669 | 1.14412 | 0.44971 | 0.17165 | 0.00000 | 2.07281 | 0.00000 | 1.77841 | 0.00000 | 0.53011 | 0.74714 | 2.11215 |
| **CTC-338M12.4** | 0.00417 | 0.53188 | 1.53032 | 2.56887 | 1.01421 | 0.00000 | 0.00000 | 0.41912 | 3.36339 | 1.17302 | 1.49710 | 1.88445 |
| **CTC-436P18.1** | 0.00376 | 0.03906 | 0.02432 | 0.01186 | 0.01463 | 0.01775 | 0.02452 | 0.02541 | 0.02152 | 0.00000 | 0.01854 | 0.05987 |
| **CTC1** | 0.78843 | 2.87471 | 3.36069 | 1.03655 | 2.40603 | 0.00000 | 0.00000 | 0.00000 | 0.96037 | 2.28434 | 5.58582 | 0.74479 |
| **CTCF** | 1.88913 | 2.48929 | 3.32316 | 4.37349 | 5.81157 | 16.04718 | 13.44465 | 17.01505 | 0.61675 | 21.64774 | 4.55878 | 6.35670 |
| **CTCFL** | 6.04543 | 0.21291 | 0.10965 | 0.13138 | 0.01634 | 0.21307 | 0.00000 | 1.11346 | 0.28708 | 0.00000 | 0.34848 | 1.13674 |
| **CTD-2151A2.1** | 0.02363 | 0.05361 | 0.02020 | 0.00575 | 0.00025 | 0.04107 | 0.01270 | 0.00792 | 0.01793 | 0.03051 | 0.02078 | 0.06108 |
| **CTD-2194D22.4** | 0.00645 | 0.01840 | 0.18325 | 0.01096 | 0.01484 | 0.01026 | 0.05127 | 0.01053 | 0.04737 | 0.00000 | 0.01007 | 0.01724 |
| **CTD-2201E9.1** | 0.00435 | 0.01045 | 0.01160 | 0.00951 | 0.00960 | 0.03485 | 0.01413 | 0.01644 | 0.01486 | 0.02779 | 0.02840 | 0.01699 |
| **CTD-2201I18.1** | 2.05623 | 0.03214 | 0.03554 | 0.00654 | 0.01850 | 0.00000 | 0.00000 | 1.02417 | 0.01633 | 0.32040 | 0.00439 | 0.00000 |
| **CTD-2258A20.5** | 0.11828 | 0.01396 | 0.04302 | 0.00606 | 0.00834 | 0.02487 | 0.01406 | 0.00656 | 0.02242 | 0.03775 | 0.01096 | 0.01548 |
| **CTD-2350J17.1** | 0.01207 | 0.01880 | 0.02406 | 0.01023 | 0.01456 | 0.02444 | 0.01729 | 0.00927 | 0.01710 | 0.01022 | 0.01443 | 0.01582 |
| **CTD-3080P12.3** | 0.14173 | 0.01113 | 0.00991 | 0.03456 | 0.00000 | 0.01875 | 0.00579 | 0.00410 | 0.11793 | 0.05284 | 0.00341 | 0.01201 |
| **CTDNEP1** | 3.60288 | 2.29702 | 5.59381 | 3.16673 | 9.14991 | 0.07156 | 12.50860 | 2.58598 | 0.00000 | 22.34155 | 1.36360 | 3.40375 |
| **CTDP1** | 1.01748 | 1.47879 | 2.34606 | 1.55087 | 3.55947 | 0.00000 | 3.17194 | 1.19596 | 0.00000 | 9.80490 | 0.41654 | 1.89390 |
| **CTDSP1** | 0.43942 | 7.01979 | 0.41151 | 9.23271 | 10.91274 | 0.46718 | 4.92052 | 1.35166 | 2.17761 | 16.06744 | 2.20452 | 5.35213 |
| **CTDSP2** | 4.28766 | 1.36628 | 1.70562 | 3.22325 | 6.79766 | 4.95964 | 8.68326 | 6.39331 | 1.06614 | 10.96634 | 1.61086 | 1.36747 |
| **CTDSPL** | 5.42746 | 0.00000 | 2.85256 | 1.08727 | 2.86025 | 0.00076 | 15.19599 | 0.34905 | 2.72297 | 8.18027 | 8.70526 | 0.05694 |

| | | | | | | | | | | | | |
|---|---|---|---|---|---|---|---|---|---|---|---|---|
| **CTDSPL2** | 3.86868 | 4.04051 | 5.38551 | 1.73428 | 1.29422 | 17.86101 | 4.60329 | 11.55276 | 6.04209 | 5.80973 | 2.72122 | 9.52651 |
| **CTF1** | 0.00467 | 0.00421 | 0.01766 | 0.05229 | 0.12862 | 0.00000 | 0.06927 | 0.00845 | 0.00000 | 0.11856 | 0.00316 | 0.05293 |
| **CTGF** | 0.00000 | 2.73156 | 3.84810 | 0.58436 | 0.85730 | 2.82139 | 0.00000 | 0.00000 | 0.78721 | 2.81919 | 0.38847 | 1.41025 |
| **CTGLF12P** | 0.00518 | 0.03198 | 0.02838 | 0.00658 | 0.01356 | 0.01080 | 0.00104 | 0.01400 | 0.03913 | 0.15500 | 0.00710 | 0.03547 |
| **CTH** | 0.00000 | 1.05552 | 5.06325 | 5.71080 | 0.00000 | 12.14834 | 8.91095 | 0.16340 | 21.07996 | 1.17650 | 13.38417 | 1.78085 |
| **CTHRC1** | 0.00494 | 0.06026 | 0.14595 | 0.40666 | 0.00000 | 0.52156 | 0.44907 | 0.01050 | 0.45206 | 0.19523 | 0.30497 | 0.33309 |
| **CTIF** | 0.64262 | 0.23290 | 1.15786 | 1.16866 | 3.49850 | 0.00000 | 1.80237 | 0.55945 | 1.38043 | 1.24457 | 2.08295 | 0.05514 |
| **CTLA4** | 0.00520 | 0.03298 | 0.10725 | 0.00757 | 0.02537 | 0.00000 | 0.08993 | 0.00947 | 0.05627 | 0.04753 | 0.00692 | 0.03443 |
| **CTNNA1** | 11.07575 | 22.76102 | 20.81220 | 14.25207 | 15.08434 | 31.07758 | 32.46474 | 13.73405 | 30.00740 | 2.47123 | 27.59152 | 21.57979 |
| **CTNNA2** | 2.93465 | 0.01442 | 0.01044 | 0.10313 | 0.00000 | 0.04626 | 0.01308 | 0.59836 | 0.00000 | 0.07030 | 0.01881 | 0.03650 |
| **CTNNA3** | 3.39611 | 0.00000 | 0.02177 | 0.19777 | 0.00000 | 0.00941 | 0.00000 | 0.97375 | 0.01399 | 0.76343 | 0.05092 | 0.00000 |
| **CTNNAL1** | 0.00000 | 7.98616 | 14.84707 | 1.43732 | 3.54247 | 23.63213 | 17.76598 | 2.38492 | 12.80970 | 12.62214 | 11.91265 | 3.30259 |
| **CTNNB1** | 9.28490 | 24.09247 | 39.15320 | 10.43641 | 16.00953 | 39.38220 | 41.71249 | 14.46329 | 20.21139 | 12.67573 | 34.63511 | 20.10631 |
| **CTNNBIP1** | 1.10750 | 0.14866 | 0.13767 | 0.26841 | 0.35414 | 0.21614 | 0.30892 | 0.05320 | 0.05504 | 0.72070 | 0.00000 | 0.02428 |
| **CTNNBL1** | 1.09541 | 15.63862 | 21.94108 | 9.14094 | 11.58474 | 10.50122 | 17.79361 | 20.39422 | 11.56541 | 2.33807 | 10.53437 | 16.28255 |
| **CTNND1** | 0.91833 | 2.13812 | 2.21596 | 0.30202 | 0.00000 | 1.30411 | 0.24136 | 0.00000 | 1.05936 | 1.02824 | 1.54622 | 2.61256 |
| **CTNND2** | 0.94197 | 0.05432 | 0.03614 | 0.56800 | 0.00000 | 0.00306 | 0.02028 | 0.04711 | 0.01826 | 0.02032 | 0.01878 | 0.00876 |
| **CTNS** | 1.27601 | 15.05653 | 4.34412 | 5.71136 | 5.16491 | 0.00000 | 2.43214 | 1.55824 | 12.23097 | 1.13721 | 3.82282 | 7.92795 |
| **CTPS1** | 1.96252 | 20.91832 | 8.49834 | 10.18326 | 11.74978 | 26.22772 | 11.65163 | 5.38770 | 27.90586 | 6.07245 | 17.73457 | 14.59155 |
| **CTPS2** | 0.09394 | 0.01577 | 0.12791 | 0.32810 | 0.00000 | 0.22880 | 0.20516 | 0.01693 | 0.09332 | 0.00000 | 0.08844 | 0.01990 |
| **CTR9** | 4.26967 | 9.30044 | 8.67030 | 5.05740 | 6.12860 | 18.48590 | 11.24167 | 9.03598 | 7.62132 | 9.55757 | 7.78174 | 12.30293 |
| **CTRB1** | 0.21504 | 0.28060 | 0.00632 | 0.00000 | 0.00000 | 0.03838 | 0.00366 | 0.01761 | 0.00789 | 0.07008 | 0.00658 | 0.00000 |
| **CTRB2** | 6.28002 | 0.08292 | 0.00000 | 0.12302 | 0.16003 | 0.04311 | 0.00664 | 1.98859 | 0.00000 | 0.00000 | 0.08468 | 0.77759 |
| **CTRC** | 0.04514 | 0.01529 | 0.01502 | 0.01184 | 0.01607 | 0.01497 | 0.02061 | 0.06829 | 0.02544 | 0.01134 | 0.01474 | 0.01531 |
| **CTRL** | 0.01121 | 0.33004 | 1.71247 | 0.40546 | 0.55400 | 0.00000 | 0.60525 | 0.00354 | 2.73418 | 0.00000 | 2.13741 | 0.24193 |
| **CTSA** | 2.65219 | 24.54980 | 8.55709 | 14.65931 | 22.45302 | 9.78009 | 0.00000 | 0.44657 | 23.16650 | 0.72425 | 14.83569 | 12.13417 |
| **CTSB** | 5.98586 | 42.21123 | 19.37055 | 22.00484 | 55.43438 | 37.61873 | 6.86088 | 14.82345 | 60.53219 | 22.27153 | 45.22438 | 39.33281 |
| **CTSC** | 4.04007 | 25.07398 | 27.19542 | 27.42819 | 6.84310 | 47.52491 | 0.00000 | 1.99580 | 53.12565 | 10.05888 | 24.50170 | 21.35720 |
| **CTSD** | 2.92050 | 27.84453 | 26.55232 | 18.09324 | 33.61955 | 0.00000 | 6.27755 | 0.00000 | 48.35979 | 13.19961 | 14.57848 | 2.43611 |
| **CTSE** | 0.00465 | 0.05437 | 5.66119 | 11.27223 | 0.43333 | 0.00000 | 0.00000 | 0.00834 | 0.35680 | 0.00000 | 0.00364 | 0.00720 |
| **CTSF** | 0.00277 | 8.56150 | 0.76489 | 2.56478 | 0.84065 | 0.00000 | 0.15073 | 1.67844 | 0.00000 | 1.51474 | 0.72611 | 10.48359 |
| **CTSH** | 0.97100 | 2.99879 | 2.45656 | 0.00000 | 5.43490 | 3.84083 | 0.00000 | 0.22285 | 13.33237 | 0.78351 | 4.55889 | 1.63424 |
| **CTSK** | 0.00320 | 0.00802 | 0.09480 | 1.12425 | 0.00000 | 0.58872 | 0.00000 | 0.09352 | 0.78552 | 2.08763 | 0.00000 | 0.16155 |
| **CTSL** | 1.11487 | 34.37747 | 38.61481 | 11.68097 | 26.12591 | 51.75309 | 8.41638 | 2.53801 | 63.22295 | 22.77342 | 31.12809 | 18.10694 |
| **CTSL3P** | 0.00821 | 0.00000 | 0.34719 | 0.06979 | 0.00000 | 0.00000 | 0.00000 | 0.01323 | 0.35368 | 0.40595 | 0.37653 | 0.30013 |
| **CTSLP8** | 0.00779 | 0.01541 | 0.01634 | 0.00853 | 0.00550 | 0.04483 | 0.01593 | 0.01773 | 0.03801 | 0.03188 | 0.01975 | 0.02025 |
| **CTSO** | 0.00652 | 0.01010 | 0.01942 | 0.03180 | 0.02494 | 0.05029 | 0.01082 | 0.01288 | 0.03037 | 0.00185 | 0.01505 | 0.03083 |
| **CTSS** | 0.46066 | 0.40588 | 0.01396 | 0.00000 | 0.23629 | 0.00000 | 0.01097 | 0.86608 | 0.00000 | 0.11052 | 0.00220 | 0.08128 |
| **CTSV** | 2.42840 | 37.57157 | 44.38675 | 10.73171 | 14.36696 | 54.27639 | 25.46607 | 23.21236 | 59.72985 | 20.03855 | 26.54143 | 43.79171 |
| **CTSW** | 0.00148 | 0.33975 | 0.01395 | 0.05748 | 0.02141 | 0.03586 | 0.01153 | 0.54769 | 0.02338 | 0.00000 | 0.00749 | 0.37363 |
| **CTSZ** | 0.59741 | 0.14949 | 0.26526 | 0.04888 | 1.58660 | 0.33176 | 0.00000 | 0.06299 | 0.00000 | 1.75341 | 0.96835 | 0.04330 |
| **CTTN** | 12.44390 | 24.42721 | 24.17903 | 33.12820 | 27.78382 | 14.75172 | 41.22724 | 6.01566 | 17.90871 | 0.00000 | 34.68306 | 12.89157 |
| **CTTNBP2** | 2.24862 | 0.00000 | 2.44372 | 0.00000 | 5.63409 | 0.78562 | 2.77453 | 0.39186 | 7.09425 | 2.05205 | 5.46491 | 0.36716 |
| **CTTNBP2NL** | 9.05132 | 2.65400 | 3.20496 | 3.08639 | 4.05905 | 4.48608 | 5.14247 | 3.67814 | 2.84443 | 1.79522 | 2.53206 | 2.82620 |
| **CTU1** | 0.05309 | 0.42412 | 0.26508 | 0.16596 | 0.38843 | 0.00000 | 0.06956 | 0.02292 | 0.00000 | 0.85515 | 0.15964 | 0.18776 |
| **CTU2** | 3.84274 | 10.59869 | 7.96988 | 5.86900 | 11.16843 | 0.00000 | 4.87691 | 1.12839 | 10.43340 | 0.00000 | 9.25678 | 7.87721 |
| **CTXN1** | 0.05728 | 0.19281 | 0.01658 | 0.00899 | 0.05221 | 0.00000 | 0.06328 | 0.00000 | 0.00000 | 0.93813 | 0.00000 | 0.40732 |
| **CTXN2** | 0.00576 | 0.01793 | 0.02499 | 0.01430 | 0.00914 | 0.02686 | 0.01148 | 0.01050 | 0.01502 | 0.03125 | 0.01341 | 0.01681 |

| | | | | | | | | | | | | |
|---|---|---|---|---|---|---|---|---|---|---|---|---|
| **CUBN** | 0.61358 | 0.00324 | 0.00862 | 1.06420 | 0.00677 | 0.00465 | 0.02200 | 0.23307 | 0.01609 | 0.01425 | 0.01097 | 0.19887 |
| **CUEDC1** | 0.45547 | 0.00968 | 0.03879 | 0.02032 | 0.19821 | 0.00000 | 0.13749 | 0.00428 | 0.00000 | 0.05066 | 0.03022 | 0.04304 |
| **CUEDC2** | 0.00000 | 4.63969 | 1.70545 | 3.91901 | 2.85015 | 3.22703 | 0.49944 | 0.00000 | 2.82255 | 1.03139 | 4.79618 | 1.33006 |
| **CUL1** | 8.70189 | 16.43488 | 6.94151 | 7.95285 | 6.14148 | 13.16896 | 17.97202 | 8.75555 | 10.05070 | 4.08000 | 8.66069 | 5.65704 |
| **CUL2** | 0.00000 | 10.42431 | 9.32889 | 3.63884 | 0.00000 | 24.09077 | 10.82654 | 34.89451 | 5.48183 | 4.68393 | 7.61637 | 27.52163 |
| **CUL3** | 1.06164 | 1.96764 | 8.00415 | 1.65642 | 5.32232 | 12.98351 | 18.15304 | 5.01303 | 0.02900 | 22.85606 | 2.33758 | 2.09846 |
| **CUL4A** | 0.25066 | 9.58589 | 10.31046 | 3.14056 | 6.73477 | 14.82574 | 11.49556 | 3.79157 | 12.77378 | 7.07936 | 9.50299 | 6.58804 |
| **CUL4B** | 4.97181 | 1.34621 | 4.29389 | 2.90810 | 0.00000 | 8.15986 | 7.49469 | 1.61326 | 3.33736 | 5.48928 | 3.09116 | 5.55392 |
| **CUL5** | 20.87775 | 2.12702 | 5.08179 | 0.73671 | 1.78706 | 16.83125 | 9.61768 | 15.89169 | 6.80350 | 13.33741 | 7.93644 | 4.37912 |
| **CUL7** | 0.32535 | 0.09468 | 0.75407 | 1.23867 | 0.54810 | 0.00000 | 0.00000 | 0.04101 | 0.31581 | 1.59013 | 0.68942 | 0.35550 |
| **CUL9** | 0.29745 | 0.27635 | 0.22548 | 0.24604 | 0.00000 | 0.00000 | 0.90509 | 0.00000 | 0.00000 | 4.68083 | 0.74055 | 0.93123 |
| **CUTA** | 3.47725 | 6.34762 | 2.74599 | 4.28052 | 2.60401 | 1.91231 | 5.01302 | 1.49196 | 7.81059 | 0.00000 | 2.13577 | 9.00717 |
| **CUTC** | 2.18762 | 8.95016 | 5.32820 | 5.29563 | 1.00190 | 13.18258 | 7.69383 | 0.97990 | 3.35288 | 0.25332 | 8.99508 | 10.74290 |
| **CUX1** | 6.22545 | 1.17218 | 1.13442 | 2.03458 | 2.00481 | 0.72354 | 1.34277 | 0.85976 | 2.78737 | 8.21579 | 1.73045 | 0.63065 |
| **CUX2** | 0.98049 | 0.07339 | 0.02651 | 0.18006 | 0.00000 | 0.02426 | 0.02606 | 0.00000 | 0.00000 | 0.25282 | 0.00000 | 0.25491 |
| **CWC15** | 6.53071 | 18.43014 | 12.65456 | 8.18031 | 12.12804 | 29.53179 | 24.10538 | 19.95842 | 11.65488 | 2.24472 | 16.70894 | 26.09682 |
| **CWC22** | 0.50480 | 4.68978 | 7.76439 | 4.90511 | 0.09040 | 19.28191 | 5.32640 | 6.05251 | 6.98999 | 10.01805 | 6.97161 | 8.46616 |
| **CWC25** | 8.97341 | 7.41921 | 7.37727 | 5.37908 | 5.99987 | 5.15039 | 8.46481 | 17.34795 | 5.38673 | 8.55990 | 4.68217 | 9.08934 |
| **CWC27** | 0.35997 | 5.67338 | 4.23784 | 2.56288 | 2.44161 | 12.70622 | 6.40069 | 2.87706 | 4.06856 | 8.48170 | 3.81774 | 5.08769 |
| **CWF19L1** | 0.97987 | 11.45947 | 4.40271 | 5.96072 | 3.29345 | 9.91734 | 8.75164 | 4.95235 | 11.63164 | 0.00000 | 8.33743 | 11.60784 |
| **CWF19L2** | 7.67537 | 2.76778 | 4.11597 | 2.51946 | 3.48563 | 9.91801 | 4.02259 | 2.15494 | 3.84876 | 13.43358 | 2.43868 | 6.90031 |
| **CWH43** | 0.00610 | 0.01402 | 0.02093 | 0.01225 | 0.01982 | 0.01567 | 0.01842 | 0.01038 | 0.01370 | 0.01608 | 0.01919 | 0.02121 |
| **CX3CL1** | 0.00347 | 0.01524 | 0.16234 | 0.00989 | 0.01547 | 0.02178 | 0.00000 | 0.05379 | 0.06173 | 0.00467 | 0.01009 | 0.01520 |
| **CX3CR1** | 0.01979 | 0.06274 | 0.02001 | 0.22432 | 0.00000 | 0.00000 | 0.07960 | 0.00733 | 0.00000 | 0.28880 | 0.08412 | 0.11775 |
| **CXADR** | 0.00000 | 28.71610 | 0.00000 | 13.34731 | 18.54765 | 31.37317 | 0.00000 | 22.39471 | 18.93606 | 13.20457 | 17.25629 | 31.22127 |
| **CXADRP2** | 0.00635 | 0.01503 | 0.00974 | 0.01354 | 0.02249 | 0.01260 | 0.01801 | 0.01627 | 0.01924 | 0.05901 | 0.01407 | 0.02059 |
| **CXADRP3** | 0.00204 | 0.29176 | 0.00000 | 0.07647 | 0.23920 | 0.22329 | 0.00000 | 0.46858 | 0.16363 | 0.00000 | 0.10537 | 0.33699 |
| **CXCL12** | 0.00446 | 0.00000 | 0.18624 | 3.08753 | 0.04207 | 0.14997 | 0.03113 | 0.03585 | 0.20084 | 0.00000 | 0.00000 | 0.09431 |
| **CXCL14** | 0.42544 | 0.02183 | 0.05495 | 0.10709 | 0.00000 | 0.00000 | 0.82309 | 1.55650 | 0.00000 | 0.44713 | 0.00000 | 0.18201 |
| **CXCL16** | 2.22435 | 0.29647 | 0.07873 | 0.00000 | 0.03463 | 0.07477 | 0.14742 | 0.29223 | 0.03939 | 0.03261 | 0.01554 | 1.23923 |
| **CXCL17** | 0.08207 | 0.05276 | 0.05381 | 0.11067 | 0.01841 | 0.01139 | 0.01327 | 0.00574 | 0.01174 | 0.00709 | 0.01744 | 0.02427 |
| **CXCL2** | 0.00651 | 0.01602 | 0.03241 | 0.01226 | 0.00840 | 0.01669 | 0.00875 | 0.01033 | 0.07244 | 0.01660 | 0.04050 | 0.01594 |
| **CXCL3** | 0.05073 | 0.01166 | 0.01753 | 0.01316 | 0.01705 | 0.01280 | 0.01804 | 0.02634 | 0.02123 | 0.00487 | 0.02005 | 0.07918 |
| **CXCL5** | 0.05434 | 0.01232 | 0.03095 | 0.01312 | 0.01709 | 0.01401 | 0.03045 | 0.07144 | 0.00000 | 0.05369 | 0.06245 | 0.01380 |
| **CXCL8** | 0.00689 | 0.02189 | 0.01736 | 0.01395 | 0.01792 | 0.00659 | 0.03039 | 0.01114 | 0.01337 | 0.01263 | 0.02611 | 0.01838 |
| **CXCL9** | 0.06126 | 0.01959 | 0.01397 | 0.00923 | 0.01455 | 0.02679 | 0.01901 | 0.01124 | 0.02112 | 0.00000 | 0.01400 | 0.30240 |
| **CXCR1** | 0.05491 | 0.00347 | 0.01658 | 0.00772 | 0.01291 | 0.03405 | 0.01043 | 0.11599 | 0.01609 | 0.02437 | 0.01572 | 0.18956 |
| **CXCR2** | 0.00581 | 0.01555 | 0.01477 | 0.01781 | 0.01358 | 0.01343 | 0.01939 | 0.01034 | 0.01959 | 0.03465 | 0.01717 | 0.01998 |
| **CXCR2P1** | 0.00582 | 0.01473 | 0.01352 | 0.01188 | 0.01395 | 0.02393 | 0.02325 | 0.01160 | 0.01731 | 0.01075 | 0.01421 | 0.01771 |
| **CXCR4** | 0.72733 | 0.15437 | 0.09726 | 0.74420 | 0.37262 | 0.00000 | 0.00000 | 0.00000 | 0.00000 | 0.34887 | 0.00000 | 0.28112 |
| **CXCR5** | 0.00535 | 0.01692 | 0.06995 | 0.01109 | 0.01535 | 0.00802 | 0.14670 | 0.01041 | 0.00000 | 0.05403 | 0.01131 | 0.01504 |
| **CXCR6** | 0.00484 | 0.02339 | 3.48860 | 0.00087 | 0.15833 | 0.03315 | 0.28854 | 0.01126 | 0.00000 | 0.00000 | 0.02008 | 0.01685 |
| **CXXC1** | 0.53206 | 13.99402 | 14.75843 | 11.55316 | 11.15269 | 0.00000 | 16.17420 | 2.18506 | 21.60234 | 4.71626 | 6.98295 | 12.90812 |
| **CXXC4** | 0.82591 | 0.00448 | 0.11373 | 0.48518 | 0.00000 | 0.01791 | 0.00000 | 0.05047 | 0.02731 | 0.17319 | 0.00065 | 0.13475 |
| **CXXC5** | 0.09511 | 0.20210 | 1.13725 | 1.62782 | 1.92838 | 0.00000 | 0.50421 | 0.00000 | 0.71717 | 0.25837 | 0.13300 | 0.27843 |
| **CXorf22** | 0.35257 | 0.34546 | 0.12957 | 0.19475 | 0.00000 | 0.22279 | 0.00000 | 0.00072 | 0.20111 | 0.00000 | 0.05017 | 0.22795 |
| **CXorf23** | 3.33732 | 0.00000 | 1.22568 | 0.87022 | 0.20072 | 0.78004 | 0.70588 | 0.37603 | 0.00022 | 3.48456 | 1.34554 | 0.30039 |
| **CXorf30** | 0.02249 | 0.00009 | 0.00000 | 0.00000 | 0.00000 | 0.28612 | 0.00000 | 0.03223 | 0.00983 | 0.18631 | 0.09383 | 0.01153 |

| | | | | | | | | | | | | |
|---|---|---|---|---|---|---|---|---|---|---|---|---|
| **CXorf31** | 0.18565 | 0.01452 | 0.02539 | 0.00444 | 0.02144 | 0.00669 | 0.02772 | 0.03935 | 0.07084 | 0.05311 | 0.02945 | 0.01147 |
| **CXorf36** | 0.00643 | 0.01447 | 0.02678 | 0.01294 | 0.01569 | 0.01881 | 0.02020 | 0.01126 | 0.01464 | 0.01842 | 0.01576 | 0.01583 |
| **CXorf38** | 0.31079 | 3.87657 | 1.33088 | 1.29338 | 1.33591 | 3.77682 | 0.87933 | 2.34065 | 3.45574 | 1.02366 | 2.24301 | 2.89743 |
| **CXorf40A** | 1.75869 | 2.08104 | 0.14378 | 0.31991 | 0.76298 | 1.26522 | 0.00000 | 0.77823 | 1.22917 | 0.00000 | 0.03467 | 1.84753 |
| **CXorf40B** | 4.98085 | 5.43118 | 2.42689 | 1.53985 | 1.96482 | 5.94281 | 1.72457 | 11.08563 | 5.15307 | 0.00000 | 2.34054 | 7.17407 |
| **CXorf49+CXorf4** | 0.00633 | 0.01719 | 0.01338 | 0.01033 | 0.01489 | 0.02698 | 0.01661 | 0.01005 | 0.01658 | 0.01471 | 0.01415 | 0.02029 |
| **CXorf56** | 5.17822 | 2.87789 | 4.54745 | 1.74381 | 0.78106 | 6.80297 | 6.68199 | 8.92019 | 2.40561 | 1.22662 | 5.41531 | 7.29775 |
| **CXorf57** | 0.00702 | 0.42614 | 2.73682 | 0.50017 | 0.00000 | 0.18904 | 0.00180 | 0.03487 | 0.02984 | 0.00000 | 0.00416 | 2.08215 |
| **CXorf58** | 0.21488 | 0.00467 | 0.00527 | 0.02084 | 0.02974 | 0.07918 | 0.00627 | 0.02834 | 0.01601 | 0.01284 | 0.01145 | 0.06244 |
| **CXorf65** | 0.00000 | 3.51639 | 0.00000 | 0.00000 | 1.45864 | 1.58270 | 1.10405 | 0.00000 | 0.56557 | 0.29424 | 2.55799 | 0.87784 |
| **CXorf67** | 9.66244 | 0.00000 | 0.31821 | 0.00937 | 0.10519 | 0.00000 | 0.00000 | 12.24938 | 0.45792 | 0.72223 | 0.00000 | 3.22632 |
| **CYB561** | 0.85032 | 11.33030 | 2.74899 | 2.14476 | 4.16897 | 3.68283 | 0.53488 | 0.38121 | 2.71161 | 1.93399 | 0.88986 | 8.83298 |
| **CYB561A3** | 0.71613 | 4.51548 | 9.24771 | 1.61392 | 0.00000 | 0.95508 | 3.79627 | 0.12289 | 5.83058 | 0.00000 | 3.55582 | 4.80104 |
| **CYB561D1** | 0.00000 | 1.28342 | 0.88938 | 0.83608 | 2.43899 | 0.87868 | 0.98866 | 0.51922 | 1.14541 | 1.17593 | 0.77958 | 2.30235 |
| **CYB561D2** | 0.62984 | 10.43949 | 21.86293 | 6.25892 | 11.72541 | 2.77097 | 9.26196 | 2.13215 | 20.00712 | 0.00000 | 10.19986 | 12.69486 |
| **CYB5A** | 3.47763 | 3.23502 | 12.72947 | 2.22077 | 6.47378 | 6.18264 | 13.50191 | 0.04525 | 8.62550 | 0.03668 | 16.90981 | 0.13090 |
| **CYB5B** | 2.91576 | 17.59449 | 15.02524 | 12.29657 | 23.60628 | 34.77681 | 29.77173 | 20.67806 | 26.60036 | 0.01173 | 23.75471 | 19.09269 |
| **CYB5D1** | 4.79618 | 4.50518 | 7.14163 | 2.39955 | 0.00000 | 2.87501 | 7.01644 | 0.63453 | 8.25610 | 2.19561 | 5.45480 | 3.59411 |
| **CYB5D2** | 1.19290 | 2.18344 | 1.98044 | 1.36988 | 1.96933 | 0.00000 | 0.33519 | 0.70012 | 1.44020 | 1.19782 | 1.77453 | 3.59660 |
| **CYB5R1** | 0.00262 | 4.51710 | 19.15908 | 0.02060 | 5.22829 | 5.96069 | 1.70846 | 1.52515 | 17.24928 | 0.00000 | 0.49627 | 1.80372 |
| **CYB5R2** | 0.11151 | 1.31135 | 2.50337 | 0.00021 | 3.42358 | 1.95270 | 0.63511 | 0.09409 | 4.06781 | 0.00000 | 1.48590 | 0.32318 |
| **CYB5R3** | 5.28363 | 14.72552 | 13.86356 | 11.46350 | 19.17959 | 0.00000 | 6.81554 | 0.92942 | 24.22209 | 8.00389 | 14.55610 | 4.56689 |
| **CYB5R4** | 0.02154 | 3.66204 | 6.24493 | 0.04344 | 0.00000 | 11.84685 | 6.06190 | 2.00698 | 6.43330 | 0.09259 | 2.56726 | 5.66920 |
| **CYB5RL** | 0.61638 | 3.48453 | 0.53207 | 0.07156 | 1.27568 | 0.96611 | 0.02635 | 0.20860 | 1.04942 | 0.87115 | 1.27925 | 2.30608 |
| **CYBA** | 0.00517 | 0.29992 | 0.79461 | 0.52623 | 0.46106 | 0.00000 | 0.58496 | 0.01014 | 2.01849 | 0.01797 | 0.28585 | 0.00128 |
| **CYBB** | 0.01296 | 0.01403 | 0.01366 | 0.01213 | 0.01305 | 0.02055 | 0.02038 | 0.01195 | 0.01447 | 0.02197 | 0.01490 | 0.01650 |
| **CYBRD1** | 0.10798 | 0.28439 | 0.06032 | 0.00000 | 0.00000 | 0.94382 | 0.03849 | 0.03418 | 0.71971 | 0.00781 | 1.52676 | 0.77090 |
| **CYC1** | 0.12547 | 19.24814 | 24.78150 | 24.05763 | 29.90546 | 0.43742 | 25.79108 | 0.80786 | 40.71257 | 7.35154 | 40.13419 | 3.70139 |
| **CYCS** | 2.27308 | 25.67361 | 18.39825 | 13.63117 | 16.80793 | 66.77376 | 32.96330 | 28.01263 | 33.28155 | 21.49823 | 26.15482 | 37.36556 |
| **CYCSP52** | 0.00621 | 0.01337 | 0.08216 | 0.09478 | 0.21548 | 0.19909 | 0.00000 | 0.02128 | 0.00000 | 0.12718 | 0.00264 | 0.08845 |
| **CYFIP1** | 3.03609 | 11.36638 | 10.86285 | 4.39015 | 8.13022 | 9.28155 | 10.59149 | 1.29046 | 10.53547 | 0.00000 | 10.55913 | 4.57248 |
| **CYFIP2** | 3.04540 | 4.43382 | 0.53641 | 0.51735 | 2.60883 | 0.61569 | 0.00000 | 0.30951 | 0.17316 | 0.61634 | 0.00000 | 0.00000 |
| **CYGB** | 0.00316 | 0.77386 | 0.07882 | 0.00218 | 0.09976 | 0.00002 | 0.06793 | 0.43686 | 0.00000 | 0.00000 | 0.02098 | 2.35000 |
| **CYHR1** | 2.29088 | 3.72221 | 0.69800 | 0.40448 | 3.60407 | 1.36391 | 0.59330 | 3.83153 | 1.60494 | 4.04042 | 0.81636 | 2.02896 |
| **CYLC2** | 0.00619 | 0.01578 | 0.01629 | 0.01487 | 0.01484 | 0.01956 | 0.01653 | 0.01047 | 0.01748 | 0.01737 | 0.01452 | 0.01608 |
| **CYLD** | 4.71145 | 0.62697 | 0.04556 | 0.00000 | 0.52166 | 2.43135 | 0.19937 | 2.49856 | 0.00000 | 1.42184 | 0.00000 | 1.77204 |
| **CYMP** | 0.00441 | 0.01725 | 0.07598 | 0.02903 | 0.02132 | 0.00000 | 0.00996 | 0.05344 | 0.02417 | 0.01161 | 0.01606 | 0.01557 |
| **CYP11A1** | 0.16465 | 0.00000 | 29.98557 | 0.00000 | 3.02433 | 0.07227 | 22.40253 | 0.00000 | 8.73825 | 0.00000 | 8.97968 | 1.33077 |
| **CYP17A1** | 0.00599 | 0.02180 | 0.04047 | 0.02174 | 0.01923 | 0.00000 | 0.01704 | 0.00858 | 0.01818 | 0.01095 | 0.02230 | 0.07717 |
| **CYP19A1** | 0.00000 | 0.00000 | 31.06005 | 0.00000 | 0.03940 | 0.15819 | 0.00000 | 0.12379 | 3.47353 | 0.18255 | 0.00000 | 0.01344 |
| **CYP1A1** | 0.00000 | 13.66636 | 4.39279 | 8.51693 | 0.00000 | 2.85459 | 7.49707 | 0.17293 | 8.93629 | 0.51113 | 0.00000 | 0.00000 |
| **CYP1A2** | 0.00779 | 0.04092 | 0.00872 | 0.03830 | 0.00000 | 0.02003 | 0.14376 | 0.01159 | 0.00282 | 0.03838 | 0.02679 | 0.01158 |
| **CYP1B1** | 0.19865 | 31.30451 | 1.40852 | 15.95238 | 0.00000 | 2.04101 | 2.18322 | 0.00000 | 0.00000 | 7.36692 | 0.00000 | 0.23071 |
| **CYP1B1-AS1** | 0.00812 | 0.46806 | 0.01208 | 0.01908 | 0.00000 | 0.02301 | 0.04334 | 0.01073 | 0.00000 | 0.07253 | 0.00605 | 0.00000 |
| **CYP20A1** | 0.62675 | 4.72881 | 2.60491 | 6.51295 | 4.89661 | 7.10916 | 0.00000 | 0.10648 | 5.32677 | 10.70584 | 3.39360 | 1.61812 |
| **CYP21A1P** | 0.00875 | 0.01415 | 0.02650 | 0.00992 | 0.01234 | 0.01915 | 0.01956 | 0.00944 | 0.03471 | 0.01555 | 0.01319 | 0.02247 |
| **CYP21A2.uid-84** | 0.00597 | 0.01226 | 0.04190 | 0.00927 | 0.01128 | 0.01386 | 0.04868 | 0.00872 | 0.01024 | 0.02041 | 0.02465 | 0.04848 |
| **CYP24A1** | 1.75247 | 0.01657 | 0.01447 | 0.01274 | 0.01674 | 0.00460 | 0.02951 | 0.00268 | 0.01570 | 0.00001 | 0.02517 | 0.03909 |

| | | | | | | | | | | | | |
|---|---|---|---|---|---|---|---|---|---|---|---|---|
| **CYP26A1** | 0.00666 | 20.55845 | 0.00000 | 0.00000 | 18.57041 | 0.23128 | 0.00000 | 0.58586 | 0.00000 | 0.00230 | 5.08410 | 0.00000 |
| **CYP26B1** | 0.20048 | 0.00000 | 0.08515 | 0.00485 | 0.00018 | 0.01757 | 0.00042 | 0.08708 | 0.03479 | 0.42056 | 0.00000 | 0.20349 |
| **CYP26C1** | 0.03575 | 0.03493 | 0.01915 | 0.01192 | 0.02275 | 0.01048 | 0.03076 | 0.00773 | 0.01791 | 0.00045 | 0.01983 | 0.01486 |
| **CYP27A1** | 0.04844 | 0.00000 | 0.02182 | 0.41259 | 0.00000 | 0.02625 | 0.00000 | 0.03703 | 0.12698 | 0.06613 | 0.11914 | 0.04378 |
| **CYP27B1** | 0.47579 | 0.22837 | 0.20869 | 0.00000 | 0.63685 | 0.00000 | 0.00000 | 0.08786 | 1.56764 | 2.08456 | 0.00000 | 1.74911 |
| **CYP27C1** | 0.02405 | 0.33766 | 0.30279 | 0.50319 | 0.08125 | 0.00192 | 0.00000 | 0.00103 | 0.22578 | 0.05492 | 0.00853 | 1.25009 |
| **CYP2A6** | 0.00597 | 0.01411 | 0.01926 | 0.01270 | 0.01918 | 0.01198 | 0.01784 | 0.01060 | 0.00341 | 0.02596 | 0.06098 | 0.01654 |
| **CYP2A7** | 0.00608 | 0.01509 | 0.01350 | 0.02013 | 0.01773 | 0.00964 | 0.02176 | 0.01049 | 0.00769 | 0.03043 | 0.03117 | 0.01605 |
| **CYP2B6** | 0.00619 | 0.00778 | 0.31534 | 0.39463 | 0.00000 | 0.02954 | 0.06959 | 0.01130 | 0.01093 | 0.01631 | 0.01057 | 0.01434 |
| **CYP2B7P** | 0.05283 | 0.01266 | 0.04424 | 0.05480 | 0.03472 | 0.01230 | 0.02100 | 0.00634 | 0.10922 | 0.00444 | 0.00265 | 0.01943 |
| **CYP2C18** | 0.99926 | 0.00000 | 1.97497 | 0.00000 | 0.31471 | 0.30158 | 0.25269 | 2.19179 | 0.13878 | 0.73701 | 0.43727 | 0.05245 |
| **CYP2C19** | 0.00226 | 0.00000 | 0.03564 | 0.02256 | 0.05254 | 0.03303 | 0.02489 | 0.04054 | 0.04158 | 0.19200 | 0.03149 | 0.09490 |
| **CYP2C8** | 0.00371 | 0.06180 | 0.11563 | 0.06398 | 0.07527 | 0.00000 | 0.00767 | 0.00761 | 0.17133 | 0.17143 | 0.04916 | 0.06036 |
| **CYP2C9** | 0.00377 | 0.00948 | 0.32757 | 0.02019 | 0.00000 | 0.01813 | 0.16342 | 0.01483 | 0.03135 | 0.03287 | 0.06001 | 0.07443 |
| **CYP2D7P** | 0.07336 | 0.00908 | 0.05451 | 0.00000 | 0.09372 | 0.07285 | 0.00000 | 0.00000 | 0.19628 | 0.33941 | 0.09308 | 0.13385 |
| **20.00000** | 0.00634 | 4.38375 | 0.06572 | 4.86039 | 0.00000 | 0.57080 | 0.00000 | 0.00000 | 0.00000 | 0.12422 | 0.91073 | 3.03211 |
| **CYP2F1** | 0.05374 | 0.22401 | 0.01090 | 0.52934 | 0.00000 | 0.07657 | 0.19689 | 0.01168 | 0.03467 | 0.00000 | 0.02025 | 0.16394 |
| **CYP2G1P** | 0.00636 | 0.01771 | 0.02714 | 0.01683 | 0.02475 | 0.00000 | 0.01463 | 0.01062 | 0.04274 | 0.01701 | 0.01258 | 0.01728 |
| **CYP2J2** | 0.00488 | 0.03619 | 0.00318 | 0.00831 | 0.02755 | 0.09506 | 0.01352 | 0.01852 | 0.00000 | 0.10473 | 0.00448 | 0.05521 |
| **CYP2R1** | 1.67392 | 0.00000 | 0.00000 | 0.23134 | 0.00000 | 0.11262 | 0.18608 | 1.98768 | 0.01110 | 0.00000 | 0.07581 | 0.19341 |
| **CYP2S1** | 1.51920 | 39.09915 | 2.15512 | 24.98835 | 37.22718 | 0.00000 | 0.21590 | 0.60877 | 30.37757 | 8.50743 | 8.97579 | 13.62635 |
| **CYP2U1** | 0.00233 | 0.03146 | 0.04556 | 0.20385 | 0.00484 | 0.04128 | 0.00000 | 0.16392 | 0.01133 | 0.03089 | 0.01445 | 0.00757 |
| **CYP2W1** | 0.01445 | 0.16745 | 0.02811 | 0.00474 | 0.00544 | 0.00739 | 0.01310 | 0.01840 | 0.03036 | 0.03774 | 0.01455 | 0.00926 |
| **CYP39A1** | 0.04214 | 0.03215 | 0.17192 | 0.01888 | 0.00760 | 0.03936 | 0.00000 | 0.02678 | 0.00562 | 0.03423 | 0.01163 | 0.00923 |
| **CYP3A4** | 0.00533 | 0.01205 | 0.01849 | 0.03463 | 0.01792 | 0.02704 | 0.00909 | 0.00937 | 0.01916 | 0.04603 | 0.01233 | 0.02843 |
| **CYP3A43** | 0.00340 | 0.01756 | 0.02246 | 0.01304 | 0.01894 | 0.01078 | 0.01819 | 0.02542 | 0.02469 | 0.00763 | 0.01768 | 0.03162 |
| **CYP3A5** | 0.00018 | 0.01783 | 0.33926 | 0.29409 | 0.00000 | 0.44494 | 0.00000 | 0.15018 | 0.25934 | 1.05036 | 0.00941 | 0.16284 |
| **CYP3A7** | 0.00614 | 0.01369 | 0.15827 | 0.00578 | 0.00000 | 0.01676 | 0.09875 | 0.01050 | 0.06486 | 0.01022 | 0.08037 | 0.01629 |
| **CYP3A7-CYP3A** | 0.00599 | 0.01454 | 0.05744 | 0.01418 | 0.00825 | 0.01379 | 0.00941 | 0.00992 | 0.05582 | 0.03105 | 0.01964 | 0.02191 |
| **CYP46A1** | 0.55973 | 0.01426 | 0.01725 | 0.01141 | 0.05072 | 0.00000 | 0.02274 | 0.02926 | 0.02074 | 0.00000 | 0.01375 | 0.03963 |
| **CYP4B1** | 0.30330 | 0.02838 | 0.02407 | 0.00445 | 0.00428 | 0.02378 | 0.02382 | 0.32932 | 0.00431 | 0.04535 | 0.01876 | 0.00724 |
| **CYP4F11** | 0.01990 | 0.01825 | 0.03134 | 0.03300 | 0.01850 | 0.00000 | 0.01620 | 0.00905 | 0.02660 | 0.00643 | 0.02070 | 0.01788 |
| **CYP4F12** | 0.09858 | 0.01693 | 0.03093 | 0.01032 | 0.02287 | 0.02192 | 0.01330 | 0.01150 | 0.01801 | 0.01327 | 0.01232 | 0.03793 |
| **CYP4F2** | 0.08008 | 0.01020 | 0.02106 | 0.01143 | 0.00000 | 0.02761 | 0.01340 | 0.00512 | 0.04229 | 0.07971 | 0.01290 | 0.01235 |
| **CYP4F22** | 0.05008 | 0.08231 | 0.02744 | 0.11269 | 0.00000 | 0.00000 | 0.02842 | 0.00370 | 0.00000 | 0.07979 | 0.03582 | 0.17450 |
| **CYP4F24P** | 0.00196 | 0.02553 | 0.01605 | 0.01464 | 0.01570 | 0.00752 | 0.01992 | 0.10525 | 0.02280 | 0.01814 | 0.01526 | 0.03662 |
| **CYP4F3** | 0.11816 | 0.00972 | 0.01191 | 0.00512 | 0.00912 | 0.06154 | 0.03284 | 0.02302 | 0.00001 | 0.02687 | 0.01122 | 0.05365 |
| **CYP4F35P** | 0.00432 | 0.00311 | 0.00000 | 0.29034 | 0.00000 | 0.03391 | 0.00000 | 0.00745 | 0.01663 | 0.23379 | 0.00970 | 0.07806 |
| **CYP4F8** | 0.00686 | 0.01753 | 0.08247 | 0.01047 | 0.04087 | 0.02094 | 0.02140 | 0.01009 | 0.00000 | 0.01147 | 0.02647 | 0.01898 |
| **CYP4V2** | 0.41773 | 0.14470 | 0.00863 | 0.00124 | 0.02252 | 0.09683 | 0.06357 | 0.10357 | 0.04454 | 0.00000 | 0.08142 | 0.06099 |
| **CYP4X1** | 0.20912 | 0.00622 | 0.03172 | 0.28697 | 0.00000 | 0.08758 | 0.00000 | 0.14026 | 0.01918 | 0.03455 | 0.00488 | 0.14158 |
| **CYP51A1** | 0.08062 | 0.00000 | 5.87174 | 13.45983 | 24.06261 | 18.99558 | 7.76520 | 0.97110 | 30.74494 | 9.93159 | 22.08092 | 1.01313 |
| **CYP51A1-AS1** | 0.06447 | 0.10002 | 0.50378 | 0.10467 | 0.21168 | 0.00000 | 0.18151 | 0.27760 | 0.44095 | 0.00000 | 0.04746 | 0.20923 |
| **CYP7A1** | 0.00357 | 0.00245 | 0.00000 | 0.00065 | 0.00000 | 0.03767 | 0.06855 | 0.00928 | 0.00000 | 0.20291 | 0.00266 | 0.02368 |
| **CYP7B1** | 0.16826 | 0.02742 | 0.00000 | 0.09512 | 0.30813 | 0.00392 | 0.13309 | 0.03742 | 0.12933 | 0.08548 | 0.00000 | 0.00758 |
| **CYP8B1** | 0.00752 | 0.00000 | 0.13324 | 0.00000 | 0.32070 | 0.00000 | 0.00000 | 0.01236 | 0.00000 | 0.83549 | 0.41463 | 0.02328 |
| **CYR61** | 0.15231 | 1.41547 | 5.30358 | 0.18285 | 1.59398 | 0.00000 | 0.93843 | 0.03775 | 5.06272 | 0.00000 | 1.49476 | 0.00000 |
| **CYS1** | 0.00557 | 0.01180 | 0.02413 | 0.00795 | 0.06174 | 0.00000 | 0.00818 | 0.00968 | 0.01871 | 0.04065 | 0.05302 | 0.03431 |

| | | | | | | | | | | | | |
|---|---|---|---|---|---|---|---|---|---|---|---|---|
| **CYSRT1** | 0.18551 | 0.10462 | 0.03850 | 0.07441 | 0.14081 | 0.00000 | 0.04709 | 0.00392 | 0.10202 | 0.12526 | 0.00000 | 0.07018 |
| **CYSTM1** | 6.91887 | 15.72684 | 4.74458 | 4.19308 | 20.42335 | 12.89460 | 10.82644 | 4.80062 | 3.62583 | 0.08933 | 11.65605 | 5.37880 |
| **CYTH1** | 1.19239 | 2.48183 | 2.45984 | 0.23621 | 6.25897 | 0.00000 | 2.96693 | 3.85278 | 1.55846 | 0.00000 | 1.48748 | 9.30841 |
| **CYTH2** | 2.33249 | 8.13470 | 9.52027 | 7.15122 | 11.58457 | 0.00000 | 6.63814 | 7.88695 | 10.28046 | 1.94967 | 6.69783 | 7.05475 |
| **CYTH3** | 1.92417 | 3.53792 | 2.54085 | 0.00000 | 6.38295 | 2.32364 | 3.64651 | 4.57244 | 2.55336 | 9.82519 | 1.05760 | 6.59179 |
| **CYTH4** | 0.00559 | 0.01207 | 0.08364 | 0.02805 | 0.00805 | 0.02068 | 0.00729 | 0.01693 | 0.00615 | 0.03670 | 0.02584 | 0.01648 |
| **CYTIP** | 0.16538 | 0.00699 | 0.02107 | 0.00112 | 0.12473 | 0.11758 | 0.00000 | 0.05392 | 0.02787 | 0.01894 | 0.00000 | 0.06115 |
| **CYYR1** | 0.05118 | 0.01203 | 0.03075 | 0.02513 | 0.00000 | 0.03423 | 0.01080 | 0.01488 | 0.00632 | 0.05000 | 0.01997 | 0.01378 |
| **D2HGDH** | 0.06488 | 0.27462 | 0.23604 | 0.00000 | 0.01842 | 0.00000 | 0.00000 | 0.76826 | 0.20928 | 0.38550 | 0.21937 | 0.26828 |
| **DAAM1** | 13.04253 | 0.21741 | 4.38255 | 2.00752 | 4.70972 | 16.31342 | 4.90388 | 5.02191 | 0.00000 | 15.70939 | 6.40149 | 2.89731 |
| **DAAM2** | 4.34274 | 0.00000 | 0.17250 | 0.00000 | 0.00000 | 3.00847 | 0.87199 | 0.81391 | 0.00000 | 5.22452 | 0.00000 | 0.60957 |
| **DAB1** | 2.33895 | 0.00879 | 0.05137 | 0.32061 | 0.05830 | 0.00017 | 0.04759 | 0.27155 | 0.04843 | 0.00088 | 0.01063 | 0.01172 |
| **DAB2** | 0.00000 | 5.13261 | 14.03507 | 0.00000 | 58.35174 | 21.69025 | 38.38575 | 4.40569 | 38.34182 | 10.79307 | 44.03551 | 18.69013 |
| **DAB2IP** | 2.54955 | 0.68813 | 0.21335 | 0.96557 | 2.12926 | 0.54528 | 0.28245 | 0.47144 | 1.29858 | 4.10927 | 1.52287 | 0.00000 |
| **DACH1** | 0.40524 | 0.04644 | 0.42371 | 0.12510 | 0.23977 | 0.00182 | 0.17853 | 0.04062 | 0.00000 | 0.00000 | 0.01299 | 0.09826 |
| **DACH2** | 0.00388 | 0.00950 | 0.04922 | 0.11060 | 0.00305 | 0.03113 | 0.00704 | 0.05095 | 0.01127 | 0.03116 | 0.01820 | 0.02970 |
| **DACT1** | 0.11735 | 0.00000 | 1.87489 | 2.30461 | 0.25333 | 1.42389 | 1.71742 | 0.00000 | 0.37684 | 0.78131 | 0.00000 | 0.00000 |
| **DACT2** | 0.00385 | 0.00000 | 0.18154 | 0.11319 | 1.86933 | 0.00000 | 0.41533 | 0.00926 | 0.45462 | 0.08866 | 0.50097 | 0.00917 |
| **DACT3** | 0.10613 | 0.01621 | 0.02358 | 0.01054 | 0.01401 | 0.02190 | 0.05480 | 0.02453 | 0.02729 | 0.00000 | 0.01529 | 0.03063 |
| **DACT3-AS1** | 0.10735 | 0.00998 | 0.03400 | 0.00855 | 0.00742 | 0.01965 | 0.01908 | 0.00744 | 0.03335 | 0.03891 | 0.01760 | 0.04741 |
| **DAD1** | 1.76819 | 15.76745 | 24.16732 | 13.29113 | 23.61035 | 39.65440 | 14.87402 | 2.87726 | 35.60170 | 14.71255 | 22.82917 | 28.10311 |
| **DAG1** | 3.54416 | 16.36345 | 24.58188 | 6.02325 | 37.23155 | 6.94624 | 5.89436 | 4.30045 | 46.03538 | 28.65186 | 27.61235 | 8.11395 |
| **DAGLA** | 0.38309 | 0.78402 | 0.24234 | 1.46401 | 0.48734 | 0.00000 | 0.00000 | 0.00000 | 0.10810 | 0.00616 | 0.00000 | 0.20873 |
| **DAGLB** | 2.89007 | 8.47197 | 5.78652 | 6.70951 | 3.18013 | 0.00000 | 0.00000 | 8.60957 | 13.65530 | 0.90493 | 1.73579 | 11.00752 |
| **DAK** | 0.00102 | 3.18062 | 0.01724 | 0.77119 | 2.37620 | 0.00000 | 0.93088 | 0.03544 | 1.83790 | 1.19952 | 3.03778 | 1.22457 |
| **DALRD3** | 0.84939 | 6.86145 | 4.57831 | 6.19482 | 4.41822 | 0.00000 | 1.60781 | 0.00000 | 5.32161 | 0.00000 | 5.60491 | 2.35492 |
| **DANCR** | 1.40106 | 24.21992 | 22.95723 | 10.57122 | 19.83620 | 21.63581 | 41.91860 | 7.91148 | 44.19174 | 18.40483 | 28.34073 | 35.81246 |
| **DAND5** | 2.41607 | 0.00990 | 0.03437 | 0.01194 | 0.01753 | 0.00000 | 0.03036 | 0.20874 | 0.00000 | 0.00712 | 0.06344 | 0.02013 |
| **DAO** | 0.00453 | 0.01348 | 0.22676 | 0.00849 | 0.00000 | 0.02379 | 0.06601 | 0.03234 | 0.03730 | 0.02310 | 0.04668 | 0.01328 |
| **DAP** | 4.17686 | 6.32719 | 7.97475 | 3.07895 | 3.76785 | 1.76710 | 3.79088 | 0.00000 | 8.40772 | 0.00000 | 7.04854 | 3.44304 |
| **DAP3** | 4.07415 | 15.12590 | 19.73808 | 14.02620 | 10.21594 | 27.66910 | 19.85053 | 10.86701 | 14.35296 | 6.70063 | 19.07576 | 19.42859 |
| **DAPK1** | 0.00000 | 7.11775 | 0.00000 | 3.09807 | 7.57994 | 4.35506 | 3.70635 | 0.16252 | 7.21528 | 5.33449 | 6.36096 | 2.40485 |
| **DAPK2** | 0.10369 | 0.15621 | 0.01138 | 0.10560 | 0.19468 | 0.00000 | 0.04377 | 0.00622 | 0.00140 | 0.01680 | 0.01586 | 0.01249 |
| **DAPK3** | 1.71659 | 7.15841 | 7.92804 | 3.17940 | 7.31317 | 0.00000 | 1.39586 | 1.75351 | 5.08470 | 6.80013 | 3.71499 | 3.29724 |
| **DAPP1** | 0.69940 | 0.13576 | 0.00000 | 0.00786 | 0.21528 | 0.39237 | 0.00000 | 0.00000 | 0.00000 | 0.02957 | 0.01865 | 0.00000 |
| **DARS** | 2.28833 | 13.63754 | 15.30759 | 8.50876 | 14.52556 | 36.31575 | 27.47747 | 1.97239 | 10.88402 | 13.81972 | 24.42444 | 9.78702 |
| **DARS2** | 0.90534 | 8.09576 | 6.69683 | 4.36600 | 3.37505 | 22.06365 | 5.05408 | 0.79349 | 7.36822 | 0.00464 | 11.37637 | 3.89562 |
| **DAW1** | 3.86207 | 0.00346 | 0.00113 | 0.05048 | 0.01521 | 0.00000 | 0.00579 | 0.42579 | 0.03910 | 0.09142 | 0.09647 | 0.03743 |
| **DAXX** | 0.35833 | 22.27361 | 17.83684 | 17.45567 | 17.50409 | 0.00001 | 27.84528 | 13.14178 | 22.54920 | 4.11452 | 17.36881 | 28.64724 |
| **DAZAP1** | 0.59551 | 3.07137 | 7.20035 | 5.13362 | 11.77989 | 0.00006 | 13.54181 | 0.29964 | 1.00947 | 18.12776 | 2.99865 | 1.94328 |
| **DAZAP2** | 5.78466 | 7.03247 | 3.92971 | 3.34059 | 1.04671 | 14.29802 | 7.79350 | 19.39217 | 3.07616 | 3.77004 | 6.32613 | 14.51032 |
| **DAZL** | 5.97685 | 0.28294 | 1.06216 | 0.03064 | 0.00000 | 0.58320 | 0.00000 | 0.79190 | 0.41241 | 0.00000 | 0.07550 | 4.32132 |
| **DBET** | 0.04482 | 0.07393 | 0.04960 | 0.01245 | 0.03643 | 0.03622 | 0.09124 | 0.04106 | 0.00705 | 0.08994 | 0.03776 | 0.01838 |
| **DBF4** | 2.27715 | 9.05907 | 4.58799 | 1.81438 | 3.45793 | 13.19152 | 9.28007 | 4.68580 | 9.77853 | 7.37402 | 4.34253 | 15.70283 |
| **DBF4B** | 5.34518 | 2.21337 | 1.65252 | 0.86163 | 2.47460 | 0.00000 | 3.57321 | 3.58432 | 5.09970 | 4.81687 | 2.67819 | 4.52988 |
| **DBH** | 0.70987 | 0.01110 | 0.02032 | 0.00976 | 0.01038 | 0.01391 | 0.02052 | 0.01111 | 0.02391 | 0.03235 | 0.01209 | 0.01857 |
| **DBH-AS1** | 0.13000 | 0.00565 | 0.02100 | 0.00570 | 0.00523 | 0.03152 | 0.01630 | 0.05983 | 0.01759 | 0.04991 | 0.01096 | 0.07319 |
| **DBI** | 5.04047 | 6.77569 | 23.62926 | 25.26178 | 33.54558 | 27.21792 | 43.89080 | 11.91775 | 32.60909 | 9.60791 | 31.81338 | 13.88687 |

| | | | | | | | | | | | | |
|---|---|---|---|---|---|---|---|---|---|---|---|---|
| **DBIL5P** | 0.01122 | 0.03381 | 0.18465 | 0.02542 | 0.00000 | 0.07361 | 0.03199 | 0.50219 | 0.26103 | 0.00247 | 0.01179 | 0.44129 |
| **DBIL5P2** | 0.00419 | 0.01903 | 0.00776 | 0.07519 | 0.01703 | 0.00000 | 0.05367 | 0.01423 | 0.01551 | 0.03287 | 0.02178 | 0.04433 |
| **DBN1** | 0.15822 | 11.17260 | 0.65120 | 15.78099 | 15.75965 | 0.00000 | 0.00202 | 0.21330 | 13.44221 | 5.51214 | 7.02785 | 3.31757 |
| **DBNDD1** | 0.05366 | 0.36264 | 0.29996 | 0.00000 | 1.08921 | 0.02470 | 0.38449 | 0.00986 | 0.02168 | 0.03067 | 0.31064 | 0.00000 |
| **DBNDD2** | 0.00232 | 0.41248 | 0.45563 | 0.00000 | 1.65806 | 0.29543 | 0.55868 | 0.04918 | 0.50023 | 0.00000 | 0.87603 | 0.08490 |
| **DBNL** | 0.39969 | 16.50449 | 13.15554 | 7.43069 | 14.43592 | 0.00000 | 4.50043 | 0.59526 | 5.24102 | 0.00000 | 14.51700 | 8.72383 |
| **DBP** | 0.00764 | 0.77851 | 0.03458 | 0.40009 | 0.15328 | 0.00000 | 0.01760 | 2.94741 | 0.17967 | 0.95689 | 0.00006 | 6.00309 |
| **DBR1** | 3.20789 | 6.03380 | 6.91814 | 4.27418 | 0.00000 | 16.20295 | 7.50093 | 27.13449 | 8.65833 | 0.76268 | 8.32111 | 8.09710 |
| **DBT** | 0.45425 | 2.68151 | 5.38925 | 1.47658 | 0.00002 | 10.36050 | 4.31526 | 1.70606 | 4.08725 | 2.72198 | 2.93786 | 3.60606 |
| **DBX1** | 0.03991 | 0.01395 | 0.03889 | 0.01083 | 0.01619 | 0.02136 | 0.01353 | 0.05468 | 0.01758 | 0.00883 | 0.01707 | 0.02208 |
| **DBX2** | 0.02328 | 0.01559 | 0.01661 | 0.00833 | 0.21457 | 0.00000 | 0.02486 | 0.00825 | 0.00723 | 0.01170 | 0.00885 | 0.02438 |
| **DCAF10** | 0.13496 | 0.14708 | 0.91666 | 0.41081 | 0.94494 | 1.46259 | 1.57921 | 0.04898 | 0.99398 | 1.39396 | 0.19858 | 0.01310 |
| **DCAF11** | 0.26078 | 2.58868 | 6.96156 | 7.64167 | 5.80608 | 2.26202 | 7.61089 | 0.15775 | 6.67546 | 0.00000 | 6.29755 | 1.46705 |
| **DCAF12** | 4.97300 | 8.27119 | 15.81677 | 3.18926 | 3.03885 | 15.06871 | 12.15490 | 7.24339 | 14.80819 | 0.00000 | 6.82523 | 10.38771 |
| **DCAF12L1** | 0.00172 | 0.02221 | 0.09658 | 0.06379 | 0.02905 | 0.03931 | 0.01375 | 0.24991 | 0.00000 | 0.00000 | 0.05867 | 0.06444 |
| **DCAF12L2** | 0.00608 | 0.01194 | 0.01801 | 0.06351 | 0.01742 | 0.00000 | 0.02139 | 0.00867 | 0.02700 | 0.00475 | 0.01810 | 0.08398 |
| **DCAF13** | 1.98561 | 10.08528 | 20.37932 | 6.17976 | 0.00000 | 33.16792 | 23.64388 | 21.62355 | 17.92731 | 0.76620 | 16.32114 | 20.69664 |
| **DCAF13P3** | 0.10570 | 0.80788 | 2.11287 | 0.30905 | 0.00000 | 2.22160 | 2.56505 | 3.13727 | 1.44765 | 0.68758 | 1.30459 | 2.42108 |
| **DCAF15** | 1.38056 | 0.26943 | 0.62805 | 0.44351 | 1.23246 | 0.00000 | 0.87841 | 0.16601 | 1.03423 | 2.99008 | 0.07359 | 0.77831 |
| **DCAF16** | 2.19247 | 1.30675 | 4.66855 | 0.00000 | 1.94383 | 9.53389 | 7.39501 | 6.74455 | 5.89702 | 6.68082 | 5.80873 | 6.10199 |
| **DCAF17** | 4.35231 | 4.44726 | 2.15467 | 0.00000 | 0.50389 | 10.67670 | 3.80263 | 1.36487 | 4.11219 | 3.68309 | 2.11988 | 3.17580 |
| **DCAF4** | 1.22546 | 6.25987 | 11.53958 | 8.64811 | 8.10690 | 10.13741 | 6.79889 | 9.49915 | 21.11749 | 5.75185 | 10.87347 | 6.94360 |
| **DCAF4L1** | 8.09165 | 2.10139 | 1.14206 | 0.19056 | 0.00000 | 2.19309 | 1.02207 | 12.06686 | 2.36044 | 3.02232 | 0.96137 | 13.90345 |
| **DCAF4L2** | 1.32197 | 3.02915 | 3.93248 | 0.53232 | 0.00000 | 1.51334 | 3.52712 | 2.48689 | 1.22451 | 0.00000 | 1.39236 | 5.28603 |
| **DCAF5** | 11.23557 | 0.94246 | 2.11965 | 4.67323 | 3.10284 | 3.27838 | 2.93910 | 3.66191 | 1.25519 | 6.97920 | 1.39074 | 1.05886 |
| **DCAF6** | 4.51979 | 4.52187 | 8.03826 | 2.92940 | 2.79214 | 10.90509 | 9.49705 | 2.97108 | 4.61916 | 4.92422 | 0.87033 | 3.82275 |
| **DCAF7** | 14.42980 | 10.79499 | 6.26188 | 12.24202 | 7.36892 | 9.71649 | 13.17141 | 8.54488 | 6.71455 | 2.27813 | 7.28534 | 12.38432 |
| **DCAF8** | 4.39788 | 7.79545 | 9.25069 | 7.42812 | 8.43445 | 3.36518 | 7.11754 | 3.79080 | 14.18326 | 3.64497 | 8.65974 | 8.13413 |
| **DCAF8L1** | 0.00322 | 0.15358 | 0.00000 | 0.09645 | 0.00000 | 0.13063 | 0.20656 | 0.27932 | 0.12507 | 0.00000 | 0.20414 | 0.05190 |
| **DCAF8L2** | 0.01932 | 0.02891 | 0.10858 | 0.02914 | 0.04270 | 0.03099 | 0.14523 | 0.08877 | 0.00000 | 0.00000 | 0.05040 | 0.18697 |
| **DCAKD** | 1.47171 | 3.89224 | 4.97143 | 1.56156 | 1.92370 | 0.75325 | 5.23182 | 2.80795 | 5.61519 | 2.00044 | 10.13071 | 2.54324 |
| **DCBLD1** | 0.10016 | 0.34872 | 1.55276 | 0.00015 | 1.71553 | 0.38577 | 0.00400 | 0.12234 | 2.30411 | 0.93868 | 0.77674 | 0.00157 |
| **DCBLD2** | 0.19713 | 1.10240 | 2.18888 | 0.66683 | 0.00000 | 4.44002 | 1.48452 | 0.36986 | 0.71746 | 6.19792 | 0.27607 | 0.63964 |
| **DCC** | 2.56740 | 0.42065 | 0.00000 | 0.05361 | 0.00000 | 0.04084 | 0.00960 | 0.05560 | 0.00000 | 0.12529 | 0.00830 | 0.25192 |
| **DCD** | 0.01108 | 0.00940 | 0.00286 | 0.00000 | 0.00000 | 0.06692 | 0.05284 | 0.01230 | 0.00000 | 0.11043 | 0.01539 | 0.01280 |
| **DCDC1** | 0.19727 | 0.10212 | 0.02893 | 0.00000 | 0.03762 | 0.06672 | 0.00531 | 0.00000 | 0.00000 | 0.10430 | 0.01657 | 0.32196 |
| **DCDC2** | 16.36433 | 0.29351 | 0.15329 | 0.00000 | 1.91009 | 0.48752 | 0.00000 | 4.72055 | 0.00000 | 0.05955 | 0.73130 | 0.89005 |
| **DCDC2B** | 0.03898 | 0.07363 | 0.14670 | 0.09835 | 0.15127 | 0.00000 | 0.01332 | 0.07558 | 0.15168 | 0.04152 | 0.09852 | 0.08129 |
| **DCDC2C** | 1.13186 | 0.00712 | 0.01302 | 0.01055 | 0.04669 | 0.03972 | 0.01800 | 0.03319 | 0.00756 | 0.00521 | 0.01378 | 0.02108 |
| **DCDC5** | 0.00618 | 0.01122 | 0.01070 | 0.02539 | 0.00809 | 0.04066 | 0.01572 | 0.01016 | 0.01486 | 0.01654 | 0.01365 | 0.01500 |
| **DCHS1** | 0.00620 | 0.16267 | 0.08119 | 0.92563 | 0.14487 | 0.00000 | 0.14427 | 0.00272 | 0.16265 | 0.34778 | 0.41188 | 0.16832 |
| **DCHS2** | 2.96460 | 0.00313 | 0.03313 | 0.01365 | 0.01678 | 0.00000 | 0.00983 | 0.90520 | 0.02264 | 0.02868 | 0.01406 | 0.13216 |
| **DCK** | 0.29778 | 3.48211 | 2.47171 | 0.23609 | 0.00000 | 6.89184 | 4.51249 | 2.42700 | 8.14951 | 0.00000 | 3.35895 | 3.62286 |
| **DCLK1** | 0.01024 | 0.03222 | 0.11713 | 0.29673 | 0.00000 | 0.03911 | 0.00000 | 0.00846 | 0.04543 | 0.02292 | 0.01485 | 0.02091 |
| **DCLK2** | 6.96701 | 0.02371 | 0.00000 | 0.52742 | 0.06321 | 0.08515 | 0.00000 | 0.01706 | 0.03650 | 0.06254 | 0.01259 | 0.02274 |
| **DCLK3** | 0.04506 | 0.10452 | 0.14840 | 0.00844 | 0.01210 | 0.00000 | 0.03006 | 0.00731 | 0.06388 | 0.01954 | 0.01936 | 0.00908 |
| **DCLRE1A** | 6.15858 | 2.02305 | 0.00000 | 0.00000 | 0.00032 | 7.37448 | 1.50818 | 0.77364 | 0.41272 | 1.50240 | 4.00985 | 0.11072 |
| **DCLRE1B** | 6.27153 | 7.51651 | 1.66393 | 4.54508 | 2.71723 | 6.22230 | 4.86130 | 15.85451 | 5.76048 | 0.00000 | 5.60397 | 9.57620 |

| | | | | | | | | | | | | |
|---|---|---|---|---|---|---|---|---|---|---|---|---|
| **DCLRE1C** | 0.63717 | 2.08784 | 3.66327 | 3.11886 | 0.55266 | 4.61407 | 1.52973 | 1.97609 | 9.13104 | 6.71546 | 7.76122 | 3.20034 |
| **DCP1A** | 9.38417 | 7.87039 | 15.28473 | 10.39781 | 8.49100 | 12.69573 | 19.83158 | 28.15142 | 13.17411 | 3.72243 | 9.34425 | 15.06401 |
| **DCP1B** | 0.23331 | 3.47074 | 0.00000 | 0.87150 | 0.61182 | 4.10164 | 0.25728 | 0.00000 | 0.00000 | 0.20780 | 1.10361 | 0.91322 |
| **DCP2** | 16.35120 | 10.91678 | 8.84511 | 2.94328 | 0.00036 | 28.75625 | 13.98203 | 12.46040 | 2.05032 | 14.69574 | 11.87053 | 12.49822 |
| **DCPS** | 2.74501 | 4.58582 | 8.21335 | 3.52298 | 0.96878 | 2.91925 | 10.15139 | 1.32751 | 6.21453 | 0.00000 | 7.81840 | 3.47260 |
| **DCST1** | 0.01438 | 0.04836 | 0.04031 | 0.01349 | 0.02433 | 0.00498 | 0.13398 | 0.00878 | 0.00000 | 0.04318 | 0.07454 | 0.04240 |
| **DCST2** | 0.00326 | 0.03340 | 0.04945 | 0.07999 | 0.00000 | 0.00000 | 0.16721 | 0.00824 | 0.00000 | 0.36493 | 0.03855 | 0.03280 |
| **DCT** | 5.21654 | 0.00309 | 0.00000 | 0.00292 | 0.07304 | 0.01338 | 0.05361 | 0.33577 | 0.13110 | 0.00000 | 0.00431 | 0.27041 |
| **DCTD** | 0.18271 | 2.17362 | 9.98485 | 0.86791 | 4.40336 | 9.77281 | 5.13642 | 0.44177 | 10.17142 | 0.00000 | 6.72602 | 3.01501 |
| **DCTN1** | 6.04605 | 15.70317 | 6.90374 | 16.85146 | 17.34700 | 0.00000 | 13.19223 | 0.49647 | 12.96119 | 7.92795 | 10.55795 | 12.51216 |
| **DCTN1-AS1** | 0.01113 | 0.06121 | 0.01126 | 0.05255 | 0.00159 | 0.00000 | 0.00837 | 0.00865 | 0.07310 | 0.06841 | 0.01491 | 0.07217 |
| **DCTN2** | 0.90791 | 16.60484 | 15.22035 | 17.01526 | 14.82119 | 14.60797 | 16.19299 | 0.11513 | 17.17985 | 2.11959 | 11.09531 | 7.70251 |
| **DCTN3** | 1.87309 | 8.60599 | 6.40500 | 5.06419 | 9.09115 | 8.67773 | 10.18265 | 11.83037 | 6.87355 | 0.00000 | 6.96352 | 14.19140 |
| **DCTN4** | 1.89717 | 5.64606 | 8.64476 | 0.61743 | 0.51460 | 14.80377 | 8.43514 | 1.09436 | 6.55424 | 7.54331 | 6.77066 | 4.91894 |
| **DCTN5** | 5.30041 | 18.40825 | 17.30136 | 19.76079 | 11.06210 | 20.45496 | 17.74970 | 8.85908 | 11.80122 | 0.00035 | 14.39172 | 26.11633 |
| **DCTN6** | 0.27629 | 5.88597 | 9.16777 | 1.97757 | 1.79615 | 21.41438 | 7.25563 | 0.06918 | 7.89190 | 0.77727 | 7.31422 | 4.57674 |
| **DCTPP1** | 3.84324 | 13.97757 | 14.91002 | 20.18176 | 16.27398 | 6.98169 | 32.08864 | 0.50470 | 24.91399 | 0.86679 | 19.21411 | 12.29429 |
| **DCUN1D1** | 1.96537 | 3.11473 | 3.60560 | 0.24555 | 0.00000 | 15.25135 | 7.38680 | 0.95462 | 5.24946 | 0.00001 | 5.56484 | 3.73126 |
| **DCUN1D2** | 0.19458 | 0.20336 | 1.14465 | 0.39904 | 1.37074 | 2.21521 | 2.90370 | 0.31260 | 2.82988 | 6.37532 | 2.08029 | 0.35258 |
| **DCUN1D3** | 2.13745 | 1.11341 | 1.45706 | 3.88350 | 0.60289 | 0.00000 | 1.51249 | 3.37204 | 0.42078 | 0.50571 | 0.94247 | 1.26432 |
| **DCUN1D4** | 1.22930 | 4.82075 | 0.00000 | 3.92261 | 1.72062 | 18.33385 | 3.70999 | 5.75828 | 6.09294 | 0.00000 | 4.79532 | 3.77556 |
| **DCUN1D5** | 5.59897 | 18.50637 | 14.96901 | 5.43831 | 12.50321 | 31.07262 | 24.91689 | 4.79919 | 25.32222 | 7.27484 | 12.45359 | 11.29169 |
| **DCX** | 0.55154 | 0.08200 | 0.00467 | 0.03117 | 0.00000 | 0.01054 | 0.08658 | 0.01363 | 0.00000 | 0.04779 | 0.01519 | 0.00898 |
| **DCXR** | 4.56909 | 6.47905 | 17.02820 | 15.19586 | 21.63242 | 0.00000 | 23.14946 | 1.52678 | 16.70418 | 3.20036 | 18.20064 | 0.00000 |
| **DDA1** | 4.42567 | 6.36128 | 12.38853 | 5.24929 | 5.65611 | 0.88116 | 11.31424 | 13.63256 | 10.17669 | 4.04515 | 5.59082 | 12.96504 |
| **DDAH1** | 1.46985 | 0.00004 | 0.66892 | 0.10050 | 0.07685 | 1.29966 | 0.57569 | 0.00000 | 0.35569 | 0.00000 | 0.47002 | 0.00000 |
| **DDAH2** | 0.00804 | 2.00843 | 2.76428 | 4.43515 | 2.41963 | 0.00000 | 1.49198 | 6.30212 | 3.52657 | 1.65443 | 0.54948 | 2.89249 |
| **DDB1** | 5.94510 | 22.46077 | 35.58983 | 19.00126 | 20.43094 | 8.89132 | 41.34923 | 5.80063 | 27.50801 | 6.78425 | 28.12472 | 17.38375 |
| **DDB2** | 0.90059 | 8.17924 | 1.34022 | 4.11348 | 1.21502 | 3.28684 | 0.77792 | 1.84875 | 1.83837 | 0.00000 | 0.86923 | 6.37550 |
| **DDC** | 0.00290 | 0.23859 | 0.34033 | 1.08474 | 0.00000 | 0.00533 | 0.02359 | 0.15442 | 0.01196 | 0.00274 | 0.08149 | 0.15462 |
| **DDC-AS1** | 0.00625 | 0.01557 | 0.01888 | 0.01534 | 0.01420 | 0.01245 | 0.01738 | 0.01036 | 0.01673 | 0.01446 | 0.02737 | 0.01997 |
| **DDHD1** | 2.03962 | 1.63987 | 6.32541 | 5.05883 | 1.26916 | 20.75211 | 3.54688 | 0.34454 | 2.20297 | 21.79556 | 3.70526 | 5.94032 |
| **DDHD2** | 1.37367 | 0.59832 | 2.85833 | 0.00000 | 1.56025 | 0.79386 | 2.42376 | 0.09127 | 6.80447 | 1.61995 | 2.79534 | 0.25641 |
| **DDI1** | 0.00632 | 0.01427 | 0.02004 | 0.01171 | 0.01436 | 0.02320 | 0.01682 | 0.01020 | 0.01917 | 0.01229 | 0.01406 | 0.01904 |
| **DDI2** | 0.04554 | 1.61747 | 1.02734 | 0.00000 | 0.46600 | 0.42703 | 0.32509 | 4.09887 | 0.51945 | 0.48169 | 1.58200 | 0.75852 |
| **DDIAS** | 1.43658 | 3.87889 | 6.87805 | 0.88862 | 0.29985 | 12.74440 | 4.43252 | 21.67341 | 6.66832 | 1.23697 | 1.88554 | 11.39801 |
| **DDIT3** | 6.00539 | 3.61930 | 4.76375 | 7.68105 | 2.35556 | 4.88807 | 0.95719 | 22.96151 | 2.75416 | 1.32482 | 0.00000 | 4.59988 |
| **DDIT4** | 2.07061 | 2.47383 | 11.26666 | 0.00000 | 1.04787 | 12.21984 | 29.33276 | 0.52296 | 37.91084 | 3.87731 | 5.04245 | 0.19544 |
| **DDIT4L** | 0.20362 | 5.11928 | 0.00000 | 0.00000 | 2.58059 | 7.85298 | 8.99448 | 0.00000 | 7.36172 | 0.00000 | 8.63881 | 0.77934 |
| **DDN** | 0.21583 | 0.00699 | 0.32685 | 0.00114 | 0.00000 | 0.00292 | 0.55909 | 0.00262 | 0.39615 | 0.06135 | 0.00000 | 0.02935 |
| **DDO** | 0.00617 | 0.00925 | 0.00000 | 0.24742 | 0.03325 | 0.05875 | 0.00981 | 0.00978 | 0.01565 | 0.00013 | 0.01270 | 0.01383 |
| **DDOST** | 3.22016 | 26.88181 | 28.74308 | 21.85775 | 21.30498 | 26.49879 | 6.56732 | 7.24980 | 60.23242 | 0.00000 | 27.54160 | 17.77733 |
| **DDR1** | 2.54329 | 2.57972 | 4.22295 | 1.48506 | 1.15953 | 0.77652 | 1.15331 | 0.65482 | 7.63093 | 4.96583 | 0.52639 | 4.16247 |
| **DDR2** | 0.08241 | 0.00336 | 0.06144 | 0.21563 | 0.00070 | 0.01435 | 0.00012 | 0.05302 | 0.17011 | 0.04395 | 0.00820 | 0.10944 |
| **DDRGK1** | 0.12558 | 6.41183 | 12.68608 | 2.40499 | 6.65884 | 0.00000 | 0.68508 | 0.52816 | 11.24217 | 14.66499 | 3.99452 | 0.75350 |
| **DDT** | 0.00565 | 0.01341 | 0.01633 | 0.01077 | 0.01969 | 0.02876 | 0.01708 | 0.01252 | 0.02306 | 0.01312 | 0.01319 | 0.01727 |
| **DDTL** | 0.00647 | 1.33562 | 0.53796 | 0.43283 | 0.67880 | 0.49932 | 0.00000 | 0.45639 | 1.13640 | 0.00000 | 0.71203 | 0.40073 |
| **DDX1** | 1.60532 | 22.67434 | 11.34440 | 14.36288 | 14.70438 | 41.90356 | 23.04977 | 5.91757 | 15.97240 | 19.05264 | 25.01082 | 16.31670 |

| | | | | | | | | | | | | |
|---|---|---|---|---|---|---|---|---|---|---|---|---|
| **DDX10** | 15.06292 | 4.73819 | 9.15421 | 0.40067 | 5.90722 | 20.24482 | 11.76220 | 11.39641 | 9.35379 | 12.30501 | 6.88669 | 8.91682 |
| **DDX11** | 1.34906 | 2.00330 | 1.97580 | 2.69256 | 1.41671 | 0.00000 | 1.92072 | 0.07210 | 4.69911 | 5.02791 | 0.55599 | 0.94315 |
| **DDX11-AS1** | 0.17336 | 0.27835 | 0.14353 | 0.05406 | 0.00000 | 0.10010 | 0.04562 | 0.01100 | 0.05428 | 0.00000 | 0.26063 | 0.01391 |
| **DDX11L2** | 0.00685 | 0.01022 | 0.02936 | 0.00700 | 0.07937 | 0.01323 | 0.05400 | 0.68188 | 0.09447 | 0.06545 | 0.03765 | 0.36112 |
| **DDX11L5** | 0.02320 | 0.03528 | 0.01257 | 0.00682 | 0.00931 | 0.03583 | 0.01469 | 0.01233 | 0.01504 | 0.01697 | 0.01287 | 0.03591 |
| **DDX12P** | 0.23150 | 0.36556 | 0.26800 | 0.68975 | 0.54377 | 0.00000 | 0.00000 | 0.05257 | 1.10409 | 0.64405 | 0.20939 | 0.24346 |
| **DDX17** | 4.04232 | 13.88042 | 19.23449 | 16.60846 | 12.27583 | 31.22916 | 35.50695 | 8.00346 | 18.07626 | 37.02298 | 25.72784 | 15.54976 |
| **DDX18** | 4.14908 | 17.25073 | 15.12144 | 2.48066 | 8.45199 | 35.68309 | 26.27266 | 12.74702 | 12.36694 | 23.94720 | 18.08580 | 21.53214 |
| **DDX19A** | 2.58291 | 10.70964 | 8.92949 | 5.99612 | 8.19640 | 3.79712 | 11.28583 | 1.75931 | 16.18166 | 0.00000 | 11.74972 | 6.49778 |
| **DDX19B** | 0.24296 | 3.62529 | 2.35966 | 1.36039 | 2.37493 | 4.91478 | 3.13341 | 0.27375 | 3.32626 | 0.00000 | 4.17277 | 5.14562 |
| **DDX20** | 1.59529 | 12.20803 | 5.22886 | 3.44382 | 1.62135 | 20.96832 | 9.75822 | 10.59316 | 7.87412 | 2.35225 | 6.43028 | 12.23327 |
| **DDX21** | 1.06208 | 18.89503 | 25.87522 | 11.16029 | 20.32589 | 48.67207 | 47.40965 | 29.21348 | 22.42675 | 49.91141 | 27.04247 | 32.77291 |
| **DDX23** | 1.30611 | 18.41340 | 12.60877 | 14.70970 | 11.89904 | 9.04197 | 18.03129 | 2.35865 | 17.64670 | 6.17977 | 10.37854 | 19.01693 |
| **DDX24** | 8.98120 | 19.14003 | 16.13816 | 9.40925 | 12.84049 | 17.35898 | 21.45524 | 22.06584 | 15.71413 | 10.60531 | 16.08389 | 20.92380 |
| **DDX25** | 0.00371 | 1.10862 | 0.26627 | 0.00585 | 0.00078 | 0.12443 | 0.05567 | 0.37218 | 0.00000 | 0.00000 | 0.20897 | 2.96342 |
| **DDX26B** | 0.17592 | 0.27441 | 0.09100 | 1.19152 | 0.00000 | 0.37335 | 0.30337 | 4.26943 | 0.36674 | 0.21103 | 0.61051 | 4.48269 |
| **DDX26B-AS1** | 0.00597 | 0.01832 | 0.01473 | 0.01231 | 0.01835 | 0.01545 | 0.01876 | 0.02306 | 0.01980 | 0.00889 | 0.01551 | 0.03055 |
| **DDX27** | 2.23874 | 18.30910 | 17.13116 | 11.58202 | 18.41155 | 8.20733 | 26.63397 | 10.20268 | 18.04083 | 27.00988 | 17.15829 | 20.90908 |
| **DDX28** | 3.21626 | 9.39536 | 6.37702 | 6.10601 | 8.47864 | 11.53201 | 5.48800 | 2.34026 | 8.80570 | 0.00000 | 8.73099 | 10.21454 |
| **DDX31** | 0.00000 | 8.96312 | 2.32234 | 5.20332 | 3.40494 | 3.32560 | 4.97769 | 1.04479 | 8.24629 | 4.58675 | 4.67940 | 12.19803 |
| **DDX39A** | 6.90874 | 34.34308 | 22.45025 | 21.61853 | 32.97605 | 13.76337 | 34.83613 | 35.32545 | 39.41979 | 0.00000 | 27.74824 | 38.50646 |
| **DDX39B** | 0.00549 | 0.18241 | 0.08784 | 0.15575 | 0.00000 | 0.12515 | 0.27276 | 0.19605 | 0.20078 | 0.00000 | 0.04804 | 0.12336 |
| **DDX3X** | 13.30660 | 16.97031 | 18.46793 | 12.40795 | 8.46743 | 51.01272 | 26.11049 | 25.42330 | 22.12855 | 19.93835 | 25.90851 | 26.19566 |
| **DDX3Y** | 0.92046 | 9.11202 | 12.50153 | 2.67651 | 0.01440 | 29.56816 | 18.16063 | 15.73634 | 20.87248 | 16.44560 | 3.45145 | 26.74823 |
| **DDX4** | 5.22284 | 0.07045 | 0.00507 | 0.00000 | 0.14961 | 0.24944 | 0.00000 | 4.13392 | 0.00000 | 0.00675 | 0.01058 | 2.86374 |
| **DDX41** | 0.33226 | 14.97773 | 7.95668 | 15.21604 | 17.63902 | 1.92008 | 8.29717 | 9.40047 | 15.61983 | 2.11475 | 18.75721 | 12.98335 |
| **DDX42** | 5.92354 | 5.94004 | 7.09301 | 5.44389 | 6.35397 | 11.87253 | 7.68188 | 5.93481 | 2.92910 | 22.30058 | 5.92941 | 9.42686 |
| **DDX43** | 3.18423 | 4.06727 | 1.09249 | 3.30210 | 1.04087 | 9.15299 | 0.00000 | 4.96767 | 1.83493 | 3.60407 | 0.00000 | 10.25388 |
| **DDX46** | 2.36598 | 12.33611 | 13.02437 | 8.22172 | 16.77229 | 24.76421 | 23.13177 | 7.60883 | 15.88829 | 37.67109 | 11.99406 | 18.16952 |
| **DDX47** | 3.48055 | 15.58100 | 15.14604 | 6.23524 | 4.44195 | 29.16450 | 13.94370 | 17.84005 | 14.36281 | 0.00000 | 16.92003 | 16.92748 |
| **DDX49** | 1.68120 | 14.60744 | 11.18684 | 8.31201 | 16.40434 | 0.00030 | 20.72993 | 11.47093 | 18.24802 | 0.00000 | 11.15709 | 19.36680 |
| **DDX5** | 7.35050 | 39.66361 | 25.24315 | 25.04275 | 17.40282 | 67.11871 | 35.87995 | 43.04357 | 37.54809 | 8.38232 | 37.03222 | 47.26947 |
| **DDX50** | 6.32374 | 3.29297 | 3.23734 | 1.78535 | 3.38548 | 9.51695 | 1.48968 | 3.87925 | 0.99774 | 2.15202 | 5.75280 | 2.74713 |
| **DDX51** | 1.02057 | 1.52171 | 1.56132 | 1.92326 | 4.42498 | 0.00000 | 4.91968 | 3.03621 | 2.63357 | 14.97566 | 0.62503 | 1.48986 |
| **DDX52** | 1.81618 | 10.62081 | 9.07462 | 3.37197 | 0.00000 | 27.45855 | 18.22455 | 6.92293 | 5.23896 | 16.64259 | 7.08500 | 11.62119 |
| **DDX53** | 0.00620 | 0.00220 | 0.11707 | 0.01525 | 0.00324 | 0.07613 | 0.01219 | 0.02001 | 0.02729 | 0.00241 | 0.03172 | 0.07197 |
| **DDX54** | 7.41475 | 14.35352 | 8.92650 | 9.29467 | 9.69560 | 0.00000 | 8.58051 | 4.63694 | 15.15697 | 7.94111 | 10.68096 | 9.48820 |
| **DDX55** | 2.52717 | 6.82593 | 5.53040 | 6.61545 | 6.89788 | 11.91894 | 8.03019 | 5.81380 | 9.81109 | 8.27652 | 8.29353 | 13.15510 |
| **DDX56** | 1.63508 | 30.42266 | 21.92410 | 16.70815 | 21.19229 | 9.26157 | 25.26645 | 12.19031 | 31.71614 | 0.00000 | 21.65848 | 24.65967 |
| **DDX58** | 2.62955 | 0.00000 | 1.28168 | 1.67936 | 0.31542 | 0.00294 | 0.00000 | 1.99413 | 0.00000 | 2.97497 | 0.59517 | 1.24076 |
| **DDX59** | 5.40778 | 2.20388 | 8.13791 | 2.54050 | 0.99621 | 6.60797 | 3.22346 | 5.71146 | 1.01624 | 1.00970 | 2.00580 | 6.94005 |
| **DDX6** | 0.00000 | 9.48961 | 10.96896 | 5.08753 | 4.40264 | 25.27423 | 17.66415 | 13.73531 | 4.81085 | 18.72995 | 17.14771 | 14.50129 |
| **DDX60** | 0.00609 | 0.04359 | 0.01957 | 0.05625 | 0.00000 | 0.04113 | 0.02297 | 0.03400 | 0.01372 | 0.01727 | 0.01229 | 0.04154 |
| **DDX60L** | 0.97646 | 0.05612 | 0.00005 | 0.02176 | 0.10425 | 0.17291 | 0.09622 | 0.09803 | 0.01428 | 0.00000 | 0.03062 | 0.37416 |
| **DEAF1** | 6.50597 | 0.63239 | 1.45407 | 0.81702 | 0.55512 | 1.87939 | 1.68426 | 0.75569 | 2.48566 | 4.91266 | 0.06513 | 1.94331 |
| **DEAR** | 0.00906 | 0.02140 | 0.01314 | 0.01241 | 0.01727 | 0.02038 | 0.04024 | 0.02123 | 0.01433 | 0.01180 | 0.01754 | 0.01904 |
| **DEC1** | 0.54638 | 0.02038 | 0.00936 | 0.02819 | 0.00697 | 0.02397 | 0.05987 | 0.00253 | 0.00169 | 0.01176 | 0.01343 | 0.01452 |
| **DECR1** | 0.33942 | 0.36072 | 2.38031 | 0.48489 | 0.37554 | 3.58448 | 1.20035 | 0.00000 | 1.83305 | 0.00000 | 2.86219 | 0.00000 |

| | | | | | | | | | | | | |
|---|---|---|---|---|---|---|---|---|---|---|---|---|
| **DECR2** | 0.87808 | 8.81155 | 3.39687 | 7.62768 | 16.14531 | 0.00000 | 8.72177 | 0.68440 | 13.69124 | 7.19061 | 14.24568 | 13.82305 |
| **DEDD** | 8.01048 | 8.49036 | 10.66467 | 4.67660 | 4.03108 | 10.09312 | 6.88147 | 10.65652 | 9.82584 | 0.00000 | 6.37829 | 18.74217 |
| **DEDD2** | 1.27873 | 8.27177 | 18.48448 | 4.20620 | 7.49380 | 0.00000 | 10.44225 | 0.00000 | 11.63755 | 1.01056 | 6.58650 | 3.79561 |
| **DEF6** | 0.29720 | 0.24665 | 0.40681 | 0.21598 | 0.36903 | 0.00000 | 0.80130 | 0.53316 | 0.98730 | 0.45173 | 0.00000 | 0.82801 |
| **DEF8** | 4.81583 | 14.52941 | 3.49050 | 12.47090 | 19.12527 | 7.49471 | 7.13682 | 2.94925 | 11.54101 | 0.01710 | 10.36475 | 4.18410 |
| **DEFA6** | 0.00583 | 0.00915 | 0.46550 | 0.35859 | 0.00000 | 0.02902 | 0.00000 | 0.00993 | 0.01843 | 0.02006 | 0.02129 | 0.01393 |
| **DEFB1** | 0.00558 | 0.00997 | 0.00000 | 0.48500 | 0.00000 | 0.02902 | 0.02999 | 0.00995 | 0.01267 | 0.02364 | 0.01007 | 0.01383 |
| **DEFB109P1** | 0.00629 | 0.01370 | 0.01246 | 0.01506 | 0.01176 | 0.02704 | 0.01568 | 0.00998 | 0.01603 | 0.01948 | 0.01444 | 0.01943 |
| **DEFB118** | 0.00597 | 0.04465 | 0.02127 | 0.00514 | 0.00045 | 0.10303 | 0.00000 | 0.00944 | 0.00779 | 0.07174 | 0.01041 | 0.01001 |
| **DEFB119** | 0.00482 | 0.22692 | 0.00222 | 0.01155 | 0.04524 | 0.02939 | 0.01085 | 0.01854 | 0.00000 | 0.13085 | 0.00295 | 0.02865 |
| **DEFB121** | 0.00612 | 0.03301 | 0.01067 | 0.00794 | 0.00577 | 0.01067 | 0.01121 | 0.02501 | 0.01307 | 0.07627 | 0.01554 | 0.05318 |
| **DEFB122** | 0.03268 | 4.78973 | 0.00000 | 1.16823 | 1.36091 | 0.00000 | 0.01716 | 0.21179 | 0.00000 | 0.01468 | 0.00000 | 2.83169 |
| **DEFB123** | 0.03045 | 0.02514 | 0.03495 | 0.75784 | 0.22599 | 0.00000 | 0.00965 | 0.14680 | 0.02740 | 0.08278 | 0.00279 | 0.04700 |
| **DEFB124** | 0.00146 | 0.29046 | 0.09193 | 1.25255 | 0.05780 | 0.00000 | 0.02521 | 0.17993 | 0.00000 | 0.06683 | 0.02427 | 0.27802 |
| **DEFB126** | 0.02337 | 0.03685 | 0.01343 | 0.01860 | 0.01336 | 0.02972 | 0.01977 | 0.00605 | 0.01393 | 0.00485 | 0.01932 | 0.07820 |
| **DEFB136** | 0.00610 | 0.03539 | 0.01751 | 0.00741 | 0.01157 | 0.03224 | 0.01582 | 0.01035 | 0.01431 | 0.01402 | 0.01403 | 0.01681 |
| **DEGS1** | 1.48830 | 18.38510 | 8.56204 | 8.75432 | 8.27310 | 38.46226 | 0.00000 | 6.84628 | 39.16033 | 3.89544 | 14.61763 | 8.23267 |
| **DEGS2** | 6.56561 | 1.92004 | 0.00000 | 0.00000 | 0.28068 | 0.00000 | 0.06326 | 4.92664 | 0.03448 | 0.00000 | 0.10892 | 0.00000 |
| **DEK** | 5.53630 | 15.47768 | 13.22966 | 10.59474 | 7.98330 | 41.38445 | 30.45670 | 6.98931 | 8.05705 | 33.49567 | 14.77154 | 16.79234 |
| **DENND1A** | 3.63551 | 1.68027 | 3.37458 | 2.10106 | 5.20760 | 0.78902 | 0.53089 | 0.33700 | 3.00335 | 1.32385 | 1.31682 | 0.00000 |
| **DENND1B** | 0.00000 | 0.17860 | 0.57211 | 1.87982 | 0.00000 | 1.36394 | 0.58486 | 0.01726 | 0.00000 | 1.03837 | 0.00002 | 0.11967 |
| **DENND1C** | 0.90201 | 7.09746 | 0.03693 | 0.20524 | 4.30678 | 0.00000 | 0.28344 | 0.00000 | 0.60020 | 1.58291 | 2.17327 | 3.16351 |
| **DENND2A** | 0.64989 | 1.40348 | 0.38365 | 1.93814 | 0.50217 | 0.00000 | 0.00000 | 1.42008 | 0.00000 | 0.01903 | 0.01529 | 1.10683 |
| **DENND2C** | 0.00201 | 0.05717 | 0.20065 | 5.25592 | 0.00000 | 0.30811 | 0.00000 | 0.23949 | 0.00000 | 0.12272 | 0.07969 | 0.70777 |
| **DENND2D** | 0.02240 | 1.81587 | 3.47767 | 0.00000 | 0.89516 | 1.61812 | 1.03873 | 0.00084 | 0.32324 | 0.00000 | 1.18884 | 0.51166 |
| **DENND3** | 0.23280 | 0.30839 | 0.11159 | 0.63212 | 0.05689 | 0.00000 | 0.24812 | 0.00000 | 0.00000 | 0.13830 | 0.02336 | 0.21702 |
| **DENND4A** | 1.54963 | 0.23056 | 2.19731 | 0.58334 | 0.88059 | 5.13705 | 0.00000 | 0.38371 | 0.15584 | 2.70623 | 0.50889 | 0.63199 |
| **DENND4B** | 0.15560 | 2.13037 | 0.41302 | 3.26168 | 5.55751 | 0.00000 | 0.70227 | 0.00230 | 0.00000 | 6.48018 | 1.90519 | 0.09198 |
| **DENND4C** | 0.11742 | 1.91571 | 10.90932 | 4.95041 | 0.00000 | 18.98342 | 14.81933 | 1.40986 | 1.72753 | 22.43446 | 3.03686 | 3.21308 |
| **DENND5A** | 4.70418 | 2.18862 | 6.91361 | 5.47633 | 6.18092 | 8.44272 | 8.05835 | 2.41896 | 9.09059 | 10.71469 | 2.53942 | 4.07017 |
| **DENND5B** | 1.66903 | 1.09623 | 0.00000 | 0.39460 | 0.42741 | 0.92574 | 0.31489 | 1.05642 | 0.00000 | 1.79771 | 0.00000 | 1.60483 |
| **DENND5B-AS1** | 0.04146 | 0.00000 | 0.29505 | 0.24582 | 0.10651 | 0.00000 | 0.03322 | 0.01548 | 0.00000 | 0.18244 | 0.19064 | 0.07189 |
| **DENND6A** | 0.36902 | 0.31343 | 0.36413 | 0.00000 | 0.24420 | 1.65460 | 2.12601 | 0.13529 | 0.31350 | 2.73735 | 1.45535 | 0.00000 |
| **DENND6B** | 0.04334 | 0.08583 | 0.02957 | 0.00949 | 0.00000 | 0.11983 | 0.18169 | 0.02122 | 0.00919 | 0.00000 | 0.01576 | 0.00506 |
| **DENR** | 0.00000 | 23.17189 | 11.01075 | 13.12477 | 8.11584 | 49.91077 | 24.77036 | 14.86729 | 17.49967 | 19.04181 | 15.16035 | 22.60821 |
| **DEPDC1** | 0.00000 | 3.84358 | 7.37983 | 0.37342 | 2.27169 | 11.87376 | 8.48785 | 0.16554 | 5.08707 | 0.89084 | 1.40152 | 0.08212 |
| **DEPDC1-AS1** | 0.00471 | 0.05401 | 0.07389 | 0.10203 | 0.00000 | 0.09069 | 0.06116 | 0.00919 | 0.09510 | 0.06466 | 0.04432 | 0.01264 |
| **DEPDC1B** | 3.09917 | 7.43417 | 8.80751 | 2.18259 | 0.00000 | 11.16426 | 5.23726 | 4.37281 | 6.29304 | 0.00000 | 0.00000 | 7.27831 |
| **DEPDC4** | 0.27972 | 0.45090 | 0.33504 | 0.00000 | 0.00841 | 0.57814 | 0.00045 | 0.17715 | 0.00000 | 0.28584 | 0.15335 | 0.62208 |
| **DEPDC5** | 0.79547 | 2.01447 | 3.68824 | 4.05540 | 3.02050 | 3.57879 | 0.70152 | 2.35662 | 2.59150 | 4.17281 | 3.99188 | 2.25868 |
| **DEPDC7** | 4.42263 | 0.80710 | 0.26356 | 4.86825 | 0.00000 | 1.29084 | 0.00000 | 8.27152 | 0.00000 | 0.00000 | 0.39888 | 6.00201 |
| **DEPTOR** | 0.48116 | 13.44895 | 0.00000 | 14.02087 | 0.31658 | 4.08353 | 0.00000 | 0.00000 | 0.00000 | 1.20900 | 0.29309 | 10.67865 |
| **DERA** | 2.83699 | 13.82922 | 2.00591 | 7.06952 | 5.16495 | 21.28582 | 2.06622 | 1.67118 | 0.00000 | 0.00030 | 5.32920 | 4.20480 |
| **DERL1** | 3.31013 | 10.37394 | 11.25673 | 9.09035 | 5.54777 | 17.71521 | 0.00000 | 2.72422 | 28.53394 | 5.22359 | 7.63838 | 6.99850 |
| **DERL2** | 0.00139 | 8.02902 | 8.38968 | 7.23625 | 4.85439 | 20.14976 | 1.77735 | 1.69171 | 12.11266 | 0.05287 | 5.28872 | 13.43011 |
| **DERL3** | 2.68399 | 0.26448 | 0.19337 | 0.75179 | 0.48649 | 0.00000 | 0.00000 | 8.76574 | 0.59165 | 0.62371 | 0.67811 | 0.85394 |
| **DES** | 1.70445 | 0.02331 | 0.16544 | 0.01408 | 0.30581 | 0.00000 | 0.06229 | 0.70403 | 0.00000 | 0.00000 | 0.00769 | 0.02400 |
| **DESI1** | 2.08716 | 7.49162 | 7.32516 | 4.69921 | 11.75953 | 1.26958 | 4.10120 | 0.11732 | 7.99210 | 1.19554 | 15.99692 | 2.45934 |

| | | | | | | | | | | | | |
|---|---|---|---|---|---|---|---|---|---|---|---|---|
| **DESI2** | 2.30970 | 6.04206 | 9.03408 | 5.19829 | 4.20439 | 16.42366 | 20.52284 | 2.44955 | 8.25697 | 20.02299 | 4.13459 | 4.17950 |
| **DET1** | 2.72110 | 4.16975 | 1.09444 | 3.22112 | 3.06226 | 1.84198 | 0.00000 | 0.46346 | 0.61642 | 0.00000 | 2.36215 | 1.42616 |
| **DEXI** | 0.03343 | 0.33167 | 0.38042 | 0.00000 | 1.64638 | 0.34111 | 1.80494 | 0.00000 | 0.00000 | 2.68147 | 0.49216 | 0.35427 |
| **DFFA** | 2.98022 | 12.22027 | 19.05493 | 19.01068 | 18.45518 | 6.04336 | 21.49374 | 5.54683 | 21.37810 | 0.01637 | 26.30959 | 13.51199 |
| **DFFB** | 0.76178 | 4.49715 | 2.16323 | 0.18396 | 0.29555 | 3.11349 | 2.57377 | 0.00000 | 6.42206 | 0.00000 | 5.00338 | 1.70975 |
| **DFNA5** | 7.16792 | 0.00000 | 2.39035 | 0.16420 | 0.00000 | 0.00000 | 0.49438 | 0.05024 | 0.00000 | 0.77655 | 0.23643 | 0.27279 |
| **DFNB31** | 5.70325 | 0.09057 | 0.19991 | 0.61956 | 0.07902 | 0.00000 | 0.00000 | 0.10177 | 0.31127 | 0.08970 | 0.00000 | 0.14706 |
| **DFNB59** | 0.00419 | 0.00000 | 0.04082 | 0.11582 | 0.00000 | 0.36483 | 0.00000 | 0.08389 | 0.05266 | 0.84639 | 0.00778 | 0.22553 |
| **DGAT1** | 1.04854 | 0.42252 | 2.45084 | 2.14115 | 3.64965 | 0.00000 | 0.00173 | 0.13492 | 1.45082 | 7.88555 | 0.96464 | 0.44500 |
| **DGAT2** | 0.20478 | 1.35842 | 0.00000 | 2.32497 | 0.00000 | 0.00931 | 0.00323 | 0.14300 | 0.00000 | 0.07777 | 0.00000 | 2.87465 |
| **DGCR10** | 0.00419 | 0.00949 | 0.00224 | 0.01669 | 0.00000 | 0.02139 | 0.02178 | 0.00920 | 0.08025 | 0.15761 | 0.00584 | 0.02425 |
| **DGCR11** | 0.03163 | 0.37943 | 0.32397 | 0.23270 | 0.46434 | 0.84833 | 0.00000 | 0.06108 | 0.96262 | 0.66832 | 0.09391 | 0.06395 |
| **DGCR14** | 1.69256 | 10.42807 | 3.36235 | 5.53863 | 7.23882 | 0.00000 | 4.38173 | 10.31505 | 7.63980 | 0.00000 | 5.00282 | 10.87451 |
| **DGCR2** | 1.57364 | 11.12525 | 5.19365 | 8.14536 | 9.72560 | 0.00000 | 0.00927 | 0.00000 | 13.64373 | 4.79919 | 4.91314 | 5.31831 |
| **DGCR5** | 0.00368 | 0.12857 | 0.05385 | 0.01849 | 0.14670 | 0.00041 | 0.01545 | 0.01439 | 0.00000 | 0.02549 | 0.04418 | 0.15278 |
| **DGCR6** | 0.98121 | 3.41527 | 0.00000 | 1.96559 | 10.07099 | 3.05901 | 0.85702 | 0.09952 | 0.00000 | 0.00000 | 5.16966 | 0.53990 |
| **DGCR6L** | 1.04921 | 6.17849 | 2.96291 | 4.27591 | 5.34904 | 0.00003 | 2.05843 | 0.00000 | 2.96306 | 0.00000 | 4.26081 | 5.22542 |
| **DGCR8** | 3.35678 | 4.69962 | 4.21912 | 3.51933 | 4.77965 | 2.18894 | 9.49472 | 0.56273 | 5.94155 | 8.51586 | 4.60719 | 0.36014 |
| **DGCR9** | 0.00301 | 0.02122 | 0.13406 | 0.00000 | 0.13812 | 0.08623 | 0.00353 | 0.00897 | 0.14859 | 0.20310 | 0.00000 | 0.01662 |
| **DGKA** | 0.00448 | 0.19839 | 0.78997 | 0.53303 | 0.00000 | 0.00000 | 0.05612 | 0.00851 | 0.43383 | 0.23916 | 0.00000 | 0.02762 |
| **DGKB** | 1.99065 | 0.29741 | 0.00354 | 0.00000 | 0.20661 | 0.00047 | 0.00466 | 0.21970 | 0.09316 | 0.22942 | 0.01406 | 0.05694 |
| **DGKD** | 0.02019 | 1.59964 | 4.68643 | 2.86790 | 3.06388 | 0.00000 | 4.90874 | 1.48427 | 2.98667 | 13.66160 | 0.00000 | 4.35186 |
| **DGKE** | 3.30545 | 0.35097 | 0.06256 | 0.30914 | 0.02190 | 0.11341 | 0.00000 | 0.79552 | 0.01317 | 0.00000 | 0.00000 | 1.28595 |
| **DGKG** | 0.25022 | 0.01378 | 0.04976 | 0.01677 | 0.01394 | 0.01579 | 0.01965 | 0.00338 | 0.01492 | 0.00616 | 0.01992 | 0.01570 |
| **DGKH** | 0.03041 | 0.06573 | 0.00329 | 0.04250 | 0.04938 | 0.00000 | 0.00708 | 0.08962 | 0.18978 | 0.36627 | 0.00341 | 0.22424 |
| **DGKI** | 0.94135 | 0.00730 | 0.02051 | 0.02288 | 0.01373 | 0.01462 | 0.00451 | 0.22730 | 0.03370 | 0.09210 | 0.00970 | 0.02663 |
| **DGKK** | 0.00508 | 0.03416 | 0.06432 | 0.39660 | 0.28431 | 0.06191 | 0.00000 | 0.00899 | 0.00000 | 0.03681 | 0.02641 | 0.02420 |
| **DGKQ** | 0.04030 | 0.68526 | 0.82512 | 0.27075 | 1.02186 | 0.00000 | 0.00671 | 0.00000 | 0.35181 | 1.67439 | 0.00000 | 0.52240 |
| **DGKZ** | 0.21773 | 0.91902 | 3.97485 | 1.39842 | 3.76514 | 0.00000 | 2.86166 | 0.00000 | 1.23659 | 8.76851 | 0.34638 | 0.55983 |
| **DGUOK** | 3.42087 | 6.39126 | 11.26069 | 9.31220 | 6.14276 | 13.01497 | 11.82296 | 2.87277 | 4.15238 | 2.42302 | 8.94688 | 10.81728 |
| **DGUOK-AS1** | 0.33475 | 0.17654 | 0.46280 | 0.77820 | 0.31622 | 0.38704 | 0.24754 | 0.07200 | 0.49369 | 0.00000 | 0.25740 | 0.18954 |
| **DHCR24** | 0.30755 | 13.96793 | 12.81241 | 39.46673 | 41.69402 | 18.88710 | 18.53934 | 1.09260 | 61.04812 | 22.34661 | 37.36145 | 7.05111 |
| **DHCR7** | 4.64214 | 2.87914 | 18.72310 | 17.50897 | 38.56133 | 3.15530 | 7.77736 | 0.46071 | 55.16126 | 2.95358 | 24.98361 | 0.32506 |
| **DHDDS** | 0.88216 | 12.41745 | 7.74685 | 1.26312 | 6.58367 | 7.96032 | 4.72458 | 6.49634 | 3.47968 | 4.81018 | 11.35939 | 12.80966 |
| **DHDH** | 0.00903 | 0.79697 | 0.21647 | 0.42569 | 0.00000 | 0.06889 | 0.00000 | 0.46344 | 0.09953 | 0.47824 | 0.05718 | 1.10078 |
| **DHFR** | 0.03883 | 0.83301 | 0.31028 | 0.56755 | 1.09613 | 0.59850 | 0.58282 | 0.07622 | 0.00000 | 2.60629 | 1.44452 | 0.83082 |
| **DHFRL1** | 0.34410 | 0.55518 | 0.17069 | 0.00000 | 0.00000 | 0.75952 | 0.00000 | 0.20293 | 0.07789 | 0.52691 | 0.58137 | 0.52011 |
| **DHH** | 0.24849 | 0.00000 | 0.01157 | 0.10572 | 0.22530 | 0.00000 | 0.03096 | 0.00000 | 0.07241 | 0.00000 | 0.00198 | 0.31701 |
| **DHODH** | 0.00357 | 5.15586 | 3.05654 | 2.08830 | 4.90300 | 1.57349 | 2.21015 | 5.36931 | 4.01908 | 0.00000 | 5.47293 | 5.82591 |
| **DHPS** | 0.88711 | 6.02729 | 8.23564 | 5.55686 | 2.48191 | 0.13601 | 6.19118 | 1.96547 | 7.56409 | 0.00000 | 4.76351 | 6.29115 |
| **DHRS1** | 0.07107 | 0.46763 | 1.89000 | 0.58518 | 0.32786 | 1.78321 | 0.00000 | 0.50641 | 2.00653 | 0.00000 | 1.69621 | 0.85404 |
| **DHRS11** | 0.36097 | 0.81769 | 0.26905 | 0.28163 | 0.64840 | 0.00029 | 1.39330 | 0.23024 | 0.00557 | 4.20175 | 0.41031 | 0.74085 |
| **DHRS12** | 0.02806 | 0.66632 | 0.40569 | 0.16334 | 1.29682 | 0.00000 | 0.00000 | 0.17947 | 0.67544 | 0.48769 | 0.41711 | 0.38313 |
| **DHRS13** | 0.00085 | 3.69794 | 5.53944 | 2.47768 | 3.58366 | 0.00000 | 2.99633 | 0.00000 | 6.12629 | 4.36582 | 3.32919 | 5.80860 |
| **DHRS2** | 0.01660 | 0.01591 | 0.09196 | 0.02954 | 0.01030 | 0.00029 | 0.01095 | 0.00979 | 0.07307 | 0.03793 | 0.01719 | 0.01720 |
| **DHRS3** | 0.00266 | 22.42518 | 1.01941 | 7.06259 | 12.94749 | 3.34000 | 0.00000 | 0.00000 | 0.00000 | 0.00000 | 1.75270 | 1.14172 |
| **DHRS4** | 0.02177 | 0.06137 | 0.11612 | 0.08932 | 0.11700 | 0.00000 | 0.01551 | 0.01239 | 0.01581 | 0.00000 | 0.05334 | 0.05445 |
| **DHRS4-AS1** | 0.00833 | 0.56647 | 0.09407 | 0.23219 | 0.00000 | 0.00000 | 0.40547 | 0.00137 | 0.03983 | 0.58924 | 0.30974 | 0.69389 |

| | | | | | | | | | | | | |
|---|---|---|---|---|---|---|---|---|---|---|---|---|
| **DHRS4L1** | 0.02014 | 0.01051 | 0.06755 | 0.01135 | 0.00000 | 0.03476 | 0.02347 | 0.00787 | 0.05880 | 0.04951 | 0.00864 | 0.02174 |
| **DHRS4L2** | 0.00492 | 0.02248 | 0.13641 | 0.07429 | 0.09770 | 0.00000 | 0.04223 | 0.01669 | 0.03915 | 0.01545 | 0.01493 | 0.06945 |
| **DHRS7** | 0.00000 | 2.10289 | 2.02845 | 0.10819 | 0.16804 | 7.21054 | 0.00000 | 0.80511 | 3.20740 | 1.14792 | 0.19382 | 2.71648 |
| **DHRS7B** | 0.78306 | 11.05289 | 6.90535 | 1.24285 | 2.01063 | 6.17697 | 0.23430 | 1.51476 | 5.22008 | 0.00000 | 4.36477 | 9.90369 |
| **DHTKD1** | 0.43287 | 9.09529 | 5.25082 | 0.68823 | 6.60378 | 4.26593 | 7.38413 | 0.61648 | 13.00395 | 7.11180 | 11.68540 | 5.94211 |
| **DHX15** | 1.58170 | 20.09395 | 21.44414 | 11.83985 | 11.26843 | 40.86591 | 35.70940 | 24.66496 | 30.36955 | 14.09831 | 21.99487 | 22.13536 |
| **DHX16** | 2.95021 | 10.20411 | 7.98751 | 10.95979 | 10.12591 | 2.20747 | 9.22288 | 7.28029 | 12.17524 | 0.00000 | 9.82986 | 12.87139 |
| **DHX29** | 2.52982 | 5.44237 | 6.48718 | 3.29047 | 2.42118 | 22.12716 | 9.03170 | 7.58194 | 4.41979 | 19.12058 | 8.10828 | 9.58586 |
| **DHX30** | 1.61036 | 21.23566 | 9.43535 | 13.82666 | 16.69446 | 0.01679 | 17.09753 | 1.44110 | 21.29999 | 4.69595 | 16.38008 | 14.27855 |
| **DHX32** | 0.05507 | 0.53491 | 4.28827 | 1.60916 | 4.06044 | 5.36014 | 2.68355 | 0.26277 | 0.06282 | 3.44594 | 2.04771 | 0.13577 |
| **DHX33** | 3.04113 | 1.53460 | 4.51279 | 7.17873 | 5.42230 | 3.49464 | 19.26764 | 4.58136 | 3.17490 | 27.28946 | 8.97074 | 8.11531 |
| **DHX34** | 0.11474 | 4.97209 | 3.52208 | 0.83581 | 3.76451 | 0.00000 | 3.16702 | 3.38460 | 6.56252 | 4.08075 | 2.25147 | 5.22669 |
| **DHX35** | 0.26933 | 3.73605 | 2.31700 | 2.22755 | 0.31380 | 5.73938 | 1.18769 | 0.00000 | 2.60428 | 1.32936 | 3.11298 | 3.09104 |
| **DHX36** | 0.40374 | 0.00000 | 8.52574 | 1.22233 | 3.68462 | 13.85136 | 14.62278 | 3.10923 | 0.37216 | 31.02456 | 5.12838 | 4.05309 |
| **DHX37** | 0.00000 | 6.33934 | 5.95786 | 3.98284 | 8.39461 | 0.00000 | 13.91303 | 1.55265 | 10.55819 | 13.75781 | 2.02790 | 7.81909 |
| **DHX38** | 5.62272 | 9.31567 | 10.57436 | 9.85928 | 6.11286 | 0.00000 | 11.46722 | 4.84211 | 17.22753 | 9.44729 | 5.18871 | 8.00700 |
| **DHX40** | 0.24393 | 3.13757 | 2.73848 | 4.02023 | 0.00000 | 8.52894 | 6.06247 | 7.64098 | 2.79562 | 7.25429 | 2.47129 | 5.43678 |
| **DHX57** | 0.94598 | 1.63687 | 4.21073 | 0.73098 | 0.00000 | 4.99769 | 5.87230 | 0.44511 | 4.36713 | 5.45461 | 2.52041 | 1.96589 |
| **DHX58** | 0.86320 | 1.09180 | 1.75193 | 0.00103 | 0.01112 | 0.00000 | 0.36764 | 4.89874 | 0.00000 | 0.00070 | 0.03282 | 6.00336 |
| **DHX8** | 3.55672 | 11.83550 | 6.93662 | 8.48572 | 10.91792 | 4.27949 | 6.79065 | 15.24048 | 12.40777 | 5.26881 | 8.58470 | 11.53189 |
| **DHX9** | 0.75654 | 20.13091 | 18.36006 | 15.67832 | 15.31802 | 29.46226 | 23.72877 | 18.61599 | 26.92766 | 15.79690 | 19.08886 | 24.35768 |
| **DIABLO** | 5.13346 | 20.46634 | 17.58300 | 14.71038 | 15.33962 | 28.27038 | 15.95656 | 20.51159 | 15.91607 | 0.79878 | 20.64038 | 29.32456 |
| **DIAPH1** | 0.32516 | 4.68102 | 8.70981 | 5.19102 | 15.78265 | 8.18262 | 3.32517 | 4.33974 | 10.83688 | 25.28016 | 2.14805 | 10.22000 |
| **DIAPH2** | 2.96118 | 0.12664 | 2.31450 | 1.58370 | 0.00047 | 0.62896 | 0.00000 | 0.18575 | 0.00000 | 1.20032 | 0.11384 | 0.39174 |
| **DIAPH3** | 3.76699 | 2.42185 | 5.06990 | 0.00000 | 8.51801 | 15.17420 | 7.43310 | 1.17025 | 12.93742 | 14.61122 | 8.74849 | 7.20256 |
| **DIAPH3-AS1** | 0.01673 | 0.00000 | 0.00000 | 0.04115 | 0.27237 | 0.11589 | 0.00000 | 0.04892 | 0.00000 | 0.85628 | 0.11152 | 0.14610 |
| **DIAPH3-AS2** | 0.01142 | 0.02180 | 0.01801 | 0.04690 | 0.09296 | 0.00000 | 0.01584 | 0.01591 | 0.05229 | 0.00469 | 0.06742 | 0.06457 |
| **DICER1** | 4.45015 | 1.89237 | 1.19303 | 1.71798 | 2.22706 | 11.40773 | 6.06584 | 0.42959 | 4.14020 | 18.70935 | 3.32806 | 0.92360 |
| **DICER1-AS1** | 0.00259 | 0.02782 | 0.00807 | 0.28269 | 0.05038 | 0.06447 | 0.16332 | 0.05108 | 0.00000 | 0.00000 | 0.12405 | 0.07130 |
| **DIDO1** | 2.20180 | 4.33446 | 4.09994 | 2.82514 | 5.75696 | 5.44821 | 2.37197 | 10.94157 | 6.74607 | 9.59585 | 2.93213 | 6.24020 |
| **DIEXF** | 0.00000 | 7.37565 | 5.81845 | 2.47939 | 2.03597 | 11.14048 | 13.59933 | 12.83108 | 8.26193 | 2.71221 | 8.95151 | 12.44049 |
| **DIMT1** | 0.00000 | 9.92458 | 15.98529 | 3.02030 | 2.24205 | 29.18357 | 20.24425 | 9.99663 | 20.64517 | 0.37907 | 15.04691 | 13.56988 |
| **DIO1** | 0.00384 | 0.04220 | 0.01598 | 0.00000 | 0.00224 | 0.01464 | 0.00000 | 0.03399 | 0.06298 | 0.05864 | 0.65696 | 0.03751 |
| **DIO2** | 1.11482 | 0.08121 | 0.08376 | 0.27430 | 0.00000 | 0.00601 | 0.02042 | 0.16754 | 0.44374 | 0.04816 | 0.10175 | 0.06619 |
| **DIO2-AS1** | 0.00585 | 0.01388 | 0.01048 | 0.02902 | 0.00799 | 0.02097 | 0.01803 | 0.01129 | 0.03917 | 0.02324 | 0.01250 | 0.01689 |
| **DIO3** | 0.54724 | 0.00472 | 0.03739 | 0.01013 | 0.00000 | 0.00089 | 0.00217 | 0.02266 | 0.01242 | 0.12634 | 0.20610 | 0.01218 |
| **DIP2A** | 0.22602 | 2.59719 | 3.87081 | 3.50682 | 2.92975 | 0.00000 | 1.11272 | 2.00031 | 5.37191 | 7.12052 | 3.62806 | 2.02697 |
| **DIP2A-IT1** | 0.01099 | 0.05702 | 0.08390 | 0.01186 | 0.00000 | 0.07057 | 0.03800 | 0.85645 | 0.01507 | 0.05561 | 0.07922 | 0.23861 |
| **DIP2B** | 0.24781 | 6.89133 | 16.36147 | 2.05007 | 1.64438 | 5.12539 | 9.49134 | 4.07943 | 7.56936 | 7.98265 | 7.60421 | 5.60126 |
| **DIP2C** | 5.61127 | 0.29552 | 1.73357 | 0.00000 | 0.11744 | 0.72471 | 0.82773 | 0.81547 | 0.74253 | 2.61439 | 0.10927 | 0.94329 |
| **DIRAS1** | 1.47489 | 0.06690 | 0.00517 | 0.00411 | 0.17224 | 0.00000 | 0.05789 | 0.00357 | 0.00097 | 0.07633 | 0.01472 | 0.17370 |
| **DIRAS2** | 0.07909 | 0.01772 | 0.01585 | 0.01543 | 0.01947 | 0.00567 | 0.01842 | 0.02274 | 0.03406 | 0.00260 | 0.01458 | 0.01518 |
| **DIRAS3** | 5.15588 | 0.00000 | 0.04525 | 0.04526 | 0.04516 | 0.40494 | 0.00000 | 0.67676 | 0.00777 | 0.00000 | 0.00000 | 0.06175 |
| **DIRC1** | 0.04084 | 0.01487 | 0.01492 | 0.01203 | 0.01400 | 0.01401 | 0.01965 | 0.05579 | 0.01248 | 0.01863 | 0.02699 | 0.01312 |
| **DIRC2** | 0.98025 | 1.54272 | 8.58361 | 1.27885 | 4.92235 | 13.11006 | 0.00000 | 0.13125 | 6.63110 | 13.69307 | 1.02891 | 1.41746 |
| **DIRC3** | 0.16690 | 0.18706 | 0.00000 | 0.00000 | 0.00000 | 0.00379 | 0.00190 | 0.00579 | 0.09636 | 0.27607 | 0.00396 | 0.01950 |
| **DIS3** | 0.20949 | 5.79643 | 9.17764 | 2.59764 | 3.91456 | 18.63877 | 7.44673 | 15.46558 | 12.81845 | 11.64942 | 12.37041 | 10.80867 |
| **DIS3L** | 0.00000 | 5.08488 | 2.33898 | 0.00000 | 2.50861 | 4.76152 | 6.26359 | 8.29354 | 6.15490 | 0.89845 | 7.36615 | 19.46054 |

| | | | | | | | | | | | | |
|---|---|---|---|---|---|---|---|---|---|---|---|---|
| **DIS3L2** | 0.86834 | 2.03634 | 2.24748 | 1.23836 | 3.78309 | 0.95977 | 0.20079 | 1.72352 | 4.51927 | 3.75999 | 3.25003 | 5.78516 |
| **DISC1** | 0.04463 | 0.00838 | 0.02561 | 0.00950 | 0.01673 | 0.07473 | 0.00726 | 0.01686 | 0.00630 | 0.02429 | 0.01934 | 0.04618 |
| **DISC1FP1** | 0.26094 | 0.03957 | 0.03305 | 0.02525 | 0.01769 | 0.02486 | 0.04338 | 0.04678 | 0.00000 | 0.05836 | 0.03811 | 0.03802 |
| **DISC2** | 0.00612 | 0.01548 | 0.01492 | 0.01199 | 0.00505 | 0.01208 | 0.03493 | 0.01250 | 0.00000 | 0.02953 | 0.08084 | 0.01803 |
| **DISP1** | 1.78429 | 0.41377 | 0.43724 | 0.01320 | 0.03000 | 0.43932 | 0.00000 | 0.40737 | 0.84979 | 0.00000 | 0.32173 | 0.39706 |
| **DISP2** | 0.35732 | 0.00608 | 0.16846 | 0.05722 | 0.00333 | 0.08486 | 0.00000 | 0.00131 | 0.00000 | 0.06949 | 0.10016 | 0.02115 |
| **DIXDC1** | 0.01597 | 0.00003 | 0.27298 | 0.64313 | 0.00000 | 0.39296 | 0.00000 | 0.17449 | 0.39491 | 0.56871 | 0.00000 | 2.43884 |
| **DKC1** | 1.99600 | 22.31197 | 20.83284 | 13.90291 | 22.30056 | 20.44015 | 37.17231 | 21.14236 | 23.43419 | 14.07632 | 26.79747 | 26.84366 |
| **DKFZP434A062** | 0.00418 | 0.01009 | 0.01588 | 0.00331 | 0.00000 | 0.01336 | 0.00652 | 0.01008 | 0.06747 | 0.18987 | 0.00665 | 0.01082 |
| **DKFZP434H168** | 0.00710 | 0.01542 | 0.01362 | 0.01185 | 0.01648 | 0.02034 | 0.01730 | 0.01259 | 0.02214 | 0.01023 | 0.01486 | 0.02110 |
| **DKFZP434I0714** | 0.09784 | 0.49731 | 0.27955 | 0.04601 | 0.04369 | 1.14225 | 0.00000 | 0.81365 | 0.00000 | 0.00000 | 1.18582 | 0.88905 |
| **DKFZP586I1420** | 0.16601 | 0.11949 | 2.17491 | 0.50039 | 0.86466 | 1.11198 | 3.85577 | 0.16950 | 0.31893 | 3.12925 | 0.63882 | 0.40681 |
| **DKFZp434J0226** | 0.01603 | 0.00376 | 0.04516 | 0.00893 | 0.00111 | 0.02660 | 0.00000 | 0.00611 | 0.07747 | 0.05286 | 0.01086 | 0.15874 |
| **DKFZp451B082** | 0.00606 | 0.01553 | 0.02507 | 0.01378 | 0.01282 | 0.00738 | 0.01776 | 0.01067 | 0.02004 | 0.03119 | 0.03093 | 0.01598 |
| **1.00000** | 0.02769 | 0.01115 | 0.42815 | 0.07617 | 0.00650 | 0.00000 | 0.00205 | 0.00609 | 0.27285 | 0.00098 | 0.00585 | 0.15181 |
| **3.00000** | 0.10415 | 0.17759 | 0.04878 | 0.20449 | 0.80993 | 0.07313 | 0.00000 | 0.21921 | 0.04548 | 0.08138 | 0.03923 | 0.00058 |
| **4.00000** | 0.00260 | 0.01554 | 0.02555 | 0.01160 | 0.01548 | 0.01581 | 0.02160 | 0.04136 | 0.01798 | 0.01164 | 0.01604 | 0.01559 |
| **DKKL1** | 0.03441 | 0.46236 | 0.00443 | 0.00000 | 0.00000 | 0.01626 | 0.03579 | 0.08427 | 0.16541 | 0.02920 | 0.01350 | 0.65854 |
| **DLAT** | 1.03786 | 23.05910 | 8.75332 | 13.15727 | 11.05253 | 35.47165 | 20.24891 | 0.00000 | 24.02915 | 5.05565 | 21.03811 | 12.52747 |
| **DLC1** | 1.10433 | 17.19913 | 0.00000 | 4.98175 | 15.94688 | 11.42485 | 19.50050 | 2.42841 | 14.21871 | 14.56601 | 16.55621 | 8.84160 |
| **DLD** | 3.64405 | 20.72625 | 12.21699 | 8.62048 | 3.84573 | 37.30880 | 16.39292 | 6.01751 | 21.44625 | 0.01144 | 23.69551 | 12.18031 |
| **DLEC1** | 0.72316 | 0.07216 | 0.04612 | 0.34866 | 0.23622 | 0.00000 | 0.00576 | 0.96132 | 0.20719 | 0.03604 | 0.15669 | 1.76816 |
| **DLEU1** | 2.29581 | 8.95430 | 4.48733 | 0.00000 | 3.57664 | 19.03667 | 10.17907 | 0.33563 | 8.39361 | 7.87291 | 10.00617 | 3.00741 |
| **DLEU2** | 0.79342 | 0.64344 | 0.00000 | 0.34989 | 0.29829 | 3.10466 | 0.00000 | 0.69076 | 0.00000 | 1.08862 | 0.51023 | 0.99957 |
| **DLEU2L** | 0.34319 | 0.35195 | 0.00000 | 0.18150 | 0.19275 | 1.13733 | 0.00000 | 0.16103 | 0.00006 | 0.38840 | 0.13321 | 0.43596 |
| **DLEU7** | 0.47447 | 0.02614 | 0.03765 | 0.00858 | 0.02761 | 0.00000 | 0.00227 | 5.11605 | 0.05410 | 0.00000 | 0.08803 | 0.00939 |
| **DLEU7-AS1** | 0.00492 | 0.01314 | 0.01048 | 0.01769 | 0.01108 | 0.02372 | 0.00974 | 0.01332 | 0.02215 | 0.04477 | 0.01530 | 0.01500 |
| **DLG1** | 3.85468 | 0.70966 | 3.38214 | 1.79263 | 1.40639 | 13.04300 | 3.75560 | 0.30978 | 3.81094 | 14.26288 | 1.90902 | 0.92942 |
| **DLG1-AS1** | 0.06859 | 0.03050 | 0.05485 | 0.05582 | 0.02242 | 0.05157 | 0.05521 | 0.05828 | 0.00765 | 0.02745 | 0.01605 | 0.03767 |
| **DLG2** | 1.43535 | 0.00858 | 2.83626 | 0.01299 | 0.16888 | 0.00000 | 4.00380 | 0.36951 | 3.55690 | 0.03359 | 3.73444 | 0.07572 |
| **DLG3** | 5.64667 | 6.33376 | 0.00000 | 8.50116 | 13.16764 | 7.87528 | 6.12345 | 0.21716 | 0.00000 | 6.11632 | 2.13773 | 1.49606 |
| **DLG3-AS1** | 0.00892 | 0.04214 | 0.00623 | 0.08786 | 0.07056 | 0.07545 | 0.01254 | 0.01329 | 0.04595 | 0.00010 | 0.02007 | 0.03049 |
| **DLG4** | 0.09600 | 0.08366 | 0.25703 | 0.00155 | 0.00615 | 0.00000 | 0.00000 | 0.01511 | 0.32402 | 0.45339 | 0.00001 | 0.07423 |
| **DLG5** | 1.20776 | 2.39295 | 33.17522 | 1.08355 | 5.36117 | 13.28946 | 28.15402 | 0.15574 | 15.19657 | 17.24145 | 12.35786 | 1.89471 |
| **DLG5-AS1** | 0.02371 | 0.21541 | 0.89997 | 0.00000 | 0.00000 | 0.02195 | 0.03873 | 0.02032 | 0.22257 | 0.46161 | 0.07422 | 0.20767 |
| **DLGAP1** | 0.30378 | 0.50843 | 0.06658 | 0.70223 | 0.18257 | 0.40933 | 0.00000 | 0.10440 | 0.43600 | 0.00000 | 0.00000 | 0.51684 |
| **DLGAP1-AS1** | 0.00102 | 0.68139 | 0.24087 | 0.14244 | 0.02914 | 0.26465 | 0.83745 | 0.30073 | 0.00069 | 0.00000 | 0.21763 | 3.31871 |
| **DLGAP1-AS2** | 0.00000 | 0.97120 | 0.00000 | 0.84049 | 0.11318 | 1.23940 | 0.00000 | 0.04351 | 0.00000 | 0.75372 | 0.00000 | 2.25437 |
| **DLGAP1-AS3** | 0.00569 | 0.00980 | 0.00796 | 0.04246 | 0.00000 | 0.04484 | 0.00000 | 0.00999 | 0.05963 | 0.04520 | 0.00865 | 0.01253 |
| **DLGAP1-AS4** | 0.00599 | 0.01267 | 0.01397 | 0.03187 | 0.00516 | 0.03292 | 0.01064 | 0.01028 | 0.01951 | 0.02730 | 0.01265 | 0.01428 |
| **DLGAP1-AS5** | 0.00236 | 0.00000 | 0.00000 | 0.00000 | 0.00000 | 0.04395 | 0.07620 | 0.01496 | 0.00779 | 0.21802 | 0.02233 | 0.07819 |
| **DLGAP2** | 0.40824 | 0.00000 | 0.02347 | 0.06034 | 0.15112 | 0.00000 | 0.01945 | 0.07271 | 0.01882 | 0.00000 | 0.14390 | 0.31885 |
| **DLGAP3** | 0.25430 | 0.48249 | 0.19823 | 2.63699 | 0.68599 | 0.05972 | 0.00000 | 0.00000 | 0.01220 | 0.39474 | 0.00000 | 0.20076 |
| **DLGAP4** | 1.48261 | 6.94543 | 17.87761 | 6.97031 | 18.30692 | 0.00001 | 9.45230 | 1.80030 | 13.44787 | 0.00000 | 10.50088 | 5.73945 |
| **DLGAP5** | 16.89256 | 7.56310 | 6.84238 | 7.17823 | 7.14277 | 41.00156 | 13.12473 | 12.14104 | 9.63381 | 6.06994 | 6.14981 | 8.68518 |
| **DLK1** | 0.03100 | 0.08431 | 0.03573 | 0.00204 | 0.02021 | 0.06766 | 0.00000 | 0.01224 | 0.02053 | 0.01726 | 0.00751 | 0.01155 |
| **DLK2** | 0.08336 | 0.47682 | 0.13134 | 0.10439 | 0.01757 | 0.00000 | 0.00340 | 1.07812 | 0.29648 | 0.00000 | 0.00000 | 4.60778 |
| **DLL1** | 0.00471 | 0.01666 | 0.11914 | 0.08418 | 0.02934 | 0.00000 | 0.10900 | 0.07993 | 0.00000 | 0.01896 | 0.01157 | 0.01287 |

| | | | | | | | | | | | | |
|---|---|---|---|---|---|---|---|---|---|---|---|---|
| **DLL3** | 1.45044 | 1.16865 | 0.59971 | 1.97664 | 0.00000 | 0.15508 | 0.55278 | 0.87201 | 0.78460 | 0.00000 | 0.51739 | 9.89128 |
| **DLL4** | 0.31592 | 0.17348 | 0.03395 | 0.00000 | 0.00000 | 0.01071 | 0.15020 | 0.38832 | 0.30202 | 0.12103 | 0.00000 | 0.00837 |
| **DLST** | 5.94426 | 0.78769 | 2.00689 | 1.09140 | 0.41356 | 0.00000 | 0.61175 | 2.26359 | 0.00000 | 1.10886 | 6.26109 | 1.84947 |
| **DLX1** | 0.32482 | 0.03233 | 0.11488 | 0.11767 | 0.06869 | 0.00000 | 0.00372 | 0.80227 | 0.04333 | 0.00000 | 0.01383 | 0.09453 |
| **DLX2** | 0.60973 | 0.00000 | 0.15137 | 0.21711 | 0.38778 | 0.30200 | 0.00000 | 0.29070 | 0.00000 | 0.37899 | 0.00000 | 0.06163 |
| **DLX3** | 0.00344 | 0.00000 | 26.58994 | 2.29554 | 11.28372 | 0.00001 | 28.14858 | 0.02065 | 23.61770 | 0.16177 | 12.34434 | 0.00000 |
| **DLX4** | 0.00245 | 0.24919 | 2.64803 | 1.59711 | 2.43616 | 0.01743 | 6.30526 | 0.00567 | 5.41176 | 0.22643 | 2.76403 | 0.02102 |
| **DLX5** | 0.19238 | 0.00000 | 22.96424 | 1.71568 | 5.05532 | 1.08028 | 4.58096 | 0.03858 | 0.69499 | 0.00000 | 1.46936 | 0.00000 |
| **DLX6** | 1.07525 | 1.48398 | 12.32462 | 0.67056 | 7.85466 | 1.10001 | 2.07459 | 1.03205 | 0.00000 | 0.00000 | 5.30356 | 2.73685 |
| **DLX6-AS1** | 0.00612 | 0.05984 | 0.49055 | 0.00864 | 0.44984 | 0.09135 | 0.07955 | 0.00670 | 0.00000 | 0.00000 | 0.64493 | 0.08166 |
| **DMAP1** | 0.00000 | 9.66547 | 4.54041 | 7.87227 | 8.61199 | 1.37771 | 8.43552 | 4.37199 | 7.80020 | 3.53753 | 4.17866 | 10.69231 |
| **DMBT1** | 0.21036 | 0.06497 | 0.01567 | 0.00000 | 0.00000 | 0.06332 | 0.01423 | 0.00000 | 0.00000 | 0.03190 | 0.03490 | 0.10235 |
| **DMBT1P1** | 0.00625 | 0.00620 | 0.01199 | 0.00045 | 0.01246 | 0.05473 | 0.01939 | 0.00962 | 0.00823 | 0.08955 | 0.01007 | 0.01582 |
| **DMBX1** | 0.26591 | 0.03799 | 0.12655 | 0.17399 | 0.00000 | 0.06475 | 0.00000 | 0.00721 | 0.15007 | 0.00855 | 0.00000 | 0.16244 |
| **DMC1** | 0.00199 | 1.40709 | 0.05607 | 0.00000 | 0.07437 | 0.67694 | 0.00000 | 0.05559 | 0.00000 | 0.23830 | 0.10276 | 0.44877 |
| **DMD** | 5.91078 | 1.63552 | 5.34663 | 0.82083 | 0.00000 | 5.75323 | 2.18834 | 0.73592 | 2.36672 | 3.14538 | 2.53153 | 6.14670 |
| **DMGDH** | 0.64074 | 0.00766 | 0.04991 | 0.04605 | 0.01324 | 0.00004 | 0.00259 | 0.43682 | 0.00000 | 0.23341 | 0.10485 | 0.00000 |
| **DMKN** | 2.93648 | 0.22176 | 0.49406 | 0.66190 | 0.53755 | 0.04439 | 0.14803 | 0.65861 | 0.88135 | 0.00000 | 0.53637 | 0.06222 |
| **DMP1** | 0.00512 | 0.00736 | 0.02060 | 0.01016 | 0.03044 | 0.01892 | 0.00571 | 0.00885 | 0.01638 | 0.07268 | 0.02318 | 0.04225 |
| **DMPK** | 0.15207 | 0.23053 | 1.94036 | 0.01375 | 0.67766 | 0.00000 | 1.16035 | 0.16460 | 0.95903 | 6.48831 | 0.00000 | 0.03142 |
| **DMRT1** | 0.43767 | 0.03309 | 0.01597 | 0.10591 | 0.00734 | 0.00000 | 0.05230 | 0.00000 | 0.01465 | 0.01630 | 0.01511 | 0.20083 |
| **DMRT2** | 0.06376 | 0.00000 | 0.06060 | 0.01051 | 0.00920 | 0.03543 | 0.01213 | 0.00353 | 0.00000 | 0.04462 | 0.01490 | 0.21029 |
| **DMRT3** | 0.03358 | 0.05805 | 0.01978 | 0.00647 | 0.00985 | 0.02072 | 0.02450 | 0.02630 | 0.01727 | 0.01079 | 0.01422 | 0.03745 |
| **DMRTA1** | 0.35671 | 0.06711 | 0.05340 | 0.14956 | 0.01783 | 0.03146 | 0.02938 | 0.04811 | 0.04444 | 0.00000 | 0.00926 | 0.01059 |
| **DMRTA2** | 0.02172 | 0.01166 | 0.00636 | 0.72147 | 0.07955 | 0.00000 | 0.01915 | 0.01064 | 0.03543 | 0.00671 | 0.00778 | 0.01620 |
| **DMRTB1** | 2.76957 | 0.71127 | 0.23890 | 0.55850 | 0.51870 | 0.00000 | 0.14382 | 0.67906 | 0.00000 | 0.00084 | 0.24553 | 6.84031 |
| **DMRTC2** | 1.73971 | 0.00639 | 0.01774 | 0.01276 | 0.01983 | 0.00000 | 0.02755 | 0.23355 | 0.03146 | 0.01778 | 0.01578 | 0.11695 |
| **DMTF1** | 1.36113 | 2.94317 | 3.02046 | 2.16906 | 0.34539 | 8.04699 | 2.41207 | 3.97376 | 7.99917 | 1.98877 | 3.34784 | 4.13643 |
| **DMTN** | 0.95425 | 0.01886 | 0.35178 | 0.35675 | 0.03662 | 0.00000 | 0.00000 | 0.03223 | 0.00000 | 0.30275 | 0.00401 | 0.02164 |
| **DMWD** | 1.24056 | 0.53157 | 2.33673 | 0.00000 | 2.92969 | 0.00000 | 3.02549 | 0.13477 | 1.93657 | 15.20707 | 0.00000 | 1.33110 |
| **DMXL1** | 5.74272 | 2.00144 | 2.44479 | 4.61878 | 3.51611 | 14.90797 | 2.65308 | 1.01335 | 0.38765 | 12.68364 | 2.48944 | 1.65663 |
| **DMXL2** | 1.28493 | 0.36248 | 1.66243 | 0.28725 | 0.00000 | 3.84200 | 0.81370 | 0.32243 | 0.00000 | 3.63061 | 0.33986 | 1.02355 |
| **DNA2** | 2.64936 | 3.34118 | 4.26252 | 2.89676 | 0.00000 | 12.21077 | 4.70468 | 1.79389 | 9.25844 | 2.70449 | 5.19471 | 3.99899 |
| **DNAAF1** | 0.01128 | 0.03403 | 0.01404 | 0.01415 | 0.00431 | 0.01928 | 0.02403 | 0.09271 | 0.03652 | 0.02125 | 0.01261 | 0.10742 |
| **DNAAF2** | 1.37872 | 3.17354 | 8.87862 | 2.41285 | 2.53058 | 5.65449 | 7.74469 | 0.45761 | 7.52186 | 4.53857 | 9.77757 | 1.78609 |
| **DNAAF3** | 0.14094 | 0.04165 | 0.00861 | 0.03500 | 0.00410 | 0.00000 | 0.07065 | 0.48341 | 0.06273 | 0.11399 | 0.00120 | 0.32937 |
| **DNAH1** | 0.04706 | 0.07492 | 0.08279 | 0.28074 | 0.33609 | 0.00000 | 0.25039 | 0.00410 | 0.00000 | 0.93750 | 0.12310 | 0.15084 |
| **DNAH10** | 8.32112 | 0.02609 | 0.04986 | 0.02221 | 0.02166 | 0.00000 | 0.00192 | 1.67227 | 0.03370 | 0.05212 | 0.01973 | 0.47011 |
| **DNAH11** | 9.49055 | 0.00378 | 0.11166 | 0.21540 | 0.22404 | 0.00000 | 0.04647 | 0.90503 | 0.00000 | 0.08578 | 0.01595 | 0.60794 |
| **DNAH12** | 1.82792 | 0.50943 | 0.00000 | 0.32755 | 0.02159 | 0.30606 | 0.00000 | 1.22871 | 0.00000 | 0.39760 | 0.00000 | 1.11972 |
| **DNAH14** | 0.18708 | 0.10211 | 0.97995 | 0.00000 | 0.03365 | 5.13587 | 0.00041 | 0.60845 | 3.02185 | 4.02449 | 1.66418 | 0.95166 |
| **DNAH17** | 3.67336 | 0.19784 | 0.24389 | 0.28780 | 0.66485 | 0.00000 | 0.00000 | 0.20568 | 0.36427 | 1.59606 | 0.22946 | 0.32131 |
| **DNAH17-AS1** | 0.00856 | 0.08547 | 0.05011 | 0.00370 | 0.03101 | 0.00000 | 0.00503 | 0.01005 | 0.01059 | 0.09170 | 0.02630 | 0.02002 |
| **DNAH2** | 0.36428 | 0.08340 | 0.04245 | 0.48662 | 0.15125 | 0.00000 | 0.01109 | 0.09840 | 0.00000 | 0.05101 | 0.03977 | 0.29943 |
| **DNAH3** | 4.24031 | 1.40001 | 0.00000 | 0.83130 | 1.71906 | 1.00680 | 0.83847 | 0.62427 | 0.00000 | 0.00000 | 2.32160 | 0.07141 |
| **DNAH5** | 1.00430 | 0.02312 | 0.03413 | 0.04411 | 0.00000 | 0.07492 | 0.03846 | 0.49124 | 0.00000 | 0.10058 | 0.02865 | 0.05520 |
| **DNAH6** | 0.37965 | 0.08488 | 0.10452 | 0.06370 | 0.00976 | 0.00000 | 0.00000 | 0.00934 | 0.09366 | 0.03754 | 0.00707 | 0.05100 |
| **DNAH7** | 1.14127 | 0.01843 | 0.00000 | 0.01222 | 0.00000 | 0.14874 | 0.12810 | 0.08597 | 0.00000 | 0.24503 | 0.00481 | 0.21281 |

| | | | | | | | | | | | | |
|---|---|---|---|---|---|---|---|---|---|---|---|---|
| **DNAH8** | 0.00256 | 0.00665 | 0.12257 | 0.06568 | 0.00000 | 0.01014 | 0.01947 | 0.14946 | 0.00798 | 0.13125 | 0.04960 | 0.03204 |
| **DNAH9** | 1.15602 | 0.00018 | 0.07774 | 0.40768 | 0.04307 | 0.04118 | 0.00000 | 0.44563 | 0.03177 | 0.09206 | 0.00507 | 0.03320 |
| **DNAI1** | 0.46652 | 0.00000 | 0.25915 | 0.00322 | 0.00000 | 0.37911 | 0.41254 | 0.36697 | 0.09582 | 0.00000 | 0.00000 | 0.32034 |
| **DNAI2** | 0.06795 | 0.00945 | 0.03110 | 0.00932 | 0.01430 | 0.03024 | 0.01343 | 0.00771 | 0.01661 | 0.00758 | 0.01416 | 0.05726 |
| **DNAJA1** | 14.65373 | 21.53709 | 29.69943 | 20.54066 | 19.48397 | 39.54245 | 38.26495 | 27.02384 | 39.94547 | 11.40901 | 26.56148 | 30.16932 |
| **DNAJA1P5** | 0.01985 | 0.03970 | 0.04998 | 0.01391 | 0.04226 | 0.05833 | 0.02480 | 0.04425 | 0.03652 | 0.00306 | 0.03887 | 0.03835 |
| **DNAJA2** | 7.79714 | 21.15058 | 18.28443 | 15.28913 | 8.92637 | 39.56051 | 20.50347 | 15.39199 | 26.64292 | 10.81018 | 15.91940 | 17.66877 |
| **DNAJA3** | 3.26439 | 14.49639 | 14.26689 | 6.87454 | 5.48896 | 11.31128 | 11.59974 | 2.24332 | 18.13287 | 0.00000 | 16.58001 | 7.69745 |
| **DNAJA4** | 2.89110 | 1.32354 | 1.45223 | 0.37339 | 1.96622 | 1.93508 | 1.84234 | 0.29355 | 0.05836 | 0.24723 | 1.31851 | 0.34880 |
| **DNAJB1** | 11.56130 | 23.14373 | 27.26498 | 12.74060 | 19.27680 | 9.15881 | 30.13867 | 39.20760 | 28.39235 | 0.00000 | 18.72127 | 32.21127 |
| **DNAJB11** | 3.42762 | 19.37130 | 16.79749 | 7.61454 | 10.80205 | 14.84473 | 0.00000 | 12.72962 | 26.29509 | 10.01155 | 12.81892 | 11.11527 |
| **DNAJB12** | 0.63832 | 6.38314 | 8.12037 | 6.68358 | 9.03696 | 2.32081 | 7.56068 | 5.88109 | 16.85697 | 4.62761 | 5.78253 | 11.79778 |
| **DNAJB13** | 1.07061 | 0.13764 | 0.00892 | 0.15315 | 0.00000 | 0.01260 | 0.02508 | 0.00000 | 0.03754 | 0.01743 | 0.01262 | 0.28219 |
| **DNAJB14** | 0.05202 | 0.73573 | 2.02118 | 1.84180 | 0.49512 | 7.04273 | 0.77863 | 7.42095 | 4.16137 | 5.06727 | 1.00825 | 5.02410 |
| **DNAJB2** | 1.46773 | 0.06253 | 0.45792 | 0.40981 | 1.75866 | 0.00000 | 2.28135 | 0.00000 | 0.00000 | 10.21186 | 0.08043 | 1.29076 |
| **DNAJB3** | 0.00772 | 0.01341 | 0.02455 | 0.01819 | 0.01469 | 0.02650 | 0.02426 | 0.02239 | 0.01980 | 0.00825 | 0.01814 | 0.01525 |
| **DNAJB4** | 0.45005 | 5.23786 | 2.16925 | 2.54447 | 0.00000 | 11.18411 | 0.00000 | 5.72617 | 1.24487 | 0.27963 | 1.11394 | 8.01478 |
| **DNAJB5** | 0.21638 | 0.36382 | 0.10713 | 0.32683 | 0.04642 | 0.00000 | 0.00000 | 0.00000 | 0.16412 | 0.04599 | 0.07627 | 0.21905 |
| **DNAJB5-AS1** | 0.31885 | 0.02268 | 0.02083 | 0.00698 | 0.05975 | 0.00000 | 0.01998 | 0.08156 | 0.05436 | 0.07886 | 0.01157 | 0.15009 |
| **DNAJB6** | 2.36562 | 11.14371 | 13.51448 | 11.65120 | 3.95023 | 16.37071 | 23.88626 | 13.69109 | 18.65995 | 4.17322 | 9.65584 | 8.68260 |
| **DNAJB7** | 0.02551 | 0.05096 | 0.14751 | 0.00000 | 0.13963 | 0.46717 | 0.00000 | 0.05216 | 0.17042 | 0.27008 | 0.36605 | 0.39464 |
| **DNAJB8** | 0.00607 | 0.01431 | 0.42940 | 0.09399 | 0.01403 | 0.00000 | 0.00000 | 0.01040 | 0.00000 | 0.02609 | 0.01942 | 0.01638 |
| **DNAJB8-AS1** | 0.00597 | 0.01589 | 0.17790 | 0.01596 | 0.02620 | 0.00000 | 0.00316 | 0.01057 | 0.01561 | 0.03779 | 0.01418 | 0.01592 |
| **DNAJB9** | 0.94956 | 3.04871 | 15.82364 | 0.56976 | 0.00000 | 12.18438 | 0.00000 | 12.59454 | 6.52288 | 0.00000 | 3.91078 | 9.19765 |
| **DNAJC1** | 3.46877 | 1.11576 | 3.06414 | 2.49437 | 10.97942 | 3.87850 | 7.37200 | 3.44779 | 15.54195 | 13.91588 | 8.74155 | 1.41726 |
| **DNAJC10** | 1.43875 | 7.15708 | 6.42393 | 1.50094 | 3.38960 | 16.57184 | 6.42346 | 1.67796 | 13.67968 | 4.77943 | 7.77742 | 8.21711 |
| **DNAJC11** | 0.62440 | 17.51914 | 5.55511 | 11.91263 | 2.12905 | 12.58615 | 0.00000 | 3.29392 | 17.04860 | 0.00000 | 14.88288 | 8.87940 |
| **DNAJC12** | 0.00298 | 0.08323 | 0.04435 | 0.02271 | 0.00147 | 0.18481 | 0.01293 | 0.04592 | 0.00000 | 0.00105 | 0.05376 | 0.05558 |
| **DNAJC13** | 7.42963 | 2.13268 | 1.33053 | 3.20475 | 1.02659 | 10.24781 | 2.53774 | 2.91213 | 0.00000 | 8.78912 | 1.00660 | 2.93430 |
| **DNAJC14** | 2.35928 | 9.18041 | 7.65699 | 5.13485 | 1.07298 | 3.64061 | 0.00000 | 1.57425 | 11.97275 | 0.00000 | 7.02205 | 8.27442 |
| **DNAJC15** | 3.68222 | 48.78054 | 21.02550 | 24.33421 | 62.55773 | 81.37041 | 43.35974 | 29.18427 | 48.91805 | 40.23040 | 43.30854 | 57.93872 |
| **DNAJC16** | 2.17001 | 6.22832 | 0.00000 | 3.69135 | 3.72391 | 6.73246 | 0.00000 | 0.66482 | 0.00000 | 5.82921 | 3.08085 | 4.29369 |
| **DNAJC17** | 0.75951 | 3.65975 | 4.50437 | 1.61183 | 2.57594 | 1.15733 | 5.34666 | 5.35452 | 0.92029 | 0.00000 | 4.51102 | 7.80691 |
| **DNAJC18** | 1.15641 | 0.16862 | 0.62390 | 0.65425 | 0.11733 | 0.00000 | 0.20228 | 0.24588 | 0.71930 | 0.00001 | 0.02682 | 0.19760 |
| **DNAJC19** | 1.27138 | 11.22776 | 4.33059 | 2.20735 | 6.53671 | 13.05589 | 5.03278 | 0.08046 | 7.18904 | 2.32401 | 8.86931 | 7.89356 |
| **DNAJC2** | 2.24449 | 12.27860 | 11.84261 | 4.07635 | 8.32779 | 25.98643 | 19.52347 | 4.71179 | 20.87566 | 19.19351 | 7.60951 | 11.35197 |
| **DNAJC21** | 1.97143 | 1.86280 | 2.37945 | 0.00000 | 6.07683 | 5.50030 | 8.36653 | 3.05605 | 0.00000 | 22.42293 | 2.60505 | 1.34124 |
| **DNAJC22** | 0.15372 | 0.07121 | 0.07119 | 3.09099 | 0.07098 | 0.14973 | 0.00000 | 0.09151 | 0.24911 | 0.00000 | 0.04694 | 0.09105 |
| **DNAJC24** | 0.16300 | 0.95952 | 1.96870 | 0.01934 | 0.12789 | 4.98056 | 2.22220 | 1.36235 | 3.35684 | 4.96812 | 0.45624 | 2.06672 |
| **DNAJC25** | 0.22614 | 3.40517 | 1.41829 | 5.00758 | 0.49262 | 6.49723 | 0.00000 | 2.63720 | 1.02829 | 9.19750 | 0.00000 | 1.02939 |
| **DNAJC27** | 1.00318 | 0.21439 | 0.22880 | 0.56074 | 0.69371 | 1.33375 | 0.00000 | 2.35333 | 2.24334 | 1.81158 | 0.14567 | 0.14290 |
| **DNAJC27-AS1** | 0.00736 | 0.12990 | 0.03515 | 0.00535 | 0.06850 | 0.00306 | 0.06206 | 0.00523 | 0.04713 | 0.11044 | 0.01514 | 0.31341 |
| **DNAJC28** | 0.39608 | 0.91416 | 0.26592 | 0.47283 | 0.22119 | 1.71623 | 0.00000 | 0.00000 | 0.00000 | 0.00000 | 1.29361 | 0.96775 |
| **DNAJC3** | 6.77627 | 9.26579 | 17.74592 | 13.47258 | 3.66010 | 38.14789 | 0.00000 | 9.09832 | 15.66613 | 11.82911 | 10.53631 | 6.02706 |
| **DNAJC30** | 1.02940 | 5.62275 | 3.32712 | 1.02253 | 3.78667 | 2.53500 | 1.32409 | 0.72239 | 2.91163 | 1.69687 | 4.45862 | 6.83464 |
| **DNAJC4** | 1.08701 | 0.63173 | 0.96447 | 1.79006 | 0.57056 | 0.00000 | 0.91792 | 0.17035 | 0.50305 | 0.49525 | 0.46090 | 0.35536 |
| **DNAJC5** | 0.64842 | 5.26329 | 6.99584 | 1.81436 | 12.52913 | 0.73687 | 12.32603 | 0.90904 | 4.50199 | 11.81475 | 4.94427 | 3.33021 |
| **DNAJC5B** | 0.00826 | 0.01468 | 0.02896 | 0.01775 | 0.01304 | 0.01713 | 0.01181 | 0.07070 | 0.01714 | 0.02230 | 0.01508 | 0.01258 |

| | | | | | | | | | | | | |
|---|---|---|---|---|---|---|---|---|---|---|---|---|
| **DNAJC5G** | 0.34841 | 0.01633 | 0.00000 | 0.03216 | 0.00000 | 0.12128 | 0.00000 | 0.10599 | 0.01397 | 0.20136 | 0.02373 | 0.00000 |
| **DNAJC6** | 0.45966 | 0.27949 | 0.00000 | 0.00000 | 8.74440 | 0.32038 | 5.83229 | 0.00147 | 5.05870 | 3.60758 | 9.07189 | 0.00000 |
| **DNAJC7** | 2.13577 | 11.38139 | 12.30845 | 8.63562 | 9.38503 | 21.70356 | 15.23411 | 10.37811 | 19.15103 | 0.72426 | 14.16929 | 15.89068 |
| **DNAJC8** | 5.02690 | 14.63109 | 19.77365 | 10.41561 | 9.43313 | 19.07075 | 28.99663 | 18.85768 | 13.33682 | 6.43979 | 15.76309 | 22.12248 |
| **DNAJC9** | 6.62076 | 8.04759 | 10.90606 | 5.93807 | 5.18904 | 16.98855 | 15.05676 | 8.57451 | 15.44074 | 1.21923 | 12.39443 | 16.47370 |
| **DNAJC9-AS1** | 0.42369 | 0.12956 | 0.06161 | 0.05814 | 0.09841 | 0.00000 | 0.00062 | 0.12459 | 0.21506 | 0.86053 | 0.00000 | 0.09286 |
| **DNAL1** | 1.91615 | 0.92982 | 1.43956 | 0.83212 | 1.21846 | 1.07126 | 2.23578 | 9.59854 | 0.00000 | 0.02935 | 1.71688 | 5.70143 |
| **DNAL4** | 6.01999 | 3.97677 | 0.77272 | 0.61180 | 1.05934 | 0.39137 | 1.07220 | 0.00000 | 1.05544 | 0.00000 | 1.55298 | 5.90447 |
| **DNALI1** | 0.00480 | 0.02851 | 1.34485 | 1.06043 | 0.00000 | 0.18626 | 1.27224 | 0.05706 | 0.66290 | 0.06785 | 0.88948 | 0.06738 |
| **DNASE1** | 0.00045 | 1.01069 | 0.78933 | 0.40475 | 0.69404 | 0.00000 | 0.14430 | 1.44755 | 0.42186 | 1.55671 | 0.53614 | 0.64603 |
| **DNASE1L1** | 0.00382 | 0.13104 | 0.08294 | 0.09309 | 0.00000 | 0.00000 | 0.18155 | 0.08451 | 0.00000 | 0.12318 | 0.58179 | 0.33374 |
| **DNASE1L2** | 0.00151 | 0.22681 | 0.19573 | 0.19198 | 0.00000 | 0.00000 | 0.06706 | 0.05666 | 0.03066 | 0.09645 | 0.01937 | 0.33572 |
| **DNASE1L3** | 0.01619 | 0.05014 | 0.64587 | 0.19262 | 0.00000 | 0.07804 | 0.00000 | 0.05112 | 0.50774 | 0.00000 | 0.00394 | 0.18036 |
| **DNASE2** | 0.61414 | 13.74571 | 9.92179 | 4.59242 | 14.31071 | 0.00000 | 4.61584 | 5.43518 | 16.00394 | 0.00000 | 10.82877 | 13.64834 |
| **DNASE2B** | 0.00701 | 0.00296 | 0.01475 | 0.01206 | 0.02958 | 0.00000 | 0.00655 | 0.00461 | 0.04924 | 0.02506 | 0.00884 | 0.40269 |
| **DND1** | 1.00074 | 0.97824 | 2.91494 | 7.85375 | 0.26412 | 0.00000 | 1.50664 | 1.11724 | 1.61520 | 0.69709 | 1.13960 | 0.98424 |
| **DNER** | 0.18422 | 0.00550 | 0.03254 | 0.02304 | 1.48470 | 0.21495 | 0.00000 | 0.09186 | 0.03225 | 0.01346 | 0.05136 | 0.04316 |
| **DNHD1** | 0.81844 | 0.38035 | 1.46660 | 1.39824 | 0.62508 | 0.00000 | 0.01547 | 1.00105 | 3.20900 | 5.82322 | 2.11553 | 1.70310 |
| **DNLZ** | 0.00394 | 0.02223 | 0.58842 | 0.65559 | 1.54591 | 0.00000 | 1.48624 | 0.35994 | 0.00000 | 3.89269 | 0.00000 | 0.61747 |
| **DNM1** | 0.00254 | 0.01066 | 0.06215 | 0.66273 | 0.01356 | 0.02252 | 0.02859 | 0.07694 | 0.02152 | 0.00640 | 0.01089 | 0.20434 |
| **DNM1L** | 3.35323 | 8.62883 | 13.44234 | 7.99123 | 8.62346 | 15.33661 | 18.37603 | 1.88142 | 18.91143 | 3.69214 | 13.09365 | 3.41863 |
| **DNM1P35** | 0.16235 | 0.01367 | 0.03556 | 0.01683 | 0.00365 | 0.00000 | 0.02284 | 0.00512 | 0.00000 | 0.03917 | 0.11341 | 0.01437 |
| **DNM1P41** | 0.03173 | 0.01463 | 0.01619 | 0.01201 | 0.01507 | 0.02023 | 0.01726 | 0.00764 | 0.01860 | 0.01363 | 0.01469 | 0.01556 |
| **DNM1P46** | 0.01663 | 0.02184 | 0.02156 | 0.01262 | 0.01020 | 0.01240 | 0.01186 | 0.06510 | 0.01378 | 0.04020 | 0.02611 | 0.02041 |
| **DNM2** | 1.25759 | 4.57727 | 10.45463 | 9.65037 | 7.99592 | 0.00000 | 10.80341 | 0.23960 | 7.44056 | 16.38988 | 2.81193 | 8.51862 |
| **DNM3** | 1.28771 | 0.00000 | 0.04238 | 0.55391 | 0.02075 | 0.03874 | 0.00000 | 1.76764 | 0.10138 | 0.33129 | 0.00004 | 0.04738 |
| **DNM3OS** | 0.00989 | 0.00000 | 0.06215 | 0.00932 | 0.00000 | 0.04418 | 0.00000 | 0.14954 | 0.00054 | 0.22268 | 0.08451 | 0.07511 |
| **DNMBP** | 0.94770 | 2.18542 | 0.10216 | 3.03630 | 3.87459 | 4.16869 | 0.52520 | 2.31564 | 1.86123 | 3.98916 | 1.18085 | 2.24696 |
| **DNMBP-AS1** | 0.00411 | 0.01318 | 0.00361 | 0.01885 | 0.02967 | 0.08007 | 0.04038 | 0.13320 | 0.03275 | 0.00000 | 0.11486 | 0.17360 |
| **DNMT1** | 41.03528 | 7.66033 | 29.18756 | 7.46292 | 11.00955 | 1.90299 | 23.14736 | 30.56387 | 12.42603 | 8.45942 | 5.72097 | 22.52227 |
| **DNMT3A** | 6.23360 | 22.48045 | 0.00000 | 15.16273 | 22.43656 | 3.84346 | 14.67826 | 1.13915 | 12.51806 | 10.94732 | 21.08341 | 11.49614 |
| **DNMT3B** | 5.43110 | 17.93839 | 26.09913 | 18.20314 | 32.56232 | 13.49734 | 33.46569 | 0.00000 | 29.98732 | 14.99290 | 17.76422 | 0.00000 |
| **DNMT3L** | 0.00000 | 64.86242 | 33.47188 | 44.17753 | 70.63009 | 59.56608 | 47.41771 | 0.36884 | 52.31199 | 17.15108 | 51.82860 | 18.86147 |
| **DNPEP** | 0.30238 | 1.47582 | 4.29704 | 3.37181 | 4.74041 | 0.17634 | 2.97054 | 6.18038 | 1.53536 | 1.32522 | 0.00000 | 2.26077 |
| **DNPH1** | 0.20535 | 6.02412 | 1.68567 | 6.39981 | 7.90486 | 1.22232 | 5.14604 | 0.00000 | 2.98568 | 0.00000 | 10.31873 | 1.06736 |
| **DNTT** | 0.03535 | 0.21984 | 0.01105 | 0.04754 | 0.00000 | 0.00388 | 0.03407 | 0.00654 | 0.01343 | 0.01570 | 0.01586 | 0.03292 |
| **DNTTIP1** | 0.34046 | 3.99452 | 5.02662 | 3.57080 | 5.85356 | 0.55463 | 2.71964 | 0.81383 | 1.53896 | 6.02178 | 1.68104 | 3.51507 |
| **DNTTIP2** | 2.25292 | 18.71498 | 23.72045 | 7.31618 | 10.18296 | 42.30389 | 27.45970 | 25.09432 | 21.16956 | 16.71754 | 24.27770 | 28.83642 |
| **DOC2A** | 0.00183 | 0.19374 | 0.17598 | 0.00000 | 0.04686 | 0.27026 | 0.01489 | 0.00634 | 0.00000 | 0.75063 | 0.06839 | 0.00766 |
| **DOC2B** | 0.53460 | 0.02296 | 0.01757 | 0.01363 | 0.01645 | 0.00000 | 0.04064 | 0.02263 | 0.00388 | 0.00000 | 0.03829 | 0.08905 |
| **DOCK1** | 1.30875 | 2.73498 | 1.71432 | 5.05160 | 2.59489 | 1.86452 | 0.69066 | 0.63643 | 2.82111 | 2.29727 | 0.83006 | 1.78211 |
| **DOCK10** | 1.93531 | 0.01444 | 0.01577 | 0.01270 | 0.01736 | 0.00000 | 0.01675 | 0.19932 | 0.03024 | 0.00407 | 0.01913 | 0.00287 |
| **DOCK11** | 0.51644 | 0.06867 | 0.01728 | 0.09104 | 0.00000 | 0.50993 | 0.00000 | 0.21262 | 0.00000 | 0.02544 | 0.00857 | 0.66224 |
| **DOCK2** | 3.49691 | 0.05141 | 0.02243 | 0.00261 | 0.00000 | 0.02665 | 0.00943 | 0.16576 | 0.02256 | 0.05017 | 0.04415 | 0.07038 |
| **DOCK3** | 0.18279 | 1.19018 | 0.55254 | 2.57795 | 2.67124 | 0.00000 | 0.00000 | 0.33684 | 0.21895 | 2.16114 | 0.40564 | 1.21467 |
| **DOCK4** | 1.26811 | 0.48963 | 4.99245 | 1.92824 | 1.92311 | 2.22651 | 1.83248 | 0.15336 | 1.63364 | 0.77217 | 0.40135 | 0.22529 |
| **DOCK4-AS1** | 0.03856 | 0.00632 | 0.12359 | 0.00199 | 0.00872 | 0.07194 | 0.07237 | 0.00974 | 0.02241 | 0.01253 | 0.01663 | 0.02340 |
| **DOCK5** | 0.89565 | 0.61954 | 0.27765 | 0.05382 | 0.60287 | 0.31684 | 0.01619 | 0.72785 | 0.00000 | 0.04489 | 0.12661 | 0.06108 |

| | | | | | | | | | | | | |
|---|---|---|---|---|---|---|---|---|---|---|---|---|
| **DOCK6** | 0.46997 | 4.23170 | 0.77997 | 11.55076 | 6.21114 | 0.00000 | 0.57758 | 0.00000 | 2.77343 | 3.35039 | 0.00000 | 2.04704 |
| **DOCK7** | 0.63673 | 0.92715 | 2.31122 | 0.60145 | 1.67005 | 4.52316 | 3.34465 | 0.49848 | 4.45328 | 5.39075 | 0.98472 | 0.04508 |
| **DOCK8** | 0.43228 | 0.13966 | 1.59112 | 0.59876 | 0.00000 | 0.17699 | 0.77092 | 0.12441 | 0.58179 | 0.00000 | 1.74718 | 0.24310 |
| **DOCK9** | 4.33056 | 0.00000 | 12.12479 | 0.00000 | 4.23393 | 2.10533 | 21.21860 | 0.84615 | 5.72282 | 0.00000 | 10.75891 | 0.00000 |
| **DOCK9-AS1** | 0.00596 | 0.01271 | 0.02545 | 0.01026 | 0.01322 | 0.02340 | 0.01908 | 0.01044 | 0.00556 | 0.02243 | 0.05432 | 0.01538 |
| **DOCK9-AS2** | 0.25913 | 0.02135 | 0.08649 | 0.04282 | 0.00760 | 0.04274 | 0.00000 | 0.00235 | 0.04575 | 0.01937 | 0.00883 | 0.00982 |
| **DOHH** | 0.21865 | 6.66361 | 13.19364 | 0.00000 | 0.67620 | 0.00000 | 0.00000 | 1.62494 | 5.93709 | 4.78635 | 3.29846 | 0.73201 |
| **DOK1** | 0.91067 | 0.42550 | 1.92738 | 2.21039 | 0.07934 | 0.00000 | 1.90591 | 0.14480 | 0.36532 | 0.00000 | 0.00000 | 0.79817 |
| **DOK2** | 1.94077 | 0.06344 | 0.13322 | 0.00782 | 0.01340 | 0.00000 | 0.00000 | 0.00000 | 0.79720 | 0.39047 | 0.00146 | 0.13110 |
| **DOK3** | 0.02349 | 0.29326 | 0.51634 | 0.26909 | 0.06630 | 0.00000 | 1.51452 | 0.09280 | 3.56682 | 8.26619 | 0.00000 | 2.80788 |
| **DOK4** | 0.00166 | 0.00000 | 4.20229 | 2.58312 | 1.88073 | 0.39902 | 2.75153 | 2.22075 | 3.26619 | 0.00000 | 0.87880 | 0.42905 |
| **DOK5** | 2.85567 | 0.04580 | 0.01740 | 0.00577 | 0.01353 | 0.02968 | 0.01222 | 0.13641 | 0.01749 | 0.00265 | 0.01363 | 0.07816 |
| **DOK6** | 0.64918 | 0.00616 | 0.05238 | 0.01593 | 0.01919 | 0.03948 | 0.00199 | 0.08698 | 0.00499 | 0.05961 | 0.02298 | 0.02169 |
| **DOK7** | 0.01606 | 0.02329 | 0.01867 | 0.13778 | 0.00000 | 0.00540 | 0.05123 | 0.00767 | 0.02209 | 0.11714 | 0.02881 | 0.02246 |
| **DOLK** | 0.69635 | 8.02016 | 9.84921 | 5.25806 | 9.00769 | 8.35733 | 0.00000 | 0.19126 | 7.39608 | 0.00000 | 4.50745 | 5.35461 |
| **DOLPP1** | 1.60528 | 6.54167 | 15.54581 | 6.11651 | 8.32628 | 0.00000 | 1.98023 | 0.35731 | 20.22585 | 2.46217 | 4.24576 | 5.64631 |
| **DONSON** | 0.10727 | 3.33776 | 5.42244 | 1.64355 | 0.31039 | 7.00407 | 3.45521 | 0.58395 | 2.30422 | 11.86442 | 2.99329 | 0.31487 |
| **DOPEY1** | 9.25382 | 0.00000 | 0.29438 | 0.28817 | 0.55966 | 1.16494 | 0.00000 | 2.18252 | 0.42281 | 1.49480 | 0.80883 | 0.73403 |
| **DOPEY2** | 2.93183 | 0.46047 | 1.37268 | 0.12687 | 1.33920 | 0.00151 | 2.11084 | 0.66277 | 0.97280 | 0.89083 | 0.67957 | 0.97866 |
| **DOT1L** | 8.76424 | 5.02047 | 4.21679 | 0.51803 | 17.93309 | 0.00000 | 1.32655 | 3.33156 | 12.13752 | 23.14380 | 2.50631 | 6.78746 |
| **DPAGT1** | 0.56008 | 11.58716 | 12.05691 | 10.09361 | 6.98679 | 5.38441 | 1.42094 | 0.93357 | 21.32278 | 0.00000 | 12.72774 | 8.75135 |
| **DPCD** | 0.85704 | 6.87650 | 3.47650 | 2.35499 | 2.67285 | 8.10232 | 3.96264 | 2.69714 | 2.43461 | 0.15244 | 3.12056 | 8.39829 |
| **DPCR1** | 0.33052 | 0.04486 | 0.03229 | 0.13518 | 0.04659 | 0.03168 | 0.03243 | 0.03930 | 0.01506 | 0.01231 | 0.01425 | 0.03505 |
| **DPEP1** | 1.94086 | 1.44710 | 0.00000 | 0.03900 | 0.23084 | 0.00000 | 0.00000 | 2.04580 | 0.00000 | 0.28715 | 0.07564 | 2.19229 |
| **DPEP2** | 0.00650 | 0.01693 | 0.11838 | 0.01492 | 0.03907 | 0.00000 | 0.01522 | 0.01051 | 0.04458 | 0.00030 | 0.04907 | 0.01999 |
| **DPEP3** | 0.04308 | 0.41265 | 0.01888 | 0.00000 | 0.15886 | 0.00278 | 0.02461 | 0.00091 | 0.00000 | 0.02877 | 0.09588 | 0.61569 |
| **DPF1** | 1.99660 | 0.48681 | 0.00000 | 4.43863 | 1.62576 | 0.00000 | 0.00424 | 0.28421 | 0.38261 | 0.69148 | 0.05234 | 0.08015 |
| **DPF2** | 9.68070 | 16.20674 | 13.38729 | 11.03190 | 14.58135 | 5.43422 | 14.43317 | 1.53319 | 13.46864 | 0.00000 | 16.62621 | 17.74941 |
| **DPF3** | 1.02443 | 0.33581 | 4.53728 | 0.00000 | 5.08194 | 0.86256 | 4.82178 | 0.10712 | 5.04416 | 2.40942 | 2.25224 | 0.10462 |
| **DPH1** | 0.11255 | 7.84482 | 2.54258 | 0.91207 | 0.79908 | 0.00000 | 0.00000 | 0.00000 | 2.79162 | 0.00000 | 5.75935 | 2.14292 |
| **DPH2** | 0.55794 | 21.94707 | 17.18288 | 12.09102 | 14.60054 | 5.70698 | 21.88173 | 8.97937 | 24.54705 | 0.00000 | 21.58423 | 25.22657 |
| **DPH3** | 0.54639 | 14.87833 | 13.69711 | 4.29943 | 7.93971 | 32.32888 | 13.88382 | 18.77719 | 15.85801 | 8.54477 | 8.30875 | 25.55263 |
| **DPH3P1** | 0.32491 | 0.68869 | 0.72574 | 0.17869 | 0.78857 | 2.10875 | 0.47417 | 0.96633 | 0.78729 | 0.52637 | 0.28553 | 1.86598 |
| **DPH5** | 0.00000 | 5.90649 | 1.88850 | 3.92627 | 5.27941 | 14.78952 | 6.24231 | 0.00000 | 5.15061 | 2.65422 | 5.12822 | 7.39286 |
| **DPH6** | 0.85536 | 5.55646 | 1.42739 | 1.01516 | 4.45093 | 11.04682 | 0.00000 | 0.00173 | 0.00006 | 2.88739 | 5.06041 | 6.44605 |
| **DPH6-AS1** | 0.02143 | 0.01699 | 0.01697 | 0.01762 | 0.01694 | 0.00445 | 0.04779 | 0.02001 | 0.01203 | 0.02665 | 0.02179 | 0.03139 |
| **DPH7** | 0.15480 | 8.25902 | 6.49929 | 4.95966 | 6.46110 | 1.59064 | 9.76763 | 0.90216 | 12.80936 | 2.17462 | 7.94357 | 5.16948 |
| **DPM1** | 1.38584 | 13.71644 | 14.57370 | 3.01895 | 1.56766 | 41.05877 | 9.70139 | 11.43652 | 15.78991 | 7.57137 | 13.68221 | 19.23345 |
| **DPM2** | 0.68510 | 12.67359 | 11.35297 | 6.62787 | 7.57098 | 1.56041 | 9.59193 | 7.10623 | 11.97238 | 0.00000 | 10.31868 | 17.14779 |
| **DPM3** | 0.00000 | 1.52217 | 2.55192 | 0.30408 | 0.80234 | 1.60680 | 1.86014 | 2.13458 | 3.01050 | 3.55094 | 4.00871 | 2.28561 |
| **DPP10** | 1.37558 | 0.02843 | 0.05143 | 0.00002 | 0.05990 | 0.08026 | 0.00000 | 0.20629 | 0.00000 | 0.01314 | 0.02376 | 0.00833 |
| **DPP10-AS1** | 0.00482 | 0.01026 | 0.06655 | 0.11140 | 0.06786 | 0.00000 | 0.01236 | 0.01051 | 0.04440 | 0.09428 | 0.02004 | 0.01382 |
| **DPP3** | 4.13593 | 2.14243 | 11.71214 | 8.95257 | 1.73758 | 0.00000 | 6.65153 | 2.38476 | 9.18559 | 0.00000 | 13.84249 | 2.96200 |
| **DPP4** | 0.08326 | 0.94118 | 3.35700 | 9.42149 | 0.66166 | 0.00000 | 0.00000 | 0.00000 | 0.60464 | 0.00000 | 0.36005 | 4.26198 |
| **DPP6** | 1.65423 | 0.00000 | 0.03240 | 0.00000 | 0.25972 | 0.05465 | 0.04495 | 0.52133 | 0.00000 | 0.58768 | 0.00000 | 0.01112 |
| **DPP7** | 0.48455 | 0.27391 | 0.01191 | 0.17560 | 0.00000 | 0.00000 | 0.29520 | 0.00474 | 0.00000 | 0.68478 | 0.02074 | 0.08106 |
| **DPP8** | 2.52469 | 2.42371 | 3.88981 | 2.17932 | 1.74503 | 10.28702 | 5.93569 | 2.66305 | 3.55404 | 5.11612 | 3.66147 | 4.75199 |
| **DPP9** | 0.04969 | 9.31332 | 8.28614 | 6.11757 | 10.45816 | 0.00000 | 13.01075 | 0.66696 | 14.62274 | 4.03275 | 4.35017 | 4.34079 |

| | | | | | | | | | | | | |
|---|---|---|---|---|---|---|---|---|---|---|---|---|
| **DPP9-AS1** | 0.00356 | 0.11728 | 0.11432 | 0.08964 | 0.11437 | 0.00000 | 0.00048 | 0.06482 | 0.34485 | 0.06245 | 0.01466 | 0.10222 |
| **DPPA2** | 3.23656 | 2.11251 | 4.92776 | 12.57974 | 1.96901 | 4.79541 | 0.00000 | 6.58784 | 3.78177 | 0.57792 | 2.08427 | 9.41409 |
| **DPPA2P3** | 0.05733 | 0.01786 | 0.04983 | 0.15030 | 0.01965 | 0.14479 | 0.02820 | 0.04229 | 0.04281 | 0.00058 | 0.02484 | 0.13964 |
| **DPPA3** | 16.32887 | 25.83006 | 6.41864 | 12.70017 | 22.97938 | 52.64530 | 20.12045 | 28.25453 | 28.55672 | 9.24026 | 29.92796 | 46.88225 |
| **DPPA4** | 0.64023 | 13.39743 | 14.04352 | 20.27990 | 5.32678 | 25.85013 | 12.90313 | 1.09773 | 6.98207 | 9.51528 | 10.68236 | 20.74149 |
| **DPPA5** | 34.79472 | 27.56657 | 8.15360 | 37.29868 | 6.18054 | 1.99330 | 0.00000 | 42.30029 | 0.00000 | 0.51063 | 0.00000 | 39.76287 |
| **DPRX** | 0.00936 | 1.36799 | 0.27127 | 0.12844 | 0.05232 | 0.24696 | 0.00000 | 8.60004 | 0.00000 | 0.00000 | 0.04508 | 33.00348 |
| **DPRXP4** | 0.00104 | 0.17142 | 0.04913 | 0.08392 | 0.00000 | 0.03496 | 0.02434 | 0.18334 | 0.01176 | 0.95917 | 0.02640 | 1.56103 |
| **DPY19L1** | 0.09490 | 0.00000 | 1.44256 | 3.20750 | 0.01791 | 3.78738 | 0.00000 | 0.00681 | 0.28015 | 2.76467 | 0.08812 | 0.04031 |
| **DPY19L1P1** | 0.02258 | 0.03125 | 0.15422 | 0.22225 | 0.06446 | 0.19281 | 0.19795 | 0.01694 | 0.00000 | 0.77423 | 0.02214 | 0.00000 |
| **DPY19L2** | 0.00577 | 0.04660 | 0.01747 | 0.02693 | 0.00000 | 0.05623 | 0.02502 | 0.00867 | 0.01081 | 0.02899 | 0.01174 | 0.03741 |
| **DPY19L2P1** | 0.03188 | 0.04461 | 0.04370 | 0.09083 | 0.00961 | 0.08329 | 0.01352 | 0.03686 | 0.01078 | 0.00000 | 0.01354 | 0.10219 |
| **DPY19L2P2** | 0.09056 | 0.02298 | 0.07277 | 0.02262 | 0.01109 | 0.00523 | 0.01775 | 0.02859 | 0.07087 | 0.02618 | 0.00605 | 0.12146 |
| **DPY19L2P3** | 0.00765 | 0.06687 | 0.05877 | 0.01548 | 0.02133 | 0.00629 | 0.08724 | 0.01059 | 0.00000 | 0.01767 | 0.05975 | 0.03546 |
| **DPY19L2P4** | 0.02787 | 0.01809 | 0.01590 | 0.01826 | 0.01935 | 0.00372 | 0.02021 | 0.00833 | 0.02259 | 0.00799 | 0.01537 | 0.06811 |
| **DPY19L3** | 3.14890 | 0.66271 | 1.87875 | 2.50130 | 0.64731 | 1.87631 | 0.00000 | 0.48939 | 3.41974 | 3.36004 | 1.15645 | 0.84423 |
| **DPY19L4** | 0.01745 | 0.32880 | 0.00000 | 1.11023 | 0.15665 | 6.51405 | 0.00000 | 0.28187 | 0.37174 | 4.49711 | 1.26807 | 0.26598 |
| **DPY30** | 2.15271 | 12.68884 | 12.95610 | 7.35587 | 7.98109 | 30.76366 | 19.80996 | 1.19290 | 12.39371 | 7.91655 | 13.03916 | 16.45656 |
| **DPYD** | 0.04707 | 0.02017 | 0.02264 | 0.05930 | 0.02219 | 0.04346 | 0.01883 | 0.07784 | 0.00000 | 0.01731 | 0.03188 | 0.01067 |
| **DPYS** | 0.14753 | 0.01593 | 0.01473 | 0.01704 | 0.01681 | 0.01419 | 0.02573 | 0.21158 | 0.01885 | 0.00942 | 0.01635 | 0.01642 |
| **DPYSL2** | 3.43045 | 5.68917 | 2.99535 | 8.54932 | 0.00000 | 5.07990 | 4.63976 | 5.98135 | 0.00000 | 0.00000 | 2.29873 | 8.63221 |
| **DPYSL3** | 0.00184 | 0.01510 | 1.34243 | 2.09746 | 0.00000 | 0.00000 | 0.25345 | 0.00649 | 0.35204 | 0.85966 | 0.09964 | 0.02143 |
| **DPYSL4** | 0.45459 | 0.06093 | 0.02234 | 0.00834 | 0.00671 | 0.00925 | 0.01404 | 0.23782 | 0.05129 | 0.02118 | 0.00877 | 0.40962 |
| **DPYSL5** | 0.69296 | 0.29232 | 0.23588 | 0.12724 | 0.15330 | 0.12244 | 0.33686 | 0.00000 | 0.25600 | 0.83196 | 0.78853 | 0.12259 |
| **DQX1** | 0.02200 | 0.58255 | 0.08066 | 0.72487 | 0.11126 | 0.08030 | 0.01923 | 0.05033 | 0.19223 | 0.00000 | 0.15830 | 0.08823 |
| **DR1** | 1.20925 | 11.92596 | 11.84208 | 3.61836 | 6.75947 | 17.64386 | 7.08535 | 13.66065 | 17.28968 | 0.00000 | 7.20897 | 18.05238 |
| **DRAM1** | 0.00000 | 4.54849 | 6.90924 | 0.00252 | 0.00000 | 10.43599 | 0.00000 | 6.35249 | 9.88161 | 5.80060 | 0.00000 | 11.40810 |
| **DRAM2** | 0.14479 | 13.69115 | 7.16551 | 3.51436 | 1.56279 | 26.62465 | 0.00000 | 0.48910 | 11.22220 | 9.61035 | 6.69634 | 13.01837 |
| **DRAP1** | 0.65861 | 6.79439 | 2.65053 | 7.28762 | 8.50009 | 0.00218 | 3.95528 | 4.57703 | 2.48515 | 6.04855 | 2.03356 | 4.11046 |
| **DRAXIN** | 0.00586 | 0.01804 | 0.04349 | 0.11016 | 0.02254 | 0.00000 | 0.00989 | 0.00923 | 0.02995 | 0.02614 | 0.01364 | 0.03654 |
| **DRC1** | 0.00400 | 0.02427 | 0.03587 | 0.05350 | 0.00558 | 0.00000 | 0.01070 | 0.18986 | 0.07626 | 0.13557 | 0.01121 | 0.01403 |
| **DRD1** | 0.00549 | 0.00471 | 0.01462 | 0.00370 | 0.00000 | 0.06493 | 0.10933 | 0.48091 | 0.00000 | 0.01074 | 0.01149 | 0.12425 |
| **DRD2** | 0.00522 | 0.01182 | 0.00451 | 0.23910 | 0.00182 | 0.02841 | 0.00395 | 0.01082 | 0.00657 | 0.10422 | 0.01493 | 0.01507 |
| **DRD3** | 0.00867 | 0.01521 | 0.03098 | 0.01302 | 0.01548 | 0.01319 | 0.01316 | 0.03655 | 0.01738 | 0.02004 | 0.02381 | 0.01422 |
| **DRD4** | 0.00261 | 0.00000 | 0.00000 | 0.01656 | 0.19450 | 0.00000 | 0.04859 | 0.00760 | 0.08016 | 0.67089 | 0.04786 | 0.02947 |
| **DRD5P2+LOC1(** | 0.01092 | 0.02852 | 0.01539 | 0.01181 | 0.01361 | 0.01543 | 0.02323 | 0.00971 | 0.02659 | 0.00699 | 0.01352 | 0.01941 |
| **DRG1** | 2.20956 | 16.83831 | 19.58371 | 11.96417 | 14.87784 | 27.46459 | 32.21755 | 14.09005 | 21.14395 | 5.42694 | 23.38419 | 21.45270 |
| **DRG2** | 0.82418 | 11.84478 | 12.56276 | 6.25689 | 10.01821 | 3.33012 | 11.07716 | 1.90307 | 15.58452 | 0.00000 | 11.48298 | 9.96260 |
| **DRGX** | 0.02816 | 0.01110 | 0.00813 | 0.00600 | 0.01124 | 0.02302 | 0.00002 | 0.04461 | 0.05248 | 0.06540 | 0.00443 | 0.01444 |
| **DRICH1** | 0.01476 | 0.03456 | 0.06913 | 0.00000 | 0.00000 | 0.12727 | 0.00000 | 0.00632 | 0.10573 | 0.27063 | 0.04791 | 0.06167 |
| **DROSHA** | 1.78239 | 1.80477 | 6.14965 | 2.68638 | 2.72160 | 9.75035 | 5.37823 | 0.00000 | 12.19820 | 19.74895 | 5.71883 | 0.35669 |
| **DRP2** | 0.00175 | 0.00588 | 0.03281 | 0.00761 | 0.00000 | 0.03137 | 0.07452 | 0.08867 | 0.04553 | 0.01598 | 0.06687 | 0.07528 |
| **DSC1** | 0.04896 | 0.05009 | 0.33922 | 0.00439 | 0.12313 | 0.00000 | 0.00000 | 0.10025 | 0.15857 | 0.01093 | 0.01267 | 0.04172 |
| **DSC2** | 0.00000 | 5.74837 | 29.83468 | 5.05906 | 8.26115 | 19.58665 | 0.00002 | 0.00000 | 26.05355 | 9.63644 | 7.38448 | 0.00000 |
| **DSC3** | 0.03833 | 0.07927 | 2.76309 | 0.00948 | 0.00000 | 0.00000 | 0.30803 | 0.10222 | 1.05590 | 0.24364 | 0.00000 | 0.00000 |
| **DSCAM** | 0.35075 | 0.01814 | 0.01534 | 0.00257 | 0.04334 | 0.00000 | 0.01018 | 0.03672 | 0.00000 | 0.16675 | 0.05233 | 0.01523 |
| **DSCAML1** | 0.64206 | 1.52557 | 0.00253 | 0.77429 | 0.01669 | 0.06166 | 0.03638 | 0.04341 | 0.00000 | 0.00000 | 0.26695 | 0.00191 |
| **DSCC1** | 0.58929 | 7.13638 | 5.99835 | 3.07089 | 0.00000 | 18.32748 | 5.06067 | 3.12463 | 8.48458 | 1.14222 | 5.84441 | 7.23534 |

| | | | | | | | | | | | | |
|---|---|---|---|---|---|---|---|---|---|---|---|---|
| **DSCR10** | 0.10138 | 0.01686 | 0.01190 | 0.01267 | 0.02068 | 0.00271 | 0.04708 | 0.19646 | 0.02288 | 0.02255 | 0.01217 | 0.00808 |
| **DSCR3** | 1.85046 | 5.84427 | 8.13143 | 5.00219 | 2.05003 | 5.74386 | 7.84615 | 0.66197 | 12.28652 | 0.00000 | 11.27389 | 1.08243 |
| **DSCR4** | 0.40954 | 0.80375 | 1.26740 | 0.28934 | 0.84845 | 1.26752 | 0.81698 | 0.05006 | 1.28300 | 0.00000 | 0.95161 | 0.16447 |
| **DSCR8** | 2.00740 | 2.42766 | 2.84689 | 1.84289 | 7.27576 | 7.27658 | 1.58921 | 0.48246 | 1.65922 | 1.80662 | 3.13324 | 0.00000 |
| **DSCR9** | 0.15223 | 0.21404 | 0.27963 | 0.00000 | 0.10737 | 0.21751 | 0.24246 | 0.75947 | 0.28933 | 0.01128 | 0.20549 | 0.29500 |
| **DSE** | 0.00000 | 0.00000 | 2.06841 | 1.32749 | 1.04894 | 1.44088 | 0.55142 | 1.31799 | 0.00000 | 0.74169 | 0.00000 | 0.17036 |
| **DSEL** | 0.00140 | 0.94280 | 1.85466 | 1.02449 | 0.00000 | 0.00000 | 0.12612 | 3.89480 | 0.00000 | 0.80503 | 0.00000 | 2.36454 |
| **DSG1** | 0.07669 | 0.09202 | 0.02342 | 0.00000 | 0.00000 | 0.04045 | 0.02300 | 0.00000 | 0.07737 | 0.05272 | 0.04220 | 0.43114 |
| **DSG2** | 0.00000 | 6.07257 | 7.12939 | 3.44486 | 11.11609 | 34.71826 | 2.79165 | 0.32317 | 29.30498 | 33.92825 | 13.47848 | 1.79826 |
| **DSG3** | 0.00290 | 0.00000 | 1.90628 | 0.00000 | 0.19162 | 0.43783 | 0.00000 | 0.00121 | 1.08084 | 0.38797 | 0.86084 | 0.14946 |
| **DSG4** | 0.00627 | 0.01470 | 0.04900 | 0.01036 | 0.01244 | 0.01406 | 0.01399 | 0.01034 | 0.00461 | 0.02354 | 0.03900 | 0.01797 |
| **DSN1** | 2.56238 | 9.03703 | 3.87113 | 4.44400 | 6.80498 | 16.80503 | 3.95097 | 2.17224 | 1.72122 | 0.00083 | 11.56667 | 7.13122 |
| **DSP** | 0.00000 | 7.33885 | 20.31123 | 11.67414 | 18.59850 | 29.61585 | 24.36725 | 0.57962 | 25.34767 | 28.49739 | 9.02163 | 6.92339 |
| **DSPP** | 0.00629 | 0.01637 | 0.02925 | 0.02240 | 0.01445 | 0.00471 | 0.01227 | 0.01069 | 0.03358 | 0.02503 | 0.01465 | 0.01648 |
| **DST** | 0.71024 | 1.58109 | 1.73587 | 18.13959 | 10.20480 | 13.23345 | 0.00000 | 1.16076 | 0.00000 | 17.63839 | 0.89119 | 1.44632 |
| **DSTN** | 8.07565 | 8.69686 | 16.76890 | 12.86728 | 13.12410 | 35.78860 | 28.29054 | 2.42459 | 19.38010 | 11.25905 | 18.47682 | 5.42978 |
| **DSTNP2** | 0.11146 | 1.89179 | 0.20210 | 5.76918 | 0.00000 | 3.70891 | 0.97981 | 0.67083 | 0.02760 | 1.17336 | 0.80978 | 2.32976 |
| **DSTYK** | 1.58250 | 0.58170 | 0.79677 | 1.74267 | 1.58449 | 1.11843 | 1.59934 | 0.30978 | 0.67938 | 3.37678 | 1.10825 | 1.09770 |
| **DTD1** | 1.18485 | 8.71548 | 2.43231 | 5.80148 | 5.60858 | 12.11613 | 4.33132 | 1.43062 | 3.30097 | 0.00000 | 6.02988 | 1.47342 |
| **DTD2** | 0.10027 | 0.48542 | 0.28869 | 0.51726 | 0.96199 | 2.74449 | 0.00000 | 0.00000 | 0.00000 | 0.84680 | 1.63248 | 0.71020 |
| **DTL** | 8.90424 | 12.85716 | 5.04072 | 13.68453 | 3.97235 | 25.46187 | 6.10717 | 10.57472 | 10.53878 | 3.10705 | 13.26006 | 21.61275 |
| **DTNA** | 0.00242 | 0.12943 | 0.21689 | 0.03687 | 0.00990 | 0.00953 | 0.17083 | 0.04216 | 0.03048 | 0.00007 | 0.02192 | 0.01610 |
| **DTNB** | 1.07687 | 5.53264 | 0.00000 | 6.80657 | 5.01403 | 1.23363 | 0.20050 | 1.13935 | 4.82754 | 1.92170 | 2.96020 | 2.96542 |
| **DTNBP1** | 1.73708 | 1.62677 | 1.54063 | 1.62109 | 1.24231 | 1.85976 | 2.90413 | 0.42842 | 0.00000 | 0.33371 | 0.96980 | 1.38111 |
| **DTWD1** | 0.08258 | 5.09894 | 4.45315 | 1.14418 | 0.00000 | 7.93864 | 2.22011 | 0.00000 | 6.09648 | 0.91308 | 2.03308 | 1.62202 |
| **DTWD2** | 0.88520 | 1.21836 | 0.64157 | 1.17507 | 0.00000 | 1.14220 | 0.23391 | 0.38732 | 0.13651 | 1.26670 | 0.28506 | 0.65120 |
| **DTX1** | 0.00360 | 0.00536 | 0.01210 | 0.00918 | 0.00556 | 0.03079 | 0.01447 | 0.02069 | 0.08627 | 0.03476 | 0.00173 | 0.14254 |
| **DTX2** | 5.58178 | 4.44367 | 3.31286 | 1.69766 | 3.13483 | 0.00000 | 4.48843 | 0.07672 | 2.43203 | 2.34081 | 1.73456 | 3.79015 |
| **DTX2P1-UPK3B** | 0.19274 | 0.25761 | 0.21617 | 0.07363 | 0.07318 | 0.00000 | 0.04356 | 0.01050 | 0.25043 | 0.00000 | 0.01592 | 0.27713 |
| **DTX3** | 0.03486 | 0.01112 | 0.01471 | 0.13891 | 0.00000 | 0.01483 | 0.06317 | 0.01397 | 0.00000 | 0.13921 | 0.04439 | 0.01105 |
| **DTX3L** | 0.00547 | 0.14362 | 0.05779 | 0.34121 | 0.09783 | 0.26403 | 0.02212 | 0.00922 | 0.00000 | 0.00000 | 0.01652 | 0.00334 |
| **DTX4** | 0.00542 | 0.00000 | 2.07759 | 0.16608 | 7.99702 | 0.00050 | 12.11171 | 0.00457 | 6.32861 | 3.96374 | 0.00005 | 0.00243 |
| **DTYMK** | 0.00000 | 20.82898 | 16.36408 | 12.60876 | 17.52474 | 7.36252 | 24.43695 | 6.09787 | 19.30822 | 8.06076 | 21.09341 | 20.69598 |
| **DUOX1** | 0.00448 | 0.02189 | 0.42593 | 0.00000 | 0.00000 | 0.07103 | 0.00710 | 0.00675 | 0.05164 | 0.11265 | 0.08314 | 0.11183 |
| **DUOX2** | 0.05252 | 0.01036 | 0.06637 | 0.08691 | 0.02001 | 0.04225 | 0.00000 | 0.00635 | 0.01972 | 0.04539 | 0.00858 | 0.01611 |
| **DUOXA1** | 0.02006 | 0.01249 | 0.80733 | 0.00336 | 0.04934 | 0.00000 | 0.00000 | 0.01145 | 0.00000 | 0.09849 | 0.03689 | 0.02794 |
| **DUPD1** | 0.00392 | 0.01868 | 0.01094 | 0.01325 | 0.02507 | 0.00000 | 0.03368 | 0.03784 | 0.02745 | 0.03952 | 0.01940 | 0.02302 |
| **DUS1L** | 0.21895 | 3.78395 | 3.20480 | 2.35267 | 10.38335 | 0.00000 | 7.35851 | 0.00000 | 2.96105 | 18.32846 | 4.25389 | 2.93925 |
| **DUS2** | 0.00000 | 9.58991 | 8.52655 | 6.04797 | 6.39985 | 4.96793 | 14.77980 | 1.89172 | 14.90032 | 0.00000 | 10.23364 | 8.99128 |
| **DUS3L** | 0.60241 | 17.74300 | 11.65664 | 10.41576 | 8.26650 | 0.00000 | 8.29185 | 4.39566 | 18.17503 | 0.00000 | 12.42400 | 15.17261 |
| **DUS4L** | 2.14280 | 5.58189 | 1.01186 | 0.30516 | 1.28583 | 4.31933 | 0.92852 | 0.00000 | 1.14435 | 0.00000 | 1.67464 | 4.01149 |
| **DUSP1** | 0.24174 | 9.02584 | 1.02192 | 0.00000 | 4.70088 | 6.34642 | 1.53022 | 0.04366 | 0.39893 | 1.74903 | 2.34719 | 0.40366 |
| **DUSP10** | 3.96098 | 0.08448 | 0.11126 | 1.59160 | 0.00000 | 0.10531 | 0.00000 | 0.38443 | 0.00000 | 0.10553 | 0.00860 | 0.29524 |
| **DUSP11** | 1.32123 | 6.51588 | 6.12016 | 2.31528 | 0.11439 | 21.61511 | 5.82284 | 7.08630 | 5.39500 | 0.00000 | 7.85704 | 10.62210 |
| **DUSP12** | 0.48900 | 11.44363 | 1.27649 | 5.76350 | 3.77426 | 13.58018 | 4.07889 | 18.53888 | 2.67852 | 0.00001 | 3.58724 | 14.75187 |
| **DUSP13** | 0.02204 | 0.00000 | 1.59964 | 0.00000 | 0.15849 | 0.09331 | 0.92214 | 0.00499 | 0.43733 | 0.00000 | 0.05188 | 0.18899 |
| **DUSP14** | 3.50680 | 22.37837 | 10.06326 | 4.81558 | 9.36741 | 17.53098 | 16.74414 | 10.76615 | 19.23356 | 0.00000 | 11.62856 | 25.12957 |
| **DUSP15** | 0.00298 | 0.04958 | 0.01915 | 0.04640 | 0.01430 | 0.01979 | 0.01282 | 0.04153 | 0.01675 | 0.00935 | 0.02870 | 0.07030 |

| | | | | | | | | | | | | |
|---|---|---|---|---|---|---|---|---|---|---|---|---|
| **DUSP16** | 3.28152 | 6.09790 | 0.24597 | 7.52197 | 5.70784 | 3.61052 | 1.30319 | 0.62343 | 0.00000 | 5.72770 | 2.72164 | 4.33091 |
| **DUSP18** | 5.39033 | 1.96761 | 1.04770 | 2.44049 | 2.44666 | 0.97948 | 1.98098 | 8.19708 | 1.02499 | 0.00000 | 0.98057 | 11.47512 |
| **DUSP19** | 4.09417 | 0.79731 | 0.08244 | 0.00000 | 0.00000 | 0.56880 | 0.00000 | 0.30218 | 0.00000 | 0.25841 | 0.35291 | 1.21925 |
| **DUSP2** | 0.05537 | 0.01580 | 0.06092 | 0.01795 | 0.08049 | 0.00857 | 0.01232 | 0.01227 | 0.02641 | 0.00067 | 0.01903 | 0.01623 |
| **DUSP22** | 5.96754 | 2.48684 | 0.58288 | 1.52234 | 1.75580 | 0.18883 | 2.25838 | 0.42947 | 0.00000 | 2.81500 | 0.79502 | 1.60274 |
| **DUSP23** | 0.73850 | 4.38864 | 1.30768 | 4.70600 | 4.69539 | 0.12440 | 6.00797 | 0.86563 | 3.24313 | 0.00000 | 2.16816 | 5.10418 |
| **DUSP27** | 0.00117 | 3.68655 | 0.42587 | 0.37092 | 1.67261 | 0.00000 | 0.00000 | 0.36181 | 0.00000 | 0.00003 | 0.46944 | 10.05304 |
| **DUSP28** | 0.30818 | 0.41563 | 0.50263 | 0.14929 | 1.13013 | 0.00000 | 0.51404 | 0.01041 | 0.41592 | 0.00000 | 0.54921 | 0.35799 |
| **DUSP3** | 2.58810 | 7.93622 | 5.67060 | 5.52367 | 5.69829 | 5.47290 | 4.87765 | 2.26726 | 3.20132 | 0.00000 | 2.38727 | 9.89156 |
| **DUSP4** | 0.00031 | 5.54199 | 2.44674 | 11.72849 | 4.22021 | 0.00000 | 0.85369 | 0.14008 | 0.00000 | 1.04272 | 0.00000 | 5.24325 |
| **DUSP5** | 3.85706 | 3.51793 | 17.91414 | 1.87955 | 4.07365 | 0.00000 | 8.07053 | 1.64356 | 0.00000 | 6.94370 | 0.00000 | 6.71859 |
| **DUSP5P1** | 0.00723 | 0.00000 | 0.01015 | 0.00000 | 0.00000 | 0.02711 | 0.10225 | 0.00929 | 0.14797 | 0.87465 | 0.00000 | 0.00817 |
| **DUSP6** | 2.04951 | 0.00000 | 3.04832 | 6.27031 | 10.37241 | 1.84665 | 14.53776 | 0.15577 | 18.35218 | 2.72732 | 5.11785 | 0.00000 |
| **DUSP7** | 8.66034 | 3.19314 | 0.32308 | 4.04357 | 2.75381 | 0.87972 | 2.37171 | 1.32335 | 0.86970 | 4.76077 | 0.84443 | 3.06783 |
| **DUSP8** | 1.00005 | 0.01858 | 0.38156 | 0.01920 | 0.45778 | 0.00000 | 0.01951 | 0.02573 | 0.00000 | 0.18297 | 0.02611 | 0.14073 |
| **DUSP9** | 0.00187 | 0.34208 | 0.32218 | 0.38759 | 5.98161 | 0.00000 | 0.12112 | 0.00523 | 0.00000 | 2.12283 | 0.00000 | 0.01742 |
| **DUT** | 1.51945 | 16.65317 | 14.40246 | 5.87895 | 12.39868 | 33.16060 | 18.04343 | 6.45188 | 14.29897 | 15.27785 | 20.05081 | 22.18007 |
| **DUXA** | 5.07462 | 0.00000 | 0.00095 | 0.00120 | 0.02182 | 0.01346 | 0.00251 | 36.73188 | 0.00456 | 0.01615 | 0.00000 | 31.42445 |
| **DUXAP8** | 0.02404 | 0.08423 | 0.07564 | 0.00767 | 0.00000 | 0.00929 | 0.16055 | 0.04838 | 0.00028 | 0.03494 | 0.01862 | 0.08093 |
| **DVL1** | 1.03199 | 0.53564 | 3.06319 | 0.10439 | 5.72182 | 0.00001 | 8.74745 | 0.00000 | 2.95550 | 14.12444 | 1.59572 | 0.01844 |
| **DVL2** | 0.92406 | 4.39750 | 3.96578 | 6.14035 | 3.73579 | 0.00000 | 1.87420 | 0.40880 | 4.76794 | 4.33796 | 7.08113 | 1.38732 |
| **DVL3** | 1.32315 | 2.62530 | 12.85236 | 1.96106 | 13.02920 | 3.74674 | 8.87320 | 0.00000 | 9.24652 | 20.82788 | 0.98599 | 3.89935 |
| **DXO** | 0.42594 | 1.55315 | 4.41911 | 1.62778 | 0.00000 | 0.00000 | 3.99699 | 1.26764 | 3.91196 | 0.00000 | 4.46459 | 2.04948 |
| **DYDC2** | 0.06307 | 0.10379 | 0.00417 | 0.00000 | 0.03247 | 0.08925 | 0.00942 | 0.18723 | 0.07482 | 0.00693 | 0.00003 | 0.05280 |
| **DYM** | 4.63879 | 3.25724 | 5.64216 | 3.80371 | 5.60764 | 8.05935 | 4.88505 | 0.36466 | 9.66056 | 0.00000 | 4.12711 | 0.77335 |
| **DYNAP** | 0.06911 | 0.01090 | 0.00536 | 0.00562 | 0.00659 | 0.04003 | 0.00606 | 0.00971 | 0.01516 | 0.12787 | 0.00850 | 0.04265 |
| **DYNC1H1** | 0.64242 | 8.88431 | 9.58705 | 13.33899 | 15.23640 | 9.57191 | 12.85224 | 2.04527 | 15.29410 | 23.71653 | 6.08903 | 2.42320 |
| **DYNC1I1** | 0.31332 | 0.40231 | 0.25797 | 0.12189 | 0.00592 | 0.17368 | 0.13348 | 0.11813 | 0.00000 | 0.00000 | 0.02114 | 0.07714 |
| **DYNC1I2** | 8.02398 | 5.27537 | 3.91968 | 3.42205 | 6.34857 | 9.06873 | 9.49961 | 3.40541 | 2.77882 | 12.68969 | 5.81755 | 1.60809 |
| **DYNC1LI1** | 1.15651 | 9.62959 | 8.10796 | 4.88330 | 7.17202 | 16.93777 | 13.70626 | 7.58064 | 11.26943 | 4.85591 | 5.47499 | 29.87359 |
| **DYNC1LI2** | 6.16337 | 5.66226 | 13.84621 | 7.04775 | 8.84193 | 18.52252 | 18.71537 | 1.84690 | 11.08199 | 15.96328 | 10.88604 | 14.74468 |
| **DYNC2H1** | 7.15188 | 0.02039 | 0.00000 | 0.75718 | 0.08048 | 0.52002 | 0.74841 | 2.52246 | 1.22843 | 2.78281 | 0.17553 | 1.04812 |
| **DYNC2LI1** | 1.46668 | 2.41860 | 0.58500 | 0.00000 | 0.57646 | 3.31031 | 0.00000 | 0.00000 | 0.49105 | 0.18539 | 1.62373 | 1.05674 |
| **DYNLL1** | 7.39240 | 11.47838 | 14.66899 | 8.09510 | 9.80938 | 19.57038 | 22.46967 | 14.51976 | 21.39464 | 2.74134 | 4.69750 | 18.16246 |
| **DYNLL1-AS1** | 1.79265 | 1.01282 | 1.06818 | 0.00000 | 0.00000 | 0.13754 | 2.08812 | 11.18922 | 2.29685 | 0.00000 | 0.75908 | 7.47683 |
| **DYNLL2** | 3.84503 | 0.58120 | 0.90133 | 0.28634 | 5.47798 | 0.00000 | 3.37725 | 6.50967 | 0.00000 | 11.59571 | 1.57905 | 1.45037 |
| **DYNLRB1** | 3.48520 | 10.00819 | 10.21568 | 9.19447 | 13.56079 | 7.77342 | 14.93965 | 1.00848 | 11.56937 | 4.24554 | 11.93129 | 5.45031 |
| **DYNLRB2** | 0.13537 | 0.16282 | 0.06326 | 0.00033 | 0.00165 | 0.12908 | 0.07091 | 0.05080 | 0.06919 | 0.00406 | 0.00306 | 0.05275 |
| **DYNLT1** | 0.60681 | 10.43252 | 8.83292 | 4.92851 | 3.68759 | 25.94443 | 11.03192 | 3.86922 | 5.26699 | 0.01969 | 13.48700 | 16.65002 |
| **DYNLT3** | 0.12695 | 2.01967 | 3.03115 | 3.23016 | 0.00000 | 2.34121 | 0.65680 | 0.87596 | 0.08669 | 0.00000 | 0.00000 | 3.68998 |
| **DYRK1A** | 6.05134 | 1.66009 | 3.55431 | 2.02670 | 0.00000 | 8.98868 | 5.47627 | 0.40155 | 0.00000 | 11.78850 | 3.10628 | 3.27072 |
| **DYRK1B** | 0.15917 | 0.26346 | 0.13515 | 0.29797 | 0.99266 | 0.00000 | 0.06270 | 0.04367 | 0.73370 | 1.79368 | 0.51921 | 0.66085 |
| **DYRK2** | 1.71351 | 0.66517 | 2.50310 | 0.19562 | 1.79812 | 1.25620 | 4.80265 | 2.76884 | 1.39543 | 6.41237 | 0.00000 | 2.33634 |
| **DYRK3** | 0.30884 | 0.01433 | 0.24766 | 0.00825 | 0.00000 | 0.07078 | 0.00000 | 0.06889 | 0.00000 | 0.06778 | 0.00937 | 0.06677 |
| **DYRK4** | 0.50200 | 3.41948 | 0.00000 | 0.96056 | 0.93865 | 3.61299 | 0.71922 | 0.06523 | 0.00000 | 0.00000 | 0.03436 | 2.16316 |
| **DYSF** | 4.06288 | 0.00000 | 5.14977 | 3.62693 | 13.63231 | 0.00000 | 8.55334 | 0.92820 | 10.17894 | 8.47113 | 9.71348 | 0.01931 |
| **DYTN** | 0.01086 | 0.01273 | 0.01374 | 0.01174 | 0.01554 | 0.01829 | 0.01694 | 0.00832 | 0.02092 | 0.01576 | 0.01513 | 0.03746 |
| **DYX1C1** | 0.11929 | 0.23121 | 0.05900 | 0.05608 | 0.00000 | 0.23676 | 0.06526 | 0.12188 | 0.00000 | 0.04961 | 0.08718 | 0.40138 |

| | | | | | | | | | | | | |
|---|---|---|---|---|---|---|---|---|---|---|---|---|
| **DZANK1** | 1.27217 | 0.80366 | 0.64070 | 1.63314 | 1.15989 | 0.35655 | 0.63241 | 0.16324 | 0.00000 | 0.00000 | 1.42591 | 0.54146 |
| **DZIP1** | 3.66008 | 0.15072 | 0.16296 | 0.28911 | 0.85864 | 0.00000 | 0.04107 | 0.00000 | 0.00000 | 0.62097 | 0.08270 | 0.27850 |
| **DZIP1L** | 0.11174 | 0.28156 | 0.09564 | 0.95599 | 0.00000 | 0.06379 | 0.00000 | 0.00000 | 0.00000 | 0.07972 | 0.00937 | 0.90545 |
| **DZIP3** | 2.18216 | 3.83371 | 1.57467 | 0.80475 | 2.82032 | 6.97928 | 0.71106 | 0.59613 | 1.03020 | 3.32595 | 1.75323 | 1.57102 |
| **E2F1** | 2.57208 | 4.47661 | 2.45398 | 3.22559 | 6.78749 | 0.00000 | 1.04323 | 0.04652 | 4.12523 | 7.91853 | 0.05156 | 2.31145 |
| **E2F2** | 0.00000 | 3.25471 | 2.63301 | 2.11268 | 3.52846 | 0.34224 | 1.61688 | 0.05029 | 0.97718 | 1.39496 | 0.21363 | 0.85690 |
| **E2F3** | 0.00000 | 0.56005 | 2.47506 | 0.90055 | 3.33936 | 4.73864 | 5.34257 | 0.37400 | 2.98716 | 10.56534 | 1.95729 | 1.05849 |
| **E2F4** | 1.37680 | 10.84483 | 11.78523 | 8.40049 | 11.52191 | 0.00112 | 9.82485 | 0.00000 | 12.84955 | 19.02754 | 7.69370 | 7.98280 |
| **E2F5** | 0.00440 | 0.10802 | 0.27961 | 0.00000 | 0.00000 | 0.34675 | 0.75307 | 0.04074 | 0.44990 | 0.42571 | 0.22053 | 0.05803 |
| **E2F6** | 0.00000 | 2.64438 | 2.21952 | 0.50479 | 0.81923 | 0.00000 | 1.83538 | 2.21690 | 1.48707 | 0.99286 | 1.93185 | 1.34120 |
| **E2F7** | 0.29453 | 0.49657 | 3.35211 | 0.49639 | 0.00000 | 3.23434 | 2.33683 | 0.00000 | 2.61076 | 2.44796 | 0.17411 | 0.52590 |
| **E2F8** | 0.65438 | 2.27743 | 10.00515 | 1.22930 | 2.67477 | 12.10100 | 2.49311 | 0.04473 | 7.88686 | 0.93902 | 1.89957 | 2.51834 |
| **E4F1** | 2.73960 | 13.94311 | 6.39952 | 8.42034 | 7.86927 | 0.00000 | 3.35633 | 5.97092 | 8.11309 | 9.72332 | 6.59069 | 13.32734 |
| **EAF1** | 0.46355 | 6.43915 | 17.40232 | 2.08498 | 0.00000 | 18.70633 | 18.97065 | 12.55684 | 4.96535 | 0.87655 | 10.55827 | 8.12701 |
| **EAF2** | 0.00000 | 0.26333 | 3.64277 | 0.33415 | 1.45345 | 2.32224 | 4.76832 | 0.02227 | 0.05708 | 1.25598 | 5.87687 | 0.89544 |
| **EAPP** | 0.00000 | 3.82655 | 6.29809 | 5.18180 | 2.22699 | 22.01178 | 11.05539 | 5.08993 | 4.75390 | 4.61717 | 8.20884 | 11.72552 |
| **EARS2** | 1.14155 | 13.80834 | 8.72970 | 9.19133 | 8.70799 | 6.80644 | 8.44177 | 1.52736 | 9.57710 | 0.00000 | 10.34813 | 8.80875 |
| **EBAG9** | 4.98540 | 6.53466 | 5.07662 | 1.03826 | 3.95397 | 15.79738 | 1.99406 | 0.47002 | 6.72410 | 7.26631 | 3.86762 | 4.05556 |
| **EBF1** | 5.84631 | 0.01127 | 0.02445 | 0.08487 | 0.00000 | 0.03854 | 0.01005 | 1.15005 | 0.02471 | 0.00340 | 0.00725 | 0.03092 |
| **EBF2** | 5.52150 | 0.02113 | 0.02227 | 0.01115 | 0.01206 | 0.02631 | 0.00941 | 0.93047 | 0.01407 | 0.00643 | 0.01236 | 0.01121 |
| **EBF3** | 0.07698 | 0.13934 | 0.00804 | 0.01697 | 0.01124 | 0.06838 | 0.00000 | 0.23068 | 0.01131 | 0.03004 | 0.00731 | 0.06235 |
| **EBF4** | 0.02597 | 0.00000 | 0.01180 | 0.09861 | 0.00000 | 0.04265 | 0.00000 | 0.00877 | 0.01153 | 0.16124 | 0.01338 | 0.11082 |
| **EBI3** | 0.00231 | 0.05635 | 1.05174 | 0.03440 | 0.01895 | 0.00000 | 0.00000 | 0.04966 | 0.57936 | 0.00000 | 0.00389 | 0.05655 |
| **EBLN1** | 8.66077 | 0.02200 | 0.02974 | 0.00552 | 0.00718 | 0.00832 | 0.00397 | 0.59544 | 0.11268 | 0.00041 | 0.02113 | 0.02758 |
| **EBLN2** | 0.03109 | 0.01744 | 0.18151 | 0.02261 | 0.23063 | 1.34167 | 0.00000 | 0.26145 | 1.12689 | 3.81344 | 0.00000 | 0.56232 |
| **EBLN3** | 2.35641 | 3.84619 | 4.91320 | 2.51772 | 1.37336 | 10.76838 | 5.47272 | 11.13665 | 4.38588 | 5.28088 | 3.01869 | 15.65370 |
| **EBNA1BP2** | 0.84990 | 24.69372 | 25.97398 | 12.67639 | 24.38407 | 35.02090 | 42.16329 | 27.87834 | 33.89221 | 10.15716 | 25.85169 | 32.99617 |
| **EBP** | 6.62188 | 17.83064 | 18.35460 | 24.49288 | 44.05399 | 23.03972 | 16.36231 | 11.89605 | 48.01150 | 9.75205 | 33.52403 | 13.23281 |
| **EBPL** | 0.76835 | 3.99198 | 6.70092 | 1.88016 | 3.58541 | 3.79802 | 1.83929 | 0.15447 | 9.51531 | 4.83395 | 2.40349 | 2.86128 |
| **ECD** | 0.00000 | 7.07385 | 8.57210 | 1.34202 | 1.10521 | 10.53629 | 5.63632 | 4.82498 | 7.78612 | 0.00000 | 10.15330 | 8.43955 |
| **ECE1** | 0.97344 | 5.26169 | 5.18785 | 4.59626 | 3.04700 | 0.00000 | 1.08467 | 0.73616 | 6.07081 | 8.19869 | 2.40789 | 0.47189 |
| **ECE2** | 0.15927 | 11.39340 | 13.96885 | 5.18224 | 7.95090 | 1.31745 | 15.94923 | 2.63997 | 21.44002 | 0.00000 | 9.58851 | 17.74756 |
| **ECEL1** | 0.00377 | 0.03229 | 0.08282 | 0.00661 | 0.06744 | 0.00000 | 0.02307 | 0.01638 | 0.00000 | 0.09284 | 0.01203 | 0.01277 |
| **ECEL1P2** | 0.08309 | 0.00000 | 0.80864 | 0.00000 | 1.78139 | 0.00000 | 3.93158 | 0.00962 | 0.47968 | 1.14422 | 4.65660 | 0.01143 |
| **ECH1** | 2.01948 | 11.02229 | 8.70775 | 18.45948 | 14.43139 | 6.83405 | 16.83306 | 2.75416 | 9.96675 | 1.53485 | 20.64283 | 11.21943 |
| **ECHDC1** | 0.13574 | 2.57909 | 10.73868 | 0.68563 | 0.02148 | 16.87726 | 9.82296 | 2.42886 | 10.64019 | 0.17347 | 2.83096 | 6.73659 |
| **ECHDC2** | 0.04293 | 1.78161 | 3.44385 | 0.43953 | 2.35435 | 1.07399 | 3.04601 | 0.16899 | 5.70325 | 4.58344 | 5.32946 | 1.58734 |
| **ECHDC3** | 0.00886 | 0.00015 | 0.53321 | 0.00000 | 0.85834 | 0.16413 | 0.31443 | 0.03890 | 1.76614 | 0.94221 | 0.21589 | 0.01504 |
| **ECHS1** | 4.03819 | 20.05155 | 18.32430 | 19.77594 | 33.17007 | 18.03714 | 45.58614 | 1.66450 | 36.77872 | 9.61370 | 34.76209 | 1.82896 |
| **ECI1** | 0.00000 | 7.50717 | 3.76489 | 3.76276 | 15.28651 | 0.00000 | 15.00612 | 0.06277 | 5.60383 | 16.50029 | 3.73656 | 2.14909 |
| **ECI2** | 0.82420 | 0.00000 | 0.19704 | 2.10383 | 0.00000 | 4.99343 | 8.20851 | 0.81769 | 12.03745 | 0.00000 | 3.65264 | 1.50310 |
| **ECM1** | 0.00379 | 0.06408 | 1.25468 | 0.00000 | 0.10930 | 0.84226 | 0.00000 | 0.00000 | 0.21277 | 0.26354 | 0.54821 | 0.32073 |
| **ECM2** | 0.20393 | 0.14118 | 0.28662 | 0.35632 | 0.21539 | 0.40165 | 0.00000 | 0.13533 | 0.04241 | 0.56342 | 0.13942 | 0.08408 |
| **ECSCR** | 0.45721 | 0.01403 | 0.01464 | 0.01324 | 0.01494 | 0.01350 | 0.01785 | 0.00607 | 0.02064 | 0.01125 | 0.01386 | 0.01189 |
| **ECSIT** | 2.65275 | 13.55794 | 8.08726 | 12.14976 | 15.48346 | 1.47353 | 17.00992 | 2.13472 | 19.96103 | 0.00000 | 9.24142 | 10.01439 |
| **ECT2** | 10.47614 | 7.47995 | 6.41152 | 7.38498 | 0.14513 | 31.22842 | 8.68241 | 8.61788 | 8.71692 | 10.11549 | 3.70847 | 13.07309 |
| **ECT2L** | 0.06284 | 0.03558 | 0.13069 | 0.01055 | 0.03999 | 0.04436 | 0.00328 | 0.09267 | 0.03906 | 0.58793 | 0.01118 | 0.04916 |
| **EDA** | 1.14949 | 1.08741 | 0.10055 | 0.68784 | 0.00000 | 0.00000 | 0.00000 | 0.76577 | 0.16963 | 0.36530 | 0.09501 | 1.37824 |

| | | | | | | | | | | | | |
|---|---|---|---|---|---|---|---|---|---|---|---|---|
| **EDA2R** | 0.06699 | 1.71470 | 1.59532 | 1.35524 | 0.09046 | 1.43996 | 0.00000 | 0.00000 | 3.49531 | 0.00000 | 1.21002 | 0.02287 |
| **EDAR** | 1.98505 | 0.00920 | 0.03818 | 0.01004 | 0.01452 | 0.00000 | 0.00000 | 0.00451 | 0.32401 | 0.02611 | 0.00000 | 0.04037 |
| **EDARADD** | 0.01157 | 25.89335 | 24.03856 | 23.67699 | 17.05352 | 33.72924 | 25.19462 | 8.40234 | 35.98608 | 0.00000 | 27.08432 | 23.87024 |
| **EDC3** | 1.91210 | 10.11150 | 9.13556 | 6.69329 | 6.71497 | 2.47675 | 15.78274 | 9.69166 | 10.40965 | 0.00000 | 9.71923 | 15.48370 |
| **EDC4** | 0.92374 | 9.90036 | 8.95210 | 4.75069 | 12.37213 | 0.00000 | 7.90639 | 0.04809 | 10.94256 | 15.28093 | 3.22408 | 4.46298 |
| **EDDM3A** | 0.00636 | 0.01299 | 0.04438 | 0.00947 | 0.00872 | 0.03400 | 0.03753 | 0.01033 | 0.02402 | 0.00000 | 0.02971 | 0.01654 |
| **EDEM1** | 2.44605 | 1.93663 | 5.87000 | 7.80638 | 0.00000 | 11.24793 | 2.80661 | 3.36528 | 1.99801 | 2.17969 | 1.49661 | 1.31268 |
| **EDEM2** | 3.10638 | 10.54635 | 9.20570 | 6.47572 | 10.93874 | 6.98764 | 0.00000 | 3.98054 | 14.02066 | 0.00000 | 10.29016 | 7.28061 |
| **EDEM3** | 0.34338 | 0.40629 | 2.38730 | 0.83122 | 0.00573 | 10.65921 | 0.00000 | 1.47826 | 5.23373 | 10.44406 | 2.62636 | 1.02060 |
| **EDF1** | 3.99093 | 20.67425 | 25.87392 | 20.23743 | 28.36213 | 13.48632 | 36.26811 | 4.42411 | 30.33606 | 11.30266 | 21.73603 | 14.37039 |
| **EDIL3** | 0.26711 | 0.18778 | 0.38019 | 0.00000 | 0.00000 | 0.00000 | 0.85860 | 0.26424 | 0.00000 | 0.06521 | 0.86257 | 0.00000 |
| **EDN1** | 0.27395 | 0.53808 | 0.12818 | 0.00913 | 0.00000 | 0.00000 | 0.01886 | 0.00000 | 0.01777 | 0.09039 | 0.03222 | 0.67975 |
| **EDN3** | 0.00347 | 0.02185 | 0.00723 | 0.00000 | 0.06812 | 0.00592 | 0.02417 | 0.01329 | 0.00006 | 0.18839 | 0.00813 | 0.02421 |
| **EDNRA** | 0.05580 | 0.02436 | 0.00543 | 0.21223 | 0.03149 | 0.03332 | 0.04669 | 0.01034 | 0.00000 | 0.05944 | 0.01197 | 0.01889 |
| **EDNRB** | 0.00203 | 0.00000 | 9.31817 | 0.00333 | 0.05083 | 0.04931 | 0.00000 | 0.06313 | 5.49770 | 0.00000 | 1.14381 | 0.10520 |
| **EDNRB-AS1** | 0.94913 | 0.00458 | 0.02573 | 0.02131 | 0.00595 | 0.03264 | 0.00000 | 0.29207 | 0.04070 | 0.06200 | 0.05640 | 0.02531 |
| **EDRF1** | 2.99583 | 3.21379 | 4.31642 | 0.23375 | 0.00000 | 5.18368 | 6.71542 | 4.27205 | 7.52133 | 4.28160 | 3.70966 | 3.18278 |
| **EDRF1-AS1** | 0.00282 | 0.04622 | 0.02419 | 0.02710 | 0.00000 | 0.05523 | 0.00000 | 0.04234 | 0.13427 | 0.13099 | 0.01886 | 0.03201 |
| **EEA1** | 2.92636 | 0.00000 | 6.40780 | 4.15711 | 11.66861 | 13.72363 | 1.68791 | 3.58434 | 10.62798 | 20.26720 | 5.52759 | 5.79939 |
| **EED** | 0.30286 | 5.01574 | 14.11647 | 0.77458 | 0.63412 | 13.57568 | 6.12372 | 3.01882 | 1.19319 | 6.39149 | 7.51130 | 8.98231 |
| **EEF1A1** | 5.74073 | 20.40967 | 12.85206 | 14.20849 | 13.14814 | 40.20106 | 13.97532 | 3.97947 | 29.70713 | 0.00000 | 13.91239 | 20.11317 |
| **EEF1A2** | 0.48727 | 0.17868 | 0.01634 | 0.00000 | 0.39561 | 0.00000 | 0.03422 | 0.13875 | 0.00000 | 0.39871 | 0.00654 | 0.01410 |
| **EEF1B2** | 2.40243 | 22.23339 | 16.88868 | 18.18096 | 23.78525 | 21.14305 | 20.61650 | 12.68592 | 32.43689 | 0.00000 | 17.59327 | 19.11694 |
| **EEF1D** | 5.13492 | 26.64413 | 23.49866 | 22.86003 | 39.00593 | 2.97841 | 41.17830 | 4.30392 | 41.96126 | 27.75291 | 22.12851 | 25.97387 |
| **EEF1DP3** | 0.01037 | 0.03173 | 0.01626 | 0.04989 | 0.03429 | 0.01557 | 0.02245 | 0.08735 | 0.00582 | 0.00000 | 0.05371 | 0.07244 |
| **EEF1E1.uid-815** | 0.00955 | 0.02092 | 0.07558 | 0.01098 | 0.11262 | 0.00000 | 0.10771 | 0.02473 | 0.00000 | 0.01794 | 0.01319 | 0.03644 |
| **EEF1E1.uid-816** | 1.38523 | 1.85793 | 11.76024 | 0.00000 | 0.00000 | 22.64418 | 16.25615 | 0.93267 | 12.50715 | 7.16482 | 6.44166 | 5.53516 |
| **EEF1G** | 7.05425 | 20.73015 | 23.30309 | 21.15704 | 29.83442 | 11.98570 | 36.59478 | 7.57154 | 32.22187 | 16.66100 | 20.71844 | 20.95707 |
| **EEF2** | 19.94070 | 42.22130 | 40.96044 | 40.49508 | 47.69780 | 3.43589 | 62.38873 | 16.22971 | 57.81626 | 35.66601 | 36.47029 | 49.46361 |
| **EEF2K** | 8.08222 | 2.52903 | 2.02171 | 1.79631 | 1.17122 | 0.00000 | 2.51510 | 0.59743 | 7.13703 | 3.53166 | 4.21417 | 1.76243 |
| **EEFSEC** | 0.29450 | 3.35573 | 1.83190 | 2.92207 | 3.06215 | 1.65059 | 2.17998 | 0.12887 | 2.87078 | 0.00000 | 3.76048 | 0.55002 |
| **EEPD1** | 1.67529 | 0.08140 | 1.43545 | 0.94761 | 0.00004 | 0.00000 | 0.37523 | 1.04633 | 0.00000 | 1.41327 | 0.07754 | 0.52873 |
| **EFCAB1** | 0.00613 | 0.01459 | 0.01829 | 0.01271 | 0.01400 | 0.02127 | 0.01848 | 0.01055 | 0.01649 | 0.01685 | 0.01441 | 0.01632 |
| **EFCAB10** | 2.41290 | 0.12432 | 0.04941 | 0.03053 | 0.06032 | 0.09439 | 0.04159 | 0.24174 | 0.04280 | 0.06695 | 0.03284 | 0.21078 |
| **EFCAB11** | 3.39192 | 2.84241 | 1.99936 | 2.78034 | 2.15121 | 9.21423 | 5.03917 | 2.50510 | 0.00000 | 1.17512 | 5.83988 | 5.28005 |
| **EFCAB12** | 0.13998 | 0.00213 | 0.23703 | 0.00350 | 0.01392 | 0.02133 | 0.00000 | 0.25263 | 0.01461 | 0.08273 | 0.01962 | 0.12566 |
| **EFCAB13** | 0.00807 | 0.31048 | 0.07008 | 0.56722 | 0.32630 | 0.86769 | 0.00000 | 0.65062 | 0.00012 | 1.46442 | 0.10330 | 1.22270 |
| **EFCAB14** | 6.41030 | 8.36928 | 7.72359 | 4.37202 | 4.14537 | 16.58831 | 3.58284 | 9.22536 | 12.74657 | 6.73405 | 9.13645 | 8.31660 |
| **EFCAB14-AS1** | 0.01200 | 0.00000 | 0.00346 | 0.00726 | 0.01928 | 0.09814 | 0.04201 | 0.01769 | 0.03445 | 0.19075 | 0.00000 | 0.01392 |
| **EFCAB2** | 0.67171 | 1.07936 | 1.20837 | 2.06123 | 0.93658 | 6.11952 | 0.00000 | 0.28286 | 3.47987 | 1.05810 | 0.38438 | 0.48191 |
| **EFCAB3** | 0.00591 | 0.01486 | 0.01326 | 0.01116 | 0.01702 | 0.01424 | 0.02042 | 0.01049 | 0.01952 | 0.02787 | 0.01536 | 0.01687 |
| **EFCAB4A** | 0.37597 | 0.06435 | 0.24450 | 0.44994 | 0.10707 | 0.00000 | 0.20486 | 0.10974 | 1.62872 | 0.30300 | 0.31108 | 0.00000 |
| **EFCAB4B** | 0.20532 | 0.05901 | 0.06866 | 0.01690 | 0.00000 | 0.00955 | 0.04126 | 0.00774 | 0.00000 | 0.01071 | 0.17369 | 0.01271 |
| **EFCAB5** | 0.03457 | 0.02683 | 0.05903 | 0.00229 | 0.03405 | 0.06347 | 0.03642 | 0.05895 | 0.00000 | 0.09956 | 0.12296 | 0.05736 |
| **EFCAB6** | 3.10766 | 0.46400 | 0.00032 | 0.01131 | 0.00000 | 0.08864 | 0.00000 | 0.33257 | 0.00000 | 0.09994 | 0.01987 | 0.34326 |
| **EFCAB6-AS1** | 0.00463 | 0.04856 | 0.01096 | 0.00735 | 0.00291 | 0.04221 | 0.02214 | 0.02582 | 0.01012 | 0.01999 | 0.01402 | 0.01095 |
| **EFCAB7** | 0.24824 | 0.23642 | 1.67397 | 0.00000 | 0.00000 | 1.93053 | 0.38230 | 0.60099 | 1.07396 | 0.10143 | 0.60871 | 0.71664 |
| **EFCAB9** | 0.00399 | 0.00869 | 0.00747 | 0.00629 | 0.03707 | 0.04184 | 0.03486 | 0.08547 | 0.03655 | 0.01748 | 0.00882 | 0.02319 |

| | | | | | | | | | | | | |
|---|---|---|---|---|---|---|---|---|---|---|---|---|
| EFCC1 | 0.09294 | 0.05007 | 0.04220 | 0.02481 | 0.03830 | 0.00000 | 0.00757 | 0.04267 | 0.01980 | 0.06056 | 0.01682 | 0.03266 |
| EFEMP1 | 0.00000 | 1.92215 | 0.15453 | 3.38620 | 0.00000 | 0.42777 | 0.00000 | 0.00369 | 0.00000 | 0.38482 | 0.74753 | 0.00000 |
| EFEMP2 | 0.06049 | 0.06253 | 0.10387 | 0.01960 | 0.09374 | 0.00000 | 0.01213 | 0.06416 | 0.04702 | 0.08898 | 0.01308 | 0.11346 |
| EFHB | 0.28073 | 0.04513 | 0.03849 | 0.01310 | 0.00671 | 0.00641 | 0.03071 | 0.00345 | 0.04349 | 0.07943 | 0.00671 | 0.03864 |
| EFHC1 | 0.19637 | 0.57535 | 1.13196 | 0.39031 | 0.00000 | 2.10948 | 0.00000 | 1.68680 | 2.15846 | 2.17908 | 0.72551 | 2.05582 |
| EFHC2 | 1.04125 | 0.01535 | 0.01561 | 0.02453 | 0.02758 | 0.00000 | 0.02116 | 0.19679 | 0.02606 | 0.00000 | 0.01513 | 0.00801 |
| EFHD1 | 1.63370 | 7.51393 | 1.17037 | 0.00000 | 30.06143 | 0.00000 | 6.08791 | 0.00174 | 0.00000 | 7.49280 | 8.19535 | 0.00000 |
| EFHD2 | 0.00000 | 1.74076 | 3.28023 | 2.85894 | 16.87148 | 0.00000 | 5.70621 | 0.51236 | 0.84408 | 11.51468 | 1.80285 | 0.29596 |
| EFNA1 | 0.00000 | 14.01686 | 6.97401 | 0.00000 | 21.57619 | 6.56399 | 12.71654 | 2.69998 | 43.89868 | 1.73398 | 17.56666 | 0.00000 |
| EFNA2 | 0.03462 | 0.09639 | 0.00000 | 0.00868 | 0.15006 | 0.30940 | 0.00000 | 0.10791 | 0.00000 | 0.88777 | 0.15324 | 1.54225 |
| EFNA3 | 1.05981 | 1.61831 | 0.15291 | 0.14616 | 1.77324 | 0.61841 | 0.00178 | 0.12169 | 0.00000 | 0.57602 | 0.18294 | 0.29796 |
| EFNA4 | 1.03416 | 15.93378 | 0.49036 | 2.63871 | 7.47343 | 5.23363 | 1.17283 | 1.04990 | 8.53393 | 0.57597 | 1.92277 | 14.52657 |
| EFNA5 | 1.27857 | 0.08661 | 0.15210 | 0.06657 | 0.00133 | 0.27643 | 0.09044 | 0.31182 | 0.00000 | 0.22675 | 0.21099 | 0.20672 |
| EFNB1 | 0.95778 | 0.11617 | 0.13739 | 0.43932 | 0.27089 | 0.00035 | 0.00000 | 0.47737 | 0.28754 | 0.70560 | 0.00000 | 0.39123 |
| EFNB2 | 4.48865 | 0.27766 | 1.72309 | 1.25182 | 3.02519 | 0.00000 | 0.99085 | 2.14229 | 0.63307 | 7.00690 | 0.04171 | 0.02482 |
| EFNB3 | 0.61517 | 0.32057 | 0.00000 | 0.08084 | 0.00000 | 0.00000 | 0.00216 | 1.55238 | 0.00420 | 0.71779 | 0.00000 | 5.10420 |
| EFR3A | 3.18227 | 2.12226 | 1.94175 | 1.26180 | 2.70495 | 12.07848 | 6.90732 | 0.83608 | 2.23767 | 4.56239 | 5.17425 | 1.68554 |
| EFR3B | 4.15285 | 12.89109 | 0.00000 | 8.12069 | 8.21301 | 0.00000 | 0.61805 | 1.35374 | 2.10529 | 3.25013 | 9.91306 | 9.72886 |
| EFS | 0.00215 | 0.21913 | 0.82217 | 0.00000 | 0.43446 | 0.00000 | 3.21172 | 0.11005 | 1.76266 | 2.43251 | 0.00000 | 0.16518 |
| EFTUD1 | 2.50897 | 1.01665 | 2.00934 | 6.86997 | 2.05787 | 0.60035 | 3.03044 | 1.56670 | 2.94677 | 0.46184 | 4.25301 | 1.84451 |
| EFTUD1P1 | 0.03129 | 0.01267 | 0.01432 | 0.00957 | 0.01366 | 0.03486 | 0.01926 | 0.03968 | 0.02189 | 0.00683 | 0.01418 | 0.02130 |
| EFTUD2 | 4.23909 | 25.10211 | 17.61975 | 15.31997 | 18.84153 | 7.36712 | 21.85817 | 2.76668 | 27.23336 | 3.59604 | 23.13598 | 24.76823 |
| EGF | 3.35206 | 0.00426 | 0.05812 | 0.08222 | 0.00000 | 0.00000 | 0.00483 | 0.04995 | 0.01081 | 0.25107 | 0.01034 | 0.00361 |
| EGFEM1P | 0.38486 | 0.01379 | 0.03238 | 0.01420 | 0.01921 | 0.00634 | 0.02495 | 0.08144 | 0.01046 | 0.02040 | 0.01819 | 0.01659 |
| EGFL6 | 0.13573 | 0.11217 | 0.00000 | 0.36687 | 0.00016 | 0.16773 | 0.00000 | 0.00000 | 0.00000 | 0.05384 | 0.01115 | 0.10960 |
| EGFL7 | 0.04442 | 6.27623 | 4.08091 | 4.28727 | 6.31896 | 0.00000 | 0.12252 | 0.00912 | 5.78623 | 0.00000 | 1.88486 | 2.14743 |
| EGFL8.uid-8475 | 0.00469 | 0.04875 | 0.00747 | 0.00510 | 0.02966 | 0.02898 | 0.00548 | 0.00954 | 0.00876 | 0.09905 | 0.00894 | 0.02232 |
| EGFLAM | 0.06734 | 5.27922 | 0.00000 | 7.47870 | 2.70826 | 1.34588 | 0.00000 | 0.29392 | 0.46940 | 1.43657 | 0.37500 | 3.20405 |
| EGFLAM-AS2 | 0.00621 | 0.01526 | 0.01407 | 0.01325 | 0.01644 | 0.01181 | 0.02434 | 0.00995 | 0.02073 | 0.01403 | 0.01488 | 0.02467 |
| EGFLAM-AS4 | 0.00482 | 0.00000 | 0.01486 | 0.01886 | 0.00000 | 0.02356 | 0.00991 | 0.07786 | 0.00000 | 0.12194 | 0.09843 | 0.27901 |
| EGFR | 0.00131 | 0.07673 | 0.03017 | 0.00000 | 0.34754 | 0.00000 | 0.56200 | 0.44255 | 1.54089 | 0.05179 | 0.00000 | 0.11396 |
| EGFR-AS1 | 0.00637 | 0.24759 | 0.09276 | 0.01742 | 0.03345 | 0.00000 | 0.06405 | 0.01118 | 0.00000 | 0.00000 | 0.02208 | 0.00695 |
| EGLN1 | 0.90940 | 0.84835 | 0.73202 | 0.22722 | 0.88871 | 3.66272 | 3.44735 | 0.61308 | 1.12576 | 4.99190 | 0.82881 | 1.60486 |
| EGLN2.uid-2379 | 0.02610 | 0.70943 | 0.43074 | 0.43104 | 0.13610 | 0.00000 | 0.08660 | 0.08274 | 0.59618 | 0.00000 | 0.37393 | 0.53501 |
| EGLN2.uid-2379 | 0.00632 | 0.09075 | 0.06947 | 0.11841 | 0.02948 | 0.00000 | 0.02363 | 0.01021 | 0.07002 | 0.00000 | 0.02454 | 0.02179 |
| EGLN3 | 0.00000 | 11.02673 | 17.39305 | 43.78838 | 1.72946 | 23.26295 | 9.44300 | 0.75705 | 9.76690 | 5.04635 | 1.24253 | 6.02993 |
| EGOT | 0.00590 | 0.01504 | 0.02202 | 0.01554 | 0.01048 | 0.00609 | 0.01255 | 0.01059 | 0.03068 | 0.03964 | 0.03662 | 0.01567 |
| EGR1 | 2.02270 | 0.19939 | 0.33042 | 1.19852 | 3.66867 | 0.00000 | 0.21005 | 7.73891 | 0.00000 | 2.66327 | 0.11997 | 0.53759 |
| EGR2 | 0.96308 | 0.19482 | 0.37242 | 0.73719 | 0.00000 | 0.09970 | 0.00000 | 0.90442 | 0.03630 | 0.00279 | 0.00026 | 0.07466 |
| EGR3 | 0.01457 | 0.00799 | 0.29755 | 0.00157 | 0.02680 | 0.06915 | 0.00000 | 0.01308 | 0.00000 | 0.01378 | 0.01127 | 0.03703 |
| EHBP1 | 2.87802 | 2.16968 | 0.89765 | 2.39643 | 1.17632 | 9.07066 | 0.00325 | 1.42492 | 1.60320 | 7.95955 | 1.93840 | 1.64806 |
| EHBP1L1 | 0.00550 | 0.01780 | 0.30853 | 0.06090 | 0.20574 | 0.00000 | 0.03332 | 0.01084 | 0.00000 | 0.31469 | 0.03041 | 0.01021 |
| EHD1 | 5.01908 | 8.84906 | 16.17884 | 1.70142 | 16.12125 | 0.00000 | 10.79759 | 4.75305 | 10.49250 | 8.17840 | 8.86884 | 6.07178 |
| EHD2 | 0.18473 | 0.73733 | 0.05447 | 0.00070 | 0.00000 | 0.00000 | 0.04173 | 0.00000 | 0.00000 | 0.04755 | 0.00988 | 0.85896 |
| EHD3 | 1.18962 | 0.06458 | 1.39889 | 1.74453 | 0.59541 | 0.00000 | 0.16918 | 0.27342 | 0.16000 | 0.05734 | 0.00000 | 0.76769 |
| EHD4 | 0.00000 | 14.22209 | 11.72959 | 8.44533 | 24.26512 | 1.08349 | 18.38047 | 0.81595 | 32.54794 | 0.00000 | 22.83254 | 4.27001 |
| EHF | 0.00595 | 0.01659 | 0.03916 | 0.01995 | 0.03442 | 0.00000 | 0.06171 | 0.01068 | 0.00000 | 0.02607 | 0.01680 | 0.01649 |
| EHHADH | 0.26479 | 1.85889 | 0.00000 | 1.71896 | 0.00445 | 0.50311 | 0.00000 | 0.00000 | 0.00000 | 0.00107 | 0.19687 | 0.73871 |

| | | | | | | | | | | | | |
|---|---|---|---|---|---|---|---|---|---|---|---|---|
| **EHMT1** | 6.69980 | 5.59060 | 8.19959 | 4.72014 | 7.12706 | 0.00000 | 2.13351 | 2.37070 | 9.31743 | 7.70676 | 5.60866 | 4.35900 |
| **EHMT1-IT1** | 0.15016 | 0.10970 | 0.01832 | 0.00000 | 0.00000 | 0.00000 | 0.00048 | 0.25538 | 0.17128 | 0.08563 | 0.01193 | 0.03803 |
| **EHMT2** | 1.64120 | 10.30422 | 4.56913 | 9.53834 | 10.14188 | 0.00000 | 9.12603 | 2.29685 | 3.79517 | 21.44705 | 5.03937 | 17.62383 |
| **EI24** | 3.54535 | 15.87337 | 9.36220 | 7.01216 | 15.38068 | 14.80809 | 14.23486 | 0.00000 | 12.74211 | 21.46933 | 10.04917 | 5.91428 |
| **EID1** | 2.58613 | 16.96203 | 10.36961 | 11.68471 | 6.31260 | 30.15110 | 13.36813 | 2.92791 | 4.10065 | 10.07963 | 16.54525 | 13.09558 |
| **EID2** | 0.00000 | 0.48759 | 0.56629 | 0.45392 | 0.00000 | 1.31041 | 0.94449 | 0.00000 | 0.00000 | 5.91884 | 1.39091 | 1.16846 |
| **EID2B** | 0.01331 | 0.11766 | 0.04915 | 0.22838 | 0.00000 | 0.00000 | 0.05128 | 0.05439 | 0.10292 | 0.00000 | 0.21547 | 0.86529 |
| **EID3** | 0.96718 | 0.26585 | 0.09534 | 0.98882 | 0.23680 | 0.69391 | 0.00000 | 7.31018 | 0.00000 | 2.92143 | 0.10982 | 16.67869 |
| **EIF1** | 18.72910 | 36.11857 | 31.95362 | 23.52749 | 29.41944 | 45.78848 | 41.70275 | 30.07878 | 50.91293 | 25.21180 | 26.83061 | 50.19738 |
| **EIF1AD** | 0.65787 | 9.79313 | 11.70821 | 7.82667 | 11.68386 | 11.56186 | 15.97467 | 33.74893 | 11.54609 | 5.24225 | 14.04362 | 20.42068 |
| **EIF1AX** | 0.87520 | 4.28841 | 3.13866 | 0.75003 | 3.05044 | 18.67606 | 13.28453 | 4.72798 | 1.56374 | 19.81468 | 7.52774 | 5.74473 |
| **EIF1AX-AS1** | 0.00477 | 0.01048 | 0.02745 | 0.02483 | 0.06173 | 0.05438 | 0.03473 | 0.01522 | 0.00000 | 0.00000 | 0.05620 | 0.02883 |
| **EIF1AY** | 0.95512 | 7.56860 | 9.58873 | 2.29285 | 5.37197 | 30.48013 | 14.03140 | 0.02057 | 9.48808 | 6.47268 | 6.86000 | 8.08388 |
| **EIF1B** | 7.97592 | 12.65921 | 12.63872 | 5.96522 | 5.84780 | 48.55972 | 15.43029 | 22.84871 | 20.06366 | 16.87131 | 15.29483 | 21.39789 |
| **EIF1B-AS1** | 0.26409 | 0.08799 | 0.02066 | 0.06846 | 0.02674 | 0.00763 | 0.02279 | 0.51401 | 0.03913 | 0.03379 | 0.10526 | 0.17649 |
| **EIF2A** | 7.60624 | 9.03024 | 15.74858 | 6.17861 | 2.48997 | 32.92417 | 18.75763 | 6.63591 | 20.13510 | 7.34236 | 14.53093 | 13.60037 |
| **EIF2AK1** | 3.40552 | 4.74613 | 17.60967 | 6.08566 | 7.60610 | 4.04424 | 16.01040 | 0.00000 | 6.20162 | 9.70320 | 7.07610 | 3.19869 |
| **EIF2AK2** | 3.29602 | 0.00000 | 3.24756 | 0.00000 | 0.00000 | 0.82914 | 4.48729 | 0.54687 | 1.48878 | 1.38830 | 1.44441 | 0.78585 |
| **EIF2AK3** | 0.03018 | 0.10026 | 1.76619 | 0.94625 | 0.52812 | 1.65318 | 0.00000 | 0.40025 | 2.15994 | 4.57459 | 0.21649 | 0.00000 |
| **EIF2AK4** | 2.38543 | 3.25077 | 5.38485 | 7.87900 | 16.51664 | 4.29364 | 6.33306 | 0.25406 | 0.00000 | 10.22392 | 7.49802 | 0.00000 |
| **EIF2B1** | 4.04784 | 16.52301 | 15.00891 | 6.89256 | 2.61257 | 24.95326 | 16.70837 | 7.79250 | 18.44498 | 0.00000 | 20.18970 | 20.27262 |
| **EIF2B2** | 0.58019 | 13.82214 | 20.62628 | 5.85817 | 6.40364 | 10.46912 | 18.66113 | 8.99319 | 10.76609 | 0.00000 | 11.10381 | 11.92196 |
| **EIF2B3** | 1.10161 | 11.06251 | 7.72075 | 7.25194 | 6.16449 | 14.15717 | 12.08385 | 3.51396 | 10.83895 | 0.55434 | 18.93615 | 12.07963 |
| **EIF2B4** | 1.33481 | 18.16979 | 15.83149 | 14.67124 | 19.27953 | 5.72485 | 17.53092 | 1.56174 | 12.95868 | 7.19279 | 15.17765 | 9.90290 |
| **EIF2B5** | 0.00000 | 3.74811 | 4.50013 | 5.80281 | 9.42663 | 0.92388 | 7.42561 | 1.29235 | 3.21184 | 7.19396 | 2.45183 | 2.42124 |
| **EIF2D** | 1.13434 | 6.54022 | 0.59459 | 2.69964 | 1.17021 | 2.67851 | 0.00000 | 0.05266 | 2.20221 | 0.00000 | 3.97771 | 2.97113 |
| **EIF2S1** | 2.27521 | 17.43221 | 21.56246 | 13.74864 | 12.57399 | 58.78347 | 28.30382 | 22.01803 | 29.21456 | 4.25131 | 29.39542 | 29.17343 |
| **EIF2S2** | 0.86981 | 12.09903 | 11.77839 | 7.57395 | 11.61459 | 18.42352 | 31.85761 | 10.52932 | 6.12085 | 28.65627 | 14.22358 | 17.60444 |
| **EIF2S3** | 3.25219 | 16.84684 | 15.23516 | 13.21734 | 8.14322 | 42.30896 | 40.20327 | 12.07456 | 17.05618 | 24.45952 | 30.47265 | 24.44320 |
| **EIF3A** | 3.76561 | 14.19648 | 19.26465 | 16.51070 | 22.30004 | 33.35912 | 33.15618 | 17.25569 | 26.23601 | 46.29963 | 18.30065 | 28.63111 |
| **EIF3B** | 0.85831 | 12.71179 | 17.67976 | 16.91181 | 25.15754 | 1.04115 | 38.42785 | 9.11434 | 10.72070 | 38.68220 | 8.33715 | 15.45831 |
| **EIF3D** | 1.99893 | 34.65089 | 30.11767 | 27.70321 | 33.49132 | 36.92643 | 41.61089 | 13.19374 | 41.36164 | 12.83616 | 36.38347 | 38.34403 |
| **EIF3E** | 0.37153 | 16.27951 | 20.45403 | 11.40520 | 5.59264 | 51.74717 | 26.44008 | 7.80130 | 29.48560 | 7.67964 | 22.05323 | 19.31922 |
| **EIF3F** | 6.18022 | 12.77531 | 13.34932 | 13.05120 | 14.59424 | 6.72295 | 20.61753 | 10.67231 | 18.03919 | 4.00122 | 9.39180 | 14.75277 |
| **EIF3G** | 1.56618 | 27.78139 | 28.11421 | 20.92244 | 26.14141 | 9.20309 | 25.02386 | 21.84588 | 32.24979 | 0.00000 | 19.71314 | 32.94188 |
| **EIF3H** | 6.00137 | 23.32297 | 23.79018 | 20.65761 | 21.01226 | 45.67501 | 28.25854 | 16.81101 | 33.13350 | 11.88443 | 20.13594 | 28.52504 |
| **EIF3I** | 1.40814 | 31.69093 | 30.86278 | 18.50543 | 25.06296 | 34.03991 | 37.99268 | 26.77994 | 35.23492 | 0.03903 | 29.73243 | 36.57675 |
| **EIF3IP1** | 0.01988 | 2.73982 | 3.36431 | 1.54514 | 1.71624 | 2.92718 | 3.27250 | 2.86882 | 2.07860 | 0.00000 | 3.05526 | 3.39949 |
| **EIF3J** | 2.77040 | 12.10598 | 17.11288 | 4.44642 | 15.74611 | 15.36016 | 21.73871 | 16.14916 | 14.29550 | 27.27652 | 12.94252 | 12.26322 |
| **EIF3J-AS1** | 0.01787 | 0.79523 | 2.56216 | 0.63895 | 0.48071 | 0.60329 | 1.13155 | 3.91179 | 2.40348 | 0.21558 | 1.95910 | 1.03396 |
| **EIF3K** | 7.95960 | 37.52749 | 22.98730 | 34.05305 | 36.04818 | 32.13832 | 42.94688 | 16.43384 | 32.53084 | 12.26530 | 27.67473 | 37.96605 |
| **EIF3L** | 5.88947 | 26.77212 | 25.96822 | 24.62128 | 23.70089 | 29.39166 | 41.12601 | 19.92428 | 37.24853 | 14.84239 | 27.06997 | 28.97009 |
| **EIF3M** | 2.76342 | 22.81715 | 26.02448 | 12.88618 | 14.98293 | 47.19775 | 34.34944 | 17.63996 | 31.61417 | 3.39389 | 28.31760 | 26.17198 |
| **EIF4A2** | 15.97257 | 17.92875 | 8.67909 | 13.17894 | 12.61274 | 43.69682 | 8.32816 | 16.07854 | 26.91448 | 22.11409 | 15.60909 | 21.58406 |
| **EIF4A3** | 5.06296 | 27.50079 | 29.04843 | 14.68503 | 18.54850 | 26.26129 | 26.50842 | 41.16622 | 31.96040 | 1.07190 | 30.94570 | 33.34309 |
| **EIF4B** | 3.91468 | 7.99951 | 9.43367 | 8.17286 | 2.66891 | 20.15182 | 16.32244 | 6.96968 | 10.12429 | 8.22925 | 13.62940 | 10.13465 |
| **EIF4E** | 1.31012 | 6.20953 | 9.83555 | 1.60387 | 0.00000 | 28.34365 | 17.47554 | 5.96928 | 6.60153 | 6.29297 | 11.83552 | 8.94183 |
| **EIF4E1B** | 11.73407 | 0.05206 | 0.06315 | 0.04770 | 0.05917 | 0.00000 | 0.00515 | 4.80119 | 0.03646 | 0.00000 | 0.00391 | 0.82176 |

| | | | | | | | | | | | | |
|---|---|---|---|---|---|---|---|---|---|---|---|---|
| EIF4E2 | 2.33893 | 19.52685 | 19.41662 | 15.43308 | 18.13279 | 23.68590 | 25.61277 | 10.51263 | 20.44485 | 3.57945 | 18.13409 | 25.78502 |
| EIF4E3 | 0.55218 | 0.26175 | 0.70643 | 0.51985 | 1.01941 | 1.11638 | 1.27596 | 0.15491 | 0.27730 | 0.95068 | 0.00000 | 0.00000 |
| EIF4EBP1 | 0.00000 | 18.04499 | 8.14852 | 31.51940 | 26.40067 | 16.53593 | 27.47737 | 0.65736 | 22.31410 | 7.09800 | 23.27845 | 10.23911 |
| EIF4EBP2 | 0.29322 | 9.66561 | 10.83454 | 15.54744 | 6.28521 | 22.27644 | 17.34249 | 0.25369 | 6.55259 | 17.30034 | 11.87774 | 3.48908 |
| EIF4EBP3 | 0.05793 | 1.71994 | 0.78414 | 1.24655 | 1.34945 | 0.31243 | 0.82846 | 0.06913 | 0.17849 | 0.00000 | 1.32648 | 1.85660 |
| EIF4ENIF1 | 10.78509 | 3.62279 | 1.12410 | 2.11350 | 4.37929 | 1.62700 | 4.30377 | 3.74321 | 1.28865 | 9.44441 | 2.62687 | 2.86460 |
| EIF4G1 | 4.24351 | 28.81476 | 34.41749 | 26.08823 | 36.51435 | 5.64680 | 36.23482 | 14.50660 | 39.41168 | 35.22338 | 29.45974 | 42.29509 |
| EIF4G2 | 13.31937 | 40.68582 | 47.41896 | 26.46419 | 30.03656 | 67.94704 | 57.62879 | 31.14519 | 50.37920 | 27.79406 | 44.77984 | 47.47581 |
| EIF4G3 | 2.74063 | 1.13565 | 3.40210 | 4.15036 | 8.84533 | 16.06377 | 5.97224 | 1.46245 | 5.16451 | 26.47169 | 3.82940 | 4.44594 |
| EIF4H | 1.71714 | 28.67012 | 15.06498 | 13.99574 | 21.56900 | 9.84357 | 20.71365 | 6.21316 | 20.85478 | 0.00000 | 13.90525 | 15.86923 |
| EIF5 | 9.23480 | 31.03473 | 27.62737 | 24.22407 | 28.54532 | 61.31125 | 42.75389 | 41.88961 | 36.53262 | 20.99203 | 24.90158 | 44.25004 |
| EIF5A | 2.53569 | 37.63685 | 42.92632 | 28.31820 | 37.37645 | 0.13067 | 49.19596 | 30.11198 | 38.77154 | 10.32239 | 38.77425 | 34.06056 |
| EIF5A2 | 0.00000 | 3.20465 | 1.97909 | 1.57358 | 0.00000 | 4.10775 | 3.60746 | 11.82878 | 3.03421 | 1.48355 | 0.55919 | 7.47560 |
| EIF5AL1 | 0.11110 | 5.42777 | 6.99489 | 2.09199 | 1.33725 | 5.16564 | 4.48134 | 2.59606 | 4.90317 | 0.00000 | 5.13915 | 3.01768 |
| EIF5B | 10.65686 | 14.03002 | 21.49116 | 8.87486 | 19.92479 | 31.03494 | 40.18626 | 21.29162 | 8.53050 | 56.00550 | 21.23797 | 23.03783 |
| EIF6 | 0.00000 | 31.06049 | 21.96989 | 28.07065 | 34.88553 | 22.29576 | 32.56164 | 21.83770 | 30.81194 | 0.00000 | 25.83535 | 31.63336 |
| ELAC1 | 0.00290 | 1.74570 | 0.00000 | 0.44190 | 1.64029 | 1.19868 | 0.00000 | 0.21311 | 0.00000 | 0.00125 | 0.15636 | 1.20430 |
| ELAC2 | 6.44491 | 25.16832 | 14.52198 | 15.45138 | 22.18944 | 7.85968 | 20.43294 | 12.09480 | 32.44674 | 0.00000 | 19.83639 | 23.14027 |
| ELANE | 5.47535 | 0.21333 | 0.00444 | 0.00000 | 0.02742 | 0.01558 | 0.01298 | 1.25021 | 0.00872 | 0.01641 | 0.00943 | 0.52630 |
| ELAVL1 | 6.02512 | 8.75728 | 14.37433 | 5.74734 | 8.87472 | 2.44430 | 9.48603 | 0.77217 | 12.38835 | 14.93607 | 9.37731 | 2.42437 |
| ELAVL2 | 2.63602 | 0.27974 | 0.12925 | 0.58386 | 0.00000 | 0.07137 | 0.00000 | 0.37033 | 0.00000 | 0.46521 | 0.00000 | 0.57537 |
| ELAVL3 | 0.46524 | 0.00000 | 0.06300 | 0.07146 | 0.00000 | 0.01585 | 0.03085 | 0.67619 | 0.00000 | 0.51718 | 0.00000 | 0.21767 |
| ELAVL4 | 0.10232 | 0.04440 | 0.01796 | 0.06066 | 0.03657 | 0.00452 | 0.04934 | 0.01274 | 0.00421 | 0.00000 | 0.01779 | 0.02956 |
| ELF1 | 6.24755 | 0.58529 | 6.21783 | 5.11858 | 3.00724 | 9.47469 | 9.22865 | 7.54086 | 4.20408 | 4.52838 | 2.50110 | 4.94765 |
| ELF2 | 1.12365 | 1.33141 | 4.27523 | 3.48080 | 3.72258 | 2.83416 | 5.60902 | 2.44834 | 2.65494 | 9.73180 | 3.38474 | 4.84238 |
| ELF3 | 0.00000 | 18.96846 | 5.23260 | 12.70744 | 40.46709 | 13.11961 | 8.27915 | 0.00000 | 23.45304 | 13.63167 | 15.57189 | 7.26753 |
| ELF4 | 0.03448 | 2.67632 | 3.88574 | 0.98600 | 7.97037 | 0.00038 | 6.55773 | 0.00210 | 2.19627 | 4.01478 | 1.75304 | 0.01880 |
| ELF5 | 0.00640 | 0.00939 | 0.28813 | 0.00298 | 0.07409 | 0.02947 | 0.08654 | 0.01029 | 0.00000 | 0.00631 | 0.02171 | 0.01584 |
| ELFN1 | 0.13280 | 0.06286 | 0.01121 | 0.07347 | 0.02718 | 0.00000 | 0.01584 | 0.16101 | 0.01632 | 0.01094 | 0.04176 | 0.15346 |
| ELFN1-AS1 | 0.00595 | 0.03483 | 0.01759 | 0.00823 | 0.00618 | 0.03123 | 0.01984 | 0.01035 | 0.01456 | 0.02125 | 0.01972 | 0.01336 |
| ELFN2 | 0.84214 | 0.01052 | 0.01905 | 0.00003 | 0.08096 | 0.01896 | 0.00125 | 0.00146 | 0.00000 | 0.05227 | 0.14514 | 0.00950 |
| ELK1 | 1.79235 | 3.23992 | 1.57025 | 2.94289 | 2.42757 | 0.03865 | 1.77831 | 0.61908 | 3.68802 | 0.00000 | 2.15609 | 3.43387 |
| ELK2AP | 0.01560 | 0.01733 | 0.01706 | 0.01883 | 0.01826 | 0.01297 | 0.01726 | 0.01767 | 0.02036 | 0.01136 | 0.01458 | 0.02313 |
| ELK3 | 4.63378 | 0.88329 | 0.18981 | 1.53095 | 0.00000 | 0.87104 | 0.00000 | 0.37291 | 0.20121 | 0.30497 | 0.00000 | 2.02504 |
| ELK4 | 4.77356 | 0.00000 | 0.77934 | 0.00172 | 0.20569 | 8.53784 | 2.47264 | 1.45917 | 0.25409 | 6.56963 | 1.95690 | 1.09081 |
| ELL | 1.01036 | 6.79719 | 7.15889 | 4.34759 | 7.76287 | 0.00000 | 4.23124 | 2.96710 | 3.74839 | 4.54770 | 3.24031 | 4.28601 |
| ELL2 | 0.94181 | 0.95748 | 0.60327 | 0.52453 | 1.89243 | 2.99177 | 0.52780 | 3.63897 | 0.00000 | 4.35445 | 0.86637 | 0.97600 |
| ELL3 | 0.06889 | 2.27631 | 0.79586 | 0.00000 | 0.00000 | 0.00000 | 0.00000 | 8.77721 | 1.04253 | 1.35215 | 0.13543 | 17.13941 |
| ELMO1 | 6.69069 | 0.10773 | 12.22490 | 3.22520 | 0.09322 | 3.46516 | 3.17185 | 2.50353 | 1.99560 | 1.16173 | 0.26131 | 0.00000 |
| ELMO1-AS1 | 0.00506 | 0.01307 | 0.08082 | 0.01457 | 0.00000 | 0.02272 | 0.02972 | 0.01006 | 0.03728 | 0.07984 | 0.01608 | 0.01506 |
| ELMO2 | 10.85101 | 5.48475 | 1.56701 | 4.15857 | 0.53998 | 3.41965 | 2.40423 | 4.13933 | 0.19160 | 0.00000 | 2.99745 | 5.72082 |
| ELMO3 | 0.00196 | 0.35885 | 0.63828 | 1.13096 | 0.51505 | 0.00000 | 0.52682 | 0.00570 | 0.90398 | 2.20820 | 0.47940 | 0.12579 |
| ELMOD1 | 0.32331 | 3.71119 | 0.12838 | 2.27090 | 0.52082 | 1.10798 | 0.00000 | 0.39055 | 0.17442 | 0.37992 | 0.07213 | 0.08296 |
| ELMOD2 | 0.05490 | 4.19270 | 0.72718 | 1.57441 | 0.74594 | 12.31810 | 0.00000 | 1.43038 | 8.39241 | 2.02908 | 1.10151 | 6.15493 |
| ELMOD3 | 0.05605 | 3.94946 | 1.45048 | 2.04135 | 0.45381 | 0.19296 | 1.66193 | 1.29740 | 3.92898 | 0.00000 | 1.16569 | 2.21632 |
| ELMSAN1 | 4.25468 | 7.90235 | 12.12345 | 4.04926 | 4.23906 | 13.60335 | 10.81888 | 17.19042 | 3.16840 | 9.34851 | 11.93131 | 5.45000 |
| ELN | 0.01529 | 0.01523 | 0.15011 | 0.20271 | 0.00801 | 0.00000 | 0.04568 | 0.01017 | 0.00000 | 0.02258 | 0.02234 | 0.01747 |
| ELOF1 | 6.99233 | 27.54213 | 19.71685 | 14.50143 | 18.63633 | 3.22311 | 22.87658 | 18.87985 | 28.56418 | 0.00000 | 18.67299 | 34.52538 |

| | | | | | | | | | | | | |
|---|---|---|---|---|---|---|---|---|---|---|---|---|
| **ELOVL1** | 0.61434 | 8.08455 | 21.92338 | 9.08553 | 19.15910 | 6.87150 | 17.03278 | 2.93733 | 32.48106 | 0.00000 | 15.91598 | 7.95126 |
| **ELOVL2** | 1.20381 | 0.11204 | 0.00000 | 0.00000 | 0.00000 | 0.00008 | 0.00000 | 0.37416 | 0.52585 | 0.46301 | 0.07559 | 0.09683 |
| **ELOVL2-AS1** | 0.07457 | 0.05010 | 0.02611 | 0.01287 | 0.01613 | 0.00000 | 0.02088 | 0.10088 | 0.03559 | 0.00000 | 0.01564 | 0.06891 |
| **ELOVL3** | 2.51982 | 1.34662 | 0.50611 | 0.18595 | 0.33293 | 0.59434 | 0.00000 | 3.91959 | 0.47985 | 0.00000 | 0.00000 | 3.14972 |
| **ELOVL4** | 4.63759 | 0.46191 | 1.02819 | 2.99253 | 0.00000 | 0.96075 | 0.32446 | 7.39644 | 0.88881 | 0.00000 | 0.00000 | 0.73082 |
| **ELOVL5** | 11.45429 | 4.28790 | 19.37073 | 4.65302 | 5.96482 | 21.44785 | 1.31991 | 4.18648 | 31.27282 | 0.00081 | 7.06398 | 4.01367 |
| **ELOVL6** | 1.64077 | 0.99539 | 2.22003 | 3.70339 | 21.27523 | 13.09640 | 8.28722 | 1.07075 | 35.33056 | 8.43574 | 16.31516 | 4.74705 |
| **ELOVL7** | 0.09254 | 0.94897 | 0.07637 | 1.18372 | 0.00000 | 0.01755 | 0.04388 | 0.55876 | 0.00000 | 0.00650 | 0.16668 | 5.23005 |
| **ELP2** | 0.00518 | 11.53161 | 7.20974 | 3.66699 | 4.17929 | 18.81150 | 14.51159 | 1.98360 | 12.44986 | 0.00000 | 10.41661 | 10.17067 |
| **ELP3** | 8.21616 | 9.18285 | 4.51614 | 4.73978 | 5.46586 | 13.63100 | 5.49084 | 4.63429 | 8.86889 | 3.95546 | 7.23798 | 9.67348 |
| **ELP4** | 0.30242 | 3.44747 | 2.50985 | 0.90477 | 0.68756 | 8.39449 | 1.53456 | 2.73529 | 2.95814 | 0.00000 | 3.10724 | 1.88652 |
| **ELP5** | 1.61142 | 16.81463 | 8.99428 | 6.13967 | 9.66804 | 11.52776 | 8.52126 | 8.48563 | 16.07923 | 6.04366 | 15.02111 | 13.57548 |
| **ELP6** | 0.00836 | 7.22665 | 7.27548 | 6.00106 | 3.37252 | 6.59070 | 9.62832 | 9.00463 | 3.12444 | 0.00000 | 8.28654 | 10.86265 |
| **ELSPBP1** | 0.03467 | 0.00986 | 0.04811 | 0.01430 | 0.01515 | 0.00081 | 0.01467 | 0.07149 | 0.00449 | 0.03196 | 0.03905 | 0.08583 |
| **ELTD1** | 0.00530 | 0.01876 | 0.02018 | 0.03325 | 0.01690 | 0.00000 | 0.02281 | 0.03389 | 0.02656 | 0.00484 | 0.03678 | 0.02166 |
| **EMB** | 0.06984 | 0.36251 | 0.43953 | 0.29335 | 0.03881 | 0.06861 | 0.00000 | 0.15419 | 0.05390 | 0.00000 | 0.00476 | 0.08709 |
| **EMBP1** | 0.07842 | 0.01817 | 0.01011 | 0.03164 | 0.02690 | 0.00488 | 0.10477 | 0.00748 | 0.00000 | 0.00000 | 0.01496 | 0.02081 |
| **EMC1** | 0.00000 | 8.51738 | 10.34326 | 3.98614 | 8.58012 | 7.42894 | 0.00000 | 0.72916 | 25.53849 | 7.17582 | 9.95131 | 7.34803 |
| **EMC10** | 0.04794 | 1.39450 | 1.45223 | 3.67837 | 6.26910 | 0.00003 | 3.20466 | 0.20307 | 0.82513 | 18.80055 | 0.00000 | 1.64592 |
| **EMC2** | 1.38815 | 8.90417 | 4.38479 | 3.14721 | 0.14254 | 20.84220 | 2.40530 | 0.99601 | 3.04529 | 4.00021 | 3.86921 | 9.73537 |
| **EMC3** | 3.42105 | 18.38124 | 12.92001 | 7.27852 | 11.15067 | 19.14216 | 0.00000 | 17.10966 | 28.05953 | 3.10674 | 10.65265 | 25.44548 |
| **EMC3-AS1** | 1.43456 | 0.71025 | 3.15780 | 0.53422 | 2.39935 | 0.00000 | 2.88820 | 5.31987 | 2.09729 | 9.97687 | 1.89415 | 4.51329 |
| **EMC4** | 0.02505 | 19.36011 | 22.58822 | 11.70670 | 12.53073 | 30.55253 | 17.91952 | 21.56717 | 30.73399 | 0.14082 | 15.73990 | 28.65235 |
| **EMC6** | 0.29455 | 5.45192 | 4.28232 | 4.00996 | 4.14386 | 1.67449 | 2.95709 | 1.00709 | 5.98453 | 0.00000 | 5.11965 | 5.50966 |
| **EMC7** | 1.70682 | 10.30500 | 24.56226 | 6.59289 | 6.77455 | 24.16535 | 0.00000 | 8.92816 | 30.70308 | 4.34901 | 10.77778 | 19.58545 |
| **EMC8** | 3.90263 | 18.79803 | 13.67047 | 10.47197 | 12.90903 | 0.00000 | 12.09026 | 5.09027 | 13.12476 | 11.27450 | 10.63872 | 7.80224 |
| **EMC9** | 0.00000 | 2.87007 | 2.84773 | 2.02311 | 4.39111 | 1.20483 | 2.34426 | 1.64347 | 3.45350 | 2.29257 | 1.10744 | 4.75838 |
| **EMCN** | 0.00602 | 0.01335 | 0.02798 | 0.01276 | 0.01116 | 0.02273 | 0.01515 | 0.01330 | 0.00877 | 0.02607 | 0.02310 | 0.01494 |
| **EMD** | 0.30617 | 4.39424 | 3.45835 | 4.19241 | 6.94310 | 0.00000 | 5.87811 | 1.25686 | 12.41423 | 1.26647 | 7.23929 | 3.51149 |
| **EME1** | 0.66035 | 9.36694 | 0.58226 | 2.34691 | 1.61230 | 4.65250 | 0.00000 | 0.36172 | 1.33068 | 0.00000 | 2.85662 | 6.35079 |
| **EME2** | 0.00617 | 0.06150 | 0.09239 | 0.13794 | 0.05453 | 0.00000 | 0.01292 | 0.01099 | 0.11196 | 0.22106 | 0.01617 | 0.01138 |
| **EMG1** | 2.00038 | 19.94194 | 26.24144 | 12.21467 | 16.98391 | 22.69367 | 26.35569 | 10.67299 | 12.10875 | 7.06464 | 18.01543 | 24.29351 |
| **EMID1** | 0.00376 | 0.05649 | 0.17637 | 0.14154 | 0.04449 | 0.00000 | 0.00186 | 0.00870 | 0.00000 | 0.15196 | 0.01305 | 0.04370 |
| **EMILIN1** | 0.12234 | 0.01506 | 0.03839 | 0.02809 | 0.00409 | 0.01897 | 0.00942 | 0.00476 | 0.02959 | 0.08170 | 0.01121 | 0.01632 |
| **EMILIN2** | 7.11223 | 1.36166 | 1.16931 | 0.00000 | 4.64466 | 2.06123 | 5.53453 | 1.90847 | 7.30239 | 10.36629 | 3.44067 | 0.00000 |
| **EMILIN3** | 0.00415 | 0.02086 | 0.05911 | 0.04737 | 0.04553 | 0.00000 | 0.00483 | 0.03736 | 0.03745 | 0.00449 | 0.01754 | 0.32474 |
| **EML1** | 0.42534 | 0.04976 | 0.03836 | 0.08849 | 0.04135 | 0.08954 | 0.00000 | 0.07921 | 0.00000 | 0.14556 | 0.04904 | 0.04969 |
| **EML2** | 0.16463 | 4.33704 | 0.46921 | 0.44782 | 0.51119 | 1.57398 | 0.00000 | 0.12296 | 0.47517 | 0.56263 | 3.09324 | 1.28733 |
| **EML3** | 0.07020 | 0.14651 | 1.52236 | 1.68603 | 5.96055 | 0.00000 | 0.09597 | 0.99329 | 4.35477 | 5.62940 | 5.14355 | 0.51037 |
| **EML4** | 3.59587 | 4.97801 | 4.03549 | 5.46474 | 5.82239 | 7.76707 | 4.45172 | 6.62728 | 5.10615 | 15.99955 | 4.30938 | 5.54863 |
| **EML5** | 4.11803 | 0.00000 | 0.06349 | 0.00000 | 0.00000 | 0.32392 | 1.27619 | 1.01284 | 0.71792 | 1.76538 | 0.08607 | 0.11931 |
| **EML6** | 1.42579 | 1.24186 | 0.00000 | 0.52044 | 0.00000 | 0.00000 | 0.28061 | 0.00000 | 0.10291 | 0.99534 | 0.00000 | 1.93998 |
| **EMP1** | 0.00671 | 0.01675 | 0.01573 | 0.01791 | 0.01630 | 0.00000 | 0.01635 | 0.01074 | 0.03047 | 0.02559 | 0.04786 | 0.01687 |
| **EMP2** | 1.64044 | 5.06207 | 1.84592 | 0.00000 | 14.12414 | 4.44109 | 26.30972 | 0.28956 | 52.72448 | 8.98928 | 28.42808 | 0.87333 |
| **EMP3** | 0.00232 | 0.14009 | 0.10011 | 0.48068 | 0.05825 | 0.00000 | 0.05554 | 0.45658 | 0.12385 | 0.00000 | 0.01218 | 2.29263 |
| **EMR1** | 0.71941 | 0.22874 | 0.02315 | 0.00000 | 0.00000 | 0.00000 | 0.07523 | 1.10713 | 0.05046 | 0.20606 | 0.00000 | 0.11373 |
| **EMR2** | 0.09848 | 0.35673 | 0.08759 | 0.01311 | 0.00000 | 0.00021 | 0.75697 | 0.09020 | 2.53488 | 0.00000 | 0.44322 | 0.06381 |
| **EMR3** | 0.01847 | 0.01419 | 0.01911 | 0.01067 | 0.01050 | 0.01755 | 0.03054 | 0.09366 | 0.00735 | 0.03536 | 0.01721 | 0.01216 |

| | | | | | | | | | | | | |
|---|---|---|---|---|---|---|---|---|---|---|---|---|
| **EMR4P** | 0.00617 | 0.01503 | 0.02627 | 0.01821 | 0.02140 | 0.00000 | 0.01792 | 0.00992 | 0.02216 | 0.00823 | 0.02727 | 0.04687 |
| **EMX1** | 7.27667 | 0.00354 | 0.07014 | 0.02452 | 0.00317 | 0.03037 | 0.08438 | 0.67427 | 0.00367 | 0.00036 | 0.02955 | 0.09610 |
| **EMX2** | 0.01249 | 0.01440 | 0.01170 | 0.00401 | 0.01078 | 0.02777 | 0.03442 | 0.02053 | 0.00297 | 0.07944 | 0.02824 | 0.01426 |
| **EMX2OS** | 0.00551 | 0.00499 | 0.00000 | 0.00856 | 0.00000 | 0.05185 | 0.08091 | 0.00934 | 0.00000 | 0.11189 | 0.00526 | 0.01303 |
| **EN1** | 0.67433 | 0.25907 | 0.00000 | 0.00000 | 0.00000 | 0.30808 | 0.68645 | 1.03023 | 0.00000 | 1.39907 | 0.00000 | 1.12441 |
| **EN2** | 0.37500 | 0.00141 | 0.00000 | 0.35123 | 0.03952 | 0.07382 | 0.00622 | 0.17031 | 0.00000 | 0.09583 | 0.00297 | 0.10697 |
| **ENAH** | 3.88450 | 1.92399 | 11.09256 | 10.86249 | 0.00000 | 34.79495 | 32.46122 | 3.68431 | 0.00000 | 33.88540 | 3.08069 | 12.85607 |
| **ENAM** | 0.00238 | 0.00905 | 0.07491 | 0.00950 | 0.00997 | 0.02090 | 0.01025 | 0.13931 | 0.00000 | 0.02835 | 0.03663 | 0.06324 |
| **ENC1** | 0.00423 | 0.01325 | 1.45708 | 1.31919 | 0.00000 | 0.00000 | 0.37146 | 0.10269 | 0.08095 | 0.53434 | 0.08075 | 0.00770 |
| **ENDOD1** | 0.63725 | 3.60523 | 0.00000 | 0.00000 | 0.64221 | 0.66408 | 0.00000 | 0.08656 | 2.40156 | 1.63203 | 0.03363 | 2.14472 |
| **ENDOG** | 0.11729 | 0.40455 | 0.88945 | 0.92070 | 1.63323 | 0.00000 | 0.46986 | 0.09716 | 1.00113 | 0.00000 | 0.91142 | 0.13714 |
| **ENDOU** | 0.00210 | 0.00000 | 3.64224 | 0.02281 | 1.39870 | 0.43527 | 0.00000 | 0.06300 | 0.09813 | 0.00128 | 0.00000 | 0.00329 |
| **ENDOV** | 0.39675 | 9.06006 | 1.91037 | 1.59073 | 1.00542 | 3.76315 | 1.82916 | 0.53414 | 1.72533 | 1.19919 | 2.32345 | 8.35369 |
| **ENG** | 10.59940 | 0.01681 | 1.69729 | 0.29366 | 0.05136 | 0.00000 | 0.08191 | 0.45468 | 0.00000 | 0.01220 | 0.05145 | 0.39162 |
| **ENGASE** | 1.36529 | 0.52144 | 0.77966 | 0.06079 | 0.94324 | 0.00000 | 0.40742 | 0.03852 | 0.16311 | 0.00000 | 0.03031 | 0.10314 |
| **ENHO** | 0.25609 | 0.21521 | 0.15370 | 0.54258 | 0.00000 | 0.00000 | 0.22770 | 0.00000 | 0.00000 | 0.27666 | 0.06181 | 0.17070 |
| **ENKD1** | 0.23998 | 1.00180 | 0.88009 | 0.48059 | 1.73357 | 0.00001 | 0.96699 | 0.00000 | 0.00307 | 1.85612 | 0.22288 | 0.59551 |
| **ENKUR** | 0.07853 | 0.06766 | 0.10869 | 0.00000 | 0.00000 | 0.02528 | 0.03946 | 0.00686 | 0.00000 | 0.06398 | 0.10924 | 0.03408 |
| **ENO1** | 0.00000 | 63.19578 | 58.71879 | 56.53931 | 65.17580 | 62.23276 | 82.62258 | 30.36051 | 77.89664 | 23.63549 | 62.44717 | 61.80979 |
| **ENO1-AS1** | 0.06690 | 0.03885 | 0.18171 | 0.00000 | 0.10292 | 0.05610 | 0.00000 | 0.03346 | 0.07273 | 0.05897 | 0.00939 | 0.12043 |
| **ENO2** | 1.95640 | 16.30304 | 0.66386 | 31.22954 | 2.24299 | 2.53419 | 0.00000 | 12.36963 | 0.00000 | 2.36695 | 0.68732 | 24.96662 |
| **ENO3** | 0.04146 | 0.52439 | 0.46027 | 2.83339 | 1.25092 | 0.19878 | 1.20486 | 1.38232 | 0.00000 | 2.18966 | 2.94316 | 2.31221 |
| **ENO4** | 0.03492 | 0.08375 | 0.18729 | 0.00000 | 0.00000 | 0.03089 | 0.28930 | 0.08443 | 0.87302 | 0.59757 | 0.00000 | 0.50495 |
| **ENOPH1** | 1.85002 | 10.76122 | 13.91387 | 6.35680 | 3.35995 | 33.94471 | 15.59305 | 1.69474 | 21.80745 | 0.00480 | 17.71754 | 11.46333 |
| **ENOSF1** | 0.40037 | 8.78310 | 1.39590 | 4.38100 | 4.69880 | 6.73940 | 2.28597 | 1.41483 | 5.93443 | 6.11560 | 8.99559 | 2.65792 |
| **ENOX1** | 1.10863 | 0.13877 | 0.02118 | 0.00000 | 0.00000 | 0.02252 | 0.00000 | 0.45103 | 0.06270 | 0.10147 | 0.02881 | 0.26815 |
| **ENOX2** | 1.44837 | 2.14563 | 2.90728 | 1.66881 | 2.34923 | 5.29160 | 4.19112 | 0.24509 | 2.74147 | 1.85271 | 3.50902 | 1.03345 |
| **ENPEP** | 0.00000 | 0.00000 | 3.55549 | 0.00000 | 22.83135 | 0.66476 | 6.24078 | 0.00000 | 31.46255 | 0.92301 | 13.68250 | 0.00000 |
| **ENPP1** | 0.90937 | 0.00000 | 1.14687 | 5.07719 | 2.12502 | 1.82690 | 0.00000 | 0.42755 | 2.60986 | 4.31092 | 5.58731 | 0.00000 |
| **ENPP2** | 0.92208 | 0.00000 | 0.17451 | 0.86104 | 0.00000 | 0.00000 | 0.00000 | 5.17501 | 0.02046 | 0.06122 | 0.07143 | 0.00000 |
| **ENPP3** | 0.05062 | 0.01497 | 0.00995 | 0.01069 | 0.01243 | 0.02184 | 0.01646 | 0.03492 | 0.04841 | 0.00851 | 0.01099 | 0.01362 |
| **ENPP4** | 0.00138 | 0.54792 | 1.20302 | 0.02780 | 0.00000 | 0.64153 | 0.00000 | 0.10855 | 0.26703 | 0.00000 | 0.19149 | 0.19961 |
| **ENPP5** | 0.00255 | 0.02416 | 0.13792 | 0.00195 | 0.01146 | 0.06854 | 0.01450 | 0.06419 | 0.50191 | 0.00000 | 0.12941 | 0.00897 |
| **ENPP6** | 0.00200 | 0.00995 | 0.03432 | 0.01896 | 0.02921 | 0.00000 | 0.02626 | 0.05045 | 0.01419 | 0.02649 | 0.01536 | 0.09719 |
| **ENSA** | 3.94745 | 10.88680 | 14.49425 | 13.60780 | 12.58040 | 14.19338 | 20.08144 | 7.01976 | 10.71191 | 2.73580 | 14.00594 | 18.54366 |
| **ENTHD1** | 0.06460 | 0.01573 | 0.02497 | 0.01575 | 0.01691 | 0.00729 | 0.15329 | 0.00631 | 0.00000 | 0.00000 | 0.01193 | 0.03523 |
| **ENTHD2** | 0.28261 | 2.02129 | 0.38482 | 0.69959 | 3.43834 | 0.00000 | 2.83915 | 0.69699 | 0.07198 | 7.95626 | 0.86412 | 1.43008 |
| **ENTPD1** | 0.00000 | 0.88997 | 10.96395 | 0.00000 | 11.09107 | 9.40354 | 16.81770 | 0.50002 | 13.24965 | 8.61910 | 15.53104 | 6.65446 |
| **ENTPD1-AS1** | 0.00387 | 0.15840 | 0.09537 | 0.07022 | 0.22246 | 0.40218 | 0.36630 | 0.82073 | 0.00000 | 0.00000 | 0.12943 | 0.36889 |
| **ENTPD2** | 0.33797 | 0.01348 | 0.01623 | 0.00098 | 0.03083 | 0.08386 | 0.00000 | 0.01418 | 0.09898 | 0.02219 | 0.00302 | 0.06472 |
| **ENTPD3** | 0.00628 | 0.01473 | 0.00758 | 0.01022 | 0.00927 | 0.01928 | 0.01955 | 0.01041 | 0.07174 | 0.01824 | 0.00747 | 0.01576 |
| **ENTPD3-AS1** | 0.07088 | 0.14591 | 0.04659 | 0.12965 | 0.24559 | 0.65241 | 0.00000 | 0.04728 | 0.00000 | 0.74338 | 0.34829 | 0.29440 |
| **ENTPD4** | 0.64085 | 3.36862 | 2.82406 | 1.62935 | 3.52472 | 7.09818 | 0.00000 | 0.72950 | 5.34768 | 1.25343 | 0.00031 | 5.85820 |
| **ENTPD5** | 1.00965 | 1.00689 | 0.00000 | 0.00000 | 0.11651 | 0.37739 | 0.00000 | 0.05365 | 0.02869 | 0.44220 | 0.45239 | 0.40653 |
| **ENTPD6** | 0.34571 | 14.60961 | 5.61616 | 15.38557 | 13.64066 | 0.00000 | 4.09118 | 0.00000 | 13.41653 | 3.88495 | 4.73809 | 1.04297 |
| **ENTPD7** | 1.22401 | 1.44359 | 2.53457 | 1.11009 | 0.00000 | 4.07915 | 2.61276 | 0.00000 | 2.18013 | 0.00000 | 1.40448 | 0.86815 |
| **ENTPD8** | 0.00400 | 0.16008 | 0.28098 | 0.00000 | 0.07118 | 0.00000 | 0.03223 | 0.00971 | 0.45458 | 0.24076 | 0.30800 | 0.00319 |
| **ENY2** | 12.75817 | 13.06012 | 11.84523 | 10.24705 | 9.39280 | 34.21859 | 25.65266 | 18.41785 | 14.57054 | 15.07650 | 16.92173 | 22.90841 |

| | | | | | | | | | | | | |
|---|---|---|---|---|---|---|---|---|---|---|---|---|
| **EOGT** | 2.98700 | 3.06121 | 1.96227 | 0.08829 | 0.00000 | 5.55149 | 0.00000 | 1.26685 | 2.71157 | 1.47682 | 1.41585 | 1.68572 |
| **EOMES** | 0.51425 | 0.31972 | 0.26861 | 0.42193 | 0.00000 | 0.00000 | 0.00000 | 0.04781 | 0.00000 | 0.84263 | 0.00000 | 0.00000 |
| **EP300** | 8.32485 | 1.57056 | 10.16088 | 10.65188 | 11.93599 | 13.18549 | 20.74241 | 9.27054 | 0.11074 | 34.22593 | 4.14905 | 16.40855 |
| **EP300-AS1** | 0.09001 | 0.65262 | 0.29481 | 0.00000 | 0.16077 | 0.00000 | 0.03105 | 0.20145 | 0.00000 | 0.20967 | 0.04395 | 0.45416 |
| **EP400** | 5.25224 | 2.31280 | 3.57249 | 1.96646 | 7.10119 | 6.21050 | 6.03322 | 2.30207 | 5.17437 | 21.62238 | 2.24304 | 3.76876 |
| **EP400NL** | 0.09613 | 0.24412 | 0.29612 | 0.07703 | 0.29646 | 0.33573 | 0.14854 | 0.05540 | 0.48928 | 0.24738 | 0.28406 | 0.05105 |
| **EPAS1** | 8.46925 | 2.41194 | 6.75211 | 0.00000 | 2.36651 | 0.00000 | 2.28295 | 1.72084 | 0.52540 | 5.92420 | 0.00000 | 3.63428 |
| **EPB41** | 1.91837 | 0.96701 | 1.07429 | 0.00000 | 1.68381 | 2.16814 | 0.00002 | 2.21359 | 2.83295 | 1.06741 | 0.89202 | 0.88041 |
| **EPB41L1** | 0.55498 | 2.80306 | 2.19197 | 6.84253 | 5.51977 | 0.36364 | 1.88100 | 0.00000 | 4.05508 | 8.74075 | 12.26740 | 0.54730 |
| **EPB41L2** | 10.81376 | 4.17177 | 2.04085 | 12.81033 | 1.44351 | 7.10227 | 0.00000 | 7.02575 | 0.97635 | 3.75751 | 0.95522 | 9.80048 |
| **EPB41L3** | 1.47075 | 0.00000 | 6.72861 | 0.62003 | 2.04397 | 4.05059 | 12.74501 | 0.20556 | 7.23720 | 3.82474 | 0.00000 | 0.37438 |
| **EPB41L4A** | 3.68845 | 3.09873 | 0.42160 | 2.36510 | 1.47980 | 1.74169 | 0.00000 | 1.96279 | 4.11740 | 7.81132 | 0.90700 | 2.45728 |
| **EPB41L4A-AS1** | 1.65169 | 3.40267 | 6.31146 | 6.95256 | 5.98574 | 7.11703 | 9.94449 | 3.41901 | 12.13284 | 8.34776 | 0.79248 | 6.80829 |
| **EPB41L4A-AS2** | 0.01818 | 0.05187 | 0.04265 | 0.07344 | 0.01281 | 0.01483 | 0.02446 | 0.00466 | 0.01382 | 0.01311 | 0.02707 | 0.11765 |
| **EPB41L4B** | 2.24592 | 1.84735 | 3.96956 | 0.18136 | 0.33591 | 1.18023 | 2.01880 | 0.18430 | 0.33190 | 0.42652 | 0.41860 | 0.87968 |
| **EPB41L5** | 6.22024 | 10.50807 | 0.00000 | 7.72563 | 13.15586 | 31.15364 | 3.74804 | 8.60271 | 16.29733 | 0.00000 | 10.76868 | 12.89110 |
| **EPB42** | 0.00591 | 0.01375 | 0.01806 | 0.06195 | 0.04200 | 0.00000 | 0.05797 | 0.03133 | 0.00918 | 0.00000 | 0.02388 | 0.02412 |
| **EPC1** | 5.57330 | 2.55081 | 1.62228 | 3.18332 | 1.87444 | 3.22571 | 4.72439 | 4.10316 | 0.00000 | 4.94848 | 2.91531 | 5.88895 |
| **EPC2** | 0.83213 | 0.88212 | 1.43613 | 0.49440 | 0.00000 | 4.24570 | 3.58529 | 1.65666 | 0.35816 | 5.92066 | 0.73551 | 1.49031 |
| **EPCAM** | 3.35733 | 9.04998 | 21.08001 | 19.71222 | 32.99232 | 30.32028 | 12.73848 | 3.23277 | 74.41791 | 3.15971 | 26.81383 | 0.00000 |
| **EPDR1** | 0.00198 | 0.01665 | 0.08993 | 0.03304 | 0.00607 | 0.03662 | 0.00000 | 0.07039 | 0.01843 | 0.03866 | 0.00949 | 0.04704 |
| **EPG5** | 2.37509 | 0.30067 | 0.93275 | 0.00000 | 1.43176 | 6.19060 | 0.85405 | 0.02443 | 0.00000 | 4.57806 | 0.87212 | 1.14249 |
| **EPGN** | 0.00442 | 0.00375 | 0.00263 | 0.00441 | 0.11898 | 0.01439 | 0.01937 | 0.01016 | 0.04131 | 0.12588 | 0.12479 | 0.01199 |
| **EPHA1** | 0.24188 | 0.45771 | 0.43879 | 4.74417 | 2.33531 | 0.00000 | 0.00274 | 0.48628 | 5.28587 | 0.00000 | 1.41572 | 0.96318 |
| **EPHA1-AS1** | 0.15203 | 0.04848 | 0.00504 | 0.07609 | 0.00000 | 0.06750 | 0.08359 | 0.86016 | 0.00000 | 0.26215 | 0.09431 | 0.17857 |
| **EPHA10** | 0.50569 | 0.54069 | 0.09588 | 0.54890 | 0.04128 | 0.11388 | 0.00000 | 0.19216 | 0.00000 | 0.14490 | 0.00009 | 0.65708 |
| **EPHA2** | 0.59234 | 0.66179 | 1.81216 | 0.00000 | 1.01348 | 0.00000 | 0.00000 | 0.00000 | 0.31865 | 0.59920 | 0.57178 | 1.46527 |
| **EPHA3** | 0.00585 | 0.05752 | 0.04175 | 0.00000 | 0.02139 | 0.05786 | 0.01119 | 0.00952 | 0.01345 | 0.02853 | 0.00813 | 0.02154 |
| **EPHA4** | 5.65822 | 0.00000 | 0.07220 | 15.52799 | 0.38325 | 0.08902 | 0.15547 | 1.54858 | 0.00000 | 0.00001 | 0.10963 | 0.48034 |
| **EPHA5** | 0.00269 | 0.01698 | 0.01382 | 0.02344 | 0.00091 | 0.05552 | 0.00929 | 0.03369 | 0.01367 | 0.05324 | 0.01147 | 0.00958 |
| **EPHA6** | 0.01385 | 0.00041 | 0.01441 | 0.03771 | 0.08111 | 0.09143 | 0.00144 | 0.06433 | 0.00000 | 0.05134 | 0.00930 | 0.01770 |
| **EPHA7** | 0.23447 | 0.21676 | 0.00000 | 2.26648 | 0.00000 | 0.20113 | 0.00000 | 0.85229 | 0.00000 | 0.14208 | 0.00000 | 0.27005 |
| **EPHA8** | 0.60618 | 0.09042 | 0.01655 | 0.02753 | 0.04309 | 0.04946 | 0.00000 | 0.01131 | 0.00000 | 0.63430 | 0.01477 | 0.43836 |
| **EPHB1** | 4.83427 | 1.82887 | 0.05628 | 0.38943 | 0.17801 | 0.36615 | 0.00000 | 0.96687 | 0.00000 | 0.01713 | 0.05680 | 1.10552 |
| **EPHB2** | 0.33775 | 0.40185 | 0.00000 | 2.85838 | 1.24331 | 0.00000 | 0.13352 | 0.00645 | 0.45455 | 1.22539 | 0.55255 | 0.00000 |
| **EPHB3** | 0.04970 | 0.00000 | 2.20352 | 0.38371 | 0.04685 | 0.02161 | 0.18872 | 0.07596 | 0.06880 | 0.73908 | 0.00000 | 0.00136 |
| **EPHB4** | 0.00000 | 5.19233 | 15.69167 | 9.14001 | 2.15922 | 0.00000 | 5.91842 | 0.57607 | 5.57066 | 5.03111 | 0.26670 | 4.85680 |
| **EPHB6** | 0.00155 | 0.15041 | 0.00000 | 0.30767 | 0.04145 | 0.01763 | 0.04645 | 0.07567 | 0.03840 | 0.04420 | 0.00468 | 0.12228 |
| **EPHX1** | 0.77506 | 0.74712 | 0.75219 | 0.39130 | 0.50876 | 0.00000 | 0.00000 | 0.62928 | 0.00000 | 0.67142 | 0.01585 | 2.05855 |
| **EPHX2** | 0.34643 | 0.11728 | 0.21911 | 0.47911 | 0.09648 | 0.00000 | 0.00006 | 0.02106 | 0.03950 | 0.00315 | 0.25271 | 0.09982 |
| **EPHX3** | 0.48878 | 4.19809 | 2.07466 | 2.89656 | 0.60911 | 0.00000 | 0.00000 | 0.14506 | 3.18565 | 0.00000 | 1.11963 | 2.08154 |
| **EPHX4** | 0.02303 | 0.10935 | 0.13263 | 0.12333 | 0.08079 | 0.42710 | 0.40453 | 1.72001 | 0.00000 | 0.70292 | 0.03719 | 0.21365 |
| **EPM2A** | 0.00201 | 0.04940 | 0.33586 | 0.06103 | 0.19460 | 0.00000 | 0.15990 | 0.06470 | 0.00000 | 0.00000 | 0.09869 | 0.19692 |
| **EPM2AIP1** | 1.67896 | 0.10215 | 1.34024 | 1.26857 | 0.93197 | 7.87241 | 0.00000 | 7.82903 | 0.00000 | 6.05861 | 3.81421 | 2.06294 |
| **EPN1** | 0.96638 | 0.34495 | 3.26365 | 1.02756 | 4.63134 | 0.00000 | 2.85878 | 0.22995 | 0.00000 | 12.92862 | 0.65548 | 0.81310 |
| **EPN2** | 5.59270 | 2.66824 | 3.21908 | 3.99765 | 3.62988 | 0.00000 | 0.60688 | 3.55731 | 2.49142 | 2.53000 | 1.67063 | 7.84872 |
| **EPN2-AS1** | 0.00332 | 0.02457 | 0.00709 | 0.04701 | 0.00000 | 0.06122 | 0.00539 | 0.01761 | 0.00538 | 0.04424 | 0.01707 | 0.03459 |
| **EPN2-IT1** | 0.34043 | 0.03135 | 0.19012 | 0.03561 | 0.05766 | 0.03416 | 0.07217 | 5.60972 | 0.23104 | 0.00000 | 0.00328 | 1.06072 |

| | | | | | | | | | | | | |
|---|---|---|---|---|---|---|---|---|---|---|---|---|
| **EPN3** | 0.00201 | 0.00524 | 0.45663 | 0.00000 | 0.19818 | 0.00000 | 0.27452 | 0.03929 | 0.00000 | 0.10948 | 0.03181 | 0.01042 |
| **EPO** | 0.01199 | 0.07103 | 2.27172 | 0.18982 | 0.00000 | 0.00000 | 0.21926 | 0.02722 | 0.00000 | 0.04486 | 0.01392 | 0.06550 |
| **EPOR** | 0.49888 | 1.00997 | 0.52359 | 1.10396 | 0.10263 | 0.00000 | 0.15855 | 1.70509 | 0.53116 | 0.54197 | 0.07544 | 1.47867 |
| **EPPIN** | 0.00367 | 0.01351 | 0.03392 | 0.00187 | 0.08015 | 0.08785 | 0.00000 | 0.02029 | 0.10372 | 0.00968 | 0.00000 | 0.01268 |
| **EPPIN-WFDC6** | 0.00586 | 0.00787 | 0.01431 | 0.01465 | 0.00638 | 0.02134 | 0.00918 | 0.00820 | 0.01256 | 0.03532 | 0.05842 | 0.05881 |
| **EPPK1** | 0.02036 | 6.74584 | 1.62304 | 3.91008 | 9.97036 | 1.44528 | 1.11348 | 0.03982 | 10.84616 | 15.74756 | 2.97116 | 0.45955 |
| **EPRS** | 4.29890 | 11.55238 | 18.04917 | 6.15142 | 10.19556 | 29.31305 | 25.46414 | 2.19676 | 19.46552 | 25.98437 | 25.54715 | 15.06505 |
| **EPS15** | 5.17150 | 3.98510 | 8.73931 | 3.98027 | 0.43211 | 16.34116 | 7.42269 | 5.69070 | 6.64799 | 5.77852 | 4.32111 | 3.11324 |
| **EPS15L1** | 6.46888 | 3.68460 | 7.19761 | 4.61682 | 8.52163 | 0.00000 | 7.61611 | 0.83022 | 9.94380 | 3.09045 | 10.09620 | 2.12926 |
| **EPS8** | 0.35368 | 0.19820 | 3.89454 | 0.47718 | 0.00015 | 3.63705 | 1.20446 | 0.43721 | 3.04435 | 0.07874 | 0.00000 | 0.34410 |
| **EPS8L1** | 2.18385 | 0.00000 | 0.65949 | 0.65066 | 0.01848 | 0.00000 | 0.15836 | 7.12109 | 0.00000 | 0.41709 | 0.01369 | 0.13803 |
| **EPS8L2** | 0.00000 | 4.20745 | 1.81173 | 1.14585 | 18.58749 | 0.00000 | 0.25276 | 0.09593 | 4.95804 | 3.38912 | 10.74206 | 0.00000 |
| **EPS8L3** | 0.02744 | 0.01578 | 0.15962 | 0.01196 | 0.01454 | 0.01696 | 0.02620 | 0.04993 | 0.02101 | 0.00745 | 0.02339 | 0.01353 |
| **EPSTI1** | 8.17275 | 0.00000 | 0.62451 | 2.25158 | 5.08985 | 2.69229 | 2.42491 | 0.21728 | 0.00917 | 0.00000 | 3.70554 | 0.06317 |
| **EPT1** | 2.65060 | 8.05312 | 6.47672 | 5.88468 | 6.72202 | 30.41252 | 0.00000 | 3.79041 | 22.51064 | 22.24131 | 12.81600 | 7.44718 |
| **EPX** | 0.00844 | 0.03066 | 0.03085 | 0.05764 | 0.00740 | 0.01814 | 0.01190 | 0.01523 | 0.01102 | 0.02384 | 0.04514 | 0.02341 |
| **EQTN** | 0.00482 | 0.01344 | 0.01989 | 0.07543 | 0.06457 | 0.06961 | 0.00000 | 0.00970 | 0.11968 | 0.05907 | 0.02043 | 0.00919 |
| **ERAL1** | 2.53594 | 27.77027 | 8.89326 | 14.14192 | 18.67808 | 11.89387 | 12.73923 | 3.05541 | 12.28587 | 0.00000 | 16.82981 | 21.17753 |
| **ERAP1** | 0.01690 | 0.00000 | 0.00384 | 0.00000 | 0.26259 | 0.04967 | 0.12741 | 0.01587 | 0.00000 | 0.65301 | 0.05989 | 0.00197 |
| **ERAP2** | 0.00405 | 0.01053 | 0.01612 | 0.01533 | 0.00720 | 0.00717 | 0.02754 | 0.01363 | 0.00357 | 0.04477 | 0.06197 | 0.06565 |
| **ERAS** | 0.00511 | 0.00804 | 0.01862 | 0.01575 | 0.02903 | 0.00000 | 0.02313 | 0.01349 | 0.03085 | 0.00000 | 0.02065 | 0.17806 |
| **ERBB2** | 0.00000 | 2.96478 | 3.98291 | 0.91981 | 5.80897 | 0.00000 | 0.74704 | 0.46669 | 5.17232 | 4.30519 | 3.92581 | 0.08113 |
| **ERBB2IP** | 1.39179 | 4.78368 | 4.74622 | 6.37690 | 0.85554 | 26.76657 | 4.37039 | 1.62505 | 0.06462 | 28.28723 | 2.39134 | 4.76249 |
| **ERBB3** | 0.27564 | 7.45355 | 11.47720 | 4.74045 | 8.23878 | 6.90040 | 0.02307 | 1.18362 | 4.34048 | 14.84008 | 1.88651 | 10.80774 |
| **ERBB4** | 12.50522 | 0.00000 | 0.00000 | 3.27685 | 0.48012 | 0.46527 | 0.09192 | 1.75670 | 0.00000 | 0.27133 | 0.14401 | 0.78141 |
| **ERC1** | 3.69041 | 0.18124 | 5.01337 | 1.52143 | 8.28494 | 4.10421 | 6.76229 | 3.60950 | 5.46017 | 19.16845 | 3.82832 | 3.76836 |
| **ERC2** | 3.48450 | 0.49792 | 0.00000 | 0.06357 | 0.09396 | 0.00000 | 0.06164 | 0.04978 | 0.73619 | 1.18703 | 0.00000 | 0.45453 |
| **ERC2-IT1** | 0.00639 | 0.01551 | 0.01385 | 0.01180 | 0.01931 | 0.00731 | 0.02004 | 0.01186 | 0.03222 | 0.02686 | 0.02030 | 0.01609 |
| **ERCC1** | 1.04576 | 6.31891 | 1.33073 | 2.87171 | 10.15984 | 4.84440 | 0.00000 | 1.77917 | 0.40203 | 0.01307 | 1.69077 | 5.75281 |
| **ERCC2** | 7.26531 | 5.12935 | 2.39324 | 3.34197 | 7.38230 | 0.00000 | 2.12322 | 0.32742 | 3.29564 | 0.00000 | 5.74550 | 3.49985 |
| **ERCC3** | 3.10908 | 14.05553 | 10.85542 | 9.19529 | 3.92812 | 12.29598 | 13.88493 | 0.57074 | 17.22731 | 0.00000 | 13.52457 | 9.83039 |
| **ERCC4** | 2.46834 | 1.71985 | 2.53801 | 1.20842 | 0.38539 | 5.01312 | 2.13777 | 0.57102 | 2.51388 | 4.68883 | 1.84408 | 3.25875 |
| **ERCC5** | 0.61958 | 0.32694 | 0.35447 | 0.17582 | 0.29817 | 0.85621 | 0.00000 | 0.31195 | 0.46041 | 0.00000 | 0.16168 | 0.35021 |
| **ERCC6** | 3.11785 | 1.06796 | 1.20821 | 0.71201 | 1.38626 | 3.21893 | 2.02742 | 0.98195 | 1.84378 | 1.31830 | 1.41473 | 2.47185 |
| **ERCC6L** | 3.36404 | 4.68721 | 2.94737 | 3.23733 | 0.54759 | 9.93953 | 1.91645 | 4.32897 | 6.77666 | 0.00000 | 5.36814 | 7.22809 |
| **ERCC6L2** | 2.15294 | 0.55655 | 2.17553 | 0.51719 | 0.44226 | 5.74294 | 4.63272 | 0.72844 | 3.33391 | 2.85016 | 1.77480 | 1.14933 |
| **ERCC8** | 0.01445 | 2.81580 | 3.03486 | 0.53757 | 1.48479 | 4.60768 | 1.76883 | 0.80380 | 4.01249 | 0.00000 | 2.26227 | 2.10662 |
| **EREG** | 0.00738 | 0.02110 | 0.02459 | 0.01169 | 0.02031 | 0.01231 | 0.02358 | 0.01030 | 0.01739 | 0.01252 | 0.01604 | 0.01858 |
| **ERF** | 1.15812 | 1.12496 | 0.79242 | 1.88177 | 12.26945 | 0.00000 | 2.65507 | 0.00000 | 1.93526 | 15.66109 | 0.00000 | 3.07266 |
| **ERG** | 1.52151 | 0.07701 | 0.02493 | 0.00000 | 0.01515 | 0.01052 | 0.01164 | 0.00000 | 0.01103 | 0.06757 | 0.01723 | 0.13164 |
| **ERGIC1** | 0.04923 | 1.25478 | 0.07128 | 0.11995 | 2.67815 | 0.00000 | 0.14780 | 0.09929 | 1.28870 | 3.49719 | 10.56198 | 0.56790 |
| **ERGIC2** | 1.79854 | 5.02395 | 12.61012 | 1.57572 | 0.18345 | 36.04034 | 1.29397 | 5.95189 | 16.14409 | 9.10813 | 7.48325 | 9.88279 |
| **ERGIC3** | 1.51104 | 22.68226 | 20.76063 | 16.89195 | 18.85277 | 14.92367 | 11.26952 | 6.67364 | 28.76460 | 7.80482 | 14.80923 | 17.81295 |
| **ERH** | 7.63337 | 11.15430 | 23.22300 | 14.00952 | 12.74574 | 34.73791 | 37.48236 | 13.78797 | 20.79254 | 15.53246 | 30.92241 | 35.74376 |
| **ERI1** | 4.02925 | 3.93501 | 8.18472 | 3.72473 | 1.45149 | 27.89590 | 7.62238 | 12.84196 | 9.59982 | 7.19332 | 8.11387 | 11.03434 |
| **ERI2** | 1.07028 | 4.84327 | 2.01206 | 0.34601 | 0.95571 | 9.57286 | 0.00000 | 1.04555 | 3.08085 | 0.00000 | 2.23664 | 3.21588 |
| **ERI3** | 0.72891 | 6.55046 | 8.08765 | 8.32268 | 10.77830 | 1.04237 | 12.46201 | 0.05432 | 10.89688 | 0.00000 | 8.25577 | 5.86775 |
| **ERI3-IT1** | 0.04691 | 0.01792 | 0.09320 | 0.05683 | 0.02453 | 0.00000 | 0.00732 | 0.05402 | 0.05667 | 0.04732 | 0.08066 | 0.08498 |

| | | | | | | | | | | | | |
|---|---|---|---|---|---|---|---|---|---|---|---|---|
| **ERICH1** | 1.82775 | 7.55998 | 5.06139 | 3.94959 | 13.61742 | 6.01798 | 7.80540 | 1.49979 | 6.71727 | 13.82378 | 4.90587 | 3.03955 |
| **ERICH1-AS1** | 0.44516 | 0.01202 | 0.26397 | 0.05606 | 0.45083 | 0.00000 | 0.00468 | 0.08600 | 0.32770 | 0.00000 | 0.12529 | 0.04591 |
| **ERICH2** | 0.32816 | 0.75496 | 0.00181 | 1.25449 | 0.25601 | 1.11059 | 0.00000 | 0.00000 | 0.34429 | 0.00000 | 0.36705 | 1.25373 |
| **ERICH3** | 0.00364 | 0.08329 | 0.22057 | 0.05466 | 0.00000 | 0.36489 | 0.00000 | 0.00656 | 0.00000 | 0.14700 | 0.02821 | 0.00000 |
| **ERICH3-AS1** | 0.00645 | 0.01890 | 0.04072 | 0.03771 | 0.01764 | 0.00427 | 0.05427 | 0.01062 | 0.00548 | 0.00000 | 0.01720 | 0.01825 |
| **ERICH4** | 0.04658 | 0.01132 | 0.01671 | 0.00906 | 0.00888 | 0.02417 | 0.01364 | 0.00738 | 0.01636 | 0.04397 | 0.01273 | 0.02721 |
| **ERICH5** | 0.03595 | 0.51158 | 1.40747 | 3.49496 | 0.00000 | 2.21553 | 1.76546 | 0.48505 | 0.00000 | 0.40329 | 0.78230 | 0.45906 |
| **ERICH6** | 0.02817 | 0.02576 | 0.01223 | 0.02656 | 0.01308 | 0.01312 | 0.01523 | 0.01263 | 0.02559 | 0.02771 | 0.01593 | 0.02873 |
| **ERICH6-AS1** | 0.04322 | 0.05841 | 0.00000 | 0.22017 | 0.00000 | 0.10683 | 0.05362 | 0.03695 | 0.11751 | 0.18547 | 0.01308 | 0.10846 |
| **ERICH6B** | 0.10840 | 0.01449 | 0.01115 | 0.02179 | 0.02773 | 0.00000 | 0.03070 | 0.00757 | 0.01872 | 0.06224 | 0.01904 | 0.01322 |
| **ERLEC1** | 2.79711 | 8.41738 | 9.14146 | 2.88883 | 2.80607 | 17.48191 | 0.93680 | 2.82080 | 12.31690 | 10.10756 | 6.38129 | 6.84786 |
| **ERLIN1** | 4.67763 | 12.18514 | 10.05962 | 3.87579 | 6.65455 | 16.60928 | 0.00000 | 3.64169 | 23.39383 | 3.92092 | 13.12983 | 12.46369 |
| **ERLIN2** | 0.00000 | 1.74321 | 2.76212 | 0.00000 | 1.13466 | 6.79634 | 0.00000 | 0.66459 | 0.00000 | 11.35412 | 3.37412 | 4.29978 |
| **ERMAP** | 0.00120 | 0.00000 | 0.32893 | 0.18728 | 0.15178 | 1.10232 | 0.16732 | 1.89929 | 0.50057 | 0.00000 | 0.70863 | 1.39387 |
| **ERMARD** | 1.65306 | 2.55832 | 0.61657 | 0.70307 | 0.95109 | 5.43531 | 0.54438 | 0.98596 | 2.02767 | 1.70044 | 2.71088 | 2.28072 |
| **ERMN** | 0.00568 | 0.11156 | 0.01949 | 0.03322 | 0.00000 | 0.01230 | 0.85113 | 0.01159 | 0.04018 | 0.00000 | 0.26397 | 0.01926 |
| **ERMP1** | 1.51823 | 1.28795 | 0.97807 | 0.00000 | 3.23644 | 5.33534 | 0.00000 | 0.00000 | 3.32191 | 8.79474 | 2.16605 | 0.38320 |
| **1.00000** | 5.04822 | 0.13018 | 1.07119 | 0.08508 | 0.74361 | 0.16370 | 0.11167 | 0.66509 | 0.10991 | 0.00000 | 0.00000 | 0.53205 |
| **2.00000** | 0.00650 | 0.00341 | 0.07547 | 0.19140 | 0.05163 | 0.01578 | 0.07392 | 0.01322 | 0.00000 | 0.07229 | 0.01951 | 0.03395 |
| **ERO1L** | 1.35270 | 0.00000 | 2.70341 | 0.00000 | 0.59796 | 6.69218 | 3.37437 | 0.88530 | 2.98907 | 5.17904 | 0.90110 | 0.00000 |
| **ERO1LB** | 5.02986 | 0.30393 | 0.37518 | 0.63757 | 0.00000 | 0.32807 | 0.00000 | 1.79316 | 0.51008 | 1.51252 | 0.00000 | 0.02152 |
| **ERP27** | 0.00572 | 0.00000 | 2.10259 | 0.00101 | 0.00000 | 0.03871 | 0.53083 | 0.04761 | 0.58851 | 0.00000 | 0.47457 | 0.01062 |
| **ERP29** | 3.47852 | 23.40826 | 26.63128 | 21.78516 | 25.37146 | 15.93186 | 19.11008 | 2.10929 | 51.25319 | 24.24664 | 29.06166 | 18.48190 |
| **ERP44** | 3.99727 | 12.57156 | 11.45841 | 9.79491 | 7.64381 | 24.67497 | 0.00000 | 7.80101 | 25.14714 | 0.00000 | 10.06884 | 8.86834 |
| **ERRFI1** | 1.26618 | 1.17352 | 31.38009 | 11.15324 | 5.22225 | 2.59157 | 24.45183 | 6.24307 | 15.93808 | 0.00042 | 6.49138 | 5.42400 |
| **ERV3-1** | 0.09549 | 0.00000 | 1.46115 | 0.27043 | 0.06337 | 0.62496 | 0.00000 | 0.40083 | 2.48573 | 0.21650 | 0.00000 | 0.35716 |
| **ERVFRD-1** | 0.70379 | 0.08737 | 7.71477 | 0.00264 | 0.10067 | 0.00000 | 0.15904 | 0.03008 | 0.51374 | 0.00001 | 0.00000 | 0.12411 |
| **ERVK13-1** | 0.16838 | 0.33851 | 0.31403 | 0.36661 | 0.33068 | 0.00000 | 0.00000 | 3.03805 | 0.55384 | 0.90503 | 0.08220 | 0.48362 |
| **ERVMER34-1** | 0.00000 | 1.57597 | 9.26693 | 0.00000 | 0.00000 | 0.00784 | 4.93572 | 0.06696 | 9.91787 | 0.00000 | 3.51325 | 0.76702 |
| **ERVV-1** | 0.00713 | 0.00405 | 7.92563 | 0.00278 | 0.03268 | 0.00000 | 0.00000 | 0.00862 | 0.31503 | 0.05768 | 0.18643 | 0.04536 |
| **ERVV-2** | 0.06882 | 0.00000 | 10.47766 | 0.00292 | 1.07882 | 0.03646 | 0.00000 | 0.00446 | 0.02377 | 0.00559 | 3.80862 | 0.03665 |
| **ESAM** | 0.07816 | 0.21260 | 0.00000 | 4.91123 | 0.18963 | 0.72044 | 0.00000 | 0.07496 | 0.09789 | 0.00000 | 0.00000 | 0.14239 |
| **ESCO1** | 1.61430 | 2.41969 | 6.23057 | 10.01215 | 0.00000 | 19.77707 | 3.13903 | 0.23875 | 3.76639 | 10.42785 | 3.83949 | 6.62435 |
| **ESCO2** | 9.60388 | 1.62275 | 2.26394 | 0.80399 | 0.00141 | 11.47499 | 2.90658 | 7.68755 | 3.11209 | 2.59927 | 3.45341 | 7.52827 |
| **ESD** | 0.73369 | 0.74599 | 6.13856 | 3.03628 | 0.16005 | 15.26548 | 9.19869 | 0.00000 | 8.93554 | 0.75223 | 10.83507 | 0.21377 |
| **ESF1** | 3.79054 | 4.69407 | 6.43264 | 2.57385 | 0.85788 | 25.59286 | 11.83479 | 10.29477 | 3.61988 | 33.51679 | 12.67294 | 17.58535 |
| **ESPL1** | 4.51856 | 8.00971 | 5.21132 | 4.64269 | 9.10868 | 0.02072 | 9.89862 | 0.50739 | 14.35721 | 11.64730 | 5.76423 | 17.95436 |
| **ESPN** | 0.10732 | 0.07537 | 0.00043 | 0.02709 | 0.00000 | 0.00000 | 0.03886 | 0.00730 | 0.09901 | 0.45574 | 0.02241 | 0.06283 |
| **ESPNL** | 0.00614 | 0.01168 | 0.00679 | 0.01761 | 0.01954 | 0.03796 | 0.03459 | 0.02349 | 0.03304 | 0.01758 | 0.00899 | 0.01403 |
| **ESPNP** | 0.00979 | 0.02002 | 0.01620 | 0.01361 | 0.02101 | 0.00672 | 0.03097 | 0.01300 | 0.01973 | 0.01811 | 0.01392 | 0.01602 |
| **ESR1** | 0.70239 | 0.00000 | 0.12347 | 0.00830 | 0.00432 | 0.00000 | 0.05348 | 0.00000 | 0.00000 | 0.33255 | 0.00600 | 0.30256 |
| **ESR2** | 1.47813 | 0.31435 | 0.30936 | 0.16498 | 0.03486 | 0.24547 | 0.00000 | 0.01129 | 0.59660 | 0.94952 | 0.23072 | 0.34736 |
| **ESRG** | 0.08342 | 3.88502 | 4.33424 | 16.72536 | 6.06786 | 7.80189 | 3.29460 | 0.00000 | 0.00005 | 8.70702 | 2.58721 | 0.00000 |
| **ESRP1** | 10.63754 | 15.79396 | 8.27214 | 12.12268 | 13.64454 | 29.71770 | 16.53111 | 5.83363 | 12.60199 | 7.61546 | 6.54774 | 15.79525 |
| **ESRP2** | 0.00000 | 8.77768 | 10.39258 | 4.16508 | 12.30654 | 0.00000 | 2.07993 | 1.59309 | 10.16627 | 4.18528 | 10.12208 | 2.94967 |
| **ESRRA** | 0.00686 | 6.21250 | 4.45196 | 5.68619 | 11.06292 | 0.00000 | 5.78043 | 0.08575 | 8.11492 | 9.10323 | 3.66467 | 0.67093 |
| **ESRRB** | 0.00000 | 11.27813 | 0.00000 | 0.70578 | 14.03169 | 0.52107 | 0.00000 | 0.00000 | 0.00000 | 4.37304 | 3.45666 | 7.24533 |
| **ESRRG** | 1.35652 | 2.28592 | 9.57852 | 0.00000 | 3.46645 | 0.00000 | 0.40614 | 0.02887 | 0.06718 | 2.60302 | 1.98578 | 0.00000 |

| | | | | | | | | | | | | |
|---|---|---|---|---|---|---|---|---|---|---|---|---|
| **ESX1** | 0.00500 | 0.00507 | 0.02493 | 0.01919 | 0.00634 | 0.02529 | 0.00464 | 0.00834 | 0.01240 | 0.08631 | 0.01202 | 0.06734 |
| **ESYT1** | 0.35597 | 6.06405 | 6.79008 | 5.47586 | 1.62967 | 4.08049 | 0.00000 | 3.87809 | 11.62408 | 1.82672 | 0.00000 | 20.43651 |
| **ESYT2** | 1.84629 | 2.65120 | 8.38404 | 2.69473 | 10.52892 | 7.21308 | 3.29687 | 2.70300 | 11.79214 | 9.36547 | 2.44614 | 2.77944 |
| **ESYT3** | 1.85046 | 0.25652 | 0.45095 | 0.03297 | 0.00000 | 0.00000 | 0.02228 | 0.82134 | 0.00000 | 0.11451 | 0.03352 | 0.17167 |
| **ETAA1** | 1.25521 | 0.73327 | 2.11664 | 0.58495 | 0.00000 | 6.67093 | 0.00066 | 2.20029 | 1.75238 | 1.37492 | 0.60959 | 1.69020 |
| **ETF1** | 0.21010 | 18.36253 | 20.00038 | 8.79402 | 14.18016 | 32.60955 | 28.75881 | 22.57509 | 18.20744 | 18.60369 | 24.04188 | 20.70671 |
| **ETFA** | 13.65991 | 5.81127 | 15.74856 | 8.31420 | 6.67237 | 36.71730 | 18.75683 | 8.57827 | 21.45852 | 1.64524 | 19.31859 | 3.68521 |
| **ETFB** | 0.00000 | 8.78439 | 8.21563 | 10.28368 | 11.91869 | 0.05711 | 9.76715 | 1.02461 | 3.51658 | 2.04585 | 10.32839 | 2.66423 |
| **ETFDH** | 0.96569 | 1.86138 | 3.34524 | 2.46153 | 0.45165 | 1.86642 | 6.25243 | 5.80034 | 5.75293 | 0.00000 | 5.24286 | 2.14514 |
| **ETHE1** | 4.17126 | 13.38770 | 1.45729 | 9.90679 | 13.15435 | 7.96253 | 0.00000 | 1.08685 | 4.12226 | 0.00000 | 2.18382 | 1.83879 |
| **ETNK1** | 0.09647 | 1.62214 | 5.18979 | 1.20641 | 0.56963 | 18.47447 | 4.27383 | 9.61074 | 13.57013 | 0.93893 | 5.68893 | 3.62120 |
| **ETNK2** | 0.02400 | 1.61575 | 1.92350 | 4.26120 | 0.94858 | 0.00000 | 0.90999 | 0.15253 | 2.56793 | 0.14496 | 0.26617 | 0.57563 |
| **ETNPPL** | 1.31613 | 11.98529 | 0.00000 | 0.01019 | 0.47135 | 0.00000 | 0.01947 | 0.07706 | 0.00000 | 0.00000 | 0.00000 | 27.36363 |
| **ETS1** | 1.16115 | 0.06944 | 0.08574 | 0.23153 | 0.00000 | 0.45474 | 0.09757 | 0.11889 | 0.12489 | 0.01161 | 0.00000 | 0.82740 |
| **ETS2** | 0.32194 | 1.68562 | 4.67451 | 0.44387 | 1.60842 | 0.00000 | 2.92887 | 0.07993 | 3.33310 | 0.63744 | 0.57239 | 0.59136 |
| **ETV1** | 0.59062 | 0.00000 | 0.22603 | 6.62410 | 0.03586 | 0.14425 | 0.00000 | 0.07492 | 0.00000 | 0.01875 | 0.05583 | 0.00939 |
| **ETV2** | 0.00217 | 0.04795 | 0.12884 | 0.18713 | 0.06785 | 0.00000 | 0.00899 | 0.28866 | 0.17975 | 0.20720 | 0.07431 | 0.00192 |
| **ETV3** | 1.71397 | 0.88831 | 0.15890 | 0.15618 | 4.46666 | 2.14467 | 0.00000 | 1.95941 | 0.80656 | 2.40562 | 0.46071 | 1.69535 |
| **ETV3L** | 0.04566 | 0.01934 | 0.02415 | 0.01200 | 0.01724 | 0.01104 | 0.01758 | 0.04833 | 0.01990 | 0.01098 | 0.01514 | 0.02279 |
| **ETV4** | 0.89160 | 0.19229 | 3.52833 | 20.76706 | 0.23651 | 1.01106 | 0.00000 | 3.56096 | 0.55188 | 0.00000 | 0.00066 | 5.40194 |
| **ETV5** | 4.32103 | 0.92882 | 1.57313 | 5.21247 | 0.37010 | 1.02654 | 0.43494 | 7.33648 | 0.00000 | 0.00012 | 0.37236 | 2.11928 |
| **ETV6** | 1.00373 | 1.15103 | 1.12327 | 3.00983 | 0.66225 | 1.02097 | 4.75733 | 0.06404 | 0.96540 | 3.60297 | 0.00000 | 0.87832 |
| **ETV7** | 0.23658 | 0.01344 | 0.05097 | 0.01361 | 0.01002 | 0.01707 | 0.03712 | 0.00249 | 0.03633 | 0.01752 | 0.01146 | 0.01392 |
| **EVA1A** | 0.03350 | 0.01173 | 0.02070 | 0.00896 | 0.00675 | 0.02186 | 0.03441 | 0.02151 | 0.00000 | 0.06891 | 0.01359 | 0.04027 |
| **EVA1B** | 0.01528 | 0.18051 | 0.54126 | 0.47685 | 0.38546 | 0.00000 | 0.30734 | 0.58674 | 0.00000 | 0.53981 | 0.08119 | 0.12423 |
| **EVA1C** | 0.00539 | 0.01387 | 0.04609 | 0.02833 | 0.01317 | 0.01543 | 0.02274 | 0.01281 | 0.00563 | 0.02041 | 0.01742 | 0.01604 |
| **EVC** | 0.58246 | 0.08918 | 0.06040 | 0.14854 | 0.00000 | 0.00000 | 0.09859 | 0.00251 | 0.00000 | 0.37021 | 0.05758 | 0.12455 |
| **EVC2** | 1.73494 | 0.00045 | 0.04092 | 0.58621 | 0.00000 | 0.07530 | 0.01308 | 0.12005 | 0.10091 | 0.05911 | 0.02976 | 0.06039 |
| **EVI2A** | 0.01503 | 0.01793 | 0.31754 | 0.29125 | 0.00000 | 0.11070 | 0.00000 | 0.00851 | 0.23501 | 0.73290 | 0.00602 | 0.10317 |
| **EVI2B** | 0.00570 | 0.04439 | 0.13422 | 0.04806 | 0.02770 | 0.05172 | 0.06053 | 0.00997 | 0.24411 | 0.00490 | 0.04449 | 0.01490 |
| **EVI5** | 1.13510 | 0.68537 | 5.68557 | 0.00000 | 0.00000 | 8.23801 | 9.73383 | 1.32132 | 1.58688 | 14.61249 | 4.46198 | 3.09534 |
| **EVI5L** | 0.67543 | 0.01253 | 0.14673 | 0.00951 | 0.04254 | 0.00000 | 0.13053 | 0.02568 | 0.00000 | 0.00000 | 0.09264 | 0.04123 |
| **EVL** | 0.33770 | 0.12308 | 0.07373 | 0.13865 | 0.67915 | 0.10671 | 0.76366 | 0.00780 | 0.43783 | 2.66981 | 0.11824 | 0.00797 |
| **EVPL** | 0.33609 | 0.02481 | 0.60749 | 0.00000 | 0.63818 | 0.00000 | 0.31845 | 0.09771 | 0.00000 | 0.13487 | 0.34648 | 0.02187 |
| **EVPLL** | 0.01256 | 0.05276 | 0.02074 | 0.07593 | 0.01140 | 0.00000 | 0.02312 | 0.00873 | 0.01635 | 0.01789 | 0.01995 | 0.12093 |
| **EVX2** | 0.00574 | 0.01439 | 0.01437 | 0.01239 | 0.01632 | 0.01793 | 0.01887 | 0.02932 | 0.01816 | 0.01116 | 0.01517 | 0.02224 |
| **EWSR1** | 2.79515 | 32.04305 | 27.42845 | 28.92815 | 23.65467 | 16.21178 | 37.06866 | 19.68433 | 33.81462 | 3.17700 | 27.03314 | 29.37629 |
| **EXD1** | 0.29471 | 0.22833 | 0.02757 | 0.08676 | 0.00000 | 0.01610 | 0.01463 | 0.94088 | 0.00000 | 0.11014 | 0.08964 | 0.42379 |
| **EXD2** | 1.52902 | 2.90831 | 5.07885 | 0.88946 | 0.04954 | 8.72272 | 0.00000 | 0.93458 | 13.32895 | 1.06558 | 3.64378 | 5.74447 |
| **EXD3** | 3.65911 | 0.50089 | 1.03836 | 0.21273 | 0.14698 | 0.13077 | 0.03428 | 0.17625 | 0.50180 | 0.00000 | 0.65512 | 0.79393 |
| **EXO1** | 0.51060 | 10.34892 | 15.95229 | 4.18363 | 5.77625 | 20.18791 | 4.75113 | 7.54843 | 5.80014 | 1.23372 | 5.56453 | 9.58012 |
| **EXO5** | 3.30391 | 2.69554 | 0.28807 | 1.31294 | 0.43920 | 1.41425 | 1.39350 | 7.12920 | 4.35102 | 0.04013 | 2.53609 | 6.35446 |
| **EXOC1** | 1.33393 | 4.96715 | 2.84250 | 2.66695 | 1.17615 | 6.85743 | 5.73954 | 3.28366 | 0.59603 | 4.98638 | 4.54481 | 3.89081 |
| **EXOC2** | 1.93114 | 2.40285 | 0.67686 | 4.24990 | 1.47703 | 5.28247 | 3.02207 | 0.55781 | 3.48476 | 0.44436 | 1.76579 | 1.01109 |
| **EXOC3** | 2.79037 | 2.17475 | 1.93165 | 2.13553 | 4.90636 | 1.17342 | 4.82179 | 1.51454 | 3.07719 | 5.04094 | 6.86491 | 2.32980 |
| **EXOC3L1** | 0.03483 | 0.01270 | 0.12868 | 1.77985 | 0.36646 | 0.00037 | 0.01264 | 0.00996 | 0.00000 | 0.00000 | 0.18723 | 0.03214 |
| **EXOC3L2** | 7.14510 | 0.04235 | 0.05879 | 0.00000 | 0.13623 | 0.00000 | 0.00280 | 1.32635 | 0.03003 | 0.00000 | 0.00458 | 1.22239 |
| **EXOC3L4** | 0.00644 | 0.01943 | 0.02413 | 0.01958 | 0.01798 | 0.00000 | 0.03519 | 0.01160 | 0.01539 | 0.01569 | 0.02407 | 0.01745 |

| | | | | | | | | | | | | |
|---|---|---|---|---|---|---|---|---|---|---|---|---|
| **EXOC4** | 4.49740 | 1.83134 | 3.66198 | 2.35521 | 1.44236 | 6.15575 | 0.83441 | 0.33946 | 1.99403 | 5.32491 | 3.50396 | 0.67283 |
| **EXOC5** | 4.37023 | 1.72050 | 8.07457 | 3.58585 | 0.00000 | 25.76090 | 11.28211 | 4.84258 | 5.82258 | 12.84411 | 5.96179 | 3.62394 |
| **EXOC6** | 0.47272 | 2.63806 | 1.26315 | 0.00000 | 0.58292 | 4.67250 | 2.90107 | 0.63491 | 1.48588 | 2.04906 | 2.56072 | 1.57831 |
| **EXOC6B** | 0.11396 | 0.29628 | 2.22362 | 0.42237 | 0.60701 | 1.94592 | 0.03717 | 0.79562 | 0.66444 | 1.27414 | 1.14791 | 1.20301 |
| **EXOC7** | 0.71564 | 4.33423 | 7.84194 | 6.45476 | 7.39759 | 0.00000 | 6.62567 | 0.16528 | 5.95594 | 2.53563 | 7.08925 | 2.15347 |
| **EXOC8** | 0.00000 | 1.57606 | 5.41342 | 0.00000 | 0.00000 | 4.57550 | 0.56278 | 2.60735 | 1.39800 | 3.65586 | 1.44550 | 0.96734 |
| **EXOG** | 0.32774 | 0.30382 | 2.40410 | 0.97470 | 0.69447 | 3.23874 | 0.89263 | 0.00000 | 0.42841 | 0.00000 | 0.67386 | 0.31343 |
| **EXOSC1** | 0.20486 | 13.36610 | 10.58697 | 6.31411 | 6.37026 | 16.37964 | 15.45201 | 6.22482 | 11.16810 | 0.84779 | 9.23967 | 17.18733 |
| **EXOSC10** | 3.33805 | 21.48094 | 15.43083 | 10.46820 | 10.74196 | 20.07717 | 16.90132 | 15.70627 | 25.24043 | 0.00000 | 18.67983 | 19.79841 |
| **EXOSC2** | 0.00000 | 10.14188 | 5.05527 | 6.12324 | 8.23189 | 13.53212 | 14.62328 | 3.90401 | 15.40491 | 3.89277 | 13.78482 | 14.86736 |
| **EXOSC3** | 9.22208 | 16.25969 | 15.62917 | 5.65750 | 7.36557 | 27.09493 | 18.00840 | 17.95898 | 16.79423 | 0.00000 | 18.22115 | 21.58831 |
| **EXOSC4** | 0.26181 | 17.50891 | 17.44410 | 12.49744 | 22.16315 | 0.15817 | 30.05338 | 5.85711 | 16.48604 | 1.99276 | 19.07214 | 25.89731 |
| **EXOSC5** | 0.29783 | 16.98879 | 13.67685 | 8.31911 | 10.98411 | 4.03121 | 17.93533 | 1.30115 | 27.79467 | 0.00000 | 15.75364 | 24.85596 |
| **EXOSC6** | 0.08684 | 6.59827 | 2.26481 | 3.68548 | 2.38304 | 3.85727 | 3.10307 | 4.00325 | 0.00000 | 10.65545 | 2.41085 | 5.32913 |
| **EXOSC7** | 6.61418 | 14.49252 | 12.58005 | 11.10248 | 14.39733 | 11.19159 | 17.08444 | 5.57409 | 12.06138 | 0.00000 | 16.04384 | 14.81383 |
| **EXOSC8** | 0.89378 | 16.90553 | 12.85480 | 12.80926 | 15.56458 | 38.86356 | 19.74275 | 2.31497 | 14.64167 | 3.65972 | 22.94733 | 13.97961 |
| **EXOSC9** | 1.48285 | 8.16975 | 13.87071 | 4.56658 | 6.02601 | 23.44303 | 18.37758 | 6.25786 | 13.49289 | 2.40111 | 11.45550 | 16.75070 |
| **EXPH5** | 0.00000 | 0.83276 | 4.93871 | 0.00000 | 6.54959 | 15.17277 | 1.61069 | 0.00000 | 12.61786 | 9.74393 | 5.99252 | 0.00000 |
| **EXT1** | 5.12822 | 0.54987 | 0.54467 | 10.00065 | 2.79109 | 0.53543 | 1.33654 | 1.11500 | 1.65048 | 4.12384 | 0.92326 | 1.74466 |
| **EXT2** | 2.30922 | 10.05394 | 4.84800 | 0.45866 | 6.06976 | 11.97374 | 0.00000 | 2.55261 | 13.53114 | 1.45385 | 5.25152 | 1.85088 |
| **EXTL1** | 0.30613 | 0.04015 | 0.04294 | 0.02605 | 0.24327 | 0.00000 | 0.01323 | 0.03967 | 0.05213 | 0.11749 | 0.05047 | 0.00442 |
| **EXTL2** | 0.00000 | 0.96891 | 8.68853 | 0.90665 | 0.00000 | 6.39162 | 4.05397 | 1.19218 | 8.13612 | 4.01662 | 0.89474 | 3.90343 |
| **EXTL3** | 7.72304 | 7.30186 | 2.67211 | 4.84251 | 2.51375 | 1.48684 | 1.79104 | 3.35003 | 4.96625 | 6.62264 | 0.69781 | 15.04804 |
| **EYA1** | 2.78491 | 0.00436 | 0.00000 | 0.35609 | 0.08388 | 0.13444 | 0.02787 | 0.56518 | 0.00000 | 0.13124 | 0.04755 | 0.05714 |
| **EYA2** | 2.21831 | 0.09770 | 0.03123 | 0.00700 | 0.00310 | 0.06621 | 0.00000 | 0.93619 | 0.07156 | 0.00000 | 0.00442 | 0.00755 |
| **EYA3** | 2.38584 | 6.62509 | 13.73893 | 8.27329 | 3.41674 | 6.38590 | 14.52961 | 17.92137 | 12.54958 | 1.66000 | 9.73257 | 8.31480 |
| **EYA4** | 0.15428 | 0.01475 | 0.05966 | 0.07605 | 0.00000 | 0.10736 | 0.00000 | 0.16616 | 0.00000 | 0.23851 | 0.00000 | 0.02436 |
| **EYA4-AS1** | 0.86394 | 0.01179 | 0.01284 | 0.00946 | 0.02284 | 0.01923 | 0.01413 | 0.13075 | 0.02311 | 0.00708 | 0.01375 | 0.01056 |
| **EYS** | 0.77691 | 0.04020 | 0.00000 | 0.02371 | 0.00000 | 0.07931 | 0.44938 | 0.27187 | 0.14244 | 0.79339 | 0.00000 | 0.32008 |
| **EZH1** | 0.49627 | 0.46722 | 1.72634 | 0.45635 | 0.00199 | 0.28586 | 0.30202 | 0.00000 | 4.06628 | 0.00000 | 1.43134 | 1.31751 |
| **EZH2** | 1.31085 | 15.85584 | 10.60894 | 2.94484 | 9.10710 | 14.37134 | 7.65870 | 8.13468 | 15.70066 | 8.99666 | 6.52553 | 11.82065 |
| **EZR** | 11.67409 | 27.43465 | 30.83917 | 26.21005 | 50.97980 | 47.82383 | 53.89558 | 9.34613 | 53.06740 | 29.87414 | 48.64813 | 19.37002 |
| **EZR-AS1** | 0.00568 | 0.01795 | 0.03780 | 0.01872 | 0.03033 | 0.08750 | 0.04142 | 0.01596 | 0.05934 | 0.02718 | 0.07860 | 0.03690 |
| **F10** | 0.49273 | 0.01376 | 0.01366 | 0.01492 | 0.01397 | 0.00702 | 0.01926 | 0.10641 | 0.02498 | 0.02083 | 0.01558 | 0.01026 |
| **F11** | 0.42299 | 0.02328 | 0.01824 | 0.00766 | 0.02263 | 0.01732 | 0.02197 | 0.00698 | 0.01881 | 0.01824 | 0.01055 | 0.01227 |
| **F11-AS1** | 0.07582 | 0.02506 | 0.03179 | 0.02374 | 0.01221 | 0.00000 | 0.01033 | 0.02569 | 0.10539 | 0.09705 | 0.01989 | 0.03632 |
| **F11R** | 0.00000 | 37.47814 | 34.88052 | 25.17648 | 25.77294 | 34.31246 | 4.96067 | 28.33763 | 53.46337 | 24.48156 | 24.88290 | 44.70745 |
| **F12** | 0.03987 | 0.07204 | 0.45909 | 0.03014 | 0.03057 | 0.00000 | 0.02463 | 0.06413 | 0.10861 | 0.00000 | 0.00396 | 0.01970 |
| **F13A1** | 1.37246 | 0.01395 | 0.01463 | 0.01320 | 0.01562 | 0.00273 | 0.02692 | 0.69507 | 0.01446 | 0.01595 | 0.01367 | 0.00000 |
| **F13B** | 0.00839 | 0.01610 | 0.04520 | 0.02024 | 0.01924 | 0.01063 | 0.00942 | 0.01150 | 0.08827 | 0.00000 | 0.00654 | 0.03747 |
| **F2** | 2.39330 | 0.04463 | 0.41737 | 0.16529 | 0.00000 | 0.00000 | 0.00000 | 0.45287 | 0.01886 | 0.09697 | 0.02546 | 0.01568 |
| **F2R** | 0.00144 | 2.61581 | 0.44978 | 0.50582 | 0.00000 | 0.68356 | 0.23844 | 0.03085 | 1.80481 | 0.06836 | 0.00000 | 1.40643 |
| **F2RL1** | 0.00000 | 24.37244 | 1.58552 | 20.99085 | 0.85358 | 34.67843 | 0.00000 | 5.94477 | 38.58790 | 7.17529 | 4.36853 | 28.37474 |
| **F2RL2** | 0.00284 | 0.00000 | 0.00000 | 0.00000 | 0.56349 | 0.00000 | 0.15439 | 0.00600 | 0.00000 | 1.09593 | 0.25574 | 0.06725 |
| **F2RL3** | 0.00700 | 0.01756 | 0.02413 | 0.01395 | 0.04260 | 0.00000 | 0.04188 | 0.01542 | 0.01796 | 0.03442 | 0.01838 | 0.02145 |
| **F3** | 0.00000 | 2.38637 | 6.23181 | 10.10151 | 0.00000 | 5.24577 | 3.77149 | 0.04534 | 8.70214 | 0.34587 | 17.11286 | 2.63101 |
| **F5** | 0.00583 | 0.00000 | 0.07125 | 0.00355 | 0.41714 | 0.07537 | 0.00472 | 0.00949 | 0.13308 | 0.00000 | 0.24017 | 0.01274 |
| **F8** | 0.05265 | 0.22401 | 0.08685 | 0.45970 | 0.00000 | 0.00000 | 0.09649 | 0.05732 | 0.00000 | 0.01309 | 0.26419 | 0.28128 |

| | | | | | | | | | | | | |
|---|---|---|---|---|---|---|---|---|---|---|---|---|
| **F8A1.uid-13426** | 0.04415 | 0.04708 | 0.05792 | 0.00769 | 0.00000 | 0.02439 | 0.05655 | 0.00418 | 0.00001 | 0.09398 | 0.01418 | 0.04018 |
| **FA2H** | 0.02864 | 11.54583 | 0.28076 | 1.21734 | 5.00208 | 0.00000 | 0.00000 | 0.05623 | 0.00000 | 2.10991 | 0.80010 | 4.17944 |
| **FAAH** | 0.05961 | 1.45297 | 0.73467 | 1.77212 | 2.18820 | 0.00000 | 0.00000 | 0.33102 | 0.92930 | 0.75339 | 0.30117 | 1.66165 |
| **FAAH2** | 0.01463 | 0.04750 | 0.00000 | 0.59782 | 0.01052 | 0.01141 | 0.01269 | 0.00799 | 0.03874 | 0.04764 | 0.02188 | 0.01942 |
| **FAAHP1** | 0.00443 | 0.02236 | 0.03888 | 0.05682 | 0.00000 | 0.00020 | 0.18318 | 0.00728 | 0.00000 | 0.13005 | 0.04469 | 0.08948 |
| **FABP1** | 0.00589 | 0.01432 | 0.01947 | 0.01807 | 0.01666 | 0.01441 | 0.03469 | 0.01054 | 0.01135 | 0.03063 | 0.01505 | 0.01544 |
| **FABP12** | 0.01956 | 0.00998 | 0.01634 | 0.00530 | 0.01946 | 0.05277 | 0.02189 | 0.02112 | 0.02084 | 0.01503 | 0.01384 | 0.01431 |
| **FABP2** | 0.03864 | 0.01247 | 0.01306 | 0.00867 | 0.00000 | 0.01763 | 0.19748 | 0.00401 | 0.01684 | 0.06151 | 0.02190 | 0.10480 |
| **FABP3** | 0.00000 | 0.00000 | 0.00000 | 0.76252 | 21.29950 | 0.12974 | 31.88454 | 0.19775 | 62.62808 | 1.58793 | 48.74944 | 0.00000 |
| **FABP4** | 0.00300 | 0.02714 | 0.06182 | 0.02283 | 0.00974 | 0.02103 | 0.01079 | 0.09834 | 0.07547 | 0.00206 | 0.00789 | 0.01219 |
| **FABP5** | 0.07201 | 2.59690 | 10.99154 | 12.13301 | 6.94351 | 4.19484 | 6.79120 | 0.02636 | 13.68101 | 0.00000 | 3.89524 | 0.20345 |
| **FABP5P3** | 0.05864 | 0.01727 | 0.24003 | 0.20567 | 0.00000 | 0.27651 | 0.16788 | 0.00365 | 0.26244 | 0.07938 | 0.04464 | 0.01910 |
| **FABP6** | 0.00202 | 0.16917 | 0.00000 | 0.50966 | 0.02894 | 0.33837 | 0.10697 | 0.22391 | 0.05350 | 0.00000 | 0.00658 | 0.01848 |
| **FABP7** | 0.10921 | 0.00000 | 3.05634 | 0.00000 | 1.28703 | 0.84261 | 5.65719 | 0.00000 | 5.70706 | 0.00713 | 4.97192 | 0.00000 |
| **FADD** | 0.79948 | 5.75140 | 7.79911 | 4.78085 | 6.47350 | 0.00000 | 6.40121 | 0.00000 | 9.84498 | 0.14196 | 8.95112 | 8.73402 |
| **FADS1** | 1.74122 | 0.00000 | 17.16870 | 0.00000 | 0.00000 | 0.00000 | 19.29475 | 0.65349 | 27.49860 | 0.00000 | 4.86431 | 1.69752 |
| **FADS2** | 0.00001 | 0.00000 | 8.12999 | 0.00000 | 0.00000 | 0.00000 | 1.30406 | 0.02447 | 23.69927 | 1.48171 | 5.73919 | 0.26195 |
| **FADS3** | 4.94891 | 1.13281 | 7.18636 | 0.64786 | 0.82610 | 0.00000 | 1.01755 | 0.57565 | 5.87794 | 4.69188 | 0.00000 | 0.67189 |
| **FADS6** | 0.00629 | 0.01428 | 0.03329 | 0.00713 | 0.00506 | 0.02656 | 0.02151 | 0.02314 | 0.01668 | 0.05745 | 0.01307 | 0.01187 |
| **FAF1** | 0.56262 | 4.40367 | 7.25857 | 7.64976 | 8.42731 | 7.19111 | 7.18705 | 0.00073 | 3.32816 | 12.93957 | 1.42091 | 4.53530 |
| **FAF2** | 0.71137 | 11.20364 | 11.89464 | 7.49237 | 4.17519 | 20.47958 | 2.80747 | 12.30169 | 17.18882 | 11.66006 | 9.99812 | 18.80018 |
| **FAH** | 0.00000 | 0.47777 | 5.81658 | 0.00000 | 1.32712 | 0.89472 | 2.37358 | 0.04935 | 2.70669 | 0.48249 | 2.34005 | 0.17547 |
| **FAHD1** | 0.80794 | 4.22767 | 1.32996 | 4.60917 | 3.55653 | 4.99057 | 2.20463 | 0.17486 | 1.16628 | 1.59646 | 4.42955 | 0.40825 |
| **FAHD2A** | 0.75320 | 1.62897 | 0.96759 | 0.94083 | 1.88918 | 1.87538 | 0.28007 | 1.28708 | 1.26660 | 0.00000 | 1.75521 | 1.16249 |
| **FAHD2B** | 0.08440 | 0.56747 | 0.17499 | 0.29023 | 0.65993 | 0.17177 | 0.04817 | 0.21019 | 0.08985 | 0.00000 | 0.16035 | 0.27552 |
| **FAHD2CP** | 0.00556 | 0.07247 | 0.00065 | 0.00146 | 0.00000 | 0.09017 | 0.00000 | 0.00583 | 0.04564 | 0.05074 | 0.01035 | 0.11916 |
| **FAIM** | 0.22820 | 1.17275 | 3.36309 | 0.33946 | 0.82983 | 4.95499 | 6.19233 | 0.07002 | 2.73189 | 0.82324 | 2.11976 | 0.00000 |
| **FAIM2** | 2.66271 | 0.08314 | 0.01434 | 0.01465 | 0.01142 | 0.00000 | 0.01424 | 0.04462 | 0.03509 | 0.00000 | 0.01185 | 0.04585 |
| **FAIM3** | 0.06491 | 0.01526 | 0.02329 | 0.02089 | 0.01122 | 0.00844 | 0.03126 | 0.00626 | 0.00000 | 0.01604 | 0.05129 | 0.01574 |
| **FAM101B** | 0.38316 | 0.84116 | 0.00000 | 0.87623 | 24.08238 | 0.04536 | 13.56232 | 0.00000 | 0.00000 | 13.69693 | 18.38247 | 0.00000 |
| **FAM102A** | 6.11912 | 4.25182 | 4.40437 | 2.96417 | 5.30137 | 0.00598 | 3.62969 | 0.17965 | 2.36831 | 5.80879 | 0.41637 | 2.74244 |
| **FAM102B** | 0.68932 | 4.65552 | 0.97056 | 10.65113 | 0.00000 | 8.78653 | 4.76389 | 0.13435 | 0.00000 | 5.37043 | 0.00000 | 4.58446 |
| **FAM103A1** | 0.49658 | 4.29457 | 2.19732 | 2.65791 | 1.94923 | 5.50531 | 4.02398 | 4.00030 | 0.00000 | 0.62893 | 3.30224 | 5.21040 |
| **FAM104A** | 6.38651 | 13.16019 | 20.93208 | 5.09669 | 8.27188 | 14.52338 | 11.81129 | 24.67303 | 12.80408 | 0.00011 | 7.69856 | 20.60564 |
| **FAM104B** | 0.06611 | 4.23122 | 3.06873 | 2.33590 | 2.13648 | 6.81364 | 2.13597 | 1.78596 | 2.54336 | 0.00000 | 3.41005 | 10.42533 |
| **FAM105A** | 0.56601 | 1.32193 | 0.05685 | 0.21667 | 0.00000 | 0.49792 | 0.00000 | 3.21799 | 0.00010 | 0.64017 | 0.00000 | 0.33055 |
| **FAM106A** | 0.00955 | 0.00869 | 0.00368 | 0.00571 | 0.01169 | 0.00807 | 0.00295 | 0.00709 | 0.10246 | 0.11830 | 0.00867 | 0.05826 |
| **FAM106CP** | 0.00565 | 0.01002 | 0.02210 | 0.01003 | 0.02447 | 0.01929 | 0.01263 | 0.00920 | 0.01292 | 0.03043 | 0.01626 | 0.04593 |
| **FAM107A** | 0.16782 | 0.09803 | 0.02617 | 0.04405 | 0.00000 | 0.02146 | 0.02141 | 0.00913 | 0.00614 | 0.06984 | 0.03545 | 0.04968 |
| **FAM107B** | 2.22570 | 3.17098 | 0.00000 | 4.83762 | 5.02038 | 11.31677 | 6.81591 | 0.41165 | 0.46954 | 12.21882 | 1.25985 | 0.71124 |
| **FAM109A** | 0.45965 | 1.67145 | 0.84722 | 2.81475 | 1.48467 | 0.00000 | 1.46897 | 0.35966 | 1.02421 | 3.24270 | 0.00000 | 1.49729 |
| **FAM109B** | 0.16854 | 0.45261 | 0.05695 | 1.07994 | 0.29091 | 2.19253 | 0.31550 | 0.33421 | 0.00000 | 0.00000 | 0.57675 | 3.17400 |
| **FAM110A** | 2.65064 | 0.39725 | 16.63701 | 0.00000 | 5.82341 | 0.00000 | 6.17353 | 0.26335 | 5.44354 | 1.71185 | 1.69202 | 0.06833 |
| **FAM110B** | 0.00000 | 1.01976 | 1.77678 | 0.00000 | 1.49373 | 2.09790 | 3.81841 | 0.00000 | 7.68606 | 1.71348 | 5.99796 | 0.23108 |
| **FAM110C** | 0.00517 | 0.01125 | 0.10742 | 0.00093 | 0.00437 | 0.03075 | 0.00000 | 0.01011 | 0.02920 | 0.08197 | 0.00620 | 0.01179 |
| **FAM110D** | 0.00329 | 0.00919 | 0.01465 | 0.01567 | 0.07983 | 0.00000 | 0.04472 | 0.73916 | 0.00000 | 0.00000 | 0.01266 | 0.39748 |
| **FAM111A** | 0.04768 | 0.01235 | 0.01723 | 0.06833 | 0.01218 | 0.00864 | 0.04328 | 0.05397 | 0.00586 | 0.00000 | 0.03865 | 0.07254 |
| **FAM111B** | 0.43121 | 0.00000 | 0.85864 | 0.65437 | 1.67703 | 0.28345 | 0.00000 | 0.00000 | 0.00000 | 0.26752 | 1.83874 | 0.57424 |

| | | | | | | | | | | | | |
|---|---|---|---|---|---|---|---|---|---|---|---|---|
| **FAM114A1** | 0.00692 | 0.06177 | 0.04337 | 0.00833 | 0.01635 | 0.00000 | 0.00810 | 0.05006 | 0.04427 | 0.18628 | 0.00376 | 0.00585 |
| **FAM114A2** | 1.75884 | 4.98643 | 2.97576 | 0.00001 | 0.00000 | 11.39646 | 2.23145 | 3.99998 | 1.32783 | 1.37940 | 5.17703 | 5.42847 |
| **FAM115A** | 0.11045 | 0.12348 | 5.92458 | 0.12251 | 0.32050 | 0.03140 | 0.80248 | 0.13096 | 0.33492 | 0.05263 | 1.07400 | 0.20444 |
| **FAM115C** | 0.01883 | 0.01313 | 0.15733 | 0.00473 | 0.08481 | 0.00001 | 0.00550 | 0.00847 | 0.07934 | 0.04988 | 0.02312 | 0.02026 |
| **FAM117A** | 2.65920 | 0.46930 | 0.93909 | 0.12955 | 0.95098 | 0.27527 | 1.30959 | 0.00075 | 0.46902 | 1.27752 | 0.00000 | 2.25995 |
| **FAM117B** | 0.91889 | 4.19638 | 0.00000 | 4.09139 | 0.00000 | 10.65169 | 1.89652 | 11.18314 | 0.60965 | 8.22081 | 1.23879 | 8.98225 |
| **FAM118A** | 4.07125 | 6.52833 | 10.88343 | 0.56941 | 12.20763 | 5.77969 | 5.31410 | 2.21112 | 14.91396 | 6.62828 | 11.64894 | 7.07561 |
| **FAM118B** | 3.25479 | 9.12333 | 10.22968 | 5.98669 | 2.81166 | 11.91926 | 15.24399 | 10.93585 | 8.12520 | 0.00000 | 9.71945 | 9.90055 |
| **FAM120A** | 1.21901 | 1.08914 | 7.04177 | 4.02446 | 9.80624 | 10.88304 | 19.06082 | 0.16464 | 0.00830 | 33.63400 | 5.49227 | 2.20650 |
| **FAM120AOS** | 2.07102 | 3.98144 | 5.13832 | 1.59244 | 6.26326 | 9.23931 | 3.97802 | 0.53272 | 0.73504 | 11.27169 | 4.71424 | 2.89514 |
| **FAM120B** | 3.45959 | 2.57521 | 0.97574 | 1.31387 | 1.63949 | 0.00000 | 0.01242 | 1.73244 | 3.30454 | 0.00000 | 1.55358 | 3.42823 |
| **FAM120C** | 0.93838 | 0.09547 | 2.02451 | 1.61996 | 0.27262 | 0.67289 | 1.96299 | 0.52604 | 2.83280 | 0.38416 | 2.50587 | 0.76152 |
| **FAM122A** | 5.72381 | 3.75358 | 4.41659 | 0.00000 | 1.05416 | 3.29901 | 1.01803 | 2.58791 | 1.68852 | 4.02396 | 2.73659 | 1.23387 |
| **FAM122B** | 1.69909 | 0.00000 | 2.42240 | 2.89683 | 0.00003 | 7.36711 | 3.24462 | 0.31750 | 1.16821 | 6.03524 | 1.56971 | 0.21377 |
| **FAM122C** | 0.18844 | 6.03765 | 0.00000 | 4.80401 | 0.00000 | 3.45214 | 0.00000 | 2.38906 | 0.00000 | 0.07500 | 0.11293 | 6.09497 |
| **FAM124A** | 0.49708 | 2.59430 | 0.07411 | 4.17214 | 0.35618 | 0.09141 | 0.15055 | 0.00000 | 0.00000 | 0.53988 | 0.00000 | 1.41822 |
| **FAM124B** | 0.09361 | 0.18127 | 0.25645 | 0.21417 | 0.06535 | 0.00000 | 0.00000 | 0.00000 | 0.00000 | 0.29873 | 0.00000 | 0.43198 |
| **FAM126A** | 0.82460 | 9.92012 | 3.90012 | 5.33563 | 4.65757 | 22.91525 | 1.63883 | 3.26093 | 0.82131 | 7.87369 | 6.58450 | 4.93440 |
| **FAM126B** | 1.65340 | 2.25371 | 0.84005 | 6.39732 | 0.00000 | 17.10763 | 5.46679 | 11.26810 | 0.37984 | 8.12479 | 4.38481 | 3.16580 |
| **FAM127A** | 0.40693 | 1.58542 | 5.93927 | 8.33436 | 5.17453 | 0.01356 | 9.42180 | 2.64808 | 8.81045 | 0.00000 | 5.99362 | 1.52623 |
| **FAM127B** | 0.00000 | 2.08720 | 4.48207 | 3.96221 | 1.73726 | 1.00664 | 4.81200 | 2.87768 | 8.60647 | 0.20357 | 3.28349 | 3.59138 |
| **FAM127C** | 0.28038 | 0.44016 | 0.78047 | 1.20769 | 0.00000 | 0.00000 | 0.00298 | 0.35344 | 1.05790 | 0.52652 | 0.24287 | 0.76491 |
| **FAM129A** | 0.62478 | 0.01383 | 0.08348 | 0.05431 | 0.00587 | 0.00487 | 0.03045 | 0.09779 | 0.00000 | 0.03990 | 0.03623 | 0.04241 |
| **FAM129B** | 1.14019 | 0.01716 | 2.48893 | 0.00000 | 0.00000 | 0.00000 | 0.90515 | 0.28471 | 0.81013 | 1.26792 | 0.01496 | 0.12655 |
| **FAM129C** | 0.00714 | 0.02634 | 0.02800 | 0.00611 | 0.01865 | 0.04189 | 0.01544 | 0.00731 | 0.01342 | 0.01089 | 0.01683 | 0.08517 |
| **FAM131A** | 0.99403 | 0.63891 | 4.12615 | 0.80986 | 0.15224 | 0.00000 | 0.00000 | 0.00000 | 0.70152 | 0.60369 | 0.10565 | 0.26722 |
| **FAM131B** | 0.00667 | 0.02158 | 0.03735 | 0.01173 | 0.00755 | 0.03565 | 0.01961 | 0.01036 | 0.00684 | 0.02573 | 0.01376 | 0.01420 |
| **FAM131C** | 0.00647 | 0.01446 | 0.01561 | 0.01066 | 0.01356 | 0.02375 | 0.01472 | 0.01069 | 0.01547 | 0.02481 | 0.01424 | 0.02018 |
| **FAM132A** | 0.00175 | 1.04870 | 0.10200 | 0.53914 | 0.83329 | 0.00000 | 0.33709 | 0.13250 | 0.00000 | 0.00000 | 0.08587 | 0.00420 |
| **FAM132B** | 0.01097 | 0.05597 | 0.05013 | 0.07684 | 0.00000 | 0.02995 | 0.04865 | 0.00733 | 0.00600 | 0.06043 | 0.01719 | 0.09205 |
| **FAM133A** | 0.00000 | 0.07698 | 1.66696 | 0.21150 | 0.00000 | 0.71109 | 1.79693 | 0.34485 | 0.00000 | 1.64552 | 0.59325 | 1.59559 |
| **FAM133B** | 0.52666 | 0.21517 | 0.08220 | 0.22110 | 0.00000 | 0.12263 | 0.34703 | 0.22469 | 0.24632 | 0.76823 | 0.22830 | 0.16677 |
| **FAM133CP** | 0.15514 | 0.14804 | 0.00000 | 0.02565 | 0.03987 | 0.34415 | 0.24939 | 0.30859 | 0.07202 | 0.63822 | 0.02088 | 0.17756 |
| **FAM134A** | 1.46976 | 3.65304 | 4.48966 | 4.63308 | 5.07074 | 1.06373 | 10.18783 | 0.72473 | 0.00066 | 17.96507 | 2.92653 | 4.36637 |
| **FAM134B** | 0.05187 | 0.00000 | 0.30352 | 0.00000 | 0.25569 | 0.31391 | 0.00000 | 0.00374 | 0.76396 | 0.40471 | 0.06625 | 0.04189 |
| **FAM134C** | 1.60552 | 6.21141 | 4.97625 | 3.50855 | 4.12847 | 5.24281 | 1.39094 | 1.52400 | 13.62482 | 1.89386 | 1.41468 | 7.35119 |
| **FAM135A** | 1.94623 | 1.81428 | 2.32138 | 2.73615 | 0.00000 | 11.78322 | 3.37637 | 2.77960 | 9.31109 | 5.72699 | 1.87418 | 1.58745 |
| **FAM135B** | 0.74855 | 0.02701 | 0.01505 | 0.01227 | 0.00892 | 0.00750 | 0.03864 | 0.01287 | 0.00364 | 0.04169 | 0.02743 | 0.00813 |
| **FAM136A** | 4.82783 | 24.66174 | 21.82185 | 11.94821 | 20.46166 | 29.09053 | 38.51921 | 5.11518 | 27.57191 | 3.54015 | 30.14117 | 23.13047 |
| **FAM13A** | 16.26116 | 0.11612 | 0.29123 | 0.01539 | 0.05485 | 0.00000 | 0.20987 | 1.76850 | 0.38036 | 0.45925 | 0.13068 | 1.17429 |
| **FAM13A-AS1** | 0.00256 | 0.04400 | 0.20456 | 0.00000 | 0.02582 | 0.16773 | 0.00000 | 0.02067 | 0.18682 | 0.16003 | 0.21057 | 0.00887 |
| **FAM13B** | 0.07934 | 0.61821 | 1.60975 | 2.23242 | 0.00000 | 2.09906 | 1.75577 | 4.56328 | 2.18500 | 5.04773 | 3.52973 | 4.05777 |
| **FAM13C** | 0.04702 | 0.02225 | 0.02902 | 0.16805 | 0.00003 | 0.04825 | 0.00804 | 0.03222 | 0.01329 | 0.01038 | 0.01171 | 0.01039 |
| **FAM149A** | 1.16325 | 0.02173 | 0.00802 | 0.05353 | 0.00000 | 0.03048 | 0.03659 | 0.00601 | 0.00000 | 0.05803 | 0.02204 | 0.13937 |
| **FAM149B1** | 0.27220 | 0.06054 | 0.24620 | 0.28779 | 0.21469 | 2.80896 | 0.78998 | 0.19586 | 0.39385 | 3.51919 | 0.59137 | 0.58394 |
| **FAM150A** | 0.03121 | 0.00635 | 0.01406 | 0.13334 | 0.05056 | 0.05891 | 0.04191 | 0.01991 | 0.02887 | 0.00000 | 0.01005 | 0.02047 |
| **FAM150B** | 0.00945 | 0.01602 | 0.01314 | 0.00206 | 0.02764 | 0.02642 | 0.00874 | 0.03611 | 0.14595 | 0.05084 | 0.01390 | 0.04092 |
| **FAM151A** | 0.00000 | 47.59365 | 0.00000 | 20.94269 | 10.75898 | 0.00000 | 0.00000 | 22.08674 | 0.00000 | 3.45320 | 1.79309 | 51.10931 |

| | | | | | | | | | | | | |
|---|---|---|---|---|---|---|---|---|---|---|---|---|
| **FAM151B** | 0.10956 | 0.20168 | 0.59780 | 0.00000 | 0.08217 | 0.37030 | 0.57322 | 0.13059 | 0.78014 | 0.42705 | 0.50079 | 0.00000 |
| **FAM153A** | 0.01294 | 0.01594 | 0.01693 | 0.01463 | 0.02130 | 0.00979 | 0.02801 | 0.01179 | 0.02276 | 0.00423 | 0.01499 | 0.02251 |
| **FAM153B** | 0.00655 | 0.01135 | 0.01205 | 0.00944 | 0.01113 | 0.02962 | 0.01343 | 0.01257 | 0.01459 | 0.03036 | 0.01377 | 0.03174 |
| **FAM154A** | 6.71321 | 0.02622 | 0.03139 | 0.06575 | 0.07423 | 0.00000 | 0.17681 | 0.78913 | 0.04154 | 0.00000 | 0.00000 | 0.29659 |
| **FAM154B** | 0.00965 | 0.08598 | 0.07559 | 0.18749 | 0.00000 | 0.00158 | 0.07728 | 0.62239 | 0.17198 | 0.00000 | 0.04789 | 0.22638 |
| **FAM155A** | 0.00298 | 0.03159 | 0.03894 | 0.23598 | 0.05150 | 0.00000 | 0.01405 | 0.00839 | 0.00000 | 0.36731 | 0.00000 | 0.02432 |
| **FAM155A-IT1** | 0.00222 | 0.00959 | 0.06129 | 0.03678 | 0.02726 | 0.06200 | 0.01771 | 0.03013 | 0.00000 | 0.07141 | 0.00796 | 0.02394 |
| **FAM155B** | 0.00000 | 0.25648 | 0.14338 | 0.12910 | 0.18113 | 0.90290 | 0.41632 | 0.00510 | 0.00000 | 1.59636 | 0.00000 | 0.00000 |
| **FAM156A.uid-12** | 0.00563 | 0.01332 | 0.06560 | 0.01438 | 0.00528 | 0.04679 | 0.01076 | 0.00971 | 0.07282 | 0.03858 | 0.01981 | 0.01917 |
| **FAM157A** | 0.02327 | 0.02374 | 0.07727 | 0.07249 | 0.01529 | 0.01906 | 0.01016 | 0.03444 | 0.03856 | 0.00000 | 0.02628 | 0.03821 |
| **FAM157B** | 0.00696 | 0.01496 | 0.01329 | 0.01186 | 0.01501 | 0.01910 | 0.01934 | 0.01203 | 0.02562 | 0.01155 | 0.01260 | 0.01708 |
| **FAM159A** | 0.00932 | 0.02648 | 0.04899 | 0.01245 | 0.04712 | 0.00000 | 0.01713 | 0.00734 | 0.02458 | 0.00000 | 0.01417 | 0.10615 |
| **FAM159B** | 0.03464 | 0.03159 | 0.23441 | 0.12272 | 0.00000 | 0.06068 | 0.08421 | 0.02167 | 0.00000 | 0.07319 | 0.03555 | 0.00907 |
| **FAM160A1** | 2.07451 | 2.12654 | 0.00000 | 3.26423 | 0.04909 | 2.69451 | 0.00000 | 1.67859 | 0.12902 | 0.83219 | 0.07108 | 1.28871 |
| **FAM160A2** | 3.86942 | 0.20533 | 14.39257 | 0.00000 | 1.60805 | 0.95492 | 4.55737 | 0.96306 | 4.63764 | 4.71664 | 0.33812 | 3.36571 |
| **FAM160B1** | 0.25906 | 2.42919 | 4.62679 | 0.00000 | 2.60370 | 13.01641 | 6.47321 | 0.09804 | 0.80953 | 7.54627 | 2.62062 | 1.05838 |
| **FAM160B2** | 0.58701 | 3.06805 | 1.89400 | 1.68899 | 2.98394 | 0.00000 | 0.44166 | 0.30311 | 1.05197 | 3.14839 | 0.38500 | 3.59749 |
| **FAM161A** | 2.90993 | 1.70139 | 2.09461 | 1.04524 | 0.00000 | 2.04014 | 0.00000 | 0.49017 | 2.16048 | 1.54020 | 0.76762 | 2.00343 |
| **FAM161B** | 1.80999 | 0.55042 | 0.51180 | 0.00000 | 0.21161 | 0.34929 | 0.00000 | 1.19689 | 0.81870 | 0.00000 | 0.23781 | 0.86986 |
| **FAM162A** | 0.00000 | 11.26930 | 8.37941 | 13.25595 | 9.02833 | 21.19153 | 18.31730 | 0.45823 | 16.25174 | 8.85697 | 19.18984 | 8.49103 |
| **FAM162B** | 0.00000 | 2.32588 | 0.59977 | 1.05055 | 0.18561 | 0.22629 | 0.00000 | 6.03877 | 0.00000 | 0.28991 | 0.13846 | 15.98353 |
| **FAM163A** | 0.00615 | 0.01205 | 0.02323 | 0.00720 | 0.02978 | 0.04238 | 0.01051 | 0.01012 | 0.01837 | 0.01323 | 0.01457 | 0.01518 |
| **FAM163B** | 0.25418 | 0.79957 | 0.00000 | 0.15352 | 0.00100 | 0.10628 | 0.00283 | 0.00141 | 0.00063 | 0.01467 | 0.00222 | 0.00000 |
| **FAM166A** | 0.00254 | 0.01796 | 0.02811 | 0.01014 | 0.05245 | 0.01389 | 0.00795 | 0.04347 | 0.07587 | 0.00000 | 0.00976 | 0.01905 |
| **FAM166B** | 0.04225 | 0.04538 | 0.01607 | 0.01375 | 0.01960 | 0.00792 | 0.02190 | 0.00235 | 0.02402 | 0.00000 | 0.01565 | 0.12059 |
| **FAM167A** | 2.57869 | 0.05034 | 0.00616 | 0.05063 | 0.00000 | 0.08205 | 0.00000 | 0.07202 | 0.02068 | 0.01542 | 0.00410 | 0.00000 |
| **FAM167B** | 0.08354 | 0.01218 | 0.01346 | 0.01095 | 0.07395 | 0.00000 | 0.01274 | 0.00873 | 0.03135 | 0.02902 | 0.01059 | 0.02278 |
| **FAM168A** | 1.87600 | 1.94016 | 0.48468 | 4.29784 | 3.95233 | 3.56035 | 2.46923 | 1.59100 | 0.76511 | 3.95379 | 0.51486 | 1.09124 |
| **FAM168B** | 3.66706 | 5.92850 | 9.42509 | 3.69886 | 9.39438 | 11.61428 | 17.30976 | 4.57763 | 8.25687 | 13.81994 | 7.17791 | 8.07929 |
| **FAM169A** | 4.07400 | 4.41571 | 3.18762 | 6.88542 | 1.11513 | 14.26601 | 3.28228 | 1.84617 | 1.37212 | 4.76606 | 0.46692 | 3.29187 |
| **FAM169B** | 0.39567 | 0.01268 | 0.03278 | 0.01107 | 0.00872 | 0.01205 | 0.01727 | 0.28781 | 0.00326 | 0.02643 | 0.04260 | 0.00419 |
| **FAM171A1** | 0.00102 | 2.68291 | 0.36480 | 2.33432 | 1.14628 | 0.00000 | 0.10524 | 1.04775 | 0.00000 | 0.95440 | 0.00000 | 1.98175 |
| **FAM171A2** | 0.05038 | 0.06528 | 0.01902 | 0.00241 | 0.06355 | 0.00000 | 0.01437 | 0.00495 | 0.00748 | 0.05415 | 0.02834 | 0.06267 |
| **FAM171B** | 0.61029 | 0.01831 | 0.02722 | 0.02138 | 0.01815 | 0.00000 | 0.02173 | 0.23090 | 0.01265 | 0.04509 | 0.01601 | 0.04712 |
| **FAM172A** | 2.15477 | 1.21936 | 1.57762 | 0.40988 | 0.53565 | 6.55282 | 2.10018 | 0.48022 | 1.86856 | 3.57446 | 1.09437 | 3.88136 |
| **FAM172BP** | 5.91694 | 0.02341 | 0.05361 | 0.00298 | 0.01313 | 0.02909 | 0.00038 | 0.28573 | 0.01464 | 0.01511 | 0.01562 | 0.00000 |
| **FAM173A** | 0.00000 | 3.99064 | 2.49408 | 0.81002 | 3.98759 | 0.00000 | 2.00405 | 0.02302 | 6.96547 | 3.08162 | 5.68135 | 0.24849 |
| **FAM173B** | 0.14620 | 1.06363 | 1.32435 | 0.68485 | 0.61314 | 3.88044 | 0.00000 | 0.95909 | 0.15778 | 0.29059 | 1.52272 | 3.77012 |
| **FAM174A** | 0.14064 | 2.99390 | 2.88222 | 0.27737 | 0.60951 | 4.24281 | 0.00000 | 1.44060 | 3.19448 | 0.00000 | 1.78263 | 0.90309 |
| **FAM174B** | 3.59956 | 0.52808 | 0.14620 | 0.45563 | 0.00000 | 0.29757 | 0.09388 | 2.06244 | 0.00995 | 0.00000 | 0.01010 | 0.39559 |
| **FAM175A** | 0.38772 | 1.45020 | 0.40710 | 0.81959 | 0.00000 | 3.67226 | 0.00000 | 0.31785 | 2.72212 | 1.76314 | 2.85743 | 3.19546 |
| **FAM175B** | 4.94818 | 7.70586 | 1.58102 | 2.78625 | 3.39980 | 14.50366 | 3.24531 | 5.36052 | 2.64137 | 0.00000 | 4.84731 | 11.49572 |
| **FAM177A1** | 0.30399 | 0.49593 | 5.89918 | 4.92537 | 4.96177 | 8.81730 | 8.22555 | 9.19703 | 3.85512 | 13.27888 | 6.70257 | 3.60471 |
| **FAM177B** | 0.00601 | 0.01276 | 0.01738 | 0.01043 | 0.01088 | 0.02350 | 0.01739 | 0.01041 | 0.00940 | 0.02082 | 0.03184 | 0.02042 |
| **FAM178A** | 2.47057 | 3.21677 | 2.41895 | 3.20910 | 0.83816 | 9.20319 | 3.11344 | 1.00698 | 4.28775 | 5.79643 | 4.25587 | 4.64562 |
| **FAM178B** | 0.08570 | 0.43969 | 0.77419 | 0.00101 | 0.49963 | 0.00008 | 0.32001 | 0.86796 | 0.39939 | 0.00000 | 0.25098 | 2.43964 |
| **FAM179A** | 2.35677 | 0.02738 | 0.05734 | 0.02338 | 0.00284 | 0.01710 | 0.02273 | 0.26551 | 0.01696 | 0.02558 | 0.01096 | 0.01553 |
| **FAM179B** | 0.69129 | 0.14389 | 0.68692 | 0.82373 | 1.56337 | 2.19144 | 0.00000 | 0.16524 | 3.84105 | 2.66406 | 0.00000 | 2.78569 |

| | | | | | | | | | | | | |
|---|---|---|---|---|---|---|---|---|---|---|---|---|
| **FAM180A** | 0.23168 | 0.01700 | 0.30384 | 0.01303 | 0.02096 | 0.00000 | 0.07436 | 0.00340 | 0.04693 | 0.01198 | 0.01011 | 0.01390 |
| **FAM180B** | 0.30092 | 0.02903 | 0.06576 | 0.00000 | 0.04291 | 0.00000 | 0.00224 | 0.01525 | 0.12039 | 0.17914 | 0.00000 | 0.07834 |
| **FAM181B** | 0.00607 | 0.01508 | 0.01200 | 0.11719 | 0.05750 | 0.00000 | 0.01818 | 0.01039 | 0.02264 | 0.00823 | 0.01430 | 0.02162 |
| **FAM182A** | 0.00536 | 0.02571 | 0.03224 | 0.10057 | 0.00000 | 0.02011 | 0.03424 | 0.00963 | 0.03078 | 0.03657 | 0.00782 | 0.03132 |
| **FAM182B** | 0.00894 | 0.13626 | 0.06249 | 0.23918 | 0.00000 | 0.04128 | 0.01576 | 0.00871 | 0.00958 | 0.00001 | 0.14571 | 0.02151 |
| **FAM183A** | 0.07145 | 0.00049 | 0.00530 | 0.00252 | 0.01087 | 0.11000 | 0.01402 | 0.01785 | 0.00000 | 0.05558 | 0.08965 | 0.02336 |
| **FAM183B** | 0.04962 | 0.05901 | 0.04559 | 0.11574 | 0.00000 | 0.05485 | 0.00000 | 0.00216 | 0.00787 | 0.10718 | 0.00826 | 0.11196 |
| **FAM183CP** | 0.06931 | 0.01456 | 0.01442 | 0.01144 | 0.00906 | 0.03374 | 0.01004 | 0.00551 | 0.05576 | 0.00121 | 0.01595 | 0.01474 |
| **FAM184A** | 0.70790 | 0.00000 | 2.45078 | 0.00000 | 0.00000 | 3.35456 | 5.56181 | 0.00000 | 3.24606 | 7.43288 | 0.75671 | 0.00000 |
| **FAM184B** | 2.42446 | 0.42001 | 0.31484 | 0.24177 | 0.33994 | 0.00000 | 0.12232 | 0.65527 | 0.84803 | 1.16883 | 0.32367 | 0.18752 |
| **FAM185A** | 0.30091 | 0.73394 | 0.61410 | 0.32107 | 0.54069 | 0.50519 | 0.00000 | 0.02990 | 1.63266 | 0.12164 | 0.97996 | 0.15311 |
| **FAM186A** | 0.03669 | 0.00824 | 0.03254 | 0.08598 | 0.00000 | 0.00000 | 0.01635 | 0.00738 | 0.00000 | 0.10535 | 0.09753 | 0.02883 |
| **FAM186B** | 0.03841 | 0.08765 | 0.10632 | 0.04084 | 0.06001 | 0.00000 | 0.03910 | 0.01446 | 0.19607 | 0.23310 | 0.08833 | 0.25268 |
| **FAM188A** | 3.88622 | 8.12555 | 3.70379 | 0.79492 | 0.04010 | 8.42037 | 2.48170 | 8.88137 | 5.83860 | 0.00000 | 3.33886 | 8.41377 |
| **FAM188B** | 0.10901 | 0.07359 | 0.04191 | 0.00425 | 0.04662 | 0.04660 | 0.04478 | 0.00166 | 0.10864 | 0.00000 | 0.10114 | 0.09237 |
| **FAM189A1** | 0.02786 | 0.03909 | 0.03852 | 0.10399 | 0.02318 | 0.00000 | 0.04421 | 0.02872 | 0.00000 | 0.04900 | 0.01251 | 0.02351 |
| **FAM189A2** | 6.16558 | 0.00061 | 0.33985 | 0.00000 | 0.00000 | 0.00000 | 0.21249 | 0.22856 | 0.19360 | 0.02537 | 0.49176 | 0.03351 |
| **FAM189B** | 0.25470 | 0.00000 | 0.59958 | 2.00370 | 0.81575 | 0.23977 | 4.09989 | 0.00793 | 2.20992 | 3.54273 | 0.52918 | 0.02083 |
| **FAM192A** | 3.12261 | 14.17156 | 5.20391 | 8.52927 | 9.96596 | 14.00263 | 8.63775 | 3.51010 | 6.86758 | 3.75858 | 8.89535 | 12.79153 |
| **FAM193A** | 6.71495 | 2.92745 | 1.57024 | 3.91779 | 7.09912 | 2.03212 | 4.51825 | 0.98001 | 3.63225 | 13.91921 | 1.54961 | 3.28193 |
| **FAM193B** | 0.16123 | 1.16036 | 2.42640 | 6.94603 | 5.24398 | 0.00000 | 2.12273 | 0.08851 | 2.79305 | 16.02448 | 0.17858 | 4.21999 |
| **FAM195A** | 0.00000 | 15.67424 | 0.24486 | 5.37560 | 15.15260 | 0.00000 | 5.03288 | 0.87025 | 7.83066 | 8.51562 | 8.55515 | 16.09910 |
| **FAM195B** | 1.61646 | 0.28975 | 10.70342 | 5.73923 | 1.41680 | 0.00000 | 8.45443 | 0.22618 | 7.04856 | 0.00000 | 5.84949 | 1.83142 |
| **FAM196A** | 0.00546 | 0.00000 | 0.00000 | 0.65129 | 0.18941 | 0.33224 | 0.00000 | 0.00812 | 0.32917 | 1.71719 | 0.00000 | 0.22357 |
| **FAM198A** | 0.00835 | 0.00000 | 0.11159 | 0.02112 | 0.18158 | 0.08002 | 0.00000 | 0.00546 | 0.00000 | 0.02198 | 0.09287 | 0.15336 |
| **FAM198B** | 0.27442 | 0.01560 | 2.01030 | 0.72993 | 0.00000 | 0.27129 | 0.37289 | 0.00199 | 0.00000 | 0.10006 | 0.00000 | 0.00517 |
| **FAM199X** | 1.94179 | 1.46339 | 1.99345 | 2.30587 | 2.32437 | 7.80653 | 1.88156 | 0.32475 | 3.05931 | 5.17321 | 0.95860 | 0.15405 |
| **FAM19A1** | 0.13824 | 0.01531 | 0.02457 | 0.01489 | 0.01623 | 0.01019 | 0.01677 | 0.00947 | 0.01824 | 0.01757 | 0.01480 | 0.01430 |
| **FAM19A2** | 1.92140 | 0.01373 | 0.14769 | 0.01747 | 0.01118 | 0.00001 | 0.05795 | 0.00512 | 0.00000 | 0.07086 | 0.00992 | 0.00886 |
| **FAM19A3** | 0.03305 | 1.08288 | 0.02542 | 0.00000 | 0.00000 | 0.00000 | 0.00163 | 2.19702 | 0.03672 | 0.04425 | 0.00053 | 6.86547 |
| **FAM19A4** | 5.65596 | 0.00000 | 0.07532 | 0.06209 | 0.00000 | 0.20960 | 0.31090 | 0.47066 | 0.04402 | 0.00000 | 0.00000 | 1.16064 |
| **FAM19A5** | 1.53023 | 0.02656 | 0.02884 | 0.06939 | 0.27374 | 0.09117 | 0.00812 | 0.06610 | 0.00000 | 0.10341 | 0.02080 | 0.00000 |
| **FAM200A** | 0.00089 | 3.97304 | 0.00000 | 1.44014 | 0.34897 | 3.94579 | 0.81156 | 3.37865 | 0.96948 | 0.00000 | 1.20939 | 5.93827 |
| **FAM200B** | 1.60221 | 2.92072 | 4.67705 | 2.87174 | 2.10274 | 13.91193 | 3.04834 | 4.88852 | 2.09514 | 4.46265 | 5.07217 | 5.29081 |
| **FAM204A** | 1.11141 | 3.35749 | 4.38354 | 0.61009 | 1.58322 | 15.27375 | 6.24366 | 3.30274 | 5.93380 | 6.77286 | 5.24146 | 7.24393 |
| **FAM205B** | 0.00629 | 0.01370 | 0.01428 | 0.01502 | 0.01033 | 0.01811 | 0.02379 | 0.01044 | 0.00812 | 0.01278 | 0.05346 | 0.01668 |
| **FAM206A** | 0.81911 | 7.47928 | 5.73043 | 1.38388 | 0.00000 | 10.58701 | 8.63584 | 0.14418 | 3.30275 | 0.00000 | 4.30787 | 9.51347 |
| **FAM207A** | 0.02226 | 1.83129 | 1.02889 | 1.07914 | 3.96767 | 0.00000 | 1.71745 | 0.41136 | 0.00000 | 7.58669 | 1.40837 | 1.32928 |
| **FAM208A** | 5.64060 | 2.15625 | 1.16178 | 1.27374 | 0.01518 | 11.87815 | 0.95646 | 3.79302 | 0.00000 | 9.83124 | 0.91331 | 4.82153 |
| **FAM208B** | 12.90948 | 1.56755 | 3.32903 | 2.70037 | 0.62765 | 16.25724 | 5.14860 | 3.19869 | 6.30346 | 12.21904 | 2.84917 | 0.00000 |
| **FAM209A** | 0.00336 | 0.01755 | 0.04129 | 0.05020 | 0.02813 | 0.00751 | 0.04857 | 0.01830 | 0.00000 | 0.07249 | 0.01076 | 0.03974 |
| **FAM209B** | 0.03202 | 0.19774 | 0.09827 | 0.15070 | 0.20202 | 0.12677 | 0.04818 | 0.00000 | 0.00000 | 0.09278 | 0.02153 | 0.29445 |
| **FAM20A** | 0.11953 | 0.00366 | 0.08759 | 1.16780 | 0.00858 | 0.00000 | 0.00000 | 0.01452 | 0.10278 | 0.25312 | 0.09781 | 0.02160 |
| **FAM20B** | 0.19870 | 2.04981 | 1.96435 | 3.43144 | 4.96236 | 9.15466 | 7.31497 | 0.00000 | 0.57200 | 16.00274 | 2.94459 | 0.03963 |
| **FAM20C** | 0.06149 | 0.01157 | 0.04949 | 0.02022 | 0.01808 | 0.04144 | 0.00604 | 0.00744 | 0.00179 | 0.03439 | 0.01524 | 0.02118 |
| **FAM210A** | 3.16142 | 12.39290 | 10.93841 | 2.41125 | 6.55514 | 14.16858 | 12.47503 | 10.72206 | 14.26024 | 3.00921 | 15.01400 | 5.56668 |
| **FAM210B** | 0.71524 | 0.74791 | 0.96357 | 0.00000 | 0.18744 | 1.12455 | 0.00000 | 0.00000 | 0.00001 | 1.07450 | 0.00323 | 0.02430 |
| **FAM212A** | 0.06959 | 0.15111 | 0.04267 | 0.00004 | 0.00404 | 0.00783 | 0.01575 | 0.07118 | 0.02774 | 0.04250 | 0.01354 | 0.22288 |

| | | | | | | | | | | | | |
|---|---|---|---|---|---|---|---|---|---|---|---|---|
| **FAM212B** | 0.04533 | 1.96896 | 0.03142 | 0.11364 | 0.00000 | 0.09198 | 0.00000 | 0.00000 | 0.00000 | 0.10224 | 0.00000 | 1.04546 |
| **FAM212B-AS1** | 0.30446 | 0.73655 | 0.00312 | 0.00000 | 0.00000 | 0.06893 | 0.00000 | 0.12961 | 0.00000 | 0.04692 | 0.01200 | 1.53938 |
| **FAM213A** | 0.32781 | 3.91616 | 2.04622 | 6.50666 | 7.68987 | 20.97475 | 9.65612 | 0.00000 | 30.49985 | 9.98243 | 21.95655 | 4.21502 |
| **FAM213B** | 0.21757 | 0.42940 | 0.21655 | 0.10148 | 0.00000 | 0.20651 | 0.00000 | 0.15644 | 0.20389 | 0.13546 | 0.00456 | 0.23227 |
| **FAM214A** | 5.20748 | 1.43870 | 8.39418 | 1.77125 | 0.08364 | 4.46188 | 0.00000 | 8.09437 | 0.17397 | 2.03984 | 0.00000 | 7.89412 |
| **FAM214B** | 1.44488 | 0.53402 | 0.28589 | 2.87590 | 0.37203 | 0.00000 | 0.20557 | 3.31961 | 0.77202 | 0.38443 | 0.05711 | 5.45480 |
| **FAM215A** | 0.28557 | 0.01489 | 0.01579 | 0.01415 | 0.01599 | 0.01352 | 0.01764 | 0.00579 | 0.02636 | 0.00467 | 0.01325 | 0.02346 |
| **FAM216A** | 0.01910 | 0.34280 | 3.01425 | 0.75704 | 0.15782 | 2.72091 | 5.79090 | 0.52573 | 0.39611 | 0.00005 | 3.19758 | 1.59112 |
| **FAM216B** | 0.00552 | 0.01501 | 0.01478 | 0.01120 | 0.01517 | 0.01888 | 0.01736 | 0.01298 | 0.02161 | 0.01768 | 0.01445 | 0.01572 |
| **FAM217A** | 0.02807 | 0.01484 | 0.01476 | 0.01085 | 0.01250 | 0.02246 | 0.01854 | 0.00900 | 0.01657 | 0.01647 | 0.02288 | 0.01534 |
| **FAM217B** | 0.01045 | 1.49445 | 1.39243 | 2.08787 | 0.00000 | 6.39026 | 7.60423 | 0.00000 | 1.96568 | 1.06154 | 2.57781 | 0.09578 |
| **FAM218A** | 0.02053 | 0.11150 | 0.23007 | 0.10633 | 0.00565 | 0.87095 | 0.00000 | 3.82471 | 0.12670 | 0.00000 | 0.23351 | 1.28162 |
| **FAM219A** | 0.92835 | 0.15353 | 7.14286 | 0.66868 | 0.78051 | 0.00000 | 3.54171 | 0.00000 | 1.28237 | 2.55624 | 0.27458 | 0.02307 |
| **FAM219B** | 0.00000 | 2.76779 | 5.35925 | 4.50094 | 2.42887 | 1.68650 | 1.50452 | 0.41079 | 4.86083 | 4.92192 | 2.80104 | 2.63388 |
| **FAM21A** | 0.26976 | 0.32590 | 0.23965 | 0.07943 | 0.40824 | 0.42643 | 0.21938 | 0.15989 | 0.85949 | 0.10743 | 0.23844 | 0.23449 |
| **FAM21C** | 0.30926 | 0.51558 | 0.36051 | 0.30273 | 0.32162 | 0.71047 | 0.41919 | 0.15135 | 1.24081 | 0.32462 | 0.65350 | 0.38330 |
| **FAM21EP** | 0.00604 | 0.02411 | 0.06376 | 0.02417 | 0.02117 | 0.02644 | 0.00900 | 0.01009 | 0.04092 | 0.00004 | 0.04338 | 0.05526 |
| **FAM220A** | 18.33519 | 3.08350 | 1.35452 | 1.50674 | 1.12861 | 0.61436 | 0.50743 | 1.91929 | 2.43269 | 1.61845 | 3.15579 | 0.39641 |
| **FAM221A** | 0.33028 | 0.32540 | 0.20280 | 5.54603 | 0.12420 | 1.07320 | 0.00036 | 0.00000 | 0.13332 | 0.00000 | 0.53984 | 0.35941 |
| **FAM221B** | 0.00617 | 0.01325 | 0.02689 | 0.03695 | 0.02018 | 0.02054 | 0.03026 | 0.00994 | 0.00121 | 0.01035 | 0.03719 | 0.02199 |
| **FAM222A** | 1.25071 | 0.00000 | 1.30905 | 0.75158 | 0.34271 | 0.00000 | 2.29961 | 0.27325 | 0.13177 | 2.80836 | 0.14504 | 0.68159 |
| **FAM222A-AS1** | 0.45261 | 0.00473 | 0.01720 | 0.00076 | 0.00000 | 0.04397 | 0.01981 | 0.18160 | 0.00000 | 0.17898 | 0.00520 | 0.09500 |
| **FAM222B** | 14.55760 | 9.40745 | 10.65320 | 14.10263 | 14.56244 | 3.24474 | 16.55489 | 3.21041 | 8.71080 | 10.36445 | 10.48176 | 10.94711 |
| **FAM223A+FAM2** | 0.00616 | 0.01268 | 0.02063 | 0.00984 | 0.01292 | 0.02923 | 0.01498 | 0.01036 | 0.01985 | 0.01848 | 0.01772 | 0.01526 |
| **FAM223A+FAM2** | 0.01388 | 0.01222 | 0.02743 | 0.01329 | 0.00997 | 0.03427 | 0.01715 | 0.00936 | 0.01657 | 0.02766 | 0.01158 | 0.01498 |
| **FAM225A** | 0.00618 | 0.00875 | 0.19096 | 0.01318 | 0.01577 | 0.04131 | 0.02703 | 0.00923 | 0.00000 | 0.00000 | 0.01482 | 0.04999 |
| **FAM225B** | 0.00435 | 0.00368 | 0.04228 | 0.00173 | 0.00000 | 0.03698 | 0.04974 | 0.00959 | 0.00000 | 0.16960 | 0.00612 | 0.02333 |
| **FAM226A+FAM2** | 0.00413 | 0.00408 | 0.00000 | 0.03375 | 0.00000 | 0.06054 | 0.00371 | 0.01061 | 0.00785 | 0.16154 | 0.00929 | 0.00825 |
| **FAM227A** | 0.52618 | 0.10203 | 0.22648 | 0.34602 | 0.00000 | 0.00000 | 0.00580 | 0.04051 | 0.37348 | 0.37812 | 0.00000 | 0.05847 |
| **FAM227B** | 0.44405 | 0.36986 | 0.16582 | 0.37469 | 0.00000 | 0.38337 | 0.00000 | 0.02908 | 0.07936 | 0.17710 | 0.14103 | 0.63884 |
| **FAM228A** | 4.53557 | 0.28194 | 0.04423 | 0.27461 | 0.21875 | 0.00000 | 0.06822 | 0.48278 | 0.00000 | 0.00000 | 0.05080 | 1.31553 |
| **FAM228B** | 0.07437 | 0.01738 | 0.09771 | 0.00003 | 0.04608 | 0.10565 | 0.00000 | 0.02096 | 0.00000 | 0.10961 | 0.09485 | 0.03285 |
| **FAM229A** | 0.00177 | 0.00000 | 0.06126 | 0.03742 | 0.00000 | 0.04675 | 0.28056 | 0.03211 | 0.07604 | 0.24947 | 0.01778 | 0.03185 |
| **FAM229B** | 0.36704 | 1.39205 | 0.00000 | 0.02200 | 0.52619 | 2.06481 | 0.53613 | 0.00000 | 0.00000 | 0.32305 | 0.54687 | 1.06230 |
| **FAM230B** | 0.00622 | 0.01475 | 0.01509 | 0.02016 | 0.01387 | 0.01380 | 0.01796 | 0.01011 | 0.01711 | 0.01928 | 0.01814 | 0.02225 |
| **FAM24A** | 0.00597 | 0.01805 | 0.01468 | 0.01061 | 0.01268 | 0.01880 | 0.02606 | 0.01052 | 0.01465 | 0.02450 | 0.01547 | 0.01506 |
| **FAM24B** | 1.35118 | 0.38340 | 2.21068 | 0.55816 | 0.00000 | 1.44830 | 1.25956 | 0.31334 | 0.07518 | 2.14806 | 0.56361 | 0.20766 |
| **FAM24B-CUZD1** | 0.00629 | 0.01455 | 0.01869 | 0.01131 | 0.01414 | 0.02144 | 0.01867 | 0.01038 | 0.02143 | 0.01116 | 0.01459 | 0.01776 |
| **FAM25A** | 0.00973 | 0.01548 | 0.02107 | 0.01151 | 0.01531 | 0.01739 | 0.01588 | 0.02535 | 0.01927 | 0.01401 | 0.01545 | 0.01519 |
| **FAM26D** | 1.31440 | 0.07338 | 0.02919 | 0.02878 | 0.25892 | 0.06570 | 0.00000 | 0.00000 | 0.00000 | 0.00327 | 0.22000 | 0.16742 |
| **FAM26E** | 0.00350 | 0.02271 | 0.09847 | 0.01032 | 0.09215 | 0.00000 | 0.00157 | 0.04628 | 0.06576 | 0.00000 | 0.24727 | 0.03002 |
| **FAM26F** | 0.00562 | 0.05175 | 0.12607 | 0.29620 | 0.09549 | 0.23909 | 0.00000 | 0.00300 | 0.35202 | 0.08177 | 0.09627 | 0.16760 |
| **FAM32A** | 3.43574 | 20.06103 | 19.37202 | 14.79409 | 14.78785 | 14.67114 | 32.67642 | 35.26305 | 23.17413 | 3.35379 | 19.12416 | 34.99566 |
| **FAM35A** | 0.01902 | 1.56093 | 1.12261 | 1.91197 | 0.00000 | 3.31063 | 3.18581 | 1.08301 | 2.63860 | 4.01146 | 1.19608 | 1.84121 |
| **FAM35BP** | 0.00599 | 0.01540 | 0.01228 | 0.01033 | 0.00999 | 0.02866 | 0.01998 | 0.01028 | 0.01436 | 0.02432 | 0.01384 | 0.01576 |
| **FAM35DP** | 0.00615 | 0.01718 | 0.01339 | 0.01409 | 0.01192 | 0.03555 | 0.01581 | 0.01027 | 0.01863 | 0.01367 | 0.01311 | 0.01542 |
| **FAM3A** | 0.34986 | 2.62466 | 1.55281 | 3.12633 | 2.78352 | 0.00000 | 0.41553 | 0.00000 | 1.98084 | 1.14638 | 2.77437 | 1.14498 |
| **FAM3B** | 0.26796 | 1.12284 | 0.20876 | 0.00247 | 0.04860 | 0.02686 | 0.00000 | 0.00000 | 0.08219 | 0.00000 | 0.05080 | 0.82877 |

| | | | | | | | | | | | | |
|---|---|---|---|---|---|---|---|---|---|---|---|---|
| **FAM3C** | 0.03097 | 0.02724 | 2.47056 | 0.00000 | 1.07699 | 3.95516 | 0.98538 | 0.07602 | 3.92065 | 5.62125 | 1.44630 | 0.00000 |
| **FAM41C** | 0.34666 | 0.01954 | 0.01254 | 0.01548 | 0.00959 | 0.00007 | 0.18485 | 0.00682 | 0.00000 | 0.02869 | 0.01202 | 0.02484 |
| **FAM43A** | 1.16614 | 0.00704 | 0.10704 | 0.01619 | 0.00000 | 0.02770 | 0.09124 | 0.20487 | 0.00000 | 0.04445 | 0.00818 | 0.01310 |
| **FAM43B** | 0.10466 | 0.04802 | 0.00844 | 0.00000 | 0.00600 | 0.03695 | 0.00113 | 0.00143 | 0.00740 | 0.13629 | 0.00651 | 0.10709 |
| **FAM45A** | 0.03975 | 0.11435 | 0.03139 | 0.07076 | 0.09098 | 0.10433 | 0.01438 | 0.03459 | 0.06392 | 0.00000 | 0.01417 | 0.11012 |
| **FAM45B.uid-12** | 0.30533 | 2.75591 | 0.26489 | 2.33334 | 2.25942 | 5.88936 | 0.00000 | 0.05273 | 0.59279 | 0.00000 | 0.91296 | 1.18191 |
| **FAM45B.uid-14** | 0.00376 | 0.01424 | 0.01061 | 0.00353 | 0.01087 | 0.06309 | 0.01098 | 0.01467 | 0.01723 | 0.08809 | 0.00803 | 0.01110 |
| **FAM46A** | 0.38597 | 2.41843 | 13.10630 | 0.00000 | 3.57470 | 8.44213 | 3.99839 | 3.75284 | 6.36112 | 7.69725 | 7.41928 | 2.67095 |
| **FAM46B** | 7.56070 | 13.52250 | 2.35206 | 11.67242 | 13.09308 | 0.00000 | 4.00864 | 2.21005 | 0.00000 | 8.40081 | 2.76075 | 0.42829 |
| **FAM46C** | 25.83770 | 0.96117 | 0.44876 | 0.09534 | 0.00000 | 0.81518 | 0.00000 | 11.41019 | 0.00000 | 0.95338 | 0.00000 | 3.57827 |
| **FAM46D** | 0.00283 | 0.13261 | 0.06945 | 0.00000 | 0.00421 | 0.03141 | 0.00693 | 0.13588 | 0.01859 | 0.02453 | 0.01374 | 0.19199 |
| **FAM47E** | 0.00568 | 0.12351 | 0.01158 | 0.12111 | 0.00000 | 0.05043 | 0.00000 | 0.01072 | 0.00674 | 0.04860 | 0.01487 | 0.01773 |
| **FAM49A** | 0.00250 | 0.17507 | 13.66306 | 0.00427 | 0.05552 | 0.96953 | 0.00000 | 0.16822 | 0.30180 | 0.00000 | 0.12473 | 0.23042 |
| **FAM49B** | 4.39249 | 11.65722 | 19.20153 | 6.05275 | 0.01587 | 29.50196 | 13.38552 | 1.26005 | 17.43025 | 4.74759 | 9.97916 | 10.68070 |
| **FAM50A** | 1.07354 | 12.30202 | 11.36296 | 8.86798 | 15.28079 | 4.59887 | 14.98558 | 0.14407 | 7.96822 | 9.10430 | 13.44253 | 5.47223 |
| **FAM50B** | 0.02853 | 0.09355 | 0.28925 | 0.01328 | 0.11714 | 0.00000 | 0.03150 | 0.11168 | 0.78559 | 0.86100 | 0.00903 | 0.82526 |
| **FAM53A** | 0.06572 | 0.07249 | 0.02489 | 0.05266 | 0.16671 | 0.00000 | 0.00000 | 0.06274 | 0.31424 | 0.36701 | 0.00000 | 0.06340 |
| **FAM53B** | 2.04899 | 2.92738 | 1.25740 | 1.23228 | 5.61832 | 0.00000 | 2.43719 | 6.14920 | 4.81887 | 7.29067 | 1.23926 | 2.08768 |
| **FAM53B-AS1** | 0.03682 | 0.00162 | 0.02699 | 0.00621 | 0.35002 | 0.00000 | 0.00285 | 0.25577 | 0.04780 | 0.22795 | 0.00429 | 0.06993 |
| **FAM53C** | 6.92473 | 5.13486 | 6.44133 | 9.49193 | 7.88896 | 1.80012 | 14.79142 | 9.02250 | 12.58542 | 3.05259 | 4.63264 | 14.80882 |
| **FAM57A** | 0.05610 | 0.43711 | 0.89528 | 0.00000 | 2.09067 | 0.05760 | 2.51264 | 0.25237 | 2.06788 | 3.05779 | 1.62650 | 0.07746 |
| **FAM57B** | 0.00625 | 0.02339 | 0.03134 | 0.01200 | 0.01654 | 0.02406 | 0.01551 | 0.01042 | 0.01681 | 0.01030 | 0.01602 | 0.01570 |
| **FAM58A** | 2.51270 | 1.88486 | 0.84895 | 1.38158 | 2.96267 | 0.00000 | 3.68992 | 5.39126 | 1.18561 | 3.21144 | 0.18375 | 6.01955 |
| **FAM60A** | 2.43499 | 7.62069 | 10.35826 | 7.33549 | 6.79107 | 15.42760 | 12.54484 | 7.77798 | 14.30752 | 0.00000 | 8.89443 | 8.92578 |
| **FAM63A** | 0.18267 | 0.14853 | 0.34445 | 0.42122 | 0.00000 | 0.00000 | 0.06256 | 0.18565 | 0.28885 | 0.10224 | 0.26857 | 0.06925 |
| **FAM63B** | 2.41132 | 6.97890 | 5.56686 | 11.40968 | 6.90444 | 6.08422 | 1.17775 | 3.00501 | 12.80167 | 17.47462 | 9.35468 | 7.03449 |
| **FAM64A** | 0.00148 | 0.13572 | 0.11190 | 6.28902 | 0.00000 | 0.00000 | 0.36394 | 1.63000 | 0.00068 | 0.13515 | 0.01396 | 0.77332 |
| **FAM65A** | 0.02460 | 0.11329 | 4.47529 | 0.00050 | 0.21391 | 0.13970 | 0.29566 | 0.02861 | 0.84254 | 0.00000 | 0.00000 | 0.61545 |
| **FAM65B** | 1.66107 | 0.73419 | 4.10477 | 0.00000 | 0.90738 | 0.16279 | 2.44548 | 0.73704 | 0.00000 | 1.07329 | 0.00000 | 0.00000 |
| **FAM65C** | 1.06434 | 0.04656 | 0.00000 | 0.00000 | 0.00000 | 0.29188 | 0.00000 | 0.12072 | 0.28383 | 0.64003 | 0.00000 | 0.07893 |
| **FAM66A** | 0.00616 | 0.01481 | 0.01363 | 0.01375 | 0.01670 | 0.02465 | 0.01763 | 0.01026 | 0.01525 | 0.01256 | 0.01787 | 0.01883 |
| **FAM66B** | 0.01933 | 0.01927 | 0.01431 | 0.01204 | 0.01483 | 0.01591 | 0.02165 | 0.01195 | 0.01764 | 0.01395 | 0.01526 | 0.01537 |
| **FAM66C** | 0.11936 | 0.05834 | 0.01528 | 0.34409 | 0.00000 | 0.00741 | 0.02324 | 0.02255 | 0.00711 | 0.03801 | 0.01370 | 0.17839 |
| **FAM66D** | 0.00479 | 0.06933 | 0.01520 | 0.03066 | 0.00000 | 0.01692 | 0.03203 | 0.03268 | 0.00532 | 0.03516 | 0.01622 | 0.16430 |
| **FAM69A** | 0.26441 | 1.64591 | 0.42127 | 0.35220 | 0.30103 | 0.69609 | 0.00000 | 0.51613 | 1.23677 | 0.09850 | 0.28993 | 0.81055 |
| **FAM69B** | 1.54923 | 0.89122 | 1.50326 | 3.90378 | 3.97263 | 0.00000 | 2.98515 | 0.15315 | 1.13957 | 4.01848 | 0.00000 | 0.24474 |
| **FAM69C** | 0.00624 | 0.01611 | 0.01864 | 0.02869 | 0.02561 | 0.01232 | 0.05889 | 0.00977 | 0.00890 | 0.00000 | 0.01807 | 0.03306 |
| **FAM71A** | 0.00874 | 0.01711 | 0.01184 | 0.01886 | 0.01817 | 0.01165 | 0.04980 | 0.00988 | 0.01248 | 0.00841 | 0.01404 | 0.01683 |
| **FAM71C** | 0.21790 | 0.01299 | 0.02896 | 0.01144 | 0.01361 | 0.01937 | 0.02035 | 0.03589 | 0.01487 | 0.01195 | 0.01556 | 0.01962 |
| **FAM71E1** | 3.21445 | 2.88806 | 0.77506 | 3.62776 | 3.64426 | 0.00000 | 2.06937 | 1.00938 | 2.20370 | 1.10634 | 2.41398 | 2.27236 |
| **FAM71E2** | 0.45072 | 0.01742 | 0.02096 | 0.01681 | 0.02382 | 0.00004 | 0.07265 | 0.01230 | 0.00103 | 0.00003 | 0.01640 | 0.01395 |
| **FAM71F1** | 0.00944 | 0.02735 | 0.16241 | 0.12405 | 0.05025 | 0.00000 | 0.03779 | 0.00840 | 0.02991 | 0.07258 | 0.01326 | 0.05476 |
| **FAM71F2** | 0.00432 | 0.13001 | 0.00045 | 0.25837 | 0.00000 | 0.00000 | 0.33137 | 0.02277 | 0.00000 | 1.62492 | 0.27826 | 0.23124 |
| **FAM72B** | 0.24986 | 0.84861 | 0.43578 | 0.23737 | 0.75663 | 0.45413 | 0.18771 | 0.10546 | 0.48803 | 0.01345 | 0.25276 | 1.12143 |
| **FAM73A** | 0.99928 | 2.25035 | 5.98903 | 0.00000 | 0.98664 | 11.04268 | 0.00004 | 0.31089 | 9.53798 | 12.64158 | 2.30151 | 2.00074 |
| **FAM73B** | 0.01924 | 3.72315 | 2.33578 | 2.67501 | 4.47885 | 0.00000 | 0.02689 | 0.35745 | 6.04142 | 4.59612 | 0.58715 | 2.07783 |
| **FAM76A** | 2.43415 | 5.60453 | 13.15278 | 0.94794 | 0.00000 | 16.57344 | 11.32176 | 14.21046 | 0.00000 | 4.34864 | 15.58346 | 12.19428 |
| **FAM76B** | 0.52459 | 2.27029 | 0.00000 | 0.92211 | 0.00000 | 12.38504 | 3.08063 | 2.86184 | 1.43003 | 10.20489 | 1.42843 | 2.82550 |

| | | | | | | | | | | | | |
|---|---|---|---|---|---|---|---|---|---|---|---|---|
| **FAM78A** | 0.67820 | 0.06076 | 0.12330 | 0.00000 | 1.22027 | 0.00000 | 2.33632 | 0.00000 | 1.10541 | 0.29913 | 0.00006 | 0.14361 |
| **FAM78B** | 0.00184 | 0.09811 | 0.00000 | 0.31634 | 0.07470 | 0.00000 | 0.00096 | 0.17779 | 0.10109 | 0.43248 | 0.00000 | 0.01292 |
| **FAM81A** | 0.79515 | 0.01665 | 0.29445 | 0.00667 | 0.08127 | 0.12831 | 0.29790 | 0.12203 | 0.00000 | 0.00000 | 0.08193 | 0.00000 |
| **FAM81B** | 1.98353 | 0.00972 | 0.01058 | 0.01059 | 0.01619 | 0.00141 | 0.03469 | 0.18794 | 0.02219 | 0.01180 | 0.02163 | 0.04233 |
| **FAM83A** | 0.00272 | 0.00000 | 0.10792 | 0.06343 | 0.00000 | 0.00000 | 0.00003 | 0.14243 | 0.06837 | 0.39129 | 0.00883 | 0.64113 |
| **FAM83A-AS1** | 0.01089 | 0.01410 | 0.02001 | 0.01635 | 0.01699 | 0.01656 | 0.02737 | 0.06434 | 0.02900 | 0.00000 | 0.01287 | 0.04361 |
| **FAM83B** | 0.00000 | 0.00000 | 3.22983 | 0.00000 | 0.43637 | 9.07623 | 7.86791 | 0.07224 | 3.80207 | 1.16103 | 0.88066 | 0.00000 |
| **FAM83C** | 0.00562 | 0.01736 | 0.05007 | 0.00829 | 0.03401 | 0.00896 | 0.01374 | 0.01055 | 0.01047 | 0.04693 | 0.01429 | 0.01604 |
| **FAM83C-AS1** | 0.01244 | 0.05321 | 0.09182 | 0.01725 | 0.02678 | 0.01835 | 0.01013 | 0.00891 | 0.02663 | 0.02182 | 0.03021 | 0.02051 |
| **FAM83D** | 0.00000 | 13.53196 | 10.85582 | 6.43015 | 18.91457 | 16.55046 | 8.22992 | 8.24205 | 17.93950 | 5.95491 | 16.02789 | 6.59196 |
| **FAM83E** | 2.90927 | 0.00000 | 0.21018 | 0.00000 | 0.00000 | 0.00000 | 0.48181 | 0.31183 | 0.11361 | 0.75939 | 0.00000 | 0.03391 |
| **FAM83F** | 0.00561 | 0.33166 | 1.87023 | 1.18768 | 0.91900 | 0.00000 | 0.90955 | 0.01623 | 0.10843 | 1.44617 | 0.18573 | 0.00000 |
| **FAM83G** | 0.01114 | 1.09780 | 3.15546 | 0.21264 | 1.22581 | 0.00000 | 0.00000 | 0.03430 | 2.44957 | 0.03784 | 0.27617 | 0.03089 |
| **FAM83H** | 0.00000 | 1.26348 | 3.30861 | 1.68006 | 8.55577 | 0.56930 | 7.70235 | 0.00000 | 0.38060 | 15.02343 | 1.35379 | 0.10236 |
| **FAM83H-AS1** | 0.00137 | 0.73488 | 6.32740 | 0.00000 | 1.97784 | 0.79627 | 3.94160 | 0.00000 | 3.59962 | 1.46844 | 1.08358 | 0.13631 |
| **FAM84A** | 0.00711 | 0.08097 | 0.28292 | 0.00000 | 0.06209 | 0.01642 | 0.00000 | 0.01031 | 0.00000 | 0.13268 | 0.01277 | 0.01077 |
| **FAM84B** | 4.59150 | 15.69529 | 0.00000 | 7.12952 | 8.84532 | 12.10138 | 6.63128 | 0.59718 | 5.35704 | 12.17685 | 4.71751 | 2.32493 |
| **FAM86A** | 0.18811 | 2.75794 | 1.27286 | 1.04761 | 3.31663 | 0.66579 | 0.02750 | 0.00000 | 1.40759 | 0.00000 | 1.44467 | 2.33743 |
| **FAM86B1** | 0.19712 | 0.09279 | 0.00582 | 0.00266 | 0.13362 | 0.00971 | 0.00837 | 0.01456 | 0.02638 | 0.06271 | 0.00753 | 0.19375 |
| **FAM86B2** | 0.04919 | 0.02537 | 0.02289 | 0.01502 | 0.01862 | 0.00722 | 0.01744 | 0.00616 | 0.02877 | 0.01203 | 0.01480 | 0.04365 |
| **FAM86B3P** | 0.00121 | 0.14648 | 0.09573 | 0.00291 | 0.00000 | 0.00000 | 0.05965 | 0.06292 | 0.34815 | 0.30892 | 0.12137 | 0.08240 |
| **FAM86C1** | 0.40658 | 1.81794 | 0.29085 | 0.34432 | 0.95150 | 1.54888 | 0.14348 | 0.15771 | 0.00000 | 0.00000 | 0.86848 | 0.95195 |
| **FAM86C2P** | 0.04262 | 0.39221 | 0.03472 | 0.09746 | 0.00356 | 0.21002 | 0.00000 | 0.11114 | 0.08400 | 0.11000 | 0.10622 | 0.59291 |
| **FAM86DP** | 0.05316 | 1.38157 | 0.40214 | 0.06760 | 0.31299 | 0.18941 | 0.00000 | 0.09494 | 0.37008 | 0.42730 | 0.08963 | 0.13954 |
| **FAM86EP** | 0.36350 | 1.12921 | 0.15981 | 0.97845 | 0.56632 | 0.00000 | 0.00816 | 0.04455 | 0.35031 | 0.00000 | 0.19870 | 1.28342 |
| **FAM86FP** | 0.05251 | 0.07121 | 0.01804 | 0.09591 | 0.00000 | 0.07523 | 0.04283 | 0.05766 | 0.00000 | 0.13884 | 0.01728 | 0.07233 |
| **FAM86HP** | 0.02336 | 0.03452 | 0.02954 | 0.04700 | 0.00000 | 0.01012 | 0.01119 | 0.03531 | 0.00181 | 0.10965 | 0.17881 | 0.06660 |
| **FAM86JP** | 0.34912 | 0.31748 | 0.16127 | 0.02101 | 0.00818 | 0.04736 | 0.00000 | 0.00000 | 0.16293 | 0.00000 | 0.13117 | 0.46717 |
| **FAM87A** | 0.10059 | 0.01523 | 0.02305 | 0.02202 | 0.00000 | 0.01060 | 0.02587 | 0.05477 | 0.00000 | 0.08617 | 0.06893 | 0.01147 |
| **FAM87B** | 0.00765 | 0.01643 | 0.02933 | 0.03819 | 0.01983 | 0.00000 | 0.02660 | 0.14633 | 0.04391 | 0.00000 | 0.01311 | 0.03649 |
| **FAM89A** | 0.37543 | 0.02444 | 8.07718 | 1.19893 | 0.00000 | 0.40677 | 2.03256 | 2.19682 | 0.00000 | 0.68386 | 0.00000 | 0.90473 |
| **FAM89B** | 0.78064 | 0.29534 | 2.28391 | 0.16694 | 1.11019 | 0.01932 | 2.96717 | 5.50184 | 1.31250 | 4.22788 | 0.00000 | 0.93214 |
| **FAM8A1** | 0.23561 | 0.10560 | 3.02106 | 0.00000 | 0.00000 | 5.50928 | 2.26430 | 0.88798 | 2.24280 | 6.58057 | 1.08816 | 0.28388 |
| **FAM90A1** | 0.41537 | 0.02657 | 0.00290 | 0.17025 | 0.00000 | 0.05911 | 0.00000 | 0.64968 | 0.05021 | 0.02290 | 0.00153 | 0.00000 |
| **FAM90A25P** | 0.02952 | 0.01475 | 0.01424 | 0.01221 | 0.01550 | 0.01696 | 0.01801 | 0.02725 | 0.01870 | 0.01368 | 0.01489 | 0.01714 |
| **FAM90A27P** | 0.66574 | 0.01261 | 0.01273 | 0.01002 | 0.01317 | 0.01702 | 0.01503 | 0.29298 | 0.01924 | 0.01617 | 0.01309 | 0.00997 |
| **FAM90A2P** | 0.00660 | 0.01542 | 0.01885 | 0.01190 | 0.01591 | 0.01742 | 0.01829 | 0.01441 | 0.01723 | 0.01258 | 0.01526 | 0.01604 |
| **FAM91A1** | 0.00000 | 3.12486 | 4.99736 | 0.00000 | 0.05933 | 11.47789 | 6.59537 | 1.57788 | 3.61275 | 9.12509 | 3.32834 | 3.50651 |
| **FAM92A1** | 0.09112 | 0.53963 | 0.30350 | 1.75961 | 0.00000 | 2.54181 | 0.67612 | 0.04241 | 0.00000 | 1.17565 | 0.63600 | 0.17523 |
| **FAM92A1P2** | 0.00389 | 0.00736 | 0.01535 | 0.02550 | 0.00474 | 0.04009 | 0.00200 | 0.04085 | 0.02643 | 0.06677 | 0.01698 | 0.02900 |
| **FAM92B** | 0.00670 | 0.01434 | 0.01870 | 0.01726 | 0.00788 | 0.00939 | 0.02828 | 0.01041 | 0.00000 | 0.01965 | 0.07779 | 0.01679 |
| **FAM95A** | 0.00505 | 0.02078 | 0.01452 | 0.09790 | 0.15583 | 0.00667 | 0.03703 | 0.01365 | 0.00000 | 0.00000 | 0.01142 | 0.13336 |
| **FAM95C** | 0.00438 | 0.01858 | 0.06260 | 0.11207 | 0.01234 | 0.00000 | 0.00633 | 0.02783 | 0.02410 | 0.06204 | 0.01162 | 0.03355 |
| **FAM96A** | 2.63569 | 15.31668 | 10.31233 | 11.14556 | 6.22620 | 37.60570 | 18.35290 | 1.36698 | 13.47426 | 3.71812 | 15.85387 | 13.70478 |
| **FAM96B** | 7.30831 | 27.00653 | 16.14889 | 21.13897 | 26.99433 | 16.66617 | 28.74840 | 12.73022 | 23.69288 | 4.14523 | 27.47073 | 22.58335 |
| **FAM98A** | 1.23614 | 8.21605 | 8.91593 | 5.35867 | 9.18265 | 16.68315 | 15.69764 | 0.94074 | 5.22290 | 12.36050 | 14.41961 | 12.97999 |
| **FAM98B** | 0.00000 | 0.00000 | 4.96188 | 0.62618 | 0.00000 | 3.95501 | 7.29368 | 0.10825 | 0.00000 | 10.29217 | 1.08931 | 0.38831 |
| **FAM98C** | 0.01978 | 1.58154 | 0.41752 | 0.07215 | 0.33624 | 0.00000 | 0.04477 | 0.07987 | 0.27360 | 1.42554 | 0.37350 | 0.99441 |

| | | | | | | | | | | | | |
|---|---|---|---|---|---|---|---|---|---|---|---|---|
| **FAM9A** | 0.37547 | 0.00392 | 0.02265 | 0.00463 | 0.00000 | 0.01527 | 0.20030 | 1.64974 | 0.00000 | 0.10517 | 0.00425 | 0.07228 |
| **FAM9B** | 0.13369 | 0.13666 | 0.01572 | 0.00000 | 0.00000 | 0.39780 | 0.18002 | 0.56493 | 0.00000 | 0.02784 | 0.00000 | 0.00000 |
| **FAM9C** | 0.75449 | 0.00000 | 0.32519 | 0.02185 | 0.83472 | 0.67963 | 0.40311 | 0.05785 | 1.76479 | 0.94681 | 2.08831 | 0.00000 |
| **FAN1** | 0.00000 | 3.45073 | 0.00000 | 1.04678 | 0.18743 | 3.27635 | 0.00000 | 0.03144 | 0.00000 | 1.73193 | 3.81600 | 3.41488 |
| **FANCA** | 1.22860 | 6.36454 | 1.62130 | 11.77863 | 6.82702 | 0.00000 | 1.08065 | 1.61988 | 1.46632 | 2.66861 | 2.68710 | 2.65280 |
| **FANCB** | 0.00001 | 0.00000 | 3.01786 | 1.30183 | 0.00000 | 3.83399 | 2.53156 | 3.59127 | 2.50961 | 3.51896 | 2.91980 | 2.17779 |
| **FANCC** | 0.75191 | 1.54971 | 0.88517 | 0.00000 | 1.44583 | 1.18401 | 1.52085 | 0.50820 | 2.23184 | 0.34663 | 1.01505 | 3.25722 |
| **FANCD2** | 3.40333 | 9.12770 | 5.30070 | 8.64907 | 4.75404 | 13.56809 | 6.71344 | 4.00561 | 2.88357 | 12.21887 | 11.54913 | 8.92590 |
| **FANCD2OS** | 0.01568 | 0.04704 | 0.02692 | 0.03301 | 0.09457 | 0.00000 | 0.00991 | 0.01431 | 0.02860 | 0.17366 | 0.04451 | 0.04249 |
| **FANCE** | 1.19557 | 4.27113 | 5.61510 | 2.95796 | 3.17864 | 2.13326 | 5.35574 | 1.25442 | 2.45668 | 6.64621 | 1.02952 | 16.43720 |
| **FANCF** | 1.48059 | 2.60292 | 2.06136 | 0.12433 | 1.06465 | 3.20815 | 1.20027 | 1.00772 | 0.63887 | 0.35010 | 2.15498 | 3.99161 |
| **FANCG** | 0.35587 | 2.29321 | 7.40255 | 6.91387 | 3.23049 | 2.75442 | 2.61763 | 0.00000 | 7.05839 | 1.31479 | 5.70240 | 1.19274 |
| **FANCI** | 6.95365 | 6.08792 | 9.44153 | 3.77420 | 0.92978 | 16.96321 | 9.58535 | 6.50032 | 9.28453 | 4.81388 | 8.69268 | 6.24234 |
| **FANCL** | 0.83518 | 1.89284 | 3.94953 | 1.56188 | 0.00000 | 4.36447 | 2.04887 | 1.09791 | 1.24872 | 1.68238 | 3.67398 | 2.76869 |
| **FANCM** | 2.83703 | 0.29223 | 1.83904 | 1.38421 | 1.10495 | 7.28750 | 0.00000 | 6.32675 | 3.87652 | 5.36300 | 1.16472 | 3.18649 |
| **FANK1** | 0.01908 | 0.85300 | 0.48853 | 0.11248 | 0.52443 | 0.73576 | 0.09573 | 0.04057 | 0.49607 | 0.00000 | 0.12802 | 1.31066 |
| **FAP** | 0.00686 | 0.01639 | 0.02740 | 0.01416 | 0.01754 | 0.02120 | 0.04102 | 0.01017 | 0.00000 | 0.01469 | 0.02843 | 0.01642 |
| **FAR1** | 0.72519 | 0.00000 | 7.04857 | 2.65413 | 3.43217 | 13.14412 | 13.84236 | 9.21613 | 11.56518 | 2.60176 | 14.02518 | 3.92353 |
| **FAR2** | 1.76192 | 0.01590 | 0.24358 | 0.00000 | 0.02142 | 0.41600 | 0.00000 | 0.30388 | 0.20240 | 0.14831 | 0.00000 | 0.00000 |
| **FAR2P1** | 0.00195 | 0.00000 | 0.13632 | 0.00000 | 0.00000 | 0.34539 | 0.00051 | 0.26108 | 0.00000 | 0.22963 | 0.00000 | 0.72539 |
| **FAR2P2** | 0.00594 | 0.03411 | 0.05988 | 0.01891 | 0.02293 | 0.00000 | 0.01140 | 0.01169 | 0.01775 | 0.01851 | 0.02218 | 0.04335 |
| **FARP1** | 4.47368 | 0.73540 | 0.00000 | 2.49424 | 8.58633 | 4.07901 | 3.08673 | 0.12432 | 3.61635 | 5.65007 | 5.13919 | 0.00000 |
| **FARP2** | 0.03938 | 0.77116 | 1.66634 | 0.86422 | 1.64193 | 0.00000 | 0.39453 | 3.39627 | 2.29872 | 1.26866 | 0.66420 | 0.87265 |
| **FARS2** | 1.42969 | 6.06781 | 2.79692 | 4.07112 | 2.45846 | 2.22917 | 1.96984 | 1.46064 | 4.45809 | 0.00000 | 4.41089 | 4.88622 |
| **FARSA** | 1.57939 | 28.94845 | 26.34359 | 17.72461 | 30.31260 | 1.67726 | 34.47862 | 0.66540 | 25.70532 | 0.00000 | 28.68723 | 16.04370 |
| **FARSB** | 2.55864 | 17.71899 | 9.95344 | 2.86383 | 6.32162 | 32.39397 | 9.16122 | 2.44278 | 11.93931 | 0.00000 | 17.50780 | 12.36771 |
| **FAS** | 0.03547 | 0.20555 | 0.00394 | 0.00000 | 0.13295 | 0.07082 | 0.00122 | 0.00512 | 0.02634 | 0.00733 | 0.01131 | 0.08152 |
| **FAS-AS1** | 0.03837 | 0.01266 | 0.03011 | 0.01614 | 0.01415 | 0.01896 | 0.01390 | 0.00705 | 0.01620 | 0.02113 | 0.01425 | 0.02219 |
| **FASLG** | 0.00624 | 0.00996 | 0.62036 | 0.00326 | 0.01019 | 0.05173 | 0.00000 | 0.00988 | 0.01169 | 0.01069 | 0.01024 | 0.01388 |
| **FASN** | 0.21900 | 1.81937 | 14.91187 | 4.37529 | 26.10264 | 0.00000 | 17.33379 | 0.05938 | 32.86665 | 31.40608 | 9.80610 | 0.31752 |
| **FASTK** | 0.97192 | 11.85875 | 6.03185 | 1.39090 | 3.42276 | 0.00000 | 1.98362 | 0.00000 | 8.54538 | 2.86003 | 6.58922 | 7.81438 |
| **FASTKD1** | 2.62110 | 16.89912 | 0.00000 | 6.20954 | 3.85689 | 35.14753 | 9.00581 | 1.37102 | 13.37861 | 9.61212 | 6.44043 | 10.70552 |
| **FASTKD2** | 1.62726 | 12.63927 | 5.52486 | 5.87397 | 3.78038 | 27.26502 | 9.51699 | 7.26342 | 13.14309 | 2.18217 | 11.97303 | 12.08198 |
| **FASTKD3** | 0.79506 | 5.15525 | 3.61697 | 1.32408 | 0.00000 | 9.65682 | 6.26895 | 3.76350 | 5.24700 | 0.00000 | 6.73365 | 6.66972 |
| **FASTKD5** | 0.00000 | 11.00084 | 7.52650 | 7.93665 | 2.18526 | 15.42273 | 12.34570 | 18.10940 | 16.07368 | 0.83055 | 8.01549 | 16.47112 |
| **FAT1** | 6.09995 | 0.00000 | 0.98383 | 0.00000 | 7.40260 | 5.55395 | 0.00000 | 4.40459 | 8.92103 | 10.73312 | 1.97918 | 1.57373 |
| **FAT2** | 1.63375 | 0.00000 | 0.49575 | 0.25801 | 0.11172 | 0.00000 | 0.17573 | 0.89079 | 0.64636 | 0.29758 | 0.02059 | 0.11894 |
| **FAT3** | 1.47500 | 0.03316 | 0.00384 | 0.03207 | 0.00000 | 0.00878 | 0.00240 | 0.00159 | 0.00909 | 0.39227 | 0.01002 | 0.03561 |
| **FAT4** | 0.00564 | 0.01439 | 0.02823 | 0.02551 | 0.03141 | 0.00927 | 0.01820 | 0.00928 | 0.01144 | 0.03050 | 0.02969 | 0.04149 |
| **FAU** | 9.34285 | 39.81171 | 37.23187 | 29.32491 | 47.88702 | 31.90941 | 53.65232 | 32.10549 | 48.44979 | 24.84079 | 37.16923 | 48.69620 |
| **FAXC** | 0.00702 | 1.24342 | 0.41679 | 0.36172 | 0.00000 | 0.07084 | 0.25648 | 0.16640 | 0.06278 | 0.57727 | 0.00000 | 1.15095 |
| **FAXDC2** | 0.46138 | 0.01927 | 0.05942 | 0.03965 | 0.06108 | 0.00000 | 0.00000 | 0.11210 | 0.36378 | 0.60095 | 0.09228 | 0.00148 |
| **FBF1** | 0.01450 | 0.53226 | 0.64980 | 0.06046 | 0.81259 | 0.00000 | 0.00000 | 0.00971 | 1.05972 | 0.00225 | 0.98287 | 0.56469 |
| **FBL** | 3.47550 | 19.82150 | 19.59149 | 13.47468 | 30.30565 | 0.09853 | 39.10215 | 16.72248 | 23.85550 | 27.93972 | 25.78595 | 29.07603 |
| **FBLIM1** | 0.22093 | 0.00000 | 0.87654 | 1.23102 | 0.00000 | 0.13830 | 1.49927 | 0.00000 | 0.11115 | 0.00000 | 0.24415 | 0.04445 |
| **FBLL1** | 0.00266 | 0.00000 | 0.20460 | 0.00469 | 0.10518 | 0.00000 | 0.30875 | 0.01574 | 0.00000 | 0.53449 | 0.00000 | 0.01980 |
| **FBLN1** | 0.00136 | 0.88244 | 18.05130 | 4.49647 | 7.62799 | 0.00000 | 0.05172 | 0.48416 | 8.99522 | 5.77690 | 13.40339 | 0.57639 |
| **FBLN2** | 0.99481 | 0.00377 | 2.49131 | 0.00423 | 2.21129 | 0.00000 | 0.00000 | 0.22878 | 1.87878 | 0.00028 | 0.03658 | 0.13920 |

| | | | | | | | | | | | | |
|---|---|---|---|---|---|---|---|---|---|---|---|---|
| **FBLN5** | 4.37355 | 0.17136 | 0.00000 | 0.00011 | 0.06102 | 0.47704 | 0.00000 | 1.24250 | 0.00000 | 0.00047 | 0.05404 | 0.10718 |
| **FBLN7** | 9.25662 | 0.00000 | 0.00000 | 0.00000 | 0.05952 | 0.05234 | 0.00000 | 0.72604 | 0.00000 | 0.19177 | 0.04162 | 0.17695 |
| **FBN1** | 0.48205 | 0.00000 | 0.13504 | 0.18589 | 0.00000 | 0.00000 | 0.22443 | 0.04043 | 0.30533 | 0.18517 | 0.06600 | 0.16867 |
| **FBN2** | 0.27843 | 0.01759 | 2.40934 | 0.24070 | 0.00000 | 0.17468 | 0.76996 | 0.05796 | 0.36747 | 0.04492 | 0.00000 | 0.05796 |
| **FBN3** | 0.10123 | 0.04299 | 0.05064 | 0.01369 | 0.01248 | 0.00000 | 0.00797 | 0.01328 | 0.00000 | 0.09365 | 0.06723 | 0.01186 |
| **FBP1** | 0.31348 | 0.00898 | 1.48699 | 17.44961 | 0.50852 | 0.00000 | 0.00000 | 0.47494 | 0.00000 | 0.64163 | 0.00000 | 0.26708 |
| **FBP2** | 0.70561 | 0.03901 | 0.08004 | 0.00367 | 0.00000 | 0.06885 | 0.02072 | 0.12285 | 0.10057 | 0.02196 | 0.00690 | 0.34481 |
| **FBRS** | 3.76574 | 1.02581 | 8.40065 | 1.34947 | 3.46639 | 0.00000 | 3.90712 | 0.60555 | 4.21397 | 12.23444 | 0.70771 | 3.27936 |
| **FBRSL1** | 0.56731 | 0.00000 | 0.00000 | 0.00000 | 2.34497 | 0.27741 | 1.76209 | 0.00000 | 0.00000 | 7.04542 | 0.21360 | 0.00000 |
| **FBXL12** | 4.41079 | 14.65659 | 5.69455 | 9.28699 | 8.30253 | 1.31265 | 7.06902 | 11.26109 | 9.23841 | 0.00000 | 6.07108 | 13.62700 |
| **FBXL13** | 0.81456 | 0.02912 | 0.17571 | 0.00524 | 0.02574 | 0.07403 | 0.00061 | 0.02870 | 0.00000 | 0.04991 | 0.08093 | 0.12036 |
| **FBXL14** | 0.31020 | 0.04091 | 0.00000 | 0.39474 | 0.14165 | 0.00579 | 0.54513 | 0.14170 | 0.00000 | 1.08479 | 0.07546 | 0.08864 |
| **FBXL15** | 1.16036 | 8.74816 | 7.16565 | 9.71729 | 10.64330 | 0.00000 | 7.55777 | 0.56819 | 7.47378 | 1.70855 | 7.66611 | 3.56708 |
| **FBXL16** | 0.00595 | 0.00065 | 0.33271 | 0.00000 | 0.19232 | 0.00000 | 0.14550 | 0.01270 | 0.00000 | 0.29252 | 0.07696 | 0.03453 |
| **FBXL17** | 0.27358 | 0.00646 | 0.20226 | 0.08709 | 0.00000 | 0.14733 | 0.68094 | 0.03643 | 0.08024 | 0.36057 | 0.00000 | 0.00335 |
| **FBXL18** | 2.86448 | 1.85251 | 4.22203 | 0.49376 | 12.24326 | 0.00000 | 10.26946 | 0.98577 | 11.01788 | 11.04310 | 5.24124 | 2.72744 |
| **FBXL19** | 0.64680 | 0.00000 | 0.13097 | 0.36765 | 0.56307 | 0.00000 | 0.29886 | 0.00000 | 0.00000 | 4.69066 | 0.00000 | 0.20461 |
| **FBXL19-AS1** | 0.01264 | 0.00000 | 0.82259 | 0.29027 | 0.98700 | 0.00000 | 0.00000 | 3.32727 | 1.79007 | 2.71002 | 0.79593 | 0.48077 |
| **FBXL2** | 1.65968 | 0.00000 | 0.11068 | 0.00000 | 0.00000 | 0.37762 | 0.00000 | 1.12915 | 0.03936 | 0.76145 | 0.18567 | 1.04305 |
| **FBXL20** | 3.26268 | 27.62723 | 0.86947 | 12.99982 | 10.26754 | 8.59414 | 0.00000 | 12.07882 | 0.00000 | 6.61465 | 4.24567 | 23.55887 |
| **FBXL21** | 0.80250 | 0.75562 | 0.49432 | 0.00000 | 0.00000 | 0.10411 | 0.63438 | 0.01227 | 5.42451 | 0.00148 | 0.29014 | 0.08469 |
| **FBXL22** | 1.42570 | 0.02766 | 0.00896 | 0.01146 | 0.00608 | 0.01143 | 0.01869 | 0.15974 | 0.01967 | 0.04657 | 0.02246 | 0.07186 |
| **FBXL3** | 1.46801 | 1.41564 | 0.91959 | 0.02229 | 0.00000 | 7.43803 | 2.61998 | 3.89262 | 0.22656 | 6.29966 | 0.26897 | 2.05530 |
| **FBXL4** | 0.06617 | 1.10968 | 1.10828 | 1.29726 | 0.00000 | 4.57383 | 0.58437 | 0.26273 | 1.15028 | 1.12912 | 0.63138 | 1.04488 |
| **FBXL5** | 8.04271 | 2.07273 | 3.31230 | 3.69830 | 0.00000 | 15.29026 | 4.37506 | 5.47486 | 2.32510 | 0.89666 | 9.40154 | 3.35664 |
| **FBXL6** | 1.28602 | 4.00495 | 1.46031 | 1.23200 | 6.72795 | 0.00000 | 0.17341 | 0.14746 | 2.01109 | 6.43599 | 1.49930 | 1.16548 |
| **FBXL7** | 1.96399 | 0.03245 | 0.00330 | 0.02032 | 0.01240 | 0.10032 | 0.03884 | 0.01731 | 0.12090 | 0.00039 | 0.00000 | 0.00422 |
| **FBXL8** | 0.00452 | 0.06418 | 0.08513 | 0.06066 | 0.06887 | 0.04535 | 0.00000 | 0.00557 | 0.13750 | 0.13039 | 0.06635 | 0.11889 |
| **FBXO10** | 3.71158 | 1.22704 | 0.77182 | 1.14300 | 1.09154 | 0.00000 | 0.05456 | 0.14897 | 2.04232 | 1.64211 | 0.53043 | 1.68357 |
| **FBXO11** | 1.53569 | 0.99783 | 8.59159 | 0.00000 | 5.38352 | 11.10352 | 8.86688 | 3.34483 | 3.34663 | 15.15109 | 3.09297 | 0.44491 |
| **FBXO15** | 1.17484 | 0.50143 | 0.18584 | 0.00000 | 0.00000 | 0.00000 | 0.01372 | 0.41412 | 0.00120 | 0.02631 | 0.05309 | 0.05057 |
| **FBXO16** | 0.10045 | 0.21740 | 0.00000 | 0.04766 | 0.11918 | 0.00000 | 0.04662 | 0.26251 | 0.00000 | 0.13775 | 0.13679 | 0.71146 |
| **FBXO17** | 0.03341 | 1.53082 | 0.37090 | 1.81852 | 0.20407 | 0.00017 | 0.36721 | 0.00259 | 0.65211 | 0.00000 | 0.53070 | 2.12076 |
| **FBXO18** | 4.24881 | 2.66231 | 3.76970 | 1.89718 | 4.89481 | 1.13367 | 8.19058 | 0.59636 | 4.44909 | 10.62439 | 1.54438 | 2.63430 |
| **FBXO2** | 0.00156 | 0.17947 | 0.00582 | 0.02217 | 0.00000 | 0.10509 | 0.00000 | 0.53799 | 0.00000 | 0.15423 | 0.02010 | 0.00000 |
| **FBXO21** | 0.00000 | 6.84766 | 8.20503 | 6.02529 | 4.24495 | 13.09460 | 17.62481 | 1.85617 | 9.19678 | 0.00000 | 8.74126 | 5.06028 |
| **FBXO22** | 0.09528 | 6.88496 | 12.84716 | 4.70127 | 3.24589 | 24.62978 | 3.43531 | 0.51544 | 7.08665 | 0.00000 | 10.64494 | 7.09980 |
| **FBXO22-AS1** | 0.00626 | 0.01830 | 0.02121 | 0.01560 | 0.01890 | 0.01436 | 0.02054 | 0.01048 | 0.01721 | 0.00807 | 0.02758 | 0.01850 |
| **FBXO24** | 0.00362 | 0.34860 | 0.17072 | 0.08467 | 0.00000 | 0.00000 | 0.00043 | 0.03968 | 0.19953 | 0.11369 | 0.06751 | 0.21667 |
| **FBXO25** | 2.49790 | 4.89182 | 2.94310 | 1.06923 | 6.98512 | 5.12490 | 5.49193 | 0.00000 | 1.71611 | 2.52802 | 4.19452 | 2.55513 |
| **FBXO27** | 4.15352 | 0.14919 | 0.15787 | 0.28495 | 0.00000 | 0.18181 | 0.23461 | 0.08322 | 0.00008 | 0.01109 | 0.00620 | 0.90697 |
| **FBXO28** | 1.37990 | 10.42626 | 4.58878 | 4.51299 | 0.02888 | 24.18871 | 11.40025 | 14.18452 | 4.85624 | 5.47096 | 8.25435 | 8.81501 |
| **FBXO3** | 0.40542 | 2.16068 | 2.23871 | 0.58035 | 0.00383 | 8.63737 | 2.73049 | 0.84294 | 0.62167 | 0.00000 | 2.90742 | 1.36805 |
| **FBXO30** | 4.22934 | 0.97071 | 6.55517 | 0.00000 | 0.00000 | 6.12382 | 4.47512 | 2.22535 | 2.77248 | 3.71879 | 2.36717 | 0.68426 |
| **FBXO31** | 0.06707 | 0.75085 | 0.60989 | 2.82720 | 1.59924 | 0.00000 | 2.16216 | 4.59085 | 1.49075 | 6.64605 | 1.33281 | 3.87136 |
| **FBXO32** | 0.27145 | 2.63367 | 6.26986 | 0.00000 | 0.79861 | 0.94099 | 0.39052 | 0.59947 | 0.00000 | 1.16277 | 0.71185 | 3.11842 |
| **FBXO33** | 1.35591 | 0.00000 | 2.35278 | 0.86498 | 1.59030 | 3.29470 | 5.74616 | 6.41787 | 0.34047 | 7.44006 | 0.70322 | 1.17374 |
| **FBXO34** | 3.99198 | 3.03035 | 2.58950 | 3.42574 | 4.35637 | 6.72426 | 0.00000 | 0.13556 | 7.69682 | 1.37653 | 6.44739 | 3.99495 |

| | | | | | | | | | | | | |
|---|---|---|---|---|---|---|---|---|---|---|---|---|
| **FBXO36** | 0.22765 | 7.24550 | 1.44678 | 2.44355 | 1.24692 | 6.67671 | 0.00000 | 0.41604 | 0.00000 | 0.00000 | 3.23847 | 10.67847 |
| **FBXO38** | 2.40694 | 3.21154 | 3.07601 | 1.63285 | 0.00000 | 3.61173 | 2.27314 | 0.86485 | 4.08840 | 2.48909 | 1.95689 | 2.83663 |
| **FBXO4** | 0.05153 | 0.17148 | 0.24182 | 0.00000 | 0.05158 | 0.34728 | 0.00000 | 0.05813 | 0.87877 | 0.85179 | 0.70172 | 0.40298 |
| **FBXO40** | 0.00507 | 0.00432 | 0.02455 | 0.00963 | 0.00000 | 0.01509 | 0.19782 | 0.03482 | 0.00000 | 0.08754 | 0.01749 | 0.14024 |
| **FBXO41** | 0.56292 | 0.30902 | 0.00000 | 0.54643 | 0.00000 | 0.00207 | 0.02060 | 0.00000 | 0.22659 | 0.21990 | 0.00000 | 0.19238 |
| **FBXO42** | 0.53987 | 3.29915 | 3.64837 | 3.78651 | 3.02097 | 1.52632 | 1.21873 | 1.57173 | 5.62248 | 0.00000 | 3.51986 | 2.87820 |
| **FBXO43** | 2.05763 | 0.02135 | 0.03113 | 0.02702 | 0.00000 | 0.00000 | 0.01160 | 0.00000 | 0.00017 | 0.03153 | 0.12897 | 0.37185 |
| **FBXO44** | 0.01508 | 0.86361 | 0.12691 | 0.36658 | 0.10716 | 0.00000 | 0.03924 | 0.79320 | 0.00399 | 0.00000 | 0.01418 | 0.10403 |
| **FBXO45** | 5.18645 | 1.08883 | 2.87567 | 4.31197 | 3.34097 | 15.58413 | 7.98591 | 8.94020 | 0.00021 | 18.00001 | 4.17636 | 4.25695 |
| **FBXO46** | 0.00000 | 1.49713 | 3.24437 | 0.98783 | 2.85993 | 0.00175 | 3.17657 | 2.58307 | 0.00000 | 7.01584 | 1.78612 | 2.17767 |
| **FBXO47** | 0.18050 | 0.02965 | 0.01381 | 0.00581 | 0.00799 | 0.02504 | 0.02845 | 0.27675 | 0.01196 | 0.02499 | 0.01240 | 0.05418 |
| **FBXO48** | 0.40435 | 0.58983 | 0.19553 | 0.00000 | 0.13538 | 1.58857 | 0.11815 | 0.17425 | 0.00000 | 0.57535 | 0.33545 | 1.27990 |
| **FBXO5** | 5.21702 | 7.43829 | 2.09119 | 5.90400 | 0.82120 | 23.99173 | 7.53558 | 27.26821 | 11.31197 | 2.18306 | 11.07684 | 25.06883 |
| **FBXO6** | 0.00096 | 7.87938 | 0.00000 | 0.44258 | 1.43795 | 1.97534 | 0.00000 | 5.84762 | 0.00000 | 0.00171 | 0.34316 | 1.24488 |
| **FBXO7** | 3.99796 | 19.64530 | 13.75690 | 14.44756 | 12.38805 | 19.51013 | 17.89911 | 4.17571 | 16.71119 | 3.47774 | 13.06241 | 14.43265 |
| **FBXO8** | 1.53240 | 2.63984 | 2.03087 | 4.00324 | 0.48850 | 10.75339 | 0.00000 | 3.17337 | 1.02506 | 0.38347 | 1.25551 | 3.83236 |
| **FBXO9** | 0.89186 | 1.51881 | 10.97111 | 5.92973 | 5.45362 | 6.28103 | 5.28835 | 0.16283 | 6.30893 | 8.60998 | 8.32748 | 3.21660 |
| **FBXW10** | 0.00453 | 0.01178 | 0.03065 | 0.02066 | 0.02446 | 0.00000 | 0.02311 | 0.02209 | 0.03559 | 0.00002 | 0.01671 | 0.15694 |
| **FBXW11** | 3.31788 | 6.31504 | 2.37677 | 3.19292 | 2.15363 | 5.51716 | 4.15336 | 3.07913 | 7.22519 | 7.43889 | 8.05164 | 7.55644 |
| **FBXW12** | 7.48611 | 0.00000 | 0.02691 | 0.00613 | 0.03423 | 0.01651 | 0.00244 | 4.07661 | 0.01490 | 0.03044 | 0.05232 | 0.09585 |
| **FBXW2** | 6.60559 | 7.24477 | 5.69043 | 4.59403 | 2.19945 | 16.79629 | 9.08351 | 5.40731 | 8.53296 | 5.98672 | 9.73214 | 10.17254 |
| **FBXW4** | 1.09553 | 0.56637 | 0.43551 | 2.32412 | 1.97505 | 1.12863 | 1.29397 | 0.00000 | 0.82303 | 2.22766 | 1.01041 | 0.43887 |
| **FBXW4P1** | 0.13761 | 0.00000 | 0.39837 | 0.27991 | 0.58110 | 0.00000 | 0.15013 | 0.60358 | 0.14352 | 1.63546 | 0.23907 | 0.39247 |
| **FBXW5** | 0.26576 | 8.94990 | 11.97758 | 12.46350 | 12.78196 | 0.00000 | 12.89163 | 0.41493 | 11.65894 | 7.20162 | 16.72448 | 5.36896 |
| **FBXW7** | 1.55286 | 0.25455 | 3.47381 | 2.16480 | 3.73163 | 6.63143 | 6.35404 | 16.60203 | 0.00000 | 15.58930 | 2.95727 | 3.03630 |
| **FBXW8** | 0.04134 | 1.06483 | 0.19500 | 0.00000 | 0.55614 | 0.77176 | 0.36977 | 0.27305 | 0.84195 | 0.00000 | 0.81132 | 0.59545 |
| **FBXW9** | 1.73981 | 2.04764 | 2.53844 | 0.73862 | 2.50602 | 0.00000 | 3.24877 | 0.51329 | 3.66354 | 4.74332 | 1.39211 | 2.52542 |
| **FCAR** | 0.00374 | 0.04424 | 0.02468 | 0.00905 | 0.00790 | 0.00704 | 0.00756 | 0.01108 | 0.01885 | 0.14923 | 0.01114 | 0.03906 |
| **FCER1G** | 0.00363 | 0.02606 | 0.13359 | 2.15591 | 0.00000 | 0.02586 | 0.36877 | 0.00573 | 0.07814 | 0.30914 | 0.08941 | 0.08127 |
| **FCER2** | 0.00594 | 0.01008 | 0.02082 | 0.02683 | 0.43095 | 0.00567 | 0.24229 | 0.01033 | 0.00000 | 0.00000 | 0.10107 | 0.01743 |
| **FCF1** | 1.17810 | 1.65905 | 3.50230 | 0.78285 | 2.16704 | 7.44997 | 7.59424 | 3.10020 | 0.85424 | 9.96445 | 4.53914 | 6.92749 |
| **FCGBP** | 0.08886 | 0.15959 | 1.87593 | 3.64686 | 0.00000 | 0.00000 | 0.26523 | 0.02662 | 0.42942 | 0.55253 | 0.72658 | 0.18083 |
| **FCGR1A** | 0.11808 | 0.01385 | 0.01982 | 0.01984 | 0.00946 | 0.02199 | 0.01234 | 0.01545 | 0.02841 | 0.02527 | 0.01536 | 0.02746 |
| **FCGR1B** | 0.00751 | 0.01226 | 0.02436 | 0.03315 | 0.00783 | 0.02366 | 0.01584 | 0.01924 | 0.01022 | 0.03430 | 0.01532 | 0.01855 |
| **FCGR1C** | 0.01007 | 0.00973 | 0.01405 | 0.02352 | 0.01273 | 0.04455 | 0.01364 | 0.01631 | 0.01359 | 0.01492 | 0.01325 | 0.01837 |
| **FCGR2A** | 0.05521 | 0.01522 | 0.01226 | 0.01509 | 0.00994 | 0.03410 | 0.01487 | 0.01650 | 0.01600 | 0.04386 | 0.01247 | 0.06954 |
| **FCGR2B** | 0.00404 | 0.01701 | 0.01460 | 0.06495 | 0.01736 | 0.00784 | 0.01672 | 0.02290 | 0.04845 | 0.00000 | 0.01582 | 0.01778 |
| **FCGR2C** | 0.00594 | 0.01451 | 0.02049 | 0.01878 | 0.01235 | 0.01535 | 0.01510 | 0.01052 | 0.02728 | 0.02895 | 0.01329 | 0.01569 |
| **FCGR3B** | 0.00627 | 0.01402 | 0.01680 | 0.01507 | 0.01437 | 0.01657 | 0.01758 | 0.01054 | 0.01311 | 0.01062 | 0.05148 | 0.01689 |
| **FCGRT** | 0.65068 | 0.61540 | 0.20108 | 1.52953 | 0.23181 | 0.00000 | 0.06150 | 0.04239 | 0.12196 | 0.55638 | 0.07542 | 0.79060 |
| **FCHO1** | 3.57091 | 0.00000 | 0.13738 | 0.31230 | 0.23130 | 0.00000 | 0.31747 | 0.24923 | 0.32031 | 0.12570 | 0.00000 | 0.50112 |
| **FCHO2** | 0.13651 | 0.31385 | 3.55089 | 0.00000 | 0.00000 | 5.88407 | 3.05908 | 0.88803 | 0.00000 | 2.44141 | 1.20040 | 0.86824 |
| **FCHSD1** | 0.60429 | 0.14881 | 0.81127 | 1.03385 | 0.00000 | 0.17986 | 0.00000 | 0.09845 | 1.13348 | 0.00000 | 0.00254 | 0.22241 |
| **FCHSD2** | 1.01906 | 0.51242 | 0.82700 | 0.57106 | 0.61880 | 1.67145 | 0.61473 | 0.09507 | 1.01089 | 1.38171 | 0.43224 | 0.22389 |
| **FCN2** | 0.01891 | 0.01600 | 0.02508 | 0.01261 | 0.01645 | 0.01561 | 0.01823 | 0.04668 | 0.01760 | 0.00935 | 0.01545 | 0.01400 |
| **FCRL1** | 0.00612 | 0.01446 | 0.01413 | 0.01172 | 0.01407 | 0.01928 | 0.01999 | 0.01048 | 0.01816 | 0.02013 | 0.01515 | 0.01580 |
| **FCRL2** | 0.00617 | 0.01384 | 0.06999 | 0.00887 | 0.01071 | 0.02654 | 0.03451 | 0.01040 | 0.02163 | 0.01243 | 0.01090 | 0.01578 |
| **FCRL5** | 0.00240 | 0.01200 | 0.04045 | 0.00928 | 0.01479 | 0.00585 | 0.01687 | 0.07456 | 0.01690 | 0.05938 | 0.02109 | 0.04803 |

| | | | | | | | | | | | | |
|---|---|---|---|---|---|---|---|---|---|---|---|---|
| **FCRL6** | 0.00662 | 0.01773 | 0.05781 | 0.01655 | 0.02324 | 0.00000 | 0.01955 | 0.01060 | 0.01358 | 0.00794 | 0.02948 | 0.01809 |
| **FCRLB** | 0.00285 | 0.07286 | 0.10075 | 0.00850 | 0.01436 | 0.05010 | 0.02347 | 0.03346 | 0.00000 | 0.00000 | 0.16340 | 0.01118 |
| **FDCSP** | 0.00724 | 0.01348 | 0.01903 | 0.00821 | 0.00958 | 0.03157 | 0.01788 | 0.01466 | 0.03582 | 0.01350 | 0.01284 | 0.01481 |
| **FDFT1** | 12.57273 | 1.86520 | 13.06316 | 18.78482 | 39.92284 | 17.12975 | 28.38900 | 4.40676 | 39.94418 | 4.10089 | 24.48703 | 2.57436 |
| **FDPS** | 0.00672 | 6.56937 | 15.20445 | 15.38662 | 26.34781 | 13.30849 | 22.59725 | 0.47099 | 36.87709 | 0.00000 | 34.17437 | 4.96446 |
| **FDPSP2** | 0.02545 | 0.00000 | 0.09892 | 0.02602 | 0.02086 | 0.00000 | 0.00173 | 0.02016 | 0.05529 | 0.29994 | 0.30980 | 0.10924 |
| **FDX1** | 0.10541 | 2.60723 | 7.78886 | 2.08324 | 2.85090 | 5.48570 | 5.62051 | 0.05674 | 3.30464 | 7.36722 | 4.02755 | 0.26337 |
| **FDX1L** | 0.30381 | 7.13813 | 6.75429 | 3.33126 | 4.82798 | 0.00000 | 5.49401 | 3.39021 | 4.77862 | 4.95489 | 3.30899 | 8.58604 |
| **FDXACB1** | 0.43398 | 3.16298 | 1.30982 | 0.00560 | 0.00000 | 2.58178 | 1.09626 | 1.08286 | 1.78232 | 0.00000 | 2.55280 | 2.77950 |
| **FDXR** | 0.74464 | 23.71003 | 6.88829 | 7.24177 | 12.81889 | 0.00000 | 5.65291 | 0.25025 | 7.36214 | 2.44104 | 6.87298 | 4.71912 |
| **FECH** | 0.42244 | 3.78749 | 3.06791 | 0.00000 | 2.54757 | 2.13434 | 5.05661 | 0.07463 | 6.19094 | 0.00000 | 7.20437 | 0.80207 |
| **FEM1A** | 0.72032 | 4.40998 | 6.14908 | 6.43837 | 5.52962 | 3.43937 | 12.62784 | 0.65102 | 7.14682 | 6.39942 | 7.05767 | 7.82234 |
| **FEM1B** | 0.63955 | 1.49754 | 5.51820 | 4.72804 | 0.39363 | 11.45077 | 2.66752 | 1.48497 | 1.23332 | 12.77881 | 1.71928 | 2.09121 |
| **FEM1C** | 2.30709 | 3.06425 | 1.45299 | 0.00000 | 0.00000 | 6.42114 | 1.84733 | 1.92331 | 1.91480 | 4.27112 | 2.50763 | 2.83591 |
| **FEN1** | 4.65162 | 15.31247 | 19.73652 | 16.66452 | 15.87902 | 15.43602 | 26.10007 | 1.72333 | 25.65691 | 0.00000 | 21.90799 | 17.48263 |
| **FENDRR** | 0.00577 | 0.02161 | 0.03459 | 0.01156 | 0.01323 | 0.01183 | 0.00994 | 0.01057 | 0.02325 | 0.03952 | 0.01252 | 0.01491 |
| **FER** | 2.93528 | 2.31960 | 1.45396 | 0.98070 | 3.89420 | 5.54883 | 0.00000 | 0.70253 | 1.46623 | 4.08756 | 2.41993 | 1.76893 |
| **FER1L4** | 0.69825 | 0.04552 | 0.44261 | 0.00216 | 0.07789 | 0.00000 | 0.06966 | 0.10960 | 0.00000 | 0.03238 | 0.01283 | 0.02925 |
| **FER1L5** | 0.06108 | 0.01092 | 0.17924 | 0.03822 | 0.06070 | 0.00000 | 0.00449 | 0.01515 | 0.00000 | 0.37393 | 0.05825 | 0.04572 |
| **FER1L6** | 1.56125 | 0.00638 | 0.00221 | 0.00196 | 0.03900 | 0.00000 | 0.00933 | 0.11241 | 0.03116 | 0.28447 | 0.00000 | 0.00224 |
| **FER1L6-AS2** | 0.50809 | 0.03811 | 0.06313 | 0.01162 | 0.03508 | 0.02588 | 0.06823 | 0.00279 | 0.06060 | 0.00000 | 0.03506 | 0.09595 |
| **FERMT1** | 0.80019 | 0.00000 | 1.92547 | 0.19696 | 0.02316 | 0.11191 | 0.00000 | 0.00000 | 0.26530 | 0.01473 | 0.08540 | 0.00468 |
| **FERMT2** | 4.34293 | 7.22075 | 2.03589 | 17.58069 | 7.37140 | 22.50618 | 4.41684 | 2.88070 | 0.00000 | 14.74299 | 5.60038 | 15.16687 |
| **FERMT3** | 0.00309 | 0.06524 | 0.37368 | 1.01138 | 1.69633 | 0.00000 | 0.29172 | 0.00000 | 0.78392 | 3.51752 | 0.00000 | 0.55259 |
| **FES** | 0.12730 | 3.38737 | 0.81713 | 6.88197 | 5.70780 | 0.00000 | 1.05521 | 0.00000 | 0.68855 | 0.00000 | 1.64559 | 0.49716 |
| **FETUB** | 0.03963 | 0.01272 | 0.06573 | 0.01257 | 0.01889 | 0.03836 | 0.00596 | 0.00874 | 0.00480 | 0.03060 | 0.02083 | 0.02124 |
| **FEV** | 0.04779 | 0.02550 | 0.01554 | 0.01385 | 0.00550 | 0.02929 | 0.05034 | 0.00900 | 0.02744 | 0.01795 | 0.00728 | 0.02150 |
| **FEZ1** | 0.25644 | 0.52429 | 0.18076 | 0.02449 | 0.00382 | 0.23891 | 0.00000 | 0.08480 | 0.00000 | 0.00402 | 0.04923 | 2.21124 |
| **FEZ2** | 13.61209 | 7.13433 | 0.00000 | 2.34157 | 13.69626 | 28.64264 | 7.14389 | 6.03010 | 0.00000 | 34.83229 | 24.65907 | 9.69163 |
| **FEZF1** | 0.00290 | 0.01076 | 0.01361 | 0.00895 | 0.01919 | 0.03523 | 0.01657 | 0.02979 | 0.01503 | 0.00978 | 0.01328 | 0.05393 |
| **FEZF1-AS1** | 0.00217 | 0.01570 | 0.01151 | 0.00625 | 0.03232 | 0.04382 | 0.01323 | 0.14379 | 0.01089 | 0.01747 | 0.01095 | 0.02234 |
| **FEZF2** | 1.62621 | 0.00448 | 0.01105 | 0.05124 | 0.10862 | 0.02843 | 0.01816 | 0.04496 | 0.01185 | 0.00000 | 0.01020 | 0.06427 |
| **FFAR2** | 0.01445 | 0.01359 | 0.00736 | 0.00857 | 0.00986 | 0.01987 | 0.07987 | 0.00912 | 0.01028 | 0.03397 | 0.01096 | 0.01496 |
| **FFAR3** | 0.00899 | 0.01515 | 0.01563 | 0.01461 | 0.01483 | 0.01669 | 0.02484 | 0.00974 | 0.01686 | 0.01402 | 0.01514 | 0.01632 |
| **FFAR4** | 0.00383 | 0.03968 | 0.03773 | 0.00734 | 0.00510 | 0.04949 | 0.02521 | 0.06359 | 0.05525 | 0.00000 | 0.03904 | 0.01683 |
| **FGA** | 0.01259 | 0.00078 | 0.28668 | 0.42633 | 0.00978 | 0.00000 | 0.00000 | 0.01213 | 0.00000 | 0.26090 | 0.15384 | 0.01911 |
| **FGB** | 0.00600 | 0.01722 | 0.05645 | 0.08048 | 0.01094 | 0.00000 | 0.01764 | 0.01232 | 0.08265 | 0.00935 | 0.00805 | 0.01938 |
| **FGD1** | 0.06003 | 0.42379 | 0.30471 | 1.87372 | 0.00000 | 0.00000 | 0.24884 | 0.12771 | 0.66095 | 1.32403 | 0.11604 | 3.08568 |
| **FGD2** | 0.00677 | 0.03074 | 0.00514 | 0.18793 | 0.02357 | 0.00000 | 0.00535 | 0.01032 | 0.05488 | 0.03826 | 0.00602 | 0.01465 |
| **FGD3** | 0.67624 | 0.76547 | 0.75616 | 0.07115 | 0.22339 | 0.00000 | 0.91187 | 0.19450 | 0.00000 | 0.00000 | 0.71142 | 0.70989 |
| **FGD4** | 1.40224 | 1.68537 | 0.00000 | 2.47363 | 2.15354 | 4.88207 | 0.00000 | 1.21612 | 1.47903 | 8.82521 | 0.08954 | 3.85254 |
| **FGD5** | 1.04159 | 0.01639 | 0.01862 | 0.02338 | 0.00622 | 0.02262 | 0.04080 | 0.11301 | 0.00892 | 0.05978 | 0.02353 | 0.02473 |
| **FGD5-AS1** | 0.31884 | 5.95388 | 5.12509 | 2.80637 | 3.51348 | 15.02267 | 10.56351 | 7.99246 | 0.04923 | 16.16560 | 6.01409 | 9.07498 |
| **FGD5P1** | 0.21516 | 0.00892 | 0.04869 | 0.00836 | 0.01046 | 0.01099 | 0.00382 | 0.02970 | 0.02050 | 0.06301 | 0.01147 | 0.05417 |
| **FGD6** | 2.84144 | 0.65655 | 8.18099 | 9.39653 | 0.00000 | 4.87831 | 5.05811 | 4.11735 | 3.26039 | 3.57230 | 0.65189 | 3.24885 |
| **FGF1** | 0.03312 | 0.01681 | 0.00197 | 0.48204 | 0.00000 | 0.02241 | 0.02463 | 0.01160 | 0.01433 | 0.01162 | 0.01678 | 0.01584 |
| **FGF10** | 0.01131 | 0.01452 | 0.01833 | 0.01203 | 0.01210 | 0.02367 | 0.03600 | 0.01015 | 0.03200 | 0.00002 | 0.01591 | 0.02248 |
| **FGF10-AS1** | 0.00636 | 0.01518 | 0.01480 | 0.01243 | 0.01412 | 0.02088 | 0.01906 | 0.01047 | 0.02097 | 0.01046 | 0.02229 | 0.01668 |

| | | | | | | | | | | | | |
|---|---|---|---|---|---|---|---|---|---|---|---|---|
| **FGF11** | 0.04743 | 0.05464 | 0.52688 | 0.45846 | 0.00000 | 0.00000 | 0.94222 | 0.01871 | 0.60175 | 0.09220 | 0.00000 | 0.44243 |
| **FGF12** | 0.09855 | 0.55334 | 0.53544 | 0.09617 | 0.72793 | 0.00000 | 0.14769 | 0.00655 | 0.00000 | 0.62569 | 0.07986 | 0.05945 |
| **FGF13** | 0.21286 | 7.30871 | 0.00000 | 0.40427 | 1.38472 | 2.56901 | 0.21168 | 0.32057 | 0.00000 | 1.70150 | 4.16288 | 2.30996 |
| **FGF13-AS1** | 0.00363 | 0.07637 | 0.00000 | 0.00000 | 0.09509 | 0.18897 | 0.00000 | 0.00594 | 0.00000 | 0.10948 | 0.13273 | 0.06583 |
| **FGF14** | 11.20187 | 0.14929 | 0.00130 | 0.08247 | 0.00160 | 0.00453 | 0.01282 | 1.11994 | 0.04715 | 0.00000 | 0.00276 | 0.00000 |
| **FGF14-AS2** | 0.40878 | 0.32123 | 0.13737 | 0.06738 | 0.01265 | 0.49427 | 0.00000 | 0.28165 | 0.18281 | 0.00000 | 0.24668 | 0.52878 |
| **FGF14-IT1** | 0.22096 | 0.10033 | 0.00849 | 0.00000 | 0.00000 | 0.04254 | 0.01475 | 0.01005 | 0.00432 | 0.05192 | 0.03017 | 0.07304 |
| **FGF16** | 0.00633 | 0.01446 | 0.02743 | 0.01082 | 0.01395 | 0.02378 | 0.01383 | 0.01044 | 0.02169 | 0.01142 | 0.01362 | 0.01606 |
| **FGF17** | 0.01183 | 0.02787 | 0.04766 | 0.01148 | 0.01061 | 0.00259 | 0.02144 | 0.00767 | 0.05361 | 0.06031 | 0.01138 | 0.06863 |
| **FGF18** | 0.00810 | 0.24867 | 0.89781 | 0.82971 | 1.28309 | 0.44568 | 0.29177 | 0.65901 | 0.00000 | 1.28937 | 0.14167 | 0.09129 |
| **FGF19** | 0.00448 | 0.01095 | 0.05953 | 0.02800 | 0.00515 | 0.04138 | 0.00892 | 0.02261 | 0.02220 | 0.01295 | 0.01275 | 0.01410 |
| **FGF2** | 0.00985 | 0.02589 | 0.30138 | 0.42967 | 0.00000 | 0.02293 | 0.02993 | 0.01203 | 0.02832 | 0.02114 | 0.14016 | 0.14286 |
| **FGF20** | 0.00935 | 0.00912 | 0.06654 | 0.00984 | 0.01836 | 0.00607 | 0.00635 | 0.03776 | 0.03051 | 0.02038 | 0.01257 | 0.19122 |
| **FGF21** | 0.00498 | 0.01529 | 0.06492 | 0.00000 | 0.03324 | 0.04008 | 0.29632 | 0.01072 | 0.01991 | 0.04594 | 0.02063 | 0.01373 |
| **FGF22** | 0.00624 | 0.01797 | 0.02375 | 0.00983 | 0.03134 | 0.01172 | 0.01849 | 0.01056 | 0.02608 | 0.01317 | 0.01430 | 0.01667 |
| **FGF23** | 0.01187 | 0.02258 | 0.00000 | 0.00000 | 0.00000 | 0.56897 | 0.03068 | 0.07202 | 0.00000 | 1.26612 | 0.00000 | 0.09782 |
| **FGF4** | 0.00571 | 0.00915 | 0.06055 | 0.89602 | 0.01505 | 0.06017 | 0.01095 | 0.01015 | 0.03992 | 0.00000 | 0.01036 | 0.01427 |
| **FGF5** | 0.00564 | 0.01910 | 0.01847 | 0.01318 | 0.02661 | 0.00982 | 0.01945 | 0.01423 | 0.02095 | 0.00668 | 0.01404 | 0.02924 |
| **FGF6** | 0.02063 | 0.01647 | 0.01198 | 0.01927 | 0.00808 | 0.02918 | 0.01356 | 0.01966 | 0.01790 | 0.02353 | 0.01365 | 0.01447 |
| **FGF7** | 0.01720 | 0.01074 | 0.01503 | 0.00796 | 0.00527 | 0.04490 | 0.02386 | 0.06601 | 0.02416 | 0.02759 | 0.00858 | 0.01474 |
| **FGF8** | 0.05225 | 0.49262 | 0.13161 | 0.56490 | 0.00000 | 0.00000 | 0.08482 | 0.00129 | 0.13268 | 0.43350 | 0.02434 | 0.25647 |
| **FGF9** | 3.99030 | 0.00318 | 0.05790 | 0.02546 | 0.10367 | 0.06113 | 0.00000 | 0.18766 | 0.10148 | 0.01698 | 0.00214 | 0.03728 |
| **FGFBP1** | 0.00310 | 0.00227 | 0.00000 | 0.27432 | 0.00000 | 0.01079 | 0.00000 | 0.00937 | 2.43419 | 0.24532 | 0.00000 | 0.00722 |
| **FGFBP2** | 0.00633 | 0.75309 | 0.00000 | 0.00053 | 1.24112 | 0.12333 | 0.12035 | 0.00800 | 0.00000 | 0.00000 | 0.54587 | 0.03282 |
| **FGFBP3** | 0.77997 | 0.00000 | 0.04689 | 0.02454 | 0.10301 | 0.33933 | 0.07053 | 0.00000 | 0.00000 | 0.00000 | 0.00259 | 0.12944 |
| **FGFR1** | 3.66591 | 5.81505 | 7.94727 | 24.35783 | 2.24333 | 1.82849 | 0.00000 | 0.60901 | 0.00000 | 7.96991 | 0.00000 | 2.51915 |
| **FGFR1OP** | 8.65506 | 12.87391 | 6.21583 | 4.47361 | 3.69403 | 29.31935 | 9.26575 | 4.98969 | 9.38658 | 2.86708 | 7.00404 | 3.95432 |
| **FGFR1OP2** | 3.47958 | 2.49742 | 5.43044 | 2.88003 | 1.81562 | 14.91021 | 4.13883 | 13.62845 | 7.31337 | 3.77430 | 3.42739 | 4.93678 |
| **FGFR2** | 0.47669 | 9.26205 | 2.01906 | 11.75490 | 0.37840 | 8.67200 | 1.14652 | 0.90998 | 0.00000 | 11.87564 | 0.00000 | 0.00000 |
| **FGFR3** | 0.10711 | 3.60268 | 0.00000 | 2.72257 | 14.96301 | 0.00000 | 2.08902 | 0.00000 | 1.94692 | 26.98142 | 1.26050 | 6.29315 |
| **FGFR4** | 0.00000 | 2.60821 | 11.03954 | 10.00658 | 19.81800 | 0.00000 | 5.41195 | 0.11660 | 34.63549 | 8.36397 | 9.28527 | 1.03213 |
| **FGFRL1** | 2.16964 | 0.20059 | 0.20329 | 0.14112 | 1.03281 | 0.00000 | 0.59106 | 0.61643 | 0.15798 | 1.29168 | 0.16159 | 0.53809 |
| **FGG** | 0.00696 | 0.14321 | 0.05531 | 0.18782 | 0.00400 | 0.06460 | 0.00000 | 0.00965 | 0.01146 | 0.02189 | 0.00701 | 0.00600 |
| **FGGY** | 3.06510 | 0.79639 | 0.14206 | 0.22537 | 0.17984 | 0.27747 | 0.12078 | 1.17107 | 0.00000 | 0.61770 | 0.29325 | 0.63641 |
| **FGL1** | 0.00597 | 0.01436 | 0.06605 | 0.02754 | 0.01418 | 0.01262 | 0.01347 | 0.01073 | 0.00744 | 0.02360 | 0.01758 | 0.01606 |
| **FGL2** | 0.00619 | 0.01585 | 0.01368 | 0.01481 | 0.01783 | 0.01485 | 0.01446 | 0.01043 | 0.03566 | 0.02099 | 0.01171 | 0.01739 |
| **FGR** | 0.00509 | 0.01154 | 0.00658 | 0.00539 | 0.00000 | 0.02388 | 0.09758 | 0.01028 | 0.00000 | 0.07944 | 0.01651 | 0.01231 |
| **FH** | 4.89832 | 11.84981 | 7.28517 | 8.53540 | 10.97290 | 19.78521 | 11.55280 | 4.51695 | 13.33796 | 1.74769 | 18.99929 | 2.36341 |
| **FHAD1** | 0.35831 | 0.04630 | 0.04511 | 0.00560 | 0.02564 | 0.02905 | 0.04217 | 0.14352 | 0.02408 | 0.00010 | 0.01979 | 0.03777 |
| **FHDC1** | 0.04077 | 0.22259 | 16.84068 | 1.97694 | 1.35691 | 0.00000 | 1.23755 | 0.00000 | 1.26758 | 4.04534 | 0.00000 | 0.11526 |
| **FHIT** | 2.49166 | 0.15998 | 0.18741 | 0.10005 | 0.52950 | 0.38734 | 0.07151 | 0.30807 | 0.39285 | 0.00000 | 0.80784 | 0.07141 |
| **FHL1** | 0.00877 | 0.17403 | 0.13442 | 0.40618 | 0.00000 | 0.10258 | 0.14071 | 0.44690 | 0.00000 | 0.02714 | 0.28115 | 0.48191 |
| **FHL2** | 0.04195 | 4.55921 | 11.38800 | 0.00000 | 18.21450 | 10.81575 | 6.13509 | 0.00000 | 12.20571 | 0.00000 | 10.11217 | 1.32327 |
| **FHL3** | 0.23252 | 5.09241 | 2.03069 | 3.14879 | 3.62778 | 0.00000 | 1.18061 | 0.00000 | 5.57636 | 0.77807 | 3.39985 | 1.03040 |
| **FHL5** | 0.04092 | 0.01618 | 0.01676 | 0.01356 | 0.01845 | 0.00673 | 0.01745 | 0.06372 | 0.02276 | 0.01310 | 0.02313 | 0.01494 |
| **FHOD1** | 1.13030 | 0.00000 | 2.56119 | 0.21957 | 3.85923 | 0.00000 | 1.22480 | 0.41253 | 1.38097 | 8.68176 | 0.76057 | 2.69432 |
| **FHOD3** | 5.92313 | 0.03818 | 0.00504 | 0.01758 | 0.00944 | 0.07173 | 0.00000 | 0.57562 | 0.07507 | 0.00000 | 0.01299 | 0.18880 |
| **FIBCD1** | 1.67057 | 0.02909 | 0.33219 | 0.12683 | 0.30302 | 0.00000 | 0.00000 | 0.00925 | 0.70758 | 0.00000 | 0.00462 | 0.00713 |

| | | | | | | | | | | | | |
|---|---|---|---|---|---|---|---|---|---|---|---|---|
| **FIBP** | 0.64494 | 7.98229 | 7.98215 | 12.73442 | 9.79643 | 2.82815 | 12.54232 | 3.58841 | 9.65028 | 0.00000 | 10.37890 | 8.31893 |
| **FICD** | 0.02557 | 0.46418 | 0.32152 | 0.50484 | 0.06931 | 0.89724 | 0.15722 | 0.15599 | 0.00000 | 0.19811 | 0.51240 | 0.20976 |
| **FIG4** | 0.87617 | 0.51044 | 1.63162 | 1.18327 | 0.00000 | 2.81182 | 0.00000 | 0.47691 | 0.32560 | 0.00000 | 1.26234 | 0.27688 |
| **FIGLA** | 5.98554 | 0.00378 | 0.00842 | 0.02025 | 0.05396 | 0.00000 | 0.01955 | 1.01040 | 0.01603 | 0.00000 | 0.01045 | 0.25239 |
| **FIGN** | 2.58213 | 0.26312 | 7.99590 | 0.00000 | 3.19264 | 1.56070 | 1.84839 | 0.02410 | 7.06483 | 3.47859 | 2.08390 | 1.02212 |
| **FIGNL1** | 5.63712 | 7.39763 | 2.74362 | 2.65126 | 0.00000 | 10.57996 | 5.31770 | 7.81607 | 2.10590 | 0.94490 | 9.59892 | 7.80649 |
| **FIGNL2** | 0.04570 | 0.00000 | 1.96804 | 0.95662 | 0.00000 | 0.00000 | 3.15986 | 0.01606 | 1.10580 | 1.05984 | 0.00000 | 0.15457 |
| **FILIP1** | 0.00000 | 3.01305 | 0.79119 | 12.38266 | 9.32979 | 3.75809 | 5.43651 | 0.31710 | 2.05074 | 8.50416 | 7.99818 | 3.75112 |
| **FILIP1L** | 0.13937 | 0.00000 | 0.54385 | 0.07691 | 0.36472 | 0.18428 | 0.00000 | 0.00132 | 0.00000 | 0.77275 | 0.02703 | 0.15009 |
| **FIP1L1** | 5.12383 | 17.74794 | 24.65533 | 9.59640 | 11.79358 | 23.65744 | 16.97039 | 4.79230 | 12.94739 | 8.63946 | 13.52176 | 18.59891 |
| **FIRRE** | 0.09959 | 0.34833 | 0.03527 | 0.18335 | 0.27478 | 0.00000 | 0.02330 | 0.04667 | 0.67189 | 0.06919 | 0.09005 | 0.33402 |
| **FIS1** | 1.41311 | 14.83088 | 12.94397 | 15.89995 | 18.33729 | 0.00218 | 14.26191 | 11.79390 | 6.80040 | 0.00000 | 10.64015 | 10.48082 |
| **FITM1** | 0.00392 | 0.02378 | 0.09701 | 0.07706 | 0.08389 | 0.00000 | 0.00032 | 0.00976 | 0.04336 | 0.18517 | 0.00697 | 0.01415 |
| **FITM2** | 0.00084 | 0.82554 | 0.64942 | 0.31129 | 0.02087 | 1.59298 | 0.36485 | 0.72002 | 1.00128 | 0.00000 | 0.55552 | 3.50006 |
| **FIZ1** | 0.15731 | 0.18140 | 0.21565 | 0.20667 | 0.62387 | 0.00000 | 0.11122 | 0.00000 | 0.00000 | 1.66543 | 0.01739 | 0.21731 |
| **FJX1** | 0.03307 | 0.00493 | 0.51916 | 0.48765 | 0.04291 | 0.00000 | 0.14608 | 0.00461 | 0.00059 | 0.58223 | 0.00003 | 0.02299 |
| **FKBP10** | 0.01228 | 0.00680 | 0.10671 | 0.00232 | 0.05231 | 0.02903 | 0.03816 | 0.01008 | 0.00000 | 0.07796 | 0.01094 | 0.02445 |
| **FKBP11** | 0.00000 | 19.59154 | 5.26474 | 10.64970 | 0.00001 | 2.32377 | 2.22266 | 0.06279 | 7.34675 | 3.91878 | 2.97412 | 8.48927 |
| **FKBP14** | 0.02655 | 0.57549 | 3.37998 | 0.73845 | 1.20152 | 1.88235 | 1.54673 | 0.11181 | 4.88637 | 4.02837 | 1.31219 | 1.82547 |
| **FKBP15** | 2.20952 | 5.09905 | 5.10010 | 1.39754 | 3.68303 | 1.95259 | 4.39141 | 1.74259 | 6.01447 | 1.77800 | 4.64636 | 3.96408 |
| **FKBP1A** | 6.92989 | 8.48748 | 19.62988 | 17.99092 | 8.34168 | 25.10538 | 28.30038 | 0.41531 | 16.20075 | 0.00000 | 14.54166 | 10.93916 |
| **FKBP1AP1** | 0.86228 | 0.54617 | 2.08855 | 1.68661 | 0.59724 | 3.87266 | 1.18362 | 0.03248 | 3.15048 | 0.01800 | 1.20014 | 1.05255 |
| **FKBP1B** | 0.00000 | 0.85440 | 0.27154 | 1.38757 | 0.83422 | 0.00000 | 1.12737 | 0.55184 | 0.54960 | 2.03854 | 0.35105 | 3.64855 |
| **FKBP2** | 1.33750 | 11.10566 | 13.66269 | 10.81117 | 14.00595 | 0.00000 | 9.45071 | 7.01396 | 15.98119 | 11.92552 | 10.33052 | 11.37503 |
| **FKBP3** | 0.00000 | 1.47743 | 11.47847 | 6.51939 | 4.27895 | 41.60706 | 21.69247 | 0.86055 | 11.84391 | 16.44033 | 17.42788 | 5.04972 |
| **FKBP4** | 5.99314 | 24.37243 | 14.49641 | 24.45577 | 32.17676 | 13.22647 | 28.80313 | 12.42051 | 28.25980 | 12.46434 | 15.23451 | 15.06903 |
| **FKBP5** | 1.03779 | 2.13386 | 1.21351 | 1.21236 | 2.39366 | 6.83856 | 1.19453 | 1.66469 | 5.92214 | 2.61692 | 2.81324 | 2.47368 |
| **FKBP6** | 2.68288 | 0.88637 | 0.00000 | 0.00000 | 0.00000 | 0.00000 | 0.00512 | 0.90363 | 0.01601 | 0.30597 | 0.00000 | 3.82768 |
| **FKBP7** | 0.00462 | 0.04054 | 0.28632 | 0.15694 | 0.14557 | 0.01751 | 0.04437 | 0.02523 | 0.67109 | 0.00000 | 0.00000 | 0.52933 |
| **FKBP8** | 0.10976 | 0.87014 | 3.21394 | 2.22042 | 3.28865 | 0.00000 | 2.64187 | 0.04578 | 0.40464 | 4.22196 | 0.78981 | 1.10242 |
| **FKBP9** | 0.00433 | 0.32846 | 0.00000 | 1.15645 | 0.65218 | 0.66801 | 0.47866 | 0.01768 | 0.85172 | 0.57183 | 0.97972 | 0.14318 |
| **FKBP9P1** | 0.02372 | 0.00714 | 0.06736 | 0.05474 | 0.00000 | 0.04514 | 0.03861 | 0.01106 | 0.00000 | 0.04331 | 0.03369 | 0.01585 |
| **FKBPL** | 0.00141 | 11.34379 | 4.48977 | 2.34785 | 6.01497 | 0.00000 | 6.12735 | 7.81105 | 4.06307 | 0.00000 | 4.01906 | 19.90669 |
| **FKRP** | 0.00085 | 1.74608 | 0.90654 | 0.93605 | 1.63452 | 0.05124 | 0.00008 | 0.00000 | 3.34121 | 1.57454 | 0.89753 | 2.04887 |
| **FKSG29** | 0.00564 | 0.01432 | 0.02761 | 0.00833 | 0.00675 | 0.02462 | 0.01185 | 0.00957 | 0.02214 | 0.04340 | 0.01598 | 0.02400 |
| **FKTN** | 0.10072 | 2.12453 | 0.08859 | 0.00000 | 0.00000 | 2.87855 | 0.00000 | 0.00000 | 1.62072 | 0.58729 | 0.32906 | 2.39528 |
| **FLAD1** | 1.85623 | 19.87573 | 7.59926 | 17.41551 | 15.50977 | 3.75657 | 12.84416 | 4.40167 | 19.36346 | 0.00028 | 20.56640 | 19.02145 |
| **FLCN** | 1.92227 | 1.51492 | 4.42934 | 3.12490 | 1.85977 | 2.03353 | 2.45369 | 2.91776 | 2.40474 | 2.32627 | 2.66659 | 2.14221 |
| **FLG** | 0.00105 | 0.03435 | 0.00292 | 0.37813 | 0.11152 | 0.00000 | 0.06816 | 0.09527 | 0.02007 | 0.45049 | 0.07801 | 0.06391 |
| **FLG-AS1** | 0.00457 | 0.00000 | 0.05377 | 0.32631 | 0.38072 | 0.00000 | 0.17524 | 0.03075 | 0.09655 | 0.00000 | 0.27522 | 0.23274 |
| **FLG2** | 0.00189 | 0.00000 | 0.00000 | 0.07311 | 0.00000 | 0.02507 | 0.01572 | 0.00869 | 0.01021 | 1.28601 | 0.00000 | 0.03028 |
| **FLI1** | 0.34689 | 0.02204 | 0.01369 | 0.02301 | 0.04494 | 0.00000 | 0.03175 | 0.08422 | 0.01000 | 0.04161 | 0.01446 | 0.02754 |
| **FLII** | 4.05791 | 15.91286 | 7.63071 | 12.46562 | 15.23148 | 1.16257 | 7.47431 | 0.37680 | 14.10247 | 0.00000 | 9.53358 | 4.01030 |
| **FLJ10038** | 0.00762 | 0.00000 | 0.47188 | 0.37851 | 0.47544 | 0.00000 | 0.25994 | 0.12853 | 0.00000 | 2.92338 | 0.20638 | 0.62876 |
| **FLJ12825** | 0.01751 | 0.06128 | 0.00426 | 0.02565 | 0.00552 | 0.02056 | 0.00653 | 0.01476 | 0.15067 | 0.11078 | 0.02920 | 0.13718 |
| **FLJ13224** | 0.00218 | 0.06898 | 0.23373 | 0.09984 | 0.02545 | 0.00000 | 0.00000 | 0.26670 | 0.11125 | 0.04871 | 0.01031 | 0.04808 |
| **FLJ16171** | 0.02424 | 0.01941 | 0.01391 | 0.02910 | 0.01250 | 0.03792 | 0.00949 | 0.01999 | 0.00717 | 0.06418 | 0.01013 | 0.00966 |
| **FLJ16779** | 0.00624 | 0.01501 | 0.10497 | 0.01105 | 0.00918 | 0.00000 | 0.01185 | 0.01043 | 0.05665 | 0.01991 | 0.11252 | 0.01831 |

| | | | | | | | | | | | | |
|---|---|---|---|---|---|---|---|---|---|---|---|---|
| **FLJ20021** | 0.36884 | 0.05164 | 0.11711 | 0.41463 | 0.09451 | 0.00000 | 0.01806 | 0.04738 | 0.00000 | 0.00000 | 0.09206 | 0.04130 |
| **FLJ20518.uid-5** | 0.10282 | 0.01432 | 0.01460 | 0.01230 | 0.01400 | 0.01633 | 0.01610 | 0.02505 | 0.01975 | 0.01609 | 0.01730 | 0.01487 |
| **FLJ21408** | 0.00816 | 0.05432 | 0.02373 | 0.01222 | 0.01488 | 0.02093 | 0.02226 | 0.00944 | 0.03013 | 0.00000 | 0.02872 | 0.13004 |
| **FLJ22184** | 1.19884 | 0.00987 | 0.04988 | 0.01892 | 0.02374 | 0.00000 | 0.06997 | 0.01142 | 0.00000 | 0.02094 | 0.01224 | 0.05019 |
| **FLJ22447** | 0.01730 | 0.00988 | 0.18795 | 0.18910 | 0.00000 | 0.02736 | 0.00429 | 0.01088 | 0.01793 | 0.02566 | 0.01449 | 0.01415 |
| **FLJ22763** | 0.01377 | 0.02828 | 0.00459 | 0.40355 | 0.01614 | 0.00000 | 0.02183 | 0.02746 | 0.01571 | 0.01813 | 0.01531 | 0.08968 |
| **FLJ23867** | 0.00180 | 0.02435 | 0.36010 | 0.01601 | 0.03611 | 0.00000 | 0.01244 | 0.09249 | 0.00000 | 0.09022 | 0.02547 | 0.11072 |
| **FLJ25758** | 0.67528 | 0.07247 | 0.04086 | 0.15650 | 0.00000 | 0.10260 | 0.00000 | 2.29574 | 0.00000 | 0.01873 | 0.00815 | 2.32435 |
| **FLJ26850** | 0.29833 | 0.11052 | 0.00802 | 0.16948 | 0.00755 | 0.00000 | 0.09277 | 0.14059 | 0.00108 | 0.07311 | 0.01238 | 0.00000 |
| **FLJ30403** | 0.15506 | 0.04890 | 0.18728 | 0.01606 | 0.00000 | 0.23126 | 0.11280 | 0.20919 | 0.14736 | 0.00853 | 0.05642 | 0.13220 |
| **FLJ30679** | 0.14459 | 0.05064 | 0.01925 | 0.00148 | 0.03933 | 0.00000 | 0.03408 | 0.05674 | 0.12589 | 0.04280 | 0.00950 | 0.06028 |
| **FLJ31104** | 0.00226 | 0.03793 | 0.09094 | 0.12316 | 0.01486 | 0.01082 | 0.05867 | 0.13850 | 0.00000 | 0.05830 | 0.01877 | 0.05188 |
| **FLJ31306** | 0.30762 | 1.27543 | 2.46357 | 2.23308 | 0.81528 | 8.78600 | 3.30367 | 8.37669 | 3.19612 | 9.01901 | 2.11135 | 4.59328 |
| **FLJ31356** | 0.00531 | 0.01152 | 0.05069 | 0.00932 | 0.02286 | 0.01919 | 0.00736 | 0.01001 | 0.01180 | 0.05939 | 0.01438 | 0.01983 |
| **FLJ32255** | 0.81420 | 0.02881 | 0.00000 | 0.15253 | 0.00000 | 0.01906 | 0.00000 | 0.32625 | 0.00410 | 0.34983 | 0.00000 | 0.37188 |
| **FLJ33360** | 0.00964 | 0.00107 | 0.00000 | 0.00000 | 0.03473 | 0.09897 | 0.04668 | 0.01218 | 0.08000 | 0.03517 | 0.07251 | 0.06933 |
| **FLJ33534** | 0.27914 | 0.05535 | 0.04458 | 0.00000 | 0.03161 | 0.07797 | 0.03376 | 0.13955 | 0.00003 | 0.00765 | 0.00917 | 0.12789 |
| **FLJ34503** | 0.03409 | 0.00924 | 0.00788 | 0.02534 | 0.00205 | 0.02005 | 0.00251 | 0.02332 | 0.05369 | 0.05451 | 0.06325 | 0.01253 |
| **FLJ35934** | 0.00880 | 0.01758 | 0.01673 | 0.01121 | 0.01231 | 0.01136 | 0.01529 | 0.00936 | 0.01559 | 0.04749 | 0.01511 | 0.03730 |
| **FLJ36000** | 0.47394 | 0.02513 | 0.02446 | 0.03719 | 0.00000 | 0.02331 | 0.04282 | 0.00598 | 0.00339 | 0.07671 | 0.01095 | 0.06930 |
| **FLJ36777** | 0.00508 | 0.01862 | 0.01704 | 0.01433 | 0.01836 | 0.00333 | 0.02162 | 0.01457 | 0.02174 | 0.01034 | 0.02985 | 0.02261 |
| **FLJ37035** | 0.00461 | 0.05362 | 0.02208 | 0.00324 | 0.00372 | 0.01582 | 0.01148 | 0.03310 | 0.06045 | 0.07106 | 0.00588 | 0.10869 |
| **FLJ37201** | 0.23730 | 0.02776 | 0.03181 | 0.00180 | 0.01845 | 0.02346 | 0.01327 | 0.08479 | 0.00000 | 0.09225 | 0.13822 | 0.09057 |
| **FLJ37453** | 0.16178 | 0.82502 | 0.32060 | 0.51644 | 0.19932 | 0.63853 | 0.43196 | 0.58615 | 0.00000 | 1.32552 | 0.40967 | 1.17470 |
| **FLJ37505** | 0.00614 | 0.01287 | 0.01120 | 0.01828 | 0.05100 | 0.04448 | 0.03279 | 0.01014 | 0.00278 | 0.01151 | 0.01150 | 0.01428 |
| **FLJ39080** | 0.03058 | 0.17584 | 0.01280 | 0.06434 | 0.00496 | 0.05238 | 0.01210 | 0.00835 | 0.01172 | 0.02090 | 0.04900 | 0.11072 |
| **FLJ40194** | 0.01256 | 0.00000 | 0.01771 | 0.04514 | 0.05001 | 0.04507 | 0.00000 | 0.16110 | 0.00000 | 0.29364 | 0.00000 | 0.05933 |
| **FLJ40288** | 0.00450 | 0.01457 | 0.02665 | 0.01337 | 0.01405 | 0.01126 | 0.01544 | 0.04130 | 0.02018 | 0.02593 | 0.01646 | 0.02103 |
| **FLJ41200** | 0.00683 | 0.01537 | 0.01702 | 0.01174 | 0.01521 | 0.02070 | 0.01718 | 0.01107 | 0.01880 | 0.01221 | 0.01485 | 0.01609 |
| **FLJ41278** | 0.00746 | 0.00911 | 0.05531 | 0.01981 | 0.01739 | 0.05008 | 0.01586 | 0.01189 | 0.00000 | 0.02451 | 0.01496 | 0.02044 |
| **FLJ41941** | 0.00660 | 0.01399 | 0.00987 | 0.00962 | 0.01401 | 0.02500 | 0.00479 | 0.01027 | 0.07184 | 0.00709 | 0.01157 | 0.01673 |
| **FLJ42102** | 0.00674 | 0.01305 | 0.01070 | 0.00783 | 0.00994 | 0.02365 | 0.03588 | 0.01063 | 0.00266 | 0.05183 | 0.01724 | 0.01393 |
| **FLJ42351** | 0.00606 | 0.01420 | 0.02232 | 0.00906 | 0.01983 | 0.02321 | 0.01390 | 0.01021 | 0.01652 | 0.01893 | 0.01464 | 0.01880 |
| **FLJ42393** | 0.00206 | 0.03488 | 0.01753 | 0.78604 | 1.31362 | 0.00548 | 0.00000 | 0.06766 | 0.53507 | 1.13393 | 0.00000 | 0.81549 |
| **FLJ42627** | 1.41567 | 0.01312 | 0.13906 | 0.09247 | 0.03679 | 0.00000 | 0.03178 | 0.43081 | 0.04586 | 0.33398 | 0.00000 | 0.02087 |
| **FLJ42969** | 0.00643 | 0.01504 | 0.25839 | 0.00322 | 0.05651 | 0.02578 | 0.00001 | 0.00981 | 0.44802 | 0.00000 | 0.12280 | 0.01739 |
| **FLJ43681** | 0.01683 | 0.00057 | 0.02247 | 0.01483 | 0.00000 | 0.10491 | 0.00000 | 0.02017 | 0.02860 | 0.06068 | 0.00732 | 0.04051 |
| **FLJ43879** | 0.00632 | 0.01827 | 0.01482 | 0.01300 | 0.01647 | 0.01532 | 0.01865 | 0.01052 | 0.02165 | 0.01129 | 0.01498 | 0.01685 |
| **FLJ44087** | 0.03644 | 0.01660 | 0.01667 | 0.00918 | 0.02397 | 0.02039 | 0.01416 | 0.01232 | 0.01665 | 0.02147 | 0.01747 | 0.01411 |
| **FLJ44313** | 0.13241 | 0.04547 | 0.02071 | 0.00292 | 0.00648 | 0.04876 | 0.01023 | 0.00858 | 0.00058 | 0.06946 | 0.01022 | 0.14760 |
| **FLJ44511** | 0.12499 | 0.00409 | 0.03914 | 0.00000 | 0.05492 | 0.02463 | 0.13616 | 0.00315 | 0.00000 | 0.07775 | 0.32009 | 0.02257 |
| **FLJ44635** | 1.92479 | 0.03418 | 1.13242 | 0.02388 | 0.39707 | 8.89380 | 4.18536 | 0.59926 | 0.00000 | 13.00128 | 0.17930 | 2.97068 |
| **FLJ45079** | 0.02987 | 0.02195 | 0.03211 | 0.01533 | 0.03550 | 0.00000 | 0.02865 | 0.01212 | 0.01459 | 0.00411 | 0.01565 | 0.04845 |
| **FLJ45513** | 0.00341 | 0.11449 | 0.46959 | 0.49617 | 0.19890 | 0.00000 | 0.13366 | 0.16239 | 0.34486 | 0.09455 | 0.31913 | 0.62172 |
| **FLJ46066** | 0.00563 | 0.01432 | 0.04187 | 0.00701 | 0.01055 | 0.02971 | 0.02834 | 0.01048 | 0.01220 | 0.03457 | 0.02483 | 0.01519 |
| **FLJ46284** | 0.00833 | 0.01780 | 0.10189 | 0.01840 | 0.01626 | 0.00000 | 0.00149 | 0.01099 | 0.12201 | 0.00523 | 0.00629 | 0.01937 |
| **FLJ46906** | 0.00964 | 0.38864 | 0.83659 | 0.65436 | 0.90415 | 0.22420 | 0.47968 | 0.14527 | 0.43364 | 0.14216 | 0.58188 | 0.02828 |
| **FLNA** | 1.67081 | 8.71037 | 8.22928 | 7.57532 | 15.55845 | 0.00000 | 8.19122 | 0.64530 | 18.09313 | 15.25764 | 2.15816 | 0.00000 |

| | | | | | | | | | | | | |
|---|---|---|---|---|---|---|---|---|---|---|---|---|
| **FLNB** | 4.12492 | 15.57721 | 8.89082 | 12.17475 | 39.04458 | 6.13624 | 7.90691 | 0.46841 | 45.78770 | 29.49630 | 11.34222 | 3.43853 |
| **FLNC** | 1.30542 | 0.29855 | 2.07834 | 2.30172 | 3.36075 | 0.00000 | 0.29780 | 0.00559 | 0.42092 | 0.52929 | 0.79350 | 0.03725 |
| **FLOT1** | 10.59392 | 3.73927 | 3.23698 | 8.08802 | 3.57901 | 0.00000 | 3.16422 | 8.41089 | 3.21352 | 4.26769 | 4.00289 | 3.53173 |
| **FLOT2** | 0.80600 | 9.46368 | 6.34713 | 7.91087 | 8.78760 | 0.00000 | 8.29910 | 0.12842 | 8.30562 | 5.06702 | 9.08359 | 1.17936 |
| **FLRT1** | 0.00473 | 0.08904 | 0.20600 | 0.19890 | 0.00000 | 0.00000 | 0.00000 | 0.01434 | 0.18735 | 0.03563 | 0.04027 | 0.05343 |
| **FLRT2** | 0.07029 | 0.00485 | 0.25105 | 0.23870 | 0.00041 | 0.06355 | 0.00000 | 0.10981 | 0.00000 | 0.01547 | 0.04990 | 0.01125 |
| **FLRT3** | 0.05363 | 0.12816 | 1.29918 | 4.98461 | 0.05574 | 0.00000 | 0.00000 | 0.12495 | 0.00000 | 0.02736 | 0.10404 | 0.00961 |
| **FLT1** | 0.63952 | 0.06906 | 0.04912 | 0.01435 | 0.05663 | 0.03292 | 0.02198 | 0.00663 | 0.04559 | 0.00000 | 0.01761 | 0.50000 |
| **FLT3** | 0.00308 | 0.25160 | 0.00678 | 0.02428 | 0.00000 | 0.05246 | 0.00000 | 0.32979 | 0.01946 | 0.03543 | 0.20309 | 0.03782 |
| **FLT3LG** | 0.44381 | 0.23322 | 0.19435 | 0.00982 | 0.38570 | 0.00000 | 0.24413 | 0.90651 | 0.22164 | 0.75722 | 0.24786 | 1.04889 |
| **FLT4** | 0.00341 | 1.85380 | 0.01943 | 0.00669 | 0.44108 | 0.00000 | 0.02319 | 0.00751 | 0.00000 | 0.09088 | 0.04608 | 0.29192 |
| **FLVCR1** | 0.03527 | 0.03326 | 3.19280 | 5.20695 | 2.90593 | 5.39118 | 2.32718 | 0.17232 | 5.07152 | 12.41293 | 1.74456 | 0.00000 |
| **FLVCR1-AS1** | 0.19255 | 0.17920 | 0.66363 | 1.25016 | 1.72907 | 1.11880 | 0.08573 | 0.25650 | 0.00000 | 0.00000 | 0.83489 | 0.37630 |
| **FLVCR2** | 4.11017 | 0.99507 | 7.84597 | 0.16770 | 0.50338 | 0.68029 | 0.00000 | 1.70349 | 7.42061 | 0.00000 | 3.20185 | 1.72987 |
| **FLYWCH1** | 0.34497 | 2.24180 | 1.76010 | 1.51315 | 2.02332 | 0.00000 | 1.82009 | 0.44748 | 1.51537 | 3.86954 | 0.00000 | 1.46707 |
| **FLYWCH2** | 1.38035 | 5.73714 | 2.28649 | 2.76512 | 5.00927 | 0.00000 | 3.33557 | 1.77458 | 4.40043 | 0.00058 | 2.21249 | 5.08363 |
| **FMN1** | 8.59781 | 0.28852 | 1.23978 | 1.74287 | 0.00000 | 0.00000 | 0.04694 | 0.70090 | 0.00000 | 0.71519 | 0.05600 | 1.29358 |
| **FMN2** | 1.31754 | 0.16346 | 0.00639 | 0.19765 | 0.00006 | 0.17609 | 0.00000 | 0.09050 | 0.00000 | 0.56419 | 0.00000 | 0.21269 |
| **FMNL1** | 0.78410 | 0.05332 | 0.42159 | 0.02986 | 0.06045 | 0.00000 | 0.27021 | 0.00000 | 0.09953 | 0.00000 | 0.00000 | 0.19659 |
| **FMNL2** | 2.72673 | 0.99356 | 9.41860 | 0.67938 | 1.58268 | 6.77179 | 1.91060 | 0.00000 | 0.60451 | 6.85524 | 0.68113 | 1.08851 |
| **FMNL3** | 0.09189 | 0.13666 | 0.04446 | 0.65193 | 0.02662 | 0.05388 | 0.00000 | 0.49858 | 0.07086 | 0.93104 | 0.11425 | 0.27571 |
| **FMO1** | 0.00649 | 0.14963 | 0.03037 | 0.00486 | 0.07062 | 0.01494 | 0.02046 | 0.01082 | 0.00506 | 0.01713 | 0.01485 | 0.00707 |
| **FMO2** | 2.01604 | 0.21869 | 0.01036 | 0.00379 | 0.15490 | 0.00000 | 0.02437 | 0.53389 | 0.00000 | 0.02309 | 0.01868 | 0.81168 |
| **FMO3** | 0.02874 | 0.01447 | 0.01562 | 0.01484 | 0.02676 | 0.00030 | 0.02186 | 0.02581 | 0.01996 | 0.01034 | 0.01664 | 0.09514 |
| **FMO4** | 0.04434 | 0.16768 | 0.19026 | 0.00000 | 0.00000 | 0.00000 | 0.16115 | 0.06528 | 0.01425 | 0.03293 | 0.10093 | 0.12905 |
| **FMO5** | 0.15653 | 0.32114 | 0.07620 | 2.81572 | 0.00000 | 0.00001 | 0.02298 | 0.16146 | 0.01753 | 0.00000 | 0.29912 | 0.48126 |
| **FMO6P** | 0.00597 | 0.01407 | 0.01747 | 0.01335 | 0.01295 | 0.01702 | 0.01512 | 0.01001 | 0.01502 | 0.02989 | 0.01513 | 0.02310 |
| **FMO9P** | 0.00349 | 0.00513 | 0.05293 | 0.00318 | 0.00000 | 0.02878 | 0.00857 | 0.00970 | 0.00000 | 0.31208 | 0.00272 | 0.00771 |
| **FMOD** | 0.01093 | 0.00872 | 0.02061 | 0.02176 | 0.01643 | 0.02502 | 0.01835 | 0.00814 | 0.01381 | 0.00607 | 0.01899 | 0.07204 |
| **FMR1** | 1.00845 | 0.56093 | 1.83751 | 0.39820 | 0.43199 | 7.85172 | 3.08183 | 2.99476 | 0.00000 | 10.74017 | 0.38131 | 1.39109 |
| **FMR1-AS1** | 0.02927 | 0.06723 | 0.09022 | 0.04098 | 0.02333 | 0.09935 | 0.00000 | 0.03386 | 0.08271 | 0.04889 | 0.02698 | 0.13197 |
| **FMR1NB** | 0.00000 | 10.90872 | 0.23650 | 0.00000 | 3.96801 | 9.84760 | 1.40530 | 4.77698 | 4.97063 | 1.89482 | 3.42222 | 10.11200 |
| **FN1** | 16.17037 | 0.46008 | 8.63930 | 8.77615 | 0.00000 | 0.00000 | 0.00000 | 11.10818 | 0.00000 | 2.89141 | 0.78105 | 1.42657 |
| **FN3K** | 0.67655 | 0.19115 | 0.16768 | 0.08371 | 3.40702 | 0.54990 | 0.00000 | 0.04316 | 0.26097 | 0.48445 | 0.18475 | 0.21812 |
| **FN3KRP** | 2.47009 | 11.08222 | 2.07779 | 4.55311 | 7.82332 | 4.28365 | 3.72118 | 0.31664 | 3.73126 | 0.20574 | 3.37107 | 4.92204 |
| **FNBP1** | 3.70151 | 2.86262 | 1.87575 | 1.08462 | 6.29068 | 2.77827 | 2.92310 | 6.88329 | 4.45553 | 8.57330 | 0.00000 | 15.87973 |
| **FNBP1L** | 6.18462 | 8.63968 | 4.03732 | 1.37105 | 7.13426 | 28.90157 | 10.43646 | 0.00000 | 20.74594 | 17.05082 | 13.97689 | 6.98329 |
| **FNBP4** | 2.12368 | 4.02589 | 9.52390 | 8.32681 | 9.18162 | 6.17823 | 4.66218 | 5.62466 | 16.34509 | 27.72646 | 3.53982 | 4.67500 |
| **FNDC1** | 0.00564 | 0.01517 | 0.02037 | 0.01843 | 0.01977 | 0.01182 | 0.04387 | 0.01182 | 0.00505 | 0.02020 | 0.01555 | 0.01622 |
| **FNDC3A** | 0.82583 | 0.02881 | 5.07402 | 0.66704 | 1.01806 | 9.83888 | 2.49658 | 0.00000 | 2.45495 | 5.02517 | 3.65163 | 0.83321 |
| **FNDC3B** | 8.10112 | 0.49241 | 6.88427 | 2.46479 | 2.28149 | 8.91945 | 6.07884 | 1.23522 | 2.56350 | 6.84020 | 6.50273 | 0.00000 |
| **FNDC4** | 0.57860 | 0.17941 | 0.20243 | 5.85574 | 0.00000 | 0.00000 | 0.00641 | 0.45531 | 0.00000 | 0.19014 | 0.00326 | 0.10115 |
| **FNDC5** | 0.12246 | 0.09603 | 0.00000 | 0.05806 | 0.00000 | 0.07310 | 0.00642 | 0.13613 | 0.05467 | 0.16269 | 0.00492 | 0.18759 |
| **FNDC7** | 0.32436 | 0.01403 | 0.00394 | 0.02233 | 0.00852 | 0.01768 | 0.02368 | 0.11731 | 0.09930 | 0.00138 | 0.04312 | 0.01123 |
| **FNDC8** | 0.00994 | 0.15033 | 0.02638 | 0.09170 | 0.03547 | 0.03685 | 0.00955 | 0.02170 | 0.64014 | 0.11806 | 0.04976 | 0.15675 |
| **FNIP1** | 0.00000 | 1.42578 | 2.22049 | 1.53819 | 0.00000 | 8.39556 | 2.94136 | 2.75630 | 2.93795 | 2.83393 | 3.71836 | 1.25077 |
| **FNIP2** | 7.37176 | 8.33460 | 0.00000 | 1.18858 | 2.89060 | 9.27131 | 3.00680 | 0.83631 | 0.07743 | 13.06080 | 0.79542 | 2.43308 |
| **FNTA** | 2.02179 | 4.69683 | 6.63128 | 6.97653 | 3.72671 | 7.70821 | 11.25063 | 0.92642 | 1.56659 | 9.09410 | 3.87928 | 1.21140 |

| | | | | | | | | | | | | |
|---|---|---|---|---|---|---|---|---|---|---|---|---|
| **FNTB** | 0.00117 | 1.00272 | 0.32750 | 0.71611 | 0.01488 | 0.81780 | 0.00000 | 0.12451 | 0.93415 | 0.00000 | 0.93708 | 0.41943 |
| **FOCAD** | 1.38956 | 2.91658 | 1.62663 | 1.40705 | 2.56764 | 4.54293 | 3.92322 | 0.58662 | 2.53409 | 4.18027 | 2.13884 | 1.57385 |
| **FOCAD-AS1** | 0.17567 | 0.01253 | 0.03062 | 0.01545 | 0.01764 | 0.00844 | 0.02220 | 0.02419 | 0.04688 | 0.00000 | 0.00908 | 0.04500 |
| **FOLH1** | 0.43253 | 0.04973 | 0.01002 | 0.00000 | 0.10650 | 0.03011 | 0.00000 | 0.24765 | 0.00000 | 0.18155 | 0.02252 | 0.15380 |
| **FOLH1B** | 0.02704 | 0.01522 | 0.01553 | 0.01193 | 0.01160 | 0.02708 | 0.01450 | 0.07104 | 0.01919 | 0.01378 | 0.02863 | 0.01818 |
| **FOLR1** | 0.00000 | 11.73464 | 13.31887 | 2.09952 | 25.22601 | 6.90289 | 9.90826 | 1.27003 | 53.62740 | 2.34941 | 26.90817 | 0.00000 |
| **FOLR2** | 0.00173 | 0.08652 | 0.02263 | 0.00000 | 0.00028 | 0.05108 | 0.01346 | 0.05857 | 0.16694 | 0.03395 | 0.02529 | 0.00537 |
| **FOLR3** | 0.08048 | 0.19716 | 0.05881 | 0.68918 | 0.63007 | 0.00000 | 0.00693 | 0.09505 | 0.00000 | 0.14295 | 0.27210 | 0.03652 |
| **FOLR4** | 5.16776 | 0.01936 | 0.01781 | 0.00623 | 0.02001 | 0.00516 | 0.01317 | 1.14892 | 0.02105 | 0.00000 | 0.01102 | 0.01842 |
| **FOPNL** | 2.41021 | 7.36662 | 8.77038 | 1.25274 | 0.00154 | 16.25037 | 7.49319 | 1.71711 | 7.27031 | 0.44145 | 5.24766 | 9.93246 |
| **FOS** | 5.76661 | 1.47561 | 4.81797 | 3.22554 | 14.50543 | 2.77271 | 2.11467 | 7.55918 | 4.39880 | 0.00000 | 1.04843 | 4.44676 |
| **FOSB** | 1.32903 | 0.00000 | 4.35694 | 0.00000 | 0.80865 | 0.00000 | 0.97273 | 0.12559 | 0.00000 | 1.32802 | 0.00000 | 0.00117 |
| **FOSL1** | 0.10420 | 0.00678 | 4.50649 | 0.36752 | 0.00000 | 0.37699 | 1.27376 | 0.01178 | 0.51024 | 0.00000 | 0.00000 | 0.00027 |
| **FOSL2** | 0.96914 | 3.37411 | 0.23575 | 1.88497 | 3.15613 | 0.00000 | 1.10199 | 0.00000 | 0.00000 | 5.63942 | 0.97442 | 1.04677 |
| **FOXA1** | 0.03186 | 0.00736 | 0.00000 | 0.56988 | 0.00000 | 0.19307 | 0.00000 | 0.00304 | 0.01924 | 0.02237 | 0.00267 | 0.03508 |
| **FOXA2** | 0.01594 | 0.00222 | 0.25312 | 4.06221 | 0.00000 | 0.02926 | 0.00000 | 0.01187 | 0.00000 | 0.16119 | 0.00299 | 0.02881 |
| **FOXA3** | 0.12834 | 0.00446 | 0.58324 | 0.45136 | 0.01163 | 0.00000 | 0.00000 | 0.00000 | 0.05872 | 0.28542 | 0.00000 | 0.18419 |
| **FOXB1** | 0.00637 | 0.02561 | 0.02932 | 0.01895 | 0.02175 | 0.01288 | 0.03211 | 0.01061 | 0.01291 | 0.00474 | 0.01573 | 0.01631 |
| **FOXB2** | 0.00401 | 0.01409 | 0.01826 | 0.00965 | 0.01270 | 0.01901 | 0.01629 | 0.02828 | 0.01833 | 0.03257 | 0.01413 | 0.01410 |
| **FOXC1** | 0.21884 | 0.00672 | 0.07042 | 0.39799 | 0.27382 | 0.00000 | 0.54029 | 0.16067 | 0.00000 | 0.22920 | 0.00000 | 0.02196 |
| **FOXC2** | 0.00563 | 0.01492 | 0.01428 | 0.01215 | 0.01933 | 0.02047 | 0.01872 | 0.01718 | 0.01839 | 0.01115 | 0.01491 | 0.01572 |
| **FOXD1** | 0.04599 | 0.00000 | 0.08025 | 0.00000 | 0.00000 | 0.06166 | 0.41338 | 0.29265 | 0.00000 | 0.81537 | 0.00000 | 0.00423 |
| **FOXD2** | 0.08515 | 0.02012 | 0.00444 | 0.10405 | 0.00000 | 0.00000 | 0.20056 | 2.16744 | 0.05899 | 0.13694 | 0.01640 | 0.32970 |
| **FOXD2-AS1** | 0.00289 | 0.42441 | 0.02815 | 0.00000 | 0.00000 | 0.18161 | 0.01447 | 2.65856 | 0.43667 | 0.75977 | 0.02091 | 1.90263 |
| **FOXD3** | 0.07601 | 0.06358 | 0.01992 | 1.00728 | 0.00000 | 0.06311 | 0.00000 | 0.09400 | 0.00000 | 0.27105 | 0.01852 | 0.60318 |
| **FOXD3-AS1** | 0.03055 | 0.36974 | 0.40119 | 4.43863 | 0.03443 | 0.31287 | 0.10004 | 0.97667 | 0.00000 | 0.02440 | 0.02801 | 0.91373 |
| **FOXD4** | 0.02244 | 0.02088 | 0.02075 | 0.01223 | 0.01368 | 0.00535 | 0.01309 | 0.01258 | 0.03694 | 0.09202 | 0.00843 | 0.08218 |
| **FOXD4L1** | 0.00402 | 0.01907 | 0.04097 | 0.01222 | 0.01404 | 0.01737 | 0.01601 | 0.03178 | 0.02799 | 0.01319 | 0.01185 | 0.01528 |
| **FOXE1** | 0.11854 | 0.70891 | 0.06060 | 0.00000 | 0.86149 | 0.00000 | 0.00299 | 0.00000 | 0.00000 | 1.04854 | 0.65152 | 0.26654 |
| **FOXE3** | 0.07142 | 0.01026 | 0.02228 | 0.00247 | 0.00759 | 0.09263 | 0.02725 | 0.00770 | 0.00138 | 0.02824 | 0.02249 | 0.03283 |
| **FOXF1** | 0.04701 | 0.03291 | 0.01284 | 0.03423 | 0.07389 | 0.00000 | 0.03339 | 0.03011 | 0.02043 | 0.02156 | 0.01217 | 0.03377 |
| **FOXF2** | 0.01496 | 0.01034 | 0.06189 | 0.10455 | 0.00770 | 0.01605 | 0.01145 | 0.01950 | 0.00568 | 0.06275 | 0.01611 | 0.01365 |
| **FOXG1** | 0.90452 | 0.00000 | 0.09647 | 0.00000 | 0.00000 | 0.04069 | 0.58455 | 0.00000 | 0.02676 | 1.09809 | 0.00000 | 0.41649 |
| **FOXH1** | 0.00475 | 19.15124 | 0.00160 | 14.09183 | 9.95525 | 0.00000 | 0.00312 | 0.27445 | 5.50320 | 0.00000 | 6.96719 | 1.24337 |
| **FOXI2** | 0.01452 | 0.01731 | 0.00886 | 0.01213 | 0.00577 | 0.04684 | 0.01214 | 0.01509 | 0.01213 | 0.03557 | 0.01227 | 0.01363 |
| **FOXI3** | 0.20849 | 1.94449 | 0.91204 | 0.00000 | 0.78971 | 0.02687 | 0.20008 | 0.03591 | 0.00000 | 0.00000 | 0.49402 | 2.90640 |
| **FOXJ1** | 0.02254 | 0.01341 | 0.02717 | 0.05583 | 0.00778 | 0.01445 | 0.00959 | 0.09234 | 0.03475 | 0.01623 | 0.02117 | 0.02187 |
| **FOXJ2** | 0.34773 | 0.00000 | 0.21550 | 0.70167 | 0.85286 | 1.27640 | 3.01823 | 0.86293 | 0.01941 | 2.47890 | 1.84398 | 2.66542 |
| **FOXJ3** | 5.74186 | 2.45153 | 5.42991 | 3.36595 | 0.95346 | 4.64371 | 2.36496 | 8.48253 | 5.99425 | 10.85135 | 2.32472 | 3.91740 |
| **FOXK1** | 2.39094 | 10.20417 | 2.37837 | 3.70452 | 4.79457 | 0.00001 | 9.95497 | 0.13924 | 6.87593 | 10.31546 | 7.56748 | 3.79163 |
| **FOXK2** | 6.25919 | 4.13215 | 5.60426 | 5.34664 | 9.08369 | 3.78851 | 15.26509 | 2.60749 | 0.52436 | 18.63266 | 6.95365 | 4.03787 |
| **FOXL2** | 0.00584 | 0.01680 | 0.07792 | 0.01504 | 0.02824 | 0.00022 | 0.04521 | 0.01151 | 0.00000 | 0.01558 | 0.05542 | 0.01789 |
| **FOXL2NB** | 0.04737 | 0.01597 | 0.02980 | 0.01335 | 0.02950 | 0.00747 | 0.03471 | 0.00682 | 0.02479 | 0.01105 | 0.01225 | 0.02300 |
| **FOXM1** | 3.68565 | 6.69858 | 7.27601 | 8.87234 | 12.61297 | 4.30777 | 19.42412 | 0.00000 | 3.54153 | 18.28480 | 4.24466 | 8.85770 |
| **FOXN1** | 0.00618 | 0.01069 | 0.01663 | 0.00775 | 0.03208 | 0.04702 | 0.02881 | 0.01029 | 0.01187 | 0.00485 | 0.02178 | 0.01506 |
| **FOXN2** | 1.16860 | 1.54589 | 2.27978 | 2.77952 | 0.64006 | 13.89089 | 5.24684 | 18.72037 | 1.66676 | 9.58650 | 6.14610 | 3.54609 |
| **FOXN3** | 1.32772 | 3.15561 | 1.16989 | 6.84425 | 14.81152 | 5.00035 | 9.11324 | 0.11895 | 1.76880 | 17.20573 | 4.04475 | 0.00000 |
| **FOXN3-AS1** | 0.00514 | 0.31283 | 0.04277 | 0.06848 | 0.09998 | 0.00000 | 0.00033 | 0.01089 | 0.03267 | 0.22625 | 0.01792 | 0.00336 |

| | | | | | | | | | | | | |
|---|---|---|---|---|---|---|---|---|---|---|---|---|
| **FOXN3-AS2** | 0.00618 | 0.01202 | 0.01197 | 0.08930 | 0.00649 | 0.01917 | 0.00594 | 0.01073 | 0.04538 | 0.01238 | 0.01976 | 0.01715 |
| **FOXN4** | 0.15337 | 0.38270 | 0.00000 | 0.00000 | 1.99366 | 0.00000 | 0.05894 | 0.00149 | 0.00000 | 0.69014 | 0.29746 | 0.02942 |
| **FOXO1** | 2.68057 | 4.75010 | 1.07150 | 4.72112 | 1.39806 | 5.77614 | 0.67206 | 1.40177 | 0.00000 | 16.10745 | 0.00000 | 4.21061 |
| **FOXO3** | 4.33251 | 0.31708 | 0.24011 | 0.95498 | 1.19917 | 0.84179 | 2.13896 | 0.90794 | 0.07869 | 4.20542 | 0.00000 | 3.23302 |
| **FOXO3B** | 0.33017 | 0.00000 | 0.75653 | 0.00000 | 0.00000 | 0.00000 | 1.08292 | 0.49021 | 0.41678 | 1.50350 | 0.15572 | 0.21459 |
| **FOXO4** | 0.18323 | 0.00000 | 0.21643 | 0.46852 | 1.99604 | 0.00000 | 2.11654 | 0.02328 | 0.42144 | 1.90064 | 1.18929 | 0.37733 |
| **FOXO6** | 0.02430 | 0.15533 | 3.49723 | 0.00000 | 1.13404 | 0.00000 | 0.52309 | 0.00555 | 0.00000 | 2.34147 | 0.00000 | 0.06600 |
| **FOXP1** | 0.96420 | 0.27562 | 1.13156 | 7.21797 | 5.00654 | 0.90687 | 0.25326 | 0.00000 | 0.00000 | 2.46839 | 0.39087 | 0.00000 |
| **FOXP2** | 0.04775 | 0.02756 | 0.05066 | 0.01110 | 0.01670 | 0.07896 | 0.07741 | 0.06671 | 0.00000 | 0.00001 | 0.07134 | 0.03626 |
| **FOXP3** | 0.07826 | 0.02192 | 0.00563 | 0.04079 | 0.05062 | 0.00000 | 0.02567 | 0.00558 | 0.02023 | 0.18492 | 0.02036 | 0.03019 |
| **FOXP4** | 1.73216 | 0.86749 | 0.27888 | 1.21910 | 3.40688 | 0.00000 | 1.59503 | 0.12539 | 1.10154 | 6.11664 | 0.00000 | 0.79844 |
| **FOXQ1** | 0.18634 | 0.01432 | 0.13436 | 0.08567 | 0.00311 | 0.02055 | 0.00000 | 0.00304 | 0.01050 | 0.05902 | 0.01314 | 0.01576 |
| **FOXR1** | 4.33745 | 28.25614 | 0.00000 | 8.56626 | 19.13187 | 0.00000 | 1.55871 | 38.20349 | 0.00000 | 0.31137 | 7.79907 | 38.28977 |
| **FOXR2** | 0.00647 | 0.00383 | 0.01529 | 0.29492 | 0.00304 | 0.00698 | 0.05699 | 0.03578 | 0.02040 | 0.00998 | 0.01395 | 0.21651 |
| **FOXRED1** | 0.05538 | 16.75702 | 7.02014 | 12.36447 | 15.62379 | 4.61124 | 9.09980 | 0.00000 | 17.56644 | 2.02885 | 12.21862 | 5.36805 |
| **FOXRED2** | 1.50187 | 2.95136 | 0.22810 | 1.53251 | 0.00000 | 0.00000 | 4.86715 | 0.93406 | 2.59390 | 1.81164 | 3.71721 | 4.42174 |
| **FPGS** | 1.31897 | 9.19128 | 8.22178 | 3.96409 | 6.21632 | 0.00000 | 3.28084 | 1.81697 | 10.15452 | 0.00000 | 5.04421 | 5.98926 |
| **FPGT** | 0.42366 | 1.92707 | 1.93841 | 0.00267 | 0.00000 | 10.84963 | 1.36122 | 0.06761 | 1.91846 | 0.00000 | 2.51954 | 3.23341 |
| **FPR3** | 0.16632 | 0.01351 | 0.02135 | 0.03059 | 0.05241 | 0.00000 | 0.01810 | 0.04040 | 0.02233 | 0.00000 | 0.01395 | 0.09539 |
| **FRA10AC1** | 0.60938 | 2.18744 | 0.54729 | 0.62640 | 0.73698 | 6.48951 | 1.85899 | 0.21066 | 0.58620 | 6.59272 | 1.80499 | 3.55051 |
| **FRAS1** | 0.00115 | 0.49276 | 1.77212 | 7.98977 | 0.60267 | 1.84260 | 1.43906 | 0.18715 | 3.31383 | 2.30958 | 0.63740 | 0.00000 |
| **FRAT1** | 0.92998 | 0.04792 | 0.33373 | 0.69102 | 0.27217 | 0.00000 | 0.31191 | 0.00939 | 0.00000 | 0.63377 | 0.00000 | 0.46167 |
| **FRAT2** | 2.69827 | 8.28914 | 2.46721 | 11.08444 | 5.97210 | 6.04745 | 7.96958 | 9.62324 | 0.00000 | 30.87146 | 0.00000 | 25.77890 |
| **FREM1** | 0.18209 | 0.01377 | 0.09086 | 0.02295 | 0.08444 | 0.00000 | 0.02370 | 0.00000 | 0.00000 | 0.04225 | 0.02152 | 0.12298 |
| **FREM2** | 0.00552 | 0.00000 | 2.52224 | 0.00000 | 0.72324 | 0.08646 | 0.13534 | 0.53271 | 2.04167 | 1.79874 | 0.00063 | 0.14024 |
| **FREM3** | 0.01471 | 0.01094 | 0.03780 | 0.01319 | 0.00399 | 0.01654 | 0.00253 | 0.00912 | 0.03536 | 0.10023 | 0.00725 | 0.01681 |
| **FRG1** | 7.21972 | 7.87516 | 6.42362 | 6.38484 | 13.69154 | 7.96262 | 15.86553 | 15.12051 | 1.17593 | 9.41063 | 6.34578 | 14.10516 |
| **FRG1B** | 2.47189 | 1.88366 | 1.78207 | 3.54286 | 4.66227 | 3.48244 | 4.04138 | 2.63663 | 0.69984 | 4.91625 | 2.69725 | 3.25878 |
| **FRG2.uid-6569** | 0.95805 | 0.01434 | 0.02210 | 0.01288 | 0.01784 | 0.00195 | 0.01336 | 0.14748 | 0.03004 | 0.01748 | 0.01201 | 0.01986 |
| **FRG2B** | 0.92115 | 0.02686 | 0.02455 | 0.01367 | 0.01732 | 0.00129 | 0.02108 | 0.05629 | 0.02038 | 0.00406 | 0.01559 | 0.01223 |
| **FRK** | 0.03291 | 0.21434 | 0.36970 | 0.02445 | 0.00000 | 0.42096 | 0.16406 | 0.07014 | 0.11402 | 0.00000 | 0.00548 | 0.05100 |
| **FRMD1** | 0.00626 | 0.03544 | 0.18648 | 0.02589 | 0.05740 | 0.00000 | 0.05486 | 0.01073 | 0.00000 | 0.00000 | 0.03171 | 0.01644 |
| **FRMD3** | 9.76495 | 0.31173 | 0.15008 | 0.24366 | 0.00000 | 0.01504 | 0.37996 | 2.74741 | 0.11814 | 0.21015 | 0.00000 | 0.53061 |
| **FRMD4A** | 1.93677 | 0.00000 | 7.23212 | 0.00000 | 7.65315 | 0.00000 | 6.54194 | 0.62053 | 2.75787 | 3.97957 | 4.14804 | 0.13324 |
| **FRMD4B** | 3.72231 | 3.88247 | 2.88750 | 0.00000 | 9.12162 | 13.42289 | 13.37535 | 3.87335 | 5.87179 | 9.72628 | 17.85360 | 11.25593 |
| **FRMD5** | 0.59819 | 0.00386 | 0.19191 | 0.06427 | 0.14000 | 0.70920 | 0.00000 | 0.67258 | 0.70676 | 2.00549 | 0.89854 | 0.87068 |
| **FRMD6** | 0.93092 | 0.38692 | 10.17489 | 12.17334 | 2.42894 | 12.52512 | 6.47172 | 1.63679 | 4.70289 | 5.15099 | 2.72877 | 2.04991 |
| **FRMD6-AS1** | 0.00000 | 0.18855 | 1.12847 | 0.36771 | 0.02508 | 0.64779 | 0.00000 | 0.06591 | 1.05676 | 0.36570 | 0.04832 | 0.20300 |
| **FRMD6-AS2** | 0.00537 | 0.00857 | 0.03993 | 0.00283 | 0.00434 | 0.06035 | 0.04527 | 0.01088 | 0.07222 | 0.01739 | 0.00925 | 0.01942 |
| **FRMD8** | 0.56477 | 11.75345 | 7.17746 | 3.44393 | 5.55371 | 0.00000 | 5.68052 | 1.28016 | 10.96577 | 3.82586 | 3.39065 | 5.44417 |
| **FRMD8P1** | 0.00460 | 0.61080 | 0.34780 | 0.11156 | 0.01339 | 0.00000 | 0.00486 | 0.03431 | 0.31947 | 0.00000 | 0.10938 | 0.10291 |
| **FRMPD1** | 0.06098 | 0.01205 | 0.02321 | 0.01307 | 0.01342 | 0.00000 | 0.03037 | 0.00591 | 0.00969 | 0.04445 | 0.04206 | 0.08955 |
| **FRMPD3** | 0.00725 | 1.28825 | 0.00000 | 0.20721 | 0.38038 | 0.15488 | 0.00165 | 0.00280 | 0.00000 | 0.17964 | 0.03381 | 1.11359 |
| **FRMPD4** | 0.05618 | 0.01240 | 0.01483 | 0.03350 | 0.01612 | 0.01376 | 0.02968 | 0.01578 | 0.02061 | 0.00759 | 0.01703 | 0.02960 |
| **FRRS1** | 0.22858 | 0.21237 | 1.01650 | 0.00000 | 2.08167 | 1.63891 | 0.00000 | 0.19866 | 1.36065 | 0.40759 | 0.35745 | 0.27149 |
| **FRS2** | 7.52634 | 1.63955 | 4.74096 | 0.65895 | 0.00000 | 10.38360 | 9.17750 | 15.06689 | 8.58488 | 7.10276 | 0.75324 | 4.53823 |
| **FRS3** | 0.80732 | 0.19942 | 0.08084 | 0.24787 | 0.00733 | 0.00000 | 0.00000 | 0.10677 | 0.44381 | 0.48798 | 0.00000 | 0.28208 |
| **FRY** | 0.00612 | 0.00805 | 0.25892 | 0.92924 | 0.12277 | 0.00000 | 0.00000 | 0.09194 | 0.00000 | 0.25501 | 0.00569 | 0.06559 |

| | | | | | | | | | | | | |
|---|---|---|---|---|---|---|---|---|---|---|---|---|
| **FRY-AS1** | 0.01479 | 0.01538 | 0.00000 | 0.30072 | 0.04287 | 0.08809 | 0.02261 | 0.08964 | 0.05130 | 0.00697 | 0.01729 | 0.19830 |
| **FRYL** | 1.15065 | 0.00000 | 3.96703 | 2.57387 | 0.53264 | 9.15573 | 4.20229 | 0.72409 | 2.29275 | 14.53927 | 1.03932 | 1.73164 |
| **FRZB** | 0.00546 | 0.00584 | 0.50030 | 2.91311 | 0.00000 | 0.01984 | 0.00000 | 0.04096 | 0.13413 | 0.00590 | 0.02173 | 0.01363 |
| **FSCN1** | 0.79116 | 1.00171 | 12.17569 | 11.56611 | 10.33510 | 0.00000 | 28.18436 | 0.81813 | 23.12030 | 10.90386 | 2.26328 | 3.36397 |
| **FSCN2** | 0.03164 | 0.01434 | 0.01714 | 0.00639 | 0.02950 | 0.01562 | 0.00347 | 0.06302 | 0.07639 | 0.02754 | 0.00601 | 0.01395 |
| **FSCN3** | 0.00743 | 0.01076 | 0.04813 | 0.07729 | 0.00845 | 0.02183 | 0.00000 | 0.00979 | 0.02755 | 0.07350 | 0.01881 | 0.01965 |
| **FSD1** | 0.38790 | 0.10362 | 1.53651 | 0.62152 | 0.00000 | 0.00000 | 0.26050 | 0.00304 | 0.18916 | 0.00000 | 0.02413 | 0.05630 |
| **FSD1L** | 1.83328 | 0.00000 | 0.29871 | 0.08985 | 0.05737 | 0.29384 | 0.11371 | 0.03206 | 0.40589 | 1.48018 | 0.32955 | 0.22104 |
| **FSD2** | 0.13518 | 0.00000 | 0.46772 | 0.00000 | 0.11501 | 1.89213 | 0.00000 | 0.70874 | 1.44818 | 0.02010 | 0.28897 | 0.82983 |
| **FSHR** | 0.00765 | 0.03762 | 0.01410 | 0.01462 | 0.02497 | 0.00665 | 0.00335 | 0.01115 | 0.03786 | 0.00840 | 0.27894 | 0.01989 |
| **FSIP1** | 0.00205 | 0.08448 | 0.15241 | 0.00688 | 0.05519 | 0.00000 | 0.02805 | 0.69893 | 0.17245 | 0.00000 | 0.03403 | 0.39708 |
| **FSIP2** | 0.81729 | 0.03974 | 0.01660 | 0.02220 | 0.00000 | 0.40279 | 0.00000 | 0.07266 | 0.11169 | 0.40999 | 0.00000 | 0.27962 |
| **FST** | 0.15114 | 0.04139 | 13.44597 | 1.48732 | 0.00000 | 0.31473 | 0.00000 | 0.14710 | 0.00000 | 0.02653 | 0.11840 | 0.00000 |
| **FSTL1** | 0.15193 | 0.00000 | 2.95944 | 1.69306 | 1.56921 | 1.82026 | 0.57829 | 0.04481 | 6.46446 | 0.00000 | 9.70259 | 0.00000 |
| **FSTL3** | 0.12264 | 0.56327 | 0.23460 | 0.29404 | 0.05783 | 0.00000 | 0.20052 | 0.21913 | 0.55364 | 0.33134 | 0.26355 | 2.62265 |
| **FSTL4** | 4.49898 | 0.07473 | 0.01670 | 0.02301 | 0.00000 | 0.01617 | 0.01018 | 0.33806 | 0.06263 | 0.06153 | 0.01089 | 0.18276 |
| **FSTL5** | 0.61176 | 0.04008 | 0.00000 | 0.00000 | 0.00000 | 0.07912 | 0.01768 | 0.04082 | 0.02559 | 0.16439 | 0.01459 | 0.01705 |
| **FTCD** | 4.25793 | 0.00899 | 0.02954 | 0.00653 | 0.00275 | 0.03651 | 0.00602 | 0.70811 | 0.00864 | 0.02748 | 0.01710 | 0.07420 |
| **FTCDNL1** | 0.03173 | 0.09422 | 0.01190 | 0.00962 | 0.14100 | 0.03381 | 0.00137 | 0.04102 | 0.09831 | 0.00000 | 0.03805 | 0.01181 |
| **FTH1** | 3.67394 | 27.50976 | 22.57574 | 36.77177 | 32.78847 | 28.40820 | 32.03683 | 8.81822 | 35.58837 | 2.97530 | 14.07481 | 26.77501 |
| **FTH1P3** | 0.01601 | 0.13938 | 0.12444 | 0.56215 | 0.25071 | 0.00000 | 0.00239 | 2.70090 | 0.24452 | 1.59962 | 0.14293 | 0.45914 |
| **FTHL17** | 0.47667 | 0.19670 | 0.23786 | 0.32566 | 0.02057 | 0.47505 | 0.08155 | 0.20031 | 0.07806 | 0.00000 | 0.24822 | 0.08681 |
| **FTL** | 25.36373 | 11.27720 | 53.95218 | 26.45055 | 33.98451 | 29.31127 | 58.26015 | 22.70329 | 62.68831 | 11.48806 | 31.56455 | 28.66359 |
| **FTMT** | 0.07759 | 0.00621 | 0.18654 | 0.04260 | 0.00000 | 0.05149 | 0.00000 | 0.02545 | 0.00000 | 0.06368 | 0.01349 | 0.00954 |
| **FTO** | 0.13448 | 2.33205 | 3.20461 | 4.16991 | 0.79480 | 2.53939 | 3.34572 | 3.38838 | 2.95410 | 0.00000 | 4.81601 | 1.76702 |
| **FTO-IT1** | 0.00176 | 0.06338 | 0.11555 | 0.06732 | 0.10147 | 0.47614 | 0.35985 | 0.47777 | 0.00000 | 0.02923 | 0.10541 | 0.50066 |
| **FTSJ1** | 0.50636 | 4.65891 | 5.94677 | 9.99925 | 5.89317 | 4.26208 | 7.06205 | 0.35978 | 15.47801 | 0.00000 | 12.29158 | 6.52600 |
| **FTSJ2** | 2.98572 | 18.27169 | 11.62705 | 10.56821 | 9.62235 | 21.64249 | 13.52136 | 7.26621 | 14.70110 | 0.00000 | 15.24122 | 19.35488 |
| **FTSJ3** | 0.82139 | 17.49043 | 23.12547 | 14.01388 | 19.92957 | 6.48983 | 31.55900 | 15.75529 | 20.35517 | 10.90949 | 19.54620 | 24.95580 |
| **FTX** | 0.01833 | 0.03054 | 0.28147 | 0.33111 | 0.00279 | 1.04664 | 0.11559 | 0.03614 | 1.36703 | 0.42914 | 0.09699 | 0.44068 |
| **FUBP1** | 2.68131 | 13.02619 | 14.87262 | 10.77995 | 11.92282 | 29.65175 | 21.84062 | 14.31517 | 18.95968 | 15.66982 | 7.53103 | 25.43368 |
| **FUBP3** | 2.57875 | 3.59330 | 4.67292 | 3.06967 | 2.09640 | 11.46964 | 10.70508 | 3.00374 | 6.10972 | 10.78248 | 2.62152 | 3.41012 |
| **FUCA1** | 1.83410 | 16.70197 | 2.07676 | 10.92994 | 9.25252 | 16.56832 | 0.00000 | 5.33256 | 3.69243 | 0.00000 | 3.96152 | 9.73981 |
| **FUCA2** | 0.00092 | 0.34312 | 9.48959 | 3.23453 | 4.90313 | 17.66429 | 0.00000 | 0.16119 | 34.24066 | 0.04427 | 11.46130 | 0.00000 |
| **FUK** | 0.00000 | 1.07817 | 0.00000 | 0.73980 | 0.70453 | 0.00000 | 0.27661 | 0.02844 | 1.02876 | 2.34516 | 0.38244 | 0.45748 |
| **FUNDC1** | 1.08176 | 11.14046 | 0.39869 | 6.50893 | 5.56147 | 20.01536 | 4.07967 | 5.40172 | 1.14951 | 4.32417 | 6.16599 | 10.67548 |
| **FUNDC2** | 0.09273 | 5.49738 | 8.23700 | 5.18043 | 6.05521 | 11.27504 | 11.73632 | 2.64733 | 8.15194 | 0.00000 | 9.41071 | 9.03496 |
| **FUNDC2P2** | 0.40716 | 0.11781 | 0.18925 | 0.06004 | 0.14075 | 0.47318 | 0.21263 | 0.18185 | 0.23777 | 0.00000 | 0.26123 | 0.25666 |
| **FUOM** | 0.00042 | 4.76502 | 1.18502 | 6.03577 | 15.14511 | 0.00000 | 4.02531 | 0.00362 | 3.52383 | 0.63278 | 12.34483 | 0.00000 |
| **FURIN** | 8.60494 | 9.44732 | 16.77220 | 8.16916 | 29.26822 | 0.00000 | 0.28885 | 1.41393 | 27.02636 | 20.78176 | 6.31794 | 8.95150 |
| **FUS** | 4.20336 | 31.59034 | 27.63236 | 24.48500 | 22.71731 | 0.00503 | 36.67423 | 22.89765 | 36.71917 | 18.25465 | 29.85642 | 37.75600 |
| **FUT1** | 0.00937 | 0.05493 | 0.00000 | 1.34393 | 6.00123 | 0.14577 | 0.81791 | 0.00618 | 5.13878 | 0.00000 | 2.01532 | 0.00454 |
| **FUT10** | 3.58125 | 1.98230 | 1.37418 | 0.40591 | 0.00000 | 2.42416 | 1.05039 | 0.60207 | 2.61747 | 0.00000 | 2.43482 | 1.37571 |
| **FUT11** | 0.08904 | 0.28037 | 1.71024 | 0.34989 | 0.00000 | 0.12084 | 0.37695 | 0.02104 | 4.15760 | 0.00000 | 0.49994 | 0.25122 |
| **FUT2** | 1.44363 | 0.20085 | 0.00000 | 0.15829 | 0.00000 | 0.06782 | 0.00296 | 0.05998 | 0.00000 | 0.12373 | 0.00298 | 0.00000 |
| **FUT3** | 0.00418 | 1.34102 | 0.00000 | 0.00000 | 0.00000 | 0.00000 | 0.02085 | 1.33112 | 0.04245 | 0.09692 | 0.00000 | 18.36483 |
| **FUT4** | 0.00000 | 0.00000 | 3.18306 | 0.00000 | 5.51307 | 3.20309 | 4.89533 | 8.47431 | 13.12525 | 10.66607 | 2.52865 | 10.02216 |
| **FUT5** | 1.36354 | 0.02896 | 0.04001 | 0.04645 | 0.02485 | 0.00000 | 0.01067 | 0.35360 | 0.04159 | 0.00000 | 0.01051 | 0.05267 |

| | | | | | | | | | | | | |
|---|---|---|---|---|---|---|---|---|---|---|---|---|
| **FUT6** | 0.02740 | 0.19914 | 0.14403 | 0.07093 | 0.18881 | 0.00000 | 0.02357 | 0.07386 | 0.04189 | 0.00000 | 0.05191 | 0.39998 |
| **FUT8** | 5.00940 | 1.33638 | 10.29233 | 0.68771 | 5.16731 | 12.44803 | 7.49431 | 1.42184 | 22.25270 | 7.04427 | 7.87845 | 0.00000 |
| **FUT8-AS1** | 0.06950 | 0.03572 | 0.06119 | 0.07874 | 0.00000 | 0.06158 | 0.00000 | 0.02720 | 0.08420 | 0.01310 | 0.09751 | 0.04783 |
| **FUT9** | 0.25925 | 0.21928 | 0.00729 | 0.00000 | 0.22361 | 0.44638 | 0.00000 | 0.41381 | 0.00000 | 0.01518 | 0.10326 | 0.00000 |
| **FUZ** | 0.62349 | 0.84690 | 2.40393 | 2.22406 | 0.87911 | 0.00000 | 0.84768 | 0.00000 | 3.25342 | 0.00000 | 2.40142 | 1.42713 |
| **FXN** | 0.53958 | 15.44189 | 5.06650 | 12.07742 | 5.23888 | 16.70307 | 11.69076 | 0.07245 | 9.99265 | 4.94052 | 12.29334 | 8.62197 |
| **FXR1** | 2.81683 | 7.76375 | 2.67055 | 12.49956 | 3.14565 | 24.28209 | 18.47725 | 8.44133 | 8.78867 | 20.23986 | 13.19132 | 14.38404 |
| **FXR2** | 0.99216 | 3.25970 | 8.46155 | 2.72744 | 3.51524 | 0.00000 | 7.34583 | 1.41250 | 3.44341 | 12.40784 | 2.38199 | 4.65388 |
| **FXYD1** | 0.00612 | 0.01604 | 0.02422 | 0.01915 | 0.01851 | 0.00697 | 0.02009 | 0.02150 | 0.02081 | 0.01720 | 0.01534 | 0.03307 |
| **FXYD3** | 0.04197 | 0.48742 | 0.08221 | 0.00000 | 0.31694 | 0.61177 | 0.09428 | 0.01052 | 0.21175 | 0.17856 | 0.22339 | 0.04878 |
| **FXYD4** | 0.00000 | 22.09289 | 0.00000 | 1.10107 | 10.54990 | 1.37800 | 5.00599 | 0.69088 | 10.51821 | 6.25704 | 10.69380 | 7.76682 |
| **FXYD5** | 0.00477 | 0.00000 | 11.53155 | 2.26284 | 0.12345 | 1.27313 | 2.81489 | 0.00318 | 8.42027 | 0.00000 | 0.56221 | 0.06020 |
| **FXYD6** | 0.01369 | 0.00992 | 0.00922 | 0.13633 | 0.03497 | 0.04285 | 0.02681 | 0.00993 | 0.02375 | 0.00000 | 0.01081 | 0.01524 |
| **FXYD7** | 0.00507 | 0.00859 | 0.04220 | 0.06368 | 0.05889 | 0.00000 | 0.00209 | 0.01011 | 0.06537 | 0.21191 | 0.08476 | 0.01699 |
| **FYB** | 0.00433 | 0.00000 | 23.86487 | 0.00000 | 0.00000 | 0.03686 | 17.65390 | 0.00940 | 0.44184 | 0.00000 | 0.00000 | 0.01517 |
| **FYCO1** | 0.09468 | 0.00000 | 0.64700 | 1.05277 | 0.70268 | 0.31749 | 0.03336 | 0.12637 | 2.11390 | 1.37892 | 1.75053 | 0.48370 |
| **FYN** | 4.32443 | 1.59797 | 1.09798 | 2.34162 | 0.47068 | 0.60747 | 0.00000 | 1.25577 | 0.00000 | 1.30790 | 0.00000 | 0.50070 |
| **FYTTD1** | 1.26588 | 6.55073 | 0.36449 | 1.09480 | 4.81496 | 15.33195 | 0.00000 | 7.16368 | 1.02515 | 1.48468 | 0.76552 | 9.67200 |
| **FZD1** | 0.65888 | 0.12812 | 0.09315 | 0.00000 | 0.13280 | 0.04410 | 0.01145 | 0.01453 | 0.15901 | 0.09834 | 0.03397 | 0.23899 |
| **FZD10** | 0.02003 | 0.02459 | 1.13100 | 0.30954 | 0.05220 | 0.00311 | 0.00000 | 0.00870 | 0.00000 | 0.01849 | 0.01461 | 0.22205 |
| **FZD10-AS1** | 0.03833 | 0.01730 | 0.04570 | 0.05921 | 0.00000 | 0.01252 | 0.00038 | 0.01256 | 0.01824 | 0.26870 | 0.00374 | 0.01438 |
| **FZD2** | 0.00598 | 0.01818 | 0.02177 | 0.00968 | 0.03569 | 0.00238 | 0.04185 | 0.01062 | 0.02051 | 0.03002 | 0.01563 | 0.01630 |
| **FZD3** | 5.63845 | 4.75196 | 2.78071 | 0.00000 | 0.40981 | 7.99103 | 2.37199 | 0.37451 | 7.60492 | 8.85386 | 2.86579 | 6.19037 |
| **FZD4** | 0.47007 | 1.96826 | 0.00000 | 1.33182 | 4.04048 | 3.42674 | 0.53208 | 4.68970 | 0.00004 | 3.28461 | 0.40789 | 0.96049 |
| **FZD5** | 0.37992 | 2.00115 | 16.58429 | 16.97114 | 2.67996 | 11.15653 | 10.76752 | 1.63855 | 16.51205 | 15.54834 | 9.83127 | 0.49010 |
| **FZD6** | 1.00520 | 0.00005 | 2.36029 | 0.00270 | 0.00000 | 0.51166 | 0.36277 | 0.25490 | 5.84195 | 0.00000 | 0.00880 | 0.00000 |
| **FZD7** | 0.61580 | 0.32940 | 0.41980 | 1.65246 | 3.78888 | 0.00035 | 0.00000 | 0.74178 | 0.00000 | 1.21141 | 0.00000 | 1.51116 |
| **FZD8** | 0.00219 | 0.02075 | 0.20925 | 0.34803 | 0.07431 | 0.04648 | 0.01459 | 0.00890 | 0.00000 | 0.48793 | 0.00000 | 0.02389 |
| **FZD9** | 0.05809 | 0.00397 | 0.07682 | 0.21757 | 0.00000 | 0.02515 | 0.00674 | 0.01187 | 0.00000 | 0.11335 | 0.01443 | 0.03395 |
| **FZR1** | 1.75542 | 2.41196 | 7.18547 | 2.00638 | 4.26439 | 0.00000 | 4.55407 | 0.36292 | 0.94843 | 9.95659 | 1.80774 | 2.61106 |
| **G0S2** | 0.00645 | 0.02051 | 0.01193 | 0.03244 | 0.01466 | 0.01680 | 0.05121 | 0.00946 | 0.04742 | 0.00000 | 0.01945 | 0.03224 |
| **G2E3** | 0.09589 | 1.74536 | 8.04665 | 2.62833 | 2.10146 | 21.67603 | 13.42487 | 6.29926 | 7.92836 | 11.62308 | 6.16342 | 7.49347 |
| **G3BP1** | 6.70462 | 27.76017 | 24.73609 | 20.13298 | 22.32906 | 49.96932 | 35.18189 | 6.59261 | 32.64817 | 0.36980 | 38.90275 | 23.06718 |
| **G3BP2** | 4.31875 | 14.92611 | 5.33340 | 14.50632 | 8.39069 | 43.55379 | 21.07083 | 6.84448 | 11.38641 | 15.29547 | 15.50392 | 34.26482 |
| **G6PC** | 0.00414 | 0.02620 | 0.00000 | 0.00000 | 0.01195 | 0.02105 | 0.00984 | 0.00791 | 0.00000 | 0.40897 | 0.05409 | 0.04436 |
| **G6PC2** | 0.00232 | 0.01570 | 0.02002 | 0.03558 | 0.01568 | 0.00325 | 0.02180 | 0.04958 | 0.01370 | 0.02187 | 0.02894 | 0.01513 |
| **G6PC3** | 0.68445 | 5.83269 | 8.36226 | 9.94968 | 7.10542 | 0.00000 | 0.00000 | 0.07190 | 11.93750 | 0.00657 | 6.37971 | 1.76774 |
| **G6PD** | 2.19045 | 4.51164 | 6.69256 | 6.05911 | 8.25418 | 0.00000 | 8.61481 | 0.40278 | 14.99720 | 0.00000 | 9.85109 | 5.00669 |
| **GAA** | 1.12877 | 1.51274 | 1.50893 | 0.32409 | 0.82979 | 0.00000 | 0.00000 | 0.00000 | 1.67898 | 1.02108 | 1.69810 | 1.21594 |
| **GAB1** | 21.19790 | 3.30852 | 5.22300 | 2.47385 | 6.76883 | 8.19207 | 3.47084 | 2.49777 | 0.00000 | 4.48579 | 3.84225 | 1.53606 |
| **GAB2** | 9.35616 | 1.50192 | 12.58364 | 0.00000 | 16.21728 | 1.76227 | 11.93084 | 0.20554 | 10.96305 | 10.17853 | 15.03433 | 0.25505 |
| **GAB3** | 0.00231 | 0.02911 | 0.04487 | 0.07011 | 0.03225 | 0.10675 | 0.00000 | 0.08262 | 0.00000 | 0.06089 | 0.02213 | 0.04232 |
| **GAB4** | 0.02004 | 0.01305 | 0.01363 | 0.01328 | 0.01449 | 0.00974 | 0.01586 | 0.00895 | 0.01883 | 0.03157 | 0.01503 | 0.05693 |
| **GABARAP** | 2.84560 | 4.35436 | 19.43348 | 12.83535 | 14.38660 | 12.28862 | 21.65333 | 2.95760 | 19.60655 | 6.99601 | 16.08028 | 3.35516 |
| **GABARAPL1** | 0.56073 | 32.55609 | 18.10349 | 23.76712 | 9.37698 | 13.83478 | 9.40393 | 7.00623 | 9.32703 | 4.65747 | 0.49443 | 52.21710 |
| **GABARAPL2** | 6.60629 | 17.53398 | 10.59872 | 5.93143 | 10.31072 | 30.77068 | 13.55448 | 21.71443 | 10.36946 | 0.00000 | 8.77496 | 18.14505 |
| **GABARAPL3** | 0.00336 | 0.05751 | 0.03402 | 0.12665 | 0.01697 | 0.05126 | 0.01562 | 0.03478 | 0.03061 | 0.00000 | 0.04011 | 0.20651 |
| **GABBR1** | 0.04002 | 0.05955 | 0.24518 | 3.74759 | 0.09464 | 0.00000 | 0.20625 | 0.00307 | 0.13884 | 2.19907 | 0.19390 | 0.34728 |

| | | | | | | | | | | | | |
|---|---|---|---|---|---|---|---|---|---|---|---|---|
| **GABBR2** | 5.55970 | 0.00000 | 0.44070 | 0.00000 | 3.38143 | 0.00000 | 3.32557 | 0.58328 | 1.94685 | 1.94594 | 3.84528 | 0.00000 |
| **GABPA** | 1.31926 | 4.39643 | 4.31417 | 1.40204 | 0.00000 | 7.91707 | 5.64397 | 13.04349 | 1.52245 | 4.84202 | 3.42710 | 5.93938 |
| **GABPB1** | 3.70633 | 11.84015 | 7.91908 | 2.04515 | 6.50428 | 17.28439 | 8.45427 | 4.03869 | 10.03026 | 0.11928 | 7.97032 | 7.87363 |
| **GABPB1-AS1** | 0.00004 | 1.02860 | 2.93109 | 0.76305 | 0.62255 | 0.67562 | 0.02500 | 0.88724 | 3.31920 | 6.65040 | 1.62683 | 1.78891 |
| **GABPB2** | 0.37499 | 1.20083 | 0.12516 | 0.72819 | 0.68721 | 1.04995 | 0.00000 | 1.18791 | 1.20408 | 0.31898 | 0.00570 | 1.18949 |
| **GABRA1** | 0.00259 | 0.00737 | 0.04137 | 0.01317 | 0.00000 | 0.06791 | 0.01700 | 0.20966 | 0.00307 | 0.03291 | 0.02004 | 0.00739 |
| **GABRA2** | 0.26405 | 0.00509 | 0.02659 | 0.00685 | 0.04175 | 0.16783 | 0.00000 | 0.08537 | 0.01367 | 0.00656 | 0.00464 | 0.04348 |
| **GABRA3** | 0.00793 | 0.22339 | 0.03479 | 0.04058 | 0.00000 | 0.00008 | 0.01148 | 0.07041 | 0.00000 | 0.05368 | 0.21437 | 0.13541 |
| **GABRA4** | 0.00573 | 0.01920 | 0.00630 | 0.01284 | 0.01288 | 0.00000 | 0.14778 | 0.01064 | 0.00000 | 0.03843 | 0.01025 | 0.01551 |
| **GABRA5** | 0.21667 | 0.44591 | 0.73646 | 0.00110 | 1.10063 | 0.04352 | 0.41398 | 0.11462 | 0.00000 | 0.00000 | 0.47864 | 0.00000 |
| **GABRA6** | 0.00267 | 0.03248 | 0.00889 | 0.01319 | 0.01757 | 0.00090 | 0.14191 | 0.11597 | 0.00000 | 0.00000 | 0.01448 | 0.02986 |
| **GABRB1** | 0.00468 | 0.00496 | 0.06208 | 0.00010 | 0.00000 | 0.03759 | 0.00039 | 0.04580 | 0.01153 | 0.38237 | 0.00015 | 0.00348 |
| **GABRB2** | 0.00214 | 0.01649 | 0.10735 | 0.02449 | 0.11457 | 0.00000 | 0.00303 | 0.03193 | 0.29277 | 0.39356 | 0.00000 | 0.01066 |
| **GABRB3** | 0.01253 | 3.92594 | 4.61771 | 1.15368 | 3.87090 | 10.93206 | 0.00000 | 1.23677 | 2.57014 | 4.20752 | 0.40995 | 0.70449 |
| **GABRD** | 0.11227 | 0.49933 | 0.00000 | 0.17901 | 0.00000 | 0.01227 | 0.00000 | 0.00593 | 0.00000 | 0.17189 | 0.02987 | 1.07272 |
| **GABRE** | 0.00243 | 0.05511 | 1.59306 | 0.04506 | 0.00000 | 0.00051 | 0.00000 | 0.00967 | 0.00000 | 0.59798 | 0.00256 | 0.00597 |
| **GABRG1** | 0.04005 | 0.00919 | 0.03809 | 0.01787 | 0.00000 | 0.05604 | 0.54786 | 0.00889 | 0.00594 | 0.00000 | 0.00000 | 0.02924 |
| **GABRG2** | 0.08190 | 0.05280 | 0.00000 | 0.25128 | 0.02556 | 0.47878 | 0.17723 | 0.00000 | 0.00000 | 0.09244 | 0.03913 | 0.06496 |
| **GABRG3** | 0.36689 | 0.03093 | 0.06985 | 0.00000 | 0.11404 | 0.04452 | 0.07043 | 0.15885 | 0.00000 | 0.06497 | 0.01407 | 0.05398 |
| **GABRP** | 1.02315 | 0.01691 | 0.02695 | 0.00974 | 0.01298 | 0.02095 | 0.01318 | 0.10914 | 0.01482 | 0.01351 | 0.01978 | 0.09030 |
| **GABRQ** | 0.05153 | 0.00990 | 0.00527 | 0.57036 | 0.00000 | 0.01797 | 0.00000 | 1.09589 | 0.01702 | 0.09965 | 0.02836 | 0.07109 |
| **GABRR1** | 0.00478 | 0.49492 | 0.00450 | 0.00258 | 0.00000 | 0.09617 | 0.00000 | 0.01179 | 0.29855 | 0.00502 | 0.11794 | 0.03809 |
| **GABRR2** | 0.00697 | 0.00841 | 0.10058 | 0.01862 | 0.00000 | 0.04954 | 0.03384 | 0.03004 | 0.00000 | 0.06241 | 0.02126 | 0.00992 |
| **GABRR3** | 8.51041 | 0.01107 | 0.00932 | 0.00716 | 0.01965 | 0.00368 | 0.01776 | 0.45817 | 0.02312 | 0.00000 | 0.01914 | 0.00000 |
| **GAD1** | 1.03285 | 0.62614 | 0.00000 | 2.40501 | 0.27519 | 0.00000 | 0.08364 | 0.36228 | 0.47590 | 0.29274 | 0.00000 | 0.48403 |
| **GAD2** | 0.00619 | 0.01822 | 0.03577 | 0.16159 | 0.00003 | 0.05660 | 0.00659 | 0.01535 | 0.00958 | 0.02162 | 0.01344 | 0.01607 |
| **GADD45A** | 2.54992 | 2.45338 | 4.08417 | 0.50731 | 0.97580 | 3.81780 | 4.59710 | 3.48531 | 0.66774 | 10.57187 | 0.00000 | 0.00000 |
| **GADD45B** | 0.00592 | 0.29439 | 1.44304 | 0.00295 | 1.77792 | 0.01057 | 1.04024 | 18.35455 | 0.00000 | 0.00000 | 3.01069 | 0.45623 |
| **GADD45G** | 0.00211 | 0.22647 | 4.33110 | 1.31722 | 3.81522 | 0.00000 | 1.26483 | 0.00690 | 0.53034 | 0.65903 | 0.23624 | 0.00000 |
| **GADD45GIP1** | 0.24254 | 15.67002 | 12.14140 | 10.90834 | 13.77481 | 0.00000 | 10.95396 | 0.92685 | 6.15880 | 23.94846 | 11.23671 | 8.51352 |
| **GADL1** | 0.20339 | 0.02126 | 0.05015 | 0.15426 | 0.00570 | 0.07270 | 0.00000 | 0.00555 | 0.01414 | 0.03370 | 0.00933 | 0.05416 |
| **GAGE1** | 0.01735 | 0.00000 | 0.00000 | 0.00075 | 0.33837 | 0.96461 | 0.00000 | 0.00108 | 0.14601 | 0.00000 | 0.03337 | 2.10978 |
| **GAGE10** | 0.00627 | 0.01611 | 0.01243 | 0.01021 | 0.02772 | 0.02254 | 0.02296 | 0.01068 | 0.02087 | 0.01361 | 0.01730 | 0.02133 |
| **GAGE12J** | 0.00478 | 0.20035 | 0.00000 | 0.00091 | 0.09027 | 0.63396 | 0.78654 | 0.01902 | 0.26087 | 0.00000 | 0.00000 | 0.12271 |
| **GAGE2A.uid-12** | 0.00183 | 0.15729 | 0.02095 | 0.00000 | 0.00000 | 0.20547 | 1.27245 | 0.03746 | 0.04704 | 0.01147 | 0.00000 | 0.00109 |
| **GAGE2B** | 0.00533 | 0.09805 | 0.00552 | 0.00000 | 0.00404 | 0.08927 | 0.00000 | 0.01014 | 0.08811 | 0.02948 | 0.01873 | 0.00599 |
| **GAK** | 3.77479 | 10.35580 | 7.51467 | 8.98522 | 14.08496 | 0.00000 | 6.98874 | 0.73542 | 14.20449 | 17.53392 | 6.42111 | 3.77461 |
| **GAL** | 0.01854 | 5.12069 | 2.87893 | 0.25856 | 3.06197 | 0.58410 | 0.00024 | 0.34631 | 3.47421 | 4.13657 | 0.00018 | 1.89237 |
| **GAL3ST1** | 0.20855 | 0.00665 | 0.12672 | 0.41526 | 0.02462 | 0.00000 | 0.01019 | 0.00299 | 0.00057 | 0.09849 | 0.00963 | 0.01200 |
| **GAL3ST3** | 0.05723 | 0.00251 | 0.06978 | 0.01241 | 0.03384 | 0.06203 | 0.03982 | 0.07036 | 0.05699 | 0.00000 | 0.00806 | 0.41904 |
| **GAL3ST4** | 0.03428 | 0.00000 | 0.20137 | 0.46257 | 0.37116 | 0.00000 | 0.30293 | 0.07251 | 0.89959 | 0.69255 | 0.00000 | 0.14670 |
| **GALC** | 2.56421 | 0.42247 | 0.43192 | 1.23985 | 0.00002 | 1.49809 | 0.00000 | 2.37666 | 1.14192 | 0.00000 | 0.46470 | 0.58705 |
| **GALE** | 0.00000 | 8.29319 | 3.27366 | 6.90568 | 9.46412 | 7.67191 | 1.10545 | 0.04914 | 8.01275 | 0.80333 | 13.07912 | 1.94081 |
| **GALK1** | 0.10852 | 31.99340 | 2.91614 | 0.64486 | 15.45320 | 2.21770 | 4.59783 | 0.00000 | 1.17862 | 0.00000 | 14.12826 | 13.59103 |
| **GALK2** | 0.85315 | 2.32169 | 3.78663 | 4.32244 | 0.89736 | 7.59688 | 1.05506 | 1.05986 | 3.46780 | 0.00000 | 2.13432 | 1.37670 |
| **GALM** | 0.00000 | 7.58095 | 0.00000 | 5.42323 | 6.65841 | 1.56403 | 6.06461 | 5.55815 | 3.50648 | 3.87066 | 10.53703 | 10.48480 |
| **GALNS** | 0.16979 | 0.60991 | 3.00610 | 0.23937 | 0.99811 | 0.00000 | 0.00000 | 0.05072 | 7.23089 | 0.12249 | 0.95409 | 0.31087 |
| **GALNT1** | 1.51772 | 1.34126 | 2.09096 | 0.00000 | 3.14418 | 8.30941 | 4.61687 | 0.89148 | 0.00080 | 14.26322 | 2.94221 | 1.88948 |

| | | | | | | | | | | | | |
|---|---|---|---|---|---|---|---|---|---|---|---|---|
| GALNT10 | 0.00001 | 0.00000 | 0.00000 | 0.65432 | 15.68184 | 0.00000 | 4.28698 | 0.22899 | 8.35082 | 8.70970 | 15.98500 | 0.23821 |
| GALNT11 | 1.38862 | 2.42525 | 7.28480 | 0.32106 | 1.23166 | 2.67924 | 5.89803 | 2.52757 | 23.63685 | 3.78941 | 16.00638 | 1.85390 |
| GALNT12 | 0.11496 | 0.55761 | 0.00001 | 0.28929 | 0.56414 | 0.35195 | 0.00000 | 0.07277 | 0.00000 | 0.25300 | 0.00000 | 0.00000 |
| GALNT13 | 1.84150 | 0.09713 | 0.10440 | 0.05358 | 0.00000 | 0.02508 | 0.12451 | 0.00000 | 0.01756 | 0.00000 | 0.01185 | 0.03790 |
| GALNT14 | 0.53322 | 0.02111 | 0.74553 | 0.00000 | 0.00000 | 0.00000 | 0.01682 | 0.56331 | 1.06818 | 0.13272 | 1.66778 | 0.14275 |
| GALNT15 | 0.64479 | 0.00944 | 0.01566 | 0.00902 | 0.01165 | 0.02554 | 0.03981 | 0.03242 | 0.00918 | 0.00897 | 0.01326 | 0.03045 |
| GALNT16 | 0.00193 | 0.10103 | 0.16778 | 0.00431 | 0.08219 | 0.02069 | 0.00324 | 0.06835 | 0.10358 | 0.00000 | 0.01955 | 0.00832 |
| GALNT18 | 1.31014 | 0.34161 | 0.90670 | 0.64532 | 0.04955 | 0.00000 | 0.00000 | 0.17661 | 0.42964 | 0.00000 | 0.08693 | 0.00000 |
| GALNT2 | 0.51528 | 5.09659 | 7.62947 | 3.09183 | 5.62498 | 0.00000 | 0.05886 | 0.89328 | 6.17431 | 3.43930 | 0.56490 | 1.93414 |
| GALNT3 | 1.85954 | 0.33536 | 2.08027 | 0.02778 | 0.00000 | 0.76504 | 0.00325 | 0.10400 | 2.44925 | 0.98809 | 1.45485 | 0.43626 |
| GALNT4 | 0.00618 | 0.02402 | 0.04112 | 0.01388 | 0.06500 | 0.04651 | 0.00405 | 0.01017 | 0.16369 | 0.00000 | 0.03975 | 0.01450 |
| GALNT5 | 0.90498 | 0.06825 | 0.24565 | 0.01366 | 0.00703 | 0.00000 | 0.00000 | 0.10959 | 0.02799 | 0.05650 | 0.11063 | 0.33377 |
| GALNT6 | 0.21071 | 5.23734 | 0.01538 | 7.95337 | 13.37548 | 4.77080 | 0.00000 | 0.31688 | 13.49889 | 11.16851 | 7.20374 | 7.40348 |
| GALNT7 | 0.05830 | 0.73728 | 1.17066 | 3.83033 | 0.00000 | 0.61564 | 0.00000 | 1.23850 | 5.33897 | 3.72799 | 1.00865 | 0.16078 |
| GALNT8 | 0.00184 | 0.86047 | 0.01630 | 1.42036 | 0.01506 | 0.31849 | 0.16504 | 0.24692 | 0.28620 | 0.00000 | 0.12138 | 0.69935 |
| GALNT9 | 0.13806 | 0.01335 | 0.03327 | 0.01338 | 0.00940 | 0.02248 | 0.01170 | 0.06475 | 0.01186 | 0.03217 | 0.01471 | 0.01632 |
| GALNTL5 | 0.01258 | 0.01434 | 0.01354 | 0.01162 | 0.01095 | 0.01672 | 0.01963 | 0.02279 | 0.01705 | 0.02057 | 0.03328 | 0.01547 |
| GALNTL6 | 0.38855 | 0.12768 | 0.01301 | 0.00376 | 0.00673 | 0.00162 | 0.03520 | 0.43940 | 0.01660 | 0.02168 | 0.01387 | 0.00353 |
| GALR1 | 0.00593 | 0.09801 | 0.01455 | 0.00935 | 0.03618 | 0.01896 | 0.01797 | 0.01202 | 0.02853 | 0.00000 | 0.01538 | 0.02635 |
| GALR2 | 0.05673 | 0.07186 | 1.02994 | 0.00174 | 0.09085 | 0.21208 | 0.05270 | 0.00421 | 0.29486 | 0.00000 | 0.01239 | 0.01528 |
| GALR3 | 0.00650 | 0.02153 | 0.03006 | 0.01412 | 0.02924 | 0.01029 | 0.01870 | 0.01018 | 0.02694 | 0.00178 | 0.01695 | 0.02177 |
| GALT | 0.78640 | 2.81249 | 6.99366 | 4.51666 | 2.71310 | 0.70711 | 3.66700 | 0.87681 | 1.29624 | 0.43487 | 6.86920 | 1.49239 |
| GAMT | 0.00764 | 1.70410 | 0.43992 | 1.57709 | 1.16734 | 0.00000 | 0.36255 | 0.00052 | 0.42437 | 0.77776 | 1.71751 | 0.26320 |
| GAN | 1.16385 | 0.82478 | 3.09375 | 0.00000 | 2.46983 | 3.79665 | 1.82702 | 0.56229 | 0.85313 | 3.13508 | 1.41903 | 1.04048 |
| GANAB | 1.33460 | 14.77392 | 19.87436 | 12.72814 | 16.19017 | 7.22392 | 2.59786 | 0.33294 | 40.47506 | 10.45975 | 18.72622 | 14.53447 |
| GANC | 0.90205 | 1.16679 | 0.56593 | 0.24149 | 0.01563 | 1.48157 | 0.00000 | 1.63159 | 1.00860 | 0.00000 | 0.84546 | 1.62178 |
| GAP43 | 0.00244 | 0.01475 | 0.25757 | 0.22854 | 0.12807 | 0.03895 | 0.00407 | 0.22254 | 0.00000 | 0.00000 | 0.08051 | 0.12452 |
| GAPDH | 4.56964 | 55.30538 | 57.13079 | 45.01174 | 64.64247 | 50.37055 | 80.12284 | 8.38206 | 83.97675 | 27.94895 | 56.13983 | 33.85909 |
| GAPDHS | 0.00622 | 0.09168 | 0.11362 | 0.06406 | 0.18027 | 0.00000 | 0.02455 | 0.00608 | 0.13689 | 0.18513 | 0.17564 | 0.16077 |
| GAPVD1 | 3.81740 | 6.54054 | 13.13856 | 7.55657 | 7.15423 | 16.07515 | 16.54402 | 12.19682 | 10.52840 | 11.40078 | 13.13531 | 10.79946 |
| GAR1 | 0.06809 | 4.12536 | 11.58041 | 3.99163 | 10.13500 | 2.95061 | 18.40725 | 10.08674 | 9.53485 | 9.45536 | 9.59835 | 16.54003 |
| GAREM | 0.80406 | 0.40089 | 1.20040 | 2.52474 | 0.00000 | 0.52461 | 1.93622 | 0.13407 | 0.00000 | 2.34616 | 0.00000 | 0.55370 |
| GAREML | 1.55443 | 0.21423 | 0.06226 | 0.15659 | 0.25381 | 0.00000 | 0.06053 | 0.00000 | 0.15189 | 0.92390 | 0.00000 | 0.99200 |
| GARNL3 | 0.04960 | 0.00000 | 0.00000 | 0.06139 | 0.00000 | 0.04980 | 0.08963 | 0.25811 | 0.12479 | 0.13772 | 0.00000 | 0.00721 |
| GARS | 8.80002 | 13.35874 | 16.79948 | 20.78844 | 10.74717 | 15.48059 | 36.91383 | 5.19959 | 28.36511 | 5.40607 | 23.56140 | 9.81908 |
| GART | 0.34742 | 25.05901 | 18.84034 | 10.77215 | 14.34266 | 32.61178 | 24.48056 | 4.83994 | 30.23077 | 3.83512 | 33.78122 | 18.14548 |
| GAS1 | 0.00292 | 0.05502 | 0.00000 | 0.00000 | 0.19770 | 0.07537 | 0.14044 | 0.01025 | 0.00000 | 0.62301 | 0.00000 | 0.00000 |
| GAS2 | 0.00905 | 0.01438 | 0.03688 | 0.01344 | 0.01300 | 0.00000 | 0.01020 | 0.07322 | 0.02629 | 0.10102 | 0.05431 | 0.02903 |
| GAS2L1 | 0.00300 | 1.52423 | 0.10033 | 1.78114 | 2.82038 | 0.00000 | 0.18842 | 0.22871 | 1.31581 | 6.52426 | 0.02245 | 0.67994 |
| GAS2L1P2 | 0.00975 | 0.01523 | 0.01384 | 0.00923 | 0.01282 | 0.01933 | 0.03508 | 0.03846 | 0.03755 | 0.01574 | 0.00857 | 0.01453 |
| GAS2L2 | 0.00597 | 0.01557 | 0.01275 | 0.01185 | 0.01444 | 0.01073 | 0.02554 | 0.01046 | 0.02007 | 0.02945 | 0.01814 | 0.01705 |
| GAS2L3 | 0.62891 | 0.17255 | 4.98712 | 0.00000 | 1.14492 | 0.00000 | 0.25212 | 0.00103 | 4.86410 | 0.01683 | 0.00001 | 0.02558 |
| GAS5 | 3.46914 | 7.84508 | 7.52316 | 14.41100 | 15.32278 | 16.62567 | 11.35784 | 11.97658 | 15.80433 | 23.99165 | 2.96565 | 12.18726 |
| GAS5-AS1 | 0.01732 | 0.00000 | 0.13021 | 0.02633 | 0.00000 | 0.12679 | 0.10203 | 0.04600 | 0.03290 | 0.39547 | 0.06468 | 0.08160 |
| GAS6 | 0.10904 | 13.96642 | 5.50861 | 9.04739 | 15.83083 | 3.06404 | 2.21013 | 0.02202 | 4.76682 | 12.16799 | 7.69608 | 0.28259 |
| GAS6-AS1 | 0.06670 | 0.02237 | 0.00963 | 0.06032 | 0.06095 | 0.00000 | 0.00595 | 0.00344 | 0.05439 | 0.10317 | 0.00899 | 0.09983 |
| GAS6-AS2 | 0.01693 | 1.52365 | 0.03267 | 0.00000 | 0.00000 | 2.01513 | 0.00000 | 0.00497 | 0.00000 | 0.38522 | 0.00058 | 0.37426 |
| GAS7 | 5.51021 | 2.02385 | 0.31014 | 6.68666 | 15.97301 | 1.36848 | 5.28686 | 1.37610 | 0.00000 | 8.84632 | 5.03387 | 1.58967 |

| | | | | | | | | | | | | |
|---|---|---|---|---|---|---|---|---|---|---|---|---|
| **GAS8** | 0.81550 | 0.44201 | 0.39504 | 0.73524 | 0.00000 | 0.66083 | 0.05797 | 0.47878 | 0.23226 | 0.00000 | 0.71214 | 0.32170 |
| **GAST** | 0.09108 | 0.00733 | 12.31160 | 0.00159 | 0.00000 | 0.00631 | 0.24596 | 0.79052 | 0.55168 | 0.04955 | 0.00000 | 0.01059 |
| **GATA1** | 0.00612 | 0.01524 | 0.01460 | 0.01132 | 0.01474 | 0.01453 | 0.02007 | 0.01048 | 0.02851 | 0.02217 | 0.01244 | 0.01650 |
| **GATA2** | 1.61383 | 0.00000 | 48.41752 | 1.54695 | 52.53754 | 0.00000 | 45.10682 | 0.08081 | 35.27207 | 5.80374 | 23.31471 | 0.00000 |
| **GATA3** | 0.00000 | 15.09007 | 35.36693 | 0.00000 | 49.97756 | 3.91394 | 42.68319 | 0.58447 | 17.60652 | 30.71740 | 26.58636 | 0.00000 |
| **GATA3-AS1** | 0.59119 | 0.81062 | 2.83359 | 0.00000 | 0.57148 | 0.04659 | 1.22169 | 3.01124 | 0.27356 | 0.58035 | 0.79915 | 0.00000 |
| **GATA4** | 1.04548 | 0.00396 | 2.13312 | 7.61264 | 0.01829 | 0.00000 | 0.00919 | 0.22811 | 0.06990 | 0.00000 | 0.01755 | 0.06167 |
| **GATA5** | 0.00342 | 0.00693 | 0.11757 | 0.29034 | 0.10700 | 0.00000 | 0.14558 | 0.02148 | 0.00000 | 0.03933 | 0.00611 | 0.01474 |
| **GATA6** | 0.00000 | 20.71449 | 0.00000 | 7.70863 | 13.77163 | 18.09069 | 6.70503 | 8.56588 | 0.00000 | 27.55913 | 8.90807 | 28.09065 |
| **GATA6-AS1** | 0.00000 | 3.30414 | 0.00000 | 2.02003 | 0.67777 | 3.29514 | 0.02710 | 1.65091 | 1.82148 | 1.72978 | 2.09508 | 6.50350 |
| **GATAD1** | 0.00445 | 0.14930 | 3.49164 | 1.36049 | 3.06668 | 1.46970 | 7.67271 | 0.00000 | 0.02737 | 6.89291 | 1.50050 | 0.00000 |
| **GATAD2A** | 4.53833 | 5.98788 | 14.37713 | 8.04455 | 13.42713 | 2.50942 | 23.37451 | 3.56606 | 13.53632 | 17.69381 | 9.33205 | 9.31708 |
| **GATAD2B** | 3.76971 | 0.62274 | 0.47812 | 1.42693 | 1.27872 | 2.48870 | 2.03435 | 3.89185 | 2.12402 | 6.33891 | 1.92991 | 2.02388 |
| **GATC** | 0.97595 | 5.63006 | 7.50245 | 2.58987 | 5.33294 | 14.44618 | 8.34365 | 8.85772 | 10.14053 | 3.04219 | 5.63549 | 7.73239 |
| **GATM** | 0.68651 | 0.22480 | 0.59011 | 0.32114 | 0.00000 | 0.07467 | 0.28734 | 0.26740 | 0.26251 | 0.13205 | 0.70575 | 0.59748 |
| **GATS** | 2.30233 | 0.89658 | 1.04348 | 1.23199 | 1.34186 | 0.00000 | 0.57378 | 0.50754 | 0.53706 | 0.00000 | 0.98213 | 0.40515 |
| **GATSL2** | 0.00644 | 0.01490 | 0.01458 | 0.01266 | 0.01456 | 0.02489 | 0.02165 | 0.01070 | 0.02382 | 0.01229 | 0.01586 | 0.01758 |
| **GATSL3** | 0.45072 | 0.89840 | 3.71515 | 0.43556 | 1.01410 | 0.00000 | 0.63320 | 0.00000 | 0.00000 | 0.00000 | 0.63214 | 0.81381 |
| **GBA** | 0.05834 | 4.32457 | 3.48158 | 1.95854 | 2.78135 | 1.92457 | 0.00000 | 0.04706 | 6.74383 | 0.00000 | 3.52931 | 1.96596 |
| **GBA2** | 0.29305 | 2.52203 | 4.81805 | 5.97537 | 3.88368 | 0.00000 | 4.81883 | 0.48928 | 7.04972 | 5.40343 | 2.13448 | 2.32409 |
| **GBA3** | 0.00270 | 0.01148 | 0.03619 | 0.29387 | 0.00738 | 0.00393 | 0.01143 | 0.19727 | 0.03777 | 0.04941 | 0.01250 | 0.09216 |
| **GBAP1** | 0.04667 | 0.20090 | 0.15357 | 0.15594 | 0.00000 | 0.02801 | 0.06474 | 0.03298 | 0.09198 | 0.13992 | 0.22412 | 0.04890 |
| **GBAS** | 0.91021 | 2.38067 | 3.80442 | 5.92270 | 1.99647 | 8.15798 | 3.33159 | 0.48731 | 2.04117 | 0.00000 | 7.71749 | 0.40890 |
| **GBE1** | 0.51496 | 1.81266 | 2.44270 | 1.59963 | 0.17976 | 4.24771 | 0.62786 | 0.97773 | 0.22108 | 0.61602 | 1.59598 | 0.96272 |
| **GBF1** | 6.75829 | 7.43846 | 6.44332 | 7.23611 | 9.47660 | 0.00000 | 5.64892 | 0.52447 | 6.28141 | 7.39454 | 4.43681 | 6.30233 |
| **GBGT1** | 0.00233 | 0.00000 | 0.23876 | 0.00000 | 0.26060 | 0.00000 | 0.61300 | 0.00721 | 3.24568 | 0.39922 | 4.40161 | 0.00000 |
| **GBP1** | 0.00300 | 0.04308 | 0.10613 | 0.39869 | 0.02863 | 0.11096 | 0.00274 | 0.02833 | 0.00000 | 0.00226 | 0.01125 | 0.39364 |
| **GBP1P1** | 0.00348 | 0.18285 | 0.01376 | 0.00337 | 0.07285 | 0.08860 | 0.01806 | 0.05255 | 0.00000 | 0.07762 | 0.00081 | 0.22557 |
| **GBP2** | 0.00441 | 0.03976 | 0.13050 | 0.09103 | 0.07103 | 0.00000 | 0.04312 | 0.31066 | 0.00000 | 0.10568 | 0.00817 | 0.32657 |
| **GBP3** | 0.00587 | 0.00986 | 0.05076 | 0.00663 | 0.02161 | 0.03360 | 0.01934 | 0.00975 | 0.00000 | 0.01256 | 0.05816 | 0.02956 |
| **GBP4** | 0.00756 | 0.01535 | 0.53658 | 0.03445 | 0.01715 | 0.00004 | 0.00000 | 0.00978 | 0.00000 | 0.03184 | 0.01796 | 0.01978 |
| **GBP5** | 0.00543 | 0.00957 | 0.15781 | 0.00634 | 0.00000 | 0.05066 | 0.10799 | 0.01112 | 0.01379 | 0.01109 | 0.01091 | 0.01937 |
| **GBP6** | 0.24802 | 0.01473 | 0.01635 | 0.01673 | 0.02821 | 0.01185 | 0.02298 | 0.09196 | 0.02116 | 0.00315 | 0.01454 | 0.01007 |
| **GBP7** | 0.00630 | 0.02546 | 0.02826 | 0.01208 | 0.01562 | 0.01163 | 0.01929 | 0.01062 | 0.01498 | 0.01337 | 0.01757 | 0.01559 |
| **GBX1** | 0.34376 | 0.01493 | 0.01213 | 0.03767 | 0.02998 | 0.00000 | 0.04233 | 0.02320 | 0.01660 | 0.00000 | 0.01278 | 0.01313 |
| **GBX2** | 1.97036 | 0.15337 | 0.10222 | 1.28292 | 0.18829 | 0.00000 | 0.03389 | 1.61170 | 0.10023 | 0.06996 | 0.02388 | 1.87328 |
| **GC** | 0.07932 | 0.00774 | 0.01003 | 0.08437 | 0.00491 | 0.03330 | 0.01055 | 0.00390 | 0.02187 | 0.01648 | 0.01196 | 0.04883 |
| **GCA** | 2.74395 | 1.66705 | 2.69951 | 2.88524 | 0.00000 | 6.02641 | 2.03760 | 0.30389 | 3.17884 | 1.02168 | 1.07429 | 0.26649 |
| **GCAT** | 5.17762 | 6.59238 | 3.83901 | 5.88021 | 4.56407 | 0.90509 | 5.05077 | 1.68799 | 3.39547 | 0.00000 | 5.28249 | 7.67142 |
| **GCC1** | 3.79491 | 7.20025 | 4.33794 | 2.93529 | 5.94334 | 3.36946 | 7.10382 | 3.23076 | 8.30242 | 0.00000 | 3.56641 | 7.49932 |
| **GCC2** | 5.66834 | 2.49964 | 7.22473 | 3.20275 | 0.00000 | 20.78440 | 6.20279 | 3.41060 | 6.96962 | 17.45363 | 4.71403 | 3.98773 |
| **GCDH** | 0.81930 | 7.94729 | 6.96741 | 5.72612 | 7.51034 | 1.11549 | 9.89698 | 0.91255 | 7.77023 | 0.00000 | 13.47444 | 9.04944 |
| **GCFC2** | 1.72043 | 5.79625 | 2.86413 | 0.78633 | 0.73598 | 9.75389 | 1.21182 | 1.91704 | 1.12583 | 5.54362 | 3.42847 | 3.18294 |
| **GCG** | 0.00812 | 0.03179 | 0.02192 | 0.01156 | 0.02635 | 0.00471 | 0.02012 | 0.03032 | 0.02040 | 0.00714 | 0.01548 | 0.01465 |
| **GCGR** | 0.02523 | 0.17394 | 0.00040 | 1.19844 | 0.00000 | 0.00000 | 0.16946 | 0.36191 | 0.00000 | 0.07907 | 0.00481 | 0.03018 |
| **GCH1** | 4.88864 | 0.00000 | 2.03545 | 0.00000 | 0.00000 | 3.62948 | 20.92182 | 0.04547 | 18.68638 | 10.29714 | 13.54252 | 0.13384 |
| **GCHFR** | 1.54246 | 10.37017 | 1.59993 | 6.78225 | 3.65171 | 0.00000 | 0.45987 | 2.71324 | 0.52988 | 1.74645 | 1.72499 | 18.50032 |
| **GCK** | 0.00184 | 0.14049 | 0.00000 | 0.44988 | 0.61065 | 0.04808 | 0.05402 | 1.48131 | 0.00000 | 0.00000 | 0.14652 | 1.45942 |

| | | | | | | | | | | | | |
|---|---|---|---|---|---|---|---|---|---|---|---|---|
| GCKR | 0.01210 | 0.00812 | 0.19979 | 1.11325 | 0.00000 | 0.00000 | 0.00000 | 0.10832 | 0.01781 | 0.15759 | 0.01921 | 0.04685 |
| GCLC | 1.84198 | 0.28603 | 4.52818 | 0.16394 | 1.24920 | 9.89070 | 8.10825 | 0.00000 | 1.80052 | 5.79743 | 0.76584 | 0.00000 |
| GCLM | 0.25615 | 2.65448 | 10.86809 | 0.00000 | 6.73836 | 4.01037 | 11.54705 | 0.00000 | 2.65589 | 8.05129 | 5.39221 | 0.00000 |
| GCM1 | 0.01779 | 0.00000 | 61.31131 | 13.02771 | 0.00000 | 0.78273 | 22.18211 | 0.19453 | 4.94247 | 0.00000 | 0.00000 | 0.00000 |
| GCM2 | 0.23723 | 0.00445 | 0.03601 | 0.05519 | 0.00000 | 0.06695 | 0.51653 | 0.01659 | 0.00000 | 0.08886 | 0.08494 | 0.32789 |
| GCN1L1 | 2.55081 | 14.08486 | 12.64792 | 11.85983 | 18.33891 | 5.91162 | 16.47789 | 2.17285 | 18.25553 | 31.63783 | 9.44857 | 12.11497 |
| GCNT1 | 1.99554 | 0.00000 | 2.06568 | 3.76347 | 0.90241 | 1.67782 | 0.00000 | 6.68014 | 2.58187 | 0.00000 | 3.65787 | 1.22822 |
| GCNT2 | 0.03170 | 0.32941 | 6.25537 | 1.40303 | 3.73636 | 1.81136 | 4.51151 | 0.00156 | 5.55295 | 2.57629 | 4.13010 | 0.00000 |
| GCNT3 | 12.64351 | 0.48937 | 0.36330 | 2.30441 | 0.03740 | 0.00000 | 0.00000 | 3.33438 | 0.00000 | 0.02762 | 0.00392 | 2.70895 |
| GCNT4 | 0.00186 | 0.01121 | 0.01268 | 0.01195 | 0.01508 | 0.02893 | 0.01574 | 0.35569 | 0.01905 | 0.01221 | 0.01383 | 0.01938 |
| GCNT7 | 0.00896 | 0.00000 | 0.20995 | 0.04198 | 0.05113 | 0.00000 | 0.00012 | 0.20436 | 0.09960 | 0.95665 | 0.10441 | 0.11841 |
| GCOM1.uid-190 | 0.06008 | 0.01080 | 0.01442 | 0.01252 | 0.00845 | 0.03372 | 0.02217 | 0.00866 | 0.01580 | 0.01737 | 0.01671 | 0.02111 |
| GCSAM | 0.00234 | 0.11430 | 0.07923 | 0.04004 | 0.19916 | 0.32735 | 0.00000 | 0.00683 | 0.20889 | 0.16659 | 0.06207 | 0.00000 |
| GCSAML | 0.04183 | 0.01968 | 0.02102 | 0.01719 | 0.05845 | 0.00009 | 0.03132 | 0.02665 | 0.03411 | 0.00000 | 0.01957 | 0.09381 |
| GCSAML-AS1 | 0.00586 | 0.20793 | 0.00848 | 0.00806 | 0.01763 | 0.05596 | 0.01496 | 0.01034 | 0.00933 | 0.01077 | 0.00907 | 0.00810 |
| GCSH | 0.00000 | 1.96505 | 1.39524 | 1.23126 | 2.03386 | 2.43113 | 2.79188 | 0.21704 | 2.10001 | 1.67016 | 3.06041 | 1.19386 |
| GCSHP3 | 0.06851 | 0.18368 | 0.28035 | 0.13824 | 1.11253 | 0.05734 | 2.09895 | 0.00615 | 0.00002 | 3.32565 | 0.54178 | 0.24510 |
| GDA | 1.57333 | 0.00000 | 0.63651 | 0.00000 | 0.11724 | 0.66263 | 0.00000 | 0.38556 | 0.56119 | 0.71161 | 0.86491 | 0.23377 |
| GDAP1 | 15.32392 | 0.37924 | 1.36903 | 0.26262 | 0.00000 | 1.28504 | 0.25407 | 7.14497 | 1.25984 | 0.00000 | 0.01240 | 0.81076 |
| GDAP1L1 | 0.01074 | 0.01857 | 0.02326 | 0.01590 | 0.02608 | 0.00000 | 0.03085 | 0.01192 | 0.01835 | 0.02571 | 0.01463 | 0.02697 |
| GDAP2 | 1.03406 | 1.68766 | 6.28851 | 2.15879 | 0.00269 | 11.52095 | 2.93776 | 8.69897 | 4.30159 | 2.11245 | 3.39364 | 3.04250 |
| GDE1 | 0.34771 | 25.42612 | 11.43488 | 2.29867 | 20.40426 | 23.48848 | 4.84626 | 0.00000 | 19.31664 | 11.20071 | 11.42603 | 28.03314 |
| GDF11 | 0.01544 | 0.23624 | 1.72252 | 1.73239 | 0.95695 | 0.00000 | 3.27794 | 1.58570 | 0.00000 | 6.20337 | 0.36770 | 1.21995 |
| GDF15 | 0.60024 | 6.48254 | 11.99252 | 1.26167 | 12.41942 | 0.00000 | 0.00000 | 0.03044 | 6.29930 | 1.58282 | 0.00000 | 0.00000 |
| GDF3 | 0.15567 | 0.02530 | 6.01533 | 33.23197 | 0.00000 | 1.64309 | 0.00000 | 0.20243 | 1.74077 | 0.00000 | 0.28359 | 0.55187 |
| GDF5 | 0.00605 | 0.01362 | 0.02162 | 0.01964 | 0.01766 | 0.02697 | 0.01491 | 0.01121 | 0.01749 | 0.00788 | 0.01634 | 0.01670 |
| GDF6 | 0.11829 | 0.01176 | 0.04962 | 0.05786 | 0.01656 | 0.00000 | 0.06501 | 0.00874 | 0.00000 | 0.04884 | 0.01277 | 0.02166 |
| GDF7 | 0.00587 | 0.02424 | 0.01847 | 0.01546 | 0.01635 | 0.01388 | 0.02171 | 0.01161 | 0.02272 | 0.00947 | 0.01811 | 0.01882 |
| GDF9 | 27.63908 | 1.36052 | 1.22840 | 1.03514 | 2.06378 | 0.37537 | 1.49106 | 34.74759 | 1.35293 | 0.00000 | 1.84375 | 3.83529 |
| GDI1 | 5.15068 | 1.99112 | 3.48918 | 2.43973 | 3.91578 | 0.00000 | 3.80797 | 2.00400 | 2.54753 | 7.09083 | 1.93544 | 4.59049 |
| GDI2 | 18.30317 | 22.30447 | 21.39284 | 15.52152 | 18.17081 | 49.92908 | 33.29970 | 13.48500 | 29.36460 | 15.49638 | 26.40769 | 23.08556 |
| GDNF | 0.12109 | 0.01566 | 0.01509 | 0.00997 | 0.01538 | 0.02163 | 0.02363 | 0.11722 | 0.01475 | 0.02260 | 0.01397 | 0.01400 |
| GDNF-AS1 | 0.00590 | 0.01540 | 0.01288 | 0.01208 | 0.01236 | 0.01450 | 0.06582 | 0.01047 | 0.00000 | 0.02909 | 0.01529 | 0.01578 |
| GDPD1 | 0.00000 | 5.56138 | 0.00000 | 5.47966 | 0.00001 | 2.30518 | 0.00000 | 0.08754 | 1.89487 | 2.75559 | 0.00000 | 14.99264 |
| GDPD2 | 0.00074 | 12.45502 | 0.00000 | 7.90563 | 8.04953 | 1.10217 | 0.00000 | 0.00010 | 0.00000 | 0.15599 | 0.00000 | 0.05963 |
| GDPD3 | 0.01657 | 0.00000 | 0.00000 | 0.02090 | 2.83086 | 0.00000 | 0.12766 | 0.04848 | 7.83934 | 2.64508 | 6.77867 | 0.07819 |
| GDPD4 | 0.29961 | 0.02218 | 0.02360 | 0.08047 | 0.02032 | 0.00781 | 0.02287 | 0.02737 | 0.01102 | 0.02018 | 0.03139 | 0.13086 |
| GDPD5 | 0.20568 | 1.39038 | 1.54145 | 1.21713 | 9.61984 | 0.00000 | 0.77409 | 0.19588 | 1.53508 | 3.08785 | 0.57151 | 0.38248 |
| GDPGP1 | 0.00674 | 0.97276 | 0.46703 | 0.00000 | 0.02687 | 0.16724 | 0.07952 | 0.00855 | 0.35485 | 0.10707 | 0.47483 | 0.37458 |
| GEM | 0.85826 | 0.46840 | 0.00877 | 0.99105 | 0.02084 | 0.32402 | 0.00000 | 0.57923 | 0.00000 | 0.00810 | 0.00000 | 7.73035 |
| GEMIN2 | 3.63449 | 4.57923 | 3.21208 | 5.44937 | 1.32571 | 14.69935 | 2.31968 | 0.63784 | 3.30667 | 2.89050 | 6.03837 | 5.93831 |
| GEMIN4 | 0.00780 | 17.00064 | 9.98931 | 7.74681 | 9.10124 | 0.00029 | 11.61114 | 1.98143 | 19.80806 | 0.00000 | 14.37392 | 15.93596 |
| GEMIN5 | 19.17742 | 4.15531 | 2.19450 | 4.63942 | 3.79206 | 4.26586 | 9.02192 | 5.65497 | 5.45066 | 9.25798 | 9.54693 | 5.53432 |
| GEMIN6 | 0.00000 | 10.64579 | 7.52971 | 3.23832 | 4.99856 | 11.58865 | 14.41859 | 6.13867 | 10.73430 | 5.71446 | 13.81657 | 17.83864 |
| GEMIN7 | 1.31497 | 13.99131 | 2.71514 | 8.31541 | 7.32889 | 15.06759 | 5.97813 | 1.19766 | 3.27719 | 0.00000 | 4.20011 | 11.54091 |
| GEMIN8 | 0.46726 | 1.80200 | 3.75082 | 3.14914 | 2.31209 | 3.51518 | 3.24843 | 1.08469 | 1.93839 | 0.00000 | 5.40285 | 2.95563 |
| GEMIN8P4 | 0.00687 | 1.55441 | 6.06075 | 0.00000 | 0.24886 | 0.78021 | 0.51966 | 0.94885 | 1.54334 | 0.00000 | 0.16107 | 5.95377 |
| GEN1 | 0.08421 | 1.33261 | 1.23636 | 0.00000 | 0.87814 | 8.49810 | 1.65051 | 0.90172 | 3.09207 | 8.37698 | 2.13242 | 3.18364 |

| | | | | | | | | | | | | |
|---|---|---|---|---|---|---|---|---|---|---|---|---|
| **GET4** | 2.56093 | 12.68681 | 13.19148 | 8.47848 | 17.63847 | 0.00000 | 16.58547 | 7.14841 | 11.69070 | 16.90047 | 9.08925 | 3.53967 |
| **GFAP** | 0.01852 | 0.00925 | 0.03195 | 0.01305 | 0.07265 | 0.00018 | 0.02298 | 0.02243 | 0.06491 | 0.06969 | 0.02267 | 0.04123 |
| **GFER** | 0.00344 | 6.81069 | 6.03960 | 2.07715 | 8.32470 | 0.00000 | 5.26897 | 0.05344 | 0.00000 | 9.36218 | 4.77122 | 2.44513 |
| **GFI1** | 0.31914 | 0.62043 | 0.01869 | 0.00000 | 0.12795 | 0.29387 | 0.00000 | 0.01183 | 0.00000 | 0.04894 | 0.00000 | 0.00708 |
| **GFI1B** | 0.00611 | 0.01473 | 0.01061 | 0.06869 | 0.01655 | 0.01958 | 0.01633 | 0.01043 | 0.01840 | 0.01432 | 0.01533 | 0.01582 |
| **GFM1** | 3.34182 | 10.27882 | 9.44946 | 4.63506 | 1.93230 | 26.40147 | 5.02232 | 6.11300 | 16.21951 | 4.81962 | 16.09316 | 11.45578 |
| **GFM2** | 4.26527 | 5.17039 | 4.15888 | 5.22904 | 0.15127 | 12.18506 | 2.05403 | 6.91336 | 9.33219 | 0.00000 | 6.80674 | 2.59711 |
| **GFOD1** | 0.04803 | 0.31437 | 0.06185 | 0.12384 | 0.00000 | 0.02064 | 0.00023 | 0.60617 | 0.00000 | 0.25850 | 0.06140 | 0.08369 |
| **GFOD2** | 0.19000 | 3.51256 | 8.52792 | 2.03735 | 3.78440 | 0.00000 | 4.84805 | 5.71944 | 10.36059 | 0.00000 | 4.35530 | 7.48238 |
| **GFPT1** | 10.05662 | 7.31712 | 11.68961 | 2.11618 | 6.19026 | 24.33336 | 10.23084 | 3.79429 | 7.01795 | 5.21171 | 7.49287 | 2.87952 |
| **GFPT2** | 0.00859 | 0.03327 | 1.92931 | 0.63184 | 0.00000 | 0.19080 | 0.48856 | 1.96545 | 0.00000 | 1.24640 | 0.83071 | 1.00318 |
| **GFRA1** | 0.66835 | 0.00152 | 0.01842 | 0.00830 | 0.03007 | 0.01972 | 0.01221 | 0.12658 | 0.00000 | 0.01590 | 0.13729 | 0.11258 |
| **GFRA2** | 7.11635 | 0.20340 | 0.05510 | 0.06426 | 0.00000 | 0.00000 | 0.00000 | 0.61068 | 0.03072 | 0.18429 | 0.00000 | 0.14299 |
| **GFRA3** | 0.01012 | 0.05658 | 0.09973 | 0.09193 | 0.00000 | 0.07766 | 0.00000 | 0.29829 | 0.00000 | 0.34242 | 0.04514 | 0.00000 |
| **GFRAL** | 0.00239 | 0.01347 | 0.01760 | 0.01120 | 0.01396 | 0.02632 | 0.01736 | 0.04546 | 0.02005 | 0.00718 | 0.01413 | 0.02732 |
| **GFY** | 1.18873 | 0.01506 | 0.01113 | 0.07405 | 0.00911 | 0.05864 | 0.01662 | 0.20552 | 0.02314 | 0.00197 | 0.00784 | 0.03100 |
| **GGA1** | 1.70941 | 1.71561 | 4.28189 | 6.71935 | 9.23516 | 0.00000 | 5.79600 | 0.23664 | 9.13996 | 10.88509 | 4.35876 | 1.81784 |
| **GGA2** | 2.30224 | 12.75019 | 5.08895 | 10.79312 | 16.74192 | 1.99178 | 10.13125 | 2.22093 | 10.54377 | 18.64247 | 7.40699 | 4.83047 |
| **GGA3** | 4.82576 | 4.17699 | 3.88927 | 5.55015 | 8.78899 | 2.58849 | 3.02855 | 0.59530 | 3.22578 | 6.94549 | 8.88241 | 10.01052 |
| **GGACT** | 8.49398 | 0.48969 | 0.00294 | 0.17626 | 1.04883 | 0.08536 | 0.00000 | 0.42803 | 0.27302 | 0.08214 | 0.19469 | 0.10615 |
| **GGCT** | 2.67689 | 2.64500 | 13.02919 | 1.59870 | 3.37264 | 15.95278 | 23.72143 | 0.13015 | 14.63601 | 5.43974 | 12.03565 | 1.55929 |
| **GGCX** | 0.70479 | 2.59122 | 9.57231 | 2.14211 | 5.06562 | 3.87733 | 1.31406 | 0.91226 | 9.88852 | 2.21356 | 5.32664 | 0.00000 |
| **GGH** | 2.90078 | 4.37451 | 7.97277 | 0.24772 | 2.26835 | 17.24693 | 0.00000 | 0.92327 | 9.45919 | 3.53792 | 6.74192 | 4.29751 |
| **GGN** | 0.09700 | 0.03005 | 0.00000 | 0.10918 | 0.15508 | 0.00000 | 0.00059 | 0.05023 | 0.14238 | 0.10724 | 0.01399 | 0.01262 |
| **GGNBP1** | 0.01592 | 0.01710 | 0.02109 | 0.00811 | 0.00511 | 0.03399 | 0.00937 | 0.00943 | 0.01168 | 0.05685 | 0.01293 | 0.01720 |
| **GGNBP2** | 6.43299 | 11.36135 | 9.32942 | 9.08848 | 11.35892 | 12.41941 | 12.54896 | 17.72617 | 12.17136 | 21.81683 | 10.99337 | 7.37326 |
| **GGPS1** | 0.48669 | 3.47251 | 7.91077 | 3.18678 | 1.75528 | 12.45786 | 8.76346 | 6.99598 | 6.44688 | 0.00000 | 5.62597 | 8.49535 |
| **GGT1** | 0.08964 | 0.18301 | 0.96686 | 1.21876 | 0.03340 | 0.00000 | 0.00000 | 0.20332 | 0.03371 | 0.16791 | 0.01869 | 0.31186 |
| **GGT3P** | 0.01155 | 0.01372 | 0.02073 | 0.02139 | 0.01221 | 0.02260 | 0.01520 | 0.00968 | 0.01796 | 0.01691 | 0.01551 | 0.01724 |
| **GGT5** | 0.00370 | 0.05600 | 0.54219 | 0.00000 | 0.00000 | 0.08300 | 0.85705 | 0.80155 | 0.06128 | 0.23376 | 0.19978 | 0.37576 |
| **GGT6** | 1.33441 | 0.00479 | 0.02794 | 0.00805 | 0.00216 | 0.02493 | 0.01501 | 0.03046 | 0.00000 | 0.04975 | 0.06511 | 0.03099 |
| **GGT7** | 1.54235 | 0.34929 | 0.82198 | 1.18375 | 1.05700 | 0.00000 | 0.00000 | 0.31103 | 1.81072 | 0.25729 | 0.53736 | 0.88043 |
| **GGT8P** | 0.00717 | 0.01589 | 0.02242 | 0.01856 | 0.01725 | 0.00656 | 0.01615 | 0.01118 | 0.02159 | 0.01594 | 0.02799 | 0.01991 |
| **GGTA1P** | 0.04790 | 0.06863 | 0.04189 | 0.03404 | 0.01466 | 0.00307 | 0.01288 | 0.02709 | 0.01194 | 0.03988 | 0.01558 | 0.03449 |
| **GGTLC1** | 0.00611 | 0.03646 | 0.00972 | 0.00230 | 0.03637 | 0.00338 | 0.01052 | 0.00566 | 0.02010 | 0.09272 | 0.00992 | 0.48146 |
| **GGTLC2** | 0.00626 | 0.01269 | 0.01497 | 0.01099 | 0.01132 | 0.02567 | 0.01813 | 0.01131 | 0.01361 | 0.02631 | 0.01432 | 0.02237 |
| **GH1** | 0.00302 | 0.01700 | 0.01619 | 0.01365 | 0.01813 | 0.01055 | 0.02086 | 0.09738 | 0.01914 | 0.00747 | 0.01566 | 0.02433 |
| **GH2** | 0.11335 | 0.01497 | 0.02962 | 0.01123 | 0.01659 | 0.01007 | 0.01260 | 0.11032 | 0.02926 | 0.01197 | 0.01239 | 0.01611 |
| **GHDC** | 0.00245 | 0.06412 | 0.19635 | 0.49066 | 0.79086 | 0.00000 | 0.14276 | 0.00780 | 0.70035 | 0.55418 | 0.47815 | 0.00778 |
| **GHITM** | 2.19371 | 30.78836 | 30.79409 | 29.85074 | 22.61401 | 67.80978 | 43.09748 | 21.53150 | 38.41461 | 6.59864 | 38.71536 | 29.80921 |
| **GHR** | 0.00722 | 0.00000 | 0.16329 | 0.01668 | 0.00000 | 0.01896 | 0.17818 | 0.01051 | 0.79280 | 0.24456 | 0.21145 | 0.00591 |
| **GHRH** | 0.04209 | 0.01594 | 0.01386 | 0.01762 | 0.00848 | 0.02899 | 0.01620 | 0.01234 | 0.01414 | 0.02268 | 0.01690 | 0.01332 |
| **GHRHR** | 0.78208 | 0.01610 | 0.01404 | 0.01181 | 0.01417 | 0.01348 | 0.01559 | 0.05946 | 0.02527 | 0.00740 | 0.01712 | 0.00749 |
| **GHRL** | 0.00848 | 0.02253 | 0.09730 | 0.04545 | 0.01735 | 0.00000 | 0.00585 | 0.05803 | 0.28358 | 0.00000 | 0.07685 | 0.05348 |
| **GHRLOS** | 0.00000 | 0.22246 | 0.54172 | 0.13116 | 0.28278 | 1.32554 | 0.75553 | 0.23269 | 0.00000 | 3.86240 | 0.33023 | 0.42741 |
| **GID4** | 3.91772 | 1.19141 | 1.11220 | 2.53327 | 2.08206 | 3.23423 | 3.11663 | 0.00000 | 1.99809 | 3.62379 | 1.81548 | 1.94920 |
| **GID8** | 4.32956 | 10.75703 | 10.38010 | 8.93533 | 11.93568 | 8.94498 | 17.74043 | 3.58083 | 12.66592 | 7.23976 | 7.35900 | 8.27735 |
| **GIF** | 0.00722 | 0.07052 | 0.03161 | 0.01654 | 0.00000 | 0.06518 | 0.00000 | 0.00990 | 0.01690 | 0.07211 | 0.01552 | 0.01294 |

| | | | | | | | | | | | | |
|---|---|---|---|---|---|---|---|---|---|---|---|---|
| **GIGYF1** | 1.73910 | 2.92187 | 6.92760 | 9.17409 | 1.86050 | 0.00000 | 4.70648 | 1.25196 | 6.39334 | 20.84524 | 10.41877 | 3.43522 |
| **GIGYF2** | 6.07878 | 7.42399 | 9.00451 | 6.30417 | 10.61421 | 9.49830 | 11.64307 | 4.27723 | 10.46783 | 23.50952 | 8.41866 | 10.43741 |
| **GIMAP1** | 0.01196 | 0.01407 | 0.04701 | 0.01024 | 0.01582 | 0.01283 | 0.02926 | 0.04290 | 0.01033 | 0.02114 | 0.01566 | 0.03942 |
| **GIMAP6** | 0.00550 | 0.00000 | 0.01100 | 0.00428 | 0.06062 | 0.03617 | 0.00069 | 0.00623 | 0.01766 | 0.04187 | 0.09258 | 0.20238 |
| **GIMAP7** | 0.10289 | 0.09609 | 0.02173 | 0.10838 | 0.04378 | 0.00000 | 0.00459 | 0.12438 | 0.02217 | 0.05338 | 0.00917 | 0.04126 |
| **GIN1** | 0.06054 | 2.01801 | 1.25928 | 0.00000 | 0.93058 | 3.37552 | 0.00000 | 2.46774 | 1.31359 | 0.08126 | 1.12132 | 1.98005 |
| **GINM1** | 0.40949 | 3.95133 | 2.98831 | 0.33583 | 2.56971 | 7.32713 | 0.00000 | 0.66773 | 4.93254 | 0.82457 | 1.26283 | 2.98313 |
| **GINS1** | 2.41990 | 10.43585 | 6.09119 | 6.77742 | 6.18190 | 29.66084 | 13.59550 | 2.04710 | 11.77314 | 9.11356 | 12.29238 | 7.88427 |
| **GINS2** | 10.71423 | 22.39778 | 7.86082 | 5.21493 | 20.90398 | 25.35755 | 9.01394 | 6.01066 | 11.37025 | 1.53804 | 17.11877 | 14.29146 |
| **GINS3** | 2.76825 | 25.96400 | 0.82179 | 16.11332 | 3.66935 | 19.63256 | 1.75904 | 15.30131 | 10.97413 | 3.72995 | 3.98992 | 27.25510 |
| **GINS4** | 0.89109 | 7.56170 | 4.43073 | 6.33602 | 5.19546 | 10.65116 | 4.24161 | 6.23007 | 5.48179 | 3.93608 | 5.93668 | 9.61716 |
| **GIP** | 0.59889 | 1.92517 | 1.66630 | 0.10449 | 0.00000 | 0.25728 | 0.25518 | 13.23628 | 0.61644 | 0.00000 | 0.34312 | 10.20972 |
| **GIPC1** | 0.99874 | 1.19493 | 0.57014 | 1.38196 | 5.28142 | 0.00000 | 8.56787 | 0.14910 | 1.18863 | 7.19638 | 0.58005 | 0.57076 |
| **GIPC2** | 0.04334 | 1.82779 | 0.00000 | 1.48219 | 1.30591 | 1.66016 | 1.40672 | 0.82081 | 0.00000 | 1.46815 | 1.77803 | 0.97543 |
| **GIPC3** | 0.00387 | 0.01196 | 0.01892 | 0.01297 | 0.02008 | 0.00969 | 0.01841 | 0.01949 | 0.01871 | 0.01726 | 0.01645 | 0.06716 |
| **GIPR** | 0.11449 | 0.23115 | 0.03957 | 0.00000 | 0.00000 | 0.01025 | 0.00898 | 0.07424 | 0.01934 | 0.09577 | 0.04445 | 0.33339 |
| **GIT1** | 3.85616 | 6.80366 | 1.98519 | 3.73646 | 7.18673 | 0.00000 | 5.57952 | 0.00000 | 1.38464 | 8.92320 | 2.03390 | 3.66825 |
| **GIT2** | 19.39031 | 8.39501 | 0.84574 | 2.06553 | 1.39403 | 12.11730 | 3.31771 | 8.16627 | 1.05006 | 4.79432 | 7.32629 | 5.98957 |
| **GJA1** | 0.34254 | 8.69230 | 11.66105 | 16.67208 | 3.53450 | 49.61203 | 7.04433 | 2.09972 | 28.92681 | 66.86498 | 20.35237 | 26.72508 |
| **GJA10** | 0.00342 | 0.20495 | 0.05266 | 0.29507 | 0.00000 | 0.03644 | 0.03508 | 0.03038 | 0.05702 | 0.00000 | 0.00484 | 0.33519 |
| **GJA3** | 0.12463 | 0.01198 | 0.02035 | 0.00857 | 0.01110 | 0.02332 | 0.01921 | 0.20432 | 0.02951 | 0.01559 | 0.01238 | 0.02694 |
| **GJA4** | 0.02347 | 0.00549 | 0.09416 | 0.06021 | 0.00849 | 0.00000 | 0.00363 | 0.04838 | 0.00000 | 0.00972 | 0.26360 | 0.10277 |
| **GJA5** | 0.00291 | 0.00000 | 30.98321 | 1.91773 | 0.00000 | 0.31258 | 0.00000 | 0.00270 | 1.83437 | 0.24977 | 0.97433 | 0.00387 |
| **GJA8** | 0.00516 | 0.00572 | 0.01789 | 0.00695 | 0.00974 | 0.01851 | 0.00978 | 0.00777 | 0.00000 | 0.07399 | 0.05424 | 0.08301 |
| **GJA9-MYCBP.ui** | 0.20967 | 0.03682 | 0.06688 | 0.08252 | 0.05982 | 0.00000 | 0.00935 | 0.04798 | 0.05356 | 0.10244 | 0.01516 | 0.06601 |
| **GJB1** | 0.05527 | 0.00924 | 0.03752 | 0.40191 | 0.00000 | 0.00846 | 0.09299 | 0.13016 | 0.02492 | 0.02589 | 0.01197 | 0.02538 |
| **GJB2** | 0.05044 | 0.01346 | 0.16040 | 0.01197 | 0.02401 | 0.00947 | 0.00374 | 0.01070 | 0.00440 | 0.01826 | 0.01811 | 0.02470 |
| **GJB3** | 0.01016 | 0.00000 | 0.49892 | 0.08923 | 0.17549 | 0.07051 | 0.00000 | 0.04620 | 0.33112 | 0.00080 | 0.11127 | 0.03007 |
| **GJB4** | 0.00305 | 0.23722 | 0.14907 | 0.34035 | 0.00000 | 0.09973 | 0.00000 | 0.14355 | 0.00000 | 0.02171 | 0.00842 | 0.00295 |
| **GJB5** | 0.00279 | 0.25260 | 0.17423 | 0.58804 | 0.00000 | 0.23214 | 0.08428 | 0.02313 | 0.07671 | 0.13302 | 0.00000 | 0.00000 |
| **GJB6** | 0.00622 | 0.00992 | 0.03844 | 0.00917 | 0.00611 | 0.04425 | 0.01435 | 0.01008 | 0.00002 | 0.03760 | 0.01891 | 0.01310 |
| **GJB7** | 0.00120 | 0.61200 | 0.40027 | 0.00293 | 0.00000 | 1.50195 | 0.97749 | 0.04539 | 0.70676 | 0.00000 | 0.41148 | 0.52529 |
| **GJC1** | 6.79296 | 0.00000 | 1.98906 | 0.05292 | 2.82796 | 2.02908 | 6.07220 | 4.41443 | 11.30854 | 15.97541 | 8.07780 | 1.18433 |
| **GJC2** | 0.00481 | 0.00992 | 0.01541 | 0.05290 | 0.04745 | 0.00390 | 0.00040 | 0.01121 | 0.07401 | 0.09161 | 0.00384 | 0.01414 |
| **GJC3** | 0.00646 | 0.01806 | 0.02081 | 0.02742 | 0.01424 | 0.01290 | 0.01674 | 0.01055 | 0.02769 | 0.00615 | 0.01543 | 0.01730 |
| **GJD3** | 0.06410 | 0.05099 | 0.09488 | 0.01736 | 0.01166 | 0.00002 | 0.03943 | 0.00647 | 0.04552 | 0.09060 | 0.04901 | 0.00940 |
| **GJD4** | 0.00928 | 0.03531 | 0.02289 | 0.00296 | 0.09182 | 0.00000 | 0.00171 | 0.03016 | 0.15480 | 0.08255 | 0.00676 | 0.09918 |
| **GK** | 0.00158 | 0.13884 | 0.01906 | 1.50654 | 0.00000 | 0.23234 | 0.00000 | 0.03841 | 0.01155 | 0.25201 | 0.02151 | 0.20550 |
| **GK2** | 0.00624 | 0.01789 | 0.01913 | 0.01258 | 0.01342 | 0.01448 | 0.02089 | 0.01055 | 0.01109 | 0.01398 | 0.03217 | 0.01626 |
| **GK3P** | 0.00756 | 0.04741 | 0.02553 | 0.17052 | 0.00369 | 0.08986 | 0.00000 | 0.00928 | 0.02445 | 0.01564 | 0.01275 | 0.01981 |
| **GK5** | 0.25772 | 1.70498 | 2.00066 | 0.00000 | 2.52710 | 5.51126 | 2.47637 | 0.00000 | 0.00000 | 9.78666 | 4.81408 | 2.35799 |
| **GKAP1** | 1.48429 | 2.65569 | 0.00000 | 0.22352 | 3.03833 | 0.00000 | 0.28059 | 0.75656 | 0.00000 | 5.57878 | 0.00000 | 1.54164 |
| **GLA** | 0.69132 | 3.66418 | 17.98803 | 4.51458 | 12.41237 | 16.34673 | 8.59147 | 11.98917 | 40.54708 | 0.15656 | 11.06791 | 5.58634 |
| **GLB1** | 4.39527 | 6.85709 | 4.90390 | 9.53398 | 6.97281 | 11.49363 | 5.69355 | 5.04669 | 34.15291 | 0.12321 | 15.18353 | 3.01976 |
| **GLB1L** | 2.18678 | 0.27960 | 0.31268 | 0.17131 | 0.00000 | 0.00000 | 0.40504 | 0.99154 | 0.00000 | 0.55403 | 0.70384 | 1.01411 |
| **GLB1L2** | 1.91695 | 0.01708 | 0.20723 | 0.01164 | 0.00296 | 0.01395 | 0.11528 | 0.21443 | 0.35767 | 0.01530 | 0.00000 | 0.00855 |
| **GLB1L3** | 0.05778 | 0.25642 | 0.75805 | 0.00000 | 0.00000 | 1.03761 | 0.11070 | 0.22203 | 1.50286 | 1.42859 | 0.59280 | 0.48122 |
| **GLCCI1** | 2.79829 | 0.08920 | 0.17010 | 0.34963 | 0.41242 | 0.17151 | 1.80364 | 0.73770 | 0.00000 | 3.27141 | 0.08409 | 0.08115 |

| | | | | | | | | | | | | |
|---|---|---|---|---|---|---|---|---|---|---|---|---|
| **GLCE** | 0.63664 | 0.55477 | 0.76322 | 0.00000 | 0.00270 | 1.75907 | 0.00000 | 2.98101 | 1.10720 | 1.01768 | 0.41033 | 1.26955 |
| **GLDC** | 0.00563 | 0.00000 | 8.36975 | 12.87152 | 11.68355 | 5.78241 | 11.84307 | 0.03641 | 7.17484 | 8.49501 | 3.03681 | 0.00000 |
| **GLDN** | 1.52360 | 0.00588 | 1.77198 | 0.47340 | 0.01522 | 0.00000 | 0.00000 | 0.03983 | 0.30219 | 0.00349 | 0.00097 | 0.07020 |
| **GLE1** | 1.90783 | 8.09508 | 5.86055 | 2.68638 | 5.80982 | 3.88231 | 7.31606 | 6.86972 | 7.20030 | 0.00000 | 6.90896 | 8.45729 |
| **GLG1** | 2.63306 | 11.72558 | 8.40136 | 9.26268 | 15.14254 | 6.71454 | 1.26498 | 9.32496 | 19.58993 | 23.62517 | 9.33638 | 12.51414 |
| **GLI1** | 0.00418 | 0.09136 | 0.11800 | 0.03104 | 0.03902 | 0.00000 | 0.16022 | 0.00869 | 0.06393 | 0.16638 | 0.01369 | 0.03661 |
| **GLI2** | 0.00295 | 0.00113 | 0.32561 | 0.06461 | 0.00000 | 0.01895 | 0.00000 | 0.01173 | 0.00000 | 1.04396 | 0.00000 | 0.00455 |
| **GLI3** | 0.05259 | 0.00000 | 0.02917 | 0.25834 | 0.00000 | 0.00000 | 1.07351 | 0.07476 | 0.29680 | 0.07342 | 0.30236 | 0.03026 |
| **GLI4** | 0.12518 | 0.36122 | 0.00000 | 0.18800 | 0.02438 | 0.00000 | 0.00131 | 0.00000 | 0.29420 | 0.98607 | 0.07910 | 0.25675 |
| **GLIDR** | 0.00502 | 0.06311 | 0.04610 | 0.01508 | 0.04105 | 0.08868 | 0.02889 | 0.01207 | 0.11096 | 0.00000 | 0.03102 | 0.12494 |
| **GLIPR1** | 2.49566 | 0.11653 | 1.35353 | 0.10170 | 0.00000 | 0.58512 | 0.47911 | 0.30567 | 0.30625 | 0.00000 | 0.00000 | 0.22176 |
| **GLIPR1L1** | 0.02529 | 0.01510 | 0.04517 | 0.01433 | 0.01501 | 0.01763 | 0.01252 | 0.01053 | 0.03426 | 0.00574 | 0.02442 | 0.01637 |
| **GLIPR1L2** | 0.00765 | 0.01137 | 0.04259 | 0.02200 | 0.00920 | 0.03088 | 0.00787 | 0.01050 | 0.02943 | 0.02236 | 0.01734 | 0.01610 |
| **GLIPR2** | 0.00000 | 29.58609 | 0.00000 | 5.73347 | 25.84475 | 13.71435 | 12.78940 | 0.16976 | 10.21953 | 10.75882 | 28.39520 | 12.68120 |
| **GLIS1** | 0.01504 | 0.00909 | 0.03532 | 0.00437 | 0.00671 | 0.03995 | 0.00015 | 0.01931 | 0.02257 | 0.07900 | 0.01665 | 0.01396 |
| **GLIS2** | 0.65017 | 0.58243 | 0.33672 | 0.75877 | 0.93698 | 0.00000 | 0.00000 | 0.01814 | 0.86479 | 1.23610 | 0.00000 | 0.95190 |
| **GLIS3** | 0.06091 | 0.00000 | 0.43790 | 0.00000 | 0.15435 | 0.00000 | 0.89016 | 0.00508 | 0.95625 | 0.52242 | 0.84873 | 0.23603 |
| **GLIS3-AS1** | 0.14432 | 0.01798 | 0.01976 | 0.00935 | 0.02263 | 0.02196 | 0.01526 | 0.00748 | 0.03080 | 0.00455 | 0.01574 | 0.02027 |
| **GLMN** | 0.09929 | 3.03063 | 5.27242 | 0.69725 | 0.00000 | 13.35150 | 8.21952 | 3.55491 | 4.96333 | 3.58078 | 5.04293 | 5.68285 |
| **GLO1** | 2.85786 | 8.22428 | 19.72967 | 7.44070 | 4.78299 | 44.34070 | 30.07139 | 4.54969 | 28.31490 | 8.74263 | 22.45827 | 24.93225 |
| **GLOD4** | 6.70158 | 18.16467 | 11.72972 | 8.71838 | 10.58286 | 26.85064 | 17.84319 | 1.33098 | 24.27941 | 4.67750 | 19.29642 | 15.48759 |
| **GLP2R** | 0.00459 | 0.01571 | 0.01487 | 0.01193 | 0.01760 | 0.01694 | 0.01891 | 0.14554 | 0.01759 | 0.01239 | 0.01502 | 0.01350 |
| **GLRA1** | 0.00430 | 0.04536 | 0.00907 | 0.01842 | 0.02059 | 0.00000 | 0.00689 | 0.00901 | 0.05738 | 0.11416 | 0.00815 | 0.03997 |
| **GLRA2** | 0.59571 | 0.02870 | 0.07922 | 0.03308 | 0.03147 | 0.00000 | 0.06850 | 0.13651 | 0.00000 | 0.00000 | 0.07488 | 0.05493 |
| **GLRA3** | 0.00214 | 0.04845 | 0.03941 | 0.00565 | 0.03029 | 0.01511 | 0.01697 | 0.05779 | 0.00000 | 0.01384 | 0.07059 | 0.01081 |
| **GLRA4** | 0.02439 | 0.02313 | 0.06166 | 0.00508 | 0.13122 | 0.02060 | 0.05208 | 0.10277 | 0.17513 | 0.00000 | 0.02778 | 0.02582 |
| **GLRB** | 0.01652 | 0.19206 | 0.03184 | 0.00000 | 0.01740 | 0.08232 | 0.00000 | 0.00595 | 0.00000 | 0.02640 | 0.05789 | 0.24415 |
| **GLRX** | 3.33074 | 0.00000 | 0.00000 | 2.05681 | 0.10062 | 0.00460 | 1.00258 | 0.28740 | 5.94731 | 0.00000 | 18.52350 | 0.00000 |
| **GLRX2** | 0.11565 | 3.09673 | 5.75364 | 1.71846 | 1.92433 | 11.94735 | 7.00367 | 0.58781 | 3.41691 | 0.92164 | 3.52313 | 4.24600 |
| **GLRX3** | 1.11816 | 12.41582 | 21.49657 | 7.49696 | 5.40578 | 32.00469 | 24.07818 | 7.07138 | 21.59589 | 0.00000 | 16.48067 | 14.45004 |
| **GLRX5** | 0.88739 | 4.80829 | 11.25921 | 4.92585 | 10.53514 | 9.18722 | 23.14595 | 3.86365 | 8.20120 | 26.05152 | 8.55487 | 3.96212 |
| **GLS** | 0.00000 | 0.00000 | 10.30029 | 1.18654 | 0.00000 | 3.74316 | 10.07394 | 5.79323 | 3.76780 | 12.17061 | 1.84960 | 1.10707 |
| **GLS2** | 0.40962 | 0.55050 | 0.07325 | 1.78987 | 0.12290 | 0.00000 | 0.00771 | 0.34793 | 0.04818 | 0.00118 | 0.06955 | 0.05873 |
| **GLT1D1** | 0.50183 | 0.26274 | 0.03791 | 1.83884 | 2.52375 | 0.01626 | 0.30039 | 0.03747 | 0.22613 | 0.00000 | 0.96280 | 0.00000 |
| **GLT8D1** | 1.74617 | 4.70844 | 5.10602 | 1.38521 | 0.00000 | 4.16147 | 0.00000 | 0.94547 | 4.49784 | 4.69762 | 2.64407 | 0.86458 |
| **GLT8D2** | 0.03282 | 0.00000 | 0.08817 | 0.01719 | 0.00000 | 0.04605 | 0.15196 | 0.00576 | 0.14021 | 0.12571 | 0.01721 | 0.09198 |
| **GLTP** | 0.14962 | 0.15835 | 4.10530 | 0.00000 | 7.42278 | 5.67785 | 2.80653 | 2.14627 | 0.00000 | 1.63544 | 5.97095 | 0.36549 |
| **GLTPD2** | 0.07395 | 0.71249 | 0.00000 | 0.20494 | 0.65067 | 0.00000 | 0.00179 | 0.17293 | 0.21915 | 0.75879 | 0.03559 | 0.24705 |
| **GLTSCR1** | 1.42370 | 0.12228 | 1.09780 | 0.36334 | 0.47405 | 0.00000 | 0.24689 | 0.00000 | 0.78720 | 5.49237 | 0.00000 | 0.96196 |
| **GLTSCR1L** | 0.85899 | 0.31912 | 1.04749 | 0.57646 | 0.00000 | 1.71179 | 0.78754 | 0.00963 | 1.35773 | 2.43752 | 0.99814 | 0.04156 |
| **GLTSCR2** | 3.14976 | 3.35089 | 5.06142 | 2.93999 | 8.73208 | 0.00000 | 9.57853 | 4.27187 | 0.00000 | 24.94102 | 1.46172 | 4.50859 |
| **GLUD1** | 0.53378 | 11.96906 | 2.09126 | 10.16842 | 4.80556 | 23.03189 | 4.75871 | 3.03352 | 0.00000 | 24.42451 | 3.54359 | 12.97282 |
| **GLUD1P3** | 0.30321 | 0.00000 | 0.39857 | 0.00000 | 0.29473 | 0.00000 | 0.00000 | 0.00000 | 0.58498 | 1.90730 | 0.00000 | 0.43719 |
| **GLUD2** | 0.12054 | 3.79714 | 0.75227 | 1.98457 | 0.56360 | 3.40834 | 0.00004 | 0.72130 | 1.34648 | 0.00000 | 0.68379 | 2.66613 |
| **GLUL** | 0.76981 | 36.55964 | 17.08608 | 32.83014 | 10.57769 | 31.80815 | 31.80296 | 40.56538 | 39.34131 | 0.00000 | 17.29040 | 42.78482 |
| **GLYAT** | 0.00628 | 0.01265 | 0.10344 | 0.00533 | 0.01113 | 0.03798 | 0.01303 | 0.01019 | 0.01865 | 0.00954 | 0.01206 | 0.01511 |
| **GLYATL1** | 7.07346 | 0.00000 | 0.01073 | 0.01185 | 0.01515 | 0.00000 | 0.00910 | 1.29427 | 0.00000 | 0.07078 | 0.07476 | 0.04981 |
| **GLYATL3** | 0.28672 | 0.01082 | 0.01491 | 0.01256 | 0.01452 | 0.00999 | 0.01614 | 0.00449 | 0.01956 | 0.02585 | 0.01430 | 0.04029 |

| | | | | | | | | | | | | |
|---|---|---|---|---|---|---|---|---|---|---|---|---|
| **GLYCTK** | 0.00473 | 2.14280 | 1.04751 | 2.08180 | 1.37720 | 0.23944 | 0.64979 | 0.07070 | 0.36747 | 0.85937 | 2.10724 | 1.89169 |
| **GLYCTK-AS1** | 0.00593 | 0.06469 | 0.04405 | 0.01629 | 0.00270 | 0.00342 | 0.00610 | 0.01008 | 0.01782 | 0.03314 | 0.06188 | 0.02513 |
| **GLYR1** | 1.61028 | 4.02590 | 8.19398 | 4.13877 | 8.73311 | 0.00000 | 11.13410 | 2.39822 | 8.45282 | 15.99959 | 9.47938 | 5.59228 |
| **GM2A** | 3.21950 | 0.78311 | 6.17092 | 0.00000 | 28.00811 | 0.00000 | 6.38279 | 2.06035 | 23.61979 | 10.81690 | 16.70210 | 1.14779 |
| **GMCL1** | 1.60380 | 1.85652 | 2.87293 | 1.82994 | 0.57434 | 1.07135 | 1.25888 | 1.46886 | 4.29054 | 0.00000 | 4.01035 | 2.20503 |
| **GMCL1P1** | 0.11161 | 0.09070 | 0.23667 | 0.14032 | 0.00000 | 0.24206 | 0.07951 | 0.13302 | 0.30033 | 0.00000 | 0.38133 | 0.17121 |
| **GMDS** | 1.65190 | 2.52666 | 4.21289 | 2.24790 | 3.16093 | 0.26923 | 1.25770 | 0.80148 | 2.49376 | 0.00000 | 4.69465 | 0.46338 |
| **GMDS-AS1** | 0.09548 | 0.02324 | 0.02118 | 0.00429 | 0.00952 | 0.03509 | 0.00585 | 0.16232 | 0.02737 | 0.03455 | 0.02281 | 0.08758 |
| **GMEB1** | 1.92986 | 2.76958 | 10.48383 | 4.19555 | 2.75972 | 4.76711 | 6.28783 | 3.48865 | 4.66097 | 0.00000 | 4.17088 | 3.59278 |
| **GMEB2** | 0.72809 | 1.92450 | 3.83211 | 2.59798 | 0.90175 | 0.00000 | 2.26541 | 0.63108 | 1.62588 | 6.54131 | 0.68553 | 3.42035 |
| **GMFB** | 7.04813 | 5.76193 | 0.00000 | 10.76013 | 6.49798 | 25.25393 | 0.00000 | 8.90132 | 0.00000 | 0.75261 | 1.58394 | 11.06204 |
| **GMFG** | 0.00600 | 0.03186 | 0.11680 | 0.07433 | 0.06310 | 0.02721 | 0.06242 | 0.03173 | 0.00000 | 0.00000 | 0.05884 | 0.08344 |
| **GMIP** | 0.00000 | 1.33925 | 0.59308 | 0.53558 | 0.00000 | 0.00000 | 0.95236 | 0.05194 | 0.16166 | 2.00446 | 0.51759 | 0.70285 |
| **GML** | 1.17585 | 0.01406 | 0.02051 | 0.01262 | 0.01675 | 0.00516 | 0.04203 | 0.18205 | 0.00565 | 0.00753 | 0.01547 | 0.00517 |
| **GMNC** | 0.06011 | 0.00693 | 0.01220 | 0.01827 | 0.02615 | 0.03722 | 0.02239 | 0.00559 | 0.01025 | 0.00000 | 0.04803 | 0.08379 |
| **GMNN** | 6.04398 | 17.33510 | 18.04411 | 11.81398 | 10.72962 | 43.61110 | 24.30697 | 21.65664 | 15.98858 | 0.00000 | 19.57290 | 22.05525 |
| **GMPPA** | 0.50971 | 9.61819 | 8.02803 | 6.03710 | 6.91350 | 0.12705 | 6.86956 | 0.87809 | 4.54404 | 1.98425 | 9.58921 | 4.32799 |
| **GMPPB** | 0.57842 | 4.99015 | 10.40351 | 3.50755 | 4.97215 | 0.00000 | 1.16672 | 0.41686 | 2.54570 | 0.18196 | 8.03792 | 1.22105 |
| **GMPR** | 0.74164 | 0.10150 | 0.04783 | 0.30709 | 0.00163 | 0.06025 | 0.00043 | 0.19742 | 0.00601 | 0.00896 | 0.01288 | 0.22959 |
| **GMPR2** | 1.13705 | 3.39974 | 10.10656 | 5.41601 | 0.00000 | 7.61637 | 10.51568 | 3.52949 | 9.56028 | 0.00000 | 12.04961 | 7.10262 |
| **GMPS** | 0.48866 | 15.50722 | 10.64403 | 3.81447 | 2.34821 | 31.68980 | 11.69290 | 8.89188 | 19.25765 | 0.00000 | 15.59906 | 23.59572 |
| **GNA11** | 4.84048 | 4.21416 | 2.53371 | 3.00207 | 9.16170 | 0.16328 | 5.19286 | 1.59932 | 0.02416 | 20.79203 | 0.73937 | 8.45213 |
| **GNA12** | 8.38756 | 1.39784 | 29.30782 | 1.15508 | 0.00000 | 2.38261 | 23.68288 | 0.64642 | 3.90562 | 0.00000 | 0.00000 | 3.35618 |
| **GNA13** | 10.51170 | 8.62626 | 6.75685 | 10.68585 | 0.00000 | 24.19149 | 10.35658 | 7.98876 | 1.03057 | 13.96770 | 2.91556 | 8.67945 |
| **GNA14** | 7.53053 | 0.04376 | 0.00000 | 0.30448 | 0.00000 | 0.12104 | 0.00000 | 0.35008 | 0.11868 | 0.00000 | 0.00000 | 0.33968 |
| **GNA15** | 0.00868 | 0.00594 | 0.05879 | 0.06847 | 0.00000 | 0.03347 | 0.00778 | 0.01065 | 0.00000 | 0.10161 | 0.01901 | 0.01120 |
| **GNAI1** | 8.03649 | 1.76008 | 7.00416 | 0.00000 | 0.00563 | 3.62365 | 0.18843 | 3.87927 | 0.00000 | 0.00000 | 0.00000 | 1.27981 |
| **GNAI2** | 2.55147 | 1.13062 | 1.38169 | 8.69287 | 7.33419 | 0.00000 | 5.93018 | 4.45069 | 0.00000 | 11.92668 | 0.00000 | 2.51611 |
| **GNAI3** | 4.91338 | 14.77605 | 20.23359 | 7.11567 | 1.42176 | 39.10440 | 23.02663 | 22.15631 | 18.18348 | 0.00000 | 14.77693 | 16.47327 |
| **GNAL** | 0.00000 | 0.94023 | 1.52212 | 0.28365 | 0.89460 | 1.00048 | 1.30945 | 3.10950 | 1.08973 | 1.21119 | 0.98527 | 2.66839 |
| **GNAO1** | 0.88244 | 0.02188 | 0.03833 | 0.02017 | 0.03564 | 0.00000 | 0.19082 | 0.11740 | 0.00000 | 0.22479 | 0.00000 | 0.27218 |
| **GNAQ** | 3.12468 | 0.17436 | 0.62313 | 1.81740 | 1.54112 | 9.73885 | 1.90397 | 0.34647 | 0.00000 | 9.58822 | 0.54305 | 0.98184 |
| **GNAS** | 4.21973 | 2.89039 | 19.13336 | 19.60905 | 28.39139 | 11.81192 | 31.94542 | 1.45494 | 15.41132 | 27.12401 | 10.59609 | 2.12654 |
| **GNAS-AS1** | 0.05212 | 0.15611 | 0.28376 | 0.19870 | 0.12427 | 0.00000 | 0.05679 | 0.01391 | 0.00000 | 0.34062 | 0.16770 | 0.20910 |
| **GNAT1** | 0.00585 | 0.01246 | 0.02122 | 0.01128 | 0.00964 | 0.01441 | 0.02430 | 0.01039 | 0.00000 | 0.03487 | 0.07253 | 0.01653 |
| **GNAT2** | 2.21240 | 0.02700 | 0.23325 | 0.00421 | 0.08159 | 0.00000 | 0.37467 | 0.67530 | 0.22888 | 0.00000 | 0.47299 | 0.27338 |
| **GNAT3** | 0.02032 | 0.01415 | 0.01894 | 0.00676 | 0.01357 | 0.02309 | 0.01836 | 0.00972 | 0.00645 | 0.04949 | 0.02306 | 0.01520 |
| **GNAZ** | 0.25673 | 0.02606 | 0.01256 | 0.12234 | 0.05515 | 0.00000 | 0.04076 | 0.00430 | 0.01912 | 0.00000 | 0.01532 | 0.04135 |
| **GNB1** | 8.29239 | 9.92625 | 9.25899 | 3.62003 | 14.89298 | 12.69926 | 21.92193 | 0.32883 | 6.51802 | 35.98431 | 5.30396 | 13.61000 |
| **GNB1L** | 0.05663 | 5.58689 | 5.51504 | 3.30103 | 6.55921 | 0.00000 | 7.16785 | 0.00000 | 11.40148 | 0.97862 | 8.29129 | 4.51821 |
| **GNB2** | 3.66532 | 6.25101 | 9.00367 | 7.09498 | 15.26870 | 0.00000 | 15.14709 | 1.33090 | 0.09246 | 26.56181 | 2.16493 | 5.30816 |
| **GNB2L1** | 3.20680 | 46.57005 | 43.75791 | 41.48503 | 51.64976 | 35.48502 | 69.19680 | 28.44987 | 60.36027 | 32.82628 | 41.18973 | 60.47911 |
| **GNB3** | 0.19993 | 0.19973 | 0.15009 | 0.25188 | 0.00000 | 0.33023 | 0.07508 | 0.05645 | 0.28494 | 0.00000 | 0.34074 | 0.27839 |
| **GNB4** | 3.68823 | 3.50722 | 8.88373 | 0.00000 | 0.00000 | 2.21267 | 8.09377 | 2.17820 | 0.14677 | 1.00074 | 1.73044 | 5.53297 |
| **GNB5** | 1.43484 | 1.52885 | 2.27791 | 2.88118 | 0.17987 | 3.41144 | 1.75804 | 0.80200 | 0.77919 | 0.00000 | 0.52189 | 2.00582 |
| **GNE** | 0.39968 | 0.00000 | 7.63467 | 0.01412 | 0.40873 | 0.00000 | 1.02496 | 0.07104 | 0.00000 | 1.26183 | 0.25558 | 0.49632 |
| **GNG10** | 0.03167 | 2.83561 | 0.12510 | 0.47632 | 2.29523 | 1.32232 | 0.37800 | 0.00000 | 1.54599 | 0.00000 | 1.18709 | 0.34148 |
| **GNG11** | 0.49837 | 0.02059 | 0.05729 | 0.02012 | 0.00416 | 0.02198 | 0.01264 | 0.72745 | 0.00854 | 0.03276 | 0.01383 | 0.00310 |

| | | | | | | | | | | | | |
|---|---|---|---|---|---|---|---|---|---|---|---|---|
| **GNG12** | 7.10148 | 0.16538 | 10.16056 | 0.14984 | 0.00000 | 3.80667 | 2.87029 | 3.27563 | 0.00519 | 1.97819 | 1.60891 | 1.44487 |
| **GNG12-AS1** | 0.36181 | 0.00462 | 0.12919 | 0.01144 | 0.14238 | 0.02805 | 0.00000 | 0.04086 | 0.01975 | 0.04606 | 0.04592 | 0.05347 |
| **GNG13** | 0.04344 | 0.01552 | 0.01098 | 0.01156 | 0.01501 | 0.00464 | 0.02067 | 0.01755 | 0.03351 | 0.04747 | 0.01819 | 0.01682 |
| **GNG2** | 4.39678 | 0.00725 | 1.34489 | 0.43151 | 0.00000 | 0.00000 | 0.00000 | 0.65189 | 0.10416 | 0.05145 | 0.00037 | 0.05460 |
| **GNG3** | 0.00162 | 0.26819 | 0.08192 | 0.13401 | 0.14067 | 0.19512 | 0.07493 | 2.19369 | 0.26442 | 0.35808 | 0.00000 | 0.40824 |
| **GNG4** | 0.60373 | 0.95614 | 0.17007 | 0.00120 | 0.44175 | 0.00000 | 0.00443 | 0.00000 | 0.08671 | 0.00000 | 0.06664 | 0.12237 |
| **GNG5** | 5.14663 | 17.85651 | 15.63835 | 9.61191 | 15.37762 | 30.03502 | 19.93923 | 12.29338 | 17.54246 | 5.85090 | 17.31769 | 32.38146 |
| **GNG7** | 0.39742 | 0.87164 | 0.02067 | 0.22622 | 0.09127 | 0.04481 | 0.00615 | 0.00758 | 0.00000 | 0.01271 | 0.00714 | 0.61666 |
| **GNG8** | 0.00352 | 0.01700 | 0.01437 | 0.01231 | 0.01545 | 0.01650 | 0.01920 | 0.03906 | 0.01812 | 0.01337 | 0.01526 | 0.01474 |
| **GNGT1** | 0.21584 | 0.02830 | 0.01270 | 0.04716 | 0.00006 | 0.03530 | 0.00505 | 0.09180 | 0.04393 | 0.04030 | 0.00736 | 0.00509 |
| **GNGT2** | 0.03732 | 0.05199 | 0.03220 | 0.02113 | 0.24036 | 0.00001 | 0.01398 | 0.39484 | 0.01821 | 0.00000 | 0.04540 | 0.14231 |
| **GNL1** | 1.48559 | 14.52385 | 18.19024 | 13.26505 | 21.60118 | 0.00000 | 21.24372 | 10.10674 | 18.28163 | 0.00000 | 14.67763 | 12.10221 |
| **GNL2** | 0.78946 | 19.26771 | 15.20782 | 12.27636 | 10.30710 | 27.54613 | 20.58501 | 14.95164 | 25.11549 | 0.00000 | 18.94123 | 22.77196 |
| **GNL3** | 6.55743 | 22.28781 | 21.85001 | 15.39933 | 17.92153 | 40.19613 | 29.37695 | 11.51753 | 40.08378 | 4.91769 | 28.01272 | 30.35788 |
| **GNL3L** | 0.12400 | 1.24013 | 7.03819 | 3.06834 | 5.86374 | 3.27425 | 9.98009 | 3.19451 | 7.52865 | 17.42089 | 9.93414 | 7.46933 |
| **GNLY** | 0.00599 | 0.13064 | 0.01418 | 0.00508 | 0.00000 | 0.00000 | 0.00858 | 0.00476 | 0.01528 | 0.10812 | 0.03406 | 0.32778 |
| **GNMT** | 0.00885 | 0.00302 | 0.06771 | 0.19872 | 0.07094 | 0.13789 | 0.00000 | 0.06985 | 0.05075 | 0.03793 | 0.02085 | 0.02984 |
| **GNN** | 0.88419 | 0.09014 | 0.07145 | 0.00000 | 0.01942 | 0.00000 | 0.00000 | 0.03882 | 0.02695 | 0.40477 | 0.06395 | 0.18167 |
| **GNPAT** | 6.01136 | 2.59166 | 6.95017 | 3.81372 | 0.00000 | 8.96052 | 8.93372 | 2.43557 | 5.83532 | 0.53654 | 5.23219 | 2.61139 |
| **GNPDA1** | 3.41408 | 15.17212 | 3.69195 | 9.12578 | 20.53991 | 19.83886 | 22.99675 | 0.00000 | 25.66307 | 3.73259 | 23.85092 | 11.66347 |
| **GNPDA2** | 6.37982 | 0.91974 | 3.73456 | 0.02922 | 0.00000 | 2.39384 | 2.72880 | 2.68485 | 2.06665 | 0.00000 | 1.06188 | 2.56698 |
| **GNPNAT1** | 0.00000 | 4.66144 | 4.27770 | 7.17242 | 4.76687 | 30.29797 | 15.40736 | 8.05898 | 0.82798 | 17.65195 | 20.26927 | 8.56327 |
| **GNPTAB** | 5.91441 | 3.13213 | 0.47403 | 2.38600 | 4.60083 | 14.28917 | 0.00000 | 3.40020 | 0.00000 | 13.20831 | 1.27715 | 6.29498 |
| **GNPTG** | 0.51834 | 6.69637 | 5.82942 | 4.28828 | 3.98277 | 0.00000 | 0.00000 | 0.77232 | 8.59715 | 1.25248 | 7.05809 | 2.32856 |
| **GNRH1** | 0.28018 | 0.00000 | 0.18210 | 0.00000 | 0.00000 | 0.18655 | 0.57135 | 0.67294 | 0.16915 | 0.26847 | 0.24830 | 0.70663 |
| **GNRH2** | 0.00140 | 0.12146 | 0.00000 | 0.49802 | 0.00000 | 0.04355 | 0.00000 | 0.04079 | 0.00000 | 0.40454 | 0.00389 | 0.00088 |
| **GNRHR** | 0.01720 | 0.01682 | 0.00839 | 0.00432 | 0.08395 | 0.00000 | 0.00878 | 0.01165 | 0.24455 | 0.01888 | 0.04694 | 0.02956 |
| **GNRHR2** | 0.00740 | 0.06869 | 0.04631 | 0.01076 | 0.02868 | 0.01236 | 0.07335 | 0.02798 | 0.02422 | 0.00000 | 0.02182 | 0.01604 |
| **GNS** | 14.26117 | 6.26187 | 5.64348 | 1.06087 | 3.57582 | 8.95270 | 0.00000 | 12.67491 | 19.62186 | 3.82846 | 1.47776 | 4.05535 |
| **GOLGA1** | 3.21864 | 2.95967 | 1.19027 | 1.64018 | 2.30963 | 5.11219 | 2.12207 | 0.30509 | 0.70127 | 4.03079 | 3.35181 | 1.82980 |
| **GOLGA2** | 1.74907 | 1.43264 | 6.93074 | 4.09075 | 6.13517 | 2.07577 | 4.99256 | 2.32279 | 3.58057 | 9.56461 | 2.73358 | 3.87556 |
| **GOLGA2P5** | 2.48243 | 0.07603 | 0.51413 | 0.29708 | 0.01212 | 0.00000 | 0.00000 | 0.41914 | 0.69513 | 0.67215 | 0.18275 | 0.13551 |
| **GOLGA2P6** | 0.00429 | 0.00635 | 0.01497 | 0.00473 | 0.00000 | 0.04946 | 0.05015 | 0.01138 | 0.03606 | 0.09077 | 0.00297 | 0.01267 |
| **GOLGA2P7** | 0.00418 | 0.05300 | 0.04234 | 0.00549 | 0.17475 | 0.00000 | 0.08561 | 0.00947 | 0.00000 | 0.31652 | 0.00479 | 0.02089 |
| **GOLGA2P9** | 0.21703 | 0.01933 | 0.01294 | 0.00995 | 0.00991 | 0.02836 | 0.02431 | 0.16263 | 0.01867 | 0.01187 | 0.01936 | 0.00935 |
| **GOLGA3** | 10.12203 | 8.08470 | 8.74718 | 4.74411 | 4.51061 | 5.84137 | 11.04448 | 13.35656 | 7.83241 | 16.78814 | 9.19248 | 7.62952 |
| **GOLGA4** | 5.65912 | 0.83481 | 13.28162 | 7.40623 | 13.20543 | 30.59690 | 4.95023 | 4.62159 | 20.13784 | 56.06445 | 8.84114 | 8.08645 |
| **GOLGA5** | 4.80620 | 7.09467 | 8.20994 | 3.66533 | 1.98483 | 14.33238 | 6.76088 | 2.35290 | 6.01677 | 0.40617 | 6.14027 | 9.39654 |
| **GOLGA6A** | 0.01625 | 0.01467 | 0.01258 | 0.03225 | 0.01344 | 0.01107 | 0.01769 | 0.00871 | 0.01822 | 0.02034 | 0.01538 | 0.01646 |
| **GOLGA6B** | 0.05582 | 0.01453 | 0.01390 | 0.01287 | 0.01494 | 0.01904 | 0.01663 | 0.01478 | 0.02331 | 0.01140 | 0.01450 | 0.01503 |
| **GOLGA6C** | 0.00973 | 0.01448 | 0.02056 | 0.01493 | 0.01386 | 0.01952 | 0.01659 | 0.01179 | 0.01614 | 0.01651 | 0.01539 | 0.01576 |
| **GOLGA6D** | 0.03990 | 0.01739 | 0.01546 | 0.01254 | 0.01513 | 0.01706 | 0.01802 | 0.00693 | 0.01857 | 0.01272 | 0.01485 | 0.01550 |
| **GOLGA6L10** | 0.00691 | 0.01398 | 0.01375 | 0.00971 | 0.01213 | 0.02232 | 0.01761 | 0.01233 | 0.01871 | 0.02958 | 0.01335 | 0.01772 |
| **GOLGA6L4** | 0.00598 | 0.01932 | 0.02035 | 0.01055 | 0.01372 | 0.01615 | 0.01660 | 0.01022 | 0.01470 | 0.02285 | 0.01650 | 0.02039 |
| **GOLGA6L5P** | 0.00746 | 0.01784 | 0.01723 | 0.01534 | 0.03706 | 0.01422 | 0.01445 | 0.00988 | 0.02608 | 0.00724 | 0.01646 | 0.02456 |
| **GOLGA6L7P** | 0.06902 | 0.01542 | 0.00912 | 0.01831 | 0.01179 | 0.02831 | 0.00573 | 0.02608 | 0.01884 | 0.06248 | 0.01172 | 0.02059 |
| **GOLGA6L9.uid-** | 0.00618 | 0.01359 | 0.01361 | 0.01034 | 0.01875 | 0.02619 | 0.01638 | 0.01040 | 0.01755 | 0.01361 | 0.01406 | 0.01704 |
| **GOLGA6L9.uid-** | 0.00638 | 0.01564 | 0.01195 | 0.02310 | 0.01584 | 0.02976 | 0.01732 | 0.01034 | 0.02225 | 0.00566 | 0.01379 | 0.01604 |

| | | | | | | | | | | | | |
|---|---|---|---|---|---|---|---|---|---|---|---|---|
| GOLGA7 | 8.16444 | 21.93902 | 24.90482 | 7.66689 | 0.00001 | 31.49787 | 13.66232 | 3.68203 | 8.61726 | 6.32073 | 8.53173 | 19.88125 |
| GOLGA7B | 0.12619 | 2.51634 | 0.53232 | 1.73837 | 0.00000 | 0.06445 | 0.03640 | 0.20606 | 0.00000 | 0.03239 | 0.00337 | 1.70065 |
| GOLGA8A | 0.00000 | 0.32845 | 0.35546 | 0.08760 | 0.00000 | 4.51896 | 0.39060 | 0.21175 | 3.52078 | 11.63187 | 0.42705 | 0.88250 |
| GOLGA8B | 0.00000 | 0.27557 | 0.00000 | 0.90761 | 0.22423 | 0.53584 | 0.00000 | 0.01154 | 1.82686 | 4.45166 | 0.37115 | 0.82553 |
| GOLGA8H | 0.17094 | 0.03039 | 0.01074 | 0.01424 | 0.01220 | 0.01406 | 0.02187 | 0.02463 | 0.02642 | 0.01871 | 0.01395 | 0.01382 |
| GOLGA8J | 0.01375 | 0.01439 | 0.01568 | 0.01175 | 0.01340 | 0.01884 | 0.01883 | 0.01048 | 0.01304 | 0.01544 | 0.02529 | 0.01668 |
| GOLGA8M | 0.01751 | 0.01803 | 0.01439 | 0.01100 | 0.01296 | 0.02352 | 0.01866 | 0.02916 | 0.01581 | 0.01679 | 0.01459 | 0.01385 |
| GOLGA8N | 0.06560 | 0.01110 | 0.01200 | 0.00776 | 0.02455 | 0.02427 | 0.00866 | 0.01164 | 0.02175 | 0.03502 | 0.01405 | 0.02380 |
| GOLGA8S | 0.00957 | 0.01421 | 0.01737 | 0.01896 | 0.01548 | 0.01903 | 0.01532 | 0.01186 | 0.02294 | 0.01537 | 0.01487 | 0.01628 |
| GOLGB1 | 0.00000 | 0.27700 | 7.22786 | 7.32934 | 6.52100 | 16.90419 | 0.35651 | 6.21150 | 14.36045 | 31.83026 | 1.65831 | 4.44085 |
| GOLIM4 | 0.26609 | 0.20300 | 0.85060 | 2.60678 | 0.00000 | 0.69981 | 1.19376 | 0.22309 | 0.50575 | 0.99748 | 0.97159 | 0.35962 |
| GOLM1 | 13.29353 | 1.15645 | 2.46524 | 1.59031 | 0.00000 | 3.87098 | 1.86928 | 8.47256 | 0.00018 | 6.49178 | 0.11208 | 0.37111 |
| GOLPH3 | 4.15185 | 7.22964 | 14.64022 | 2.83856 | 0.00000 | 26.74060 | 18.76121 | 2.26924 | 5.11243 | 25.64941 | 5.78841 | 2.68433 |
| GOLPH3L | 3.42724 | 1.35281 | 1.28329 | 3.81128 | 0.83465 | 6.98847 | 3.60213 | 0.10061 | 2.85574 | 4.93955 | 3.07019 | 0.85615 |
| GOLT1A | 0.27118 | 0.14782 | 0.15518 | 0.02216 | 0.00393 | 0.00000 | 0.18482 | 0.08610 | 0.03276 | 0.00000 | 0.01584 | 0.02192 |
| GOLT1B | 3.18175 | 6.50309 | 10.01063 | 0.05256 | 0.00000 | 22.96525 | 0.00000 | 9.97517 | 15.69508 | 4.67904 | 4.75799 | 13.49429 |
| GON4L | 4.24278 | 6.01475 | 6.49361 | 7.63508 | 4.42463 | 7.16667 | 1.47039 | 5.60210 | 8.39077 | 11.62352 | 4.03656 | 8.88716 |
| GOPC | 1.07984 | 3.29438 | 11.01259 | 1.16243 | 4.26217 | 5.30014 | 5.90022 | 1.48876 | 3.55425 | 8.31727 | 3.82325 | 1.63325 |
| GORAB | 0.00000 | 1.07128 | 0.71321 | 0.56178 | 0.33507 | 3.03175 | 0.90850 | 0.81471 | 2.01074 | 0.67563 | 1.07998 | 2.24083 |
| GORASP1 | 1.64994 | 10.99578 | 2.74534 | 5.61959 | 10.60381 | 2.53415 | 4.38099 | 0.09210 | 7.41864 | 0.00000 | 6.78814 | 3.48860 |
| GORASP2 | 5.19147 | 14.23010 | 15.62703 | 7.67193 | 7.72832 | 26.06633 | 14.30209 | 9.08201 | 19.52679 | 0.00000 | 17.45109 | 18.42261 |
| GOSR1 | 0.30662 | 8.91206 | 6.64904 | 6.02455 | 3.18428 | 22.26130 | 9.75933 | 3.09560 | 5.65706 | 11.04825 | 9.21110 | 16.69978 |
| GOSR2 | 5.14199 | 11.04632 | 8.22647 | 4.98972 | 9.60577 | 10.23583 | 5.07190 | 11.97939 | 9.93020 | 2.76557 | 10.57499 | 17.57728 |
| GOT1 | 3.03988 | 14.85332 | 18.93537 | 6.71834 | 11.42850 | 25.85865 | 18.02675 | 2.25751 | 21.42822 | 0.00000 | 19.55823 | 3.51748 |
| GOT1L1 | 0.00547 | 0.01375 | 0.02984 | 0.01029 | 0.01389 | 0.02144 | 0.01272 | 0.01117 | 0.01469 | 0.03964 | 0.01380 | 0.02028 |
| GOT2 | 10.53117 | 18.81441 | 14.28092 | 16.54915 | 20.15120 | 20.37750 | 28.26283 | 0.75179 | 29.15850 | 0.00000 | 23.98178 | 7.15201 |
| GP1BA | 0.22964 | 0.04856 | 0.11418 | 0.04078 | 0.00000 | 0.02549 | 0.03813 | 0.07256 | 0.05014 | 0.27064 | 0.16120 | 0.16217 |
| GP2 | 0.00620 | 0.01451 | 0.01053 | 0.07920 | 0.05497 | 0.00368 | 0.01231 | 0.00832 | 0.02608 | 0.01741 | 0.01536 | 0.05801 |
| GP5 | 0.00594 | 0.06830 | 0.01144 | 0.07210 | 0.00000 | 0.04734 | 0.03932 | 0.00967 | 0.00218 | 0.01260 | 0.01284 | 0.01895 |
| GP6 | 0.00471 | 0.16391 | 0.14576 | 0.11595 | 0.13766 | 0.00000 | 0.00123 | 0.00696 | 0.21382 | 0.53772 | 0.00000 | 0.19700 |
| GPA33 | 0.00606 | 0.09280 | 0.08496 | 0.00097 | 0.03079 | 0.11632 | 0.01782 | 0.00621 | 0.21507 | 0.00000 | 0.00000 | 0.09261 |
| GPAA1 | 0.54213 | 17.09115 | 12.26619 | 15.36441 | 21.79788 | 0.00000 | 0.70424 | 0.00000 | 18.66848 | 2.99865 | 15.95527 | 7.63786 |
| GPALPP1 | 2.04815 | 1.19265 | 2.07034 | 0.39674 | 0.00000 | 3.09351 | 1.49917 | 0.70967 | 0.73454 | 0.00000 | 1.85821 | 2.10762 |
| GPAM | 1.69462 | 4.92875 | 1.59233 | 0.00000 | 3.27040 | 7.09407 | 3.57332 | 2.15154 | 8.00684 | 9.17100 | 4.65688 | 5.57080 |
| GPANK1 | 5.55050 | 19.74791 | 14.56537 | 12.12634 | 15.95190 | 0.27618 | 15.00396 | 6.17969 | 9.10618 | 5.26399 | 9.81480 | 25.66978 |
| GPAT2 | 0.28025 | 0.01977 | 0.02192 | 0.25765 | 0.00000 | 0.06284 | 0.04055 | 0.06291 | 0.04436 | 0.00000 | 0.04417 | 0.10977 |
| GPATCH1 | 1.66721 | 4.50904 | 0.51774 | 1.95614 | 3.51632 | 8.36596 | 2.67920 | 0.45371 | 3.42167 | 4.01427 | 3.52208 | 4.24767 |
| GPATCH11 | 1.34322 | 1.31170 | 1.93895 | 0.11109 | 1.23750 | 5.78016 | 2.16334 | 11.24953 | 0.00000 | 2.96927 | 3.01752 | 6.26454 |
| GPATCH2 | 2.15874 | 3.85865 | 2.79466 | 2.80089 | 4.60078 | 9.66096 | 1.18527 | 7.48397 | 2.92611 | 8.82520 | 1.61184 | 9.07813 |
| GPATCH2L | 1.19796 | 1.27733 | 3.81187 | 1.29097 | 1.20175 | 4.21770 | 3.85171 | 10.47317 | 5.23659 | 5.91957 | 1.98622 | 3.13485 |
| GPATCH3 | 1.50648 | 7.08193 | 12.63474 | 9.04962 | 8.36995 | 0.00000 | 13.81038 | 23.90727 | 8.58906 | 0.00000 | 7.75664 | 27.38665 |
| GPATCH4 | 1.48686 | 14.26430 | 8.60777 | 9.13011 | 20.90239 | 17.02496 | 22.62964 | 9.93215 | 9.44672 | 19.13030 | 15.87068 | 24.19548 |
| GPATCH8 | 4.37246 | 5.90487 | 8.14813 | 7.93757 | 6.11585 | 10.51434 | 6.55958 | 2.04595 | 16.64224 | 15.43540 | 7.80979 | 9.53697 |
| GPBAR1 | 0.35947 | 0.05475 | 0.10998 | 0.16524 | 0.09444 | 0.02218 | 0.01744 | 0.11401 | 0.04195 | 0.00000 | 0.01267 | 0.06218 |
| GPBP1 | 5.89633 | 13.37351 | 13.90349 | 2.36536 | 6.45009 | 35.08830 | 14.43043 | 20.42937 | 8.63011 | 16.57108 | 8.55016 | 25.39522 |
| GPBP1L1 | 2.76396 | 11.20991 | 9.30381 | 9.47107 | 6.40949 | 16.47709 | 10.31549 | 16.23581 | 15.37289 | 12.40598 | 4.87047 | 12.39431 |
| GPC1 | 0.99037 | 0.28529 | 2.84454 | 0.00000 | 4.46821 | 0.00000 | 0.00014 | 0.20839 | 2.30687 | 6.75368 | 1.87367 | 0.08273 |
| GPC2 | 0.19740 | 0.01738 | 0.06733 | 0.07890 | 0.00000 | 0.00000 | 0.14430 | 0.01707 | 0.28817 | 0.17544 | 0.13951 | 0.01530 |

| | | | | | | | | | | | | |
|---|---|---|---|---|---|---|---|---|---|---|---|---|
| GPC3 | 0.61008 | 0.74764 | 1.35029 | 11.23098 | 0.01518 | 0.00000 | 0.00000 | 1.60873 | 1.10578 | 0.00000 | 0.29981 | 0.69118 |
| GPC4 | 0.00189 | 0.00011 | 1.38524 | 4.49953 | 0.00000 | 0.27406 | 0.00000 | 0.25876 | 0.00000 | 0.04141 | 0.01014 | 0.04055 |
| GPC5 | 0.33388 | 0.01086 | 0.35843 | 0.01494 | 0.01054 | 0.00000 | 0.00000 | 0.64064 | 0.04607 | 0.02486 | 0.04619 | 0.02980 |
| GPC5-AS1 | 0.51633 | 0.01521 | 0.01260 | 0.00915 | 0.01028 | 0.02411 | 0.01506 | 0.09032 | 0.01809 | 0.01525 | 0.01262 | 0.07508 |
| GPC6 | 1.85216 | 0.05399 | 0.21150 | 0.22100 | 0.00000 | 0.01320 | 0.00000 | 0.38598 | 0.00000 | 0.14415 | 0.02713 | 0.03958 |
| GPC6-AS1 | 0.13739 | 0.01259 | 0.01432 | 0.01196 | 0.01417 | 0.01813 | 0.01587 | 0.06769 | 0.02066 | 0.01388 | 0.01418 | 0.02507 |
| GPC6-AS2 | 0.00855 | 0.01505 | 0.01407 | 0.01207 | 0.01506 | 0.01870 | 0.01807 | 0.04117 | 0.01781 | 0.01427 | 0.01499 | 0.01413 |
| GPCPD1 | 2.25413 | 22.55604 | 0.00084 | 6.52954 | 7.41589 | 13.72721 | 0.00000 | 4.27725 | 0.00000 | 9.85720 | 8.10546 | 13.30363 |
| GPD1 | 1.01533 | 0.02667 | 0.01445 | 0.00353 | 0.00000 | 0.00000 | 0.00634 | 0.17053 | 0.16412 | 0.11344 | 0.00000 | 0.30657 |
| GPD1L | 1.97823 | 4.46478 | 3.25149 | 0.91332 | 3.26602 | 7.32906 | 1.19023 | 1.20652 | 0.00000 | 0.41515 | 6.31784 | 0.05446 |
| GPD2 | 6.97503 | 15.09076 | 0.00000 | 5.35490 | 18.29589 | 30.00719 | 0.00000 | 1.81882 | 0.00000 | 11.21233 | 8.60032 | 2.18775 |
| GPER1 | 0.00154 | 0.00216 | 0.12507 | 0.00324 | 0.08738 | 0.00000 | 0.00000 | 0.13785 | 0.01685 | 0.30545 | 0.00980 | 0.06273 |
| GPHA2 | 3.62124 | 0.97970 | 0.00000 | 0.00000 | 0.28919 | 0.00000 | 0.46119 | 2.22028 | 0.00000 | 0.00000 | 0.00000 | 0.15018 |
| GPHB5 | 0.00950 | 0.01415 | 0.01192 | 0.00632 | 0.00535 | 0.03493 | 0.00990 | 0.03490 | 0.01699 | 0.04777 | 0.01165 | 0.03321 |
| GPHN | 0.42139 | 1.14196 | 1.24784 | 3.02382 | 2.51807 | 3.71995 | 0.59391 | 0.68229 | 3.89420 | 1.94509 | 0.74459 | 0.78922 |
| GPI | 4.95910 | 26.37035 | 29.79742 | 24.97444 | 29.11834 | 10.11841 | 47.98020 | 0.00000 | 41.27290 | 11.90106 | 32.76782 | 9.00120 |
| GPIHBP1 | 0.00625 | 0.01617 | 0.01865 | 0.01292 | 0.01624 | 0.01659 | 0.01805 | 0.01052 | 0.01838 | 0.01491 | 0.01488 | 0.01618 |
| GPKOW | 4.25044 | 9.78174 | 0.89437 | 3.89680 | 6.31107 | 6.39822 | 2.95100 | 3.15477 | 3.50456 | 0.00000 | 3.88988 | 10.84286 |
| GPLD1 | 4.67655 | 0.26426 | 0.84828 | 0.00000 | 1.99827 | 0.53518 | 0.10607 | 2.03447 | 0.53954 | 0.75599 | 0.00000 | 0.26964 |
| GPM6A | 0.63098 | 0.42852 | 0.00000 | 0.16681 | 0.04094 | 0.95912 | 0.00000 | 0.00567 | 0.00000 | 0.71128 | 0.00000 | 0.00000 |
| GPM6B | 3.36452 | 0.00264 | 1.04612 | 1.18517 | 0.00000 | 0.88209 | 0.00000 | 1.25892 | 0.51097 | 0.57338 | 0.57864 | 0.45320 |
| GPN1 | 1.48541 | 17.84733 | 9.82510 | 9.63347 | 8.69732 | 14.02093 | 12.31923 | 9.27356 | 12.21520 | 0.00000 | 12.49205 | 16.99944 |
| GPN2 | 0.84131 | 8.04599 | 7.20148 | 5.67870 | 3.80566 | 0.79125 | 4.41532 | 3.56366 | 5.86207 | 0.00000 | 6.18439 | 9.09273 |
| GPN3 | 2.01226 | 10.63736 | 10.03821 | 3.38712 | 2.04918 | 24.83216 | 11.82735 | 12.44973 | 10.73270 | 0.08314 | 8.75884 | 16.07208 |
| GPNMB | 0.33958 | 0.73564 | 0.29060 | 0.92836 | 0.66595 | 0.00001 | 0.01212 | 1.01308 | 0.12440 | 0.00000 | 0.31002 | 3.40478 |
| GPR1 | 0.93286 | 0.25427 | 0.09234 | 0.08093 | 0.00153 | 0.00000 | 0.00000 | 0.00221 | 0.25806 | 0.16354 | 0.00000 | 0.02687 |
| GPR1-AS | 0.00000 | 1.42958 | 0.60281 | 2.66264 | 2.36371 | 0.00000 | 0.43311 | 0.04436 | 2.30053 | 2.67026 | 0.23041 | 1.28327 |
| GPR107 | 4.33340 | 2.09043 | 6.47416 | 3.72528 | 4.79539 | 6.49477 | 2.87696 | 0.60850 | 7.50836 | 11.89417 | 3.43877 | 6.06251 |
| GPR108 | 3.66476 | 9.20225 | 8.62414 | 14.19950 | 8.25367 | 0.00000 | 0.54091 | 0.00000 | 16.76498 | 0.11027 | 7.36599 | 5.26638 |
| GPR110 | 0.00479 | 0.01248 | 0.12130 | 0.00922 | 0.00936 | 0.01629 | 0.00188 | 0.01186 | 0.00512 | 0.07638 | 0.02247 | 0.01327 |
| GPR111 | 0.00623 | 0.00748 | 0.11237 | 0.49187 | 0.00192 | 0.04185 | 0.00000 | 0.00941 | 0.02748 | 0.00651 | 0.01220 | 0.02206 |
| GPR112 | 0.01000 | 0.01398 | 0.03389 | 0.00873 | 0.05611 | 0.00000 | 0.01704 | 0.01199 | 0.01396 | 0.05245 | 0.01497 | 0.01505 |
| GPR113 | 0.30241 | 0.25363 | 0.20982 | 0.14497 | 0.00000 | 0.00000 | 0.00000 | 0.52920 | 0.36500 | 0.53172 | 0.46132 | 0.20467 |
| GPR114 | 0.01191 | 0.00511 | 0.92572 | 0.00000 | 0.16752 | 0.00000 | 0.63863 | 0.00623 | 2.48578 | 1.06776 | 0.38310 | 0.00463 |
| GPR115 | 0.01653 | 0.00876 | 0.30165 | 0.47796 | 0.02905 | 0.02680 | 0.01127 | 0.00821 | 0.02496 | 0.00002 | 0.01730 | 0.01504 |
| GPR116 | 0.01109 | 0.00418 | 0.08844 | 0.00011 | 0.00000 | 0.05054 | 0.05788 | 0.01157 | 0.10022 | 0.05690 | 0.00000 | 0.04364 |
| GPR119 | 0.06080 | 0.01640 | 0.01346 | 0.01051 | 0.01271 | 0.01932 | 0.01602 | 0.07979 | 0.01755 | 0.02314 | 0.01410 | 0.01125 |
| GPR12 | 0.00672 | 0.04765 | 0.00881 | 0.36753 | 0.14889 | 0.05012 | 0.00000 | 0.00889 | 0.00000 | 0.07330 | 0.01099 | 0.07752 |
| GPR123 | 0.13891 | 0.01675 | 0.01501 | 0.02102 | 0.00371 | 0.03303 | 0.00979 | 0.05347 | 0.01486 | 0.04152 | 0.01216 | 0.00816 |
| GPR124 | 0.00630 | 0.04320 | 0.05014 | 0.00000 | 0.18209 | 0.00000 | 0.03902 | 0.05167 | 0.11895 | 0.25521 | 0.00000 | 0.08903 |
| GPR125 | 2.23594 | 0.88895 | 1.22023 | 2.46009 | 0.54284 | 2.22646 | 2.21852 | 0.48299 | 0.72990 | 2.95384 | 0.00000 | 0.00863 |
| GPR126 | 1.13279 | 0.00000 | 1.86066 | 0.00000 | 6.36350 | 0.24079 | 1.08742 | 0.29175 | 2.54348 | 0.14349 | 3.55744 | 0.00000 |
| GPR128 | 0.00179 | 0.01580 | 0.01044 | 0.01150 | 0.06988 | 0.00000 | 0.00576 | 0.14453 | 0.07834 | 0.01181 | 0.01212 | 0.01456 |
| GPR132 | 0.05499 | 0.01971 | 0.07102 | 0.03473 | 0.08060 | 0.00000 | 0.01968 | 0.00676 | 0.00000 | 0.08278 | 0.01631 | 0.01236 |
| GPR133 | 2.95974 | 1.63434 | 0.00000 | 1.71062 | 0.63217 | 0.15150 | 0.00000 | 0.18183 | 0.00000 | 0.51305 | 0.13382 | 0.55131 |
| GPR135 | 0.00238 | 0.00000 | 0.12701 | 0.05859 | 0.00000 | 0.06981 | 0.04175 | 0.04988 | 0.00000 | 0.09799 | 0.05141 | 0.04584 |
| GPR137 | 1.25403 | 0.92874 | 1.85886 | 1.78058 | 1.49061 | 0.00000 | 1.13065 | 0.02223 | 0.99280 | 3.31560 | 0.00000 | 1.43993 |
| GPR137B | 2.27105 | 0.02315 | 0.10068 | 0.09169 | 0.10452 | 0.00000 | 0.01126 | 0.06780 | 0.00450 | 0.02976 | 0.01665 | 0.03490 |

| | | | | | | | | | | | | |
|---|---|---|---|---|---|---|---|---|---|---|---|---|
| **GPR137C** | 0.45154 | 0.00000 | 0.13176 | 0.00000 | 0.00000 | 0.41727 | 0.18771 | 0.00000 | 0.03952 | 0.30561 | 0.13022 | 0.00000 |
| **GPR142** | 0.00563 | 0.01215 | 0.04777 | 0.00590 | 0.01274 | 0.02702 | 0.01207 | 0.01028 | 0.00707 | 0.04682 | 0.01530 | 0.01427 |
| **GPR143** | 8.02385 | 0.42570 | 0.07698 | 0.02787 | 0.00000 | 0.00000 | 0.02309 | 6.83726 | 0.14032 | 0.07588 | 0.11815 | 1.82834 |
| **GPR146** | 0.41399 | 0.34565 | 0.81546 | 0.37641 | 0.00000 | 0.00000 | 0.00000 | 0.07785 | 0.00000 | 0.57220 | 0.00000 | 0.05429 |
| **GPR151** | 0.00459 | 0.00803 | 0.09233 | 0.01326 | 0.00650 | 0.00915 | 0.01386 | 0.01273 | 0.00000 | 0.09996 | 0.03041 | 0.04502 |
| **GPR153** | 0.28229 | 0.67214 | 0.12306 | 0.94428 | 0.91708 | 0.00000 | 0.00000 | 0.16915 | 0.00000 | 0.47241 | 0.00000 | 0.23212 |
| **GPR155** | 0.01638 | 0.00000 | 0.11114 | 0.00000 | 0.03697 | 0.00000 | 0.00000 | 12.70323 | 2.57878 | 0.11229 | 3.22840 | 6.67500 |
| **GPR156** | 0.41416 | 0.00585 | 0.23643 | 0.00000 | 0.00000 | 0.05886 | 0.10895 | 0.15223 | 0.00000 | 0.19638 | 0.00000 | 0.15002 |
| **GPR157** | 0.01513 | 0.08430 | 0.36181 | 0.00799 | 0.22272 | 0.00000 | 0.00406 | 0.40713 | 0.46927 | 0.00000 | 0.14564 | 0.09536 |
| **GPR158** | 2.59407 | 0.05264 | 0.00508 | 0.00412 | 0.00000 | 0.00000 | 0.04196 | 0.45403 | 0.00000 | 0.09558 | 0.13046 | 0.08091 |
| **GPR160** | 1.62071 | 13.39189 | 0.00000 | 12.99573 | 0.75579 | 11.04896 | 0.00000 | 0.21335 | 0.00000 | 3.81887 | 2.31792 | 6.76893 |
| **GPR161** | 0.00176 | 1.49159 | 0.93561 | 2.39828 | 0.21609 | 0.00000 | 0.00000 | 0.50236 | 0.74747 | 0.49486 | 0.21520 | 3.31644 |
| **GPR162** | 0.00561 | 0.01634 | 0.01547 | 0.01909 | 0.01842 | 0.01387 | 0.02321 | 0.01127 | 0.02045 | 0.00633 | 0.01508 | 0.03366 |
| **GPR17** | 0.02299 | 0.01041 | 0.00940 | 0.00707 | 0.06533 | 0.00153 | 0.03875 | 0.00802 | 0.00000 | 0.07022 | 0.02152 | 0.01681 |
| **GPR171** | 0.03819 | 0.00000 | 2.19442 | 0.00000 | 0.17450 | 0.00000 | 0.00000 | 0.20732 | 0.38595 | 1.74092 | 0.09302 | 0.17656 |
| **GPR176** | 0.09012 | 2.30265 | 0.20423 | 13.20752 | 3.66358 | 0.00000 | 0.08689 | 0.05887 | 0.00000 | 0.00000 | 0.00000 | 1.50811 |
| **GPR179** | 0.00307 | 0.04485 | 0.07864 | 0.00000 | 0.07184 | 0.00000 | 0.00000 | 0.00365 | 0.26212 | 0.68930 | 0.00000 | 0.20256 |
| **GPR18** | 0.00252 | 0.05602 | 0.13958 | 0.19733 | 0.26059 | 0.41126 | 0.00000 | 0.00000 | 0.18708 | 2.44645 | 0.00072 | 0.30218 |
| **GPR180** | 0.00092 | 0.23080 | 0.77325 | 0.79664 | 0.24875 | 2.90927 | 0.00012 | 0.68779 | 1.68385 | 2.50025 | 1.70420 | 0.60461 |
| **GPR182** | 0.01305 | 0.01239 | 0.01007 | 0.05485 | 0.00931 | 0.01843 | 0.01386 | 0.01088 | 0.02044 | 0.02712 | 0.01321 | 0.01655 |
| **GPR183** | 0.00817 | 0.18117 | 0.12556 | 0.28662 | 0.05973 | 0.40733 | 0.00000 | 0.00000 | 0.10574 | 0.34073 | 0.21740 | 0.25797 |
| **GPR19** | 0.31306 | 0.25960 | 0.81753 | 0.78063 | 0.25842 | 1.00756 | 0.00000 | 0.23806 | 0.50910 | 0.00000 | 0.26001 | 0.45087 |
| **GPR21** | 0.00558 | 0.00681 | 0.01625 | 0.10678 | 0.00000 | 0.04950 | 0.00856 | 0.01061 | 0.01172 | 0.03026 | 0.01701 | 0.02336 |
| **GPR22** | 0.02357 | 0.26800 | 0.14022 | 0.61879 | 0.00000 | 1.43903 | 0.00000 | 0.21026 | 0.32741 | 0.31689 | 0.19421 | 0.80331 |
| **GPR25** | 0.00572 | 0.01229 | 0.01218 | 0.00254 | 0.03013 | 0.08179 | 0.00680 | 0.01067 | 0.00383 | 0.01839 | 0.02055 | 0.01358 |
| **GPR26** | 0.00592 | 0.01443 | 0.02995 | 0.02424 | 0.01077 | 0.01121 | 0.03186 | 0.01057 | 0.00689 | 0.03449 | 0.02192 | 0.01552 |
| **GPR27** | 0.00269 | 0.12067 | 0.00000 | 0.06801 | 0.00000 | 0.07415 | 0.00000 | 0.02003 | 0.02378 | 0.11068 | 0.02515 | 0.01763 |
| **GPR3** | 1.40027 | 0.01426 | 0.08146 | 0.34196 | 0.16466 | 0.00000 | 0.00000 | 0.14736 | 0.17714 | 0.14558 | 0.00030 | 0.27220 |
| **GPR32** | 0.00578 | 0.01739 | 3.32802 | 0.00735 | 0.01594 | 0.00000 | 0.00000 | 0.01030 | 0.00000 | 0.01835 | 0.02000 | 0.01515 |
| **GPR34** | 0.00348 | 0.00712 | 0.00000 | 0.15020 | 0.00000 | 0.08134 | 0.00000 | 0.00823 | 0.05634 | 0.17442 | 0.04930 | 0.02248 |
| **GPR35** | 0.00694 | 0.00807 | 0.08690 | 0.06335 | 0.02228 | 0.00541 | 0.00567 | 0.00677 | 0.15077 | 0.00000 | 0.01853 | 0.11173 |
| **GPR37** | 4.54075 | 0.00000 | 1.15486 | 0.00000 | 0.16810 | 0.32153 | 1.27485 | 0.97074 | 6.06179 | 4.32779 | 0.00000 | 0.10116 |
| **GPR37L1** | 0.02286 | 1.06395 | 0.04981 | 0.00000 | 0.63070 | 0.00000 | 0.00234 | 2.67160 | 0.63998 | 0.04943 | 0.24725 | 3.86607 |
| **GPR39** | 5.44021 | 0.01307 | 0.02847 | 0.19665 | 0.00000 | 0.00000 | 0.01455 | 0.59691 | 0.01003 | 0.03877 | 0.01256 | 0.00000 |
| **GPR4** | 0.00320 | 0.03943 | 0.09358 | 0.03120 | 0.00000 | 0.24789 | 0.04798 | 0.13361 | 0.00000 | 0.04022 | 0.00273 | 0.78075 |
| **GPR45** | 0.01093 | 0.03919 | 0.08773 | 0.04895 | 0.04506 | 0.00000 | 0.04853 | 0.00727 | 0.01897 | 0.00000 | 0.06221 | 0.15421 |
| **GPR50** | 0.00416 | 0.01015 | 0.11203 | 0.10724 | 0.00461 | 0.01271 | 0.00546 | 0.67622 | 0.01455 | 0.02352 | 0.01682 | 0.01408 |
| **GPR52** | 0.00595 | 0.01535 | 0.04815 | 0.00930 | 0.01035 | 0.01269 | 0.00561 | 0.01023 | 0.05681 | 0.02624 | 0.01405 | 0.02739 |
| **GPR55** | 0.00383 | 0.04084 | 0.01876 | 0.13327 | 0.00000 | 0.03688 | 0.01102 | 0.02423 | 0.02641 | 0.00921 | 0.01301 | 0.03698 |
| **GPR56** | 0.53360 | 0.00000 | 5.08103 | 0.03884 | 0.16233 | 0.01595 | 0.17472 | 0.10575 | 1.73510 | 0.08752 | 0.00000 | 0.01866 |
| **GPR61** | 0.00601 | 0.01490 | 0.01395 | 0.01619 | 0.01343 | 0.01380 | 0.01562 | 0.01048 | 0.01874 | 0.02796 | 0.02517 | 0.01587 |
| **GPR62** | 0.06420 | 0.00820 | 0.02357 | 0.01266 | 0.00970 | 0.04488 | 0.01027 | 0.00495 | 0.01692 | 0.01335 | 0.01300 | 0.03526 |
| **GPR63** | 0.37084 | 0.53084 | 0.58357 | 0.00000 | 0.00000 | 0.21581 | 0.43985 | 0.07580 | 0.31499 | 0.14127 | 0.08243 | 0.77722 |
| **GPR64** | 0.15357 | 0.05970 | 0.07032 | 0.21700 | 0.00534 | 0.09960 | 0.00000 | 0.36689 | 0.00000 | 0.03983 | 0.03935 | 0.34466 |
| **GPR65** | 0.00618 | 0.01579 | 0.01671 | 0.01544 | 0.01519 | 0.01691 | 0.01801 | 0.01051 | 0.01737 | 0.01713 | 0.01749 | 0.01610 |
| **GPR68** | 0.06735 | 0.00758 | 0.06259 | 0.15775 | 0.00000 | 0.00000 | 0.00003 | 0.00325 | 0.01868 | 0.18131 | 0.03606 | 0.10751 |
| **GPR75** | 0.01824 | 0.21148 | 0.45169 | 0.69956 | 0.00000 | 0.04505 | 0.48019 | 0.05681 | 1.31217 | 0.51255 | 0.21731 | 0.21199 |
| **GPR78** | 1.29808 | 0.01854 | 4.81985 | 0.00000 | 0.00000 | 0.00000 | 0.12850 | 0.04110 | 0.00000 | 0.57634 | 0.10728 | 0.00789 |

| | | | | | | | | | | | | |
|---|---|---|---|---|---|---|---|---|---|---|---|---|
| GPR82 | 0.00148 | 0.00000 | 0.00000 | 0.15027 | 0.00000 | 0.18610 | 0.00000 | 0.04827 | 0.00000 | 0.11276 | 0.01845 | 0.00242 |
| GPR83 | 0.00615 | 0.25200 | 0.64768 | 0.00000 | 0.00000 | 0.00000 | 0.07760 | 0.97612 | 0.45406 | 0.03916 | 0.04706 | 0.55085 |
| GPR84 | 0.00617 | 0.01432 | 0.02581 | 0.01228 | 0.01070 | 0.01953 | 0.02585 | 0.01061 | 0.02287 | 0.02194 | 0.01393 | 0.01570 |
| GPR85 | 0.00428 | 0.02421 | 0.03845 | 0.06885 | 0.00000 | 0.06793 | 0.03469 | 0.01284 | 0.00000 | 0.28983 | 0.00461 | 0.00692 |
| GPR87 | 0.00315 | 0.00000 | 0.91249 | 0.03777 | 0.00000 | 0.09008 | 0.00000 | 0.00725 | 0.21046 | 0.23552 | 0.24660 | 0.02754 |
| GPR89A | 0.15752 | 0.38542 | 1.39409 | 0.22500 | 0.86602 | 1.92467 | 0.00000 | 0.77079 | 3.01930 | 2.33292 | 0.55768 | 0.64486 |
| GPR89B | 0.03880 | 0.18412 | 0.42653 | 0.37753 | 0.26213 | 0.27271 | 0.00000 | 0.07095 | 0.85982 | 0.00000 | 0.14606 | 0.14400 |
| GPR97 | 0.00575 | 0.01362 | 0.01058 | 0.01601 | 0.00483 | 0.01794 | 0.00442 | 0.01026 | 0.08337 | 0.05309 | 0.00680 | 0.01428 |
| GPR98 | 0.89876 | 1.03913 | 0.42434 | 1.54714 | 0.22494 | 0.28107 | 0.20804 | 0.76781 | 0.24925 | 0.37134 | 0.00000 | 0.18134 |
| GPRASP1 | 0.00452 | 0.02929 | 0.00989 | 0.01577 | 0.07886 | 0.01688 | 0.00531 | 0.01216 | 0.02400 | 0.08040 | 0.00492 | 0.02745 |
| GPRASP2 | 0.01262 | 0.01796 | 0.01452 | 0.01199 | 0.01806 | 0.01524 | 0.01762 | 0.00944 | 0.01994 | 0.01198 | 0.01806 | 0.01685 |
| GPRC5A | 0.05278 | 8.63136 | 0.00000 | 0.00000 | 46.28979 | 0.00002 | 0.00219 | 0.03824 | 19.62819 | 2.58824 | 21.71049 | 0.32854 |
| GPRC5B | 0.00219 | 1.85802 | 3.62072 | 18.70875 | 0.00000 | 0.00159 | 0.00000 | 0.28817 | 0.52207 | 0.01244 | 0.00000 | 4.01586 |
| GPRC5C | 0.29766 | 0.39925 | 2.55033 | 17.93107 | 1.96292 | 2.65695 | 0.00016 | 0.00000 | 0.39388 | 0.00000 | 0.00000 | 4.61159 |
| GPRC5D | 0.00184 | 0.02508 | 0.24090 | 0.05421 | 0.45982 | 0.00000 | 0.29459 | 0.19251 | 0.35279 | 0.00000 | 0.72947 | 0.79510 |
| GPRC6A | 0.00186 | 0.01898 | 0.03019 | 0.01696 | 0.00521 | 0.03782 | 0.03952 | 0.08385 | 0.00996 | 0.02377 | 0.01265 | 0.12164 |
| GPRIN1 | 0.00255 | 0.03279 | 0.94848 | 0.11984 | 0.04153 | 0.00000 | 0.88520 | 0.00333 | 0.00000 | 1.08754 | 0.00000 | 0.40324 |
| GPRIN3 | 0.00599 | 0.01019 | 0.00671 | 0.06778 | 0.00839 | 0.03739 | 0.01246 | 0.01011 | 0.00793 | 0.02963 | 0.02892 | 0.01385 |
| GPS1 | 2.71675 | 21.73728 | 19.80593 | 17.82698 | 21.33560 | 1.36892 | 24.93964 | 3.30517 | 27.72519 | 0.00000 | 19.68515 | 18.46201 |
| GPS2 | 3.65639 | 11.77250 | 11.98582 | 10.61671 | 15.70146 | 1.09383 | 14.47099 | 5.66544 | 8.36071 | 17.18197 | 13.41142 | 11.97113 |
| GPSM1 | 0.14311 | 0.02169 | 0.01152 | 0.03679 | 0.14345 | 0.00457 | 0.03001 | 0.00203 | 0.02489 | 0.18161 | 0.01772 | 0.04399 |
| GPSM2 | 1.00366 | 0.87528 | 3.17278 | 0.08249 | 0.17493 | 1.21116 | 0.52112 | 0.17908 | 0.08701 | 0.12217 | 0.36076 | 0.24777 |
| GPSM3 | 0.02746 | 1.05267 | 0.14294 | 1.81801 | 1.40964 | 0.00000 | 0.37337 | 0.29178 | 0.49160 | 0.00000 | 0.00003 | 2.85495 |
| GPT | 0.20343 | 0.01441 | 0.03807 | 0.05011 | 0.01589 | 0.00000 | 0.00156 | 0.00180 | 0.09029 | 0.03491 | 0.03517 | 0.03395 |
| GPT2 | 4.49864 | 0.75474 | 5.88326 | 2.70031 | 3.09424 | 0.13127 | 11.24802 | 0.05269 | 3.14858 | 11.67130 | 4.82931 | 0.62078 |
| GPX1 | 9.96846 | 24.28284 | 12.32956 | 19.40453 | 35.83923 | 12.12416 | 10.69608 | 7.03585 | 10.13227 | 10.79974 | 10.44171 | 23.52239 |
| GPX2 | 0.00238 | 0.00000 | 9.49151 | 59.51510 | 1.59576 | 0.00000 | 0.20557 | 0.04402 | 1.47943 | 0.60532 | 0.00000 | 0.01983 |
| GPX3 | 0.00550 | 0.00000 | 3.18671 | 1.06979 | 0.00000 | 0.93970 | 1.22960 | 0.12196 | 0.99046 | 0.00000 | 0.00000 | 0.02305 |
| GPX4 | 1.24006 | 21.50370 | 24.83688 | 20.97558 | 35.02666 | 1.34292 | 31.91755 | 6.25595 | 32.44877 | 8.22708 | 21.26757 | 31.26105 |
| GPX6 | 0.23718 | 0.26441 | 0.01187 | 0.03621 | 0.00081 | 0.00709 | 0.02834 | 0.00000 | 0.06064 | 0.00000 | 0.03104 | 0.09948 |
| GPX7 | 0.00848 | 0.01659 | 0.16660 | 0.29812 | 0.00000 | 0.06869 | 0.00000 | 0.00426 | 0.00000 | 0.02272 | 0.02718 | 0.25486 |
| GPX8 | 0.03210 | 0.42443 | 0.23456 | 0.00000 | 0.00000 | 0.00000 | 1.52120 | 0.07467 | 1.77856 | 0.13349 | 0.24996 | 0.22366 |
| GRAMD1A | 2.22859 | 0.73396 | 2.43676 | 7.02673 | 0.00000 | 0.00000 | 0.53699 | 0.00950 | 1.41831 | 0.28895 | 0.30460 | 0.00000 |
| GRAMD1B | 0.00151 | 0.00000 | 0.08109 | 0.13577 | 0.00000 | 0.00000 | 0.01059 | 3.77295 | 0.00000 | 0.39068 | 0.03678 | 2.52029 |
| GRAMD1C | 0.50560 | 0.65902 | 0.18413 | 0.67499 | 0.59146 | 0.61038 | 0.00000 | 0.50754 | 0.00000 | 0.02132 | 0.00000 | 0.57297 |
| GRAMD2 | 0.90472 | 0.72071 | 20.25099 | 0.43635 | 3.78970 | 0.54430 | 12.99547 | 0.00000 | 0.56991 | 0.00000 | 5.44646 | 0.00000 |
| GRAMD3 | 0.43592 | 0.00000 | 15.52104 | 0.04310 | 1.78767 | 0.03558 | 1.20636 | 0.00000 | 5.14870 | 0.41006 | 2.71282 | 0.17342 |
| GRAMD4 | 0.59036 | 0.26415 | 0.16697 | 0.09691 | 0.47177 | 0.27219 | 0.00000 | 0.00000 | 0.83943 | 1.64041 | 0.61617 | 0.00296 |
| GRAP | 0.01832 | 0.34865 | 0.00449 | 0.00000 | 0.23626 | 0.00000 | 0.05999 | 0.04089 | 0.00000 | 0.03601 | 0.22258 | 0.01707 |
| GRAP2 | 0.54148 | 0.01093 | 0.03339 | 0.01224 | 0.01429 | 0.00001 | 0.06344 | 0.00827 | 0.00766 | 0.03134 | 0.01428 | 0.05111 |
| GRASP | 0.15055 | 0.01945 | 0.09614 | 0.07217 | 0.14636 | 0.00000 | 0.12075 | 0.03641 | 0.00000 | 0.29990 | 0.00000 | 0.08114 |
| GRB10 | 2.01592 | 4.52262 | 0.00000 | 4.77943 | 11.18368 | 5.27965 | 12.18836 | 0.88626 | 4.30724 | 23.61375 | 5.44401 | 3.85761 |
| GRB14 | 0.00000 | 0.48248 | 0.50991 | 0.00000 | 0.47155 | 0.62675 | 1.17005 | 0.10098 | 0.00075 | 0.49403 | 0.11446 | 0.24699 |
| GRB2 | 1.84637 | 19.86559 | 28.15138 | 17.26867 | 20.80352 | 11.14828 | 37.88686 | 33.29376 | 22.89331 | 7.38741 | 26.33517 | 19.17800 |
| GRB7 | 7.14383 | 6.61694 | 2.47163 | 11.81494 | 8.95258 | 0.00000 | 0.00044 | 0.11485 | 3.83829 | 4.63786 | 1.40792 | 0.91887 |
| GREB1 | 1.37157 | 0.11210 | 0.06933 | 0.00000 | 0.40009 | 0.00000 | 0.12058 | 0.09382 | 0.76565 | 0.64030 | 0.16750 | 0.03208 |
| GREB1L | 0.37849 | 1.06760 | 4.90949 | 0.00000 | 4.70372 | 0.00000 | 3.62011 | 0.25314 | 11.73515 | 3.16543 | 5.57571 | 0.60965 |
| GREM2 | 0.00498 | 0.00000 | 34.21013 | 0.74573 | 4.17926 | 0.82266 | 4.24144 | 0.00000 | 0.38882 | 0.00000 | 2.84353 | 0.00000 |

| | | | | | | | | | | | | |
|---|---|---|---|---|---|---|---|---|---|---|---|---|
| **GRHL1** | 3.60325 | 1.88401 | 24.35569 | 0.00000 | 9.75220 | 4.85478 | 20.71746 | 0.37419 | 18.90377 | 1.21053 | 10.76861 | 0.00000 |
| **GRHL2** | 4.95573 | 5.72831 | 4.22946 | 0.00000 | 14.01265 | 6.19046 | 11.69303 | 0.78351 | 4.58990 | 3.01440 | 14.31083 | 0.00000 |
| **GRHL3** | 5.40481 | 0.07865 | 19.23583 | 0.66340 | 18.94233 | 2.85401 | 1.01342 | 0.37389 | 2.57734 | 0.00000 | 0.00000 | 0.00000 |
| **GRHPR** | 3.23473 | 3.77170 | 10.30383 | 2.92169 | 10.15574 | 3.61154 | 14.84555 | 1.19668 | 18.25662 | 0.00000 | 19.80723 | 1.09668 |
| **GRIA1** | 1.42302 | 0.00676 | 0.00377 | 0.08221 | 0.00000 | 0.00575 | 0.00087 | 0.15894 | 0.01990 | 0.49690 | 0.00007 | 0.00594 |
| **GRIA2** | 0.00521 | 0.02041 | 0.01767 | 0.01174 | 0.02324 | 0.02152 | 0.01758 | 0.01379 | 0.01940 | 0.00739 | 0.01432 | 0.06215 |
| **GRIA3** | 0.08924 | 0.00780 | 0.08302 | 0.00934 | 0.00677 | 0.02049 | 0.01099 | 0.05085 | 0.02291 | 0.04378 | 0.08516 | 0.09332 |
| **GRIA4** | 0.02702 | 0.02622 | 0.05626 | 0.02203 | 0.01115 | 0.05396 | 0.00394 | 0.01334 | 0.02331 | 0.07268 | 0.02423 | 0.02332 |
| **GRID1** | 0.05892 | 0.08080 | 0.03258 | 0.03291 | 0.01962 | 0.00000 | 0.09138 | 0.12380 | 0.00594 | 0.00000 | 0.04017 | 0.13597 |
| **GRID2** | 0.30015 | 0.04374 | 0.01695 | 0.02872 | 0.01878 | 0.12736 | 0.03400 | 0.02773 | 0.00000 | 0.00000 | 0.10325 | 0.02425 |
| **GRID2IP** | 0.53019 | 0.01516 | 0.05849 | 0.02730 | 0.06391 | 0.00000 | 0.00000 | 0.09513 | 0.05557 | 0.07633 | 0.04758 | 0.01382 |
| **GRIFIN** | 0.00615 | 0.01426 | 0.01753 | 0.01825 | 0.01194 | 0.01600 | 0.01758 | 0.01056 | 0.01335 | 0.01690 | 0.03392 | 0.01635 |
| **GRIK1** | 0.03647 | 0.00000 | 0.11210 | 0.14400 | 0.00000 | 0.00000 | 0.14655 | 0.00576 | 0.36957 | 0.30502 | 0.38747 | 0.01125 |
| **GRIK1-AS1** | 0.06361 | 0.01352 | 0.02096 | 0.00880 | 0.00976 | 0.02206 | 0.01460 | 0.11018 | 0.02973 | 0.03014 | 0.00992 | 0.00897 |
| **GRIK1-AS2** | 0.03731 | 0.02408 | 0.01375 | 0.03636 | 0.02375 | 0.02284 | 0.09429 | 0.01435 | 0.11383 | 0.00000 | 0.00536 | 0.01549 |
| **GRIK2** | 1.75043 | 0.01534 | 0.03743 | 0.00789 | 0.00708 | 0.00228 | 0.02308 | 0.16930 | 0.00878 | 0.06797 | 0.01626 | 0.11584 |
| **GRIK3** | 0.01250 | 0.00510 | 0.08856 | 0.05178 | 0.00000 | 0.03151 | 0.00000 | 0.02989 | 0.00000 | 0.18361 | 0.00937 | 0.00757 |
| **GRIK4** | 1.95511 | 0.02936 | 0.07753 | 0.00401 | 0.01409 | 0.01114 | 0.00000 | 0.48988 | 0.01920 | 0.04323 | 0.01010 | 0.02549 |
| **GRIK5** | 0.08546 | 0.01857 | 0.02129 | 0.01470 | 0.00000 | 0.07044 | 0.02023 | 0.00209 | 0.00000 | 0.11067 | 0.02157 | 0.13955 |
| **GRIN1** | 0.01006 | 0.01514 | 0.07795 | 0.02499 | 0.02050 | 0.01057 | 0.01979 | 0.01509 | 0.02312 | 0.00586 | 0.01546 | 0.02327 |
| **GRIN2A** | 2.06441 | 0.00000 | 0.00000 | 0.27407 | 0.58237 | 0.28550 | 0.02817 | 0.20497 | 0.00000 | 0.17399 | 0.11573 | 0.00880 |
| **GRIN2B** | 2.79654 | 0.01786 | 0.01697 | 0.01584 | 0.03567 | 0.00066 | 0.03980 | 0.50787 | 0.01220 | 0.00000 | 0.01289 | 0.00778 |
| **GRIN2C** | 0.00091 | 0.10393 | 0.08386 | 0.04467 | 0.04318 | 0.05234 | 0.00585 | 0.06829 | 0.00000 | 0.31241 | 0.22581 | 0.05511 |
| **GRIN2D** | 0.00824 | 0.07158 | 0.33014 | 0.11360 | 1.00291 | 0.00000 | 0.74082 | 0.48355 | 0.00000 | 3.13119 | 0.34691 | 0.83986 |
| **GRIN3A** | 0.00230 | 0.07520 | 0.03253 | 0.02562 | 0.07587 | 0.00000 | 0.15464 | 0.07522 | 0.00000 | 0.03538 | 0.12249 | 0.03995 |
| **GRIN3B** | 0.02421 | 0.01525 | 0.01049 | 0.00987 | 0.01157 | 0.01637 | 0.02123 | 0.00862 | 0.02118 | 0.12055 | 0.01484 | 0.01870 |
| **GRINA** | 1.40085 | 0.59587 | 9.59962 | 2.77367 | 3.03967 | 0.00000 | 3.48453 | 0.24590 | 2.66186 | 9.63917 | 1.15269 | 1.51359 |
| **GRIP1** | 0.70994 | 0.00000 | 0.28807 | 0.44174 | 0.04465 | 1.35474 | 0.23194 | 0.27472 | 0.00000 | 0.64314 | 0.00000 | 0.05713 |
| **GRIP2** | 0.00349 | 0.21898 | 0.04269 | 0.94146 | 0.00000 | 0.00000 | 0.04118 | 0.02222 | 0.00446 | 0.55510 | 0.26174 | 0.00000 |
| **GRIPAP1** | 3.27800 | 2.59979 | 2.73606 | 5.25889 | 4.80385 | 0.00000 | 6.07566 | 0.00000 | 3.21262 | 4.91274 | 4.64540 | 3.46740 |
| **GRK1** | 0.13956 | 0.01311 | 0.02174 | 0.01152 | 0.01173 | 0.01879 | 0.02313 | 0.00622 | 0.00946 | 0.01886 | 0.03208 | 0.01440 |
| **GRK4** | 0.07533 | 0.04750 | 0.09535 | 0.04801 | 0.00000 | 0.04045 | 0.12814 | 0.25470 | 0.00000 | 0.18915 | 0.05308 | 0.02505 |
| **GRK5** | 2.84307 | 1.35517 | 0.14122 | 3.20290 | 0.00000 | 0.23587 | 0.13995 | 0.92649 | 0.00000 | 1.28129 | 0.00000 | 0.62960 |
| **GRK6** | 0.00058 | 0.81213 | 3.31983 | 2.03580 | 4.62358 | 0.00000 | 5.11724 | 0.00357 | 3.14375 | 5.60077 | 1.61486 | 0.00194 |
| **GRK7** | 0.28061 | 0.01380 | 0.01219 | 0.01309 | 0.02965 | 0.00000 | 0.01598 | 0.19152 | 0.10072 | 0.00000 | 0.00716 | 0.03514 |
| **GRM1** | 0.00581 | 0.01765 | 0.03203 | 0.01134 | 0.01307 | 0.01467 | 0.01672 | 0.01040 | 0.01159 | 0.02991 | 0.01890 | 0.01932 |
| **GRM2** | 0.07745 | 0.22542 | 0.02504 | 0.23780 | 0.00000 | 0.09783 | 0.09070 | 0.00000 | 0.00642 | 0.00000 | 0.06124 | 0.23634 |
| **GRM3** | 0.00495 | 0.00699 | 0.01280 | 0.00003 | 0.01657 | 0.04015 | 0.00363 | 0.00956 | 0.00000 | 0.17423 | 0.05166 | 0.00956 |
| **GRM4** | 1.50331 | 0.15724 | 0.28659 | 0.30230 | 2.63901 | 0.00000 | 0.00000 | 0.01937 | 0.00000 | 0.10377 | 0.03132 | 0.18910 |
| **GRM5** | 0.29177 | 0.01419 | 0.03845 | 0.01432 | 0.01478 | 0.00000 | 0.02852 | 0.05208 | 0.00639 | 0.01237 | 0.04666 | 0.03192 |
| **GRM6** | 0.01302 | 0.01305 | 0.02870 | 0.01110 | 0.01903 | 0.00000 | 0.00534 | 0.01073 | 0.02904 | 0.06074 | 0.08018 | 0.03001 |
| **GRM7** | 2.19842 | 0.00184 | 0.00920 | 0.00714 | 0.01516 | 0.02480 | 0.01948 | 0.37435 | 0.02386 | 0.01411 | 0.01109 | 0.16707 |
| **GRM8** | 2.06748 | 0.02260 | 0.07929 | 0.02614 | 0.02501 | 0.02641 | 0.01236 | 0.14919 | 0.00302 | 0.01409 | 0.01718 | 0.10133 |
| **GRN** | 6.98413 | 34.95939 | 6.75755 | 18.25042 | 17.96137 | 0.00000 | 0.00000 | 5.49738 | 16.47667 | 5.27407 | 7.89872 | 37.93994 |
| **GRP** | 0.01074 | 0.21693 | 0.00839 | 0.00000 | 0.08374 | 0.01862 | 0.02563 | 0.00358 | 0.00364 | 0.03821 | 0.01071 | 0.29553 |
| **GRPEL1** | 6.63393 | 27.15681 | 23.69678 | 15.33100 | 25.24806 | 27.20153 | 32.48501 | 18.36760 | 31.46663 | 0.00002 | 25.83833 | 27.95502 |
| **GRPEL2** | 1.50495 | 7.07246 | 11.41284 | 2.88089 | 1.25190 | 16.23523 | 12.99933 | 10.06980 | 11.94039 | 3.99548 | 11.55680 | 13.63395 |
| **GRPR** | 1.19400 | 0.00826 | 0.14688 | 0.32880 | 0.02174 | 0.00000 | 0.01457 | 0.00001 | 0.00000 | 0.03000 | 0.07163 | 0.23790 |

| | | | | | | | | | | | | |
|---|---|---|---|---|---|---|---|---|---|---|---|---|
| **GRSF1** | 2.47496 | 32.95577 | 19.15815 | 12.99982 | 15.83894 | 36.63587 | 18.39647 | 8.23066 | 31.97788 | 10.18289 | 14.51316 | 32.94235 |
| **GRTP1** | 4.97313 | 5.65151 | 2.11537 | 2.57688 | 5.00790 | 4.58472 | 2.60096 | 1.37058 | 5.25442 | 0.00000 | 1.87993 | 0.89175 |
| **GRTP1-AS1** | 0.81702 | 0.00867 | 0.01327 | 0.09384 | 0.08252 | 0.05835 | 0.03383 | 0.16559 | 0.02549 | 0.00077 | 0.05258 | 0.12464 |
| **GRWD1** | 0.67786 | 26.83400 | 21.80427 | 16.33461 | 21.92069 | 1.95851 | 28.23293 | 18.44873 | 28.69014 | 0.00000 | 19.17648 | 27.84434 |
| **GRXCR2** | 0.00746 | 0.02261 | 0.01424 | 0.01145 | 0.01209 | 0.01518 | 0.01763 | 0.01081 | 0.01635 | 0.01629 | 0.02997 | 0.01549 |
| **GS1-259H13.2** | 0.00183 | 0.06702 | 0.14989 | 0.00000 | 0.15275 | 0.00000 | 0.02471 | 0.06413 | 0.11569 | 0.05823 | 0.03693 | 0.03658 |
| **GS1-279B7.1** | 0.00671 | 0.01700 | 0.01201 | 0.01115 | 0.02183 | 0.00681 | 0.01297 | 0.01047 | 0.09519 | 0.00681 | 0.00421 | 0.01739 |
| **GS1-600G8.3** | 0.00439 | 0.01503 | 0.01663 | 0.16910 | 0.00000 | 0.00000 | 0.01155 | 0.00968 | 0.05212 | 0.12423 | 0.05438 | 0.02174 |
| **GSAP** | 2.78499 | 0.01588 | 0.03544 | 0.03869 | 0.05711 | 0.03803 | 0.07755 | 0.17248 | 0.00081 | 0.00024 | 0.00914 | 0.07517 |
| **GSC** | 2.11161 | 1.58128 | 0.00000 | 3.69662 | 0.00000 | 0.00110 | 0.00000 | 1.06030 | 0.00000 | 0.22021 | 0.03436 | 0.25330 |
| **GSDMA** | 0.00871 | 0.13152 | 0.19254 | 0.00616 | 0.04015 | 0.16477 | 0.00000 | 0.00601 | 0.00000 | 0.07744 | 0.01178 | 0.07457 |
| **GSDMB** | 0.00088 | 0.49522 | 0.19159 | 0.18472 | 0.00000 | 0.40033 | 0.16184 | 0.07768 | 1.19772 | 2.39275 | 0.07439 | 0.58125 |
| **GSDMC** | 0.03193 | 0.01269 | 0.01387 | 0.02401 | 0.01072 | 0.02556 | 0.02336 | 0.01023 | 0.01259 | 0.02255 | 0.02253 | 0.01493 |
| **GSDMD** | 0.00357 | 0.03077 | 0.10962 | 1.91642 | 0.00000 | 0.00023 | 0.05762 | 0.00878 | 0.08506 | 0.16641 | 0.02903 | 0.01364 |
| **GSE1** | 9.61865 | 7.17618 | 21.38379 | 4.35277 | 14.42511 | 12.75073 | 16.02746 | 1.00478 | 13.84083 | 18.13146 | 4.59652 | 0.04232 |
| **GSG1** | 0.01202 | 0.54430 | 0.00000 | 0.27808 | 0.00000 | 0.03559 | 0.00000 | 0.33905 | 0.04233 | 0.07074 | 0.00149 | 0.10906 |
| **GSG1L** | 0.00750 | 0.01400 | 0.05090 | 0.01264 | 0.01162 | 0.02145 | 0.04498 | 0.01024 | 0.00212 | 0.02708 | 0.01529 | 0.01520 |
| **GSG2** | 1.27045 | 5.15366 | 4.85477 | 2.12697 | 1.80647 | 2.92748 | 0.45624 | 1.09412 | 5.21937 | 0.00000 | 5.15638 | 5.25584 |
| **GSK3A** | 4.06574 | 0.00000 | 1.55984 | 0.91372 | 4.38085 | 0.00000 | 4.51540 | 0.00000 | 0.20679 | 16.05319 | 0.00000 | 2.64003 |
| **GSK3B** | 1.51603 | 0.60178 | 13.70740 | 1.97711 | 4.37098 | 5.78576 | 13.33125 | 0.73269 | 1.97671 | 15.03290 | 0.73260 | 1.69413 |
| **GSKIP** | 5.69635 | 10.44224 | 16.52226 | 4.13517 | 0.67481 | 35.04778 | 15.54219 | 19.59940 | 9.58684 | 0.00000 | 7.64166 | 7.36004 |
| **GSN** | 2.00454 | 3.95183 | 0.67656 | 17.07832 | 0.03155 | 1.79913 | 0.00000 | 0.10937 | 0.59007 | 0.00000 | 0.23838 | 1.69504 |
| **GSN-AS1** | 0.46574 | 0.04101 | 0.05159 | 0.35462 | 0.00000 | 0.17384 | 0.00000 | 0.04980 | 0.00000 | 0.17367 | 0.00000 | 0.27578 |
| **GSPT1** | 6.09580 | 7.65169 | 16.14694 | 6.90693 | 16.00171 | 19.85017 | 32.31189 | 12.21900 | 8.68324 | 32.26121 | 14.59080 | 5.80505 |
| **GSPT2** | 0.09204 | 0.42817 | 0.42962 | 0.22363 | 0.07746 | 1.18916 | 0.68698 | 0.24467 | 0.75497 | 0.23083 | 0.25440 | 0.57518 |
| **GSR** | 1.34822 | 1.52189 | 7.21119 | 4.55801 | 3.55619 | 0.57546 | 11.37621 | 0.31509 | 0.00000 | 7.12649 | 0.83004 | 0.55493 |
| **GSS** | 0.48374 | 4.56629 | 4.85497 | 3.59095 | 0.20304 | 5.36293 | 2.15422 | 0.00000 | 4.36561 | 0.00000 | 5.71270 | 4.15060 |
| **GSTA1** | 0.00664 | 0.01040 | 0.03608 | 0.03066 | 0.01633 | 0.05036 | 0.01242 | 0.01027 | 0.00662 | 0.00807 | 0.02001 | 0.01795 |
| **GSTA2** | 0.00393 | 0.01822 | 0.00505 | 0.55328 | 0.10406 | 0.14850 | 0.00000 | 0.00860 | 0.00000 | 0.08000 | 0.00397 | 0.00267 |
| **GSTA3** | 0.05417 | 0.05262 | 1.66093 | 0.50749 | 0.82419 | 1.38851 | 0.00000 | 0.00000 | 0.39977 | 0.00000 | 1.42144 | 0.00177 |
| **GSTA4** | 0.00073 | 0.10181 | 2.15187 | 2.06493 | 0.00000 | 3.94640 | 4.09204 | 0.09397 | 3.19330 | 0.00000 | 4.47271 | 0.13182 |
| **GSTA5** | 0.00776 | 0.01590 | 0.02019 | 0.01224 | 0.01648 | 0.01365 | 0.01883 | 0.01034 | 0.02401 | 0.01249 | 0.01431 | 0.01663 |
| **GSTCD** | 2.93314 | 0.93824 | 2.30070 | 1.12754 | 0.74160 | 8.67306 | 3.36935 | 2.68011 | 1.45650 | 5.84439 | 4.22563 | 4.58752 |
| **GSTK1** | 0.68922 | 7.53275 | 8.84178 | 4.91904 | 2.72186 | 5.68553 | 4.92625 | 0.20377 | 15.41178 | 0.71015 | 16.30935 | 3.91014 |
| **GSTM1** | 0.00277 | 0.01592 | 0.23520 | 0.01740 | 0.00000 | 0.01462 | 0.06258 | 0.04170 | 0.00000 | 0.01066 | 0.34356 | 0.04395 |
| **GSTM2** | 0.08230 | 0.00915 | 0.11059 | 0.05922 | 0.00000 | 0.03800 | 0.00000 | 0.00943 | 0.07965 | 0.01752 | 0.01185 | 0.01695 |
| **GSTM2P1** | 0.00402 | 0.01452 | 0.40854 | 0.10248 | 0.00000 | 0.16751 | 0.04291 | 0.06568 | 0.00001 | 0.00224 | 0.12967 | 0.00975 |
| **GSTM3** | 1.80939 | 0.00000 | 5.01758 | 0.00000 | 0.00000 | 0.49256 | 1.71974 | 8.43334 | 2.53040 | 0.00000 | 2.21362 | 2.77680 |
| **GSTM4** | 0.00883 | 0.60809 | 1.71762 | 1.94139 | 0.00000 | 0.47902 | 1.35633 | 0.00224 | 0.52965 | 0.06710 | 1.80755 | 0.00000 |
| **GSTM5** | 0.00692 | 0.00895 | 0.34736 | 0.01920 | 0.00000 | 0.04986 | 0.00000 | 0.03799 | 0.00000 | 0.02828 | 0.01909 | 0.01136 |
| **GSTO1** | 0.00000 | 8.27398 | 5.35149 | 21.82656 | 9.87642 | 18.77728 | 4.48586 | 0.00000 | 1.85393 | 5.79511 | 10.94760 | 0.41017 |
| **GSTO2** | 0.00681 | 5.80439 | 0.00000 | 1.09519 | 1.41527 | 0.93974 | 0.00876 | 0.00000 | 0.00000 | 0.00000 | 0.74077 | 5.96874 |
| **GSTP1** | 0.65617 | 42.52865 | 35.37019 | 57.61743 | 72.04072 | 47.28967 | 46.43570 | 1.64297 | 42.77671 | 15.28240 | 42.42159 | 12.38736 |
| **GSTT1** | 0.00415 | 0.00287 | 8.70435 | 0.00050 | 0.00000 | 0.00402 | 5.53002 | 0.03166 | 3.37776 | 0.00001 | 2.32009 | 0.08046 |
| **GSTTP2** | 0.00614 | 0.01634 | 0.06453 | 0.01006 | 0.01675 | 0.01644 | 0.01946 | 0.01501 | 0.01703 | 0.01129 | 0.01615 | 0.01681 |
| **GSTZ1** | 0.03315 | 13.17260 | 2.50508 | 6.94226 | 12.05067 | 2.29595 | 2.45725 | 0.00000 | 4.44845 | 1.50437 | 5.79690 | 4.50102 |
| **GSX2** | 0.00476 | 0.01550 | 0.03545 | 0.01990 | 0.01381 | 0.01817 | 0.01436 | 0.01632 | 0.01239 | 0.01882 | 0.01693 | 0.01572 |
| **GTDC1** | 0.78474 | 3.67957 | 2.31759 | 2.78508 | 0.94953 | 5.99529 | 3.93615 | 3.65346 | 6.46053 | 1.63721 | 3.52152 | 2.26361 |

| | | | | | | | | | | | | |
|---|---|---|---|---|---|---|---|---|---|---|---|---|
| **GTF2A1** | 2.14328 | 1.17971 | 3.85408 | 0.01456 | 0.00000 | 11.10280 | 5.45366 | 4.24029 | 0.00225 | 15.62843 | 1.89909 | 4.75535 |
| **GTF2A1L** | 0.14358 | 0.01802 | 0.02068 | 0.00988 | 0.02088 | 0.01928 | 0.01617 | 0.28079 | 0.01799 | 0.00922 | 0.01730 | 0.01681 |
| **GTF2A2** | 4.29190 | 19.92131 | 22.26738 | 5.07087 | 18.94372 | 66.20683 | 29.92179 | 27.29781 | 24.06148 | 24.09177 | 32.75749 | 30.59544 |
| **GTF2B** | 5.67358 | 13.40430 | 9.87766 | 5.42980 | 2.42823 | 29.45762 | 9.31771 | 10.65427 | 16.20420 | 0.20199 | 5.31074 | 15.35120 |
| **GTF2E1** | 0.94577 | 9.59032 | 3.26141 | 3.34161 | 0.00000 | 19.08787 | 9.33438 | 9.89066 | 6.10867 | 0.00000 | 6.62611 | 10.81329 |
| **GTF2E2** | 0.01961 | 2.97630 | 4.11957 | 4.41689 | 5.93612 | 5.95065 | 5.96347 | 6.05475 | 1.53248 | 8.97666 | 1.99313 | 3.00330 |
| **GTF2F1** | 8.24250 | 14.58330 | 11.95954 | 13.15969 | 16.68259 | 0.79545 | 21.19565 | 11.44805 | 10.54182 | 13.44131 | 8.07428 | 14.73972 |
| **GTF2F2** | 5.19967 | 9.86473 | 8.29929 | 4.66382 | 5.12123 | 24.51140 | 13.66889 | 17.91761 | 10.26343 | 8.38694 | 7.82712 | 11.48172 |
| **GTF2H1** | 1.38203 | 7.44762 | 12.78300 | 4.79233 | 2.67239 | 22.97975 | 10.90204 | 16.34507 | 11.75223 | 0.00000 | 13.31828 | 9.52717 |
| **GTF2H2** | 0.03913 | 0.34769 | 0.70135 | 0.09762 | 0.00000 | 1.41508 | 0.26903 | 0.00000 | 0.43571 | 0.00234 | 0.79317 | 0.38727 |
| **GTF2H2B** | 0.00893 | 0.05554 | 0.16756 | 0.01764 | 0.01673 | 0.00458 | 0.03017 | 0.00928 | 0.04655 | 0.04850 | 0.02636 | 0.02517 |
| **GTF2H2C** | 0.00581 | 0.00904 | 0.01410 | 0.00574 | 0.04815 | 0.00000 | 0.02405 | 0.00962 | 0.02076 | 0.17816 | 0.01466 | 0.01437 |
| **GTF2H2C-2** | 0.04385 | 0.07931 | 0.07790 | 0.09029 | 0.15687 | 0.23910 | 0.00000 | 0.00192 | 0.21712 | 0.17742 | 0.32161 | 0.17521 |
| **GTF2H3** | 0.12617 | 5.64965 | 2.89045 | 1.25003 | 1.34062 | 14.13087 | 0.56133 | 0.08377 | 3.33389 | 3.00114 | 5.46993 | 3.16513 |
| **GTF2H4** | 0.42687 | 2.13639 | 1.03929 | 4.45094 | 0.99922 | 0.00000 | 0.67341 | 0.00000 | 3.84749 | 0.00000 | 4.04190 | 2.99261 |
| **GTF2H5** | 1.11402 | 18.47566 | 5.18671 | 5.04154 | 11.62774 | 26.41800 | 8.13347 | 0.43174 | 5.37681 | 10.90182 | 15.74157 | 12.23401 |
| **GTF2I** | 0.81428 | 1.73331 | 0.94973 | 1.88613 | 1.31743 | 0.32890 | 0.45991 | 0.10834 | 1.32285 | 1.09712 | 0.51200 | 0.37923 |
| **GTF2IP1.uid-92** | 0.00783 | 0.01511 | 0.01704 | 0.01193 | 0.02569 | 0.01502 | 0.01722 | 0.01030 | 0.01723 | 0.01971 | 0.01594 | 0.01611 |
| **GTF2IRD1** | 0.15018 | 2.52177 | 0.13292 | 1.32261 | 0.00000 | 0.00219 | 0.19672 | 0.43033 | 0.27010 | 0.00000 | 0.00000 | 2.67844 |
| **GTF2IRD1P1** | 0.10954 | 0.08343 | 0.02226 | 0.00277 | 0.01200 | 0.05548 | 0.01715 | 0.00459 | 0.01198 | 0.02547 | 0.01228 | 0.03686 |
| **GTF2IRD2** | 0.10094 | 0.04149 | 0.00921 | 0.01052 | 0.00406 | 0.04572 | 0.03190 | 0.10146 | 0.02343 | 0.01715 | 0.00807 | 0.02119 |
| **GTF2IRD2+GTF** | 0.06707 | 0.02753 | 0.01476 | 0.00877 | 0.02064 | 0.02766 | 0.02014 | 0.00951 | 0.01679 | 0.00878 | 0.01568 | 0.01817 |
| **GTF3A** | 0.23388 | 19.25409 | 13.45440 | 15.23898 | 18.65187 | 22.84351 | 29.88681 | 1.36912 | 32.24446 | 10.06068 | 30.36310 | 23.17631 |
| **GTF3C1** | 3.69705 | 11.31424 | 5.62134 | 15.92067 | 7.42916 | 0.00000 | 5.88473 | 4.19875 | 15.76599 | 13.26553 | 2.46878 | 12.97868 |
| **GTF3C2** | 4.87701 | 24.66544 | 14.28996 | 13.46600 | 13.75864 | 3.98666 | 17.66390 | 5.68815 | 21.41556 | 5.83674 | 15.24053 | 19.68572 |
| **GTF3C2-AS1** | 0.02367 | 0.17829 | 0.06857 | 0.16580 | 0.09811 | 0.00000 | 0.04310 | 0.06597 | 0.23326 | 0.16892 | 0.07855 | 0.11133 |
| **GTF3C3** | 0.57243 | 9.92205 | 3.05198 | 2.70285 | 2.54948 | 21.18665 | 7.24047 | 2.30526 | 3.61118 | 8.59880 | 8.43787 | 8.75725 |
| **GTF3C4** | 1.16281 | 2.16558 | 2.74027 | 3.10485 | 0.72389 | 12.10251 | 3.17845 | 2.07027 | 3.74897 | 7.80604 | 1.58520 | 7.47046 |
| **GTF3C5** | 3.92147 | 23.55957 | 18.63710 | 12.13716 | 14.02551 | 4.03005 | 21.57789 | 3.86411 | 23.94462 | 0.00000 | 12.32561 | 22.75316 |
| **GTF3C6** | 0.24171 | 9.01556 | 9.79998 | 9.76988 | 2.28764 | 30.32001 | 16.83889 | 5.90529 | 11.53612 | 7.54352 | 11.74553 | 22.14292 |
| **GTPBP1** | 1.48940 | 3.85251 | 4.03417 | 4.58435 | 5.24856 | 0.00000 | 2.85536 | 0.87088 | 5.34448 | 5.54674 | 0.93076 | 2.07758 |
| **GTPBP10** | 4.35236 | 5.96480 | 5.60446 | 0.46630 | 1.84639 | 14.64139 | 4.95016 | 0.93460 | 8.37643 | 3.07556 | 5.05584 | 6.27625 |
| **GTPBP2** | 1.43106 | 1.14866 | 2.36815 | 3.59587 | 1.71935 | 0.00000 | 2.22601 | 0.21846 | 2.37965 | 0.56570 | 0.97037 | 1.64802 |
| **GTPBP3** | 1.75349 | 0.00000 | 0.60714 | 2.55127 | 3.04834 | 0.00000 | 2.32925 | 1.23809 | 12.65001 | 0.00000 | 5.65601 | 7.74956 |
| **GTPBP4** | 1.05418 | 18.42154 | 17.36753 | 8.70679 | 13.80913 | 30.41525 | 25.97074 | 24.72706 | 24.96285 | 12.11916 | 20.50523 | 26.25116 |
| **GTPBP8** | 0.64844 | 6.81037 | 0.69956 | 4.78096 | 4.84379 | 11.68383 | 5.85844 | 1.96446 | 4.44462 | 1.09451 | 7.12077 | 10.30158 |
| **GTSE1** | 6.06576 | 10.95261 | 6.91851 | 9.62598 | 13.53390 | 0.33043 | 17.16801 | 2.07813 | 15.52112 | 9.56016 | 7.96615 | 9.73895 |
| **GTSE1-AS1** | 0.00372 | 0.36720 | 0.07427 | 0.01367 | 0.00000 | 0.00000 | 0.12002 | 0.02394 | 0.01687 | 0.00000 | 0.24449 | 0.74062 |
| **GTSF1** | 6.35639 | 13.23918 | 16.28236 | 12.15956 | 15.68507 | 29.86422 | 26.70266 | 5.81951 | 19.42179 | 0.77186 | 22.34946 | 11.22113 |
| **GTSF1L** | 0.35391 | 0.05986 | 0.18743 | 0.11154 | 0.00000 | 0.05974 | 0.00000 | 0.00764 | 0.00000 | 0.03506 | 0.05158 | 0.00337 |
| **GUCA1A** | 0.01046 | 1.21740 | 0.41883 | 3.20135 | 0.00000 | 0.41516 | 0.00000 | 0.14514 | 0.11774 | 0.00000 | 0.15879 | 1.48840 |
| **GUCA1B** | 0.00111 | 0.09942 | 0.06168 | 0.15519 | 0.04274 | 0.22287 | 0.02946 | 0.09231 | 0.06900 | 0.10519 | 0.06351 | 0.27512 |
| **GUCA1C** | 0.01138 | 0.01349 | 0.01873 | 0.01148 | 0.01049 | 0.02140 | 0.05199 | 0.00974 | 0.00096 | 0.02970 | 0.01510 | 0.01476 |
| **GUCA2A** | 0.00386 | 0.00407 | 0.65658 | 0.54026 | 0.00000 | 0.08280 | 0.00000 | 0.00905 | 0.00000 | 0.10467 | 0.06256 | 0.00640 |
| **GUCD1** | 10.04498 | 12.83325 | 4.68789 | 9.07779 | 11.56022 | 2.03430 | 10.80180 | 10.36801 | 10.76531 | 4.32702 | 5.91024 | 10.24760 |
| **GUCY1A2** | 3.55698 | 0.02519 | 0.29861 | 0.00790 | 0.00000 | 0.00000 | 0.00000 | 0.21353 | 0.06498 | 0.08629 | 0.01502 | 0.01490 |
| **GUCY1A3** | 0.70030 | 0.01557 | 0.00983 | 0.02934 | 0.00629 | 0.00000 | 0.01059 | 0.05491 | 0.01517 | 0.08687 | 0.03233 | 0.01212 |
| **GUCY1B2** | 0.10083 | 0.08565 | 0.00164 | 0.00000 | 0.00000 | 0.04617 | 0.00000 | 0.12518 | 0.04164 | 0.10679 | 0.00284 | 0.00652 |

| | | | | | | | | | | | | |
|---|---|---|---|---|---|---|---|---|---|---|---|---|
| **GUCY1B3** | 0.00699 | 0.00539 | 0.02379 | 0.02975 | 0.06041 | 0.05771 | 0.03466 | 0.00914 | 0.01568 | 0.00000 | 0.02761 | 0.02282 |
| **GUCY2C** | 0.23075 | 0.39780 | 0.02391 | 0.31001 | 0.00000 | 0.01054 | 0.00146 | 0.04412 | 0.13686 | 0.02912 | 0.06800 | 0.38606 |
| **GUCY2D** | 0.11586 | 0.00809 | 0.03658 | 0.01552 | 0.01590 | 0.00000 | 0.00807 | 0.05290 | 0.03160 | 0.04312 | 0.04083 | 0.08539 |
| **GUCY2EP** | 0.00835 | 0.01145 | 0.01540 | 0.20432 | 0.01330 | 0.01453 | 0.02519 | 0.00958 | 0.01083 | 0.02517 | 0.02594 | 0.01546 |
| **GUCY2F** | 0.00990 | 0.01511 | 0.01278 | 0.01036 | 0.02500 | 0.00000 | 0.08640 | 0.04051 | 0.00000 | 0.05688 | 0.01029 | 0.01601 |
| **GUCY2GP** | 0.04837 | 0.02031 | 0.01601 | 0.01474 | 0.01424 | 0.01011 | 0.01753 | 0.00625 | 0.02080 | 0.01902 | 0.01966 | 0.01725 |
| **GUF1** | 0.00000 | 12.07091 | 6.34692 | 3.25108 | 5.32052 | 28.96283 | 8.17687 | 5.85097 | 13.64136 | 8.10560 | 10.27304 | 12.18532 |
| **GUK1** | 0.00000 | 12.51362 | 17.20164 | 8.58507 | 15.42424 | 0.00000 | 16.07618 | 1.57795 | 19.75454 | 12.62813 | 19.21298 | 3.76600 |
| **GULP1** | 0.00000 | 0.00633 | 6.89669 | 0.00000 | 0.18602 | 9.94260 | 5.29614 | 0.03264 | 2.16489 | 3.88926 | 3.49806 | 2.24001 |
| **GUSB** | 0.52616 | 14.05352 | 4.25759 | 4.92091 | 7.71259 | 0.00000 | 2.35516 | 2.38862 | 11.33107 | 0.02153 | 5.97000 | 8.54712 |
| **GUSBP1** | 0.33937 | 1.18137 | 1.95979 | 0.01505 | 1.23949 | 1.68206 | 1.14994 | 14.17817 | 4.18012 | 1.23998 | 0.72728 | 3.31340 |
| **GUSBP10** | 0.17228 | 0.07137 | 0.00865 | 0.05202 | 0.00362 | 0.03528 | 0.01527 | 0.15189 | 0.01971 | 0.00171 | 0.02317 | 0.03160 |
| **GUSBP11** | 0.00001 | 1.13851 | 1.49262 | 1.66116 | 1.29867 | 0.00000 | 0.00116 | 1.05134 | 6.68044 | 3.92727 | 1.24872 | 1.33153 |
| **GUSBP2** | 0.02590 | 0.01373 | 0.03713 | 0.01886 | 0.01159 | 0.03675 | 0.00839 | 0.01120 | 0.07750 | 0.01771 | 0.03204 | 0.01689 |
| **GUSBP3.uid-67** | 0.00622 | 0.01421 | 0.01500 | 0.01324 | 0.01544 | 0.02016 | 0.01855 | 0.01043 | 0.01659 | 0.01511 | 0.01524 | 0.01704 |
| **GUSBP4** | 0.14141 | 0.15057 | 0.10775 | 0.19303 | 0.14008 | 0.40360 | 0.00000 | 2.21689 | 0.53938 | 0.18935 | 0.00003 | 0.66164 |
| **GUSBP5** | 0.07236 | 0.33196 | 0.00707 | 0.00000 | 0.00000 | 0.12109 | 0.04052 | 0.08801 | 0.02833 | 0.07015 | 0.03399 | 0.26392 |
| **GVINP1** | 0.34104 | 0.14391 | 0.39681 | 0.96441 | 0.13527 | 0.00000 | 0.00000 | 0.22524 | 1.16329 | 0.21967 | 0.49032 | 0.72461 |
| **GXYLT1** | 0.00000 | 4.44941 | 3.10437 | 1.89850 | 0.00000 | 13.57046 | 0.00000 | 0.45262 | 2.62966 | 16.92009 | 1.20350 | 1.43814 |
| **GXYLT2** | 0.08085 | 0.01293 | 0.05160 | 0.09333 | 0.08970 | 0.02620 | 0.07316 | 0.00409 | 0.00000 | 0.15262 | 0.00915 | 0.00942 |
| **GYG1** | 16.44233 | 9.41822 | 3.93984 | 9.26680 | 2.51578 | 13.26838 | 10.17938 | 1.80054 | 9.72979 | 2.73921 | 4.58731 | 4.39769 |
| **GYG2** | 0.00325 | 0.17315 | 0.47985 | 3.54477 | 0.00000 | 1.01995 | 0.93547 | 0.61203 | 0.78677 | 0.00000 | 0.00000 | 0.82079 |
| **GYG2P1** | 0.00785 | 0.01049 | 0.09093 | 0.34595 | 0.07713 | 0.00022 | 0.02155 | 0.01130 | 0.08805 | 0.00000 | 0.27105 | 0.02103 |
| **GYLTL1B** | 0.00000 | 8.32570 | 6.87289 | 0.00000 | 31.56013 | 0.07294 | 14.32126 | 0.00000 | 27.58732 | 16.29873 | 14.46908 | 0.00000 |
| **GYPA** | 0.00610 | 0.01505 | 0.01579 | 0.01224 | 0.01654 | 0.01491 | 0.03543 | 0.01054 | 0.00968 | 0.02001 | 0.01494 | 0.01609 |
| **GYPC** | 0.00527 | 0.00807 | 0.39793 | 5.70932 | 0.00000 | 0.02146 | 0.01629 | 0.00855 | 0.01259 | 0.01187 | 0.02204 | 0.01129 |
| **GYPE** | 0.00568 | 0.01107 | 0.02999 | 0.00840 | 0.00712 | 0.03558 | 0.01830 | 0.01027 | 0.01133 | 0.03960 | 0.01245 | 0.01355 |
| **GYS1** | 3.20050 | 10.54280 | 11.71101 | 4.65665 | 17.24081 | 0.00000 | 14.44686 | 0.93291 | 17.00943 | 10.18328 | 18.26668 | 4.55820 |
| **GYS2** | 0.03561 | 0.02272 | 0.03767 | 0.01307 | 0.00664 | 0.01615 | 0.03493 | 0.00874 | 0.16778 | 0.02127 | 0.01082 | 0.02046 |
| **GZF1** | 0.78356 | 1.02708 | 1.96797 | 0.74353 | 0.28080 | 4.56874 | 2.35511 | 0.54330 | 2.23641 | 2.67264 | 2.22745 | 0.93294 |
| **GZMM** | 0.01580 | 0.00841 | 0.05149 | 0.15005 | 0.00838 | 0.00000 | 0.00827 | 0.00406 | 0.13965 | 0.00672 | 0.04106 | 0.35862 |
| **H19** | 0.09385 | 0.00000 | 23.75403 | 0.04072 | 0.00000 | 0.07171 | 9.69151 | 0.00181 | 0.19114 | 0.13073 | 0.00000 | 0.00447 |
| **H1F0** | 10.86048 | 14.34600 | 14.57938 | 15.55389 | 3.58731 | 19.95715 | 13.96052 | 5.68661 | 11.39678 | 1.20496 | 4.40124 | 23.91736 |
| **H1FNT** | 0.00420 | 0.04362 | 0.01385 | 0.00204 | 0.04713 | 0.01335 | 0.01076 | 2.46878 | 0.01332 | 0.02094 | 0.00952 | 0.22773 |
| **H1FOO** | 32.26483 | 0.04947 | 0.00060 | 0.00000 | 0.17485 | 0.00000 | 0.00372 | 12.92145 | 0.00613 | 0.00039 | 0.00000 | 0.98850 |
| **H1FX** | 0.00268 | 0.86512 | 0.76031 | 0.31870 | 1.46077 | 0.00000 | 2.09271 | 0.00029 | 0.00000 | 10.18873 | 0.00000 | 0.96230 |
| **H1FX-AS1** | 0.05350 | 0.04287 | 0.06859 | 0.05017 | 0.07901 | 0.00000 | 0.06270 | 0.00204 | 0.32862 | 0.45550 | 0.00000 | 0.12724 |
| **H2AFJ** | 0.00000 | 3.30795 | 16.17679 | 3.02699 | 0.00080 | 3.47687 | 10.54818 | 1.45475 | 6.99216 | 4.42059 | 2.69615 | 2.45172 |
| **H2AFV** | 0.93555 | 17.66581 | 6.66182 | 8.12560 | 12.63564 | 13.48967 | 10.24060 | 15.20458 | 4.32138 | 15.86401 | 7.49052 | 15.53985 |
| **H2AFX** | 3.16729 | 6.77048 | 12.59775 | 3.58254 | 11.33242 | 0.00000 | 12.06189 | 1.51025 | 3.49403 | 26.84246 | 0.00000 | 5.34498 |
| **H2AFY** | 0.00000 | 6.48344 | 9.70679 | 11.03771 | 13.17660 | 15.84102 | 20.38823 | 0.53875 | 0.00000 | 24.86081 | 8.70471 | 4.53527 |
| **H2AFY2** | 1.96533 | 2.56936 | 0.80837 | 4.73164 | 1.30550 | 0.21544 | 0.00000 | 0.07414 | 0.00052 | 0.00742 | 0.91723 | 0.00000 |
| **H2AFZ** | 7.95170 | 32.53763 | 35.07481 | 30.68770 | 28.14200 | 66.74158 | 63.42608 | 48.21771 | 46.52732 | 16.26984 | 42.86024 | 55.75447 |
| **H2BFM** | 0.00707 | 0.04810 | 0.01289 | 0.00277 | 0.01139 | 0.04326 | 0.00000 | 0.00948 | 0.03519 | 0.03521 | 0.00820 | 0.05043 |
| **H2BFWT** | 0.00607 | 0.07252 | 0.02198 | 0.04635 | 0.00000 | 0.03039 | 0.01856 | 0.00974 | 0.01162 | 0.02141 | 0.01534 | 0.08723 |
| **H3F3A** | 0.20255 | 1.13950 | 1.33088 | 1.16246 | 0.87176 | 0.95361 | 0.80002 | 0.97662 | 1.67460 | 0.00000 | 0.96818 | 1.83603 |
| **H3F3B** | 23.40669 | 39.43607 | 42.57362 | 27.58436 | 33.93904 | 69.22529 | 56.54586 | 50.14858 | 40.64962 | 19.25435 | 37.94680 | 51.63428 |
| **H3F3C** | 6.50444 | 7.66811 | 8.60150 | 5.09940 | 6.62918 | 10.39529 | 8.80219 | 23.01136 | 5.90184 | 0.00000 | 5.10761 | 11.10440 |

| | | | | | | | | | | | | |
|---|---|---|---|---|---|---|---|---|---|---|---|---|
| **H6PD** | 0.44729 | 0.05377 | 0.10261 | 0.00002 | 0.35276 | 0.00000 | 0.00880 | 0.14747 | 0.16754 | 0.11962 | 0.01495 | 0.18873 |
| **HAAO** | 0.00474 | 0.00575 | 0.14578 | 0.14223 | 0.09805 | 0.00000 | 0.00000 | 0.01037 | 0.05387 | 0.08613 | 0.01060 | 0.03315 |
| **HABP2** | 14.39585 | 0.00000 | 0.22377 | 1.42438 | 0.08694 | 0.00013 | 0.00000 | 2.31811 | 0.00326 | 0.00944 | 0.01033 | 0.15911 |
| **HABP4** | 1.31224 | 0.09182 | 0.34013 | 0.39526 | 0.64648 | 0.00000 | 0.02778 | 0.34046 | 0.51107 | 0.32490 | 0.00921 | 0.31375 |
| **HACE1** | 1.51358 | 0.81050 | 1.74768 | 0.41044 | 0.02531 | 2.46553 | 0.52284 | 1.96083 | 1.71987 | 0.39317 | 0.84349 | 0.53616 |
| **HACL1** | 0.57536 | 5.83562 | 8.56071 | 1.16689 | 1.53111 | 14.78088 | 8.13784 | 1.80583 | 9.26723 | 0.00000 | 6.08224 | 6.71338 |
| **HADH** | 2.37533 | 2.88512 | 5.50031 | 6.09441 | 3.21799 | 7.71020 | 14.70044 | 1.10269 | 4.03895 | 0.00000 | 9.85127 | 0.75331 |
| **HADHA** | 2.94230 | 8.23606 | 7.21560 | 10.41240 | 12.59757 | 14.97864 | 11.83027 | 1.03814 | 18.29154 | 3.83947 | 12.08587 | 9.47731 |
| **HADHB** | 1.51355 | 9.91907 | 2.84894 | 7.63645 | 5.87416 | 17.53381 | 6.82399 | 0.00000 | 9.86835 | 0.74633 | 11.31401 | 4.32015 |
| **HAGH** | 8.07359 | 1.67912 | 2.67560 | 3.86034 | 3.89689 | 0.17094 | 2.79749 | 3.08680 | 2.20855 | 0.00000 | 2.79383 | 0.10312 |
| **HAGHL** | 0.00102 | 0.10096 | 0.36372 | 0.00000 | 0.00000 | 0.00000 | 0.53454 | 0.02281 | 0.06385 | 1.23158 | 0.15467 | 0.02295 |
| **HAGLR** | 0.94973 | 0.00000 | 0.00000 | 1.66940 | 0.06921 | 0.00000 | 0.00093 | 0.00000 | 0.06360 | 0.14263 | 0.00769 | 1.58582 |
| **HAGLROS** | 0.05823 | 0.00852 | 0.07669 | 0.17784 | 0.00967 | 0.01272 | 0.01360 | 0.00546 | 0.01573 | 0.01070 | 0.01722 | 0.10042 |
| **HAL** | 3.03426 | 0.00287 | 0.07162 | 0.01450 | 0.01192 | 0.07461 | 0.13003 | 0.92066 | 0.02939 | 0.00000 | 0.00477 | 0.03714 |
| **HAMP** | 0.07655 | 0.01395 | 0.16550 | 0.04544 | 0.00954 | 0.01749 | 0.00000 | 0.00495 | 0.13516 | 0.00572 | 0.01043 | 0.01519 |
| **HAND1** | 0.34835 | 0.00000 | 2.45893 | 2.92568 | 14.84134 | 0.00000 | 15.70502 | 0.23268 | 16.52130 | 0.00738 | 0.00000 | 0.00287 |
| **HAND2** | 0.01072 | 0.01518 | 0.01081 | 0.01105 | 0.01276 | 0.01350 | 0.02496 | 0.25171 | 0.00685 | 0.06390 | 0.01405 | 0.02123 |
| **HAO1** | 0.00893 | 0.01402 | 0.01412 | 0.01341 | 0.01146 | 0.01993 | 0.01812 | 0.03421 | 0.01489 | 0.02476 | 0.02023 | 0.01394 |
| **HAP1** | 11.81423 | 0.42748 | 3.62279 | 0.00000 | 0.00000 | 0.00000 | 0.61514 | 1.86075 | 0.00000 | 1.82792 | 0.08115 | 1.44560 |
| **HAPLN1** | 0.02353 | 0.00000 | 14.25745 | 0.00000 | 0.00000 | 0.00000 | 18.48023 | 0.00371 | 40.22921 | 0.30980 | 10.27602 | 0.00000 |
| **HAPLN2** | 0.00626 | 0.01413 | 0.01672 | 0.00809 | 0.01928 | 0.02570 | 0.01936 | 0.01073 | 0.00000 | 0.02833 | 0.05321 | 0.01671 |
| **HAPLN3** | 0.80388 | 1.01018 | 0.01344 | 0.50605 | 0.00000 | 0.00000 | 0.00000 | 0.76354 | 0.06369 | 0.05783 | 0.00000 | 0.33489 |
| **HAPLN4** | 0.00406 | 0.00004 | 0.01276 | 0.04383 | 0.00000 | 0.06628 | 0.03408 | 0.00953 | 0.01724 | 0.15806 | 0.00000 | 0.00812 |
| **HAR1A** | 0.00972 | 0.01645 | 0.03033 | 0.01038 | 0.01563 | 0.01888 | 0.01177 | 0.01034 | 0.01730 | 0.02191 | 0.02166 | 0.01740 |
| **HARBI1** | 0.42009 | 1.84452 | 0.79852 | 1.43526 | 0.65067 | 2.59281 | 0.00010 | 1.41941 | 0.88879 | 0.00000 | 2.64829 | 3.61664 |
| **HARS** | 1.34503 | 18.76729 | 19.07230 | 9.83252 | 16.28588 | 19.14761 | 21.01967 | 10.93266 | 20.79723 | 0.00000 | 23.34558 | 17.67528 |
| **HARS2** | 0.00358 | 12.27292 | 4.19586 | 3.59536 | 7.63673 | 8.74796 | 2.55877 | 2.62539 | 7.01510 | 0.00000 | 7.86881 | 2.73404 |
| **HAS1** | 0.22330 | 0.02686 | 0.01230 | 0.00891 | 0.01667 | 0.01370 | 0.02124 | 0.02400 | 0.02239 | 0.01535 | 0.01164 | 0.09671 |
| **HAS2** | 1.94568 | 0.00000 | 0.12676 | 13.57275 | 0.00000 | 0.05250 | 0.00000 | 0.11319 | 0.16701 | 0.55702 | 0.06169 | 0.00000 |
| **HAS2-AS1** | 0.04361 | 0.00913 | 0.02756 | 0.44054 | 0.00000 | 0.04076 | 0.02044 | 0.00740 | 0.02704 | 0.01701 | 0.02646 | 0.01594 |
| **HAS3** | 2.65350 | 0.50770 | 2.02416 | 0.00000 | 1.08549 | 0.00000 | 0.30752 | 0.09247 | 0.00000 | 1.01463 | 2.93577 | 2.65946 |
| **HAT1** | 0.55690 | 16.98437 | 11.96876 | 13.73652 | 0.00000 | 41.53294 | 11.47528 | 17.98534 | 7.25456 | 0.00067 | 10.60118 | 30.61318 |
| **HAUS1** | 0.24874 | 3.85125 | 6.42317 | 6.93373 | 1.41386 | 19.27962 | 16.23252 | 0.00000 | 8.48930 | 1.48599 | 8.89855 | 1.74626 |
| **HAUS2** | 0.00000 | 6.26766 | 11.19914 | 1.64193 | 3.70032 | 20.01887 | 8.92900 | 3.79476 | 8.02927 | 6.29093 | 7.12698 | 13.29510 |
| **HAUS3** | 0.52341 | 3.65087 | 2.59321 | 0.71365 | 0.00000 | 8.13792 | 4.35460 | 9.92557 | 1.42790 | 3.53685 | 3.22502 | 6.27371 |
| **HAUS4** | 9.67943 | 6.30328 | 6.25498 | 6.15787 | 4.37298 | 11.39085 | 7.00117 | 3.12152 | 5.47582 | 0.00000 | 12.23676 | 0.51492 |
| **HAUS5** | 1.19696 | 8.75893 | 2.66462 | 5.23515 | 6.48350 | 0.00000 | 2.25739 | 0.32488 | 5.00452 | 1.39402 | 3.79016 | 5.56936 |
| **HAUS6** | 0.00000 | 1.70415 | 4.57501 | 0.49860 | 0.00000 | 10.69502 | 8.13864 | 2.54375 | 4.32242 | 8.23069 | 2.58471 | 6.26477 |
| **HAUS7** | 1.02320 | 5.55255 | 5.03845 | 4.28006 | 9.01365 | 0.00049 | 10.20736 | 1.40986 | 9.32738 | 0.00000 | 7.96611 | 5.58421 |
| **HAUS8** | 1.23885 | 5.44781 | 6.73126 | 2.74937 | 4.56145 | 5.21454 | 5.54452 | 2.10087 | 5.15608 | 2.83648 | 3.17056 | 5.45840 |
| **HAVCR1** | 0.00000 | 21.38561 | 6.76144 | 6.71232 | 16.40074 | 15.20650 | 0.00000 | 0.95974 | 29.70675 | 0.50933 | 34.83472 | 15.68139 |
| **HAVCR1P1** | 0.00266 | 2.66127 | 0.46230 | 0.64337 | 1.33548 | 1.78142 | 0.00000 | 0.22787 | 2.74874 | 0.00000 | 6.03923 | 1.60933 |
| **HAVCR2** | 0.00000 | 1.30945 | 0.59942 | 0.00000 | 1.34440 | 0.35702 | 0.21550 | 0.00000 | 0.50678 | 1.74333 | 2.35652 | 1.29651 |
| **HAX1** | 0.57030 | 22.00500 | 7.69255 | 12.63175 | 18.33656 | 10.81282 | 13.01419 | 0.30239 | 17.79155 | 2.52987 | 12.74350 | 21.74184 |
| **HBA1** | 0.00205 | 1.23961 | 0.04883 | 0.00000 | 0.30851 | 0.00000 | 0.13319 | 0.22074 | 0.20238 | 0.25601 | 0.16926 | 1.25906 |
| **HBA2** | 0.00745 | 0.91813 | 0.05014 | 0.00000 | 0.14829 | 0.00000 | 0.01351 | 0.40170 | 0.00000 | 0.48623 | 0.06716 | 1.14888 |
| **HBB** | 0.00205 | 0.00689 | 0.02703 | 0.01419 | 0.01000 | 0.07435 | 0.00785 | 0.04444 | 0.00000 | 0.02688 | 0.07103 | 0.03865 |
| **HBBP1** | 0.00622 | 0.01331 | 0.01631 | 0.01096 | 0.01623 | 0.02418 | 0.01593 | 0.01040 | 0.01426 | 0.01480 | 0.02724 | 0.01590 |

| | | | | | | | | | | | | |
|---|---|---|---|---|---|---|---|---|---|---|---|---|
| **HBD** | 0.00535 | 0.00390 | 0.04190 | 0.00440 | 0.01494 | 0.06830 | 0.02739 | 0.01057 | 0.00000 | 0.04639 | 0.02908 | 0.05548 |
| **HBE1** | 0.14210 | 0.00000 | 0.07768 | 0.05187 | 0.00000 | 0.66933 | 0.52253 | 0.04719 | 0.12043 | 1.67305 | 0.92460 | 0.00000 |
| **HBEGF** | 0.26578 | 0.12040 | 1.98451 | 0.07820 | 0.03585 | 0.00004 | 0.64288 | 3.74717 | 0.40479 | 0.00000 | 0.53226 | 1.47940 |
| **HBG1** | 0.00734 | 0.01223 | 0.01332 | 0.01415 | 0.01212 | 0.02755 | 0.01865 | 0.01889 | 0.01832 | 0.01282 | 0.01461 | 0.02584 |
| **HBG2** | 0.00408 | 0.01158 | 0.01998 | 0.02284 | 0.00829 | 0.01244 | 0.00955 | 0.01882 | 0.02703 | 0.06461 | 0.02072 | 0.02150 |
| **HBM** | 1.12264 | 0.41055 | 0.00225 | 0.00216 | 0.09502 | 0.00000 | 0.17223 | 0.57559 | 0.05606 | 0.00000 | 0.06626 | 0.14991 |
| **HBP1** | 3.32361 | 6.81562 | 7.48810 | 4.84729 | 0.75150 | 8.50636 | 3.39949 | 12.74077 | 5.00832 | 0.00000 | 2.87142 | 9.90888 |
| **HBQ1** | 0.09026 | 0.00659 | 0.15157 | 0.00582 | 0.00000 | 0.00729 | 0.32174 | 0.36310 | 0.23014 | 0.00000 | 0.35958 | 0.00737 |
| **HBS1L** | 2.26978 | 8.09869 | 9.42515 | 5.07901 | 0.00000 | 18.60609 | 8.08994 | 12.65345 | 12.65288 | 3.13467 | 10.69294 | 9.70082 |
| **HBZ** | 0.42608 | 0.00698 | 0.45693 | 0.05623 | 0.29248 | 0.00000 | 0.00482 | 0.34328 | 0.00000 | 0.00000 | 0.21400 | 0.05369 |
| **HCAR1** | 0.00740 | 0.01930 | 0.05731 | 0.20935 | 0.00001 | 0.04495 | 0.01978 | 0.01757 | 0.00458 | 0.02080 | 0.01767 | 0.02894 |
| **HCAR2** | 0.00127 | 0.45314 | 0.00000 | 0.25731 | 0.83633 | 0.00000 | 0.00046 | 0.18394 | 0.75599 | 0.11443 | 0.18538 | 0.13056 |
| **HCAR3** | 0.11131 | 0.00983 | 0.00000 | 0.08059 | 0.51037 | 0.19080 | 0.01354 | 0.08577 | 0.04731 | 0.31402 | 0.00000 | 0.02879 |
| **HCCS** | 0.71896 | 13.42724 | 8.82971 | 6.73989 | 5.45578 | 20.72067 | 15.12975 | 11.02926 | 14.80481 | 0.00000 | 15.03210 | 11.78603 |
| **HCFC1** | 6.90722 | 14.04307 | 12.00078 | 12.76249 | 18.73209 | 0.00000 | 21.20539 | 0.00145 | 13.80091 | 31.96499 | 12.09087 | 17.14414 |
| **HCFC1-AS1** | 0.00556 | 0.01091 | 0.04494 | 0.00747 | 0.05041 | 0.01470 | 0.00645 | 0.00970 | 0.01005 | 0.04777 | 0.01170 | 0.02400 |
| **HCFC1R1** | 0.49068 | 5.78251 | 4.74390 | 4.84665 | 6.70846 | 0.00000 | 5.44463 | 0.65461 | 5.81777 | 2.83300 | 6.73885 | 2.79608 |
| **HCFC2** | 1.79179 | 0.88173 | 0.96474 | 0.00000 | 0.00000 | 3.53587 | 0.00000 | 2.95874 | 0.00000 | 1.27615 | 0.91038 | 1.32037 |
| **HCG11** | 0.01095 | 0.53140 | 0.87906 | 0.00000 | 0.00000 | 0.00000 | 0.30262 | 0.52099 | 0.28589 | 1.10353 | 0.79098 | 0.55099 |
| **HCG14** | 0.02855 | 0.02134 | 0.00825 | 0.01876 | 0.06986 | 0.00000 | 0.09015 | 0.14572 | 0.00000 | 0.18827 | 0.00686 | 0.01617 |
| **HCG17** | 0.11413 | 0.11625 | 0.12319 | 0.03479 | 0.04563 | 0.00000 | 0.01113 | 1.13200 | 0.11941 | 0.00863 | 0.03787 | 0.15665 |
| **HCG18** | 4.81235 | 2.87713 | 5.25056 | 1.40961 | 0.18711 | 5.27132 | 2.79292 | 7.99925 | 4.73876 | 2.87702 | 3.50634 | 5.89736 |
| **HCG2040054** | 0.00374 | 0.01828 | 0.04809 | 0.00956 | 0.02822 | 0.01368 | 0.01133 | 0.02109 | 0.04108 | 0.01104 | 0.01518 | 0.03355 |
| **HCG22** | 0.00331 | 0.01111 | 0.01242 | 0.07914 | 0.01200 | 0.01601 | 0.01900 | 0.09263 | 0.01845 | 0.00041 | 0.01828 | 0.07321 |
| **HCG23** | 0.01252 | 0.01365 | 0.01362 | 0.01284 | 0.01584 | 0.01879 | 0.01876 | 0.05774 | 0.01785 | 0.01638 | 0.01521 | 0.04782 |
| **HCG25** | 0.03821 | 0.04120 | 0.03876 | 0.17092 | 0.17201 | 0.09437 | 0.00087 | 0.04486 | 0.13969 | 0.42372 | 0.01653 | 0.03942 |
| **HCG26** | 0.05270 | 0.01120 | 0.00853 | 0.03816 | 0.00276 | 0.00641 | 0.00390 | 0.08337 | 0.02179 | 0.13581 | 0.01006 | 0.01534 |
| **HCG27** | 0.00172 | 0.00869 | 0.01930 | 0.01787 | 0.01136 | 0.01068 | 0.01347 | 0.07074 | 0.01648 | 0.03811 | 0.01578 | 0.15179 |
| **HCG4** | 0.01310 | 0.01446 | 0.02122 | 0.01879 | 0.01388 | 0.01922 | 0.02134 | 0.00942 | 0.01615 | 0.01401 | 0.01467 | 0.01778 |
| **HCG4B** | 0.00338 | 0.04453 | 0.03021 | 0.01270 | 0.06986 | 0.00000 | 0.01507 | 0.02579 | 0.01888 | 0.01859 | 0.01969 | 0.02088 |
| **HCG8** | 0.26942 | 0.12316 | 0.00902 | 0.08220 | 0.00000 | 0.16149 | 0.00000 | 0.00000 | 0.07345 | 0.05348 | 0.02530 | 0.16304 |
| **HCG9** | 0.00426 | 0.00749 | 0.01596 | 0.01340 | 0.03493 | 0.06232 | 0.00850 | 0.01680 | 0.00997 | 0.02143 | 0.01728 | 0.04995 |
| **HCK** | 0.00437 | 0.00689 | 0.11516 | 0.12889 | 0.00000 | 0.03040 | 0.00000 | 0.29050 | 0.00000 | 0.06106 | 0.01881 | 0.01345 |
| **HCLS1** | 0.00081 | 0.25482 | 1.56669 | 7.06189 | 4.25996 | 3.63918 | 0.55990 | 0.26668 | 0.86147 | 0.00000 | 2.08256 | 0.12503 |
| **HCN1** | 0.00057 | 0.36009 | 0.00467 | 0.00000 | 0.48468 | 0.00000 | 0.22222 | 0.12696 | 0.02573 | 0.85014 | 0.19462 | 0.15301 |
| **HCN2** | 0.47169 | 0.03317 | 0.00621 | 0.01101 | 0.08885 | 0.00000 | 0.03008 | 0.03468 | 0.06234 | 0.15810 | 0.00354 | 0.09589 |
| **HCN3** | 0.00201 | 0.24167 | 0.07839 | 0.25426 | 0.00000 | 0.00000 | 0.00000 | 0.10961 | 0.41883 | 0.27019 | 0.03330 | 0.23663 |
| **HCN4** | 2.13306 | 0.00000 | 0.00000 | 0.05984 | 0.00000 | 0.02772 | 0.00000 | 0.00000 | 0.00751 | 0.23713 | 0.00201 | 0.12732 |
| **HCP5** | 0.01559 | 0.00000 | 0.16985 | 0.07509 | 0.00000 | 0.00000 | 0.00000 | 0.06198 | 0.03691 | 0.21791 | 0.01062 | 0.54056 |
| **HCST** | 0.00246 | 0.03198 | 0.07441 | 0.02440 | 0.02042 | 0.00964 | 0.00512 | 0.08304 | 0.02299 | 0.08198 | 0.01721 | 0.02604 |
| **HDAC1** | 1.24844 | 15.37850 | 21.95081 | 18.34431 | 10.05251 | 15.15254 | 29.87586 | 15.16207 | 24.26823 | 0.00000 | 24.46189 | 28.37618 |
| **HDAC10** | 0.05565 | 0.03111 | 0.13652 | 0.18788 | 0.25022 | 0.00000 | 0.00479 | 0.01196 | 0.42899 | 0.72910 | 0.42343 | 0.02272 |
| **HDAC11** | 0.86708 | 0.42262 | 0.22380 | 0.13006 | 0.29423 | 0.00000 | 0.39716 | 0.33274 | 0.00801 | 0.00000 | 0.05990 | 0.39735 |
| **HDAC11-AS1** | 0.27903 | 0.02016 | 0.01736 | 0.01256 | 0.00670 | 0.01296 | 0.01824 | 0.03087 | 0.02230 | 0.07338 | 0.01382 | 0.07055 |
| **HDAC2** | 3.27731 | 18.79334 | 23.43565 | 15.17073 | 12.00637 | 44.82543 | 28.20007 | 21.45166 | 21.62028 | 15.52003 | 19.26633 | 23.34766 |
| **HDAC3** | 1.16094 | 9.64529 | 11.08809 | 3.83255 | 4.63848 | 2.16927 | 7.45323 | 2.30721 | 11.20908 | 0.00000 | 6.02625 | 7.33813 |
| **HDAC4** | 0.97394 | 0.31657 | 0.59618 | 0.83231 | 1.26322 | 0.00000 | 0.48968 | 0.00223 | 0.00000 | 0.76249 | 0.13560 | 0.18052 |
| **HDAC5** | 0.47184 | 3.37174 | 11.12001 | 4.52506 | 2.74965 | 0.00000 | 0.61827 | 0.18586 | 6.27117 | 3.67751 | 1.14227 | 3.89687 |

| | | | | | | | | | | | | |
|---|---|---|---|---|---|---|---|---|---|---|---|---|
| **HDAC6** | 0.42063 | 2.72460 | 3.75887 | 4.35987 | 4.71316 | 0.00000 | 0.00444 | 0.47408 | 2.93792 | 0.00000 | 2.92159 | 1.98301 |
| **HDAC7** | 0.13201 | 0.33610 | 0.42340 | 3.36971 | 1.43205 | 0.00000 | 0.43054 | 0.00000 | 0.80245 | 1.25654 | 0.00000 | 0.26113 |
| **HDAC8** | 0.36275 | 2.70227 | 0.97910 | 4.10590 | 0.69045 | 6.89067 | 1.23904 | 0.68246 | 2.39223 | 0.00000 | 2.81464 | 8.27748 |
| **HDAC9** | 10.12193 | 0.00459 | 0.02903 | 0.28413 | 0.00716 | 0.06604 | 0.00000 | 0.82366 | 0.03954 | 0.00000 | 0.02760 | 0.22112 |
| **HDC** | 14.72577 | 0.00756 | 0.11070 | 0.01339 | 0.00698 | 0.01686 | 0.01810 | 6.26917 | 0.00327 | 0.00000 | 0.00683 | 0.28420 |
| **HDDC2** | 0.23022 | 20.29270 | 13.39798 | 11.70088 | 15.64693 | 25.27799 | 20.61118 | 1.96031 | 15.16822 | 9.29694 | 19.37592 | 18.35733 |
| **HDDC3** | 0.35908 | 9.57330 | 1.03536 | 4.54281 | 11.54489 | 4.80334 | 0.49280 | 0.00000 | 4.30410 | 0.52496 | 11.78410 | 1.41050 |
| **HDGF** | 2.62146 | 4.14443 | 9.93762 | 8.92564 | 15.30858 | 6.98801 | 31.69235 | 0.00000 | 6.98451 | 35.47104 | 9.18789 | 5.17995 |
| **HDGFL1** | 0.26656 | 0.08758 | 0.00566 | 0.00000 | 0.28543 | 0.11925 | 0.00000 | 0.02608 | 0.00000 | 0.22351 | 0.00000 | 0.87251 |
| **HDGFRP2** | 1.94014 | 0.64388 | 1.17288 | 0.49113 | 1.86534 | 0.00000 | 0.74365 | 1.31516 | 0.00000 | 5.41077 | 0.12945 | 0.49131 |
| **HDGFRP3** | 1.21586 | 1.23659 | 0.25617 | 2.59617 | 1.62063 | 1.82886 | 0.24047 | 0.00000 | 0.00000 | 1.78275 | 0.00000 | 0.24507 |
| **HDHD1** | 1.05590 | 7.94180 | 5.38712 | 1.51971 | 2.79027 | 10.65348 | 6.58625 | 0.00000 | 2.56304 | 0.00000 | 9.87109 | 4.02474 |
| **HDHD2** | 1.77870 | 3.81184 | 2.46154 | 0.94565 | 0.92343 | 8.02892 | 0.00000 | 2.85530 | 3.26749 | 0.00000 | 5.72728 | 4.21245 |
| **HDHD3** | 3.72163 | 8.60545 | 9.91799 | 5.45531 | 10.13750 | 2.15154 | 8.21485 | 5.52575 | 9.92832 | 0.00000 | 9.12257 | 10.44567 |
| **HDLBP** | 10.44769 | 28.52018 | 30.53724 | 20.41842 | 31.40236 | 15.08028 | 35.61528 | 4.66488 | 30.11087 | 17.85793 | 23.95629 | 23.00701 |
| **HDX** | 0.00332 | 0.00736 | 0.03184 | 0.08350 | 0.00000 | 0.08377 | 0.03911 | 0.02662 | 0.15434 | 0.00694 | 0.00000 | 0.02424 |
| **HEATR1** | 7.86043 | 6.84732 | 6.21836 | 0.96386 | 3.54834 | 15.60735 | 7.49771 | 9.38055 | 15.22857 | 3.50388 | 6.18267 | 16.99376 |
| **HEATR2** | 1.37518 | 7.12982 | 4.02983 | 2.97668 | 8.06052 | 1.19220 | 12.84281 | 0.86354 | 4.37006 | 10.34068 | 6.55357 | 4.15032 |
| **HEATR3** | 0.14333 | 2.20418 | 1.44635 | 0.56560 | 0.66339 | 4.15331 | 0.00000 | 0.91102 | 1.87781 | 2.29140 | 2.02663 | 2.10001 |
| **HEATR4** | 0.17074 | 0.00000 | 0.05220 | 0.22569 | 0.00000 | 0.15536 | 0.00000 | 0.08165 | 0.00000 | 0.56274 | 0.00000 | 0.06957 |
| **HEATR5A** | 1.72765 | 2.70411 | 0.00000 | 3.88404 | 8.85767 | 18.05499 | 9.74528 | 0.63958 | 4.95102 | 8.41349 | 11.17939 | 2.43311 |
| **HEATR5B** | 2.27307 | 3.61428 | 4.64276 | 1.46665 | 0.00000 | 7.49199 | 0.83510 | 0.79550 | 3.79997 | 7.17180 | 7.92339 | 4.05589 |
| **HEATR6** | 0.00464 | 2.06670 | 3.46421 | 1.86093 | 2.56495 | 3.96110 | 1.55130 | 0.00000 | 4.57785 | 0.00000 | 3.33705 | 2.67549 |
| **HEATR9** | 0.00451 | 0.03863 | 0.01292 | 0.01455 | 0.02171 | 0.02940 | 0.00912 | 0.12202 | 0.01951 | 0.03291 | 0.01086 | 0.45210 |
| **HEBP1** | 0.29610 | 0.74351 | 0.42139 | 2.46747 | 3.97830 | 2.56428 | 0.00000 | 0.02996 | 3.22931 | 0.00000 | 5.38878 | 0.88076 |
| **HEBP2** | 0.00000 | 0.00000 | 4.48306 | 0.00000 | 4.14079 | 3.83318 | 11.38274 | 0.63692 | 0.76258 | 12.19858 | 3.24741 | 0.58699 |
| **HECA** | 0.59639 | 0.29273 | 1.94054 | 0.00000 | 2.38472 | 3.75858 | 6.29179 | 2.28611 | 0.00000 | 5.48633 | 0.00000 | 1.34795 |
| **HECTD1** | 3.10665 | 0.00016 | 9.32876 | 6.62255 | 3.94052 | 22.23776 | 4.01154 | 1.45721 | 11.74344 | 25.57595 | 5.35809 | 5.23749 |
| **HECTD2** | 3.01782 | 0.56101 | 0.43149 | 0.00000 | 0.00000 | 0.96958 | 0.26881 | 0.07332 | 0.28713 | 0.08901 | 0.00000 | 0.50014 |
| **HECTD3** | 2.49932 | 14.83973 | 7.25182 | 8.81954 | 8.37951 | 0.00000 | 3.25169 | 2.32349 | 10.10582 | 6.62402 | 9.35008 | 6.31289 |
| **HECTD4** | 2.54086 | 1.17976 | 1.34342 | 3.25317 | 3.25479 | 0.00000 | 2.56651 | 0.02902 | 3.37831 | 6.82988 | 2.47864 | 0.59453 |
| **HECW1** | 0.05781 | 0.06226 | 0.04102 | 0.02646 | 0.00000 | 0.03285 | 0.00751 | 0.03828 | 0.01471 | 0.03085 | 0.02471 | 0.06519 |
| **HECW2** | 1.15949 | 0.06129 | 0.00000 | 0.00000 | 0.15961 | 0.29753 | 0.88458 | 0.37915 | 1.09127 | 0.00000 | 0.68410 | 0.03217 |
| **HEG1** | 0.00000 | 0.14496 | 0.45283 | 0.34614 | 3.34998 | 2.52191 | 2.06821 | 0.00000 | 2.80259 | 1.28321 | 0.64329 | 0.37052 |
| **HEIH** | 0.63491 | 0.07743 | 1.37621 | 0.10158 | 0.05884 | 0.00000 | 1.24926 | 0.24428 | 0.61307 | 0.00000 | 0.18727 | 0.53283 |
| **HELB** | 0.00251 | 0.42877 | 1.93454 | 0.29436 | 0.25628 | 1.72272 | 0.39117 | 3.41140 | 2.35719 | 0.40157 | 0.36599 | 4.06749 |
| **HELLS** | 2.41044 | 1.30903 | 6.97102 | 6.62736 | 3.51712 | 10.99107 | 9.13021 | 2.53551 | 8.89799 | 24.45829 | 7.28900 | 5.85037 |
| **HELQ** | 1.37090 | 3.79902 | 2.41239 | 3.22045 | 1.46284 | 8.10046 | 0.00000 | 2.56871 | 3.18868 | 0.74335 | 2.84657 | 6.54731 |
| **HELT** | 0.05251 | 0.01462 | 0.02298 | 0.01503 | 0.01671 | 0.01434 | 0.04090 | 0.00841 | 0.00974 | 0.00219 | 0.01687 | 0.02945 |
| **HELZ** | 12.13077 | 4.98325 | 5.70991 | 6.96709 | 3.53043 | 16.53913 | 9.28179 | 5.06810 | 10.80737 | 20.39598 | 8.87621 | 5.79732 |
| **HELZ2** | 0.02248 | 0.00483 | 0.10931 | 0.05352 | 0.12241 | 0.00000 | 0.10014 | 0.03091 | 0.00091 | 0.12740 | 0.02496 | 0.02944 |
| **HEMGN** | 0.00417 | 0.00703 | 0.03083 | 0.00682 | 0.02796 | 0.02456 | 0.02212 | 0.01288 | 0.00501 | 0.03687 | 0.01676 | 0.05469 |
| **HEMK1** | 0.29890 | 3.19044 | 0.29407 | 3.42722 | 4.18748 | 0.00000 | 0.38982 | 0.00000 | 1.36034 | 3.46465 | 4.08208 | 0.42607 |
| **HENMT1** | 14.19752 | 16.62987 | 4.26695 | 6.89697 | 3.71720 | 13.32231 | 5.53588 | 14.35770 | 0.22423 | 0.53161 | 11.88000 | 7.94999 |
| **HEPACAM** | 0.03378 | 0.00942 | 0.01328 | 0.02399 | 0.03476 | 0.05523 | 0.01296 | 0.00965 | 0.01318 | 0.01359 | 0.01345 | 0.01404 |
| **HEPACAM2** | 0.00576 | 0.05216 | 0.01662 | 0.01161 | 0.01661 | 0.01358 | 0.01918 | 0.01370 | 0.02775 | 0.00000 | 0.02777 | 0.02278 |
| **HEPH** | 0.13125 | 0.00000 | 0.00000 | 0.25885 | 0.08620 | 0.00000 | 0.05842 | 0.00000 | 0.53580 | 0.00000 | 0.02157 | 0.71649 |
| **HEPHL1** | 0.90001 | 0.02181 | 0.02669 | 0.00000 | 0.00560 | 0.03657 | 0.06428 | 0.22043 | 0.03317 | 0.06831 | 0.07754 | 0.00834 |

| | | | | | | | | | | | | |
|---|---|---|---|---|---|---|---|---|---|---|---|---|
| **HERC1** | 4.30833 | 0.66511 | 3.03959 | 3.92214 | 4.07928 | 2.99273 | 2.09135 | 0.74180 | 2.18194 | 7.60134 | 2.48322 | 0.00000 |
| **HERC2** | 1.23502 | 2.71303 | 0.45871 | 4.81355 | 6.68862 | 1.89823 | 0.88857 | 1.44350 | 0.06751 | 14.47349 | 3.37459 | 2.64811 |
| **HERC2P10** | 0.04434 | 0.02219 | 0.00697 | 0.07933 | 0.00000 | 0.02160 | 0.00000 | 0.15251 | 0.08476 | 0.07100 | 0.02983 | 0.08070 |
| **HERC2P2** | 0.00000 | 0.15665 | 0.04846 | 0.33011 | 0.52860 | 1.00623 | 0.00000 | 0.08010 | 0.13881 | 0.96904 | 0.11645 | 0.35015 |
| **HERC2P3** | 0.01109 | 0.07163 | 0.00613 | 0.08946 | 0.13665 | 0.00000 | 0.00863 | 0.01276 | 0.06653 | 0.11475 | 0.07599 | 0.03773 |
| **HERC2P4** | 0.08171 | 0.02282 | 0.02901 | 0.00000 | 0.10287 | 0.01789 | 0.07621 | 0.01408 | 0.03452 | 0.06486 | 0.03745 | 0.06866 |
| **HERC2P9** | 0.00343 | 0.23386 | 0.02381 | 0.13827 | 0.07929 | 0.34126 | 0.00000 | 0.06217 | 0.11141 | 0.36378 | 0.29318 | 0.16041 |
| **HERC3** | 7.69733 | 0.96431 | 1.66252 | 0.71533 | 0.00002 | 6.77095 | 1.85416 | 1.23111 | 7.21928 | 1.47153 | 1.25375 | 0.98633 |
| **HERC4** | 0.99937 | 1.26305 | 3.21491 | 0.00000 | 0.00000 | 15.60186 | 5.96345 | 1.34664 | 7.44425 | 6.87747 | 4.54831 | 4.57420 |
| **HERC5** | 0.73242 | 1.79250 | 6.16069 | 0.85381 | 0.93757 | 1.37010 | 10.65941 | 3.08285 | 0.31941 | 2.68467 | 4.16739 | 6.14635 |
| **HERC6** | 15.02761 | 0.06720 | 0.08712 | 0.00000 | 0.55855 | 0.00000 | 0.04554 | 5.55702 | 0.00000 | 0.00000 | 0.12442 | 0.33719 |
| **HERPUD1** | 1.14672 | 4.07455 | 19.03041 | 2.31696 | 1.87435 | 11.98749 | 12.96862 | 1.84756 | 42.81411 | 2.67773 | 14.92589 | 3.68439 |
| **HERPUD2** | 1.19839 | 0.71162 | 2.60234 | 0.23137 | 1.28545 | 0.33930 | 2.25767 | 0.57020 | 0.00000 | 2.16644 | 1.93227 | 1.06519 |
| **HES1** | 0.55029 | 0.00000 | 2.71868 | 0.00000 | 4.13106 | 0.22600 | 4.18827 | 1.18131 | 0.02963 | 2.42351 | 0.46214 | 0.17480 |
| **HES2** | 0.59761 | 5.22111 | 1.18825 | 0.00000 | 0.00000 | 0.00000 | 0.00000 | 1.47398 | 0.25014 | 1.32004 | 0.00000 | 4.08875 |
| **HES4** | 0.00769 | 0.64039 | 0.00000 | 0.00000 | 1.13956 | 0.00000 | 0.00853 | 0.00850 | 0.18064 | 0.05046 | 0.09848 | 0.03120 |
| **HES5** | 0.00384 | 0.01499 | 0.03914 | 0.00880 | 0.02094 | 0.01539 | 0.02165 | 0.02182 | 0.01475 | 0.02527 | 0.01280 | 0.01541 |
| **HES6** | 3.33805 | 0.14951 | 0.11687 | 0.69346 | 2.89705 | 0.00000 | 3.81195 | 0.19018 | 1.78263 | 5.42581 | 0.52543 | 0.00000 |
| **HES7** | 0.07968 | 0.00000 | 0.05032 | 0.00490 | 0.00000 | 0.01364 | 0.18166 | 0.03256 | 0.00000 | 0.17919 | 0.02345 | 0.19379 |
| **HESX1** | 0.00258 | 0.06368 | 0.61115 | 3.81199 | 0.10964 | 0.46108 | 0.00000 | 0.85733 | 0.00000 | 0.00000 | 0.05179 | 0.82228 |
| **HEXA** | 2.51855 | 7.12170 | 11.51638 | 7.60044 | 7.22124 | 3.38957 | 0.00000 | 1.97496 | 23.02033 | 0.75267 | 7.75888 | 2.79296 |
| **HEXA-AS1** | 0.14934 | 0.03506 | 0.23445 | 0.04161 | 0.10200 | 0.12266 | 0.00000 | 0.00162 | 0.00000 | 0.04965 | 0.06777 | 0.11125 |
| **HEXB** | 3.04227 | 8.33688 | 12.13057 | 4.47750 | 10.35833 | 16.60051 | 0.00000 | 2.07087 | 32.43653 | 3.55100 | 11.29823 | 0.85687 |
| **HEXDC** | 0.22596 | 0.20450 | 0.19138 | 0.20862 | 0.24558 | 0.00000 | 0.41757 | 0.04506 | 0.15239 | 1.79829 | 0.30365 | 0.20115 |
| **HEXIM1** | 1.73471 | 2.23813 | 25.97785 | 4.16434 | 4.31188 | 3.12516 | 8.80474 | 21.13741 | 5.24158 | 6.97532 | 7.16244 | 6.90522 |
| **HEXIM2** | 2.40146 | 7.87789 | 3.89726 | 5.00071 | 2.70553 | 0.06474 | 2.42134 | 0.58210 | 0.07808 | 2.09937 | 5.32977 | 8.84655 |
| **HEY1** | 0.00985 | 0.10281 | 1.59964 | 0.63909 | 0.28197 | 0.00000 | 0.00000 | 4.03913 | 0.00000 | 0.00012 | 0.01340 | 1.96743 |
| **HEY2** | 0.22223 | 3.48064 | 0.09348 | 2.67271 | 2.14588 | 0.00000 | 0.00000 | 0.69351 | 0.00000 | 0.53212 | 0.00000 | 2.26818 |
| **HEYL** | 0.00933 | 0.38486 | 0.16288 | 0.00000 | 0.09582 | 0.00000 | 0.09690 | 0.00990 | 0.02263 | 0.04862 | 0.04549 | 0.89631 |
| **HFE** | 0.00270 | 0.01667 | 0.09424 | 0.07863 | 0.01518 | 0.00000 | 0.01810 | 0.04610 | 0.02851 | 0.00000 | 0.04409 | 0.02017 |
| **HFE2** | 0.01272 | 0.00000 | 0.19476 | 0.01846 | 0.05544 | 0.01612 | 0.01742 | 0.07151 | 0.00000 | 0.01616 | 0.02034 | 1.06982 |
| **HFM1** | 0.00810 | 0.26286 | 0.00000 | 0.00404 | 0.00000 | 0.12528 | 0.00000 | 0.00661 | 0.15391 | 0.02524 | 0.01776 | 0.08708 |
| **HGC6.3** | 0.04168 | 0.01743 | 0.05268 | 0.02169 | 0.02065 | 0.00000 | 0.01516 | 0.00809 | 0.01480 | 0.01717 | 0.01755 | 0.01644 |
| **HGD** | 0.03311 | 0.00954 | 0.14678 | 0.39658 | 0.00000 | 0.09437 | 0.00000 | 0.00638 | 0.00000 | 0.03956 | 0.00693 | 0.03230 |
| **HGF** | 0.01344 | 0.01358 | 0.02434 | 0.00929 | 0.01125 | 0.02332 | 0.01512 | 0.01040 | 0.01780 | 0.02853 | 0.01348 | 0.01463 |
| **HGFAC** | 0.00641 | 0.01473 | 0.01439 | 0.01255 | 0.01390 | 0.01766 | 0.03455 | 0.01246 | 0.01504 | 0.01888 | 0.01407 | 0.01610 |
| **HGH1.uid-11150** | 0.01045 | 0.08883 | 0.02654 | 0.02079 | 0.04290 | 0.02604 | 0.01191 | 0.01258 | 0.06295 | 0.00000 | 0.02035 | 0.02007 |
| **HGH1.uid-11154** | 0.00816 | 0.01329 | 0.01541 | 0.01355 | 0.01953 | 0.01133 | 0.01949 | 0.00940 | 0.01996 | 0.01002 | 0.01554 | 0.04446 |
| **HGS** | 2.67780 | 11.85280 | 17.35316 | 9.51729 | 5.19663 | 0.00000 | 7.15381 | 6.94634 | 13.25923 | 7.17796 | 10.05551 | 9.08757 |
| **HGSNAT** | 0.36683 | 1.87824 | 0.48392 | 1.98928 | 0.39277 | 1.95993 | 0.00000 | 0.80994 | 1.20539 | 1.65588 | 0.41976 | 0.00000 |
| **HHAT** | 4.72306 | 0.43016 | 0.28151 | 0.25066 | 0.00000 | 0.23674 | 0.00000 | 1.67656 | 0.00000 | 0.00891 | 0.02990 | 0.35774 |
| **HHATL** | 0.05980 | 0.01355 | 0.01361 | 0.01532 | 0.00989 | 0.01998 | 0.00753 | 0.04841 | 0.02699 | 0.07670 | 0.01003 | 0.00973 |
| **HHEX** | 2.65136 | 0.20917 | 0.21706 | 1.47383 | 0.00000 | 0.20793 | 0.00000 | 0.28555 | 0.04507 | 0.04568 | 0.00927 | 0.26013 |
| **HHIP** | 0.00483 | 0.00895 | 0.02204 | 0.00167 | 0.02691 | 0.03514 | 0.02977 | 0.01119 | 0.04462 | 0.06540 | 0.01717 | 0.01521 |
| **HHIP-AS1** | 0.00593 | 0.01254 | 0.02260 | 0.03042 | 0.00625 | 0.02459 | 0.01571 | 0.01056 | 0.04748 | 0.02775 | 0.00865 | 0.01489 |
| **HHIPL1** | 0.06349 | 0.00000 | 0.06746 | 0.00517 | 0.19978 | 0.01365 | 0.00425 | 0.00569 | 0.00635 | 0.06377 | 0.00979 | 0.17278 |
| **HHIPL2** | 0.00148 | 0.28966 | 0.05373 | 0.00930 | 0.00000 | 0.00000 | 0.00705 | 0.19956 | 0.03796 | 0.15291 | 0.08680 | 0.10258 |
| **HHLA1** | 1.70581 | 0.02844 | 0.60282 | 1.01311 | 0.00735 | 0.00000 | 0.04034 | 0.00000 | 0.00942 | 0.00000 | 0.01432 | 0.04554 |

| | | | | | | | | | | | | |
|---|---|---|---|---|---|---|---|---|---|---|---|---|
| **HHLA2** | 13.76727 | 0.02040 | 0.00423 | 0.06464 | 0.03640 | 0.01448 | 0.00000 | 6.61322 | 0.00590 | 0.02787 | 0.01238 | 0.69312 |
| **HHLA3** | 0.34889 | 0.72076 | 1.20799 | 0.89090 | 1.31026 | 1.45042 | 1.50165 | 0.61498 | 1.29354 | 1.39283 | 0.68764 | 1.15681 |
| **HIAT1** | 6.45327 | 6.76811 | 9.74422 | 6.23954 | 0.17067 | 19.40076 | 0.00000 | 9.46243 | 19.78534 | 3.92801 | 9.47386 | 6.00838 |
| **HIATL1** | 2.32098 | 3.14229 | 4.48328 | 2.92240 | 2.55812 | 21.19132 | 0.00000 | 10.47756 | 6.81872 | 19.82912 | 2.09771 | 6.13293 |
| **HIATL2** | 0.00836 | 0.07490 | 0.41140 | 0.06848 | 0.23776 | 0.00000 | 0.01886 | 0.73805 | 0.24429 | 0.00000 | 0.10924 | 0.23807 |
| **HIBADH** | 1.50013 | 2.88467 | 3.33537 | 2.97176 | 2.34882 | 10.21837 | 0.00000 | 0.00000 | 2.24220 | 0.01044 | 6.64394 | 1.40217 |
| **HIBCH** | 1.92669 | 1.79640 | 6.65389 | 3.33857 | 0.55848 | 6.25660 | 5.82383 | 2.68926 | 3.03312 | 0.07421 | 2.74972 | 2.66253 |
| **HIC1** | 0.29893 | 0.02543 | 0.03812 | 0.00873 | 0.00931 | 0.02254 | 0.01243 | 0.03934 | 0.03042 | 0.01993 | 0.01473 | 0.01715 |
| **HIC2** | 9.95224 | 14.80105 | 22.09677 | 9.71846 | 34.88235 | 6.78490 | 36.77755 | 1.32433 | 20.54723 | 56.98633 | 17.83125 | 10.69178 |
| **HID1** | 2.86344 | 0.18257 | 0.41204 | 0.06939 | 0.53285 | 0.00000 | 0.13570 | 0.00000 | 1.16762 | 0.00000 | 0.22591 | 0.32876 |
| **HIF1A** | 4.61241 | 1.29454 | 14.12998 | 4.95404 | 0.06832 | 17.08561 | 3.13661 | 1.82706 | 13.48950 | 0.00000 | 4.82851 | 4.23555 |
| **HIF1AN** | 1.83798 | 12.35927 | 3.32975 | 7.60194 | 5.98235 | 9.37688 | 9.38850 | 1.97095 | 6.34102 | 5.00747 | 11.27109 | 15.31338 |
| **HIF3A** | 0.59014 | 0.12678 | 0.00065 | 0.00000 | 0.09081 | 0.00000 | 0.00000 | 0.02847 | 0.10795 | 0.41341 | 0.00000 | 0.14074 |
| **HIGD1A** | 4.62719 | 15.79119 | 17.18524 | 11.50653 | 10.43062 | 54.98164 | 23.12970 | 2.53204 | 17.12866 | 7.58712 | 22.23700 | 30.06477 |
| **HIGD1B** | 0.03565 | 0.05788 | 0.08479 | 0.08559 | 0.00000 | 0.00000 | 0.00836 | 0.00645 | 0.08561 | 0.15899 | 0.05517 | 0.14054 |
| **HIGD1C** | 0.00553 | 0.01043 | 0.02219 | 0.00499 | 0.00000 | 0.00253 | 0.10485 | 0.06212 | 0.09082 | 0.06156 | 0.26365 | 0.01104 |
| **HIGD2A** | 6.74416 | 14.50537 | 12.62623 | 16.09865 | 12.71257 | 12.52243 | 19.56171 | 7.01133 | 16.77738 | 3.34558 | 15.61922 | 15.01940 |
| **HIGD2B** | 0.00745 | 0.02033 | 0.03314 | 0.00964 | 0.01786 | 0.04248 | 0.05400 | 0.04728 | 0.00144 | 0.01195 | 0.03905 | 0.11604 |
| **HILPDA** | 1.98927 | 21.06633 | 9.21098 | 3.19948 | 2.52608 | 6.52845 | 13.79249 | 2.77947 | 9.79232 | 10.24351 | 11.06286 | 24.69590 |
| **HINFP** | 1.19192 | 6.15233 | 4.53848 | 3.30054 | 2.03612 | 3.61679 | 4.07369 | 5.78869 | 10.23119 | 0.00000 | 3.68851 | 8.81743 |
| **HINT1** | 16.16343 | 5.75067 | 29.67442 | 14.81950 | 13.90047 | 19.03762 | 47.83849 | 23.95324 | 42.74931 | 1.41933 | 28.00295 | 0.44068 |
| **HINT2** | 1.02071 | 2.35103 | 0.43959 | 2.62623 | 3.01981 | 0.01977 | 1.39354 | 0.45598 | 2.63230 | 0.84785 | 4.72161 | 3.17517 |
| **HINT3** | 0.00000 | 5.12426 | 6.71383 | 0.81421 | 0.71567 | 11.21498 | 6.45824 | 0.00000 | 3.23217 | 3.79502 | 5.07381 | 4.00832 |
| **HIP1** | 1.69265 | 0.12089 | 0.00000 | 0.00000 | 13.50367 | 1.59291 | 10.27537 | 0.04441 | 9.61855 | 13.30763 | 13.82677 | 0.48464 |
| **HIP1R** | 4.22998 | 6.48446 | 4.53874 | 6.87166 | 4.89575 | 0.00000 | 1.27181 | 2.14270 | 12.20858 | 0.00000 | 0.46489 | 2.66369 |
| **HIPK1** | 4.96924 | 2.90240 | 2.50239 | 3.49690 | 3.21090 | 15.01149 | 11.10562 | 3.06041 | 2.46309 | 17.52331 | 4.14009 | 9.08933 |
| **HIPK1-AS1** | 0.00922 | 0.04594 | 0.05952 | 0.01221 | 0.03524 | 0.05580 | 0.01613 | 0.03179 | 0.00504 | 0.00428 | 0.04428 | 0.16732 |
| **HIPK2** | 7.14428 | 2.07890 | 0.12413 | 1.06817 | 2.19259 | 0.65799 | 0.00000 | 0.21140 | 0.00000 | 0.25224 | 0.00000 | 1.01073 |
| **HIPK3** | 5.26133 | 1.91480 | 1.50068 | 2.42686 | 2.51107 | 12.59500 | 9.36012 | 36.09678 | 1.20364 | 9.40663 | 1.50114 | 6.72223 |
| **HIPK4** | 1.79276 | 0.06756 | 0.02262 | 0.00220 | 0.00415 | 0.01189 | 0.01326 | 0.48257 | 0.00000 | 0.02890 | 0.05720 | 1.00336 |
| **HIRA** | 1.83323 | 1.33149 | 1.82328 | 2.88045 | 3.48368 | 0.00000 | 4.01264 | 0.28817 | 1.29205 | 6.41823 | 2.88828 | 1.39814 |
| **HIRIP3** | 7.90496 | 3.28185 | 1.99995 | 4.50418 | 3.86504 | 0.00000 | 5.77029 | 5.20564 | 0.47365 | 2.06680 | 4.25317 | 4.62488 |
| **HIST1H1A** | 26.72902 | 0.59808 | 4.54141 | 0.21833 | 1.78387 | 12.85721 | 2.77170 | 3.90524 | 8.22191 | 5.63314 | 3.17716 | 6.99132 |
| **HIST1H1B** | 0.09118 | 0.18642 | 2.56286 | 1.37679 | 2.78064 | 2.56541 | 0.00005 | 0.51049 | 3.03835 | 2.25322 | 0.18218 | 1.79307 |
| **HIST1H1C** | 0.92504 | 0.00000 | 6.48088 | 0.11571 | 2.08422 | 4.33988 | 1.37238 | 2.23421 | 3.55100 | 4.05634 | 1.01175 | 6.02484 |
| **HIST1H1D** | 0.05292 | 0.00000 | 3.58224 | 0.42718 | 2.50722 | 4.88723 | 2.59442 | 0.00000 | 6.34927 | 2.49841 | 0.00017 | 6.13331 |
| **HIST1H1E** | 2.08085 | 0.00000 | 1.43665 | 0.12595 | 2.22569 | 3.03291 | 0.00001 | 0.67274 | 3.46947 | 2.14640 | 1.65060 | 4.29730 |
| **HIST1H1T** | 0.00374 | 0.11414 | 0.28880 | 0.00004 | 0.00000 | 0.03105 | 0.01113 | 0.25692 | 0.39162 | 0.00792 | 0.00000 | 0.06141 |
| **HIST1H2AA** | 2.18689 | 0.59412 | 2.27087 | 0.00000 | 0.89684 | 0.00000 | 2.25530 | 0.41143 | 0.92087 | 0.00000 | 0.00000 | 2.86364 |
| **HIST1H2AB** | 0.30923 | 0.00883 | 0.70248 | 0.09273 | 0.13020 | 0.00000 | 0.01547 | 0.41298 | 0.10862 | 0.00000 | 0.00035 | 0.14600 |
| **HIST1H2AC** | 0.78680 | 0.26897 | 3.30645 | 0.99309 | 0.43896 | 0.00000 | 0.00000 | 11.18296 | 1.11628 | 0.00000 | 0.00000 | 1.78634 |
| **HIST1H2AD** | 0.00352 | 0.03017 | 0.07758 | 0.02335 | 0.06160 | 0.09645 | 0.00000 | 0.04921 | 0.00000 | 0.03192 | 0.00883 | 0.03411 |
| **HIST1H2AE** | 0.19943 | 0.00000 | 1.51015 | 0.19665 | 0.46580 | 0.21604 | 0.25385 | 0.55159 | 0.12950 | 0.00001 | 0.00000 | 0.99461 |
| **HIST1H2AG** | 1.22542 | 0.16099 | 2.10343 | 0.12870 | 0.21975 | 0.10107 | 0.00000 | 1.68808 | 0.00000 | 0.18694 | 0.00201 | 0.12827 |
| **HIST1H2AH** | 4.56452 | 0.00000 | 1.79998 | 0.15853 | 3.92116 | 0.00000 | 0.00000 | 0.46077 | 2.27648 | 1.91098 | 0.20604 | 2.04397 |
| **HIST1H2AI** | 0.00457 | 0.00000 | 1.49457 | 0.08867 | 0.00000 | 0.25260 | 0.00000 | 0.86793 | 0.00000 | 0.08155 | 0.00391 | 0.07917 |
| **HIST1H2AJ** | 0.09074 | 0.20672 | 2.08183 | 0.01426 | 1.03208 | 0.71641 | 0.00000 | 0.08139 | 1.56897 | 0.02071 | 2.54766 | 1.55751 |
| **HIST1H2AK** | 0.03545 | 0.00691 | 0.44350 | 0.00178 | 0.18555 | 0.00000 | 0.00000 | 0.02737 | 0.02745 | 0.08140 | 0.02176 | 0.09890 |

| | | | | | | | | | | | | |
|---|---|---|---|---|---|---|---|---|---|---|---|---|
| **HIST1H2AL** | 0.25547 | 0.07859 | 1.96993 | 0.12579 | 0.39816 | 0.00000 | 0.00000 | 0.81729 | 0.00000 | 0.55945 | 0.10899 | 0.43973 |
| **HIST1H2AM** | 2.41111 | 0.00000 | 3.04533 | 0.66768 | 2.17750 | 0.70669 | 0.00000 | 8.93750 | 2.77582 | 0.00000 | 1.93373 | 3.58702 |
| **HIST1H2APS1** | 0.04882 | 0.00896 | 0.01339 | 0.00749 | 0.00910 | 0.00703 | 0.02213 | 0.12485 | 0.01489 | 0.11112 | 0.01541 | 0.03011 |
| **HIST1H2BA** | 0.09804 | 0.02541 | 0.45459 | 0.00427 | 0.10810 | 0.00274 | 0.05870 | 0.00365 | 0.00000 | 0.04195 | 0.00650 | 0.04246 |
| **HIST1H2BB** | 0.00927 | 0.00377 | 0.90820 | 0.01711 | 0.00000 | 0.02660 | 0.00000 | 0.85693 | 0.00000 | 0.22919 | 0.02675 | 0.09379 |
| **HIST1H2BC** | 1.94645 | 0.17091 | 0.84192 | 0.24668 | 0.33777 | 0.00000 | 0.00000 | 13.22185 | 0.16065 | 0.09320 | 0.00000 | 2.47390 |
| **HIST1H2BD** | 0.72966 | 0.04005 | 0.94024 | 0.21569 | 0.33850 | 0.08868 | 0.29664 | 7.95062 | 0.08091 | 0.00000 | 0.10054 | 1.91392 |
| **HIST1H2BE** | 0.11619 | 0.00000 | 0.41889 | 0.00000 | 0.08667 | 0.20858 | 0.00000 | 0.26444 | 0.09176 | 0.55165 | 0.02572 | 0.76350 |
| **HIST1H2BF** | 0.00165 | 0.00000 | 0.54578 | 0.00000 | 0.24849 | 0.16815 | 0.00000 | 0.12036 | 0.01423 | 0.50133 | 0.06874 | 0.43258 |
| **HIST1H2BG** | 0.00737 | 0.07133 | 1.84165 | 0.00000 | 0.22871 | 0.00000 | 0.00000 | 9.00975 | 0.38043 | 0.08473 | 0.00000 | 1.40810 |
| **HIST1H2BH** | 0.04450 | 0.13090 | 3.14689 | 0.28097 | 0.39866 | 0.37306 | 0.00000 | 5.67965 | 0.55778 | 0.06465 | 0.09158 | 1.58215 |
| **HIST1H2BI** | 0.07869 | 0.00741 | 0.24401 | 0.01289 | 0.08446 | 0.00000 | 0.00592 | 0.00990 | 0.15254 | 0.35149 | 0.00000 | 0.07071 |
| **HIST1H2BJ** | 1.48512 | 0.04615 | 2.21292 | 0.35301 | 0.37062 | 0.06799 | 0.10195 | 3.15982 | 0.00019 | 0.25873 | 0.06043 | 0.43900 |
| **HIST1H2BK** | 2.99944 | 0.23707 | 8.62126 | 2.41364 | 0.46138 | 2.57858 | 0.95260 | 11.41602 | 1.36054 | 0.78765 | 0.00000 | 7.68967 |
| **HIST1H2BL** | 0.01141 | 0.00000 | 0.42471 | 0.10289 | 0.00000 | 0.02652 | 0.04787 | 0.61867 | 0.06348 | 0.10935 | 0.01717 | 0.13542 |
| **HIST1H2BM** | 0.00147 | 0.03866 | 1.22500 | 0.13704 | 0.19338 | 0.00000 | 0.00000 | 0.06836 | 0.08463 | 0.22539 | 0.02374 | 0.07090 |
| **HIST1H2BN** | 0.41472 | 0.00000 | 0.28650 | 0.01864 | 0.00470 | 0.00000 | 0.00000 | 0.21084 | 0.06490 | 0.51372 | 0.02222 | 0.22800 |
| **HIST1H2BO** | 0.01203 | 0.00007 | 0.51348 | 0.02264 | 0.06347 | 0.12536 | 0.00000 | 1.81182 | 0.04697 | 0.24085 | 0.00000 | 0.03616 |
| **HIST1H3A** | 2.04837 | 0.02389 | 1.15203 | 0.12128 | 0.00000 | 0.00000 | 0.00001 | 4.08401 | 0.40576 | 0.42998 | 0.01623 | 0.08036 |
| **HIST1H3B** | 0.30041 | 0.00028 | 0.70371 | 0.07614 | 0.31792 | 0.43653 | 0.00000 | 0.49741 | 0.03146 | 1.80383 | 0.02097 | 0.36758 |
| **HIST1H3C** | 0.29671 | 0.00000 | 1.09964 | 0.00000 | 1.87063 | 0.49434 | 0.00000 | 2.85942 | 0.63000 | 3.73537 | 0.00000 | 1.62461 |
| **HIST1H3D** | 0.00184 | 0.09273 | 1.76908 | 0.53123 | 0.46805 | 0.00000 | 0.28366 | 1.67550 | 0.66024 | 0.00000 | 0.01666 | 0.71781 |
| **HIST1H3E** | 0.04488 | 0.00308 | 0.03774 | 0.01399 | 0.02011 | 0.15983 | 0.09834 | 0.09536 | 0.18196 | 0.13323 | 0.00000 | 0.07525 |
| **HIST1H3F** | 0.10967 | 0.05535 | 0.48102 | 0.08551 | 0.05484 | 0.00000 | 0.06783 | 0.25854 | 0.04355 | 0.00000 | 0.02451 | 0.12933 |
| **HIST1H3G** | 4.41229 | 0.00000 | 2.72241 | 0.50765 | 0.46771 | 0.00000 | 0.07267 | 6.36141 | 0.58073 | 1.89533 | 0.00000 | 0.48625 |
| **HIST1H3H** | 0.01721 | 0.01780 | 1.21591 | 0.13037 | 0.50891 | 0.13907 | 0.11884 | 1.36885 | 0.19051 | 0.00000 | 0.00777 | 0.36029 |
| **HIST1H3I** | 0.15396 | 0.21727 | 1.20887 | 0.50379 | 0.00000 | 0.17215 | 0.34332 | 0.04731 | 0.70359 | 0.00000 | 0.99244 | 0.88492 |
| **HIST1H3J** | 0.42943 | 0.00000 | 1.51405 | 0.17020 | 1.31009 | 0.00000 | 0.00000 | 0.39847 | 0.71531 | 1.33143 | 0.00000 | 1.44228 |
| **HIST1H4A** | 0.06087 | 0.02982 | 0.23425 | 0.00000 | 0.02880 | 0.00000 | 0.00000 | 0.00000 | 0.06988 | 0.13682 | 0.02741 | 0.13937 |
| **HIST1H4B** | 0.17329 | 0.02826 | 0.70164 | 0.25714 | 0.17280 | 0.00000 | 0.03299 | 0.02792 | 0.00000 | 0.06120 | 0.04584 | 0.04854 |
| **HIST1H4C** | 0.61310 | 19.85339 | 38.56508 | 27.22627 | 43.39941 | 35.17542 | 72.23813 | 0.40742 | 37.61796 | 31.94000 | 45.91617 | 43.93321 |
| **HIST1H4D** | 0.06971 | 0.00134 | 0.49278 | 0.18599 | 0.00000 | 0.28395 | 0.00000 | 0.06549 | 0.03571 | 0.24978 | 0.03840 | 0.12593 |
| **HIST1H4E** | 0.07790 | 0.00333 | 0.40457 | 0.33467 | 0.52545 | 0.69127 | 0.00000 | 0.10033 | 0.14579 | 0.69015 | 0.00000 | 0.33145 |
| **HIST1H4F** | 0.03775 | 0.00000 | 0.12969 | 0.15773 | 0.03801 | 0.23216 | 0.00000 | 0.14405 | 0.00000 | 0.55768 | 0.00000 | 0.08889 |
| **HIST1H4H** | 1.53541 | 0.00000 | 0.59655 | 0.10246 | 0.00000 | 0.22017 | 0.00000 | 4.03035 | 0.00000 | 0.70564 | 0.00000 | 0.10700 |
| **HIST1H4I** | 0.03449 | 0.47418 | 0.60262 | 0.84412 | 0.00645 | 0.00000 | 0.33127 | 0.12601 | 1.12728 | 0.00000 | 0.18167 | 0.10796 |
| **HIST1H4J** | 0.08231 | 0.01366 | 0.44427 | 0.13131 | 0.04449 | 0.06277 | 0.00000 | 0.14899 | 0.02090 | 0.15448 | 0.07556 | 0.12336 |
| **HIST1H4K** | 0.02006 | 0.01807 | 0.03909 | 0.02174 | 0.01592 | 0.02802 | 0.02659 | 0.09612 | 0.02206 | 0.01249 | 0.01347 | 0.04025 |
| **HIST1H4L** | 0.00179 | 0.00186 | 0.45513 | 0.00344 | 0.00000 | 0.10062 | 0.00000 | 0.01522 | 0.00000 | 0.37416 | 0.03092 | 0.08942 |
| **HIST2H2AB** | 0.04534 | 0.01999 | 0.37239 | 0.09373 | 0.00000 | 0.06200 | 0.00000 | 0.03019 | 0.04587 | 0.12853 | 0.01370 | 0.06591 |
| **HIST2H2AC** | 0.20094 | 0.00000 | 1.02442 | 0.00000 | 0.82253 | 0.25718 | 0.56545 | 0.00000 | 0.26019 | 1.37721 | 0.00000 | 0.59416 |
| **HIST2H2BA** | 0.13065 | 0.01261 | 0.02875 | 0.02904 | 0.02982 | 0.00000 | 0.00793 | 0.26766 | 0.02822 | 0.04387 | 0.01277 | 0.23215 |
| **HIST2H2BC** | 0.07153 | 0.00560 | 0.12251 | 0.00211 | 0.02169 | 0.15200 | 0.00000 | 0.05850 | 0.01679 | 0.14846 | 0.01412 | 0.04532 |
| **HIST2H2BE** | 0.85141 | 0.00000 | 1.18485 | 0.58189 | 0.00000 | 0.28177 | 0.00000 | 7.62999 | 0.00000 | 0.46682 | 0.00000 | 1.15133 |
| **HIST2H2BF** | 0.43795 | 0.01116 | 0.20673 | 0.02673 | 0.15365 | 0.00000 | 0.00208 | 0.10024 | 0.00000 | 0.05359 | 0.01563 | 0.06855 |
| **HIST2H3D** | 0.00427 | 0.01334 | 0.05947 | 0.04534 | 0.01292 | 0.02173 | 0.01202 | 0.01752 | 0.06039 | 0.06396 | 0.00504 | 0.02238 |
| **HIST3H2A** | 0.02325 | 0.00000 | 4.47456 | 0.26022 | 0.11061 | 0.70786 | 0.00000 | 0.14242 | 1.51569 | 0.80792 | 0.36720 | 0.00000 |
| **HIST3H2BB** | 0.00529 | 0.01267 | 0.21037 | 0.03858 | 0.18721 | 0.00000 | 0.10451 | 0.03062 | 0.00000 | 0.00000 | 0.02381 | 0.02191 |

| | | | | | | | | | | | | |
|---|---|---|---|---|---|---|---|---|---|---|---|---|
| HIST3H3 | 0.00705 | 0.01108 | 0.34201 | 0.04536 | 0.00000 | 0.00747 | 0.15014 | 0.02081 | 0.00000 | 0.19692 | 0.00640 | 0.01301 |
| HIST4H4 | 0.00767 | 0.00762 | 0.24128 | 0.00329 | 0.02218 | 0.13579 | 0.06116 | 0.10322 | 0.18047 | 0.00000 | 0.00229 | 0.09601 |
| HIVEP1 | 7.44304 | 0.49965 | 4.72267 | 0.44376 | 3.00126 | 3.10658 | 3.74280 | 0.14489 | 9.13396 | 7.19161 | 2.19656 | 2.48422 |
| HIVEP2 | 0.00278 | 0.00000 | 0.00000 | 0.00000 | 1.69572 | 0.18088 | 0.70097 | 0.09721 | 0.27446 | 1.40667 | 0.76569 | 0.00000 |
| HIVEP3 | 0.40364 | 0.18661 | 0.04394 | 0.20904 | 0.16257 | 0.00000 | 0.00221 | 0.00000 | 0.00000 | 0.31216 | 0.24520 | 0.18210 |
| HJURP | 8.73984 | 24.22390 | 6.85418 | 1.69423 | 7.48870 | 15.01166 | 9.78918 | 16.67986 | 15.03687 | 0.00000 | 6.31613 | 29.29418 |
| HK1 | 0.00000 | 13.82840 | 19.83712 | 11.01346 | 14.31651 | 0.00000 | 14.34990 | 3.44593 | 6.77583 | 0.00000 | 5.34431 | 17.61748 |
| HK2 | 3.04143 | 27.21719 | 2.11120 | 10.84874 | 18.79199 | 21.29578 | 12.15294 | 17.54108 | 8.78906 | 15.68395 | 17.86628 | 28.61429 |
| HK3 | 0.00408 | 0.01422 | 0.04611 | 0.01926 | 0.01694 | 0.00002 | 0.02839 | 0.01379 | 0.01094 | 0.06095 | 0.02094 | 0.03585 |
| HKDC1 | 0.00250 | 2.93616 | 0.11901 | 0.00000 | 0.08115 | 0.06151 | 0.00000 | 0.03212 | 0.00000 | 0.06077 | 0.00000 | 4.22565 |
| HKR1 | 0.21100 | 10.43822 | 1.91805 | 6.17454 | 2.03786 | 4.05533 | 1.28651 | 6.40874 | 4.61792 | 1.64549 | 2.34552 | 12.29495 |
| HLA-A | 0.02088 | 0.05363 | 0.21367 | 1.55544 | 0.00000 | 0.09695 | 0.00000 | 0.35280 | 0.07618 | 0.00000 | 0.00762 | 0.18432 |
| HLA-B | 0.00538 | 0.41281 | 0.18160 | 1.05844 | 0.00000 | 0.18861 | 0.31957 | 0.09070 | 0.00231 | 0.04985 | 0.00000 | 1.71232 |
| HLA-C | 0.17557 | 0.28197 | 3.98867 | 6.14471 | 0.80391 | 0.00000 | 1.02198 | 0.12800 | 5.71813 | 0.00000 | 0.14822 | 1.36833 |
| HLA-DMA | 0.00516 | 0.26157 | 0.57164 | 0.21341 | 0.00000 | 0.08902 | 0.17309 | 0.02443 | 0.00799 | 0.00000 | 0.00000 | 0.18715 |
| HLA-DOA | 0.04810 | 0.01683 | 0.01633 | 0.06550 | 0.01500 | 0.00000 | 0.03074 | 0.01241 | 0.02080 | 0.00745 | 0.01819 | 0.01750 |
| HLA-DOB | 0.00702 | 0.01196 | 0.01714 | 0.00696 | 0.09211 | 0.00003 | 0.07294 | 0.01034 | 0.00000 | 0.04414 | 0.00920 | 0.01698 |
| HLA-DPA1 | 0.00537 | 0.01993 | 0.24030 | 0.02036 | 0.05568 | 0.06579 | 0.00000 | 0.01089 | 0.00000 | 0.16540 | 0.00751 | 0.12009 |
| HLA-DPB1 | 0.08250 | 0.03197 | 0.24894 | 0.46683 | 0.00000 | 0.03066 | 0.17521 | 0.02117 | 0.01976 | 0.00000 | 0.01440 | 0.08438 |
| HLA-DPB2 | 0.05869 | 0.00919 | 0.01694 | 0.18368 | 0.00298 | 0.03761 | 0.01693 | 0.00883 | 0.02810 | 0.00929 | 0.01286 | 0.01912 |
| HLA-DQA2 | 0.00548 | 0.00628 | 0.00883 | 0.00534 | 0.01413 | 0.04726 | 0.01361 | 0.00862 | 0.00477 | 0.05074 | 0.01209 | 0.03864 |
| HLA-DQB1 | 0.02044 | 0.24800 | 0.02295 | 0.00000 | 0.00000 | 0.07555 | 0.00000 | 0.52855 | 0.00000 | 0.05565 | 0.00362 | 0.80343 |
| HLA-DQB2 | 0.00780 | 0.09149 | 0.00712 | 0.00000 | 0.00000 | 0.09120 | 0.00000 | 0.20567 | 0.00408 | 0.04078 | 0.00696 | 0.13522 |
| HLA-DRA | 0.00416 | 0.00790 | 0.00822 | 0.00820 | 0.00764 | 0.05554 | 0.02842 | 0.01824 | 0.02471 | 0.00442 | 0.00967 | 0.03494 |
| HLA-DRB1 | 0.00101 | 0.00849 | 2.06208 | 0.12701 | 0.00000 | 0.00000 | 2.55246 | 0.24465 | 0.00000 | 0.00000 | 2.20817 | 0.20919 |
| HLA-DRB5 | 0.00135 | 0.00747 | 0.32179 | 0.23474 | 0.00757 | 0.00000 | 0.44725 | 0.55314 | 0.01009 | 0.00000 | 0.09760 | 0.12234 |
| HLA-DRB6 | 0.02219 | 0.00107 | 0.41114 | 0.01215 | 0.12540 | 0.01267 | 0.60261 | 0.03234 | 0.20942 | 0.00000 | 0.20569 | 0.04646 |
| HLA-E | 0.15899 | 9.62089 | 8.97442 | 15.80994 | 7.41526 | 4.10713 | 4.20630 | 3.83467 | 16.32661 | 3.37246 | 0.00000 | 3.55057 |
| HLA-F | 0.00106 | 15.40291 | 0.00000 | 11.12320 | 11.79226 | 0.00000 | 0.50144 | 0.97538 | 0.29475 | 0.00000 | 0.85414 | 3.20781 |
| HLA-F-AS1 | 0.00166 | 0.61625 | 0.02363 | 0.27184 | 0.33303 | 0.00000 | 0.02022 | 0.12718 | 0.08504 | 0.00000 | 0.03533 | 0.17999 |
| HLA-G | 0.06713 | 0.05787 | 0.09094 | 0.48968 | 0.00000 | 0.01959 | 0.05720 | 0.04236 | 0.34760 | 0.00009 | 0.00000 | 0.18649 |
| HLA-H | 0.00731 | 0.01540 | 0.02661 | 0.03679 | 0.00216 | 0.03865 | 0.01483 | 0.02470 | 0.02462 | 0.04850 | 0.01031 | 0.06534 |
| HLA-J | 0.02411 | 0.04573 | 0.03057 | 0.10759 | 0.03196 | 0.04484 | 0.00735 | 0.01261 | 0.04421 | 0.02883 | 0.02601 | 0.16616 |
| HLA-L | 0.03156 | 0.01823 | 0.22066 | 0.11378 | 0.17584 | 0.00000 | 0.02570 | 0.03989 | 0.00000 | 0.30902 | 0.04453 | 0.05214 |
| HLCS | 0.54534 | 1.78913 | 1.74676 | 2.24786 | 0.34396 | 0.95017 | 1.69560 | 0.03867 | 1.27661 | 2.40674 | 4.87203 | 1.05086 |
| HLF | 0.14247 | 0.01525 | 0.05069 | 0.02440 | 0.02720 | 0.00055 | 0.02910 | 0.06569 | 0.01459 | 0.00000 | 0.03071 | 0.06539 |
| HLTF | 0.18791 | 1.85170 | 0.00000 | 7.40600 | 0.47635 | 9.99690 | 0.00000 | 2.67105 | 0.00000 | 3.24191 | 1.33082 | 4.27064 |
| HLX | 0.00179 | 0.02393 | 0.44915 | 0.01471 | 0.00000 | 0.01595 | 0.07703 | 0.05183 | 0.19359 | 0.06910 | 0.00000 | 0.07644 |
| HLX-AS1 | 0.00636 | 0.01593 | 0.02797 | 0.01703 | 0.02106 | 0.01450 | 0.01452 | 0.01050 | 0.02084 | 0.01133 | 0.01494 | 0.01660 |
| HM13 | 3.02435 | 22.95604 | 22.32345 | 20.73968 | 22.19736 | 0.00068 | 8.31921 | 4.67638 | 42.02335 | 6.83150 | 18.47106 | 19.51279 |
| HM13-AS1 | 0.02654 | 0.04076 | 0.06972 | 0.08299 | 0.06546 | 0.04468 | 0.00000 | 0.06966 | 0.05097 | 0.05881 | 0.00787 | 0.11221 |
| HMBOX1 | 2.70760 | 0.82542 | 2.79767 | 5.89973 | 0.40469 | 1.57813 | 0.03178 | 0.98883 | 4.91567 | 6.40173 | 2.06346 | 1.36826 |
| HMBS | 0.00000 | 13.47828 | 7.47537 | 6.77995 | 6.07472 | 4.46201 | 8.78505 | 0.00000 | 2.84180 | 0.07504 | 19.69904 | 4.53318 |
| HMCES | 4.98563 | 15.37660 | 13.36227 | 6.01437 | 6.51864 | 13.79642 | 17.62402 | 2.87555 | 8.64325 | 0.00000 | 14.52342 | 9.79535 |
| HMCN1 | 6.17538 | 0.62896 | 0.13437 | 0.57690 | 0.00000 | 0.00000 | 0.01861 | 1.60678 | 0.01186 | 0.00724 | 0.01502 | 0.59593 |
| HMCN2 | 0.00734 | 0.16087 | 0.19751 | 0.00000 | 0.37272 | 0.00000 | 0.01142 | 0.00709 | 0.00000 | 0.21924 | 0.15305 | 0.14457 |
| HMG20A | 3.19759 | 6.29698 | 4.12562 | 4.12414 | 0.00000 | 7.90894 | 8.21364 | 0.28614 | 2.65683 | 0.00000 | 5.79017 | 4.13770 |
| HMG20B | 1.51374 | 5.17784 | 10.68439 | 5.05896 | 10.74192 | 0.00000 | 5.43295 | 3.03263 | 6.01746 | 8.53544 | 4.90645 | 1.72390 |

| | | | | | | | | | | | | |
|---|---|---|---|---|---|---|---|---|---|---|---|---|
| **HMGA1** | 10.46469 | 37.05457 | 39.54784 | 31.92377 | 51.34523 | 2.25067 | 54.98450 | 5.43263 | 51.99926 | 30.40048 | 29.55096 | 37.17975 |
| **HMGA1P7** | 0.00776 | 0.00816 | 0.10234 | 0.02769 | 0.00681 | 0.05215 | 0.00578 | 0.01005 | 0.00000 | 0.02809 | 0.02376 | 0.01510 |
| **HMGA2** | 0.00890 | 0.00660 | 0.62053 | 0.25581 | 0.00000 | 0.01595 | 0.00000 | 0.14090 | 0.00000 | 0.05159 | 0.02062 | 0.03445 |
| **HMGB1** | 2.74626 | 10.97979 | 6.81310 | 7.01469 | 7.99676 | 33.41615 | 22.68182 | 4.10282 | 7.07072 | 15.38342 | 12.97223 | 20.17542 |
| **HMGB2** | 9.77228 | 28.64479 | 20.76380 | 13.16078 | 17.94556 | 55.17674 | 35.40278 | 15.05845 | 25.23218 | 9.55855 | 14.39910 | 34.98427 |
| **HMGB3** | 10.88344 | 23.44320 | 5.04868 | 19.46626 | 21.03792 | 36.19853 | 15.36040 | 18.33534 | 5.82603 | 9.82863 | 12.47906 | 19.06856 |
| **HMGB3P1** | 0.00842 | 0.02493 | 0.01316 | 0.00809 | 0.02371 | 0.02427 | 0.04036 | 0.01257 | 0.00062 | 0.05143 | 0.01908 | 0.01395 |
| **HMGCL** | 1.43164 | 3.59667 | 0.61782 | 2.16726 | 1.65199 | 0.50629 | 0.42333 | 0.63147 | 0.92703 | 0.00000 | 1.83486 | 1.49399 |
| **HMGCLL1** | 0.48513 | 0.01593 | 0.01614 | 0.01331 | 0.01685 | 0.00332 | 0.02006 | 0.06939 | 0.02149 | 0.00614 | 0.02401 | 0.02517 |
| **HMGCR** | 2.33913 | 7.89897 | 11.51755 | 30.01777 | 30.23773 | 37.39220 | 0.00000 | 7.52002 | 35.96711 | 6.85904 | 17.51307 | 5.12537 |
| **HMGCS1** | 0.00000 | 0.00000 | 9.41877 | 12.86858 | 32.42806 | 18.73313 | 44.68464 | 4.37244 | 9.64304 | 9.82792 | 37.21161 | 1.24227 |
| **HMGCS2** | 0.00617 | 0.01170 | 0.02582 | 0.00729 | 0.00441 | 0.03916 | 0.01764 | 0.01030 | 0.06825 | 0.01923 | 0.00256 | 0.01434 |
| **HMGN1** | 1.29152 | 14.22560 | 10.62927 | 13.00183 | 12.12459 | 23.54706 | 22.78946 | 0.00000 | 17.79964 | 8.46012 | 16.01284 | 13.95304 |
| **HMGN2** | 4.49422 | 13.23398 | 3.59258 | 9.71626 | 9.91110 | 9.25446 | 8.69797 | 6.55233 | 9.13722 | 0.00000 | 11.26646 | 13.40633 |
| **HMGN2P46** | 0.52505 | 0.35134 | 0.01437 | 0.00000 | 0.02185 | 0.03918 | 0.04698 | 0.89872 | 0.00000 | 0.36137 | 0.00000 | 0.28901 |
| **HMGN3** | 0.00000 | 0.97720 | 5.23214 | 6.67536 | 0.00000 | 5.22407 | 6.60410 | 0.09206 | 3.76740 | 0.63624 | 3.30956 | 0.00000 |
| **HMGN3-AS1** | 0.00215 | 0.01008 | 0.20383 | 0.23415 | 0.22941 | 0.10451 | 0.00000 | 0.00797 | 0.09397 | 0.51902 | 0.08696 | 0.00024 |
| **HMGN4** | 1.65300 | 8.63405 | 13.97223 | 4.69826 | 0.00000 | 22.88229 | 17.18903 | 12.81809 | 10.20057 | 0.00000 | 6.70192 | 19.77751 |
| **HMGN5** | 0.21043 | 0.45211 | 0.10299 | 1.22540 | 1.29759 | 3.04896 | 2.46197 | 0.00379 | 0.00000 | 3.18870 | 1.27368 | 0.32750 |
| **HMGXB3** | 4.63580 | 8.94482 | 4.40112 | 4.65419 | 5.75624 | 0.00000 | 4.20322 | 5.58902 | 12.67967 | 4.88019 | 2.49303 | 9.42309 |
| **HMGXB4** | 9.38480 | 3.78526 | 1.49251 | 6.45815 | 2.05286 | 12.07775 | 3.73317 | 2.35616 | 5.72992 | 11.33299 | 2.72901 | 7.11326 |
| **HMHA1** | 8.63697 | 6.53272 | 13.85311 | 1.94025 | 7.37659 | 0.00000 | 6.55373 | 1.71552 | 7.29492 | 3.64581 | 1.52234 | 0.00000 |
| **HMMR** | 4.19324 | 12.95205 | 7.84259 | 1.25380 | 9.89947 | 36.48909 | 0.00000 | 5.19121 | 7.63025 | 15.85264 | 4.49466 | 20.28829 |
| **HMMR-AS1** | 0.09372 | 0.51795 | 0.11189 | 0.14126 | 0.28721 | 0.80862 | 0.00000 | 0.18130 | 0.22339 | 0.57586 | 0.07634 | 0.65378 |
| **HMOX1** | 0.11303 | 10.59090 | 42.52757 | 3.36220 | 26.98308 | 9.57439 | 24.01897 | 0.89696 | 37.56841 | 2.60107 | 8.10720 | 4.49546 |
| **HMOX2** | 2.11054 | 16.96312 | 21.11559 | 9.14425 | 17.24566 | 6.05658 | 24.03085 | 10.99361 | 22.24953 | 0.00000 | 26.25177 | 10.60830 |
| **HMP19** | 0.00548 | 0.01819 | 0.03928 | 0.00732 | 0.02689 | 0.00000 | 0.00000 | 0.00973 | 0.10282 | 0.06292 | 0.00323 | 0.03035 |
| **HMX1** | 0.15399 | 0.00802 | 0.01848 | 0.04218 | 0.06906 | 0.02003 | 0.01591 | 0.00266 | 0.01627 | 0.02895 | 0.01338 | 0.02975 |
| **HMX2** | 0.03359 | 0.02765 | 0.03787 | 0.01243 | 0.01867 | 0.01491 | 0.01928 | 0.00771 | 0.01021 | 0.01969 | 0.01591 | 0.02595 |
| **HMX3** | 0.00308 | 0.01151 | 0.04884 | 0.02611 | 0.00697 | 0.02453 | 0.02339 | 0.06461 | 0.00530 | 0.02778 | 0.03803 | 0.01720 |
| **HN1** | 5.32403 | 41.54236 | 28.67833 | 21.64228 | 45.65381 | 35.65547 | 44.10865 | 12.87457 | 43.83088 | 20.29751 | 24.72275 | 54.08974 |
| **HN1L** | 1.53708 | 16.04357 | 24.28929 | 12.10408 | 20.17347 | 11.26880 | 34.47542 | 0.26931 | 15.19485 | 6.86363 | 32.59183 | 9.78906 |
| **HNF1A** | 8.24910 | 0.55788 | 0.21786 | 0.43778 | 0.00000 | 0.00000 | 0.07333 | 0.55293 | 0.69028 | 0.03387 | 0.00000 | 0.43539 |
| **HNF1A-AS1** | 0.11129 | 0.00106 | 0.00937 | 1.13572 | 0.04603 | 0.01972 | 0.00000 | 0.07279 | 0.02532 | 0.10640 | 0.01000 | 0.00797 |
| **HNF1B** | 0.01690 | 0.26606 | 0.33003 | 13.03032 | 1.25340 | 0.05979 | 0.00202 | 0.00616 | 0.00000 | 0.52658 | 0.00000 | 0.00000 |
| **HNF4A** | 0.00000 | 8.80009 | 0.19663 | 15.99038 | 2.47917 | 2.79872 | 0.62450 | 1.61840 | 0.00000 | 4.70574 | 0.37966 | 21.88187 |
| **HNF4A-AS1** | 0.06440 | 0.00520 | 0.02400 | 0.02083 | 0.01309 | 0.01235 | 0.01976 | 0.00813 | 0.01266 | 0.03578 | 0.01485 | 0.19978 |
| **HNF4G** | 0.00568 | 0.01977 | 0.01238 | 0.49503 | 0.00000 | 0.05031 | 0.00800 | 0.00924 | 0.01155 | 0.01445 | 0.03997 | 0.02908 |
| **HNMT** | 0.11557 | 0.15538 | 0.00000 | 0.40923 | 0.06896 | 0.09324 | 0.01837 | 0.03322 | 0.02520 | 0.05476 | 0.09197 | 0.18287 |
| **HNRNPA0** | 1.03200 | 8.46109 | 22.33153 | 6.72731 | 6.17014 | 0.00000 | 16.31825 | 2.24468 | 11.86889 | 20.11512 | 18.71993 | 8.90781 |
| **HNRNPA1** | 0.65379 | 4.28755 | 5.91369 | 3.34351 | 0.00000 | 22.25737 | 18.69105 | 4.10645 | 3.27426 | 15.30195 | 5.73813 | 11.16756 |
| **HNRNPA1L2** | 1.62573 | 1.74580 | 0.00000 | 0.30755 | 1.30998 | 2.25908 | 0.24729 | 2.22649 | 0.35660 | 0.00000 | 0.53022 | 2.60586 |
| **HNRNPA1P33** | 0.01107 | 0.04877 | 0.06453 | 0.00000 | 1.04682 | 0.40290 | 1.28579 | 0.02058 | 0.09093 | 0.29547 | 0.61106 | 0.04780 |
| **HNRNPA2B1** | 9.39978 | 32.49392 | 30.67075 | 27.20088 | 23.68094 | 69.31961 | 51.33927 | 26.22441 | 32.86054 | 44.97274 | 27.92057 | 35.04240 |
| **HNRNPA3** | 0.00000 | 1.05150 | 2.12660 | 2.15935 | 2.54316 | 12.43214 | 14.74344 | 2.28242 | 0.00000 | 25.19625 | 4.80987 | 3.19402 |
| **HNRNPA3P1** | 0.00549 | 0.00584 | 0.14794 | 0.03646 | 0.00000 | 0.07018 | 0.00000 | 0.02311 | 0.00000 | 0.08365 | 0.03562 | 0.01898 |
| **HNRNPAB** | 0.32353 | 8.93893 | 12.28851 | 7.21371 | 16.84135 | 20.55223 | 34.94640 | 16.94521 | 4.53960 | 54.45159 | 7.90593 | 13.23624 |
| **HNRNPC** | 6.57793 | 29.90448 | 26.35417 | 21.34597 | 21.87345 | 51.38418 | 37.79097 | 25.64257 | 37.44123 | 10.67726 | 32.72810 | 36.62776 |

| | | | | | | | | | | | | |
|---|---|---|---|---|---|---|---|---|---|---|---|---|
| **HNRNPCL1** | 0.76601 | 0.00832 | 0.01103 | 0.00941 | 0.01599 | 0.00000 | 0.01254 | 6.11922 | 0.01738 | 0.02065 | 0.01206 | 0.03867 |
| **HNRNPCL2** | 4.04122 | 0.00000 | 0.00918 | 0.00043 | 0.03983 | 0.08812 | 0.00001 | 20.11412 | 0.05950 | 0.00000 | 0.01903 | 0.54047 |
| **HNRNPD** | 3.03201 | 6.19643 | 16.68135 | 7.93981 | 11.12845 | 34.01522 | 41.27536 | 6.20300 | 0.00000 | 55.95566 | 17.41731 | 17.59625 |
| **HNRNPDL** | 6.11067 | 27.38867 | 19.24301 | 18.49541 | 11.86014 | 49.99184 | 24.53166 | 20.52153 | 30.53415 | 25.82166 | 23.57125 | 35.05008 |
| **HNRNPF** | 11.16761 | 45.70741 | 31.76121 | 31.27611 | 28.40686 | 54.57563 | 48.30874 | 34.67830 | 46.32893 | 9.11791 | 43.93724 | 46.96848 |
| **HNRNPH1** | 3.37727 | 17.81852 | 28.94367 | 19.60334 | 9.85563 | 63.11940 | 38.44983 | 32.47005 | 25.44647 | 34.62574 | 29.82785 | 34.91551 |
| **HNRNPH2** | 0.05364 | 0.84501 | 0.53065 | 0.15014 | 0.00364 | 2.25454 | 0.40084 | 1.16083 | 1.08254 | 0.00000 | 1.96484 | 0.94732 |
| **HNRNPH3** | 1.21390 | 21.47309 | 16.27303 | 9.36540 | 14.10898 | 32.91221 | 23.90444 | 6.04952 | 18.22112 | 7.19542 | 12.88639 | 28.40125 |
| **HNRNPK** | 4.10391 | 29.23627 | 25.25629 | 21.81965 | 15.69767 | 35.63639 | 37.05606 | 21.62740 | 27.90594 | 5.26355 | 36.95252 | 30.61186 |
| **HNRNPKP3** | 0.03892 | 0.11553 | 0.48349 | 0.13118 | 0.00000 | 0.43113 | 0.00000 | 0.22198 | 0.06925 | 0.06217 | 0.18086 | 0.13828 |
| **HNRNPL** | 0.00000 | 5.80562 | 10.25486 | 5.19972 | 5.63359 | 10.57097 | 27.97735 | 13.57991 | 0.00000 | 33.93806 | 10.49163 | 11.98765 |
| **HNRNPLL** | 0.00000 | 3.29345 | 3.31470 | 1.70097 | 1.58300 | 8.59417 | 2.08108 | 1.33117 | 3.68932 | 6.66754 | 1.88080 | 1.39040 |
| **HNRNPM** | 3.24940 | 30.99172 | 30.31688 | 29.60351 | 28.67446 | 19.10410 | 52.42093 | 24.69532 | 28.93645 | 27.63736 | 34.25972 | 33.62431 |
| **HNRNPR** | 2.24234 | 21.52998 | 26.59773 | 13.01048 | 17.77347 | 47.63018 | 28.41944 | 13.49719 | 27.05486 | 10.18832 | 21.88157 | 25.22375 |
| **HNRNPU** | 3.22326 | 9.82225 | 22.40618 | 10.52691 | 17.69120 | 40.46637 | 37.23830 | 21.75695 | 7.83551 | 52.24865 | 17.57787 | 20.95204 |
| **HNRNPU-AS1** | 0.07360 | 0.00000 | 0.84218 | 1.08182 | 1.42968 | 4.49734 | 0.00000 | 1.18906 | 3.23284 | 15.53999 | 0.66538 | 0.45491 |
| **HNRNPUL1** | 3.62285 | 3.33175 | 8.96872 | 7.16446 | 12.71014 | 1.42840 | 16.79988 | 3.96450 | 2.86968 | 24.74445 | 7.68851 | 15.26204 |
| **HNRNPUL2** | 0.50004 | 0.25084 | 0.49963 | 0.45893 | 1.11229 | 0.51513 | 3.48492 | 0.37140 | 0.00000 | 8.22950 | 0.00000 | 0.60409 |
| **HOGA1** | 0.01374 | 0.01550 | 0.02889 | 0.01210 | 0.00823 | 0.00597 | 0.01350 | 0.01104 | 0.00000 | 0.01932 | 0.12821 | 0.01725 |
| **HOMER1** | 1.71435 | 0.00000 | 2.66683 | 0.61717 | 1.30206 | 3.00780 | 2.56772 | 0.00000 | 1.06933 | 7.01299 | 0.15160 | 0.30783 |
| **HOMER2** | 7.11013 | 3.86290 | 0.12440 | 1.97701 | 1.38486 | 0.68537 | 0.00000 | 1.75264 | 0.00000 | 1.11027 | 0.00000 | 0.78722 |
| **HOMER3** | 0.10309 | 0.49291 | 0.36315 | 0.41019 | 0.23144 | 0.00000 | 0.05543 | 0.06549 | 0.17183 | 0.33119 | 0.00000 | 0.97521 |
| **HOMEZ** | 0.07944 | 0.40584 | 1.32087 | 0.02251 | 1.54778 | 0.21787 | 0.00000 | 0.03610 | 0.63237 | 0.00000 | 0.73038 | 0.29046 |
| **HOOK1** | 1.35547 | 0.00000 | 0.00000 | 0.00000 | 3.50879 | 2.87594 | 8.43593 | 0.41473 | 0.69587 | 6.42906 | 4.40703 | 0.14825 |
| **HOOK2** | 0.68348 | 3.10542 | 5.04030 | 3.37282 | 0.90659 | 0.00000 | 2.59195 | 0.19052 | 5.86796 | 0.60329 | 6.77480 | 0.82722 |
| **HOOK3** | 2.61755 | 0.83037 | 4.31082 | 2.60222 | 2.57360 | 16.17447 | 3.94329 | 2.36284 | 0.00000 | 26.75447 | 3.49900 | 2.36456 |
| **HOPX** | 0.00115 | 0.02924 | 14.38463 | 0.01317 | 0.20149 | 0.00000 | 1.32800 | 0.05975 | 0.00000 | 0.00000 | 0.00063 | 0.03621 |
| **HORMAD1** | 0.11401 | 5.34134 | 0.00000 | 0.95921 | 0.00001 | 6.16358 | 0.00000 | 0.51067 | 1.84423 | 2.24197 | 0.71260 | 3.55765 |
| **HORMAD2** | 0.11657 | 0.19488 | 0.01839 | 0.10606 | 0.00000 | 0.04289 | 0.06359 | 0.03430 | 0.00000 | 0.03789 | 0.00796 | 0.39289 |
| **HORMAD2-AS1** | 0.01723 | 10.35170 | 0.00000 | 16.05029 | 3.25338 | 2.82795 | 0.00001 | 0.01054 | 0.00000 | 1.43893 | 1.25894 | 0.06590 |
| **HOTAIR** | 0.00491 | 0.01298 | 0.01378 | 0.01029 | 0.00900 | 0.01389 | 0.01360 | 0.01143 | 0.02373 | 0.04654 | 0.01917 | 0.06309 |
| **HOTAIRM1** | 0.42080 | 0.14556 | 0.49608 | 0.32050 | 0.00000 | 0.31667 | 0.43682 | 0.30263 | 0.34438 | 0.02487 | 0.00000 | 2.56045 |
| **HOTTIP** | 0.00587 | 0.01490 | 0.01397 | 0.01627 | 0.01238 | 0.01236 | 0.02209 | 0.01046 | 0.02165 | 0.03455 | 0.01556 | 0.01603 |
| **HOXA-AS2** | 0.01661 | 0.01749 | 0.01334 | 0.03003 | 0.02078 | 0.02211 | 0.03162 | 0.02345 | 0.01085 | 0.00057 | 0.01689 | 0.01679 |
| **HOXA-AS3** | 0.04067 | 0.00965 | 0.01093 | 0.01123 | 0.02259 | 0.03583 | 0.01268 | 0.00987 | 0.01018 | 0.03354 | 0.01314 | 0.01300 |
| **HOXA1** | 0.47321 | 0.48446 | 0.65248 | 1.48209 | 0.00130 | 0.42288 | 0.94869 | 0.10972 | 0.00000 | 0.00000 | 0.02772 | 2.30002 |
| **HOXA10** | 0.03202 | 0.00443 | 0.01907 | 0.28553 | 0.01741 | 0.02504 | 0.00014 | 0.19236 | 0.00000 | 0.19373 | 0.00622 | 0.12129 |
| **HOXA11** | 0.00286 | 0.00909 | 0.04159 | 0.01955 | 0.00617 | 0.00870 | 0.00003 | 0.12839 | 0.01031 | 0.23434 | 0.02096 | 0.02207 |
| **HOXA11-AS** | 0.08156 | 0.00800 | 0.01302 | 0.14426 | 0.00000 | 0.01344 | 0.00221 | 0.00776 | 0.02064 | 0.12676 | 0.01084 | 0.03041 |
| **HOXA13** | 0.10229 | 0.05593 | 0.04718 | 0.01010 | 0.01938 | 0.01180 | 0.02405 | 0.00322 | 0.00901 | 0.00133 | 0.01881 | 0.08991 |
| **HOXA2** | 0.05593 | 0.01178 | 0.07283 | 0.22612 | 0.02667 | 0.00000 | 0.04922 | 0.00454 | 0.00000 | 0.00000 | 0.01624 | 0.07479 |
| **HOXA3** | 0.34383 | 0.05614 | 0.00258 | 0.24970 | 0.00000 | 0.00000 | 0.00767 | 0.02379 | 0.00502 | 0.37360 | 0.00718 | 0.20691 |
| **HOXA4** | 0.63677 | 0.05390 | 0.05836 | 0.00501 | 0.03460 | 0.02619 | 0.00132 | 0.00215 | 0.05205 | 0.01449 | 0.01240 | 0.04635 |
| **HOXA5** | 0.00525 | 0.01462 | 0.02713 | 0.00696 | 0.00000 | 0.03269 | 0.02543 | 0.00616 | 0.01974 | 0.06624 | 0.01084 | 0.12857 |
| **HOXA7** | 3.90773 | 0.11376 | 0.04299 | 0.31847 | 0.12874 | 0.06151 | 0.00000 | 0.24781 | 0.06078 | 0.00761 | 0.02380 | 0.14005 |
| **HOXA9** | 0.33284 | 0.00904 | 0.00654 | 0.07941 | 0.01677 | 0.04236 | 0.01406 | 0.03548 | 0.01847 | 0.00012 | 0.01388 | 0.01862 |
| **HOXB-AS1** | 0.01911 | 0.00767 | 0.08242 | 0.04208 | 0.03881 | 0.05954 | 0.00001 | 0.01478 | 0.09131 | 0.01520 | 0.05394 | 0.03574 |
| **HOXB-AS3** | 0.08832 | 0.03222 | 0.00000 | 0.00750 | 0.00000 | 0.02431 | 0.18106 | 0.00464 | 0.34522 | 0.00000 | 0.00000 | 0.04738 |

| | | | | | | | | | | | | |
|---|---|---|---|---|---|---|---|---|---|---|---|---|
| **HOXB1** | 0.00597 | 0.00858 | 0.01249 | 0.29350 | 0.00258 | 0.03478 | 0.02156 | 0.01000 | 0.02497 | 0.00918 | 0.02216 | 0.01501 |
| **HOXB13** | 0.46026 | 0.07942 | 0.00000 | 0.27824 | 0.02261 | 0.00000 | 0.25576 | 0.51864 | 0.16302 | 0.00000 | 0.14075 | 0.87369 |
| **HOXB2** | 0.05620 | 0.02456 | 0.85460 | 0.66383 | 0.49743 | 0.00998 | 0.33082 | 0.03343 | 0.05552 | 0.34187 | 0.14783 | 0.04228 |
| **HOXB3** | 0.00266 | 0.00000 | 0.44883 | 0.33866 | 0.09193 | 0.06654 | 0.35196 | 0.00712 | 0.98273 | 0.62631 | 0.00000 | 0.06442 |
| **HOXB4** | 0.00267 | 0.00000 | 0.03951 | 0.00000 | 0.00000 | 0.19671 | 2.35059 | 0.07772 | 0.00020 | 2.39003 | 0.91552 | 0.37122 |
| **HOXB5** | 0.03291 | 0.00000 | 0.02557 | 0.04897 | 0.03763 | 0.07362 | 0.03180 | 0.06422 | 0.15825 | 0.00037 | 0.01190 | 0.30455 |
| **HOXB6** | 0.61018 | 0.00115 | 0.09135 | 0.02244 | 0.03322 | 0.09097 | 0.00035 | 0.32394 | 0.11730 | 0.00000 | 0.00901 | 0.18277 |
| **HOXB7** | 0.12763 | 0.84530 | 0.00000 | 2.61877 | 1.07506 | 2.09685 | 2.74823 | 0.13809 | 0.00359 | 1.07907 | 1.05832 | 2.16862 |
| **HOXB8** | 0.01691 | 0.02163 | 0.00593 | 0.02358 | 0.00267 | 0.07022 | 0.00977 | 0.01237 | 0.00843 | 0.02291 | 0.01089 | 0.24877 |
| **HOXB9** | 0.11291 | 0.17237 | 0.00000 | 0.00150 | 0.70795 | 0.38205 | 0.00000 | 0.08161 | 0.05227 | 0.00000 | 0.00000 | 1.25153 |
| **HOXC-AS1** | 0.00606 | 0.04943 | 0.02226 | 0.00689 | 0.01394 | 0.00515 | 0.04836 | 0.01011 | 0.08808 | 0.01861 | 0.03542 | 0.03772 |
| **HOXC-AS3** | 0.00498 | 0.01331 | 0.00786 | 0.07960 | 0.01270 | 0.00673 | 0.01688 | 0.01429 | 0.04146 | 0.01937 | 0.00986 | 0.02300 |
| **HOXC10** | 1.08453 | 0.17917 | 0.00000 | 0.15386 | 1.00651 | 0.00000 | 0.00000 | 1.73583 | 0.00000 | 0.33741 | 0.09191 | 1.05521 |
| **HOXC11** | 0.10546 | 0.05187 | 0.01090 | 0.00401 | 0.01384 | 0.06221 | 0.01276 | 0.05771 | 0.00905 | 0.01579 | 0.01146 | 0.10293 |
| **HOXC12** | 0.09161 | 0.01320 | 0.01444 | 0.01275 | 0.02612 | 0.01886 | 0.01482 | 0.00724 | 0.02321 | 0.00358 | 0.02942 | 0.02536 |
| **HOXC13** | 0.29235 | 0.00000 | 0.00000 | 0.36303 | 2.74030 | 0.00000 | 3.17695 | 0.08595 | 0.00000 | 2.60853 | 0.33975 | 0.27105 |
| **HOXC13-AS** | 0.00582 | 0.00000 | 0.04243 | 0.10286 | 0.99415 | 0.06652 | 0.75993 | 0.00531 | 0.04947 | 0.00000 | 0.24674 | 0.14634 |
| **HOXC4** | 0.01417 | 0.01809 | 0.12755 | 0.00000 | 0.00000 | 0.27738 | 0.00002 | 0.00772 | 0.00000 | 0.05030 | 0.00818 | 0.00249 |
| **HOXC5** | 0.00639 | 0.17486 | 0.00000 | 0.15532 | 0.00000 | 0.07941 | 0.00000 | 0.00326 | 0.05339 | 0.08006 | 0.00000 | 0.24778 |
| **HOXC6** | 0.01413 | 0.02685 | 0.02872 | 0.03503 | 0.00442 | 0.00008 | 0.05164 | 0.04518 | 0.00000 | 0.05812 | 0.08353 | 0.01385 |
| **HOXC8** | 0.04987 | 0.02201 | 0.02016 | 0.02567 | 0.03517 | 0.00000 | 0.02661 | 0.00734 | 0.03632 | 0.00280 | 0.01268 | 0.01960 |
| **HOXC9** | 0.27072 | 1.96785 | 0.29065 | 0.34024 | 1.13255 | 1.25458 | 0.36915 | 0.47967 | 1.96150 | 0.34046 | 0.00000 | 2.22818 |
| **HOXD1** | 0.14495 | 0.00211 | 0.14092 | 2.11303 | 0.01448 | 0.00000 | 0.00478 | 0.77217 | 0.11032 | 0.00000 | 0.00220 | 0.48000 |
| **HOXD11** | 0.08232 | 0.01240 | 0.01333 | 0.01478 | 0.02256 | 0.00301 | 0.01810 | 0.00389 | 0.01964 | 0.03310 | 0.01524 | 0.04630 |
| **HOXD13** | 0.00777 | 0.00000 | 0.00000 | 0.98686 | 0.00000 | 0.21731 | 0.48505 | 0.00068 | 0.00000 | 0.27788 | 0.00000 | 0.26871 |
| **HOXD3** | 0.00575 | 0.01532 | 0.03204 | 0.01556 | 0.04787 | 0.00000 | 0.01995 | 0.00976 | 0.01709 | 0.01694 | 0.01430 | 0.04197 |
| **HOXD4** | 0.00456 | 0.01308 | 0.00728 | 0.10762 | 0.02488 | 0.02106 | 0.01636 | 0.02180 | 0.02224 | 0.00227 | 0.01711 | 0.03682 |
| **HOXD8** | 0.68303 | 0.00000 | 0.02319 | 0.00000 | 0.18466 | 0.08449 | 0.00000 | 0.01519 | 0.00000 | 0.20067 | 0.00000 | 0.30550 |
| **HOXD9** | 0.01574 | 0.14140 | 0.00000 | 0.00953 | 0.18976 | 0.13323 | 0.00000 | 0.00397 | 0.00000 | 0.15619 | 0.00000 | 0.09871 |
| **HP** | 0.00588 | 0.01078 | 0.00607 | 0.07844 | 0.00309 | 0.05987 | 0.01008 | 0.01013 | 0.01537 | 0.01095 | 0.01203 | 0.01959 |
| **HP08942** | 0.00626 | 0.02972 | 0.02083 | 0.00976 | 0.02304 | 0.04289 | 0.02100 | 0.01015 | 0.04377 | 0.00000 | 0.01349 | 0.02160 |
| **HP09025** | 0.05140 | 0.01501 | 0.02502 | 0.01715 | 0.01508 | 0.00886 | 0.02447 | 0.00572 | 0.00744 | 0.02354 | 0.03270 | 0.01558 |
| **HP09053** | 0.03915 | 0.20894 | 0.03360 | 0.18665 | 0.01650 | 0.17421 | 0.00000 | 0.37595 | 0.24917 | 0.07024 | 0.21183 | 0.59033 |
| **HP1BP3** | 1.51428 | 6.85119 | 7.51060 | 6.04246 | 4.22566 | 13.36086 | 7.45285 | 4.01623 | 8.16393 | 20.22673 | 7.40559 | 9.03571 |
| **HPCA** | 0.10393 | 0.00226 | 0.02528 | 0.03323 | 0.00000 | 0.04993 | 0.02343 | 0.01469 | 0.00217 | 0.07769 | 0.01263 | 0.11944 |
| **HPCAL1** | 0.00106 | 4.53859 | 4.09086 | 0.00000 | 4.91619 | 0.00000 | 1.99239 | 0.02893 | 3.55659 | 4.92138 | 2.59063 | 0.10286 |
| **HPCAL4** | 0.00261 | 0.00496 | 0.05934 | 0.01227 | 0.00000 | 0.03268 | 0.01209 | 0.39824 | 0.00000 | 0.04736 | 0.04031 | 0.08112 |
| **HPD** | 0.17695 | 0.15186 | 2.41035 | 0.45703 | 0.02433 | 0.00000 | 0.00000 | 0.02723 | 0.03352 | 0.00000 | 0.04425 | 0.11600 |
| **HPDL** | 0.00452 | 19.06349 | 7.83437 | 11.01046 | 10.74705 | 3.81685 | 4.35503 | 0.00000 | 17.00645 | 0.00000 | 11.85355 | 17.62796 |
| **HPGD** | 0.00000 | 7.61675 | 8.73281 | 14.03522 | 16.63498 | 23.09495 | 2.81193 | 0.00000 | 2.38970 | 1.98210 | 17.29555 | 0.00000 |
| **HPGDS** | 0.00822 | 0.00492 | 0.00576 | 0.01979 | 0.01551 | 0.08347 | 0.00653 | 0.00952 | 0.00469 | 0.02728 | 0.01477 | 0.07020 |
| **HPN** | 0.00208 | 0.01081 | 0.01829 | 0.07673 | 0.00000 | 0.01798 | 0.03670 | 0.05273 | 0.01344 | 0.04117 | 0.01472 | 0.01908 |
| **HPN-AS1** | 0.00591 | 0.01415 | 0.20739 | 0.01126 | 0.02754 | 0.01586 | 0.00000 | 0.01200 | 0.00000 | 0.05191 | 0.03592 | 0.01980 |
| **HPR** | 0.00212 | 0.00049 | 0.01481 | 0.00000 | 0.05239 | 0.01688 | 0.08460 | 0.04835 | 0.00000 | 0.45162 | 0.00000 | 0.00693 |
| **HPRT1** | 6.69080 | 11.24649 | 1.46820 | 0.00000 | 7.48425 | 31.49924 | 3.79096 | 3.01686 | 10.76807 | 2.40963 | 10.98148 | 10.22149 |
| **HPS1** | 0.03844 | 8.19099 | 6.91949 | 5.09477 | 5.27657 | 2.37135 | 6.45963 | 0.93340 | 11.27420 | 0.00000 | 5.60623 | 6.45505 |
| **HPS3** | 1.45002 | 1.36069 | 2.02379 | 0.79995 | 0.35874 | 4.28900 | 2.71038 | 0.49411 | 1.54832 | 0.00000 | 3.56409 | 0.20543 |
| **HPS4** | 5.52221 | 5.39047 | 1.74870 | 3.68538 | 0.00000 | 5.57526 | 2.10016 | 2.26045 | 2.68200 | 3.52487 | 3.00814 | 5.83593 |

| | | | | | | | | | | | | |
|---|---|---|---|---|---|---|---|---|---|---|---|---|
| **HPS5** | 5.06373 | 1.94819 | 3.87371 | 0.24484 | 7.51015 | 7.27845 | 2.09654 | 4.92652 | 1.65507 | 1.58592 | 3.55979 | 4.35585 |
| **HPS6** | 0.32413 | 2.14330 | 1.24211 | 3.03776 | 2.41809 | 0.47597 | 2.77587 | 0.00000 | 0.09208 | 4.74881 | 1.39778 | 3.14936 |
| **HPSE** | 0.55449 | 0.02906 | 0.09867 | 2.07729 | 1.48539 | 0.30834 | 0.00000 | 0.20362 | 0.74973 | 0.00000 | 0.28215 | 0.27295 |
| **HPSE2** | 3.63993 | 0.04870 | 0.01358 | 0.07594 | 0.01358 | 0.00000 | 0.00544 | 0.11717 | 0.03748 | 0.01838 | 0.01054 | 0.00681 |
| **HPX** | 0.00274 | 0.00000 | 0.10194 | 0.34538 | 0.77175 | 0.25740 | 0.00000 | 0.18122 | 0.04166 | 0.00000 | 0.12708 | 0.08247 |
| **HR** | 0.02063 | 0.00000 | 0.00404 | 0.00954 | 0.00000 | 0.04819 | 0.03313 | 0.03505 | 0.00000 | 0.26337 | 0.01828 | 0.08515 |
| **HRAS** | 0.59158 | 1.51339 | 5.66941 | 2.16614 | 6.59994 | 0.00000 | 11.90025 | 1.69084 | 0.71816 | 7.55277 | 1.39320 | 0.16371 |
| **HRASLS** | 0.20746 | 0.04689 | 0.21604 | 0.00435 | 0.02036 | 0.05954 | 0.00000 | 0.07679 | 0.00000 | 0.04970 | 0.02241 | 0.08486 |
| **HRASLS2** | 0.05590 | 0.01653 | 0.08077 | 0.00800 | 0.00698 | 0.02826 | 0.02012 | 0.02426 | 0.00000 | 0.03433 | 0.02096 | 0.01866 |
| **HRASLS5** | 0.93532 | 13.85939 | 0.00000 | 1.17072 | 1.31145 | 6.64572 | 0.00000 | 0.18806 | 0.00000 | 1.17612 | 2.04420 | 21.17198 |
| **HRC** | 0.00213 | 0.00278 | 0.01662 | 0.00000 | 0.01315 | 0.03236 | 0.00000 | 0.10140 | 0.01903 | 0.69989 | 0.00000 | 0.00333 |
| **HRCT1** | 0.01550 | 0.01689 | 0.01716 | 0.01853 | 0.01517 | 0.01244 | 0.01939 | 0.01356 | 0.02009 | 0.01329 | 0.01498 | 0.02064 |
| **HRG** | 0.65250 | 0.00000 | 0.00307 | 0.19438 | 0.28394 | 0.33917 | 0.00000 | 0.13270 | 0.00000 | 0.01190 | 0.00000 | 0.08924 |
| **HRH1** | 0.87610 | 0.00000 | 0.88178 | 0.00000 | 0.63459 | 0.87621 | 0.00000 | 2.04294 | 2.16057 | 0.00703 | 0.00000 | 0.90811 |
| **HRH2** | 5.80922 | 0.00000 | 0.09840 | 0.00000 | 0.00000 | 0.00000 | 0.11610 | 0.91360 | 0.09892 | 0.37353 | 0.13589 | 0.05207 |
| **HRH3** | 0.00677 | 0.05067 | 0.00864 | 0.00340 | 0.04034 | 0.00000 | 0.00453 | 0.00922 | 0.01895 | 0.21508 | 0.00815 | 0.02096 |
| **HRH4** | 0.00490 | 0.03872 | 0.03505 | 0.01233 | 0.00824 | 0.01358 | 0.01540 | 0.10400 | 0.02087 | 0.04624 | 0.01719 | 0.01197 |
| **HRK** | 0.00565 | 0.04508 | 0.02336 | 0.24997 | 0.19988 | 0.05826 | 0.01160 | 0.01018 | 0.00000 | 0.09507 | 0.00000 | 0.15647 |
| **HRNR** | 0.00185 | 0.44773 | 0.30173 | 0.00000 | 0.12049 | 0.00000 | 0.00000 | 0.04135 | 0.24876 | 0.18007 | 0.16788 | 0.73518 |
| **HRSP12** | 0.07400 | 6.76611 | 8.02877 | 5.12706 | 1.41344 | 15.03471 | 13.01492 | 0.32348 | 4.94875 | 5.48442 | 15.45877 | 2.31103 |
| **HS1BP3** | 0.60460 | 3.75790 | 1.23796 | 0.78134 | 1.88228 | 0.00000 | 1.67758 | 0.00000 | 0.00075 | 0.00000 | 2.49407 | 1.53675 |
| **HS1BP3-IT1** | 0.00323 | 0.01418 | 0.01415 | 0.01030 | 0.01147 | 0.02327 | 0.03669 | 0.14339 | 0.01262 | 0.01801 | 0.01362 | 0.01000 |
| **HS2ST1** | 0.65606 | 2.48994 | 1.54964 | 1.36074 | 0.00008 | 6.31562 | 1.26046 | 0.94490 | 3.44624 | 9.07663 | 0.95028 | 0.82362 |
| **HS3ST1** | 0.07663 | 0.50495 | 0.05042 | 0.99286 | 0.00000 | 0.00000 | 0.06873 | 0.04582 | 0.00002 | 0.40110 | 0.12261 | 0.06315 |
| **HS3ST2** | 0.05786 | 0.00538 | 0.00321 | 0.12541 | 0.00000 | 0.07705 | 0.00000 | 0.00598 | 0.01032 | 0.02688 | 0.01077 | 0.06753 |
| **HS3ST3A1** | 0.01985 | 0.00000 | 0.13339 | 0.67730 | 0.13263 | 0.07237 | 0.33657 | 0.00554 | 0.73556 | 0.61998 | 0.12038 | 0.00000 |
| **HS3ST3B1** | 0.03632 | 0.03958 | 1.41265 | 0.04066 | 0.00001 | 0.00799 | 0.00000 | 0.03593 | 0.31897 | 0.01613 | 0.00046 | 0.05510 |
| **HS3ST4** | 0.00561 | 0.08377 | 0.14471 | 0.37104 | 0.00000 | 0.05524 | 0.00000 | 0.01011 | 0.00000 | 0.04076 | 0.04077 | 0.00470 |
| **HS3ST5** | 0.16375 | 0.75260 | 0.00000 | 0.00741 | 0.46688 | 0.15285 | 0.00000 | 0.01767 | 0.00000 | 0.00246 | 0.10332 | 0.00000 |
| **HS3ST6** | 0.61244 | 0.04306 | 0.01258 | 0.02184 | 0.00798 | 0.00012 | 0.01461 | 0.09170 | 0.01985 | 0.05106 | 0.01228 | 0.05573 |
| **HS6ST1** | 0.00839 | 5.41873 | 0.00000 | 6.00797 | 9.56457 | 0.00000 | 0.59785 | 0.00000 | 0.00000 | 5.76375 | 1.47952 | 2.12974 |
| **HS6ST2** | 0.00873 | 0.05344 | 3.24625 | 0.00000 | 0.00000 | 0.01505 | 0.17718 | 0.00780 | 9.79978 | 0.01933 | 0.98127 | 0.01177 |
| **HS6ST3** | 0.10188 | 0.06803 | 0.03364 | 0.01576 | 0.06690 | 0.00465 | 0.05280 | 0.00768 | 0.00179 | 0.00897 | 0.00971 | 0.08779 |
| **HSBP1** | 3.72704 | 16.62330 | 21.14983 | 13.52771 | 17.66600 | 35.37803 | 32.74807 | 12.13956 | 23.61192 | 18.01050 | 23.28834 | 15.69822 |
| **HSBP1L1** | 0.00000 | 0.47681 | 0.89995 | 0.15107 | 0.00000 | 0.00000 | 1.42324 | 0.29073 | 0.00000 | 1.68388 | 2.99133 | 0.86103 |
| **HSCB** | 0.21725 | 3.76333 | 5.08416 | 2.90388 | 1.72685 | 4.63404 | 5.61449 | 1.59327 | 3.52307 | 3.68417 | 3.14626 | 3.21185 |
| **HSD11B1L** | 0.39740 | 0.13337 | 0.78064 | 0.28197 | 0.00000 | 0.00000 | 0.18894 | 0.37004 | 0.81571 | 0.94986 | 0.52438 | 0.13491 |
| **HSD11B2** | 0.01173 | 0.84986 | 0.97327 | 0.29668 | 0.38496 | 0.00000 | 0.21593 | 0.00774 | 0.94658 | 0.00000 | 0.00639 | 0.04901 |
| **HSD17B1** | 0.00070 | 0.46651 | 4.95996 | 1.43959 | 8.49455 | 0.00000 | 9.28864 | 0.00000 | 12.57236 | 9.12152 | 3.93313 | 0.35536 |
| **HSD17B10** | 1.64592 | 16.64624 | 16.44249 | 17.17852 | 20.23712 | 10.50454 | 25.53968 | 3.62643 | 20.96632 | 0.00000 | 19.24854 | 12.77754 |
| **HSD17B11** | 0.15541 | 1.09388 | 11.91834 | 3.71088 | 0.00000 | 4.37894 | 0.44758 | 0.00000 | 23.64525 | 1.90841 | 0.00000 | 2.52453 |
| **HSD17B12** | 1.05458 | 0.00000 | 12.90461 | 6.47670 | 10.67221 | 11.11700 | 0.00700 | 2.18435 | 29.04980 | 5.11827 | 8.72205 | 0.46031 |
| **HSD17B13** | 0.01228 | 0.01431 | 0.01879 | 0.01266 | 0.00812 | 0.01902 | 0.05150 | 0.01739 | 0.01899 | 0.03038 | 0.01164 | 0.01417 |
| **HSD17B14** | 0.00212 | 0.07130 | 0.14793 | 1.31206 | 0.13628 | 0.00000 | 0.19024 | 0.66299 | 0.17577 | 0.07152 | 0.00584 | 0.30869 |
| **HSD17B2** | 0.00577 | 0.01059 | 0.12383 | 0.45104 | 0.01149 | 0.00000 | 0.01432 | 0.00988 | 0.00407 | 0.01272 | 0.01999 | 0.03198 |
| **HSD17B3** | 0.01328 | 1.10152 | 0.06536 | 0.24919 | 0.00000 | 0.02312 | 0.03615 | 0.14292 | 0.00000 | 0.12551 | 0.00684 | 0.08069 |
| **HSD17B4** | 1.76916 | 0.77888 | 8.06262 | 8.00298 | 0.00000 | 11.05797 | 11.72179 | 0.37375 | 15.13657 | 0.00000 | 13.00673 | 0.00000 |
| **HSD17B6** | 0.09082 | 0.91712 | 0.67403 | 0.31413 | 0.00000 | 0.52363 | 0.63072 | 0.11527 | 0.06381 | 0.65184 | 0.38958 | 0.71894 |

| | | | | | | | | | | | | |
|---|---|---|---|---|---|---|---|---|---|---|---|---|
| **HSD17B7** | 0.18121 | 0.00000 | 2.76424 | 2.18928 | 6.82349 | 0.33523 | 1.82842 | 0.05474 | 4.42422 | 0.74952 | 2.97757 | 0.01651 |
| **HSD17B7P2** | 0.03246 | 0.00643 | 0.25285 | 0.02848 | 0.12278 | 0.16929 | 0.00206 | 0.03173 | 0.47277 | 0.01122 | 0.15747 | 0.00820 |
| **HSD17B8** | 0.22005 | 1.80833 | 0.57790 | 0.33225 | 0.69153 | 0.41599 | 0.00000 | 0.00000 | 0.00000 | 0.24609 | 1.06052 | 0.12959 |
| **HSD3B1** | 0.00731 | 0.00005 | 8.84840 | 0.00000 | 0.00000 | 0.05582 | 5.96446 | 0.00785 | 2.69882 | 0.00000 | 0.00000 | 0.01235 |
| **HSD3B2** | 0.00690 | 0.01536 | 0.01584 | 0.01738 | 0.01951 | 0.02055 | 0.02609 | 0.01031 | 0.01616 | 0.00573 | 0.01582 | 0.01659 |
| **HSD3B7** | 0.09796 | 0.00000 | 0.14776 | 0.17917 | 0.00000 | 0.12934 | 0.12689 | 0.00000 | 0.58353 | 0.06611 | 0.00000 | 0.55534 |
| **HSD3BP4** | 0.00466 | 0.01107 | 0.12040 | 0.00818 | 0.00000 | 0.00212 | 0.00000 | 0.01029 | 0.01437 | 0.13236 | 0.13367 | 0.01298 |
| **HSDL1** | 5.30199 | 3.08269 | 1.16569 | 1.97316 | 0.00000 | 0.05011 | 1.02977 | 0.77239 | 1.02976 | 0.00000 | 0.94937 | 3.46623 |
| **HSDL2** | 0.51566 | 4.65018 | 4.63372 | 3.57565 | 0.77360 | 9.01769 | 9.38869 | 0.00000 | 8.78144 | 2.60603 | 4.19435 | 0.00000 |
| **HSF1** | 0.56012 | 8.14154 | 15.61789 | 7.38608 | 8.91400 | 0.00000 | 5.53006 | 2.53075 | 12.97889 | 2.34687 | 7.81276 | 5.05716 |
| **HSF2** | 0.06100 | 2.64803 | 3.02603 | 0.00000 | 0.00000 | 7.64733 | 2.07805 | 1.48528 | 8.60914 | 0.53758 | 3.04017 | 2.73687 |
| **HSF2BP** | 13.39653 | 1.10609 | 0.46904 | 0.18147 | 0.99016 | 1.56982 | 0.00000 | 4.14931 | 0.95641 | 0.51980 | 0.02827 | 1.99517 |
| **HSF4** | 0.10506 | 0.12073 | 0.12535 | 0.00837 | 0.09831 | 0.00000 | 0.02404 | 0.02793 | 0.04384 | 0.22950 | 0.00000 | 0.01793 |
| **HSF5** | 0.73328 | 0.11539 | 0.06246 | 0.13723 | 0.00000 | 0.00049 | 0.01126 | 0.95033 | 0.02188 | 0.00318 | 0.05003 | 0.60120 |
| **HSH2D** | 0.13746 | 0.62032 | 1.44990 | 1.22902 | 2.71373 | 0.00037 | 2.86083 | 0.09405 | 0.00000 | 0.77645 | 0.73177 | 0.77484 |
| **HSP90AA1** | 14.88389 | 30.08193 | 49.48326 | 37.03059 | 50.59159 | 71.15496 | 78.63820 | 20.08247 | 45.53705 | 67.79153 | 49.14039 | 24.58485 |
| **HSP90AB1** | 21.37172 | 49.14216 | 49.93953 | 37.27631 | 45.95729 | 61.52209 | 74.20933 | 11.76706 | 59.18956 | 36.22013 | 51.65073 | 31.11206 |
| **HSP90AB4P** | 0.02515 | 0.00000 | 0.82488 | 0.23948 | 0.00000 | 0.27031 | 0.03041 | 0.39229 | 0.00000 | 2.73704 | 0.85844 | 0.00695 |
| **HSP90B1** | 2.72741 | 6.72261 | 23.40651 | 21.74504 | 17.14822 | 49.76932 | 5.36199 | 3.28759 | 50.04788 | 29.77252 | 22.74211 | 0.70560 |
| **HSP90B2P** | 0.24354 | 0.28480 | 5.10380 | 5.02699 | 3.00875 | 9.55669 | 0.00000 | 0.52850 | 11.50042 | 5.61604 | 4.92676 | 0.00000 |
| **HSPA12A** | 1.01926 | 0.14593 | 0.40262 | 0.43270 | 0.22447 | 0.07004 | 0.01201 | 0.00617 | 0.00000 | 0.00000 | 0.01610 | 0.01791 |
| **HSPA12B** | 0.80785 | 0.00657 | 0.02139 | 0.25474 | 0.14911 | 0.00000 | 0.02717 | 0.07960 | 0.13475 | 0.32903 | 0.00000 | 0.21457 |
| **HSPA13** | 0.75481 | 4.24215 | 9.59060 | 2.15933 | 0.00000 | 9.83913 | 0.00000 | 2.96123 | 13.69366 | 2.01595 | 1.18150 | 4.13047 |
| **HSPA14** | 1.91262 | 19.19566 | 13.96591 | 9.45623 | 10.45224 | 33.31544 | 18.41258 | 17.64239 | 20.96608 | 9.26067 | 14.99564 | 17.57395 |
| **HSPA1A** | 2.29453 | 1.29067 | 5.25169 | 1.25505 | 2.18472 | 0.00000 | 1.95795 | 2.25996 | 1.70002 | 0.34862 | 0.15274 | 0.26273 |
| **HSPA1B** | 2.18863 | 1.31579 | 16.62299 | 1.99752 | 2.21020 | 0.11971 | 10.33202 | 1.36091 | 4.47768 | 2.59301 | 1.88591 | 0.55917 |
| **HSPA1L** | 0.91868 | 0.47830 | 1.97432 | 0.00000 | 0.00000 | 0.76903 | 0.00002 | 0.43179 | 0.00000 | 0.63112 | 0.05424 | 0.00000 |
| **HSPA2** | 1.40674 | 0.38121 | 9.81132 | 1.25859 | 0.00000 | 9.74082 | 5.84747 | 12.85808 | 1.60201 | 0.00000 | 2.36658 | 4.85221 |
| **HSPA4** | 0.81842 | 24.43955 | 24.57637 | 14.09541 | 15.34281 | 41.16496 | 34.26808 | 5.89323 | 25.82450 | 9.27911 | 28.33336 | 17.71101 |
| **HSPA4L** | 4.30021 | 3.31912 | 0.00000 | 0.00000 | 0.73832 | 8.86493 | 0.00000 | 1.96552 | 0.00000 | 3.02072 | 2.95368 | 0.86785 |
| **HSPA5** | 4.70909 | 21.68089 | 37.21003 | 21.68475 | 26.03772 | 45.18885 | 4.26669 | 44.93994 | 62.72752 | 15.75396 | 29.01436 | 28.55382 |
| **HSPA6** | 0.00179 | 0.00631 | 1.45850 | 0.00913 | 0.00166 | 0.12014 | 0.00000 | 0.10130 | 0.00000 | 0.01776 | 0.00882 | 0.06742 |
| **HSPA7** | 0.02475 | 0.02268 | 0.02223 | 0.01432 | 0.03347 | 0.00000 | 0.02786 | 0.02213 | 0.02012 | 0.00000 | 0.01468 | 0.04230 |
| **HSPA8** | 23.94154 | 61.03732 | 50.74862 | 40.26184 | 55.01117 | 76.40450 | 59.70622 | 32.59651 | 70.43448 | 20.76457 | 60.12990 | 58.84671 |
| **HSPA9** | 4.66766 | 32.43567 | 25.13139 | 24.07891 | 23.80349 | 53.44482 | 42.82742 | 10.85394 | 43.49226 | 22.75370 | 33.13768 | 32.09058 |
| **HSPB1** | 12.83259 | 5.77638 | 6.52035 | 7.57604 | 1.81837 | 0.75108 | 0.93473 | 16.80873 | 0.00000 | 1.12560 | 0.00000 | 11.87466 |
| **HSPB11** | 2.49768 | 2.37998 | 1.00377 | 0.00000 | 9.19967 | 7.89653 | 7.04975 | 6.06590 | 0.00000 | 0.99970 | 19.39951 | 8.04385 |
| **HSPB2** | 0.00559 | 0.01783 | 0.03630 | 0.01518 | 0.00991 | 0.03569 | 0.01344 | 0.00971 | 0.02083 | 0.01556 | 0.01434 | 0.03812 |
| **HSPB2-C11orf5** | 0.00936 | 0.01864 | 0.02199 | 0.01360 | 0.01677 | 0.01585 | 0.02632 | 0.01011 | 0.01775 | 0.01575 | 0.01841 | 0.01567 |
| **HSPB3** | 0.01610 | 0.10502 | 0.05775 | 0.25337 | 0.16290 | 0.00000 | 0.01383 | 0.03601 | 0.00000 | 0.01789 | 0.72693 | 0.09373 |
| **HSPB6** | 0.49281 | 0.01132 | 0.04355 | 0.01638 | 0.01088 | 0.01291 | 0.02013 | 0.17728 | 0.02029 | 0.00507 | 0.03393 | 0.02924 |
| **HSPB7** | 0.00590 | 0.00369 | 0.13542 | 0.00867 | 0.08202 | 0.07168 | 0.13092 | 0.00976 | 0.00000 | 0.00040 | 0.13880 | 0.01343 |
| **HSPB8** | 0.00395 | 0.00000 | 25.95082 | 2.27655 | 15.89280 | 0.03343 | 4.72204 | 0.00651 | 0.64337 | 0.00000 | 0.00000 | 0.03326 |
| **HSPB9** | 0.23540 | 0.07588 | 0.37668 | 0.15424 | 0.03771 | 0.00000 | 0.20095 | 0.01693 | 0.17902 | 0.07415 | 0.09599 | 0.10912 |
| **HSPBAP1** | 0.94828 | 1.86501 | 5.96276 | 2.68895 | 3.08166 | 3.85233 | 4.07309 | 0.72134 | 3.06683 | 1.55399 | 0.68662 | 2.75010 |
| **HSPBP1** | 1.01644 | 16.65178 | 19.01778 | 7.94551 | 11.09921 | 0.00000 | 14.84286 | 6.31540 | 20.48036 | 1.66831 | 12.85826 | 15.58347 |
| **HSPD1** | 1.86384 | 31.37188 | 28.51741 | 21.67041 | 30.83247 | 50.90395 | 45.38894 | 0.24424 | 40.88266 | 23.82566 | 28.85966 | 5.05659 |
| **HSPE1** | 0.58144 | 11.41078 | 20.58799 | 10.68891 | 14.83747 | 36.78446 | 36.65845 | 0.29680 | 14.95815 | 22.85361 | 14.68043 | 1.14886 |

| | | | | | | | | | | | | |
|---|---|---|---|---|---|---|---|---|---|---|---|---|
| **HSPG2** | 0.01779 | 0.09795 | 0.73540 | 0.62014 | 0.01452 | 0.00000 | 0.00000 | 0.00727 | 2.11544 | 0.00000 | 0.00624 | 0.07301 |
| **HSPH1** | 1.07030 | 6.23732 | 18.30262 | 4.14457 | 7.92446 | 14.36960 | 26.67962 | 8.90091 | 10.45599 | 26.37322 | 4.25582 | 2.12630 |
| **HTATIP2** | 0.99209 | 0.00000 | 22.18758 | 1.51634 | 6.50383 | 9.30858 | 20.12564 | 0.00000 | 18.99871 | 0.00000 | 15.12929 | 0.00000 |
| **HTATSF1** | 6.06077 | 7.04308 | 10.15325 | 2.07848 | 1.98665 | 7.41118 | 7.02057 | 6.10042 | 6.58176 | 10.87420 | 7.72826 | 5.31058 |
| **HTATSF1P2** | 0.00109 | 0.00000 | 0.07628 | 0.31762 | 0.16476 | 0.52502 | 0.06316 | 0.03330 | 0.00000 | 1.12339 | 0.42168 | 0.01219 |
| **HTR1D** | 0.00336 | 0.20893 | 0.30347 | 0.27433 | 0.00000 | 0.00000 | 0.07153 | 0.01090 | 0.00000 | 0.20963 | 0.04907 | 0.01030 |
| **HTR1F** | 0.00571 | 0.04155 | 0.01065 | 0.06422 | 0.01154 | 0.00000 | 0.02017 | 0.01005 | 0.01901 | 0.02181 | 0.01584 | 0.03037 |
| **HTR2A** | 1.00279 | 0.01449 | 0.05275 | 0.01661 | 0.01747 | 0.00000 | 0.01946 | 0.12454 | 0.01098 | 0.00817 | 0.03590 | 0.00949 |
| **HTR2A-AS1** | 0.51733 | 0.01568 | 0.01338 | 0.01106 | 0.01382 | 0.01498 | 0.02049 | 0.07372 | 0.02605 | 0.00802 | 0.01274 | 0.00804 |
| **HTR2B** | 0.02271 | 0.06542 | 0.27194 | 0.00155 | 0.20339 | 0.19370 | 0.00000 | 0.38132 | 0.00593 | 1.36885 | 0.11755 | 0.08848 |
| **HTR2C** | 0.02563 | 0.44087 | 0.00921 | 0.09364 | 0.16048 | 0.00000 | 0.02809 | 0.01687 | 0.02462 | 0.00000 | 0.02703 | 0.05300 |
| **HTR3A** | 0.00609 | 0.01350 | 0.03550 | 0.02657 | 0.01311 | 0.02023 | 0.01656 | 0.01038 | 0.00687 | 0.02420 | 0.02306 | 0.01542 |
| **HTR3B** | 0.02470 | 0.01882 | 0.09699 | 0.09019 | 0.03272 | 0.00000 | 0.09752 | 0.04036 | 0.00000 | 0.02160 | 0.01775 | 0.02482 |
| **HTR3C** | 0.70064 | 0.13675 | 0.00642 | 0.01180 | 0.00000 | 0.00490 | 0.00996 | 0.02918 | 0.01998 | 0.09387 | 0.01369 | 0.12417 |
| **HTR3D** | 0.00674 | 0.05437 | 0.01228 | 0.01710 | 0.00860 | 0.00228 | 0.01454 | 0.04166 | 0.02374 | 0.04105 | 0.01233 | 0.63969 |
| **HTR3E** | 0.00203 | 0.00000 | 0.01844 | 0.00997 | 0.01509 | 0.02226 | 0.01821 | 0.05445 | 0.01601 | 0.00810 | 0.01540 | 0.58856 |
| **HTR5A** | 0.77362 | 0.02241 | 0.00759 | 0.15017 | 0.00000 | 0.01760 | 0.01908 | 0.00132 | 0.00574 | 0.03345 | 0.02782 | 0.03174 |
| **HTR5AOS** | 0.00666 | 0.13534 | 0.01067 | 0.07386 | 0.00874 | 0.00000 | 0.02027 | 0.01082 | 0.01635 | 0.01060 | 0.02688 | 0.01024 |
| **HTR6** | 0.00602 | 0.02110 | 0.02287 | 0.01393 | 0.00819 | 0.00000 | 0.02049 | 0.01143 | 0.00000 | 0.01770 | 0.11167 | 0.01706 |
| **HTR7** | 0.10221 | 0.05892 | 0.00768 | 0.07575 | 0.03678 | 0.03910 | 0.05201 | 0.00274 | 0.00487 | 0.00000 | 0.01373 | 0.03173 |
| **HTR7P1** | 0.02129 | 0.06755 | 0.03207 | 0.01161 | 0.01327 | 0.02684 | 0.01454 | 0.44545 | 0.04345 | 0.00188 | 0.05426 | 0.15837 |
| **HTRA1** | 0.00530 | 0.01459 | 0.01006 | 0.00271 | 0.05978 | 0.03628 | 0.00976 | 0.01026 | 0.01312 | 0.05174 | 0.02256 | 0.01190 |
| **HTRA2** | 1.17473 | 7.87934 | 7.81179 | 5.32660 | 11.46854 | 2.07654 | 4.23850 | 0.75163 | 14.74165 | 0.00000 | 7.32616 | 5.07778 |
| **HTRA3** | 0.00610 | 0.00996 | 0.06640 | 0.00108 | 0.01407 | 0.02916 | 0.01424 | 0.01007 | 0.04223 | 0.09737 | 0.01383 | 0.01114 |
| **HTRA4** | 0.00628 | 0.01558 | 1.24526 | 0.01198 | 0.00783 | 0.00000 | 0.01776 | 0.01294 | 0.05700 | 0.03217 | 0.04323 | 0.01665 |
| **HTT** | 3.53973 | 2.68695 | 2.79355 | 10.26877 | 11.68227 | 0.00000 | 3.22032 | 1.04630 | 4.78123 | 11.67579 | 0.94026 | 3.89750 |
| **HULC** | 0.00474 | 0.01018 | 0.01368 | 0.02573 | 0.01695 | 0.02793 | 0.01790 | 0.01378 | 0.01397 | 0.02196 | 0.01529 | 0.03680 |
| **HUNK** | 0.98016 | 3.64652 | 9.86522 | 0.00000 | 2.91442 | 0.60330 | 8.78239 | 0.00000 | 3.33776 | 3.79239 | 1.04713 | 0.38235 |
| **HUS1** | 0.00139 | 11.19470 | 7.34101 | 13.71636 | 3.83078 | 16.71762 | 7.91527 | 2.63439 | 11.28627 | 0.00000 | 5.23357 | 12.93745 |
| **HUS1B** | 0.00365 | 0.40484 | 0.26499 | 0.01162 | 0.00000 | 0.00000 | 0.40364 | 0.01416 | 0.00000 | 0.03386 | 0.21246 | 0.53571 |
| **HUWE1** | 2.73770 | 14.02807 | 10.90321 | 22.85624 | 34.25293 | 26.50164 | 26.10606 | 3.35622 | 18.00207 | 49.80485 | 20.71479 | 20.64803 |
| **HVCN1** | 1.23569 | 1.73621 | 0.22157 | 0.41737 | 0.00000 | 0.80096 | 0.00000 | 2.14525 | 0.18763 | 0.00000 | 0.00004 | 4.98080 |
| **HYAL1** | 0.09335 | 0.88371 | 0.67358 | 2.54773 | 0.00048 | 0.98938 | 0.00000 | 0.07611 | 0.26962 | 1.21163 | 0.06761 | 1.16016 |
| **HYAL2** | 0.68147 | 10.57647 | 5.59853 | 5.40031 | 5.46773 | 0.00000 | 2.00506 | 1.08736 | 9.97892 | 2.19362 | 4.12530 | 6.25814 |
| **HYAL3** | 0.14905 | 0.25748 | 1.48700 | 0.42733 | 1.03816 | 0.00000 | 1.13919 | 0.20587 | 1.26957 | 0.00000 | 0.62562 | 1.40977 |
| **HYAL4** | 0.00318 | 2.37682 | 0.64207 | 4.45008 | 0.05383 | 4.79118 | 0.00000 | 0.16958 | 0.56703 | 0.00000 | 2.67895 | 0.18049 |
| **HYALP1** | 0.06092 | 0.20818 | 0.03536 | 0.27401 | 0.03678 | 0.09518 | 0.00000 | 0.01180 | 0.00271 | 0.07947 | 0.03906 | 0.04222 |
| **HYDIN** | 0.44741 | 0.29052 | 0.03491 | 0.12329 | 0.00000 | 0.00000 | 0.00026 | 0.32461 | 0.00000 | 0.27646 | 0.05281 | 0.55561 |
| **HYKK** | 0.17056 | 1.34875 | 0.04403 | 0.94107 | 0.44542 | 0.56229 | 0.00000 | 0.16544 | 0.00000 | 0.00000 | 0.35742 | 0.48689 |
| **HYLS1** | 4.57603 | 15.14700 | 1.45812 | 3.05794 | 2.52431 | 12.96092 | 4.26492 | 15.57599 | 3.07703 | 0.00512 | 5.61965 | 31.89848 |
| **HYMAI** | 0.06646 | 0.01092 | 0.01570 | 0.00548 | 0.00544 | 0.04797 | 0.00876 | 0.00492 | 0.00999 | 0.03602 | 0.01236 | 0.04791 |
| **HYOU1** | 3.14914 | 22.64769 | 14.03710 | 21.79001 | 20.41322 | 8.93276 | 2.59525 | 3.09026 | 25.48100 | 37.30326 | 3.73163 | 16.82780 |
| **HYPK** | 0.02775 | 0.06071 | 0.05214 | 0.01983 | 0.02031 | 0.04087 | 0.01498 | 0.02795 | 0.04221 | 0.01210 | 0.01689 | 0.06407 |
| **HYPM** | 0.01754 | 0.01039 | 0.23673 | 0.02470 | 0.01307 | 0.00000 | 0.05462 | 0.22054 | 0.03088 | 0.03013 | 0.02053 | 0.12854 |
| **IAH1** | 8.31819 | 0.48834 | 1.00374 | 3.33891 | 2.53753 | 2.74310 | 0.49759 | 1.37341 | 1.38008 | 0.00000 | 6.67349 | 0.37022 |
| **IAPP** | 0.01903 | 0.00871 | 0.01008 | 0.01929 | 0.02729 | 0.04717 | 0.01069 | 0.01083 | 0.01805 | 0.00772 | 0.01688 | 0.03761 |
| **IARS** | 0.47936 | 14.07318 | 16.25798 | 13.01977 | 4.85790 | 35.49617 | 21.17702 | 3.21632 | 29.12850 | 9.73840 | 27.98451 | 12.64666 |
| **IARS2** | 3.05356 | 15.91376 | 13.93440 | 6.20913 | 5.84767 | 27.49153 | 6.02083 | 1.28231 | 26.13926 | 12.79016 | 15.07790 | 8.48606 |

| | | | | | | | | | | | | |
|---|---|---|---|---|---|---|---|---|---|---|---|---|
| **IBA57** | 0.00052 | 2.24970 | 1.51521 | 1.16961 | 4.04375 | 0.00000 | 3.29763 | 0.24606 | 0.81456 | 2.87865 | 2.96890 | 2.01370 |
| **IBA57-AS1** | 0.06884 | 0.00792 | 0.00952 | 0.00000 | 0.26266 | 0.00000 | 0.00079 | 0.00625 | 0.04148 | 0.13060 | 0.00274 | 0.01828 |
| **IBSP** | 0.00615 | 0.01394 | 0.01395 | 0.00886 | 0.02142 | 0.03197 | 0.01591 | 0.01011 | 0.01418 | 0.01424 | 0.01379 | 0.01820 |
| **IBTK** | 0.00000 | 5.55223 | 0.00000 | 2.13079 | 7.92923 | 18.87783 | 0.81570 | 0.13133 | 6.14986 | 11.09478 | 12.17676 | 4.00134 |
| **ICA1** | 2.20016 | 0.80589 | 0.45812 | 0.12537 | 1.89453 | 0.78461 | 0.54560 | 0.57738 | 0.05012 | 0.45217 | 0.26750 | 0.00000 |
| **ICA1L** | 1.97499 | 0.72998 | 0.92064 | 0.00000 | 1.29949 | 3.54127 | 2.11199 | 0.59112 | 2.07327 | 4.88686 | 2.61651 | 0.97026 |
| **ICAM1** | 1.59316 | 0.00638 | 0.93890 | 0.03084 | 0.00723 | 0.00000 | 0.00388 | 1.02847 | 0.15612 | 0.00000 | 0.04201 | 1.24057 |
| **ICAM2** | 0.01238 | 0.09928 | 0.03460 | 0.01101 | 0.00192 | 0.00647 | 0.02306 | 0.09655 | 0.02942 | 0.03553 | 0.01487 | 0.00597 |
| **ICAM3** | 0.93065 | 0.29812 | 2.20595 | 1.30325 | 0.00000 | 0.00000 | 3.57160 | 0.68884 | 2.11363 | 0.53754 | 2.49566 | 2.12744 |
| **ICAM4** | 0.00456 | 0.01277 | 0.09008 | 0.07642 | 0.00000 | 0.02181 | 0.08249 | 0.04124 | 0.05590 | 0.02427 | 0.00696 | 0.01542 |
| **ICAM5** | 0.43580 | 0.02295 | 0.01395 | 0.02286 | 0.03100 | 0.00000 | 0.08917 | 0.28760 | 0.00000 | 0.05845 | 0.00928 | 0.00598 |
| **ICE1** | 8.67488 | 1.38339 | 2.55748 | 2.56981 | 1.17564 | 9.88435 | 5.64623 | 2.61546 | 4.44334 | 21.92978 | 3.43534 | 4.62585 |
| **ICE2** | 4.24226 | 3.86098 | 4.30718 | 2.44097 | 2.48409 | 16.37874 | 3.57355 | 4.02800 | 2.96920 | 3.57598 | 4.41373 | 5.40697 |
| **ICK** | 2.03215 | 2.36374 | 3.69304 | 1.29594 | 5.07485 | 8.10845 | 8.52951 | 0.04558 | 4.94067 | 7.35977 | 3.73128 | 0.72331 |
| **ICMT** | 5.16752 | 25.39838 | 17.27380 | 13.12694 | 16.94941 | 24.31797 | 8.76493 | 1.82882 | 27.20196 | 8.34581 | 21.39886 | 18.30405 |
| **ICOSLG** | 0.00000 | 5.46727 | 0.40687 | 1.12950 | 1.34328 | 0.00000 | 0.00051 | 0.10143 | 1.24529 | 4.43408 | 0.00003 | 4.99198 |
| **ICT1** | 0.28452 | 10.53121 | 3.15507 | 9.22479 | 10.11614 | 19.56513 | 6.68292 | 2.38446 | 9.66251 | 3.07355 | 11.82137 | 15.39562 |
| **ID1** | 0.87418 | 0.00000 | 4.44896 | 28.69226 | 21.99845 | 7.36935 | 2.05269 | 2.18953 | 0.00000 | 0.00000 | 0.00000 | 16.38518 |
| **ID2** | 2.70015 | 0.00000 | 16.66516 | 7.76232 | 9.90724 | 4.04375 | 3.85883 | 5.89674 | 0.35177 | 0.03185 | 0.00000 | 0.63753 |
| **ID3** | 0.84931 | 0.00000 | 15.29396 | 30.62132 | 20.69568 | 0.82134 | 7.87714 | 1.06213 | 1.57411 | 0.00000 | 0.00000 | 0.57149 |
| **ID4** | 1.28384 | 0.34420 | 2.46793 | 2.08621 | 1.83273 | 0.67531 | 0.00000 | 2.61908 | 0.00000 | 0.64113 | 0.00000 | 0.71544 |
| **IDE** | 5.65109 | 6.40583 | 7.77644 | 4.30206 | 4.25795 | 19.21587 | 13.41269 | 4.09911 | 10.19011 | 5.45198 | 11.44926 | 5.46937 |
| **IDH1** | 5.15473 | 27.22271 | 5.31044 | 34.07128 | 22.89812 | 53.28879 | 26.77064 | 0.78494 | 25.88990 | 11.66893 | 38.22199 | 7.25440 |
| **IDH1-AS1** | 0.00727 | 0.18971 | 0.04281 | 0.08175 | 0.09444 | 0.22696 | 0.02680 | 0.03551 | 0.00000 | 0.00000 | 0.26863 | 0.11171 |
| **IDH2** | 7.47849 | 1.50434 | 8.29004 | 1.25788 | 0.00000 | 2.02003 | 3.10086 | 6.32384 | 4.43739 | 0.00000 | 10.58861 | 0.10888 |
| **IDH3A** | 1.70260 | 16.52429 | 15.25303 | 4.77849 | 13.26013 | 23.93158 | 12.56831 | 2.15124 | 10.41106 | 0.44492 | 20.40844 | 6.93920 |
| **IDH3B** | 0.60547 | 25.40720 | 21.42483 | 19.80636 | 23.71827 | 15.59016 | 24.57003 | 4.38018 | 27.80268 | 1.99437 | 20.92128 | 19.55687 |
| **IDH3G** | 2.98528 | 4.67048 | 7.22458 | 10.14494 | 10.00106 | 4.54528 | 6.76332 | 2.34686 | 10.64463 | 0.00000 | 14.00999 | 4.63961 |
| **IDI1** | 1.30316 | 2.81741 | 13.24369 | 13.41907 | 29.13181 | 37.16667 | 29.92186 | 11.31958 | 24.02958 | 3.40853 | 27.36649 | 7.50300 |
| **IDI2** | 0.00175 | 0.00000 | 0.82491 | 0.00000 | 0.54186 | 0.00000 | 1.41851 | 0.20657 | 0.15936 | 3.19441 | 0.40628 | 0.33967 |
| **IDI2-AS1** | 0.00841 | 0.00000 | 0.39854 | 0.26226 | 0.24258 | 2.15821 | 0.00001 | 0.43152 | 1.12206 | 0.00000 | 0.81547 | 0.20871 |
| **IDNK** | 0.19304 | 0.04807 | 0.20209 | 0.58946 | 0.31037 | 0.00000 | 0.12212 | 0.08760 | 0.10287 | 1.28889 | 0.08171 | 0.27184 |
| **IDO1** | 0.00246 | 0.01238 | 0.01520 | 0.07647 | 0.00689 | 0.00712 | 0.01542 | 0.18406 | 0.01192 | 0.02111 | 0.01912 | 1.14932 |
| **IDO2** | 0.01383 | 0.00586 | 0.01320 | 0.01241 | 0.02677 | 0.02145 | 0.00847 | 0.00640 | 0.02260 | 0.03450 | 0.01373 | 0.09507 |
| **IDS** | 4.06759 | 0.79816 | 1.11688 | 0.58094 | 0.00000 | 0.69956 | 0.29681 | 0.63695 | 0.07011 | 0.00000 | 0.00000 | 1.67465 |
| **IDUA** | 0.02332 | 0.01391 | 0.01601 | 0.00830 | 0.01143 | 0.01932 | 0.01571 | 0.00845 | 0.03148 | 0.04087 | 0.01628 | 0.01402 |
| **IER2** | 5.67599 | 18.44404 | 21.90362 | 13.68571 | 19.94698 | 0.00000 | 9.75204 | 0.60847 | 16.74844 | 7.24205 | 10.93187 | 9.87547 |
| **IER3** | 7.22728 | 11.64181 | 7.48184 | 0.87987 | 0.00000 | 0.00000 | 1.98393 | 10.89692 | 2.05317 | 0.59099 | 0.00000 | 15.59414 |
| **IER3IP1** | 7.16901 | 9.21058 | 12.24937 | 4.33997 | 2.69068 | 25.74512 | 11.02260 | 9.58321 | 17.60628 | 2.42528 | 8.16543 | 17.32338 |
| **IER5** | 2.06319 | 0.34024 | 1.46713 | 3.46888 | 0.32630 | 0.00000 | 0.16211 | 5.77545 | 0.00000 | 3.71253 | 0.00000 | 9.65282 |
| **IER5L** | 0.18364 | 0.04933 | 0.46535 | 0.00000 | 0.16540 | 0.00000 | 0.03336 | 0.15741 | 0.00000 | 1.01524 | 0.00000 | 0.00000 |
| **IFFO1** | 0.24122 | 0.31379 | 0.32113 | 0.64390 | 0.00000 | 0.00000 | 0.00000 | 0.90045 | 0.27769 | 0.98962 | 0.24962 | 0.48083 |
| **IFFO2** | 0.30647 | 0.03618 | 0.74258 | 0.07662 | 0.28581 | 0.00000 | 0.05442 | 0.44341 | 0.00000 | 0.12935 | 0.07081 | 0.12890 |
| **IFI16** | 0.00188 | 0.57719 | 0.75733 | 21.54185 | 0.47073 | 0.71141 | 0.00000 | 0.03234 | 0.00000 | 0.00178 | 0.00046 | 0.24439 |
| **IFI27** | 0.00700 | 0.28468 | 0.09319 | 0.55819 | 0.29739 | 0.10925 | 0.00000 | 0.14630 | 0.19673 | 0.40603 | 0.00564 | 0.01950 |
| **IFI27L1** | 0.56414 | 0.62621 | 1.14668 | 1.26231 | 0.45094 | 0.28107 | 1.00892 | 1.52637 | 1.45179 | 2.31691 | 1.12693 | 0.92973 |
| **IFI27L2** | 0.07890 | 0.01142 | 0.05939 | 0.02706 | 0.03124 | 0.02916 | 0.00923 | 0.09961 | 0.04428 | 0.03150 | 0.00693 | 0.03051 |
| **IFI30** | 1.10730 | 32.23448 | 1.66276 | 0.01108 | 29.63801 | 5.81387 | 0.00000 | 4.70621 | 26.48681 | 3.35614 | 8.64201 | 24.72983 |

| | | | | | | | | | | | | |
|---|---|---|---|---|---|---|---|---|---|---|---|---|
| **IFI35** | 0.08910 | 0.00000 | 0.04722 | 0.28566 | 0.19604 | 0.38901 | 0.08128 | 0.36924 | 0.03384 | 0.00000 | 0.01977 | 0.07408 |
| **IFI44L** | 0.05903 | 0.01163 | 0.01835 | 0.01278 | 0.00835 | 0.02082 | 0.00971 | 0.02068 | 0.02181 | 0.03590 | 0.02156 | 0.02081 |
| **IFI6** | 0.02246 | 0.00000 | 4.84768 | 0.00000 | 0.00000 | 0.16925 | 0.61019 | 0.02415 | 0.81705 | 0.16830 | 0.34267 | 0.01740 |
| **IFIH1** | 0.01270 | 0.01055 | 0.01504 | 0.02022 | 0.00767 | 0.04829 | 0.01296 | 0.00912 | 0.01706 | 0.01833 | 0.01154 | 0.01975 |
| **IFIT1** | 0.00570 | 0.00855 | 0.00694 | 0.00337 | 0.00000 | 0.06755 | 0.06732 | 0.00991 | 0.02388 | 0.02277 | 0.00438 | 0.01355 |
| **IFIT3** | 0.00960 | 0.01788 | 0.03810 | 0.02044 | 0.02026 | 0.00046 | 0.02294 | 0.00981 | 0.02407 | 0.01413 | 0.01502 | 0.01741 |
| **IFIT5** | 0.00261 | 0.00726 | 0.10292 | 0.34183 | 0.00856 | 0.00000 | 0.16213 | 0.47520 | 0.00000 | 0.01770 | 0.01522 | 0.12032 |
| **IFITM1** | 0.00009 | 0.13524 | 19.30526 | 40.35955 | 1.21877 | 2.61320 | 2.93258 | 0.30224 | 14.37473 | 0.00000 | 5.09940 | 0.00000 |
| **IFITM10** | 0.00384 | 0.00000 | 0.42961 | 0.04287 | 0.00960 | 0.35625 | 0.00205 | 0.00586 | 0.09658 | 0.60628 | 0.08358 | 0.00000 |
| **IFITM2** | 0.00000 | 2.36867 | 3.32533 | 7.13481 | 14.49294 | 0.69278 | 20.54075 | 0.00000 | 2.26411 | 3.33980 | 28.38743 | 0.00000 |
| **IFITM3** | 0.00412 | 0.45396 | 19.06033 | 30.62750 | 28.53530 | 0.15220 | 6.93845 | 0.00200 | 4.93076 | 0.01485 | 15.46247 | 0.00556 |
| **IFITM4P** | 0.00594 | 0.01334 | 0.01207 | 0.02146 | 0.01681 | 0.02110 | 0.00966 | 0.01060 | 0.03513 | 0.02547 | 0.01908 | 0.01573 |
| **IFITM5** | 0.00613 | 0.00908 | 0.00529 | 0.05256 | 0.03826 | 0.05643 | 0.00626 | 0.01032 | 0.03694 | 0.00355 | 0.00874 | 0.01485 |
| **IFNA22P** | 0.06672 | 0.01464 | 0.01459 | 0.01229 | 0.01451 | 0.01727 | 0.01598 | 0.00700 | 0.02082 | 0.01597 | 0.01438 | 0.01493 |
| **IFNAR1** | 0.37314 | 3.28280 | 0.45443 | 2.12879 | 2.93316 | 6.35230 | 0.00000 | 0.23117 | 8.26755 | 1.57802 | 5.57941 | 3.39589 |
| **IFNAR2** | 3.25591 | 0.88722 | 0.00000 | 0.20185 | 3.19630 | 3.13878 | 0.24695 | 2.18430 | 2.31256 | 1.63657 | 3.50197 | 0.48240 |
| **IFNE** | 0.01912 | 0.01362 | 0.01464 | 0.01055 | 0.00969 | 0.02158 | 0.02265 | 0.03882 | 0.01479 | 0.01731 | 0.03554 | 0.01350 |
| **IFNGR1** | 0.00363 | 6.23693 | 14.19646 | 3.44134 | 0.92992 | 12.51417 | 0.00000 | 3.18174 | 14.22566 | 0.00000 | 5.34536 | 7.69032 |
| **IFNGR2** | 2.00188 | 1.46751 | 1.25727 | 0.87652 | 0.28492 | 0.00000 | 0.00000 | 1.02465 | 1.64811 | 0.00000 | 0.58771 | 0.56272 |
| **IFNK** | 0.00582 | 0.01196 | 0.01335 | 0.00664 | 0.05448 | 0.02822 | 0.01365 | 0.01041 | 0.01347 | 0.02850 | 0.01288 | 0.01472 |
| **IFNL1** | 0.00606 | 0.01016 | 0.04013 | 0.00656 | 0.02476 | 0.02105 | 0.05352 | 0.00834 | 0.02591 | 0.01194 | 0.03874 | 0.05563 |
| **IFNL4** | 0.00641 | 0.01472 | 0.01865 | 0.01892 | 0.01700 | 0.01604 | 0.01681 | 0.01045 | 0.03999 | 0.00694 | 0.02370 | 0.01695 |
| **IFNLR1** | 0.00021 | 1.08494 | 0.18261 | 0.24935 | 0.00000 | 0.19250 | 0.00000 | 0.06361 | 0.00000 | 0.46175 | 0.03383 | 1.37225 |
| **IFNW1** | 0.07433 | 0.01056 | 0.01888 | 0.00933 | 0.01128 | 0.02702 | 0.01409 | 0.00644 | 0.01400 | 0.02267 | 0.01442 | 0.03748 |
| **IFRD1** | 2.42575 | 16.32443 | 29.40832 | 6.23457 | 9.41700 | 14.29887 | 10.98507 | 12.11635 | 9.92489 | 0.00000 | 6.65541 | 9.36920 |
| **IFRD2** | 0.17614 | 19.03725 | 14.00128 | 13.01008 | 8.50938 | 3.76458 | 15.61153 | 3.33661 | 27.18556 | 0.00000 | 21.01682 | 16.92160 |
| **IFT122** | 0.82229 | 2.62192 | 1.04172 | 2.31332 | 1.63123 | 1.59750 | 0.85932 | 0.87602 | 0.60207 | 1.00218 | 1.78116 | 2.57810 |
| **IFT140** | 2.02261 | 1.01033 | 0.62694 | 2.09764 | 0.49991 | 0.00000 | 0.00000 | 0.20297 | 1.37609 | 1.14461 | 0.20680 | 0.74142 |
| **IFT172** | 1.59858 | 1.98004 | 0.87484 | 7.92365 | 1.61144 | 0.57513 | 0.43230 | 1.32341 | 1.32243 | 0.47554 | 0.58453 | 5.04846 |
| **IFT20** | 0.59510 | 4.13279 | 4.42078 | 2.09500 | 2.78245 | 4.85033 | 4.97692 | 6.60438 | 3.56710 | 0.00000 | 2.48261 | 7.82089 |
| **IFT22** | 5.47530 | 14.31807 | 0.49364 | 3.69731 | 1.19203 | 9.51802 | 1.52831 | 7.07008 | 2.25886 | 0.00000 | 3.17365 | 16.99298 |
| **IFT27** | 1.21570 | 0.36515 | 1.44013 | 0.20025 | 0.48687 | 0.65321 | 0.46183 | 0.25177 | 0.00000 | 0.44745 | 0.96808 | 0.54703 |
| **IFT43** | 0.11955 | 2.61613 | 1.19978 | 4.57832 | 1.95947 | 1.08084 | 0.40151 | 0.00000 | 1.12578 | 0.27780 | 1.07948 | 0.37597 |
| **IFT46** | 2.99015 | 5.32750 | 0.00000 | 1.27322 | 5.48258 | 0.85281 | 0.00000 | 2.76429 | 0.00000 | 0.66586 | 1.13985 | 8.30740 |
| **IFT52** | 2.65037 | 4.40864 | 0.68127 | 3.45559 | 0.43279 | 10.36544 | 3.35269 | 1.48176 | 1.13760 | 0.00000 | 2.74755 | 3.92818 |
| **IFT57** | 2.41469 | 0.00000 | 2.74747 | 2.51748 | 0.00000 | 3.26817 | 2.22728 | 1.44831 | 1.46342 | 0.00000 | 2.88301 | 0.94477 |
| **IFT74** | 0.95456 | 0.00000 | 2.00966 | 0.81029 | 0.00000 | 1.94635 | 2.64294 | 0.44626 | 1.03835 | 0.84945 | 0.83509 | 0.25245 |
| **IFT74-AS1** | 0.00494 | 0.01685 | 0.02815 | 0.01226 | 0.01701 | 0.01228 | 0.01801 | 0.01623 | 0.03121 | 0.00439 | 0.01959 | 0.01724 |
| **IFT80** | 0.00000 | 0.23926 | 1.25332 | 0.38145 | 0.00000 | 5.27484 | 0.00000 | 2.55459 | 0.00000 | 1.67903 | 0.87727 | 1.30345 |
| **IFT81** | 0.29679 | 1.24491 | 0.47335 | 0.94764 | 0.00000 | 0.36199 | 0.23097 | 0.62806 | 0.43487 | 1.22973 | 0.02373 | 1.95834 |
| **IFT88** | 0.89015 | 1.12961 | 1.98150 | 0.48455 | 1.67814 | 1.39661 | 1.09988 | 0.95953 | 0.01225 | 0.00062 | 0.61717 | 2.33651 |
| **IGBP1** | 2.91711 | 7.42593 | 7.33253 | 7.33984 | 5.55134 | 12.17212 | 13.73059 | 3.16899 | 4.84197 | 4.03702 | 7.06542 | 9.09520 |
| **IGBP1P1** | 0.17083 | 0.16453 | 0.26782 | 0.16104 | 0.08308 | 0.82263 | 0.29790 | 0.14829 | 0.14877 | 0.40734 | 0.27867 | 0.29174 |
| **IGDCC3** | 3.14477 | 0.00000 | 1.03917 | 0.00000 | 0.23554 | 0.00000 | 0.37504 | 0.23658 | 2.80763 | 1.88903 | 0.07831 | 0.06029 |
| **IGDCC4** | 0.18323 | 0.09302 | 0.00697 | 0.00000 | 0.00000 | 0.00283 | 0.06157 | 0.02374 | 0.15843 | 0.08088 | 0.06464 | 0.00983 |
| **IGF1** | 0.02275 | 0.00000 | 1.07021 | 4.82922 | 0.07988 | 0.00000 | 0.00000 | 0.06108 | 0.23935 | 1.18312 | 0.00000 | 0.09965 |
| **IGF1R** | 9.00025 | 3.89890 | 2.00518 | 0.74009 | 2.24255 | 5.61878 | 0.00000 | 6.12262 | 0.00000 | 5.95442 | 0.00000 | 3.87133 |
| **IGF2.uid-15072** | 0.03586 | 0.01653 | 0.07932 | 0.01568 | 0.02175 | 0.00353 | 0.01179 | 0.01263 | 0.01151 | 0.01552 | 0.01718 | 0.02967 |

| | | | | | | | | | | | | |
|---|---|---|---|---|---|---|---|---|---|---|---|---|
| **IGF2.uid-15081** | 0.01601 | 0.01423 | 0.04566 | 0.01066 | 0.01512 | 0.01916 | 0.01128 | 0.01349 | 0.01471 | 0.01814 | 0.01553 | 0.01799 |
| **IGF2BP1** | 0.00000 | 23.72940 | 24.99411 | 19.85859 | 31.67525 | 14.57449 | 38.06554 | 9.63942 | 32.26496 | 18.18208 | 26.15495 | 30.33181 |
| **IGF2BP2** | 1.01388 | 0.56330 | 2.71093 | 2.33842 | 6.69266 | 1.25313 | 5.67409 | 1.94312 | 2.68734 | 12.48999 | 0.75090 | 3.48001 |
| **IGF2BP3** | 5.32230 | 0.00000 | 12.39044 | 3.80908 | 3.60057 | 5.49987 | 12.19117 | 0.25632 | 5.13192 | 7.87418 | 6.76611 | 1.30337 |
| **IGF2R** | 1.96186 | 2.19911 | 4.34367 | 1.98562 | 4.49353 | 0.00000 | 0.00000 | 1.01496 | 5.13408 | 15.89580 | 1.28270 | 2.54105 |
| **IGFALS** | 0.55949 | 0.01429 | 0.03061 | 0.01419 | 0.13602 | 0.00056 | 0.04864 | 0.01168 | 0.00000 | 0.00000 | 0.01432 | 0.01235 |
| **IGFBP1** | 0.89821 | 0.03243 | 0.01325 | 0.02145 | 0.00961 | 0.02024 | 0.01921 | 0.31160 | 0.02102 | 0.00000 | 0.01307 | 0.01557 |
| **IGFBP2** | 1.41366 | 0.49548 | 2.52874 | 5.81276 | 0.73268 | 0.00000 | 0.57633 | 1.13781 | 0.46819 | 1.81916 | 0.03021 | 0.31906 |
| **IGFBP3** | 7.73311 | 0.08840 | 1.43176 | 0.00000 | 0.00000 | 0.00000 | 0.00000 | 1.40874 | 0.40203 | 0.24312 | 0.00000 | 0.06322 |
| **IGFBP4** | 0.00471 | 0.00732 | 0.21903 | 0.07652 | 0.00000 | 0.01783 | 0.00000 | 0.01207 | 0.02628 | 0.14778 | 0.00969 | 0.01112 |
| **IGFBP5** | 0.16643 | 0.04672 | 0.02246 | 0.01685 | 0.01979 | 0.00947 | 0.01660 | 0.06793 | 0.02324 | 0.01661 | 0.01274 | 0.05912 |
| **IGFBP6** | 1.48576 | 0.38538 | 0.24253 | 0.88604 | 0.00000 | 0.00000 | 0.00000 | 2.01042 | 0.85283 | 0.10875 | 0.42295 | 0.09484 |
| **IGFBP7** | 0.02731 | 0.00502 | 0.00000 | 0.00298 | 0.00236 | 0.01811 | 0.00000 | 0.00864 | 0.09061 | 0.15681 | 0.04643 | 0.04092 |
| **IGFBP7-AS1** | 0.00206 | 0.14110 | 0.19863 | 0.49493 | 0.00000 | 0.02801 | 0.00000 | 0.06114 | 0.00532 | 0.03038 | 0.01513 | 0.16270 |
| **IGFBPL1** | 1.97098 | 0.55755 | 0.21046 | 0.57532 | 0.00000 | 0.17761 | 0.00000 | 1.92847 | 0.03462 | 0.14878 | 0.00000 | 0.33417 |
| **IGFL2** | 0.06950 | 0.01445 | 0.04894 | 0.02048 | 0.02341 | 0.00000 | 0.01480 | 0.00244 | 0.02959 | 0.00476 | 0.03113 | 0.08564 |
| **IGFL3** | 0.00588 | 0.01280 | 0.01349 | 0.01190 | 0.01454 | 0.01820 | 0.01744 | 0.00916 | 0.01824 | 0.01848 | 0.01695 | 0.04885 |
| **IGFL4** | 0.01502 | 0.06519 | 0.01104 | 0.00345 | 0.00000 | 0.03386 | 0.01247 | 0.00650 | 0.01188 | 0.04193 | 0.01270 | 0.04138 |
| **IGFLR1** | 0.15249 | 0.48751 | 0.67979 | 2.61624 | 0.27518 | 0.48179 | 0.00000 | 0.04000 | 0.51827 | 0.00000 | 1.33503 | 0.19020 |
| **IGFN1** | 0.01759 | 0.01653 | 0.07395 | 0.03201 | 0.01816 | 0.00000 | 0.01847 | 0.01215 | 0.00000 | 0.01888 | 0.05706 | 0.02038 |
| **IGHMBP2** | 1.61850 | 4.61759 | 4.62250 | 4.68000 | 4.78004 | 0.00000 | 1.67678 | 2.25537 | 7.57586 | 6.74770 | 1.40749 | 6.76029 |
| **IGIP** | 0.02673 | 0.00000 | 0.10471 | 0.01650 | 0.00000 | 0.30060 | 0.29292 | 0.01916 | 0.01504 | 0.46109 | 0.22468 | 0.03410 |
| **IGJ** | 0.00742 | 0.01306 | 0.01334 | 0.01115 | 0.01391 | 0.02347 | 0.01923 | 0.16851 | 0.01747 | 0.01129 | 0.01509 | 0.01707 |
| **IGLL1** | 0.04974 | 0.01963 | 0.04880 | 0.00734 | 0.01137 | 0.03113 | 0.05835 | 0.00943 | 0.10114 | 0.00321 | 0.00000 | 0.01998 |
| **IGLL3P** | 1.30143 | 0.07990 | 0.02023 | 0.13084 | 0.16283 | 0.00000 | 0.02342 | 0.17941 | 0.09683 | 0.23757 | 0.02939 | 0.25917 |
| **IGLL5** | 0.01062 | 0.01608 | 0.03471 | 0.01249 | 0.01191 | 0.02425 | 0.01221 | 0.00919 | 0.01002 | 0.02403 | 0.01668 | 0.01573 |
| **IGLON5** | 0.00742 | 0.01414 | 0.01622 | 0.01176 | 0.01789 | 0.01732 | 0.02131 | 0.01424 | 0.01369 | 0.01959 | 0.02775 | 0.02112 |
| **IGSF1** | 0.02985 | 0.01781 | 0.09155 | 2.92886 | 0.00000 | 0.05110 | 0.25063 | 0.01382 | 0.00000 | 0.53136 | 0.00000 | 0.16489 |
| **IGSF10** | 5.08232 | 0.16066 | 0.11039 | 0.00000 | 0.47494 | 0.23899 | 0.00000 | 2.74280 | 0.21400 | 0.19153 | 0.11300 | 0.13785 |
| **IGSF11** | 1.78305 | 0.04300 | 0.06545 | 0.00093 | 0.07392 | 0.00000 | 0.01717 | 0.00817 | 0.05204 | 0.06363 | 0.27801 | 0.17631 |
| **IGSF21** | 0.00619 | 0.01785 | 0.01440 | 0.01205 | 0.01475 | 0.01984 | 0.02085 | 0.01042 | 0.01574 | 0.01448 | 0.01481 | 0.01700 |
| **IGSF22** | 0.02494 | 0.02858 | 0.13716 | 0.19580 | 0.00000 | 0.00000 | 0.00178 | 0.01995 | 0.02197 | 0.03865 | 0.06643 | 0.17559 |
| **IGSF23** | 0.00594 | 0.01026 | 0.00803 | 0.09301 | 0.00719 | 0.02850 | 0.01064 | 0.00725 | 0.01620 | 0.02524 | 0.01659 | 0.09678 |
| **IGSF3** | 0.07093 | 0.26254 | 3.68886 | 7.90998 | 6.59567 | 2.50665 | 0.68712 | 0.34128 | 9.70978 | 6.11398 | 0.29927 | 0.00000 |
| **IGSF5** | 0.49605 | 0.00755 | 0.02742 | 0.01365 | 0.00716 | 0.02103 | 0.02105 | 0.05356 | 0.00716 | 0.04410 | 0.01473 | 0.05338 |
| **IGSF6** | 0.01976 | 0.00724 | 0.01303 | 0.06077 | 0.00000 | 0.04776 | 0.02359 | 0.02041 | 0.02301 | 0.07403 | 0.00922 | 0.02739 |
| **IGSF8** | 0.00128 | 0.76345 | 3.46145 | 1.15752 | 0.00675 | 0.00000 | 0.00000 | 0.44113 | 3.14744 | 0.00000 | 0.41987 | 0.58455 |
| **IGSF9** | 3.04797 | 1.43260 | 0.00000 | 0.51086 | 1.88712 | 0.00000 | 0.34329 | 0.32526 | 3.78887 | 1.35594 | 0.09616 | 1.97769 |
| **IGSF9B** | 0.19353 | 0.06599 | 0.00322 | 0.11480 | 0.24338 | 0.00000 | 0.00109 | 0.00173 | 0.04388 | 0.34577 | 0.01195 | 0.05405 |
| **IHH** | 0.00739 | 0.01458 | 0.23714 | 0.06089 | 0.10317 | 0.00000 | 0.00177 | 0.01015 | 0.01005 | 0.01580 | 0.01985 | 0.01678 |
| **IK** | 3.19604 | 11.69450 | 7.08570 | 7.59665 | 9.94073 | 21.15222 | 16.36695 | 7.52505 | 8.36649 | 8.96413 | 11.94363 | 8.97709 |
| **IKBIP** | 0.67591 | 0.94585 | 0.43034 | 0.53734 | 0.16771 | 0.69830 | 0.00000 | 0.31807 | 0.34187 | 2.17483 | 0.00000 | 0.35626 |
| **IKBKAP** | 0.93075 | 10.06525 | 9.11552 | 3.97064 | 4.16403 | 10.22142 | 14.37982 | 0.00000 | 6.87477 | 4.73240 | 9.18599 | 5.17315 |
| **IKBKB** | 1.36504 | 0.85785 | 3.53855 | 2.01504 | 2.55399 | 0.00000 | 2.26939 | 1.17560 | 4.70196 | 5.01631 | 0.97055 | 1.72700 |
| **IKBKE** | 0.12236 | 0.16700 | 0.01721 | 0.02748 | 0.22299 | 0.00000 | 0.04473 | 0.01558 | 0.00000 | 0.00000 | 0.00962 | 0.04121 |
| **IKBKG** | 0.02406 | 0.51995 | 0.55048 | 1.01544 | 0.49826 | 0.00000 | 0.03487 | 0.27980 | 0.78796 | 0.00000 | 0.59592 | 0.77627 |
| **IKZF1** | 0.00471 | 0.00120 | 0.01304 | 0.00394 | 0.22085 | 0.00000 | 0.03368 | 0.01142 | 0.00000 | 0.32984 | 0.00000 | 0.01587 |
| **IKZF2** | 0.04499 | 0.03994 | 0.02219 | 0.20003 | 0.24273 | 0.00000 | 0.00522 | 0.01873 | 0.00000 | 0.15426 | 0.01626 | 0.01081 |

| | | | | | | | | | | | | |
|---|---|---|---|---|---|---|---|---|---|---|---|---|
| **IKZF3** | 0.00185 | 0.02908 | 0.16952 | 0.11431 | 0.00000 | 0.09319 | 0.00000 | 0.11207 | 0.00000 | 0.04118 | 0.02984 | 0.25898 |
| **IKZF4** | 0.50942 | 0.13976 | 0.33328 | 0.97216 | 0.30646 | 1.93597 | 0.22927 | 0.00000 | 0.46280 | 3.84063 | 0.14755 | 0.97111 |
| **IKZF5** | 0.80180 | 3.67316 | 8.17068 | 0.83127 | 1.05861 | 13.21758 | 4.90497 | 11.45754 | 5.68727 | 4.67809 | 4.83417 | 4.60442 |
| **IL10** | 0.00606 | 0.01316 | 0.01715 | 0.01192 | 0.01291 | 0.02816 | 0.01589 | 0.00997 | 0.01550 | 0.01423 | 0.01469 | 0.02322 |
| **IL10RA** | 0.00646 | 0.01209 | 0.00530 | 0.01043 | 0.00613 | 0.05883 | 0.01009 | 0.00994 | 0.10349 | 0.00000 | 0.01272 | 0.01427 |
| **IL10RB** | 0.56588 | 8.53173 | 1.55895 | 0.12099 | 2.61935 | 3.54818 | 0.00000 | 0.12095 | 5.98965 | 0.00000 | 4.05342 | 3.09121 |
| **IL10RB-AS1** | 0.16978 | 0.03881 | 0.09170 | 0.01125 | 0.08933 | 0.00170 | 0.04610 | 0.02091 | 0.02849 | 0.00000 | 0.03392 | 0.08117 |
| **IL11** | 0.11375 | 0.02026 | 0.01923 | 0.02194 | 0.03539 | 0.00009 | 0.02640 | 0.12493 | 0.02035 | 0.00853 | 0.01559 | 0.02823 |
| **IL11RA** | 0.00109 | 0.67338 | 4.00562 | 1.58782 | 0.21838 | 0.00000 | 0.00000 | 0.00447 | 2.84244 | 1.01247 | 1.39136 | 0.44484 |
| **IL12A** | 0.00583 | 0.06472 | 0.21008 | 0.00127 | 0.05328 | 0.10699 | 0.00000 | 0.00941 | 0.09529 | 0.00000 | 0.01078 | 0.02055 |
| **IL12A-AS1** | 0.01216 | 0.01308 | 0.03244 | 0.00356 | 0.00365 | 0.07273 | 0.00000 | 0.00852 | 0.03794 | 0.04031 | 0.02220 | 0.01327 |
| **IL12B** | 0.04585 | 0.00772 | 0.00447 | 0.03413 | 0.00000 | 0.02138 | 0.00212 | 0.04566 | 0.02031 | 0.12438 | 0.00851 | 0.01390 |
| **IL12RB1** | 0.03158 | 0.01352 | 0.04808 | 0.02013 | 0.00889 | 0.00791 | 0.01199 | 0.04111 | 0.00000 | 0.01784 | 0.14109 | 0.09146 |
| **IL12RB2** | 0.00251 | 0.00000 | 0.02255 | 0.04602 | 0.09212 | 0.00000 | 0.08156 | 0.00793 | 0.00000 | 0.78247 | 0.01071 | 0.11051 |
| **IL13RA1** | 3.15859 | 0.39246 | 0.43033 | 0.73736 | 0.00000 | 0.26231 | 0.00000 | 1.09561 | 0.05402 | 0.04969 | 0.07160 | 0.65344 |
| **IL13RA2** | 0.00177 | 0.12233 | 0.00192 | 0.10825 | 0.00000 | 0.03338 | 0.00836 | 0.21429 | 0.00922 | 0.02161 | 0.00963 | 3.04094 |
| **IL15** | 0.00514 | 0.08467 | 0.32689 | 0.94367 | 0.00000 | 0.42591 | 0.10740 | 0.00630 | 0.02064 | 0.00000 | 0.00125 | 0.00356 |
| **IL15RA** | 0.12530 | 0.01888 | 0.02139 | 0.02686 | 0.07056 | 0.01118 | 0.02211 | 0.00908 | 0.00266 | 0.04465 | 0.01014 | 0.01398 |
| **IL16** | 2.49369 | 0.19918 | 0.02264 | 0.00000 | 0.00000 | 0.01878 | 0.00000 | 0.07280 | 0.21208 | 0.08393 | 0.00001 | 0.35467 |
| **IL17B** | 4.83283 | 2.02349 | 0.00000 | 0.00000 | 0.02783 | 0.00000 | 0.02787 | 11.40171 | 0.00000 | 0.02519 | 0.00000 | 6.70702 |
| **IL17C** | 1.21775 | 0.00777 | 0.02150 | 0.01071 | 0.01927 | 0.00950 | 0.01365 | 0.04132 | 0.03207 | 0.00000 | 0.01626 | 0.12657 |
| **IL17D** | 0.00367 | 0.08411 | 0.52795 | 0.00000 | 0.00000 | 0.30225 | 0.25720 | 0.03003 | 0.27460 | 0.40247 | 0.01337 | 0.09262 |
| **IL17F** | 0.05449 | 0.01656 | 0.00717 | 0.00595 | 0.01255 | 0.01598 | 0.01994 | 0.00843 | 0.02992 | 0.07766 | 0.01228 | 0.01191 |
| **IL17RA** | 1.02345 | 1.31118 | 1.07560 | 0.43245 | 1.82753 | 0.00000 | 0.84292 | 0.30334 | 0.00000 | 5.33415 | 0.08732 | 0.76831 |
| **IL17RB** | 0.01356 | 0.27440 | 0.86709 | 3.17764 | 1.58314 | 1.07620 | 0.91114 | 0.00000 | 0.70539 | 0.44538 | 1.07755 | 0.40318 |
| **IL17RC** | 0.10229 | 0.10926 | 0.01872 | 0.53625 | 0.14944 | 0.00000 | 0.00850 | 0.00000 | 0.21012 | 0.00352 | 0.00924 | 0.81415 |
| **IL17RD** | 2.31202 | 0.78285 | 1.33335 | 1.05127 | 0.01550 | 0.24927 | 0.00000 | 0.47522 | 0.06816 | 0.01969 | 0.00002 | 0.52857 |
| **IL17RE** | 0.00304 | 0.00000 | 0.37943 | 0.19703 | 0.00000 | 0.12976 | 0.07020 | 0.02723 | 0.95362 | 0.02007 | 1.76034 | 0.01332 |
| **IL17REL** | 0.03098 | 0.01531 | 0.04779 | 0.01238 | 0.02123 | 0.00000 | 0.06328 | 0.00898 | 0.00000 | 0.04478 | 0.02357 | 0.02059 |
| **IL18** | 0.00736 | 0.01368 | 0.00000 | 0.09065 | 0.04525 | 0.04221 | 0.02198 | 0.05269 | 0.17392 | 0.04692 | 0.02795 | 0.04393 |
| **IL18BP** | 0.16359 | 0.33348 | 0.43924 | 0.46302 | 0.49516 | 0.00000 | 0.00000 | 0.12945 | 2.64635 | 1.98047 | 0.00039 | 0.57637 |
| **IL18R1** | 0.00598 | 0.01039 | 0.03147 | 0.00878 | 0.00000 | 0.01944 | 0.01303 | 0.01023 | 0.00000 | 0.02851 | 0.14862 | 0.01758 |
| **IL19** | 0.00499 | 0.01171 | 0.01062 | 0.16798 | 0.04082 | 0.00008 | 0.02889 | 0.01425 | 0.01013 | 0.02038 | 0.01495 | 0.01756 |
| **IL1A** | 0.00130 | 0.00000 | 0.04040 | 0.00133 | 0.00000 | 0.05678 | 0.02737 | 0.14264 | 0.00000 | 0.80881 | 0.10164 | 1.23200 |
| **IL1B** | 0.32614 | 0.01108 | 0.00000 | 0.08368 | 0.00031 | 0.04235 | 0.00567 | 0.00056 | 0.07135 | 0.03955 | 0.00620 | 0.02399 |
| **IL1F10** | 0.00639 | 0.01608 | 0.09354 | 0.01104 | 0.01604 | 0.00000 | 0.00822 | 0.01057 | 0.00018 | 0.01793 | 0.09590 | 0.01741 |
| **IL1R1** | 0.00108 | 0.06862 | 6.85293 | 0.00000 | 0.10273 | 0.22544 | 0.00011 | 0.01419 | 0.21271 | 0.21229 | 0.00000 | 0.05000 |
| **IL1R2** | 0.00665 | 0.01716 | 0.45436 | 0.01538 | 0.02293 | 0.00000 | 0.00000 | 0.01044 | 0.00000 | 0.00522 | 0.06299 | 0.01781 |
| **IL1RAP** | 1.44176 | 0.93518 | 1.11069 | 1.78989 | 0.00000 | 0.41971 | 0.00000 | 0.53148 | 0.00000 | 0.23255 | 0.00000 | 1.29788 |
| **IL1RAPL1** | 2.11317 | 0.05600 | 0.03608 | 0.00248 | 0.03860 | 0.00844 | 0.01263 | 0.45671 | 0.00000 | 0.00000 | 0.10224 | 0.00000 |
| **IL1RAPL2** | 0.63726 | 0.00925 | 0.01585 | 0.00494 | 0.06317 | 0.01130 | 0.01316 | 0.19961 | 0.01517 | 0.04369 | 0.01549 | 0.01094 |
| **IL1RL1** | 0.02542 | 0.01785 | 0.00947 | 0.01796 | 0.02362 | 0.00002 | 0.02055 | 0.05607 | 0.03866 | 0.06002 | 0.00966 | 0.01108 |
| **IL1RL2** | 0.00501 | 0.00887 | 0.05730 | 0.00577 | 0.14560 | 0.00000 | 0.01375 | 0.01053 | 0.00000 | 0.08510 | 0.00771 | 0.01437 |
| **IL1RN** | 0.13469 | 0.01225 | 0.04459 | 0.02218 | 0.12835 | 0.00000 | 0.04037 | 0.76758 | 0.00000 | 0.00000 | 0.02996 | 0.08385 |
| **IL20RA** | 0.26157 | 0.01684 | 0.01404 | 0.01201 | 0.01829 | 0.00000 | 0.07193 | 0.01639 | 0.00000 | 0.01503 | 0.07421 | 0.01277 |
| **IL20RB** | 0.00065 | 0.54813 | 1.36033 | 0.68190 | 0.12829 | 0.15613 | 0.17156 | 1.21173 | 1.04635 | 0.35432 | 0.40189 | 0.41076 |
| **IL21-AS1** | 0.02379 | 0.00005 | 0.01946 | 0.05611 | 0.00000 | 0.01592 | 0.00000 | 0.00393 | 0.00913 | 0.24492 | 0.00735 | 0.08444 |
| **IL21R** | 0.00520 | 0.00887 | 0.02722 | 0.01080 | 0.00747 | 0.00061 | 0.04034 | 0.01196 | 0.16548 | 0.05243 | 0.02851 | 0.07271 |

| | | | | | | | | | | | | |
|---|---|---|---|---|---|---|---|---|---|---|---|---|
| **IL21R-AS1** | 0.00324 | 0.00329 | 0.00688 | 0.00402 | 0.00131 | 0.01545 | 0.00007 | 0.00852 | 0.02450 | 0.40444 | 0.00473 | 0.02594 |
| **IL22RA1** | 0.00533 | 0.19528 | 0.13133 | 0.09152 | 0.00000 | 0.00000 | 0.00000 | 0.00278 | 0.01726 | 0.17749 | 0.01417 | 0.47208 |
| **IL22RA2** | 0.02515 | 0.01951 | 0.02888 | 0.01314 | 0.02257 | 0.00887 | 0.01788 | 0.02346 | 0.02150 | 0.00760 | 0.01422 | 0.01548 |
| **IL23A** | 0.51303 | 0.11847 | 0.14614 | 1.98266 | 0.00000 | 0.05561 | 0.09172 | 0.05860 | 0.04460 | 0.00000 | 0.05552 | 0.31101 |
| **IL23R** | 4.23972 | 0.00000 | 0.01620 | 0.00380 | 0.02663 | 0.00000 | 0.02052 | 0.28705 | 0.01916 | 0.06745 | 0.02164 | 0.97696 |
| **IL27** | 0.07901 | 0.03657 | 0.03712 | 0.05835 | 0.00751 | 0.06788 | 0.04245 | 0.00549 | 0.01563 | 0.01044 | 0.02379 | 0.03848 |
| **IL27RA** | 0.01153 | 0.00164 | 0.00694 | 0.00244 | 0.00000 | 0.06604 | 0.00000 | 0.22352 | 0.27288 | 0.02233 | 0.00000 | 0.10161 |
| **IL2RA** | 1.35212 | 0.01125 | 0.01446 | 0.01228 | 0.01859 | 0.00026 | 0.01812 | 0.19196 | 0.02369 | 0.00769 | 0.01393 | 0.06589 |
| **IL2RB** | 0.05279 | 0.00437 | 3.72225 | 0.00139 | 0.33380 | 0.00000 | 0.00000 | 0.03782 | 1.05209 | 0.01623 | 0.35770 | 0.01240 |
| **IL2RG** | 0.00126 | 0.69646 | 0.00000 | 0.00000 | 0.27689 | 0.11387 | 0.00000 | 0.10311 | 0.00011 | 0.28255 | 0.00000 | 0.14153 |
| **IL3** | 0.00600 | 0.02874 | 0.01765 | 0.01233 | 0.01997 | 0.01175 | 0.01962 | 0.01051 | 0.01901 | 0.00656 | 0.01873 | 0.02643 |
| **IL31RA** | 0.29686 | 0.19500 | 0.00842 | 0.00923 | 0.00000 | 0.00000 | 0.02759 | 0.27758 | 0.02400 | 0.08732 | 0.01980 | 0.07573 |
| **IL32** | 0.00221 | 0.00000 | 1.77715 | 2.72141 | 0.23552 | 0.00000 | 3.12276 | 0.02222 | 0.75558 | 0.07864 | 3.82988 | 0.00078 |
| **IL34** | 0.56039 | 4.19495 | 0.00000 | 2.15712 | 3.86909 | 0.00000 | 0.84516 | 0.05817 | 3.03623 | 3.61218 | 9.57053 | 4.73989 |
| **IL36B** | 0.00656 | 0.07790 | 0.09012 | 0.07998 | 0.00860 | 0.02596 | 0.03884 | 0.01046 | 0.03050 | 0.00000 | 0.01549 | 0.01126 |
| **IL36G** | 0.00580 | 0.01018 | 0.05493 | 0.08982 | 0.01076 | 0.02374 | 0.00942 | 0.01110 | 0.00000 | 0.00838 | 0.07585 | 0.02299 |
| **IL36RN** | 0.00582 | 0.00659 | 3.19521 | 0.00191 | 0.01078 | 0.00994 | 1.01895 | 0.00977 | 0.00000 | 0.00000 | 0.30248 | 0.01514 |
| **IL37** | 0.00629 | 0.09346 | 0.01525 | 0.04130 | 0.01073 | 0.02184 | 0.01290 | 0.01037 | 0.03116 | 0.01108 | 0.01419 | 0.01313 |
| **IL4** | 0.72521 | 0.01782 | 0.01693 | 0.01245 | 0.02521 | 0.00271 | 0.02174 | 0.00222 | 0.02645 | 0.00216 | 0.01376 | 0.01636 |
| **IL4I1** | 0.12131 | 0.23344 | 0.01920 | 0.03983 | 0.00756 | 0.00000 | 0.11991 | 0.12240 | 0.12742 | 0.00000 | 0.11792 | 0.33554 |
| **IL4R** | 0.21409 | 0.70647 | 0.22531 | 2.02936 | 0.18278 | 0.00000 | 0.25908 | 0.00000 | 0.07363 | 0.83255 | 0.19472 | 0.07507 |
| **IL5** | 0.02850 | 0.00207 | 0.00605 | 0.01604 | 0.03610 | 0.05279 | 0.01276 | 0.00338 | 0.00069 | 0.05890 | 0.01062 | 0.09445 |
| **IL5RA** | 0.47577 | 0.00453 | 0.00655 | 0.09935 | 0.00835 | 0.01337 | 0.02329 | 0.04624 | 0.00859 | 0.15777 | 0.00988 | 0.00660 |
| **IL6** | 0.02446 | 0.71381 | 11.03308 | 0.00000 | 1.53537 | 1.06901 | 0.45550 | 0.01534 | 0.67683 | 0.00041 | 1.91018 | 1.12986 |
| **IL6R** | 0.14566 | 0.85573 | 3.38942 | 39.34260 | 0.04299 | 0.00000 | 0.00000 | 2.60400 | 0.00000 | 0.98901 | 0.33177 | 3.80065 |
| **IL6ST** | 0.05081 | 2.06848 | 5.86264 | 14.04688 | 0.00000 | 18.71972 | 0.00000 | 0.93988 | 3.91893 | 16.20008 | 1.37218 | 1.21550 |
| **IL9** | 0.07188 | 0.03363 | 0.01389 | 0.00933 | 0.06334 | 0.00539 | 0.04188 | 0.00502 | 0.03742 | 0.00000 | 0.00616 | 0.01979 |
| **ILDR1** | 0.07382 | 0.04890 | 0.21139 | 0.00512 | 0.00088 | 0.11620 | 0.00000 | 0.00390 | 0.00000 | 0.02258 | 0.03754 | 0.05466 |
| **ILDR2** | 2.50538 | 0.05072 | 0.05724 | 0.12394 | 0.00000 | 0.00000 | 0.00000 | 0.08367 | 0.01569 | 0.33954 | 0.00005 | 0.00999 |
| **ILF2** | 9.32602 | 35.32183 | 29.78431 | 22.82445 | 21.46383 | 49.24342 | 44.43270 | 33.01068 | 41.92790 | 8.16784 | 33.77745 | 36.26070 |
| **ILF3** | 8.14621 | 23.89626 | 26.28378 | 22.99377 | 31.62457 | 13.49095 | 37.60962 | 13.49444 | 38.38199 | 41.80362 | 26.15968 | 37.21964 |
| **ILF3-AS1** | 0.50736 | 1.53819 | 1.95311 | 1.05123 | 1.00217 | 0.37020 | 1.09829 | 1.30207 | 0.39236 | 2.64282 | 1.30729 | 4.24073 |
| **ILK** | 1.04124 | 10.48680 | 10.40284 | 13.42078 | 11.91105 | 6.17187 | 9.95383 | 4.57206 | 7.17592 | 0.00000 | 15.93208 | 5.37917 |
| **ILKAP** | 5.15103 | 8.44830 | 4.95306 | 7.35172 | 7.33190 | 3.05205 | 8.83937 | 1.96703 | 12.07938 | 4.55538 | 9.67322 | 8.03291 |
| **ILVBL** | 0.03153 | 5.51155 | 6.51728 | 3.91687 | 5.23493 | 0.00000 | 1.82474 | 0.00000 | 14.47147 | 2.45305 | 11.99111 | 3.08338 |
| **IMMP1L** | 0.99374 | 2.57453 | 2.63617 | 0.57454 | 0.91953 | 5.51397 | 2.65426 | 1.11079 | 1.31692 | 0.00000 | 1.12781 | 3.71680 |
| **IMMP2L** | 0.64139 | 0.89851 | 0.92748 | 0.94606 | 0.36444 | 2.99279 | 1.38224 | 0.14543 | 0.45292 | 0.72727 | 1.34587 | 1.06535 |
| **IMMT** | 9.70607 | 20.60411 | 13.44144 | 15.18501 | 16.56916 | 26.51284 | 19.62478 | 10.68433 | 24.45329 | 12.51253 | 23.14425 | 18.22667 |
| **IMP3** | 3.75512 | 20.31205 | 23.69633 | 15.39509 | 24.91578 | 18.84244 | 27.89653 | 24.14201 | 32.42115 | 4.93221 | 22.00211 | 33.60594 |
| **IMP4** | 1.73177 | 25.80368 | 20.00444 | 18.58753 | 29.56319 | 5.03031 | 31.37896 | 22.60597 | 25.68198 | 6.34887 | 26.62367 | 30.62545 |
| **IMPA1** | 5.90714 | 3.81292 | 3.86169 | 1.15410 | 0.00000 | 7.80154 | 5.41990 | 3.12298 | 4.40735 | 2.26570 | 3.46458 | 5.33132 |
| **IMPA2** | 0.62628 | 3.38298 | 8.21532 | 1.97252 | 0.00000 | 5.93556 | 18.23898 | 0.03547 | 2.09309 | 10.26097 | 1.20192 | 0.95054 |
| **IMPACT** | 0.28401 | 6.95309 | 3.22520 | 0.00000 | 2.21969 | 15.24667 | 4.68577 | 1.55419 | 0.00642 | 5.39229 | 7.90173 | 3.90884 |
| **IMPAD1** | 0.00000 | 3.27097 | 1.80169 | 12.27322 | 3.13498 | 23.51401 | 4.41569 | 1.82627 | 12.39550 | 37.57591 | 1.31923 | 1.89754 |
| **IMPDH1** | 1.47667 | 40.08776 | 9.87936 | 5.87817 | 14.96447 | 0.00000 | 14.67110 | 11.14666 | 19.70860 | 0.00000 | 7.14395 | 33.31394 |
| **IMPDH2** | 6.08032 | 32.93475 | 35.31743 | 24.05856 | 37.08244 | 21.70529 | 54.72438 | 10.35400 | 55.91618 | 10.45672 | 40.60294 | 27.02044 |
| **IMPG1** | 0.55940 | 0.00675 | 0.01151 | 0.00872 | 0.00000 | 0.03599 | 0.00058 | 0.00186 | 0.16918 | 0.09530 | 0.05336 | 0.02296 |
| **IMPG2** | 0.06870 | 0.03569 | 0.04270 | 0.03713 | 0.00000 | 0.03017 | 0.03532 | 0.00593 | 0.06670 | 0.07216 | 0.04192 | 0.01345 |

| | | | | | | | | | | | | |
|---|---|---|---|---|---|---|---|---|---|---|---|---|
| **INA** | 0.61713 | 0.00000 | 4.80403 | 0.00000 | 0.53966 | 0.00000 | 15.00421 | 0.00317 | 12.68899 | 1.02783 | 10.12666 | 0.00000 |
| **INADL** | 1.06899 | 6.07804 | 1.28235 | 0.20535 | 4.01410 | 11.26614 | 0.49227 | 0.00000 | 9.23076 | 13.48863 | 5.66049 | 5.64394 |
| **INCA1** | 0.67577 | 0.01853 | 0.13134 | 0.40787 | 0.05150 | 0.23906 | 0.00000 | 0.05844 | 0.39581 | 0.01700 | 0.10245 | 0.16659 |
| **INCENP** | 5.31023 | 0.00000 | 1.91160 | 0.36417 | 4.44840 | 0.00000 | 1.34905 | 0.46612 | 4.11537 | 10.58074 | 0.00170 | 1.94956 |
| **INE1** | 0.00751 | 0.00000 | 0.03622 | 0.23688 | 0.00000 | 0.01410 | 0.00064 | 0.04719 | 0.19482 | 0.15311 | 0.42576 | 0.27933 |
| **INF2** | 1.04075 | 3.61876 | 9.72569 | 1.30241 | 13.05978 | 0.00000 | 4.62120 | 0.00000 | 13.75257 | 2.92170 | 2.82200 | 0.37890 |
| **ING1** | 1.15793 | 1.45119 | 3.93949 | 2.05133 | 2.73916 | 1.52407 | 8.24817 | 6.96031 | 1.22876 | 4.11161 | 1.43363 | 2.46363 |
| **ING2** | 0.61395 | 2.51663 | 3.15148 | 1.78383 | 2.32868 | 3.27922 | 3.88734 | 9.62201 | 0.00821 | 8.66600 | 1.91156 | 8.25813 |
| **ING3** | 11.67148 | 6.83120 | 4.56560 | 3.75563 | 1.92394 | 11.63872 | 6.12265 | 9.12359 | 3.12335 | 1.09788 | 8.32954 | 7.43801 |
| **ING4** | 0.20168 | 2.11094 | 1.43036 | 0.98370 | 0.00886 | 0.72383 | 0.16647 | 1.52204 | 2.67818 | 0.01287 | 0.54572 | 3.50613 |
| **ING5** | 0.54994 | 2.80158 | 4.72982 | 1.25364 | 3.45424 | 0.00000 | 3.48477 | 0.29403 | 2.53519 | 7.14006 | 2.60735 | 2.08901 |
| **INGX** | 0.01610 | 0.02437 | 0.01116 | 0.01538 | 0.01222 | 0.01869 | 0.01306 | 0.03609 | 0.01970 | 0.03684 | 0.01263 | 0.02835 |
| **INHA** | 0.00601 | 0.01046 | 0.16996 | 0.00333 | 0.11857 | 0.04153 | 0.13095 | 0.01025 | 0.00000 | 0.00000 | 0.00742 | 0.01492 |
| **INHBA** | 0.00988 | 0.00538 | 0.00000 | 0.03947 | 0.00000 | 0.11965 | 0.12097 | 0.00966 | 0.01364 | 0.00763 | 0.00546 | 0.01055 |
| **INHBA-AS1** | 0.00625 | 0.01501 | 0.02090 | 0.01361 | 0.01476 | 0.01766 | 0.02046 | 0.01049 | 0.01487 | 0.01464 | 0.01678 | 0.01630 |
| **INHBB** | 0.00296 | 0.01694 | 0.03659 | 0.05531 | 0.02345 | 0.01269 | 0.01366 | 0.14988 | 0.01670 | 0.01024 | 0.01445 | 0.01214 |
| **INHBC** | 0.00586 | 0.01902 | 0.01909 | 0.02006 | 0.00607 | 0.05519 | 0.01545 | 0.01027 | 0.01231 | 0.01368 | 0.02549 | 0.01789 |
| **INHBE** | 0.00772 | 0.02338 | 0.01468 | 0.01456 | 0.01286 | 0.00000 | 0.24677 | 0.01112 | 0.00000 | 0.03488 | 0.02337 | 0.02186 |
| **INIP** | 1.80715 | 2.94720 | 3.53657 | 1.55550 | 1.99890 | 8.77039 | 6.21313 | 3.06848 | 4.18462 | 0.14752 | 2.16587 | 5.60845 |
| **INMT** | 0.00310 | 0.05461 | 0.03152 | 0.00000 | 0.00000 | 0.07788 | 0.11833 | 0.01422 | 0.00000 | 0.10217 | 0.06599 | 0.01265 |
| **INO80** | 1.53925 | 3.51944 | 5.46912 | 3.60076 | 4.19176 | 2.78459 | 8.31126 | 7.05369 | 3.78716 | 17.67812 | 2.48426 | 5.07851 |
| **INO80B** | 0.04919 | 0.01323 | 0.08807 | 0.02251 | 0.29466 | 0.00000 | 0.37618 | 0.08861 | 0.00000 | 0.60019 | 0.00928 | 0.32322 |
| **INO80C** | 0.93634 | 4.57916 | 2.01553 | 1.70707 | 1.17933 | 8.52202 | 3.98470 | 13.22208 | 5.16795 | 0.00000 | 3.13519 | 16.73223 |
| **INO80D** | 3.23246 | 2.51888 | 3.51917 | 4.40946 | 0.00000 | 10.26496 | 2.70937 | 2.77607 | 3.14818 | 16.29799 | 3.96172 | 6.98241 |
| **INO80E** | 0.37127 | 0.86248 | 2.01284 | 1.50152 | 2.90324 | 0.00000 | 1.61470 | 0.00000 | 2.20415 | 1.72170 | 1.94464 | 1.30284 |
| **INPP1** | 0.86561 | 0.00000 | 0.00558 | 0.15846 | 0.00000 | 0.01950 | 0.09504 | 0.06060 | 0.00000 | 0.16338 | 0.01864 | 0.04395 |
| **INPP4A** | 1.03151 | 1.47615 | 2.34279 | 0.79285 | 1.06580 | 3.54751 | 1.05337 | 0.00000 | 1.41322 | 1.28571 | 0.49658 | 0.56163 |
| **INPP4B** | 0.71715 | 0.01516 | 0.00363 | 0.01694 | 0.00000 | 0.03002 | 0.02234 | 0.24411 | 0.04865 | 0.07901 | 0.00454 | 0.01667 |
| **INPP5A** | 0.67668 | 2.01228 | 0.52609 | 0.85737 | 4.42295 | 1.53306 | 2.77692 | 0.41159 | 0.00000 | 3.56264 | 0.44643 | 0.39912 |
| **INPP5B** | 0.80516 | 4.65944 | 1.91529 | 4.52431 | 1.01305 | 3.14649 | 0.91023 | 1.61917 | 2.69084 | 2.32174 | 2.79941 | 3.71720 |
| **INPP5D** | 0.86677 | 11.64115 | 0.53836 | 6.47540 | 8.73755 | 0.00000 | 0.00000 | 0.00000 | 0.00000 | 4.75671 | 9.11561 | 3.91139 |
| **INPP5E** | 1.68160 | 0.93030 | 0.19259 | 2.01876 | 2.28799 | 0.00000 | 0.39875 | 0.00000 | 1.90036 | 3.49896 | 0.00000 | 0.48921 |
| **INPP5F** | 10.49492 | 12.29828 | 5.41416 | 7.69683 | 8.40022 | 23.98227 | 5.27572 | 8.88890 | 4.71005 | 23.71238 | 0.47323 | 13.13951 |
| **INPP5J** | 0.00402 | 0.05206 | 0.00000 | 0.98628 | 0.32079 | 0.00000 | 0.05873 | 2.60297 | 0.78368 | 0.28219 | 0.07157 | 2.24124 |
| **INPP5K** | 3.19502 | 10.09590 | 6.21238 | 6.29267 | 5.46653 | 0.00000 | 8.90824 | 5.51718 | 6.21749 | 2.75004 | 5.56441 | 9.07259 |
| **INPPL1** | 0.01225 | 2.39638 | 2.12839 | 5.72070 | 6.25361 | 0.00000 | 7.39394 | 0.00000 | 5.29044 | 15.73546 | 1.31326 | 0.25980 |
| **INSC** | 1.06396 | 0.02900 | 0.05508 | 0.02514 | 0.00000 | 0.00000 | 0.10043 | 0.02182 | 0.10672 | 0.02717 | 0.12583 | 0.00751 |
| **INSIG1** | 2.40482 | 4.24152 | 8.19264 | 8.78080 | 22.25295 | 13.75730 | 0.03632 | 6.39845 | 25.73541 | 7.37804 | 10.54974 | 5.02215 |
| **INSIG2** | 0.66066 | 1.09875 | 3.40331 | 0.62063 | 0.00000 | 7.87719 | 0.00000 | 0.98033 | 3.61115 | 2.28887 | 1.74012 | 2.65149 |
| **INSL3** | 0.00245 | 0.01813 | 0.01347 | 0.01441 | 0.01754 | 0.00656 | 0.07525 | 0.04505 | 0.00458 | 0.00586 | 0.01611 | 0.02433 |
| **INSL4** | 0.00516 | 0.01212 | 0.05997 | 0.00000 | 0.00000 | 0.00000 | 0.12247 | 0.00834 | 2.56612 | 0.04906 | 0.25404 | 0.05100 |
| **INSL6** | 0.22425 | 0.11826 | 0.09792 | 0.00135 | 0.03855 | 0.11473 | 0.00000 | 0.00000 | 0.06711 | 0.00000 | 0.00446 | 0.66613 |
| **INSM1** | 0.24602 | 2.77225 | 0.00000 | 0.43279 | 0.09623 | 0.09188 | 0.00000 | 0.00000 | 0.00000 | 0.10878 | 0.00000 | 0.87725 |
| **INSM2** | 0.19637 | 0.01278 | 0.01831 | 0.00924 | 0.01853 | 0.02463 | 0.01713 | 0.00983 | 0.01615 | 0.01513 | 0.01368 | 0.01247 |
| **INSR** | 0.10407 | 5.27066 | 0.66995 | 9.04908 | 3.28189 | 7.19656 | 0.72801 | 6.58030 | 3.63857 | 7.00188 | 3.37644 | 4.24820 |
| **INSRR** | 0.00561 | 0.01643 | 0.03528 | 0.01628 | 0.08994 | 0.00000 | 0.02634 | 0.01124 | 0.00696 | 0.03272 | 0.01113 | 0.02164 |
| **INTS1** | 1.53870 | 3.97050 | 6.62382 | 6.85770 | 8.48913 | 0.00000 | 3.39149 | 2.30870 | 6.12733 | 17.48688 | 1.28840 | 4.95281 |
| **INTS10** | 3.09304 | 6.98648 | 9.56674 | 4.49038 | 1.37213 | 15.51060 | 6.49735 | 1.91134 | 13.29691 | 0.00000 | 9.37420 | 6.04357 |

| | | | | | | | | | | | | |
|---|---|---|---|---|---|---|---|---|---|---|---|---|
| INTS12 | 1.36682 | 4.36081 | 3.98121 | 0.37832 | 1.09142 | 11.43796 | 4.78582 | 5.58559 | 5.17039 | 0.00000 | 3.95585 | 10.60673 |
| INTS2 | 0.00000 | 2.06098 | 2.97380 | 0.61790 | 0.00000 | 9.19527 | 2.78925 | 2.69969 | 2.56690 | 5.60170 | 3.05344 | 2.38791 |
| INTS3 | 2.08588 | 4.04316 | 4.64955 | 5.05812 | 6.21413 | 0.16266 | 5.60537 | 0.15793 | 8.89819 | 6.29603 | 3.45097 | 3.62463 |
| INTS4 | 0.90467 | 5.11404 | 5.61911 | 3.79589 | 4.58064 | 4.25082 | 4.70212 | 4.98026 | 6.67930 | 1.14675 | 2.74598 | 5.71890 |
| INTS4L2 | 0.00393 | 0.01283 | 0.01334 | 0.01003 | 0.01563 | 0.03123 | 0.01669 | 0.02444 | 0.01535 | 0.01095 | 0.01668 | 0.02516 |
| INTS5 | 1.99720 | 3.61174 | 3.10708 | 3.85967 | 3.76507 | 0.26723 | 2.03510 | 0.50228 | 3.28992 | 0.00000 | 4.81967 | 3.54409 |
| INTS6 | 2.78987 | 0.34853 | 1.52624 | 1.72234 | 2.67953 | 8.35063 | 3.09769 | 3.19801 | 0.10515 | 10.20053 | 0.40030 | 1.65909 |
| INTS6-AS1 | 0.00611 | 0.11234 | 0.09127 | 0.00000 | 0.00000 | 0.25912 | 0.00000 | 0.18275 | 0.00000 | 0.20965 | 0.08739 | 0.25741 |
| INTS7 | 1.49310 | 12.74830 | 5.25040 | 10.40455 | 4.31846 | 19.98484 | 6.38868 | 3.28635 | 9.36894 | 0.00000 | 10.35703 | 13.98043 |
| INTS8 | 7.15432 | 8.40579 | 2.57103 | 4.16813 | 5.44539 | 13.88024 | 2.68849 | 2.76163 | 3.76295 | 6.93293 | 2.44523 | 2.43151 |
| INTS9 | 7.28996 | 9.08885 | 5.49438 | 4.08294 | 1.90891 | 2.76167 | 3.19396 | 0.99976 | 8.18499 | 0.00000 | 6.40280 | 7.77804 |
| INTU | 0.00000 | 0.09185 | 0.52822 | 0.00000 | 0.53731 | 2.11460 | 0.00000 | 0.32769 | 1.32962 | 2.54419 | 0.01799 | 1.29374 |
| INVS | 0.51147 | 2.01467 | 1.21944 | 1.46993 | 0.97681 | 1.56643 | 0.00000 | 0.47399 | 1.24625 | 0.22586 | 1.66709 | 0.66196 |
| IP6K1 | 2.21340 | 8.90811 | 8.08539 | 2.40347 | 7.34311 | 0.01845 | 9.09590 | 0.51623 | 8.52684 | 5.33349 | 4.05482 | 3.48877 |
| IP6K2 | 0.88185 | 18.03730 | 15.36212 | 18.36706 | 16.65810 | 7.84403 | 24.17328 | 7.25444 | 22.01920 | 0.00000 | 12.32434 | 18.45654 |
| IP6K3 | 0.00315 | 0.00423 | 0.15369 | 0.00228 | 0.00000 | 0.02334 | 0.00000 | 0.00956 | 0.01356 | 0.41836 | 0.00001 | 0.00693 |
| IPCEF1 | 1.27809 | 0.07915 | 0.02662 | 0.01442 | 0.00316 | 0.00277 | 0.02192 | 0.22550 | 0.00274 | 0.01843 | 0.05301 | 0.00000 |
| IPMK | 0.08994 | 1.93950 | 0.30793 | 2.70519 | 0.00000 | 6.66884 | 2.01460 | 0.30844 | 0.00000 | 7.11926 | 0.00000 | 2.40831 |
| IPO11 | 0.00190 | 8.64714 | 3.15037 | 1.23694 | 0.00000 | 11.71461 | 0.00000 | 6.10000 | 9.99371 | 0.00000 | 4.64342 | 7.46810 |
| IPO13 | 0.78872 | 2.45673 | 3.43133 | 2.51853 | 5.03545 | 0.00000 | 4.79923 | 0.25366 | 4.41274 | 6.14903 | 2.61217 | 1.38301 |
| IPO4 | 1.52602 | 12.81985 | 13.32182 | 11.10196 | 9.25452 | 0.14403 | 15.49611 | 1.08351 | 20.05901 | 0.00000 | 16.39630 | 15.86299 |
| IPO5 | 2.24106 | 23.63285 | 14.18203 | 12.58847 | 16.72908 | 46.93204 | 32.06686 | 2.65847 | 24.79704 | 23.60012 | 24.66820 | 19.69036 |
| IPO5P1 | 1.09456 | 3.48473 | 3.45291 | 0.81657 | 0.85425 | 3.61417 | 0.00000 | 6.93301 | 2.04248 | 4.04971 | 1.91272 | 4.27686 |
| IPO7 | 1.04377 | 0.00000 | 6.61207 | 1.62217 | 0.00000 | 20.01991 | 25.83023 | 2.26963 | 0.18475 | 35.41493 | 10.24047 | 5.93603 |
| IPO8 | 20.94228 | 2.17093 | 2.10235 | 0.72035 | 0.00000 | 8.85079 | 6.68778 | 3.00535 | 0.29785 | 5.34455 | 2.82688 | 0.80334 |
| IPO9 | 2.72324 | 6.06140 | 11.89666 | 10.19421 | 7.40247 | 13.00258 | 8.81768 | 1.81229 | 13.13751 | 9.72007 | 13.54676 | 7.08766 |
| IPO9-AS1 | 0.00874 | 0.08500 | 0.03121 | 0.01384 | 0.02997 | 0.03737 | 0.03618 | 0.01139 | 0.02945 | 0.00597 | 0.05521 | 0.02488 |
| IPP | 0.42702 | 3.21944 | 1.19396 | 3.81548 | 2.54978 | 3.04099 | 0.85319 | 2.36937 | 0.00000 | 1.09966 | 2.63324 | 4.52600 |
| IPPK | 0.66227 | 5.33448 | 7.68385 | 2.52505 | 3.73038 | 1.98507 | 1.29397 | 5.32461 | 8.70275 | 0.00000 | 4.03036 | 2.52901 |
| IPW | 0.00826 | 1.04816 | 0.60564 | 2.41591 | 0.93945 | 3.88467 | 0.00000 | 0.05096 | 0.00000 | 4.61992 | 0.00000 | 1.28698 |
| IQCA1 | 6.65242 | 0.00000 | 0.51352 | 0.00000 | 0.13613 | 0.81309 | 0.50641 | 1.98951 | 0.51032 | 1.23960 | 0.83127 | 0.00000 |
| IQCB1 | 2.21358 | 3.71313 | 3.04001 | 0.67223 | 2.57079 | 10.93727 | 5.14156 | 0.31327 | 6.98199 | 5.14674 | 7.62567 | 2.61046 |
| IQCC | 0.04582 | 2.49247 | 1.09157 | 0.34892 | 0.47429 | 2.10885 | 0.59611 | 0.42546 | 0.81000 | 1.14902 | 1.89454 | 2.10009 |
| IQCD | 1.89718 | 0.28825 | 1.46436 | 0.00000 | 0.79345 | 0.46752 | 0.11247 | 1.75259 | 0.00000 | 0.28973 | 0.27351 | 0.40139 |
| IQCE | 0.82485 | 2.23899 | 0.94455 | 1.32949 | 1.68858 | 0.00000 | 0.47368 | 0.42714 | 0.00000 | 1.52029 | 0.03878 | 2.39291 |
| IQCF1 | 0.01541 | 0.10954 | 0.00000 | 0.15401 | 0.01268 | 0.05950 | 0.00000 | 0.04418 | 0.13452 | 0.02081 | 0.04407 | 0.04703 |
| IQCF2 | 0.01741 | 0.01331 | 0.01627 | 0.01384 | 0.01801 | 0.01308 | 0.02009 | 0.03975 | 0.01972 | 0.00456 | 0.01575 | 0.06133 |
| IQCF3 | 0.00610 | 0.02472 | 0.01477 | 0.01735 | 0.01554 | 0.01917 | 0.01879 | 0.01058 | 0.01619 | 0.01661 | 0.01624 | 0.01482 |
| IQCF4 | 0.00653 | 0.01320 | 0.01339 | 0.00995 | 0.02786 | 0.01376 | 0.02504 | 0.04524 | 0.00676 | 0.02529 | 0.02938 | 0.01435 |
| IQCF5 | 0.00640 | 0.02191 | 0.00322 | 0.03728 | 0.01134 | 0.09615 | 0.00433 | 0.00891 | 0.00567 | 0.01779 | 0.01487 | 0.01654 |
| IQCG | 1.33075 | 2.03650 | 2.69610 | 0.94764 | 0.00000 | 0.00267 | 2.98215 | 1.02978 | 1.15666 | 0.54267 | 6.62736 | 1.67119 |
| IQCH | 0.66735 | 0.12168 | 0.23499 | 0.24226 | 0.00000 | 0.18094 | 0.00000 | 0.08416 | 0.11207 | 0.00000 | 0.14793 | 0.34580 |
| IQCH-AS1 | 0.64889 | 1.01001 | 0.95310 | 0.00000 | 1.34748 | 1.18069 | 1.17553 | 0.00000 | 1.25377 | 2.38604 | 2.29189 | 0.10595 |
| IQCJ-SCHIP1-A | 0.02692 | 0.00884 | 0.05999 | 0.00276 | 0.01216 | 0.02660 | 0.00754 | 0.04797 | 0.00000 | 0.17324 | 0.00805 | 0.19114 |
| IQCJ.uid-4645 | 0.27366 | 0.01608 | 0.01181 | 0.05451 | 0.04048 | 0.01045 | 0.03783 | 0.05216 | 0.00708 | 0.00000 | 0.01662 | 0.01104 |
| IQCJ.uid-4646 | 0.02728 | 0.01558 | 0.01554 | 0.01268 | 0.01689 | 0.01634 | 0.01808 | 0.01344 | 0.01860 | 0.01265 | 0.01507 | 0.01541 |
| IQCK | 1.35607 | 3.37497 | 1.73168 | 1.84401 | 1.08860 | 0.02849 | 1.45909 | 1.99781 | 1.12247 | 1.16517 | 1.05057 | 3.88863 |
| IQGAP1 | 11.40453 | 9.95163 | 16.63729 | 2.05877 | 15.14510 | 33.44778 | 0.00000 | 8.76094 | 28.27788 | 25.73921 | 12.59073 | 7.61225 |

| | | | | | | | | | | | | |
|---|---|---|---|---|---|---|---|---|---|---|---|---|
| **IQGAP2** | 10.20886 | 1.91798 | 4.09763 | 0.00000 | 24.31243 | 12.69845 | 1.78063 | 6.33274 | 15.45194 | 35.12526 | 10.93158 | 0.00000 |
| **IQGAP3** | 0.61274 | 8.67311 | 0.00000 | 13.06429 | 6.97031 | 6.07471 | 0.00000 | 0.91377 | 2.11820 | 15.04861 | 5.41162 | 3.72000 |
| **IQSEC1** | 5.28327 | 1.35722 | 1.91779 | 0.00009 | 7.25452 | 1.30015 | 4.12176 | 0.98676 | 2.42132 | 10.55955 | 3.79841 | 0.44770 |
| **IQSEC2** | 0.02202 | 0.18255 | 0.04417 | 0.21189 | 0.00000 | 0.08260 | 0.00000 | 0.01102 | 0.00000 | 0.16099 | 0.00757 | 0.00442 |
| **IQSEC3** | 0.07100 | 0.01074 | 0.03038 | 0.01423 | 0.01558 | 0.00139 | 0.01440 | 0.00328 | 0.02256 | 0.02217 | 0.03123 | 0.08218 |
| **IQUB** | 7.30008 | 0.22777 | 0.12943 | 0.00000 | 0.00250 | 0.00000 | 0.16375 | 1.89618 | 0.05372 | 0.01398 | 0.00000 | 0.19182 |
| **IRAK1** | 0.40930 | 1.67130 | 3.41384 | 2.32737 | 0.00000 | 0.00000 | 10.15601 | 0.00000 | 5.87145 | 5.35997 | 1.22860 | 2.02574 |
| **IRAK1BP1** | 2.08733 | 0.33231 | 0.00000 | 0.40314 | 0.19103 | 0.98430 | 0.15072 | 0.03725 | 0.03990 | 0.43921 | 0.85496 | 0.48864 |
| **IRAK2** | 1.23497 | 0.05899 | 2.44639 | 0.02250 | 0.00000 | 0.00000 | 1.80825 | 0.13610 | 0.58759 | 0.53121 | 0.00000 | 0.02283 |
| **IRAK3** | 0.00249 | 0.01263 | 0.12254 | 0.01747 | 0.02003 | 0.00000 | 0.00952 | 0.34070 | 0.11540 | 0.00000 | 0.00715 | 0.85027 |
| **IRAK4** | 0.00000 | 0.91673 | 1.56169 | 0.44988 | 1.20101 | 4.04820 | 0.00000 | 0.00000 | 0.69826 | 0.87329 | 1.32975 | 0.50707 |
| **IREB2** | 1.49814 | 6.45189 | 11.59912 | 1.25976 | 2.96464 | 21.48731 | 14.14420 | 1.80617 | 6.00241 | 15.81181 | 8.64881 | 5.77007 |
| **IRF1** | 1.75204 | 1.86145 | 0.25748 | 1.96014 | 1.74929 | 0.00000 | 0.82219 | 3.06113 | 1.87102 | 0.29042 | 1.56701 | 2.90090 |
| **IRF2** | 0.00629 | 0.01010 | 0.17765 | 0.13694 | 0.06749 | 0.04459 | 0.00000 | 0.00976 | 0.27227 | 0.00000 | 0.00864 | 0.01455 |
| **IRF2BP1** | 0.55239 | 0.21285 | 1.20644 | 0.30060 | 1.23125 | 0.00000 | 0.99575 | 0.14944 | 0.00000 | 5.40440 | 0.28224 | 1.37773 |
| **IRF2BP2** | 0.49427 | 0.96475 | 1.94247 | 2.26029 | 4.37308 | 5.56947 | 8.37681 | 0.45463 | 0.90036 | 14.73227 | 1.62140 | 1.00389 |
| **IRF2BPL** | 2.73576 | 2.62502 | 2.68573 | 1.61617 | 0.00007 | 3.80776 | 8.59093 | 0.24825 | 0.00000 | 10.84665 | 6.99645 | 3.08323 |
| **IRF3** | 0.02453 | 9.29995 | 13.70926 | 9.81688 | 14.60686 | 0.00000 | 21.31176 | 0.66351 | 15.98998 | 2.42974 | 8.75800 | 7.07924 |
| **IRF4** | 1.22702 | 0.00000 | 0.02670 | 0.02981 | 0.00000 | 0.00000 | 0.00898 | 9.10105 | 0.00000 | 0.18389 | 0.11327 | 0.00000 |
| **IRF5** | 0.12007 | 0.04321 | 0.11275 | 0.04356 | 0.07136 | 0.01450 | 0.05162 | 0.03759 | 0.00578 | 0.00697 | 0.01032 | 0.01798 |
| **IRF6** | 6.50223 | 1.56714 | 3.51081 | 0.00000 | 1.42111 | 0.02578 | 7.02390 | 0.07277 | 1.62174 | 0.00000 | 0.85282 | 0.55701 |
| **IRF7** | 0.01031 | 0.68412 | 2.33616 | 2.22540 | 1.71271 | 0.00000 | 0.71176 | 0.20528 | 0.53410 | 0.00000 | 0.66839 | 0.60815 |
| **IRF8** | 1.80588 | 0.01325 | 0.65057 | 0.26682 | 0.00000 | 0.00000 | 0.00053 | 0.55738 | 0.00000 | 0.02058 | 0.07775 | 1.17374 |
| **IRF9** | 0.57073 | 0.22739 | 0.66001 | 2.16458 | 0.66119 | 0.31362 | 0.67563 | 0.16613 | 2.66261 | 4.63380 | 0.00000 | 0.44572 |
| **IRGM** | 1.36509 | 0.01851 | 0.04242 | 0.00000 | 0.04560 | 0.02408 | 0.00000 | 0.02513 | 0.00324 | 0.15266 | 0.00849 | 0.01589 |
| **IRGQ** | 1.51590 | 9.71590 | 2.07516 | 1.51252 | 0.00000 | 7.93969 | 5.95816 | 4.42809 | 1.29585 | 15.63502 | 2.25810 | 25.97205 |
| **IRS1** | 0.49439 | 0.00000 | 3.34522 | 0.00000 | 1.02496 | 0.49098 | 3.60385 | 0.00000 | 1.14362 | 0.85194 | 0.21118 | 0.15156 |
| **IRS2** | 0.05901 | 0.00000 | 0.36801 | 0.16382 | 0.00000 | 0.03103 | 0.74333 | 0.00906 | 0.00000 | 0.57038 | 0.00000 | 0.89715 |
| **IRS4** | 0.00570 | 0.01148 | 0.09530 | 0.02159 | 0.01530 | 0.00000 | 0.00444 | 0.00734 | 0.00713 | 0.07095 | 0.01608 | 0.08103 |
| **IRX2** | 0.66218 | 0.00000 | 6.75180 | 0.00000 | 0.34445 | 0.09001 | 15.08915 | 0.54475 | 0.00362 | 6.97692 | 0.00000 | 0.00000 |
| **IRX3** | 0.00190 | 0.05909 | 0.00000 | 0.76951 | 0.57297 | 0.00000 | 0.00000 | 1.00549 | 0.00000 | 0.68604 | 0.77381 | 1.41774 |
| **IRX4** | 0.02559 | 0.00000 | 5.00546 | 0.00000 | 0.09093 | 0.00620 | 1.14433 | 0.22130 | 0.60547 | 1.17770 | 0.00000 | 0.00000 |
| **IRX5** | 0.14461 | 0.00000 | 0.03847 | 0.00859 | 0.01881 | 0.00000 | 0.04912 | 1.80923 | 0.03159 | 0.03415 | 0.01126 | 0.97488 |
| **IRX6** | 0.03810 | 0.01490 | 0.01283 | 0.05086 | 0.02416 | 0.00000 | 0.03200 | 0.00835 | 0.02196 | 0.00771 | 0.01600 | 0.02558 |
| **ISCA1** | 0.00000 | 8.76257 | 4.61734 | 4.24940 | 1.44842 | 16.40265 | 8.26115 | 5.43956 | 9.24238 | 1.15775 | 5.88513 | 9.55823 |
| **ISCA2** | 0.53261 | 9.24334 | 5.66423 | 7.17927 | 10.30588 | 14.25195 | 7.32367 | 0.71988 | 7.03546 | 2.54719 | 9.33743 | 9.54827 |
| **ISCU** | 5.59726 | 22.47959 | 12.09710 | 6.92552 | 11.16674 | 39.46792 | 16.74712 | 7.52866 | 15.86130 | 9.31274 | 15.74070 | 20.51933 |
| **ISG15** | 0.00452 | 0.16879 | 0.41348 | 0.93519 | 0.44457 | 0.00000 | 0.72851 | 0.13010 | 0.16862 | 0.00000 | 0.08733 | 0.05303 |
| **ISG20** | 0.32902 | 0.07996 | 0.63977 | 0.30178 | 0.72625 | 0.00000 | 0.16328 | 0.00000 | 0.00000 | 0.67106 | 0.00000 | 0.13955 |
| **ISG20L2** | 0.00000 | 7.99503 | 11.33031 | 4.85735 | 6.28637 | 3.37604 | 12.62337 | 7.65151 | 13.81313 | 0.00000 | 8.83332 | 14.71329 |
| **ISL1** | 1.19646 | 0.33099 | 0.54710 | 1.72222 | 0.00000 | 0.00000 | 0.06830 | 1.37733 | 0.42534 | 0.00000 | 0.00000 | 0.84813 |
| **ISL2** | 0.00325 | 0.00484 | 0.07146 | 0.02295 | 0.02907 | 0.01780 | 0.03893 | 3.60086 | 0.00378 | 0.00000 | 0.01197 | 1.05960 |
| **ISLR2** | 0.00274 | 0.01964 | 0.04219 | 0.01434 | 0.02044 | 0.00897 | 0.01555 | 0.14354 | 0.01265 | 0.02096 | 0.01652 | 0.01643 |
| **ISM1** | 0.01417 | 0.05179 | 0.01531 | 0.19957 | 0.00232 | 0.15975 | 0.04029 | 0.02161 | 0.00000 | 0.03603 | 0.00929 | 0.02903 |
| **ISM1-AS1** | 0.00550 | 0.04007 | 0.02541 | 0.04076 | 0.00366 | 0.02839 | 0.00304 | 0.00995 | 0.03354 | 0.03628 | 0.02198 | 0.02705 |
| **ISM2** | 0.00172 | 0.75103 | 1.46138 | 0.48551 | 0.20889 | 0.29624 | 0.00000 | 0.00334 | 1.17251 | 0.45570 | 0.27235 | 0.18855 |
| **ISOC1** | 3.12611 | 12.97876 | 15.61719 | 0.00001 | 0.00000 | 19.28250 | 17.75804 | 0.94129 | 15.11375 | 0.00000 | 9.74118 | 15.43063 |
| **ISOC2** | 0.70776 | 2.98257 | 1.65559 | 12.51661 | 7.83531 | 0.00000 | 7.80431 | 0.14652 | 7.32236 | 0.00000 | 3.90404 | 2.72678 |

| | | | | | | | | | | | | |
|---|---|---|---|---|---|---|---|---|---|---|---|---|
| **ISPD** | 3.33679 | 0.00000 | 0.00000 | 0.00000 | 0.28782 | 1.17620 | 1.32268 | 0.65013 | 0.10747 | 2.03461 | 0.80925 | 0.11310 |
| **ISPD-AS1** | 0.32692 | 0.00454 | 0.00000 | 0.00000 | 0.03062 | 0.05368 | 0.00480 | 0.05507 | 0.01874 | 0.18537 | 0.03115 | 0.00333 |
| **IST1** | 1.83253 | 21.36899 | 17.76127 | 11.51287 | 14.05219 | 18.00366 | 22.13395 | 21.19709 | 25.15811 | 2.64532 | 15.92675 | 21.35609 |
| **ISX** | 0.00593 | 0.00727 | 0.03879 | 0.31739 | 0.00000 | 0.03568 | 0.00000 | 0.00984 | 0.02347 | 0.02853 | 0.01148 | 0.01374 |
| **ISY1** | 0.30771 | 4.76213 | 6.36263 | 2.06908 | 3.59506 | 2.77062 | 9.72793 | 12.46766 | 5.87362 | 4.85908 | 4.61503 | 6.93249 |
| **ISYNA1** | 1.18824 | 2.34007 | 17.08465 | 6.99585 | 6.90174 | 0.00198 | 11.14571 | 0.46757 | 11.74432 | 0.00000 | 4.95753 | 0.00000 |
| **ITCH** | 3.05980 | 5.59914 | 8.07449 | 4.82756 | 3.91573 | 20.53905 | 1.99638 | 3.69836 | 5.54445 | 6.98153 | 6.88002 | 9.59850 |
| **ITFG1** | 0.09509 | 5.90189 | 8.15087 | 3.42613 | 3.31346 | 10.61123 | 0.00000 | 0.12277 | 2.76458 | 3.82533 | 5.63496 | 4.99896 |
| **ITFG2** | 0.23553 | 2.03935 | 2.41725 | 3.41392 | 3.83269 | 0.00000 | 0.00000 | 1.24687 | 4.75569 | 1.73863 | 1.63714 | 4.38108 |
| **ITFG3** | 3.05166 | 21.39674 | 1.74150 | 17.68044 | 5.71781 | 0.00000 | 0.50237 | 7.49213 | 2.49109 | 7.40371 | 2.67187 | 23.96537 |
| **ITGA1** | 0.02201 | 0.01004 | 2.78409 | 1.37119 | 0.00977 | 0.00000 | 0.00000 | 0.01066 | 0.00000 | 0.00996 | 0.04533 | 0.01780 |
| **ITGA10** | 1.13794 | 0.02109 | 0.04520 | 0.00407 | 0.03520 | 0.00000 | 0.01131 | 0.00000 | 0.19780 | 0.30581 | 0.09343 | 0.69741 |
| **ITGA11** | 0.86575 | 0.02369 | 0.01399 | 0.15329 | 0.06256 | 0.00000 | 0.00885 | 0.40820 | 0.01719 | 0.04309 | 0.01672 | 0.00870 |
| **ITGA2** | 0.21386 | 0.00000 | 0.31944 | 3.17796 | 0.00000 | 0.01520 | 0.06501 | 0.15587 | 0.09436 | 0.05962 | 0.07178 | 0.00573 |
| **ITGA2B** | 0.08340 | 0.69156 | 0.40973 | 0.07943 | 0.19898 | 0.00000 | 0.65334 | 0.00000 | 1.23937 | 1.06526 | 0.00000 | 0.18995 |
| **ITGA3** | 0.58411 | 0.21796 | 0.01088 | 0.37192 | 0.16477 | 0.00000 | 0.00000 | 0.00000 | 0.20389 | 0.55923 | 0.00000 | 0.05199 |
| **ITGA4** | 4.38131 | 0.00137 | 0.01430 | 0.03086 | 0.00000 | 0.01387 | 0.00000 | 0.98311 | 0.03361 | 0.05273 | 0.01958 | 0.11940 |
| **ITGA5** | 0.25597 | 1.75672 | 14.85457 | 0.02297 | 0.00000 | 0.28533 | 0.70930 | 0.54149 | 16.46406 | 1.03268 | 0.00000 | 18.90864 |
| **ITGA6** | 3.12661 | 2.14667 | 2.04346 | 8.55557 | 1.31847 | 12.28317 | 0.00000 | 1.10868 | 4.16518 | 10.99033 | 1.47426 | 0.43406 |
| **ITGA7** | 0.00126 | 0.03563 | 0.24009 | 0.40401 | 0.02904 | 0.02039 | 0.15667 | 0.02582 | 0.04225 | 0.43675 | 0.21617 | 0.16527 |
| **ITGA8** | 0.12076 | 0.04129 | 0.01592 | 0.04010 | 0.03301 | 0.00000 | 0.05083 | 0.19344 | 0.00000 | 0.00287 | 0.05298 | 0.00975 |
| **ITGA9** | 0.86505 | 0.30302 | 0.08753 | 0.04049 | 0.00000 | 0.00000 | 0.00639 | 0.32561 | 0.06468 | 0.02554 | 0.03946 | 0.01445 |
| **ITGA9-AS1** | 0.07010 | 0.00680 | 0.05540 | 0.01950 | 0.03269 | 0.00887 | 0.01422 | 0.03154 | 0.06082 | 0.00000 | 0.08560 | 0.09782 |
| **ITGAD** | 0.02678 | 0.00981 | 0.01730 | 0.00793 | 0.01943 | 0.03856 | 0.01451 | 0.00945 | 0.01575 | 0.03484 | 0.01189 | 0.01649 |
| **ITGAE** | 3.46796 | 3.20895 | 7.65310 | 4.80325 | 4.33706 | 9.50619 | 9.98612 | 0.91777 | 5.75346 | 4.25784 | 4.64005 | 4.47722 |
| **ITGAL** | 0.02979 | 0.23630 | 0.02046 | 0.54981 | 0.00000 | 0.02467 | 0.00000 | 1.36358 | 0.02312 | 0.06141 | 0.01576 | 0.20381 |
| **ITGAM** | 0.01608 | 3.00101 | 0.04484 | 7.65600 | 0.39768 | 0.00000 | 0.00004 | 0.14438 | 0.00000 | 0.66068 | 0.00000 | 0.94615 |
| **ITGAV** | 2.51052 | 0.00000 | 1.03614 | 2.45285 | 0.00000 | 1.68255 | 0.00000 | 1.63904 | 0.50379 | 1.62913 | 0.08279 | 0.00000 |
| **ITGAX** | 0.00429 | 0.06191 | 0.01573 | 0.52948 | 0.03447 | 0.00000 | 0.00277 | 0.25548 | 0.00000 | 0.81355 | 0.03332 | 0.41006 |
| **ITGB1** | 0.91949 | 4.27252 | 8.74265 | 3.61042 | 2.85097 | 16.38523 | 0.00000 | 5.30365 | 16.57786 | 2.53429 | 6.69940 | 3.42598 |
| **ITGB1BP1** | 1.41311 | 7.02188 | 9.98692 | 6.47761 | 9.90298 | 19.22122 | 17.36366 | 0.11407 | 13.23553 | 2.85258 | 13.18655 | 9.56741 |
| **ITGB1BP2** | 0.00094 | 0.25006 | 0.00000 | 0.22650 | 0.00000 | 0.00656 | 0.37671 | 0.34732 | 0.01980 | 0.58038 | 0.16602 | 0.59025 |
| **ITGB2** | 0.01007 | 0.00115 | 0.01291 | 0.05085 | 0.06072 | 0.12873 | 0.00000 | 0.00863 | 0.02947 | 0.06913 | 0.00000 | 0.01663 |
| **ITGB3** | 0.00451 | 0.40407 | 0.30129 | 0.25330 | 0.00000 | 0.00000 | 0.00000 | 0.00747 | 0.00000 | 0.17218 | 0.00365 | 0.64066 |
| **ITGB3BP** | 1.90363 | 5.22140 | 3.69700 | 0.00000 | 0.00000 | 15.79828 | 6.21394 | 1.00392 | 5.01271 | 5.30102 | 5.00485 | 4.53775 |
| **ITGB4** | 0.10625 | 0.00000 | 3.67721 | 0.00000 | 0.00000 | 0.00000 | 2.47997 | 0.01659 | 1.48639 | 0.08033 | 2.17178 | 0.04671 |
| **ITGB5** | 1.28298 | 3.36285 | 8.99689 | 6.18357 | 8.76481 | 0.34392 | 7.50226 | 0.00000 | 4.23085 | 13.80854 | 1.14114 | 2.82040 |
| **ITGB6** | 0.01242 | 0.00636 | 0.01178 | 0.08408 | 0.00000 | 0.06907 | 0.00654 | 0.00821 | 0.00694 | 0.02329 | 0.01201 | 0.01244 |
| **ITGB7** | 0.00906 | 0.25486 | 0.00000 | 0.00000 | 0.34099 | 0.00000 | 0.06906 | 0.00000 | 0.00000 | 0.59163 | 0.00000 | 1.44669 |
| **ITGB8** | 0.01202 | 0.00000 | 0.10531 | 0.28897 | 0.00000 | 0.13476 | 0.02377 | 0.27626 | 0.00001 | 0.06848 | 0.01599 | 0.23662 |
| **ITGBL1** | 1.92966 | 0.02451 | 0.09088 | 0.04719 | 0.15459 | 0.00000 | 0.00000 | 0.00101 | 0.13475 | 0.32938 | 0.05838 | 0.03640 |
| **ITIH1** | 0.00565 | 0.01328 | 0.01235 | 0.02695 | 0.00741 | 0.01569 | 0.02840 | 0.01041 | 0.01223 | 0.04357 | 0.02109 | 0.01548 |
| **ITIH2** | 0.58232 | 0.04042 | 0.01802 | 0.00541 | 0.14694 | 0.00000 | 0.00741 | 0.04674 | 0.08207 | 0.09132 | 0.00768 | 0.09571 |
| **ITIH3** | 0.00604 | 0.01271 | 0.01285 | 0.01102 | 0.01466 | 0.02458 | 0.02121 | 0.01015 | 0.01568 | 0.02172 | 0.01343 | 0.01882 |
| **ITIH4** | 0.01091 | 0.35290 | 0.11996 | 0.11629 | 0.00000 | 0.00000 | 0.00000 | 0.00201 | 0.08037 | 0.76364 | 0.00000 | 0.20811 |
| **ITIH4-AS1** | 0.01626 | 0.01501 | 0.01599 | 0.01150 | 0.01387 | 0.01638 | 0.01607 | 0.01021 | 0.02346 | 0.02502 | 0.01291 | 0.01654 |
| **ITIH5** | 4.02551 | 0.35939 | 1.32172 | 2.77199 | 0.76963 | 0.00000 | 0.23316 | 0.53818 | 0.11064 | 0.00000 | 0.39030 | 0.17965 |
| **ITIH6** | 0.01180 | 0.10116 | 0.02823 | 0.00947 | 0.00000 | 0.01986 | 0.02347 | 0.00754 | 0.00000 | 0.02436 | 0.05477 | 0.10789 |

| | | | | | | | | | | | | |
|---|---|---|---|---|---|---|---|---|---|---|---|---|
| **ITK** | 0.00466 | 0.00332 | 0.04886 | 0.02696 | 0.00532 | 0.04823 | 0.01128 | 0.01178 | 0.01581 | 0.00433 | 0.01969 | 0.17504 |
| **ITLN1** | 0.31432 | 0.21901 | 0.00000 | 0.00000 | 0.00474 | 0.26683 | 0.02276 | 0.23618 | 0.00000 | 0.04847 | 0.00000 | 0.02020 |
| **ITLN2** | 0.17111 | 2.33292 | 0.00000 | 2.96790 | 0.00814 | 0.93664 | 0.00000 | 0.20709 | 0.00000 | 0.00134 | 0.46508 | 1.46697 |
| **ITM2A** | 0.00152 | 0.06374 | 0.00000 | 0.00000 | 1.04676 | 0.23245 | 0.07202 | 0.00541 | 0.37269 | 0.34828 | 1.16602 | 0.00000 |
| **ITM2B** | 1.13377 | 13.62009 | 21.67231 | 6.00720 | 8.52677 | 44.38331 | 0.00000 | 1.73167 | 33.90991 | 17.92608 | 17.94267 | 10.88896 |
| **ITM2C** | 7.42192 | 38.12411 | 4.36581 | 20.63551 | 13.02593 | 0.00000 | 0.00000 | 4.49375 | 8.92471 | 2.84860 | 3.26718 | 11.84851 |
| **ITPA** | 1.52155 | 15.06035 | 14.62103 | 9.61686 | 13.83975 | 8.90584 | 20.94151 | 0.42945 | 16.88289 | 4.82763 | 14.71989 | 18.85543 |
| **ITPK1** | 4.05053 | 11.90889 | 2.93940 | 5.62195 | 9.58688 | 0.00000 | 8.11742 | 0.00000 | 4.43474 | 14.55744 | 1.90750 | 3.87543 |
| **ITPK1-AS1** | 0.00297 | 0.04715 | 0.06227 | 0.12700 | 0.68501 | 0.00000 | 0.00000 | 0.00283 | 0.07528 | 0.83197 | 0.00000 | 0.29010 |
| **ITPKA** | 0.12870 | 0.02866 | 0.16691 | 0.42001 | 0.08035 | 0.00000 | 0.00000 | 1.31710 | 0.07832 | 0.35389 | 0.00000 | 1.35035 |
| **ITPKB** | 0.00618 | 0.01959 | 0.01685 | 0.07021 | 0.00000 | 0.01133 | 0.02090 | 0.01048 | 0.00000 | 0.02951 | 0.07416 | 0.01563 |
| **ITPKC** | 0.00000 | 3.38377 | 6.31494 | 0.92152 | 4.18495 | 0.00003 | 5.64353 | 0.40227 | 0.74833 | 3.94108 | 0.90704 | 3.88082 |
| **ITPR1** | 4.79472 | 0.35201 | 0.39153 | 1.17990 | 0.00000 | 0.00019 | 0.55797 | 0.37285 | 2.17120 | 1.81550 | 0.19798 | 0.50006 |
| **ITPR1-AS1** | 0.01253 | 0.14955 | 0.01296 | 0.15713 | 0.00000 | 0.03006 | 0.00973 | 0.01187 | 0.00231 | 0.05204 | 0.18424 | 0.01657 |
| **ITPR2** | 0.34756 | 0.00000 | 0.73920 | 2.49477 | 0.49111 | 2.06087 | 0.18937 | 0.00000 | 0.00000 | 1.11549 | 0.00000 | 0.09448 |
| **ITPR3** | 2.53913 | 2.73188 | 0.00000 | 9.81721 | 9.89683 | 0.00000 | 4.59322 | 0.00000 | 11.07617 | 7.05408 | 2.13327 | 0.20679 |
| **ITPRIP** | 1.51452 | 0.09147 | 0.18095 | 1.27642 | 1.55402 | 0.00000 | 0.50875 | 1.73304 | 1.22335 | 0.23919 | 1.16703 | 1.11052 |
| **ITPRIPL1** | 0.00604 | 0.00000 | 0.80607 | 0.33808 | 0.00000 | 0.00000 | 0.59462 | 0.00281 | 1.71360 | 0.00000 | 1.25045 | 0.58826 |
| **ITPRIPL2** | 0.93474 | 0.00000 | 0.60058 | 0.00000 | 0.00000 | 0.40516 | 1.48313 | 0.60621 | 0.00000 | 3.04153 | 3.09923 | 0.03855 |
| **ITSN1** | 13.95851 | 1.53457 | 14.73860 | 9.40869 | 23.22200 | 11.81746 | 23.11252 | 8.19803 | 21.63858 | 22.34280 | 15.85169 | 9.50604 |
| **ITSN2** | 2.68471 | 1.67164 | 4.33833 | 0.91139 | 7.44856 | 7.14895 | 5.35993 | 0.94660 | 8.53153 | 20.89659 | 3.73189 | 2.10753 |
| **IVD** | 2.26792 | 1.98092 | 1.60198 | 6.11472 | 2.98024 | 0.07641 | 5.41647 | 1.02176 | 3.04771 | 0.91908 | 6.81941 | 1.40197 |
| **IVL** | 0.01887 | 0.01581 | 0.02267 | 0.01134 | 0.01710 | 0.01052 | 0.02529 | 0.01015 | 0.01075 | 0.03063 | 0.01606 | 0.01572 |
| **IVNS1ABP** | 1.53068 | 2.32747 | 16.46663 | 3.20350 | 0.00000 | 4.97912 | 8.32662 | 3.27908 | 18.25777 | 0.00000 | 18.50943 | 4.70268 |
| **IWS1** | 14.65068 | 11.91269 | 11.98630 | 10.99186 | 12.22967 | 17.12946 | 15.45185 | 10.53744 | 10.87976 | 12.73726 | 9.53574 | 16.81955 |
| **IYD** | 0.00836 | 0.01528 | 0.01614 | 0.01592 | 0.02032 | 0.00375 | 0.02223 | 0.04575 | 0.02683 | 0.00000 | 0.01573 | 0.09048 |
| **IZUMO1** | 0.00197 | 0.00441 | 0.00854 | 0.02789 | 0.00000 | 0.02130 | 0.18005 | 0.03822 | 0.10154 | 0.15257 | 0.00000 | 0.01410 |
| **IZUMO2** | 0.00607 | 0.02643 | 0.01050 | 0.00781 | 0.00856 | 0.02873 | 0.01843 | 0.01040 | 0.02586 | 0.02087 | 0.01294 | 0.04799 |
| **IZUMO4** | 0.00313 | 0.18353 | 0.04847 | 0.00490 | 0.00428 | 0.00000 | 0.00599 | 0.53840 | 0.25883 | 0.00000 | 0.13725 | 0.15759 |
| **JADE1** | 1.18953 | 5.23314 | 19.90748 | 6.12202 | 4.44578 | 8.92213 | 17.04471 | 0.46188 | 8.95362 | 12.54989 | 10.65443 | 3.83464 |
| **JADE2** | 0.17771 | 0.14558 | 0.00000 | 9.59191 | 7.03711 | 0.18069 | 0.91065 | 0.00000 | 0.77128 | 0.72237 | 0.16662 | 0.00000 |
| **JADE3** | 2.32625 | 2.76195 | 5.55031 | 1.03277 | 3.27852 | 7.71670 | 5.75080 | 1.02608 | 3.09851 | 0.00000 | 2.53217 | 1.45050 |
| **JAG1** | 11.66793 | 0.44630 | 2.08869 | 2.81492 | 0.00000 | 0.00000 | 0.00000 | 5.46329 | 0.46181 | 1.19277 | 0.00902 | 3.68236 |
| **JAG2** | 2.01237 | 6.30243 | 0.56345 | 1.54484 | 8.68570 | 0.00000 | 0.04212 | 0.00000 | 0.00000 | 5.24108 | 0.14672 | 0.85582 |
| **JAGN1** | 2.30743 | 16.19495 | 11.87973 | 8.79774 | 11.10283 | 22.45951 | 0.00003 | 2.81015 | 34.97633 | 4.71317 | 13.17801 | 23.88944 |
| **JAK1** | 3.65307 | 4.23414 | 1.41862 | 4.28155 | 2.07159 | 5.03171 | 0.82133 | 0.53818 | 6.63778 | 5.49743 | 2.29967 | 1.33819 |
| **JAK2** | 1.42550 | 0.03377 | 0.13369 | 0.50875 | 0.00000 | 0.61912 | 0.34511 | 0.44279 | 2.14797 | 0.56991 | 0.00000 | 0.40407 |
| **JAK3** | 0.00421 | 0.02933 | 0.05886 | 0.38689 | 0.00000 | 0.00337 | 0.01092 | 0.02238 | 0.00578 | 0.13004 | 0.01170 | 0.42027 |
| **JAKMIP1** | 0.71515 | 0.00000 | 0.00000 | 0.00000 | 0.00000 | 0.26886 | 0.12685 | 0.10325 | 0.00000 | 0.65178 | 0.03808 | 0.02724 |
| **JAKMIP2** | 0.00301 | 0.00000 | 0.00002 | 0.00000 | 0.00000 | 0.08545 | 0.23499 | 0.01797 | 0.00000 | 0.10113 | 0.02316 | 0.00751 |
| **JAKMIP2-AS1** | 0.00625 | 0.01617 | 0.01538 | 0.01023 | 0.02955 | 0.00358 | 0.09168 | 0.01059 | 0.02943 | 0.01534 | 0.01297 | 0.01728 |
| **JAKMIP3** | 0.69263 | 0.00737 | 0.02912 | 0.01504 | 0.09387 | 0.01662 | 0.02271 | 0.04698 | 0.00724 | 0.01942 | 0.01244 | 0.02903 |
| **JAM2** | 0.73092 | 0.08717 | 0.34834 | 0.08699 | 0.00000 | 0.00000 | 0.16879 | 0.62402 | 0.00000 | 0.30245 | 0.06236 | 0.11258 |
| **JAM3** | 4.85554 | 8.23479 | 11.22204 | 2.86360 | 1.38104 | 11.98397 | 0.00073 | 4.06994 | 2.99111 | 7.18075 | 1.45980 | 2.65873 |
| **JARID2** | 13.21688 | 5.16347 | 7.58384 | 15.26590 | 4.78131 | 7.04918 | 14.05936 | 5.11311 | 13.87348 | 26.80229 | 2.13366 | 7.95877 |
| **JARID2-AS1** | 0.00339 | 0.03679 | 0.04031 | 0.00789 | 0.02603 | 0.00000 | 0.00666 | 0.11617 | 0.05340 | 0.02879 | 0.01823 | 0.04319 |
| **JAZF1** | 7.91619 | 1.46637 | 0.54155 | 0.57138 | 1.20687 | 0.38164 | 0.00000 | 1.19316 | 0.20307 | 0.13740 | 0.37063 | 1.19300 |
| **JAZF1-AS1** | 0.02690 | 0.04134 | 0.02891 | 0.02729 | 0.02887 | 0.00000 | 0.02480 | 0.02219 | 0.01363 | 0.01349 | 0.01711 | 0.04197 |

| | | | | | | | | | | | | |
|---|---|---|---|---|---|---|---|---|---|---|---|---|
| JDP2 | 0.58541 | 0.00000 | 1.92240 | 0.00000 | 9.54685 | 0.54450 | 15.11974 | 0.28135 | 0.65271 | 13.13093 | 8.61704 | 0.47759 |
| JHDM1D-AS1 | 0.66383 | 1.21836 | 0.34335 | 0.49085 | 0.11572 | 0.13207 | 0.00000 | 0.05968 | 0.08636 | 0.20759 | 0.23954 | 2.36661 |
| JKAMP | 0.03320 | 4.69508 | 12.56882 | 2.80967 | 0.00230 | 28.55197 | 0.00000 | 9.68346 | 22.74090 | 0.00000 | 10.51295 | 17.78196 |
| JMJD1C | 1.26912 | 0.44104 | 4.27500 | 5.77168 | 0.54227 | 8.69156 | 1.08503 | 1.13144 | 4.95309 | 13.16267 | 1.85761 | 1.36440 |
| JMJD1C-AS1 | 0.00806 | 0.06082 | 0.02873 | 0.00192 | 0.00913 | 0.04522 | 0.00000 | 0.05730 | 0.01631 | 0.06433 | 0.01148 | 0.04718 |
| JMJD4 | 0.30560 | 5.81388 | 3.37896 | 2.63113 | 3.58174 | 0.00000 | 1.30812 | 0.24105 | 2.39744 | 0.00000 | 3.92895 | 4.75371 |
| JMJD6 | 1.47630 | 16.91413 | 17.27777 | 10.87690 | 16.95471 | 4.43189 | 21.91599 | 7.92031 | 23.18450 | 0.00000 | 12.93306 | 11.67152 |
| JMJD7 | 0.88580 | 0.61902 | 1.06452 | 0.49768 | 0.84910 | 0.00000 | 0.00144 | 0.21255 | 0.37311 | 0.25290 | 0.53520 | 0.22420 |
| JMJD8 | 0.77383 | 2.93725 | 3.07191 | 1.77911 | 4.46192 | 0.00000 | 2.97353 | 0.78432 | 4.27025 | 5.07319 | 1.45060 | 1.05190 |
| JMY | 8.42354 | 5.12751 | 0.37031 | 6.38796 | 0.00000 | 7.69526 | 5.00257 | 1.21824 | 0.00000 | 21.05350 | 0.00001 | 20.78484 |
| JOSD1 | 0.42425 | 1.32434 | 3.67583 | 1.27680 | 0.00000 | 3.06151 | 4.84745 | 0.23622 | 1.63394 | 8.48593 | 1.67349 | 1.91457 |
| JOSD2 | 0.54307 | 0.77880 | 6.78054 | 1.02147 | 3.18267 | 0.00000 | 3.95894 | 0.22291 | 2.02089 | 0.00025 | 2.03542 | 1.55313 |
| JPH1 | 0.39641 | 0.35281 | 0.00000 | 0.29416 | 0.09456 | 0.32860 | 0.02561 | 0.01753 | 0.00000 | 0.31083 | 0.01778 | 0.60709 |
| JPH2 | 0.36249 | 0.00278 | 0.04365 | 0.00390 | 0.00283 | 0.02244 | 0.00012 | 0.06227 | 0.01620 | 0.05275 | 0.25847 | 0.00925 |
| JPH3 | 3.75845 | 0.63318 | 0.11938 | 0.11965 | 0.24087 | 0.00000 | 0.09828 | 0.39416 | 0.04970 | 0.67082 | 0.00691 | 0.28166 |
| JPH4 | 0.00596 | 0.01176 | 0.06065 | 0.04801 | 0.00000 | 0.01739 | 0.01446 | 0.01007 | 0.02119 | 0.06173 | 0.05230 | 0.01444 |
| JPX | 0.00000 | 0.87286 | 0.71209 | 1.76206 | 2.58650 | 2.85601 | 0.91535 | 0.21486 | 1.14053 | 1.52312 | 2.26021 | 1.99882 |
| JRK | 0.00041 | 0.71058 | 0.55082 | 0.01901 | 0.36774 | 0.00000 | 0.42961 | 0.32944 | 0.00000 | 3.99929 | 2.12054 | 1.17394 |
| JRKL | 0.00000 | 1.25957 | 0.87780 | 0.00000 | 0.53845 | 1.13658 | 0.67156 | 1.24708 | 0.90834 | 0.11045 | 0.60568 | 3.67463 |
| JSRP1 | 0.00468 | 0.00988 | 0.06814 | 0.00463 | 0.00000 | 0.00000 | 0.00096 | 0.01120 | 0.01497 | 0.26386 | 0.08180 | 0.01088 |
| JTB | 0.30392 | 21.54743 | 18.77527 | 5.63459 | 23.00177 | 0.00005 | 17.29461 | 0.26641 | 24.98578 | 24.46168 | 15.66054 | 10.03016 |
| JUN | 0.02215 | 0.00000 | 0.35636 | 0.60366 | 1.35687 | 0.00000 | 0.18588 | 0.18823 | 0.00000 | 0.70584 | 0.18186 | 1.12639 |
| JUNB | 1.06342 | 1.31559 | 9.21831 | 1.93192 | 5.27249 | 0.00000 | 0.00051 | 1.10359 | 3.54625 | 5.31913 | 0.76840 | 2.29436 |
| JUND | 0.23844 | 0.08739 | 4.14774 | 0.00000 | 6.74942 | 0.24919 | 5.79452 | 0.21120 | 0.00000 | 19.16584 | 1.21806 | 1.25169 |
| JUP | 2.84864 | 27.56162 | 27.65038 | 11.71576 | 50.50884 | 6.39220 | 26.74921 | 0.00000 | 51.02141 | 3.72425 | 19.02121 | 0.00000 |
| KAAG1 | 0.01900 | 0.01380 | 0.09016 | 0.02229 | 0.01744 | 0.03221 | 0.02277 | 0.03263 | 0.00000 | 0.01105 | 0.05116 | 0.08879 |
| KAL1 | 0.64197 | 0.13510 | 0.12920 | 0.00000 | 0.14004 | 0.00000 | 0.00000 | 0.12130 | 0.19798 | 0.13906 | 0.00000 | 0.07369 |
| KALRN | 12.63519 | 0.00000 | 7.51831 | 2.99672 | 0.27165 | 2.38291 | 0.70753 | 2.08130 | 4.62822 | 0.00026 | 0.31962 | 0.16402 |
| KANK1 | 2.07458 | 11.04768 | 9.80079 | 6.75451 | 5.04968 | 2.77616 | 4.30813 | 0.00000 | 8.47721 | 4.28327 | 7.35155 | 6.35307 |
| KANK2 | 1.43470 | 3.78000 | 0.75910 | 4.46806 | 0.32175 | 1.28137 | 1.95965 | 0.35879 | 0.30447 | 2.59848 | 1.33408 | 2.08787 |
| KANK3 | 0.04543 | 1.03596 | 0.16714 | 0.36426 | 0.07989 | 0.32113 | 0.17351 | 0.09791 | 0.92540 | 0.77135 | 0.00000 | 0.49896 |
| KANK4 | 0.00093 | 0.58668 | 6.32050 | 0.00000 | 5.91688 | 0.00000 | 16.22716 | 0.00000 | 2.45504 | 4.70305 | 2.05586 | 0.11829 |
| KANSL1 | 3.26563 | 0.16686 | 1.22161 | 3.41829 | 1.31568 | 6.07485 | 3.91285 | 1.72632 | 1.15352 | 17.08137 | 0.51560 | 3.38139 |
| KANSL1-AS1 | 0.00641 | 0.09485 | 0.00000 | 0.14189 | 0.00000 | 0.52343 | 0.00000 | 0.00026 | 0.20976 | 0.53397 | 0.39209 | 0.10073 |
| KANSL1L | 0.00809 | 0.15495 | 0.30984 | 0.28906 | 0.00462 | 1.51273 | 0.05003 | 0.39568 | 0.67113 | 2.10148 | 0.45831 | 0.73861 |
| KANSL2 | 3.89704 | 15.23375 | 7.99979 | 11.32465 | 3.28931 | 18.66292 | 8.97217 | 17.92797 | 14.17690 | 0.00005 | 10.66518 | 16.08771 |
| KANSL3 | 2.67032 | 7.38958 | 6.30190 | 7.20932 | 7.95158 | 0.35108 | 9.24381 | 4.52078 | 6.70574 | 7.10376 | 6.39779 | 8.40236 |
| KANTR | 0.01776 | 0.01783 | 1.42371 | 0.20953 | 0.23202 | 0.11465 | 0.00000 | 0.09813 | 0.55916 | 0.10581 | 0.79168 | 0.31082 |
| KARS | 4.91147 | 33.86511 | 26.05125 | 12.82121 | 23.19040 | 36.65776 | 37.17453 | 8.05787 | 35.29513 | 7.54530 | 26.40521 | 33.96619 |
| KAT2A | 1.21521 | 4.64148 | 2.75612 | 4.51379 | 7.15247 | 0.00000 | 8.31796 | 0.00000 | 1.31805 | 11.25747 | 3.35330 | 3.10744 |
| KAT2B | 0.00000 | 0.65206 | 3.46685 | 0.00000 | 0.00000 | 1.79735 | 3.65656 | 0.01957 | 0.00000 | 4.31803 | 0.00000 | 3.86671 |
| KAT5 | 0.73904 | 3.37189 | 2.99938 | 2.70186 | 2.67062 | 0.00000 | 2.29360 | 5.75177 | 1.22395 | 0.00000 | 3.14218 | 2.48941 |
| KAT6A | 3.23201 | 1.11721 | 3.44805 | 3.17358 | 2.85034 | 4.89011 | 4.74986 | 2.15175 | 0.92919 | 12.92897 | 2.32654 | 3.25305 |
| KAT6B | 1.00373 | 2.15752 | 0.49407 | 6.74140 | 5.02200 | 4.38872 | 6.56525 | 3.65304 | 0.71958 | 18.19316 | 3.03086 | 7.29921 |
| KAT7 | 4.51789 | 5.43746 | 4.45141 | 13.57259 | 1.89823 | 8.26242 | 6.80978 | 8.24144 | 2.62763 | 0.00000 | 6.82798 | 10.81707 |
| KAT8 | 1.58709 | 3.04159 | 4.09634 | 1.53688 | 1.88839 | 0.05691 | 3.44727 | 0.32144 | 2.62191 | 3.25077 | 0.67544 | 2.66089 |
| KATNA1 | 1.89358 | 8.75762 | 9.24043 | 3.36686 | 4.07508 | 16.49634 | 12.07137 | 1.70401 | 8.60956 | 0.00000 | 10.23982 | 6.93613 |
| KATNAL1 | 13.87805 | 3.84193 | 0.00000 | 2.50522 | 1.90186 | 2.47140 | 0.14141 | 1.45802 | 0.28414 | 1.63435 | 0.72510 | 3.09368 |

| | | | | | | | | | | | | |
|---|---|---|---|---|---|---|---|---|---|---|---|---|
| **KATNAL2** | 0.01334 | 0.00971 | 0.06656 | 0.13913 | 0.00776 | 0.09604 | 0.00000 | 0.09846 | 0.09989 | 0.26161 | 0.01472 | 0.03035 |
| **KATNB1** | 5.73502 | 6.22726 | 1.77560 | 4.63823 | 4.42523 | 0.00000 | 1.40830 | 1.44837 | 6.25616 | 0.00000 | 4.25404 | 2.13446 |
| **KATNBL1** | 0.21637 | 1.46286 | 14.10855 | 0.00000 | 0.00002 | 6.88469 | 4.66016 | 0.50648 | 1.93413 | 0.77026 | 1.78681 | 0.63764 |
| **KATNBL1P6** | 0.15813 | 0.57392 | 6.54743 | 0.00088 | 0.00000 | 2.74373 | 0.90997 | 0.13190 | 0.87048 | 0.00000 | 0.77146 | 0.25424 |
| **KAZALD1** | 0.39794 | 0.04923 | 0.12546 | 0.01512 | 0.00952 | 0.00883 | 0.00310 | 0.00381 | 0.01900 | 0.01218 | 0.01677 | 0.00917 |
| **KAZN** | 0.00000 | 4.74959 | 1.42365 | 4.01011 | 4.89763 | 0.04369 | 0.75038 | 0.00412 | 0.00000 | 4.77491 | 0.29258 | 0.00000 |
| **KBTBD11** | 0.06706 | 0.04092 | 0.09542 | 0.00000 | 0.00000 | 0.15313 | 0.12161 | 0.01333 | 0.05038 | 0.13293 | 0.00000 | 0.02071 |
| **KBTBD12** | 0.00638 | 0.01417 | 0.02750 | 0.01032 | 0.02149 | 0.01077 | 0.00631 | 0.01034 | 0.04493 | 0.03588 | 0.01713 | 0.01539 |
| **KBTBD13** | 0.02530 | 0.01481 | 0.01204 | 0.01863 | 0.01094 | 0.00512 | 0.05532 | 0.00979 | 0.00151 | 0.04859 | 0.03435 | 0.01456 |
| **KBTBD2** | 2.68257 | 0.20145 | 6.82373 | 6.02018 | 0.00000 | 1.42412 | 5.56271 | 4.63922 | 0.02095 | 0.00000 | 1.10153 | 4.29753 |
| **KBTBD3** | 0.05654 | 0.17229 | 0.67003 | 0.28406 | 0.29329 | 0.28550 | 0.00000 | 0.10088 | 0.82480 | 0.48978 | 0.17685 | 0.36736 |
| **KBTBD4** | 0.00038 | 1.38053 | 6.69719 | 2.12199 | 0.00000 | 0.62362 | 3.52844 | 4.35060 | 4.68824 | 0.00000 | 3.52575 | 2.34076 |
| **KBTBD6** | 1.36068 | 2.04547 | 2.34709 | 0.49768 | 0.00000 | 3.41890 | 7.58986 | 12.30880 | 4.25359 | 1.26893 | 6.28498 | 5.69463 |
| **KBTBD7** | 1.43614 | 1.00954 | 0.31155 | 1.00705 | 0.00000 | 2.63112 | 2.67924 | 2.10070 | 2.89705 | 0.00000 | 2.97716 | 2.45947 |
| **KBTBD8** | 1.38236 | 0.72159 | 3.58151 | 1.43360 | 0.00000 | 5.77969 | 3.38935 | 6.13510 | 2.17323 | 0.22572 | 2.67565 | 1.11716 |
| **KC6** | 0.00456 | 0.06192 | 0.11107 | 0.05650 | 0.00000 | 0.03029 | 0.00000 | 0.00954 | 0.03678 | 0.11536 | 0.00466 | 0.01862 |
| **KCMF1** | 2.79525 | 1.53958 | 7.10192 | 4.47383 | 10.58902 | 6.52849 | 16.24216 | 0.21154 | 2.04923 | 14.72860 | 1.39107 | 0.00000 |
| **KCNA1** | 1.27795 | 0.00854 | 0.04502 | 0.01920 | 0.00457 | 0.04200 | 0.01436 | 0.32388 | 0.00303 | 0.01238 | 0.01740 | 0.01668 |
| **KCNA2** | 0.00197 | 0.01356 | 0.01584 | 0.01138 | 0.01134 | 0.02740 | 0.01636 | 0.06214 | 0.01385 | 0.01635 | 0.01647 | 0.04539 |
| **KCNA3** | 0.01114 | 0.00460 | 0.00000 | 0.06653 | 0.00000 | 0.01462 | 0.00000 | 0.01028 | 0.01285 | 0.31350 | 0.00732 | 0.01024 |
| **KCNA4** | 0.17533 | 0.03941 | 0.18070 | 0.00481 | 0.01037 | 0.01838 | 0.00000 | 0.00935 | 0.01295 | 0.01922 | 0.01509 | 0.01045 |
| **KCNA6** | 0.00497 | 0.35062 | 0.06602 | 0.51458 | 0.06685 | 0.00000 | 0.29785 | 0.00903 | 0.07353 | 0.06752 | 0.02600 | 0.05198 |
| **KCNA7** | 0.07299 | 0.00000 | 0.08337 | 0.01751 | 0.00000 | 0.01289 | 0.25052 | 0.60957 | 0.00000 | 0.68810 | 0.00000 | 0.02311 |
| **KCNAB1** | 0.05999 | 0.00000 | 0.13804 | 0.03812 | 0.00000 | 0.21190 | 0.23535 | 0.00000 | 1.01251 | 1.74944 | 0.04913 | 0.27740 |
| **KCNAB2** | 0.00582 | 0.22676 | 0.04164 | 0.23578 | 0.02340 | 0.00000 | 0.02301 | 0.00826 | 0.12617 | 0.00000 | 0.08748 | 0.07602 |
| **KCNAB3** | 0.23268 | 0.03664 | 0.41509 | 0.20265 | 0.00000 | 0.12420 | 0.00000 | 0.00000 | 0.28053 | 0.21140 | 0.00000 | 0.21225 |
| **KCNB1** | 0.00266 | 0.00296 | 0.00154 | 0.00404 | 0.00188 | 0.02233 | 0.00711 | 0.11706 | 0.01382 | 0.27717 | 0.00757 | 0.07465 |
| **KCNB2** | 0.01055 | 0.07651 | 0.08218 | 0.01674 | 0.04263 | 0.00000 | 0.01605 | 0.01128 | 0.02460 | 0.00759 | 0.01002 | 0.01504 |
| **KCNC1** | 0.00578 | 0.01167 | 0.00742 | 0.02134 | 0.03419 | 0.02377 | 0.02130 | 0.01033 | 0.04394 | 0.03182 | 0.00600 | 0.01455 |
| **KCNC2** | 0.36423 | 0.00709 | 0.00813 | 0.00384 | 0.00596 | 0.04279 | 0.00321 | 0.06056 | 0.01581 | 0.06106 | 0.01688 | 0.02430 |
| **KCNC3** | 0.07968 | 0.00000 | 0.28795 | 0.00000 | 1.10109 | 0.00000 | 0.41489 | 0.16053 | 1.34699 | 2.74680 | 0.30751 | 0.20539 |
| **KCNC4** | 3.54043 | 0.97030 | 0.00000 | 1.92181 | 2.13879 | 0.00000 | 0.03724 | 0.67498 | 0.00000 | 2.32674 | 0.00000 | 3.53227 |
| **KCNC4-AS1** | 0.34369 | 0.10891 | 0.01265 | 0.04387 | 0.01200 | 0.00321 | 0.01410 | 0.11305 | 0.03734 | 0.00000 | 0.01110 | 0.20811 |
| **KCND1** | 0.00295 | 0.47869 | 0.00941 | 0.40597 | 0.00767 | 0.00039 | 0.01941 | 0.00277 | 2.31036 | 0.84090 | 0.00000 | 0.21487 |
| **KCND2** | 0.70863 | 0.01210 | 0.06034 | 0.11133 | 0.01011 | 0.02746 | 0.00000 | 0.00286 | 0.01848 | 0.03329 | 0.01454 | 0.03305 |
| **KCND3** | 0.03247 | 0.00000 | 0.00804 | 0.00972 | 0.02649 | 0.03765 | 0.00947 | 0.03334 | 0.01190 | 0.04458 | 0.03033 | 0.35719 |
| **KCNE1** | 0.17525 | 0.00915 | 0.02184 | 0.01970 | 0.00576 | 0.00000 | 0.02196 | 0.00648 | 0.00000 | 0.04717 | 0.10141 | 0.06618 |
| **KCNE1L** | 0.09140 | 0.15682 | 0.08112 | 0.01003 | 0.07773 | 0.00000 | 0.06100 | 0.02430 | 0.04581 | 0.00000 | 0.03970 | 0.28808 |
| **KCNE2** | 0.34530 | 0.17243 | 0.00000 | 0.00283 | 0.01988 | 0.12741 | 0.00000 | 0.14268 | 0.06951 | 0.00000 | 0.21177 | 0.21864 |
| **KCNE3** | 0.13826 | 0.10711 | 3.87360 | 0.16905 | 0.00000 | 0.04515 | 0.00000 | 0.06001 | 0.66087 | 0.00000 | 0.27910 | 0.01182 |
| **KCNE4** | 0.00785 | 0.00020 | 0.00277 | 0.01046 | 0.00466 | 0.11251 | 0.00000 | 0.01257 | 0.03523 | 0.01967 | 0.04223 | 0.06806 |
| **KCNF1** | 0.01701 | 0.00799 | 0.00878 | 0.09472 | 0.00550 | 0.00050 | 0.02787 | 0.02978 | 0.00912 | 0.04368 | 0.02460 | 0.19586 |
| **KCNG1** | 0.00502 | 0.24143 | 0.00197 | 0.00000 | 0.04713 | 0.03442 | 0.04399 | 0.00610 | 0.00000 | 0.08822 | 0.01056 | 0.11050 |
| **KCNG2** | 0.33321 | 0.02106 | 0.06805 | 0.02903 | 0.00880 | 0.00000 | 0.00537 | 0.04971 | 0.03743 | 0.03340 | 0.03146 | 0.02974 |
| **KCNG3** | 0.02208 | 0.02899 | 0.09783 | 0.35499 | 0.08562 | 0.00000 | 0.01323 | 0.14526 | 0.09605 | 0.29550 | 0.00000 | 0.03924 |
| **KCNG4** | 0.35076 | 0.01454 | 0.01296 | 0.02046 | 0.02230 | 0.01651 | 0.02716 | 0.00492 | 0.01532 | 0.00180 | 0.01558 | 0.01262 |
| **KCNH1** | 1.32138 | 0.17655 | 0.00000 | 0.00000 | 0.35376 | 0.00000 | 0.00069 | 0.26320 | 0.00764 | 1.00660 | 0.40505 | 0.00000 |
| **KCNH2** | 0.15893 | 0.01193 | 0.00000 | 0.80591 | 0.44934 | 0.19377 | 0.00000 | 0.00399 | 0.05562 | 0.50943 | 0.03089 | 0.00000 |

| | | | | | | | | | | | | |
|---|---|---|---|---|---|---|---|---|---|---|---|---|
| **KCNH3** | 0.55082 | 0.01712 | 0.00000 | 0.01192 | 0.00000 | 0.03407 | 0.00416 | 0.10248 | 0.00000 | 0.53702 | 0.03121 | 0.03022 |
| **KCNH4** | 0.01028 | 0.00000 | 0.08629 | 0.03030 | 0.00000 | 0.02904 | 0.00739 | 0.00762 | 0.00000 | 0.11282 | 0.05728 | 0.15288 |
| **KCNH5** | 0.00437 | 0.21694 | 0.40225 | 0.13068 | 0.02542 | 0.38327 | 0.00000 | 1.87082 | 1.12151 | 0.00000 | 0.28512 | 1.96305 |
| **KCNH6** | 0.02395 | 0.01963 | 0.02654 | 0.29074 | 0.01173 | 0.00000 | 0.01733 | 0.01424 | 0.00695 | 0.02636 | 0.02904 | 0.01695 |
| **KCNH7** | 0.66048 | 0.02100 | 0.02248 | 0.03016 | 0.02530 | 0.00000 | 0.03803 | 0.01528 | 0.02430 | 0.00000 | 0.06319 | 0.04104 |
| **KCNH8** | 1.37615 | 0.00000 | 0.03464 | 0.11939 | 0.03329 | 0.00000 | 0.03797 | 0.10832 | 0.00000 | 0.61953 | 0.01298 | 0.16596 |
| **KCNIP2** | 0.00600 | 0.22328 | 0.31140 | 0.22159 | 0.00000 | 0.04171 | 0.05090 | 0.34201 | 0.00000 | 0.07970 | 0.37433 | 0.21559 |
| **KCNIP2-AS1** | 0.51801 | 0.00000 | 0.07518 | 0.00196 | 0.00501 | 0.00000 | 0.00000 | 0.98335 | 0.07067 | 0.49473 | 0.05802 | 0.21238 |
| **KCNIP3** | 0.00938 | 0.00872 | 0.02128 | 0.00498 | 0.03353 | 0.03258 | 0.02197 | 0.00969 | 0.00000 | 0.06097 | 0.01139 | 0.01326 |
| **KCNIP4** | 0.79061 | 0.14346 | 0.05592 | 0.02490 | 0.00000 | 0.11739 | 0.26292 | 0.00000 | 0.08559 | 0.12506 | 0.04264 | 0.15159 |
| **KCNIP4-IT1** | 0.00631 | 0.01538 | 0.01577 | 0.01548 | 0.01717 | 0.01458 | 0.02045 | 0.01054 | 0.01882 | 0.01240 | 0.01482 | 0.01668 |
| **KCNJ1** | 0.00627 | 0.01372 | 0.05119 | 0.01388 | 0.01402 | 0.02382 | 0.00744 | 0.01050 | 0.02869 | 0.01332 | 0.01363 | 0.01606 |
| **KCNJ10** | 0.00574 | 0.00714 | 0.02800 | 0.01178 | 0.00831 | 0.01586 | 0.01426 | 0.00683 | 0.00563 | 0.04863 | 0.02737 | 0.10864 |
| **KCNJ11** | 0.00750 | 0.12040 | 0.05945 | 0.00000 | 0.00588 | 0.05209 | 0.02532 | 0.00761 | 0.09866 | 0.03682 | 0.00598 | 0.03728 |
| **KCNJ12** | 0.00592 | 0.01944 | 0.02557 | 0.00997 | 0.01218 | 0.02469 | 0.01731 | 0.01085 | 0.00912 | 0.03169 | 0.04024 | 0.02173 |
| **KCNJ13** | 0.00161 | 0.00000 | 1.41175 | 1.12109 | 0.00000 | 0.23368 | 0.33480 | 0.15024 | 1.08825 | 1.87112 | 2.50459 | 0.14319 |
| **KCNJ14** | 0.09335 | 0.00000 | 0.11865 | 0.10960 | 0.24600 | 0.00000 | 0.17286 | 0.00000 | 1.03532 | 2.58444 | 0.00000 | 0.31608 |
| **KCNJ15** | 0.02930 | 0.17661 | 0.04359 | 0.00000 | 0.12047 | 0.02141 | 0.14686 | 0.01030 | 0.00000 | 0.03300 | 0.03362 | 0.00551 |
| **KCNJ16** | 0.08790 | 0.00351 | 0.28797 | 1.53772 | 0.00000 | 0.03812 | 0.01293 | 0.12303 | 0.00000 | 0.04119 | 0.00967 | 0.00736 |
| **KCNJ2** | 0.44579 | 0.00000 | 0.13625 | 0.01657 | 0.00000 | 0.11771 | 0.04219 | 0.00000 | 0.00000 | 0.29324 | 0.02064 | 0.01831 |
| **KCNJ3** | 0.02060 | 0.09872 | 0.00000 | 0.00000 | 0.00000 | 0.31636 | 1.26019 | 0.23465 | 0.00000 | 1.34055 | 0.00000 | 0.26405 |
| **KCNJ4** | 0.15770 | 0.01160 | 0.01742 | 0.02039 | 0.01502 | 0.00000 | 0.02720 | 0.00580 | 0.00000 | 0.03940 | 0.04490 | 0.03640 |
| **KCNJ5** | 0.00766 | 0.01485 | 0.02395 | 0.01148 | 0.01249 | 0.01651 | 0.04892 | 0.01003 | 0.00518 | 0.01816 | 0.02272 | 0.01606 |
| **KCNJ6** | 0.65874 | 0.11068 | 0.14203 | 0.01093 | 0.01077 | 0.01072 | 0.10190 | 0.05107 | 0.00010 | 0.00713 | 0.01554 | 0.02190 |
| **KCNJ8** | 0.27768 | 0.00977 | 0.02658 | 0.26568 | 0.02966 | 0.06256 | 0.07987 | 0.16717 | 0.03384 | 0.00000 | 0.00826 | 0.02198 |
| **KCNJ9** | 0.00629 | 0.01405 | 0.01364 | 0.01104 | 0.01610 | 0.02002 | 0.01329 | 0.01048 | 0.04592 | 0.01780 | 0.01398 | 0.01581 |
| **KCNK1** | 0.00000 | 0.00000 | 0.43341 | 0.60484 | 10.42355 | 13.93700 | 3.35988 | 0.53467 | 5.46706 | 10.95537 | 1.44223 | 3.06697 |
| **KCNK10** | 0.09688 | 0.01829 | 0.02109 | 0.01029 | 0.00032 | 0.06510 | 0.01138 | 0.11238 | 0.00419 | 0.02648 | 0.01830 | 0.11501 |
| **KCNK12** | 0.47686 | 0.05247 | 0.21725 | 0.45497 | 0.06779 | 0.00000 | 0.00000 | 0.05955 | 0.02939 | 0.03075 | 0.01276 | 0.01753 |
| **KCNK13** | 0.10485 | 0.08820 | 0.02129 | 0.56621 | 0.17740 | 0.00000 | 0.02397 | 0.05952 | 0.01122 | 0.00000 | 0.03405 | 0.06413 |
| **KCNK15** | 0.15669 | 0.02257 | 0.04173 | 0.01817 | 0.02875 | 0.00000 | 0.02021 | 0.01481 | 0.01571 | 0.00000 | 0.01596 | 0.03946 |
| **KCNK16** | 0.85713 | 0.02154 | 0.01853 | 0.01146 | 0.02990 | 0.00000 | 0.01944 | 0.07167 | 0.02967 | 0.00475 | 0.01145 | 0.01795 |
| **KCNK17** | 0.16384 | 0.01503 | 0.01404 | 0.01241 | 0.01804 | 0.00953 | 0.01570 | 0.05124 | 0.02492 | 0.01418 | 0.01604 | 0.01374 |
| **KCNK18** | 0.98063 | 0.01526 | 0.01343 | 0.01485 | 0.01695 | 0.01147 | 0.01817 | 0.00270 | 0.02396 | 0.00316 | 0.01282 | 0.01192 |
| **KCNK2** | 0.01544 | 0.07670 | 0.04487 | 0.04803 | 0.18472 | 0.07165 | 0.05844 | 0.05049 | 0.03395 | 0.07874 | 0.05639 | 0.00281 |
| **KCNK3** | 0.00561 | 0.08271 | 0.01322 | 0.00156 | 0.00000 | 0.02422 | 0.01137 | 0.01133 | 0.02019 | 0.10618 | 0.00832 | 0.00761 |
| **KCNK4** | 0.00533 | 0.02849 | 0.03656 | 0.01258 | 0.09428 | 0.00000 | 0.01001 | 0.00995 | 0.04801 | 0.01312 | 0.01077 | 0.16444 |
| **KCNK5** | 3.79491 | 0.63242 | 0.80109 | 2.98131 | 0.00000 | 0.00000 | 0.20288 | 2.35315 | 1.17657 | 1.25851 | 0.00000 | 7.46991 |
| **KCNK6** | 0.00100 | 4.48140 | 0.61440 | 2.89485 | 1.09259 | 0.00000 | 0.06229 | 0.00000 | 1.00242 | 7.41912 | 0.50531 | 1.30173 |
| **KCNK7** | 0.03008 | 0.02853 | 0.06816 | 0.00715 | 0.01161 | 0.01669 | 0.01894 | 0.00919 | 0.11973 | 0.00000 | 0.00244 | 0.03357 |
| **KCNK9** | 11.37990 | 0.04545 | 0.03939 | 0.01453 | 0.03868 | 0.00000 | 0.19065 | 2.20458 | 0.00000 | 0.00000 | 0.02790 | 0.16961 |
| **KCNMA1** | 0.09894 | 0.01417 | 0.12623 | 0.36105 | 0.00000 | 0.05575 | 0.03957 | 0.13080 | 0.00000 | 0.11438 | 0.01712 | 0.04945 |
| **KCNMB2** | 0.13385 | 0.00000 | 0.00000 | 0.08926 | 0.12701 | 0.02240 | 0.00000 | 0.00626 | 0.11811 | 0.21994 | 0.07082 | 0.04619 |
| **KCNMB3** | 0.00127 | 0.01232 | 0.14576 | 0.25446 | 0.31418 | 0.15659 | 0.20334 | 0.08038 | 0.04994 | 0.88081 | 0.09339 | 0.44398 |
| **KCNMB4** | 0.00000 | 0.64082 | 0.00000 | 2.81140 | 0.56211 | 1.15592 | 0.89863 | 1.20933 | 0.00000 | 3.72412 | 0.88965 | 1.98677 |
| **KCNN1** | 0.10184 | 1.02465 | 0.00376 | 0.58604 | 0.83426 | 0.00000 | 0.00065 | 0.01279 | 0.00000 | 0.43106 | 0.10299 | 0.00758 |
| **KCNN2** | 2.51522 | 0.55026 | 0.00000 | 0.53039 | 0.00000 | 0.07498 | 0.00000 | 0.13019 | 0.33893 | 0.43990 | 0.00000 | 0.77659 |
| **KCNN3** | 0.27845 | 0.02280 | 0.02980 | 0.34343 | 0.07855 | 0.08017 | 0.01177 | 0.01342 | 0.03962 | 0.00000 | 0.05474 | 0.03161 |

| | | | | | | | | | | | | |
|---|---|---|---|---|---|---|---|---|---|---|---|---|
| **KCNN4** | 0.01001 | 0.00000 | 10.21320 | 0.00000 | 0.00000 | 0.00000 | 1.17322 | 0.00681 | 6.17501 | 2.38927 | 3.71673 | 0.00000 |
| **KCNQ1** | 0.06859 | 0.01319 | 0.03997 | 0.03450 | 0.01233 | 0.01207 | 0.01672 | 0.03903 | 0.00637 | 0.02721 | 0.02934 | 0.02530 |
| **KCNQ1OT1** | 0.08869 | 0.00000 | 3.62769 | 2.65386 | 4.08081 | 1.55328 | 0.16785 | 1.35766 | 4.40108 | 10.49681 | 0.33101 | 0.17080 |
| **KCNQ2** | 0.22569 | 0.55877 | 0.07480 | 0.23572 | 0.17783 | 0.06763 | 0.00000 | 0.09605 | 0.00000 | 0.32823 | 0.00000 | 0.13171 |
| **KCNQ3** | 0.19992 | 0.02803 | 0.02182 | 0.01265 | 0.04303 | 0.00912 | 0.01882 | 0.03981 | 0.00588 | 0.05210 | 0.03001 | 0.01281 |
| **KCNQ4** | 0.01782 | 1.44753 | 0.05650 | 0.00000 | 0.13768 | 0.32396 | 0.12978 | 0.00000 | 1.84348 | 1.89118 | 0.79357 | 0.33230 |
| **KCNQ5** | 0.05442 | 0.00000 | 0.00000 | 0.00000 | 0.00000 | 0.32477 | 0.00000 | 0.00000 | 0.34592 | 0.69654 | 0.09011 | 0.36678 |
| **KCNQ5-AS1** | 0.00607 | 0.01099 | 0.01111 | 0.03853 | 0.00420 | 0.03072 | 0.00639 | 0.01020 | 0.04471 | 0.02714 | 0.01370 | 0.01464 |
| **KCNQ5-IT1** | 0.01765 | 0.04349 | 0.00528 | 0.00336 | 0.01067 | 0.10014 | 0.02735 | 0.00369 | 0.36954 | 0.00000 | 0.10687 | 0.21100 |
| **KCNRG** | 0.00204 | 0.00000 | 0.19172 | 0.49450 | 0.54029 | 0.02273 | 0.50540 | 0.22462 | 0.00000 | 1.44264 | 0.40088 | 0.71245 |
| **KCNS1** | 0.08866 | 0.02448 | 0.13954 | 0.04615 | 0.14137 | 0.00000 | 0.00000 | 0.00000 | 0.54613 | 0.07783 | 0.00000 | 0.36607 |
| **KCNS3** | 0.43360 | 0.52716 | 2.09920 | 2.38027 | 0.00000 | 0.67795 | 0.00000 | 0.07333 | 0.37933 | 0.00000 | 0.00000 | 0.12580 |
| **KCNT1** | 0.32118 | 0.00540 | 0.02399 | 0.01448 | 0.07467 | 0.00759 | 0.02974 | 0.07100 | 0.00000 | 0.04021 | 0.01231 | 0.09899 |
| **KCNT2** | 0.00219 | 0.00910 | 0.00000 | 0.08248 | 0.00000 | 0.03340 | 0.00000 | 0.02546 | 0.02013 | 0.20865 | 0.00288 | 0.05404 |
| **KCNU1** | 0.00605 | 0.01362 | 0.03249 | 0.01399 | 0.01176 | 0.02170 | 0.01819 | 0.01037 | 0.01028 | 0.02504 | 0.01564 | 0.01574 |
| **KCNV1** | 0.01097 | 0.19074 | 0.00542 | 0.00118 | 0.00000 | 0.00475 | 0.91427 | 0.00776 | 4.76003 | 0.00000 | 0.00000 | 0.09389 |
| **KCNV2** | 0.21700 | 0.71291 | 0.22719 | 1.32349 | 0.46300 | 0.00000 | 0.00049 | 0.08465 | 1.10579 | 0.00000 | 0.08585 | 2.29071 |
| **KCP** | 0.22279 | 0.04606 | 0.01688 | 0.18673 | 0.19838 | 0.05060 | 0.00000 | 0.05094 | 0.01056 | 0.32219 | 0.00000 | 0.20481 |
| **KCTD1** | 0.05497 | 1.20985 | 0.17945 | 0.61939 | 3.95637 | 2.33970 | 2.01355 | 0.05607 | 0.30253 | 4.69366 | 1.35576 | 0.20520 |
| **KCTD10** | 8.62004 | 9.26821 | 13.90677 | 6.09705 | 2.15535 | 12.99539 | 14.13155 | 6.87263 | 13.86454 | 0.00000 | 9.24346 | 12.89185 |
| **KCTD11** | 0.02667 | 0.09397 | 0.81942 | 0.03347 | 0.25422 | 0.00044 | 0.04407 | 0.04192 | 0.00002 | 0.00000 | 1.34056 | 0.34971 |
| **KCTD12** | 0.09121 | 0.00000 | 0.93940 | 0.00000 | 0.48056 | 1.96460 | 7.11667 | 0.03460 | 2.09217 | 2.79116 | 0.00000 | 0.10810 |
| **KCTD13** | 3.49667 | 2.60106 | 5.01032 | 0.17159 | 0.61964 | 0.00000 | 0.00010 | 0.86325 | 3.85858 | 4.86109 | 1.45048 | 0.74242 |
| **KCTD14.uid-159** | 0.02127 | 0.01014 | 0.00776 | 0.01471 | 0.03941 | 0.04859 | 0.01361 | 0.01687 | 0.04267 | 0.00000 | 0.01125 | 0.02465 |
| **KCTD15** | 0.26993 | 5.42957 | 0.00000 | 3.54789 | 9.93248 | 0.00000 | 6.24910 | 0.00000 | 4.26228 | 7.10735 | 1.70872 | 3.51678 |
| **KCTD16** | 0.10826 | 0.00030 | 0.02296 | 0.01420 | 0.00000 | 0.04875 | 0.00341 | 0.07553 | 0.00000 | 0.66834 | 0.00000 | 0.00215 |
| **KCTD17** | 0.74875 | 0.17016 | 0.07573 | 0.05296 | 0.00000 | 0.04537 | 0.00000 | 0.05542 | 0.00000 | 0.36502 | 0.02371 | 0.14154 |
| **KCTD18** | 0.01669 | 0.87097 | 0.23510 | 0.28458 | 0.00000 | 1.42528 | 0.00000 | 0.27871 | 0.15528 | 0.00000 | 0.29150 | 0.38812 |
| **KCTD19** | 0.01361 | 0.05544 | 0.01837 | 0.01453 | 0.00773 | 0.00574 | 0.01803 | 0.00577 | 0.01992 | 0.01415 | 0.03502 | 0.20364 |
| **KCTD2** | 3.25835 | 1.76168 | 2.02941 | 3.63209 | 3.20230 | 3.12487 | 7.33761 | 1.15744 | 0.01403 | 14.10942 | 0.57158 | 4.21001 |
| **KCTD20** | 2.76553 | 1.55961 | 7.59162 | 0.00000 | 2.64331 | 4.71320 | 14.31846 | 1.16830 | 5.65561 | 0.00000 | 5.78886 | 2.42919 |
| **KCTD21** | 2.62641 | 0.34990 | 0.38410 | 0.28861 | 0.14634 | 1.09348 | 0.16712 | 0.14155 | 0.67826 | 0.17607 | 0.24390 | 0.16674 |
| **KCTD21-AS1** | 0.01169 | 0.09356 | 0.29560 | 0.08164 | 0.00146 | 0.03158 | 0.00000 | 0.03050 | 0.12861 | 0.01413 | 0.20276 | 0.18368 |
| **KCTD3** | 0.54779 | 0.77127 | 0.00000 | 0.00000 | 3.08288 | 13.44026 | 2.76689 | 0.96006 | 0.00000 | 14.87028 | 1.59506 | 2.45419 |
| **KCTD4** | 0.18819 | 0.10279 | 0.02676 | 0.01729 | 0.00770 | 0.00022 | 0.00235 | 0.00858 | 0.06421 | 0.05767 | 0.03281 | 0.01480 |
| **KCTD5** | 4.93048 | 5.00023 | 8.49065 | 3.25796 | 2.83049 | 0.72438 | 4.40494 | 4.99091 | 5.23378 | 8.12822 | 1.11972 | 3.85756 |
| **KCTD6** | 0.02035 | 3.10957 | 0.00000 | 3.39910 | 0.52436 | 6.86766 | 0.00000 | 2.19763 | 0.00000 | 0.73524 | 1.88614 | 5.70679 |
| **KCTD7** | 0.99979 | 1.79962 | 0.34315 | 3.04257 | 0.00000 | 0.00000 | 0.04135 | 0.04659 | 0.89883 | 0.52807 | 0.31631 | 1.68515 |
| **KCTD8** | 0.01506 | 0.06136 | 0.11428 | 0.02186 | 0.06890 | 0.00000 | 0.00312 | 0.00937 | 0.00000 | 0.15153 | 0.00587 | 0.01031 |
| **KCTD9** | 0.13828 | 1.17233 | 2.47609 | 0.30346 | 0.02478 | 5.99356 | 6.67346 | 4.22060 | 0.00006 | 3.67470 | 1.28032 | 0.93333 |
| **KDELC1** | 0.23267 | 0.17529 | 1.46659 | 1.51248 | 0.00000 | 2.09029 | 0.00000 | 1.25101 | 1.48339 | 0.18254 | 1.33671 | 0.05977 |
| **KDELC2** | 0.00000 | 0.00000 | 1.96839 | 0.00000 | 0.66314 | 2.78556 | 0.00000 | 0.10451 | 3.72506 | 2.02944 | 1.74608 | 0.48354 |
| **KDELR1** | 1.75591 | 7.30994 | 11.02891 | 10.89442 | 14.87385 | 5.52040 | 0.62800 | 1.17275 | 21.34442 | 10.43940 | 4.18666 | 4.62253 |
| **KDELR2** | 7.47972 | 14.99831 | 18.91058 | 9.85540 | 10.42415 | 40.00199 | 7.11985 | 3.72181 | 40.18140 | 9.77016 | 17.29866 | 14.54117 |
| **KDELR3** | 0.44080 | 0.20631 | 0.23274 | 0.00259 | 0.00000 | 0.19961 | 0.00000 | 0.01561 | 0.00000 | 0.27816 | 0.20447 | 0.19162 |
| **KDF1** | 1.83369 | 5.97589 | 3.26642 | 5.56275 | 5.47412 | 1.01784 | 2.56605 | 0.00000 | 6.60567 | 0.00000 | 5.42088 | 5.23069 |
| **KDM1A** | 7.21319 | 4.49879 | 15.79876 | 5.21284 | 7.20117 | 14.11343 | 23.63878 | 5.02465 | 3.32973 | 21.96301 | 2.55864 | 3.85094 |
| **KDM1B** | 11.38300 | 0.94970 | 0.70068 | 0.00000 | 0.40416 | 1.80913 | 0.56695 | 0.51810 | 0.00347 | 1.84697 | 0.32058 | 1.63865 |

| | | | | | | | | | | | | |
|---|---|---|---|---|---|---|---|---|---|---|---|---|
| **KDM2A** | 2.60541 | 3.14681 | 8.11353 | 7.10205 | 7.60181 | 1.50518 | 10.88349 | 2.55996 | 7.72884 | 16.91820 | 6.38025 | 4.00395 |
| **KDM2B** | 1.47716 | 2.77860 | 4.56918 | 3.55006 | 3.72358 | 5.87123 | 7.44800 | 2.50178 | 2.52744 | 13.63078 | 3.51301 | 2.85519 |
| **KDM3A** | 9.12158 | 3.31573 | 2.25781 | 3.18512 | 0.33305 | 8.13419 | 2.36066 | 7.57477 | 2.15820 | 4.30557 | 0.00000 | 7.76862 |
| **KDM3B** | 2.23391 | 2.49803 | 2.61288 | 5.12250 | 5.54464 | 8.60082 | 7.57453 | 0.48820 | 2.64653 | 8.82570 | 1.57048 | 1.95378 |
| **KDM4A** | 4.79306 | 1.75144 | 3.47894 | 5.85505 | 2.15917 | 2.52599 | 7.94498 | 10.61255 | 3.84621 | 0.00000 | 3.35019 | 9.60387 |
| **KDM4A-AS1** | 0.10927 | 0.11920 | 0.33306 | 0.11858 | 0.02123 | 0.00000 | 0.00000 | 0.28724 | 0.71054 | 0.26760 | 0.18772 | 0.41032 |
| **KDM4B** | 0.18043 | 0.00000 | 0.45965 | 0.26201 | 2.28069 | 0.00000 | 2.42060 | 0.00384 | 0.22607 | 3.56383 | 0.51621 | 0.04431 |
| **KDM4C** | 1.18209 | 5.28352 | 2.71288 | 7.37220 | 1.11549 | 8.48274 | 0.00000 | 5.43814 | 4.47977 | 6.81595 | 2.16656 | 3.43979 |
| **KDM4D** | 2.35616 | 0.63420 | 0.40061 | 0.29033 | 0.00000 | 0.00000 | 0.00000 | 1.03594 | 1.25118 | 0.16887 | 0.48560 | 3.23302 |
| **KDM4E** | 2.75280 | 0.60318 | 0.01665 | 0.43620 | 0.00210 | 0.00000 | 0.00368 | 3.85928 | 0.03348 | 0.00213 | 0.02090 | 2.62629 |
| **KDM5A** | 9.39695 | 5.41179 | 5.61724 | 7.66564 | 10.71404 | 10.70092 | 10.68536 | 11.41596 | 6.70795 | 22.58737 | 4.92511 | 9.29920 |
| **KDM5B** | 1.99182 | 20.04488 | 6.82156 | 7.76000 | 17.96644 | 30.28300 | 19.07076 | 34.14079 | 19.07357 | 32.50213 | 14.40645 | 22.32053 |
| **KDM5C** | 3.04478 | 0.89694 | 3.70378 | 2.32430 | 2.49750 | 0.00000 | 0.29188 | 2.13472 | 4.41867 | 7.39253 | 0.46787 | 1.94549 |
| **KDM5D** | 0.00000 | 6.58952 | 1.24729 | 3.11865 | 4.49210 | 6.91475 | 1.56060 | 0.00000 | 2.51791 | 4.03523 | 1.06227 | 6.29580 |
| **KDM6A** | 1.50173 | 3.03623 | 3.30270 | 3.00332 | 0.80084 | 8.21518 | 0.00000 | 1.15819 | 3.17811 | 5.55213 | 1.36056 | 2.77489 |
| **KDM6B** | 6.02559 | 0.91230 | 7.95100 | 5.47285 | 3.98230 | 0.00000 | 5.46235 | 0.33809 | 3.37774 | 11.59717 | 0.00000 | 0.30913 |
| **KDM7A** | 8.44985 | 1.80684 | 2.00658 | 6.18966 | 0.01237 | 6.76859 | 0.27854 | 2.44583 | 0.00000 | 7.53063 | 0.00000 | 6.55590 |
| **KDM8** | 0.05164 | 1.35055 | 0.50377 | 0.30448 | 0.00000 | 0.00000 | 1.16622 | 0.20016 | 0.47219 | 0.00000 | 1.08172 | 1.18256 |
| **KDR** | 0.00960 | 0.00511 | 0.10221 | 0.00791 | 0.14680 | 0.01442 | 0.03888 | 0.00965 | 0.02983 | 0.05016 | 0.01427 | 0.06450 |
| **KDSR** | 0.34650 | 7.74758 | 9.31723 | 0.76234 | 2.84677 | 14.35086 | 0.00000 | 0.12517 | 17.25960 | 7.39470 | 5.06864 | 3.37917 |
| **KEAP1** | 1.04833 | 11.00472 | 13.52708 | 10.01428 | 8.08121 | 0.12191 | 18.89447 | 2.34553 | 16.41777 | 0.00000 | 12.36731 | 6.19687 |
| **KEL** | 0.00668 | 0.02020 | 0.05325 | 0.07079 | 0.01689 | 0.00000 | 0.00827 | 0.01053 | 0.06096 | 0.00282 | 0.01031 | 0.01776 |
| **KGFLP2** | 0.00710 | 0.02321 | 0.01464 | 0.01029 | 0.01215 | 0.03345 | 0.01249 | 0.00998 | 0.03538 | 0.01214 | 0.01430 | 0.03517 |
| **KHDC1** | 2.62208 | 2.65375 | 3.90389 | 0.76334 | 2.67953 | 2.23884 | 1.00150 | 17.02704 | 7.29028 | 0.64329 | 0.17901 | 4.31030 |
| **KHDC1L** | 9.80883 | 63.78550 | 12.37323 | 64.51263 | 78.15628 | 48.76801 | 35.13168 | 59.52585 | 31.42811 | 30.20544 | 47.87274 | 73.01000 |
| **KHDC3L** | 27.90375 | 0.08481 | 2.85211 | 5.85427 | 0.00246 | 0.00000 | 0.00000 | 22.54503 | 0.12222 | 0.00000 | 0.00000 | 3.73297 |
| **KHDRBS1** | 0.71588 | 2.35844 | 15.23439 | 7.35876 | 11.36866 | 26.09457 | 29.49192 | 4.47410 | 1.13332 | 34.07911 | 13.22342 | 6.98708 |
| **KHDRBS2** | 0.01703 | 0.01179 | 0.01466 | 0.00821 | 0.03084 | 0.02952 | 0.01773 | 0.01844 | 0.01546 | 0.01431 | 0.02041 | 0.01844 |
| **KHDRBS3** | 0.99878 | 0.00000 | 6.08201 | 0.00000 | 17.18507 | 5.66061 | 16.98824 | 0.44000 | 0.00000 | 22.87114 | 6.66647 | 0.24835 |
| **KHK** | 2.18230 | 0.47576 | 1.26001 | 0.38015 | 0.00000 | 0.00000 | 1.13236 | 0.72924 | 0.78969 | 0.00003 | 3.91547 | 0.36398 |
| **KHNYN** | 0.08773 | 0.15182 | 1.52335 | 1.76180 | 1.06468 | 1.91745 | 1.80076 | 0.14172 | 0.79192 | 2.49496 | 0.99266 | 0.23555 |
| **KHSRP** | 2.63143 | 2.57548 | 5.86108 | 5.94935 | 14.99464 | 0.00000 | 23.58709 | 2.72311 | 0.00001 | 45.23074 | 5.63676 | 6.67286 |
| **KIAA0020** | 1.97545 | 12.02119 | 15.68582 | 7.17113 | 10.84435 | 30.43414 | 21.06752 | 19.34410 | 20.87802 | 20.04563 | 18.13454 | 25.91060 |
| **KIAA0040** | 1.11663 | 2.35825 | 0.70359 | 0.00000 | 3.44075 | 1.93048 | 0.00000 | 2.18023 | 1.01728 | 2.25304 | 1.61826 | 3.47233 |
| **KIAA0087** | 0.01001 | 0.01456 | 0.01169 | 0.01127 | 0.01305 | 0.01756 | 0.02235 | 0.01009 | 0.02489 | 0.02565 | 0.01225 | 0.01562 |
| **KIAA0100** | 3.50459 | 5.11213 | 9.65119 | 9.10622 | 11.08921 | 2.01851 | 10.53790 | 1.91781 | 8.47262 | 26.88193 | 5.61550 | 8.26018 |
| **KIAA0101** | 0.34174 | 0.00000 | 15.20186 | 0.00000 | 0.35583 | 8.42795 | 25.80123 | 0.25162 | 14.59742 | 1.79084 | 16.78172 | 0.00000 |
| **KIAA0125** | 0.06346 | 0.01995 | 0.07291 | 0.01394 | 0.02542 | 0.00243 | 0.02196 | 0.00733 | 0.00366 | 0.03559 | 0.01573 | 0.01634 |
| **KIAA0141** | 0.33124 | 2.29999 | 3.25209 | 4.38035 | 1.48414 | 2.24740 | 4.68736 | 0.33885 | 5.50166 | 1.00205 | 3.55761 | 0.98759 |
| **KIAA0195** | 1.55936 | 1.74261 | 4.42244 | 1.81043 | 3.67644 | 0.00000 | 0.10080 | 0.03447 | 6.67083 | 9.98312 | 1.49362 | 1.90599 |
| **KIAA0196** | 1.02964 | 4.94183 | 3.13330 | 3.10672 | 1.96406 | 16.99804 | 0.00000 | 2.74467 | 12.36826 | 4.75415 | 9.57419 | 3.02700 |
| **KIAA0226** | 2.33825 | 1.94734 | 2.26260 | 4.95275 | 1.63302 | 2.94344 | 3.67848 | 5.36377 | 0.53164 | 4.40862 | 3.17441 | 2.50277 |
| **KIAA0226L** | 0.02100 | 0.53546 | 0.03691 | 0.07900 | 0.00687 | 0.51792 | 0.12792 | 0.56586 | 0.00000 | 0.00000 | 0.20202 | 0.73384 |
| **KIAA0232** | 1.72427 | 2.21477 | 0.00000 | 2.29092 | 1.96378 | 15.13651 | 1.22781 | 0.33751 | 4.13820 | 12.54328 | 2.51491 | 1.51620 |
| **KIAA0247** | 13.56233 | 2.89091 | 0.82981 | 1.48095 | 1.93532 | 3.35401 | 0.00000 | 3.43387 | 2.54726 | 1.93223 | 0.65015 | 2.23473 |
| **KIAA0319** | 2.44838 | 0.06576 | 0.58924 | 0.00705 | 0.00611 | 0.00000 | 0.00000 | 0.53098 | 0.05993 | 0.01271 | 0.05731 | 1.04699 |
| **KIAA0319L** | 0.76178 | 2.77007 | 3.25831 | 4.22963 | 6.38229 | 0.00000 | 0.00000 | 0.26025 | 6.58579 | 5.08248 | 3.80098 | 1.45981 |
| **KIAA0355** | 3.08211 | 1.08290 | 3.36331 | 0.87537 | 0.74289 | 0.96895 | 1.50547 | 0.39057 | 2.49928 | 2.28742 | 1.66021 | 2.43194 |

| | | | | | | | | | | | | |
|---|---|---|---|---|---|---|---|---|---|---|---|---|
| **KIAA0368** | 1.35751 | 1.95023 | 8.64743 | 8.70884 | 8.20471 | 12.68020 | 14.34084 | 0.56481 | 7.13631 | 11.36134 | 8.64590 | 0.73495 |
| **KIAA0391** | 0.26603 | 7.73141 | 5.10375 | 4.38811 | 4.79741 | 16.18624 | 2.42144 | 2.41940 | 5.60521 | 4.28065 | 8.54066 | 14.15691 |
| **KIAA0430** | 3.06154 | 1.07036 | 4.16710 | 0.11993 | 1.26298 | 6.47359 | 3.75920 | 3.09929 | 0.99081 | 6.34979 | 1.06088 | 3.21330 |
| **KIAA0513** | 1.19319 | 0.98574 | 0.38730 | 0.47773 | 0.00496 | 0.11305 | 0.42077 | 0.37363 | 0.00000 | 0.00000 | 0.41597 | 0.71528 |
| **KIAA0556** | 1.02156 | 1.88566 | 1.81582 | 0.92410 | 1.15036 | 0.88082 | 0.88601 | 0.73563 | 1.94700 | 0.68590 | 0.55265 | 2.29955 |
| **KIAA0586** | 1.81159 | 0.18886 | 1.26797 | 1.45240 | 1.01656 | 6.17466 | 1.14875 | 1.05401 | 2.48778 | 4.77715 | 2.59322 | 2.48047 |
| **KIAA0753** | 3.36164 | 1.81490 | 0.80205 | 1.25389 | 0.38291 | 1.01272 | 2.05560 | 0.06453 | 2.81206 | 0.55327 | 1.51622 | 2.77409 |
| **KIAA0754** | 0.07857 | 0.00000 | 0.00000 | 0.46332 | 0.21864 | 0.00000 | 0.66930 | 0.02426 | 0.15435 | 1.30862 | 0.00000 | 0.20767 |
| **KIAA0825** | 1.35221 | 0.04134 | 0.18120 | 0.00000 | 0.00000 | 0.49147 | 0.61645 | 0.47279 | 0.00000 | 1.17631 | 0.01974 | 0.24027 |
| **KIAA0895** | 0.20257 | 0.03417 | 0.68054 | 0.01354 | 0.00167 | 0.00000 | 0.54417 | 0.00000 | 0.77326 | 0.11829 | 0.58405 | 0.06431 |
| **KIAA0895L** | 0.14005 | 0.49498 | 0.62386 | 0.22162 | 0.21631 | 0.44768 | 0.05276 | 0.00000 | 0.53644 | 0.72994 | 0.56484 | 0.17392 |
| **KIAA0907** | 0.58125 | 1.16827 | 4.30623 | 2.92508 | 0.00000 | 6.46581 | 3.85029 | 9.51485 | 7.71010 | 10.52040 | 5.67637 | 3.91533 |
| **KIAA0922** | 10.19958 | 4.58500 | 0.00000 | 1.43268 | 3.49538 | 0.00000 | 0.00000 | 4.47165 | 5.76709 | 3.08974 | 5.84187 | 0.45081 |
| **KIAA0930** | 1.95550 | 0.76199 | 1.13365 | 1.97906 | 10.04942 | 0.00000 | 5.13748 | 0.51642 | 0.00000 | 6.02743 | 1.27801 | 0.27594 |
| **KIAA1024** | 0.34840 | 0.13838 | 0.48163 | 0.36798 | 0.08052 | 0.00000 | 0.00000 | 0.00000 | 0.24726 | 0.70927 | 0.15390 | 0.08180 |
| **KIAA1033** | 4.57218 | 0.91534 | 3.28323 | 2.13977 | 0.27200 | 12.68456 | 7.07522 | 5.10271 | 8.78244 | 9.77229 | 6.06886 | 3.81960 |
| **KIAA1045** | 0.19349 | 0.55472 | 0.00812 | 1.00378 | 0.01987 | 0.00000 | 0.02531 | 0.00000 | 0.18435 | 0.26026 | 0.00000 | 0.25758 |
| **KIAA1107** | 0.28438 | 0.36163 | 0.09252 | 0.37860 | 0.07427 | 0.81088 | 0.00000 | 0.02994 | 0.00000 | 0.45654 | 0.26631 | 0.88376 |
| **KIAA1109** | 4.71986 | 0.01553 | 0.36936 | 0.02929 | 0.47591 | 4.75108 | 0.00000 | 2.01600 | 3.24947 | 9.32416 | 0.41655 | 0.89109 |
| **KIAA1143** | 0.57859 | 2.47039 | 7.00145 | 0.29539 | 0.00000 | 12.32280 | 8.97267 | 2.01591 | 6.24127 | 6.25413 | 2.27240 | 6.18769 |
| **KIAA1147** | 9.04932 | 0.61308 | 0.67823 | 0.00000 | 0.10526 | 0.20429 | 0.25446 | 0.33312 | 0.26357 | 0.00000 | 1.14922 | 0.75842 |
| **KIAA1161** | 0.24441 | 1.07858 | 2.66882 | 0.18869 | 7.02106 | 0.00000 | 0.00000 | 0.03609 | 4.50974 | 0.99670 | 7.72805 | 0.19974 |
| **KIAA1191** | 5.36766 | 7.33002 | 4.52105 | 6.28549 | 2.87925 | 12.22083 | 9.30269 | 24.88721 | 3.57829 | 8.83571 | 4.33743 | 18.03291 |
| **KIAA1210** | 0.44862 | 0.00892 | 0.02805 | 0.00950 | 0.00510 | 0.02348 | 0.00627 | 0.20106 | 0.00337 | 0.04904 | 0.05812 | 0.00387 |
| **KIAA1211** | 0.53777 | 0.01984 | 0.46306 | 0.00000 | 0.00000 | 1.75298 | 2.27677 | 0.12176 | 0.83681 | 1.47469 | 0.00000 | 0.27406 |
| **KIAA1211L** | 0.02449 | 0.01365 | 0.01198 | 0.00824 | 0.00803 | 0.05078 | 0.03310 | 0.01459 | 0.00446 | 0.04008 | 0.04131 | 0.02444 |
| **KIAA1217** | 0.10463 | 0.00000 | 25.98272 | 6.93678 | 1.74268 | 1.29689 | 11.26888 | 0.08049 | 3.06923 | 1.13906 | 0.00000 | 0.00000 |
| **KIAA1244** | 2.33020 | 0.00000 | 0.27230 | 0.16848 | 0.00000 | 0.10712 | 0.09344 | 0.75473 | 0.02314 | 1.21758 | 0.02288 | 0.13795 |
| **KIAA1257** | 4.39273 | 0.60635 | 0.07757 | 0.00000 | 0.00310 | 0.61999 | 0.00000 | 0.63106 | 0.00000 | 0.11379 | 0.00000 | 1.27974 |
| **KIAA1279** | 2.37018 | 8.08278 | 0.17014 | 4.18080 | 2.15595 | 12.60329 | 0.00000 | 0.26048 | 3.38163 | 2.43604 | 7.51411 | 0.44259 |
| **KIAA1324** | 0.00248 | 0.06452 | 0.85119 | 0.00068 | 0.07490 | 0.13139 | 0.11071 | 0.05537 | 0.09529 | 0.00000 | 0.00194 | 0.36127 |
| **KIAA1324L** | 2.37117 | 0.85734 | 0.48490 | 0.00869 | 0.24096 | 0.23505 | 0.13584 | 1.31333 | 0.42009 | 0.00000 | 0.00000 | 0.15954 |
| **KIAA1328** | 0.31111 | 0.28335 | 0.38738 | 0.16993 | 0.77295 | 1.25060 | 0.47405 | 0.25251 | 0.00000 | 0.00000 | 0.18525 | 0.25223 |
| **KIAA1377** | 0.16212 | 0.00000 | 0.67122 | 0.00000 | 0.49184 | 0.11961 | 1.09560 | 0.20355 | 0.25438 | 0.97562 | 0.15599 | 0.40099 |
| **KIAA1407** | 0.84732 | 0.21961 | 0.21786 | 0.01194 | 0.04720 | 0.12296 | 0.03145 | 0.28512 | 0.07505 | 0.00000 | 0.03086 | 0.25682 |
| **KIAA1429** | 1.03827 | 5.36505 | 5.01945 | 2.50185 | 2.42432 | 13.61276 | 8.10259 | 4.71562 | 6.01266 | 16.43755 | 6.99446 | 8.55987 |
| **KIAA1430** | 0.61293 | 1.46582 | 0.94664 | 1.18022 | 1.17810 | 0.99469 | 0.00000 | 17.24589 | 0.59771 | 0.57282 | 0.00000 | 13.36644 |
| **KIAA1432** | 1.87358 | 0.60315 | 1.70735 | 0.65060 | 0.00000 | 3.72153 | 2.67214 | 0.03720 | 4.39970 | 3.92089 | 1.19580 | 1.40836 |
| **KIAA1456** | 0.09153 | 0.02169 | 0.02315 | 0.03315 | 0.03818 | 0.00000 | 0.02683 | 0.07464 | 0.00771 | 0.10107 | 0.08862 | 0.01549 |
| **KIAA1462** | 0.07791 | 0.00000 | 2.50661 | 1.86068 | 6.17821 | 0.18946 | 6.60196 | 0.30335 | 3.79123 | 4.25099 | 7.39831 | 0.29939 |
| **KIAA1467** | 2.12745 | 1.98238 | 1.96084 | 0.26956 | 1.58672 | 0.72729 | 1.49065 | 2.19219 | 4.07633 | 0.88928 | 3.64299 | 4.88702 |
| **KIAA1468** | 0.09347 | 1.44193 | 2.80996 | 1.06628 | 1.81297 | 5.18987 | 2.95265 | 0.79487 | 3.52350 | 0.96356 | 2.52398 | 1.10522 |
| **KIAA1522** | 1.36763 | 1.80855 | 3.91937 | 2.50945 | 0.59849 | 0.50553 | 2.70109 | 0.28487 | 0.42090 | 4.38254 | 0.00000 | 5.73615 |
| **KIAA1524** | 4.42689 | 0.53900 | 5.25118 | 1.35014 | 2.87272 | 10.31534 | 9.48205 | 4.89649 | 6.78654 | 5.53794 | 8.40564 | 7.31307 |
| **KIAA1549** | 0.85183 | 0.86944 | 3.03640 | 0.00000 | 2.15577 | 0.13822 | 1.14332 | 0.07502 | 0.93680 | 1.47184 | 0.04519 | 0.00000 |
| **KIAA1549L** | 5.03241 | 0.02689 | 0.27528 | 0.70112 | 0.13050 | 0.00000 | 0.00000 | 0.49374 | 0.00000 | 0.61607 | 0.00000 | 0.10265 |
| **KIAA1551** | 9.53033 | 2.80550 | 0.00000 | 14.84290 | 4.21011 | 46.19046 | 0.20522 | 5.37952 | 13.62317 | 32.61439 | 4.90259 | 24.90437 |
| **KIAA1586** | 0.01125 | 0.00000 | 1.71763 | 1.06846 | 0.00000 | 0.93253 | 0.91464 | 0.43659 | 0.64462 | 0.66151 | 1.07992 | 3.37524 |

| | | | | | | | | | | | | |
|---|---|---|---|---|---|---|---|---|---|---|---|---|
| **KIAA1598** | 5.37439 | 4.80437 | 8.54630 | 3.33530 | 0.00000 | 16.97646 | 13.35181 | 1.75694 | 19.27403 | 8.81243 | 6.44855 | 13.66122 |
| **KIAA1614** | 0.00518 | 0.06998 | 0.06057 | 0.00000 | 0.04019 | 0.00000 | 0.03248 | 0.00712 | 0.08184 | 0.13617 | 0.06641 | 0.13192 |
| **KIAA1644** | 0.13014 | 0.30628 | 0.00754 | 0.00000 | 0.04678 | 0.17403 | 0.00000 | 0.00520 | 0.00000 | 0.04398 | 0.00000 | 0.12452 |
| **KIAA1656** | 0.00504 | 0.04945 | 0.18436 | 0.07090 | 0.05077 | 0.00000 | 0.00223 | 0.14642 | 0.08444 | 1.13835 | 0.11927 | 0.12270 |
| **KIAA1671** | 4.27881 | 5.45101 | 1.55286 | 5.20239 | 5.16352 | 0.45986 | 0.00000 | 0.00000 | 0.00000 | 3.95119 | 1.95616 | 1.35346 |
| **KIAA1683** | 0.00320 | 0.00487 | 0.00000 | 0.00046 | 0.00202 | 0.02120 | 0.00465 | 0.00947 | 0.02111 | 0.45280 | 0.00000 | 0.00724 |
| **KIAA1715** | 1.60142 | 2.25198 | 2.55576 | 0.21816 | 0.76772 | 5.09817 | 0.00000 | 3.93131 | 3.64649 | 4.34723 | 1.29978 | 2.99318 |
| **KIAA1731** | 2.83089 | 0.76523 | 2.81029 | 0.68702 | 2.42831 | 11.08782 | 1.47403 | 2.33635 | 3.01609 | 23.42243 | 0.19268 | 3.39818 |
| **KIAA1751** | 1.09760 | 0.07770 | 0.08395 | 0.02214 | 0.00000 | 0.00000 | 0.00000 | 0.34490 | 0.02998 | 0.21775 | 0.00137 | 0.15824 |
| **KIAA1755** | 0.00315 | 0.01320 | 0.05265 | 0.01523 | 0.01399 | 0.02042 | 0.01069 | 0.04928 | 0.01303 | 0.02944 | 0.01479 | 0.01384 |
| **KIAA1804** | 0.00125 | 0.09213 | 0.18768 | 1.08687 | 0.42272 | 1.03357 | 0.05102 | 0.48557 | 0.00000 | 0.84810 | 0.21513 | 1.72841 |
| **KIAA1841** | 1.07323 | 1.22737 | 0.82291 | 0.75346 | 0.00000 | 0.62699 | 0.72157 | 0.23617 | 0.17615 | 0.08594 | 0.28002 | 0.54686 |
| **KIAA1875** | 0.02908 | 0.05083 | 0.09546 | 0.15191 | 0.00000 | 0.02729 | 0.00000 | 0.00706 | 0.23635 | 0.10042 | 0.27536 | 0.06263 |
| **KIAA1919** | 0.44252 | 2.78031 | 4.40332 | 0.87561 | 0.80127 | 6.37230 | 0.00000 | 2.98615 | 4.20663 | 5.20396 | 2.63684 | 5.21272 |
| **KIAA1958** | 1.27257 | 1.62201 | 2.07787 | 1.52330 | 2.23602 | 3.32571 | 1.10140 | 0.14027 | 0.99758 | 2.42978 | 3.00764 | 0.23187 |
| **KIAA2013** | 0.93773 | 4.01125 | 2.23191 | 5.91844 | 10.64955 | 0.00000 | 3.78632 | 0.40666 | 6.48772 | 17.79074 | 0.97012 | 4.98254 |
| **KIAA2018** | 1.03673 | 0.37795 | 4.05935 | 4.76017 | 1.81987 | 5.54209 | 4.82884 | 1.53365 | 0.00743 | 10.42353 | 1.23176 | 0.94594 |
| **KIAA2022** | 0.00336 | 0.01004 | 0.05691 | 0.01952 | 0.00272 | 0.08541 | 0.03313 | 0.04196 | 0.00030 | 0.02455 | 0.01617 | 0.01458 |
| **KIAA2026** | 2.85271 | 0.00000 | 0.28897 | 1.83022 | 0.42000 | 5.32367 | 1.04114 | 1.12267 | 0.79963 | 9.03575 | 0.00000 | 2.28561 |
| **KIDINS220** | 2.52012 | 1.90245 | 2.59825 | 0.14455 | 2.10916 | 5.65266 | 0.00000 | 8.47904 | 4.35455 | 4.71479 | 0.56325 | 2.74653 |
| **KIF11** | 1.99701 | 6.56578 | 3.26063 | 4.73489 | 0.00000 | 26.91830 | 9.12269 | 3.30963 | 9.51988 | 9.73811 | 6.07783 | 6.47964 |
| **KIF12** | 0.01087 | 0.38699 | 0.07288 | 0.27129 | 0.35550 | 0.00000 | 0.04104 | 0.39067 | 0.00000 | 0.00000 | 0.04657 | 1.15417 |
| **KIF13A** | 0.34207 | 0.33615 | 2.47361 | 0.23128 | 0.00000 | 1.64582 | 2.36950 | 0.45561 | 0.00000 | 4.40966 | 0.86136 | 0.88177 |
| **KIF13B** | 5.57396 | 0.09044 | 0.00000 | 0.66141 | 2.11501 | 0.03768 | 1.70652 | 0.31232 | 3.46843 | 4.02668 | 0.95583 | 0.70707 |
| **KIF14** | 0.92231 | 0.00000 | 11.81910 | 0.33942 | 0.00000 | 5.87326 | 5.12033 | 0.98456 | 5.64036 | 9.11450 | 1.41296 | 7.54392 |
| **KIF15** | 2.51402 | 0.93483 | 3.17061 | 2.22492 | 2.41205 | 10.35664 | 4.12685 | 2.93228 | 8.62872 | 7.49809 | 3.81619 | 5.86553 |
| **KIF16B** | 1.32249 | 2.79264 | 3.02245 | 1.42355 | 1.87957 | 4.27738 | 4.03160 | 0.26792 | 3.24800 | 6.13775 | 3.76580 | 2.87382 |
| **KIF17** | 0.00637 | 1.20031 | 0.11704 | 1.40573 | 2.24445 | 0.07358 | 0.28975 | 0.06288 | 0.46296 | 0.91996 | 1.73356 | 0.54056 |
| **KIF18A** | 1.03705 | 3.40107 | 0.78004 | 0.64312 | 0.01182 | 11.70790 | 2.77662 | 7.59132 | 5.67351 | 0.00005 | 2.36050 | 7.33329 |
| **KIF18B** | 1.35148 | 5.14353 | 3.87554 | 2.99015 | 5.22036 | 0.00000 | 8.37252 | 0.52249 | 8.38580 | 6.86471 | 2.71166 | 3.37780 |
| **KIF19** | 0.00807 | 0.18986 | 0.04184 | 0.00000 | 0.05066 | 0.07621 | 0.00000 | 0.12903 | 0.00597 | 0.01636 | 0.00910 | 0.26320 |
| **KIF1A** | 4.29542 | 0.09113 | 0.10212 | 0.00000 | 0.33686 | 0.02276 | 0.03228 | 0.36198 | 0.00000 | 0.15058 | 0.07521 | 0.09799 |
| **KIF1B** | 3.96216 | 3.08270 | 3.95598 | 3.50166 | 3.79683 | 9.08118 | 7.97321 | 6.70749 | 1.60597 | 14.32419 | 2.10551 | 4.19863 |
| **KIF1C** | 1.01933 | 2.38694 | 3.38304 | 10.10239 | 10.31056 | 0.00000 | 6.72993 | 0.00000 | 2.16873 | 13.39347 | 1.17122 | 0.95119 |
| **KIF20A** | 1.00096 | 23.84413 | 10.40154 | 7.01026 | 7.68979 | 22.27317 | 15.31580 | 3.75668 | 23.25948 | 0.00000 | 11.71543 | 20.39419 |
| **KIF20B** | 7.95134 | 2.20316 | 9.46706 | 2.34089 | 1.57689 | 27.64134 | 11.93438 | 11.28922 | 6.16484 | 26.19780 | 5.22903 | 11.61348 |
| **KIF21A** | 0.92453 | 0.00000 | 14.18336 | 5.84262 | 8.52305 | 2.57721 | 11.97715 | 0.16257 | 1.96172 | 6.24519 | 9.53481 | 0.00000 |
| **KIF21B** | 1.94600 | 0.03598 | 0.09854 | 0.14828 | 0.05891 | 0.00000 | 0.00000 | 0.01501 | 0.15813 | 0.02663 | 0.00850 | 0.62708 |
| **KIF22** | 1.34042 | 23.59528 | 19.39731 | 18.94542 | 17.72737 | 5.46683 | 26.64882 | 7.06118 | 27.26743 | 0.00000 | 22.36812 | 19.20528 |
| **KIF23** | 5.76463 | 18.13384 | 8.06093 | 6.71266 | 9.23271 | 28.05612 | 11.86214 | 22.44252 | 16.89030 | 4.23644 | 9.16436 | 22.40016 |
| **KIF24** | 0.00811 | 1.31682 | 1.35193 | 0.38567 | 0.78649 | 0.00000 | 0.51948 | 1.54002 | 2.79306 | 0.26785 | 1.63677 | 1.73038 |
| **KIF25** | 0.00346 | 0.01657 | 0.01701 | 0.01390 | 0.01269 | 0.00171 | 0.03023 | 0.10163 | 0.00000 | 0.01534 | 0.07566 | 0.01328 |
| **KIF26A** | 0.01744 | 1.35815 | 0.00000 | 0.44668 | 2.62054 | 0.00000 | 0.20827 | 0.00342 | 0.00000 | 5.45011 | 0.22733 | 0.59699 |
| **KIF26B** | 6.12690 | 0.15929 | 0.03308 | 0.08605 | 0.00000 | 0.00000 | 0.01941 | 0.04059 | 0.04744 | 0.04727 | 0.01474 | 0.36631 |
| **KIF27** | 0.64272 | 0.68765 | 0.82428 | 0.43367 | 0.34077 | 0.45008 | 0.25237 | 3.44495 | 0.18062 | 0.89594 | 0.41293 | 0.72590 |
| **KIF2A** | 0.38503 | 1.06968 | 5.13599 | 0.47102 | 1.39641 | 8.54512 | 0.00000 | 2.77214 | 1.91769 | 5.79728 | 1.46365 | 1.09715 |
| **KIF2C** | 8.31170 | 19.43952 | 16.72526 | 10.60578 | 12.20132 | 7.71380 | 23.27379 | 27.37902 | 24.43487 | 0.00000 | 12.88255 | 20.00837 |
| **KIF3A** | 9.30863 | 0.49269 | 2.62068 | 0.81913 | 0.00000 | 4.89596 | 6.42186 | 4.67975 | 2.91154 | 5.16564 | 5.23020 | 2.90130 |

| | | | | | | | | | | | | |
|---|---|---|---|---|---|---|---|---|---|---|---|---|
| **KIF3B** | 1.94828 | 3.75102 | 2.84612 | 3.27988 | 1.40063 | 7.51035 | 9.22070 | 4.07350 | 0.50429 | 11.65847 | 2.19490 | 5.70293 |
| **KIF3C** | 3.95133 | 1.42132 | 1.19634 | 1.74019 | 1.30122 | 0.76470 | 0.22795 | 0.99540 | 0.50548 | 2.42271 | 0.27924 | 0.97134 |
| **KIF4A** | 8.84104 | 6.15489 | 2.64010 | 3.66308 | 3.89737 | 11.51118 | 4.94330 | 4.00204 | 4.07278 | 5.54922 | 2.42871 | 10.59970 |
| **KIF4B** | 2.24933 | 0.80269 | 0.71902 | 0.37575 | 0.49625 | 2.93693 | 0.00000 | 0.65100 | 1.37748 | 0.00000 | 0.42439 | 2.32137 |
| **KIF5A** | 0.31785 | 0.09294 | 0.10358 | 1.20901 | 0.00004 | 0.05098 | 0.02841 | 0.00000 | 0.05803 | 0.00000 | 0.00168 | 0.13667 |
| **KIF5B** | 5.21317 | 16.33375 | 14.08805 | 13.40845 | 12.42933 | 43.79650 | 25.21660 | 8.25544 | 25.88239 | 24.45673 | 19.56404 | 21.55569 |
| **KIF5C** | 1.44606 | 0.02874 | 0.41425 | 1.78679 | 0.08353 | 0.00000 | 0.41331 | 0.94707 | 0.00000 | 0.00427 | 0.00819 | 0.30605 |
| **KIF6** | 0.36230 | 0.10880 | 0.04331 | 0.03227 | 0.01735 | 0.00000 | 0.10086 | 0.00903 | 0.02261 | 0.09168 | 0.01019 | 0.06941 |
| **KIF7** | 0.82733 | 0.54302 | 0.09429 | 0.72870 | 0.27751 | 0.00000 | 0.14372 | 0.00236 | 0.72319 | 0.38245 | 0.06304 | 0.02917 |
| **KIF9** | 1.37910 | 7.81263 | 0.00000 | 4.70455 | 2.96785 | 3.99576 | 2.31659 | 0.74189 | 1.93936 | 2.01359 | 0.94416 | 6.97234 |
| **KIF9-AS1** | 0.12639 | 1.04705 | 0.01936 | 0.28033 | 0.34048 | 0.42714 | 0.00000 | 1.13279 | 0.02592 | 0.58974 | 0.16833 | 0.75634 |
| **KIFAP3** | 1.04130 | 2.84577 | 0.31335 | 10.03375 | 1.85904 | 13.09897 | 2.19354 | 1.96738 | 1.81164 | 2.26053 | 1.81490 | 2.25706 |
| **KIFC1** | 1.94358 | 6.64682 | 10.63304 | 8.20575 | 6.47097 | 0.00000 | 6.82911 | 2.92336 | 12.51040 | 0.00000 | 17.65219 | 14.91975 |
| **KIFC2** | 0.64330 | 0.16682 | 0.31718 | 0.00000 | 0.68436 | 0.00000 | 0.01419 | 0.00000 | 0.00167 | 1.47572 | 0.14583 | 0.07359 |
| **KIFC3** | 2.47308 | 0.24819 | 1.11945 | 1.38229 | 0.36910 | 0.00000 | 0.47664 | 0.16488 | 0.26959 | 0.00000 | 0.00000 | 2.25899 |
| **KIN** | 1.46421 | 5.42879 | 6.63844 | 1.82170 | 1.85796 | 7.95114 | 7.39350 | 8.77095 | 3.86271 | 11.73321 | 3.68236 | 6.43041 |
| **KIR2DL4.uid-24** | 0.00510 | 0.01044 | 0.02148 | 0.01708 | 0.00007 | 0.01827 | 0.00797 | 0.01031 | 0.00000 | 0.08894 | 0.04535 | 0.01462 |
| **KIR2DS4** | 0.00493 | 0.01318 | 0.01547 | 0.00830 | 0.00932 | 0.01055 | 0.01965 | 0.01042 | 0.01973 | 0.10123 | 0.01048 | 0.01291 |
| **KIR3DL1** | 0.00530 | 0.01042 | 0.01601 | 0.02098 | 0.00517 | 0.01826 | 0.01273 | 0.01160 | 0.01139 | 0.05894 | 0.02452 | 0.02333 |
| **KIR3DL2** | 0.00446 | 0.00754 | 0.01134 | 0.00823 | 0.00590 | 0.04637 | 0.04092 | 0.01368 | 0.02399 | 0.01041 | 0.01699 | 0.05703 |
| **KIR3DL3** | 0.00622 | 0.01526 | 0.01530 | 0.01247 | 0.01570 | 0.01490 | 0.02412 | 0.01053 | 0.01289 | 0.01460 | 0.02569 | 0.01653 |
| **KIR3DX1** | 0.00572 | 0.01453 | 0.02527 | 0.01007 | 0.01180 | 0.01723 | 0.01529 | 0.01041 | 0.01349 | 0.04224 | 0.01475 | 0.01522 |
| **KIRREL** | 0.02448 | 0.10272 | 0.04555 | 0.14179 | 0.00000 | 0.00899 | 0.00768 | 0.01569 | 0.00289 | 0.10818 | 0.01807 | 0.02485 |
| **KIRREL2** | 0.00852 | 0.01582 | 0.09659 | 0.01470 | 0.00000 | 0.02749 | 0.08382 | 0.06170 | 0.08637 | 0.01239 | 0.00583 | 0.02217 |
| **KIRREL3** | 0.16713 | 0.02794 | 0.06001 | 0.16714 | 0.09996 | 0.00000 | 0.05590 | 0.00099 | 0.11361 | 0.00000 | 0.02030 | 0.07210 |
| **KIRREL3-AS3** | 0.04798 | 0.01819 | 0.01486 | 0.01421 | 0.02266 | 0.00608 | 0.02386 | 0.02187 | 0.02264 | 0.00837 | 0.01393 | 0.02326 |
| **KIT** | 8.26804 | 8.84725 | 0.00000 | 8.87560 | 0.28929 | 0.32928 | 0.00000 | 2.39692 | 0.00000 | 0.13338 | 1.15300 | 1.20893 |
| **KITLG** | 0.32372 | 2.20577 | 0.00000 | 0.00000 | 0.00000 | 0.86952 | 1.61787 | 5.93722 | 0.80269 | 2.11419 | 1.07588 | 13.65304 |
| **KIZ** | 0.27551 | 2.28520 | 1.70447 | 1.76671 | 1.35678 | 4.55463 | 2.06838 | 1.61070 | 2.47198 | 0.00000 | 0.00000 | 3.45592 |
| **KL** | 0.00617 | 0.01481 | 0.03012 | 0.01072 | 0.01616 | 0.02160 | 0.01109 | 0.01045 | 0.01866 | 0.01937 | 0.01420 | 0.01555 |
| **KLB** | 0.00205 | 0.00000 | 1.23769 | 1.90863 | 0.40138 | 0.94980 | 0.00000 | 0.52770 | 0.54166 | 0.73744 | 0.31623 | 0.00000 |
| **KLC1** | 3.06634 | 6.50446 | 8.71396 | 2.77801 | 7.49601 | 1.42743 | 1.88455 | 1.93253 | 2.29863 | 8.41180 | 5.27344 | 1.35387 |
| **KLC2** | 2.95712 | 1.20717 | 2.22241 | 1.18838 | 1.27006 | 0.00000 | 0.00000 | 0.17600 | 2.94165 | 2.16506 | 0.43407 | 1.75820 |
| **KLC3** | 0.43955 | 0.58699 | 0.51697 | 0.25519 | 0.28985 | 0.00000 | 0.34782 | 0.01792 | 0.30833 | 0.00000 | 0.45215 | 0.09917 |
| **KLC4** | 0.20650 | 1.90131 | 3.43739 | 1.96616 | 1.58117 | 0.00000 | 2.57989 | 1.89152 | 2.47412 | 0.00000 | 4.35000 | 1.20702 |
| **KLF1** | 0.07442 | 0.04515 | 0.11535 | 0.06232 | 0.03420 | 0.00298 | 0.17223 | 0.00827 | 0.06276 | 0.00000 | 0.01951 | 0.15798 |
| **KLF10** | 0.01075 | 7.46270 | 7.40834 | 1.27753 | 1.78353 | 16.13901 | 6.83846 | 15.45656 | 8.61630 | 0.00000 | 3.01177 | 2.76507 |
| **KLF11** | 6.42151 | 5.94388 | 0.00000 | 1.65359 | 4.69087 | 1.96668 | 0.00000 | 6.69231 | 0.00000 | 3.15844 | 0.00000 | 7.48913 |
| **KLF12** | 3.17003 | 0.06200 | 0.00000 | 0.01570 | 0.02100 | 0.18013 | 0.00000 | 0.62786 | 0.27664 | 0.00073 | 0.14302 | 2.26436 |
| **KLF13** | 0.00000 | 1.55739 | 0.25283 | 0.00009 | 2.54028 | 1.45634 | 3.40756 | 0.37131 | 0.00000 | 9.50123 | 0.17098 | 0.53688 |
| **KLF14** | 0.03504 | 0.01761 | 0.00980 | 0.00889 | 0.10371 | 0.02621 | 0.02845 | 0.03111 | 0.00000 | 0.04283 | 0.01245 | 0.04066 |
| **KLF15** | 0.03077 | 0.11662 | 0.22871 | 0.09726 | 0.06419 | 0.07270 | 0.39409 | 0.06806 | 0.04563 | 0.85677 | 0.08589 | 0.01578 |
| **KLF16** | 2.87420 | 0.60244 | 0.89153 | 2.27736 | 6.30459 | 0.00000 | 3.61745 | 2.02945 | 1.83065 | 17.04556 | 0.35556 | 3.78086 |
| **KLF17** | 4.92874 | 0.66462 | 0.38379 | 5.98586 | 0.00000 | 0.00000 | 0.06250 | 40.64151 | 0.00000 | 0.05288 | 0.00000 | 20.57089 |
| **KLF2** | 2.80079 | 0.00644 | 0.00084 | 0.02860 | 0.00000 | 0.01022 | 0.04684 | 0.00190 | 0.04380 | 0.23064 | 0.00000 | 0.06295 |
| **KLF3** | 0.00000 | 8.34697 | 0.00000 | 2.92236 | 3.98173 | 16.45847 | 4.87743 | 6.90533 | 0.00000 | 10.36294 | 0.75970 | 8.05612 |
| **KLF3-AS1** | 0.00582 | 0.01402 | 0.00455 | 0.05867 | 0.00087 | 0.06716 | 0.01850 | 0.00914 | 0.01545 | 0.01932 | 0.01709 | 0.02430 |
| **KLF4** | 1.66827 | 25.62646 | 9.60529 | 27.01133 | 15.45637 | 4.86032 | 0.00000 | 3.62701 | 0.00000 | 5.12999 | 0.00000 | 21.92159 |

| | | | | | | | | | | | | |
|---|---|---|---|---|---|---|---|---|---|---|---|---|
| **KLF5** | 2.42154 | 6.34026 | 0.00000 | 0.00000 | 16.79705 | 18.11990 | 16.01697 | 9.25421 | 0.00000 | 43.25751 | 16.10327 | 16.06637 |
| **KLF6** | 0.14772 | 0.00000 | 3.69872 | 13.74953 | 5.99271 | 4.07219 | 13.13444 | 0.00000 | 6.67587 | 6.32088 | 10.30403 | 0.14716 |
| **KLF7** | 0.63789 | 0.11163 | 0.09271 | 2.16115 | 0.33016 | 0.00000 | 0.01037 | 0.01849 | 0.20275 | 1.07181 | 0.00000 | 0.11610 |
| **KLF8** | 0.00226 | 6.53741 | 0.83665 | 3.96848 | 2.73124 | 3.75172 | 0.00000 | 0.00000 | 2.82632 | 0.00000 | 1.63568 | 3.99310 |
| **KLF9** | 0.00000 | 0.75428 | 0.15094 | 2.28202 | 0.00000 | 3.14638 | 0.00000 | 3.98663 | 0.00000 | 8.29786 | 0.00000 | 4.19678 |
| **KLHDC1** | 0.04575 | 0.01100 | 0.03381 | 0.04422 | 0.00702 | 0.03343 | 0.01207 | 0.03881 | 0.00724 | 0.05242 | 0.01221 | 0.02369 |
| **KLHDC10** | 1.82301 | 0.00024 | 1.48096 | 0.46616 | 0.70965 | 1.68245 | 4.98474 | 0.00000 | 0.03993 | 4.79603 | 2.38975 | 0.35537 |
| **KLHDC2** | 2.24209 | 5.21547 | 3.47071 | 4.51808 | 2.97576 | 4.95028 | 1.37437 | 0.98877 | 1.25044 | 3.59759 | 2.67053 | 4.79617 |
| **KLHDC3** | 13.88171 | 21.92381 | 15.15741 | 18.84203 | 14.35808 | 4.34679 | 17.67590 | 3.63415 | 30.03169 | 0.00000 | 16.94723 | 20.75371 |
| **KLHDC4** | 0.57230 | 7.10410 | 4.40478 | 3.20162 | 6.00368 | 0.00000 | 5.23974 | 1.52238 | 5.05641 | 2.95066 | 5.35902 | 2.61973 |
| **KLHDC7A** | 0.48347 | 0.01477 | 0.04198 | 0.01644 | 0.00995 | 0.02281 | 0.01790 | 0.00275 | 0.02139 | 0.01134 | 0.01624 | 0.01334 |
| **KLHDC7B** | 0.00574 | 0.01798 | 0.02260 | 0.01330 | 0.01263 | 0.01409 | 0.02682 | 0.01054 | 0.00919 | 0.03744 | 0.01565 | 0.01471 |
| **KLHDC8A** | 0.00653 | 0.01219 | 0.01673 | 0.02195 | 0.00433 | 0.01858 | 0.02484 | 0.01034 | 0.00000 | 0.02296 | 0.07686 | 0.01602 |
| **KLHDC8B** | 0.00266 | 1.96352 | 1.19985 | 1.96524 | 0.00000 | 0.00000 | 1.20397 | 0.29557 | 1.98207 | 0.00000 | 1.26317 | 0.74544 |
| **KLHDC9** | 0.14654 | 1.15724 | 0.05715 | 0.22990 | 0.00000 | 0.04586 | 0.00000 | 0.00053 | 0.00000 | 0.28397 | 0.10640 | 0.05628 |
| **KLHL1** | 0.21530 | 0.01543 | 0.02369 | 0.01970 | 0.01286 | 0.02803 | 0.02418 | 0.01378 | 0.02922 | 0.00578 | 0.01498 | 0.06743 |
| **KLHL10** | 0.01993 | 0.04465 | 0.07313 | 0.07718 | 0.01353 | 0.04198 | 0.00000 | 0.00863 | 0.01422 | 0.03343 | 0.01039 | 0.04460 |
| **KLHL11** | 7.38637 | 3.59898 | 3.03460 | 0.52890 | 1.83202 | 1.42526 | 0.00000 | 11.12820 | 3.94729 | 0.00000 | 0.00000 | 7.04567 |
| **KLHL12** | 5.04989 | 6.55203 | 2.65248 | 2.86954 | 0.59893 | 10.22189 | 2.41159 | 1.72699 | 3.24238 | 0.00000 | 4.07802 | 4.28873 |
| **KLHL13** | 0.00729 | 0.09489 | 0.82354 | 0.00484 | 0.00000 | 0.24301 | 0.15623 | 0.00520 | 0.00000 | 0.01881 | 0.05624 | 0.20896 |
| **KLHL14** | 0.02252 | 0.73343 | 0.00149 | 0.00022 | 0.00000 | 0.24943 | 0.00000 | 0.08404 | 0.03254 | 0.29671 | 0.00000 | 0.09630 |
| **KLHL15** | 3.38596 | 1.37336 | 2.29248 | 0.00000 | 1.05436 | 1.94476 | 2.40844 | 13.46086 | 0.74676 | 2.33183 | 2.43630 | 7.35209 |
| **KLHL17** | 0.10773 | 0.76764 | 0.31657 | 1.22515 | 1.78489 | 0.00000 | 0.50041 | 0.64865 | 2.60741 | 6.79191 | 0.63869 | 1.70978 |
| **KLHL18** | 7.71541 | 18.59941 | 5.83497 | 24.00235 | 9.09454 | 3.86528 | 8.89460 | 5.66662 | 7.22278 | 2.44657 | 7.00484 | 19.76670 |
| **KLHL2** | 0.00446 | 0.00000 | 1.34180 | 0.00000 | 0.48277 | 4.55065 | 4.07730 | 0.46932 | 0.04282 | 2.99810 | 1.75921 | 0.70920 |
| **KLHL20** | 1.41839 | 0.69680 | 2.98648 | 2.54323 | 0.24813 | 7.72268 | 1.51247 | 4.41268 | 1.71249 | 2.88049 | 1.78931 | 4.25935 |
| **KLHL21** | 1.97718 | 6.38354 | 9.29265 | 2.76249 | 6.53055 | 0.00000 | 13.07965 | 5.25896 | 16.92715 | 8.33253 | 5.86461 | 3.37054 |
| **KLHL22** | 0.02387 | 0.97719 | 1.93931 | 1.54242 | 0.54977 | 0.00000 | 0.45821 | 0.57660 | 0.68732 | 2.22254 | 2.42669 | 0.44572 |
| **KLHL23** | 0.12440 | 0.07660 | 0.18350 | 0.01137 | 0.00000 | 0.00003 | 0.98803 | 0.01652 | 0.64920 | 0.00000 | 1.04214 | 0.02159 |
| **KLHL24** | 1.21465 | 2.18350 | 2.76354 | 2.75173 | 0.00000 | 8.85233 | 6.20598 | 6.55719 | 2.90494 | 6.23195 | 0.00000 | 5.08980 |
| **KLHL25** | 0.27011 | 0.91160 | 3.52051 | 1.01263 | 2.81068 | 0.00000 | 1.28180 | 0.68384 | 1.99551 | 0.00000 | 0.61631 | 1.01925 |
| **KLHL26** | 2.89940 | 1.34607 | 0.19288 | 0.00000 | 0.19102 | 0.31869 | 0.10786 | 0.00000 | 0.85927 | 0.00000 | 0.06770 | 1.04709 |
| **KLHL28** | 1.50268 | 0.53651 | 3.01467 | 1.04413 | 0.05837 | 4.50554 | 1.91386 | 3.00138 | 2.65842 | 0.41705 | 2.21430 | 3.38308 |
| **KLHL29** | 0.18890 | 0.03788 | 0.26998 | 0.18661 | 0.06878 | 0.00000 | 0.72853 | 0.03463 | 0.16894 | 0.37593 | 0.03188 | 0.05623 |
| **KLHL3** | 0.14763 | 0.02594 | 0.11198 | 2.08225 | 0.00000 | 0.21810 | 0.00000 | 0.13834 | 0.08103 | 0.00000 | 0.07167 | 0.08914 |
| **KLHL30** | 0.08354 | 0.01863 | 0.01709 | 0.00667 | 0.08816 | 0.00000 | 0.00336 | 0.00571 | 0.03938 | 0.05022 | 0.02208 | 0.01605 |
| **KLHL31** | 0.00865 | 0.24420 | 0.00000 | 0.08835 | 0.43248 | 0.49178 | 0.00000 | 0.00533 | 0.00000 | 0.04651 | 0.02286 | 0.00000 |
| **KLHL32** | 5.42177 | 0.16760 | 0.01026 | 0.00000 | 0.00000 | 0.06896 | 0.00000 | 0.52089 | 0.07095 | 0.11912 | 0.00937 | 0.00000 |
| **KLHL33** | 0.00528 | 0.01269 | 0.05975 | 0.01648 | 0.00771 | 0.02163 | 0.00870 | 0.01121 | 0.00453 | 0.06150 | 0.01822 | 0.01350 |
| **KLHL34** | 0.00787 | 0.01441 | 0.02565 | 0.01086 | 0.01484 | 0.01921 | 0.01316 | 0.01060 | 0.02027 | 0.01836 | 0.01455 | 0.01705 |
| **KLHL35** | 1.66749 | 1.86406 | 0.01242 | 0.16699 | 0.00000 | 0.13201 | 0.00000 | 0.04103 | 0.00000 | 0.01024 | 0.03771 | 1.90552 |
| **KLHL36** | 1.08624 | 0.99959 | 1.44377 | 0.85090 | 1.16994 | 0.00000 | 0.46262 | 0.00000 | 0.59942 | 0.18181 | 0.01757 | 0.12713 |
| **KLHL40** | 0.00555 | 0.01796 | 0.01483 | 0.01788 | 0.01908 | 0.00275 | 0.01893 | 0.01008 | 0.02250 | 0.02820 | 0.01470 | 0.03446 |
| **KLHL41** | 0.17223 | 0.08845 | 0.00070 | 0.00688 | 0.23625 | 0.05033 | 0.00000 | 0.14430 | 0.17227 | 0.00000 | 0.00145 | 0.14108 |
| **KLHL42** | 0.46790 | 2.07087 | 2.47828 | 3.64683 | 2.77848 | 2.90598 | 6.61376 | 0.33263 | 1.36452 | 11.24447 | 1.34991 | 1.16114 |
| **KLHL5** | 0.06850 | 0.00000 | 0.00000 | 0.55860 | 2.74145 | 1.33959 | 1.87644 | 0.00000 | 0.00000 | 2.66226 | 1.07954 | 0.00000 |
| **KLHL6** | 0.01247 | 0.01092 | 0.06818 | 0.10552 | 0.00000 | 0.01894 | 0.01829 | 0.01172 | 0.00304 | 0.02316 | 0.05231 | 0.02653 |
| **KLHL7** | 10.71617 | 8.08094 | 9.83755 | 2.26068 | 4.16373 | 15.88957 | 7.53884 | 5.68989 | 6.85216 | 0.00000 | 7.36366 | 11.05879 |

| | | | | | | | | | | | | |
|---|---|---|---|---|---|---|---|---|---|---|---|---|
| **KLHL7-AS1** | 0.12412 | 0.37741 | 0.11460 | 0.05257 | 0.05329 | 0.08505 | 0.00000 | 0.00308 | 0.00000 | 0.00000 | 0.33714 | 0.31829 |
| **KLHL8** | 1.48459 | 0.59153 | 2.36045 | 0.99048 | 0.00000 | 2.84945 | 3.85615 | 1.63546 | 0.76839 | 2.23098 | 2.38875 | 0.99583 |
| **KLHL9** | 0.59213 | 3.36635 | 2.31330 | 0.00000 | 0.00000 | 6.53524 | 1.03446 | 1.74057 | 1.74700 | 2.56577 | 3.35113 | 4.85514 |
| **KLK1** | 0.00408 | 0.01034 | 0.00644 | 0.15977 | 0.18936 | 0.00000 | 0.05028 | 0.03762 | 0.00000 | 0.00000 | 0.01083 | 0.12904 |
| **KLK10** | 0.00294 | 0.06054 | 0.00000 | 0.00000 | 0.08770 | 0.02554 | 0.00000 | 2.81041 | 0.00000 | 0.15972 | 0.05204 | 1.96041 |
| **KLK11** | 0.03046 | 0.00000 | 0.00000 | 0.00000 | 0.13416 | 0.05622 | 0.00000 | 17.67219 | 0.00000 | 0.01389 | 0.00000 | 5.18449 |
| **KLK12** | 0.00205 | 0.01542 | 0.01312 | 0.02412 | 0.02258 | 0.01503 | 0.01817 | 0.11198 | 0.01897 | 0.00932 | 0.01545 | 0.01918 |
| **KLK13** | 0.09874 | 0.13589 | 0.00000 | 3.35385 | 0.28985 | 0.16178 | 0.00413 | 0.00000 | 0.00000 | 0.00239 | 0.01274 | 0.13060 |
| **KLK14** | 0.01136 | 0.02827 | 0.02023 | 0.01091 | 0.03098 | 0.00000 | 0.01552 | 0.01742 | 0.05765 | 0.02170 | 0.02843 | 0.03618 |
| **KLK15** | 0.01890 | 0.10920 | 0.01487 | 0.00347 | 0.06747 | 0.01886 | 0.03154 | 0.08463 | 0.00838 | 0.00000 | 0.01061 | 0.20718 |
| **KLK2** | 0.05648 | 0.11427 | 0.01288 | 0.00000 | 0.00000 | 0.03146 | 0.18510 | 0.03349 | 0.03066 | 0.03581 | 0.02506 | 0.50404 |
| **KLK4** | 0.00325 | 0.00752 | 0.01617 | 0.01370 | 0.01810 | 0.00666 | 0.01705 | 0.22634 | 0.02221 | 0.01708 | 0.01515 | 0.17549 |
| **KLK5** | 0.00821 | 0.13284 | 0.00595 | 0.00000 | 0.04413 | 0.04802 | 0.02497 | 0.00900 | 0.00239 | 0.04994 | 0.00812 | 0.01222 |
| **KLK6** | 0.00523 | 0.01202 | 0.02835 | 1.62366 | 0.00000 | 0.01952 | 0.00843 | 0.00931 | 0.00871 | 0.03405 | 0.02831 | 0.01299 |
| **KLK7** | 0.03743 | 1.04895 | 0.28936 | 0.00778 | 0.64232 | 0.00000 | 0.00000 | 0.00000 | 2.83994 | 0.39355 | 0.00000 | 0.64917 |
| **KLK8** | 0.01014 | 5.09124 | 0.69668 | 1.66708 | 1.64664 | 1.13784 | 0.05208 | 1.10421 | 5.10328 | 0.00000 | 1.22384 | 7.67048 |
| **KLK9** | 0.00343 | 0.04188 | 0.05636 | 0.01683 | 0.02900 | 0.00000 | 0.02367 | 0.07821 | 0.03013 | 0.01696 | 0.01454 | 0.01591 |
| **KLKB1** | 0.03879 | 0.01602 | 0.04959 | 0.01888 | 0.01582 | 0.01222 | 0.02089 | 0.00586 | 0.01384 | 0.00759 | 0.01704 | 0.02434 |
| **KLKP1** | 0.00476 | 0.01191 | 0.01958 | 0.01648 | 0.02762 | 0.00032 | 0.02849 | 0.08600 | 0.01182 | 0.03312 | 0.01406 | 0.26473 |
| **KLLN** | 0.00998 | 0.33142 | 0.19903 | 0.01915 | 0.00000 | 0.00073 | 0.06739 | 0.47865 | 0.00000 | 0.08641 | 0.19035 | 0.18978 |
| **KLRAP1** | 0.00000 | 0.00809 | 0.00000 | 0.00000 | 0.00000 | 0.85316 | 0.06624 | 0.17065 | 0.00000 | 1.22578 | 0.00008 | 0.06659 |
| **KLRB1** | 0.16003 | 0.03208 | 0.06840 | 0.04125 | 0.00540 | 0.01910 | 0.00541 | 0.26955 | 0.00478 | 0.03974 | 0.03611 | 0.28196 |
| **KLRC1** | 0.01013 | 0.00995 | 0.05877 | 0.00555 | 0.00533 | 0.04379 | 0.05038 | 0.00895 | 0.05857 | 0.00000 | 0.01074 | 0.04306 |
| **KLRC2** | 0.00656 | 0.01437 | 0.02787 | 0.00793 | 0.01272 | 0.02935 | 0.01990 | 0.01027 | 0.05667 | 0.00209 | 0.01067 | 0.01625 |
| **KLRD1** | 0.03371 | 0.00000 | 0.07496 | 0.02025 | 0.00056 | 0.18599 | 0.00000 | 0.09316 | 0.00000 | 0.07479 | 0.00325 | 0.06855 |
| **KLRF1** | 0.12072 | 0.01300 | 0.01604 | 0.01136 | 0.02121 | 0.02242 | 0.01401 | 0.00445 | 0.02841 | 0.00393 | 0.01519 | 0.04716 |
| **KLRG1** | 0.34098 | 0.11933 | 0.56602 | 0.28031 | 0.15305 | 0.00003 | 0.00000 | 1.40050 | 0.61665 | 0.00000 | 0.10839 | 0.97468 |
| **KLRG2** | 0.00000 | 12.88490 | 3.81276 | 4.35967 | 9.56675 | 0.00000 | 0.32168 | 0.08698 | 2.07106 | 9.98959 | 3.24639 | 1.18803 |
| **KLRK1** | 0.01468 | 0.01528 | 0.01349 | 0.01161 | 0.01339 | 0.01503 | 0.02170 | 0.00860 | 0.01177 | 0.02266 | 0.02922 | 0.01574 |
| **KMO** | 0.00629 | 0.00864 | 0.05839 | 0.00000 | 0.00000 | 0.09876 | 0.27466 | 0.00932 | 0.00021 | 0.02038 | 0.00246 | 0.01243 |
| **KMT2A** | 2.18540 | 0.91247 | 0.00000 | 0.41139 | 3.33900 | 6.62014 | 1.46924 | 0.72039 | 6.02131 | 11.23287 | 0.27420 | 0.30576 |
| **KMT2B** | 4.55470 | 1.11341 | 1.79768 | 3.55769 | 6.33555 | 0.00000 | 4.29866 | 0.51085 | 1.73576 | 22.13769 | 0.46754 | 2.31932 |
| **KMT2C** | 18.51671 | 5.52608 | 6.11742 | 9.21496 | 10.74085 | 18.55661 | 6.94983 | 12.75956 | 5.38609 | 42.37292 | 5.99337 | 14.07841 |
| **KMT2D** | 7.49433 | 1.81863 | 4.35288 | 2.66214 | 8.81749 | 0.00000 | 6.14557 | 0.71632 | 9.60109 | 32.48359 | 1.24417 | 4.41213 |
| **KMT2E** | 5.29122 | 2.87056 | 0.81710 | 4.41674 | 6.26798 | 9.43534 | 0.03289 | 2.19633 | 4.82924 | 22.76047 | 3.28134 | 4.39855 |
| **KMT2E-AS1** | 0.36437 | 1.25161 | 1.21737 | 0.81555 | 1.74923 | 0.00221 | 1.47477 | 0.35095 | 0.75764 | 1.88533 | 2.57632 | 1.52963 |
| **KNDC1** | 0.64262 | 0.01266 | 0.31754 | 0.13876 | 0.04329 | 0.00000 | 0.00217 | 0.00362 | 0.00000 | 0.02113 | 0.01534 | 0.01177 |
| **KNG1** | 0.06772 | 2.03386 | 0.00660 | 0.00000 | 0.30501 | 0.10004 | 0.07366 | 0.00094 | 0.00000 | 1.52764 | 0.05591 | 0.71465 |
| **KNOP1** | 6.09408 | 13.65869 | 8.28790 | 5.64811 | 12.28196 | 12.89222 | 17.64082 | 5.75379 | 8.67530 | 22.95056 | 15.36797 | 19.39608 |
| **KNSTRN** | 5.44045 | 19.95850 | 18.09186 | 6.93890 | 12.94471 | 38.69927 | 23.33700 | 9.91205 | 16.10450 | 0.00000 | 18.55448 | 34.07807 |
| **KNTC1** | 2.34939 | 2.15752 | 1.28877 | 0.70766 | 0.02993 | 12.94983 | 0.00000 | 2.70120 | 4.37809 | 10.59163 | 1.11715 | 1.99542 |
| **KPNA1** | 0.28012 | 8.75530 | 6.59671 | 8.68882 | 2.25414 | 22.03237 | 11.50507 | 10.07647 | 13.55084 | 4.06686 | 8.91703 | 14.55219 |
| **KPNA2** | 11.57569 | 39.83558 | 27.37001 | 18.68862 | 23.08095 | 57.84501 | 39.02905 | 45.83473 | 42.10970 | 0.08744 | 31.54925 | 39.03142 |
| **KPNA3** | 7.03471 | 12.88175 | 9.61556 | 7.66626 | 17.99788 | 44.68781 | 12.55587 | 6.07890 | 5.51044 | 13.56243 | 12.36437 | 19.03511 |
| **KPNA4** | 0.00000 | 2.42626 | 6.40571 | 2.34252 | 5.16469 | 14.73912 | 17.07617 | 1.99949 | 0.01148 | 18.37434 | 4.60645 | 3.13894 |
| **KPNA5** | 0.13903 | 0.20449 | 0.53642 | 0.14136 | 0.66520 | 1.48736 | 0.14729 | 0.12803 | 2.10895 | 0.38421 | 0.64761 | 0.87628 |
| **KPNA6** | 5.94438 | 16.75507 | 16.06785 | 13.34488 | 9.06430 | 20.51329 | 33.72029 | 28.20869 | 20.79595 | 17.09219 | 22.51507 | 31.77653 |
| **KPNA7** | 28.77550 | 0.00922 | 0.01616 | 0.01065 | 0.05008 | 0.00000 | 0.01744 | 21.79891 | 0.00000 | 0.01083 | 0.00454 | 0.26017 |

| | | | | | | | | | | | | |
|---|---|---|---|---|---|---|---|---|---|---|---|---|
| **KPNB1** | 4.12706 | 12.14212 | 19.77853 | 17.98045 | 25.34001 | 24.83205 | 45.74274 | 13.82721 | 15.86046 | 50.45769 | 21.40875 | 17.75202 |
| **KPRP** | 0.00647 | 0.01401 | 0.01529 | 0.01147 | 0.01827 | 0.01899 | 0.02177 | 0.01040 | 0.01027 | 0.01442 | 0.02901 | 0.01628 |
| **KPTN** | 0.00000 | 4.63704 | 5.05562 | 2.35536 | 2.94620 | 1.88720 | 5.29984 | 10.74867 | 4.80556 | 2.02415 | 2.21085 | 8.86603 |
| **KRAS** | 0.00000 | 12.99190 | 6.13136 | 3.48611 | 3.02634 | 36.85130 | 23.30736 | 13.29684 | 5.50123 | 33.91348 | 5.48386 | 17.44160 |
| **KRBA1** | 0.00039 | 0.83018 | 1.22735 | 1.09990 | 0.26826 | 0.00000 | 0.23234 | 0.18640 | 2.78040 | 0.44367 | 0.27761 | 0.64173 |
| **KRBA2** | 0.82992 | 0.72548 | 0.02455 | 0.55182 | 0.04984 | 0.12828 | 0.00000 | 0.11175 | 0.06351 | 0.24341 | 0.00643 | 0.69687 |
| **KRBOX1** | 0.43528 | 2.59490 | 0.00000 | 1.88426 | 4.51471 | 6.89516 | 0.00000 | 1.62983 | 0.00000 | 0.92968 | 0.00000 | 9.34431 |
| **KRBOX1-AS1** | 0.02201 | 0.02700 | 0.04796 | 0.01318 | 0.01577 | 0.03415 | 0.01354 | 0.04204 | 0.01242 | 0.00500 | 0.01462 | 0.07096 |
| **KRBOX4** | 0.15609 | 1.41918 | 0.30173 | 0.73259 | 0.83137 | 2.50381 | 0.49077 | 2.59830 | 0.74258 | 1.18320 | 1.15782 | 2.92597 |
| **KRCC1** | 0.21203 | 0.23308 | 0.02797 | 0.20514 | 0.00000 | 0.89101 | 0.00000 | 0.00000 | 0.00000 | 1.09988 | 0.00000 | 0.15047 |
| **KREMEN1** | 0.01940 | 0.78968 | 3.54742 | 3.37167 | 3.05378 | 0.11118 | 0.11456 | 0.01603 | 1.82898 | 0.85739 | 0.00000 | 0.00000 |
| **KREMEN2** | 0.12043 | 0.49536 | 0.71867 | 0.33208 | 0.17514 | 0.00000 | 0.13339 | 0.05000 | 0.43906 | 1.58137 | 0.11969 | 0.03573 |
| **KRI1** | 1.47175 | 8.10877 | 6.38563 | 8.66152 | 11.58635 | 0.00000 | 5.33346 | 4.38354 | 20.04005 | 11.89325 | 6.95560 | 9.35835 |
| **KRIT1** | 0.02047 | 5.84158 | 5.65656 | 1.85161 | 0.00000 | 11.11161 | 5.97369 | 2.02513 | 3.66690 | 11.91255 | 5.84240 | 5.66377 |
| **KRR1** | 4.14157 | 4.23229 | 12.76233 | 1.26655 | 0.00000 | 17.10690 | 11.98685 | 7.62297 | 10.19194 | 7.72629 | 4.36047 | 9.86993 |
| **KRT1** | 0.00616 | 0.01592 | 0.01856 | 0.01296 | 0.02711 | 0.02247 | 0.06287 | 0.01021 | 0.00568 | 0.01721 | 0.03339 | 0.01908 |
| **KRT10** | 0.18846 | 5.22432 | 5.39612 | 0.00000 | 0.80788 | 6.57844 | 6.73389 | 4.99402 | 6.94180 | 10.09155 | 1.98260 | 7.76556 |
| **KRT12** | 0.00234 | 0.01643 | 0.02485 | 0.01839 | 0.01265 | 0.01220 | 0.04876 | 0.03760 | 0.00000 | 0.04857 | 0.02274 | 0.01844 |
| **KRT13** | 0.05387 | 0.00641 | 0.02968 | 0.02380 | 0.00000 | 0.03368 | 0.03674 | 0.00653 | 0.04304 | 0.03351 | 0.03368 | 0.03261 |
| **KRT14** | 0.00288 | 0.01838 | 1.25482 | 0.01154 | 0.02301 | 0.00000 | 0.00000 | 0.03845 | 0.00000 | 0.01057 | 0.01808 | 0.01556 |
| **KRT15** | 0.13336 | 0.00786 | 0.02620 | 0.01594 | 0.00158 | 0.03705 | 0.01783 | 0.23810 | 0.44172 | 0.00000 | 0.02934 | 0.00985 |
| **KRT16** | 0.06393 | 0.01185 | 0.39688 | 0.01938 | 0.00000 | 0.03669 | 0.03742 | 0.00818 | 0.06243 | 0.04968 | 0.00000 | 0.02141 |
| **KRT16P1** | 0.00958 | 0.00764 | 0.00386 | 0.09500 | 0.00000 | 0.02813 | 0.00707 | 0.01376 | 0.02299 | 0.08770 | 0.01286 | 0.02386 |
| **KRT16P2** | 0.01259 | 0.01295 | 0.11434 | 0.04149 | 0.01165 | 0.02481 | 0.03167 | 0.00927 | 0.00000 | 0.02039 | 0.01371 | 0.01646 |
| **KRT16P3** | 0.00600 | 0.01249 | 0.05415 | 0.17770 | 0.00946 | 0.00891 | 0.00719 | 0.01015 | 0.02425 | 0.02026 | 0.01447 | 0.02129 |
| **KRT17** | 0.03949 | 0.20262 | 0.15494 | 0.00000 | 0.00000 | 0.09089 | 0.12130 | 0.28820 | 0.00000 | 0.11522 | 0.00936 | 0.00000 |
| **KRT17P5** | 0.00621 | 0.01494 | 0.01725 | 0.01146 | 0.01669 | 0.01674 | 0.01898 | 0.01044 | 0.01738 | 0.01710 | 0.01517 | 0.01682 |
| **KRT18** | 0.00000 | 49.90849 | 48.71700 | 25.37493 | 85.38589 | 35.65455 | 70.93171 | 0.00000 | 71.00601 | 6.57738 | 62.36769 | 6.47593 |
| **KRT18P55** | 0.44388 | 0.00850 | 0.12510 | 0.00000 | 0.71215 | 0.00000 | 0.54187 | 0.00000 | 0.00000 | 0.46548 | 0.22779 | 0.13631 |
| **KRT19** | 0.06946 | 0.47516 | 35.59159 | 1.49100 | 6.20473 | 1.69089 | 71.84632 | 0.20243 | 78.52199 | 2.36146 | 35.29523 | 0.00000 |
| **KRT19P2** | 0.00511 | 0.01108 | 0.03642 | 0.01064 | 0.01507 | 0.00871 | 0.04596 | 0.00959 | 0.02568 | 0.05400 | 0.12058 | 0.02754 |
| **KRT2** | 0.00646 | 0.01557 | 0.01486 | 0.01676 | 0.01507 | 0.01549 | 0.02380 | 0.01043 | 0.02757 | 0.01438 | 0.01358 | 0.01633 |
| **KRT20** | 0.04827 | 0.01743 | 0.06192 | 0.01624 | 0.02226 | 0.00008 | 0.00713 | 0.03999 | 0.02706 | 0.01854 | 0.01464 | 0.01411 |
| **KRT222** | 0.00681 | 0.00780 | 0.01447 | 0.01465 | 0.10747 | 0.00000 | 0.02839 | 0.52382 | 0.00973 | 0.00413 | 0.01227 | 0.09674 |
| **KRT23** | 0.10717 | 0.00937 | 25.30844 | 1.38386 | 0.00000 | 0.00531 | 0.86962 | 0.04959 | 0.70446 | 0.00000 | 0.00000 | 0.02626 |
| **KRT24** | 0.00576 | 0.01580 | 0.01709 | 0.01050 | 0.01620 | 0.02400 | 0.02818 | 0.01067 | 0.00702 | 0.02559 | 0.01438 | 0.01744 |
| **KRT25** | 0.00573 | 0.01430 | 0.01426 | 0.00963 | 0.01021 | 0.01687 | 0.03617 | 0.01004 | 0.00374 | 0.05043 | 0.01525 | 0.01810 |
| **KRT26** | 0.00811 | 0.00906 | 0.04137 | 0.02456 | 0.00542 | 0.04178 | 0.00741 | 0.01045 | 0.00000 | 0.03618 | 0.02639 | 0.01364 |
| **KRT27** | 0.08793 | 0.02020 | 0.01593 | 0.01286 | 0.02156 | 0.00000 | 0.02040 | 0.00311 | 0.02523 | 0.00000 | 0.01410 | 0.52056 |
| **KRT3** | 0.00608 | 0.01352 | 0.02035 | 0.01327 | 0.01324 | 0.02522 | 0.02680 | 0.01042 | 0.01278 | 0.01760 | 0.01805 | 0.01551 |
| **KRT32** | 0.00614 | 0.01464 | 0.01755 | 0.00921 | 0.01129 | 0.02785 | 0.01461 | 0.01037 | 0.02254 | 0.01940 | 0.01265 | 0.01512 |
| **KRT33A** | 0.00541 | 0.01484 | 0.06107 | 0.00989 | 0.01524 | 0.02247 | 0.02265 | 0.01381 | 0.01000 | 0.00868 | 0.01609 | 0.01648 |
| **KRT35** | 0.00612 | 0.01505 | 0.01697 | 0.01189 | 0.01362 | 0.01450 | 0.02981 | 0.01053 | 0.00914 | 0.01808 | 0.02829 | 0.01630 |
| **KRT36** | 0.20160 | 0.01454 | 0.01731 | 0.01204 | 0.01562 | 0.01299 | 0.01732 | 0.16689 | 0.02042 | 0.01277 | 0.01450 | 0.00967 |
| **KRT4** | 0.00796 | 0.00889 | 0.07627 | 0.02315 | 0.01444 | 0.04311 | 0.01486 | 0.01035 | 0.00000 | 0.02898 | 0.01876 | 0.01351 |
| **KRT40** | 0.00626 | 0.01605 | 0.01053 | 0.00887 | 0.01042 | 0.03229 | 0.08459 | 0.01042 | 0.00426 | 0.00500 | 0.01160 | 0.01550 |
| **KRT42P** | 0.00533 | 0.01384 | 0.12340 | 0.00921 | 0.01724 | 0.01897 | 0.00000 | 0.01350 | 0.01566 | 0.02531 | 0.01418 | 0.01524 |
| **KRT5** | 0.00508 | 0.01110 | 0.05574 | 0.03006 | 0.00000 | 0.02791 | 0.01451 | 0.01664 | 0.00000 | 0.16500 | 0.02198 | 0.00939 |

| | | | | | | | | | | | | |
|---|---|---|---|---|---|---|---|---|---|---|---|---|
| **KRT6A** | 0.00453 | 0.01371 | 0.01742 | 0.01482 | 0.02398 | 0.02671 | 0.02720 | 0.01654 | 0.01307 | 0.02898 | 0.01334 | 0.01421 |
| **KRT6B** | 0.00478 | 0.01093 | 0.01119 | 0.19544 | 0.00000 | 0.01889 | 0.01686 | 0.01200 | 0.00826 | 0.06294 | 0.01583 | 0.04227 |
| **KRT7** | 0.30217 | 2.30360 | 29.53259 | 2.02382 | 8.84093 | 0.00000 | 6.10117 | 0.05476 | 2.73139 | 0.00000 | 2.11349 | 0.00000 |
| **KRT71** | 0.00367 | 0.01304 | 0.04448 | 0.00545 | 0.00257 | 0.02355 | 0.05842 | 0.01996 | 0.03201 | 0.03616 | 0.01713 | 0.01703 |
| **KRT72** | 0.00692 | 0.00751 | 0.03804 | 0.02365 | 0.00085 | 0.05687 | 0.01069 | 0.01363 | 0.00298 | 0.03394 | 0.02159 | 0.01237 |
| **KRT73** | 0.00403 | 0.00956 | 0.04122 | 0.02538 | 0.00000 | 0.04029 | 0.01699 | 0.03247 | 0.00000 | 0.04284 | 0.01799 | 0.01445 |
| **KRT75** | 0.00539 | 0.01416 | 0.01075 | 0.18199 | 0.00839 | 0.04063 | 0.01989 | 0.01319 | 0.04757 | 0.00000 | 0.03437 | 0.01472 |
| **KRT76** | 0.00618 | 0.01750 | 0.01675 | 0.02016 | 0.03446 | 0.00115 | 0.03499 | 0.01069 | 0.01171 | 0.01485 | 0.01473 | 0.01732 |
| **KRT77** | 0.04570 | 0.01675 | 0.01869 | 0.01390 | 0.02596 | 0.00268 | 0.02400 | 0.00892 | 0.02678 | 0.00957 | 0.03784 | 0.01678 |
| **KRT78** | 0.00560 | 0.00977 | 0.01423 | 0.04055 | 0.12791 | 0.00000 | 0.02190 | 0.01053 | 0.00599 | 0.03597 | 0.01052 | 0.01694 |
| **KRT79** | 0.00669 | 0.02082 | 0.01382 | 0.00836 | 0.01131 | 0.02555 | 0.01293 | 0.00991 | 0.02587 | 0.02999 | 0.01597 | 0.01717 |
| **KRT8** | 0.16388 | 49.40641 | 47.39392 | 32.22750 | 75.37808 | 46.29007 | 73.81843 | 0.00000 | 74.92123 | 6.43286 | 62.81633 | 14.92413 |
| **KRT80** | 0.39903 | 0.02097 | 4.54455 | 0.00267 | 0.01532 | 0.00000 | 0.00000 | 0.07422 | 0.00000 | 0.10424 | 0.04788 | 0.00847 |
| **KRT81** | 0.01025 | 0.01624 | 0.22220 | 0.01162 | 0.01479 | 0.01308 | 0.05005 | 0.05129 | 0.00000 | 0.01019 | 0.01645 | 0.01389 |
| **KRT82** | 0.00633 | 0.03307 | 0.04187 | 0.05929 | 0.01123 | 0.00000 | 0.00671 | 0.01066 | 0.03860 | 0.01636 | 0.01325 | 0.01538 |
| **KRT84** | 0.00604 | 0.02193 | 0.02324 | 0.01480 | 0.02872 | 0.00000 | 0.03820 | 0.01061 | 0.01040 | 0.01873 | 0.06530 | 0.01843 |
| **KRT85** | 0.00580 | 0.01246 | 0.01347 | 0.02175 | 0.00773 | 0.02388 | 0.01817 | 0.01339 | 0.01097 | 0.02125 | 0.03500 | 0.01550 |
| **KRT86** | 0.00647 | 0.02026 | 0.52080 | 0.00848 | 0.03163 | 0.00000 | 0.08469 | 0.01051 | 0.00000 | 0.00000 | 0.01606 | 0.01786 |
| **KRT8P41** | 0.00679 | 0.09441 | 0.13645 | 0.05285 | 0.34661 | 0.09120 | 0.20072 | 0.00607 | 0.11075 | 0.00000 | 0.41223 | 0.11873 |
| **KRT9** | 0.00201 | 0.01123 | 0.03180 | 0.00967 | 0.08876 | 0.00000 | 0.08309 | 0.06103 | 0.00000 | 0.03166 | 0.01047 | 0.08875 |
| **KRTAP10-10** | 0.00775 | 0.01576 | 0.03077 | 0.01234 | 0.01585 | 0.01278 | 0.01649 | 0.01110 | 0.01620 | 0.01721 | 0.02066 | 0.01629 |
| **KRTAP10-2** | 0.05638 | 0.01444 | 0.01778 | 0.01175 | 0.01422 | 0.01705 | 0.01670 | 0.00584 | 0.01705 | 0.01392 | 0.02104 | 0.01647 |
| **KRTAP10-6** | 0.01203 | 0.01489 | 0.01534 | 0.01217 | 0.01535 | 0.01617 | 0.01862 | 0.01155 | 0.01967 | 0.01601 | 0.01454 | 0.01814 |
| **KRTAP11-1** | 0.00169 | 0.01306 | 0.04178 | 0.00841 | 0.01409 | 0.03079 | 0.01371 | 0.09701 | 0.02416 | 0.00141 | 0.01233 | 0.02443 |
| **KRTAP19-1** | 0.00719 | 0.01701 | 0.25343 | 0.01516 | 0.02266 | 0.00000 | 0.00314 | 0.01035 | 0.00000 | 0.01003 | 0.03875 | 0.01766 |
| **KRTAP19-8** | 0.08962 | 0.01462 | 0.03353 | 0.01621 | 0.01393 | 0.01623 | 0.02290 | 0.02841 | 0.04162 | 0.00000 | 0.00990 | 0.02552 |
| **KRTAP22-2** | 0.00583 | 0.01638 | 0.01316 | 0.01066 | 0.01165 | 0.01924 | 0.01600 | 0.00873 | 0.01573 | 0.02705 | 0.01547 | 0.05071 |
| **KRTAP4-1** | 0.00414 | 0.30811 | 0.00169 | 0.00000 | 0.00000 | 0.02995 | 0.00000 | 0.01059 | 0.00328 | 0.15888 | 0.00342 | 0.00012 |
| **KRTAP5-10** | 0.00610 | 0.01626 | 0.01760 | 0.03083 | 0.02839 | 0.00282 | 0.02408 | 0.00931 | 0.02387 | 0.00000 | 0.01582 | 0.05541 |
| **KRTAP5-7** | 0.00570 | 0.03806 | 0.01139 | 0.00610 | 0.00001 | 0.05754 | 0.00762 | 0.00979 | 0.01147 | 0.03344 | 0.01111 | 0.01420 |
| **KRTAP5-9** | 0.00369 | 0.01195 | 0.02345 | 0.01756 | 0.03621 | 0.00000 | 0.05054 | 0.01281 | 0.00000 | 0.08844 | 0.01173 | 0.26511 |
| **KRTAP5-AS1** | 0.01672 | 0.01361 | 0.01850 | 0.01012 | 0.01757 | 0.02763 | 0.02274 | 0.01086 | 0.01492 | 0.00749 | 0.01432 | 0.02340 |
| **KRTAP7-1** | 0.00582 | 0.02111 | 0.01365 | 0.00717 | 0.00658 | 0.03520 | 0.03905 | 0.01112 | 0.00147 | 0.01844 | 0.02880 | 0.01486 |
| **KRTCAP2** | 1.67838 | 16.00558 | 7.63835 | 6.61500 | 14.74594 | 8.43799 | 2.25102 | 1.72531 | 18.88862 | 13.91880 | 9.73410 | 12.12706 |
| **KRTCAP3** | 0.00162 | 6.96890 | 4.45451 | 10.16473 | 4.67722 | 3.61789 | 0.81539 | 0.51344 | 2.16931 | 0.00000 | 2.67468 | 2.54728 |
| **KRTDAP** | 0.00587 | 0.01868 | 0.01147 | 0.01358 | 0.01045 | 0.01487 | 0.02020 | 0.01053 | 0.02630 | 0.03643 | 0.01303 | 0.01467 |
| **KSR1** | 2.59791 | 0.18420 | 0.96749 | 0.47884 | 0.80133 | 0.00000 | 0.88434 | 0.00000 | 2.50237 | 0.60018 | 0.13489 | 0.11056 |
| **KSR2** | 3.36415 | 0.06448 | 0.16576 | 0.17730 | 0.00000 | 0.00000 | 0.07732 | 2.11433 | 0.19166 | 0.03431 | 0.00000 | 0.05890 |
| **KTN1** | 8.86442 | 2.04227 | 11.37788 | 8.47523 | 11.38412 | 38.74518 | 1.87812 | 3.62581 | 23.09908 | 51.85759 | 15.18394 | 5.75594 |
| **KTN1-AS1** | 0.54710 | 0.35929 | 0.58560 | 0.00696 | 0.79326 | 1.73885 | 0.33730 | 0.04091 | 0.00000 | 0.00000 | 0.91188 | 0.84349 |
| **KU-MEL-3** | 0.00229 | 0.01734 | 0.06865 | 0.02015 | 0.02234 | 0.00341 | 0.04396 | 0.05228 | 0.00000 | 0.00725 | 0.03683 | 0.01614 |
| **KXD1** | 2.61980 | 26.33839 | 17.01075 | 18.94968 | 21.70752 | 6.23814 | 24.35048 | 0.55532 | 28.22028 | 0.00000 | 19.66963 | 16.71518 |
| **KY** | 0.11572 | 0.03415 | 0.08005 | 0.00714 | 0.02452 | 0.00452 | 0.00975 | 0.05065 | 0.01121 | 0.04402 | 0.01572 | 0.04956 |
| **KYNU** | 0.20470 | 0.13505 | 2.30365 | 0.00322 | 0.09426 | 0.15499 | 0.00000 | 0.07355 | 0.41434 | 0.00000 | 0.00000 | 0.14232 |
| **L1CAM** | 0.09987 | 0.00000 | 0.00000 | 0.00000 | 0.36819 | 0.00000 | 0.04219 | 0.04623 | 0.36597 | 0.53355 | 0.29495 | 0.03570 |
| **L1TD1** | 0.00000 | 3.67120 | 24.97762 | 30.02517 | 9.25109 | 38.15260 | 26.84291 | 2.92975 | 0.55433 | 29.73232 | 4.55575 | 30.84070 |
| **L2HGDH** | 0.01923 | 0.63169 | 4.28355 | 3.04783 | 2.16240 | 5.97648 | 3.54267 | 0.66377 | 5.30598 | 0.86767 | 2.60705 | 2.50805 |
| **L3HYPDH** | 0.74971 | 0.77570 | 5.84414 | 0.00000 | 0.00000 | 4.78536 | 6.88405 | 0.67618 | 7.53450 | 0.00000 | 9.94617 | 0.95035 |

| | | | | | | | | | | | | |
|---|---|---|---|---|---|---|---|---|---|---|---|---|
| **L3MBTL1** | 0.00366 | 0.15478 | 0.04120 | 0.03311 | 0.04351 | 0.06976 | 0.20466 | 0.03637 | 0.41487 | 0.38856 | 0.00000 | 0.11450 |
| **L3MBTL2** | 0.61757 | 7.24709 | 8.30086 | 5.29509 | 3.90830 | 3.37921 | 6.70968 | 0.79796 | 12.54849 | 4.88264 | 8.46987 | 5.32269 |
| **L3MBTL3** | 0.16654 | 0.00000 | 0.00000 | 1.01266 | 0.69468 | 0.21679 | 0.96940 | 0.15218 | 0.69652 | 2.82355 | 0.37307 | 0.21044 |
| **L3MBTL4** | 0.00594 | 0.00962 | 0.00789 | 0.01688 | 0.00663 | 0.02826 | 0.03518 | 0.01070 | 0.00707 | 0.04622 | 0.01278 | 0.02496 |
| **LACC1** | 1.51601 | 0.00000 | 1.81400 | 0.00000 | 0.28173 | 0.02366 | 0.74611 | 3.99836 | 0.00003 | 0.03901 | 0.49501 | 0.54753 |
| **LACE1** | 0.04763 | 2.69704 | 0.39834 | 0.50245 | 1.13702 | 2.66235 | 1.13048 | 0.25003 | 0.27839 | 0.14500 | 3.46609 | 0.98225 |
| **LACTB** | 0.00134 | 0.86437 | 0.83907 | 6.86879 | 2.80967 | 2.15169 | 1.16781 | 0.74737 | 0.00001 | 2.47452 | 0.74471 | 0.00000 |
| **LACTB2** | 0.00937 | 1.99577 | 1.02463 | 1.89457 | 3.08985 | 9.25869 | 0.78784 | 0.00003 | 0.00000 | 0.93969 | 2.61849 | 0.00000 |
| **LAD1** | 19.28914 | 0.00000 | 6.83347 | 0.55640 | 15.67106 | 1.07863 | 8.63140 | 2.13340 | 11.18304 | 0.00000 | 4.53183 | 0.45957 |
| **LAG3** | 0.12329 | 0.00514 | 0.02356 | 0.01071 | 0.07561 | 0.00000 | 0.01859 | 0.01677 | 0.02181 | 0.08892 | 0.02699 | 0.06260 |
| **LAGE3** | 2.69232 | 3.99805 | 6.64957 | 4.40091 | 6.86935 | 0.00000 | 5.60661 | 1.60627 | 3.60417 | 7.06058 | 6.88908 | 0.42216 |
| **LAIR1** | 0.00647 | 0.06712 | 0.01093 | 0.00825 | 0.05360 | 0.03029 | 0.02795 | 0.00654 | 0.01920 | 0.00001 | 0.02622 | 0.14425 |
| **LAIR2** | 0.00573 | 0.01111 | 0.04116 | 0.00646 | 0.00407 | 0.01928 | 0.03927 | 0.01194 | 0.00000 | 0.01816 | 0.17177 | 0.01994 |
| **LALBA** | 0.00475 | 0.28301 | 0.00818 | 0.00281 | 0.18401 | 0.02106 | 0.02072 | 0.01031 | 0.00457 | 0.00718 | 0.06229 | 0.80680 |
| **LAMA1** | 0.00000 | 16.84921 | 17.06817 | 11.12594 | 30.27922 | 28.45492 | 2.72490 | 0.28134 | 60.86490 | 49.15993 | 21.68236 | 7.92019 |
| **LAMA2** | 2.25846 | 0.00000 | 0.11999 | 0.05846 | 0.00003 | 0.18131 | 0.00000 | 0.17792 | 0.13984 | 0.69250 | 0.04088 | 0.04212 |
| **LAMA3** | 0.00266 | 0.63185 | 0.02276 | 0.29360 | 0.00000 | 0.12397 | 0.00000 | 0.17499 | 0.10031 | 0.09720 | 0.07945 | 0.82436 |
| **LAMA4** | 0.94531 | 0.64005 | 0.02341 | 10.05181 | 0.00000 | 0.00000 | 0.00000 | 0.36538 | 0.00000 | 0.33537 | 0.00000 | 1.12138 |
| **LAMA5** | 2.82749 | 3.31205 | 5.70173 | 4.67015 | 17.59319 | 0.00000 | 0.75587 | 0.27368 | 21.74976 | 28.43795 | 2.94124 | 0.00000 |
| **LAMA5-AS1** | 0.00804 | 0.03926 | 0.02927 | 0.00961 | 0.04185 | 0.00000 | 0.01704 | 0.01052 | 0.03035 | 0.03284 | 0.01256 | 0.01765 |
| **LAMB1** | 5.60171 | 5.18076 | 13.88353 | 15.86776 | 28.44709 | 20.65239 | 9.72822 | 4.68326 | 44.67397 | 23.11510 | 23.46818 | 1.58459 |
| **LAMB2** | 0.00000 | 0.60867 | 0.51059 | 0.00000 | 0.93268 | 0.00000 | 0.15851 | 0.04008 | 7.22398 | 1.41714 | 1.42890 | 0.15708 |
| **LAMB2P1** | 0.02640 | 0.01013 | 0.02988 | 0.00262 | 0.00000 | 0.02302 | 0.05149 | 0.00703 | 0.02920 | 0.10573 | 0.00375 | 0.02213 |
| **LAMB3** | 0.08077 | 0.01528 | 0.53984 | 0.00000 | 0.94333 | 0.01999 | 0.37985 | 0.06316 | 5.16852 | 0.00797 | 1.18517 | 0.06904 |
| **LAMB4** | 0.65794 | 0.04376 | 0.04945 | 0.20069 | 0.01865 | 0.00000 | 0.05398 | 0.00235 | 0.00000 | 0.18326 | 0.26704 | 0.01009 |
| **LAMC1** | 1.36481 | 6.10688 | 11.13928 | 10.11615 | 29.09716 | 27.12131 | 2.72559 | 3.12783 | 43.54469 | 47.82039 | 18.75875 | 2.20266 |
| **LAMC2** | 0.00074 | 1.02647 | 1.45880 | 4.98438 | 0.73344 | 1.11828 | 0.00000 | 0.10577 | 2.41118 | 0.00000 | 0.00049 | 0.00000 |
| **LAMC3** | 0.16381 | 1.75800 | 0.26971 | 0.19721 | 1.42794 | 0.00000 | 0.00000 | 0.05189 | 0.00000 | 2.64298 | 0.75651 | 0.00000 |
| **LAMP1** | 2.28427 | 1.23753 | 8.29620 | 0.74774 | 8.58556 | 1.92517 | 9.07995 | 1.48359 | 0.49958 | 35.24239 | 2.03609 | 2.21756 |
| **LAMP2** | 2.71787 | 3.34919 | 16.16881 | 2.53482 | 13.69713 | 37.39368 | 0.00000 | 5.82797 | 48.98868 | 11.94114 | 29.87956 | 1.67971 |
| **LAMP3** | 1.12526 | 0.22707 | 0.00000 | 0.00000 | 0.02185 | 0.08398 | 0.04090 | 0.62777 | 0.01894 | 0.02766 | 0.03475 | 0.15802 |
| **LAMP5** | 0.00610 | 0.01155 | 0.03748 | 0.01826 | 0.01756 | 0.01342 | 0.01674 | 0.00832 | 0.01505 | 0.00699 | 0.01652 | 0.08066 |
| **LAMP5-AS1** | 0.00516 | 0.01269 | 0.00874 | 0.00809 | 0.01072 | 0.01432 | 0.01293 | 0.01042 | 0.02241 | 0.07979 | 0.01384 | 0.01340 |
| **LAMTOR1** | 0.72189 | 24.28598 | 20.76653 | 14.08351 | 24.93854 | 11.01491 | 24.42772 | 1.02478 | 25.25142 | 5.12242 | 21.85814 | 9.60212 |
| **LAMTOR2** | 3.79780 | 21.20493 | 9.65485 | 18.80110 | 17.52969 | 12.44195 | 21.61083 | 2.67413 | 10.70637 | 12.82772 | 19.57107 | 11.49766 |
| **LAMTOR3** | 5.09685 | 7.76265 | 12.73648 | 2.10239 | 1.97423 | 23.05016 | 10.89437 | 12.78893 | 8.86521 | 0.00000 | 14.15571 | 19.84836 |
| **LAMTOR4** | 0.67640 | 15.80005 | 15.32609 | 17.82111 | 30.57671 | 0.87988 | 37.97660 | 9.61520 | 13.43993 | 7.79839 | 25.75498 | 14.41899 |
| **LAMTOR5** | 4.24022 | 19.47473 | 11.39820 | 13.78905 | 19.83496 | 37.35362 | 25.92285 | 17.50084 | 20.25033 | 8.28218 | 23.05390 | 27.32887 |
| **LAMTOR5-AS1** | 0.08426 | 0.16445 | 0.11109 | 0.00178 | 0.51682 | 0.47530 | 0.02635 | 0.05919 | 0.00000 | 0.00000 | 0.29431 | 0.64212 |
| **LANCL1** | 0.74758 | 1.73059 | 1.09586 | 0.23738 | 0.00000 | 2.41454 | 0.00000 | 0.01221 | 0.00000 | 0.47222 | 0.79905 | 0.21167 |
| **LANCL1-AS1** | 0.00505 | 0.03648 | 0.01257 | 0.00332 | 0.01139 | 0.05415 | 0.00793 | 0.00901 | 0.00000 | 0.07920 | 0.09286 | 0.02117 |
| **LANCL2** | 0.69762 | 7.37370 | 1.93147 | 2.91954 | 3.82945 | 2.57080 | 5.34229 | 0.19109 | 2.61557 | 1.44217 | 1.88927 | 2.49347 |
| **LANCL3** | 0.00546 | 0.01057 | 0.05906 | 0.14455 | 0.00495 | 0.02436 | 0.01216 | 0.01112 | 0.01287 | 0.03078 | 0.01446 | 0.01482 |
| **LAP3** | 0.15148 | 7.25562 | 12.50786 | 6.70034 | 3.25930 | 13.26293 | 10.08416 | 7.82427 | 8.38904 | 1.34653 | 10.85063 | 6.97684 |
| **LAPTM4A** | 8.73104 | 10.85781 | 15.98153 | 15.21641 | 2.78770 | 39.43603 | 0.00000 | 20.18691 | 30.82198 | 12.44788 | 14.56146 | 28.81523 |
| **LAPTM4B** | 5.81893 | 13.39225 | 5.36570 | 14.25483 | 6.71225 | 21.09001 | 0.00000 | 2.92994 | 5.53366 | 22.61191 | 0.24993 | 14.63850 |
| **LAPTM5** | 0.03997 | 2.04901 | 0.38611 | 0.06198 | 1.70836 | 1.62072 | 0.00000 | 0.00632 | 0.19627 | 0.37457 | 0.00000 | 0.72132 |
| **LARGE** | 0.72338 | 0.57560 | 0.47331 | 0.79909 | 1.67049 | 1.08733 | 1.20852 | 0.17899 | 0.10931 | 3.07698 | 0.00000 | 0.00000 |

| | | | | | | | | | | | | |
|---|---|---|---|---|---|---|---|---|---|---|---|---|
| **LARGE-AS1** | 0.15816 | 0.00640 | 0.07907 | 0.05915 | 0.27898 | 0.00000 | 0.00113 | 0.02675 | 0.03931 | 0.81762 | 0.02589 | 0.02245 |
| **LARP1** | 13.60573 | 12.41088 | 17.79624 | 16.65838 | 39.41970 | 20.85003 | 42.72039 | 1.92993 | 18.34807 | 44.08650 | 23.90214 | 4.91842 |
| **LARP1B** | 2.68464 | 10.13399 | 7.24023 | 5.81006 | 3.04048 | 10.97137 | 8.57262 | 12.83874 | 0.82913 | 17.05012 | 2.20402 | 28.20412 |
| **LARP4** | 6.05063 | 12.73723 | 16.91837 | 3.36427 | 5.73025 | 36.72860 | 17.32561 | 18.33995 | 11.79299 | 18.00832 | 11.97309 | 15.64160 |
| **LARP4B** | 1.78600 | 10.41217 | 10.58767 | 9.94089 | 10.09241 | 13.87909 | 18.83448 | 19.47642 | 7.00775 | 9.04414 | 13.98908 | 6.27243 |
| **LARP6** | 0.00000 | 2.61664 | 0.00000 | 8.96839 | 3.44693 | 0.00000 | 0.02411 | 0.04395 | 0.00000 | 3.00623 | 0.58895 | 0.84612 |
| **LARP7** | 7.30552 | 10.76820 | 5.84191 | 18.01704 | 4.72341 | 34.07528 | 12.56475 | 2.83610 | 3.01946 | 26.99544 | 15.47052 | 20.33697 |
| **LARS** | 3.17913 | 9.60997 | 16.77725 | 8.05713 | 13.15377 | 32.53164 | 21.55281 | 2.02544 | 19.69435 | 23.51665 | 22.00236 | 1.41010 |
| **LARS2** | 6.44748 | 9.63084 | 3.48480 | 7.01041 | 11.04879 | 0.00008 | 7.60001 | 1.67539 | 14.24964 | 0.00022 | 4.33214 | 5.40306 |
| **LARS2-AS1** | 0.01149 | 0.04052 | 0.18480 | 0.00000 | 0.07825 | 0.35468 | 0.00000 | 0.00550 | 0.06464 | 1.04570 | 0.09765 | 0.05914 |
| **LAS1L** | 2.43387 | 15.12560 | 11.87546 | 10.51224 | 11.98505 | 6.12828 | 18.65931 | 4.23652 | 8.50615 | 0.00000 | 23.12101 | 19.24323 |
| **LASP1** | 1.69398 | 35.86553 | 6.43819 | 16.17400 | 32.52189 | 11.75631 | 12.89718 | 1.91813 | 7.23220 | 19.74523 | 7.54632 | 14.23048 |
| **LAT** | 0.08183 | 0.20758 | 0.28118 | 0.20739 | 0.00000 | 0.13050 | 0.12979 | 0.03711 | 1.80664 | 1.82542 | 0.00000 | 0.36996 |
| **LAT2** | 0.10432 | 0.33960 | 0.05223 | 0.03633 | 0.06534 | 0.36299 | 0.13802 | 0.16285 | 0.36489 | 0.55516 | 0.00000 | 0.79934 |
| **LATS1** | 1.90585 | 2.06081 | 4.46581 | 1.91480 | 0.00000 | 4.84193 | 2.32922 | 8.65551 | 2.77779 | 5.16930 | 1.13987 | 5.76128 |
| **LATS2** | 0.36277 | 1.34277 | 1.39857 | 0.00000 | 7.60582 | 9.24278 | 7.30738 | 1.36797 | 0.00000 | 16.04732 | 5.13916 | 6.01208 |
| **LAX1** | 0.00982 | 0.05975 | 0.02599 | 0.03617 | 0.00000 | 0.00000 | 0.00109 | 0.17129 | 0.04895 | 0.31010 | 0.01116 | 0.37661 |
| **LAYN** | 0.37345 | 2.80797 | 0.00000 | 0.00000 | 7.19159 | 0.00000 | 0.06362 | 0.00000 | 2.62723 | 1.04580 | 4.58448 | 0.81917 |
| **LBH** | 2.39847 | 0.35445 | 2.52582 | 24.09445 | 0.00000 | 0.00000 | 0.00000 | 5.37628 | 0.00000 | 1.00705 | 0.10567 | 1.49385 |
| **LBP** | 0.00364 | 0.08944 | 0.00561 | 0.13618 | 0.08055 | 0.00000 | 0.06354 | 0.01154 | 0.00000 | 0.35072 | 0.00000 | 0.13662 |
| **LBR** | 2.61508 | 10.05607 | 10.82888 | 4.64234 | 7.38622 | 38.91472 | 0.00000 | 7.63577 | 27.36301 | 11.69032 | 9.17807 | 9.04454 |
| **LBX1** | 0.00620 | 0.01368 | 0.01920 | 0.01588 | 0.01634 | 0.02324 | 0.01679 | 0.01042 | 0.01465 | 0.01648 | 0.01538 | 0.01581 |
| **LBX1-AS1** | 0.00636 | 0.01151 | 0.03401 | 0.02474 | 0.01010 | 0.04238 | 0.02125 | 0.01031 | 0.01227 | 0.00799 | 0.01626 | 0.01535 |
| **LBX2** | 0.00483 | 0.02495 | 0.13010 | 0.05679 | 0.23128 | 0.00000 | 0.02690 | 0.01032 | 0.12119 | 0.06658 | 0.03653 | 0.01288 |
| **LBX2-AS1** | 0.00022 | 0.64583 | 1.12178 | 0.15547 | 0.28200 | 0.21498 | 0.02115 | 0.01939 | 0.32896 | 0.43301 | 0.09597 | 0.00000 |
| **LCA5** | 0.84432 | 0.00000 | 1.15477 | 0.11024 | 0.00000 | 1.11394 | 1.36221 | 0.11937 | 0.20986 | 0.00001 | 0.08882 | 0.31270 |
| **LCA5L** | 0.55366 | 0.39810 | 0.01363 | 0.16443 | 0.00000 | 0.00000 | 0.00457 | 0.14306 | 0.14053 | 0.09019 | 0.15220 | 1.01500 |
| **LCAT** | 0.00826 | 0.19390 | 0.90865 | 0.23165 | 1.04265 | 0.00000 | 0.25480 | 0.01301 | 0.97317 | 2.10590 | 0.00000 | 0.08549 |
| **LCK** | 0.00281 | 0.00852 | 0.36757 | 0.10187 | 0.00000 | 0.00513 | 0.10816 | 0.03185 | 0.07296 | 0.04344 | 0.02446 | 0.08376 |
| **LCLAT1** | 0.31459 | 2.58413 | 0.38890 | 3.51812 | 0.65635 | 12.98794 | 0.00000 | 2.02521 | 6.85202 | 4.15706 | 0.69967 | 5.97903 |
| **LCMT1** | 9.91303 | 7.65772 | 9.15962 | 4.61259 | 7.51569 | 10.92112 | 7.87414 | 3.89007 | 12.79635 | 0.00006 | 7.64397 | 5.37328 |
| **LCMT2** | 0.09893 | 2.15837 | 3.40679 | 0.75447 | 1.15545 | 2.52395 | 0.57093 | 1.64859 | 2.68400 | 0.52877 | 3.72932 | 4.15898 |
| **LCN1** | 0.01865 | 0.01640 | 0.01371 | 0.01340 | 0.01564 | 0.02089 | 0.01675 | 0.01009 | 0.01640 | 0.02025 | 0.01408 | 0.01598 |
| **LCN12** | 0.00354 | 0.01581 | 0.00312 | 0.00001 | 0.04142 | 0.00902 | 0.00057 | 0.00983 | 0.01622 | 0.33065 | 0.00073 | 0.00883 |
| **LCN15** | 0.00239 | 0.01606 | 0.01347 | 0.01062 | 0.02847 | 0.01687 | 0.01330 | 0.05440 | 0.05048 | 0.00302 | 0.01126 | 0.01664 |
| **LCN2** | 0.00560 | 0.00579 | 0.00000 | 0.00000 | 0.00000 | 0.09122 | 0.11606 | 0.00953 | 0.00605 | 0.02939 | 0.00469 | 0.01274 |
| **LCNL1** | 0.00632 | 0.01601 | 0.02623 | 0.01678 | 0.01692 | 0.01489 | 0.01671 | 0.01040 | 0.01675 | 0.01046 | 0.02035 | 0.01876 |
| **LCOR** | 6.22895 | 1.23122 | 1.70648 | 2.10545 | 0.00000 | 5.51108 | 0.00000 | 12.10912 | 3.58669 | 6.69581 | 0.57280 | 2.87583 |
| **LCORL** | 2.87355 | 0.89122 | 1.83762 | 1.16291 | 0.00000 | 5.25782 | 2.66767 | 1.86297 | 1.00898 | 2.92900 | 0.84019 | 1.26231 |
| **LCP1** | 0.00000 | 0.00000 | 35.50797 | 37.65415 | 10.37574 | 7.38709 | 28.68241 | 0.17338 | 11.15934 | 0.00000 | 5.25592 | 0.00000 |
| **LCP2** | 0.56543 | 0.02217 | 0.01738 | 0.01341 | 0.02305 | 0.00085 | 0.06186 | 0.01017 | 0.00072 | 0.00582 | 0.01686 | 0.01532 |
| **LCT** | 0.00186 | 0.00000 | 0.09256 | 0.00000 | 0.00000 | 0.18417 | 0.00000 | 0.09060 | 0.00948 | 0.16037 | 0.00303 | 0.03126 |
| **LCTL** | 0.00809 | 0.08102 | 0.00000 | 0.14078 | 0.00000 | 0.55604 | 0.00000 | 0.11596 | 0.14512 | 0.51287 | 0.39527 | 0.16408 |
| **LDB1** | 1.81644 | 2.75516 | 2.30260 | 5.32377 | 10.50445 | 0.00022 | 3.87572 | 0.50667 | 3.01618 | 13.68252 | 1.67862 | 2.22666 |
| **LDB2** | 0.12717 | 0.00724 | 0.08497 | 0.08112 | 0.03433 | 0.03619 | 0.00000 | 0.03036 | 0.01406 | 0.03162 | 0.01714 | 0.02668 |
| **LDB3** | 0.01630 | 0.00000 | 0.06254 | 0.04548 | 0.15065 | 0.19777 | 0.04657 | 0.01796 | 0.18452 | 0.00000 | 0.00000 | 0.12684 |
| **LDHA** | 2.28440 | 54.58513 | 42.04341 | 29.57036 | 24.86769 | 55.14033 | 62.87781 | 19.49982 | 59.71028 | 12.05046 | 49.57051 | 72.85639 |
| **LDHAL6A** | 8.47828 | 0.00000 | 0.10057 | 0.12349 | 0.00000 | 0.00000 | 0.13047 | 2.34288 | 0.02065 | 0.44817 | 0.18115 | 0.07421 |

| | | | | | | | | | | | | |
|---|---|---|---|---|---|---|---|---|---|---|---|---|
| **LDHAL6B** | 0.01490 | 0.00798 | 0.21829 | 0.04007 | 0.00265 | 0.04689 | 0.00000 | 0.05098 | 0.00000 | 0.03044 | 0.02065 | 0.01298 |
| **LDHB** | 0.00000 | 47.01866 | 39.93092 | 35.96338 | 38.52857 | 75.92052 | 61.89451 | 4.31497 | 55.11680 | 26.09863 | 49.82372 | 42.64455 |
| **LDHC** | 1.56377 | 0.09368 | 0.01302 | 0.06986 | 0.16568 | 0.00000 | 0.09900 | 1.19771 | 0.00000 | 0.00000 | 0.00768 | 0.53628 |
| **LDHD** | 0.35999 | 1.86318 | 0.72671 | 2.38841 | 0.83852 | 0.00000 | 0.00000 | 0.00000 | 0.00000 | 0.00000 | 0.34302 | 0.10446 |
| **LDLR** | 3.00282 | 0.00000 | 6.43422 | 20.14721 | 25.94503 | 0.00000 | 4.07613 | 0.27818 | 26.03796 | 3.67467 | 6.63748 | 0.03674 |
| **LDLRAD1** | 0.00752 | 0.04205 | 0.03936 | 0.02189 | 0.02728 | 0.00000 | 0.01808 | 0.01284 | 0.01558 | 0.01681 | 0.01915 | 0.01656 |
| **LDLRAD2** | 0.00816 | 0.01948 | 0.04892 | 0.01304 | 0.01274 | 0.00609 | 0.00921 | 0.01083 | 0.03997 | 0.01891 | 0.02965 | 0.01625 |
| **LDLRAD3** | 2.27485 | 1.90922 | 1.20439 | 0.93480 | 0.82368 | 0.00000 | 0.00000 | 0.56818 | 2.23672 | 0.00000 | 0.27378 | 0.59477 |
| **LDLRAD4** | 0.04390 | 0.00000 | 0.02393 | 0.01378 | 0.15145 | 0.00000 | 0.00108 | 0.08064 | 0.22461 | 0.34247 | 0.00000 | 1.00429 |
| **LDLRAP1** | 5.03532 | 12.82785 | 1.02558 | 6.46447 | 1.04177 | 0.00000 | 0.00000 | 0.00000 | 2.45706 | 3.59113 | 0.07589 | 6.78941 |
| **LDOC1** | 3.23632 | 0.23574 | 1.92412 | 0.16283 | 0.00000 | 2.75934 | 3.57676 | 0.35858 | 7.68369 | 1.52924 | 10.44121 | 0.23979 |
| **LDOC1L** | 0.45941 | 0.40381 | 0.23264 | 0.00000 | 0.22924 | 0.48283 | 1.31370 | 0.00000 | 0.00000 | 0.02774 | 1.10219 | 0.04475 |
| **LEAP2** | 0.00275 | 0.13446 | 0.34912 | 0.23292 | 0.20327 | 0.47368 | 0.24507 | 0.95940 | 0.77852 | 0.49718 | 0.03306 | 0.75698 |
| **LECT1** | 0.00836 | 0.00012 | 0.16664 | 0.11101 | 0.00000 | 0.11293 | 0.00883 | 0.08182 | 0.00000 | 0.15658 | 0.00563 | 0.01993 |
| **LECT2** | 0.01143 | 0.01434 | 0.00415 | 0.00000 | 0.00000 | 0.00526 | 0.33645 | 0.01102 | 0.39605 | 0.26118 | 0.00000 | 0.01972 |
| **LEF1** | 7.71499 | 0.01437 | 0.01007 | 0.00504 | 0.00755 | 0.03198 | 0.02564 | 0.32317 | 0.02562 | 0.00000 | 0.00558 | 0.08899 |
| **LEF1-AS1** | 0.35929 | 0.00940 | 0.01653 | 0.01359 | 0.01750 | 0.00934 | 0.02154 | 0.01735 | 0.02015 | 0.00454 | 0.01503 | 0.10416 |
| **LEFTY1** | 0.02022 | 0.02354 | 3.25310 | 11.23528 | 0.00000 | 0.00000 | 0.00000 | 0.00243 | 0.00000 | 0.04732 | 0.01506 | 0.10918 |
| **LEFTY2** | 0.06427 | 0.00610 | 9.68526 | 2.59634 | 0.00000 | 0.00000 | 0.00000 | 0.00535 | 0.09452 | 0.03490 | 0.01138 | 0.01044 |
| **LEKR1** | 0.05771 | 0.05052 | 0.12370 | 0.00152 | 0.02714 | 0.05851 | 0.00000 | 0.01815 | 0.11632 | 0.01893 | 0.00958 | 0.09561 |
| **LEMD1** | 0.00906 | 0.01063 | 0.13884 | 0.07962 | 0.01003 | 0.03584 | 0.05799 | 0.01674 | 0.01591 | 0.00000 | 0.00994 | 0.01544 |
| **LEMD1-AS1** | 0.00609 | 0.01258 | 0.02723 | 0.01233 | 0.02032 | 0.00000 | 0.01374 | 0.01056 | 0.00000 | 0.02619 | 0.15211 | 0.01731 |
| **LEMD2** | 0.47612 | 1.81182 | 2.27517 | 2.15857 | 3.60249 | 0.00000 | 0.00005 | 0.00000 | 3.24212 | 5.01513 | 0.49565 | 0.50780 |
| **LEMD3** | 1.20806 | 1.39886 | 0.70061 | 1.57755 | 1.59755 | 6.13981 | 0.00000 | 2.74603 | 0.61191 | 6.91107 | 0.50602 | 1.33273 |
| **LENEP** | 0.02573 | 0.01137 | 0.03951 | 0.01867 | 0.01000 | 0.02499 | 0.03263 | 0.00543 | 0.01744 | 0.03713 | 0.01376 | 0.05545 |
| **LENG1** | 0.08076 | 0.78347 | 3.89715 | 3.90104 | 6.31648 | 0.96020 | 4.81279 | 3.71852 | 3.79989 | 0.00000 | 3.24907 | 3.60234 |
| **LENG8** | 2.08416 | 1.80530 | 2.41083 | 1.04587 | 0.82659 | 0.00000 | 0.00000 | 3.28204 | 3.32601 | 10.70350 | 1.65890 | 2.78066 |
| **LENG9** | 0.02934 | 0.01411 | 0.14894 | 0.12062 | 0.00000 | 0.02212 | 0.00887 | 0.00981 | 0.06759 | 0.05497 | 0.12692 | 0.01516 |
| **LEO1** | 0.04540 | 9.39115 | 8.84179 | 6.53027 | 0.69553 | 19.83602 | 11.21041 | 7.73875 | 9.27397 | 5.15253 | 10.37043 | 14.28669 |
| **LEP** | 0.00194 | 0.00000 | 1.24260 | 0.00000 | 0.00000 | 0.00000 | 0.80504 | 0.00770 | 1.90633 | 0.84978 | 0.62485 | 0.00151 |
| **LEPR** | 0.00380 | 0.29236 | 0.91815 | 0.01793 | 0.00000 | 0.79779 | 1.55740 | 0.15288 | 3.55810 | 0.00000 | 0.00000 | 0.09904 |
| **LEPRE1** | 0.69768 | 8.43985 | 4.54857 | 14.49737 | 0.93122 | 0.00000 | 0.00000 | 0.36297 | 11.21888 | 7.44457 | 6.84950 | 5.93263 |
| **LEPREL1** | 0.53822 | 0.01854 | 1.96932 | 1.14553 | 0.00000 | 0.00000 | 0.00000 | 0.03151 | 0.00000 | 0.07919 | 0.03141 | 0.04526 |
| **LEPREL2** | 0.00978 | 0.01491 | 0.03362 | 0.02833 | 0.00901 | 0.01466 | 0.01727 | 0.01526 | 0.02888 | 0.02074 | 0.01428 | 0.04370 |
| **LEPREL4** | 0.00825 | 0.29972 | 0.39107 | 0.01380 | 0.09722 | 0.27977 | 0.00000 | 0.03388 | 0.19532 | 0.37763 | 0.03696 | 0.04264 |
| **LEPROT** | 1.83613 | 3.69628 | 4.74258 | 1.04966 | 1.39962 | 17.48278 | 0.00000 | 0.38323 | 5.30626 | 8.15234 | 6.07557 | 5.34036 |
| **LEPROTL1** | 1.97873 | 8.99267 | 13.01473 | 4.17653 | 1.64728 | 18.76837 | 4.80021 | 1.19094 | 24.15459 | 4.90582 | 10.47721 | 7.02320 |
| **LETM1** | 1.60935 | 25.74320 | 9.97173 | 10.77165 | 22.34132 | 9.58433 | 20.04451 | 0.32381 | 11.62194 | 17.48571 | 15.79437 | 14.89879 |
| **LETM2** | 0.32159 | 0.16613 | 0.54951 | 0.28544 | 0.00000 | 0.13820 | 0.00000 | 0.96029 | 0.17057 | 0.00000 | 0.00828 | 0.35212 |
| **LETMD1** | 1.36075 | 2.72706 | 1.25671 | 1.46979 | 4.13655 | 4.88618 | 2.89580 | 0.36622 | 9.20931 | 7.77029 | 6.75411 | 2.36540 |
| **LEUTX** | 8.17317 | 0.01000 | 0.00407 | 0.00449 | 0.01410 | 0.00000 | 0.00551 | 23.49490 | 0.01831 | 0.00131 | 0.00537 | 0.00000 |
| **LFNG** | 0.61248 | 0.62624 | 0.09849 | 0.75314 | 0.13181 | 0.00000 | 0.00000 | 0.45576 | 0.00000 | 0.72928 | 0.00000 | 1.01174 |
| **LGALS1** | 11.60128 | 1.22268 | 4.75050 | 15.52881 | 0.83094 | 0.00000 | 6.85680 | 18.53625 | 6.57376 | 0.00000 | 0.66608 | 0.41784 |
| **LGALS12** | 1.35018 | 0.16084 | 0.40956 | 4.50286 | 0.22722 | 0.00000 | 0.03311 | 0.24306 | 0.03196 | 0.00000 | 0.02689 | 0.59491 |
| **LGALS13** | 0.00462 | 0.01271 | 0.71420 | 0.00929 | 0.00977 | 0.00000 | 0.15971 | 0.01033 | 0.00000 | 0.10859 | 0.00464 | 0.01218 |
| **LGALS14** | 0.00682 | 0.01811 | 0.04000 | 0.01012 | 0.06795 | 0.00004 | 0.02514 | 0.01054 | 0.03209 | 0.00000 | 0.00973 | 0.01875 |
| **LGALS16** | 0.00463 | 0.00788 | 17.26678 | 0.00000 | 0.00000 | 0.01636 | 0.73127 | 0.00954 | 0.00000 | 0.02610 | 0.00253 | 0.02630 |
| **LGALS17A** | 0.00601 | 0.01302 | 0.13007 | 0.00711 | 0.01018 | 0.02532 | 0.00000 | 0.01026 | 0.02466 | 0.03205 | 0.01149 | 0.01445 |

| | | | | | | | | | | | | |
|---|---|---|---|---|---|---|---|---|---|---|---|---|
| **LGALS2** | 0.09289 | 0.00463 | 0.15717 | 11.98810 | 0.02330 | 0.05703 | 0.11184 | 0.40815 | 0.02040 | 0.00000 | 0.03502 | 0.00802 |
| **LGALS3** | 0.00249 | 0.61053 | 35.18302 | 0.00002 | 0.00000 | 0.03453 | 15.33717 | 0.06684 | 1.60351 | 0.00000 | 1.16038 | 0.25828 |
| **LGALS3BP** | 0.13483 | 0.04775 | 3.04097 | 0.72133 | 0.00000 | 0.20261 | 0.00000 | 0.00394 | 0.00000 | 0.00808 | 0.02299 | 0.00582 |
| **LGALS4** | 1.26455 | 3.57304 | 0.00000 | 0.72824 | 1.32101 | 0.00000 | 0.04830 | 1.20120 | 0.00000 | 0.00000 | 0.01550 | 6.43322 |
| **LGALS7** | 0.00576 | 0.01538 | 0.01442 | 0.01082 | 0.02249 | 0.01689 | 0.01724 | 0.01369 | 0.01627 | 0.01939 | 0.01628 | 0.02433 |
| **LGALS7B** | 0.00692 | 0.03589 | 0.01047 | 0.06473 | 0.00875 | 0.03658 | 0.01398 | 0.03209 | 0.01813 | 0.01818 | 0.01654 | 0.03727 |
| **LGALS8** | 7.09202 | 0.46142 | 0.30304 | 0.11434 | 0.12917 | 1.27112 | 0.00000 | 2.03809 | 0.09841 | 2.01451 | 0.29844 | 0.45046 |
| **LGALS8-AS1** | 0.02072 | 0.01371 | 0.05658 | 0.01523 | 0.01337 | 0.02668 | 0.01393 | 0.01263 | 0.00730 | 0.01312 | 0.01613 | 0.01490 |
| **LGALS9** | 0.00831 | 0.00927 | 0.27977 | 0.00635 | 0.07827 | 0.00000 | 0.79609 | 0.00982 | 0.26289 | 0.39969 | 0.00000 | 0.00619 |
| **LGALS9C** | 0.00601 | 0.01427 | 0.01329 | 0.01379 | 0.01217 | 0.01991 | 0.03077 | 0.01039 | 0.01279 | 0.02437 | 0.01390 | 0.01707 |
| **LGALSL** | 0.33572 | 0.34721 | 2.32874 | 0.27823 | 0.00000 | 1.40125 | 1.33350 | 0.11425 | 0.00512 | 1.03887 | 0.18167 | 0.22842 |
| **LGI1** | 0.00295 | 0.01502 | 0.05410 | 0.12897 | 0.01217 | 0.00000 | 0.01283 | 0.10712 | 0.00879 | 0.03783 | 0.01738 | 0.01103 |
| **LGI2** | 0.11954 | 0.02981 | 0.12288 | 0.04174 | 0.01814 | 0.08538 | 0.00000 | 0.01340 | 0.00000 | 0.01713 | 0.01405 | 0.02162 |
| **LGI3** | 0.02140 | 0.00873 | 0.00188 | 0.00405 | 0.00000 | 0.01628 | 0.00225 | 0.01170 | 0.02021 | 0.21849 | 0.02573 | 0.00951 |
| **LGI4** | 0.00537 | 0.01389 | 0.03105 | 0.01437 | 0.00813 | 0.01500 | 0.09347 | 0.01133 | 0.00000 | 0.03891 | 0.01474 | 0.01465 |
| **LGMN** | 3.75804 | 6.70626 | 19.51272 | 5.12327 | 3.85476 | 10.21916 | 1.17137 | 25.94141 | 23.02371 | 0.00142 | 7.65924 | 30.13867 |
| **LGR4** | 0.13744 | 0.28192 | 14.17238 | 1.16741 | 4.66532 | 18.05523 | 0.00000 | 0.53992 | 7.29317 | 12.69464 | 1.46585 | 0.25450 |
| **LGR5** | 0.32504 | 0.00483 | 0.36987 | 0.69164 | 0.00000 | 0.00000 | 0.00000 | 0.16977 | 0.00851 | 0.12112 | 0.01094 | 0.00828 |
| **LGR6** | 0.18126 | 0.19784 | 0.29095 | 1.26156 | 0.00000 | 0.00000 | 0.00000 | 0.00164 | 0.00000 | 0.17802 | 0.00386 | 0.01968 |
| **LGSN** | 0.01553 | 0.12892 | 0.00310 | 0.01269 | 0.00000 | 0.25937 | 0.00000 | 0.00137 | 1.11140 | 0.02939 | 0.22761 | 0.12573 |
| **LHB** | 0.00541 | 0.03391 | 0.23430 | 0.00917 | 0.05778 | 0.00000 | 0.00160 | 1.56261 | 0.04614 | 0.03936 | 0.00892 | 1.05892 |
| **LHCGR** | 0.00585 | 0.02261 | 0.01964 | 0.01211 | 0.02228 | 0.01615 | 0.01037 | 0.01243 | 0.03251 | 0.02910 | 0.01179 | 0.02879 |
| **LHFP** | 0.40304 | 1.32145 | 1.76582 | 0.00083 | 0.55103 | 3.14792 | 0.94112 | 0.05329 | 2.25699 | 0.00000 | 1.68627 | 0.15379 |
| **LHFPL1** | 0.00568 | 0.02914 | 0.02884 | 0.01472 | 0.02552 | 0.02321 | 0.00981 | 0.00907 | 0.01465 | 0.03300 | 0.01259 | 0.03410 |
| **LHFPL2** | 1.17594 | 0.34732 | 2.21554 | 0.00000 | 0.25489 | 2.15797 | 4.82571 | 0.30714 | 4.56406 | 3.64567 | 0.00000 | 0.17314 |
| **LHFPL3** | 0.35376 | 0.62336 | 0.19138 | 0.78640 | 0.00000 | 0.22325 | 0.23238 | 0.48124 | 0.27649 | 0.36765 | 0.41513 | 1.22208 |
| **LHFPL3-AS1** | 0.00500 | 0.04792 | 0.06484 | 0.06686 | 0.00000 | 0.00361 | 0.00005 | 0.03126 | 0.06396 | 0.05133 | 0.06791 | 0.08400 |
| **LHFPL3-AS2** | 1.10441 | 1.49625 | 1.10085 | 1.27113 | 0.91155 | 1.87802 | 0.70169 | 3.65206 | 1.39820 | 0.12466 | 0.35446 | 5.10099 |
| **LHFPL4** | 0.00287 | 0.76010 | 0.16469 | 0.94978 | 0.14572 | 0.00000 | 0.23599 | 0.02145 | 0.00000 | 0.79748 | 0.01014 | 2.49701 |
| **LHFPL5** | 0.00562 | 0.03778 | 0.00485 | 0.11125 | 0.06154 | 0.16524 | 0.00000 | 0.00977 | 0.01010 | 0.00856 | 0.00782 | 0.00724 |
| **LHPP** | 0.67957 | 1.21321 | 0.55333 | 1.36872 | 3.38348 | 0.67217 | 0.25816 | 0.10864 | 0.00000 | 1.52803 | 2.28575 | 0.20536 |
| **LHX1** | 0.02247 | 0.00600 | 0.00762 | 0.58001 | 0.00000 | 0.01173 | 0.03850 | 0.02664 | 0.00000 | 0.07054 | 0.01155 | 0.05724 |
| **LHX2** | 1.98036 | 0.11337 | 0.01117 | 0.16084 | 0.00000 | 0.00000 | 0.00000 | 1.99975 | 0.03719 | 0.16944 | 0.00000 | 2.75951 |
| **LHX3** | 0.00623 | 0.01781 | 0.01844 | 0.01651 | 0.01581 | 0.01140 | 0.01958 | 0.01057 | 0.02420 | 0.01736 | 0.01461 | 0.01625 |
| **LHX4** | 0.84795 | 0.57251 | 0.00000 | 0.00000 | 1.20742 | 0.31044 | 0.31768 | 0.08490 | 0.00000 | 0.00000 | 1.11631 | 0.00000 |
| **LHX4-AS1** | 0.02652 | 0.08043 | 0.26707 | 0.01762 | 0.03187 | 0.43104 | 0.10393 | 0.00035 | 0.15292 | 0.00000 | 0.44749 | 0.15502 |
| **LHX5** | 0.09092 | 0.01434 | 0.01511 | 0.01092 | 0.01635 | 0.02125 | 0.01661 | 0.00526 | 0.01750 | 0.01545 | 0.01388 | 0.01435 |
| **LHX6** | 0.67344 | 0.08385 | 0.10356 | 0.11018 | 0.00000 | 0.38467 | 0.00000 | 0.18636 | 0.04427 | 0.00900 | 0.00000 | 0.56179 |
| **LHX8** | 2.21052 | 0.04740 | 0.05358 | 0.12344 | 0.04957 | 0.04976 | 0.00332 | 0.06496 | 0.00000 | 0.01966 | 0.01856 | 0.03405 |
| **LHX9** | 0.11497 | 0.01228 | 0.01439 | 0.04280 | 0.00000 | 0.00851 | 0.02663 | 0.00682 | 0.00000 | 0.03322 | 0.08471 | 0.03078 |
| **LIAS** | 0.48422 | 7.34114 | 3.19470 | 1.03191 | 1.84217 | 10.66661 | 2.71424 | 4.14279 | 5.26165 | 2.57142 | 4.78490 | 13.08611 |
| **LIF** | 0.00972 | 1.76148 | 0.14312 | 0.67948 | 0.00000 | 0.00000 | 0.00000 | 0.00820 | 0.12774 | 0.15422 | 0.00362 | 0.02547 |
| **LIFR** | 2.65493 | 0.00000 | 1.95583 | 0.00000 | 0.00654 | 4.35631 | 0.00000 | 1.30389 | 0.00000 | 9.77424 | 0.90214 | 6.16745 |
| **LIFR-AS1** | 0.17119 | 0.15254 | 0.11708 | 0.01185 | 0.02231 | 0.01673 | 0.03474 | 0.02181 | 0.00661 | 0.00000 | 0.05783 | 0.12050 |
| **LIG1** | 3.54677 | 9.23168 | 3.43003 | 9.01097 | 5.72875 | 4.16490 | 2.73043 | 0.68061 | 9.43616 | 4.09012 | 9.89362 | 5.20937 |
| **LIG3** | 1.61417 | 11.39292 | 9.17350 | 7.27393 | 3.47599 | 4.79860 | 5.38570 | 0.98067 | 13.93485 | 0.00000 | 14.07512 | 5.70305 |
| **LIG4** | 0.20713 | 1.40769 | 2.40262 | 0.00000 | 0.06365 | 5.25800 | 2.46621 | 0.44321 | 4.16762 | 0.00136 | 0.87443 | 1.60657 |
| **LILRA1** | 0.02198 | 0.01167 | 0.01463 | 0.00894 | 0.00806 | 0.02645 | 0.01303 | 0.01779 | 0.01709 | 0.04565 | 0.01333 | 0.02268 |

| | | | | | | | | | | | | |
|---|---|---|---|---|---|---|---|---|---|---|---|---|
| **LILRA2** | 0.00498 | 0.01040 | 0.01243 | 0.02833 | 0.00179 | 0.01828 | 0.00344 | 0.01014 | 0.02089 | 0.10544 | 0.01068 | 0.01329 |
| **LILRA3** | 0.00682 | 0.00000 | 0.03121 | 0.01354 | 0.04532 | 0.00000 | 0.03062 | 0.00396 | 0.01493 | 0.00000 | 0.01721 | 0.56890 |
| **LILRA4** | 0.77657 | 0.22399 | 0.03006 | 0.00418 | 0.00286 | 0.00000 | 0.02000 | 0.03985 | 0.04414 | 0.01844 | 0.00899 | 0.00000 |
| **LILRA5** | 0.00574 | 0.01485 | 0.01956 | 0.02963 | 0.02279 | 0.00000 | 0.07459 | 0.01068 | 0.00000 | 0.03670 | 0.01480 | 0.01635 |
| **LILRA6** | 0.00226 | 0.01420 | 0.07263 | 0.08109 | 0.15951 | 0.00000 | 0.03244 | 0.10042 | 0.06649 | 0.00000 | 0.01107 | 0.01641 |
| **LILRB1** | 0.00523 | 0.01393 | 0.00828 | 0.00813 | 0.00889 | 0.01209 | 0.01565 | 0.00942 | 0.01245 | 0.13099 | 0.01456 | 0.01963 |
| **LILRB2** | 0.01357 | 0.01561 | 0.01475 | 0.01244 | 0.02287 | 0.00681 | 0.05255 | 0.01006 | 0.00113 | 0.02717 | 0.01551 | 0.01615 |
| **LILRB3** | 0.04021 | 0.11807 | 0.00491 | 0.44979 | 0.46077 | 0.00000 | 0.00243 | 0.06349 | 0.26319 | 0.34872 | 0.00000 | 0.07758 |
| **LILRB4** | 0.12359 | 0.01380 | 0.01705 | 0.01315 | 0.01203 | 0.02072 | 0.01672 | 0.00579 | 0.01658 | 0.01355 | 0.02173 | 0.02257 |
| **LILRB5** | 0.01095 | 0.03963 | 0.01737 | 0.00489 | 0.05913 | 0.02319 | 0.03496 | 0.00996 | 0.02051 | 0.00544 | 0.00790 | 0.01742 |
| **LILRP2** | 0.00544 | 0.01026 | 0.02936 | 0.00715 | 0.01031 | 0.03672 | 0.01083 | 0.00946 | 0.00561 | 0.04031 | 0.01894 | 0.02903 |
| **LIM2** | 0.00508 | 0.06168 | 0.09933 | 1.02716 | 0.00000 | 0.22556 | 0.00000 | 0.11611 | 0.02169 | 0.01220 | 0.00711 | 0.84235 |
| **LIMA1** | 24.64952 | 0.91500 | 16.82238 | 2.05104 | 17.03218 | 10.20293 | 8.52026 | 7.95517 | 22.79373 | 5.33088 | 20.17126 | 17.47426 |
| **LIMCH1** | 0.10607 | 0.12634 | 0.23665 | 5.66674 | 0.01577 | 0.00344 | 0.00013 | 0.05749 | 0.02232 | 0.21094 | 0.00144 | 0.10943 |
| **LIMD1** | 0.49559 | 0.21874 | 1.77477 | 0.03962 | 2.95866 | 3.67241 | 2.32435 | 0.05224 | 3.06351 | 7.64027 | 1.13064 | 0.43022 |
| **LIMD1-AS1** | 0.11953 | 0.00828 | 0.01206 | 0.00423 | 0.01240 | 0.03457 | 0.00000 | 0.00666 | 0.08484 | 0.10468 | 0.00269 | 0.03395 |
| **LIMD2** | 1.26710 | 0.19792 | 0.79445 | 1.34718 | 0.00000 | 0.00000 | 0.14119 | 0.33368 | 0.00000 | 0.52477 | 0.00000 | 0.32941 |
| **LIME1** | 0.00135 | 0.39930 | 0.47850 | 0.34980 | 0.00000 | 0.00000 | 0.47376 | 0.01478 | 1.04876 | 3.23793 | 0.18314 | 0.18712 |
| **LIMK1** | 5.29435 | 2.89802 | 0.59490 | 2.08559 | 3.58732 | 0.36611 | 1.62653 | 0.00000 | 0.00000 | 1.76139 | 0.00000 | 1.76091 |
| **LIMK2** | 1.47004 | 8.96147 | 12.86166 | 4.28655 | 26.16298 | 3.62043 | 18.01387 | 0.37811 | 16.10697 | 0.00000 | 11.65337 | 7.41978 |
| **LIMS1** | 0.00000 | 2.98320 | 5.72564 | 0.53939 | 0.77376 | 9.80047 | 10.06676 | 0.62739 | 1.18234 | 12.97099 | 2.74556 | 0.49248 |
| **LIMS2** | 3.50575 | 0.53280 | 0.03804 | 0.05465 | 0.20445 | 0.00000 | 0.00180 | 0.00000 | 0.15688 | 0.00000 | 0.26915 | 0.12914 |
| **LIN28A** | 0.15145 | 24.35774 | 13.82062 | 15.56921 | 21.09668 | 12.05156 | 27.73482 | 5.67824 | 27.49817 | 19.05145 | 26.53831 | 35.45745 |
| **LIN28B** | 0.34273 | 5.88865 | 14.58875 | 1.22971 | 1.26724 | 16.48891 | 10.88134 | 1.24340 | 4.93568 | 7.37136 | 3.20465 | 6.88300 |
| **LIN37** | 0.40389 | 4.57958 | 6.45428 | 1.69551 | 7.60629 | 2.03741 | 2.64088 | 0.76630 | 4.34773 | 0.00000 | 2.96394 | 2.05404 |
| **LIN52** | 1.94150 | 2.89370 | 3.40227 | 1.44478 | 0.08328 | 4.76029 | 0.00000 | 5.33958 | 3.33050 | 0.00000 | 1.70956 | 4.88844 |
| **LIN54** | 1.83106 | 1.54801 | 5.59484 | 1.30763 | 0.00000 | 7.94778 | 5.13536 | 4.85904 | 1.95498 | 3.81686 | 2.68289 | 1.35120 |
| **LIN7A** | 0.00589 | 0.01339 | 0.05444 | 0.01180 | 0.01443 | 0.01982 | 0.01361 | 0.00995 | 0.01002 | 0.01749 | 0.01659 | 0.03006 |
| **LIN7B** | 0.13829 | 0.34183 | 0.20043 | 0.00082 | 0.39578 | 0.72860 | 0.28946 | 0.08596 | 0.35125 | 0.00000 | 0.06607 | 0.37549 |
| **LIN7C** | 1.87156 | 7.50074 | 5.24114 | 1.72500 | 0.00000 | 33.02773 | 9.95103 | 6.59106 | 9.44170 | 6.88602 | 9.32896 | 6.63468 |
| **LIN9** | 1.23630 | 3.49239 | 2.72138 | 1.31837 | 0.00000 | 11.81503 | 2.88985 | 4.32649 | 2.29121 | 0.62035 | 3.90059 | 5.61114 |
| **LINC-PINT.uid-1** | 0.04914 | 0.00968 | 0.01950 | 0.00464 | 0.02491 | 0.06191 | 0.00656 | 0.00570 | 0.01548 | 0.01414 | 0.01850 | 0.02823 |
| **LINC-PINT.uid-9** | 0.82023 | 0.26529 | 0.44118 | 0.56961 | 0.48745 | 1.10162 | 0.00000 | 0.17178 | 0.00000 | 0.51764 | 0.25980 | 0.07626 |
| **LINC-ROR** | 0.05265 | 0.05239 | 0.62894 | 1.57946 | 0.00000 | 0.00000 | 0.00000 | 0.07677 | 0.27848 | 0.07841 | 0.00280 | 0.02209 |
| **LINC00028** | 0.00559 | 0.07255 | 0.01762 | 0.01665 | 0.00935 | 0.00070 | 0.02429 | 0.00918 | 0.01450 | 0.02385 | 0.02000 | 0.05416 |
| **LINC00029** | 0.13315 | 0.02653 | 0.04143 | 0.00518 | 0.07831 | 0.01407 | 0.05041 | 0.01115 | 0.00000 | 0.00000 | 0.01095 | 0.15273 |
| **LINC00032** | 0.59522 | 0.00150 | 0.01328 | 0.19785 | 0.00000 | 0.06050 | 0.02053 | 0.19550 | 0.00000 | 0.05910 | 0.01691 | 0.03751 |
| **LINC00035** | 0.04906 | 0.01718 | 0.04691 | 0.02574 | 0.01564 | 0.01636 | 0.00574 | 0.00800 | 0.09151 | 0.00000 | 0.00556 | 0.03108 |
| **LINC00051** | 0.00628 | 0.01487 | 0.01639 | 0.01281 | 0.01603 | 0.01875 | 0.01935 | 0.01048 | 0.01657 | 0.01366 | 0.01508 | 0.01628 |
| **LINC00052** | 0.09899 | 0.10419 | 0.00363 | 0.00000 | 0.23667 | 0.00000 | 0.00984 | 0.00000 | 0.03342 | 0.17744 | 0.00000 | 0.29223 |
| **LINC00086** | 0.00521 | 0.05543 | 0.00000 | 1.01358 | 0.03035 | 0.02230 | 0.00000 | 0.00355 | 0.01755 | 0.14206 | 0.00971 | 0.25243 |
| **LINC00087** | 0.00603 | 0.00249 | 0.00883 | 0.11182 | 0.11479 | 0.08684 | 0.00000 | 0.39001 | 0.00000 | 0.03529 | 0.00647 | 0.08431 |
| **LINC00092** | 0.02305 | 0.01454 | 0.01413 | 0.01294 | 0.01421 | 0.01995 | 0.01833 | 0.01968 | 0.01774 | 0.01461 | 0.01496 | 0.01566 |
| **LINC00094** | 3.26149 | 2.42796 | 1.03439 | 2.47286 | 0.49470 | 0.22536 | 0.82838 | 0.09850 | 0.38966 | 0.00000 | 0.94011 | 1.57659 |
| **LINC00113** | 0.01017 | 0.00846 | 0.00615 | 0.00618 | 0.00832 | 0.07294 | 0.04767 | 0.01433 | 0.01696 | 0.01102 | 0.01142 | 0.01248 |
| **LINC00115** | 0.00292 | 0.06714 | 0.14958 | 0.14909 | 0.28226 | 0.29103 | 0.05189 | 0.50676 | 0.23704 | 0.00001 | 0.19850 | 0.25744 |
| **LINC00116** | 0.15662 | 8.01866 | 1.06783 | 3.59053 | 10.35164 | 2.36982 | 4.41438 | 0.66911 | 1.67503 | 5.20827 | 5.63144 | 4.27371 |
| **LINC00152** | 0.00048 | 0.04466 | 2.48050 | 0.23720 | 0.98253 | 0.22505 | 1.81090 | 0.00386 | 1.32324 | 0.69207 | 0.54179 | 0.00000 |

| | | | | | | | | | | | | |
|---|---|---|---|---|---|---|---|---|---|---|---|---|
| **LINC00158** | 0.04158 | 0.00841 | 0.01471 | 0.00857 | 0.01362 | 0.03252 | 0.05795 | 0.03996 | 0.00000 | 0.02506 | 0.01325 | 0.04392 |
| **LINC00159** | 0.00656 | 0.01596 | 0.01655 | 0.01253 | 0.01783 | 0.01589 | 0.01667 | 0.04279 | 0.01836 | 0.01103 | 0.02385 | 0.01541 |
| **LINC00162** | 0.33055 | 0.00594 | 0.01359 | 0.00000 | 0.07362 | 0.05695 | 0.01369 | 0.00138 | 0.00175 | 0.07911 | 0.00325 | 0.00742 |
| **LINC00167** | 0.63202 | 0.02362 | 0.03217 | 0.00000 | 0.02171 | 0.03418 | 0.00000 | 0.00604 | 0.00000 | 0.41103 | 0.05460 | 0.02848 |
| **LINC00173** | 0.00432 | 0.05502 | 0.00622 | 0.10382 | 0.00000 | 0.10118 | 0.11208 | 0.01103 | 0.00000 | 0.03916 | 0.04122 | 0.02358 |
| **LINC00174** | 0.00456 | 0.58226 | 0.50056 | 0.96895 | 0.47805 | 0.00000 | 0.00662 | 0.14138 | 0.11367 | 0.68664 | 0.52938 | 1.14808 |
| **LINC00176** | 0.03276 | 0.03311 | 0.27368 | 0.01656 | 0.01769 | 0.00000 | 0.00837 | 0.32710 | 0.00000 | 0.45609 | 0.02316 | 0.16610 |
| **LINC00189** | 0.52764 | 0.69541 | 0.00000 | 0.64429 | 0.43342 | 0.14604 | 0.07635 | 0.15221 | 0.00000 | 0.00000 | 0.33152 | 0.19704 |
| **LINC00200** | 0.00391 | 0.46501 | 0.53912 | 0.07818 | 0.01241 | 0.52693 | 0.00000 | 0.08247 | 0.38405 | 0.00000 | 0.40324 | 1.03131 |
| **LINC00202-1** | 0.00231 | 0.01921 | 0.03752 | 0.06516 | 0.06072 | 0.00000 | 0.00698 | 0.06933 | 0.02788 | 0.10094 | 0.03329 | 0.03188 |
| **LINC00202-2** | 0.00881 | 0.23677 | 0.07639 | 0.00000 | 0.00000 | 0.01806 | 0.21893 | 0.00354 | 0.20662 | 0.06859 | 0.00000 | 0.31772 |
| **LINC00221** | 4.84039 | 3.26541 | 0.00000 | 0.65763 | 3.10013 | 5.02495 | 3.72581 | 2.68382 | 2.95272 | 0.00000 | 2.48626 | 7.35012 |
| **LINC00226** | 0.36201 | 0.02596 | 0.04229 | 0.01985 | 0.01336 | 0.01034 | 0.01263 | 0.24741 | 0.01548 | 0.01260 | 0.01512 | 0.04706 |
| **LINC00229** | 0.20335 | 0.01303 | 0.01287 | 0.01024 | 0.01171 | 0.02201 | 0.01489 | 0.00302 | 0.01775 | 0.02323 | 0.01356 | 0.01555 |
| **LINC00235** | 0.00597 | 0.02533 | 0.03394 | 0.02381 | 0.01746 | 0.01137 | 0.01937 | 0.00975 | 0.00000 | 0.00676 | 0.07914 | 0.03324 |
| **LINC00238** | 0.00499 | 0.02177 | 0.00995 | 0.00720 | 0.02765 | 0.00404 | 0.02720 | 0.01071 | 0.02143 | 0.08175 | 0.01962 | 0.01375 |
| **LINC00240** | 0.06120 | 0.04058 | 0.14044 | 0.00000 | 0.00000 | 0.06567 | 0.10191 | 0.05729 | 0.00765 | 0.03436 | 0.03173 | 0.06344 |
| **LINC00242** | 0.02082 | 0.01243 | 0.09034 | 0.16668 | 0.00205 | 0.02698 | 0.01597 | 0.02925 | 0.00618 | 0.01272 | 0.01631 | 0.02937 |
| **LINC00260** | 0.06253 | 0.05499 | 0.23423 | 0.17194 | 0.62200 | 0.00000 | 0.16667 | 0.00224 | 0.00000 | 0.43559 | 0.12660 | 0.03360 |
| **LINC00261** | 0.00255 | 0.00049 | 1.46407 | 8.83741 | 0.08217 | 0.00000 | 0.00000 | 0.00763 | 0.00000 | 0.27994 | 0.00063 | 0.00650 |
| **LINC00263** | 0.29217 | 0.13553 | 2.73069 | 1.72420 | 0.33927 | 0.07312 | 3.30390 | 0.08504 | 2.61688 | 0.40557 | 0.33949 | 1.50787 |
| **LINC00264** | 0.00599 | 0.01332 | 0.02532 | 0.00990 | 0.01642 | 0.02582 | 0.01473 | 0.01079 | 0.01723 | 0.01440 | 0.01577 | 0.02159 |
| **LINC00265** | 0.03186 | 0.00000 | 0.31838 | 0.01457 | 0.04731 | 0.08636 | 0.02544 | 0.00584 | 0.07395 | 0.29280 | 0.09050 | 0.04871 |
| **LINC00266-3.ui** | 0.00409 | 0.01856 | 0.01391 | 0.01089 | 0.01781 | 0.02126 | 0.01819 | 0.02973 | 0.01727 | 0.01299 | 0.01527 | 0.01904 |
| **LINC00269** | 0.01003 | 0.01567 | 0.01963 | 0.01355 | 0.01762 | 0.00923 | 0.01744 | 0.06496 | 0.02197 | 0.01110 | 0.01529 | 0.03557 |
| **LINC00271** | 0.00689 | 0.01621 | 0.02392 | 0.01697 | 0.05088 | 0.00000 | 0.02634 | 0.01036 | 0.01810 | 0.00000 | 0.02311 | 0.02825 |
| **LINC00273** | 0.01085 | 0.20794 | 0.33522 | 0.08793 | 0.27118 | 0.00000 | 0.00000 | 0.00602 | 0.55326 | 0.63846 | 0.13332 | 0.05938 |
| **LINC00276** | 0.44874 | 0.01505 | 0.02765 | 0.01204 | 0.02470 | 0.00444 | 0.01742 | 0.04746 | 0.01684 | 0.01390 | 0.01571 | 0.04767 |
| **LINC00277** | 0.40579 | 0.01307 | 0.02114 | 0.01384 | 0.00869 | 0.01090 | 0.01546 | 0.00460 | 0.00963 | 0.01564 | 0.05888 | 0.01232 |
| **LINC00278** | 0.00585 | 0.00000 | 0.28198 | 0.01529 | 0.10319 | 0.05377 | 0.16433 | 0.00977 | 0.14046 | 0.00000 | 0.15172 | 0.01443 |
| **LINC00282** | 0.12660 | 0.06781 | 0.19431 | 0.05103 | 0.00000 | 0.00000 | 0.04160 | 0.01703 | 0.14105 | 0.00000 | 0.17107 | 0.04916 |
| **LINC00284** | 0.07199 | 0.06462 | 0.12017 | 0.01567 | 0.00000 | 0.03446 | 0.00386 | 0.00928 | 0.00000 | 0.03361 | 0.01586 | 0.01906 |
| **LINC00293** | 0.00597 | 0.01468 | 0.01835 | 0.01202 | 0.02316 | 0.02377 | 0.02945 | 0.01144 | 0.00901 | 0.00877 | 0.01462 | 0.01742 |
| **LINC00294** | 0.24641 | 0.04483 | 0.47702 | 0.63939 | 0.00000 | 0.63809 | 1.27572 | 0.07830 | 0.25260 | 0.00000 | 0.42994 | 0.25600 |
| **LINC00298** | 0.19238 | 0.01378 | 0.01620 | 0.01072 | 0.01311 | 0.02108 | 0.02018 | 0.14812 | 0.02546 | 0.01065 | 0.01240 | 0.00830 |
| **LINC00299** | 0.25783 | 0.01007 | 0.01224 | 0.00987 | 0.01109 | 0.02396 | 0.02128 | 0.00798 | 0.01408 | 0.02367 | 0.01334 | 0.02841 |
| **LINC00301** | 0.06836 | 0.01538 | 0.01343 | 0.01322 | 0.01508 | 0.02020 | 0.01608 | 0.02762 | 0.01852 | 0.01652 | 0.01378 | 0.02837 |
| **LINC00303** | 0.00432 | 0.01111 | 0.02327 | 0.09709 | 0.00634 | 0.02217 | 0.02738 | 0.05052 | 0.01137 | 0.05181 | 0.01847 | 0.04849 |
| **LINC00305** | 0.19315 | 0.01465 | 0.01509 | 0.01199 | 0.01485 | 0.01558 | 0.01795 | 0.02562 | 0.01934 | 0.01209 | 0.01603 | 0.01223 |
| **LINC00308** | 0.01075 | 0.01576 | 0.04856 | 0.00784 | 0.03705 | 0.00000 | 0.00405 | 0.01876 | 0.06133 | 0.03480 | 0.00714 | 0.02104 |
| **LINC00309** | 0.00748 | 0.00000 | 0.00886 | 0.25333 | 0.00000 | 0.11385 | 0.01185 | 0.00841 | 0.00000 | 0.14002 | 0.00190 | 0.00483 |
| **LINC00310** | 0.00941 | 0.03601 | 0.07832 | 0.00371 | 0.00000 | 0.05357 | 0.03439 | 0.02151 | 0.02347 | 0.02234 | 0.16746 | 0.05807 |
| **LINC00311** | 0.03573 | 0.03603 | 0.01734 | 0.04636 | 0.05205 | 0.09785 | 0.00000 | 0.00483 | 0.01401 | 0.06873 | 0.00210 | 0.03302 |
| **LINC00312** | 0.03588 | 0.15487 | 0.01922 | 0.02561 | 0.00000 | 0.00586 | 0.01484 | 0.00687 | 0.00000 | 0.21719 | 0.05584 | 0.02916 |
| **LINC00313** | 0.02792 | 0.01658 | 0.05000 | 0.00126 | 0.14310 | 0.00000 | 0.00180 | 0.00947 | 0.02418 | 0.17023 | 0.00058 | 0.02786 |
| **LINC00319** | 0.04029 | 0.02531 | 0.01582 | 0.01020 | 0.02871 | 0.00689 | 0.02593 | 0.00839 | 0.01592 | 0.03459 | 0.01589 | 0.01401 |
| **LINC00320** | 0.01076 | 0.02325 | 0.06707 | 0.02233 | 0.02257 | 0.00000 | 0.01558 | 0.01147 | 0.01880 | 0.00633 | 0.01822 | 0.01943 |
| **LINC00324** | 1.01859 | 0.00000 | 0.13533 | 0.20276 | 0.09830 | 0.00000 | 0.02275 | 8.28441 | 0.12977 | 0.00000 | 0.10075 | 0.58599 |

| | | | | | | | | | | | | |
|---|---|---|---|---|---|---|---|---|---|---|---|---|
| **LINC00327** | 0.00532 | 0.04721 | 0.12103 | 0.01005 | 0.01534 | 0.00000 | 0.00775 | 0.01051 | 0.01259 | 0.05755 | 0.01488 | 0.01905 |
| **LINC00330** | 0.00908 | 1.28387 | 0.00000 | 0.12523 | 0.00000 | 0.02837 | 0.00000 | 0.00000 | 0.00000 | 0.23154 | 0.00000 | 1.36903 |
| **LINC00333** | 0.00615 | 0.01320 | 0.04238 | 0.02011 | 0.01509 | 0.02488 | 0.00881 | 0.01030 | 0.02114 | 0.01971 | 0.01481 | 0.01606 |
| **LINC00336** | 0.00786 | 0.00708 | 0.07370 | 0.11797 | 0.01395 | 0.00000 | 0.00168 | 0.00676 | 0.00876 | 0.11463 | 0.01102 | 0.14260 |
| **LINC00337** | 0.05943 | 4.03376 | 1.24786 | 1.78739 | 3.29442 | 0.00000 | 0.52044 | 1.83190 | 0.68243 | 0.12367 | 1.11053 | 6.68828 |
| **LINC00339** | 0.61867 | 3.06315 | 0.00015 | 2.97826 | 0.00000 | 0.56283 | 0.08187 | 7.02424 | 0.00078 | 0.18615 | 0.40680 | 3.13720 |
| **LINC00341** | 0.00216 | 0.00000 | 0.04313 | 0.00000 | 0.00000 | 0.00296 | 0.48217 | 0.00802 | 0.46517 | 0.96277 | 1.02579 | 0.00232 |
| **LINC00342** | 0.00226 | 0.01023 | 0.00178 | 0.08468 | 0.08919 | 0.00000 | 0.18121 | 0.02976 | 0.00000 | 0.19162 | 0.02618 | 0.14881 |
| **LINC00346** | 0.00483 | 0.02847 | 0.11992 | 0.01816 | 0.00850 | 0.00473 | 0.04236 | 0.02274 | 0.09454 | 0.01021 | 0.00878 | 0.02009 |
| **LINC00347** | 0.34438 | 0.00824 | 0.01413 | 0.01072 | 0.01588 | 0.02188 | 0.01817 | 0.29827 | 0.01600 | 0.00293 | 0.02022 | 0.07963 |
| **LINC00348** | 0.00512 | 0.02684 | 0.01983 | 0.00567 | 0.01669 | 0.00979 | 0.03365 | 0.01360 | 0.02961 | 0.06758 | 0.01795 | 0.01259 |
| **LINC00354** | 0.23947 | 0.01423 | 0.01402 | 0.01174 | 0.01492 | 0.01728 | 0.01771 | 0.02721 | 0.02009 | 0.01032 | 0.01398 | 0.01221 |
| **LINC00366** | 0.32556 | 0.01328 | 0.02003 | 0.01087 | 0.01444 | 0.01698 | 0.01435 | 0.05318 | 0.02144 | 0.01229 | 0.01348 | 0.01372 |
| **LINC00367** | 0.00614 | 0.01491 | 0.16189 | 0.00985 | 0.06031 | 0.00000 | 0.00973 | 0.01074 | 0.06594 | 0.00000 | 0.02834 | 0.03383 |
| **LINC00368** | 0.00834 | 0.01410 | 0.01152 | 0.00960 | 0.00982 | 0.01692 | 0.08648 | 0.01107 | 0.00000 | 0.05498 | 0.02595 | 0.01698 |
| **LINC00371** | 0.06410 | 0.51239 | 0.00000 | 0.67251 | 0.94121 | 0.09894 | 0.00000 | 0.02597 | 0.00000 | 0.00398 | 0.65801 | 0.18217 |
| **LINC00378** | 0.32272 | 0.01886 | 0.02633 | 0.01423 | 0.01729 | 0.00983 | 0.02202 | 0.01111 | 0.01707 | 0.00494 | 0.01579 | 0.01577 |
| **LINC00379** | 0.00338 | 0.01505 | 1.57059 | 0.01118 | 0.00000 | 0.01265 | 1.99194 | 0.04161 | 0.08152 | 0.00000 | 0.00000 | 0.01255 |
| **LINC00381** | 0.00688 | 0.02902 | 0.01282 | 0.01089 | 0.00516 | 0.03420 | 0.01408 | 0.00966 | 0.01445 | 0.02762 | 0.02073 | 0.02128 |
| **LINC00392** | 0.01775 | 0.06135 | 0.05160 | 0.22596 | 0.09011 | 0.00073 | 0.04612 | 0.02376 | 0.00257 | 0.08199 | 0.00874 | 0.16976 |
| **LINC00400** | 0.03333 | 0.01229 | 0.08811 | 0.03342 | 0.04760 | 0.00000 | 0.09210 | 0.05476 | 0.04357 | 0.01259 | 0.02224 | 0.04210 |
| **LINC00408** | 0.00210 | 0.02174 | 0.04759 | 0.01588 | 0.02640 | 0.00000 | 0.01583 | 0.15888 | 0.02033 | 0.00000 | 0.01670 | 0.04107 |
| **LINC00412** | 0.00453 | 0.03753 | 0.00567 | 0.01210 | 0.00206 | 0.04343 | 0.00000 | 0.01321 | 0.06185 | 0.02715 | 0.02790 | 0.06143 |
| **LINC00417** | 0.00642 | 0.01712 | 0.01370 | 0.02855 | 0.08421 | 0.00000 | 0.01706 | 0.01034 | 0.02991 | 0.00000 | 0.01581 | 0.02290 |
| **LINC00423** | 0.01590 | 0.01501 | 0.02003 | 0.01360 | 0.01171 | 0.00034 | 0.02173 | 0.01237 | 0.00000 | 0.01974 | 0.10160 | 0.01906 |
| **LINC00426** | 0.02087 | 0.09566 | 0.08273 | 0.09724 | 0.08878 | 0.00000 | 0.00224 | 0.15659 | 0.02256 | 0.02014 | 0.01410 | 0.57096 |
| **LINC00441** | 0.00348 | 0.01803 | 0.07565 | 0.01070 | 0.00000 | 0.00000 | 0.07525 | 0.02292 | 0.00000 | 0.09067 | 0.05431 | 0.03559 |
| **LINC00442** | 0.00588 | 0.01695 | 0.01187 | 0.01080 | 0.01740 | 0.01168 | 0.03549 | 0.01044 | 0.01322 | 0.03154 | 0.01439 | 0.01673 |
| **LINC00443** | 0.01855 | 0.02071 | 0.01544 | 0.01184 | 0.01568 | 0.01456 | 0.02533 | 0.06040 | 0.01747 | 0.00970 | 0.01510 | 0.01330 |
| **LINC00446** | 0.26018 | 0.01501 | 0.01463 | 0.01276 | 0.01686 | 0.01220 | 0.01880 | 0.01242 | 0.02164 | 0.00775 | 0.01420 | 0.01356 |
| **LINC00448** | 0.77724 | 0.01456 | 0.01415 | 0.01220 | 0.01689 | 0.00818 | 0.01864 | 0.11025 | 0.02225 | 0.00686 | 0.01385 | 0.00638 |
| **LINC00452** | 0.01332 | 0.06086 | 0.03768 | 0.01337 | 0.04525 | 0.00000 | 0.03176 | 0.00972 | 0.00575 | 0.01115 | 0.01491 | 0.02261 |
| **LINC00457** | 0.00683 | 0.01295 | 0.01168 | 0.06651 | 0.02203 | 0.02059 | 0.01955 | 0.01422 | 0.00004 | 0.04099 | 0.09786 | 0.01455 |
| **LINC00458** | 0.37235 | 0.02922 | 8.41908 | 0.01269 | 0.00000 | 0.00000 | 0.55515 | 0.01381 | 0.00000 | 0.02269 | 0.00494 | 0.03876 |
| **LINC00461** | 0.00665 | 0.06341 | 0.00223 | 0.88011 | 0.08585 | 0.00188 | 0.02172 | 0.02578 | 0.00938 | 0.04298 | 0.00830 | 0.01490 |
| **LINC00466** | 0.31262 | 0.04033 | 0.01056 | 0.00921 | 0.00000 | 0.05588 | 0.01137 | 0.00976 | 0.01805 | 0.02831 | 0.00898 | 0.05137 |
| **LINC00467** | 1.19816 | 0.68582 | 0.53269 | 0.58883 | 0.12209 | 0.38353 | 0.00000 | 0.34645 | 0.00000 | 0.05865 | 0.08426 | 2.59209 |
| **LINC00470** | 0.00000 | 0.03895 | 1.56076 | 0.00297 | 0.17324 | 0.58337 | 1.81219 | 0.16050 | 1.00913 | 1.93977 | 0.40754 | 0.16231 |
| **LINC00471** | 0.20428 | 0.38337 | 0.02406 | 0.04804 | 0.00000 | 0.11436 | 0.00000 | 1.09159 | 0.06286 | 0.12125 | 0.40010 | 0.82099 |
| **LINC00472** | 0.01486 | 0.10482 | 0.03776 | 0.13523 | 0.00651 | 0.00000 | 0.00960 | 0.01077 | 0.02808 | 0.07064 | 0.01989 | 0.01360 |
| **LINC00475** | 0.01390 | 0.08900 | 0.01777 | 0.01855 | 0.00186 | 0.04426 | 0.00606 | 0.01268 | 0.00000 | 0.06299 | 0.05304 | 0.01286 |
| **LINC00476.uid-** | 0.42625 | 0.20326 | 0.47871 | 0.00544 | 0.60095 | 0.43132 | 0.95316 | 0.48810 | 0.07347 | 0.00000 | 0.38176 | 0.49804 |
| **LINC00476.uid-** | 0.05665 | 0.09235 | 0.00000 | 0.00000 | 0.25092 | 0.00000 | 0.00743 | 0.00392 | 0.20601 | 0.11214 | 0.09771 | 0.06027 |
| **LINC00477** | 0.00183 | 0.05414 | 0.04657 | 0.01855 | 0.04058 | 0.00960 | 0.04931 | 0.07792 | 0.02512 | 0.04615 | 0.02869 | 0.06369 |
| **LINC00478** | 0.34163 | 0.01702 | 0.02408 | 0.01049 | 0.01120 | 0.00819 | 0.07284 | 0.00191 | 0.00000 | 0.03535 | 0.01975 | 0.04344 |
| **LINC00479** | 0.15224 | 1.56017 | 0.03993 | 0.72727 | 0.00000 | 0.13157 | 1.21133 | 0.00613 | 0.48600 | 0.03000 | 2.15484 | 0.00000 |
| **LINC00482** | 0.00541 | 0.01539 | 0.09040 | 0.00812 | 0.00578 | 0.01628 | 0.10164 | 0.01068 | 0.07440 | 0.04324 | 0.07239 | 0.02157 |
| **LINC00483** | 0.66734 | 0.01845 | 0.03213 | 0.01292 | 0.03322 | 0.00000 | 0.01047 | 0.22422 | 0.02282 | 0.02633 | 0.01401 | 0.03185 |

| | | | | | | | | | | | | |
|---|---|---|---|---|---|---|---|---|---|---|---|---|
| **LINC00485** | 0.00449 | 0.00774 | 0.03003 | 0.12858 | 0.00000 | 0.01222 | 0.00000 | 0.01012 | 0.02355 | 0.14198 | 0.00811 | 0.01443 |
| **LINC00486** | 0.04336 | 0.00879 | 0.00885 | 0.03737 | 0.00513 | 0.05299 | 0.01154 | 0.03703 | 0.01371 | 0.01723 | 0.01327 | 0.01602 |
| **LINC00488** | 0.00674 | 0.01711 | 0.01566 | 0.04756 | 0.01603 | 0.00001 | 0.02258 | 0.02316 | 0.01820 | 0.01749 | 0.02175 | 0.01717 |
| **LINC00491** | 0.20582 | 0.14608 | 0.05080 | 0.06650 | 0.00000 | 0.29810 | 0.07053 | 0.15615 | 0.24530 | 0.72524 | 0.24551 | 0.08520 |
| **LINC00492** | 0.01117 | 0.01465 | 0.02546 | 0.00873 | 0.00937 | 0.02910 | 0.00798 | 0.00973 | 0.01645 | 0.03758 | 0.01287 | 0.02059 |
| **LINC00493** | 0.00002 | 9.97131 | 13.55993 | 7.12024 | 6.75548 | 20.28594 | 16.79270 | 2.80239 | 14.05970 | 7.32725 | 16.35376 | 12.93168 |
| **LINC00494** | 0.03130 | 0.00508 | 0.01220 | 0.00636 | 0.00988 | 0.06211 | 0.01297 | 0.00672 | 0.00477 | 0.01375 | 0.02488 | 0.04608 |
| **LINC00501** | 0.38117 | 0.56289 | 0.41152 | 0.17893 | 0.03659 | 0.75679 | 0.00000 | 0.58814 | 0.00000 | 0.50652 | 0.01687 | 1.24332 |
| **LINC00506** | 0.00427 | 0.00824 | 0.02036 | 0.00695 | 0.01951 | 0.02114 | 0.01410 | 0.01400 | 0.01280 | 0.05010 | 0.01316 | 0.05462 |
| **LINC00511** | 0.18950 | 0.17413 | 0.00307 | 0.26165 | 0.26686 | 0.00000 | 0.01779 | 0.00000 | 0.00000 | 0.05520 | 0.00292 | 0.20665 |
| **LINC00514** | 0.00697 | 0.04490 | 0.02848 | 0.01922 | 0.05374 | 0.03005 | 0.03531 | 0.01502 | 0.00000 | 0.00524 | 0.03764 | 0.05604 |
| **LINC00515** | 0.00547 | 0.00560 | 0.02075 | 0.02980 | 0.00000 | 0.06165 | 0.00000 | 0.00898 | 0.01993 | 0.04924 | 0.01981 | 0.03852 |
| **LINC00518** | 0.02851 | 0.42964 | 0.06550 | 0.00000 | 0.40560 | 0.00000 | 0.24496 | 0.01408 | 0.00000 | 0.08177 | 0.02428 | 0.00000 |
| **LINC00521** | 0.00601 | 0.01382 | 0.02104 | 0.01362 | 0.01589 | 0.01907 | 0.02454 | 0.01046 | 0.01453 | 0.02375 | 0.01363 | 0.01559 |
| **LINC00525** | 0.00607 | 0.01338 | 0.01989 | 0.01772 | 0.01535 | 0.02258 | 0.01493 | 0.00935 | 0.01940 | 0.00965 | 0.02239 | 0.04164 |
| **LINC00526** | 0.00224 | 0.53764 | 0.23755 | 0.40876 | 0.00000 | 0.64516 | 0.00000 | 0.61412 | 0.27157 | 0.00926 | 0.16069 | 0.27601 |
| **LINC00528** | 0.04103 | 0.01843 | 0.03798 | 0.01498 | 0.02158 | 0.00503 | 0.05303 | 0.04935 | 0.00078 | 0.00562 | 0.01808 | 0.01428 |
| **LINC00534** | 0.01279 | 0.00619 | 0.00886 | 0.01009 | 0.03142 | 0.02383 | 0.00959 | 0.01249 | 0.00000 | 0.07051 | 0.07245 | 0.01304 |
| **LINC00535** | 0.08675 | 0.04396 | 0.03101 | 0.02818 | 0.01171 | 0.01132 | 0.01609 | 0.07151 | 0.01541 | 0.01512 | 0.01750 | 0.04433 |
| **LINC00536** | 0.03909 | 0.01427 | 0.01529 | 0.01148 | 0.01924 | 0.02058 | 0.01492 | 0.02720 | 0.02153 | 0.01280 | 0.01610 | 0.01398 |
| **LINC00539** | 0.07520 | 0.04231 | 0.00963 | 0.04161 | 0.02702 | 0.05088 | 0.00000 | 0.05238 | 0.00921 | 0.09358 | 0.10384 | 0.05899 |
| **LINC00540** | 0.01283 | 1.78842 | 2.05831 | 0.00000 | 0.00629 | 1.97625 | 1.46284 | 0.26657 | 2.00153 | 0.00000 | 1.07611 | 2.64318 |
| **LINC00545** | 0.00686 | 0.01523 | 0.00920 | 0.13581 | 0.01155 | 0.01149 | 0.01592 | 0.01008 | 0.01217 | 0.02186 | 0.03584 | 0.01686 |
| **LINC00547** | 0.00368 | 0.01690 | 0.00799 | 0.01190 | 0.01306 | 0.01000 | 0.15097 | 0.02367 | 0.00000 | 0.00263 | 0.02523 | 0.01728 |
| **LINC00548** | 0.00654 | 0.14154 | 0.00170 | 0.08378 | 0.00000 | 0.04979 | 0.00000 | 0.00985 | 0.00883 | 0.07030 | 0.01323 | 0.00252 |
| **LINC00550** | 0.01265 | 0.01347 | 0.02036 | 0.01133 | 0.00905 | 0.01752 | 0.01445 | 0.01871 | 0.01133 | 0.04241 | 0.02624 | 0.01430 |
| **LINC00552** | 0.19294 | 0.03530 | 0.03078 | 0.00395 | 0.00992 | 0.03852 | 0.01950 | 0.00344 | 0.01668 | 0.01553 | 0.00933 | 0.00977 |
| **LINC00558** | 0.59850 | 0.03274 | 0.01321 | 0.00789 | 0.02407 | 0.01910 | 0.02438 | 0.15255 | 0.01685 | 0.00000 | 0.01224 | 0.02842 |
| **LINC00563** | 0.00580 | 0.01094 | 0.19805 | 0.00461 | 0.02385 | 0.02950 | 0.04512 | 0.01027 | 0.00000 | 0.02693 | 0.01431 | 0.01430 |
| **LINC00568** | 0.03613 | 0.01626 | 0.11894 | 0.04913 | 0.04197 | 0.00000 | 0.00884 | 0.00558 | 0.01778 | 0.03201 | 0.13262 | 0.06834 |
| **LINC00570** | 0.00782 | 0.01230 | 0.01418 | 0.01078 | 0.00801 | 0.01490 | 0.02602 | 0.03369 | 0.02248 | 0.02213 | 0.04873 | 0.04471 |
| **LINC00571** | 0.01472 | 0.01607 | 0.01045 | 0.00489 | 0.06215 | 0.03471 | 0.00589 | 0.01070 | 0.03585 | 0.01168 | 0.00578 | 0.01626 |
| **LINC00574** | 0.14627 | 0.00992 | 0.04118 | 0.00926 | 0.00000 | 0.06626 | 0.04851 | 0.00860 | 0.00000 | 0.08074 | 0.00817 | 0.05992 |
| **LINC00577** | 0.00536 | 0.03696 | 0.05265 | 0.00715 | 0.02092 | 0.02878 | 0.01038 | 0.09057 | 0.01937 | 0.00001 | 0.01524 | 0.12404 |
| **LINC00578** | 0.11616 | 0.00000 | 0.00630 | 0.05386 | 0.05827 | 0.04328 | 0.09989 | 0.00849 | 0.04391 | 0.03045 | 0.01453 | 0.24387 |
| **LINC00581** | 0.00638 | 0.01689 | 0.02373 | 0.01163 | 0.01182 | 0.00952 | 0.00626 | 0.01057 | 0.06255 | 0.01968 | 0.02689 | 0.01631 |
| **LINC00592** | 0.01677 | 0.01338 | 0.08920 | 0.00713 | 0.00978 | 0.03575 | 0.02086 | 0.00795 | 0.08497 | 0.00000 | 0.00194 | 0.01565 |
| **LINC00597** | 0.00101 | 0.05994 | 0.40451 | 0.00000 | 0.00000 | 0.32971 | 0.16240 | 0.00538 | 0.00002 | 0.76800 | 0.18896 | 0.00000 |
| **LINC00598** | 0.16031 | 0.01793 | 0.01265 | 0.11181 | 0.00273 | 0.01335 | 0.01074 | 0.02365 | 0.03241 | 0.02765 | 0.01921 | 0.01724 |
| **LINC00599** | 0.13946 | 0.02909 | 0.01750 | 0.00945 | 0.00727 | 0.03265 | 0.01495 | 0.18666 | 0.01327 | 0.03217 | 0.01564 | 0.00748 |
| **LINC00601** | 0.01477 | 0.01541 | 0.01931 | 0.01183 | 0.01556 | 0.01358 | 0.01597 | 0.30462 | 0.01659 | 0.01796 | 0.01537 | 0.00693 |
| **LINC00605** | 0.02456 | 0.00000 | 0.00000 | 0.12368 | 0.53462 | 0.05712 | 0.00000 | 0.02615 | 0.00000 | 0.26865 | 0.00000 | 0.16041 |
| **LINC00607** | 0.13906 | 0.15483 | 0.09456 | 0.13190 | 0.01842 | 0.00000 | 0.03277 | 0.03318 | 0.00000 | 0.02492 | 0.01976 | 0.01460 |
| **LINC00609** | 0.06330 | 0.01228 | 0.01361 | 0.00977 | 0.03347 | 0.01511 | 0.01618 | 0.02532 | 0.01820 | 0.02149 | 0.01404 | 0.02312 |
| **LINC00612** | 0.00338 | 0.01400 | 0.05503 | 0.00866 | 0.07288 | 0.00000 | 0.02729 | 0.04471 | 0.03256 | 0.02294 | 0.00944 | 0.02129 |
| **LINC00615** | 0.00978 | 0.01494 | 0.04153 | 0.00800 | 0.01198 | 0.01769 | 0.01107 | 0.03524 | 0.00748 | 0.03737 | 0.02112 | 0.01321 |
| **LINC00616** | 0.00735 | 0.02217 | 0.01366 | 0.03026 | 0.05449 | 0.03282 | 0.02216 | 0.03332 | 0.00887 | 0.00381 | 0.01554 | 0.03956 |
| **LINC00618** | 0.01843 | 0.00000 | 0.02049 | 0.00000 | 0.03716 | 0.21425 | 0.00000 | 0.01468 | 0.02573 | 0.18053 | 0.00000 | 0.04501 |

| | | | | | | | | | | | | |
|---|---|---|---|---|---|---|---|---|---|---|---|---|
| **LINC00620** | 0.00628 | 0.01455 | 0.07070 | 0.01182 | 0.02067 | 0.02556 | 0.00816 | 0.01035 | 0.01904 | 0.01276 | 0.01311 | 0.01566 |
| **LINC00622** | 0.03908 | 0.01058 | 0.14269 | 0.00312 | 0.00000 | 0.04128 | 0.08347 | 0.00691 | 0.12421 | 0.01278 | 0.00000 | 0.01361 |
| **LINC00624** | 0.51012 | 2.99088 | 2.96342 | 0.28715 | 1.94771 | 1.68050 | 0.00002 | 2.18391 | 4.32134 | 0.00000 | 1.02882 | 3.44798 |
| **LINC00630** | 0.58728 | 0.47623 | 0.28030 | 0.00388 | 0.00618 | 0.93228 | 0.00000 | 0.85871 | 0.72323 | 0.00000 | 0.31032 | 0.57759 |
| **LINC00632** | 0.04755 | 0.00347 | 0.02425 | 0.01595 | 0.00000 | 0.04283 | 0.00250 | 0.06985 | 0.00674 | 0.19376 | 0.03249 | 0.01079 |
| **LINC00634** | 0.00237 | 0.01423 | 0.02419 | 0.02969 | 0.00335 | 0.01803 | 0.01758 | 0.03766 | 0.01901 | 0.06343 | 0.01268 | 0.01564 |
| **LINC00635** | 0.03205 | 0.01395 | 0.05404 | 0.01153 | 0.01486 | 0.01569 | 0.00765 | 0.04732 | 0.01256 | 0.03846 | 0.01454 | 0.01177 |
| **LINC00636** | 0.00412 | 0.01960 | 0.01851 | 0.02672 | 0.01082 | 0.03127 | 0.01339 | 0.02398 | 0.00921 | 0.03123 | 0.01389 | 0.06130 |
| **LINC00637** | 0.03004 | 0.01062 | 0.00967 | 0.00626 | 0.00601 | 0.04027 | 0.00853 | 0.00921 | 0.01234 | 0.05043 | 0.01243 | 0.02690 |
| **LINC00638** | 0.00573 | 0.01599 | 0.01796 | 0.01154 | 0.03610 | 0.00998 | 0.02259 | 0.01355 | 0.00691 | 0.04514 | 0.02239 | 0.01664 |
| **LINC00639** | 0.05169 | 0.01709 | 0.05874 | 0.06110 | 0.00533 | 0.04426 | 0.02621 | 0.02436 | 0.00864 | 0.01316 | 0.01627 | 0.05097 |
| **LINC00640** | 0.01006 | 0.00201 | 0.08042 | 0.19300 | 0.12998 | 0.09175 | 0.00857 | 0.02886 | 0.00000 | 0.02553 | 0.05007 | 0.02720 |
| **LINC00641** | 0.02079 | 0.00000 | 2.48315 | 1.38180 | 0.00000 | 1.83415 | 1.37191 | 0.07710 | 3.55970 | 9.70538 | 1.01578 | 0.29418 |
| **LINC00643** | 0.05063 | 0.00000 | 12.17880 | 0.00000 | 0.00000 | 0.01133 | 1.40071 | 0.23794 | 0.04349 | 0.23075 | 0.00000 | 0.00000 |
| **LINC00645** | 0.00225 | 0.00000 | 0.00000 | 0.00000 | 0.00000 | 0.01563 | 0.03009 | 0.00677 | 0.49057 | 1.11696 | 0.34970 | 0.05438 |
| **LINC00648** | 0.19921 | 0.09959 | 0.18992 | 0.00090 | 0.00050 | 0.59496 | 0.33008 | 0.11334 | 0.49541 | 0.00496 | 0.20950 | 0.11195 |
| **LINC00649** | 0.13065 | 0.68663 | 0.10906 | 0.07282 | 0.20424 | 0.13283 | 0.02770 | 0.01133 | 0.00000 | 0.09485 | 0.35811 | 0.14715 |
| **LINC00652** | 0.00289 | 0.01697 | 0.14125 | 0.00047 | 0.00000 | 0.05777 | 0.29277 | 0.10428 | 0.02912 | 0.04292 | 0.04694 | 0.00572 |
| **LINC00654** | 0.02453 | 0.02837 | 0.04359 | 0.11621 | 0.02278 | 0.00151 | 0.00700 | 0.03501 | 0.08254 | 0.00000 | 0.02640 | 0.07201 |
| **LINC00656** | 0.00691 | 0.01639 | 0.02931 | 0.01386 | 0.02108 | 0.01132 | 0.01602 | 0.01148 | 0.02372 | 0.00849 | 0.01721 | 0.01707 |
| **LINC00657** | 0.91917 | 1.44767 | 2.48769 | 0.65750 | 0.45490 | 6.41861 | 3.38699 | 1.58629 | 0.00000 | 12.38074 | 1.66254 | 2.30096 |
| **LINC00661** | 0.00372 | 0.01383 | 0.02904 | 0.01361 | 0.01195 | 0.02071 | 0.01630 | 0.02219 | 0.00900 | 0.02118 | 0.02329 | 0.01511 |
| **LINC00662** | 0.31794 | 1.98957 | 0.48647 | 0.10831 | 0.04613 | 0.96130 | 0.51601 | 0.58093 | 1.74079 | 0.00000 | 1.09322 | 1.31911 |
| **LINC00663** | 0.37195 | 0.04024 | 0.08082 | 0.02808 | 0.00000 | 0.08399 | 0.01441 | 0.01396 | 0.00000 | 0.10013 | 0.04478 | 0.17499 |
| **LINC00664** | 0.03974 | 1.26525 | 0.00000 | 0.49394 | 0.00000 | 0.28081 | 0.00000 | 0.20725 | 0.07406 | 0.22903 | 0.02634 | 0.85271 |
| **LINC00665** | 3.34435 | 0.75590 | 2.09791 | 0.92333 | 0.00000 | 0.05200 | 5.27150 | 1.19126 | 5.28527 | 0.00000 | 1.04794 | 0.94903 |
| **LINC00667** | 0.09421 | 4.58659 | 1.76325 | 2.83901 | 4.18830 | 7.15405 | 0.00000 | 0.31956 | 2.26696 | 1.27533 | 1.72373 | 4.01484 |
| **LINC00668** | 0.00000 | 3.88971 | 13.56732 | 0.00000 | 3.04552 | 3.49399 | 12.00888 | 0.66667 | 6.50694 | 0.00000 | 17.97143 | 3.37122 |
| **LINC00669** | 0.03913 | 0.07186 | 0.30981 | 0.08065 | 0.12836 | 0.89653 | 0.00000 | 0.03263 | 0.00000 | 0.25358 | 0.26315 | 0.08846 |
| **LINC00670** | 0.40199 | 0.01273 | 0.01826 | 0.01746 | 0.01044 | 0.01945 | 0.02787 | 0.00895 | 0.01216 | 0.01249 | 0.02889 | 0.01152 |
| **LINC00671** | 0.01198 | 0.01794 | 0.02029 | 0.01252 | 0.05516 | 0.00000 | 0.02468 | 0.00866 | 0.02711 | 0.00819 | 0.01518 | 0.04265 |
| **LINC00672** | 0.04785 | 0.17975 | 0.13584 | 0.63799 | 0.00000 | 0.16122 | 0.21735 | 0.03703 | 0.00013 | 0.07516 | 0.00000 | 0.33550 |
| **LINC00673** | 0.00526 | 0.13272 | 0.05880 | 3.06586 | 0.19944 | 0.00000 | 0.00000 | 0.37528 | 0.00000 | 0.20204 | 0.00611 | 0.03562 |
| **LINC00674** | 1.20245 | 0.12963 | 0.41332 | 1.31325 | 0.08423 | 0.22676 | 0.29488 | 0.19178 | 0.49299 | 0.15640 | 0.25467 | 0.12377 |
| **LINC00676** | 0.02362 | 0.01392 | 0.01127 | 0.00977 | 0.01002 | 0.01234 | 0.03883 | 0.01923 | 0.00787 | 0.06134 | 0.01279 | 0.01264 |
| **LINC00678** | 0.00532 | 0.09471 | 0.73981 | 2.42566 | 1.94568 | 0.46560 | 0.24573 | 0.00790 | 0.00000 | 0.00000 | 0.00000 | 0.34701 |
| **LINC00682** | 0.08670 | 0.00914 | 0.02119 | 0.01601 | 0.02081 | 0.00000 | 0.01587 | 1.06278 | 0.02368 | 0.00652 | 0.01512 | 0.12517 |
| **LINC00687** | 0.00547 | 0.01414 | 0.01094 | 0.00595 | 0.01789 | 0.02987 | 0.07531 | 0.01054 | 0.00259 | 0.03423 | 0.01171 | 0.03370 |
| **LINC00689** | 0.22373 | 0.08958 | 0.05155 | 0.10057 | 0.00475 | 0.08796 | 0.00000 | 0.00000 | 0.12432 | 0.00000 | 0.07695 | 1.16528 |
| **LINC00691** | 0.00799 | 0.01719 | 0.01490 | 0.00950 | 0.01233 | 0.02306 | 0.01980 | 0.01058 | 0.01890 | 0.03316 | 0.01333 | 0.02633 |
| **LINC00692** | 0.02514 | 0.00823 | 0.00637 | 0.03536 | 0.00000 | 0.03059 | 0.00000 | 0.01123 | 0.01899 | 0.11014 | 0.00800 | 0.02591 |
| **LINC00693** | 0.24539 | 0.01481 | 0.01304 | 0.03137 | 0.02533 | 0.01275 | 0.05245 | 0.00257 | 0.00885 | 0.00000 | 0.01526 | 0.01750 |
| **LINC00696** | 0.00481 | 0.00653 | 0.00698 | 0.04668 | 0.00000 | 0.01951 | 0.00000 | 0.01068 | 0.01158 | 0.25272 | 0.00850 | 0.01660 |
| **LINC00698** | 0.15628 | 0.00882 | 0.13986 | 0.35549 | 0.00000 | 0.00691 | 0.01380 | 0.04024 | 0.03105 | 0.05947 | 0.01251 | 0.01057 |
| **LINC00701** | 0.03632 | 0.15993 | 0.01849 | 0.00186 | 0.00000 | 0.06833 | 0.15546 | 0.22668 | 0.00188 | 0.00001 | 0.00289 | 0.32055 |
| **LINC00703** | 0.00547 | 0.01525 | 0.02948 | 0.01646 | 0.03090 | 0.00005 | 0.02995 | 0.01137 | 0.00852 | 0.04538 | 0.01600 | 0.01579 |
| **LINC00707** | 1.68775 | 1.18495 | 0.00000 | 0.96641 | 0.00921 | 0.42688 | 0.00000 | 0.34781 | 0.00000 | 0.00206 | 0.04535 | 1.96245 |
| **LINC00708** | 0.00630 | 0.01464 | 0.02180 | 0.01163 | 0.01459 | 0.02074 | 0.01792 | 0.01047 | 0.02372 | 0.01232 | 0.01348 | 0.01620 |

| | | | | | | | | | | | | |
|---|---|---|---|---|---|---|---|---|---|---|---|---|
| **LINC00836** | 0.00353 | 0.02987 | 0.03309 | 0.00000 | 0.00010 | 0.04871 | 0.06902 | 0.00973 | 0.00000 | 0.23729 | 0.12113 | 0.00916 |
| **LINC00837** | 0.00459 | 0.01075 | 0.04361 | 0.02254 | 0.03009 | 0.00000 | 0.00547 | 0.03375 | 0.02620 | 0.02552 | 0.05235 | 0.05797 |
| **LINC00839** | 0.00773 | 0.08575 | 1.07576 | 0.00912 | 0.01325 | 0.00372 | 0.55030 | 2.59938 | 0.00000 | 0.00040 | 0.00410 | 0.36176 |
| **LINC00840** | 0.01900 | 0.01426 | 0.01728 | 0.01152 | 0.01449 | 0.01727 | 0.02074 | 0.00893 | 0.01824 | 0.01927 | 0.01542 | 0.01820 |
| **LINC00842** | 0.00079 | 0.34649 | 0.35358 | 0.00000 | 2.14477 | 0.25068 | 0.29891 | 0.00398 | 2.31606 | 0.29106 | 2.48605 | 0.00000 |
| **LINC00844** | 0.00744 | 0.01380 | 0.01374 | 0.00445 | 0.04573 | 0.03702 | 0.00246 | 0.01567 | 0.01705 | 0.07189 | 0.00623 | 0.01342 |
| **LINC00845** | 0.00657 | 0.01460 | 0.04050 | 0.00845 | 0.01597 | 0.02537 | 0.02153 | 0.01036 | 0.03856 | 0.00000 | 0.01050 | 0.01688 |
| **LINC00847** | 0.59513 | 0.10925 | 0.25003 | 0.04880 | 0.00000 | 0.00000 | 0.12095 | 0.15208 | 0.08827 | 0.00000 | 0.17393 | 0.50229 |
| **LINC00852** | 0.00039 | 0.46325 | 0.50720 | 0.39467 | 0.27886 | 0.12056 | 0.40992 | 0.16852 | 0.67052 | 0.31070 | 0.04777 | 0.07102 |
| **LINC00853** | 0.02666 | 0.03428 | 0.00689 | 0.01463 | 0.01016 | 0.00755 | 0.08206 | 0.03920 | 0.00000 | 0.07508 | 0.01226 | 0.02632 |
| **LINC00854** | 0.04670 | 0.01277 | 0.00513 | 0.06005 | 0.01296 | 0.03634 | 0.00890 | 0.00781 | 0.05362 | 0.12559 | 0.00016 | 0.03090 |
| **LINC00857** | 0.00509 | 0.00000 | 0.03762 | 0.12055 | 0.27673 | 0.45975 | 0.42147 | 0.00590 | 0.84551 | 0.00000 | 0.16558 | 0.01522 |
| **LINC00858** | 0.00324 | 0.27872 | 0.21293 | 0.00000 | 0.05657 | 0.35417 | 0.00000 | 0.00973 | 0.00000 | 0.02099 | 0.14780 | 0.24056 |
| **LINC00861** | 0.01380 | 0.01367 | 0.00989 | 0.00909 | 0.02909 | 0.03050 | 0.07440 | 0.02650 | 0.00346 | 0.01002 | 0.01287 | 0.01376 |
| **LINC00862** | 0.05249 | 0.00054 | 0.03651 | 0.21367 | 0.09223 | 0.04748 | 0.03232 | 0.09399 | 0.00000 | 0.04099 | 0.08064 | 0.01870 |
| **LINC00864** | 0.01246 | 0.01486 | 0.01445 | 0.01109 | 0.01433 | 0.01811 | 0.01957 | 0.01119 | 0.02849 | 0.01317 | 0.01710 | 0.01596 |
| **LINC00865** | 0.00187 | 0.01505 | 0.06545 | 0.04733 | 0.01003 | 0.01567 | 0.01030 | 0.11193 | 0.01315 | 0.02032 | 0.02704 | 0.01578 |
| **LINC00867** | 0.02666 | 0.00553 | 0.01383 | 0.00023 | 0.04001 | 0.09770 | 0.00576 | 0.00521 | 0.00000 | 0.04258 | 0.01317 | 0.04294 |
| **LINC00869.uid-** | 0.00623 | 0.01365 | 0.02222 | 0.01314 | 0.01492 | 0.02420 | 0.02304 | 0.01042 | 0.01341 | 0.01269 | 0.01735 | 0.01608 |
| **LINC00871** | 0.26706 | 0.00648 | 0.00408 | 0.06116 | 0.00000 | 0.04428 | 0.04780 | 0.02867 | 0.00001 | 0.04608 | 0.01186 | 0.10923 |
| **LINC00877** | 0.04261 | 0.01448 | 0.02593 | 0.01308 | 0.03261 | 0.00584 | 0.01400 | 0.01504 | 0.01859 | 0.02881 | 0.01426 | 0.01477 |
| **LINC00879** | 0.00629 | 0.01626 | 0.02358 | 0.00981 | 0.01016 | 0.02322 | 0.02610 | 0.01099 | 0.02148 | 0.02325 | 0.01300 | 0.02193 |
| **LINC00880** | 0.00629 | 0.01529 | 0.02041 | 0.01296 | 0.01554 | 0.01694 | 0.01771 | 0.01049 | 0.01962 | 0.01374 | 0.01481 | 0.01641 |
| **LINC00882** | 0.06721 | 0.01837 | 0.02805 | 0.02606 | 0.00767 | 0.02759 | 0.02650 | 0.08089 | 0.02669 | 0.00766 | 0.01131 | 0.09784 |
| **LINC00883** | 0.11729 | 0.18237 | 0.69331 | 0.28345 | 0.00000 | 0.00645 | 0.26362 | 0.54861 | 0.00000 | 0.02228 | 0.38526 | 0.52350 |
| **LINC00884** | 0.07344 | 0.41164 | 0.00000 | 0.00000 | 0.06876 | 0.02334 | 0.35551 | 0.11152 | 0.00000 | 0.09269 | 0.06967 | 0.63379 |
| **LINC00885** | 0.79961 | 0.02455 | 0.24755 | 0.00000 | 0.11672 | 0.03251 | 0.00000 | 0.14763 | 0.00000 | 0.03158 | 0.06017 | 0.00222 |
| **LINC00886** | 0.32315 | 0.01595 | 0.01841 | 0.02702 | 0.03389 | 0.00663 | 0.02537 | 0.01116 | 0.02685 | 0.04538 | 0.02160 | 0.02634 |
| **LINC00887** | 1.71383 | 0.08636 | 0.02330 | 0.03208 | 0.01138 | 0.04420 | 0.01016 | 0.60064 | 0.01267 | 0.00134 | 0.00866 | 0.02526 |
| **LINC00888** | 0.37709 | 0.61242 | 1.07407 | 2.84586 | 0.68430 | 4.92351 | 1.07196 | 0.00000 | 0.15212 | 1.98191 | 1.32671 | 0.30880 |
| **LINC00889** | 0.00589 | 0.02354 | 0.01285 | 0.01332 | 0.01269 | 0.00566 | 0.02967 | 0.01008 | 0.01677 | 0.03855 | 0.02160 | 0.02174 |
| **LINC00890** | 0.00598 | 0.01269 | 0.01196 | 0.01483 | 0.01105 | 0.03283 | 0.01932 | 0.00982 | 0.01073 | 0.02809 | 0.01410 | 0.02068 |
| **LINC00893** | 0.03861 | 0.03654 | 0.02313 | 0.01249 | 0.02639 | 0.00000 | 0.01384 | 0.06681 | 0.04615 | 0.00874 | 0.01407 | 0.02856 |
| **LINC00894** | 0.07432 | 0.05130 | 0.00105 | 0.00000 | 0.00000 | 0.16248 | 0.00000 | 0.08875 | 0.11669 | 0.44191 | 0.01693 | 0.06959 |
| **LINC00896** | 0.00630 | 0.05048 | 0.01413 | 0.00791 | 0.01653 | 0.01739 | 0.02231 | 0.00793 | 0.01280 | 0.01357 | 0.01679 | 0.10954 |
| **LINC00898** | 0.14229 | 0.01540 | 0.02845 | 0.01871 | 0.01031 | 0.01586 | 0.01774 | 0.00306 | 0.02073 | 0.02118 | 0.02223 | 0.01385 |
| **LINC00899** | 0.01014 | 0.01480 | 0.01714 | 0.01220 | 0.02915 | 0.00779 | 0.01663 | 0.03472 | 0.01668 | 0.02836 | 0.01746 | 0.01460 |
| **LINC00907** | 1.56658 | 0.00000 | 0.02321 | 0.08391 | 0.10423 | 0.20585 | 0.00000 | 0.21901 | 0.00000 | 0.07518 | 0.02377 | 0.57963 |
| **LINC00908** | 0.00629 | 0.01446 | 0.03361 | 0.01294 | 0.01512 | 0.02123 | 0.01726 | 0.01038 | 0.01622 | 0.01231 | 0.01623 | 0.01685 |
| **LINC00909** | 0.00233 | 0.72154 | 0.64981 | 0.00000 | 0.33415 | 3.04709 | 1.02162 | 2.04233 | 2.23017 | 1.57538 | 1.02580 | 1.78315 |
| **LINC00910** | 0.08132 | 0.28246 | 0.32466 | 0.25128 | 0.04785 | 0.00000 | 0.35564 | 0.20837 | 0.00632 | 0.32332 | 0.09857 | 0.59360 |
| **LINC00917** | 0.00706 | 0.01300 | 0.02068 | 0.02485 | 0.01161 | 0.00000 | 0.03106 | 0.00598 | 0.00358 | 0.01003 | 0.06033 | 0.33306 |
| **LINC00920** | 0.00620 | 0.01298 | 0.15822 | 0.03700 | 0.06972 | 0.00000 | 0.00969 | 0.00982 | 0.00450 | 0.00000 | 0.01747 | 0.03752 |
| **LINC00921** | 0.07418 | 0.01233 | 0.04171 | 0.02685 | 0.00000 | 0.08977 | 0.04028 | 0.16886 | 0.00000 | 0.03301 | 0.06022 | 0.05015 |
| **LINC00922** | 0.83477 | 0.00590 | 0.03659 | 0.25618 | 0.00000 | 0.00590 | 0.00596 | 0.03670 | 0.02071 | 0.03479 | 0.01362 | 0.08168 |
| **LINC00923** | 0.04495 | 0.06901 | 0.02333 | 0.00761 | 0.11398 | 0.00000 | 0.03039 | 0.02248 | 0.00000 | 0.05033 | 0.01173 | 0.05285 |
| **LINC00924** | 0.01738 | 0.01615 | 0.01466 | 0.01449 | 0.01791 | 0.01302 | 0.05311 | 0.01016 | 0.00357 | 0.00657 | 0.01610 | 0.01665 |
| **LINC00925** | 0.01386 | 1.06398 | 0.39678 | 0.00000 | 0.47292 | 0.02130 | 0.52216 | 2.13504 | 0.06063 | 0.00000 | 0.08673 | 4.29520 |

| | | | | | | | | | | | | |
|---|---|---|---|---|---|---|---|---|---|---|---|---|
| **LINC00926** | 0.02169 | 0.00541 | 0.02670 | 0.07182 | 0.00000 | 0.00465 | 0.09732 | 1.67456 | 0.25471 | 0.14814 | 0.07485 | 0.00000 |
| **LINC00927** | 0.02648 | 0.00728 | 0.00992 | 0.00262 | 0.00933 | 0.03376 | 0.00000 | 0.00939 | 0.01540 | 0.13243 | 0.00708 | 0.00985 |
| **LINC00929** | 0.00638 | 0.01598 | 0.04169 | 0.01524 | 0.01948 | 0.01334 | 0.02241 | 0.01053 | 0.01185 | 0.00820 | 0.01610 | 0.01699 |
| **LINC00933** | 0.18758 | 0.01834 | 0.02677 | 0.01553 | 0.02597 | 0.00311 | 0.05220 | 0.05692 | 0.00000 | 0.02140 | 0.01888 | 0.02949 |
| **LINC00934** | 0.00377 | 0.01040 | 0.00184 | 0.00506 | 0.00960 | 0.00000 | 0.00520 | 0.00996 | 0.03350 | 0.30549 | 0.01906 | 0.01084 |
| **LINC00935** | 0.00175 | 0.00493 | 0.00000 | 0.16189 | 0.00241 | 0.01684 | 0.00249 | 0.93290 | 0.00682 | 0.17357 | 0.00829 | 0.67584 |
| **LINC00936** | 0.53484 | 0.26801 | 0.95319 | 0.77737 | 0.75999 | 0.00000 | 0.00000 | 0.61265 | 2.29878 | 0.00024 | 1.17892 | 0.58604 |
| **LINC00937** | 0.18048 | 0.01935 | 0.01898 | 0.12612 | 0.03270 | 0.01159 | 0.02169 | 0.01180 | 0.02026 | 0.00000 | 0.01568 | 0.04409 |
| **LINC00938** | 0.14157 | 1.92212 | 1.53343 | 0.73252 | 0.00000 | 3.39128 | 0.00000 | 0.22927 | 1.56743 | 0.00424 | 1.96251 | 0.77093 |
| **LINC00940** | 0.00606 | 0.01357 | 0.01616 | 0.01074 | 0.01300 | 0.02274 | 0.01981 | 0.01038 | 0.00897 | 0.02098 | 0.02756 | 0.01611 |
| **LINC00941** | 0.16930 | 0.00613 | 0.02348 | 0.20514 | 0.00021 | 0.04695 | 0.02571 | 0.02808 | 0.00954 | 0.00141 | 0.01437 | 0.07779 |
| **LINC00942** | 0.00623 | 0.01506 | 0.04534 | 0.00935 | 0.03636 | 0.00927 | 0.02085 | 0.01054 | 0.05765 | 0.01078 | 0.02767 | 0.01686 |
| **LINC00943** | 1.58983 | 0.00550 | 0.00863 | 0.07887 | 0.01189 | 0.00000 | 0.01792 | 0.53363 | 0.00669 | 0.08102 | 0.01188 | 0.01035 |
| **LINC00948** | 0.00118 | 0.00000 | 0.04246 | 0.03667 | 0.00000 | 0.13258 | 0.00000 | 7.50965 | 0.39260 | 0.79166 | 0.60638 | 6.66265 |
| **LINC00950** | 0.00626 | 0.01671 | 0.04415 | 0.01228 | 0.01850 | 0.00000 | 0.01338 | 0.01061 | 0.01735 | 0.02112 | 0.05551 | 0.01715 |
| **LINC00951** | 0.00278 | 0.01489 | 0.01662 | 0.01141 | 0.01376 | 0.01965 | 0.01784 | 0.15814 | 0.01552 | 0.01855 | 0.01504 | 0.00969 |
| **LINC00954** | 0.27200 | 0.00917 | 0.00683 | 0.00707 | 0.00000 | 0.01775 | 0.07004 | 0.05474 | 0.00000 | 0.11168 | 0.01646 | 0.00650 |
| **LINC00955** | 0.00523 | 0.01353 | 0.05351 | 0.00711 | 0.01432 | 0.00898 | 0.00430 | 0.01054 | 0.02918 | 0.07956 | 0.00824 | 0.01370 |
| **LINC00957** | 0.05243 | 0.09547 | 0.00345 | 0.00575 | 0.00000 | 0.17032 | 0.00000 | 0.01601 | 0.01760 | 0.06272 | 0.00436 | 0.01799 |
| **LINC00958** | 1.20540 | 0.05292 | 3.56193 | 0.00000 | 0.80439 | 0.38475 | 1.46129 | 1.01901 | 2.53682 | 0.11247 | 0.34474 | 2.49175 |
| **LINC00959** | 0.74730 | 0.23051 | 0.07342 | 0.10111 | 0.00189 | 0.05633 | 0.03320 | 0.14736 | 0.00370 | 0.00000 | 0.19257 | 0.14135 |
| **LINC00960.uid-!** | 0.10545 | 0.32996 | 0.22999 | 0.10388 | 0.00000 | 0.00000 | 0.31013 | 0.00357 | 0.10328 | 0.11073 | 0.03767 | 0.05160 |
| **LINC00963** | 0.06422 | 0.00000 | 0.60816 | 0.00118 | 0.40737 | 0.17572 | 0.00000 | 0.00480 | 0.56653 | 0.00000 | 0.08709 | 0.01406 |
| **LINC00967** | 0.00172 | 0.07655 | 0.02210 | 0.03952 | 0.01378 | 0.00000 | 0.02102 | 0.12246 | 0.08508 | 0.00000 | 0.01349 | 0.12368 |
| **LINC00969** | 0.09624 | 0.32764 | 0.09279 | 0.06414 | 0.13375 | 0.13124 | 0.00000 | 0.02364 | 0.32990 | 0.32071 | 0.09604 | 0.19383 |
| **LINC00970** | 0.02339 | 0.01428 | 0.01600 | 0.01624 | 0.01165 | 0.01681 | 0.01519 | 0.03329 | 0.01713 | 0.02072 | 0.02627 | 0.01374 |
| **LINC00971** | 0.10044 | 0.02055 | 0.00777 | 0.08431 | 0.02102 | 0.00000 | 0.04732 | 0.00680 | 0.00000 | 0.07093 | 0.01340 | 0.02679 |
| **LINC00974** | 0.00539 | 0.00000 | 0.03478 | 0.00000 | 0.00000 | 0.01381 | 0.56046 | 0.00917 | 0.71574 | 0.01863 | 0.29788 | 0.20237 |
| **LINC00977** | 1.32083 | 0.01615 | 0.01447 | 0.01308 | 0.01981 | 0.00000 | 0.02110 | 0.12490 | 0.02448 | 0.00000 | 0.01404 | 0.01978 |
| **LINC00982** | 0.00540 | 0.01050 | 0.01283 | 0.01019 | 0.00960 | 0.02590 | 0.01713 | 0.00988 | 0.01320 | 0.03736 | 0.01399 | 0.04441 |
| **LINC00987** | 0.02022 | 0.03862 | 0.09203 | 0.19399 | 0.03950 | 0.24023 | 0.19196 | 0.07877 | 0.00000 | 0.12668 | 0.07191 | 0.04320 |
| **LINC00989** | 0.00677 | 0.00000 | 0.00000 | 0.68058 | 0.00000 | 0.28287 | 0.00000 | 0.00208 | 0.00000 | 0.02283 | 0.00329 | 0.17179 |
| **LINC00992** | 0.00385 | 0.09337 | 0.01169 | 0.03335 | 0.06056 | 0.08814 | 0.03384 | 0.01002 | 0.00087 | 0.08302 | 0.01592 | 0.03825 |
| **LINC00996** | 0.00369 | 0.00768 | 0.00826 | 0.00680 | 0.00728 | 0.03471 | 0.06805 | 0.01777 | 0.00000 | 0.03904 | 0.01061 | 0.04235 |
| **LINC00997** | 0.00415 | 0.00000 | 1.00477 | 0.36547 | 0.07317 | 0.03909 | 1.21422 | 0.02471 | 0.00016 | 0.29714 | 0.28086 | 0.01200 |
| **LINC00998** | 0.16617 | 1.88916 | 3.66083 | 0.02872 | 0.01011 | 7.31827 | 2.32176 | 1.91787 | 2.30592 | 0.20672 | 4.21804 | 4.52453 |
| **LINC00999** | 0.18086 | 0.09459 | 0.05633 | 0.03559 | 0.05447 | 0.00001 | 0.09522 | 0.00000 | 0.00000 | 0.10540 | 0.11641 | 0.15780 |
| **LINC01000** | 0.16306 | 1.18788 | 0.40988 | 1.43141 | 0.80374 | 0.00000 | 0.12562 | 0.04007 | 2.31888 | 6.81468 | 0.10965 | 0.90114 |
| **LINC01003** | 0.08547 | 0.41114 | 0.18521 | 0.00000 | 0.59291 | 0.00000 | 0.00000 | 0.00544 | 0.00000 | 0.49282 | 0.13612 | 0.09417 |
| **LINC01004** | 0.00000 | 0.00000 | 0.00000 | 0.35356 | 0.00203 | 1.23230 | 0.00000 | 0.05221 | 0.68608 | 3.48216 | 1.47634 | 0.37533 |
| **LINC01006** | 0.06653 | 0.14164 | 0.01343 | 0.04134 | 0.00000 | 0.02653 | 0.02941 | 0.07393 | 0.00783 | 0.01591 | 0.01519 | 0.00519 |
| **LINC01007** | 0.12508 | 0.00000 | 0.01392 | 0.00183 | 0.02947 | 0.02563 | 0.00196 | 0.00000 | 0.00000 | 0.22577 | 0.03069 | 0.10243 |
| **LINC01010** | 0.06486 | 0.28603 | 0.00996 | 0.00000 | 0.08728 | 0.03216 | 0.01469 | 0.22227 | 0.00431 | 0.00462 | 0.02318 | 0.78656 |
| **LINC01011** | 0.23258 | 0.61448 | 0.15541 | 0.36145 | 0.00000 | 0.44453 | 0.00000 | 0.00000 | 0.00000 | 0.39819 | 0.00000 | 0.17771 |
| **LINC01012** | 0.02181 | 0.23522 | 0.62989 | 0.20933 | 0.18021 | 0.13189 | 0.34512 | 0.06319 | 0.61922 | 0.00000 | 0.42886 | 0.01046 |
| **LINC01017** | 0.00698 | 0.01564 | 0.02204 | 0.00474 | 0.00737 | 0.03179 | 0.00468 | 0.01096 | 0.03635 | 0.03854 | 0.04545 | 0.01375 |
| **LINC01018** | 0.02178 | 0.02135 | 0.01279 | 0.00772 | 0.02446 | 0.01562 | 0.01986 | 0.01966 | 0.01555 | 0.03576 | 0.01830 | 0.01353 |
| **LINC01019** | 1.10248 | 0.01260 | 0.24748 | 0.00154 | 0.02307 | 0.00000 | 0.22564 | 0.16419 | 0.32649 | 0.00000 | 0.37436 | 0.00906 |

| | | | | | | | | | | | | |
|---|---|---|---|---|---|---|---|---|---|---|---|---|
| **LINC01021** | 0.52244 | 0.47533 | 1.23349 | 0.07950 | 0.25157 | 0.79611 | 0.00000 | 0.24417 | 2.27168 | 0.69255 | 0.45777 | 0.01384 |
| **LINC01023** | 0.11146 | 0.34696 | 0.20758 | 0.14721 | 0.23157 | 0.00000 | 0.00000 | 0.13631 | 0.00000 | 0.40299 | 0.22135 | 0.20452 |
| **LINC01024** | 0.42216 | 0.03047 | 0.04703 | 0.06193 | 0.16556 | 0.75118 | 0.00000 | 0.37984 | 0.43556 | 0.00000 | 0.09477 | 1.01160 |
| **LINC01028** | 0.00602 | 0.01600 | 0.02384 | 0.00887 | 0.01089 | 0.03062 | 0.01797 | 0.01022 | 0.01614 | 0.02025 | 0.01473 | 0.01605 |
| **LINC01029** | 0.00814 | 0.06392 | 0.39207 | 0.00000 | 0.00000 | 0.01394 | 0.47075 | 0.13300 | 0.35620 | 0.18535 | 0.00000 | 0.15308 |
| **LINC01030** | 0.93537 | 0.00000 | 0.15470 | 0.11398 | 0.24429 | 0.48679 | 0.00000 | 3.56871 | 0.02549 | 0.00000 | 0.10166 | 0.47403 |
| **LINC01049** | 0.02403 | 0.01594 | 0.02141 | 0.01564 | 0.01617 | 0.01366 | 0.02198 | 0.00994 | 0.03407 | 0.00608 | 0.01351 | 0.01657 |
| **LINC01056** | 0.35344 | 0.01438 | 0.01178 | 0.00532 | 0.01985 | 0.03098 | 0.00375 | 0.32536 | 0.01198 | 0.05702 | 0.01684 | 0.31422 |
| **LINC01057** | 0.02630 | 0.01966 | 0.00000 | 0.59625 | 0.19225 | 0.08429 | 0.00000 | 0.03497 | 0.21323 | 0.00000 | 0.01444 | 0.01200 |
| **LINC01060** | 0.00608 | 0.01519 | 0.05551 | 0.02749 | 0.07357 | 0.01201 | 0.18703 | 0.00967 | 0.00000 | 0.00000 | 0.03090 | 0.06607 |
| **LINC01061** | 0.04987 | 0.18829 | 0.03652 | 0.02518 | 0.17201 | 0.00000 | 0.22704 | 0.05483 | 0.06785 | 0.60156 | 0.13630 | 0.31482 |
| **LINC01081** | 0.19937 | 0.00925 | 0.01547 | 0.00986 | 0.00523 | 0.03777 | 0.00815 | 0.08580 | 0.01166 | 0.04207 | 0.01255 | 0.02363 |
| **LINC01082** | 0.05044 | 0.01202 | 0.03668 | 0.00735 | 0.00780 | 0.02488 | 0.01901 | 0.01412 | 0.00647 | 0.03516 | 0.03262 | 0.01335 |
| **LINC01087** | 0.00297 | 0.04245 | 0.00351 | 0.05772 | 0.00000 | 0.07774 | 0.00000 | 0.07546 | 0.05578 | 0.27121 | 0.02357 | 0.07839 |
| **LINC01088** | 0.13487 | 0.02618 | 0.03345 | 0.00629 | 0.00941 | 0.05720 | 0.01287 | 0.01272 | 0.09516 | 0.00000 | 0.02522 | 0.10433 |
| **LINC01089** | 1.01573 | 1.00330 | 2.37765 | 1.22942 | 0.17567 | 0.00000 | 0.87848 | 0.94303 | 1.56160 | 4.45486 | 0.97430 | 1.54499 |
| **LINC01091** | 0.28974 | 0.02314 | 0.76482 | 0.00000 | 0.39214 | 0.75862 | 0.00000 | 0.00000 | 0.79342 | 0.17178 | 0.00000 | 0.05535 |
| **LINC01094** | 0.01549 | 0.01136 | 0.01773 | 0.06552 | 0.00172 | 0.03887 | 0.00007 | 0.00861 | 0.04792 | 0.07953 | 0.05045 | 0.07009 |
| **LINC01096** | 0.00936 | 0.01691 | 0.06824 | 0.02510 | 0.02027 | 0.00000 | 0.01270 | 0.06804 | 0.01230 | 0.01873 | 0.01751 | 0.02290 |
| **LINC01097** | 0.02157 | 0.01843 | 0.01506 | 0.01887 | 0.01957 | 0.00528 | 0.02290 | 0.04186 | 0.02514 | 0.00643 | 0.01541 | 0.01641 |
| **LINC01098** | 0.73474 | 0.04773 | 0.08955 | 0.03721 | 0.04725 | 0.11608 | 0.00000 | 0.15083 | 0.00000 | 0.14459 | 0.02537 | 0.00283 |
| **LINC01099** | 0.43272 | 0.01970 | 0.01035 | 0.00000 | 0.02892 | 0.05724 | 0.05747 | 0.08771 | 0.00076 | 0.16704 | 0.00000 | 0.02950 |
| **LINC01104** | 0.00607 | 0.01357 | 0.03502 | 0.01378 | 0.01224 | 0.01939 | 0.01211 | 0.01044 | 0.01208 | 0.02703 | 0.02183 | 0.01541 |
| **LINC01105** | 0.00931 | 0.01428 | 0.02917 | 0.01227 | 0.01414 | 0.01117 | 0.01691 | 0.01093 | 0.00000 | 0.02375 | 0.06589 | 0.01623 |
| **LINC01108** | 0.00339 | 0.86176 | 0.65888 | 10.20800 | 1.35062 | 0.00000 | 0.00000 | 0.00000 | 0.00000 | 1.34230 | 0.92779 | 1.36305 |
| **LINC01111** | 0.01640 | 0.01078 | 0.04402 | 0.01994 | 0.00794 | 0.06536 | 0.02036 | 0.03018 | 0.00000 | 0.05811 | 0.01702 | 0.04456 |
| **LINC01115** | 0.07016 | 0.01433 | 0.01427 | 0.01196 | 0.01519 | 0.01873 | 0.01904 | 0.00627 | 0.01871 | 0.01115 | 0.01451 | 0.01676 |
| **LINC01116** | 0.00665 | 0.02168 | 0.01554 | 0.01513 | 0.02992 | 0.00637 | 0.02150 | 0.02417 | 0.01726 | 0.01153 | 0.01522 | 0.03124 |
| **LINC01118** | 11.78646 | 0.00306 | 0.00484 | 0.00354 | 0.00793 | 0.00000 | 0.00057 | 3.18644 | 0.02444 | 0.07330 | 0.00743 | 0.01106 |
| **LINC01119** | 0.00528 | 0.01095 | 0.03643 | 0.01014 | 0.01538 | 0.04097 | 0.01552 | 0.01266 | 0.00856 | 0.00874 | 0.03822 | 0.01871 |
| **LINC01121** | 0.01588 | 0.01349 | 0.02425 | 0.01388 | 0.01369 | 0.01609 | 0.02047 | 0.03023 | 0.01281 | 0.02248 | 0.01564 | 0.02458 |
| **LINC01122** | 0.85681 | 0.03186 | 0.01438 | 0.00711 | 0.01597 | 0.02451 | 0.05080 | 0.01792 | 0.00438 | 0.01399 | 0.01530 | 0.04298 |
| **LINC01123.uid-** | 0.01202 | 0.02533 | 0.01552 | 0.01096 | 0.03597 | 0.00000 | 0.07253 | 0.01039 | 0.07278 | 0.02713 | 0.04582 | 0.08829 |
| **LINC01123.uid-** | 0.00447 | 0.01830 | 0.04508 | 0.00048 | 0.02258 | 0.03682 | 0.00564 | 0.00948 | 0.05479 | 0.08887 | 0.00936 | 0.05587 |
| **LINC01125** | 0.25870 | 0.07171 | 0.04749 | 0.01386 | 0.02505 | 0.12343 | 0.00000 | 0.00000 | 0.09371 | 0.00000 | 0.03026 | 0.09193 |
| **LINC01126** | 0.02504 | 0.01736 | 0.01891 | 0.00867 | 0.01186 | 0.01200 | 0.00093 | 0.00763 | 0.03352 | 0.13250 | 0.00495 | 0.01481 |
| **LINC01127** | 0.00623 | 0.01393 | 0.05451 | 0.01032 | 0.02583 | 0.01596 | 0.00704 | 0.01044 | 0.02455 | 0.01924 | 0.01232 | 0.01611 |
| **LINC01128** | 6.58404 | 3.67269 | 1.16277 | 0.00000 | 1.27897 | 1.91079 | 2.44299 | 1.94990 | 4.90196 | 0.55297 | 1.81862 | 2.64979 |
| **LINC01132** | 0.00790 | 0.01496 | 0.01820 | 0.01301 | 0.01381 | 0.01387 | 0.01971 | 0.01123 | 0.00976 | 0.01751 | 0.03232 | 0.01631 |
| **LINC01133** | 0.01463 | 0.04003 | 0.00000 | 0.05942 | 0.26856 | 0.68496 | 0.00000 | 0.00336 | 0.00000 | 0.01089 | 0.01802 | 0.00000 |
| **LINC01134** | 0.09557 | 0.08236 | 0.08726 | 0.00000 | 0.03572 | 0.00000 | 0.00142 | 0.00000 | 0.15960 | 0.11635 | 0.05961 | 0.17713 |
| **LINC01136** | 0.00605 | 0.01586 | 0.02489 | 0.01517 | 0.02594 | 0.01178 | 0.01366 | 0.01047 | 0.01911 | 0.02356 | 0.01515 | 0.01669 |
| **LINC01137** | 0.00243 | 0.33043 | 0.23609 | 0.00000 | 0.00000 | 0.01579 | 0.10250 | 0.00666 | 0.16749 | 0.37862 | 0.04969 | 0.11358 |
| **LINC01138** | 0.00609 | 0.01407 | 0.01516 | 0.01235 | 0.01432 | 0.02205 | 0.01912 | 0.01391 | 0.01957 | 0.01273 | 0.01428 | 0.02268 |
| **LINC01140** | 0.04781 | 0.10071 | 0.49374 | 0.14579 | 0.04829 | 0.05218 | 0.00000 | 0.08951 | 0.09635 | 0.14368 | 0.01079 | 0.34181 |
| **LINC01141** | 0.09068 | 0.02880 | 0.05895 | 0.02685 | 0.03419 | 0.00000 | 0.01440 | 0.00560 | 0.01490 | 0.01219 | 0.01386 | 0.01700 |
| **LINC01144** | 0.00323 | 0.10263 | 0.08281 | 0.01599 | 0.01409 | 0.00000 | 0.00558 | 0.11651 | 0.09827 | 0.00000 | 0.01001 | 0.15457 |
| **LINC01150** | 0.01037 | 0.01028 | 0.01031 | 0.02124 | 0.00268 | 0.05419 | 0.04248 | 0.01735 | 0.07599 | 0.00000 | 0.00997 | 0.01442 |

| | | | | | | | | | | | | |
|---|---|---|---|---|---|---|---|---|---|---|---|---|
| **LINC01152** | 9.76721 | 0.03035 | 0.00721 | 0.00246 | 0.01080 | 0.04915 | 0.00426 | 0.99125 | 0.00431 | 0.03071 | 0.00796 | 0.28391 |
| **LINC01159** | 0.00645 | 0.01623 | 0.01781 | 0.01843 | 0.01503 | 0.00869 | 0.02857 | 0.01051 | 0.00969 | 0.01200 | 0.03794 | 0.01729 |
| **LINC01160** | 0.00000 | 7.40291 | 1.66151 | 9.71296 | 2.67917 | 2.00291 | 1.53240 | 0.07263 | 0.00000 | 1.48685 | 0.00000 | 0.95326 |
| **LINC01163** | 0.00623 | 0.01444 | 0.01900 | 0.01288 | 0.01380 | 0.02027 | 0.01663 | 0.01048 | 0.01743 | 0.01831 | 0.01529 | 0.01588 |
| **LINC01164** | 0.00450 | 0.01177 | 0.01766 | 0.05600 | 0.00759 | 0.00737 | 0.02000 | 0.02990 | 0.01648 | 0.08660 | 0.02030 | 0.07517 |
| **LINC01166** | 0.17748 | 0.13391 | 0.01135 | 0.00419 | 0.00000 | 0.01837 | 0.02061 | 0.00164 | 0.01445 | 0.02521 | 0.01324 | 0.22026 |
| **LINC01168** | 0.14865 | 0.02929 | 0.02705 | 0.00581 | 0.00995 | 0.03507 | 0.01254 | 0.00459 | 0.01875 | 0.03604 | 0.01541 | 0.09081 |
| **LINC01169** | 0.20143 | 0.03026 | 0.01487 | 0.00787 | 0.05602 | 0.05391 | 0.01683 | 0.00358 | 0.01901 | 0.00000 | 0.02000 | 0.01094 |
| **LINC01176** | 0.00326 | 0.00000 | 0.11608 | 0.00663 | 0.00000 | 0.04152 | 0.08113 | 0.00854 | 0.27732 | 0.24374 | 0.09527 | 0.02558 |
| **LINC01179** | 0.03187 | 0.01131 | 0.01250 | 0.01120 | 0.01920 | 0.03097 | 0.02087 | 0.01274 | 0.01443 | 0.01642 | 0.01411 | 0.01883 |
| **LINC01180** | 0.00631 | 0.01613 | 0.01250 | 0.01042 | 0.01311 | 0.02212 | 0.02751 | 0.01054 | 0.01835 | 0.02988 | 0.01338 | 0.02308 |
| **LINC01182** | 0.00892 | 0.01666 | 0.01595 | 0.01213 | 0.01618 | 0.02085 | 0.03031 | 0.01839 | 0.00842 | 0.01740 | 0.02399 | 0.01893 |
| **LINC01184** | 0.01333 | 0.03388 | 0.01488 | 0.15243 | 0.02395 | 0.00000 | 0.03128 | 0.01526 | 0.02307 | 0.00000 | 0.01849 | 0.06524 |
| **LINC01185** | 0.00612 | 0.01532 | 0.01728 | 0.01239 | 0.01779 | 0.01324 | 0.02251 | 0.01034 | 0.01634 | 0.01703 | 0.01706 | 0.01974 |
| **LINC01186** | 0.00766 | 0.12541 | 0.00875 | 0.00000 | 0.06307 | 0.09161 | 0.01663 | 0.00376 | 0.00000 | 0.01214 | 0.01494 | 0.33143 |
| **LINC01187** | 0.15229 | 0.01502 | 0.01538 | 0.01191 | 0.01538 | 0.01392 | 0.01753 | 0.10930 | 0.01885 | 0.01434 | 0.01459 | 0.00948 |
| **LINC01191** | 0.01822 | 0.00502 | 0.02083 | 0.03580 | 0.00594 | 0.00000 | 0.02173 | 0.01095 | 0.09450 | 0.28396 | 0.00159 | 0.04085 |
| **LINC01192** | 0.01562 | 0.01553 | 0.00999 | 0.00727 | 0.00690 | 0.01765 | 0.01086 | 0.01412 | 0.02164 | 0.10584 | 0.00978 | 0.01175 |
| **LINC01193** | 0.02839 | 0.04765 | 0.01247 | 0.00771 | 0.01030 | 0.00520 | 0.01803 | 0.67998 | 0.03178 | 0.03880 | 0.01458 | 0.03884 |
| **LINC01194** | 0.00846 | 0.14941 | 0.44676 | 0.17583 | 0.12212 | 0.20630 | 0.00000 | 0.03424 | 0.71154 | 0.31095 | 0.21343 | 0.28128 |
| **LINC01203** | 0.01985 | 0.00499 | 0.10672 | 0.03786 | 0.01033 | 0.05729 | 0.00000 | 0.00825 | 0.12070 | 0.07135 | 0.11536 | 0.04517 |
| **LINC01204** | 0.00567 | 0.02233 | 0.02316 | 0.03418 | 0.00083 | 0.04204 | 0.00874 | 0.01013 | 0.01248 | 0.03614 | 0.01187 | 0.01262 |
| **LINC01205** | 0.00400 | 0.51182 | 0.22776 | 0.99641 | 1.11085 | 0.92902 | 0.00000 | 0.00919 | 1.00122 | 1.16562 | 0.47643 | 0.17737 |
| **LINC01206** | 0.06313 | 0.00000 | 0.44068 | 0.91193 | 0.00081 | 0.00000 | 0.00000 | 0.21140 | 0.00000 | 0.00917 | 0.29143 | 0.19514 |
| **LINC01207** | 0.00229 | 0.01709 | 0.02449 | 0.01302 | 0.01728 | 0.00903 | 0.04311 | 0.05444 | 0.00867 | 0.00718 | 0.03350 | 0.01507 |
| **LINC01208** | 1.14996 | 0.01363 | 0.01327 | 0.01101 | 0.01571 | 0.00849 | 0.01706 | 0.12921 | 0.02279 | 0.01026 | 0.01462 | 0.02873 |
| **LINC01209** | 0.03384 | 0.01252 | 0.01320 | 0.00890 | 0.01885 | 0.03612 | 0.02220 | 0.06141 | 0.01258 | 0.00924 | 0.01595 | 0.03409 |
| **LINC01212** | 0.13919 | 0.04960 | 0.05468 | 0.01161 | 0.02404 | 0.00000 | 0.01412 | 0.00227 | 0.02276 | 0.00609 | 0.02361 | 0.06045 |
| **LINC01214** | 0.04463 | 0.01587 | 0.03434 | 0.01182 | 0.00579 | 0.00003 | 0.11539 | 0.00706 | 0.00000 | 0.03621 | 0.08784 | 0.01552 |
| **LINC01215** | 0.00606 | 0.01526 | 0.03300 | 0.01179 | 0.01704 | 0.01196 | 0.02154 | 0.01039 | 0.00913 | 0.02058 | 0.01693 | 0.01983 |
| **LINC01220** | 0.03203 | 0.02307 | 0.02559 | 0.01787 | 0.00964 | 0.02065 | 0.02233 | 0.01000 | 0.01354 | 0.00980 | 0.04050 | 0.04622 |
| **LINC01225** | 0.02010 | 0.01459 | 0.03849 | 0.01456 | 0.01335 | 0.01399 | 0.01629 | 0.00887 | 0.00859 | 0.02646 | 0.02258 | 0.01539 |
| **LINC01226** | 0.35694 | 0.01429 | 0.02109 | 0.01002 | 0.01814 | 0.01194 | 0.01446 | 0.42023 | 0.01752 | 0.02205 | 0.01331 | 0.00640 |
| **LINC01230** | 0.01093 | 0.04971 | 0.02299 | 0.00000 | 0.01557 | 0.05461 | 0.00761 | 0.00767 | 0.00400 | 0.05247 | 0.01112 | 0.07563 |
| **LINC01232** | 0.17896 | 0.02506 | 0.04679 | 0.00400 | 0.01458 | 0.07365 | 0.02243 | 0.00122 | 0.11155 | 0.00000 | 0.04683 | 0.04185 |
| **LINC01233** | 0.05756 | 0.02500 | 0.01606 | 0.01238 | 0.03481 | 0.00603 | 0.01993 | 0.00587 | 0.02490 | 0.00000 | 0.01251 | 0.05805 |
| **LINC01234** | 0.00705 | 3.29848 | 0.12494 | 0.00000 | 1.74973 | 0.13856 | 1.93148 | 0.00000 | 0.00000 | 0.75039 | 0.62778 | 0.32247 |
| **LINC01239** | 0.30018 | 0.01969 | 0.00441 | 0.00305 | 0.00000 | 0.03668 | 0.03071 | 0.19511 | 0.00000 | 0.12760 | 0.00821 | 0.00794 |
| **LINC01241** | 0.06556 | 0.01049 | 0.02464 | 0.01231 | 0.01806 | 0.03282 | 0.02632 | 0.00671 | 0.02208 | 0.01335 | 0.01531 | 0.02077 |
| **LINC01242** | 0.00526 | 0.00154 | 0.19076 | 0.02721 | 0.01713 | 0.05794 | 0.00000 | 0.01053 | 0.00000 | 0.03952 | 0.05108 | 0.05572 |
| **LINC01245** | 0.00599 | 0.01315 | 0.02683 | 0.03167 | 0.00896 | 0.01923 | 0.01532 | 0.01041 | 0.01005 | 0.02798 | 0.02167 | 0.01541 |
| **LINC01247** | 0.00832 | 0.00633 | 0.01246 | 0.01309 | 0.00715 | 0.03340 | 0.01449 | 0.00700 | 0.01142 | 0.03156 | 0.01960 | 0.06506 |
| **LINC01248** | 0.01179 | 0.01346 | 0.01368 | 0.01478 | 0.01267 | 0.01993 | 0.01480 | 0.00911 | 0.01918 | 0.02260 | 0.01427 | 0.02019 |
| **LINC01249** | 0.13786 | 0.13371 | 0.01432 | 0.00633 | 0.00000 | 0.00951 | 0.03167 | 0.06721 | 0.00657 | 0.02479 | 0.03274 | 0.03030 |
| **LINC01250** | 0.04646 | 0.01202 | 0.00000 | 0.15573 | 0.00000 | 0.07843 | 0.00000 | 0.01473 | 0.02439 | 0.01470 | 0.02139 | 0.02424 |
| **LINC01251** | 0.03360 | 0.00990 | 0.00908 | 0.01791 | 0.00341 | 0.04573 | 0.00216 | 0.05018 | 0.02318 | 0.04976 | 0.01078 | 0.02882 |
| **LINC01252** | 5.03107 | 0.00475 | 0.03569 | 0.09715 | 0.00494 | 0.00000 | 0.05309 | 0.10756 | 0.02036 | 0.01570 | 0.02598 | 0.09516 |
| **LINC01257** | 0.07677 | 0.01368 | 0.01324 | 0.01121 | 0.01886 | 0.01728 | 0.01465 | 0.17809 | 0.02482 | 0.01319 | 0.01375 | 0.01388 |

| | | | | | | | | | | | | |
|---|---|---|---|---|---|---|---|---|---|---|---|---|
| **LINC01258** | 0.01409 | 0.01432 | 0.01688 | 0.01527 | 0.01319 | 0.02003 | 0.01825 | 0.01597 | 0.01612 | 0.01656 | 0.01669 | 0.01519 |
| **LINC01260** | 0.11876 | 0.00000 | 0.00000 | 0.63948 | 0.41255 | 0.83881 | 0.00000 | 0.08205 | 0.36327 | 0.00000 | 0.13180 | 0.77571 |
| **LINC01263** | 0.12065 | 0.03203 | 0.04251 | 0.37821 | 0.07280 | 0.00000 | 0.02907 | 0.04421 | 0.00427 | 0.02358 | 0.02885 | 0.05259 |
| **LINC01266** | 0.00603 | 0.01653 | 0.02508 | 0.00960 | 0.01297 | 0.02100 | 0.01384 | 0.01025 | 0.01409 | 0.02399 | 0.01621 | 0.01826 |
| **LINC01268** | 0.00582 | 0.01610 | 0.03535 | 0.01922 | 0.02163 | 0.02391 | 0.01402 | 0.03652 | 0.01446 | 0.01621 | 0.01553 | 0.01511 |
| **LINC01270** | 0.00533 | 0.00541 | 0.01535 | 0.00000 | 0.05208 | 0.05751 | 0.02657 | 0.00997 | 0.01140 | 0.03994 | 0.02156 | 0.01207 |
| **LINC01272** | 0.01814 | 0.01066 | 0.03274 | 0.00763 | 0.00309 | 0.02736 | 0.00683 | 0.01136 | 0.01966 | 0.11873 | 0.00808 | 0.01451 |
| **LINC01273** | 0.04582 | 0.03440 | 0.08269 | 0.01781 | 0.05790 | 0.00000 | 0.22217 | 0.00526 | 0.00001 | 0.00000 | 0.18914 | 0.21452 |
| **LINC01277** | 0.20054 | 0.03629 | 0.01652 | 0.00918 | 0.01082 | 0.02007 | 0.01937 | 0.02157 | 0.02686 | 0.00380 | 0.01330 | 0.02945 |
| **LINC01278** | 0.00000 | 1.28697 | 2.81172 | 0.15874 | 4.93130 | 2.03047 | 2.62905 | 0.44499 | 5.17805 | 0.98869 | 2.79121 | 0.06686 |
| **LINC01279** | 0.23111 | 0.00000 | 0.05331 | 0.15075 | 0.03074 | 0.03565 | 0.01683 | 0.07014 | 0.07124 | 0.02533 | 0.03049 | 0.22775 |
| **LINC01289** | 0.00597 | 0.00904 | 0.14160 | 0.00569 | 0.01225 | 0.06781 | 0.01761 | 0.00977 | 0.00000 | 0.01119 | 0.01348 | 0.01650 |
| **LINC01296.uid-** | 0.00494 | 0.04702 | 0.02900 | 0.01073 | 0.01907 | 0.00864 | 0.07145 | 0.01242 | 0.01541 | 0.02040 | 0.01551 | 0.04214 |
| **LINC01299** | 0.00674 | 0.01390 | 0.01249 | 0.01203 | 0.01471 | 0.04441 | 0.01454 | 0.00834 | 0.01853 | 0.00721 | 0.01117 | 0.09762 |
| **LINC01300** | 0.00437 | 0.02753 | 0.03681 | 0.00000 | 0.07385 | 0.00000 | 0.01386 | 0.01022 | 0.04133 | 0.14565 | 0.01778 | 0.01192 |
| **LINC01301** | 0.08522 | 0.30774 | 0.05840 | 0.01156 | 0.00000 | 0.02265 | 0.00107 | 0.00191 | 0.00000 | 0.04354 | 0.08951 | 0.08514 |
| **LINC01305** | 0.00558 | 0.01859 | 0.02545 | 0.02675 | 0.03535 | 0.00000 | 0.01995 | 0.01363 | 0.02486 | 0.00344 | 0.01591 | 0.01823 |
| **LINC01310** | 0.02224 | 0.01521 | 0.02053 | 0.01824 | 0.01666 | 0.01271 | 0.01881 | 0.01093 | 0.01569 | 0.01545 | 0.01568 | 0.01593 |
| **LINC01311** | 0.00073 | 0.55822 | 0.50353 | 0.20915 | 0.31020 | 0.00000 | 0.00008 | 0.06021 | 1.04275 | 0.62844 | 1.15164 | 0.85502 |
| **LINC01312** | 0.34972 | 0.01208 | 0.00982 | 0.01201 | 0.01555 | 0.02197 | 0.01002 | 0.10519 | 0.04812 | 0.00575 | 0.01273 | 0.01867 |
| **LINC01314** | 0.00457 | 0.01220 | 0.16195 | 0.00379 | 0.04496 | 0.00000 | 0.01408 | 0.01291 | 0.14644 | 0.07605 | 0.00824 | 0.01387 |
| **LINC01315** | 0.01983 | 0.28964 | 0.08401 | 0.00091 | 0.00000 | 0.00034 | 0.26332 | 0.01718 | 0.80840 | 0.08556 | 0.00000 | 0.27737 |
| **LINC01330** | 0.00488 | 0.01105 | 0.00733 | 0.00582 | 0.00556 | 0.01815 | 0.00135 | 0.01022 | 0.03700 | 0.11238 | 0.01273 | 0.01290 |
| **LINC01333** | 0.00612 | 0.01963 | 0.01424 | 0.01104 | 0.04888 | 0.00002 | 0.01341 | 0.01033 | 0.03255 | 0.01844 | 0.01026 | 0.02364 |
| **LINC01338** | 0.00438 | 0.03266 | 0.00702 | 0.00826 | 0.01479 | 0.03340 | 0.01709 | 0.01128 | 0.01059 | 0.08073 | 0.00997 | 0.05176 |
| **LINC01341** | 0.00754 | 0.01322 | 0.01525 | 0.01631 | 0.00655 | 0.03542 | 0.02347 | 0.03935 | 0.02787 | 0.00952 | 0.03054 | 0.01752 |
| **LINC01342** | 0.00807 | 0.01376 | 0.01314 | 0.01102 | 0.01650 | 0.02405 | 0.01850 | 0.01012 | 0.01870 | 0.01251 | 0.01697 | 0.01639 |
| **LINC01344** | 0.18002 | 0.04984 | 0.02037 | 0.03031 | 0.01373 | 0.00086 | 0.01722 | 0.08600 | 0.01993 | 0.01831 | 0.01565 | 0.01778 |
| **LINC01347** | 0.04435 | 0.01824 | 0.09599 | 0.02910 | 0.00826 | 0.04841 | 0.00000 | 0.03304 | 0.00905 | 0.04376 | 0.08248 | 0.08994 |
| **LINC01351** | 0.00308 | 0.01403 | 0.00884 | 0.10915 | 0.00635 | 0.02546 | 0.01600 | 0.03374 | 0.01691 | 0.00924 | 0.01524 | 0.01477 |
| **LINC01352** | 0.00481 | 0.01022 | 0.05987 | 0.01024 | 0.00000 | 0.01831 | 0.04003 | 0.01039 | 0.02343 | 0.10821 | 0.03430 | 0.01200 |
| **LINC01354** | 0.00232 | 0.01300 | 0.02148 | 0.01455 | 0.01847 | 0.00822 | 0.02844 | 0.13618 | 0.00000 | 0.00108 | 0.07879 | 0.12155 |
| **LINC01355** | 0.00000 | 0.00000 | 0.93972 | 0.29668 | 0.00000 | 0.91150 | 0.00000 | 0.25464 | 1.16879 | 1.49431 | 1.29423 | 0.05985 |
| **LINC01356** | 0.01456 | 0.34401 | 12.55146 | 21.52697 | 2.16943 | 0.00013 | 9.30850 | 0.03624 | 0.85415 | 0.00000 | 2.45277 | 0.43199 |
| **LINC01359** | 0.05597 | 0.03353 | 0.01289 | 0.01345 | 0.00783 | 0.02073 | 0.02122 | 0.04156 | 0.02319 | 0.00975 | 0.02777 | 0.01136 |
| **LINC01360** | 0.00595 | 0.01313 | 0.01593 | 0.02432 | 0.00692 | 0.03883 | 0.02661 | 0.00907 | 0.02621 | 0.01007 | 0.01165 | 0.03886 |
| **LINC01361** | 0.00643 | 0.01599 | 0.01585 | 0.01113 | 0.01147 | 0.01141 | 0.01489 | 0.01047 | 0.05397 | 0.01131 | 0.02767 | 0.01696 |
| **LINC01363** | 0.01093 | 0.00521 | 0.08760 | 0.00448 | 0.01033 | 0.09699 | 0.01573 | 0.06990 | 0.02620 | 0.00000 | 0.10479 | 0.04801 |
| **LINC01364** | 0.00568 | 0.02456 | 0.03872 | 0.02506 | 0.00030 | 0.03106 | 0.00570 | 0.01048 | 0.02938 | 0.03831 | 0.01767 | 0.01267 |
| **LINC01365** | 0.00515 | 0.01473 | 0.00901 | 0.00786 | 0.00732 | 0.02563 | 0.00686 | 0.01050 | 0.02605 | 0.05921 | 0.01008 | 0.01991 |
| **LINC01372** | 0.07862 | 0.02432 | 0.01615 | 0.12893 | 0.00868 | 0.00000 | 0.05916 | 0.09088 | 0.03287 | 0.08292 | 0.05155 | 0.02733 |
| **LINC01391** | 0.00838 | 0.01866 | 0.04124 | 0.01033 | 0.00984 | 0.01675 | 0.01677 | 0.00999 | 0.00000 | 0.00826 | 0.08156 | 0.01667 |
| **LINC01393** | 0.00487 | 0.08721 | 0.00481 | 0.00425 | 0.00000 | 0.07480 | 0.00000 | 0.00890 | 0.01460 | 0.06467 | 0.01023 | 0.03887 |
| **LINC01395** | 0.00650 | 0.01494 | 0.01726 | 0.05334 | 0.01713 | 0.00078 | 0.00860 | 0.00919 | 0.09905 | 0.00000 | 0.05466 | 0.04681 |
| **LINC01405** | 0.03140 | 0.00492 | 0.22337 | 0.10312 | 0.17422 | 0.00000 | 0.08557 | 0.05538 | 0.00000 | 0.11217 | 0.14156 | 0.00911 |
| **LINC01410** | 0.06265 | 0.00393 | 0.03571 | 0.00862 | 0.02723 | 0.14450 | 0.03624 | 0.00609 | 0.00000 | 0.15505 | 0.04227 | 0.03331 |
| **LINC01412** | 0.02846 | 0.01494 | 0.01495 | 0.01194 | 0.01488 | 0.01712 | 0.01788 | 0.02639 | 0.01783 | 0.01690 | 0.01499 | 0.01454 |
| **LINC01413** | 0.03820 | 0.16553 | 0.24314 | 0.01904 | 0.17924 | 0.00000 | 0.02038 | 0.00900 | 0.02241 | 0.02122 | 0.11546 | 0.02580 |

| | | | | | | | | | | | | |
|---|---|---|---|---|---|---|---|---|---|---|---|---|
| **LINC01419** | 0.00583 | 0.00480 | 0.00000 | 0.03910 | 0.00000 | 0.30676 | 0.22604 | 0.01236 | 0.00000 | 0.02527 | 0.00000 | 0.28214 |
| **LINC01420** | 0.00319 | 0.99287 | 1.11763 | 0.55348 | 0.10616 | 1.74706 | 1.07879 | 0.17790 | 0.00000 | 0.22449 | 0.52108 | 0.34863 |
| **LINC01422** | 0.00580 | 0.01766 | 0.02850 | 0.03260 | 0.00028 | 0.05046 | 0.01280 | 0.08052 | 0.00689 | 0.03978 | 0.01506 | 0.11459 |
| **LINC01424** | 0.00700 | 0.08713 | 0.05706 | 0.02619 | 0.00732 | 0.00000 | 0.01641 | 0.00836 | 0.07746 | 0.02687 | 0.03373 | 0.07599 |
| **LINC01425** | 0.16745 | 0.01757 | 0.01464 | 0.01037 | 0.02683 | 0.00000 | 0.01722 | 0.01921 | 0.02094 | 0.02997 | 0.03620 | 0.14016 |
| **LINC01426** | 0.00819 | 0.22723 | 0.13478 | 0.48297 | 0.25367 | 0.40489 | 0.08503 | 0.00451 | 0.00000 | 0.00001 | 0.00320 | 0.00971 |
| **LINC01428** | 0.14116 | 0.01644 | 0.06390 | 0.05566 | 0.01430 | 0.01251 | 0.01104 | 0.00385 | 0.01732 | 0.01381 | 0.01413 | 0.01430 |
| **LINC01429** | 0.00574 | 0.01779 | 0.02435 | 0.02938 | 0.00891 | 0.02202 | 0.02067 | 0.01021 | 0.01332 | 0.03661 | 0.01736 | 0.01874 |
| **LINC01431** | 0.00438 | 0.04557 | 0.11714 | 0.00000 | 0.00910 | 0.08850 | 0.00000 | 0.10437 | 0.11848 | 0.03212 | 0.00512 | 0.16415 |
| **LINC01432** | 0.00709 | 0.01451 | 0.01116 | 0.01024 | 0.00955 | 0.02149 | 0.01188 | 0.06691 | 0.02082 | 0.04052 | 0.01384 | 0.01087 |
| **LINC01433** | 0.10258 | 0.05362 | 0.31403 | 0.08270 | 0.11244 | 0.00000 | 0.00000 | 0.07169 | 0.06455 | 0.42732 | 0.16152 | 0.03416 |
| **LINC01436** | 0.01136 | 0.01060 | 0.01770 | 0.00714 | 0.00913 | 0.04608 | 0.02687 | 0.01223 | 0.01919 | 0.01394 | 0.01073 | 0.01699 |
| **LINC01441** | 0.00616 | 0.02070 | 0.02013 | 0.01112 | 0.01137 | 0.01936 | 0.03629 | 0.01057 | 0.01240 | 0.01555 | 0.01816 | 0.01533 |
| **LINC01443** | 0.00622 | 0.01684 | 0.01822 | 0.01026 | 0.01608 | 0.02294 | 0.01536 | 0.01037 | 0.02159 | 0.01818 | 0.01429 | 0.01600 |
| **LINC01446** | 0.01635 | 0.22808 | 0.60956 | 0.13806 | 0.21305 | 0.06871 | 0.06021 | 0.02292 | 0.08817 | 0.00000 | 0.29008 | 0.08798 |
| **LINC01447** | 0.15141 | 0.05629 | 1.79767 | 0.00493 | 0.03960 | 0.00000 | 0.00000 | 0.15805 | 0.00000 | 0.05085 | 0.01947 | 0.02613 |
| **LINC01448** | 0.02843 | 0.05447 | 0.01553 | 0.00000 | 0.09001 | 0.04693 | 0.01291 | 0.12119 | 0.00000 | 0.07378 | 0.00902 | 0.09974 |
| **LINC01452** | 0.00505 | 0.01706 | 0.00998 | 0.02348 | 0.01444 | 0.02964 | 0.01454 | 0.01499 | 0.01185 | 0.03783 | 0.01236 | 0.01369 |
| **LINCR-0001** | 0.01400 | 0.01586 | 0.02714 | 0.05876 | 0.01491 | 0.01513 | 0.02231 | 0.04109 | 0.03681 | 0.01467 | 0.01579 | 0.01983 |
| **LINCR-0002** | 0.00639 | 0.01624 | 0.02552 | 0.01835 | 0.02819 | 0.01582 | 0.02587 | 0.01049 | 0.02243 | 0.00432 | 0.01628 | 0.01694 |
| **LINCR-0003** | 0.09533 | 0.01372 | 0.03046 | 0.01555 | 0.01256 | 0.01779 | 0.01494 | 0.14593 | 0.01263 | 0.02070 | 0.01560 | 0.01015 |
| **LINGO1** | 0.45653 | 0.07386 | 0.16389 | 0.05458 | 0.50349 | 0.00000 | 0.00000 | 0.01300 | 0.00000 | 0.57463 | 0.41973 | 0.01030 |
| **LINGO2** | 5.53561 | 0.13432 | 0.01717 | 0.00210 | 0.04279 | 0.08172 | 0.00000 | 1.65811 | 0.00786 | 0.00000 | 0.00907 | 0.19291 |
| **LINGO3** | 0.00629 | 0.01431 | 0.01657 | 0.01115 | 0.02137 | 0.01903 | 0.01688 | 0.01347 | 0.01710 | 0.01601 | 0.01630 | 0.01970 |
| **LINGO4** | 0.00692 | 0.00912 | 0.01376 | 0.08253 | 0.01636 | 0.02295 | 0.02283 | 0.01554 | 0.01464 | 0.02856 | 0.01946 | 0.05326 |
| **LINS** | 0.48899 | 0.93202 | 2.98887 | 0.27221 | 0.80098 | 4.55688 | 0.00000 | 4.27759 | 3.81123 | 1.48367 | 0.62145 | 3.53303 |
| **LIPA** | 1.72005 | 0.21838 | 1.08919 | 0.22428 | 0.00000 | 0.13152 | 0.00000 | 0.54389 | 0.22718 | 0.00000 | 0.10536 | 0.92155 |
| **LIPC** | 0.00540 | 0.00000 | 0.04440 | 0.69621 | 0.60805 | 0.09323 | 0.00000 | 0.00848 | 0.27057 | 0.01973 | 0.38774 | 0.00866 |
| **LIPE** | 0.81326 | 0.03362 | 0.02363 | 0.04904 | 0.00000 | 0.02843 | 0.01819 | 0.00000 | 0.00477 | 0.07811 | 0.06899 | 0.09299 |
| **LIPE-AS1** | 1.50946 | 0.20505 | 0.05721 | 0.11469 | 0.00000 | 0.00782 | 0.01611 | 0.31633 | 0.06078 | 0.01313 | 0.05584 | 0.31998 |
| **LIPF** | 0.00473 | 0.00992 | 0.00641 | 0.00654 | 0.00332 | 0.02130 | 0.00236 | 0.01051 | 0.02537 | 0.10296 | 0.02547 | 0.01625 |
| **LIPG** | 4.24487 | 0.04452 | 5.32134 | 0.98379 | 0.00000 | 1.06674 | 1.70240 | 1.68459 | 0.36558 | 0.00000 | 0.03445 | 0.00000 |
| **LIPH** | 0.00000 | 3.15331 | 1.49297 | 0.00000 | 10.19474 | 3.33239 | 0.00000 | 0.01450 | 0.00000 | 0.80280 | 2.93663 | 0.00000 |
| **LIPI** | 0.00294 | 0.03271 | 0.01368 | 0.00254 | 0.01329 | 0.02411 | 0.00200 | 0.02118 | 0.02991 | 0.10271 | 0.00766 | 0.02535 |
| **LIPJ** | 0.00566 | 0.01216 | 0.01231 | 0.00912 | 0.00993 | 0.02506 | 0.02893 | 0.01022 | 0.00789 | 0.03768 | 0.01378 | 0.01867 |
| **LIPM** | 0.92937 | 0.01321 | 0.01619 | 0.02132 | 0.01582 | 0.00452 | 0.01757 | 0.19046 | 0.02416 | 0.00615 | 0.01368 | 0.01537 |
| **LIPT1** | 0.01719 | 3.66666 | 0.21073 | 0.00000 | 0.00000 | 6.61606 | 1.81875 | 1.26659 | 0.54597 | 1.61667 | 3.69989 | 6.30642 |
| **LIPT2** | 0.13929 | 1.06139 | 0.78340 | 0.99940 | 0.00000 | 0.20323 | 0.36533 | 0.84320 | 0.68910 | 0.00000 | 1.21112 | 0.59265 |
| **LITAF** | 0.47109 | 0.19129 | 1.35719 | 0.19116 | 0.07579 | 0.00000 | 0.46976 | 0.11586 | 0.00283 | 0.00000 | 0.08930 | 1.45894 |
| **LIX1** | 1.51026 | 0.10444 | 0.01762 | 0.00000 | 0.00000 | 0.00000 | 0.02149 | 0.00840 | 0.04875 | 0.14086 | 0.11770 | 0.07036 |
| **LIX1L** | 0.02911 | 0.00000 | 0.13851 | 0.93429 | 0.00000 | 0.09946 | 0.00000 | 0.00425 | 0.00024 | 0.28871 | 0.02123 | 0.15254 |
| **LL22NC01-81G9** | 0.00526 | 0.01593 | 0.01756 | 0.02515 | 0.01960 | 0.00703 | 0.01939 | 0.01379 | 0.01957 | 0.01961 | 0.01627 | 0.01717 |
| **LLGL1** | 0.08602 | 1.41061 | 0.73072 | 0.50107 | 0.39010 | 0.00000 | 1.47412 | 0.00000 | 0.52153 | 1.09965 | 1.45702 | 0.92996 |
| **LLGL2** | 3.14501 | 7.13786 | 0.00000 | 5.65268 | 16.87502 | 0.00000 | 8.30519 | 2.57074 | 19.11561 | 22.20656 | 12.08193 | 5.35799 |
| **LLPH** | 0.00000 | 4.97428 | 7.42902 | 3.35174 | 3.24146 | 18.52804 | 14.61589 | 10.31878 | 6.61558 | 5.63881 | 6.36025 | 13.67794 |
| **LMAN1** | 6.33726 | 12.39247 | 15.25395 | 2.20508 | 1.95402 | 32.73328 | 0.00000 | 5.40282 | 24.54106 | 24.15751 | 7.12779 | 12.90820 |
| **LMAN2** | 9.03215 | 24.03399 | 24.71898 | 24.47024 | 29.62888 | 12.47347 | 12.34709 | 13.47666 | 35.13972 | 9.86530 | 20.76094 | 22.51738 |
| **LMAN2L** | 0.00000 | 7.07113 | 5.76132 | 2.96707 | 4.28010 | 6.95449 | 0.00303 | 1.12770 | 13.06883 | 2.12370 | 2.46002 | 7.17118 |

| | | | | | | | | | | | | |
|---|---|---|---|---|---|---|---|---|---|---|---|---|
| **LMBR1** | 3.55677 | 4.91170 | 2.52421 | 0.80654 | 0.00000 | 10.87493 | 0.00000 | 1.28937 | 5.03920 | 6.74327 | 1.41340 | 0.00000 |
| **LMBR1L** | 2.09156 | 12.66721 | 1.02228 | 11.48461 | 4.45567 | 0.00000 | 2.95369 | 13.14145 | 15.05010 | 11.77635 | 14.95684 | 31.99953 |
| **LMBRD1** | 5.60007 | 5.50018 | 2.84936 | 8.13283 | 0.01128 | 21.13360 | 0.00000 | 1.44207 | 11.20756 | 5.95660 | 0.94329 | 4.59531 |
| **LMBRD2** | 1.42026 | 0.64683 | 1.32532 | 1.68308 | 0.86823 | 1.04115 | 0.00000 | 0.37345 | 0.84943 | 1.17796 | 1.62725 | 0.41380 |
| **LMCD1** | 0.33957 | 0.68229 | 0.23482 | 0.13767 | 0.00000 | 0.00000 | 0.00000 | 0.39676 | 0.09050 | 0.18089 | 0.00000 | 2.12632 |
| **LMCD1-AS1** | 0.14754 | 0.00791 | 0.10174 | 0.04104 | 0.02230 | 0.07240 | 0.00000 | 0.00388 | 0.01195 | 0.04695 | 0.00665 | 0.08234 |
| **LMF1** | 0.37603 | 0.99946 | 0.56246 | 0.25043 | 1.34911 | 0.00000 | 0.00165 | 0.37812 | 0.37419 | 0.22660 | 1.29002 | 0.86674 |
| **LMF1-AS1** | 0.02577 | 0.01120 | 0.04029 | 0.00574 | 0.01495 | 0.04180 | 0.00059 | 0.00855 | 0.02191 | 0.02019 | 0.04878 | 0.02608 |
| **LMF2** | 2.23408 | 4.00338 | 4.34440 | 3.05840 | 4.04536 | 0.00000 | 0.30151 | 0.08062 | 10.22597 | 2.87732 | 4.02026 | 2.03412 |
| **LMLN** | 1.78744 | 0.71985 | 0.44650 | 0.88733 | 0.00000 | 0.96201 | 0.00000 | 0.61573 | 0.29105 | 0.80161 | 0.57051 | 2.05869 |
| **LMLN-AS1** | 0.00581 | 0.01359 | 0.01269 | 0.01061 | 0.01839 | 0.02697 | 0.02103 | 0.01113 | 0.01494 | 0.01484 | 0.01884 | 0.01952 |
| **LMNA** | 2.56091 | 15.86252 | 24.69069 | 18.04813 | 42.94477 | 0.00023 | 38.69982 | 8.20654 | 40.63985 | 7.65377 | 39.56283 | 8.26708 |
| **LMNB1** | 2.62845 | 2.39033 | 7.86673 | 5.71729 | 2.20280 | 4.14336 | 10.93746 | 2.62588 | 3.59257 | 1.59871 | 17.95000 | 12.93972 |
| **LMNB2** | 1.82022 | 10.53560 | 17.65812 | 12.15785 | 21.24200 | 0.19451 | 28.67919 | 0.32026 | 10.55490 | 26.36857 | 9.31612 | 5.90490 |
| **LMNTD1** | 0.00810 | 0.11778 | 0.01958 | 0.00000 | 0.00000 | 0.04350 | 0.01023 | 0.00372 | 0.00745 | 0.04802 | 0.01211 | 1.59025 |
| **LMNTD2** | 0.01301 | 0.01329 | 0.10812 | 0.01368 | 0.01802 | 0.02563 | 0.01891 | 0.00916 | 0.01380 | 0.01328 | 0.02927 | 0.01659 |
| **LMO1** | 0.83709 | 0.05404 | 0.11050 | 0.00000 | 0.00000 | 0.05019 | 0.08138 | 0.04581 | 0.00000 | 0.04789 | 0.06014 | 1.35657 |
| **LMO2** | 0.00256 | 0.03631 | 1.94299 | 0.04154 | 0.00000 | 0.01041 | 0.06152 | 0.00582 | 0.00000 | 1.29038 | 0.00000 | 0.84156 |
| **LMO3** | 0.51077 | 0.02583 | 0.00376 | 0.05183 | 0.08495 | 0.08589 | 0.00000 | 0.09419 | 0.00000 | 0.11022 | 0.12482 | 0.01439 |
| **LMO4** | 1.86611 | 4.53246 | 5.81787 | 9.41320 | 12.17023 | 4.28202 | 18.96274 | 4.73682 | 0.00000 | 18.96038 | 5.29192 | 5.53434 |
| **LMO7** | 6.36056 | 0.52644 | 1.36468 | 0.12536 | 0.00000 | 1.17008 | 0.00000 | 2.12338 | 0.01698 | 1.44063 | 0.00000 | 1.75346 |
| **LMO7-AS1** | 0.42540 | 0.12548 | 0.53327 | 0.04636 | 0.17201 | 1.02565 | 0.71787 | 0.12909 | 0.19632 | 0.44789 | 0.49643 | 0.13611 |
| **LMO7DN** | 0.04520 | 0.01272 | 0.01166 | 0.00982 | 0.00990 | 0.02267 | 0.02386 | 0.01628 | 0.01164 | 0.03596 | 0.01659 | 0.01558 |
| **LMOD1** | 0.00362 | 0.00978 | 0.23370 | 0.00000 | 0.00000 | 0.00648 | 0.07457 | 0.00480 | 0.15523 | 0.31711 | 0.00000 | 0.20002 |
| **LMOD2** | 0.06161 | 0.08085 | 0.00000 | 0.00000 | 0.54154 | 0.31049 | 0.94843 | 0.04163 | 1.23371 | 0.10942 | 1.44129 | 0.00000 |
| **LMOD3** | 7.21878 | 0.42741 | 0.00000 | 0.24719 | 0.22623 | 3.40005 | 0.00000 | 0.31356 | 0.00000 | 0.79786 | 0.00000 | 0.01845 |
| **LMTK2** | 2.58924 | 3.90786 | 4.23395 | 0.54272 | 2.39939 | 3.86757 | 0.40415 | 0.85979 | 8.40818 | 8.76974 | 0.95175 | 2.65390 |
| **LMTK3** | 0.03109 | 0.02406 | 0.15895 | 0.00335 | 0.15023 | 0.00000 | 0.10415 | 0.01710 | 0.00000 | 0.60276 | 0.00000 | 0.02916 |
| **LMX1A** | 0.00390 | 0.01603 | 0.01245 | 0.01918 | 0.01195 | 0.00809 | 0.02441 | 0.11688 | 0.03427 | 0.03083 | 0.01139 | 0.01174 |
| **LMX1B** | 0.03127 | 0.18186 | 0.40040 | 0.00000 | 0.37581 | 0.00000 | 1.72261 | 0.01046 | 0.49342 | 1.70020 | 0.00000 | 0.00000 |
| **LNP1** | 0.21875 | 0.21871 | 0.34258 | 0.09622 | 0.00000 | 0.26478 | 0.31498 | 0.68819 | 0.00704 | 0.00000 | 0.05242 | 0.33136 |
| **LNPEP** | 1.15203 | 0.80165 | 0.35874 | 0.27869 | 0.00000 | 0.75116 | 0.23801 | 2.12345 | 0.31040 | 0.00000 | 0.00000 | 1.00637 |
| **LNX1** | 0.57689 | 1.46361 | 3.36635 | 0.87678 | 1.20272 | 0.75724 | 1.44640 | 0.28784 | 0.83651 | 0.96761 | 0.75921 | 0.48493 |
| **LNX1-AS2** | 0.01061 | 0.01591 | 0.02993 | 0.01199 | 0.01431 | 0.02281 | 0.01980 | 0.01233 | 0.01315 | 0.01386 | 0.01500 | 0.01686 |
| **LNX2** | 0.00101 | 1.37806 | 1.18804 | 1.01598 | 0.74274 | 2.18385 | 0.55916 | 2.09977 | 1.60473 | 3.08194 | 0.50620 | 1.20128 |
| **LOC100126784** | 0.07215 | 0.03044 | 0.01218 | 0.00946 | 0.16530 | 0.06670 | 0.00492 | 0.25029 | 0.00000 | 0.02695 | 0.00839 | 0.11952 |
| **LOC100128006** | 0.01600 | 0.01522 | 0.02648 | 0.01204 | 0.01614 | 0.01589 | 0.01508 | 0.05553 | 0.02907 | 0.01474 | 0.01272 | 0.01769 |
| **LOC100128076** | 0.00555 | 0.01275 | 0.02243 | 0.01164 | 0.03031 | 0.00460 | 0.03493 | 0.01045 | 0.00397 | 0.03211 | 0.01505 | 0.03694 |
| **LOC100128164** | 0.16059 | 0.15110 | 0.08276 | 0.17667 | 0.00000 | 0.02157 | 0.05769 | 0.27634 | 0.00000 | 0.02146 | 0.10682 | 0.60441 |
| **LOC100128176** | 0.00650 | 0.01920 | 0.01685 | 0.05644 | 0.02332 | 0.00036 | 0.02407 | 0.01056 | 0.02365 | 0.00397 | 0.02739 | 0.01854 |
| **LOC100128233** | 0.00476 | 0.01535 | 0.01953 | 0.01179 | 0.03888 | 0.02466 | 0.01733 | 0.01489 | 0.01643 | 0.01792 | 0.01967 | 0.01866 |
| **LOC100128239** | 0.29769 | 0.07454 | 0.00989 | 0.02177 | 0.00000 | 0.04389 | 0.01340 | 0.09046 | 0.01290 | 0.01632 | 0.01119 | 0.02672 |
| **LOC100128288** | 0.40815 | 0.20888 | 0.00000 | 1.24474 | 0.00000 | 0.00000 | 0.00600 | 0.21181 | 0.10995 | 0.74185 | 0.02027 | 0.61119 |
| **LOC100128361** | 0.47333 | 0.08138 | 0.26389 | 0.32795 | 0.21123 | 0.00000 | 0.51115 | 0.12921 | 0.00000 | 0.52448 | 0.00000 | 0.03706 |
| **LOC100128398** | 0.00582 | 0.08538 | 0.11651 | 0.00000 | 0.00000 | 0.00000 | 0.00414 | 0.09853 | 0.16595 | 0.75982 | 0.11723 | 0.21183 |
| **LOC100128494** | 0.00225 | 0.00000 | 0.03721 | 0.36792 | 0.20859 | 0.00000 | 0.02219 | 0.01750 | 0.04467 | 0.36852 | 0.01142 | 0.05913 |
| **LOC100128505** | 0.02140 | 0.03013 | 0.00456 | 0.00675 | 0.05006 | 0.04375 | 0.00000 | 0.01126 | 0.08652 | 0.03193 | 0.00918 | 0.05196 |
| **LOC100128531** | 0.00457 | 0.02356 | 0.02637 | 0.01434 | 0.02028 | 0.01894 | 0.02346 | 0.01332 | 0.00550 | 0.03730 | 0.01948 | 0.02450 |

| | | | | | | | | | | | | |
|---|---|---|---|---|---|---|---|---|---|---|---|---|
| **LOC100128554** | 0.10073 | 0.01329 | 0.01420 | 0.01376 | 0.01289 | 0.02021 | 0.01847 | 0.00684 | 0.01622 | 0.01160 | 0.02255 | 0.01776 |
| **LOC100128568** | 0.09686 | 0.01258 | 0.09590 | 0.07846 | 0.09269 | 0.03406 | 0.00001 | 0.03935 | 0.05888 | 0.07975 | 0.08773 | 0.16591 |
| **LOC100128573** | 0.00657 | 0.01446 | 0.04008 | 0.02039 | 0.02080 | 0.00209 | 0.01917 | 0.01137 | 0.01668 | 0.03746 | 0.01909 | 0.02691 |
| **LOC100128593** | 0.00635 | 0.01634 | 0.03128 | 0.01677 | 0.01764 | 0.01140 | 0.02259 | 0.01057 | 0.01623 | 0.00944 | 0.01644 | 0.01707 |
| **LOC100128682** | 0.00461 | 0.01849 | 0.00667 | 0.01468 | 0.02642 | 0.00020 | 0.02294 | 0.01068 | 0.02668 | 0.11762 | 0.00705 | 0.01384 |
| **LOC100128770** | 0.00401 | 0.01439 | 0.02074 | 0.00938 | 0.01156 | 0.00000 | 0.10134 | 0.01020 | 0.01224 | 0.21751 | 0.01778 | 0.01069 |
| **LOC100128885** | 0.00000 | 9.12392 | 4.00243 | 8.13317 | 11.04039 | 2.03129 | 10.20925 | 1.44717 | 8.13656 | 4.13604 | 8.68637 | 3.10698 |
| **LOC100129027** | 0.00603 | 0.01471 | 0.03006 | 0.01972 | 0.01481 | 0.01347 | 0.01809 | 0.01059 | 0.01359 | 0.02371 | 0.01993 | 0.01597 |
| **LOC100129034** | 0.00924 | 2.62677 | 2.68488 | 1.21180 | 2.72903 | 1.77537 | 3.01185 | 0.71925 | 2.31461 | 0.14262 | 2.09985 | 1.98243 |
| **LOC100129046** | 0.00470 | 0.01107 | 0.05781 | 0.01625 | 0.09411 | 0.00000 | 0.04891 | 0.01032 | 0.13308 | 0.08972 | 0.03284 | 0.01453 |
| **LOC100129083** | 0.03460 | 0.01462 | 0.01419 | 0.01220 | 0.01536 | 0.01716 | 0.01791 | 0.00808 | 0.01858 | 0.01452 | 0.01485 | 0.01724 |
| **LOC100129138** | 0.00678 | 0.11302 | 0.22030 | 0.09827 | 0.03854 | 0.09265 | 0.03554 | 0.01186 | 0.18333 | 0.00000 | 0.02693 | 0.13874 |
| **LOC100129148** | 0.02225 | 0.10200 | 0.08539 | 0.07589 | 0.15519 | 0.13571 | 0.04093 | 0.25611 | 0.13309 | 0.11414 | 0.03718 | 0.08599 |
| **LOC100129203** | 0.01304 | 0.00000 | 0.04190 | 0.45579 | 0.08112 | 0.10550 | 0.00000 | 0.54184 | 0.00000 | 0.18906 | 0.00000 | 0.17212 |
| **LOC100129216** | 0.00580 | 0.02575 | 0.01084 | 0.00556 | 0.01832 | 0.03577 | 0.00967 | 0.01600 | 0.03091 | 0.01973 | 0.01513 | 0.05199 |
| **LOC100129316** | 0.13064 | 0.00890 | 0.01807 | 0.01941 | 0.03776 | 0.05816 | 0.00000 | 0.00689 | 0.01850 | 0.03506 | 0.00813 | 0.01009 |
| **LOC100129361** | 0.00000 | 7.27964 | 10.02773 | 0.67900 | 1.19235 | 13.48759 | 7.57245 | 1.19158 | 8.18196 | 8.51769 | 6.90778 | 9.79660 |
| **LOC100129518** | 0.00275 | 0.00000 | 0.01724 | 0.08444 | 0.04676 | 0.11573 | 0.05391 | 0.02715 | 0.07033 | 0.00000 | 0.38482 | 0.01125 |
| **LOC100129520** | 0.22498 | 0.03059 | 0.09567 | 0.02234 | 0.26608 | 0.00000 | 0.11738 | 0.00000 | 0.12419 | 0.00000 | 0.73226 | 0.15638 |
| **LOC100129534** | 0.01168 | 0.27319 | 0.33219 | 0.55700 | 0.22727 | 0.00000 | 0.15218 | 0.05424 | 0.04842 | 0.27790 | 0.10546 | 0.24995 |
| **LOC100129550** | 0.13654 | 0.00914 | 1.03944 | 0.46455 | 0.00000 | 0.67183 | 0.24783 | 0.00000 | 2.72098 | 3.26864 | 0.44813 | 0.20832 |
| **LOC100129617** | 0.00517 | 0.00001 | 0.00988 | 0.00000 | 0.28965 | 0.01064 | 0.00386 | 0.00883 | 0.00000 | 0.59534 | 0.00000 | 0.00778 |
| **LOC100129620** | 0.02273 | 0.01318 | 0.01973 | 0.01287 | 0.01284 | 0.01453 | 0.02234 | 0.02706 | 0.00347 | 0.01278 | 0.04672 | 0.02659 |
| **LOC100129636** | 0.00692 | 0.00881 | 0.04135 | 0.02580 | 0.00364 | 0.04607 | 0.01100 | 0.01648 | 0.00514 | 0.03598 | 0.01623 | 0.01374 |
| **LOC100129722** | 0.00480 | 0.04040 | 0.04324 | 0.05286 | 0.00000 | 0.01341 | 0.01277 | 0.01042 | 0.00957 | 0.07677 | 0.01306 | 0.01619 |
| **LOC100129917** | 0.09420 | 0.10332 | 0.26367 | 0.20466 | 0.06442 | 0.26833 | 0.03326 | 0.14604 | 0.00000 | 0.11650 | 0.29169 | 0.54315 |
| **LOC100129924** | 0.00587 | 0.06541 | 0.09421 | 0.00261 | 0.00000 | 0.00941 | 0.70261 | 0.00952 | 0.63322 | 0.00000 | 0.00000 | 0.03151 |
| **LOC100129931** | 0.09491 | 0.00293 | 0.01403 | 0.00160 | 0.05325 | 0.00000 | 0.00252 | 0.00209 | 0.06873 | 0.26534 | 0.00000 | 0.05609 |
| **LOC100129935** | 0.01286 | 0.01478 | 7.73530 | 0.01144 | 0.01075 | 0.00000 | 0.04630 | 0.00854 | 0.00000 | 0.01864 | 0.01840 | 0.02775 |
| **LOC100129940.** | 0.00606 | 0.01372 | 0.08256 | 0.05827 | 0.00000 | 0.04130 | 0.00692 | 0.01046 | 0.00436 | 0.04320 | 0.01620 | 0.02073 |
| **LOC100129973** | 0.11009 | 0.00878 | 0.08537 | 0.07064 | 0.01307 | 0.00537 | 0.00580 | 0.03308 | 0.02006 | 0.01464 | 0.02011 | 0.12890 |
| **LOC100130075** | 0.13683 | 0.07333 | 0.57416 | 0.03597 | 0.14406 | 0.00000 | 0.06322 | 0.09559 | 0.00000 | 0.11196 | 0.04042 | 0.12589 |
| **LOC100130093** | 0.32589 | 0.87608 | 0.69650 | 0.65983 | 0.82865 | 0.00000 | 0.00208 | 0.60713 | 0.76308 | 0.00000 | 0.38599 | 0.81903 |
| **LOC100130238** | 0.19058 | 0.01446 | 0.00386 | 0.00868 | 0.00724 | 0.00588 | 0.24905 | 0.00336 | 0.00000 | 0.03663 | 0.00658 | 0.01260 |
| **LOC100130264** | 0.00573 | 0.02130 | 0.01439 | 0.01022 | 0.02341 | 0.02189 | 0.02292 | 0.01384 | 0.01634 | 0.00556 | 0.01339 | 0.03018 |
| **LOC100130298** | 0.37406 | 0.01439 | 0.11329 | 0.01088 | 0.01735 | 0.01604 | 0.00623 | 0.05607 | 0.00738 | 0.01032 | 0.01730 | 0.00964 |
| **LOC100130301** | 0.31879 | 0.01021 | 0.01440 | 0.01197 | 0.01564 | 0.01462 | 0.01788 | 0.06641 | 0.02143 | 0.00930 | 0.01394 | 0.05446 |
| **LOC100130331** | 2.33238 | 0.64261 | 0.50442 | 0.29911 | 0.07461 | 0.92129 | 0.00000 | 0.32362 | 0.20387 | 0.00000 | 0.06396 | 0.20725 |
| **LOC100130357** | 0.01867 | 0.00000 | 0.12963 | 0.08917 | 0.00000 | 0.60588 | 0.00000 | 0.03831 | 0.26845 | 0.40977 | 0.15801 | 0.48321 |
| **LOC100130370** | 0.00806 | 0.02374 | 0.00184 | 0.00000 | 0.01859 | 0.08475 | 0.00425 | 0.00884 | 0.00615 | 0.13896 | 0.00864 | 0.00621 |
| **LOC100130417** | 0.00568 | 0.01523 | 0.00990 | 0.01153 | 0.00786 | 0.01230 | 0.00863 | 0.00924 | 0.02444 | 0.07960 | 0.01271 | 0.02709 |
| **LOC100130451** | 0.00637 | 0.01514 | 0.02062 | 0.01269 | 0.01558 | 0.01956 | 0.01826 | 0.01045 | 0.01560 | 0.01234 | 0.01552 | 0.01636 |
| **LOC100130452** | 0.00407 | 0.00000 | 0.01571 | 0.01173 | 0.00000 | 0.14136 | 0.00000 | 0.02176 | 0.07659 | 0.05636 | 0.06324 | 0.03875 |
| **LOC100130476** | 0.00451 | 0.01893 | 0.41370 | 0.17045 | 0.00000 | 0.03707 | 0.03371 | 0.02142 | 0.10244 | 0.01401 | 0.00755 | 0.02520 |
| **LOC100130673** | 0.00205 | 0.00239 | 0.11273 | 0.00397 | 0.11399 | 0.15790 | 0.05555 | 0.03611 | 0.10257 | 0.00000 | 0.24625 | 0.41132 |
| **LOC100130691** | 0.00537 | 0.04268 | 0.05718 | 0.01649 | 0.11612 | 0.00000 | 0.00666 | 0.01702 | 0.04701 | 0.00000 | 0.03348 | 0.02668 |
| **LOC100130700** | 0.22193 | 0.12012 | 0.01227 | 0.09093 | 0.00687 | 0.00000 | 0.01783 | 0.12098 | 0.02743 | 0.00558 | 0.01362 | 0.01546 |
| **LOC100130705** | 0.01252 | 0.01847 | 0.11004 | 0.04506 | 0.03413 | 0.00000 | 0.00203 | 0.00932 | 0.05202 | 0.02106 | 0.05116 | 0.02571 |

| | | | | | | | | | | | | |
|---|---|---|---|---|---|---|---|---|---|---|---|---|
| **LOC100130744** | 0.07156 | 0.31350 | 0.44314 | 0.00000 | 0.11536 | 0.00000 | 0.90519 | 0.14599 | 0.00010 | 0.09603 | 0.68799 | 0.18153 |
| **LOC100130849** | 0.07572 | 0.01698 | 0.01465 | 0.01143 | 0.01482 | 0.01916 | 0.01788 | 0.01435 | 0.01837 | 0.01251 | 0.01447 | 0.01421 |
| **LOC100130872** | 0.00457 | 0.01648 | 0.00000 | 0.03525 | 0.00794 | 0.24010 | 0.00000 | 0.00994 | 0.00000 | 0.05406 | 0.01489 | 0.06059 |
| **LOC100130880** | 0.00617 | 0.03802 | 0.02631 | 0.01686 | 0.01490 | 0.00000 | 0.01442 | 0.01078 | 0.03694 | 0.02275 | 0.01587 | 0.01455 |
| **LOC100130899** | 0.00553 | 0.19481 | 0.45680 | 1.52698 | 0.49361 | 0.00000 | 1.11497 | 0.01105 | 1.28213 | 0.63820 | 0.06069 | 0.00660 |
| **LOC100130950** | 0.04346 | 0.14570 | 0.06003 | 0.12917 | 0.22458 | 0.00000 | 0.02769 | 0.10815 | 0.07380 | 0.00000 | 0.24574 | 0.55467 |
| **LOC100130954** | 0.00665 | 0.01850 | 0.07730 | 0.01769 | 0.02988 | 0.00000 | 0.01924 | 0.01064 | 0.01861 | 0.00000 | 0.01581 | 0.01854 |
| **LOC100130964** | 0.00315 | 0.01034 | 0.03594 | 0.03663 | 0.03861 | 0.00377 | 0.01956 | 0.02424 | 0.00809 | 0.03327 | 0.01571 | 0.04642 |
| **LOC100130987** | 0.11688 | 0.07990 | 0.31230 | 0.00000 | 0.21228 | 0.14293 | 0.12498 | 0.03516 | 0.32212 | 0.10353 | 0.07789 | 0.03947 |
| **LOC100130992** | 0.00725 | 0.01625 | 0.03889 | 0.01102 | 0.01567 | 0.01493 | 0.01677 | 0.01071 | 0.00831 | 0.02758 | 0.01632 | 0.01548 |
| **LOC100131257** | 0.07348 | 0.01184 | 0.01736 | 0.00000 | 0.01293 | 0.03483 | 0.03801 | 0.01448 | 0.00000 | 0.13181 | 0.02511 | 0.01856 |
| **LOC100131289** | 0.02858 | 0.01995 | 0.30651 | 0.06478 | 0.00000 | 0.30684 | 0.14785 | 0.00317 | 0.02543 | 0.00000 | 0.05564 | 0.11057 |
| **LOC100131303** | 0.00593 | 0.01390 | 0.81761 | 0.01112 | 0.07925 | 0.00000 | 0.00099 | 0.01016 | 0.03786 | 0.00779 | 0.01546 | 0.03325 |
| **LOC100131347** | 0.01199 | 0.12586 | 0.00000 | 0.01907 | 0.00282 | 0.10535 | 0.00000 | 0.00634 | 0.11565 | 0.07606 | 0.06940 | 0.05486 |
| **LOC100131496** | 0.09890 | 0.05840 | 0.23288 | 0.67892 | 0.04019 | 0.02828 | 0.09234 | 0.03597 | 0.05544 | 0.18674 | 0.08701 | 0.13816 |
| **LOC100131564** | 0.11880 | 1.35968 | 0.38182 | 0.01723 | 0.86769 | 0.75530 | 0.00000 | 0.22559 | 0.58521 | 0.00000 | 0.62449 | 1.36734 |
| **LOC100131626** | 0.00767 | 0.02525 | 0.02672 | 0.01323 | 0.01975 | 0.01908 | 0.00898 | 0.01637 | 0.04527 | 0.01680 | 0.01910 | 0.04646 |
| **LOC100131655** | 0.00152 | 0.13338 | 0.03092 | 0.02114 | 0.00000 | 0.00236 | 0.02687 | 0.07448 | 0.01958 | 0.10272 | 0.05050 | 0.02898 |
| **LOC100131691** | 0.09952 | 0.08693 | 0.01254 | 0.33557 | 0.29523 | 0.00000 | 0.07021 | 0.41834 | 0.34361 | 0.65811 | 0.07908 | 0.54448 |
| **LOC100132057.** | 0.00469 | 0.01539 | 0.01514 | 0.01209 | 0.01622 | 0.01780 | 0.01861 | 0.01949 | 0.01804 | 0.01255 | 0.01500 | 0.01682 |
| **LOC100132062-** | 0.00800 | 0.01883 | 0.01594 | 0.02018 | 0.01385 | 0.04251 | 0.00556 | 0.01855 | 0.03579 | 0.02519 | 0.01357 | 0.01383 |
| **LOC100132077** | 0.18185 | 0.03395 | 0.00000 | 0.09586 | 0.00000 | 0.13303 | 0.00000 | 0.00443 | 0.05055 | 0.10400 | 0.11410 | 0.05280 |
| **LOC100132111** | 0.01096 | 0.01835 | 0.02630 | 0.01383 | 0.00942 | 0.01256 | 0.00164 | 0.04490 | 0.06487 | 0.07120 | 0.02386 | 0.01782 |
| **LOC100132146** | 0.22928 | 0.03145 | 0.03225 | 0.04300 | 0.00976 | 0.00292 | 0.05699 | 0.00273 | 0.00000 | 0.08043 | 0.00946 | 0.00987 |
| **LOC100132215** | 0.00142 | 0.04710 | 0.03222 | 0.16247 | 0.32355 | 0.00000 | 0.00000 | 0.03602 | 0.15818 | 0.69526 | 0.22786 | 0.03209 |
| **LOC100132352** | 0.20567 | 0.19088 | 0.04099 | 0.06109 | 0.00000 | 0.86596 | 0.00006 | 0.29685 | 0.15629 | 0.39357 | 0.07493 | 0.18613 |
| **LOC100132356** | 0.20998 | 0.08236 | 0.15954 | 0.02032 | 0.00000 | 0.27270 | 0.00000 | 0.12449 | 0.08238 | 0.44217 | 0.26654 | 0.35159 |
| **LOC100132529** | 0.04536 | 0.09098 | 0.05987 | 0.03736 | 0.01641 | 0.01506 | 0.04647 | 0.01044 | 0.02630 | 0.00030 | 0.01969 | 0.07655 |
| **LOC100132735** | 0.00418 | 0.04179 | 1.19469 | 0.00000 | 0.56023 | 0.93142 | 0.40794 | 0.00431 | 1.11985 | 0.01398 | 0.45763 | 0.00000 |
| **LOC100132781** | 0.20743 | 0.00707 | 0.00585 | 0.10002 | 0.17941 | 0.00000 | 0.00531 | 0.00296 | 0.06999 | 0.26187 | 0.00080 | 0.01429 |
| **LOC100132831** | 0.01012 | 0.11257 | 0.32686 | 0.08888 | 0.13757 | 0.02728 | 0.06870 | 0.06157 | 0.36045 | 0.13519 | 0.05188 | 0.05805 |
| **LOC100132891** | 0.17577 | 0.01283 | 0.03771 | 0.00861 | 0.01125 | 0.01999 | 0.02846 | 0.00549 | 0.00945 | 0.03192 | 0.01561 | 0.01245 |
| **LOC100133077** | 0.00227 | 0.00000 | 0.01082 | 0.02146 | 0.06093 | 0.00000 | 0.03532 | 0.07542 | 0.02146 | 0.00304 | 0.01773 | 1.58573 |
| **LOC100133091** | 0.05837 | 0.02991 | 0.09622 | 0.04497 | 0.00000 | 0.03007 | 0.05299 | 0.05085 | 0.18057 | 0.20405 | 0.00765 | 0.10044 |
| **LOC100133286** | 0.34204 | 0.01147 | 0.05272 | 0.02296 | 0.01365 | 0.04001 | 0.01545 | 0.11743 | 0.05166 | 0.00010 | 0.02990 | 0.01718 |
| **LOC100133315** | 0.00256 | 0.46622 | 0.24942 | 0.22039 | 0.12295 | 0.46695 | 0.00000 | 0.64344 | 0.16663 | 0.00000 | 0.72560 | 1.07284 |
| **LOC100133331** | 0.01210 | 0.01383 | 0.00800 | 0.04301 | 0.03827 | 0.09815 | 0.00263 | 0.00637 | 0.04384 | 0.14537 | 0.01042 | 0.06760 |
| **LOC100133669** | 0.03341 | 0.03408 | 0.01958 | 0.01751 | 0.00980 | 0.01578 | 0.01810 | 0.01012 | 0.01146 | 0.01645 | 0.03414 | 0.01421 |
| **LOC100133985** | 0.00976 | 0.05823 | 0.11755 | 0.05961 | 0.02817 | 0.04110 | 0.04251 | 0.01019 | 0.10247 | 0.00000 | 0.03387 | 0.01854 |
| **LOC100134317** | 0.00302 | 0.00358 | 0.00585 | 0.12022 | 0.00295 | 0.00000 | 0.00152 | 0.01266 | 0.15643 | 0.28148 | 0.09744 | 0.03169 |
| **LOC100134368** | 0.00402 | 0.00750 | 0.00808 | 0.04618 | 0.00832 | 0.06748 | 0.00477 | 0.02173 | 0.02439 | 0.01625 | 0.01256 | 0.01304 |
| **LOC100134391** | 0.08663 | 0.01597 | 0.02901 | 0.01106 | 0.01673 | 0.01403 | 0.01770 | 0.02151 | 0.01586 | 0.01097 | 0.01551 | 0.01434 |
| **LOC100134868** | 0.00681 | 0.03958 | 0.18018 | 0.00633 | 0.05824 | 0.03485 | 0.11091 | 0.01184 | 0.22138 | 0.00000 | 0.17602 | 0.01686 |
| **LOC100144595** | 0.00755 | 0.01420 | 0.01890 | 0.01065 | 0.01268 | 0.01767 | 0.01883 | 0.01444 | 0.01509 | 0.03658 | 0.01610 | 0.01465 |
| **LOC100188947** | 0.12833 | 0.01094 | 0.01434 | 0.02804 | 0.01468 | 0.03242 | 0.01813 | 0.03136 | 0.01366 | 0.01683 | 0.01319 | 0.01705 |
| **LOC100190940** | 0.00622 | 0.03799 | 0.01816 | 0.01518 | 0.01371 | 0.00349 | 0.02098 | 0.01075 | 0.01809 | 0.01331 | 0.02979 | 0.01593 |
| **LOC100190986** | 0.00627 | 0.01481 | 0.01694 | 0.01221 | 0.01717 | 0.01750 | 0.01848 | 0.01045 | 0.01844 | 0.01376 | 0.01472 | 0.01733 |
| **LOC100240734** | 0.19211 | 0.01590 | 0.01726 | 0.01262 | 0.01816 | 0.01077 | 0.01858 | 0.00357 | 0.02197 | 0.00786 | 0.01416 | 0.02035 |

| | | | | | | | | | | | | |
|---|---|---|---|---|---|---|---|---|---|---|---|---|
| **LOC100240735** | 0.06506 | 0.00867 | 0.00346 | 0.12779 | 0.03385 | 0.02716 | 0.01460 | 0.00780 | 0.02393 | 0.01800 | 0.01195 | 0.01684 |
| **LOC100268168** | 0.17177 | 0.13826 | 0.10923 | 0.02038 | 0.00001 | 0.19691 | 0.00000 | 0.27900 | 0.02847 | 0.12473 | 0.01131 | 0.12860 |
| **LOC100270746** | 0.00222 | 0.01327 | 0.25290 | 0.03503 | 0.00000 | 0.31664 | 0.08144 | 0.02438 | 0.11981 | 0.00012 | 0.17863 | 0.14434 |
| **LOC100270804** | 0.20363 | 0.12231 | 0.07657 | 0.01351 | 0.00411 | 0.00000 | 0.08195 | 0.05259 | 0.00000 | 0.00000 | 0.28649 | 0.28864 |
| **LOC100271832** | 0.00523 | 0.01433 | 0.01031 | 0.01603 | 0.02850 | 0.00000 | 0.00404 | 0.00974 | 0.03294 | 0.16109 | 0.00517 | 0.01560 |
| **LOC100272216** | 0.01188 | 0.00553 | 0.05157 | 0.05287 | 0.02306 | 0.03027 | 0.00054 | 0.01194 | 0.06416 | 0.07998 | 0.01405 | 0.03109 |
| **LOC100272217** | 0.11535 | 0.00000 | 0.24326 | 0.00411 | 0.02386 | 0.19178 | 0.00000 | 0.06178 | 0.09177 | 0.00000 | 0.12275 | 0.71507 |
| **LOC100286922** | 0.02004 | 0.05296 | 0.03510 | 0.02127 | 0.01288 | 0.02726 | 0.03981 | 0.02407 | 0.06445 | 0.00000 | 0.12294 | 0.01508 |
| **LOC100287015** | 0.31008 | 2.14800 | 1.49269 | 0.78992 | 0.62458 | 1.33838 | 1.02737 | 1.26037 | 0.47057 | 0.22047 | 1.35197 | 4.95485 |
| **LOC100287036** | 0.02839 | 0.00000 | 0.00001 | 0.00000 | 0.05228 | 0.14795 | 0.00000 | 0.00453 | 0.01803 | 0.14631 | 0.00000 | 0.02367 |
| **LOC100287042** | 0.09600 | 0.12322 | 0.12093 | 0.00597 | 0.38193 | 0.00000 | 0.13574 | 0.00000 | 0.24115 | 0.09901 | 0.05825 | 0.19912 |
| **LOC100287072** | 0.00400 | 0.06706 | 0.00000 | 0.36660 | 0.12468 | 0.06645 | 0.19514 | 0.22329 | 0.03118 | 0.12791 | 0.03034 | 0.12696 |
| **LOC100287098** | 0.00685 | 0.04542 | 0.03621 | 0.02935 | 0.03701 | 0.00000 | 0.01013 | 0.01044 | 0.03151 | 0.03213 | 0.02437 | 0.04061 |
| **LOC100287225** | 1.35647 | 0.01196 | 0.00000 | 0.05678 | 0.06733 | 0.05790 | 0.71698 | 0.80197 | 0.00000 | 0.23629 | 0.03453 | 0.00000 |
| **LOC100287592** | 0.01636 | 0.01450 | 0.02482 | 0.01115 | 0.01538 | 0.02186 | 0.01722 | 0.00799 | 0.01752 | 0.00620 | 0.01541 | 0.03015 |
| **LOC100287632** | 0.00303 | 0.01099 | 0.02329 | 0.00824 | 0.00668 | 0.04737 | 0.03552 | 0.02563 | 0.01206 | 0.01807 | 0.01612 | 0.02674 |
| **LOC100287704** | 0.00588 | 0.01541 | 0.00890 | 0.05494 | 0.00229 | 0.03293 | 0.01267 | 0.01024 | 0.01578 | 0.02982 | 0.01519 | 0.01376 |
| **LOC100287792** | 0.03903 | 0.00000 | 0.00000 | 0.00000 | 0.00000 | 1.07171 | 0.21544 | 0.08009 | 0.00000 | 1.33834 | 0.03846 | 0.04009 |
| **LOC100287846** | 0.00558 | 0.17578 | 0.04089 | 0.05286 | 0.01480 | 0.03899 | 0.01316 | 0.01004 | 0.01205 | 0.03054 | 0.02506 | 0.02288 |
| **LOC100287944** | 0.00590 | 0.01879 | 0.03963 | 0.01960 | 0.02678 | 0.00000 | 0.01495 | 0.01194 | 0.03916 | 0.01015 | 0.01413 | 0.02174 |
| **LOC100288069** | 0.03884 | 0.01796 | 0.03648 | 0.01813 | 0.03339 | 0.02721 | 0.00582 | 0.03299 | 0.06212 | 0.11576 | 0.02451 | 0.04861 |
| **LOC100288123** | 0.00455 | 0.00509 | 0.11668 | 0.02709 | 0.06372 | 0.02780 | 0.00000 | 0.01009 | 0.03027 | 0.09664 | 0.04216 | 0.03377 |
| **LOC100288162.** | 0.00609 | 0.01571 | 0.01314 | 0.01179 | 0.01278 | 0.01860 | 0.01570 | 0.01026 | 0.03147 | 0.01833 | 0.01405 | 0.01950 |
| **LOC100288181** | 0.05840 | 0.00675 | 0.01059 | 0.00417 | 0.07205 | 0.01634 | 0.01854 | 0.14935 | 0.00216 | 0.07278 | 0.01156 | 0.01836 |
| **LOC100288637** | 0.07225 | 0.48259 | 1.06221 | 0.69442 | 0.51862 | 0.00030 | 0.89381 | 0.18095 | 2.77845 | 2.51297 | 0.60220 | 0.84394 |
| **LOC100288748** | 0.01090 | 0.19719 | 0.22875 | 0.00000 | 0.00000 | 0.33980 | 0.00000 | 0.58690 | 0.00000 | 0.47101 | 0.48179 | 0.48598 |
| **LOC100288778** | 0.02257 | 0.06165 | 0.01313 | 0.03131 | 0.09058 | 0.07833 | 0.23141 | 0.08883 | 0.05102 | 0.12978 | 0.02707 | 0.29222 |
| **LOC100288814** | 0.08717 | 0.02104 | 0.03544 | 0.00748 | 0.09910 | 0.00902 | 0.01861 | 0.00545 | 0.00796 | 0.00824 | 0.01428 | 0.01530 |
| **LOC100288842** | 0.14165 | 0.39992 | 0.22930 | 0.45147 | 0.05853 | 0.00031 | 0.42194 | 0.15785 | 0.00000 | 0.00000 | 0.43201 | 0.88146 |
| **LOC100288846** | 0.84427 | 0.09088 | 0.10827 | 0.00080 | 0.00735 | 0.19917 | 0.00000 | 0.11709 | 0.13326 | 0.12485 | 0.00004 | 0.01159 |
| **LOC100288866** | 0.00638 | 0.01291 | 0.02743 | 0.03718 | 0.02399 | 0.00850 | 0.01987 | 0.02331 | 0.00000 | 0.10312 | 0.01545 | 0.01226 |
| **LOC100288911** | 0.00130 | 0.01332 | 0.56159 | 0.18337 | 0.24959 | 0.00000 | 0.00000 | 0.09565 | 0.08898 | 0.26819 | 0.03103 | 0.01145 |
| **LOC100288974** | 0.01416 | 0.00650 | 0.00057 | 0.00000 | 0.00000 | 0.03211 | 0.02923 | 0.00494 | 0.04684 | 0.16686 | 0.01961 | 0.09884 |
| **LOC100289019** | 0.13171 | 0.37228 | 0.15558 | 0.48067 | 0.00000 | 0.40469 | 0.30748 | 0.04549 | 0.01781 | 3.26618 | 0.51341 | 0.45020 |
| **LOC100289092** | 0.03712 | 0.13724 | 0.04884 | 0.19039 | 0.10530 | 0.00000 | 0.14970 | 0.12151 | 0.14436 | 0.05846 | 0.13023 | 0.39830 |
| **LOC100289230** | 0.00758 | 0.64406 | 0.05217 | 0.00000 | 0.01412 | 0.27333 | 0.00000 | 0.26675 | 0.05938 | 0.09125 | 0.45894 | 0.59712 |
| **LOC100289333** | 0.12259 | 0.08585 | 0.02050 | 0.07996 | 0.16789 | 0.00000 | 0.05238 | 0.68520 | 0.23325 | 0.05109 | 0.00000 | 0.25886 |
| **LOC100289361** | 0.73403 | 0.33352 | 0.13557 | 0.64170 | 0.17952 | 0.35587 | 0.04800 | 0.91710 | 0.06349 | 0.00000 | 0.25299 | 0.63014 |
| **LOC100289473** | 0.00620 | 0.03182 | 0.01901 | 0.01027 | 0.02109 | 0.02305 | 0.01622 | 0.00989 | 0.01749 | 0.00830 | 0.01596 | 0.02473 |
| **LOC100289495** | 0.22653 | 0.05548 | 0.03141 | 0.01868 | 0.00000 | 0.00881 | 0.00446 | 0.10025 | 0.06890 | 0.05994 | 0.01597 | 0.02567 |
| **LOC100289511** | 0.14372 | 0.08619 | 0.15589 | 0.03289 | 0.06124 | 0.00000 | 0.00533 | 0.18040 | 0.10208 | 0.00000 | 0.06438 | 0.06800 |
| **LOC100289561** | 0.00979 | 0.03498 | 0.03811 | 0.00995 | 0.08432 | 0.00000 | 0.00869 | 0.00670 | 0.07775 | 0.27382 | 0.02575 | 0.05673 |
| **LOC100289580** | 0.22108 | 0.00976 | 0.05722 | 0.00294 | 0.02882 | 0.00000 | 0.14071 | 0.00000 | 0.10739 | 0.05903 | 0.10397 | 0.08223 |
| **LOC100289650** | 0.06308 | 0.01604 | 0.02310 | 0.01439 | 0.01571 | 0.01577 | 0.02118 | 0.01284 | 0.01892 | 0.01091 | 0.01446 | 0.02855 |
| **LOC100289656** | 0.00696 | 0.01789 | 0.01918 | 0.01660 | 0.03242 | 0.00041 | 0.01943 | 0.00873 | 0.02200 | 0.02133 | 0.01391 | 0.04812 |
| **LOC100289673** | 0.00251 | 0.01171 | 0.04819 | 0.01841 | 0.00589 | 0.02274 | 0.00795 | 0.12006 | 0.00894 | 0.06078 | 0.01385 | 0.00926 |
| **LOC100294145** | 2.61681 | 2.53037 | 2.57839 | 1.94806 | 0.00000 | 7.57816 | 0.00000 | 1.69793 | 0.00000 | 1.90785 | 2.18679 | 2.71239 |
| **LOC100294362** | 0.04751 | 0.11251 | 0.06059 | 0.16177 | 0.02476 | 0.00000 | 0.11948 | 0.48012 | 0.16565 | 0.00000 | 0.00213 | 0.15930 |

| | | | | | | | | | | | | |
|---|---|---|---|---|---|---|---|---|---|---|---|---|
| **LOC100303749** | 0.00479 | 0.03054 | 0.03993 | 0.05538 | 0.00000 | 0.09651 | 0.00000 | 0.00630 | 0.09714 | 0.54885 | 0.02028 | 0.21848 |
| **LOC100335030** | 0.35964 | 0.00000 | 0.37856 | 0.58404 | 0.00000 | 0.38044 | 0.07671 | 0.73285 | 0.00000 | 0.65150 | 0.26969 | 0.38482 |
| **LOC100379224** | 0.00754 | 0.35000 | 0.38524 | 0.03303 | 0.09749 | 0.01930 | 0.38865 | 0.14156 | 0.48524 | 0.00000 | 0.02380 | 0.87723 |
| **LOC100420587** | 0.13252 | 0.25330 | 0.15303 | 0.06481 | 0.33151 | 0.19431 | 0.00000 | 0.01364 | 0.03616 | 0.06438 | 0.13273 | 0.04206 |
| **LOC100422212** | 0.00614 | 0.01412 | 0.01451 | 0.01102 | 0.01247 | 0.02239 | 0.02112 | 0.01045 | 0.01847 | 0.01875 | 0.01506 | 0.01568 |
| **LOC100422556** | 0.05606 | 0.21619 | 0.01442 | 0.22780 | 0.05250 | 0.10680 | 0.05602 | 0.03560 | 0.04242 | 0.04984 | 0.01582 | 0.04935 |
| **LOC100499194** | 0.00712 | 0.01460 | 0.01497 | 0.01173 | 0.01366 | 0.01698 | 0.02117 | 0.01215 | 0.01624 | 0.02609 | 0.01408 | 0.01542 |
| **LOC100499484** | 0.00553 | 0.00707 | 0.01048 | 0.10021 | 0.00000 | 0.01877 | 0.00320 | 0.12213 | 0.01754 | 0.12052 | 0.01576 | 0.14144 |
| **LOC100499489** | 0.54508 | 0.02769 | 0.01426 | 0.01010 | 0.00731 | 0.01710 | 0.02096 | 0.00444 | 0.00741 | 0.01107 | 0.04825 | 0.00936 |
| **LOC100500773** | 0.02067 | 0.00000 | 0.13085 | 0.00477 | 0.00000 | 0.09752 | 0.00000 | 0.01231 | 0.04953 | 0.21638 | 0.01917 | 0.02105 |
| **LOC100505474** | 0.00569 | 0.01168 | 0.01384 | 0.01683 | 0.01032 | 0.02848 | 0.01218 | 0.00996 | 0.02004 | 0.02779 | 0.01456 | 0.02691 |
| **LOC100505549** | 0.00000 | 1.19988 | 1.59600 | 1.67287 | 0.21036 | 3.84854 | 2.94080 | 0.46095 | 1.75246 | 2.27119 | 0.64348 | 0.85763 |
| **LOC100505592** | 0.00115 | 0.14033 | 0.00789 | 0.00000 | 0.00000 | 0.05207 | 0.00945 | 0.14240 | 0.02528 | 0.14832 | 0.00000 | 0.23590 |
| **LOC100505622** | 0.00591 | 0.01339 | 0.02827 | 0.01193 | 0.07936 | 0.00000 | 0.02547 | 0.01068 | 0.00911 | 0.02932 | 0.01174 | 0.01674 |
| **LOC100505658** | 0.00229 | 0.00000 | 0.02727 | 0.04848 | 0.10883 | 0.38944 | 0.00000 | 0.05678 | 0.11314 | 0.45635 | 0.00000 | 0.09125 |
| **LOC100505666** | 0.43075 | 0.20751 | 0.14838 | 0.00012 | 0.00000 | 0.00000 | 0.22151 | 0.02909 | 0.00022 | 0.01622 | 0.39540 | 0.05364 |
| **LOC100505715** | 1.99518 | 1.22719 | 1.05503 | 0.13936 | 0.00000 | 0.35762 | 0.02738 | 0.15550 | 1.56212 | 1.38482 | 0.24326 | 0.09718 |
| **LOC100505716** | 0.01131 | 0.02193 | 0.16375 | 0.00000 | 0.00000 | 0.51093 | 0.00000 | 0.02306 | 0.00000 | 0.23377 | 0.05894 | 0.03206 |
| **LOC100505736** | 0.00816 | 0.00645 | 0.02150 | 0.01343 | 0.04750 | 0.00812 | 0.00619 | 0.06491 | 0.05907 | 0.07661 | 0.02713 | 0.13208 |
| **LOC100505782** | 0.00619 | 0.01439 | 0.02093 | 0.01180 | 0.01362 | 0.02110 | 0.01513 | 0.01043 | 0.01809 | 0.01887 | 0.01504 | 0.01574 |
| **LOC100505795** | 0.00564 | 0.01716 | 0.05221 | 0.03021 | 0.01780 | 0.00407 | 0.01020 | 0.01263 | 0.03338 | 0.04055 | 0.01190 | 0.03493 |
| **LOC100505811** | 0.00174 | 0.14982 | 0.00748 | 0.01052 | 0.00000 | 0.07141 | 0.01019 | 0.08843 | 0.00626 | 0.02592 | 0.00971 | 0.01893 |
| **LOC100505812** | 0.01258 | 0.18930 | 0.10580 | 0.00000 | 0.04186 | 0.00867 | 0.07035 | 0.00426 | 0.12658 | 0.04646 | 0.03144 | 0.16773 |
| **LOC100505817** | 1.86945 | 1.34677 | 0.00000 | 9.83703 | 0.79597 | 2.61772 | 0.00000 | 0.12744 | 0.00000 | 2.02236 | 2.93979 | 0.12625 |
| **LOC100505841** | 0.22949 | 0.01378 | 0.01403 | 0.01185 | 0.01263 | 0.01316 | 0.01642 | 0.01511 | 0.01708 | 0.01841 | 0.02684 | 0.01397 |
| **LOC100505878** | 0.74791 | 0.01579 | 0.01503 | 0.01332 | 0.02622 | 0.00133 | 0.02201 | 0.00119 | 0.02226 | 0.00407 | 0.01357 | 0.01353 |
| **LOC100505887** | 0.02400 | 0.03717 | 0.01612 | 0.01579 | 0.01902 | 0.00000 | 0.02251 | 0.02041 | 0.02105 | 0.00509 | 0.01723 | 0.06045 |
| **LOC100505912** | 0.00555 | 0.01294 | 0.08406 | 0.10108 | 0.00000 | 0.00882 | 0.08503 | 0.01019 | 0.04727 | 0.03839 | 0.00865 | 0.01745 |
| **LOC100505918** | 0.00790 | 0.00802 | 0.01172 | 0.00001 | 0.00000 | 0.17362 | 0.05326 | 0.01543 | 0.00000 | 0.17379 | 0.08057 | 0.04017 |
| **LOC100505921** | 0.01866 | 0.03394 | 0.01411 | 0.00757 | 0.01298 | 0.03081 | 0.01227 | 0.03400 | 0.02293 | 0.01656 | 0.01282 | 0.01283 |
| **LOC100505942** | 0.07737 | 0.01520 | 0.28707 | 0.00391 | 0.01001 | 0.02754 | 0.28798 | 0.00400 | 0.03022 | 0.00000 | 0.00000 | 0.02019 |
| **LOC100505978** | 0.01661 | 0.01088 | 0.05879 | 0.02735 | 0.07370 | 0.00000 | 0.01099 | 0.01439 | 0.01734 | 0.01834 | 0.02792 | 0.36368 |
| **LOC100505984** | 0.00529 | 0.03223 | 0.01268 | 0.05124 | 0.00000 | 0.01701 | 0.01429 | 0.00865 | 0.01016 | 0.07654 | 0.02875 | 0.03679 |
| **LOC100506022** | 0.00996 | 0.00816 | 0.03716 | 0.00935 | 0.00823 | 0.04642 | 0.01816 | 0.00878 | 0.00197 | 0.03998 | 0.01312 | 0.04188 |
| **LOC100506023** | 0.00421 | 0.01629 | 0.27253 | 0.27897 | 0.03530 | 0.06641 | 0.00000 | 0.30339 | 0.22717 | 0.07313 | 0.00769 | 0.12933 |
| **LOC100506071** | 0.06874 | 0.01156 | 0.11741 | 0.00600 | 0.01189 | 0.04330 | 0.04203 | 0.05719 | 0.81021 | 0.01889 | 0.16515 | 0.01343 |
| **LOC100506082** | 0.30000 | 0.50415 | 0.11292 | 0.02471 | 0.09428 | 0.09411 | 0.04511 | 0.35707 | 0.00000 | 0.00001 | 0.01095 | 0.20810 |
| **LOC100506083** | 0.00653 | 0.54967 | 0.29428 | 0.00000 | 0.44048 | 0.16064 | 0.00000 | 0.05034 | 0.14716 | 0.00377 | 0.10133 | 1.00423 |
| **LOC100506085** | 0.01162 | 0.01739 | 0.05631 | 0.07043 | 0.12918 | 0.00000 | 0.00977 | 0.03640 | 0.00000 | 0.06846 | 0.03207 | 0.02305 |
| **LOC100506100** | 0.50110 | 0.50129 | 2.84614 | 2.57363 | 0.70949 | 2.93241 | 1.04893 | 0.42799 | 2.08018 | 1.48591 | 3.27854 | 0.84965 |
| **LOC100506119** | 0.00606 | 0.01612 | 0.05679 | 0.03529 | 0.01164 | 0.00409 | 0.04012 | 0.01054 | 0.01551 | 0.02338 | 0.02049 | 0.01679 |
| **LOC100506123** | 0.00485 | 0.02734 | 0.01094 | 0.02421 | 0.01860 | 0.00550 | 0.03611 | 0.02527 | 0.04187 | 0.10432 | 0.00972 | 0.04005 |
| **LOC100506124** | 0.01495 | 0.00083 | 0.07813 | 0.02316 | 0.01117 | 0.11026 | 0.00000 | 0.03902 | 0.00000 | 0.06299 | 0.08744 | 0.11882 |
| **LOC100506125** | 0.11073 | 0.01533 | 0.02020 | 0.01233 | 0.01876 | 0.01503 | 0.01633 | 0.07275 | 0.01852 | 0.01240 | 0.01520 | 0.01437 |
| **LOC100506127** | 0.05482 | 0.23097 | 0.16236 | 0.25845 | 0.05112 | 0.07876 | 0.03771 | 0.08237 | 0.05443 | 0.00000 | 0.33172 | 0.31947 |
| **LOC100506136** | 0.18131 | 0.16300 | 0.05024 | 0.00073 | 0.00000 | 0.13624 | 0.17390 | 0.67108 | 0.01045 | 0.01548 | 0.07722 | 0.06597 |
| **LOC100506142** | 0.00195 | 0.03329 | 0.02169 | 0.09101 | 0.00000 | 0.03067 | 0.01023 | 0.04820 | 0.03266 | 0.07365 | 0.01958 | 0.11325 |
| **LOC100506159** | 0.03042 | 0.01590 | 0.01440 | 0.01250 | 0.01631 | 0.01504 | 0.02211 | 0.00936 | 0.02366 | 0.01284 | 0.01382 | 0.01720 |

| | | | | | | | | | | | | |
|---|---|---|---|---|---|---|---|---|---|---|---|---|
| **LOC100506172** | 0.00745 | 0.01481 | 0.02201 | 0.01096 | 0.01438 | 0.02078 | 0.03173 | 0.07420 | 0.00868 | 0.01259 | 0.01546 | 0.01297 |
| **LOC100506175** | 0.00388 | 0.01949 | 0.01562 | 0.01400 | 0.01478 | 0.00794 | 0.02114 | 0.06333 | 0.03000 | 0.01878 | 0.01251 | 0.03669 |
| **LOC100506178** | 0.06144 | 0.30583 | 2.56897 | 0.00000 | 0.00000 | 0.25471 | 0.71688 | 0.00000 | 1.58206 | 0.68578 | 4.34242 | 0.02021 |
| **LOC100506207** | 0.39062 | 0.07115 | 0.07423 | 0.01588 | 0.03668 | 0.04876 | 0.00000 | 0.00000 | 0.06168 | 0.00338 | 0.05434 | 0.21279 |
| **LOC100506258** | 0.14567 | 0.00948 | 0.00989 | 0.01127 | 0.00213 | 0.01700 | 0.00093 | 0.00234 | 0.02411 | 0.21233 | 0.00707 | 0.02851 |
| **LOC100506272** | 0.00999 | 0.42545 | 0.00000 | 0.09122 | 0.17845 | 0.12045 | 0.00070 | 0.00619 | 0.00000 | 0.06928 | 0.00000 | 0.07158 |
| **LOC100506281** | 0.03517 | 0.01253 | 0.04016 | 0.06217 | 0.04211 | 0.01809 | 0.06486 | 0.00653 | 0.00000 | 0.05475 | 0.01792 | 0.01660 |
| **LOC100506299** | 0.00591 | 0.07302 | 0.05202 | 0.17250 | 0.00375 | 0.00016 | 0.04755 | 0.01079 | 0.00208 | 0.01472 | 0.01824 | 0.01234 |
| **LOC100506314** | 0.28082 | 0.84689 | 0.43046 | 0.60609 | 0.73528 | 0.25397 | 1.28213 | 0.10329 | 1.18714 | 0.00000 | 2.16663 | 1.66695 |
| **LOC100506321** | 0.08040 | 0.02019 | 0.06325 | 0.06610 | 0.57212 | 0.00000 | 0.14311 | 0.14719 | 0.07758 | 0.80114 | 0.10409 | 0.32440 |
| **LOC100506325** | 0.07033 | 0.06379 | 0.13885 | 0.00026 | 0.13740 | 0.30460 | 0.00000 | 0.00023 | 0.00000 | 0.00000 | 0.07342 | 0.10662 |
| **LOC100506368** | 0.00129 | 0.05262 | 0.02550 | 0.00000 | 0.06876 | 0.12911 | 0.00000 | 0.13668 | 0.00000 | 0.02314 | 0.05562 | 0.03599 |
| **LOC100506393** | 0.00774 | 0.01532 | 0.01916 | 0.01225 | 0.01484 | 0.01913 | 0.01470 | 0.01325 | 0.02924 | 0.01045 | 0.01409 | 0.01609 |
| **LOC100506403** | 0.00846 | 0.03415 | 0.01077 | 0.00589 | 0.01249 | 0.04432 | 0.02140 | 0.02761 | 0.01330 | 0.01251 | 0.01897 | 0.01749 |
| **LOC100506422** | 0.00613 | 0.01474 | 0.01283 | 0.01788 | 0.01266 | 0.02283 | 0.01712 | 0.02355 | 0.01693 | 0.01572 | 0.01465 | 0.01795 |
| **LOC100506457** | 0.22942 | 0.00894 | 0.02021 | 0.01131 | 0.05234 | 0.00000 | 0.00941 | 0.12988 | 0.01084 | 0.05626 | 0.03367 | 0.02955 |
| **LOC100506469** | 0.00249 | 0.22079 | 0.48330 | 0.14124 | 0.13176 | 0.61076 | 0.93225 | 0.01272 | 0.48546 | 1.04156 | 0.69787 | 0.42221 |
| **LOC100506472** | 0.11043 | 0.04243 | 0.25962 | 0.19608 | 0.15234 | 0.00006 | 0.00000 | 0.00077 | 0.21018 | 0.62648 | 0.10369 | 0.05621 |
| **LOC100506474** | 0.21058 | 0.03501 | 0.01142 | 0.00599 | 0.02893 | 0.02241 | 0.04927 | 0.00332 | 0.00898 | 0.01072 | 0.00994 | 0.01244 |
| **LOC100506476** | 0.01016 | 0.02701 | 0.05098 | 0.07106 | 0.08584 | 0.00000 | 0.02645 | 0.00943 | 0.01151 | 0.07165 | 0.01786 | 0.05533 |
| **LOC100506497** | 0.00418 | 0.00817 | 0.02308 | 0.00215 | 0.26147 | 0.00000 | 0.01275 | 0.00951 | 0.00000 | 0.16598 | 0.06805 | 0.02058 |
| **LOC100506499** | 0.00423 | 0.03094 | 0.51507 | 0.00076 | 0.10298 | 0.01884 | 0.15770 | 0.07726 | 0.15353 | 0.00157 | 0.17426 | 0.09793 |
| **LOC100506548** | 0.44939 | 2.07154 | 3.66901 | 1.42252 | 1.36028 | 3.19248 | 5.79124 | 7.10896 | 7.12541 | 10.70825 | 0.00000 | 5.43584 |
| **LOC100506551** | 0.00625 | 0.14652 | 0.10853 | 0.00638 | 0.01404 | 0.00226 | 0.00369 | 0.01366 | 0.14549 | 0.00257 | 0.07060 | 0.06588 |
| **LOC100506603** | 0.02476 | 0.12790 | 0.05971 | 0.45127 | 0.31094 | 0.42849 | 0.00811 | 0.02002 | 0.08185 | 1.10255 | 0.05107 | 0.32550 |
| **LOC100506606** | 0.00717 | 0.02264 | 0.01035 | 0.09364 | 0.06057 | 0.00000 | 0.00116 | 0.14122 | 0.06078 | 0.50251 | 0.00000 | 0.25842 |
| **LOC100506639** | 0.00229 | 0.00000 | 0.14556 | 0.06434 | 0.06819 | 0.29527 | 0.00000 | 0.19949 | 0.39321 | 0.01367 | 0.23193 | 0.18750 |
| **LOC100506655** | 0.00292 | 0.00000 | 21.83704 | 0.00000 | 2.10238 | 0.02252 | 2.74955 | 0.00207 | 5.32800 | 1.89592 | 1.87701 | 0.03231 |
| **LOC100506674** | 0.00274 | 0.00000 | 0.11723 | 0.01765 | 0.00000 | 0.12936 | 0.38255 | 0.21953 | 0.00008 | 0.03696 | 0.04317 | 0.29803 |
| **LOC100506675** | 0.03394 | 0.00014 | 0.00000 | 0.00000 | 0.00000 | 1.84421 | 0.96042 | 0.10371 | 0.00000 | 2.44376 | 0.00000 | 0.17139 |
| **LOC100506682** | 0.08233 | 0.01674 | 0.01928 | 0.01129 | 0.01647 | 0.02379 | 0.01732 | 0.02026 | 0.02552 | 0.00440 | 0.01544 | 0.01518 |
| **LOC100506688** | 0.06339 | 0.41787 | 0.00000 | 0.00000 | 0.41297 | 0.54419 | 0.19035 | 0.00083 | 0.45647 | 0.01093 | 1.00057 | 0.00000 |
| **LOC100506700** | 0.00789 | 0.01952 | 0.04933 | 0.01592 | 0.00000 | 0.15842 | 0.08252 | 0.00697 | 0.21334 | 0.01172 | 0.11412 | 0.04946 |
| **LOC100506713** | 0.00427 | 0.17861 | 0.30570 | 0.06546 | 0.12130 | 0.00000 | 0.01886 | 0.03222 | 0.43320 | 0.17166 | 0.41691 | 0.40711 |
| **LOC100506725** | 0.14665 | 0.01766 | 0.02836 | 0.01181 | 0.02130 | 0.00457 | 0.03131 | 0.00707 | 0.03921 | 0.00973 | 0.01042 | 0.01762 |
| **LOC100506730** | 0.00331 | 0.00000 | 0.51460 | 0.13801 | 0.12706 | 0.13317 | 0.33068 | 0.00444 | 0.47335 | 0.32250 | 0.45551 | 0.20125 |
| **LOC100506746** | 0.19192 | 0.33815 | 0.16353 | 0.28152 | 0.19907 | 0.46254 | 0.00000 | 0.14198 | 0.00000 | 0.02233 | 0.11775 | 0.85647 |
| **LOC100506747** | 0.00235 | 0.03080 | 0.20828 | 0.01161 | 0.16053 | 0.00000 | 0.02894 | 0.13686 | 0.04206 | 0.00000 | 0.05968 | 0.08432 |
| **LOC100506787** | 0.03630 | 0.01266 | 0.02427 | 0.01091 | 0.01422 | 0.02321 | 0.01619 | 0.01560 | 0.02137 | 0.00991 | 0.01450 | 0.03444 |
| **LOC100506790** | 5.02928 | 0.37061 | 0.39303 | 1.58702 | 0.00711 | 0.00000 | 0.00000 | 7.13530 | 0.00000 | 0.03170 | 0.00376 | 4.45795 |
| **LOC100506801** | 0.00256 | 0.09820 | 0.04910 | 0.00000 | 0.10452 | 0.00000 | 0.03862 | 0.13202 | 0.00000 | 0.92981 | 0.00000 | 0.00865 |
| **LOC100506804** | 0.00514 | 0.00000 | 0.10638 | 0.02286 | 0.00000 | 0.11630 | 0.02566 | 0.29795 | 0.10319 | 0.09245 | 0.33240 | 0.23767 |
| **LOC100506834** | 0.02164 | 0.01233 | 0.01668 | 0.01014 | 0.01181 | 0.02987 | 0.02313 | 0.00830 | 0.01126 | 0.01122 | 0.03436 | 0.01602 |
| **LOC100506844** | 3.02716 | 0.13553 | 0.05826 | 0.01508 | 0.00029 | 0.03404 | 0.00215 | 1.25724 | 0.01339 | 0.02790 | 0.02044 | 0.92417 |
| **LOC100506860** | 0.00477 | 0.02558 | 0.03449 | 0.00000 | 0.16758 | 0.00000 | 0.00049 | 0.01033 | 0.11350 | 0.11401 | 0.00295 | 0.01209 |
| **LOC100506895** | 0.42224 | 0.01211 | 0.02559 | 0.00690 | 0.02996 | 0.01149 | 0.02148 | 0.10127 | 0.00722 | 0.03389 | 0.01269 | 0.04197 |
| **LOC100506985** | 0.21044 | 0.06448 | 0.11405 | 0.09043 | 0.00000 | 0.26837 | 0.12308 | 0.03656 | 0.08355 | 0.04409 | 0.26714 | 0.09488 |
| **LOC100506990** | 0.36008 | 0.29431 | 0.43190 | 0.75153 | 0.00000 | 0.56715 | 0.40739 | 1.00818 | 0.10678 | 0.00000 | 0.00000 | 1.77596 |

| | | | | | | | | | | | | |
|---|---|---|---|---|---|---|---|---|---|---|---|---|
| **LOC100507002** | 0.03392 | 1.06058 | 0.55991 | 4.74215 | 0.40893 | 0.21080 | 0.08021 | 0.00389 | 0.00000 | 0.00000 | 0.00000 | 0.15321 |
| **LOC100507006** | 0.00366 | 0.01220 | 0.01714 | 0.10749 | 0.00598 | 0.04592 | 0.02499 | 0.03086 | 0.03619 | 0.01075 | 0.00734 | 0.01754 |
| **LOC100507012** | 0.21270 | 0.02142 | 0.01943 | 0.01342 | 0.01740 | 0.00110 | 0.01898 | 0.00799 | 0.01876 | 0.00880 | 0.03085 | 0.02634 |
| **LOC100507053** | 0.79155 | 0.03658 | 0.23148 | 0.00220 | 0.04226 | 0.00000 | 0.03918 | 3.13189 | 0.09349 | 0.14246 | 0.00706 | 0.36974 |
| **LOC100507065** | 0.00620 | 0.01339 | 0.02238 | 0.00936 | 0.01275 | 0.03020 | 0.03215 | 0.01048 | 0.00973 | 0.00958 | 0.01422 | 0.01659 |
| **LOC100507073** | 0.24828 | 0.00808 | 0.00951 | 0.02437 | 0.00594 | 0.04717 | 0.01270 | 0.06895 | 0.01525 | 0.01098 | 0.01208 | 0.02316 |
| **LOC100507091** | 0.00441 | 0.03918 | 0.06711 | 0.06933 | 0.00000 | 0.00591 | 0.08201 | 0.01060 | 0.05049 | 0.13724 | 0.00000 | 0.00892 |
| **LOC100507140** | 0.01436 | 0.01598 | 0.04373 | 0.01829 | 0.01173 | 0.00852 | 0.08357 | 0.01143 | 0.00000 | 0.02056 | 0.01917 | 0.01584 |
| **LOC100507156** | 0.00572 | 0.01163 | 0.03431 | 0.01033 | 0.01211 | 0.02145 | 0.00492 | 0.00892 | 0.07080 | 0.03888 | 0.01329 | 0.03780 |
| **LOC100507175** | 0.55974 | 0.01046 | 0.01929 | 0.01136 | 0.01595 | 0.01910 | 0.01656 | 0.00262 | 0.02397 | 0.00273 | 0.01304 | 0.04092 |
| **LOC100507194** | 0.00580 | 0.01333 | 0.06345 | 0.18351 | 0.01386 | 0.01446 | 0.00393 | 0.01026 | 0.01102 | 0.02867 | 0.02324 | 0.01531 |
| **LOC100507195** | 0.91623 | 0.04592 | 0.06257 | 0.10632 | 0.00000 | 0.05864 | 0.00000 | 0.55647 | 0.00742 | 0.03549 | 0.01478 | 0.13722 |
| **LOC100507205** | 0.02217 | 0.01376 | 0.00786 | 0.01357 | 0.00700 | 0.01653 | 0.00836 | 0.01340 | 0.02739 | 0.07091 | 0.01233 | 0.01273 |
| **LOC100507217** | 0.66414 | 0.00000 | 2.98010 | 1.37779 | 1.53433 | 5.98964 | 5.99458 | 0.37057 | 0.00000 | 14.75456 | 0.99640 | 0.09965 |
| **LOC100507250** | 0.08210 | 0.38474 | 0.11212 | 0.00000 | 0.00000 | 0.27100 | 0.03453 | 0.03154 | 0.07740 | 0.04002 | 0.00332 | 0.97933 |
| **LOC100507283** | 0.10380 | 0.01764 | 0.04354 | 0.05389 | 0.00960 | 0.00000 | 0.02216 | 0.00768 | 0.06892 | 0.00659 | 0.09376 | 0.07127 |
| **LOC100507291** | 0.48385 | 0.43455 | 0.18980 | 0.00552 | 0.00000 | 0.65347 | 0.00000 | 0.03586 | 0.15505 | 0.02974 | 0.19370 | 0.47942 |
| **LOC100507316** | 0.11124 | 0.14340 | 0.01692 | 0.00379 | 0.00000 | 0.00000 | 0.02789 | 0.46240 | 0.02721 | 0.01502 | 0.08300 | 1.75568 |
| **LOC100507334** | 0.00394 | 0.01483 | 0.01425 | 0.01219 | 0.01682 | 0.01846 | 0.01860 | 0.02168 | 0.01822 | 0.01197 | 0.01504 | 0.01949 |
| **LOC100507346** | 0.00399 | 0.00637 | 0.00648 | 0.00000 | 0.01320 | 0.02987 | 0.00000 | 0.00913 | 0.04795 | 0.17711 | 0.00483 | 0.02280 |
| **LOC100507351** | 0.00509 | 0.05749 | 0.04651 | 0.08696 | 0.00002 | 0.04605 | 0.00253 | 0.00715 | 0.00000 | 0.05948 | 0.01360 | 0.07737 |
| **LOC100507373** | 0.05680 | 0.15176 | 0.16894 | 0.02368 | 0.21044 | 0.00016 | 0.07288 | 0.15041 | 0.14113 | 0.58716 | 0.08695 | 0.28756 |
| **LOC100507377** | 0.28941 | 0.31948 | 0.32471 | 0.20605 | 0.25399 | 0.66474 | 0.00000 | 0.61530 | 0.00273 | 0.00000 | 0.00378 | 0.34804 |
| **LOC100507387** | 0.00465 | 0.01431 | 0.01452 | 0.01247 | 0.01594 | 0.01726 | 0.01925 | 0.01926 | 0.01788 | 0.01232 | 0.01522 | 0.02136 |
| **LOC100507388** | 0.01089 | 0.01363 | 0.04246 | 0.01162 | 0.01280 | 0.02263 | 0.01572 | 0.02464 | 0.00742 | 0.02792 | 0.01542 | 0.01814 |
| **LOC100507391** | 0.03434 | 0.01428 | 0.03045 | 0.01091 | 0.01687 | 0.01667 | 0.01668 | 0.06424 | 0.01371 | 0.01182 | 0.01623 | 0.01768 |
| **LOC100507420.** | 0.00546 | 0.00889 | 0.24930 | 0.23675 | 0.05456 | 0.00000 | 0.00000 | 0.03040 | 0.00000 | 0.19315 | 0.00872 | 0.00899 |
| **LOC100507420.** | 0.00336 | 0.00000 | 0.04927 | 1.10351 | 1.58678 | 0.09117 | 0.00000 | 0.00666 | 0.00000 | 0.14264 | 0.02671 | 0.06607 |
| **LOC100507424** | 0.00555 | 0.00000 | 0.16609 | 0.00508 | 0.05183 | 0.21932 | 0.00000 | 0.00852 | 0.27008 | 1.10314 | 0.05658 | 0.19842 |
| **LOC100507443** | 0.00364 | 0.03935 | 0.01505 | 0.01396 | 0.01664 | 0.00730 | 0.03401 | 0.01938 | 0.00740 | 0.02285 | 0.02311 | 0.03901 |
| **LOC100507459** | 0.02221 | 0.13360 | 0.00000 | 0.01825 | 0.71395 | 0.72348 | 0.00000 | 0.07383 | 0.24701 | 0.00000 | 0.31739 | 0.32169 |
| **LOC100507462** | 0.20261 | 0.14832 | 0.31681 | 0.14574 | 0.08853 | 0.11777 | 0.15440 | 0.02377 | 0.23032 | 0.15156 | 0.35468 | 0.10469 |
| **LOC100507472** | 0.00660 | 0.02596 | 0.01122 | 0.01168 | 0.01754 | 0.00508 | 0.01584 | 0.01061 | 0.06679 | 0.00543 | 0.02464 | 0.01677 |
| **LOC100507477** | 0.00653 | 0.01750 | 0.04087 | 0.02713 | 0.01733 | 0.00856 | 0.02119 | 0.01059 | 0.04589 | 0.00000 | 0.01814 | 0.02055 |
| **LOC100507501** | 0.28473 | 0.34311 | 0.11888 | 0.37938 | 0.02096 | 0.00000 | 0.00453 | 0.07424 | 0.00000 | 0.10302 | 0.11896 | 0.10501 |
| **LOC100507506** | 0.29222 | 0.01207 | 0.01439 | 0.01205 | 0.00133 | 0.00000 | 0.02368 | 0.00705 | 0.00000 | 0.05066 | 0.12747 | 0.01230 |
| **LOC100507537** | 0.00554 | 0.01391 | 0.01417 | 0.01201 | 0.03522 | 0.01264 | 0.02553 | 0.01047 | 0.01887 | 0.04667 | 0.01340 | 0.01543 |
| **LOC100507540** | 0.35241 | 0.02361 | 0.06617 | 0.00000 | 0.00000 | 0.10515 | 0.04112 | 0.02025 | 0.02302 | 0.02217 | 0.05714 | 0.19771 |
| **LOC100507547** | 0.09831 | 0.23977 | 0.07274 | 0.07672 | 0.24305 | 0.00000 | 0.01979 | 0.06775 | 0.25978 | 0.00000 | 0.06103 | 0.37408 |
| **LOC100507557** | 0.07805 | 0.04399 | 0.14859 | 0.00000 | 0.00000 | 0.16155 | 0.06096 | 0.00701 | 0.19639 | 0.14712 | 0.00000 | 0.17962 |
| **LOC100507564** | 1.00051 | 0.17537 | 0.04081 | 0.00903 | 0.15589 | 0.00000 | 0.12564 | 0.14529 | 0.00000 | 0.00000 | 0.09620 | 0.19066 |
| **LOC100507577** | 0.33784 | 0.03059 | 0.11958 | 0.14626 | 0.00000 | 0.00000 | 0.16080 | 0.43741 | 0.09020 | 0.71759 | 0.39089 | 0.08039 |
| **LOC100507634** | 0.25922 | 0.00000 | 0.20702 | 0.00000 | 0.23691 | 0.06433 | 0.00000 | 0.00000 | 0.19669 | 0.21741 | 0.15312 | 0.24309 |
| **LOC100507639** | 0.25504 | 0.00176 | 0.33909 | 0.00156 | 0.00229 | 0.19463 | 0.23843 | 0.28490 | 0.00000 | 0.00000 | 0.00533 | 0.01712 |
| **LOC100507651** | 0.10228 | 0.03015 | 0.05369 | 0.06744 | 0.01367 | 0.16597 | 0.00000 | 0.00107 | 0.00000 | 0.22164 | 0.00000 | 0.07742 |
| **LOC100507661** | 0.00601 | 0.01211 | 0.01091 | 0.04089 | 0.00815 | 0.02587 | 0.01493 | 0.01010 | 0.02471 | 0.01704 | 0.01326 | 0.02258 |
| **LOC100508046** | 0.00566 | 0.01496 | 0.01437 | 0.01183 | 0.02068 | 0.01455 | 0.01534 | 0.01269 | 0.02311 | 0.01524 | 0.01636 | 0.01648 |
| **LOC100630918** | 0.00369 | 0.37203 | 0.28455 | 0.16114 | 0.00000 | 0.05335 | 0.00000 | 0.67768 | 0.25107 | 0.31332 | 0.08712 | 0.31622 |

| | | | | | | | | | | | | |
|---|---|---|---|---|---|---|---|---|---|---|---|---|
| **LOC100631378** | 0.20640 | 0.14774 | 0.11128 | 0.15465 | 0.00158 | 0.19738 | 0.00000 | 0.00000 | 0.06319 | 0.00000 | 0.01658 | 0.23662 |
| **LOC100652758** | 2.80862 | 0.50366 | 0.14152 | 0.55219 | 0.12032 | 0.00000 | 0.00384 | 0.04831 | 0.00000 | 0.02172 | 0.18991 | 0.18592 |
| **LOC100652768** | 0.04578 | 0.07139 | 0.08111 | 0.00002 | 0.02312 | 0.05216 | 0.00000 | 0.02579 | 0.13156 | 0.18835 | 0.03088 | 0.08941 |
| **LOC100652770** | 4.76951 | 0.03576 | 0.00000 | 0.14328 | 0.07621 | 0.02448 | 0.08648 | 0.58572 | 0.12573 | 0.29044 | 0.22102 | 0.16425 |
| **LOC100652824** | 0.99261 | 0.00958 | 0.00000 | 0.01012 | 0.00061 | 0.20453 | 0.00000 | 0.25319 | 0.02205 | 0.06556 | 0.00000 | 0.00000 |
| **LOC100652931** | 0.00409 | 0.01327 | 0.01215 | 0.01055 | 0.01163 | 0.01823 | 0.01946 | 0.03872 | 0.01742 | 0.03237 | 0.01546 | 0.02197 |
| **LOC100652999** | 0.00609 | 0.01570 | 0.01437 | 0.01064 | 0.01710 | 0.01761 | 0.02940 | 0.01052 | 0.01643 | 0.01868 | 0.01876 | 0.01588 |
| **LOC100653515** | 0.05535 | 0.00747 | 0.00425 | 0.00694 | 0.01915 | 0.01400 | 0.04583 | 0.04903 | 0.00000 | 0.13541 | 0.03272 | 0.02552 |
| **LOC100862671** | 0.01778 | 0.00918 | 7.72408 | 0.63995 | 0.27353 | 0.77576 | 3.22058 | 0.27274 | 1.50326 | 5.37543 | 2.15317 | 0.81506 |
| **LOC100996249** | 0.09898 | 1.23215 | 0.01561 | 0.09758 | 0.00000 | 0.03557 | 0.00000 | 0.00546 | 0.00000 | 0.09685 | 0.00000 | 2.20652 |
| **LOC100996251** | 0.03787 | 0.02702 | 0.04634 | 0.00000 | 0.00000 | 0.04613 | 0.01715 | 0.11093 | 0.13970 | 0.39721 | 0.00000 | 0.08981 |
| **LOC100996255** | 0.03453 | 0.00780 | 0.00948 | 0.00577 | 0.02064 | 0.02951 | 0.00786 | 0.02232 | 0.01266 | 0.10297 | 0.01366 | 0.01678 |
| **LOC100996291** | 0.07170 | 0.01445 | 0.01619 | 0.09857 | 0.00288 | 0.02728 | 0.01486 | 0.00560 | 0.01855 | 0.02091 | 0.01338 | 0.01419 |
| **LOC100996324** | 4.68882 | 0.01755 | 0.35789 | 0.16479 | 0.08897 | 0.00000 | 0.00000 | 4.76439 | 0.00000 | 0.00482 | 0.01307 | 0.94957 |
| **LOC100996348** | 0.00322 | 0.02256 | 0.08777 | 0.00272 | 0.00451 | 0.21519 | 0.03245 | 0.02448 | 0.15027 | 0.00000 | 0.00000 | 0.15348 |
| **LOC100996349** | 0.01202 | 0.00761 | 0.11427 | 0.04624 | 0.00000 | 0.04830 | 0.00000 | 0.00984 | 0.00000 | 0.03895 | 0.01925 | 0.01391 |
| **LOC100996351** | 0.00553 | 0.01201 | 0.00660 | 0.00259 | 0.07472 | 0.05942 | 0.00634 | 0.01018 | 0.01750 | 0.03053 | 0.01280 | 0.01178 |
| **LOC100996385** | 0.00879 | 0.02106 | 0.01602 | 0.01470 | 0.02717 | 0.01113 | 0.02511 | 0.01171 | 0.01768 | 0.00289 | 0.01488 | 0.02619 |
| **LOC100996437** | 0.11811 | 0.33389 | 0.22951 | 0.00128 | 0.17657 | 0.00000 | 0.11091 | 0.01940 | 0.00000 | 0.02773 | 0.31275 | 0.33697 |
| **LOC100996447** | 0.07105 | 1.00472 | 0.62241 | 0.33123 | 0.57719 | 3.56665 | 2.65888 | 0.09091 | 0.00000 | 0.00000 | 2.29919 | 0.71294 |
| **LOC100996455** | 0.00353 | 0.00000 | 0.00000 | 0.03250 | 0.06940 | 0.39576 | 0.10023 | 0.00726 | 0.44143 | 0.06987 | 0.22017 | 0.00371 |
| **LOC100996579** | 0.06735 | 0.03204 | 0.05198 | 0.16688 | 0.03310 | 0.22996 | 0.00000 | 0.00541 | 0.33318 | 0.00000 | 0.00000 | 0.03548 |
| **LOC100996590** | 0.00569 | 0.01371 | 0.01843 | 0.01163 | 0.03959 | 0.00000 | 0.02090 | 0.00940 | 0.01284 | 0.03654 | 0.01383 | 0.03825 |
| **LOC100996634** | 0.00156 | 0.12122 | 0.09832 | 0.04458 | 0.02351 | 0.08441 | 0.00930 | 0.09121 | 0.00000 | 0.11672 | 0.30307 | 0.51115 |
| **LOC100996654** | 0.00757 | 0.01252 | 0.02152 | 0.01747 | 0.01722 | 0.02539 | 0.01756 | 0.03123 | 0.01939 | 0.02681 | 0.02421 | 0.07095 |
| **LOC100996669** | 0.00598 | 0.01550 | 0.01137 | 0.01519 | 0.00745 | 0.03656 | 0.01438 | 0.01030 | 0.01505 | 0.02149 | 0.01447 | 0.01572 |
| **LOC100996693** | 0.11178 | 0.01415 | 0.02811 | 0.02375 | 0.01525 | 0.00000 | 0.11262 | 0.14341 | 0.04215 | 0.00000 | 0.02128 | 0.03262 |
| **LOC100996694** | 0.00638 | 0.00946 | 0.04578 | 0.02285 | 0.01535 | 0.01504 | 0.01241 | 0.00856 | 0.01180 | 0.02226 | 0.01647 | 0.06913 |
| **LOC101059948** | 0.13074 | 0.00315 | 0.13959 | 0.01346 | 0.00782 | 0.00000 | 0.00000 | 0.08759 | 0.05462 | 0.19926 | 0.00211 | 0.11555 |
| **LOC101060019** | 0.57210 | 0.01405 | 0.01331 | 0.01089 | 0.01219 | 0.00997 | 0.01608 | 0.09869 | 0.01601 | 0.02637 | 0.02141 | 0.01992 |
| **LOC101060091** | 0.01282 | 0.03993 | 0.13017 | 0.00669 | 0.01990 | 0.09072 | 0.00000 | 0.01906 | 0.01422 | 0.03732 | 0.00834 | 0.01604 |
| **LOC101060498** | 0.00553 | 0.06825 | 0.00382 | 0.00276 | 0.05302 | 0.00000 | 0.00250 | 0.00878 | 0.09184 | 0.07155 | 0.01764 | 0.18241 |
| **LOC101060542** | 0.00467 | 0.02020 | 0.01573 | 0.01050 | 0.00628 | 0.01719 | 0.03584 | 0.00969 | 0.00040 | 0.08798 | 0.01653 | 0.03218 |
| **LOC101060553** | 0.22577 | 0.01388 | 0.10343 | 0.00595 | 0.07075 | 0.00000 | 0.00628 | 0.04174 | 0.02349 | 0.02520 | 0.01433 | 0.02562 |
| **LOC101060691** | 0.07094 | 0.01276 | 0.36731 | 1.24055 | 0.82205 | 1.15328 | 0.00000 | 0.82980 | 3.00302 | 2.15854 | 0.57485 | 1.56161 |
| **LOC101241902** | 0.00540 | 0.01254 | 0.00994 | 0.00987 | 0.00772 | 0.01878 | 0.06199 | 0.06703 | 0.02582 | 0.03456 | 0.01310 | 0.02639 |
| **LOC101243545** | 0.00375 | 0.03230 | 0.09058 | 0.04830 | 0.00000 | 0.00000 | 0.01224 | 0.01016 | 0.02913 | 0.24636 | 0.01581 | 0.00984 |
| **LOC101409256** | 0.00777 | 0.00591 | 0.02512 | 0.01401 | 0.01219 | 0.06414 | 0.05268 | 0.00976 | 0.03555 | 0.00601 | 0.00713 | 0.07753 |
| **LOC101448202** | 0.03819 | 0.01503 | 0.01203 | 0.01304 | 0.01277 | 0.00640 | 0.08718 | 0.02244 | 0.00183 | 0.03836 | 0.01285 | 0.04049 |
| **LOC101559451** | 0.01686 | 0.24059 | 0.00000 | 0.00000 | 0.06764 | 0.24993 | 0.05060 | 0.19060 | 0.00000 | 0.13890 | 0.07432 | 0.37835 |
| **LOC101593348** | 0.82687 | 0.00310 | 0.13290 | 0.03987 | 0.00174 | 0.00000 | 0.00851 | 0.60852 | 0.01677 | 0.10004 | 0.01283 | 0.20368 |
| **LOC101805491** | 0.20688 | 0.13544 | 1.46080 | 0.03769 | 0.00000 | 0.07121 | 0.34075 | 0.41362 | 0.19589 | 0.21522 | 0.22158 | 0.14512 |
| **LOC101926889** | 0.00567 | 0.01282 | 0.02495 | 0.03844 | 0.01128 | 0.02386 | 0.01242 | 0.01140 | 0.01584 | 0.02217 | 0.01844 | 0.01769 |
| **LOC101926892** | 0.00605 | 0.00813 | 0.30675 | 0.03105 | 0.01619 | 0.12423 | 0.00000 | 0.00757 | 0.09850 | 0.00000 | 0.06517 | 0.05799 |
| **LOC101926895** | 0.08806 | 0.11240 | 0.10231 | 0.00893 | 0.00119 | 0.11000 | 0.06126 | 0.08806 | 0.05587 | 0.00000 | 0.08659 | 0.09298 |
| **LOC101926896** | 0.00631 | 0.02379 | 0.02238 | 0.01825 | 0.05341 | 0.00000 | 0.02108 | 0.01068 | 0.02413 | 0.01187 | 0.01882 | 0.01696 |
| **LOC101926905** | 0.00693 | 0.75258 | 0.01580 | 0.00000 | 0.00000 | 0.00629 | 0.00022 | 0.37893 | 0.03710 | 0.04839 | 0.02985 | 0.58385 |
| **LOC101926911** | 0.00505 | 0.02284 | 0.37472 | 0.02938 | 0.00000 | 0.29366 | 0.08195 | 0.04058 | 0.00000 | 0.07951 | 0.05075 | 0.06851 |

| | | | | | | | | | | | | |
|---|---|---|---|---|---|---|---|---|---|---|---|---|
| **LOC101926913** | 0.02893 | 0.01221 | 0.01335 | 0.02014 | 0.07662 | 0.08485 | 0.01176 | 0.02562 | 0.00000 | 0.01145 | 0.02585 | 0.01184 |
| **LOC101926928** | 0.00472 | 0.01380 | 0.01664 | 0.00933 | 0.02727 | 0.01662 | 0.01983 | 0.02853 | 0.01044 | 0.03068 | 0.01459 | 0.01482 |
| **LOC101926933** | 0.03916 | 0.00000 | 0.02646 | 0.00000 | 0.04551 | 0.13211 | 0.00000 | 0.04735 | 0.06669 | 0.14615 | 0.05334 | 0.14708 |
| **LOC101926935** | 0.00357 | 0.02035 | 0.04826 | 0.18670 | 0.00000 | 0.01698 | 0.00978 | 0.00509 | 0.08601 | 0.38365 | 0.00137 | 0.11214 |
| **LOC101926940** | 0.00621 | 0.01250 | 0.12241 | 0.00524 | 0.03534 | 0.03474 | 0.03374 | 0.01061 | 0.05861 | 0.00000 | 0.04184 | 0.01908 |
| **LOC101926941** | 0.05119 | 0.01689 | 0.00986 | 0.06230 | 0.01316 | 0.04660 | 0.01007 | 0.00896 | 0.01586 | 0.01287 | 0.01203 | 0.01397 |
| **LOC101926942** | 0.01602 | 0.01894 | 0.01408 | 0.01159 | 0.01553 | 0.01794 | 0.01955 | 0.03725 | 0.01782 | 0.01302 | 0.01553 | 0.01603 |
| **LOC101926943** | 0.02912 | 0.05729 | 0.03677 | 0.17048 | 0.04744 | 0.08901 | 0.01755 | 0.01816 | 0.10365 | 0.00000 | 0.06620 | 0.06368 |
| **LOC101926944** | 0.00636 | 0.01897 | 0.02512 | 0.01544 | 0.01262 | 0.02167 | 0.01023 | 0.01042 | 0.03230 | 0.01246 | 0.01339 | 0.01571 |
| **LOC101926950** | 0.04134 | 0.01527 | 0.01443 | 0.01256 | 0.01603 | 0.01545 | 0.01986 | 0.01196 | 0.01911 | 0.01243 | 0.01477 | 0.01588 |
| **LOC101926955** | 0.00574 | 0.01386 | 0.02049 | 0.01255 | 0.03928 | 0.00427 | 0.01878 | 0.01268 | 0.01303 | 0.04658 | 0.01258 | 0.01579 |
| **LOC101926960** | 0.00454 | 0.01858 | 0.00814 | 0.02135 | 0.00090 | 0.03469 | 0.00774 | 0.01489 | 0.01528 | 0.05738 | 0.01380 | 0.01423 |
| **LOC101926962** | 0.00569 | 0.01312 | 0.01162 | 0.04837 | 0.00874 | 0.01667 | 0.02418 | 0.01027 | 0.01490 | 0.03331 | 0.01438 | 0.02003 |
| **LOC101926963** | 0.00598 | 0.01416 | 0.02281 | 0.01207 | 0.01348 | 0.02296 | 0.00357 | 0.01036 | 0.03404 | 0.03214 | 0.01101 | 0.01462 |
| **LOC101926964** | 0.00439 | 0.01348 | 0.02339 | 0.01149 | 0.01398 | 0.01848 | 0.03395 | 0.01596 | 0.00275 | 0.03162 | 0.01551 | 0.01905 |
| **LOC101926966** | 0.00546 | 0.14022 | 0.43062 | 0.00351 | 0.10823 | 0.00842 | 0.22294 | 0.06095 | 0.00500 | 0.00000 | 0.49396 | 0.24679 |
| **LOC101926975** | 0.52305 | 0.01336 | 0.01374 | 0.01149 | 0.01489 | 0.01645 | 0.01772 | 0.00996 | 0.02087 | 0.00881 | 0.01351 | 0.01249 |
| **LOC101926987** | 0.07412 | 0.07384 | 0.17786 | 0.13859 | 0.00000 | 0.00000 | 0.00022 | 0.16827 | 0.09059 | 0.18666 | 0.01237 | 0.27348 |
| **LOC101926996** | 0.01899 | 0.03672 | 0.02827 | 0.29417 | 0.09434 | 0.00000 | 0.01740 | 0.01002 | 0.00000 | 0.33076 | 0.00281 | 0.03060 |
| **LOC101927015** | 0.02449 | 0.01494 | 0.01314 | 0.01146 | 0.02202 | 0.01612 | 0.03300 | 0.03305 | 0.01104 | 0.01636 | 0.01586 | 0.01475 |
| **LOC101927018** | 0.00998 | 0.01460 | 0.02269 | 0.04169 | 0.02080 | 0.01489 | 0.00069 | 0.00733 | 0.01908 | 0.16284 | 0.02989 | 0.04677 |
| **LOC101927021** | 0.51398 | 0.27665 | 0.12286 | 0.03275 | 0.00495 | 0.10575 | 0.00000 | 0.03596 | 0.10125 | 0.00000 | 0.03079 | 0.22215 |
| **LOC101927023** | 0.10049 | 0.02109 | 0.01329 | 0.01044 | 0.01028 | 0.01210 | 0.01395 | 0.00890 | 0.01786 | 0.02970 | 0.02878 | 0.02291 |
| **LOC101927027** | 0.42877 | 0.22387 | 0.24072 | 0.80265 | 0.00000 | 1.50172 | 0.09526 | 0.05904 | 0.00000 | 1.59635 | 0.47205 | 0.89285 |
| **LOC101927040** | 0.00454 | 0.00932 | 0.00978 | 0.00789 | 0.04482 | 0.04546 | 0.00611 | 0.04435 | 0.02873 | 0.01681 | 0.01306 | 0.02359 |
| **LOC101927043** | 0.01865 | 0.00354 | 0.00000 | 0.00000 | 0.02904 | 0.11506 | 0.00000 | 0.07682 | 0.26509 | 0.11714 | 0.00000 | 0.00457 |
| **LOC101927045** | 0.10937 | 0.14963 | 0.39094 | 0.00114 | 0.00000 | 0.98159 | 0.22216 | 0.00000 | 0.04784 | 0.00000 | 0.29907 | 0.28774 |
| **LOC101927049** | 0.00931 | 0.06181 | 0.01334 | 0.00803 | 0.02443 | 0.01236 | 0.01199 | 0.01577 | 0.00785 | 0.04686 | 0.07219 | 0.01114 |
| **LOC101927051** | 0.01982 | 0.01648 | 0.05553 | 0.02270 | 0.03485 | 0.00000 | 0.00000 | 0.01097 | 0.11617 | 0.05609 | 0.00403 | 0.01411 |
| **LOC101927053** | 0.01241 | 0.01686 | 0.02601 | 0.01183 | 0.02510 | 0.00000 | 0.07132 | 0.02884 | 0.00000 | 0.02444 | 0.02805 | 0.01779 |
| **LOC101927055** | 0.01080 | 0.00329 | 0.01582 | 0.00045 | 0.08185 | 0.06636 | 0.00528 | 0.00878 | 0.00165 | 0.04432 | 0.01187 | 0.01146 |
| **LOC101927069** | 0.01748 | 0.10705 | 0.00000 | 0.00000 | 0.40672 | 0.19652 | 0.00000 | 0.00000 | 0.00000 | 1.07280 | 0.24627 | 0.22983 |
| **LOC101927081** | 0.02751 | 0.00000 | 0.06768 | 0.27542 | 0.00000 | 0.16298 | 0.00000 | 0.00364 | 0.16454 | 0.44263 | 0.03836 | 0.15587 |
| **LOC101927087** | 0.08845 | 0.01404 | 0.02183 | 0.01021 | 0.01315 | 0.02313 | 0.02091 | 0.07595 | 0.02488 | 0.00950 | 0.01459 | 0.01153 |
| **LOC101927100** | 0.04676 | 0.01369 | 0.13729 | 0.02475 | 0.02599 | 0.10466 | 0.03597 | 0.18479 | 0.05983 | 0.05722 | 0.05957 | 0.30156 |
| **LOC101927102** | 0.02755 | 0.01686 | 0.01381 | 0.01941 | 0.00505 | 0.03100 | 0.01473 | 0.01119 | 0.01234 | 0.03737 | 0.02949 | 0.01540 |
| **LOC101927112** | 0.01895 | 0.01512 | 0.01515 | 0.02373 | 0.01093 | 0.02861 | 0.02903 | 0.00747 | 0.00940 | 0.01825 | 0.01356 | 0.02068 |
| **LOC101927123** | 0.00414 | 0.05895 | 0.03861 | 0.00000 | 0.00000 | 0.06962 | 0.00000 | 0.00987 | 0.01154 | 0.13113 | 0.00063 | 0.00383 |
| **LOC101927124** | 0.17971 | 0.05950 | 0.00000 | 0.12330 | 0.01664 | 0.57601 | 0.00000 | 0.00000 | 0.00000 | 0.00473 | 0.14472 | 0.06632 |
| **LOC101927129** | 0.31894 | 0.03668 | 0.03121 | 0.01789 | 0.15118 | 0.18795 | 0.00000 | 0.00000 | 0.00000 | 0.04017 | 0.00000 | 0.21973 |
| **LOC101927131** | 0.04507 | 0.01411 | 0.03026 | 0.01163 | 0.02222 | 0.02962 | 0.02126 | 0.00643 | 0.01609 | 0.00905 | 0.01999 | 0.01661 |
| **LOC101927132** | 0.38843 | 0.00588 | 0.13995 | 0.00715 | 0.02790 | 0.03879 | 0.00000 | 0.01007 | 0.02478 | 0.00396 | 0.01525 | 0.09857 |
| **LOC101927134** | 0.09713 | 0.06680 | 0.01274 | 0.03013 | 0.00924 | 0.00789 | 0.01751 | 0.01190 | 0.01914 | 0.01309 | 0.02121 | 0.04518 |
| **LOC101927136** | 0.00615 | 0.01306 | 0.04792 | 0.00895 | 0.01907 | 0.03399 | 0.01580 | 0.01025 | 0.00646 | 0.01277 | 0.02168 | 0.01628 |
| **LOC101927142** | 0.00650 | 0.01159 | 0.01227 | 0.00803 | 0.00988 | 0.02922 | 0.02040 | 0.00936 | 0.00876 | 0.02850 | 0.04401 | 0.02946 |
| **LOC101927143** | 0.02543 | 0.06259 | 0.05599 | 0.03971 | 0.03349 | 0.25514 | 0.00000 | 0.02424 | 0.07784 | 0.00000 | 0.01130 | 0.02815 |
| **LOC101927151** | 0.00000 | 2.13294 | 1.74420 | 0.16607 | 0.32143 | 0.41818 | 1.77992 | 3.22476 | 2.71753 | 0.65023 | 2.06045 | 3.96160 |
| **LOC101927153** | 0.01913 | 0.01478 | 0.01901 | 0.00679 | 0.00500 | 0.02503 | 0.01328 | 0.02398 | 0.03493 | 0.04877 | 0.00889 | 0.03227 |

| | | | | | | | | | | | | |
|---|---|---|---|---|---|---|---|---|---|---|---|---|
| **LOC101927164** | 0.00186 | 0.04419 | 0.01884 | 0.02371 | 0.00000 | 0.01556 | 0.01963 | 0.04157 | 0.08467 | 0.07190 | 0.05305 | 0.03898 |
| **LOC101927166** | 0.05216 | 0.01386 | 0.02123 | 0.01866 | 0.03645 | 0.00791 | 0.02537 | 0.01486 | 0.02943 | 0.00000 | 0.01887 | 0.02516 |
| **LOC101927178** | 0.00355 | 0.00235 | 0.04404 | 0.03879 | 0.02827 | 0.06394 | 0.05643 | 0.12134 | 0.03912 | 0.23225 | 0.05598 | 0.07013 |
| **LOC101927181** | 0.17329 | 0.16172 | 0.04104 | 1.05141 | 0.34967 | 0.00000 | 0.00254 | 0.00000 | 0.00000 | 0.42782 | 0.00805 | 0.16015 |
| **LOC101927190** | 0.00364 | 0.00000 | 0.00000 | 0.04354 | 0.00000 | 0.02915 | 0.33362 | 0.00946 | 0.00000 | 0.36471 | 0.06488 | 0.02331 |
| **LOC101927196** | 0.04068 | 0.02644 | 0.01037 | 0.01577 | 0.01010 | 0.06092 | 0.02689 | 0.03020 | 0.01685 | 0.02338 | 0.01626 | 0.02282 |
| **LOC101927204** | 0.00585 | 0.01224 | 0.02373 | 0.00995 | 0.09173 | 0.00000 | 0.02103 | 0.01078 | 0.00831 | 0.03216 | 0.01166 | 0.02035 |
| **LOC101927207** | 0.01230 | 0.02260 | 0.00641 | 0.03309 | 0.00000 | 0.05960 | 0.00807 | 0.01070 | 0.01136 | 0.03640 | 0.01011 | 0.02191 |
| **LOC101927211** | 0.00706 | 0.00437 | 0.05485 | 0.02972 | 0.07256 | 0.00000 | 0.00677 | 0.05580 | 0.04077 | 0.20178 | 0.00492 | 0.08788 |
| **LOC101927217** | 0.13263 | 0.03055 | 0.03085 | 0.02482 | 0.02181 | 0.01746 | 0.01143 | 0.19122 | 0.02520 | 0.01099 | 0.01324 | 0.00708 |
| **LOC101927230** | 0.00476 | 0.16336 | 0.00000 | 0.30293 | 0.22051 | 0.11304 | 0.00000 | 0.00248 | 0.00428 | 0.17698 | 0.00069 | 0.35361 |
| **LOC101927237** | 0.01473 | 0.03994 | 0.00315 | 0.00186 | 0.00312 | 0.04507 | 0.00000 | 0.24908 | 0.07581 | 0.05767 | 0.00859 | 0.03647 |
| **LOC101927239** | 0.20910 | 0.41360 | 0.04961 | 0.02853 | 0.00488 | 0.10802 | 0.07710 | 0.02016 | 0.01215 | 0.00000 | 0.23142 | 0.25659 |
| **LOC101927243** | 0.14540 | 0.01815 | 0.01456 | 0.01065 | 0.01302 | 0.01942 | 0.02975 | 0.03170 | 0.01316 | 0.01431 | 0.01391 | 0.01247 |
| **LOC101927248** | 0.12493 | 0.01412 | 0.01311 | 0.03619 | 0.02580 | 0.00000 | 0.01325 | 0.51037 | 0.03100 | 0.02167 | 0.01165 | 0.07828 |
| **LOC101927257** | 0.09007 | 0.00667 | 0.13447 | 0.11006 | 0.32437 | 0.00000 | 0.03970 | 0.06544 | 0.00000 | 0.58763 | 0.01385 | 0.15130 |
| **LOC101927258** | 1.57328 | 0.00000 | 0.00000 | 0.04670 | 0.01337 | 0.09276 | 0.00000 | 0.10226 | 0.05887 | 0.02050 | 0.00558 | 0.17522 |
| **LOC101927267** | 0.46515 | 0.21093 | 0.01600 | 0.16920 | 0.11861 | 0.06234 | 0.00024 | 0.27685 | 0.29290 | 0.16174 | 0.06845 | 0.60432 |
| **LOC101927273** | 0.00415 | 0.00000 | 0.04999 | 0.00115 | 0.35304 | 0.03900 | 0.00000 | 0.00606 | 0.15171 | 0.23213 | 0.14834 | 0.08454 |
| **LOC101927274** | 0.01286 | 0.01239 | 0.02059 | 0.01985 | 0.01254 | 0.01132 | 0.02806 | 0.02465 | 0.00705 | 0.02462 | 0.02428 | 0.19747 |
| **LOC101927275** | 0.01535 | 0.01703 | 0.25017 | 0.05569 | 0.11732 | 0.06611 | 0.00884 | 0.22345 | 0.00000 | 0.02766 | 0.18676 | 0.04730 |
| **LOC101927280** | 4.13080 | 0.00939 | 0.01221 | 0.00946 | 0.01890 | 0.00000 | 0.02659 | 0.95844 | 0.01743 | 0.00000 | 0.01317 | 0.03325 |
| **LOC101927282** | 0.00788 | 0.00985 | 0.00998 | 0.01084 | 0.00000 | 0.04732 | 0.13866 | 0.06133 | 0.00011 | 0.01574 | 0.00916 | 0.01735 |
| **LOC101927284** | 0.06923 | 0.00000 | 0.00000 | 0.00000 | 0.00000 | 0.63319 | 0.56651 | 0.17317 | 0.00000 | 1.37247 | 0.00000 | 0.01486 |
| **LOC101927285** | 0.18589 | 0.01348 | 0.06905 | 0.01575 | 0.08709 | 0.15537 | 0.00000 | 0.03080 | 0.00000 | 0.03389 | 0.12919 | 0.00276 |
| **LOC101927286** | 0.53887 | 0.01445 | 0.01561 | 0.01203 | 0.01573 | 0.01063 | 0.02177 | 0.06963 | 0.03219 | 0.00001 | 0.01726 | 0.01249 |
| **LOC101927292** | 0.00639 | 0.05046 | 0.02498 | 0.01322 | 0.01400 | 0.00000 | 0.01697 | 0.01079 | 0.01642 | 0.01689 | 0.03102 | 0.01422 |
| **LOC101927296** | 0.00596 | 0.01418 | 0.05985 | 0.05110 | 0.01460 | 0.00767 | 0.01186 | 0.01051 | 0.01121 | 0.02800 | 0.01640 | 0.01619 |
| **LOC101927314** | 0.00622 | 0.01465 | 0.01913 | 0.01810 | 0.01341 | 0.01985 | 0.02141 | 0.01054 | 0.01909 | 0.01292 | 0.01404 | 0.01622 |
| **LOC101927318** | 0.00594 | 0.01846 | 0.01339 | 0.00910 | 0.01106 | 0.02364 | 0.03730 | 0.01045 | 0.01264 | 0.02377 | 0.01266 | 0.01462 |
| **LOC101927322** | 0.20139 | 0.00725 | 0.21371 | 0.00044 | 0.03201 | 0.06135 | 0.00000 | 0.30593 | 0.00000 | 0.04177 | 0.03147 | 0.11404 |
| **LOC101927323** | 0.01525 | 0.01710 | 0.01642 | 0.01153 | 0.01515 | 0.01642 | 0.01769 | 0.01116 | 0.01643 | 0.01729 | 0.01845 | 0.01546 |
| **LOC101927324** | 0.00614 | 0.02403 | 0.01618 | 0.00970 | 0.02994 | 0.02772 | 0.01767 | 0.01092 | 0.01415 | 0.00000 | 0.04682 | 0.01928 |
| **LOC101927332** | 0.02040 | 0.01469 | 0.01312 | 0.01384 | 0.01360 | 0.01697 | 0.02615 | 0.20885 | 0.01708 | 0.01735 | 0.01372 | 0.00915 |
| **LOC101927347** | 0.01232 | 0.00813 | 0.05576 | 0.01335 | 0.00440 | 0.04939 | 0.00000 | 0.00900 | 0.03241 | 0.05391 | 0.01080 | 0.01795 |
| **LOC101927354** | 0.08493 | 0.00000 | 0.05075 | 0.00301 | 0.32952 | 0.06014 | 0.00093 | 0.03918 | 0.20779 | 0.07928 | 0.15556 | 0.05173 |
| **LOC101927356** | 0.00547 | 0.01260 | 0.00925 | 0.01839 | 0.00071 | 0.03787 | 0.00747 | 0.01061 | 0.01238 | 0.06315 | 0.01226 | 0.02706 |
| **LOC101927364** | 0.01568 | 0.00712 | 0.01083 | 0.00978 | 0.00856 | 0.02330 | 0.01111 | 0.02356 | 0.01329 | 0.05416 | 0.01409 | 0.06181 |
| **LOC101927374** | 0.01212 | 0.01373 | 0.04091 | 0.00875 | 0.01182 | 0.02005 | 0.01287 | 0.01224 | 0.00725 | 0.03499 | 0.01823 | 0.01465 |
| **LOC101927378** | 0.00789 | 0.01072 | 0.01990 | 0.01743 | 0.02276 | 0.00584 | 0.01493 | 0.01022 | 0.00000 | 0.02567 | 0.15288 | 0.01680 |
| **LOC101927379** | 0.00284 | 0.17438 | 0.00000 | 0.00000 | 1.09378 | 0.02092 | 0.09588 | 0.04192 | 0.30634 | 0.15282 | 1.03237 | 0.10829 |
| **LOC101927380** | 0.00946 | 0.00878 | 0.05052 | 0.12853 | 0.00000 | 0.09201 | 0.02029 | 0.03243 | 0.09006 | 0.48764 | 0.07642 | 0.19521 |
| **LOC101927391** | 0.34366 | 0.00000 | 0.07324 | 0.00125 | 0.70075 | 0.69271 | 0.00000 | 0.00000 | 0.00000 | 0.00000 | 0.27766 | 0.02167 |
| **LOC101927394** | 0.46760 | 0.01500 | 0.01487 | 0.01232 | 0.01698 | 0.01007 | 0.01889 | 0.08526 | 0.02193 | 0.00706 | 0.01402 | 0.00906 |
| **LOC101927396** | 0.06416 | 0.01661 | 0.01213 | 0.01981 | 0.01610 | 0.03952 | 0.00943 | 0.01326 | 0.03019 | 0.01042 | 0.01048 | 0.03974 |
| **LOC101927411** | 0.00600 | 0.01413 | 0.02534 | 0.00931 | 0.01312 | 0.02123 | 0.01753 | 0.01040 | 0.01170 | 0.02660 | 0.01515 | 0.01572 |
| **LOC101927415** | 0.13009 | 0.15442 | 0.28974 | 0.16580 | 0.00000 | 0.20432 | 0.00112 | 0.01004 | 0.00000 | 0.24630 | 0.21163 | 0.10509 |
| **LOC101927416** | 0.00617 | 0.02036 | 0.02997 | 0.01261 | 0.03060 | 0.01154 | 0.01690 | 0.01050 | 0.01354 | 0.01181 | 0.02088 | 0.01962 |

| | | | | | | | | | | | | |
|---|---|---|---|---|---|---|---|---|---|---|---|---|
| **LOC101927418** | 0.21025 | 0.04986 | 0.08435 | 0.02020 | 0.01223 | 0.01162 | 0.12022 | 1.56890 | 0.00719 | 0.00000 | 0.01105 | 0.48245 |
| **LOC101927419** | 0.00584 | 0.01423 | 0.01433 | 0.01166 | 0.02142 | 0.01537 | 0.02223 | 0.01050 | 0.01407 | 0.03371 | 0.01423 | 0.01567 |
| **LOC101927421** | 1.21028 | 0.01558 | 0.02148 | 0.01243 | 0.02023 | 0.00087 | 0.01598 | 0.16369 | 0.01980 | 0.00594 | 0.02752 | 0.04014 |
| **LOC101927437** | 0.00899 | 0.01053 | 0.01110 | 0.00915 | 0.05328 | 0.01500 | 0.03579 | 0.01510 | 0.00039 | 0.04120 | 0.01821 | 0.01546 |
| **LOC101927446** | 0.00427 | 0.01548 | 0.02959 | 0.01524 | 0.01581 | 0.01603 | 0.01635 | 0.02057 | 0.01766 | 0.01030 | 0.01624 | 0.01634 |
| **LOC101927457** | 0.13405 | 0.01677 | 0.01108 | 0.01396 | 0.01851 | 0.00000 | 0.02099 | 0.34153 | 0.05177 | 0.00241 | 0.01064 | 0.04698 |
| **LOC101927472** | 0.06380 | 0.01835 | 0.01466 | 0.01460 | 0.02777 | 0.01634 | 0.01758 | 0.00530 | 0.01980 | 0.00978 | 0.01424 | 0.01838 |
| **LOC101927476** | 0.04639 | 0.01722 | 0.02267 | 0.01804 | 0.04809 | 0.01609 | 0.10854 | 0.12036 | 0.00000 | 0.00590 | 0.01894 | 0.02409 |
| **LOC101927477** | 0.00630 | 0.01447 | 0.01600 | 0.01061 | 0.01298 | 0.02675 | 0.02179 | 0.01041 | 0.03164 | 0.00772 | 0.01087 | 0.01761 |
| **LOC101927478** | 0.00339 | 0.02612 | 0.05641 | 0.00998 | 0.01905 | 0.02893 | 0.01744 | 0.02265 | 0.02223 | 0.00913 | 0.01450 | 0.04095 |
| **LOC101927482** | 0.00138 | 0.05056 | 0.16968 | 0.00000 | 0.00000 | 0.20408 | 2.19598 | 0.00120 | 2.16253 | 1.45856 | 0.67396 | 0.15743 |
| **LOC101927492** | 0.04232 | 0.87146 | 0.34827 | 0.59232 | 0.44897 | 0.66891 | 0.98527 | 0.11720 | 0.69806 | 0.00000 | 1.34118 | 0.36907 |
| **LOC101927495** | 0.00638 | 0.01763 | 0.02157 | 0.01447 | 0.04363 | 0.00548 | 0.02249 | 0.01061 | 0.01571 | 0.00800 | 0.01523 | 0.01782 |
| **LOC101927497** | 0.00798 | 0.06709 | 0.08246 | 0.00000 | 0.07082 | 0.01866 | 0.25632 | 0.01011 | 0.13711 | 0.03525 | 0.12110 | 0.00902 |
| **LOC101927501** | 0.00480 | 0.18332 | 0.92904 | 0.00000 | 0.00000 | 0.03233 | 0.66275 | 0.01427 | 1.18516 | 0.02268 | 0.00008 | 0.00473 |
| **LOC101927502** | 0.00193 | 0.01985 | 0.02632 | 0.00638 | 0.06111 | 0.00000 | 0.00611 | 0.20667 | 0.03917 | 0.06323 | 0.01589 | 0.01106 |
| **LOC101927523** | 0.00256 | 0.01251 | 0.01538 | 0.01236 | 0.01500 | 0.01150 | 0.01856 | 0.26363 | 0.01976 | 0.02566 | 0.01406 | 0.04392 |
| **LOC101927539** | 0.04528 | 0.01498 | 0.01603 | 0.01416 | 0.01546 | 0.01670 | 0.01738 | 0.02187 | 0.02138 | 0.01121 | 0.01535 | 0.01455 |
| **LOC101927549** | 0.03482 | 0.01439 | 0.01305 | 0.01123 | 0.01377 | 0.02067 | 0.01841 | 0.01251 | 0.02380 | 0.01461 | 0.01259 | 0.01546 |
| **LOC101927550** | 0.20706 | 0.29226 | 0.10020 | 0.10904 | 0.32753 | 0.00000 | 0.25643 | 0.14004 | 0.25337 | 1.30249 | 0.03324 | 0.63896 |
| **LOC101927557** | 0.00898 | 0.01616 | 0.02700 | 0.01235 | 0.01968 | 0.01333 | 0.01076 | 0.01003 | 0.06090 | 0.00030 | 0.01002 | 0.02069 |
| **LOC101927571** | 0.03696 | 0.05521 | 0.00108 | 0.00214 | 0.10426 | 0.30155 | 0.00000 | 0.29985 | 0.00000 | 0.01099 | 0.10389 | 0.06790 |
| **LOC101927572** | 0.15121 | 0.06737 | 0.16802 | 0.08499 | 0.05825 | 0.12312 | 0.00643 | 0.00346 | 0.32662 | 0.00000 | 0.50321 | 0.07582 |
| **LOC101927575** | 0.81133 | 0.01014 | 0.01874 | 0.01337 | 0.01454 | 0.00664 | 0.01497 | 0.07485 | 0.01931 | 0.02187 | 0.01441 | 0.05300 |
| **LOC101927580** | 0.00113 | 0.00188 | 0.00000 | 0.33113 | 0.00000 | 0.24227 | 0.00000 | 0.07073 | 0.00000 | 0.24647 | 0.00000 | 0.41325 |
| **LOC101927583** | 0.00608 | 0.01290 | 0.01635 | 0.01215 | 0.01137 | 0.03100 | 0.01888 | 0.01044 | 0.01576 | 0.01820 | 0.01328 | 0.01510 |
| **LOC101927587** | 0.00708 | 0.01703 | 0.01867 | 0.01611 | 0.03536 | 0.00000 | 0.03421 | 0.01411 | 0.02071 | 0.01410 | 0.01447 | 0.01842 |
| **LOC101927592** | 0.15494 | 0.01021 | 0.00689 | 0.00000 | 0.00000 | 0.04462 | 0.02533 | 0.24989 | 0.00284 | 0.17389 | 0.00393 | 0.00000 |
| **LOC101927599** | 0.26911 | 1.08543 | 0.82359 | 0.04954 | 0.23125 | 1.48538 | 0.44283 | 0.46085 | 1.55821 | 0.07889 | 0.66408 | 1.81932 |
| **LOC101927604** | 0.00771 | 0.02014 | 0.01854 | 0.01059 | 0.02589 | 0.00770 | 0.00980 | 0.02485 | 0.04622 | 0.01278 | 0.01357 | 0.02045 |
| **LOC101927620** | 0.00364 | 0.06356 | 0.10596 | 0.00460 | 0.00000 | 0.07567 | 0.02633 | 0.00947 | 0.00000 | 0.18390 | 0.02761 | 0.00240 |
| **LOC101927621** | 0.15062 | 0.19862 | 0.46391 | 0.06297 | 0.00585 | 0.00000 | 0.57049 | 0.14819 | 0.57124 | 0.00000 | 0.63264 | 0.30477 |
| **LOC101927623** | 0.00537 | 0.02519 | 0.01544 | 0.01653 | 0.03070 | 0.00173 | 0.01921 | 0.01229 | 0.02187 | 0.01915 | 0.01652 | 0.02781 |
| **LOC101927630** | 0.11928 | 0.01258 | 0.01220 | 0.00805 | 0.01944 | 0.03692 | 0.01460 | 0.01919 | 0.01415 | 0.01627 | 0.01231 | 0.01377 |
| **LOC101927631** | 0.00000 | 0.25836 | 0.02495 | 0.06306 | 0.07262 | 0.40168 | 0.00000 | 0.06989 | 0.08844 | 0.46895 | 0.01199 | 0.03810 |
| **LOC101927641** | 0.00639 | 0.01066 | 0.01638 | 0.01423 | 0.03427 | 0.00000 | 0.02348 | 0.37581 | 0.01862 | 0.00332 | 0.01492 | 0.13486 |
| **LOC101927653** | 0.00622 | 0.01542 | 0.01866 | 0.01428 | 0.01354 | 0.01714 | 0.01610 | 0.01042 | 0.02339 | 0.01627 | 0.02045 | 0.01701 |
| **LOC101927661** | 0.02746 | 0.01418 | 0.01821 | 0.01120 | 0.01392 | 0.01323 | 0.01430 | 0.01520 | 0.01755 | 0.02874 | 0.01491 | 0.05072 |
| **LOC101927666** | 0.05051 | 0.01279 | 0.05901 | 0.02276 | 0.02498 | 0.01547 | 0.02372 | 0.01358 | 0.00000 | 0.02404 | 0.05851 | 0.01413 |
| **LOC101927667** | 0.07380 | 0.06750 | 0.08347 | 0.05708 | 0.00000 | 0.03673 | 0.02570 | 0.08185 | 0.02357 | 0.31552 | 0.24274 | 0.18882 |
| **LOC101927668** | 0.25919 | 0.00628 | 0.23848 | 0.46415 | 0.00135 | 0.00017 | 0.01539 | 0.09744 | 0.03919 | 0.01610 | 0.01757 | 0.08096 |
| **LOC101927688** | 0.00656 | 0.02568 | 0.01156 | 0.01330 | 0.02023 | 0.02374 | 0.01613 | 0.00939 | 0.02853 | 0.01675 | 0.01291 | 0.04066 |
| **LOC101927692** | 0.00727 | 0.01433 | 0.01404 | 0.00988 | 0.01398 | 0.02757 | 0.01904 | 0.01416 | 0.03340 | 0.01459 | 0.01131 | 0.02253 |
| **LOC101927697** | 0.03492 | 0.01485 | 0.01453 | 0.01511 | 0.01460 | 0.01458 | 0.02010 | 0.00819 | 0.01650 | 0.01783 | 0.01883 | 0.01557 |
| **LOC101927698** | 0.00984 | 0.02719 | 0.03846 | 0.00937 | 0.01031 | 0.02829 | 0.03398 | 0.00995 | 0.03270 | 0.00376 | 0.02932 | 0.01482 |
| **LOC101927700** | 0.22180 | 0.09304 | 0.03992 | 0.00000 | 0.09934 | 0.00000 | 0.00158 | 0.00079 | 0.25290 | 0.39975 | 0.00000 | 0.03039 |
| **LOC101927701** | 0.13454 | 0.01362 | 0.01415 | 0.01334 | 0.01377 | 0.01678 | 0.01646 | 0.02973 | 0.01885 | 0.01847 | 0.01422 | 0.01548 |
| **LOC101927718** | 0.00717 | 0.03390 | 0.09206 | 0.09991 | 0.01636 | 0.05500 | 0.00586 | 0.00683 | 0.00000 | 0.01179 | 0.20547 | 0.12470 |

| | | | | | | | | | | | | |
|---|---|---|---|---|---|---|---|---|---|---|---|---|
| **LOC101927720** | 0.71169 | 1.59795 | 0.00000 | 0.62670 | 0.46588 | 2.93821 | 0.67398 | 0.18591 | 0.63820 | 0.16559 | 0.00000 | 2.18077 |
| **LOC101927722** | 3.12992 | 0.01380 | 0.00000 | 0.00000 | 0.00059 | 0.02932 | 0.00051 | 0.20268 | 0.02015 | 0.22240 | 0.00030 | 0.04902 |
| **LOC101927728** | 0.00797 | 0.06983 | 0.04598 | 0.03183 | 0.15183 | 0.06120 | 0.00000 | 0.00736 | 0.00000 | 0.25333 | 0.02595 | 0.15425 |
| **LOC101927730** | 0.01388 | 0.01757 | 0.18853 | 0.01468 | 0.00428 | 0.04453 | 0.00000 | 0.00845 | 0.00664 | 0.06140 | 0.01199 | 0.01978 |
| **LOC101927740** | 0.01807 | 0.01481 | 0.02410 | 0.01014 | 0.01336 | 0.01864 | 0.01332 | 0.02004 | 0.01549 | 0.02749 | 0.01584 | 0.01514 |
| **LOC101927746** | 0.00000 | 7.64914 | 0.64072 | 4.77339 | 0.00000 | 7.33375 | 2.08126 | 4.87603 | 3.25728 | 2.31639 | 4.77617 | 0.82602 |
| **LOC101927750** | 0.00622 | 0.10116 | 0.13786 | 0.00000 | 0.10112 | 0.00000 | 0.06129 | 0.01513 | 0.00000 | 0.17717 | 0.07623 | 0.11464 |
| **LOC101927755** | 0.03049 | 0.45106 | 0.07034 | 0.00000 | 0.02109 | 0.00094 | 0.11479 | 1.08996 | 0.21117 | 0.39396 | 0.51015 | 2.68517 |
| **LOC101927762** | 0.01709 | 0.01596 | 0.01517 | 0.01054 | 0.02875 | 0.02712 | 0.01701 | 0.00955 | 0.01890 | 0.03661 | 0.01304 | 0.04025 |
| **LOC101927770** | 0.14790 | 0.02248 | 0.10546 | 0.01219 | 0.16693 | 0.00000 | 0.00000 | 0.72724 | 0.00000 | 0.60769 | 0.01077 | 0.05886 |
| **LOC101927780** | 0.00586 | 0.01615 | 0.06850 | 0.01076 | 0.02280 | 0.00010 | 0.00834 | 0.01056 | 0.03407 | 0.04487 | 0.01044 | 0.01567 |
| **LOC101927795** | 0.09956 | 0.01914 | 0.01369 | 0.02080 | 0.00919 | 0.02543 | 0.01388 | 0.04288 | 0.01763 | 0.01944 | 0.01359 | 0.02664 |
| **LOC101927797** | 0.00147 | 0.01979 | 0.75862 | 0.17520 | 0.30274 | 0.56991 | 0.00000 | 0.07016 | 1.02354 | 0.00151 | 0.53011 | 0.08264 |
| **LOC101927811** | 0.41083 | 0.11099 | 0.13623 | 0.01734 | 0.00000 | 0.04801 | 0.00001 | 0.09788 | 0.00000 | 0.08588 | 0.00893 | 0.10522 |
| **LOC101927814** | 0.00358 | 0.02273 | 0.03092 | 0.01570 | 0.01992 | 0.00814 | 0.01820 | 0.05357 | 0.04055 | 0.00511 | 0.01108 | 0.01745 |
| **LOC101927817** | 0.04695 | 0.00000 | 0.07416 | 0.18811 | 0.13743 | 0.00000 | 0.00000 | 0.00000 | 0.22126 | 0.60561 | 0.10498 | 0.25428 |
| **LOC101927829** | 0.00615 | 0.01439 | 0.01842 | 0.01233 | 0.01303 | 0.01938 | 0.01707 | 0.01047 | 0.01376 | 0.01882 | 0.02311 | 0.01593 |
| **LOC101927839** | 0.00641 | 0.01640 | 0.06366 | 0.01163 | 0.01939 | 0.01444 | 0.01413 | 0.01050 | 0.01099 | 0.00802 | 0.01864 | 0.01722 |
| **LOC101927843** | 0.16818 | 0.02757 | 0.02868 | 0.01248 | 0.03864 | 0.00000 | 0.00714 | 0.00605 | 0.05747 | 0.02656 | 0.00949 | 0.01329 |
| **LOC101927844** | 0.00716 | 0.02342 | 0.01248 | 0.01405 | 0.09137 | 0.00000 | 0.02462 | 0.02782 | 0.03274 | 0.03264 | 0.00537 | 0.01487 |
| **LOC101927854** | 0.30744 | 0.03445 | 0.45762 | 0.05808 | 0.08898 | 0.25555 | 0.00921 | 0.02703 | 0.00000 | 0.02796 | 0.00027 | 0.01308 |
| **LOC101927865** | 0.17702 | 0.06286 | 0.02477 | 0.02080 | 0.00661 | 0.01307 | 0.01221 | 0.04440 | 0.02213 | 0.03580 | 0.04595 | 0.02223 |
| **LOC101927870** | 1.48748 | 0.01311 | 0.01450 | 0.01295 | 0.01985 | 0.00150 | 0.01943 | 0.01290 | 0.02675 | 0.00000 | 0.01651 | 0.03582 |
| **LOC101927873** | 1.81866 | 1.21864 | 0.03414 | 0.33326 | 0.14556 | 0.08022 | 0.00148 | 0.65784 | 0.00000 | 0.00000 | 0.09393 | 0.82418 |
| **LOC101927881** | 0.00527 | 0.05528 | 0.01047 | 0.03246 | 0.00769 | 0.07128 | 0.01096 | 0.01437 | 0.00933 | 0.01032 | 0.01629 | 0.05054 |
| **LOC101927901** | 0.32795 | 0.01215 | 0.02728 | 0.07292 | 0.02936 | 0.00000 | 0.02806 | 0.11546 | 0.03621 | 0.00931 | 0.01037 | 0.01623 |
| **LOC101927905** | 0.01771 | 0.01930 | 0.01334 | 0.03653 | 0.00903 | 0.02021 | 0.01524 | 0.02159 | 0.01942 | 0.01553 | 0.02000 | 0.02757 |
| **LOC101927907** | 0.10253 | 0.01256 | 0.01648 | 0.00925 | 0.06861 | 0.01939 | 0.00991 | 0.13069 | 0.01856 | 0.02292 | 0.01320 | 0.02819 |
| **LOC101927911** | 0.00629 | 0.00857 | 0.04191 | 0.01188 | 0.03058 | 0.00000 | 0.01938 | 0.00906 | 0.00000 | 0.02170 | 0.07116 | 0.10587 |
| **LOC101927914** | 0.00799 | 0.01144 | 0.00402 | 0.23154 | 0.00000 | 0.00748 | 0.01433 | 0.01292 | 0.01924 | 0.05046 | 0.01608 | 0.01460 |
| **LOC101927919** | 0.01537 | 0.10812 | 0.84720 | 0.03301 | 0.32010 | 0.00000 | 0.20721 | 0.21474 | 0.24834 | 0.00000 | 0.17422 | 0.38094 |
| **LOC101927921** | 1.98439 | 0.01113 | 0.01398 | 0.01148 | 0.06943 | 0.00000 | 0.02185 | 0.67820 | 0.01331 | 0.00000 | 0.03181 | 0.05506 |
| **LOC101927922** | 0.08835 | 0.01562 | 0.02657 | 0.01689 | 0.01688 | 0.01066 | 0.01666 | 0.00995 | 0.01787 | 0.01199 | 0.01562 | 0.01517 |
| **LOC101927924** | 0.00621 | 0.01522 | 0.01287 | 0.01324 | 0.01441 | 0.01513 | 0.04367 | 0.01049 | 0.00678 | 0.01183 | 0.02422 | 0.01719 |
| **LOC101927931** | 0.10518 | 0.01327 | 0.04194 | 0.00859 | 0.01260 | 0.02515 | 0.01908 | 0.19733 | 0.02154 | 0.01060 | 0.01153 | 0.01030 |
| **LOC101927934** | 0.04278 | 0.09107 | 0.01464 | 0.00000 | 0.24137 | 0.05484 | 0.02607 | 0.00864 | 0.00000 | 0.03824 | 0.17418 | 0.00380 |
| **LOC101927948** | 0.00417 | 0.00443 | 0.02174 | 0.00000 | 0.14545 | 0.04165 | 0.11528 | 0.00961 | 0.07914 | 0.13345 | 0.03860 | 0.01141 |
| **LOC101927950** | 0.00561 | 0.01648 | 0.00637 | 0.09612 | 0.16615 | 0.00000 | 0.01443 | 0.01045 | 0.03444 | 0.03365 | 0.01665 | 0.01576 |
| **LOC101927954** | 0.06720 | 0.03620 | 0.02388 | 0.03277 | 0.01791 | 0.00000 | 0.01811 | 0.05486 | 0.02721 | 0.02244 | 0.03034 | 0.02508 |
| **LOC101927964** | 0.00612 | 0.02306 | 0.01335 | 0.02350 | 0.01115 | 0.01986 | 0.01927 | 0.01047 | 0.00902 | 0.01088 | 0.04538 | 0.01712 |
| **LOC101927967** | 0.01365 | 0.01635 | 0.03973 | 0.00521 | 0.00562 | 0.03352 | 0.02128 | 0.01188 | 0.05435 | 0.04500 | 0.00958 | 0.01524 |
| **LOC101928001** | 0.00450 | 0.01643 | 0.02581 | 0.00000 | 0.01838 | 0.06046 | 0.04484 | 0.01151 | 0.00000 | 0.10115 | 0.02660 | 0.01426 |
| **LOC101928002** | 0.02354 | 0.01231 | 0.08573 | 0.00753 | 0.02653 | 0.00000 | 0.02045 | 0.01703 | 0.00575 | 0.12504 | 0.00892 | 0.01210 |
| **LOC101928008** | 0.00558 | 0.00793 | 0.03681 | 0.00284 | 0.06092 | 0.04601 | 0.00000 | 0.00990 | 0.06357 | 0.03834 | 0.00242 | 0.03552 |
| **LOC101928012** | 0.07389 | 0.56902 | 0.00000 | 0.42055 | 1.07738 | 0.82953 | 0.00000 | 0.01554 | 0.00000 | 0.09267 | 0.16228 | 0.90115 |
| **LOC101928020** | 0.26452 | 0.00000 | 0.00298 | 0.10830 | 0.14735 | 0.00000 | 0.16042 | 0.37804 | 0.05978 | 0.44439 | 0.31280 | 0.17385 |
| **LOC101928035** | 0.02868 | 0.04595 | 0.01015 | 0.00000 | 0.11473 | 0.03846 | 0.03051 | 0.00938 | 0.01016 | 0.03871 | 0.03461 | 0.00882 |
| **LOC101928036** | 0.00654 | 0.03019 | 0.06473 | 0.01077 | 0.01296 | 0.05701 | 0.00940 | 0.00997 | 0.01464 | 0.00963 | 0.01102 | 0.01253 |

| | | | | | | | | | | | | |
|---|---|---|---|---|---|---|---|---|---|---|---|---|
| **LOC101928043** | 0.01422 | 0.12597 | 0.00922 | 0.00000 | 0.02996 | 0.08322 | 0.00779 | 0.33130 | 0.00000 | 0.01399 | 0.01351 | 0.25077 |
| **LOC101928053** | 0.10767 | 0.07490 | 0.24793 | 0.00000 | 0.00000 | 0.00000 | 0.00000 | 2.26371 | 0.16731 | 0.27767 | 0.02462 | 0.24617 |
| **LOC101928062** | 0.00446 | 0.39288 | 0.13828 | 0.30451 | 0.01327 | 1.95358 | 0.00000 | 2.33559 | 1.23515 | 0.00000 | 0.23276 | 2.52217 |
| **LOC101928063** | 0.04688 | 0.04241 | 0.00330 | 0.01162 | 0.00000 | 0.07191 | 0.01082 | 0.00368 | 0.00000 | 0.06791 | 0.02489 | 0.02834 |
| **LOC101928068** | 0.18908 | 0.03981 | 0.12549 | 0.23033 | 0.06939 | 0.00000 | 0.00003 | 0.03493 | 0.29782 | 0.12940 | 0.02386 | 0.14986 |
| **LOC101928069** | 0.00419 | 0.01221 | 0.10061 | 0.01391 | 0.01950 | 0.06839 | 0.00000 | 0.01584 | 0.05141 | 0.01025 | 0.08122 | 0.05223 |
| **LOC101928075** | 0.00973 | 0.01614 | 0.01730 | 0.01180 | 0.02112 | 0.01870 | 0.02171 | 0.02249 | 0.03579 | 0.00795 | 0.01544 | 0.01905 |
| **LOC101928093** | 0.00837 | 0.02002 | 0.01574 | 0.01161 | 0.01697 | 0.01537 | 0.02603 | 0.01338 | 0.01938 | 0.01112 | 0.01716 | 0.01601 |
| **LOC101928099** | 0.42925 | 0.12134 | 0.47557 | 0.45031 | 0.00000 | 0.00000 | 0.45342 | 1.70714 | 0.54637 | 0.00000 | 0.33915 | 4.27171 |
| **LOC101928100** | 0.34614 | 0.00000 | 0.14238 | 0.00000 | 0.59674 | 0.43760 | 0.04763 | 0.11400 | 0.00002 | 0.22854 | 0.31793 | 0.17546 |
| **LOC101928103** | 0.14247 | 0.94282 | 0.00124 | 0.23164 | 0.00000 | 0.29726 | 0.00000 | 0.00605 | 0.00000 | 0.17374 | 0.00000 | 0.02774 |
| **LOC101928104** | 0.00610 | 0.01513 | 0.01639 | 0.00863 | 0.01711 | 0.02486 | 0.01285 | 0.01037 | 0.04021 | 0.01921 | 0.01612 | 0.01600 |
| **LOC101928105** | 0.00873 | 0.01120 | 0.02582 | 0.00623 | 0.00000 | 0.02515 | 0.19936 | 0.02780 | 0.00000 | 0.03104 | 0.01263 | 0.03666 |
| **LOC101928113** | 0.00792 | 0.01548 | 0.01391 | 0.01688 | 0.02520 | 0.00895 | 0.01746 | 0.01032 | 0.02171 | 0.01891 | 0.01395 | 0.01676 |
| **LOC101928122** | 0.89477 | 0.20985 | 0.00175 | 0.01660 | 0.15504 | 0.02463 | 0.02586 | 0.26193 | 0.00000 | 0.07793 | 0.06129 | 0.41960 |
| **LOC101928131** | 0.00186 | 0.01388 | 0.01243 | 0.01131 | 0.01453 | 0.02160 | 0.04367 | 0.09173 | 0.00943 | 0.00895 | 0.01748 | 0.01801 |
| **LOC101928135** | 0.08427 | 0.01647 | 0.01961 | 0.01549 | 0.01311 | 0.01514 | 0.01633 | 0.00552 | 0.01779 | 0.02117 | 0.01462 | 0.01455 |
| **LOC101928137** | 0.03509 | 0.01139 | 0.05672 | 0.01487 | 0.05714 | 0.03268 | 0.02061 | 0.02086 | 0.02247 | 0.00000 | 0.02964 | 0.04272 |
| **LOC101928150** | 0.01357 | 0.01150 | 0.02498 | 0.02448 | 0.00000 | 0.07428 | 0.01569 | 0.00916 | 0.00000 | 0.11742 | 0.01380 | 0.02107 |
| **LOC101928163** | 0.00911 | 0.03331 | 0.02065 | 0.00402 | 0.02802 | 0.01690 | 0.04099 | 0.01189 | 0.00972 | 0.07179 | 0.00992 | 0.01554 |
| **LOC101928167** | 0.00627 | 0.04451 | 0.00855 | 0.01006 | 0.01519 | 0.00667 | 0.02214 | 0.00979 | 0.09389 | 0.00000 | 0.01500 | 0.04263 |
| **LOC101928168** | 0.04680 | 0.00034 | 0.00000 | 0.00000 | 0.00243 | 0.95485 | 0.00000 | 0.03951 | 0.00000 | 1.36064 | 0.08014 | 0.13337 |
| **LOC101928174** | 0.01993 | 0.01513 | 0.05958 | 0.00966 | 0.01143 | 0.00613 | 0.03760 | 0.01175 | 0.00000 | 0.04665 | 0.02923 | 0.01736 |
| **LOC101928203** | 0.25889 | 0.01756 | 0.03573 | 0.03061 | 0.01612 | 0.07195 | 0.06656 | 0.10209 | 0.03463 | 0.01411 | 0.01040 | 0.20981 |
| **LOC101928222** | 0.03519 | 0.01266 | 0.02472 | 0.01271 | 0.01157 | 0.02183 | 0.01059 | 0.00869 | 0.02551 | 0.02774 | 0.01207 | 0.02587 |
| **LOC101928227** | 0.02914 | 0.01640 | 0.01248 | 0.00918 | 0.02166 | 0.02227 | 0.02566 | 0.00816 | 0.00984 | 0.02449 | 0.01341 | 0.01502 |
| **LOC101928243** | 0.04096 | 0.26711 | 0.36805 | 0.19711 | 0.27773 | 0.29740 | 0.08425 | 0.03310 | 0.41265 | 0.00000 | 0.68552 | 0.20054 |
| **LOC101928244** | 0.00962 | 0.01542 | 0.01794 | 0.01346 | 0.01272 | 0.02190 | 0.02605 | 0.01150 | 0.02525 | 0.00916 | 0.01630 | 0.01606 |
| **LOC101928254** | 0.02632 | 0.31329 | 0.11052 | 2.56144 | 0.00000 | 0.03912 | 0.00000 | 0.02808 | 0.00000 | 0.23274 | 0.01140 | 0.06617 |
| **LOC101928259** | 0.00105 | 0.05497 | 0.00036 | 0.06997 | 0.00539 | 0.00000 | 0.00067 | 0.07217 | 0.08680 | 0.62347 | 0.07994 | 0.54645 |
| **LOC101928266** | 0.11026 | 0.01236 | 0.02930 | 0.00847 | 0.00299 | 0.01941 | 0.03862 | 0.00187 | 0.00000 | 0.11135 | 0.01157 | 0.03693 |
| **LOC101928269** | 0.02455 | 0.01139 | 0.00901 | 0.00590 | 0.00000 | 0.04007 | 0.05989 | 0.04819 | 0.01701 | 0.03074 | 0.01067 | 0.01086 |
| **LOC101928272** | 0.32415 | 0.03262 | 0.10107 | 0.04662 | 0.00000 | 0.01242 | 0.00000 | 0.01105 | 0.01807 | 0.14938 | 0.01213 | 0.16737 |
| **LOC101928283** | 0.01732 | 0.09048 | 0.02973 | 0.19378 | 0.09266 | 0.00000 | 0.02242 | 0.01013 | 0.00000 | 0.03770 | 0.05412 | 0.01277 |
| **LOC101928290** | 0.04995 | 0.04636 | 0.07981 | 0.01995 | 0.01001 | 0.05946 | 0.00000 | 0.11836 | 0.08934 | 0.00000 | 0.03377 | 0.10423 |
| **LOC101928292** | 0.00621 | 0.01525 | 0.02208 | 0.01310 | 0.01258 | 0.01261 | 0.01648 | 0.01053 | 0.01194 | 0.01709 | 0.04359 | 0.01655 |
| **LOC101928295** | 0.23959 | 0.00000 | 0.04276 | 0.04136 | 0.00602 | 0.09925 | 0.01642 | 0.00000 | 0.00000 | 0.36789 | 0.03803 | 0.67858 |
| **LOC101928303** | 0.00569 | 0.00896 | 0.37777 | 0.00461 | 0.32337 | 0.04071 | 0.08963 | 0.00680 | 0.01848 | 0.02985 | 0.06495 | 0.12733 |
| **LOC101928335** | 0.00339 | 0.00775 | 0.01264 | 0.01021 | 0.01063 | 0.03879 | 0.01884 | 0.02358 | 0.01643 | 0.00973 | 0.01418 | 0.05970 |
| **LOC101928336** | 0.09917 | 0.01243 | 0.04455 | 0.03002 | 0.01590 | 0.01895 | 0.01260 | 0.00753 | 0.00000 | 0.06318 | 0.04950 | 0.05203 |
| **LOC101928358** | 0.05032 | 0.04829 | 0.02088 | 0.02543 | 0.01587 | 0.00686 | 0.02323 | 0.02040 | 0.02430 | 0.00572 | 0.01414 | 0.06828 |
| **LOC101928363** | 0.00654 | 0.00000 | 0.14230 | 0.09049 | 0.38673 | 0.00000 | 0.00676 | 0.01368 | 0.20964 | 0.35135 | 0.13883 | 0.02609 |
| **LOC101928371** | 0.11761 | 0.01107 | 0.13638 | 0.00000 | 0.01476 | 0.06958 | 0.00000 | 0.33486 | 0.15487 | 0.15910 | 0.09146 | 0.05794 |
| **LOC101928372** | 0.01492 | 0.62677 | 0.00000 | 0.60796 | 0.02112 | 0.12845 | 0.00000 | 0.42106 | 0.00000 | 0.22549 | 0.05428 | 1.59800 |
| **LOC101928378** | 0.00243 | 0.03379 | 0.59051 | 0.40497 | 0.31541 | 0.00000 | 0.00000 | 0.04585 | 1.14783 | 2.43720 | 0.00000 | 0.04983 |
| **LOC101928386** | 0.00714 | 0.01715 | 0.01608 | 0.01197 | 0.02738 | 0.05432 | 0.00068 | 0.01722 | 0.03227 | 0.10216 | 0.06946 | 0.01067 |
| **LOC101928388** | 0.00933 | 0.13519 | 0.00000 | 0.00036 | 0.14609 | 0.07510 | 0.10736 | 0.00663 | 0.32883 | 0.00000 | 0.00000 | 0.06705 |
| **LOC101928401** | 0.04088 | 0.04208 | 0.03569 | 0.01283 | 0.00000 | 0.02712 | 0.02406 | 0.02186 | 0.00000 | 0.01998 | 0.06772 | 0.09829 |

| | | | | | | | | | | | | |
|---|---|---|---|---|---|---|---|---|---|---|---|---|
| **LOC101928404** | 0.01078 | 0.00385 | 0.08064 | 0.04611 | 0.01214 | 0.01952 | 0.03051 | 0.00425 | 0.00356 | 0.00000 | 0.01712 | 0.46450 |
| **LOC101928414** | 0.09425 | 0.03428 | 0.03161 | 0.01277 | 0.02154 | 0.00587 | 0.04821 | 0.01202 | 0.01798 | 0.00000 | 0.01490 | 0.11034 |
| **LOC101928437** | 0.12309 | 0.01631 | 0.05195 | 0.03489 | 0.01765 | 0.01005 | 0.02370 | 0.03067 | 0.01262 | 0.00898 | 0.02076 | 0.01550 |
| **LOC101928438** | 0.04483 | 0.01986 | 0.01207 | 0.01225 | 0.01154 | 0.00969 | 0.02495 | 0.01860 | 0.00977 | 0.04832 | 0.02186 | 0.04961 |
| **LOC101928440** | 0.03398 | 0.01481 | 0.01411 | 0.01188 | 0.01497 | 0.01927 | 0.01784 | 0.02213 | 0.01842 | 0.01296 | 0.01472 | 0.01454 |
| **LOC101928443** | 0.00571 | 0.01221 | 0.04100 | 0.00819 | 0.05264 | 0.00185 | 0.02306 | 0.00986 | 0.00967 | 0.03890 | 0.01488 | 0.04021 |
| **LOC101928445** | 0.03809 | 0.21334 | 0.08682 | 0.16181 | 0.06110 | 0.00000 | 0.06514 | 0.07797 | 0.06759 | 0.19953 | 0.09908 | 0.12546 |
| **LOC101928446** | 0.03999 | 0.01258 | 0.01403 | 0.01270 | 0.01391 | 0.01512 | 0.01731 | 0.03533 | 0.01511 | 0.04726 | 0.01833 | 0.01349 |
| **LOC101928453** | 0.00640 | 0.01654 | 0.04262 | 0.01597 | 0.01908 | 0.00952 | 0.01714 | 0.01058 | 0.01840 | 0.00960 | 0.01660 | 0.01717 |
| **LOC101928460** | 0.02047 | 0.11812 | 0.02122 | 0.00000 | 0.03854 | 0.00000 | 0.00702 | 0.00930 | 0.11850 | 0.06478 | 0.02175 | 0.04107 |
| **LOC101928470** | 0.01066 | 0.02327 | 0.01380 | 0.00997 | 0.01097 | 0.02732 | 0.01573 | 0.00912 | 0.01667 | 0.01779 | 0.01390 | 0.01611 |
| **LOC101928489** | 0.16648 | 0.01270 | 0.01763 | 0.03194 | 0.00961 | 0.01067 | 0.01879 | 0.03679 | 0.01161 | 0.03927 | 0.01515 | 0.02705 |
| **LOC101928491** | 0.00544 | 0.02110 | 0.00901 | 0.01265 | 0.02151 | 0.02292 | 0.03618 | 0.02866 | 0.02227 | 0.02457 | 0.06301 | 0.01846 |
| **LOC101928495** | 0.01123 | 0.04141 | 0.02962 | 0.02639 | 0.06119 | 0.03802 | 0.00940 | 0.00922 | 0.00000 | 0.02378 | 0.04302 | 0.01299 |
| **LOC101928496** | 0.03171 | 0.01363 | 0.02808 | 0.01108 | 0.01365 | 0.02611 | 0.01483 | 0.02373 | 0.01228 | 0.02934 | 0.01450 | 0.01596 |
| **LOC101928505** | 0.01177 | 0.06094 | 0.01133 | 0.02600 | 0.00000 | 0.04949 | 0.02937 | 0.01288 | 0.00552 | 0.01263 | 0.01499 | 0.01321 |
| **LOC101928514** | 0.09839 | 0.01651 | 0.01447 | 0.00537 | 0.00000 | 0.02631 | 0.02256 | 0.27197 | 0.03322 | 0.06012 | 0.02051 | 0.03974 |
| **LOC101928516** | 0.00897 | 0.01467 | 0.01571 | 0.01193 | 0.01404 | 0.01830 | 0.01851 | 0.01922 | 0.01428 | 0.01659 | 0.02142 | 0.01551 |
| **LOC101928517** | 0.75434 | 0.02114 | 0.01411 | 0.01230 | 0.01690 | 0.00240 | 0.01656 | 0.34194 | 0.02179 | 0.01748 | 0.01346 | 0.00755 |
| **LOC101928519** | 0.62740 | 0.01403 | 0.01435 | 0.01224 | 0.01649 | 0.01172 | 0.01885 | 0.01725 | 0.02266 | 0.00500 | 0.01377 | 0.01214 |
| **LOC101928523** | 0.08557 | 0.14170 | 0.02440 | 0.00000 | 0.00000 | 0.04419 | 0.06264 | 1.31035 | 0.00000 | 0.04378 | 0.00771 | 0.61968 |
| **LOC101928525** | 0.01673 | 0.91169 | 0.01871 | 0.45226 | 0.42680 | 0.27962 | 0.04321 | 0.00804 | 0.53535 | 0.00000 | 0.33473 | 0.50445 |
| **LOC101928530** | 0.00204 | 0.04388 | 0.15305 | 0.02033 | 0.01348 | 0.00000 | 0.00692 | 0.10484 | 0.09914 | 0.00000 | 0.02646 | 0.23437 |
| **LOC101928535** | 0.43846 | 0.02864 | 0.00626 | 0.00412 | 0.00060 | 0.04480 | 0.05944 | 0.00000 | 0.02568 | 0.01838 | 0.00797 | 0.08740 |
| **LOC101928537** | 0.62947 | 0.35748 | 0.00000 | 0.22994 | 0.00000 | 0.05321 | 0.00000 | 1.18186 | 0.00155 | 0.05741 | 0.01293 | 0.73142 |
| **LOC101928539** | 0.00582 | 0.01446 | 0.10238 | 0.01100 | 0.01813 | 0.00965 | 0.00867 | 0.01353 | 0.03105 | 0.02320 | 0.02935 | 0.02229 |
| **LOC101928545** | 0.12579 | 0.02490 | 0.02361 | 0.07559 | 0.01865 | 0.00216 | 0.02900 | 0.28208 | 0.03552 | 0.00000 | 0.02852 | 0.10010 |
| **LOC101928550** | 0.02200 | 0.01373 | 0.03218 | 0.01266 | 0.01979 | 0.01641 | 0.01909 | 0.01070 | 0.02958 | 0.01540 | 0.01503 | 0.01790 |
| **LOC101928559** | 5.40275 | 0.02086 | 0.01058 | 0.00372 | 0.01727 | 0.01655 | 0.01993 | 2.02254 | 0.01765 | 0.00000 | 0.01032 | 0.01898 |
| **LOC101928564** | 0.02557 | 0.03029 | 0.01659 | 0.02898 | 0.00860 | 0.00743 | 0.04495 | 0.00776 | 0.00542 | 0.09088 | 0.01154 | 0.01733 |
| **LOC101928567** | 0.19022 | 0.15449 | 0.02453 | 0.00000 | 0.00000 | 0.14656 | 0.03207 | 0.00044 | 0.10251 | 0.16106 | 0.01128 | 0.00882 |
| **LOC101928571** | 0.03212 | 0.09685 | 0.02600 | 0.02406 | 0.00658 | 0.02656 | 0.00791 | 0.05092 | 0.03847 | 0.01117 | 0.00927 | 0.22853 |
| **LOC101928580** | 0.01104 | 0.01418 | 0.01647 | 0.01329 | 0.01731 | 0.01365 | 0.01800 | 0.07278 | 0.02995 | 0.02256 | 0.01410 | 0.06005 |
| **LOC101928597.** | 0.04587 | 0.02570 | 0.02012 | 0.01912 | 0.01522 | 0.01017 | 0.03256 | 0.02718 | 0.04362 | 0.00623 | 0.01460 | 0.01723 |
| **LOC101928663** | 0.00104 | 0.05233 | 0.00634 | 0.11133 | 0.03791 | 0.39248 | 0.15127 | 0.06541 | 0.02475 | 0.21049 | 0.10387 | 0.14145 |
| **LOC101928674** | 0.00403 | 0.03727 | 0.20770 | 0.35072 | 0.28812 | 0.00000 | 0.20447 | 0.00871 | 0.00000 | 0.20381 | 0.01873 | 0.02570 |
| **LOC101928682** | 0.00142 | 0.00703 | 0.00000 | 0.00501 | 0.00000 | 0.01715 | 0.09089 | 0.06096 | 0.00000 | 0.22946 | 0.01213 | 0.01512 |
| **LOC101928694** | 2.50172 | 0.01380 | 0.01356 | 0.01015 | 0.03707 | 0.00000 | 0.01965 | 0.03392 | 0.02081 | 0.00379 | 0.01930 | 0.01689 |
| **LOC101928696** | 0.00628 | 0.01460 | 0.02138 | 0.01709 | 0.01553 | 0.02245 | 0.03551 | 0.01025 | 0.01169 | 0.00881 | 0.01855 | 0.01887 |
| **LOC101928700** | 0.07660 | 0.01231 | 0.02455 | 0.01706 | 0.01389 | 0.01068 | 0.01921 | 0.01425 | 0.00927 | 0.01870 | 0.03221 | 0.02801 |
| **LOC101928708** | 0.00629 | 0.01511 | 0.01835 | 0.01336 | 0.01434 | 0.01754 | 0.01700 | 0.01050 | 0.02018 | 0.01361 | 0.01783 | 0.01642 |
| **LOC101928710** | 0.10283 | 0.00893 | 0.11654 | 0.01942 | 0.00625 | 0.00000 | 0.10713 | 0.05728 | 0.00000 | 0.18052 | 0.03317 | 0.04114 |
| **LOC101928720** | 0.05116 | 0.01817 | 0.04845 | 0.01687 | 0.00932 | 0.02110 | 0.06654 | 0.01775 | 0.00000 | 0.02557 | 0.01183 | 0.01317 |
| **LOC101928731** | 0.00341 | 0.00350 | 0.04511 | 0.00002 | 0.03358 | 0.02365 | 0.00000 | 0.00772 | 0.00000 | 0.30185 | 0.10786 | 0.03921 |
| **LOC101928736** | 0.00352 | 0.07764 | 0.09778 | 0.14498 | 0.65611 | 0.00000 | 0.07097 | 10.11321 | 0.23516 | 0.00000 | 0.09312 | 5.83337 |
| **LOC101928737** | 0.02903 | 0.02261 | 0.01881 | 0.01092 | 0.04932 | 0.00000 | 0.02313 | 0.02629 | 0.01629 | 0.02265 | 0.03032 | 0.03865 |
| **LOC101928738** | 0.00781 | 0.01264 | 0.02808 | 0.01139 | 0.01534 | 0.01808 | 0.02355 | 0.01103 | 0.02064 | 0.01144 | 0.01564 | 0.03865 |
| **LOC101928766** | 0.00651 | 0.01967 | 0.03268 | 0.02233 | 0.03298 | 0.00073 | 0.02599 | 0.01057 | 0.01861 | 0.00000 | 0.01812 | 0.02023 |

| | | | | | | | | | | | | |
|---|---|---|---|---|---|---|---|---|---|---|---|---|
| **LOC101928767** | 0.01864 | 0.01978 | 0.01339 | 0.05709 | 0.00000 | 0.01776 | 0.00000 | 0.17167 | 0.02101 | 0.42156 | 0.00000 | 0.02844 |
| **LOC101928769** | 0.66455 | 0.03562 | 0.01209 | 0.00784 | 0.01436 | 0.00000 | 0.01610 | 3.34061 | 0.01824 | 0.01091 | 0.01352 | 0.00000 |
| **LOC101928775** | 1.41788 | 0.09735 | 0.02567 | 0.01312 | 0.01629 | 0.27021 | 0.00000 | 0.06691 | 0.00000 | 0.02661 | 0.04143 | 0.31715 |
| **LOC101928778** | 0.06970 | 0.02068 | 0.01700 | 0.01290 | 0.01821 | 0.00000 | 0.05026 | 0.01490 | 0.00096 | 0.04418 | 0.01548 | 0.01546 |
| **LOC101928782** | 0.20129 | 0.03441 | 0.01121 | 0.00744 | 0.00433 | 0.03858 | 0.01226 | 0.00242 | 0.01519 | 0.02395 | 0.01158 | 0.00980 |
| **LOC101928784** | 0.91633 | 0.34662 | 0.15155 | 0.00000 | 0.51436 | 0.23813 | 0.02648 | 0.23510 | 0.17247 | 0.11108 | 0.00863 | 0.72937 |
| **LOC101928786** | 0.00594 | 0.00724 | 0.03884 | 0.04129 | 0.03321 | 0.00000 | 0.01559 | 0.00514 | 0.01086 | 0.03571 | 0.01584 | 0.31863 |
| **LOC101928794** | 0.19340 | 0.04851 | 0.42232 | 0.01794 | 0.10470 | 0.00000 | 0.00436 | 0.00299 | 0.00000 | 0.00000 | 0.15688 | 0.06848 |
| **LOC101928796** | 0.14919 | 0.01006 | 0.11063 | 0.02114 | 0.10982 | 0.00000 | 0.00625 | 0.07823 | 0.02416 | 0.00000 | 0.08683 | 0.03307 |
| **LOC101928812** | 0.00583 | 0.01299 | 0.02623 | 0.01820 | 0.01762 | 0.00489 | 0.01872 | 0.01030 | 0.01799 | 0.01052 | 0.02594 | 0.06313 |
| **LOC101928823** | 1.11977 | 0.05011 | 0.00737 | 0.07009 | 0.00952 | 0.00784 | 0.01585 | 0.19507 | 0.03705 | 0.00000 | 0.00936 | 0.08976 |
| **LOC101928837** | 0.00588 | 0.01149 | 0.03494 | 0.02435 | 0.00170 | 0.03289 | 0.01584 | 0.01022 | 0.05087 | 0.02821 | 0.01571 | 0.01423 |
| **LOC101928844** | 0.12800 | 0.00000 | 0.00000 | 0.00518 | 0.03380 | 0.00000 | 0.00254 | 1.16875 | 0.01888 | 0.17540 | 0.00000 | 3.68137 |
| **LOC101928847** | 0.00624 | 0.01466 | 0.01348 | 0.01118 | 0.01628 | 0.02126 | 0.02668 | 0.01037 | 0.01295 | 0.01391 | 0.01578 | 0.01719 |
| **LOC101928855** | 0.00054 | 0.27863 | 0.06865 | 0.16726 | 0.00000 | 0.00000 | 0.00000 | 0.04182 | 0.03530 | 1.70321 | 0.44924 | 0.23821 |
| **LOC101928861** | 0.00563 | 0.00633 | 0.05898 | 0.00192 | 0.13305 | 0.02983 | 0.01304 | 0.01194 | 0.03327 | 0.03048 | 0.02964 | 0.02396 |
| **LOC101928865** | 0.00374 | 0.04236 | 0.20678 | 0.00000 | 0.02203 | 0.00000 | 0.01390 | 0.00929 | 0.13025 | 0.24625 | 0.06548 | 0.06293 |
| **LOC101928869** | 0.00614 | 0.01201 | 0.05532 | 0.00563 | 0.03008 | 0.03072 | 0.02339 | 0.01020 | 0.00963 | 0.01324 | 0.01445 | 0.01796 |
| **LOC101928882** | 0.00568 | 0.01287 | 0.01221 | 0.01716 | 0.01399 | 0.02554 | 0.01891 | 0.04758 | 0.01724 | 0.02327 | 0.01319 | 0.01305 |
| **LOC101928886** | 0.00582 | 0.01513 | 0.06452 | 0.00669 | 0.01234 | 0.03959 | 0.00000 | 0.01213 | 0.01675 | 0.03008 | 0.01101 | 0.02410 |
| **LOC101928891** | 0.00607 | 0.02134 | 0.01793 | 0.01911 | 0.03544 | 0.00948 | 0.01715 | 0.01389 | 0.02801 | 0.00330 | 0.01814 | 0.02367 |
| **LOC101928894** | 0.00195 | 1.10400 | 0.00000 | 0.00000 | 0.27071 | 0.00228 | 1.02996 | 0.17159 | 0.12743 | 0.06131 | 0.63901 | 0.00000 |
| **LOC101928896** | 0.01828 | 0.00000 | 0.09812 | 0.01669 | 0.09286 | 0.00000 | 0.00016 | 0.03647 | 0.09596 | 0.28201 | 0.00310 | 0.10000 |
| **LOC101928909** | 0.00735 | 0.00000 | 0.15979 | 0.01209 | 0.26152 | 0.00737 | 0.00000 | 0.00733 | 0.21942 | 0.58900 | 0.01049 | 0.11014 |
| **LOC101928911** | 0.00498 | 0.01592 | 0.01282 | 0.01369 | 0.01385 | 0.02424 | 0.01866 | 0.01842 | 0.01757 | 0.01405 | 0.01532 | 0.03212 |
| **LOC101928913** | 0.00316 | 0.04297 | 0.05870 | 0.03657 | 0.00000 | 0.38051 | 0.00000 | 0.04354 | 0.00000 | 0.02013 | 0.01522 | 0.03808 |
| **LOC101928936** | 0.00600 | 0.01813 | 0.01699 | 0.01327 | 0.01981 | 0.01340 | 0.01844 | 0.01129 | 0.01791 | 0.01280 | 0.01965 | 0.01799 |
| **LOC101928940** | 0.03394 | 0.06216 | 0.02421 | 0.02009 | 0.03779 | 0.00000 | 0.03257 | 0.04194 | 0.03674 | 0.01864 | 0.01302 | 0.01810 |
| **LOC101928943** | 0.01437 | 0.01227 | 0.01239 | 0.01246 | 0.01082 | 0.01584 | 0.01982 | 0.00938 | 0.04426 | 0.05951 | 0.01243 | 0.01368 |
| **LOC101928944** | 0.19164 | 0.01404 | 0.01624 | 0.01257 | 0.01621 | 0.01418 | 0.02179 | 0.00925 | 0.01745 | 0.01105 | 0.01453 | 0.01795 |
| **LOC101928989** | 0.24967 | 0.01529 | 0.01012 | 0.00867 | 0.01320 | 0.04181 | 0.05942 | 0.05752 | 0.00747 | 0.00000 | 0.01078 | 0.07961 |
| **LOC101928992** | 0.00528 | 0.01477 | 0.01566 | 0.01504 | 0.02036 | 0.01361 | 0.01760 | 0.01374 | 0.01707 | 0.01929 | 0.01483 | 0.01617 |
| **LOC101928994** | 0.02913 | 0.01626 | 0.14290 | 0.00972 | 0.01940 | 0.00637 | 0.01561 | 0.02550 | 0.00556 | 0.00624 | 0.04072 | 0.02097 |
| **LOC101929010** | 0.00267 | 0.00926 | 0.00927 | 0.03644 | 0.00000 | 0.02074 | 0.00156 | 0.05659 | 0.01809 | 0.12215 | 0.00925 | 0.00983 |
| **LOC101929025** | 0.00602 | 0.00630 | 0.01459 | 0.02587 | 0.02692 | 0.07100 | 0.01260 | 0.01009 | 0.01311 | 0.00720 | 0.01228 | 0.01373 |
| **LOC101929057** | 0.00448 | 0.04584 | 0.10673 | 0.08636 | 0.18748 | 0.00000 | 0.02590 | 0.10586 | 0.10195 | 0.24196 | 0.19492 | 0.12839 |
| **LOC101929059** | 0.01291 | 0.27042 | 0.08266 | 0.00556 | 0.14056 | 0.00000 | 0.04461 | 0.10465 | 0.37591 | 0.00000 | 0.00166 | 0.73545 |
| **LOC101929076** | 3.00923 | 0.01836 | 0.02240 | 0.00819 | 0.01074 | 0.01956 | 0.02655 | 0.05764 | 0.01113 | 0.00000 | 0.03940 | 0.07221 |
| **LOC101929089** | 0.01750 | 0.03003 | 0.01897 | 0.01589 | 0.01023 | 0.01678 | 0.00175 | 0.00817 | 0.04456 | 0.07978 | 0.01768 | 0.10643 |
| **LOC101929099** | 0.02072 | 0.02628 | 0.03279 | 0.03417 | 0.02078 | 0.02677 | 0.02330 | 0.02152 | 0.02972 | 0.00000 | 0.01557 | 0.07107 |
| **LOC101929106** | 0.03263 | 0.01485 | 0.01716 | 0.08268 | 0.00310 | 0.00000 | 0.01130 | 0.00932 | 0.01371 | 0.16125 | 0.04632 | 0.08168 |
| **LOC101929122** | 0.13581 | 0.01285 | 0.01380 | 0.01206 | 0.01540 | 0.01107 | 0.01649 | 0.35459 | 0.01935 | 0.01716 | 0.01469 | 0.03344 |
| **LOC101929124** | 0.03536 | 0.01266 | 0.02075 | 0.01746 | 0.01365 | 0.04801 | 0.01030 | 0.08679 | 0.00000 | 0.02370 | 0.06209 | 0.03438 |
| **LOC101929125** | 0.00861 | 0.00294 | 0.07816 | 0.01709 | 0.00000 | 0.01726 | 0.03391 | 0.10232 | 0.02909 | 0.15956 | 0.02015 | 0.12915 |
| **LOC101929140** | 0.02348 | 0.01701 | 0.13648 | 0.00942 | 0.04602 | 0.19982 | 0.12917 | 0.00299 | 0.29863 | 0.11116 | 0.19944 | 0.15893 |
| **LOC101929144** | 0.01199 | 0.00811 | 0.02225 | 0.02676 | 0.00209 | 0.02284 | 0.00710 | 0.03604 | 0.01288 | 0.10803 | 0.01041 | 0.01698 |
| **LOC101929153** | 0.00478 | 0.02417 | 0.05690 | 0.00632 | 0.00000 | 0.00000 | 0.02420 | 0.00950 | 0.07812 | 0.08774 | 0.24060 | 0.03087 |
| **LOC101929162** | 0.03975 | 0.03545 | 0.06235 | 0.01609 | 0.12735 | 0.09512 | 0.01449 | 0.02140 | 0.22973 | 0.30836 | 0.05870 | 0.05800 |

| | | | | | | | | | | | | |
|---|---|---|---|---|---|---|---|---|---|---|---|---|
| **LOC101929173** | 0.00195 | 0.01494 | 0.01249 | 0.01126 | 0.01493 | 0.01459 | 0.00948 | 0.17623 | 0.03193 | 0.02714 | 0.01520 | 0.00882 |
| **LOC101929181** | 0.00741 | 0.01873 | 0.03360 | 0.01709 | 0.02201 | 0.00023 | 0.06333 | 0.01700 | 0.00247 | 0.01133 | 0.03540 | 0.01780 |
| **LOC101929196** | 0.00429 | 0.01972 | 0.01014 | 0.01396 | 0.03853 | 0.04636 | 0.00000 | 0.10479 | 0.06965 | 0.01572 | 0.01150 | 0.05671 |
| **LOC101929218** | 0.00232 | 0.01847 | 0.01942 | 0.01550 | 0.01218 | 0.01372 | 0.01789 | 0.05939 | 0.01688 | 0.02397 | 0.01628 | 0.03578 |
| **LOC101929224** | 0.00489 | 0.02344 | 0.03368 | 0.01119 | 0.01131 | 0.00296 | 0.02502 | 0.02255 | 0.02253 | 0.04187 | 0.03370 | 0.05248 |
| **LOC101929225** | 0.00097 | 0.00000 | 0.07637 | 0.09453 | 0.11012 | 0.00000 | 0.12707 | 0.04682 | 0.26964 | 1.01313 | 0.01742 | 0.05307 |
| **LOC101929231** | 0.18149 | 0.05183 | 0.18802 | 0.09094 | 0.13412 | 0.11699 | 0.00711 | 0.45464 | 0.01606 | 0.00000 | 0.03269 | 0.07600 |
| **LOC101929234** | 0.06708 | 0.01876 | 0.03251 | 0.02575 | 0.04794 | 0.00000 | 0.04218 | 0.04957 | 0.03980 | 0.01381 | 0.01334 | 0.02491 |
| **LOC101929259** | 0.00599 | 0.01189 | 0.01877 | 0.01038 | 0.00000 | 0.07389 | 0.03339 | 0.01283 | 0.00000 | 0.05943 | 0.03156 | 0.01355 |
| **LOC101929260** | 0.00570 | 0.01244 | 0.03671 | 0.00659 | 0.00938 | 0.02677 | 0.02369 | 0.01034 | 0.00229 | 0.03930 | 0.02213 | 0.01411 |
| **LOC101929279** | 0.00579 | 0.01535 | 0.00866 | 0.01031 | 0.01428 | 0.02207 | 0.01936 | 0.01741 | 0.03913 | 0.02043 | 0.01175 | 0.01478 |
| **LOC101929282** | 0.00683 | 0.01528 | 0.01467 | 0.01323 | 0.01369 | 0.01647 | 0.01282 | 0.02367 | 0.02969 | 0.01856 | 0.01587 | 0.01502 |
| **LOC101929284** | 0.03917 | 0.01581 | 0.01403 | 0.01161 | 0.01459 | 0.02014 | 0.01764 | 0.01247 | 0.01847 | 0.01298 | 0.01452 | 0.01487 |
| **LOC101929295** | 0.32815 | 0.01581 | 0.06227 | 0.00774 | 0.08382 | 0.00000 | 0.01316 | 0.02733 | 0.01625 | 0.00587 | 0.01844 | 0.01304 |
| **LOC101929302** | 0.01908 | 0.01237 | 0.01243 | 0.02614 | 0.01593 | 0.02252 | 0.02358 | 0.00998 | 0.00483 | 0.01898 | 0.03876 | 0.01519 |
| **LOC101929319** | 0.09807 | 0.01425 | 0.01421 | 0.01160 | 0.01437 | 0.01731 | 0.01708 | 0.02500 | 0.02112 | 0.01606 | 0.01372 | 0.01766 |
| **LOC101929335** | 0.01932 | 0.00352 | 0.04487 | 0.02479 | 0.00000 | 0.01665 | 0.02244 | 0.00833 | 0.00000 | 0.07712 | 0.29236 | 0.01804 |
| **LOC101929340** | 0.00178 | 0.25894 | 0.11221 | 0.09082 | 0.00000 | 0.13913 | 0.08813 | 0.20913 | 0.02965 | 0.10766 | 0.02679 | 0.11091 |
| **LOC101929344** | 0.00538 | 0.06172 | 0.11878 | 1.83119 | 0.00000 | 0.02908 | 0.04244 | 0.00498 | 0.00000 | 0.01964 | 0.01355 | 0.14329 |
| **LOC101929372** | 0.00200 | 0.07926 | 0.07573 | 0.15986 | 0.00000 | 0.00872 | 0.02807 | 0.03500 | 0.12043 | 0.13985 | 0.01044 | 0.10676 |
| **LOC101929374** | 0.00484 | 0.23638 | 0.00849 | 0.02073 | 0.00000 | 0.02458 | 0.00645 | 0.00610 | 0.02388 | 0.07384 | 0.00687 | 0.17067 |
| **LOC101929380** | 0.15624 | 0.01582 | 0.05866 | 0.04440 | 0.01592 | 0.00381 | 0.01196 | 0.09876 | 0.02950 | 0.01223 | 0.01617 | 0.01643 |
| **LOC101929397** | 0.00302 | 0.37531 | 0.34598 | 0.00000 | 0.10524 | 0.00000 | 0.00000 | 3.26793 | 0.12973 | 0.61248 | 0.00000 | 1.98862 |
| **LOC101929412** | 1.97417 | 0.01107 | 0.01279 | 0.01141 | 0.00974 | 0.00463 | 0.03262 | 0.00365 | 0.03837 | 0.00000 | 0.04931 | 0.01424 |
| **LOC101929413** | 0.00846 | 0.01473 | 0.01127 | 0.04775 | 0.02078 | 0.01084 | 0.02085 | 0.01110 | 0.01951 | 0.01418 | 0.01531 | 0.02834 |
| **LOC101929420** | 0.02678 | 0.03821 | 0.02506 | 0.01476 | 0.02313 | 0.01111 | 0.01879 | 0.00868 | 0.01557 | 0.01405 | 0.01606 | 0.01962 |
| **LOC101929427** | 0.00590 | 0.00689 | 0.00470 | 0.02559 | 0.00002 | 0.07718 | 0.00000 | 0.00974 | 0.02871 | 0.02527 | 0.01052 | 0.01197 |
| **LOC101929431** | 0.00568 | 0.06250 | 0.06571 | 0.00647 | 0.00890 | 0.00000 | 0.00122 | 0.00922 | 0.04236 | 0.04089 | 0.01231 | 0.04860 |
| **LOC101929454** | 0.43119 | 0.01049 | 0.01311 | 0.01131 | 0.01280 | 0.02067 | 0.01729 | 0.28864 | 0.01883 | 0.01249 | 0.01309 | 0.01882 |
| **LOC101929486** | 0.04377 | 0.02879 | 0.01941 | 0.01442 | 0.01410 | 0.00872 | 0.01497 | 0.01169 | 0.02404 | 0.01715 | 0.01535 | 0.01448 |
| **LOC101929497** | 0.00834 | 0.02355 | 0.01666 | 0.01737 | 0.04577 | 0.00082 | 0.02939 | 0.01097 | 0.02335 | 0.00000 | 0.03219 | 0.01775 |
| **LOC101929505** | 0.02284 | 0.03469 | 0.01237 | 0.01391 | 0.01433 | 0.00004 | 0.01140 | 0.00831 | 0.08478 | 0.00333 | 0.01824 | 0.02153 |
| **LOC101929524** | 5.19784 | 0.00898 | 0.00802 | 0.00799 | 0.01344 | 0.03096 | 0.01404 | 0.65091 | 0.01963 | 0.00000 | 0.03333 | 0.13069 |
| **LOC101929526** | 0.00628 | 0.01509 | 0.01405 | 0.01264 | 0.01589 | 0.01858 | 0.01843 | 0.01048 | 0.01793 | 0.01287 | 0.01884 | 0.01634 |
| **LOC101929549** | 0.00000 | 1.00135 | 0.00000 | 2.74539 | 0.00000 | 0.28210 | 0.04070 | 0.03707 | 0.01799 | 0.29913 | 0.00000 | 1.04963 |
| **LOC101929555** | 0.00586 | 0.01369 | 0.03256 | 0.01408 | 0.01370 | 0.01938 | 0.01668 | 0.01945 | 0.01432 | 0.01866 | 0.01540 | 0.02030 |
| **LOC101929563** | 0.00260 | 0.03845 | 0.05362 | 0.04430 | 0.03690 | 0.02042 | 0.01892 | 0.03861 | 0.00000 | 0.10641 | 0.01013 | 0.01574 |
| **LOC101929567** | 0.00612 | 0.01233 | 0.00640 | 0.00839 | 0.00846 | 0.02005 | 0.00731 | 0.01043 | 0.03831 | 0.07427 | 0.00889 | 0.01381 |
| **LOC101929570** | 0.00944 | 0.01407 | 0.02740 | 0.07058 | 0.01157 | 0.00000 | 0.01310 | 0.02021 | 0.01665 | 0.05671 | 0.01422 | 0.01448 |
| **LOC101929574** | 0.01366 | 0.01869 | 0.01625 | 0.02770 | 0.08786 | 0.05268 | 0.02262 | 0.00587 | 0.00000 | 0.00000 | 0.01484 | 0.18133 |
| **LOC101929584** | 0.00638 | 0.01690 | 0.01117 | 0.01285 | 0.03189 | 0.01072 | 0.01192 | 0.01052 | 0.06192 | 0.01095 | 0.01312 | 0.01672 |
| **LOC101929591** | 0.00383 | 0.00904 | 0.00467 | 0.00533 | 0.00000 | 0.04064 | 0.00000 | 0.01171 | 0.03924 | 0.21249 | 0.01483 | 0.01377 |
| **LOC101929596** | 0.02295 | 0.02763 | 0.12242 | 0.00633 | 0.01722 | 0.01006 | 0.00049 | 0.02992 | 0.03754 | 0.04020 | 0.02244 | 0.01150 |
| **LOC101929607** | 0.00631 | 0.01382 | 0.01978 | 0.05779 | 0.00734 | 0.01077 | 0.01257 | 0.02030 | 0.01348 | 0.04266 | 0.01673 | 0.01432 |
| **LOC101929613** | 0.00315 | 0.01323 | 0.04783 | 0.00974 | 0.00916 | 0.03506 | 0.02264 | 0.02828 | 0.04813 | 0.00051 | 0.00936 | 0.01859 |
| **LOC101929631** | 0.01597 | 0.01380 | 0.02057 | 0.01113 | 0.01411 | 0.02126 | 0.01396 | 0.01587 | 0.01921 | 0.01595 | 0.01575 | 0.01805 |
| **LOC101929633** | 0.00315 | 0.11917 | 0.13401 | 0.00000 | 0.22497 | 0.14392 | 0.00000 | 0.03203 | 0.00000 | 0.00940 | 0.00272 | 0.00267 |
| **LOC101929645** | 0.01107 | 0.00348 | 0.05932 | 0.00048 | 0.00000 | 0.01612 | 0.00000 | 0.02520 | 0.09184 | 0.35390 | 0.03909 | 0.00689 |

| | | | | | | | | | | | | |
|---|---|---|---|---|---|---|---|---|---|---|---|---|
| **LOC101929653** | 0.19612 | 0.02698 | 0.00750 | 0.00854 | 0.00832 | 0.00216 | 0.11231 | 0.14315 | 0.00000 | 0.04773 | 0.01538 | 0.01033 |
| **LOC101929655** | 0.72192 | 0.03078 | 0.11148 | 0.00214 | 0.00000 | 0.00000 | 0.00986 | 0.06990 | 0.08652 | 0.08942 | 0.06612 | 0.08798 |
| **LOC101929657** | 0.01193 | 0.01932 | 0.12721 | 0.21727 | 0.03828 | 0.00000 | 0.04977 | 0.01219 | 0.03969 | 0.02271 | 0.02408 | 0.01600 |
| **LOC101929679** | 0.04732 | 0.02891 | 0.01327 | 0.08655 | 0.26688 | 0.00000 | 0.00655 | 0.01374 | 0.04488 | 0.01273 | 0.05312 | 0.03852 |
| **LOC101929680** | 0.01749 | 0.01440 | 0.01797 | 0.01310 | 0.01623 | 0.01078 | 0.02516 | 0.01871 | 0.01408 | 0.02679 | 0.01496 | 0.02515 |
| **LOC101929681** | 0.01063 | 0.01275 | 0.01988 | 0.00931 | 0.01113 | 0.02584 | 0.02487 | 0.01177 | 0.00851 | 0.03261 | 0.01536 | 0.01420 |
| **LOC101929684** | 0.00635 | 0.01477 | 0.03776 | 0.01050 | 0.01380 | 0.02122 | 0.01927 | 0.01045 | 0.01708 | 0.00855 | 0.02258 | 0.01656 |
| **LOC101929690** | 0.00748 | 0.01371 | 0.01876 | 0.01504 | 0.00305 | 0.02254 | 0.01508 | 0.02251 | 0.00000 | 0.02530 | 0.08703 | 0.01524 |
| **LOC101929694** | 0.00917 | 0.02180 | 0.02813 | 0.03055 | 0.00000 | 0.00000 | 0.00811 | 0.03433 | 0.05577 | 0.44176 | 0.03446 | 0.00918 |
| **LOC101929698** | 0.03418 | 0.01211 | 0.02177 | 0.03123 | 0.00977 | 0.03778 | 0.00868 | 0.01804 | 0.02277 | 0.01350 | 0.01592 | 0.01923 |
| **LOC101929715** | 1.47938 | 0.01881 | 0.08563 | 0.03798 | 0.00555 | 0.00000 | 0.00000 | 1.71626 | 0.02363 | 0.17279 | 0.00608 | 0.08403 |
| **LOC101929717** | 0.01677 | 0.01456 | 0.01743 | 0.01546 | 0.01969 | 0.01704 | 0.02304 | 0.01706 | 0.01560 | 0.01093 | 0.01525 | 0.01561 |
| **LOC101929718** | 0.02282 | 0.03474 | 0.11716 | 0.35773 | 0.04558 | 0.35699 | 0.00000 | 0.09671 | 0.29839 | 0.36078 | 0.30240 | 0.23260 |
| **LOC101929723** | 0.34590 | 0.01986 | 0.01726 | 0.02109 | 0.00547 | 0.02540 | 0.00529 | 0.00046 | 0.02737 | 0.15538 | 0.00496 | 0.03544 |
| **LOC101929733** | 0.03780 | 0.02727 | 0.03262 | 0.01246 | 0.01180 | 0.01949 | 0.01480 | 0.03954 | 0.01241 | 0.01541 | 0.01644 | 0.10671 |
| **LOC101929743** | 0.01665 | 0.08063 | 0.00000 | 0.75717 | 0.41578 | 0.00297 | 0.00071 | 0.00000 | 0.00000 | 0.38388 | 0.13261 | 0.44636 |
| **LOC101929754** | 0.06629 | 0.01328 | 0.01316 | 0.01080 | 0.01228 | 0.02265 | 0.01550 | 0.00715 | 0.01729 | 0.02117 | 0.01416 | 0.02602 |
| **LOC101929760** | 0.00595 | 0.01342 | 0.01226 | 0.01092 | 0.01016 | 0.02148 | 0.00793 | 0.01042 | 0.03485 | 0.03082 | 0.02585 | 0.01498 |
| **LOC101929765** | 0.00659 | 0.01574 | 0.01523 | 0.00893 | 0.00708 | 0.02074 | 0.01212 | 0.00969 | 0.01092 | 0.05227 | 0.02300 | 0.03828 |
| **LOC101929767** | 0.00052 | 0.14057 | 0.94354 | 4.17502 | 0.00000 | 0.12409 | 0.51442 | 0.01380 | 0.87378 | 0.80552 | 0.42956 | 0.34557 |
| **LOC101930452** | 0.49759 | 0.02920 | 0.10306 | 0.00000 | 0.00096 | 0.18623 | 0.00000 | 0.02484 | 0.29591 | 1.05268 | 0.00000 | 0.05729 |
| **LOC102060414** | 0.03906 | 0.01232 | 0.00870 | 0.06389 | 0.00601 | 0.00487 | 0.04171 | 0.05117 | 0.02337 | 0.07943 | 0.01333 | 0.02257 |
| **LOC102288414** | 0.11607 | 2.50987 | 3.78086 | 2.84508 | 4.23427 | 2.27883 | 4.07939 | 3.04851 | 5.05200 | 0.00000 | 2.63681 | 6.48562 |
| **LOC102467073** | 0.00201 | 0.06719 | 0.12411 | 0.01215 | 0.06733 | 0.18358 | 0.00000 | 0.10622 | 0.06890 | 0.00595 | 0.02545 | 0.32127 |
| **LOC102467213** | 0.12173 | 0.62245 | 0.10941 | 0.06115 | 0.00000 | 0.00000 | 0.01250 | 0.01074 | 0.03175 | 0.10778 | 0.01242 | 0.08228 |
| **LOC102467217** | 0.00278 | 0.01218 | 0.01225 | 0.00980 | 0.01082 | 0.02648 | 0.01414 | 0.03111 | 0.01456 | 0.03108 | 0.01441 | 0.02532 |
| **LOC102467223** | 0.23856 | 0.02454 | 0.03473 | 0.02558 | 0.00961 | 0.00000 | 0.02507 | 0.24777 | 0.00000 | 0.01338 | 0.06562 | 0.01608 |
| **LOC102467224** | 0.13354 | 0.01298 | 0.01196 | 0.03105 | 0.01201 | 0.01396 | 0.01730 | 0.04832 | 0.01808 | 0.01859 | 0.01468 | 0.01548 |
| **LOC102467225** | 0.12426 | 0.01418 | 0.01419 | 0.01254 | 0.01629 | 0.01247 | 0.02189 | 0.00464 | 0.01861 | 0.01390 | 0.01458 | 0.02103 |
| **LOC102467226** | 0.24804 | 0.01623 | 0.18465 | 0.01081 | 0.02294 | 0.00357 | 0.00639 | 0.00352 | 0.00616 | 0.00920 | 0.01856 | 0.01410 |
| **LOC102467655** | 0.01107 | 0.01484 | 0.01546 | 0.01176 | 0.01517 | 0.01785 | 0.01787 | 0.01418 | 0.01933 | 0.01531 | 0.01474 | 0.01654 |
| **LOC102503427** | 0.02802 | 0.01331 | 0.02375 | 0.00903 | 0.02190 | 0.01897 | 0.01994 | 0.03973 | 0.00825 | 0.02988 | 0.01447 | 0.01288 |
| **LOC102546226** | 0.00662 | 0.06847 | 0.01928 | 0.01151 | 0.02968 | 0.00000 | 0.01571 | 0.00756 | 0.05783 | 0.00000 | 0.06553 | 0.09760 |
| **LOC102546229** | 0.03980 | 0.82212 | 0.01809 | 0.34124 | 0.88599 | 0.19475 | 0.21968 | 0.07477 | 0.23213 | 0.00000 | 0.30622 | 0.33996 |
| **LOC102546294** | 0.00661 | 0.03004 | 0.13147 | 0.01379 | 0.01651 | 0.00000 | 0.00146 | 0.22698 | 0.02465 | 0.02336 | 0.04682 | 0.13075 |
| **LOC102546298** | 0.00481 | 0.09898 | 0.04386 | 0.07795 | 0.09242 | 0.00000 | 0.01836 | 0.13564 | 0.16401 | 0.00000 | 0.03724 | 0.01499 |
| **LOC102577426** | 0.00920 | 0.01189 | 0.01735 | 0.01736 | 0.02079 | 0.02972 | 0.01326 | 0.07661 | 0.04413 | 0.00105 | 0.01578 | 0.06104 |
| **LOC102606465** | 0.04169 | 0.08592 | 0.00000 | 0.00000 | 0.36041 | 0.76759 | 0.00000 | 0.08562 | 0.18773 | 0.10372 | 0.25492 | 0.24666 |
| **LOC102659288** | 1.53271 | 0.00697 | 0.03284 | 0.00985 | 0.02072 | 0.00000 | 0.04556 | 0.91748 | 0.00549 | 0.00000 | 0.01525 | 0.20409 |
| **LOC102723344** | 0.19370 | 0.01428 | 0.02863 | 0.01066 | 0.01483 | 0.01551 | 0.01698 | 0.01717 | 0.01604 | 0.01667 | 0.01471 | 0.01298 |
| **LOC102723354** | 0.01616 | 0.93007 | 0.19588 | 0.25662 | 0.03548 | 0.10461 | 0.00000 | 0.14129 | 0.00000 | 0.28133 | 0.00000 | 1.02730 |
| **LOC102723373** | 0.03991 | 0.02237 | 0.08636 | 0.00625 | 0.02975 | 0.03689 | 0.00689 | 0.00706 | 0.02847 | 0.00181 | 0.01348 | 0.01393 |
| **LOC102723385** | 0.01095 | 0.12967 | 0.01389 | 0.15985 | 0.08325 | 0.00168 | 0.15815 | 0.03275 | 0.00332 | 0.24609 | 0.06729 | 0.15180 |
| **LOC102723439** | 0.05534 | 0.00000 | 0.13300 | 0.00803 | 0.26787 | 0.01769 | 0.00000 | 0.02592 | 0.15786 | 0.30383 | 0.14035 | 0.00956 |
| **LOC102723483** | 0.00857 | 0.00434 | 0.11600 | 0.00122 | 0.05034 | 0.02135 | 0.00000 | 0.06466 | 0.01003 | 0.10053 | 0.01820 | 0.07114 |
| **LOC102723505** | 0.00395 | 0.01593 | 0.01416 | 0.01377 | 0.01626 | 0.01894 | 0.01807 | 0.07474 | 0.01892 | 0.01239 | 0.01438 | 0.01458 |
| **LOC102723510** | 0.07209 | 0.24041 | 0.21344 | 0.00000 | 0.04816 | 0.07736 | 0.00000 | 0.05795 | 0.39914 | 0.02926 | 0.01749 | 0.58833 |
| **LOC102723513** | 0.00561 | 0.02356 | 0.01989 | 0.01288 | 0.01585 | 0.01134 | 0.01772 | 0.01301 | 0.01698 | 0.01742 | 0.01550 | 0.03119 |

| | | | | | | | | | | | | |
|---|---|---|---|---|---|---|---|---|---|---|---|---|
| **LOC102723517** | 0.37358 | 0.03074 | 0.01112 | 0.00563 | 0.00000 | 0.05808 | 0.00000 | 0.23151 | 0.02828 | 0.03102 | 0.01027 | 0.00500 |
| **LOC102723544** | 0.07767 | 0.01383 | 0.01541 | 0.01105 | 0.01436 | 0.02598 | 0.02183 | 0.00715 | 0.02388 | 0.00524 | 0.01409 | 0.01626 |
| **LOC102723562** | 0.00728 | 0.01390 | 0.01271 | 0.01023 | 0.00906 | 0.01804 | 0.01130 | 0.02019 | 0.02766 | 0.04130 | 0.01543 | 0.02965 |
| **LOC102723631** | 0.01705 | 0.01155 | 0.01312 | 0.00955 | 0.01854 | 0.03030 | 0.01746 | 0.01765 | 0.01796 | 0.00780 | 0.01299 | 0.10680 |
| **LOC102723641** | 0.00604 | 0.01483 | 0.01266 | 0.01070 | 0.01233 | 0.02133 | 0.01800 | 0.01039 | 0.01878 | 0.02401 | 0.01569 | 0.01589 |
| **LOC102723703** | 0.00898 | 0.21196 | 0.18836 | 0.50752 | 0.75862 | 0.00000 | 0.00000 | 0.01932 | 0.28599 | 0.59147 | 0.00000 | 0.30583 |
| **LOC102723769** | 0.01121 | 0.00603 | 0.03201 | 0.01987 | 0.00266 | 0.06888 | 0.01104 | 0.00941 | 0.01782 | 0.16553 | 0.00757 | 0.02807 |
| **LOC102723809** | 0.00539 | 0.02170 | 0.01206 | 0.02768 | 0.00000 | 0.02614 | 0.02966 | 0.01021 | 0.03360 | 0.04746 | 0.01865 | 0.01576 |
| **LOC102723824** | 0.07802 | 0.01467 | 0.02844 | 0.01016 | 0.02038 | 0.01492 | 0.01813 | 0.00812 | 0.01407 | 0.01859 | 0.01569 | 0.01485 |
| **LOC102723854** | 0.78180 | 0.00000 | 0.27320 | 0.00089 | 0.24293 | 0.06695 | 0.27159 | 0.04921 | 0.76099 | 0.00000 | 0.00000 | 0.03330 |
| **LOC102723885** | 0.13840 | 0.47081 | 0.30131 | 0.25653 | 0.00000 | 0.17109 | 0.76427 | 0.04839 | 0.15170 | 0.00000 | 0.56867 | 0.08932 |
| **LOC102723927** | 0.00377 | 0.01465 | 0.02466 | 0.01123 | 0.01408 | 0.02163 | 0.01639 | 0.03858 | 0.01529 | 0.01324 | 0.01740 | 0.01498 |
| **LOC102724009** | 0.00894 | 0.11413 | 0.03696 | 0.08608 | 0.04523 | 0.00000 | 0.01029 | 0.00870 | 0.05840 | 0.09314 | 0.01495 | 0.03753 |
| **LOC102724020** | 0.00641 | 0.08268 | 0.03064 | 0.00365 | 0.00163 | 0.03624 | 0.00994 | 0.02109 | 0.03052 | 0.01738 | 0.01162 | 0.03718 |
| **LOC102724034.** | 0.01417 | 0.01569 | 0.01659 | 0.01341 | 0.01709 | 0.01587 | 0.01913 | 0.01040 | 0.01745 | 0.01238 | 0.01512 | 0.01767 |
| **LOC102724050** | 0.01276 | 0.00000 | 0.06538 | 0.06818 | 0.08282 | 0.00000 | 0.05361 | 0.06029 | 0.03758 | 0.68715 | 0.01312 | 0.60989 |
| **LOC102724058** | 0.02590 | 0.00492 | 0.00437 | 0.09040 | 0.00000 | 0.01077 | 0.00796 | 0.00565 | 0.01132 | 0.10470 | 0.01365 | 0.06076 |
| **LOC102724084** | 0.04128 | 0.02425 | 0.01688 | 0.01153 | 0.01270 | 0.01786 | 0.01742 | 0.01294 | 0.01707 | 0.01727 | 0.01532 | 0.01386 |
| **LOC102724094** | 0.00260 | 0.01426 | 0.01790 | 0.01177 | 0.01596 | 0.01556 | 0.01674 | 0.41601 | 0.01220 | 0.01440 | 0.02178 | 0.22044 |
| **LOC102724153** | 1.52209 | 0.00000 | 0.02655 | 0.03931 | 0.11965 | 0.03833 | 0.07138 | 0.36067 | 0.04444 | 0.01073 | 0.02117 | 1.60822 |
| **LOC102724163** | 0.00626 | 0.01464 | 0.01579 | 0.01797 | 0.01322 | 0.02170 | 0.01652 | 0.01044 | 0.01789 | 0.01409 | 0.01472 | 0.01600 |
| **LOC102724246** | 0.01148 | 0.01530 | 0.03574 | 0.01180 | 0.02069 | 0.00873 | 0.02439 | 0.01941 | 0.01800 | 0.00707 | 0.01488 | 0.03068 |
| **LOC102724301** | 0.01693 | 0.04765 | 0.29037 | 0.00457 | 0.05039 | 0.00128 | 0.03275 | 0.09073 | 0.05476 | 0.00000 | 0.02649 | 0.12995 |
| **LOC102724316** | 0.99959 | 2.21151 | 2.28835 | 2.56160 | 2.76332 | 3.18244 | 4.96071 | 0.01637 | 4.12107 | 0.00000 | 3.40641 | 0.65515 |
| **LOC102724321** | 0.01201 | 0.01534 | 0.01023 | 0.00915 | 0.00838 | 0.01955 | 0.01056 | 0.01898 | 0.02149 | 0.05237 | 0.01394 | 0.01573 |
| **LOC102724323** | 0.00644 | 0.02500 | 0.00950 | 0.08734 | 0.01979 | 0.01870 | 0.01733 | 0.01046 | 0.03459 | 0.00000 | 0.01311 | 0.01607 |
| **LOC102724467** | 0.03127 | 0.01548 | 0.06840 | 0.03961 | 0.02809 | 0.00000 | 0.00385 | 0.00607 | 0.06189 | 0.07040 | 0.02228 | 0.05243 |
| **LOC102724484** | 0.00494 | 0.01163 | 0.02591 | 0.01009 | 0.00052 | 0.01689 | 0.07382 | 0.00975 | 0.00000 | 0.07349 | 0.02982 | 0.02312 |
| **LOC102724508** | 0.07141 | 0.02048 | 0.01820 | 0.03011 | 0.05005 | 0.00000 | 0.04553 | 0.06412 | 0.01888 | 0.00000 | 0.01354 | 0.09573 |
| **LOC102724555** | 0.01530 | 0.01424 | 0.01348 | 0.01268 | 0.01501 | 0.01929 | 0.01761 | 0.03366 | 0.03052 | 0.00578 | 0.01291 | 0.03048 |
| **LOC102724589** | 0.00639 | 0.01438 | 0.01475 | 0.03882 | 0.02143 | 0.01516 | 0.02206 | 0.01044 | 0.01960 | 0.00552 | 0.02585 | 0.01729 |
| **LOC102724596** | 0.05890 | 0.01146 | 0.01794 | 0.01446 | 0.08158 | 0.00000 | 0.01605 | 0.00723 | 0.03064 | 0.03317 | 0.01017 | 0.06224 |
| **LOC102724601** | 0.00828 | 0.09958 | 0.05244 | 0.00000 | 0.00746 | 0.09998 | 0.00000 | 0.00450 | 0.00000 | 0.07631 | 0.00276 | 0.17684 |
| **LOC102724663** | 0.00296 | 0.01273 | 0.01330 | 0.01088 | 0.01277 | 0.02881 | 0.01695 | 0.03292 | 0.01635 | 0.01274 | 0.01446 | 0.02440 |
| **LOC102724691** | 0.01236 | 0.03107 | 0.02968 | 0.02063 | 0.07538 | 0.04481 | 0.00668 | 0.01388 | 0.02014 | 0.00352 | 0.03328 | 0.02313 |
| **LOC102724719** | 0.00589 | 0.01488 | 0.01519 | 0.06053 | 0.01380 | 0.00543 | 0.01610 | 0.01326 | 0.02059 | 0.02536 | 0.01578 | 0.01816 |
| **LOC102724784** | 0.00369 | 0.00552 | 0.00239 | 0.03980 | 0.23559 | 0.00000 | 0.00346 | 0.00625 | 0.00000 | 0.36450 | 0.08485 | 0.11929 |
| **LOC102724814** | 0.00239 | 0.03557 | 0.46732 | 0.75630 | 0.00000 | 0.02971 | 0.00061 | 0.03421 | 1.22859 | 0.31458 | 0.18665 | 0.06556 |
| **LOC102724849** | 0.01349 | 0.01531 | 0.07509 | 0.05276 | 0.01275 | 0.01058 | 0.02645 | 0.06928 | 0.00325 | 0.01421 | 0.01835 | 0.01312 |
| **LOC102724890** | 0.04422 | 0.01433 | 0.01452 | 0.01205 | 0.01533 | 0.01816 | 0.02131 | 0.06829 | 0.01850 | 0.00985 | 0.01460 | 0.01955 |
| **LOC102724927** | 0.05184 | 0.02228 | 0.01842 | 0.00264 | 0.03269 | 0.01918 | 0.01547 | 0.02491 | 0.00069 | 0.12797 | 0.02681 | 0.01522 |
| **LOC102724933** | 0.00619 | 0.01516 | 0.01581 | 0.02104 | 0.01375 | 0.01276 | 0.01698 | 0.01042 | 0.01800 | 0.01698 | 0.02257 | 0.01793 |
| **LOC102724957** | 0.16719 | 0.01461 | 0.01653 | 0.01173 | 0.01529 | 0.01650 | 0.01684 | 0.00866 | 0.02090 | 0.01150 | 0.01393 | 0.01368 |
| **LOC102800447** | 0.02054 | 0.01199 | 0.01600 | 0.01375 | 0.01493 | 0.01021 | 0.01916 | 0.00760 | 0.01481 | 0.01447 | 0.02611 | 0.10906 |
| **LOC103091866** | 0.00999 | 0.01120 | 0.01327 | 0.01106 | 0.01176 | 0.02616 | 0.03107 | 0.00866 | 0.00969 | 0.02049 | 0.01512 | 0.03548 |
| **LOC103344931** | 0.09873 | 0.43154 | 0.16791 | 0.14770 | 0.00000 | 0.00000 | 0.01253 | 0.00523 | 0.64320 | 0.38685 | 0.41953 | 0.02576 |
| **LOC113230** | 0.60251 | 0.00681 | 0.00468 | 0.00000 | 0.05494 | 0.05211 | 0.00000 | 0.60476 | 0.00000 | 0.33000 | 0.03282 | 0.17944 |
| **LOC143666** | 0.06383 | 0.01573 | 0.08571 | 0.00312 | 0.00618 | 0.02834 | 0.02423 | 0.00576 | 0.08040 | 0.03444 | 0.02663 | 0.05136 |

| | | | | | | | | | | | | |
|---|---|---|---|---|---|---|---|---|---|---|---|---|
| **LOC145474** | 0.16391 | 0.00000 | 0.15297 | 0.00000 | 0.16585 | 0.00000 | 0.00000 | 0.28712 | 0.31534 | 3.20746 | 0.00000 | 0.25299 |
| **LOC145783** | 0.26819 | 0.17927 | 0.27714 | 0.52164 | 0.07325 | 0.00000 | 0.16203 | 0.22167 | 0.49498 | 0.00000 | 0.32873 | 0.25115 |
| **LOC145837** | 1.47431 | 0.02007 | 0.05061 | 0.00000 | 0.20960 | 0.01635 | 0.00007 | 0.01617 | 0.00358 | 0.21990 | 0.01744 | 0.12143 |
| **LOC145845** | 0.00783 | 0.01368 | 0.01872 | 0.01140 | 0.01512 | 0.01803 | 0.01710 | 0.01160 | 0.02067 | 0.01797 | 0.01482 | 0.02172 |
| **LOC146513** | 0.00567 | 0.01465 | 0.02125 | 0.01452 | 0.05112 | 0.00000 | 0.02706 | 0.01048 | 0.01655 | 0.05051 | 0.02460 | 0.02218 |
| **LOC146880** | 0.01776 | 0.21907 | 0.49099 | 0.43910 | 0.07788 | 1.02184 | 0.51325 | 0.12274 | 0.00003 | 3.14450 | 0.04973 | 0.17986 |
| **LOC148413** | 0.36001 | 9.20437 | 3.87984 | 6.29112 | 2.86960 | 0.00000 | 5.90574 | 5.93287 | 3.70077 | 0.00000 | 7.20591 | 10.97389 |
| **LOC148696** | 0.00197 | 0.00000 | 0.01195 | 0.00000 | 0.00677 | 0.03094 | 0.00000 | 0.74461 | 0.02643 | 0.26025 | 0.00000 | 0.05072 |
| **LOC148709** | 0.88267 | 0.29652 | 1.07829 | 0.00000 | 0.63081 | 0.44413 | 0.19789 | 0.21722 | 1.07099 | 0.36701 | 0.71643 | 0.03079 |
| **LOC150776** | 0.52194 | 2.09164 | 0.68516 | 2.10587 | 1.92932 | 0.00000 | 1.09749 | 1.30450 | 3.95627 | 3.43491 | 0.54695 | 2.98659 |
| **LOC151121** | 0.00218 | 0.01321 | 0.08504 | 0.01842 | 0.00831 | 0.00002 | 0.08280 | 0.06499 | 0.02680 | 0.03844 | 0.01670 | 0.05204 |
| **LOC151174** | 0.00117 | 0.19512 | 0.00063 | 0.00000 | 0.00000 | 0.00000 | 0.06785 | 0.04529 | 0.00000 | 0.52923 | 0.69025 | 0.02127 |
| **LOC151475** | 0.00642 | 0.01817 | 0.16574 | 0.01328 | 0.03550 | 0.02149 | 0.00595 | 0.01042 | 0.01695 | 0.00304 | 0.02591 | 0.01628 |
| **LOC152225** | 0.55381 | 0.01222 | 0.02446 | 0.00835 | 0.00861 | 0.02456 | 0.01148 | 0.00167 | 0.01715 | 0.03334 | 0.01299 | 0.01655 |
| **LOC152578** | 0.00675 | 0.03782 | 0.01838 | 0.00372 | 0.01848 | 0.05891 | 0.02854 | 0.00890 | 0.01848 | 0.01673 | 0.01209 | 0.02829 |
| **LOC153684** | 0.00525 | 0.09210 | 0.01058 | 0.01010 | 0.08770 | 0.04418 | 0.01649 | 0.06030 | 0.00278 | 0.02575 | 0.01305 | 0.35527 |
| **LOC154761** | 0.00749 | 0.01385 | 0.02343 | 0.01148 | 0.01677 | 0.02281 | 0.01741 | 0.01059 | 0.01532 | 0.01261 | 0.01533 | 0.01978 |
| **LOC154872** | 0.05136 | 0.01111 | 0.01296 | 0.00680 | 0.07030 | 0.00001 | 0.02628 | 0.00730 | 0.00415 | 0.06980 | 0.02101 | 0.01398 |
| **LOC155060** | 0.03192 | 0.12014 | 0.18654 | 0.26917 | 0.12471 | 0.24616 | 0.02502 | 0.00664 | 0.06158 | 0.17730 | 0.05243 | 0.12096 |
| **LOC157273** | 0.00663 | 0.01047 | 0.01845 | 0.02025 | 0.00134 | 0.03775 | 0.00773 | 0.01282 | 0.04208 | 0.02605 | 0.02328 | 0.01832 |
| **LOC158434** | 0.01156 | 0.01031 | 0.01517 | 0.01209 | 0.01948 | 0.01852 | 0.01990 | 0.06784 | 0.02189 | 0.01344 | 0.01266 | 0.04967 |
| **LOC158435** | 0.00532 | 0.01117 | 0.00575 | 0.02380 | 0.00249 | 0.02143 | 0.01829 | 0.01028 | 0.03234 | 0.06806 | 0.01060 | 0.01385 |
| **LOC158960** | 0.07857 | 0.51118 | 0.22802 | 0.35737 | 0.94724 | 0.20734 | 1.06919 | 0.15577 | 0.00000 | 0.25712 | 0.90922 | 0.33314 |
| **LOC1720** | 0.00625 | 0.01544 | 0.01223 | 0.01092 | 0.01257 | 0.02354 | 0.01408 | 0.01027 | 0.04547 | 0.01828 | 0.01082 | 0.01881 |
| **LOC200726** | 0.01852 | 0.01321 | 0.02589 | 0.01704 | 0.00000 | 0.02798 | 0.02418 | 0.02164 | 0.00000 | 0.09308 | 0.01731 | 0.02022 |
| **LOC202181** | 0.00254 | 0.03079 | 0.15522 | 0.51268 | 0.41045 | 0.01875 | 0.00000 | 0.13170 | 0.39009 | 0.24287 | 0.25447 | 0.16968 |
| **LOC220729** | 0.35103 | 1.87867 | 0.65514 | 1.29378 | 1.47723 | 3.49229 | 0.00000 | 1.98143 | 2.14019 | 0.00000 | 0.99533 | 1.85427 |
| **LOC253044** | 0.00642 | 0.01293 | 0.04154 | 0.04839 | 0.04453 | 0.01363 | 0.01313 | 0.01046 | 0.01612 | 0.00202 | 0.01544 | 0.01760 |
| **LOC253573** | 0.00575 | 0.01958 | 0.01910 | 0.00830 | 0.01151 | 0.02393 | 0.01170 | 0.00956 | 0.01869 | 0.03329 | 0.02708 | 0.02510 |
| **LOC254028** | 0.00773 | 0.00811 | 0.14273 | 0.00487 | 0.00793 | 0.03842 | 0.04473 | 0.01155 | 0.00121 | 0.01223 | 0.01387 | 0.06088 |
| **LOC254896** | 0.00594 | 0.02372 | 0.03158 | 0.01439 | 0.00914 | 0.02211 | 0.01234 | 0.01038 | 0.02073 | 0.02915 | 0.01211 | 0.01523 |
| **LOC255187** | 0.00479 | 0.01646 | 0.14291 | 0.00336 | 0.00575 | 0.02544 | 0.00000 | 0.01015 | 0.02517 | 0.07276 | 0.00898 | 0.04787 |
| **LOC255654** | 0.02650 | 0.01144 | 0.01333 | 0.01176 | 0.01212 | 0.01773 | 0.01513 | 0.10562 | 0.01880 | 0.03101 | 0.01398 | 0.02760 |
| **LOC256880** | 0.05280 | 0.23027 | 0.14597 | 0.00000 | 0.00384 | 0.21256 | 0.00000 | 6.31977 | 0.26819 | 0.00420 | 0.03437 | 0.51430 |
| **LOC257396** | 0.83548 | 0.10213 | 0.26233 | 0.01345 | 0.01838 | 0.16656 | 0.00000 | 0.00000 | 0.00000 | 0.00000 | 0.30414 | 0.42807 |
| **LOC283038** | 0.01764 | 0.08930 | 0.03236 | 0.00000 | 0.00444 | 0.09995 | 0.02323 | 0.04119 | 0.01142 | 0.07874 | 0.04295 | 0.17653 |
| **LOC283045** | 0.00270 | 0.01323 | 0.01552 | 0.01067 | 0.01358 | 0.01721 | 0.01683 | 0.03931 | 0.01980 | 0.02892 | 0.01373 | 0.02612 |
| **LOC283070** | 0.18717 | 0.00997 | 0.12877 | 0.07544 | 0.09699 | 0.00000 | 0.01710 | 0.04118 | 0.00000 | 0.02154 | 0.01595 | 0.04996 |
| **LOC283177** | 0.00323 | 0.01333 | 0.01125 | 0.01182 | 0.01050 | 0.01949 | 0.01827 | 0.02469 | 0.01366 | 0.04397 | 0.01589 | 0.01571 |
| **LOC283194** | 0.00839 | 0.01587 | 0.02059 | 0.01211 | 0.01810 | 0.00474 | 0.01567 | 0.00956 | 0.01271 | 0.02324 | 0.06434 | 0.02551 |
| **LOC283214** | 0.00624 | 0.01458 | 0.01533 | 0.01096 | 0.01399 | 0.02115 | 0.03099 | 0.01040 | 0.01841 | 0.01241 | 0.01330 | 0.01681 |
| **LOC283299** | 10.00495 | 0.02329 | 0.37775 | 0.00929 | 0.09205 | 0.00000 | 0.00000 | 4.11760 | 0.06853 | 0.26419 | 0.00000 | 0.00000 |
| **LOC283335** | 0.00884 | 0.04319 | 0.12525 | 0.00000 | 0.31079 | 0.07486 | 0.00000 | 0.00590 | 0.12699 | 0.16451 | 0.28621 | 0.09030 |
| **LOC283352** | 0.39709 | 0.16019 | 0.00592 | 0.00000 | 0.24215 | 0.02413 | 0.00977 | 0.00056 | 0.00000 | 0.05553 | 0.03390 | 0.04640 |
| **LOC283440** | 0.10417 | 0.01471 | 0.03198 | 0.01073 | 0.01566 | 0.01642 | 0.01731 | 0.00366 | 0.01263 | 0.00944 | 0.02232 | 0.01520 |
| **LOC283575** | 0.06407 | 0.01455 | 0.01686 | 0.01511 | 0.01299 | 0.01010 | 0.01438 | 0.05005 | 0.01941 | 0.04972 | 0.02054 | 0.01249 |
| **LOC283585** | 0.01418 | 0.01277 | 0.01219 | 0.01022 | 0.01108 | 0.02377 | 0.01588 | 0.03226 | 0.02070 | 0.02858 | 0.01495 | 0.01703 |
| **LOC283683** | 0.11326 | 0.08494 | 0.14455 | 0.19508 | 0.04445 | 0.07274 | 0.00000 | 0.03075 | 0.00000 | 0.04623 | 0.04617 | 0.03824 |

| | | | | | | | | | | | | |
|---|---|---|---|---|---|---|---|---|---|---|---|---|
| **LOC283692** | 0.07346 | 0.29159 | 0.33590 | 0.13119 | 0.23346 | 0.38186 | 0.40394 | 0.48155 | 0.27545 | 0.09044 | 0.10427 | 0.33288 |
| **LOC283693** | 0.02579 | 0.07850 | 0.05118 | 0.00000 | 0.00000 | 0.06744 | 0.00000 | 0.11683 | 0.02168 | 0.22425 | 0.01871 | 0.08731 |
| **LOC283710** | 0.00753 | 0.01462 | 0.01413 | 0.00988 | 0.01282 | 0.00789 | 0.02479 | 0.01553 | 0.01919 | 0.05812 | 0.01643 | 0.01405 |
| **LOC283731** | 0.00580 | 0.01367 | 0.03757 | 0.01034 | 0.01696 | 0.01804 | 0.03933 | 0.01041 | 0.00010 | 0.03344 | 0.01570 | 0.01545 |
| **LOC283856** | 0.00285 | 0.00862 | 0.00480 | 0.06755 | 0.00000 | 0.04188 | 0.00639 | 0.02551 | 0.01236 | 0.05214 | 0.01168 | 0.02350 |
| **LOC283914** | 0.09683 | 0.01543 | 0.02237 | 0.01160 | 0.01555 | 0.01651 | 0.01889 | 0.02398 | 0.02088 | 0.00964 | 0.01481 | 0.01422 |
| **LOC283922** | 0.15501 | 0.22280 | 0.07265 | 0.03640 | 0.02085 | 0.00000 | 0.00812 | 0.07377 | 0.18685 | 0.00000 | 0.17497 | 0.34630 |
| **LOC284009** | 0.22933 | 0.04600 | 0.04342 | 0.20022 | 0.10019 | 0.00000 | 0.00438 | 0.05173 | 0.08842 | 0.19507 | 0.07474 | 0.30177 |
| **LOC284023** | 0.00269 | 0.32049 | 0.12344 | 0.38801 | 0.12251 | 0.00000 | 0.09407 | 2.83196 | 0.28857 | 0.78014 | 0.19535 | 0.90602 |
| **LOC284294** | 0.65753 | 0.03774 | 0.02083 | 0.00576 | 0.01890 | 0.01060 | 0.03232 | 0.05057 | 0.00661 | 0.04401 | 0.01297 | 0.01168 |
| **LOC284344** | 0.00557 | 0.03418 | 0.24179 | 0.00796 | 0.01543 | 0.00000 | 0.02439 | 0.01064 | 0.00000 | 0.04934 | 0.01583 | 0.01327 |
| **LOC284379** | 0.04119 | 0.01812 | 3.04931 | 0.16714 | 0.00366 | 0.03419 | 0.04696 | 0.06320 | 0.00015 | 0.00192 | 0.10057 | 0.02270 |
| **LOC284395** | 0.00738 | 0.01412 | 0.01700 | 0.01169 | 0.01386 | 0.01870 | 0.01840 | 0.01021 | 0.01535 | 0.01583 | 0.02165 | 0.02012 |
| **LOC284412** | 0.00155 | 0.01604 | 0.01147 | 0.13093 | 0.00000 | 0.00000 | 0.00108 | 0.03308 | 0.31108 | 0.41983 | 0.05598 | 0.04087 |
| **LOC284454** | 0.08227 | 0.03170 | 4.80894 | 0.17387 | 3.31850 | 0.00000 | 0.42573 | 0.02934 | 1.74193 | 0.35129 | 0.26368 | 0.12493 |
| **LOC284578** | 0.36615 | 0.02795 | 0.01210 | 0.11287 | 0.01723 | 0.00000 | 0.01603 | 0.00000 | 0.05254 | 0.00000 | 0.01087 | 0.16007 |
| **LOC284581** | 0.08536 | 0.44675 | 0.34883 | 0.21781 | 0.00000 | 0.00000 | 0.00572 | 0.00000 | 0.03485 | 0.28214 | 0.12682 | 0.50687 |
| **LOC284632** | 0.00788 | 0.03881 | 0.01556 | 0.00538 | 0.00681 | 0.04523 | 0.01455 | 0.01127 | 0.01328 | 0.01528 | 0.01262 | 0.01547 |
| **LOC284648** | 0.18664 | 0.01528 | 0.04980 | 0.01999 | 0.02704 | 0.01624 | 0.03168 | 0.00635 | 0.00181 | 0.00227 | 0.01810 | 0.01390 |
| **LOC284798** | 0.03762 | 0.01109 | 0.00908 | 0.02059 | 0.00654 | 0.06026 | 0.00982 | 0.01913 | 0.01353 | 0.01595 | 0.01142 | 0.01567 |
| **LOC284837** | 0.01199 | 0.05608 | 0.07436 | 0.00000 | 0.08590 | 0.00000 | 0.00000 | 0.00055 | 0.24238 | 1.60914 | 0.00000 | 0.23550 |
| **LOC284865** | 0.04157 | 0.01096 | 0.01520 | 0.01723 | 0.01087 | 0.01933 | 0.02576 | 0.00659 | 0.01078 | 0.01967 | 0.03548 | 0.03307 |
| **LOC284889** | 0.12580 | 13.23339 | 10.62434 | 23.34153 | 19.20651 | 0.00039 | 18.90429 | 3.62657 | 15.50069 | 0.00000 | 8.82912 | 10.93164 |
| **LOC284930** | 0.49816 | 0.14916 | 0.00000 | 0.45776 | 0.00000 | 0.26365 | 0.00000 | 0.15336 | 0.01974 | 0.10070 | 0.03493 | 0.21391 |
| **LOC284950** | 0.50381 | 0.01464 | 0.01502 | 0.01504 | 0.01952 | 0.00888 | 0.01957 | 0.00278 | 0.02313 | 0.00376 | 0.01432 | 0.01969 |
| **LOC285000** | 0.00612 | 0.01333 | 0.01047 | 0.01105 | 0.00817 | 0.03450 | 0.03761 | 0.01030 | 0.02366 | 0.01377 | 0.01192 | 0.01500 |
| **LOC285074** | 0.16277 | 0.01328 | 0.06396 | 0.09250 | 0.02590 | 0.06658 | 0.00335 | 0.18209 | 0.10196 | 0.34163 | 0.03045 | 0.03225 |
| **LOC285484** | 0.01569 | 0.06492 | 0.07265 | 0.01726 | 0.00799 | 0.00808 | 0.03063 | 0.01069 | 0.00880 | 0.02338 | 0.01777 | 0.01200 |
| **LOC285593** | 0.00635 | 0.01701 | 0.05003 | 0.01524 | 0.03171 | 0.00016 | 0.01333 | 0.00951 | 0.02296 | 0.00637 | 0.01574 | 0.03775 |
| **LOC285626** | 0.00668 | 0.06947 | 0.02310 | 0.01191 | 0.01426 | 0.00000 | 0.02002 | 0.01095 | 0.01929 | 0.02024 | 0.01584 | 0.01959 |
| **LOC285629** | 0.00646 | 0.01404 | 0.01728 | 0.01127 | 0.00834 | 0.01179 | 0.03662 | 0.01033 | 0.00000 | 0.04041 | 0.04276 | 0.01522 |
| **LOC285692** | 0.00614 | 0.01456 | 0.02512 | 0.01076 | 0.00973 | 0.01380 | 0.01820 | 0.01049 | 0.00970 | 0.01958 | 0.05724 | 0.01646 |
| **LOC285696** | 0.00357 | 0.06065 | 0.00000 | 0.00000 | 0.15090 | 0.16206 | 0.00000 | 0.00814 | 0.03865 | 0.14210 | 0.29531 | 0.01430 |
| **LOC285740** | 0.01908 | 0.01487 | 0.05124 | 0.02901 | 0.00552 | 0.00000 | 0.00959 | 0.03247 | 0.00000 | 0.04028 | 0.12901 | 0.01458 |
| **LOC285762** | 0.02802 | 0.01386 | 0.01390 | 0.01105 | 0.01996 | 0.01981 | 0.01707 | 0.01468 | 0.01718 | 0.01663 | 0.01398 | 0.01683 |
| **LOC285768** | 0.00661 | 0.01580 | 0.02655 | 0.01489 | 0.01548 | 0.01635 | 0.01246 | 0.01040 | 0.04199 | 0.00286 | 0.01644 | 0.01787 |
| **LOC285819** | 0.00547 | 0.01165 | 0.01159 | 0.00896 | 0.01683 | 0.02004 | 0.03671 | 0.01066 | 0.01333 | 0.04242 | 0.02805 | 0.01463 |
| **LOC285847** | 0.02800 | 0.01480 | 0.02259 | 0.01182 | 0.00701 | 0.03550 | 0.02502 | 0.01236 | 0.01801 | 0.01638 | 0.02188 | 0.02272 |
| **LOC285889** | 4.50061 | 0.00327 | 0.00749 | 0.00475 | 0.12208 | 0.00000 | 0.00292 | 0.45325 | 0.03484 | 0.03396 | 0.00349 | 0.03986 |
| **LOC286059** | 0.36845 | 0.01365 | 0.02670 | 0.01069 | 0.01544 | 0.01881 | 0.01102 | 0.00096 | 0.04056 | 0.00914 | 0.00986 | 0.01524 |
| **LOC286083** | 0.00211 | 0.01248 | 0.01487 | 0.00651 | 0.24491 | 0.02138 | 0.01699 | 0.04757 | 0.08287 | 0.03240 | 0.00532 | 0.05681 |
| **LOC286189** | 0.15031 | 0.01143 | 0.01489 | 0.01270 | 0.01399 | 0.02435 | 0.01687 | 0.00417 | 0.01863 | 0.00872 | 0.01396 | 0.02867 |
| **LOC286190** | 0.00414 | 0.00000 | 0.05586 | 0.05041 | 0.06960 | 0.17666 | 0.00000 | 0.00819 | 0.00000 | 0.40598 | 0.01443 | 0.00623 |
| **LOC286238** | 0.00422 | 0.01419 | 0.01242 | 0.01446 | 0.01309 | 0.03183 | 0.01432 | 0.02940 | 0.02024 | 0.01505 | 0.01327 | 0.05032 |
| **LOC286359** | 0.00538 | 0.02147 | 0.01367 | 0.01010 | 0.02001 | 0.02078 | 0.01838 | 0.07855 | 0.01543 | 0.02273 | 0.01414 | 0.03487 |
| **LOC286367** | 0.00582 | 0.00945 | 0.06499 | 0.09933 | 0.00000 | 0.01565 | 0.00700 | 0.00969 | 0.02000 | 0.03103 | 0.07309 | 0.02461 |
| **LOC286370** | 0.00659 | 0.00695 | 0.09614 | 0.00071 | 0.11271 | 0.00000 | 0.00930 | 0.01005 | 0.00000 | 0.10326 | 0.06827 | 0.01519 |
| **LOC286437** | 0.00105 | 0.00555 | 0.17142 | 0.00456 | 0.00000 | 0.00000 | 0.03129 | 1.10254 | 0.19002 | 0.81007 | 0.51831 | 0.06410 |

| | | | | | | | | | | | | |
|---|---|---|---|---|---|---|---|---|---|---|---|---|
| **LOC338963** | 0.01590 | 0.01714 | 0.04351 | 0.00903 | 0.01477 | 0.02151 | 0.01506 | 0.07647 | 0.01048 | 0.01258 | 0.01624 | 0.01260 |
| **LOC339059** | 0.00620 | 0.01776 | 0.01306 | 0.00662 | 0.02780 | 0.04014 | 0.01686 | 0.01134 | 0.02744 | 0.02178 | 0.01033 | 0.01963 |
| **LOC339166** | 2.19176 | 0.01794 | 0.01481 | 0.03653 | 0.01392 | 0.01794 | 0.02852 | 0.32041 | 0.07425 | 0.00000 | 0.00782 | 0.06897 |
| **LOC339298** | 0.00521 | 0.01573 | 0.00904 | 0.00728 | 0.01714 | 0.01316 | 0.01529 | 0.01035 | 0.02131 | 0.08557 | 0.00906 | 0.01382 |
| **LOC339529** | 0.03121 | 0.01715 | 0.04872 | 0.01119 | 0.01754 | 0.01293 | 0.01564 | 0.00894 | 0.03406 | 0.00149 | 0.02138 | 0.01690 |
| **LOC339539** | 0.04672 | 0.01628 | 0.04527 | 0.03401 | 0.00000 | 0.09524 | 0.03414 | 0.04376 | 0.00000 | 0.08228 | 0.08322 | 0.01137 |
| **LOC339568** | 0.00619 | 0.01934 | 0.01413 | 0.01097 | 0.01294 | 0.02263 | 0.01673 | 0.01052 | 0.01950 | 0.01684 | 0.01665 | 0.01520 |
| **LOC339593** | 0.00582 | 0.01230 | 0.02018 | 0.01784 | 0.00830 | 0.02897 | 0.02143 | 0.01052 | 0.01260 | 0.02795 | 0.01302 | 0.01464 |
| **LOC339622** | 0.00596 | 0.01128 | 0.01282 | 0.01667 | 0.03064 | 0.03000 | 0.01876 | 0.01056 | 0.01751 | 0.01860 | 0.01841 | 0.01948 |
| **LOC339666** | 0.02007 | 0.00205 | 0.03302 | 0.01351 | 0.05462 | 0.00000 | 0.07125 | 0.01001 | 0.03151 | 0.31253 | 0.00000 | 0.05288 |
| **LOC339803** | 0.84944 | 3.30777 | 3.86857 | 1.81732 | 1.85402 | 4.21154 | 1.78572 | 0.12773 | 2.51309 | 1.27888 | 4.22866 | 2.81010 |
| **LOC339807** | 0.00600 | 0.00562 | 0.02330 | 0.00270 | 0.11580 | 0.05368 | 0.01699 | 0.01004 | 0.00000 | 0.00838 | 0.01668 | 0.01457 |
| **LOC339862** | 0.04933 | 0.08443 | 0.08640 | 0.00000 | 0.08383 | 0.29052 | 0.04085 | 0.01708 | 0.00000 | 0.03632 | 0.01577 | 0.08742 |
| **LOC339874** | 0.01067 | 0.06722 | 0.05938 | 0.10846 | 0.00000 | 0.00000 | 0.10688 | 0.12326 | 0.00000 | 0.17617 | 0.03388 | 0.15169 |
| **LOC339975** | 0.02425 | 0.09610 | 0.11684 | 0.00000 | 0.10065 | 0.00000 | 0.00043 | 0.00614 | 0.25071 | 0.08932 | 0.01190 | 0.12831 |
| **LOC340017** | 0.01745 | 0.01646 | 0.02241 | 0.01153 | 0.01434 | 0.01789 | 0.01553 | 0.01217 | 0.01736 | 0.01746 | 0.01491 | 0.01526 |
| **LOC340074** | 0.00492 | 0.01120 | 0.01925 | 0.00545 | 0.00464 | 0.02061 | 0.00064 | 0.01024 | 0.03295 | 0.10370 | 0.01164 | 0.01214 |
| **LOC340113** | 0.01119 | 0.01671 | 0.01235 | 0.01164 | 0.01279 | 0.00516 | 0.00879 | 0.07766 | 0.07311 | 0.02919 | 0.00640 | 0.01203 |
| **LOC340357** | 0.11435 | 0.01495 | 0.01423 | 0.01199 | 0.01550 | 0.01658 | 0.01787 | 0.03694 | 0.01967 | 0.01153 | 0.01439 | 0.01324 |
| **LOC341056** | 0.00476 | 0.86377 | 0.94408 | 0.30257 | 0.12759 | 1.04130 | 1.24141 | 0.48296 | 1.10747 | 0.00000 | 0.79108 | 0.38982 |
| **LOC344887** | 0.00996 | 0.05992 | 0.02688 | 0.06335 | 0.01455 | 0.00000 | 0.01385 | 0.09583 | 0.02372 | 0.01092 | 0.01611 | 0.01065 |
| **LOC344967** | 0.05801 | 1.68097 | 0.98734 | 0.36278 | 1.19167 | 0.40805 | 0.86539 | 0.14472 | 1.68044 | 0.00000 | 0.66261 | 0.37888 |
| **LOC349160** | 0.06321 | 0.03091 | 0.88929 | 0.03598 | 0.00000 | 0.12404 | 0.00000 | 0.04343 | 0.00000 | 0.15864 | 0.00781 | 0.01606 |
| **LOC374443** | 2.63386 | 0.51022 | 1.53209 | 0.06742 | 1.07460 | 0.23279 | 0.37499 | 0.56111 | 0.93369 | 0.00000 | 0.47937 | 1.28153 |
| **LOC375196** | 0.06806 | 0.09032 | 0.04397 | 0.02720 | 0.06943 | 0.15068 | 0.00000 | 0.48729 | 0.01063 | 0.03843 | 0.01531 | 1.03848 |
| **LOC387810** | 0.02379 | 0.01277 | 0.00891 | 0.04839 | 0.00796 | 0.01870 | 0.01016 | 0.01084 | 0.04000 | 0.02817 | 0.01593 | 0.01540 |
| **LOC388436+LO** | 0.00278 | 0.02153 | 0.02572 | 0.02136 | 0.01605 | 0.01310 | 0.02124 | 0.03730 | 0.01442 | 0.00860 | 0.01816 | 0.06159 |
| **LOC388553** | 0.00219 | 0.34584 | 0.20284 | 0.02455 | 0.78625 | 0.00000 | 0.01604 | 0.19269 | 0.18868 | 0.00000 | 0.31246 | 0.74809 |
| **LOC388780** | 0.02889 | 0.25149 | 0.11548 | 0.00000 | 0.00000 | 0.08088 | 0.04178 | 0.14925 | 0.06903 | 0.00953 | 0.00000 | 0.29405 |
| **LOC388813** | 0.06613 | 0.01491 | 0.01761 | 0.01311 | 0.01444 | 0.01614 | 0.01721 | 0.29169 | 0.01880 | 0.01495 | 0.01461 | 0.00644 |
| **LOC388882** | 0.01936 | 0.11187 | 0.01268 | 0.06203 | 0.00178 | 0.01289 | 0.01049 | 0.04010 | 0.03086 | 0.01315 | 0.01325 | 0.03287 |
| **LOC388942** | 0.00606 | 0.01541 | 0.01667 | 0.01210 | 0.01579 | 0.01654 | 0.01816 | 0.04765 | 0.01822 | 0.01384 | 0.01504 | 0.01435 |
| **LOC389033** | 0.00549 | 0.03176 | 0.01161 | 0.00360 | 0.00185 | 0.04517 | 0.01845 | 0.00996 | 0.00867 | 0.03456 | 0.03147 | 0.01740 |
| **LOC389247** | 0.02253 | 0.22402 | 0.07784 | 0.01121 | 0.09871 | 0.18010 | 0.05111 | 0.50631 | 0.16233 | 0.00000 | 0.06267 | 0.26884 |
| **LOC389332** | 0.00426 | 0.04636 | 0.15778 | 0.06019 | 0.84194 | 0.08261 | 0.00000 | 0.01394 | 0.00000 | 0.05919 | 0.00427 | 0.00843 |
| **LOC389641** | 0.00692 | 0.08189 | 0.04354 | 0.00000 | 0.21757 | 0.00000 | 0.01320 | 0.01082 | 0.00000 | 0.08778 | 0.00831 | 0.00951 |
| **LOC389705** | 1.03915 | 0.00976 | 0.01725 | 0.00846 | 0.01704 | 0.02015 | 0.01705 | 0.23239 | 0.01936 | 0.03215 | 0.01293 | 0.01843 |
| **LOC389765** | 0.10837 | 0.05218 | 0.01211 | 0.11625 | 0.00000 | 0.13432 | 0.00001 | 0.01176 | 0.05534 | 0.24709 | 0.08659 | 0.14964 |
| **LOC389895** | 0.14633 | 0.01478 | 0.01332 | 0.01012 | 0.00871 | 0.01072 | 0.01665 | 0.45266 | 0.00534 | 0.03107 | 0.04831 | 0.13933 |
| **LOC389906** | 0.01532 | 0.00000 | 0.06015 | 0.09686 | 0.49269 | 0.28648 | 0.56542 | 0.17017 | 0.04026 | 1.94951 | 0.25429 | 0.25240 |
| **LOC391003.uid-** | 2.07038 | 1.07516 | 0.00000 | 0.00000 | 0.36068 | 0.00000 | 0.00000 | 3.48216 | 0.00000 | 0.11903 | 0.05009 | 4.49654 |
| **LOC391322** | 0.00911 | 0.06051 | 0.10917 | 0.06966 | 0.16135 | 0.11957 | 0.00000 | 0.08320 | 0.08950 | 0.66935 | 0.41610 | 0.06307 |
| **LOC392196** | 0.02552 | 0.01445 | 0.02894 | 0.01822 | 0.01248 | 0.02900 | 0.01124 | 0.00831 | 0.01746 | 0.01171 | 0.01482 | 0.01700 |
| **LOC392232** | 0.00798 | 0.01363 | 0.01301 | 0.00979 | 0.01951 | 0.02597 | 0.03208 | 0.01532 | 0.00913 | 0.02121 | 0.01465 | 0.01456 |
| **LOC392364** | 0.00565 | 0.01415 | 0.02038 | 0.00818 | 0.00769 | 0.03053 | 0.00575 | 0.01007 | 0.02495 | 0.04536 | 0.01158 | 0.01600 |
| **LOC399715** | 1.40770 | 0.21383 | 0.01760 | 0.34110 | 0.00000 | 0.00000 | 0.04034 | 1.55722 | 0.00457 | 0.10610 | 0.02319 | 0.34354 |
| **LOC399815** | 0.10036 | 0.73982 | 1.13048 | 0.00000 | 0.00000 | 1.84987 | 2.22350 | 0.00000 | 1.16586 | 0.75074 | 1.50426 | 0.41643 |
| **LOC399886** | 0.00634 | 0.01682 | 0.01319 | 0.01478 | 0.01618 | 0.01691 | 0.02544 | 0.01305 | 0.01642 | 0.01264 | 0.01553 | 0.01630 |

| | | | | | | | | | | | | |
|---|---|---|---|---|---|---|---|---|---|---|---|---|
| **LOC400043** | 0.00708 | 0.85715 | 0.00000 | 5.41051 | 0.67021 | 0.75119 | 0.25120 | 0.00000 | 0.01560 | 0.20536 | 0.00000 | 0.89976 |
| **LOC400511** | 0.00544 | 0.01178 | 0.02831 | 0.01741 | 0.01296 | 0.01429 | 0.03887 | 0.01027 | 0.00000 | 0.06610 | 0.01502 | 0.01505 |
| **LOC400548** | 0.00874 | 0.01577 | 0.01574 | 0.01638 | 0.01443 | 0.00848 | 0.02078 | 0.01084 | 0.00791 | 0.02295 | 0.05749 | 0.01838 |
| **LOC400620** | 0.21505 | 0.01386 | 0.02975 | 0.01487 | 0.01460 | 0.01401 | 0.01527 | 0.05420 | 0.01510 | 0.01548 | 0.01552 | 0.01302 |
| **LOC400655** | 0.43222 | 0.73045 | 3.30477 | 0.25342 | 1.21010 | 5.72727 | 1.34676 | 0.00000 | 5.58477 | 0.00000 | 1.02403 | 1.00323 |
| **LOC400661** | 0.09680 | 0.01504 | 0.01013 | 0.00593 | 0.00043 | 0.03425 | 0.00637 | 0.06629 | 0.00925 | 0.09634 | 0.01120 | 0.03567 |
| **LOC400684** | 0.01328 | 0.21977 | 0.16492 | 0.00119 | 0.04430 | 0.03992 | 0.10398 | 0.02033 | 0.01700 | 0.00000 | 0.03453 | 0.33694 |
| **LOC400685** | 0.00609 | 0.01700 | 0.02276 | 0.01474 | 0.02677 | 0.00624 | 0.02046 | 0.01062 | 0.01433 | 0.02264 | 0.01488 | 0.01647 |
| **LOC400736+PR** | 0.07357 | 0.01265 | 0.01302 | 0.00979 | 0.02683 | 0.01570 | 0.02524 | 0.03564 | 0.01036 | 0.02439 | 0.01359 | 0.01656 |
| **LOC400736+PR** | 0.66396 | 0.03539 | 0.01323 | 0.00875 | 0.01612 | 0.00000 | 0.01629 | 1.63735 | 0.01991 | 0.01177 | 0.01413 | 0.00853 |
| **LOC400794** | 0.00494 | 0.03753 | 0.15428 | 0.06133 | 0.00091 | 0.01464 | 0.00893 | 0.01319 | 0.00253 | 0.05348 | 0.01893 | 0.01437 |
| **LOC400863** | 0.00320 | 0.15259 | 0.02537 | 0.00356 | 0.00728 | 0.01036 | 0.02597 | 0.02414 | 0.19515 | 0.00031 | 0.00000 | 0.16532 |
| **LOC400867** | 0.80405 | 0.01003 | 0.01565 | 0.00581 | 0.12734 | 0.00000 | 0.05559 | 0.00330 | 0.00000 | 0.05428 | 0.00607 | 0.01538 |
| **LOC400891** | 0.05553 | 0.04605 | 0.01175 | 0.05991 | 0.01301 | 0.00000 | 0.01015 | 0.00742 | 0.11287 | 0.07143 | 0.03521 | 0.01237 |
| **LOC400927** | 0.04437 | 0.10471 | 0.05557 | 0.09431 | 0.06758 | 0.00000 | 0.03190 | 0.15557 | 0.06001 | 0.00000 | 0.03432 | 0.19596 |
| **LOC400958** | 0.00633 | 0.01335 | 0.02235 | 0.03143 | 0.04584 | 0.00002 | 0.04880 | 0.00871 | 0.00558 | 0.00227 | 0.01672 | 0.06056 |
| **LOC400997** | 0.00208 | 0.00000 | 0.00000 | 0.00000 | 0.00779 | 0.10027 | 0.00000 | 0.00824 | 0.29973 | 0.52333 | 0.05705 | 0.00133 |
| **LOC401010** | 0.31947 | 2.38674 | 1.47414 | 1.22588 | 1.61745 | 0.00000 | 1.34621 | 0.82499 | 1.94424 | 0.41733 | 1.85350 | 1.92629 |
| **LOC401052** | 0.00564 | 0.00222 | 0.00591 | 0.09569 | 0.13683 | 0.00000 | 0.05895 | 0.03683 | 0.04896 | 0.15950 | 0.01392 | 0.03645 |
| **LOC401127** | 0.52039 | 2.18633 | 0.80833 | 1.11862 | 1.24319 | 1.73559 | 0.00000 | 0.44985 | 1.26086 | 0.00000 | 1.36066 | 1.51089 |
| **LOC401134** | 0.01297 | 0.01727 | 0.04552 | 0.01865 | 0.02889 | 0.00000 | 0.01894 | 0.01170 | 0.01316 | 0.00778 | 0.01720 | 0.01767 |
| **LOC401177** | 0.04276 | 0.01482 | 0.01615 | 0.01399 | 0.01876 | 0.00988 | 0.01504 | 0.00669 | 0.02087 | 0.02142 | 0.02759 | 0.02427 |
| **LOC401242** | 0.00121 | 0.01990 | 1.14612 | 0.02833 | 0.00000 | 0.27484 | 0.00000 | 0.10308 | 0.00000 | 0.14253 | 0.00075 | 0.03699 |
| **LOC401286** | 0.00627 | 0.01472 | 0.06110 | 0.02274 | 0.01404 | 0.02058 | 0.01288 | 0.01046 | 0.01454 | 0.01183 | 0.01485 | 0.01615 |
| **LOC401312** | 0.00586 | 0.73017 | 0.57310 | 0.00000 | 0.46959 | 0.00000 | 0.00000 | 0.00048 | 0.70817 | 0.37648 | 0.81945 | 0.63890 |
| **LOC401320** | 0.12606 | 0.33106 | 0.36125 | 0.03609 | 0.26314 | 0.00000 | 0.08526 | 0.04715 | 0.17509 | 0.15401 | 0.18433 | 0.65262 |
| **LOC401324** | 0.00874 | 0.00919 | 0.00595 | 0.00500 | 0.00053 | 0.03111 | 0.01577 | 0.01664 | 0.01488 | 0.10580 | 0.01187 | 0.01502 |
| **LOC401463** | 0.01004 | 0.02834 | 0.02347 | 0.00448 | 0.01013 | 0.01872 | 0.02589 | 0.01059 | 0.00897 | 0.06183 | 0.01929 | 0.08888 |
| **LOC401557** | 0.17073 | 0.01817 | 0.02361 | 0.00890 | 0.01084 | 0.01622 | 0.01280 | 0.00350 | 0.01836 | 0.04085 | 0.01381 | 0.01370 |
| **LOC402160** | 0.04345 | 0.01491 | 0.01512 | 0.01207 | 0.01546 | 0.01839 | 0.01839 | 0.02530 | 0.01834 | 0.01515 | 0.01695 | 0.01451 |
| **LOC403323** | 0.06722 | 0.06781 | 0.29631 | 0.01746 | 0.00000 | 0.00000 | 0.10979 | 0.00958 | 0.00000 | 0.13019 | 0.28132 | 0.01035 |
| **LOC407835** | 0.12239 | 0.29134 | 0.77904 | 0.57899 | 1.05694 | 0.00000 | 1.61383 | 0.29699 | 0.00001 | 1.74903 | 0.39688 | 0.24994 |
| **LOC414300** | 0.05706 | 0.01149 | 0.04363 | 0.00658 | 0.00848 | 0.01681 | 0.00026 | 0.08663 | 0.02100 | 0.07772 | 0.01308 | 0.02362 |
| **LOC439933** | 0.00917 | 0.01487 | 0.01557 | 0.01088 | 0.01310 | 0.01809 | 0.02483 | 0.01012 | 0.01470 | 0.02685 | 0.01401 | 0.01783 |
| **LOC439994** | 0.22118 | 0.14406 | 0.10306 | 0.08326 | 0.07731 | 0.00000 | 0.03682 | 0.22553 | 0.14044 | 0.06022 | 0.05100 | 0.21957 |
| **LOC440028** | 0.62399 | 1.04995 | 0.03736 | 0.00000 | 0.13987 | 0.00000 | 0.09369 | 0.00000 | 0.00000 | 0.09476 | 0.13596 | 1.04867 |
| **LOC440040** | 0.08910 | 0.01317 | 0.01408 | 0.01345 | 0.01755 | 0.01278 | 0.02853 | 0.04032 | 0.01709 | 0.00491 | 0.01506 | 0.04027 |
| **LOC440173** | 0.06899 | 0.01198 | 0.06852 | 0.04698 | 0.01628 | 0.03172 | 0.00000 | 0.07774 | 0.04259 | 0.03643 | 0.00687 | 0.10787 |
| **LOC440300** | 0.02059 | 0.01313 | 0.01179 | 0.00893 | 0.01971 | 0.03113 | 0.01486 | 0.00959 | 0.02039 | 0.02057 | 0.01162 | 0.01550 |
| **LOC440311** | 0.28030 | 0.48023 | 0.93573 | 0.42267 | 1.32503 | 0.00000 | 0.70343 | 0.43573 | 0.00000 | 3.07037 | 0.11578 | 0.41021 |
| **LOC440416** | 0.00129 | 0.23049 | 1.05403 | 0.38978 | 0.00000 | 0.06540 | 2.40974 | 0.69337 | 0.24241 | 1.08056 | 0.06073 | 1.09675 |
| **LOC440434** | 0.01201 | 0.38322 | 0.37435 | 0.68091 | 0.00000 | 2.18100 | 0.00000 | 0.94298 | 1.79646 | 1.61655 | 0.66439 | 0.95581 |
| **LOC440446** | 0.01216 | 0.12471 | 0.00840 | 0.01035 | 0.00424 | 0.03874 | 0.00708 | 0.02140 | 0.05440 | 0.01655 | 0.00650 | 0.00580 |
| **LOC440461** | 0.00479 | 0.01103 | 0.04314 | 0.00470 | 0.00000 | 0.02307 | 0.04124 | 0.01024 | 0.01511 | 0.10011 | 0.01431 | 0.01183 |
| **LOC440600** | 0.16253 | 0.51594 | 0.03628 | 0.12705 | 0.35288 | 0.00000 | 0.45699 | 0.14996 | 0.35101 | 0.00000 | 0.24900 | 0.83612 |
| **LOC440602** | 0.00510 | 0.01462 | 0.01713 | 0.01547 | 0.01282 | 0.01827 | 0.01548 | 0.01481 | 0.02714 | 0.01738 | 0.01321 | 0.01593 |
| **LOC440700** | 0.00449 | 0.01479 | 0.01782 | 0.01079 | 0.00947 | 0.01136 | 0.04967 | 0.12275 | 0.01965 | 0.02451 | 0.02601 | 0.02588 |
| **LOC440896** | 0.00616 | 0.02252 | 0.01515 | 0.01598 | 0.02787 | 0.00303 | 0.02696 | 0.01158 | 0.02232 | 0.00570 | 0.01709 | 0.01732 |

| | | | | | | | | | | | | |
|---|---|---|---|---|---|---|---|---|---|---|---|---|
| **LOC440910** | 0.03334 | 0.00409 | 0.01482 | 0.00000 | 0.02561 | 0.01897 | 0.00319 | 0.02276 | 0.03022 | 0.30846 | 0.00925 | 0.04458 |
| **LOC441081** | 0.01875 | 0.01430 | 0.01853 | 0.01190 | 0.01447 | 0.02151 | 0.01545 | 0.01169 | 0.01704 | 0.02161 | 0.01464 | 0.01488 |
| **LOC441155** | 0.00756 | 0.01403 | 0.01479 | 0.01228 | 0.01651 | 0.01891 | 0.01811 | 0.01134 | 0.02083 | 0.01474 | 0.01406 | 0.01922 |
| **LOC441204** | 0.00229 | 0.01447 | 0.03144 | 0.01252 | 0.01143 | 0.01522 | 0.02214 | 0.17679 | 0.02262 | 0.03226 | 0.01583 | 0.00828 |
| **LOC441242** | 0.01050 | 0.23571 | 0.31593 | 0.24696 | 0.45424 | 0.00000 | 0.00000 | 0.05151 | 0.09466 | 0.01764 | 0.24459 | 0.10900 |
| **LOC441454** | 0.02885 | 0.01213 | 0.01572 | 0.03271 | 0.02747 | 0.08111 | 0.09721 | 0.02315 | 0.02539 | 0.12240 | 0.05145 | 0.09841 |
| **LOC441455** | 0.93202 | 0.32500 | 0.15815 | 0.49139 | 0.01980 | 0.00000 | 0.01802 | 0.70117 | 0.15239 | 0.10671 | 0.04757 | 2.54016 |
| **LOC441461** | 0.01745 | 0.10159 | 0.14328 | 0.00619 | 0.00422 | 0.00000 | 0.04352 | 0.03989 | 0.01769 | 0.00459 | 0.05911 | 0.02599 |
| **LOC441601** | 1.00073 | 2.27838 | 3.10408 | 0.32713 | 2.15976 | 5.55725 | 2.91225 | 1.99653 | 2.68912 | 0.00000 | 5.29234 | 2.76161 |
| **LOC441666** | 0.00556 | 0.10094 | 1.23825 | 0.08920 | 0.28631 | 0.00115 | 0.00000 | 0.05460 | 1.76929 | 0.06416 | 0.11167 | 0.16450 |
| **LOC442028** | 0.00277 | 0.01591 | 0.01159 | 0.15560 | 0.01029 | 0.00000 | 0.01507 | 0.11081 | 0.04193 | 0.03798 | 0.01858 | 0.16011 |
| **LOC442132** | 0.05878 | 0.01584 | 0.02548 | 0.01423 | 0.02485 | 0.00559 | 0.02545 | 0.03560 | 0.01729 | 0.01622 | 0.01404 | 0.02007 |
| **LOC442497** | 0.03276 | 0.00794 | 0.01696 | 0.00426 | 0.00000 | 0.01315 | 0.06933 | 0.00691 | 0.00000 | 0.26365 | 0.00137 | 0.00882 |
| **LOC493754** | 0.41888 | 5.55141 | 3.48509 | 5.34966 | 4.00906 | 0.00000 | 0.50253 | 18.31873 | 5.20147 | 4.48086 | 1.59233 | 7.52763 |
| **LOC494127** | 0.15054 | 0.80115 | 0.49391 | 0.48279 | 0.32022 | 0.76596 | 0.28314 | 0.37575 | 0.21378 | 0.00000 | 0.38520 | 0.45940 |
| **LOC494141** | 0.01452 | 0.01212 | 0.01380 | 0.01086 | 0.01252 | 0.03759 | 0.02357 | 0.01119 | 0.01643 | 0.00748 | 0.01743 | 0.03761 |
| **LOC54944** | 0.08935 | 0.01248 | 0.03348 | 0.03073 | 0.01550 | 0.00000 | 0.01831 | 0.00787 | 0.16330 | 0.16398 | 0.00000 | 0.09542 |
| **LOC553103** | 0.10221 | 0.10327 | 0.01499 | 0.02315 | 0.01477 | 0.08656 | 0.00000 | 0.00150 | 0.03047 | 0.02698 | 0.08806 | 0.08363 |
| **LOC554206** | 0.13048 | 0.14916 | 0.23694 | 0.00219 | 0.00258 | 0.03431 | 0.00830 | 0.00731 | 0.10532 | 0.00000 | 0.07749 | 0.04280 |
| **LOC574538** | 0.00665 | 0.01398 | 0.01292 | 0.01264 | 0.01343 | 0.02170 | 0.02314 | 0.01131 | 0.01626 | 0.01620 | 0.01433 | 0.01725 |
| **LOC613037** | 0.00410 | 0.01051 | 0.01269 | 0.08725 | 0.00000 | 0.05183 | 0.01497 | 0.02357 | 0.03076 | 0.12955 | 0.01794 | 0.01341 |
| **LOC63930** | 0.01274 | 0.03951 | 0.03204 | 0.01670 | 0.02590 | 0.00000 | 0.02230 | 0.01697 | 0.01697 | 0.00000 | 0.01829 | 0.06255 |
| **LOC641367** | 0.32453 | 0.16065 | 0.15808 | 0.30178 | 0.20073 | 0.45801 | 0.00000 | 0.13709 | 0.02912 | 0.23633 | 0.00371 | 0.44870 |
| **LOC641515** | 0.00208 | 0.05642 | 0.00714 | 0.06428 | 0.23282 | 0.28902 | 0.00000 | 0.02477 | 0.00000 | 0.03943 | 0.06520 | 0.00092 |
| **LOC641746** | 0.28726 | 0.52920 | 0.43568 | 0.08501 | 0.35548 | 0.43013 | 0.00014 | 0.59334 | 0.05978 | 1.63901 | 0.12986 | 0.55290 |
| **LOC642236** | 0.11233 | 0.12448 | 0.09670 | 0.14803 | 0.06636 | 0.85219 | 0.13215 | 0.22097 | 0.00000 | 1.10846 | 0.05254 | 0.20398 |
| **LOC642361** | 0.26341 | 0.14800 | 0.01645 | 0.02205 | 0.06890 | 0.18995 | 0.00000 | 0.44212 | 0.06410 | 0.09579 | 0.00703 | 0.22131 |
| **LOC642366** | 0.02170 | 0.00000 | 0.02849 | 0.00001 | 0.00000 | 0.05863 | 0.00000 | 0.06314 | 0.00000 | 0.11960 | 0.10100 | 0.07006 |
| **LOC642757** | 0.01036 | 0.00930 | 0.07450 | 0.02755 | 0.06745 | 0.04392 | 0.02459 | 0.01664 | 0.00000 | 0.41717 | 0.00000 | 0.02137 |
| **LOC642846** | 0.09066 | 0.38248 | 0.49063 | 0.28166 | 0.19397 | 0.00000 | 0.00000 | 0.04772 | 0.62007 | 0.90133 | 0.14859 | 0.08285 |
| **LOC642852** | 0.83824 | 1.12183 | 1.35313 | 0.39305 | 0.00000 | 0.23149 | 0.60217 | 0.00000 | 1.20797 | 1.51457 | 0.86891 | 1.47872 |
| **LOC642943** | 0.04464 | 0.01553 | 0.02511 | 0.01510 | 0.01465 | 0.00647 | 0.07341 | 0.00988 | 0.00000 | 0.03654 | 0.01349 | 0.01549 |
| **LOC643072** | 0.03242 | 0.04625 | 0.09157 | 0.03151 | 0.05968 | 0.00000 | 0.01816 | 0.01103 | 0.08486 | 0.00000 | 0.01979 | 0.12636 |
| **LOC643201** | 0.11808 | 0.27163 | 0.30546 | 0.27221 | 0.00000 | 0.04756 | 0.41752 | 0.09147 | 0.00000 | 0.88300 | 0.00000 | 0.60429 |
| **LOC643339** | 0.16739 | 0.15089 | 0.02884 | 0.00158 | 0.00451 | 0.14664 | 0.00000 | 0.04343 | 0.03192 | 0.03061 | 0.10773 | 0.25143 |
| **LOC643355** | 0.08087 | 0.03080 | 0.01332 | 0.00000 | 0.04280 | 0.04845 | 0.18239 | 0.15253 | 0.01348 | 0.36112 | 0.02298 | 0.02226 |
| **LOC643387** | 0.04730 | 0.08384 | 0.33479 | 0.08696 | 0.00000 | 0.35561 | 0.36117 | 0.18354 | 0.00000 | 0.10227 | 0.22710 | 0.20080 |
| **LOC643406** | 0.09463 | 0.01815 | 0.01584 | 0.01319 | 0.01837 | 0.00722 | 0.01936 | 0.00644 | 0.02163 | 0.01093 | 0.01482 | 0.02261 |
| **LOC643441** | 0.02207 | 0.00000 | 1.24734 | 0.00007 | 0.60155 | 0.31331 | 0.00000 | 0.00000 | 2.36519 | 0.00000 | 4.12467 | 0.51075 |
| **LOC643486** | 0.00527 | 0.03359 | 0.01519 | 0.06615 | 0.00680 | 0.12092 | 0.00000 | 0.29019 | 0.03343 | 0.17041 | 0.01117 | 0.09677 |
| **LOC643623** | 1.86204 | 3.11965 | 0.03877 | 0.71525 | 0.20441 | 0.00000 | 0.00000 | 0.28437 | 0.00000 | 0.04799 | 0.01396 | 2.36556 |
| **LOC643711** | 0.23576 | 0.01409 | 0.01615 | 0.01877 | 0.02824 | 0.01326 | 0.02239 | 0.07291 | 0.01561 | 0.01328 | 0.01480 | 0.01454 |
| **LOC643770** | 0.01129 | 0.31339 | 0.00000 | 0.00000 | 0.03930 | 0.09823 | 0.01412 | 0.00616 | 0.01911 | 0.22665 | 0.04869 | 0.10220 |
| **LOC643802** | 0.00721 | 0.01520 | 0.01409 | 0.01063 | 0.02757 | 0.02153 | 0.01927 | 0.01130 | 0.01736 | 0.01064 | 0.01583 | 0.01604 |
| **LOC643923** | 0.00401 | 1.23426 | 0.01872 | 0.00850 | 0.06060 | 0.08163 | 0.04627 | 0.07622 | 0.01233 | 0.01191 | 0.06454 | 0.58732 |
| **LOC644050** | 0.00606 | 0.00759 | 0.02501 | 0.16544 | 0.00000 | 0.02382 | 0.00000 | 0.01082 | 0.06997 | 0.01159 | 0.07123 | 0.01743 |
| **LOC644145** | 0.03407 | 0.01387 | 0.01287 | 0.02332 | 0.01187 | 0.01694 | 0.01554 | 0.01035 | 0.02047 | 0.02174 | 0.01453 | 0.01539 |
| **LOC644172.uid-** | 0.00348 | 0.01647 | 0.00779 | 0.05365 | 0.00567 | 0.00000 | 0.01339 | 0.02414 | 0.09643 | 0.11730 | 0.05617 | 0.01434 |

| | | | | | | | | | | | | |
|---|---|---|---|---|---|---|---|---|---|---|---|---|
| **LOC644172.uid-** | 0.01369 | 0.06065 | 0.00852 | 0.07163 | 0.03768 | 0.00000 | 0.01457 | 0.02576 | 0.06008 | 0.06361 | 0.02970 | 0.03142 |
| **LOC644189** | 0.02149 | 0.00000 | 0.11158 | 0.04126 | 0.04925 | 0.08084 | 0.02412 | 0.05378 | 0.08526 | 0.18220 | 0.10015 | 0.03580 |
| **LOC644554** | 0.00823 | 0.00766 | 0.03244 | 0.08863 | 0.00043 | 0.15174 | 0.05123 | 0.14441 | 0.00000 | 0.00000 | 0.25878 | 0.08301 |
| **LOC644656** | 0.29155 | 0.47277 | 1.09816 | 0.25222 | 0.00000 | 1.21487 | 0.00000 | 0.57658 | 1.30695 | 0.35611 | 0.43934 | 0.78468 |
| **LOC644669** | 0.00621 | 0.01548 | 0.01638 | 0.01217 | 0.01529 | 0.01744 | 0.02080 | 0.01050 | 0.01557 | 0.01667 | 0.01532 | 0.01612 |
| **LOC644762** | 0.20143 | 0.30719 | 0.43999 | 0.06050 | 0.12829 | 2.12050 | 0.92697 | 0.06123 | 1.04561 | 0.11467 | 0.76668 | 0.17185 |
| **LOC644838** | 0.00535 | 0.02537 | 0.01496 | 0.01061 | 0.00849 | 0.01940 | 0.02260 | 0.01494 | 0.00012 | 0.02865 | 0.04664 | 0.05058 |
| **LOC644919** | 0.12846 | 0.10980 | 0.07856 | 0.25649 | 0.21871 | 0.62167 | 0.12182 | 0.02058 | 0.18471 | 0.03771 | 0.05225 | 0.07722 |
| **LOC644936** | 0.51157 | 17.90111 | 12.91301 | 7.13038 | 19.12672 | 9.38972 | 12.35201 | 1.00702 | 17.93328 | 0.00000 | 18.93335 | 2.60088 |
| **LOC644961** | 1.34454 | 10.80641 | 7.88479 | 7.17268 | 5.81769 | 5.34464 | 3.85836 | 5.44469 | 6.24101 | 0.00000 | 1.12985 | 4.78402 |
| **LOC645166** | 0.23187 | 0.48730 | 0.36628 | 0.17831 | 0.48268 | 0.25054 | 0.35599 | 0.18459 | 0.51835 | 0.00000 | 0.20774 | 0.85212 |
| **LOC645354+LO** | 0.26604 | 0.01435 | 0.01139 | 0.02113 | 0.01882 | 0.01451 | 0.01376 | 1.24407 | 0.02018 | 0.01082 | 0.01351 | 0.00000 |
| **LOC645354+LO** | 0.65034 | 0.01545 | 0.01340 | 0.01173 | 0.01824 | 0.01150 | 0.01666 | 0.54792 | 0.02017 | 0.00910 | 0.01377 | 0.00000 |
| **LOC645513** | 0.00396 | 0.54576 | 0.37504 | 0.52734 | 0.00000 | 1.21252 | 0.17285 | 1.05128 | 0.69230 | 0.00000 | 0.25545 | 0.99322 |
| **LOC645752** | 0.29141 | 0.01445 | 0.01423 | 0.02354 | 0.01228 | 0.00651 | 0.03139 | 0.04468 | 0.01209 | 0.04473 | 0.01376 | 0.01555 |
| **LOC645949** | 0.30260 | 0.01589 | 0.02228 | 0.01886 | 0.01832 | 0.00349 | 0.01801 | 0.08156 | 0.02067 | 0.01020 | 0.01911 | 0.00986 |
| **LOC646214** | 0.63262 | 0.62730 | 1.30942 | 0.79190 | 0.38504 | 1.96334 | 0.33632 | 1.49651 | 1.91374 | 1.03826 | 1.20807 | 0.41457 |
| **LOC646471** | 0.00657 | 0.24320 | 0.21772 | 0.64929 | 0.16626 | 0.00000 | 0.05044 | 0.07311 | 0.56433 | 0.39134 | 0.07628 | 0.18138 |
| **LOC646522** | 0.03025 | 0.01703 | 0.03056 | 0.01154 | 0.02070 | 0.01103 | 0.01777 | 0.02787 | 0.01517 | 0.01039 | 0.01597 | 0.01754 |
| **LOC646626** | 0.05939 | 0.04664 | 0.15269 | 0.01945 | 0.07273 | 0.10138 | 0.24875 | 0.00471 | 0.00000 | 0.12005 | 0.06880 | 0.11336 |
| **LOC646719** | 0.33747 | 0.38887 | 0.97687 | 0.46036 | 0.18663 | 0.00000 | 0.00000 | 2.28447 | 0.93013 | 0.83663 | 0.37705 | 1.53808 |
| **LOC646736** | 0.00742 | 0.01412 | 0.02559 | 0.01147 | 0.01424 | 0.02086 | 0.01878 | 0.01118 | 0.01713 | 0.00954 | 0.01909 | 0.02034 |
| **LOC646762** | 0.87699 | 0.57804 | 1.96715 | 0.25897 | 0.00000 | 1.45791 | 6.50219 | 0.00000 | 2.02725 | 1.46039 | 1.77190 | 0.28812 |
| **LOC646813** | 0.00547 | 0.01367 | 0.00901 | 0.00857 | 0.00927 | 0.01897 | 0.00641 | 0.01234 | 0.03155 | 0.06546 | 0.01007 | 0.01750 |
| **LOC646862** | 0.00743 | 0.01664 | 0.01579 | 0.01195 | 0.01342 | 0.02376 | 0.02029 | 0.01027 | 0.02331 | 0.01164 | 0.01510 | 0.01578 |
| **LOC646903** | 0.01297 | 0.02177 | 0.01988 | 0.00000 | 0.00042 | 0.08927 | 0.01708 | 0.03332 | 0.13468 | 0.03773 | 0.06022 | 0.08501 |
| **LOC647859** | 0.00584 | 0.05954 | 0.05235 | 0.01505 | 0.01870 | 0.01089 | 0.00767 | 0.01176 | 0.02489 | 0.02587 | 0.01673 | 0.01165 |
| **LOC648691** | 0.01016 | 0.05899 | 0.01708 | 0.01162 | 0.01821 | 0.00539 | 0.02040 | 0.09289 | 0.01986 | 0.01189 | 0.01631 | 0.01885 |
| **LOC648987** | 0.22688 | 0.01490 | 0.13776 | 0.53932 | 0.19261 | 0.41669 | 0.00000 | 0.09518 | 0.00000 | 1.20052 | 0.07273 | 0.98688 |
| **LOC649330** | 0.36712 | 0.01277 | 0.01340 | 0.01128 | 0.01749 | 0.00609 | 0.01680 | 1.29759 | 0.01891 | 0.01151 | 0.01414 | 0.01824 |
| **LOC649352** | 0.00627 | 0.00842 | 0.02000 | 0.05406 | 0.00000 | 0.02532 | 0.01236 | 0.00908 | 0.00240 | 0.07885 | 0.01431 | 0.02701 |
| **LOC650226** | 0.75539 | 1.57111 | 3.10089 | 0.68523 | 2.25821 | 2.18368 | 0.00000 | 1.06742 | 8.10140 | 15.88929 | 0.03196 | 2.98248 |
| **LOC650368** | 0.27329 | 0.22475 | 0.00082 | 0.37354 | 0.09367 | 0.00000 | 0.13738 | 0.14798 | 0.00635 | 0.21805 | 0.03849 | 0.23648 |
| **LOC652276** | 0.28240 | 0.21782 | 1.93532 | 1.18238 | 0.83016 | 0.00000 | 1.18910 | 0.06600 | 1.24537 | 1.23727 | 0.29502 | 0.88477 |
| **LOC653160** | 0.01508 | 0.01737 | 0.06004 | 0.01693 | 0.00763 | 0.03607 | 0.01653 | 0.01067 | 0.01404 | 0.01862 | 0.01599 | 0.01559 |
| **LOC653513** | 0.02442 | 0.04928 | 0.08880 | 0.10874 | 0.01488 | 0.03401 | 0.00626 | 0.49651 | 0.02187 | 0.02961 | 0.02492 | 0.09671 |
| **LOC653653** | 0.02750 | 0.07992 | 0.09896 | 0.25327 | 0.00000 | 0.14377 | 0.12760 | 0.11239 | 0.05682 | 0.00000 | 0.05311 | 0.16036 |
| **LOC653712** | 0.16967 | 0.00838 | 0.00000 | 0.30301 | 0.22052 | 0.00000 | 0.00055 | 0.01640 | 0.09749 | 0.59302 | 0.14739 | 0.07920 |
| **LOC653786** | 0.04149 | 0.32378 | 0.03827 | 0.14185 | 0.14368 | 0.00000 | 0.02757 | 0.07237 | 0.01436 | 0.00000 | 0.01905 | 0.14711 |
| **LOC654342** | 0.50263 | 2.20002 | 1.39167 | 0.65317 | 1.28150 | 0.61617 | 2.39831 | 0.41699 | 1.80501 | 0.00000 | 1.10463 | 3.33600 |
| **LOC654841** | 0.03318 | 0.12971 | 0.13599 | 0.03122 | 0.00000 | 0.06505 | 0.00000 | 0.03010 | 0.03040 | 0.05207 | 0.05805 | 0.09893 |
| **LOC727896** | 0.04976 | 1.92106 | 3.40347 | 0.61020 | 0.85675 | 4.31204 | 3.70403 | 2.15525 | 3.47727 | 0.99843 | 3.34131 | 2.24449 |
| **LOC727924** | 0.00911 | 0.02549 | 0.01445 | 0.00945 | 0.01373 | 0.02826 | 0.01769 | 0.06288 | 0.01706 | 0.01076 | 0.01334 | 0.01245 |
| **LOC728024** | 0.97523 | 0.39514 | 0.52200 | 0.49650 | 0.10641 | 0.41837 | 0.53680 | 1.71496 | 0.56978 | 0.00000 | 0.89473 | 1.39406 |
| **LOC728084** | 0.00844 | 0.00969 | 0.00000 | 0.00588 | 0.00000 | 0.02882 | 0.53048 | 0.03110 | 0.49411 | 0.00000 | 0.00000 | 0.01330 |
| **LOC728175** | 0.00571 | 0.01396 | 0.01819 | 0.01095 | 0.01488 | 0.01437 | 0.02977 | 0.01065 | 0.00687 | 0.03662 | 0.01748 | 0.01719 |
| **LOC728323** | 1.10141 | 0.23936 | 0.44534 | 0.11839 | 1.42030 | 0.00000 | 0.07588 | 0.00000 | 0.67920 | 1.96900 | 0.18932 | 0.28000 |
| **LOC728392** | 0.03566 | 0.01719 | 0.13365 | 0.32233 | 0.00000 | 0.00000 | 0.02738 | 0.00207 | 0.08698 | 0.31167 | 0.00000 | 0.41406 |

| | | | | | | | | | | | | |
|---|---|---|---|---|---|---|---|---|---|---|---|---|
| **LOC728554** | 1.23652 | 1.80608 | 1.36644 | 2.02320 | 2.27296 | 2.57662 | 2.61361 | 2.12463 | 1.45735 | 0.00000 | 1.84440 | 3.17856 |
| **LOC728613** | 0.64667 | 0.41728 | 0.17716 | 0.45008 | 1.01752 | 0.64292 | 0.00001 | 1.55815 | 0.43875 | 0.00011 | 0.26266 | 0.15577 |
| **LOC728637** | 0.14803 | 0.34331 | 0.00000 | 0.00000 | 0.50846 | 0.00002 | 0.28944 | 4.70833 | 0.49551 | 0.26976 | 2.92243 | 5.77210 |
| **LOC728730** | 1.74868 | 0.43775 | 0.10376 | 0.09491 | 0.38460 | 0.12820 | 0.00000 | 0.06176 | 0.00000 | 0.24787 | 0.24097 | 0.27012 |
| **LOC728739** | 0.23932 | 0.18419 | 0.12081 | 0.09120 | 0.08125 | 0.22560 | 0.04624 | 0.15615 | 0.43475 | 0.00000 | 0.07869 | 0.05476 |
| **LOC728743** | 0.00843 | 2.78608 | 0.57306 | 1.00842 | 0.37785 | 0.00000 | 0.18490 | 0.04239 | 0.00000 | 1.52881 | 0.03158 | 1.19592 |
| **LOC728752** | 0.00959 | 0.37752 | 0.50023 | 0.19163 | 0.04128 | 0.00000 | 0.04285 | 2.84293 | 0.15141 | 0.05484 | 0.09732 | 1.06063 |
| **LOC728819** | 0.13874 | 1.30985 | 0.00000 | 0.00000 | 0.14023 | 0.13570 | 0.00000 | 0.00000 | 0.00000 | 0.15163 | 0.02272 | 1.19389 |
| **LOC728989** | 0.01079 | 0.01196 | 0.01919 | 0.01754 | 0.03025 | 0.01200 | 0.02558 | 0.01732 | 0.02230 | 0.02032 | 0.01944 | 0.02926 |
| **LOC729080** | 0.00775 | 0.03573 | 0.01964 | 0.03176 | 0.05060 | 0.10538 | 0.00986 | 0.00880 | 0.12880 | 0.00000 | 0.04850 | 0.07506 |
| **LOC729083** | 0.03383 | 0.00000 | 0.04495 | 0.00000 | 0.07670 | 0.02921 | 0.01827 | 0.20019 | 0.04938 | 0.27149 | 0.00000 | 0.62601 |
| **LOC729218** | 0.12010 | 0.58808 | 0.10222 | 0.24504 | 0.40905 | 1.14617 | 0.00000 | 0.11019 | 0.55665 | 1.28754 | 0.00000 | 0.58936 |
| **LOC729296** | 0.03358 | 0.01196 | 0.01466 | 0.01309 | 0.01592 | 0.01977 | 0.01840 | 0.00906 | 0.01888 | 0.00749 | 0.01563 | 0.04310 |
| **LOC729506** | 0.02327 | 0.01569 | 0.05060 | 0.01801 | 0.02963 | 0.00000 | 0.01874 | 0.03779 | 0.01870 | 0.00005 | 0.01780 | 0.04574 |
| **LOC729603** | 0.00126 | 0.00000 | 0.49739 | 0.24582 | 0.08143 | 0.03544 | 0.00000 | 0.16740 | 0.22852 | 1.42964 | 0.12769 | 0.21098 |
| **LOC729609** | 0.00334 | 0.01240 | 0.11441 | 0.02127 | 0.00000 | 0.04220 | 0.02734 | 0.02668 | 0.00000 | 0.05779 | 0.03173 | 0.01537 |
| **LOC729683** | 0.00720 | 0.07007 | 0.01555 | 0.02856 | 0.01520 | 0.00096 | 0.01894 | 0.02009 | 0.02372 | 0.01167 | 0.01878 | 0.01396 |
| **LOC729732** | 0.04147 | 0.02686 | 0.05012 | 0.01826 | 0.00000 | 0.04521 | 0.00000 | 0.03808 | 0.05791 | 0.05287 | 0.00800 | 0.03388 |
| **LOC729737** | 0.00815 | 0.01360 | 0.06644 | 0.05247 | 0.03666 | 0.00000 | 0.00985 | 0.11237 | 0.06966 | 0.10175 | 0.03124 | 0.14608 |
| **LOC729739** | 0.00267 | 0.00000 | 0.97011 | 0.01752 | 0.21723 | 0.27012 | 0.13294 | 0.16735 | 0.29861 | 0.00000 | 0.16964 | 1.19576 |
| **LOC729930** | 0.11194 | 0.12513 | 0.00000 | 0.03422 | 0.12886 | 0.33187 | 0.03041 | 0.10047 | 0.02614 | 0.02460 | 0.17277 | 0.02095 |
| **LOC729966** | 0.00296 | 0.01462 | 0.03522 | 0.02116 | 0.02322 | 0.01220 | 0.01766 | 0.03085 | 0.01906 | 0.00498 | 0.01558 | 0.02950 |
| **LOC729970** | 0.00846 | 0.03529 | 0.01178 | 0.11587 | 0.06404 | 0.15043 | 0.00185 | 0.00583 | 0.00000 | 0.05639 | 0.11459 | 0.08114 |
| **LOC730101** | 0.02426 | 0.00000 | 2.15689 | 0.38287 | 0.79220 | 5.75410 | 1.81150 | 0.06342 | 2.96691 | 0.05584 | 2.21079 | 0.00000 |
| **LOC730102** | 0.01227 | 0.00000 | 0.07310 | 0.01518 | 0.00117 | 0.18586 | 0.00000 | 0.03098 | 0.33928 | 0.00000 | 0.15377 | 0.04790 |
| **LOC730183** | 0.17747 | 0.39964 | 0.27782 | 0.00003 | 0.21682 | 0.15843 | 0.02180 | 0.02096 | 0.24623 | 0.84484 | 0.29894 | 0.66688 |
| **LOC730668** | 0.00665 | 0.01839 | 0.07709 | 0.01969 | 0.02899 | 0.00000 | 0.01598 | 0.01064 | 0.01830 | 0.00000 | 0.01725 | 0.01867 |
| **LOC731157** | 0.00698 | 0.06270 | 0.12619 | 0.04578 | 0.05649 | 0.33253 | 0.00000 | 0.00390 | 0.00000 | 0.02632 | 0.01251 | 0.13763 |
| **LOC731424** | 0.09365 | 0.00421 | 0.18524 | 0.04338 | 0.02919 | 0.00000 | 0.00000 | 0.01674 | 0.10070 | 0.26148 | 0.00000 | 0.01828 |
| **LOC80154.uid-1** | 0.00768 | 0.01469 | 0.01749 | 0.01322 | 0.01665 | 0.01947 | 0.01793 | 0.01214 | 0.01879 | 0.01427 | 0.01457 | 0.01684 |
| **LOC81691** | 1.21688 | 2.12268 | 0.49597 | 0.66439 | 1.33046 | 1.29133 | 0.00000 | 0.63008 | 0.65016 | 0.00000 | 0.61987 | 2.66080 |
| **LOC90246** | 0.00838 | 0.00969 | 0.08389 | 0.03432 | 0.11045 | 0.00000 | 0.08396 | 0.01434 | 0.00000 | 0.00000 | 0.20028 | 0.02541 |
| **LOC90784** | 0.00131 | 0.00000 | 1.85213 | 1.47145 | 1.88596 | 0.70859 | 3.60640 | 0.71335 | 1.13485 | 6.42932 | 1.49341 | 1.83648 |
| **LOC90834** | 0.12464 | 0.03618 | 0.09600 | 0.47784 | 0.21644 | 0.00000 | 0.00000 | 0.00249 | 0.40148 | 0.67862 | 0.00000 | 0.06856 |
| **LOC91450** | 0.00475 | 0.01188 | 0.06355 | 0.00593 | 0.00000 | 0.01439 | 0.09243 | 0.01036 | 0.00498 | 0.10609 | 0.01351 | 0.01222 |
| **LOC93432** | 0.86614 | 0.00000 | 0.06031 | 0.00000 | 0.00000 | 0.07719 | 0.08927 | 0.64866 | 0.00000 | 0.05747 | 0.01418 | 0.22847 |
| **LOC93622** | 0.35875 | 0.33053 | 0.41917 | 0.87385 | 2.04500 | 0.00675 | 3.51716 | 0.02544 | 0.05525 | 4.78785 | 0.40643 | 0.48379 |
| **LOH12CR1** | 6.55653 | 2.64697 | 1.07073 | 0.95299 | 1.51846 | 2.82441 | 0.00000 | 5.65306 | 1.77612 | 0.00000 | 1.97513 | 4.12423 |
| **LOH12CR2** | 0.13655 | 0.02700 | 0.17940 | 0.00592 | 0.02599 | 0.00000 | 0.00000 | 0.01104 | 0.06792 | 0.03020 | 0.01864 | 0.44471 |
| **LONP1** | 0.49608 | 8.72852 | 10.83083 | 14.28363 | 18.72585 | 0.02130 | 24.35232 | 2.35352 | 9.71335 | 21.62564 | 5.95221 | 4.36564 |
| **LONP2** | 5.33569 | 6.01816 | 4.07436 | 3.36989 | 5.81066 | 8.25203 | 0.00000 | 3.25630 | 3.27912 | 2.38615 | 8.84181 | 2.66587 |
| **LONRF1** | 0.98353 | 0.04343 | 0.00000 | 1.45035 | 2.76805 | 2.29411 | 1.93270 | 0.01267 | 0.63592 | 3.40645 | 0.74886 | 0.00000 |
| **LONRF2** | 0.49460 | 0.00990 | 0.09279 | 0.02598 | 0.03986 | 0.00000 | 0.07813 | 0.00379 | 0.00346 | 0.08916 | 0.03976 | 0.01243 |
| **LONRF3** | 0.21061 | 0.06558 | 0.02633 | 0.15941 | 0.00000 | 0.00071 | 0.00560 | 0.11126 | 0.01006 | 0.10186 | 0.01286 | 0.20182 |
| **LOR** | 0.00530 | 0.01448 | 0.00870 | 0.01354 | 0.01297 | 0.01186 | 0.10731 | 0.02001 | 0.00061 | 0.01838 | 0.01362 | 0.07634 |
| **LOX** | 0.03202 | 0.00000 | 0.04658 | 0.00000 | 0.13786 | 0.07158 | 0.18472 | 0.29517 | 0.35093 | 0.15916 | 0.00000 | 0.08538 |
| **LOXHD1** | 2.43876 | 0.01624 | 0.01446 | 0.00979 | 0.03087 | 0.00000 | 0.11469 | 0.77267 | 0.00000 | 0.12879 | 0.00428 | 0.05001 |
| **LOXL1** | 0.29295 | 0.29182 | 0.04856 | 0.00000 | 0.06748 | 0.03791 | 0.00489 | 0.02255 | 0.00000 | 0.16197 | 0.00000 | 0.09005 |

| | | | | | | | | | | | | |
|---|---|---|---|---|---|---|---|---|---|---|---|---|
| LOXL1-AS1 | 0.04447 | 0.31789 | 0.02391 | 0.03934 | 0.01675 | 0.04469 | 0.07593 | 0.01098 | 0.02420 | 0.00000 | 0.00490 | 0.38494 |
| LOXL2 | 0.01394 | 3.47391 | 0.67723 | 0.50672 | 0.66092 | 0.00000 | 0.00000 | 1.22337 | 0.69623 | 0.83365 | 0.00000 | 2.98161 |
| LOXL3 | 0.43253 | 0.49784 | 0.60947 | 0.63784 | 0.50026 | 0.04894 | 0.58787 | 0.07655 | 0.49377 | 0.32786 | 0.32584 | 0.15744 |
| LOXL4 | 0.00704 | 0.01035 | 0.13167 | 0.07876 | 0.00000 | 0.01682 | 0.00000 | 0.00982 | 0.03749 | 0.06381 | 0.02801 | 0.02370 |
| LPA | 0.01337 | 0.01085 | 0.01768 | 0.00638 | 0.00445 | 0.02563 | 0.02406 | 0.95374 | 0.02061 | 0.05145 | 0.01714 | 0.00464 |
| LPAL2 | 0.00623 | 0.02644 | 0.02306 | 0.02219 | 0.01033 | 0.01490 | 0.01499 | 1.03257 | 0.00937 | 0.03671 | 0.01516 | 0.00000 |
| LPAR1 | 0.94350 | 2.19563 | 0.07179 | 2.24473 | 0.00000 | 0.00000 | 0.01009 | 0.58812 | 0.00000 | 0.03524 | 0.16993 | 2.31897 |
| LPAR2 | 0.00000 | 13.44695 | 4.66401 | 8.00430 | 4.60565 | 0.00000 | 2.76707 | 0.92613 | 7.92751 | 5.59477 | 6.88737 | 13.84286 |
| LPAR3 | 0.01706 | 0.53937 | 0.01674 | 0.00000 | 0.01359 | 0.01580 | 0.00000 | 0.06121 | 0.12004 | 0.05389 | 0.00796 | 0.10620 |
| LPAR4 | 0.00525 | 0.01724 | 0.01411 | 0.12622 | 0.00000 | 0.05023 | 0.02630 | 0.06374 | 0.00000 | 0.02724 | 0.15327 | 0.04776 |
| LPAR5 | 0.01697 | 0.28073 | 0.00000 | 0.00036 | 0.55981 | 0.02069 | 0.00083 | 0.05266 | 0.89496 | 0.00157 | 0.03320 | 0.00367 |
| LPAR6 | 0.00332 | 0.00375 | 0.07781 | 0.23681 | 0.00000 | 0.06023 | 0.00000 | 0.05487 | 0.20778 | 0.11965 | 0.01781 | 0.02248 |
| LPCAT1 | 4.89125 | 7.89069 | 4.80860 | 5.26871 | 8.22834 | 0.00003 | 0.07183 | 0.63333 | 6.56947 | 6.67053 | 1.73830 | 0.13230 |
| LPCAT2 | 0.36797 | 0.86486 | 0.81708 | 0.00000 | 0.00000 | 0.77798 | 0.31485 | 1.08652 | 0.73863 | 0.48579 | 0.00000 | 1.75663 |
| LPCAT3 | 0.25261 | 37.55345 | 7.58001 | 12.56222 | 23.87599 | 16.38859 | 0.00000 | 0.11355 | 24.36957 | 3.24710 | 27.51214 | 26.09474 |
| LPCAT4 | 12.93658 | 3.05107 | 0.44786 | 7.36729 | 6.40301 | 1.37699 | 0.54908 | 1.79613 | 3.76937 | 10.16629 | 3.67524 | 2.82275 |
| LPGAT1 | 0.18399 | 3.66332 | 0.00000 | 0.00000 | 3.72821 | 11.08473 | 0.00000 | 0.35551 | 0.46658 | 7.62088 | 0.00000 | 2.47838 |
| LPHN1 | 3.81114 | 1.43025 | 0.21458 | 1.47076 | 1.56834 | 0.00000 | 0.40941 | 0.48562 | 0.71231 | 8.69756 | 0.00000 | 4.35313 |
| LPHN2 | 0.08342 | 0.00000 | 0.00000 | 0.00000 | 0.00000 | 1.53431 | 4.53830 | 0.16469 | 16.15826 | 3.73058 | 7.69457 | 0.00000 |
| LPHN3 | 0.67959 | 0.60082 | 7.64370 | 0.00000 | 0.50509 | 0.10397 | 0.17866 | 1.85811 | 0.00000 | 0.03901 | 0.42428 | 4.14687 |
| LPHN3-AS1 | 0.02481 | 0.01012 | 0.09495 | 0.00422 | 0.00870 | 0.03837 | 0.03441 | 0.01715 | 0.00524 | 0.02878 | 0.01561 | 0.01815 |
| LPIN1 | 0.20925 | 7.33959 | 2.66510 | 13.60772 | 11.70918 | 13.72635 | 8.26081 | 0.60489 | 7.84705 | 20.59726 | 4.82872 | 3.86095 |
| LPIN2 | 7.38557 | 13.93484 | 0.00000 | 4.28195 | 7.60542 | 12.54817 | 1.57944 | 1.89595 | 0.00000 | 14.23668 | 8.24574 | 9.06409 |
| LPIN3 | 0.63707 | 2.06626 | 1.10089 | 2.11105 | 0.69424 | 1.76231 | 0.25617 | 1.68527 | 1.29474 | 1.28908 | 0.94023 | 0.67444 |
| LPL | 0.00905 | 0.00758 | 0.00918 | 0.00440 | 0.00000 | 0.03159 | 0.01078 | 0.01975 | 0.03560 | 0.13699 | 0.00320 | 0.01156 |
| LPO | 0.11261 | 0.01127 | 0.02796 | 0.02116 | 0.00973 | 0.01522 | 0.01177 | 0.00836 | 0.01413 | 0.03361 | 0.02419 | 0.02316 |
| LPP | 0.54686 | 2.99027 | 5.01545 | 8.37840 | 7.57270 | 10.19408 | 4.94080 | 0.00000 | 3.64254 | 16.40119 | 4.51087 | 6.50678 |
| LPP-AS1 | 0.00579 | 0.01707 | 0.02635 | 0.01534 | 0.01497 | 0.01224 | 0.00879 | 0.01033 | 0.03756 | 0.02674 | 0.01170 | 0.02536 |
| LPP-AS2 | 0.33865 | 0.09962 | 0.05227 | 0.39492 | 0.36618 | 0.00000 | 0.00000 | 0.27234 | 0.30405 | 0.27621 | 0.15452 | 1.43709 |
| LPPR1 | 0.90069 | 0.03143 | 0.01707 | 0.00000 | 0.00000 | 0.08104 | 0.03357 | 0.37334 | 0.00083 | 0.05569 | 0.00758 | 0.01595 |
| LPPR2 | 2.35812 | 1.10985 | 0.67098 | 2.02791 | 0.41188 | 0.00000 | 0.25427 | 0.00000 | 1.06709 | 0.00000 | 0.07022 | 0.17536 |
| LPPR3 | 0.00626 | 0.09380 | 0.03730 | 0.00055 | 0.00612 | 0.00000 | 0.01208 | 0.01336 | 0.19957 | 0.80848 | 0.01402 | 0.01884 |
| LPPR5 | 0.14318 | 0.01129 | 0.01078 | 0.00889 | 0.00726 | 0.02028 | 0.00740 | 0.00501 | 0.02610 | 0.05729 | 0.01228 | 0.01509 |
| LPXN | 0.01934 | 0.27388 | 0.13912 | 0.36915 | 0.05336 | 0.20318 | 0.02480 | 0.20719 | 0.68283 | 0.53231 | 0.13038 | 0.19160 |
| LRAT | 4.71449 | 0.09556 | 0.53501 | 0.47626 | 0.00000 | 0.00000 | 0.00000 | 1.89383 | 0.11084 | 0.20488 | 0.00000 | 0.14341 |
| LRBA | 0.25936 | 0.75785 | 2.36602 | 4.65890 | 3.06378 | 5.21154 | 2.20431 | 0.00189 | 3.98713 | 7.61741 | 3.16235 | 2.36893 |
| LRCH1 | 2.30576 | 2.77205 | 0.12101 | 3.31729 | 0.69148 | 7.68861 | 1.16603 | 0.00000 | 3.62299 | 10.44179 | 2.43058 | 4.14788 |
| LRCH2 | 0.00920 | 0.01081 | 0.05497 | 0.00749 | 0.00831 | 0.04350 | 0.06081 | 0.01081 | 0.00798 | 0.01307 | 0.01569 | 0.02789 |
| LRCH3 | 3.02985 | 2.32146 | 1.01134 | 1.81399 | 0.00000 | 1.85154 | 0.39533 | 1.39621 | 3.57364 | 0.00000 | 1.61271 | 3.51840 |
| LRCH4 | 1.55184 | 1.84825 | 1.66961 | 0.74312 | 3.80597 | 0.00000 | 0.08717 | 0.31204 | 1.70132 | 2.98181 | 0.38418 | 0.57960 |
| LRFN1 | 0.00594 | 0.00076 | 0.46302 | 0.00000 | 0.54809 | 0.00000 | 0.67325 | 0.00851 | 0.03246 | 1.54157 | 0.10130 | 0.00451 |
| LRFN2 | 0.00637 | 0.01966 | 0.02094 | 0.02153 | 0.02415 | 0.00000 | 0.06366 | 0.01160 | 0.02011 | 0.00000 | 0.01628 | 0.01834 |
| LRFN3 | 0.07407 | 0.37940 | 0.26242 | 0.18767 | 0.00000 | 0.00000 | 0.00000 | 0.00000 | 0.00000 | 0.46029 | 0.02732 | 0.49242 |
| LRFN4 | 0.00071 | 1.90656 | 0.09708 | 0.00000 | 0.00000 | 0.00000 | 0.32566 | 0.00481 | 0.63346 | 4.10212 | 0.37344 | 0.44857 |
| LRFN5 | 0.06025 | 0.00269 | 0.63406 | 0.21163 | 0.00000 | 0.09894 | 0.00000 | 0.00508 | 0.08165 | 0.00000 | 0.02969 | 0.01468 |
| LRG1 | 0.00969 | 0.83301 | 0.08534 | 0.59469 | 0.79945 | 0.34548 | 0.00000 | 0.00489 | 0.15624 | 0.00000 | 0.21247 | 0.08921 |
| LRGUK | 0.63474 | 0.16362 | 0.00000 | 0.06382 | 0.09517 | 0.03115 | 0.00555 | 0.00308 | 0.07134 | 0.19012 | 0.00000 | 0.16181 |
| LRIF1 | 0.92090 | 3.44980 | 7.41117 | 1.36161 | 0.00000 | 12.32838 | 5.97833 | 5.63553 | 4.13171 | 2.35535 | 2.04421 | 5.84217 |

| | | | | | | | | | | | | |
|---|---|---|---|---|---|---|---|---|---|---|---|---|
| **LRIG1** | 5.99652 | 0.24763 | 0.61401 | 4.00401 | 0.00000 | 0.00001 | 0.54782 | 2.22630 | 0.40172 | 0.58960 | 0.17366 | 0.74529 |
| **LRIG2** | 2.54282 | 5.96346 | 1.11319 | 1.55021 | 1.67508 | 3.53958 | 0.00000 | 6.69719 | 6.29843 | 3.18474 | 0.00360 | 4.39993 |
| **LRIG3** | 0.43927 | 18.43163 | 3.12829 | 0.00000 | 6.53686 | 12.09170 | 0.00000 | 0.53224 | 10.44784 | 5.33705 | 5.60966 | 7.53592 |
| **LRIT2** | 0.00733 | 0.01963 | 0.01597 | 0.06951 | 0.02560 | 0.00674 | 0.03665 | 0.01031 | 0.00368 | 0.00012 | 0.03993 | 0.01749 |
| **LRIT3** | 0.09269 | 0.00000 | 0.11357 | 0.06356 | 0.00207 | 0.00000 | 0.00000 | 0.21944 | 0.12922 | 0.35299 | 0.11802 | 1.02480 |
| **LRMP** | 2.80401 | 0.02117 | 0.03370 | 0.16060 | 0.00000 | 0.03690 | 0.00667 | 0.72430 | 0.00000 | 0.11941 | 0.00922 | 0.18018 |
| **LRP1** | 0.31833 | 2.58391 | 5.77363 | 2.04872 | 11.42622 | 0.00000 | 0.00000 | 0.17111 | 7.56747 | 9.69499 | 0.62285 | 2.12268 |
| **LRP10** | 0.57027 | 0.82092 | 7.64567 | 4.05009 | 4.32260 | 0.00000 | 0.35503 | 1.09948 | 5.54473 | 6.67992 | 0.14580 | 1.83490 |
| **LRP11** | 0.05131 | 0.02597 | 0.05344 | 0.10306 | 0.16093 | 0.00000 | 0.05432 | 0.00369 | 0.00000 | 0.34400 | 0.09964 | 0.13392 |
| **LRP12** | 0.00000 | 1.34676 | 0.62326 | 0.00000 | 0.03097 | 2.54237 | 0.00000 | 3.26722 | 1.27076 | 0.92245 | 0.11447 | 1.91221 |
| **LRP1B** | 3.44087 | 0.00000 | 0.09599 | 0.02959 | 0.15054 | 0.00000 | 0.21417 | 0.00000 | 0.00000 | 0.01937 | 0.26927 | 0.17094 |
| **LRP2** | 0.00000 | 0.00000 | 16.32853 | 1.23589 | 37.85572 | 1.31802 | 23.09071 | 0.07993 | 62.78496 | 23.97129 | 31.96900 | 0.00000 |
| **LRP2BP** | 0.47847 | 0.06730 | 0.20507 | 0.61605 | 0.39268 | 0.00000 | 0.09523 | 0.65865 | 0.27466 | 0.02256 | 0.00000 | 0.17538 |
| **LRP3** | 0.66649 | 0.36262 | 0.04903 | 0.18768 | 0.00000 | 0.04726 | 0.06559 | 0.00000 | 0.01699 | 0.19765 | 0.00000 | 0.18114 |
| **LRP4** | 0.14388 | 3.30427 | 0.91697 | 11.41426 | 0.00021 | 0.00000 | 0.00000 | 2.08680 | 0.00000 | 2.28929 | 0.00000 | 5.35151 |
| **LRP4-AS1** | 0.27125 | 0.02481 | 0.03136 | 0.02866 | 0.09829 | 0.00000 | 0.03255 | 0.12828 | 0.14082 | 0.15570 | 0.00008 | 0.14303 |
| **LRP5** | 3.72548 | 0.65095 | 0.18473 | 0.02810 | 1.14003 | 0.00000 | 0.00000 | 1.32667 | 0.71277 | 0.26848 | 0.16156 | 0.34377 |
| **LRP5L** | 0.00396 | 0.22320 | 0.39309 | 0.13315 | 0.81914 | 0.00000 | 0.13311 | 0.00700 | 1.14039 | 0.10985 | 0.12489 | 0.08978 |
| **LRP6** | 12.99393 | 1.23712 | 3.64420 | 2.35869 | 1.94510 | 9.19774 | 2.34334 | 4.05315 | 8.29658 | 16.14051 | 2.59720 | 2.69711 |
| **LRP8** | 0.00959 | 0.00000 | 0.69921 | 1.64741 | 0.71379 | 1.45039 | 1.39856 | 0.01334 | 1.26388 | 1.39650 | 0.33549 | 0.00000 |
| **LRPAP1** | 1.46803 | 34.52835 | 23.77278 | 22.57439 | 47.57232 | 22.30193 | 3.59994 | 10.03824 | 57.53550 | 11.53079 | 28.61380 | 20.65998 |
| **LRPPRC** | 0.35441 | 25.14113 | 15.64465 | 10.54748 | 9.41390 | 60.78798 | 28.31935 | 4.09286 | 28.68586 | 27.35598 | 20.00424 | 28.13002 |
| **LRR1** | 2.32167 | 7.04919 | 6.32491 | 7.85294 | 6.25923 | 23.93579 | 16.56980 | 5.12105 | 11.20639 | 2.97816 | 12.61342 | 19.84282 |
| **LRRC1** | 1.66052 | 0.82773 | 3.91054 | 0.59860 | 7.70925 | 7.34298 | 5.52971 | 0.82589 | 0.66233 | 4.82791 | 3.74158 | 0.00000 |
| **LRRC10** | 0.00690 | 0.01281 | 0.01259 | 0.00868 | 0.00855 | 0.01968 | 0.02325 | 0.01079 | 0.02129 | 0.04917 | 0.01788 | 0.01578 |
| **LRRC10B** | 0.00249 | 0.01447 | 0.02868 | 0.03154 | 0.01183 | 0.01833 | 0.01867 | 0.03912 | 0.01927 | 0.01830 | 0.01581 | 0.01470 |
| **LRRC14** | 0.14544 | 5.82169 | 4.90573 | 7.86043 | 4.12612 | 0.00000 | 5.95240 | 1.32626 | 5.89848 | 1.85554 | 4.97493 | 5.15814 |
| **LRRC15** | 0.02965 | 0.01272 | 0.04446 | 0.00465 | 0.00607 | 0.03447 | 0.02087 | 0.00743 | 0.00007 | 0.04788 | 0.01434 | 0.01298 |
| **LRRC16A** | 17.23556 | 5.66135 | 2.81733 | 10.95092 | 1.13407 | 8.21308 | 8.82329 | 11.16846 | 1.03693 | 1.67082 | 9.62089 | 6.51764 |
| **LRRC16B** | 0.03084 | 0.01543 | 0.01655 | 0.01408 | 0.01021 | 0.02693 | 0.01171 | 0.03932 | 0.02505 | 0.04229 | 0.00969 | 0.01659 |
| **LRRC17** | 7.77301 | 0.04293 | 0.00000 | 0.01747 | 0.10206 | 0.09188 | 0.01445 | 1.13889 | 0.00000 | 0.00000 | 0.02451 | 0.05498 |
| **LRRC19** | 0.11418 | 0.00000 | 0.09530 | 0.06219 | 0.02129 | 0.11863 | 0.00000 | 0.00346 | 0.18583 | 0.08376 | 0.00000 | 0.08899 |
| **LRRC2** | 0.00425 | 1.45104 | 0.10879 | 0.58366 | 0.66691 | 0.00000 | 0.01033 | 0.00776 | 0.00000 | 0.08624 | 0.45313 | 0.07746 |
| **LRRC2-AS1** | 0.01626 | 0.08966 | 0.02060 | 0.00813 | 0.02930 | 0.00000 | 0.01759 | 0.00961 | 0.01924 | 0.02746 | 0.01396 | 0.01206 |
| **LRRC20** | 0.01440 | 0.71505 | 0.15928 | 0.30264 | 0.00000 | 0.05792 | 0.00000 | 0.22403 | 0.03207 | 0.02515 | 0.01292 | 0.73070 |
| **LRRC23** | 0.32177 | 0.91152 | 0.96693 | 0.39706 | 0.31952 | 0.00000 | 0.00104 | 0.13730 | 0.00000 | 0.10711 | 0.54150 | 1.74896 |
| **LRRC25** | 0.00477 | 0.01084 | 0.02777 | 0.02949 | 0.12442 | 0.00000 | 0.01400 | 0.01039 | 0.00000 | 0.10006 | 0.01737 | 0.01414 |
| **LRRC26** | 0.06372 | 0.04492 | 0.04453 | 0.01064 | 0.00970 | 0.02583 | 0.00578 | 0.00627 | 0.03461 | 0.01178 | 0.01597 | 0.01284 |
| **LRRC27** | 0.62341 | 1.00941 | 0.16708 | 0.00000 | 0.14299 | 0.29126 | 0.00000 | 0.07221 | 0.98511 | 0.20983 | 0.00000 | 0.34674 |
| **LRRC28** | 3.52040 | 2.94925 | 3.22229 | 2.96898 | 0.74219 | 3.12609 | 1.31497 | 0.22878 | 1.59074 | 0.00000 | 4.00591 | 2.63176 |
| **LRRC29** | 0.00920 | 0.06059 | 0.03181 | 0.01018 | 0.02313 | 0.00181 | 0.02254 | 0.02749 | 0.03072 | 0.00000 | 0.02255 | 0.07597 |
| **LRRC3** | 0.05639 | 0.82382 | 0.12384 | 0.00000 | 0.08845 | 0.07762 | 0.00808 | 0.00000 | 0.00000 | 0.21554 | 0.07708 | 0.34518 |
| **LRRC3-AS1** | 0.22239 | 0.01723 | 0.01001 | 0.15663 | 0.02110 | 0.00000 | 0.02096 | 0.59796 | 0.02553 | 0.00000 | 0.01628 | 0.02217 |
| **LRRC31** | 0.01812 | 0.02989 | 0.02200 | 0.03456 | 0.02659 | 0.00713 | 0.02868 | 0.00895 | 0.01884 | 0.00000 | 0.01517 | 0.04499 |
| **LRRC32** | 0.14721 | 1.51724 | 0.17812 | 2.40051 | 0.29137 | 0.05888 | 0.00000 | 0.00000 | 0.00000 | 0.20801 | 0.00000 | 0.31005 |
| **LRRC34** | 0.08640 | 0.89019 | 0.58671 | 0.53921 | 0.00000 | 0.00001 | 0.05337 | 0.29084 | 0.39337 | 0.41259 | 0.31289 | 0.67548 |
| **LRRC36** | 0.00174 | 0.13393 | 0.02396 | 0.00214 | 0.00823 | 0.02247 | 0.01524 | 0.07539 | 0.01513 | 0.00994 | 0.01411 | 0.24039 |
| **LRRC37A** | 0.01482 | 0.01142 | 0.01729 | 0.01905 | 0.03415 | 0.03357 | 0.00993 | 0.02135 | 0.06048 | 0.01367 | 0.00876 | 0.02953 |

| | | | | | | | | | | | | |
|---|---|---|---|---|---|---|---|---|---|---|---|---|
| **LRRC37A11P** | 0.01250 | 0.87949 | 0.21356 | 0.38911 | 0.42719 | 0.87754 | 0.00000 | 0.04243 | 0.29298 | 0.10414 | 0.09509 | 1.53079 |
| **LRRC37A2** | 0.01959 | 0.08614 | 0.04056 | 0.06709 | 0.09533 | 0.03917 | 0.00000 | 0.06752 | 0.24235 | 0.13757 | 0.01702 | 0.13576 |
| **LRRC37A3** | 0.09946 | 0.03818 | 0.00000 | 0.48066 | 0.00000 | 0.35398 | 0.04102 | 0.13061 | 0.13708 | 0.39195 | 0.07433 | 0.07316 |
| **LRRC37A4P** | 0.06471 | 0.01811 | 0.01477 | 0.00126 | 0.00000 | 0.05440 | 0.02954 | 0.01048 | 0.00386 | 0.09555 | 0.00880 | 0.50264 |
| **LRRC37A6P** | 0.01103 | 0.02704 | 0.10975 | 0.05805 | 0.13646 | 0.11873 | 0.15208 | 0.02161 | 0.17607 | 0.07843 | 0.01223 | 0.09626 |
| **LRRC37B** | 0.05102 | 0.00000 | 0.19375 | 0.00000 | 0.00157 | 0.48339 | 0.00000 | 0.02275 | 0.65493 | 0.53124 | 0.02042 | 0.00000 |
| **LRRC37BP1** | 0.01456 | 0.65184 | 0.14132 | 3.58171 | 0.12587 | 0.21082 | 0.19711 | 3.03361 | 0.09025 | 2.68547 | 0.11217 | 3.05600 |
| **LRRC38** | 0.54023 | 0.09826 | 0.04257 | 0.06152 | 0.00000 | 0.02946 | 0.00000 | 3.27908 | 0.00000 | 0.14267 | 0.00467 | 0.19359 |
| **LRRC39** | 0.00945 | 0.05213 | 0.06602 | 0.01516 | 0.00000 | 0.17539 | 0.02750 | 0.08040 | 0.00000 | 0.25668 | 0.06531 | 0.02585 |
| **LRRC3B** | 0.17939 | 0.02422 | 0.04409 | 0.02728 | 0.00469 | 0.03771 | 0.00676 | 0.02237 | 0.01878 | 0.02067 | 0.01118 | 0.00961 |
| **LRRC4** | 2.79020 | 0.67621 | 0.00000 | 0.00000 | 0.55452 | 1.19622 | 0.00000 | 0.28882 | 0.00000 | 2.43691 | 0.38554 | 2.22240 |
| **LRRC40** | 2.12901 | 2.63935 | 2.78293 | 1.51938 | 0.00000 | 10.68343 | 2.62602 | 1.34206 | 1.19093 | 1.19481 | 6.35054 | 1.06305 |
| **LRRC41** | 1.15632 | 3.61772 | 13.47109 | 15.87012 | 4.72694 | 2.78865 | 3.37216 | 3.90696 | 11.76988 | 0.00000 | 6.14694 | 2.79839 |
| **LRRC42** | 7.25047 | 5.58189 | 2.01064 | 2.58095 | 9.00928 | 8.49666 | 5.74189 | 1.24364 | 5.08402 | 4.21741 | 5.79397 | 10.28214 |
| **LRRC43** | 0.12128 | 0.08432 | 0.05149 | 0.14210 | 0.00000 | 0.03785 | 0.00590 | 0.05206 | 0.04981 | 0.01979 | 0.02246 | 0.08970 |
| **LRRC45** | 0.53526 | 0.70041 | 2.34658 | 0.87849 | 3.24656 | 0.00000 | 0.82084 | 0.00208 | 0.77208 | 2.24048 | 2.83061 | 0.05385 |
| **LRRC46** | 0.56675 | 0.28572 | 0.10236 | 0.18805 | 0.25151 | 0.00000 | 0.11712 | 0.00000 | 0.00948 | 0.00000 | 0.07801 | 0.60888 |
| **LRRC47** | 1.75216 | 7.35498 | 4.46545 | 11.75919 | 12.06537 | 5.55435 | 11.31325 | 2.84328 | 0.75672 | 15.83546 | 4.69889 | 6.26803 |
| **LRRC48** | 0.50416 | 0.28912 | 0.17431 | 0.43349 | 0.02615 | 0.25188 | 0.08897 | 0.06500 | 0.28030 | 0.00000 | 0.36688 | 0.20991 |
| **LRRC49** | 1.12370 | 0.45115 | 0.00000 | 1.01383 | 0.00000 | 0.41931 | 0.00038 | 0.98047 | 0.00000 | 0.24890 | 0.00000 | 0.00598 |
| **LRRC4B** | 1.12661 | 0.05181 | 0.01311 | 0.09792 | 0.08796 | 0.00000 | 0.01214 | 0.04681 | 0.00000 | 0.19977 | 0.02280 | 0.00321 |
| **LRRC4C** | 0.00343 | 0.01147 | 0.02424 | 0.00980 | 0.00993 | 0.03061 | 0.00730 | 0.03991 | 0.01604 | 0.03776 | 0.01686 | 0.01361 |
| **LRRC56** | 0.07084 | 0.00000 | 0.14408 | 0.00001 | 0.04253 | 0.08202 | 0.00000 | 0.06841 | 0.42223 | 0.33677 | 0.03482 | 0.07239 |
| **LRRC57** | 0.05822 | 5.17959 | 5.71234 | 1.89535 | 0.27994 | 6.48210 | 1.60848 | 5.42414 | 2.97562 | 0.00000 | 3.93978 | 9.97440 |
| **LRRC58** | 0.00000 | 2.01193 | 19.64638 | 1.31313 | 0.00000 | 6.72467 | 18.86510 | 0.00000 | 6.25391 | 3.71404 | 6.11046 | 0.00000 |
| **LRRC59** | 6.20430 | 14.77065 | 17.17407 | 14.17197 | 21.54309 | 11.31168 | 17.08499 | 10.80693 | 22.79127 | 0.03366 | 14.45809 | 17.38201 |
| **LRRC6** | 0.82675 | 0.35708 | 0.01981 | 0.00000 | 0.00000 | 0.58015 | 1.02641 | 0.28457 | 0.00000 | 1.48227 | 0.45033 | 0.69915 |
| **LRRC61** | 0.71922 | 9.42963 | 12.69200 | 2.59170 | 6.35458 | 0.00000 | 4.24789 | 0.00000 | 12.26472 | 4.16062 | 5.64554 | 0.35973 |
| **LRRC63** | 0.01285 | 0.06000 | 0.16753 | 0.10150 | 0.00803 | 0.07727 | 0.01039 | 0.17103 | 0.00722 | 0.00002 | 0.01151 | 0.26663 |
| **LRRC66** | 0.00537 | 0.01504 | 0.07426 | 0.05379 | 0.02262 | 0.00748 | 0.02087 | 0.01573 | 0.00004 | 0.05688 | 0.01521 | 0.03795 |
| **LRRC69** | 0.04955 | 0.48007 | 0.15534 | 0.00000 | 0.35074 | 1.15937 | 0.00000 | 0.12499 | 0.33852 | 0.26674 | 0.36872 | 0.80731 |
| **LRRC7** | 2.36207 | 0.02622 | 0.01352 | 0.00562 | 0.04999 | 0.02010 | 0.00427 | 0.99604 | 0.00359 | 0.06005 | 0.01884 | 0.22735 |
| **LRRC70** | 0.05170 | 0.10985 | 0.21202 | 0.14051 | 0.00000 | 0.06734 | 0.01350 | 0.29954 | 0.00000 | 0.42082 | 0.01546 | 0.08169 |
| **LRRC71** | 0.00602 | 0.01564 | 0.01707 | 0.00929 | 0.02781 | 0.01206 | 0.03085 | 0.00920 | 0.02875 | 0.02751 | 0.01332 | 0.03864 |
| **LRRC72** | 0.00467 | 0.01446 | 0.01655 | 0.01095 | 0.01412 | 0.02091 | 0.01726 | 0.01669 | 0.01739 | 0.01745 | 0.01528 | 0.01559 |
| **LRRC73** | 0.17502 | 0.02216 | 0.00908 | 0.00575 | 0.00000 | 0.00000 | 0.03753 | 0.00156 | 0.02430 | 0.14497 | 0.05168 | 0.05671 |
| **LRRC74** | 0.00878 | 0.01264 | 0.01092 | 0.01562 | 0.00347 | 0.03525 | 0.00886 | 0.01835 | 0.02094 | 0.04311 | 0.01270 | 0.03091 |
| **LRRC75A** | 0.12063 | 0.04351 | 0.41052 | 0.22435 | 0.41973 | 0.00000 | 0.01787 | 0.10416 | 0.00000 | 1.23092 | 0.53365 | 0.06767 |
| **LRRC75A-AS1** | 9.25867 | 34.22370 | 31.26937 | 37.35334 | 36.11934 | 51.30878 | 52.75488 | 20.19458 | 40.09102 | 38.42003 | 33.97234 | 43.51971 |
| **LRRC75B** | 0.00619 | 0.01546 | 0.05447 | 0.16866 | 0.03726 | 0.00000 | 0.01081 | 0.01089 | 0.06907 | 0.00000 | 0.00962 | 0.02294 |
| **LRRC8A** | 6.46000 | 2.58827 | 6.72979 | 6.10839 | 5.89966 | 4.62752 | 10.78373 | 6.34389 | 9.07983 | 17.29108 | 0.86292 | 6.21783 |
| **LRRC8B** | 3.72998 | 0.21981 | 7.60378 | 0.00000 | 2.86747 | 2.07384 | 6.04945 | 0.11934 | 3.70824 | 3.59355 | 0.00000 | 0.00000 |
| **LRRC8C** | 1.06253 | 1.00489 | 0.73093 | 0.37887 | 0.00000 | 0.00000 | 0.01566 | 0.19171 | 0.00000 | 0.02273 | 0.01250 | 0.96328 |
| **LRRC8D** | 3.87770 | 4.14081 | 10.87816 | 2.15035 | 2.82733 | 10.06113 | 3.24724 | 0.83823 | 6.06500 | 2.51066 | 1.75800 | 0.04297 |
| **LRRC8E** | 1.75549 | 0.40132 | 0.40636 | 0.45129 | 0.00000 | 0.00000 | 0.00000 | 0.13989 | 0.85183 | 0.39290 | 0.00000 | 0.06053 |
| **LRRC9** | 0.00917 | 0.00525 | 0.10275 | 0.01069 | 0.02189 | 0.24933 | 0.11407 | 0.00510 | 0.06876 | 0.00000 | 0.00000 | 0.07118 |
| **LRRCC1** | 0.00000 | 2.16092 | 0.16899 | 0.09883 | 2.43742 | 9.89072 | 0.00000 | 0.06174 | 0.00000 | 5.86303 | 1.87034 | 3.54493 |
| **LRRD1** | 0.03190 | 0.06086 | 0.04421 | 0.00902 | 0.01228 | 0.00243 | 0.04265 | 0.08607 | 0.00639 | 0.03301 | 0.01601 | 0.02624 |

| | | | | | | | | | | | | |
|---|---|---|---|---|---|---|---|---|---|---|---|---|
| **LRRFIP1** | 3.15896 | 3.20478 | 10.07345 | 1.65759 | 15.32677 | 10.02572 | 40.19733 | 1.66464 | 15.94157 | 36.45265 | 20.85010 | 0.28855 |
| **LRRFIP2** | 4.57993 | 7.13230 | 6.41498 | 2.88813 | 2.16185 | 10.09269 | 4.58877 | 4.82945 | 5.53600 | 4.09376 | 4.32618 | 6.98989 |
| **LRRIQ1** | 0.11493 | 0.09059 | 0.00485 | 0.07022 | 0.09930 | 0.00000 | 0.01921 | 0.17471 | 0.00007 | 0.10475 | 0.02574 | 0.07634 |
| **LRRIQ3** | 0.11556 | 0.45650 | 0.10726 | 0.01603 | 0.00000 | 0.00000 | 0.00000 | 0.16329 | 0.03228 | 0.72933 | 0.09275 | 1.70976 |
| **LRRIQ4** | 0.00597 | 0.01333 | 0.03634 | 0.01345 | 0.00562 | 0.01380 | 0.02017 | 0.00955 | 0.00007 | 0.03996 | 0.05160 | 0.05128 |
| **LRRK1** | 3.16786 | 0.00137 | 0.71804 | 0.24071 | 0.17808 | 0.00000 | 0.00000 | 0.17655 | 0.00000 | 0.17355 | 0.00688 | 0.03367 |
| **LRRK2** | 0.00426 | 0.11269 | 0.00653 | 0.70786 | 0.00000 | 0.00000 | 0.07432 | 0.00634 | 0.00000 | 0.13214 | 0.01906 | 0.11260 |
| **LRRN1** | 0.53367 | 0.00957 | 0.83496 | 0.92634 | 0.00000 | 0.02496 | 0.00000 | 0.04354 | 0.18089 | 0.15271 | 0.00000 | 0.16169 |
| **LRRN2** | 0.00584 | 0.01635 | 0.04117 | 0.02713 | 0.02311 | 0.00718 | 0.01838 | 0.00940 | 0.02808 | 0.00695 | 0.06829 | 0.05676 |
| **LRRN3** | 0.00327 | 0.08633 | 0.00103 | 0.00000 | 0.15334 | 0.00000 | 0.00143 | 0.00718 | 0.14494 | 0.26042 | 0.02480 | 0.08494 |
| **LRRN4** | 1.23043 | 0.00248 | 0.00241 | 0.28388 | 0.00000 | 0.00000 | 0.00205 | 0.18872 | 0.02495 | 0.12723 | 0.02275 | 0.05227 |
| **LRRN4CL** | 0.00495 | 0.00596 | 0.00273 | 0.00679 | 0.00000 | 0.01123 | 0.01197 | 0.00901 | 0.04979 | 0.16698 | 0.03890 | 0.04775 |
| **LRRTM1** | 0.00629 | 0.02543 | 0.49451 | 0.16508 | 0.01162 | 0.00000 | 0.00000 | 0.00816 | 0.00000 | 0.02323 | 0.02783 | 0.12672 |
| **LRRTM2** | 0.00458 | 0.00000 | 0.42976 | 0.32445 | 0.03645 | 0.00000 | 0.00000 | 0.02015 | 0.31827 | 2.55939 | 0.15886 | 0.15447 |
| **LRRTM3** | 0.01698 | 0.01023 | 0.02653 | 0.01218 | 0.00518 | 0.02986 | 0.02841 | 0.03765 | 0.00890 | 0.07429 | 0.01560 | 0.01023 |
| **LRRTM4** | 11.50355 | 0.26650 | 0.12241 | 0.02489 | 0.05669 | 0.20760 | 0.03090 | 11.21099 | 0.00000 | 0.00000 | 0.24800 | 0.00000 |
| **LRSAM1** | 0.29346 | 2.05868 | 2.50836 | 2.05762 | 0.98035 | 0.00000 | 1.57437 | 0.21515 | 4.90737 | 0.89807 | 2.31071 | 1.83209 |
| **LRTM1** | 0.01396 | 0.05994 | 0.03322 | 0.09383 | 0.10730 | 0.18109 | 0.00000 | 0.00628 | 0.00000 | 0.11503 | 0.42408 | 0.00000 |
| **LRTM2** | 0.00679 | 0.37295 | 0.00000 | 0.01370 | 0.00000 | 0.10039 | 0.00000 | 0.01092 | 0.03320 | 0.03425 | 0.00008 | 0.09224 |
| **LRTOMT** | 1.94723 | 1.77596 | 0.98732 | 0.57565 | 1.31814 | 0.16330 | 1.01882 | 2.72486 | 0.59958 | 0.00000 | 0.53374 | 4.58649 |
| **LRWD1** | 4.48819 | 9.49605 | 7.89660 | 4.76734 | 2.13932 | 0.00000 | 5.68473 | 0.83571 | 12.69040 | 0.00000 | 7.73219 | 1.59229 |
| **LSAMP** | 0.08300 | 0.03681 | 0.17048 | 0.20384 | 0.10045 | 0.07554 | 0.00000 | 0.00649 | 0.00000 | 0.05577 | 0.04133 | 0.02081 |
| **LSAMP-AS1** | 0.00756 | 0.20199 | 0.00663 | 0.13495 | 0.01131 | 0.13927 | 0.00000 | 0.00344 | 0.16911 | 0.00000 | 0.00438 | 0.28258 |
| **LSG1** | 0.79263 | 11.07480 | 10.57728 | 6.30673 | 8.37027 | 11.95032 | 16.89951 | 16.15256 | 11.18974 | 7.64857 | 7.71201 | 17.00659 |
| **1.00000** | 0.00602 | 15.65916 | 12.06458 | 8.34316 | 9.17500 | 30.46115 | 14.41470 | 17.91176 | 13.41030 | 0.00000 | 9.68933 | 25.54581 |
| **10.00000** | 4.23294 | 11.65862 | 9.56493 | 8.26240 | 9.68457 | 9.32351 | 15.79931 | 11.73844 | 15.88629 | 0.00000 | 11.49455 | 16.88240 |
| **11.00000** | 1.87816 | 0.34374 | 0.24705 | 0.00000 | 0.58415 | 0.31397 | 0.00000 | 0.12586 | 0.08905 | 0.86544 | 0.00001 | 0.53102 |
| **12.00000** | 0.00923 | 1.57431 | 1.86954 | 0.87460 | 0.52833 | 4.41090 | 2.54037 | 1.92215 | 2.78634 | 0.00000 | 2.20707 | 2.34029 |
| **LSM14A** | 4.70209 | 14.48724 | 12.01784 | 10.09132 | 10.33608 | 22.53095 | 25.62809 | 16.14158 | 13.04585 | 17.15782 | 15.50605 | 16.66523 |
| **LSM14B** | 1.58039 | 0.09051 | 0.06652 | 0.49868 | 1.01101 | 0.21430 | 1.00765 | 0.28270 | 0.00000 | 5.47861 | 0.30831 | 0.44538 |
| **2.00000** | 2.79509 | 19.33566 | 19.64995 | 12.50527 | 16.46967 | 8.78869 | 27.24072 | 4.47494 | 25.92228 | 3.40726 | 17.31250 | 21.49595 |
| **3.00000** | 1.46596 | 12.34897 | 17.34730 | 9.50101 | 12.39271 | 39.49162 | 27.13685 | 3.72434 | 11.88322 | 14.90781 | 19.61767 | 18.67218 |
| **4.00000** | 1.15107 | 27.68489 | 21.79934 | 17.21000 | 33.85941 | 11.00515 | 27.93937 | 0.00000 | 31.81290 | 9.68731 | 25.57125 | 18.52376 |
| **5.00000** | 0.52298 | 14.13721 | 8.92511 | 6.29932 | 12.13446 | 45.02489 | 20.67812 | 0.95725 | 13.16673 | 14.23203 | 18.62745 | 9.98277 |
| **6.00000** | 0.84610 | 7.88594 | 5.58005 | 3.76622 | 5.63507 | 26.62518 | 8.85131 | 0.00000 | 5.59895 | 7.07356 | 10.77010 | 7.03065 |
| **7.00000** | 6.69432 | 11.31911 | 15.83930 | 13.89104 | 15.33302 | 0.07025 | 27.77059 | 11.35826 | 17.84511 | 8.70787 | 15.48094 | 9.81076 |
| **8.00000** | 1.54063 | 12.08772 | 12.89382 | 1.33821 | 5.24039 | 27.39107 | 14.30907 | 0.94004 | 13.70936 | 10.94488 | 9.58595 | 12.86364 |
| **LSMEM1** | 0.09263 | 0.13343 | 0.28051 | 0.21687 | 0.08226 | 0.00000 | 0.00000 | 0.71716 | 0.72117 | 0.46075 | 0.00000 | 0.20025 |
| **LSMEM2** | 0.00552 | 0.08796 | 0.03445 | 0.01773 | 0.00470 | 0.00000 | 0.00319 | 0.01047 | 0.14554 | 0.11282 | 0.00558 | 0.00865 |
| **LSP1** | 0.00387 | 0.01616 | 0.13392 | 0.00814 | 0.01660 | 0.01905 | 0.00703 | 0.02370 | 0.01500 | 0.01115 | 0.02447 | 0.01589 |
| **LSP1P3** | 0.04299 | 0.17203 | 0.10124 | 0.11875 | 0.07215 | 0.02492 | 0.10057 | 0.04381 | 0.12739 | 0.00000 | 0.01946 | 0.31492 |
| **LSR** | 0.93916 | 6.37359 | 6.44103 | 3.64094 | 5.74810 | 0.00000 | 0.22687 | 0.35582 | 11.06889 | 10.38936 | 3.63745 | 4.55727 |
| **LSS** | 2.22240 | 3.46818 | 4.52671 | 10.22235 | 21.70691 | 0.00000 | 9.26285 | 2.51708 | 19.33228 | 7.52494 | 19.58889 | 2.16881 |
| **LST1** | 0.06534 | 0.02280 | 0.01367 | 0.01113 | 0.02006 | 0.00857 | 0.01484 | 0.00562 | 0.02110 | 0.02495 | 0.01564 | 0.08640 |
| **LTA** | 0.00915 | 0.38132 | 0.11500 | 0.00773 | 0.10613 | 0.00425 | 0.00536 | 0.01390 | 0.00000 | 0.01112 | 0.18740 | 0.05037 |
| **LTA4H** | 8.68074 | 5.01888 | 4.29690 | 8.74229 | 4.11506 | 12.29508 | 15.47866 | 3.73661 | 14.66240 | 0.00000 | 13.05239 | 0.00000 |
| **LTB4R** | 0.66158 | 0.25271 | 1.40910 | 0.34456 | 0.16013 | 0.00000 | 0.00000 | 0.02398 | 2.48295 | 3.85596 | 0.76783 | 1.43549 |
| **LTB4R2** | 0.06352 | 0.02881 | 0.06349 | 0.00000 | 0.04325 | 0.02721 | 0.11211 | 0.04881 | 0.00000 | 0.87707 | 0.04983 | 0.06317 |

| | | | | | | | | | | | | |
|---|---|---|---|---|---|---|---|---|---|---|---|---|
| **LTBP1** | 3.91686 | 0.00000 | 0.05442 | 0.92357 | 3.27687 | 0.51869 | 0.31823 | 1.34742 | 0.00000 | 2.95998 | 0.22993 | 0.00000 |
| **LTBP2** | 1.61382 | 0.28212 | 0.18218 | 0.29047 | 0.04436 | 0.00000 | 0.00000 | 0.07283 | 0.10922 | 0.11712 | 0.19744 | 0.22914 |
| **LTBP3** | 0.38556 | 1.29455 | 1.65654 | 0.00000 | 1.65699 | 0.00000 | 1.21683 | 0.79604 | 5.52877 | 0.00000 | 1.17839 | 0.67970 |
| **LTBP4** | 0.86275 | 4.55815 | 0.98078 | 5.66575 | 3.47388 | 0.00000 | 0.20975 | 3.55394 | 2.37090 | 2.36586 | 0.01781 | 4.18620 |
| **LTBR** | 0.00040 | 0.55132 | 3.16277 | 3.09695 | 3.69815 | 0.00000 | 2.45935 | 0.01931 | 1.95952 | 0.25508 | 0.49248 | 0.02730 |
| **LTC4S** | 0.00381 | 0.00933 | 0.00423 | 0.02199 | 0.00000 | 0.04912 | 0.01310 | 0.01602 | 0.03926 | 0.05241 | 0.03082 | 0.02175 |
| **LTF** | 0.00565 | 0.00906 | 0.03427 | 0.08893 | 0.00022 | 0.03230 | 0.03704 | 0.06935 | 0.02183 | 0.02626 | 0.02826 | 0.01111 |
| **LTK** | 0.33073 | 0.01522 | 0.18949 | 0.21640 | 0.00000 | 0.04890 | 0.16539 | 0.00199 | 0.06276 | 0.08018 | 0.00000 | 0.11100 |
| **LTN1** | 3.34367 | 1.69244 | 5.69717 | 3.35232 | 3.51068 | 20.55566 | 4.51183 | 2.73805 | 5.61846 | 8.43632 | 2.47303 | 3.52126 |
| **LTV1** | 5.48100 | 14.02443 | 10.52105 | 4.07007 | 3.31860 | 36.30264 | 14.25011 | 14.25875 | 16.49651 | 4.89350 | 17.13422 | 17.05510 |
| **LUC7L** | 0.49351 | 10.97127 | 7.04010 | 9.65888 | 7.76012 | 8.83603 | 14.27555 | 10.69997 | 15.89450 | 10.30504 | 7.94841 | 9.93475 |
| **LUC7L2.uid-939** | 0.00584 | 0.02506 | 0.04783 | 0.02551 | 0.00000 | 0.02457 | 0.03201 | 0.12837 | 0.02516 | 0.03303 | 0.03281 | 0.11098 |
| **LUC7L2.uid-939** | 0.18416 | 2.46389 | 1.53978 | 1.34267 | 1.46551 | 0.00000 | 0.70914 | 4.15728 | 3.05924 | 0.00000 | 1.36451 | 2.16110 |
| **LUC7L3** | 3.42962 | 8.94666 | 9.88904 | 8.59605 | 8.20273 | 31.43694 | 26.53394 | 16.24624 | 14.52364 | 39.10227 | 13.57673 | 20.43679 |
| **LUCAT1** | 0.00273 | 0.18776 | 0.09374 | 0.13028 | 0.95574 | 0.13367 | 0.00000 | 0.00196 | 0.00000 | 0.51116 | 0.25522 | 0.18998 |
| **LURAP1** | 0.08723 | 0.18119 | 0.15049 | 0.35526 | 1.69841 | 0.10566 | 0.00000 | 0.13029 | 0.00000 | 0.54592 | 1.75880 | 0.17631 |
| **LURAP1L** | 0.00215 | 0.01496 | 2.68813 | 0.01663 | 0.00431 | 0.01296 | 0.00000 | 0.04050 | 0.11450 | 0.03862 | 0.00873 | 0.01309 |
| **LUZP1** | 7.09636 | 9.66017 | 8.37991 | 7.11569 | 27.23291 | 12.84972 | 11.86497 | 16.67014 | 11.57794 | 20.64472 | 14.85384 | 17.63209 |
| **LUZP2** | 0.71790 | 0.01456 | 0.01430 | 0.01899 | 0.01552 | 0.00022 | 0.01608 | 0.00327 | 0.02673 | 0.01914 | 0.01531 | 0.04732 |
| **LUZP4** | 0.00208 | 0.00000 | 0.04120 | 0.26418 | 0.00000 | 0.19278 | 0.00000 | 0.02544 | 0.00000 | 0.80842 | 0.04229 | 4.24880 |
| **LUZP6+MTPN** | 4.68789 | 3.79589 | 16.08796 | 4.23985 | 7.25427 | 14.68630 | 23.40235 | 8.35653 | 1.74480 | 20.91143 | 11.65251 | 12.81478 |
| **LXN** | 0.00652 | 0.08601 | 1.31902 | 0.00000 | 0.00000 | 0.10625 | 1.25902 | 0.21991 | 0.39300 | 0.20143 | 0.00000 | 0.00000 |
| **LY6E** | 0.15635 | 7.73554 | 5.06615 | 9.07316 | 19.50078 | 0.00000 | 2.16675 | 0.25338 | 16.28851 | 5.38433 | 19.16875 | 2.40878 |
| **LY6G5B** | 0.41197 | 1.46782 | 0.55699 | 0.04005 | 0.82904 | 0.00000 | 0.00000 | 1.31144 | 3.10836 | 1.56181 | 0.12512 | 1.94424 |
| **LY6G5C** | 0.05583 | 0.00000 | 0.17739 | 0.02253 | 0.69586 | 0.00000 | 0.05014 | 0.00000 | 0.05788 | 0.70140 | 0.00000 | 0.15009 |
| **LY6G6C** | 0.00359 | 0.00000 | 3.34020 | 0.00000 | 1.06113 | 0.04324 | 0.00000 | 0.00795 | 0.50531 | 0.10992 | 1.04842 | 0.00982 |
| **LY6G6D** | 0.00257 | 0.01648 | 0.01165 | 0.01196 | 0.01375 | 0.01814 | 0.03395 | 0.04737 | 0.04681 | 0.00093 | 0.01121 | 0.01496 |
| **LY6H** | 0.00795 | 0.02471 | 0.01648 | 0.01106 | 0.01966 | 0.01947 | 0.02185 | 0.01176 | 0.01957 | 0.00629 | 0.01356 | 0.01992 |
| **LY6K** | 0.08233 | 0.01394 | 0.01403 | 0.01251 | 0.01858 | 0.01496 | 0.02098 | 0.01158 | 0.02198 | 0.00912 | 0.01403 | 0.04402 |
| **LY75** | 0.00286 | 0.01170 | 0.09655 | 0.05677 | 0.00361 | 0.03030 | 0.00001 | 0.04176 | 0.02887 | 0.02066 | 0.01134 | 0.01341 |
| **LY86** | 0.00891 | 0.01798 | 0.04709 | 0.01660 | 0.02829 | 0.00000 | 0.01575 | 0.02076 | 0.02001 | 0.00326 | 0.01604 | 0.01985 |
| **LY86-AS1** | 0.01818 | 0.02560 | 0.04602 | 0.02423 | 0.03268 | 0.00000 | 0.02563 | 0.02408 | 0.01613 | 0.00000 | 0.02506 | 0.02946 |
| **LY9** | 0.00658 | 0.09871 | 1.42103 | 0.08449 | 1.42632 | 0.00000 | 0.01462 | 0.00481 | 0.00000 | 0.00000 | 0.38673 | 0.29963 |
| **LY96** | 0.00440 | 0.01045 | 0.00786 | 0.09389 | 0.01604 | 0.03568 | 0.02503 | 0.01802 | 0.00988 | 0.00359 | 0.03406 | 0.02743 |
| **LYAR** | 3.26801 | 12.29574 | 12.27004 | 2.85801 | 8.14249 | 27.31446 | 21.84403 | 15.19033 | 19.62761 | 14.13617 | 14.79020 | 20.18392 |
| **LYG1** | 0.03627 | 0.04741 | 0.28649 | 0.22479 | 0.00000 | 0.00084 | 0.94834 | 0.80669 | 0.00000 | 0.78329 | 0.03463 | 0.34389 |
| **LYG2** | 0.00159 | 0.38172 | 0.01321 | 0.04780 | 0.00000 | 0.23304 | 0.00000 | 0.21462 | 0.19717 | 0.20556 | 0.08941 | 0.85000 |
| **LYL1** | 0.00413 | 0.00855 | 0.00740 | 0.03601 | 0.01344 | 0.01739 | 0.02830 | 0.01006 | 0.00000 | 0.17442 | 0.01069 | 0.01088 |
| **LYN** | 0.00000 | 2.58878 | 33.01471 | 0.00000 | 20.96546 | 14.56838 | 31.48314 | 0.00000 | 13.88105 | 13.65220 | 9.18040 | 0.00000 |
| **LYNX1** | 0.17332 | 0.01574 | 0.01914 | 0.00884 | 0.16957 | 0.00000 | 0.01799 | 0.27635 | 0.00000 | 0.05625 | 0.10471 | 0.00658 |
| **LYPD1** | 3.08444 | 0.00205 | 0.25735 | 0.40887 | 0.00100 | 0.00000 | 0.00000 | 0.95780 | 0.00784 | 0.04207 | 0.02007 | 0.07846 |
| **LYPD2** | 0.00578 | 0.00264 | 0.00647 | 0.00453 | 0.01622 | 0.00000 | 0.00355 | 0.00973 | 0.00000 | 0.08335 | 3.19364 | 0.01374 |
| **LYPD3** | 0.00230 | 0.00000 | 14.15653 | 0.52395 | 12.02625 | 0.00000 | 0.00000 | 0.57588 | 5.02547 | 0.00000 | 2.16931 | 0.64354 |
| **LYPD4** | 4.39855 | 0.00920 | 0.00000 | 0.01729 | 0.01447 | 0.11881 | 0.00000 | 0.49151 | 0.01100 | 0.03545 | 0.00941 | 0.17815 |
| **LYPD5** | 0.02272 | 0.08393 | 4.97368 | 0.00457 | 0.00701 | 0.00924 | 0.18566 | 0.03286 | 0.32282 | 0.00000 | 0.05891 | 0.30662 |
| **LYPD6** | 1.81237 | 0.00495 | 0.10366 | 0.03692 | 0.05957 | 0.14727 | 0.00000 | 0.00000 | 0.19731 | 0.00000 | 0.05387 | 0.02778 |
| **LYPD6B** | 0.21098 | 1.48016 | 0.23599 | 0.18536 | 0.17255 | 1.81404 | 0.00000 | 0.02421 | 3.28829 | 0.91840 | 0.00000 | 0.12980 |
| **LYPLA1** | 0.00000 | 10.71242 | 7.22830 | 10.38743 | 1.19699 | 47.71270 | 10.81012 | 2.80386 | 9.10568 | 13.38874 | 12.99552 | 9.31277 |

| | | | | | | | | | | | | |
|---|---|---|---|---|---|---|---|---|---|---|---|---|
| **LYPLA2** | 0.93543 | 9.97001 | 15.19876 | 9.94557 | 11.48669 | 0.00000 | 16.81268 | 0.00000 | 18.87805 | 0.00000 | 12.72227 | 3.99789 |
| **LYPLA2P2** | 0.05837 | 0.53725 | 1.00010 | 0.65200 | 0.25061 | 0.22860 | 0.47052 | 0.00327 | 1.51026 | 0.00000 | 0.76001 | 0.17338 |
| **LYPLAL1** | 0.63552 | 2.75421 | 6.64573 | 1.68041 | 2.27823 | 11.98299 | 4.11648 | 0.20629 | 2.07012 | 0.00000 | 6.05425 | 3.32662 |
| **LYPLAL1-AS1** | 0.06034 | 0.01119 | 0.01515 | 0.01190 | 0.07092 | 0.00000 | 0.01941 | 0.01060 | 0.01420 | 0.02107 | 0.02110 | 0.03236 |
| **LYRM1** | 0.94525 | 6.26554 | 3.25207 | 3.08029 | 0.66046 | 7.12794 | 1.08978 | 2.53363 | 1.61825 | 4.64183 | 5.39104 | 8.89004 |
| **LYRM2** | 0.46292 | 2.89971 | 8.07083 | 1.26453 | 5.39048 | 13.27345 | 6.38908 | 1.43243 | 0.40896 | 5.64528 | 4.88965 | 6.97253 |
| **LYRM4** | 0.69679 | 8.02774 | 9.71797 | 6.37356 | 5.85194 | 14.75309 | 14.93407 | 1.02591 | 15.89034 | 4.98391 | 8.11268 | 12.67795 |
| **LYRM5** | 1.11516 | 0.35600 | 0.46957 | 0.47390 | 0.50023 | 4.21129 | 0.00000 | 0.14422 | 0.00000 | 0.83630 | 2.51168 | 0.12146 |
| **LYRM7** | 0.10975 | 1.83627 | 4.32069 | 0.59378 | 1.51732 | 10.99318 | 5.58799 | 0.23031 | 3.69979 | 10.29003 | 10.26719 | 1.90156 |
| **LYRM9** | 0.67798 | 0.11115 | 0.10865 | 0.00504 | 0.00093 | 0.02003 | 0.07907 | 0.17911 | 0.00000 | 0.00894 | 0.33339 | 0.08475 |
| **LYSMD1** | 0.72328 | 2.07523 | 5.58036 | 0.42172 | 0.42162 | 3.25198 | 2.90437 | 11.91262 | 2.23568 | 4.22137 | 2.97562 | 3.50369 |
| **LYSMD2** | 0.00267 | 0.20580 | 0.15385 | 0.00323 | 0.00022 | 0.03834 | 0.00000 | 0.03810 | 0.03546 | 0.02210 | 0.02579 | 0.06845 |
| **LYSMD3** | 0.19052 | 3.34298 | 3.96543 | 0.00000 | 0.00000 | 13.62832 | 2.88864 | 7.18589 | 4.74129 | 1.53979 | 2.74586 | 3.56092 |
| **LYSMD4** | 0.48623 | 2.82075 | 1.20125 | 2.92973 | 3.15861 | 0.13961 | 1.03855 | 2.15684 | 1.62997 | 0.00000 | 1.41297 | 3.44383 |
| **LYST** | 2.00113 | 0.71044 | 0.40665 | 0.65667 | 0.00000 | 0.87580 | 0.65777 | 0.29192 | 0.00514 | 0.08413 | 0.38431 | 1.44284 |
| **LYVE1** | 0.00719 | 0.01163 | 0.01024 | 0.05161 | 0.00669 | 0.03562 | 0.01465 | 0.01509 | 0.01367 | 0.01883 | 0.02400 | 0.01415 |
| **LYZ** | 0.40110 | 0.07203 | 0.00000 | 2.64275 | 0.00000 | 0.15200 | 0.01541 | 0.56653 | 0.00000 | 0.02907 | 0.00677 | 0.00000 |
| **LYZL1** | 0.00583 | 0.01718 | 0.02161 | 0.01639 | 0.01635 | 0.00469 | 0.01627 | 0.01004 | 0.02281 | 0.02289 | 0.01517 | 0.03222 |
| **LYZL6** | 0.00612 | 0.01464 | 0.02086 | 0.01055 | 0.01460 | 0.01994 | 0.01631 | 0.01033 | 0.01459 | 0.02046 | 0.01523 | 0.01753 |
| **LZIC** | 1.37232 | 5.44129 | 4.51110 | 2.51753 | 3.95734 | 14.93457 | 4.88670 | 7.63013 | 6.21064 | 6.18123 | 5.74027 | 13.16496 |
| **LZTFL1** | 0.22978 | 1.34181 | 0.37830 | 0.51682 | 0.00000 | 0.72389 | 0.91358 | 1.66166 | 1.01611 | 0.00000 | 0.41496 | 3.90797 |
| **LZTR1** | 0.19618 | 1.05408 | 2.62508 | 0.78191 | 0.00001 | 0.00000 | 3.83669 | 0.12117 | 2.88031 | 6.97564 | 1.85689 | 0.39162 |
| **LZTS1** | 0.00338 | 0.01387 | 0.68395 | 4.51889 | 0.00000 | 0.03984 | 0.02187 | 0.00927 | 0.00000 | 0.10595 | 0.01281 | 0.00896 |
| **LZTS1-AS1** | 0.00504 | 0.00373 | 0.03200 | 0.11003 | 0.00000 | 0.06469 | 0.03178 | 0.00984 | 0.00006 | 0.05990 | 0.00857 | 0.01113 |
| **LZTS2** | 0.21371 | 1.12479 | 4.19033 | 2.18023 | 0.60214 | 0.00000 | 0.49479 | 0.05150 | 2.73015 | 2.23147 | 0.99485 | 0.17349 |
| **LZTS3** | 0.09464 | 0.12676 | 0.02447 | 0.74006 | 0.00000 | 0.00000 | 0.00252 | 0.04321 | 0.16115 | 0.81290 | 0.00000 | 1.47976 |
| **M1AP** | 0.99004 | 4.28829 | 0.65548 | 0.00000 | 0.77790 | 0.02165 | 0.47190 | 0.50188 | 0.70778 | 0.00000 | 2.95570 | 8.04932 |
| **M6PR** | 2.83846 | 4.78032 | 13.13239 | 4.15329 | 1.16804 | 14.73145 | 2.77473 | 1.29995 | 23.31690 | 0.00000 | 7.95230 | 8.31127 |
| **MAATS1** | 0.09842 | 0.18014 | 0.09190 | 0.22614 | 0.00124 | 0.00949 | 0.00227 | 0.18893 | 0.23762 | 0.00000 | 0.00098 | 0.00468 |
| **MAB21L1** | 0.16385 | 0.01331 | 0.01222 | 0.00823 | 0.00972 | 0.02271 | 0.01426 | 0.07102 | 0.05155 | 0.01876 | 0.01107 | 0.01060 |
| **MAB21L2** | 0.06104 | 0.00826 | 0.01336 | 0.07203 | 0.01635 | 0.00000 | 0.01797 | 0.03515 | 0.02187 | 0.14109 | 0.01203 | 0.05392 |
| **MAB21L3** | 0.00229 | 0.00000 | 1.86709 | 0.08240 | 0.00000 | 0.07258 | 0.28986 | 0.01612 | 0.00000 | 0.13157 | 0.12109 | 0.01322 |
| **MACC1** | 6.65950 | 0.04741 | 0.01891 | 0.02556 | 0.03246 | 0.27619 | 0.00000 | 0.58890 | 0.00000 | 0.24804 | 0.01150 | 0.00000 |
| **MACF1** | 1.09231 | 3.38889 | 3.18225 | 11.13504 | 6.37251 | 5.77530 | 1.31997 | 1.60254 | 8.56417 | 17.37570 | 1.27443 | 8.44108 |
| **MACROD1** | 0.14673 | 0.27046 | 0.03900 | 0.84847 | 1.42259 | 0.00000 | 0.16649 | 0.01674 | 0.00000 | 0.92985 | 0.13553 | 0.00420 |
| **MACROD2** | 0.35307 | 0.06641 | 0.06210 | 0.00065 | 0.05007 | 0.05952 | 0.03836 | 0.03237 | 0.05103 | 0.01310 | 0.08492 | 0.08704 |
| **MACROD2-AS1** | 0.01039 | 0.00269 | 0.00208 | 0.00146 | 0.00934 | 0.13736 | 0.02878 | 0.00793 | 0.00000 | 0.01496 | 0.02544 | 0.01014 |
| **MACROD2-IT1** | 0.01567 | 0.01520 | 0.01310 | 0.02351 | 0.01896 | 0.01489 | 0.01785 | 0.01149 | 0.02022 | 0.01232 | 0.01542 | 0.01624 |
| **MAD1L1** | 2.02685 | 10.56281 | 8.03694 | 9.17982 | 8.39935 | 0.00000 | 8.36111 | 6.92917 | 9.22667 | 0.00000 | 9.66499 | 5.65573 |
| **MAD2L1** | 9.18765 | 11.21772 | 13.20467 | 4.85941 | 2.78245 | 41.93220 | 24.02392 | 11.78526 | 19.69687 | 3.62869 | 14.46971 | 20.02781 |
| **MAD2L1BP** | 4.28209 | 18.49081 | 5.88230 | 10.82104 | 9.11283 | 15.86408 | 13.01468 | 36.72594 | 15.14975 | 0.00000 | 13.66165 | 24.85928 |
| **MAD2L2** | 2.46260 | 11.30216 | 3.96233 | 13.76670 | 8.75724 | 0.45581 | 2.99735 | 8.04779 | 5.53330 | 0.00000 | 1.81986 | 10.80098 |
| **MADCAM1** | 0.11446 | 0.01443 | 0.01136 | 0.00867 | 0.01001 | 0.01922 | 0.03950 | 0.00630 | 0.00702 | 0.04107 | 0.01532 | 0.01467 |
| **MADD** | 8.01577 | 0.59690 | 2.32211 | 2.14913 | 4.85058 | 0.00000 | 2.72768 | 0.00000 | 5.02670 | 3.69522 | 1.12117 | 1.29192 |
| **MAEA** | 6.02114 | 19.73752 | 11.48248 | 9.23287 | 17.13206 | 14.72037 | 19.26252 | 2.54962 | 22.64863 | 2.67470 | 17.75691 | 14.82126 |
| **MAEL** | 3.58311 | 12.21309 | 0.11088 | 3.31678 | 2.10219 | 4.16043 | 0.00000 | 3.25417 | 0.00000 | 0.30437 | 0.00000 | 17.98019 |
| **MAF** | 0.00617 | 0.00000 | 0.21217 | 0.55531 | 0.11207 | 0.06519 | 0.00000 | 0.16708 | 0.00000 | 0.53400 | 0.00000 | 0.00000 |
| **1.00000** | 1.04947 | 14.62907 | 12.17636 | 9.55385 | 16.96772 | 0.00000 | 8.93164 | 0.59295 | 16.93449 | 17.02013 | 4.75216 | 4.65681 |

| | | | | | | | | | | | | |
|---|---|---|---|---|---|---|---|---|---|---|---|---|
| **MAFA** | 0.14542 | 0.05263 | 0.01315 | 0.00000 | 0.81049 | 0.00000 | 0.15447 | 0.00000 | 0.00000 | 1.68074 | 0.23422 | 0.29596 |
| **MAFB** | 0.18800 | 0.00519 | 0.17683 | 0.02509 | 0.12546 | 0.00000 | 0.00000 | 0.04516 | 0.01154 | 0.37400 | 0.00426 | 0.00491 |
| **MAFF** | 0.61425 | 1.25860 | 5.98614 | 1.05637 | 0.74906 | 0.00000 | 0.00000 | 0.82331 | 0.23874 | 0.94843 | 0.00000 | 0.99702 |
| **MAFG** | 1.99980 | 2.32102 | 11.81854 | 3.80193 | 6.89011 | 0.00000 | 5.78635 | 1.31589 | 8.54795 | 16.05636 | 1.98534 | 1.62194 |
| **MAFG-AS1** | 0.41322 | 0.01899 | 0.73442 | 0.00710 | 0.00000 | 0.00000 | 0.36329 | 0.11230 | 0.00000 | 1.06906 | 0.01717 | 0.08729 |
| **MAFK** | 0.53339 | 0.16225 | 1.23866 | 0.27512 | 0.98447 | 0.00000 | 0.52282 | 0.00100 | 0.73134 | 0.00000 | 0.00000 | 0.06451 |
| **MAG** | 0.00466 | 5.12796 | 0.58502 | 0.00000 | 3.32328 | 0.00000 | 0.08308 | 0.01808 | 0.00000 | 0.10028 | 0.00000 | 0.72213 |
| **MAGEA1** | 0.00137 | 0.35733 | 0.14252 | 0.00582 | 0.15988 | 0.23583 | 0.01183 | 1.23730 | 0.00000 | 0.00000 | 0.02344 | 0.20052 |
| **MAGEA10** | 0.00167 | 0.25101 | 0.01835 | 0.29057 | 0.13282 | 0.08111 | 0.82225 | 0.52063 | 0.11386 | 0.00000 | 0.47417 | 2.08538 |
| **MAGEA10-MAG** | 0.01124 | 0.01287 | 0.01211 | 0.00980 | 0.00877 | 0.02948 | 0.01298 | 0.01032 | 0.01398 | 0.03377 | 0.01471 | 0.03052 |
| **MAGEA11** | 0.00448 | 0.01154 | 0.23727 | 0.01510 | 0.00785 | 0.01303 | 0.00549 | 0.87925 | 0.66238 | 0.00000 | 0.07922 | 0.03099 |
| **MAGEA12** | 0.00399 | 0.00000 | 0.12937 | 0.01262 | 0.00903 | 0.00000 | 0.22661 | 11.29341 | 0.02262 | 0.00000 | 0.00360 | 0.77000 |
| **MAGEA2.uid-13** | 0.00632 | 0.01720 | 0.01879 | 0.01405 | 0.01766 | 0.01093 | 0.01938 | 0.02077 | 0.02074 | 0.01012 | 0.01551 | 0.01957 |
| **MAGEA3** | 0.00430 | 0.00268 | 0.08932 | 0.01333 | 0.02569 | 0.00000 | 0.02357 | 4.64872 | 0.05958 | 0.00000 | 0.00850 | 0.40434 |
| **MAGEA4** | 0.03083 | 12.42846 | 0.04634 | 0.00000 | 0.00000 | 3.23714 | 7.99490 | 12.16731 | 0.00000 | 2.36767 | 3.47111 | 22.27014 |
| **MAGEA6** | 0.01190 | 0.01041 | 0.03681 | 0.02157 | 0.02433 | 0.00000 | 0.05139 | 3.08847 | 0.01223 | 0.00000 | 0.01355 | 0.15107 |
| **MAGEA8** | 0.02621 | 0.01243 | 0.70697 | 0.05028 | 0.02958 | 0.00000 | 0.00549 | 0.17179 | 0.03523 | 0.00000 | 0.01952 | 0.08237 |
| **MAGEA8-AS1** | 0.00632 | 0.02016 | 0.01360 | 0.01637 | 0.01435 | 0.02140 | 0.02887 | 0.01041 | 0.02246 | 0.00826 | 0.01670 | 0.01675 |
| **MAGEB1** | 0.00094 | 1.95018 | 1.17622 | 0.42099 | 0.00000 | 0.52883 | 1.29055 | 0.71945 | 0.07904 | 0.00000 | 0.84649 | 6.45085 |
| **MAGEB10** | 0.00626 | 0.13042 | 0.03449 | 0.00003 | 0.00087 | 0.01431 | 0.01053 | 0.26214 | 0.00828 | 0.02158 | 0.01455 | 1.77538 |
| **MAGEB16** | 0.15234 | 0.00324 | 0.07403 | 0.00357 | 0.06240 | 0.10722 | 0.00000 | 0.00280 | 0.01741 | 0.00000 | 0.05769 | 0.01304 |
| **MAGEB17** | 0.00383 | 0.03602 | 0.02497 | 0.14579 | 0.00401 | 0.00701 | 0.01611 | 0.02729 | 0.02507 | 0.01429 | 0.02644 | 0.18405 |
| **MAGEB2** | 0.00000 | 6.84494 | 1.89152 | 0.00268 | 1.48849 | 8.13585 | 5.05963 | 4.65600 | 0.22695 | 1.63993 | 3.40352 | 21.73749 |
| **MAGEB3** | 0.00178 | 0.04610 | 0.02052 | 0.00715 | 0.01551 | 0.00088 | 0.01680 | 0.51271 | 0.01642 | 0.00401 | 0.01520 | 2.76855 |
| **MAGEB4** | 0.00532 | 0.01070 | 0.00942 | 0.00621 | 0.01126 | 0.03112 | 0.00899 | 0.01022 | 0.01924 | 0.05894 | 0.01040 | 0.01439 |
| **MAGEC1** | 0.00044 | 0.08820 | 0.00000 | 0.33729 | 0.00000 | 0.33868 | 0.81634 | 0.00443 | 0.00000 | 1.72659 | 0.21545 | 0.00000 |
| **MAGEC2** | 0.06762 | 0.01477 | 0.03506 | 0.01001 | 0.01131 | 0.01248 | 0.05185 | 0.00717 | 0.07263 | 0.00000 | 0.01493 | 0.02828 |
| **MAGEC3** | 0.00553 | 0.00940 | 0.00683 | 0.03653 | 0.00000 | 0.03619 | 0.04323 | 0.01076 | 0.02164 | 0.03641 | 0.02738 | 0.01474 |
| **MAGED1** | 0.93724 | 6.30212 | 19.16443 | 0.94985 | 3.62993 | 2.86852 | 13.06098 | 0.16410 | 14.30235 | 0.00000 | 3.10694 | 3.00360 |
| **MAGED2** | 0.02368 | 1.97939 | 1.48882 | 10.28640 | 1.28330 | 0.03176 | 1.90713 | 0.00000 | 2.31591 | 0.00000 | 1.15954 | 1.97556 |
| **MAGEE1** | 0.07375 | 0.00938 | 0.00107 | 0.00568 | 0.00000 | 0.02081 | 0.08351 | 0.02097 | 0.00000 | 0.17958 | 0.00553 | 0.02333 |
| **MAGEF1** | 0.00000 | 0.81327 | 4.49180 | 0.65045 | 1.59925 | 1.05832 | 3.24511 | 0.00000 | 0.06844 | 4.80963 | 1.51502 | 0.14124 |
| **MAGEH1** | 0.11922 | 0.31314 | 0.19217 | 0.07377 | 0.03329 | 0.00548 | 0.24772 | 0.23619 | 0.20078 | 0.00000 | 0.30717 | 0.81815 |
| **MAGI1** | 1.85441 | 3.16708 | 6.03917 | 1.76892 | 7.15251 | 0.00000 | 1.28292 | 0.32856 | 0.00000 | 15.05394 | 1.77009 | 2.38878 |
| **MAGI1-AS1** | 0.00320 | 0.04671 | 0.12590 | 0.07916 | 0.22860 | 0.00000 | 0.00000 | 0.00464 | 0.04154 | 0.59896 | 0.10850 | 0.13981 |
| **MAGI2** | 1.49135 | 0.00239 | 0.01713 | 0.09713 | 0.01418 | 0.00000 | 0.02385 | 0.06947 | 0.00463 | 0.10839 | 0.01356 | 0.07748 |
| **MAGI2-AS3** | 6.78354 | 0.01638 | 0.02358 | 0.01074 | 0.02000 | 0.00000 | 0.02855 | 0.15791 | 0.01644 | 0.01792 | 0.01608 | 0.06345 |
| **MAGI3** | 0.09940 | 0.00000 | 6.32328 | 0.71400 | 2.94589 | 3.35481 | 7.02076 | 0.24259 | 2.18435 | 8.44942 | 6.49110 | 0.16944 |
| **MAGIX** | 0.00830 | 0.03368 | 0.25371 | 0.26417 | 0.14776 | 0.00000 | 0.53080 | 0.09984 | 0.02004 | 0.31154 | 0.24145 | 0.80105 |
| **MAGOH** | 8.23773 | 4.46887 | 11.49298 | 6.22456 | 3.77119 | 12.78996 | 14.21071 | 8.29816 | 5.03173 | 7.00511 | 15.79898 | 15.73520 |
| **MAGOH2** | 1.11324 | 0.78227 | 1.33441 | 0.85806 | 0.19900 | 1.75216 | 1.56097 | 0.87166 | 0.61548 | 0.32658 | 2.00730 | 2.00199 |
| **MAGOHB** | 0.47570 | 10.83130 | 5.78071 | 2.30373 | 3.91883 | 19.03492 | 9.10481 | 1.05013 | 4.01459 | 4.89063 | 6.30070 | 11.05929 |
| **MAGT1** | 0.68916 | 2.31425 | 3.29580 | 3.92033 | 5.43427 | 18.06035 | 0.00000 | 5.62001 | 8.79961 | 6.06335 | 3.46992 | 5.38279 |
| **MAK** | 2.23284 | 0.41094 | 0.06014 | 0.70627 | 0.22732 | 1.58097 | 0.00000 | 10.10194 | 1.00819 | 1.44019 | 0.64040 | 2.61611 |
| **MAK16** | 1.07758 | 8.43994 | 10.09176 | 1.23412 | 0.00000 | 25.39842 | 15.13069 | 9.20208 | 12.05328 | 7.67586 | 11.83796 | 8.95936 |
| **MAL** | 0.06154 | 0.98389 | 0.00000 | 0.83178 | 0.17414 | 0.01577 | 0.00000 | 0.01050 | 0.00000 | 0.27994 | 0.00000 | 0.00000 |
| **MAL2** | 1.14046 | 0.48599 | 0.97661 | 0.00000 | 0.05493 | 1.80626 | 0.79594 | 0.00000 | 0.00000 | 3.49313 | 0.33435 | 0.14272 |
| **MALAT1** | 6.14736 | 9.38512 | 45.38802 | 23.32114 | 36.00372 | 105.12378 | 27.42660 | 14.09510 | 39.59459 | 89.60696 | 38.95732 | 43.96128 |

| | | | | | | | | | | | | |
|---|---|---|---|---|---|---|---|---|---|---|---|---|
| **MALL** | 0.00547 | 0.01702 | 0.01522 | 0.01101 | 0.01594 | 0.02801 | 0.02197 | 0.01396 | 0.02369 | 0.00810 | 0.01215 | 0.01872 |
| **MALRD1** | 2.84559 | 0.00936 | 0.01801 | 0.01701 | 0.00000 | 0.02470 | 0.04029 | 0.40137 | 0.00946 | 0.05980 | 0.01219 | 0.07335 |
| **MALSU1** | 1.85698 | 16.42556 | 7.69987 | 7.99584 | 14.52744 | 17.19751 | 15.95829 | 8.81956 | 17.21443 | 6.92230 | 11.08398 | 22.35420 |
| **MALT1** | 2.34172 | 0.00000 | 1.22959 | 0.78997 | 2.38637 | 3.01583 | 2.24895 | 0.13294 | 0.10019 | 4.49142 | 0.00000 | 0.13929 |
| **MAMDC2** | 0.13849 | 0.01781 | 0.02219 | 0.00001 | 0.06417 | 0.30895 | 0.00000 | 0.13647 | 0.00000 | 0.00721 | 0.01559 | 0.09734 |
| **MAMDC2-AS1** | 0.00442 | 0.01631 | 0.01609 | 0.01433 | 0.01325 | 0.00772 | 0.01817 | 0.07154 | 0.01926 | 0.01484 | 0.03935 | 0.01461 |
| **MAMDC4** | 0.00155 | 0.46290 | 0.71985 | 0.22334 | 0.03367 | 0.00000 | 0.00000 | 0.00851 | 0.69542 | 1.52342 | 0.06441 | 0.12789 |
| **MAML1** | 0.23900 | 4.19015 | 3.08191 | 3.83246 | 2.33584 | 7.26321 | 7.01910 | 0.49872 | 7.19569 | 9.05507 | 2.21597 | 3.40831 |
| **MAML2** | 0.35409 | 0.00469 | 0.26120 | 0.03214 | 0.01348 | 0.07225 | 0.00000 | 0.02158 | 0.00000 | 0.08407 | 0.03455 | 0.06740 |
| **MAML3** | 0.22731 | 0.26044 | 1.58523 | 0.61753 | 1.18230 | 0.00000 | 1.02410 | 0.04474 | 0.00000 | 1.06540 | 0.69493 | 0.05440 |
| **MAMLD1** | 0.00648 | 0.05298 | 0.01715 | 0.00416 | 0.00727 | 0.04086 | 0.00821 | 0.01046 | 0.02161 | 0.02600 | 0.01246 | 0.01169 |
| **MAMSTR** | 0.52025 | 0.03072 | 0.08334 | 0.36719 | 0.05901 | 0.00000 | 0.03381 | 0.04275 | 0.04387 | 0.00000 | 0.14179 | 0.10469 |
| **MAN1A1** | 2.37612 | 0.00000 | 0.00000 | 0.00000 | 0.00000 | 3.28066 | 0.00000 | 0.13142 | 2.73424 | 3.76459 | 0.00000 | 0.35442 |
| **MAN1A2** | 4.72608 | 0.05704 | 9.63253 | 3.85944 | 5.87518 | 10.42382 | 3.98896 | 1.74250 | 4.02979 | 18.56018 | 3.17455 | 2.66950 |
| **MAN1B1** | 0.15580 | 8.78286 | 4.13854 | 5.89249 | 6.62577 | 0.00000 | 0.00002 | 0.92937 | 6.91682 | 4.29858 | 8.36409 | 4.35432 |
| **MAN1B1-AS1** | 0.00807 | 0.08440 | 0.12938 | 0.00000 | 0.08449 | 0.00000 | 0.08161 | 0.00695 | 0.00000 | 0.14659 | 0.02561 | 0.08473 |
| **MAN1C1** | 0.06645 | 2.22825 | 0.59569 | 9.09932 | 0.00000 | 0.27641 | 0.00000 | 0.19140 | 0.00000 | 0.99185 | 0.00000 | 0.44630 |
| **MAN2A1** | 0.95581 | 0.17560 | 7.24680 | 1.49052 | 2.75729 | 14.79793 | 0.93013 | 0.57823 | 10.66325 | 20.80287 | 2.97513 | 4.18034 |
| **MAN2A2** | 0.00507 | 2.69581 | 6.08651 | 0.80543 | 5.26971 | 1.47151 | 0.00000 | 0.16528 | 3.19935 | 5.72627 | 3.43584 | 0.13821 |
| **MAN2B1** | 0.24452 | 3.90856 | 2.92700 | 3.43184 | 3.50459 | 0.00000 | 0.74914 | 0.02628 | 6.60950 | 1.71082 | 3.46072 | 0.69001 |
| **MAN2B2** | 0.00491 | 1.36477 | 1.43670 | 1.55803 | 1.53975 | 0.00000 | 0.01478 | 0.00000 | 5.99996 | 1.48136 | 0.59750 | 0.58083 |
| **MAN2C1** | 0.14333 | 1.88287 | 1.37308 | 2.51543 | 2.41970 | 0.00000 | 0.00000 | 0.00000 | 2.10235 | 3.12424 | 3.49077 | 1.23663 |
| **MANBA** | 0.23354 | 0.00000 | 0.96545 | 0.75740 | 3.55593 | 0.62725 | 0.94219 | 0.00000 | 13.83312 | 4.29533 | 4.30605 | 0.00000 |
| **MANBAL** | 0.00000 | 8.16048 | 11.76757 | 6.54167 | 2.53758 | 3.27402 | 8.46824 | 4.21137 | 14.27613 | 0.00000 | 9.23103 | 14.81600 |
| **MANEA** | 0.00079 | 1.14734 | 2.33163 | 2.21778 | 0.42253 | 11.75600 | 0.00000 | 3.23501 | 5.10465 | 0.00000 | 0.65920 | 7.78937 |
| **MANEA-AS1** | 0.00069 | 0.12833 | 0.62869 | 0.46247 | 0.00000 | 0.00000 | 0.00000 | 0.13457 | 0.27265 | 0.45118 | 0.13337 | 0.36617 |
| **MANEAL** | 0.05882 | 0.39984 | 0.56658 | 0.00000 | 0.29322 | 0.95451 | 1.78859 | 1.85211 | 0.05873 | 2.25645 | 0.00000 | 1.67589 |
| **MANF** | 2.52389 | 0.55014 | 9.74596 | 3.10181 | 4.60365 | 3.78455 | 7.80234 | 4.29230 | 2.22475 | 20.55740 | 6.21932 | 1.04538 |
| **MANSC1** | 0.98295 | 0.00000 | 0.94775 | 0.09176 | 0.62519 | 1.59122 | 0.00000 | 0.02428 | 4.75486 | 0.51392 | 1.53186 | 0.07089 |
| **MANSC4** | 0.00180 | 0.01136 | 0.01293 | 0.01689 | 0.01698 | 0.00000 | 0.03883 | 0.10455 | 0.00978 | 0.01997 | 0.01992 | 0.43144 |
| **MAOA** | 0.24375 | 0.00000 | 1.41008 | 0.12689 | 0.53695 | 0.40437 | 0.18572 | 0.00000 | 0.68605 | 0.60530 | 0.34708 | 0.30897 |
| **MAOB** | 0.00731 | 0.01198 | 0.21821 | 0.01270 | 0.00793 | 0.03267 | 0.00000 | 0.01023 | 0.00000 | 0.02013 | 0.02372 | 0.03386 |
| **MAP10** | 0.11597 | 0.15611 | 0.22543 | 0.16724 | 0.00720 | 0.19011 | 0.00000 | 0.00000 | 0.00000 | 0.00000 | 0.11960 | 0.17734 |
| **MAP1A** | 1.14204 | 0.00000 | 0.08025 | 0.57145 | 0.85390 | 0.00000 | 0.69124 | 2.24223 | 0.24724 | 0.45548 | 0.00000 | 0.40468 |
| **MAP1B** | 0.26469 | 0.42030 | 0.84541 | 4.36364 | 0.14006 | 1.28711 | 0.14639 | 4.63251 | 0.00095 | 0.92673 | 0.20046 | 7.49801 |
| **MAP1LC3A** | 0.72547 | 0.90845 | 1.87700 | 0.66846 | 0.22917 | 0.00000 | 0.00000 | 1.07764 | 0.75980 | 0.39285 | 0.11358 | 1.47243 |
| **MAP1LC3B** | 2.89524 | 11.96072 | 14.79723 | 4.76915 | 3.49179 | 19.57897 | 19.51785 | 28.61360 | 14.40905 | 5.12048 | 7.80361 | 31.48165 |
| **MAP1LC3B2** | 0.62654 | 3.97746 | 3.45038 | 0.65958 | 1.09657 | 5.70462 | 3.30143 | 12.09527 | 3.57735 | 0.00000 | 1.77524 | 10.28513 |
| **MAP1LC3C** | 0.40280 | 0.86124 | 0.00000 | 1.23257 | 0.01083 | 0.30364 | 0.00000 | 0.00065 | 0.03658 | 0.00000 | 0.01581 | 0.04445 |
| **MAP1S** | 3.56631 | 10.65045 | 2.95040 | 7.04512 | 10.20977 | 0.00000 | 3.17887 | 0.83975 | 10.46374 | 0.00000 | 4.64315 | 15.34866 |
| **MAP2** | 0.13649 | 0.01050 | 0.03559 | 0.04382 | 0.02408 | 0.04500 | 0.02377 | 0.37405 | 0.00000 | 0.00205 | 0.03478 | 0.04626 |
| **MAP2K1** | 7.06406 | 0.11671 | 7.81484 | 0.00000 | 4.59200 | 5.92779 | 11.06749 | 1.31610 | 0.45926 | 11.54884 | 2.99513 | 0.35262 |
| **MAP2K2** | 1.02985 | 2.48439 | 4.19626 | 3.85164 | 8.39334 | 0.00000 | 13.90758 | 0.37062 | 0.52337 | 18.78669 | 2.07329 | 2.66612 |
| **MAP2K3** | 3.93663 | 3.93996 | 7.02239 | 5.73869 | 3.01518 | 0.00000 | 5.44911 | 0.37933 | 5.59796 | 1.31584 | 0.87962 | 0.24813 |
| **MAP2K4** | 1.37854 | 1.38228 | 3.39710 | 5.95092 | 1.65154 | 7.61034 | 5.60264 | 0.27806 | 1.66088 | 8.49662 | 0.69730 | 1.62875 |
| **MAP2K4P1** | 0.03473 | 0.18970 | 0.15821 | 0.22887 | 0.00567 | 0.23253 | 0.06777 | 0.17682 | 0.32564 | 0.04417 | 0.01521 | 0.18904 |
| **MAP2K5** | 0.49642 | 1.15521 | 2.01323 | 1.23227 | 1.86145 | 1.63170 | 1.58017 | 0.03414 | 3.39067 | 0.94894 | 0.86772 | 0.17728 |
| **MAP2K6** | 1.21913 | 0.72712 | 0.09660 | 5.30575 | 0.00000 | 0.87111 | 0.39744 | 0.84274 | 0.88694 | 0.00000 | 2.66301 | 0.48781 |

| | | | | | | | | | | | | |
|---|---|---|---|---|---|---|---|---|---|---|---|---|
| **MAP2K7** | 1.05087 | 0.24555 | 1.67052 | 1.83323 | 2.73122 | 0.00000 | 5.61788 | 0.38177 | 0.13308 | 12.19585 | 1.30031 | 1.82085 |
| **MAP3K1** | 0.00000 | 1.46205 | 1.88137 | 3.39393 | 0.23089 | 6.28946 | 0.48341 | 1.10249 | 0.00000 | 5.04760 | 0.00659 | 2.61060 |
| **MAP3K10** | 0.25144 | 0.00233 | 0.04531 | 0.01154 | 0.13302 | 0.00000 | 0.03346 | 0.00268 | 0.04572 | 0.61705 | 0.00292 | 0.02009 |
| **MAP3K11** | 0.85262 | 1.11055 | 2.92293 | 1.32607 | 3.19936 | 0.00000 | 3.13537 | 0.00000 | 1.45816 | 7.45387 | 0.32838 | 0.88921 |
| **MAP3K12** | 0.42725 | 0.00000 | 0.08286 | 0.37855 | 0.29715 | 0.00000 | 0.00000 | 0.07175 | 0.41753 | 1.00021 | 0.02231 | 0.46913 |
| **MAP3K13** | 0.74060 | 4.22445 | 2.89986 | 1.53424 | 1.57294 | 8.86410 | 2.59006 | 4.58293 | 3.25694 | 6.85465 | 3.68661 | 14.20701 |
| **MAP3K14** | 0.30564 | 0.98940 | 0.14606 | 0.21782 | 0.52587 | 0.62734 | 0.53849 | 0.21978 | 0.00000 | 0.31203 | 0.47210 | 0.34387 |
| **MAP3K14-AS1** | 0.11127 | 0.00461 | 0.04915 | 0.01132 | 0.10800 | 0.04445 | 0.03197 | 0.00324 | 0.00000 | 0.00000 | 0.06865 | 0.04463 |
| **MAP3K15** | 0.95561 | 0.07718 | 0.00514 | 0.09246 | 0.04848 | 0.00000 | 0.00515 | 0.00000 | 0.00000 | 0.37713 | 0.40349 | 0.06634 |
| **MAP3K19** | 0.00539 | 0.05304 | 0.01296 | 0.27675 | 0.00440 | 0.04553 | 0.00003 | 0.00777 | 0.00000 | 0.19708 | 0.00144 | 0.06572 |
| **MAP3K2** | 0.00000 | 1.32850 | 2.02358 | 3.34867 | 1.33121 | 11.69727 | 5.09102 | 0.55367 | 0.49194 | 15.30337 | 2.13663 | 1.51567 |
| **MAP3K3** | 2.24393 | 0.51580 | 0.71037 | 3.52894 | 0.00000 | 0.00000 | 0.43681 | 2.10501 | 0.16379 | 0.92729 | 0.11306 | 1.17087 |
| **MAP3K4** | 3.81398 | 7.61850 | 8.68100 | 4.65699 | 1.85365 | 9.37132 | 7.89438 | 2.03468 | 0.00000 | 7.78893 | 2.45155 | 5.37640 |
| **MAP3K5** | 0.71948 | 2.22359 | 0.00000 | 1.45970 | 2.41467 | 0.64967 | 0.00000 | 0.06926 | 0.00000 | 1.18339 | 0.11056 | 0.93827 |
| **MAP3K6** | 0.75233 | 1.39283 | 0.41010 | 0.61958 | 0.59377 | 0.46961 | 0.00000 | 0.05825 | 0.16276 | 0.84518 | 0.00097 | 0.74024 |
| **MAP3K7** | 0.22486 | 7.19533 | 7.21263 | 5.55645 | 2.17120 | 11.43512 | 10.41609 | 7.24335 | 9.72872 | 1.46017 | 8.03402 | 5.68360 |
| **MAP3K7CL** | 11.46802 | 0.00000 | 0.02515 | 0.00940 | 0.00789 | 0.07922 | 0.00271 | 4.43819 | 0.00000 | 0.10205 | 0.04462 | 0.22046 |
| **MAP3K8** | 0.01659 | 2.15433 | 9.23494 | 0.90839 | 0.50852 | 8.00146 | 3.00239 | 1.12762 | 2.86564 | 0.00000 | 0.71552 | 2.48525 |
| **MAP3K9** | 0.95542 | 0.55319 | 2.90383 | 2.21110 | 2.69969 | 0.00000 | 1.55599 | 1.19923 | 0.34688 | 5.13387 | 0.93742 | 1.41353 |
| **MAP4** | 7.67027 | 24.07694 | 9.48829 | 20.14426 | 40.74077 | 9.73814 | 19.14018 | 1.80152 | 21.66432 | 19.19990 | 26.69078 | 18.12585 |
| **MAP4K1** | 0.93272 | 0.19249 | 0.51516 | 5.49603 | 0.58355 | 0.00000 | 0.30966 | 0.56133 | 0.00000 | 1.33660 | 0.00000 | 0.30016 |
| **MAP4K2** | 0.00255 | 2.92972 | 1.88471 | 2.74775 | 2.62606 | 0.00000 | 1.57942 | 2.00852 | 1.14048 | 1.05790 | 1.53824 | 9.13537 |
| **MAP4K3** | 3.73758 | 1.54539 | 2.17088 | 2.04567 | 3.09055 | 9.83962 | 0.94841 | 1.26933 | 1.31687 | 12.26817 | 0.29974 | 1.23995 |
| **MAP4K4** | 3.43953 | 5.72372 | 8.79435 | 4.70106 | 9.04464 | 9.90903 | 8.25239 | 0.45341 | 0.00000 | 20.34761 | 0.00000 | 2.11031 |
| **MAP4K5** | 0.31960 | 0.00000 | 4.43004 | 0.00000 | 1.46157 | 6.10516 | 5.42224 | 0.63362 | 1.48012 | 7.09747 | 1.08367 | 1.79991 |
| **MAP6** | 0.00435 | 0.00507 | 0.13072 | 0.31386 | 0.00246 | 0.08515 | 0.00000 | 0.01408 | 0.02201 | 0.04534 | 0.02661 | 0.02059 |
| **MAP6D1** | 0.55873 | 0.56283 | 0.06464 | 0.00000 | 0.18490 | 0.22795 | 0.75301 | 0.18646 | 0.65312 | 0.68240 | 0.00000 | 0.48929 |
| **MAP7** | 3.47846 | 1.35065 | 2.93115 | 1.62290 | 3.61712 | 10.38665 | 0.00000 | 0.32633 | 9.71202 | 10.15391 | 4.34670 | 1.21402 |
| **MAP7D1** | 1.63142 | 0.89034 | 3.15082 | 2.26034 | 0.34255 | 0.12373 | 4.87694 | 2.61380 | 0.00000 | 3.48098 | 0.00000 | 0.69013 |
| **MAP7D2** | 6.31679 | 3.22811 | 0.00000 | 0.52300 | 1.41751 | 1.16535 | 0.37842 | 1.79227 | 0.90165 | 0.00000 | 0.94545 | 3.00386 |
| **MAP7D3** | 0.43473 | 5.56812 | 3.68081 | 2.48036 | 0.12612 | 8.74069 | 0.96122 | 1.68099 | 0.43966 | 6.16245 | 1.12717 | 7.53378 |
| **MAP9** | 1.61440 | 0.23685 | 0.69806 | 1.78300 | 0.00000 | 2.45287 | 1.53038 | 2.29770 | 0.00000 | 0.98016 | 0.64159 | 1.25443 |
| **MAPK1** | 0.83236 | 6.45236 | 2.08648 | 7.50884 | 3.17262 | 18.75640 | 6.77121 | 5.00760 | 0.22045 | 15.79188 | 2.82171 | 15.00857 |
| **MAPK10** | 1.49668 | 0.00630 | 0.02769 | 0.01728 | 0.01590 | 0.00779 | 0.01935 | 0.14901 | 0.01444 | 0.01609 | 0.01422 | 0.09727 |
| **MAPK11** | 0.00608 | 0.01567 | 0.05368 | 0.02499 | 0.01299 | 0.02002 | 0.02026 | 0.01027 | 0.03578 | 0.00483 | 0.01263 | 0.03525 |
| **MAPK12** | 0.95807 | 0.53020 | 0.38119 | 0.84937 | 0.34651 | 0.00000 | 0.00000 | 0.06841 | 0.00000 | 0.62999 | 0.01965 | 0.38824 |
| **MAPK13** | 0.28918 | 3.40088 | 0.00000 | 0.59856 | 0.23348 | 1.93202 | 0.00000 | 0.48396 | 0.32976 | 1.55594 | 0.64626 | 5.40902 |
| **MAPK14** | 0.00000 | 4.55321 | 3.84977 | 0.35104 | 11.86212 | 10.94457 | 10.58969 | 0.77321 | 5.96318 | 17.46452 | 7.04768 | 3.44142 |
| **MAPK15** | 0.06606 | 0.02913 | 0.01212 | 0.01967 | 0.00000 | 0.01642 | 0.02441 | 0.08138 | 0.19052 | 0.13231 | 0.01558 | 0.02490 |
| **MAPK1IP1L** | 0.75253 | 14.98971 | 17.06627 | 13.97371 | 9.70872 | 42.24206 | 23.99885 | 13.95857 | 19.24227 | 7.51637 | 23.70519 | 16.04972 |
| **MAPK3** | 0.00041 | 6.89248 | 0.00000 | 4.97828 | 19.81281 | 0.00000 | 5.64547 | 0.59720 | 2.02009 | 7.47376 | 6.99610 | 3.59251 |
| **MAPK4** | 2.74451 | 0.05904 | 0.52475 | 0.46434 | 0.00533 | 0.02037 | 0.14346 | 0.00308 | 0.00000 | 0.01246 | 0.01997 | 0.00243 |
| **MAPK6** | 4.23826 | 7.73544 | 9.10932 | 3.34631 | 1.80977 | 15.36543 | 10.81934 | 12.99384 | 7.55646 | 4.09849 | 3.93644 | 14.96644 |
| **MAPK7** | 1.76313 | 1.93745 | 1.21878 | 2.69196 | 0.62336 | 0.00000 | 0.00000 | 0.44224 | 1.90311 | 1.52725 | 1.62082 | 1.52800 |
| **MAPK8** | 10.71843 | 0.00000 | 1.02466 | 0.59097 | 1.12430 | 8.76916 | 2.78708 | 4.04658 | 7.58437 | 6.14029 | 2.22024 | 4.35651 |
| **MAPK8IP1** | 0.53363 | 0.63235 | 0.03332 | 0.00000 | 0.48976 | 0.15034 | 1.01208 | 0.00000 | 0.40009 | 2.23931 | 0.00000 | 0.19413 |
| **MAPK8IP2** | 0.07919 | 0.44105 | 0.00000 | 0.35012 | 0.58675 | 0.00000 | 0.05870 | 0.16116 | 0.00000 | 0.61896 | 0.00000 | 0.18398 |
| **MAPK8IP3** | 1.50948 | 0.86111 | 1.76766 | 1.93797 | 0.00000 | 0.00000 | 1.49013 | 0.20731 | 3.53952 | 1.95098 | 1.52132 | 0.00000 |

| | | | | | | | | | | | | |
|---|---|---|---|---|---|---|---|---|---|---|---|---|
| **MAPK9** | 1.51417 | 1.22036 | 3.69697 | 1.42290 | 5.96513 | 6.82377 | 9.19841 | 0.17691 | 0.42435 | 7.86488 | 4.24904 | 0.02544 |
| **MAPKAP1** | 3.77558 | 6.73273 | 5.76185 | 9.30438 | 9.16274 | 4.92711 | 8.96379 | 0.65145 | 2.95880 | 16.63011 | 2.66078 | 2.34640 |
| **MAPKAPK2** | 5.20544 | 0.21459 | 0.41780 | 1.20979 | 2.06297 | 0.00000 | 2.02086 | 1.32553 | 0.00000 | 8.50219 | 0.00000 | 0.47674 |
| **MAPKAPK3** | 0.01521 | 0.15624 | 3.64457 | 1.91818 | 1.90027 | 0.00000 | 2.56050 | 0.00865 | 2.70280 | 0.35280 | 1.98134 | 0.17179 |
| **MAPKAPK5** | 1.29558 | 4.79467 | 2.90507 | 3.33335 | 5.35580 | 3.63714 | 7.54162 | 0.67508 | 2.18678 | 7.57454 | 1.30586 | 3.31489 |
| **MAPKAPK5-AS** | 2.27891 | 3.14588 | 3.96726 | 1.39736 | 0.00000 | 4.95995 | 8.43684 | 6.90901 | 6.53802 | 0.00000 | 5.88240 | 7.53669 |
| **MAPKBP1** | 3.36011 | 0.89124 | 3.38361 | 1.57080 | 0.88816 | 0.00000 | 0.67151 | 1.31987 | 1.97164 | 4.33299 | 0.34851 | 6.54405 |
| **MAPRE1** | 5.98277 | 17.77109 | 18.36117 | 10.33786 | 13.71586 | 35.46502 | 22.32167 | 16.63323 | 24.60130 | 3.85598 | 23.52417 | 20.85709 |
| **MAPRE2** | 3.45430 | 7.00605 | 0.50915 | 0.63986 | 0.76699 | 2.47353 | 1.01545 | 4.88890 | 1.56554 | 0.00000 | 0.00000 | 4.45771 |
| **MAPRE3** | 1.54824 | 0.19812 | 0.61579 | 0.01169 | 0.02075 | 0.13309 | 0.18736 | 0.10527 | 0.00748 | 0.00000 | 0.10720 | 0.03885 |
| **MAPT** | 1.59846 | 1.05985 | 1.89012 | 0.10110 | 0.42860 | 0.15083 | 0.00000 | 0.07014 | 0.00000 | 0.21938 | 0.03389 | 0.60212 |
| **MAPT-IT1** | 0.00626 | 0.07044 | 0.01685 | 0.01032 | 0.02269 | 0.01593 | 0.01562 | 0.01074 | 0.02117 | 0.01108 | 0.01216 | 0.01302 |
| **MARC1** | 0.04680 | 0.54575 | 4.17924 | 0.00000 | 0.55410 | 1.07076 | 0.83132 | 0.07500 | 2.88141 | 0.37956 | 0.00000 | 0.00000 |
| **MARC2** | 0.78010 | 0.00312 | 2.61570 | 0.03294 | 0.28386 | 0.05677 | 0.00000 | 0.02887 | 0.00000 | 0.02547 | 0.00402 | 0.00872 |
| **MARCH1** | 1.54751 | 0.00000 | 0.00000 | 0.00000 | 0.00000 | 0.76268 | 0.00000 | 0.35955 | 1.36046 | 0.97706 | 0.82209 | 0.97626 |
| **MARCH10** | 0.16648 | 0.03197 | 0.08227 | 0.02771 | 0.02904 | 0.00000 | 0.04146 | 0.00468 | 0.00676 | 0.00000 | 0.02018 | 0.03535 |
| **MARCH11** | 0.05870 | 0.16004 | 0.01149 | 0.02405 | 0.02091 | 0.02122 | 0.01194 | 0.00716 | 0.03695 | 0.00000 | 0.01197 | 0.04922 |
| **MARCH2** | 0.30363 | 0.40988 | 0.60257 | 0.44336 | 0.55069 | 0.00000 | 0.24668 | 0.19153 | 0.00000 | 0.00000 | 0.46054 | 2.85804 |
| **MARCH3** | 0.00000 | 2.38965 | 0.65666 | 1.64941 | 0.72130 | 1.78477 | 0.16158 | 2.38359 | 0.59213 | 0.88999 | 0.22493 | 0.28076 |
| **MARCH4** | 1.96424 | 0.01288 | 0.01385 | 0.02813 | 0.04022 | 0.00000 | 0.07723 | 0.06518 | 0.00000 | 0.00000 | 0.02563 | 0.00613 |
| **MARCH5** | 3.60870 | 3.70825 | 3.70512 | 2.47719 | 2.19965 | 11.59463 | 3.66197 | 3.40455 | 11.80924 | 0.14120 | 2.46450 | 4.03468 |
| **MARCH6** | 3.74353 | 5.80391 | 12.31537 | 3.27237 | 0.00930 | 27.08299 | 0.00000 | 1.80184 | 26.99313 | 13.75894 | 12.73780 | 2.38232 |
| **MARCH7** | 4.97686 | 10.39483 | 8.74288 | 4.29513 | 1.47759 | 31.78697 | 9.72996 | 16.13381 | 13.45924 | 9.51192 | 12.36766 | 14.67789 |
| **MARCH8** | 2.62548 | 8.39393 | 0.00000 | 7.19540 | 9.35439 | 2.63553 | 6.02141 | 9.38886 | 6.26783 | 12.08149 | 3.26631 | 21.54777 |
| **MARCH9** | 0.01745 | 1.69589 | 0.24601 | 1.29494 | 1.68987 | 0.00000 | 0.88343 | 0.00000 | 0.09505 | 2.17600 | 0.18097 | 0.74129 |
| **MARCKS** | 0.00000 | 7.57372 | 0.00000 | 9.31426 | 7.09909 | 16.20711 | 2.06245 | 0.14840 | 0.00000 | 27.70276 | 8.01814 | 1.56321 |
| **MARCKSL1** | 0.63985 | 10.68428 | 14.45603 | 10.60415 | 8.70091 | 4.16599 | 29.84463 | 1.27767 | 14.21479 | 15.07479 | 2.56372 | 19.06339 |
| **MARCO** | 0.00854 | 0.04029 | 0.04234 | 0.00958 | 0.08999 | 0.01223 | 0.03795 | 0.01007 | 0.00859 | 0.00000 | 0.02358 | 0.01983 |
| **MARK1** | 0.00000 | 1.69766 | 1.24467 | 1.00322 | 0.01859 | 4.85760 | 1.04943 | 0.03684 | 0.00000 | 6.19416 | 0.00000 | 1.53640 |
| **MARK2** | 0.90711 | 1.25679 | 5.03981 | 1.25767 | 6.14141 | 0.00000 | 4.11842 | 0.36720 | 0.00000 | 18.19519 | 1.59888 | 2.92407 |
| **MARK2P9** | 0.02855 | 0.00000 | 0.01812 | 0.17794 | 0.07394 | 0.00000 | 0.01130 | 0.25398 | 0.14429 | 0.44247 | 0.05880 | 0.21767 |
| **MARK3** | 6.85016 | 1.58053 | 4.89919 | 4.63733 | 7.40794 | 3.65069 | 10.86848 | 4.31307 | 2.36716 | 20.25707 | 1.03905 | 2.26209 |
| **MARK4** | 3.16142 | 0.35346 | 1.74048 | 0.41286 | 0.08864 | 0.00000 | 1.03397 | 0.18343 | 0.00000 | 4.17432 | 0.22673 | 0.27108 |
| **MARS** | 4.38218 | 39.13402 | 16.21164 | 27.20909 | 16.15362 | 22.02986 | 22.72788 | 14.80415 | 41.18030 | 0.00002 | 24.20852 | 27.95223 |
| **MARS2** | 0.00000 | 8.48254 | 2.86339 | 2.43687 | 2.94172 | 5.17342 | 2.06774 | 3.86593 | 9.31215 | 0.93335 | 7.53348 | 4.36541 |
| **MARVELD1** | 0.14167 | 1.99512 | 0.24828 | 0.81631 | 0.15485 | 0.11269 | 0.32127 | 1.17791 | 0.00000 | 0.00000 | 0.10517 | 1.52874 |
| **MARVELD2** | 0.33996 | 3.10271 | 4.02779 | 1.30168 | 2.49497 | 5.21516 | 0.00000 | 0.00000 | 4.48385 | 7.20740 | 3.93242 | 1.97933 |
| **MARVELD3** | 0.00000 | 0.07745 | 1.52617 | 0.00000 | 2.90249 | 1.10766 | 2.80390 | 0.01837 | 1.47154 | 9.28250 | 0.63306 | 1.03341 |
| **MAS1** | 0.03438 | 0.00793 | 0.05745 | 0.00905 | 0.01031 | 0.06323 | 0.01339 | 0.05381 | 0.03126 | 0.00000 | 0.01901 | 0.01867 |
| **MASP1** | 0.00776 | 0.01925 | 0.03816 | 0.06246 | 0.05132 | 0.00000 | 0.03252 | 0.11087 | 0.00101 | 0.01268 | 0.02258 | 0.01692 |
| **MASP2** | 0.00202 | 0.12781 | 0.26202 | 0.05862 | 0.22165 | 0.00378 | 1.95289 | 0.67057 | 1.55006 | 4.34971 | 0.34824 | 0.86794 |
| **MAST1** | 0.00481 | 0.03618 | 0.23047 | 0.13899 | 0.20648 | 0.00000 | 0.00434 | 0.09413 | 0.24396 | 0.04170 | 0.05524 | 0.39198 |
| **MAST2** | 1.86501 | 10.54652 | 1.99845 | 10.66780 | 8.81755 | 0.00000 | 5.28631 | 2.60312 | 4.76869 | 13.52576 | 0.00000 | 12.84142 |
| **MAST3** | 2.21137 | 1.69153 | 0.31327 | 0.08163 | 1.00559 | 0.00000 | 0.27623 | 8.35546 | 1.02554 | 1.30950 | 0.10163 | 8.44735 |
| **MAST4** | 6.11669 | 0.93153 | 1.54142 | 0.77803 | 1.58577 | 0.35594 | 0.24038 | 0.85849 | 0.00000 | 1.48264 | 0.12565 | 0.61343 |
| **MASTL** | 2.37037 | 6.49914 | 5.43821 | 6.36037 | 3.67154 | 27.35890 | 0.00000 | 20.95074 | 9.47707 | 6.40812 | 8.85981 | 11.38149 |
| **MAT1A** | 1.37139 | 0.02336 | 0.01619 | 0.00425 | 0.02958 | 0.00000 | 0.05060 | 0.20328 | 0.00000 | 0.11164 | 0.03634 | 0.00850 |
| **MAT2A** | 3.35858 | 21.37921 | 13.66830 | 8.07477 | 6.13568 | 27.04959 | 9.88239 | 19.96525 | 30.80993 | 0.00000 | 18.11608 | 17.90682 |

| | | | | | | | | | | | | |
|---|---|---|---|---|---|---|---|---|---|---|---|---|
| **MAT2B** | 5.16854 | 10.78882 | 9.68511 | 6.78191 | 0.00000 | 20.26197 | 14.45191 | 3.87237 | 8.37537 | 0.00000 | 6.89592 | 8.85983 |
| **MATK** | 1.20354 | 0.21705 | 0.03237 | 0.11894 | 1.00625 | 0.00000 | 0.03004 | 0.00438 | 0.00000 | 0.04254 | 0.08981 | 0.00614 |
| **MATN1** | 0.12507 | 0.01672 | 0.03064 | 0.04098 | 0.01272 | 0.00000 | 0.14651 | 0.09862 | 0.00128 | 0.00670 | 0.02207 | 0.01164 |
| **MATN1-AS1** | 0.02670 | 0.01718 | 0.06842 | 0.01401 | 0.01942 | 0.00681 | 0.04664 | 0.01891 | 0.00790 | 0.00714 | 0.02072 | 0.01649 |
| **MATN2** | 0.59911 | 1.00952 | 0.00000 | 0.00000 | 0.00000 | 0.38380 | 0.00000 | 0.59068 | 0.00000 | 0.51711 | 0.06807 | 0.49479 |
| **MATN3** | 0.03044 | 0.01260 | 0.09007 | 0.00000 | 0.20766 | 0.00004 | 0.00000 | 0.08903 | 0.37205 | 0.88684 | 0.04185 | 0.00152 |
| **MATN4** | 0.01073 | 1.08673 | 0.00000 | 0.00000 | 0.06217 | 0.00000 | 0.00098 | 0.00121 | 0.04180 | 0.53322 | 0.02487 | 0.33515 |
| **MATR3** | 0.47984 | 16.87971 | 14.15345 | 12.14334 | 5.71761 | 42.54203 | 26.37172 | 24.94026 | 20.23172 | 17.53834 | 23.88642 | 22.65469 |
| **MAU2** | 0.33952 | 5.12445 | 5.56745 | 5.22492 | 5.14202 | 0.00000 | 7.23174 | 0.00000 | 8.22257 | 6.93320 | 4.57219 | 4.75928 |
| **MAVS** | 1.02413 | 7.18501 | 5.69779 | 4.03362 | 7.67232 | 0.00000 | 5.63463 | 0.61459 | 3.00394 | 13.91977 | 8.38707 | 7.90304 |
| **MAX** | 8.68003 | 10.19158 | 9.79585 | 8.00109 | 16.32881 | 20.02768 | 15.90622 | 7.63142 | 12.52935 | 10.23695 | 18.13963 | 23.36053 |
| **MAZ** | 1.99051 | 10.77676 | 16.81740 | 13.60598 | 23.93863 | 0.00000 | 27.57635 | 1.70657 | 15.64479 | 33.76478 | 10.20981 | 9.30498 |
| **MB** | 0.00850 | 0.60992 | 0.00000 | 0.15675 | 0.87512 | 0.11716 | 0.00012 | 0.00795 | 0.00000 | 0.01391 | 0.00000 | 0.00009 |
| **MB21D1** | 0.00578 | 0.01169 | 0.23874 | 0.10454 | 0.05254 | 0.00000 | 0.00000 | 0.61217 | 0.06801 | 0.02753 | 0.04436 | 0.02834 |
| **MB21D2** | 2.43695 | 0.82670 | 0.81001 | 0.46186 | 0.87369 | 0.90760 | 0.93737 | 0.20458 | 1.19149 | 0.67528 | 0.64757 | 0.87871 |
| **MBD1** | 0.00000 | 7.08641 | 9.17187 | 4.25201 | 5.77109 | 1.47106 | 8.27789 | 4.63075 | 9.49538 | 6.82094 | 5.72733 | 6.32306 |
| **MBD2** | 0.13333 | 1.31494 | 1.49495 | 0.66686 | 2.48470 | 1.85078 | 4.21323 | 0.58166 | 0.00010 | 11.26355 | 1.33984 | 0.79553 |
| **MBD3** | 0.03261 | 10.86119 | 10.59271 | 7.92253 | 18.05730 | 0.00000 | 23.38506 | 0.15634 | 7.65132 | 35.61898 | 2.06339 | 6.27253 |
| **MBD3L2** | 6.63549 | 0.00310 | 0.00716 | 0.00633 | 0.05793 | 0.00000 | 0.01039 | 6.76419 | 0.01478 | 0.00000 | 0.00980 | 0.11076 |
| **MBD3L3** | 3.84913 | 0.00749 | 0.00804 | 0.00663 | 0.01535 | 0.00000 | 0.01104 | 9.39843 | 0.01639 | 0.00398 | 0.01390 | 0.00000 |
| **MBD3L4** | 0.01660 | 0.01509 | 0.01418 | 0.01210 | 0.01532 | 0.01804 | 0.01808 | 0.01947 | 0.01830 | 0.01393 | 0.01491 | 0.01527 |
| **MBD3L5** | 1.63437 | 0.01153 | 0.01252 | 0.01073 | 0.01809 | 0.00000 | 0.01650 | 1.82477 | 0.02140 | 0.00407 | 0.01326 | 0.00000 |
| **MBD4** | 7.60557 | 7.96191 | 14.04124 | 8.04292 | 3.85840 | 17.26455 | 12.52581 | 9.25952 | 8.54092 | 7.54798 | 6.73202 | 12.43080 |
| **MBD5** | 0.08486 | 0.35696 | 0.34950 | 0.00096 | 0.38780 | 1.01032 | 0.00000 | 0.75811 | 1.10019 | 0.00000 | 0.00000 | 0.57803 |
| **MBD6** | 2.43554 | 0.49300 | 2.58737 | 1.48917 | 2.39964 | 0.00000 | 0.52669 | 0.19780 | 1.87636 | 7.50307 | 0.22649 | 0.96391 |
| **MBIP** | 2.65777 | 2.06280 | 3.06682 | 2.60089 | 0.00000 | 12.79468 | 2.73261 | 14.08751 | 1.61385 | 0.00000 | 2.03802 | 11.22305 |
| **MBL1P** | 0.00687 | 0.03066 | 0.01344 | 0.01196 | 0.00000 | 0.00817 | 0.01815 | 0.01675 | 0.06809 | 0.18667 | 0.04628 | 0.03368 |
| **MBLAC1** | 0.65934 | 0.09911 | 0.08144 | 0.00751 | 0.82937 | 0.00000 | 0.26914 | 0.23699 | 0.00000 | 0.19846 | 0.00981 | 0.35374 |
| **MBLAC2** | 0.09684 | 0.05956 | 0.24714 | 0.45566 | 0.00000 | 0.77254 | 0.09073 | 0.11645 | 0.00000 | 0.25733 | 1.19049 | 0.37117 |
| **MBNL1** | 2.42576 | 0.43900 | 3.54874 | 0.00000 | 0.00000 | 5.84825 | 6.96167 | 4.61607 | 0.65639 | 4.74201 | 1.00503 | 5.47920 |
| **MBNL1-AS1** | 0.04377 | 0.05842 | 0.01411 | 0.00100 | 0.00000 | 0.05787 | 0.00000 | 0.00744 | 0.00000 | 0.24709 | 0.00034 | 0.00404 |
| **MBNL2** | 0.55178 | 0.00000 | 7.48313 | 0.00000 | 0.00000 | 0.34542 | 0.69227 | 0.00437 | 0.00000 | 0.14041 | 0.19273 | 0.00307 |
| **MBNL3** | 0.00000 | 1.72632 | 13.59765 | 0.51104 | 2.69450 | 8.68045 | 12.56897 | 0.00000 | 0.00000 | 7.94960 | 4.25433 | 1.02059 |
| **MBOAT1** | 0.21127 | 2.46510 | 0.73277 | 1.03844 | 4.96456 | 6.93013 | 0.00000 | 0.22349 | 0.00000 | 2.68722 | 2.47412 | 0.38500 |
| **MBOAT2** | 1.71190 | 0.11007 | 0.00000 | 1.95348 | 1.68649 | 4.85241 | 0.83315 | 0.56649 | 4.59367 | 5.84583 | 0.33018 | 0.92317 |
| **MBOAT4** | 0.03834 | 0.23397 | 0.13180 | 0.10725 | 0.08188 | 0.00000 | 0.30208 | 0.10381 | 0.00000 | 0.00000 | 0.25911 | 0.04106 |
| **MBOAT7** | 0.00214 | 3.77913 | 13.89678 | 7.54197 | 9.04347 | 0.00000 | 7.27119 | 0.00000 | 22.98656 | 21.04359 | 1.67272 | 2.43781 |
| **MBP** | 1.65528 | 0.40280 | 1.02504 | 0.02279 | 1.02060 | 0.00000 | 0.55346 | 0.10381 | 0.61977 | 1.64140 | 1.93278 | 0.00000 |
| **MBTD1** | 0.61352 | 0.00000 | 0.57198 | 0.09882 | 0.00000 | 3.72599 | 2.19472 | 0.69282 | 0.00000 | 10.11916 | 0.52117 | 2.38905 |
| **MBTPS1** | 4.34363 | 5.44409 | 5.19880 | 4.46211 | 2.10356 | 3.75909 | 3.16928 | 1.66050 | 15.19182 | 5.23682 | 9.38103 | 1.47347 |
| **MBTPS2** | 0.83532 | 0.57109 | 1.21803 | 0.33044 | 0.08874 | 2.67798 | 0.00000 | 0.09935 | 2.63122 | 0.53552 | 2.91869 | 0.00000 |
| **MC1R** | 0.00325 | 0.37848 | 0.54817 | 0.81057 | 0.00000 | 0.00000 | 0.41981 | 0.05295 | 2.38932 | 4.02631 | 0.00000 | 0.26758 |
| **MC2R** | 0.01051 | 0.19151 | 0.01675 | 0.00365 | 0.00346 | 0.01525 | 0.01899 | 0.01227 | 0.02157 | 0.00814 | 0.01534 | 0.02726 |
| **MC5R** | 0.03100 | 0.03736 | 0.01804 | 0.00948 | 0.00000 | 0.00000 | 0.03617 | 0.00816 | 0.00000 | 0.05251 | 0.10198 | 0.02719 |
| **MCAM** | 2.78615 | 4.55825 | 0.77022 | 4.28697 | 1.21326 | 0.00000 | 0.00000 | 0.53225 | 1.77030 | 3.28185 | 0.00000 | 5.69370 |
| **MCAT** | 0.47288 | 8.82344 | 8.19308 | 1.48176 | 4.38265 | 0.00000 | 4.56528 | 0.94084 | 6.51114 | 6.37206 | 11.42537 | 4.15658 |
| **MCC** | 1.96993 | 2.89902 | 0.49221 | 1.96604 | 0.00000 | 0.88518 | 1.19451 | 1.28588 | 0.68577 | 0.76774 | 1.64646 | 1.97495 |
| **MCCC1** | 1.38615 | 0.00000 | 3.22816 | 0.00000 | 0.00000 | 0.74920 | 3.94247 | 0.03543 | 1.89881 | 0.00000 | 4.43529 | 0.12121 |

| | | | | | | | | | | | | |
|---|---|---|---|---|---|---|---|---|---|---|---|---|
| MCCC2 | 2.08501 | 5.03396 | 14.13167 | 5.75246 | 6.68121 | 16.43451 | 15.21707 | 0.13402 | 18.59534 | 2.87928 | 14.81761 | 2.35667 |
| MCEE | 0.30080 | 0.19544 | 5.08564 | 0.00000 | 0.76148 | 5.18315 | 5.01450 | 0.43719 | 3.79871 | 2.76809 | 5.32031 | 0.37679 |
| MCEMP1 | 0.01920 | 0.01527 | 0.02471 | 0.01021 | 0.01718 | 0.01376 | 0.01492 | 0.01092 | 0.03116 | 0.02038 | 0.01285 | 0.01568 |
| 2.00000 | 0.00674 | 0.01077 | 0.29089 | 0.02907 | 0.00000 | 0.06214 | 0.00000 | 0.01859 | 0.00000 | 0.09066 | 0.02528 | 0.01449 |
| MCF2L | 0.04820 | 0.01657 | 0.04169 | 0.04911 | 0.05700 | 0.00000 | 0.02334 | 0.00957 | 0.08384 | 0.00000 | 0.08940 | 0.02771 |
| MCF2L-AS1 | 0.01409 | 0.63577 | 0.01052 | 0.46483 | 0.26118 | 0.00000 | 0.00790 | 0.00751 | 0.10003 | 0.69202 | 0.87348 | 0.01155 |
| MCF2L2 | 0.04044 | 0.01393 | 0.04020 | 0.01560 | 0.04083 | 0.00037 | 0.01276 | 0.07910 | 0.02478 | 0.07932 | 0.02616 | 0.01041 |
| MCFD2 | 0.13468 | 0.00000 | 18.41388 | 2.27898 | 0.37253 | 18.66591 | 13.36945 | 1.56641 | 27.15943 | 11.91857 | 15.71244 | 3.88168 |
| MCHR1 | 0.13817 | 0.02778 | 0.05098 | 0.06734 | 0.01655 | 0.00000 | 0.01549 | 0.00842 | 0.04992 | 0.00557 | 0.01236 | 0.14400 |
| MCHR2 | 0.07427 | 0.01848 | 0.00183 | 0.00365 | 0.00019 | 0.02637 | 0.00000 | 0.00320 | 0.09426 | 0.06465 | 0.00649 | 0.07196 |
| MCHR2-AS1 | 0.01319 | 0.01633 | 0.01865 | 0.01441 | 0.01169 | 0.00542 | 0.01329 | 0.00957 | 0.02497 | 0.01338 | 0.06399 | 0.01714 |
| MCIDAS | 0.00430 | 0.44247 | 0.00473 | 0.22418 | 0.00000 | 0.00017 | 0.02041 | 0.15185 | 0.04940 | 0.04233 | 0.06808 | 0.61056 |
| MCL1 | 0.66674 | 13.15281 | 14.24157 | 19.68678 | 4.34261 | 29.85167 | 16.85639 | 17.08021 | 2.78856 | 26.85186 | 6.72172 | 27.24649 |
| MCM10 | 3.82138 | 14.45522 | 9.91007 | 10.34468 | 11.61417 | 22.23439 | 8.87417 | 6.62642 | 14.07956 | 10.29281 | 12.30708 | 19.56661 |
| MCM2 | 8.63513 | 29.32107 | 16.57889 | 17.20775 | 17.73199 | 1.83205 | 19.97971 | 3.46706 | 26.05007 | 6.92722 | 23.30618 | 22.16507 |
| MCM3 | 13.16395 | 21.59178 | 18.78162 | 27.74945 | 15.94327 | 22.44656 | 29.77496 | 11.20865 | 25.65548 | 4.28279 | 28.23201 | 24.64081 |
| MCM3AP | 4.45130 | 11.32977 | 8.24465 | 6.37360 | 8.91272 | 0.10588 | 16.94757 | 0.88346 | 9.10121 | 19.16451 | 9.00753 | 6.68882 |
| MCM3AP-AS1 | 0.11525 | 0.39594 | 0.40159 | 0.56983 | 0.74662 | 1.60151 | 0.00000 | 0.29694 | 1.84674 | 1.88050 | 0.28544 | 0.97797 |
| MCM4 | 3.10511 | 28.26416 | 18.08363 | 21.36899 | 24.99449 | 33.06491 | 22.99409 | 13.53600 | 31.13142 | 15.93841 | 31.57889 | 22.97399 |
| MCM5 | 11.22111 | 31.23098 | 15.62596 | 23.82500 | 29.08554 | 11.95423 | 15.30576 | 2.44798 | 27.82969 | 4.35830 | 22.95012 | 32.50653 |
| MCM6 | 2.04563 | 18.65947 | 10.71321 | 12.79812 | 10.33746 | 30.88847 | 12.89338 | 7.06217 | 11.69229 | 9.36911 | 19.11134 | 21.10040 |
| MCM7 | 8.23003 | 37.08116 | 27.86639 | 28.00250 | 28.51999 | 4.80417 | 34.32744 | 16.06549 | 41.86846 | 8.18852 | 30.30271 | 29.67771 |
| MCM8 | 0.00000 | 1.97999 | 3.78421 | 0.78496 | 2.72014 | 7.27010 | 3.02184 | 1.33648 | 2.40142 | 16.05015 | 2.58383 | 1.85157 |
| MCM9 | 0.55066 | 1.17432 | 2.46795 | 0.81063 | 0.00000 | 4.56847 | 1.86976 | 2.44518 | 3.93922 | 0.07939 | 2.88662 | 1.41193 |
| MCMBP | 3.73503 | 6.03715 | 2.28260 | 5.94061 | 5.51652 | 15.66497 | 10.09201 | 0.56300 | 2.44413 | 13.99673 | 5.23472 | 5.28176 |
| MCMDC2 | 0.05117 | 0.18003 | 0.08210 | 0.01233 | 0.00000 | 0.07743 | 0.05660 | 0.26478 | 0.29905 | 0.59381 | 0.00000 | 0.98975 |
| MCOLN1 | 2.90502 | 15.51427 | 8.52055 | 4.30633 | 19.14997 | 0.00000 | 2.73923 | 1.03034 | 28.95340 | 0.05459 | 12.84865 | 16.28016 |
| MCOLN2 | 1.07972 | 0.07522 | 0.20553 | 0.00000 | 0.00000 | 0.00000 | 0.00000 | 0.20765 | 0.26473 | 0.50805 | 0.00000 | 0.01462 |
| MCOLN3 | 0.01027 | 0.03419 | 0.08618 | 0.00637 | 0.00093 | 0.05669 | 0.00000 | 0.14728 | 0.00000 | 0.04184 | 0.01274 | 0.04776 |
| MCPH1 | 2.13887 | 8.10924 | 6.10241 | 1.23951 | 3.80377 | 18.55307 | 0.00000 | 5.97614 | 4.89767 | 5.77557 | 4.66896 | 12.07189 |
| MCRS1 | 5.98078 | 18.90848 | 14.15463 | 15.61322 | 21.16085 | 6.37921 | 13.11964 | 7.09341 | 22.51085 | 0.00000 | 12.13256 | 22.04616 |
| MCTP1 | 4.24410 | 0.00000 | 0.00000 | 0.00000 | 0.01923 | 0.49429 | 0.00000 | 0.86171 | 0.10271 | 1.03631 | 0.00000 | 0.25808 |
| MCTP2 | 3.15419 | 0.04875 | 0.12359 | 0.01973 | 0.00000 | 0.00747 | 0.00000 | 0.40840 | 0.00930 | 0.27174 | 0.00716 | 0.00000 |
| MCTS1 | 2.51639 | 7.74884 | 8.02607 | 7.27969 | 10.10444 | 27.57107 | 10.72079 | 2.47010 | 9.02903 | 5.76882 | 13.56157 | 9.51786 |
| MCTS2P | 0.14947 | 1.27730 | 2.09641 | 1.20212 | 0.17575 | 4.27422 | 0.70408 | 0.05293 | 2.72740 | 0.00000 | 1.34230 | 0.99992 |
| MCU | 0.93885 | 3.16832 | 3.05872 | 5.60999 | 1.73967 | 2.67479 | 2.24859 | 0.57379 | 3.18929 | 0.02380 | 3.05758 | 1.42151 |
| MCUR1 | 0.77044 | 8.20222 | 12.50975 | 9.44770 | 9.61593 | 9.79660 | 10.42809 | 0.00000 | 4.69270 | 10.52215 | 6.24762 | 0.73337 |
| 1.00000 | 0.95521 | 7.21279 | 3.74973 | 2.98555 | 12.70737 | 1.01261 | 5.18750 | 2.14295 | 3.90533 | 7.79364 | 2.45044 | 7.35291 |
| MDFI | 0.00495 | 0.00000 | 8.67718 | 2.03554 | 0.37881 | 0.00000 | 0.47161 | 0.00740 | 0.00000 | 0.19067 | 0.03115 | 0.02028 |
| MDFIC | 0.00395 | 0.00662 | 0.03361 | 0.00000 | 0.05046 | 0.22692 | 0.18047 | 0.01450 | 0.00000 | 0.21416 | 0.00000 | 0.00945 |
| MDGA1 | 0.05880 | 0.00867 | 0.02523 | 0.05852 | 0.00933 | 0.00000 | 0.02453 | 0.00279 | 0.00000 | 0.01334 | 0.13299 | 0.12681 |
| MDGA2 | 1.67946 | 0.00411 | 0.63555 | 1.14948 | 0.00000 | 0.00086 | 0.11269 | 0.58772 | 0.00000 | 0.12754 | 0.01814 | 0.25609 |
| MDH1 | 3.59154 | 23.75590 | 18.53294 | 21.00879 | 17.74766 | 52.25145 | 25.58115 | 0.52465 | 25.54720 | 4.80496 | 34.06151 | 14.61452 |
| MDH1B | 0.09932 | 0.19594 | 0.00000 | 0.30606 | 0.07343 | 0.27879 | 0.00104 | 0.21028 | 0.00000 | 0.00006 | 0.22706 | 0.41354 |
| MDH2 | 6.25515 | 25.85699 | 32.55696 | 35.98345 | 42.87118 | 25.19049 | 50.01466 | 4.86972 | 52.71343 | 0.00005 | 43.35286 | 5.84750 |
| MDK | 1.51958 | 13.66731 | 18.89707 | 25.00368 | 17.79771 | 12.58642 | 8.24400 | 0.64777 | 27.43092 | 3.95199 | 2.84482 | 5.30526 |
| MDM1 | 1.70360 | 1.08752 | 2.27187 | 0.66541 | 0.00000 | 2.87642 | 1.97384 | 0.97456 | 2.01891 | 0.48986 | 1.20961 | 2.67830 |
| MDM2 | 2.50884 | 12.10786 | 24.79911 | 3.66521 | 1.81081 | 38.37475 | 17.00235 | 20.03373 | 5.01866 | 10.14127 | 11.25727 | 8.47976 |

| | | | | | | | | | | | | |
|---|---|---|---|---|---|---|---|---|---|---|---|---|
| **MDM4** | 8.67779 | 7.17811 | 7.07257 | 5.71036 | 7.11871 | 29.47635 | 4.19906 | 10.76909 | 13.03488 | 25.09993 | 15.37286 | 14.16771 |
| **MDN1** | 0.00000 | 4.53545 | 5.65834 | 10.34550 | 11.86275 | 12.50043 | 11.15981 | 4.26160 | 10.06588 | 35.85171 | 8.73411 | 12.57625 |
| **MDP1** | 0.01383 | 0.37680 | 0.02845 | 1.02671 | 0.15024 | 0.31592 | 0.05404 | 0.01197 | 0.13997 | 0.00000 | 0.15306 | 0.33037 |
| **MDS2** | 0.01952 | 0.01932 | 0.01547 | 0.01862 | 0.01775 | 0.01320 | 0.01293 | 0.00884 | 0.02882 | 0.01630 | 0.01416 | 0.01680 |
| **ME1** | 1.05511 | 0.00000 | 1.71877 | 0.15219 | 0.00000 | 0.18585 | 1.31297 | 0.10132 | 0.26495 | 0.00000 | 0.28287 | 0.06214 |
| **ME2** | 0.00000 | 0.05977 | 6.65212 | 0.00000 | 0.00000 | 4.74343 | 7.53891 | 0.02631 | 5.34780 | 4.60817 | 3.26422 | 0.06190 |
| **ME3** | 0.89966 | 0.04190 | 0.10916 | 0.08253 | 0.00000 | 0.00000 | 0.00720 | 0.58403 | 0.00002 | 0.14657 | 0.14009 | 0.12303 |
| **MEA1** | 1.64883 | 14.82176 | 17.27149 | 13.44026 | 12.64215 | 19.65836 | 18.46572 | 9.32541 | 14.08936 | 0.00000 | 17.27190 | 16.53389 |
| **MEAF6** | 2.41350 | 9.79970 | 5.34211 | 7.67828 | 2.46574 | 22.04547 | 8.80570 | 5.03552 | 2.19413 | 6.24899 | 8.31425 | 11.45281 |
| **MECOM** | 1.20646 | 0.00000 | 0.81950 | 0.06123 | 0.66250 | 0.00000 | 0.00000 | 0.27745 | 0.21726 | 1.07366 | 0.00000 | 1.16537 |
| **MECP2** | 4.42221 | 0.60596 | 0.93316 | 1.87873 | 1.31020 | 0.00000 | 3.07768 | 0.99184 | 1.68979 | 3.81337 | 1.04228 | 1.24749 |
| **MECR** | 1.47991 | 10.63108 | 6.73250 | 7.27654 | 15.27847 | 1.66392 | 7.36391 | 0.43196 | 14.27822 | 0.03767 | 12.57746 | 4.69149 |
| **MED1** | 1.94128 | 2.21636 | 9.24163 | 4.58028 | 2.97720 | 8.86290 | 11.79723 | 4.45474 | 8.98436 | 20.10216 | 9.35161 | 9.27258 |
| **MED10** | 2.92303 | 13.28194 | 17.54694 | 2.95991 | 5.89612 | 14.15894 | 18.47270 | 17.61719 | 10.50321 | 2.05163 | 5.68065 | 14.19068 |
| **MED11** | 2.04142 | 5.61534 | 4.62191 | 3.35294 | 1.73438 | 6.29698 | 1.98243 | 2.71547 | 4.47618 | 0.00000 | 5.52446 | 11.00986 |
| **MED12** | 0.66346 | 6.25295 | 0.98494 | 2.15904 | 5.21970 | 1.85006 | 3.60564 | 0.06672 | 6.04318 | 7.14467 | 1.80780 | 5.10030 |
| **MED12L** | 4.46943 | 1.34970 | 22.84820 | 0.00484 | 3.08617 | 4.53430 | 10.43542 | 4.16444 | 3.80763 | 8.00639 | 7.04267 | 3.39757 |
| **MED13** | 8.05760 | 3.43303 | 12.19895 | 5.04511 | 0.00355 | 27.91694 | 10.30477 | 7.38216 | 4.88736 | 38.03326 | 2.99851 | 7.55386 |
| **MED13L** | 5.26951 | 3.81424 | 0.00000 | 5.28265 | 10.27896 | 19.31844 | 8.83219 | 2.94978 | 3.96826 | 28.04269 | 3.57894 | 3.74800 |
| **MED14** | 6.32634 | 1.87231 | 4.45851 | 1.24210 | 1.08183 | 6.58571 | 2.87685 | 4.44366 | 1.54177 | 4.38741 | 1.38487 | 3.36179 |
| **MED14OS** | 0.12127 | 0.03038 | 0.10830 | 0.04445 | 0.12668 | 0.03964 | 0.00000 | 0.07289 | 0.02617 | 0.10233 | 0.02194 | 0.09338 |
| **MED15** | 8.53369 | 15.96347 | 10.31283 | 15.82873 | 12.32006 | 0.00000 | 8.22186 | 5.17719 | 8.71915 | 15.84215 | 5.02922 | 22.99916 |
| **MED15P9** | 0.04474 | 0.04357 | 0.02310 | 0.01558 | 0.00986 | 0.00849 | 0.01593 | 0.30827 | 0.01558 | 0.02351 | 0.01594 | 0.02607 |
| **MED16** | 9.26508 | 9.95612 | 7.08752 | 7.81349 | 6.98516 | 0.00000 | 6.43582 | 1.97583 | 13.55080 | 0.00000 | 5.54636 | 8.30070 |
| **MED17** | 0.00000 | 6.05843 | 6.27451 | 0.45261 | 0.23214 | 7.20428 | 5.00743 | 7.47100 | 12.71693 | 6.69541 | 3.76077 | 4.87054 |
| **MED18** | 0.10013 | 5.24782 | 5.23061 | 3.62253 | 4.66723 | 3.88643 | 8.95797 | 0.62276 | 3.99294 | 0.00000 | 8.97454 | 11.97510 |
| **MED19** | 6.20965 | 11.58865 | 9.06621 | 7.84492 | 8.54989 | 10.63796 | 10.08425 | 22.69830 | 13.12280 | 6.41959 | 6.41975 | 20.64309 |
| **MED20** | 6.00428 | 9.07695 | 3.11680 | 4.45338 | 4.24413 | 12.85188 | 2.68381 | 4.28359 | 6.59975 | 0.00000 | 5.80275 | 8.96917 |
| **MED21** | 4.34846 | 4.23174 | 8.58368 | 3.30819 | 3.76429 | 21.28575 | 14.61265 | 12.33001 | 9.43875 | 0.22671 | 6.31170 | 9.76952 |
| **MED22** | 1.07541 | 3.83576 | 8.42354 | 4.35589 | 2.58424 | 0.00000 | 4.87656 | 1.01824 | 10.78698 | 3.92151 | 6.93080 | 4.97428 |
| **MED23** | 3.84652 | 3.36506 | 1.03922 | 0.82637 | 0.02123 | 5.97596 | 0.59511 | 0.92494 | 3.42726 | 5.07100 | 1.48358 | 3.96758 |
| **MED24** | 1.96110 | 8.52421 | 8.57963 | 7.85606 | 8.93118 | 0.00000 | 11.93611 | 2.51806 | 11.69025 | 4.47694 | 8.84776 | 6.36068 |
| **MED25** | 3.20082 | 1.52190 | 2.76640 | 1.65250 | 2.53242 | 0.00000 | 3.34115 | 0.13248 | 1.96898 | 4.11848 | 0.88982 | 2.48320 |
| **MED26** | 2.33319 | 0.21970 | 1.18561 | 1.03861 | 1.57624 | 0.70902 | 1.82450 | 2.34425 | 0.00000 | 9.22124 | 0.00000 | 0.68832 |
| **MED27** | 1.73321 | 7.50847 | 5.26507 | 4.68296 | 6.12223 | 0.80650 | 6.15118 | 2.87875 | 5.10421 | 0.00000 | 4.17200 | 5.17746 |
| **MED28** | 0.00000 | 11.66160 | 12.88645 | 7.96135 | 6.26302 | 26.18865 | 11.25197 | 12.12062 | 16.03039 | 0.00000 | 12.32625 | 18.71662 |
| **MED29** | 1.37896 | 10.89422 | 7.44696 | 10.98052 | 9.33325 | 0.00113 | 9.01113 | 4.87477 | 8.38001 | 2.15165 | 12.51207 | 14.91407 |
| **MED30** | 2.67521 | 4.77087 | 5.52735 | 2.22360 | 2.45714 | 10.06007 | 5.02729 | 1.76948 | 5.26689 | 0.00000 | 4.65193 | 5.52171 |
| **MED31** | 0.81461 | 1.81676 | 5.09549 | 0.00000 | 0.55221 | 4.41129 | 2.90380 | 15.69027 | 5.11169 | 0.08569 | 2.32997 | 6.68262 |
| **MED4** | 1.55796 | 5.36335 | 9.27820 | 2.81649 | 0.00000 | 22.77102 | 9.20247 | 13.19864 | 4.36815 | 10.58501 | 7.90886 | 14.94640 |
| **MED4-AS1** | 0.03129 | 0.02381 | 0.06886 | 0.05526 | 0.04081 | 0.00942 | 0.00021 | 0.02644 | 0.11092 | 0.05809 | 0.06944 | 0.10827 |
| **MED6** | 2.01104 | 6.16295 | 10.78510 | 3.74284 | 2.19242 | 23.28793 | 9.43318 | 20.63742 | 9.79253 | 2.14458 | 8.44352 | 23.64069 |
| **MED7** | 0.00000 | 7.37661 | 4.83838 | 2.26199 | 0.00000 | 14.75505 | 3.61838 | 11.22703 | 4.10758 | 0.75754 | 6.31330 | 9.97958 |
| **MED8** | 4.65990 | 9.49446 | 13.74953 | 3.39651 | 6.85980 | 14.20558 | 8.41127 | 6.17464 | 11.32818 | 0.00000 | 5.79937 | 13.61804 |
| **MED9** | 0.44978 | 11.55451 | 2.34352 | 6.13519 | 0.83426 | 6.85159 | 2.50807 | 1.73208 | 7.82311 | 0.00000 | 3.92094 | 9.31506 |
| **MEF2A** | 0.97453 | 2.66003 | 2.40441 | 3.53895 | 2.39718 | 5.59324 | 0.40634 | 1.49536 | 2.13931 | 3.48984 | 1.62188 | 3.10257 |
| **MEF2B** | 0.00633 | 0.01453 | 0.02031 | 0.01292 | 0.01501 | 0.02172 | 0.02062 | 0.01042 | 0.01585 | 0.01227 | 0.01812 | 0.01623 |
| **MEF2BNB.uid-2** | 1.16141 | 0.31862 | 1.20860 | 0.39372 | 0.00000 | 0.40166 | 1.40381 | 0.61215 | 0.33012 | 0.68057 | 0.56578 | 1.33312 |

| | | | | | | | | | | | | |
|---|---|---|---|---|---|---|---|---|---|---|---|---|
| **MEF2BNB.uid-2** | 0.18190 | 3.30461 | 1.95091 | 1.13951 | 1.35331 | 1.73220 | 1.27858 | 0.00000 | 0.00000 | 0.00000 | 2.67600 | 4.33221 |
| **MEF2C** | 0.02466 | 0.01215 | 0.01245 | 0.00876 | 0.01240 | 0.03603 | 0.06560 | 0.00992 | 0.00479 | 0.00516 | 0.01199 | 0.01640 |
| **MEF2C-AS1** | 0.00827 | 0.01240 | 0.01657 | 0.01066 | 0.01356 | 0.02536 | 0.01703 | 0.01046 | 0.01583 | 0.01330 | 0.01716 | 0.02974 |
| **MEF2D** | 2.25581 | 2.27929 | 0.56659 | 2.00355 | 1.88534 | 0.00000 | 1.98445 | 1.30287 | 0.61840 | 3.70036 | 0.90744 | 2.16689 |
| **MEFV** | 0.00295 | 0.00880 | 0.00400 | 0.02298 | 0.00000 | 0.02667 | 0.00922 | 0.02168 | 0.02165 | 0.11332 | 0.01026 | 0.01234 |
| **MEG3** | 0.27879 | 0.05975 | 3.34645 | 12.82861 | 0.05625 | 0.00000 | 0.00000 | 0.01186 | 0.00000 | 0.60141 | 0.03590 | 0.00320 |
| **MEG8** | 0.00406 | 0.01582 | 0.02330 | 0.24124 | 0.00520 | 0.00718 | 0.01277 | 0.03190 | 0.02259 | 0.01925 | 0.01573 | 0.01489 |
| **MEG9** | 0.06334 | 0.00893 | 0.07143 | 1.20505 | 0.00000 | 0.00000 | 0.01900 | 0.00582 | 0.00712 | 0.04116 | 0.01528 | 0.01625 |
| **MEGF10** | 0.81249 | 0.00811 | 0.01454 | 0.06790 | 0.00732 | 0.00031 | 0.01679 | 0.42279 | 0.02091 | 0.06475 | 0.01072 | 0.06198 |
| **MEGF11** | 0.58636 | 0.05261 | 0.18815 | 0.10939 | 0.22633 | 0.00000 | 0.00000 | 1.35817 | 0.58316 | 1.02916 | 0.18113 | 0.16028 |
| **MEGF6** | 0.00868 | 0.20549 | 0.02844 | 0.00000 | 0.00000 | 0.00889 | 0.04659 | 0.01042 | 0.00000 | 0.23141 | 0.00887 | 0.00109 |
| **MEGF8** | 0.00278 | 0.55479 | 0.24568 | 1.36140 | 4.57888 | 0.00000 | 0.00101 | 0.05346 | 1.40804 | 4.66115 | 0.69851 | 0.70520 |
| **MEGF9** | 0.40280 | 0.37797 | 0.49312 | 0.10160 | 0.08785 | 0.83645 | 0.13194 | 0.08518 | 0.00368 | 0.47183 | 0.00000 | 0.00000 |
| **MEI1** | 0.11329 | 2.68153 | 0.76905 | 2.32673 | 2.70150 | 0.76686 | 0.00032 | 1.47566 | 0.00000 | 0.06835 | 1.69632 | 4.34375 |
| **MEI4** | 0.91569 | 0.09283 | 0.07912 | 0.04687 | 0.02665 | 0.02416 | 0.19551 | 0.00000 | 0.00000 | 0.00000 | 0.14393 | 0.09336 |
| **MEIG1** | 0.00231 | 0.26406 | 0.00000 | 0.00000 | 0.14439 | 0.13034 | 0.00000 | 0.06520 | 0.05182 | 0.00731 | 0.00019 | 0.10332 |
| **MEIOB** | 0.05353 | 0.04814 | 0.01700 | 0.14921 | 0.12944 | 0.00022 | 0.00887 | 0.01427 | 0.00000 | 0.08751 | 0.01850 | 1.44922 |
| **MEIS1** | 0.08923 | 0.85896 | 0.56538 | 1.37909 | 0.54519 | 0.36072 | 1.37864 | 0.29235 | 1.25565 | 1.02019 | 1.57746 | 1.26015 |
| **MEIS1-AS3** | 0.03041 | 0.01048 | 0.00892 | 0.00566 | 0.01118 | 0.02480 | 0.01014 | 0.00871 | 0.02565 | 0.07778 | 0.00985 | 0.01208 |
| **MEIS2** | 0.00356 | 0.00000 | 0.42157 | 1.41065 | 0.28682 | 0.00896 | 0.90979 | 0.00792 | 0.49931 | 0.42603 | 0.00000 | 0.07707 |
| **MEIS3** | 0.71389 | 0.04255 | 0.08341 | 0.06545 | 0.56780 | 0.00000 | 0.04088 | 0.17270 | 0.04111 | 0.00000 | 0.05281 | 0.09913 |
| **MEIS3P1** | 0.10931 | 0.01367 | 0.07724 | 0.11309 | 0.06581 | 0.00002 | 0.17154 | 0.02294 | 0.02806 | 0.02785 | 0.01565 | 0.05608 |
| **MELK** | 7.55081 | 14.87217 | 13.14094 | 9.84470 | 4.28760 | 37.08500 | 12.71726 | 7.85402 | 15.43394 | 0.00587 | 10.69154 | 11.87014 |
| **MEMO1** | 0.17456 | 2.82369 | 2.48758 | 0.84938 | 1.10941 | 4.76577 | 0.67358 | 0.10403 | 3.54595 | 1.36456 | 1.21539 | 2.08858 |
| **MEN1** | 0.55004 | 1.13756 | 1.50327 | 0.77215 | 4.19122 | 0.00227 | 5.54620 | 0.00000 | 0.56780 | 9.89330 | 1.06964 | 2.56683 |
| **MEOX2** | 0.00620 | 0.01443 | 0.02157 | 0.01206 | 0.01431 | 0.02050 | 0.01780 | 0.01047 | 0.01600 | 0.01691 | 0.01518 | 0.01593 |
| **MEP1A** | 0.00567 | 0.00916 | 0.01253 | 0.10618 | 0.00000 | 0.02956 | 0.01709 | 0.01016 | 0.00000 | 0.01856 | 0.07349 | 0.02204 |
| **MEP1B** | 0.03517 | 0.01947 | 0.00000 | 0.00849 | 0.00000 | 0.04061 | 0.00000 | 0.37439 | 0.08326 | 0.11505 | 0.00000 | 0.01037 |
| **MEPCE** | 7.39997 | 5.35864 | 2.39551 | 5.22725 | 6.68550 | 2.49096 | 12.09121 | 1.20123 | 0.00081 | 17.89510 | 2.77313 | 5.98715 |
| **MEPE** | 0.00606 | 0.00619 | 0.01387 | 0.18086 | 0.02233 | 0.00664 | 0.02291 | 0.00563 | 0.01916 | 0.01884 | 0.01706 | 0.21561 |
| **MERTK** | 1.59435 | 8.23692 | 0.52856 | 10.03331 | 2.64596 | 10.60233 | 0.00000 | 0.50471 | 14.75213 | 5.76064 | 3.38758 | 8.38771 |
| **MESDC1** | 0.60029 | 0.00000 | 0.35637 | 2.29752 | 0.64056 | 0.34013 | 0.87817 | 1.83113 | 0.00000 | 3.31948 | 0.00000 | 0.64179 |
| **MESDC2** | 3.54460 | 13.78902 | 10.65995 | 9.21850 | 29.46425 | 32.64309 | 5.22545 | 10.25383 | 31.56187 | 21.65287 | 14.17032 | 14.88648 |
| **MESP1** | 0.74069 | 0.01673 | 0.03293 | 0.02402 | 0.53843 | 0.00000 | 0.08465 | 0.04490 | 0.76839 | 0.29012 | 0.93308 | 0.00091 |
| **MESP2** | 0.00229 | 0.03802 | 0.00000 | 0.67970 | 0.35392 | 0.00000 | 0.03824 | 0.00901 | 0.00000 | 0.58560 | 0.36127 | 0.06093 |
| **MEST** | 8.42177 | 25.51646 | 4.96661 | 12.71984 | 9.97140 | 27.17639 | 0.00000 | 2.22733 | 12.72050 | 0.00000 | 15.60471 | 15.74244 |
| **MESTIT1** | 0.00398 | 0.04652 | 0.01530 | 0.01006 | 0.01533 | 0.01229 | 0.00931 | 0.02563 | 0.02482 | 0.04797 | 0.01807 | 0.02441 |
| **MET** | 0.00000 | 1.93351 | 1.84487 | 0.78617 | 0.00000 | 2.05844 | 1.41673 | 0.13473 | 2.29351 | 4.18167 | 0.10545 | 1.26687 |
| **METAP1** | 1.58224 | 7.53474 | 14.22677 | 7.74479 | 9.61957 | 30.54365 | 20.23427 | 4.81594 | 17.67511 | 5.02724 | 14.16542 | 7.84165 |
| **METAP1D** | 0.02112 | 2.81497 | 1.25662 | 0.52061 | 0.33225 | 2.40554 | 0.00000 | 0.00000 | 1.80084 | 0.07068 | 1.96296 | 0.87415 |
| **METAP2** | 9.46130 | 16.62531 | 11.62077 | 7.88084 | 11.90928 | 35.07943 | 27.87696 | 16.84118 | 13.28271 | 30.49412 | 15.19495 | 21.38565 |
| **METRN** | 0.26854 | 0.12224 | 0.00437 | 0.48116 | 0.17621 | 0.00000 | 0.10576 | 0.00000 | 0.00000 | 0.48768 | 0.00000 | 0.16332 |
| **METRNL** | 0.51249 | 0.08707 | 2.72517 | 0.00000 | 5.70037 | 0.00000 | 3.25218 | 0.40151 | 0.00000 | 8.51319 | 0.00000 | 0.49069 |
| **METTL1** | 0.22495 | 17.49301 | 6.93168 | 3.07379 | 9.82911 | 8.50303 | 10.94322 | 3.74650 | 6.46805 | 0.55843 | 7.05471 | 17.99176 |
| **METTL10** | 0.00000 | 2.51141 | 0.00000 | 1.39861 | 3.26138 | 13.77423 | 0.00000 | 9.50260 | 0.00052 | 2.81804 | 4.46599 | 2.60392 |
| **METTL11B** | 0.01369 | 0.01322 | 0.05012 | 0.03101 | 0.02017 | 0.02790 | 0.06385 | 0.03216 | 0.03176 | 0.00000 | 0.00806 | 0.02592 |
| **METTL12** | 0.04593 | 2.20231 | 5.06622 | 2.03323 | 4.30692 | 0.32443 | 9.73643 | 3.17950 | 3.68987 | 2.73211 | 6.33259 | 3.96854 |
| **METTL13** | 1.01032 | 13.86966 | 12.34502 | 7.34286 | 11.02001 | 3.06946 | 19.97926 | 6.65746 | 16.80958 | 0.00000 | 14.04264 | 14.56853 |

| | | | | | | | | | | | | |
|---|---|---|---|---|---|---|---|---|---|---|---|---|
| **METTL14** | 2.04816 | 4.05754 | 3.87347 | 1.99262 | 1.16600 | 11.70619 | 4.05823 | 3.62784 | 6.90954 | 0.96502 | 4.27613 | 7.47756 |
| **METTL15** | 0.32894 | 1.26477 | 0.00029 | 0.69964 | 1.62352 | 3.85040 | 1.01008 | 0.00000 | 1.39454 | 0.19384 | 1.28883 | 0.85569 |
| **METTL16** | 1.05219 | 9.72171 | 8.98522 | 10.63493 | 7.01791 | 3.42420 | 10.97794 | 6.80992 | 6.43618 | 12.14529 | 7.70484 | 11.57223 |
| **METTL17** | 1.34334 | 3.18284 | 4.33664 | 6.94707 | 1.30747 | 5.35242 | 2.06693 | 1.41413 | 13.96340 | 0.00000 | 4.16771 | 7.14086 |
| **METTL18** | 0.02582 | 3.27620 | 0.81501 | 0.42048 | 0.00148 | 6.86654 | 2.26300 | 1.00573 | 2.31131 | 1.17483 | 1.60950 | 6.20669 |
| **METTL20** | 0.00695 | 0.15022 | 0.28749 | 0.00000 | 0.00000 | 0.35317 | 0.41501 | 2.02779 | 0.00151 | 0.45488 | 0.16577 | 2.52965 |
| **METTL21A** | 3.02389 | 4.47055 | 6.62489 | 0.32712 | 2.86718 | 1.79171 | 3.13994 | 0.29984 | 2.18849 | 3.00997 | 5.92637 | 5.66000 |
| **METTL21B** | 0.03242 | 0.15637 | 0.54344 | 0.29751 | 0.00000 | 0.00000 | 0.00000 | 0.17636 | 0.79574 | 0.00598 | 0.61772 | 1.19924 |
| **METTL21C** | 0.01951 | 0.00793 | 0.10251 | 0.05830 | 0.00000 | 0.05728 | 0.00432 | 0.04766 | 0.12635 | 0.00859 | 0.00324 | 0.01118 |
| **METTL21EP** | 0.03678 | 0.01387 | 0.02289 | 0.01000 | 0.01395 | 0.02362 | 0.01900 | 0.00754 | 0.01885 | 0.01537 | 0.01409 | 0.01507 |
| **METTL22** | 0.74928 | 5.96750 | 3.43121 | 1.52567 | 1.61895 | 0.00000 | 2.27472 | 3.01412 | 3.87939 | 0.00000 | 2.81226 | 8.23324 |
| **METTL23** | 0.42645 | 7.10713 | 4.19811 | 3.89309 | 4.49039 | 5.16440 | 0.92087 | 0.00000 | 0.00000 | 0.86079 | 4.87160 | 1.43788 |
| **METTL25** | 0.11993 | 0.21355 | 1.43498 | 0.91656 | 0.00000 | 0.77654 | 1.15119 | 0.28308 | 1.60384 | 0.82363 | 2.25216 | 0.23021 |
| **METTL2A** | 0.00000 | 6.18238 | 5.05337 | 2.93956 | 5.54292 | 5.22138 | 4.31487 | 0.49135 | 9.44023 | 3.43642 | 4.36791 | 3.74656 |
| **METTL2B** | 0.08939 | 4.84110 | 4.26382 | 2.17966 | 4.02798 | 4.90798 | 2.85664 | 0.67286 | 6.79956 | 0.82587 | 5.31405 | 3.03563 |
| **METTL3** | 0.00271 | 1.73673 | 10.58715 | 1.29758 | 1.38775 | 7.20698 | 1.92664 | 1.16106 | 15.61006 | 0.00000 | 7.48179 | 3.14651 |
| **METTL4** | 0.01440 | 2.78537 | 6.95461 | 1.54091 | 0.00000 | 10.59081 | 5.28865 | 5.31953 | 7.74070 | 5.03217 | 2.20077 | 9.33564 |
| **METTL5** | 0.28712 | 16.82448 | 10.70522 | 6.35819 | 6.51735 | 27.91721 | 11.06179 | 0.00000 | 17.15673 | 0.00006 | 17.44393 | 6.60957 |
| **METTL6** | 0.23102 | 3.71740 | 3.37093 | 0.00000 | 1.25962 | 8.81208 | 3.17593 | 1.05651 | 1.76181 | 3.78559 | 2.23316 | 5.74503 |
| **METTL7A** | 0.10754 | 0.00000 | 0.23443 | 0.96802 | 0.00000 | 0.00000 | 0.05058 | 0.01683 | 0.00622 | 0.30845 | 0.30318 | 0.04626 |
| **METTL7B** | 0.00538 | 0.10676 | 0.10385 | 0.09074 | 0.01457 | 0.03549 | 0.02299 | 0.00771 | 0.00000 | 0.06696 | 0.01131 | 0.06978 |
| **METTL8** | 0.35041 | 5.46276 | 4.20786 | 1.36037 | 1.30300 | 9.31076 | 7.80027 | 0.00000 | 9.37573 | 6.35202 | 8.61758 | 2.40138 |
| **METTL9** | 7.59268 | 8.71719 | 10.39574 | 6.50014 | 0.83863 | 9.98640 | 6.90291 | 3.35127 | 5.34108 | 11.83418 | 2.27861 | 4.32107 |
| **MEX3A** | 1.29937 | 0.19485 | 0.80729 | 0.05919 | 0.00000 | 0.00000 | 5.29643 | 0.39441 | 1.09973 | 8.23386 | 0.41965 | 0.16823 |
| **MEX3B** | 0.51992 | 0.11904 | 0.00015 | 0.00000 | 0.00000 | 0.07909 | 0.51180 | 2.24553 | 0.00000 | 1.10466 | 0.00000 | 0.55001 |
| **MEX3C** | 4.43143 | 0.00000 | 2.76425 | 1.91772 | 0.00000 | 6.38392 | 13.34254 | 2.91137 | 4.33218 | 10.45194 | 1.40244 | 0.68951 |
| **MEX3D** | 0.38732 | 0.12991 | 0.13791 | 0.00000 | 0.00000 | 0.44544 | 2.29472 | 0.07957 | 0.00000 | 4.40788 | 0.00000 | 0.14191 |
| **MFAP1** | 6.21156 | 13.17915 | 16.21601 | 4.22994 | 8.77463 | 22.63361 | 14.01299 | 29.38428 | 8.98097 | 7.22883 | 14.42292 | 23.26113 |
| **MFAP2** | 1.21006 | 0.12524 | 1.05256 | 0.37960 | 0.63328 | 0.00000 | 0.00000 | 0.29442 | 0.86291 | 0.00000 | 0.14433 | 0.06436 |
| **MFAP3** | 0.00005 | 2.24608 | 3.94204 | 0.00000 | 0.00232 | 12.34115 | 0.00000 | 6.01588 | 8.23215 | 2.81386 | 3.06785 | 14.02750 |
| **MFAP3L** | 3.84651 | 1.99636 | 0.00000 | 4.50344 | 6.05069 | 6.84581 | 0.00000 | 3.78610 | 0.00000 | 3.79041 | 4.47034 | 2.43643 |
| **MFAP4** | 0.36870 | 0.03332 | 0.00000 | 0.03872 | 0.03072 | 0.05842 | 0.00000 | 0.00000 | 0.04896 | 0.06092 | 0.00638 | 0.09472 |
| **MFAP5** | 0.07080 | 1.05566 | 0.49191 | 0.00000 | 0.72211 | 0.46487 | 0.00000 | 0.00000 | 0.00000 | 0.05217 | 0.34832 | 1.10182 |
| **MFF** | 2.38602 | 3.67133 | 3.64972 | 2.24760 | 1.27288 | 16.84412 | 7.81371 | 0.62057 | 9.82994 | 1.75791 | 7.46470 | 1.90955 |
| **MFGE8** | 0.18337 | 0.89849 | 0.39867 | 3.32542 | 1.02250 | 0.00000 | 0.03237 | 0.22064 | 0.11368 | 0.32506 | 0.14021 | 0.84692 |
| **MFHAS1** | 3.00965 | 0.91301 | 1.87510 | 3.71818 | 4.74605 | 0.52070 | 5.28026 | 0.00000 | 1.34673 | 3.83245 | 2.85843 | 0.57559 |
| **MFI2** | 0.00303 | 0.17574 | 0.08686 | 0.27618 | 0.07010 | 0.00000 | 0.04546 | 0.08806 | 0.15271 | 0.41424 | 0.00000 | 0.17773 |
| **MFI2-AS1** | 0.18012 | 0.01265 | 0.02760 | 0.01232 | 0.01912 | 0.00820 | 0.01976 | 0.01770 | 0.01498 | 0.09562 | 0.03702 | 0.09856 |
| **MFN1** | 5.17543 | 30.23383 | 1.71477 | 0.00000 | 10.88074 | 36.15571 | 0.00000 | 21.15217 | 4.96109 | 5.34347 | 7.06354 | 36.26313 |
| **MFN2** | 4.64244 | 30.19709 | 7.34396 | 7.65337 | 28.17398 | 7.91668 | 15.84923 | 2.01393 | 14.65270 | 19.99894 | 16.10770 | 19.15859 |
| **MFNG** | 0.00355 | 0.15224 | 0.03505 | 0.00000 | 0.04762 | 0.01454 | 0.06726 | 0.01062 | 0.08185 | 0.24846 | 0.00000 | 0.00074 |
| **MFSD1** | 1.13328 | 5.97180 | 14.51366 | 1.58745 | 4.73909 | 16.66567 | 0.00000 | 0.38046 | 12.39195 | 0.00000 | 10.00729 | 3.99805 |
| **MFSD10** | 11.30548 | 3.45200 | 10.46388 | 1.73889 | 0.49652 | 0.00000 | 2.33260 | 3.26987 | 20.75123 | 4.61628 | 5.56115 | 2.06977 |
| **MFSD11** | 2.48139 | 4.75287 | 3.53849 | 2.84255 | 1.33035 | 10.85746 | 0.00000 | 7.43748 | 13.57491 | 4.83328 | 4.82946 | 5.73657 |
| **MFSD12** | 1.06322 | 0.76780 | 6.15610 | 0.39050 | 3.41749 | 0.00000 | 0.83992 | 0.00000 | 3.46624 | 4.38274 | 0.96701 | 0.79610 |
| **MFSD2A** | 3.99225 | 0.00000 | 0.52835 | 0.37239 | 0.30729 | 0.00000 | 0.00000 | 26.97247 | 0.00000 | 0.00207 | 0.00000 | 30.38683 |
| **MFSD2B** | 0.00444 | 0.02389 | 0.06915 | 0.00849 | 0.01362 | 0.00889 | 0.02866 | 0.09205 | 0.01958 | 0.02918 | 0.01318 | 0.04735 |
| **MFSD3** | 0.17943 | 0.42915 | 1.81472 | 0.97610 | 2.52438 | 0.00000 | 0.00000 | 0.00228 | 1.23937 | 1.29629 | 4.23136 | 0.01094 |

| | | | | | | | | | | | | |
|---|---|---|---|---|---|---|---|---|---|---|---|---|
| **MFSD4** | 2.62029 | 0.00000 | 0.09512 | 0.09378 | 0.07028 | 0.00000 | 0.00000 | 0.32377 | 0.09592 | 0.16088 | 0.00088 | 0.89737 |
| **MFSD5** | 0.94920 | 10.62685 | 13.66519 | 3.22680 | 6.80897 | 0.00000 | 3.54257 | 3.65907 | 10.47391 | 12.36515 | 4.53999 | 6.45037 |
| **MFSD6** | 0.07799 | 0.04775 | 1.05381 | 0.11523 | 0.13934 | 0.00234 | 0.01937 | 0.00390 | 0.00000 | 0.12037 | 0.08233 | 0.06392 |
| **MFSD6L** | 0.00401 | 0.01256 | 0.01128 | 0.01331 | 0.01758 | 0.02963 | 0.01608 | 0.08888 | 0.02312 | 0.00973 | 0.02071 | 0.01265 |
| **MFSD7** | 0.00266 | 0.04568 | 0.22505 | 0.24543 | 0.68299 | 0.00000 | 0.00000 | 0.00743 | 0.00300 | 0.36742 | 0.16121 | 0.03685 |
| **MFSD8** | 0.25496 | 1.92695 | 5.24367 | 3.17004 | 1.14057 | 8.66057 | 0.00000 | 0.64571 | 8.88507 | 1.76418 | 4.08630 | 5.17109 |
| **MFSD9** | 0.83900 | 3.34884 | 5.30707 | 1.77667 | 2.32519 | 3.05729 | 0.33448 | 0.59584 | 7.07457 | 0.00000 | 3.12206 | 3.24479 |
| **MGA** | 6.54517 | 1.50394 | 7.08388 | 4.87159 | 1.53972 | 21.53622 | 8.15608 | 8.58887 | 3.57657 | 23.67298 | 6.84360 | 13.41448 |
| **MGAM** | 0.16809 | 0.11038 | 0.09422 | 0.62665 | 0.07251 | 0.00000 | 0.00638 | 0.02859 | 0.00000 | 0.25524 | 0.01701 | 0.07435 |
| **MGARP** | 0.13477 | 0.00352 | 0.02398 | 0.18714 | 0.00000 | 0.30040 | 0.07449 | 0.11290 | 0.01827 | 0.00223 | 0.08725 | 0.08547 |
| **MGAT1** | 0.85729 | 16.63982 | 26.93309 | 11.57392 | 10.92926 | 0.00000 | 8.68436 | 4.13835 | 34.40674 | 5.55732 | 4.82659 | 11.97637 |
| **MGAT2** | 2.78054 | 2.54907 | 1.13155 | 4.27127 | 1.66407 | 6.34451 | 1.69206 | 2.62133 | 5.75154 | 0.00000 | 4.34143 | 6.70745 |
| **MGAT3** | 0.01091 | 0.48928 | 1.38337 | 0.14671 | 1.14749 | 0.00000 | 0.01140 | 0.00726 | 0.77502 | 0.71909 | 0.18400 | 0.00000 |
| **MGAT4A** | 3.44765 | 1.05318 | 8.86519 | 0.00947 | 0.03112 | 1.07525 | 0.00000 | 1.49508 | 2.35138 | 1.30404 | 2.38521 | 0.01644 |
| **MGAT4B** | 1.44750 | 0.43520 | 0.42088 | 1.02607 | 1.90382 | 0.11944 | 0.37184 | 0.00000 | 0.32844 | 5.86556 | 0.19807 | 0.00000 |
| **MGAT4C** | 0.10973 | 0.01792 | 0.01959 | 0.01800 | 0.10840 | 0.00010 | 0.02394 | 0.03619 | 0.07560 | 0.00000 | 0.00878 | 0.01791 |
| **MGAT5** | 8.42010 | 0.37371 | 2.00701 | 1.01428 | 0.74320 | 2.17613 | 2.39970 | 1.97450 | 2.70321 | 5.57624 | 0.04638 | 0.22189 |
| **MGAT5B** | 6.28645 | 0.08494 | 0.02251 | 0.01530 | 0.02149 | 0.07689 | 0.00000 | 0.34959 | 0.03146 | 0.02589 | 0.00008 | 0.20072 |
| **MGC12916** | 0.10167 | 0.01241 | 1.67639 | 0.02493 | 0.00000 | 0.02233 | 0.65092 | 0.12205 | 0.08364 | 0.00000 | 0.00000 | 0.01218 |
| **MGC15885** | 0.00765 | 0.00678 | 0.01266 | 0.05446 | 0.09992 | 0.01094 | 0.02837 | 0.00940 | 0.03154 | 0.12160 | 0.02503 | 0.01206 |
| **MGC16025** | 0.00611 | 0.01479 | 0.06031 | 0.18271 | 0.01368 | 0.00000 | 0.01090 | 0.01050 | 0.01634 | 0.01970 | 0.01668 | 0.01689 |
| **MGC16142** | 0.05242 | 0.10554 | 0.21683 | 0.19675 | 0.04963 | 0.06928 | 0.10300 | 0.44808 | 0.03789 | 0.32279 | 0.03629 | 0.26518 |
| **MGC16275** | 0.93059 | 0.05136 | 0.27544 | 0.70135 | 0.00000 | 0.00000 | 1.37312 | 0.39160 | 0.00000 | 0.74790 | 0.23374 | 0.11202 |
| **MGC27345** | 0.51099 | 2.34586 | 0.43592 | 1.05900 | 2.05150 | 0.18977 | 5.52231 | 0.57925 | 2.81957 | 2.92404 | 3.65107 | 2.45190 |
| **MGC27382** | 0.01656 | 0.01299 | 0.01584 | 0.01799 | 0.01919 | 0.00285 | 0.02055 | 0.08655 | 0.02149 | 0.00705 | 0.01861 | 0.15187 |
| **MGC2889** | 0.00642 | 0.01531 | 0.18712 | 0.00861 | 0.01540 | 0.01759 | 0.00313 | 0.01035 | 0.00000 | 0.01564 | 0.02489 | 0.01593 |
| **MGC32805** | 0.01440 | 0.00917 | 0.00000 | 0.54796 | 0.06171 | 0.07728 | 0.00000 | 0.00820 | 0.01523 | 0.09674 | 0.02762 | 0.08955 |
| **MGC34796** | 0.00141 | 0.04029 | 0.00000 | 0.00000 | 0.00000 | 0.04401 | 0.06509 | 0.13222 | 0.05146 | 0.15274 | 0.01619 | 0.03602 |
| **MGC45800** | 0.01474 | 0.00826 | 2.55502 | 0.01385 | 0.00000 | 0.00541 | 0.12042 | 0.00873 | 0.02201 | 0.09866 | 0.00797 | 0.05136 |
| **MGC45922** | 0.00786 | 0.12121 | 0.01345 | 0.00000 | 0.01590 | 0.06851 | 0.00000 | 0.06980 | 0.04758 | 0.02099 | 0.01681 | 0.08299 |
| **MGC57346** | 0.90947 | 1.04676 | 0.29276 | 0.60242 | 0.59634 | 1.12142 | 0.84611 | 0.44213 | 0.61496 | 0.54584 | 0.67801 | 1.06876 |
| **MGC72080** | 2.13506 | 0.97947 | 0.91288 | 0.77373 | 1.16387 | 0.21680 | 1.33098 | 0.07909 | 0.88287 | 0.58682 | 1.45049 | 0.15685 |
| **MGEA5** | 1.70755 | 4.07115 | 3.49183 | 4.16991 | 0.58568 | 18.89671 | 9.27141 | 6.36196 | 5.16726 | 30.43359 | 0.00000 | 8.12584 |
| **MGLL** | 0.00851 | 0.18798 | 4.25927 | 1.75107 | 4.19596 | 0.00000 | 3.93727 | 0.00000 | 3.29023 | 0.56807 | 10.79612 | 0.16140 |
| **MGME1** | 2.82375 | 6.00291 | 7.27842 | 8.41033 | 1.11892 | 20.80584 | 6.20358 | 4.34930 | 5.22387 | 0.09846 | 6.76787 | 12.05295 |
| **MGMT** | 2.80981 | 1.64597 | 0.61572 | 2.12182 | 2.96498 | 0.92313 | 2.23024 | 0.82258 | 0.00000 | 0.31201 | 1.75035 | 0.19635 |
| **MGP** | 0.10198 | 0.01277 | 0.01860 | 0.02523 | 0.01096 | 0.01051 | 0.01662 | 0.03795 | 0.02236 | 0.03262 | 0.01335 | 0.02622 |
| **MGRN1** | 13.30066 | 1.61130 | 2.68720 | 1.60971 | 3.88749 | 0.00000 | 3.42342 | 0.93319 | 3.37506 | 2.90044 | 2.76959 | 0.25467 |
| **MGST1** | 0.01971 | 0.00000 | 9.82651 | 4.89447 | 12.45292 | 14.35505 | 9.65571 | 0.02528 | 21.41595 | 1.06633 | 29.78051 | 0.12064 |
| **MGST2** | 5.68111 | 10.15739 | 2.23232 | 5.51344 | 3.82111 | 6.60414 | 0.00000 | 2.05148 | 5.62336 | 1.04366 | 1.73380 | 6.34014 |
| **MGST3** | 1.26201 | 6.66216 | 27.92199 | 0.47470 | 11.22585 | 18.36456 | 22.44417 | 4.75355 | 37.59661 | 0.43386 | 22.86318 | 7.53550 |
| **MIA** | 0.00483 | 0.01521 | 0.05017 | 0.01444 | 0.01410 | 0.01376 | 0.02002 | 0.01940 | 0.01558 | 0.01612 | 0.02026 | 0.05991 |
| **MIA3** | 4.41847 | 5.28145 | 4.19765 | 5.51299 | 4.83052 | 11.96155 | 0.00000 | 5.54180 | 7.73775 | 8.50088 | 3.03497 | 3.70558 |
| **MIAT** | 0.27440 | 0.02203 | 0.08007 | 3.54459 | 0.01204 | 0.00000 | 0.42942 | 0.21668 | 0.00000 | 0.08203 | 0.04831 | 0.11782 |
| **MIATNB** | 0.00441 | 0.27285 | 0.05698 | 0.18914 | 0.00654 | 0.00000 | 0.08811 | 0.59314 | 0.00934 | 0.00000 | 0.10159 | 0.31799 |
| **MIB1** | 4.16604 | 0.17468 | 2.20509 | 0.00000 | 0.56524 | 5.79015 | 4.53883 | 0.55530 | 0.00000 | 9.90889 | 1.12450 | 1.31493 |
| **MIB2** | 0.34058 | 0.36545 | 0.70769 | 0.32099 | 1.86546 | 0.00000 | 0.01001 | 2.70314 | 0.45054 | 5.42942 | 0.01732 | 0.40226 |
| **MICA** | 0.00431 | 0.54375 | 0.13662 | 1.34521 | 0.00000 | 0.00000 | 0.00000 | 0.00066 | 0.34919 | 0.49232 | 0.00000 | 2.65765 |

| | | | | | | | | | | | | |
|---|---|---|---|---|---|---|---|---|---|---|---|---|
| **MICAL1** | 0.43564 | 2.93243 | 1.02604 | 2.79185 | 1.94619 | 0.00000 | 0.00000 | 0.32769 | 1.15188 | 4.54792 | 0.53521 | 2.07966 |
| **MICAL2** | 4.00923 | 1.50309 | 5.20122 | 0.95156 | 6.04090 | 1.46804 | 3.68911 | 0.49925 | 3.07687 | 1.81106 | 3.58992 | 0.00000 |
| **MICAL3** | 3.45583 | 3.17109 | 3.09155 | 2.17246 | 3.08732 | 0.00000 | 0.76143 | 2.14339 | 3.64328 | 7.01208 | 0.00000 | 2.30153 |
| **MICALCL** | 3.92027 | 0.00046 | 0.21521 | 0.00000 | 0.49912 | 1.16895 | 0.00000 | 0.06350 | 0.00000 | 1.01231 | 0.11990 | 0.05016 |
| **MICALL1** | 0.22814 | 2.41505 | 6.94540 | 2.98046 | 9.71597 | 0.00000 | 8.29630 | 1.27287 | 0.00000 | 5.18793 | 1.62677 | 0.90574 |
| **MICALL2** | 1.43503 | 0.47191 | 2.41276 | 0.49444 | 0.63225 | 0.00000 | 0.02913 | 0.06862 | 0.25541 | 2.00723 | 0.00000 | 0.15170 |
| **MICB** | 0.00194 | 3.98675 | 0.12407 | 4.99526 | 0.77427 | 0.48834 | 0.00000 | 0.42187 | 0.48559 | 0.52230 | 0.62917 | 3.28555 |
| **MICU1** | 3.40109 | 4.30633 | 1.95053 | 5.20876 | 6.38322 | 5.92295 | 4.06212 | 9.71819 | 2.49880 | 0.75355 | 2.20909 | 2.96187 |
| **MICU2** | 1.46434 | 4.77296 | 9.39094 | 3.21182 | 0.99012 | 20.91867 | 5.39348 | 2.11988 | 8.60022 | 3.05477 | 9.20275 | 4.08449 |
| **MICU3** | 0.00000 | 0.00000 | 0.90536 | 0.00000 | 0.08625 | 0.46745 | 0.21299 | 0.03750 | 0.91318 | 1.06961 | 0.13746 | 0.04484 |
| **MID1** | 2.98332 | 1.16086 | 12.50154 | 6.99483 | 0.00000 | 2.37685 | 12.10684 | 0.08351 | 10.87777 | 0.00000 | 1.08913 | 4.80437 |
| **MID1IP1** | 0.11323 | 0.00000 | 1.24012 | 0.73356 | 1.32296 | 0.01411 | 4.35959 | 0.00142 | 0.42588 | 4.69554 | 0.53338 | 0.00442 |
| **MID1IP1-AS1** | 0.04907 | 0.00605 | 0.03195 | 0.00153 | 0.00000 | 0.03621 | 0.01022 | 0.00553 | 0.03802 | 0.17321 | 0.00391 | 0.00842 |
| **MID2** | 0.03577 | 0.11762 | 0.03479 | 0.10822 | 0.00004 | 0.00000 | 0.01592 | 0.01116 | 0.01623 | 0.01431 | 0.02703 | 0.02103 |
| **MIDN** | 2.44137 | 7.06097 | 8.45753 | 4.19419 | 18.10749 | 0.00000 | 11.48872 | 0.88411 | 1.61560 | 34.57857 | 3.31874 | 3.74642 |
| **MIEF1** | 1.42195 | 17.86822 | 14.94145 | 15.42291 | 10.38284 | 15.96855 | 22.47287 | 5.73297 | 17.46842 | 7.43318 | 19.36639 | 18.30345 |
| **MIEF2** | 0.06099 | 2.56953 | 1.12489 | 1.53330 | 1.66729 | 0.00000 | 0.20649 | 0.01368 | 1.40630 | 0.00000 | 2.16857 | 1.35593 |
| **MIEN1** | 0.00000 | 16.62045 | 8.59431 | 7.62444 | 16.79634 | 8.54098 | 9.92308 | 3.65852 | 13.37476 | 0.00000 | 16.94943 | 9.47333 |
| **MIER1** | 2.11595 | 2.11068 | 12.28159 | 3.27378 | 0.00000 | 17.01013 | 10.83181 | 8.48746 | 4.38867 | 9.96578 | 7.79719 | 6.81338 |
| **MIER2** | 1.19966 | 2.62291 | 2.26508 | 2.31151 | 1.13402 | 0.00000 | 0.00000 | 0.00000 | 3.87721 | 4.55330 | 1.12205 | 1.31487 |
| **MIER3** | 0.42915 | 1.35099 | 8.99707 | 0.00000 | 0.00000 | 6.12551 | 7.14720 | 2.91865 | 1.35770 | 5.90619 | 3.34373 | 4.73916 |
| **MIF4GD** | 2.13072 | 2.28805 | 0.89329 | 1.27255 | 2.39018 | 0.05350 | 0.42960 | 0.47527 | 1.04807 | 0.00000 | 1.78615 | 0.20323 |
| **MIIP** | 3.97520 | 33.22595 | 8.60134 | 18.94636 | 34.80119 | 0.00000 | 21.31648 | 3.31845 | 25.84626 | 8.52111 | 19.63729 | 21.33374 |
| **MILR1** | 0.00827 | 0.00509 | 0.00646 | 0.19473 | 0.00000 | 0.05363 | 0.01019 | 0.03615 | 0.00747 | 0.02849 | 0.01282 | 0.03684 |
| **MIMT1** | 0.00714 | 0.01289 | 0.01985 | 0.01170 | 0.02782 | 0.03074 | 0.01861 | 0.01632 | 0.02249 | 0.00568 | 0.01489 | 0.10540 |
| **MINA** | 6.25767 | 6.35881 | 2.34962 | 0.00000 | 0.34770 | 0.96463 | 1.91741 | 1.27739 | 4.70063 | 0.00000 | 4.66559 | 6.38718 |
| **MINK1** | 1.07626 | 1.73320 | 1.99463 | 1.21339 | 2.22229 | 0.00000 | 0.00000 | 0.20155 | 3.82531 | 4.62975 | 2.51928 | 1.20485 |
| **MINOS1** | 0.69709 | 6.04131 | 14.03168 | 8.63437 | 8.10474 | 19.95855 | 20.90627 | 0.41188 | 8.72554 | 13.63734 | 18.37843 | 3.30216 |
| **MINOS1P1** | 0.00059 | 0.00000 | 0.51925 | 0.33767 | 0.00000 | 0.00000 | 0.03564 | 0.03535 | 0.98614 | 2.84559 | 0.72910 | 0.44362 |
| **MINPP1** | 0.00154 | 6.41807 | 3.66951 | 1.39222 | 0.21173 | 5.96686 | 0.00000 | 2.34838 | 8.15152 | 0.00000 | 3.91616 | 4.52200 |
| **MIOS** | 1.94449 | 3.07766 | 2.30574 | 0.00000 | 4.00905 | 6.96028 | 2.84485 | 0.60196 | 4.52228 | 0.80413 | 4.93709 | 2.31905 |
| **MIOX** | 0.13217 | 0.01575 | 0.02091 | 0.01470 | 0.03044 | 0.00000 | 0.00973 | 0.00315 | 0.05941 | 0.00610 | 0.01684 | 0.04568 |
| **MIP** | 0.07231 | 0.00000 | 0.27650 | 0.13143 | 0.02680 | 0.02134 | 0.11665 | 0.29311 | 0.04831 | 0.00000 | 0.02845 | 0.66447 |
| **MIPEP** | 1.56020 | 6.68434 | 1.27953 | 2.96873 | 4.33934 | 9.80808 | 2.93252 | 0.80848 | 4.55879 | 1.95414 | 9.89610 | 3.33488 |
| **MIPEPP3** | 0.44937 | 0.41828 | 0.00969 | 0.00492 | 0.36189 | 1.02622 | 0.23893 | 1.70740 | 0.35394 | 0.00000 | 0.49095 | 1.77001 |
| **MIPOL1** | 0.23645 | 0.22098 | 0.54156 | 0.00000 | 0.00000 | 1.20051 | 0.49349 | 0.00000 | 0.00000 | 1.05468 | 0.33913 | 0.50628 |
| **MIR100HG** | 0.24395 | 0.05147 | 0.00994 | 0.01747 | 0.00000 | 0.05647 | 0.01314 | 0.03365 | 0.00542 | 0.02056 | 0.03171 | 0.00512 |
| **MIR103A2** | 0.00803 | 0.08430 | 0.05074 | 0.06188 | 0.06381 | 0.11116 | 0.09252 | 0.07739 | 0.13361 | 0.11746 | 0.13632 | 0.09985 |
| **MIR106B** | 0.00528 | 0.02581 | 0.00817 | 0.03567 | 0.01127 | 0.01784 | 0.01523 | 0.00879 | 0.04317 | 0.04790 | 0.00719 | 0.05945 |
| **MIR1204** | 0.03478 | 0.04428 | 0.13898 | 0.09773 | 0.29260 | 0.09019 | 0.01051 | 0.06467 | 0.20650 | 0.18375 | 0.00014 | 0.03658 |
| **MIR1206** | 0.00645 | 0.01601 | 0.01408 | 0.01294 | 0.01701 | 0.01498 | 0.02296 | 0.01051 | 0.02983 | 0.00662 | 0.01607 | 0.01700 |
| **MIR1207** | 0.00627 | 0.01247 | 0.01588 | 0.01570 | 0.01105 | 0.02832 | 0.02864 | 0.01028 | 0.00418 | 0.02113 | 0.03029 | 0.01514 |
| **MIR1226** | 0.01370 | 0.18875 | 0.10863 | 0.15493 | 0.24688 | 0.25996 | 0.24667 | 0.00932 | 0.13486 | 0.33753 | 0.16213 | 0.20153 |
| **MIR1227** | 0.00321 | 0.04216 | 0.02816 | 0.02633 | 0.00139 | 0.04230 | 0.00481 | 0.01636 | 0.05465 | 0.15539 | 0.02431 | 0.03900 |
| **MIR1228** | 0.00586 | 0.01894 | 0.02026 | 0.01231 | 0.01343 | 0.02605 | 0.01275 | 0.01085 | 0.03065 | 0.01920 | 0.01697 | 0.01576 |
| **MIR1229** | 0.02125 | 0.02304 | 0.01788 | 0.01223 | 0.02481 | 0.00804 | 0.00976 | 0.00932 | 0.03945 | 0.03121 | 0.01241 | 0.01456 |
| **MIR1231** | 0.01041 | 0.01634 | 0.01407 | 0.03577 | 0.01496 | 0.00622 | 0.02736 | 0.01021 | 0.02771 | 0.01226 | 0.01567 | 0.01727 |
| **MIR1234** | 0.00622 | 0.03085 | 0.02367 | 0.02583 | 0.04438 | 0.00000 | 0.01665 | 0.01138 | 0.05993 | 0.00351 | 0.02285 | 0.01900 |

| | | | | | | | | | | | | |
|---|---|---|---|---|---|---|---|---|---|---|---|---|
| **MIR1236** | 0.01241 | 0.01595 | 0.02735 | 0.01717 | 0.02147 | 0.01442 | 0.00432 | 0.05390 | 0.05334 | 0.01928 | 0.01361 | 0.02200 |
| **MIR1237** | 0.00617 | 0.01648 | 0.01509 | 0.01009 | 0.02430 | 0.01946 | 0.01743 | 0.01040 | 0.02445 | 0.01522 | 0.01399 | 0.01665 |
| **MIR1238** | 0.00476 | 0.03497 | 0.02327 | 0.04204 | 0.02593 | 0.01252 | 0.03024 | 0.03271 | 0.04769 | 0.00000 | 0.01570 | 0.04760 |
| **MIR124-2HG** | 0.01184 | 0.01174 | 0.03110 | 0.01069 | 0.02190 | 0.01946 | 0.01724 | 0.02648 | 0.01636 | 0.00825 | 0.01545 | 0.07385 |
| **MIR1244-1+MIR** | 0.00602 | 0.01448 | 0.01367 | 0.01137 | 0.01613 | 0.02115 | 0.01854 | 0.01113 | 0.01708 | 0.01528 | 0.01483 | 0.01679 |
| **MIR1244-1+MIR** | 0.00694 | 0.02258 | 0.01488 | 0.01275 | 0.01668 | 0.01685 | 0.01884 | 0.01064 | 0.01871 | 0.01101 | 0.01531 | 0.01648 |
| **MIR1248** | 0.00627 | 0.01488 | 0.01358 | 0.01147 | 0.01445 | 0.02276 | 0.01960 | 0.01038 | 0.01919 | 0.01254 | 0.01398 | 0.01694 |
| **MIR1272** | 0.00510 | 0.01242 | 0.01436 | 0.00625 | 0.03680 | 0.02777 | 0.00379 | 0.00944 | 0.01468 | 0.08666 | 0.01109 | 0.02418 |
| **MIR128-1** | 0.00621 | 0.01444 | 0.02072 | 0.01232 | 0.01293 | 0.02458 | 0.02202 | 0.01043 | 0.01592 | 0.01411 | 0.01378 | 0.01577 |
| **MIR1281** | 0.00865 | 0.01589 | 0.01461 | 0.01114 | 0.01730 | 0.02143 | 0.01776 | 0.01056 | 0.01756 | 0.01269 | 0.01441 | 0.01675 |
| **MIR1283-1** | 0.00621 | 0.01664 | 0.02346 | 0.01164 | 0.02128 | 0.01417 | 0.03318 | 0.01014 | 0.01775 | 0.01428 | 0.01322 | 0.02236 |
| **MIR1283-2** | 0.00612 | 0.01436 | 0.01898 | 0.01212 | 0.01390 | 0.01754 | 0.01420 | 0.01031 | 0.02500 | 0.02017 | 0.01840 | 0.01918 |
| **MIR1291** | 0.01033 | 0.01379 | 0.01571 | 0.01239 | 0.01751 | 0.02508 | 0.01493 | 0.01185 | 0.02696 | 0.01651 | 0.01339 | 0.01586 |
| **MIR1292** | 0.06386 | 0.39022 | 0.25439 | 0.24086 | 0.41433 | 0.00000 | 0.03990 | 0.16754 | 0.57538 | 0.00000 | 0.12570 | 0.73729 |
| **MIR1296** | 0.01167 | 0.00959 | 0.01750 | 0.03215 | 0.01037 | 0.03340 | 0.00584 | 0.01046 | 0.02520 | 0.02864 | 0.01551 | 0.03731 |
| **MIR1297** | 0.00623 | 0.01417 | 0.01678 | 0.01172 | 0.01484 | 0.02048 | 0.01687 | 0.01034 | 0.01849 | 0.01391 | 0.01469 | 0.01941 |
| **MIR1304** | 0.00154 | 0.04430 | 0.07516 | 0.05729 | 0.34083 | 0.00050 | 0.00005 | 0.02757 | 0.15155 | 0.59257 | 0.00000 | 0.08420 |
| **MIR130A** | 0.00524 | 0.01876 | 0.01303 | 0.02936 | 0.01937 | 0.01061 | 0.01095 | 0.01137 | 0.03906 | 0.03645 | 0.02491 | 0.02233 |
| **MIR130B** | 0.00572 | 0.01393 | 0.02326 | 0.00693 | 0.01155 | 0.04288 | 0.01888 | 0.01059 | 0.02231 | 0.01309 | 0.01407 | 0.02127 |
| **MIR1322** | 0.00618 | 0.01895 | 0.01319 | 0.01272 | 0.01597 | 0.03228 | 0.01284 | 0.01011 | 0.01933 | 0.01263 | 0.02347 | 0.01851 |
| **MIR1323** | 0.00615 | 0.01470 | 0.05221 | 0.00441 | 0.01478 | 0.02876 | 0.02769 | 0.01313 | 0.06798 | 0.04400 | 0.02049 | 0.02469 |
| **MIR133A1HG** | 0.01325 | 0.01274 | 0.09864 | 0.03502 | 0.04891 | 0.04936 | 0.00000 | 0.03184 | 0.17451 | 0.07230 | 0.02819 | 0.07248 |
| **MIR1343** | 0.00346 | 0.01765 | 0.01234 | 0.01669 | 0.01027 | 0.02557 | 0.01786 | 0.02306 | 0.01745 | 0.01578 | 0.01929 | 0.02428 |
| **MIR137HG** | 0.04552 | 0.01473 | 0.01405 | 0.01274 | 0.01491 | 0.01821 | 0.02308 | 0.00897 | 0.01788 | 0.01349 | 0.01430 | 0.01521 |
| **MIR138-2** | 0.00604 | 0.01443 | 0.02369 | 0.01286 | 0.01311 | 0.02049 | 0.01508 | 0.01002 | 0.01361 | 0.02100 | 0.01557 | 0.02388 |
| **MIR141** | 0.00531 | 0.02099 | 0.01291 | 0.01919 | 0.01117 | 0.02353 | 0.02196 | 0.01266 | 0.02147 | 0.01708 | 0.01938 | 0.02019 |
| **MIR1469** | 0.02278 | 0.01484 | 0.02670 | 0.01051 | 0.01490 | 0.01921 | 0.01704 | 0.00877 | 0.01763 | 0.01631 | 0.01453 | 0.01929 |
| **MIR146B** | 0.00590 | 0.01256 | 0.01063 | 0.08322 | 0.04357 | 0.00000 | 0.02094 | 0.01058 | 0.01350 | 0.02392 | 0.01533 | 0.01729 |
| **MIR148B** | 0.00623 | 0.02004 | 0.01513 | 0.01489 | 0.01443 | 0.01493 | 0.01560 | 0.01056 | 0.02368 | 0.01648 | 0.01515 | 0.01626 |
| **MIR149** | 0.00626 | 0.01485 | 0.01717 | 0.01213 | 0.01562 | 0.01950 | 0.01648 | 0.01046 | 0.01831 | 0.01517 | 0.01512 | 0.01610 |
| **MIR150** | 0.00553 | 0.01522 | 0.01412 | 0.01213 | 0.01542 | 0.01815 | 0.01894 | 0.04743 | 0.01804 | 0.01319 | 0.01492 | 0.01407 |
| **MIR1539** | 0.01340 | 0.01700 | 0.01723 | 0.01358 | 0.01932 | 0.00825 | 0.03406 | 0.00908 | 0.02767 | 0.00000 | 0.01402 | 0.02825 |
| **MIR155HG** | 0.00276 | 0.03523 | 0.01971 | 0.11337 | 0.08943 | 0.00000 | 0.03213 | 0.03298 | 0.00590 | 0.00765 | 0.01468 | 0.02431 |
| **MIR15A** | 0.00897 | 0.01486 | 0.01544 | 0.01305 | 0.01364 | 0.02177 | 0.01793 | 0.01087 | 0.01745 | 0.01541 | 0.01459 | 0.01572 |
| **MIR15B** | 0.00587 | 0.01430 | 0.01772 | 0.00526 | 0.00533 | 0.04623 | 0.01806 | 0.00980 | 0.02847 | 0.02497 | 0.00803 | 0.01756 |
| **MIR16-2** | 0.04308 | 0.00064 | 0.02706 | 0.02970 | 0.06496 | 0.15300 | 0.00000 | 0.03321 | 0.05417 | 0.20929 | 0.01338 | 0.06345 |
| **MIR17HG** | 0.25008 | 0.13161 | 5.14650 | 1.28810 | 0.00000 | 24.79309 | 0.00000 | 0.00000 | 5.43800 | 20.64546 | 5.18355 | 0.95092 |
| **MIR181A2** | 0.00562 | 0.00968 | 0.01570 | 0.01022 | 0.08964 | 0.01145 | 0.00562 | 0.01065 | 0.06300 | 0.04027 | 0.01706 | 0.02540 |
| **MIR181A2HG** | 0.01151 | 0.06087 | 0.13401 | 0.04959 | 0.00000 | 0.06337 | 0.00000 | 0.05833 | 0.00000 | 0.49396 | 0.02287 | 0.00025 |
| **MIR181B2** | 0.00622 | 0.01385 | 0.02821 | 0.01126 | 0.02244 | 0.02142 | 0.01261 | 0.01034 | 0.02283 | 0.01453 | 0.01514 | 0.01717 |
| **MIR186** | 0.00422 | 0.01220 | 0.00381 | 0.00174 | 0.01044 | 0.02551 | 0.00599 | 0.00976 | 0.01981 | 0.15778 | 0.01624 | 0.01341 |
| **MIR188** | 0.00742 | 0.02037 | 0.02152 | 0.01207 | 0.03558 | 0.01657 | 0.01827 | 0.01079 | 0.01620 | 0.01046 | 0.01747 | 0.01927 |
| **MIR1909** | 0.00421 | 0.02979 | 0.01760 | 0.00818 | 0.03158 | 0.00000 | 0.00210 | 0.01025 | 0.04433 | 0.16496 | 0.00861 | 0.01454 |
| **MIR191** | 0.00524 | 0.01484 | 0.03220 | 0.01889 | 0.01248 | 0.00609 | 0.02159 | 0.01093 | 0.03569 | 0.05503 | 0.01727 | 0.01745 |
| **MIR1910** | 0.00619 | 0.01498 | 0.01442 | 0.01481 | 0.01660 | 0.01878 | 0.01653 | 0.01083 | 0.02198 | 0.01291 | 0.01557 | 0.01864 |
| **MIR1914** | 0.00307 | 0.03680 | 0.02781 | 0.03476 | 0.03982 | 0.00000 | 0.00633 | 0.19763 | 0.11653 | 0.04418 | 0.00493 | 0.01836 |
| **MIR192** | 0.00491 | 0.16799 | 0.00897 | 0.01796 | 0.00000 | 0.00364 | 0.00448 | 0.01209 | 0.02612 | 0.14421 | 0.00705 | 0.32006 |
| **MIR193A** | 0.00623 | 0.01875 | 0.01724 | 0.01022 | 0.01740 | 0.02156 | 0.01377 | 0.01046 | 0.02251 | 0.01518 | 0.01374 | 0.01540 |

| | | | | | | | | | | | | |
|---|---|---|---|---|---|---|---|---|---|---|---|---|
| **MIR194-1** | 0.00599 | 0.01993 | 0.01484 | 0.01678 | 0.00226 | 0.04217 | 0.01045 | 0.00986 | 0.01219 | 0.03430 | 0.02579 | 0.01622 |
| **MIR197** | 0.02224 | 0.02261 | 0.01750 | 0.02386 | 0.02909 | 0.00000 | 0.03596 | 0.01656 | 0.03279 | 0.02167 | 0.02195 | 0.02319 |
| **MIR1972-1+MIR** | 0.00633 | 0.01454 | 0.01537 | 0.01129 | 0.01896 | 0.01886 | 0.01516 | 0.01048 | 0.02711 | 0.01363 | 0.01457 | 0.01626 |
| **MIR1973** | 0.00692 | 0.00230 | 0.03483 | 0.03194 | 0.25822 | 0.49971 | 0.00000 | 0.01915 | 0.25167 | 0.45818 | 0.00000 | 0.10902 |
| **MIR200C** | 0.00614 | 0.01490 | 0.01719 | 0.01564 | 0.01541 | 0.01852 | 0.01722 | 0.01047 | 0.01576 | 0.01933 | 0.01528 | 0.01587 |
| **MIR202HG** | 0.00511 | 0.01008 | 0.02617 | 0.01236 | 0.00000 | 0.02856 | 0.04797 | 0.01039 | 0.00000 | 0.05659 | 0.03655 | 0.01699 |
| **MIR205** | 0.00610 | 0.01483 | 0.04908 | 0.01479 | 0.01361 | 0.01784 | 0.04906 | 0.01052 | 0.02271 | 0.01428 | 0.01528 | 0.01619 |
| **MIR205HG** | 0.00394 | 0.00000 | 3.65810 | 0.35335 | 0.00000 | 0.04961 | 3.07991 | 0.00707 | 2.50177 | 0.19597 | 1.10036 | 0.00000 |
| **MIR210HG** | 0.03259 | 0.00000 | 0.58965 | 0.01692 | 0.00350 | 0.02724 | 0.01475 | 0.05464 | 0.33914 | 0.38557 | 0.15599 | 0.00469 |
| **MIR2116** | 0.00649 | 0.01622 | 0.01660 | 0.01417 | 0.03244 | 0.01070 | 0.01698 | 0.01051 | 0.02734 | 0.00653 | 0.01403 | 0.01772 |
| **MIR2117** | 0.00580 | 0.00791 | 0.00841 | 0.01715 | 0.08641 | 0.00000 | 0.01244 | 0.00879 | 0.02464 | 0.06838 | 0.01305 | 0.06761 |
| **MIR215** | 0.00619 | 0.02354 | 0.01540 | 0.02352 | 0.00423 | 0.03628 | 0.00799 | 0.01000 | 0.01899 | 0.03115 | 0.01702 | 0.01540 |
| **MIR219A1** | 0.00628 | 0.01472 | 0.01999 | 0.01151 | 0.01626 | 0.02226 | 0.01860 | 0.01046 | 0.01540 | 0.01196 | 0.01532 | 0.01611 |
| **MIR221** | 0.00586 | 0.01128 | 0.01126 | 0.01287 | 0.01291 | 0.02861 | 0.01613 | 0.00942 | 0.01785 | 0.02668 | 0.01346 | 0.02937 |
| **MIR222** | 0.00629 | 0.01535 | 0.01838 | 0.01319 | 0.01466 | 0.01976 | 0.01704 | 0.01047 | 0.01861 | 0.01297 | 0.01467 | 0.01618 |
| **MIR2276** | 0.00616 | 0.01422 | 0.01444 | 0.01920 | 0.01231 | 0.02277 | 0.01501 | 0.01029 | 0.01961 | 0.01967 | 0.01416 | 0.01722 |
| **MIR22HG** | 3.65496 | 2.29837 | 8.13439 | 3.18881 | 7.11167 | 5.48051 | 6.26070 | 1.09159 | 5.92555 | 2.44292 | 0.00000 | 4.88469 |
| **MIR23A** | 0.00626 | 0.01511 | 0.01868 | 0.01168 | 0.01583 | 0.01730 | 0.01777 | 0.01049 | 0.01735 | 0.01462 | 0.01698 | 0.01632 |
| **MIR23B** | 0.00452 | 0.01144 | 0.00357 | 0.00521 | 0.00917 | 0.02554 | 0.00972 | 0.01005 | 0.03261 | 0.13962 | 0.00964 | 0.01094 |
| **MIR24-2** | 0.00612 | 0.01478 | 0.02857 | 0.01087 | 0.01426 | 0.01806 | 0.01677 | 0.01047 | 0.01183 | 0.02131 | 0.01791 | 0.01580 |
| **MIR25** | 0.00539 | 0.20591 | 0.10702 | 0.20383 | 0.30498 | 0.00000 | 0.00000 | 0.03768 | 0.39016 | 0.17602 | 0.07028 | 0.26280 |
| **MIR26A1** | 0.00618 | 0.01512 | 0.01592 | 0.01846 | 0.01163 | 0.02087 | 0.01359 | 0.01040 | 0.03552 | 0.01679 | 0.02417 | 0.01585 |
| **MIR26A2** | 0.00461 | 0.04010 | 0.01657 | 0.01153 | 0.01491 | 0.01648 | 0.01882 | 0.03057 | 0.02020 | 0.00660 | 0.01756 | 0.02271 |
| **MIR26B** | 0.01781 | 0.02025 | 0.01213 | 0.01743 | 0.02073 | 0.01960 | 0.01493 | 0.00800 | 0.02869 | 0.01756 | 0.01131 | 0.01594 |
| **MIR27A** | 0.00630 | 0.01507 | 0.01702 | 0.01182 | 0.01513 | 0.01868 | 0.01854 | 0.01048 | 0.01767 | 0.01278 | 0.01689 | 0.01635 |
| **MIR27B** | 0.00500 | 0.02559 | 0.00493 | 0.00103 | 0.01553 | 0.03088 | 0.00399 | 0.00948 | 0.01838 | 0.14223 | 0.04812 | 0.01287 |
| **MIR29B1** | 0.00896 | 0.00903 | 0.11727 | 0.03854 | 0.00000 | 0.04748 | 0.00000 | 0.01023 | 0.00000 | 0.03641 | 0.01989 | 0.01281 |
| **MIR300** | 0.00608 | 0.01490 | 0.01120 | 0.04324 | 0.01253 | 0.01314 | 0.01817 | 0.01024 | 0.01741 | 0.01944 | 0.01571 | 0.01943 |
| **MIR301A** | 0.00612 | 0.02083 | 0.01548 | 0.01246 | 0.04306 | 0.00259 | 0.01943 | 0.01102 | 0.02447 | 0.00715 | 0.01551 | 0.02535 |
| **MIR302A** | 0.00287 | 0.02389 | 0.01188 | 0.01545 | 0.01708 | 0.06993 | 0.00000 | 0.01299 | 0.00000 | 0.17485 | 0.00613 | 0.05751 |
| **MIR302B** | 0.00713 | 0.02176 | 0.01869 | 0.02813 | 0.00794 | 0.09423 | 0.00000 | 0.01263 | 0.00000 | 0.21458 | 0.00165 | 0.12825 |
| **MIR302C** | 0.00321 | 0.03758 | 0.02758 | 0.02694 | 0.00046 | 0.05444 | 0.00000 | 0.00991 | 0.00000 | 0.16630 | 0.02101 | 0.06966 |
| **MIR302D** | 0.02890 | 0.04650 | 0.00768 | 0.02254 | 0.00695 | 0.07912 | 0.00000 | 0.00896 | 0.00000 | 0.15317 | 0.01577 | 0.06909 |
| **MIR3064** | 0.00155 | 0.24387 | 0.04480 | 0.13646 | 0.06114 | 0.42030 | 0.00000 | 1.25060 | 0.23746 | 0.00000 | 0.05267 | 0.38807 |
| **MIR3074** | 0.00616 | 0.01871 | 0.01897 | 0.01197 | 0.01981 | 0.01264 | 0.01852 | 0.01538 | 0.02200 | 0.01097 | 0.01796 | 0.01730 |
| **MIR30A** | 0.00457 | 0.00149 | 0.00483 | 0.01924 | 0.09606 | 0.03297 | 0.00380 | 0.00882 | 0.00000 | 0.12323 | 0.00964 | 0.02770 |
| **MIR30C1** | 0.00593 | 0.01245 | 0.01225 | 0.00940 | 0.01335 | 0.03009 | 0.01350 | 0.00978 | 0.01481 | 0.02686 | 0.01540 | 0.02174 |
| **MIR3115** | 0.00642 | 0.01448 | 0.01796 | 0.01112 | 0.01402 | 0.02080 | 0.01610 | 0.01192 | 0.01662 | 0.01811 | 0.01501 | 0.01710 |
| **MIR3124** | 0.00406 | 0.01710 | 0.01534 | 0.01326 | 0.02935 | 0.03027 | 0.01290 | 0.01713 | 0.01181 | 0.02820 | 0.01306 | 0.02147 |
| **MIR3127** | 0.00624 | 0.01874 | 0.02002 | 0.01160 | 0.01388 | 0.01732 | 0.01774 | 0.01047 | 0.01964 | 0.01527 | 0.01572 | 0.01624 |
| **MIR3129** | 0.00766 | 0.01576 | 0.01540 | 0.01176 | 0.01436 | 0.02090 | 0.01895 | 0.01029 | 0.01798 | 0.01382 | 0.01448 | 0.01600 |
| **MIR3130-1+MIR** | 0.00601 | 0.01496 | 0.01422 | 0.01172 | 0.01682 | 0.02005 | 0.01745 | 0.01349 | 0.01924 | 0.01247 | 0.01640 | 0.01602 |
| **MIR3134** | 0.00580 | 0.01465 | 0.01531 | 0.01108 | 0.04272 | 0.00010 | 0.02252 | 0.01063 | 0.01924 | 0.03955 | 0.01252 | 0.01622 |
| **MIR3140** | 0.00606 | 0.01408 | 0.01646 | 0.01154 | 0.01941 | 0.02088 | 0.01572 | 0.01067 | 0.01858 | 0.01634 | 0.01698 | 0.01732 |
| **MIR3143** | 0.00494 | 0.01600 | 0.05912 | 0.00738 | 0.01521 | 0.02138 | 0.00910 | 0.01445 | 0.01996 | 0.02535 | 0.01211 | 0.01506 |
| **MIR3153** | 0.00629 | 0.01411 | 0.01805 | 0.01153 | 0.01355 | 0.02429 | 0.01639 | 0.01040 | 0.02695 | 0.01230 | 0.01444 | 0.01597 |
| **MIR3173** | 0.00611 | 0.01359 | 0.01391 | 0.01138 | 0.01339 | 0.01872 | 0.01618 | 0.01129 | 0.02234 | 0.02101 | 0.01567 | 0.02291 |
| **MIR3175** | 0.01136 | 0.01514 | 0.01538 | 0.01301 | 0.01437 | 0.01166 | 0.01956 | 0.01303 | 0.01622 | 0.01895 | 0.03299 | 0.01690 |

| | | | | | | | | | | | | |
|---|---|---|---|---|---|---|---|---|---|---|---|---|
| **MIR3176** | 0.00597 | 0.02976 | 0.03494 | 0.01441 | 0.02106 | 0.00000 | 0.02048 | 0.01068 | 0.04673 | 0.02981 | 0.01721 | 0.01514 |
| **MIR3178** | 0.00691 | 0.01575 | 0.02457 | 0.01129 | 0.01665 | 0.01784 | 0.01478 | 0.01049 | 0.03240 | 0.01216 | 0.01155 | 0.01792 |
| **MIR3188** | 0.00566 | 0.01452 | 0.02712 | 0.01047 | 0.01455 | 0.01175 | 0.01902 | 0.01057 | 0.03722 | 0.04458 | 0.00962 | 0.01485 |
| **MIR3189** | 0.00604 | 0.01372 | 0.01264 | 0.00918 | 0.02821 | 0.03144 | 0.01582 | 0.01034 | 0.02471 | 0.01819 | 0.01268 | 0.01485 |
| **MIR31HG** | 0.00731 | 0.06121 | 0.12331 | 0.09994 | 0.03362 | 0.00211 | 0.01387 | 0.07256 | 0.00943 | 0.00000 | 0.04944 | 0.02421 |
| **MIR32** | 0.00632 | 0.01683 | 0.01411 | 0.01262 | 0.01521 | 0.01711 | 0.02087 | 0.01046 | 0.02631 | 0.01093 | 0.01376 | 0.01698 |
| **MIR320A** | 0.04048 | 0.01716 | 0.02120 | 0.01530 | 0.01858 | 0.01691 | 0.01877 | 0.02463 | 0.02824 | 0.00647 | 0.01680 | 0.01921 |
| **MIR320B2** | 0.00600 | 0.00695 | 0.01153 | 0.00422 | 0.02952 | 0.02792 | 0.01110 | 0.01065 | 0.00000 | 0.08892 | 0.03047 | 0.01391 |
| **MIR320C1** | 0.00384 | 0.01708 | 0.01556 | 0.01546 | 0.01750 | 0.01906 | 0.01776 | 0.02312 | 0.02081 | 0.00992 | 0.01497 | 0.01807 |
| **MIR320D1** | 0.00565 | 0.01479 | 0.01682 | 0.01466 | 0.01400 | 0.01451 | 0.01684 | 0.01159 | 0.02359 | 0.02602 | 0.01575 | 0.02531 |
| **MIR320D2** | 0.00563 | 0.01366 | 0.00966 | 0.02506 | 0.00751 | 0.01772 | 0.02774 | 0.01044 | 0.01884 | 0.04407 | 0.02263 | 0.01488 |
| **MIR320E** | 0.01083 | 0.01045 | 0.02115 | 0.01612 | 0.01272 | 0.02717 | 0.00953 | 0.01573 | 0.03340 | 0.05764 | 0.01121 | 0.01869 |
| **MIR324** | 0.00839 | 0.02532 | 0.01970 | 0.02427 | 0.02803 | 0.00000 | 0.00787 | 0.01030 | 0.06821 | 0.02015 | 0.01777 | 0.01775 |
| **MIR326** | 0.00612 | 0.01531 | 0.01886 | 0.01525 | 0.01248 | 0.02260 | 0.01768 | 0.01037 | 0.01666 | 0.01950 | 0.01424 | 0.01609 |
| **MIR328** | 0.00622 | 0.01642 | 0.01441 | 0.01208 | 0.01396 | 0.02322 | 0.01773 | 0.01044 | 0.01763 | 0.01533 | 0.01446 | 0.01569 |
| **MIR330** | 0.00837 | 0.02856 | 0.01432 | 0.01281 | 0.03068 | 0.02227 | 0.00900 | 0.01006 | 0.03928 | 0.02105 | 0.01132 | 0.02225 |
| **MIR331** | 0.00694 | 0.01801 | 0.02364 | 0.01664 | 0.02081 | 0.02006 | 0.01752 | 0.01347 | 0.03062 | 0.00049 | 0.01904 | 0.01836 |
| **MIR345** | 0.00594 | 0.01718 | 0.01074 | 0.04964 | 0.01837 | 0.03646 | 0.01201 | 0.01025 | 0.01401 | 0.02467 | 0.01196 | 0.01941 |
| **MIR34A** | 0.00596 | 0.01510 | 0.02281 | 0.01172 | 0.01040 | 0.03649 | 0.02117 | 0.01078 | 0.01626 | 0.01087 | 0.01377 | 0.01913 |
| **MIR34C** | 0.00635 | 0.01526 | 0.01389 | 0.01155 | 0.01665 | 0.01833 | 0.01714 | 0.01046 | 0.02382 | 0.01387 | 0.01349 | 0.01630 |
| **MIR3607** | 0.00400 | 0.00542 | 0.00255 | 0.00000 | 0.03923 | 0.14292 | 0.00000 | 0.00787 | 0.05947 | 0.13351 | 0.03825 | 0.02080 |
| **MIR3609** | 0.03519 | 0.01897 | 0.02337 | 0.01418 | 0.02147 | 0.01586 | 0.00806 | 0.01836 | 0.03128 | 0.02007 | 0.01249 | 0.01564 |
| **MIR3620** | 0.25484 | 0.76390 | 0.52283 | 0.53664 | 0.48535 | 0.54606 | 0.29904 | 0.36269 | 0.96115 | 0.00000 | 0.45031 | 0.43507 |
| **MIR3646** | 0.02890 | 0.00571 | 0.01018 | 0.00837 | 0.01168 | 0.04254 | 0.00321 | 0.04278 | 0.01257 | 0.07260 | 0.00936 | 0.04831 |
| **MIR3648** | 0.00568 | 0.00549 | 0.05762 | 0.00089 | 0.11687 | 0.00000 | 0.00000 | 0.00871 | 0.03459 | 0.55500 | 0.00000 | 0.05956 |
| **MIR3651** | 0.00963 | 0.01332 | 0.01724 | 0.03155 | 0.01821 | 0.02017 | 0.01239 | 0.01942 | 0.03060 | 0.03273 | 0.03355 | 0.02271 |
| **MIR3656** | 0.00652 | 0.01799 | 0.01337 | 0.00992 | 0.01965 | 0.01850 | 0.01485 | 0.01202 | 0.01893 | 0.02301 | 0.01521 | 0.01745 |
| **MIR365A** | 0.00627 | 0.01399 | 0.01597 | 0.01256 | 0.01815 | 0.02276 | 0.01800 | 0.01047 | 0.01827 | 0.01221 | 0.01444 | 0.01611 |
| **MIR3660** | 0.00559 | 0.01314 | 0.01353 | 0.01319 | 0.00931 | 0.01773 | 0.02353 | 0.01029 | 0.01433 | 0.04924 | 0.01561 | 0.01578 |
| **MIR3661** | 0.05322 | 0.51103 | 0.39960 | 0.06107 | 0.05148 | 0.00000 | 0.03843 | 0.43470 | 0.39305 | 0.00000 | 0.25532 | 0.12917 |
| **MIR3662** | 0.00544 | 0.01304 | 0.01807 | 0.00518 | 0.02336 | 0.03177 | 0.00497 | 0.01609 | 0.02775 | 0.03970 | 0.01406 | 0.01736 |
| **MIR3668** | 0.00659 | 0.01701 | 0.01456 | 0.01177 | 0.01911 | 0.01843 | 0.01624 | 0.01014 | 0.01974 | 0.01646 | 0.01478 | 0.01918 |
| **MIR367** | 0.00531 | 0.05426 | 0.00000 | 0.23454 | 0.00000 | 0.07574 | 0.00000 | 0.00762 | 0.00000 | 0.14782 | 0.02043 | 0.13772 |
| **MIR3671** | 0.00507 | 0.02073 | 0.01109 | 0.05102 | 0.00000 | 0.04140 | 0.00694 | 0.01000 | 0.03190 | 0.04744 | 0.03960 | 0.02073 |
| **MIR3675.uid-91** | 0.00616 | 0.01290 | 0.01270 | 0.00884 | 0.01961 | 0.02616 | 0.01554 | 0.01038 | 0.02925 | 0.01750 | 0.01135 | 0.01551 |
| **MIR3675.uid-91** | 0.00640 | 0.01519 | 0.01353 | 0.01178 | 0.01681 | 0.02005 | 0.02078 | 0.01047 | 0.03006 | 0.00814 | 0.01168 | 0.01654 |
| **MIR3679** | 0.00581 | 0.02240 | 0.03866 | 0.03044 | 0.03010 | 0.00000 | 0.00806 | 0.00978 | 0.07857 | 0.03434 | 0.01680 | 0.02913 |
| **MIR3682** | 0.00329 | 0.00000 | 0.09807 | 0.00638 | 0.00000 | 0.19120 | 0.16097 | 0.02703 | 0.08265 | 0.19418 | 0.00762 | 0.02386 |
| **MIR3685** | 0.00637 | 0.01801 | 0.01213 | 0.03507 | 0.00000 | 0.06391 | 0.00857 | 0.00992 | 0.05356 | 0.04370 | 0.00919 | 0.01521 |
| **MIR3687** | 0.00912 | 0.01052 | 0.01419 | 0.01155 | 0.06244 | 0.00000 | 0.01086 | 0.01106 | 0.03119 | 0.11839 | 0.00416 | 0.01915 |
| **MIR371A** | 0.00600 | 0.01298 | 0.01226 | 0.01315 | 0.01180 | 0.02572 | 0.01624 | 0.01018 | 0.02353 | 0.02331 | 0.01408 | 0.01775 |
| **MIR371B** | 0.00611 | 0.01567 | 0.01334 | 0.00976 | 0.01414 | 0.02916 | 0.01784 | 0.01039 | 0.01433 | 0.01594 | 0.02189 | 0.01586 |
| **MIR372** | 0.00313 | 0.11427 | 0.00635 | 0.30704 | 0.32748 | 0.13584 | 0.00000 | 0.10261 | 0.38896 | 0.96660 | 0.42980 | 0.35070 |
| **MIR373** | 0.01259 | 0.23447 | 0.11782 | 0.01057 | 0.14208 | 0.28569 | 0.00000 | 0.02604 | 0.22341 | 0.31471 | 0.21110 | 0.23187 |
| **MIR378J** | 0.00501 | 0.01148 | 0.01609 | 0.00469 | 0.03041 | 0.01934 | 0.00318 | 0.01603 | 0.02862 | 0.07227 | 0.00993 | 0.01863 |
| **MIR3911** | 0.09606 | 0.02565 | 0.02492 | 0.00553 | 0.01765 | 0.03764 | 0.00695 | 0.01248 | 0.03079 | 0.01593 | 0.01143 | 0.03786 |
| **MIR3916** | 0.00434 | 0.02878 | 0.06296 | 0.09782 | 0.07684 | 0.05480 | 0.04273 | 0.16892 | 0.30991 | 0.02240 | 0.01408 | 0.07124 |
| **MIR3917** | 0.00572 | 2.85923 | 0.48363 | 2.85240 | 1.52438 | 3.38126 | 0.83234 | 0.20465 | 1.10563 | 0.00000 | 3.47617 | 1.88583 |

| | | | | | | | | | | | | |
|---|---|---|---|---|---|---|---|---|---|---|---|---|
| **MIR3928** | 0.00551 | 0.01795 | 0.02311 | 0.01079 | 0.01317 | 0.01889 | 0.00973 | 0.01224 | 0.02903 | 0.02590 | 0.01198 | 0.01523 |
| **MIR3939** | 0.00542 | 0.02810 | 0.02915 | 0.02854 | 0.02594 | 0.00000 | 0.00515 | 0.00884 | 0.06424 | 0.04258 | 0.08679 | 0.05413 |
| **MIR3940** | 0.01817 | 0.02492 | 0.05450 | 0.01093 | 0.05632 | 0.00000 | 0.00686 | 0.01592 | 0.05027 | 0.05009 | 0.03134 | 0.03310 |
| **MIR3941** | 0.02094 | 0.01628 | 0.01103 | 0.07722 | 0.02756 | 0.02066 | 0.01700 | 0.01055 | 0.02007 | 0.02268 | 0.01757 | 0.02221 |
| **MIR3942** | 0.00449 | 0.01117 | 0.03045 | 0.01882 | 0.00161 | 0.06029 | 0.00474 | 0.01085 | 0.02865 | 0.04780 | 0.01685 | 0.05098 |
| **MIR3944** | 0.00650 | 0.04921 | 0.03191 | 0.01668 | 0.03525 | 0.00000 | 0.01077 | 0.01101 | 0.09147 | 0.00230 | 0.01587 | 0.01672 |
| **MIR3978** | 0.01220 | 0.01585 | 0.01339 | 0.01932 | 0.01504 | 0.01941 | 0.01499 | 0.01143 | 0.02420 | 0.01586 | 0.01464 | 0.01818 |
| **MIR421** | 0.00570 | 0.01193 | 0.02023 | 0.01312 | 0.01526 | 0.02128 | 0.01192 | 0.00995 | 0.04607 | 0.03498 | 0.02052 | 0.02269 |
| **MIR423** | 0.00531 | 0.01610 | 0.02015 | 0.01152 | 0.01925 | 0.02305 | 0.01613 | 0.01361 | 0.03010 | 0.00664 | 0.01335 | 0.02183 |
| **MIR425** | 0.00524 | 0.01408 | 0.03740 | 0.02359 | 0.01179 | 0.00682 | 0.02628 | 0.01081 | 0.03245 | 0.06050 | 0.01512 | 0.01487 |
| **MIR4257** | 0.00483 | 0.01104 | 0.02226 | 0.00386 | 0.00651 | 0.01344 | 0.02604 | 0.01046 | 0.04202 | 0.08902 | 0.04129 | 0.01749 |
| **MIR4258** | 0.01817 | 0.03395 | 0.02113 | 0.04355 | 0.06802 | 0.00000 | 0.03233 | 0.03134 | 0.06691 | 0.04163 | 0.02174 | 0.01885 |
| **MIR4263** | 0.00576 | 0.02154 | 0.02137 | 0.00707 | 0.02343 | 0.04960 | 0.02987 | 0.00907 | 0.00000 | 0.02379 | 0.04137 | 0.03546 |
| **MIR4269** | 0.00714 | 0.02562 | 0.00506 | 0.00050 | 0.00000 | 0.04263 | 0.00100 | 0.00937 | 0.01282 | 0.12109 | 0.00852 | 0.01445 |
| **MIR4276** | 0.00627 | 0.01414 | 0.01298 | 0.01123 | 0.01387 | 0.02099 | 0.01327 | 0.01043 | 0.02447 | 0.01423 | 0.02505 | 0.01613 |
| **MIR4278** | 0.00503 | 0.01429 | 0.01066 | 0.00651 | 0.02553 | 0.01723 | 0.01302 | 0.01019 | 0.01639 | 0.07482 | 0.01048 | 0.01685 |
| **MIR4279** | 0.00388 | 0.00995 | 0.00648 | 0.00908 | 0.01348 | 0.00550 | 0.00908 | 0.01336 | 0.04023 | 0.10805 | 0.03074 | 0.03176 |
| **MIR4284** | 0.00463 | 0.01681 | 0.01316 | 0.01193 | 0.05029 | 0.00000 | 0.04672 | 0.00984 | 0.00000 | 0.13880 | 0.03933 | 0.01884 |
| **MIR4292** | 0.00586 | 0.02388 | 0.01668 | 0.02128 | 0.03628 | 0.00000 | 0.04643 | 0.01074 | 0.01344 | 0.02583 | 0.01409 | 0.01711 |
| **MIR4295** | 0.01089 | 0.01826 | 0.02096 | 0.01270 | 0.01242 | 0.01665 | 0.01298 | 0.01099 | 0.02503 | 0.01886 | 0.01879 | 0.01565 |
| **MIR4296** | 0.00613 | 0.01513 | 0.01427 | 0.01228 | 0.01385 | 0.01727 | 0.02074 | 0.01046 | 0.02130 | 0.01945 | 0.01492 | 0.01722 |
| **MIR4306** | 0.00835 | 0.01316 | 0.01131 | 0.01940 | 0.00670 | 0.04294 | 0.01306 | 0.00955 | 0.01263 | 0.01827 | 0.01702 | 0.01410 |
| **MIR4307** | 0.00507 | 0.01218 | 0.01929 | 0.01533 | 0.02115 | 0.03260 | 0.00613 | 0.01364 | 0.04022 | 0.02118 | 0.01010 | 0.01966 |
| **MIR4308** | 0.00625 | 0.01424 | 0.01396 | 0.01187 | 0.01442 | 0.02024 | 0.01320 | 0.01048 | 0.02721 | 0.01722 | 0.01617 | 0.01595 |
| **MIR4312** | 0.00580 | 0.01637 | 0.01851 | 0.01251 | 0.01256 | 0.02234 | 0.02173 | 0.01004 | 0.01507 | 0.01502 | 0.01427 | 0.03132 |
| **MIR4324** | 0.05743 | 0.01193 | 0.02279 | 0.01087 | 0.01713 | 0.02621 | 0.01064 | 0.01194 | 0.01943 | 0.02780 | 0.01517 | 0.02756 |
| **MIR4329** | 0.00634 | 0.01537 | 0.01575 | 0.01091 | 0.01308 | 0.02353 | 0.01506 | 0.01033 | 0.01647 | 0.02153 | 0.01567 | 0.01837 |
| **MIR4427** | 0.00552 | 0.01206 | 0.01738 | 0.00694 | 0.02548 | 0.02020 | 0.01136 | 0.00980 | 0.01740 | 0.04680 | 0.01338 | 0.02312 |
| **MIR4432** | 0.00622 | 0.01446 | 0.01346 | 0.01153 | 0.01358 | 0.01808 | 0.01639 | 0.01000 | 0.01670 | 0.02359 | 0.01786 | 0.02109 |
| **MIR4434** | 0.00363 | 0.01898 | 0.00967 | 0.01087 | 0.01119 | 0.03401 | 0.00837 | 0.01856 | 0.02554 | 0.03561 | 0.01393 | 0.03017 |
| **MIR4435-1HG** | 0.00045 | 0.00000 | 3.80645 | 0.41986 | 1.22044 | 0.49641 | 3.09044 | 0.00095 | 0.35535 | 0.58008 | 1.18419 | 0.00000 |
| **MIR4442** | 0.02635 | 0.01910 | 0.01794 | 0.01816 | 0.01588 | 0.01482 | 0.02262 | 0.00764 | 0.02932 | 0.00781 | 0.01481 | 0.01602 |
| **MIR4444-1+MIR** | 0.00595 | 0.01497 | 0.01380 | 0.01186 | 0.01647 | 0.01892 | 0.01764 | 0.01251 | 0.02034 | 0.01344 | 0.01468 | 0.01675 |
| **MIR4449** | 0.02275 | 0.02149 | 0.02507 | 0.00713 | 0.05701 | 0.00037 | 0.00816 | 0.01072 | 0.07750 | 0.09026 | 0.01086 | 0.05868 |
| **MIR4450** | 0.00635 | 0.01928 | 0.02123 | 0.01580 | 0.05392 | 0.00000 | 0.01716 | 0.01010 | 0.04329 | 0.00124 | 0.02156 | 0.02828 |
| **MIR4451** | 0.00624 | 0.01396 | 0.01482 | 0.01285 | 0.01287 | 0.02325 | 0.01565 | 0.01045 | 0.01928 | 0.01575 | 0.02090 | 0.01583 |
| **MIR4458HG** | 0.00000 | 0.61612 | 0.22296 | 0.00000 | 0.05406 | 0.92882 | 0.86289 | 0.07213 | 1.40210 | 0.54687 | 0.41850 | 1.02291 |
| **MIR4461** | 0.18567 | 0.05015 | 0.00703 | 0.00161 | 0.21504 | 0.10772 | 0.00000 | 0.23322 | 0.03096 | 0.29124 | 0.05604 | 0.13581 |
| **MIR4469** | 0.00621 | 0.01798 | 0.01684 | 0.01325 | 0.01647 | 0.01599 | 0.01887 | 0.01026 | 0.01811 | 0.00970 | 0.01579 | 0.02471 |
| **MIR4472-2** | 0.00522 | 0.00870 | 0.02616 | 0.01375 | 0.00309 | 0.08147 | 0.03611 | 0.01089 | 0.00870 | 0.03016 | 0.00726 | 0.01682 |
| **MIR4479** | 0.00797 | 0.01550 | 0.01730 | 0.01216 | 0.01666 | 0.01633 | 0.01743 | 0.01222 | 0.02098 | 0.01096 | 0.01450 | 0.01805 |
| **MIR4482** | 0.00507 | 0.01868 | 0.03387 | 0.02373 | 0.03969 | 0.00000 | 0.02366 | 0.01531 | 0.03236 | 0.03607 | 0.02941 | 0.02391 |
| **MIR4489** | 0.00638 | 0.01616 | 0.01877 | 0.01152 | 0.01496 | 0.01119 | 0.01313 | 0.01058 | 0.03675 | 0.02053 | 0.01204 | 0.01656 |
| **MIR4497** | 0.00632 | 0.01523 | 0.01379 | 0.01271 | 0.01550 | 0.01822 | 0.01886 | 0.01053 | 0.02202 | 0.01135 | 0.01482 | 0.01647 |
| **MIR4500HG** | 2.30443 | 0.01023 | 0.01110 | 0.00699 | 0.02164 | 0.01436 | 0.01447 | 0.08317 | 0.01637 | 0.02627 | 0.01463 | 0.06359 |
| **MIR4505** | 0.00628 | 0.01663 | 0.01804 | 0.00973 | 0.01728 | 0.02514 | 0.01652 | 0.01036 | 0.01504 | 0.01499 | 0.01451 | 0.01575 |
| **MIR4510** | 0.00471 | 0.01063 | 0.01285 | 0.01029 | 0.01199 | 0.03064 | 0.01780 | 0.02544 | 0.01451 | 0.01579 | 0.01446 | 0.03163 |
| **MIR4517** | 0.00801 | 0.01963 | 0.06366 | 0.01176 | 0.03811 | 0.02296 | 0.00000 | 0.03550 | 0.03963 | 0.07338 | 0.06160 | 0.05801 |

| | | | | | | | | | | | | |
|---|---|---|---|---|---|---|---|---|---|---|---|---|
| **MIR4519** | 0.00553 | 0.02001 | 0.01329 | 0.00674 | 0.01864 | 0.02923 | 0.01058 | 0.02234 | 0.02394 | 0.02717 | 0.01569 | 0.03151 |
| **MIR4522** | 0.00873 | 0.01490 | 0.01887 | 0.01025 | 0.01305 | 0.02426 | 0.01622 | 0.01126 | 0.01456 | 0.01902 | 0.01481 | 0.01718 |
| **MIR454** | 0.00883 | 0.01089 | 0.02097 | 0.00735 | 0.01471 | 0.02967 | 0.01139 | 0.00988 | 0.02554 | 0.02792 | 0.01818 | 0.02971 |
| **MIR4632** | 0.03150 | 0.01512 | 0.01491 | 0.01277 | 0.01571 | 0.01736 | 0.01871 | 0.00735 | 0.01897 | 0.01118 | 0.01491 | 0.01599 |
| **MIR4633** | 0.00491 | 0.01523 | 0.01317 | 0.01152 | 0.01402 | 0.01879 | 0.01652 | 0.01591 | 0.02196 | 0.01832 | 0.01433 | 0.01696 |
| **MIR4639** | 0.05112 | 0.01463 | 0.02089 | 0.01276 | 0.02258 | 0.03271 | 0.01481 | 0.01406 | 0.04188 | 0.02511 | 0.01378 | 0.02483 |
| **MIR4640** | 0.02877 | 0.02342 | 0.05624 | 0.01409 | 0.03097 | 0.00000 | 0.01034 | 0.01061 | 0.05435 | 0.00000 | 0.01436 | 0.02375 |
| **MIR4641** | 0.01459 | 0.01662 | 0.01750 | 0.01021 | 0.03253 | 0.01218 | 0.01660 | 0.00924 | 0.02355 | 0.01837 | 0.01359 | 0.01846 |
| **MIR4644** | 0.00520 | 0.01262 | 0.01336 | 0.01114 | 0.01311 | 0.02690 | 0.01741 | 0.01504 | 0.01750 | 0.01121 | 0.01859 | 0.02409 |
| **MIR4646** | 0.00623 | 0.01569 | 0.01538 | 0.01161 | 0.01877 | 0.01488 | 0.01791 | 0.01039 | 0.02031 | 0.01674 | 0.01489 | 0.01762 |
| **MIR4648** | 0.00624 | 0.01832 | 0.01574 | 0.01690 | 0.01863 | 0.01541 | 0.01821 | 0.01035 | 0.01951 | 0.01034 | 0.01532 | 0.02029 |
| **MIR4651** | 0.01190 | 0.01567 | 0.02302 | 0.01240 | 0.01453 | 0.01821 | 0.01478 | 0.00973 | 0.02200 | 0.01367 | 0.01596 | 0.01668 |
| **MIR4652** | 0.00551 | 0.01229 | 0.01553 | 0.00866 | 0.01930 | 0.01751 | 0.01855 | 0.01030 | 0.04388 | 0.05141 | 0.00872 | 0.01560 |
| **MIR4653** | 0.00876 | 0.01822 | 0.03112 | 0.01613 | 0.02217 | 0.00837 | 0.01794 | 0.01062 | 0.03791 | 0.00483 | 0.01578 | 0.02023 |
| **MIR4656** | 0.00636 | 0.02089 | 0.01583 | 0.01612 | 0.01536 | 0.01054 | 0.01579 | 0.01054 | 0.03201 | 0.01373 | 0.01366 | 0.01645 |
| **MIR4658** | 0.01137 | 0.01550 | 0.01385 | 0.03128 | 0.01685 | 0.01253 | 0.01734 | 0.00926 | 0.02182 | 0.01110 | 0.01937 | 0.01677 |
| **MIR4660** | 0.00622 | 0.01356 | 0.02251 | 0.01316 | 0.01861 | 0.02185 | 0.01364 | 0.01049 | 0.02207 | 0.01657 | 0.01343 | 0.01590 |
| **MIR4664** | 0.00606 | 0.01491 | 0.01928 | 0.01433 | 0.01664 | 0.01342 | 0.03874 | 0.01060 | 0.00614 | 0.01858 | 0.03126 | 0.01631 |
| **MIR4667** | 0.04658 | 0.02385 | 0.02631 | 0.01322 | 0.03468 | 0.00208 | 0.02961 | 0.01262 | 0.01881 | 0.03526 | 0.01525 | 0.05041 |
| **MIR4668** | 0.01520 | 0.16167 | 0.00000 | 0.04827 | 0.63104 | 0.24841 | 0.00000 | 0.00740 | 0.08431 | 0.13403 | 0.17154 | 0.01387 |
| **MIR4670** | 0.00565 | 0.01254 | 0.00941 | 0.01896 | 0.02440 | 0.01741 | 0.00458 | 0.01021 | 0.02409 | 0.06593 | 0.01526 | 0.01412 |
| **MIR4676** | 0.00620 | 0.01409 | 0.02157 | 0.02101 | 0.01328 | 0.02271 | 0.01377 | 0.01043 | 0.01997 | 0.01581 | 0.01536 | 0.01586 |
| **MIR4677** | 0.00604 | 0.01669 | 0.01535 | 0.01337 | 0.01614 | 0.01895 | 0.01885 | 0.01122 | 0.01946 | 0.01182 | 0.01451 | 0.01684 |
| **MIR4685** | 0.00764 | 0.03561 | 0.03691 | 0.01704 | 0.01772 | 0.00087 | 0.03191 | 0.01498 | 0.02228 | 0.00705 | 0.01819 | 0.02427 |
| **MIR4690** | 0.00862 | 0.03104 | 0.01583 | 0.02046 | 0.02302 | 0.00888 | 0.00723 | 0.01349 | 0.06927 | 0.02987 | 0.03467 | 0.01498 |
| **MIR4691** | 0.00589 | 0.07226 | 0.04272 | 0.05752 | 0.06071 | 0.00000 | 0.00101 | 0.00850 | 0.20554 | 0.02022 | 0.06136 | 0.06099 |
| **MIR4695** | 0.00634 | 0.01563 | 0.02865 | 0.01398 | 0.01667 | 0.01411 | 0.01687 | 0.01056 | 0.01716 | 0.01339 | 0.01542 | 0.01742 |
| **MIR4697HG** | 0.00340 | 0.00000 | 0.02141 | 0.00053 | 0.03066 | 0.26684 | 0.00000 | 0.00407 | 0.11027 | 0.29872 | 0.11248 | 0.06428 |
| **MIR4706** | 0.00616 | 0.01588 | 0.01448 | 0.01223 | 0.01453 | 0.02205 | 0.02218 | 0.01030 | 0.01732 | 0.01060 | 0.01458 | 0.02056 |
| **MIR4712** | 0.00532 | 0.03644 | 0.05180 | 0.02990 | 0.02010 | 0.02883 | 0.00003 | 0.00985 | 0.02776 | 0.06157 | 0.04109 | 0.01737 |
| **MIR4720** | 0.00601 | 0.01585 | 0.02823 | 0.01235 | 0.02323 | 0.02916 | 0.01960 | 0.01202 | 0.04968 | 0.00000 | 0.04831 | 0.02010 |
| **MIR4721** | 0.00569 | 0.50885 | 0.38977 | 0.30831 | 0.46727 | 0.18571 | 0.24437 | 0.03935 | 0.71991 | 0.00000 | 0.31547 | 0.32938 |
| **MIR4722** | 0.00689 | 0.03097 | 0.01543 | 0.01172 | 0.01683 | 0.00000 | 0.01389 | 0.01053 | 0.05540 | 0.04539 | 0.01876 | 0.02279 |
| **MIR4723** | 0.00502 | 0.17416 | 0.08416 | 0.06014 | 0.05103 | 0.25958 | 0.10976 | 0.12210 | 0.18368 | 0.02412 | 0.16048 | 0.15247 |
| **MIR4725** | 0.00604 | 0.01515 | 0.01422 | 0.01239 | 0.02373 | 0.01173 | 0.02052 | 0.01055 | 0.01904 | 0.02377 | 0.01535 | 0.01606 |
| **MIR4727** | 0.01882 | 0.01897 | 0.02353 | 0.01329 | 0.02191 | 0.01478 | 0.01605 | 0.01027 | 0.02498 | 0.01514 | 0.01214 | 0.01809 |
| **MIR4728** | 0.00580 | 0.01625 | 0.01369 | 0.01164 | 0.01682 | 0.02764 | 0.01956 | 0.01463 | 0.01916 | 0.01097 | 0.01919 | 0.01521 |
| **MIR4729** | 0.00619 | 0.01505 | 0.01330 | 0.01008 | 0.01916 | 0.01961 | 0.01529 | 0.00985 | 0.01673 | 0.02504 | 0.01537 | 0.02281 |
| **MIR4737** | 0.00587 | 0.05109 | 0.02725 | 0.03897 | 0.00616 | 0.05636 | 0.01187 | 0.00905 | 0.03754 | 0.00944 | 0.01544 | 0.04425 |
| **MIR4738** | 0.00629 | 0.01475 | 0.02433 | 0.01190 | 0.01509 | 0.02033 | 0.01671 | 0.01048 | 0.01580 | 0.01268 | 0.01540 | 0.01628 |
| **MIR4742** | 0.00595 | 0.01453 | 0.01400 | 0.01172 | 0.02364 | 0.02007 | 0.01639 | 0.01143 | 0.01835 | 0.01471 | 0.01515 | 0.01607 |
| **MIR4743** | 0.01939 | 0.01509 | 0.01543 | 0.01820 | 0.01847 | 0.01695 | 0.01481 | 0.01423 | 0.03033 | 0.01234 | 0.01302 | 0.01653 |
| **MIR4745** | 0.00674 | 0.01837 | 0.01238 | 0.01084 | 0.01279 | 0.02517 | 0.01440 | 0.01014 | 0.02452 | 0.01809 | 0.02267 | 0.01614 |
| **MIR4746** | 0.04559 | 0.03091 | 0.02042 | 0.01984 | 0.08485 | 0.00000 | 0.04178 | 0.04142 | 0.07980 | 0.20519 | 0.00664 | 0.13894 |
| **MIR4747** | 0.00625 | 0.01390 | 0.01893 | 0.01074 | 0.02561 | 0.01901 | 0.01711 | 0.01045 | 0.01906 | 0.01323 | 0.01746 | 0.01649 |
| **MIR4748** | 0.00617 | 0.01725 | 0.01999 | 0.01620 | 0.01383 | 0.00537 | 0.01718 | 0.01046 | 0.02634 | 0.02008 | 0.03233 | 0.02147 |
| **MIR4749** | 0.02233 | 0.03275 | 0.06889 | 0.02886 | 0.04154 | 0.00000 | 0.04514 | 0.01083 | 0.04663 | 0.01717 | 0.03323 | 0.02402 |
| **MIR4750** | 0.03364 | 0.02078 | 0.07856 | 0.02338 | 0.04283 | 0.00000 | 0.02070 | 0.01094 | 0.04043 | 0.02277 | 0.01553 | 0.02667 |

| | | | | | | | | | | | | |
|---|---|---|---|---|---|---|---|---|---|---|---|---|
| **MIR4754** | 0.00508 | 0.02006 | 0.02513 | 0.05509 | 0.03364 | 0.00222 | 0.01212 | 0.01261 | 0.06580 | 0.03490 | 0.02824 | 0.02866 |
| **MIR4755** | 0.00806 | 0.01396 | 0.01479 | 0.01993 | 0.11220 | 0.01003 | 0.01593 | 0.03464 | 0.03506 | 0.00000 | 0.01327 | 0.01988 |
| **MIR4762** | 0.00772 | 0.01895 | 0.01453 | 0.01185 | 0.01736 | 0.01569 | 0.01649 | 0.01030 | 0.02101 | 0.02000 | 0.01591 | 0.02031 |
| **MIR4766** | 0.01565 | 0.01041 | 0.01967 | 0.01380 | 0.00857 | 0.04778 | 0.01075 | 0.01214 | 0.01959 | 0.02602 | 0.01525 | 0.01544 |
| **MIR4767** | 0.00649 | 0.01524 | 0.01420 | 0.01524 | 0.01752 | 0.01928 | 0.02306 | 0.01043 | 0.01594 | 0.00973 | 0.01876 | 0.01644 |
| **MIR4774** | 0.00620 | 0.01670 | 0.01474 | 0.01282 | 0.01076 | 0.02853 | 0.01578 | 0.01040 | 0.01785 | 0.01471 | 0.02144 | 0.01518 |
| **MIR4781** | 0.00982 | 0.01560 | 0.01839 | 0.02249 | 0.01564 | 0.01136 | 0.01780 | 0.01383 | 0.01946 | 0.01560 | 0.01563 | 0.01690 |
| **MIR4784** | 0.00886 | 0.01519 | 0.01269 | 0.01141 | 0.01517 | 0.01751 | 0.01838 | 0.01027 | 0.02460 | 0.03491 | 0.01617 | 0.02330 |
| **MIR4785** | 0.00511 | 0.02117 | 0.01490 | 0.01228 | 0.02026 | 0.01553 | 0.01958 | 0.01597 | 0.01973 | 0.00927 | 0.01544 | 0.01936 |
| **MIR4786** | 0.00287 | 0.02106 | 0.01678 | 0.01566 | 0.16603 | 0.00000 | 0.01896 | 0.02178 | 0.00533 | 0.10476 | 0.01616 | 0.01605 |
| **MIR4787** | 0.00631 | 0.01634 | 0.01820 | 0.01615 | 0.02004 | 0.01508 | 0.01925 | 0.01051 | 0.02160 | 0.01162 | 0.01418 | 0.01650 |
| **MIR4793** | 0.00598 | 0.01747 | 0.01311 | 0.01068 | 0.01388 | 0.01970 | 0.02150 | 0.01046 | 0.02404 | 0.02527 | 0.01528 | 0.01532 |
| **MIR4794** | 0.00625 | 0.01465 | 0.01582 | 0.01293 | 0.02412 | 0.01620 | 0.01722 | 0.01033 | 0.01647 | 0.02348 | 0.01379 | 0.01678 |
| **MIR4802** | 0.00640 | 0.01937 | 0.01844 | 0.01580 | 0.02616 | 0.00109 | 0.02717 | 0.01062 | 0.02919 | 0.00000 | 0.01898 | 0.02361 |
| **MIR491** | 0.00621 | 0.01862 | 0.01360 | 0.01266 | 0.01501 | 0.03722 | 0.02178 | 0.01007 | 0.05582 | 0.01090 | 0.00559 | 0.01717 |
| **MIR492** | 0.00881 | 0.00000 | 1.09305 | 0.00000 | 0.00000 | 0.01527 | 3.26021 | 0.01107 | 3.90595 | 0.00000 | 1.55093 | 0.01769 |
| **MIR497HG** | 0.07435 | 0.14623 | 0.21931 | 0.16660 | 0.08545 | 0.00000 | 0.12135 | 0.16829 | 0.28518 | 0.03838 | 0.07399 | 0.22980 |
| **MIR498** | 0.00593 | 0.02327 | 0.06200 | 0.01785 | 0.01242 | 0.04712 | 0.00000 | 0.00934 | 0.08250 | 0.04930 | 0.01858 | 0.01881 |
| **MIR5000** | 0.00552 | 0.01698 | 0.01083 | 0.00800 | 0.00390 | 0.04267 | 0.00000 | 0.00941 | 0.04672 | 0.05158 | 0.01050 | 0.02355 |
| **MIR5001** | 0.01188 | 0.03470 | 0.02111 | 0.02497 | 0.02135 | 0.01800 | 0.02061 | 0.01671 | 0.04163 | 0.00001 | 0.02965 | 0.05051 |
| **MIR5006** | 0.04812 | 0.01274 | 0.01662 | 0.00958 | 0.01971 | 0.02482 | 0.01071 | 0.01211 | 0.02882 | 0.01142 | 0.01662 | 0.01867 |
| **MIR5010** | 0.00605 | 0.02038 | 0.02405 | 0.04051 | 0.05903 | 0.00000 | 0.01674 | 0.01054 | 0.01804 | 0.02304 | 0.01327 | 0.01807 |
| **MIR503HG** | 0.00295 | 0.00000 | 0.00000 | 0.00000 | 0.03997 | 0.13277 | 0.10576 | 0.02156 | 0.00000 | 0.10615 | 0.02331 | 0.09780 |
| **MIR5047** | 0.02636 | 0.30248 | 0.05610 | 0.23321 | 0.48845 | 0.53627 | 0.00000 | 1.43063 | 0.53197 | 0.00000 | 0.00000 | 1.29420 |
| **MIR5087** | 0.00415 | 0.00702 | 0.02166 | 0.01074 | 0.00555 | 0.05705 | 0.02369 | 0.01763 | 0.01153 | 0.00895 | 0.03747 | 0.03666 |
| **MIR5088** | 0.06811 | 0.14433 | 0.24647 | 0.18638 | 0.16241 | 0.28938 | 0.24689 | 0.08453 | 0.75893 | 0.00000 | 0.29803 | 0.28949 |
| **MIR5090** | 0.02317 | 0.01754 | 0.02335 | 0.01493 | 0.01796 | 0.01187 | 0.02115 | 0.00847 | 0.02090 | 0.00409 | 0.01613 | 0.03248 |
| **MIR5091** | 0.01115 | 0.01588 | 0.01183 | 0.03345 | 0.01034 | 0.01022 | 0.02086 | 0.00983 | 0.01595 | 0.02894 | 0.03575 | 0.05196 |
| **MIR5094** | 0.00588 | 0.01837 | 0.01464 | 0.01297 | 0.00949 | 0.02107 | 0.01619 | 0.00980 | 0.02600 | 0.02225 | 0.02707 | 0.02507 |
| **MIR516A1** | 0.00629 | 0.01684 | 0.01957 | 0.01066 | 0.01603 | 0.02672 | 0.01833 | 0.01007 | 0.03662 | 0.00652 | 0.01307 | 0.02068 |
| **MIR516A2** | 0.00629 | 0.01421 | 0.01891 | 0.01557 | 0.01454 | 0.02481 | 0.01663 | 0.01045 | 0.02225 | 0.01323 | 0.01531 | 0.01736 |
| **MIR516B1** | 0.00608 | 0.01619 | 0.01527 | 0.01077 | 0.01965 | 0.01552 | 0.01459 | 0.01012 | 0.03265 | 0.01527 | 0.01557 | 0.02362 |
| **MIR516B2** | 0.00612 | 0.01466 | 0.01961 | 0.05204 | 0.01413 | 0.01206 | 0.02481 | 0.01065 | 0.01774 | 0.01726 | 0.01868 | 0.01906 |
| **MIR517A** | 0.00600 | 0.02427 | 0.03586 | 0.00802 | 0.02674 | 0.03217 | 0.00793 | 0.01013 | 0.04353 | 0.01776 | 0.03905 | 0.01886 |
| **MIR517B** | 0.00740 | 0.01438 | 0.03288 | 0.01723 | 0.00768 | 0.02776 | 0.00124 | 0.01027 | 0.06533 | 0.05777 | 0.00992 | 0.02017 |
| **MIR517C** | 0.00337 | 0.01213 | 0.01868 | 0.01377 | 0.00724 | 0.05345 | 0.01442 | 0.02000 | 0.02273 | 0.03342 | 0.01589 | 0.03910 |
| **MIR5187** | 0.00880 | 0.01807 | 0.02252 | 0.01588 | 0.01499 | 0.01434 | 0.01681 | 0.00989 | 0.02434 | 0.01172 | 0.01503 | 0.01638 |
| **MIR5188** | 0.00645 | 0.01768 | 0.04199 | 0.00872 | 0.04143 | 0.00999 | 0.01089 | 0.01020 | 0.01764 | 0.03709 | 0.01158 | 0.01665 |
| **MIR518A1** | 0.00603 | 0.01547 | 0.02489 | 0.01798 | 0.00960 | 0.03835 | 0.02857 | 0.01025 | 0.01726 | 0.01097 | 0.01235 | 0.01852 |
| **MIR518A2** | 0.00613 | 0.01491 | 0.01733 | 0.00961 | 0.02308 | 0.02158 | 0.01771 | 0.01025 | 0.01484 | 0.01598 | 0.01789 | 0.01867 |
| **MIR518B** | 0.00630 | 0.01486 | 0.01791 | 0.01352 | 0.01481 | 0.01884 | 0.01642 | 0.01045 | 0.02155 | 0.01212 | 0.01463 | 0.01718 |
| **MIR518D** | 0.00645 | 0.01798 | 0.01701 | 0.01014 | 0.01324 | 0.01863 | 0.01213 | 0.01026 | 0.04689 | 0.01686 | 0.01127 | 0.01710 |
| **MIR518E** | 0.00623 | 0.01452 | 0.01845 | 0.01130 | 0.01718 | 0.01984 | 0.01716 | 0.01034 | 0.02071 | 0.01355 | 0.01471 | 0.01827 |
| **MIR518F** | 0.00614 | 0.01457 | 0.02165 | 0.01136 | 0.01757 | 0.02466 | 0.01283 | 0.01041 | 0.02064 | 0.01865 | 0.01455 | 0.01550 |
| **MIR5196** | 0.00625 | 0.02355 | 0.03461 | 0.01658 | 0.04275 | 0.00000 | 0.01564 | 0.01048 | 0.01612 | 0.01864 | 0.01754 | 0.02112 |
| **MIR519A1** | 0.00779 | 0.01500 | 0.01877 | 0.01398 | 0.01864 | 0.01516 | 0.01417 | 0.01079 | 0.02318 | 0.01817 | 0.01767 | 0.01613 |
| **MIR519A2** | 0.00522 | 0.01404 | 0.05676 | 0.01575 | 0.01634 | 0.02017 | 0.01707 | 0.01339 | 0.03867 | 0.03073 | 0.02933 | 0.01840 |
| **MIR519B** | 0.00601 | 0.01711 | 0.03556 | 0.01209 | 0.01926 | 0.01696 | 0.02491 | 0.01054 | 0.03266 | 0.01504 | 0.01335 | 0.01983 |

| | | | | | | | | | | | | |
|---|---|---|---|---|---|---|---|---|---|---|---|---|
| **MIR519C** | 0.00650 | 0.01665 | 0.01596 | 0.01258 | 0.01463 | 0.01861 | 0.01531 | 0.01054 | 0.02939 | 0.01581 | 0.01778 | 0.01874 |
| **MIR519D** | 0.00627 | 0.01480 | 0.01966 | 0.01861 | 0.01226 | 0.01878 | 0.01283 | 0.01041 | 0.03315 | 0.01715 | 0.01255 | 0.01639 |
| **MIR519E** | 0.00638 | 0.01487 | 0.01364 | 0.01224 | 0.01414 | 0.02096 | 0.01697 | 0.01040 | 0.03052 | 0.00976 | 0.01191 | 0.01707 |
| **MIR520A** | 0.00611 | 0.01507 | 0.02603 | 0.01335 | 0.01838 | 0.01256 | 0.02543 | 0.01060 | 0.03328 | 0.03043 | 0.01942 | 0.01557 |
| **MIR520C** | 0.00645 | 0.01314 | 0.01551 | 0.00519 | 0.01564 | 0.04433 | 0.01168 | 0.00918 | 0.06847 | 0.00885 | 0.00446 | 0.03049 |
| **MIR520D** | 0.00533 | 0.01648 | 0.13415 | 0.05799 | 0.04443 | 0.05372 | 0.00000 | 0.01429 | 0.13648 | 0.15657 | 0.00706 | 0.05835 |
| **MIR520E** | 0.00590 | 0.00875 | 0.02500 | 0.00865 | 0.01535 | 0.05186 | 0.00859 | 0.01095 | 0.04287 | 0.03340 | 0.01532 | 0.01334 |
| **MIR520F** | 0.00787 | 0.01382 | 0.13698 | 0.04764 | 0.03076 | 0.03263 | 0.00492 | 0.01409 | 0.15605 | 0.02998 | 0.02955 | 0.03256 |
| **MIR520G** | 0.00543 | 0.01471 | 0.03241 | 0.00794 | 0.01210 | 0.02806 | 0.02804 | 0.01002 | 0.00959 | 0.04078 | 0.02193 | 0.01937 |
| **MIR520H** | 0.00568 | 0.01362 | 0.03738 | 0.00689 | 0.01655 | 0.02745 | 0.01011 | 0.00970 | 0.02458 | 0.03158 | 0.02079 | 0.02489 |
| **MIR521-1** | 0.00609 | 0.01489 | 0.02078 | 0.01144 | 0.01589 | 0.01739 | 0.01568 | 0.01106 | 0.02235 | 0.01334 | 0.01656 | 0.01819 |
| **MIR521-2** | 0.00627 | 0.01305 | 0.02063 | 0.02336 | 0.01516 | 0.02589 | 0.01440 | 0.01027 | 0.04001 | 0.01213 | 0.01367 | 0.01699 |
| **MIR522** | 0.00616 | 0.01397 | 0.01816 | 0.01037 | 0.01987 | 0.01935 | 0.01652 | 0.01019 | 0.02647 | 0.01454 | 0.01431 | 0.02245 |
| **MIR523** | 0.00610 | 0.01415 | 0.01554 | 0.01442 | 0.01266 | 0.02410 | 0.01505 | 0.01013 | 0.02108 | 0.01901 | 0.01490 | 0.01959 |
| **MIR524** | 0.00571 | 0.01266 | 0.01432 | 0.00631 | 0.04235 | 0.02432 | 0.01682 | 0.01040 | 0.01471 | 0.03369 | 0.01335 | 0.01443 |
| **MIR525** | 0.00560 | 0.01597 | 0.02113 | 0.01100 | 0.01680 | 0.00709 | 0.03210 | 0.01056 | 0.01074 | 0.04296 | 0.01641 | 0.01709 |
| **MIR526A1** | 0.00497 | 0.04691 | 0.02373 | 0.00770 | 0.00000 | 0.04130 | 0.00000 | 0.00968 | 0.07680 | 0.07126 | 0.00467 | 0.01666 |
| **MIR526A2** | 0.00591 | 0.01398 | 0.03523 | 0.01296 | 0.05184 | 0.01581 | 0.00405 | 0.01036 | 0.05454 | 0.03078 | 0.02706 | 0.01535 |
| **MIR526B** | 0.00611 | 0.01388 | 0.02649 | 0.01096 | 0.01406 | 0.02202 | 0.01982 | 0.01037 | 0.01395 | 0.02070 | 0.01595 | 0.01615 |
| **MIR527** | 0.00600 | 0.01463 | 0.01678 | 0.01108 | 0.01668 | 0.01743 | 0.01630 | 0.01091 | 0.03016 | 0.01429 | 0.01307 | 0.02050 |
| **MIR545** | 0.00632 | 0.01524 | 0.01403 | 0.01330 | 0.01694 | 0.01996 | 0.01984 | 0.01034 | 0.01734 | 0.01016 | 0.01916 | 0.01791 |
| **MIR548AC** | 0.00614 | 0.01380 | 0.02130 | 0.00930 | 0.01550 | 0.02957 | 0.01799 | 0.01032 | 0.01448 | 0.01309 | 0.01781 | 0.01804 |
| **MIR548AE2** | 0.00848 | 0.01495 | 0.01435 | 0.01180 | 0.01528 | 0.01873 | 0.01791 | 0.01438 | 0.01965 | 0.01477 | 0.01458 | 0.01609 |
| **MIR548AZ** | 0.00608 | 0.01555 | 0.02158 | 0.01280 | 0.01319 | 0.01647 | 0.02558 | 0.01048 | 0.01915 | 0.02148 | 0.01408 | 0.01617 |
| **MIR548F2** | 0.00611 | 0.01389 | 0.01554 | 0.01245 | 0.01298 | 0.02066 | 0.01641 | 0.01044 | 0.03096 | 0.02079 | 0.01741 | 0.01570 |
| **MIR548G** | 0.00621 | 0.01400 | 0.01584 | 0.01222 | 0.01435 | 0.02244 | 0.01806 | 0.01019 | 0.01679 | 0.01217 | 0.01783 | 0.02023 |
| **MIR548H4** | 0.00620 | 0.01443 | 0.01385 | 0.01100 | 0.01483 | 0.02192 | 0.01968 | 0.01046 | 0.01862 | 0.01560 | 0.01556 | 0.01589 |
| **MIR548J** | 0.00587 | 0.01554 | 0.01302 | 0.01393 | 0.01341 | 0.02336 | 0.01131 | 0.01005 | 0.02750 | 0.02574 | 0.01325 | 0.02117 |
| **MIR548K** | 0.00627 | 0.01529 | 0.01495 | 0.01860 | 0.01760 | 0.01369 | 0.01569 | 0.01055 | 0.02250 | 0.01589 | 0.01695 | 0.01647 |
| **MIR548Y** | 0.00623 | 0.01354 | 0.03135 | 0.00959 | 0.01503 | 0.02380 | 0.00651 | 0.00990 | 0.03592 | 0.01469 | 0.01503 | 0.02276 |
| **MIR553** | 0.00604 | 0.01135 | 0.01455 | 0.00892 | 0.02295 | 0.03271 | 0.01604 | 0.01031 | 0.01805 | 0.01946 | 0.01378 | 0.01497 |
| **MIR554** | 0.02494 | 0.01642 | 0.05072 | 0.00926 | 0.01323 | 0.02444 | 0.00731 | 0.17342 | 0.00677 | 0.03060 | 0.04763 | 0.03134 |
| **MIR555** | 0.01441 | 0.01110 | 0.02687 | 0.01505 | 0.01804 | 0.03192 | 0.02037 | 0.01418 | 0.02295 | 0.02391 | 0.02176 | 0.01588 |
| **MIR558** | 0.00617 | 0.01341 | 0.02025 | 0.00681 | 0.00872 | 0.04941 | 0.00909 | 0.00972 | 0.01530 | 0.02888 | 0.01874 | 0.01842 |
| **MIR5581** | 0.00506 | 0.02315 | 0.01582 | 0.01881 | 0.01793 | 0.01229 | 0.02165 | 0.01543 | 0.01855 | 0.01039 | 0.01725 | 0.01723 |
| **MIR5586** | 0.00576 | 0.01585 | 0.01891 | 0.01053 | 0.01733 | 0.01861 | 0.01557 | 0.01190 | 0.02057 | 0.01617 | 0.01440 | 0.01613 |
| **MIR559** | 0.00626 | 0.01685 | 0.01456 | 0.01386 | 0.01465 | 0.01989 | 0.01817 | 0.01050 | 0.01866 | 0.01339 | 0.01492 | 0.01596 |
| **MIR561** | 0.00565 | 0.01132 | 0.03375 | 0.00801 | 0.01397 | 0.03358 | 0.00949 | 0.00998 | 0.02421 | 0.03708 | 0.01175 | 0.01853 |
| **MIR564** | 0.01104 | 0.01536 | 0.02785 | 0.02017 | 0.01318 | 0.02115 | 0.01476 | 0.00964 | 0.02551 | 0.01004 | 0.01836 | 0.02684 |
| **MIR567** | 0.00562 | 0.01424 | 0.01444 | 0.00871 | 0.01649 | 0.01864 | 0.02965 | 0.01041 | 0.01508 | 0.05072 | 0.01443 | 0.01435 |
| **MIR5684** | 0.00564 | 0.01545 | 0.01478 | 0.01449 | 0.01529 | 0.01651 | 0.02058 | 0.01282 | 0.02181 | 0.01148 | 0.01488 | 0.01654 |
| **MIR5687** | 0.00536 | 0.03764 | 0.00000 | 0.11185 | 0.00000 | 0.09248 | 0.00000 | 0.00752 | 0.03477 | 0.03074 | 0.00681 | 0.06809 |
| **MIR570** | 0.01594 | 0.01392 | 0.02510 | 0.00007 | 0.04652 | 0.04718 | 0.00541 | 0.00856 | 0.14217 | 0.16865 | 0.00085 | 0.03962 |
| **MIR5703** | 0.00611 | 0.02448 | 0.01851 | 0.01597 | 0.01911 | 0.01403 | 0.01803 | 0.01163 | 0.02010 | 0.01249 | 0.01525 | 0.01553 |
| **MIR572** | 0.00785 | 0.02370 | 0.02068 | 0.01778 | 0.03679 | 0.00965 | 0.02081 | 0.01106 | 0.03673 | 0.00422 | 0.01978 | 0.02084 |
| **MIR573** | 0.02590 | 0.02223 | 0.02755 | 0.01454 | 0.02157 | 0.00954 | 0.09439 | 0.03755 | 0.03191 | 0.00016 | 0.04584 | 0.02816 |
| **MIR581** | 0.00605 | 0.01413 | 0.01516 | 0.01270 | 0.01123 | 0.02305 | 0.01544 | 0.01006 | 0.01711 | 0.02066 | 0.01902 | 0.02079 |
| **MIR589** | 0.01242 | 0.05373 | 0.01891 | 0.04004 | 0.10160 | 0.00000 | 0.02536 | 0.03210 | 0.06612 | 0.06131 | 0.05900 | 0.06272 |

| | | | | | | | | | | | | |
|---|---|---|---|---|---|---|---|---|---|---|---|---|
| **MIR590** | 0.00950 | 0.03534 | 0.01444 | 0.01167 | 0.01233 | 0.02328 | 0.01273 | 0.00939 | 0.03258 | 0.00666 | 0.01456 | 0.01772 |
| **MIR593** | 0.00586 | 0.01547 | 0.01674 | 0.00999 | 0.01039 | 0.01719 | 0.00997 | 0.00970 | 0.03253 | 0.03689 | 0.01136 | 0.02481 |
| **MIR597** | 0.00653 | 0.01402 | 0.01337 | 0.01135 | 0.01172 | 0.02541 | 0.00883 | 0.01037 | 0.04863 | 0.00911 | 0.01361 | 0.01612 |
| **MIR600HG** | 0.00041 | 1.02646 | 1.19235 | 0.13683 | 1.13900 | 0.00000 | 2.42792 | 0.20080 | 1.60615 | 1.96008 | 0.96932 | 0.28243 |
| **MIR603** | 0.00570 | 0.01172 | 0.06538 | 0.02730 | 0.00722 | 0.02953 | 0.00448 | 0.01042 | 0.01700 | 0.03719 | 0.01392 | 0.01420 |
| **MIR610** | 0.00671 | 0.01480 | 0.01679 | 0.01053 | 0.01659 | 0.01830 | 0.01155 | 0.01037 | 0.03427 | 0.01699 | 0.01312 | 0.01971 |
| **MIR611** | 0.04577 | 0.35015 | 0.43762 | 0.44224 | 0.85930 | 0.00120 | 0.52769 | 0.07077 | 0.35926 | 0.00000 | 0.32816 | 0.26529 |
| **MIR612** | 0.00609 | 0.01396 | 0.02334 | 0.03595 | 0.01181 | 0.01678 | 0.01162 | 0.01041 | 0.02106 | 0.01837 | 0.01975 | 0.01843 |
| **MIR6124** | 0.00668 | 0.02688 | 0.03034 | 0.02049 | 0.04737 | 0.00000 | 0.01161 | 0.01065 | 0.05390 | 0.00000 | 0.01652 | 0.01730 |
| **MIR6126** | 0.02374 | 0.05230 | 0.08169 | 0.04401 | 0.01498 | 0.00112 | 0.02857 | 0.01354 | 0.11341 | 0.00000 | 0.03340 | 0.04203 |
| **MIR6132** | 0.00621 | 0.01527 | 0.01935 | 0.01075 | 0.01328 | 0.02355 | 0.01645 | 0.01012 | 0.01883 | 0.01380 | 0.01561 | 0.02543 |
| **MIR614** | 0.00284 | 0.33939 | 0.00000 | 0.00000 | 1.41585 | 0.00000 | 0.00166 | 0.01380 | 0.50837 | 0.15391 | 0.83172 | 0.00571 |
| **MIR6165.uid-22** | 0.00861 | 0.01641 | 0.01456 | 0.01407 | 0.01583 | 0.01090 | 0.02286 | 0.03511 | 0.01970 | 0.01662 | 0.01569 | 0.01516 |
| **MIR6165.uid-77** | 0.00596 | 0.01341 | 0.01214 | 0.00984 | 0.01971 | 0.02527 | 0.01335 | 0.01120 | 0.02713 | 0.01404 | 0.01575 | 0.01722 |
| **MIR622** | 0.00638 | 0.02354 | 0.02363 | 0.00825 | 0.04191 | 0.03026 | 0.03318 | 0.01027 | 0.07046 | 0.00000 | 0.05003 | 0.01755 |
| **MIR623** | 0.00600 | 0.02376 | 0.02428 | 0.00916 | 0.01216 | 0.01743 | 0.01652 | 0.01067 | 0.01593 | 0.03310 | 0.01614 | 0.01955 |
| **MIR624** | 0.03431 | 0.01246 | 0.01205 | 0.03871 | 0.01258 | 0.01086 | 0.02050 | 0.00640 | 0.02359 | 0.04101 | 0.02472 | 0.02741 |
| **MIR628** | 0.00624 | 0.01362 | 0.01371 | 0.01220 | 0.01434 | 0.02104 | 0.01715 | 0.01001 | 0.01732 | 0.01746 | 0.01481 | 0.02136 |
| **MIR630** | 0.00666 | 0.01331 | 0.02008 | 0.00864 | 0.02003 | 0.01272 | 0.03641 | 0.01284 | 0.02157 | 0.05851 | 0.02118 | 0.01397 |
| **MIR631** | 0.00609 | 0.01521 | 0.01725 | 0.03254 | 0.01086 | 0.02200 | 0.01194 | 0.00977 | 0.04340 | 0.01713 | 0.01984 | 0.02499 |
| **MIR632** | 0.00302 | 0.01308 | 0.01489 | 0.00812 | 0.01888 | 0.02175 | 0.02157 | 0.06539 | 0.00858 | 0.06156 | 0.01813 | 0.01629 |
| **MIR635** | 0.00739 | 0.01736 | 0.01740 | 0.01029 | 0.02735 | 0.02189 | 0.03188 | 0.01128 | 0.00751 | 0.00369 | 0.01961 | 0.02003 |
| **MIR636** | 0.00618 | 0.01804 | 0.01170 | 0.07101 | 0.00797 | 0.02881 | 0.01484 | 0.01476 | 0.02100 | 0.00596 | 0.01373 | 0.01728 |
| **MIR637** | 0.00630 | 0.02533 | 0.04371 | 0.01478 | 0.02781 | 0.00438 | 0.01153 | 0.01018 | 0.03963 | 0.00744 | 0.01785 | 0.02614 |
| **MIR639** | 0.00714 | 0.01870 | 0.02526 | 0.01109 | 0.01756 | 0.02311 | 0.03144 | 0.01033 | 0.00897 | 0.01611 | 0.01665 | 0.02131 |
| **MIR644A** | 0.01036 | 0.01072 | 0.00946 | 0.01551 | 0.01191 | 0.09260 | 0.00000 | 0.01064 | 0.04741 | 0.06067 | 0.01352 | 0.01925 |
| **MIR645** | 0.00778 | 0.01445 | 0.01559 | 0.01067 | 0.01608 | 0.02247 | 0.01957 | 0.01103 | 0.02111 | 0.01386 | 0.01497 | 0.01585 |
| **MIR646HG** | 1.42736 | 0.16499 | 0.10365 | 0.22357 | 0.00000 | 0.00000 | 0.30402 | 0.21171 | 0.00000 | 0.08325 | 0.13573 | 0.17808 |
| **MIR647** | 0.00337 | 0.00278 | 0.01642 | 0.02672 | 0.11619 | 0.00000 | 0.00155 | 0.00774 | 0.03889 | 0.31177 | 0.00306 | 0.05606 |
| **MIR6505** | 0.00608 | 0.03758 | 0.02643 | 0.02545 | 0.02324 | 0.01190 | 0.02355 | 0.01465 | 0.06395 | 0.00000 | 0.02826 | 0.03828 |
| **MIR6514** | 0.00505 | 0.01519 | 0.01106 | 0.04225 | 0.01632 | 0.01489 | 0.07891 | 0.02356 | 0.00473 | 0.01458 | 0.02608 | 0.03672 |
| **MIR6515** | 0.03719 | 0.27129 | 0.54424 | 0.42906 | 0.54742 | 0.00000 | 0.05507 | 0.07924 | 1.32389 | 0.00000 | 0.33939 | 0.16971 |
| **MIR658** | 0.00589 | 0.01678 | 0.01886 | 0.01366 | 0.01401 | 0.01833 | 0.01645 | 0.01187 | 0.02625 | 0.01174 | 0.01376 | 0.01615 |
| **MIR659** | 0.00608 | 0.01477 | 0.01884 | 0.02327 | 0.01399 | 0.01950 | 0.02697 | 0.01044 | 0.02194 | 0.01744 | 0.01472 | 0.01646 |
| **MIR663A** | 0.00535 | 0.00000 | 0.00383 | 0.00000 | 0.28171 | 0.00000 | 0.00111 | 0.00617 | 0.11703 | 0.81477 | 0.00000 | 0.06282 |
| **MIR663AHG** | 0.00111 | 0.16733 | 0.77869 | 0.13849 | 1.18753 | 0.00000 | 0.00000 | 0.03339 | 0.66182 | 1.55464 | 0.00000 | 0.09981 |
| **MIR663B** | 0.00601 | 0.01454 | 0.01741 | 0.01103 | 0.01585 | 0.01802 | 0.01487 | 0.01026 | 0.02074 | 0.02566 | 0.01433 | 0.01855 |
| **MIR6716** | 0.00690 | 0.02215 | 0.01553 | 0.01221 | 0.01898 | 0.01359 | 0.01931 | 0.01287 | 0.02813 | 0.01477 | 0.01689 | 0.01695 |
| **MIR6717** | 0.00607 | 0.01352 | 0.01241 | 0.01426 | 0.01477 | 0.01700 | 0.01264 | 0.01039 | 0.01896 | 0.03162 | 0.02626 | 0.01550 |
| **MIR6719** | 0.00571 | 0.01053 | 0.01661 | 0.01465 | 0.04681 | 0.00000 | 0.00395 | 0.00973 | 0.06774 | 0.09631 | 0.02310 | 0.03009 |
| **MIR6721** | 0.01035 | 0.01465 | 0.01917 | 0.01162 | 0.01731 | 0.01714 | 0.01511 | 0.01118 | 0.02179 | 0.01598 | 0.01737 | 0.02035 |
| **MIR6723** | 0.10010 | 0.08811 | 0.00253 | 0.09350 | 0.22354 | 0.00000 | 0.01188 | 0.06842 | 0.15066 | 0.19408 | 0.04506 | 0.08894 |
| **MIR6727** | 0.00967 | 0.01634 | 0.02169 | 0.03660 | 0.02403 | 0.00000 | 0.03614 | 0.01747 | 0.01184 | 0.09178 | 0.03009 | 0.02140 |
| **MIR6728** | 0.00547 | 0.03702 | 0.03380 | 0.01743 | 0.02829 | 0.00000 | 0.02338 | 0.01374 | 0.03922 | 0.00000 | 0.03107 | 0.04103 |
| **MIR6729** | 0.00811 | 0.06990 | 0.03430 | 0.03018 | 0.10500 | 0.00000 | 0.01041 | 0.02003 | 0.12150 | 0.00000 | 0.01372 | 0.07065 |
| **MIR6730** | 0.00628 | 0.02024 | 0.01366 | 0.01684 | 0.01687 | 0.02013 | 0.01782 | 0.01045 | 0.01958 | 0.00823 | 0.01564 | 0.01885 |
| **MIR6732** | 0.01351 | 0.01518 | 0.03474 | 0.02750 | 0.05339 | 0.03569 | 0.01079 | 0.00933 | 0.04410 | 0.00000 | 0.01016 | 0.02371 |
| **MIR6733** | 0.00289 | 0.01468 | 0.03128 | 0.05398 | 0.08627 | 0.02636 | 0.00000 | 0.04090 | 0.05029 | 0.10728 | 0.01734 | 0.03163 |

| | | | | | | | | | | | | |
|---|---|---|---|---|---|---|---|---|---|---|---|---|
| **MIR6734** | 0.00509 | 0.01910 | 0.05936 | 0.00212 | 0.04485 | 0.05651 | 0.01024 | 0.01006 | 0.03166 | 0.05021 | 0.05717 | 0.01127 |
| **MIR6735** | 0.00788 | 0.01579 | 0.01454 | 0.01582 | 0.01477 | 0.02897 | 0.01846 | 0.01037 | 0.02152 | 0.01510 | 0.01870 | 0.02052 |
| **MIR6736** | 0.01260 | 0.03995 | 0.03185 | 0.02864 | 0.00000 | 0.03657 | 0.02468 | 0.01567 | 0.01703 | 0.01422 | 0.03787 | 0.04450 |
| **MIR6737** | 0.00582 | 0.01795 | 0.01912 | 0.01021 | 0.01379 | 0.02699 | 0.01816 | 0.01222 | 0.02351 | 0.01022 | 0.01420 | 0.01534 |
| **MIR6739** | 0.02611 | 0.04045 | 0.08782 | 0.04666 | 0.01471 | 0.03592 | 0.02367 | 0.01666 | 0.06290 | 0.00137 | 0.08623 | 0.04977 |
| **MIR6740** | 0.00916 | 0.00050 | 0.00400 | 0.05337 | 0.05304 | 0.09402 | 0.00000 | 0.00960 | 0.06474 | 0.11403 | 0.04513 | 0.02259 |
| **MIR6741** | 0.00638 | 0.01513 | 0.01823 | 0.01582 | 0.01728 | 0.01669 | 0.01531 | 0.01047 | 0.02910 | 0.01079 | 0.01685 | 0.01659 |
| **MIR6742** | 0.00647 | 0.02017 | 0.03035 | 0.01877 | 0.02890 | 0.00984 | 0.01823 | 0.01059 | 0.03129 | 0.00534 | 0.01546 | 0.01732 |
| **MIR6743** | 0.05117 | 0.09717 | 0.11667 | 0.05100 | 0.06589 | 0.00000 | 0.01866 | 0.13004 | 0.10087 | 0.00000 | 0.03807 | 0.09366 |
| **MIR6745** | 0.02472 | 0.01344 | 0.01227 | 0.01554 | 0.01561 | 0.02167 | 0.01981 | 0.01623 | 0.01947 | 0.02049 | 0.01372 | 0.01715 |
| **MIR6746** | 0.00603 | 0.01346 | 0.01974 | 0.01542 | 0.02028 | 0.01658 | 0.01219 | 0.01030 | 0.02832 | 0.02574 | 0.01340 | 0.01810 |
| **MIR6747** | 0.00619 | 0.01161 | 0.03122 | 0.01204 | 0.02582 | 0.05151 | 0.01049 | 0.00966 | 0.03353 | 0.00665 | 0.01383 | 0.02006 |
| **MIR6748** | 0.04463 | 0.13242 | 0.07030 | 0.15392 | 0.26587 | 0.20937 | 0.00000 | 0.06787 | 0.15086 | 0.00000 | 0.16665 | 0.31228 |
| **MIR6749** | 0.00565 | 0.01463 | 0.01500 | 0.01576 | 0.00958 | 0.02083 | 0.01734 | 0.01004 | 0.02096 | 0.03772 | 0.01341 | 0.02149 |
| **MIR6750** | 0.00596 | 0.01510 | 0.01423 | 0.01022 | 0.01756 | 0.02815 | 0.01554 | 0.00885 | 0.01689 | 0.02086 | 0.01459 | 0.04795 |
| **MIR6751** | 0.00624 | 0.01504 | 0.02588 | 0.01135 | 0.02006 | 0.01614 | 0.01689 | 0.01049 | 0.01678 | 0.01603 | 0.01552 | 0.01627 |
| **MIR6752** | 0.07348 | 0.05559 | 0.04006 | 0.07899 | 0.05396 | 0.00047 | 0.09034 | 0.10923 | 0.02373 | 0.03844 | 0.03236 | 0.15954 |
| **MIR6753** | 0.00653 | 0.03641 | 0.04618 | 0.01444 | 0.01991 | 0.00000 | 0.00498 | 0.01020 | 0.05164 | 0.01600 | 0.02048 | 0.02102 |
| **MIR6754** | 0.01396 | 0.02866 | 0.03715 | 0.03864 | 0.02060 | 0.04283 | 0.00963 | 0.00804 | 0.10021 | 0.00000 | 0.03100 | 0.03258 |
| **MIR6758** | 0.03015 | 1.24881 | 0.17713 | 0.52862 | 0.00000 | 0.56993 | 0.18251 | 0.21674 | 0.87602 | 0.00000 | 0.39278 | 0.46131 |
| **MIR6759** | 0.00539 | 0.01456 | 0.01314 | 0.01470 | 0.01075 | 0.01706 | 0.02418 | 0.01223 | 0.01944 | 0.07823 | 0.01097 | 0.02623 |
| **MIR6761** | 0.00866 | 0.04610 | 0.06647 | 0.54124 | 0.13891 | 0.07996 | 0.07443 | 0.01044 | 0.04781 | 0.00000 | 0.08755 | 0.01096 |
| **MIR6762** | 0.01399 | 0.01749 | 0.01578 | 0.01090 | 0.01960 | 0.02671 | 0.01081 | 0.00911 | 0.02109 | 0.02017 | 0.03221 | 0.01759 |
| **MIR6763** | 0.00863 | 0.01303 | 0.01424 | 0.00979 | 0.01662 | 0.02753 | 0.01721 | 0.00969 | 0.01808 | 0.01481 | 0.01739 | 0.01559 |
| **MIR6764** | 0.02422 | 0.04104 | 0.06984 | 0.12782 | 0.13502 | 0.00000 | 0.07117 | 0.21565 | 0.08426 | 0.69918 | 0.00000 | 0.14613 |
| **MIR6765** | 0.01198 | 0.01985 | 0.01590 | 0.00827 | 0.01893 | 0.01788 | 0.01914 | 0.00900 | 0.02209 | 0.03450 | 0.01692 | 0.01407 |
| **MIR6767** | 0.01021 | 0.21188 | 0.05550 | 0.06439 | 0.04647 | 0.00000 | 0.01989 | 0.02171 | 0.17340 | 0.00000 | 0.08507 | 0.10586 |
| **MIR6768** | 0.01329 | 0.05970 | 0.04518 | 0.03505 | 0.03254 | 0.00000 | 0.06656 | 0.01209 | 0.04480 | 0.00000 | 0.08612 | 0.03844 |
| **MIR6769B** | 0.00790 | 0.01423 | 0.01452 | 0.01468 | 0.02121 | 0.02243 | 0.01934 | 0.01124 | 0.01458 | 0.01399 | 0.01440 | 0.01697 |
| **MIR6773** | 0.00632 | 0.01554 | 0.01613 | 0.01097 | 0.01409 | 0.02657 | 0.01417 | 0.01039 | 0.03005 | 0.01033 | 0.01897 | 0.01577 |
| **MIR6774** | 0.01074 | 0.01622 | 0.01809 | 0.01449 | 0.01209 | 0.02486 | 0.01693 | 0.01555 | 0.01630 | 0.01331 | 0.01607 | 0.02204 |
| **MIR6775** | 0.00489 | 0.00804 | 0.01654 | 0.00297 | 0.17822 | 0.00000 | 0.01788 | 0.00999 | 0.04731 | 0.12841 | 0.01495 | 0.01558 |
| **MIR6776** | 0.00625 | 0.02835 | 0.01473 | 0.01298 | 0.02161 | 0.00082 | 0.01498 | 0.01062 | 0.03758 | 0.01426 | 0.02329 | 0.01986 |
| **MIR6777** | 0.00583 | 0.00375 | 0.00995 | 0.00002 | 0.03656 | 0.02764 | 0.02293 | 0.00918 | 0.04544 | 0.25185 | 0.00040 | 0.00843 |
| **MIR6778** | 0.00496 | 0.01681 | 0.02549 | 0.03100 | 0.03546 | 0.00428 | 0.01889 | 0.01470 | 0.01940 | 0.01444 | 0.03093 | 0.01929 |
| **MIR6779** | 0.00670 | 0.22586 | 0.01953 | 0.06494 | 0.03832 | 0.00000 | 0.02293 | 0.02045 | 0.05947 | 0.00000 | 0.03772 | 0.03319 |
| **MIR6780B** | 0.01786 | 0.01787 | 0.02085 | 0.04146 | 0.02869 | 0.01708 | 0.01683 | 0.00900 | 0.04876 | 0.01673 | 0.02643 | 0.01569 |
| **MIR6781** | 0.00355 | 0.01391 | 0.07979 | 0.03960 | 0.00807 | 0.04183 | 0.01082 | 0.05474 | 0.05276 | 0.02411 | 0.04583 | 0.04302 |
| **MIR6783** | 0.00573 | 0.01942 | 0.02661 | 0.02241 | 0.00849 | 0.00724 | 0.01883 | 0.01018 | 0.00949 | 0.02949 | 0.06751 | 0.02253 |
| **MIR6784** | 0.00567 | 0.02589 | 0.01262 | 0.01872 | 0.00930 | 0.01561 | 0.02147 | 0.01013 | 0.02160 | 0.03911 | 0.02446 | 0.01883 |
| **MIR6785** | 0.00546 | 0.01344 | 0.05025 | 0.02601 | 0.05098 | 0.00000 | 0.02019 | 0.00997 | 0.05758 | 0.09628 | 0.01431 | 0.03047 |
| **MIR6786** | 0.03300 | 0.10113 | 0.10279 | 0.02961 | 0.01562 | 0.02922 | 0.03880 | 0.01123 | 0.07760 | 0.00004 | 0.05829 | 0.04585 |
| **MIR6787** | 0.00621 | 0.01487 | 0.03080 | 0.00958 | 0.01303 | 0.01557 | 0.02644 | 0.01052 | 0.03661 | 0.01231 | 0.03634 | 0.01650 |
| **MIR6789** | 0.00593 | 0.03266 | 0.04499 | 0.01988 | 0.05592 | 0.00000 | 0.00897 | 0.01296 | 0.07388 | 0.00000 | 0.03904 | 0.02251 |
| **MIR6790** | 0.16700 | 0.12086 | 0.19388 | 0.24592 | 0.35061 | 0.00000 | 0.27478 | 0.10778 | 0.12470 | 0.56471 | 0.13400 | 0.31075 |
| **MIR6791** | 0.00662 | 0.01778 | 0.02021 | 0.04282 | 0.03959 | 0.00000 | 0.01728 | 0.01060 | 0.05236 | 0.00000 | 0.01278 | 0.01881 |
| **MIR6792** | 0.01011 | 0.01767 | 0.01817 | 0.01318 | 0.02916 | 0.00256 | 0.02367 | 0.01105 | 0.02441 | 0.01094 | 0.01497 | 0.01752 |
| **MIR6793** | 0.00561 | 0.01563 | 0.01598 | 0.01532 | 0.01695 | 0.01332 | 0.02397 | 0.01959 | 0.02096 | 0.01256 | 0.01438 | 0.01763 |

| | | | | | | | | | | | | |
|---|---|---|---|---|---|---|---|---|---|---|---|---|
| **MIR6794** | 0.00652 | 0.01471 | 0.01303 | 0.01208 | 0.01806 | 0.01560 | 0.01912 | 0.01028 | 0.02136 | 0.02074 | 0.01523 | 0.02224 |
| **MIR6796** | 0.02986 | 0.28503 | 0.32074 | 0.02425 | 0.09064 | 0.00000 | 0.00223 | 0.02378 | 0.35348 | 0.00000 | 0.07186 | 0.24745 |
| **MIR6797** | 2.46319 | 5.26650 | 4.99095 | 4.29566 | 8.35622 | 1.29118 | 8.75912 | 3.01509 | 8.22153 | 8.52470 | 2.19570 | 7.18628 |
| **MIR6798** | 0.01046 | 0.18742 | 0.11232 | 0.08846 | 0.23174 | 0.00006 | 0.12243 | 0.02312 | 0.24838 | 0.00000 | 0.22391 | 0.11383 |
| **MIR6799** | 0.00970 | 0.03609 | 0.04726 | 0.04454 | 0.25414 | 0.00000 | 0.01979 | 0.01059 | 0.12380 | 0.00916 | 0.00612 | 0.04932 |
| **MIR6800** | 0.00527 | 0.01440 | 0.01419 | 0.01363 | 0.01425 | 0.02001 | 0.01812 | 0.01660 | 0.02173 | 0.01293 | 0.01392 | 0.01948 |
| **MIR6801** | 0.00827 | 0.02983 | 0.01932 | 0.01273 | 0.02127 | 0.02706 | 0.03420 | 0.01281 | 0.01598 | 0.00495 | 0.01584 | 0.01719 |
| **MIR6802** | 0.00622 | 0.01776 | 0.01804 | 0.01625 | 0.01504 | 0.01481 | 0.01822 | 0.01057 | 0.02699 | 0.01682 | 0.01355 | 0.01604 |
| **MIR6803** | 0.00613 | 0.01119 | 0.00975 | 0.01783 | 0.02211 | 0.03421 | 0.02199 | 0.01039 | 0.02555 | 0.08243 | 0.01040 | 0.01766 |
| **MIR6806** | 0.00809 | 0.02376 | 0.01627 | 0.01159 | 0.01329 | 0.01752 | 0.02235 | 0.03252 | 0.01126 | 0.01261 | 0.02972 | 0.01766 |
| **MIR6807** | 0.13461 | 0.41176 | 0.16701 | 0.78534 | 1.90191 | 0.00000 | 1.94916 | 0.22339 | 0.00000 | 6.20731 | 0.11305 | 1.23385 |
| **MIR6810** | 0.00585 | 0.03505 | 0.02298 | 0.01051 | 0.01749 | 0.01083 | 0.01126 | 0.00954 | 0.02609 | 0.01724 | 0.03761 | 0.04040 |
| **MIR6812** | 0.00444 | 0.01395 | 0.01828 | 0.00918 | 0.01678 | 0.02201 | 0.01071 | 0.01323 | 0.05641 | 0.04993 | 0.01575 | 0.03430 |
| **MIR6816** | 0.00601 | 0.01837 | 0.02076 | 0.01441 | 0.01785 | 0.01314 | 0.01615 | 0.01000 | 0.02159 | 0.03278 | 0.01896 | 0.02074 |
| **MIR6817** | 0.01237 | 0.05131 | 0.11164 | 0.02283 | 0.11549 | 0.00000 | 0.05536 | 0.02575 | 0.08816 | 0.00000 | 0.07996 | 0.07305 |
| **MIR6818** | 0.02919 | 0.02159 | 0.02153 | 0.01515 | 0.01706 | 0.01440 | 0.01930 | 0.02180 | 0.01930 | 0.01373 | 0.01724 | 0.02320 |
| **MIR6820** | 0.07574 | 0.71466 | 0.86492 | 0.50957 | 0.80998 | 0.55034 | 0.80174 | 1.44386 | 0.55822 | 0.58132 | 0.49861 | 1.54801 |
| **MIR6821** | 0.07902 | 0.04962 | 0.14100 | 0.03371 | 0.14287 | 0.00000 | 0.02785 | 0.07246 | 0.13035 | 0.02168 | 0.01587 | 0.04974 |
| **MIR6822** | 0.00623 | 0.01492 | 0.01403 | 0.01261 | 0.01537 | 0.02157 | 0.01776 | 0.01038 | 0.02003 | 0.01347 | 0.01444 | 0.01737 |
| **MIR6825** | 0.00606 | 0.01508 | 0.02204 | 0.01351 | 0.01805 | 0.02099 | 0.01376 | 0.01042 | 0.02920 | 0.02306 | 0.01283 | 0.01524 |
| **MIR6826** | 0.05162 | 0.10760 | 0.19560 | 0.13248 | 0.09882 | 0.02218 | 0.07053 | 0.03596 | 0.13707 | 0.00000 | 0.22881 | 0.07056 |
| **MIR6828** | 0.00594 | 0.01455 | 0.01433 | 0.01260 | 0.01600 | 0.01598 | 0.01856 | 0.01153 | 0.01839 | 0.01418 | 0.01520 | 0.02253 |
| **MIR6831** | 0.00587 | 0.01883 | 0.01374 | 0.03804 | 0.01938 | 0.01875 | 0.00930 | 0.01091 | 0.04839 | 0.01691 | 0.01752 | 0.02945 |
| **MIR6832** | 0.00879 | 0.01970 | 0.03529 | 0.03247 | 0.02586 | 0.00000 | 0.01596 | 0.00932 | 0.05902 | 0.00420 | 0.01514 | 0.03359 |
| **MIR6833** | 0.00496 | 0.02293 | 0.02928 | 0.03403 | 0.00769 | 0.02491 | 0.03445 | 0.01969 | 0.03348 | 0.02642 | 0.06320 | 0.05373 |
| **MIR6834** | 0.04405 | 0.07890 | 0.14834 | 0.02532 | 0.09165 | 0.10535 | 0.00000 | 0.03350 | 0.09585 | 0.06028 | 0.24605 | 0.15141 |
| **MIR6835** | 0.00644 | 0.03662 | 0.06928 | 0.03626 | 0.07461 | 0.00000 | 0.01521 | 0.01554 | 0.08047 | 0.00000 | 0.04516 | 0.05973 |
| **MIR6836** | 0.01667 | 0.02709 | 0.01555 | 0.01590 | 0.01933 | 0.00468 | 0.02836 | 0.01980 | 0.01492 | 0.02017 | 0.01716 | 0.02817 |
| **MIR6837** | 0.01436 | 0.26375 | 0.12638 | 0.07162 | 0.10008 | 0.05374 | 0.01111 | 0.00973 | 0.06241 | 0.00000 | 0.18145 | 0.10119 |
| **MIR6839** | 0.14260 | 0.05028 | 0.03182 | 0.01861 | 0.01340 | 0.03562 | 0.00903 | 0.03136 | 0.04389 | 0.00939 | 0.01712 | 0.05281 |
| **MIR6840** | 0.00417 | 0.04102 | 0.04368 | 0.02953 | 0.01797 | 0.00000 | 0.00415 | 0.04589 | 0.07077 | 0.02828 | 0.04498 | 0.03286 |
| **MIR6842** | 0.00989 | 0.01720 | 0.01526 | 0.01824 | 0.01501 | 0.02504 | 0.01493 | 0.01143 | 0.01210 | 0.03333 | 0.01495 | 0.01661 |
| **MIR6845** | 0.00567 | 0.01601 | 0.01602 | 0.01273 | 0.03699 | 0.00557 | 0.01625 | 0.00991 | 0.01659 | 0.03023 | 0.01614 | 0.03161 |
| **MIR6847** | 0.01042 | 0.11247 | 0.10510 | 0.16052 | 0.18212 | 0.01361 | 0.16722 | 0.04121 | 0.22796 | 0.00000 | 0.12391 | 0.35160 |
| **MIR6848** | 0.00619 | 0.01381 | 0.02044 | 0.00984 | 0.02804 | 0.01892 | 0.01071 | 0.01024 | 0.02527 | 0.01953 | 0.01244 | 0.01841 |
| **MIR6849** | 0.00569 | 0.02852 | 0.01500 | 0.01031 | 0.00791 | 0.03391 | 0.01276 | 0.00984 | 0.01779 | 0.04384 | 0.01150 | 0.01749 |
| **MIR6850** | 0.96092 | 5.19585 | 3.40075 | 6.00247 | 4.53771 | 3.26187 | 4.01483 | 2.54687 | 8.11698 | 0.00000 | 3.92196 | 4.64638 |
| **MIR6851** | 0.00623 | 0.01500 | 0.01402 | 0.01201 | 0.01805 | 0.02214 | 0.01324 | 0.01046 | 0.02487 | 0.01687 | 0.01351 | 0.01642 |
| **MIR6853** | 0.00775 | 0.16824 | 0.03922 | 0.10978 | 0.10600 | 0.09561 | 0.02425 | 0.08256 | 0.06600 | 0.00000 | 0.07269 | 0.19877 |
| **MIR6855** | 0.01191 | 0.03071 | 0.01474 | 0.01131 | 0.01924 | 0.01316 | 0.02027 | 0.00982 | 0.02455 | 0.00786 | 0.01539 | 0.01565 |
| **MIR6856** | 0.02466 | 0.04102 | 0.02563 | 0.02230 | 0.06650 | 0.04026 | 0.03282 | 0.03676 | 0.08676 | 0.00000 | 0.02579 | 0.03848 |
| **MIR6857** | 0.03744 | 0.01796 | 0.05278 | 0.07031 | 0.06156 | 0.08482 | 0.00712 | 0.02132 | 0.10334 | 0.02988 | 0.03824 | 0.04597 |
| **MIR6858** | 0.00557 | 0.02263 | 0.03556 | 0.01582 | 0.03360 | 0.00000 | 0.01268 | 0.01374 | 0.04055 | 0.02572 | 0.01779 | 0.01609 |
| **MIR6861** | 0.00568 | 0.01684 | 0.01190 | 0.01587 | 0.00850 | 0.01588 | 0.01462 | 0.01051 | 0.02813 | 0.04520 | 0.01478 | 0.01431 |
| **MIR6863** | 0.00488 | 0.03386 | 0.01018 | 0.02759 | 0.05838 | 0.00000 | 0.01724 | 0.01018 | 0.04142 | 0.09002 | 0.08135 | 0.01769 |
| **MIR6865** | 0.12935 | 0.01429 | 0.01747 | 0.01176 | 0.01436 | 0.01572 | 0.01430 | 0.02300 | 0.02549 | 0.01541 | 0.01340 | 0.01677 |
| **MIR6866** | 0.03649 | 0.06278 | 0.03057 | 0.02119 | 0.00543 | 0.03289 | 0.02127 | 0.00696 | 0.08598 | 0.00318 | 0.03206 | 0.03325 |
| **MIR6869** | 0.15799 | 0.21333 | 0.55979 | 0.42098 | 0.21120 | 0.18106 | 0.28634 | 0.02188 | 0.58083 | 0.00000 | 0.20087 | 0.21468 |

| | | | | | | | | | | | | |
|---|---|---|---|---|---|---|---|---|---|---|---|---|
| **MIR6871** | 0.00745 | 0.02822 | 0.01956 | 0.01351 | 0.01525 | 0.01584 | 0.01661 | 0.00998 | 0.02652 | 0.03247 | 0.03609 | 0.01466 |
| **MIR6873** | 0.00540 | 0.01534 | 0.01997 | 0.01563 | 0.02470 | 0.00000 | 0.00112 | 0.01070 | 0.09751 | 0.07511 | 0.00621 | 0.01787 |
| **MIR6875** | 0.00633 | 0.01587 | 0.07377 | 0.01170 | 0.01818 | 0.00988 | 0.01828 | 0.01052 | 0.03720 | 0.00968 | 0.03170 | 0.01710 |
| **MIR6876** | 0.02120 | 0.01448 | 0.01391 | 0.01324 | 0.01575 | 0.02373 | 0.01556 | 0.01286 | 0.01734 | 0.01892 | 0.01419 | 0.01513 |
| **MIR6877** | 0.05440 | 0.20279 | 0.10945 | 0.05480 | 0.05726 | 0.03839 | 0.08937 | 0.02455 | 0.25153 | 0.00000 | 0.08417 | 0.16144 |
| **MIR6878** | 0.01070 | 0.19471 | 0.02730 | 0.04413 | 0.02010 | 0.07531 | 0.00111 | 0.02548 | 0.13246 | 0.00000 | 0.05198 | 0.08412 |
| **MIR6879** | 0.01752 | 0.41752 | 0.17161 | 0.26719 | 0.31221 | 0.02743 | 0.35320 | 0.01657 | 0.33658 | 0.00000 | 0.31500 | 0.13623 |
| **MIR6884** | 0.00604 | 0.01576 | 0.01589 | 0.02162 | 0.01771 | 0.01237 | 0.01893 | 0.01264 | 0.01890 | 0.01281 | 0.01537 | 0.02461 |
| **MIR6886** | 0.04593 | 0.01014 | 0.04395 | 0.11834 | 0.21435 | 0.00000 | 0.00384 | 0.01150 | 0.17885 | 0.02803 | 0.03004 | 0.01679 |
| **MIR6887** | 0.00619 | 0.02712 | 0.01901 | 0.00483 | 0.01566 | 0.05816 | 0.00851 | 0.01065 | 0.03019 | 0.02777 | 0.01681 | 0.01835 |
| **MIR6888** | 0.00668 | 0.01577 | 0.01545 | 0.01547 | 0.01442 | 0.01644 | 0.01667 | 0.01108 | 0.02045 | 0.01864 | 0.01436 | 0.01663 |
| **MIR6889** | 0.00798 | 0.01603 | 0.01396 | 0.01493 | 0.01407 | 0.01954 | 0.01297 | 0.01063 | 0.02452 | 0.01988 | 0.01494 | 0.01834 |
| **MIR6890** | 0.00532 | 0.05393 | 0.01864 | 0.01938 | 0.01870 | 0.00000 | 0.00370 | 0.01052 | 0.05897 | 0.06368 | 0.01913 | 0.01526 |
| **MIR6892** | 0.01739 | 0.07093 | 0.04876 | 0.01948 | 0.05959 | 0.00000 | 0.02075 | 0.01162 | 0.07347 | 0.00000 | 0.02859 | 0.02305 |
| **MIR6893** | 0.00623 | 0.01524 | 0.01426 | 0.01182 | 0.01419 | 0.01753 | 0.01784 | 0.01030 | 0.02475 | 0.01576 | 0.01407 | 0.01880 |
| **MIR6894** | 0.00614 | 0.01382 | 0.01554 | 0.01076 | 0.01405 | 0.02466 | 0.01638 | 0.01020 | 0.01609 | 0.01701 | 0.01560 | 0.01819 |
| **MIR7-1** | 0.01141 | 0.01123 | 0.01378 | 0.00763 | 0.00614 | 0.10235 | 0.00000 | 0.01037 | 0.08285 | 0.07312 | 0.02058 | 0.02359 |
| **MIR7-3HG** | 0.38874 | 0.02668 | 0.00993 | 0.00191 | 0.00135 | 0.11523 | 0.01304 | 0.51856 | 0.00000 | 0.03101 | 0.00890 | 0.00285 |
| **MIR7107** | 0.00697 | 0.01677 | 0.01373 | 0.01074 | 0.01594 | 0.02816 | 0.01695 | 0.01138 | 0.01629 | 0.01247 | 0.01391 | 0.02138 |
| **MIR7108** | 0.01034 | 0.05119 | 0.05861 | 0.01873 | 0.02316 | 0.01942 | 0.03661 | 0.01216 | 0.01735 | 0.03210 | 0.03720 | 0.01917 |
| **MIR7109** | 0.00788 | 0.03139 | 0.02509 | 0.03621 | 0.02249 | 0.00000 | 0.01364 | 0.01637 | 0.06507 | 0.00874 | 0.01986 | 0.02070 |
| **MIR711** | 0.00640 | 0.01677 | 0.01529 | 0.01400 | 0.01660 | 0.01388 | 0.01924 | 0.01055 | 0.02379 | 0.00944 | 0.01504 | 0.01684 |
| **MIR7110** | 0.00341 | 0.00000 | 0.04048 | 0.00605 | 0.11968 | 0.06701 | 0.02263 | 0.01113 | 0.05771 | 0.17598 | 0.00488 | 0.02143 |
| **MIR7111** | 0.00944 | 0.14280 | 0.08040 | 0.13370 | 0.24398 | 0.06380 | 0.00000 | 0.02646 | 0.20733 | 0.26560 | 0.00879 | 0.06784 |
| **MIR7113** | 0.01048 | 0.20556 | 0.12268 | 0.07534 | 0.08491 | 0.00947 | 0.23049 | 0.02996 | 0.26494 | 0.08976 | 0.05530 | 0.10533 |
| **MIR7114** | 0.00601 | 0.01285 | 0.01403 | 0.01149 | 0.01283 | 0.02474 | 0.01529 | 0.01557 | 0.01893 | 0.01670 | 0.01442 | 0.02432 |
| **MIR7161** | 0.01050 | 0.01362 | 0.01407 | 0.00845 | 0.00621 | 0.02298 | 0.01122 | 0.01724 | 0.02997 | 0.04995 | 0.01552 | 0.02275 |
| **MIR744** | 0.00617 | 0.01385 | 0.02097 | 0.02914 | 0.01242 | 0.01817 | 0.01436 | 0.01043 | 0.02458 | 0.01865 | 0.01695 | 0.01654 |
| **MIR761** | 0.00600 | 0.01436 | 0.01294 | 0.01572 | 0.01746 | 0.02075 | 0.01766 | 0.01068 | 0.01788 | 0.02230 | 0.01519 | 0.01831 |
| **MIR7703** | 0.00281 | 0.01838 | 0.01233 | 0.01966 | 0.02172 | 0.02169 | 0.03647 | 0.04393 | 0.02215 | 0.02630 | 0.02419 | 0.03623 |
| **MIR7846** | 0.00629 | 0.01467 | 0.02122 | 0.01162 | 0.01485 | 0.02056 | 0.01578 | 0.01045 | 0.01769 | 0.01395 | 0.01504 | 0.01615 |
| **MIR7851.uid-20** | 0.00610 | 0.01769 | 0.01641 | 0.01987 | 0.01732 | 0.01076 | 0.02038 | 0.01052 | 0.02181 | 0.02037 | 0.01890 | 0.01676 |
| **MIR7851.uid-46** | 0.00528 | 0.01546 | 0.01484 | 0.01349 | 0.03199 | 0.00662 | 0.02060 | 0.01375 | 0.01898 | 0.01652 | 0.01398 | 0.01889 |
| **MIR7854** | 0.00570 | 0.01339 | 0.01493 | 0.01746 | 0.00771 | 0.02502 | 0.00287 | 0.01015 | 0.03951 | 0.04322 | 0.01448 | 0.01695 |
| **MIR7856** | 0.00617 | 0.01436 | 0.01791 | 0.01085 | 0.01283 | 0.02101 | 0.01372 | 0.01044 | 0.02668 | 0.01957 | 0.01258 | 0.01570 |
| **MIR7974** | 0.00625 | 0.01524 | 0.01736 | 0.01267 | 0.01498 | 0.01784 | 0.01781 | 0.01045 | 0.01914 | 0.01400 | 0.01543 | 0.01730 |
| **MIR8055** | 0.02960 | 0.01686 | 0.02048 | 0.01687 | 0.00924 | 0.02233 | 0.01451 | 0.00815 | 0.03040 | 0.01306 | 0.03074 | 0.02695 |
| **MIR8069.uid-25** | 0.01037 | 0.01461 | 0.01362 | 0.01142 | 0.01392 | 0.02375 | 0.01734 | 0.01665 | 0.01858 | 0.01232 | 0.01463 | 0.01852 |
| **MIR8072** | 0.01041 | 0.21892 | 0.22333 | 0.08959 | 0.12838 | 0.32516 | 0.10383 | 0.55040 | 0.48744 | 0.00000 | 0.17682 | 0.31136 |
| **MIR8076** | 0.00669 | 0.01904 | 0.01438 | 0.01293 | 0.01481 | 0.01793 | 0.01851 | 0.01058 | 0.01958 | 0.01038 | 0.01627 | 0.02265 |
| **MIR8079** | 0.00611 | 0.01484 | 0.00929 | 0.01069 | 0.01161 | 0.01767 | 0.05148 | 0.01048 | 0.03033 | 0.01716 | 0.01345 | 0.01592 |
| **MIR8085** | 0.00629 | 0.01517 | 0.01868 | 0.01724 | 0.01407 | 0.01740 | 0.01566 | 0.01053 | 0.02454 | 0.01270 | 0.01452 | 0.01650 |
| **MIR877** | 0.00233 | 0.12305 | 0.08981 | 0.05743 | 0.08007 | 0.00435 | 0.12906 | 0.15830 | 0.26420 | 0.00000 | 0.05803 | 0.29734 |
| **MIR9-1** | 0.00617 | 0.01501 | 0.01494 | 0.01270 | 0.01747 | 0.01642 | 0.01936 | 0.01067 | 0.01890 | 0.00947 | 0.01681 | 0.02350 |
| **MIR93** | 0.00473 | 0.03349 | 0.00934 | 0.04430 | 0.01835 | 0.00180 | 0.01938 | 0.00881 | 0.04786 | 0.08716 | 0.01162 | 0.05034 |
| **MIR933** | 0.01037 | 0.01637 | 0.01511 | 0.01379 | 0.01856 | 0.01096 | 0.02109 | 0.01303 | 0.02452 | 0.01127 | 0.01445 | 0.01655 |
| **MIR934** | 0.00631 | 0.01496 | 0.01781 | 0.01130 | 0.01570 | 0.02069 | 0.01999 | 0.01047 | 0.01854 | 0.01102 | 0.01572 | 0.01639 |
| **MIR937** | 0.00634 | 0.01580 | 0.01575 | 0.01334 | 0.01889 | 0.01412 | 0.01768 | 0.01052 | 0.02328 | 0.01201 | 0.01497 | 0.01670 |

| | | | | | | | | | | | | |
|---|---|---|---|---|---|---|---|---|---|---|---|---|
| **MIR939** | 0.00595 | 0.01345 | 0.01880 | 0.05774 | 0.05345 | 0.00001 | 0.03123 | 0.00997 | 0.01042 | 0.13320 | 0.02994 | 0.02595 |
| **MIR940** | 0.00656 | 0.03465 | 0.02181 | 0.03407 | 0.01395 | 0.00497 | 0.03680 | 0.01015 | 0.01965 | 0.00864 | 0.02073 | 0.02772 |
| **MIR943** | 0.00592 | 0.01240 | 0.01952 | 0.01627 | 0.01611 | 0.02942 | 0.01811 | 0.01030 | 0.01650 | 0.02576 | 0.01367 | 0.01603 |
| **MIR98** | 0.00494 | 0.02050 | 0.01245 | 0.06110 | 0.07440 | 0.00002 | 0.01454 | 0.00972 | 0.06245 | 0.05666 | 0.02440 | 0.03210 |
| **MIRLET7A1** | 0.00564 | 0.01238 | 0.01802 | 0.00908 | 0.00840 | 0.02475 | 0.01111 | 0.00982 | 0.02350 | 0.04011 | 0.02214 | 0.02396 |
| **MIRLET7BHG** | 0.01560 | 0.02096 | 0.03062 | 0.01725 | 0.02438 | 0.00113 | 0.03513 | 0.00951 | 0.01159 | 0.01862 | 0.02784 | 0.01745 |
| **MIRLET7D** | 0.00904 | 0.01813 | 0.00154 | 0.00928 | 0.05160 | 0.11733 | 0.00000 | 0.00868 | 0.02853 | 0.09330 | 0.11016 | 0.05910 |
| **MIRLET7DHG** | 0.02836 | 0.19962 | 0.13036 | 0.46148 | 0.09499 | 0.26655 | 0.00000 | 0.13760 | 0.71844 | 0.66208 | 0.38112 | 0.44764 |
| **MIRLET7F1** | 0.00600 | 0.03177 | 0.02255 | 0.01348 | 0.01952 | 0.04360 | 0.00000 | 0.00889 | 0.08937 | 0.01307 | 0.01825 | 0.03894 |
| **MIRLET7F2** | 0.00690 | 0.01484 | 0.01447 | 0.01249 | 0.01558 | 0.01831 | 0.01670 | 0.01047 | 0.02206 | 0.01421 | 0.01509 | 0.01791 |
| **MIRLET7G** | 0.00612 | 0.01399 | 0.02011 | 0.01140 | 0.01170 | 0.02215 | 0.01055 | 0.01024 | 0.03844 | 0.02356 | 0.01137 | 0.01784 |
| **MIRLET7I** | 0.00877 | 0.01170 | 0.06905 | 0.00812 | 0.00865 | 0.05019 | 0.00000 | 0.00978 | 0.01895 | 0.06957 | 0.01547 | 0.01686 |
| **MIS12** | 0.74224 | 4.02446 | 4.26126 | 3.45479 | 2.25435 | 9.59299 | 4.95685 | 1.89183 | 7.36377 | 0.00000 | 4.25370 | 6.84803 |
| **MIS18A** | 7.39996 | 18.06871 | 5.61318 | 7.35336 | 9.21088 | 17.24723 | 20.45241 | 6.99765 | 5.66542 | 9.81820 | 10.82350 | 16.85501 |
| **MIS18BP1** | 3.39620 | 0.12044 | 4.61550 | 2.07980 | 0.00000 | 27.61086 | 6.70053 | 7.68999 | 10.15794 | 21.78326 | 3.98473 | 7.69896 |
| **MISP** | 0.47594 | 6.10469 | 0.00000 | 0.00000 | 7.27341 | 0.00000 | 3.29603 | 0.54796 | 5.45499 | 0.82574 | 5.85450 | 1.86599 |
| **MITD1** | 0.71781 | 2.67098 | 6.07348 | 0.31391 | 0.17021 | 8.92826 | 3.23969 | 0.03042 | 4.72961 | 3.47126 | 6.79116 | 3.62533 |
| **MITF** | 4.69307 | 0.75024 | 3.38338 | 0.00000 | 0.98073 | 2.15947 | 1.63094 | 1.06798 | 2.87367 | 2.47049 | 2.18066 | 0.72573 |
| **MIXL1** | 6.64661 | 0.04537 | 0.83354 | 0.74187 | 0.00000 | 0.00000 | 0.30993 | 0.54849 | 0.00000 | 1.08324 | 0.01703 | 0.21045 |
| **MKI67** | 0.62548 | 8.26638 | 13.09917 | 6.92273 | 23.51927 | 20.28930 | 10.31366 | 9.97889 | 17.59659 | 48.38566 | 2.67548 | 16.83581 |
| **MKKS** | 0.00000 | 17.00986 | 1.99079 | 5.75000 | 12.86525 | 37.66720 | 6.53795 | 0.00000 | 13.32173 | 11.48454 | 16.60732 | 20.08826 |
| **MKL1** | 5.96772 | 5.61742 | 4.92757 | 3.83435 | 4.46859 | 0.00000 | 1.84922 | 1.59623 | 5.54652 | 5.59595 | 2.66383 | 1.93723 |
| **MKL2** | 2.76907 | 0.98899 | 2.11539 | 0.33141 | 0.58305 | 3.84821 | 3.66442 | 0.23861 | 7.16738 | 2.44147 | 2.48257 | 1.25035 |
| **MKLN1** | 3.71860 | 4.03088 | 3.14279 | 5.33580 | 2.20290 | 18.80203 | 6.45636 | 2.58172 | 8.57647 | 11.95264 | 10.30909 | 4.03105 |
| **MKNK1** | 1.75802 | 10.28141 | 3.70194 | 5.01636 | 4.57114 | 9.15870 | 1.89028 | 1.47816 | 6.17658 | 0.00113 | 4.98119 | 6.45579 |
| **MKNK1-AS1** | 0.05304 | 0.02057 | 0.01903 | 0.07885 | 0.00000 | 0.06456 | 0.00369 | 0.00623 | 0.00000 | 0.11208 | 0.10190 | 0.06751 |
| **MKNK2** | 5.97631 | 1.07859 | 5.51228 | 2.55489 | 7.72963 | 0.00000 | 16.05829 | 0.82659 | 8.23252 | 18.71637 | 1.70478 | 2.01469 |
| **MKRN1** | 4.61617 | 16.03820 | 4.65768 | 18.96527 | 7.00664 | 13.05306 | 11.03063 | 20.19282 | 1.14965 | 24.28649 | 4.64695 | 23.41215 |
| **MKRN2** | 2.50468 | 8.75888 | 4.52984 | 2.72101 | 2.33250 | 12.19163 | 1.52163 | 12.33954 | 6.94001 | 0.00000 | 8.81703 | 8.94583 |
| **MKRN2OS** | 0.00540 | 0.03302 | 0.08158 | 0.14364 | 0.03843 | 0.00000 | 0.00773 | 0.00802 | 0.00032 | 0.05958 | 0.16555 | 0.06187 |
| **MKRN3** | 0.02010 | 2.52879 | 3.80674 | 0.45181 | 0.00000 | 0.37972 | 1.09133 | 1.04840 | 2.46939 | 0.00000 | 6.41761 | 3.50815 |
| **MKRN7P** | 0.02883 | 0.00646 | 0.06843 | 0.01660 | 0.00199 | 0.06649 | 0.00955 | 0.01840 | 0.00000 | 0.05632 | 0.01412 | 0.04586 |
| **MKRN9P** | 0.24027 | 0.74137 | 0.74754 | 0.57885 | 0.00000 | 0.19051 | 0.02968 | 1.17474 | 0.19633 | 0.49705 | 0.44944 | 0.76732 |
| **MKS1** | 1.92007 | 9.24403 | 1.19868 | 4.03363 | 0.63902 | 6.04542 | 1.52642 | 0.79999 | 1.58194 | 0.00000 | 5.53281 | 6.88158 |
| **MKX** | 0.02072 | 0.02540 | 0.01812 | 0.13707 | 0.00101 | 0.08860 | 0.00422 | 0.00662 | 0.00000 | 0.08242 | 0.00619 | 0.02172 |
| **MLANA** | 0.00417 | 0.02178 | 0.00961 | 0.01613 | 0.01056 | 0.02150 | 0.00719 | 0.01609 | 0.03512 | 0.05887 | 0.01314 | 0.04075 |
| **MLC1** | 0.00526 | 0.00594 | 0.00449 | 0.13051 | 0.00992 | 0.04574 | 0.01051 | 0.00967 | 0.01345 | 0.05279 | 0.01271 | 0.01520 |
| **MLEC** | 0.14573 | 8.96878 | 7.24834 | 13.35151 | 15.32361 | 4.49817 | 6.86530 | 2.78732 | 10.17111 | 30.80026 | 3.97894 | 5.66838 |
| **1.00000** | 6.77190 | 2.52353 | 1.54886 | 0.01611 | 0.00000 | 3.16224 | 0.83851 | 10.91771 | 0.18485 | 0.00000 | 0.42621 | 13.40326 |
| **2.00000** | 1.47223 | 20.95977 | 24.60862 | 23.38208 | 22.15063 | 14.72050 | 27.85711 | 18.65704 | 29.92037 | 0.00000 | 20.87451 | 21.54840 |
| **MLH1** | 0.44631 | 5.08201 | 4.46673 | 2.88945 | 1.75535 | 10.84060 | 1.83722 | 4.00859 | 6.25383 | 1.87425 | 3.24479 | 7.15958 |
| **MLH3** | 0.00000 | 0.81315 | 4.11715 | 0.53079 | 0.92023 | 3.14633 | 0.01180 | 0.33882 | 5.40081 | 2.84377 | 2.51894 | 1.49281 |
| **MLIP** | 0.10359 | 0.01631 | 0.00632 | 0.22174 | 0.01821 | 0.03038 | 0.01273 | 0.02721 | 0.01879 | 0.00322 | 0.01523 | 0.01214 |
| **MLIP-IT1** | 0.00881 | 0.01043 | 0.01294 | 0.00755 | 0.01620 | 0.04648 | 0.01155 | 0.00719 | 0.01079 | 0.02324 | 0.01280 | 0.09900 |
| **MLK7-AS1** | 0.41085 | 0.08199 | 2.03804 | 0.00000 | 0.00000 | 0.23073 | 0.00008 | 0.87987 | 1.14753 | 0.38401 | 0.00000 | 0.17613 |
| **MLKL** | 0.18540 | 0.05155 | 0.16295 | 0.37474 | 0.00000 | 0.07491 | 0.00000 | 0.19458 | 0.01981 | 0.00746 | 0.01353 | 0.07185 |
| **MLLT1** | 1.55459 | 2.03118 | 11.16860 | 3.04356 | 2.85588 | 0.00000 | 7.43208 | 0.07658 | 4.70430 | 11.02971 | 0.49010 | 2.80693 |
| **MLLT10** | 1.94638 | 0.83361 | 5.92422 | 2.55801 | 2.14954 | 11.66475 | 3.99286 | 3.33145 | 4.48796 | 21.37612 | 2.06393 | 4.17525 |

| | | | | | | | | | | | | |
|---|---|---|---|---|---|---|---|---|---|---|---|---|
| **MLLT10P1** | 0.01689 | 0.11914 | 0.09130 | 0.00028 | 0.00000 | 0.54032 | 0.00000 | 0.14788 | 0.25193 | 0.32148 | 0.01611 | 0.24464 |
| **MLLT11** | 2.61009 | 0.31632 | 19.00772 | 0.30529 | 0.00000 | 0.61034 | 2.72905 | 1.29376 | 0.40272 | 0.00000 | 0.17431 | 1.48299 |
| **MLLT3** | 2.92805 | 0.64669 | 0.22090 | 2.28489 | 0.00000 | 3.30203 | 0.00000 | 0.11289 | 0.00000 | 2.04955 | 0.45862 | 0.00000 |
| **MLLT4** | 3.07525 | 0.41201 | 3.35592 | 0.82763 | 5.95265 | 11.95163 | 10.53519 | 1.59726 | 0.74697 | 29.79450 | 2.09335 | 2.25201 |
| **MLLT4-AS1** | 0.00423 | 0.04640 | 0.34507 | 0.10896 | 0.00426 | 0.18702 | 0.00000 | 0.41545 | 0.19968 | 0.00000 | 0.56577 | 0.26918 |
| **MLLT6** | 0.85431 | 0.90897 | 0.56621 | 2.99858 | 2.85545 | 0.23927 | 0.12545 | 0.00675 | 0.80468 | 1.11813 | 0.50752 | 0.60111 |
| **MLN** | 0.00607 | 0.01882 | 0.00497 | 0.30360 | 0.03567 | 0.00809 | 0.02182 | 0.01035 | 0.03028 | 0.00857 | 0.01417 | 0.01623 |
| **MLNR** | 0.00510 | 0.00666 | 0.01029 | 0.06773 | 0.13686 | 0.00000 | 0.00986 | 0.30423 | 0.04087 | 0.00000 | 0.02034 | 0.56922 |
| **MLPH** | 0.52348 | 0.01237 | 0.32225 | 0.10714 | 0.00974 | 0.00000 | 0.04734 | 0.01702 | 0.01442 | 0.00051 | 0.02498 | 0.04707 |
| **MLST8** | 0.67660 | 13.08342 | 9.79293 | 10.15374 | 10.33808 | 0.00000 | 12.57579 | 0.30256 | 19.97760 | 1.58651 | 12.50583 | 5.26996 |
| **MLX** | 7.96668 | 11.08775 | 17.62456 | 8.84298 | 3.79296 | 9.93972 | 6.79323 | 4.88106 | 13.30764 | 0.00000 | 12.10758 | 14.36970 |
| **MLXIP** | 4.09310 | 9.06561 | 0.00000 | 7.32292 | 10.78248 | 0.72842 | 6.19740 | 0.17557 | 0.02250 | 15.97259 | 3.12473 | 5.94807 |
| **MLXIPL** | 2.97398 | 0.11055 | 0.00000 | 0.00000 | 0.34239 | 0.00000 | 0.20899 | 0.12790 | 0.00000 | 0.86459 | 0.06991 | 0.01007 |
| **MLYCD** | 0.78416 | 0.22291 | 0.19117 | 0.92592 | 1.89585 | 0.37589 | 1.93565 | 0.50328 | 0.00280 | 0.00013 | 0.29179 | 0.13965 |
| **MMAA** | 1.60075 | 0.50196 | 0.19251 | 0.79517 | 0.00629 | 1.69143 | 0.02821 | 0.70622 | 0.46177 | 0.00000 | 0.41692 | 5.43025 |
| **MMAB** | 0.00000 | 0.00000 | 4.22911 | 3.95059 | 12.65873 | 3.20057 | 8.41082 | 0.03308 | 5.03741 | 1.57426 | 10.39242 | 0.30999 |
| **MMACHC** | 0.45420 | 3.52858 | 1.50734 | 1.14811 | 1.68945 | 0.76116 | 2.45353 | 2.26117 | 2.21298 | 1.43110 | 5.27819 | 8.16646 |
| **MMADHC** | 0.00000 | 18.11363 | 20.26966 | 6.52412 | 6.43080 | 42.62667 | 26.71496 | 21.20783 | 24.38508 | 3.96577 | 18.82220 | 21.02443 |
| **MMD** | 0.27163 | 0.37738 | 2.77303 | 2.35514 | 0.00000 | 1.22906 | 1.35416 | 0.66017 | 2.84108 | 0.05140 | 0.34649 | 0.52814 |
| **MME** | 0.10017 | 0.10164 | 2.77341 | 0.00000 | 0.09733 | 0.09976 | 0.05762 | 0.08211 | 0.00000 | 0.00512 | 0.10257 | 0.01409 |
| **MMEL1** | 0.00719 | 0.00983 | 0.01720 | 0.00859 | 0.02907 | 0.02874 | 0.02925 | 0.00936 | 0.00467 | 0.03235 | 0.01271 | 0.02245 |
| **MMGT1** | 7.12679 | 7.37262 | 2.12180 | 0.00000 | 0.00000 | 16.72920 | 2.85038 | 4.42854 | 11.27986 | 6.46276 | 4.08064 | 10.19208 |
| **MMP11** | 0.14219 | 0.36485 | 0.09988 | 0.05567 | 0.20691 | 0.00000 | 0.06149 | 0.02242 | 0.20288 | 0.00000 | 0.00834 | 0.00175 |
| **MMP14** | 0.00111 | 0.00000 | 2.79498 | 0.04709 | 0.10473 | 0.00000 | 0.14059 | 0.04505 | 0.92860 | 0.85378 | 0.13694 | 0.14599 |
| **MMP15** | 0.13079 | 0.39755 | 1.00606 | 1.39761 | 3.12821 | 0.00000 | 0.00000 | 0.26832 | 0.00000 | 1.79437 | 0.00000 | 0.64337 |
| **MMP16** | 0.03407 | 0.00000 | 0.09452 | 0.00000 | 0.09084 | 0.05212 | 0.11679 | 0.00821 | 0.19389 | 0.07043 | 0.06694 | 0.01639 |
| **MMP17** | 0.00235 | 0.18765 | 0.94857 | 0.08805 | 0.69133 | 0.00000 | 0.00000 | 0.01572 | 1.02136 | 0.30549 | 0.44283 | 0.01761 |
| **MMP19** | 0.26707 | 0.02235 | 3.35169 | 0.00000 | 0.00000 | 0.04690 | 0.00000 | 0.00000 | 1.07370 | 0.06471 | 0.00000 | 0.07887 |
| **MMP2** | 0.44214 | 0.72568 | 0.10056 | 5.93510 | 0.00000 | 0.27333 | 0.00000 | 0.00000 | 0.00000 | 0.02253 | 0.00000 | 0.15563 |
| **MMP20** | 0.00579 | 0.01065 | 0.03162 | 0.01738 | 0.01412 | 0.03266 | 0.03336 | 0.01025 | 0.02911 | 0.02776 | 0.01522 | 0.01423 |
| **MMP21** | 0.01092 | 0.00718 | 0.02683 | 0.00652 | 0.00879 | 0.03897 | 0.00000 | 0.01211 | 0.06793 | 0.02712 | 0.02560 | 0.04098 |
| **MMP24** | 0.53066 | 0.72842 | 0.28797 | 0.81455 | 0.61809 | 0.00000 | 0.00000 | 0.03569 | 0.00000 | 0.28273 | 0.06620 | 1.81583 |
| **MMP24-AS1** | 0.32055 | 0.29320 | 0.13875 | 0.97739 | 1.41126 | 0.00000 | 0.39824 | 0.14339 | 0.00000 | 2.39822 | 0.00000 | 1.27540 |
| **MMP25** | 0.02228 | 0.09670 | 0.06294 | 0.07180 | 0.02910 | 0.00000 | 0.00000 | 0.00874 | 0.25538 | 0.13716 | 0.15881 | 0.08149 |
| **MMP28** | 0.19745 | 0.01604 | 0.04454 | 0.01135 | 0.01920 | 0.00508 | 0.02787 | 0.07597 | 0.01161 | 0.01335 | 0.01541 | 0.01239 |
| **MMP8** | 0.00467 | 0.01470 | 0.02997 | 0.00953 | 0.01487 | 0.02561 | 0.02946 | 0.01424 | 0.00004 | 0.06495 | 0.01578 | 0.01641 |
| **MMP9** | 0.15829 | 0.03186 | 0.11111 | 1.01424 | 0.00000 | 0.00000 | 0.03601 | 0.38835 | 0.02584 | 0.03752 | 0.01504 | 0.06492 |
| **MMRN1** | 0.01495 | 0.01348 | 0.01674 | 0.01060 | 0.04107 | 0.00133 | 0.04550 | 0.01982 | 0.00000 | 0.04679 | 0.01304 | 0.01671 |
| **MMRN2** | 0.00397 | 0.11896 | 0.23614 | 0.07062 | 0.00000 | 0.02130 | 0.31109 | 0.00218 | 0.12147 | 0.35029 | 0.04455 | 0.74783 |
| **MMS19** | 3.58495 | 11.23212 | 7.72247 | 12.27510 | 5.54240 | 3.81234 | 3.84934 | 1.75933 | 11.73592 | 6.45577 | 7.93297 | 6.11341 |
| **MMS22L** | 2.41594 | 4.76474 | 4.66387 | 0.95597 | 0.00000 | 13.23226 | 5.82742 | 2.60053 | 5.73505 | 4.75668 | 4.17475 | 3.87664 |
| **MN1** | 0.09470 | 0.01121 | 0.10657 | 0.28282 | 0.01795 | 0.00000 | 0.03844 | 0.00729 | 0.09173 | 0.00000 | 0.01569 | 0.08460 |
| **MNAT1** | 2.19963 | 2.87204 | 1.72587 | 1.00691 | 4.99012 | 15.78889 | 4.41110 | 0.33378 | 3.93859 | 5.69374 | 4.88854 | 6.34016 |
| **MND1** | 2.71169 | 5.97319 | 0.00000 | 1.75077 | 2.26529 | 12.52897 | 1.53038 | 0.44700 | 3.55059 | 4.92245 | 3.66428 | 3.07474 |
| **MNDA** | 0.00602 | 0.01317 | 0.00615 | 0.18741 | 0.00000 | 0.03644 | 0.01267 | 0.01010 | 0.01523 | 0.01468 | 0.01710 | 0.01429 |
| **MNS1** | 0.76140 | 0.87451 | 0.00000 | 0.16930 | 0.51336 | 1.38904 | 0.05458 | 1.44742 | 0.00000 | 1.43943 | 0.31946 | 3.20413 |
| **MNT** | 2.13187 | 0.00434 | 0.44111 | 0.79376 | 0.65694 | 0.00000 | 1.20956 | 0.09570 | 1.00965 | 7.07073 | 0.00000 | 0.41097 |
| **MNX1** | 0.02034 | 0.03372 | 0.00118 | 0.03648 | 0.29220 | 0.00000 | 0.10860 | 0.00719 | 0.09375 | 0.55216 | 0.00000 | 0.00872 |

| | | | | | | | | | | | | |
|---|---|---|---|---|---|---|---|---|---|---|---|---|
| **MNX1-AS1** | 0.02126 | 0.02192 | 0.02662 | 0.00002 | 0.24325 | 0.06860 | 0.00308 | 0.00982 | 0.00000 | 0.11801 | 0.01633 | 0.00716 |
| **MOAP1** | 0.00205 | 4.12353 | 3.55271 | 0.75146 | 0.00000 | 2.58754 | 4.64103 | 5.43492 | 3.93295 | 0.00000 | 2.58152 | 6.83283 |
| **MOB1A** | 1.92327 | 7.23251 | 7.81039 | 3.98057 | 0.55920 | 28.45177 | 7.52936 | 4.65348 | 13.33013 | 0.00000 | 5.73509 | 7.47585 |
| **MOB1B** | 1.93218 | 2.19260 | 5.29844 | 0.00000 | 0.00000 | 9.27997 | 11.31577 | 6.10459 | 3.97507 | 9.60996 | 3.58304 | 1.87553 |
| **MOB2** | 0.33381 | 3.56337 | 5.97289 | 0.79970 | 2.67937 | 0.00000 | 0.95000 | 1.39581 | 1.30489 | 6.93266 | 1.14140 | 1.78631 |
| **MOB3A** | 1.05858 | 3.85449 | 0.69461 | 1.52127 | 2.56474 | 0.15203 | 0.76631 | 1.15540 | 2.11400 | 0.00000 | 0.24063 | 2.77623 |
| **MOB3B** | 3.53584 | 8.86187 | 10.36425 | 11.19086 | 22.16714 | 11.15443 | 23.95718 | 0.00000 | 19.79391 | 1.90068 | 22.68920 | 0.00000 |
| **MOB3C** | 0.03024 | 0.00785 | 0.20737 | 0.02279 | 0.00000 | 0.11944 | 0.00000 | 0.00508 | 0.00000 | 0.02136 | 0.01107 | 0.07791 |
| **MOB4.uid-3968** | 0.60218 | 1.69757 | 1.66400 | 0.82355 | 0.45658 | 1.98568 | 0.00000 | 0.45358 | 1.20607 | 0.00000 | 0.72628 | 1.59809 |
| **MOB4.uid-3970** | 0.01291 | 0.01683 | 0.02164 | 0.02050 | 0.02093 | 0.01303 | 0.02719 | 0.01530 | 0.02003 | 0.00128 | 0.02248 | 0.02122 |
| **MOBP** | 0.00426 | 0.20879 | 0.17021 | 0.58444 | 0.00000 | 0.00000 | 0.00000 | 0.20284 | 0.00000 | 0.44586 | 0.00041 | 0.48234 |
| **MOCOS** | 0.94989 | 0.00000 | 0.38169 | 0.22483 | 0.32687 | 0.00000 | 0.58214 | 0.02945 | 0.54285 | 0.48492 | 0.74887 | 0.02178 |
| **MOCS1** | 0.07118 | 1.59489 | 2.44759 | 1.06609 | 1.64171 | 1.40196 | 0.00009 | 0.02191 | 4.62441 | 0.00000 | 1.38875 | 0.06333 |
| **MOCS2** | 1.73652 | 6.16503 | 8.79819 | 5.01738 | 1.01063 | 13.11187 | 7.25000 | 0.13244 | 2.17899 | 0.04690 | 8.00331 | 4.91902 |
| **MOCS3** | 0.63750 | 5.53325 | 3.93636 | 3.13412 | 1.35549 | 6.34561 | 1.82118 | 1.14770 | 3.87795 | 0.00000 | 3.34947 | 7.01282 |
| **MOG** | 0.00389 | 0.00000 | 0.24266 | 0.37936 | 0.00000 | 0.00000 | 0.00000 | 0.00520 | 0.04670 | 0.26334 | 0.10470 | 0.16352 |
| **MOGAT1** | 0.00225 | 0.01212 | 0.02404 | 0.02331 | 0.01391 | 0.01580 | 0.02008 | 0.05268 | 0.01250 | 0.01993 | 0.03301 | 0.02823 |
| **MOGAT2** | 0.00634 | 0.02282 | 0.00309 | 0.09156 | 0.00139 | 0.05862 | 0.00000 | 0.00932 | 0.01084 | 0.06011 | 0.01049 | 0.01962 |
| **MOGAT3** | 0.01391 | 0.44897 | 0.00002 | 1.08686 | 0.65307 | 0.12690 | 0.00000 | 0.01001 | 0.00000 | 0.02823 | 0.03496 | 0.00000 |
| **MOGS** | 0.04024 | 3.22286 | 8.07214 | 3.08835 | 7.72186 | 0.00000 | 0.00000 | 0.15494 | 10.42266 | 14.48774 | 4.80381 | 5.21493 |
| **MOK** | 4.61615 | 1.12644 | 0.87966 | 0.00000 | 0.00000 | 0.18939 | 1.19259 | 7.44838 | 0.22843 | 0.52084 | 0.36121 | 3.62258 |
| **MON1A** | 0.09695 | 4.44996 | 2.33006 | 3.91060 | 3.63894 | 0.00000 | 5.18806 | 1.11318 | 5.35810 | 2.04443 | 2.60408 | 6.72895 |
| **MON1B** | 2.41447 | 6.17622 | 10.64917 | 3.18093 | 7.08367 | 0.62267 | 8.97642 | 2.68469 | 11.77840 | 0.00000 | 4.00894 | 5.08709 |
| **MON2** | 2.60224 | 0.58474 | 5.03915 | 0.06902 | 1.01539 | 6.38709 | 1.45114 | 4.06559 | 3.91146 | 4.14691 | 3.70306 | 0.93924 |
| **MORC1** | 0.24262 | 0.21174 | 0.22863 | 0.05505 | 0.00000 | 0.04642 | 0.00000 | 0.08637 | 0.00000 | 0.33406 | 0.00583 | 0.10385 |
| **MORC2** | 1.55552 | 1.95761 | 3.64907 | 1.95113 | 1.98438 | 1.86929 | 7.75026 | 0.94442 | 5.16511 | 8.09954 | 1.26754 | 4.95861 |
| **MORC2-AS1** | 1.23447 | 1.89500 | 7.56198 | 7.13193 | 1.75100 | 0.90996 | 11.56314 | 1.18623 | 2.54652 | 3.97187 | 7.71746 | 4.52285 |
| **MORC3** | 0.22425 | 3.86895 | 5.12906 | 2.70263 | 0.00023 | 12.74856 | 5.94040 | 8.22185 | 6.93693 | 4.71934 | 4.15649 | 7.74707 |
| **MORC4** | 0.56329 | 0.86655 | 4.45198 | 0.96535 | 6.31642 | 5.63490 | 5.20294 | 0.00000 | 0.00036 | 7.35802 | 4.05043 | 0.33308 |
| **MORF4L1** | 3.85616 | 7.06703 | 12.00410 | 10.02051 | 5.44665 | 26.13094 | 21.78627 | 1.81771 | 4.05532 | 15.63217 | 12.26931 | 10.08198 |
| **MORF4L2** | 1.79171 | 10.83056 | 24.87175 | 4.45371 | 13.10222 | 45.63182 | 28.75444 | 6.11764 | 26.20185 | 3.18265 | 20.48724 | 12.94186 |
| **MORF4L2-AS1** | 0.00554 | 0.04128 | 0.17374 | 0.00000 | 0.00000 | 0.00000 | 0.00060 | 0.03783 | 0.66420 | 0.27462 | 0.24012 | 0.04923 |
| **MORN1** | 0.05153 | 0.37728 | 0.12734 | 0.13598 | 0.23760 | 0.00000 | 0.00287 | 0.01913 | 0.11950 | 0.00000 | 0.56772 | 0.78622 |
| **MORN2** | 3.57806 | 5.36272 | 2.38692 | 0.00000 | 0.00000 | 10.64894 | 2.43253 | 1.57159 | 2.74200 | 4.49529 | 1.70759 | 12.17988 |
| **MORN3** | 0.33008 | 0.00000 | 0.84830 | 0.00059 | 0.42682 | 0.06058 | 0.34398 | 0.37577 | 0.23126 | 0.00000 | 0.13765 | 1.27149 |
| **MORN4** | 0.75216 | 0.32948 | 0.14629 | 0.04758 | 0.00000 | 0.00000 | 0.00422 | 0.09120 | 0.06876 | 0.04831 | 0.18945 | 0.20457 |
| **MORN5** | 0.00933 | 0.01889 | 0.04066 | 0.01009 | 0.07613 | 0.00000 | 0.01879 | 0.01091 | 0.01491 | 0.01345 | 0.01169 | 0.01888 |
| **MOS** | 2.53541 | 0.02095 | 0.02317 | 0.00750 | 0.01279 | 0.01705 | 0.01700 | 0.06129 | 0.01722 | 0.00630 | 0.01203 | 0.05749 |
| **MOSPD1** | 0.00000 | 4.66953 | 1.11088 | 2.18955 | 0.70694 | 19.45054 | 5.17529 | 18.17216 | 0.00334 | 3.78553 | 3.74848 | 18.94168 |
| **MOSPD2** | 1.61908 | 0.07242 | 3.29343 | 0.07175 | 0.21984 | 7.04263 | 0.88146 | 2.56195 | 1.78454 | 0.91729 | 1.82739 | 1.33788 |
| **MOSPD3** | 0.24416 | 5.84565 | 2.40924 | 1.38666 | 0.67421 | 0.00000 | 0.82803 | 0.69365 | 0.67388 | 1.73175 | 1.02009 | 3.14984 |
| **MOV10** | 2.02302 | 8.29154 | 2.20658 | 6.17558 | 3.23340 | 2.45568 | 3.29645 | 0.18448 | 8.72997 | 7.31506 | 4.41933 | 1.56418 |
| **MOV10L1** | 0.11293 | 0.04433 | 0.07204 | 0.00581 | 0.04618 | 0.00408 | 0.09227 | 1.05614 | 0.00961 | 0.05788 | 0.00517 | 0.58158 |
| **MOXD1** | 0.00975 | 0.02011 | 0.01512 | 0.01216 | 0.00945 | 0.01233 | 0.01539 | 0.03379 | 0.01202 | 0.01695 | 0.05672 | 0.01426 |
| **MPC1** | 2.61716 | 1.58646 | 4.95934 | 6.42642 | 2.50806 | 8.53065 | 5.75889 | 1.14030 | 6.90131 | 0.84256 | 5.89117 | 0.05676 |
| **MPC1L** | 0.39490 | 0.58909 | 0.00004 | 0.00000 | 0.12015 | 0.11110 | 0.00000 | 0.00000 | 0.00000 | 0.05021 | 0.00000 | 2.42738 |
| **MPC2** | 3.29725 | 5.49622 | 0.00000 | 6.88875 | 0.51124 | 9.59985 | 0.00000 | 2.37234 | 0.00000 | 4.45827 | 6.14226 | 3.10757 |
| **MPDU1** | 0.00000 | 15.91322 | 20.59444 | 12.62228 | 15.24749 | 0.00000 | 9.23784 | 1.02125 | 30.35359 | 9.49869 | 13.89920 | 13.20354 |

| | | | | | | | | | | | | |
|---|---|---|---|---|---|---|---|---|---|---|---|---|
| **MPDZ** | 0.39522 | 0.00000 | 1.45874 | 2.22302 | 0.32924 | 2.86020 | 3.99174 | 0.17574 | 1.32036 | 3.53210 | 1.90265 | 2.85690 |
| **MPEG1** | 0.00658 | 0.01389 | 0.01475 | 0.01263 | 0.01546 | 0.02571 | 0.01919 | 0.01162 | 0.01868 | 0.01183 | 0.02246 | 0.01643 |
| **MPG** | 8.25260 | 1.05433 | 5.06151 | 4.38694 | 4.51668 | 0.00000 | 2.90802 | 4.05968 | 8.83610 | 0.00000 | 8.20180 | 0.00000 |
| **MPHOSPH10** | 0.98297 | 10.46538 | 7.59496 | 2.71397 | 4.47942 | 30.47627 | 14.52759 | 14.20390 | 4.80820 | 18.50780 | 8.87121 | 12.91647 |
| **MPHOSPH6** | 2.11353 | 8.04209 | 9.12798 | 2.16880 | 3.85300 | 21.73156 | 11.61110 | 2.76445 | 11.51882 | 5.84358 | 9.60606 | 11.51087 |
| **MPHOSPH8** | 6.60268 | 3.21411 | 9.11252 | 5.43850 | 8.07095 | 9.24824 | 13.86554 | 10.20095 | 2.91333 | 23.41554 | 7.89105 | 6.95428 |
| **MPHOSPH9** | 1.37764 | 3.41238 | 0.18848 | 1.88155 | 0.00000 | 7.31260 | 0.00000 | 2.44062 | 3.82731 | 0.81643 | 2.39192 | 4.09627 |
| **MPI** | 0.00000 | 9.11185 | 5.73672 | 7.53784 | 4.92477 | 4.03665 | 3.86138 | 0.48170 | 3.77029 | 3.38431 | 15.14405 | 7.82825 |
| **MPL** | 0.11160 | 0.10755 | 0.16204 | 0.05164 | 0.00168 | 0.02980 | 0.00685 | 0.02611 | 0.04186 | 0.13870 | 0.17568 | 0.10641 |
| **MPLKIP** | 2.51942 | 8.23570 | 10.29280 | 3.52604 | 4.06390 | 8.23515 | 5.29839 | 7.49273 | 10.15147 | 2.74466 | 10.75998 | 16.20240 |
| **MPND** | 1.05365 | 0.29524 | 0.33090 | 0.42045 | 1.51422 | 0.00000 | 0.75693 | 0.29504 | 0.00000 | 2.69866 | 0.25800 | 0.47276 |
| **MPO** | 0.48048 | 0.01267 | 0.42000 | 0.00478 | 0.03367 | 0.00000 | 0.02186 | 0.00423 | 0.78673 | 0.00000 | 0.00000 | 0.01912 |
| **MPP1** | 1.09120 | 1.10738 | 32.01596 | 2.06228 | 1.34309 | 4.10237 | 18.81318 | 0.14979 | 18.83354 | 0.00000 | 19.68312 | 0.19332 |
| **MPP2** | 6.58225 | 0.25316 | 0.28948 | 0.32862 | 0.00000 | 0.09700 | 0.00000 | 0.83417 | 0.04872 | 0.09099 | 0.00288 | 0.23737 |
| **MPP3** | 0.42843 | 0.26619 | 0.06735 | 0.08960 | 0.21422 | 0.00000 | 0.00000 | 1.28222 | 0.08691 | 0.20374 | 0.00000 | 0.97858 |
| **MPP4** | 0.59691 | 0.00440 | 0.02916 | 0.02254 | 0.01228 | 0.00000 | 0.02366 | 0.00383 | 0.01635 | 0.04857 | 0.02501 | 0.13276 |
| **MPP5** | 3.11277 | 0.89207 | 4.94086 | 0.89003 | 2.31782 | 5.72183 | 7.16234 | 12.23679 | 3.05776 | 9.50298 | 4.00803 | 6.89545 |
| **MPP6** | 0.58055 | 0.00000 | 7.57537 | 0.56769 | 0.00000 | 4.61418 | 9.01334 | 0.27571 | 0.76028 | 15.73012 | 0.36635 | 0.82405 |
| **MPP7** | 1.75109 | 1.63924 | 9.48460 | 3.07239 | 1.52289 | 3.10697 | 3.13810 | 0.60150 | 0.00000 | 0.00000 | 1.50736 | 5.21885 |
| **MPPE1** | 0.34336 | 1.58350 | 0.51903 | 0.49954 | 0.69199 | 0.00000 | 0.14537 | 0.64740 | 2.55411 | 0.00000 | 2.56171 | 1.79419 |
| **MPPED1** | 0.05912 | 0.00935 | 0.02151 | 0.00652 | 0.00844 | 0.02925 | 0.00768 | 0.00656 | 0.00799 | 0.07242 | 0.01251 | 0.02041 |
| **MPPED2** | 0.02751 | 0.06947 | 0.16927 | 0.03344 | 0.00000 | 0.03448 | 0.00000 | 0.00715 | 0.00000 | 0.03079 | 0.02149 | 0.02653 |
| **MPRIP** | 1.65431 | 0.99331 | 3.91842 | 2.24724 | 6.08404 | 1.29349 | 6.18290 | 0.02850 | 5.58747 | 13.50761 | 3.84843 | 0.57837 |
| **MPST** | 0.03732 | 7.52887 | 7.69861 | 0.42750 | 1.94873 | 0.00000 | 17.46457 | 0.00000 | 10.08252 | 9.37203 | 8.05442 | 3.15795 |
| **MPV17** | 1.39375 | 10.10418 | 8.92933 | 3.47454 | 8.77687 | 7.42510 | 7.74760 | 1.57939 | 9.00959 | 0.00000 | 15.23230 | 8.78301 |
| **MPV17L** | 0.00631 | 0.59689 | 0.06643 | 0.14110 | 0.19980 | 0.22654 | 0.04919 | 0.00000 | 0.00000 | 0.28028 | 3.41864 | 0.88894 |
| **MPV17L2** | 2.33104 | 17.10897 | 7.95806 | 14.22835 | 30.51042 | 0.00194 | 14.46069 | 3.93751 | 23.00442 | 0.00000 | 13.26015 | 17.36680 |
| **MPZ** | 0.02261 | 0.00000 | 0.17676 | 0.11391 | 0.07595 | 0.02923 | 0.17304 | 0.02114 | 0.04246 | 0.01714 | 0.22722 | 0.03866 |
| **MPZL1** | 0.94255 | 21.65285 | 28.45186 | 0.00000 | 29.15578 | 25.93618 | 10.75821 | 3.82506 | 43.85817 | 18.34488 | 25.73048 | 9.27700 |
| **MPZL2** | 0.50149 | 0.05654 | 0.23582 | 0.30848 | 0.00000 | 0.24652 | 0.00000 | 0.11880 | 0.00000 | 0.01402 | 0.00529 | 0.14196 |
| **MPZL3** | 0.38924 | 0.67613 | 0.65321 | 0.01125 | 0.00058 | 0.14631 | 0.00000 | 1.95159 | 1.55744 | 0.00000 | 0.11479 | 0.64613 |
| **MR1** | 0.00381 | 0.12779 | 0.00000 | 0.00000 | 0.16345 | 0.10820 | 0.00000 | 0.20535 | 0.00000 | 0.61297 | 0.00000 | 0.02098 |
| **MRAP** | 0.01054 | 0.01749 | 0.01442 | 0.00724 | 0.02649 | 0.02969 | 0.01101 | 0.01664 | 0.02212 | 0.03043 | 0.03822 | 0.02751 |
| **MRAP2** | 0.06588 | 0.21442 | 1.24536 | 2.86019 | 0.00000 | 0.00000 | 0.00000 | 0.01118 | 0.92263 | 0.20707 | 0.00000 | 0.08864 |
| **MRAS** | 2.01662 | 0.36783 | 0.53474 | 0.85610 | 0.00000 | 0.00000 | 0.22895 | 0.54064 | 0.11272 | 0.05879 | 0.00000 | 0.79715 |
| **MRC1** | 0.12058 | 0.01346 | 0.01559 | 0.01321 | 0.01505 | 0.01770 | 0.01598 | 0.09883 | 0.02296 | 0.01148 | 0.01547 | 0.01701 |
| **MRC2** | 0.23071 | 0.00327 | 0.53090 | 1.74455 | 0.00000 | 0.00000 | 0.00000 | 0.17474 | 0.25486 | 0.07756 | 0.01246 | 0.06692 |
| **MRE11A** | 3.42630 | 3.05850 | 4.58885 | 1.56848 | 1.32332 | 10.33131 | 3.58021 | 1.09018 | 6.77860 | 6.25437 | 6.99566 | 3.67933 |
| **MREG** | 0.03911 | 0.77975 | 4.48218 | 0.00000 | 0.00000 | 2.28852 | 4.12366 | 0.00000 | 0.14308 | 1.10617 | 2.07351 | 0.28320 |
| **MRFAP1** | 7.76036 | 29.18363 | 26.34593 | 21.91849 | 27.90742 | 49.50349 | 44.38213 | 10.16498 | 38.82261 | 9.55892 | 31.10290 | 36.74670 |
| **MRFAP1L1** | 0.92351 | 8.31430 | 10.25388 | 6.03756 | 4.48844 | 29.29408 | 18.49437 | 2.93241 | 15.17575 | 0.75089 | 11.62076 | 18.43680 |
| **MRGBP** | 1.06962 | 0.95345 | 2.76284 | 0.96237 | 2.25472 | 0.00000 | 5.55659 | 0.45641 | 1.14723 | 5.61406 | 1.28210 | 0.10704 |
| **MRGPRD** | 0.00635 | 0.03031 | 0.01917 | 0.00802 | 0.08501 | 0.00494 | 0.01662 | 0.01070 | 0.02137 | 0.00933 | 0.01775 | 0.01611 |
| **MRGPRF** | 0.00583 | 0.03056 | 0.06548 | 0.08913 | 0.01110 | 0.11818 | 0.04583 | 0.00944 | 0.00000 | 0.00000 | 0.10908 | 0.01090 |
| **MRGPRX1** | 0.00664 | 0.00000 | 22.93502 | 0.00218 | 0.55367 | 0.14516 | 0.00000 | 0.00640 | 4.98611 | 0.00000 | 3.17185 | 0.00937 |
| **MRGPRX2** | 0.00688 | 0.01254 | 0.53997 | 0.13096 | 0.00394 | 0.02849 | 0.00191 | 0.03387 | 0.00000 | 0.00773 | 0.03706 | 0.01796 |
| **MRGPRX3** | 0.00209 | 0.01699 | 0.14564 | 0.01308 | 0.21754 | 0.00000 | 0.01790 | 0.12633 | 0.00000 | 0.00434 | 0.14860 | 0.31674 |
| **MRGPRX4** | 0.00634 | 0.01524 | 0.02077 | 0.01214 | 0.01726 | 0.01691 | 0.01460 | 0.01052 | 0.02502 | 0.01361 | 0.01506 | 0.01637 |

| | | | | | | | | | | | | |
|---|---|---|---|---|---|---|---|---|---|---|---|---|
| **MRI1** | 0.20653 | 1.81058 | 5.80067 | 3.14054 | 2.27701 | 0.00000 | 4.10196 | 0.00000 | 4.96598 | 2.24189 | 6.54328 | 3.01666 |
| **MRM1** | 0.00282 | 5.37995 | 4.72603 | 1.53006 | 2.11267 | 0.00000 | 6.39800 | 0.05928 | 2.65556 | 0.77878 | 9.37833 | 8.17598 |
| **MRO** | 0.74742 | 0.00821 | 0.01302 | 0.00803 | 0.01365 | 0.01904 | 0.04262 | 0.00686 | 0.00907 | 0.03373 | 0.01533 | 0.05190 |
| **MROH1** | 0.51844 | 2.06488 | 1.87260 | 1.93355 | 2.27989 | 0.02843 | 1.31903 | 11.62273 | 1.54277 | 0.94309 | 2.71221 | 1.47434 |
| **MROH2A** | 0.32636 | 0.01253 | 0.05626 | 0.01272 | 0.01364 | 0.00762 | 0.00906 | 0.00276 | 0.01798 | 0.04296 | 0.01556 | 0.02564 |
| **MROH2B** | 0.00588 | 0.01611 | 0.01501 | 0.01520 | 0.00808 | 0.00631 | 0.02801 | 0.01059 | 0.00000 | 0.03190 | 0.07757 | 0.01606 |
| **MROH5** | 0.00616 | 0.01164 | 0.01040 | 0.12839 | 0.16028 | 0.00000 | 0.05484 | 0.01069 | 0.00000 | 0.00000 | 0.06879 | 0.01830 |
| **MROH6** | 0.00081 | 0.00000 | 1.65566 | 0.00000 | 0.00000 | 0.00123 | 4.58252 | 0.04349 | 1.87801 | 8.65816 | 0.00000 | 0.38033 |
| **MROH7** | 0.08763 | 0.31906 | 0.00997 | 0.01051 | 0.01293 | 0.00293 | 0.01438 | 0.00000 | 0.01592 | 0.03168 | 0.01116 | 0.72002 |
| **MROH7-TTC4** | 0.00359 | 0.01420 | 0.01800 | 0.01358 | 0.01280 | 0.01552 | 0.01522 | 0.08705 | 0.01652 | 0.03077 | 0.01462 | 0.03678 |
| **MROH8** | 0.21859 | 0.57840 | 0.05928 | 0.17225 | 0.00000 | 0.49384 | 0.23845 | 0.25178 | 0.00000 | 0.02804 | 0.06303 | 0.92342 |
| **MROH9** | 0.00621 | 0.01782 | 0.02281 | 0.01140 | 0.01121 | 0.02693 | 0.01418 | 0.01037 | 0.01738 | 0.01414 | 0.01945 | 0.01557 |
| **MRPL1** | 0.00000 | 10.24948 | 0.00000 | 3.33443 | 5.80843 | 23.83814 | 4.73837 | 0.87086 | 1.14189 | 11.25388 | 9.60418 | 16.10838 |
| **MRPL10** | 2.50904 | 25.93919 | 10.20785 | 11.79010 | 19.09871 | 22.70697 | 9.89744 | 0.82917 | 16.38140 | 0.00000 | 15.99075 | 11.93114 |
| **MRPL11** | 1.55916 | 21.75743 | 16.85696 | 16.98089 | 23.25271 | 22.54980 | 27.17831 | 0.58354 | 25.48830 | 7.20344 | 20.47104 | 18.62590 |
| **MRPL12** | 0.00000 | 39.43426 | 23.54851 | 24.00274 | 35.97322 | 0.00000 | 22.41808 | 11.40426 | 37.53765 | 11.00285 | 26.84063 | 30.86551 |
| **MRPL13** | 0.85059 | 14.65041 | 14.90082 | 10.65894 | 9.76138 | 46.64763 | 18.80737 | 4.74505 | 18.22936 | 0.00006 | 22.70268 | 15.28206 |
| **MRPL14** | 2.96117 | 27.97161 | 29.08260 | 24.14299 | 33.08220 | 18.32654 | 44.85834 | 11.03126 | 28.72709 | 8.38246 | 33.46305 | 30.71057 |
| **MRPL15** | 0.70720 | 26.86040 | 13.70175 | 18.77663 | 20.83789 | 50.55432 | 20.22237 | 3.73106 | 23.49855 | 0.00379 | 9.98266 | 28.64819 |
| **MRPL16** | 0.00000 | 21.75515 | 9.83974 | 13.77926 | 10.95484 | 23.92478 | 24.72184 | 20.42612 | 23.05548 | 3.94110 | 23.89434 | 23.53701 |
| **MRPL17** | 0.03245 | 21.91722 | 15.51033 | 17.91514 | 19.40873 | 18.20918 | 23.07987 | 5.80189 | 18.81078 | 3.99259 | 20.33007 | 23.09862 |
| **MRPL18** | 10.54889 | 25.17012 | 13.13727 | 26.20552 | 19.50105 | 40.48279 | 19.96837 | 13.35784 | 19.24930 | 5.43013 | 19.67479 | 23.84673 |
| **MRPL19** | 3.57466 | 13.11941 | 6.58671 | 9.23102 | 6.94829 | 32.61436 | 16.66824 | 10.59573 | 10.57267 | 8.56464 | 15.64507 | 15.69726 |
| **MRPL2** | 6.95292 | 23.51978 | 11.54047 | 12.18158 | 23.41172 | 10.66823 | 20.22202 | 6.86836 | 12.91347 | 0.00000 | 23.67334 | 17.61570 |
| **MRPL20** | 3.42989 | 26.37647 | 20.05191 | 17.01485 | 22.54829 | 25.33864 | 29.57344 | 19.88553 | 22.38739 | 11.18942 | 22.79228 | 29.08600 |
| **MRPL21** | 0.21966 | 22.31897 | 13.04404 | 16.09338 | 24.02380 | 19.60179 | 24.71657 | 9.16333 | 18.39125 | 4.35154 | 18.19178 | 24.98807 |
| **MRPL22** | 3.10001 | 13.96771 | 13.62872 | 10.94785 | 10.49735 | 31.46268 | 23.44127 | 8.15161 | 15.07402 | 3.36966 | 19.33200 | 20.41500 |
| **MRPL23** | 0.03160 | 7.45281 | 3.71374 | 7.35281 | 12.63783 | 0.00000 | 5.15846 | 0.21121 | 6.76792 | 1.42501 | 6.03762 | 2.35114 |
| **MRPL23-AS1** | 0.56005 | 0.02118 | 0.03757 | 0.01403 | 0.01489 | 0.00443 | 0.01581 | 0.00000 | 0.01641 | 0.01307 | 0.02672 | 0.03211 |
| **MRPL24** | 1.99893 | 14.23162 | 7.46982 | 5.89187 | 14.27222 | 7.83177 | 12.56804 | 2.54132 | 7.82896 | 3.57711 | 17.99715 | 12.55280 |
| **MRPL27** | 0.39116 | 22.58219 | 21.54154 | 13.59944 | 18.35796 | 18.72473 | 33.67150 | 3.84808 | 13.70240 | 12.44585 | 19.51024 | 23.73325 |
| **MRPL28** | 0.00000 | 23.98017 | 14.42133 | 15.73481 | 28.54781 | 3.68477 | 32.86505 | 2.44576 | 16.71600 | 7.21952 | 23.65183 | 24.84592 |
| **MRPL3** | 2.33960 | 22.44133 | 16.10638 | 8.44485 | 12.16036 | 45.64981 | 29.20125 | 4.45641 | 27.26760 | 0.51794 | 23.04535 | 25.29662 |
| **MRPL30** | 1.41974 | 6.06027 | 5.70135 | 1.20424 | 3.91964 | 18.61468 | 8.94622 | 2.93663 | 8.99254 | 6.14243 | 6.90356 | 11.36556 |
| **MRPL32** | 4.19547 | 11.09334 | 16.68735 | 8.46966 | 8.61080 | 32.13538 | 16.09432 | 4.75309 | 21.02930 | 0.00000 | 14.03908 | 17.72318 |
| **MRPL33** | 0.15807 | 10.18671 | 14.92442 | 6.16105 | 7.09824 | 31.15047 | 23.74057 | 1.70071 | 11.94995 | 19.25775 | 18.85417 | 14.76676 |
| **MRPL34** | 1.31010 | 20.80503 | 15.99217 | 18.96732 | 21.77488 | 21.22226 | 19.37864 | 3.73166 | 20.65593 | 8.19710 | 23.39701 | 22.65575 |
| **MRPL35** | 1.18846 | 16.71177 | 2.96598 | 5.56718 | 8.68327 | 27.67061 | 9.56497 | 5.06876 | 12.71325 | 6.30618 | 15.54705 | 16.98991 |
| **MRPL36** | 1.13616 | 12.43706 | 13.37970 | 5.84729 | 8.93414 | 18.31572 | 16.89371 | 0.15066 | 15.53201 | 2.93221 | 16.39015 | 16.38715 |
| **MRPL37** | 6.36641 | 36.21328 | 18.53181 | 20.55306 | 34.00027 | 30.80675 | 32.56799 | 4.15509 | 25.34041 | 13.90600 | 27.54469 | 31.64692 |
| **MRPL38** | 0.73887 | 23.54885 | 19.18783 | 19.92047 | 28.10094 | 2.88780 | 26.03443 | 1.57805 | 29.67965 | 3.30792 | 23.71913 | 18.57368 |
| **MRPL39** | 0.49520 | 18.69109 | 11.66431 | 7.17191 | 6.12977 | 41.17859 | 11.11089 | 2.53823 | 9.06795 | 0.00000 | 14.33180 | 13.96798 |
| **MRPL4** | 6.02963 | 26.39241 | 28.40711 | 19.53585 | 35.08140 | 0.18994 | 30.26689 | 4.23035 | 26.97444 | 8.88026 | 24.61637 | 20.89593 |
| **MRPL40** | 1.60369 | 17.60598 | 5.21676 | 7.81384 | 17.28599 | 21.63926 | 9.65809 | 2.94787 | 14.24391 | 1.06571 | 14.67018 | 12.91861 |
| **MRPL41** | 1.98042 | 23.40647 | 10.57936 | 12.54743 | 26.17472 | 0.00000 | 10.86734 | 12.48086 | 18.90720 | 14.16195 | 16.02449 | 18.72753 |
| **MRPL42** | 0.00000 | 9.71560 | 17.69944 | 4.93744 | 2.88250 | 45.58290 | 21.97232 | 2.56068 | 21.93568 | 3.30964 | 18.63330 | 10.03501 |
| **MRPL42P5** | 0.00719 | 0.00350 | 0.14148 | 0.03298 | 0.00000 | 0.06429 | 0.00000 | 0.01074 | 0.00000 | 0.08970 | 0.02708 | 0.01213 |
| **MRPL43** | 3.49736 | 19.16865 | 13.88272 | 12.71670 | 11.44550 | 13.81821 | 15.44980 | 19.76689 | 17.53325 | 4.00437 | 12.88294 | 23.18072 |

| | | | | | | | | | | | | |
|---|---|---|---|---|---|---|---|---|---|---|---|---|
| **MRPL44** | 6.80858 | 16.73633 | 12.40888 | 8.47146 | 7.07125 | 31.56253 | 20.30587 | 10.46299 | 21.04566 | 0.77967 | 13.96011 | 20.34279 |
| **MRPL45** | 0.76990 | 17.45662 | 13.21459 | 8.00638 | 9.87595 | 25.35164 | 23.04954 | 4.23294 | 14.60048 | 3.77277 | 16.78221 | 17.47380 |
| **MRPL45P2** | 0.00079 | 0.26206 | 0.34037 | 0.21407 | 0.48589 | 0.44199 | 0.16478 | 0.42604 | 0.77570 | 1.58563 | 0.17567 | 0.80188 |
| **MRPL46** | 0.21992 | 17.96317 | 11.89874 | 7.38334 | 20.03000 | 23.95467 | 15.20193 | 6.13773 | 12.00378 | 4.87185 | 18.20419 | 16.64855 |
| **MRPL47** | 0.49295 | 13.78681 | 14.74895 | 9.03257 | 6.70151 | 38.45609 | 23.59816 | 3.51442 | 17.21861 | 14.15405 | 15.15942 | 16.23770 |
| **MRPL48** | 0.00000 | 14.59139 | 2.20735 | 9.46625 | 11.53657 | 28.45627 | 8.25017 | 3.91073 | 5.41299 | 0.71459 | 11.22226 | 18.49003 |
| **MRPL49** | 2.77843 | 21.70961 | 20.83238 | 14.50387 | 20.98980 | 14.59829 | 22.33638 | 10.92136 | 28.70216 | 0.00000 | 22.44159 | 17.49636 |
| **MRPL50** | 0.52191 | 12.14024 | 14.88804 | 5.47085 | 4.37176 | 39.45216 | 16.77887 | 11.95324 | 14.55225 | 6.22231 | 16.58690 | 21.54937 |
| **MRPL51** | 1.53491 | 25.78596 | 18.16103 | 19.05908 | 24.28252 | 35.16200 | 30.05745 | 5.94168 | 15.85001 | 16.08280 | 20.57885 | 26.96993 |
| **MRPL52** | 0.95725 | 11.04949 | 17.83278 | 9.59043 | 24.87174 | 19.74606 | 26.17061 | 3.10259 | 29.92041 | 10.16872 | 19.26874 | 20.95251 |
| **MRPL53** | 3.82016 | 8.75612 | 7.86510 | 9.21847 | 16.61733 | 5.80943 | 9.56180 | 6.47236 | 7.65343 | 8.45473 | 8.69687 | 12.70652 |
| **MRPL54** | 0.10596 | 10.32771 | 5.46027 | 6.64502 | 11.84582 | 4.74853 | 13.41651 | 0.79499 | 7.91309 | 3.34760 | 8.59419 | 11.85579 |
| **MRPL55** | 5.82804 | 25.21728 | 14.20758 | 9.73298 | 25.54231 | 2.88467 | 22.91261 | 10.28843 | 17.77946 | 3.26841 | 19.32901 | 22.86445 |
| **MRPL57** | 0.36690 | 20.53546 | 17.56784 | 17.60710 | 21.95321 | 22.23909 | 20.36052 | 10.37649 | 16.94480 | 14.42375 | 24.24608 | 21.86858 |
| **MRPL9** | 1.31447 | 24.28632 | 11.78805 | 16.16524 | 16.07047 | 27.16425 | 17.86215 | 1.51846 | 14.47842 | 4.91304 | 19.67972 | 27.46476 |
| **MRPS10** | 0.76597 | 7.57959 | 4.67481 | 3.45249 | 4.32197 | 18.70494 | 7.30650 | 1.33437 | 6.24595 | 3.54133 | 7.89143 | 7.21863 |
| **MRPS11** | 0.70914 | 21.13954 | 20.46182 | 15.15840 | 22.27208 | 18.54307 | 27.89062 | 9.04179 | 21.08518 | 0.00000 | 27.77978 | 24.11162 |
| **MRPS12** | 0.72922 | 27.01911 | 22.21665 | 23.96085 | 34.49175 | 0.03539 | 39.02119 | 6.05696 | 34.92324 | 1.46915 | 31.92249 | 24.34024 |
| **MRPS14** | 1.96142 | 8.95663 | 10.60702 | 7.20942 | 9.40316 | 28.32805 | 12.13175 | 7.64578 | 10.09038 | 5.06712 | 16.36162 | 10.02791 |
| **MRPS15** | 1.06335 | 22.83155 | 27.07201 | 19.47871 | 26.79686 | 26.87002 | 33.28710 | 7.11254 | 33.65502 | 0.88099 | 21.38251 | 26.74727 |
| **MRPS16** | 3.75859 | 24.02630 | 16.28408 | 17.78467 | 23.25296 | 22.56356 | 25.68466 | 5.13680 | 20.92811 | 17.76676 | 29.19405 | 19.79054 |
| **MRPS17** | 0.00000 | 14.86375 | 11.04588 | 3.27840 | 7.23182 | 18.80824 | 25.69834 | 1.84579 | 14.49393 | 9.70299 | 17.63662 | 12.53943 |
| **MRPS18A** | 0.31498 | 23.08200 | 17.80976 | 20.01629 | 22.36155 | 10.20236 | 28.53425 | 4.50687 | 17.85607 | 2.46700 | 22.47275 | 21.91215 |
| **MRPS18B** | 2.08463 | 23.75435 | 18.54446 | 13.18039 | 16.23564 | 28.50254 | 23.72221 | 9.27040 | 16.59556 | 2.08807 | 25.03323 | 22.71807 |
| **MRPS18C** | 0.13863 | 9.20909 | 9.02367 | 4.72525 | 9.25552 | 18.56433 | 13.53796 | 3.88073 | 9.35804 | 0.00000 | 12.30188 | 19.21191 |
| **MRPS2** | 0.84849 | 26.50880 | 1.32113 | 17.87578 | 25.12797 | 6.05368 | 10.89691 | 0.39608 | 17.13186 | 0.00000 | 12.90129 | 19.53626 |
| **MRPS21** | 2.80281 | 17.06625 | 19.76487 | 10.09635 | 14.50247 | 19.79800 | 28.19694 | 12.23017 | 21.00501 | 15.47961 | 25.75586 | 29.67986 |
| **MRPS22** | 0.56055 | 10.50793 | 11.59882 | 6.32627 | 3.15833 | 27.36758 | 14.24838 | 4.79079 | 12.09259 | 0.00000 | 12.65375 | 14.14405 |
| **MRPS23** | 2.16232 | 22.62922 | 11.15936 | 26.30117 | 9.60470 | 36.98838 | 17.64623 | 19.62148 | 15.35280 | 6.27329 | 17.80500 | 31.35775 |
| **MRPS24** | 0.04992 | 2.71116 | 1.02702 | 1.18833 | 1.32317 | 0.00000 | 0.85221 | 0.56130 | 1.64883 | 0.00000 | 1.21286 | 1.71914 |
| **MRPS25** | 6.24592 | 20.80785 | 14.65024 | 21.31562 | 23.27674 | 24.55663 | 24.54117 | 9.69492 | 18.70214 | 9.77825 | 22.63574 | 22.26121 |
| **MRPS26** | 6.27043 | 31.84770 | 13.20065 | 22.59440 | 26.29895 | 0.00054 | 21.62128 | 2.66917 | 25.18336 | 4.43455 | 20.25204 | 10.96106 |
| **MRPS27** | 0.80668 | 13.07322 | 13.42578 | 7.27631 | 9.94346 | 13.32888 | 16.70149 | 0.68662 | 18.64916 | 0.00000 | 16.41087 | 10.10418 |
| **MRPS28** | 0.70688 | 12.02747 | 11.73445 | 7.80527 | 8.26375 | 23.06454 | 15.74846 | 6.80172 | 12.64114 | 0.00921 | 10.44484 | 18.99184 |
| **MRPS30** | 0.00000 | 17.71323 | 10.58998 | 10.56424 | 11.07074 | 26.65205 | 15.13234 | 8.57291 | 20.70463 | 3.39083 | 14.84639 | 16.05616 |
| **MRPS31** | 1.21797 | 12.81156 | 5.55237 | 6.58777 | 10.13863 | 21.25300 | 8.95185 | 0.94040 | 3.53224 | 5.70679 | 12.06751 | 12.05310 |
| **MRPS31P5** | 0.00000 | 0.71781 | 0.68499 | 0.80041 | 0.45171 | 2.55436 | 0.23262 | 0.45025 | 1.10946 | 1.94565 | 0.19837 | 1.01649 |
| **MRPS33** | 3.16752 | 15.84618 | 3.48487 | 4.60918 | 9.55165 | 23.73513 | 5.03337 | 0.64165 | 6.86319 | 7.56410 | 11.68354 | 18.05560 |
| **MRPS34** | 0.73816 | 26.52313 | 21.07554 | 17.92912 | 29.03148 | 2.20922 | 33.30142 | 7.16657 | 31.64225 | 8.81916 | 25.81169 | 20.69398 |
| **MRPS35** | 1.33690 | 14.94938 | 9.25285 | 12.44233 | 12.52474 | 41.01164 | 19.70126 | 5.39382 | 19.24673 | 4.68256 | 17.07099 | 12.94566 |
| **MRPS36** | 0.14419 | 8.32512 | 8.72831 | 7.88155 | 4.28115 | 20.14170 | 11.59233 | 0.01530 | 6.80604 | 7.63931 | 7.85871 | 2.49022 |
| **MRPS5** | 2.38066 | 19.04821 | 10.56187 | 18.31345 | 20.78751 | 18.03158 | 27.12892 | 1.54246 | 14.79702 | 20.30136 | 23.24350 | 16.32240 |
| **MRPS6** | 3.12600 | 11.53196 | 13.87610 | 4.42148 | 3.71656 | 17.87852 | 16.64091 | 1.17192 | 19.16746 | 1.96890 | 9.53424 | 2.97199 |
| **MRPS7** | 2.98186 | 26.13249 | 22.83803 | 18.80980 | 19.87564 | 28.05680 | 33.19197 | 16.69731 | 27.10715 | 3.45664 | 32.84458 | 29.49975 |
| **MRPS9** | 1.39197 | 14.35054 | 9.44591 | 6.89550 | 6.43407 | 29.20726 | 16.43385 | 1.88879 | 16.31547 | 4.64076 | 18.75458 | 10.19830 |
| **MRRF** | 4.48420 | 11.52866 | 5.49001 | 5.53373 | 8.78940 | 13.23524 | 7.77183 | 2.26412 | 10.96105 | 0.00000 | 11.42406 | 12.54762 |
| **MRS2** | 0.35926 | 25.97353 | 8.47730 | 24.18975 | 6.87793 | 36.14124 | 15.64347 | 4.53502 | 12.70679 | 1.33820 | 12.82742 | 32.03441 |
| **MRS2P2** | 0.04258 | 0.88303 | 0.59635 | 1.09462 | 0.00000 | 1.98339 | 0.08150 | 0.52573 | 0.97287 | 0.01366 | 0.53505 | 1.78351 |

| | | | | | | | | | | | | |
|---|---|---|---|---|---|---|---|---|---|---|---|---|
| **MRTO4** | 1.60650 | 17.32777 | 14.94936 | 9.26199 | 20.22443 | 6.75823 | 31.87042 | 16.58219 | 21.75288 | 9.85759 | 15.04452 | 24.60432 |
| **MRVI1** | 7.18234 | 0.24552 | 0.04981 | 0.05029 | 0.32998 | 0.23435 | 0.10765 | 0.24203 | 0.00000 | 0.00000 | 0.40715 | 0.15711 |
| **MRVI1-AS1** | 0.87449 | 0.02943 | 0.03359 | 0.02251 | 0.00000 | 0.00000 | 0.01759 | 0.53586 | 0.01886 | 0.05995 | 0.08176 | 0.03781 |
| **MS4A1** | 0.00317 | 0.00000 | 0.00000 | 0.00000 | 0.08629 | 0.06512 | 0.13460 | 0.00496 | 0.00000 | 0.29295 | 0.00875 | 0.14457 |
| **MS4A10** | 0.00608 | 0.02325 | 0.02389 | 0.01176 | 0.00582 | 0.00000 | 0.01727 | 0.01053 | 0.00000 | 0.02477 | 0.12539 | 0.01774 |
| **MS4A12** | 0.13017 | 0.59644 | 0.00000 | 0.33213 | 0.04452 | 0.09184 | 0.00000 | 0.00000 | 0.00000 | 0.02313 | 0.00008 | 0.48426 |
| **MS4A13** | 0.03591 | 0.02069 | 0.00000 | 0.00000 | 0.05718 | 0.10148 | 0.00000 | 0.00300 | 0.08636 | 0.13412 | 0.00000 | 0.19364 |
| **MS4A14** | 0.00537 | 0.01383 | 0.05062 | 0.00902 | 0.00745 | 0.01248 | 0.01535 | 0.00872 | 0.05215 | 0.05202 | 0.02294 | 0.04912 |
| **MS4A15** | 0.01404 | 1.84974 | 1.83828 | 2.60469 | 7.46680 | 0.53003 | 0.10219 | 0.00229 | 0.00000 | 0.00000 | 2.31611 | 0.00047 |
| **MS4A2** | 0.00408 | 0.00171 | 0.02727 | 0.01667 | 0.00000 | 0.02207 | 0.00000 | 0.00689 | 0.13693 | 0.21468 | 0.00000 | 0.10884 |
| **MS4A3** | 0.37538 | 0.08552 | 0.07116 | 0.00000 | 0.00000 | 0.02368 | 0.00656 | 0.16306 | 0.36795 | 0.01236 | 0.00000 | 0.28795 |
| **MS4A4A** | 0.05158 | 0.00944 | 0.03863 | 0.05095 | 0.00246 | 0.03621 | 0.01781 | 0.00713 | 0.05051 | 0.02598 | 0.00519 | 0.02240 |
| **MS4A5** | 0.00577 | 0.01331 | 0.01228 | 0.01383 | 0.00898 | 0.02940 | 0.01662 | 0.00848 | 0.01663 | 0.02586 | 0.01341 | 0.05624 |
| **MS4A6A** | 0.00735 | 0.01549 | 0.01698 | 0.01145 | 0.02315 | 0.01010 | 0.02384 | 0.00958 | 0.00703 | 0.01612 | 0.04298 | 0.02500 |
| **MS4A6E** | 0.01044 | 0.01477 | 0.03819 | 0.00994 | 0.01525 | 0.02112 | 0.01390 | 0.00984 | 0.01692 | 0.01123 | 0.01835 | 0.01957 |
| **MS4A7** | 0.00246 | 0.02609 | 0.06933 | 0.00272 | 0.01295 | 0.03493 | 0.00000 | 0.06327 | 0.04550 | 0.02850 | 0.02907 | 0.10598 |
| **MS4A8** | 0.00184 | 0.00752 | 0.02158 | 0.01624 | 0.00930 | 0.09207 | 0.00000 | 0.31008 | 0.03717 | 0.03540 | 0.02039 | 0.19118 |
| **MSANTD1** | 0.01679 | 0.00558 | 0.00166 | 0.19216 | 0.00000 | 0.00000 | 0.02590 | 0.01031 | 0.00401 | 0.22522 | 0.03954 | 0.03369 |
| **MSANTD2** | 0.00000 | 0.54754 | 1.13893 | 0.80155 | 0.00000 | 1.94891 | 1.25961 | 1.47502 | 0.70618 | 2.01821 | 0.08746 | 0.16660 |
| **MSANTD3** | 0.47391 | 0.62612 | 1.91895 | 0.17641 | 0.00000 | 0.58293 | 2.81088 | 0.00000 | 0.00000 | 2.12940 | 0.54025 | 0.26119 |
| **MSANTD4** | 0.23892 | 0.18227 | 0.90008 | 1.46718 | 0.00000 | 2.05123 | 2.50960 | 0.16042 | 2.41138 | 0.19883 | 3.30731 | 0.73674 |
| **MSC** | 0.01123 | 0.01727 | 0.01496 | 0.11946 | 0.01678 | 0.01895 | 0.02214 | 0.01020 | 0.02789 | 0.00000 | 0.01386 | 0.02431 |
| **MSH2** | 6.11612 | 19.59291 | 5.81838 | 11.37439 | 0.56478 | 35.69986 | 9.81186 | 21.43318 | 11.65143 | 9.56381 | 16.91542 | 18.83307 |
| **MSH3** | 6.59954 | 1.28088 | 1.18302 | 2.03813 | 1.52018 | 5.16578 | 4.08573 | 1.84222 | 1.37533 | 9.00708 | 2.88577 | 2.54341 |
| **MSH4** | 0.00259 | 0.00191 | 0.00000 | 0.00000 | 0.17989 | 0.08027 | 0.02054 | 0.00714 | 0.00000 | 0.33602 | 0.13923 | 0.06264 |
| **MSH5-SAPCD1** | 0.00408 | 0.05420 | 0.08289 | 0.09186 | 0.00000 | 0.00000 | 0.00121 | 0.00939 | 0.05779 | 0.11439 | 0.06120 | 0.03460 |
| **MSH5.uid-7870** | 0.03910 | 0.02581 | 0.06741 | 0.01475 | 0.11502 | 0.00000 | 0.01076 | 0.01377 | 0.08747 | 0.00000 | 0.07455 | 0.02312 |
| **MSH5.uid-7872** | 0.00652 | 0.02652 | 0.02060 | 0.01641 | 0.02399 | 0.00876 | 0.01820 | 0.01341 | 0.03675 | 0.01384 | 0.02170 | 0.01975 |
| **MSH6** | 2.96628 | 24.95035 | 4.46256 | 30.68411 | 5.25658 | 19.39696 | 7.02685 | 15.88026 | 11.58059 | 15.50406 | 10.36022 | 25.49885 |
| **MSI1** | 0.00000 | 2.55374 | 1.31061 | 2.45293 | 2.40339 | 0.00000 | 5.26565 | 0.49624 | 0.93745 | 0.83366 | 0.76629 | 0.85207 |
| **MSI2** | 6.51153 | 0.15362 | 2.31482 | 0.00000 | 0.00000 | 0.79138 | 1.53932 | 0.12637 | 0.46926 | 2.88788 | 0.16589 | 0.14056 |
| **MSL1** | 2.34757 | 1.87607 | 2.65910 | 1.14904 | 0.00000 | 7.87580 | 6.78126 | 2.82142 | 2.73800 | 10.93180 | 2.94131 | 2.14828 |
| **MSL2** | 9.50433 | 1.37299 | 0.96551 | 0.56562 | 2.01551 | 4.56972 | 3.49853 | 9.19413 | 1.25183 | 7.66205 | 1.70447 | 3.86047 |
| **MSL3** | 7.76212 | 3.23536 | 2.55267 | 2.85336 | 1.85159 | 4.38548 | 5.60113 | 5.50484 | 4.91345 | 2.58141 | 2.79703 | 4.53547 |
| **MSL3P1** | 0.14332 | 5.42588 | 1.39250 | 0.07215 | 0.00000 | 3.07141 | 1.13401 | 2.31541 | 3.21011 | 0.00000 | 0.83010 | 11.63883 |
| **MSLN** | 0.00686 | 0.01296 | 0.05228 | 0.20553 | 0.04218 | 0.00000 | 0.00329 | 0.01022 | 0.12854 | 0.00064 | 0.00649 | 0.01828 |
| **MSMB** | 0.02482 | 0.39386 | 0.13432 | 0.59574 | 0.44820 | 0.41516 | 0.00000 | 0.97030 | 0.00000 | 0.12140 | 0.12767 | 0.48594 |
| **MSMO1** | 0.02343 | 0.00000 | 15.99792 | 17.51401 | 36.03766 | 38.35867 | 0.00000 | 0.41742 | 47.67125 | 6.52424 | 22.60301 | 13.43593 |
| **MSMP** | 0.24047 | 0.24986 | 0.08409 | 0.06747 | 0.37526 | 0.00000 | 0.07453 | 0.00000 | 0.27669 | 0.11174 | 0.07098 | 0.17893 |
| **MSN** | 0.32283 | 7.82285 | 1.24725 | 5.72421 | 0.79512 | 3.21628 | 0.70837 | 0.06718 | 0.00000 | 1.86519 | 0.23286 | 0.84405 |
| **MSR1** | 0.37963 | 0.02725 | 0.07943 | 0.25883 | 0.00000 | 0.10059 | 0.00000 | 0.48861 | 0.00000 | 0.20096 | 0.00045 | 0.16831 |
| **MSRA** | 7.96788 | 0.10745 | 0.19890 | 0.08132 | 0.26284 | 0.00000 | 0.00001 | 1.19300 | 0.29276 | 0.00000 | 0.22768 | 0.60575 |
| **MSRB1** | 0.90687 | 1.51148 | 14.19697 | 4.80569 | 1.77849 | 0.00000 | 4.54814 | 3.22968 | 3.03127 | 0.00000 | 1.64728 | 6.50341 |
| **MSRB2** | 0.36775 | 0.14154 | 0.14968 | 0.12121 | 0.06764 | 0.00000 | 0.00000 | 0.13170 | 0.04398 | 0.39620 | 0.00936 | 0.00000 |
| **MSRB3** | 0.27296 | 0.41665 | 0.03630 | 0.00000 | 0.02387 | 0.01841 | 0.06655 | 0.04787 | 0.00000 | 0.08066 | 0.01542 | 0.87246 |
| **MSS51** | 0.00096 | 0.11305 | 0.05504 | 0.02459 | 0.00000 | 0.06585 | 0.05423 | 0.03787 | 0.00000 | 0.66932 | 0.47972 | 0.21045 |
| **MST1** | 0.00858 | 0.09076 | 1.10333 | 0.38336 | 0.38846 | 0.06175 | 0.68644 | 0.04995 | 0.51859 | 0.21210 | 0.39292 | 0.07682 |
| **MST1L** | 0.00807 | 0.00772 | 0.06612 | 0.33981 | 0.00000 | 0.02702 | 0.01501 | 0.01440 | 0.02722 | 0.04183 | 0.02891 | 0.02629 |

| | | | | | | | | | | | | |
|---|---|---|---|---|---|---|---|---|---|---|---|---|
| **MST1P2** | 0.02243 | 0.06821 | 0.12947 | 0.19620 | 0.00000 | 0.02230 | 0.00000 | 0.01211 | 0.05586 | 0.06263 | 0.01650 | 0.02137 |
| **MST1R** | 0.03790 | 0.26812 | 0.00618 | 0.22621 | 0.00105 | 0.02903 | 0.00000 | 0.00373 | 0.05053 | 0.43138 | 0.00000 | 0.03837 |
| **MST4** | 0.00000 | 2.19677 | 4.86605 | 0.00000 | 0.00000 | 3.04353 | 6.79910 | 0.00234 | 5.35377 | 1.10290 | 4.49880 | 0.12893 |
| **MSTO1** | 0.09008 | 0.60837 | 0.80240 | 3.75551 | 1.86784 | 0.00000 | 1.05594 | 0.12419 | 0.77383 | 0.82626 | 2.08045 | 1.05661 |
| **MSTO2P.uid-19** | 0.00939 | 0.02772 | 0.06910 | 0.02784 | 0.02820 | 0.04985 | 0.01464 | 0.02463 | 0.05988 | 0.00508 | 0.05610 | 0.08863 |
| **MSX1** | 0.90872 | 0.00000 | 0.00000 | 0.31453 | 0.38429 | 0.20866 | 0.88739 | 0.83331 | 0.00000 | 1.49417 | 0.00000 | 0.09015 |
| **MSX2** | 0.61207 | 6.19473 | 31.73274 | 4.31172 | 13.04955 | 0.00000 | 24.31977 | 0.84680 | 0.63609 | 11.31886 | 4.22643 | 1.90559 |
| **MSX2P1** | 0.17117 | 0.06435 | 0.53036 | 0.05795 | 0.23083 | 0.00000 | 0.39451 | 0.01603 | 0.00000 | 0.25733 | 0.01843 | 0.00969 |
| **MT1A** | 0.08668 | 0.30511 | 0.08335 | 0.24930 | 0.00000 | 0.09982 | 0.00000 | 0.20691 | 0.11921 | 0.09092 | 0.03383 | 0.03227 |
| **MT1B** | 0.00733 | 0.00587 | 0.97711 | 0.03514 | 0.00000 | 0.06670 | 0.00000 | 0.23534 | 0.00000 | 0.02696 | 0.01318 | 0.00881 |
| **MT1DP** | 0.00213 | 0.01121 | 0.21516 | 0.16653 | 0.00000 | 0.09879 | 0.00000 | 0.11129 | 0.00000 | 0.03316 | 0.00948 | 0.00785 |
| **MT1E** | 4.06874 | 3.10912 | 0.00000 | 1.28610 | 0.42045 | 0.01361 | 0.00000 | 0.27752 | 0.00000 | 0.11612 | 0.00000 | 0.40176 |
| **MT1F** | 1.32308 | 0.95256 | 0.08672 | 4.03397 | 0.83686 | 0.14578 | 0.00000 | 0.10787 | 0.00000 | 0.06947 | 0.00000 | 0.15819 |
| **MT1G** | 3.58234 | 6.37159 | 0.00000 | 16.00561 | 3.93652 | 0.00000 | 0.00000 | 0.92697 | 0.00000 | 1.15338 | 0.00000 | 1.21860 |
| **MT1H** | 0.33029 | 2.40031 | 0.00000 | 9.07307 | 1.96380 | 0.02865 | 0.00000 | 0.08053 | 0.00000 | 0.17002 | 0.00000 | 0.72224 |
| **MT1HL1** | 0.06724 | 0.39043 | 0.00000 | 2.88590 | 0.35709 | 0.12287 | 0.00000 | 0.13494 | 0.00000 | 0.16371 | 0.00000 | 0.14646 |
| **MT1IP** | 0.00717 | 0.02007 | 0.01229 | 0.06622 | 0.05993 | 0.00453 | 0.02412 | 0.01006 | 0.01839 | 0.00000 | 0.01512 | 0.02241 |
| **MT1JP** | 0.00608 | 0.01376 | 0.01438 | 0.01391 | 0.01470 | 0.02122 | 0.01845 | 0.01026 | 0.01561 | 0.01966 | 0.01455 | 0.01827 |
| **MT1L** | 0.04185 | 0.63175 | 0.20887 | 0.22591 | 0.33464 | 0.00000 | 0.00000 | 0.02364 | 0.00000 | 0.16605 | 0.00000 | 0.28779 |
| **MT1M** | 0.12286 | 0.78931 | 0.00000 | 0.22116 | 0.46849 | 0.03561 | 0.00000 | 0.23242 | 0.00000 | 0.15699 | 0.00000 | 0.26532 |
| **MT1X** | 11.47962 | 28.53536 | 1.59298 | 19.15485 | 8.46201 | 0.00000 | 0.00000 | 9.78756 | 0.00000 | 0.36305 | 0.00000 | 12.11574 |
| **MT2A** | 4.11405 | 7.59251 | 0.10975 | 4.87343 | 0.74795 | 0.97471 | 0.00000 | 1.30115 | 0.00000 | 0.01324 | 0.00000 | 1.76190 |
| **MT3** | 0.25385 | 0.02507 | 0.01633 | 0.01170 | 0.01508 | 0.01341 | 0.03680 | 0.01075 | 0.00800 | 0.02025 | 0.02324 | 0.01571 |
| **MT4** | 0.00463 | 0.01265 | 0.02127 | 0.01207 | 0.01107 | 0.03133 | 0.01461 | 0.01656 | 0.02044 | 0.01693 | 0.01269 | 0.01497 |
| **MTA1** | 1.13399 | 0.00000 | 0.60258 | 0.00000 | 1.46272 | 0.00000 | 1.50337 | 0.00000 | 0.00000 | 3.47968 | 0.08510 | 0.27330 |
| **MTA2** | 0.51117 | 8.02477 | 10.31970 | 5.73621 | 10.99456 | 0.00000 | 10.15902 | 15.83003 | 10.95360 | 13.83547 | 7.28214 | 3.96416 |
| **MTA3** | 3.10844 | 3.83921 | 4.74745 | 8.82534 | 6.36959 | 10.43398 | 6.61409 | 2.10412 | 2.11832 | 14.76124 | 0.59433 | 4.84872 |
| **MTAP** | 0.81085 | 0.00000 | 16.05268 | 1.02743 | 1.51769 | 7.30338 | 11.27303 | 0.00000 | 9.98544 | 1.31359 | 12.78696 | 0.00000 |
| **MTBP** | 0.61349 | 1.36508 | 1.52294 | 1.04345 | 0.00137 | 3.92863 | 0.66797 | 1.48750 | 2.18682 | 0.00000 | 1.30621 | 3.19748 |
| **MTCH1** | 1.90424 | 13.30294 | 13.17351 | 17.30898 | 12.75887 | 0.00000 | 17.98627 | 6.21534 | 11.95743 | 16.81840 | 8.75386 | 11.43567 |
| **MTCH2** | 2.71262 | 19.41840 | 22.84213 | 16.49105 | 16.21273 | 29.66018 | 26.24802 | 3.64254 | 27.12793 | 1.47665 | 28.34323 | 16.29950 |
| **MTCL1** | 0.22089 | 0.54114 | 0.27047 | 3.75308 | 0.24473 | 0.00000 | 0.00003 | 0.47465 | 0.00000 | 1.44254 | 0.00000 | 4.45378 |
| **MTCL1-AS1** | 0.00315 | 0.00605 | 0.01987 | 0.01465 | 0.00665 | 0.02073 | 0.01308 | 0.03456 | 0.00000 | 0.03328 | 0.08327 | 0.07132 |
| **MTCP1** | 0.06469 | 0.11809 | 0.16397 | 0.04036 | 0.08531 | 0.00000 | 0.01888 | 0.18065 | 0.62028 | 0.00000 | 0.00000 | 0.11889 |
| **MTDH** | 2.74027 | 6.95783 | 9.77236 | 0.00000 | 11.69772 | 9.16001 | 10.50825 | 6.77193 | 14.06644 | 41.30867 | 11.87507 | 6.82395 |
| **MTERF** | 0.00303 | 7.37918 | 4.28126 | 0.14701 | 0.00000 | 10.10818 | 4.88068 | 5.97895 | 2.74171 | 0.00000 | 5.80789 | 9.82784 |
| **MTERFD1** | 4.46167 | 9.14489 | 2.57221 | 2.26828 | 0.00000 | 20.69664 | 3.86674 | 2.08501 | 3.39778 | 0.00000 | 8.87717 | 12.19861 |
| **MTERFD2** | 0.22598 | 9.47319 | 5.82169 | 4.98502 | 6.24064 | 7.87228 | 6.63748 | 2.39402 | 8.45454 | 1.28554 | 8.10257 | 12.46513 |
| **MTERFD3** | 0.00046 | 0.22881 | 0.41945 | 0.06863 | 0.71234 | 2.31687 | 0.63133 | 1.89234 | 1.24467 | 0.05629 | 0.47663 | 0.77236 |
| **MTF1** | 3.48388 | 4.11868 | 6.01509 | 3.92309 | 3.76017 | 1.36645 | 7.16594 | 7.55729 | 8.68149 | 4.73974 | 3.51054 | 5.23514 |
| **MTF2** | 6.50268 | 7.81095 | 7.57661 | 7.58797 | 5.50872 | 21.40557 | 18.82875 | 8.84716 | 7.75246 | 14.54892 | 10.82302 | 11.89819 |
| **MTFMT** | 0.39761 | 5.99910 | 3.15215 | 1.52739 | 2.42207 | 6.96248 | 3.66191 | 0.00000 | 1.39755 | 0.00000 | 5.99337 | 5.40856 |
| **MTFP1** | 0.00357 | 12.56429 | 4.12843 | 14.83038 | 19.96665 | 3.76077 | 14.45725 | 0.00000 | 13.48962 | 0.00000 | 14.68110 | 3.94326 |
| **MTFR1** | 4.99148 | 3.45787 | 2.00475 | 5.70546 | 0.00000 | 12.13877 | 6.05167 | 4.77094 | 0.28205 | 10.53247 | 1.20099 | 6.07135 |
| **MTFR1L** | 0.97264 | 10.56022 | 4.72146 | 5.12523 | 5.45889 | 8.32849 | 3.66350 | 3.23071 | 6.22513 | 3.15862 | 8.47014 | 5.80185 |
| **MTFR2** | 2.90359 | 5.89632 | 1.74845 | 1.80275 | 2.60821 | 15.84251 | 2.94177 | 4.28496 | 5.66279 | 2.50719 | 3.05374 | 6.78042 |
| **MTG1** | 0.00000 | 14.99011 | 9.98195 | 11.45197 | 14.82368 | 3.30806 | 8.44892 | 2.54913 | 17.03472 | 5.05775 | 16.66221 | 10.77721 |
| **MTG2** | 2.56101 | 6.06120 | 3.16054 | 3.85987 | 3.86564 | 0.00000 | 2.58990 | 3.80480 | 11.21013 | 0.00000 | 4.83730 | 10.09319 |

| | | | | | | | | | | | | |
|---|---|---|---|---|---|---|---|---|---|---|---|---|
| **MTHFD1** | 2.51713 | 9.56579 | 14.99413 | 14.70415 | 6.32075 | 16.96183 | 22.95368 | 0.45449 | 26.95658 | 0.44240 | 23.43992 | 6.83334 |
| **MTHFD1L** | 0.05132 | 0.99973 | 4.60033 | 1.55421 | 6.67308 | 9.73390 | 18.05479 | 0.00000 | 14.28188 | 18.40240 | 4.74534 | 0.00000 |
| **MTHFD2** | 0.00000 | 10.31180 | 11.40746 | 8.30766 | 7.27012 | 34.21317 | 33.63351 | 5.91287 | 35.97893 | 11.62307 | 16.48161 | 8.22216 |
| **MTHFD2L** | 0.03675 | 0.59223 | 1.39304 | 0.00000 | 0.58768 | 0.54523 | 0.36371 | 0.05282 | 0.72649 | 0.00000 | 0.92765 | 0.42853 |
| **MTHFD2P1** | 0.00543 | 0.04388 | 0.03952 | 0.02914 | 0.00527 | 0.10003 | 0.06251 | 0.02725 | 0.22256 | 0.00000 | 0.13030 | 0.03968 |
| **MTHFR** | 1.20599 | 0.94392 | 2.07778 | 1.56282 | 1.66457 | 0.35821 | 2.22877 | 0.00000 | 1.75773 | 3.71773 | 0.63400 | 0.36439 |
| **MTHFS.uid-196** | 0.00727 | 0.02000 | 0.03468 | 0.01186 | 0.04940 | 0.00381 | 0.02179 | 0.01352 | 0.02536 | 0.01637 | 0.01790 | 0.04215 |
| **MTHFS.uid-196** | 0.03302 | 1.23745 | 0.50367 | 1.50668 | 1.84088 | 1.02044 | 0.51278 | 0.04910 | 1.59158 | 0.00000 | 0.56385 | 0.40039 |
| **MTHFSD** | 0.35426 | 2.95678 | 0.70305 | 0.00975 | 0.55741 | 0.73954 | 0.24838 | 0.00000 | 1.63122 | 1.45015 | 0.98391 | 3.04696 |
| **MTIF2** | 0.18068 | 17.29720 | 6.06236 | 6.98157 | 9.44290 | 32.88319 | 9.71897 | 3.44965 | 9.29576 | 11.24389 | 11.69103 | 16.98298 |
| **MTIF3** | 1.54574 | 4.95766 | 4.66013 | 4.01977 | 7.60621 | 13.10271 | 11.64579 | 2.06820 | 7.07617 | 1.28156 | 10.14988 | 11.59679 |
| **5.00000** | 2.47572 | 0.05101 | 0.72659 | 0.30789 | 0.00000 | 0.07705 | 0.00006 | 0.05082 | 0.60230 | 0.17230 | 0.26281 | 0.02540 |
| **MTM1** | 3.42083 | 2.37808 | 0.63033 | 2.61464 | 0.00000 | 2.12623 | 1.50190 | 0.35239 | 0.94571 | 0.00000 | 2.54009 | 0.40348 |
| **MTMR1** | 3.75242 | 0.00000 | 0.00000 | 0.22837 | 1.49545 | 1.69162 | 1.02099 | 0.73561 | 0.00000 | 3.01020 | 0.96881 | 0.00789 |
| **MTMR10** | 1.65357 | 0.02718 | 5.89713 | 1.08583 | 3.16028 | 4.59911 | 5.83961 | 0.03614 | 0.47038 | 3.91652 | 3.04295 | 0.35059 |
| **MTMR11** | 0.02060 | 0.02048 | 0.04706 | 1.10914 | 0.00000 | 0.00000 | 0.00288 | 0.00650 | 0.13911 | 0.47211 | 0.00000 | 0.09632 |
| **MTMR12** | 0.94216 | 5.57494 | 3.05192 | 3.50041 | 0.85691 | 8.20014 | 5.11853 | 1.81919 | 0.59058 | 7.43724 | 0.35002 | 7.54343 |
| **MTMR14** | 1.58255 | 27.10405 | 14.23192 | 22.79245 | 6.27811 | 0.33844 | 13.43073 | 12.45022 | 8.22162 | 0.00000 | 6.00282 | 41.90896 |
| **MTMR2** | 2.84550 | 8.18366 | 10.70813 | 4.06368 | 4.51452 | 24.41879 | 10.49975 | 1.83113 | 10.43406 | 0.00000 | 10.67703 | 0.93172 |
| **MTMR3** | 19.34678 | 2.46169 | 3.02658 | 2.56938 | 0.50871 | 1.21799 | 0.00000 | 4.85163 | 5.20766 | 5.38916 | 0.27158 | 4.66267 |
| **MTMR4** | 1.77049 | 1.50456 | 3.51591 | 2.83584 | 0.12221 | 6.87113 | 2.92002 | 0.35386 | 6.47457 | 6.57826 | 2.12026 | 2.67064 |
| **MTMR6** | 0.98243 | 4.43572 | 3.97222 | 0.32531 | 2.96841 | 13.14894 | 4.40699 | 0.13162 | 3.20948 | 1.45544 | 3.61854 | 1.32241 |
| **MTMR7** | 0.90232 | 0.08607 | 3.51843 | 0.00000 | 0.10342 | 1.43079 | 1.63668 | 0.20926 | 3.96386 | 0.87921 | 2.68272 | 0.03865 |
| **MTMR8** | 0.43327 | 0.07337 | 0.00000 | 0.47557 | 0.00000 | 0.40177 | 0.00000 | 0.52858 | 0.08952 | 0.00000 | 0.20083 | 0.18813 |
| **MTMR9** | 1.10925 | 1.67693 | 2.28578 | 2.72125 | 0.54983 | 9.42726 | 5.26764 | 1.93450 | 2.18599 | 9.80515 | 2.47215 | 2.92125 |
| **MTMR9LP** | 0.03710 | 0.03574 | 0.09727 | 0.12638 | 0.13488 | 0.00000 | 0.01501 | 0.00679 | 0.29219 | 0.02320 | 0.00734 | 0.28899 |
| **MTNR1A** | 0.00504 | 0.03008 | 0.00777 | 0.00000 | 0.12065 | 0.00000 | 0.00049 | 0.01410 | 0.14732 | 0.05690 | 0.00000 | 0.03590 |
| **MTNR1B** | 0.01660 | 0.04420 | 0.03996 | 0.03000 | 0.02900 | 0.00000 | 0.01671 | 0.03333 | 0.01628 | 0.01020 | 0.01688 | 0.01561 |
| **MTO1** | 0.84232 | 8.35318 | 5.25220 | 3.52956 | 3.16336 | 18.83766 | 7.15106 | 5.85373 | 8.77008 | 0.66253 | 6.81909 | 6.31637 |
| **MTOR** | 9.10540 | 10.83646 | 7.54045 | 14.13389 | 11.07466 | 8.95730 | 6.34793 | 1.08405 | 7.42413 | 18.49996 | 6.20603 | 3.40637 |
| **MTOR-AS1** | 0.01168 | 0.01542 | 0.01737 | 0.02881 | 0.01564 | 0.00000 | 0.00638 | 0.00905 | 0.01856 | 0.19840 | 0.04135 | 0.04399 |
| **MTPAP** | 4.73131 | 17.95127 | 5.02737 | 9.03158 | 3.66306 | 29.55230 | 9.66730 | 16.30764 | 13.07475 | 4.96901 | 15.56060 | 18.99164 |
| **MTR** | 4.60760 | 1.41172 | 0.44369 | 1.04647 | 2.37462 | 11.18118 | 1.65069 | 1.09783 | 1.03977 | 8.74996 | 5.96887 | 2.31425 |
| **MTRF1** | 0.47497 | 3.66340 | 4.19010 | 2.97586 | 1.84340 | 8.46682 | 7.52532 | 0.22985 | 6.24958 | 0.09484 | 5.65543 | 2.74391 |
| **MTRF1L** | 0.28714 | 1.90236 | 1.91866 | 1.69521 | 0.91730 | 0.96136 | 1.47113 | 1.61624 | 2.00905 | 0.55800 | 1.16586 | 2.90231 |
| **MTRNR2L1** | 1.68772 | 0.00000 | 0.00000 | 30.54623 | 60.82979 | 48.92446 | 17.13951 | 10.69962 | 13.21705 | 0.00000 | 0.00000 | 47.53490 |
| **MTRNR2L10** | 0.10362 | 0.69125 | 0.00000 | 0.00658 | 0.68180 | 5.21900 | 0.00000 | 0.32417 | 3.23595 | 4.11442 | 0.00000 | 0.12244 |
| **MTRNR2L2** | 0.22677 | 0.39842 | 0.00000 | 0.28358 | 1.64199 | 2.72290 | 0.00000 | 0.35056 | 1.71200 | 3.63309 | 0.00000 | 0.52326 |
| **MTRNR2L3** | 0.14138 | 0.04036 | 0.02672 | 0.21794 | 0.56609 | 0.51478 | 0.00000 | 0.07413 | 0.42106 | 0.65348 | 0.00000 | 0.38542 |
| **MTRNR2L4** | 0.01832 | 0.01173 | 0.02196 | 0.02208 | 0.01329 | 0.16424 | 0.00000 | 0.00555 | 0.02141 | 0.24108 | 0.00009 | 0.05443 |
| **MTRNR2L5** | 0.03088 | 0.00000 | 0.23605 | 0.27321 | 0.09258 | 0.68858 | 0.00000 | 0.02240 | 0.05331 | 0.75915 | 0.00000 | 0.03406 |
| **MTRNR2L6** | 0.06058 | 0.06626 | 1.13127 | 0.22078 | 0.00428 | 2.05940 | 0.00000 | 0.06924 | 0.09699 | 1.95307 | 0.00000 | 0.07094 |
| **MTRNR2L7** | 0.01854 | 0.01618 | 0.02318 | 0.00589 | 0.03021 | 0.17012 | 0.00000 | 0.03641 | 0.01697 | 0.28474 | 0.00189 | 0.03037 |
| **MTRNR2L8** | 0.35531 | 0.02033 | 0.08327 | 0.09157 | 0.93859 | 1.28502 | 0.00000 | 0.30029 | 0.72198 | 2.40471 | 0.00000 | 0.46626 |
| **MTRNR2L9** | 0.09603 | 0.07135 | 0.00648 | 0.15673 | 0.48767 | 0.39049 | 0.00000 | 0.04571 | 0.45177 | 0.94226 | 0.00000 | 0.21613 |
| **MTRR** | 2.24437 | 3.04474 | 3.83163 | 3.18746 | 0.78186 | 11.42456 | 3.85160 | 5.84660 | 5.23610 | 0.09746 | 7.54667 | 0.63458 |
| **MTSS1** | 4.39798 | 0.70904 | 0.49089 | 2.01596 | 0.00000 | 0.69167 | 0.00000 | 0.00000 | 0.48425 | 1.21821 | 0.12628 | 1.22788 |
| **MTSS1L** | 1.61840 | 0.37375 | 0.00000 | 0.18357 | 0.18604 | 0.00000 | 0.04807 | 0.05570 | 0.34086 | 1.14759 | 0.26002 | 0.06561 |

| | | | | | | | | | | | | |
|---|---|---|---|---|---|---|---|---|---|---|---|---|
| **MTTP** | 0.00445 | 0.00938 | 0.18213 | 0.92636 | 0.00000 | 0.03740 | 0.00000 | 0.06407 | 0.05885 | 0.00306 | 0.00658 | 0.08595 |
| **MTURN** | 0.07310 | 0.15351 | 0.13062 | 0.37772 | 0.11931 | 0.94308 | 0.00000 | 0.05149 | 0.00000 | 0.93357 | 0.00000 | 0.26303 |
| **MTUS1** | 10.95185 | 0.00000 | 7.34381 | 1.44435 | 1.66132 | 6.48886 | 4.38599 | 0.31170 | 3.99662 | 5.18014 | 1.96416 | 1.55052 |
| **MTUS2** | 4.93522 | 0.97653 | 0.00000 | 0.00000 | 1.24380 | 0.00000 | 1.37862 | 0.00000 | 1.29273 | 2.03759 | 5.41664 | 0.36744 |
| **MTVR2** | 0.27355 | 0.02044 | 0.00561 | 0.02078 | 0.00630 | 0.01783 | 0.11613 | 0.00149 | 0.00000 | 0.02029 | 0.00994 | 0.01214 |
| **MTX1** | 0.01154 | 7.75777 | 6.26140 | 3.70449 | 5.45219 | 3.42766 | 3.13818 | 0.18253 | 6.80268 | 0.00000 | 5.66624 | 5.24214 |
| **MTX2** | 0.95592 | 13.45076 | 7.96921 | 4.02689 | 2.91340 | 28.84492 | 6.86938 | 0.45688 | 10.72007 | 2.18325 | 11.09441 | 8.64635 |
| **MTX3** | 1.34011 | 2.28167 | 3.12611 | 0.00000 | 0.00000 | 9.69657 | 4.52690 | 7.03262 | 3.08127 | 6.58242 | 1.41983 | 4.51202 |
| **MUC1** | 0.03370 | 0.00179 | 0.12920 | 0.17094 | 0.00000 | 0.06971 | 0.00235 | 0.05161 | 0.17841 | 0.08228 | 0.02103 | 0.00610 |
| **MUC12** | 0.08312 | 0.02175 | 0.01107 | 0.00157 | 0.05162 | 0.04376 | 0.01102 | 1.44695 | 0.00000 | 0.01951 | 0.20565 | 0.48958 |
| **MUC13** | 0.08257 | 0.00957 | 0.02027 | 0.26439 | 0.00000 | 0.02670 | 0.01134 | 0.00505 | 0.00000 | 0.02445 | 0.04761 | 0.01323 |
| **MUC15** | 0.06142 | 0.00000 | 23.61409 | 0.02032 | 1.75396 | 0.11170 | 0.00000 | 0.00590 | 3.89802 | 0.00000 | 0.01987 | 0.00891 |
| **MUC16** | 3.12354 | 0.06434 | 0.23724 | 0.00000 | 0.30414 | 0.00000 | 0.10530 | 0.00000 | 0.00000 | 0.60141 | 0.23248 | 2.31183 |
| **MUC17** | 0.00918 | 0.01645 | 0.02496 | 0.01518 | 0.01962 | 0.01388 | 0.01378 | 0.06835 | 0.02388 | 0.01648 | 0.01468 | 0.01272 |
| **MUC19** | 0.00600 | 0.00000 | 0.00109 | 0.01542 | 0.00000 | 0.01826 | 0.21052 | 0.07661 | 0.00000 | 0.47825 | 0.00000 | 0.00791 |
| **MUC2** | 0.09479 | 0.00204 | 0.02309 | 0.00915 | 0.03543 | 0.00000 | 0.03946 | 0.23366 | 0.00000 | 0.16581 | 0.05675 | 0.05464 |
| **MUC20** | 2.40499 | 0.07118 | 0.05669 | 0.02845 | 0.00000 | 0.00000 | 0.20928 | 0.85992 | 0.10525 | 0.66671 | 0.11577 | 0.04834 |
| **MUC21** | 0.00214 | 0.22249 | 0.01328 | 0.42145 | 0.01578 | 0.08032 | 0.05359 | 0.35859 | 0.00000 | 0.00000 | 0.01432 | 0.92627 |
| **MUC22** | 0.03529 | 0.01682 | 0.01174 | 0.01146 | 0.01241 | 0.00080 | 0.10114 | 0.09470 | 0.00000 | 0.04635 | 0.01361 | 0.03379 |
| **MUC3A** | 4.38920 | 0.09669 | 0.33168 | 0.02924 | 1.05947 | 0.00000 | 0.09941 | 1.70810 | 0.00000 | 1.08068 | 0.44688 | 1.69334 |
| **MUC4** | 0.13215 | 0.21861 | 1.36373 | 2.87908 | 0.00000 | 0.00000 | 0.20405 | 0.00618 | 0.12840 | 0.32878 | 0.00000 | 0.00000 |
| **MUC5B** | 0.12230 | 0.43781 | 0.00000 | 0.04705 | 0.19506 | 0.02557 | 0.00000 | 0.00000 | 0.00886 | 0.25215 | 0.02055 | 0.55689 |
| **MUC6** | 0.00272 | 0.00267 | 0.09169 | 0.00368 | 0.07035 | 0.00000 | 0.00033 | 0.00880 | 0.00241 | 0.97787 | 0.00000 | 0.00718 |
| **MUCL1** | 0.00626 | 0.04434 | 0.18165 | 0.01655 | 0.05554 | 0.00000 | 0.02128 | 0.01084 | 0.00000 | 0.01392 | 0.02462 | 0.02269 |
| **MUL1** | 1.42084 | 20.62230 | 4.46535 | 4.89625 | 8.67336 | 7.34151 | 4.37639 | 3.06198 | 12.93601 | 1.46927 | 9.48062 | 14.59524 |
| **MUM1** | 0.25762 | 6.87695 | 3.10861 | 5.38847 | 4.19715 | 0.00000 | 0.77035 | 1.03597 | 3.58791 | 8.34857 | 4.96668 | 3.13401 |
| **MUM1L1** | 0.06040 | 0.01240 | 0.04292 | 0.01591 | 0.01348 | 0.01017 | 0.01014 | 0.01804 | 0.01336 | 0.02463 | 0.02332 | 0.03976 |
| **MURC** | 0.00277 | 0.26433 | 0.43305 | 0.00000 | 0.00000 | 0.00000 | 0.16617 | 0.00562 | 0.00000 | 0.51729 | 0.14121 | 0.48662 |
| **MUS81** | 1.53048 | 12.94245 | 5.80681 | 6.69073 | 8.54958 | 0.00000 | 0.92213 | 0.08741 | 7.09515 | 0.00000 | 3.96941 | 4.75281 |
| **MUSK** | 0.06839 | 0.00984 | 0.02302 | 0.00836 | 0.02873 | 0.03680 | 0.00817 | 0.00560 | 0.01425 | 0.02931 | 0.01091 | 0.01452 |
| **MUSTN1** | 0.00463 | 0.08438 | 0.05700 | 0.02100 | 0.04297 | 0.00979 | 0.05610 | 0.00954 | 0.01727 | 0.06442 | 0.03910 | 0.03541 |
| **MUT** | 1.52738 | 0.85521 | 3.36763 | 1.55215 | 0.06643 | 10.28775 | 0.00621 | 1.39262 | 3.48390 | 0.56820 | 3.65198 | 2.22833 |
| **MUTYH** | 0.29155 | 4.30783 | 1.03744 | 3.94666 | 0.94010 | 1.19689 | 1.22478 | 0.13565 | 1.57723 | 0.65217 | 1.51390 | 5.87723 |
| **MVB12A** | 0.14162 | 3.18796 | 4.89448 | 5.37463 | 5.38212 | 0.00000 | 4.30530 | 1.93644 | 6.57838 | 0.00000 | 5.39462 | 5.61914 |
| **MVB12B** | 0.68438 | 1.83348 | 0.11265 | 0.23733 | 0.09267 | 0.00000 | 0.00000 | 0.01503 | 0.00000 | 0.57519 | 0.47437 | 0.21438 |
| **MVD** | 0.35264 | 3.97207 | 7.55487 | 11.34422 | 27.17758 | 0.00000 | 12.19363 | 0.44683 | 18.48591 | 11.28975 | 25.41527 | 0.82800 |
| **MVK** | 1.33598 | 0.00000 | 3.05546 | 4.43494 | 14.20963 | 0.00000 | 4.08047 | 0.00000 | 10.46599 | 1.08779 | 3.16579 | 2.13516 |
| **MVP** | 14.22260 | 0.39069 | 5.03714 | 4.78282 | 1.87877 | 0.00000 | 1.04053 | 8.38953 | 2.22719 | 1.34740 | 1.87928 | 2.68019 |
| **MX1** | 0.00000 | 22.51300 | 0.00000 | 31.18837 | 19.28799 | 3.40224 | 0.00000 | 0.00000 | 0.00000 | 2.91667 | 16.89934 | 2.54953 |
| **MX2** | 0.92851 | 0.00048 | 0.10465 | 0.05267 | 0.00000 | 0.17228 | 0.00000 | 0.05937 | 0.11918 | 0.00000 | 0.02782 | 0.04957 |
| **MXD1** | 10.35505 | 0.22722 | 16.95434 | 0.62688 | 3.22801 | 3.49569 | 1.29290 | 1.42681 | 1.84058 | 0.96509 | 1.93203 | 0.00000 |
| **MXD3** | 0.01141 | 0.86366 | 0.72536 | 2.02031 | 1.09201 | 0.00000 | 0.29316 | 1.60183 | 1.83340 | 2.00141 | 0.50778 | 0.34338 |
| **MXD4** | 5.84101 | 0.30413 | 0.50741 | 2.32340 | 1.15158 | 0.00000 | 0.42737 | 0.20264 | 0.68489 | 4.27475 | 0.00000 | 0.20583 |
| **MXI1** | 0.69476 | 0.53671 | 0.00000 | 1.87261 | 2.19361 | 2.50542 | 0.44107 | 0.01689 | 0.11223 | 2.53343 | 0.10293 | 0.88761 |
| **MXRA5** | 0.01297 | 0.01910 | 0.04675 | 0.03251 | 0.00646 | 0.05932 | 0.00000 | 0.00892 | 0.00000 | 0.06925 | 0.01008 | 0.00983 |
| **MXRA7** | 0.00000 | 1.83636 | 0.59532 | 0.00000 | 4.54183 | 2.48260 | 0.57904 | 0.01181 | 0.53047 | 2.23224 | 1.33545 | 0.01883 |
| **MXRA8** | 0.04229 | 0.00000 | 10.30397 | 2.03617 | 0.94475 | 0.00000 | 1.01640 | 0.01242 | 14.71621 | 3.23567 | 1.18617 | 0.12871 |
| **MYADM** | 5.99705 | 30.46587 | 3.78959 | 8.30030 | 38.72554 | 0.00000 | 0.73449 | 8.42763 | 17.47213 | 14.80924 | 26.49724 | 20.21213 |

| | | | | | | | | | | | | |
|---|---|---|---|---|---|---|---|---|---|---|---|---|
| **MYADML** | 0.07913 | 0.11256 | 0.00000 | 0.11425 | 0.03598 | 0.67170 | 0.00000 | 0.00166 | 0.00000 | 0.47088 | 0.05034 | 0.00000 |
| **MYADML2** | 0.02275 | 0.01413 | 0.01514 | 0.01194 | 0.01294 | 0.01957 | 0.01725 | 0.01087 | 0.01206 | 0.03063 | 0.02040 | 0.01443 |
| **MYB** | 0.40542 | 0.41084 | 0.02541 | 0.66321 | 0.20321 | 0.23266 | 0.00000 | 0.99924 | 0.00000 | 0.00407 | 0.00745 | 0.53367 |
| **MYBBP1A** | 4.70989 | 13.75304 | 10.70289 | 4.54814 | 5.24663 | 0.00000 | 3.16259 | 0.44119 | 27.91747 | 3.82262 | 13.53099 | 11.80874 |
| **MYBL1** | 1.05312 | 0.00000 | 0.00000 | 0.03346 | 0.14165 | 0.60635 | 0.31877 | 0.06593 | 0.00000 | 1.28117 | 0.09851 | 0.27928 |
| **MYBL2** | 6.61858 | 31.08348 | 5.66294 | 31.19671 | 55.57915 | 17.56129 | 34.21978 | 8.71058 | 12.48644 | 46.99940 | 12.84301 | 30.13498 |
| **MYBPC1** | 0.00371 | 0.01276 | 0.01478 | 0.01258 | 0.02084 | 0.00960 | 0.02001 | 0.85853 | 0.01861 | 0.00582 | 0.01614 | 0.04839 |
| **MYBPC2** | 0.54075 | 0.02575 | 0.05375 | 0.25844 | 0.01761 | 0.00000 | 0.08561 | 0.15292 | 0.02726 | 0.00000 | 0.01535 | 0.02305 |
| **MYBPC3** | 0.05181 | 0.01095 | 0.10952 | 0.01834 | 0.01467 | 0.01454 | 0.00619 | 0.00656 | 0.00576 | 0.09328 | 0.03572 | 0.01231 |
| **MYBPH** | 0.00974 | 0.02831 | 0.00544 | 0.00305 | 0.10152 | 0.00000 | 0.15447 | 0.00936 | 0.02881 | 0.01950 | 0.09225 | 0.01809 |
| **MYBPHL** | 0.27691 | 0.06881 | 0.00970 | 0.00419 | 0.07326 | 0.01717 | 0.00057 | 0.00156 | 0.30679 | 0.00000 | 0.03994 | 0.01897 |
| **MYC** | 0.49367 | 13.00949 | 10.06193 | 0.00000 | 17.69871 | 9.95707 | 17.35714 | 38.56595 | 28.22677 | 0.00000 | 34.38453 | 30.29214 |
| **MYCBP** | 0.15235 | 0.76694 | 0.36343 | 0.08427 | 0.58378 | 0.37123 | 0.31912 | 0.31199 | 0.50710 | 0.00000 | 0.50099 | 1.26286 |
| **MYCBP2** | 0.76469 | 0.26699 | 5.41116 | 3.15965 | 3.10899 | 10.79606 | 9.08720 | 0.20062 | 2.01141 | 19.67583 | 2.73229 | 3.20474 |
| **MYCBP2-AS1** | 0.00567 | 0.01685 | 0.01011 | 0.00887 | 0.02705 | 0.02804 | 0.01425 | 0.01038 | 0.03457 | 0.03615 | 0.01289 | 0.01344 |
| **MYCBPAP** | 0.01235 | 0.00863 | 0.01072 | 0.00980 | 0.00013 | 0.02769 | 0.00157 | 0.01305 | 0.02403 | 0.12063 | 0.02153 | 0.01063 |
| **MYCL** | 0.20789 | 7.81252 | 0.31130 | 8.59308 | 1.57572 | 0.00000 | 1.28072 | 0.00000 | 1.08997 | 3.05007 | 0.21792 | 4.40404 |
| **MYCN** | 0.00000 | 4.90970 | 4.26019 | 4.95942 | 5.91052 | 13.33090 | 2.92002 | 1.54518 | 0.00000 | 28.56110 | 0.70481 | 5.17232 |
| **MYCNOS** | 0.04178 | 1.41265 | 0.11678 | 0.39417 | 0.63763 | 0.18817 | 0.00000 | 0.08163 | 0.22979 | 0.90410 | 0.09933 | 1.30639 |
| **MYCT1** | 0.08960 | 0.72235 | 0.70682 | 0.48369 | 5.82715 | 1.62849 | 5.85184 | 0.00000 | 14.17081 | 0.00000 | 3.48076 | 0.00000 |
| **MYD88** | 0.17272 | 1.28011 | 5.18939 | 6.56642 | 0.00000 | 1.03388 | 4.26188 | 0.00000 | 6.78316 | 0.00000 | 3.20925 | 0.01871 |
| **MYEF2** | 0.63506 | 0.07371 | 0.37327 | 0.00000 | 0.39039 | 0.00000 | 0.07569 | 0.27834 | 0.00000 | 0.90307 | 0.05501 | 0.00000 |
| **MYEOV2** | 0.82815 | 13.25243 | 8.64980 | 7.31419 | 16.12430 | 10.58327 | 23.16941 | 6.87780 | 11.82058 | 22.23515 | 11.20367 | 18.28655 |
| **MYF6** | 0.00673 | 0.01326 | 0.03124 | 0.01165 | 0.01070 | 0.02508 | 0.01146 | 0.01375 | 0.01820 | 0.02580 | 0.01349 | 0.01488 |
| **MYH1** | 0.01573 | 0.14888 | 0.07864 | 0.00000 | 0.28158 | 0.07360 | 0.00000 | 0.02850 | 0.04852 | 0.03029 | 0.17917 | 0.13730 |
| **MYH10** | 3.14169 | 6.08170 | 12.74796 | 13.88685 | 21.17316 | 19.15840 | 19.34539 | 2.44769 | 34.52843 | 23.35871 | 22.15516 | 0.14311 |
| **MYH11** | 2.84653 | 0.09010 | 0.13003 | 0.12155 | 0.00000 | 0.04757 | 0.22703 | 0.20451 | 0.04853 | 0.35898 | 0.00001 | 0.22350 |
| **MYH13** | 0.29403 | 0.19728 | 0.15677 | 0.13168 | 0.00000 | 0.23086 | 0.06862 | 0.17225 | 0.00000 | 0.47140 | 0.08084 | 0.20784 |
| **MYH14** | 0.16075 | 0.13432 | 0.00026 | 0.10733 | 0.37172 | 0.10274 | 0.09258 | 0.00678 | 0.00000 | 0.39099 | 0.03475 | 0.03607 |
| **MYH15** | 4.69137 | 0.00000 | 0.14434 | 0.21705 | 0.00000 | 0.00000 | 0.04077 | 1.91208 | 0.20414 | 0.17980 | 0.00001 | 0.16911 |
| **MYH16** | 0.19279 | 0.07781 | 0.01167 | 0.01060 | 0.00982 | 0.03434 | 0.01513 | 0.03566 | 0.01660 | 0.01223 | 0.02859 | 0.01007 |
| **MYH2** | 0.00241 | 0.02682 | 0.00378 | 0.00253 | 0.02596 | 0.18834 | 0.00000 | 0.02575 | 0.05460 | 0.00544 | 0.10274 | 0.09032 |
| **MYH3** | 0.00341 | 0.00532 | 0.27461 | 0.49291 | 0.13921 | 0.23209 | 0.01213 | 0.35891 | 0.46634 | 1.33834 | 0.14317 | 0.58519 |
| **MYH4** | 0.00475 | 0.11376 | 0.02264 | 0.02431 | 0.17861 | 0.02338 | 0.00000 | 0.02677 | 0.01861 | 0.35343 | 0.05143 | 0.10399 |
| **MYH6** | 0.00733 | 0.01754 | 0.02380 | 0.01554 | 0.01485 | 0.00463 | 0.01844 | 0.05677 | 0.02041 | 0.02302 | 0.02651 | 0.02111 |
| **MYH7** | 0.41817 | 0.00175 | 0.01974 | 0.01337 | 0.01579 | 0.00000 | 0.01438 | 0.10377 | 0.02083 | 0.04318 | 0.01574 | 0.26960 |
| **MYH7B** | 0.04773 | 0.24369 | 0.03694 | 0.67587 | 0.18170 | 0.00836 | 0.00000 | 0.00199 | 0.01669 | 0.41188 | 0.02538 | 0.08189 |
| **MYH8** | 0.04472 | 0.06738 | 0.03992 | 0.00000 | 0.12596 | 0.10924 | 0.00000 | 0.00514 | 0.01096 | 0.36574 | 0.08129 | 0.25178 |
| **MYH9** | 0.36450 | 20.20303 | 9.76869 | 12.96805 | 37.53052 | 2.39640 | 25.27903 | 1.72640 | 24.75256 | 29.47659 | 14.36436 | 14.59965 |
| **MYL1** | 0.16258 | 0.01672 | 0.06956 | 0.01164 | 0.01664 | 0.00810 | 0.00991 | 0.10485 | 0.00909 | 0.01261 | 0.03110 | 0.01107 |
| **MYL10** | 0.00238 | 0.00596 | 0.13407 | 0.02133 | 0.00000 | 0.06436 | 0.00000 | 0.07525 | 0.00000 | 0.03650 | 0.01564 | 0.01912 |
| **MYL12A** | 1.53760 | 17.62417 | 23.41785 | 13.96241 | 11.35097 | 31.56176 | 25.53406 | 3.70667 | 22.33514 | 2.04047 | 21.59537 | 9.58719 |
| **MYL12B** | 2.43842 | 17.68556 | 26.54351 | 11.14164 | 22.31975 | 39.12223 | 34.79661 | 9.84044 | 24.73055 | 0.00000 | 28.71752 | 13.70733 |
| **MYL2** | 0.00584 | 0.01625 | 0.01604 | 0.01548 | 0.02315 | 0.00173 | 0.02499 | 0.05438 | 0.02137 | 0.00430 | 0.01668 | 0.04122 |
| **MYL3** | 0.01550 | 0.00761 | 0.14730 | 0.61336 | 0.00000 | 0.05353 | 0.04154 | 0.01704 | 0.00358 | 0.00977 | 0.02423 | 0.01200 |
| **MYL4** | 0.43467 | 0.52857 | 1.45744 | 4.83157 | 0.32437 | 0.00000 | 0.00000 | 0.24509 | 0.33042 | 0.00000 | 0.06899 | 0.01903 |
| **MYL5** | 0.00000 | 1.39577 | 0.00000 | 1.06392 | 3.79772 | 0.88454 | 0.57045 | 0.01457 | 0.00000 | 1.73817 | 1.38738 | 0.11873 |
| **MYL6** | 0.82167 | 36.26798 | 32.23169 | 32.12727 | 43.12661 | 37.43106 | 47.97626 | 9.50218 | 24.52288 | 24.93767 | 31.01282 | 37.49823 |

| | | | | | | | | | | | | |
|---|---|---|---|---|---|---|---|---|---|---|---|---|
| **MYL6B** | 0.83048 | 4.22860 | 8.43310 | 7.35814 | 5.64758 | 0.03928 | 7.49279 | 0.07326 | 8.54568 | 0.00000 | 4.12517 | 3.14398 |
| **MYL7** | 0.00540 | 0.59443 | 0.02915 | 0.00000 | 2.06302 | 0.01556 | 0.15883 | 0.01006 | 0.00000 | 0.00783 | 0.66890 | 0.00457 |
| **MYL9** | 0.00087 | 0.00000 | 1.62847 | 0.00000 | 0.00000 | 0.05272 | 1.48625 | 0.20587 | 0.37207 | 0.28215 | 0.53165 | 0.00000 |
| **MYLIP** | 5.39225 | 0.00000 | 0.41953 | 0.00000 | 3.46819 | 4.87806 | 3.34147 | 2.27457 | 3.17227 | 12.17197 | 0.57935 | 0.06453 |
| **MYLK** | 2.16793 | 0.24481 | 0.55623 | 0.00000 | 0.00000 | 0.95895 | 0.00000 | 0.26612 | 0.00000 | 0.55904 | 0.33459 | 0.93338 |
| **MYLK-AS1** | 0.00381 | 0.17117 | 0.34032 | 0.08703 | 0.00000 | 0.48119 | 0.00000 | 0.13024 | 0.08938 | 0.35670 | 0.44523 | 0.70940 |
| **MYLK2** | 0.19386 | 0.16115 | 0.15478 | 0.00000 | 0.36118 | 0.00000 | 0.42733 | 0.26483 | 0.12690 | 0.38737 | 0.00000 | 0.45998 |
| **MYLK3** | 1.05684 | 0.00000 | 0.40657 | 0.10358 | 0.00000 | 0.00000 | 0.26072 | 1.63836 | 0.10755 | 0.58960 | 0.06203 | 0.00000 |
| **MYLK4** | 0.02007 | 0.02647 | 0.01611 | 0.00928 | 0.00521 | 0.02585 | 0.01294 | 0.00867 | 0.01845 | 0.04130 | 0.01381 | 0.01352 |
| **MYLPF** | 0.00429 | 1.86925 | 0.91551 | 0.00000 | 3.78707 | 0.13264 | 2.94750 | 0.05350 | 1.50040 | 4.03232 | 12.08073 | 0.63769 |
| **MYNN** | 1.94376 | 3.73056 | 3.26883 | 1.65387 | 3.28422 | 13.49504 | 1.73918 | 5.40835 | 4.21603 | 3.48448 | 4.47515 | 6.67332 |
| **MYO10** | 5.66217 | 4.52011 | 28.59356 | 3.38066 | 3.77234 | 0.00037 | 9.47249 | 3.73519 | 0.96158 | 6.48456 | 0.33266 | 7.17499 |
| **MYO15A** | 0.56482 | 0.00000 | 0.01298 | 0.00859 | 0.13210 | 0.00000 | 0.00971 | 0.00000 | 0.00000 | 0.31711 | 0.19565 | 0.04611 |
| **MYO15B** | 0.12002 | 0.11689 | 0.05808 | 0.15888 | 0.00000 | 0.00000 | 0.62903 | 0.00943 | 0.02330 | 0.58330 | 0.36160 | 0.03178 |
| **MYO16** | 3.36019 | 0.08995 | 0.04444 | 0.00001 | 0.06265 | 0.00000 | 0.07133 | 0.10595 | 0.00978 | 0.01927 | 0.01131 | 0.31037 |
| **MYO18A** | 3.13256 | 0.33643 | 0.52540 | 3.73434 | 0.11683 | 0.00000 | 0.08860 | 0.17631 | 0.50080 | 0.39414 | 0.17572 | 0.43393 |
| **MYO18B** | 0.49977 | 0.04784 | 0.55386 | 0.00000 | 0.28721 | 0.00000 | 0.00000 | 0.03203 | 0.00000 | 0.49488 | 0.01607 | 0.07172 |
| **MYO19** | 2.90438 | 12.15099 | 11.61758 | 11.53239 | 10.74978 | 1.60149 | 8.83240 | 1.00631 | 24.75664 | 2.59605 | 14.57337 | 8.71932 |
| **MYO1A** | 0.00511 | 0.00496 | 0.02149 | 0.05072 | 0.04730 | 0.00476 | 0.00491 | 0.00571 | 0.04339 | 0.08832 | 0.01432 | 0.17440 |
| **MYO1B** | 0.70900 | 10.24024 | 6.48250 | 8.00391 | 2.70635 | 22.66623 | 13.93404 | 2.45302 | 6.60854 | 26.14533 | 1.91349 | 8.18712 |
| **MYO1C** | 1.32380 | 7.57283 | 7.49452 | 5.36975 | 10.76881 | 0.00000 | 11.79283 | 0.88171 | 13.10437 | 7.18480 | 8.00506 | 8.51123 |
| **MYO1D** | 0.04622 | 0.34154 | 2.76865 | 1.66338 | 0.45207 | 1.92095 | 0.62964 | 0.00000 | 0.44999 | 0.00004 | 0.23170 | 0.40938 |
| **MYO1E** | 3.38327 | 4.19582 | 19.03116 | 5.66567 | 5.90033 | 0.00000 | 4.67344 | 2.36045 | 0.60356 | 0.66758 | 1.66067 | 6.12954 |
| **MYO1F** | 0.38544 | 0.13012 | 0.11268 | 0.04025 | 0.05114 | 0.00000 | 0.02444 | 0.09304 | 0.04530 | 0.23511 | 0.05471 | 0.13998 |
| **MYO1G** | 0.00547 | 0.18158 | 0.03151 | 1.48559 | 0.74450 | 0.09405 | 0.00000 | 0.00475 | 0.00000 | 0.02187 | 0.02758 | 0.23060 |
| **MYO1H** | 3.51260 | 0.17371 | 0.20278 | 0.07330 | 0.11707 | 0.16063 | 0.00000 | 0.26630 | 0.00000 | 0.17703 | 0.00000 | 0.28665 |
| **MYO3A** | 0.04472 | 0.04892 | 0.00000 | 0.18728 | 0.03086 | 0.04807 | 0.00000 | 0.04286 | 0.10624 | 0.08846 | 0.03484 | 0.02107 |
| **MYO3B** | 1.69018 | 0.07664 | 0.03001 | 0.50862 | 0.18579 | 0.00491 | 0.02966 | 0.17542 | 0.00000 | 0.00000 | 0.03170 | 0.05564 |
| **MYO5A** | 0.71817 | 4.62108 | 5.94967 | 4.26070 | 4.35938 | 9.96219 | 0.00000 | 4.83823 | 1.89696 | 6.90748 | 0.82471 | 18.71334 |
| **MYO5B** | 4.95912 | 0.51120 | 1.80447 | 3.67938 | 6.81377 | 5.71377 | 6.37878 | 1.37280 | 1.55116 | 3.60430 | 2.94568 | 0.00000 |
| **MYO5C** | 0.00000 | 1.12739 | 3.26683 | 1.52985 | 0.84302 | 2.98126 | 0.44901 | 1.10943 | 0.00000 | 1.94051 | 0.99427 | 4.04595 |
| **MYO6** | 6.47569 | 0.62161 | 6.64334 | 0.00000 | 1.84282 | 18.90265 | 13.38998 | 3.19374 | 17.95292 | 19.57395 | 13.31519 | 1.06692 |
| **MYO7A** | 2.81391 | 0.00000 | 3.92841 | 0.00972 | 0.08394 | 0.00011 | 1.13720 | 0.06844 | 5.46014 | 0.00000 | 2.21898 | 0.15769 |
| **MYO7B** | 2.21986 | 0.00000 | 0.05556 | 0.02291 | 0.02446 | 0.00000 | 0.03247 | 0.20350 | 0.00000 | 0.05625 | 0.01371 | 0.20197 |
| **MYO9A** | 6.20510 | 4.56265 | 3.29187 | 12.15469 | 3.77711 | 8.22892 | 0.00000 | 2.52366 | 1.38614 | 7.92221 | 0.72394 | 2.00156 |
| **MYO9B** | 0.92885 | 0.21927 | 7.75803 | 0.99354 | 4.28619 | 0.00000 | 3.37360 | 0.29893 | 1.31014 | 16.49238 | 1.40737 | 0.43080 |
| **MYOC** | 0.29404 | 0.01177 | 0.02617 | 0.05882 | 0.02387 | 0.00190 | 0.00637 | 0.02729 | 0.02248 | 0.01810 | 0.05150 | 0.04564 |
| **MYOCD** | 1.14488 | 0.01136 | 0.09919 | 0.01279 | 0.01050 | 0.01479 | 0.05723 | 0.01167 | 0.00000 | 0.01657 | 0.02209 | 0.01386 |
| **MYOD1** | 0.02311 | 0.03681 | 0.00589 | 0.01541 | 0.03944 | 0.00000 | 0.01370 | 0.36887 | 0.25723 | 0.02630 | 0.00047 | 0.32305 |
| **MYOF** | 4.72993 | 7.04976 | 6.80757 | 1.29023 | 24.84471 | 19.32780 | 18.97099 | 1.37205 | 16.90310 | 10.73337 | 12.27543 | 0.00000 |
| **MYOG** | 0.00793 | 0.01589 | 0.01413 | 0.01285 | 0.01717 | 0.01848 | 0.02148 | 0.01119 | 0.01558 | 0.01535 | 0.01721 | 0.01607 |
| **MYOM1** | 0.00505 | 0.79377 | 0.18171 | 0.00000 | 0.43641 | 0.12209 | 0.00000 | 0.04383 | 0.00000 | 0.34413 | 0.24925 | 0.23913 |
| **MYOM2** | 0.17533 | 0.92876 | 0.00040 | 0.05299 | 0.00192 | 0.33547 | 0.08215 | 0.10379 | 0.00566 | 0.00000 | 0.14481 | 1.59156 |
| **MYOM3** | 0.02041 | 0.07328 | 0.01174 | 0.10765 | 0.00692 | 0.05186 | 0.01483 | 0.11618 | 0.01006 | 0.00394 | 0.02086 | 0.19285 |
| **MYOT** | 0.20178 | 0.14188 | 0.07800 | 0.06245 | 0.00000 | 0.08550 | 0.00000 | 0.00000 | 0.00000 | 0.09018 | 0.18845 | 0.28874 |
| **MYOZ1** | 0.00890 | 0.00000 | 0.00000 | 0.08194 | 0.21270 | 0.00663 | 0.16136 | 0.00744 | 0.00000 | 0.29518 | 0.11654 | 0.11620 |
| **MYOZ2** | 0.00782 | 0.01162 | 0.01125 | 0.07391 | 0.00513 | 0.02689 | 0.01415 | 0.01008 | 0.01484 | 0.01849 | 0.01563 | 0.02343 |
| **MYOZ3** | 0.02450 | 0.08086 | 0.09019 | 0.04511 | 0.00611 | 0.20344 | 0.00000 | 0.00000 | 0.00000 | 0.02849 | 0.00355 | 0.43632 |

| | | | | | | | | | | | | |
|---|---|---|---|---|---|---|---|---|---|---|---|---|
| **MYPN** | 0.00157 | 0.11548 | 0.01105 | 0.00443 | 0.00000 | 0.02667 | 0.04844 | 0.22007 | 0.00389 | 0.02245 | 0.01290 | 0.07647 |
| **MYPOP** | 0.20489 | 0.12044 | 0.97128 | 0.28623 | 0.51531 | 0.00000 | 1.52305 | 0.00000 | 0.00000 | 3.18628 | 0.22594 | 0.42616 |
| **MYRF** | 0.00307 | 0.03358 | 0.02019 | 2.07792 | 0.65172 | 0.00000 | 0.00000 | 0.09501 | 0.75380 | 0.62131 | 0.22023 | 0.19285 |
| **MYRIP** | 2.50827 | 0.03147 | 0.08071 | 0.90404 | 0.00438 | 0.26169 | 0.00000 | 0.07534 | 0.00000 | 0.00000 | 0.28456 | 0.02766 |
| **MYSM1** | 0.70049 | 1.12973 | 2.16391 | 1.81028 | 0.00000 | 12.78090 | 0.00000 | 1.66370 | 3.75644 | 16.02952 | 1.15652 | 2.12767 |
| **MYT1** | 0.24207 | 0.11668 | 0.07297 | 0.02281 | 0.00119 | 0.00000 | 0.00666 | 0.39691 | 0.01101 | 0.03605 | 0.04175 | 1.66466 |
| **MYT1L** | 0.22100 | 0.00001 | 0.14440 | 0.00000 | 0.00000 | 0.00000 | 0.03435 | 0.11655 | 0.06076 | 0.48144 | 0.19663 | 0.91352 |
| **MYZAP** | 0.69315 | 0.00224 | 0.08973 | 0.09987 | 0.10081 | 0.07512 | 0.00295 | 0.08830 | 0.09658 | 0.00000 | 0.01196 | 0.00731 |
| **MZB1** | 0.00638 | 0.01020 | 0.10726 | 0.00681 | 0.00434 | 0.05124 | 0.01837 | 0.01005 | 0.10674 | 0.00000 | 0.00941 | 0.01523 |
| **MZF1** | 0.00071 | 0.00000 | 1.93524 | 0.34574 | 0.00000 | 0.00000 | 0.65855 | 0.17867 | 3.90571 | 3.98428 | 0.97620 | 1.29200 |
| **MZT1** | 0.66173 | 6.29091 | 8.64058 | 1.56285 | 0.45924 | 35.37419 | 14.36084 | 7.67153 | 9.97712 | 0.00000 | 9.07321 | 14.45849 |
| **MZT2A** | 0.37184 | 1.76059 | 2.37356 | 1.01109 | 1.45432 | 0.00000 | 5.50769 | 0.26800 | 1.92937 | 6.07404 | 1.97639 | 0.60815 |
| **MZT2B** | 0.12045 | 0.45479 | 1.62007 | 2.44706 | 3.27476 | 0.00000 | 5.82830 | 0.37079 | 0.60904 | 7.93172 | 0.85859 | 0.56059 |
| **N4BP1** | 2.35662 | 1.37700 | 1.27681 | 1.30703 | 1.32861 | 4.42759 | 2.76490 | 0.20829 | 1.06110 | 8.45904 | 1.00797 | 1.67836 |
| **N4BP2** | 2.91004 | 4.83493 | 0.00000 | 10.41913 | 9.11119 | 24.56126 | 0.00000 | 1.54144 | 0.97236 | 21.41568 | 5.56448 | 16.42799 |
| **N4BP2L1** | 0.01579 | 0.02131 | 0.06417 | 0.01060 | 0.00000 | 0.08157 | 0.15890 | 0.01431 | 0.08350 | 0.00000 | 0.00000 | 0.10668 |
| **N4BP2L2** | 5.01914 | 5.03669 | 8.49305 | 3.52980 | 0.00000 | 14.94237 | 4.88091 | 5.40936 | 12.10348 | 8.89573 | 6.58137 | 14.19807 |
| **N4BP2L2-IT2** | 0.09552 | 0.00000 | 0.06579 | 0.00307 | 0.00000 | 0.59217 | 0.00000 | 0.01712 | 0.77337 | 1.59769 | 0.00000 | 0.24065 |
| **N4BP3** | 0.30276 | 4.91735 | 1.76323 | 6.07476 | 2.39901 | 0.00000 | 0.86674 | 0.01002 | 2.47713 | 0.61604 | 1.76514 | 0.02299 |
| **N6AMT1** | 0.00187 | 5.72118 | 2.21157 | 0.53626 | 0.89739 | 3.78558 | 1.05562 | 0.72218 | 2.33445 | 0.00000 | 3.62208 | 2.82093 |
| **N6AMT2** | 0.00930 | 2.49776 | 0.97094 | 0.00000 | 0.00000 | 2.69129 | 2.55399 | 0.64761 | 0.28467 | 0.00000 | 3.41043 | 5.22114 |
| **NAA10** | 0.00000 | 12.88932 | 15.42822 | 12.48322 | 18.10395 | 5.18189 | 24.12365 | 3.83962 | 24.47805 | 0.00000 | 18.02374 | 9.42887 |
| **NAA11** | 2.94497 | 1.83521 | 4.80863 | 3.49724 | 2.83002 | 13.28522 | 2.42941 | 3.60635 | 9.01490 | 0.00000 | 2.50600 | 5.29563 |
| **NAA15** | 2.88122 | 11.41308 | 20.50170 | 9.21019 | 7.83799 | 32.33563 | 31.04958 | 8.84223 | 19.37028 | 18.97804 | 19.19456 | 19.33461 |
| **NAA16** | 1.23152 | 0.38338 | 2.88884 | 3.69827 | 0.00000 | 7.27721 | 2.61794 | 0.50857 | 3.35589 | 6.50450 | 0.81374 | 0.96809 |
| **NAA20** | 1.62473 | 14.69227 | 13.30011 | 5.08450 | 0.80678 | 32.18541 | 12.62595 | 2.81900 | 13.47259 | 6.41534 | 12.93979 | 16.89456 |
| **NAA25** | 0.00000 | 4.17592 | 10.48687 | 3.57961 | 0.00000 | 28.18498 | 8.25105 | 3.57226 | 15.86019 | 17.03083 | 8.33047 | 8.68454 |
| **NAA30** | 5.00928 | 0.54739 | 1.27465 | 2.71055 | 0.51951 | 8.79899 | 4.44167 | 1.39071 | 0.00000 | 14.48849 | 1.93468 | 1.66346 |
| **NAA35** | 8.00428 | 4.22968 | 6.82965 | 3.25673 | 1.72887 | 10.40408 | 9.04729 | 5.64541 | 4.25575 | 6.67009 | 6.96520 | 6.89327 |
| **NAA38** | 3.53875 | 14.79995 | 17.86942 | 14.30487 | 22.16397 | 0.00170 | 27.08430 | 4.35758 | 19.86559 | 8.82416 | 15.71787 | 13.93470 |
| **NAA40** | 1.32172 | 7.30831 | 5.24189 | 3.62918 | 5.64146 | 2.75649 | 3.05190 | 4.11471 | 5.80140 | 1.19179 | 3.15617 | 10.57941 |
| **NAA50** | 5.58892 | 23.71608 | 23.51497 | 13.16369 | 11.19612 | 57.81803 | 38.40787 | 15.95478 | 25.43257 | 11.34123 | 24.92748 | 25.73823 |
| **NAA60** | 5.90089 | 8.28558 | 10.69791 | 10.55830 | 0.00000 | 0.00000 | 10.46161 | 4.70233 | 15.68005 | 2.01729 | 5.18537 | 10.07112 |
| **NAAA** | 0.23774 | 0.22998 | 0.10266 | 0.29509 | 1.73760 | 0.52765 | 0.00000 | 0.06883 | 1.40059 | 0.10414 | 1.41958 | 1.03333 |
| **NAALAD2** | 0.65625 | 0.52970 | 0.00000 | 0.00000 | 0.00000 | 3.53144 | 10.16577 | 2.07125 | 11.73133 | 2.49757 | 5.95782 | 1.44218 |
| **NAALADL1** | 0.10123 | 0.00954 | 0.01666 | 0.01011 | 0.00979 | 0.03378 | 0.01708 | 0.00158 | 0.01686 | 0.01546 | 0.01270 | 0.07543 |
| **NAALADL2** | 0.55286 | 0.00193 | 0.08391 | 0.03485 | 0.00520 | 0.11228 | 0.00000 | 0.10489 | 0.00000 | 0.01826 | 0.04729 | 0.02008 |
| **NAALADL2-AS3** | 0.00939 | 0.01298 | 0.02062 | 0.01831 | 0.01044 | 0.02220 | 0.01439 | 0.01143 | 0.01483 | 0.03035 | 0.01460 | 0.01492 |
| **NAB1** | 1.19722 | 2.27327 | 1.27341 | 0.00000 | 1.10932 | 3.04962 | 1.97992 | 0.15992 | 0.68933 | 4.68240 | 2.32036 | 1.52197 |
| **NAB2** | 0.00107 | 0.06299 | 0.98148 | 0.00623 | 0.00000 | 0.00000 | 0.40362 | 0.11280 | 0.47245 | 0.22283 | 0.00000 | 0.12578 |
| **NABP1** | 1.80339 | 4.76880 | 33.35068 | 2.90879 | 0.00000 | 18.68037 | 15.28647 | 23.14279 | 10.62782 | 10.59786 | 10.98850 | 12.30972 |
| **NABP2** | 1.78635 | 2.32106 | 6.56363 | 1.47185 | 0.00000 | 0.17737 | 9.02346 | 2.49635 | 8.23693 | 2.52464 | 3.95708 | 5.56587 |
| **NACA** | 7.76651 | 27.72277 | 30.43872 | 22.43610 | 27.93725 | 50.34298 | 45.01871 | 22.28043 | 39.97412 | 23.10634 | 32.97426 | 38.35271 |
| **NACA2** | 1.70117 | 8.72871 | 10.98314 | 8.68524 | 10.10079 | 19.19008 | 19.56181 | 6.69726 | 12.80534 | 7.09743 | 11.59972 | 13.51015 |
| **NACAD** | 0.13089 | 0.08697 | 0.11170 | 1.05115 | 0.00000 | 0.00000 | 0.21832 | 0.03565 | 0.15975 | 1.66290 | 0.00000 | 0.18703 |
| **NACAP1** | 0.17419 | 1.25796 | 2.88846 | 1.62451 | 1.15867 | 3.05769 | 2.18050 | 1.11986 | 2.07576 | 0.00740 | 1.67709 | 2.00288 |
| **NACC1** | 0.63645 | 0.60617 | 1.54026 | 0.80246 | 3.38087 | 0.00000 | 3.91668 | 0.00000 | 0.28035 | 13.72711 | 0.75682 | 0.87952 |
| **NACC2** | 0.25740 | 2.31784 | 0.00000 | 0.00000 | 2.19217 | 0.95769 | 0.00000 | 0.00509 | 0.00000 | 3.73589 | 0.37197 | 0.00000 |

| | | | | | | | | | | | | |
|---|---|---|---|---|---|---|---|---|---|---|---|---|
| **NADK** | 0.09433 | 3.05819 | 7.72442 | 11.93557 | 7.37775 | 0.00000 | 4.76031 | 0.00000 | 5.32547 | 2.78298 | 6.26929 | 1.19535 |
| **NADK2** | 0.27496 | 2.35116 | 0.42781 | 0.43112 | 0.00000 | 3.10909 | 1.35935 | 0.21070 | 0.05918 | 2.72690 | 0.93396 | 0.00000 |
| **NADSYN1** | 0.88465 | 4.38427 | 5.17988 | 4.49706 | 6.77796 | 0.17131 | 5.14178 | 0.07271 | 9.90611 | 1.52300 | 3.73200 | 2.08415 |
| **NAE1** | 5.13633 | 13.42455 | 12.17129 | 4.61819 | 7.33103 | 39.55294 | 14.47173 | 6.31872 | 17.58411 | 2.00265 | 18.91231 | 12.87932 |
| **NAF1** | 0.82296 | 1.87920 | 2.47166 | 0.85967 | 3.10925 | 0.94135 | 4.79238 | 7.73447 | 2.29616 | 1.52342 | 2.69848 | 6.59544 |
| **NAGA** | 1.17052 | 3.65936 | 0.20914 | 8.46038 | 19.89080 | 5.23123 | 0.00000 | 1.00072 | 0.00000 | 2.68485 | 8.19597 | 1.20214 |
| **NAGK** | 1.63286 | 8.38645 | 14.10368 | 6.68752 | 4.72703 | 2.19617 | 9.38993 | 1.06753 | 10.18167 | 0.00000 | 8.31328 | 6.51632 |
| **NAGLU** | 0.00448 | 0.83108 | 0.76993 | 1.11289 | 0.80319 | 0.00000 | 0.00041 | 0.00061 | 0.00000 | 2.28197 | 0.70942 | 0.14314 |
| **NAGPA** | 1.98916 | 6.60426 | 8.04230 | 1.42512 | 2.66991 | 0.00000 | 1.85756 | 0.19893 | 12.29740 | 7.48185 | 4.19324 | 2.61745 |
| **NAGPA-AS1** | 0.02839 | 0.11978 | 0.12707 | 0.01629 | 0.00000 | 0.00000 | 0.00000 | 0.00688 | 0.16301 | 0.03071 | 0.17833 | 0.09411 |
| **NAGS** | 0.32794 | 0.01564 | 0.33758 | 0.16488 | 0.17239 | 0.00000 | 0.00350 | 0.13603 | 0.01431 | 0.00000 | 0.03270 | 0.17352 |
| **NAIF1** | 1.35643 | 0.38811 | 1.79611 | 0.55653 | 1.18957 | 0.19774 | 1.00817 | 1.05993 | 0.37415 | 1.41231 | 0.60484 | 1.55111 |
| **NAIP** | 0.00623 | 0.01284 | 0.02972 | 0.03516 | 0.00256 | 0.02848 | 0.01221 | 0.08712 | 0.01153 | 0.07175 | 0.01616 | 0.01470 |
| **NALCN** | 0.65344 | 0.00944 | 0.01010 | 0.00938 | 0.00778 | 0.00809 | 0.03509 | 0.08815 | 0.00010 | 0.12528 | 0.01067 | 0.01497 |
| **NALCN-AS1** | 0.11240 | 0.01542 | 0.01880 | 0.01755 | 0.00453 | 0.04022 | 0.00815 | 0.06040 | 0.01409 | 0.02860 | 0.01627 | 0.01592 |
| **NAMA** | 0.00642 | 0.01766 | 0.03249 | 0.01372 | 0.02019 | 0.00000 | 0.01596 | 0.01063 | 0.02990 | 0.01287 | 0.03447 | 0.01769 |
| **NAMPT** | 1.49304 | 4.33318 | 3.20190 | 2.09863 | 0.00000 | 9.37292 | 5.49933 | 3.48676 | 3.65657 | 1.03835 | 3.38679 | 6.13751 |
| **NANOG** | 0.67903 | 0.59207 | 0.04994 | 8.83515 | 0.00000 | 0.24556 | 0.08407 | 2.15740 | 0.00000 | 0.06730 | 0.00000 | 2.60020 |
| **NANOGNB** | 2.31271 | 8.06713 | 0.00000 | 0.48823 | 6.96002 | 0.00000 | 0.00000 | 13.61873 | 0.00000 | 0.56864 | 0.39406 | 20.85131 |
| **NANOS1** | 5.52771 | 0.37326 | 0.00000 | 0.47774 | 3.87449 | 2.73637 | 1.55330 | 0.00000 | 0.00000 | 5.66195 | 0.78166 | 0.55718 |
| **NANOS3** | 0.14479 | 0.06240 | 0.04036 | 0.05912 | 0.00937 | 0.02784 | 0.00901 | 0.00341 | 0.02010 | 0.00579 | 0.01611 | 0.01001 |
| **NANP** | 0.59363 | 3.26071 | 3.77318 | 0.43581 | 0.00482 | 8.14376 | 2.10331 | 2.26621 | 6.42887 | 2.39128 | 6.65426 | 4.12984 |
| **NANS** | 0.00000 | 7.13441 | 11.71740 | 8.80173 | 13.83973 | 8.12674 | 19.98809 | 18.68806 | 15.13001 | 0.00000 | 16.79829 | 11.76344 |
| **NAP1L1** | 10.26373 | 12.36924 | 14.72658 | 10.47377 | 9.22760 | 43.63258 | 38.70629 | 9.31205 | 22.57321 | 39.77171 | 12.85973 | 12.14098 |
| **NAP1L2** | 0.05477 | 0.00063 | 0.17750 | 0.17030 | 0.00000 | 0.08595 | 0.00000 | 0.03535 | 0.00000 | 0.05032 | 0.01818 | 0.05174 |
| **NAP1L3** | 0.37610 | 0.18021 | 0.11837 | 0.01237 | 0.00000 | 0.08452 | 0.02926 | 0.00000 | 0.00824 | 0.00256 | 0.02062 | 0.34123 |
| **NAP1L4** | 7.31182 | 18.85638 | 22.38081 | 20.34069 | 22.08747 | 11.94022 | 30.38597 | 14.84830 | 18.95509 | 9.62551 | 21.82340 | 15.71924 |
| **NAP1L5** | 0.31526 | 0.65387 | 0.31622 | 0.00639 | 0.02443 | 0.43639 | 0.06207 | 0.68973 | 0.01114 | 0.00000 | 0.20271 | 0.43468 |
| **NAP1L6** | 0.00820 | 0.02264 | 0.03497 | 0.14667 | 0.07467 | 0.00000 | 0.10421 | 0.00992 | 0.00000 | 0.00000 | 0.06016 | 0.01823 |
| **NAPA** | 6.02928 | 14.62665 | 19.23878 | 14.15744 | 18.39128 | 5.26028 | 17.99996 | 10.66486 | 20.27731 | 0.00000 | 13.73561 | 13.87173 |
| **NAPA-AS1** | 0.01502 | 0.00990 | 0.16531 | 0.12554 | 0.31416 | 0.00000 | 0.00091 | 0.08335 | 0.09323 | 0.13665 | 0.09316 | 0.17360 |
| **NAPB** | 0.38791 | 1.84135 | 0.36886 | 0.36223 | 0.00000 | 5.22419 | 0.00000 | 1.10787 | 3.56188 | 3.85525 | 2.90327 | 2.52552 |
| **NAPEPLD** | 0.03759 | 0.16121 | 1.02284 | 0.53825 | 0.48969 | 1.20134 | 1.26868 | 0.03172 | 0.36750 | 0.06811 | 0.72214 | 0.62679 |
| **NAPG** | 0.56116 | 8.48693 | 4.54530 | 4.91536 | 5.49613 | 17.32814 | 9.13239 | 2.89866 | 8.94817 | 8.88345 | 5.55565 | 5.21220 |
| **NAPRT1** | 0.00000 | 5.78036 | 1.55847 | 4.24965 | 6.12870 | 0.00000 | 0.39284 | 0.09586 | 1.39077 | 8.00356 | 1.60079 | 1.50615 |
| **NAPSA** | 0.02441 | 0.12049 | 0.03744 | 0.10540 | 0.00473 | 0.01552 | 0.04236 | 0.00232 | 0.09773 | 0.09295 | 0.00000 | 0.27249 |
| **NAPSB** | 0.10910 | 0.02044 | 0.01224 | 0.00923 | 0.02198 | 0.01327 | 0.00941 | 0.20756 | 0.08186 | 0.00443 | 0.00910 | 0.03497 |
| **NARF** | 5.24931 | 21.61459 | 13.97728 | 12.13914 | 13.73217 | 14.93648 | 22.36537 | 12.70805 | 26.66109 | 0.07909 | 12.96007 | 22.27958 |
| **NARFL** | 0.54160 | 7.49231 | 9.46082 | 6.33033 | 5.28111 | 0.00000 | 5.79621 | 0.00000 | 10.51925 | 0.00000 | 7.56147 | 6.65135 |
| **NARS** | 0.66238 | 27.00289 | 21.68423 | 16.94198 | 11.67208 | 52.02704 | 35.16616 | 9.36255 | 33.84506 | 18.66046 | 34.94532 | 21.87135 |
| **NARS2** | 2.77734 | 7.88289 | 7.82335 | 3.75980 | 0.71118 | 14.20284 | 7.43166 | 3.54809 | 11.81667 | 0.00004 | 6.87603 | 7.37506 |
| **NASP** | 12.26622 | 30.11999 | 21.61021 | 22.77992 | 27.08580 | 38.02259 | 35.01394 | 14.67796 | 32.91706 | 26.54616 | 19.53619 | 21.91492 |
| **NAT1** | 0.79380 | 0.00000 | 2.18427 | 1.23852 | 0.66110 | 0.92329 | 3.33853 | 0.05249 | 2.44297 | 0.61318 | 1.54319 | 0.00000 |
| **NAT10** | 2.91684 | 12.99579 | 9.88024 | 8.43034 | 11.72132 | 8.72267 | 16.39912 | 5.10446 | 10.69168 | 9.47814 | 13.65830 | 12.18511 |
| **NAT14** | 0.60632 | 0.59133 | 0.14792 | 0.49173 | 0.88385 | 0.00000 | 0.65052 | 0.00000 | 0.00000 | 2.19081 | 0.40691 | 0.33508 |
| **NAT16** | 0.00697 | 0.30239 | 0.09249 | 0.02030 | 0.09471 | 0.00000 | 0.00368 | 0.00967 | 0.07873 | 0.00000 | 0.01008 | 0.04973 |
| **NAT2** | 0.62481 | 0.01335 | 0.01236 | 0.01746 | 0.00837 | 0.02823 | 0.03292 | 0.04773 | 0.05544 | 0.00000 | 0.00289 | 0.00982 |
| **NAT6** | 0.79409 | 1.57537 | 0.84131 | 2.49546 | 1.32996 | 0.00000 | 0.70231 | 0.85910 | 0.61087 | 0.00000 | 2.28815 | 1.70501 |

| | | | | | | | | | | | | |
|---|---|---|---|---|---|---|---|---|---|---|---|---|
| **NAT8** | 0.01764 | 0.01680 | 0.01615 | 0.01320 | 0.01541 | 0.00796 | 0.02481 | 0.01112 | 0.02471 | 0.01061 | 0.03120 | 0.01687 |
| **NAT8L** | 0.15340 | 0.10095 | 0.27832 | 0.00000 | 0.73808 | 0.00000 | 0.00000 | 0.01649 | 0.17158 | 0.83588 | 0.01460 | 0.00346 |
| **NAT9** | 0.79094 | 6.69210 | 10.18536 | 6.47126 | 8.18532 | 0.00000 | 11.64515 | 5.46426 | 12.69448 | 0.00000 | 7.57593 | 10.13187 |
| **NATD1** | 1.78947 | 4.43575 | 14.91822 | 0.91108 | 2.20554 | 0.00000 | 2.47128 | 0.09707 | 2.37775 | 2.95391 | 0.00000 | 11.12930 |
| **NAV1** | 0.22078 | 0.27862 | 6.74058 | 4.62834 | 0.29934 | 0.92788 | 3.15957 | 0.00000 | 0.00000 | 1.91706 | 0.26454 | 0.21025 |
| **NAV2** | 5.86661 | 1.26754 | 1.39712 | 2.10972 | 9.23058 | 1.16219 | 1.95638 | 2.02656 | 0.00000 | 2.02742 | 2.23255 | 1.21174 |
| **NAV2-AS2** | 0.01513 | 0.01630 | 0.03834 | 0.01397 | 0.03610 | 0.01079 | 0.01453 | 0.00976 | 0.04546 | 0.00332 | 0.01803 | 0.02512 |
| **NAV2-AS4** | 0.00732 | 0.01310 | 0.23991 | 0.00625 | 0.26188 | 0.00000 | 0.00261 | 0.01010 | 0.00000 | 0.01521 | 0.01284 | 0.01676 |
| **NAV3** | 4.71448 | 0.00242 | 0.01891 | 0.01001 | 0.01490 | 0.02594 | 0.00787 | 0.09634 | 0.01265 | 0.02486 | 0.01193 | 0.08596 |
| **NBAS** | 2.54859 | 11.86846 | 1.16955 | 12.78294 | 11.48956 | 5.99757 | 0.00000 | 1.36837 | 2.82738 | 8.70928 | 2.15490 | 3.48718 |
| **NBEA** | 0.32532 | 0.02241 | 0.30802 | 0.09475 | 0.00000 | 0.13237 | 0.00000 | 0.70569 | 0.00000 | 0.88804 | 0.05318 | 0.13368 |
| **NBEAL1** | 2.14107 | 0.96368 | 1.83131 | 0.61001 | 0.09604 | 6.51577 | 0.00000 | 0.49191 | 1.59968 | 5.68788 | 2.51390 | 2.38431 |
| **NBEAL2** | 0.15023 | 1.21961 | 0.52250 | 2.86547 | 3.53069 | 0.00000 | 0.00271 | 0.00000 | 2.10568 | 4.33636 | 0.64208 | 0.05559 |
| **NBEAP1** | 0.18306 | 0.01838 | 0.05597 | 0.00000 | 0.01357 | 0.09445 | 0.02360 | 1.33713 | 0.01141 | 0.03002 | 0.00938 | 0.29311 |
| **NBN** | 2.29949 | 4.62246 | 2.16403 | 0.94062 | 0.18421 | 20.08204 | 3.05559 | 0.13428 | 9.40510 | 3.85270 | 3.84871 | 5.03236 |
| **NBPF1** | 2.61976 | 1.90808 | 1.84242 | 0.69389 | 0.80516 | 0.00000 | 2.01724 | 1.67793 | 3.88920 | 2.69841 | 1.11689 | 3.94801 |
| **NBPF10** | 0.01107 | 0.00000 | 0.55118 | 0.01598 | 0.06997 | 0.03720 | 0.33199 | 0.01428 | 0.00000 | 0.35401 | 0.03355 | 0.06143 |
| **NBPF11.uid-434** | 0.00602 | 0.01646 | 0.01402 | 0.02098 | 0.01641 | 0.01158 | 0.01992 | 0.01076 | 0.01889 | 0.02549 | 0.01426 | 0.01884 |
| **NBPF12** | 0.00701 | 0.03397 | 0.02699 | 0.26884 | 0.02697 | 0.00000 | 0.01271 | 0.04656 | 0.07223 | 0.02380 | 0.06242 | 0.08339 |
| **NBPF13P** | 0.00761 | 0.01459 | 0.01588 | 0.02620 | 0.01370 | 0.01553 | 0.05146 | 0.01377 | 0.03138 | 0.00916 | 0.02226 | 0.01856 |
| **NBPF15** | 0.08126 | 0.85306 | 0.25329 | 0.80666 | 0.00000 | 0.09485 | 0.12028 | 0.14486 | 1.01185 | 0.09940 | 0.39449 | 0.77852 |
| **NBPF22P** | 0.00500 | 0.00435 | 0.01946 | 0.00590 | 0.00000 | 0.06812 | 0.01607 | 0.00971 | 0.04206 | 0.06126 | 0.03559 | 0.01263 |
| **NBPF25P.uid-43** | 0.01292 | 0.02232 | 0.04052 | 0.05685 | 0.02480 | 0.00000 | 0.00449 | 0.01342 | 0.05529 | 0.05491 | 0.03756 | 0.02330 |
| **NBPF25P.uid-43** | 0.01148 | 0.01734 | 0.02221 | 0.01097 | 0.02157 | 0.01152 | 0.01936 | 0.01196 | 0.02900 | 0.00849 | 0.02725 | 0.01672 |
| **NBPF3** | 0.22330 | 1.04145 | 0.79622 | 1.32497 | 0.62284 | 1.18164 | 0.00000 | 0.24589 | 0.00000 | 0.00000 | 1.37816 | 1.30283 |
| **NBPF4** | 0.00461 | 0.01377 | 0.02113 | 0.01177 | 0.01571 | 0.01528 | 0.06440 | 0.01597 | 0.00940 | 0.01658 | 0.03227 | 0.02159 |
| **NBPF6** | 0.00555 | 0.01677 | 0.02795 | 0.01639 | 0.02415 | 0.00043 | 0.02093 | 0.02041 | 0.02793 | 0.01211 | 0.02843 | 0.02324 |
| **NBPF8.uid-74** | 0.03504 | 0.02035 | 0.02957 | 0.03772 | 0.02861 | 0.08195 | 0.00000 | 0.00824 | 0.07494 | 0.01202 | 0.01515 | 0.05027 |
| **NBPF8.uid-77** | 0.00599 | 0.01983 | 0.01473 | 0.01560 | 0.00843 | 0.06002 | 0.01116 | 0.00981 | 0.02844 | 0.01280 | 0.01646 | 0.01794 |
| **NBPF9.uid-78** | 0.00633 | 0.01282 | 0.35992 | 0.00000 | 0.00000 | 0.04638 | 0.12437 | 0.00914 | 0.00320 | 0.12663 | 0.02388 | 0.02025 |
| **NBR1** | 2.32552 | 7.09330 | 2.15328 | 10.59963 | 0.18428 | 9.72317 | 1.36685 | 0.57316 | 5.01359 | 1.30826 | 2.11513 | 4.74755 |
| **NBR2** | 0.34533 | 0.87769 | 0.69093 | 0.83823 | 0.87148 | 0.14492 | 1.12055 | 1.00208 | 0.42735 | 0.00000 | 0.78312 | 1.52718 |
| **NCALD** | 4.38526 | 0.16332 | 0.30409 | 0.14587 | 0.00294 | 0.89116 | 0.00000 | 0.21894 | 0.00000 | 0.00000 | 0.82082 | 0.00000 |
| **NCAM1** | 0.15828 | 0.00000 | 0.06888 | 0.00000 | 0.01597 | 0.46054 | 0.05931 | 0.26067 | 0.00000 | 0.55503 | 0.00000 | 0.16211 |
| **NCAM2** | 0.01225 | 1.01051 | 0.25734 | 0.00000 | 0.06220 | 0.24982 | 0.00000 | 0.02098 | 0.08231 | 0.03618 | 0.24126 | 0.13065 |
| **NCAN** | 0.00197 | 0.01664 | 0.01639 | 0.01301 | 0.08857 | 0.00691 | 0.06001 | 0.08927 | 0.00000 | 0.00000 | 0.03986 | 0.01386 |
| **NCAPD2** | 2.24997 | 15.55245 | 8.96249 | 9.41579 | 13.96666 | 9.71150 | 14.80600 | 8.30080 | 12.76190 | 6.28817 | 6.26970 | 16.22888 |
| **NCAPD3** | 3.79870 | 5.19463 | 6.57705 | 4.66636 | 2.32888 | 5.64313 | 6.58772 | 0.84878 | 7.64907 | 6.66479 | 4.88121 | 4.22319 |
| **NCAPG** | 1.68151 | 6.08028 | 8.08192 | 2.83380 | 2.85532 | 25.45143 | 14.67663 | 8.54095 | 16.05149 | 0.00000 | 10.39062 | 12.51119 |
| **NCAPG2** | 4.26456 | 7.83573 | 6.26535 | 5.45692 | 0.89893 | 11.93466 | 4.86307 | 2.11098 | 10.33123 | 2.20958 | 7.65056 | 3.97733 |
| **NCAPH** | 7.49950 | 7.94884 | 6.86582 | 6.78155 | 6.39265 | 11.88690 | 11.86266 | 2.45787 | 12.40914 | 2.40682 | 10.42644 | 5.92095 |
| **NCAPH2** | 1.97146 | 15.68701 | 11.87533 | 11.12736 | 15.94863 | 0.00000 | 17.06139 | 4.86636 | 17.13516 | 3.96718 | 15.53659 | 11.84459 |
| **NCBP1** | 1.29558 | 9.43742 | 8.29336 | 4.89596 | 5.04583 | 26.97521 | 12.00067 | 10.45002 | 14.17214 | 9.19913 | 12.30250 | 7.00356 |
| **NCBP2** | 5.63604 | 18.36603 | 13.75795 | 24.55491 | 8.33993 | 35.77748 | 22.02440 | 22.25270 | 19.50731 | 11.51849 | 16.77420 | 26.20828 |
| **NCBP2-AS2** | 1.78009 | 10.89863 | 6.65051 | 17.07832 | 8.20628 | 9.13181 | 7.15027 | 24.69160 | 7.52031 | 4.43812 | 10.25496 | 17.43790 |
| **NCCRP1** | 0.02234 | 0.22427 | 3.20043 | 2.29368 | 0.00000 | 0.37287 | 0.66554 | 0.00135 | 0.28403 | 0.00036 | 0.00000 | 0.08544 |
| **NCDN** | 0.45660 | 3.36610 | 3.50915 | 3.30146 | 3.36863 | 0.00000 | 4.03730 | 0.00000 | 5.19484 | 3.21230 | 2.23613 | 5.53580 |
| **NCEH1** | 2.99559 | 0.80663 | 0.02739 | 4.34180 | 2.48404 | 16.53585 | 0.00000 | 1.15558 | 0.95253 | 4.85005 | 4.56687 | 3.95050 |

| | | | | | | | | | | | | |
|---|---|---|---|---|---|---|---|---|---|---|---|---|
| **NCF1** | 0.23045 | 0.01114 | 0.01381 | 0.01416 | 0.01767 | 0.01470 | 0.01646 | 0.23212 | 0.01841 | 0.01357 | 0.01363 | 0.15626 |
| **NCF1B** | 0.18327 | 0.01455 | 0.02738 | 0.01102 | 0.03887 | 0.00207 | 0.01919 | 0.09023 | 0.00273 | 0.03335 | 0.04767 | 0.25610 |
| **NCF1C** | 0.07878 | 0.05291 | 0.01502 | 0.00877 | 0.01781 | 0.00571 | 0.01679 | 0.19114 | 0.02025 | 0.01460 | 0.01451 | 0.27955 |
| **NCF2** | 1.36446 | 0.53307 | 0.05001 | 0.56215 | 0.00000 | 0.00000 | 0.17211 | 0.14473 | 0.00000 | 0.37657 | 1.15360 | 0.07083 |
| **NCF4** | 0.00619 | 0.02419 | 0.12710 | 0.00578 | 0.00920 | 0.02261 | 0.03089 | 0.01049 | 0.04137 | 0.01151 | 0.00760 | 0.01476 |
| **NCK1** | 0.00005 | 5.35912 | 5.40045 | 2.64406 | 0.00000 | 16.27822 | 5.59617 | 8.49956 | 8.82648 | 0.19209 | 4.76748 | 11.86076 |
| **NCK1-AS1** | 0.80924 | 0.89572 | 1.05910 | 0.08405 | 0.49032 | 0.13902 | 0.55142 | 6.51810 | 0.03688 | 0.00000 | 0.22981 | 1.56345 |
| **NCK2** | 4.87561 | 1.54764 | 7.97648 | 3.05644 | 5.86336 | 3.10199 | 15.11188 | 0.50155 | 10.25195 | 3.75842 | 5.70713 | 1.61787 |
| **NCKAP1** | 2.19901 | 2.74282 | 8.34669 | 3.28157 | 2.91218 | 28.06465 | 17.38187 | 1.51179 | 1.10813 | 25.83243 | 5.39889 | 7.44256 |
| **NCKAP1L** | 0.01511 | 0.03261 | 0.05447 | 0.10670 | 0.00000 | 0.01513 | 0.02186 | 0.00538 | 0.00000 | 0.06615 | 0.02006 | 0.12297 |
| **NCKAP5** | 0.29139 | 0.04935 | 0.01120 | 0.00869 | 0.00475 | 0.04250 | 0.01508 | 0.05600 | 0.01826 | 0.00786 | 0.01073 | 0.05149 |
| **NCKAP5L** | 3.53404 | 0.07774 | 0.59096 | 2.77950 | 0.58681 | 0.00000 | 1.50921 | 0.00000 | 0.34638 | 1.43156 | 0.00000 | 0.73322 |
| **NCKIPSD** | 1.56816 | 4.57196 | 3.04982 | 4.07345 | 5.30301 | 0.00000 | 2.97435 | 0.00000 | 7.49316 | 0.95508 | 6.29755 | 1.13904 |
| **NCL** | 8.04101 | 46.48540 | 41.35309 | 31.63629 | 44.55911 | 56.69873 | 64.92962 | 33.73891 | 55.54507 | 43.21891 | 48.95687 | 54.94854 |
| **NCLN** | 0.99579 | 17.21290 | 14.09359 | 9.94343 | 18.63927 | 0.00000 | 4.07075 | 0.57858 | 33.92174 | 19.97950 | 10.32169 | 13.22163 |
| **NCMAP** | 0.29374 | 0.03937 | 0.64208 | 0.05852 | 2.06109 | 0.00024 | 0.39858 | 0.04299 | 0.00000 | 0.00000 | 3.30154 | 0.14907 |
| **NCOA1** | 4.51965 | 3.56930 | 0.59262 | 7.49763 | 3.94960 | 6.12667 | 1.71296 | 3.60670 | 0.00000 | 9.98208 | 0.37102 | 5.49712 |
| **NCOA2** | 7.72469 | 2.00248 | 3.65457 | 4.57623 | 2.79883 | 1.04911 | 0.12884 | 1.14234 | 0.88530 | 5.34115 | 1.66108 | 1.70464 |
| **NCOA3** | 9.75906 | 8.75788 | 12.07821 | 15.24297 | 5.07644 | 21.54530 | 8.04613 | 7.31756 | 8.69900 | 15.75812 | 7.30446 | 5.16694 |
| **NCOA4** | 1.84944 | 27.89626 | 19.53768 | 18.14280 | 21.51935 | 61.91846 | 24.20797 | 10.53575 | 23.88069 | 12.05380 | 34.79425 | 23.20887 |
| **NCOA5** | 0.88189 | 5.19214 | 1.72644 | 4.56391 | 4.39219 | 3.83269 | 7.00215 | 0.31755 | 6.97070 | 3.45066 | 2.03687 | 3.51547 |
| **NCOA6** | 12.46531 | 2.48406 | 4.60653 | 4.90842 | 8.68135 | 4.97966 | 8.78922 | 3.82179 | 4.62247 | 15.82250 | 2.98429 | 4.26662 |
| **NCOA7** | 0.33908 | 0.28264 | 1.20193 | 2.47962 | 0.43355 | 7.08511 | 2.58996 | 0.00000 | 0.00000 | 8.05306 | 2.51591 | 1.25519 |
| **NCOR1** | 11.93933 | 10.08877 | 7.92027 | 13.08480 | 8.75372 | 22.54960 | 9.88395 | 14.78615 | 17.28168 | 28.03457 | 13.56375 | 14.15919 |
| **NCOR1P1** | 0.01025 | 0.01074 | 0.04269 | 0.01350 | 0.01069 | 0.02884 | 0.02097 | 0.51732 | 0.00498 | 0.01752 | 0.01923 | 0.05955 |
| **NCOR2** | 2.17555 | 0.00000 | 4.69957 | 0.00000 | 5.01912 | 0.00000 | 5.11821 | 0.19614 | 1.25368 | 8.31503 | 0.00000 | 0.35633 |
| **NCR1** | 0.27623 | 0.93334 | 0.00000 | 0.51231 | 0.57103 | 0.15606 | 0.01600 | 0.27357 | 0.05431 | 0.36731 | 0.30010 | 0.13204 |
| **NCR3** | 0.00648 | 0.01617 | 0.00909 | 0.02107 | 0.00588 | 0.04487 | 0.00000 | 0.01008 | 0.11113 | 0.00902 | 0.01031 | 0.01412 |
| **NCR3LG1** | 0.00743 | 0.00000 | 0.00611 | 0.12079 | 0.02948 | 0.00126 | 0.00583 | 0.06985 | 0.03770 | 0.24427 | 0.00374 | 1.16165 |
| **NCRNA00185** | 0.00685 | 0.00000 | 0.45144 | 0.13407 | 0.20244 | 0.00000 | 0.38758 | 0.00772 | 0.21400 | 0.64174 | 0.33310 | 0.00428 |
| **NCRUPAR** | 0.00629 | 0.01508 | 0.01279 | 0.01399 | 0.01671 | 0.01641 | 0.01938 | 0.01034 | 0.02953 | 0.00788 | 0.01399 | 0.02141 |
| **NCS1** | 0.16719 | 0.11380 | 0.12787 | 0.06439 | 0.04337 | 0.00000 | 0.00434 | 0.00267 | 0.03921 | 0.16440 | 0.00266 | 0.03382 |
| **NCSTN** | 1.08414 | 15.51115 | 11.91808 | 8.90555 | 8.31719 | 10.28171 | 0.00000 | 3.58874 | 17.57083 | 0.00000 | 8.61708 | 12.61370 |
| **NDC1** | 5.32030 | 15.74926 | 6.24323 | 5.92617 | 5.02974 | 34.45747 | 0.00000 | 0.00000 | 29.18467 | 21.27952 | 11.46681 | 14.23712 |
| **NDC80** | 0.61481 | 9.97332 | 6.37447 | 5.06203 | 1.75914 | 27.30167 | 11.33120 | 1.24470 | 9.48908 | 2.00640 | 4.83096 | 10.37403 |
| **NDE1** | 8.75972 | 2.00834 | 2.47676 | 0.76786 | 1.50279 | 0.61036 | 2.31542 | 2.92384 | 2.19254 | 0.00000 | 0.20345 | 1.84465 |
| **NDEL1** | 4.28766 | 5.07809 | 8.48555 | 4.10362 | 2.67107 | 0.16050 | 2.83006 | 0.33933 | 9.99900 | 4.32579 | 6.02424 | 1.81280 |
| **NDFIP1** | 2.50977 | 9.73892 | 0.17409 | 16.83658 | 5.20321 | 28.27571 | 4.48007 | 3.32910 | 21.51540 | 10.82396 | 7.13273 | 7.21399 |
| **NDFIP2** | 4.69456 | 7.66116 | 19.16058 | 1.92577 | 10.18866 | 39.87760 | 0.00000 | 4.27342 | 21.88764 | 0.00000 | 9.99844 | 10.89126 |
| **NDFIP2-AS1** | 0.01093 | 0.01382 | 0.01256 | 0.00880 | 0.01545 | 0.03953 | 0.01500 | 0.00964 | 0.01466 | 0.01204 | 0.01345 | 0.02688 |
| **NDN** | 0.24171 | 1.08672 | 3.61375 | 1.12681 | 0.00000 | 0.00000 | 4.47050 | 0.42134 | 0.92578 | 2.89013 | 1.76271 | 0.89273 |
| **NDNF** | 0.00722 | 0.02111 | 0.05881 | 0.01963 | 0.02212 | 0.00000 | 0.00759 | 0.16636 | 0.01763 | 0.23947 | 0.01077 | 0.03034 |
| **NDNL2** | 0.88077 | 1.26439 | 1.57007 | 0.40808 | 0.71883 | 1.16373 | 1.71936 | 0.00000 | 0.09022 | 5.49914 | 0.11509 | 1.32543 |
| **NDOR1** | 1.06509 | 4.48479 | 4.12280 | 2.65996 | 8.25928 | 0.00000 | 2.78657 | 0.00000 | 6.30497 | 17.63693 | 1.21325 | 4.56567 |
| **NDRG1** | 1.85638 | 1.05028 | 1.48533 | 0.64668 | 6.28366 | 1.37085 | 0.79266 | 0.07078 | 0.96852 | 1.56541 | 0.00000 | 0.00000 |
| **NDRG2** | 0.00556 | 0.00000 | 1.05838 | 6.31673 | 14.32808 | 0.11383 | 2.26697 | 0.11676 | 0.00000 | 0.00000 | 2.00614 | 0.18370 |
| **NDRG3** | 7.07505 | 11.52014 | 7.88766 | 5.12416 | 7.46224 | 9.78080 | 8.33505 | 1.65638 | 6.98375 | 2.63891 | 10.72890 | 11.83021 |
| **NDRG4** | 4.69060 | 0.13318 | 0.23108 | 0.01695 | 0.23536 | 0.00000 | 0.00000 | 0.31518 | 0.54451 | 0.00822 | 0.20515 | 0.38220 |

| | | | | | | | | | | | | |
|---|---|---|---|---|---|---|---|---|---|---|---|---|
| **NDST1** | 1.65140 | 7.43257 | 3.58533 | 5.04297 | 9.83115 | 0.00000 | 2.08126 | 2.00339 | 10.96094 | 10.04550 | 10.18716 | 3.07954 |
| **NDST2** | 0.74574 | 0.88800 | 1.00083 | 1.29482 | 0.38872 | 0.99929 | 0.00000 | 0.27420 | 0.72671 | 3.59200 | 0.06373 | 0.63870 |
| **NDST3** | 0.81119 | 0.02274 | 0.04182 | 0.01401 | 0.07572 | 0.01849 | 0.00770 | 0.18676 | 0.03208 | 0.08925 | 0.06097 | 0.00245 |
| **NDST4** | 0.00619 | 0.01603 | 0.03736 | 0.05598 | 0.01715 | 0.01211 | 0.05420 | 0.01059 | 0.00066 | 0.00535 | 0.02202 | 0.01729 |
| **NDUFA1** | 0.49894 | 16.46661 | 10.93239 | 17.73186 | 17.10375 | 36.47403 | 30.41015 | 10.79246 | 11.83927 | 22.76553 | 30.45273 | 24.76253 |
| **NDUFA10** | 4.27755 | 20.06351 | 13.26025 | 14.63322 | 17.12813 | 18.58667 | 25.36912 | 2.33727 | 27.39015 | 7.26498 | 18.73194 | 13.44847 |
| **NDUFA11** | 2.57730 | 19.72854 | 22.09896 | 21.41633 | 27.42514 | 7.79542 | 38.26608 | 5.82482 | 28.23100 | 9.58210 | 24.72413 | 14.78257 |
| **NDUFA12** | 0.88816 | 14.41711 | 9.93867 | 21.94467 | 16.22257 | 37.62875 | 20.97374 | 1.26314 | 9.04752 | 8.26309 | 19.55581 | 15.04383 |
| **NDUFA13** | 7.97804 | 19.08147 | 25.57813 | 29.81686 | 31.57105 | 13.21459 | 38.03243 | 19.99783 | 18.31184 | 12.78105 | 25.11852 | 22.62980 |
| **NDUFA2** | 0.78944 | 16.93840 | 10.73062 | 19.32928 | 22.83325 | 24.69681 | 28.20742 | 15.91565 | 22.35355 | 16.14248 | 26.19319 | 22.63147 |
| **NDUFA3** | 1.44624 | 17.39533 | 14.54718 | 17.95997 | 27.59449 | 2.62685 | 30.84232 | 3.19411 | 14.26662 | 13.62271 | 20.37048 | 13.79392 |
| **NDUFA4** | 1.85178 | 19.29322 | 26.02743 | 21.95800 | 22.81746 | 54.75794 | 36.60983 | 9.36985 | 35.67507 | 20.46747 | 33.00756 | 24.93680 |
| **NDUFA4L2** | 0.00739 | 0.01110 | 0.27517 | 0.08269 | 0.00000 | 0.05070 | 0.00000 | 0.01655 | 0.00000 | 0.03168 | 0.01759 | 0.02049 |
| **NDUFA5** | 0.00000 | 9.00710 | 4.52922 | 2.96523 | 0.00000 | 34.85505 | 13.32242 | 0.99611 | 9.34502 | 5.98792 | 11.75602 | 4.28734 |
| **NDUFA6** | 10.64558 | 23.15389 | 22.51198 | 27.00604 | 32.73831 | 37.69593 | 35.54825 | 21.14004 | 25.26041 | 23.77371 | 34.04607 | 22.27038 |
| **NDUFA6-AS1** | 0.05828 | 0.52162 | 0.08768 | 0.02129 | 0.21734 | 0.10284 | 0.00000 | 2.16005 | 0.22854 | 0.00000 | 0.08232 | 2.39835 |
| **NDUFA7** | 2.09843 | 13.47897 | 5.36772 | 11.27958 | 12.59559 | 7.66134 | 9.90159 | 1.42958 | 9.42343 | 2.04758 | 14.56903 | 13.96305 |
| **NDUFA8** | 1.76319 | 22.96412 | 15.46688 | 19.88380 | 22.77609 | 28.22104 | 26.51413 | 0.00000 | 17.94976 | 2.54780 | 26.26590 | 14.14851 |
| **NDUFA9** | 0.62259 | 23.13115 | 13.90666 | 17.39938 | 19.92934 | 27.77814 | 23.67176 | 3.69780 | 22.52291 | 3.98572 | 19.10008 | 17.85426 |
| **NDUFAB1** | 6.94566 | 41.70316 | 22.39477 | 39.36536 | 38.56135 | 55.17610 | 39.10487 | 15.24754 | 24.76903 | 27.90031 | 32.41459 | 38.30154 |
| **NDUFAF1** | 0.93146 | 6.07497 | 6.19979 | 3.99788 | 5.77189 | 17.64789 | 0.00000 | 3.22319 | 5.46348 | 0.00000 | 7.98492 | 8.55537 |
| **NDUFAF2** | 0.55336 | 1.73800 | 3.50966 | 0.00000 | 1.78709 | 4.63225 | 5.66142 | 0.76561 | 3.55916 | 4.31282 | 2.41200 | 2.90871 |
| **NDUFAF3** | 0.59277 | 7.77022 | 3.56837 | 5.82186 | 12.17499 | 4.99944 | 0.54616 | 0.20148 | 2.10814 | 0.26410 | 4.68985 | 1.26697 |
| **NDUFAF4** | 1.57314 | 15.58896 | 9.88289 | 6.98252 | 9.23011 | 35.62680 | 18.93080 | 5.44184 | 12.97139 | 5.90065 | 14.38189 | 14.25861 |
| **NDUFAF4P1** | 0.05207 | 0.32048 | 0.16479 | 0.03142 | 0.09969 | 1.30890 | 0.21678 | 0.32862 | 0.26860 | 0.72634 | 0.68505 | 0.52991 |
| **NDUFAF5** | 0.09306 | 4.69303 | 1.89136 | 5.62768 | 1.58610 | 9.95689 | 4.89508 | 4.20333 | 5.65916 | 0.00880 | 3.34811 | 4.08607 |
| **NDUFAF6** | 0.11674 | 4.74333 | 3.84642 | 2.88851 | 1.41449 | 8.20022 | 4.87315 | 0.00000 | 7.87474 | 2.02981 | 7.83530 | 0.53793 |
| **NDUFAF7** | 0.90727 | 6.23524 | 5.70214 | 12.77600 | 2.09922 | 16.63974 | 5.11388 | 12.24466 | 8.21916 | 2.36754 | 5.93264 | 8.54625 |
| **NDUFB1** | 1.40203 | 7.58142 | 8.15391 | 8.70890 | 10.72327 | 20.97904 | 15.43665 | 0.51644 | 4.12854 | 21.37005 | 16.03229 | 6.71610 |
| **NDUFB10** | 3.32541 | 20.36455 | 19.18000 | 24.49131 | 28.42847 | 14.97436 | 34.48964 | 14.38708 | 35.64949 | 12.82270 | 30.56425 | 9.73892 |
| **NDUFB11** | 0.91819 | 17.69617 | 14.41325 | 16.16977 | 28.61371 | 13.39562 | 19.31494 | 9.68607 | 26.47237 | 12.67502 | 19.20571 | 21.96387 |
| **NDUFB2** | 5.11470 | 22.07095 | 19.17709 | 22.39496 | 31.29969 | 22.20610 | 39.77054 | 6.97261 | 18.55336 | 13.42910 | 30.61998 | 12.08843 |
| **NDUFB2-AS1** | 0.55909 | 2.03653 | 0.45335 | 0.30969 | 1.32988 | 0.00000 | 0.28376 | 0.11060 | 0.47862 | 1.38536 | 0.50918 | 0.80603 |
| **NDUFB3** | 0.00000 | 12.98752 | 6.26845 | 8.70259 | 9.76430 | 35.88489 | 9.42963 | 1.66935 | 3.80335 | 5.68133 | 17.06396 | 15.29893 |
| **NDUFB4** | 0.20863 | 10.91409 | 15.05248 | 11.66412 | 13.82640 | 29.39791 | 21.62321 | 2.55458 | 19.66688 | 9.02132 | 18.88911 | 11.71302 |
| **NDUFB5** | 5.50493 | 13.00497 | 10.06712 | 4.93019 | 8.02236 | 41.67493 | 6.40770 | 0.87756 | 10.19512 | 5.81484 | 17.81723 | 10.47197 |
| **NDUFB6** | 1.36346 | 15.62837 | 10.78519 | 13.66447 | 12.47755 | 42.20181 | 21.63009 | 9.27045 | 16.95671 | 11.20468 | 20.46959 | 20.03633 |
| **NDUFB7** | 1.55670 | 24.94414 | 13.89435 | 21.82055 | 33.68215 | 3.75870 | 35.02303 | 7.77474 | 24.89178 | 16.91717 | 24.03958 | 22.61349 |
| **NDUFB8** | 8.28435 | 21.90016 | 20.97040 | 26.55191 | 27.37409 | 19.57118 | 37.37094 | 15.01266 | 25.46370 | 3.63059 | 30.50884 | 15.43977 |
| **NDUFB9** | 6.32065 | 28.94429 | 21.99565 | 30.05656 | 36.00572 | 42.03156 | 21.71822 | 14.43044 | 44.51790 | 25.86966 | 30.34173 | 27.90760 |
| **NDUFC1** | 2.16238 | 10.90051 | 15.56188 | 14.71173 | 19.63299 | 26.06109 | 29.92312 | 3.15125 | 18.48930 | 8.98542 | 25.54589 | 13.19038 |
| **NDUFC2** | 0.00000 | 8.43565 | 6.32252 | 5.76467 | 6.18403 | 39.56095 | 7.68498 | 1.13264 | 12.24471 | 21.36028 | 9.68851 | 8.49479 |
| **NDUFS1** | 0.85783 | 13.14848 | 11.87065 | 12.57635 | 7.38814 | 31.02994 | 12.78759 | 3.44856 | 16.04996 | 1.37353 | 22.33174 | 8.46413 |
| **NDUFS2** | 4.05445 | 21.42954 | 18.34657 | 18.15405 | 17.00752 | 19.69643 | 15.37133 | 4.85926 | 16.32447 | 0.00000 | 28.83818 | 16.47332 |
| **NDUFS3** | 3.57159 | 21.20673 | 23.18487 | 20.23821 | 24.61038 | 16.02205 | 32.95719 | 5.58402 | 30.52139 | 3.32642 | 21.18936 | 14.28978 |
| **NDUFS4** | 0.04386 | 7.54402 | 8.75354 | 5.05708 | 6.09908 | 23.09550 | 7.93474 | 1.10491 | 9.49857 | 5.57635 | 11.55957 | 5.70996 |
| **NDUFS5** | 1.12324 | 27.52060 | 21.07643 | 24.23856 | 32.25758 | 40.88085 | 41.18115 | 0.00000 | 27.32920 | 25.02897 | 30.70451 | 21.99376 |
| **NDUFS6** | 0.68395 | 23.28750 | 18.82859 | 21.53389 | 32.25537 | 22.09666 | 41.48434 | 0.58786 | 22.99906 | 21.20531 | 25.19178 | 13.48651 |

| | | | | | | | | | | | | |
|---|---|---|---|---|---|---|---|---|---|---|---|---|
| NDUFS7 | 0.32718 | 16.36328 | 9.10783 | 17.62074 | 25.74970 | 3.52956 | 22.03934 | 1.30612 | 21.49077 | 0.00000 | 22.01018 | 6.24808 |
| NDUFS8 | 0.05959 | 18.65049 | 17.00630 | 18.79917 | 30.99955 | 2.12135 | 29.55262 | 1.81577 | 22.31789 | 2.27987 | 24.06081 | 13.28309 |
| NDUFV1 | 0.00000 | 31.45849 | 14.74375 | 38.84063 | 44.40185 | 10.92437 | 32.08185 | 9.32873 | 24.72410 | 6.29983 | 34.30996 | 20.04813 |
| NDUFV2 | 0.27876 | 15.27941 | 13.70424 | 11.14200 | 11.34591 | 29.03655 | 19.21819 | 2.35000 | 14.21962 | 6.04173 | 20.92713 | 14.69247 |
| NDUFV3 | 2.08321 | 12.70292 | 4.88720 | 7.21290 | 19.62810 | 9.28601 | 6.19557 | 0.23385 | 10.48503 | 3.03645 | 17.70029 | 8.72485 |
| NEAT1 | 0.94744 | 0.77183 | 3.84645 | 0.35953 | 0.88508 | 4.23025 | 0.00000 | 0.00000 | 0.00000 | 9.96479 | 0.42258 | 0.05828 |
| NEB | 2.71633 | 0.03830 | 1.48789 | 0.83619 | 0.00000 | 0.72388 | 0.00000 | 0.55367 | 0.76163 | 3.57572 | 0.03756 | 0.94769 |
| NEBL | 3.26624 | 0.00000 | 0.23505 | 0.00000 | 0.72892 | 0.13505 | 0.67268 | 0.28737 | 0.10369 | 0.52907 | 0.00000 | 0.24759 |
| NEBL-AS1 | 0.00365 | 0.01465 | 0.01618 | 0.01152 | 0.01482 | 0.02121 | 0.01691 | 0.02843 | 0.01780 | 0.01486 | 0.01485 | 0.01471 |
| NECAB1 | 0.15705 | 0.00000 | 0.12048 | 0.11482 | 0.00000 | 0.40971 | 0.21105 | 0.08524 | 0.55239 | 0.00757 | 0.00817 | 0.32509 |
| NECAB2 | 0.09092 | 0.12556 | 0.01749 | 0.00000 | 0.00000 | 0.01987 | 0.01712 | 0.21030 | 0.00626 | 0.04160 | 0.03606 | 0.02949 |
| NECAB3 | 0.86628 | 1.97412 | 4.14830 | 2.15450 | 6.68920 | 0.00000 | 2.03396 | 0.02172 | 0.00000 | 6.64291 | 1.41132 | 0.31965 |
| NECAP1 | 2.34995 | 9.68616 | 3.63305 | 3.96303 | 1.14217 | 9.71798 | 3.09668 | 2.53816 | 4.75603 | 1.15571 | 7.44118 | 6.41756 |
| NECAP2 | 7.96416 | 11.20995 | 4.44208 | 6.21758 | 7.18644 | 4.91645 | 9.23659 | 2.24624 | 10.09968 | 0.00000 | 6.42703 | 11.79644 |
| NEDD1 | 8.34742 | 2.89031 | 7.16913 | 0.47040 | 0.00000 | 12.91759 | 9.40419 | 4.63339 | 6.88524 | 0.37907 | 4.81759 | 5.85006 |
| NEDD4 | 0.00000 | 1.01837 | 0.69404 | 1.67771 | 0.00000 | 2.68482 | 1.03888 | 0.73119 | 0.00000 | 0.91665 | 0.37089 | 2.07960 |
| NEDD4L | 5.10844 | 0.79161 | 5.78428 | 3.62532 | 4.46229 | 2.25642 | 0.35905 | 0.17011 | 0.00000 | 2.72127 | 0.00000 | 0.21304 |
| NEDD8 | 2.64464 | 8.77398 | 16.84492 | 14.62774 | 16.88742 | 24.92590 | 30.12201 | 2.41452 | 12.72288 | 17.10756 | 20.50509 | 19.79575 |
| NEDD9 | 0.07576 | 2.75690 | 1.76184 | 0.29734 | 0.98359 | 1.08185 | 1.06786 | 0.23426 | 0.52399 | 0.00000 | 0.23232 | 1.41705 |
| NEFH | 5.49794 | 9.17147 | 0.00000 | 10.13444 | 7.66587 | 3.17888 | 5.84944 | 7.22242 | 15.55494 | 11.49003 | 2.07082 | 9.72764 |
| NEFL | 3.36520 | 12.81170 | 0.00049 | 4.73579 | 0.60023 | 0.00001 | 0.00463 | 1.99401 | 0.00000 | 0.08994 | 0.00000 | 1.11681 |
| NEFM | 6.67589 | 0.33746 | 0.15704 | 0.00000 | 0.05259 | 0.51597 | 0.44617 | 9.69260 | 0.00000 | 0.61531 | 0.00000 | 2.01612 |
| NEGR1 | 0.73366 | 0.02580 | 0.00000 | 0.02994 | 0.02078 | 0.23333 | 0.00000 | 0.00392 | 0.00000 | 0.09522 | 0.06829 | 0.05020 |
| NEIL1 | 0.21625 | 0.06394 | 0.55813 | 1.66849 | 0.00000 | 0.00000 | 0.16773 | 0.00684 | 0.74989 | 0.37753 | 0.16814 | 0.10792 |
| NEIL2 | 1.21224 | 8.97427 | 0.00000 | 4.29602 | 5.70776 | 0.00000 | 1.53185 | 2.27083 | 0.91446 | 4.17746 | 3.98939 | 5.92297 |
| NEIL3 | 0.78147 | 0.79703 | 2.63234 | 1.39100 | 0.54020 | 2.21714 | 2.79686 | 0.39170 | 3.12378 | 0.00000 | 3.02069 | 0.64336 |
| NEK1 | 13.30892 | 2.12248 | 1.05694 | 1.95353 | 0.98989 | 13.47941 | 0.00000 | 5.26335 | 2.57486 | 10.72367 | 2.21200 | 3.52547 |
| NEK10 | 0.63651 | 0.04469 | 0.09654 | 0.00669 | 0.00000 | 0.00000 | 0.03648 | 0.01054 | 0.00000 | 0.04476 | 0.06427 | 0.02683 |
| NEK11 | 0.18222 | 0.75309 | 0.39175 | 0.00000 | 0.11857 | 0.37429 | 0.10516 | 0.29541 | 0.00000 | 0.20262 | 0.06551 | 1.46986 |
| NEK2 | 2.54023 | 10.02675 | 0.32130 | 6.13075 | 2.40437 | 18.44251 | 3.60876 | 7.11656 | 3.65872 | 1.41626 | 0.98072 | 11.57782 |
| NEK3 | 0.68984 | 3.39514 | 0.74087 | 4.90316 | 4.37509 | 3.80702 | 0.93795 | 0.07606 | 0.11760 | 0.00000 | 0.91280 | 1.53756 |
| NEK4 | 4.03668 | 3.99339 | 0.00000 | 1.64191 | 1.46841 | 9.00725 | 0.82768 | 0.17950 | 0.00000 | 2.88345 | 2.67140 | 3.92721 |
| NEK5 | 0.54755 | 0.14044 | 0.13857 | 0.07743 | 0.00809 | 0.37087 | 0.00000 | 0.01418 | 0.00000 | 0.05480 | 0.04168 | 0.24267 |
| NEK6 | 1.15012 | 14.39432 | 7.65754 | 2.56411 | 13.42307 | 0.00002 | 2.24047 | 0.00000 | 9.12564 | 0.19402 | 17.50763 | 0.70022 |
| NEK7 | 1.12140 | 0.19738 | 3.63646 | 0.00000 | 0.00000 | 13.14655 | 2.88974 | 1.26556 | 0.00000 | 4.46808 | 0.58234 | 1.59815 |
| NEK8 | 0.26525 | 0.80656 | 0.92729 | 1.65948 | 1.08070 | 0.00000 | 0.39908 | 0.07976 | 2.29267 | 1.10451 | 0.88598 | 0.47463 |
| NEK9 | 0.09474 | 2.32514 | 4.91281 | 2.06363 | 3.74463 | 6.25166 | 4.85401 | 0.27927 | 6.46842 | 6.09398 | 8.67088 | 2.18399 |
| NELFA | 5.83844 | 10.89732 | 9.58344 | 6.04643 | 5.43179 | 3.13657 | 10.04167 | 2.10523 | 12.86422 | 1.38322 | 6.54105 | 10.98935 |
| NELFB | 0.43025 | 7.01613 | 3.56456 | 4.11879 | 7.46330 | 0.00000 | 8.54627 | 0.85565 | 2.13163 | 16.85676 | 2.45871 | 9.30647 |
| NELFCD | 0.00000 | 13.24055 | 15.72853 | 12.89779 | 8.69921 | 10.31562 | 14.07347 | 6.66380 | 24.54655 | 0.00000 | 13.55383 | 17.66054 |
| NELFE | 1.30956 | 23.98281 | 16.98481 | 15.84872 | 22.06136 | 12.43692 | 20.62243 | 23.56783 | 13.46274 | 3.11014 | 18.31182 | 21.11772 |
| NELL1 | 2.43280 | 0.10140 | 0.01592 | 0.02146 | 0.00705 | 0.00000 | 0.01020 | 2.74331 | 0.03086 | 0.01265 | 0.02348 | 0.08907 |
| NELL2 | 0.52417 | 2.23608 | 0.00000 | 0.00000 | 0.05580 | 0.94917 | 0.00000 | 0.55821 | 0.00000 | 0.00711 | 0.00000 | 2.21499 |
| NEMF | 3.09988 | 1.42448 | 4.49471 | 5.12340 | 8.33945 | 23.14106 | 7.82201 | 3.20589 | 8.71124 | 23.87607 | 6.27784 | 2.63928 |
| NENF | 0.60106 | 0.54588 | 3.84131 | 0.00000 | 12.81852 | 1.62885 | 10.67356 | 0.04516 | 12.94009 | 8.37747 | 4.66076 | 0.31604 |
| NEO1 | 13.39651 | 1.38011 | 0.24727 | 2.79806 | 0.17427 | 2.81616 | 0.37764 | 1.27914 | 0.74977 | 4.56637 | 0.00000 | 4.36778 |
| NES | 0.33627 | 0.00000 | 0.00299 | 0.72341 | 0.00000 | 0.00000 | 0.00477 | 0.37707 | 0.03017 | 0.12680 | 0.02054 | 0.65280 |
| NET1 | 2.68254 | 4.70087 | 35.23678 | 0.16159 | 3.91891 | 8.97557 | 20.26447 | 5.35962 | 7.12033 | 2.16773 | 4.44101 | 3.39889 |

| | | | | | | | | | | | | |
|---|---|---|---|---|---|---|---|---|---|---|---|---|
| **NETO1** | 0.00379 | 0.01444 | 0.01848 | 0.04134 | 0.00002 | 0.06711 | 0.00000 | 0.05642 | 0.02008 | 0.02724 | 0.02064 | 0.01010 |
| **NETO2** | 2.74761 | 2.69075 | 2.01537 | 3.05061 | 0.00000 | 5.05124 | 2.28279 | 0.13669 | 0.00000 | 7.48354 | 0.00000 | 2.70704 |
| **NEU1** | 0.81432 | 14.52203 | 18.47537 | 14.49471 | 11.81568 | 7.56699 | 0.83629 | 7.19696 | 18.25834 | 0.00000 | 9.62749 | 23.71352 |
| **NEU2** | 0.04995 | 0.01571 | 0.00568 | 0.01234 | 0.01041 | 0.00469 | 0.17394 | 0.05691 | 0.00000 | 0.03708 | 0.00831 | 0.01130 |
| **NEU3** | 0.09323 | 0.49457 | 0.10187 | 0.00687 | 0.02514 | 0.00000 | 0.03841 | 0.17024 | 0.51701 | 0.00000 | 0.25495 | 0.64790 |
| **NEU4** | 0.00856 | 0.01340 | 0.02465 | 0.01060 | 0.00575 | 0.01377 | 0.01547 | 0.00998 | 0.00000 | 0.03674 | 0.07362 | 0.01532 |
| **NEURL1** | 0.78582 | 0.80813 | 6.43524 | 0.52722 | 0.06353 | 0.00000 | 0.77956 | 0.46077 | 0.00294 | 0.00000 | 0.00000 | 4.73147 |
| **NEURL1B** | 0.41620 | 0.00162 | 0.00781 | 0.01881 | 0.00231 | 0.05722 | 0.00606 | 0.01672 | 0.02533 | 0.07753 | 0.01535 | 0.03840 |
| **NEURL2** | 0.00239 | 1.64186 | 0.30772 | 3.40730 | 2.45573 | 0.00000 | 0.67230 | 0.00302 | 0.07712 | 1.61082 | 0.10583 | 0.06444 |
| **NEURL3** | 0.00418 | 0.01675 | 0.01833 | 0.03988 | 0.01347 | 0.00509 | 0.02649 | 0.02015 | 0.12702 | 0.01257 | 0.00798 | 0.02505 |
| **NEURL4** | 0.03201 | 2.26764 | 1.26422 | 3.63063 | 4.95857 | 0.00000 | 1.70706 | 0.00000 | 2.61982 | 9.39752 | 1.59125 | 2.68680 |
| **NEUROD1** | 0.01432 | 0.00542 | 0.03992 | 0.02501 | 0.00000 | 0.05533 | 0.04901 | 0.14453 | 0.00000 | 0.08470 | 0.02827 | 0.01075 |
| **NEUROD2** | 0.04689 | 0.00404 | 0.00000 | 0.07659 | 0.00277 | 0.12772 | 0.05177 | 0.02306 | 0.00000 | 0.21091 | 0.03529 | 0.02516 |
| **NEUROD4** | 0.03113 | 0.01198 | 0.01171 | 0.01053 | 0.01083 | 0.02997 | 0.01272 | 0.01174 | 0.01523 | 0.04103 | 0.01275 | 0.02625 |
| **NEUROD6** | 0.03612 | 0.01464 | 0.01984 | 0.01132 | 0.01689 | 0.01751 | 0.01583 | 0.00889 | 0.01514 | 0.02344 | 0.01510 | 0.01484 |
| **NEUROG1** | 0.25965 | 0.02600 | 0.06421 | 0.09889 | 0.00758 | 0.00011 | 0.18909 | 0.01762 | 0.00000 | 0.00000 | 0.47425 | 0.73401 |
| **NEUROG2** | 0.08150 | 0.01091 | 0.01069 | 0.03288 | 0.01916 | 0.02919 | 0.01369 | 0.06882 | 0.01765 | 0.01587 | 0.01374 | 0.01854 |
| **NEUROG3** | 0.02161 | 0.01598 | 0.01772 | 0.01337 | 0.04949 | 0.00000 | 0.00911 | 0.01482 | 0.04833 | 0.02753 | 0.03413 | 0.06223 |
| **NEXN** | 0.23022 | 1.58879 | 0.00000 | 1.23857 | 0.38199 | 1.79740 | 0.00000 | 0.06310 | 0.00000 | 2.51816 | 0.00000 | 0.11729 |
| **NEXN-AS1** | 0.28106 | 0.05436 | 0.00317 | 0.05086 | 0.00000 | 0.06660 | 0.01891 | 0.08057 | 0.01958 | 0.28859 | 0.01722 | 0.02969 |
| **NF1** | 7.85742 | 0.00000 | 5.29444 | 0.20286 | 0.80082 | 10.38365 | 5.11750 | 1.27866 | 7.25045 | 15.11196 | 2.43686 | 1.99615 |
| **NF1P2.uid-1918** | 0.01788 | 0.01536 | 0.01414 | 0.01223 | 0.01591 | 0.01729 | 0.01811 | 0.02584 | 0.01982 | 0.01223 | 0.01499 | 0.01533 |
| **NF2** | 0.55778 | 23.25893 | 2.01400 | 27.31648 | 23.59210 | 2.63738 | 15.86416 | 15.71550 | 15.08755 | 5.20030 | 27.25969 | 24.18336 |
| **NFAM1** | 0.12689 | 0.01256 | 0.01602 | 0.03162 | 0.00606 | 0.02201 | 0.01338 | 0.00715 | 0.02133 | 0.04906 | 0.01205 | 0.02092 |
| **NFASC** | 0.81527 | 0.05329 | 0.05822 | 0.00473 | 0.01565 | 0.00000 | 0.01291 | 0.17802 | 0.00000 | 0.17664 | 0.05490 | 0.07957 |
| **NFAT5** | 0.16210 | 0.64195 | 5.76513 | 1.61093 | 1.78579 | 5.28724 | 2.51487 | 3.07054 | 4.66665 | 9.13596 | 3.04934 | 0.07303 |
| **NFATC1** | 5.67937 | 0.48433 | 0.36135 | 0.06937 | 0.01293 | 0.00000 | 0.15978 | 1.45080 | 0.04443 | 0.00000 | 0.01013 | 1.15331 |
| **NFATC2** | 1.37832 | 0.00000 | 0.07007 | 0.00000 | 0.42150 | 0.55161 | 1.43022 | 0.02668 | 0.16382 | 0.14160 | 1.90444 | 0.34663 |
| **NFATC2IP** | 1.65479 | 4.71665 | 5.25546 | 5.13076 | 8.82636 | 4.74701 | 7.88858 | 8.25413 | 5.89239 | 21.79527 | 4.39706 | 10.05850 |
| **NFATC3** | 4.64081 | 3.91519 | 9.71484 | 4.79811 | 3.96638 | 4.85006 | 12.51283 | 1.17648 | 5.22993 | 12.21434 | 10.50525 | 4.84601 |
| **NFATC4** | 0.00523 | 0.00997 | 0.01646 | 0.17488 | 0.05087 | 0.02512 | 0.03342 | 0.01497 | 0.02884 | 0.00000 | 0.02204 | 0.02380 |
| **NFE2** | 0.17146 | 0.13277 | 2.94924 | 0.00055 | 0.00000 | 0.03864 | 0.39913 | 0.00000 | 0.58897 | 0.00000 | 0.13534 | 0.18616 |
| **NFE2L1** | 2.44495 | 1.41676 | 4.10561 | 3.31367 | 10.00116 | 4.84804 | 2.08740 | 0.00000 | 3.22320 | 22.88929 | 3.63431 | 2.27091 |
| **NFE2L2** | 1.51117 | 9.44915 | 11.20350 | 0.00000 | 7.67570 | 21.83633 | 15.58516 | 0.15951 | 10.62824 | 14.17317 | 11.32528 | 2.20440 |
| **NFE2L3** | 0.42818 | 3.63342 | 23.83831 | 0.00000 | 3.85339 | 18.67695 | 4.19662 | 0.69260 | 10.01578 | 17.97041 | 0.00000 | 0.50557 |
| **NFIA** | 0.09864 | 0.14964 | 0.16528 | 0.16583 | 0.14727 | 0.00000 | 0.00000 | 0.12341 | 0.08975 | 0.28653 | 0.00692 | 0.12391 |
| **NFIB** | 0.53943 | 0.08564 | 0.12887 | 0.20662 | 0.00000 | 0.15416 | 0.00000 | 0.02284 | 0.00000 | 0.15772 | 0.00900 | 0.09704 |
| **NFIC** | 2.14776 | 0.30137 | 0.61703 | 0.22779 | 0.48855 | 0.44874 | 1.10013 | 0.23941 | 0.78451 | 1.24730 | 0.31132 | 0.87480 |
| **NFIL3** | 0.32261 | 0.63958 | 5.09214 | 0.00000 | 4.34975 | 2.26993 | 7.66601 | 0.59283 | 0.00000 | 8.32900 | 0.00000 | 0.94346 |
| **NFIX** | 0.25243 | 0.01865 | 0.00747 | 0.06104 | 0.02740 | 0.01693 | 0.01657 | 0.02486 | 0.07445 | 0.04569 | 0.00645 | 0.01205 |
| **NFKB1** | 1.10311 | 0.00000 | 1.82297 | 0.92472 | 0.38375 | 0.00000 | 7.74650 | 0.01378 | 1.28688 | 0.94267 | 2.65767 | 0.00000 |
| **NFKB2** | 2.88996 | 2.30250 | 1.06701 | 8.68511 | 12.78452 | 0.00000 | 1.41842 | 0.26892 | 0.00000 | 11.56122 | 0.00000 | 1.51454 |
| **NFKBIA** | 19.89270 | 13.83474 | 5.44937 | 21.03486 | 23.91781 | 11.71960 | 3.11708 | 11.25287 | 3.87834 | 14.74768 | 9.53199 | 21.50219 |
| **NFKBIB** | 0.34797 | 2.02600 | 5.49557 | 4.61809 | 4.77636 | 0.00000 | 4.61270 | 0.62850 | 7.27008 | 5.90451 | 4.42838 | 5.47124 |
| **NFKBID** | 0.00273 | 0.05494 | 0.06769 | 0.35124 | 0.00000 | 0.00000 | 0.00000 | 0.01864 | 0.52126 | 0.08924 | 0.27037 | 0.04293 |
| **NFKBIE** | 0.23534 | 1.03279 | 2.11490 | 0.55231 | 1.39677 | 0.09138 | 4.31635 | 0.01807 | 0.20394 | 0.00000 | 0.16057 | 0.19666 |
| **NFKBIL1** | 0.14530 | 8.57639 | 3.48438 | 0.41234 | 4.10521 | 0.00000 | 0.00045 | 0.31442 | 1.45176 | 2.74502 | 2.39493 | 2.88029 |
| **NFKBIZ** | 0.00372 | 0.53895 | 1.28067 | 0.00000 | 0.26510 | 0.62317 | 0.14098 | 0.26980 | 2.56831 | 0.05709 | 1.04825 | 0.43580 |

| | | | | | | | | | | | | |
|---|---|---|---|---|---|---|---|---|---|---|---|---|
| **NFRKB** | 4.88426 | 8.64429 | 4.51333 | 8.73452 | 6.58338 | 3.04603 | 14.81833 | 5.04989 | 13.49476 | 10.68053 | 9.07213 | 9.34178 |
| **NFS1** | 2.10764 | 7.66171 | 8.68415 | 5.39139 | 5.17038 | 9.75532 | 8.61783 | 0.93730 | 15.64819 | 0.00000 | 13.53897 | 7.21818 |
| **NFU1** | 0.71298 | 2.77568 | 6.55808 | 1.41970 | 2.16475 | 15.36809 | 1.35516 | 2.08113 | 3.33614 | 0.00000 | 6.09428 | 3.66132 |
| **NFX1** | 1.93794 | 12.93334 | 5.40519 | 2.67697 | 5.31972 | 16.32040 | 6.04007 | 7.62155 | 5.60514 | 10.15024 | 9.85473 | 9.03919 |
| **NFXL1** | 2.30526 | 8.83257 | 6.23728 | 5.00169 | 1.41675 | 13.36635 | 3.29611 | 0.00000 | 10.61699 | 6.69873 | 9.75889 | 5.46455 |
| **NFYA** | 0.21501 | 4.16576 | 3.96175 | 3.46449 | 3.34112 | 7.63659 | 2.02287 | 21.55363 | 3.18018 | 4.31047 | 3.96228 | 7.64126 |
| **NFYB** | 2.93781 | 11.00894 | 8.19139 | 2.92530 | 2.49249 | 37.06516 | 10.38186 | 7.70002 | 6.20746 | 10.19939 | 10.24112 | 2.54167 |
| **NFYC** | 3.60456 | 16.95511 | 6.75434 | 8.48440 | 11.09560 | 9.62099 | 13.52938 | 7.44009 | 10.41407 | 0.00000 | 6.66419 | 12.88798 |
| **NFYC-AS1** | 0.00153 | 0.23320 | 0.12454 | 0.94014 | 0.10735 | 0.12291 | 0.00000 | 0.25562 | 0.73425 | 1.22713 | 0.94192 | 0.00000 |
| **NGB** | 0.28312 | 0.07029 | 0.01383 | 0.00257 | 0.00000 | 0.01327 | 0.01129 | 0.00000 | 0.00772 | 0.05818 | 0.03813 | 0.60140 |
| **NGDN** | 5.11491 | 14.60547 | 12.07431 | 7.38980 | 7.69531 | 29.24939 | 11.60907 | 23.86770 | 14.98551 | 1.06836 | 10.83468 | 26.72817 |
| **NGEF** | 0.36118 | 5.11056 | 0.00000 | 4.54311 | 0.27175 | 0.00000 | 0.00344 | 0.00000 | 0.00000 | 0.24507 | 0.00000 | 2.12644 |
| **NGF** | 0.00586 | 0.00863 | 0.00924 | 0.17552 | 0.00000 | 0.04003 | 0.00000 | 0.00996 | 0.03362 | 0.03606 | 0.00824 | 0.01332 |
| **NGFR** | 0.62868 | 0.02188 | 0.05218 | 0.72571 | 0.00000 | 0.05941 | 0.00000 | 1.12267 | 0.06089 | 0.12581 | 0.00000 | 0.34252 |
| **NGFRAP1** | 0.00000 | 0.00005 | 20.69639 | 2.50436 | 0.00000 | 7.21212 | 33.40066 | 0.75217 | 14.75529 | 2.85151 | 18.31676 | 6.40934 |
| **NGLY1** | 0.69766 | 7.09093 | 9.08311 | 2.41957 | 4.18452 | 16.41142 | 0.00000 | 2.29213 | 7.06343 | 0.00000 | 11.76907 | 7.37998 |
| **NGRN** | 3.88651 | 21.73832 | 17.84948 | 12.57710 | 20.46390 | 20.30170 | 18.09250 | 14.38531 | 19.40884 | 1.44510 | 15.65059 | 28.87566 |
| **NHEG1** | 0.04646 | 0.01490 | 0.01365 | 0.01223 | 0.01732 | 0.01310 | 0.02515 | 0.01230 | 0.01566 | 0.01855 | 0.01489 | 0.01649 |
| **NHEJ1** | 0.36766 | 3.76683 | 1.72393 | 3.69381 | 2.74715 | 6.51248 | 1.74260 | 1.11674 | 6.21607 | 0.48510 | 3.19248 | 2.26010 |
| **NHLH1** | 0.00360 | 0.00000 | 0.40134 | 0.00000 | 0.37007 | 0.00000 | 0.00000 | 0.00524 | 0.10870 | 0.26650 | 0.00227 | 0.19314 |
| **NHLH2** | 0.03627 | 0.02986 | 0.03583 | 0.00779 | 0.01026 | 0.01958 | 0.03512 | 0.00901 | 0.00406 | 0.02811 | 0.01510 | 0.03138 |
| **NHLRC1** | 0.00789 | 0.26283 | 0.02869 | 0.00501 | 0.00687 | 0.01663 | 0.01816 | 0.00413 | 0.02187 | 0.00000 | 0.06007 | 0.49120 |
| **NHLRC2** | 4.79287 | 2.52433 | 1.72716 | 0.50395 | 2.07279 | 7.65442 | 1.94078 | 0.90368 | 2.73627 | 0.64519 | 2.79228 | 3.14360 |
| **NHLRC3** | 5.88161 | 0.52723 | 3.75852 | 0.77504 | 1.64147 | 0.99127 | 0.52396 | 1.84945 | 5.26397 | 2.37199 | 3.86295 | 1.46167 |
| **NHLRC4** | 0.25403 | 0.02635 | 0.01733 | 0.01160 | 0.01357 | 0.00000 | 0.02715 | 0.00348 | 0.03614 | 0.01642 | 0.05338 | 0.01336 |
| **NHP2** | 1.85649 | 23.24715 | 17.74346 | 19.25293 | 21.95152 | 13.65859 | 29.71651 | 28.00391 | 25.74599 | 2.11426 | 21.40849 | 35.18335 |
| **NHP2L1** | 4.82199 | 30.41236 | 34.78480 | 23.27523 | 29.46816 | 29.40811 | 43.71487 | 34.03502 | 44.01513 | 12.80765 | 31.49660 | 38.91994 |
| **NHS** | 0.00313 | 0.12002 | 0.13019 | 1.57115 | 0.00144 | 1.23646 | 0.00000 | 0.08312 | 0.00000 | 0.23361 | 0.00000 | 0.04665 |
| **NHSL1** | 5.65192 | 2.21239 | 3.14024 | 5.09968 | 3.26978 | 12.37681 | 6.98697 | 1.46795 | 6.84773 | 8.64107 | 7.58069 | 0.42568 |
| **NHSL2** | 0.00732 | 0.00000 | 0.00000 | 0.46291 | 0.02965 | 0.17559 | 0.00000 | 0.00445 | 0.00000 | 0.02550 | 0.00216 | 0.08494 |
| **NICN1** | 0.23246 | 1.27814 | 0.97492 | 0.74886 | 0.79530 | 0.00000 | 0.00000 | 1.46767 | 0.00000 | 0.00000 | 2.36356 | 1.78151 |
| **NID1** | 0.00000 | 0.75020 | 8.94384 | 24.07968 | 12.84847 | 15.25697 | 1.98880 | 0.00000 | 33.43929 | 8.97583 | 12.16858 | 0.00000 |
| **NID2** | 6.11954 | 0.00000 | 1.58004 | 7.78337 | 0.00000 | 0.03613 | 0.12018 | 0.85402 | 0.40753 | 0.05717 | 0.00551 | 1.10853 |
| **NIF3L1** | 0.46266 | 10.28092 | 4.82821 | 6.37093 | 3.46046 | 6.93611 | 4.34003 | 6.53290 | 6.04730 | 0.00000 | 10.49039 | 9.51446 |
| **NIFK** | 2.82028 | 14.51781 | 12.76421 | 6.28528 | 8.09698 | 38.09621 | 18.23056 | 14.22604 | 20.73318 | 12.00450 | 15.13585 | 18.11413 |
| **NIFK-AS1** | 0.01446 | 0.90497 | 0.32538 | 0.28273 | 0.78600 | 0.64107 | 0.46553 | 0.65555 | 0.83247 | 1.30508 | 0.72080 | 1.70566 |
| **NIM1K** | 0.00070 | 0.00000 | 0.00000 | 0.00000 | 1.99168 | 0.01737 | 1.68603 | 0.09294 | 3.05932 | 0.00000 | 11.15075 | 0.12940 |
| **NIN** | 1.08600 | 0.47094 | 1.03338 | 0.93255 | 1.53442 | 3.96604 | 0.00000 | 0.35531 | 0.00000 | 3.10384 | 0.50570 | 1.03057 |
| **NINJ1** | 1.55660 | 2.45203 | 2.89455 | 0.00250 | 0.69317 | 0.00418 | 0.00114 | 0.25664 | 1.62134 | 0.00000 | 0.30472 | 1.39172 |
| **NINJ2** | 0.00837 | 2.46540 | 0.29517 | 0.10696 | 0.05016 | 0.00095 | 0.03791 | 0.09298 | 0.00000 | 0.00000 | 0.45791 | 0.49773 |
| **NINL** | 0.00000 | 2.85838 | 0.00000 | 6.41886 | 2.11941 | 1.33492 | 0.80466 | 0.90024 | 2.00948 | 6.86136 | 1.66525 | 1.60729 |
| **NIP7** | 0.00000 | 18.81801 | 15.79917 | 7.25617 | 10.18842 | 21.98883 | 16.57821 | 24.84745 | 11.19056 | 4.79101 | 13.18449 | 23.88782 |
| **NIPA1** | 2.64199 | 0.06083 | 0.51919 | 3.07520 | 2.22938 | 0.00000 | 0.42706 | 0.09765 | 1.44113 | 3.04266 | 0.01877 | 0.00000 |
| **NIPA2** | 2.54341 | 17.50652 | 24.29955 | 10.23089 | 9.68352 | 24.75428 | 12.23141 | 10.80015 | 30.34958 | 10.79509 | 13.86203 | 13.20956 |
| **NIPAL1** | 0.00000 | 3.62220 | 0.83136 | 0.22924 | 4.47491 | 2.66948 | 0.00000 | 0.12817 | 5.07932 | 0.79571 | 5.71495 | 10.55952 |
| **NIPAL2** | 0.00252 | 0.05196 | 0.00018 | 0.02393 | 0.10775 | 0.05925 | 0.02666 | 0.03191 | 0.00972 | 0.08655 | 0.02243 | 0.07860 |
| **NIPAL3** | 5.24415 | 1.25575 | 2.05888 | 0.64222 | 0.54763 | 1.54207 | 1.65857 | 0.82534 | 0.95643 | 0.57670 | 3.01932 | 0.37549 |
| **NIPAL4** | 0.00811 | 0.01422 | 0.01805 | 0.01255 | 0.01433 | 0.01983 | 0.03341 | 0.01906 | 0.00766 | 0.01446 | 0.02424 | 0.01529 |

| | | | | | | | | | | | | |
|---|---|---|---|---|---|---|---|---|---|---|---|---|
| **NIPBL** | 5.91598 | 1.94946 | 7.86683 | 5.39802 | 7.78254 | 19.44819 | 10.71295 | 7.78811 | 2.57156 | 26.83051 | 3.51711 | 6.66779 |
| **NIPSNAP1** | 0.82502 | 3.17874 | 0.46654 | 3.86168 | 3.34678 | 2.32936 | 2.61354 | 1.82940 | 0.32155 | 0.00000 | 5.69001 | 3.39168 |
| **NIPSNAP3A** | 0.03416 | 1.96654 | 1.32067 | 0.66338 | 0.00000 | 5.03522 | 0.83829 | 0.08152 | 0.52637 | 0.28793 | 2.83860 | 1.10681 |
| **NIPSNAP3B** | 0.00614 | 0.00822 | 0.06419 | 0.16702 | 0.02590 | 0.12203 | 0.06714 | 0.01048 | 0.00000 | 0.01310 | 0.07030 | 0.02538 |
| **NISCH** | 3.23389 | 4.67816 | 4.50850 | 4.01260 | 4.94383 | 1.66279 | 6.06265 | 1.20459 | 7.64500 | 7.50905 | 3.31890 | 3.73682 |
| **NIT1** | 0.63807 | 4.30491 | 6.61454 | 3.61931 | 2.67330 | 4.25496 | 1.78709 | 0.31326 | 2.18415 | 0.00000 | 3.45132 | 2.50629 |
| **NIT2** | 0.17065 | 10.51878 | 6.04555 | 6.93039 | 8.14465 | 20.69781 | 6.12645 | 0.35937 | 7.65177 | 1.29734 | 13.54000 | 1.40512 |
| **NKAIN1** | 0.00856 | 0.13949 | 0.14472 | 0.09420 | 0.00000 | 0.00973 | 0.13219 | 0.05439 | 0.03616 | 0.11246 | 0.00000 | 0.09618 |
| **NKAIN2** | 0.25743 | 0.00578 | 0.04494 | 0.00818 | 0.01198 | 0.03732 | 0.02129 | 0.00000 | 0.00009 | 0.00930 | 0.02790 | 0.12809 |
| **NKAIN4** | 0.00364 | 0.00064 | 0.16668 | 0.00000 | 0.00000 | 0.00000 | 0.16566 | 0.01056 | 0.50608 | 0.31491 | 0.50709 | 0.00721 |
| **NKAP** | 0.46421 | 6.44373 | 0.25567 | 3.82267 | 6.91530 | 7.90603 | 3.87326 | 1.68173 | 1.98917 | 9.42597 | 5.61327 | 12.41729 |
| **NKAPL** | 0.55799 | 0.07592 | 0.18822 | 0.00374 | 0.11320 | 0.25340 | 0.30737 | 0.29072 | 0.00005 | 0.00005 | 0.14498 | 0.49495 |
| **NKAPP1** | 0.00189 | 0.53825 | 0.00000 | 0.85643 | 0.40017 | 0.13991 | 0.00000 | 0.38010 | 0.00000 | 0.58709 | 0.25409 | 1.16269 |
| **NKD1** | 2.17920 | 0.41547 | 0.94591 | 0.02358 | 0.35894 | 0.17324 | 0.00000 | 1.34492 | 0.00000 | 0.06774 | 0.00000 | 1.37261 |
| **NKD2** | 4.72648 | 2.56089 | 0.95267 | 0.00000 | 14.57385 | 0.00000 | 4.91392 | 0.32661 | 2.61870 | 11.03494 | 1.46835 | 0.06067 |
| **NKG7** | 0.01438 | 0.02050 | 0.03254 | 0.01271 | 0.00528 | 0.03152 | 0.01167 | 0.00928 | 0.01850 | 0.01475 | 0.03837 | 0.01556 |
| **NKIRAS1** | 0.74280 | 1.54988 | 2.81847 | 1.42916 | 0.00000 | 5.87253 | 2.94548 | 17.23180 | 1.50620 | 3.22883 | 2.54090 | 5.00060 |
| **NKIRAS2** | 2.99750 | 12.57391 | 6.04932 | 6.70619 | 10.44184 | 9.43627 | 10.56728 | 3.06553 | 7.42751 | 0.00000 | 8.06037 | 20.58650 |
| **NKPD1** | 0.02517 | 0.00297 | 0.00151 | 0.00108 | 0.00177 | 0.02587 | 0.00100 | 0.00747 | 0.01566 | 0.42829 | 0.00000 | 0.02067 |
| **NKRF** | 0.40797 | 0.25375 | 1.60269 | 0.26661 | 1.16779 | 7.65303 | 6.30850 | 1.74786 | 0.52232 | 12.15780 | 3.14568 | 1.21104 |
| **NKTR** | 6.58275 | 3.51132 | 4.32976 | 3.87347 | 2.63487 | 17.06522 | 4.17188 | 7.23526 | 8.42336 | 35.92048 | 8.71857 | 8.64108 |
| **NKX1-1** | 0.01326 | 0.01075 | 0.01020 | 0.00908 | 0.01205 | 0.02862 | 0.00920 | 0.01924 | 0.01593 | 0.05903 | 0.01249 | 0.01214 |
| **NKX1-2** | 0.00656 | 0.01493 | 0.01742 | 0.01088 | 0.01995 | 0.02093 | 0.01584 | 0.01123 | 0.01828 | 0.01343 | 0.01531 | 0.01603 |
| **NKX2-1** | 0.01520 | 0.01404 | 0.01437 | 0.01109 | 0.01450 | 0.02302 | 0.01792 | 0.05546 | 0.01808 | 0.01317 | 0.01663 | 0.01746 |
| **NKX2-2** | 0.01724 | 0.00530 | 0.00533 | 0.21723 | 0.09653 | 0.03053 | 0.00000 | 0.23291 | 0.00000 | 0.11535 | 0.00565 | 0.12230 |
| **NKX2-3** | 0.00267 | 0.06519 | 0.00000 | 0.07108 | 0.05626 | 0.00000 | 0.00060 | 0.04115 | 0.21217 | 0.21653 | 0.01495 | 0.03381 |
| **NKX2-4** | 0.01488 | 0.03932 | 0.00316 | 0.10368 | 0.00000 | 0.03119 | 0.01127 | 0.01003 | 0.01529 | 0.04918 | 0.00992 | 0.01724 |
| **NKX2-5** | 0.00170 | 0.00000 | 0.01708 | 0.01223 | 0.03722 | 0.00000 | 0.03214 | 0.89124 | 0.00288 | 0.02413 | 0.01244 | 2.04222 |
| **NKX2-8** | 0.03501 | 0.00453 | 0.00643 | 0.30453 | 0.00000 | 0.00975 | 0.01850 | 0.00501 | 0.01870 | 0.03090 | 0.01532 | 0.12757 |
| **NKX3-1** | 8.31797 | 0.45083 | 0.23841 | 0.00000 | 0.00000 | 1.25007 | 0.63082 | 2.43153 | 0.00000 | 2.54502 | 0.13590 | 0.41012 |
| **NKX3-2** | 0.17747 | 0.01000 | 0.01795 | 0.00904 | 0.02498 | 0.02011 | 0.01222 | 0.12691 | 0.01457 | 0.03762 | 0.01477 | 0.01901 |
| **NKX6-1** | 0.00800 | 0.03435 | 0.01419 | 0.01053 | 0.01302 | 0.01747 | 0.01777 | 0.05582 | 0.01814 | 0.01411 | 0.01549 | 0.01269 |
| **NKX6-2** | 0.00346 | 0.00570 | 0.20678 | 0.07066 | 0.00000 | 0.04763 | 0.00000 | 0.62662 | 0.00000 | 0.04915 | 0.01491 | 0.00000 |
| **NKX6-3** | 0.00633 | 0.01548 | 0.01731 | 0.01558 | 0.01556 | 0.01687 | 0.02270 | 0.01050 | 0.01462 | 0.01003 | 0.01859 | 0.01668 |
| **NLE1** | 0.00114 | 9.87430 | 3.51245 | 4.08666 | 8.42786 | 0.00000 | 6.97520 | 10.92827 | 9.95848 | 0.00000 | 8.16918 | 9.07901 |
| **NLGN1** | 1.89765 | 0.01176 | 0.01726 | 0.01560 | 0.00000 | 0.03608 | 0.00000 | 0.32640 | 0.00000 | 0.61487 | 0.00000 | 0.10177 |
| **NLGN2** | 1.19318 | 0.09894 | 0.40878 | 0.78348 | 0.13425 | 0.00000 | 0.17952 | 0.13479 | 0.89702 | 2.71528 | 0.04848 | 0.03630 |
| **NLGN3** | 0.14126 | 0.00000 | 0.18808 | 0.03822 | 0.06538 | 0.00000 | 0.20373 | 0.02826 | 0.28197 | 0.03933 | 0.02651 | 0.25075 |
| **NLGN4X** | 1.08232 | 0.61466 | 0.40958 | 0.00062 | 0.00000 | 1.47337 | 1.60101 | 0.00000 | 0.00000 | 2.38614 | 1.61776 | 0.00000 |
| **NLGN4Y** | 0.01792 | 0.05837 | 0.01898 | 0.01494 | 0.02069 | 0.00000 | 0.02318 | 0.01117 | 0.02104 | 0.00405 | 0.01671 | 0.01776 |
| **NLK** | 2.01874 | 1.80841 | 2.83921 | 1.09327 | 1.60099 | 3.74974 | 2.13844 | 0.75608 | 0.00000 | 5.27739 | 0.47772 | 1.82171 |
| **NLN** | 1.70273 | 6.72086 | 6.34942 | 1.50889 | 2.13189 | 14.00759 | 15.01755 | 0.72274 | 13.66475 | 1.64999 | 11.73588 | 2.85609 |
| **NLRC3** | 0.07873 | 0.02828 | 0.28873 | 0.33749 | 0.00000 | 0.01282 | 0.00000 | 0.00000 | 0.06508 | 0.38802 | 0.00061 | 0.28079 |
| **NLRC4** | 0.00874 | 0.03581 | 0.14066 | 1.02324 | 0.00000 | 0.00000 | 0.04800 | 0.40793 | 0.28019 | 0.74551 | 0.00773 | 1.14146 |
| **NLRC5** | 0.00550 | 0.02224 | 0.03373 | 0.08235 | 0.00000 | 0.00000 | 0.02786 | 0.02812 | 0.00000 | 0.16467 | 0.03312 | 0.00948 |
| **NLRP1** | 0.41921 | 0.00000 | 0.88051 | 0.01023 | 0.00595 | 0.00000 | 0.00000 | 0.05244 | 0.00000 | 0.18265 | 0.02421 | 0.40058 |
| **NLRP10** | 0.00623 | 0.01604 | 0.08470 | 0.00640 | 0.01965 | 0.02353 | 0.05504 | 0.01037 | 0.00000 | 0.00850 | 0.01304 | 0.01661 |
| **NLRP11** | 25.64629 | 0.02509 | 0.00000 | 0.02387 | 0.01337 | 0.04990 | 0.01743 | 13.32954 | 0.00000 | 0.10469 | 0.00000 | 0.22513 |

| | | | | | | | | | | | | |
|---|---|---|---|---|---|---|---|---|---|---|---|---|
| **NLRP12** | 2.24995 | 6.42997 | 1.47448 | 6.42470 | 3.88462 | 2.11414 | 0.00000 | 0.19381 | 0.44444 | 1.44581 | 2.71067 | 1.29722 |
| **NLRP13** | 28.14064 | 0.13061 | 0.00320 | 0.02281 | 0.07638 | 0.00000 | 0.00000 | 17.16102 | 0.00583 | 0.01074 | 0.00714 | 0.73242 |
| **NLRP14** | 2.04226 | 0.00699 | 0.21709 | 0.00062 | 0.04002 | 0.24641 | 0.05821 | 0.72436 | 0.00000 | 0.00000 | 0.00845 | 0.00241 |
| **NLRP2** | 14.03540 | 54.53806 | 35.62166 | 42.53230 | 52.47206 | 55.98519 | 43.65750 | 4.46928 | 49.85882 | 14.87631 | 35.11807 | 42.11836 |
| **NLRP3** | 0.14053 | 0.01144 | 0.04287 | 0.04071 | 0.00000 | 0.03426 | 0.04600 | 0.00443 | 0.00000 | 0.04735 | 0.01272 | 0.01648 |
| **NLRP4** | 36.54431 | 0.13299 | 0.15856 | 0.25419 | 0.54813 | 0.00000 | 0.04524 | 18.81202 | 0.00000 | 0.00000 | 0.06068 | 1.33880 |
| **NLRP5** | 13.64298 | 0.08605 | 0.00000 | 0.00000 | 0.30339 | 0.03596 | 0.00870 | 3.94170 | 0.00000 | 0.00881 | 0.00000 | 0.30520 |
| **NLRP6** | 0.00641 | 0.01476 | 0.03044 | 0.01338 | 0.01537 | 0.02484 | 0.02222 | 0.01041 | 0.01418 | 0.00661 | 0.01503 | 0.01644 |
| **NLRP7** | 7.92245 | 57.66949 | 22.14889 | 46.97417 | 55.20012 | 48.31376 | 47.37085 | 2.46451 | 37.37272 | 30.21245 | 50.53169 | 23.10015 |
| **NLRP8** | 8.63239 | 0.00482 | 0.00943 | 0.03170 | 0.01023 | 0.00000 | 0.00873 | 3.29523 | 0.02032 | 0.00804 | 0.01200 | 0.00000 |
| **NLRP9** | 25.96718 | 0.00000 | 0.04619 | 2.88637 | 0.21023 | 0.32595 | 0.00000 | 7.51911 | 0.00000 | 0.94291 | 0.00000 | 1.03096 |
| **NLRX1** | 0.05785 | 0.30788 | 0.48023 | 0.00161 | 0.18136 | 0.00000 | 0.08145 | 0.08328 | 0.00000 | 0.00000 | 0.12146 | 0.07938 |
| **NMB** | 0.00000 | 1.47533 | 1.34178 | 0.00000 | 0.00000 | 0.16980 | 1.73791 | 0.07301 | 0.96587 | 1.18532 | 0.14231 | 0.88816 |
| **NMBR** | 0.01683 | 0.01494 | 0.01487 | 0.01570 | 0.01204 | 0.01218 | 0.01983 | 0.01090 | 0.01568 | 0.02188 | 0.02968 | 0.01640 |
| **NMD3** | 2.11756 | 9.44014 | 5.07598 | 0.65427 | 2.68528 | 21.74540 | 3.75225 | 4.20447 | 5.86737 | 2.82726 | 9.37032 | 6.78258 |
| **NME1** | 0.74387 | 16.34375 | 20.57291 | 9.05098 | 18.51445 | 26.47611 | 41.36152 | 0.71332 | 26.41145 | 16.60528 | 25.36117 | 14.67672 |
| **NME1-NME2** | 0.01764 | 0.09372 | 0.21481 | 0.01584 | 0.06480 | 0.06707 | 0.13620 | 0.03445 | 0.21203 | 0.00866 | 0.07006 | 0.11339 |
| **NME2.uid-22041** | 0.40060 | 6.23828 | 5.63751 | 5.92800 | 6.88243 | 1.03793 | 5.59589 | 0.63838 | 11.64237 | 0.00000 | 6.40259 | 4.27604 |
| **NME2.uid-22044** | 0.03915 | 0.95521 | 0.24742 | 1.11867 | 1.40941 | 0.00000 | 0.39611 | 0.01033 | 1.11657 | 0.00000 | 0.51549 | 1.54893 |
| **NME2.uid-22045** | 0.00757 | 0.01826 | 0.04620 | 0.08423 | 0.01539 | 0.02335 | 0.06578 | 0.01872 | 0.02083 | 0.00468 | 0.03535 | 0.03031 |
| **NME2.uid-22046** | 0.23365 | 3.00881 | 1.87919 | 3.05224 | 4.17545 | 0.00015 | 2.09155 | 0.24038 | 5.37178 | 0.00000 | 2.33701 | 1.77401 |
| **NME3** | 0.00285 | 1.26824 | 2.60296 | 0.56811 | 1.15710 | 0.00000 | 0.81429 | 0.28576 | 3.12404 | 3.03561 | 2.57057 | 0.27976 |
| **NME4** | 0.98247 | 4.65312 | 7.26893 | 0.51192 | 0.00000 | 0.00000 | 5.17945 | 0.01887 | 2.06175 | 2.94334 | 3.96145 | 1.42213 |
| **NME5** | 0.00338 | 0.16212 | 0.00858 | 0.00000 | 0.00000 | 0.00000 | 0.36014 | 0.24286 | 0.00000 | 0.24173 | 0.29926 | 0.92348 |
| **NME6** | 0.87904 | 7.02136 | 5.17520 | 3.76628 | 5.50327 | 7.84671 | 6.47374 | 3.29225 | 8.40685 | 0.00000 | 8.71238 | 10.33198 |
| **NME7** | 1.17456 | 0.82211 | 2.32330 | 1.52336 | 0.00000 | 2.46491 | 3.58684 | 1.90767 | 1.33969 | 0.00000 | 1.99856 | 1.92996 |
| **NME8** | 0.00626 | 0.01239 | 0.03567 | 0.00856 | 0.00703 | 0.03916 | 0.06690 | 0.01028 | 0.01844 | 0.00537 | 0.00879 | 0.01534 |
| **NME9** | 0.00217 | 0.00000 | 0.00000 | 0.20077 | 0.01712 | 0.15109 | 0.00000 | 0.01513 | 0.11212 | 1.70340 | 0.16693 | 0.13648 |
| **NMI** | 0.00145 | 0.00000 | 4.33726 | 0.38836 | 3.90936 | 1.50827 | 9.35888 | 0.15016 | 0.78057 | 0.38607 | 6.83085 | 0.00000 |
| **NMNAT1** | 1.11683 | 3.33445 | 1.98761 | 1.81812 | 0.85050 | 1.89511 | 0.41961 | 9.68242 | 0.50539 | 0.00000 | 0.98849 | 3.77392 |
| **NMNAT2** | 0.00503 | 0.46005 | 2.21464 | 2.36307 | 0.00000 | 0.11518 | 0.00000 | 0.05642 | 0.00000 | 0.29021 | 0.23336 | 0.59942 |
| **NMNAT3** | 0.43524 | 0.12012 | 0.02508 | 0.00000 | 0.01804 | 0.03275 | 0.00882 | 0.00368 | 0.00778 | 0.03657 | 0.01482 | 0.02606 |
| **NMRAL1** | 0.69816 | 4.86179 | 8.53084 | 5.35744 | 6.55018 | 0.05633 | 7.99766 | 0.00000 | 5.51351 | 0.00000 | 12.60062 | 5.39490 |
| **NMRK1** | 0.04224 | 0.61716 | 0.06347 | 0.35452 | 0.02936 | 0.96419 | 1.30983 | 0.02795 | 0.00223 | 0.00000 | 0.54172 | 0.43241 |
| **NMRK2** | 1.25987 | 1.71377 | 0.00000 | 1.72316 | 9.78163 | 0.00000 | 1.94527 | 1.66656 | 3.78168 | 0.00000 | 1.43597 | 1.96764 |
| **NMT1** | 0.24156 | 14.30889 | 16.18574 | 12.29944 | 18.92375 | 2.42129 | 38.07584 | 11.87063 | 5.23715 | 5.71065 | 23.71073 | 22.22105 |
| **NMT2** | 0.00000 | 2.96152 | 0.00000 | 7.38635 | 1.41791 | 1.79597 | 0.74616 | 13.50543 | 0.22019 | 1.02781 | 0.29261 | 4.07789 |
| **NMU** | 0.01768 | 14.85905 | 0.00000 | 2.87037 | 18.29550 | 2.45538 | 0.00000 | 0.00000 | 0.00000 | 0.74702 | 6.79378 | 2.23206 |
| **NMUR1** | 0.01932 | 0.54705 | 0.00000 | 0.57047 | 0.00000 | 0.03259 | 0.00000 | 0.00540 | 0.00000 | 0.31093 | 0.00000 | 0.05427 |
| **NMUR2** | 0.16127 | 0.02563 | 0.12685 | 0.48508 | 0.00000 | 0.04758 | 0.00000 | 0.00723 | 0.08322 | 0.01904 | 0.00000 | 0.03436 |
| **NNAT** | 0.07433 | 0.17978 | 1.64418 | 0.45558 | 0.00000 | 0.04456 | 1.00377 | 0.00435 | 0.17526 | 0.00000 | 0.00832 | 0.11496 |
| **NNMT** | 0.00619 | 0.00871 | 0.27040 | 0.31439 | 0.00000 | 0.00686 | 0.43848 | 0.01022 | 0.42899 | 0.00000 | 0.00000 | 0.01584 |
| **NNT** | 1.06813 | 0.00001 | 0.53492 | 9.35050 | 2.62746 | 4.77017 | 0.00000 | 0.56689 | 0.00000 | 0.09385 | 3.70099 | 0.01667 |
| **NNT-AS1** | 0.00000 | 0.40737 | 1.75522 | 2.61179 | 1.47897 | 4.57195 | 1.70855 | 1.60268 | 0.00000 | 2.20237 | 1.93917 | 3.89005 |
| **NOA1** | 2.42588 | 14.35379 | 8.43939 | 6.61032 | 7.09224 | 3.99686 | 8.83928 | 2.55189 | 15.58272 | 5.60725 | 7.23602 | 9.25345 |
| **NOB1** | 2.36486 | 18.08341 | 10.89750 | 9.77409 | 12.63719 | 7.23349 | 17.84334 | 8.81915 | 21.09917 | 0.00121 | 14.42856 | 22.17554 |
| **NOBOX** | 0.74141 | 0.01286 | 0.02021 | 0.01086 | 0.01053 | 0.01047 | 0.01315 | 0.02289 | 0.01563 | 0.04873 | 0.01307 | 0.01861 |
| **NOC2L** | 0.49301 | 20.63018 | 13.44398 | 10.93304 | 13.35361 | 0.00000 | 11.48434 | 5.89886 | 23.34311 | 0.00000 | 17.11723 | 14.28142 |

| | | | | | | | | | | | | |
|---|---|---|---|---|---|---|---|---|---|---|---|---|
| **NOC3L** | 5.19706 | 3.72725 | 5.99946 | 0.00000 | 0.00021 | 21.49268 | 9.07474 | 6.02249 | 7.28682 | 18.37800 | 10.48962 | 9.18868 |
| **NOC4L** | 0.56627 | 13.89374 | 10.07236 | 5.83085 | 11.75882 | 0.00000 | 6.17688 | 3.90049 | 10.84037 | 5.86057 | 9.55001 | 13.37579 |
| **NOD1** | 4.81588 | 0.69696 | 0.27191 | 0.46157 | 0.00000 | 0.00000 | 0.31739 | 0.50417 | 0.54252 | 0.78845 | 0.28892 | 1.17672 |
| **NOD2** | 0.00662 | 0.00177 | 0.00450 | 0.00262 | 0.02728 | 0.03557 | 0.00000 | 0.05816 | 0.01799 | 0.31524 | 0.00394 | 0.01628 |
| **NODAL** | 0.41732 | 0.93001 | 5.79084 | 12.69953 | 0.00000 | 0.00000 | 0.00000 | 0.01051 | 0.26500 | 0.25173 | 0.04576 | 0.14232 |
| **NOG** | 0.03935 | 0.00000 | 0.48120 | 1.18837 | 0.00000 | 0.39934 | 0.12716 | 0.03927 | 0.00000 | 1.31829 | 0.01464 | 0.16725 |
| **NOL10** | 3.00615 | 10.63575 | 11.66550 | 3.52372 | 5.83632 | 23.08308 | 12.56446 | 12.05909 | 14.39969 | 7.34511 | 14.06082 | 13.26025 |
| **NOL11** | 1.70957 | 10.31732 | 11.09063 | 6.57094 | 3.11207 | 20.98556 | 15.35260 | 23.67650 | 16.17712 | 8.81964 | 12.81104 | 14.77245 |
| **NOL12** | 3.78113 | 14.59107 | 7.59647 | 6.25225 | 11.05201 | 6.63914 | 12.66227 | 19.79817 | 12.46383 | 0.00000 | 8.21875 | 17.75274 |
| **NOL3** | 0.00243 | 1.26332 | 0.40334 | 0.29391 | 1.04784 | 0.00000 | 0.00064 | 0.01028 | 0.90722 | 0.71145 | 0.17681 | 0.62799 |
| **NOL4** | 0.64050 | 0.00697 | 0.00368 | 0.00026 | 0.02428 | 0.02108 | 0.00000 | 0.00000 | 0.03252 | 0.16039 | 0.01755 | 0.06414 |
| **NOL4L** | 4.23723 | 1.45244 | 0.25983 | 2.34549 | 0.31929 | 0.73661 | 0.67606 | 0.54809 | 0.62943 | 2.21090 | 0.31763 | 0.63763 |
| **NOL6** | 0.63119 | 14.79587 | 10.38072 | 7.50718 | 11.13385 | 1.18416 | 10.63882 | 3.60014 | 24.72614 | 5.35473 | 7.54731 | 18.01395 |
| **NOL7** | 0.73515 | 14.46811 | 16.96261 | 5.55705 | 10.50045 | 19.12954 | 5.54947 | 6.92236 | 20.81116 | 11.65190 | 14.79245 | 13.19933 |
| **NOL8** | 5.26117 | 6.42333 | 3.68097 | 1.41066 | 2.52985 | 21.21582 | 10.04380 | 8.81390 | 7.63475 | 18.54468 | 8.43345 | 9.45209 |
| **NOL9** | 0.39012 | 6.12143 | 8.37767 | 4.76786 | 5.06645 | 4.75467 | 16.02304 | 14.43647 | 1.49448 | 14.87900 | 11.08843 | 15.38770 |
| **NOLC1** | 3.25323 | 34.26856 | 25.91502 | 13.90350 | 24.76326 | 31.94407 | 36.76873 | 31.47101 | 35.21942 | 19.98137 | 31.28317 | 43.10141 |
| **NOM1** | 0.71736 | 2.12611 | 2.16971 | 2.19214 | 3.58316 | 0.00154 | 2.87952 | 4.84839 | 1.08611 | 11.36368 | 2.80668 | 1.41889 |
| **NOMO1** | 0.00070 | 1.42410 | 1.74469 | 0.83893 | 2.01444 | 0.70525 | 0.53250 | 0.22445 | 2.20182 | 5.00321 | 0.98235 | 0.52156 |
| **NOMO2** | 0.02069 | 0.21221 | 0.25593 | 0.12581 | 0.08487 | 0.00234 | 0.00164 | 0.04018 | 0.47712 | 0.00000 | 0.12802 | 0.06069 |
| **NOMO3** | 0.00647 | 0.07442 | 0.07111 | 0.08769 | 0.05554 | 0.07021 | 0.00000 | 0.01890 | 0.20680 | 0.00000 | 0.17890 | 0.02772 |
| **NONO** | 0.00000 | 22.82156 | 15.75650 | 20.19959 | 19.46747 | 24.66942 | 35.96546 | 13.08192 | 28.30271 | 7.01095 | 21.29330 | 21.83057 |
| **NOP10** | 5.60874 | 24.05555 | 30.01234 | 20.32568 | 27.77902 | 27.50805 | 38.94575 | 25.03892 | 26.55637 | 11.04246 | 30.90723 | 41.18599 |
| **NOP14** | 0.00009 | 3.26655 | 8.43967 | 1.98244 | 6.42936 | 0.00000 | 3.86670 | 2.41583 | 8.76177 | 11.70839 | 3.13013 | 3.12639 |
| **NOP14-AS1** | 0.71921 | 1.00376 | 0.23263 | 0.66324 | 0.98490 | 1.55050 | 0.30103 | 0.11008 | 0.27626 | 0.23667 | 0.18874 | 0.33722 |
| **NOP16** | 1.88864 | 27.74418 | 17.42297 | 20.20818 | 29.81202 | 25.44384 | 32.65249 | 19.96986 | 31.77109 | 5.71484 | 32.14053 | 28.35555 |
| **NOP2** | 2.06711 | 23.57218 | 19.68776 | 15.44564 | 19.83315 | 6.14149 | 21.68982 | 14.32565 | 33.04370 | 3.96653 | 21.84510 | 29.69097 |
| **NOP56** | 5.56358 | 27.13779 | 21.00471 | 21.77989 | 33.29561 | 30.79460 | 30.95951 | 16.09815 | 42.00513 | 39.03759 | 20.97297 | 40.43030 |
| **NOP58** | 2.14875 | 22.31401 | 20.41932 | 11.65489 | 15.29585 | 44.36023 | 32.33356 | 30.38205 | 24.54974 | 34.29388 | 27.82268 | 22.92754 |
| **NOP9** | 0.60093 | 0.84480 | 3.24137 | 3.98821 | 5.99280 | 0.00000 | 5.88194 | 3.21771 | 0.00765 | 4.49022 | 2.27454 | 1.88237 |
| **NOS1** | 0.00219 | 0.00409 | 0.04392 | 0.04946 | 0.00000 | 0.00001 | 0.05452 | 1.16562 | 0.02353 | 0.10419 | 0.01797 | 0.28082 |
| **NOS1AP** | 6.79453 | 0.37054 | 0.00000 | 0.57048 | 0.38775 | 0.00000 | 0.28281 | 0.45494 | 1.23898 | 1.94339 | 2.76703 | 0.31043 |
| **NOS2** | 0.09905 | 0.00911 | 8.04263 | 0.00551 | 0.00341 | 0.00002 | 0.04248 | 0.13143 | 4.10169 | 0.01333 | 0.00000 | 0.02523 |
| **NOS3** | 0.03112 | 1.24823 | 0.45986 | 0.72562 | 0.33341 | 0.00000 | 0.06208 | 0.58227 | 0.00000 | 0.10370 | 0.19593 | 2.00260 |
| **NOSIP** | 2.87924 | 17.52564 | 18.43496 | 14.42715 | 22.86396 | 3.75355 | 33.65350 | 27.77025 | 15.03222 | 1.97658 | 16.97925 | 21.76338 |
| **NOSTRIN** | 1.26296 | 0.00000 | 0.30376 | 1.20908 | 0.00000 | 0.30883 | 0.16667 | 0.04400 | 0.25197 | 0.24593 | 0.06442 | 0.17360 |
| **NOTCH1** | 0.04709 | 2.47477 | 0.00000 | 0.99676 | 0.18815 | 0.00000 | 0.00022 | 0.21373 | 0.00000 | 2.17229 | 0.00000 | 0.79704 |
| **NOTCH2** | 0.16937 | 0.88605 | 0.27437 | 0.18034 | 0.89248 | 1.39138 | 0.22293 | 0.02515 | 1.15657 | 1.02237 | 0.00000 | 0.19159 |
| **NOTCH2NL** | 0.59913 | 0.77298 | 1.70757 | 0.87736 | 1.11465 | 0.00000 | 0.07810 | 0.37989 | 0.27280 | 0.34552 | 0.37865 | 0.27812 |
| **NOTCH3** | 0.01102 | 0.00000 | 0.58972 | 0.26033 | 0.06047 | 0.13820 | 0.00000 | 0.00278 | 0.26912 | 0.83600 | 0.00001 | 0.06498 |
| **NOTCH4** | 0.40589 | 0.04129 | 0.12370 | 0.01018 | 0.10853 | 0.00000 | 0.00805 | 0.00000 | 0.08034 | 0.06040 | 0.08451 | 0.06903 |
| **NOTO** | 0.00950 | 0.40827 | 0.00941 | 0.00000 | 0.00000 | 0.00304 | 0.01667 | 0.00150 | 0.00501 | 0.03854 | 0.02199 | 1.51347 |
| **NOTUM** | 0.01658 | 0.00364 | 1.58672 | 0.01775 | 0.00907 | 0.11986 | 0.00000 | 0.13001 | 0.07169 | 0.06997 | 0.02606 | 0.01029 |
| **NOVA1** | 0.59460 | 1.57376 | 0.00000 | 0.76131 | 0.00000 | 8.91062 | 0.00000 | 0.00000 | 0.00000 | 5.15356 | 0.54100 | 0.00000 |
| **NOVA2** | 0.00627 | 0.01827 | 0.01725 | 0.01272 | 0.01674 | 0.01070 | 0.01817 | 0.01124 | 0.02129 | 0.01434 | 0.01846 | 0.02389 |
| **NOX1** | 0.00298 | 0.00863 | 0.01280 | 0.06201 | 0.00000 | 0.05721 | 0.31260 | 0.03788 | 0.00000 | 0.51802 | 0.07864 | 0.01344 |
| **NOX3** | 0.00605 | 0.01353 | 0.01541 | 0.01375 | 0.01235 | 0.02558 | 0.01531 | 0.01027 | 0.01515 | 0.02486 | 0.01473 | 0.01630 |
| **NOX4** | 0.39139 | 0.02889 | 0.03036 | 0.00000 | 0.00000 | 0.13434 | 0.06144 | 0.64764 | 0.00000 | 0.05060 | 0.01969 | 0.06906 |

| | | | | | | | | | | | | |
|---|---|---|---|---|---|---|---|---|---|---|---|---|
| **NOX5** | 1.99862 | 0.01095 | 0.01133 | 0.02339 | 0.01359 | 0.01150 | 0.01636 | 0.32244 | 0.02293 | 0.00000 | 0.01413 | 0.07761 |
| **NOXA1** | 0.00614 | 0.01485 | 0.04747 | 0.01474 | 0.01696 | 0.01288 | 0.01865 | 0.01046 | 0.00595 | 0.02901 | 0.01863 | 0.01563 |
| **NOXO1** | 0.15800 | 0.00000 | 0.32522 | 0.00000 | 0.43622 | 0.00000 | 0.21466 | 0.07192 | 0.34070 | 0.81647 | 0.09837 | 0.05755 |
| **NOXRED1** | 0.00207 | 0.12123 | 0.17635 | 0.10989 | 0.08412 | 0.00005 | 0.00059 | 0.08587 | 0.19899 | 0.00270 | 0.11615 | 0.48454 |
| **NPAP1** | 0.03677 | 0.00512 | 0.00102 | 0.00538 | 0.00000 | 0.00000 | 0.00711 | 0.01413 | 0.00000 | 0.74141 | 0.09608 | 0.00667 |
| **NPAS1** | 0.91930 | 0.62672 | 0.00000 | 0.12601 | 0.51294 | 0.00000 | 0.00000 | 0.26112 | 0.11430 | 0.34158 | 0.00000 | 0.29519 |
| **NPAS2** | 0.53386 | 0.51365 | 0.17538 | 0.19812 | 0.47814 | 0.00000 | 0.00000 | 0.11306 | 0.00000 | 0.21963 | 0.01443 | 0.13972 |
| **NPAS3** | 2.01367 | 0.00463 | 0.10692 | 0.10424 | 0.00000 | 0.02164 | 0.00000 | 0.77348 | 0.01239 | 0.15558 | 0.00892 | 0.00000 |
| **NPAS4** | 0.00521 | 0.01510 | 0.07472 | 0.02229 | 0.01852 | 0.00197 | 0.02976 | 0.01018 | 0.02534 | 0.01025 | 0.02227 | 0.12223 |
| **NPAT** | 5.97566 | 1.56060 | 4.89344 | 1.34472 | 1.04427 | 10.76750 | 2.38988 | 6.84910 | 7.63696 | 10.23469 | 2.30591 | 5.22483 |
| **NPB** | 0.00756 | 0.38057 | 0.86882 | 0.00000 | 0.05650 | 0.00000 | 0.00000 | 0.16925 | 0.00000 | 0.06423 | 0.04355 | 0.35574 |
| **NPBWR1** | 0.00436 | 0.00828 | 0.03841 | 0.01783 | 0.00949 | 0.01518 | 0.00554 | 0.02946 | 0.04053 | 0.00000 | 0.58355 | 0.01637 |
| **NPC1** | 11.68293 | 4.16614 | 8.83942 | 3.64365 | 10.33690 | 0.00214 | 0.78262 | 1.73743 | 15.07922 | 0.00000 | 8.45883 | 3.33131 |
| **NPC1L1** | 0.00496 | 0.00750 | 0.04399 | 1.31558 | 0.01399 | 0.00415 | 0.00037 | 0.00974 | 0.12696 | 0.05987 | 0.01632 | 0.01339 |
| **NPC2** | 4.46397 | 15.28605 | 6.93008 | 3.13535 | 28.33646 | 24.57035 | 7.17319 | 2.08833 | 42.96369 | 12.08388 | 21.68439 | 15.40784 |
| **NPDC1** | 0.37054 | 0.11743 | 0.15833 | 0.15972 | 0.68243 | 0.00000 | 0.32486 | 0.04388 | 0.00000 | 0.11795 | 0.22222 | 0.00246 |
| **NPEPL1.uid-246** | 0.00598 | 0.00994 | 0.01142 | 0.00896 | 0.03338 | 0.03716 | 0.01236 | 0.01002 | 0.01780 | 0.01910 | 0.01584 | 0.01891 |
| **NPEPL1.uid-246** | 0.04905 | 0.05599 | 0.01905 | 0.07818 | 0.02817 | 0.03684 | 0.03750 | 0.00714 | 0.02834 | 0.00667 | 0.05299 | 0.01198 |
| **NPEPPS** | 3.83051 | 7.98308 | 10.44402 | 18.38617 | 11.01071 | 25.33749 | 17.33611 | 6.28640 | 11.10783 | 20.06669 | 11.22318 | 7.09769 |
| **NPFF** | 0.00262 | 0.11833 | 0.09479 | 0.15537 | 0.05636 | 0.09363 | 0.00002 | 0.08345 | 0.10144 | 0.61042 | 0.09394 | 0.06078 |
| **NPFFR2** | 0.19712 | 1.13282 | 0.88380 | 0.00000 | 0.00053 | 1.95181 | 0.31575 | 0.03234 | 1.96475 | 0.11701 | 0.64039 | 0.08719 |
| **NPHP1** | 0.31398 | 1.33405 | 0.06697 | 0.12731 | 0.00000 | 0.63809 | 0.12732 | 1.85172 | 0.00000 | 0.01887 | 0.45921 | 2.47701 |
| **NPHP3** | 0.00084 | 0.00000 | 0.05684 | 0.13353 | 0.00000 | 0.30272 | 0.27944 | 0.04332 | 0.00000 | 0.63342 | 0.03302 | 0.03160 |
| **NPHP3-ACAD11** | 0.00721 | 0.05171 | 0.00000 | 0.28831 | 0.02515 | 0.08790 | 0.00000 | 0.01113 | 0.06044 | 0.07126 | 0.00085 | 0.02324 |
| **NPHP3-AS1** | 0.05463 | 0.04921 | 0.01905 | 0.01534 | 0.01122 | 0.01675 | 0.02737 | 0.04392 | 0.01547 | 0.00107 | 0.01607 | 0.04968 |
| **NPHP4** | 5.35010 | 1.00912 | 0.24270 | 0.78028 | 0.88899 | 0.59191 | 0.28473 | 0.26793 | 0.00000 | 1.63110 | 0.00000 | 0.67522 |
| **NPHS1** | 0.01071 | 0.02579 | 0.00808 | 0.04027 | 0.00000 | 0.02533 | 0.04160 | 0.00729 | 0.02897 | 0.12102 | 0.01785 | 0.05263 |
| **NPHS2** | 0.00649 | 0.01512 | 0.01604 | 0.01327 | 0.01498 | 0.01334 | 0.02111 | 0.01051 | 0.01213 | 0.01487 | 0.03260 | 0.01665 |
| **NPIPA1** | 0.01841 | 0.07494 | 0.02547 | 0.06677 | 0.06952 | 0.00000 | 0.09119 | 0.08364 | 0.01804 | 0.71615 | 0.02726 | 0.03543 |
| **NPIPA5** | 0.00484 | 0.07610 | 0.02195 | 0.11920 | 0.03155 | 0.03111 | 0.03704 | 0.04198 | 0.14752 | 0.22599 | 0.01388 | 0.08727 |
| **NPIPB3** | 0.00677 | 0.21505 | 0.12923 | 0.26252 | 0.32220 | 0.00000 | 0.00000 | 0.11562 | 0.57586 | 0.07095 | 0.16576 | 0.23961 |
| **NPIPB5** | 0.00310 | 0.19714 | 0.12918 | 0.22940 | 0.22164 | 0.40641 | 0.00000 | 0.14434 | 0.59502 | 0.75213 | 0.21959 | 0.22427 |
| **NPIPB6** | 0.00668 | 0.09919 | 0.03115 | 0.05977 | 0.03809 | 0.00000 | 0.00108 | 0.09615 | 0.07893 | 0.17419 | 0.00595 | 0.35097 |
| **NPIPB9** | 0.01710 | 0.04071 | 0.02599 | 0.05108 | 0.00834 | 0.00368 | 0.02109 | 0.00920 | 0.02852 | 0.05196 | 0.01531 | 0.02234 |
| **NPL** | 0.44555 | 0.65595 | 0.70685 | 1.70300 | 0.34538 | 1.27806 | 0.42404 | 0.58952 | 0.19062 | 0.00000 | 0.27923 | 3.15599 |
| **NPLOC4** | 1.21314 | 10.45939 | 11.59926 | 9.00544 | 16.46262 | 1.50683 | 17.57295 | 3.20422 | 5.67178 | 19.11863 | 4.98675 | 7.64176 |
| **NPM1** | 6.11100 | 31.08352 | 23.75083 | 21.80278 | 21.18491 | 50.90626 | 42.13736 | 19.37497 | 40.23526 | 19.89336 | 27.27510 | 38.24729 |
| **NPM2** | 11.71091 | 0.05976 | 0.00000 | 0.13242 | 0.02326 | 0.00000 | 0.14537 | 3.37083 | 0.00000 | 0.28447 | 0.07521 | 0.79320 |
| **NPM3** | 14.07708 | 11.75907 | 13.49945 | 9.46205 | 4.59403 | 0.00000 | 20.91350 | 4.08746 | 15.27179 | 0.00000 | 22.16414 | 5.10635 |
| **NPNT** | 0.09255 | 0.00000 | 0.91820 | 0.20921 | 0.00000 | 0.00000 | 1.42609 | 0.00000 | 4.87900 | 0.29851 | 0.08804 | 0.09460 |
| **NPPA** | 0.01016 | 0.00000 | 0.33651 | 0.20252 | 0.20425 | 0.00000 | 0.00000 | 0.02979 | 0.84044 | 0.91135 | 0.51817 | 0.02206 |
| **NPPA-AS1** | 0.00620 | 0.00000 | 0.27764 | 0.11411 | 0.00523 | 0.00000 | 0.00473 | 0.03889 | 0.88138 | 0.32037 | 0.21491 | 0.04112 |
| **NPPB** | 0.01513 | 0.36255 | 0.00000 | 0.00000 | 3.66900 | 0.00000 | 0.37736 | 0.00861 | 1.58874 | 0.34258 | 0.66349 | 0.10851 |
| **NPPC** | 0.18532 | 0.23875 | 0.05214 | 0.00000 | 0.03215 | 0.00000 | 0.00001 | 0.33193 | 0.01569 | 0.13737 | 0.00342 | 0.24449 |
| **1.00000** | 0.35011 | 0.02034 | 0.62937 | 0.00000 | 0.00000 | 0.00000 | 0.16887 | 0.07571 | 0.00000 | 0.26917 | 0.07477 | 0.28644 |
| **2.00000** | 0.10900 | 0.09237 | 0.09476 | 0.05905 | 0.02432 | 0.09656 | 0.00000 | 0.01695 | 0.03483 | 0.01714 | 0.00304 | 0.20555 |
| **3.00000** | 0.01402 | 0.00407 | 0.02491 | 0.00230 | 0.00000 | 0.09806 | 0.02932 | 0.45561 | 0.04577 | 0.03093 | 0.00294 | 0.02306 |
| **NPRL2** | 1.88491 | 7.12625 | 6.55808 | 6.77230 | 3.14377 | 2.31119 | 7.55451 | 1.65259 | 8.28026 | 0.00000 | 10.28515 | 5.35430 |

| | | | | | | | | | | | | |
|---|---|---|---|---|---|---|---|---|---|---|---|---|
| **NPRL3** | 2.82709 | 3.79393 | 7.03495 | 2.95746 | 3.61143 | 0.00000 | 7.94104 | 0.51040 | 6.63825 | 9.28549 | 5.31270 | 8.42187 |
| **NPSR1** | 0.02097 | 0.07331 | 0.06733 | 0.12284 | 0.00000 | 0.02664 | 0.00000 | 0.01017 | 0.01422 | 0.04676 | 0.01162 | 0.00758 |
| **NPSR1-AS1** | 0.46375 | 0.02036 | 0.03746 | 0.09380 | 0.01665 | 0.00000 | 0.02040 | 0.09785 | 0.00868 | 0.01310 | 0.04473 | 0.09445 |
| **NPTN** | 4.62033 | 3.76331 | 11.66045 | 5.34205 | 2.61676 | 22.89849 | 12.34592 | 7.87014 | 6.00807 | 17.46299 | 10.68423 | 7.64893 |
| **NPTN-IT1** | 0.00926 | 0.30433 | 0.46116 | 0.62346 | 0.65493 | 0.00001 | 0.00000 | 0.35125 | 1.60218 | 2.77952 | 0.58051 | 0.40503 |
| **NPTX1** | 0.11068 | 5.98356 | 0.01685 | 0.00000 | 6.57041 | 0.29148 | 0.00000 | 0.00041 | 0.00000 | 3.21106 | 0.28725 | 0.00000 |
| **NPTX2** | 2.25578 | 0.00641 | 0.09509 | 0.00422 | 0.00763 | 0.00000 | 0.01737 | 2.07040 | 0.08447 | 0.02223 | 0.02344 | 1.23049 |
| **NPTXR** | 0.00520 | 0.27635 | 0.03259 | 0.31158 | 0.11628 | 0.00000 | 0.00585 | 0.01155 | 0.00000 | 0.06967 | 0.06568 | 0.00434 |
| **NPW** | 0.14046 | 0.25687 | 0.06453 | 0.00257 | 0.12415 | 0.00000 | 0.04126 | 0.09669 | 0.20026 | 0.02717 | 0.00000 | 0.24924 |
| **NPY** | 0.00174 | 0.00713 | 0.04988 | 0.09259 | 0.00154 | 0.00737 | 0.04431 | 0.04057 | 0.00000 | 0.09654 | 0.01075 | 0.08462 |
| **NPY1R** | 0.01237 | 0.24628 | 0.22548 | 0.00000 | 0.21556 | 0.00000 | 0.00000 | 0.05432 | 0.17677 | 0.05868 | 0.04031 | 0.16172 |
| **NPY4R** | 0.03964 | 0.01730 | 0.02144 | 0.01216 | 0.04310 | 0.00777 | 0.01456 | 0.00708 | 0.04203 | 0.00000 | 0.01445 | 0.01725 |
| **NQO1** | 0.09272 | 3.18597 | 5.09836 | 5.12159 | 12.36242 | 8.10106 | 7.70068 | 0.32741 | 8.06858 | 0.18525 | 5.45033 | 0.00000 |
| **NQO2** | 0.00000 | 6.97362 | 7.75274 | 13.08540 | 12.63747 | 13.42168 | 0.00000 | 1.62518 | 0.00003 | 1.49418 | 1.98736 | 2.09030 |
| **NR0B1** | 0.00921 | 0.01526 | 0.39712 | 0.03110 | 0.00750 | 0.03936 | 0.00000 | 0.00774 | 0.00000 | 0.02109 | 0.01775 | 0.06599 |
| **NR0B2** | 0.00161 | 0.00562 | 0.01200 | 0.01967 | 0.01099 | 0.00000 | 0.00826 | 0.15969 | 0.03917 | 0.03897 | 0.05524 | 0.17045 |
| **NR1D1** | 0.18195 | 0.18499 | 0.15651 | 0.51843 | 0.00000 | 0.02436 | 0.39753 | 0.17567 | 0.28033 | 0.28234 | 0.19896 | 1.10953 |
| **NR1D2** | 0.00000 | 0.95263 | 0.15481 | 1.38789 | 0.00000 | 4.55022 | 0.91215 | 3.31409 | 0.00000 | 6.80394 | 0.00771 | 3.67587 |
| **NR1H2** | 4.64568 | 19.04230 | 8.01740 | 21.20423 | 9.55844 | 0.00000 | 5.53502 | 1.89893 | 18.02471 | 0.00000 | 3.36290 | 18.22250 |
| **NR1H3** | 0.73106 | 6.88481 | 5.22469 | 0.10904 | 4.13025 | 2.21716 | 0.83342 | 0.00000 | 3.00901 | 0.19257 | 1.15429 | 3.30179 |
| **NR1H4** | 2.44446 | 0.00987 | 0.75162 | 0.01370 | 0.00000 | 0.00043 | 0.24187 | 0.00878 | 0.13657 | 0.00000 | 0.00000 | 0.03458 |
| **NR1I2** | 0.00509 | 0.11344 | 0.12747 | 0.00078 | 0.00223 | 0.03102 | 0.00000 | 0.01146 | 0.00000 | 0.04763 | 0.05719 | 0.00899 |
| **NR1I3** | 0.00732 | 0.02458 | 0.07423 | 0.03611 | 0.02057 | 0.00077 | 0.02743 | 0.01078 | 0.06175 | 0.01158 | 0.01439 | 0.02570 |
| **NR2C1** | 2.26949 | 9.79883 | 2.70749 | 8.45003 | 1.88703 | 23.22460 | 0.00000 | 7.47202 | 8.70129 | 10.27671 | 3.69417 | 15.19344 |
| **NR2C2** | 0.00000 | 1.18487 | 0.70141 | 2.78297 | 1.10385 | 6.92287 | 0.00097 | 2.04896 | 5.34389 | 6.87870 | 1.44524 | 2.38513 |
| **NR2C2AP** | 1.75608 | 3.58398 | 6.74299 | 3.86436 | 3.69201 | 1.37063 | 7.37373 | 2.17878 | 8.59213 | 0.00000 | 6.26252 | 7.21611 |
| **NR2E1** | 7.96865 | 0.11322 | 0.00176 | 0.00000 | 0.02455 | 0.02381 | 0.00747 | 0.06254 | 0.00000 | 0.03069 | 0.12191 | 2.98916 |
| **NR2E3** | 0.00513 | 0.00337 | 0.12029 | 0.08729 | 0.04733 | 0.01574 | 0.04519 | 0.01032 | 0.07019 | 0.08562 | 0.02072 | 0.08637 |
| **NR2F1** | 0.09793 | 0.18681 | 0.00308 | 0.02512 | 0.12236 | 0.00000 | 0.00052 | 0.03468 | 0.05743 | 0.29183 | 0.01508 | 0.02088 |
| **NR2F1-AS1** | 0.45496 | 0.16928 | 0.08331 | 0.24528 | 0.10705 | 0.10282 | 0.00000 | 0.18266 | 0.00000 | 0.01008 | 0.00323 | 0.04951 |
| **NR2F2** | 1.17902 | 0.00000 | 33.81799 | 0.00000 | 3.48993 | 0.31799 | 30.19355 | 0.03589 | 3.94799 | 1.29342 | 2.49379 | 0.00000 |
| **NR2F2-AS1** | 0.00839 | 0.00861 | 0.83920 | 0.00400 | 0.01133 | 0.05450 | 0.00000 | 0.02688 | 0.00000 | 0.04059 | 0.05829 | 0.01014 |
| **NR2F6** | 0.00000 | 2.17736 | 11.80105 | 10.79303 | 19.73786 | 0.00000 | 25.37719 | 0.38429 | 8.99496 | 22.04993 | 3.27196 | 0.23986 |
| **NR3C1** | 2.28624 | 0.03588 | 0.00000 | 0.90358 | 0.00000 | 0.63047 | 0.06137 | 0.11865 | 0.00000 | 0.65521 | 0.14320 | 0.59523 |
| **NR3C2** | 9.70206 | 0.00000 | 0.03909 | 0.13812 | 0.42992 | 0.00000 | 0.00000 | 0.83548 | 0.04951 | 1.39750 | 0.10468 | 0.45320 |
| **NR4A1** | 0.18092 | 0.56042 | 2.27712 | 0.35069 | 2.22924 | 0.00000 | 0.06295 | 0.00000 | 0.00000 | 2.73423 | 0.00000 | 0.90436 |
| **NR4A2** | 0.08483 | 1.10883 | 0.59425 | 0.63272 | 1.15564 | 1.40208 | 0.16687 | 1.68209 | 0.20499 | 0.97987 | 0.00000 | 3.85283 |
| **NR4A3** | 0.28144 | 0.01361 | 0.22173 | 0.07828 | 0.02955 | 0.00226 | 0.03621 | 0.41969 | 0.00000 | 0.04253 | 0.04129 | 0.06301 |
| **NR5A1** | 0.00715 | 0.01024 | 0.14390 | 0.01472 | 0.01044 | 0.01930 | 0.00000 | 0.03512 | 0.00972 | 0.07958 | 0.03575 | 0.01804 |
| **NR5A2** | 0.09845 | 3.49957 | 0.00000 | 3.44251 | 0.16399 | 1.57100 | 0.00000 | 3.51893 | 1.82324 | 3.15588 | 0.50875 | 4.07093 |
| **NR6A1** | 4.19469 | 7.28903 | 37.96768 | 16.33134 | 10.28120 | 12.46444 | 52.25826 | 7.76222 | 18.60683 | 15.48693 | 26.82434 | 4.53779 |
| **NRADDP** | 0.00998 | 0.00280 | 0.08573 | 0.05385 | 0.18331 | 0.00000 | 0.00064 | 0.00965 | 0.05645 | 0.09780 | 0.04826 | 0.02286 |
| **NRAP** | 7.06757 | 0.17707 | 0.08626 | 0.17471 | 0.00000 | 0.03425 | 0.00000 | 0.94832 | 0.00000 | 0.01724 | 0.03182 | 0.50665 |
| **NRARP** | 0.06098 | 0.08230 | 0.21001 | 0.69153 | 0.02273 | 0.00000 | 0.12566 | 0.10252 | 0.00000 | 1.09320 | 0.00000 | 0.00000 |
| **NRAS** | 4.35603 | 11.36521 | 8.89573 | 5.01788 | 1.70997 | 30.39916 | 13.01043 | 8.52899 | 10.22759 | 3.91231 | 7.26870 | 16.19847 |
| **NRBF2** | 1.36585 | 13.77467 | 6.35556 | 5.89551 | 8.88892 | 23.89937 | 6.85416 | 19.01109 | 5.77985 | 0.83834 | 5.78969 | 23.99043 |
| **NRBP1** | 6.36251 | 24.43326 | 14.86416 | 30.83055 | 17.61931 | 6.13995 | 24.01005 | 17.40587 | 23.82651 | 0.00000 | 14.65609 | 22.76546 |
| **NRBP2** | 0.33875 | 2.59690 | 0.00000 | 6.17755 | 0.78912 | 0.03236 | 0.00000 | 0.38609 | 0.07043 | 3.45376 | 0.05087 | 1.88855 |

| | | | | | | | | | | | | |
|---|---|---|---|---|---|---|---|---|---|---|---|---|
| **NRCAM** | 1.24874 | 0.77707 | 0.00000 | 2.63911 | 0.17925 | 0.00000 | 0.08606 | 0.49645 | 0.01745 | 0.00998 | 0.06115 | 0.14551 |
| **NRD1** | 0.00000 | 14.78335 | 14.98039 | 21.74192 | 16.91604 | 25.83618 | 19.75642 | 2.23663 | 18.41956 | 8.76449 | 14.30199 | 13.34713 |
| **NRDE2** | 2.23148 | 2.95240 | 3.65768 | 1.29271 | 1.02531 | 2.64527 | 4.68453 | 3.49549 | 7.60691 | 4.88227 | 1.92023 | 3.93571 |
| **NREP** | 2.57115 | 0.04625 | 0.70350 | 1.60678 | 0.00000 | 0.05795 | 0.55511 | 0.35654 | 0.26583 | 0.35452 | 0.00000 | 0.06754 |
| **NREP-AS1** | 0.05570 | 0.01522 | 0.01867 | 0.01173 | 0.01704 | 0.01864 | 0.01682 | 0.00865 | 0.02050 | 0.00937 | 0.01417 | 0.01553 |
| **NRF1** | 0.68805 | 5.96242 | 3.83759 | 7.07222 | 5.12255 | 1.52236 | 4.61609 | 17.80137 | 4.04694 | 3.26183 | 4.06970 | 9.03240 |
| **NRG1** | 2.78132 | 0.25982 | 0.04512 | 1.37717 | 0.00000 | 0.00000 | 0.08739 | 0.72046 | 0.00000 | 0.58158 | 0.00000 | 0.63478 |
| **NRG1-IT1** | 0.07390 | 0.01021 | 0.01152 | 0.00605 | 0.02869 | 0.07364 | 0.01186 | 0.00614 | 0.02214 | 0.00296 | 0.01238 | 0.09492 |
| **NRG2** | 0.01054 | 0.03929 | 0.08418 | 0.40265 | 0.00000 | 0.01071 | 0.04195 | 0.02710 | 0.01102 | 0.05051 | 0.01788 | 0.02601 |
| **NRG3** | 0.01228 | 0.01369 | 0.01119 | 0.00785 | 0.01577 | 0.03672 | 0.01399 | 0.01544 | 0.01929 | 0.01914 | 0.01590 | 0.01417 |
| **NRG4** | 2.35341 | 0.00000 | 0.00000 | 0.12586 | 0.00000 | 0.74134 | 0.39417 | 0.02381 | 0.27685 | 2.29105 | 0.55358 | 0.58784 |
| **NRGN** | 0.01030 | 0.01118 | 0.00000 | 2.41178 | 0.64118 | 0.00000 | 0.06416 | 0.12772 | 0.00000 | 0.69445 | 0.00000 | 0.08998 |
| **NRIP1** | 1.83695 | 0.00000 | 5.40347 | 0.00000 | 0.14801 | 0.18620 | 0.78911 | 0.42195 | 0.00000 | 1.36991 | 0.00000 | 0.05949 |
| **NRIP2** | 0.00504 | 0.00299 | 0.31573 | 0.01004 | 0.20537 | 0.00000 | 0.05138 | 0.02303 | 0.08495 | 0.70233 | 0.17122 | 0.46978 |
| **NRIP3** | 2.20875 | 9.78979 | 0.00000 | 9.96851 | 9.92356 | 7.78145 | 0.31389 | 3.85267 | 0.00000 | 0.17378 | 3.87168 | 5.19940 |
| **NRK** | 0.17636 | 0.12932 | 0.11990 | 0.18899 | 0.03574 | 0.13752 | 0.13060 | 0.00000 | 0.00000 | 0.00000 | 0.00000 | 0.30862 |
| **NRL** | 0.08009 | 0.03183 | 0.34244 | 0.17300 | 0.02550 | 0.00000 | 0.00000 | 0.00128 | 0.06236 | 0.03365 | 0.06379 | 0.18669 |
| **NRM** | 0.37821 | 1.92548 | 1.33211 | 0.75629 | 0.93789 | 3.28631 | 0.00000 | 0.87070 | 2.59735 | 1.20558 | 0.52225 | 0.97056 |
| **NRN1** | 0.61794 | 0.36360 | 0.19046 | 3.29664 | 0.61410 | 1.29341 | 0.00000 | 0.08855 | 0.00000 | 0.00000 | 0.96812 | 0.00000 |
| **NRN1L** | 0.01134 | 0.00000 | 1.53524 | 0.00000 | 0.00000 | 0.00079 | 2.78504 | 0.00603 | 3.23546 | 0.06771 | 1.18352 | 0.03618 |
| **NRON** | 0.00563 | 0.01606 | 0.02657 | 0.00894 | 0.01145 | 0.01827 | 0.01854 | 0.01132 | 0.00706 | 0.03463 | 0.02164 | 0.01554 |
| **NRP1** | 0.00430 | 0.00000 | 23.89954 | 0.08657 | 0.00000 | 0.03606 | 0.81254 | 0.10325 | 0.02900 | 0.00000 | 0.34777 | 0.01257 |
| **NRP2** | 0.01362 | 0.24442 | 0.56332 | 0.19834 | 0.00000 | 0.00000 | 0.00000 | 0.00751 | 0.00000 | 0.64535 | 0.00134 | 0.09452 |
| **NRROS** | 0.00773 | 0.01126 | 0.01239 | 0.12021 | 0.00454 | 0.03785 | 0.01106 | 0.01164 | 0.01461 | 0.02119 | 0.01455 | 0.01412 |
| **NRSN1** | 0.00747 | 0.01444 | 0.01487 | 0.01149 | 0.01992 | 0.02030 | 0.02304 | 0.02103 | 0.01340 | 0.01397 | 0.01501 | 0.01673 |
| **NRSN2** | 0.67329 | 3.23747 | 2.96711 | 6.46030 | 1.22879 | 2.09010 | 0.60418 | 0.02320 | 4.18129 | 0.00000 | 5.15409 | 0.07715 |
| **NRTN** | 0.01668 | 0.02423 | 0.05743 | 0.01168 | 0.06597 | 0.00000 | 0.00013 | 0.03036 | 0.03711 | 0.19298 | 0.00941 | 0.04997 |
| **NRXN1** | 0.70275 | 0.00417 | 0.06682 | 0.00275 | 0.12017 | 0.05842 | 0.00000 | 0.36625 | 0.00000 | 0.08594 | 0.00000 | 0.06968 |
| **NRXN2** | 0.24490 | 0.81148 | 0.31812 | 0.35527 | 2.11643 | 0.00000 | 0.14410 | 0.00000 | 0.00000 | 2.87117 | 1.00906 | 0.59892 |
| **NRXN3** | 1.51020 | 0.00000 | 2.39864 | 0.00000 | 0.18632 | 0.00001 | 1.18096 | 0.36977 | 2.12391 | 0.72001 | 0.00000 | 0.03888 |
| **NS3BP** | 0.05426 | 0.30862 | 0.01021 | 0.17571 | 0.69807 | 0.00000 | 0.00302 | 0.00000 | 0.79396 | 0.25123 | 0.47517 | 0.36642 |
| **NSA2** | 0.20388 | 6.57958 | 4.41029 | 3.66071 | 3.77252 | 21.52398 | 12.79250 | 10.36303 | 8.16486 | 13.38915 | 7.23994 | 12.78945 |
| **NSD1** | 3.67649 | 8.87545 | 5.37703 | 11.20711 | 10.78228 | 14.83673 | 7.31850 | 5.55337 | 18.00653 | 35.28251 | 6.69621 | 14.83409 |
| **NSDHL** | 2.50839 | 6.60104 | 8.88984 | 8.61828 | 17.85399 | 5.82682 | 8.18359 | 7.15533 | 20.55484 | 0.00000 | 11.82326 | 7.69180 |
| **NSF** | 2.12542 | 6.49676 | 4.45379 | 4.43183 | 2.27700 | 10.72495 | 5.41971 | 1.21556 | 5.36532 | 0.38117 | 5.76042 | 4.68475 |
| **NSFL1C** | 4.14052 | 16.09356 | 10.71878 | 14.51725 | 18.01093 | 8.87450 | 15.25425 | 6.56833 | 11.33713 | 0.20831 | 14.72682 | 14.43703 |
| **NSG1** | 14.45258 | 2.20555 | 0.76019 | 0.00000 | 1.40185 | 0.87001 | 0.00000 | 3.17413 | 0.49557 | 0.01171 | 0.68866 | 6.20049 |
| **NSL1** | 2.21952 | 8.24056 | 3.20375 | 9.41856 | 0.91908 | 22.21586 | 4.94497 | 1.32602 | 6.68637 | 6.13359 | 9.55499 | 7.20776 |
| **NSMAF** | 0.14984 | 8.22007 | 6.70667 | 11.18596 | 0.00000 | 13.52043 | 7.18565 | 2.07618 | 3.89385 | 5.81058 | 2.65104 | 3.85233 |
| **NSMCE1** | 0.40481 | 6.71481 | 3.87499 | 1.29171 | 3.21807 | 5.02861 | 4.76040 | 0.79947 | 0.00105 | 2.09999 | 6.62262 | 4.64068 |
| **NSMCE2** | 1.55575 | 4.41910 | 3.52582 | 1.25155 | 3.96364 | 15.72780 | 7.42566 | 4.87019 | 8.29462 | 1.94599 | 5.43720 | 10.43637 |
| **NSMCE4A** | 11.72561 | 12.15389 | 6.67692 | 3.15426 | 6.74701 | 12.94858 | 2.47321 | 6.79161 | 7.80801 | 7.31398 | 12.39790 | 3.02934 |
| **NSMF** | 1.62152 | 1.81846 | 0.56715 | 0.30080 | 2.76040 | 0.00000 | 0.00322 | 1.45116 | 0.66813 | 3.42556 | 0.00000 | 1.61296 |
| **NSRP1** | 0.25511 | 4.62425 | 7.42762 | 4.61989 | 4.60834 | 16.16820 | 12.78556 | 6.50038 | 1.02706 | 20.93326 | 9.30101 | 8.70175 |
| **NSUN2** | 3.64104 | 18.77666 | 17.60667 | 5.68412 | 7.62640 | 13.10650 | 16.16775 | 3.44409 | 24.19352 | 5.59929 | 17.95944 | 11.27317 |
| **NSUN3** | 0.07402 | 0.48686 | 0.51413 | 0.58002 | 0.00000 | 2.69111 | 1.14115 | 2.23317 | 0.46967 | 0.00000 | 0.35050 | 0.96478 |
| **NSUN4** | 1.64088 | 8.56574 | 2.80413 | 6.61317 | 8.31485 | 8.94122 | 5.88139 | 3.26810 | 7.97788 | 4.73426 | 7.51959 | 9.71668 |
| **NSUN5** | 1.48297 | 25.87277 | 8.30408 | 11.22562 | 17.58788 | 0.00000 | 14.05846 | 2.67560 | 15.62551 | 0.00000 | 9.05770 | 17.92257 |

| | | | | | | | | | | | | |
|---|---|---|---|---|---|---|---|---|---|---|---|---|
| **NSUN5P1** | 0.18380 | 1.09450 | 1.27991 | 0.57990 | 0.48984 | 0.73065 | 1.39482 | 0.09145 | 2.75587 | 1.32582 | 1.38296 | 0.66717 |
| **NSUN5P2** | 0.00931 | 0.02280 | 0.01956 | 0.01314 | 0.01722 | 0.01202 | 0.01188 | 0.01138 | 0.04518 | 0.01028 | 0.01165 | 0.01735 |
| **NSUN6** | 4.06262 | 2.22821 | 2.68459 | 2.55253 | 0.84917 | 0.67408 | 1.30409 | 1.46346 | 0.87495 | 0.58551 | 1.88569 | 2.84626 |
| **NSUN7** | 0.24058 | 3.59149 | 0.17100 | 0.00000 | 0.85272 | 3.42955 | 0.00000 | 0.00000 | 3.19953 | 3.70338 | 0.53555 | 3.69987 |
| **NT5C** | 0.14869 | 7.89751 | 8.28705 | 10.11660 | 6.64215 | 1.55807 | 9.48472 | 2.18136 | 6.97063 | 2.40543 | 7.00971 | 5.25514 |
| **NT5C1B** | 0.01094 | 0.01933 | 0.01462 | 0.01396 | 0.01219 | 0.01559 | 0.01197 | 0.01608 | 0.02790 | 0.01744 | 0.02279 | 0.02007 |
| **NT5C2** | 2.45363 | 5.06698 | 5.21043 | 2.76932 | 2.19959 | 12.80350 | 12.88399 | 4.73201 | 12.97251 | 0.00000 | 9.52676 | 6.18741 |
| **NT5C3A** | 1.02770 | 1.62299 | 3.17381 | 1.36932 | 2.57697 | 9.44130 | 2.66292 | 0.70251 | 9.27097 | 3.30472 | 1.45965 | 1.88042 |
| **NT5C3B** | 0.30776 | 6.52463 | 9.22984 | 3.46358 | 3.45525 | 4.34473 | 10.62685 | 0.18305 | 4.20250 | 6.64292 | 4.53642 | 2.43188 |
| **NT5DC1** | 0.23622 | 6.95393 | 0.87497 | 2.51910 | 0.00000 | 13.21341 | 2.03020 | 0.00000 | 0.48902 | 1.47834 | 3.81395 | 1.00741 |
| **NT5DC2** | 0.08268 | 0.00000 | 3.73628 | 1.81086 | 0.00006 | 0.00000 | 7.17439 | 0.03209 | 1.50306 | 3.81939 | 0.48162 | 0.00521 |
| **NT5DC3** | 2.22489 | 4.61484 | 3.14465 | 3.29319 | 0.70808 | 6.14573 | 6.76652 | 16.59966 | 7.39933 | 4.76581 | 15.84283 | 5.00520 |
| **NT5E** | 0.00568 | 0.32629 | 0.01322 | 0.91188 | 0.24242 | 0.04788 | 0.00000 | 0.00970 | 0.04037 | 0.01884 | 0.35430 | 0.00429 |
| **NT5M** | 0.51121 | 0.14493 | 0.01110 | 0.31054 | 0.23397 | 0.00000 | 0.07851 | 0.00000 | 0.00000 | 0.00000 | 0.05569 | 0.06201 |
| **NTAN1** | 1.28275 | 2.19393 | 1.23698 | 0.80176 | 0.62205 | 2.24991 | 0.00000 | 0.85744 | 2.49767 | 0.54842 | 1.20467 | 0.63849 |
| **NTF3** | 0.09496 | 0.01094 | 0.07396 | 0.44264 | 0.00000 | 0.00906 | 0.03748 | 0.00770 | 0.00699 | 0.02849 | 0.02267 | 0.01547 |
| **NTF4** | 0.04004 | 0.04462 | 0.28161 | 0.01138 | 0.00676 | 0.00015 | 0.13822 | 0.11222 | 0.01139 | 0.01323 | 0.01373 | 0.38169 |
| **NTHL1** | 3.70604 | 11.83015 | 9.17682 | 8.28851 | 6.00015 | 0.00000 | 10.13231 | 2.81456 | 10.62978 | 1.58682 | 13.75102 | 5.73010 |
| **NTM** | 0.21066 | 0.01380 | 0.01098 | 0.01359 | 0.01297 | 0.05768 | 0.01543 | 0.00601 | 0.03698 | 0.00538 | 0.00821 | 0.01093 |
| **NTMT1** | 0.68265 | 17.85284 | 13.18508 | 7.91054 | 17.44748 | 8.59611 | 15.04946 | 11.56965 | 17.82639 | 0.00000 | 17.07112 | 18.06344 |
| **NTN1** | 1.29385 | 0.06307 | 0.07842 | 0.03312 | 0.05390 | 0.09550 | 0.00000 | 0.02423 | 0.00000 | 0.16585 | 0.00560 | 0.01361 |
| **NTN3** | 0.00626 | 0.01627 | 0.01615 | 0.01371 | 0.02461 | 0.00000 | 0.02929 | 0.01011 | 0.01728 | 0.02155 | 0.02765 | 0.02395 |
| **NTN4** | 0.66831 | 0.92549 | 0.00000 | 1.23157 | 0.40833 | 0.26789 | 0.00000 | 0.00000 | 0.00000 | 0.11446 | 0.04434 | 0.73250 |
| **NTN5** | 0.00593 | 0.01257 | 0.01279 | 0.01578 | 0.01320 | 0.01375 | 0.02205 | 0.00951 | 0.01424 | 0.03047 | 0.01520 | 0.03472 |
| **NTNG1** | 0.08189 | 0.13532 | 0.00763 | 1.30378 | 0.06169 | 0.00446 | 0.00439 | 0.00000 | 0.01223 | 0.03565 | 0.00603 | 2.72452 |
| **NTNG2** | 0.15084 | 0.01725 | 0.04823 | 0.01553 | 0.02586 | 0.00000 | 0.01400 | 0.00419 | 0.02115 | 0.00538 | 0.01526 | 0.01648 |
| **NTPCR** | 0.67510 | 10.91367 | 5.44171 | 3.93927 | 11.45525 | 8.88941 | 6.30091 | 0.47876 | 7.69360 | 0.00000 | 7.90942 | 5.24047 |
| **NTRK1** | 0.00582 | 0.06993 | 0.02867 | 0.00722 | 0.06949 | 0.00511 | 0.03577 | 0.01101 | 0.00750 | 0.00000 | 0.01611 | 0.03070 |
| **NTRK2** | 2.88345 | 0.00000 | 0.19454 | 0.00000 | 0.04274 | 0.00000 | 0.31068 | 0.07346 | 0.82137 | 0.63165 | 0.15317 | 0.03621 |
| **NTRK3** | 5.40340 | 0.26310 | 0.00000 | 0.74896 | 0.41152 | 0.29579 | 0.25838 | 0.22745 | 0.09847 | 0.04297 | 0.00000 | 0.01148 |
| **NTS** | 0.00610 | 0.02624 | 0.39051 | 0.02044 | 0.16771 | 0.00000 | 0.00000 | 0.01081 | 0.02451 | 0.00627 | 0.01126 | 0.02163 |
| **NTSR1** | 0.01419 | 0.01285 | 0.01805 | 0.01374 | 0.02351 | 0.00163 | 0.03624 | 0.00755 | 0.01438 | 0.01971 | 0.01461 | 0.05105 |
| **NTSR2** | 0.10797 | 0.02421 | 0.01273 | 0.00999 | 0.01424 | 0.01085 | 0.01665 | 0.00000 | 0.02691 | 0.00393 | 0.01239 | 1.10243 |
| **NUAK1** | 0.05680 | 0.02625 | 0.08629 | 0.63512 | 0.00000 | 0.60342 | 0.41581 | 0.00000 | 0.00000 | 0.79159 | 0.00000 | 0.22052 |
| **NUAK2** | 0.41454 | 2.28999 | 10.52615 | 0.00000 | 10.21895 | 0.00286 | 7.56869 | 0.00156 | 9.32272 | 0.00000 | 5.82417 | 0.02757 |
| **NUB1** | 3.00450 | 7.74706 | 4.40208 | 2.02343 | 2.31161 | 10.88397 | 8.60473 | 0.00000 | 6.55141 | 0.00000 | 4.01844 | 2.89382 |
| **NUBP1** | 0.00360 | 4.72450 | 5.21978 | 3.66459 | 2.77580 | 5.00917 | 9.68365 | 0.58180 | 6.39836 | 0.00000 | 7.00154 | 5.43849 |
| **NUBP2** | 0.85782 | 16.52236 | 12.03856 | 11.23144 | 18.86072 | 0.00000 | 14.31145 | 4.83894 | 18.80889 | 2.81102 | 17.71833 | 15.01247 |
| **NUBPL** | 0.06651 | 2.66456 | 1.56811 | 3.44380 | 1.19106 | 9.45497 | 4.28130 | 0.36632 | 4.26143 | 4.56977 | 3.21646 | 2.63774 |
| **NUCB1** | 0.97240 | 27.21974 | 10.58060 | 21.92291 | 32.43525 | 0.00000 | 1.24064 | 3.33839 | 19.89194 | 11.73915 | 7.55299 | 10.19872 |
| **NUCB1-AS1** | 0.03872 | 0.47682 | 0.10928 | 0.32475 | 0.30214 | 0.00000 | 0.00539 | 0.01966 | 0.28332 | 0.00107 | 0.13681 | 0.10295 |
| **NUCB2** | 1.56354 | 0.26931 | 9.50608 | 0.40584 | 0.32090 | 1.51707 | 0.00000 | 0.76401 | 0.63499 | 0.38235 | 0.22918 | 2.83552 |
| **NUCKS1** | 2.56303 | 4.68590 | 1.99665 | 10.76755 | 13.43362 | 19.94367 | 18.39298 | 3.56284 | 0.00000 | 34.96166 | 12.78016 | 5.11003 |
| **NUDC** | 0.61522 | 27.36190 | 26.95499 | 24.88840 | 32.64579 | 16.35628 | 47.85589 | 21.81656 | 29.88243 | 5.63089 | 30.37721 | 24.24861 |
| **NUDCD1** | 3.41129 | 7.97897 | 7.25453 | 5.57448 | 0.89125 | 21.75827 | 13.31467 | 14.75664 | 15.04307 | 0.00000 | 12.62343 | 9.26182 |
| **NUDCD2** | 3.59482 | 10.78174 | 9.05834 | 6.98001 | 4.58603 | 28.96496 | 12.60378 | 3.42830 | 12.54324 | 2.18310 | 11.44350 | 11.89171 |
| **NUDCD3** | 2.69634 | 9.30597 | 6.01805 | 6.90790 | 6.19374 | 1.87523 | 7.51425 | 0.29411 | 8.95642 | 3.18414 | 6.04180 | 2.49338 |
| **NUDT1** | 0.43946 | 14.75869 | 4.33239 | 9.47505 | 12.80449 | 5.03223 | 7.98045 | 1.30613 | 1.44967 | 4.93129 | 6.38350 | 16.92393 |

| | | | | | | | | | | | | |
|---|---|---|---|---|---|---|---|---|---|---|---|---|
| **NUDT10** | 0.04502 | 0.01792 | 0.02112 | 0.03562 | 0.02044 | 0.00055 | 0.02595 | 0.06761 | 0.03728 | 0.00291 | 0.01315 | 0.02123 |
| **NUDT11** | 0.80211 | 0.05071 | 0.72120 | 0.57987 | 0.01305 | 0.20415 | 0.27635 | 0.26554 | 0.13400 | 0.00000 | 0.00000 | 0.48717 |
| **NUDT12** | 0.00768 | 0.07223 | 0.37681 | 0.22765 | 0.00000 | 0.44653 | 0.04867 | 0.07590 | 0.21456 | 0.00000 | 0.21263 | 0.35794 |
| **NUDT13** | 0.05709 | 0.46995 | 0.42577 | 0.49678 | 0.00000 | 0.34124 | 0.00000 | 0.49452 | 0.47302 | 0.68774 | 0.55966 | 1.46364 |
| **NUDT14** | 0.80288 | 1.45619 | 1.23104 | 0.57226 | 3.24625 | 0.28950 | 0.06997 | 0.51344 | 0.85314 | 0.00000 | 1.07947 | 0.46531 |
| **NUDT15** | 2.26600 | 4.13179 | 9.86886 | 7.36141 | 0.00000 | 20.03733 | 14.42085 | 3.90306 | 13.10271 | 1.24316 | 10.57273 | 6.91743 |
| **NUDT16** | 3.16524 | 1.13194 | 4.53147 | 2.89571 | 4.70425 | 2.05758 | 4.89024 | 0.33572 | 4.70448 | 1.64449 | 3.28317 | 0.15367 |
| **NUDT16L1** | 0.78996 | 11.14908 | 10.26758 | 8.76588 | 10.41756 | 0.00000 | 11.05252 | 8.78315 | 9.08312 | 0.00000 | 7.55023 | 9.87534 |
| **NUDT16P1** | 0.97518 | 0.25389 | 0.60674 | 1.84081 | 0.26167 | 2.46068 | 0.00000 | 0.00642 | 0.21450 | 0.00000 | 1.21542 | 0.57343 |
| **NUDT17** | 0.02656 | 0.27225 | 0.14140 | 0.23757 | 0.04768 | 0.00000 | 0.00000 | 0.10178 | 0.48518 | 0.22826 | 0.32793 | 0.47217 |
| **NUDT18** | 0.01033 | 0.00000 | 0.11371 | 0.00613 | 0.29130 | 0.06238 | 0.00000 | 0.00738 | 0.17579 | 0.37698 | 0.00035 | 0.07101 |
| **NUDT19** | 0.00466 | 0.00000 | 1.34165 | 0.11951 | 0.00000 | 0.50050 | 1.69301 | 0.07301 | 0.92216 | 0.00000 | 2.26144 | 0.30039 |
| **NUDT2** | 0.72801 | 3.83706 | 4.53430 | 1.82981 | 0.86404 | 8.19650 | 4.65301 | 5.40616 | 3.22505 | 0.00000 | 4.16674 | 5.83279 |
| **NUDT21** | 7.33772 | 15.83271 | 16.97807 | 6.67972 | 8.79771 | 33.88491 | 18.96224 | 6.36429 | 15.34356 | 6.65025 | 16.11812 | 18.17330 |
| **NUDT22** | 0.08704 | 3.20168 | 6.06748 | 5.77756 | 4.91499 | 0.00000 | 2.88901 | 0.02012 | 5.18629 | 0.00000 | 4.79690 | 2.11496 |
| **NUDT3** | 0.07748 | 0.13141 | 0.25905 | 0.05257 | 0.13203 | 0.29638 | 0.07990 | 0.16411 | 0.16412 | 0.00365 | 0.05120 | 0.17121 |
| **NUDT4** | 0.00000 | 0.88063 | 0.14605 | 1.45276 | 0.61560 | 1.72102 | 2.20405 | 0.43802 | 0.00000 | 4.29234 | 0.69251 | 1.77101 |
| **NUDT5** | 2.36028 | 15.22569 | 20.61607 | 12.71838 | 13.84863 | 26.89665 | 24.51828 | 5.92430 | 22.89276 | 0.00000 | 18.66718 | 17.57744 |
| **NUDT6** | 0.14088 | 1.37548 | 0.24232 | 0.63436 | 0.24408 | 0.96252 | 0.49893 | 0.21835 | 0.00000 | 0.00000 | 0.32505 | 4.01547 |
| **NUDT7** | 2.49306 | 1.12064 | 0.08802 | 0.03118 | 0.00146 | 1.05524 | 0.37447 | 0.01867 | 0.00000 | 0.00000 | 0.56062 | 0.70985 |
| **NUDT8** | 0.01175 | 0.51080 | 0.08487 | 0.91645 | 1.44073 | 0.00000 | 0.00095 | 0.07100 | 0.64531 | 0.90567 | 0.93186 | 0.29688 |
| **NUDT9** | 0.56692 | 3.26986 | 4.57104 | 3.08445 | 2.46152 | 9.02638 | 0.00000 | 0.18288 | 14.69120 | 3.08485 | 8.09414 | 8.01214 |
| **NUDT9P1** | 0.00522 | 0.01838 | 0.03967 | 0.01335 | 0.00552 | 0.04816 | 0.01924 | 0.00971 | 0.04378 | 0.03481 | 0.04545 | 0.07057 |
| **NUF2** | 4.47872 | 4.19934 | 4.29320 | 2.65188 | 0.50909 | 16.85069 | 6.04658 | 1.51297 | 4.06249 | 0.00000 | 4.15089 | 6.61099 |
| **NUFIP1** | 2.81732 | 8.12392 | 4.15661 | 2.29940 | 5.47016 | 4.88472 | 6.27099 | 15.11016 | 5.56150 | 4.32369 | 3.86007 | 12.37028 |
| **NUFIP2** | 5.50371 | 4.88815 | 11.02621 | 1.20657 | 4.92807 | 23.17553 | 24.62204 | 17.19547 | 5.36407 | 33.98505 | 13.88323 | 14.50491 |
| **NUGGC** | 0.00511 | 0.01195 | 0.08269 | 0.04670 | 0.00090 | 0.01262 | 0.01157 | 0.01019 | 0.03893 | 0.07653 | 0.02090 | 0.01573 |
| **NUMA1** | 3.52530 | 5.09948 | 7.43754 | 8.37032 | 7.82053 | 0.00000 | 7.07168 | 1.45688 | 12.73327 | 13.53557 | 0.17268 | 3.02711 |
| **NUMB** | 10.16001 | 5.90440 | 7.31617 | 12.26664 | 5.27823 | 9.93358 | 2.56922 | 6.11162 | 8.62326 | 0.89986 | 3.33567 | 3.08975 |
| **NUMBL** | 0.07200 | 0.63460 | 0.89472 | 0.29465 | 0.32608 | 0.00000 | 0.76813 | 0.00474 | 0.71718 | 2.07792 | 0.03438 | 0.40375 |
| **NUP107** | 2.41485 | 11.34471 | 8.73675 | 6.63915 | 2.64827 | 29.02664 | 12.63996 | 3.00304 | 14.73019 | 6.58013 | 8.97843 | 14.38203 |
| **NUP133** | 10.56705 | 7.04577 | 7.13880 | 3.84250 | 3.13198 | 19.46550 | 9.70793 | 6.37149 | 6.16758 | 3.84782 | 8.80986 | 7.19974 |
| **NUP153** | 7.94612 | 9.83489 | 16.58149 | 4.44984 | 3.16021 | 27.37511 | 15.74863 | 20.61013 | 7.76998 | 25.99408 | 4.82628 | 11.37646 |
| **NUP155** | 1.62656 | 8.64513 | 6.67168 | 1.57122 | 1.71704 | 19.56930 | 3.38467 | 9.56878 | 14.31591 | 6.12757 | 5.35656 | 13.37026 |
| **NUP160** | 4.52387 | 13.50860 | 16.50193 | 8.33288 | 4.40121 | 22.76224 | 19.00011 | 5.90216 | 23.70018 | 0.68422 | 14.07671 | 13.54717 |
| **NUP188** | 0.75780 | 14.03202 | 13.97262 | 13.21015 | 10.68178 | 0.00070 | 18.46338 | 3.39620 | 23.09181 | 0.00000 | 13.63337 | 13.67442 |
| **NUP205** | 4.22901 | 11.42961 | 7.47931 | 8.61133 | 5.17836 | 10.80269 | 13.81939 | 0.00000 | 10.02022 | 8.44820 | 7.23444 | 11.02517 |
| **NUP210** | 0.04995 | 0.53831 | 0.45152 | 0.23871 | 1.03568 | 0.00000 | 0.00000 | 0.20810 | 0.00000 | 0.01171 | 0.55383 | 0.00279 |
| **NUP210L** | 0.05996 | 0.03151 | 0.07258 | 0.07752 | 0.11163 | 0.00000 | 0.02363 | 0.08551 | 0.03100 | 0.34200 | 0.11054 | 0.21369 |
| **NUP214** | 4.73377 | 8.20334 | 8.19817 | 6.96993 | 10.39232 | 1.71825 | 7.59113 | 2.97819 | 8.65729 | 10.66489 | 6.92265 | 11.42396 |
| **NUP35** | 2.82356 | 7.89070 | 6.60612 | 2.80525 | 2.89681 | 24.05996 | 8.12905 | 15.02236 | 8.08809 | 1.88798 | 7.86413 | 14.46487 |
| **NUP37** | 2.42670 | 9.32118 | 5.56147 | 1.49481 | 5.20162 | 14.52215 | 3.59999 | 0.59002 | 5.07276 | 2.28812 | 7.32316 | 8.33191 |
| **NUP43** | 2.37252 | 6.73396 | 6.66403 | 6.44626 | 3.86137 | 9.26026 | 21.33152 | 7.49082 | 1.14789 | 7.46099 | 12.94837 | 8.58823 |
| **NUP50** | 3.62833 | 14.52548 | 13.62496 | 7.67183 | 8.41701 | 28.29574 | 26.93027 | 13.57466 | 15.11837 | 16.10936 | 13.44562 | 16.38522 |
| **NUP50-AS1** | 0.37322 | 2.37744 | 2.35472 | 1.83895 | 2.53208 | 0.00000 | 0.22516 | 1.33888 | 1.42259 | 0.86613 | 3.09811 | 3.66204 |
| **NUP54** | 4.86498 | 10.01373 | 10.18481 | 9.10140 | 3.20078 | 29.76373 | 11.37970 | 12.49546 | 11.62665 | 0.01199 | 8.79166 | 16.16039 |
| **NUP62** | 6.47164 | 22.82150 | 21.37565 | 16.12689 | 20.04607 | 9.01036 | 35.94145 | 14.18727 | 24.04501 | 8.99704 | 20.37708 | 30.26453 |
| **NUP62CL** | 0.00250 | 0.09610 | 0.19194 | 0.07755 | 0.54471 | 1.88186 | 0.00000 | 0.15140 | 0.15385 | 0.00000 | 0.29582 | 0.00000 |

| | | | | | | | | | | | | |
|---|---|---|---|---|---|---|---|---|---|---|---|---|
| **NUP85** | 4.64457 | 13.44031 | 13.12987 | 9.61921 | 11.07245 | 9.15976 | 16.31738 | 5.89462 | 19.86550 | 3.66537 | 12.93321 | 13.79265 |
| **NUP88** | 3.64070 | 12.99274 | 10.85269 | 8.14332 | 6.69686 | 16.99646 | 10.92823 | 3.13435 | 15.23362 | 0.00000 | 15.45360 | 12.35624 |
| **NUP93** | 10.58308 | 22.15034 | 11.38428 | 8.58618 | 11.45838 | 18.75570 | 20.17998 | 4.38685 | 20.05580 | 0.00000 | 15.52432 | 9.85701 |
| **NUP98** | 8.14047 | 14.83333 | 11.64678 | 9.45144 | 16.79433 | 13.04572 | 16.93660 | 13.71868 | 19.92573 | 4.41391 | 16.03271 | 28.78827 |
| **NUPL1** | 6.20798 | 6.41071 | 8.67505 | 1.39555 | 1.43696 | 18.47105 | 8.04973 | 5.96928 | 7.30301 | 4.84575 | 6.64277 | 7.68106 |
| **NUPL2** | 0.82129 | 5.03075 | 3.89995 | 1.24752 | 0.26035 | 11.38121 | 6.78856 | 6.96555 | 7.83626 | 0.00000 | 9.11013 | 6.80781 |
| **NUPR1** | 0.08339 | 0.00000 | 0.95024 | 0.80802 | 0.79589 | 0.53529 | 0.74809 | 0.20632 | 0.00000 | 0.02529 | 0.00000 | 0.00000 |
| **NUPR1L** | 0.02164 | 0.00000 | 0.07867 | 0.02238 | 0.01461 | 0.06629 | 0.04465 | 0.00996 | 0.00000 | 0.13640 | 0.09677 | 0.03776 |
| **NUS1** | 0.35576 | 0.26876 | 4.79545 | 0.00000 | 0.31880 | 6.51980 | 4.28415 | 1.05538 | 3.36402 | 7.57292 | 1.74092 | 1.03177 |
| **NUSAP1** | 5.61548 | 29.24486 | 20.53150 | 19.63888 | 14.78734 | 30.03118 | 24.91299 | 35.54728 | 19.55444 | 6.98730 | 11.32836 | 27.97872 |
| **NUTF2** | 3.01753 | 16.94716 | 18.10086 | 19.69593 | 19.58276 | 10.07446 | 30.08298 | 4.31939 | 21.63617 | 0.00000 | 22.25484 | 14.96606 |
| **NUTM1** | 0.03841 | 0.58252 | 0.11726 | 0.07465 | 0.10900 | 0.00000 | 0.00000 | 0.00234 | 0.20490 | 0.45599 | 0.37640 | 1.07901 |
| **NUTM2A-AS1** | 0.00077 | 2.01383 | 0.74686 | 0.55906 | 1.55519 | 2.48357 | 1.38717 | 2.41218 | 1.37169 | 0.00044 | 1.33151 | 2.12676 |
| **NUTM2D** | 0.00487 | 0.01714 | 0.01941 | 0.00543 | 0.03910 | 0.03657 | 0.00744 | 0.01050 | 0.00770 | 0.07804 | 0.02403 | 0.01277 |
| **NUTM2F** | 0.00633 | 0.01484 | 0.02450 | 0.02208 | 0.01754 | 0.01580 | 0.01347 | 0.01042 | 0.02503 | 0.01096 | 0.01482 | 0.01975 |
| **NUTM2G** | 0.06070 | 0.06295 | 0.04337 | 0.12897 | 0.14266 | 0.00000 | 0.02229 | 0.35730 | 0.15019 | 0.32558 | 0.03340 | 0.04984 |
| **NVL** | 0.77276 | 10.87421 | 3.89322 | 4.17327 | 1.00060 | 14.53670 | 6.31550 | 2.32249 | 7.82527 | 2.09322 | 9.68217 | 14.58693 |
| **NWD1** | 0.12248 | 0.04103 | 0.06500 | 0.01160 | 0.04562 | 0.06031 | 0.00703 | 0.00409 | 0.01678 | 0.00909 | 0.01570 | 0.03051 |
| **NWD2** | 0.03005 | 0.01553 | 0.09748 | 0.00024 | 0.12882 | 0.00000 | 0.00080 | 0.03198 | 0.06727 | 0.05617 | 0.02202 | 0.02082 |
| **NXF1** | 9.62170 | 10.50103 | 17.29272 | 13.22927 | 15.50500 | 11.39633 | 23.23545 | 23.47178 | 31.13487 | 2.91385 | 16.14587 | 18.83038 |
| **NXF2+NXF2B.ui** | 0.00405 | 0.01431 | 0.01320 | 0.00498 | 0.00000 | 0.01452 | 0.06101 | 0.01597 | 0.05723 | 0.14156 | 0.00406 | 0.01222 |
| **NXF2+NXF2B.ui** | 0.00767 | 0.02579 | 0.02362 | 0.00080 | 0.19517 | 0.08703 | 0.03666 | 0.01447 | 0.00000 | 0.03409 | 0.03247 | 0.02658 |
| **NXF3** | 0.00548 | 0.08866 | 0.04537 | 0.00167 | 0.00000 | 0.01253 | 0.00707 | 0.00995 | 0.00429 | 0.05430 | 0.05747 | 0.02354 |
| **NXN** | 0.99669 | 1.71505 | 3.37074 | 0.00000 | 0.00000 | 0.72609 | 1.29399 | 0.21773 | 0.00000 | 1.83978 | 0.48688 | 0.71045 |
| **NXNL1** | 0.03112 | 0.01604 | 0.01484 | 0.01298 | 0.01290 | 0.01655 | 0.01694 | 0.00789 | 0.02226 | 0.01947 | 0.01834 | 0.01806 |
| **NXNL2** | 0.00949 | 0.01226 | 0.35259 | 0.01229 | 0.05360 | 0.05629 | 0.00000 | 0.00900 | 0.23959 | 0.00000 | 0.04990 | 0.02096 |
| **NXPE3** | 0.26737 | 0.63040 | 2.26666 | 0.06309 | 0.00000 | 1.34201 | 1.00196 | 0.36439 | 0.09928 | 0.00000 | 0.00000 | 0.60282 |
| **NXPE4** | 0.00610 | 0.01537 | 0.01544 | 0.01124 | 0.01159 | 0.02145 | 0.01597 | 0.01059 | 0.04191 | 0.01615 | 0.01470 | 0.01754 |
| **NXPH1** | 0.66615 | 0.02330 | 0.03369 | 0.00048 | 0.00000 | 0.04405 | 0.01047 | 0.00000 | 0.00000 | 0.07833 | 0.01002 | 0.04793 |
| **NXPH2** | 0.00522 | 0.46939 | 0.00000 | 0.00000 | 0.00003 | 0.21590 | 0.00000 | 0.00358 | 0.00000 | 0.07649 | 0.00432 | 0.19925 |
| **NXPH3** | 0.19119 | 0.00584 | 0.08699 | 0.07729 | 0.00000 | 0.00000 | 0.30683 | 0.01543 | 0.03629 | 0.06946 | 0.04467 | 0.24798 |
| **NXPH4** | 2.27659 | 0.01127 | 0.01381 | 0.01112 | 0.01286 | 0.02860 | 0.06365 | 0.02372 | 0.01517 | 0.03196 | 0.00515 | 0.08418 |
| **NXT1** | 2.31226 | 9.57215 | 9.73330 | 6.90973 | 9.95115 | 5.30511 | 14.29067 | 8.84575 | 14.11568 | 0.00000 | 10.30771 | 17.94187 |
| **NXT2** | 1.55343 | 3.42199 | 0.71006 | 0.14520 | 0.00000 | 7.86680 | 1.87594 | 1.91059 | 0.20212 | 0.73546 | 1.65153 | 9.67923 |
| **NYAP1** | 0.01953 | 0.02611 | 0.02496 | 0.02216 | 0.01858 | 0.00907 | 0.04308 | 0.05087 | 0.00000 | 0.08428 | 0.02150 | 0.01006 |
| **NYAP2** | 1.25344 | 0.01321 | 0.01413 | 0.01250 | 0.02216 | 0.00033 | 0.00916 | 0.05095 | 0.01958 | 0.07065 | 0.01360 | 0.07686 |
| **NYNRIN** | 0.15967 | 0.00000 | 0.32815 | 0.00000 | 0.73575 | 0.00000 | 0.00000 | 0.00000 | 0.47882 | 0.33742 | 0.51084 | 0.21085 |
| **NYX** | 0.02235 | 0.00850 | 0.01105 | 0.01029 | 0.04557 | 0.12556 | 0.00000 | 0.00569 | 0.00000 | 0.03993 | 0.00524 | 0.22094 |
| **OACYLP** | 0.53404 | 0.03251 | 0.02404 | 0.05640 | 0.01370 | 0.00000 | 0.01841 | 0.00222 | 0.02576 | 0.00000 | 0.01362 | 0.03293 |
| **OAF** | 0.86702 | 0.00000 | 0.80314 | 0.00000 | 1.27128 | 0.01302 | 0.73071 | 0.20679 | 1.14057 | 1.33154 | 1.67452 | 0.00000 |
| **OARD1** | 4.92885 | 6.69609 | 6.08713 | 5.61846 | 3.31453 | 15.55636 | 3.76662 | 7.45171 | 4.74624 | 0.20723 | 5.02376 | 13.42330 |
| **OAS1** | 0.01671 | 0.00929 | 0.12670 | 0.35356 | 0.00000 | 0.03072 | 0.00000 | 0.03338 | 0.03663 | 0.00965 | 0.01607 | 0.01418 |
| **OAS2** | 0.00323 | 0.00395 | 0.00524 | 0.00000 | 0.00000 | 0.01564 | 0.07037 | 0.00957 | 0.04138 | 0.42758 | 0.00000 | 0.00757 |
| **OAS3** | 1.65157 | 0.00993 | 0.03054 | 0.01382 | 0.07741 | 0.00000 | 0.03669 | 0.05167 | 0.00022 | 0.02041 | 0.01860 | 0.09761 |
| **OASL** | 0.00143 | 0.09727 | 0.02561 | 0.00988 | 0.02905 | 0.00000 | 0.03539 | 0.48704 | 0.00000 | 0.02992 | 0.01290 | 3.34873 |
| **OAT** | 11.51924 | 17.47888 | 8.82831 | 6.48495 | 29.45142 | 43.82112 | 9.02080 | 8.16627 | 10.72626 | 1.49847 | 34.46078 | 2.59010 |
| **OAZ1** | 11.54794 | 42.28523 | 45.04793 | 43.64194 | 56.63370 | 35.13858 | 67.62141 | 28.91998 | 54.89863 | 20.37270 | 45.48528 | 43.76363 |
| **OAZ2** | 0.84408 | 2.85010 | 7.85389 | 5.66838 | 0.00678 | 0.07961 | 15.90574 | 9.49639 | 10.48757 | 1.29744 | 1.88678 | 5.49986 |

| | | | | | | | | | | | | |
|---|---|---|---|---|---|---|---|---|---|---|---|---|
| **OAZ3** | 0.03670 | 0.84611 | 1.21900 | 0.63891 | 0.10237 | 0.86101 | 0.22359 | 0.59677 | 0.75405 | 0.00000 | 0.00549 | 2.02994 |
| **OBFC1** | 0.50760 | 4.27713 | 1.67793 | 3.79600 | 1.18824 | 2.65266 | 2.18871 | 2.25136 | 1.41238 | 0.00000 | 4.29336 | 2.47520 |
| **OBSCN** | 0.00092 | 0.00000 | 0.09706 | 0.08746 | 0.00000 | 0.00000 | 0.04792 | 0.11630 | 0.12057 | 1.39582 | 0.02088 | 0.06911 |
| **OBSL1** | 0.77604 | 0.30718 | 0.68386 | 0.11019 | 0.81933 | 0.00000 | 0.30972 | 1.88374 | 0.00000 | 1.02666 | 0.00000 | 0.10721 |
| **OC90** | 0.00421 | 0.00239 | 0.14868 | 1.04242 | 0.00000 | 0.00000 | 0.07430 | 0.00787 | 0.01919 | 0.08251 | 0.01796 | 0.07015 |
| **OCA2** | 2.52221 | 1.35142 | 0.00121 | 0.14581 | 0.86044 | 0.02293 | 0.00028 | 1.15317 | 0.00000 | 0.01242 | 0.00000 | 2.64463 |
| **OCEL1** | 0.06305 | 2.02338 | 4.29635 | 3.46475 | 2.59477 | 0.00000 | 2.76897 | 0.00000 | 1.87077 | 2.83788 | 2.84025 | 1.03321 |
| **OCIAD1** | 2.60255 | 10.73911 | 10.44899 | 5.21131 | 2.36935 | 41.87517 | 5.46840 | 2.07291 | 24.52440 | 19.17694 | 9.07597 | 15.29451 |
| **OCIAD2** | 0.24256 | 0.06260 | 2.01450 | 0.56134 | 0.00000 | 0.27961 | 0.26078 | 0.03938 | 0.02908 | 0.00000 | 0.10015 | 0.01524 |
| **OCLM** | 0.00274 | 0.16999 | 0.21289 | 0.02604 | 0.00000 | 0.00000 | 0.00000 | 0.00782 | 0.30438 | 0.29589 | 0.20040 | 0.05732 |
| **OCLN** | 0.00099 | 7.63475 | 0.00000 | 5.51824 | 2.68051 | 10.20031 | 0.00000 | 0.45718 | 9.98984 | 0.00000 | 7.57058 | 0.25929 |
| **OCM** | 0.00374 | 0.01734 | 0.01781 | 0.01013 | 0.02855 | 0.01612 | 0.03454 | 0.02185 | 0.00171 | 0.02801 | 0.01418 | 0.03247 |
| **OCM2** | 0.06544 | 0.01408 | 0.01523 | 0.01401 | 0.01886 | 0.00878 | 0.02152 | 0.01846 | 0.02019 | 0.00673 | 0.01550 | 0.03198 |
| **OCRL** | 5.11933 | 2.99309 | 0.00000 | 0.02484 | 1.51984 | 4.02106 | 0.00000 | 0.00214 | 0.50183 | 2.04400 | 0.82689 | 2.15451 |
| **ODAM** | 0.00731 | 0.00000 | 10.46305 | 0.00000 | 0.00000 | 0.01264 | 17.65688 | 0.00626 | 13.02659 | 0.01175 | 9.43828 | 0.00418 |
| **ODC1** | 17.96778 | 32.84699 | 22.02860 | 0.00000 | 40.74030 | 36.57227 | 40.33537 | 17.96486 | 37.06104 | 38.61638 | 15.26025 | 34.71956 |
| **ODF2** | 4.55395 | 7.08998 | 5.68232 | 6.36508 | 5.69021 | 7.47985 | 4.29751 | 6.50913 | 9.22734 | 0.00000 | 4.29048 | 5.73578 |
| **ODF2L** | 0.68241 | 0.03717 | 5.19332 | 0.00879 | 0.00000 | 3.46853 | 2.63877 | 0.00000 | 4.99447 | 0.88774 | 0.98938 | 0.43226 |
| **ODF3** | 0.00751 | 0.00989 | 0.00582 | 0.00639 | 0.02534 | 0.01447 | 0.02244 | 0.01035 | 0.02759 | 0.10223 | 0.00934 | 0.01388 |
| **ODF3B** | 0.00650 | 0.01521 | 0.03100 | 0.00911 | 0.01165 | 0.02359 | 0.03168 | 0.01041 | 0.05056 | 0.00000 | 0.01563 | 0.01694 |
| **ODF3L1** | 0.00897 | 0.13567 | 0.00621 | 0.00000 | 0.00000 | 0.06581 | 0.00360 | 0.08009 | 0.00838 | 0.04581 | 0.00758 | 0.05600 |
| **ODF3L2** | 1.18003 | 0.05783 | 0.01204 | 0.00858 | 0.02647 | 0.00000 | 0.02655 | 0.00000 | 0.07147 | 0.00000 | 0.00797 | 0.04868 |
| **ODF4** | 0.00358 | 0.02784 | 0.01447 | 0.00434 | 0.06664 | 0.03631 | 0.01980 | 0.01884 | 0.01169 | 0.00000 | 0.02358 | 0.15642 |
| **OFD1** | 10.74105 | 5.25595 | 6.16329 | 2.04843 | 4.97597 | 12.49240 | 6.24130 | 6.71000 | 5.22064 | 14.12841 | 4.03422 | 6.06109 |
| **OGDH** | 1.98971 | 13.47140 | 14.11876 | 11.21306 | 13.84342 | 0.00000 | 14.16264 | 2.24055 | 17.61598 | 0.00000 | 20.91769 | 3.46197 |
| **OGDHL** | 0.53148 | 0.52719 | 0.00000 | 1.28107 | 0.00000 | 0.00000 | 0.20020 | 0.25146 | 0.38146 | 1.80063 | 0.00000 | 0.00000 |
| **OGFOD1** | 8.30808 | 13.52506 | 11.03155 | 7.18960 | 7.74017 | 12.45063 | 17.49974 | 9.14527 | 9.80179 | 3.21681 | 14.25364 | 10.31127 |
| **OGFOD2** | 0.84324 | 6.63525 | 6.42065 | 3.70935 | 3.12963 | 0.00000 | 4.92898 | 4.97641 | 2.85851 | 0.00000 | 6.01626 | 11.18103 |
| **OGFOD3** | 0.17714 | 3.19413 | 1.04931 | 1.77573 | 3.74218 | 0.00000 | 0.76436 | 0.00000 | 2.24616 | 2.31424 | 2.42491 | 1.02034 |
| **OGFR** | 0.06926 | 0.00000 | 1.39701 | 0.71311 | 2.50758 | 0.00000 | 3.16361 | 0.05346 | 0.00000 | 6.02295 | 0.06167 | 0.00000 |
| **OGFR-AS1** | 0.08372 | 0.01800 | 0.03009 | 0.02566 | 0.02364 | 0.00000 | 0.02760 | 0.01480 | 0.05049 | 0.00000 | 0.01045 | 0.02361 |
| **OGFRL1** | 0.01127 | 1.74175 | 0.00000 | 0.93421 | 0.64629 | 0.99983 | 0.00000 | 0.00000 | 0.00000 | 0.79728 | 0.00000 | 0.56889 |
| **OGFRP1** | 0.00485 | 0.00821 | 0.04197 | 0.02862 | 0.06357 | 0.04093 | 0.01696 | 0.12738 | 0.02337 | 0.00916 | 0.02305 | 0.03861 |
| **OGG1** | 1.86528 | 2.41318 | 2.04815 | 3.32159 | 0.73104 | 0.00000 | 2.28911 | 1.89279 | 2.55311 | 2.94609 | 1.74730 | 6.00322 |
| **OGN** | 0.03517 | 0.01166 | 0.09962 | 0.10942 | 0.00000 | 0.33770 | 0.00000 | 0.07368 | 0.15532 | 0.64384 | 0.00974 | 0.04263 |
| **OGT** | 5.39132 | 6.62719 | 2.05726 | 9.40516 | 7.43052 | 20.73162 | 3.70686 | 2.40379 | 11.72799 | 12.87234 | 15.81611 | 11.86277 |
| **OIP5** | 2.59994 | 15.73520 | 13.27832 | 10.80643 | 4.28318 | 26.53200 | 11.83796 | 6.07692 | 13.71400 | 0.00000 | 8.65886 | 14.99030 |
| **OIP5-AS1** | 0.28796 | 10.41991 | 3.79545 | 8.37898 | 6.70035 | 17.47922 | 3.02192 | 0.71047 | 0.00000 | 2.59083 | 7.29899 | 8.38725 |
| **OIT3** | 0.00200 | 0.00000 | 0.00000 | 0.20530 | 0.00000 | 0.03663 | 0.13910 | 0.06303 | 0.01105 | 0.00000 | 0.02429 | 0.29519 |
| **OLA1** | 0.31017 | 3.81632 | 10.33999 | 5.11266 | 4.97798 | 16.22358 | 18.59591 | 0.61535 | 10.75252 | 2.60634 | 8.34033 | 2.87626 |
| **OLAH** | 0.01541 | 5.55140 | 0.02982 | 0.28397 | 0.00000 | 0.08024 | 0.00000 | 0.01571 | 0.00000 | 0.38911 | 0.00000 | 1.15530 |
| **OLFM1** | 0.58127 | 0.27200 | 0.00546 | 0.00000 | 0.01431 | 0.04514 | 0.00000 | 0.18514 | 0.00511 | 0.10774 | 0.00215 | 0.56732 |
| **OLFM2** | 1.46297 | 0.17740 | 0.01884 | 0.00550 | 0.13249 | 0.02290 | 0.01480 | 0.08439 | 0.08094 | 0.00000 | 0.00847 | 0.09937 |
| **OLFM3** | 0.00571 | 0.03383 | 0.01804 | 0.00859 | 0.07390 | 0.01075 | 0.04228 | 0.01249 | 0.00416 | 0.00418 | 0.02465 | 0.03043 |
| **OLFM4** | 0.48100 | 0.05929 | 0.06671 | 0.00000 | 0.00000 | 0.01025 | 0.00635 | 0.25620 | 0.02844 | 0.13159 | 0.00448 | 0.02588 |
| **OLFML1** | 4.76895 | 2.49261 | 0.00000 | 0.00000 | 8.24934 | 0.00000 | 2.56213 | 0.97258 | 7.09329 | 1.37977 | 10.85880 | 0.47736 |
| **OLFML2A** | 0.16340 | 0.18857 | 0.03205 | 0.01015 | 0.00000 | 0.00000 | 0.01363 | 0.00164 | 0.00000 | 0.10244 | 0.05401 | 0.05611 |
| **OLFML2B** | 0.00513 | 0.01335 | 0.04393 | 0.00611 | 0.03033 | 0.00000 | 0.16666 | 0.01056 | 0.00000 | 0.08158 | 0.00547 | 0.01386 |

| | | | | | | | | | | | | |
|---|---|---|---|---|---|---|---|---|---|---|---|---|
| **OLFML3** | 0.00243 | 0.00000 | 0.33877 | 1.83889 | 0.17618 | 0.00000 | 0.00000 | 0.01214 | 0.00000 | 0.59928 | 1.87397 | 0.00574 |
| **OLIG1** | 0.65490 | 0.01091 | 0.02657 | 0.00775 | 0.00888 | 0.04052 | 0.00005 | 0.00165 | 0.02694 | 0.07169 | 0.00583 | 0.01415 |
| **OLIG3** | 0.00473 | 0.01022 | 0.00692 | 0.00762 | 0.00543 | 0.01984 | 0.01930 | 0.01025 | 0.02643 | 0.11440 | 0.01105 | 0.01195 |
| **OLR1** | 0.00000 | 2.41002 | 4.44283 | 0.00000 | 8.23593 | 9.52074 | 14.88414 | 0.00000 | 12.73373 | 0.76737 | 6.75674 | 0.58063 |
| **OMA1** | 0.00000 | 5.79846 | 0.51517 | 0.59790 | 1.01604 | 17.50445 | 0.00000 | 0.60532 | 1.30994 | 2.31191 | 6.74640 | 4.29093 |
| **OMD** | 0.07928 | 0.00000 | 0.09399 | 0.05877 | 0.22569 | 0.00000 | 0.00000 | 0.03522 | 0.00000 | 0.68435 | 0.57121 | 0.04754 |
| **OMG** | 0.02713 | 0.10375 | 0.00931 | 0.06533 | 0.00000 | 0.13791 | 0.00000 | 0.00724 | 0.12224 | 0.10628 | 0.00467 | 0.05455 |
| **OMP** | 0.00620 | 0.01770 | 0.01665 | 0.02188 | 0.01152 | 0.02470 | 0.01604 | 0.01036 | 0.02266 | 0.01875 | 0.01272 | 0.01511 |
| **ONECUT1** | 0.03634 | 0.03281 | 0.02176 | 0.00622 | 0.00580 | 0.02832 | 0.03003 | 0.00823 | 0.01287 | 0.02777 | 0.02351 | 0.02194 |
| **ONECUT2** | 0.00607 | 0.06151 | 0.00912 | 0.00000 | 0.09639 | 0.15093 | 0.00153 | 0.03831 | 0.00000 | 0.56511 | 0.20226 | 0.00001 |
| **OOEP** | 25.92979 | 14.02078 | 0.33327 | 4.10740 | 5.39573 | 3.69113 | 0.00000 | 23.78401 | 0.00000 | 2.39127 | 0.69852 | 16.11741 |
| **OOSP2** | 36.31396 | 0.06789 | 0.00000 | 0.00000 | 0.03677 | 0.14851 | 0.00000 | 16.48957 | 0.00000 | 0.00513 | 0.00000 | 2.26111 |
| **OPA1** | 5.83252 | 5.59569 | 2.20810 | 2.64299 | 0.81760 | 27.23998 | 8.60419 | 9.18687 | 9.47525 | 10.60605 | 6.97171 | 4.38149 |
| **OPA1-AS1** | 0.02990 | 0.04570 | 0.04807 | 0.02804 | 0.03547 | 0.09576 | 0.00011 | 0.13543 | 0.08289 | 0.01879 | 0.04666 | 0.02169 |
| **OPA3** | 1.39292 | 5.78847 | 7.43820 | 4.93496 | 6.96419 | 0.00013 | 10.97647 | 3.40217 | 8.25232 | 3.35831 | 7.10076 | 5.61670 |
| **OPALIN** | 1.47196 | 0.01096 | 0.01396 | 0.00998 | 0.01223 | 0.02202 | 0.02589 | 0.09718 | 0.02626 | 0.00000 | 0.00994 | 0.02116 |
| **OPCML** | 0.07676 | 0.01592 | 0.01698 | 0.01451 | 0.01601 | 0.01107 | 0.02424 | 0.02241 | 0.02508 | 0.01421 | 0.01311 | 0.01423 |
| **OPHN1** | 0.07473 | 0.00000 | 0.03552 | 0.20572 | 0.00000 | 0.16162 | 0.04470 | 0.02681 | 0.33342 | 0.09324 | 0.00000 | 0.29714 |
| **OPLAH** | 0.02759 | 0.00480 | 0.00000 | 0.40651 | 1.16140 | 0.00000 | 0.11806 | 0.04042 | 0.40054 | 0.34331 | 0.48511 | 0.00463 |
| **OPN1SW** | 0.00416 | 0.05133 | 0.02912 | 0.01692 | 0.00707 | 0.00000 | 0.01970 | 0.21398 | 0.01333 | 0.02017 | 0.05090 | 0.03595 |
| **OPN3** | 0.00893 | 0.02311 | 4.94282 | 0.00000 | 0.27065 | 0.13831 | 0.12273 | 0.09004 | 0.00000 | 0.42752 | 0.18577 | 0.25003 |
| **OPN4** | 0.00619 | 0.01446 | 0.01441 | 0.01053 | 0.01588 | 0.02104 | 0.01321 | 0.01043 | 0.02604 | 0.01899 | 0.01343 | 0.01570 |
| **OPN5** | 0.00548 | 0.02572 | 0.02718 | 0.00899 | 0.00762 | 0.01881 | 0.01068 | 0.01048 | 0.00873 | 0.05613 | 0.01645 | 0.01248 |
| **OPRD1** | 0.00556 | 0.00000 | 0.01053 | 0.00522 | 0.06241 | 0.03151 | 0.00774 | 0.00704 | 0.01671 | 0.02906 | 0.01701 | 0.25596 |
| **OPRK1** | 0.00481 | 0.01209 | 0.02668 | 0.00000 | 0.00000 | 0.01079 | 0.12027 | 0.01229 | 0.11422 | 0.09046 | 0.03778 | 0.01254 |
| **OPRL1** | 0.62602 | 0.04935 | 0.03012 | 0.05595 | 0.00000 | 0.01732 | 0.02725 | 0.00677 | 0.00040 | 0.02249 | 0.04177 | 0.02560 |
| **OPRM1** | 0.47710 | 0.00979 | 0.00351 | 0.01135 | 0.03181 | 0.05102 | 0.00041 | 0.06902 | 0.01307 | 0.36752 | 0.00327 | 0.00896 |
| **OPTC** | 0.00514 | 0.01225 | 0.04961 | 0.01673 | 0.00675 | 0.01151 | 0.04041 | 0.01049 | 0.00015 | 0.08349 | 0.01706 | 0.01348 |
| **OPTN** | 0.49248 | 3.28888 | 1.74218 | 0.85491 | 1.31534 | 3.06719 | 1.04387 | 11.69070 | 0.34381 | 0.57043 | 0.83675 | 4.86895 |
| **OR10A2** | 0.00586 | 0.01269 | 0.01747 | 0.00936 | 0.02177 | 0.01368 | 0.00999 | 0.00982 | 0.03592 | 0.03152 | 0.02055 | 0.02725 |
| **OR10A4** | 0.00379 | 0.00912 | 0.02657 | 0.00000 | 0.00041 | 0.02068 | 0.00000 | 0.02055 | 0.09765 | 0.47491 | 0.00000 | 0.01791 |
| **OR10A5** | 0.00690 | 0.01154 | 0.01958 | 0.00848 | 0.01014 | 0.02650 | 0.05498 | 0.00960 | 0.04265 | 0.00669 | 0.01250 | 0.03968 |
| **OR10AD1** | 0.00620 | 0.01726 | 0.02912 | 0.01461 | 0.01835 | 0.00342 | 0.01588 | 0.01061 | 0.02523 | 0.02127 | 0.02959 | 0.01676 |
| **OR10G7** | 0.00621 | 0.01559 | 0.01668 | 0.01395 | 0.01322 | 0.02281 | 0.01684 | 0.01043 | 0.01760 | 0.01588 | 0.01443 | 0.01578 |
| **OR11A1** | 0.00616 | 0.01398 | 0.02680 | 0.01043 | 0.02102 | 0.01526 | 0.02691 | 0.01046 | 0.01122 | 0.02554 | 0.01854 | 0.01592 |
| **OR11H4** | 0.01038 | 0.02247 | 0.01850 | 0.01383 | 0.03279 | 0.00000 | 0.02787 | 0.04758 | 0.01620 | 0.00000 | 0.01992 | 0.04592 |
| **OR11H6** | 0.00835 | 0.01276 | 0.01364 | 0.01175 | 0.01422 | 0.01589 | 0.01700 | 0.05120 | 0.01719 | 0.02322 | 0.01505 | 0.03910 |
| **OR13A1** | 0.00730 | 0.08159 | 0.00959 | 0.00261 | 0.00015 | 0.04094 | 0.03084 | 0.00750 | 0.00656 | 0.02053 | 0.01574 | 0.07536 |
| **OR13C2** | 0.00408 | 0.01263 | 0.01165 | 0.01908 | 0.00815 | 0.01763 | 0.00844 | 0.01690 | 0.02259 | 0.05825 | 0.01180 | 0.01442 |
| **OR13C5** | 0.02308 | 0.01618 | 0.01411 | 0.01919 | 0.02567 | 0.01022 | 0.02435 | 0.00754 | 0.02064 | 0.00984 | 0.01631 | 0.01668 |
| **OR13F1** | 0.01548 | 0.01259 | 0.01477 | 0.01438 | 0.01045 | 0.02545 | 0.01380 | 0.05267 | 0.02123 | 0.02084 | 0.01443 | 0.01963 |
| **OR1D5** | 0.00610 | 0.01502 | 0.01261 | 0.01261 | 0.01157 | 0.01110 | 0.02078 | 0.01058 | 0.02581 | 0.02513 | 0.03078 | 0.01610 |
| **OR1E1** | 0.01657 | 0.01492 | 0.02779 | 0.01029 | 0.01436 | 0.01332 | 0.02245 | 0.00933 | 0.01978 | 0.02206 | 0.02249 | 0.01881 |
| **OR1E2** | 0.29574 | 0.01699 | 0.02025 | 0.01677 | 0.01457 | 0.00172 | 0.03646 | 0.02004 | 0.06086 | 0.00000 | 0.01861 | 0.01321 |
| **OR1F1** | 0.00196 | 0.01534 | 0.02848 | 0.01430 | 0.01552 | 0.01497 | 0.01595 | 0.07434 | 0.03148 | 0.01013 | 0.01317 | 0.02634 |
| **OR1F2P** | 0.09121 | 0.01638 | 0.01400 | 0.01201 | 0.02169 | 0.00917 | 0.06417 | 0.06243 | 0.00000 | 0.00833 | 0.01668 | 0.01461 |
| **OR1J1** | 0.00599 | 0.01144 | 0.01438 | 0.01005 | 0.01025 | 0.01141 | 0.01494 | 0.01047 | 0.00000 | 0.02734 | 0.13478 | 0.01637 |
| **OR1J4** | 0.00750 | 0.01806 | 0.05579 | 0.01707 | 0.02255 | 0.00001 | 0.01269 | 0.05566 | 0.02279 | 0.01604 | 0.01558 | 0.01546 |

| | | | | | | | | | | | | |
|---|---|---|---|---|---|---|---|---|---|---|---|---|
| **OR1L8** | 0.08187 | 0.01118 | 0.01465 | 0.01958 | 0.01923 | 0.02110 | 0.01269 | 0.00584 | 0.01621 | 0.02607 | 0.01874 | 0.01669 |
| **OR1N1** | 2.01142 | 0.16669 | 0.17586 | 0.00066 | 0.06150 | 0.00018 | 0.00175 | 0.08336 | 0.10702 | 0.00000 | 0.02090 | 0.12592 |
| **OR1N2** | 0.00517 | 0.01322 | 0.00299 | 0.00865 | 0.00000 | 0.01783 | 0.16859 | 0.00998 | 0.00000 | 0.05516 | 0.00845 | 0.02280 |
| **OR2A5** | 0.00611 | 0.01350 | 0.01379 | 0.01333 | 0.01482 | 0.02267 | 0.02278 | 0.01014 | 0.01963 | 0.00746 | 0.01439 | 0.02621 |
| **OR2AG1** | 0.01426 | 0.01972 | 0.01147 | 0.01148 | 0.01745 | 0.02744 | 0.00528 | 0.00861 | 0.05038 | 0.01484 | 0.00968 | 0.01986 |
| **OR2AG2** | 0.19080 | 0.01798 | 0.01650 | 0.01452 | 0.00323 | 0.01875 | 0.01043 | 0.01201 | 0.08523 | 0.02161 | 0.04989 | 0.04139 |
| **OR2B2** | 0.22167 | 0.00921 | 0.01168 | 0.01939 | 0.03404 | 0.03415 | 0.01207 | 0.18528 | 0.01553 | 0.01425 | 0.01290 | 0.00595 |
| **OR2B6** | 0.01084 | 0.00980 | 0.13948 | 0.02906 | 0.00000 | 0.03553 | 0.00000 | 0.60928 | 0.11539 | 0.08305 | 0.00940 | 0.09077 |
| **OR2C1** | 0.00576 | 0.01009 | 0.01249 | 0.00823 | 0.01080 | 0.03691 | 0.01342 | 0.00956 | 0.00827 | 0.02213 | 0.03484 | 0.02896 |
| **OR2C3** | 0.02952 | 0.03602 | 0.02828 | 0.01519 | 0.01545 | 0.03445 | 0.01128 | 0.00526 | 0.00810 | 0.02696 | 0.01273 | 0.04505 |
| **OR2D2** | 0.00492 | 0.00730 | 0.05932 | 0.04573 | 0.00000 | 0.03224 | 0.01410 | 0.00982 | 0.00000 | 0.07454 | 0.06781 | 0.01721 |
| **OR2D3** | 0.00529 | 0.00000 | 0.00000 | 0.03484 | 0.09528 | 0.11443 | 0.03421 | 0.00551 | 0.00000 | 0.05547 | 0.06139 | 0.11380 |
| **OR2F1** | 0.00633 | 0.01483 | 0.10606 | 0.00810 | 0.01582 | 0.02038 | 0.00564 | 0.01038 | 0.01733 | 0.01517 | 0.01656 | 0.01588 |
| **OR2H1** | 0.00519 | 0.01353 | 0.02916 | 0.01025 | 0.01150 | 0.02470 | 0.01211 | 0.01269 | 0.01767 | 0.03126 | 0.01323 | 0.01655 |
| **OR2H2** | 0.00645 | 0.01621 | 0.16664 | 0.01277 | 0.03509 | 0.00826 | 0.00486 | 0.01048 | 0.00760 | 0.00989 | 0.02022 | 0.01698 |
| **OR2K2** | 0.00394 | 0.01486 | 0.01459 | 0.01163 | 0.01474 | 0.01859 | 0.01755 | 0.13034 | 0.01699 | 0.01581 | 0.01512 | 0.01166 |
| **OR2L13** | 0.10676 | 0.01997 | 0.02391 | 0.02041 | 0.02165 | 0.01322 | 0.01462 | 0.02762 | 0.03452 | 0.00447 | 0.01634 | 0.01977 |
| **OR2M3** | 0.25212 | 0.01457 | 0.01475 | 0.01552 | 0.01560 | 0.01281 | 0.01826 | 0.00178 | 0.02064 | 0.01024 | 0.01433 | 0.01384 |
| **OR2M7** | 0.00933 | 0.01729 | 0.02944 | 0.01854 | 0.01965 | 0.00458 | 0.01813 | 0.01709 | 0.01734 | 0.01245 | 0.01801 | 0.01660 |
| **OR2S2** | 0.02104 | 0.01569 | 0.01441 | 0.01216 | 0.01270 | 0.01830 | 0.01827 | 0.01269 | 0.01499 | 0.02073 | 0.02133 | 0.01490 |
| **OR2T2** | 0.01238 | 0.01842 | 0.01768 | 0.01578 | 0.01697 | 0.00795 | 0.02631 | 0.00919 | 0.00949 | 0.00455 | 0.05022 | 0.02780 |
| **OR2T27** | 0.01216 | 0.01933 | 0.01512 | 0.01199 | 0.01639 | 0.00969 | 0.03614 | 0.01478 | 0.00711 | 0.03801 | 0.01529 | 0.04520 |
| **OR2T29** | 0.00676 | 0.01409 | 0.01433 | 0.01206 | 0.01469 | 0.02347 | 0.02216 | 0.01157 | 0.01395 | 0.00866 | 0.01874 | 0.02654 |
| **OR2T4** | 0.74382 | 0.01111 | 0.01358 | 0.00901 | 0.02675 | 0.01055 | 0.01780 | 0.12171 | 0.01388 | 0.01780 | 0.02379 | 0.04374 |
| **OR2V1** | 0.00768 | 0.01056 | 0.05233 | 0.00891 | 0.00646 | 0.04624 | 0.01701 | 0.08532 | 0.00478 | 0.02398 | 0.01672 | 0.01086 |
| **OR2W3** | 0.00607 | 0.07847 | 0.05765 | 0.02086 | 0.05057 | 0.07903 | 0.01554 | 0.01000 | 0.00000 | 0.00000 | 0.01809 | 0.00959 |
| **OR3A1** | 0.44311 | 0.10778 | 0.02946 | 0.00956 | 0.04446 | 0.16397 | 0.01140 | 0.04346 | 0.00000 | 0.03426 | 0.12164 | 0.11456 |
| **OR3A2** | 0.13070 | 0.22599 | 0.00000 | 0.01414 | 0.02109 | 0.14776 | 0.00002 | 0.00000 | 0.13386 | 0.00000 | 0.06029 | 0.11738 |
| **OR3A3** | 0.00628 | 0.03010 | 0.00909 | 0.01200 | 0.01047 | 0.00323 | 0.04289 | 0.01014 | 0.07836 | 0.00734 | 0.01622 | 0.02532 |
| **OR3A4P** | 0.00580 | 0.01256 | 0.05910 | 0.00722 | 0.00923 | 0.02659 | 0.00867 | 0.00966 | 0.02045 | 0.02991 | 0.01428 | 0.02563 |
| **OR4C12** | 0.00627 | 0.01497 | 0.01738 | 0.01259 | 0.01588 | 0.01986 | 0.01876 | 0.01048 | 0.01763 | 0.01337 | 0.01459 | 0.01619 |
| **OR4D1** | 0.00582 | 0.01086 | 0.02650 | 0.00705 | 0.01024 | 0.03529 | 0.02420 | 0.01185 | 0.01204 | 0.02153 | 0.01242 | 0.02352 |
| **OR4D10** | 0.03604 | 0.01538 | 0.01636 | 0.00991 | 0.01968 | 0.01982 | 0.01974 | 0.01011 | 0.01731 | 0.01835 | 0.01415 | 0.01517 |
| **OR4F13P** | 0.05553 | 0.01272 | 0.03008 | 0.01310 | 0.00716 | 0.01505 | 0.01445 | 0.00772 | 0.00000 | 0.03200 | 0.05762 | 0.01491 |
| **OR4F16+OR4F2** | 0.00617 | 0.01472 | 0.01332 | 0.01035 | 0.01683 | 0.02206 | 0.01764 | 0.01036 | 0.02295 | 0.01638 | 0.01291 | 0.01739 |
| **OR4F21** | 0.00623 | 0.01779 | 0.02178 | 0.01344 | 0.01419 | 0.01843 | 0.01750 | 0.01051 | 0.01631 | 0.01501 | 0.01574 | 0.01579 |
| **OR4F4** | 0.00572 | 0.01327 | 0.01089 | 0.01199 | 0.01263 | 0.01889 | 0.01343 | 0.01042 | 0.02126 | 0.04297 | 0.01291 | 0.01471 |
| **OR4K14** | 0.00621 | 0.01583 | 0.01562 | 0.01301 | 0.01600 | 0.01246 | 0.02189 | 0.01055 | 0.01965 | 0.01782 | 0.01596 | 0.01641 |
| **OR4K2** | 0.00644 | 0.01571 | 0.01550 | 0.01311 | 0.01396 | 0.00000 | 0.05231 | 0.01054 | 0.00000 | 0.02590 | 0.06804 | 0.01696 |
| **OR51B2** | 0.00612 | 0.01712 | 0.01476 | 0.00301 | 0.00639 | 0.12927 | 0.00000 | 0.00934 | 0.00967 | 0.02127 | 0.03925 | 0.00885 |
| **OR51B4** | 0.08860 | 0.00742 | 0.00000 | 0.19169 | 0.00000 | 0.04235 | 0.08109 | 0.00464 | 0.03503 | 0.07624 | 0.01979 | 0.01367 |
| **OR51B5** | 2.23587 | 0.21576 | 0.00000 | 0.00000 | 1.16342 | 0.31796 | 1.51948 | 0.39746 | 0.83832 | 1.24040 | 1.87976 | 0.78771 |
| **OR51B6** | 0.00602 | 0.00000 | 0.04888 | 0.03323 | 0.00000 | 0.16271 | 0.02539 | 0.03228 | 0.00000 | 0.18630 | 0.11321 | 0.00415 |
| **OR51D1** | 0.02932 | 0.01485 | 0.01399 | 0.01255 | 0.01489 | 0.01811 | 0.01772 | 0.02306 | 0.01824 | 0.01498 | 0.01480 | 0.01464 |
| **OR51E1** | 1.42842 | 0.01039 | 0.01164 | 0.04046 | 0.02272 | 0.00000 | 0.02718 | 0.01476 | 0.01945 | 0.00000 | 0.01491 | 0.17010 |
| **OR51E2** | 0.03555 | 0.01645 | 0.04850 | 0.01718 | 0.02847 | 0.00255 | 0.01766 | 0.00751 | 0.01503 | 0.00720 | 0.02404 | 0.01722 |
| **OR51I1** | 0.02564 | 0.00000 | 0.00000 | 0.00000 | 0.14234 | 0.03154 | 0.00537 | 0.00248 | 0.28158 | 0.29879 | 0.24109 | 0.09744 |
| **OR51I2** | 0.36040 | 0.00556 | 0.01163 | 0.00532 | 0.05831 | 0.03685 | 0.04921 | 0.00000 | 0.00924 | 0.01039 | 0.01425 | 0.04375 |

| | | | | | | | | | | | | |
|---|---|---|---|---|---|---|---|---|---|---|---|---|
| **OR51M1** | 0.00521 | 0.01174 | 0.00752 | 0.00914 | 0.00000 | 0.01693 | 0.02045 | 0.01034 | 0.04529 | 0.07372 | 0.04859 | 0.01338 |
| **OR51Q1** | 0.01195 | 0.00581 | 0.01853 | 0.01395 | 0.06916 | 0.03473 | 0.00652 | 0.01034 | 0.10321 | 0.00001 | 0.10922 | 0.02049 |
| **OR52D1** | 0.00162 | 0.00000 | 0.08825 | 0.00000 | 0.06555 | 0.20649 | 0.08046 | 0.08693 | 0.00000 | 0.26538 | 0.21767 | 0.04717 |
| **OR52E4** | 0.00149 | 0.01221 | 0.06806 | 0.07696 | 0.00000 | 0.02012 | 0.00000 | 0.04357 | 0.24524 | 0.29988 | 0.00000 | 0.01251 |
| **OR52E6** | 0.21799 | 0.01629 | 0.02101 | 0.10669 | 0.00671 | 0.00000 | 0.00318 | 0.00519 | 0.22926 | 0.14306 | 0.02749 | 0.02067 |
| **OR52E8** | 0.00671 | 0.01650 | 0.03149 | 0.01912 | 0.02633 | 0.00000 | 0.00613 | 0.01078 | 0.13452 | 0.02947 | 0.00162 | 0.01759 |
| **OR52K1** | 0.00592 | 0.01115 | 0.00961 | 0.04650 | 0.01047 | 0.02688 | 0.01606 | 0.00988 | 0.01639 | 0.02282 | 0.01886 | 0.02113 |
| **OR52K2** | 0.22560 | 0.00861 | 0.01254 | 0.03493 | 0.00024 | 0.04943 | 0.00821 | 0.00000 | 0.01289 | 0.02807 | 0.01152 | 0.04070 |
| **OR52N1** | 0.00390 | 0.01482 | 0.11521 | 0.00003 | 0.01468 | 0.07642 | 0.00000 | 0.00970 | 0.01412 | 0.14371 | 0.05533 | 0.00635 |
| **OR52N2** | 0.00698 | 0.00466 | 0.05326 | 0.02207 | 0.04352 | 0.03980 | 0.01394 | 0.00948 | 0.00618 | 0.07377 | 0.01346 | 0.01656 |
| **OR52N4** | 0.00289 | 0.00000 | 0.10652 | 0.00000 | 0.12752 | 0.11996 | 0.00444 | 0.00804 | 0.19833 | 0.29816 | 0.27312 | 0.00805 |
| **OR52N5** | 0.00388 | 0.00297 | 0.08865 | 0.22155 | 0.00000 | 0.01582 | 0.03887 | 0.00945 | 0.00000 | 0.18801 | 0.26477 | 0.01898 |
| **OR56A1** | 0.00629 | 0.01391 | 0.01606 | 0.01113 | 0.01107 | 0.02505 | 0.01863 | 0.00966 | 0.03090 | 0.01309 | 0.01231 | 0.02898 |
| **OR56A3** | 0.00659 | 0.01681 | 0.10656 | 0.01340 | 0.01622 | 0.00000 | 0.00700 | 0.00986 | 0.11401 | 0.00000 | 0.00532 | 0.05221 |
| **OR56A5** | 0.00833 | 0.01098 | 0.01670 | 0.01317 | 0.01415 | 0.01736 | 0.02784 | 0.00949 | 0.01395 | 0.01792 | 0.01459 | 0.05521 |
| **OR56B1** | 0.00608 | 0.01528 | 0.03873 | 0.03315 | 0.01003 | 0.02023 | 0.01393 | 0.01043 | 0.01504 | 0.02087 | 0.01551 | 0.01543 |
| **OR56B4** | 0.02260 | 0.01182 | 0.01702 | 0.00966 | 0.01103 | 0.02849 | 0.05613 | 0.00649 | 0.03283 | 0.00000 | 0.00730 | 0.06182 |
| **OR5A1** | 0.17128 | 0.01451 | 0.01782 | 0.01130 | 0.01225 | 0.00575 | 0.05459 | 0.11705 | 0.00000 | 0.03992 | 0.02180 | 0.00840 |
| **OR5AU1** | 0.00559 | 0.03909 | 0.01969 | 0.03479 | 0.00000 | 0.31817 | 0.00000 | 0.00623 | 0.00000 | 0.53140 | 0.05806 | 0.00000 |
| **OR5D18** | 0.00624 | 0.01671 | 0.03626 | 0.02202 | 0.01986 | 0.00409 | 0.01837 | 0.01026 | 0.01652 | 0.01119 | 0.01659 | 0.02361 |
| **OR5E1P** | 0.25875 | 0.01075 | 0.03227 | 0.00908 | 0.01059 | 0.03027 | 0.01377 | 0.02514 | 0.03302 | 0.00506 | 0.01982 | 0.01875 |
| **OR5P2** | 0.01726 | 0.01122 | 0.02927 | 0.01350 | 0.01628 | 0.02386 | 0.02868 | 0.03042 | 0.02066 | 0.00207 | 0.01498 | 0.04906 |
| **OR5P3** | 0.44805 | 0.01394 | 0.05271 | 0.03160 | 0.01878 | 0.01418 | 0.01101 | 0.01204 | 0.02834 | 0.00000 | 0.02032 | 0.01979 |
| **OR5W2** | 0.00621 | 0.01598 | 0.01315 | 0.01368 | 0.01382 | 0.02254 | 0.01728 | 0.01042 | 0.01922 | 0.01553 | 0.01415 | 0.01606 |
| **OR6A2** | 0.02249 | 0.00740 | 0.01498 | 0.01450 | 0.06668 | 0.00749 | 0.00172 | 0.01031 | 0.04473 | 0.06482 | 0.06341 | 0.02145 |
| **OR6C2** | 0.01281 | 0.00914 | 0.01290 | 0.01623 | 0.01202 | 0.03568 | 0.01217 | 0.03719 | 0.03147 | 0.00572 | 0.01555 | 0.05763 |
| **OR6C68** | 0.00606 | 0.02402 | 0.01383 | 0.01102 | 0.01254 | 0.02282 | 0.01790 | 0.01038 | 0.01798 | 0.01576 | 0.01443 | 0.01765 |
| **OR6C70** | 0.00313 | 0.06555 | 0.01572 | 0.01076 | 0.03051 | 0.00589 | 0.02426 | 0.03009 | 0.02185 | 0.00000 | 0.01443 | 0.08312 |
| **OR6F1** | 0.00629 | 0.01396 | 0.02145 | 0.01208 | 0.01583 | 0.01830 | 0.01916 | 0.01118 | 0.02002 | 0.00810 | 0.01558 | 0.02732 |
| **OR6S1** | 0.14243 | 0.01287 | 0.01362 | 0.01244 | 0.01955 | 0.02977 | 0.01946 | 0.02374 | 0.01564 | 0.00016 | 0.02174 | 0.03245 |
| **OR6V1** | 0.04780 | 0.01457 | 0.03615 | 0.03659 | 0.01924 | 0.00852 | 0.01039 | 0.09724 | 0.01879 | 0.02252 | 0.01509 | 0.01925 |
| **OR6W1P** | 0.04141 | 0.01441 | 0.04393 | 0.01511 | 0.01153 | 0.00950 | 0.00457 | 0.09199 | 0.01977 | 0.06819 | 0.01525 | 0.01474 |
| **OR7C1** | 0.00584 | 0.01518 | 0.01905 | 0.01114 | 0.01390 | 0.01968 | 0.02785 | 0.01134 | 0.00784 | 0.01966 | 0.01589 | 0.01569 |
| **OR7D2** | 0.03839 | 0.01301 | 0.00397 | 0.01875 | 0.00000 | 0.03234 | 0.11822 | 0.03090 | 0.00519 | 0.11169 | 0.00653 | 0.00822 |
| **OR7E12P** | 0.50125 | 0.00112 | 0.06253 | 0.00000 | 0.00000 | 0.03761 | 0.00571 | 0.18938 | 0.00000 | 0.40448 | 0.02774 | 0.08892 |
| **OR7E14P** | 0.04380 | 0.02090 | 0.02178 | 0.00000 | 0.09566 | 0.02352 | 0.05872 | 0.04527 | 0.15331 | 0.41662 | 0.00512 | 0.03752 |
| **OR7E156P** | 0.00554 | 0.00800 | 0.00523 | 0.15445 | 0.00000 | 0.08265 | 0.00000 | 0.00962 | 0.00756 | 0.03181 | 0.01028 | 0.01062 |
| **OR7E24** | 0.35452 | 0.01063 | 0.00698 | 0.00724 | 0.01008 | 0.01251 | 0.00256 | 0.07646 | 0.03733 | 0.09113 | 0.00649 | 0.00801 |
| **OR7E2P** | 0.00642 | 0.02429 | 0.01615 | 0.02125 | 0.02096 | 0.00919 | 0.01837 | 0.01061 | 0.02131 | 0.00783 | 0.01728 | 0.01643 |
| **OR7E47P** | 0.01132 | 0.00735 | 0.01064 | 0.00431 | 0.25001 | 0.00000 | 0.01139 | 0.01926 | 0.01560 | 0.24178 | 0.00000 | 0.05362 |
| **OR7E5P** | 0.03726 | 0.28840 | 0.00972 | 0.27882 | 0.00000 | 0.10049 | 0.00000 | 0.00551 | 0.00000 | 0.03084 | 0.02155 | 0.03675 |
| **OR7E91P** | 0.01087 | 0.01122 | 0.61555 | 0.01829 | 0.00000 | 0.05573 | 0.18458 | 0.01151 | 0.16190 | 0.00000 | 0.00000 | 0.01391 |
| **OR8A1** | 0.00559 | 0.13861 | 0.04039 | 0.05749 | 0.00000 | 0.00540 | 0.05273 | 0.01115 | 0.03960 | 0.03922 | 0.01016 | 0.00664 |
| **OR8B12** | 0.00315 | 0.01243 | 0.01470 | 0.09872 | 0.00955 | 0.01815 | 0.02598 | 0.02756 | 0.01468 | 0.01149 | 0.01824 | 0.01613 |
| **OR8B2** | 0.00527 | 0.01506 | 0.04995 | 0.00492 | 0.00871 | 0.02853 | 0.03084 | 0.01028 | 0.03980 | 0.06092 | 0.00457 | 0.01235 |
| **OR8B3** | 0.00586 | 0.01439 | 0.05149 | 0.00862 | 0.01606 | 0.01538 | 0.01735 | 0.01048 | 0.00986 | 0.03404 | 0.01605 | 0.01519 |
| **OR8B4** | 0.00777 | 0.01657 | 0.06089 | 0.02227 | 0.00781 | 0.04427 | 0.00142 | 0.00960 | 0.02240 | 0.03385 | 0.01507 | 0.01269 |
| **OR8B8** | 0.00624 | 0.01580 | 0.01418 | 0.01869 | 0.01569 | 0.01611 | 0.01552 | 0.01054 | 0.02622 | 0.01509 | 0.01447 | 0.01825 |

| | | | | | | | | | | | | |
|---|---|---|---|---|---|---|---|---|---|---|---|---|
| **OR8D1** | 0.00766 | 0.04321 | 0.01402 | 0.02796 | 0.00000 | 0.10760 | 0.00114 | 0.00871 | 0.00729 | 0.05405 | 0.00898 | 0.01307 |
| **OR8D2** | 0.00397 | 0.00623 | 0.00844 | 0.05674 | 0.00343 | 0.05101 | 0.01216 | 0.00966 | 0.04262 | 0.15919 | 0.00647 | 0.00836 |
| **OR8G1** | 0.00704 | 0.03798 | 0.03085 | 0.04755 | 0.00022 | 0.04106 | 0.04093 | 0.01658 | 0.02872 | 0.01183 | 0.02034 | 0.01179 |
| **OR8G2** | 0.03638 | 0.01172 | 0.03772 | 0.03752 | 0.01619 | 0.01802 | 0.00984 | 0.01002 | 0.02417 | 0.03844 | 0.01588 | 0.03644 |
| **OR8G5** | 1.09532 | 0.04857 | 0.00961 | 0.24697 | 0.01303 | 0.00000 | 0.00745 | 0.12387 | 0.08832 | 0.00000 | 0.02226 | 0.11917 |
| **OR8S1** | 0.01087 | 0.00473 | 0.01761 | 0.00185 | 0.05537 | 0.00000 | 0.00873 | 0.00878 | 0.10632 | 0.10191 | 0.26231 | 0.01356 |
| **OR9A2** | 0.00607 | 0.01299 | 0.02046 | 0.01482 | 0.00574 | 0.02423 | 0.03185 | 0.01031 | 0.00000 | 0.03277 | 0.03614 | 0.01466 |
| **OR9Q1** | 0.03941 | 0.01381 | 0.01239 | 0.00998 | 0.01592 | 0.02392 | 0.02411 | 0.05410 | 0.01195 | 0.02302 | 0.01401 | 0.01916 |
| **ORAI1** | 1.13169 | 4.21968 | 2.84475 | 8.30175 | 8.77485 | 0.00000 | 2.82654 | 0.25527 | 0.00000 | 4.93182 | 1.64236 | 1.71729 |
| **ORAI2** | 0.03211 | 0.18564 | 0.04245 | 0.01590 | 0.12427 | 0.00000 | 0.10815 | 0.02054 | 0.28715 | 1.58677 | 0.15630 | 0.45686 |
| **ORAI3** | 0.00335 | 0.00585 | 0.00000 | 0.40345 | 0.00000 | 0.06164 | 0.06703 | 0.00143 | 0.09179 | 0.97288 | 0.03400 | 0.31398 |
| **ORAOV1** | 2.29703 | 3.55298 | 2.15997 | 1.98634 | 3.69428 | 0.00000 | 2.75570 | 0.59653 | 6.27056 | 0.00000 | 1.26763 | 5.52206 |
| **ORC1** | 1.41307 | 12.85833 | 10.03979 | 11.00516 | 8.69333 | 12.89702 | 11.14345 | 7.76071 | 17.20943 | 0.00000 | 11.13095 | 15.37879 |
| **ORC2** | 3.02640 | 5.38818 | 7.91979 | 5.41165 | 3.13716 | 13.76249 | 4.60122 | 3.08266 | 9.42571 | 3.59512 | 5.64359 | 10.15188 |
| **ORC3** | 0.80466 | 5.22283 | 5.60954 | 2.16910 | 0.00000 | 16.09639 | 5.27262 | 2.66969 | 6.57982 | 0.58883 | 8.15336 | 5.63280 |
| **ORC4** | 0.77578 | 6.33581 | 6.53508 | 5.52455 | 0.00850 | 24.04590 | 13.58707 | 8.35328 | 13.87651 | 2.01087 | 9.45131 | 11.55276 |
| **ORC5** | 3.26024 | 11.29521 | 5.99469 | 2.55393 | 2.50128 | 18.99900 | 2.33439 | 3.66534 | 6.86245 | 0.03966 | 6.97800 | 8.49333 |
| **ORC6** | 2.73101 | 13.83265 | 10.74190 | 14.06489 | 2.76111 | 20.89326 | 12.77280 | 7.12621 | 14.29482 | 11.01431 | 7.05774 | 7.62433 |
| **ORMDL1** | 4.92855 | 8.20160 | 16.12693 | 3.67559 | 0.64511 | 23.78804 | 8.06399 | 2.70357 | 19.69919 | 2.25902 | 12.75405 | 14.42457 |
| **ORMDL2** | 4.97321 | 13.49911 | 14.90176 | 7.63488 | 13.85711 | 15.61485 | 0.00000 | 14.60523 | 11.55374 | 4.50430 | 9.48342 | 18.26025 |
| **ORMDL3** | 4.74485 | 1.36540 | 1.96082 | 1.19187 | 2.16027 | 0.00000 | 1.54995 | 2.35721 | 4.49534 | 4.81289 | 0.69633 | 3.69657 |
| **OS9** | 1.12396 | 12.78022 | 14.09353 | 11.37288 | 13.33974 | 7.15203 | 0.15316 | 3.00310 | 24.32959 | 9.05350 | 9.68527 | 11.64710 |
| **OSBP** | 8.46190 | 2.48313 | 5.13919 | 3.36376 | 5.13458 | 11.83478 | 10.43936 | 3.77948 | 0.34274 | 17.56471 | 3.06226 | 1.01451 |
| **OSBP2** | 1.07525 | 2.35805 | 0.07465 | 0.71663 | 0.08516 | 0.00000 | 0.12442 | 0.18692 | 0.20685 | 0.62777 | 0.00686 | 0.46628 |
| **OSBPL10** | 16.78723 | 0.06702 | 2.22160 | 1.40593 | 0.00857 | 2.78281 | 1.16209 | 4.92639 | 0.34313 | 1.61573 | 0.47508 | 0.00000 |
| **OSBPL10-AS1** | 0.00479 | 0.01008 | 0.02099 | 0.05495 | 0.00314 | 0.01405 | 0.00594 | 0.01039 | 0.02424 | 0.10757 | 0.01271 | 0.01284 |
| **OSBPL11** | 1.22831 | 4.60738 | 10.53705 | 5.24300 | 2.03560 | 20.98004 | 1.54134 | 2.05972 | 6.23658 | 11.16700 | 7.26096 | 14.95090 |
| **OSBPL1A** | 2.87249 | 0.62260 | 0.00000 | 0.00000 | 1.43853 | 2.46125 | 2.76466 | 0.80283 | 1.45268 | 1.51658 | 4.65804 | 0.27681 |
| **OSBPL2** | 2.94526 | 5.68208 | 7.19568 | 3.49493 | 5.62794 | 3.52209 | 5.95468 | 2.07369 | 11.27224 | 0.00000 | 5.08019 | 2.47814 |
| **OSBPL3** | 1.06067 | 2.48507 | 3.36535 | 2.75965 | 4.14595 | 6.53655 | 2.42091 | 0.00000 | 3.88856 | 6.00794 | 2.86552 | 2.26574 |
| **OSBPL5** | 1.98361 | 0.23981 | 0.02347 | 0.00000 | 0.00000 | 0.00000 | 0.00009 | 0.11534 | 0.00000 | 0.20064 | 0.21175 | 0.00000 |
| **OSBPL6** | 0.00544 | 12.74133 | 0.00000 | 0.00000 | 11.02167 | 4.74964 | 3.27911 | 11.53159 | 3.94195 | 0.00000 | 7.84950 | 6.79178 |
| **OSBPL7** | 0.14676 | 0.41149 | 0.06588 | 0.00000 | 0.57125 | 0.15193 | 0.00000 | 0.06364 | 0.00000 | 0.07399 | 0.02133 | 0.14448 |
| **OSBPL8** | 6.46028 | 0.92090 | 7.69581 | 2.99354 | 3.97428 | 15.07650 | 6.65265 | 4.23430 | 5.15720 | 16.20233 | 4.52540 | 2.67572 |
| **OSBPL9** | 6.52865 | 5.10532 | 17.68714 | 2.53671 | 4.69048 | 22.02054 | 10.06152 | 2.24328 | 13.75863 | 0.49299 | 9.13282 | 3.35609 |
| **OSCAR** | 0.47314 | 0.10652 | 0.10124 | 0.11778 | 0.12468 | 0.00000 | 0.16988 | 0.21177 | 0.15725 | 0.01413 | 0.39898 | 0.24347 |
| **OSCP1** | 1.16446 | 0.27357 | 0.20380 | 0.02100 | 0.00000 | 0.42260 | 0.00000 | 0.29019 | 0.00000 | 0.08905 | 0.11998 | 1.46574 |
| **OSER1** | 3.65588 | 13.08787 | 17.29085 | 6.31633 | 6.34760 | 19.76286 | 18.31567 | 30.06301 | 14.27545 | 0.00000 | 12.81058 | 24.41824 |
| **OSER1-AS1** | 0.40701 | 1.22404 | 1.47301 | 0.55579 | 0.00000 | 0.00000 | 0.59913 | 0.38434 | 0.59314 | 0.00000 | 1.03334 | 3.58443 |
| **OSGEP** | 0.26821 | 11.30484 | 0.99423 | 9.16413 | 6.35996 | 21.05075 | 5.82471 | 1.80939 | 11.33818 | 0.57348 | 5.60516 | 16.80585 |
| **OSGEPL1** | 0.43176 | 4.65759 | 1.48830 | 0.27150 | 0.10866 | 7.17836 | 1.42723 | 0.11244 | 0.39186 | 3.68060 | 4.21995 | 5.94305 |
| **OSGEPL1-AS1** | 0.00530 | 0.06250 | 0.15114 | 0.00000 | 0.02317 | 0.06848 | 0.18143 | 0.00668 | 0.00000 | 0.01663 | 0.07705 | 0.16969 |
| **OSGIN1** | 0.00546 | 0.27828 | 0.25584 | 0.09572 | 0.00000 | 0.00000 | 0.79391 | 0.01131 | 0.16618 | 0.08222 | 0.00000 | 0.01180 |
| **OSGIN2** | 0.00000 | 0.93728 | 1.07719 | 1.55534 | 0.69429 | 3.54324 | 5.37146 | 0.00005 | 2.94912 | 0.27559 | 2.15866 | 0.10353 |
| **OSM** | 0.00605 | 0.01611 | 0.04912 | 0.01734 | 0.01581 | 0.00714 | 0.03897 | 0.01060 | 0.01373 | 0.02159 | 0.01390 | 0.01648 |
| **OSMR** | 1.22357 | 0.20777 | 0.68277 | 2.51495 | 0.27013 | 0.00000 | 0.00000 | 0.49549 | 0.28421 | 0.22031 | 0.04901 | 0.37718 |
| **OSMR-AS1** | 0.01578 | 0.02020 | 0.03263 | 0.01695 | 0.01547 | 0.01433 | 0.01450 | 0.01388 | 0.01824 | 0.01783 | 0.01545 | 0.02290 |
| **OSR1** | 0.02959 | 0.00000 | 0.01769 | 0.02167 | 0.03054 | 0.00000 | 0.02197 | 0.11093 | 0.01748 | 0.00000 | 0.01659 | 1.32435 |

| | | | | | | | | | | | | |
|---|---|---|---|---|---|---|---|---|---|---|---|---|
| **OSR2** | 6.08576 | 5.36449 | 0.55695 | 2.42100 | 0.73681 | 0.41178 | 1.70989 | 1.98116 | 0.00009 | 0.00000 | 0.62846 | 14.04278 |
| **OST4** | 2.53931 | 21.77680 | 17.62278 | 17.81895 | 23.60375 | 19.78499 | 17.16569 | 5.03207 | 20.08639 | 0.07221 | 19.36849 | 22.91018 |
| **OSTC** | 1.50448 | 10.82036 | 16.48019 | 8.18918 | 5.78404 | 33.99945 | 9.45585 | 4.95191 | 25.51253 | 6.47244 | 11.16125 | 20.10216 |
| **OSTCP1** | 0.08357 | 0.60609 | 0.92944 | 0.61491 | 0.10684 | 2.26629 | 0.00000 | 0.21148 | 1.41223 | 0.00000 | 0.57102 | 1.25101 |
| **OSTF1** | 3.66104 | 1.85948 | 4.91341 | 1.56643 | 3.49196 | 9.12335 | 5.19353 | 2.99323 | 4.87712 | 0.00000 | 4.33589 | 2.61070 |
| **OSTM1** | 0.41396 | 2.56998 | 6.91103 | 0.00000 | 0.30831 | 6.68854 | 0.00000 | 1.92148 | 8.62895 | 5.25364 | 6.20797 | 3.24243 |
| **OTC** | 0.00210 | 0.00807 | 0.01241 | 0.00732 | 0.07432 | 0.02312 | 0.01150 | 0.04524 | 0.00542 | 0.05074 | 0.01988 | 0.02373 |
| **OTOA** | 0.54718 | 0.04442 | 0.01565 | 0.01005 | 0.01670 | 0.00000 | 0.02555 | 0.02312 | 0.02623 | 0.00982 | 0.01694 | 0.03976 |
| **OTOF** | 0.00225 | 0.00000 | 0.00000 | 0.12577 | 0.11729 | 0.01243 | 0.00000 | 0.00800 | 0.10857 | 1.16372 | 0.00000 | 0.00714 |
| **OTOG** | 0.00443 | 0.01360 | 0.05102 | 0.01907 | 0.00116 | 0.07638 | 0.00939 | 0.01694 | 0.02728 | 0.03637 | 0.01076 | 0.03419 |
| **OTOGL** | 1.22561 | 0.00431 | 0.35118 | 0.01562 | 0.02858 | 0.00000 | 0.00868 | 0.18529 | 0.00000 | 0.05677 | 0.11367 | 0.10587 |
| **OTOP3** | 0.07113 | 0.27414 | 0.01676 | 0.01115 | 0.01911 | 0.00000 | 0.01833 | 0.00733 | 0.01729 | 0.02912 | 0.01131 | 0.00388 |
| **OTP** | 0.00581 | 0.01420 | 0.02622 | 0.02211 | 0.01751 | 0.01904 | 0.01726 | 0.01101 | 0.01498 | 0.01363 | 0.01524 | 0.03927 |
| **OTUB1** | 5.80253 | 19.32372 | 24.39292 | 21.93078 | 23.35941 | 5.28711 | 25.53818 | 11.55879 | 32.17558 | 0.00000 | 21.46883 | 23.40917 |
| **OTUB2** | 0.01773 | 0.00000 | 0.34421 | 0.07508 | 0.00000 | 0.05286 | 0.68869 | 0.02310 | 0.09919 | 0.11031 | 0.46327 | 0.28315 |
| **OTUD1** | 0.20037 | 0.96925 | 0.00000 | 1.89862 | 0.00000 | 3.02456 | 0.77212 | 0.26815 | 0.00000 | 2.59701 | 0.12452 | 1.10553 |
| **OTUD3** | 7.25348 | 0.82438 | 4.44333 | 0.07311 | 0.19011 | 3.48426 | 2.76015 | 1.68617 | 2.64277 | 2.86158 | 2.09172 | 1.33485 |
| **OTUD4** | 6.01014 | 0.48422 | 2.00482 | 2.38131 | 2.32919 | 5.26502 | 6.94341 | 0.96978 | 0.00010 | 12.05404 | 1.68524 | 4.55183 |
| **OTUD5** | 5.16228 | 2.65752 | 2.56587 | 2.95154 | 8.68863 | 0.00000 | 6.38950 | 2.28429 | 1.90504 | 13.55346 | 0.57443 | 3.65604 |
| **OTUD6A** | 0.01912 | 0.00000 | 0.18937 | 0.14650 | 0.01196 | 0.00000 | 0.03681 | 1.02539 | 0.00525 | 0.00000 | 0.02454 | 0.91070 |
| **OTUD6B** | 0.86016 | 2.54634 | 4.59792 | 0.28413 | 0.00000 | 18.73105 | 6.40820 | 0.52613 | 4.82721 | 4.89513 | 3.73947 | 4.70101 |
| **OTUD6B-AS1** | 0.76848 | 4.16210 | 5.57272 | 0.00000 | 1.16653 | 15.73334 | 2.60833 | 1.87919 | 3.49353 | 8.42017 | 3.45291 | 11.45524 |
| **OTUD7A** | 0.01669 | 0.00767 | 0.03177 | 0.01108 | 0.11469 | 0.00000 | 0.03923 | 0.07585 | 0.00000 | 0.03160 | 0.02107 | 0.18301 |
| **OTUD7B** | 1.61288 | 0.29591 | 1.68070 | 0.30163 | 0.30076 | 2.75875 | 2.85788 | 0.00000 | 0.00323 | 2.79142 | 0.55046 | 0.43484 |
| **OTULIN** | 1.97856 | 0.88280 | 3.45907 | 1.63094 | 3.59820 | 3.44602 | 5.21966 | 1.25149 | 0.90284 | 6.43074 | 1.06263 | 1.33388 |
| **OTX1** | 1.16327 | 0.97407 | 0.00000 | 1.54549 | 1.06138 | 0.00000 | 0.01475 | 3.62920 | 1.13562 | 1.13079 | 1.66134 | 7.04929 |
| **OTX2** | 36.40720 | 0.29282 | 0.00000 | 12.44777 | 0.00000 | 0.60137 | 0.06026 | 11.71163 | 0.00000 | 0.00000 | 0.31458 | 4.08106 |
| **OTX2-AS1** | 0.00877 | 0.01260 | 0.01711 | 0.01989 | 0.01082 | 0.03008 | 0.06377 | 0.01252 | 0.01179 | 0.00732 | 0.01357 | 0.01747 |
| **OVCA2** | 0.00571 | 2.40251 | 1.51909 | 1.15511 | 1.68140 | 0.00000 | 0.20956 | 0.22445 | 1.60769 | 0.00000 | 1.09062 | 1.63875 |
| **OVCH1** | 1.54286 | 0.01091 | 0.00470 | 0.00319 | 0.02725 | 0.01503 | 0.12973 | 0.49453 | 0.00510 | 0.05221 | 0.02783 | 0.00260 |
| **OVCH1-AS1** | 0.04831 | 0.09328 | 0.06784 | 0.00062 | 0.06858 | 0.19695 | 0.06105 | 0.11053 | 0.00560 | 0.00000 | 0.10713 | 0.09272 |
| **OVCH2** | 1.02525 | 0.00939 | 0.08298 | 0.01330 | 0.00458 | 0.02803 | 0.01796 | 0.03163 | 0.06047 | 0.00000 | 0.05271 | 0.01275 |
| **OVGP1** | 0.33866 | 0.04481 | 1.96940 | 1.18429 | 0.65187 | 0.00000 | 0.70938 | 0.36755 | 1.56126 | 1.81395 | 0.00000 | 0.13548 |
| **OVOL1** | 6.54391 | 6.74083 | 33.79265 | 5.96775 | 20.78811 | 3.38874 | 7.43769 | 7.16113 | 10.93825 | 0.00000 | 0.00000 | 0.00000 |
| **OVOL1-AS1** | 0.00909 | 0.01734 | 0.05217 | 0.00874 | 0.02017 | 0.01400 | 0.05787 | 0.01079 | 0.00000 | 0.04812 | 0.01489 | 0.01460 |
| **OVOL2** | 0.37789 | 5.34112 | 0.00000 | 2.87105 | 3.56791 | 0.00000 | 1.03195 | 1.54091 | 0.00000 | 5.36737 | 1.21366 | 4.83534 |
| **OVOL3** | 0.01951 | 0.02124 | 0.03803 | 0.07777 | 0.00556 | 0.00000 | 0.01736 | 0.07973 | 0.15381 | 0.11035 | 0.02826 | 0.18753 |
| **OXA1L** | 1.55598 | 14.07748 | 9.11108 | 17.88343 | 16.31694 | 23.58493 | 13.38023 | 5.07851 | 26.27040 | 11.67960 | 25.50874 | 18.69384 |
| **OXCT1** | 0.29532 | 2.21705 | 0.55594 | 0.58525 | 0.25762 | 1.86023 | 0.00000 | 0.48960 | 0.20009 | 0.00000 | 0.00000 | 3.29646 |
| **OXCT2** | 0.02349 | 0.01384 | 0.02187 | 0.01047 | 0.04022 | 0.00831 | 0.01691 | 0.00807 | 0.01751 | 0.03527 | 0.01296 | 0.02109 |
| **OXER1** | 0.00563 | 0.01156 | 0.02674 | 0.05460 | 0.01126 | 0.01995 | 0.00661 | 0.01032 | 0.02036 | 0.04940 | 0.01242 | 0.01414 |
| **OXGR1** | 0.00392 | 0.00000 | 0.04231 | 0.00000 | 0.00000 | 0.17386 | 0.04248 | 0.09682 | 0.00000 | 0.10265 | 0.08292 | 0.00498 |
| **OXLD1** | 0.09625 | 2.65005 | 1.83142 | 1.09091 | 1.14945 | 0.00101 | 0.42599 | 0.00000 | 1.46844 | 1.57475 | 3.22960 | 3.44457 |
| **OXNAD1** | 2.07658 | 4.81246 | 2.58578 | 4.35922 | 3.32249 | 15.93585 | 2.10086 | 2.44266 | 3.09324 | 0.00000 | 5.20637 | 4.95564 |
| **OXR1** | 0.90007 | 0.85178 | 3.08889 | 0.00000 | 0.49473 | 2.47524 | 1.00079 | 0.14568 | 2.30835 | 7.07634 | 1.35690 | 0.09386 |
| **OXSM** | 0.35192 | 8.78527 | 3.38512 | 0.33007 | 0.00000 | 8.85913 | 2.21476 | 0.57978 | 3.68714 | 0.00000 | 5.72429 | 12.33562 |
| **OXSR1** | 1.38104 | 3.65329 | 5.26299 | 1.95854 | 4.27776 | 11.83274 | 7.51216 | 2.32112 | 6.06395 | 14.36258 | 3.31036 | 10.33380 |
| **OXT** | 9.53548 | 2.26411 | 0.00000 | 21.61211 | 13.23080 | 0.00000 | 0.33840 | 7.31934 | 0.00000 | 3.18907 | 1.64361 | 6.23020 |

| | | | | | | | | | | | | |
|---|---|---|---|---|---|---|---|---|---|---|---|---|
| **OXTR** | 0.00685 | 0.18275 | 0.04048 | 0.14603 | 0.11965 | 0.62011 | 0.20083 | 0.04183 | 0.14888 | 1.61913 | 0.27639 | 0.00035 |
| **P2RX1** | 0.59344 | 0.01254 | 0.02515 | 0.01731 | 0.01165 | 0.01908 | 0.01173 | 0.00104 | 0.02862 | 0.00938 | 0.01203 | 0.01117 |
| **P2RX2** | 0.00947 | 0.01455 | 0.02264 | 0.00693 | 0.00245 | 0.02867 | 0.02224 | 0.00963 | 0.00000 | 0.05267 | 0.03489 | 0.03354 |
| **P2RX3** | 0.00444 | 0.01917 | 0.04525 | 0.00917 | 0.01666 | 0.02506 | 0.01145 | 0.26318 | 0.01703 | 0.00247 | 0.01529 | 0.03161 |
| **P2RX4** | 1.30997 | 2.61304 | 1.89330 | 0.33246 | 0.96792 | 3.35941 | 0.00000 | 0.41652 | 3.62453 | 0.79511 | 2.81204 | 0.00000 |
| **P2RX5** | 0.72908 | 0.20876 | 0.05660 | 0.19536 | 0.00440 | 0.00000 | 0.00000 | 1.48503 | 0.15691 | 0.70769 | 0.00000 | 0.23414 |
| **P2RX5-TAX1BP3** | 0.23462 | 0.69892 | 0.70754 | 1.13113 | 0.66074 | 0.00000 | 0.38986 | 0.14305 | 0.98951 | 0.72494 | 0.49321 | 0.93406 |
| **P2RX6** | 0.00421 | 0.03234 | 0.00284 | 0.00167 | 0.00000 | 0.00000 | 0.00508 | 0.00763 | 0.28016 | 0.11951 | 0.21551 | 0.08883 |
| **P2RX7** | 0.02486 | 0.00677 | 0.03039 | 0.01081 | 0.00002 | 0.02249 | 0.11722 | 0.00626 | 0.00000 | 0.04112 | 0.04123 | 0.10744 |
| **P2RY1** | 0.03834 | 0.09985 | 0.13550 | 0.12977 | 0.00000 | 0.00000 | 0.00000 | 0.33909 | 0.11600 | 0.06995 | 0.01246 | 0.13324 |
| **P2RY11** | 0.00172 | 0.16193 | 0.16107 | 0.05168 | 0.15089 | 0.00000 | 0.40168 | 0.06543 | 0.02815 | 0.67758 | 0.04104 | 0.17925 |
| **P2RY12** | 0.00321 | 0.00717 | 0.98590 | 0.13322 | 0.03425 | 0.00000 | 0.00000 | 0.00705 | 0.16542 | 0.40049 | 0.04435 | 0.08266 |
| **P2RY13** | 0.00220 | 0.00000 | 1.41164 | 0.20798 | 0.03053 | 0.00000 | 0.45378 | 0.00626 | 0.00000 | 1.06831 | 0.19702 | 0.03641 |
| **P2RY14** | 0.00239 | 0.00000 | 2.15353 | 0.00000 | 0.00000 | 0.00000 | 0.00000 | 0.03294 | 1.37929 | 1.41868 | 0.47880 | 0.25609 |
| **P2RY2** | 0.00523 | 0.00000 | 1.43960 | 0.02540 | 1.13413 | 0.11111 | 0.98212 | 0.00857 | 0.26678 | 0.00000 | 0.24705 | 0.00810 |
| **P2RY6** | 0.00881 | 0.63449 | 22.80714 | 2.58543 | 11.63223 | 3.74140 | 6.00596 | 0.00000 | 19.04957 | 0.00000 | 0.55196 | 0.00000 |
| **P4HA1** | 2.49536 | 2.74272 | 2.95139 | 9.56342 | 0.00000 | 1.02000 | 0.00000 | 2.88982 | 9.89394 | 2.40708 | 4.16692 | 9.16707 |
| **P4HA2** | 0.00000 | 5.62466 | 1.31500 | 0.94725 | 2.21018 | 4.90379 | 0.00000 | 0.55814 | 2.13353 | 1.08856 | 1.26623 | 1.51132 |
| **P4HA2-AS1** | 0.00310 | 0.06011 | 0.01193 | 0.02763 | 0.00000 | 0.25984 | 0.00000 | 0.01098 | 0.00000 | 0.11509 | 0.03254 | 0.05480 |
| **P4HA3** | 0.00402 | 0.34474 | 0.03397 | 0.06784 | 0.00000 | 0.00449 | 0.01305 | 0.10575 | 0.02443 | 0.02599 | 0.01926 | 0.37669 |
| **P4HB** | 6.44210 | 31.76013 | 33.84615 | 25.94419 | 35.49608 | 13.21893 | 15.66044 | 6.15104 | 67.46297 | 22.49532 | 25.49147 | 27.05459 |
| **P4HTM** | 0.00610 | 0.06007 | 0.42588 | 0.02280 | 0.00000 | 0.00037 | 0.16991 | 0.01134 | 0.14390 | 0.00000 | 0.15679 | 0.02016 |
| **PA2G4** | 3.68458 | 26.13057 | 29.20855 | 18.93841 | 29.11613 | 23.68412 | 45.34287 | 11.34051 | 32.56905 | 25.82818 | 28.99996 | 30.01547 |
| **PA2G4P4** | 0.17927 | 4.16242 | 4.40641 | 2.20712 | 3.58801 | 5.62095 | 9.67465 | 1.41338 | 5.93803 | 4.37694 | 5.01871 | 4.76539 |
| **PAAF1** | 0.00103 | 7.66357 | 0.69332 | 4.01110 | 4.60355 | 7.09325 | 0.00000 | 2.02890 | 0.42424 | 0.00000 | 5.57359 | 3.49226 |
| **PABPC1** | 7.00537 | 24.76373 | 28.91955 | 30.96160 | 47.54349 | 42.04960 | 55.20456 | 15.64480 | 36.56511 | 66.13014 | 33.15213 | 35.43825 |
| **PABPC1L** | 3.12605 | 0.15452 | 0.77746 | 1.18582 | 0.00000 | 0.00000 | 2.26366 | 0.22304 | 0.48964 | 3.04278 | 0.63504 | 0.17187 |
| **PABPC1P2** | 0.04856 | 0.02364 | 0.02721 | 0.03404 | 0.12175 | 0.13567 | 0.16918 | 0.03312 | 0.00000 | 0.23785 | 0.09049 | 0.12524 |
| **PABPC3** | 1.42784 | 2.42609 | 6.14029 | 7.07985 | 13.61413 | 12.26642 | 15.91297 | 1.47165 | 5.93618 | 29.30401 | 6.00974 | 6.77242 |
| **PABPC4** | 5.23435 | 15.61876 | 3.68806 | 2.79142 | 24.72313 | 18.49116 | 27.75467 | 2.25051 | 14.02296 | 44.75592 | 10.28817 | 20.87942 |
| **PABPC4L** | 0.14430 | 0.00000 | 0.03118 | 0.03927 | 0.00000 | 0.00000 | 0.02720 | 0.00287 | 0.01141 | 0.25507 | 0.06568 | 0.17306 |
| **PABPN1** | 0.09985 | 0.16973 | 0.80427 | 0.17585 | 0.79167 | 0.00000 | 0.30266 | 0.29395 | 0.38057 | 1.18611 | 0.00000 | 0.57719 |
| **PABPN1L** | 19.58602 | 0.03881 | 0.00127 | 0.00170 | 0.03813 | 0.00000 | 0.00642 | 13.07569 | 0.01840 | 0.00000 | 0.03227 | 0.47871 |
| **PACRG** | 0.09453 | 0.05959 | 0.05284 | 0.00418 | 0.05992 | 0.00000 | 0.03334 | 0.02091 | 0.00000 | 0.08287 | 0.01662 | 0.00982 |
| **PACRGL** | 1.29690 | 3.46150 | 0.73760 | 0.27926 | 0.00000 | 5.77860 | 0.00000 | 0.81426 | 3.00550 | 0.00000 | 0.11530 | 3.96799 |
| **PACS1** | 2.62207 | 0.29615 | 0.37805 | 0.20695 | 1.85293 | 0.00000 | 1.66266 | 0.02568 | 1.14622 | 2.24368 | 0.85822 | 0.20002 |
| **PACS2** | 2.51269 | 0.98563 | 0.56488 | 0.24044 | 2.10350 | 0.00000 | 0.64241 | 0.30635 | 0.70577 | 0.76159 | 0.19867 | 0.21483 |
| **PACSIN1** | 0.04691 | 0.54734 | 0.08632 | 0.37371 | 0.83497 | 0.00000 | 0.00000 | 0.05113 | 1.17808 | 0.08220 | 0.00334 | 0.03972 |
| **PACSIN2** | 2.93052 | 15.91663 | 13.23480 | 10.57835 | 14.60419 | 2.80586 | 13.15645 | 7.36769 | 12.85454 | 0.00000 | 14.86012 | 6.94707 |
| **PACSIN3** | 0.71324 | 6.56938 | 6.29682 | 9.73104 | 19.90838 | 0.00000 | 4.41493 | 0.48987 | 2.11744 | 3.28935 | 12.24345 | 3.24886 |
| **PADI1** | 0.03660 | 0.00386 | 0.05083 | 0.02408 | 0.00667 | 0.02308 | 0.01483 | 0.05407 | 0.00000 | 0.02687 | 0.04606 | 0.11901 |
| **PADI2** | 0.04312 | 0.09184 | 0.00000 | 0.00224 | 0.05738 | 0.02399 | 0.18536 | 0.00762 | 0.06192 | 0.00000 | 0.24746 | 0.08096 |
| **PADI3** | 0.02182 | 0.01661 | 0.02799 | 0.12550 | 0.00765 | 0.00551 | 0.02860 | 0.10263 | 0.07940 | 0.01058 | 0.01396 | 0.01187 |
| **PADI4** | 0.00608 | 0.01272 | 0.01263 | 0.01373 | 0.00495 | 0.03002 | 0.02808 | 0.01049 | 0.04788 | 0.01436 | 0.02032 | 0.01561 |
| **PADI6** | 18.77367 | 0.16108 | 0.01143 | 0.00000 | 0.00057 | 0.07608 | 0.00000 | 9.65285 | 0.00000 | 0.03742 | 0.01231 | 0.75287 |
| **PAEP** | 0.00607 | 0.01428 | 0.18174 | 0.01055 | 0.01589 | 0.01814 | 0.00711 | 0.01048 | 0.00000 | 0.02143 | 0.01888 | 0.01647 |
| **PAF1** | 4.32028 | 16.17731 | 9.06211 | 13.52467 | 13.74935 | 1.85177 | 17.71551 | 6.40151 | 12.04901 | 0.00000 | 12.01339 | 17.08477 |
| **PAFAH1B1** | 1.81250 | 9.54570 | 11.19839 | 5.56410 | 8.00959 | 17.20049 | 18.68771 | 4.44121 | 4.01722 | 16.77516 | 6.36287 | 9.09687 |

| | | | | | | | | | | | | |
|---|---|---|---|---|---|---|---|---|---|---|---|---|
| **PAFAH1B2** | 6.59512 | 16.34277 | 14.46398 | 12.93698 | 8.96759 | 38.27289 | 21.04013 | 32.23820 | 10.17559 | 14.30021 | 20.40452 | 21.57383 |
| **PAFAH1B3** | 0.33619 | 9.88216 | 7.11930 | 11.64722 | 10.75266 | 0.53653 | 18.68629 | 4.65915 | 14.79857 | 0.00000 | 11.96665 | 12.07694 |
| **PAFAH2** | 0.95080 | 12.40001 | 3.20478 | 1.72013 | 3.53998 | 10.21800 | 2.26644 | 3.21959 | 3.30890 | 0.56360 | 5.59434 | 8.21695 |
| **PAG1** | 5.98663 | 0.26267 | 0.38583 | 0.60037 | 0.09036 | 0.00000 | 0.00278 | 0.79703 | 0.03218 | 0.55439 | 0.00000 | 0.13311 |
| **PAGE1** | 0.24836 | 0.63646 | 0.00000 | 0.48902 | 2.44973 | 0.00000 | 0.64208 | 0.01012 | 1.50296 | 0.00000 | 2.24241 | 0.02195 |
| **PAGE2** | 0.00000 | 0.00000 | 1.33612 | 0.43990 | 0.00000 | 3.45821 | 3.77739 | 0.06719 | 0.00000 | 1.88580 | 0.36675 | 0.70205 |
| **PAGE2B** | 0.07133 | 1.52828 | 0.30377 | 0.00000 | 0.44651 | 0.09205 | 0.77874 | 0.03976 | 0.93388 | 0.00000 | 0.06393 | 7.02631 |
| **PAGE4** | 0.21944 | 0.01405 | 13.95560 | 0.00237 | 0.00000 | 0.03400 | 0.53242 | 0.11836 | 0.15970 | 0.00000 | 0.03646 | 0.03237 |
| **PAGE5** | 0.00091 | 3.97915 | 0.00000 | 0.00000 | 0.00000 | 2.41651 | 1.68761 | 0.01326 | 0.84073 | 0.04805 | 0.00000 | 4.30408 |
| **PAGR1** | 5.26700 | 15.66320 | 21.40601 | 6.25986 | 7.27961 | 0.00000 | 6.18089 | 0.82927 | 12.51927 | 14.31927 | 14.70964 | 4.79124 |
| **PAH** | 0.00594 | 0.01316 | 0.05729 | 0.01404 | 0.01138 | 0.02046 | 0.00920 | 0.01034 | 0.00712 | 0.02915 | 0.02214 | 0.01819 |
| **PAICS** | 0.89412 | 19.02086 | 21.49455 | 11.28518 | 9.60441 | 35.68726 | 36.02052 | 0.68371 | 28.31232 | 11.36178 | 29.91006 | 4.12249 |
| **PAIP1** | 18.53923 | 8.85357 | 3.25286 | 2.11305 | 4.84039 | 25.31379 | 0.00000 | 4.78837 | 1.27251 | 4.32479 | 5.78671 | 0.64107 |
| **PAIP2** | 6.31053 | 14.91869 | 18.50906 | 11.96698 | 3.84020 | 43.30533 | 25.74343 | 15.69035 | 17.58080 | 8.13179 | 13.93637 | 25.51179 |
| **PAIP2B** | 7.41044 | 0.09940 | 4.29554 | 1.04676 | 0.19892 | 1.15352 | 3.38342 | 0.92841 | 0.00000 | 0.00000 | 4.76186 | 0.27404 |
| **PAK1** | 4.36221 | 1.31769 | 2.04471 | 9.37553 | 3.11988 | 7.88569 | 2.27512 | 0.20767 | 1.10295 | 2.20315 | 0.00000 | 1.04938 |
| **PAK1IP1** | 6.27575 | 15.34382 | 15.23316 | 7.97614 | 6.44879 | 30.57614 | 17.34485 | 12.15212 | 15.85267 | 7.32821 | 17.96778 | 21.77008 |
| **PAK2** | 5.56226 | 3.34656 | 7.91035 | 5.80457 | 4.35045 | 19.52553 | 14.05596 | 0.74462 | 10.16268 | 19.03481 | 9.55057 | 3.75634 |
| **PAK3** | 0.04321 | 0.01491 | 2.64475 | 0.04063 | 0.02738 | 0.01070 | 0.11976 | 0.01023 | 0.00000 | 0.01044 | 0.01416 | 0.02492 |
| **PAK4** | 2.29114 | 2.44279 | 9.45099 | 2.35385 | 6.28878 | 0.00000 | 9.09778 | 0.22258 | 2.20756 | 15.12692 | 0.00000 | 2.86407 |
| **PAK6** | 0.32785 | 1.07961 | 2.17809 | 0.18196 | 0.73394 | 0.41177 | 1.65742 | 0.00374 | 3.55218 | 0.00000 | 0.28562 | 0.11586 |
| **PAK7** | 0.58346 | 0.02571 | 0.00495 | 0.00255 | 0.01746 | 0.07459 | 0.00000 | 0.04380 | 0.04233 | 0.01226 | 0.01655 | 0.00494 |
| **PALB2** | 5.05991 | 0.65960 | 0.99461 | 1.20170 | 1.04919 | 5.83070 | 2.42224 | 1.24244 | 1.39442 | 2.97054 | 3.33172 | 3.32807 |
| **PALD1** | 0.05043 | 0.00451 | 0.10601 | 0.09890 | 0.00000 | 0.05408 | 0.00000 | 0.11027 | 0.02345 | 0.02557 | 0.03370 | 0.03375 |
| **PALLD** | 1.91853 | 1.29528 | 8.43971 | 0.00000 | 11.49974 | 4.73485 | 29.38421 | 0.00000 | 10.96471 | 18.78192 | 15.21973 | 1.77717 |
| **PALM** | 0.00616 | 0.11902 | 0.01177 | 0.02261 | 0.01579 | 0.00000 | 0.03112 | 0.42277 | 0.00000 | 0.28415 | 0.00000 | 2.64821 |
| **PALM2** | 0.34749 | 0.01732 | 0.03216 | 0.02549 | 0.03284 | 0.00000 | 0.04148 | 0.01011 | 0.02765 | 0.00000 | 0.01814 | 0.01798 |
| **PALM3** | 0.93780 | 0.00000 | 1.58096 | 0.00000 | 0.16821 | 0.00000 | 1.70443 | 0.34169 | 0.29542 | 0.61882 | 0.00000 | 1.23388 |
| **PALMD** | 0.01216 | 0.00000 | 0.00000 | 0.00000 | 0.00000 | 0.14737 | 0.03321 | 0.00697 | 0.05860 | 0.10386 | 0.00000 | 0.02158 |
| **PAM** | 0.12066 | 3.95477 | 0.65566 | 0.00000 | 0.52596 | 1.71955 | 0.00000 | 0.02082 | 0.00000 | 0.85154 | 0.87652 | 0.08868 |
| **PAM16** | 0.22731 | 0.90991 | 0.42104 | 0.37616 | 0.78444 | 0.00000 | 0.22977 | 1.30520 | 0.60806 | 0.51792 | 0.76835 | 1.17971 |
| **PAMR1** | 2.79679 | 1.81578 | 0.00000 | 4.53746 | 0.73061 | 0.08093 | 0.00000 | 0.56814 | 0.00000 | 0.00535 | 0.00000 | 0.05309 |
| **PAN2** | 4.31752 | 1.04337 | 0.72468 | 2.34687 | 0.45484 | 0.87890 | 0.00000 | 0.93256 | 0.34805 | 1.22699 | 0.59412 | 0.63758 |
| **PAN3** | 3.69469 | 3.13426 | 1.14999 | 9.47778 | 0.00000 | 26.84089 | 5.62744 | 15.87200 | 1.25818 | 6.34187 | 9.60533 | 6.33199 |
| **PAN3-AS1** | 0.15901 | 0.02532 | 0.11854 | 0.01631 | 0.26222 | 0.23958 | 0.00000 | 0.01408 | 0.41790 | 0.00000 | 0.21403 | 0.26706 |
| **PANDAR** | 0.00205 | 1.14687 | 2.60369 | 1.31382 | 0.30940 | 2.01399 | 1.02713 | 1.46202 | 1.19575 | 0.00000 | 1.84886 | 1.01454 |
| **PANK1** | 0.30271 | 0.00000 | 2.98345 | 2.28370 | 0.28832 | 0.60960 | 3.30601 | 0.02926 | 0.65763 | 0.00000 | 0.18527 | 0.00000 |
| **PANK2** | 0.47859 | 0.22427 | 2.49325 | 2.56245 | 2.66799 | 6.42792 | 6.26692 | 1.51847 | 2.13109 | 6.54920 | 1.50746 | 0.83110 |
| **PANK3** | 2.80165 | 4.52789 | 1.17309 | 4.97250 | 2.65699 | 17.82868 | 3.93206 | 0.00000 | 1.70078 | 10.33249 | 1.87107 | 5.19048 |
| **PANK4** | 0.13216 | 1.89389 | 3.26065 | 2.72819 | 0.96315 | 1.15068 | 3.56341 | 0.23135 | 1.13805 | 0.47597 | 5.54925 | 0.90732 |
| **PANO** | 0.00413 | 0.01222 | 0.00291 | 0.00075 | 0.05607 | 0.00000 | 0.00564 | 0.01020 | 0.05896 | 0.34154 | 0.00000 | 0.01739 |
| **PANX1** | 4.45268 | 2.73827 | 3.23087 | 0.00000 | 1.31158 | 1.50242 | 0.00000 | 6.14819 | 4.77954 | 6.18218 | 0.72613 | 5.43070 |
| **PANX2** | 0.00903 | 0.22020 | 0.25457 | 0.11753 | 0.52261 | 0.00000 | 0.22288 | 0.01310 | 0.00000 | 0.22512 | 0.00000 | 0.05042 |
| **PANX3** | 0.00573 | 0.01065 | 0.01636 | 0.07583 | 0.00139 | 0.03645 | 0.01650 | 0.01011 | 0.01149 | 0.03554 | 0.01235 | 0.01476 |
| **PAOX** | 1.43481 | 0.55571 | 0.19840 | 0.28023 | 1.51072 | 0.00000 | 0.16851 | 0.25781 | 0.00000 | 0.16849 | 0.19833 | 0.12059 |
| **PAPD4** | 3.68819 | 4.26568 | 5.82205 | 3.24496 | 1.38915 | 15.57899 | 4.04681 | 5.35412 | 7.05942 | 2.32284 | 7.58223 | 4.46430 |
| **PAPD5** | 3.42337 | 1.21257 | 4.86789 | 3.41090 | 3.34438 | 9.96308 | 14.23253 | 7.67111 | 0.00000 | 26.73827 | 3.65139 | 4.43517 |
| **PAPD7** | 15.49000 | 2.63121 | 2.53514 | 1.23101 | 2.87593 | 5.00935 | 1.04560 | 1.86399 | 7.36709 | 12.67232 | 0.36977 | 1.95272 |

| | | | | | | | | | | | | |
|---|---|---|---|---|---|---|---|---|---|---|---|---|
| **PAPL** | 0.03657 | 0.04265 | 0.00959 | 0.14324 | 0.00000 | 0.01182 | 0.02627 | 0.00450 | 0.02599 | 0.06802 | 0.00954 | 0.15679 |
| **PAPLN** | 0.24791 | 0.18431 | 0.32074 | 0.56991 | 0.15223 | 0.31494 | 0.00000 | 0.35960 | 0.03543 | 0.44087 | 0.00000 | 0.09422 |
| **PAPOLA** | 5.27123 | 18.28135 | 56.17488 | 4.19077 | 11.23724 | 38.67667 | 35.54733 | 19.00790 | 20.29499 | 27.89740 | 18.45012 | 29.93210 |
| **PAPOLB** | 0.11686 | 0.52304 | 5.05832 | 0.01108 | 0.14857 | 1.32989 | 0.00000 | 1.05402 | 0.43559 | 0.19602 | 0.77199 | 1.19057 |
| **PAPOLG** | 4.10185 | 3.89379 | 0.92344 | 2.26727 | 0.00000 | 12.25037 | 2.03555 | 5.85965 | 2.44081 | 3.63643 | 2.32124 | 4.18866 |
| **PAPPA** | 0.13485 | 0.24381 | 0.00000 | 0.12034 | 0.32972 | 0.09520 | 0.00000 | 0.00000 | 0.00000 | 0.58334 | 0.08772 | 0.06947 |
| **PAPPA2** | 0.31098 | 0.00358 | 0.02245 | 0.07628 | 0.00000 | 0.02283 | 0.00837 | 0.00000 | 0.01944 | 0.15673 | 0.02389 | 0.05829 |
| **PAPSS1** | 3.63245 | 3.87284 | 6.34104 | 7.78063 | 0.00000 | 6.23246 | 3.63736 | 1.69710 | 6.53963 | 0.00000 | 5.23153 | 2.84523 |
| **PAPSS2** | 0.39286 | 0.25825 | 0.24680 | 1.74754 | 0.00000 | 0.17708 | 0.00000 | 0.16497 | 0.04355 | 0.15297 | 0.02266 | 0.23162 |
| **PAQR3** | 0.00000 | 0.18567 | 2.27746 | 2.16524 | 0.76009 | 13.86501 | 2.49519 | 0.00000 | 0.00000 | 6.95810 | 0.73496 | 0.73427 |
| **PAQR4** | 0.14071 | 1.45672 | 1.08614 | 0.73642 | 0.45852 | 0.00000 | 0.00005 | 0.00000 | 0.92980 | 1.71397 | 0.17107 | 0.14862 |
| **PAQR5** | 1.22668 | 0.76795 | 0.05673 | 0.19121 | 0.00000 | 0.46323 | 0.00000 | 0.95921 | 0.00000 | 0.21622 | 0.09501 | 0.79553 |
| **PAQR6** | 0.00043 | 0.26120 | 0.48434 | 0.19411 | 0.05529 | 0.00000 | 0.00000 | 0.04954 | 0.26965 | 1.80004 | 0.00000 | 0.72351 |
| **PAQR7** | 4.62339 | 0.66287 | 4.20326 | 0.79458 | 0.57058 | 0.00000 | 0.47398 | 6.05423 | 0.00000 | 1.87300 | 2.79807 | 0.48945 |
| **PAQR8** | 0.00094 | 0.02727 | 0.60298 | 2.62575 | 0.20585 | 0.19732 | 0.24863 | 0.09264 | 0.00000 | 0.32667 | 0.15774 | 0.37476 |
| **PAQR9** | 0.03891 | 0.01425 | 0.02722 | 0.01074 | 0.01437 | 0.02349 | 0.02083 | 0.01059 | 0.01318 | 0.01002 | 0.01553 | 0.01510 |
| **PARD3** | 6.39365 | 3.53989 | 3.74740 | 14.21355 | 10.17725 | 7.73300 | 9.66712 | 2.86070 | 0.13673 | 16.63959 | 0.75188 | 6.67551 |
| **PARD3-AS1** | 0.01018 | 0.01885 | 0.01217 | 0.02062 | 0.01415 | 0.03226 | 0.01906 | 0.13041 | 0.04414 | 0.00000 | 0.01600 | 0.04509 |
| **PARD3B** | 2.48112 | 0.00000 | 0.00000 | 0.00000 | 1.01706 | 0.00000 | 0.20492 | 0.42999 | 0.47177 | 1.28008 | 0.00000 | 0.22057 |
| **PARD6A** | 0.07132 | 0.02870 | 0.99628 | 0.00000 | 0.34616 | 0.00000 | 0.28604 | 0.21569 | 0.00000 | 0.07711 | 1.17053 | 0.00298 |
| **PARD6B** | 0.16508 | 9.54435 | 11.86624 | 1.82354 | 0.00000 | 37.55629 | 16.84176 | 18.31719 | 6.49384 | 15.35738 | 22.71109 | 11.65335 |
| **PARD6G** | 2.43414 | 0.78265 | 0.11572 | 0.00000 | 0.00000 | 0.42500 | 0.23610 | 0.93319 | 0.14375 | 0.01058 | 0.00000 | 0.59048 |
| **PARD6G-AS1** | 0.16839 | 0.01178 | 0.07248 | 0.00810 | 0.00000 | 0.05981 | 0.00000 | 0.01307 | 0.04355 | 0.04442 | 0.06578 | 0.08605 |
| **PARG** | 0.86986 | 2.85071 | 1.10549 | 1.45688 | 0.00000 | 6.93503 | 3.63809 | 1.03371 | 2.42971 | 2.12161 | 2.61748 | 4.95296 |
| **PARK2** | 0.26626 | 2.51182 | 0.00000 | 2.12665 | 3.37988 | 0.00000 | 0.00603 | 1.18880 | 0.00000 | 0.00000 | 0.23395 | 5.62577 |
| **PARK7** | 3.80848 | 20.78333 | 26.01034 | 20.35438 | 25.29273 | 36.17568 | 36.04583 | 8.91877 | 31.02045 | 2.45819 | 30.66000 | 17.01307 |
| **PARL** | 18.01288 | 11.68691 | 10.80363 | 6.99190 | 6.54436 | 15.69454 | 11.98629 | 24.33986 | 10.68256 | 6.30855 | 8.76244 | 12.77270 |
| **PARM1** | 0.01895 | 1.29882 | 0.07250 | 1.89983 | 0.05515 | 0.38720 | 0.00000 | 0.04986 | 0.00000 | 0.00965 | 0.01543 | 0.95083 |
| **PARN** | 4.79981 | 16.31929 | 6.53782 | 12.87798 | 7.60596 | 19.03200 | 9.29245 | 7.05866 | 9.60467 | 4.93263 | 8.54183 | 12.92059 |
| **PARP1** | 11.07487 | 23.39491 | 16.65648 | 29.89083 | 16.69971 | 16.51636 | 33.10103 | 2.71638 | 25.48223 | 10.98554 | 22.79577 | 19.34161 |
| **PARP10** | 0.00946 | 0.00728 | 0.15322 | 0.05177 | 0.00713 | 0.00000 | 0.00522 | 0.00535 | 0.00000 | 0.02913 | 0.07663 | 0.13218 |
| **PARP11** | 0.96887 | 0.59696 | 0.53302 | 0.53587 | 0.22098 | 2.22487 | 0.00000 | 2.89115 | 1.23471 | 0.00000 | 0.07788 | 1.97534 |
| **PARP12** | 7.06442 | 0.67431 | 0.00000 | 1.10610 | 0.73862 | 0.42133 | 0.00000 | 0.08254 | 0.00000 | 0.00472 | 0.00000 | 0.07422 |
| **PARP14** | 0.03888 | 0.18587 | 0.05468 | 0.90277 | 0.09268 | 0.00000 | 0.00164 | 0.17426 | 0.10591 | 0.16091 | 0.00122 | 0.44122 |
| **PARP15** | 0.01232 | 0.01265 | 0.06292 | 0.02910 | 0.00686 | 0.02797 | 0.01467 | 0.01196 | 0.00735 | 0.03199 | 0.03233 | 0.01382 |
| **PARP16** | 5.91831 | 2.25503 | 0.56346 | 3.95156 | 0.00000 | 0.45899 | 0.69407 | 4.47934 | 0.00000 | 0.06943 | 0.07840 | 1.87723 |
| **PARP2** | 2.48131 | 11.38693 | 8.49469 | 4.57221 | 7.28420 | 30.97060 | 7.08700 | 6.62715 | 17.10305 | 1.73875 | 13.26168 | 22.69647 |
| **PARP3** | 0.02261 | 0.05331 | 0.13855 | 0.13494 | 0.09412 | 0.07911 | 0.00000 | 0.00944 | 0.00603 | 0.03240 | 0.02634 | 0.02789 |
| **PARP4** | 0.16124 | 1.90220 | 0.87626 | 1.16016 | 0.02911 | 3.46977 | 0.00000 | 0.00000 | 0.00017 | 0.69205 | 0.00000 | 0.97307 |
| **PARP6** | 0.43617 | 0.89871 | 1.30320 | 2.98128 | 2.09617 | 0.83400 | 1.34138 | 0.29316 | 2.79475 | 5.26489 | 0.77945 | 0.94764 |
| **PARP8** | 0.27116 | 0.04516 | 0.39677 | 0.19744 | 0.00000 | 0.08391 | 0.07875 | 0.36819 | 1.06465 | 0.00000 | 0.00000 | 0.57784 |
| **PARP9** | 0.16869 | 0.20697 | 0.01505 | 1.37220 | 0.03263 | 0.17728 | 0.00000 | 0.04018 | 0.00000 | 0.02288 | 0.00698 | 0.00000 |
| **PARPBP** | 3.12843 | 3.19647 | 4.61199 | 2.08332 | 0.71532 | 16.56346 | 5.38761 | 4.53037 | 6.18690 | 0.00000 | 6.48321 | 6.85384 |
| **PARS2** | 2.52243 | 4.92196 | 1.52109 | 0.80579 | 1.18075 | 5.83426 | 0.00000 | 1.03672 | 1.59950 | 0.00000 | 2.48254 | 5.97861 |
| **PART1** | 0.22161 | 0.00773 | 0.07764 | 0.01585 | 0.00000 | 0.03618 | 0.06708 | 0.16934 | 0.07923 | 0.04421 | 0.03252 | 0.02529 |
| **PARVA** | 1.24544 | 9.11697 | 13.65389 | 3.35786 | 6.76153 | 12.34039 | 11.11623 | 2.73338 | 5.06570 | 0.00000 | 7.97081 | 10.32180 |
| **PARVB** | 0.24783 | 0.17953 | 0.08880 | 0.30708 | 0.03327 | 0.00000 | 0.04747 | 0.09768 | 0.26615 | 0.05413 | 0.26769 | 0.09545 |
| **PARVG** | 3.12900 | 0.01298 | 0.02435 | 0.01173 | 0.01817 | 0.00166 | 0.01801 | 0.13684 | 0.02009 | 0.00000 | 0.01424 | 0.06552 |

| | | | | | | | | | | | | |
|---|---|---|---|---|---|---|---|---|---|---|---|---|
| **PASD1** | 0.00221 | 0.06263 | 0.01285 | 0.07615 | 0.01229 | 0.00000 | 0.01959 | 0.10431 | 0.00671 | 0.02354 | 0.04514 | 0.56854 |
| **PASK** | 1.92683 | 6.22491 | 4.56986 | 2.68297 | 2.17902 | 1.21888 | 3.40056 | 0.00000 | 4.48679 | 5.78848 | 5.31715 | 1.60942 |
| **PATE1** | 0.00538 | 0.01638 | 0.04071 | 0.00727 | 0.01537 | 0.01344 | 0.01135 | 0.00955 | 0.02215 | 0.05786 | 0.01115 | 0.03623 |
| **PATE2** | 0.00426 | 0.32895 | 0.17248 | 0.00000 | 0.21523 | 0.00000 | 0.04183 | 0.00502 | 0.05733 | 0.23804 | 0.00000 | 0.15965 |
| **PATE3** | 0.00420 | 0.01008 | 0.01150 | 0.03870 | 0.02937 | 0.05827 | 0.02236 | 0.00782 | 0.00114 | 0.09545 | 0.02633 | 0.05369 |
| **PATE4** | 0.00599 | 0.01756 | 0.01647 | 0.01461 | 0.01555 | 0.00639 | 0.02155 | 0.00963 | 0.01939 | 0.01864 | 0.02082 | 0.03905 |
| **PATL1** | 4.44833 | 0.75541 | 2.24506 | 3.27325 | 5.59370 | 10.10881 | 7.13696 | 2.67765 | 4.71758 | 14.27295 | 3.11600 | 4.46763 |
| **PATL2** | 13.33327 | 0.00565 | 0.00000 | 0.00000 | 0.01207 | 0.09640 | 0.00013 | 3.92091 | 0.00013 | 0.43142 | 0.05606 | 0.12959 |
| **PATZ1** | 0.06828 | 0.79338 | 0.69151 | 1.02186 | 0.93079 | 0.10168 | 4.12256 | 0.00000 | 0.01359 | 4.23766 | 1.33243 | 1.78149 |
| **PAWR** | 0.59321 | 0.00000 | 3.55006 | 0.00156 | 3.92143 | 5.04105 | 6.43606 | 0.00000 | 0.00000 | 12.19825 | 1.22408 | 0.46115 |
| **PAX1** | 0.00618 | 0.01439 | 0.01844 | 0.01318 | 0.01260 | 0.01886 | 0.01999 | 0.01049 | 0.01246 | 0.01659 | 0.02681 | 0.01613 |
| **PAX2** | 0.00441 | 0.02567 | 0.01318 | 0.01232 | 0.03527 | 0.00094 | 0.07777 | 0.04327 | 0.00000 | 0.05520 | 0.02212 | 0.02153 |
| **PAX3** | 0.02834 | 0.00467 | 0.02156 | 0.01211 | 0.00507 | 0.03968 | 0.02692 | 0.00423 | 0.14413 | 0.00000 | 0.01195 | 0.13472 |
| **PAX4** | 0.00612 | 0.01449 | 0.08505 | 0.00741 | 0.01450 | 0.02664 | 0.00731 | 0.00968 | 0.00587 | 0.01504 | 0.01773 | 0.02618 |
| **PAX5** | 0.05966 | 0.10203 | 0.28041 | 0.08328 | 0.00000 | 0.04382 | 0.00000 | 0.01443 | 0.00000 | 0.43982 | 0.00000 | 0.21335 |
| **PAX6** | 0.26052 | 0.15253 | 0.30572 | 0.33973 | 0.00000 | 0.20397 | 0.00000 | 0.66869 | 0.00000 | 1.19829 | 0.29803 | 0.30546 |
| **PAX7** | 0.00908 | 0.01552 | 0.02075 | 0.03046 | 0.02630 | 0.00883 | 0.04220 | 0.01023 | 0.02024 | 0.00000 | 0.01205 | 0.09072 |
| **PAX8** | 1.45533 | 0.28088 | 0.23385 | 0.07762 | 0.00000 | 0.00000 | 0.05408 | 0.00000 | 0.22379 | 0.02188 | 0.30295 | 0.51962 |
| **PAX8-AS1.uid-3** | 0.32734 | 1.31835 | 2.20000 | 1.77897 | 2.54524 | 1.02041 | 1.53378 | 0.00000 | 2.92418 | 0.09462 | 1.54441 | 0.71435 |
| **PAX8-AS1.uid-3** | 0.00000 | 5.16879 | 4.29931 | 1.75383 | 1.99330 | 2.70852 | 4.33628 | 0.28869 | 2.84298 | 1.09227 | 7.06619 | 2.43716 |
| **PAX9** | 2.81913 | 0.01906 | 0.24898 | 0.00027 | 0.06684 | 0.00000 | 0.02750 | 0.52030 | 0.70189 | 0.00000 | 0.02729 | 1.79916 |
| **PAXBP1** | 1.50349 | 1.68038 | 3.34730 | 0.13471 | 2.33389 | 12.89123 | 4.94628 | 6.69683 | 0.31307 | 16.82621 | 0.81846 | 2.24724 |
| **PAXBP1-AS1** | 0.09127 | 0.01576 | 0.25759 | 0.00000 | 0.00000 | 0.18410 | 0.00000 | 0.27393 | 0.05173 | 0.67990 | 0.02384 | 0.18335 |
| **PAXIP1** | 6.07406 | 5.59373 | 1.08038 | 6.66918 | 0.51139 | 5.19453 | 4.77998 | 0.00000 | 3.41016 | 9.06113 | 0.86132 | 4.78355 |
| **PAXIP1-AS1** | 0.23252 | 1.93407 | 1.49068 | 0.81478 | 0.00000 | 0.53330 | 0.00000 | 0.01677 | 0.77858 | 1.10248 | 1.96014 | 0.96268 |
| **PAXIP1OS** | 0.31520 | 0.29021 | 0.04971 | 0.46305 | 0.09226 | 0.27862 | 0.00115 | 0.04387 | 0.22324 | 0.81971 | 0.10044 | 0.21217 |
| **PBDC1** | 0.00000 | 5.45833 | 6.46313 | 8.28020 | 15.80011 | 17.71536 | 21.44559 | 4.85750 | 12.94788 | 7.10066 | 13.45257 | 16.67897 |
| **PBK** | 0.56993 | 7.51168 | 0.72411 | 2.05125 | 0.00000 | 23.66987 | 4.52443 | 0.72374 | 3.00544 | 4.55830 | 4.53387 | 7.16151 |
| **PBLD** | 0.10901 | 0.61824 | 0.07461 | 0.15424 | 0.00000 | 0.54028 | 0.00000 | 0.18694 | 0.00000 | 0.18410 | 0.00000 | 0.27577 |
| **PBOV1** | 0.00518 | 0.01598 | 0.01788 | 0.01316 | 0.01512 | 0.02145 | 0.02338 | 0.01421 | 0.02107 | 0.01443 | 0.01407 | 0.01560 |
| **PBRM1** | 3.55362 | 0.00000 | 7.93234 | 1.83507 | 4.40419 | 13.11173 | 13.49630 | 1.11549 | 4.36081 | 23.43959 | 4.10711 | 2.80040 |
| **PBX1** | 3.39499 | 0.00000 | 0.00000 | 3.87028 | 1.45176 | 0.67669 | 0.03850 | 0.45069 | 0.00000 | 0.75530 | 0.09687 | 0.12672 |
| **PBX2** | 0.10332 | 0.17485 | 0.65746 | 1.12292 | 0.99126 | 0.00000 | 0.76698 | 0.00889 | 0.05532 | 3.40964 | 0.65917 | 0.45463 |
| **PBX3** | 1.81655 | 2.70281 | 0.12797 | 7.68714 | 1.17825 | 1.79176 | 0.50335 | 2.38083 | 0.00000 | 2.72690 | 0.00000 | 3.80362 |
| **PBX4** | 0.00357 | 3.06727 | 0.54567 | 2.19835 | 0.14448 | 0.00027 | 0.63187 | 0.00609 | 1.79606 | 1.50107 | 0.76640 | 4.53263 |
| **PBXIP1** | 0.05784 | 0.50827 | 0.92071 | 0.97495 | 1.75015 | 0.00000 | 0.30013 | 0.19226 | 1.95439 | 1.49019 | 1.07343 | 0.07211 |
| **PC** | 0.12813 | 1.39937 | 0.97700 | 2.50910 | 0.95911 | 0.00000 | 2.05852 | 0.73033 | 5.18056 | 1.60500 | 0.96216 | 1.62793 |
| **PCA3** | 0.01777 | 0.00836 | 0.02134 | 0.11316 | 0.05968 | 0.00000 | 0.03897 | 0.00785 | 0.00304 | 0.09261 | 0.01105 | 0.05375 |
| **PCAT1** | 0.00593 | 0.00000 | 0.00000 | 0.00016 | 0.00756 | 0.03460 | 0.00230 | 0.00980 | 0.00000 | 0.40524 | 0.04324 | 0.02342 |
| **PCAT18** | 0.02297 | 0.05296 | 0.08554 | 0.10055 | 0.02217 | 0.00000 | 0.09501 | 0.00688 | 0.01997 | 0.10327 | 0.01008 | 0.05886 |
| **PCAT19** | 0.01073 | 0.01222 | 0.03055 | 0.01767 | 0.01617 | 0.03820 | 0.05482 | 0.06380 | 0.00974 | 0.02050 | 0.01018 | 0.05437 |
| **PCAT2** | 0.01170 | 0.01759 | 0.01563 | 0.01039 | 0.01731 | 0.03439 | 0.01345 | 0.02165 | 0.01961 | 0.01152 | 0.01264 | 0.01723 |
| **PCAT4** | 0.00617 | 0.00000 | 0.00000 | 0.43808 | 0.08563 | 0.10159 | 0.00498 | 0.81849 | 0.00000 | 0.00000 | 0.02067 | 0.93510 |
| **PCAT6** | 0.06948 | 0.33540 | 0.54532 | 0.00000 | 0.82140 | 0.09102 | 0.44083 | 1.66248 | 0.65721 | 0.06215 | 1.37964 | 0.29536 |
| **PCBD1** | 0.17066 | 1.91328 | 2.83077 | 6.42290 | 3.25859 | 1.82879 | 7.54431 | 0.00000 | 6.99850 | 0.00041 | 9.27924 | 0.00000 |
| **PCBD2** | 0.02244 | 3.57372 | 0.00000 | 2.06739 | 3.22626 | 7.06272 | 0.00000 | 0.00000 | 0.44123 | 0.77749 | 4.10259 | 1.97429 |
| **PCBP1** | 13.41596 | 20.24510 | 26.65060 | 12.72916 | 26.71939 | 22.39191 | 35.09421 | 5.49890 | 13.77715 | 44.75497 | 18.26081 | 20.32422 |
| **PCBP1-AS1** | 0.12818 | 0.45090 | 0.35898 | 0.01566 | 0.14517 | 0.21036 | 0.00168 | 0.50570 | 0.56788 | 0.13420 | 0.45156 | 0.46066 |

| | | | | | | | | | | | | |
|---|---|---|---|---|---|---|---|---|---|---|---|---|
| **PCBP2** | 9.18799 | 5.40933 | 8.72589 | 9.11676 | 13.58849 | 21.16892 | 29.72103 | 6.84322 | 6.93197 | 48.33278 | 9.29686 | 9.88216 |
| **PCBP2-OT1** | 0.20066 | 0.10281 | 0.16859 | 0.15075 | 0.34106 | 0.00000 | 0.24786 | 0.05839 | 0.22275 | 1.33456 | 0.08849 | 0.12762 |
| **PCBP3** | 0.19799 | 0.06704 | 0.00000 | 0.17261 | 0.17954 | 0.26603 | 0.01265 | 0.12321 | 0.00000 | 0.05479 | 0.00000 | 0.05772 |
| **PCBP4** | 0.85866 | 0.54662 | 0.68253 | 2.01681 | 0.43651 | 0.00000 | 0.35997 | 1.15284 | 0.51762 | 0.32557 | 0.10759 | 2.35666 |
| **PCCA** | 1.22712 | 0.11992 | 2.13708 | 3.42188 | 1.15599 | 2.74926 | 0.00000 | 0.49372 | 5.30487 | 0.00000 | 1.05892 | 0.03296 |
| **PCCA-AS1** | 0.00579 | 0.01657 | 0.01974 | 0.02370 | 0.00253 | 0.03490 | 0.00769 | 0.00973 | 0.04604 | 0.03035 | 0.03743 | 0.02218 |
| **PCCB** | 4.85697 | 10.86743 | 9.37610 | 10.62520 | 7.69738 | 13.27812 | 7.56403 | 2.69555 | 14.63357 | 0.00000 | 12.86361 | 5.89664 |
| **PCDH1** | 0.34295 | 7.04380 | 0.00000 | 1.80898 | 19.48990 | 0.00000 | 0.00906 | 0.25018 | 7.51535 | 10.98412 | 6.82016 | 0.84612 |
| **PCDH10** | 0.09876 | 0.01063 | 0.47570 | 0.02028 | 0.00000 | 0.01751 | 0.06879 | 0.21824 | 0.00616 | 0.02041 | 0.05079 | 0.04571 |
| **PCDH11X** | 4.61778 | 0.01586 | 0.02956 | 0.00806 | 0.01698 | 0.02166 | 0.01935 | 0.96113 | 0.01307 | 0.00000 | 0.02467 | 0.01280 |
| **PCDH11Y** | 0.00513 | 0.01491 | 0.00348 | 0.00559 | 0.03445 | 0.05405 | 0.02005 | 0.18010 | 0.09441 | 0.00000 | 0.01518 | 0.05509 |
| **PCDH12** | 0.00845 | 0.01496 | 0.06941 | 0.14198 | 0.25279 | 0.00000 | 0.16934 | 0.06967 | 0.02243 | 0.01346 | 0.04148 | 0.05020 |
| **PCDH15** | 5.03098 | 3.24948 | 0.00000 | 0.62386 | 0.39346 | 0.74066 | 0.00000 | 1.38703 | 0.00000 | 0.00790 | 0.00000 | 0.13201 |
| **PCDH17** | 0.00166 | 0.00000 | 0.07904 | 0.46665 | 0.27034 | 0.27882 | 0.97286 | 0.00357 | 1.37666 | 1.21416 | 0.12821 | 0.00000 |
| **PCDH18** | 0.00639 | 0.05911 | 0.02950 | 0.14195 | 0.00399 | 0.01623 | 0.02881 | 0.01041 | 0.02418 | 0.00000 | 0.01791 | 0.01480 |
| **PCDH19** | 0.61271 | 0.01177 | 0.05520 | 0.42044 | 0.00000 | 0.05348 | 0.00000 | 0.39368 | 0.00000 | 0.28884 | 0.00000 | 0.04941 |
| **PCDH20** | 0.00181 | 0.02464 | 0.06637 | 0.07215 | 0.03108 | 0.00000 | 0.01092 | 0.40433 | 0.00690 | 0.06377 | 0.01648 | 0.18736 |
| **PCDH7** | 0.13127 | 0.02148 | 0.00000 | 0.22711 | 0.06480 | 0.08172 | 0.01436 | 0.01106 | 0.00619 | 0.00145 | 0.02363 | 0.08607 |
| **PCDH8** | 0.50218 | 0.06417 | 0.03074 | 0.47465 | 0.00000 | 0.17405 | 0.00000 | 0.50126 | 0.00000 | 0.04774 | 0.00495 | 0.16838 |
| **PCDH9** | 1.50404 | 0.26362 | 0.05334 | 0.00000 | 0.03380 | 0.17095 | 0.10083 | 2.54900 | 0.00000 | 0.11147 | 0.10022 | 0.22817 |
| **PCDH9-AS2** | 0.01393 | 0.01115 | 0.00727 | 0.00904 | 0.00452 | 0.02419 | 0.00555 | 0.00984 | 0.02513 | 0.08259 | 0.01186 | 0.01408 |
| **PCDH9-AS3** | 0.00592 | 0.01085 | 0.01390 | 0.01118 | 0.01135 | 0.02559 | 0.01397 | 0.00907 | 0.01819 | 0.02378 | 0.01420 | 0.03829 |
| **PCDHA1** | 0.00354 | 0.07409 | 0.00717 | 0.00000 | 0.05295 | 0.03249 | 0.01415 | 0.16479 | 0.05944 | 0.01061 | 0.00435 | 0.39988 |
| **PCDHA10** | 0.00502 | 0.01471 | 0.00785 | 0.00361 | 0.00605 | 0.07499 | 0.03390 | 0.00733 | 0.04470 | 0.00000 | 0.02339 | 0.51253 |
| **PCDHA11** | 0.00515 | 0.03216 | 0.00687 | 0.00615 | 0.02138 | 0.00315 | 0.10553 | 0.00959 | 0.00000 | 0.06483 | 0.01014 | 0.05748 |
| **PCDHA12** | 0.00614 | 0.01197 | 0.03506 | 0.00060 | 0.05027 | 0.02400 | 0.00000 | 0.01134 | 0.14598 | 0.04191 | 0.00001 | 0.02942 |
| **PCDHA13** | 0.00569 | 0.00677 | 0.01504 | 0.01514 | 0.01979 | 0.06798 | 0.00917 | 0.02925 | 0.02442 | 0.00192 | 0.02489 | 0.03102 |
| **PCDHA2** | 0.00529 | 0.01387 | 0.02557 | 0.01497 | 0.01431 | 0.02198 | 0.02621 | 0.01322 | 0.01740 | 0.00709 | 0.01447 | 0.02732 |
| **PCDHA3** | 0.00190 | 0.00629 | 0.03711 | 0.01095 | 0.00608 | 0.03072 | 0.08690 | 0.09067 | 0.00091 | 0.00000 | 0.07227 | 0.10923 |
| **PCDHA4** | 0.00659 | 0.02050 | 0.01121 | 0.02235 | 0.02976 | 0.00000 | 0.01716 | 0.01078 | 0.06457 | 0.00000 | 0.04554 | 0.01804 |
| **PCDHA5** | 0.00660 | 0.00942 | 0.02017 | 0.01441 | 0.01928 | 0.00000 | 0.01958 | 0.00773 | 0.04310 | 0.01339 | 0.01482 | 0.13584 |
| **PCDHA7** | 0.00314 | 0.00545 | 0.00367 | 0.06477 | 0.00000 | 0.01415 | 0.00137 | 0.01248 | 0.02282 | 0.35067 | 0.00000 | 0.01787 |
| **PCDHA9** | 0.00901 | 0.01720 | 0.01320 | 0.00468 | 0.00875 | 0.04538 | 0.02006 | 0.01102 | 0.00971 | 0.06458 | 0.01711 | 0.02065 |
| **PCDHAC1** | 0.01712 | 0.00624 | 0.11318 | 0.02353 | 0.00837 | 0.02427 | 0.00403 | 0.00661 | 0.00000 | 0.03255 | 0.03620 | 0.10291 |
| **PCDHAC2** | 0.00275 | 0.05675 | 0.07467 | 0.00869 | 0.01071 | 0.01115 | 0.01134 | 0.06386 | 0.01343 | 0.02161 | 0.01727 | 0.01274 |
| **PCDHB1** | 0.00154 | 0.10465 | 0.01417 | 0.00143 | 0.00339 | 0.02772 | 0.00623 | 0.09301 | 0.02428 | 0.02549 | 0.02667 | 0.10138 |
| **PCDHB11** | 0.00683 | 0.01753 | 0.02913 | 0.02358 | 0.00518 | 0.03879 | 0.01806 | 0.01029 | 0.01632 | 0.03344 | 0.01316 | 0.01593 |
| **PCDHB12** | 0.00410 | 0.02641 | 0.01511 | 0.00974 | 0.00693 | 0.02323 | 0.01562 | 0.04284 | 0.01581 | 0.03084 | 0.03311 | 0.05677 |
| **PCDHB13** | 0.00544 | 0.00955 | 0.01576 | 0.01116 | 0.01100 | 0.02918 | 0.01632 | 0.01254 | 0.01803 | 0.01965 | 0.01328 | 0.05804 |
| **PCDHB14** | 0.00573 | 0.02032 | 0.02361 | 0.01554 | 0.02791 | 0.00198 | 0.03048 | 0.01610 | 0.01705 | 0.01208 | 0.01622 | 0.02006 |
| **PCDHB16** | 0.03209 | 0.05037 | 0.01366 | 0.00878 | 0.00739 | 0.02259 | 0.01379 | 0.00432 | 0.01775 | 0.02430 | 0.01266 | 0.09866 |
| **PCDHB18** | 0.07766 | 0.01139 | 0.01651 | 0.01300 | 0.01905 | 0.01474 | 0.02417 | 0.00263 | 0.02427 | 0.00000 | 0.01427 | 0.06950 |
| **PCDHB19P** | 0.00392 | 0.01213 | 0.02976 | 0.00233 | 0.00167 | 0.04878 | 0.00569 | 0.45039 | 0.00404 | 0.05130 | 0.01212 | 0.42442 |
| **PCDHB2** | 0.09779 | 1.36724 | 0.09658 | 0.00004 | 0.00000 | 0.05536 | 0.00822 | 0.11314 | 0.08950 | 0.00628 | 0.02708 | 2.80142 |
| **PCDHB3** | 0.00138 | 0.47649 | 0.42724 | 0.00000 | 0.00307 | 0.20724 | 0.00000 | 0.28360 | 0.00000 | 0.00938 | 0.08321 | 2.20754 |
| **PCDHB4** | 0.00625 | 0.00335 | 0.01718 | 0.00793 | 0.02429 | 0.02259 | 0.00388 | 0.00663 | 0.03858 | 0.03003 | 0.01740 | 0.18000 |
| **PCDHB5** | 0.02870 | 0.61315 | 0.00264 | 0.00000 | 0.00000 | 0.01117 | 0.01280 | 0.21418 | 0.02944 | 0.05146 | 0.00909 | 1.68670 |
| **PCDHB6** | 0.01639 | 0.02546 | 0.01586 | 0.01228 | 0.01952 | 0.01446 | 0.02040 | 0.04162 | 0.01870 | 0.00681 | 0.01545 | 0.02614 |

| | | | | | | | | | | | | |
|---|---|---|---|---|---|---|---|---|---|---|---|---|
| **PCDHB7** | 0.00650 | 0.01740 | 0.01345 | 0.01059 | 0.01272 | 0.02745 | 0.01682 | 0.01020 | 0.01732 | 0.01152 | 0.01452 | 0.04550 |
| **PCDHB9** | 0.12534 | 0.00582 | 0.04255 | 0.01286 | 0.00909 | 0.03916 | 0.01006 | 0.00051 | 0.00744 | 0.01580 | 0.01520 | 0.12202 |
| **PCDHGA1** | 0.00269 | 0.01123 | 0.04966 | 0.00631 | 0.00675 | 0.04642 | 0.03765 | 0.04353 | 0.00329 | 0.01785 | 0.01142 | 0.01192 |
| **PCDHGA10** | 0.00227 | 0.17062 | 0.09005 | 0.04270 | 0.03072 | 0.00000 | 0.00631 | 0.55848 | 0.02568 | 0.00000 | 0.05215 | 0.57477 |
| **PCDHGA11** | 0.00153 | 0.01968 | 0.00000 | 0.05446 | 0.18974 | 0.05398 | 0.00000 | 0.59794 | 0.02374 | 0.03545 | 0.06257 | 0.27244 |
| **PCDHGA12** | 0.06139 | 0.00044 | 0.04799 | 0.01364 | 0.00888 | 0.01069 | 0.00281 | 1.63005 | 0.01674 | 0.03272 | 0.01858 | 1.03145 |
| **PCDHGA2** | 0.00665 | 0.07918 | 0.01411 | 0.00480 | 0.01203 | 0.01768 | 0.02043 | 0.00487 | 0.01476 | 0.00423 | 0.01557 | 0.65989 |
| **PCDHGA3** | 0.00310 | 0.01995 | 0.03426 | 0.02250 | 0.00097 | 0.04890 | 0.01084 | 0.07723 | 0.00000 | 0.02284 | 0.02123 | 0.21207 |
| **PCDHGA4** | 0.00132 | 0.10554 | 0.01026 | 0.00000 | 0.00546 | 0.05411 | 0.00770 | 0.15711 | 0.01206 | 0.02672 | 0.00863 | 0.12318 |
| **PCDHGA5** | 0.00536 | 0.01439 | 0.01443 | 0.01571 | 0.01549 | 0.01207 | 0.02546 | 0.01198 | 0.01502 | 0.01821 | 0.01557 | 0.03511 |
| **PCDHGA7** | 0.00366 | 0.03337 | 0.03035 | 0.01688 | 0.00012 | 0.07051 | 0.00000 | 0.03228 | 0.00000 | 0.03176 | 0.03447 | 0.15323 |
| **PCDHGA8** | 0.00486 | 0.00805 | 0.02240 | 0.01548 | 0.00098 | 0.04741 | 0.02369 | 0.01046 | 0.01550 | 0.02310 | 0.02760 | 0.07909 |
| **PCDHGA9** | 0.00300 | 0.01813 | 0.13140 | 0.05172 | 0.00000 | 0.02690 | 0.00983 | 0.12992 | 0.00000 | 0.04418 | 0.01787 | 0.05862 |
| **PCDHGB1** | 0.00314 | 0.07651 | 0.00919 | 0.00747 | 0.01140 | 0.01058 | 0.02218 | 0.03321 | 0.08956 | 0.00000 | 0.02424 | 0.01187 |
| **PCDHGB2** | 0.00605 | 0.02264 | 0.01242 | 0.01067 | 0.01239 | 0.01612 | 0.02093 | 0.01006 | 0.02125 | 0.02183 | 0.01395 | 0.02933 |
| **PCDHGB4** | 0.00677 | 0.08508 | 0.01992 | 0.00302 | 0.00673 | 0.04437 | 0.01658 | 0.00893 | 0.02089 | 0.00636 | 0.02685 | 0.08278 |
| **PCDHGB5** | 0.00627 | 0.00840 | 0.01658 | 0.01155 | 0.01005 | 0.01002 | 0.03076 | 0.00840 | 0.00481 | 0.02814 | 0.05838 | 0.11990 |
| **PCDHGB6** | 0.00169 | 0.00716 | 0.01545 | 0.01494 | 0.02045 | 0.01038 | 0.01937 | 0.11770 | 0.02872 | 0.00000 | 0.01624 | 0.18868 |
| **PCDHGB7** | 0.00877 | 0.00506 | 0.04801 | 0.01217 | 0.01928 | 0.01023 | 0.01141 | 0.12783 | 0.02448 | 0.00261 | 0.01599 | 0.25035 |
| **PCDHGB8P** | 0.05701 | 0.00405 | 0.01229 | 0.01063 | 0.01278 | 0.02327 | 0.01489 | 0.55486 | 0.01512 | 0.01610 | 0.01454 | 0.22654 |
| **PCDHGC3** | 0.00312 | 0.03974 | 0.19098 | 0.02570 | 0.01357 | 0.02279 | 0.00100 | 0.02896 | 0.10880 | 0.00000 | 0.00856 | 0.04221 |
| **PCDHGC4** | 0.00637 | 0.00915 | 0.02031 | 0.01479 | 0.02147 | 0.02101 | 0.01309 | 0.00953 | 0.03879 | 0.00000 | 0.02355 | 0.08389 |
| **PCDP1** | 0.17419 | 0.03555 | 0.03136 | 0.04188 | 0.11165 | 0.00000 | 0.03582 | 0.14028 | 0.00000 | 0.11765 | 0.00795 | 0.06725 |
| **PCED1A** | 0.14172 | 0.22985 | 0.67363 | 0.19940 | 0.00000 | 0.00000 | 0.27860 | 0.04005 | 0.00000 | 0.59932 | 0.59796 | 0.53972 |
| **PCED1B** | 0.28926 | 0.00000 | 0.59925 | 6.09698 | 0.49642 | 0.00000 | 0.19053 | 0.31345 | 0.13347 | 0.00000 | 0.48303 | 2.85704 |
| **PCED1B-AS1** | 0.00576 | 0.00997 | 0.02151 | 0.02878 | 0.01808 | 0.00000 | 0.01601 | 0.01034 | 0.02171 | 0.02595 | 0.05435 | 0.11528 |
| **PCF11** | 5.50826 | 1.98811 | 7.96844 | 3.98089 | 2.24045 | 9.29170 | 6.20155 | 5.86224 | 7.91359 | 17.96796 | 4.11180 | 9.26908 |
| **PCGF1** | 6.06223 | 3.34142 | 4.75467 | 1.73515 | 3.84397 | 2.09984 | 3.59869 | 5.24052 | 9.26292 | 0.00000 | 1.15133 | 7.08253 |
| **PCGF2** | 0.08136 | 0.04652 | 0.55615 | 1.37493 | 0.78944 | 0.00000 | 0.91781 | 0.00315 | 0.00356 | 0.81438 | 0.12888 | 0.01064 |
| **PCGF3** | 9.57865 | 3.46590 | 8.44940 | 2.93089 | 8.45354 | 5.99727 | 7.19542 | 7.08178 | 17.87616 | 14.18069 | 1.24605 | 4.69246 |
| **PCGF5** | 4.57047 | 8.66902 | 0.47025 | 0.00000 | 5.47114 | 7.05747 | 1.49169 | 1.44940 | 1.68739 | 4.36949 | 4.49098 | 6.83866 |
| **PCGF6** | 1.72604 | 6.12656 | 5.06524 | 2.22809 | 0.22069 | 10.28885 | 3.78230 | 4.55872 | 6.69126 | 2.88182 | 3.68074 | 13.85232 |
| **PCID2** | 5.96024 | 9.54391 | 5.38722 | 5.99735 | 3.86813 | 18.25080 | 5.22761 | 1.23130 | 7.48272 | 0.00000 | 12.74103 | 7.74715 |
| **PCIF1** | 4.67381 | 1.98349 | 3.18906 | 2.99754 | 0.49537 | 0.00000 | 0.58505 | 1.05714 | 5.91336 | 0.00000 | 2.89082 | 2.31402 |
| **PCK1** | 0.00600 | 0.01864 | 0.01437 | 0.02473 | 0.00753 | 0.02337 | 0.01515 | 0.01044 | 0.01150 | 0.02125 | 0.03129 | 0.01489 |
| **PCK2** | 0.00513 | 0.00000 | 0.76256 | 1.39423 | 0.00000 | 0.09997 | 0.46836 | 0.00861 | 0.86030 | 0.01125 | 0.21466 | 0.00871 |
| **PCLO** | 4.65704 | 0.10794 | 0.11921 | 1.33240 | 0.43875 | 0.00000 | 0.21551 | 1.01949 | 0.00000 | 0.64426 | 0.01357 | 1.58913 |
| **PCM1** | 6.76098 | 5.87575 | 5.85993 | 6.42515 | 6.57904 | 17.40018 | 9.39728 | 4.54951 | 8.37936 | 21.76544 | 4.36266 | 10.30650 |
| **PCMT1** | 6.75998 | 26.25346 | 18.56794 | 19.24228 | 16.32919 | 55.25592 | 10.68568 | 18.89823 | 44.04932 | 9.93926 | 22.18973 | 24.02197 |
| **PCMTD1** | 5.35243 | 1.33798 | 0.77112 | 0.07456 | 0.00000 | 7.96918 | 3.08717 | 0.90226 | 1.16888 | 5.74976 | 3.47714 | 0.96223 |
| **PCMTD2** | 0.56358 | 5.61593 | 0.00000 | 5.84732 | 0.00000 | 21.17602 | 0.00000 | 2.20753 | 0.00000 | 7.50154 | 2.64983 | 8.82513 |
| **PCNA** | 16.35817 | 18.27425 | 15.35015 | 6.61294 | 1.04713 | 53.84011 | 22.85865 | 15.57051 | 18.04851 | 18.60553 | 19.19301 | 24.85912 |
| **PCNAP1** | 0.02669 | 0.00941 | 0.58415 | 0.03582 | 0.00000 | 0.13519 | 0.00000 | 0.06530 | 0.00000 | 0.03012 | 0.02755 | 0.01631 |
| **PCNP** | 3.28331 | 7.25173 | 9.40026 | 7.67752 | 0.00006 | 37.76826 | 13.65989 | 2.84065 | 8.19997 | 9.83408 | 11.26070 | 9.88717 |
| **PCNT** | 2.32546 | 4.53567 | 5.65037 | 9.85802 | 12.50447 | 0.00000 | 3.78383 | 2.18217 | 18.16003 | 17.90279 | 2.69375 | 10.50651 |
| **PCNX** | 0.03353 | 0.37108 | 1.93854 | 1.36249 | 1.38964 | 4.78449 | 0.23347 | 2.52136 | 2.53948 | 8.63864 | 1.20261 | 4.48701 |
| **PCNXL2** | 0.09101 | 0.21384 | 0.01802 | 1.05589 | 0.21820 | 0.00000 | 0.00182 | 0.04974 | 0.11866 | 0.12990 | 0.03690 | 0.72232 |
| **PCNXL3** | 0.62748 | 3.37714 | 5.61001 | 5.86413 | 6.11762 | 0.00000 | 2.94978 | 0.36423 | 10.76172 | 14.04059 | 3.60731 | 3.54795 |

| | | | | | | | | | | | | |
|---|---|---|---|---|---|---|---|---|---|---|---|---|
| **PCNXL4** | 3.13095 | 1.07973 | 4.47695 | 0.70711 | 1.31779 | 5.96884 | 0.00000 | 0.26171 | 6.94411 | 0.41250 | 0.99277 | 2.78058 |
| **PCOLCE** | 0.07008 | 0.11132 | 1.05695 | 0.73802 | 0.00000 | 0.17454 | 0.00000 | 0.09953 | 0.10143 | 0.08189 | 0.15434 | 0.23305 |
| **PCOLCE-AS1** | 0.02023 | 0.03451 | 0.02417 | 0.01894 | 0.01122 | 0.00703 | 0.03533 | 0.01313 | 0.00980 | 0.00816 | 0.03306 | 0.02095 |
| **PCOLCE2** | 1.87081 | 1.47653 | 0.48048 | 0.00882 | 0.00000 | 0.32553 | 0.00000 | 0.28164 | 0.22073 | 0.00425 | 0.00000 | 1.30540 |
| **PCP2** | 0.03601 | 0.01807 | 0.04606 | 0.01122 | 0.01417 | 0.00253 | 0.02449 | 0.00870 | 0.00478 | 0.03181 | 0.06708 | 0.04443 |
| **PCP4** | 0.00832 | 0.00000 | 0.04851 | 0.02419 | 0.00086 | 0.08389 | 0.00000 | 0.06259 | 0.13757 | 0.01871 | 0.28428 | 0.08882 |
| **PCP4L1** | 32.11290 | 0.06449 | 1.32737 | 5.17880 | 0.00000 | 0.53693 | 0.00000 | 14.84719 | 0.00000 | 0.00000 | 0.03600 | 3.27340 |
| **PCSK1** | 0.07253 | 0.02346 | 0.11301 | 0.39184 | 0.00000 | 0.09602 | 0.00000 | 0.00080 | 0.02552 | 0.07380 | 0.00850 | 0.07265 |
| **PCSK1N** | 0.30345 | 0.15922 | 0.49415 | 0.87211 | 0.14029 | 0.00000 | 0.17605 | 0.18162 | 0.02142 | 0.18745 | 0.04259 | 0.00000 |
| **PCSK2** | 0.02136 | 0.00399 | 0.00707 | 0.04135 | 0.00448 | 0.03976 | 0.00837 | 0.22472 | 0.00761 | 0.07855 | 0.04787 | 0.06043 |
| **PCSK4** | 0.47214 | 0.37175 | 0.14673 | 0.34422 | 0.04038 | 0.00000 | 0.00000 | 0.00000 | 0.10405 | 0.19535 | 0.00027 | 0.31031 |
| **PCSK5** | 5.92407 | 0.27249 | 0.38311 | 3.24609 | 0.32487 | 0.46996 | 0.52834 | 2.74836 | 0.00000 | 2.32845 | 0.28823 | 0.55008 |
| **PCSK6** | 0.48575 | 0.00000 | 1.50738 | 0.58590 | 3.03316 | 0.15207 | 1.53287 | 0.06149 | 2.55074 | 1.16693 | 1.68322 | 0.16822 |
| **PCSK7** | 0.00000 | 4.36147 | 3.30674 | 8.85476 | 5.52556 | 0.00000 | 2.62101 | 2.73026 | 8.10257 | 1.14169 | 2.87017 | 3.69659 |
| **PCSK9** | 0.03346 | 0.43979 | 0.00922 | 2.38574 | 0.54486 | 0.00000 | 0.03720 | 0.23358 | 0.07382 | 0.00000 | 0.35893 | 1.12734 |
| **PCTP** | 1.66161 | 2.04064 | 5.68361 | 3.17317 | 0.00000 | 4.25560 | 5.64385 | 0.19917 | 1.08336 | 0.00000 | 1.75097 | 0.18261 |
| **PCYOX1** | 1.80208 | 17.66531 | 5.54560 | 10.15295 | 5.96555 | 24.51189 | 0.00000 | 6.39884 | 11.51730 | 17.49843 | 8.41667 | 13.31708 |
| **PCYOX1L** | 1.34344 | 0.18365 | 0.46360 | 0.17075 | 0.00000 | 0.00000 | 0.00000 | 0.40504 | 0.33534 | 0.09242 | 0.28626 | 0.08449 |
| **PCYT1A** | 2.05253 | 3.72122 | 9.54087 | 2.96759 | 1.53592 | 7.76480 | 7.49304 | 2.08400 | 5.22488 | 0.00725 | 5.62781 | 4.94542 |
| **PCYT1B** | 1.25311 | 0.01434 | 0.14486 | 0.10487 | 0.01484 | 0.00000 | 0.00000 | 0.07247 | 0.04493 | 0.01879 | 0.01081 | 0.00842 |
| **PCYT2** | 0.02662 | 5.58954 | 4.31839 | 5.03546 | 26.22965 | 0.00000 | 6.86383 | 0.22511 | 5.22278 | 11.09226 | 17.50502 | 0.24691 |
| **PDAP1** | 4.22190 | 15.32555 | 13.93134 | 10.97231 | 14.29082 | 0.00000 | 16.19313 | 3.13540 | 17.41899 | 9.14053 | 10.34349 | 14.26037 |
| **PDC** | 0.00626 | 0.01416 | 0.02651 | 0.01298 | 0.00887 | 0.01435 | 0.02781 | 0.01047 | 0.01553 | 0.01234 | 0.06491 | 0.01683 |
| **PDCD1** | 0.00405 | 0.03059 | 0.01411 | 0.07850 | 0.04062 | 0.00000 | 0.03528 | 0.01639 | 0.01154 | 0.04016 | 0.01906 | 0.02897 |
| **PDCD10** | 1.39417 | 6.85322 | 10.06528 | 6.93761 | 5.71871 | 37.27207 | 14.41648 | 6.88874 | 8.29228 | 2.96556 | 10.96117 | 10.69019 |
| **PDCD11** | 0.01053 | 12.66964 | 13.63122 | 10.34691 | 20.73682 | 9.34384 | 20.11408 | 12.48527 | 30.37816 | 30.14481 | 10.27676 | 20.36102 |
| **PDCD1LG2** | 0.02594 | 0.00755 | 0.01091 | 0.00596 | 0.00313 | 0.03555 | 0.00782 | 0.00769 | 0.02352 | 0.08503 | 0.00896 | 0.02623 |
| **PDCD2** | 1.78436 | 5.88046 | 7.46946 | 1.97183 | 3.98754 | 13.67619 | 14.47172 | 0.00000 | 14.60445 | 8.68064 | 5.40419 | 3.77643 |
| **PDCD2L** | 0.17951 | 14.23184 | 7.93781 | 3.27582 | 5.44483 | 10.53174 | 6.87219 | 6.59416 | 8.25016 | 0.00000 | 7.74196 | 18.61653 |
| **PDCD4** | 0.65381 | 0.00000 | 1.14377 | 3.75509 | 10.79046 | 14.46015 | 1.53345 | 1.67306 | 0.00000 | 18.07253 | 0.00000 | 1.71410 |
| **PDCD4-AS1** | 0.00594 | 0.00493 | 0.01656 | 0.02140 | 0.01317 | 0.03039 | 0.07460 | 0.02772 | 0.00000 | 0.02591 | 0.02168 | 0.10494 |
| **PDCD5** | 5.10989 | 12.21999 | 10.62823 | 5.74228 | 14.64003 | 32.73473 | 17.93374 | 0.18248 | 12.17345 | 10.22182 | 14.39105 | 4.35608 |
| **PDCD6** | 0.77077 | 22.53168 | 15.38671 | 10.46331 | 18.66893 | 29.91560 | 24.69536 | 6.60464 | 20.51278 | 3.84643 | 21.89236 | 22.71909 |
| **PDCD6IP** | 3.48918 | 13.73008 | 14.95819 | 9.13188 | 7.36590 | 22.13719 | 14.79630 | 2.32249 | 16.08363 | 3.98740 | 14.37108 | 8.50513 |
| **PDCD6IPP2** | 0.14066 | 0.25701 | 0.02926 | 0.11677 | 0.00000 | 0.00927 | 0.12995 | 0.11935 | 0.00000 | 0.04541 | 0.26116 | 0.04892 |
| **PDCD7** | 3.80595 | 1.75200 | 1.92872 | 1.61809 | 4.63074 | 5.12939 | 4.73479 | 8.41918 | 0.00000 | 13.28452 | 2.92860 | 5.30445 |
| **PDCL** | 5.33319 | 10.02551 | 9.40294 | 12.91191 | 3.76626 | 14.39492 | 10.32320 | 21.80751 | 12.61608 | 0.00000 | 7.41544 | 19.35951 |
| **PDCL2** | 0.10854 | 5.36831 | 0.00000 | 1.55223 | 2.20294 | 4.20056 | 0.00000 | 0.03433 | 0.00000 | 1.19648 | 1.59193 | 1.61850 |
| **PDCL3** | 6.96030 | 42.84697 | 1.72006 | 2.96090 | 40.81805 | 35.29943 | 4.50911 | 31.06692 | 0.00000 | 4.76609 | 21.72508 | 43.99476 |
| **PDCL3P4** | 0.14847 | 1.46050 | 0.00000 | 2.16111 | 2.55347 | 0.00645 | 0.01158 | 0.77607 | 0.24002 | 2.23561 | 1.16385 | 1.13858 |
| **PDDC1** | 0.46611 | 1.93834 | 3.66858 | 1.44003 | 1.78102 | 0.00000 | 4.13309 | 0.19079 | 4.24333 | 4.58282 | 3.85187 | 2.95740 |
| **PDE10A** | 0.40112 | 0.00341 | 0.00230 | 0.02173 | 0.01107 | 0.04301 | 0.00000 | 0.00344 | 0.06725 | 0.04576 | 0.01576 | 0.07021 |
| **PDE11A** | 1.28807 | 0.00000 | 0.08337 | 0.04985 | 0.04753 | 0.05377 | 0.00000 | 1.53703 | 0.01297 | 0.00931 | 0.02192 | 1.68534 |
| **PDE12** | 2.39366 | 9.90523 | 5.59503 | 3.71553 | 3.09453 | 17.03137 | 3.95748 | 6.85730 | 8.98556 | 0.00000 | 11.03465 | 4.84576 |
| **PDE1A** | 0.60639 | 0.10831 | 0.00000 | 0.00000 | 0.30133 | 0.00000 | 0.00143 | 0.16513 | 0.45104 | 0.40135 | 0.00000 | 0.00000 |
| **PDE1B** | 0.02420 | 0.01314 | 0.08554 | 0.06086 | 0.00580 | 0.00956 | 0.05942 | 0.01818 | 0.00256 | 0.02639 | 0.03480 | 0.02768 |
| **PDE1C** | 0.04099 | 0.01146 | 0.11223 | 0.16005 | 0.08608 | 0.00000 | 0.00209 | 0.47445 | 0.07564 | 0.02882 | 0.02336 | 0.00523 |
| **PDE2A** | 0.07098 | 0.15818 | 0.01128 | 0.00000 | 1.31780 | 0.00000 | 0.19369 | 0.02031 | 0.18421 | 0.23165 | 0.12606 | 0.26850 |

| | | | | | | | | | | | | |
|---|---|---|---|---|---|---|---|---|---|---|---|---|
| **PDE3A** | 4.25863 | 0.00618 | 0.10182 | 0.28300 | 0.07799 | 0.00000 | 0.02494 | 0.49982 | 0.00000 | 0.02313 | 0.01381 | 0.04643 |
| **PDE3B** | 0.01541 | 0.09948 | 0.66826 | 0.00000 | 0.00000 | 0.00147 | 1.02780 | 0.09346 | 0.63201 | 0.52365 | 0.03295 | 0.01662 |
| **PDE4A** | 0.25032 | 0.53358 | 0.50419 | 0.73279 | 0.05008 | 0.00000 | 0.00000 | 0.55075 | 0.60079 | 1.09463 | 0.00000 | 2.94101 |
| **PDE4B** | 0.00815 | 0.05088 | 0.02376 | 0.00511 | 0.00000 | 0.03493 | 0.01372 | 0.09761 | 0.01838 | 0.04494 | 0.01515 | 0.01216 |
| **PDE4C** | 0.00600 | 0.52489 | 0.07091 | 0.21207 | 0.00000 | 0.00000 | 0.00138 | 0.30695 | 0.13902 | 0.28710 | 0.00000 | 0.10246 |
| **PDE4D** | 0.92720 | 0.00000 | 3.60045 | 2.41274 | 0.00000 | 1.13393 | 0.00000 | 0.12712 | 0.73303 | 2.38891 | 0.39256 | 0.52680 |
| **PDE4DIP** | 1.26136 | 4.97950 | 6.35009 | 7.69519 | 1.55424 | 2.98544 | 7.03734 | 4.01954 | 14.26778 | 2.32448 | 11.27648 | 5.99880 |
| **PDE5A** | 0.34151 | 0.10156 | 0.11531 | 1.21602 | 0.03648 | 1.61089 | 0.00000 | 0.38460 | 0.37586 | 1.67142 | 0.17283 | 0.36227 |
| **PDE6A** | 0.00375 | 0.01450 | 0.00868 | 0.14035 | 0.21270 | 0.00000 | 0.03815 | 0.02637 | 0.00000 | 0.19784 | 0.05468 | 0.01813 |
| **PDE6B** | 2.15024 | 0.14037 | 0.16181 | 0.19530 | 0.04204 | 0.00000 | 0.00000 | 1.49962 | 0.04434 | 0.09725 | 0.20140 | 0.01604 |
| **PDE6C** | 3.29288 | 0.00823 | 0.03866 | 0.00764 | 0.02116 | 0.00000 | 0.01206 | 1.86316 | 0.01908 | 0.02242 | 0.05335 | 0.00000 |
| **PDE6D** | 0.26818 | 6.30080 | 3.48325 | 4.53580 | 4.18561 | 14.07017 | 4.76517 | 1.68946 | 3.04876 | 0.00060 | 5.08508 | 6.20566 |
| **PDE6G** | 0.00003 | 2.36765 | 0.07000 | 1.61468 | 0.46374 | 0.17514 | 0.00000 | 0.05280 | 0.00000 | 0.28049 | 0.25182 | 0.10580 |
| **PDE7A** | 2.95584 | 0.00761 | 3.48548 | 2.11412 | 0.49360 | 3.48313 | 4.30445 | 0.17367 | 0.19529 | 7.01440 | 0.13088 | 0.25512 |
| **PDE7B** | 0.08691 | 0.01455 | 0.03840 | 0.08943 | 0.00486 | 0.00000 | 0.04929 | 0.00661 | 0.00000 | 0.15775 | 0.02008 | 0.01684 |
| **PDE8A** | 1.83093 | 1.03733 | 3.02205 | 1.57465 | 1.70066 | 2.89717 | 2.20168 | 0.03567 | 3.49015 | 4.37245 | 0.69544 | 0.32968 |
| **PDE8B** | 20.75787 | 0.25994 | 0.00000 | 0.17866 | 0.11591 | 0.16230 | 0.00810 | 3.55209 | 0.08881 | 0.22304 | 0.03409 | 0.87196 |
| **PDE9A** | 2.91572 | 0.09404 | 0.00811 | 0.01416 | 0.03957 | 0.09686 | 0.04654 | 0.34907 | 0.00000 | 0.00000 | 0.00908 | 0.00450 |
| **PDF** | 0.07504 | 1.99979 | 0.77640 | 0.62303 | 0.66756 | 0.00000 | 0.00020 | 0.16675 | 1.05641 | 0.26092 | 0.84814 | 0.53308 |
| **PDGFA** | 2.18710 | 11.97392 | 0.00000 | 0.02174 | 22.56556 | 7.32929 | 25.42994 | 1.54414 | 9.40586 | 17.18872 | 18.82226 | 1.64789 |
| **PDGFB** | 0.00634 | 0.02251 | 0.18239 | 0.01565 | 0.02788 | 0.00051 | 0.15215 | 0.01062 | 0.00000 | 0.00000 | 0.01614 | 0.01738 |
| **PDGFC** | 0.00222 | 0.00000 | 0.11848 | 0.08002 | 0.00000 | 0.18212 | 0.15947 | 0.29191 | 0.00000 | 0.18759 | 0.00000 | 0.16608 |
| **PDGFD** | 0.02863 | 0.58036 | 0.13180 | 0.00398 | 0.13041 | 0.04386 | 0.06237 | 0.33395 | 0.12697 | 0.00000 | 0.00000 | 0.00000 |
| **PDGFRA** | 0.59006 | 0.00106 | 1.76777 | 36.21918 | 0.00000 | 0.17753 | 0.00000 | 1.06509 | 0.60518 | 0.10737 | 0.04401 | 0.00000 |
| **PDGFRB** | 0.00484 | 0.00764 | 0.04097 | 0.02434 | 0.03177 | 0.00000 | 0.04800 | 0.01342 | 0.02067 | 0.09121 | 0.08541 | 0.02865 |
| **PDGFRL** | 6.03826 | 0.01357 | 0.53865 | 0.35758 | 0.00000 | 0.01625 | 0.03017 | 1.73763 | 0.60560 | 0.00000 | 0.14510 | 0.20531 |
| **PDHA1** | 3.32017 | 24.63630 | 10.32994 | 24.65909 | 26.46307 | 36.00757 | 32.40903 | 4.66775 | 33.86448 | 7.90631 | 43.14372 | 23.92021 |
| **PDHA2** | 0.09756 | 0.11563 | 0.07687 | 0.21607 | 0.03295 | 0.58030 | 0.09210 | 0.08639 | 0.18558 | 0.17951 | 0.64643 | 0.18977 |
| **PDHB** | 0.00000 | 21.72994 | 14.97956 | 22.80982 | 14.83218 | 36.31098 | 19.81544 | 6.72451 | 20.11693 | 0.00000 | 28.27270 | 19.14509 |
| **PDHX** | 2.49936 | 9.31980 | 8.99828 | 9.62108 | 3.68418 | 27.22764 | 10.56564 | 1.84528 | 13.27744 | 0.00000 | 14.10959 | 8.66582 |
| **PDIA2** | 0.00688 | 0.02889 | 0.03468 | 0.05020 | 0.04087 | 0.00559 | 0.01471 | 0.01063 | 0.01903 | 0.00870 | 0.02232 | 0.01655 |
| **PDIA3** | 3.39273 | 7.48842 | 26.79444 | 4.39915 | 6.44338 | 35.45160 | 1.63713 | 12.71166 | 44.33048 | 11.85301 | 22.86353 | 11.17880 |
| **PDIA3P1** | 1.12494 | 2.53842 | 13.09239 | 2.20889 | 2.61845 | 14.97997 | 0.00000 | 5.78689 | 21.15739 | 0.00000 | 11.88650 | 4.59151 |
| **PDIA4** | 2.46816 | 14.62419 | 18.87978 | 14.25728 | 12.36613 | 8.63657 | 1.97586 | 5.35198 | 27.37243 | 4.50183 | 14.49102 | 8.51301 |
| **PDIA5** | 0.00202 | 0.22061 | 1.98191 | 0.03329 | 2.63743 | 0.99631 | 0.00039 | 0.75693 | 2.36831 | 6.64493 | 2.94310 | 0.23237 |
| **PDIA6** | 2.98665 | 14.70423 | 24.79682 | 12.31771 | 13.36231 | 24.52262 | 0.00000 | 5.43712 | 45.94191 | 0.00000 | 21.85777 | 7.63994 |
| **PDIK1L** | 0.91326 | 0.40480 | 0.29096 | 0.00000 | 0.21373 | 1.59432 | 0.53417 | 0.44750 | 0.00000 | 1.01467 | 0.64921 | 0.57044 |
| **PDK1** | 6.20990 | 0.38048 | 0.91233 | 3.73186 | 0.00000 | 1.01009 | 0.69889 | 1.17025 | 0.00270 | 0.00000 | 0.00000 | 1.38367 |
| **PDK2** | 3.01233 | 0.21121 | 0.02622 | 0.01191 | 0.21855 | 0.00000 | 0.09244 | 0.45302 | 0.00000 | 0.00000 | 0.01434 | 0.11614 |
| **PDK3** | 0.31036 | 1.21622 | 2.13383 | 1.91777 | 0.00757 | 1.32606 | 0.15112 | 2.68958 | 0.09228 | 0.00000 | 0.18967 | 2.72351 |
| **PDK4** | 0.00827 | 0.03151 | 0.01075 | 0.03264 | 0.01644 | 0.00426 | 0.03155 | 0.01053 | 0.03857 | 0.00000 | 0.01271 | 0.02001 |
| **PDLIM1** | 7.85475 | 21.05817 | 0.72398 | 3.31510 | 24.64882 | 15.98678 | 26.07508 | 0.37350 | 36.49164 | 12.71304 | 27.58741 | 11.88317 |
| **PDLIM2** | 5.01954 | 0.36183 | 15.59216 | 0.00000 | 0.23255 | 0.00000 | 1.14502 | 0.38970 | 0.60954 | 0.00000 | 0.12589 | 0.43390 |
| **PDLIM3** | 0.42413 | 0.00225 | 1.21866 | 0.00850 | 0.02109 | 0.00022 | 0.00000 | 0.12284 | 0.00000 | 0.03054 | 0.01335 | 0.47380 |
| **PDLIM4** | 0.00340 | 0.21499 | 0.11209 | 0.39555 | 0.19058 | 0.25704 | 0.00000 | 0.01350 | 0.04142 | 0.05419 | 0.01216 | 0.03788 |
| **PDLIM5** | 0.31454 | 4.37387 | 5.19681 | 7.25366 | 7.10235 | 7.75520 | 6.45178 | 6.45111 | 4.63378 | 0.00000 | 0.86689 | 7.89029 |
| **PDLIM7** | 0.00133 | 7.81176 | 4.71105 | 8.68219 | 7.63587 | 0.00000 | 3.56464 | 3.27009 | 5.20805 | 0.00000 | 2.62259 | 6.11904 |
| **PDP1** | 1.25568 | 1.85353 | 0.00010 | 3.66548 | 0.00000 | 4.33054 | 1.47613 | 10.09351 | 1.00953 | 2.10663 | 1.89068 | 4.38925 |

| | | | | | | | | | | | | |
|---|---|---|---|---|---|---|---|---|---|---|---|---|
| PDP2 | 0.41665 | 0.52633 | 2.04288 | 0.00000 | 4.36585 | 3.42591 | 9.08663 | 1.23152 | 0.41853 | 13.19677 | 14.47656 | 1.09959 |
| PDPK1 | 3.19436 | 3.43770 | 7.17987 | 3.21028 | 6.99937 | 3.37835 | 6.24796 | 3.01144 | 3.26440 | 14.79068 | 3.93957 | 1.07646 |
| PDPN | 10.71397 | 8.15783 | 4.19732 | 16.43450 | 17.24720 | 0.00000 | 0.00000 | 4.29799 | 0.00000 | 0.00109 | 6.62717 | 0.67961 |
| PDPR | 5.06210 | 1.88198 | 0.31705 | 2.63730 | 0.00000 | 0.76674 | 3.86485 | 1.78667 | 2.94598 | 9.68985 | 3.35676 | 9.76114 |
| PDRG1 | 5.19981 | 16.98665 | 9.82898 | 5.37484 | 9.99592 | 1.81136 | 18.11204 | 18.38884 | 10.01521 | 0.00000 | 6.45707 | 23.87040 |
| PDS5A | 1.13275 | 5.09270 | 5.85912 | 6.78090 | 3.15451 | 25.75660 | 13.34722 | 3.32620 | 5.04296 | 27.41010 | 4.77019 | 11.21094 |
| PDS5B | 0.00000 | 2.81317 | 4.26795 | 3.54389 | 0.83876 | 20.97217 | 13.23285 | 8.14020 | 0.87505 | 18.54288 | 6.46608 | 9.61085 |
| PDSS1 | 0.00000 | 3.97630 | 2.53483 | 1.33383 | 2.89226 | 3.50615 | 0.98686 | 0.24925 | 6.52541 | 2.90731 | 2.72832 | 0.73203 |
| PDSS2 | 0.39336 | 3.58212 | 2.16100 | 2.83825 | 2.59528 | 6.83147 | 4.02887 | 0.21791 | 3.57622 | 0.88272 | 3.72356 | 3.32121 |
| PDX1 | 0.06638 | 0.00282 | 0.01494 | 0.00209 | 0.01302 | 0.01732 | 0.01075 | 0.03086 | 0.01370 | 0.25881 | 0.00116 | 0.04115 |
| PDX1-AS1 | 0.01684 | 0.01503 | 0.02370 | 0.01239 | 0.00818 | 0.00693 | 0.01425 | 0.01069 | 0.00999 | 0.02010 | 0.09116 | 0.01999 |
| PDXDC1 | 11.23761 | 6.21210 | 10.78714 | 4.54696 | 7.01800 | 10.35537 | 17.66008 | 2.77993 | 12.42328 | 11.32964 | 10.38149 | 1.35849 |
| PDXDC2P | 0.15591 | 0.29303 | 0.81600 | 1.16397 | 0.15116 | 1.19955 | 0.00000 | 0.44904 | 1.03406 | 1.63695 | 0.04194 | 0.32902 |
| PDXK | 0.86234 | 6.58356 | 2.61004 | 7.68944 | 29.62105 | 0.00000 | 21.60619 | 0.00000 | 13.04172 | 22.68273 | 10.25492 | 0.79000 |
| PDXP | 2.76619 | 1.02430 | 0.24391 | 2.03525 | 2.24694 | 0.40079 | 3.28618 | 1.17652 | 0.50616 | 6.53054 | 0.63894 | 3.08212 |
| PDZD11 | 0.65688 | 6.77428 | 4.37351 | 5.76789 | 6.46359 | 12.94917 | 4.93872 | 3.89615 | 3.12482 | 0.00000 | 6.36927 | 6.49793 |
| PDZD2 | 2.91863 | 1.83621 | 0.37934 | 0.46345 | 1.32582 | 1.57486 | 0.45268 | 0.22135 | 2.01583 | 3.66469 | 3.84200 | 1.68265 |
| PDZD3 | 0.00522 | 0.05064 | 0.02436 | 0.00729 | 0.00685 | 0.03703 | 0.00541 | 0.00827 | 0.02231 | 0.04697 | 0.01544 | 0.06151 |
| PDZD4 | 0.12522 | 0.02926 | 0.07703 | 0.02818 | 0.02348 | 0.00000 | 0.00567 | 0.18294 | 0.04934 | 0.00264 | 0.01131 | 0.00945 |
| PDZD7 | 2.67672 | 0.00398 | 0.09049 | 0.02404 | 0.00619 | 0.05626 | 0.00000 | 0.04137 | 0.06977 | 0.00061 | 0.00482 | 0.09148 |
| PDZD8 | 1.60200 | 0.00000 | 0.77182 | 1.54146 | 0.00000 | 2.10027 | 1.92234 | 2.79151 | 0.00000 | 5.92671 | 0.28119 | 2.32568 |
| PDZD9 | 0.00643 | 0.01672 | 0.02543 | 0.00640 | 0.00793 | 0.03214 | 0.00000 | 0.01405 | 0.12915 | 0.00556 | 0.01027 | 0.01581 |
| PDZK1 | 0.39584 | 0.01714 | 0.11182 | 0.11809 | 0.31547 | 0.41533 | 0.00000 | 0.24339 | 0.00000 | 0.00000 | 0.34855 | 0.07150 |
| PDZK1IP1 | 0.02551 | 0.01258 | 0.01258 | 0.01105 | 0.02298 | 0.03412 | 0.01625 | 0.03428 | 0.01544 | 0.01174 | 0.01457 | 0.01325 |
| PDZRN3 | 0.96573 | 0.01909 | 0.36189 | 1.79434 | 0.04756 | 0.26128 | 0.00000 | 0.04173 | 0.04337 | 0.00000 | 0.00471 | 0.00000 |
| PDZRN3-AS1 | 0.56406 | 0.25984 | 0.01878 | 0.00210 | 0.00000 | 0.01284 | 0.01477 | 0.00361 | 0.01935 | 0.01705 | 0.01040 | 0.00184 |
| PDZRN4 | 0.38007 | 0.00678 | 0.22980 | 0.02400 | 0.00000 | 0.03631 | 0.00000 | 0.00148 | 0.00000 | 0.06816 | 0.01345 | 0.00887 |
| PEA15 | 1.05448 | 2.81228 | 9.18423 | 2.93110 | 9.88586 | 0.00000 | 12.55705 | 2.91377 | 2.63986 | 9.80589 | 3.61072 | 3.40145 |
| PEAK1 | 4.90777 | 0.53403 | 8.00985 | 0.00000 | 2.01094 | 1.93583 | 4.24889 | 1.09355 | 6.03327 | 9.17624 | 2.55191 | 0.22040 |
| PEAR1 | 0.00696 | 0.03546 | 0.04581 | 0.01920 | 0.01291 | 0.00000 | 0.01596 | 0.01158 | 0.04423 | 0.04297 | 0.13902 | 0.01966 |
| PEBP1 | 6.17006 | 16.91172 | 27.57837 | 23.75762 | 13.60465 | 11.19596 | 50.11514 | 1.57807 | 35.39611 | 10.01704 | 32.54929 | 0.49370 |
| PEBP4 | 1.04454 | 0.01484 | 0.01728 | 0.00889 | 0.01609 | 0.02027 | 0.01590 | 0.52502 | 0.02039 | 0.00334 | 0.01261 | 0.00669 |
| PECAM1 | 0.69498 | 0.00000 | 0.12067 | 0.76536 | 0.00000 | 0.05507 | 0.00000 | 0.60261 | 0.00000 | 0.57010 | 0.00000 | 0.15833 |
| PECR | 3.55776 | 1.40710 | 1.48268 | 0.66159 | 0.13348 | 2.85453 | 2.33312 | 0.46031 | 4.04857 | 0.00068 | 4.61311 | 0.03551 |
| PEF1 | 1.47665 | 17.92372 | 11.73078 | 6.83301 | 10.57194 | 9.61177 | 14.46001 | 1.22491 | 14.60727 | 0.00000 | 10.27644 | 15.07590 |
| PEG10 | 0.31820 | 0.00000 | 31.04408 | 0.59557 | 0.00000 | 0.00000 | 23.67331 | 0.16278 | 3.33770 | 2.44268 | 4.11929 | 0.00066 |
| PEG3 | 0.11528 | 0.39456 | 3.12422 | 0.55273 | 0.00000 | 0.93111 | 2.11935 | 0.52184 | 0.00004 | 2.22744 | 0.46188 | 1.34078 |
| PELI1 | 2.55687 | 2.47473 | 1.61227 | 11.08550 | 0.00000 | 9.19235 | 3.08718 | 6.87062 | 3.16908 | 1.67008 | 3.50093 | 2.88585 |
| PELI2 | 1.68182 | 0.56477 | 0.19295 | 2.15848 | 0.12249 | 0.55843 | 0.00000 | 0.52880 | 0.00000 | 0.47862 | 0.00000 | 0.79835 |
| PELI3 | 0.67428 | 0.46449 | 0.12790 | 0.47337 | 1.32445 | 0.00000 | 0.74247 | 0.02895 | 0.00000 | 0.34478 | 0.11763 | 0.69127 |
| PELO | 2.64645 | 7.16532 | 29.30849 | 3.81415 | 0.00000 | 9.69836 | 17.39668 | 0.90968 | 17.37658 | 0.00000 | 8.04722 | 5.51823 |
| PELP1 | 0.56928 | 2.01462 | 3.56848 | 3.01310 | 7.11661 | 0.00000 | 11.97821 | 0.00023 | 0.17745 | 29.76593 | 1.94370 | 5.75758 |
| PEMT | 0.01751 | 18.35149 | 2.06653 | 15.50309 | 19.82575 | 2.51598 | 0.66682 | 0.01262 | 7.28510 | 0.01911 | 6.20438 | 0.43637 |
| PENK | 0.00223 | 0.00000 | 0.23884 | 0.00645 | 0.00279 | 0.02640 | 0.00659 | 0.26500 | 0.26968 | 0.00000 | 0.07248 | 0.81071 |
| PEPD | 0.40113 | 17.88713 | 9.58107 | 8.53347 | 9.72049 | 0.39755 | 11.24532 | 0.00000 | 18.13346 | 0.00000 | 10.66305 | 5.84180 |
| PER1 | 1.55482 | 0.00000 | 1.09956 | 1.38445 | 0.00000 | 0.00000 | 3.88726 | 0.76605 | 2.28247 | 8.37594 | 0.84091 | 1.51717 |
| PER2 | 4.35521 | 3.03088 | 1.60978 | 0.98757 | 0.00000 | 2.79777 | 7.01233 | 2.69094 | 0.15199 | 5.20837 | 5.49920 | 4.35066 |
| PER3 | 2.58478 | 6.26344 | 2.04938 | 0.00000 | 0.60104 | 4.17312 | 3.27084 | 0.91137 | 1.46200 | 0.00000 | 1.20800 | 4.78174 |

| | | | | | | | | | | | | |
|---|---|---|---|---|---|---|---|---|---|---|---|---|
| **PERM1** | 0.01934 | 0.01456 | 0.03146 | 0.00383 | 0.03980 | 0.02972 | 0.00000 | 0.00792 | 0.03031 | 0.05996 | 0.00810 | 0.02184 |
| **PERP** | 0.18025 | 11.68271 | 20.62474 | 0.04347 | 16.51007 | 35.53789 | 6.34272 | 1.46734 | 38.95998 | 15.41537 | 16.48455 | 1.54515 |
| **1.00000** | 3.13297 | 10.34068 | 8.78981 | 7.73639 | 10.45181 | 0.00000 | 11.69941 | 4.40793 | 20.42094 | 7.35813 | 12.75338 | 17.30477 |
| **PET100** | 0.49853 | 4.09357 | 3.16226 | 2.93890 | 3.99925 | 3.08685 | 7.29315 | 2.92591 | 1.78635 | 5.72934 | 4.44240 | 7.98066 |
| **PET112** | 0.00000 | 8.83363 | 8.35751 | 8.54267 | 6.77256 | 8.80059 | 7.77800 | 1.54866 | 13.15174 | 0.00000 | 14.44803 | 4.67651 |
| **PET117** | 0.12959 | 3.44738 | 0.00000 | 2.72915 | 5.73451 | 4.96767 | 1.93966 | 0.54335 | 0.35734 | 3.59417 | 2.40542 | 3.10428 |
| **PEX1** | 1.20387 | 4.33642 | 0.13618 | 2.54606 | 0.51173 | 6.76514 | 0.00000 | 0.70595 | 1.42853 | 3.87903 | 0.86462 | 0.88109 |
| **PEX10** | 0.24065 | 2.59354 | 1.33779 | 0.82905 | 1.42513 | 0.00000 | 0.00567 | 0.00000 | 2.28611 | 0.00000 | 1.79516 | 0.56799 |
| **PEX11A** | 1.31553 | 2.95596 | 0.00000 | 0.00000 | 4.22218 | 2.66171 | 3.00437 | 1.28566 | 7.47330 | 0.00000 | 10.76682 | 2.95900 |
| **PEX11B** | 2.92860 | 4.22486 | 0.59049 | 1.66719 | 1.79921 | 2.50712 | 1.67023 | 1.99233 | 1.94876 | 0.47027 | 2.48997 | 3.78224 |
| **PEX11G** | 0.02800 | 0.25314 | 0.20138 | 0.00000 | 0.07478 | 0.00000 | 0.09346 | 0.13409 | 0.08820 | 0.02824 | 0.77742 | 0.71851 |
| **PEX12** | 2.07260 | 4.37736 | 0.00000 | 0.69010 | 1.23700 | 4.55011 | 0.00000 | 5.12796 | 2.19995 | 0.00000 | 0.99191 | 3.37807 |
| **PEX13** | 2.62181 | 3.84704 | 4.20149 | 4.36293 | 0.23304 | 19.51349 | 0.00000 | 5.56157 | 6.16558 | 0.00000 | 6.17669 | 5.69598 |
| **PEX14** | 1.89260 | 2.58181 | 2.09000 | 2.25685 | 2.15181 | 0.00000 | 1.29625 | 0.42482 | 3.86368 | 2.88070 | 2.15960 | 2.48273 |
| **PEX16** | 0.48598 | 14.50238 | 7.88148 | 4.99412 | 16.42314 | 0.91944 | 8.62755 | 0.38686 | 8.43087 | 0.00000 | 8.76430 | 7.24466 |
| **PEX19** | 1.71456 | 11.28475 | 10.47092 | 6.08661 | 4.50862 | 20.39327 | 7.97850 | 1.88528 | 8.76608 | 0.00000 | 9.79948 | 10.18318 |
| **PEX2** | 1.36748 | 4.81796 | 5.83095 | 0.22468 | 0.00006 | 13.87875 | 3.30611 | 5.95004 | 6.61497 | 0.07475 | 4.06821 | 8.83237 |
| **PEX26** | 0.92984 | 2.16947 | 2.24539 | 0.81580 | 1.72003 | 0.00000 | 1.09730 | 0.00000 | 2.94734 | 4.40229 | 2.84220 | 0.55313 |
| **PEX3** | 0.67859 | 2.79732 | 3.22957 | 0.77671 | 1.04518 | 20.89107 | 4.74219 | 0.68012 | 11.24649 | 0.00000 | 15.29921 | 14.82818 |
| **PEX5** | 2.36266 | 2.46310 | 0.00000 | 4.08708 | 1.32632 | 1.82143 | 3.03212 | 0.36718 | 0.00000 | 5.91789 | 7.12017 | 5.14536 |
| **PEX5L** | 0.18845 | 0.00124 | 0.02812 | 0.05897 | 0.00000 | 0.00884 | 0.00302 | 0.04734 | 0.03581 | 0.41808 | 0.00613 | 0.07392 |
| **PEX6** | 0.28574 | 0.36818 | 1.84166 | 1.15004 | 1.46076 | 0.00000 | 2.88783 | 0.00000 | 0.09137 | 1.21824 | 0.84268 | 0.40923 |
| **PEX7** | 0.07779 | 0.77570 | 1.12192 | 0.00000 | 0.00000 | 3.69876 | 0.58785 | 0.00000 | 0.00000 | 0.57578 | 0.62055 | 0.00000 |
| **PFAS** | 0.00000 | 11.41003 | 9.12099 | 8.88760 | 8.39962 | 2.13805 | 12.49168 | 0.66089 | 14.04529 | 13.11287 | 11.83261 | 8.94876 |
| **PFDN1** | 2.65673 | 13.88856 | 18.34600 | 10.27586 | 12.06470 | 17.58122 | 21.72397 | 11.04843 | 17.90401 | 3.93335 | 12.68306 | 30.74783 |
| **PFDN2** | 4.43805 | 13.76707 | 24.00429 | 10.72677 | 20.76499 | 19.75714 | 28.32382 | 18.52061 | 19.81219 | 11.39672 | 22.56103 | 24.18581 |
| **PFDN4** | 1.59693 | 6.44452 | 9.76411 | 4.53474 | 1.86586 | 31.97647 | 17.80513 | 1.54755 | 6.97605 | 14.67192 | 9.55067 | 6.54525 |
| **PFDN5** | 7.43347 | 19.90429 | 12.26857 | 20.36664 | 26.12641 | 20.96542 | 22.04756 | 14.63881 | 14.27767 | 20.71534 | 19.70410 | 28.52975 |
| **PFDN6** | 2.24312 | 2.37832 | 3.58387 | 2.60725 | 2.54723 | 0.00000 | 2.61418 | 1.97389 | 2.43279 | 1.16296 | 10.64324 | 5.66392 |
| **PFKFB1** | 0.00596 | 0.02429 | 0.04186 | 0.00670 | 0.00812 | 0.05898 | 0.00000 | 0.01221 | 0.11695 | 0.00000 | 0.00804 | 0.04634 |
| **PFKFB2** | 3.89093 | 1.73095 | 0.99557 | 0.00000 | 2.27674 | 4.70518 | 4.47739 | 1.53486 | 0.34490 | 2.83678 | 5.06683 | 1.49755 |
| **PFKFB3** | 4.63026 | 14.14116 | 4.57905 | 4.52830 | 0.00000 | 0.00001 | 1.28615 | 19.04677 | 1.32230 | 0.52328 | 0.11361 | 32.07873 |
| **PFKFB4** | 0.10144 | 8.42294 | 9.46618 | 3.78685 | 8.07299 | 1.54948 | 7.07974 | 0.03491 | 19.07833 | 1.61661 | 8.38765 | 0.00000 |
| **PFKL** | 0.58859 | 26.91302 | 11.36281 | 21.43843 | 27.25443 | 0.00000 | 8.92320 | 0.02004 | 26.12956 | 6.63274 | 26.04703 | 3.03392 |
| **PFKM** | 0.23542 | 3.63543 | 2.50329 | 2.67998 | 0.00000 | 2.35856 | 3.46247 | 0.33607 | 3.61685 | 0.00000 | 3.19039 | 4.93891 |
| **PFKP** | 5.35357 | 40.19726 | 26.49334 | 29.92004 | 23.34171 | 10.39433 | 25.67304 | 6.01326 | 45.74565 | 6.34987 | 9.19340 | 18.03394 |
| **PFN1** | 11.46039 | 47.03702 | 43.24607 | 47.40557 | 52.29398 | 35.66809 | 65.99966 | 37.41928 | 55.24021 | 23.54677 | 50.38343 | 50.86023 |
| **PFN1P2** | 0.00274 | 0.00838 | 0.01385 | 0.23118 | 0.05988 | 0.01112 | 0.02261 | 0.02463 | 0.02531 | 0.09680 | 0.03001 | 0.06250 |
| **PFN2** | 5.33200 | 4.67301 | 18.69697 | 5.29926 | 8.34897 | 26.22195 | 20.28159 | 0.20774 | 10.15491 | 0.00000 | 11.13042 | 3.83077 |
| **PFN3** | 0.00282 | 0.01942 | 0.01162 | 0.00865 | 0.01437 | 0.03827 | 0.01255 | 0.05367 | 0.01149 | 0.02438 | 0.01839 | 0.02045 |
| **PFN4** | 0.00460 | 0.02821 | 0.01397 | 0.03691 | 0.01188 | 0.00000 | 0.02238 | 0.01316 | 0.01708 | 0.01838 | 0.03784 | 0.06447 |
| **PGA3** | 0.10546 | 0.01450 | 0.02895 | 0.00767 | 0.01146 | 0.04797 | 0.00710 | 0.04128 | 0.01120 | 0.02386 | 0.01263 | 0.03687 |
| **PGA5** | 0.00745 | 0.00850 | 0.12180 | 0.02427 | 0.00832 | 0.04833 | 0.00000 | 0.00987 | 0.03701 | 0.01434 | 0.01241 | 0.02322 |
| **PGAM1** | 0.00000 | 6.41972 | 7.28642 | 8.62188 | 0.04646 | 21.47721 | 8.75190 | 4.05460 | 12.34218 | 0.00000 | 8.85385 | 7.19132 |
| **PGAM1P5** | 0.35748 | 0.05403 | 0.10317 | 0.09614 | 0.03457 | 0.07488 | 0.04669 | 0.01929 | 0.03795 | 0.00000 | 0.14512 | 0.07158 |
| **PGAM2** | 0.00814 | 0.00055 | 0.66105 | 0.05185 | 0.00000 | 0.09483 | 0.00000 | 0.01346 | 0.00000 | 0.04410 | 0.01643 | 0.06814 |
| **PGAM4** | 0.03769 | 0.21147 | 0.11646 | 0.08714 | 0.10975 | 0.12638 | 0.24883 | 0.09306 | 0.19440 | 0.00000 | 0.25332 | 0.11429 |
| **PGAM5** | 2.41951 | 4.22893 | 7.40516 | 3.47222 | 9.16589 | 4.67741 | 18.61360 | 0.43508 | 2.31812 | 17.11637 | 7.33073 | 1.74071 |

| | | | | | | | | | | | | |
|---|---|---|---|---|---|---|---|---|---|---|---|---|
| **PGAP1** | 1.14753 | 0.75743 | 3.00399 | 0.13287 | 0.00000 | 0.91674 | 0.00000 | 0.89220 | 2.83452 | 2.49924 | 1.24013 | 0.10743 |
| **PGAP2** | 3.13844 | 16.01442 | 8.37720 | 3.98389 | 9.54943 | 4.19418 | 8.00617 | 2.27409 | 13.87296 | 0.00000 | 13.35664 | 18.38796 |
| **PGAP3** | 0.34295 | 0.42467 | 0.15335 | 0.02624 | 0.55834 | 0.00000 | 0.00008 | 0.56033 | 0.53586 | 0.00700 | 0.00711 | 0.73601 |
| **PGBD1** | 0.00000 | 2.26572 | 0.00000 | 1.19693 | 1.78842 | 1.90420 | 0.00000 | 1.03213 | 0.00000 | 0.81181 | 1.14521 | 4.24654 |
| **PGBD2** | 0.65012 | 1.29313 | 2.35118 | 0.25228 | 0.03971 | 3.48329 | 1.62082 | 3.04548 | 4.85370 | 0.00000 | 1.18003 | 6.79473 |
| **PGBD4** | 0.00395 | 0.00000 | 0.84568 | 0.37852 | 0.01509 | 0.50900 | 0.00000 | 0.36138 | 0.33285 | 0.00001 | 0.52901 | 0.47696 |
| **PGBD5** | 0.23636 | 0.41399 | 0.10316 | 0.08018 | 0.26861 | 0.11999 | 0.09003 | 0.00000 | 0.01058 | 0.00974 | 0.00000 | 0.16357 |
| **PGC** | 0.00816 | 0.00000 | 8.71301 | 0.00126 | 0.03465 | 0.03625 | 0.81756 | 0.03443 | 0.71054 | 0.00000 | 0.00000 | 0.10782 |
| **PGCP1** | 0.00615 | 0.01573 | 0.01376 | 0.01679 | 0.01493 | 0.01249 | 0.01617 | 0.00954 | 0.04108 | 0.00479 | 0.01436 | 0.04603 |
| **PGD** | 7.93161 | 20.69339 | 16.15309 | 25.10672 | 22.26581 | 12.97950 | 26.08132 | 2.22103 | 32.98508 | 3.66738 | 15.57663 | 13.07640 |
| **PGF** | 1.04205 | 2.07326 | 72.48514 | 0.00000 | 1.05972 | 1.54016 | 18.43520 | 0.00000 | 13.51365 | 0.00000 | 0.00000 | 0.85559 |
| **PGGT1B** | 1.36620 | 1.61157 | 5.14734 | 2.35279 | 0.00000 | 11.30237 | 6.06970 | 3.88499 | 0.67628 | 0.00009 | 5.04482 | 4.38255 |
| **1.00000** | 3.86097 | 11.53373 | 28.52074 | 23.71672 | 20.45658 | 46.48306 | 41.35969 | 2.23442 | 41.92203 | 0.00000 | 36.54834 | 11.88889 |
| **2.00000** | 0.00795 | 0.00628 | 0.14832 | 0.02411 | 0.00000 | 0.08990 | 0.01329 | 0.00970 | 0.00000 | 0.02463 | 0.05072 | 0.01384 |
| **PGLS** | 0.06943 | 6.64826 | 3.90380 | 7.50667 | 7.87173 | 0.00000 | 5.40581 | 0.00000 | 8.24184 | 1.79321 | 6.98627 | 2.42156 |
| **PGLYRP1** | 0.03612 | 0.02755 | 0.02245 | 0.00886 | 0.11871 | 0.00000 | 0.02467 | 0.01074 | 0.07353 | 0.00000 | 0.01707 | 0.04244 |
| **PGLYRP2** | 0.00461 | 0.00621 | 0.01425 | 0.01417 | 0.01186 | 0.01389 | 0.00895 | 0.08230 | 0.02504 | 0.06056 | 0.01292 | 0.18389 |
| **PGLYRP4** | 0.00688 | 0.09034 | 0.02440 | 0.00371 | 0.00000 | 0.03258 | 0.00999 | 0.01502 | 0.01023 | 0.02230 | 0.05016 | 0.01484 |
| **PGM1** | 0.62847 | 8.61974 | 3.26937 | 11.96890 | 8.65135 | 8.62277 | 0.65689 | 0.06862 | 8.07547 | 0.00000 | 4.09401 | 0.00000 |
| **PGM2** | 0.64985 | 16.76105 | 4.26067 | 16.30307 | 7.86732 | 25.50777 | 4.30577 | 0.04024 | 8.84324 | 0.03569 | 10.08750 | 3.74288 |
| **PGM2L1** | 0.47099 | 4.66721 | 1.58678 | 7.88723 | 0.00000 | 3.19430 | 4.54457 | 2.05711 | 0.06106 | 6.45840 | 2.10063 | 8.50066 |
| **PGM3** | 1.40526 | 7.00214 | 13.92813 | 1.17992 | 3.08010 | 26.93631 | 11.04680 | 8.84444 | 11.18932 | 0.00000 | 11.71012 | 7.06895 |
| **PGM5** | 0.23259 | 0.00794 | 0.00343 | 0.04456 | 0.00000 | 0.00563 | 0.00173 | 0.06417 | 0.02166 | 0.16463 | 0.02178 | 0.00917 |
| **PGM5P2** | 0.08678 | 0.01550 | 0.01470 | 0.01007 | 0.09267 | 0.00000 | 0.02578 | 0.06451 | 0.01492 | 0.09734 | 0.01580 | 0.03046 |
| **PGP** | 0.00000 | 3.86682 | 7.62142 | 6.63386 | 14.26524 | 0.00000 | 11.97020 | 0.35915 | 1.53135 | 29.62298 | 3.91657 | 2.26337 |
| **PGPEP1** | 4.87593 | 0.58603 | 0.84935 | 0.00000 | 6.18759 | 1.73629 | 0.00000 | 0.12150 | 0.00000 | 1.05724 | 2.09162 | 0.90427 |
| **PGPEP1L** | 0.06416 | 0.01345 | 0.15426 | 0.01095 | 0.01570 | 0.00209 | 0.00522 | 0.04686 | 0.00002 | 0.05600 | 0.01832 | 0.05233 |
| **PGR** | 0.00733 | 0.01203 | 0.01473 | 0.01730 | 0.13027 | 0.00000 | 0.04544 | 0.04178 | 0.01322 | 0.00000 | 0.01234 | 0.04021 |
| **PGRMC1** | 0.00000 | 10.67479 | 4.58755 | 1.53266 | 0.63726 | 27.93802 | 0.00000 | 1.30828 | 18.87759 | 11.11641 | 9.07757 | 2.23690 |
| **PGRMC2** | 1.23330 | 0.04818 | 10.77905 | 1.29062 | 1.65326 | 5.19834 | 5.62308 | 2.24061 | 0.51456 | 16.20721 | 0.72888 | 3.20981 |
| **PGS1** | 1.09715 | 4.43014 | 6.21257 | 1.34971 | 2.05913 | 2.73897 | 3.01184 | 1.75808 | 9.05345 | 0.00000 | 5.92072 | 1.57686 |
| **PHACTR1** | 0.81319 | 0.30175 | 0.07955 | 0.89580 | 0.00000 | 0.00152 | 0.53461 | 0.46273 | 0.37655 | 0.00000 | 0.04446 | 0.59951 |
| **PHACTR2** | 0.06636 | 3.57635 | 5.34880 | 6.96658 | 11.60238 | 5.44199 | 12.03631 | 0.00000 | 4.65357 | 6.53576 | 8.86149 | 0.00000 |
| **PHACTR3** | 8.34325 | 0.05331 | 0.01049 | 0.00262 | 0.03447 | 0.02817 | 0.00772 | 2.73650 | 0.01032 | 0.00385 | 0.01147 | 0.40042 |
| **PHACTR4** | 1.33975 | 3.69325 | 6.02703 | 4.67998 | 0.00000 | 2.31050 | 3.52139 | 7.62798 | 8.79520 | 9.20332 | 3.66962 | 4.89713 |
| **PHAX** | 0.50157 | 14.07092 | 11.28895 | 8.29042 | 3.74946 | 28.57201 | 21.45145 | 22.66416 | 8.61770 | 6.55242 | 14.46972 | 23.08964 |
| **PHB** | 2.30971 | 32.77695 | 17.46080 | 24.23219 | 32.27240 | 18.69898 | 41.12115 | 11.98085 | 28.64912 | 6.96165 | 31.45166 | 32.04296 |
| **PHB2** | 7.34201 | 40.35924 | 32.35934 | 30.93957 | 36.34233 | 33.39238 | 37.80935 | 10.77339 | 46.51147 | 12.17527 | 37.42580 | 31.27784 |
| **PHC1** | 0.17069 | 0.09290 | 0.81815 | 0.64691 | 0.02579 | 0.02108 | 0.38906 | 0.04126 | 0.13199 | 0.00001 | 0.04017 | 0.22355 |
| **PHC2** | 1.49642 | 1.19253 | 3.90065 | 1.59045 | 3.95975 | 0.00000 | 12.05413 | 0.03973 | 3.03189 | 7.33323 | 1.40900 | 0.34715 |
| **PHC3** | 1.78158 | 1.93726 | 5.78172 | 2.02210 | 1.01698 | 7.70257 | 0.84326 | 4.55689 | 5.33098 | 8.73571 | 3.99067 | 6.42085 |
| **PHEX** | 7.71698 | 0.00000 | 0.00076 | 0.00000 | 0.02293 | 0.00000 | 0.00000 | 0.72237 | 0.07238 | 0.46266 | 0.00000 | 0.14008 |
| **PHF1** | 1.47278 | 1.01231 | 1.83075 | 5.00239 | 1.22253 | 0.00000 | 1.04355 | 0.25111 | 0.20418 | 1.76078 | 0.50528 | 0.65206 |
| **PHF10** | 0.20116 | 0.01386 | 0.56841 | 0.27953 | 0.52814 | 1.26896 | 0.84224 | 0.01448 | 0.00000 | 1.54413 | 0.30947 | 0.26893 |
| **PHF11** | 0.00196 | 0.80423 | 1.15113 | 1.26906 | 0.51808 | 1.68166 | 0.52613 | 0.00451 | 0.74702 | 0.20992 | 0.97285 | 0.07685 |
| **PHF12** | 3.67597 | 1.81644 | 1.37395 | 3.17021 | 3.75641 | 0.80038 | 1.94538 | 3.15213 | 1.92917 | 12.44478 | 1.64451 | 3.16145 |
| **PHF13** | 7.13967 | 3.03872 | 3.20343 | 4.49414 | 3.42284 | 0.00000 | 2.64167 | 8.26749 | 2.20770 | 6.09903 | 1.03996 | 2.98460 |
| **PHF14** | 4.42106 | 0.50934 | 5.61007 | 1.78132 | 4.28452 | 8.53663 | 14.08004 | 2.38619 | 2.28215 | 24.81719 | 3.06264 | 3.71708 |

| | | | | | | | | | | | | |
|---|---|---|---|---|---|---|---|---|---|---|---|---|
| **PHF19** | 0.03976 | 0.03967 | 0.30618 | 1.33680 | 0.01160 | 0.03341 | 0.07246 | 0.12589 | 0.00000 | 0.00000 | 0.08817 | 0.03222 |
| **PHF2** | 1.02968 | 0.34663 | 0.96242 | 0.61535 | 0.59463 | 2.23700 | 0.00000 | 0.00000 | 0.00000 | 2.12982 | 0.42155 | 0.00000 |
| **PHF20** | 2.84994 | 4.16197 | 2.94658 | 3.19018 | 2.05733 | 9.24687 | 5.03322 | 1.61927 | 5.31100 | 6.62133 | 2.96000 | 6.46966 |
| **PHF20L1** | 0.41171 | 2.18886 | 2.98987 | 1.02848 | 1.36675 | 6.55272 | 5.06405 | 2.73212 | 0.00035 | 13.56878 | 1.81962 | 2.16034 |
| **PHF21A** | 4.42515 | 0.65214 | 0.01641 | 1.17041 | 1.25164 | 1.61374 | 0.85926 | 0.00000 | 1.95723 | 4.57720 | 1.99008 | 0.37920 |
| **PHF21B** | 0.00048 | 0.30104 | 0.25166 | 0.40903 | 0.46886 | 0.43773 | 0.00000 | 0.08115 | 0.00000 | 2.92899 | 0.22535 | 0.35072 |
| **PHF23** | 1.92890 | 3.93871 | 8.85678 | 4.62713 | 6.68831 | 3.33980 | 10.50025 | 7.90459 | 2.20640 | 24.22377 | 3.21506 | 9.39138 |
| **PHF3** | 4.56369 | 6.11078 | 6.38054 | 4.03340 | 2.91218 | 19.54583 | 8.08634 | 5.83437 | 10.18174 | 23.64001 | 4.91749 | 12.32853 |
| **PHF5A** | 6.61244 | 11.23207 | 17.67594 | 8.12312 | 7.43814 | 15.63445 | 18.17625 | 11.64614 | 20.24646 | 6.10097 | 15.82497 | 23.85091 |
| **PHF6** | 6.13060 | 3.14758 | 1.03577 | 0.77262 | 0.00000 | 6.83010 | 3.66194 | 4.63146 | 4.24167 | 4.07066 | 2.95662 | 5.69627 |
| **PHF7** | 1.87477 | 2.04613 | 0.74438 | 0.46461 | 0.11524 | 0.00000 | 0.68175 | 1.27503 | 2.23707 | 2.99868 | 2.08642 | 1.82034 |
| **PHF8** | 14.91284 | 3.13945 | 1.74434 | 2.20313 | 4.29709 | 4.90047 | 3.55916 | 2.21277 | 5.83212 | 6.44951 | 2.78027 | 10.17804 |
| **PHGDH** | 0.00706 | 0.00000 | 9.19050 | 9.90198 | 11.55211 | 1.43329 | 26.86246 | 0.00011 | 18.16749 | 1.43491 | 11.77111 | 1.39886 |
| **PHGR1** | 0.00645 | 0.01630 | 0.01804 | 0.01344 | 0.02274 | 0.00373 | 0.01895 | 0.01063 | 0.01169 | 0.00852 | 0.06168 | 0.01785 |
| **PHIP** | 1.55220 | 3.85912 | 4.55130 | 6.97881 | 4.90143 | 20.61159 | 10.53008 | 1.71316 | 9.01039 | 30.39557 | 0.56717 | 4.06431 |
| **PHKA1** | 0.00000 | 5.44447 | 0.95171 | 9.02578 | 5.27780 | 2.85837 | 9.44161 | 0.06853 | 11.95142 | 7.80094 | 6.52386 | 0.00000 |
| **PHKA2** | 0.84106 | 0.00000 | 0.00000 | 2.09210 | 0.20162 | 0.84226 | 0.95490 | 0.20416 | 0.23020 | 3.29246 | 1.07747 | 1.79502 |
| **PHKA2-AS1** | 0.74520 | 0.00000 | 0.06147 | 1.67428 | 0.27817 | 0.00000 | 0.02212 | 0.00841 | 0.00000 | 0.66295 | 0.32639 | 0.80324 |
| **PHKB** | 2.93584 | 6.13892 | 4.52100 | 3.52015 | 4.61454 | 15.28911 | 3.93262 | 0.46350 | 4.16345 | 5.09784 | 6.58033 | 3.56922 |
| **PHKG1** | 0.03193 | 0.21727 | 0.08496 | 0.03976 | 0.13136 | 0.00000 | 0.00000 | 0.29893 | 0.26562 | 0.18001 | 0.05060 | 0.14767 |
| **PHKG2** | 0.81241 | 7.69202 | 28.47238 | 8.88151 | 11.45012 | 0.00000 | 14.77436 | 2.27187 | 10.11062 | 8.45833 | 7.81590 | 6.02907 |
| **PHLDA1** | 0.00479 | 1.23392 | 1.32946 | 21.72165 | 0.61555 | 1.04559 | 0.48300 | 0.17262 | 0.00000 | 0.72442 | 0.00000 | 0.00000 |
| **PHLDA2** | 0.00074 | 7.75830 | 14.79474 | 7.10914 | 10.43793 | 0.00000 | 4.64245 | 16.76950 | 4.15485 | 0.00150 | 2.85025 | 22.31469 |
| **PHLDA3** | 0.46651 | 4.18173 | 5.19759 | 3.38325 | 4.90135 | 0.00233 | 5.13424 | 0.00000 | 0.62417 | 9.31116 | 0.00000 | 0.62407 |
| **PHLDB1** | 0.51514 | 1.95958 | 0.37119 | 2.99222 | 3.86585 | 0.49132 | 2.33992 | 0.13561 | 4.15054 | 5.48554 | 2.18752 | 0.50860 |
| **PHLDB2** | 1.30220 | 0.26509 | 7.65612 | 6.22491 | 0.00369 | 0.64243 | 1.17819 | 0.00000 | 0.00000 | 1.13761 | 0.12129 | 1.59870 |
| **PHLDB3** | 0.33594 | 8.38765 | 14.01263 | 1.60097 | 2.92853 | 0.00000 | 0.30925 | 0.00000 | 1.95156 | 0.00000 | 0.00000 | 7.82917 |
| **PHLPP1** | 2.59321 | 0.75290 | 7.20331 | 4.70661 | 5.81243 | 2.68492 | 6.70029 | 1.17026 | 0.06530 | 8.28342 | 1.02024 | 3.51357 |
| **PHLPP2** | 0.08677 | 2.99482 | 1.47223 | 0.31577 | 1.49958 | 1.73543 | 3.60766 | 1.56674 | 0.00000 | 7.42679 | 0.61356 | 1.68539 |
| **PHOSPHO1** | 2.58250 | 27.30176 | 0.17447 | 2.18882 | 44.69683 | 5.80813 | 0.00000 | 19.47443 | 5.08039 | 0.00000 | 3.37531 | 45.07954 |
| **PHOSPHO2** | 0.57838 | 0.14211 | 0.71767 | 0.31431 | 0.00000 | 1.77993 | 1.22404 | 0.86302 | 0.37144 | 0.41389 | 1.89043 | 1.19536 |
| **PHOX2A** | 0.02345 | 0.19428 | 0.00486 | 0.00157 | 0.16148 | 0.00000 | 0.03550 | 0.00935 | 0.00000 | 0.13855 | 0.02932 | 0.06225 |
| **PHOX2B** | 0.02034 | 0.00507 | 0.01433 | 0.01053 | 0.05002 | 0.00000 | 0.01192 | 1.04684 | 0.01173 | 0.04894 | 0.01536 | 0.19345 |
| **PHPT1** | 1.68798 | 9.47386 | 12.24733 | 15.90552 | 17.65895 | 3.54948 | 18.59286 | 4.04689 | 17.02139 | 5.43214 | 13.19422 | 6.38819 |
| **PHRF1** | 5.02677 | 2.03138 | 4.49069 | 3.10406 | 4.94995 | 0.00000 | 1.35656 | 3.62645 | 4.12082 | 11.78549 | 2.30059 | 1.95216 |
| **PHTF1** | 0.72001 | 0.90308 | 6.96446 | 0.76191 | 0.69640 | 2.92745 | 5.34395 | 2.35996 | 0.89796 | 3.04474 | 0.80460 | 0.28806 |
| **PHTF2** | 7.06594 | 2.46906 | 7.23369 | 0.93602 | 1.30001 | 11.33640 | 0.00000 | 5.81496 | 14.21788 | 3.46315 | 3.66099 | 1.64036 |
| **PHYH** | 0.01391 | 0.02495 | 0.64160 | 0.28382 | 0.08653 | 0.32177 | 0.19339 | 1.05080 | 0.00000 | 0.00000 | 0.06449 | 0.24465 |
| **PHYHD1** | 0.89243 | 1.81495 | 0.19629 | 4.39479 | 1.27484 | 1.30065 | 0.91685 | 0.34823 | 1.47116 | 0.31756 | 1.77771 | 0.85795 |
| **PHYHIP** | 0.00447 | 0.13249 | 0.64828 | 0.00000 | 0.00000 | 0.00006 | 0.02943 | 0.16713 | 0.05950 | 0.05055 | 0.07571 | 0.81329 |
| **PHYHIPL** | 0.32866 | 0.01263 | 0.51210 | 0.03613 | 0.09768 | 0.00000 | 0.02882 | 0.00000 | 0.25046 | 0.00000 | 0.00000 | 0.20954 |
| **PHYKPL** | 0.02928 | 1.93936 | 2.01711 | 2.04335 | 1.16206 | 0.00000 | 0.00000 | 0.95448 | 3.25786 | 1.22166 | 2.58974 | 1.50885 |
| **PI15** | 0.00292 | 0.02462 | 0.12402 | 0.04193 | 0.00000 | 0.26912 | 0.00000 | 0.01687 | 0.06397 | 0.01240 | 0.00087 | 0.05537 |
| **PI16** | 0.03627 | 0.40394 | 0.06263 | 2.96663 | 0.04135 | 0.00000 | 0.01188 | 0.00343 | 0.00000 | 0.01746 | 0.02496 | 0.07358 |
| **PI4K2A** | 0.00000 | 1.39180 | 4.07434 | 4.87035 | 5.12082 | 0.51636 | 6.75803 | 1.66642 | 1.22406 | 5.25649 | 0.99771 | 5.12157 |
| **PI4K2B** | 1.03016 | 0.00009 | 0.80310 | 1.36302 | 0.00000 | 2.92676 | 0.00000 | 0.96963 | 0.00000 | 2.38096 | 1.71438 | 0.00000 |
| **PI4KA** | 0.14965 | 1.31515 | 3.85940 | 3.19687 | 6.17577 | 0.59968 | 3.15803 | 0.43982 | 5.28148 | 4.84095 | 3.69434 | 0.65568 |
| **PI4KAP1** | 0.00207 | 0.82016 | 3.40660 | 1.12942 | 2.72146 | 0.00000 | 0.00000 | 0.04241 | 3.78227 | 4.41959 | 2.23677 | 0.41977 |

| | | | | | | | | | | | | |
|---|---|---|---|---|---|---|---|---|---|---|---|---|
| **PI4KAP2** | 0.07337 | 0.94057 | 1.11438 | 0.51453 | 2.28127 | 0.00000 | 0.02862 | 0.06421 | 3.88691 | 1.86666 | 1.05405 | 0.48928 |
| **PI4KB** | 7.64941 | 15.26454 | 7.21075 | 6.91854 | 8.00386 | 0.05704 | 8.00806 | 3.96156 | 9.05274 | 0.00000 | 4.60342 | 15.05767 |
| **PIANP** | 0.00670 | 0.00504 | 0.05515 | 0.02538 | 0.00448 | 0.02222 | 0.00570 | 0.99490 | 0.01071 | 0.04519 | 0.02267 | 0.10840 |
| **PIAS1** | 11.07138 | 6.50914 | 3.85203 | 5.00467 | 2.27279 | 19.36774 | 5.82952 | 14.01161 | 7.30544 | 2.29988 | 8.29965 | 10.62498 |
| **PIAS2** | 3.71197 | 3.90491 | 2.10268 | 3.89100 | 0.00000 | 8.59013 | 0.00000 | 4.95282 | 1.79669 | 8.29425 | 1.78043 | 3.26768 |
| **PIAS3** | 0.47200 | 2.79686 | 1.54808 | 2.06691 | 0.00000 | 0.00000 | 2.09704 | 0.96703 | 2.52443 | 1.55594 | 0.57313 | 4.63519 |
| **PIAS4** | 4.97186 | 10.93445 | 8.91041 | 7.62905 | 7.75069 | 0.00000 | 5.76718 | 3.83270 | 7.73113 | 3.73400 | 9.65441 | 11.48810 |
| **PIBF1** | 1.20988 | 1.20602 | 0.59632 | 0.00000 | 0.97296 | 10.26303 | 0.00000 | 1.57324 | 6.62214 | 4.35336 | 9.32820 | 1.85301 |
| **PICALM** | 3.41112 | 3.20501 | 11.14988 | 2.09351 | 0.00078 | 17.40002 | 15.62065 | 5.86927 | 0.00000 | 16.75998 | 1.84826 | 7.10882 |
| **PICK1** | 0.00000 | 1.66743 | 1.51418 | 0.74516 | 0.94303 | 0.00000 | 0.27016 | 0.16015 | 0.00000 | 3.35766 | 0.61000 | 0.21353 |
| **PID1** | 1.01377 | 1.08407 | 14.97374 | 0.00000 | 1.87147 | 1.06722 | 3.84090 | 0.68380 | 2.38662 | 0.00000 | 0.24167 | 0.93351 |
| **PIDD1** | 0.35059 | 1.66595 | 3.47728 | 2.64817 | 4.86284 | 0.00000 | 2.21225 | 0.00000 | 8.13417 | 4.83421 | 0.90121 | 0.32506 |
| **PIEZO1** | 3.57730 | 1.40841 | 1.79616 | 0.00000 | 6.37931 | 0.00000 | 1.76309 | 0.24001 | 11.39993 | 14.33540 | 2.85225 | 0.94249 |
| **PIEZO2** | 0.00659 | 0.01746 | 0.11883 | 0.01879 | 0.18283 | 0.00011 | 0.00114 | 0.06634 | 0.00000 | 0.13877 | 0.10936 | 0.11188 |
| **PIF1** | 0.00000 | 7.27623 | 3.50801 | 2.71426 | 4.10695 | 0.00219 | 0.41014 | 0.41617 | 5.76684 | 7.62392 | 1.88917 | 2.68361 |
| **PIFO** | 0.40288 | 1.29831 | 1.70667 | 3.60302 | 0.00000 | 1.51896 | 0.17132 | 0.00000 | 0.00000 | 0.03666 | 0.00000 | 1.03175 |
| **PIGA** | 3.81817 | 2.66253 | 3.86433 | 1.53659 | 1.53235 | 8.97274 | 2.31582 | 5.25039 | 1.17278 | 0.00000 | 4.37422 | 3.53690 |
| **PIGB** | 0.37578 | 3.04374 | 2.44901 | 1.58215 | 1.00040 | 4.14494 | 0.89043 | 2.61797 | 3.13041 | 0.00000 | 2.40401 | 4.41682 |
| **PIGC** | 0.00050 | 7.25252 | 4.57237 | 3.03641 | 1.42010 | 10.65475 | 2.23458 | 0.53050 | 6.92217 | 0.00000 | 5.33982 | 8.40966 |
| **PIGF** | 0.94062 | 1.18474 | 2.43470 | 0.67577 | 0.36690 | 4.58318 | 0.00000 | 1.59185 | 4.14406 | 0.00000 | 2.26078 | 4.11449 |
| **PIGG** | 0.00000 | 5.84796 | 7.40097 | 5.18931 | 5.89184 | 2.42505 | 0.15493 | 7.39605 | 21.79926 | 4.56464 | 7.13474 | 5.99815 |
| **PIGH** | 2.08199 | 1.13532 | 8.27778 | 0.24655 | 1.01668 | 10.60819 | 0.00000 | 6.37123 | 10.78077 | 0.00000 | 4.32894 | 9.32205 |
| **PIGK** | 0.15473 | 7.49369 | 2.71064 | 0.66744 | 1.74018 | 14.83462 | 0.00000 | 0.64630 | 5.27227 | 2.69015 | 3.63697 | 4.44672 |
| **PIGL** | 0.00335 | 6.36989 | 2.81163 | 2.83976 | 2.33301 | 1.13161 | 0.19274 | 0.42805 | 9.25171 | 0.81211 | 1.90980 | 5.97244 |
| **PIGM** | 1.04349 | 5.04031 | 0.81756 | 0.00000 | 3.03570 | 2.85996 | 0.00000 | 0.03739 | 0.73234 | 1.65891 | 2.85900 | 4.03069 |
| **PIGN** | 0.28258 | 2.81155 | 3.27025 | 0.79970 | 0.03124 | 4.81913 | 0.00000 | 1.18630 | 6.51280 | 0.51840 | 2.72625 | 1.18411 |
| **PIGO** | 0.25567 | 11.96152 | 2.71617 | 6.92777 | 8.45793 | 0.00000 | 0.00000 | 0.00000 | 6.89418 | 0.30340 | 7.09061 | 5.09150 |
| **PIGP** | 0.72914 | 3.71941 | 1.48944 | 0.00000 | 0.01205 | 6.33689 | 0.00000 | 0.00000 | 1.94699 | 2.61822 | 4.07769 | 3.24586 |
| **PIGQ** | 0.45424 | 2.57210 | 1.54176 | 1.37109 | 1.55663 | 0.00000 | 0.00002 | 0.00000 | 1.83867 | 1.93913 | 3.11082 | 0.37292 |
| **PIGR** | 0.00483 | 0.01459 | 0.12225 | 0.00697 | 0.01687 | 0.01784 | 0.00731 | 0.00975 | 0.00000 | 0.09362 | 0.03514 | 0.01829 |
| **PIGS** | 1.18950 | 11.49667 | 10.10730 | 6.24213 | 8.88440 | 1.91717 | 4.25239 | 1.18490 | 12.92081 | 1.83834 | 10.30110 | 8.93586 |
| **PIGT** | 0.80976 | 12.22126 | 12.67043 | 8.42525 | 16.36092 | 0.06247 | 0.48978 | 5.66205 | 29.47325 | 0.00000 | 9.22104 | 14.06573 |
| **PIGU** | 6.74291 | 6.18236 | 2.56462 | 3.03903 | 5.98778 | 0.00000 | 0.00000 | 0.55213 | 7.69225 | 2.46197 | 3.61476 | 2.59927 |
| **PIGV** | 2.04489 | 2.58833 | 1.03617 | 0.86345 | 0.27472 | 1.50415 | 0.00000 | 0.97878 | 2.22174 | 0.00000 | 1.69739 | 3.66096 |
| **PIGW** | 1.15170 | 6.10567 | 4.91620 | 2.59876 | 0.36217 | 10.77236 | 1.67184 | 2.05218 | 9.02978 | 0.00000 | 8.53993 | 9.93558 |
| **PIGX** | 0.95445 | 0.35177 | 0.43515 | 0.53618 | 1.80995 | 1.99355 | 0.43062 | 0.00000 | 0.00000 | 5.10100 | 0.73962 | 0.44589 |
| **PIGY+PYURF** | 0.87670 | 5.84562 | 6.74122 | 2.85580 | 2.88490 | 21.75762 | 12.32302 | 5.94963 | 17.08702 | 0.00000 | 11.74540 | 14.32746 |
| **PIGZ** | 0.35362 | 0.00606 | 0.01426 | 0.10818 | 0.05390 | 0.00000 | 0.03310 | 0.17235 | 0.00000 | 0.12227 | 0.01148 | 0.15046 |
| **PIH1D1** | 0.29036 | 5.77876 | 15.60311 | 14.39074 | 6.23856 | 2.04201 | 17.78797 | 7.08125 | 15.68387 | 0.00000 | 9.32058 | 10.12558 |
| **PIH1D2** | 0.19492 | 0.27201 | 1.10825 | 0.00761 | 0.00001 | 0.49552 | 0.00000 | 0.07157 | 0.68482 | 0.39908 | 0.01095 | 0.68187 |
| **PIH1D3** | 0.00579 | 0.01578 | 0.01773 | 0.01109 | 0.01030 | 0.01474 | 0.03685 | 0.01058 | 0.00000 | 0.03192 | 0.05179 | 0.01520 |
| **PIK3AP1** | 0.00093 | 1.13794 | 0.19707 | 0.51101 | 0.00000 | 0.00000 | 0.00000 | 0.70437 | 0.00000 | 0.71428 | 0.11834 | 5.32384 |
| **PIK3C2A** | 10.48432 | 0.21092 | 13.87583 | 4.42255 | 1.43586 | 12.99691 | 13.06487 | 3.11146 | 9.77889 | 13.25239 | 4.64577 | 5.23123 |
| **PIK3C2B** | 4.03747 | 0.00000 | 1.06272 | 0.00000 | 0.50242 | 0.22285 | 2.11594 | 0.48235 | 2.44431 | 2.76135 | 0.46463 | 0.89047 |
| **PIK3C2G** | 0.01199 | 0.00403 | 0.03511 | 0.00043 | 0.00400 | 0.05057 | 0.00000 | 0.03240 | 0.01946 | 0.32920 | 0.01637 | 0.00386 |
| **PIK3C3** | 1.27986 | 3.93891 | 5.56484 | 2.37332 | 0.49749 | 11.41903 | 4.64724 | 13.60513 | 7.58785 | 1.53403 | 2.21037 | 8.79276 |
| **PIK3CA** | 1.63133 | 0.11391 | 0.85754 | 0.75460 | 0.83019 | 2.17206 | 0.03804 | 1.93959 | 0.00000 | 1.73155 | 0.42446 | 1.31955 |
| **PIK3CB** | 0.21094 | 3.98012 | 11.47001 | 0.00000 | 5.27171 | 14.91603 | 8.81122 | 2.76226 | 0.00000 | 8.29198 | 4.32846 | 7.10772 |

| | | | | | | | | | | | | |
|---|---|---|---|---|---|---|---|---|---|---|---|---|
| **PIK3CD** | 0.63926 | 1.79103 | 0.00000 | 0.50851 | 0.60329 | 0.00000 | 0.21153 | 0.02896 | 0.00000 | 1.39350 | 0.19713 | 1.12377 |
| **PIK3CG** | 1.34622 | 0.04049 | 0.02004 | 0.07561 | 0.03183 | 0.00000 | 0.09010 | 0.00000 | 0.07894 | 0.00000 | 0.03676 | 0.20706 |
| **PIK3IP1** | 0.05001 | 0.24062 | 0.26745 | 0.04359 | 0.05676 | 0.88228 | 0.00000 | 0.42294 | 0.53586 | 0.00000 | 0.00000 | 1.95450 |
| **PIK3R1** | 3.85016 | 0.57897 | 1.38308 | 3.26115 | 0.00000 | 2.08415 | 0.37258 | 0.92061 | 0.23469 | 1.24932 | 0.00000 | 0.33033 |
| **PIK3R2** | 0.81499 | 0.37879 | 2.11437 | 1.05915 | 1.96764 | 0.00000 | 1.59365 | 0.04198 | 0.00000 | 6.34473 | 0.68821 | 0.33653 |
| **PIK3R3** | 2.74824 | 0.00000 | 2.73475 | 4.86922 | 0.00000 | 3.66707 | 8.39779 | 8.72736 | 9.74014 | 0.00000 | 2.15421 | 2.99588 |
| **PIK3R4** | 0.00000 | 6.55776 | 3.74587 | 3.01575 | 2.63692 | 10.28237 | 0.00000 | 12.02641 | 5.99628 | 9.74678 | 4.43729 | 11.65309 |
| **PIK3R5** | 0.00590 | 0.06304 | 0.11732 | 0.87399 | 0.00000 | 0.01654 | 0.00034 | 0.02298 | 0.00000 | 0.02475 | 0.03463 | 0.05931 |
| **PIK3R6** | 0.00399 | 0.01223 | 0.02406 | 0.04670 | 0.00184 | 0.08912 | 0.00000 | 0.13306 | 0.01745 | 0.02396 | 0.01129 | 0.00562 |
| **PIKFYVE** | 5.48631 | 0.50336 | 0.97895 | 2.58995 | 1.17426 | 8.64785 | 0.88557 | 2.43583 | 3.05445 | 11.79284 | 1.00057 | 2.12683 |
| **PILRA** | 0.08429 | 0.26123 | 0.06407 | 0.00616 | 0.11022 | 0.00000 | 0.07155 | 0.02180 | 0.24435 | 0.00000 | 0.00000 | 0.14617 |
| **PILRB** | 0.00542 | 0.12029 | 0.17159 | 0.09962 | 0.03367 | 0.00000 | 0.00000 | 0.18950 | 0.22111 | 0.12175 | 0.02063 | 0.07388 |
| **PIM1** | 5.25988 | 17.24676 | 17.05449 | 2.33786 | 0.00000 | 2.50412 | 14.40068 | 21.45407 | 18.69822 | 6.51674 | 1.66831 | 36.58781 |
| **PIM2** | 0.34789 | 4.19738 | 2.63045 | 22.75886 | 4.96142 | 0.03650 | 6.52645 | 0.98148 | 9.25452 | 0.00000 | 5.75271 | 1.07057 |
| **PIM3** | 2.61451 | 1.41700 | 6.21893 | 1.11672 | 8.12086 | 0.00000 | 7.15391 | 4.19090 | 3.45476 | 12.96397 | 1.58976 | 1.45335 |
| **PIN1** | 2.22243 | 17.83124 | 10.83924 | 8.99731 | 13.45926 | 0.00000 | 6.95334 | 1.27190 | 15.42327 | 0.00000 | 14.68847 | 9.58662 |
| **PIN1P1** | 0.10127 | 0.88577 | 0.49496 | 0.42486 | 0.51880 | 0.02673 | 0.28504 | 0.04550 | 0.42187 | 0.00000 | 0.87587 | 0.31447 |
| **PIN4** | 0.00000 | 8.26562 | 2.24283 | 5.74472 | 3.88144 | 7.61580 | 8.94440 | 2.35345 | 3.23179 | 7.25399 | 7.93550 | 9.87015 |
| **PIN4P1** | 0.00000 | 3.08267 | 0.88627 | 2.53927 | 1.35083 | 2.15617 | 3.08989 | 0.98950 | 0.74101 | 2.77815 | 2.77484 | 2.93977 |
| **PINK1** | 0.18897 | 3.58177 | 0.05126 | 2.82804 | 5.14392 | 0.00000 | 0.07831 | 0.00000 | 0.55597 | 2.16596 | 0.41755 | 2.87713 |
| **PINK1-AS** | 0.00445 | 0.65174 | 0.26952 | 0.89376 | 1.23824 | 0.00000 | 0.35864 | 0.00857 | 0.34735 | 0.62896 | 0.32307 | 0.71175 |
| **PINLYP** | 0.00287 | 0.22650 | 0.61850 | 0.27169 | 0.52954 | 0.53641 | 0.42517 | 0.02293 | 0.00000 | 0.00000 | 0.48794 | 0.19160 |
| **PINX1** | 2.70978 | 13.11574 | 6.80412 | 4.81576 | 11.60854 | 5.46514 | 9.70680 | 8.22018 | 10.22921 | 0.05718 | 8.32170 | 15.19510 |
| **PIP4K2A** | 8.56236 | 5.28767 | 3.47545 | 4.00979 | 4.60242 | 15.85664 | 18.71365 | 3.93619 | 5.52317 | 9.82839 | 13.84385 | 1.82315 |
| **PIP4K2B** | 1.37831 | 2.89163 | 2.92077 | 3.85778 | 2.04827 | 0.00664 | 6.44867 | 1.27044 | 5.18364 | 5.69752 | 3.15969 | 2.70224 |
| **PIP4K2C** | 7.81412 | 12.38527 | 9.33403 | 6.98419 | 6.45777 | 13.38890 | 10.69161 | 4.83888 | 15.05309 | 0.00000 | 7.31657 | 9.15253 |
| **PIP5K1A** | 0.68632 | 6.47970 | 13.78740 | 6.61748 | 5.64866 | 10.44862 | 9.67563 | 14.41291 | 12.46179 | 0.00000 | 8.38049 | 9.78565 |
| **PIP5K1B** | 1.34299 | 13.13379 | 0.00000 | 0.00000 | 9.20092 | 0.00000 | 4.23957 | 0.30979 | 3.36844 | 2.92290 | 10.83521 | 10.10878 |
| **PIP5K1C** | 1.70259 | 1.72420 | 2.16214 | 0.37880 | 4.22413 | 0.00000 | 0.87488 | 0.16505 | 4.85890 | 7.23645 | 1.19072 | 1.16885 |
| **PIP5K1P1** | 0.00344 | 0.13100 | 0.29760 | 0.06251 | 0.12556 | 0.29425 | 0.07271 | 0.32877 | 0.30070 | 0.00000 | 0.18125 | 0.19690 |
| **PIP5KL1** | 0.11983 | 0.10176 | 0.15005 | 0.01182 | 0.01153 | 0.03921 | 0.07298 | 0.10633 | 0.00603 | 0.01345 | 0.02985 | 0.07993 |
| **PIPOX** | 0.00577 | 0.12007 | 0.72385 | 0.33662 | 0.00000 | 0.00000 | 0.00000 | 0.00594 | 0.02735 | 0.01391 | 0.11290 | 0.23013 |
| **PIPSL** | 0.94508 | 8.67622 | 11.87773 | 6.53500 | 7.11084 | 7.70403 | 13.43051 | 4.78294 | 6.67088 | 0.00000 | 12.92758 | 8.62503 |
| **PIR** | 0.00406 | 0.28694 | 1.81201 | 0.31795 | 3.15982 | 1.18143 | 2.82483 | 0.00122 | 1.60849 | 0.01734 | 2.33174 | 0.00000 |
| **PIR-FIGF** | 0.00606 | 0.01603 | 0.04736 | 0.00949 | 0.04025 | 0.00325 | 0.02257 | 0.01028 | 0.02715 | 0.01054 | 0.04163 | 0.02645 |
| **PISD** | 1.50339 | 11.67561 | 10.02634 | 7.98971 | 7.54086 | 9.52134 | 8.56372 | 3.92997 | 16.05626 | 0.00000 | 10.07331 | 5.65769 |
| **PISRT1** | 0.00588 | 0.01398 | 0.01100 | 0.08316 | 0.01190 | 0.00356 | 0.02335 | 0.01053 | 0.02122 | 0.02854 | 0.02071 | 0.01715 |
| **PITHD1** | 8.22550 | 21.27389 | 19.62276 | 7.35843 | 9.88998 | 13.11234 | 17.36053 | 11.65836 | 18.80880 | 12.40031 | 17.26087 | 12.81340 |
| **PITPNA** | 3.35468 | 2.89514 | 5.74715 | 0.95356 | 6.76380 | 4.07310 | 10.46420 | 1.80559 | 2.76570 | 13.72337 | 1.22878 | 3.73809 |
| **PITPNA-AS1** | 0.80390 | 3.45762 | 0.90576 | 0.67222 | 1.05458 | 2.41075 | 0.01579 | 0.41169 | 1.91173 | 3.04699 | 3.27868 | 8.97347 |
| **PITPNB** | 8.37180 | 13.43254 | 21.78765 | 13.36510 | 4.53953 | 30.22309 | 19.08972 | 15.68401 | 17.34768 | 4.56592 | 12.72859 | 10.47096 |
| **PITPNC1** | 0.15009 | 1.96560 | 0.64528 | 5.61422 | 11.26375 | 1.95548 | 7.72202 | 0.24512 | 2.11084 | 12.89164 | 6.90865 | 0.26310 |
| **PITPNM1** | 0.53734 | 0.00201 | 0.17657 | 1.38021 | 0.41175 | 0.00000 | 0.58927 | 0.00004 | 0.34646 | 0.89742 | 0.16197 | 0.72943 |
| **PITPNM2** | 0.01407 | 0.08335 | 1.01262 | 2.52654 | 0.09749 | 0.00000 | 0.30500 | 0.00814 | 0.43306 | 0.10415 | 0.00000 | 0.03942 |
| **PITPNM3** | 0.21176 | 0.04676 | 0.01111 | 0.03880 | 0.12337 | 0.00000 | 0.00487 | 0.00974 | 0.14829 | 0.09739 | 0.03192 | 0.09615 |
| **PITRM1** | 13.16148 | 5.67704 | 5.67131 | 4.60417 | 6.54758 | 6.63702 | 4.56369 | 5.34072 | 16.33885 | 0.00000 | 6.20327 | 6.56696 |
| **PITRM1-AS1** | 0.49235 | 0.06993 | 0.43504 | 0.43990 | 0.20685 | 0.71270 | 0.58529 | 0.10321 | 0.52927 | 0.95389 | 0.00000 | 0.21955 |
| **PITX1** | 0.64854 | 2.23056 | 1.34518 | 5.05498 | 3.12473 | 0.00000 | 1.72052 | 1.03868 | 0.45915 | 6.30033 | 0.28506 | 1.83593 |

| | | | | | | | | | | | | |
|---|---|---|---|---|---|---|---|---|---|---|---|---|
| PITX2 | 0.46860 | 0.02919 | 2.35908 | 6.91694 | 0.22315 | 0.14622 | 0.69504 | 0.06148 | 0.00000 | 0.83961 | 0.25386 | 0.81539 |
| PITX3 | 0.01409 | 0.34753 | 0.01271 | 0.08405 | 0.02789 | 0.01259 | 0.00000 | 0.00246 | 0.01217 | 0.32036 | 0.00000 | 0.30381 |
| PIWIL1 | 0.95233 | 0.01183 | 0.01205 | 0.07658 | 0.01211 | 0.06594 | 0.00741 | 0.46782 | 0.01128 | 0.01645 | 0.00880 | 0.01756 |
| PIWIL2 | 1.10485 | 2.80081 | 0.31745 | 0.04624 | 0.18510 | 0.00000 | 0.39807 | 2.10979 | 0.02322 | 0.25447 | 0.00000 | 6.45032 |
| PIWIL3 | 4.12146 | 0.00000 | 0.14003 | 0.00000 | 0.98099 | 0.00000 | 0.24831 | 0.90688 | 0.88731 | 1.92078 | 4.04955 | 0.22323 |
| PIWIL4 | 0.09157 | 0.00000 | 0.34919 | 0.00691 | 0.38424 | 0.00503 | 0.80545 | 0.28508 | 0.80205 | 0.00000 | 0.96689 | 0.15989 |
| PJA1 | 0.24323 | 2.41821 | 2.07747 | 1.35144 | 0.00000 | 0.26704 | 2.28866 | 0.51376 | 4.72805 | 0.00000 | 0.87544 | 2.89978 |
| PJA2 | 4.07149 | 5.16393 | 4.41306 | 5.01471 | 0.00000 | 12.78691 | 2.13024 | 0.37680 | 0.00000 | 7.30581 | 1.99269 | 5.42068 |
| PKD1 | 0.66388 | 1.65727 | 0.18288 | 0.80801 | 0.66143 | 0.00000 | 0.00000 | 0.28083 | 1.45959 | 2.22342 | 0.07467 | 0.76409 |
| PKD1L1 | 0.03696 | 0.64629 | 0.14303 | 2.01746 | 0.16777 | 0.18753 | 0.00000 | 0.18464 | 0.20549 | 0.60822 | 0.00000 | 1.33646 |
| PKD1L2 | 0.01830 | 1.38586 | 0.38183 | 0.00000 | 1.39753 | 0.00000 | 1.53078 | 0.15592 | 0.00007 | 1.23203 | 0.59557 | 0.33317 |
| PKD1L3 | 6.38887 | 0.00271 | 0.01451 | 0.00783 | 0.01439 | 0.00000 | 0.00354 | 1.79752 | 0.04286 | 0.01108 | 0.02189 | 0.29795 |
| PKD1P1 | 0.01172 | 0.03919 | 0.05978 | 0.07645 | 0.02959 | 0.00000 | 0.08801 | 0.03506 | 0.03567 | 0.09997 | 0.06023 | 0.03866 |
| PKD2 | 0.02828 | 0.00015 | 0.00000 | 0.75933 | 0.26572 | 0.22699 | 0.00000 | 0.00792 | 0.28282 | 0.08365 | 0.00000 | 0.00625 |
| PKD2L1 | 0.02160 | 0.01660 | 0.01987 | 0.03000 | 0.01906 | 0.00196 | 0.03355 | 0.12900 | 0.01307 | 0.03152 | 0.02495 | 0.01968 |
| PKD2L2 | 0.00668 | 0.00591 | 0.05768 | 0.01232 | 0.00000 | 0.07444 | 0.04340 | 0.01309 | 0.00000 | 0.02594 | 0.06241 | 0.08235 |
| PKDCC | 0.05327 | 0.29169 | 0.07659 | 1.07679 | 0.00000 | 0.00000 | 0.00000 | 0.00094 | 0.00754 | 1.38923 | 0.00000 | 0.05682 |
| PKDREJ | 0.01313 | 0.01104 | 0.01757 | 0.00857 | 0.01894 | 0.02847 | 0.01840 | 0.00982 | 0.01248 | 0.02222 | 0.01352 | 0.02328 |
| PKHD1 | 1.78203 | 0.49462 | 0.09637 | 1.26852 | 0.00000 | 0.20254 | 0.06284 | 1.12971 | 0.00000 | 0.00000 | 0.07650 | 1.63453 |
| PKHD1L1 | 0.00217 | 0.08250 | 0.02509 | 0.00000 | 0.05829 | 0.00378 | 0.04397 | 0.77379 | 0.00000 | 0.05889 | 0.02165 | 0.76789 |
| PKI55 | 0.00301 | 0.17676 | 0.16979 | 0.05861 | 0.00000 | 0.00000 | 1.21034 | 0.00368 | 0.03685 | 0.67366 | 0.17789 | 0.19382 |
| PKIA | 0.00238 | 0.01478 | 10.45645 | 0.04406 | 0.00000 | 0.01977 | 1.81393 | 1.96268 | 0.07010 | 0.01152 | 0.00000 | 0.00454 |
| PKIB | 0.23791 | 0.47612 | 5.34597 | 0.00000 | 1.62063 | 10.01986 | 13.66735 | 1.33992 | 13.99552 | 1.12343 | 2.29555 | 1.02251 |
| PKIG | 0.26858 | 3.33991 | 5.26171 | 4.71504 | 2.54015 | 0.87730 | 3.09121 | 0.12228 | 1.69943 | 0.35138 | 2.05580 | 0.64173 |
| PKLR | 0.07285 | 0.01359 | 0.25024 | 0.03575 | 0.00000 | 0.02101 | 0.00000 | 0.18704 | 0.00000 | 0.07912 | 0.08185 | 0.00401 |
| PKM | 3.82002 | 24.99518 | 47.25841 | 34.50603 | 36.69097 | 3.50339 | 59.08667 | 12.00477 | 54.97874 | 6.29546 | 36.22569 | 24.22248 |
| PKMYT1 | 1.07939 | 13.04615 | 9.10861 | 6.80943 | 9.26737 | 0.00000 | 0.23941 | 0.00000 | 20.12310 | 0.00000 | 10.38398 | 7.57151 |
| PKN1 | 1.87225 | 1.58685 | 0.16207 | 0.47483 | 2.33910 | 0.00000 | 0.79633 | 0.88495 | 1.22589 | 1.93576 | 1.06034 | 1.29018 |
| PKN2 | 2.71662 | 2.87477 | 8.27460 | 5.43962 | 2.50962 | 24.95399 | 8.54780 | 7.27700 | 11.76482 | 21.92473 | 3.57303 | 9.72247 |
| PKN2-AS1 | 0.03373 | 0.01132 | 0.02105 | 0.01178 | 0.01549 | 0.01911 | 0.02867 | 0.02334 | 0.00000 | 0.02445 | 0.06233 | 0.03116 |
| PKN3 | 0.25311 | 0.62027 | 1.94090 | 0.97379 | 1.99663 | 0.00000 | 1.06359 | 0.56479 | 1.36432 | 2.13746 | 2.02320 | 0.29148 |
| PKNOX1 | 3.23835 | 6.02384 | 3.86521 | 3.61783 | 3.73489 | 12.25053 | 6.65591 | 8.28691 | 5.86588 | 7.20414 | 6.92525 | 10.48194 |
| PKNOX2 | 2.20902 | 0.00894 | 0.02579 | 0.01369 | 0.01226 | 0.00000 | 0.01535 | 0.30624 | 0.03361 | 0.02543 | 0.01287 | 0.08804 |
| PKP1 | 0.00539 | 0.01122 | 0.04954 | 0.00932 | 0.01005 | 0.02544 | 0.00956 | 0.01030 | 0.10867 | 0.05224 | 0.02596 | 0.01347 |
| PKP2 | 0.72572 | 6.34432 | 6.18071 | 7.64343 | 14.16129 | 5.08763 | 17.88586 | 0.01767 | 18.88164 | 0.00000 | 15.24739 | 0.00000 |
| PKP3 | 0.00026 | 0.02179 | 12.68229 | 0.00000 | 4.60929 | 0.00000 | 10.00709 | 0.00499 | 8.72543 | 6.13717 | 12.30727 | 0.00000 |
| PKP4 | 1.22810 | 1.79027 | 1.97491 | 0.00000 | 5.11077 | 2.47518 | 4.83501 | 0.08272 | 0.00000 | 5.75246 | 1.88811 | 0.00000 |
| PLA1A | 0.03500 | 0.00444 | 0.15805 | 0.01655 | 0.00000 | 0.05214 | 0.00000 | 0.00763 | 0.06148 | 0.05618 | 0.00988 | 0.02389 |
| PLA2G10 | 0.00155 | 0.09122 | 0.56421 | 1.64737 | 0.96541 | 0.12350 | 0.00000 | 0.03341 | 0.17707 | 0.12729 | 0.33927 | 0.01264 |
| PLA2G12A | 0.00239 | 3.67371 | 4.45999 | 3.99139 | 4.24841 | 8.30364 | 0.00000 | 3.30820 | 12.71683 | 2.45237 | 5.48873 | 4.30032 |
| PLA2G12B | 0.23766 | 0.00000 | 0.01143 | 0.58666 | 0.00000 | 0.04558 | 0.01056 | 0.06701 | 0.25695 | 0.00000 | 0.22542 | 0.01366 |
| PLA2G15 | 0.00000 | 7.60848 | 13.25166 | 3.26220 | 3.04827 | 0.00000 | 0.00000 | 5.64264 | 9.03130 | 0.68025 | 2.20200 | 13.84398 |
| PLA2G16 | 13.08129 | 2.53094 | 11.27155 | 10.89216 | 2.68157 | 6.77858 | 3.91294 | 10.91267 | 6.79114 | 0.00000 | 3.48147 | 1.49297 |
| PLA2G1B | 0.00891 | 0.00768 | 0.27675 | 0.03069 | 0.00000 | 0.05241 | 0.00000 | 0.01011 | 0.00000 | 0.03489 | 0.01712 | 0.01345 |
| PLA2G2A | 0.00478 | 0.00919 | 0.39335 | 0.01658 | 0.02472 | 0.00000 | 0.00239 | 0.75515 | 0.00000 | 0.00900 | 0.02907 | 0.09366 |
| PLA2G2F | 0.00617 | 0.01621 | 0.03822 | 0.01542 | 0.02106 | 0.00510 | 0.02142 | 0.01060 | 0.00534 | 0.01939 | 0.03996 | 0.01682 |
| PLA2G3 | 0.00607 | 0.01070 | 0.00973 | 0.08978 | 0.29601 | 0.00222 | 0.10355 | 0.01042 | 0.00000 | 0.00000 | 0.00927 | 0.01784 |
| PLA2G4A | 0.09349 | 0.01338 | 0.00000 | 0.04042 | 0.06352 | 0.01370 | 0.18300 | 0.01967 | 0.16188 | 0.00000 | 0.00000 | 0.03338 |

| | | | | | | | | | | | | |
|---|---|---|---|---|---|---|---|---|---|---|---|---|
| **PLA2G4B** | 0.00639 | 0.02367 | 0.01474 | 0.01203 | 0.01595 | 0.01465 | 0.01899 | 0.01036 | 0.01835 | 0.01622 | 0.01685 | 0.01757 |
| **PLA2G4C** | 3.66820 | 0.01934 | 0.00310 | 0.00063 | 0.02291 | 0.00000 | 0.01742 | 1.87987 | 0.01012 | 0.09473 | 0.00862 | 0.68905 |
| **PLA2G4D** | 0.00701 | 0.01528 | 0.05571 | 0.01491 | 0.01241 | 0.04903 | 0.01235 | 0.01194 | 0.01938 | 0.00518 | 0.01059 | 0.03487 |
| **PLA2G4E** | 0.05326 | 0.06555 | 0.08970 | 0.00000 | 0.07384 | 0.00000 | 0.03151 | 0.00438 | 0.02999 | 0.23325 | 0.03498 | 0.01797 |
| **PLA2G4F** | 0.24770 | 4.15513 | 0.20431 | 5.01599 | 0.24710 | 0.00000 | 0.00000 | 0.00000 | 0.00000 | 0.54351 | 1.55165 | 2.69480 |
| **PLA2G5** | 0.00589 | 0.01650 | 0.01618 | 0.01349 | 0.01812 | 0.01200 | 0.02710 | 0.00918 | 0.01819 | 0.00002 | 0.01644 | 0.08321 |
| **PLA2G6** | 0.09788 | 0.49604 | 0.08728 | 0.47014 | 0.39985 | 0.00000 | 0.10314 | 0.14289 | 0.59100 | 0.17304 | 0.40428 | 0.32575 |
| **PLA2G7** | 0.00534 | 0.01593 | 0.03131 | 0.01075 | 0.02301 | 0.01748 | 0.01124 | 0.01380 | 0.02157 | 0.02657 | 0.01340 | 0.02518 |
| **PLA2R1** | 0.12739 | 0.09293 | 0.05536 | 0.25427 | 0.26339 | 0.13386 | 0.00000 | 0.00162 | 0.04122 | 0.00000 | 0.07269 | 0.00834 |
| **PLAA** | 3.71914 | 8.84907 | 15.07920 | 4.73050 | 1.81015 | 22.66702 | 16.41736 | 3.18011 | 17.10241 | 2.03195 | 8.81163 | 4.18806 |
| **PLAC1** | 0.32776 | 0.15088 | 0.00245 | 0.02227 | 0.01696 | 0.07004 | 0.00000 | 0.23782 | 0.83470 | 0.00000 | 0.16950 | 0.17680 |
| **PLAC4** | 0.00611 | 0.01273 | 0.67113 | 0.00334 | 0.00000 | 0.01882 | 0.21972 | 0.01008 | 0.00000 | 0.05845 | 0.00513 | 0.01227 |
| **PLAC8** | 0.00017 | 0.00000 | 0.00000 | 0.53619 | 10.23256 | 6.61312 | 5.66870 | 0.00000 | 2.55032 | 0.02914 | 11.43448 | 0.85850 |
| **PLAC8L1** | 0.10855 | 0.00000 | 0.07090 | 0.06817 | 0.00000 | 0.20325 | 0.83895 | 0.00046 | 0.18451 | 1.16686 | 0.46282 | 0.00000 |
| **PLAC9** | 0.04021 | 0.11329 | 0.42215 | 0.00089 | 0.20369 | 0.00464 | 0.07643 | 0.05586 | 0.05185 | 0.00000 | 0.00352 | 0.15735 |
| **PLAG1** | 3.35315 | 0.81231 | 0.21406 | 0.00000 | 0.27924 | 2.01737 | 1.00580 | 4.69934 | 0.00666 | 0.15041 | 0.00000 | 1.05317 |
| **PLAGL1** | 0.97209 | 0.00334 | 0.07672 | 0.00608 | 0.01138 | 0.03127 | 0.06657 | 0.27886 | 0.00000 | 0.02893 | 0.06581 | 0.07495 |
| **PLAGL2** | 9.42185 | 3.44401 | 0.00000 | 0.56900 | 5.84585 | 1.79878 | 2.98662 | 1.36155 | 4.73405 | 4.99923 | 2.37926 | 4.93238 |
| **PLAT** | 0.00750 | 0.11999 | 0.03788 | 0.06054 | 0.00000 | 0.00000 | 0.00138 | 0.04303 | 0.22888 | 0.00823 | 0.09032 | 0.05990 |
| **PLAU** | 0.00829 | 0.00000 | 0.20739 | 4.31646 | 0.54274 | 0.25867 | 0.16121 | 0.00297 | 0.11816 | 0.12482 | 0.05849 | 0.33206 |
| **PLAUR** | 0.00118 | 2.58902 | 0.14183 | 0.42139 | 0.90254 | 1.69705 | 0.00000 | 0.51952 | 0.44127 | 0.00000 | 0.24990 | 1.80291 |
| **PLB1** | 0.81461 | 0.14167 | 0.46338 | 0.08219 | 0.08057 | 0.00000 | 0.03203 | 0.48992 | 0.00000 | 0.14249 | 0.04886 | 0.12823 |
| **PLBD1** | 0.00000 | 17.19524 | 7.45488 | 8.01480 | 3.06126 | 12.44749 | 0.00000 | 2.54511 | 15.90053 | 3.47266 | 6.68571 | 16.60364 |
| **PLBD2** | 0.00138 | 1.11368 | 1.79484 | 1.79829 | 5.02839 | 0.00000 | 0.00000 | 0.04672 | 3.61857 | 1.30594 | 0.71460 | 0.72748 |
| **PLCB1** | 2.65960 | 0.64683 | 0.03050 | 1.59721 | 0.00566 | 1.67424 | 0.24940 | 1.34156 | 0.44858 | 2.12435 | 0.00000 | 1.77828 |
| **PLCB2** | 0.00873 | 0.05962 | 0.07632 | 0.00606 | 0.02097 | 0.00407 | 0.03316 | 0.00822 | 0.04735 | 0.02388 | 0.00733 | 0.15150 |
| **PLCB3** | 0.00082 | 2.14234 | 0.29816 | 1.74649 | 2.77872 | 0.00000 | 0.96818 | 1.27314 | 2.20624 | 10.17146 | 0.54560 | 3.45316 |
| **PLCB4** | 0.13916 | 0.68520 | 0.00000 | 4.97434 | 0.80156 | 0.11364 | 0.00000 | 0.01824 | 0.00000 | 0.51877 | 0.00000 | 0.08743 |
| **PLCD1** | 0.00282 | 12.12709 | 0.00000 | 10.52903 | 3.52431 | 1.09689 | 0.00000 | 0.45059 | 0.00000 | 0.16058 | 0.45148 | 9.51463 |
| **PLCD3** | 0.87258 | 4.38992 | 1.73389 | 0.00000 | 9.42786 | 0.00000 | 6.99949 | 0.62337 | 6.21145 | 9.86169 | 8.16525 | 2.53678 |
| **PLCD4** | 0.13761 | 0.39776 | 0.03729 | 0.32039 | 1.03944 | 0.00000 | 0.01470 | 0.08043 | 0.71211 | 1.10956 | 0.00732 | 0.97039 |
| **PLCE1** | 6.62466 | 2.67203 | 0.00000 | 2.46807 | 9.66622 | 1.82464 | 0.81487 | 1.32902 | 0.00000 | 4.09884 | 3.04313 | 0.44644 |
| **PLCE1-AS1** | 0.10802 | 0.04612 | 0.05001 | 0.00089 | 0.10963 | 0.08566 | 0.01069 | 0.02472 | 0.00415 | 0.03948 | 0.02518 | 0.01090 |
| **PLCG1** | 0.47996 | 1.27816 | 1.17145 | 1.69660 | 2.35212 | 0.00000 | 2.34375 | 0.01953 | 2.97351 | 5.89925 | 7.26876 | 0.41363 |
| **PLCG1-AS1** | 0.20534 | 0.01682 | 0.35879 | 0.23845 | 0.04797 | 0.19913 | 0.02582 | 0.16392 | 0.34876 | 1.34705 | 0.00212 | 0.14404 |
| **PLCG2** | 1.83409 | 1.35085 | 0.00000 | 2.41907 | 0.68174 | 0.45463 | 0.06880 | 0.23858 | 0.68170 | 2.14462 | 0.67777 | 1.00478 |
| **PLCH1** | 1.12017 | 0.11089 | 0.00000 | 0.48084 | 0.00000 | 0.02146 | 0.05666 | 0.72149 | 0.00000 | 0.06797 | 0.03810 | 0.22268 |
| **PLCH2** | 0.55186 | 0.01237 | 0.01326 | 0.02234 | 0.02719 | 0.02281 | 0.03878 | 0.00090 | 0.00842 | 0.00000 | 0.01539 | 0.01767 |
| **PLCL1** | 8.10746 | 0.31272 | 0.00342 | 0.15998 | 0.00000 | 0.03048 | 0.00490 | 0.83402 | 0.01120 | 0.01386 | 0.00712 | 0.01824 |
| **PLCL2** | 0.01732 | 0.20088 | 0.19830 | 0.23222 | 0.00000 | 0.00000 | 0.00903 | 0.34480 | 0.00000 | 0.05203 | 0.11190 | 0.87027 |
| **PLCXD2** | 0.00575 | 0.12783 | 0.01660 | 0.00000 | 0.00851 | 0.02889 | 0.00804 | 0.00999 | 0.07501 | 0.03492 | 0.02295 | 0.01187 |
| **PLCXD3** | 0.01155 | 0.03137 | 0.04507 | 0.00888 | 0.01497 | 0.00449 | 0.06687 | 0.02342 | 0.00000 | 0.04801 | 0.01201 | 0.02468 |
| **PLCZ1** | 0.00977 | 0.01133 | 0.01556 | 0.02054 | 0.01397 | 0.03882 | 0.01869 | 0.11203 | 0.01562 | 0.01133 | 0.01238 | 0.01204 |
| **PLD1** | 1.88370 | 0.02578 | 0.65544 | 0.24101 | 0.62079 | 1.70771 | 1.25566 | 0.28706 | 0.36408 | 1.42339 | 0.06725 | 0.09969 |
| **PLD2** | 0.02555 | 9.75669 | 5.26556 | 1.61967 | 4.86747 | 0.00000 | 2.76988 | 3.55608 | 11.90958 | 2.30627 | 4.73235 | 15.50517 |
| **PLD3** | 0.85036 | 13.54273 | 15.04640 | 2.63008 | 6.69432 | 0.00000 | 0.58448 | 1.98340 | 20.68780 | 0.00000 | 4.79019 | 14.09812 |
| **PLD4** | 0.00546 | 0.00807 | 0.02132 | 0.01744 | 0.01203 | 0.03244 | 0.03715 | 0.01044 | 0.00253 | 0.01021 | 0.01934 | 0.06548 |
| **PLD5** | 0.16935 | 0.01121 | 0.05171 | 0.02417 | 0.03169 | 0.00000 | 0.09018 | 0.00352 | 0.00000 | 0.06855 | 0.03493 | 0.08638 |

| | | | | | | | | | | | | |
|---|---|---|---|---|---|---|---|---|---|---|---|---|
| **PLD6** | 1.24415 | 0.25435 | 0.76971 | 0.00000 | 0.00000 | 0.00000 | 0.00000 | 0.12212 | 2.16089 | 0.31085 | 2.27159 | 1.29990 |
| **PLEC** | 0.00000 | 11.93817 | 14.32281 | 13.01433 | 28.57882 | 0.00000 | 0.00000 | 0.17009 | 23.10250 | 31.46318 | 0.00000 | 1.55878 |
| **PLEK** | 0.00526 | 0.02979 | 0.01807 | 0.02531 | 0.00000 | 0.02641 | 0.01622 | 0.00876 | 0.01558 | 0.05644 | 0.01264 | 0.05256 |
| **PLEK2** | 16.42905 | 0.14737 | 1.26700 | 0.00000 | 0.24890 | 0.00000 | 1.86933 | 6.09303 | 0.72981 | 0.73557 | 1.88978 | 0.51425 |
| **PLEKHA1** | 1.10396 | 5.93963 | 0.00000 | 0.38906 | 7.74073 | 17.14002 | 9.08542 | 0.37923 | 4.41683 | 10.37667 | 10.78759 | 4.86960 |
| **PLEKHA2** | 0.00451 | 0.17940 | 0.02520 | 0.00000 | 0.39598 | 0.00000 | 0.11617 | 0.02023 | 0.00000 | 0.58445 | 0.27533 | 0.24023 |
| **PLEKHA3** | 0.57841 | 0.00618 | 0.00000 | 1.36465 | 0.61745 | 2.60931 | 1.61680 | 0.01688 | 0.00000 | 4.37527 | 0.00000 | 1.88677 |
| **PLEKHA4** | 2.59189 | 0.62758 | 1.96827 | 6.50550 | 1.79913 | 0.00000 | 0.04126 | 0.07968 | 0.60665 | 1.03020 | 0.00000 | 0.23645 |
| **PLEKHA5** | 3.93502 | 7.44918 | 0.80174 | 10.99882 | 0.51646 | 9.73887 | 0.00000 | 4.69597 | 0.07877 | 10.84055 | 0.00000 | 14.26336 |
| **PLEKHA6** | 0.21797 | 0.27011 | 8.68871 | 0.00000 | 2.19923 | 0.00072 | 10.96335 | 0.00788 | 2.41961 | 0.00000 | 0.00000 | 0.19997 |
| **PLEKHA7** | 4.48863 | 3.56080 | 0.22012 | 4.53546 | 3.97827 | 2.52433 | 0.28128 | 0.00000 | 4.76798 | 1.98657 | 0.24406 | 2.95213 |
| **PLEKHA8** | 0.22757 | 1.96276 | 1.36949 | 1.13417 | 1.03458 | 5.61192 | 1.90717 | 0.00000 | 1.91668 | 8.58037 | 1.32596 | 0.76253 |
| **PLEKHA8P1** | 1.09146 | 0.20165 | 1.47134 | 0.33097 | 0.00000 | 1.31332 | 2.93981 | 0.22456 | 0.94606 | 1.22208 | 1.10067 | 0.61912 |
| **PLEKHB1** | 8.46062 | 0.90302 | 0.00833 | 0.12832 | 0.27152 | 0.00000 | 0.00000 | 2.76589 | 0.00000 | 0.05117 | 0.03132 | 2.54223 |
| **PLEKHB2** | 8.34494 | 31.29576 | 24.74494 | 25.30142 | 33.75082 | 42.04504 | 33.21114 | 24.29981 | 33.49847 | 7.21316 | 33.28609 | 29.58221 |
| **PLEKHD1** | 0.00545 | 0.00358 | 0.02776 | 0.07597 | 0.10010 | 0.07133 | 0.00000 | 0.04052 | 0.01097 | 0.06811 | 0.02362 | 0.01263 |
| **PLEKHF1** | 0.00554 | 0.00053 | 21.26766 | 0.00000 | 0.30877 | 0.00000 | 3.83563 | 0.00658 | 2.24850 | 0.68608 | 4.90130 | 0.00000 |
| **PLEKHF2** | 3.45719 | 3.35573 | 3.01671 | 0.28374 | 1.76597 | 14.34368 | 3.06465 | 0.20879 | 7.64425 | 1.08266 | 8.30117 | 0.00000 |
| **PLEKHG1** | 6.18752 | 0.44438 | 0.03661 | 0.00000 | 0.00000 | 0.00000 | 0.26094 | 0.57395 | 0.00974 | 0.00555 | 0.02267 | 0.10750 |
| **PLEKHG2** | 0.12506 | 0.29324 | 2.56366 | 1.07919 | 0.09026 | 0.00000 | 0.89427 | 0.02836 | 0.67635 | 2.69057 | 0.43532 | 0.73544 |
| **PLEKHG3** | 1.94633 | 1.63230 | 0.83355 | 1.78702 | 3.99553 | 0.00000 | 2.14437 | 0.02360 | 2.47319 | 5.25030 | 1.14455 | 1.24312 |
| **PLEKHG4** | 0.00153 | 0.75764 | 0.06975 | 0.10555 | 0.03252 | 0.00000 | 0.00439 | 0.05962 | 0.09699 | 0.13450 | 0.06366 | 0.13595 |
| **PLEKHG4B** | 0.11060 | 0.12569 | 1.51112 | 0.01886 | 0.25140 | 0.00000 | 0.74438 | 0.08141 | 2.93958 | 0.26742 | 3.16580 | 0.41387 |
| **PLEKHG5** | 1.06695 | 0.57043 | 0.02767 | 1.20255 | 0.68757 | 0.00000 | 0.04147 | 0.06905 | 0.00000 | 0.08675 | 0.00000 | 0.38135 |
| **PLEKHG6** | 0.00122 | 0.85340 | 1.94912 | 0.97217 | 0.01559 | 0.15062 | 0.00000 | 0.00100 | 0.26270 | 1.60776 | 0.00000 | 0.19580 |
| **PLEKHG7** | 0.01151 | 0.00000 | 0.02064 | 0.06112 | 0.05845 | 0.00491 | 0.11686 | 0.01029 | 0.00938 | 0.14285 | 0.17291 | 0.02987 |
| **PLEKHH1** | 0.05144 | 0.15493 | 0.49339 | 2.29291 | 0.59819 | 1.88835 | 0.37154 | 0.00000 | 0.71422 | 1.81053 | 2.74898 | 0.58849 |
| **PLEKHH2** | 0.35915 | 0.48359 | 0.00935 | 0.00000 | 0.00000 | 0.00000 | 0.00000 | 0.28147 | 0.00000 | 1.52617 | 0.09217 | 0.07282 |
| **PLEKHH3** | 1.13281 | 1.30699 | 2.51275 | 0.59734 | 0.80266 | 0.00000 | 0.86269 | 0.99334 | 1.99839 | 3.98204 | 0.00000 | 0.39280 |
| **PLEKHJ1** | 2.53197 | 20.01369 | 12.09371 | 16.02128 | 16.71698 | 0.00000 | 16.62427 | 0.48193 | 19.41948 | 0.00050 | 20.96640 | 11.30991 |
| **PLEKHM1** | 0.71426 | 0.35704 | 1.99138 | 2.04090 | 1.37526 | 0.00000 | 1.29238 | 1.44767 | 2.58783 | 2.62039 | 0.11098 | 2.78726 |
| **PLEKHM1P** | 0.03934 | 0.54630 | 0.78770 | 0.30860 | 0.45725 | 0.00000 | 0.00000 | 1.14263 | 0.74033 | 0.36200 | 0.41682 | 0.55153 |
| **PLEKHM2** | 5.95775 | 3.44763 | 3.23832 | 4.40774 | 10.44506 | 0.00000 | 6.35401 | 1.00724 | 4.21593 | 10.87795 | 1.63891 | 2.97696 |
| **PLEKHM3** | 1.30971 | 1.45323 | 4.01903 | 0.84155 | 0.54535 | 0.84538 | 0.82376 | 1.38206 | 1.95632 | 0.00000 | 1.43105 | 3.38113 |
| **PLEKHN1** | 0.05007 | 0.13072 | 0.16181 | 0.00000 | 0.02525 | 0.00000 | 0.07889 | 0.03565 | 0.00000 | 0.08334 | 0.03578 | 0.04456 |
| **PLEKHO1** | 1.31105 | 0.26962 | 0.19362 | 0.29048 | 0.01730 | 0.00000 | 0.68933 | 0.00000 | 0.00000 | 0.96556 | 0.00000 | 4.78816 |
| **PLEKHO2** | 0.12067 | 1.31540 | 5.39334 | 2.07101 | 3.98873 | 0.00000 | 3.16156 | 2.71416 | 0.00000 | 1.73835 | 5.26498 | 0.00000 |
| **PLEKHS1** | 0.00519 | 0.02857 | 0.00487 | 0.11668 | 0.00000 | 0.01420 | 0.01147 | 0.01030 | 0.02015 | 0.06354 | 0.01315 | 0.01317 |
| **PLET1** | 0.00501 | 0.00764 | 0.04814 | 0.00190 | 0.00865 | 0.03135 | 0.01352 | 0.00855 | 0.10543 | 0.07105 | 0.00868 | 0.03886 |
| **PLG** | 0.04437 | 0.00739 | 0.03869 | 0.00576 | 0.00211 | 0.04175 | 0.03212 | 0.01015 | 0.00974 | 0.04349 | 0.01157 | 0.03637 |
| **PLGLA** | 0.01171 | 0.01508 | 0.01401 | 0.01630 | 0.01364 | 0.01719 | 0.01490 | 0.02388 | 0.01998 | 0.02266 | 0.01416 | 0.01550 |
| **PLGRKT** | 2.37420 | 3.97498 | 4.02787 | 1.17423 | 2.40554 | 9.19677 | 2.11469 | 2.39655 | 1.29578 | 0.00000 | 5.68169 | 1.38084 |
| **PLIN1** | 0.00345 | 0.00907 | 0.02300 | 0.00000 | 0.00000 | 0.02576 | 0.00000 | 0.00926 | 0.22447 | 0.19354 | 0.13276 | 0.02676 |
| **PLIN2** | 0.00000 | 34.60679 | 51.91120 | 5.90962 | 13.42617 | 23.36846 | 42.78099 | 14.30661 | 33.33494 | 0.17039 | 12.70002 | 40.75155 |
| **PLIN3** | 2.23976 | 11.33480 | 8.05616 | 18.24798 | 14.12947 | 0.88432 | 14.17268 | 12.48497 | 11.53849 | 0.00000 | 13.33441 | 8.29145 |
| **PLIN4** | 0.00548 | 0.25984 | 0.11892 | 0.18065 | 0.11677 | 0.00000 | 0.00000 | 0.18207 | 0.34725 | 0.34338 | 0.36064 | 0.05081 |
| **PLIN5** | 0.85679 | 0.02051 | 0.01841 | 0.03077 | 0.02359 | 0.00000 | 0.00323 | 0.22768 | 0.03486 | 0.22914 | 0.03454 | 0.02016 |
| **PLK1** | 6.27701 | 44.88954 | 14.68076 | 24.54951 | 32.22165 | 14.83252 | 29.51546 | 21.28968 | 36.14839 | 0.00000 | 16.08985 | 36.07551 |

| | | | | | | | | | | | | |
|---|---|---|---|---|---|---|---|---|---|---|---|---|
| **PLK2** | 0.56386 | 3.07909 | 13.33079 | 0.35917 | 0.00000 | 2.83968 | 0.00000 | 4.92615 | 0.00000 | 1.93404 | 0.41410 | 2.61126 |
| **PLK3** | 2.53331 | 6.81652 | 3.86018 | 1.00950 | 4.18745 | 0.00000 | 0.00000 | 1.63340 | 2.41029 | 3.69325 | 0.00000 | 3.65821 |
| **PLK4** | 0.00000 | 1.81757 | 5.03609 | 3.20185 | 0.00000 | 12.87982 | 3.83894 | 5.01688 | 6.95507 | 1.17052 | 5.01004 | 4.97177 |
| **PLK5** | 0.20636 | 0.01517 | 0.04392 | 0.01302 | 0.02880 | 0.00000 | 0.01128 | 0.00301 | 0.03012 | 0.01082 | 0.07119 | 0.03909 |
| **PLLP** | 0.36264 | 20.01365 | 0.00000 | 10.40428 | 1.09468 | 0.44175 | 0.00028 | 1.26986 | 0.00000 | 0.20296 | 0.27229 | 9.70319 |
| **PLN** | 0.22297 | 0.00064 | 0.06522 | 0.04977 | 0.02904 | 0.05179 | 0.00000 | 0.05310 | 0.02649 | 0.06251 | 0.03180 | 0.08200 |
| **PLOD1** | 0.08977 | 8.82709 | 5.31355 | 12.47089 | 9.66336 | 0.00000 | 5.55592 | 0.35343 | 18.77371 | 4.84189 | 8.51355 | 1.62814 |
| **PLOD2** | 0.93579 | 0.40442 | 4.81029 | 11.67416 | 0.00000 | 4.39529 | 0.00000 | 0.75665 | 5.14925 | 4.17567 | 0.00000 | 1.15486 |
| **PLOD3** | 0.35131 | 4.38533 | 5.01540 | 8.23387 | 7.68433 | 0.00000 | 0.00000 | 1.41822 | 14.90780 | 2.93881 | 3.42982 | 0.79155 |
| **PLP1** | 0.00556 | 0.23664 | 0.39417 | 0.30470 | 0.14814 | 0.07698 | 0.00000 | 0.00640 | 0.01861 | 0.00687 | 0.02668 | 0.18005 |
| **PLP2** | 0.10368 | 0.13200 | 0.81745 | 0.60277 | 0.12839 | 0.43504 | 0.56875 | 0.01837 | 0.99097 | 0.18236 | 0.00000 | 0.06728 |
| **PLRG1** | 1.99960 | 15.53094 | 15.71193 | 8.21371 | 8.28371 | 26.31230 | 19.65366 | 13.71101 | 18.90845 | 0.00000 | 15.80305 | 17.60246 |
| **PLS1** | 10.42206 | 1.99519 | 2.71610 | 0.02254 | 0.31112 | 4.44494 | 2.31538 | 3.72604 | 1.60016 | 0.00000 | 0.85285 | 1.39169 |
| **PLS3** | 0.00000 | 10.87821 | 18.08415 | 1.60797 | 9.81170 | 32.44573 | 23.59537 | 0.76208 | 28.14475 | 2.40757 | 23.39665 | 8.96272 |
| **PLS3-AS1** | 0.00395 | 0.00000 | 0.11663 | 0.22940 | 0.15733 | 0.00000 | 0.00000 | 0.00745 | 0.21388 | 0.16084 | 0.12472 | 0.06265 |
| **PLSCR1** | 0.02364 | 3.63844 | 11.96834 | 13.60036 | 0.00000 | 14.32166 | 5.04126 | 0.00000 | 3.05067 | 0.00000 | 1.44802 | 0.00000 |
| **PLSCR2** | 0.03156 | 0.02179 | 0.09748 | 0.06332 | 0.05996 | 0.03702 | 0.05504 | 0.00688 | 0.00893 | 0.05617 | 0.01863 | 0.02128 |
| **PLSCR3** | 0.00518 | 0.03003 | 0.03210 | 0.03071 | 0.02601 | 0.00221 | 0.02037 | 0.01526 | 0.03905 | 0.00000 | 0.01678 | 0.03161 |
| **PLSCR4** | 0.42548 | 1.40811 | 0.15536 | 2.82245 | 0.00000 | 0.07550 | 0.00000 | 1.33031 | 0.00000 | 0.00393 | 0.04052 | 1.59205 |
| **PLTP** | 0.00000 | 11.18398 | 13.72767 | 12.50979 | 16.22830 | 0.00000 | 0.00000 | 0.41197 | 40.52868 | 0.59008 | 7.41991 | 3.39589 |
| **PLVAP** | 0.91488 | 0.00880 | 0.00866 | 0.00499 | 0.02119 | 0.00000 | 0.00935 | 8.15593 | 0.01373 | 0.01363 | 0.00753 | 4.30722 |
| **PLXDC1** | 0.51656 | 0.26764 | 0.04444 | 0.06376 | 0.01038 | 0.11937 | 0.04145 | 0.26882 | 0.00000 | 0.08676 | 0.07221 | 0.09594 |
| **PLXDC2** | 0.30094 | 0.15331 | 9.35932 | 0.25032 | 0.00000 | 0.99242 | 3.88841 | 0.13023 | 6.00601 | 0.22394 | 0.49876 | 0.00000 |
| **PLXNA1** | 2.29667 | 1.50284 | 9.37101 | 0.00000 | 6.06478 | 0.00000 | 1.45895 | 0.10997 | 19.70487 | 18.59109 | 2.89639 | 1.34705 |
| **PLXNA2** | 0.00000 | 2.94230 | 16.07114 | 0.59483 | 5.55225 | 0.00000 | 1.64449 | 0.21790 | 2.30421 | 6.27011 | 0.00000 | 0.03576 |
| **PLXNA3** | 0.91859 | 0.16832 | 0.76717 | 0.76782 | 0.78245 | 0.00000 | 0.00000 | 0.11975 | 3.42223 | 4.36380 | 0.77232 | 0.38695 |
| **PLXNA4** | 3.63782 | 0.04381 | 0.02154 | 0.11372 | 0.00000 | 0.00000 | 0.00154 | 6.45174 | 0.00000 | 0.10624 | 0.04746 | 0.03049 |
| **PLXNB1** | 0.00000 | 0.00051 | 6.48670 | 0.00000 | 1.49648 | 0.48381 | 4.72793 | 0.00000 | 7.39955 | 2.03652 | 6.67002 | 0.00000 |
| **PLXNB2** | 0.22694 | 12.02102 | 4.22799 | 6.49254 | 5.92056 | 1.78385 | 0.00000 | 0.00000 | 4.65188 | 4.94236 | 0.00000 | 1.69308 |
| **PLXNB3** | 2.16285 | 0.00857 | 0.03905 | 0.01237 | 0.00880 | 0.00051 | 0.00001 | 1.01310 | 0.04713 | 0.06912 | 0.01012 | 0.00000 |
| **PLXNC1** | 2.62714 | 0.16615 | 0.14392 | 0.19623 | 0.00000 | 0.00000 | 0.00488 | 0.37691 | 0.73238 | 1.41806 | 0.02842 | 0.20835 |
| **PLXND1** | 0.00000 | 3.59111 | 10.66440 | 9.84936 | 8.10601 | 0.00000 | 0.34927 | 0.34098 | 4.12600 | 13.81716 | 1.50190 | 1.00545 |
| **PM20D1** | 0.00387 | 0.00374 | 0.05851 | 0.09902 | 0.00000 | 0.01594 | 0.01025 | 1.63664 | 0.00000 | 0.07924 | 0.06822 | 0.11961 |
| **PM20D2** | 0.24066 | 0.14109 | 0.75653 | 0.70198 | 0.00000 | 0.13367 | 10.14212 | 0.78224 | 1.02322 | 2.34061 | 0.88330 | 0.00000 |
| **PMAIP1** | 0.00000 | 3.67160 | 2.21870 | 14.08951 | 0.00000 | 4.71427 | 0.00000 | 17.60702 | 0.00000 | 2.70761 | 0.00000 | 4.11459 |
| **PMCH** | 0.00177 | 0.01167 | 0.02294 | 0.04288 | 0.02765 | 0.03927 | 0.05146 | 0.05792 | 0.06529 | 0.21457 | 0.00324 | 0.02325 |
| **PMCHL2** | 0.00594 | 0.01513 | 0.01685 | 0.01119 | 0.01603 | 0.02120 | 0.02168 | 0.01175 | 0.01758 | 0.01168 | 0.01439 | 0.01640 |
| **PMEL** | 0.04268 | 0.20472 | 7.47548 | 0.57046 | 1.08391 | 0.00000 | 2.14057 | 0.01485 | 6.40244 | 0.00000 | 1.62408 | 0.83977 |
| **PMEPA1** | 0.05174 | 0.01561 | 0.93513 | 0.71306 | 0.02469 | 0.00000 | 0.00000 | 0.00828 | 0.00000 | 0.04960 | 0.06051 | 0.01651 |
| **PMF1** | 0.33490 | 5.17661 | 6.38991 | 6.17881 | 9.91009 | 1.14587 | 13.01390 | 0.55586 | 7.16864 | 1.13464 | 8.47489 | 0.44917 |
| **PMFBP1** | 0.08445 | 0.05499 | 0.18034 | 0.24942 | 0.02155 | 0.00000 | 0.00167 | 0.00237 | 0.08661 | 0.16785 | 0.00251 | 0.05646 |
| **PML** | 0.24307 | 1.61988 | 3.78185 | 1.68249 | 2.03270 | 0.00000 | 3.43850 | 0.97696 | 0.47055 | 0.55149 | 1.01914 | 1.81294 |
| **PMM1** | 3.59952 | 4.61105 | 6.07317 | 6.35560 | 0.00000 | 1.16396 | 6.41521 | 10.21358 | 3.71676 | 1.59216 | 2.87980 | 5.36466 |
| **PMM2** | 3.57001 | 22.20100 | 13.40832 | 12.60877 | 18.06568 | 3.99661 | 18.50149 | 8.24383 | 24.24006 | 2.63406 | 12.87722 | 25.82142 |
| **PMP2** | 0.04354 | 0.02312 | 0.05551 | 0.00817 | 0.03048 | 0.00000 | 0.00089 | 0.00773 | 0.12175 | 0.01251 | 0.00332 | 0.02491 |
| **PMP22** | 0.00000 | 8.79004 | 0.53252 | 6.05022 | 4.62868 | 1.23719 | 0.00000 | 0.18509 | 0.00000 | 0.42376 | 1.35965 | 0.58312 |
| **PMPCA** | 0.50140 | 17.92104 | 15.13567 | 12.55002 | 14.27878 | 2.63561 | 20.06010 | 8.47439 | 19.13103 | 0.00000 | 19.79431 | 17.00675 |
| **PMPCB** | 0.28609 | 17.18515 | 10.99113 | 7.38904 | 4.03749 | 28.19242 | 9.45065 | 2.50226 | 14.82247 | 4.01957 | 14.89534 | 13.87954 |

| | | | | | | | | | | | | |
|---|---|---|---|---|---|---|---|---|---|---|---|---|
| **PMS1** | 2.30176 | 4.18964 | 3.18112 | 1.11218 | 0.00000 | 9.85249 | 2.28670 | 0.00000 | 8.31805 | 7.28656 | 8.73038 | 2.86975 |
| **PMS2** | 0.58472 | 1.64116 | 0.88009 | 0.73304 | 0.93446 | 2.18048 | 0.00000 | 0.54019 | 0.39133 | 0.22588 | 1.64363 | 0.79701 |
| **PMS2CL** | 0.00726 | 0.12382 | 0.06663 | 0.04844 | 0.21518 | 0.21161 | 0.05030 | 0.09202 | 0.00000 | 0.09264 | 0.22350 | 0.20849 |
| **PMS2L2** | 0.03438 | 0.02308 | 0.01673 | 0.01513 | 0.02840 | 0.01425 | 0.01569 | 0.01556 | 0.01974 | 0.01501 | 0.01648 | 0.03705 |
| **PMS2P1** | 0.43404 | 1.21308 | 0.21333 | 0.99329 | 0.91267 | 0.61821 | 0.17999 | 1.12497 | 0.11710 | 0.09616 | 0.55680 | 1.50583 |
| **PMS2P3** | 0.00171 | 0.05269 | 0.02885 | 0.09591 | 0.04983 | 0.00000 | 0.02120 | 0.02083 | 0.13008 | 0.37380 | 0.04934 | 0.05446 |
| **PMS2P4** | 0.04168 | 0.75720 | 0.00000 | 0.02203 | 0.22539 | 0.37576 | 0.00000 | 0.14146 | 0.01435 | 0.10472 | 0.00000 | 0.57729 |
| **PMS2P5.uid-92!** | 0.05863 | 0.16261 | 0.11507 | 0.08705 | 0.12104 | 0.22735 | 0.03089 | 0.45048 | 0.00000 | 0.00000 | 0.08187 | 0.37806 |
| **PMS2P9** | 0.02996 | 0.02288 | 0.01410 | 0.01688 | 0.05316 | 0.03709 | 0.00682 | 0.01552 | 0.10314 | 0.00000 | 0.00838 | 0.09831 |
| **PMVK** | 0.18753 | 4.73007 | 4.36225 | 6.61672 | 11.79851 | 0.00000 | 6.18156 | 0.45076 | 3.29351 | 3.36182 | 8.77019 | 2.09437 |
| **PNCK** | 0.05127 | 0.04268 | 0.05197 | 0.06771 | 0.00748 | 0.00000 | 0.00876 | 0.00312 | 0.00000 | 0.15448 | 0.12581 | 0.06953 |
| **PNISR** | 3.90911 | 9.97091 | 11.09328 | 10.40920 | 3.27437 | 16.94545 | 19.07342 | 21.50936 | 12.77091 | 31.16277 | 8.95832 | 18.81301 |
| **PNKD** | 1.59766 | 16.70304 | 4.50927 | 7.98285 | 6.88797 | 1.28847 | 10.74111 | 9.62323 | 10.65651 | 5.43561 | 11.69044 | 18.51640 |
| **PNKP** | 0.06389 | 0.84814 | 2.87700 | 4.30626 | 2.49198 | 0.00000 | 0.07549 | 0.12444 | 2.13852 | 3.15238 | 1.79085 | 0.33110 |
| **PNLDC1** | 5.96184 | 0.86926 | 0.03253 | 1.20540 | 1.60530 | 0.84118 | 0.00000 | 3.01835 | 0.00000 | 0.00000 | 0.29685 | 1.40878 |
| **PNLIPRP1** | 0.04045 | 0.03287 | 0.02463 | 0.01147 | 0.01502 | 0.00839 | 0.01828 | 0.01772 | 0.01813 | 0.01069 | 0.01824 | 0.01382 |
| **PNMA1** | 3.98566 | 18.58765 | 7.65601 | 10.40238 | 7.23049 | 9.61472 | 6.09628 | 9.21520 | 2.31599 | 2.91296 | 14.54594 | 27.16133 |
| **PNMA2** | 0.02878 | 1.22539 | 0.23996 | 4.11636 | 0.00000 | 0.13640 | 0.00000 | 0.28839 | 0.00000 | 0.00000 | 0.17362 | 4.43430 |
| **PNMA3** | 0.00642 | 0.01249 | 0.04652 | 0.01532 | 0.02663 | 0.00061 | 0.01304 | 0.00763 | 0.02085 | 0.01123 | 0.01646 | 0.11324 |
| **PNMA5** | 0.03151 | 5.20139 | 0.00377 | 0.00000 | 0.23555 | 0.04757 | 0.00000 | 0.02736 | 0.00416 | 0.01335 | 0.02707 | 9.76420 |
| **PNMAL1** | 1.52811 | 2.59039 | 0.00000 | 0.74160 | 0.21976 | 0.31744 | 0.00000 | 0.00000 | 1.38709 | 0.00000 | 0.46650 | 1.34509 |
| **PNMAL2** | 0.25075 | 0.01563 | 0.04321 | 0.01287 | 0.01780 | 0.01374 | 0.01949 | 0.04440 | 0.01839 | 0.00000 | 0.01457 | 0.03441 |
| **PNMT** | 0.00565 | 0.00457 | 0.00675 | 0.03558 | 0.01976 | 0.00000 | 0.07150 | 0.00869 | 0.24812 | 0.01867 | 0.19104 | 0.06990 |
| **PNN** | 9.95262 | 8.77164 | 13.21829 | 10.67233 | 13.04383 | 32.02686 | 18.51022 | 23.81871 | 28.98171 | 38.80215 | 16.57697 | 20.15682 |
| **PNO1** | 0.30992 | 14.20169 | 15.76540 | 7.59409 | 7.53151 | 26.83029 | 21.46514 | 13.80668 | 20.08963 | 4.99983 | 12.02432 | 23.35520 |
| **PNOC** | 0.01494 | 0.01418 | 0.02467 | 0.01220 | 0.01296 | 0.02128 | 0.01575 | 0.07696 | 0.01653 | 0.01975 | 0.01448 | 0.01177 |
| **PNP** | 5.43873 | 0.00000 | 22.89796 | 17.51693 | 0.00000 | 5.96021 | 26.03544 | 18.88261 | 28.74561 | 0.00000 | 0.00000 | 5.34783 |
| **PNPLA1** | 0.00464 | 0.03230 | 0.03633 | 0.01065 | 0.00623 | 0.00919 | 0.00570 | 0.01179 | 0.07184 | 0.05549 | 0.04028 | 0.03209 |
| **PNPLA2** | 9.12742 | 1.45323 | 3.37116 | 0.78536 | 2.75524 | 0.00000 | 1.32406 | 1.81549 | 3.79396 | 7.82299 | 0.00000 | 0.95973 |
| **PNPLA3** | 0.53913 | 0.40662 | 1.18328 | 1.34137 | 1.18417 | 0.00000 | 3.39501 | 0.00000 | 3.00567 | 0.63513 | 0.95626 | 0.82317 |
| **PNPLA4** | 3.43160 | 3.76359 | 1.03861 | 2.32069 | 5.13798 | 8.10220 | 1.71850 | 2.13556 | 2.88273 | 0.00000 | 4.96319 | 0.72865 |
| **PNPLA5** | 0.00621 | 0.01599 | 0.01793 | 0.01086 | 0.01356 | 0.01833 | 0.02431 | 0.01066 | 0.03117 | 0.01083 | 0.01188 | 0.01742 |
| **PNPLA6** | 0.46732 | 5.30556 | 6.60066 | 6.66366 | 10.25097 | 0.00000 | 1.75969 | 0.00000 | 14.81768 | 10.47856 | 2.39819 | 0.89839 |
| **PNPLA7** | 0.00393 | 0.30336 | 0.14129 | 0.05354 | 0.15736 | 0.00000 | 0.00264 | 0.17590 | 0.28815 | 0.40194 | 0.00825 | 0.03422 |
| **PNPLA8** | 2.06859 | 1.71865 | 6.80893 | 5.11966 | 3.13181 | 19.47235 | 6.34863 | 6.27720 | 9.51024 | 3.36612 | 8.40843 | 4.78111 |
| **PNPO** | 0.21919 | 3.44251 | 0.12568 | 3.83498 | 3.40815 | 2.11267 | 0.66310 | 0.09036 | 0.00000 | 1.87265 | 3.70899 | 1.27322 |
| **PNPT1** | 3.23063 | 27.36078 | 4.18100 | 12.18678 | 7.38093 | 46.91253 | 0.00000 | 3.10665 | 16.28493 | 9.06965 | 15.79689 | 18.51050 |
| **PNRC1** | 3.68653 | 1.70435 | 0.00000 | 9.30372 | 0.00000 | 5.76505 | 1.73209 | 12.83905 | 0.00000 | 5.99834 | 0.00000 | 9.68416 |
| **PNRC2** | 1.48089 | 8.75067 | 7.43188 | 4.11936 | 1.53268 | 26.65608 | 13.66975 | 12.10293 | 10.00290 | 5.52107 | 7.38119 | 13.47903 |
| **POC1A** | 0.37164 | 13.86171 | 3.89970 | 5.50247 | 6.08138 | 0.02221 | 2.42605 | 0.10171 | 3.96593 | 0.00000 | 6.79423 | 5.61870 |
| **POC1B** | 2.11758 | 1.10113 | 0.32021 | 4.93544 | 0.00541 | 7.78090 | 2.83850 | 0.25069 | 4.04793 | 0.00000 | 3.13406 | 0.00000 |
| **POC5** | 0.23219 | 3.93819 | 5.32552 | 1.30584 | 0.42304 | 16.61160 | 0.02523 | 4.25557 | 5.93534 | 0.00000 | 2.55307 | 4.33967 |
| **PODN** | 0.00629 | 0.02638 | 0.01457 | 0.00854 | 0.25463 | 0.00614 | 0.02318 | 0.01052 | 0.00824 | 0.00724 | 0.00908 | 0.01685 |
| **PODNL1** | 0.03951 | 0.06007 | 0.00767 | 0.12916 | 0.00000 | 0.04578 | 0.00000 | 0.06162 | 0.05397 | 0.31458 | 0.02464 | 0.04107 |
| **PODXL** | 2.43204 | 3.70185 | 2.04666 | 7.90025 | 0.00000 | 0.00000 | 0.76355 | 0.15509 | 1.65796 | 3.35823 | 0.28864 | 2.90322 |
| **PODXL2** | 0.00263 | 0.10875 | 0.02765 | 0.00000 | 0.19026 | 0.00000 | 0.00009 | 0.08801 | 0.26100 | 0.68090 | 0.17270 | 0.01287 |
| **POF1B** | 0.04561 | 0.08379 | 0.01630 | 0.00000 | 0.00295 | 0.08955 | 0.00000 | 0.00648 | 0.01176 | 0.07348 | 0.00302 | 0.02599 |
| **POFUT1** | 0.68628 | 9.00840 | 1.56398 | 3.23955 | 6.09762 | 5.75235 | 1.79762 | 0.44899 | 9.63546 | 1.63704 | 5.60534 | 5.08221 |

| | | | | | | | | | | | | |
|---|---|---|---|---|---|---|---|---|---|---|---|---|
| **POFUT2** | 1.61674 | 0.80264 | 1.44073 | 1.30549 | 0.54164 | 0.00000 | 0.27624 | 1.19813 | 2.05927 | 1.33223 | 0.03767 | 1.97239 |
| **POGK** | 0.00000 | 3.25629 | 0.67455 | 1.58503 | 0.00000 | 4.86098 | 2.75343 | 0.00000 | 0.36602 | 5.01974 | 1.17712 | 3.61578 |
| **POGLUT1** | 0.00000 | 0.50705 | 0.97294 | 2.02301 | 1.00982 | 0.07702 | 0.34863 | 0.02766 | 3.45776 | 1.22800 | 2.10682 | 0.00000 |
| **POGZ** | 3.60086 | 0.97845 | 2.93623 | 5.46229 | 2.75017 | 4.26241 | 5.43008 | 0.84835 | 0.00001 | 11.35490 | 2.73041 | 4.62749 |
| **POLA1** | 6.43595 | 2.15347 | 2.60193 | 3.14751 | 1.87728 | 7.88883 | 2.85625 | 1.88294 | 2.78551 | 7.78986 | 4.17580 | 2.08531 |
| **POLA2** | 1.71498 | 7.62452 | 6.75573 | 6.78964 | 6.14900 | 2.59356 | 6.44280 | 0.21126 | 3.71410 | 0.00000 | 9.87163 | 4.38874 |
| **POLB** | 6.11707 | 7.41004 | 5.90497 | 0.30783 | 2.73407 | 14.31406 | 5.26741 | 0.99900 | 8.49087 | 0.61863 | 2.58204 | 9.72194 |
| **POLD1** | 1.92020 | 17.58379 | 8.65313 | 19.35002 | 10.09114 | 0.00000 | 8.71180 | 0.08882 | 15.52486 | 11.31037 | 9.29679 | 10.29769 |
| **POLD2** | 0.60237 | 17.95514 | 7.66039 | 11.36365 | 19.72332 | 2.53744 | 11.87830 | 0.55728 | 17.85478 | 0.00000 | 19.11186 | 8.29511 |
| **POLD3** | 11.22580 | 8.00469 | 5.67796 | 5.24212 | 3.16995 | 19.15808 | 6.13193 | 8.78327 | 6.70878 | 9.77203 | 8.66346 | 16.85823 |
| **POLD4** | 0.00900 | 1.95764 | 1.06600 | 5.17321 | 2.03232 | 0.00000 | 1.35924 | 0.13485 | 0.15247 | 0.45748 | 0.81762 | 2.60953 |
| **POLDIP2** | 4.58016 | 26.13473 | 17.30453 | 11.28995 | 22.95612 | 0.48035 | 19.42762 | 3.80320 | 16.66638 | 6.98106 | 17.42152 | 10.31742 |
| **POLDIP3** | 5.83471 | 21.39156 | 18.00290 | 16.09947 | 20.58004 | 3.28056 | 24.13613 | 16.49741 | 23.16430 | 0.12091 | 14.97450 | 19.04912 |
| **POLE** | 3.14331 | 7.37437 | 7.05593 | 7.21922 | 12.31404 | 0.00000 | 9.76468 | 1.81756 | 9.72981 | 17.33476 | 4.46412 | 5.08998 |
| **POLE2** | 0.31588 | 3.16226 | 2.77743 | 0.19526 | 2.12723 | 15.57976 | 0.02028 | 0.00000 | 3.65850 | 2.62921 | 5.63630 | 2.23895 |
| **POLE3** | 5.64248 | 14.67252 | 10.71942 | 4.88320 | 10.74652 | 26.01430 | 26.52840 | 25.98844 | 17.80715 | 11.39885 | 10.79377 | 20.30236 |
| **POLE4** | 1.68245 | 4.23723 | 6.48805 | 1.84299 | 4.06391 | 0.02551 | 6.03182 | 1.88798 | 4.20491 | 1.64241 | 6.29854 | 5.61346 |
| **POLG** | 0.82935 | 4.85569 | 3.37887 | 3.61660 | 4.61807 | 0.00000 | 2.05294 | 2.70289 | 5.66626 | 6.39879 | 1.92731 | 4.08214 |
| **POLG2** | 1.49663 | 3.15221 | 2.64703 | 1.83381 | 1.50830 | 5.60971 | 2.41934 | 0.47387 | 3.96850 | 0.32898 | 3.50334 | 2.27690 |
| **POLH** | 0.82482 | 7.86476 | 3.23931 | 4.10032 | 4.36854 | 4.78126 | 0.55167 | 11.29106 | 4.87570 | 7.43513 | 3.34243 | 8.76139 |
| **POLI** | 0.42907 | 2.76290 | 3.19722 | 1.19831 | 0.00087 | 5.76690 | 0.00000 | 2.32689 | 2.41044 | 0.00000 | 2.12785 | 2.83431 |
| **POLK** | 3.51079 | 0.84207 | 2.06352 | 0.66241 | 0.00000 | 4.53761 | 1.83195 | 0.72224 | 3.21243 | 5.15554 | 2.04616 | 0.23864 |
| **POLL** | 2.81799 | 4.61115 | 2.62014 | 1.89742 | 3.07278 | 0.00000 | 2.43226 | 0.02197 | 4.11976 | 4.96661 | 2.08463 | 3.33859 |
| **POLM** | 0.52859 | 4.53250 | 2.80635 | 2.61974 | 5.77553 | 0.00000 | 2.18677 | 0.09634 | 4.37364 | 0.00000 | 2.00633 | 2.68979 |
| **POLN** | 0.08362 | 0.07899 | 0.10262 | 0.17613 | 0.21300 | 0.00000 | 0.00267 | 0.30676 | 0.48364 | 0.12220 | 0.39256 | 0.31120 |
| **POLQ** | 1.61768 | 1.10762 | 0.00000 | 1.97522 | 3.32484 | 3.84826 | 0.00000 | 2.67573 | 0.87813 | 3.60202 | 2.41413 | 5.07308 |
| **POLR1A** | 0.83014 | 7.48031 | 5.51740 | 1.75455 | 7.18362 | 0.00112 | 4.83977 | 5.47216 | 8.16267 | 4.92631 | 5.02240 | 4.69064 |
| **POLR1B** | 4.01149 | 9.72753 | 7.84685 | 3.68283 | 4.39089 | 13.39087 | 11.73973 | 1.87929 | 18.09075 | 0.00000 | 17.68559 | 4.31998 |
| **POLR1C** | 3.02520 | 16.84265 | 12.08299 | 9.13050 | 14.56453 | 12.82009 | 18.19956 | 23.14184 | 22.19427 | 0.00000 | 14.06176 | 23.75151 |
| **POLR1D** | 7.55259 | 16.25793 | 19.59329 | 11.51245 | 18.29061 | 30.38116 | 30.23269 | 12.43545 | 30.01188 | 13.00336 | 14.67400 | 26.27658 |
| **POLR1E** | 3.00680 | 4.14866 | 3.92834 | 2.23700 | 7.35737 | 5.88875 | 7.13281 | 6.67984 | 5.42190 | 4.39767 | 4.76884 | 8.26428 |
| **POLR2A** | 8.72109 | 13.63244 | 15.79383 | 14.36753 | 18.30733 | 0.00405 | 25.63206 | 7.41541 | 24.98187 | 20.08019 | 20.49975 | 19.97666 |
| **POLR2B** | 0.00000 | 12.16749 | 13.83876 | 9.08152 | 4.86451 | 31.10306 | 17.24393 | 13.11768 | 20.92603 | 15.28898 | 12.39990 | 16.21148 |
| **POLR2C** | 5.53595 | 12.61886 | 13.37734 | 8.88181 | 9.49709 | 17.16592 | 12.16029 | 22.50827 | 15.58051 | 0.00000 | 8.11321 | 11.60195 |
| **POLR2D** | 2.86832 | 16.87128 | 14.27089 | 7.25674 | 11.83726 | 22.25516 | 18.89218 | 5.12348 | 19.53934 | 0.00000 | 17.05908 | 17.16501 |
| **POLR2E** | 3.96348 | 35.29217 | 31.72222 | 27.39275 | 39.04314 | 7.59934 | 43.92723 | 26.07605 | 44.91343 | 0.00089 | 32.32829 | 39.57209 |
| **POLR2F** | 5.90373 | 27.89336 | 26.57537 | 19.07711 | 28.54374 | 19.37059 | 38.30469 | 27.76546 | 23.49862 | 6.83807 | 21.61050 | 37.15952 |
| **POLR2G** | 0.25104 | 14.55044 | 21.59447 | 14.40472 | 18.57794 | 18.29336 | 26.65780 | 5.13843 | 18.82797 | 0.00000 | 16.22438 | 15.92676 |
| **POLR2H** | 5.80667 | 23.02358 | 26.77963 | 14.61643 | 18.20166 | 17.38002 | 37.27662 | 1.37618 | 24.04278 | 9.22117 | 21.06205 | 8.34916 |
| **POLR2I** | 0.65707 | 20.39977 | 10.29421 | 21.52785 | 31.06874 | 8.94840 | 24.55689 | 5.59481 | 13.39532 | 7.69794 | 17.09250 | 18.67582 |
| **POLR2J** | 8.42615 | 14.19566 | 6.63981 | 7.15445 | 14.59716 | 3.09344 | 13.41021 | 2.72284 | 10.18037 | 8.79272 | 9.28408 | 12.34066 |
| **POLR2J2** | 0.03386 | 0.10185 | 0.14251 | 0.05185 | 0.03373 | 0.20583 | 0.00000 | 0.02886 | 0.13863 | 0.03519 | 0.02645 | 0.02168 |
| **POLR2J3** | 0.04130 | 0.15036 | 0.06515 | 0.17885 | 0.33631 | 0.30050 | 0.16377 | 0.01261 | 0.00000 | 0.07339 | 0.05261 | 0.09453 |
| **POLR2J4** | 0.37667 | 0.94617 | 0.47624 | 0.22590 | 0.31970 | 0.00206 | 0.36057 | 0.06284 | 1.35444 | 0.00078 | 0.46615 | 1.16822 |
| **POLR2K** | 1.88017 | 12.58236 | 14.44354 | 3.11009 | 0.00000 | 45.43654 | 20.86040 | 8.07508 | 12.28785 | 13.65689 | 14.64402 | 20.94546 |
| **POLR2L** | 0.89085 | 30.46021 | 25.86779 | 24.75325 | 39.16759 | 7.42827 | 49.18277 | 22.60784 | 37.72725 | 13.49602 | 32.54228 | 39.67644 |
| **POLR2M** | 0.35325 | 0.36855 | 0.39054 | 0.14789 | 0.45446 | 0.03355 | 0.12321 | 0.38505 | 0.40027 | 0.07459 | 0.12231 | 0.42357 |
| **POLR3A** | 0.81027 | 6.88570 | 8.94219 | 4.44678 | 5.06913 | 1.52191 | 10.09971 | 8.40994 | 9.50705 | 7.85931 | 8.50340 | 8.33427 |

| | | | | | | | | | | | | |
|---|---|---|---|---|---|---|---|---|---|---|---|---|
| **POLR3B** | 1.41883 | 6.64841 | 3.19776 | 0.93109 | 4.28593 | 8.24148 | 5.77635 | 3.15574 | 3.82335 | 2.81761 | 6.15427 | 6.58773 |
| **POLR3C** | 0.01591 | 17.61131 | 1.91218 | 9.01570 | 14.66135 | 17.05644 | 6.53011 | 12.78599 | 5.64783 | 0.00000 | 7.30721 | 18.62543 |
| **POLR3D** | 0.72642 | 7.14643 | 9.38179 | 1.09291 | 6.33999 | 0.00000 | 2.72044 | 4.14098 | 8.19212 | 4.08438 | 5.09380 | 6.37345 |
| **POLR3E** | 3.31119 | 10.04105 | 12.24230 | 9.34848 | 8.22152 | 3.69368 | 7.91069 | 2.58110 | 21.22029 | 5.43599 | 13.91648 | 8.17462 |
| **POLR3F** | 0.00011 | 4.40521 | 3.13713 | 1.99497 | 0.00018 | 15.10808 | 5.82114 | 4.85720 | 6.55352 | 0.05940 | 5.62422 | 7.17707 |
| **POLR3G** | 2.82254 | 3.88366 | 0.98451 | 0.00000 | 1.52410 | 13.34640 | 3.42220 | 1.04314 | 4.35730 | 7.74127 | 7.64547 | 12.71718 |
| **POLR3GL** | 4.14857 | 3.31587 | 0.04256 | 0.76748 | 0.00000 | 0.46437 | 0.00000 | 3.26757 | 0.00000 | 1.62863 | 0.00000 | 1.96235 |
| **POLR3H** | 1.47243 | 16.55538 | 12.66119 | 5.20566 | 11.21440 | 0.00000 | 13.08735 | 14.30510 | 22.52069 | 0.00000 | 18.15946 | 14.36425 |
| **POLR3K** | 0.35345 | 18.12380 | 17.14839 | 13.01494 | 24.12443 | 11.39308 | 24.73603 | 28.54824 | 25.22160 | 0.00241 | 23.70719 | 19.14841 |
| **POLRMT** | 0.11248 | 8.87992 | 3.26861 | 2.15237 | 4.88801 | 0.00000 | 2.94057 | 1.18021 | 3.12775 | 14.93368 | 2.21100 | 10.32071 |
| **POM121** | 4.78344 | 2.30123 | 2.81296 | 1.47704 | 4.35485 | 0.00000 | 3.50836 | 1.46655 | 5.65898 | 8.52719 | 1.50682 | 3.25901 |
| **POM121C** | 2.96508 | 1.37674 | 2.49616 | 2.55863 | 4.54118 | 0.00000 | 4.34796 | 1.70076 | 5.16325 | 12.34922 | 1.61691 | 3.66880 |
| **POM121L10P** | 0.00389 | 0.00694 | 0.02642 | 0.16692 | 0.13948 | 0.00000 | 0.03949 | 0.01486 | 0.00000 | 0.08077 | 0.00732 | 0.01692 |
| **POM121L1P** | 0.00564 | 0.01434 | 0.01426 | 0.01272 | 0.01225 | 0.01910 | 0.02234 | 0.01165 | 0.01155 | 0.02404 | 0.02342 | 0.01606 |
| **POM121L2** | 0.00487 | 0.01143 | 0.01793 | 0.00000 | 0.00000 | 0.12097 | 0.00000 | 0.00968 | 0.06345 | 0.06471 | 0.00000 | 0.01360 |
| **POM121L8P** | 0.00652 | 0.01453 | 0.01488 | 0.01426 | 0.01704 | 0.01685 | 0.01950 | 0.01169 | 0.01926 | 0.01342 | 0.01566 | 0.01815 |
| **POM121L9P** | 0.01242 | 0.01296 | 0.02334 | 0.01968 | 0.01834 | 0.00658 | 0.04337 | 0.01139 | 0.00304 | 0.02312 | 0.01799 | 0.05323 |
| **POMC** | 0.78213 | 3.53178 | 1.49315 | 0.97509 | 12.30257 | 0.00000 | 0.00000 | 5.98708 | 1.73197 | 3.82856 | 15.33609 | 4.32585 |
| **POMGNT1** | 2.33206 | 4.57412 | 6.82087 | 0.86197 | 6.01743 | 1.10391 | 3.65272 | 0.46763 | 4.69605 | 12.51743 | 6.29231 | 3.98595 |
| **POMGNT2** | 0.07338 | 6.84026 | 2.12675 | 0.48319 | 1.85974 | 2.32548 | 1.09350 | 0.00000 | 2.79331 | 6.32085 | 2.64071 | 3.17285 |
| **POMK** | 4.43371 | 2.95868 | 1.38568 | 1.31938 | 1.07520 | 0.00000 | 1.30380 | 3.55909 | 2.23399 | 3.30446 | 4.52928 | 2.82328 |
| **POMP** | 1.48315 | 20.29620 | 33.00832 | 15.52038 | 16.06111 | 58.26877 | 52.48044 | 20.11367 | 17.53708 | 25.81631 | 28.76948 | 37.72292 |
| **POMT1** | 1.91014 | 4.13159 | 8.26933 | 5.01043 | 3.86524 | 0.00000 | 1.39418 | 0.40270 | 9.86322 | 4.55590 | 4.60549 | 1.48727 |
| **POMT2** | 0.21468 | 1.52328 | 1.55467 | 2.03056 | 4.07254 | 0.00000 | 0.36804 | 0.45339 | 7.32606 | 12.91848 | 2.78290 | 2.60679 |
| **POMZP3** | 0.20518 | 0.25442 | 0.29833 | 0.13280 | 0.22309 | 0.00000 | 0.25357 | 0.75523 | 0.27925 | 0.25393 | 0.04322 | 0.78534 |
| **PON1** | 0.26529 | 0.05881 | 0.03198 | 0.02629 | 0.01622 | 0.00000 | 0.00708 | 0.10523 | 0.06080 | 0.01289 | 0.03512 | 0.06393 |
| **PON2** | 3.63491 | 1.27296 | 1.70258 | 0.28671 | 0.00000 | 1.13026 | 0.00000 | 2.87728 | 2.26086 | 0.25086 | 0.01192 | 0.00000 |
| **PON3** | 0.07504 | 0.00808 | 0.01047 | 0.01574 | 0.01262 | 0.04262 | 0.02601 | 0.28088 | 0.02369 | 0.00819 | 0.01588 | 0.03418 |
| **POP1** | 0.82989 | 3.86123 | 5.03453 | 3.96157 | 4.02498 | 5.91409 | 5.96399 | 2.48468 | 4.28842 | 8.72432 | 6.07170 | 8.94608 |
| **POP4** | 1.50273 | 14.46800 | 13.03777 | 4.34893 | 10.12236 | 20.32283 | 18.55878 | 3.98149 | 8.05272 | 2.58180 | 14.99707 | 20.82670 |
| **POP5** | 1.00477 | 10.42434 | 11.52610 | 5.35751 | 4.65825 | 10.49224 | 13.60500 | 5.96037 | 10.20000 | 0.96546 | 13.85521 | 15.12383 |
| **POP7** | 3.67744 | 30.64807 | 24.97086 | 15.55484 | 21.08765 | 14.66667 | 32.29069 | 16.74923 | 33.79492 | 5.64009 | 22.50767 | 33.07453 |
| **POPDC2** | 0.00637 | 0.05332 | 0.04067 | 0.16197 | 0.00000 | 0.00577 | 0.07300 | 0.01478 | 0.04474 | 0.04009 | 0.02534 | 0.05907 |
| **POPDC3** | 0.24205 | 0.27365 | 0.06839 | 0.02233 | 0.00000 | 0.21690 | 0.08726 | 0.05593 | 0.00000 | 0.07869 | 0.05250 | 0.00000 |
| **POR** | 3.36447 | 17.13771 | 18.79928 | 6.02538 | 22.52122 | 0.00000 | 0.75141 | 2.48313 | 40.49311 | 5.84688 | 11.44912 | 8.93451 |
| **PORCN** | 0.59446 | 0.51450 | 0.31266 | 3.18631 | 6.17542 | 0.92188 | 0.00000 | 0.00000 | 0.49230 | 0.00000 | 1.23811 | 0.00000 |
| **POSTN** | 0.00588 | 0.01060 | 0.01339 | 0.00981 | 0.01090 | 0.03249 | 0.02356 | 0.09722 | 0.01470 | 0.01432 | 0.01364 | 0.02672 |
| **POT1** | 2.36457 | 4.23106 | 1.88999 | 0.35707 | 0.24868 | 6.58400 | 0.99634 | 2.83085 | 1.93265 | 0.00000 | 1.35779 | 3.02332 |
| **POTEA** | 0.00561 | 0.01299 | 0.01384 | 0.01404 | 0.01195 | 0.01559 | 0.01889 | 0.01123 | 0.01163 | 0.02691 | 0.02673 | 0.02414 |
| **POTEC** | 0.01614 | 0.04340 | 0.01853 | 0.00891 | 0.02689 | 0.00387 | 0.02252 | 0.05308 | 0.01843 | 0.00696 | 0.05096 | 0.03611 |
| **POTEE** | 0.00993 | 0.04951 | 0.23073 | 0.10967 | 0.03553 | 0.00000 | 0.00000 | 0.08571 | 0.09040 | 0.36278 | 0.00316 | 0.44887 |
| **POTEF** | 0.17098 | 0.04006 | 1.27351 | 0.11493 | 0.00000 | 0.22447 | 0.00000 | 0.33473 | 0.00000 | 0.14842 | 0.02689 | 0.43426 |
| **POTEG** | 0.01015 | 0.00756 | 0.01328 | 0.01006 | 0.00924 | 0.04391 | 0.02149 | 0.02534 | 0.03341 | 0.03083 | 0.01713 | 0.06235 |
| **POTEH** | 0.00705 | 0.01048 | 0.02949 | 0.01452 | 0.01563 | 0.01083 | 0.01910 | 0.06918 | 0.04928 | 0.09236 | 0.00716 | 0.05507 |
| **POTEI** | 0.00861 | 0.00914 | 0.22013 | 0.05541 | 0.00000 | 0.05302 | 0.00000 | 0.01628 | 0.00000 | 0.10095 | 0.01221 | 0.04378 |
| **POTEJ** | 0.00598 | 0.02298 | 0.01899 | 0.01849 | 0.01686 | 0.03812 | 0.01360 | 0.01073 | 0.02339 | 0.00986 | 0.02425 | 0.01986 |
| **POTEKP** | 0.00750 | 0.01876 | 0.02561 | 0.03912 | 0.00188 | 0.03745 | 0.04167 | 0.01744 | 0.00823 | 0.08052 | 0.01210 | 0.08706 |
| **POTEM** | 0.00579 | 0.00967 | 0.01729 | 0.00829 | 0.00755 | 0.02842 | 0.00748 | 0.02153 | 0.03988 | 0.04724 | 0.02862 | 0.02474 |

| | | | | | | | | | | | | |
|---|---|---|---|---|---|---|---|---|---|---|---|---|
| **POU2AF1** | 0.00536 | 0.05079 | 0.01296 | 0.00402 | 0.12936 | 0.04216 | 0.02530 | 0.01256 | 0.01944 | 0.00000 | 0.00573 | 0.03718 |
| **POU2F1** | 6.37747 | 6.16263 | 8.72713 | 4.57076 | 12.68933 | 6.53873 | 13.88762 | 10.23141 | 5.12565 | 22.69433 | 4.04598 | 5.40906 |
| **POU2F2** | 0.01462 | 0.28460 | 0.88705 | 0.00000 | 0.07175 | 0.00000 | 0.00000 | 0.18861 | 0.28187 | 1.57855 | 0.00000 | 0.07689 |
| **POU2F3** | 0.00480 | 0.07142 | 1.62229 | 0.12688 | 0.13171 | 0.00000 | 0.02466 | 0.02254 | 0.00000 | 0.06575 | 0.03504 | 0.02447 |
| **POU3F1** | 0.12050 | 0.07853 | 0.08905 | 0.24985 | 0.00000 | 0.01520 | 0.01038 | 0.20175 | 0.01785 | 0.11668 | 0.01290 | 0.03928 |
| **POU3F2** | 0.25551 | 0.09486 | 0.00191 | 0.00000 | 0.00000 | 0.01197 | 0.01188 | 0.00686 | 0.01043 | 0.30452 | 0.01076 | 0.01805 |
| **POU4F1** | 0.62321 | 0.36857 | 0.00000 | 0.00000 | 0.24614 | 0.21347 | 1.91843 | 0.06176 | 0.00461 | 1.95842 | 0.00000 | 0.18916 |
| **POU4F2** | 0.00688 | 0.02057 | 0.00785 | 0.01350 | 0.00045 | 0.11314 | 0.02728 | 0.03231 | 0.00268 | 0.01884 | 0.00719 | 0.01289 |
| **POU5F1** | 0.18588 | 19.95913 | 1.53194 | 23.10598 | 13.38610 | 4.88413 | 0.00000 | 1.38256 | 0.00000 | 1.05520 | 3.57490 | 18.22553 |
| **POU5F1B** | 0.18605 | 0.75223 | 0.00001 | 0.90521 | 0.39434 | 0.28649 | 0.00000 | 0.01071 | 0.00000 | 0.39633 | 0.00000 | 0.50800 |
| **POU5F1P3** | 0.03822 | 5.82086 | 0.18348 | 7.58767 | 2.96730 | 1.07396 | 0.00000 | 0.25322 | 0.00000 | 0.06672 | 0.70464 | 4.88730 |
| **POU5F1P4** | 0.02757 | 2.41661 | 0.14795 | 3.16729 | 1.08137 | 0.53477 | 0.00000 | 0.07764 | 0.00000 | 0.00000 | 0.39881 | 1.84503 |
| **POU5F2** | 0.00873 | 0.00436 | 0.16481 | 0.03001 | 0.00000 | 0.02831 | 0.00000 | 0.00950 | 0.00000 | 0.08248 | 0.08710 | 0.02166 |
| **POU6F1** | 0.00186 | 0.79053 | 0.43197 | 0.45088 | 0.94789 | 0.15995 | 0.00000 | 0.00127 | 0.03979 | 0.69895 | 0.08060 | 0.16803 |
| **POU6F2** | 0.19525 | 0.12633 | 0.10493 | 0.00000 | 0.31281 | 0.06319 | 0.00000 | 0.01004 | 0.17733 | 0.31036 | 0.09643 | 0.07048 |
| **PP12613** | 0.00722 | 0.01946 | 0.16914 | 0.00901 | 0.03569 | 0.06311 | 0.01420 | 0.00919 | 0.00000 | 0.13292 | 0.00419 | 0.00780 |
| **PP14571** | 0.00396 | 0.00000 | 0.41808 | 0.02022 | 2.14063 | 0.00204 | 0.76735 | 0.01114 | 0.65592 | 0.02512 | 4.56883 | 0.02872 |
| **PP2D1** | 0.01207 | 0.00258 | 0.00141 | 0.00000 | 0.13065 | 0.13074 | 0.00000 | 0.09577 | 0.09252 | 0.14735 | 0.00666 | 0.11347 |
| **PP7080** | 0.00000 | 7.18053 | 1.31533 | 0.28947 | 6.79350 | 3.47362 | 0.00000 | 0.63800 | 0.00000 | 8.09291 | 2.51802 | 2.37042 |
| **PPA1** | 0.36910 | 20.99051 | 18.56986 | 14.49784 | 4.70145 | 37.21659 | 27.49936 | 6.78215 | 23.69435 | 7.20735 | 23.24048 | 19.20660 |
| **PPA2** | 0.95612 | 4.20602 | 9.75596 | 8.88551 | 4.95187 | 18.86165 | 17.31256 | 0.43892 | 12.35689 | 0.00000 | 15.88168 | 2.97753 |
| **PPAN** | 0.00024 | 0.59229 | 1.08326 | 0.13439 | 0.78564 | 0.00000 | 0.43850 | 0.46926 | 0.00000 | 4.43117 | 0.02227 | 0.59058 |
| **PPAP2A** | 0.00000 | 13.83397 | 0.93469 | 9.19320 | 3.38641 | 0.00000 | 0.00000 | 0.22266 | 0.00000 | 2.48312 | 0.55125 | 8.89938 |
| **PPAP2B** | 0.42201 | 2.20897 | 21.04305 | 6.74965 | 9.12695 | 5.38166 | 5.56842 | 0.00000 | 28.71148 | 4.02343 | 1.04027 | 0.00000 |
| **PPAP2C** | 0.16762 | 7.62795 | 5.07387 | 1.72771 | 0.00000 | 0.00000 | 0.00000 | 12.11325 | 0.41357 | 0.29811 | 0.00000 | 33.01182 |
| **PPAPDC1A** | 1.15867 | 0.04569 | 0.00000 | 0.25056 | 1.10801 | 0.12390 | 0.00000 | 0.46405 | 0.00000 | 0.00242 | 0.00000 | 0.10215 |
| **PPAPDC1B** | 0.01454 | 0.05294 | 6.25378 | 2.04341 | 0.00000 | 1.07279 | 0.00000 | 0.12357 | 2.39754 | 0.00002 | 0.62290 | 0.26693 |
| **PPAPDC2** | 0.96638 | 2.29431 | 1.56878 | 0.01098 | 0.88528 | 2.35396 | 0.00000 | 0.02313 | 0.00000 | 0.00000 | 1.89159 | 1.60529 |
| **PPAPDC3** | 0.81671 | 0.00400 | 0.00000 | 0.05894 | 0.00001 | 0.32025 | 0.00000 | 0.15477 | 0.11115 | 0.01472 | 0.36656 | 0.00000 |
| **PPARA** | 0.23231 | 0.90206 | 0.00000 | 0.00000 | 1.32069 | 0.45943 | 0.81138 | 0.00000 | 0.00001 | 2.05314 | 0.43398 | 0.48061 |
| **PPARD** | 2.98423 | 3.30532 | 9.82247 | 3.02929 | 5.68432 | 0.00000 | 7.76598 | 4.42815 | 2.57843 | 6.73888 | 2.27893 | 2.57438 |
| **PPARG** | 0.01031 | 0.00000 | 14.80790 | 1.13823 | 4.10327 | 3.00011 | 9.55215 | 0.26007 | 8.30043 | 0.00000 | 1.81416 | 0.27857 |
| **PPARGC1A** | 0.35197 | 0.01790 | 0.00069 | 0.10866 | 0.09844 | 0.00000 | 0.00435 | 0.01371 | 0.01904 | 0.27854 | 0.01010 | 0.00786 |
| **PPARGC1B** | 2.06468 | 10.99235 | 0.44818 | 8.48279 | 7.76225 | 7.30943 | 0.39216 | 2.53628 | 6.00322 | 14.23358 | 3.23342 | 12.76898 |
| **PPAT** | 0.48568 | 6.20394 | 9.07917 | 3.58326 | 2.03046 | 20.69325 | 10.60486 | 7.35095 | 13.28409 | 11.91913 | 10.13740 | 6.81338 |
| **PPCDC** | 0.59563 | 3.40900 | 2.25631 | 1.55132 | 2.99324 | 1.94987 | 3.41733 | 0.10791 | 0.27308 | 0.00000 | 2.98937 | 2.63769 |
| **PPCS** | 1.01153 | 7.82792 | 6.20356 | 6.26555 | 6.53237 | 18.86820 | 10.31198 | 0.00000 | 9.99560 | 5.14924 | 14.46241 | 4.27346 |
| **PPDPF** | 2.81989 | 0.00892 | 0.69344 | 0.33731 | 4.71644 | 0.00000 | 1.00434 | 0.00064 | 0.24591 | 6.37189 | 0.00000 | 1.22905 |
| **PPEF1** | 2.97286 | 0.00327 | 0.00314 | 0.28604 | 0.00409 | 0.00000 | 0.01254 | 0.94379 | 0.01991 | 0.01133 | 0.01201 | 0.19063 |
| **PPEF2** | 0.02752 | 0.02995 | 0.01134 | 0.00665 | 0.01217 | 0.00000 | 0.02216 | 0.27416 | 0.01355 | 0.07343 | 0.01436 | 0.67201 |
| **PPFIA1** | 8.15569 | 4.43016 | 10.58190 | 5.26729 | 6.67972 | 10.08486 | 12.12094 | 2.40436 | 3.30853 | 27.68518 | 5.28697 | 4.50914 |
| **PPFIA2** | 1.62156 | 0.01418 | 0.02055 | 0.00770 | 0.02075 | 0.00000 | 0.13577 | 0.81102 | 0.00000 | 0.01554 | 0.01182 | 0.04833 |
| **PPFIA3** | 0.40882 | 0.22666 | 0.79696 | 0.28130 | 0.43257 | 0.00000 | 0.25141 | 0.91837 | 2.11491 | 0.55621 | 0.42787 | 0.31664 |
| **PPFIA4** | 0.01395 | 0.01543 | 0.02394 | 0.07224 | 0.00000 | 0.00000 | 0.06639 | 0.04461 | 0.02654 | 0.56095 | 0.01323 | 0.41759 |
| **PPFIBP1** | 1.09180 | 1.45650 | 3.57066 | 3.56274 | 1.39476 | 6.28050 | 1.22651 | 1.30260 | 1.88102 | 4.89681 | 2.71102 | 3.80575 |
| **PPFIBP2** | 4.31949 | 9.61159 | 4.28443 | 11.31506 | 14.82091 | 2.76777 | 9.21200 | 0.00000 | 11.37567 | 7.94620 | 13.48370 | 6.45431 |
| **PPHLN1** | 5.93499 | 11.91358 | 9.78266 | 10.44284 | 9.91238 | 28.55384 | 16.35623 | 11.12789 | 9.04162 | 11.06813 | 10.01575 | 17.65574 |
| **PPIA** | 5.57134 | 22.03587 | 10.68461 | 15.88283 | 19.95234 | 31.17894 | 17.37567 | 4.43847 | 31.28623 | 0.00000 | 11.44990 | 16.49584 |

| | | | | | | | | | | | | |
|---|---|---|---|---|---|---|---|---|---|---|---|---|
| **PPIAL4B+PPIAL** | 0.00581 | 0.01589 | 0.01480 | 0.01548 | 0.02113 | 0.01455 | 0.01975 | 0.01200 | 0.01780 | 0.01229 | 0.01496 | 0.01659 |
| **PPIAL4G** | 0.00759 | 0.01315 | 0.01268 | 0.00973 | 0.01981 | 0.03040 | 0.03472 | 0.01244 | 0.02902 | 0.03754 | 0.01128 | 0.02015 |
| **PPIAP30** | 0.00773 | 0.01213 | 0.01047 | 0.01008 | 0.02032 | 0.05889 | 0.01082 | 2.09441 | 0.01646 | 0.09946 | 0.01518 | 0.00369 |
| **PPIB** | 0.91487 | 10.68668 | 24.69411 | 9.45493 | 10.91260 | 21.01741 | 0.00000 | 2.79489 | 37.89668 | 0.00000 | 13.62839 | 10.30884 |
| **PPIC** | 2.00964 | 11.15629 | 1.39950 | 7.50610 | 14.11214 | 12.45337 | 0.00000 | 1.86345 | 2.93777 | 3.88909 | 12.92475 | 0.31038 |
| **PPID** | 0.56430 | 14.54337 | 18.40551 | 5.54578 | 6.89334 | 29.83032 | 27.15178 | 4.85271 | 20.37243 | 4.63523 | 18.61592 | 8.88623 |
| **PPIE** | 1.88952 | 14.18876 | 8.24060 | 8.05772 | 12.49115 | 12.28565 | 11.85471 | 9.08988 | 12.08188 | 5.05020 | 9.27671 | 12.27696 |
| **PPIEL** | 0.11222 | 0.01369 | 0.12845 | 0.53807 | 0.54363 | 0.00000 | 0.03035 | 0.06863 | 0.34192 | 0.32017 | 0.21616 | 0.19485 |
| **PPIF** | 4.62968 | 32.01532 | 9.86953 | 13.65655 | 25.59613 | 25.51950 | 42.23391 | 7.64628 | 41.93886 | 8.41459 | 21.30760 | 19.80658 |
| **PPIG** | 9.03015 | 8.25888 | 10.38143 | 7.46037 | 9.59137 | 30.39827 | 22.87651 | 20.48645 | 3.26368 | 39.05874 | 10.69612 | 15.13038 |
| **PPIH** | 1.47905 | 12.33554 | 7.28175 | 7.62483 | 14.58994 | 11.37227 | 15.96677 | 0.03422 | 13.85619 | 6.37015 | 7.45877 | 6.81698 |
| **PPIL1** | 3.39255 | 19.16718 | 13.72189 | 5.22119 | 8.04534 | 33.91083 | 19.09652 | 3.92867 | 20.15297 | 0.00000 | 11.19351 | 21.46944 |
| **PPIL2** | 3.27990 | 10.86858 | 4.76859 | 6.86942 | 5.77312 | 0.90596 | 4.67724 | 0.12637 | 6.66038 | 0.00000 | 8.69813 | 5.72105 |
| **PPIL3** | 1.59936 | 6.42143 | 5.26412 | 3.09617 | 4.20689 | 20.61985 | 2.92448 | 8.65436 | 6.58017 | 3.00650 | 7.28405 | 11.33239 |
| **PPIL4** | 5.02076 | 5.73421 | 6.06161 | 1.34077 | 0.00111 | 27.80140 | 6.73579 | 1.19309 | 3.97721 | 12.48433 | 8.91381 | 5.03700 |
| **PPIL6** | 0.18161 | 0.66852 | 0.14625 | 0.00000 | 0.02825 | 0.05578 | 0.00000 | 0.07300 | 0.12700 | 0.00537 | 0.16834 | 0.06411 |
| **PPIP5K1** | 0.00299 | 0.14400 | 0.35005 | 0.00142 | 0.25187 | 0.05345 | 0.41795 | 0.77130 | 0.00000 | 0.00381 | 0.16716 | 2.19537 |
| **PPIP5K2** | 2.30701 | 1.74436 | 7.14254 | 0.00000 | 0.35194 | 13.20129 | 9.15518 | 0.34717 | 6.64067 | 8.69194 | 2.53365 | 1.66413 |
| **PPL** | 6.80174 | 0.04468 | 0.73092 | 0.00000 | 0.02885 | 0.39498 | 0.72711 | 0.42931 | 2.77473 | 0.88856 | 1.01676 | 0.26109 |
| **PPM1A** | 4.05970 | 0.39476 | 6.22591 | 1.95145 | 1.88264 | 13.56829 | 10.07595 | 3.20111 | 0.00003 | 11.15328 | 1.58770 | 6.76554 |
| **PPM1B** | 3.19828 | 1.15424 | 23.37530 | 12.46118 | 0.00000 | 13.07561 | 12.62564 | 2.23814 | 3.43083 | 20.00874 | 0.27950 | 2.30198 |
| **PPM1D** | 2.07859 | 11.84024 | 6.92830 | 3.87349 | 1.34598 | 7.55027 | 2.70865 | 12.34183 | 9.59263 | 0.00000 | 3.26302 | 9.13032 |
| **PPM1E** | 1.44163 | 0.13858 | 0.00000 | 0.64533 | 0.00000 | 0.45926 | 0.38174 | 0.65232 | 0.46402 | 0.82154 | 0.00000 | 6.88639 |
| **PPM1F** | 1.07373 | 5.54165 | 4.42543 | 3.35737 | 4.57881 | 0.00000 | 2.10419 | 0.10787 | 3.83127 | 3.70895 | 7.66832 | 2.23092 |
| **PPM1G** | 8.92249 | 29.67240 | 24.01486 | 25.00437 | 25.66435 | 12.63095 | 29.78619 | 5.53709 | 30.87643 | 18.26478 | 18.91735 | 29.25289 |
| **PPM1H** | 1.85426 | 0.44551 | 0.76062 | 3.14483 | 0.76850 | 1.21429 | 3.77319 | 0.30958 | 0.69549 | 0.00000 | 0.87942 | 0.00000 |
| **PPM1J** | 0.01041 | 2.11160 | 0.31844 | 3.31920 | 0.31601 | 0.32906 | 0.00000 | 0.00000 | 0.93647 | 0.00000 | 0.89618 | 0.49767 |
| **PPM1K** | 0.11137 | 0.10795 | 0.75171 | 0.81057 | 0.00000 | 0.90702 | 0.00000 | 0.27450 | 0.43810 | 1.51790 | 0.00235 | 1.10628 |
| **PPM1L** | 0.12177 | 0.13024 | 0.37467 | 0.00165 | 0.01790 | 0.19219 | 0.00000 | 0.00530 | 0.20323 | 0.00000 | 0.00000 | 0.09659 |
| **PPM1M** | 0.03851 | 0.18881 | 0.17071 | 0.18651 | 0.00000 | 0.00000 | 0.00000 | 1.74469 | 0.66960 | 1.73388 | 0.38542 | 0.26630 |
| **PPM1N** | 0.62243 | 2.45210 | 8.26793 | 0.00000 | 0.00000 | 1.17010 | 2.24028 | 1.43929 | 1.08466 | 3.28844 | 0.20317 | 8.67852 |
| **PPME1** | 3.17199 | 4.40381 | 16.81635 | 2.57410 | 14.71940 | 4.37514 | 23.18564 | 5.50086 | 12.82637 | 5.18564 | 26.32984 | 3.53968 |
| **PPOX** | 0.16789 | 0.90671 | 0.43316 | 3.13951 | 0.00000 | 0.00000 | 1.13555 | 0.57401 | 0.28243 | 0.00000 | 2.44484 | 0.58055 |
| **PPP1CA** | 1.38095 | 24.75505 | 22.65347 | 23.40152 | 22.76087 | 20.49833 | 26.98312 | 15.70122 | 26.51560 | 0.00000 | 26.11775 | 19.98511 |
| **PPP1CB** | 5.75152 | 10.97345 | 38.74728 | 8.25956 | 0.00000 | 55.16095 | 36.67572 | 4.59765 | 8.02366 | 15.42665 | 26.07588 | 8.57728 |
| **PPP1CC** | 7.18707 | 9.39794 | 12.34875 | 8.11723 | 8.84968 | 37.78091 | 30.17441 | 14.17495 | 4.66042 | 54.76481 | 8.33505 | 12.90776 |
| **PPP1R10** | 9.61740 | 7.32342 | 16.73715 | 12.21473 | 14.40158 | 0.09311 | 14.81753 | 4.60634 | 16.72788 | 15.93912 | 8.55632 | 18.77283 |
| **PPP1R11** | 2.87574 | 8.43010 | 9.19690 | 7.54970 | 7.49751 | 10.57198 | 10.71771 | 0.65729 | 9.41852 | 0.00000 | 10.91353 | 16.63808 |
| **PPP1R12A** | 8.81911 | 4.82742 | 12.62917 | 8.14398 | 9.78769 | 23.71640 | 17.86515 | 14.21220 | 4.20873 | 24.17210 | 13.20805 | 8.19641 |
| **PPP1R12B** | 1.61346 | 0.41433 | 1.41258 | 7.58476 | 1.68141 | 1.01825 | 0.00001 | 1.53121 | 2.93279 | 4.09836 | 0.62449 | 1.22986 |
| **PPP1R12C** | 0.39716 | 0.00000 | 0.62731 | 0.32611 | 1.83707 | 0.00000 | 0.44023 | 0.23505 | 1.13629 | 6.86411 | 0.20373 | 0.56329 |
| **PPP1R13B** | 9.23835 | 1.84979 | 0.60309 | 1.32785 | 2.05614 | 0.00000 | 0.26284 | 2.09774 | 0.00000 | 2.21552 | 0.20211 | 6.18417 |
| **PPP1R13L** | 0.00000 | 0.24953 | 4.31267 | 0.68051 | 7.05391 | 0.00000 | 6.73976 | 0.06523 | 2.98849 | 2.81253 | 0.93686 | 0.49116 |
| **PPP1R14A** | 0.00000 | 39.62532 | 12.53254 | 16.08136 | 38.01264 | 2.88639 | 17.29161 | 1.25103 | 0.11045 | 13.63767 | 21.99335 | 11.31224 |
| **PPP1R14B** | 0.04961 | 2.09763 | 1.11105 | 1.40135 | 5.63543 | 0.47464 | 5.01067 | 0.36998 | 0.03250 | 12.70907 | 0.51005 | 2.89703 |
| **PPP1R14C** | 3.87819 | 0.27634 | 0.00000 | 0.11177 | 0.03795 | 0.09413 | 0.00000 | 0.36700 | 0.00000 | 0.07098 | 0.00000 | 0.06113 |
| **PPP1R14D** | 0.00414 | 0.22346 | 1.55046 | 0.10717 | 1.39917 | 0.00000 | 0.05299 | 0.02325 | 0.30440 | 0.00000 | 0.57376 | 0.58934 |
| **PPP1R15A** | 6.04705 | 19.26239 | 13.28791 | 13.42513 | 6.70630 | 0.00000 | 5.26466 | 22.39499 | 10.13026 | 4.08820 | 1.80246 | 27.04203 |

| | | | | | | | | | | | | |
|---|---|---|---|---|---|---|---|---|---|---|---|---|
| **PPP1R15B** | 1.70006 | 5.00447 | 9.87953 | 0.00000 | 1.10475 | 22.74782 | 0.00000 | 5.64047 | 14.00558 | 14.37631 | 4.03836 | 5.62540 |
| **PPP1R16A** | 0.25406 | 2.89040 | 1.06472 | 2.45832 | 3.49661 | 0.00000 | 1.42274 | 0.29411 | 6.87159 | 5.72271 | 2.48323 | 0.68229 |
| **PPP1R16B** | 0.02112 | 0.93842 | 0.23125 | 1.11975 | 0.00000 | 0.00000 | 0.00000 | 0.00596 | 0.00000 | 0.14968 | 0.04347 | 0.26516 |
| **PPP1R17** | 0.02430 | 0.00624 | 0.06902 | 0.03830 | 0.00155 | 0.08887 | 0.00000 | 0.01179 | 0.00944 | 0.01656 | 0.01072 | 0.01167 |
| **PPP1R18** | 0.69348 | 0.32957 | 0.51519 | 1.23701 | 0.18640 | 0.00000 | 0.93899 | 0.18305 | 1.12507 | 2.00761 | 0.13904 | 0.48066 |
| **PPP1R1A** | 0.00331 | 0.00000 | 0.19627 | 0.20677 | 0.14593 | 0.11451 | 0.10814 | 0.01122 | 0.00000 | 0.21572 | 0.02484 | 0.00258 |
| **PPP1R1B** | 0.26334 | 0.09122 | 0.01810 | 0.14085 | 0.00240 | 0.07030 | 0.00004 | 0.12948 | 0.05911 | 0.01463 | 0.00000 | 0.02387 |
| **PPP1R1C** | 0.05816 | 0.32307 | 0.06062 | 0.00000 | 0.10393 | 0.55453 | 0.00000 | 0.10895 | 0.00000 | 0.08564 | 0.00554 | 0.18187 |
| **PPP1R2** | 1.99406 | 9.27596 | 5.87214 | 0.86766 | 0.00000 | 19.18724 | 13.29345 | 7.42523 | 17.90473 | 8.95747 | 13.07044 | 12.54793 |
| **PPP1R21** | 0.00000 | 0.04963 | 1.26819 | 1.78634 | 3.79675 | 4.04079 | 1.28462 | 0.49145 | 2.98644 | 6.85019 | 3.15537 | 0.55152 |
| **PPP1R26** | 0.00767 | 2.82230 | 0.04370 | 0.99173 | 0.53005 | 0.00000 | 0.00000 | 0.00000 | 0.00000 | 0.64634 | 0.00000 | 2.68470 |
| **PPP1R26-AS1** | 0.03533 | 0.22663 | 0.00280 | 0.32803 | 0.14487 | 0.09428 | 0.00000 | 0.00580 | 0.00000 | 0.01824 | 0.00692 | 0.36839 |
| **PPP1R27** | 0.06515 | 0.02983 | 0.17307 | 0.01753 | 0.11370 | 0.00000 | 0.00361 | 0.03089 | 0.08609 | 0.01249 | 0.00634 | 0.05796 |
| **PPP1R2P3** | 0.27302 | 0.51946 | 0.24210 | 0.06081 | 0.02736 | 1.01600 | 0.28491 | 0.53386 | 1.45635 | 0.37095 | 0.85132 | 0.98315 |
| **PPP1R2P9** | 0.10107 | 0.08699 | 0.58677 | 0.08291 | 0.00000 | 0.84669 | 0.00000 | 0.02385 | 0.88030 | 0.57110 | 0.00000 | 0.46005 |
| **PPP1R32** | 0.02977 | 0.68202 | 0.21568 | 0.07616 | 0.16779 | 0.05011 | 0.02827 | 0.00404 | 0.02975 | 0.00000 | 0.16094 | 0.07868 |
| **PPP1R35** | 5.18224 | 0.91435 | 1.36339 | 0.10783 | 1.23492 | 0.00000 | 1.52575 | 3.35057 | 1.68677 | 2.44231 | 1.97287 | 0.10395 |
| **PPP1R36** | 0.15875 | 0.04525 | 0.02941 | 0.03634 | 0.00396 | 0.14859 | 0.00000 | 0.00000 | 0.03722 | 0.00000 | 0.03904 | 0.05643 |
| **PPP1R37** | 0.38445 | 0.20454 | 0.32643 | 0.07562 | 0.89860 | 0.00000 | 0.72179 | 0.52177 | 0.00000 | 4.96637 | 0.00000 | 0.42771 |
| **PPP1R3A** | 5.12144 | 0.00844 | 0.00170 | 0.01428 | 0.00507 | 0.01412 | 0.04098 | 0.02365 | 0.24715 | 0.00000 | 0.00970 | 0.00444 |
| **PPP1R3B** | 0.04516 | 10.96954 | 0.23633 | 10.76793 | 7.43689 | 13.55006 | 4.50107 | 17.00669 | 3.97302 | 6.00595 | 3.96129 | 29.99605 |
| **PPP1R3C** | 3.09575 | 0.00291 | 0.08059 | 0.01387 | 0.01446 | 0.00002 | 0.01529 | 0.59587 | 0.04807 | 0.00364 | 0.00971 | 0.27738 |
| **PPP1R3D** | 2.66243 | 0.00000 | 0.12759 | 0.00000 | 0.25283 | 0.03998 | 0.44514 | 0.00000 | 0.08854 | 0.11283 | 0.05869 | 0.11531 |
| **PPP1R3E** | 0.54305 | 0.00000 | 0.52989 | 0.00191 | 0.00865 | 0.03103 | 0.20804 | 1.36637 | 0.00000 | 0.60345 | 0.27239 | 0.25145 |
| **PPP1R3F** | 0.23259 | 0.00312 | 0.03179 | 0.23627 | 0.00985 | 0.08354 | 0.00000 | 0.07218 | 0.14231 | 0.00000 | 0.00001 | 0.08146 |
| **PPP1R3G** | 0.02153 | 0.01967 | 0.09541 | 0.00786 | 0.00000 | 0.22517 | 0.02340 | 0.05155 | 0.00000 | 0.02215 | 0.01125 | 0.48120 |
| **PPP1R42** | 0.01459 | 0.04218 | 0.01283 | 0.00000 | 0.00000 | 0.06670 | 0.02441 | 0.03344 | 0.00299 | 0.05114 | 0.00910 | 0.01098 |
| **PPP1R7** | 2.73398 | 8.57775 | 9.61649 | 8.06032 | 16.96635 | 9.49446 | 17.21594 | 0.72966 | 8.18482 | 1.06414 | 16.89693 | 8.25913 |
| **PPP1R8** | 0.59319 | 2.57788 | 3.74186 | 3.12385 | 0.00000 | 4.90234 | 7.15844 | 5.27327 | 0.00000 | 11.90246 | 17.47212 | 10.94970 |
| **PPP1R9A** | 0.76333 | 5.83325 | 0.98032 | 1.79404 | 7.17403 | 4.93395 | 1.64907 | 0.00000 | 1.86545 | 10.66598 | 5.17609 | 1.44944 |
| **PPP1R9B** | 0.41325 | 0.24503 | 1.49065 | 0.28706 | 0.54714 | 0.00000 | 0.48721 | 0.17545 | 0.30474 | 3.33136 | 0.00000 | 0.08065 |
| **PPP2CA** | 4.92538 | 24.40752 | 28.86325 | 7.28665 | 9.43279 | 26.69477 | 24.47222 | 19.79842 | 18.61250 | 20.92824 | 20.91271 | 15.03634 |
| **PPP2CB** | 0.00000 | 6.04537 | 6.96485 | 4.56534 | 4.47239 | 15.49305 | 14.56056 | 3.49064 | 3.67360 | 14.46409 | 0.83343 | 7.47311 |
| **PPP2R1A** | 10.89520 | 23.83727 | 23.86844 | 24.89047 | 24.76060 | 8.20402 | 28.88058 | 5.43270 | 33.07838 | 0.05374 | 20.10473 | 14.32697 |
| **PPP2R1B** | 4.71869 | 9.07015 | 11.27131 | 0.79562 | 0.48430 | 22.41847 | 7.88708 | 4.87718 | 9.57610 | 1.01033 | 7.02673 | 6.65101 |
| **PPP2R2A** | 3.22065 | 1.65577 | 7.08121 | 3.37978 | 5.19278 | 13.70664 | 16.84616 | 3.24010 | 5.08976 | 18.27415 | 3.93204 | 8.04167 |
| **PPP2R2B** | 2.76303 | 0.00882 | 0.10547 | 1.36099 | 0.01210 | 0.00000 | 0.01090 | 0.06530 | 0.02319 | 0.04542 | 0.02408 | 0.03154 |
| **PPP2R2C** | 1.63647 | 0.03810 | 0.01207 | 0.00000 | 0.05501 | 0.00000 | 0.00000 | 0.01926 | 0.04360 | 0.25789 | 0.01266 | 0.00238 |
| **PPP2R2D** | 0.00000 | 2.10001 | 2.35906 | 3.66516 | 5.31022 | 6.12110 | 5.22117 | 2.39451 | 0.08225 | 14.61410 | 1.30462 | 0.97221 |
| **PPP2R3A** | 0.08967 | 0.12666 | 9.02928 | 0.65972 | 1.00869 | 3.32229 | 4.10506 | 1.59809 | 5.38092 | 3.50047 | 2.40724 | 1.09401 |
| **PPP2R3C** | 3.66492 | 4.36022 | 7.00457 | 2.54413 | 5.98058 | 19.69688 | 6.94131 | 5.29573 | 5.63773 | 9.72179 | 5.86934 | 18.44758 |
| **PPP2R4** | 2.09899 | 4.34864 | 12.26961 | 7.57337 | 9.07717 | 0.00000 | 13.53942 | 0.81616 | 9.29423 | 8.72418 | 6.36818 | 4.07656 |
| **PPP2R5A** | 0.00000 | 2.78688 | 0.73845 | 3.92941 | 3.30420 | 19.07019 | 15.92562 | 0.81541 | 2.58413 | 31.92362 | 3.26107 | 8.08289 |
| **PPP2R5B** | 0.19867 | 2.41083 | 0.34206 | 0.43242 | 0.00000 | 0.00000 | 0.00000 | 0.35714 | 0.32165 | 1.26929 | 0.15079 | 2.65320 |
| **PPP2R5C** | 6.68687 | 6.37057 | 8.15889 | 4.31060 | 4.69398 | 21.32130 | 11.83253 | 2.55767 | 8.05018 | 1.14122 | 10.24424 | 6.48326 |
| **PPP2R5D** | 4.28956 | 9.62809 | 9.83458 | 6.94101 | 6.07108 | 1.60466 | 11.58841 | 1.93437 | 10.84556 | 0.00000 | 8.46768 | 5.46965 |
| **PPP2R5E** | 1.22689 | 0.00000 | 8.23850 | 3.09453 | 4.63421 | 11.07070 | 17.97226 | 3.09785 | 0.00052 | 20.17548 | 1.34646 | 2.90535 |
| **PPP3CA** | 4.58589 | 0.76437 | 1.87476 | 0.00000 | 0.00000 | 2.66643 | 1.46125 | 3.61117 | 0.00000 | 1.88518 | 0.16814 | 1.71460 |

| | | | | | | | | | | | | |
|---|---|---|---|---|---|---|---|---|---|---|---|---|
| **PPP3CB** | 2.10198 | 0.18375 | 2.10062 | 0.79106 | 0.80469 | 3.20277 | 3.60401 | 0.04515 | 1.18749 | 4.63278 | 0.52763 | 0.99101 |
| **PPP3CC** | 0.69143 | 1.62777 | 1.73692 | 0.00000 | 2.07226 | 0.65755 | 0.96266 | 0.04806 | 1.05684 | 0.76201 | 0.68701 | 0.12928 |
| **PPP3R1** | 2.87962 | 0.36977 | 1.29547 | 1.36731 | 0.00000 | 3.12111 | 2.28715 | 2.36988 | 0.00000 | 8.29801 | 3.83992 | 1.65531 |
| **PPP3R2** | 0.01926 | 0.01344 | 0.01855 | 0.01277 | 0.00948 | 0.01935 | 0.01727 | 0.01244 | 0.01069 | 0.02853 | 0.03122 | 0.01460 |
| **PPP4C** | 4.29199 | 14.23146 | 19.37690 | 11.38048 | 17.45411 | 0.00000 | 18.00528 | 10.57721 | 9.99382 | 25.03077 | 13.03155 | 8.64489 |
| **PPP4R1** | 4.86509 | 4.35008 | 7.98589 | 1.87786 | 1.56478 | 11.31883 | 13.24372 | 0.69183 | 7.34197 | 3.73071 | 4.91090 | 2.29727 |
| **PPP4R1L** | 0.02593 | 0.41605 | 0.37968 | 0.47817 | 0.03419 | 0.01516 | 0.00510 | 1.18536 | 0.42459 | 0.00000 | 0.09298 | 1.16313 |
| **PPP4R2** | 1.58702 | 0.01226 | 3.25158 | 0.14548 | 1.25123 | 4.54229 | 3.24235 | 3.22758 | 0.00000 | 11.50556 | 0.57679 | 1.69061 |
| **PPP4R4** | 3.38251 | 0.03663 | 0.00925 | 0.00000 | 0.00000 | 0.03914 | 0.00000 | 1.12754 | 0.02030 | 0.06699 | 0.06448 | 0.04173 |
| **PPP5C** | 0.13001 | 15.48693 | 12.83656 | 9.55059 | 6.57451 | 0.00000 | 7.69718 | 0.00000 | 18.68396 | 0.00000 | 11.25518 | 7.25817 |
| **PPP5D1** | 0.00219 | 0.16757 | 0.28206 | 0.10336 | 0.03899 | 0.00000 | 0.06702 | 0.05275 | 0.04670 | 0.00000 | 0.11140 | 0.11173 |
| **PPP6C** | 10.66755 | 11.21679 | 12.68399 | 6.26723 | 4.75153 | 25.20905 | 15.04532 | 6.41089 | 13.00126 | 3.85888 | 8.58234 | 14.39830 |
| **PPP6R1** | 2.95087 | 5.94763 | 7.97213 | 4.64925 | 9.07288 | 0.00000 | 14.12537 | 0.00000 | 5.07178 | 17.71197 | 1.57615 | 5.49812 |
| **PPP6R2** | 0.79089 | 2.98204 | 4.28735 | 3.04073 | 4.14847 | 0.00000 | 1.63679 | 0.08284 | 4.20401 | 6.00005 | 4.75200 | 3.66756 |
| **PPP6R3** | 3.15341 | 9.03628 | 8.86906 | 10.30123 | 6.28408 | 29.33635 | 19.24178 | 10.57599 | 7.95978 | 15.55014 | 12.49809 | 13.18230 |
| **PPRC1** | 11.25873 | 14.49995 | 8.80462 | 8.71696 | 15.12216 | 0.00000 | 11.20755 | 6.25722 | 10.90147 | 27.42566 | 10.61100 | 22.01401 |
| **PPT1** | 1.09046 | 21.37650 | 12.31878 | 0.00000 | 20.23554 | 39.59895 | 14.51846 | 5.28809 | 70.34295 | 5.93477 | 38.86416 | 15.09477 |
| **PPT2** | 4.09357 | 17.71639 | 1.65404 | 4.76375 | 20.48974 | 11.59718 | 0.00000 | 0.94016 | 5.85413 | 7.42612 | 2.04869 | 10.85681 |
| **PPT2-EGFL8** | 0.02749 | 0.79051 | 0.01322 | 0.33073 | 0.21066 | 0.00000 | 0.00278 | 0.00957 | 0.34800 | 0.00000 | 0.03475 | 0.75945 |
| **PPTC7** | 1.60130 | 0.56205 | 1.79216 | 1.19033 | 1.48487 | 5.68278 | 0.31650 | 5.69770 | 0.00000 | 8.06871 | 0.89390 | 2.02181 |
| **PPWD1** | 3.46841 | 8.90644 | 10.43129 | 3.54080 | 1.69637 | 15.80586 | 8.85941 | 11.39232 | 9.34768 | 0.01049 | 8.55136 | 10.94158 |
| **PPY** | 0.77737 | 0.00776 | 0.25793 | 0.01444 | 0.03086 | 0.01405 | 0.00599 | 0.03040 | 0.04338 | 0.00000 | 0.08239 | 0.06239 |
| **PPY2** | 0.00713 | 0.14614 | 0.11734 | 0.01125 | 0.02498 | 0.01616 | 0.01282 | 0.00772 | 0.02841 | 0.00000 | 0.01976 | 0.08785 |
| **PQBP1** | 0.60878 | 12.10378 | 8.99778 | 11.15349 | 16.16042 | 5.30778 | 15.48952 | 23.43030 | 16.30574 | 4.25886 | 10.03925 | 28.18547 |
| **PQLC1** | 0.00434 | 6.32990 | 0.74374 | 7.18566 | 2.86299 | 0.00000 | 2.39247 | 1.63455 | 0.00000 | 5.73985 | 0.13349 | 6.77646 |
| **PQLC2** | 1.35111 | 3.16387 | 7.89566 | 0.85656 | 3.20820 | 0.00000 | 3.16619 | 0.00000 | 15.42940 | 0.00000 | 5.52353 | 2.55033 |
| **PQLC3** | 4.18462 | 13.06993 | 1.15032 | 16.46224 | 8.99015 | 21.10040 | 0.00000 | 1.84762 | 0.00000 | 1.65703 | 6.96322 | 6.01843 |
| **PRAC1** | 0.00608 | 0.01306 | 0.02445 | 0.01332 | 0.01229 | 0.03105 | 0.01996 | 0.01034 | 0.02058 | 0.01696 | 0.01498 | 0.01569 |
| **PRAC2** | 0.02610 | 0.03558 | 0.01161 | 0.01608 | 0.01350 | 0.03477 | 0.00904 | 0.00491 | 0.01208 | 0.06407 | 0.00977 | 0.05452 |
| **PRADC1** | 4.38068 | 5.87969 | 2.51673 | 4.67472 | 8.76123 | 0.00000 | 1.55950 | 1.26146 | 5.25146 | 0.35010 | 8.45183 | 0.68256 |
| **PRAF2** | 0.87988 | 0.69744 | 1.05607 | 0.62568 | 0.00000 | 0.00000 | 0.41031 | 1.18819 | 1.58251 | 0.00000 | 2.72467 | 3.59156 |
| **PRAM1** | 0.01705 | 0.00809 | 0.02753 | 0.01092 | 0.01233 | 0.04780 | 0.02640 | 0.12635 | 0.00143 | 0.03351 | 0.01426 | 0.08195 |
| **PRAME** | 0.48111 | 4.24115 | 0.00000 | 0.00000 | 0.31182 | 0.79592 | 0.00000 | 10.14434 | 0.00000 | 0.10878 | 0.69600 | 8.98177 |
| **PRAMEF1** | 5.98358 | 0.00387 | 0.01158 | 0.00691 | 0.04883 | 0.00000 | 0.01421 | 6.94150 | 0.00990 | 0.00000 | 0.01194 | 0.10031 |
| **PRAMEF11** | 3.88437 | 0.00000 | 0.00720 | 0.00470 | 0.01882 | 0.00000 | 0.01100 | 16.90386 | 0.00419 | 0.00393 | 0.04118 | 0.44494 |
| **PRAMEF12** | 7.03757 | 0.00000 | 0.00000 | 0.00011 | 0.02554 | 0.00000 | 0.00000 | 22.99444 | 0.15062 | 0.00000 | 0.45501 | 1.67240 |
| **PRAMEF15.uid-** | 0.00567 | 0.01439 | 0.01425 | 0.01209 | 0.01527 | 0.02037 | 0.01859 | 0.02179 | 0.01799 | 0.01060 | 0.01513 | 0.02106 |
| **PRAMEF17.uid-** | 0.00973 | 0.22470 | 0.01045 | 0.00000 | 0.00343 | 0.00709 | 0.01559 | 1.82644 | 0.01279 | 0.01375 | 0.01172 | 1.15763 |
| **PRAMEF19.uid-** | 0.00874 | 2.87774 | 0.02598 | 0.00369 | 1.05011 | 0.07471 | 0.43986 | 7.21452 | 0.00000 | 0.00000 | 2.28159 | 3.95256 |
| **PRAMEF19.uid-** | 0.25386 | 0.29730 | 0.01059 | 0.00293 | 0.05767 | 0.00000 | 0.00874 | 1.18451 | 0.00000 | 0.00000 | 0.33998 | 0.55580 |
| **PRAMEF2** | 3.88019 | 0.03740 | 0.00669 | 0.01148 | 0.02027 | 0.00000 | 0.00865 | 11.45014 | 0.02061 | 0.00000 | 0.00870 | 0.00000 |
| **PRAMEF20.uid-** | 0.26642 | 0.02933 | 0.01489 | 0.01387 | 0.03073 | 0.00803 | 0.01967 | 0.09802 | 0.02351 | 0.00009 | 0.01220 | 0.01739 |
| **PRAMEF20.uid-** | 0.01587 | 0.30654 | 0.00575 | 0.00000 | 0.00333 | 0.04440 | 0.00806 | 2.95599 | 0.00465 | 0.02082 | 0.00843 | 0.39685 |
| **PRAMEF4** | 0.29034 | 0.01180 | 0.00829 | 0.00709 | 0.01825 | 0.00000 | 0.01035 | 17.72063 | 0.01740 | 0.01190 | 0.00939 | 0.27203 |
| **PRAMEF5.uid-8** | 0.00238 | 0.01056 | 0.01098 | 0.01112 | 0.01508 | 0.00883 | 0.01274 | 3.35827 | 0.01459 | 0.02646 | 0.01394 | 0.01034 |
| **PRAMEF6.uid-8** | 0.01081 | 0.00688 | 0.01008 | 0.01339 | 0.03391 | 0.00000 | 0.01328 | 7.45627 | 0.01357 | 0.01423 | 0.01834 | 0.16309 |
| **PRAMEF7+PRA** | 0.33559 | 0.01263 | 0.01550 | 0.01684 | 0.01549 | 0.00326 | 0.02337 | 0.79907 | 0.00943 | 0.00509 | 0.03986 | 0.04534 |
| **PRAP1** | 4.61174 | 26.90194 | 0.00000 | 5.91436 | 38.54498 | 6.47517 | 0.00000 | 5.87952 | 3.68837 | 8.10368 | 21.98600 | 0.00000 |

| | | | | | | | | | | | | |
|---|---|---|---|---|---|---|---|---|---|---|---|---|
| **PRB3** | 0.00225 | 0.34316 | 0.02774 | 0.00820 | 0.00228 | 0.03122 | 0.01917 | 0.05616 | 0.08612 | 0.00490 | 0.00826 | 0.44483 |
| **PRB4** | 0.00608 | 0.01607 | 0.01710 | 0.01210 | 0.01533 | 0.02023 | 0.01865 | 0.01025 | 0.01744 | 0.00876 | 0.01529 | 0.02816 |
| **PRC1** | 10.60487 | 13.44886 | 14.99963 | 3.85576 | 13.39282 | 19.14843 | 13.02611 | 7.36778 | 19.98910 | 6.64899 | 8.90874 | 11.75684 |
| **PRC1-AS1** | 0.06838 | 0.00000 | 0.13472 | 0.08726 | 0.01651 | 0.10474 | 0.00000 | 0.00803 | 0.00694 | 0.36155 | 0.06150 | 0.08214 |
| **PRCC** | 3.94233 | 9.76885 | 12.45782 | 8.98273 | 12.32349 | 1.30961 | 22.90582 | 5.22970 | 17.61907 | 10.97202 | 7.02189 | 13.86403 |
| **PRCD** | 0.01380 | 0.08653 | 0.01736 | 0.00862 | 0.01162 | 0.00331 | 0.02296 | 0.03335 | 0.01284 | 0.01389 | 0.03384 | 0.02530 |
| **PRCP** | 0.00000 | 3.19788 | 4.58049 | 15.79535 | 3.18786 | 8.55339 | 0.00000 | 0.00000 | 8.53431 | 0.25909 | 1.03152 | 0.00000 |
| **PRDM1** | 0.00833 | 0.42648 | 0.20136 | 5.10933 | 0.02144 | 1.14208 | 0.00000 | 0.00327 | 0.00000 | 0.63411 | 0.00000 | 0.00000 |
| **PRDM10** | 11.74890 | 1.11075 | 1.74562 | 0.36360 | 0.67643 | 3.08330 | 0.43665 | 1.50231 | 3.01042 | 3.11941 | 1.68583 | 0.14373 |
| **PRDM11** | 0.00667 | 0.00002 | 0.68369 | 1.23406 | 0.31675 | 0.00000 | 0.06698 | 0.01505 | 0.00000 | 0.00000 | 0.06882 | 0.08917 |
| **PRDM12** | 0.00801 | 0.01773 | 0.02710 | 0.00523 | 0.00796 | 0.04777 | 0.01180 | 0.01213 | 0.03816 | 0.01470 | 0.00992 | 0.02347 |
| **PRDM13** | 0.09819 | 0.03069 | 0.01528 | 0.05455 | 0.00986 | 0.11422 | 0.00000 | 0.00734 | 0.07873 | 0.11268 | 0.00091 | 0.01116 |
| **PRDM14** | 0.00126 | 5.91082 | 0.94544 | 14.25130 | 0.11409 | 0.00000 | 0.00000 | 1.20156 | 0.00000 | 0.00000 | 0.10869 | 7.34080 |
| **PRDM15** | 0.88732 | 1.15384 | 1.07768 | 0.00817 | 0.00000 | 2.97707 | 0.25104 | 0.10923 | 2.29011 | 4.92640 | 1.57353 | 1.68473 |
| **PRDM16** | 1.70411 | 0.01271 | 0.05425 | 0.01719 | 0.01196 | 0.00000 | 0.02026 | 0.09228 | 0.01195 | 0.01954 | 0.03387 | 0.01595 |
| **PRDM2** | 7.15795 | 1.31971 | 8.39710 | 0.54591 | 3.51641 | 5.85242 | 7.38034 | 4.05236 | 4.17440 | 17.38498 | 3.61972 | 4.41565 |
| **PRDM4** | 6.53150 | 0.51964 | 1.08419 | 0.96997 | 2.46190 | 0.89935 | 6.36741 | 0.86102 | 3.22280 | 5.59673 | 1.27328 | 0.60839 |
| **PRDM5** | 0.15278 | 0.03766 | 0.69319 | 0.02343 | 0.03240 | 0.00000 | 0.06208 | 0.01089 | 0.00000 | 0.00000 | 0.01531 | 0.03262 |
| **PRDM6** | 0.46303 | 0.50047 | 0.24805 | 0.00000 | 0.02445 | 0.25394 | 0.00000 | 0.00169 | 0.00000 | 0.07009 | 0.02676 | 0.05869 |
| **PRDM7** | 0.01110 | 0.01142 | 0.01339 | 0.00719 | 0.00580 | 0.01775 | 0.00819 | 0.03111 | 0.02076 | 0.10277 | 0.01239 | 0.01112 |
| **PRDM8** | 0.21330 | 0.01536 | 0.00907 | 0.00642 | 0.00000 | 0.02454 | 0.00688 | 0.10468 | 0.00000 | 0.08605 | 0.05243 | 0.09102 |
| **PRDM9** | 0.01962 | 0.01273 | 0.50384 | 0.16246 | 0.00000 | 0.00986 | 0.03316 | 0.09743 | 0.00000 | 0.06041 | 0.01149 | 0.04354 |
| **PRDX1** | 4.13744 | 62.38085 | 45.57386 | 45.25500 | 64.12873 | 85.98911 | 64.02409 | 16.05121 | 58.80067 | 40.67864 | 52.76378 | 44.10275 |
| **PRDX2** | 3.15498 | 15.52223 | 30.11716 | 26.09926 | 23.30326 | 5.43954 | 51.97063 | 8.58713 | 29.57807 | 0.00000 | 34.76656 | 5.16217 |
| **PRDX3** | 5.09284 | 26.01495 | 18.63987 | 26.34228 | 12.59445 | 56.55893 | 36.78292 | 6.90880 | 27.52721 | 4.11724 | 31.87271 | 21.36457 |
| **PRDX4** | 0.00000 | 17.55245 | 7.33278 | 8.72820 | 6.01065 | 30.12363 | 5.27301 | 0.16774 | 34.11868 | 4.40088 | 7.50241 | 9.21343 |
| **PRDX5** | 4.50033 | 14.60238 | 21.65000 | 28.97709 | 26.89908 | 4.03261 | 40.82982 | 9.56809 | 25.72272 | 5.74710 | 28.94347 | 16.41350 |
| **PRDX6** | 0.80035 | 12.70193 | 25.58675 | 20.18878 | 16.64070 | 28.17048 | 29.39867 | 2.96064 | 28.51254 | 6.78186 | 19.03700 | 3.96211 |
| **PREB** | 4.11917 | 24.13887 | 19.67468 | 9.23111 | 16.28476 | 0.00000 | 6.71066 | 1.06439 | 18.37842 | 4.31856 | 15.69218 | 9.04680 |
| **PRELID1** | 0.45565 | 14.80758 | 15.73884 | 9.49021 | 20.33956 | 2.66766 | 4.69186 | 3.51938 | 23.25287 | 0.00000 | 19.66642 | 8.99900 |
| **PRELID2** | 12.43862 | 0.00000 | 0.19035 | 0.48681 | 0.64061 | 0.00000 | 0.00096 | 1.01775 | 0.24191 | 0.02314 | 2.29598 | 0.02159 |
| **PRELP** | 0.11037 | 0.08151 | 0.18656 | 0.08190 | 0.00000 | 0.00000 | 0.00000 | 0.00197 | 0.12143 | 0.19597 | 0.02872 | 0.07478 |
| **PREP** | 4.06450 | 4.65159 | 6.28836 | 2.14790 | 2.77846 | 5.94589 | 0.00070 | 1.04745 | 5.25852 | 0.00000 | 4.70574 | 0.96577 |
| **PREPL** | 11.21327 | 6.11583 | 2.84165 | 0.76702 | 2.53468 | 13.26706 | 3.56894 | 5.70125 | 2.39924 | 4.79173 | 7.21291 | 8.65284 |
| **PREX1** | 4.84756 | 1.62651 | 1.87354 | 1.62266 | 1.63398 | 0.00000 | 0.42985 | 0.00000 | 3.41284 | 2.73986 | 0.92671 | 4.11253 |
| **PREX2** | 0.01161 | 0.01589 | 0.01589 | 0.00964 | 0.02305 | 0.03075 | 0.02604 | 0.13385 | 0.00000 | 0.01555 | 0.06337 | 0.00968 |
| **PRF1** | 0.00639 | 0.01311 | 0.00628 | 0.33149 | 0.06052 | 0.00592 | 0.02689 | 0.01025 | 0.01713 | 0.00000 | 0.01478 | 0.01749 |
| **PRG2** | 0.00968 | 0.39285 | 0.09364 | 0.00000 | 0.00623 | 0.08068 | 0.00000 | 0.00139 | 0.00000 | 0.03143 | 0.00733 | 0.58934 |
| **PRG3** | 0.21407 | 0.01279 | 0.01707 | 0.01241 | 0.01582 | 0.01263 | 0.01658 | 0.78191 | 0.01769 | 0.01552 | 0.01525 | 0.00317 |
| **PRG4** | 0.11560 | 0.02718 | 0.10758 | 0.01738 | 0.00000 | 0.12326 | 0.00000 | 0.12363 | 0.07293 | 0.39737 | 0.00000 | 0.00137 |
| **PRH1** | 0.00630 | 0.02787 | 0.15369 | 0.00369 | 0.00999 | 0.07135 | 0.01922 | 0.00927 | 0.09303 | 0.01587 | 0.00223 | 0.02211 |
| **PRH1-PRR4** | 0.00701 | 0.01061 | 0.04582 | 0.04293 | 0.00012 | 0.06115 | 0.00691 | 0.00945 | 0.02759 | 0.03066 | 0.02344 | 0.02743 |
| **PRH2** | 0.06987 | 0.00498 | 0.23737 | 0.00070 | 0.29549 | 0.00000 | 0.00000 | 0.00610 | 0.25285 | 0.26032 | 0.00000 | 0.01771 |
| **PRICKLE1** | 0.20053 | 4.21571 | 0.69536 | 8.75050 | 0.00000 | 0.00000 | 0.00000 | 0.35212 | 0.00000 | 1.64832 | 0.03166 | 1.04527 |
| **PRICKLE2** | 0.28725 | 0.05520 | 0.31340 | 0.01018 | 0.01991 | 0.00000 | 0.00000 | 0.15256 | 0.00000 | 0.05874 | 0.01659 | 0.00686 |
| **PRICKLE2-AS3** | 0.00451 | 0.01472 | 0.02081 | 0.01037 | 0.01726 | 0.02438 | 0.01311 | 0.01520 | 0.01625 | 0.03300 | 0.01468 | 0.01865 |
| **PRICKLE3** | 0.00865 | 0.61333 | 0.26756 | 0.56828 | 2.34632 | 0.00000 | 0.48232 | 0.04684 | 3.00337 | 0.00000 | 0.82994 | 0.82983 |
| **PRICKLE4** | 0.05665 | 0.60262 | 1.25372 | 0.01558 | 0.38591 | 0.75800 | 0.00000 | 1.15518 | 2.06246 | 0.07436 | 0.19250 | 0.23328 |

| | | | | | | | | | | | | |
|---|---|---|---|---|---|---|---|---|---|---|---|---|
| **PRIM1** | 0.22770 | 10.50419 | 6.79913 | 4.55106 | 7.72909 | 26.63445 | 3.73346 | 0.00000 | 2.67370 | 1.78914 | 18.57993 | 10.56361 |
| **PRIM2** | 5.38152 | 3.16323 | 4.51154 | 1.24025 | 0.58924 | 11.70517 | 3.68624 | 0.83663 | 6.36836 | 0.00000 | 0.12737 | 3.95762 |
| **PRIMA1** | 0.07875 | 0.17202 | 0.01892 | 0.00000 | 0.02624 | 0.04106 | 0.01457 | 0.00728 | 0.04736 | 0.07034 | 0.01520 | 0.07257 |
| **PRIMPOL** | 0.50411 | 0.50848 | 0.34892 | 0.25666 | 0.02166 | 1.72296 | 0.51016 | 0.43228 | 0.00000 | 0.00000 | 1.21885 | 0.41064 |
| **PRKAA1** | 0.26361 | 4.94656 | 3.05802 | 2.26654 | 0.00000 | 12.29405 | 2.52696 | 8.86005 | 2.08736 | 3.41485 | 2.12290 | 7.75763 |
| **PRKAA2** | 0.13423 | 0.00000 | 0.94665 | 0.00000 | 0.00000 | 0.00000 | 4.10037 | 0.07166 | 1.34403 | 0.88318 | 2.46009 | 0.36498 |
| **PRKAB1** | 1.16329 | 13.42029 | 9.31833 | 4.25946 | 0.74045 | 11.60886 | 8.41825 | 4.35480 | 11.05217 | 2.94056 | 2.78907 | 4.15598 |
| **PRKAB2** | 0.00000 | 3.41319 | 1.14974 | 7.39236 | 2.78911 | 4.05412 | 1.26278 | 2.30047 | 0.00000 | 3.32521 | 0.80388 | 11.02595 |
| **PRKACA** | 1.29319 | 0.43894 | 0.93668 | 0.88175 | 2.27575 | 0.00000 | 1.56096 | 0.00000 | 0.00000 | 4.22680 | 0.89540 | 0.81525 |
| **PRKACB** | 0.09461 | 0.36572 | 0.81264 | 0.32692 | 0.12995 | 0.53155 | 0.59354 | 0.10701 | 0.22968 | 0.39993 | 0.40618 | 0.00000 |
| **PRKACG** | 0.02334 | 0.01383 | 0.01489 | 0.01381 | 0.02303 | 0.05157 | 0.01364 | 0.00929 | 0.02528 | 0.01939 | 0.01663 | 0.02984 |
| **PRKAG1** | 3.41426 | 11.50716 | 10.15351 | 5.57928 | 4.86525 | 11.32897 | 8.36300 | 8.35835 | 11.88533 | 0.00000 | 13.94593 | 16.52576 |
| **PRKAG2** | 0.42814 | 1.09420 | 0.75477 | 0.56684 | 0.00000 | 0.16306 | 0.57453 | 0.17959 | 0.00000 | 0.39484 | 0.04274 | 0.13323 |
| **PRKAG2-AS1** | 0.00428 | 0.02936 | 0.01884 | 0.01037 | 0.02467 | 0.02051 | 0.02737 | 0.19647 | 0.01207 | 0.00317 | 0.01399 | 0.01134 |
| **PRKAG3** | 0.03335 | 0.01596 | 0.05185 | 0.01340 | 0.02492 | 0.00017 | 0.01485 | 0.02682 | 0.01068 | 0.03924 | 0.01557 | 0.01392 |
| **PRKAR1A** | 4.19222 | 9.07692 | 9.84227 | 9.25273 | 7.07503 | 22.54732 | 10.34223 | 7.16380 | 10.58981 | 0.23939 | 11.62970 | 7.74181 |
| **PRKAR1B** | 1.05001 | 1.67027 | 2.17876 | 0.36403 | 3.27769 | 0.65871 | 6.82700 | 0.28258 | 5.21806 | 0.00000 | 7.90079 | 0.23911 |
| **PRKAR2A** | 5.96071 | 1.06223 | 2.31747 | 1.65548 | 3.05042 | 0.00000 | 0.91734 | 4.28191 | 4.20570 | 0.00000 | 5.31475 | 1.90357 |
| **PRKAR2A-AS1** | 0.30248 | 0.01508 | 0.07308 | 0.03009 | 0.01912 | 0.00000 | 0.03767 | 0.00536 | 0.00000 | 0.06038 | 0.24818 | 0.04046 |
| **PRKAR2B** | 0.89412 | 0.60354 | 0.00000 | 0.00000 | 0.19168 | 0.55510 | 1.10602 | 0.27708 | 0.52430 | 1.04192 | 0.00000 | 0.88088 |
| **PRKCA** | 0.06248 | 0.76211 | 0.46795 | 0.34551 | 2.60429 | 1.88460 | 0.32515 | 0.38127 | 0.00773 | 4.03343 | 0.84106 | 0.46588 |
| **PRKCB** | 0.00049 | 0.70326 | 0.00000 | 0.00000 | 0.01517 | 0.08696 | 0.00000 | 0.21892 | 0.00000 | 0.50879 | 0.00220 | 0.23924 |
| **PRKCD** | 0.57423 | 2.61517 | 3.28468 | 5.02144 | 4.16484 | 2.08170 | 4.34819 | 0.36678 | 0.00002 | 4.40994 | 0.36229 | 0.15210 |
| **PRKCDBP** | 0.75199 | 0.82938 | 0.91633 | 0.75541 | 0.00000 | 0.00000 | 0.79488 | 2.42924 | 0.05943 | 0.00000 | 0.66609 | 0.97407 |
| **PRKCE** | 0.95755 | 0.00000 | 0.19524 | 0.01212 | 0.01832 | 0.00000 | 0.00000 | 0.31529 | 0.04014 | 0.16153 | 0.00177 | 0.18453 |
| **PRKCG** | 8.33257 | 0.20516 | 0.01551 | 0.30606 | 0.18340 | 0.00000 | 0.00000 | 3.53075 | 0.12225 | 0.08567 | 0.00000 | 0.91432 |
| **PRKCH** | 0.48127 | 0.00287 | 10.86843 | 0.00000 | 0.00000 | 0.03399 | 0.83065 | 0.00555 | 0.61890 | 0.04991 | 0.00000 | 0.01132 |
| **PRKCI** | 0.75369 | 3.16710 | 6.16704 | 1.85081 | 3.19496 | 9.46360 | 8.76513 | 0.28758 | 2.55409 | 9.41535 | 5.52477 | 2.19763 |
| **PRKCQ** | 2.66032 | 2.50298 | 0.17254 | 0.16360 | 0.24001 | 0.00000 | 0.42877 | 0.29660 | 0.00000 | 0.00000 | 0.00000 | 1.48305 |
| **PRKCQ-AS1** | 0.16984 | 1.66190 | 0.28904 | 0.00000 | 0.17935 | 1.06046 | 0.00000 | 0.08336 | 0.00942 | 0.71160 | 0.00000 | 1.34008 |
| **PRKCSH** | 0.83683 | 7.59718 | 9.33989 | 6.09576 | 4.48551 | 0.00000 | 0.64827 | 0.90000 | 22.24141 | 12.07548 | 6.12647 | 7.56905 |
| **PRKCZ** | 1.32370 | 0.53868 | 2.07831 | 0.10968 | 0.82739 | 0.38919 | 0.46735 | 0.09817 | 0.38347 | 1.22222 | 0.18223 | 0.09100 |
| **PRKD1** | 0.93172 | 0.04349 | 0.01420 | 0.37025 | 0.21193 | 0.00000 | 0.02370 | 0.02160 | 0.00002 | 0.03369 | 0.00578 | 0.20288 |
| **PRKD2** | 5.24481 | 3.59663 | 4.20039 | 1.80974 | 2.23071 | 0.00000 | 4.93735 | 15.71215 | 4.08371 | 6.00109 | 0.31077 | 19.42081 |
| **PRKD3** | 1.18207 | 1.13374 | 1.59648 | 2.64601 | 0.00000 | 7.52308 | 3.37204 | 1.70422 | 1.65787 | 12.27363 | 1.61547 | 3.34926 |
| **PRKDC** | 0.00000 | 16.93747 | 17.64353 | 13.35363 | 18.92247 | 40.99298 | 21.60236 | 3.56219 | 30.35140 | 42.88926 | 27.17387 | 23.79466 |
| **PRKG1** | 2.08742 | 0.00839 | 0.19410 | 0.00675 | 0.05284 | 0.00000 | 0.10075 | 0.23315 | 0.00517 | 0.02329 | 0.01249 | 0.24893 |
| **PRKG2** | 1.41543 | 0.26279 | 0.06510 | 0.06405 | 0.00179 | 0.39201 | 0.00000 | 0.13757 | 0.00000 | 0.00945 | 0.00000 | 0.05798 |
| **PRKRA** | 3.70878 | 5.50913 | 6.33391 | 6.53927 | 0.35798 | 12.87112 | 5.51189 | 3.24827 | 4.60374 | 2.07373 | 9.00624 | 5.88075 |
| **PRKRIP1** | 2.36704 | 1.65772 | 0.62260 | 0.46014 | 1.71616 | 0.18578 | 0.59073 | 1.41264 | 0.67589 | 0.00000 | 0.79268 | 2.92337 |
| **PRKRIR** | 0.30350 | 1.83363 | 1.09589 | 0.36141 | 0.23179 | 7.51589 | 4.21172 | 3.79617 | 0.00000 | 8.68033 | 0.07957 | 1.58260 |
| **PRKX** | 0.23800 | 1.28863 | 0.78759 | 0.52570 | 1.63696 | 2.35729 | 5.65568 | 0.34624 | 0.32912 | 7.67067 | 1.50242 | 0.50147 |
| **PRKX-AS1** | 0.00590 | 0.02057 | 0.01457 | 0.01511 | 0.01096 | 0.00976 | 0.02396 | 0.01040 | 0.01796 | 0.02995 | 0.02375 | 0.01784 |
| **PRKXP1** | 0.00682 | 0.00000 | 0.95269 | 0.28577 | 0.47372 | 0.09900 | 0.03882 | 1.44532 | 0.40451 | 1.77898 | 0.09030 | 0.95229 |
| **PRKY** | 0.00000 | 0.75780 | 1.07041 | 1.00733 | 0.86836 | 1.93193 | 0.84585 | 0.03818 | 3.15299 | 6.87881 | 0.21528 | 0.00000 |
| **PRL** | 0.11824 | 0.01507 | 0.02223 | 0.01160 | 0.01593 | 0.01547 | 0.01734 | 0.03233 | 0.01920 | 0.01024 | 0.01451 | 0.01315 |
| **PRLHR** | 0.01078 | 0.00000 | 0.04266 | 0.17308 | 0.00000 | 0.06744 | 0.00538 | 0.00250 | 0.00000 | 0.04015 | 0.01498 | 0.26290 |
| **PRLR** | 0.00409 | 0.00000 | 3.76120 | 0.66511 | 0.06461 | 0.15355 | 1.58944 | 0.15651 | 1.39030 | 0.15532 | 0.00000 | 0.00000 |

| | | | | | | | | | | | | |
|---|---|---|---|---|---|---|---|---|---|---|---|---|
| **PRMT1** | 7.94804 | 18.01174 | 25.70252 | 21.82967 | 26.67357 | 5.45994 | 31.29045 | 10.31201 | 39.87491 | 0.00000 | 23.99496 | 23.42208 |
| **PRMT2** | 6.87117 | 6.68695 | 2.87822 | 3.72839 | 2.89311 | 0.23604 | 7.49701 | 1.39667 | 3.06893 | 0.02175 | 3.76387 | 1.96674 |
| **PRMT3** | 0.00066 | 8.95210 | 7.77517 | 2.35732 | 1.58028 | 22.56730 | 2.14155 | 0.55967 | 7.12814 | 0.00000 | 12.49480 | 5.90086 |
| **PRMT5** | 0.38433 | 13.31337 | 13.45104 | 10.13846 | 3.70608 | 25.86329 | 12.35829 | 2.98561 | 24.13678 | 0.00000 | 23.44750 | 16.04416 |
| **PRMT5-AS1** | 0.00279 | 0.12860 | 0.06373 | 0.09552 | 0.11827 | 0.00000 | 0.12394 | 0.08664 | 0.28156 | 0.07931 | 0.10417 | 0.11636 |
| **PRMT6** | 0.19712 | 4.60081 | 4.78869 | 2.44493 | 2.05050 | 10.15737 | 5.52589 | 0.31326 | 5.45579 | 0.00000 | 7.97856 | 4.68182 |
| **PRMT7** | 0.18866 | 13.79107 | 8.28674 | 8.17798 | 6.71990 | 0.00000 | 10.16173 | 3.52508 | 13.45454 | 0.00000 | 12.97007 | 6.69362 |
| **PRMT8** | 0.00548 | 0.01080 | 0.05509 | 0.01008 | 0.01507 | 0.03959 | 0.01582 | 0.01309 | 0.00000 | 0.02439 | 0.01844 | 0.01611 |
| **PRMT9** | 1.42026 | 2.17464 | 3.38268 | 1.86564 | 0.58372 | 6.58190 | 0.00006 | 0.49692 | 1.63306 | 0.90818 | 2.30053 | 1.22133 |
| **PRNCR1** | 0.25833 | 0.85084 | 0.53631 | 0.37682 | 0.96356 | 1.35825 | 0.92408 | 1.08631 | 0.58907 | 2.23308 | 1.17953 | 0.61410 |
| **PRND** | 0.00225 | 0.01642 | 0.02216 | 0.01872 | 0.02124 | 0.00776 | 0.02202 | 0.05166 | 0.01769 | 0.01218 | 0.01577 | 0.01559 |
| **PRNP** | 0.01663 | 9.99332 | 0.91229 | 3.77247 | 1.55149 | 0.00003 | 0.00000 | 3.49800 | 3.66517 | 2.27415 | 2.91487 | 13.29624 |
| **PROB1** | 0.00665 | 0.01463 | 0.05562 | 0.01595 | 0.01316 | 0.03154 | 0.00535 | 0.01018 | 0.07756 | 0.00000 | 0.01693 | 0.01644 |
| **PROC** | 0.09375 | 0.01164 | 0.02682 | 0.52868 | 0.00459 | 0.01517 | 0.01401 | 0.00471 | 0.01992 | 0.00802 | 0.01506 | 0.01988 |
| **PROCA1** | 1.05802 | 0.57471 | 0.32216 | 1.42067 | 0.16862 | 0.26366 | 0.67480 | 1.35983 | 0.00804 | 0.00000 | 0.19277 | 0.73810 |
| **PROCR** | 0.07412 | 7.47355 | 10.84158 | 14.42045 | 6.39866 | 7.83062 | 3.85202 | 2.98102 | 7.61453 | 0.00001 | 2.75955 | 5.87713 |
| **PRODH** | 0.00001 | 5.34854 | 0.00000 | 6.06405 | 8.16645 | 0.00000 | 1.41834 | 0.01303 | 9.58276 | 7.63270 | 18.55994 | 1.67990 |
| **PRODH2** | 0.00533 | 0.01279 | 0.02489 | 0.02436 | 0.01517 | 0.02509 | 0.01803 | 0.01031 | 0.00559 | 0.05943 | 0.01365 | 0.01302 |
| **PROK1** | 0.05761 | 0.03576 | 0.01384 | 0.01114 | 0.02519 | 0.00000 | 0.01368 | 0.00698 | 0.02445 | 0.03875 | 0.02543 | 0.05110 |
| **PROK2** | 0.06019 | 0.04911 | 0.01393 | 0.01169 | 0.00951 | 0.01451 | 0.01769 | 0.01217 | 0.01936 | 0.01715 | 0.01652 | 0.01121 |
| **PROKR1** | 0.00579 | 0.01011 | 0.01445 | 0.00964 | 0.01478 | 0.03887 | 0.02489 | 0.00938 | 0.00875 | 0.01572 | 0.01780 | 0.03449 |
| **PROKR2** | 0.00626 | 0.01409 | 0.00979 | 0.09314 | 0.07089 | 0.00000 | 0.02979 | 0.01067 | 0.02486 | 0.00000 | 0.01601 | 0.01866 |
| **PROL1** | 0.07027 | 0.01530 | 0.01771 | 0.01219 | 0.01608 | 0.01516 | 0.02005 | 0.00770 | 0.01903 | 0.01137 | 0.01513 | 0.01519 |
| **PROM1** | 0.00000 | 16.03856 | 5.14912 | 16.95242 | 10.76137 | 36.44250 | 4.93600 | 0.90030 | 51.38507 | 17.77017 | 24.62219 | 6.38040 |
| **PROM2** | 0.00487 | 0.00238 | 0.13481 | 0.11658 | 0.26761 | 0.14848 | 0.00000 | 0.00845 | 0.00000 | 0.07038 | 0.00000 | 0.00543 |
| **PRORSD1P** | 0.00000 | 1.05583 | 0.11967 | 1.60104 | 0.00000 | 0.17494 | 0.00000 | 0.25680 | 0.00000 | 1.73195 | 0.67144 | 3.03739 |
| **PRORY** | 0.00547 | 0.01332 | 0.01618 | 0.01386 | 0.00996 | 0.01235 | 0.08950 | 0.01065 | 0.00000 | 0.03502 | 0.01311 | 0.02459 |
| **PROS1** | 0.00987 | 0.00000 | 0.66386 | 4.90688 | 0.00000 | 1.36897 | 0.00000 | 0.00138 | 0.55176 | 0.00000 | 0.03126 | 0.00249 |
| **PROSC** | 1.45158 | 6.45279 | 9.04709 | 3.19648 | 8.19979 | 10.80730 | 15.13255 | 1.28924 | 10.56931 | 0.00000 | 11.38629 | 9.59182 |
| **PROSER1** | 28.44075 | 8.11783 | 21.17512 | 2.86579 | 11.72854 | 20.83140 | 21.42060 | 6.78034 | 9.65625 | 16.97489 | 10.68196 | 3.95353 |
| **PROSER2** | 0.42328 | 0.91865 | 0.21350 | 0.07787 | 2.04236 | 0.03584 | 2.34212 | 0.06742 | 0.00000 | 5.15089 | 0.29638 | 1.38532 |
| **PROSER2-AS1** | 0.94383 | 3.39497 | 4.95916 | 0.82715 | 6.72848 | 0.11053 | 2.03944 | 1.98659 | 4.20049 | 0.00000 | 1.49035 | 7.61615 |
| **PROSER3** | 1.46718 | 0.31980 | 0.13300 | 0.27928 | 0.40014 | 0.00000 | 0.07620 | 0.20112 | 0.57320 | 0.00792 | 0.11566 | 0.99976 |
| **PROX1** | 0.35612 | 0.00000 | 0.12257 | 1.04260 | 0.00000 | 0.07378 | 0.00000 | 0.12518 | 0.00000 | 0.19435 | 0.05444 | 0.10582 |
| **PROX2** | 0.23129 | 0.04719 | 0.04820 | 0.02411 | 0.07050 | 0.16310 | 0.00000 | 0.07739 | 0.38071 | 0.00000 | 0.00000 | 0.10424 |
| **PROZ** | 0.00257 | 0.01229 | 0.07798 | 0.00046 | 0.02490 | 0.00000 | 0.15093 | 0.00711 | 0.32611 | 0.60924 | 0.00219 | 0.04722 |
| **PRPF18** | 2.43913 | 4.25591 | 2.55930 | 2.64439 | 0.77603 | 14.93597 | 4.59967 | 11.73410 | 4.49419 | 2.88191 | 2.63315 | 5.19570 |
| **PRPF19** | 7.60154 | 0.00000 | 2.50693 | 0.81644 | 14.88803 | 0.00000 | 13.99583 | 1.47498 | 2.94926 | 24.81372 | 0.00000 | 10.37515 |
| **PRPF3** | 2.15473 | 12.70826 | 6.93577 | 4.83945 | 7.22374 | 11.94653 | 10.83607 | 11.42883 | 16.22117 | 4.56430 | 12.29827 | 12.52375 |
| **PRPF31** | 1.98844 | 22.58254 | 14.85005 | 14.88956 | 16.08526 | 1.18577 | 20.82144 | 8.94714 | 24.76503 | 0.00000 | 18.97864 | 26.08238 |
| **PRPF38A** | 4.20212 | 15.97527 | 11.05501 | 6.69689 | 6.84496 | 24.30174 | 11.56189 | 22.71232 | 10.79419 | 10.36041 | 11.84609 | 23.35059 |
| **PRPF38B** | 4.85427 | 8.69512 | 8.96128 | 9.11046 | 9.89170 | 17.53686 | 19.89710 | 15.82073 | 13.44204 | 32.37214 | 10.39248 | 10.72007 |
| **PRPF39** | 0.00000 | 0.13705 | 3.46337 | 1.65842 | 1.15769 | 10.88245 | 2.81877 | 6.33277 | 4.70152 | 5.86945 | 3.93442 | 2.73171 |
| **PRPF4** | 4.87659 | 16.29029 | 12.40160 | 8.74878 | 11.68095 | 4.08835 | 17.42292 | 7.04892 | 16.68957 | 1.32050 | 15.68937 | 16.24745 |
| **PRPF40A** | 10.06183 | 16.74526 | 25.08510 | 12.64283 | 12.55648 | 44.30255 | 33.00784 | 22.17473 | 20.46173 | 32.17979 | 20.73147 | 29.09233 |
| **PRPF40B** | 0.05440 | 1.02749 | 0.68063 | 0.50179 | 0.45810 | 0.00000 | 0.00272 | 0.00067 | 0.40195 | 0.00000 | 0.68383 | 0.77380 |
| **PRPF4B** | 3.91035 | 4.53191 | 7.07147 | 4.14021 | 6.79668 | 20.20033 | 11.95036 | 8.69946 | 9.84121 | 35.54913 | 12.01039 | 8.24781 |
| **PRPF6** | 5.53911 | 16.53027 | 13.54214 | 11.83411 | 13.52260 | 0.02259 | 21.12666 | 5.32967 | 7.48669 | 5.05451 | 15.67441 | 16.56667 |

| | | | | | | | | | | | | |
|---|---|---|---|---|---|---|---|---|---|---|---|---|
| **PRPF8** | 6.37901 | 27.73857 | 16.83000 | 22.11845 | 25.80800 | 12.66069 | 28.59503 | 16.05265 | 22.65740 | 21.00434 | 14.32011 | 26.58868 |
| **PRPH** | 0.02405 | 1.64211 | 0.00748 | 0.40413 | 0.05258 | 0.00000 | 0.00921 | 5.33708 | 0.00000 | 0.12927 | 0.00339 | 5.93595 |
| **PRPH2** | 0.42466 | 0.20505 | 0.04309 | 0.10009 | 0.00000 | 0.00000 | 0.00000 | 1.14716 | 0.13143 | 0.32632 | 0.14986 | 0.80817 |
| **PRPS1** | 0.56788 | 20.62495 | 0.97978 | 0.00000 | 15.17134 | 15.99449 | 11.41476 | 4.17780 | 8.32500 | 8.34551 | 22.14845 | 13.19296 |
| **PRPS1L1** | 0.00494 | 0.97095 | 0.43736 | 0.00000 | 0.18782 | 1.69961 | 0.00002 | 0.17831 | 0.00000 | 0.11399 | 1.25532 | 0.56185 |
| **PRPS2** | 5.11213 | 4.34877 | 10.98723 | 1.85794 | 3.23570 | 20.24612 | 18.16719 | 2.99280 | 11.05192 | 0.00000 | 28.92258 | 5.44415 |
| **PRPSAP1** | 2.79898 | 11.65636 | 0.00000 | 10.46139 | 15.74652 | 12.24586 | 1.63929 | 8.34170 | 2.31551 | 0.18523 | 6.82195 | 10.28338 |
| **PRPSAP2** | 0.70447 | 7.03468 | 4.52022 | 5.27778 | 5.82224 | 15.33466 | 3.95289 | 4.19500 | 12.17217 | 0.00000 | 7.69084 | 11.96909 |
| **PRR11** | 1.10338 | 4.88001 | 3.78814 | 4.69409 | 6.94270 | 8.21810 | 11.87903 | 0.93301 | 3.26874 | 18.73072 | 2.30449 | 7.10704 |
| **PRR12** | 1.35196 | 0.05874 | 0.65705 | 0.37651 | 1.35360 | 0.00000 | 0.58253 | 0.00000 | 0.59900 | 4.31089 | 0.00000 | 0.09750 |
| **PRR13** | 2.82351 | 35.25309 | 8.42055 | 25.06394 | 25.10246 | 27.97781 | 9.50545 | 13.58161 | 2.47351 | 8.45800 | 10.65771 | 33.56295 |
| **PRR14** | 4.54094 | 1.87047 | 4.18682 | 1.86223 | 2.50265 | 0.00000 | 2.89773 | 3.18353 | 4.66532 | 6.82104 | 0.31260 | 3.94646 |
| **PRR14L** | 10.08771 | 2.45440 | 9.40139 | 7.05957 | 7.40598 | 8.12845 | 9.62467 | 5.38339 | 10.79342 | 16.65920 | 7.49609 | 10.05547 |
| **PRR15** | 13.58825 | 0.00000 | 0.08666 | 1.41585 | 0.24126 | 0.00000 | 0.01802 | 0.73149 | 0.00000 | 0.04394 | 0.15421 | 0.24745 |
| **PRR15L** | 0.00318 | 2.03335 | 0.00000 | 0.00000 | 1.73753 | 0.08639 | 0.00000 | 0.00630 | 0.00000 | 0.10105 | 1.34023 | 0.06912 |
| **PRR16** | 0.55834 | 0.04439 | 0.00000 | 0.16356 | 0.00000 | 0.01884 | 0.00000 | 0.13190 | 0.00923 | 0.29181 | 0.00000 | 0.16507 |
| **PRR18** | 0.00610 | 0.01472 | 0.01714 | 0.01079 | 0.01271 | 0.02242 | 0.02258 | 0.01042 | 0.01395 | 0.02369 | 0.01458 | 0.01525 |
| **PRR19** | 0.61349 | 2.36559 | 1.52605 | 0.18758 | 1.39825 | 0.00000 | 5.12515 | 0.47511 | 2.22146 | 0.00000 | 3.57135 | 6.17650 |
| **PRR22** | 0.01581 | 0.13672 | 0.61976 | 0.24138 | 0.73587 | 0.00000 | 0.25186 | 0.02940 | 1.44333 | 4.37616 | 0.18112 | 0.73086 |
| **PRR23A** | 1.15592 | 0.19757 | 0.18830 | 0.29384 | 0.56544 | 0.24002 | 0.54593 | 1.50616 | 0.02185 | 0.44831 | 0.23669 | 1.08596 |
| **PRR23B** | 0.59311 | 0.62608 | 0.06665 | 0.00000 | 0.00000 | 0.08884 | 0.20296 | 1.03775 | 0.21337 | 0.33031 | 0.00000 | 0.85690 |
| **PRR23C** | 1.40592 | 0.07143 | 0.09620 | 0.00000 | 0.03131 | 0.04376 | 0.22621 | 0.65967 | 0.03145 | 0.04764 | 0.00000 | 0.90458 |
| **PRR24** | 0.00445 | 0.02757 | 0.45663 | 0.09679 | 0.50904 | 0.00000 | 0.85785 | 0.24882 | 0.00000 | 1.97824 | 0.16375 | 0.04927 |
| **PRR26** | 0.00801 | 0.00000 | 0.35894 | 0.00000 | 0.32808 | 0.00000 | 0.00000 | 0.01857 | 0.06730 | 0.41946 | 0.18162 | 0.21605 |
| **PRR27** | 0.03543 | 0.00395 | 0.00586 | 0.00350 | 0.00496 | 0.01154 | 0.00319 | 0.00896 | 0.02680 | 0.53546 | 0.00000 | 0.01065 |
| **PRR29** | 0.05074 | 0.01352 | 0.05710 | 0.00767 | 0.01228 | 0.01976 | 0.00829 | 0.00551 | 0.00827 | 0.03590 | 0.01579 | 0.01401 |
| **PRR3** | 0.77337 | 1.15189 | 2.90195 | 1.42769 | 0.65900 | 1.28856 | 4.14004 | 0.89464 | 3.12686 | 0.00000 | 1.05188 | 4.84813 |
| **PRR30** | 0.01398 | 0.01501 | 0.01254 | 0.03993 | 0.01622 | 0.00394 | 0.00762 | 0.03373 | 0.03044 | 0.04451 | 0.02853 | 0.01335 |
| **PRR32** | 0.06528 | 0.01611 | 0.01420 | 0.01181 | 0.01626 | 0.01722 | 0.01773 | 0.10101 | 0.01974 | 0.01038 | 0.01431 | 0.01224 |
| **PRR34** | 0.00903 | 0.01527 | 0.02804 | 0.01344 | 0.02259 | 0.01393 | 0.01554 | 0.01271 | 0.01467 | 0.01546 | 0.01788 | 0.01823 |
| **PRR35** | 0.03885 | 0.01238 | 0.03089 | 0.03884 | 0.01789 | 0.01272 | 0.01174 | 0.00792 | 0.01785 | 0.05716 | 0.01461 | 0.01346 |
| **PRR4** | 0.00623 | 0.01464 | 0.02184 | 0.01262 | 0.01547 | 0.01738 | 0.01259 | 0.01041 | 0.02820 | 0.01822 | 0.01426 | 0.01648 |
| **PRR5** | 1.40661 | 5.08900 | 9.99075 | 0.00000 | 18.04361 | 0.00000 | 15.95284 | 0.09913 | 8.42337 | 18.50112 | 4.76174 | 1.30096 |
| **PRR5L** | 5.88023 | 0.00762 | 0.00850 | 0.00975 | 0.01750 | 0.00063 | 0.03531 | 1.31189 | 0.02377 | 0.00000 | 0.02903 | 0.00000 |
| **PRR7** | 0.53104 | 0.12833 | 0.31121 | 0.06678 | 0.12905 | 0.00000 | 0.34752 | 0.26519 | 0.26090 | 0.09889 | 0.10141 | 0.02367 |
| **PRR7-AS1** | 0.06422 | 0.02371 | 0.04675 | 0.07697 | 0.02812 | 0.00000 | 0.01036 | 0.00490 | 0.04664 | 0.01296 | 0.03709 | 0.01770 |
| **PRR9** | 0.00844 | 0.00298 | 11.10161 | 0.00000 | 0.00000 | 0.03297 | 1.88916 | 0.00687 | 0.00573 | 0.01841 | 0.00000 | 0.06235 |
| **PRRC1** | 5.38086 | 9.33538 | 8.23963 | 4.65180 | 2.29715 | 13.41819 | 10.06456 | 0.20721 | 9.58108 | 3.21351 | 6.43477 | 7.55178 |
| **PRRC2A** | 4.13752 | 2.96008 | 6.50881 | 5.26291 | 12.72325 | 0.00000 | 6.97528 | 0.00000 | 14.12881 | 24.37242 | 1.95931 | 8.67931 |
| **PRRC2B** | 4.82673 | 9.24619 | 8.32053 | 17.56848 | 16.61983 | 6.39139 | 13.64733 | 3.43433 | 14.19375 | 35.88860 | 4.51239 | 10.20804 |
| **PRRC2C** | 5.55455 | 12.96162 | 21.95545 | 13.08836 | 21.81031 | 31.69179 | 31.07632 | 14.67100 | 17.78215 | 62.42055 | 16.07985 | 19.60683 |
| **PRRG1** | 0.00009 | 1.11254 | 1.88582 | 0.80563 | 0.00000 | 2.49030 | 0.00000 | 2.47623 | 3.72560 | 2.01723 | 0.25937 | 5.65544 |
| **PRRG2** | 0.85644 | 0.00000 | 1.64707 | 0.05351 | 0.32993 | 0.00000 | 0.42744 | 0.13890 | 0.18735 | 0.57136 | 0.19490 | 0.47322 |
| **PRRG4** | 0.34643 | 0.00000 | 0.62479 | 0.00000 | 0.29236 | 0.17007 | 0.00000 | 0.42064 | 0.00000 | 0.26941 | 0.00000 | 0.36422 |
| **PRRT1** | 0.08725 | 0.31416 | 0.10190 | 0.00856 | 0.18594 | 0.00000 | 0.04478 | 0.00000 | 0.00000 | 0.00000 | 0.03404 | 2.66413 |
| **PRRT2** | 0.09738 | 0.00003 | 0.22230 | 0.40270 | 0.00000 | 0.70193 | 0.37507 | 0.23343 | 0.00000 | 1.59865 | 0.00000 | 0.09585 |
| **PRRT3** | 0.35742 | 0.43783 | 0.12175 | 0.64381 | 0.93409 | 0.00000 | 0.00289 | 0.11616 | 0.43572 | 1.19815 | 0.27612 | 0.75837 |
| **PRRT3-AS1** | 0.00824 | 0.01016 | 0.00000 | 0.06211 | 0.00000 | 0.03768 | 0.11861 | 0.00834 | 0.07319 | 0.30618 | 0.05340 | 0.03613 |

| | | | | | | | | | | | | |
|---|---|---|---|---|---|---|---|---|---|---|---|---|
| **PRRT4** | 0.33836 | 0.01335 | 0.03282 | 0.03369 | 0.00000 | 0.05574 | 0.00000 | 0.00084 | 0.00408 | 0.05373 | 0.01179 | 0.02774 |
| **PRRX1** | 0.00463 | 0.00840 | 0.01779 | 0.00887 | 0.01121 | 0.03423 | 0.01835 | 0.02615 | 0.01651 | 0.01686 | 0.01416 | 0.05302 |
| **PRRX2** | 0.00146 | 0.04968 | 0.06674 | 0.29333 | 0.06133 | 0.01146 | 0.01601 | 0.21850 | 0.00000 | 0.07043 | 0.01718 | 0.00847 |
| **PRSS1** | 0.22943 | 0.04452 | 0.00545 | 0.19216 | 0.00000 | 0.00000 | 0.08898 | 0.56972 | 0.00000 | 0.05473 | 0.12628 | 0.13050 |
| **PRSS12** | 0.00114 | 0.00000 | 0.00000 | 2.22890 | 0.00000 | 0.63473 | 6.08212 | 0.00198 | 4.40772 | 1.48079 | 0.00000 | 0.00000 |
| **PRSS16** | 2.10035 | 1.27671 | 2.61789 | 0.05148 | 2.10712 | 0.46537 | 1.02902 | 1.10312 | 1.42645 | 0.62535 | 0.51386 | 0.62661 |
| **PRSS21** | 0.00216 | 0.60012 | 0.23841 | 0.45760 | 0.06459 | 0.00000 | 0.00365 | 0.17432 | 0.80842 | 0.00000 | 0.65468 | 0.64066 |
| **PRSS22** | 0.01591 | 0.05049 | 0.66950 | 0.00673 | 0.10935 | 0.00000 | 0.00000 | 0.02264 | 0.07902 | 0.05271 | 0.04659 | 0.04006 |
| **PRSS23** | 0.20775 | 42.48851 | 0.00000 | 17.47053 | 34.41178 | 21.71063 | 0.00000 | 0.63629 | 0.31266 | 6.58262 | 14.51821 | 25.69468 |
| **PRSS27** | 0.23658 | 0.06235 | 0.12920 | 0.00558 | 0.03197 | 0.01035 | 0.00924 | 0.00293 | 0.01958 | 0.00001 | 0.02700 | 0.01351 |
| **PRSS3** | 0.29371 | 0.38415 | 0.00000 | 7.12089 | 1.07202 | 0.03704 | 0.00551 | 0.32880 | 0.00000 | 0.00000 | 0.00000 | 0.10893 |
| **PRSS30P** | 0.00993 | 0.03675 | 0.15284 | 0.00690 | 0.00000 | 0.00000 | 0.14215 | 0.20381 | 0.76299 | 0.00000 | 0.35582 | 0.14293 |
| **PRSS33** | 0.03908 | 0.01594 | 0.01623 | 0.14511 | 0.01719 | 0.00515 | 0.01387 | 0.01862 | 0.02143 | 0.01408 | 0.01449 | 0.03678 |
| **PRSS35** | 0.00623 | 0.25581 | 0.03144 | 0.20684 | 0.00000 | 0.07373 | 0.00659 | 0.00914 | 0.01245 | 0.00552 | 0.02373 | 0.02568 |
| **PRSS36** | 0.01049 | 0.18249 | 0.12111 | 0.00513 | 0.12456 | 0.00000 | 0.01773 | 0.04710 | 0.15225 | 0.04673 | 0.30633 | 1.20733 |
| **PRSS37** | 0.00529 | 0.01290 | 0.01494 | 0.01964 | 0.00000 | 0.01025 | 0.05765 | 0.01065 | 0.12378 | 0.05000 | 0.02195 | 0.01834 |
| **PRSS38** | 0.00370 | 0.01747 | 0.01505 | 0.01223 | 0.01055 | 0.00704 | 0.01975 | 0.03482 | 0.01270 | 0.02814 | 0.06460 | 0.01431 |
| **PRSS3P2** | 0.10647 | 0.03735 | 0.01458 | 0.02327 | 0.02117 | 0.00335 | 0.02627 | 0.29564 | 0.01839 | 0.00643 | 0.01515 | 0.04494 |
| **PRSS41** | 0.00189 | 2.24200 | 0.00000 | 0.40930 | 0.32822 | 0.08388 | 0.00000 | 0.61477 | 0.00000 | 0.02057 | 0.17115 | 1.81368 |
| **PRSS45** | 0.00535 | 0.06736 | 0.00917 | 0.00912 | 0.00948 | 0.05278 | 0.01055 | 0.01048 | 0.00704 | 0.03425 | 0.01346 | 0.02738 |
| **PRSS48** | 0.00418 | 0.00716 | 0.07213 | 0.22956 | 0.08038 | 0.00083 | 0.03999 | 0.00907 | 0.00000 | 0.16188 | 0.20440 | 0.02557 |
| **PRSS50** | 0.00389 | 0.75534 | 0.00968 | 0.00000 | 0.18388 | 0.00000 | 0.00076 | 0.00820 | 0.03028 | 0.23695 | 0.04572 | 0.24936 |
| **PRSS53** | 0.23714 | 0.19320 | 0.00000 | 0.00000 | 0.00000 | 0.00000 | 0.03541 | 0.16122 | 1.31219 | 1.25504 | 0.25694 | 1.05696 |
| **PRSS55** | 0.00391 | 0.00819 | 0.00637 | 0.00300 | 0.00396 | 0.01662 | 0.01077 | 0.00996 | 0.02649 | 0.23561 | 0.00430 | 0.00994 |
| **PRSS56** | 0.00621 | 0.01528 | 0.04698 | 0.00946 | 0.01509 | 0.01609 | 0.01254 | 0.01048 | 0.01736 | 0.01855 | 0.01563 | 0.01606 |
| **PRSS58** | 0.00346 | 0.00000 | 0.00000 | 0.00000 | 0.07022 | 0.02019 | 0.00918 | 1.26032 | 0.00000 | 0.38280 | 0.00000 | 0.27545 |
| **PRSS8** | 0.00054 | 10.11257 | 5.98937 | 0.80996 | 39.81445 | 0.00000 | 15.28013 | 0.00188 | 77.37810 | 4.69696 | 28.36228 | 0.00000 |
| **PRTFDC1** | 0.07900 | 0.00000 | 0.00000 | 0.16127 | 0.46824 | 3.44093 | 0.00000 | 0.00000 | 1.46902 | 0.00000 | 1.66965 | 0.16439 |
| **PRTG** | 7.53367 | 0.00000 | 6.44411 | 0.00000 | 0.00000 | 2.68676 | 6.33679 | 3.07031 | 10.32184 | 11.42470 | 0.48328 | 0.00000 |
| **PRTN3** | 0.00199 | 0.00823 | 0.01265 | 0.00933 | 0.01316 | 0.02284 | 0.01651 | 0.20737 | 0.01460 | 0.02002 | 0.01540 | 0.27957 |
| **PRUNE** | 3.18511 | 3.94158 | 0.79803 | 2.33770 | 1.33277 | 6.66040 | 1.86954 | 0.21108 | 1.29378 | 0.00000 | 4.07648 | 5.82231 |
| **PRUNE2** | 3.67183 | 0.48455 | 0.12690 | 7.51090 | 0.14080 | 0.28308 | 0.00000 | 0.31067 | 0.00000 | 1.13769 | 0.04884 | 0.77463 |
| **PRX** | 0.46646 | 0.08043 | 0.07504 | 0.00335 | 0.19649 | 0.00000 | 0.06112 | 0.82077 | 0.00000 | 0.41033 | 0.00000 | 0.36246 |
| **PSAP** | 2.59723 | 60.33450 | 40.37852 | 44.78103 | 60.58388 | 63.25955 | 14.34598 | 18.87193 | 87.75069 | 35.38132 | 48.25303 | 42.13797 |
| **PSAPL1** | 0.00646 | 0.09872 | 0.00593 | 0.03421 | 0.01489 | 0.06976 | 0.00838 | 0.00976 | 0.01338 | 0.00776 | 0.00857 | 0.01234 |
| **PSAT1** | 0.03184 | 0.00000 | 13.46153 | 2.22560 | 0.00000 | 24.40584 | 38.83085 | 0.00000 | 34.54278 | 2.67349 | 22.56823 | 1.61776 |
| **PSCA** | 0.00674 | 0.01295 | 0.01176 | 0.01123 | 0.01353 | 0.00000 | 0.00596 | 0.00771 | 0.20738 | 0.00248 | 0.00863 | 0.09349 |
| **PSD** | 0.29621 | 0.00293 | 0.00323 | 0.16769 | 0.12799 | 0.00000 | 0.02678 | 0.00101 | 0.00000 | 0.74646 | 0.00000 | 0.02339 |
| **PSD2** | 0.00603 | 0.01352 | 0.01922 | 0.03454 | 0.01052 | 0.02334 | 0.01634 | 0.01042 | 0.01584 | 0.01655 | 0.01456 | 0.01888 |
| **PSD3** | 9.41004 | 0.44780 | 0.02411 | 0.50214 | 0.68070 | 0.84412 | 1.33033 | 2.16372 | 3.48555 | 2.24261 | 0.00000 | 1.13026 |
| **PSD4** | 0.00058 | 0.24422 | 1.80268 | 0.06452 | 1.13930 | 0.00000 | 0.00000 | 0.30531 | 0.00000 | 0.33473 | 1.37442 | 1.85011 |
| **PSEN1** | 3.00551 | 9.95934 | 11.64661 | 11.08370 | 5.32234 | 19.06885 | 0.00000 | 4.22657 | 21.53468 | 3.61955 | 2.22388 | 12.58989 |
| **PSEN2** | 7.89922 | 0.33803 | 0.95506 | 1.85947 | 0.27072 | 0.00000 | 0.39840 | 0.43370 | 1.18935 | 0.00000 | 0.43323 | 0.00000 |
| **PSENEN** | 2.35362 | 11.13298 | 13.47346 | 11.26100 | 13.17747 | 10.16618 | 4.31303 | 0.35925 | 19.83128 | 2.62943 | 9.53884 | 13.97275 |
| **PSG1** | 0.00619 | 0.01562 | 0.02063 | 0.01022 | 0.01498 | 0.01997 | 0.02071 | 0.01041 | 0.01646 | 0.01696 | 0.01433 | 0.01652 |
| **PSG10P** | 0.10098 | 0.01526 | 0.02293 | 0.00884 | 0.01645 | 0.02395 | 0.01832 | 0.01891 | 0.01417 | 0.01710 | 0.01371 | 0.01974 |
| **PSG11** | 0.00659 | 0.01688 | 0.02147 | 0.01320 | 0.01746 | 0.01114 | 0.04227 | 0.01052 | 0.04055 | 0.00001 | 0.01100 | 0.01769 |
| **PSG2** | 0.00535 | 0.01483 | 0.20737 | 0.01020 | 0.02342 | 0.00000 | 0.04742 | 0.01048 | 0.00000 | 0.06523 | 0.01064 | 0.01453 |

| | | | | | | | | | | | | |
|---|---|---|---|---|---|---|---|---|---|---|---|---|
| **PSG3** | 0.00846 | 0.01544 | 0.01549 | 0.01181 | 0.01392 | 0.01818 | 0.01513 | 0.01006 | 0.03585 | 0.01210 | 0.01191 | 0.01911 |
| **PSG4** | 0.00628 | 0.01441 | 0.53704 | 0.00716 | 0.02156 | 0.01683 | 0.00000 | 0.01021 | 0.00000 | 0.01496 | 0.01720 | 0.01665 |
| **PSG5** | 0.00601 | 0.01271 | 0.06753 | 0.00582 | 0.00994 | 0.03941 | 0.04458 | 0.01034 | 0.00001 | 0.01214 | 0.01280 | 0.01576 |
| **PSG6** | 0.00605 | 0.01251 | 0.01701 | 0.00980 | 0.01483 | 0.02799 | 0.01291 | 0.01037 | 0.02548 | 0.02219 | 0.01440 | 0.01510 |
| **PSG9** | 0.00616 | 0.01444 | 0.05407 | 0.01162 | 0.01377 | 0.02208 | 0.02338 | 0.01049 | 0.01034 | 0.01410 | 0.01470 | 0.01589 |
| **PSIP1** | 5.94374 | 0.71494 | 7.20896 | 4.03028 | 0.00000 | 9.32316 | 12.28467 | 2.92715 | 0.00017 | 19.05844 | 5.68892 | 0.38591 |
| **PSKH1** | 0.22384 | 2.61794 | 0.54589 | 1.73184 | 2.22777 | 0.02237 | 0.88742 | 0.36124 | 0.00000 | 0.00000 | 1.13991 | 1.34622 |
| **PSMA1** | 0.27793 | 21.27285 | 28.70238 | 15.20156 | 14.02225 | 61.95344 | 33.80830 | 1.42918 | 29.85535 | 6.91637 | 29.61178 | 23.58168 |
| **PSMA2** | 9.65921 | 22.13792 | 32.00800 | 17.17691 | 15.51823 | 55.15516 | 37.13205 | 9.14721 | 30.21596 | 8.38112 | 27.47558 | 25.44080 |
| **PSMA3** | 2.99847 | 11.65307 | 27.21642 | 11.83642 | 9.36207 | 61.19954 | 32.44066 | 11.93138 | 28.80723 | 8.55241 | 28.33649 | 24.45272 |
| **PSMA4** | 3.25171 | 24.55888 | 33.85491 | 14.20208 | 15.39704 | 59.24617 | 40.67368 | 3.19782 | 22.61233 | 20.85130 | 30.11669 | 22.08298 |
| **PSMA5** | 12.42409 | 23.91753 | 34.78243 | 15.93214 | 19.69989 | 42.39020 | 44.16433 | 10.83346 | 32.44804 | 0.00148 | 30.05919 | 20.91557 |
| **PSMA6** | 1.77913 | 13.24584 | 23.77201 | 16.24202 | 16.14197 | 48.42056 | 30.55594 | 19.34263 | 23.21918 | 9.45510 | 23.47266 | 22.41596 |
| **PSMA7** | 9.85055 | 28.44877 | 40.06578 | 25.96005 | 40.78090 | 28.57468 | 52.57033 | 16.98368 | 39.59398 | 19.17048 | 34.40908 | 20.63002 |
| **PSMA8** | 0.01698 | 0.26725 | 0.70168 | 0.00000 | 0.15210 | 0.39010 | 0.02037 | 0.04386 | 0.00000 | 0.08811 | 0.21152 | 0.12772 |
| **PSMB1** | 3.25140 | 33.45430 | 30.86166 | 22.94373 | 33.89166 | 49.15988 | 44.26607 | 18.90129 | 36.63591 | 9.69264 | 35.68304 | 32.40797 |
| **PSMB10** | 0.00083 | 4.08297 | 7.37370 | 5.91947 | 5.27805 | 0.00000 | 13.59850 | 0.00000 | 5.98737 | 1.40068 | 14.44737 | 1.73654 |
| **PSMB11** | 0.01049 | 0.01379 | 0.03396 | 0.00961 | 0.01244 | 0.01893 | 0.01461 | 0.01483 | 0.01235 | 0.03125 | 0.02239 | 0.01449 |
| **PSMB2** | 3.81320 | 25.48424 | 34.89918 | 17.21949 | 20.13272 | 40.01562 | 42.57103 | 16.29387 | 32.64423 | 10.38077 | 27.71598 | 34.14775 |
| **PSMB3** | 7.38505 | 28.67758 | 36.09216 | 26.12135 | 32.87127 | 29.19292 | 51.24240 | 17.06587 | 30.19195 | 8.42978 | 34.15571 | 28.01239 |
| **PSMB4** | 4.83378 | 27.19562 | 36.78417 | 24.48168 | 27.26578 | 27.85817 | 48.06545 | 15.19677 | 25.94269 | 15.49317 | 30.57425 | 30.10879 |
| **PSMB5** | 1.44236 | 22.17799 | 44.44857 | 25.30802 | 28.35323 | 41.65525 | 54.84578 | 7.04101 | 39.63815 | 1.12120 | 38.84155 | 34.82748 |
| **PSMB6** | 9.75298 | 35.57170 | 33.14875 | 27.46743 | 32.45073 | 26.55794 | 50.29214 | 16.80161 | 34.01116 | 6.06671 | 32.43801 | 42.34018 |
| **PSMB7** | 0.32520 | 25.92774 | 29.59323 | 13.49186 | 14.59711 | 45.34199 | 32.52522 | 13.06821 | 31.00783 | 0.00000 | 25.56205 | 28.34464 |
| **PSMB8** | 0.18049 | 0.00000 | 0.00997 | 0.00454 | 0.19982 | 0.13227 | 0.01199 | 0.30823 | 0.00000 | 0.02340 | 0.00730 | 0.59724 |
| **PSMB9** | 0.02537 | 0.01422 | 0.01978 | 0.04681 | 0.12593 | 0.00000 | 0.01928 | 0.08353 | 0.01133 | 0.00655 | 0.01096 | 1.25875 |
| **PSMC1** | 0.10856 | 0.37446 | 0.84304 | 0.30726 | 1.04202 | 0.02083 | 0.96510 | 0.22539 | 0.18367 | 0.73109 | 0.37079 | 0.58402 |
| **PSMC2** | 1.34259 | 22.50341 | 26.87657 | 9.94040 | 10.95281 | 40.74328 | 31.08140 | 5.74338 | 28.08170 | 0.00000 | 23.81488 | 14.96679 |
| **PSMC3** | 0.00000 | 23.43838 | 28.90784 | 23.90738 | 30.12218 | 10.02194 | 40.05894 | 9.75029 | 24.39625 | 0.00000 | 28.66297 | 23.06973 |
| **PSMC3IP** | 0.03999 | 1.64715 | 3.36719 | 1.88980 | 1.66193 | 3.66568 | 2.59281 | 5.78325 | 2.95025 | 0.21498 | 2.93719 | 3.31575 |
| **PSMC4** | 6.79242 | 31.77845 | 36.87272 | 27.53340 | 22.19069 | 28.62098 | 38.10300 | 30.10181 | 31.66713 | 0.00000 | 26.89570 | 30.35237 |
| **PSMC5** | 4.92553 | 24.55536 | 32.26844 | 22.74424 | 26.41317 | 23.81220 | 37.59955 | 11.50573 | 25.99743 | 7.60784 | 29.06552 | 26.97857 |
| **PSMC6** | 0.04599 | 4.36906 | 14.12606 | 2.88313 | 1.26028 | 29.08497 | 19.08315 | 1.04195 | 10.27293 | 2.59503 | 10.92818 | 5.74650 |
| **PSMD1** | 6.02959 | 23.58574 | 30.58731 | 15.81469 | 19.10994 | 33.77885 | 42.71450 | 12.15776 | 25.06688 | 15.28533 | 27.13489 | 17.77361 |
| **PSMD10** | 5.38708 | 9.24334 | 7.88360 | 4.24530 | 3.49730 | 18.93064 | 10.90537 | 2.52618 | 4.08171 | 0.00000 | 16.25163 | 10.80035 |
| **PSMD11** | 10.66286 | 24.82605 | 34.78411 | 18.89070 | 21.53737 | 41.77493 | 49.61268 | 6.83025 | 30.48732 | 10.63993 | 29.66339 | 19.11402 |
| **PSMD12** | 0.58978 | 16.71745 | 21.37050 | 10.95717 | 7.79524 | 59.97074 | 24.29346 | 12.49089 | 20.39848 | 5.39092 | 27.00110 | 19.19872 |
| **PSMD13** | 2.37649 | 27.05303 | 31.11254 | 12.39594 | 19.75752 | 15.52136 | 38.08578 | 7.48852 | 27.73540 | 3.83269 | 17.76413 | 19.79727 |
| **PSMD14** | 1.41321 | 13.63340 | 34.60894 | 12.41024 | 15.05840 | 35.50289 | 45.76027 | 11.95148 | 29.50467 | 2.99554 | 24.43361 | 13.90922 |
| **PSMD2** | 11.52903 | 26.29439 | 35.06699 | 20.89089 | 26.14259 | 4.53092 | 32.80791 | 17.78558 | 24.72616 | 14.62644 | 28.50526 | 21.10938 |
| **PSMD3** | 3.91944 | 14.71323 | 17.56576 | 7.61899 | 17.51011 | 0.00000 | 14.69310 | 1.68616 | 17.94419 | 15.09888 | 8.75904 | 6.37831 |
| **PSMD4** | 3.84820 | 24.88872 | 27.73700 | 18.56538 | 23.57003 | 19.84460 | 36.14668 | 11.52721 | 23.08611 | 0.00000 | 29.35487 | 24.42830 |
| **PSMD5** | 0.01642 | 0.99538 | 8.81156 | 4.12774 | 0.00000 | 6.37504 | 9.54229 | 1.19322 | 9.46012 | 0.00000 | 5.83130 | 0.96754 |
| **PSMD5-AS1** | 0.08188 | 0.13131 | 1.23376 | 1.16370 | 1.11075 | 0.00000 | 0.16844 | 0.26639 | 1.55599 | 0.68221 | 0.23612 | 0.22346 |
| **PSMD6** | 2.71414 | 20.97349 | 29.69193 | 9.74367 | 9.17126 | 45.99206 | 27.39099 | 9.04945 | 23.21610 | 1.42269 | 18.89243 | 21.45067 |
| **PSMD6-AS2** | 0.01327 | 0.00293 | 0.10104 | 0.65925 | 0.00000 | 0.45886 | 0.00000 | 0.15829 | 0.88083 | 1.43835 | 1.57245 | 0.41915 |
| **PSMD7** | 2.20050 | 22.35861 | 22.61279 | 16.03153 | 29.95629 | 29.39891 | 35.96571 | 8.69669 | 23.77962 | 18.26767 | 26.02190 | 15.89388 |
| **PSMD8** | 2.42182 | 27.37800 | 29.85291 | 22.59200 | 28.85686 | 20.31287 | 36.28033 | 17.39244 | 32.15198 | 0.00018 | 24.00302 | 31.16601 |

| | | | | | | | | | | | | |
|---|---|---|---|---|---|---|---|---|---|---|---|---|
| **PSMD9** | 11.51479 | 21.26257 | 16.77892 | 14.69074 | 12.29929 | 16.68957 | 20.04921 | 21.99444 | 17.48827 | 0.00000 | 16.93828 | 24.40750 |
| **PSME1** | 0.90004 | 0.70338 | 3.95031 | 6.93264 | 1.00229 | 2.93285 | 3.84864 | 1.31530 | 1.29019 | 0.00000 | 1.42412 | 0.88706 |
| **PSME2** | 0.00000 | 0.55231 | 3.23379 | 8.98026 | 4.58150 | 6.06540 | 7.05381 | 0.96842 | 2.78113 | 2.67450 | 4.13464 | 3.06392 |
| **PSME3** | 10.14712 | 26.33944 | 21.98970 | 17.99567 | 22.91691 | 32.06985 | 26.55248 | 14.06560 | 32.97937 | 4.46386 | 27.04115 | 24.51348 |
| **PSME4** | 0.65236 | 5.29094 | 9.87632 | 15.90965 | 2.62922 | 26.11114 | 16.25861 | 4.43789 | 0.04685 | 33.67940 | 4.76024 | 12.86906 |
| **PSMF1** | 2.05525 | 17.86929 | 10.60212 | 8.66951 | 12.92573 | 17.87716 | 13.38493 | 5.47932 | 20.73661 | 0.00000 | 11.88595 | 18.55345 |
| **PSMG1** | 11.46220 | 16.52751 | 12.23792 | 10.62875 | 4.56968 | 29.13854 | 20.29917 | 5.98859 | 15.20177 | 1.29520 | 14.71517 | 8.91319 |
| **PSMG2** | 1.42041 | 13.61965 | 10.32735 | 7.35540 | 5.71589 | 27.12705 | 16.23330 | 0.24796 | 17.40363 | 2.15290 | 10.05197 | 15.77931 |
| **PSMG3** | 1.40953 | 17.94636 | 10.38397 | 16.50342 | 15.14317 | 6.76381 | 13.86072 | 2.15131 | 14.12783 | 6.70962 | 16.58940 | 13.60842 |
| **PSMG3-AS1** | 0.60864 | 0.68646 | 0.16439 | 0.99408 | 0.12667 | 0.00000 | 0.00775 | 0.00424 | 0.09271 | 0.06489 | 0.20384 | 0.15574 |
| **PSMG4** | 3.80130 | 10.52519 | 1.78115 | 2.90372 | 2.96609 | 0.27396 | 4.98003 | 0.61133 | 2.43816 | 3.20416 | 7.43143 | 6.56383 |
| **PSORS1C1** | 0.00586 | 0.43113 | 0.00655 | 0.29716 | 0.00000 | 0.06201 | 0.00000 | 0.01220 | 0.00003 | 0.04213 | 0.00998 | 0.81981 |
| **PSORS1C2** | 0.01118 | 7.49312 | 0.65591 | 2.25859 | 0.00000 | 0.00361 | 0.00000 | 0.36726 | 0.00000 | 0.02294 | 0.00201 | 20.26719 |
| **PSORS1C3** | 0.00517 | 0.09585 | 0.02027 | 0.00000 | 0.00000 | 0.02336 | 0.02794 | 0.01038 | 0.00173 | 0.05257 | 0.01626 | 0.02981 |
| **PSPC1** | 1.39239 | 22.49771 | 17.84853 | 7.66434 | 14.23616 | 19.85530 | 14.34220 | 21.70811 | 16.15978 | 2.12425 | 18.22256 | 26.88818 |
| **PSPH** | 0.57051 | 2.71706 | 5.70459 | 2.07255 | 0.00000 | 3.73358 | 10.93771 | 0.15175 | 8.87164 | 0.00000 | 6.42916 | 1.48509 |
| **PSPN** | 0.03758 | 0.04194 | 0.04245 | 0.02040 | 0.00919 | 0.00000 | 0.04446 | 0.02145 | 0.11763 | 0.00174 | 0.02486 | 0.02299 |
| **PSRC1** | 0.27350 | 11.42533 | 4.75345 | 19.20633 | 2.56263 | 3.10255 | 6.52781 | 0.55085 | 6.08617 | 0.00000 | 1.64317 | 4.61511 |
| **PSTK** | 0.00000 | 1.33262 | 1.26101 | 0.00000 | 0.70985 | 4.36584 | 2.28662 | 0.20190 | 2.11228 | 1.29192 | 1.06710 | 3.54242 |
| **PSTPIP1** | 2.69380 | 0.08456 | 0.02614 | 0.09397 | 0.12975 | 0.00000 | 0.08224 | 0.24868 | 0.00000 | 0.00000 | 0.00603 | 0.00441 |
| **PSTPIP2** | 0.02812 | 0.00558 | 0.00000 | 0.00000 | 4.58064 | 1.34098 | 0.00000 | 0.10050 | 0.00000 | 0.28107 | 0.89825 | 0.00262 |
| **PTAFR** | 0.21612 | 0.06249 | 0.76279 | 0.19923 | 0.15240 | 0.07422 | 0.27069 | 1.84223 | 0.00000 | 1.10059 | 0.45386 | 0.66152 |
| **PTAR1** | 0.00000 | 1.57765 | 1.08702 | 3.83063 | 0.68675 | 16.16514 | 0.96477 | 8.41413 | 3.72619 | 15.86226 | 1.26877 | 6.86025 |
| **PTBP1** | 7.53895 | 28.54834 | 28.06330 | 20.38424 | 36.97740 | 9.85695 | 32.34139 | 11.38702 | 38.85130 | 34.56279 | 19.01377 | 32.90592 |
| **PTBP2** | 1.86524 | 0.00016 | 2.24245 | 3.82568 | 0.00000 | 5.71884 | 0.00000 | 4.93376 | 2.82133 | 3.71238 | 1.73562 | 0.69363 |
| **PTBP3** | 0.00000 | 3.41101 | 11.33093 | 7.27476 | 7.91521 | 23.20490 | 24.89532 | 1.40196 | 10.54935 | 16.26030 | 9.14483 | 0.00000 |
| **PTCD1** | 0.01137 | 0.60953 | 0.06033 | 0.34970 | 0.42920 | 0.00000 | 0.00054 | 0.03488 | 0.33905 | 0.09824 | 0.06649 | 0.47487 |
| **PTCD2** | 0.19930 | 2.98139 | 2.26620 | 0.76102 | 0.21680 | 5.75906 | 0.00000 | 0.19127 | 2.28709 | 0.00000 | 4.01602 | 1.41951 |
| **PTCD3** | 2.47464 | 13.73313 | 11.26134 | 11.14070 | 8.89271 | 26.17473 | 12.25001 | 7.45366 | 18.93530 | 15.73908 | 17.72178 | 12.31591 |
| **PTCH1** | 0.02677 | 0.52650 | 1.75878 | 0.00000 | 0.63313 | 0.26729 | 0.32631 | 0.12133 | 0.33311 | 2.35728 | 0.10163 | 0.80366 |
| **PTCH2** | 0.00797 | 0.33021 | 0.14778 | 0.35616 | 0.24533 | 0.00000 | 0.00000 | 0.06472 | 0.32788 | 0.39409 | 0.00166 | 0.71834 |
| **PTCHD1** | 0.01088 | 0.32603 | 0.04636 | 0.65882 | 0.14218 | 0.00000 | 0.01469 | 0.01403 | 0.00000 | 0.27621 | 0.07099 | 0.11751 |
| **PTCHD1-AS** | 0.01953 | 0.03662 | 0.01440 | 0.02320 | 0.00883 | 0.05400 | 0.01067 | 0.01452 | 0.01332 | 0.01370 | 0.01248 | 0.01920 |
| **PTCHD2** | 0.35813 | 0.09459 | 0.12317 | 0.01550 | 0.02517 | 0.00000 | 0.08786 | 0.21873 | 0.47539 | 0.08145 | 0.00000 | 0.14124 |
| **PTCHD3** | 0.00680 | 0.01410 | 0.08453 | 0.00675 | 0.01011 | 0.02589 | 0.00372 | 0.03998 | 0.02375 | 0.02316 | 0.01560 | 0.06796 |
| **PTCHD4** | 0.00503 | 0.00498 | 0.02945 | 0.00000 | 0.03975 | 0.07590 | 0.00000 | 0.00964 | 0.10377 | 0.05685 | 0.01317 | 0.00979 |
| **PTCRA** | 0.00372 | 0.01470 | 0.01337 | 0.06619 | 0.01393 | 0.02421 | 0.01545 | 0.02450 | 0.01885 | 0.00631 | 0.01928 | 0.01566 |
| **PTCSC3** | 0.00582 | 0.01449 | 0.01661 | 0.01238 | 0.01339 | 0.01701 | 0.01451 | 0.01163 | 0.02154 | 0.02463 | 0.01428 | 0.01638 |
| **PTDSS1** | 7.00800 | 20.26833 | 16.13305 | 7.91573 | 7.00974 | 16.32180 | 0.00000 | 1.11925 | 35.55609 | 12.16865 | 9.78457 | 13.86262 |
| **PTDSS2** | 2.67959 | 4.36744 | 5.41529 | 4.23303 | 9.29871 | 0.00000 | 4.82023 | 0.25657 | 10.04722 | 21.96707 | 2.06357 | 0.68660 |
| **PTEN** | 3.87644 | 4.36852 | 2.01280 | 2.26430 | 0.00000 | 15.74381 | 9.46186 | 7.51099 | 0.00000 | 5.99591 | 4.90690 | 4.15021 |
| **PTENP1** | 0.30170 | 0.54445 | 0.17570 | 0.09255 | 0.39353 | 2.04158 | 0.63839 | 0.14619 | 0.00000 | 1.35011 | 0.10754 | 0.55245 |
| **PTENP1-AS** | 0.06086 | 0.18719 | 0.00853 | 0.02703 | 0.00000 | 0.03871 | 0.00730 | 0.10658 | 0.01245 | 0.02371 | 0.01660 | 0.36128 |
| **PTER** | 0.00000 | 7.25235 | 2.30334 | 0.00000 | 1.70590 | 6.33369 | 6.47102 | 0.80829 | 3.96080 | 0.47696 | 1.95781 | 2.03558 |
| **PTF1A** | 0.01235 | 0.01355 | 0.01149 | 0.01127 | 0.01064 | 0.02972 | 0.01387 | 0.02930 | 0.02545 | 0.01899 | 0.01150 | 0.01597 |
| **PTGDR** | 0.14529 | 0.00144 | 0.01029 | 0.00214 | 0.00519 | 0.11639 | 0.00557 | 0.01460 | 0.00000 | 0.03889 | 0.00939 | 0.02914 |
| **PTGDR2** | 0.36649 | 0.16009 | 0.00419 | 0.13419 | 0.00000 | 0.00000 | 0.11842 | 0.00955 | 0.00692 | 0.06453 | 0.02719 | 0.19824 |
| **PTGDS** | 0.03237 | 0.04295 | 0.47790 | 0.00351 | 0.01394 | 0.00000 | 0.02429 | 0.00988 | 0.37224 | 0.00000 | 0.56655 | 0.05740 |

| | | | | | | | | | | | | |
|---|---|---|---|---|---|---|---|---|---|---|---|---|
| **PTGER1** | 0.00562 | 0.01320 | 0.02819 | 0.00752 | 0.00929 | 0.02214 | 0.01765 | 0.01039 | 0.01364 | 0.04504 | 0.01333 | 0.01415 |
| **PTGER2** | 0.29469 | 0.01488 | 0.01777 | 0.01468 | 0.01769 | 0.00798 | 0.01584 | 0.00875 | 0.03815 | 0.00000 | 0.01323 | 0.02011 |
| **PTGER3** | 0.24193 | 0.00460 | 0.03620 | 0.01189 | 0.01367 | 0.00000 | 0.00000 | 2.17144 | 0.05621 | 0.15039 | 0.02146 | 0.00614 |
| **PTGER4** | 0.00612 | 0.48982 | 0.01172 | 0.00000 | 0.03326 | 0.00000 | 0.00237 | 0.99645 | 0.01873 | 0.13195 | 0.02337 | 0.33134 |
| **PTGER4P2-CDK** | 0.04849 | 0.02508 | 0.02929 | 0.01739 | 0.01348 | 0.01125 | 0.01541 | 0.04888 | 0.01992 | 0.01652 | 0.01861 | 0.02002 |
| **PTGES** | 0.00000 | 41.77353 | 7.43257 | 0.00000 | 60.78157 | 2.45260 | 12.62553 | 0.00000 | 34.71509 | 10.76269 | 42.07584 | 1.54835 |
| **PTGES2** | 0.00000 | 9.21379 | 1.91261 | 7.37168 | 13.48712 | 0.00000 | 2.63479 | 0.23510 | 0.00142 | 3.63572 | 4.44781 | 8.40246 |
| **PTGES2-AS1** | 0.00462 | 0.00942 | 0.00523 | 0.01343 | 0.01627 | 0.01466 | 0.01432 | 0.00963 | 0.03360 | 0.13039 | 0.00895 | 0.01809 |
| **PTGES3** | 9.86023 | 22.67452 | 32.86901 | 15.84155 | 16.99059 | 61.75544 | 47.36591 | 11.02939 | 34.46713 | 28.48112 | 27.13777 | 20.33495 |
| **PTGES3L** | 0.02191 | 0.46401 | 0.16492 | 0.03919 | 0.00399 | 0.29045 | 0.06973 | 0.01450 | 0.20157 | 0.00000 | 0.01374 | 0.11333 |
| **PTGES3L-AARS** | 0.00612 | 0.02839 | 0.01629 | 0.01463 | 0.00950 | 0.01293 | 0.01057 | 0.01052 | 0.03895 | 0.02399 | 0.01492 | 0.01776 |
| **PTGFR** | 0.00173 | 0.01505 | 0.01566 | 0.01464 | 0.01534 | 0.01687 | 0.01816 | 0.13279 | 0.01763 | 0.01401 | 0.01498 | 0.01180 |
| **PTGFRN** | 2.04036 | 5.60514 | 0.00000 | 4.67440 | 4.31179 | 1.53952 | 0.00000 | 1.72966 | 0.00000 | 1.15333 | 1.29151 | 2.18191 |
| **PTGIR** | 0.00584 | 0.01488 | 0.01821 | 0.01423 | 0.01714 | 0.01604 | 0.01814 | 0.01178 | 0.01500 | 0.01386 | 0.02337 | 0.01652 |
| **PTGIS** | 0.05071 | 0.00000 | 0.14290 | 0.00246 | 0.16704 | 0.32576 | 0.00000 | 0.01488 | 0.00000 | 0.00927 | 0.00438 | 0.02532 |
| **PTGR1** | 0.00192 | 6.57924 | 4.70305 | 28.24264 | 25.49039 | 6.73315 | 7.29791 | 0.54355 | 7.24362 | 0.00000 | 15.06019 | 0.00000 |
| **PTGR2** | 0.01911 | 1.81659 | 0.89358 | 0.16948 | 0.05220 | 1.10732 | 0.00000 | 0.30527 | 0.00000 | 0.49785 | 1.62207 | 1.33682 |
| **PTGS1** | 0.44778 | 0.07654 | 0.00028 | 0.00100 | 0.00077 | 0.15271 | 0.00000 | 0.00311 | 0.00000 | 0.07275 | 0.00159 | 0.01018 |
| **PTGS2** | 0.02372 | 0.01437 | 0.00000 | 0.03768 | 0.01104 | 0.02746 | 0.00699 | 0.05013 | 0.06320 | 0.09971 | 0.01932 | 0.01127 |
| **PTH1R** | 1.21056 | 0.69978 | 0.11482 | 1.10382 | 0.14895 | 0.00000 | 0.01541 | 0.26638 | 0.00000 | 0.02992 | 0.00354 | 0.22755 |
| **PTH2** | 0.14044 | 0.01704 | 0.01382 | 0.00993 | 0.01566 | 0.01715 | 0.01501 | 0.04397 | 0.01863 | 0.02332 | 0.01356 | 0.03560 |
| **PTH2R** | 0.34612 | 0.00927 | 0.05766 | 0.00920 | 0.00625 | 0.02142 | 0.03934 | 0.08263 | 0.05604 | 0.02266 | 0.00907 | 0.03087 |
| **PTHLH** | 0.09671 | 0.04777 | 0.00000 | 0.76498 | 1.40012 | 1.08526 | 0.00000 | 0.00000 | 0.02219 | 0.00000 | 0.23948 | 0.30146 |
| **PTK2** | 2.21789 | 4.83236 | 11.11571 | 4.10784 | 2.17417 | 10.57340 | 9.37542 | 1.30000 | 11.66070 | 3.06084 | 7.86661 | 3.79467 |
| **PTK2B** | 5.62131 | 0.22430 | 0.92989 | 3.30664 | 1.08399 | 0.00000 | 0.22303 | 0.24717 | 0.00000 | 1.31781 | 0.11525 | 0.37041 |
| **PTK6** | 0.00400 | 0.10365 | 0.04038 | 0.13593 | 0.02324 | 0.00000 | 0.03248 | 0.01909 | 0.03352 | 0.04660 | 0.01285 | 0.00889 |
| **PTK7** | 0.13128 | 2.29322 | 2.22408 | 6.74910 | 2.04205 | 0.00000 | 0.02946 | 0.18040 | 10.79937 | 3.42998 | 0.00000 | 2.20151 |
| **PTMA** | 7.32992 | 28.80093 | 22.48407 | 28.89329 | 40.52299 | 32.97455 | 51.55052 | 22.24988 | 33.16042 | 40.03403 | 26.87507 | 33.76803 |
| **PTMS** | 0.16757 | 0.55687 | 1.50633 | 6.51830 | 6.39315 | 0.00000 | 5.77445 | 0.00000 | 0.00027 | 9.23355 | 0.00000 | 0.80890 |
| **PTN** | 7.03392 | 0.00000 | 44.65561 | 0.00000 | 9.08871 | 2.05752 | 32.68000 | 3.68377 | 35.43783 | 0.54454 | 10.88107 | 0.00000 |
| **PTOV1** | 3.04884 | 1.61611 | 6.34622 | 5.63877 | 9.31839 | 0.00000 | 13.21583 | 0.55561 | 0.00274 | 15.42832 | 3.01349 | 0.60910 |
| **PTOV1-AS1** | 0.20583 | 0.04028 | 1.05922 | 0.00000 | 0.62680 | 2.09079 | 1.14105 | 0.00000 | 1.17021 | 1.14431 | 0.13997 | 0.44754 |
| **PTP4A1** | 9.82262 | 30.03133 | 25.96915 | 5.68660 | 14.93568 | 76.87671 | 33.18790 | 36.58412 | 31.25129 | 19.20135 | 31.73117 | 33.67733 |
| **PTP4A2** | 3.18023 | 4.15252 | 10.82261 | 6.68291 | 9.31373 | 13.63121 | 27.41400 | 5.73171 | 2.28140 | 23.27828 | 6.02217 | 3.22255 |
| **PTP4A3** | 0.17153 | 0.04182 | 0.00945 | 0.24584 | 0.00000 | 0.04106 | 0.00723 | 0.02290 | 0.00626 | 0.03045 | 0.01546 | 0.12724 |
| **PTPDC1** | 0.06552 | 0.27242 | 0.24922 | 0.08453 | 0.20353 | 0.00000 | 0.13709 | 0.03047 | 0.32772 | 1.37424 | 0.11879 | 0.08033 |
| **PTPLA** | 0.01716 | 0.23757 | 0.70646 | 1.12892 | 1.08388 | 1.69836 | 0.00000 | 0.03286 | 0.21313 | 0.19506 | 0.90712 | 0.00000 |
| **PTPLAD1** | 0.00000 | 27.76923 | 27.66828 | 6.37377 | 7.00835 | 38.17509 | 8.89980 | 5.31039 | 46.94749 | 17.08194 | 18.49013 | 31.21100 |
| **PTPLAD2** | 0.02775 | 0.57622 | 0.20913 | 0.05407 | 0.10600 | 0.12180 | 0.04496 | 0.05748 | 0.02465 | 0.00006 | 0.06817 | 0.20557 |
| **PTPLB** | 0.29089 | 0.70213 | 3.16196 | 0.00000 | 0.60078 | 1.25084 | 0.00000 | 0.36858 | 3.57566 | 0.65564 | 1.20068 | 0.51343 |
| **PTPMT1** | 1.35126 | 9.31264 | 8.55252 | 2.72769 | 8.05789 | 1.07060 | 5.91601 | 1.74651 | 15.48544 | 0.00010 | 11.89970 | 3.77601 |
| **PTPN1** | 2.65129 | 3.89564 | 10.83357 | 3.50699 | 5.18962 | 5.68911 | 6.03751 | 4.52966 | 4.98874 | 1.68902 | 6.82941 | 5.76565 |
| **PTPN11** | 0.92280 | 3.66061 | 7.07421 | 3.55594 | 3.42021 | 17.87626 | 10.54723 | 3.14673 | 7.22090 | 9.39383 | 7.18654 | 1.92101 |
| **PTPN12** | 2.17312 | 3.02981 | 7.17922 | 8.10614 | 0.00000 | 11.17154 | 7.20417 | 3.38515 | 0.00000 | 10.87753 | 0.00000 | 5.36243 |
| **PTPN13** | 4.21084 | 0.02705 | 0.00000 | 0.09820 | 0.18397 | 0.09415 | 0.21398 | 0.63827 | 0.12376 | 0.07913 | 0.11337 | 0.46359 |
| **PTPN14** | 3.69853 | 2.11145 | 11.32619 | 0.00000 | 13.78369 | 3.01664 | 12.70938 | 1.63058 | 3.77355 | 16.13375 | 12.42310 | 2.90319 |
| **PTPN18** | 0.05081 | 0.00000 | 1.93693 | 0.06320 | 0.00000 | 0.02789 | 1.83618 | 0.01831 | 0.93011 | 2.67182 | 1.22852 | 0.02162 |
| **PTPN2** | 3.46967 | 13.20255 | 4.36005 | 11.26367 | 2.56838 | 23.13378 | 5.42969 | 11.25047 | 5.51525 | 5.61773 | 6.68475 | 11.45562 |

| | | | | | | | | | | | | |
|---|---|---|---|---|---|---|---|---|---|---|---|---|
| PTPN20B.uid-13 | 0.00627 | 0.01539 | 0.02454 | 0.01168 | 0.01568 | 0.02716 | 0.04071 | 0.01034 | 0.02641 | 0.00083 | 0.01449 | 0.02054 |
| PTPN20B.uid-14 | 0.00858 | 0.01560 | 0.01721 | 0.01682 | 0.01486 | 0.02464 | 0.02217 | 0.01042 | 0.01994 | 0.00640 | 0.01595 | 0.02220 |
| PTPN21 | 6.21797 | 0.00000 | 3.71429 | 0.00000 | 6.81854 | 2.01922 | 2.83618 | 3.76658 | 2.46128 | 3.40644 | 2.26727 | 5.36323 |
| PTPN22 | 0.33555 | 0.01514 | 0.00491 | 0.01370 | 0.00000 | 0.06721 | 0.00846 | 0.99853 | 0.03830 | 0.01312 | 0.09416 | 0.24669 |
| PTPN23 | 0.82292 | 1.25160 | 2.84557 | 3.28468 | 4.70839 | 0.00000 | 2.27755 | 0.47701 | 1.39126 | 8.63974 | 0.89492 | 0.92876 |
| PTPN3 | 11.34232 | 3.50108 | 0.80345 | 1.68513 | 2.67917 | 5.94623 | 3.94015 | 1.67651 | 2.57525 | 1.33397 | 1.68703 | 0.75461 |
| PTPN4 | 0.25766 | 0.24125 | 2.96458 | 0.79701 | 0.00000 | 3.11128 | 3.02538 | 0.10189 | 0.56774 | 5.96656 | 0.84301 | 0.06506 |
| PTPN5 | 3.79450 | 2.17151 | 0.90841 | 3.43052 | 0.19288 | 0.00000 | 0.00000 | 0.01246 | 0.00000 | 0.00000 | 0.04890 | 0.64999 |
| PTPN6 | 3.02208 | 0.48611 | 2.61708 | 1.97590 | 0.00000 | 0.00000 | 2.17283 | 0.13280 | 2.43606 | 0.42836 | 1.97816 | 0.03394 |
| PTPN7 | 0.00262 | 0.01365 | 0.01850 | 0.09013 | 0.01834 | 0.01778 | 0.01193 | 0.03602 | 0.01903 | 0.02765 | 0.02475 | 0.01623 |
| PTPN9 | 1.65010 | 3.17296 | 0.91794 | 5.33422 | 1.62951 | 5.07838 | 0.22651 | 4.51865 | 1.15686 | 5.15523 | 2.36487 | 7.24137 |
| PTPRA | 2.90647 | 2.50278 | 1.44129 | 5.41443 | 3.54759 | 1.54639 | 0.76409 | 1.29490 | 2.40775 | 7.37062 | 1.07642 | 3.00538 |
| PTPRB | 0.27687 | 1.30275 | 0.01867 | 0.37273 | 0.37632 | 1.03509 | 0.00000 | 0.29302 | 0.00487 | 1.23273 | 0.27469 | 1.75815 |
| PTPRC | 0.00512 | 0.09418 | 0.03250 | 0.00000 | 0.00000 | 0.03217 | 0.08130 | 0.01023 | 0.00867 | 0.06335 | 0.01206 | 0.00949 |
| PTPRCAP | 0.02098 | 0.06651 | 0.14510 | 0.48713 | 0.23946 | 0.00000 | 0.15669 | 0.00731 | 0.04745 | 0.69085 | 0.00000 | 0.40062 |
| PTPRD | 3.38498 | 0.11374 | 0.73810 | 0.64032 | 0.00000 | 0.00000 | 0.00000 | 0.83945 | 0.00000 | 0.07490 | 0.04188 | 0.00000 |
| PTPRD-AS1 | 0.51889 | 0.00325 | 0.00000 | 0.04314 | 0.00000 | 0.09862 | 0.02024 | 0.00000 | 0.00000 | 0.10952 | 0.00546 | 0.07492 |
| PTPRD-AS2 | 0.00614 | 0.01539 | 0.01171 | 0.02167 | 0.01486 | 0.01328 | 0.06212 | 0.01058 | 0.00292 | 0.01309 | 0.01567 | 0.01657 |
| PTPRE | 0.38393 | 0.01619 | 7.50778 | 0.00000 | 0.01217 | 0.08459 | 0.45689 | 0.19842 | 0.02272 | 0.07232 | 0.00000 | 0.00851 |
| PTPRF | 0.78944 | 6.44671 | 12.60697 | 9.39424 | 23.79689 | 7.49531 | 11.29877 | 0.02694 | 24.47127 | 24.60919 | 9.42929 | 1.66092 |
| PTPRG | 3.49487 | 0.78762 | 0.44667 | 3.42798 | 0.56182 | 1.93620 | 0.00000 | 1.29574 | 0.32534 | 1.76171 | 0.12246 | 0.55340 |
| PTPRG-AS1 | 1.14642 | 5.33001 | 0.70971 | 0.12786 | 2.52798 | 2.62962 | 0.00000 | 0.16338 | 0.00000 | 0.42851 | 0.94707 | 4.27810 |
| PTPRH | 3.27417 | 0.01417 | 0.01518 | 0.01258 | 0.01251 | 0.00000 | 0.01129 | 1.01089 | 0.02439 | 0.01862 | 0.01164 | 0.00000 |
| PTPRJ | 2.12494 | 2.51711 | 0.55511 | 1.48279 | 1.34871 | 0.00000 | 0.00000 | 1.08608 | 0.00000 | 2.69197 | 0.00026 | 3.19002 |
| PTPRK | 6.21595 | 0.00000 | 1.27944 | 1.77193 | 0.52486 | 0.19283 | 0.00000 | 1.01557 | 4.18962 | 0.70432 | 1.92048 | 0.00000 |
| PTPRM | 0.99713 | 0.31477 | 0.49389 | 0.07392 | 0.04703 | 0.60611 | 0.00000 | 0.26084 | 0.14968 | 0.00000 | 0.00000 | 0.00003 |
| PTPRN | 0.00566 | 0.02081 | 0.01756 | 0.00617 | 0.01617 | 0.02121 | 0.02203 | 0.01033 | 0.01855 | 0.04019 | 0.01685 | 0.01507 |
| PTPRN2 | 2.69181 | 0.13808 | 0.00231 | 0.00000 | 0.00000 | 0.10409 | 0.00000 | 0.49958 | 0.00000 | 0.06019 | 0.00149 | 0.08370 |
| PTPRO | 0.10274 | 0.16582 | 0.24112 | 0.02125 | 0.03182 | 0.00000 | 0.01099 | 0.06608 | 0.19090 | 0.04928 | 0.03789 | 0.17537 |
| PTPRQ | 0.09590 | 0.01421 | 0.00512 | 0.00839 | 0.00828 | 0.00519 | 0.18639 | 0.01931 | 0.00000 | 0.05520 | 0.01038 | 0.01395 |
| PTPRR | 1.09345 | 0.04160 | 0.01487 | 0.10220 | 0.04707 | 0.43218 | 0.00000 | 0.00000 | 0.00000 | 0.02183 | 0.02866 | 0.11890 |
| PTPRS | 1.98428 | 3.85342 | 0.40210 | 3.01452 | 2.35678 | 0.00000 | 0.18104 | 2.14864 | 0.00000 | 2.26099 | 0.73733 | 5.08492 |
| PTPRT | 0.93013 | 0.04547 | 0.12886 | 0.07453 | 0.01864 | 0.00721 | 0.02768 | 0.05127 | 0.00000 | 0.07969 | 0.01259 | 0.00786 |
| PTPRU | 0.00001 | 6.87567 | 7.98375 | 6.95306 | 19.16114 | 3.19853 | 3.19689 | 0.05577 | 8.76528 | 11.84104 | 4.09868 | 0.00000 |
| PTPRVP | 0.01238 | 0.02050 | 0.03471 | 0.01277 | 0.00000 | 0.00000 | 0.00513 | 0.00949 | 0.00000 | 0.07103 | 0.18527 | 0.02281 |
| PTPRZ1 | 0.55268 | 0.06799 | 0.16485 | 1.21815 | 0.09756 | 0.00000 | 0.00550 | 0.02421 | 0.00000 | 0.09313 | 0.04813 | 0.00483 |
| PTRF | 0.08336 | 8.40780 | 0.88311 | 1.89517 | 10.52208 | 0.00000 | 0.47913 | 0.69243 | 0.00000 | 2.11309 | 0.87574 | 2.20284 |
| PTRH1 | 0.00295 | 2.29697 | 1.06477 | 1.70714 | 2.58385 | 0.00000 | 0.04163 | 0.79163 | 0.94086 | 0.24214 | 1.95014 | 2.77370 |
| PTRH2 | 1.83425 | 8.58383 | 17.68795 | 6.01849 | 10.75633 | 14.54747 | 10.66855 | 11.97237 | 26.24568 | 4.14021 | 17.20987 | 17.45779 |
| PTRHD1 | 5.31279 | 4.99324 | 5.44028 | 5.32840 | 5.44188 | 4.97197 | 10.62968 | 5.03249 | 10.61125 | 1.25186 | 9.92485 | 6.32917 |
| PTS | 0.64049 | 10.78634 | 13.00279 | 1.66476 | 2.14307 | 25.78293 | 19.15019 | 0.95581 | 17.65474 | 3.29648 | 12.27465 | 10.99144 |
| PTTG1 | 24.56686 | 17.48562 | 13.22006 | 11.46560 | 13.16131 | 26.06010 | 28.40233 | 43.72178 | 12.36253 | 0.01555 | 8.60539 | 33.23979 |
| PTTG1IP | 10.21384 | 26.72596 | 20.24109 | 19.55389 | 26.52088 | 29.45452 | 7.34670 | 3.77268 | 52.41685 | 16.38265 | 19.15484 | 13.47291 |
| PTTG2 | 3.33201 | 0.34152 | 0.30052 | 0.36700 | 0.32886 | 0.69601 | 0.99146 | 7.16070 | 0.18527 | 0.00000 | 0.15691 | 1.29217 |
| PTTG3P | 12.90240 | 3.36404 | 2.43434 | 2.16486 | 1.55599 | 7.13973 | 5.47260 | 22.93456 | 2.33659 | 0.00000 | 1.13860 | 9.27742 |
| PTX4 | 0.00586 | 0.01548 | 0.01218 | 0.01235 | 0.00814 | 0.01439 | 0.02267 | 0.01054 | 0.02419 | 0.03161 | 0.03304 | 0.01537 |
| PUF60 | 2.18313 | 36.14340 | 24.33020 | 26.99386 | 26.42069 | 12.88896 | 35.18134 | 24.48444 | 34.08470 | 0.00000 | 32.02239 | 41.55875 |
| PUM1 | 0.00049 | 5.99508 | 13.84152 | 11.87697 | 14.78561 | 18.93722 | 22.18345 | 4.09992 | 18.71583 | 18.76950 | 15.21837 | 11.72594 |

| | | | | | | | | | | | | |
|---|---|---|---|---|---|---|---|---|---|---|---|---|
| **PUM2** | 5.60092 | 1.34368 | 9.27781 | 3.73162 | 0.00021 | 22.02030 | 11.03713 | 8.60885 | 7.63350 | 19.95388 | 3.86233 | 7.01795 |
| **PURA** | 0.02845 | 0.00000 | 2.30585 | 0.86695 | 2.60718 | 1.36373 | 8.86711 | 0.00000 | 0.00000 | 15.79781 | 1.21268 | 1.02000 |
| **PURB** | 1.00094 | 0.00000 | 2.46577 | 0.00000 | 0.75323 | 2.71696 | 0.66825 | 1.88897 | 0.00000 | 7.74165 | 1.40049 | 4.10356 |
| **PURG** | 0.12218 | 0.00928 | 0.19953 | 0.00987 | 0.00460 | 0.03616 | 0.00000 | 0.01326 | 0.00000 | 0.03986 | 0.01705 | 0.01553 |
| **PUS1** | 1.20571 | 8.49835 | 5.05745 | 5.04344 | 9.60133 | 0.00000 | 14.02158 | 1.02120 | 5.68615 | 24.06925 | 3.61270 | 3.79582 |
| **PUS10** | 0.13462 | 0.95516 | 2.01189 | 0.40939 | 0.10615 | 2.08871 | 0.00000 | 0.91812 | 0.00000 | 0.00000 | 0.39530 | 1.29043 |
| **PUS3** | 0.00000 | 3.27192 | 7.05217 | 1.44718 | 0.55183 | 14.80814 | 5.32046 | 2.20094 | 6.84542 | 3.98148 | 7.50939 | 9.31645 |
| **PUS7** | 1.68983 | 6.74740 | 7.66559 | 1.22860 | 2.70383 | 10.37504 | 8.19022 | 0.14006 | 21.69113 | 0.00000 | 11.32056 | 3.76040 |
| **PUS7L** | 0.00000 | 2.00519 | 5.70535 | 0.64889 | 0.00000 | 9.26418 | 6.26628 | 0.38173 | 0.76533 | 4.82298 | 2.93325 | 2.24524 |
| **PUSL1** | 0.13434 | 8.32730 | 8.37909 | 7.53235 | 13.38435 | 0.00000 | 8.48457 | 0.01270 | 9.09979 | 0.96364 | 11.09124 | 0.78408 |
| **PVALB** | 0.00606 | 0.02055 | 0.01748 | 0.01249 | 0.01082 | 0.02296 | 0.01582 | 0.00993 | 0.01583 | 0.02226 | 0.01435 | 0.02132 |
| **PVR** | 0.97217 | 7.90842 | 2.06407 | 10.87687 | 3.68278 | 1.63596 | 1.47248 | 0.41055 | 8.93992 | 3.92641 | 4.67183 | 3.48772 |
| **PVRIG** | 0.01758 | 0.11301 | 0.19109 | 0.09332 | 0.11289 | 0.00000 | 0.00231 | 0.02127 | 0.00000 | 0.42830 | 0.20773 | 0.02789 |
| **PVRL1** | 0.73405 | 1.49634 | 4.60539 | 0.99093 | 2.90768 | 0.00000 | 8.06017 | 0.60835 | 10.48412 | 9.92954 | 0.00000 | 0.26403 |
| **PVRL2** | 0.00000 | 11.77537 | 8.03987 | 14.83940 | 29.61398 | 0.00000 | 0.00006 | 0.67183 | 13.83520 | 19.57993 | 10.61775 | 9.51564 |
| **PVRL3** | 1.69557 | 0.00000 | 4.28504 | 0.57161 | 0.00000 | 1.22969 | 0.73571 | 1.84507 | 0.00000 | 2.20105 | 0.00000 | 1.10783 |
| **PVRL3-AS1** | 0.00288 | 0.01252 | 0.01514 | 0.00714 | 0.03988 | 0.00000 | 0.05909 | 0.14458 | 0.00000 | 0.11207 | 0.00686 | 0.01904 |
| **PVRL4** | 0.00000 | 5.54327 | 16.43767 | 6.67649 | 2.82750 | 9.63345 | 0.00001 | 0.06661 | 6.50889 | 0.46324 | 0.00000 | 6.56229 |
| **PVT1** | 0.16178 | 1.13377 | 2.39393 | 0.13819 | 3.67384 | 1.42762 | 0.87493 | 0.36179 | 5.14136 | 5.54982 | 1.08743 | 2.07030 |
| **PWAR1** | 0.00347 | 0.00000 | 0.00404 | 0.00106 | 0.03565 | 0.01841 | 0.00536 | 0.00697 | 0.00471 | 0.29478 | 0.02317 | 0.07237 |
| **PWAR5** | 0.26341 | 0.23571 | 0.55523 | 0.46664 | 0.41022 | 0.46636 | 0.00000 | 0.25970 | 0.00000 | 0.93082 | 0.10827 | 0.41010 |
| **PWARSN** | 0.62413 | 1.50102 | 1.91678 | 6.03065 | 3.50179 | 2.69373 | 0.00000 | 0.26136 | 0.42539 | 7.16916 | 0.00000 | 2.74314 |
| **PWP1** | 9.28137 | 11.66395 | 18.62271 | 4.61428 | 12.48604 | 21.47714 | 26.85285 | 15.57410 | 21.79530 | 9.23879 | 24.90500 | 23.55416 |
| **PWP2** | 2.94337 | 15.55379 | 15.22892 | 12.72432 | 12.30770 | 1.87297 | 8.90087 | 4.90358 | 20.95782 | 0.00000 | 16.52660 | 11.76033 |
| **PWRN1** | 0.08742 | 0.01466 | 0.03148 | 0.02063 | 0.02450 | 0.00623 | 0.01154 | 0.04796 | 0.02152 | 0.03578 | 0.01806 | 0.02202 |
| **PWRN2** | 0.00461 | 0.01318 | 0.03144 | 0.01207 | 0.01626 | 0.02051 | 0.02587 | 0.01657 | 0.00360 | 0.03009 | 0.01575 | 0.01497 |
| **PWWP2A** | 0.00000 | 0.42119 | 0.50655 | 1.79849 | 0.57792 | 2.77969 | 3.23863 | 2.76227 | 0.00193 | 13.15967 | 3.21533 | 2.12936 |
| **PWWP2B** | 0.00069 | 0.00000 | 7.61474 | 0.00000 | 5.49001 | 0.00000 | 11.00954 | 0.00361 | 8.77857 | 1.18773 | 3.73455 | 0.00000 |
| **PXDC1** | 0.00278 | 0.01822 | 0.55646 | 0.00297 | 0.12446 | 0.00000 | 0.00009 | 0.08268 | 0.01101 | 0.07181 | 0.00973 | 0.01777 |
| **PXDN** | 3.49637 | 0.00039 | 2.17604 | 8.91079 | 0.00000 | 0.00000 | 0.00000 | 0.04988 | 0.00000 | 0.80121 | 0.00000 | 0.05823 |
| **PXDNL** | 0.10565 | 0.02938 | 0.01380 | 0.01359 | 0.00000 | 0.00104 | 0.17345 | 0.04926 | 0.01723 | 0.02224 | 0.03201 | 0.05561 |
| **PXK** | 12.59219 | 1.26229 | 0.00000 | 3.02990 | 1.45094 | 3.57819 | 0.17659 | 1.20159 | 0.30177 | 1.61668 | 1.32912 | 0.40419 |
| **PXMP2** | 0.35835 | 0.58253 | 1.13888 | 0.16832 | 2.63639 | 2.75964 | 0.00000 | 0.08688 | 1.49163 | 1.41746 | 1.70238 | 0.65425 |
| **PXMP4** | 0.00000 | 4.97024 | 1.17658 | 4.72251 | 6.76551 | 2.38515 | 1.98608 | 3.30459 | 1.67061 | 2.40909 | 3.94740 | 6.67745 |
| **PXN** | 1.42000 | 10.06411 | 11.13124 | 1.86326 | 6.87793 | 1.91751 | 12.82904 | 0.81545 | 13.28467 | 11.10600 | 10.59444 | 8.91142 |
| **PXN-AS1** | 0.30346 | 2.02107 | 5.68220 | 0.79964 | 1.59292 | 1.51182 | 5.32053 | 0.85852 | 4.28896 | 0.00000 | 5.35534 | 3.10593 |
| **PXT1** | 0.19709 | 0.23467 | 0.03752 | 0.00000 | 0.05960 | 0.21098 | 0.00000 | 0.23868 | 0.04249 | 0.00556 | 0.06933 | 0.03210 |
| **PXYLP1** | 5.62390 | 1.99674 | 0.00000 | 0.62165 | 0.89365 | 1.15005 | 0.00000 | 1.35461 | 0.00000 | 0.76548 | 1.35956 | 0.23109 |
| **PYCARD** | 0.03621 | 0.35789 | 0.32349 | 0.71842 | 0.81371 | 0.00000 | 0.01751 | 0.00759 | 0.00000 | 0.05351 | 0.01307 | 0.02640 |
| **PYCARDOS** | 0.32392 | 0.13857 | 0.02236 | 0.00378 | 0.01210 | 0.00000 | 0.00000 | 0.30438 | 0.04275 | 0.59065 | 0.01316 | 0.13806 |
| **PYCR1** | 0.46064 | 7.60262 | 1.65983 | 4.69633 | 7.61232 | 0.00000 | 15.36559 | 0.01927 | 13.06085 | 0.00000 | 11.43254 | 10.59709 |
| **PYCR2** | 1.73636 | 6.81722 | 9.09604 | 4.58762 | 2.87461 | 0.00000 | 8.71396 | 0.80405 | 4.37458 | 0.00000 | 8.56795 | 4.61892 |
| **PYCRL** | 0.00158 | 4.17856 | 2.01568 | 1.68987 | 3.61474 | 0.00000 | 0.17355 | 0.00072 | 1.49629 | 0.33546 | 2.35425 | 3.33600 |
| **PYDC1** | 0.00211 | 0.08358 | 0.01998 | 0.23476 | 0.49251 | 0.00000 | 0.05388 | 0.22691 | 0.00000 | 0.04426 | 0.00412 | 0.01079 |
| **PYDC2** | 0.00576 | 0.01654 | 0.04089 | 0.00759 | 0.01310 | 0.03432 | 0.00794 | 0.01008 | 0.00864 | 0.04093 | 0.01599 | 0.01532 |
| **PYGB** | 0.74550 | 38.90729 | 4.13141 | 45.73043 | 31.20699 | 0.00000 | 4.05364 | 1.04890 | 5.84818 | 13.37696 | 13.25736 | 18.81645 |
| **PYGL** | 0.00000 | 7.44861 | 10.32220 | 9.41809 | 3.07801 | 9.86037 | 10.43934 | 0.00641 | 4.55718 | 1.02390 | 12.27427 | 1.69385 |
| **PYGM** | 0.14475 | 0.00000 | 0.01515 | 0.11031 | 0.16660 | 0.00000 | 0.02469 | 0.40323 | 0.25403 | 0.98321 | 0.02732 | 0.25632 |

| | | | | | | | | | | | | |
|---|---|---|---|---|---|---|---|---|---|---|---|---|
| **PYGO1** | 0.04178 | 0.62414 | 2.02639 | 0.00000 | 1.12959 | 2.55847 | 1.58769 | 0.64950 | 1.33957 | 1.86519 | 1.79637 | 1.49292 |
| **PYGO2** | 2.94811 | 4.83928 | 5.95116 | 2.09427 | 5.43221 | 0.00000 | 6.73516 | 0.00000 | 6.10088 | 6.07540 | 3.41263 | 4.48741 |
| **PYHIN1** | 0.00601 | 0.01070 | 0.02376 | 0.04647 | 0.00859 | 0.04240 | 0.01040 | 0.01008 | 0.01299 | 0.02567 | 0.01317 | 0.01500 |
| **PYROXD1** | 0.30458 | 0.38519 | 1.66771 | 0.10652 | 0.08868 | 3.16129 | 1.40843 | 1.07354 | 3.67883 | 0.00000 | 0.58636 | 2.43457 |
| **PYROXD2** | 0.00424 | 0.10366 | 0.15194 | 0.00000 | 0.09024 | 0.00000 | 0.06671 | 0.00939 | 0.00000 | 0.11253 | 0.05610 | 0.02958 |
| **PYY** | 0.08157 | 0.00309 | 0.09488 | 0.15872 | 0.00865 | 0.04862 | 0.04796 | 0.19730 | 0.00633 | 0.00563 | 0.01393 | 1.09858 |
| **PYY2** | 0.05321 | 2.36184 | 1.70059 | 3.43858 | 0.50162 | 0.00000 | 0.81429 | 1.16369 | 0.00000 | 1.98540 | 0.02961 | 1.67905 |
| **PZP** | 0.00543 | 0.01231 | 0.03628 | 0.00539 | 0.00000 | 0.03735 | 0.11886 | 0.01003 | 0.02147 | 0.03230 | 0.01646 | 0.01997 |
| **QARS** | 4.95732 | 28.28909 | 21.49508 | 30.98049 | 34.69907 | 16.55504 | 30.81717 | 0.99525 | 36.10756 | 14.92468 | 32.38977 | 3.16027 |
| **QDPR** | 5.70035 | 14.31626 | 0.54302 | 5.18970 | 10.27500 | 2.53104 | 2.31181 | 1.03903 | 4.93547 | 0.88980 | 9.30861 | 3.13907 |
| **QKI** | 4.35986 | 0.00000 | 8.75282 | 2.01295 | 8.64755 | 21.47125 | 18.55081 | 3.58236 | 0.00000 | 23.22404 | 9.25102 | 3.19079 |
| **QPCT** | 0.00584 | 0.65980 | 1.08767 | 0.93113 | 0.00000 | 0.13030 | 0.00000 | 0.00350 | 0.00000 | 0.02916 | 0.00941 | 0.77428 |
| **QPCTL** | 0.19204 | 1.38384 | 0.96979 | 1.89876 | 2.49349 | 0.00000 | 0.03037 | 0.37264 | 4.31701 | 0.86231 | 1.90242 | 2.69739 |
| **QPRT** | 0.00120 | 0.00000 | 11.37852 | 3.41228 | 6.51448 | 0.00000 | 17.29398 | 0.06955 | 25.37492 | 0.00000 | 18.22987 | 0.33336 |
| **QRFP** | 0.00581 | 0.02253 | 0.03937 | 0.00716 | 0.00000 | 0.08108 | 0.06701 | 0.00883 | 0.07312 | 0.00741 | 0.00360 | 0.04244 |
| **QRFPR** | 0.00590 | 0.01903 | 0.05141 | 0.05225 | 0.04615 | 0.00000 | 0.01779 | 0.01099 | 0.02107 | 0.00000 | 0.01632 | 0.03662 |
| **QRICH1** | 7.28263 | 12.67689 | 15.34802 | 7.01593 | 6.21850 | 7.97603 | 8.63735 | 11.94892 | 13.87375 | 10.12840 | 11.18689 | 11.38638 |
| **QRICH2** | 0.02399 | 1.80936 | 0.26026 | 0.45272 | 0.63174 | 0.00000 | 0.32394 | 0.43282 | 0.45546 | 1.67888 | 0.72013 | 1.82096 |
| **QRSL1** | 1.09039 | 8.59442 | 6.81117 | 6.13087 | 5.41267 | 18.09158 | 8.39539 | 2.87310 | 9.67305 | 0.00000 | 10.06149 | 7.58206 |
| **QSER1** | 0.07394 | 0.00000 | 2.54298 | 0.75818 | 0.00000 | 1.87448 | 4.78016 | 0.12798 | 1.20265 | 1.84576 | 0.89848 | 0.20713 |
| **QSOX1** | 0.85738 | 4.61094 | 9.74765 | 2.10227 | 4.10058 | 0.00000 | 0.00000 | 0.29945 | 3.56080 | 2.10439 | 0.00000 | 2.92259 |
| **QSOX2** | 1.11592 | 0.00000 | 0.86874 | 0.00000 | 0.24033 | 2.40106 | 0.02059 | 0.37586 | 2.40063 | 7.96762 | 0.00000 | 0.27485 |
| **QTRT1** | 1.60325 | 4.94656 | 9.93262 | 8.46244 | 4.87385 | 0.00000 | 4.61784 | 5.43888 | 10.61749 | 2.85173 | 10.55614 | 7.64884 |
| **QTRTD1** | 3.52740 | 8.40519 | 8.51456 | 7.10498 | 4.20335 | 10.53028 | 12.12692 | 7.73842 | 9.88108 | 5.45606 | 8.72938 | 14.63482 |
| **R3HCC1** | 0.12439 | 9.81578 | 0.22673 | 2.91619 | 6.45251 | 0.00000 | 1.13513 | 0.41484 | 3.48983 | 0.00000 | 2.17287 | 3.83381 |
| **R3HCC1L** | 0.42694 | 6.72636 | 4.66026 | 4.43227 | 1.99720 | 5.41767 | 2.74742 | 12.46652 | 7.09912 | 2.96735 | 2.31045 | 8.33461 |
| **R3HDM1** | 2.33963 | 2.40229 | 3.03479 | 3.01631 | 1.10122 | 6.96347 | 4.93708 | 1.85953 | 2.20561 | 9.50245 | 5.53179 | 4.36298 |
| **R3HDM2** | 2.15547 | 1.36230 | 2.91794 | 3.67569 | 5.33020 | 5.90592 | 4.51401 | 2.83724 | 0.00000 | 5.22292 | 7.81761 | 1.48267 |
| **R3HDM4** | 14.60112 | 2.55585 | 4.60274 | 2.18647 | 5.98228 | 0.00000 | 2.52383 | 0.18427 | 6.11068 | 10.13572 | 1.10693 | 2.83746 |
| **R3HDML** | 0.00211 | 0.07817 | 0.01144 | 0.05351 | 0.05659 | 0.00000 | 0.02745 | 1.70533 | 0.00000 | 0.07227 | 0.00631 | 1.67084 |
| **RAB10** | 4.77138 | 19.18712 | 17.94248 | 9.52365 | 9.89256 | 22.20726 | 16.83198 | 16.78132 | 25.74155 | 5.24321 | 17.92934 | 16.56120 |
| **RAB11A** | 6.94792 | 16.95817 | 30.24009 | 14.80025 | 25.14208 | 53.31966 | 28.88385 | 20.39678 | 23.98527 | 12.67140 | 26.86497 | 26.95347 |
| **RAB11B** | 0.10815 | 3.02062 | 6.23449 | 1.77770 | 2.16961 | 0.00000 | 1.19552 | 0.86983 | 2.54831 | 7.03322 | 1.92695 | 2.01106 |
| **RAB11B-AS1** | 0.02147 | 0.28970 | 0.15025 | 0.09424 | 0.32092 | 0.39379 | 0.00000 | 0.22682 | 0.08998 | 0.00000 | 0.10478 | 0.20251 |
| **RAB11FIP1** | 0.00000 | 5.35910 | 22.17119 | 1.74978 | 17.65807 | 5.59085 | 31.90120 | 3.07607 | 16.46020 | 18.46238 | 27.86438 | 1.99296 |
| **RAB11FIP2** | 0.75434 | 0.00000 | 2.60018 | 0.44991 | 0.00000 | 8.12870 | 5.93073 | 2.01088 | 0.00000 | 13.44486 | 2.00391 | 2.01812 |
| **RAB11FIP3** | 0.58471 | 0.10341 | 0.20449 | 0.13100 | 1.94687 | 0.00000 | 3.35667 | 0.14390 | 0.00000 | 6.97200 | 0.00009 | 0.58366 |
| **RAB11FIP4** | 1.28375 | 5.71795 | 15.72477 | 0.31610 | 19.66020 | 5.19293 | 18.78887 | 0.06720 | 15.83248 | 13.24684 | 18.05444 | 0.10533 |
| **RAB11FIP5** | 0.78292 | 0.05134 | 3.33356 | 0.00000 | 0.00000 | 0.00000 | 3.03636 | 0.00000 | 5.72441 | 4.19946 | 0.48762 | 0.16396 |
| **RAB12** | 0.00000 | 0.00000 | 0.22241 | 0.73893 | 0.11068 | 2.30827 | 1.23014 | 3.00072 | 0.00000 | 5.85666 | 1.02870 | 1.02838 |
| **RAB13** | 0.56182 | 1.17486 | 1.07010 | 0.94437 | 1.13188 | 0.00000 | 1.39712 | 0.18879 | 0.57820 | 2.17910 | 0.22517 | 0.73770 |
| **RAB14** | 0.73328 | 6.45491 | 12.24113 | 5.69870 | 10.24047 | 23.86886 | 14.96216 | 2.38066 | 12.26346 | 10.32883 | 11.25773 | 9.31827 |
| **RAB15** | 0.42673 | 12.92625 | 2.44221 | 32.58425 | 19.94870 | 10.71805 | 1.53632 | 0.04510 | 0.00000 | 15.39684 | 1.00111 | 8.09342 |
| **RAB17** | 0.03035 | 0.01303 | 2.04596 | 0.00000 | 0.00000 | 0.00000 | 4.39208 | 0.03621 | 0.00000 | 0.01828 | 2.41311 | 0.00851 |
| **RAB18** | 1.76911 | 18.06884 | 11.81006 | 4.50387 | 9.18366 | 57.07788 | 15.92382 | 0.00000 | 5.72424 | 6.53488 | 24.47788 | 10.42806 |
| **RAB19** | 0.07825 | 0.28068 | 0.07655 | 0.13226 | 0.07738 | 0.00000 | 0.08776 | 0.01459 | 0.00000 | 0.00000 | 0.06781 | 0.05469 |
| **RAB1A** | 6.17879 | 20.99815 | 23.45454 | 16.49740 | 13.86733 | 50.22092 | 27.71282 | 22.98976 | 26.02987 | 3.49419 | 21.14344 | 28.13526 |
| **RAB1B** | 1.97975 | 21.71059 | 14.19948 | 14.72228 | 16.41357 | 0.00132 | 17.84475 | 3.57575 | 27.37290 | 0.00000 | 12.44818 | 15.73104 |

| | | | | | | | | | | | | |
|---|---|---|---|---|---|---|---|---|---|---|---|---|
| **RAB20** | 2.34569 | 1.21630 | 11.44217 | 0.00000 | 6.79380 | 2.78220 | 8.03045 | 0.67974 | 8.21126 | 0.00000 | 8.41238 | 1.11863 |
| **RAB21** | 1.25930 | 0.22091 | 3.26003 | 0.27272 | 0.56628 | 9.23302 | 5.80145 | 2.67211 | 0.00000 | 9.41043 | 1.28898 | 3.17269 |
| **RAB22A** | 3.66220 | 1.24346 | 2.60755 | 0.61311 | 0.00000 | 5.12307 | 1.83978 | 3.02315 | 1.23520 | 3.02585 | 0.00506 | 2.67549 |
| **RAB23** | 1.06792 | 0.75335 | 2.29684 | 2.59778 | 0.00000 | 9.37387 | 5.45627 | 0.75283 | 0.00000 | 4.93464 | 0.45817 | 4.00087 |
| **RAB24** | 0.00089 | 4.65753 | 9.26359 | 4.02188 | 5.77793 | 0.00000 | 2.29900 | 4.02016 | 3.14530 | 0.00000 | 4.81199 | 5.71122 |
| **RAB25** | 1.95032 | 3.48586 | 9.06598 | 2.40121 | 46.72124 | 6.51586 | 9.36077 | 0.12778 | 21.65754 | 0.60381 | 22.74123 | 0.00000 |
| **RAB26** | 0.42371 | 0.00242 | 0.56318 | 0.19996 | 0.70739 | 0.00000 | 0.00000 | 0.00472 | 1.12423 | 0.48069 | 0.13931 | 0.14001 |
| **RAB27A** | 2.20672 | 1.73643 | 2.20514 | 2.09312 | 0.05652 | 1.12427 | 0.70710 | 0.87373 | 0.00000 | 0.00000 | 0.97336 | 0.57869 |
| **RAB27B** | 0.61647 | 0.01417 | 0.00538 | 0.04238 | 0.00344 | 0.00000 | 0.01288 | 0.30413 | 0.05331 | 0.07474 | 0.01283 | 0.05723 |
| **RAB28** | 2.25092 | 2.14074 | 1.92396 | 1.20782 | 0.00000 | 7.17830 | 2.60117 | 1.12830 | 1.67247 | 0.70234 | 1.97404 | 3.18135 |
| **RAB29** | 4.69482 | 0.78807 | 2.83488 | 1.93618 | 0.00000 | 0.00000 | 4.04389 | 0.00000 | 0.80707 | 0.00000 | 2.46977 | 20.30797 |
| **RAB2A** | 2.48765 | 17.83579 | 21.11403 | 7.99096 | 10.61775 | 19.98215 | 9.88081 | 6.18154 | 14.06055 | 15.84593 | 12.91686 | 8.08468 |
| **RAB2B** | 1.77214 | 0.29285 | 2.76871 | 1.04102 | 0.78174 | 5.13409 | 2.64494 | 3.34255 | 1.04573 | 4.45654 | 2.72723 | 6.51842 |
| **RAB30** | 1.90677 | 2.45236 | 1.80410 | 0.00000 | 0.97190 | 5.94906 | 2.41046 | 0.13981 | 0.80906 | 2.36726 | 1.98159 | 8.22616 |
| **RAB30-AS1** | 0.06781 | 2.91879 | 2.71434 | 0.24515 | 1.19652 | 3.79194 | 1.28760 | 0.00000 | 0.00000 | 2.42195 | 2.25487 | 11.56367 |
| **RAB31** | 0.04595 | 0.00000 | 23.10818 | 0.25949 | 17.70395 | 0.94418 | 27.68632 | 0.02490 | 8.72336 | 0.25874 | 14.95291 | 0.00000 |
| **RAB32** | 0.00692 | 0.30490 | 0.05803 | 0.02056 | 0.13418 | 0.00000 | 0.18507 | 0.01342 | 0.03858 | 0.00000 | 0.02685 | 0.01201 |
| **RAB33A** | 0.00619 | 0.01121 | 0.01933 | 0.01791 | 0.01574 | 0.04766 | 0.02651 | 0.01113 | 0.02914 | 0.00000 | 0.01037 | 0.02354 |
| **RAB33B** | 1.12238 | 0.66435 | 1.90712 | 0.34636 | 0.00000 | 3.20309 | 3.92599 | 4.59645 | 0.74259 | 2.89151 | 1.85205 | 3.02314 |
| **RAB34** | 0.20808 | 0.00000 | 10.53298 | 1.47021 | 0.00000 | 0.00000 | 7.26475 | 0.18549 | 3.80481 | 0.00000 | 1.13404 | 0.08747 |
| **RAB35** | 2.31088 | 11.23766 | 10.57640 | 9.45762 | 16.92973 | 0.00000 | 15.85738 | 8.63332 | 15.02823 | 0.00002 | 9.47999 | 7.03954 |
| **RAB36** | 10.63738 | 0.00000 | 2.18982 | 0.00000 | 0.22804 | 0.00000 | 1.03655 | 1.04150 | 4.77740 | 0.04468 | 1.69893 | 0.23100 |
| **RAB37** | 0.00984 | 0.55937 | 0.00241 | 0.13783 | 0.00000 | 0.00000 | 0.10504 | 0.12296 | 0.14788 | 0.04986 | 1.82579 | 0.14387 |
| **RAB38** | 0.13221 | 0.15129 | 1.11171 | 0.00384 | 0.00000 | 0.31322 | 0.16277 | 0.03641 | 0.55293 | 0.00000 | 0.09352 | 0.04968 |
| **RAB39A** | 0.00206 | 0.09261 | 1.51664 | 0.17273 | 0.00000 | 1.47235 | 0.42787 | 1.25548 | 1.58933 | 0.00000 | 1.18263 | 0.71321 |
| **RAB39B** | 0.02423 | 0.00000 | 0.99387 | 0.18139 | 0.00000 | 0.03843 | 0.83008 | 0.30489 | 0.52191 | 0.00000 | 0.00000 | 0.34301 |
| **RAB3A** | 3.37119 | 0.62613 | 0.07329 | 0.09382 | 0.69129 | 0.00000 | 0.00860 | 0.55495 | 0.09257 | 0.52600 | 0.11273 | 0.18119 |
| **RAB3B** | 0.00000 | 0.00000 | 3.96360 | 28.51777 | 13.43528 | 1.21860 | 7.05275 | 0.07949 | 9.51037 | 9.88604 | 10.76878 | 0.00000 |
| **RAB3C** | 0.13943 | 0.17211 | 0.11096 | 0.31631 | 0.01903 | 0.00000 | 0.00942 | 0.00905 | 0.02546 | 0.03843 | 0.01434 | 0.07202 |
| **RAB3D** | 1.68537 | 9.11181 | 0.00000 | 5.55082 | 1.36916 | 4.73526 | 1.56368 | 7.22402 | 0.73959 | 0.39427 | 0.09732 | 7.46134 |
| **RAB3GAP1** | 3.56071 | 3.36883 | 3.77996 | 4.82705 | 0.00000 | 8.64985 | 6.86958 | 0.19537 | 3.34244 | 4.28021 | 2.18290 | 4.10913 |
| **RAB3GAP2** | 5.84712 | 2.81119 | 3.07662 | 0.77939 | 0.40286 | 8.88725 | 3.13473 | 0.64490 | 3.21417 | 6.80175 | 3.07591 | 1.70936 |
| **RAB3IL1** | 0.21199 | 0.26338 | 0.92546 | 0.31607 | 0.32212 | 0.00000 | 0.45289 | 0.01617 | 0.68619 | 1.17266 | 0.00000 | 0.52070 |
| **RAB3IP** | 0.00000 | 10.03746 | 2.88259 | 9.10637 | 6.82883 | 19.75262 | 5.00289 | 4.94309 | 0.00000 | 11.65793 | 5.92308 | 6.32754 |
| **RAB40A** | 0.01384 | 0.06205 | 0.06199 | 0.03164 | 0.00000 | 0.00626 | 0.18065 | 0.00792 | 0.21084 | 0.00000 | 0.04620 | 0.05875 |
| **RAB40AL** | 0.00330 | 0.00902 | 0.03869 | 0.01402 | 0.02459 | 0.00000 | 0.00123 | 0.00826 | 0.05833 | 0.33349 | 0.03482 | 0.03447 |
| **RAB40B** | 0.32605 | 1.07278 | 0.00000 | 0.52578 | 0.12787 | 0.21696 | 0.94740 | 0.96571 | 0.95681 | 0.48539 | 0.22256 | 1.30877 |
| **RAB40C** | 1.16901 | 2.97613 | 2.15556 | 0.00000 | 2.95524 | 0.00000 | 2.46290 | 0.00000 | 2.11550 | 2.61299 | 0.86319 | 0.68979 |
| **RAB41** | 0.00539 | 0.03761 | 0.00412 | 0.10682 | 0.08814 | 0.03805 | 0.00582 | 0.03773 | 0.08063 | 0.00000 | 0.01879 | 0.04826 |
| **RAB42** | 0.51910 | 1.30985 | 0.39867 | 0.44986 | 0.18238 | 0.08134 | 0.08933 | 0.14130 | 0.00000 | 0.21829 | 0.36479 | 2.79718 |
| **RAB43** | 0.05386 | 2.02184 | 0.07973 | 0.06180 | 0.30657 | 0.00000 | 0.00408 | 0.16359 | 0.13293 | 0.02452 | 0.04659 | 1.02423 |
| **RAB44** | 0.01048 | 0.01072 | 0.01583 | 0.00799 | 0.01845 | 0.03173 | 0.02112 | 0.00961 | 0.00923 | 0.02503 | 0.01675 | 0.01910 |
| **RAB4A** | 0.94191 | 1.25328 | 3.61165 | 3.78020 | 4.12044 | 2.18421 | 5.61553 | 0.38595 | 5.34372 | 2.60547 | 4.22469 | 1.88209 |
| **RAB5A** | 0.71874 | 10.30345 | 6.49092 | 5.71615 | 1.72560 | 25.68147 | 8.80969 | 14.19703 | 5.14284 | 9.94446 | 4.75789 | 20.97201 |
| **RAB5B** | 9.53695 | 1.25753 | 4.59515 | 3.32163 | 4.15452 | 4.74479 | 6.45288 | 0.86852 | 3.28639 | 6.25671 | 2.49036 | 2.16758 |
| **RAB5C** | 7.52237 | 25.44249 | 20.11163 | 15.81156 | 24.26615 | 3.93016 | 20.83100 | 11.01560 | 33.69255 | 0.00000 | 23.36475 | 24.57890 |
| **RAB6A** | 1.46186 | 21.82060 | 16.06098 | 11.41541 | 4.52296 | 29.93416 | 13.62211 | 20.27237 | 16.86376 | 0.61401 | 13.22614 | 21.81037 |
| **RAB6B** | 2.57735 | 1.35068 | 0.44788 | 0.72827 | 0.00000 | 0.00000 | 0.02737 | 0.10667 | 0.78702 | 0.01002 | 0.24774 | 0.67806 |

| | | | | | | | | | | | | |
|---|---|---|---|---|---|---|---|---|---|---|---|---|
| **RAB6C** | 0.02615 | 0.05301 | 0.04393 | 0.03403 | 0.01034 | 0.04559 | 0.01588 | 0.05416 | 0.02132 | 0.01393 | 0.02886 | 0.03357 |
| **RAB6C-AS1** | 0.02027 | 0.00913 | 0.01628 | 0.01748 | 0.00749 | 0.03234 | 0.00246 | 0.00927 | 0.02629 | 0.03903 | 0.03445 | 0.05006 |
| **RAB7A** | 6.39404 | 25.55572 | 24.27243 | 22.42434 | 18.44501 | 33.28939 | 30.33339 | 13.06427 | 29.28439 | 2.99201 | 24.77477 | 27.30342 |
| **RAB8A** | 3.29247 | 12.00260 | 17.67216 | 16.62357 | 25.74282 | 11.08604 | 32.36038 | 4.72617 | 26.20713 | 0.00000 | 25.96975 | 10.18343 |
| **RAB8B** | 2.80280 | 1.51663 | 10.47440 | 2.05448 | 0.00000 | 6.53183 | 3.85187 | 5.38900 | 0.00000 | 0.00000 | 1.43965 | 3.33776 |
| **RAB9A** | 0.00000 | 5.95467 | 9.80052 | 0.97699 | 1.82654 | 16.22342 | 9.95151 | 3.66368 | 5.22055 | 0.00000 | 9.25380 | 14.62426 |
| **RAB9B** | 0.00475 | 0.00000 | 0.00000 | 0.10169 | 0.00000 | 0.13143 | 0.18482 | 0.00723 | 0.01799 | 0.07085 | 0.00000 | 0.03700 |
| **RAB9BP1** | 0.00832 | 0.00826 | 0.03814 | 0.00357 | 0.00000 | 0.09321 | 0.00000 | 0.00863 | 0.09767 | 0.06892 | 0.00087 | 0.03453 |
| **RABAC1** | 6.17456 | 13.67734 | 6.66491 | 18.27560 | 13.70032 | 4.56909 | 2.78933 | 4.25400 | 13.22879 | 6.88192 | 4.93666 | 12.00388 |
| **RABEP1** | 7.46750 | 8.74045 | 8.07808 | 1.77942 | 8.99570 | 11.54446 | 5.54301 | 7.63516 | 10.17471 | 10.90628 | 9.89273 | 16.35166 |
| **RABEP2** | 0.00260 | 0.87489 | 1.86681 | 1.36909 | 1.48237 | 0.00000 | 2.09188 | 1.60677 | 2.64318 | 0.59435 | 1.51507 | 3.06793 |
| **RABEPK** | 0.36012 | 13.01420 | 15.82392 | 7.49012 | 11.55118 | 5.66588 | 23.93844 | 1.44321 | 22.63624 | 0.00000 | 10.37775 | 8.62177 |
| **RABGAP1** | 5.87927 | 0.79644 | 2.14699 | 1.80190 | 0.77320 | 5.33294 | 1.73578 | 1.44989 | 1.26037 | 8.05429 | 1.70489 | 0.36100 |
| **RABGAP1L** | 2.68157 | 0.08700 | 1.16046 | 0.00000 | 0.00064 | 5.22578 | 0.78792 | 0.49354 | 2.34122 | 1.01776 | 0.05185 | 2.54161 |
| **RABGEF1** | 3.40476 | 14.79438 | 8.48200 | 4.73230 | 2.44896 | 13.46853 | 4.37039 | 27.85534 | 6.56985 | 0.00000 | 8.37582 | 19.25062 |
| **RABGGTA** | 0.45300 | 2.72135 | 8.11840 | 5.51308 | 3.54029 | 0.01512 | 4.80018 | 0.08693 | 10.96020 | 0.00000 | 5.46331 | 2.08515 |
| **RABGGTB** | 2.70777 | 13.85055 | 9.09857 | 9.02866 | 8.27516 | 39.70148 | 21.00432 | 2.55114 | 25.46487 | 27.17025 | 6.60890 | 16.16783 |
| **RABIF** | 2.52571 | 11.38764 | 10.15488 | 9.22097 | 11.00397 | 14.98124 | 15.10918 | 12.08173 | 5.34409 | 1.89202 | 12.08604 | 19.86096 |
| **RABL2A** | 0.12942 | 0.50700 | 0.20473 | 0.01479 | 0.00941 | 0.22481 | 0.09651 | 0.14508 | 0.37357 | 0.00593 | 0.23755 | 0.26879 |
| **RABL2B** | 1.32427 | 1.47893 | 0.61957 | 0.72307 | 1.10422 | 0.03756 | 0.82359 | 0.04297 | 1.05014 | 0.00000 | 1.75212 | 1.41308 |
| **RABL3** | 0.48043 | 1.74688 | 5.19365 | 0.76693 | 0.35130 | 10.39837 | 5.54444 | 0.79649 | 1.34511 | 1.45515 | 3.98791 | 4.60444 |
| **RABL6** | 0.22498 | 0.00000 | 3.20363 | 1.24069 | 6.66447 | 0.00000 | 8.90618 | 0.07206 | 0.00000 | 19.84366 | 0.67712 | 0.26787 |
| **RAC1** | 3.84954 | 6.99253 | 12.28368 | 5.87206 | 9.55580 | 5.69992 | 29.39288 | 3.06785 | 5.26838 | 28.45099 | 5.87909 | 2.21231 |
| **RAC2** | 5.52419 | 1.28540 | 0.00000 | 0.75100 | 0.97983 | 0.00000 | 0.01768 | 0.69679 | 0.00000 | 0.18657 | 0.38088 | 0.01031 |
| **RAC3** | 0.10929 | 0.19763 | 2.88945 | 0.24405 | 0.57372 | 0.00000 | 0.34176 | 0.01737 | 0.73342 | 0.01147 | 0.67999 | 0.00265 |
| **RACGAP1** | 8.55542 | 8.74416 | 9.52823 | 2.66022 | 5.76295 | 19.86973 | 9.90085 | 20.82051 | 14.52942 | 1.32897 | 13.33187 | 11.85021 |
| **RACGAP1P** | 0.67553 | 0.53651 | 0.53542 | 0.06490 | 0.21665 | 1.39257 | 0.00000 | 2.87265 | 1.22508 | 0.00000 | 0.73055 | 0.73987 |
| **RAD1** | 2.77308 | 8.88751 | 5.53925 | 3.58075 | 5.51183 | 24.22803 | 8.53953 | 8.39315 | 9.11501 | 4.35452 | 7.39970 | 10.38964 |
| **RAD17** | 2.80244 | 4.55626 | 2.32205 | 1.46322 | 1.16559 | 15.87565 | 2.04769 | 3.70333 | 5.77350 | 3.29863 | 4.94903 | 7.04474 |
| **RAD18** | 1.86682 | 4.24474 | 4.48172 | 2.28726 | 0.00159 | 10.72950 | 1.28203 | 3.26642 | 6.89414 | 7.88684 | 4.18334 | 6.87780 |
| **RAD21** | 4.61660 | 4.92217 | 12.15082 | 6.58967 | 5.33628 | 34.09314 | 20.98909 | 10.39244 | 4.21314 | 33.31064 | 7.98616 | 14.26545 |
| **RAD21-AS1** | 0.00583 | 0.04196 | 0.04499 | 0.01091 | 0.08760 | 0.02055 | 0.00514 | 0.03861 | 0.16227 | 0.00000 | 0.00586 | 0.08574 |
| **RAD21L1** | 0.00418 | 0.09862 | 0.01469 | 0.00449 | 0.00122 | 0.03283 | 0.00000 | 0.57719 | 0.01629 | 0.08219 | 0.01187 | 0.02114 |
| **RAD23A** | 1.77932 | 16.88944 | 21.81388 | 16.92532 | 20.61387 | 0.49377 | 25.28091 | 2.00886 | 18.11875 | 5.56264 | 19.52291 | 18.31494 |
| **RAD23B** | 0.25023 | 0.00000 | 6.91127 | 0.00000 | 4.25776 | 9.35692 | 15.90563 | 0.47089 | 0.00000 | 31.09254 | 20.79487 | 4.01893 |
| **RAD50** | 7.95943 | 2.01956 | 3.82567 | 2.76301 | 1.11728 | 11.39027 | 6.54874 | 5.14631 | 0.58656 | 17.84911 | 11.34397 | 4.44234 |
| **RAD51** | 2.05338 | 24.18960 | 10.95499 | 21.83806 | 18.26965 | 26.26329 | 8.42949 | 10.37331 | 11.08057 | 5.29869 | 16.75305 | 19.68892 |
| **RAD51-AS1** | 0.13537 | 0.58474 | 0.98233 | 0.98149 | 1.10683 | 0.00000 | 0.00000 | 0.97958 | 0.56702 | 1.61785 | 0.49826 | 1.58315 |
| **RAD51AP1** | 3.90547 | 5.56252 | 5.54601 | 1.76434 | 0.65720 | 20.61300 | 6.46787 | 6.28308 | 4.83842 | 10.53424 | 6.26377 | 9.55539 |
| **RAD51B** | 1.66685 | 0.35890 | 0.38813 | 0.63915 | 0.72628 | 1.11849 | 0.38843 | 0.34805 | 0.00000 | 0.00000 | 1.73605 | 0.97007 |
| **RAD51C** | 2.74105 | 10.46183 | 8.71249 | 2.81781 | 2.58835 | 18.14543 | 9.72026 | 5.41317 | 12.01660 | 2.80419 | 11.17566 | 14.44064 |
| **RAD51D** | 0.20063 | 3.32527 | 0.95622 | 1.95899 | 0.70307 | 3.01280 | 0.00000 | 0.16958 | 2.01470 | 1.30697 | 1.81956 | 1.93872 |
| **RAD51L3-RFFL** | 0.00589 | 0.01344 | 0.01724 | 0.01594 | 0.01188 | 0.02529 | 0.01786 | 0.01029 | 0.03331 | 0.01878 | 0.01154 | 0.02104 |
| **RAD52** | 0.00898 | 4.20897 | 0.00000 | 0.77717 | 0.00000 | 5.31460 | 0.00000 | 4.71817 | 1.73160 | 2.32751 | 3.12548 | 4.59006 |
| **RAD54B** | 1.04265 | 2.12596 | 1.84034 | 1.45325 | 0.00000 | 11.64304 | 1.81013 | 1.78966 | 2.36635 | 1.34318 | 1.83832 | 3.99400 |
| **RAD54L** | 1.27357 | 8.81109 | 5.22293 | 10.11950 | 13.17395 | 0.53071 | 1.04453 | 9.03263 | 13.12843 | 8.25297 | 5.14168 | 5.83079 |
| **RAD54L2** | 7.19954 | 5.00436 | 1.22685 | 5.89088 | 7.29528 | 0.00000 | 4.71087 | 1.54346 | 4.04009 | 10.45056 | 3.04778 | 6.74791 |
| **RAD9A** | 0.83476 | 1.00284 | 1.53029 | 1.17010 | 2.77596 | 0.00000 | 0.48708 | 0.18541 | 2.12243 | 3.03248 | 0.82704 | 0.89242 |

| | | | | | | | | | | | | |
|---|---|---|---|---|---|---|---|---|---|---|---|---|
| **RAD9B** | 0.22925 | 0.19642 | 0.00000 | 0.02542 | 0.03719 | 0.19698 | 0.00000 | 0.12639 | 0.08113 | 0.25657 | 0.00000 | 0.52633 |
| **RADIL** | 3.18704 | 0.50846 | 0.05513 | 0.74129 | 0.55906 | 0.15850 | 0.08347 | 0.24787 | 0.06406 | 0.03891 | 0.00000 | 0.53495 |
| **RAE1** | 11.84312 | 19.24345 | 16.47557 | 7.41719 | 10.03135 | 20.56424 | 19.16701 | 11.13307 | 17.63145 | 0.00037 | 12.03109 | 23.48638 |
| **RAET1E** | 0.00339 | 0.04713 | 0.00000 | 0.08046 | 0.00000 | 0.04866 | 0.00000 | 0.00939 | 0.00000 | 0.34896 | 0.00013 | 0.00443 |
| **RAET1G** | 0.00842 | 0.01793 | 0.01642 | 0.01757 | 0.02432 | 0.00212 | 0.03031 | 0.01443 | 0.02489 | 0.00000 | 0.01577 | 0.03531 |
| **RAET1K** | 0.14980 | 0.01269 | 0.02377 | 0.00984 | 0.01321 | 0.03031 | 0.02544 | 0.03187 | 0.01433 | 0.01123 | 0.01818 | 0.01217 |
| **RAF1** | 0.85681 | 20.38418 | 17.07415 | 15.04925 | 9.78769 | 21.97723 | 14.61753 | 10.71720 | 27.64937 | 0.43559 | 12.66676 | 17.59319 |
| **RAG1** | 0.00323 | 0.05491 | 0.00000 | 0.09332 | 0.00000 | 0.23147 | 0.20455 | 0.12885 | 0.07668 | 0.01584 | 0.00389 | 0.11965 |
| **RAG2** | 0.06242 | 0.01219 | 0.00666 | 0.00654 | 0.01042 | 0.07408 | 0.00679 | 0.00371 | 0.03218 | 0.01291 | 0.00774 | 0.05096 |
| **RAI1** | 0.41013 | 0.03298 | 0.24393 | 0.09150 | 0.00000 | 0.02258 | 0.00000 | 0.05271 | 0.00000 | 0.58521 | 0.01562 | 0.06416 |
| **RAI14** | 4.11963 | 0.68605 | 9.94771 | 2.66575 | 13.74249 | 6.64902 | 13.14741 | 0.81866 | 12.18512 | 10.26538 | 11.21704 | 0.00000 |
| **RAI2** | 0.03500 | 0.17672 | 0.00000 | 0.07709 | 0.00000 | 0.09170 | 0.00025 | 0.00556 | 0.00000 | 0.01020 | 0.20523 | 0.04805 |
| **RALA** | 7.45461 | 7.62015 | 11.69006 | 10.01006 | 9.22171 | 12.31578 | 20.66786 | 1.43179 | 11.78247 | 4.64322 | 10.52702 | 6.63023 |
| **RALB** | 6.09044 | 0.80342 | 2.05613 | 1.16439 | 0.65983 | 5.62880 | 3.60820 | 1.23313 | 1.18631 | 0.00000 | 1.11119 | 0.64341 |
| **RALBP1** | 4.59295 | 6.61380 | 22.60730 | 5.81374 | 20.32890 | 12.68644 | 36.03692 | 7.43135 | 15.68817 | 25.09288 | 14.34950 | 8.24613 |
| **RALGAPA1** | 0.00475 | 0.00114 | 0.14441 | 0.21581 | 0.00007 | 0.30581 | 0.12967 | 0.00712 | 0.10497 | 0.62147 | 0.00000 | 0.08041 |
| **RALGAPA1P** | 0.00583 | 0.01389 | 0.01216 | 0.01074 | 0.01782 | 0.01435 | 0.01712 | 0.01004 | 0.02063 | 0.03083 | 0.01441 | 0.02242 |
| **RALGAPA2** | 0.47307 | 0.20062 | 6.94672 | 0.01475 | 6.42601 | 4.17371 | 7.07934 | 0.27580 | 9.79026 | 10.69081 | 5.33122 | 0.16434 |
| **RALGAPB** | 5.54181 | 2.26832 | 2.55889 | 5.98136 | 2.80614 | 8.92742 | 3.64807 | 1.97350 | 0.00000 | 12.36350 | 1.68414 | 3.51848 |
| **RALGDS** | 0.99122 | 0.21690 | 0.73363 | 0.13569 | 2.60100 | 0.00000 | 1.05435 | 0.03675 | 2.48680 | 1.99691 | 1.86630 | 0.07142 |
| **RALGPS1** | 1.35520 | 0.50040 | 0.44142 | 1.17593 | 2.26762 | 1.47412 | 1.00271 | 0.01182 | 1.58128 | 1.08078 | 1.29392 | 0.00000 |
| **RALGPS2** | 1.33022 | 3.64808 | 0.00000 | 3.24760 | 2.94051 | 10.64669 | 0.00000 | 0.96398 | 0.00000 | 6.82027 | 0.00000 | 2.81518 |
| **RALY** | 5.34846 | 9.25803 | 14.71758 | 4.64969 | 8.61627 | 0.00000 | 7.47573 | 8.28797 | 7.85025 | 16.20402 | 7.24685 | 6.66518 |
| **RALY-AS1** | 0.31282 | 0.54820 | 0.56643 | 0.04210 | 0.51566 | 0.55249 | 0.37564 | 1.06363 | 0.09297 | 0.73741 | 0.48707 | 1.57456 |
| **RALYL** | 0.78319 | 0.01899 | 0.01587 | 0.00876 | 0.03547 | 0.00779 | 0.05342 | 0.00316 | 0.01029 | 0.00000 | 0.01373 | 0.04960 |
| **RAMP1** | 0.12701 | 1.07935 | 0.00000 | 0.67087 | 0.75245 | 0.00000 | 0.00000 | 0.97497 | 0.00000 | 0.45896 | 0.11759 | 1.87439 |
| **RAMP2** | 0.00443 | 0.01517 | 0.01872 | 0.00991 | 0.00546 | 0.01822 | 0.00120 | 0.00931 | 0.03999 | 0.09511 | 0.00697 | 0.05249 |
| **RAMP2-AS1** | 0.03136 | 0.01562 | 0.03725 | 0.04355 | 0.01664 | 0.01731 | 0.01179 | 0.06753 | 0.02414 | 0.02054 | 0.01316 | 0.01334 |
| **RAN** | 9.80813 | 34.84328 | 32.85751 | 26.70354 | 28.57280 | 58.22623 | 47.70800 | 27.42171 | 41.15990 | 12.33202 | 31.02343 | 39.35139 |
| **RANBP1** | 0.66219 | 15.64885 | 15.21118 | 9.31026 | 13.07577 | 27.34378 | 26.57386 | 15.33232 | 11.99613 | 17.71213 | 14.27259 | 11.77348 |
| **RANBP10** | 0.00841 | 3.02268 | 4.33459 | 1.56870 | 4.26990 | 0.43154 | 2.04652 | 1.65052 | 3.83305 | 4.29457 | 2.34084 | 2.27114 |
| **RANBP17** | 0.35891 | 1.31506 | 1.06932 | 1.10353 | 1.07703 | 2.26861 | 0.00000 | 0.38518 | 2.13344 | 0.00014 | 0.72299 | 1.13581 |
| **RANBP2** | 3.42713 | 11.17141 | 7.15612 | 5.29337 | 2.98473 | 37.82472 | 2.68493 | 6.79314 | 6.71256 | 44.84513 | 3.16603 | 10.92330 |
| **RANBP3** | 2.72229 | 12.23677 | 11.85981 | 9.67675 | 10.33034 | 0.00000 | 7.34356 | 4.54837 | 12.63029 | 0.00000 | 11.73016 | 11.43626 |
| **RANBP6** | 0.20456 | 2.22669 | 0.57519 | 0.17475 | 0.00000 | 6.34455 | 0.00000 | 0.11158 | 0.68444 | 1.71680 | 1.99227 | 3.09378 |
| **RANBP9** | 4.93554 | 0.61249 | 7.07781 | 2.28430 | 4.93808 | 9.84682 | 4.92701 | 1.32109 | 0.06565 | 11.28768 | 0.46544 | 1.21613 |
| **RANGAP1** | 5.93019 | 32.29551 | 19.20929 | 18.18074 | 29.78623 | 0.00000 | 36.90149 | 0.00000 | 25.04985 | 30.25045 | 13.53409 | 29.50965 |
| **RANGRF** | 0.02032 | 0.94011 | 4.48230 | 1.89191 | 1.59289 | 2.49414 | 7.89517 | 0.33481 | 3.58719 | 2.60196 | 6.20140 | 0.56791 |
| **RAP1A** | 0.32871 | 10.31136 | 2.07197 | 5.77780 | 0.26371 | 15.53638 | 2.24113 | 1.05672 | 0.00000 | 13.01001 | 0.81616 | 1.87644 |
| **RAP1B** | 2.78839 | 6.29902 | 5.93818 | 3.11549 | 0.43041 | 16.30667 | 3.67809 | 14.01660 | 3.75058 | 5.16380 | 4.06184 | 12.82428 |
| **RAP1GAP** | 0.60664 | 0.26596 | 0.26523 | 0.00000 | 0.09495 | 0.00000 | 0.05185 | 0.00000 | 0.20733 | 0.28860 | 0.00000 | 0.14979 |
| **RAP1GAP2** | 1.74958 | 1.19590 | 0.43499 | 1.61472 | 4.24538 | 0.00000 | 0.12843 | 0.12973 | 0.00000 | 0.45360 | 1.10091 | 0.64010 |
| **RAP1GDS1** | 0.85605 | 3.99853 | 9.62914 | 1.92889 | 0.00000 | 15.33448 | 15.32979 | 10.29400 | 17.97160 | 0.00095 | 13.71053 | 9.49909 |
| **RAP2A** | 0.81200 | 0.00000 | 2.75736 | 0.00000 | 0.00000 | 7.22017 | 6.13774 | 3.18500 | 0.00000 | 10.30907 | 0.60401 | 2.42645 |
| **RAP2B** | 1.65074 | 17.45317 | 11.56050 | 12.75077 | 5.58177 | 33.03162 | 18.51945 | 5.62295 | 9.35942 | 13.96461 | 16.19187 | 26.56374 |
| **RAP2C** | 1.63263 | 0.31596 | 3.07588 | 1.61372 | 0.00000 | 11.45165 | 2.51450 | 0.29377 | 0.19410 | 3.08343 | 2.33793 | 0.34605 |
| **RAP2C-AS1** | 0.00548 | 0.09862 | 0.03973 | 0.00352 | 0.08938 | 0.03055 | 0.03613 | 0.03999 | 0.00000 | 0.00000 | 0.03525 | 0.01677 |
| **RAPGEF1** | 0.51110 | 0.24075 | 12.02585 | 6.20457 | 7.03534 | 0.00000 | 8.95626 | 0.15137 | 4.34301 | 6.86793 | 3.44700 | 0.22195 |

| | | | | | | | | | | | | |
|---|---|---|---|---|---|---|---|---|---|---|---|---|
| **RAPGEF2** | 1.84112 | 3.47136 | 11.59296 | 2.11290 | 17.55709 | 12.64910 | 9.06103 | 2.29624 | 4.70869 | 12.48199 | 6.61559 | 2.19005 |
| **RAPGEF3** | 0.13431 | 0.03035 | 0.04043 | 0.00553 | 0.00000 | 0.03136 | 0.16316 | 0.01626 | 0.16507 | 0.00000 | 0.03127 | 0.05310 |
| **RAPGEF4** | 0.00681 | 0.42256 | 0.09878 | 0.04486 | 0.00000 | 0.06214 | 0.00000 | 0.08931 | 0.01890 | 0.03344 | 0.04216 | 0.06355 |
| **RAPGEF4-AS1** | 0.00593 | 0.01228 | 0.01585 | 0.01108 | 0.01194 | 0.02690 | 0.01983 | 0.01011 | 0.01677 | 0.01676 | 0.01443 | 0.02523 |
| **RAPGEF5** | 0.15974 | 0.03843 | 0.02565 | 0.05769 | 0.10975 | 0.00000 | 0.04863 | 0.02806 | 0.00000 | 0.05388 | 0.01092 | 0.02096 |
| **RAPGEF6** | 4.71632 | 0.85557 | 2.85602 | 0.86008 | 0.76217 | 8.95146 | 4.07929 | 0.60356 | 5.05797 | 8.90889 | 4.21088 | 0.77750 |
| **RAPGEFL1** | 0.02525 | 0.08587 | 0.44643 | 0.34221 | 0.65420 | 0.00000 | 0.00023 | 0.00000 | 1.75145 | 1.24011 | 0.31976 | 1.17112 |
| **RAPH1** | 3.03451 | 2.95478 | 2.77260 | 2.61391 | 1.26845 | 6.77266 | 3.29840 | 7.68836 | 0.00000 | 8.23107 | 2.12298 | 4.48398 |
| **RAPSN** | 0.00474 | 0.00345 | 0.00370 | 0.18660 | 0.03185 | 0.00000 | 0.40699 | 0.00667 | 0.18231 | 0.17198 | 0.00000 | 0.08188 |
| **RARA** | 0.68705 | 0.38673 | 2.17467 | 0.00000 | 0.50419 | 0.00000 | 0.52046 | 0.63474 | 0.93904 | 1.09588 | 0.00191 | 0.56350 |
| **RARA-AS1** | 0.00672 | 0.01739 | 0.06940 | 0.02654 | 0.01628 | 0.00000 | 0.03576 | 0.07699 | 0.01224 | 0.01622 | 0.02456 | 0.01421 |
| **RARB** | 3.07255 | 0.40563 | 0.00000 | 0.00000 | 0.33954 | 0.07450 | 0.00001 | 0.18823 | 0.02548 | 0.37180 | 0.03016 | 0.38583 |
| **RARG** | 0.10356 | 2.67056 | 0.76774 | 0.11533 | 0.00000 | 0.00000 | 1.15884 | 0.23951 | 0.19627 | 0.00000 | 0.71125 | 1.30976 |
| **RARRES1** | 0.00228 | 0.00000 | 1.06737 | 0.00000 | 0.94445 | 0.07423 | 0.26545 | 0.04232 | 0.70532 | 0.53262 | 2.28567 | 0.02812 |
| **RARRES2** | 10.71241 | 1.24474 | 0.00537 | 0.00000 | 0.30314 | 0.00000 | 0.00107 | 13.17638 | 0.00000 | 0.00485 | 0.00830 | 3.90876 |
| **RARRES3** | 0.00619 | 0.01874 | 0.04712 | 0.02966 | 0.03593 | 0.01340 | 0.02279 | 0.00964 | 0.01034 | 0.00000 | 0.01812 | 0.07160 |
| **RARS** | 1.71677 | 21.48819 | 17.67250 | 4.39461 | 6.66021 | 37.26305 | 26.39713 | 3.64744 | 21.99090 | 10.63874 | 23.43473 | 17.35453 |
| **RARS2** | 2.25161 | 8.20482 | 8.81965 | 5.25500 | 2.21630 | 19.24254 | 8.70343 | 8.10978 | 9.00792 | 0.65430 | 4.88717 | 12.70491 |
| **RASA1** | 2.04522 | 0.79029 | 6.56171 | 0.00000 | 0.17309 | 6.30586 | 2.31859 | 0.43375 | 0.00000 | 8.73604 | 0.55300 | 1.47098 |
| **RASA2** | 0.00005 | 0.27339 | 2.55710 | 0.68360 | 0.00001 | 2.12734 | 0.00000 | 1.74793 | 3.11711 | 1.12715 | 1.30143 | 1.72525 |
| **RASA3** | 7.73800 | 0.31118 | 0.08049 | 0.02292 | 0.00000 | 0.00000 | 0.00000 | 0.24845 | 0.00000 | 0.04932 | 0.10054 | 0.07903 |
| **RASA4** | 0.02670 | 0.01949 | 0.01436 | 0.01043 | 0.00720 | 0.04000 | 0.01424 | 0.00711 | 0.01457 | 0.02817 | 0.01314 | 0.01350 |
| **RASA4B.uid-99** | 0.00640 | 0.01655 | 0.02175 | 0.01505 | 0.02951 | 0.01192 | 0.01867 | 0.01031 | 0.02224 | 0.00578 | 0.02006 | 0.02007 |
| **RASA4CP** | 0.53246 | 0.01331 | 0.01741 | 0.01241 | 0.02607 | 0.02497 | 0.01721 | 0.01332 | 0.00990 | 0.00631 | 0.03158 | 0.03530 |
| **RASAL1** | 0.16843 | 0.01574 | 0.09101 | 0.03034 | 0.04455 | 0.00000 | 0.00733 | 0.01587 | 0.08377 | 0.00000 | 0.02191 | 0.01495 |
| **RASAL2** | 0.74345 | 0.14997 | 5.59864 | 1.17976 | 0.17534 | 2.05699 | 1.22322 | 0.12509 | 0.83412 | 3.52313 | 0.00000 | 0.20703 |
| **RASAL2-AS1** | 0.01459 | 0.01150 | 0.08059 | 0.00000 | 0.05949 | 0.02341 | 0.00000 | 0.01079 | 0.04964 | 0.11481 | 0.03338 | 0.01866 |
| **RASAL3** | 0.55031 | 0.02160 | 0.02831 | 0.00944 | 0.02292 | 0.01232 | 0.01968 | 0.02195 | 0.02358 | 0.00000 | 0.01717 | 0.05192 |
| **RASD1** | 0.03582 | 7.82881 | 0.29441 | 9.25213 | 0.32893 | 0.30189 | 0.00000 | 0.82218 | 0.00000 | 0.21895 | 0.00000 | 0.51484 |
| **RASEF** | 0.06846 | 0.00000 | 0.22920 | 0.00000 | 0.74994 | 0.05429 | 0.10469 | 0.00552 | 0.27320 | 0.17436 | 0.00000 | 0.02929 |
| **RASGEF1A** | 0.49049 | 10.58153 | 0.00000 | 6.72190 | 6.44161 | 0.89875 | 0.00000 | 2.83654 | 0.00000 | 1.76697 | 0.82269 | 4.52000 |
| **RASGEF1B** | 0.06688 | 0.01141 | 0.13539 | 0.32593 | 0.07907 | 0.04249 | 0.02153 | 0.00752 | 0.04647 | 0.00000 | 0.01454 | 0.10373 |
| **RASGEF1C** | 0.04305 | 0.06448 | 0.05387 | 0.01134 | 0.01440 | 0.00000 | 0.04180 | 0.02332 | 0.01240 | 0.00000 | 0.03597 | 0.19516 |
| **RASGRF1** | 2.25418 | 0.00000 | 0.16922 | 0.00000 | 0.39054 | 0.00000 | 0.99264 | 0.16788 | 2.08972 | 0.08472 | 1.40601 | 0.06118 |
| **RASGRF2** | 2.61874 | 0.75477 | 0.43000 | 1.92428 | 0.17670 | 0.22599 | 1.16649 | 0.37772 | 0.00000 | 0.67023 | 0.00000 | 0.00000 |
| **RASGRF2-AS1** | 0.00587 | 0.13553 | 0.07259 | 0.00462 | 0.00617 | 0.00545 | 0.01628 | 0.01100 | 0.01193 | 0.03702 | 0.01875 | 0.00726 |
| **RASGRP1** | 0.95080 | 0.01144 | 0.46874 | 0.14455 | 0.00000 | 0.00974 | 0.18178 | 0.22859 | 0.15454 | 0.61808 | 0.26873 | 0.07846 |
| **RASGRP2** | 0.12582 | 6.26936 | 2.07499 | 2.84216 | 13.28673 | 0.00000 | 6.78342 | 0.00000 | 18.57269 | 5.92170 | 14.90834 | 3.04907 |
| **RASGRP3** | 2.57162 | 0.00000 | 0.17172 | 0.10802 | 0.05089 | 0.00000 | 0.05503 | 0.12613 | 0.00000 | 0.27796 | 0.18115 | 0.09305 |
| **RASGRP4** | 0.18220 | 0.47917 | 0.00000 | 0.06514 | 0.13379 | 0.05294 | 0.00000 | 0.03830 | 0.00290 | 0.12490 | 0.03554 | 0.40036 |
| **RASIP1** | 0.00082 | 0.07570 | 0.37043 | 0.00000 | 0.12603 | 0.00000 | 0.89871 | 0.22822 | 0.60285 | 1.96201 | 0.00000 | 1.21395 |
| **RASL10A** | 0.25240 | 0.01802 | 0.00000 | 0.19701 | 0.15425 | 0.06855 | 0.11950 | 0.19323 | 0.02890 | 1.91973 | 0.00000 | 0.03794 |
| **RASL11A** | 5.56420 | 2.17825 | 0.11067 | 0.46667 | 0.22083 | 0.62323 | 0.44269 | 0.73892 | 0.84733 | 0.00000 | 0.33626 | 0.81582 |
| **RASL11B** | 0.18037 | 1.40922 | 0.85931 | 0.22771 | 0.00000 | 0.65279 | 2.31955 | 1.41773 | 2.04214 | 0.00000 | 0.14240 | 3.06075 |
| **RASL12** | 0.00588 | 0.04056 | 0.01619 | 0.01760 | 0.00445 | 0.02678 | 0.01478 | 0.01048 | 0.01299 | 0.02957 | 0.01474 | 0.01184 |
| **RASSF1** | 0.36478 | 5.62841 | 10.20039 | 5.02799 | 3.90253 | 1.10042 | 10.52440 | 1.16261 | 10.70487 | 0.00000 | 8.41675 | 11.31390 |
| **RASSF1-AS1** | 0.01368 | 0.04842 | 0.09278 | 0.02711 | 0.01254 | 0.05281 | 0.00000 | 0.03488 | 0.00000 | 0.09904 | 0.06409 | 0.06790 |
| **RASSF10** | 0.00845 | 0.01359 | 0.06564 | 0.04287 | 0.01022 | 0.00484 | 0.01910 | 0.02038 | 0.00668 | 0.05485 | 0.01731 | 0.03056 |

| | | | | | | | | | | | | |
|---|---|---|---|---|---|---|---|---|---|---|---|---|
| **RASSF2** | 0.00000 | 0.00000 | 1.72846 | 0.44634 | 2.02220 | 0.97025 | 6.78418 | 0.02276 | 6.27530 | 0.37916 | 6.59523 | 0.09485 |
| **RASSF3** | 6.26021 | 0.57652 | 4.71970 | 1.28563 | 0.00000 | 4.68514 | 10.15591 | 2.85249 | 0.00000 | 6.61883 | 4.28019 | 1.46952 |
| **RASSF4** | 0.00115 | 0.07512 | 0.54134 | 0.50603 | 0.45322 | 0.05307 | 1.75703 | 0.05994 | 1.04846 | 0.03284 | 4.24876 | 0.00908 |
| **RASSF5** | 1.78050 | 0.04286 | 0.06730 | 0.01137 | 0.00673 | 0.00000 | 0.05681 | 0.05107 | 0.00224 | 0.03776 | 0.02079 | 0.08238 |
| **RASSF6** | 0.00625 | 0.05481 | 0.04789 | 0.98708 | 0.00000 | 0.07819 | 0.00000 | 0.00898 | 0.12562 | 0.00000 | 0.00114 | 0.00910 |
| **RASSF7** | 0.00060 | 8.20474 | 16.04105 | 4.31552 | 9.63775 | 0.00000 | 7.61159 | 0.00000 | 13.61302 | 3.42317 | 5.80914 | 3.02486 |
| **RASSF8** | 3.17492 | 0.27349 | 1.27178 | 6.65716 | 0.00000 | 7.10645 | 4.80764 | 1.63079 | 0.00000 | 9.61981 | 0.80575 | 8.49785 |
| **RASSF8-AS1** | 0.10424 | 0.38854 | 0.25667 | 0.79703 | 0.76104 | 0.15078 | 0.00000 | 1.24149 | 0.07811 | 0.79547 | 0.15447 | 2.38599 |
| **RASSF9** | 0.00605 | 0.01410 | 0.03075 | 0.00951 | 0.01328 | 0.01740 | 0.04764 | 0.00908 | 0.00000 | 0.02029 | 0.01623 | 0.09611 |
| **RAVER1** | 1.50115 | 4.32157 | 8.20001 | 4.00833 | 5.40370 | 0.00000 | 8.63899 | 0.34616 | 7.98317 | 14.38808 | 2.05224 | 5.36858 |
| **RAVER2** | 1.40527 | 0.00000 | 0.00000 | 0.83403 | 0.00000 | 0.74618 | 0.31747 | 0.03807 | 0.00000 | 0.09275 | 0.00000 | 0.03202 |
| **RAX** | 0.11095 | 0.01357 | 0.02155 | 0.01902 | 0.01649 | 0.01148 | 0.02138 | 0.00483 | 0.01569 | 0.01018 | 0.01562 | 0.02575 |
| **RAX2** | 1.48791 | 0.01236 | 0.01651 | 0.01116 | 0.01452 | 0.00911 | 0.01912 | 0.00068 | 0.01993 | 0.02466 | 0.01390 | 0.01163 |
| **RB1** | 0.70460 | 0.93963 | 1.65694 | 0.00000 | 0.00000 | 3.76636 | 2.27076 | 0.70691 | 3.28070 | 1.83070 | 1.46786 | 1.39710 |
| **RB1CC1** | 4.62578 | 5.49068 | 7.74806 | 4.59614 | 2.08930 | 30.96676 | 11.97186 | 24.74418 | 6.52903 | 30.28787 | 6.95121 | 35.61979 |
| **RBAK** | 0.00000 | 0.00000 | 0.68411 | 0.76569 | 0.00006 | 5.64445 | 0.00001 | 0.00583 | 4.81012 | 5.78542 | 0.78746 | 4.15069 |
| **RBAKDN** | 0.01409 | 0.02046 | 0.01903 | 0.01577 | 0.02341 | 0.00011 | 0.02197 | 0.05301 | 0.02683 | 0.00000 | 0.01575 | 0.05899 |
| **RBBP4** | 5.06018 | 9.96051 | 8.74583 | 14.16557 | 4.37268 | 16.10950 | 8.24019 | 2.90934 | 19.49972 | 10.55822 | 13.47478 | 11.48309 |
| **RBBP5** | 4.73531 | 5.60756 | 6.65599 | 8.88408 | 4.83403 | 7.97385 | 9.74239 | 16.77149 | 9.09769 | 2.04400 | 6.97610 | 8.06872 |
| **RBBP6** | 9.18783 | 3.07652 | 4.58509 | 3.74241 | 6.66962 | 16.37992 | 11.64108 | 11.93363 | 4.74526 | 45.54370 | 3.26212 | 9.03630 |
| **RBBP7** | 11.96814 | 20.57916 | 25.96704 | 9.76639 | 0.00000 | 35.34601 | 28.63603 | 15.07659 | 18.86895 | 0.00000 | 18.92759 | 20.36982 |
| **RBBP8** | 3.03435 | 2.59353 | 2.15298 | 0.44917 | 0.13589 | 11.89291 | 3.00734 | 1.75857 | 3.72847 | 10.76837 | 8.67127 | 7.33274 |
| **RBBP8NL** | 0.08715 | 0.00000 | 1.88918 | 0.00000 | 0.33574 | 0.00000 | 1.66791 | 0.02582 | 2.83472 | 0.51275 | 0.00000 | 0.00070 |
| **RBBP9** | 0.00000 | 6.95609 | 0.80753 | 3.05995 | 0.13127 | 5.30735 | 1.45865 | 0.62293 | 0.22654 | 3.09635 | 1.58322 | 8.17524 |
| **RBCK1** | 3.20678 | 5.42206 | 5.07011 | 3.07200 | 8.29965 | 0.76656 | 4.53918 | 0.39247 | 5.35879 | 0.48955 | 3.66409 | 3.11501 |
| **RBFA** | 0.63633 | 6.72675 | 7.96203 | 5.01849 | 3.38991 | 2.70590 | 8.96010 | 2.35900 | 10.62630 | 0.00000 | 8.95991 | 4.86862 |
| **RBFADN** | 0.00422 | 0.05613 | 0.00827 | 0.00676 | 0.01691 | 0.07360 | 0.00000 | 0.01782 | 0.15595 | 0.02533 | 0.01163 | 0.09790 |
| **RBFOX1** | 1.42060 | 0.00002 | 0.00000 | 0.05314 | 0.17241 | 0.16236 | 0.00000 | 0.20517 | 0.00398 | 0.16182 | 0.00388 | 0.01426 |
| **RBFOX2** | 3.63471 | 0.78739 | 5.06485 | 8.69568 | 2.48554 | 0.61547 | 14.07537 | 0.15563 | 6.75685 | 12.00004 | 2.47249 | 1.48965 |
| **RBFOX3** | 0.56116 | 0.07644 | 0.01845 | 0.00047 | 0.00646 | 0.03307 | 0.01518 | 0.00413 | 0.01315 | 0.04072 | 0.02650 | 0.00520 |
| **RBKS** | 0.00339 | 1.22869 | 0.05655 | 0.20447 | 1.57873 | 0.56150 | 0.00000 | 0.02790 | 0.00000 | 0.22134 | 1.13775 | 1.11735 |
| **RBL1** | 0.45147 | 1.01596 | 0.76832 | 1.16834 | 1.47323 | 4.58444 | 1.54467 | 1.31471 | 1.22069 | 2.48767 | 0.90895 | 2.60191 |
| **RBL2** | 1.50846 | 2.26240 | 0.49167 | 0.26568 | 1.00363 | 1.34932 | 0.00000 | 1.13162 | 0.37327 | 2.20006 | 2.26963 | 1.05262 |
| **RBM10** | 1.14574 | 12.16319 | 12.32086 | 9.21196 | 14.14346 | 0.00000 | 15.86453 | 2.81471 | 10.72910 | 10.35478 | 9.83875 | 7.81523 |
| **RBM11** | 0.52879 | 0.16916 | 0.54478 | 0.00000 | 0.00000 | 0.42574 | 0.28757 | 0.00000 | 0.12925 | 0.04576 | 0.05927 | 0.23051 |
| **RBM12** | 1.73969 | 10.88035 | 10.96673 | 8.45276 | 6.56438 | 12.22402 | 17.40662 | 3.02870 | 8.74965 | 17.19018 | 16.36898 | 13.81582 |
| **RBM12B** | 0.17477 | 1.22360 | 2.21242 | 0.25767 | 0.11790 | 8.41793 | 3.31950 | 3.12206 | 6.04143 | 5.05241 | 2.27434 | 7.43169 |
| **RBM12B-AS1** | 0.01342 | 0.13878 | 0.26316 | 0.13338 | 0.16033 | 0.00000 | 0.00000 | 0.12153 | 0.93123 | 0.01167 | 0.32429 | 0.33218 |
| **RBM14** | 6.06370 | 22.64273 | 16.30166 | 17.42564 | 15.21794 | 1.96539 | 23.19101 | 9.25801 | 25.55999 | 3.94799 | 21.17403 | 25.39760 |
| **RBM15** | 3.31354 | 4.26743 | 3.57881 | 4.42312 | 5.51300 | 2.77819 | 8.76999 | 9.86645 | 7.68268 | 1.38169 | 3.48309 | 12.24801 |
| **RBM15B** | 2.83499 | 1.35844 | 7.66770 | 2.41799 | 3.52275 | 4.49381 | 10.81609 | 0.00000 | 3.98500 | 10.45602 | 2.28442 | 5.83266 |
| **RBM17** | 10.12404 | 21.15391 | 14.60704 | 11.86387 | 16.66797 | 19.32661 | 20.98352 | 9.36383 | 17.89430 | 21.07021 | 12.88399 | 20.43847 |
| **RBM18** | 4.27950 | 8.06833 | 8.59155 | 4.72477 | 4.29609 | 18.70780 | 9.92299 | 8.43838 | 10.07998 | 0.00000 | 9.40332 | 9.52808 |
| **RBM19** | 0.52041 | 11.18626 | 12.39468 | 6.94772 | 8.11109 | 0.00000 | 10.19410 | 9.65088 | 13.52341 | 3.00106 | 9.75345 | 13.57144 |
| **RBM20** | 0.02970 | 0.10104 | 0.13526 | 0.12044 | 0.01955 | 0.02852 | 0.07923 | 0.01000 | 0.00000 | 0.00000 | 0.03450 | 0.01744 |
| **RBM22** | 2.47001 | 11.43932 | 12.32141 | 6.09655 | 4.79711 | 16.25645 | 15.03798 | 8.22250 | 9.93448 | 5.21199 | 11.80742 | 13.73236 |
| **RBM23** | 1.50756 | 9.04290 | 12.36167 | 8.94215 | 11.75236 | 11.18806 | 8.82388 | 7.37316 | 10.11607 | 0.00000 | 9.00735 | 8.18610 |
| **RBM24** | 0.19381 | 0.38671 | 0.00000 | 0.01408 | 0.00000 | 0.17909 | 0.00000 | 0.58525 | 0.00000 | 0.02604 | 0.00533 | 2.66201 |

| | | | | | | | | | | | | |
|---|---|---|---|---|---|---|---|---|---|---|---|---|
| **RBM25** | 4.93651 | 11.14972 | 9.72003 | 15.53775 | 18.23974 | 25.09627 | 30.36144 | 25.63133 | 13.89214 | 45.26068 | 16.23726 | 22.72208 |
| **RBM26** | 1.56971 | 0.00000 | 3.08802 | 0.54659 | 1.42288 | 11.12937 | 5.82919 | 1.96128 | 3.07091 | 21.50436 | 1.97090 | 2.76937 |
| **RBM26-AS1** | 0.15328 | 0.06792 | 0.00000 | 0.27234 | 0.24160 | 0.59709 | 0.00000 | 0.00000 | 0.03995 | 0.00108 | 0.17529 | 0.03073 |
| **RBM27** | 7.62774 | 3.01618 | 9.93625 | 5.60456 | 3.77814 | 13.88338 | 13.89762 | 5.30245 | 6.76965 | 20.51324 | 8.05761 | 8.91177 |
| **RBM28** | 2.43704 | 12.50153 | 4.56755 | 5.57750 | 10.11272 | 9.40483 | 9.94054 | 9.78281 | 10.36375 | 9.44203 | 8.94352 | 14.18436 |
| **RBM3** | 5.53025 | 18.20179 | 17.09218 | 16.28391 | 19.01411 | 45.37579 | 31.23818 | 8.61987 | 30.47388 | 7.73684 | 24.53564 | 24.64739 |
| **RBM33** | 9.53437 | 3.95230 | 4.68110 | 1.57797 | 5.94524 | 6.71347 | 6.61496 | 1.32724 | 8.09138 | 15.50637 | 2.55352 | 7.60613 |
| **RBM34** | 3.39522 | 14.51577 | 11.93541 | 6.08477 | 10.24073 | 27.95531 | 17.79761 | 14.22178 | 10.47222 | 5.10063 | 11.15475 | 20.30776 |
| **RBM38** | 8.79906 | 7.65944 | 6.54047 | 6.44455 | 18.20035 | 0.00000 | 18.67496 | 1.22812 | 5.85851 | 21.82863 | 4.72833 | 8.36968 |
| **RBM39** | 8.10524 | 20.61981 | 14.33048 | 16.48070 | 18.91876 | 37.54714 | 27.42519 | 32.16248 | 25.18505 | 25.73874 | 14.73441 | 24.87384 |
| **RBM4** | 1.22916 | 7.52341 | 4.76903 | 6.93922 | 4.60554 | 7.11493 | 3.37451 | 8.67927 | 9.80271 | 0.00000 | 6.59709 | 8.14571 |
| **RBM41** | 1.64094 | 0.00000 | 0.00000 | 1.97328 | 0.46672 | 7.68699 | 0.00000 | 0.00007 | 3.14661 | 6.27250 | 4.56350 | 0.08987 |
| **RBM42** | 4.63461 | 21.60896 | 17.73398 | 8.38781 | 19.22796 | 0.00000 | 13.49448 | 9.71918 | 20.13490 | 2.70973 | 14.55846 | 19.15871 |
| **RBM43** | 0.00617 | 0.02477 | 0.02902 | 0.01742 | 0.04846 | 0.01980 | 0.04312 | 0.01051 | 0.02159 | 0.00489 | 0.01236 | 0.01715 |
| **RBM44** | 0.54702 | 0.11303 | 0.00000 | 0.00000 | 0.21223 | 0.53715 | 0.00000 | 3.88980 | 0.00000 | 0.16703 | 0.16765 | 4.46332 |
| **RBM45** | 0.00000 | 3.59619 | 0.17907 | 0.59291 | 1.58824 | 3.37500 | 2.72792 | 1.60186 | 0.00000 | 0.94300 | 3.53793 | 0.99818 |
| **RBM46** | 5.92427 | 0.00000 | 2.04358 | 0.00000 | 0.00000 | 1.03381 | 2.14956 | 5.57451 | 1.08948 | 0.00000 | 0.24339 | 1.40379 |
| **RBM47** | 0.00000 | 25.21858 | 17.17832 | 6.46847 | 16.91266 | 38.24138 | 38.49146 | 0.16087 | 24.45550 | 26.73709 | 20.05137 | 10.14320 |
| **RBM48** | 1.28984 | 6.47542 | 4.11334 | 1.13651 | 0.00000 | 8.90690 | 4.34682 | 15.36631 | 0.39607 | 0.00000 | 3.82943 | 11.48906 |
| **RBM4B** | 3.12753 | 2.45999 | 5.05567 | 2.41082 | 1.31419 | 1.14925 | 4.91061 | 3.95863 | 7.99464 | 0.00000 | 4.64912 | 8.00713 |
| **RBM5** | 0.00000 | 11.04136 | 7.14198 | 11.76210 | 7.00968 | 7.50726 | 4.00598 | 12.54919 | 13.02122 | 13.69228 | 5.81550 | 12.68003 |
| **RBM5-AS1** | 0.00283 | 0.07259 | 0.06990 | 0.64022 | 0.55054 | 0.00005 | 0.02714 | 0.21119 | 0.49710 | 1.56358 | 0.03457 | 0.53801 |
| **RBM6** | 2.98298 | 7.14064 | 4.28448 | 10.77099 | 8.66571 | 10.78598 | 9.71043 | 11.14226 | 16.23286 | 13.92392 | 9.68876 | 9.36349 |
| **RBM7** | 1.24399 | 5.22498 | 11.45107 | 3.70685 | 0.00000 | 19.12086 | 12.59520 | 10.41217 | 11.95667 | 0.00000 | 7.54742 | 6.21696 |
| **RBM8A** | 2.93672 | 16.52991 | 13.69107 | 17.99855 | 12.65344 | 26.45741 | 26.60983 | 25.13001 | 12.16471 | 13.23826 | 15.48760 | 30.79837 |
| **RBMS1** | 0.00000 | 3.05626 | 5.87527 | 4.42306 | 1.24359 | 11.42102 | 8.60287 | 0.33273 | 0.00000 | 18.02456 | 1.83701 | 1.86577 |
| **RBMS2** | 3.04955 | 0.16110 | 0.76701 | 0.17615 | 0.68690 | 0.00000 | 0.68601 | 0.41153 | 0.00000 | 0.90467 | 0.08966 | 0.33848 |
| **RBMS3** | 0.48766 | 0.04251 | 0.03033 | 0.02053 | 0.00000 | 0.03135 | 0.00042 | 0.46699 | 0.01236 | 0.11265 | 0.00636 | 0.01178 |
| **RBMX** | 5.69088 | 17.29668 | 10.72510 | 16.47199 | 21.49104 | 24.53348 | 34.19715 | 19.17929 | 22.53173 | 23.89532 | 17.35214 | 22.44128 |
| **RBMX2** | 9.74240 | 4.61294 | 2.10172 | 1.35220 | 2.17515 | 5.49538 | 7.62648 | 6.86225 | 0.00754 | 7.25774 | 3.39972 | 5.01460 |
| **RBMXL1** | 0.63172 | 5.99548 | 8.70216 | 3.67376 | 3.90453 | 17.03081 | 9.50921 | 2.83212 | 7.29938 | 7.38848 | 6.30667 | 9.15574 |
| **RBMXL2** | 0.00136 | 0.00006 | 0.40619 | 0.00000 | 0.10711 | 0.07376 | 0.31443 | 0.06073 | 0.09405 | 0.10109 | 0.02538 | 0.02051 |
| **RBMXL3** | 0.00578 | 0.06056 | 0.21417 | 0.01198 | 0.30047 | 0.00000 | 0.00337 | 0.25573 | 0.15407 | 0.00000 | 0.02266 | 0.73621 |
| **RBMY1F+RBMY** | 0.01450 | 0.01491 | 0.01166 | 0.01186 | 0.01660 | 0.01540 | 0.02080 | 0.02239 | 0.05139 | 0.00157 | 0.01005 | 0.03268 |
| **RBMY2EP** | 0.00792 | 0.02858 | 0.01497 | 0.01233 | 0.01619 | 0.00925 | 0.02126 | 0.01052 | 0.01877 | 0.01176 | 0.01805 | 0.01864 |
| **RBMY2FP** | 0.01593 | 0.20475 | 0.21163 | 0.00000 | 0.04715 | 0.04511 | 0.12895 | 0.54992 | 0.15725 | 0.10401 | 0.00000 | 2.60493 |
| **RBP1** | 1.06787 | 0.27681 | 0.31609 | 1.69620 | 1.51426 | 0.00000 | 1.92875 | 1.31866 | 0.67086 | 0.00000 | 1.34198 | 0.36896 |
| **RBP2** | 0.00598 | 0.01268 | 0.10464 | 0.15893 | 0.00474 | 0.01498 | 0.01176 | 0.01040 | 0.00631 | 0.02187 | 0.01883 | 0.01585 |
| **RBP3** | 0.00440 | 0.00929 | 0.02937 | 0.05411 | 0.00000 | 0.01177 | 0.01789 | 0.01005 | 0.00470 | 0.16660 | 0.00895 | 0.01241 |
| **RBP4** | 0.07068 | 0.04379 | 0.03466 | 0.01146 | 0.02712 | 0.02233 | 0.01405 | 0.03122 | 0.03934 | 0.00000 | 0.02251 | 0.04349 |
| **RBP5** | 0.05203 | 0.01726 | 0.03222 | 0.11493 | 0.01162 | 0.00000 | 0.01566 | 0.01173 | 0.02567 | 0.00858 | 0.02232 | 0.01650 |
| **RBP7** | 0.02093 | 7.23504 | 0.00000 | 0.00000 | 4.55027 | 0.86518 | 0.25884 | 2.34442 | 0.00000 | 0.00000 | 0.00000 | 29.46362 |
| **RBPJ** | 3.47231 | 11.44444 | 16.52510 | 6.63720 | 8.68606 | 27.69145 | 21.92300 | 9.40781 | 11.00513 | 22.50171 | 12.38582 | 9.32259 |
| **RBPJL** | 0.00599 | 0.92165 | 0.00315 | 0.00000 | 0.00000 | 0.00011 | 0.03483 | 0.00235 | 0.00000 | 0.10525 | 0.01152 | 2.09640 |
| **RBPMS** | 1.85511 | 1.09771 | 4.30559 | 4.30042 | 8.40410 | 2.01943 | 12.44168 | 0.14744 | 3.04772 | 21.56388 | 1.80113 | 1.55199 |
| **RBPMS-AS1** | 0.29151 | 0.45287 | 0.51544 | 0.12363 | 0.46974 | 0.00000 | 0.15343 | 0.00000 | 0.40928 | 0.00000 | 0.26013 | 0.43725 |
| **RBPMS2** | 9.11204 | 13.76474 | 5.95473 | 9.81969 | 22.07374 | 10.50453 | 2.65593 | 2.59398 | 0.00000 | 23.44854 | 5.12054 | 13.69210 |
| **RBX1** | 5.01797 | 13.58462 | 19.47569 | 15.36889 | 14.31434 | 30.33106 | 30.14095 | 11.48263 | 11.77503 | 11.35194 | 19.86508 | 23.38695 |

| | | | | | | | | | | | | |
|---|---|---|---|---|---|---|---|---|---|---|---|---|
| **RC3H1** | 1.90607 | 2.07803 | 4.35039 | 3.01160 | 0.00000 | 14.94842 | 5.73855 | 14.47498 | 5.32909 | 12.38224 | 8.64162 | 2.49360 |
| **RC3H2** | 1.13380 | 1.89829 | 4.38077 | 4.92693 | 1.05877 | 5.42832 | 3.40425 | 3.85010 | 0.00000 | 6.16596 | 2.72990 | 2.43037 |
| **RCAN1** | 1.06103 | 0.27777 | 0.52949 | 2.09358 | 0.00000 | 1.87987 | 1.15240 | 0.33216 | 0.00000 | 2.84934 | 0.38623 | 0.30727 |
| **RCAN2** | 1.91407 | 0.01321 | 0.01287 | 0.01227 | 0.02295 | 0.00057 | 0.03790 | 0.00000 | 0.01403 | 0.00000 | 0.01436 | 0.02850 |
| **RCAN3** | 1.45462 | 1.02283 | 0.35455 | 0.42518 | 2.08659 | 3.59717 | 0.00417 | 0.74484 | 0.39084 | 0.95934 | 1.37802 | 1.71591 |
| **RCAN3AS** | 0.03693 | 0.01539 | 0.01441 | 0.00858 | 0.01383 | 0.02743 | 0.02690 | 0.01041 | 0.01852 | 0.01800 | 0.01445 | 0.02101 |
| **RCBTB1** | 0.57996 | 20.14903 | 0.00161 | 20.30333 | 2.75380 | 13.82326 | 0.00000 | 2.80985 | 0.00000 | 3.98976 | 1.20899 | 7.42407 |
| **RCBTB2** | 2.04164 | 0.00243 | 0.22509 | 0.25048 | 0.00000 | 0.20422 | 0.00000 | 0.00000 | 0.26680 | 0.00000 | 0.05755 | 0.14789 |
| **RCC1** | 3.11418 | 26.10591 | 26.49038 | 18.71458 | 21.66013 | 9.59962 | 33.84934 | 5.60537 | 45.66778 | 0.00000 | 31.94915 | 23.34444 |
| **RCC2** | 5.27409 | 9.51110 | 6.74385 | 18.47642 | 16.28037 | 12.56840 | 26.71100 | 2.02937 | 6.28718 | 28.76311 | 10.54092 | 13.52167 |
| **RCCD1** | 2.15956 | 1.82475 | 2.55090 | 0.99594 | 1.25923 | 1.79541 | 1.49002 | 0.44567 | 2.58591 | 4.13622 | 1.93673 | 0.89915 |
| **RCE1** | 0.35155 | 13.15826 | 9.20454 | 5.55826 | 8.16233 | 0.00000 | 0.32211 | 7.07149 | 15.91706 | 6.73871 | 10.18786 | 6.12655 |
| **RCHY1** | 0.22975 | 6.39490 | 4.00351 | 4.91545 | 1.35423 | 26.37089 | 4.06775 | 1.89731 | 4.29141 | 3.90636 | 10.52371 | 15.24735 |
| **RCL1** | 0.72247 | 13.66623 | 10.32436 | 0.13457 | 6.33462 | 19.68859 | 10.27338 | 7.85985 | 19.95065 | 0.00000 | 9.32832 | 15.69981 |
| **RCN1** | 0.00000 | 1.73937 | 12.45890 | 14.72949 | 11.78528 | 4.47605 | 5.06323 | 0.00000 | 23.20982 | 1.27908 | 6.55389 | 0.00000 |
| **RCN2** | 0.22951 | 7.02462 | 23.63337 | 5.68680 | 2.47508 | 14.79307 | 2.92171 | 0.01463 | 19.93530 | 11.17707 | 7.93313 | 0.00000 |
| **RCN3** | 0.00666 | 1.57416 | 0.20711 | 0.33556 | 0.07251 | 0.00000 | 0.02875 | 0.95968 | 0.14074 | 0.25306 | 0.00000 | 2.30323 |
| **RCOR1** | 5.23518 | 0.82312 | 7.24665 | 0.41066 | 3.40537 | 12.39445 | 6.28939 | 4.56366 | 1.63209 | 20.47144 | 2.59231 | 4.22429 |
| **RCOR2** | 0.00432 | 0.00000 | 0.22176 | 0.30570 | 0.53429 | 0.00000 | 1.05160 | 0.01438 | 0.03557 | 2.92040 | 0.00000 | 0.17748 |
| **RCOR3** | 1.75279 | 4.56828 | 0.54965 | 7.94210 | 0.51483 | 12.23926 | 0.00000 | 1.42834 | 1.43754 | 5.53698 | 2.48704 | 4.14605 |
| **RCSD1** | 0.00448 | 0.39843 | 0.00000 | 1.14827 | 0.00000 | 0.06120 | 0.00000 | 0.00756 | 0.00000 | 0.05703 | 0.00820 | 0.02346 |
| **RCVRN** | 0.03893 | 0.00492 | 0.05433 | 0.00000 | 0.05952 | 0.05910 | 0.00000 | 0.00970 | 0.00836 | 0.10646 | 0.00838 | 0.00966 |
| **RD3** | 0.00525 | 0.02901 | 0.00331 | 0.06263 | 0.00000 | 0.05863 | 0.00000 | 0.00797 | 0.01646 | 0.04982 | 0.00996 | 0.04422 |
| **RDH10** | 0.96893 | 0.24567 | 3.92349 | 0.17860 | 0.00000 | 10.98494 | 3.02139 | 7.12439 | 8.31699 | 11.03932 | 0.76302 | 3.04696 |
| **RDH11** | 0.00000 | 4.84699 | 16.86901 | 12.50835 | 16.92413 | 30.75735 | 8.76973 | 0.00000 | 44.66652 | 9.43798 | 15.23621 | 5.03788 |
| **RDH12** | 0.00589 | 0.01614 | 0.07367 | 0.04514 | 0.02917 | 0.01410 | 0.04744 | 0.00947 | 0.04716 | 0.01164 | 0.02197 | 0.04198 |
| **RDH13** | 1.67334 | 13.15230 | 5.27070 | 3.32852 | 13.22069 | 0.00000 | 10.67016 | 2.86285 | 13.38023 | 6.32635 | 9.83555 | 4.99556 |
| **RDH14** | 0.00453 | 0.37380 | 0.42892 | 0.22208 | 0.38816 | 0.00000 | 0.00653 | 0.01333 | 0.60280 | 0.00000 | 0.35320 | 0.11829 |
| **RDH16** | 0.01158 | 0.04793 | 0.06069 | 0.04805 | 0.04022 | 0.07666 | 0.00000 | 0.01107 | 0.03535 | 0.00454 | 0.01541 | 0.03814 |
| **RDH5** | 0.00342 | 0.02682 | 0.03887 | 0.07197 | 0.00411 | 0.00744 | 0.01888 | 0.02238 | 0.09775 | 0.01893 | 0.01639 | 0.04620 |
| **RDM1** | 0.00000 | 1.06431 | 0.13599 | 0.00000 | 1.17621 | 0.39485 | 0.28558 | 0.36254 | 0.00000 | 0.69985 | 2.59155 | 0.74956 |
| **RDX** | 8.48231 | 7.56282 | 13.78228 | 7.52015 | 13.97963 | 33.88030 | 18.46083 | 6.54672 | 8.53198 | 23.51531 | 16.05111 | 8.28113 |
| **REC114** | 4.24189 | 0.02670 | 0.00610 | 0.10133 | 0.00954 | 0.04719 | 0.03845 | 0.53533 | 0.01860 | 0.03541 | 0.01194 | 0.26229 |
| **REC8** | 0.00155 | 0.00000 | 1.83280 | 3.10195 | 0.19532 | 0.00000 | 0.00000 | 0.02022 | 0.99281 | 1.84385 | 0.25053 | 0.29137 |
| **RECK** | 0.11768 | 0.70845 | 0.13590 | 0.20794 | 0.95973 | 0.44969 | 0.00000 | 0.07315 | 0.14373 | 0.00000 | 0.75162 | 0.34513 |
| **RECQL** | 0.17311 | 3.72792 | 4.17408 | 1.99858 | 0.00000 | 7.74862 | 3.98364 | 1.81346 | 2.70907 | 0.00000 | 4.52489 | 3.55460 |
| **RECQL4** | 1.60250 | 3.83228 | 6.57428 | 7.02052 | 9.24468 | 0.00000 | 5.40919 | 1.33430 | 7.77132 | 22.80469 | 3.91060 | 2.33329 |
| **RECQL5** | 2.97864 | 5.06729 | 2.20305 | 1.13888 | 2.14216 | 0.00000 | 1.65574 | 0.81808 | 3.21553 | 1.60660 | 1.67690 | 5.52862 |
| **REEP1** | 0.38701 | 12.55847 | 24.19635 | 1.55051 | 9.89892 | 9.07664 | 10.15110 | 0.00000 | 29.20165 | 6.25877 | 19.43606 | 0.00000 |
| **REEP2** | 3.44667 | 0.13432 | 0.00645 | 0.07745 | 0.00627 | 0.00000 | 0.01890 | 0.39547 | 0.07733 | 0.00000 | 0.01073 | 0.20761 |
| **REEP3** | 0.00000 | 2.08324 | 5.34996 | 1.54078 | 0.68523 | 9.39010 | 2.73215 | 0.23347 | 9.47904 | 9.14185 | 4.25920 | 0.34035 |
| **REEP4** | 3.20232 | 17.97606 | 9.25932 | 8.03299 | 10.28760 | 0.00000 | 2.90685 | 0.73345 | 24.85965 | 1.87526 | 5.49475 | 14.73271 |
| **REEP5** | 0.00000 | 18.12807 | 18.66743 | 18.77377 | 14.61482 | 34.29072 | 0.00000 | 4.87554 | 41.64210 | 14.22188 | 21.17289 | 6.24148 |
| **REEP6** | 0.72118 | 2.85180 | 2.09057 | 8.41027 | 3.49319 | 1.46454 | 0.00000 | 1.03215 | 1.60444 | 0.00000 | 0.36597 | 0.20953 |
| **REG1A** | 0.00777 | 0.09312 | 0.08601 | 0.37336 | 0.01807 | 0.05997 | 0.00441 | 0.08660 | 0.00159 | 0.00790 | 0.01576 | 0.02458 |
| **REG1P** | 0.08620 | 0.01694 | 0.07692 | 0.02562 | 0.02184 | 0.00000 | 0.01216 | 0.02635 | 0.01197 | 0.01028 | 0.02291 | 0.01543 |
| **REL** | 2.21152 | 1.19208 | 2.23955 | 2.27094 | 1.56554 | 2.27820 | 0.43951 | 0.58450 | 2.49410 | 7.53268 | 0.59497 | 1.82043 |
| **RELA** | 3.44101 | 5.61055 | 5.44614 | 4.80728 | 5.57782 | 0.15266 | 8.05719 | 0.09857 | 6.00631 | 4.63942 | 2.53034 | 5.93555 |

| | | | | | | | | | | | | |
|---|---|---|---|---|---|---|---|---|---|---|---|---|
| **RELB** | 1.46951 | 0.64126 | 0.33405 | 0.94844 | 0.00000 | 0.00000 | 0.35764 | 3.35875 | 0.00000 | 0.97994 | 0.00000 | 0.76393 |
| **RELL1** | 0.00000 | 18.08389 | 9.28034 | 9.16019 | 5.84882 | 46.23815 | 0.00000 | 4.88993 | 18.90781 | 4.27392 | 18.67919 | 11.10751 |
| **RELL2** | 0.37497 | 0.14791 | 1.04762 | 0.02375 | 0.06146 | 0.00000 | 0.00000 | 0.11613 | 0.14125 | 0.00000 | 0.01443 | 0.06306 |
| **RELN** | 0.87501 | 0.02219 | 0.00990 | 0.01063 | 0.04777 | 0.00019 | 0.10741 | 0.00468 | 0.00000 | 0.04686 | 0.01609 | 0.01005 |
| **RELT** | 0.58208 | 0.70170 | 6.18678 | 0.00000 | 1.25603 | 0.00000 | 0.19607 | 0.00381 | 2.05902 | 0.75612 | 0.91563 | 0.11657 |
| **REM1** | 0.20477 | 2.59711 | 0.03254 | 1.54615 | 0.00000 | 0.00462 | 0.00199 | 0.75906 | 0.01170 | 0.01847 | 0.00122 | 1.13597 |
| **REM2** | 0.06498 | 0.03688 | 0.08751 | 0.05378 | 0.08500 | 0.00000 | 0.00000 | 0.03899 | 0.09410 | 0.25715 | 0.07260 | 0.06245 |
| **REN** | 0.00627 | 0.01384 | 0.01341 | 0.01459 | 0.02949 | 0.02508 | 0.01882 | 0.01041 | 0.01702 | 0.00984 | 0.01508 | 0.01597 |
| **RENBP** | 0.01541 | 0.17420 | 0.45820 | 0.33287 | 0.00000 | 0.12044 | 0.00000 | 0.09239 | 0.00000 | 0.11363 | 0.01579 | 0.16911 |
| **REP15** | 0.00520 | 0.34115 | 0.06838 | 0.01628 | 0.12161 | 0.00000 | 0.01166 | 0.06773 | 0.16229 | 0.33683 | 0.07606 | 0.74754 |
| **REPIN1** | 0.12530 | 2.27631 | 2.87476 | 0.65616 | 6.03838 | 0.00000 | 6.50289 | 0.00256 | 1.92920 | 6.95358 | 0.48859 | 0.00000 |
| **REPS1** | 0.00000 | 6.30941 | 2.98353 | 3.95852 | 6.34373 | 10.26367 | 9.07599 | 1.62718 | 2.03768 | 14.45182 | 0.76462 | 5.81567 |
| **REPS2** | 0.83230 | 0.92909 | 0.11475 | 2.20786 | 0.00000 | 0.22938 | 0.00000 | 0.36515 | 0.00000 | 0.20241 | 0.02052 | 0.24713 |
| **RER1** | 3.48497 | 26.04045 | 19.19465 | 11.23537 | 21.83710 | 20.26174 | 7.41246 | 2.65741 | 32.08850 | 9.94925 | 21.02219 | 20.92176 |
| **RERE** | 6.66519 | 3.58064 | 2.95077 | 6.64964 | 7.75170 | 1.70498 | 6.18665 | 1.45497 | 4.43128 | 18.65846 | 1.07618 | 1.58746 |
| **REREP3** | 0.48994 | 0.01263 | 0.05880 | 0.08986 | 0.05691 | 0.00000 | 0.02006 | 0.12366 | 0.02824 | 0.03721 | 0.00766 | 0.05531 |
| **RERG** | 0.11992 | 0.39237 | 1.31199 | 0.06702 | 1.05441 | 3.01189 | 1.20639 | 0.07171 | 0.72359 | 0.00000 | 0.76173 | 0.09103 |
| **RERG-AS1** | 0.86682 | 0.01221 | 0.01284 | 0.01264 | 0.01575 | 0.02081 | 0.01822 | 0.00368 | 0.02123 | 0.00367 | 0.01605 | 0.01366 |
| **RERGL** | 1.14747 | 0.00403 | 0.09987 | 0.11952 | 0.01862 | 0.07357 | 0.00000 | 0.15043 | 0.00561 | 0.01109 | 0.01374 | 0.00284 |
| **RESP18** | 0.18599 | 0.01550 | 0.02197 | 0.02165 | 0.01531 | 0.00689 | 0.01069 | 0.27001 | 0.02238 | 0.02963 | 0.01377 | 0.01136 |
| **REST** | 2.65447 | 7.86587 | 9.37635 | 17.39489 | 9.51437 | 28.31513 | 15.14998 | 12.97410 | 1.22254 | 33.25633 | 3.74715 | 24.03495 |
| **RET** | 0.00147 | 9.51552 | 0.00000 | 0.00218 | 3.02770 | 0.02464 | 0.00047 | 0.02280 | 0.00000 | 0.84051 | 0.47752 | 0.44257 |
| **RETSAT** | 1.03266 | 3.15672 | 2.45110 | 1.17129 | 0.00000 | 0.77893 | 0.00000 | 1.24127 | 6.09214 | 2.22714 | 4.29144 | 3.55355 |
| **REV1** | 6.16330 | 0.57615 | 3.37971 | 1.29249 | 1.80882 | 4.89252 | 3.12192 | 0.86261 | 2.20479 | 14.56446 | 0.01777 | 1.23572 |
| **REV3L** | 2.60493 | 0.00000 | 4.05811 | 0.36131 | 1.26202 | 14.99774 | 2.11758 | 5.21826 | 7.10136 | 21.49200 | 5.74983 | 3.75414 |
| **REXO1** | 3.63818 | 0.34509 | 2.57276 | 0.97249 | 2.69867 | 0.00000 | 3.83755 | 0.11714 | 1.19507 | 14.80246 | 0.36278 | 0.51314 |
| **REXO2** | 9.39863 | 8.59917 | 20.59132 | 7.29435 | 4.75461 | 16.00813 | 20.47116 | 4.53722 | 15.23648 | 2.40313 | 12.06567 | 3.52549 |
| **REXO4** | 0.01143 | 6.31489 | 7.00853 | 4.35564 | 9.51391 | 0.00000 | 14.18970 | 4.14671 | 0.47585 | 14.13709 | 4.00954 | 7.53782 |
| **RFC1** | 5.45297 | 5.17969 | 6.39470 | 6.77083 | 1.27168 | 20.90448 | 6.88464 | 9.14168 | 6.15207 | 12.96667 | 3.72174 | 11.38458 |
| **RFC2** | 0.10708 | 18.26805 | 8.62789 | 8.44108 | 6.69773 | 16.68486 | 13.87506 | 11.24794 | 9.62907 | 0.00000 | 20.34228 | 12.96979 |
| **RFC3** | 0.37387 | 11.26670 | 11.32477 | 2.46763 | 2.60115 | 28.40820 | 18.78650 | 17.28402 | 11.58930 | 7.69277 | 16.92575 | 23.06075 |
| **RFC4** | 0.00000 | 17.71241 | 8.88343 | 5.72712 | 1.20160 | 27.12121 | 11.30941 | 7.49354 | 9.06773 | 1.12216 | 12.88641 | 12.33183 |
| **RFC5** | 0.03068 | 10.92764 | 9.26602 | 7.51324 | 3.77471 | 28.39809 | 9.35582 | 4.27133 | 13.86047 | 0.00053 | 14.50497 | 5.37901 |
| **RFESD** | 0.94008 | 0.16407 | 0.37737 | 0.00720 | 0.09606 | 1.29832 | 0.00000 | 0.18610 | 0.00000 | 0.00000 | 0.77396 | 0.99680 |
| **RFFL** | 0.60164 | 0.66157 | 0.28911 | 0.46396 | 0.07433 | 0.59593 | 0.00000 | 0.27408 | 0.43424 | 0.00000 | 0.44707 | 0.00113 |
| **RFK** | 2.65644 | 8.24004 | 7.32764 | 0.00000 | 19.93557 | 36.02724 | 15.35590 | 26.60289 | 21.73382 | 10.94922 | 26.30793 | 25.60557 |
| **RFNG** | 2.29503 | 1.04021 | 1.59487 | 1.49588 | 2.59022 | 0.00000 | 1.44391 | 0.00000 | 1.38850 | 9.11378 | 2.11294 | 0.64899 |
| **RFPL1** | 1.41695 | 0.00890 | 0.01075 | 0.00985 | 0.01773 | 0.00000 | 0.01324 | 3.92823 | 0.02294 | 0.00742 | 0.01271 | 0.00000 |
| **RFPL1S** | 0.36612 | 0.00000 | 0.00000 | 0.08352 | 0.00000 | 0.15876 | 0.14085 | 0.56269 | 0.00000 | 0.24968 | 0.00435 | 0.00000 |
| **RFPL2** | 7.38320 | 0.02492 | 0.00848 | 0.00260 | 0.00914 | 0.00000 | 0.01234 | 16.04489 | 0.01775 | 0.00908 | 0.00824 | 0.00000 |
| **RFPL3** | 0.02563 | 0.03269 | 0.01293 | 0.01366 | 0.01147 | 0.01054 | 0.02032 | 1.07990 | 0.01952 | 0.02059 | 0.01331 | 0.00303 |
| **RFPL3S** | 0.20690 | 0.09705 | 0.13395 | 0.00000 | 0.18573 | 0.19753 | 0.03260 | 0.00000 | 0.26890 | 0.66014 | 0.18285 | 0.25062 |
| **RFPL4A** | 3.12383 | 0.00889 | 0.00921 | 0.01566 | 0.01686 | 0.00000 | 0.01146 | 7.11092 | 0.01931 | 0.00879 | 0.01004 | 0.00000 |
| **RFPL4AL1** | 2.65767 | 0.00932 | 0.01004 | 0.01076 | 0.01845 | 0.00000 | 0.01556 | 4.12226 | 0.02048 | 0.00000 | 0.01442 | 0.00000 |
| **RFPL4B** | 6.20519 | 3.25377 | 15.72208 | 0.00004 | 0.00000 | 7.71961 | 6.01857 | 19.72500 | 4.40644 | 0.00000 | 4.75771 | 12.64969 |
| **RFT1** | 0.79843 | 13.71034 | 3.93848 | 4.07099 | 5.23919 | 4.21247 | 0.00000 | 3.15468 | 7.22791 | 0.63826 | 7.00762 | 7.06986 |
| **RFTN1** | 0.89228 | 0.03405 | 0.13936 | 0.11017 | 0.00520 | 0.15309 | 0.00000 | 0.23046 | 0.33112 | 0.37402 | 0.14565 | 0.03536 |
| **RFTN2** | 0.00144 | 1.19729 | 0.04257 | 0.50599 | 0.00000 | 0.34843 | 0.00000 | 0.61658 | 0.02647 | 0.20959 | 0.13338 | 1.82855 |

| | | | | | | | | | | | | |
|---|---|---|---|---|---|---|---|---|---|---|---|---|
| **RFWD2** | 0.20652 | 1.75955 | 4.20540 | 3.58821 | 4.05771 | 5.40407 | 4.64003 | 0.49128 | 0.27897 | 10.47871 | 3.15989 | 1.11264 |
| **RFWD3** | 4.13211 | 14.81928 | 8.02191 | 10.33230 | 11.13090 | 11.52568 | 11.25985 | 15.85964 | 12.81968 | 9.59506 | 8.29302 | 18.32709 |
| **RFX1** | 1.07632 | 0.00000 | 0.27594 | 0.35339 | 1.67580 | 0.00000 | 0.81382 | 0.39799 | 0.93031 | 6.19002 | 0.70354 | 1.36226 |
| **RFX2** | 0.52507 | 1.07553 | 0.41913 | 0.48903 | 0.71240 | 0.55107 | 0.00000 | 0.11268 | 0.98826 | 0.80488 | 0.00000 | 0.86021 |
| **RFX3** | 1.32559 | 0.06726 | 1.23806 | 0.00000 | 0.00000 | 1.20213 | 0.00003 | 0.03414 | 1.85489 | 1.59618 | 0.22441 | 0.21866 |
| **RFX4** | 0.25989 | 0.02017 | 0.06531 | 0.02384 | 0.01901 | 0.00000 | 0.00380 | 0.24597 | 0.03173 | 0.02345 | 0.06291 | 0.25699 |
| **RFX5** | 0.00000 | 0.12976 | 0.67880 | 3.09720 | 0.41040 | 2.55959 | 0.00000 | 0.18583 | 1.44897 | 1.10960 | 0.48335 | 0.00000 |
| **RFX6** | 0.00580 | 0.01561 | 0.06289 | 0.01641 | 0.04527 | 0.00000 | 0.02042 | 0.01063 | 0.00741 | 0.02792 | 0.02700 | 0.02020 |
| **RFX7** | 1.22231 | 0.07221 | 1.18072 | 0.00000 | 0.00000 | 3.19441 | 1.96034 | 1.00093 | 1.58670 | 3.41672 | 0.34561 | 0.46900 |
| **RFX8** | 0.00939 | 0.01885 | 0.01669 | 0.01640 | 0.01094 | 0.00049 | 0.02250 | 0.01790 | 0.00497 | 0.02014 | 0.06807 | 0.04175 |
| **RFXANK** | 1.05744 | 2.11522 | 4.14912 | 4.97698 | 3.36275 | 0.00000 | 9.46489 | 0.44908 | 4.45470 | 0.96924 | 4.95573 | 2.24313 |
| **RFXAP** | 0.00026 | 0.05788 | 0.82957 | 0.21019 | 0.23108 | 0.37776 | 1.19380 | 0.01847 | 0.00000 | 2.50101 | 0.53847 | 0.03913 |
| **RGAG1** | 0.00567 | 0.01095 | 0.20929 | 0.57023 | 0.00000 | 0.00832 | 0.04500 | 0.01008 | 0.00000 | 0.02882 | 0.04803 | 0.01496 |
| **RGAG4** | 0.00561 | 0.00543 | 0.02954 | 0.00498 | 0.00469 | 0.04077 | 0.00167 | 0.00854 | 0.00489 | 0.07684 | 0.02385 | 0.11820 |
| **RGCC** | 0.00000 | 4.40660 | 0.00000 | 3.23184 | 7.69755 | 2.07379 | 0.94183 | 0.10734 | 0.68664 | 6.39394 | 7.59798 | 0.46093 |
| **RGL1** | 0.00476 | 0.00000 | 5.77003 | 0.00000 | 0.81397 | 0.00000 | 1.71692 | 0.25597 | 0.00000 | 0.74811 | 0.39018 | 0.06895 |
| **RGL2** | 0.89832 | 0.00000 | 13.07711 | 1.67930 | 6.93238 | 0.33913 | 12.43410 | 0.67366 | 7.81916 | 4.51218 | 2.02438 | 2.07838 |
| **RGL3** | 0.12506 | 0.41373 | 0.48408 | 0.48293 | 0.00000 | 0.00000 | 0.00000 | 0.21095 | 0.56727 | 0.27729 | 0.00000 | 1.18780 |
| **RGL4** | 0.08419 | 0.08047 | 0.00000 | 0.15017 | 0.15482 | 0.00000 | 0.00120 | 0.00872 | 0.75643 | 0.36386 | 0.00000 | 0.04478 |
| **RGMA** | 3.68841 | 4.05218 | 0.11400 | 0.00000 | 9.26363 | 0.00000 | 0.70134 | 0.00000 | 2.33652 | 4.66201 | 1.20826 | 0.27058 |
| **RGMB** | 2.12775 | 1.87968 | 2.57308 | 0.96420 | 1.79129 | 1.37106 | 1.17570 | 2.69537 | 5.53838 | 2.42417 | 0.69738 | 1.70073 |
| **RGMB-AS1** | 0.44943 | 0.10023 | 0.06184 | 0.05229 | 0.09379 | 0.00000 | 0.07362 | 0.06732 | 0.00000 | 0.00000 | 0.01759 | 0.12465 |
| **RGN** | 0.00591 | 0.07292 | 0.07650 | 0.06015 | 0.01267 | 0.06342 | 0.14659 | 0.00744 | 0.28709 | 0.00000 | 0.26274 | 0.12670 |
| **RGP1** | 0.67969 | 1.02995 | 0.58840 | 0.39871 | 0.64994 | 0.00187 | 0.00000 | 0.20721 | 1.81214 | 0.00000 | 1.09555 | 1.01321 |
| **RGPD1.uid-2613** | 0.00554 | 0.01142 | 0.01671 | 0.01887 | 0.00987 | 0.05650 | 0.03361 | 0.01004 | 0.03142 | 0.03144 | 0.01408 | 0.01273 |
| **RGPD1.uid-2860** | 0.00614 | 0.01150 | 0.01863 | 0.00718 | 0.00908 | 0.04393 | 0.02884 | 0.00998 | 0.03368 | 0.00750 | 0.01817 | 0.01968 |
| **RGPD2.uid-2611** | 0.00532 | 0.02506 | 0.02730 | 0.00998 | 0.05669 | 0.03741 | 0.00332 | 0.03084 | 0.04995 | 0.01435 | 0.00961 | 0.03002 |
| **RGPD2.uid-2863** | 0.00515 | 0.02756 | 0.01213 | 0.02555 | 0.04023 | 0.02417 | 0.02623 | 0.01344 | 0.00604 | 0.03553 | 0.01247 | 0.01580 |
| **RGPD3** | 0.06488 | 0.01851 | 0.00976 | 0.00688 | 0.02039 | 0.02917 | 0.02892 | 0.15011 | 0.02045 | 0.03225 | 0.01335 | 0.02981 |
| **RGPD4** | 0.08133 | 0.01188 | 0.02773 | 0.07221 | 0.02583 | 0.09928 | 0.00000 | 0.16007 | 0.04996 | 0.03518 | 0.00308 | 0.04838 |
| **RGPD8** | 0.00561 | 0.01866 | 0.00950 | 0.01490 | 0.01337 | 0.02539 | 0.01076 | 0.01037 | 0.02100 | 0.04366 | 0.01294 | 0.01352 |
| **RGS10** | 0.00000 | 11.15940 | 0.00000 | 7.20724 | 5.84129 | 8.45367 | 0.83756 | 2.47525 | 1.28390 | 5.71048 | 3.63486 | 10.06524 |
| **RGS11** | 0.10627 | 0.21563 | 0.01379 | 0.01268 | 0.06025 | 0.00000 | 0.01242 | 0.00949 | 0.00498 | 0.08888 | 0.01203 | 0.01602 |
| **RGS12** | 0.01110 | 0.34651 | 1.01044 | 0.52737 | 0.20315 | 0.23100 | 0.51473 | 0.24116 | 0.36911 | 0.00000 | 0.00000 | 0.33118 |
| **RGS13** | 0.00031 | 0.00000 | 0.00000 | 0.00000 | 1.86361 | 0.12719 | 8.01016 | 0.00283 | 5.40728 | 0.23530 | 19.18724 | 0.00000 |
| **RGS14** | 2.94450 | 0.65746 | 1.35604 | 1.58011 | 0.28265 | 0.00078 | 0.44384 | 1.68528 | 0.81785 | 0.00000 | 0.00000 | 2.02874 |
| **RGS16** | 3.28991 | 9.19826 | 28.88737 | 11.80827 | 0.00000 | 4.29570 | 22.78933 | 3.84742 | 1.93121 | 1.95481 | 0.00000 | 0.04350 |
| **RGS17** | 0.05910 | 0.29759 | 0.00054 | 0.00000 | 0.15376 | 0.40824 | 0.00000 | 0.03248 | 0.05449 | 0.10411 | 0.00299 | 1.16775 |
| **RGS18** | 0.23292 | 0.01427 | 0.01686 | 0.01099 | 0.01405 | 0.01481 | 0.01513 | 0.09785 | 0.01912 | 0.01918 | 0.01409 | 0.00895 |
| **RGS19** | 0.07680 | 1.39535 | 2.14920 | 2.51280 | 0.70647 | 0.00000 | 1.61551 | 0.00000 | 3.19312 | 0.00000 | 2.80094 | 0.90339 |
| **RGS2** | 26.28470 | 9.82700 | 1.32190 | 3.33963 | 3.89570 | 14.01669 | 0.00000 | 15.49228 | 0.00000 | 3.43321 | 1.41770 | 32.95801 |
| **RGS20** | 0.00431 | 0.06713 | 0.13526 | 0.00807 | 0.00000 | 0.14948 | 0.00000 | 0.25496 | 0.00000 | 0.11280 | 0.00302 | 0.14392 |
| **RGS22** | 1.20458 | 0.00575 | 0.02444 | 0.01183 | 0.01221 | 0.03049 | 0.01602 | 0.05854 | 0.02231 | 0.00687 | 0.01160 | 0.08931 |
| **RGS3** | 0.07530 | 0.07642 | 4.47591 | 0.21143 | 0.65676 | 0.00000 | 1.27497 | 0.01277 | 5.17462 | 0.00000 | 0.79338 | 0.04560 |
| **RGS4** | 0.00534 | 0.01747 | 0.02519 | 0.02711 | 0.01809 | 0.00108 | 0.01953 | 0.01516 | 0.01538 | 0.01042 | 0.06729 | 0.02854 |
| **RGS5** | 0.11757 | 0.73900 | 0.82117 | 5.67279 | 0.00000 | 1.56106 | 0.00000 | 0.14381 | 0.58564 | 0.00000 | 0.42988 | 1.81725 |
| **RGS6** | 0.13453 | 0.00652 | 0.02468 | 0.00613 | 0.00000 | 0.02855 | 0.00000 | 0.03529 | 0.06656 | 0.48220 | 0.02626 | 0.00284 |
| **RGS7** | 1.44805 | 0.27306 | 0.08115 | 0.03856 | 0.00037 | 0.21309 | 0.10845 | 0.14761 | 0.00000 | 0.00000 | 0.01288 | 0.28993 |

| | | | | | | | | | | | | |
|---|---|---|---|---|---|---|---|---|---|---|---|---|
| **RGS7BP** | 0.00243 | 0.01307 | 0.03401 | 0.18372 | 0.00000 | 0.00951 | 0.04221 | 0.03793 | 0.00000 | 0.15903 | 0.02785 | 0.04187 |
| **RGS8** | 0.14874 | 0.13310 | 0.04124 | 0.00860 | 0.00588 | 0.00000 | 0.00441 | 0.53403 | 0.03758 | 0.01443 | 0.01595 | 0.00314 |
| **RGS9** | 0.00744 | 0.03276 | 0.04495 | 0.31993 | 0.03168 | 0.01444 | 0.01193 | 0.01952 | 0.00000 | 0.11252 | 0.04590 | 0.01857 |
| **RGS9BP** | 0.00480 | 0.02771 | 0.04873 | 0.00339 | 0.01108 | 0.01635 | 0.00237 | 0.01121 | 0.03561 | 0.09348 | 0.00971 | 0.01469 |
| **RGSL1** | 0.02738 | 0.07068 | 0.01809 | 0.00652 | 0.01144 | 0.01170 | 0.01703 | 0.10975 | 0.01608 | 0.01611 | 0.01595 | 0.13334 |
| **RHAG** | 0.02950 | 0.01544 | 0.01243 | 0.01383 | 0.01685 | 0.01320 | 0.02049 | 0.06919 | 0.02753 | 0.01511 | 0.01224 | 0.01340 |
| **RHBDD1** | 9.38463 | 3.29831 | 0.41037 | 0.85211 | 1.53716 | 2.88912 | 0.00204 | 1.01106 | 2.91168 | 1.43911 | 1.21244 | 3.22315 |
| **RHBDD2** | 1.57810 | 6.62230 | 7.27276 | 6.54348 | 13.37850 | 0.00000 | 3.79651 | 1.82181 | 17.34790 | 0.00000 | 6.99898 | 4.67624 |
| **RHBDD3** | 0.59684 | 4.09505 | 12.43612 | 5.07037 | 6.14427 | 0.00000 | 7.03571 | 0.00000 | 16.04188 | 6.07344 | 5.62034 | 4.86509 |
| **RHBDF1** | 0.40715 | 1.50315 | 1.80535 | 1.27187 | 3.30723 | 0.00000 | 0.00000 | 0.00000 | 1.88145 | 3.38408 | 0.56567 | 0.16363 |
| **RHBDF2** | 2.63918 | 4.64853 | 1.30983 | 0.00000 | 0.09429 | 0.00000 | 0.00000 | 0.28089 | 1.97683 | 0.36104 | 1.34634 | 1.31765 |
| **RHBDL1** | 0.00272 | 0.00000 | 0.00000 | 0.30675 | 0.18968 | 0.00000 | 0.00362 | 0.03920 | 0.34913 | 1.47467 | 0.00000 | 0.07669 |
| **RHBDL2** | 0.23965 | 0.10677 | 0.13932 | 0.04742 | 0.09958 | 0.04270 | 0.00017 | 0.10848 | 0.06969 | 0.06223 | 0.00010 | 0.51749 |
| **RHBDL3** | 0.15887 | 0.03174 | 0.05673 | 0.09847 | 0.01350 | 0.00000 | 0.01444 | 0.00782 | 0.01221 | 0.06662 | 0.01696 | 0.02224 |
| **RHBG** | 0.21812 | 0.03779 | 0.16979 | 0.00000 | 0.06995 | 0.02126 | 0.05485 | 0.19479 | 0.00000 | 0.13876 | 0.02886 | 0.85686 |
| **RHCE** | 0.02611 | 0.02036 | 0.02123 | 0.01665 | 0.01169 | 0.02215 | 0.01423 | 0.05398 | 0.01843 | 0.02256 | 0.01391 | 0.01498 |
| **RHCG** | 0.00357 | 0.01696 | 0.01302 | 0.02169 | 0.01371 | 0.01491 | 0.01648 | 0.02667 | 0.05587 | 0.00000 | 0.01899 | 0.01674 |
| **RHD** | 0.00879 | 0.01638 | 0.06655 | 0.09084 | 0.05566 | 0.00000 | 0.01398 | 0.01449 | 0.10100 | 0.05468 | 0.00384 | 0.05255 |
| **RHEB** | 1.08256 | 9.54080 | 10.83741 | 3.34325 | 4.38623 | 18.01727 | 14.09420 | 2.75830 | 9.54094 | 0.69491 | 3.94128 | 5.22330 |
| **RHEBL1** | 3.07483 | 1.47788 | 1.02610 | 0.78510 | 0.33320 | 0.00000 | 2.75146 | 17.09930 | 0.49537 | 0.38654 | 0.00000 | 12.90542 |
| **RHNO1** | 1.41379 | 10.54540 | 3.39228 | 1.71750 | 1.14202 | 14.37091 | 2.38164 | 0.00000 | 5.36769 | 0.00000 | 5.42994 | 10.38107 |
| **RHO** | 0.00312 | 0.00308 | 0.04830 | 0.00000 | 0.00000 | 0.02650 | 0.00000 | 0.00939 | 0.01673 | 0.47885 | 0.00000 | 0.00647 |
| **RHOA** | 12.57167 | 34.41327 | 28.27201 | 22.56968 | 27.00995 | 53.28509 | 42.72287 | 28.34151 | 41.16537 | 11.83595 | 32.36373 | 38.86727 |
| **RHOB** | 0.19990 | 0.54865 | 2.54588 | 0.01711 | 6.38870 | 5.93098 | 1.07252 | 0.74541 | 0.00000 | 12.42297 | 0.31424 | 11.22098 |
| **RHOBTB1** | 0.26360 | 0.00000 | 31.57290 | 0.53288 | 0.00000 | 0.00000 | 6.82137 | 0.56150 | 8.30388 | 0.14456 | 2.12437 | 0.00039 |
| **RHOBTB2** | 2.18208 | 0.00000 | 1.48662 | 0.00279 | 0.00000 | 0.16911 | 7.73888 | 0.23982 | 4.84304 | 5.12624 | 3.57803 | 0.40774 |
| **RHOBTB3** | 4.75391 | 0.00000 | 0.00001 | 8.18651 | 3.33215 | 0.87448 | 1.14978 | 0.26731 | 1.90448 | 1.60272 | 2.80971 | 0.21981 |
| **RHOC** | 3.35558 | 38.72559 | 23.47607 | 26.83538 | 44.72443 | 21.97526 | 37.44538 | 0.28527 | 35.51030 | 0.22931 | 26.47174 | 2.48378 |
| **RHOD** | 0.01486 | 0.08541 | 1.73288 | 0.08078 | 0.00000 | 0.02330 | 0.30068 | 0.97465 | 1.16424 | 0.00000 | 0.15372 | 0.21250 |
| **RHOF** | 0.00659 | 0.56069 | 0.26527 | 0.49781 | 0.66766 | 0.00000 | 0.29760 | 0.11882 | 0.07188 | 0.00000 | 0.38368 | 0.22828 |
| **RHOG** | 3.38816 | 13.51815 | 14.41568 | 12.94376 | 16.41546 | 1.43841 | 16.94021 | 0.00000 | 15.84130 | 0.00000 | 7.50901 | 8.39472 |
| **RHOH** | 0.00761 | 0.03080 | 0.04221 | 0.10528 | 0.16157 | 0.00000 | 0.02836 | 0.01067 | 0.00000 | 0.03964 | 0.24277 | 0.01338 |
| **RHOJ** | 0.17693 | 0.01444 | 0.06240 | 0.01095 | 0.01758 | 0.00967 | 0.01234 | 0.00313 | 0.01135 | 0.02232 | 0.02230 | 0.01621 |
| **RHOQ** | 0.20174 | 0.15974 | 0.38311 | 0.37980 | 0.08061 | 0.55235 | 0.00000 | 0.01739 | 0.00000 | 1.25358 | 0.16592 | 0.50185 |
| **RHOT1** | 1.77882 | 1.39200 | 0.73178 | 0.39008 | 0.90912 | 6.96158 | 0.61584 | 0.07260 | 1.45548 | 6.29912 | 1.01338 | 0.00000 |
| **RHOT2** | 0.00000 | 19.04922 | 11.73398 | 15.73222 | 17.49534 | 0.00000 | 10.32623 | 2.45760 | 21.13905 | 16.05163 | 16.62682 | 6.16637 |
| **RHOU** | 0.00000 | 0.00000 | 7.87976 | 0.00000 | 6.38706 | 5.04241 | 20.37639 | 0.00000 | 6.90980 | 8.63742 | 7.94753 | 0.00000 |
| **RHOV** | 0.00050 | 8.66613 | 2.25626 | 0.15856 | 6.29691 | 0.00000 | 4.21948 | 0.00414 | 0.00000 | 8.84814 | 0.80508 | 0.00000 |
| **RHOXF1** | 0.00175 | 0.06318 | 0.00000 | 0.02970 | 0.00000 | 0.00004 | 0.04647 | 0.09082 | 0.09854 | 0.20254 | 0.03328 | 1.15879 |
| **RHPN1** | 0.14638 | 0.10645 | 0.17985 | 0.02475 | 0.02460 | 0.00000 | 0.00575 | 0.01824 | 0.10091 | 0.03451 | 0.05656 | 0.00545 |
| **RHPN1-AS1** | 0.39403 | 0.12211 | 0.05267 | 0.00714 | 0.00000 | 0.01250 | 0.02562 | 0.00250 | 0.06630 | 0.00602 | 0.12340 | 0.01694 |
| **RHPN2** | 2.11936 | 14.22264 | 0.00000 | 0.00542 | 11.47685 | 5.34038 | 11.11487 | 1.24944 | 7.07851 | 7.72464 | 8.59473 | 8.44885 |
| **RIBC1** | 0.74084 | 0.33328 | 0.09189 | 0.46902 | 0.00000 | 0.20476 | 0.07696 | 0.70800 | 0.04315 | 0.00000 | 0.07297 | 0.27994 |
| **RIBC2** | 1.15979 | 0.58429 | 0.02287 | 0.00000 | 0.21805 | 1.41848 | 0.00000 | 0.69702 | 0.00005 | 0.66153 | 0.85129 | 2.99902 |
| **RIC3** | 0.19397 | 0.56320 | 0.01426 | 0.23473 | 0.18081 | 0.58456 | 0.00000 | 0.11349 | 0.34753 | 1.24835 | 0.52105 | 0.72708 |
| **RIC8A** | 4.73099 | 12.31199 | 10.67448 | 5.98427 | 8.59950 | 0.00000 | 11.58724 | 11.85064 | 13.59652 | 2.97212 | 7.98454 | 10.82123 |
| **RIC8B** | 0.72678 | 0.20782 | 0.83617 | 0.52442 | 0.54421 | 3.10928 | 1.74307 | 0.98245 | 1.87186 | 1.79038 | 1.87258 | 3.43949 |
| **RICTOR** | 3.11183 | 0.06987 | 2.34401 | 0.00000 | 0.00000 | 6.94666 | 4.03527 | 2.16474 | 1.81225 | 9.33913 | 1.10277 | 0.10147 |

| | | | | | | | | | | | | |
|---|---|---|---|---|---|---|---|---|---|---|---|---|
| **RIF1** | 7.20691 | 2.63231 | 8.70033 | 6.72033 | 4.79549 | 33.99270 | 12.48631 | 15.28919 | 8.76857 | 47.95035 | 11.30759 | 18.28733 |
| **RIIAD1** | 5.91300 | 0.00244 | 0.02162 | 0.00888 | 0.01516 | 0.00003 | 0.01315 | 0.60933 | 0.02048 | 0.00000 | 0.02373 | 0.34695 |
| **RILP** | 0.00177 | 0.98912 | 0.00000 | 0.93427 | 3.10473 | 0.00000 | 0.20290 | 0.24358 | 0.18618 | 2.65810 | 0.03879 | 0.13793 |
| **RILPL1** | 0.04894 | 0.21919 | 2.47215 | 0.69379 | 0.32033 | 0.02643 | 1.15916 | 0.00385 | 0.31726 | 1.52710 | 0.12865 | 0.00000 |
| **RILPL2** | 0.00378 | 0.11212 | 5.64945 | 0.55640 | 0.00000 | 0.13626 | 2.07480 | 0.18188 | 1.94536 | 0.00000 | 0.96232 | 0.08207 |
| **RIMBP2** | 1.13350 | 0.01278 | 0.00452 | 0.02802 | 0.03495 | 0.02331 | 0.00000 | 0.04599 | 0.00000 | 0.18166 | 0.03992 | 0.00253 |
| **RIMBP3** | 0.00595 | 0.01578 | 0.02622 | 0.00928 | 0.01549 | 0.01233 | 0.01896 | 0.01022 | 0.01635 | 0.04576 | 0.02543 | 0.01764 |
| **RIMKLA** | 5.54785 | 0.22611 | 0.45369 | 0.00000 | 0.08638 | 0.00000 | 0.48559 | 0.67204 | 0.18998 | 0.00000 | 0.00000 | 0.13846 |
| **RIMKLB** | 1.61920 | 10.25593 | 0.44277 | 4.36152 | 9.80786 | 12.73655 | 0.00000 | 0.42490 | 0.00000 | 2.97001 | 2.34469 | 15.47017 |
| **RIMS1** | 1.26042 | 0.00000 | 0.01084 | 0.00218 | 0.21896 | 0.00000 | 0.00909 | 0.27018 | 0.00000 | 0.66834 | 0.10208 | 0.31048 |
| **RIMS2** | 0.54790 | 0.09306 | 0.02368 | 0.19641 | 0.00000 | 0.03262 | 0.45540 | 0.28876 | 0.39090 | 0.00000 | 0.01382 | 0.01840 |
| **RIMS3** | 0.59850 | 0.12541 | 0.03972 | 0.35855 | 0.43439 | 0.15570 | 0.00000 | 0.05151 | 0.25221 | 0.17166 | 0.00000 | 0.00000 |
| **RIMS4** | 0.38402 | 0.77858 | 0.00436 | 0.01258 | 0.62748 | 0.00000 | 0.17093 | 0.03010 | 0.00000 | 0.00000 | 0.00183 | 1.52902 |
| **RIN1** | 0.00244 | 0.01602 | 0.00000 | 0.05743 | 0.03987 | 0.00000 | 0.10846 | 0.04595 | 0.31577 | 0.74190 | 0.00000 | 0.07033 |
| **RIN2** | 0.00000 | 1.12821 | 0.32639 | 3.06659 | 0.20161 | 8.04181 | 0.00000 | 0.05278 | 0.00000 | 2.65368 | 5.43858 | 0.72776 |
| **RIN3** | 0.35215 | 0.00000 | 5.41821 | 0.05437 | 1.54749 | 0.00000 | 1.05748 | 0.00223 | 2.78401 | 0.50027 | 2.66803 | 0.00000 |
| **RING1** | 0.31392 | 0.49995 | 0.93637 | 0.42163 | 1.67977 | 0.02554 | 1.90266 | 0.26656 | 0.00024 | 4.77138 | 0.28899 | 0.84286 |
| **RINL** | 0.00600 | 0.31261 | 1.34709 | 1.41609 | 0.11755 | 0.00000 | 0.41462 | 0.04629 | 0.11667 | 0.86660 | 0.86047 | 0.40997 |
| **RINT1** | 1.11692 | 7.31573 | 1.22341 | 1.99382 | 0.23512 | 15.10377 | 3.35333 | 5.45145 | 4.98829 | 0.00000 | 6.10982 | 6.78932 |
| **RIOK1** | 7.74010 | 9.99407 | 12.76861 | 1.14287 | 2.78794 | 18.10885 | 18.39271 | 7.08054 | 12.61968 | 12.56230 | 12.96186 | 15.27211 |
| **RIOK2** | 1.09831 | 11.04960 | 0.00000 | 3.09257 | 1.68861 | 23.23484 | 1.85109 | 5.94835 | 2.75333 | 5.11870 | 8.32942 | 16.55612 |
| **RIOK3** | 11.62445 | 10.59569 | 11.23094 | 1.37558 | 9.18853 | 25.79985 | 11.47521 | 9.26329 | 7.54813 | 12.88607 | 9.37607 | 11.52298 |
| **RIPK1** | 0.00000 | 1.59614 | 0.87579 | 0.58275 | 2.12043 | 1.19416 | 1.26531 | 1.67775 | 1.45163 | 1.89832 | 0.53060 | 1.46072 |
| **RIPK2** | 1.66772 | 5.52789 | 7.61647 | 0.38814 | 0.64748 | 13.03037 | 6.21061 | 0.06354 | 8.17871 | 0.00000 | 5.77038 | 2.39819 |
| **RIPK3** | 0.00649 | 0.01124 | 0.01785 | 0.00649 | 0.00818 | 0.05292 | 0.02265 | 0.01002 | 0.07209 | 0.00465 | 0.00340 | 0.01445 |
| **RIPK4** | 1.36826 | 5.69691 | 7.88076 | 6.39831 | 1.22945 | 0.00000 | 3.06739 | 0.29329 | 7.77559 | 1.34519 | 0.65076 | 1.13493 |
| **RIPPLY2** | 0.04891 | 0.01024 | 0.06423 | 0.04829 | 0.00604 | 0.03737 | 0.00699 | 0.29104 | 0.01115 | 0.01180 | 0.01555 | 0.01748 |
| **RIPPLY3** | 0.16443 | 0.01380 | 0.02958 | 0.01005 | 0.01743 | 0.01620 | 0.02783 | 0.14733 | 0.00813 | 0.02860 | 0.01946 | 0.00918 |
| **RIT1** | 0.00000 | 1.00669 | 5.10993 | 1.73668 | 0.00000 | 5.23224 | 3.93172 | 7.43487 | 2.73701 | 2.04805 | 2.17694 | 3.03678 |
| **RIT2** | 0.36377 | 0.00000 | 0.32192 | 0.04600 | 0.50332 | 0.35424 | 0.22951 | 0.06876 | 0.00000 | 0.00000 | 0.74935 | 0.03645 |
| **RITA1** | 0.30980 | 10.19812 | 6.98660 | 3.96684 | 6.40722 | 0.48483 | 6.02742 | 1.25924 | 12.15313 | 0.00000 | 10.87706 | 8.31572 |
| **RLBP1** | 0.00574 | 0.01698 | 0.02082 | 0.00964 | 0.01706 | 0.01481 | 0.02695 | 0.01145 | 0.02269 | 0.02388 | 0.01249 | 0.01548 |
| **RLF** | 7.09855 | 1.50568 | 6.25121 | 0.55437 | 2.91341 | 10.95402 | 5.20201 | 14.51724 | 11.86889 | 12.91271 | 6.30938 | 9.19774 |
| **RLIM** | 2.22486 | 3.70763 | 3.26676 | 7.55316 | 9.45417 | 12.32026 | 13.71488 | 20.22067 | 3.63573 | 27.07084 | 4.29251 | 9.42114 |
| **RLN1** | 0.51474 | 0.00494 | 0.04544 | 0.03433 | 0.00000 | 0.08102 | 0.00000 | 0.01059 | 0.02262 | 0.03058 | 0.05697 | 0.00913 |
| **RLN2** | 0.00834 | 0.03463 | 0.10168 | 0.00000 | 0.00000 | 0.06513 | 0.05433 | 0.01057 | 0.00000 | 0.05575 | 0.08182 | 0.01692 |
| **RLN3** | 0.00894 | 0.04400 | 0.02619 | 0.01932 | 0.02725 | 0.00592 | 0.02036 | 0.02207 | 0.01993 | 0.00001 | 0.01523 | 0.06865 |
| **RLTPR** | 0.28758 | 0.00000 | 0.30839 | 0.07859 | 0.14535 | 0.00000 | 0.25675 | 0.00000 | 0.00000 | 0.20347 | 0.01181 | 0.09205 |
| **RMDN1** | 0.97626 | 7.05434 | 5.03914 | 5.90809 | 6.49916 | 17.43188 | 5.06372 | 0.00000 | 2.96557 | 2.30858 | 11.47646 | 2.50349 |
| **RMDN2** | 0.00000 | 1.75520 | 0.00000 | 1.15915 | 0.00000 | 1.82653 | 0.00000 | 0.26273 | 0.00000 | 0.85543 | 0.28134 | 1.12857 |
| **RMDN2-AS1** | 0.00367 | 0.05068 | 0.01358 | 0.01431 | 0.00665 | 0.03961 | 0.01201 | 0.01682 | 0.01062 | 0.02854 | 0.01218 | 0.04073 |
| **RMDN3** | 3.72681 | 15.18113 | 13.90828 | 1.09244 | 8.73360 | 5.07484 | 3.29290 | 3.15524 | 15.51137 | 0.00000 | 13.00287 | 9.05938 |
| **RMI1** | 4.17534 | 3.75053 | 1.28488 | 3.02371 | 0.00000 | 16.37159 | 2.61273 | 9.64080 | 3.89497 | 4.35508 | 5.61303 | 5.48739 |
| **RMI2** | 1.23466 | 0.72903 | 1.37128 | 1.22031 | 0.97670 | 1.84547 | 0.47841 | 0.00844 | 0.00000 | 4.93359 | 1.40463 | 1.50988 |
| **RMND1** | 0.82474 | 19.44855 | 7.22217 | 11.03021 | 5.27460 | 25.04111 | 6.83125 | 8.90486 | 10.61072 | 3.35166 | 8.27230 | 23.00809 |
| **RMND5A** | 5.74894 | 5.34155 | 13.00016 | 6.36882 | 0.00000 | 20.62720 | 23.96380 | 16.98096 | 6.29136 | 17.20289 | 13.08532 | 7.41371 |
| **RMND5B** | 1.83311 | 5.91621 | 5.70686 | 4.79199 | 5.81615 | 0.00000 | 3.25287 | 0.20467 | 2.63650 | 0.57326 | 4.31579 | 1.71927 |
| **RMRP** | 1.20244 | 0.00000 | 0.97468 | 1.04087 | 2.48374 | 0.00000 | 0.07410 | 0.01301 | 0.00000 | 1.94678 | 0.00000 | 0.18955 |

| | | | | | | | | | | | | |
|---|---|---|---|---|---|---|---|---|---|---|---|---|
| **RNASE1** | 0.02428 | 0.00000 | 2.58990 | 6.37186 | 2.72020 | 0.00000 | 0.00000 | 0.13142 | 3.43344 | 1.46720 | 11.59933 | 0.00000 |
| **RNASE10** | 0.00902 | 0.15573 | 0.00140 | 1.13041 | 0.00000 | 0.21787 | 0.00000 | 0.20510 | 0.00000 | 0.01077 | 0.00120 | 0.61758 |
| **RNASE4** | 0.00331 | 0.00000 | 0.09787 | 0.00946 | 0.00000 | 0.09758 | 0.00000 | 0.18799 | 0.24889 | 0.00780 | 0.06250 | 1.10632 |
| **RNASE7** | 0.00778 | 0.01303 | 0.05315 | 0.00909 | 0.01214 | 0.02176 | 0.01075 | 0.01266 | 0.02772 | 0.01839 | 0.01350 | 0.03062 |
| **RNASEH1** | 1.13365 | 11.56333 | 7.32107 | 7.06007 | 10.30807 | 12.45286 | 10.25414 | 3.78218 | 9.68302 | 3.93370 | 11.60801 | 11.11697 |
| **RNASEH1-AS1** | 0.00197 | 4.18037 | 1.74686 | 2.45064 | 1.22089 | 0.40063 | 2.02952 | 3.75746 | 3.78209 | 2.00471 | 2.34542 | 3.37661 |
| **RNASEH2A** | 1.96768 | 10.70209 | 9.24822 | 8.31101 | 10.23903 | 8.99597 | 17.13795 | 2.12727 | 11.67233 | 0.00000 | 16.39321 | 10.68244 |
| **RNASEH2B** | 0.00000 | 2.40641 | 3.41810 | 1.54257 | 2.85583 | 9.60505 | 2.50401 | 0.06514 | 3.29698 | 1.19682 | 5.01588 | 2.06250 |
| **RNASEH2B-AS1** | 0.01772 | 0.03290 | 0.03411 | 0.03366 | 0.00000 | 0.02368 | 0.01552 | 0.01113 | 0.03603 | 0.02118 | 0.06674 | 0.02263 |
| **RNASEH2C** | 2.73279 | 11.31767 | 8.04583 | 11.93652 | 25.33284 | 0.01852 | 9.73568 | 7.93559 | 10.52993 | 1.59387 | 13.26991 | 17.71246 |
| **RNASEK-C17or** | 0.03234 | 0.04269 | 0.02746 | 0.03387 | 0.04487 | 0.00000 | 0.06381 | 0.01259 | 0.06512 | 0.00000 | 0.02159 | 0.05527 |
| **RNASEK.uid-21** | 0.01191 | 0.41724 | 0.97556 | 0.36568 | 0.10404 | 0.05861 | 0.95412 | 0.00582 | 1.40204 | 0.00000 | 0.95048 | 0.20416 |
| **RNASEL** | 0.00235 | 0.53085 | 1.26130 | 0.57802 | 0.00000 | 0.62530 | 2.34034 | 0.27293 | 0.00198 | 0.00000 | 0.00000 | 0.47219 |
| **RNASET2** | 2.28894 | 5.87933 | 0.98890 | 6.18477 | 4.42152 | 1.66523 | 0.43177 | 3.66644 | 1.47704 | 2.55873 | 2.02769 | 0.23661 |
| **RND1** | 15.56440 | 0.44826 | 0.09769 | 0.52881 | 0.00000 | 0.09551 | 0.00000 | 6.13764 | 0.00000 | 0.05503 | 0.00734 | 2.02971 |
| **RND2** | 0.57755 | 6.08514 | 0.52369 | 7.86962 | 3.77840 | 0.21846 | 2.38892 | 0.01552 | 0.00000 | 0.90469 | 1.86726 | 3.51857 |
| **RND3** | 0.25165 | 0.00000 | 1.03217 | 0.51323 | 0.00000 | 0.02648 | 0.04146 | 2.52022 | 0.00000 | 0.06435 | 0.00818 | 0.23291 |
| **RNF10** | 14.83609 | 18.59992 | 18.59153 | 8.32359 | 15.35186 | 14.07078 | 11.87759 | 7.25504 | 19.48653 | 12.69572 | 10.06488 | 8.69858 |
| **RNF103** | 0.00000 | 0.17241 | 5.46795 | 0.00000 | 0.00000 | 0.97734 | 2.90964 | 0.75133 | 0.64713 | 2.14779 | 0.30701 | 0.71755 |
| **RNF103-CHMP3** | 0.01360 | 0.02414 | 0.23180 | 0.00702 | 0.01828 | 0.02386 | 0.01046 | 0.02556 | 0.02809 | 0.01832 | 0.01337 | 0.04994 |
| **RNF11** | 1.04019 | 1.65776 | 9.32796 | 1.41327 | 0.89524 | 14.92850 | 6.92014 | 14.87198 | 0.42642 | 10.96706 | 1.01323 | 40.85414 |
| **RNF111** | 7.17157 | 2.99053 | 9.64325 | 1.60986 | 6.63588 | 7.71874 | 8.16683 | 3.87765 | 7.58070 | 5.50607 | 3.64588 | 7.69106 |
| **RNF112** | 0.00625 | 0.05092 | 0.02664 | 0.00801 | 0.03826 | 0.00000 | 0.04538 | 0.01126 | 0.20107 | 0.00000 | 0.00454 | 0.01758 |
| **RNF113A** | 2.03350 | 6.45329 | 2.82469 | 4.77829 | 7.15801 | 3.86612 | 6.14643 | 10.64501 | 3.61887 | 0.31464 | 7.81737 | 13.04187 |
| **RNF113B** | 0.00350 | 0.14461 | 0.32713 | 0.10903 | 0.21554 | 0.08125 | 0.04426 | 0.29092 | 0.00000 | 0.02808 | 0.09142 | 0.11333 |
| **RNF114** | 9.51989 | 20.23898 | 14.90997 | 16.99660 | 21.84919 | 23.84790 | 16.91378 | 12.36393 | 18.94753 | 0.36042 | 13.09709 | 28.20639 |
| **RNF115** | 2.18681 | 9.87956 | 0.66596 | 4.31617 | 9.83594 | 13.51489 | 0.49823 | 2.87073 | 0.74166 | 8.55360 | 3.38596 | 11.13431 |
| **RNF121** | 0.94553 | 6.74462 | 7.77284 | 6.25937 | 4.09423 | 0.00003 | 0.00000 | 5.06124 | 7.24127 | 1.45848 | 5.99710 | 7.65859 |
| **RNF122** | 7.76651 | 0.06411 | 0.03056 | 0.00000 | 0.34492 | 0.00000 | 0.09025 | 6.43903 | 0.00000 | 1.05496 | 0.03531 | 0.73378 |
| **RNF123** | 0.41640 | 3.49298 | 5.83075 | 3.59312 | 3.36471 | 0.00000 | 3.09647 | 0.25515 | 6.81184 | 2.64129 | 3.60804 | 3.17232 |
| **RNF125** | 2.94845 | 7.14253 | 0.00000 | 12.75225 | 0.77369 | 7.94258 | 4.46625 | 12.54150 | 0.00000 | 16.02554 | 11.75813 | 24.28832 |
| **RNF126** | 2.76487 | 13.58391 | 7.45853 | 4.93624 | 11.68568 | 0.00000 | 4.80431 | 2.23605 | 13.39249 | 15.25944 | 2.97468 | 10.31840 |
| **RNF126P1** | 0.14028 | 0.97746 | 0.41984 | 0.38789 | 0.57845 | 0.00000 | 0.00000 | 0.14986 | 0.82507 | 0.25983 | 0.16941 | 0.59528 |
| **RNF128** | 0.00902 | 0.22664 | 0.08933 | 0.01071 | 0.00000 | 0.03405 | 0.00000 | 0.01889 | 0.00992 | 0.08094 | 0.00624 | 0.10021 |
| **RNF13** | 1.69846 | 4.41842 | 4.91296 | 0.00000 | 2.31134 | 9.10634 | 2.80757 | 1.94804 | 9.01597 | 0.45061 | 6.32852 | 10.54222 |
| **RNF130** | 2.52637 | 0.69185 | 0.69283 | 0.34997 | 0.24753 | 0.11429 | 0.61451 | 0.55928 | 0.00000 | 0.00000 | 0.04270 | 0.45650 |
| **RNF135** | 1.31171 | 5.26968 | 2.24225 | 5.74313 | 3.71752 | 4.96408 | 2.41228 | 0.25273 | 3.69511 | 2.31195 | 3.34922 | 3.75437 |
| **RNF138** | 3.16785 | 9.89823 | 3.26731 | 4.02149 | 0.00176 | 30.31336 | 9.04467 | 18.35494 | 9.43074 | 3.95954 | 8.43577 | 15.45737 |
| **RNF138P1** | 0.34347 | 1.25546 | 0.10240 | 1.55389 | 0.57736 | 2.42273 | 0.00000 | 4.49723 | 1.34383 | 0.63083 | 0.91739 | 3.39201 |
| **RNF139** | 0.11052 | 4.40079 | 1.96682 | 2.99604 | 2.72703 | 14.52071 | 0.00000 | 2.62842 | 3.97643 | 16.49742 | 0.97144 | 2.98627 |
| **RNF139-AS1** | 0.00034 | 0.08466 | 0.11587 | 0.00000 | 0.00000 | 0.32955 | 0.00000 | 0.42912 | 0.13998 | 0.71951 | 0.31444 | 0.21801 |
| **RNF14** | 8.65782 | 9.12413 | 3.72978 | 6.26469 | 4.45201 | 19.69220 | 2.33431 | 2.37129 | 3.15927 | 0.00000 | 6.92322 | 2.65888 |
| **RNF141** | 0.00000 | 5.38974 | 0.35129 | 2.00833 | 4.50439 | 9.78487 | 0.00000 | 0.09818 | 0.00000 | 1.77688 | 2.42961 | 4.70412 |
| **RNF144A** | 0.30784 | 0.45534 | 0.37024 | 0.00578 | 0.04721 | 0.00000 | 0.23503 | 0.06142 | 0.00000 | 0.00000 | 0.03797 | 0.15678 |
| **RNF144A-AS1** | 0.08782 | 0.04933 | 0.01683 | 0.00574 | 0.00725 | 0.02389 | 0.01229 | 0.09962 | 0.01422 | 0.03403 | 0.01292 | 0.03389 |
| **RNF144B** | 0.81767 | 0.93414 | 0.41897 | 0.00004 | 0.00116 | 0.60715 | 0.07678 | 1.21810 | 0.15250 | 0.00000 | 0.04064 | 0.96499 |
| **RNF145** | 1.81036 | 2.99832 | 0.55506 | 2.49683 | 1.68031 | 7.54156 | 7.30886 | 3.18242 | 4.97340 | 13.96543 | 3.67345 | 9.99307 |
| **RNF146** | 0.69003 | 2.80755 | 1.78519 | 1.71198 | 0.54252 | 3.50632 | 0.71990 | 0.07351 | 2.47988 | 0.00000 | 1.93621 | 0.61598 |

| | | | | | | | | | | | | |
|---|---|---|---|---|---|---|---|---|---|---|---|---|
| **RNF148** | 0.09423 | 0.01178 | 0.01806 | 0.01324 | 0.02181 | 0.00821 | 0.01873 | 0.09028 | 0.01691 | 0.02203 | 0.01514 | 0.06178 |
| **RNF149** | 0.30209 | 1.63490 | 7.17003 | 1.13063 | 1.32904 | 4.60916 | 0.00000 | 0.80952 | 9.87736 | 3.95219 | 3.42931 | 2.93255 |
| **RNF150** | 0.72183 | 0.05808 | 0.03115 | 0.35344 | 0.05041 | 0.00018 | 0.01231 | 0.02765 | 0.00000 | 0.12770 | 0.01617 | 0.03820 |
| **RNF151** | 0.00232 | 0.01331 | 0.01722 | 0.03166 | 0.05445 | 0.01712 | 0.01471 | 0.06185 | 0.08030 | 0.00000 | 0.01612 | 0.03034 |
| **RNF152** | 0.28299 | 0.05215 | 0.00287 | 0.02510 | 0.00000 | 0.05255 | 0.02555 | 0.11671 | 0.00133 | 0.13815 | 0.01277 | 0.07605 |
| **RNF157** | 1.03859 | 0.01229 | 0.69729 | 0.00913 | 0.01312 | 0.00000 | 0.00675 | 0.00000 | 0.00000 | 0.01660 | 0.10131 | 0.03659 |
| **RNF157-AS1** | 0.10134 | 0.01364 | 0.05876 | 0.00813 | 0.01463 | 0.02349 | 0.00329 | 0.00471 | 0.03248 | 0.01250 | 0.01353 | 0.01445 |
| **RNF165** | 0.00181 | 0.01609 | 0.00511 | 0.03510 | 0.02412 | 0.00180 | 0.12467 | 0.14283 | 0.04508 | 0.16901 | 0.06347 | 0.17802 |
| **RNF166** | 0.66511 | 1.54854 | 0.90710 | 0.26847 | 3.08501 | 0.00000 | 1.04290 | 0.01110 | 1.18056 | 2.36659 | 0.71905 | 0.42144 |
| **RNF167** | 2.27671 | 19.56402 | 12.53164 | 13.90994 | 18.32162 | 0.00000 | 16.30439 | 4.80695 | 16.53639 | 14.07452 | 14.90135 | 20.87586 |
| **RNF168** | 5.57634 | 3.67291 | 12.26177 | 9.18748 | 6.74447 | 10.12185 | 13.17912 | 38.18153 | 5.51254 | 7.04250 | 5.77109 | 15.64685 |
| **RNF169** | 1.34160 | 0.70574 | 0.84378 | 0.14252 | 0.44662 | 1.27615 | 0.00007 | 1.02177 | 0.50264 | 2.68849 | 1.14665 | 0.20705 |
| **RNF17** | 3.42124 | 1.74869 | 0.00000 | 0.82122 | 0.12221 | 5.75070 | 0.16511 | 2.92506 | 0.00000 | 5.26472 | 2.62502 | 2.69883 |
| **RNF170** | 0.71879 | 4.73771 | 0.97628 | 1.26470 | 1.69324 | 7.69574 | 0.00000 | 0.26714 | 0.89082 | 4.57161 | 3.74254 | 3.98203 |
| **RNF180** | 0.22606 | 0.09359 | 0.00000 | 0.12532 | 0.00000 | 0.10935 | 0.00000 | 0.09951 | 0.00000 | 0.01036 | 0.09005 | 0.13228 |
| **RNF181** | 4.66380 | 13.59696 | 18.90139 | 15.29248 | 24.22608 | 8.95967 | 23.77946 | 11.25732 | 12.21358 | 9.51075 | 12.10192 | 14.04301 |
| **RNF182** | 0.50165 | 0.10830 | 0.22686 | 0.00000 | 0.00000 | 0.35294 | 0.54245 | 0.00000 | 0.18598 | 0.92097 | 0.11043 | 0.05389 |
| **RNF183** | 0.17140 | 1.19512 | 0.12989 | 0.23801 | 0.09124 | 0.00000 | 0.02688 | 0.62979 | 0.00000 | 0.12656 | 0.50494 | 0.15040 |
| **RNF185** | 1.39966 | 5.19950 | 5.36943 | 2.33495 | 2.28864 | 5.76524 | 6.46307 | 2.51572 | 4.56245 | 0.00000 | 5.50448 | 4.53080 |
| **RNF187** | 2.97075 | 2.23512 | 1.55439 | 3.95089 | 12.51954 | 0.00000 | 12.85034 | 0.88537 | 0.00000 | 19.09033 | 2.02354 | 0.95618 |
| **RNF19A** | 2.14147 | 1.42983 | 5.51022 | 6.49054 | 0.00000 | 2.17387 | 3.46172 | 2.90882 | 1.52685 | 3.09771 | 3.85520 | 1.83180 |
| **RNF19B** | 5.16798 | 6.70194 | 5.63052 | 5.40686 | 2.67760 | 8.51361 | 8.49261 | 10.78539 | 0.00000 | 10.09801 | 0.00000 | 12.85520 |
| **RNF2** | 2.11277 | 8.81094 | 8.01806 | 6.67600 | 3.51872 | 24.49103 | 13.37962 | 11.22788 | 7.25553 | 0.01768 | 10.62622 | 9.88067 |
| **RNF20** | 9.02358 | 6.68974 | 4.23371 | 6.02868 | 5.09491 | 6.63513 | 6.97817 | 5.79360 | 6.43964 | 8.24817 | 3.77907 | 10.40090 |
| **RNF207** | 0.15150 | 0.11347 | 0.45655 | 0.00000 | 0.96983 | 0.00000 | 0.00000 | 0.01814 | 2.48019 | 0.87781 | 2.29421 | 0.00500 |
| **RNF208** | 0.00580 | 0.00617 | 0.03779 | 0.20027 | 0.00000 | 0.02395 | 0.20107 | 0.03863 | 0.00000 | 0.42754 | 0.03119 | 0.01607 |
| **RNF212** | 4.11000 | 4.69557 | 0.61639 | 0.38757 | 2.18512 | 4.86248 | 1.44753 | 1.58033 | 4.79925 | 0.00000 | 4.53822 | 5.97510 |
| **RNF212B** | 0.06192 | 0.58863 | 0.50122 | 0.00725 | 0.19469 | 0.44651 | 0.22696 | 0.09582 | 0.00035 | 0.14167 | 0.58368 | 0.31423 |
| **RNF213** | 3.50875 | 1.55154 | 0.56070 | 2.64800 | 1.68194 | 0.71793 | 0.00042 | 0.84081 | 0.42690 | 3.80385 | 0.00000 | 2.22929 |
| **RNF214** | 2.71445 | 2.23105 | 0.16843 | 7.90679 | 1.26880 | 5.87727 | 0.82902 | 0.16369 | 1.73289 | 8.33726 | 2.28738 | 3.69693 |
| **RNF215** | 0.09513 | 0.11692 | 0.02721 | 0.18363 | 0.00000 | 0.14464 | 0.00000 | 0.00388 | 0.00000 | 0.58937 | 0.00000 | 0.04522 |
| **RNF216** | 5.70707 | 9.57107 | 4.38183 | 8.19453 | 9.18031 | 5.04644 | 3.83707 | 3.19489 | 8.09991 | 3.36613 | 4.65222 | 9.01865 |
| **RNF216-IT1** | 0.00457 | 0.00888 | 0.01489 | 0.00743 | 0.01202 | 0.04515 | 0.00070 | 0.00973 | 0.00000 | 0.10630 | 0.07290 | 0.01258 |
| **RNF216P1** | 4.17059 | 8.30702 | 2.73069 | 4.99186 | 4.56008 | 6.90770 | 8.57067 | 7.16798 | 7.19634 | 2.92617 | 5.47126 | 15.88684 |
| **RNF217** | 0.00526 | 0.01742 | 1.52212 | 0.00000 | 0.00000 | 0.08610 | 0.09979 | 0.00964 | 0.10523 | 0.01779 | 0.00000 | 0.00930 |
| **RNF219** | 2.39710 | 6.76137 | 2.15103 | 2.57310 | 0.00000 | 13.29861 | 3.85608 | 7.56189 | 3.51812 | 9.58316 | 1.46364 | 9.87250 |
| **RNF219-AS1** | 0.03185 | 0.01877 | 0.00271 | 0.02278 | 0.00000 | 0.06189 | 0.01716 | 0.01557 | 0.01332 | 0.06383 | 0.02813 | 0.01675 |
| **RNF220** | 2.10006 | 3.45821 | 2.93277 | 4.15866 | 3.70818 | 0.00221 | 7.19321 | 0.00000 | 2.88355 | 10.27423 | 1.54422 | 6.34812 |
| **RNF222** | 0.19984 | 0.06848 | 0.12950 | 0.00401 | 0.00000 | 0.00000 | 0.04147 | 0.00449 | 0.00000 | 0.19508 | 0.02691 | 0.00810 |
| **RNF223** | 0.06592 | 0.01571 | 0.03679 | 0.00933 | 0.01466 | 0.01567 | 0.02353 | 0.00606 | 0.03701 | 0.00841 | 0.01012 | 0.01547 |
| **RNF24** | 2.71666 | 3.92449 | 0.69001 | 11.23277 | 3.14356 | 0.00000 | 1.91825 | 1.56037 | 1.49376 | 3.75522 | 0.22223 | 1.20847 |
| **RNF25** | 1.03754 | 6.76923 | 6.64188 | 5.84845 | 2.83617 | 0.91298 | 3.85558 | 4.19921 | 5.67921 | 0.00000 | 2.92086 | 11.37675 |
| **RNF26** | 1.32637 | 23.67672 | 13.73699 | 10.41975 | 19.33601 | 0.00000 | 6.61332 | 0.00000 | 20.55110 | 0.00000 | 11.23406 | 14.84256 |
| **RNF31** | 0.65854 | 2.48491 | 1.27591 | 5.60815 | 11.66124 | 1.89947 | 0.00000 | 2.13213 | 2.36323 | 8.46110 | 0.62894 | 4.37991 |
| **RNF32** | 0.51031 | 0.67231 | 0.03806 | 0.02501 | 0.00389 | 0.09871 | 0.04658 | 0.05098 | 0.14936 | 0.29211 | 0.00757 | 0.27219 |
| **RNF34** | 19.14205 | 13.49806 | 8.31261 | 6.97833 | 5.09134 | 23.69770 | 13.44026 | 15.82921 | 15.65972 | 2.30357 | 15.16666 | 16.08333 |
| **RNF38** | 15.77261 | 5.70100 | 7.07920 | 5.88982 | 2.30616 | 24.45209 | 12.88796 | 3.63519 | 6.01422 | 11.16273 | 4.95458 | 7.32894 |
| **RNF39** | 0.01867 | 0.61506 | 0.00000 | 1.18257 | 3.61741 | 0.93962 | 0.00000 | 0.30087 | 0.27068 | 0.00429 | 2.90883 | 0.81822 |

| | | | | | | | | | | | | |
|---|---|---|---|---|---|---|---|---|---|---|---|---|
| **RNF4** | 6.65438 | 16.87876 | 10.96526 | 12.13988 | 13.55307 | 17.03330 | 15.16745 | 23.60866 | 25.53606 | 6.10885 | 11.40578 | 15.77392 |
| **RNF40** | 4.61760 | 9.64299 | 10.85971 | 8.53306 | 10.28219 | 0.00000 | 9.35111 | 0.74831 | 12.03821 | 11.21056 | 7.28408 | 5.64160 |
| **RNF41** | 3.84069 | 2.69679 | 6.95193 | 4.04758 | 4.84875 | 0.67531 | 7.61001 | 2.71594 | 11.06651 | 1.58698 | 8.00246 | 4.80430 |
| **RNF43** | 0.00265 | 0.62394 | 2.96671 | 0.00000 | 0.00000 | 0.03051 | 3.43651 | 0.09771 | 6.14137 | 0.11507 | 0.00000 | 0.00000 |
| **RNF44** | 1.19521 | 8.62080 | 5.19137 | 9.97545 | 7.36012 | 6.63094 | 10.56463 | 1.00228 | 2.67197 | 26.14167 | 0.64732 | 7.27036 |
| **RNF5** | 0.00374 | 0.04350 | 0.05148 | 0.04151 | 0.01575 | 0.02719 | 0.04658 | 0.01802 | 0.03772 | 0.00000 | 0.11516 | 0.11378 |
| **RNF5P1.uid-107** | 0.00125 | 0.46841 | 0.73482 | 1.30960 | 0.00000 | 0.31780 | 0.66546 | 0.04546 | 1.48095 | 0.02112 | 1.23981 | 1.64266 |
| **RNF6** | 0.02327 | 5.91168 | 6.97052 | 1.63898 | 0.09300 | 21.00388 | 3.92422 | 4.20205 | 9.72406 | 0.00000 | 3.46889 | 9.42799 |
| **RNF7** | 1.72132 | 9.23752 | 11.58497 | 8.46846 | 10.98903 | 14.91075 | 16.73453 | 3.96728 | 16.53085 | 3.00562 | 10.66022 | 16.05532 |
| **RNF8** | 4.84396 | 5.36259 | 4.65747 | 5.01130 | 0.00000 | 5.71108 | 11.19651 | 3.73022 | 4.67287 | 1.53784 | 4.03987 | 6.89875 |
| **RNFT1** | 0.81408 | 2.06754 | 0.49422 | 0.40190 | 0.05651 | 4.49542 | 0.00000 | 1.84503 | 2.31355 | 0.00000 | 1.36423 | 4.81721 |
| **RNFT2** | 0.00000 | 3.58826 | 0.38111 | 1.36713 | 2.93358 | 0.68652 | 0.00012 | 0.00850 | 1.08224 | 1.96967 | 1.47956 | 0.44633 |
| **RNGTT** | 0.00000 | 5.11319 | 3.30365 | 4.26586 | 2.98777 | 12.44545 | 5.01075 | 6.46931 | 7.03901 | 4.40376 | 3.06945 | 6.78697 |
| **RNH1** | 2.24319 | 21.73475 | 14.00111 | 10.01827 | 21.03967 | 0.00000 | 17.81666 | 1.74306 | 22.81082 | 0.00000 | 20.29983 | 18.90376 |
| **RNLS** | 0.20013 | 0.86955 | 0.28601 | 0.00419 | 0.81782 | 2.74885 | 0.88021 | 0.18779 | 0.77744 | 0.00000 | 0.58331 | 0.59820 |
| **RNMT** | 0.55084 | 3.36833 | 4.82019 | 1.46173 | 4.74463 | 18.92788 | 9.49719 | 10.23267 | 5.51003 | 17.14015 | 6.46025 | 8.14884 |
| **RNMTL1** | 0.79128 | 14.23378 | 11.89204 | 8.15087 | 7.65932 | 11.55844 | 17.07806 | 8.96797 | 18.90774 | 0.00000 | 15.90012 | 19.32441 |
| **RNPC3** | 1.09820 | 0.64636 | 1.42369 | 0.40901 | 0.46714 | 3.05664 | 0.58395 | 2.03538 | 4.01636 | 4.56600 | 0.60374 | 1.93188 |
| **RNPEP** | 0.35409 | 1.89625 | 1.25329 | 5.48473 | 10.43788 | 3.06171 | 0.93825 | 0.00136 | 4.21783 | 8.02186 | 6.48041 | 0.51390 |
| **RNPEPL1** | 1.16034 | 0.88868 | 1.60572 | 1.89075 | 3.45606 | 0.00000 | 1.73605 | 0.00000 | 1.46956 | 2.49562 | 1.62011 | 0.09386 |
| **RNPS1** | 3.05824 | 26.44911 | 20.45249 | 18.57425 | 20.29538 | 12.68896 | 26.80191 | 17.10422 | 29.56745 | 6.29014 | 21.61962 | 29.65102 |
| **RNU11** | 0.00869 | 0.02830 | 0.14949 | 0.00000 | 0.06050 | 0.03085 | 0.00000 | 0.02217 | 0.05941 | 0.16259 | 0.00000 | 0.05123 |
| **RNU12** | 0.00183 | 0.00000 | 0.27273 | 0.00000 | 0.03566 | 0.12535 | 0.00000 | 0.32135 | 0.16821 | 0.60089 | 0.00677 | 0.61048 |
| **RNU4ATAC** | 0.11405 | 0.00000 | 0.26324 | 0.00529 | 0.79648 | 0.00000 | 0.59515 | 0.05080 | 0.00000 | 4.17690 | 0.00000 | 0.08202 |
| **RNU5F-1** | 0.00618 | 0.01455 | 0.01638 | 0.01193 | 0.01708 | 0.01833 | 0.01718 | 0.01030 | 0.01824 | 0.01862 | 0.01707 | 0.01843 |
| **RNU6-19P** | 0.00616 | 0.01291 | 0.01355 | 0.01883 | 0.01041 | 0.02048 | 0.01162 | 0.01031 | 0.02564 | 0.03755 | 0.01219 | 0.01474 |
| **RNU6-28P.uid-1** | 0.00623 | 0.01588 | 0.01427 | 0.01161 | 0.01438 | 0.01847 | 0.01888 | 0.01046 | 0.01824 | 0.01610 | 0.01654 | 0.01657 |
| **RNU6-71P** | 0.00618 | 0.01378 | 0.01485 | 0.01601 | 0.01470 | 0.02345 | 0.01501 | 0.01043 | 0.01955 | 0.01899 | 0.01386 | 0.01566 |
| **RNU6-78P** | 0.00500 | 0.01038 | 0.00650 | 0.01454 | 0.00000 | 0.04309 | 0.01591 | 0.00990 | 0.01209 | 0.07071 | 0.01403 | 0.01647 |
| **RNU6ATAC** | 0.00799 | 0.02589 | 0.08899 | 0.00788 | 0.06723 | 0.00000 | 0.00135 | 0.03321 | 0.03881 | 0.09076 | 0.00545 | 0.02258 |
| **RNU86** | 0.05971 | 0.31358 | 0.79950 | 0.46491 | 0.52036 | 0.00000 | 0.06862 | 0.21869 | 1.56293 | 0.54228 | 0.02571 | 0.60828 |
| **RNVU1-19.uid-1** | 0.00453 | 0.02497 | 0.09315 | 0.06986 | 0.05331 | 0.00587 | 0.02613 | 0.01853 | 0.01956 | 0.00551 | 0.08030 | 0.04071 |
| **RNVU1-19.uid-1** | 0.00551 | 0.02591 | 0.12274 | 0.01219 | 0.05938 | 0.00000 | 0.01054 | 0.01109 | 0.01043 | 0.04623 | 0.01340 | 0.03071 |
| **RNVU1-8** | 0.00346 | 0.03195 | 0.10088 | 0.04501 | 0.03999 | 0.00480 | 0.08916 | 0.00930 | 0.01450 | 0.18770 | 0.00337 | 0.05493 |
| **ROBO1** | 0.74083 | 0.32016 | 1.67035 | 5.39111 | 0.00000 | 0.30810 | 0.28793 | 0.84734 | 0.00135 | 0.68767 | 0.10941 | 0.56654 |
| **ROBO2** | 1.01766 | 0.01017 | 0.26397 | 0.07758 | 0.01577 | 0.00000 | 0.01093 | 0.08989 | 0.00000 | 0.05104 | 0.01607 | 0.01142 |
| **ROBO3** | 3.14871 | 0.12067 | 0.22645 | 0.62442 | 0.00000 | 0.00000 | 0.01000 | 0.35661 | 0.17489 | 0.02646 | 0.01825 | 0.15537 |
| **ROBO4** | 1.65139 | 0.01417 | 0.02257 | 0.02733 | 0.02105 | 0.00708 | 0.01694 | 0.08000 | 0.02248 | 0.00081 | 0.02091 | 0.01870 |
| **ROCK1** | 3.78979 | 2.72196 | 6.73567 | 7.13270 | 4.84557 | 8.03672 | 10.84934 | 5.30396 | 3.22829 | 21.00582 | 3.89374 | 8.45002 |
| **ROCK1P1** | 0.00479 | 0.00811 | 0.20047 | 0.01115 | 0.00000 | 0.02959 | 0.07494 | 0.01637 | 0.00000 | 0.44816 | 0.17840 | 0.08281 |
| **ROCK2** | 0.00000 | 0.00049 | 2.94643 | 0.29109 | 2.24951 | 9.26054 | 3.94175 | 1.30436 | 0.00000 | 17.31867 | 0.54550 | 1.30730 |
| **ROGDI** | 0.75470 | 1.77312 | 7.87381 | 6.09577 | 5.45119 | 0.00000 | 2.22017 | 0.18429 | 6.64102 | 2.08653 | 6.34582 | 0.37823 |
| **ROM1** | 0.00837 | 0.00000 | 0.04559 | 0.04788 | 0.65013 | 0.26965 | 0.45425 | 0.04364 | 0.77067 | 0.00000 | 0.68377 | 0.02536 |
| **ROMO1** | 0.49553 | 20.42447 | 22.90708 | 20.35421 | 33.20411 | 11.25687 | 41.04723 | 6.70031 | 19.47362 | 20.57280 | 24.57716 | 23.66117 |
| **ROPN1** | 0.00411 | 0.04079 | 0.01704 | 0.01294 | 0.01123 | 0.00062 | 0.02225 | 0.02050 | 0.00708 | 0.01400 | 0.05717 | 0.01482 |
| **ROPN1B** | 0.00555 | 0.03987 | 0.01424 | 0.00686 | 0.01522 | 0.02944 | 0.01560 | 0.00988 | 0.01085 | 0.01056 | 0.03206 | 0.07793 |
| **ROPN1L** | 0.50080 | 0.11568 | 0.00325 | 0.09278 | 0.01765 | 0.07053 | 0.02032 | 0.15971 | 0.00944 | 0.00000 | 0.05861 | 0.35370 |
| **ROPN1L-AS1** | 0.00838 | 0.05337 | 0.05483 | 0.00651 | 0.00625 | 0.05164 | 0.00684 | 0.00929 | 0.01183 | 0.01271 | 0.01543 | 0.01489 |

| | | | | | | | | | | | | |
|---|---|---|---|---|---|---|---|---|---|---|---|---|
| **ROR1** | 0.00143 | 0.04893 | 0.10230 | 1.53918 | 0.02412 | 0.00000 | 0.00000 | 0.04175 | 0.00390 | 1.25614 | 0.00000 | 0.00083 |
| **ROR1-AS1** | 0.00582 | 0.01426 | 0.02128 | 0.02947 | 0.00743 | 0.02094 | 0.00941 | 0.01042 | 0.02412 | 0.04266 | 0.01233 | 0.01453 |
| **ROR2** | 0.01809 | 0.00813 | 0.55001 | 0.55844 | 0.09713 | 0.00000 | 0.01204 | 2.28910 | 0.00000 | 0.00000 | 0.02337 | 0.18191 |
| **RORA** | 0.62635 | 1.70247 | 0.41791 | 0.00000 | 0.00000 | 9.25348 | 0.00000 | 0.79605 | 0.00000 | 10.82989 | 1.38580 | 1.04113 |
| **RORB** | 0.52056 | 0.00848 | 0.00430 | 0.00484 | 0.01087 | 0.03754 | 0.02431 | 0.04260 | 0.03982 | 0.03360 | 0.00348 | 0.05418 |
| **RORC** | 0.00575 | 0.01524 | 0.00864 | 0.08058 | 0.00790 | 0.00896 | 0.01357 | 0.01050 | 0.02140 | 0.03661 | 0.01534 | 0.01540 |
| **ROS1** | 7.11274 | 0.01206 | 0.02720 | 0.01246 | 0.01538 | 0.00314 | 0.01662 | 1.64406 | 0.01564 | 0.00000 | 0.01253 | 0.00000 |
| **RP1** | 0.08364 | 0.00221 | 0.00598 | 0.18268 | 0.00000 | 0.03153 | 0.03347 | 0.00000 | 0.01166 | 0.02394 | 0.01137 | 0.20301 |
| **RP11-87M18.2** | 0.17327 | 0.01449 | 0.01407 | 0.01560 | 0.01624 | 0.01220 | 0.01866 | 0.00420 | 0.01994 | 0.01268 | 0.01449 | 0.01594 |
| **RP1L1** | 0.01673 | 0.05198 | 0.03356 | 0.00506 | 0.01604 | 0.01725 | 0.01484 | 0.00824 | 0.00000 | 0.02278 | 0.18940 | 0.01443 |
| **RP2** | 0.00002 | 1.33508 | 20.52653 | 0.00000 | 0.00000 | 7.60939 | 13.32329 | 5.58754 | 7.46904 | 0.00000 | 7.79507 | 1.23334 |
| **RP9** | 1.47566 | 3.78623 | 0.77923 | 2.22133 | 3.01372 | 1.36313 | 0.68617 | 0.22853 | 2.56333 | 2.71982 | 1.92050 | 3.38422 |
| **RP9P** | 0.01884 | 0.01000 | 0.03397 | 0.24360 | 0.00579 | 0.00000 | 0.06808 | 0.02710 | 0.01049 | 0.07549 | 0.01407 | 0.10140 |
| **RPA1** | 14.03330 | 12.84163 | 16.50587 | 10.42917 | 3.13257 | 15.84262 | 18.94509 | 6.97065 | 18.41341 | 1.04631 | 17.49572 | 7.38119 |
| **RPA2** | 3.56802 | 17.08178 | 18.23093 | 13.11867 | 6.97177 | 24.54997 | 22.71993 | 12.70702 | 22.41158 | 0.00000 | 17.11145 | 20.68514 |
| **RPA3** | 2.54571 | 7.32789 | 7.83349 | 7.49604 | 8.48769 | 23.52388 | 9.69364 | 4.44034 | 5.06926 | 7.00024 | 12.99244 | 10.32346 |
| **RPA3OS** | 0.96418 | 1.85967 | 3.10678 | 1.71128 | 0.51061 | 8.39239 | 1.63834 | 3.35409 | 4.12060 | 0.00000 | 3.66190 | 4.04337 |
| **RPA4** | 0.01403 | 0.00000 | 0.15747 | 0.02347 | 0.17627 | 0.21259 | 0.03423 | 0.00986 | 0.00000 | 0.40800 | 0.00822 | 0.01985 |
| **RPAIN** | 1.11778 | 6.45192 | 6.67892 | 5.17909 | 8.15171 | 21.43237 | 20.20633 | 0.15232 | 16.34542 | 5.26313 | 11.48527 | 7.25534 |
| **RPAP1** | 3.58894 | 13.33776 | 10.13914 | 7.55400 | 6.83422 | 0.00000 | 4.67497 | 0.66435 | 19.31466 | 4.94805 | 7.48975 | 14.35229 |
| **RPAP2** | 1.08795 | 4.65764 | 3.90344 | 0.88680 | 1.50913 | 9.42336 | 2.52347 | 0.04871 | 2.08259 | 1.91274 | 4.85748 | 3.20960 |
| **RPAP3** | 3.78964 | 6.59423 | 5.85368 | 2.56871 | 0.39346 | 24.16779 | 12.19861 | 2.99071 | 10.15997 | 6.35372 | 5.57336 | 3.74270 |
| **RPARP-AS1** | 0.00000 | 8.91161 | 5.36767 | 2.76781 | 3.74397 | 7.75612 | 6.39258 | 1.28312 | 1.01822 | 6.75838 | 9.76961 | 7.41330 |
| **RPE** | 1.08644 | 6.24791 | 5.43741 | 2.21959 | 0.99931 | 18.35483 | 4.88641 | 3.45722 | 8.82050 | 0.00000 | 8.01925 | 7.31456 |
| **RPE65** | 0.08267 | 0.01271 | 0.11262 | 0.00649 | 0.00929 | 0.02601 | 0.00000 | 0.00852 | 0.02397 | 0.02975 | 0.01090 | 0.01311 |
| **RPEL1** | 0.12803 | 1.16281 | 0.75139 | 0.37191 | 0.18096 | 2.19594 | 0.84394 | 0.50055 | 1.09966 | 0.00000 | 1.27375 | 0.98178 |
| **RPF1** | 1.60315 | 10.95046 | 9.91498 | 1.72395 | 4.61006 | 28.96866 | 16.31979 | 19.13748 | 7.29421 | 5.23529 | 10.85834 | 20.74562 |
| **RPF2** | 0.93146 | 9.30379 | 13.79715 | 1.92739 | 1.57969 | 32.13336 | 18.03863 | 21.75823 | 12.77592 | 12.64312 | 14.83836 | 17.43200 |
| **RPGR** | 0.00000 | 1.44970 | 0.92420 | 0.54913 | 0.00000 | 2.93331 | 3.05233 | 4.08599 | 0.21185 | 2.59547 | 0.82998 | 3.27317 |
| **RPGRIP1** | 0.41356 | 0.21132 | 0.16228 | 0.00000 | 0.23030 | 0.07009 | 0.01980 | 2.16029 | 0.00000 | 0.59024 | 0.05705 | 2.52659 |
| **RPGRIP1L** | 1.00269 | 0.90877 | 0.79395 | 1.01257 | 0.28923 | 2.51136 | 0.00000 | 1.22509 | 1.67262 | 0.00000 | 1.00352 | 2.00970 |
| **RPH3A** | 8.53691 | 0.02702 | 0.00000 | 0.00000 | 0.01342 | 0.05235 | 0.00668 | 0.30207 | 0.00767 | 0.12608 | 0.01020 | 0.51092 |
| **RPH3AL** | 1.45967 | 0.12243 | 0.00617 | 0.03999 | 0.09763 | 0.00000 | 0.02521 | 0.00000 | 0.03603 | 0.02128 | 0.08658 | 0.07502 |
| **RPIA** | 2.96069 | 1.35723 | 6.50923 | 0.94672 | 1.90381 | 0.33024 | 6.87664 | 0.58771 | 3.57218 | 5.94725 | 7.59731 | 2.73369 |
| **RPL10** | 6.52801 | 16.29949 | 10.17331 | 13.04287 | 15.56700 | 11.64722 | 14.33504 | 4.99031 | 27.81574 | 0.00000 | 15.27458 | 18.98555 |
| **RPL10A** | 4.79533 | 23.84330 | 17.70403 | 18.86313 | 23.09161 | 30.94426 | 30.58931 | 11.35254 | 31.66954 | 11.77509 | 22.87607 | 28.90148 |
| **RPL10L** | 0.61189 | 0.63106 | 1.37814 | 0.24400 | 0.08995 | 2.75893 | 0.00000 | 11.55191 | 0.00000 | 0.82895 | 0.00000 | 5.40808 |
| **RPL11** | 6.83534 | 43.14968 | 34.81215 | 35.02146 | 43.33729 | 41.23863 | 52.10042 | 15.80609 | 52.07597 | 32.08045 | 34.04889 | 48.00275 |
| **RPL12** | 3.83444 | 18.68789 | 13.29969 | 16.16308 | 26.65778 | 16.99536 | 31.47916 | 2.91022 | 20.13528 | 30.22989 | 15.23086 | 22.25921 |
| **RPL13** | 2.25939 | 36.05089 | 35.89266 | 28.84334 | 42.56889 | 13.76560 | 51.67065 | 12.06716 | 43.74500 | 32.36151 | 28.89951 | 36.46089 |
| **RPL13A** | 7.16386 | 17.77229 | 28.59361 | 23.01161 | 28.73291 | 17.70938 | 47.39563 | 13.15733 | 35.75801 | 28.49032 | 18.58034 | 23.37323 |
| **RPL13AP17** | 0.00355 | 0.00584 | 0.79847 | 0.02739 | 0.00000 | 0.06360 | 0.00000 | 0.04142 | 0.00000 | 0.03485 | 0.01315 | 0.01203 |
| **RPL13AP20** | 0.01247 | 0.10597 | 0.32273 | 0.14009 | 0.29916 | 2.71601 | 2.42795 | 0.12029 | 0.00000 | 3.08071 | 0.27485 | 0.75861 |
| **RPL13AP3** | 0.08347 | 0.05027 | 0.58637 | 0.44970 | 0.13192 | 0.23190 | 0.36413 | 0.12839 | 0.00000 | 0.26429 | 0.14105 | 0.11239 |
| **RPL13AP6** | 0.67281 | 4.37577 | 4.82930 | 3.72961 | 4.17399 | 0.03216 | 5.58918 | 1.03896 | 6.67097 | 0.00000 | 2.79913 | 4.69017 |
| **RPL13P5** | 0.00000 | 1.26960 | 0.11958 | 5.71777 | 0.21718 | 0.83998 | 0.00000 | 0.24196 | 0.12929 | 1.77569 | 0.05044 | 2.13193 |
| **RPL14** | 2.72565 | 19.85848 | 17.27494 | 11.79017 | 18.07523 | 27.29686 | 28.62576 | 6.38148 | 35.87381 | 21.69187 | 13.48251 | 25.05128 |
| **RPL15** | 13.38063 | 23.73884 | 23.81480 | 21.13504 | 22.53171 | 44.19114 | 41.54910 | 14.15507 | 36.36147 | 24.53766 | 24.91116 | 32.20521 |

| | | | | | | | | | | | | |
|---|---|---|---|---|---|---|---|---|---|---|---|---|
| **RPL17** | 0.12139 | 9.09947 | 9.96940 | 7.01281 | 6.88262 | 12.23181 | 9.24658 | 2.10644 | 16.89761 | 0.00000 | 7.76608 | 9.77448 |
| **RPL18** | 10.36136 | 40.71037 | 41.09879 | 35.21311 | 48.80081 | 28.35491 | 50.61912 | 19.21287 | 61.41337 | 29.01312 | 34.83106 | 47.82983 |
| **RPL18A** | 1.60435 | 9.31045 | 9.94981 | 10.72012 | 13.62735 | 1.12957 | 16.83874 | 5.58275 | 15.69612 | 0.99377 | 8.68818 | 13.96740 |
| **RPL19** | 4.92700 | 35.06542 | 34.30342 | 29.26304 | 39.86909 | 44.90602 | 51.20095 | 12.74994 | 52.21090 | 36.73519 | 31.26108 | 40.69816 |
| **RPL19P12** | 0.54990 | 6.28307 | 7.42674 | 7.06744 | 6.67753 | 10.15027 | 8.31085 | 1.02530 | 13.10277 | 0.00001 | 5.43340 | 7.65443 |
| **RPL21P44** | 0.15311 | 0.00356 | 0.31819 | 0.01545 | 0.19747 | 0.00041 | 0.03269 | 0.14471 | 0.00001 | 0.44972 | 0.10342 | 0.02797 |
| **RPL22** | 0.48670 | 6.39631 | 6.19427 | 4.08447 | 6.00407 | 15.29482 | 15.99523 | 1.31976 | 10.98353 | 11.80077 | 7.17821 | 8.75821 |
| **RPL22L1** | 0.54457 | 8.14732 | 6.94737 | 8.02986 | 9.08712 | 25.72513 | 8.89805 | 2.33462 | 15.62039 | 21.05316 | 9.55682 | 18.78528 |
| **RPL23** | 2.37691 | 27.87121 | 31.96940 | 27.44753 | 38.20868 | 29.42118 | 59.34792 | 18.15347 | 30.14351 | 45.06127 | 35.15064 | 39.49845 |
| **RPL23A** | 7.30157 | 16.22482 | 22.50511 | 14.98753 | 16.26450 | 27.39126 | 38.73038 | 9.87475 | 27.74820 | 21.73997 | 20.08783 | 25.57338 |
| **RPL23AP32** | 0.00019 | 0.02290 | 0.00000 | 0.00000 | 0.00000 | 0.64725 | 0.28037 | 0.27111 | 0.00000 | 1.11621 | 0.03464 | 0.48049 |
| **RPL23AP53** | 1.22417 | 0.51448 | 0.31514 | 0.01372 | 0.39404 | 0.25969 | 0.61758 | 2.87065 | 0.15663 | 0.72775 | 0.41784 | 0.62996 |
| **RPL23AP64** | 0.00363 | 0.01280 | 0.10120 | 0.02322 | 0.09820 | 0.00000 | 0.00000 | 0.01315 | 0.13223 | 0.30297 | 0.03166 | 0.05299 |
| **RPL23AP7** | 0.50774 | 1.64786 | 0.78473 | 0.49326 | 0.00000 | 0.61697 | 0.86141 | 1.00664 | 0.45729 | 0.00000 | 0.86827 | 1.18046 |
| **RPL23AP82** | 0.27930 | 2.39181 | 0.56753 | 0.86227 | 1.47716 | 1.13838 | 0.96491 | 0.94356 | 1.29670 | 0.00000 | 0.85457 | 3.17585 |
| **RPL23P8** | 0.02584 | 0.98306 | 2.05499 | 1.01810 | 0.88866 | 4.41209 | 3.52640 | 0.62680 | 1.22931 | 2.35329 | 2.41696 | 2.51765 |
| **RPL24** | 4.92572 | 17.98050 | 12.68253 | 10.87132 | 19.18041 | 33.73959 | 21.22067 | 4.97733 | 24.31548 | 24.48840 | 13.48969 | 23.22668 |
| **RPL26** | 5.24560 | 21.25547 | 19.40224 | 19.38180 | 20.29041 | 37.33843 | 35.47504 | 11.84930 | 27.39945 | 33.78341 | 22.95802 | 31.34529 |
| **RPL26L1** | 2.67833 | 8.91413 | 1.78784 | 2.61537 | 6.50573 | 8.03445 | 4.62784 | 8.70272 | 4.01670 | 0.58374 | 4.41821 | 10.68057 |
| **RPL27** | 5.52757 | 30.08874 | 34.65109 | 28.39553 | 37.50174 | 37.25877 | 57.40681 | 18.81017 | 34.53526 | 35.09551 | 37.28915 | 41.27003 |
| **RPL27A** | 4.95651 | 20.92513 | 22.64401 | 18.03785 | 29.88016 | 19.80590 | 44.65318 | 8.57957 | 27.84961 | 42.10389 | 19.56110 | 24.46288 |
| **RPL28** | 13.76453 | 28.77615 | 25.21549 | 26.20967 | 45.31718 | 5.86546 | 46.76954 | 16.35135 | 34.41391 | 44.75014 | 18.58012 | 34.90032 |
| **RPL29** | 9.04768 | 24.38048 | 21.56628 | 19.66558 | 26.11431 | 14.69484 | 34.45241 | 6.06463 | 32.52971 | 18.90437 | 22.06998 | 31.24490 |
| **RPL29P2** | 0.00558 | 0.03708 | 0.02630 | 0.00930 | 0.01014 | 0.02811 | 0.04473 | 0.01230 | 0.01910 | 0.04418 | 0.04580 | 0.03066 |
| **RPL3** | 10.83831 | 42.94264 | 33.57447 | 39.80880 | 47.26576 | 30.29891 | 50.35544 | 16.48873 | 60.17133 | 7.42971 | 39.65044 | 46.25777 |
| **RPL30** | 15.23929 | 39.97812 | 36.07732 | 29.61007 | 34.17990 | 52.54336 | 53.21766 | 25.63168 | 51.28057 | 32.13482 | 32.38863 | 47.88866 |
| **RPL31** | 6.43758 | 28.97469 | 30.05484 | 23.87200 | 31.04860 | 40.69181 | 46.21139 | 10.95860 | 39.64913 | 33.48920 | 29.27036 | 37.92375 |
| **RPL31P11** | 0.03159 | 0.48683 | 0.74237 | 0.66133 | 0.38887 | 1.61553 | 1.15486 | 0.23189 | 0.59858 | 0.95318 | 0.68698 | 0.89001 |
| **RPL32** | 5.70002 | 27.19305 | 22.69428 | 22.73677 | 32.62932 | 35.53685 | 45.73267 | 11.98132 | 31.97633 | 34.43214 | 24.58964 | 35.18865 |
| **RPL32P3** | 0.72829 | 0.65319 | 0.84745 | 0.36998 | 0.55864 | 0.74399 | 0.36090 | 1.62981 | 3.76228 | 2.55087 | 0.34110 | 1.93611 |
| **RPL34** | 0.11767 | 17.85410 | 23.17251 | 18.54430 | 24.48853 | 23.29003 | 41.31943 | 8.20257 | 26.74999 | 23.65836 | 23.87319 | 29.83480 |
| **RPL34-AS1** | 0.00370 | 0.28125 | 0.57205 | 0.06876 | 0.00000 | 0.27103 | 0.17615 | 1.30804 | 0.00000 | 0.10142 | 0.24973 | 0.70752 |
| **RPL35** | 5.26271 | 29.97089 | 26.56029 | 26.46646 | 39.77483 | 17.27930 | 48.80126 | 16.33531 | 34.15414 | 32.80819 | 26.93756 | 36.61216 |
| **RPL35A** | 4.93732 | 31.20387 | 30.64713 | 23.33253 | 31.10109 | 48.56119 | 48.08781 | 17.50449 | 40.39043 | 34.09024 | 29.97859 | 42.92834 |
| **RPL36** | 20.57091 | 43.39905 | 35.10416 | 35.27005 | 47.33422 | 25.26337 | 55.63288 | 22.52369 | 49.17920 | 40.34386 | 36.58749 | 50.25903 |
| **RPL36A.uid-130** | 0.26974 | 0.01231 | 0.04102 | 0.00000 | 0.04796 | 0.15280 | 0.11890 | 0.01743 | 0.06396 | 0.18138 | 0.02457 | 0.07971 |
| **RPL36A.uid-130** | 0.10399 | 0.00210 | 0.00000 | 0.62976 | 0.68753 | 2.04452 | 0.00000 | 0.66347 | 1.48919 | 5.94555 | 1.15511 | 1.36062 |
| **RPL36AL** | 0.00000 | 11.54401 | 15.75021 | 15.50947 | 20.68690 | 46.02526 | 28.46901 | 8.02417 | 17.93438 | 16.49190 | 21.87893 | 29.73032 |
| **RPL37** | 5.03359 | 32.95990 | 30.36782 | 25.69152 | 36.28956 | 39.29647 | 45.71610 | 17.68300 | 40.51858 | 35.42037 | 27.36928 | 43.27315 |
| **RPL37A** | 10.75914 | 33.69036 | 32.53378 | 27.95060 | 36.77388 | 40.32397 | 47.07942 | 13.14681 | 46.33775 | 47.43618 | 29.87842 | 41.15080 |
| **RPL38** | 5.83999 | 30.42727 | 31.29108 | 22.75851 | 34.12181 | 39.49944 | 55.32448 | 18.77977 | 34.78188 | 47.08838 | 30.78678 | 42.12650 |
| **RPL39** | 0.06015 | 2.40803 | 3.69540 | 1.88303 | 0.00001 | 15.27025 | 12.88188 | 1.28968 | 3.38059 | 21.18169 | 4.48635 | 6.10570 |
| **RPL39L** | 0.12311 | 6.27138 | 7.65879 | 6.90139 | 3.43816 | 10.50638 | 6.51052 | 2.43721 | 2.32269 | 5.40369 | 7.61227 | 6.53118 |
| **RPL3L** | 0.46599 | 0.04130 | 0.06324 | 0.02167 | 0.06360 | 0.00000 | 0.08151 | 0.07509 | 0.00727 | 0.00000 | 0.01175 | 0.06999 |
| **RPL4** | 3.47100 | 33.84102 | 30.54573 | 25.52049 | 32.99310 | 46.41069 | 44.47165 | 10.38489 | 39.65593 | 29.94034 | 29.69864 | 38.28197 |
| **RPL41** | 1.46412 | 7.60475 | 5.37864 | 5.11088 | 6.10840 | 11.84105 | 8.47027 | 3.12002 | 6.93476 | 1.37982 | 6.92926 | 13.72326 |
| **RPL5** | 6.81105 | 33.06351 | 33.75948 | 26.69301 | 30.05054 | 56.02214 | 52.40802 | 17.85324 | 46.93749 | 29.12438 | 37.11301 | 41.33957 |
| **RPL6** | 0.62236 | 17.68608 | 13.51004 | 10.03708 | 12.55979 | 29.23266 | 20.77999 | 9.22303 | 22.14873 | 8.15450 | 14.73529 | 23.57626 |

| | | | | | | | | | | | | |
|---|---|---|---|---|---|---|---|---|---|---|---|---|
| **RPL7** | 0.02359 | 0.25029 | 0.26437 | 0.30298 | 0.33967 | 0.72485 | 0.25819 | 0.70051 | 0.60064 | 0.78699 | 0.35669 | 0.71121 |
| **RPL7A** | 7.96992 | 18.92958 | 16.67951 | 16.46537 | 22.48117 | 17.92437 | 23.72191 | 7.05051 | 28.52746 | 16.54058 | 14.11139 | 23.64580 |
| **RPL7L1** | 0.26039 | 5.57984 | 4.56673 | 5.02126 | 9.38331 | 8.02531 | 8.55929 | 2.04174 | 5.84196 | 4.22056 | 10.49892 | 12.91844 |
| **RPL8** | 17.11884 | 50.26590 | 45.03222 | 50.19761 | 62.65040 | 40.08738 | 68.18111 | 31.34331 | 61.71047 | 37.11627 | 46.76940 | 52.75552 |
| **RPL9** | 0.24215 | 4.01181 | 5.13730 | 7.51411 | 7.31074 | 7.53402 | 19.00227 | 5.38434 | 19.47925 | 0.00000 | 3.50242 | 15.49131 |
| **RPLP0** | 13.32450 | 36.00847 | 28.55542 | 29.27417 | 35.96533 | 30.68034 | 45.27263 | 13.65027 | 52.31327 | 24.35453 | 31.02966 | 34.39320 |
| **RPLP0P2** | 0.58404 | 5.51953 | 0.47562 | 3.60763 | 1.37846 | 5.19658 | 3.79920 | 0.49302 | 0.00000 | 7.47513 | 0.34073 | 1.51012 |
| **RPLP1** | 16.08078 | 45.52870 | 47.44797 | 36.10358 | 50.06248 | 39.16144 | 62.41947 | 17.95677 | 66.25292 | 40.78990 | 36.32055 | 48.71128 |
| **RPLP2** | 12.26911 | 43.89566 | 38.30383 | 30.59810 | 48.95677 | 34.80913 | 62.80244 | 20.11631 | 56.96654 | 47.63271 | 30.12146 | 50.88149 |
| **RPN1** | 3.39381 | 33.16020 | 28.94822 | 19.90663 | 20.54228 | 28.16727 | 7.17041 | 10.52383 | 58.76675 | 8.80057 | 26.34666 | 34.71954 |
| **RPN2** | 2.66204 | 34.37996 | 27.50631 | 23.90689 | 30.97430 | 43.59132 | 8.49326 | 10.69697 | 50.88388 | 28.24820 | 24.11651 | 29.03079 |
| **RPP14** | 3.64538 | 11.08180 | 4.85750 | 8.83248 | 3.49208 | 21.17488 | 11.79352 | 9.98761 | 6.02063 | 7.98068 | 9.61278 | 18.36075 |
| **RPP21** | 0.74845 | 1.89844 | 1.17813 | 1.30130 | 2.55794 | 0.85024 | 0.96764 | 1.18021 | 1.53010 | 0.00000 | 1.16521 | 1.88322 |
| **RPP25** | 0.00000 | 15.09829 | 7.01929 | 5.05189 | 6.34213 | 0.00000 | 9.32348 | 4.23484 | 11.07105 | 15.76366 | 6.54975 | 12.14845 |
| **RPP25L** | 0.64771 | 9.09010 | 5.39085 | 2.85295 | 2.45091 | 1.95714 | 5.95344 | 0.52205 | 2.69908 | 0.00000 | 6.71909 | 12.81386 |
| **RPP30** | 0.49205 | 10.23823 | 11.00091 | 4.09522 | 4.86938 | 23.66506 | 17.47726 | 13.67572 | 9.06979 | 4.82209 | 9.93785 | 13.18921 |
| **RPP38** | 1.03950 | 12.63834 | 7.59971 | 3.61519 | 8.74636 | 13.51281 | 7.58351 | 3.26278 | 10.97341 | 6.85858 | 6.10161 | 13.59866 |
| **RPP40** | 0.00000 | 5.20689 | 6.11673 | 0.48522 | 1.23192 | 13.51901 | 5.71187 | 4.59913 | 9.70170 | 0.00000 | 8.01218 | 9.25931 |
| **RPPH1** | 0.53942 | 0.00000 | 0.60141 | 0.27248 | 0.85506 | 0.00000 | 0.72797 | 3.93420 | 0.46529 | 2.24540 | 0.43554 | 3.37419 |
| **RPRD1A** | 1.13798 | 7.40131 | 4.97136 | 6.41331 | 1.72438 | 21.65581 | 12.41359 | 10.56337 | 7.51716 | 9.05704 | 5.28738 | 9.93105 |
| **RPRD1B** | 0.00000 | 4.14353 | 4.80312 | 2.20011 | 2.43410 | 12.94310 | 5.13453 | 8.13399 | 5.64078 | 10.30812 | 6.48979 | 5.82229 |
| **RPRD2** | 3.84709 | 0.73430 | 4.13698 | 5.31583 | 4.07483 | 1.77169 | 6.68031 | 1.21476 | 2.91468 | 12.90087 | 5.48876 | 5.19672 |
| **RPRM** | 0.01299 | 0.02338 | 0.24978 | 1.83660 | 0.02219 | 0.00000 | 0.00404 | 0.00185 | 0.02834 | 0.00557 | 0.03143 | 1.09985 |
| **RPRML** | 0.00283 | 0.01110 | 0.17675 | 0.00000 | 0.00000 | 0.03903 | 0.02645 | 0.00879 | 0.01875 | 0.23881 | 0.00784 | 0.10859 |
| **RPS10** | 0.32982 | 3.94628 | 4.60983 | 3.75322 | 3.59758 | 2.09662 | 8.71185 | 5.36510 | 5.15124 | 0.00000 | 4.55441 | 6.50768 |
| **RPS10P7** | 0.00979 | 0.04489 | 0.04620 | 0.00000 | 0.01716 | 0.03672 | 0.07611 | 0.02406 | 0.04537 | 0.07704 | 0.03378 | 0.04028 |
| **RPS11** | 13.95936 | 36.42345 | 37.66632 | 30.61245 | 42.04574 | 20.92624 | 65.31716 | 24.39194 | 39.26370 | 42.24076 | 33.74496 | 45.90206 |
| **RPS12** | 8.46934 | 28.79270 | 21.87261 | 17.50412 | 25.69413 | 29.26247 | 40.17057 | 6.06124 | 38.22799 | 37.29971 | 14.23596 | 29.58921 |
| **RPS13** | 12.56677 | 31.61473 | 33.40672 | 26.15568 | 32.54776 | 48.65524 | 46.22625 | 20.08032 | 45.83867 | 28.60648 | 29.50801 | 42.05215 |
| **RPS14** | 12.81537 | 43.73752 | 44.28513 | 40.69638 | 50.32003 | 40.00170 | 68.52499 | 28.28593 | 53.87131 | 29.86225 | 46.16469 | 50.90735 |
| **RPS14P3** | 1.90164 | 8.08228 | 10.49674 | 7.95339 | 10.00352 | 7.45199 | 17.93298 | 5.52234 | 10.63757 | 7.87785 | 10.12925 | 11.38800 |
| **RPS15** | 3.88474 | 23.96886 | 22.77128 | 19.96200 | 27.77166 | 1.76676 | 38.48610 | 9.25465 | 26.94076 | 17.78271 | 19.80883 | 27.73648 |
| **RPS15A** | 2.69983 | 23.51080 | 21.09683 | 17.32474 | 22.41246 | 20.27343 | 31.18336 | 9.13079 | 32.20906 | 15.36095 | 19.37918 | 26.56506 |
| **RPS15AP10** | 0.00072 | 0.02014 | 0.00000 | 0.41221 | 0.17725 | 0.43296 | 0.00000 | 0.35389 | 0.00000 | 2.99115 | 0.04320 | 0.12343 |
| **RPS16** | 9.46179 | 39.44927 | 34.00048 | 30.42337 | 46.70813 | 30.49521 | 56.49327 | 17.25456 | 50.07614 | 38.90182 | 30.84388 | 41.87683 |
| **RPS16P5** | 0.01147 | 0.00000 | 0.19773 | 0.00068 | 0.03548 | 0.00000 | 0.00000 | 0.07027 | 0.21373 | 0.31650 | 0.01459 | 0.08198 |
| **RPS17.uid-1969** | 0.00657 | 0.03641 | 0.02223 | 0.01981 | 0.02075 | 0.05770 | 0.01861 | 0.01620 | 0.05086 | 0.00002 | 0.02534 | 0.02186 |
| **RPS17.uid-1970** | 0.00448 | 0.03274 | 0.01346 | 0.00743 | 0.01636 | 0.03697 | 0.01913 | 0.01653 | 0.01551 | 0.00965 | 0.01344 | 0.02436 |
| **RPS18** | 7.50850 | 29.40758 | 24.55861 | 27.42179 | 33.04030 | 26.02482 | 34.63288 | 14.48308 | 36.45497 | 16.38798 | 27.29785 | 37.33993 |
| **RPS18P9** | 0.58841 | 3.05300 | 2.45313 | 3.16981 | 4.37720 | 0.36409 | 3.74772 | 1.21766 | 3.91823 | 2.44392 | 2.32429 | 4.30863 |
| **RPS19** | 17.22637 | 44.97914 | 43.20529 | 37.35642 | 49.69941 | 41.61058 | 64.33710 | 29.27034 | 59.82980 | 35.25112 | 37.48297 | 51.52645 |
| **RPS19BP1** | 2.04725 | 22.99902 | 25.57898 | 17.54477 | 27.12120 | 7.30514 | 33.65260 | 6.07848 | 28.01359 | 11.61913 | 16.03265 | 23.28930 |
| **RPS2** | 3.87212 | 8.52388 | 12.78543 | 11.74769 | 23.20161 | 0.00000 | 28.46594 | 4.69227 | 10.85984 | 43.54110 | 6.64609 | 9.45268 |
| **RPS20** | 10.05837 | 31.38455 | 26.16086 | 21.01702 | 31.21023 | 36.97518 | 43.49927 | 17.10731 | 42.86353 | 29.03710 | 22.97636 | 39.63009 |
| **RPS21** | 10.54134 | 30.81673 | 25.90919 | 22.20469 | 37.94279 | 22.34542 | 44.13837 | 12.10254 | 40.22684 | 37.47237 | 22.23989 | 35.64362 |
| **RPS23** | 7.90557 | 33.79218 | 32.82088 | 23.74615 | 30.34459 | 51.35051 | 45.50624 | 11.85662 | 48.27410 | 24.02660 | 30.64369 | 41.98787 |
| **RPS24** | 9.59721 | 30.76734 | 26.81738 | 24.40614 | 31.67621 | 45.06715 | 42.54461 | 15.79512 | 36.30813 | 35.50893 | 29.00859 | 37.65527 |
| **RPS25** | 5.09895 | 27.07209 | 23.66358 | 19.34368 | 25.93597 | 49.29734 | 37.43007 | 19.50049 | 36.09768 | 16.77816 | 24.64379 | 38.90467 |

| | | | | | | | | | | | | |
|---|---|---|---|---|---|---|---|---|---|---|---|---|
| RPS26 | 0.28279 | 3.39381 | 9.64124 | 2.94738 | 1.35382 | 5.19168 | 9.31747 | 3.25652 | 8.93468 | 0.00000 | 6.39566 | 9.58174 |
| RPS26P11 | 0.00865 | 0.11528 | 0.05668 | 0.06653 | 0.11762 | 0.02790 | 0.06452 | 0.04232 | 0.11154 | 0.01026 | 0.06600 | 0.15010 |
| RPS27 | 0.30831 | 4.07471 | 1.97532 | 2.85462 | 3.49594 | 1.46274 | 3.12175 | 2.27185 | 2.60428 | 1.37900 | 2.86847 | 4.33409 |
| RPS27A | 1.63906 | 20.86522 | 18.52166 | 15.79125 | 23.48415 | 32.45199 | 32.43608 | 6.35891 | 24.18267 | 29.03980 | 16.82181 | 28.00712 |
| RPS27L | 8.28393 | 14.27539 | 10.51682 | 11.80949 | 14.23920 | 47.52966 | 6.61254 | 13.94480 | 12.35296 | 17.57224 | 9.80703 | 18.29176 |
| RPS28 | 2.09770 | 6.21225 | 4.17059 | 2.90660 | 3.88339 | 4.30690 | 9.89953 | 0.79708 | 3.65828 | 21.12709 | 4.85783 | 5.16479 |
| RPS29 | 5.89846 | 18.07549 | 20.76615 | 16.34340 | 22.81591 | 46.37040 | 38.21963 | 12.10260 | 31.60277 | 40.02112 | 21.16547 | 35.24919 |
| RPS2P32 | 0.00237 | 0.04081 | 0.31547 | 0.05214 | 0.00000 | 0.00000 | 0.12969 | 0.08133 | 0.00000 | 0.55437 | 0.11444 | 0.16858 |
| RPS3 | 14.46657 | 44.25462 | 45.70614 | 41.34709 | 52.07616 | 37.63916 | 65.43383 | 17.18350 | 61.08581 | 37.19621 | 42.30699 | 47.65517 |
| RPS3A | 0.08682 | 3.41907 | 2.78262 | 3.41180 | 1.98417 | 5.22075 | 0.69882 | 1.39662 | 8.20431 | 7.11253 | 2.66768 | 3.55198 |
| RPS4X | 9.42204 | 32.42434 | 29.22515 | 27.96357 | 32.69251 | 41.42340 | 45.25958 | 12.17023 | 41.80176 | 32.25489 | 31.41803 | 38.06275 |
| RPS4Y1 | 0.00000 | 30.05156 | 17.89366 | 17.80207 | 31.14783 | 38.19099 | 35.66163 | 7.94712 | 30.72358 | 21.76469 | 12.31217 | 29.59071 |
| RPS4Y2 | 0.14714 | 0.00552 | 0.11756 | 1.15887 | 0.00000 | 0.58464 | 0.00000 | 0.23215 | 0.00000 | 0.00884 | 0.01141 | 0.05168 |
| RPS5 | 14.47389 | 47.16600 | 39.59723 | 37.41529 | 51.41443 | 30.08760 | 60.09689 | 25.62013 | 65.91909 | 25.32186 | 39.52473 | 51.40437 |
| RPS6 | 9.92106 | 42.50619 | 37.26223 | 35.91667 | 42.35752 | 64.63940 | 58.84260 | 30.74993 | 51.52984 | 43.69709 | 41.12802 | 54.53816 |
| RPS6KA1 | 0.66646 | 4.00021 | 1.54358 | 9.26688 | 12.05690 | 0.96090 | 6.79637 | 1.39812 | 2.27098 | 7.57775 | 1.93725 | 3.83887 |
| RPS6KA2 | 1.16661 | 0.03083 | 0.19868 | 2.38901 | 0.57274 | 0.00000 | 0.00789 | 0.00208 | 0.00000 | 0.02050 | 0.08518 | 0.00968 |
| RPS6KA3 | 0.29411 | 0.30943 | 1.47476 | 0.59220 | 1.73749 | 2.64416 | 4.74454 | 2.15768 | 0.00000 | 4.66492 | 0.69634 | 2.09058 |
| RPS6KA4 | 0.01630 | 0.00221 | 1.39263 | 0.00000 | 4.78202 | 0.00000 | 6.07095 | 0.00271 | 1.38526 | 5.73626 | 0.84812 | 0.01324 |
| RPS6KA5 | 1.33821 | 1.12037 | 3.50062 | 2.67983 | 1.52467 | 4.70787 | 0.00000 | 1.90240 | 1.02796 | 4.70187 | 3.60617 | 1.32502 |
| RPS6KA6 | 1.26081 | 0.00000 | 0.72777 | 0.44575 | 0.00000 | 0.23080 | 3.50874 | 0.25095 | 0.08628 | 1.41221 | 0.41852 | 0.00000 |
| RPS6KB1 | 0.00000 | 3.34647 | 7.82327 | 2.01331 | 1.73862 | 14.80999 | 9.24699 | 2.11911 | 8.97878 | 4.73789 | 6.97749 | 3.35847 |
| RPS6KB2 | 1.67805 | 12.26138 | 9.48578 | 10.71363 | 16.35414 | 0.00000 | 10.57272 | 2.85972 | 12.38338 | 0.00000 | 13.78266 | 15.90516 |
| RPS6KC1 | 4.29541 | 3.85487 | 2.22622 | 4.22832 | 0.32098 | 7.09048 | 2.11809 | 4.34023 | 3.49031 | 1.22888 | 2.07980 | 7.67113 |
| RPS6KL1 | 2.73027 | 2.03982 | 0.00000 | 0.80967 | 0.91279 | 0.64075 | 0.00001 | 0.35017 | 0.00000 | 0.45231 | 0.45017 | 2.15554 |
| RPS7 | 0.19646 | 8.28260 | 5.52508 | 5.84588 | 9.11059 | 6.46890 | 10.85475 | 3.88258 | 10.18399 | 3.88793 | 5.43875 | 11.67548 |
| RPS7P5 | 0.00277 | 0.00557 | 1.00832 | 0.20674 | 0.00000 | 0.08512 | 0.00000 | 0.06489 | 0.00000 | 0.02754 | 0.02407 | 0.01829 |
| RPS8 | 5.74848 | 31.25901 | 30.45505 | 28.12260 | 41.53854 | 29.00763 | 53.71097 | 12.30176 | 38.50463 | 42.55282 | 31.69882 | 39.29229 |
| RPS9 | 5.72609 | 31.83542 | 26.42968 | 27.94213 | 43.80583 | 11.45005 | 44.50780 | 10.88443 | 41.25227 | 27.71749 | 24.48501 | 35.43575 |
| RPSA | 3.24744 | 15.54280 | 14.80413 | 20.33066 | 21.96587 | 11.34078 | 29.25973 | 1.53003 | 20.09129 | 18.08935 | 12.93919 | 11.02863 |
| RPSAP52 | 0.03217 | 0.02990 | 0.01513 | 0.11509 | 0.02636 | 0.08705 | 0.15044 | 0.02001 | 0.03561 | 0.03049 | 0.03657 | 0.03513 |
| RPSAP58 | 0.30920 | 3.41775 | 3.85416 | 3.46168 | 0.74559 | 9.29064 | 1.71745 | 1.72760 | 7.75318 | 0.00000 | 2.84532 | 2.84338 |
| RPSAP9 | 0.33876 | 0.02253 | 0.04060 | 0.06291 | 0.00000 | 0.06340 | 0.14715 | 0.65939 | 0.07231 | 0.05849 | 0.27527 | 0.05038 |
| RPTN | 0.04114 | 0.01336 | 0.01460 | 0.01671 | 0.01185 | 0.00000 | 0.02443 | 0.00907 | 0.01900 | 0.10231 | 0.01857 | 0.01288 |
| RPTOR | 1.63849 | 4.65104 | 1.20770 | 3.01329 | 8.09424 | 0.00000 | 2.01156 | 7.50635 | 4.17984 | 6.95037 | 1.94157 | 8.97678 |
| RPUSD1 | 0.81177 | 12.87325 | 12.17756 | 5.18504 | 11.56745 | 0.00000 | 9.91040 | 0.07996 | 15.39140 | 12.98701 | 11.27263 | 3.65287 |
| RPUSD2 | 1.01577 | 12.11142 | 7.02485 | 5.17079 | 7.66372 | 0.00000 | 6.06410 | 2.61065 | 6.62074 | 0.36207 | 9.31530 | 10.98436 |
| RPUSD3 | 0.38539 | 25.85445 | 10.00196 | 20.30090 | 23.63387 | 9.44680 | 14.32553 | 3.01142 | 20.00917 | 0.00000 | 23.01462 | 25.14460 |
| RPUSD4 | 6.26883 | 9.28314 | 7.14589 | 6.85016 | 5.51949 | 6.13548 | 15.14405 | 6.76140 | 18.76747 | 0.00000 | 5.58143 | 9.49116 |
| RQCD1 | 2.50117 | 18.56812 | 16.79166 | 13.31447 | 9.51687 | 21.38143 | 28.64167 | 14.87889 | 16.52901 | 1.17890 | 24.59821 | 17.25181 |
| RRAD | 0.21314 | 3.01982 | 0.16523 | 7.94094 | 2.19047 | 0.00000 | 0.37770 | 0.00271 | 0.00000 | 1.45879 | 1.54770 | 0.00000 |
| RRAGA | 2.90462 | 8.88231 | 13.55432 | 7.38630 | 3.24816 | 11.03580 | 4.39070 | 1.52312 | 10.46858 | 0.00000 | 9.18421 | 5.65649 |
| RRAGB | 0.86803 | 0.62910 | 0.14616 | 0.39046 | 0.00000 | 1.42392 | 0.00000 | 2.25970 | 0.68599 | 0.37707 | 0.63411 | 2.22769 |
| RRAGC | 3.19835 | 0.49677 | 7.90581 | 1.07639 | 3.19002 | 4.66045 | 9.60563 | 0.57340 | 0.00000 | 14.56376 | 1.17576 | 2.54551 |
| RRAGD | 0.05853 | 0.14472 | 0.13098 | 0.25094 | 0.09360 | 0.07699 | 0.00000 | 0.02681 | 0.05169 | 0.10553 | 0.00000 | 0.00369 |
| RRAS | 0.00638 | 0.19865 | 0.13945 | 0.00442 | 0.52354 | 0.00000 | 0.38623 | 1.19406 | 0.14527 | 0.00000 | 0.04538 | 1.73817 |
| RRAS2 | 2.17022 | 2.39590 | 2.34423 | 2.24052 | 2.96274 | 7.54972 | 5.12548 | 0.00000 | 1.88353 | 11.92919 | 0.79111 | 1.85052 |
| RRBP1 | 2.92384 | 12.97817 | 13.34320 | 15.05815 | 36.65804 | 0.00000 | 4.04144 | 6.38252 | 21.39776 | 43.50966 | 6.61074 | 8.94563 |

| | | | | | | | | | | | | |
|---|---|---|---|---|---|---|---|---|---|---|---|---|
| **RREB1** | 0.77673 | 13.40409 | 9.29857 | 4.44667 | 12.64716 | 13.83240 | 13.89204 | 12.77252 | 4.17780 | 26.69575 | 12.90993 | 16.92402 |
| **RRH** | 0.00322 | 0.02439 | 0.07791 | 0.00000 | 0.00080 | 0.03763 | 0.00000 | 0.11057 | 0.07512 | 0.38636 | 0.03306 | 0.05756 |
| **RRM1** | 1.79066 | 21.99863 | 19.65437 | 9.49263 | 7.94122 | 39.15674 | 25.40586 | 10.04382 | 17.99216 | 7.53747 | 21.72389 | 29.04194 |
| **RRM2** | 8.29352 | 28.77921 | 40.78404 | 10.31815 | 11.01584 | 47.48379 | 35.75364 | 32.17831 | 37.94145 | 8.74550 | 21.57686 | 52.41971 |
| **RRM2B** | 2.33041 | 3.51408 | 2.56337 | 1.19912 | 0.00000 | 12.62272 | 2.37566 | 7.76210 | 3.08178 | 0.89134 | 1.61801 | 3.12512 |
| **RRN3** | 2.02691 | 4.75818 | 3.31741 | 1.75724 | 0.42431 | 11.33377 | 6.02339 | 16.21060 | 4.77485 | 3.77619 | 3.92201 | 5.34889 |
| **RRN3P1** | 0.07931 | 0.02619 | 0.02470 | 0.14609 | 0.00000 | 0.09740 | 0.00000 | 0.08554 | 0.00000 | 0.04792 | 0.00841 | 0.09488 |
| **RRN3P2** | 0.10164 | 0.04452 | 0.00827 | 0.07035 | 0.02368 | 0.03447 | 0.00000 | 0.09309 | 0.05583 | 0.10100 | 0.00673 | 0.08226 |
| **RRN3P3** | 0.20102 | 0.00000 | 0.44124 | 1.23491 | 0.00000 | 0.00000 | 0.00000 | 0.06358 | 0.63117 | 3.41463 | 0.22131 | 0.43067 |
| **RRNAD1** | 0.23110 | 1.44136 | 2.14796 | 1.38898 | 1.98083 | 0.00114 | 1.55993 | 0.15115 | 1.43279 | 0.80023 | 0.83060 | 1.90741 |
| **RRP1** | 0.05310 | 20.28691 | 16.95348 | 3.40821 | 8.72896 | 0.00000 | 5.13978 | 3.72177 | 16.79177 | 8.88670 | 17.50069 | 9.42131 |
| **RRP12** | 1.19208 | 17.63547 | 14.65264 | 8.56516 | 8.74150 | 0.00000 | 15.05087 | 7.55775 | 19.17142 | 8.61961 | 12.59615 | 24.75337 |
| **RRP15** | 1.77821 | 8.17637 | 9.16335 | 2.66069 | 1.37087 | 25.82969 | 16.97714 | 9.98075 | 4.96176 | 13.35848 | 11.13971 | 11.22786 |
| **RRP1B** | 2.16105 | 12.98139 | 9.13273 | 5.52304 | 9.73767 | 7.44302 | 19.56977 | 8.50566 | 20.15405 | 16.24067 | 12.61381 | 14.35470 |
| **RRP36** | 0.00000 | 4.03401 | 6.77904 | 4.99250 | 10.05012 | 6.76362 | 10.50145 | 1.23474 | 4.80402 | 20.41454 | 2.07425 | 7.28316 |
| **RRP7A** | 0.95684 | 20.67857 | 14.79254 | 17.21563 | 31.03788 | 0.39774 | 22.10628 | 5.29605 | 24.30493 | 10.51124 | 14.61716 | 24.95011 |
| **RRP7B** | 0.13704 | 2.85233 | 1.61161 | 3.60561 | 5.82993 | 0.00000 | 1.11190 | 1.56065 | 5.24814 | 6.56156 | 3.44705 | 5.30861 |
| **RRP8** | 1.44444 | 10.13718 | 1.90095 | 1.14388 | 7.98312 | 6.19945 | 5.48970 | 7.16263 | 2.98841 | 3.45671 | 5.13729 | 8.18047 |
| **RRP9** | 0.75335 | 19.40091 | 13.59285 | 14.51027 | 16.69704 | 0.05242 | 16.35107 | 8.56137 | 16.20624 | 0.00000 | 15.33428 | 18.61376 |
| **RRS1** | 1.64885 | 4.84677 | 12.46088 | 4.77393 | 7.70961 | 2.98443 | 14.22177 | 15.98653 | 18.03755 | 1.81279 | 13.57402 | 25.80068 |
| **RRS1-AS1** | 0.00381 | 0.01253 | 0.14953 | 0.08522 | 0.02299 | 0.00000 | 0.00444 | 0.21284 | 0.14188 | 0.04862 | 0.06684 | 0.13495 |
| **RS1** | 4.69936 | 0.00945 | 0.00487 | 0.01227 | 0.01026 | 0.02185 | 0.00962 | 1.33358 | 0.02839 | 0.01863 | 0.00799 | 0.15345 |
| **RSAD1** | 0.00179 | 8.41779 | 1.75706 | 3.95695 | 5.93243 | 0.00000 | 0.00000 | 0.16253 | 2.22177 | 3.42275 | 3.38363 | 2.85212 |
| **RSAD2** | 0.00669 | 0.00687 | 0.00195 | 0.00813 | 0.01991 | 0.00000 | 0.00831 | 0.03143 | 0.02624 | 0.38247 | 0.00005 | 0.01416 |
| **RSBN1** | 1.41458 | 3.14806 | 3.47024 | 4.58500 | 4.25815 | 0.14446 | 5.76297 | 5.07808 | 5.13686 | 5.55996 | 2.51288 | 5.19711 |
| **RSBN1L** | 0.61613 | 1.24983 | 1.24320 | 0.61494 | 0.61229 | 3.05905 | 1.80155 | 2.78961 | 0.00000 | 6.18061 | 0.35447 | 1.57636 |
| **RSC1A1** | 0.38784 | 1.11510 | 0.85329 | 0.00000 | 2.10582 | 1.85692 | 1.98323 | 4.48855 | 0.10038 | 0.79602 | 1.16177 | 1.77687 |
| **RSF1** | 3.26285 | 3.04405 | 0.67849 | 6.56494 | 3.98815 | 9.99736 | 2.98615 | 8.69863 | 0.23943 | 9.91923 | 0.26309 | 5.46017 |
| **RSG1** | 0.56576 | 0.48748 | 0.10390 | 0.29668 | 0.94285 | 0.00000 | 0.22837 | 0.00000 | 0.13052 | 0.09981 | 0.24520 | 0.38152 |
| **RSL1D1** | 8.31239 | 22.80607 | 25.23106 | 14.47912 | 21.20067 | 49.60690 | 37.94634 | 21.28580 | 36.50247 | 14.81041 | 30.16301 | 34.47836 |
| **RSL24D1** | 3.77271 | 10.74711 | 14.05215 | 3.42271 | 1.25556 | 45.62882 | 23.12294 | 13.22317 | 17.25935 | 10.31654 | 13.38555 | 16.52239 |
| **RSPH1** | 0.60968 | 0.06131 | 0.00356 | 0.03168 | 0.21632 | 0.05588 | 0.01803 | 0.00630 | 0.01317 | 0.00000 | 0.00928 | 0.05867 |
| **RSPH10B+RSPI** | 0.07168 | 0.37457 | 0.00885 | 0.00000 | 0.00000 | 0.03779 | 0.01176 | 0.13866 | 0.01724 | 0.02551 | 0.00819 | 0.36291 |
| **RSPH10B+RSPI** | 0.23480 | 0.53824 | 0.00251 | 0.00000 | 0.06600 | 0.12490 | 0.00000 | 0.40491 | 0.00000 | 0.03789 | 0.00000 | 0.50771 |
| **RSPH3** | 0.33772 | 1.00770 | 0.39218 | 0.72574 | 1.07420 | 0.57281 | 0.24857 | 0.38131 | 0.44405 | 0.35562 | 1.40554 | 2.02039 |
| **RSPH4A** | 1.06109 | 0.01953 | 0.03831 | 0.08368 | 0.00000 | 0.08645 | 0.00105 | 0.08194 | 0.06813 | 0.28157 | 0.06003 | 0.04560 |
| **RSPH6A** | 0.03443 | 0.01534 | 0.01349 | 0.01448 | 0.01490 | 0.01769 | 0.01936 | 0.00758 | 0.02389 | 0.01090 | 0.01457 | 0.01646 |
| **RSPH9** | 0.21011 | 0.98838 | 0.14396 | 0.11662 | 0.14187 | 0.00000 | 0.00424 | 4.62307 | 0.50510 | 0.00000 | 0.40643 | 2.33732 |
| **RSPO1** | 0.02893 | 0.00225 | 0.01547 | 0.01446 | 0.01127 | 0.02187 | 0.05350 | 0.05923 | 0.00000 | 0.02234 | 0.01564 | 0.36387 |
| **RSPO2** | 6.60194 | 0.03805 | 0.00000 | 0.00000 | 0.10078 | 0.15394 | 0.00000 | 0.74882 | 0.01923 | 0.04438 | 0.00000 | 0.40864 |
| **RSPO3** | 0.00723 | 0.01325 | 5.69735 | 9.59282 | 0.00000 | 0.05683 | 0.27050 | 0.06870 | 0.70386 | 0.00000 | 0.00000 | 0.00312 |
| **RSPO4** | 0.00434 | 0.00856 | 0.00372 | 0.00420 | 0.00000 | 0.02474 | 0.00085 | 0.01009 | 0.02596 | 0.16134 | 0.01712 | 0.01052 |
| **RSPRY1** | 2.16752 | 10.80565 | 4.84254 | 12.01819 | 3.81693 | 11.68953 | 7.42699 | 4.00213 | 7.72545 | 0.00027 | 6.10892 | 9.32780 |
| **RSRC1** | 2.23990 | 2.14617 | 1.02696 | 2.92182 | 4.13792 | 12.28899 | 3.55238 | 1.87869 | 0.95386 | 8.64763 | 6.08135 | 6.78087 |
| **RSRC2** | 6.73490 | 13.34913 | 9.73858 | 10.44404 | 9.62948 | 33.21535 | 10.69935 | 22.24936 | 16.01770 | 18.33310 | 8.05624 | 17.11137 |
| **RSRP1** | 2.79480 | 5.71972 | 3.61392 | 2.69198 | 2.42611 | 8.96668 | 2.90788 | 29.82604 | 15.80989 | 8.44598 | 4.55244 | 6.90122 |
| **RSU1** | 5.63009 | 6.27842 | 9.38100 | 5.51444 | 2.98889 | 12.31891 | 7.80303 | 1.46495 | 9.79470 | 0.00000 | 7.22132 | 3.35455 |
| **RSU1P2** | 0.00789 | 0.01528 | 0.01567 | 0.01131 | 0.01539 | 0.02060 | 0.01948 | 0.01027 | 0.01730 | 0.01299 | 0.01866 | 0.01731 |

| | | | | | | | | | | | | |
|---|---|---|---|---|---|---|---|---|---|---|---|---|
| **RTBDN** | 0.00443 | 0.54566 | 0.13235 | 0.22463 | 0.01769 | 0.02015 | 0.12470 | 0.36295 | 0.18822 | 0.02395 | 0.02102 | 1.87578 |
| **RTCA** | 1.59206 | 6.17301 | 10.01355 | 5.71897 | 0.29874 | 20.04709 | 12.97323 | 0.20387 | 9.21088 | 0.00000 | 8.55046 | 2.00948 |
| **RTCA-AS1** | 0.02732 | 0.06885 | 0.03931 | 0.00724 | 0.01462 | 0.04400 | 0.01536 | 0.00874 | 0.01846 | 0.00000 | 0.11190 | 0.03304 |
| **RTCB** | 1.07928 | 20.98180 | 16.87252 | 12.80921 | 13.12486 | 27.33644 | 23.09010 | 9.65194 | 22.71182 | 0.00000 | 23.59706 | 21.94417 |
| **RTDR1** | 0.12869 | 0.00751 | 0.07223 | 0.01411 | 0.00296 | 0.02631 | 0.09233 | 0.05443 | 0.00000 | 0.00919 | 0.06276 | 0.06226 |
| **RTEL1** | 0.02566 | 0.69524 | 0.66222 | 0.19069 | 1.06432 | 0.00000 | 0.05439 | 0.12501 | 0.00000 | 2.99965 | 0.44474 | 0.68515 |
| **RTF1** | 5.98896 | 3.51107 | 9.19692 | 7.02025 | 8.57871 | 5.72626 | 14.23181 | 11.98511 | 3.87337 | 22.62599 | 5.05837 | 5.39657 |
| **RTFDC1** | 1.33759 | 15.89580 | 14.74364 | 8.75905 | 12.78383 | 19.10059 | 19.74023 | 14.90634 | 15.95716 | 0.58559 | 14.45874 | 19.47408 |
| **RTKN** | 0.00000 | 11.49631 | 3.25047 | 10.39582 | 15.58597 | 0.39677 | 18.21277 | 0.01667 | 17.95434 | 2.67361 | 11.47282 | 0.00000 |
| **RTKN2** | 2.66898 | 2.13293 | 5.98471 | 0.00000 | 2.74455 | 6.06976 | 6.92716 | 0.19531 | 5.27518 | 0.65406 | 4.28954 | 0.76542 |
| **RTL1** | 0.00558 | 0.01168 | 0.01291 | 0.72103 | 0.00661 | 0.00340 | 0.01335 | 0.01017 | 0.01545 | 0.02807 | 0.01492 | 0.01540 |
| **RTN1** | 0.12321 | 0.01798 | 0.02707 | 0.00665 | 0.03271 | 0.00000 | 0.00235 | 0.09912 | 0.02764 | 0.40532 | 0.00000 | 0.10711 |
| **RTN2** | 0.28936 | 0.15466 | 1.89516 | 0.21231 | 0.62826 | 0.20003 | 0.74085 | 1.24214 | 0.41237 | 2.16413 | 0.16876 | 1.13151 |
| **RTN3** | 2.52793 | 14.02732 | 13.34645 | 5.04349 | 6.04958 | 21.51145 | 4.83211 | 7.17984 | 28.51020 | 3.17636 | 12.70841 | 13.52996 |
| **RTN4** | 4.67810 | 0.00000 | 7.87731 | 9.08261 | 0.00000 | 10.87165 | 13.10842 | 2.52777 | 1.46465 | 27.04233 | 6.24245 | 1.63777 |
| **RTN4IP1** | 2.94235 | 4.53337 | 4.35720 | 2.17894 | 4.74582 | 3.34902 | 4.34167 | 2.35464 | 2.53646 | 0.00000 | 5.33879 | 4.49924 |
| **RTN4R** | 1.37203 | 0.42671 | 0.00000 | 0.00000 | 0.07102 | 0.25858 | 0.42484 | 0.17093 | 0.08134 | 0.82476 | 0.00000 | 0.02061 |
| **RTN4RL1** | 0.00494 | 0.02820 | 0.01346 | 0.16966 | 0.00549 | 0.00986 | 0.01132 | 0.01299 | 0.01778 | 0.03281 | 0.01458 | 0.02112 |
| **RTN4RL2** | 0.38144 | 0.02526 | 0.07972 | 0.03067 | 0.09011 | 0.00000 | 0.02145 | 0.06868 | 0.06140 | 0.21293 | 0.00000 | 0.03411 |
| **RTP1** | 0.91911 | 0.59363 | 0.12357 | 1.92644 | 0.00000 | 0.63290 | 0.00000 | 0.00001 | 0.00000 | 0.55242 | 0.93290 | 0.83497 |
| **RTP2** | 0.02164 | 0.02176 | 0.03744 | 0.03758 | 0.02440 | 0.00910 | 0.01771 | 0.01316 | 0.01333 | 0.00798 | 0.03623 | 0.02409 |
| **RTP3** | 0.00000 | 3.84415 | 0.00000 | 6.42746 | 5.05852 | 1.66278 | 0.57178 | 0.17513 | 0.00000 | 1.13775 | 1.42566 | 2.99607 |
| **RTP5** | 0.00605 | 0.01710 | 0.04961 | 0.01727 | 0.01744 | 0.00640 | 0.01734 | 0.01058 | 0.01217 | 0.02775 | 0.01776 | 0.01611 |
| **RTTN** | 5.04625 | 0.55628 | 2.97980 | 0.72691 | 1.79035 | 3.60105 | 2.27043 | 1.34590 | 1.27992 | 2.91566 | 1.29151 | 1.18040 |
| **RUFY1** | 1.12446 | 13.46344 | 3.02292 | 5.10143 | 4.17947 | 4.01498 | 5.07386 | 3.02840 | 2.46681 | 6.28542 | 2.29887 | 6.20809 |
| **RUFY2** | 0.72572 | 1.81585 | 0.53509 | 1.04810 | 0.44905 | 6.30237 | 0.00000 | 1.89585 | 0.00000 | 3.36606 | 0.72950 | 2.18625 |
| **RUFY3** | 1.28476 | 1.96785 | 3.48293 | 8.90605 | 0.97220 | 13.01634 | 8.66511 | 0.00000 | 0.83981 | 9.76122 | 4.10310 | 4.84997 |
| **RUFY4** | 0.00577 | 0.01381 | 0.02702 | 0.01522 | 0.01189 | 0.02012 | 0.01504 | 0.01132 | 0.01868 | 0.02647 | 0.01342 | 0.01528 |
| **RUNDC1** | 0.04688 | 3.23411 | 1.77993 | 1.55493 | 0.57049 | 2.86153 | 2.73100 | 0.65089 | 0.00000 | 7.03158 | 1.85291 | 1.92557 |
| **RUNDC3A** | 0.12579 | 0.27302 | 0.13080 | 0.13605 | 0.34137 | 0.00000 | 0.15391 | 0.16887 | 0.00000 | 0.04320 | 0.00656 | 0.09237 |
| **RUNDC3B** | 1.47551 | 0.09737 | 0.01608 | 0.00000 | 0.03302 | 0.00000 | 0.01194 | 0.18610 | 0.04365 | 0.39582 | 0.02753 | 0.11193 |
| **RUNX1** | 0.63038 | 0.96961 | 7.63467 | 3.04562 | 1.56581 | 0.00035 | 5.26151 | 0.11949 | 1.29642 | 5.00323 | 0.29152 | 0.00000 |
| **RUNX1T1** | 9.36640 | 2.49875 | 0.14722 | 6.58471 | 0.24480 | 0.10290 | 0.00000 | 0.40635 | 0.00000 | 0.83562 | 0.01220 | 3.75696 |
| **RUNX2** | 0.25584 | 0.00167 | 0.05098 | 0.05745 | 0.01871 | 0.02836 | 0.01060 | 0.96181 | 0.07195 | 0.25669 | 0.01336 | 0.00000 |
| **RUNX3** | 0.06958 | 0.00031 | 0.00362 | 0.05857 | 0.18771 | 0.00000 | 0.06750 | 0.00419 | 0.00000 | 0.06050 | 0.00468 | 0.15833 |
| **RUSC1** | 1.02041 | 10.68977 | 12.68599 | 11.39901 | 6.85751 | 0.00000 | 11.29054 | 0.57142 | 13.14221 | 0.00000 | 7.66287 | 9.54782 |
| **RUSC1-AS1** | 0.00615 | 0.04051 | 0.03675 | 0.01465 | 0.03759 | 0.00000 | 0.01610 | 0.01220 | 0.02537 | 0.00561 | 0.01472 | 0.02663 |
| **RUSC2** | 0.40582 | 0.78973 | 0.80761 | 1.03741 | 0.00000 | 0.00000 | 0.00000 | 0.45464 | 0.00000 | 0.42513 | 0.20046 | 2.26395 |
| **RUVBL1** | 2.37552 | 16.44601 | 16.77365 | 11.26857 | 7.37224 | 25.18143 | 17.11797 | 0.80137 | 20.81329 | 0.00000 | 16.16140 | 17.18821 |
| **RUVBL2** | 1.07736 | 22.28630 | 19.24678 | 19.87679 | 25.08060 | 10.86908 | 24.69607 | 4.84038 | 24.81873 | 0.00000 | 24.27028 | 16.45402 |
| **RWDD1** | 0.22556 | 4.75485 | 6.07908 | 4.78626 | 8.18149 | 15.94112 | 15.40638 | 4.56903 | 10.10403 | 12.26427 | 9.44880 | 6.87563 |
| **RWDD2A** | 0.00000 | 1.43458 | 1.58017 | 0.59466 | 0.00000 | 1.38071 | 0.65001 | 0.14443 | 0.23879 | 0.96996 | 2.11580 | 1.45929 |
| **RWDD2B** | 0.81514 | 1.50417 | 1.05237 | 1.02861 | 0.00182 | 1.41450 | 0.00000 | 0.35688 | 0.00000 | 0.00000 | 2.14101 | 0.61638 |
| **RWDD3** | 0.09790 | 0.90313 | 1.30611 | 0.05855 | 0.15486 | 5.73540 | 2.82843 | 0.39366 | 0.52075 | 1.47626 | 1.82764 | 1.89937 |
| **RWDD4** | 0.39838 | 1.20530 | 1.87190 | 0.40253 | 0.17693 | 4.42667 | 1.33392 | 0.00000 | 1.22669 | 0.65661 | 1.83356 | 2.71002 |
| **RXFP1** | 0.00590 | 0.00641 | 0.01844 | 0.08581 | 0.08993 | 0.01090 | 0.01241 | 0.00770 | 0.02135 | 0.02833 | 0.01648 | 0.11159 |
| **RXFP4** | 0.00330 | 0.00460 | 0.00000 | 0.00036 | 0.00000 | 0.02351 | 0.01762 | 0.00958 | 0.01441 | 0.38991 | 0.00000 | 0.00747 |
| **RXRA** | 0.24637 | 0.10063 | 10.79914 | 0.00000 | 1.31745 | 0.02933 | 12.20291 | 0.07091 | 1.23983 | 5.30965 | 0.85763 | 0.00000 |

| | | | | | | | | | | | | |
|---|---|---|---|---|---|---|---|---|---|---|---|---|
| **RXRB** | 0.65975 | 1.09705 | 1.53563 | 1.42661 | 0.96601 | 0.00000 | 4.86057 | 0.51706 | 1.31149 | 1.29526 | 1.00687 | 2.38373 |
| **RXRG** | 0.00606 | 0.01570 | 0.07254 | 0.00836 | 0.01307 | 0.03069 | 0.01038 | 0.00963 | 0.01468 | 0.00388 | 0.01638 | 0.03699 |
| **RYBP** | 2.06138 | 0.48281 | 11.74579 | 3.44330 | 0.00000 | 13.70313 | 23.37929 | 5.80887 | 0.00000 | 37.93846 | 3.48128 | 3.02345 |
| **RYK** | 0.45871 | 1.46649 | 3.16147 | 0.00000 | 1.51060 | 7.39719 | 2.14902 | 0.13811 | 0.63549 | 10.14807 | 0.99449 | 1.71940 |
| **RYR1** | 1.00641 | 0.66132 | 0.00856 | 0.20489 | 1.17142 | 0.00000 | 0.05888 | 0.63726 | 0.83745 | 8.15667 | 0.40229 | 2.70638 |
| **RYR2** | 1.79170 | 0.67127 | 0.22731 | 0.84050 | 0.02293 | 0.03598 | 0.00000 | 0.50900 | 0.00000 | 0.12548 | 0.09974 | 0.23879 |
| **RYR3** | 3.22177 | 0.00000 | 0.99198 | 0.02572 | 0.07554 | 0.83724 | 0.00000 | 0.54188 | 0.00000 | 1.33282 | 0.08217 | 0.11259 |
| **S100A1** | 0.00292 | 0.03932 | 0.05092 | 0.06258 | 0.00000 | 0.07537 | 0.01715 | 0.01413 | 0.00000 | 0.24635 | 0.12259 | 0.04310 |
| **S100A10** | 0.00000 | 32.53933 | 16.88769 | 14.76697 | 36.54466 | 54.65539 | 45.12418 | 0.23297 | 41.56563 | 25.08109 | 39.76472 | 10.50666 |
| **S100A11** | 0.00000 | 22.17781 | 43.98075 | 14.83491 | 22.70294 | 33.60123 | 42.56319 | 2.77589 | 29.95847 | 5.87010 | 12.04304 | 25.72332 |
| **S100A12** | 0.00637 | 0.01324 | 0.01739 | 0.01395 | 0.01785 | 0.02684 | 0.03041 | 0.01045 | 0.02455 | 0.00575 | 0.02103 | 0.01640 |
| **S100A13** | 0.00000 | 18.46726 | 14.16152 | 27.37825 | 41.86095 | 13.12462 | 39.57636 | 0.64938 | 54.17671 | 22.34273 | 34.81092 | 0.00000 |
| **S100A14** | 0.00000 | 0.00000 | 6.67172 | 41.45534 | 42.91186 | 44.25452 | 54.72118 | 0.00000 | 30.64858 | 0.10267 | 30.96769 | 0.00000 |
| **S100A16** | 0.00000 | 36.54606 | 0.00000 | 22.94235 | 64.65542 | 41.33955 | 42.94832 | 0.99598 | 47.91359 | 16.10001 | 43.75048 | 3.40763 |
| **S100A2** | 0.00216 | 0.36865 | 0.00000 | 0.31100 | 0.49729 | 0.38137 | 0.09784 | 0.00412 | 0.39802 | 0.15138 | 0.51666 | 0.07169 |
| **S100A3** | 0.00492 | 0.02728 | 0.01305 | 0.05090 | 0.00000 | 0.03923 | 0.06867 | 0.00701 | 0.00301 | 0.05553 | 0.01871 | 0.13604 |
| **S100A4** | 0.50988 | 0.00000 | 0.20774 | 0.17330 | 0.00000 | 0.03543 | 1.31969 | 0.35702 | 0.29277 | 0.00000 | 0.66703 | 0.50313 |
| **S100A5** | 0.00489 | 0.00630 | 0.04106 | 0.00871 | 0.01299 | 0.02479 | 0.01200 | 0.01212 | 0.03991 | 0.00473 | 0.00974 | 0.15265 |
| **S100A6** | 0.00000 | 0.00000 | 7.73984 | 2.52470 | 14.80757 | 3.71476 | 35.62870 | 0.00000 | 52.18661 | 5.55090 | 33.36303 | 0.00000 |
| **S100A7** | 0.00455 | 0.01449 | 0.05774 | 0.01815 | 0.00092 | 0.03584 | 0.09826 | 0.01477 | 0.01415 | 0.03332 | 0.00786 | 0.01315 |
| **S100A7A** | 0.00564 | 0.01248 | 0.01160 | 0.01035 | 0.00848 | 0.02465 | 0.01230 | 0.01036 | 0.02111 | 0.04566 | 0.01401 | 0.01403 |
| **S100A8** | 0.00662 | 0.01513 | 0.01395 | 0.01743 | 0.00284 | 0.05373 | 0.02897 | 0.01044 | 0.03433 | 0.01118 | 0.00978 | 0.01593 |
| **S100A9** | 0.00767 | 0.01423 | 0.03393 | 0.05239 | 0.62479 | 0.00000 | 0.22161 | 0.00921 | 0.00000 | 0.00000 | 0.00364 | 0.04629 |
| **S100B** | 0.08376 | 0.02023 | 0.01598 | 0.01428 | 0.02572 | 0.00000 | 0.02147 | 0.17205 | 0.02322 | 0.00240 | 0.01472 | 0.05720 |
| **S100P** | 0.00524 | 0.00000 | 11.65639 | 6.12956 | 17.09801 | 0.03438 | 14.14162 | 0.00462 | 4.76981 | 0.00000 | 2.65944 | 0.00703 |
| **S100PBP** | 2.22742 | 4.17342 | 2.57708 | 6.54407 | 5.20295 | 10.94393 | 3.91097 | 2.89594 | 3.22414 | 0.18928 | 6.24954 | 6.87288 |
| **S100Z** | 0.04451 | 0.02996 | 0.00000 | 2.56303 | 0.00000 | 0.06436 | 0.00000 | 0.23729 | 0.00000 | 0.11346 | 0.01675 | 0.67049 |
| **S1PR1** | 0.16990 | 0.01203 | 0.10079 | 0.00866 | 0.00667 | 0.04807 | 0.01313 | 0.01370 | 0.00000 | 0.02067 | 0.01465 | 0.02737 |
| **S1PR2** | 0.19150 | 0.24934 | 37.61620 | 0.00000 | 0.00000 | 1.09470 | 5.24294 | 0.14603 | 3.15630 | 0.00000 | 1.33492 | 1.00194 |
| **S1PR3** | 0.00418 | 1.69397 | 0.49430 | 2.11085 | 0.00000 | 0.69190 | 0.00000 | 0.00000 | 0.83952 | 0.43018 | 0.00239 | 0.82991 |
| **S1PR4** | 0.00602 | 0.01560 | 0.02227 | 0.01273 | 0.01383 | 0.01346 | 0.05869 | 0.01052 | 0.00281 | 0.01979 | 0.01622 | 0.01660 |
| **S1PR5** | 0.00841 | 0.31290 | 0.00532 | 0.09926 | 0.00000 | 0.00000 | 0.02713 | 0.00250 | 0.03596 | 0.26220 | 0.00587 | 0.45376 |
| **SAA1** | 1.47546 | 0.05620 | 1.37475 | 0.00116 | 0.01501 | 0.00000 | 0.11381 | 2.94324 | 0.00000 | 0.00000 | 0.02582 | 0.25271 |
| **SAA2.uid-15279** | 0.00621 | 0.01565 | 0.04035 | 0.00941 | 0.01518 | 0.01321 | 0.01504 | 0.01048 | 0.03489 | 0.02011 | 0.01170 | 0.01617 |
| **SAA2.uid-15282** | 0.01799 | 0.01135 | 0.08567 | 0.00441 | 0.00526 | 0.02392 | 0.00353 | 0.01624 | 0.01594 | 0.09331 | 0.01646 | 0.01111 |
| **SAA3P** | 0.00382 | 0.01491 | 0.03281 | 0.00934 | 0.01263 | 0.02368 | 0.01845 | 0.05773 | 0.01518 | 0.01835 | 0.01510 | 0.01535 |
| **SAAL1** | 1.43996 | 7.78692 | 2.21572 | 4.48715 | 3.47430 | 17.38635 | 4.81340 | 4.83149 | 6.96780 | 0.37333 | 9.29550 | 10.94566 |
| **SAC3D1** | 0.66003 | 2.20552 | 1.60783 | 0.86478 | 0.00000 | 0.00000 | 5.87961 | 2.09564 | 0.57775 | 4.39706 | 0.68155 | 3.62283 |
| **SACM1L** | 5.17040 | 2.47593 | 1.44872 | 2.20899 | 0.00000 | 17.10227 | 7.65525 | 0.74254 | 2.07747 | 3.89999 | 5.48650 | 1.83972 |
| **SACS** | 1.33838 | 4.05919 | 2.14038 | 0.00000 | 0.00000 | 5.49839 | 0.00006 | 2.86148 | 4.80244 | 11.45657 | 0.00000 | 7.07915 |
| **SACS-AS1** | 0.00149 | 2.03360 | 0.61969 | 0.00000 | 0.00000 | 0.01284 | 0.12776 | 0.24618 | 0.03843 | 0.00561 | 0.00000 | 1.78908 |
| **SAE1** | 1.63077 | 21.30127 | 21.70368 | 14.83773 | 16.64029 | 13.30552 | 31.44384 | 12.19204 | 28.26107 | 0.01841 | 17.50650 | 14.59929 |
| **SAFB** | 0.37927 | 21.40639 | 18.56908 | 20.20274 | 20.20585 | 0.25288 | 24.26270 | 14.58236 | 24.52699 | 2.65946 | 16.57112 | 18.46020 |
| **SAFB2** | 2.38703 | 4.19484 | 8.13590 | 7.63378 | 12.85421 | 3.70927 | 18.14068 | 5.62519 | 14.17878 | 23.13487 | 2.77436 | 13.32897 |
| **SAG** | 0.05264 | 0.01976 | 0.12233 | 0.10173 | 0.03282 | 0.00001 | 0.08031 | 0.01333 | 0.02104 | 0.13606 | 0.02108 | 0.32656 |
| **SAGE1** | 0.00615 | 0.01201 | 0.01616 | 0.01208 | 0.01355 | 0.02116 | 0.01809 | 0.01007 | 0.01061 | 0.01188 | 0.03423 | 0.04101 |
| **SALL1** | 0.02740 | 0.14349 | 0.53163 | 2.84467 | 0.09568 | 0.11075 | 0.00000 | 0.22924 | 0.00000 | 0.00337 | 0.01144 | 0.51428 |
| **SALL2** | 0.51954 | 0.00000 | 0.02637 | 0.21489 | 0.03697 | 0.01824 | 0.01288 | 0.07812 | 0.01050 | 0.00000 | 0.04627 | 0.86459 |

| | | | | | | | | | | | | |
|---|---|---|---|---|---|---|---|---|---|---|---|---|
| **SALL3** | 1.04439 | 0.84262 | 0.00000 | 2.43882 | 0.25320 | 0.29175 | 0.00000 | 0.72680 | 0.00000 | 0.46502 | 0.00000 | 0.76770 |
| **SALL4** | 4.24060 | 18.89409 | 14.55410 | 20.21847 | 27.93197 | 11.45138 | 19.63771 | 8.80493 | 22.39526 | 12.30188 | 17.56904 | 28.47344 |
| **SAMD1** | 2.48052 | 0.97905 | 1.13772 | 4.01315 | 5.42624 | 0.00000 | 3.34121 | 0.43603 | 0.57521 | 8.30987 | 0.96030 | 2.28584 |
| **SAMD10** | 0.00338 | 0.03973 | 0.13788 | 0.00000 | 0.00000 | 0.00202 | 0.22007 | 0.00951 | 0.25693 | 0.70121 | 0.11401 | 0.00860 |
| **SAMD11** | 0.01194 | 0.03100 | 0.05611 | 1.25191 | 0.02828 | 0.00063 | 0.00000 | 0.05815 | 0.09311 | 0.19666 | 0.01514 | 0.01115 |
| **SAMD12** | 4.92571 | 0.00000 | 0.59904 | 0.00000 | 0.00000 | 0.40552 | 0.43967 | 2.36639 | 0.00000 | 0.44411 | 0.90434 | 0.00000 |
| **SAMD12-AS1** | 0.26688 | 0.03155 | 0.01228 | 0.01552 | 0.01898 | 0.02803 | 0.01470 | 0.04164 | 0.02412 | 0.00540 | 0.01467 | 0.01315 |
| **SAMD13** | 0.00841 | 0.34190 | 0.00900 | 0.00000 | 0.00000 | 0.04267 | 0.21940 | 0.01011 | 0.00000 | 0.01497 | 0.11572 | 0.02328 |
| **SAMD14** | 0.31318 | 0.01248 | 0.06693 | 0.01695 | 0.01000 | 0.00246 | 0.01979 | 0.02807 | 0.00129 | 0.05555 | 0.02780 | 0.01682 |
| **SAMD15** | 0.39299 | 0.32681 | 0.10820 | 0.14256 | 0.28016 | 0.00000 | 0.00000 | 0.25770 | 0.19260 | 0.13948 | 0.08778 | 0.24829 |
| **SAMD3** | 1.35642 | 0.01322 | 0.01741 | 0.01242 | 0.01863 | 0.00008 | 0.01384 | 0.86220 | 0.02498 | 0.01260 | 0.01581 | 0.00000 |
| **SAMD4A** | 3.66885 | 0.00929 | 0.28934 | 0.12522 | 0.00000 | 0.00000 | 0.46892 | 0.00000 | 0.00000 | 0.17178 | 0.00000 | 0.04940 |
| **SAMD4B** | 1.86949 | 2.05407 | 7.39412 | 5.13107 | 9.23516 | 0.00000 | 8.36123 | 0.00000 | 3.42230 | 13.36502 | 2.22865 | 4.82481 |
| **SAMD5** | 0.04795 | 0.06323 | 0.00000 | 1.72122 | 0.00000 | 0.14923 | 0.00000 | 0.00152 | 0.00000 | 0.16592 | 0.00518 | 0.04954 |
| **SAMD7** | 0.00394 | 0.01717 | 0.04152 | 0.01124 | 0.01405 | 0.00866 | 0.02012 | 0.04217 | 0.00683 | 0.03270 | 0.01960 | 0.03204 |
| **SAMD8** | 0.21612 | 7.78971 | 15.30488 | 2.41164 | 0.00000 | 11.94304 | 4.05920 | 15.84840 | 3.85104 | 7.72787 | 2.31406 | 25.87992 |
| **SAMD9L** | 0.05389 | 0.00894 | 0.01787 | 0.02338 | 0.00621 | 0.04006 | 0.01115 | 0.00551 | 0.01266 | 0.01634 | 0.02501 | 0.03333 |
| **SAMHD1** | 10.15672 | 5.62735 | 0.38378 | 14.20992 | 2.42731 | 9.30553 | 0.00000 | 2.44069 | 2.20031 | 0.00000 | 5.41413 | 0.00000 |
| **SAMM50** | 1.48386 | 15.04793 | 9.99856 | 13.75129 | 13.65041 | 8.06478 | 18.26101 | 0.67826 | 18.44296 | 0.00000 | 12.48475 | 12.21757 |
| **SAP130** | 6.64643 | 11.02049 | 16.78865 | 6.48487 | 9.35915 | 1.75066 | 12.48329 | 3.49046 | 11.03813 | 4.84045 | 5.56306 | 6.63412 |
| **SAP18** | 10.73042 | 23.77102 | 24.66140 | 17.10905 | 18.01232 | 51.66035 | 36.11349 | 28.37814 | 28.21893 | 18.74445 | 22.96726 | 35.34060 |
| **SAP25** | 0.25894 | 1.53024 | 0.00000 | 0.20658 | 2.04304 | 0.00000 | 0.10898 | 0.02367 | 0.00000 | 2.40951 | 0.00000 | 2.06908 |
| **SAP30** | 0.22639 | 1.14327 | 0.92024 | 0.20544 | 0.00000 | 0.00000 | 1.68497 | 0.14644 | 0.00000 | 3.32672 | 0.03324 | 1.03479 |
| **SAP30BP** | 2.59356 | 17.27079 | 17.38294 | 11.31034 | 14.72362 | 9.63861 | 21.52890 | 10.72344 | 16.36139 | 2.39882 | 15.96643 | 18.06283 |
| **SAP30L** | 0.01040 | 0.00000 | 0.52555 | 0.68111 | 1.12840 | 0.10227 | 4.89823 | 0.21692 | 0.00009 | 8.67899 | 0.71854 | 0.65389 |
| **SAP30L-AS1** | 0.00087 | 0.00000 | 0.00000 | 0.00000 | 2.62256 | 0.02375 | 0.02671 | 0.01060 | 1.56585 | 2.51854 | 1.97038 | 0.01214 |
| **SAPCD2** | 0.00224 | 2.20353 | 2.09349 | 5.10664 | 25.55800 | 0.00000 | 12.03380 | 0.00208 | 6.01332 | 14.83651 | 0.88974 | 0.00000 |
| **SAR1A** | 4.70047 | 20.81402 | 22.83293 | 12.15092 | 13.03370 | 38.69956 | 32.87973 | 23.16538 | 28.47802 | 12.47857 | 24.40055 | 23.84691 |
| **SAR1B** | 0.94258 | 8.56523 | 5.04370 | 3.62781 | 5.30833 | 33.07334 | 0.00000 | 2.72512 | 11.20375 | 1.17255 | 0.00000 | 7.83674 |
| **SARDH** | 1.82771 | 0.01620 | 0.02340 | 0.02453 | 0.00000 | 0.07780 | 0.00000 | 0.41957 | 0.03563 | 0.20330 | 0.00336 | 0.02084 |
| **SARM1** | 0.09792 | 0.06430 | 0.20033 | 0.05632 | 0.06125 | 0.00000 | 0.00000 | 0.00000 | 0.19865 | 0.37884 | 0.16342 | 0.15824 |
| **SARNP** | 0.77226 | 9.61851 | 12.92874 | 10.51108 | 9.03699 | 16.14334 | 18.62908 | 11.79326 | 10.86303 | 5.92986 | 11.31999 | 14.48835 |
| **SARS** | 8.36630 | 23.17747 | 32.47569 | 15.59700 | 21.47568 | 29.26676 | 34.56523 | 17.30723 | 24.11417 | 4.30554 | 24.20546 | 21.53213 |
| **SARS2** | 0.16513 | 8.11064 | 6.96207 | 8.53662 | 8.85664 | 0.00000 | 5.77849 | 0.17599 | 7.31603 | 1.62893 | 7.57798 | 4.12328 |
| **SART1** | 1.55061 | 2.27431 | 4.64165 | 5.13628 | 9.22736 | 0.00000 | 9.80586 | 2.23108 | 1.67718 | 15.53822 | 1.05059 | 4.84081 |
| **SART3** | 4.39773 | 9.70236 | 9.46543 | 8.20788 | 8.27029 | 11.17681 | 12.91800 | 3.78575 | 10.29919 | 10.28913 | 11.13443 | 11.00468 |
| **SASH1** | 0.14466 | 4.39457 | 0.36830 | 4.78736 | 2.26832 | 5.77247 | 0.36078 | 0.06227 | 0.00000 | 6.94182 | 0.00000 | 4.93618 |
| **SASH3** | 0.00573 | 0.01568 | 0.02252 | 0.01464 | 0.01709 | 0.00977 | 0.01749 | 0.01056 | 0.01724 | 0.01609 | 0.01618 | 0.03380 |
| **SASS6** | 11.63984 | 0.35543 | 4.95861 | 0.49109 | 0.00000 | 14.10290 | 6.32914 | 4.96119 | 4.96081 | 14.89465 | 5.29150 | 1.86479 |
| **SAT1** | 0.15531 | 2.60214 | 7.50662 | 30.37853 | 0.00000 | 4.26290 | 2.85444 | 0.85736 | 2.71961 | 0.72204 | 1.24499 | 8.23685 |
| **SAT2** | 0.02439 | 0.49069 | 2.03619 | 3.05902 | 0.08877 | 0.27093 | 0.79766 | 0.71409 | 0.26281 | 0.00000 | 0.52947 | 5.40246 |
| **SATB1** | 0.50744 | 0.00000 | 2.59360 | 0.00000 | 2.51183 | 0.17741 | 11.93624 | 0.18852 | 1.37058 | 1.15783 | 3.17703 | 0.06123 |
| **SATB2** | 1.16295 | 1.90127 | 0.00000 | 0.00000 | 3.25425 | 1.75291 | 1.26737 | 8.61899 | 3.09192 | 5.43854 | 5.25069 | 3.32939 |
| **SATB2-AS1** | 0.41704 | 0.00512 | 0.01599 | 0.00950 | 0.01041 | 0.00000 | 0.00696 | 0.11409 | 0.02580 | 0.11165 | 0.01643 | 0.08960 |
| **SATL1** | 0.00620 | 0.01483 | 0.01927 | 0.01951 | 0.01869 | 0.02107 | 0.01885 | 0.01036 | 0.01459 | 0.01288 | 0.01513 | 0.01719 |
| **SAV1** | 1.91234 | 0.00018 | 0.00000 | 9.11863 | 5.12822 | 10.66691 | 7.87092 | 0.77022 | 0.00000 | 21.48166 | 6.08776 | 2.81723 |
| **SAYSD1** | 3.77343 | 2.18604 | 2.35438 | 2.12221 | 0.48128 | 2.23348 | 1.12641 | 0.73003 | 3.75078 | 0.00000 | 2.17502 | 3.45101 |
| **SBDS** | 0.94086 | 17.29893 | 13.72296 | 5.06666 | 6.75891 | 29.56970 | 12.12364 | 18.72708 | 12.09937 | 2.15477 | 13.92945 | 10.54862 |

| | | | | | | | | | | | | |
|---|---|---|---|---|---|---|---|---|---|---|---|---|
| **SBDSP1** | 1.66788 | 11.27091 | 5.56388 | 1.24542 | 2.98186 | 15.46331 | 4.44867 | 15.19619 | 6.97702 | 4.50420 | 6.06119 | 10.92108 |
| **SBF1** | 4.54340 | 1.80800 | 3.32561 | 1.95441 | 8.74414 | 0.00000 | 2.57464 | 0.09346 | 4.76258 | 12.59603 | 0.56328 | 2.17121 |
| **SBF1P1** | 0.21353 | 0.07758 | 0.22135 | 0.14951 | 0.23576 | 0.00000 | 0.01350 | 0.03273 | 0.26220 | 0.26628 | 0.04889 | 0.12554 |
| **SBF2** | 1.95580 | 0.05193 | 1.25094 | 0.15843 | 2.34718 | 3.81484 | 1.10471 | 0.13060 | 2.02551 | 3.74505 | 0.42443 | 1.81329 |
| **SBF2-AS1** | 0.00716 | 2.69181 | 0.00000 | 0.00002 | 5.90993 | 4.09443 | 0.13038 | 0.00000 | 0.00000 | 0.95856 | 3.08343 | 0.52128 |
| **SBK1** | 0.51598 | 0.21078 | 0.08750 | 0.32908 | 0.29469 | 0.00000 | 0.04128 | 0.00000 | 0.00000 | 0.11223 | 0.01935 | 0.08628 |
| **SBK3** | 1.13854 | 0.01089 | 0.01659 | 0.01491 | 0.01717 | 0.01255 | 0.01812 | 0.00000 | 0.02273 | 0.00000 | 0.01320 | 0.04611 |
| **SBNO1** | 6.88420 | 16.59600 | 21.10725 | 9.38031 | 8.49529 | 30.91954 | 26.56737 | 28.13820 | 19.57195 | 27.63280 | 23.39787 | 18.50970 |
| **SBNO2** | 1.78415 | 0.11931 | 1.00226 | 0.06960 | 0.81284 | 0.00000 | 0.22395 | 0.11052 | 0.59383 | 4.04111 | 0.00000 | 0.17684 |
| **SBSN** | 0.00169 | 0.01720 | 0.38146 | 0.01091 | 0.01494 | 0.00000 | 0.11270 | 0.07108 | 0.00000 | 0.05195 | 0.00930 | 0.02768 |
| **SBSPON** | 0.20701 | 0.00000 | 3.17880 | 1.85615 | 0.90185 | 0.00020 | 2.31945 | 0.04101 | 1.34902 | 0.00000 | 0.69233 | 0.08107 |
| **SC5D** | 1.48720 | 0.00000 | 14.55801 | 2.91796 | 7.31652 | 11.05219 | 0.00000 | 0.23080 | 40.61269 | 1.73571 | 3.43800 | 0.34068 |
| **SCAF1** | 0.51716 | 0.90492 | 2.37297 | 0.28044 | 2.32921 | 0.00000 | 0.75732 | 0.00000 | 1.34806 | 9.56830 | 1.89014 | 0.85238 |
| **SCAF11** | 2.01644 | 7.52525 | 10.16121 | 5.18494 | 11.14167 | 29.11646 | 17.60696 | 10.64862 | 0.83903 | 27.14953 | 11.58100 | 12.02780 |
| **SCAF4** | 3.35426 | 11.05590 | 9.22962 | 7.84991 | 10.64242 | 0.22711 | 9.26229 | 12.50482 | 12.97515 | 10.80772 | 10.86328 | 7.19648 |
| **SCAF8** | 2.14679 | 5.54559 | 1.81816 | 3.38161 | 2.70044 | 10.74899 | 4.07985 | 7.23229 | 0.48125 | 8.98138 | 1.29551 | 5.33669 |
| **SCAI** | 4.92835 | 1.32437 | 0.65874 | 1.12386 | 1.28761 | 2.59289 | 2.31547 | 1.34927 | 1.85802 | 1.95312 | 1.27357 | 1.90795 |
| **SCAMP1** | 4.32866 | 9.33285 | 0.00000 | 11.54373 | 2.14840 | 23.74234 | 0.00000 | 0.91496 | 0.00000 | 11.83907 | 3.82943 | 10.79672 |
| **SCAMP1-AS1** | 0.78365 | 0.53358 | 0.20437 | 0.38883 | 0.19901 | 0.77611 | 0.00000 | 0.42197 | 0.00000 | 0.59320 | 0.47782 | 0.90836 |
| **SCAMP2** | 1.63730 | 10.59520 | 6.15997 | 6.95024 | 10.50266 | 0.00000 | 0.00000 | 0.78026 | 13.37274 | 0.00000 | 6.62107 | 6.11830 |
| **SCAMP3** | 0.18091 | 9.63638 | 15.24679 | 11.39665 | 7.73014 | 5.75153 | 9.25029 | 1.11354 | 37.43898 | 1.22236 | 11.80538 | 5.74751 |
| **SCAMP4** | 0.55333 | 2.80454 | 5.55343 | 1.68265 | 9.33239 | 0.00000 | 0.00000 | 0.21699 | 11.76092 | 7.90315 | 1.82987 | 8.82055 |
| **SCAMP5** | 0.00323 | 0.03627 | 1.47882 | 2.44621 | 0.12982 | 0.00000 | 0.20783 | 0.26109 | 1.05284 | 0.43373 | 0.00000 | 1.27204 |
| **SCAND1** | 2.61878 | 7.88384 | 12.40374 | 9.15940 | 8.56077 | 0.00000 | 8.34552 | 8.24868 | 11.37675 | 2.47355 | 13.24390 | 11.62444 |
| **SCAND2P** | 0.00329 | 0.58359 | 1.86241 | 0.82503 | 1.53992 | 0.14852 | 1.88132 | 1.86425 | 0.72399 | 1.49830 | 0.78426 | 1.39302 |
| **SCAP** | 9.12533 | 9.35836 | 9.20355 | 11.76322 | 14.92915 | 0.00000 | 1.98265 | 1.82204 | 16.05542 | 12.43180 | 7.01212 | 5.66784 |
| **SCAPER** | 5.14080 | 1.42188 | 3.60942 | 3.85410 | 2.25837 | 4.28453 | 0.00000 | 0.46916 | 0.91258 | 10.59934 | 0.12401 | 0.99567 |
| **SCARA3** | 0.00135 | 0.00000 | 1.41129 | 0.00000 | 1.65708 | 0.02989 | 0.62991 | 0.00558 | 0.30671 | 0.88832 | 1.43832 | 0.02425 |
| **SCARA5** | 1.37327 | 0.00000 | 0.28391 | 0.91924 | 0.00000 | 0.13002 | 0.00000 | 0.21403 | 0.48830 | 0.06563 | 0.18646 | 0.12641 |
| **SCARB1** | 0.76297 | 0.38172 | 1.34619 | 2.02166 | 1.77702 | 0.00000 | 2.33265 | 0.09152 | 6.34647 | 0.58401 | 0.17860 | 0.07627 |
| **SCARB2** | 1.99440 | 4.40788 | 1.60805 | 1.29211 | 4.00759 | 7.74053 | 0.00000 | 0.63087 | 7.25985 | 9.27528 | 1.74707 | 1.88850 |
| **SCARF1** | 0.00988 | 0.13042 | 0.03157 | 0.05158 | 0.09796 | 0.00000 | 0.00000 | 0.00560 | 0.19299 | 0.09643 | 0.01421 | 0.13906 |
| **SCARF2** | 0.13469 | 0.05026 | 0.04774 | 0.09829 | 0.46748 | 0.00000 | 0.00037 | 0.50793 | 0.00000 | 0.21173 | 0.00000 | 0.00643 |
| **SCARNA1** | 0.15462 | 0.00000 | 0.07716 | 0.11982 | 0.16932 | 0.12082 | 0.00000 | 0.02119 | 0.05846 | 0.18596 | 0.00027 | 0.22239 |
| **SCARNA10** | 0.59164 | 0.02571 | 0.02161 | 0.00764 | 0.01567 | 0.00417 | 0.03640 | 0.03299 | 0.01304 | 0.05415 | 0.01357 | 0.04612 |
| **SCARNA11** | 0.00599 | 0.01630 | 0.04062 | 0.01083 | 0.01503 | 0.01705 | 0.03385 | 0.01094 | 0.00594 | 0.01724 | 0.01847 | 0.01704 |
| **SCARNA12** | 0.02033 | 0.60344 | 0.92849 | 1.41752 | 1.18543 | 0.00000 | 0.00000 | 0.89240 | 3.30106 | 1.10050 | 0.11190 | 0.71193 |
| **SCARNA14** | 0.02860 | 0.00855 | 0.22156 | 0.04635 | 0.03089 | 0.01027 | 0.00000 | 0.02895 | 0.00084 | 0.05850 | 0.03108 | 0.03301 |
| **SCARNA16** | 0.11756 | 0.02474 | 0.00857 | 0.09858 | 0.02096 | 0.12537 | 0.00000 | 0.06284 | 0.11442 | 0.12270 | 0.04385 | 0.06764 |
| **SCARNA17** | 0.33057 | 0.03831 | 0.06806 | 0.00122 | 0.29496 | 0.00000 | 0.02305 | 0.00169 | 0.01481 | 0.64320 | 0.00000 | 0.01001 |
| **SCARNA18** | 0.02672 | 0.05893 | 0.15164 | 0.00000 | 0.02096 | 0.03768 | 0.03910 | 0.05626 | 0.12818 | 0.08638 | 0.00000 | 0.05602 |
| **SCARNA2** | 0.68796 | 0.01161 | 0.02813 | 0.18339 | 0.34035 | 0.00000 | 0.00000 | 0.05833 | 0.06863 | 0.50981 | 0.14285 | 0.11990 |
| **SCARNA20** | 0.00431 | 0.02612 | 0.05869 | 0.01531 | 0.02156 | 0.00575 | 0.03845 | 0.02704 | 0.02626 | 0.01548 | 0.01864 | 0.04799 |
| **SCARNA21** | 0.17084 | 0.01880 | 0.01481 | 0.00917 | 0.05243 | 0.01165 | 0.01982 | 0.04844 | 0.03380 | 0.00000 | 0.01104 | 0.02385 |
| **SCARNA22** | 1.42629 | 0.40728 | 0.03311 | 0.20050 | 0.54014 | 0.00000 | 0.00000 | 3.75857 | 0.36949 | 2.34950 | 0.00000 | 0.84843 |
| **SCARNA23** | 0.00730 | 0.02046 | 0.05883 | 0.00868 | 0.00000 | 0.02741 | 0.01762 | 0.04421 | 0.00000 | 0.03340 | 0.10613 | 0.03449 |
| **SCARNA27** | 0.01559 | 0.00909 | 0.04802 | 0.02657 | 0.00991 | 0.05250 | 0.00000 | 0.00906 | 0.05609 | 0.04327 | 0.00761 | 0.02677 |
| **SCARNA3** | 0.00683 | 0.01963 | 0.04347 | 0.06010 | 0.04509 | 0.02585 | 0.00000 | 0.01156 | 0.01911 | 0.11726 | 0.01209 | 0.02824 |

| | | | | | | | | | | | | |
|---|---|---|---|---|---|---|---|---|---|---|---|---|
| **SCARNA4** | 0.00630 | 0.01009 | 0.05262 | 0.02995 | 0.01256 | 0.03220 | 0.02523 | 0.01299 | 0.02145 | 0.08396 | 0.03159 | 0.02187 |
| **SCARNA5** | 0.39395 | 0.02874 | 0.00475 | 0.00000 | 0.39897 | 0.00000 | 0.00010 | 0.23304 | 0.04512 | 0.65673 | 0.08609 | 0.20401 |
| **SCARNA6** | 0.16229 | 0.02168 | 0.02473 | 0.00992 | 0.01623 | 0.01561 | 0.01555 | 0.27046 | 0.03830 | 0.00015 | 0.04959 | 0.00839 |
| **SCARNA7** | 0.03172 | 0.02982 | 0.03077 | 0.00732 | 0.04729 | 0.00000 | 0.02422 | 0.01168 | 0.02362 | 0.04298 | 0.01562 | 0.01380 |
| **SCARNA8** | 0.02324 | 0.01589 | 0.09678 | 0.01726 | 0.04583 | 0.01628 | 0.00367 | 0.02220 | 0.06492 | 0.02168 | 0.00869 | 0.08834 |
| **SCARNA9** | 0.00747 | 0.00000 | 0.08437 | 0.00000 | 0.16745 | 0.04554 | 0.00000 | 0.01933 | 0.09696 | 0.77678 | 0.00000 | 0.01756 |
| **SCARNA9L** | 0.22650 | 0.00543 | 0.12478 | 0.06324 | 0.08515 | 0.00000 | 0.00292 | 0.02228 | 0.19270 | 0.07115 | 0.08713 | 0.12076 |
| **SCART1** | 0.00305 | 0.01264 | 0.05090 | 0.08562 | 0.19662 | 0.00000 | 0.04619 | 0.00721 | 0.00000 | 0.35045 | 0.05238 | 0.05035 |
| **SCCPDH** | 2.00277 | 0.09728 | 3.77688 | 0.72732 | 0.81391 | 3.71881 | 0.00851 | 0.03827 | 1.34220 | 0.25046 | 3.63695 | 0.02668 |
| **SCD** | 0.00000 | 10.05652 | 36.37023 | 22.84564 | 26.64798 | 34.01414 | 36.45468 | 5.47813 | 75.47877 | 34.64414 | 37.21023 | 29.64868 |
| **SCD5** | 0.00091 | 0.00000 | 0.00000 | 0.31739 | 0.00000 | 0.07203 | 0.50023 | 0.10418 | 0.00000 | 0.69654 | 0.39775 | 0.01919 |
| **SCEL** | 0.41656 | 0.01129 | 0.01889 | 0.01840 | 0.02285 | 0.02987 | 0.01521 | 0.25945 | 0.01723 | 0.00000 | 0.01392 | 0.02363 |
| **SCFD1** | 2.46881 | 2.61038 | 5.79403 | 1.99022 | 0.42643 | 26.27176 | 5.25829 | 0.57034 | 6.63904 | 2.69212 | 7.42753 | 4.54540 |
| **SCFD2** | 0.17796 | 3.62850 | 5.52092 | 2.82294 | 2.78703 | 2.28606 | 2.47483 | 2.08864 | 2.09445 | 0.77398 | 4.49358 | 2.89908 |
| **SCG2** | 1.40616 | 0.23228 | 0.07209 | 0.09766 | 0.00000 | 0.00000 | 0.01272 | 1.18669 | 0.00000 | 0.04563 | 0.03119 | 0.38623 |
| **SCG3** | 0.00528 | 0.29848 | 0.13896 | 0.37973 | 0.57834 | 0.15064 | 0.00000 | 0.00475 | 0.00000 | 0.09583 | 0.00000 | 0.30408 |
| **SCG5** | 0.09308 | 0.00975 | 0.31416 | 0.11266 | 0.06467 | 0.02473 | 0.30622 | 0.20135 | 0.21904 | 0.00000 | 0.04964 | 0.09325 |
| **SCGB1A1** | 0.00975 | 0.03252 | 0.02084 | 0.01815 | 0.04184 | 0.00000 | 0.01340 | 0.00967 | 0.06048 | 0.00000 | 0.01188 | 0.01955 |
| **SCGB1B2P** | 0.00485 | 0.04157 | 0.01806 | 0.02426 | 0.00525 | 0.04272 | 0.00001 | 0.00959 | 0.07373 | 0.04529 | 0.01116 | 0.05186 |
| **SCGB1D1** | 0.00724 | 0.57753 | 0.00000 | 0.00000 | 0.00000 | 0.31808 | 0.00000 | 0.00000 | 0.05480 | 0.10000 | 0.29963 | 0.55821 |
| **SCGB1D2** | 0.00571 | 0.09853 | 0.06798 | 0.07718 | 0.00000 | 0.01281 | 0.01928 | 0.01081 | 0.03050 | 0.02950 | 0.01396 | 0.00924 |
| **SCGB1D4** | 0.00567 | 0.04751 | 0.01196 | 0.03298 | 0.02451 | 0.00698 | 0.01613 | 0.01077 | 0.01767 | 0.03897 | 0.01297 | 0.01226 |
| **SCGB2A1** | 0.00578 | 0.01285 | 0.01361 | 0.02398 | 0.05776 | 0.00000 | 0.02159 | 0.01078 | 0.01270 | 0.02616 | 0.01548 | 0.03462 |
| **SCGB3A2** | 3.35523 | 11.78753 | 0.00000 | 9.45792 | 15.05487 | 4.07907 | 0.00000 | 1.14747 | 0.00000 | 0.00000 | 3.10959 | 14.33622 |
| **SCGN** | 0.00650 | 0.01322 | 0.01437 | 0.01146 | 0.01402 | 0.01961 | 0.03068 | 0.02856 | 0.01485 | 0.01596 | 0.01397 | 0.04772 |
| **SCHIP1.uid-546** | 0.00587 | 0.01794 | 0.12904 | 0.10328 | 0.02794 | 0.00000 | 0.02171 | 0.01143 | 0.01662 | 0.00371 | 0.02729 | 0.02380 |
| **SCHLAP1** | 0.06680 | 0.00437 | 0.10892 | 0.07370 | 0.05117 | 0.03559 | 0.07927 | 0.00416 | 0.06823 | 0.21203 | 0.03315 | 0.02065 |
| **SCIMP** | 0.00824 | 0.01086 | 0.00691 | 0.02601 | 0.02300 | 0.04866 | 0.00636 | 0.00993 | 0.02154 | 0.05118 | 0.03250 | 0.05197 |
| **SCIN** | 1.40274 | 0.46206 | 0.44164 | 0.00000 | 1.58219 | 0.06580 | 0.00000 | 0.19391 | 0.07493 | 0.04901 | 0.28873 | 0.18082 |
| **SCLT1** | 0.51719 | 1.00809 | 2.48944 | 1.54590 | 1.23645 | 3.78800 | 0.00000 | 1.27322 | 3.89248 | 0.00000 | 1.60305 | 1.67576 |
| **SCLY** | 0.01520 | 0.07714 | 0.10613 | 0.03912 | 0.01446 | 0.02229 | 0.00451 | 0.02269 | 0.20919 | 0.00000 | 0.10658 | 0.04145 |
| **SCMH1** | 0.03575 | 0.18066 | 0.79135 | 1.21581 | 0.00000 | 0.00000 | 0.00000 | 0.02701 | 0.25608 | 1.21633 | 0.69003 | 0.09073 |
| **SCML1** | 3.42799 | 0.11125 | 0.00000 | 2.11953 | 2.95335 | 1.88613 | 0.71006 | 1.29421 | 1.04606 | 0.00000 | 0.58929 | 0.30774 |
| **SCML2** | 0.73373 | 5.13631 | 0.14870 | 0.00000 | 0.00000 | 4.25619 | 1.23802 | 3.26127 | 2.79712 | 1.65553 | 0.58898 | 7.63311 |
| **SCML4** | 0.10780 | 0.02397 | 0.03885 | 0.10271 | 0.00036 | 0.05039 | 0.00000 | 0.01898 | 0.07723 | 0.20993 | 0.01573 | 0.01352 |
| **SCN10A** | 0.00546 | 0.00301 | 0.01368 | 0.00594 | 0.26867 | 0.00000 | 0.03333 | 0.00937 | 0.00000 | 0.03884 | 0.00711 | 0.11343 |
| **SCN11A** | 0.00146 | 0.01031 | 0.04409 | 0.00000 | 0.18461 | 0.22189 | 0.00000 | 0.10907 | 0.22116 | 0.17195 | 0.00000 | 0.00000 |
| **SCN1A** | 2.45024 | 0.00000 | 0.03106 | 0.00680 | 0.00486 | 0.01780 | 0.01704 | 0.08610 | 0.00331 | 0.06694 | 0.02235 | 0.49022 |
| **SCN1B** | 0.01366 | 0.10694 | 0.02461 | 0.04453 | 0.02232 | 0.08311 | 0.02125 | 0.04523 | 0.00885 | 0.00000 | 0.03278 | 0.09983 |
| **SCN2A** | 0.00000 | 0.73513 | 0.00000 | 0.22148 | 0.00038 | 0.61932 | 0.00000 | 0.00460 | 0.47424 | 1.69170 | 0.00000 | 0.07758 |
| **SCN2B** | 0.00481 | 0.01498 | 0.01336 | 0.00408 | 0.00254 | 0.04380 | 0.01240 | 0.01072 | 0.01438 | 0.06491 | 0.01053 | 0.01621 |
| **SCN3A** | 0.26733 | 0.46022 | 0.00000 | 0.28741 | 0.14995 | 0.47758 | 0.00000 | 0.07906 | 0.02301 | 1.72238 | 0.15577 | 1.40132 |
| **SCN3B** | 0.07740 | 0.19793 | 0.00000 | 0.00000 | 0.46667 | 0.00000 | 0.20855 | 0.00000 | 1.02291 | 0.55028 | 0.06435 | 0.14477 |
| **SCN4A** | 0.07428 | 0.00733 | 0.17133 | 0.01299 | 0.00137 | 0.01328 | 0.04479 | 0.00594 | 0.00000 | 0.11745 | 0.03252 | 0.03058 |
| **SCN4B** | 0.04099 | 0.02203 | 0.04909 | 0.00861 | 0.02432 | 0.00000 | 0.03731 | 0.00346 | 0.00000 | 0.08254 | 0.01976 | 0.15327 |
| **SCN5A** | 2.13672 | 0.13659 | 0.12334 | 0.06607 | 0.07823 | 0.00000 | 0.14044 | 0.27476 | 0.00000 | 0.23970 | 0.00745 | 0.00413 |
| **SCN7A** | 0.58283 | 0.00842 | 0.01395 | 0.03596 | 0.00320 | 0.00000 | 0.01192 | 0.41091 | 0.04446 | 0.10896 | 0.01303 | 0.01403 |
| **SCN8A** | 1.30495 | 0.11555 | 0.53369 | 2.74477 | 0.00000 | 0.45757 | 1.40603 | 0.38427 | 0.70870 | 1.15820 | 0.00000 | 1.47775 |

| | | | | | | | | | | | | |
|---|---|---|---|---|---|---|---|---|---|---|---|---|
| **SCN9A** | 0.97950 | 0.03045 | 0.06085 | 0.12942 | 0.03267 | 0.00000 | 0.03765 | 0.02982 | 0.02371 | 0.00000 | 0.01424 | 0.06867 |
| **SCNM1** | 0.02095 | 2.12767 | 3.19080 | 1.44613 | 1.41684 | 3.78677 | 2.67337 | 9.36029 | 2.25354 | 0.00000 | 2.35858 | 5.22960 |
| **SCNN1A** | 0.01956 | 0.11989 | 0.21289 | 8.44545 | 2.77557 | 0.00000 | 0.02291 | 0.00550 | 0.51142 | 0.65804 | 0.00000 | 0.05908 |
| **SCNN1B** | 0.10864 | 0.01983 | 0.02486 | 0.02327 | 0.02416 | 0.00990 | 0.01870 | 0.00674 | 0.02936 | 0.00608 | 0.02081 | 0.02843 |
| **SCNN1D** | 0.00570 | 0.01086 | 0.02824 | 0.02311 | 0.14951 | 0.20044 | 0.00010 | 0.00720 | 0.00898 | 0.03798 | 0.04059 | 0.10426 |
| **SCNN1G** | 0.00202 | 0.00977 | 0.00989 | 0.04025 | 0.04610 | 0.00000 | 0.04915 | 0.07758 | 0.01972 | 0.01520 | 0.01069 | 0.14766 |
| **SCO1** | 0.00413 | 13.50762 | 10.10523 | 7.73973 | 8.52249 | 22.61873 | 15.49856 | 15.92900 | 15.50871 | 0.00000 | 12.67582 | 17.56039 |
| **SCO2.uid-26107** | 0.00440 | 1.55363 | 0.66253 | 0.73436 | 1.10725 | 0.00000 | 0.94774 | 0.03860 | 0.78364 | 0.00000 | 1.46946 | 0.69010 |
| **SCOC** | 2.61977 | 4.45660 | 4.00719 | 2.34464 | 0.54573 | 24.91925 | 0.03331 | 1.56427 | 3.23375 | 5.61579 | 6.54723 | 7.13641 |
| **SCOC-AS1** | 0.04533 | 0.46732 | 0.00000 | 0.06760 | 0.31971 | 0.36561 | 0.02014 | 0.01479 | 0.00000 | 0.23359 | 0.03610 | 0.83917 |
| **SCP2** | 1.80997 | 4.88338 | 6.19677 | 4.82760 | 4.06210 | 27.50994 | 6.50560 | 0.61627 | 8.34870 | 1.92834 | 10.57077 | 5.09142 |
| **SCPEP1** | 0.00000 | 16.31831 | 0.00000 | 7.23703 | 7.71291 | 17.64213 | 0.00000 | 0.14586 | 9.64646 | 3.88669 | 9.78862 | 12.32351 |
| **SCRG1** | 0.00513 | 0.03894 | 0.01149 | 0.00341 | 0.01050 | 0.03995 | 0.00948 | 0.04639 | 0.01647 | 0.03178 | 0.01539 | 0.05804 |
| **SCRIB** | 0.40783 | 1.43657 | 1.44653 | 1.39307 | 5.59525 | 0.00000 | 0.37556 | 0.13801 | 1.02317 | 17.17452 | 0.00305 | 0.49907 |
| **SCRN1** | 0.37617 | 1.82734 | 1.03691 | 8.33272 | 0.74343 | 0.00000 | 0.39234 | 0.07676 | 0.59620 | 0.73685 | 0.05415 | 0.08198 |
| **SCRN2** | 0.38637 | 0.80148 | 4.99589 | 0.15535 | 1.38789 | 0.00000 | 0.47631 | 0.08585 | 2.06514 | 0.82204 | 1.60504 | 0.25808 |
| **SCRN3** | 0.38355 | 2.22088 | 0.85552 | 0.00000 | 0.52111 | 4.56624 | 0.00000 | 0.17844 | 0.00000 | 0.16310 | 2.02902 | 1.35533 |
| **SCRT1** | 0.19658 | 0.01055 | 0.03325 | 0.00649 | 0.00890 | 0.01255 | 0.02001 | 0.01691 | 0.00311 | 0.10874 | 0.01862 | 0.05003 |
| **SCRT2** | 0.17368 | 0.01454 | 0.02840 | 0.01259 | 0.01689 | 0.01325 | 0.01865 | 0.01316 | 0.01579 | 0.01054 | 0.02011 | 0.01567 |
| **SCTR** | 0.00278 | 0.04836 | 0.06085 | 0.01354 | 0.01347 | 0.05084 | 0.00000 | 0.02273 | 0.01612 | 0.09916 | 0.00790 | 0.00716 |
| **SCUBE1** | 1.33801 | 0.01732 | 0.01625 | 0.00926 | 0.00990 | 0.02051 | 0.01870 | 0.03418 | 0.01706 | 0.00534 | 0.03009 | 0.02521 |
| **SCUBE2** | 0.21263 | 1.09116 | 0.02885 | 0.27110 | 0.13483 | 0.00000 | 0.00000 | 0.00000 | 0.05756 | 2.02584 | 0.00000 | 0.87743 |
| **SCUBE3** | 0.00145 | 0.00002 | 2.34003 | 0.11330 | 0.01170 | 0.00000 | 0.37281 | 0.03443 | 0.86855 | 1.06809 | 0.00000 | 0.07461 |
| **SCYL1** | 3.59141 | 9.22331 | 9.28900 | 9.25475 | 6.65378 | 0.00000 | 4.21064 | 1.80895 | 14.03289 | 0.04584 | 4.38452 | 9.73830 |
| **SCYL2** | 2.22016 | 1.87082 | 6.51606 | 1.04348 | 0.00043 | 18.37633 | 6.62370 | 2.33836 | 3.36666 | 5.03966 | 2.44154 | 4.41456 |
| **SCYL3** | 0.07126 | 0.80440 | 2.55097 | 1.84206 | 0.01325 | 3.39641 | 0.69961 | 6.74599 | 1.98027 | 0.00000 | 0.92110 | 1.49038 |
| **SDAD1** | 3.85699 | 7.49681 | 6.84221 | 4.33854 | 5.14312 | 20.34072 | 12.85722 | 8.19569 | 7.85005 | 9.51176 | 8.40287 | 15.50076 |
| **SDC1** | 0.09479 | 0.14567 | 28.55765 | 0.00717 | 1.51928 | 0.00000 | 0.00000 | 0.00691 | 4.41624 | 0.00000 | 0.00000 | 0.00943 |
| **SDC2** | 0.89857 | 1.75465 | 0.40349 | 0.17855 | 0.00000 | 0.07584 | 0.04773 | 1.04968 | 0.00000 | 0.19451 | 0.00000 | 1.14480 |
| **SDC3** | 0.60736 | 0.19897 | 0.63189 | 0.01717 | 0.09850 | 0.07525 | 0.00000 | 0.00077 | 0.00000 | 0.34856 | 0.05866 | 0.04331 |
| **SDC4** | 0.00000 | 30.60168 | 25.53096 | 16.05672 | 30.93271 | 34.23708 | 9.83911 | 4.44232 | 39.84303 | 18.10607 | 25.37049 | 24.17562 |
| **SDC4P** | 0.00732 | 0.01264 | 0.08572 | 0.04959 | 0.00000 | 0.01948 | 0.00346 | 0.01759 | 0.00000 | 0.03595 | 0.07574 | 0.01445 |
| **SDCBP** | 1.80309 | 19.41145 | 18.10483 | 18.35894 | 0.00000 | 33.71428 | 17.63176 | 0.72411 | 5.93721 | 0.18589 | 7.55991 | 4.15306 |
| **SDCBP2-AS1** | 0.00293 | 0.06692 | 0.32297 | 0.25072 | 0.44040 | 0.00000 | 0.00149 | 0.17010 | 0.59182 | 1.37251 | 0.51634 | 0.63015 |
| **SDCBP2.uid-24** | 0.00595 | 0.01399 | 0.01193 | 0.08671 | 0.02389 | 0.00000 | 0.02052 | 0.01030 | 0.01670 | 0.02012 | 0.01483 | 0.02241 |
| **SDCBP2.uid-24** | 0.00538 | 0.01777 | 0.01825 | 0.01111 | 0.01493 | 0.01859 | 0.02728 | 0.01392 | 0.01675 | 0.01317 | 0.01493 | 0.01879 |
| **SDCBP2.uid-24** | 0.00620 | 0.01864 | 0.02028 | 0.01383 | 0.02072 | 0.00598 | 0.02186 | 0.01023 | 0.02524 | 0.00358 | 0.01532 | 0.03142 |
| **SDCCAG3** | 0.19585 | 1.88751 | 4.86819 | 2.47152 | 6.28468 | 1.71777 | 9.51778 | 0.04953 | 0.00013 | 17.47022 | 1.46543 | 3.30304 |
| **SDCCAG8** | 6.50961 | 1.11050 | 2.50574 | 2.28408 | 2.78495 | 2.87856 | 2.50689 | 1.69486 | 2.18017 | 0.92632 | 2.81155 | 1.50468 |
| **SDE2** | 9.31482 | 5.79089 | 9.60457 | 4.45505 | 2.32540 | 12.82117 | 6.94106 | 17.10795 | 4.92077 | 0.08402 | 7.15383 | 10.16462 |
| **SDF2** | 1.47148 | 8.74267 | 15.66492 | 9.92562 | 3.20058 | 8.49712 | 0.00869 | 3.96722 | 19.59765 | 0.00000 | 6.42663 | 12.53637 |
| **SDF2L1** | 0.69396 | 3.56957 | 22.18458 | 12.13708 | 19.77141 | 1.03043 | 19.76332 | 1.29097 | 25.38736 | 7.69236 | 8.14709 | 5.07855 |
| **SDF4** | 0.18339 | 7.11441 | 10.90324 | 1.47479 | 5.02710 | 0.00000 | 0.00053 | 1.74491 | 6.60178 | 13.85311 | 3.54606 | 2.93664 |
| **SDHA** | 2.62217 | 15.15446 | 9.48758 | 11.76159 | 12.52942 | 9.59344 | 7.22417 | 2.65515 | 18.45326 | 0.00000 | 16.80345 | 8.51618 |
| **SDHAF1** | 1.55890 | 7.32326 | 6.61920 | 1.72605 | 4.19920 | 0.00000 | 2.38329 | 1.74901 | 9.42516 | 2.13172 | 3.20635 | 4.90597 |
| **SDHAF2** | 0.00000 | 13.48729 | 15.10860 | 8.39134 | 11.76856 | 9.50479 | 20.71591 | 4.02215 | 13.80946 | 0.01843 | 9.94110 | 15.03335 |
| **SDHAP1** | 0.22093 | 0.84000 | 0.68787 | 0.93457 | 1.09383 | 1.51398 | 0.09947 | 0.87189 | 2.02147 | 0.07831 | 0.33822 | 1.14886 |
| **SDHAP2** | 0.13918 | 0.55431 | 0.17543 | 0.46765 | 0.25345 | 0.30480 | 0.00000 | 0.33934 | 1.11510 | 0.00000 | 0.29838 | 0.58952 |

| | | | | | | | | | | | | |
|---|---|---|---|---|---|---|---|---|---|---|---|---|
| **SDHAP3** | 0.00564 | 0.22789 | 0.11357 | 1.70850 | 0.71157 | 0.49060 | 0.00000 | 0.63763 | 0.36407 | 0.30824 | 0.44869 | 1.22240 |
| **SDHB** | 5.74254 | 23.00936 | 19.06226 | 15.40714 | 22.73610 | 40.28872 | 28.49725 | 7.53307 | 31.16446 | 5.80115 | 30.59702 | 20.09454 |
| **SDHC** | 1.03457 | 5.33847 | 9.22191 | 6.11945 | 5.00517 | 4.86488 | 9.82288 | 0.00000 | 6.48175 | 0.00000 | 11.94312 | 6.98981 |
| **SDHD** | 0.00000 | 12.33990 | 9.73603 | 9.38657 | 2.51947 | 42.61613 | 15.17542 | 1.59366 | 10.36316 | 0.78146 | 17.65084 | 9.55957 |
| **SDK1** | 0.35909 | 0.00000 | 0.37130 | 0.00000 | 0.25242 | 0.25393 | 0.00000 | 1.15253 | 0.79744 | 0.61543 | 0.06141 | 0.00000 |
| **SDK2** | 5.38533 | 0.05847 | 0.22725 | 0.20961 | 0.12574 | 0.00019 | 0.00000 | 1.98670 | 0.00003 | 0.39961 | 0.01300 | 0.13667 |
| **SDPR** | 0.70670 | 0.00897 | 0.07156 | 0.09858 | 0.00686 | 0.00000 | 0.01202 | 0.08339 | 0.02355 | 0.04350 | 0.01138 | 0.05123 |
| **SDR16C5** | 0.00645 | 0.43951 | 0.02606 | 0.00000 | 0.00000 | 0.03744 | 0.00743 | 0.07288 | 0.00455 | 0.04706 | 0.01761 | 0.05527 |
| **SDR39U1** | 0.03982 | 0.21529 | 2.81891 | 1.17201 | 1.13015 | 0.58283 | 1.68997 | 0.08055 | 3.14455 | 0.00000 | 3.40382 | 0.84822 |
| **SDR42E1** | 0.52884 | 1.95603 | 0.00000 | 1.15229 | 0.26434 | 0.91641 | 0.00000 | 0.23587 | 0.00000 | 0.00751 | 1.59460 | 0.84330 |
| **SDS** | 0.66252 | 0.01338 | 0.02009 | 0.01180 | 0.00145 | 0.04012 | 0.01286 | 0.12442 | 0.02248 | 0.04632 | 0.01692 | 0.04370 |
| **SDSL** | 0.13888 | 0.85053 | 0.01131 | 0.00000 | 0.10662 | 0.03846 | 0.00842 | 0.00695 | 0.00000 | 0.02537 | 0.02774 | 0.00648 |
| **SEBOX** | 0.00707 | 0.01453 | 0.00876 | 0.01708 | 0.01401 | 0.01477 | 0.00506 | 0.02945 | 0.02995 | 0.07278 | 0.01292 | 0.02452 |
| **SEC11A** | 1.78880 | 13.60373 | 26.52726 | 5.98053 | 14.25288 | 30.52086 | 10.06944 | 2.65084 | 38.05289 | 13.94258 | 12.75327 | 12.52711 |
| **SEC11C** | 0.24636 | 6.63925 | 5.85756 | 4.43615 | 5.45460 | 16.50473 | 0.00000 | 0.04592 | 13.87938 | 4.46290 | 7.82891 | 1.84065 |
| **SEC13** | 7.42562 | 18.03358 | 17.98943 | 14.61531 | 13.52692 | 14.98204 | 19.04337 | 8.96680 | 26.30183 | 0.00029 | 14.93631 | 26.08688 |
| **SEC14L1** | 10.30663 | 3.02882 | 1.83264 | 6.84626 | 2.51172 | 2.18831 | 2.42694 | 3.33079 | 0.18141 | 4.54068 | 1.52952 | 7.66256 |
| **SEC14L1P1** | 2.08865 | 0.00910 | 0.32125 | 0.34117 | 0.09668 | 0.34837 | 0.00000 | 0.86078 | 0.00000 | 0.84820 | 0.15759 | 0.42748 |
| **SEC14L2** | 0.01583 | 0.10988 | 0.36446 | 0.09181 | 0.38609 | 0.00000 | 0.00001 | 0.36855 | 0.44894 | 0.00000 | 0.50765 | 0.37898 |
| **SEC14L3** | 0.00506 | 0.01108 | 0.00980 | 0.00910 | 0.01962 | 0.02407 | 0.01270 | 0.01027 | 0.01578 | 0.06635 | 0.01100 | 0.01739 |
| **SEC14L4** | 0.00812 | 0.04231 | 0.10041 | 0.05510 | 0.15474 | 0.00000 | 0.79888 | 0.00539 | 0.00000 | 0.35700 | 0.99851 | 0.47824 |
| **SEC14L5** | 0.82510 | 0.00828 | 0.09284 | 0.01331 | 0.01977 | 0.00003 | 0.02811 | 0.04586 | 0.01632 | 0.02108 | 0.01745 | 0.10063 |
| **SEC14L6** | 0.27866 | 0.02217 | 0.01997 | 0.01225 | 0.01803 | 0.01540 | 0.01547 | 0.03783 | 0.02383 | 0.00000 | 0.01797 | 0.21915 |
| **SEC16A** | 2.57289 | 3.62835 | 5.48010 | 3.16774 | 8.97395 | 0.00000 | 7.75688 | 0.74836 | 9.35229 | 12.11375 | 3.25060 | 4.41948 |
| **SEC16B** | 0.22371 | 0.01671 | 0.01772 | 0.02544 | 0.01522 | 0.00000 | 0.02153 | 0.00118 | 0.02322 | 0.02428 | 0.01390 | 0.03532 |
| **SEC1P** | 0.07201 | 0.00250 | 0.01263 | 0.00839 | 0.00549 | 0.03631 | 0.00581 | 0.00478 | 0.00902 | 0.07394 | 0.01063 | 0.35994 |
| **SEC22A** | 7.65403 | 4.15292 | 1.30262 | 0.00000 | 0.80295 | 9.76033 | 0.00000 | 0.89530 | 3.84045 | 2.62506 | 2.87457 | 1.54935 |
| **SEC22B** | 0.48656 | 7.62141 | 10.11572 | 5.01880 | 2.16760 | 37.22011 | 11.81800 | 7.52783 | 6.23440 | 2.63063 | 12.49286 | 7.68513 |
| **SEC22C** | 2.16893 | 0.24398 | 2.96242 | 3.00406 | 0.55678 | 0.93102 | 4.26532 | 0.43137 | 3.04314 | 0.00000 | 4.23771 | 0.74567 |
| **SEC23A** | 0.73001 | 2.63743 | 3.35244 | 11.93980 | 0.00000 | 9.76437 | 3.63215 | 1.57288 | 3.59565 | 2.39613 | 4.17968 | 3.78766 |
| **SEC23B** | 3.49655 | 27.21563 | 17.63227 | 10.87339 | 9.54054 | 34.30045 | 16.69108 | 8.61155 | 16.30972 | 0.00000 | 26.52628 | 17.11590 |
| **SEC23IP** | 1.23664 | 6.00156 | 9.30060 | 1.77314 | 0.00000 | 11.39149 | 9.33148 | 2.70886 | 8.11170 | 0.00000 | 6.33908 | 5.31844 |
| **SEC24A** | 6.64158 | 2.72395 | 7.34060 | 0.96960 | 2.28986 | 16.78452 | 8.13302 | 6.11758 | 3.10717 | 8.64731 | 4.84780 | 5.78904 |
| **SEC24B** | 3.42597 | 4.98087 | 4.28786 | 1.24166 | 5.05968 | 14.33163 | 4.60306 | 3.74305 | 7.51349 | 4.67304 | 8.27606 | 6.94013 |
| **SEC24B-AS1** | 0.00214 | 0.22008 | 0.13335 | 0.04030 | 0.00000 | 0.04241 | 0.04914 | 0.25394 | 0.00000 | 0.06238 | 0.25228 | 0.31853 |
| **SEC24C** | 6.70170 | 16.26132 | 14.08031 | 9.69606 | 6.19180 | 1.82588 | 11.57487 | 3.20250 | 9.43976 | 6.23583 | 11.35508 | 15.09618 |
| **SEC24D** | 0.11749 | 0.45044 | 4.09255 | 2.29670 | 0.00000 | 1.01776 | 2.77776 | 0.27371 | 1.06840 | 0.30912 | 0.47960 | 1.19152 |
| **SEC31A** | 0.60003 | 9.47894 | 12.47066 | 11.60713 | 8.13734 | 19.30820 | 13.77798 | 5.86703 | 15.72134 | 5.16837 | 14.60029 | 9.90591 |
| **SEC31B** | 0.49055 | 0.48560 | 0.32837 | 0.52694 | 0.00000 | 0.00000 | 0.00000 | 0.00000 | 0.96289 | 1.28636 | 0.31078 | 0.26511 |
| **SEC61A1** | 6.45264 | 30.22219 | 31.68551 | 18.73820 | 29.62372 | 0.00000 | 9.70549 | 19.81153 | 51.15528 | 4.37049 | 22.97401 | 29.32814 |
| **SEC61A2** | 1.17437 | 2.96182 | 0.52880 | 2.65074 | 1.19397 | 4.77519 | 0.00000 | 6.96730 | 1.24005 | 1.72409 | 0.25371 | 10.20648 |
| **SEC61B** | 5.46921 | 13.41658 | 19.84804 | 12.95828 | 15.98957 | 23.74921 | 23.96241 | 9.11066 | 19.36301 | 4.99185 | 13.20461 | 24.14033 |
| **SEC61G** | 3.81275 | 20.44153 | 17.71042 | 16.50015 | 10.46357 | 38.76574 | 23.28151 | 11.66936 | 13.02162 | 13.12284 | 12.60804 | 21.27263 |
| **SEC62** | 3.32965 | 8.26563 | 11.00245 | 7.07639 | 2.44357 | 31.21681 | 6.06403 | 7.78702 | 17.57477 | 33.84750 | 7.92380 | 16.02185 |
| **SEC63** | 1.12138 | 5.91753 | 6.40874 | 13.72892 | 12.59603 | 30.40544 | 0.00000 | 5.48761 | 9.61586 | 33.65383 | 7.92757 | 5.50119 |
| **SECISBP2** | 7.11651 | 4.79780 | 4.93801 | 6.90980 | 5.80592 | 8.25562 | 2.53170 | 2.16971 | 6.05400 | 7.29512 | 6.73266 | 5.78894 |
| **SECISBP2L** | 7.58350 | 0.22200 | 1.35271 | 2.53621 | 2.27456 | 7.47052 | 1.72913 | 0.67557 | 5.01086 | 6.00180 | 2.58850 | 2.24280 |
| **SECTM1** | 0.01163 | 0.01534 | 0.01659 | 0.01281 | 0.01702 | 0.01246 | 0.01685 | 0.01107 | 0.02105 | 0.01924 | 0.01817 | 0.01631 |

| | | | | | | | | | | | | |
|---|---|---|---|---|---|---|---|---|---|---|---|---|
| **SEH1L** | 3.74014 | 16.94544 | 19.78970 | 11.82731 | 10.16887 | 35.62267 | 21.58048 | 11.64885 | 25.84441 | 2.46933 | 20.21389 | 17.61786 |
| **SEL1L** | 2.97148 | 1.00351 | 3.16533 | 3.36053 | 1.22565 | 15.97376 | 0.00000 | 12.47160 | 9.35613 | 9.73617 | 3.07703 | 4.34564 |
| **SEL1L2** | 0.08268 | 0.21727 | 0.02847 | 0.00303 | 0.01146 | 0.12218 | 0.00000 | 0.00249 | 0.00000 | 0.02558 | 0.01079 | 0.11300 |
| **SEL1L3** | 0.13480 | 0.47370 | 0.48440 | 0.00172 | 0.36535 | 0.00000 | 0.00000 | 0.01257 | 0.36196 | 0.00995 | 0.00590 | 0.04793 |
| **SELE** | 0.00569 | 0.01623 | 0.02256 | 0.01968 | 0.00311 | 0.04288 | 0.01086 | 0.01016 | 0.00794 | 0.03803 | 0.01339 | 0.01256 |
| **SELENBP1** | 0.00565 | 0.00000 | 0.19816 | 0.10579 | 0.04798 | 0.08942 | 0.14364 | 0.00934 | 0.37466 | 0.00000 | 0.13110 | 0.01154 |
| **SELK** | 3.98770 | 7.96931 | 21.91099 | 7.23781 | 13.10071 | 21.78304 | 10.17563 | 10.64256 | 12.57545 | 6.98510 | 7.26401 | 11.33013 |
| **SELL** | 0.00234 | 0.00000 | 0.00000 | 0.09799 | 0.00000 | 0.08662 | 0.00771 | 0.03814 | 0.00001 | 0.20845 | 0.00000 | 0.00330 |
| **SELM** | 0.00088 | 0.44524 | 2.14687 | 1.18512 | 0.79061 | 0.00000 | 5.18910 | 13.58131 | 0.39750 | 0.00000 | 0.44871 | 4.41483 |
| **SELO** | 0.09954 | 0.65140 | 2.45416 | 1.94771 | 1.53021 | 0.00000 | 2.36657 | 0.17147 | 1.80027 | 5.14892 | 1.08027 | 0.34320 |
| **SELP** | 0.11984 | 0.01343 | 0.01642 | 0.01551 | 0.01049 | 0.00784 | 0.04321 | 0.00515 | 0.00000 | 0.03965 | 0.03795 | 0.01367 |
| **SELPLG** | 0.05005 | 0.06310 | 0.24770 | 0.00068 | 0.08716 | 0.03200 | 0.56096 | 0.00802 | 0.20739 | 0.26021 | 0.01244 | 0.11888 |
| **SELT** | 0.00000 | 20.71243 | 13.62747 | 26.17643 | 10.78272 | 57.56382 | 11.72642 | 2.06492 | 25.91586 | 5.00090 | 18.72153 | 27.60806 |
| **SELV** | 0.00431 | 0.00369 | 0.11620 | 0.01757 | 0.03810 | 0.04310 | 0.00014 | 0.00776 | 0.00000 | 0.12977 | 0.01315 | 0.05256 |
| **SEMA3A** | 0.12837 | 0.23811 | 0.00000 | 0.11366 | 0.08969 | 0.03955 | 0.00000 | 0.06947 | 0.00000 | 0.54560 | 0.00000 | 0.18473 |
| **SEMA3B** | 0.00711 | 0.31569 | 0.00000 | 0.19783 | 0.14648 | 0.02502 | 0.00000 | 0.11414 | 0.00000 | 0.24459 | 0.02851 | 0.48877 |
| **SEMA3C** | 0.20519 | 0.00810 | 0.00000 | 0.00000 | 0.00000 | 0.06744 | 0.06908 | 0.06657 | 0.02233 | 0.17703 | 0.00000 | 0.00728 |
| **SEMA3D** | 0.64238 | 0.02930 | 0.01580 | 0.00000 | 0.00000 | 0.01313 | 0.01751 | 0.19218 | 0.15739 | 0.14977 | 0.00082 | 0.04007 |
| **SEMA3E** | 0.55386 | 0.45509 | 0.00162 | 0.02341 | 0.00669 | 0.17484 | 0.00000 | 0.04193 | 0.00000 | 0.06993 | 0.03896 | 0.04094 |
| **SEMA3F** | 0.00129 | 0.04908 | 0.01421 | 0.00036 | 0.33292 | 0.00000 | 0.82576 | 0.09990 | 0.03372 | 3.32951 | 0.43970 | 0.08991 |
| **SEMA3G** | 0.00607 | 0.00317 | 0.03602 | 0.00740 | 0.00000 | 0.04925 | 0.03101 | 0.00888 | 0.07258 | 0.02153 | 0.03259 | 0.09659 |
| **SEMA4A** | 0.01811 | 0.90147 | 0.00000 | 1.68728 | 0.01580 | 0.00000 | 0.00973 | 0.03038 | 0.61439 | 0.00000 | 0.19143 | 0.09296 |
| **SEMA4B** | 3.53042 | 5.39664 | 3.21241 | 5.11677 | 3.65538 | 0.00000 | 0.00000 | 1.32114 | 5.58625 | 6.10783 | 0.00000 | 5.80210 |
| **SEMA4C** | 0.00793 | 0.00000 | 11.59149 | 1.15544 | 8.05451 | 0.00000 | 1.01310 | 2.42454 | 16.84036 | 21.97567 | 2.36744 | 6.84478 |
| **SEMA4D** | 1.42253 | 0.31859 | 1.47614 | 1.97494 | 1.08372 | 0.77315 | 0.62521 | 0.75252 | 1.43716 | 4.40373 | 0.00000 | 0.31806 |
| **SEMA4F** | 0.93377 | 0.17051 | 0.15013 | 0.00000 | 0.22654 | 0.00000 | 0.25120 | 0.17622 | 0.00000 | 1.22402 | 0.72180 | 1.67366 |
| **SEMA4G** | 0.68504 | 0.00000 | 0.09712 | 3.54923 | 0.14498 | 0.37765 | 0.00000 | 0.11903 | 0.26113 | 0.53779 | 0.03660 | 0.17960 |
| **SEMA5A** | 2.01974 | 0.01013 | 0.01768 | 0.01776 | 0.01196 | 0.00000 | 0.05603 | 0.80561 | 0.05222 | 0.00000 | 0.16581 | 0.00656 |
| **SEMA5B** | 0.01364 | 0.06443 | 0.02272 | 0.00743 | 0.01601 | 0.02160 | 0.00871 | 0.03046 | 0.06266 | 0.01567 | 0.01016 | 0.01294 |
| **SEMA6A** | 0.00000 | 6.30004 | 25.51331 | 26.17559 | 19.04801 | 15.79347 | 24.96415 | 1.21028 | 48.87496 | 29.50385 | 21.49192 | 0.00000 |
| **SEMA6B** | 0.01868 | 0.02089 | 0.01795 | 0.03512 | 0.02184 | 0.01239 | 0.01900 | 0.02303 | 0.02359 | 0.03689 | 0.01458 | 0.01904 |
| **SEMA6C** | 0.05638 | 0.01023 | 0.31770 | 0.06441 | 0.01422 | 0.00000 | 0.07109 | 0.76764 | 0.16574 | 0.72482 | 0.15163 | 0.14539 |
| **SEMA6D** | 0.00000 | 0.00000 | 3.19535 | 0.00000 | 15.85214 | 1.74334 | 1.03108 | 0.00000 | 12.21034 | 3.39003 | 11.92905 | 0.00000 |
| **SEMA7A** | 0.29108 | 0.14215 | 7.60284 | 0.00000 | 0.00000 | 0.00000 | 0.71277 | 0.00000 | 0.37990 | 0.60214 | 0.07997 | 0.14628 |
| **SENCR** | 0.22746 | 0.01464 | 0.01440 | 0.01243 | 0.01643 | 0.01485 | 0.01846 | 0.00615 | 0.02052 | 0.00910 | 0.01421 | 0.01337 |
| **SENP1** | 1.00972 | 3.50273 | 3.84008 | 1.35402 | 0.00000 | 6.97246 | 4.30117 | 2.91047 | 1.15774 | 3.19896 | 2.69494 | 3.84432 |
| **SENP2** | 1.37334 | 10.91329 | 5.39226 | 8.14767 | 6.34404 | 25.27801 | 2.58413 | 1.63094 | 4.29040 | 9.17468 | 6.90730 | 12.35374 |
| **SENP3** | 0.09972 | 0.11408 | 0.08643 | 0.00000 | 0.00000 | 0.12260 | 0.42137 | 0.04334 | 0.00000 | 0.55912 | 0.10051 | 0.14457 |
| **SENP3-EIF4A1** | 0.07093 | 0.03426 | 0.10681 | 0.07133 | 0.01413 | 0.00672 | 0.07617 | 0.07943 | 0.02371 | 0.01254 | 0.02427 | 0.06025 |
| **SENP5** | 2.88347 | 12.66069 | 6.76724 | 6.83833 | 3.90250 | 15.95571 | 12.77938 | 22.46794 | 11.78814 | 12.72559 | 17.03860 | 14.12396 |
| **SENP6** | 1.43901 | 2.40612 | 4.16377 | 1.53041 | 0.00001 | 15.64120 | 5.56463 | 2.49679 | 7.45259 | 17.74654 | 3.82583 | 4.49126 |
| **SENP7** | 0.98263 | 0.00000 | 0.45229 | 0.00000 | 0.00000 | 7.30859 | 1.89894 | 1.26464 | 0.00000 | 7.26971 | 1.02000 | 0.34321 |
| **SENP8** | 1.96860 | 2.57554 | 0.73213 | 0.07580 | 0.03501 | 4.21786 | 0.00000 | 0.51220 | 0.00000 | 0.00000 | 0.14793 | 0.56331 |
| **SEP15** | 8.13441 | 18.62804 | 17.05212 | 9.86245 | 5.25207 | 45.62522 | 9.25680 | 5.66051 | 27.47716 | 14.39263 | 14.37195 | 23.12461 |
| **SEPHS1** | 1.41472 | 7.97455 | 5.75524 | 6.58199 | 7.01606 | 11.31287 | 13.87931 | 2.27986 | 8.16007 | 3.50165 | 6.69244 | 11.09712 |
| **SEPHS2** | 0.89624 | 14.73795 | 16.67264 | 18.11695 | 8.24474 | 0.00003 | 14.88113 | 0.42138 | 5.50832 | 20.59065 | 11.25222 | 5.44216 |
| **SEPN1** | 2.38442 | 0.34355 | 1.11638 | 2.80895 | 3.72003 | 0.00000 | 1.34475 | 0.22775 | 0.00000 | 4.77933 | 0.65208 | 0.28300 |
| **SEPP1** | 0.00896 | 0.00000 | 0.37800 | 3.90830 | 0.00000 | 1.11350 | 5.25322 | 0.19023 | 9.40333 | 0.48084 | 11.57210 | 0.00000 |

| | | | | | | | | | | | | |
|---|---|---|---|---|---|---|---|---|---|---|---|---|
| **SEPSECS** | 0.21745 | 1.99137 | 0.59685 | 0.99274 | 0.44340 | 8.13756 | 0.00000 | 0.15911 | 0.00000 | 2.71521 | 1.86523 | 2.25898 |
| **SEPSECS-AS1** | 0.04512 | 0.21465 | 0.21746 | 0.12909 | 0.26896 | 0.44852 | 0.05080 | 0.03451 | 0.00000 | 0.38575 | 0.34473 | 0.21486 |
| **SEPT1** | 0.32819 | 0.91987 | 0.28891 | 0.30109 | 0.30211 | 0.52101 | 0.00000 | 0.12953 | 0.00000 | 0.44857 | 0.39885 | 0.93550 |
| **SEPT10** | 2.86144 | 2.55616 | 2.20153 | 2.44392 | 0.71552 | 4.97076 | 1.23272 | 2.28595 | 0.06695 | 2.89112 | 1.38533 | 2.53846 |
| **SEPT11** | 10.66168 | 20.07562 | 15.75848 | 27.66161 | 7.79375 | 25.18982 | 11.86116 | 4.31429 | 1.34482 | 22.86203 | 3.09851 | 15.31556 |
| **SEPT12** | 0.00698 | 2.47791 | 0.01784 | 0.00000 | 1.11052 | 0.27311 | 0.00000 | 2.66846 | 0.00000 | 0.07018 | 0.00000 | 2.14255 |
| **SEPT14** | 0.00697 | 0.02521 | 0.17894 | 0.01726 | 0.00000 | 0.08996 | 0.01995 | 0.29989 | 0.00000 | 0.11042 | 0.08861 | 0.34498 |
| **SEPT2** | 5.69501 | 19.28372 | 16.71509 | 13.56796 | 13.50791 | 42.64308 | 26.65392 | 12.98262 | 23.90361 | 15.42984 | 26.94775 | 14.39043 |
| **SEPT3** | 0.04801 | 0.11453 | 0.16300 | 0.00643 | 0.00000 | 0.00000 | 0.03567 | 0.10200 | 0.00000 | 0.21021 | 0.01248 | 0.50440 |
| **SEPT4** | 0.01669 | 0.03261 | 0.04115 | 0.17173 | 0.01489 | 0.00000 | 0.00261 | 0.08187 | 0.06113 | 0.09198 | 0.00737 | 0.09543 |
| **SEPT5-GP1BB** | 0.00733 | 0.02087 | 0.01397 | 0.01489 | 0.01280 | 0.01135 | 0.01485 | 0.01156 | 0.01770 | 0.02898 | 0.02593 | 0.01515 |
| **SEPT6** | 0.86607 | 0.00020 | 1.06517 | 0.99264 | 0.00000 | 0.13873 | 0.09023 | 0.15517 | 0.05399 | 0.18438 | 0.00000 | 0.00000 |
| **SEPT7** | 2.19689 | 4.74003 | 14.50141 | 6.41705 | 0.73693 | 33.36262 | 15.46540 | 7.48693 | 5.03775 | 10.15026 | 12.73677 | 13.38158 |
| **SEPT7P2** | 0.08432 | 0.48608 | 0.64718 | 0.75200 | 1.23077 | 1.71458 | 0.13583 | 0.89432 | 2.45368 | 2.66320 | 0.55761 | 0.98769 |
| **SEPT7P9** | 0.00697 | 0.02665 | 0.01960 | 0.00772 | 0.01422 | 0.03235 | 0.01579 | 0.01040 | 0.01519 | 0.01895 | 0.02281 | 0.02044 |
| **SEPT8** | 2.27050 | 2.85349 | 0.56709 | 1.14198 | 11.13782 | 6.82342 | 2.71831 | 0.56666 | 0.00013 | 3.98614 | 9.21289 | 0.08864 |
| **SEPT9** | 5.79838 | 0.41046 | 9.01664 | 5.50306 | 2.29443 | 0.00000 | 8.41364 | 0.19463 | 2.43616 | 2.71773 | 0.79000 | 0.52518 |
| **SEPW1** | 5.37400 | 29.19640 | 8.68066 | 18.58431 | 40.89347 | 18.54321 | 11.80188 | 3.30745 | 8.04273 | 13.55965 | 12.81850 | 24.77499 |
| **SERAC1** | 1.52649 | 8.93598 | 1.33649 | 0.63111 | 2.87828 | 13.60977 | 2.24338 | 0.00000 | 2.20098 | 2.28085 | 5.90914 | 0.99948 |
| **SERBP1** | 1.76912 | 22.61571 | 20.14147 | 16.33534 | 21.13693 | 35.95104 | 40.68491 | 8.78912 | 19.06421 | 38.66870 | 25.49488 | 25.69276 |
| **SERF2** | 10.28482 | 34.14915 | 26.61258 | 24.78192 | 42.77845 | 22.49083 | 40.37624 | 22.33546 | 25.01085 | 25.37117 | 32.70027 | 44.70202 |
| **SERGEF** | 0.39727 | 1.78379 | 2.62683 | 1.50382 | 3.57102 | 2.21940 | 2.55066 | 0.31346 | 6.88932 | 0.00000 | 4.82041 | 0.38823 |
| **SERHL** | 0.03285 | 0.02133 | 0.02934 | 0.01251 | 0.02013 | 0.00757 | 0.02360 | 0.03185 | 0.03267 | 0.00000 | 0.01670 | 0.01842 |
| **SERHL2** | 2.23955 | 0.53654 | 0.19659 | 0.35938 | 0.36187 | 0.00000 | 0.39041 | 0.81901 | 0.47784 | 0.95747 | 0.53126 | 0.45493 |
| **SERINC1** | 2.42296 | 9.27846 | 9.51828 | 2.31117 | 0.00000 | 17.88992 | 0.00000 | 20.97049 | 24.85354 | 9.71328 | 5.78539 | 17.26032 |
| **SERINC2** | 2.73633 | 0.07324 | 2.80652 | 0.05256 | 0.05987 | 0.00000 | 0.00000 | 0.32602 | 0.00011 | 0.65712 | 0.72065 | 0.01749 |
| **SERINC3** | 6.71859 | 13.10679 | 16.83689 | 12.07158 | 3.76112 | 22.07607 | 0.00000 | 4.40529 | 17.55241 | 0.00000 | 10.20572 | 19.63003 |
| **SERINC4** | 0.33660 | 0.30377 | 0.42537 | 0.22288 | 0.02786 | 0.00000 | 0.03223 | 0.04917 | 0.20140 | 0.00000 | 0.22370 | 0.41506 |
| **SERINC5** | 4.70670 | 3.12976 | 11.52264 | 23.48543 | 0.00056 | 18.75202 | 3.36531 | 12.15416 | 9.15656 | 3.29931 | 6.50522 | 5.21230 |
| **SERP1** | 6.04155 | 21.24832 | 30.95902 | 12.88491 | 18.97458 | 26.42954 | 30.60534 | 4.27398 | 25.27506 | 15.16963 | 23.59140 | 17.90581 |
| **SERP2** | 0.00813 | 0.04287 | 0.07670 | 0.16764 | 0.00000 | 0.02985 | 0.00010 | 0.01186 | 0.19299 | 0.04094 | 0.01034 | 0.32431 |
| **SERPINA1** | 0.00782 | 0.02152 | 0.02756 | 0.00394 | 0.01205 | 0.09297 | 0.00000 | 0.00899 | 0.00472 | 0.02177 | 0.01224 | 0.01207 |
| **SERPINA12** | 0.00266 | 0.01489 | 0.02306 | 0.01165 | 0.01476 | 0.01973 | 0.01789 | 0.06944 | 0.01360 | 0.01580 | 0.01558 | 0.01270 |
| **SERPINA3** | 0.00704 | 0.01279 | 0.01330 | 0.01117 | 0.01477 | 0.02594 | 0.02135 | 0.00956 | 0.01885 | 0.01313 | 0.01445 | 0.02371 |
| **SERPINA5** | 0.18977 | 0.00000 | 0.01062 | 0.04748 | 0.00000 | 0.23255 | 0.28681 | 0.24836 | 0.03273 | 0.00000 | 0.00000 | 0.03135 |
| **SERPINB1** | 0.00268 | 0.10472 | 5.43904 | 2.73306 | 1.44577 | 0.35945 | 1.02744 | 0.00250 | 3.42869 | 0.06293 | 1.83943 | 0.00000 |
| **SERPINB12** | 0.00629 | 0.01466 | 0.05808 | 0.01260 | 0.01583 | 0.01948 | 0.01284 | 0.01044 | 0.00907 | 0.01333 | 0.01712 | 0.01622 |
| **SERPINB13** | 0.00757 | 0.01172 | 0.01699 | 0.01336 | 0.02506 | 0.02855 | 0.02818 | 0.01020 | 0.00247 | 0.01182 | 0.03266 | 0.01582 |
| **SERPINB2** | 0.00850 | 0.00835 | 0.06789 | 0.00467 | 0.00277 | 0.04173 | 0.01489 | 0.04434 | 0.00187 | 0.06642 | 0.02087 | 0.01919 |
| **SERPINB3** | 0.03352 | 0.01289 | 0.04596 | 0.01358 | 0.01785 | 0.02226 | 0.02121 | 0.01155 | 0.00281 | 0.01718 | 0.01929 | 0.02976 |
| **SERPINB5** | 3.14918 | 3.18528 | 0.27832 | 0.76023 | 0.00000 | 0.32676 | 0.00000 | 1.84380 | 0.00000 | 0.65718 | 0.00000 | 4.88507 |
| **SERPINB6** | 0.06408 | 17.38671 | 7.04052 | 13.60946 | 7.88965 | 3.25541 | 0.00000 | 0.33662 | 0.00000 | 0.00000 | 0.73608 | 10.68818 |
| **SERPINB7** | 0.00604 | 0.01492 | 0.02455 | 0.01215 | 0.01519 | 0.01908 | 0.01744 | 0.01138 | 0.01562 | 0.01378 | 0.01561 | 0.01625 |
| **SERPINB8** | 2.06973 | 0.57958 | 2.71682 | 0.00000 | 0.53141 | 0.03870 | 2.64242 | 0.44346 | 0.00000 | 0.00000 | 0.27364 | 0.28065 |
| **SERPINB9** | 0.64737 | 10.59761 | 51.87730 | 22.80206 | 5.61945 | 18.64716 | 20.94988 | 0.00000 | 2.75930 | 0.00000 | 1.31094 | 0.00000 |
| **SERPINB9P1** | 0.07373 | 0.04213 | 0.05317 | 0.00527 | 0.01247 | 0.02714 | 0.00960 | 0.00651 | 0.01602 | 0.01162 | 0.02273 | 0.01194 |
| **SERPINC1** | 0.00315 | 0.00000 | 0.04551 | 0.00000 | 0.16123 | 0.10662 | 0.04669 | 0.01132 | 2.92076 | 0.04682 | 2.05321 | 0.00283 |
| **SERPIND1** | 0.02919 | 0.01467 | 0.07560 | 0.05387 | 0.13554 | 0.00000 | 0.03036 | 0.00941 | 0.05242 | 0.00000 | 0.01644 | 0.05358 |

| | | | | | | | | | | | | |
|---|---|---|---|---|---|---|---|---|---|---|---|---|
| **SERPINE1** | 0.01087 | 2.51529 | 6.03799 | 3.56458 | 0.40412 | 0.00000 | 0.00000 | 0.02668 | 0.00000 | 0.12510 | 0.06426 | 1.41299 |
| **SERPINE2** | 2.50157 | 6.77448 | 5.70137 | 8.48705 | 0.00106 | 2.35026 | 0.00000 | 6.17378 | 0.05620 | 2.09856 | 0.00000 | 7.16360 |
| **SERPINE3** | 0.06133 | 0.01059 | 0.01171 | 0.00911 | 0.01111 | 0.03819 | 0.05864 | 0.04521 | 0.03894 | 0.00079 | 0.04146 | 0.01571 |
| **SERPINF1** | 8.69246 | 2.50402 | 0.10446 | 0.77519 | 0.07471 | 1.06893 | 0.23548 | 12.70211 | 0.00000 | 0.00000 | 0.49475 | 4.65065 |
| **SERPINF2** | 1.27018 | 0.01800 | 0.00732 | 0.00397 | 0.08423 | 0.00000 | 0.08493 | 0.12763 | 0.00309 | 0.04504 | 0.03610 | 0.16966 |
| **SERPING1** | 0.00701 | 0.00139 | 0.40428 | 0.65125 | 0.00896 | 0.00000 | 0.00438 | 0.00493 | 0.00844 | 0.01206 | 0.04331 | 0.24040 |
| **SERPINH1** | 0.00219 | 1.41900 | 4.07663 | 33.10442 | 4.06940 | 0.00000 | 0.00000 | 0.12215 | 1.60327 | 0.00000 | 0.53444 | 0.37985 |
| **SERPINI1** | 0.25962 | 1.70542 | 0.09336 | 0.00000 | 0.15451 | 2.27592 | 0.00000 | 0.32715 | 1.07702 | 0.27664 | 0.41968 | 0.78722 |
| **SERPINI2** | 0.05600 | 0.01218 | 0.02273 | 0.00739 | 0.03100 | 0.01165 | 0.01156 | 0.03179 | 0.01564 | 0.05808 | 0.01084 | 0.01159 |
| **SERTAD1** | 0.23766 | 6.01189 | 9.83528 | 3.20546 | 8.90413 | 0.00035 | 11.51684 | 28.68739 | 8.54019 | 0.00000 | 0.00000 | 30.04883 |
| **SERTAD2** | 1.06541 | 1.45558 | 7.66903 | 1.05021 | 8.07590 | 10.30486 | 8.41556 | 1.08542 | 4.13177 | 17.98883 | 2.79717 | 0.91201 |
| **SERTAD3** | 7.51827 | 12.81820 | 7.11639 | 11.35466 | 12.34954 | 7.39735 | 8.39015 | 30.71118 | 16.36739 | 0.00861 | 3.71383 | 32.21251 |
| **SERTAD4** | 0.08129 | 0.00000 | 5.11328 | 0.00000 | 0.00000 | 0.03837 | 4.64082 | 0.24403 | 0.56227 | 0.62111 | 0.00000 | 0.16845 |
| **SERTAD4-AS1** | 0.00592 | 0.01288 | 0.17326 | 0.02898 | 0.00000 | 0.06860 | 0.12119 | 0.01881 | 0.00000 | 0.07558 | 0.01844 | 0.08203 |
| **SERTM1** | 0.27125 | 0.01151 | 0.00000 | 0.02384 | 0.23410 | 0.11385 | 0.00000 | 0.00625 | 0.00000 | 0.08513 | 0.00000 | 0.03037 |
| **SESN1** | 4.19078 | 18.68358 | 2.65150 | 3.49323 | 0.00028 | 14.55679 | 2.01675 | 1.46532 | 0.42819 | 1.15513 | 0.65452 | 4.48083 |
| **SESN2** | 4.46820 | 15.02913 | 11.06760 | 10.03015 | 9.68517 | 0.00286 | 15.12690 | 10.97852 | 13.30243 | 7.76867 | 4.48981 | 11.10156 |
| **SESN3** | 1.41422 | 2.92969 | 0.00000 | 5.13516 | 3.11086 | 1.84936 | 0.00000 | 0.80160 | 0.00000 | 1.15584 | 0.01449 | 2.18614 |
| **SESTD1** | 2.73410 | 1.31746 | 5.24372 | 6.50719 | 0.00000 | 9.73701 | 4.25862 | 1.10617 | 0.00000 | 8.85763 | 1.62056 | 4.63013 |
| **SET** | 4.81496 | 26.41570 | 26.10656 | 20.27247 | 24.93178 | 52.02520 | 55.38332 | 34.84131 | 17.78998 | 35.11584 | 40.68961 | 30.83294 |
| **SETBP1** | 0.04235 | 0.00000 | 0.28056 | 0.23788 | 0.85206 | 0.00000 | 1.30182 | 0.02888 | 1.74010 | 1.31824 | 1.65975 | 0.32249 |
| **SETD1A** | 1.15997 | 0.88438 | 0.95621 | 0.78214 | 6.39135 | 0.00000 | 4.36137 | 1.31677 | 0.44895 | 14.94357 | 0.35092 | 2.96658 |
| **SETD1B** | 5.48986 | 1.44731 | 1.68140 | 3.96279 | 2.82626 | 4.12066 | 4.11143 | 1.90452 | 3.29524 | 19.06700 | 3.17760 | 4.42871 |
| **SETD2** | 0.00000 | 4.28201 | 3.03477 | 7.94908 | 2.05605 | 17.00766 | 2.29929 | 6.10190 | 7.78039 | 23.66209 | 3.48226 | 4.94656 |
| **SETD3** | 2.10023 | 8.96871 | 11.27765 | 5.77662 | 20.87175 | 25.08590 | 17.12053 | 2.59462 | 20.76688 | 0.00000 | 22.45755 | 7.81167 |
| **SETD4** | 2.65680 | 2.98112 | 2.99717 | 1.25392 | 1.06484 | 5.76941 | 2.90427 | 4.59045 | 5.79005 | 0.00000 | 3.29886 | 4.74558 |
| **SETD5** | 3.42381 | 1.21323 | 7.69093 | 6.71999 | 8.72071 | 11.05842 | 9.20411 | 5.91993 | 7.11549 | 27.97277 | 3.07662 | 2.96117 |
| **SETD6** | 3.35193 | 1.90048 | 4.87330 | 1.89257 | 2.68856 | 1.55504 | 4.12818 | 2.55415 | 4.92022 | 2.63439 | 4.88704 | 1.35027 |
| **SETD7** | 0.05918 | 0.30372 | 3.59170 | 0.00000 | 0.00000 | 1.04551 | 9.32504 | 0.00000 | 5.90580 | 0.25370 | 0.83955 | 0.08520 |
| **SETD8** | 1.69509 | 3.09353 | 6.00348 | 2.03378 | 6.51028 | 0.00000 | 9.97458 | 9.08367 | 0.00000 | 9.86306 | 5.22809 | 4.15271 |
| **SETD9** | 0.00000 | 2.76372 | 0.51355 | 0.47304 | 0.59002 | 4.37518 | 0.51302 | 0.00000 | 0.99784 | 1.73807 | 0.87971 | 0.62150 |
| **SETDB1** | 2.31616 | 5.65581 | 4.44097 | 4.95664 | 2.29357 | 2.06245 | 4.03141 | 0.20894 | 7.63021 | 4.97651 | 4.00075 | 4.78098 |
| **SETDB2** | 2.42432 | 1.90392 | 0.19601 | 1.21136 | 0.23384 | 3.96204 | 0.02811 | 0.59718 | 0.00000 | 1.12619 | 0.77622 | 1.56738 |
| **SETMAR** | 0.00107 | 0.75430 | 1.40605 | 1.31114 | 0.00000 | 2.41313 | 2.04347 | 4.03774 | 1.82608 | 0.00000 | 3.45999 | 0.34802 |
| **SETSIP** | 0.27575 | 0.80654 | 0.77864 | 0.82708 | 1.63640 | 4.69372 | 1.80990 | 1.02908 | 1.98795 | 0.56673 | 2.15993 | 2.00934 |
| **SETX** | 5.94625 | 1.36537 | 4.49572 | 6.12431 | 1.55794 | 13.34933 | 4.50954 | 10.55012 | 5.80237 | 16.57627 | 2.38116 | 9.75689 |
| **SEZ6** | 0.00628 | 0.07343 | 0.05404 | 0.05742 | 0.22968 | 0.00000 | 0.01385 | 0.00980 | 0.02445 | 0.00000 | 0.14685 | 0.07092 |
| **SEZ6L** | 0.12185 | 0.01896 | 0.02040 | 0.01672 | 0.02557 | 0.00000 | 0.03358 | 0.01292 | 0.01457 | 0.01798 | 0.01447 | 0.02355 |
| **SEZ6L2** | 0.01761 | 1.97168 | 0.49897 | 5.61448 | 2.73044 | 0.00000 | 1.31894 | 2.26021 | 3.53044 | 1.66691 | 0.72077 | 4.70809 |
| **SF1** | 5.48304 | 8.11612 | 12.90913 | 11.16781 | 22.60161 | 5.14186 | 26.48368 | 9.31903 | 13.26988 | 37.71567 | 7.59183 | 14.52955 |
| **SF3A1** | 6.06771 | 8.95456 | 12.76262 | 6.57401 | 12.63475 | 6.31924 | 22.42497 | 2.24909 | 14.87521 | 16.40973 | 9.89369 | 11.70974 |
| **SF3A2** | 0.70548 | 1.26630 | 5.86745 | 1.20548 | 11.13322 | 0.00000 | 5.51456 | 0.55066 | 2.39814 | 21.76576 | 2.83004 | 4.34211 |
| **SF3A3** | 7.66617 | 18.01310 | 17.08137 | 12.55623 | 12.57833 | 26.05321 | 30.12807 | 13.71494 | 20.96621 | 5.17853 | 23.26472 | 24.69714 |
| **SF3B1** | 6.14155 | 19.86159 | 21.77564 | 13.42688 | 8.50138 | 45.67682 | 26.82503 | 14.85476 | 26.53785 | 21.50919 | 23.37133 | 22.45595 |
| **SF3B2** | 6.32351 | 21.47170 | 20.10041 | 17.40683 | 28.40692 | 10.85008 | 31.44003 | 10.39774 | 25.26354 | 13.77872 | 19.53222 | 25.75578 |
| **SF3B3** | 4.55531 | 21.64998 | 15.73628 | 15.52209 | 17.31583 | 14.71111 | 24.36407 | 2.88344 | 26.29670 | 13.29315 | 24.53270 | 23.65144 |
| **SF3B4** | 3.78515 | 18.77002 | 18.57985 | 13.80581 | 18.16100 | 3.10398 | 14.90094 | 11.08371 | 26.74838 | 0.00000 | 16.35047 | 21.14872 |
| **SF3B5** | 1.92622 | 20.09913 | 13.48577 | 15.73651 | 26.17200 | 15.21307 | 27.87239 | 15.16948 | 19.47440 | 12.00258 | 18.75891 | 23.70392 |

| | | | | | | | | | | | | |
|---|---|---|---|---|---|---|---|---|---|---|---|---|
| SF3B6 | 4.45258 | 13.68800 | 18.50878 | 9.69158 | 8.64663 | 40.95545 | 24.29648 | 22.23975 | 23.42307 | 15.00217 | 21.79836 | 23.95461 |
| SFI1 | 3.13232 | 1.96646 | 0.69220 | 0.82744 | 0.89790 | 0.36459 | 0.28401 | 0.08685 | 0.92003 | 0.00000 | 0.63612 | 0.66660 |
| SFMBT1 | 2.86021 | 1.33478 | 1.12439 | 0.09342 | 0.98416 | 2.21555 | 2.79012 | 0.08169 | 3.94375 | 3.77324 | 1.00207 | 4.27024 |
| SFMBT2 | 0.20358 | 0.17695 | 0.05385 | 1.29034 | 0.28954 | 0.82745 | 0.00000 | 0.01187 | 0.35068 | 0.90383 | 0.00000 | 1.00695 |
| SFN | 0.09704 | 1.00738 | 3.03624 | 0.99157 | 1.96966 | 0.00000 | 6.53160 | 0.00233 | 3.62251 | 4.77539 | 4.14678 | 0.00000 |
| SFPQ | 0.77381 | 5.50191 | 14.64088 | 8.74795 | 18.55008 | 32.86378 | 33.97142 | 10.76076 | 4.17095 | 58.37763 | 6.49067 | 15.51950 |
| SFR1 | 3.04201 | 5.27760 | 0.30893 | 0.06639 | 1.91619 | 8.42149 | 0.43429 | 0.00000 | 0.00000 | 2.88686 | 0.99251 | 5.97053 |
| SFRP1 | 0.00782 | 0.00000 | 0.74759 | 6.30573 | 0.00000 | 0.09371 | 0.28641 | 0.03409 | 0.05500 | 0.51210 | 0.05425 | 0.00000 |
| SFRP2 | 0.00517 | 0.00577 | 1.04883 | 0.65720 | 0.17302 | 0.00000 | 0.00000 | 0.00805 | 0.00000 | 0.03246 | 0.00973 | 0.04784 |
| SFRP5 | 0.00729 | 0.00913 | 0.05924 | 0.01373 | 0.00286 | 0.02985 | 0.00733 | 0.00998 | 0.00000 | 0.02923 | 0.12236 | 0.01477 |
| SFSWAP | 2.08966 | 5.06955 | 6.34659 | 4.78115 | 6.12927 | 0.00000 | 5.55382 | 3.91005 | 7.09013 | 14.50760 | 2.57543 | 7.29806 |
| SFT2D1 | 0.76130 | 10.45705 | 11.65013 | 8.28698 | 5.93789 | 21.10634 | 6.56495 | 1.27259 | 13.87039 | 4.38271 | 8.38928 | 14.08892 |
| SFT2D2 | 1.40741 | 5.78946 | 1.15923 | 2.30613 | 6.47558 | 8.75725 | 0.00000 | 3.78914 | 5.77926 | 0.51636 | 6.39904 | 2.82384 |
| SFT2D3 | 0.05685 | 0.00000 | 0.72532 | 0.09233 | 0.16989 | 0.83268 | 0.85529 | 0.11645 | 0.38462 | 2.85631 | 0.03281 | 0.21504 |
| SFTA2 | 0.00705 | 0.00140 | 0.02253 | 0.02175 | 0.00000 | 0.08863 | 0.00000 | 0.00858 | 0.00000 | 0.03263 | 0.05011 | 0.03429 |
| SFTA3 | 0.15875 | 0.02108 | 0.01512 | 0.01159 | 0.02228 | 0.01150 | 0.01830 | 0.11676 | 0.01950 | 0.00481 | 0.02194 | 0.01100 |
| SFTPB | 0.00294 | 0.03971 | 0.02533 | 0.00119 | 0.00000 | 0.04600 | 0.00000 | 0.01134 | 0.02933 | 0.30711 | 0.00961 | 0.00719 |
| SFTPD | 0.10317 | 2.73295 | 0.14995 | 0.14665 | 0.38648 | 0.00000 | 0.06203 | 3.09758 | 0.05098 | 0.00000 | 0.00000 | 3.33035 |
| SFXN1 | 0.47288 | 7.98044 | 15.03554 | 2.25141 | 8.11492 | 27.77866 | 22.88316 | 2.38181 | 15.52873 | 2.13871 | 16.66009 | 9.56298 |
| SFXN2 | 0.00166 | 3.35001 | 2.35120 | 0.94443 | 1.60267 | 2.07600 | 2.97810 | 0.04462 | 2.43962 | 0.00000 | 4.22249 | 0.75001 |
| SFXN3 | 0.01525 | 0.15576 | 0.12789 | 0.01023 | 0.00000 | 0.05464 | 0.00459 | 0.00964 | 0.01076 | 0.01761 | 0.01127 | 0.04072 |
| SFXN4 | 1.00428 | 9.56932 | 6.05013 | 1.24101 | 10.30219 | 7.43770 | 7.30132 | 0.00582 | 15.51904 | 2.70608 | 10.97894 | 0.34405 |
| SFXN5 | 0.83959 | 2.57453 | 0.55301 | 4.13708 | 3.24754 | 0.00000 | 0.21696 | 0.01596 | 1.91745 | 1.03020 | 2.85270 | 0.58305 |
| SGCA | 0.01272 | 0.01372 | 0.03278 | 0.00839 | 0.01225 | 0.02179 | 0.01436 | 0.00951 | 0.01016 | 0.03172 | 0.01529 | 0.01468 |
| SGCB | 0.00000 | 6.63622 | 13.95520 | 4.64610 | 0.00000 | 9.96588 | 0.87505 | 1.26635 | 2.21356 | 0.60854 | 0.79903 | 11.96062 |
| SGCD | 1.73998 | 0.02680 | 0.03101 | 0.01678 | 0.02836 | 0.00000 | 0.00000 | 0.43986 | 0.08316 | 0.01554 | 0.01029 | 0.00401 |
| SGCE | 0.26594 | 0.00000 | 3.53466 | 0.00000 | 0.00000 | 0.19692 | 0.33307 | 0.04353 | 2.67664 | 0.02686 | 0.00000 | 0.00824 |
| SGCG | 0.49336 | 0.20709 | 0.02733 | 0.00110 | 0.01354 | 0.07666 | 0.05968 | 1.80934 | 0.00000 | 0.00000 | 0.00763 | 0.22270 |
| SGCZ | 1.04339 | 0.00977 | 0.16969 | 0.00365 | 0.17459 | 0.00000 | 0.01110 | 0.01857 | 0.00626 | 0.03086 | 0.00569 | 0.03171 |
| SGK1 | 2.18188 | 21.77944 | 32.81605 | 16.25135 | 17.64977 | 32.96586 | 0.00000 | 30.27979 | 0.00000 | 0.00000 | 14.39317 | 15.76441 |
| SGK2 | 0.00581 | 0.04817 | 0.00000 | 0.42834 | 0.14119 | 0.11303 | 0.00000 | 0.00914 | 0.00000 | 0.01553 | 0.00871 | 0.01828 |
| SGK223 | 0.03275 | 0.14676 | 0.00000 | 1.25761 | 0.48563 | 0.00000 | 2.39975 | 0.00000 | 0.71189 | 7.33381 | 0.73981 | 1.04442 |
| SGK3.uid-10446 | 0.00182 | 0.02745 | 0.10092 | 0.01070 | 0.02445 | 0.08737 | 0.05293 | 0.20963 | 0.09152 | 0.00000 | 0.08628 | 0.09678 |
| SGK3.uid-10450 | 0.00631 | 0.01501 | 0.02551 | 0.01669 | 0.01694 | 0.03050 | 0.02637 | 0.01004 | 0.03678 | 0.02372 | 0.01874 | 0.01614 |
| SGK494 | 0.00000 | 1.55230 | 0.46581 | 1.32432 | 0.47417 | 0.55697 | 1.73321 | 0.50850 | 3.07712 | 6.31239 | 0.63621 | 2.34483 |
| SGMS1 | 0.57573 | 3.48117 | 10.15214 | 1.58966 | 8.94414 | 8.57404 | 2.51797 | 0.55249 | 12.70511 | 5.01972 | 9.03977 | 0.00000 |
| SGMS2 | 6.79518 | 0.00502 | 0.01328 | 0.00149 | 0.06227 | 0.00000 | 0.01170 | 0.63516 | 0.01001 | 0.12372 | 0.01060 | 0.07434 |
| SGOL1 | 3.14267 | 3.51117 | 5.53547 | 1.97470 | 0.95031 | 19.13158 | 5.78682 | 7.16277 | 4.51478 | 8.48815 | 3.73831 | 6.87938 |
| SGOL1-AS1 | 0.00739 | 0.03207 | 0.03327 | 0.01084 | 0.02686 | 0.04419 | 0.00801 | 0.06145 | 0.04808 | 0.02912 | 0.01763 | 0.05059 |
| SGOL2 | 4.63257 | 5.14957 | 9.61013 | 0.71757 | 0.49116 | 23.37355 | 7.79036 | 11.86376 | 9.01563 | 16.13660 | 2.72465 | 11.37144 |
| SGPL1 | 6.28239 | 9.95789 | 11.54409 | 12.03568 | 11.13572 | 24.65407 | 9.01824 | 2.21213 | 41.71093 | 2.08506 | 21.77457 | 7.07571 |
| SGPP1 | 0.34020 | 0.00000 | 8.00033 | 0.00000 | 0.00000 | 9.61606 | 8.93077 | 4.53784 | 13.77731 | 11.95253 | 1.42833 | 3.22486 |
| SGPP2 | 0.08705 | 1.95325 | 0.00000 | 0.05969 | 0.63007 | 0.00000 | 0.04285 | 0.01012 | 0.47662 | 1.81114 | 0.48777 | 0.59105 |
| SGSH | 0.71429 | 2.41107 | 1.18667 | 1.19648 | 1.94819 | 0.00000 | 0.00000 | 0.04833 | 3.31457 | 1.24369 | 2.55238 | 0.65202 |
| SGSM1 | 0.00177 | 0.09367 | 0.00000 | 0.00949 | 0.05943 | 0.21756 | 0.00000 | 0.13957 | 0.15878 | 0.15003 | 0.00823 | 0.00001 |
| SGSM2 | 0.57226 | 0.33069 | 1.77554 | 1.93032 | 0.71869 | 0.00000 | 0.00000 | 0.00466 | 1.58930 | 4.41444 | 0.73589 | 0.28780 |
| SGSM3 | 0.58920 | 4.54850 | 5.77688 | 5.62333 | 0.58954 | 0.00000 | 1.41753 | 0.04194 | 9.04179 | 5.49220 | 5.29979 | 4.12217 |
| SGTA | 5.16915 | 25.98288 | 19.03113 | 11.56744 | 21.20524 | 0.00000 | 32.14175 | 4.46242 | 33.01656 | 0.00000 | 18.82127 | 21.77792 |

| | | | | | | | | | | | | |
|---|---|---|---|---|---|---|---|---|---|---|---|---|
| **SGTB** | 0.81670 | 1.74373 | 3.48449 | 1.23893 | 0.72378 | 5.88628 | 1.56958 | 0.72536 | 3.77460 | 1.42826 | 1.87064 | 5.54829 |
| **SH2B1** | 1.04420 | 1.17505 | 0.83128 | 1.49032 | 2.17920 | 0.00000 | 0.59763 | 0.23205 | 1.21473 | 5.53979 | 0.27109 | 1.83417 |
| **SH2B2** | 0.27670 | 0.06244 | 0.49632 | 0.00000 | 0.51381 | 0.00000 | 0.01769 | 0.00220 | 0.00000 | 0.32661 | 0.02147 | 0.02968 |
| **SH2B3** | 3.10087 | 4.17884 | 0.00000 | 4.87539 | 0.28915 | 4.18422 | 2.72919 | 2.68765 | 1.57927 | 7.97663 | 0.57682 | 1.93701 |
| **SH2D1B** | 0.52728 | 0.00000 | 0.00000 | 1.88282 | 0.01744 | 0.00000 | 0.00465 | 0.16781 | 0.40065 | 0.00000 | 0.10047 | 0.33498 |
| **SH2D2A** | 0.00697 | 0.06097 | 0.03228 | 0.00539 | 0.02568 | 0.00000 | 0.00401 | 0.00663 | 0.10604 | 0.07642 | 0.00101 | 0.15323 |
| **SH2D3A** | 0.05541 | 2.14117 | 2.45100 | 0.78387 | 2.82821 | 0.00000 | 0.85572 | 0.00000 | 2.38085 | 3.67904 | 0.34380 | 0.67367 |
| **SH2D3C** | 0.00259 | 0.01230 | 0.00555 | 0.00127 | 0.00850 | 0.09731 | 0.07689 | 0.47862 | 0.01023 | 0.01808 | 0.00379 | 0.00884 |
| **SH2D4A** | 18.31392 | 0.00000 | 24.14335 | 0.00000 | 13.89350 | 0.43543 | 27.40635 | 5.01328 | 19.18073 | 3.12318 | 17.70135 | 0.00000 |
| **SH2D4B** | 2.04604 | 0.16378 | 0.06635 | 0.73653 | 0.02560 | 0.00000 | 0.00979 | 0.86921 | 0.00000 | 0.05595 | 0.18970 | 0.18729 |
| **SH2D5** | 0.00091 | 1.18436 | 2.58348 | 0.47723 | 4.14313 | 0.00000 | 1.00315 | 0.00000 | 3.94702 | 4.14498 | 1.39152 | 0.23598 |
| **SH2D6** | 0.00822 | 0.00941 | 0.03498 | 0.02786 | 0.00181 | 0.03943 | 0.00869 | 0.01055 | 0.00394 | 0.03629 | 0.02662 | 0.01351 |
| **SH2D7** | 0.00605 | 0.03773 | 0.10687 | 0.00060 | 0.19834 | 0.07580 | 0.00000 | 0.00968 | 0.00000 | 0.00997 | 0.11895 | 0.00973 |
| **SH3BGR** | 0.79770 | 1.54035 | 0.13994 | 0.00000 | 0.13412 | 0.31267 | 0.00000 | 0.28806 | 0.00000 | 0.00000 | 0.18492 | 0.58772 |
| **SH3BGRL** | 0.00573 | 0.22213 | 0.03497 | 0.68111 | 0.06195 | 0.03919 | 0.01428 | 1.60957 | 0.00871 | 0.00000 | 0.02143 | 0.08292 |
| **SH3BGRL2** | 3.59718 | 0.00000 | 12.51130 | 0.00000 | 1.57831 | 9.47378 | 17.38321 | 0.17364 | 14.73039 | 0.67897 | 14.02082 | 0.00000 |
| **SH3BGRL3** | 0.11191 | 13.47723 | 8.82017 | 12.90493 | 15.41921 | 5.99474 | 15.42194 | 21.85217 | 16.19588 | 0.00000 | 8.33845 | 24.07869 |
| **SH3BP1** | 0.27617 | 0.00000 | 1.03462 | 0.30379 | 0.45168 | 0.00000 | 1.35283 | 0.03179 | 0.00000 | 1.88876 | 0.00000 | 0.00178 |
| **SH3BP2** | 0.50196 | 0.16357 | 1.73869 | 0.00000 | 0.00000 | 0.12732 | 1.37966 | 0.00000 | 0.38030 | 2.59116 | 0.12901 | 0.03073 |
| **SH3BP4** | 1.43294 | 8.58775 | 1.23221 | 7.94761 | 10.48751 | 6.10271 | 2.73836 | 0.00000 | 9.63929 | 3.51993 | 4.32530 | 5.56811 |
| **SH3BP5** | 0.33210 | 0.11417 | 4.36548 | 0.00146 | 0.67407 | 0.00000 | 0.00000 | 0.19092 | 0.98234 | 0.00000 | 0.51909 | 0.08394 |
| **SH3BP5-AS1** | 0.01932 | 0.15687 | 0.97854 | 0.00000 | 0.24484 | 0.23649 | 0.00000 | 0.00142 | 0.77740 | 0.98646 | 0.03414 | 0.02138 |
| **SH3BP5L** | 0.52117 | 0.61553 | 0.72371 | 0.81481 | 0.86598 | 0.00000 | 3.70911 | 2.00232 | 0.98561 | 5.90575 | 0.27726 | 3.63981 |
| **SH3D19** | 0.00000 | 0.65584 | 0.31135 | 5.08928 | 3.45204 | 2.41501 | 1.85939 | 0.70726 | 1.45218 | 1.67136 | 0.00000 | 0.00000 |
| **SH3D21** | 0.02371 | 1.14818 | 0.14513 | 0.00442 | 6.20136 | 0.56797 | 0.02682 | 0.43002 | 1.65121 | 2.02929 | 1.94254 | 0.53501 |
| **SH3GL1** | 2.65462 | 2.03671 | 8.90267 | 2.48676 | 12.98651 | 0.00000 | 13.70987 | 0.85578 | 6.25385 | 22.55425 | 4.58472 | 1.86891 |
| **SH3GL1P1** | 0.00641 | 0.03586 | 0.05289 | 0.07366 | 0.17202 | 0.00000 | 0.00053 | 0.00915 | 0.17562 | 0.08338 | 0.02134 | 0.02280 |
| **SH3GL1P2** | 0.03037 | 0.04832 | 0.04178 | 0.08997 | 0.00000 | 0.04094 | 0.00000 | 0.01089 | 0.00757 | 0.11960 | 0.01543 | 0.14716 |
| **SH3GL2** | 0.85617 | 0.05877 | 0.03700 | 0.07046 | 0.00000 | 0.34884 | 0.00000 | 0.10338 | 0.00000 | 0.02964 | 0.00000 | 0.05526 |
| **SH3GL3** | 1.03530 | 0.00000 | 2.70649 | 0.00000 | 5.85023 | 3.77890 | 9.31770 | 0.00000 | 7.77341 | 0.36015 | 3.68217 | 0.00000 |
| **SH3GLB1** | 7.61494 | 7.47706 | 10.31410 | 3.64601 | 0.18154 | 16.43639 | 6.70890 | 5.14861 | 5.83223 | 6.06493 | 5.09258 | 8.68318 |
| **SH3GLB2** | 0.97566 | 1.07743 | 1.59077 | 0.87331 | 18.96176 | 0.53764 | 3.19173 | 0.08151 | 0.00000 | 8.45873 | 3.81585 | 0.00000 |
| **SH3KBP1** | 5.89446 | 2.50635 | 8.09401 | 0.00000 | 8.70178 | 4.70411 | 11.87842 | 1.72543 | 8.42257 | 11.61598 | 12.47297 | 0.80536 |
| **SH3PXD2A** | 0.00040 | 3.30779 | 0.42402 | 5.65620 | 0.00000 | 0.00000 | 0.00000 | 0.72060 | 0.00000 | 1.49828 | 0.16781 | 5.16458 |
| **SH3PXD2A-AS1** | 0.00583 | 0.01418 | 0.00827 | 0.15119 | 0.01015 | 0.00089 | 0.02080 | 0.00867 | 0.02099 | 0.01556 | 0.01574 | 0.06688 |
| **SH3PXD2B** | 0.49527 | 0.20762 | 1.29704 | 1.19058 | 1.08876 | 0.00000 | 0.07512 | 0.09033 | 0.74732 | 0.10178 | 0.07313 | 0.79296 |
| **SH3RF1** | 0.29240 | 6.80042 | 1.96828 | 4.22291 | 6.61451 | 2.24505 | 7.41362 | 0.00000 | 1.76256 | 5.62929 | 3.28117 | 4.11663 |
| **SH3RF2** | 0.00460 | 0.26566 | 0.27935 | 1.85895 | 4.43020 | 0.25464 | 0.02321 | 0.00524 | 0.88762 | 0.00000 | 1.55459 | 0.00122 |
| **SH3RF3** | 0.00534 | 0.03780 | 0.12147 | 0.50725 | 0.00000 | 0.04391 | 0.00000 | 0.00832 | 0.00000 | 0.03624 | 0.01369 | 0.03241 |
| **SH3TC1** | 0.00121 | 1.44119 | 0.54318 | 0.00000 | 11.47728 | 0.00000 | 3.98982 | 0.00342 | 5.55300 | 3.56983 | 6.78902 | 0.00000 |
| **SH3TC2** | 0.00135 | 0.05369 | 0.91563 | 0.00864 | 0.00000 | 0.05462 | 0.28379 | 0.05517 | 0.00000 | 0.06143 | 0.09577 | 0.26059 |
| **SH3YL1** | 0.61157 | 0.44940 | 0.94006 | 0.79142 | 0.00000 | 1.24552 | 0.66246 | 0.21236 | 1.26906 | 1.42655 | 1.01175 | 0.43972 |
| **SHANK1** | 0.01232 | 0.01961 | 0.02895 | 0.01126 | 0.01237 | 0.00759 | 0.01823 | 0.00964 | 0.00439 | 0.04062 | 0.05064 | 0.01800 |
| **SHANK2** | 0.65019 | 0.83398 | 0.27955 | 1.75040 | 0.21364 | 0.48824 | 0.00000 | 0.17054 | 0.21720 | 1.43453 | 0.00000 | 0.04308 |
| **SHANK2-AS1** | 0.00673 | 0.02021 | 0.01425 | 0.01522 | 0.01250 | 0.00000 | 0.02027 | 0.01058 | 0.09993 | 0.00000 | 0.03789 | 0.01821 |
| **SHANK2-AS3** | 0.01221 | 0.00862 | 0.00099 | 0.04064 | 0.04660 | 0.02713 | 0.00000 | 0.00893 | 0.17812 | 0.01435 | 0.00282 | 0.01530 |
| **SHANK3** | 0.87959 | 0.20975 | 0.14398 | 0.17145 | 0.45556 | 0.00000 | 0.04332 | 0.00000 | 0.00000 | 0.48219 | 0.02149 | 0.21339 |
| **SHARPIN** | 0.09858 | 4.65218 | 1.63237 | 2.45017 | 7.34473 | 0.00000 | 1.86090 | 0.12031 | 0.75926 | 5.43303 | 2.55859 | 1.73226 |

| | | | | | | | | | | | | |
|---|---|---|---|---|---|---|---|---|---|---|---|---|
| **SHB** | 0.73398 | 1.83857 | 0.10789 | 6.50135 | 3.12396 | 1.05603 | 0.87140 | 0.32850 | 0.46528 | 6.28903 | 0.00000 | 2.36711 |
| **SHBG** | 0.00246 | 0.10797 | 0.00937 | 0.00000 | 0.00000 | 0.00585 | 0.00288 | 0.05650 | 0.03508 | 0.18126 | 0.01509 | 0.19401 |
| **SHC1** | 6.50139 | 12.54842 | 18.31994 | 11.20987 | 8.01123 | 0.00061 | 14.12588 | 4.51963 | 13.02083 | 0.00000 | 7.13458 | 9.70047 |
| **SHC2** | 1.15411 | 0.02012 | 0.02283 | 0.02516 | 0.02816 | 0.01432 | 0.01919 | 0.05487 | 0.01945 | 0.00000 | 0.01836 | 0.27087 |
| **SHC3** | 0.23476 | 0.01544 | 0.02959 | 0.02717 | 0.03124 | 0.00000 | 0.02630 | 0.00000 | 0.00134 | 0.00000 | 0.08133 | 0.20180 |
| **SHC4** | 0.00450 | 0.00394 | 0.06548 | 0.01303 | 0.12034 | 0.00000 | 0.05969 | 0.08455 | 0.00000 | 0.07922 | 0.00874 | 0.10970 |
| **SHCBP1** | 1.94609 | 7.70346 | 5.08977 | 5.32592 | 3.29752 | 24.07572 | 7.76775 | 7.48154 | 5.17838 | 0.07846 | 7.11912 | 7.08526 |
| **SHCBP1L** | 0.00230 | 0.00818 | 0.00209 | 0.00067 | 0.22127 | 0.07431 | 0.00000 | 0.03672 | 0.26553 | 0.03500 | 0.07614 | 0.02627 |
| **SHD** | 2.74968 | 0.34975 | 0.10155 | 1.32059 | 0.09679 | 0.00000 | 0.05133 | 0.61931 | 0.00000 | 0.56252 | 0.11777 | 0.89520 |
| **SHE** | 1.65608 | 0.00318 | 0.10581 | 0.44239 | 0.02277 | 0.11828 | 0.03067 | 0.55977 | 0.04454 | 0.00000 | 0.00168 | 0.04030 |
| **SHF** | 0.35511 | 0.17815 | 1.11117 | 0.30428 | 0.00000 | 0.00000 | 0.75264 | 0.08712 | 0.00000 | 0.19829 | 0.00314 | 0.91635 |
| **SHFM1** | 0.11318 | 15.38341 | 22.03152 | 15.00971 | 19.15774 | 30.13433 | 35.40759 | 6.52836 | 14.28171 | 13.65513 | 20.82160 | 12.52447 |
| **SHH** | 0.18018 | 0.01391 | 0.01367 | 0.01158 | 0.01505 | 0.01469 | 0.01750 | 0.28297 | 0.02022 | 0.01316 | 0.01384 | 0.00856 |
| **SHISA2** | 0.00114 | 0.00632 | 0.00000 | 0.43328 | 0.07845 | 0.35042 | 0.08664 | 0.47459 | 0.00000 | 0.08522 | 0.02523 | 0.30331 |
| **SHISA3** | 0.13758 | 0.00000 | 0.00218 | 0.96304 | 0.04573 | 0.04873 | 0.00404 | 0.00599 | 0.00653 | 0.00808 | 0.01073 | 0.77124 |
| **SHISA4** | 0.03462 | 0.06449 | 0.93168 | 0.00000 | 1.19556 | 0.00000 | 0.56522 | 0.01127 | 1.59542 | 0.27788 | 0.56395 | 0.13471 |
| **SHISA5** | 3.96640 | 41.81962 | 28.14802 | 37.60873 | 51.64120 | 13.67738 | 30.90951 | 0.42389 | 56.63887 | 5.28923 | 35.42263 | 2.21116 |
| **SHISA6** | 0.00218 | 0.01501 | 0.14819 | 0.01654 | 0.01239 | 0.01793 | 0.02026 | 0.04745 | 0.00000 | 0.02141 | 0.02059 | 0.01360 |
| **SHISA7** | 0.02030 | 0.00427 | 0.00544 | 0.05217 | 0.00000 | 0.01754 | 0.00235 | 0.03936 | 0.01232 | 0.34272 | 0.00053 | 0.00736 |
| **SHISA8** | 0.54403 | 0.01636 | 0.04943 | 0.01330 | 0.02503 | 0.00000 | 0.03263 | 0.01575 | 0.02436 | 0.01551 | 0.01295 | 0.02526 |
| **SHISA9** | 0.25874 | 0.02470 | 0.00000 | 0.06640 | 0.07183 | 0.13766 | 0.00673 | 0.01068 | 0.01075 | 0.05666 | 0.02089 | 0.01901 |
| **SHKBP1** | 0.24962 | 5.41230 | 5.80079 | 9.05815 | 5.48352 | 0.00000 | 6.02825 | 0.01051 | 8.76781 | 0.00000 | 7.72122 | 1.52769 |
| **SHMT1** | 0.20931 | 9.64731 | 14.42011 | 11.85829 | 11.09444 | 4.11409 | 14.16631 | 0.91676 | 12.99187 | 0.00000 | 18.38792 | 10.33484 |
| **SHMT2** | 5.26806 | 13.27011 | 20.44984 | 13.00105 | 15.89658 | 11.96347 | 32.41869 | 1.87513 | 27.87542 | 0.00000 | 19.34203 | 10.96140 |
| **SHOC2** | 1.31500 | 1.96023 | 4.83819 | 2.52919 | 0.00000 | 15.26094 | 6.39681 | 12.36042 | 3.24770 | 4.72304 | 5.16593 | 5.13073 |
| **SHOX2** | 0.32761 | 0.00254 | 0.05855 | 0.02630 | 0.00000 | 0.06797 | 0.03828 | 0.04455 | 0.04477 | 0.00868 | 0.00922 | 0.24072 |
| **SHPK** | 0.01382 | 22.44560 | 1.95040 | 18.31988 | 5.74409 | 5.15533 | 1.13136 | 3.25841 | 3.10484 | 0.00000 | 3.12550 | 12.32981 |
| **SHPRH** | 0.49652 | 1.03109 | 1.97935 | 1.34719 | 1.42961 | 6.56730 | 0.00000 | 2.42039 | 5.30890 | 5.88780 | 2.08932 | 2.31433 |
| **SHQ1** | 2.01586 | 6.75453 | 4.11667 | 2.87671 | 1.16005 | 10.85054 | 6.98498 | 21.34318 | 9.24894 | 0.00000 | 4.81262 | 9.65395 |
| **SHROOM1** | 0.11072 | 0.00797 | 0.15873 | 0.00000 | 0.61903 | 0.00000 | 0.23403 | 0.00275 | 0.00000 | 0.13458 | 0.28238 | 0.05638 |
| **SHROOM2** | 2.42302 | 1.31966 | 0.00000 | 0.19343 | 5.91912 | 1.11286 | 4.36639 | 0.18133 | 4.67515 | 1.16795 | 4.53297 | 0.00000 |
| **SHROOM3** | 0.34443 | 4.42072 | 1.34869 | 2.72077 | 8.77525 | 4.43934 | 2.55323 | 2.56372 | 3.62248 | 8.47828 | 6.41397 | 1.54704 |
| **SHROOM4** | 0.00862 | 0.01361 | 0.05688 | 0.11603 | 0.00797 | 0.02286 | 0.01382 | 0.01173 | 0.01700 | 0.01086 | 0.01476 | 0.01629 |
| **SI** | 0.00599 | 0.01354 | 0.02205 | 0.01020 | 0.08333 | 0.00000 | 0.02244 | 0.01064 | 0.00893 | 0.02679 | 0.01225 | 0.01689 |
| **SIAE** | 0.08503 | 4.11447 | 0.49162 | 5.34520 | 3.00953 | 6.03501 | 0.00000 | 0.00000 | 2.37690 | 0.11337 | 3.69634 | 0.00000 |
| **SIAH1** | 3.25874 | 1.83719 | 2.63615 | 4.16695 | 0.63091 | 7.99189 | 3.13424 | 21.13800 | 0.00009 | 6.07247 | 0.73144 | 10.23130 |
| **SIAH2** | 1.58131 | 0.25194 | 5.04046 | 0.00000 | 0.00000 | 0.00000 | 2.55009 | 0.11815 | 1.28660 | 1.23078 | 0.00000 | 0.11328 |
| **SIAH3** | 0.75315 | 0.12097 | 0.01600 | 0.02297 | 0.00000 | 0.06692 | 0.00000 | 0.06374 | 0.04427 | 0.10556 | 0.00259 | 0.06297 |
| **SIDT1** | 0.00592 | 0.14616 | 0.06253 | 0.10544 | 0.00000 | 0.02089 | 0.02881 | 0.01039 | 0.00000 | 0.07089 | 0.03858 | 0.00745 |
| **SIDT2** | 2.68461 | 1.15255 | 4.65159 | 0.46027 | 1.89741 | 0.98482 | 0.14139 | 1.25837 | 2.28997 | 3.03109 | 0.32851 | 3.05755 |
| **SIGIRR** | 0.11056 | 2.90992 | 5.18827 | 5.10182 | 3.92702 | 0.00000 | 3.03348 | 1.64839 | 18.02914 | 0.00000 | 10.37821 | 1.25346 |
| **SIGLEC1** | 0.00591 | 0.06015 | 0.02865 | 0.00000 | 0.00000 | 0.07191 | 0.13085 | 0.01019 | 0.00000 | 0.01661 | 0.01427 | 0.00833 |
| **SIGLEC10** | 6.52401 | 0.04204 | 0.01582 | 0.00271 | 0.01861 | 0.02929 | 0.01003 | 0.51412 | 0.01609 | 0.00000 | 0.00676 | 0.34012 |
| **SIGLEC11** | 0.00371 | 0.00414 | 0.00133 | 0.21979 | 0.08178 | 0.00000 | 0.03428 | 0.00824 | 0.00000 | 0.20486 | 0.00379 | 0.08870 |
| **SIGLEC12** | 0.00788 | 0.01370 | 0.04205 | 0.01095 | 0.01250 | 0.02474 | 0.01293 | 0.02776 | 0.01147 | 0.02028 | 0.01521 | 0.01394 |
| **SIGLEC14** | 0.00539 | 0.00546 | 0.10705 | 0.00017 | 0.00002 | 0.08556 | 0.10276 | 0.01088 | 0.23236 | 0.00633 | 0.00000 | 0.01808 |
| **SIGLEC16** | 0.00574 | 0.01507 | 0.01799 | 0.07841 | 0.01054 | 0.00000 | 0.02067 | 0.01005 | 0.02001 | 0.03832 | 0.01980 | 0.02555 |
| **SIGLEC17P** | 0.00457 | 0.01652 | 0.01457 | 0.01210 | 0.01609 | 0.01624 | 0.01978 | 0.01777 | 0.02548 | 0.01033 | 0.01400 | 0.01683 |

| | | | | | | | | | | | | |
|---|---|---|---|---|---|---|---|---|---|---|---|---|
| **SIGLEC5** | 0.00766 | 0.00944 | 0.04782 | 0.00450 | 0.00374 | 0.00000 | 0.00184 | 0.02532 | 0.42329 | 0.00800 | 0.81676 | 0.01566 |
| **SIGLEC6** | 0.00259 | 0.01781 | 0.76844 | 0.01158 | 0.01630 | 0.00324 | 0.00000 | 0.04582 | 0.00000 | 0.02135 | 0.02309 | 0.01428 |
| **SIGLEC7** | 0.20314 | 0.01340 | 0.03076 | 0.00890 | 0.01174 | 0.01605 | 0.01170 | 0.06131 | 0.01104 | 0.03206 | 0.02262 | 0.00964 |
| **SIGLEC8** | 0.00549 | 0.01309 | 0.01918 | 0.00968 | 0.00821 | 0.02217 | 0.01674 | 0.01709 | 0.01589 | 0.03192 | 0.03016 | 0.01657 |
| **SIGLEC9** | 0.00595 | 0.01387 | 0.02627 | 0.01014 | 0.01073 | 0.04495 | 0.07673 | 0.01090 | 0.01117 | 0.00001 | 0.00954 | 0.01909 |
| **SIGMAR1** | 1.19178 | 7.35138 | 12.64014 | 4.04334 | 0.36709 | 0.00589 | 4.09056 | 0.69020 | 4.28031 | 0.00000 | 8.14163 | 5.51549 |
| **SIK1** | 0.19533 | 1.93473 | 1.52597 | 0.00000 | 1.95856 | 0.98676 | 2.07864 | 0.65063 | 0.00000 | 6.26579 | 0.00000 | 0.26188 |
| **SIK2** | 0.85260 | 1.47061 | 0.53204 | 1.08245 | 1.52807 | 0.40255 | 0.49063 | 1.65282 | 1.04470 | 0.56774 | 0.52235 | 0.78283 |
| **SIK3** | 0.54018 | 0.46456 | 2.51638 | 2.06089 | 0.82684 | 0.91465 | 3.18358 | 0.28999 | 0.00000 | 7.82962 | 0.39089 | 1.74060 |
| **SIKE1** | 5.28209 | 12.07433 | 10.66328 | 8.69383 | 4.94303 | 34.14306 | 13.07832 | 15.17654 | 9.83277 | 11.05536 | 7.42905 | 17.58365 |
| **SIL1** | 0.59247 | 12.41061 | 10.35705 | 6.84996 | 11.28102 | 0.00007 | 1.77152 | 2.16152 | 18.13470 | 2.97285 | 6.29787 | 9.22846 |
| **SIM1** | 0.01178 | 0.07739 | 0.02550 | 0.00437 | 0.00991 | 0.02775 | 0.01669 | 0.10688 | 0.02053 | 0.01660 | 0.01846 | 0.00562 |
| **SIM2** | 0.00306 | 0.00674 | 0.09322 | 0.04835 | 0.13986 | 0.00000 | 0.03075 | 0.02209 | 0.00000 | 0.10150 | 0.08679 | 0.01527 |
| **SIMC1** | 0.00000 | 5.52949 | 0.59792 | 3.32352 | 0.27818 | 5.28959 | 0.00061 | 0.44637 | 0.00000 | 4.94527 | 1.08626 | 1.53547 |
| **SIN3A** | 16.73218 | 10.65980 | 4.91095 | 9.14282 | 6.00542 | 14.32697 | 4.70576 | 12.06305 | 9.89121 | 10.15491 | 4.21589 | 14.51738 |
| **SIN3B** | 1.53256 | 2.73155 | 10.14071 | 1.82072 | 7.76578 | 0.00000 | 0.00055 | 0.73913 | 5.89022 | 9.40150 | 1.08002 | 2.81136 |
| **SIPA1** | 0.12784 | 0.35997 | 1.46476 | 1.38160 | 1.25450 | 0.00000 | 0.03319 | 0.00000 | 1.55339 | 3.00290 | 0.04708 | 0.22052 |
| **SIPA1L1** | 7.10209 | 0.93763 | 5.04887 | 3.43610 | 4.71850 | 4.51493 | 5.21798 | 2.83960 | 5.38183 | 5.33975 | 3.15922 | 3.46283 |
| **SIPA1L2** | 4.84170 | 1.55951 | 1.13645 | 7.01011 | 0.62143 | 0.00000 | 0.00321 | 2.13953 | 0.00000 | 1.16500 | 0.00000 | 2.28083 |
| **SIPA1L3** | 0.71416 | 2.47227 | 1.63445 | 0.63520 | 1.66604 | 0.00000 | 2.09864 | 0.16472 | 0.99450 | 8.61587 | 2.50654 | 0.38359 |
| **SIRPA** | 1.43729 | 0.02714 | 0.67856 | 0.11918 | 0.01744 | 0.00000 | 0.00000 | 0.12339 | 0.00000 | 0.01922 | 0.04472 | 0.02503 |
| **SIRPB2** | 0.00593 | 0.01706 | 0.06054 | 0.01965 | 0.02415 | 0.00192 | 0.01445 | 0.01207 | 0.01740 | 0.00704 | 0.01761 | 0.02003 |
| **SIRT1** | 0.00000 | 0.15654 | 1.60955 | 2.25795 | 0.00000 | 12.18154 | 5.17520 | 9.77107 | 0.00002 | 12.98615 | 1.02320 | 3.11882 |
| **SIRT2** | 0.19837 | 2.45795 | 3.76465 | 1.54239 | 0.00000 | 0.56258 | 3.91910 | 0.35747 | 4.22672 | 1.11545 | 4.24391 | 1.87112 |
| **SIRT3** | 0.91276 | 2.65250 | 1.71262 | 1.33521 | 1.47060 | 0.00000 | 0.00000 | 1.28424 | 0.84665 | 0.01101 | 0.68841 | 2.46850 |
| **SIRT4** | 0.21590 | 0.33464 | 0.93301 | 0.00372 | 0.22066 | 0.00000 | 0.32438 | 0.09230 | 2.14289 | 0.00000 | 0.69498 | 1.09276 |
| **SIRT5** | 1.43838 | 3.06214 | 0.96411 | 3.68733 | 2.57723 | 3.76863 | 0.00942 | 0.81759 | 1.51942 | 0.78257 | 4.26178 | 1.04574 |
| **SIRT6** | 0.49370 | 8.70087 | 6.67542 | 2.57192 | 1.82234 | 0.00000 | 4.51152 | 0.95893 | 8.67784 | 0.00000 | 3.45428 | 7.06550 |
| **SIRT7** | 0.87222 | 2.62399 | 4.09451 | 1.28406 | 6.21451 | 0.00000 | 3.27988 | 0.65394 | 1.88608 | 14.93774 | 0.74726 | 2.09088 |
| **1.00000** | 0.00545 | 0.01369 | 0.02993 | 0.01045 | 0.02270 | 0.02129 | 0.01809 | 0.01391 | 0.01369 | 0.01013 | 0.01567 | 0.02267 |
| **SIVA1** | 9.23694 | 25.16737 | 23.45712 | 14.38864 | 30.18575 | 9.19836 | 26.27469 | 6.51952 | 24.86425 | 2.39842 | 16.86953 | 18.19616 |
| **SIX1** | 0.99017 | 0.00000 | 0.02648 | 0.67497 | 0.01623 | 0.06719 | 0.01550 | 0.16212 | 0.00509 | 0.00375 | 0.01748 | 0.15228 |
| **SIX2** | 0.81675 | 0.04855 | 0.00124 | 1.67550 | 0.00000 | 0.00000 | 0.00000 | 0.32414 | 0.00000 | 1.11268 | 0.22684 | 0.56318 |
| **SIX3** | 0.12898 | 0.00293 | 0.00000 | 0.05308 | 0.00018 | 0.13593 | 0.00000 | 0.01196 | 0.02724 | 0.02520 | 0.00715 | 0.03227 |
| **SIX3-AS1** | 0.02836 | 0.02626 | 0.01547 | 0.01080 | 0.01101 | 0.02744 | 0.01616 | 0.02981 | 0.01759 | 0.01223 | 0.01373 | 0.01225 |
| **SIX4** | 0.08839 | 0.00000 | 2.26523 | 0.00000 | 0.51719 | 2.27604 | 2.65465 | 0.87987 | 2.24440 | 1.48786 | 0.15605 | 0.84598 |
| **SIX5** | 0.49615 | 1.82085 | 1.13134 | 0.72377 | 1.29214 | 0.00000 | 1.65009 | 0.00028 | 0.50858 | 2.06429 | 1.30507 | 0.02382 |
| **SKA1** | 3.19421 | 6.10604 | 8.76514 | 3.32242 | 2.96685 | 9.45560 | 13.90117 | 5.00701 | 5.98878 | 4.61284 | 6.55290 | 11.07515 |
| **SKA2** | 1.47739 | 7.35906 | 6.69574 | 4.66234 | 4.11760 | 25.09527 | 13.49953 | 16.96900 | 3.63522 | 7.12719 | 7.47470 | 20.29423 |
| **SKA3** | 1.10096 | 16.11401 | 10.08923 | 11.02002 | 3.58944 | 26.46061 | 21.04790 | 18.46016 | 10.99840 | 6.01592 | 15.60117 | 16.60422 |
| **SKAP1** | 0.00330 | 0.01510 | 0.01477 | 0.01257 | 0.01722 | 0.01348 | 0.01948 | 0.05686 | 0.02750 | 0.00852 | 0.01564 | 0.02594 |
| **SKAP2** | 4.42378 | 4.03446 | 3.48989 | 0.95646 | 3.26427 | 14.97110 | 6.64764 | 2.30968 | 2.03291 | 6.23664 | 2.34673 | 3.13383 |
| **SKI** | 5.53354 | 0.98357 | 0.95441 | 0.20687 | 2.00263 | 0.03235 | 1.68997 | 0.00000 | 1.00898 | 7.46397 | 0.36627 | 1.07383 |
| **SKIDA1** | 0.00000 | 0.00000 | 2.95488 | 1.41086 | 0.60862 | 3.54863 | 2.56373 | 0.00827 | 0.24035 | 3.31519 | 1.27122 | 0.00000 |
| **SKIL** | 0.66715 | 2.09497 | 1.29436 | 8.57154 | 0.84262 | 20.72239 | 3.75419 | 5.85106 | 0.00000 | 4.01723 | 3.20959 | 14.38312 |
| **SKINTL** | 0.00528 | 0.01912 | 0.00912 | 0.01486 | 0.01378 | 0.01938 | 0.03552 | 0.00999 | 0.02269 | 0.04504 | 0.01440 | 0.02864 |
| **SKIV2L** | 0.02026 | 6.74094 | 6.32699 | 5.21772 | 5.09979 | 0.00000 | 2.29486 | 1.08249 | 5.70033 | 1.49227 | 6.67250 | 6.34388 |
| **SKIV2L2** | 0.42650 | 5.46224 | 9.33654 | 2.69686 | 3.24158 | 23.44465 | 13.94743 | 4.97637 | 7.59809 | 9.34965 | 7.84632 | 8.15636 |

| | | | | | | | | | | | | |
|---|---|---|---|---|---|---|---|---|---|---|---|---|
| **SKOR1** | 4.00841 | 0.03350 | 0.01288 | 0.00811 | 0.02860 | 0.00000 | 0.02765 | 0.19210 | 0.01976 | 0.00000 | 0.02026 | 0.15333 |
| **SKOR2** | 0.00342 | 0.01515 | 0.01564 | 0.01394 | 0.03109 | 0.01156 | 0.02424 | 0.04488 | 0.01920 | 0.01006 | 0.01538 | 0.01490 |
| **SKP1** | 13.15795 | 12.87546 | 23.12680 | 14.92580 | 11.01624 | 51.65414 | 29.54839 | 6.26828 | 22.58471 | 10.79781 | 23.57669 | 20.32156 |
| **SKP1P2** | 2.82730 | 1.46222 | 4.69762 | 2.06700 | 0.33517 | 13.28699 | 4.16998 | 0.72759 | 3.87320 | 0.00007 | 4.64746 | 3.18242 |
| **SKP2** | 15.61156 | 12.63611 | 18.14241 | 3.32957 | 5.70582 | 20.86383 | 20.56510 | 4.68510 | 16.23315 | 0.00000 | 29.64262 | 3.35234 |
| **SLA** | 0.00610 | 0.01508 | 0.01755 | 0.01174 | 0.01709 | 0.01962 | 0.01752 | 0.01135 | 0.02212 | 0.00923 | 0.01461 | 0.01652 |
| **SLA2** | 2.85465 | 0.06915 | 0.05919 | 0.03362 | 0.07342 | 0.05185 | 0.00000 | 0.12348 | 0.14638 | 0.18474 | 0.08637 | 0.00916 |
| **SLAIN1** | 0.23120 | 0.14598 | 0.18474 | 0.20701 | 0.12724 | 0.00000 | 0.55723 | 0.09811 | 0.00000 | 0.17976 | 0.04397 | 0.02957 |
| **SLAIN2** | 3.78734 | 6.88208 | 5.12239 | 6.26155 | 5.16811 | 9.62976 | 2.75704 | 4.37457 | 1.54014 | 15.01474 | 1.99084 | 2.65553 |
| **SLAMF1** | 0.00536 | 0.00402 | 0.02569 | 0.05718 | 0.00000 | 0.09385 | 0.00000 | 0.00842 | 0.00000 | 0.04179 | 0.01060 | 0.03885 |
| **SLAMF6** | 0.00596 | 0.01483 | 0.02412 | 0.01354 | 0.01340 | 0.01532 | 0.01857 | 0.01047 | 0.01214 | 0.02940 | 0.01590 | 0.01606 |
| **SLAMF7** | 0.00563 | 0.01475 | 0.49259 | 0.01063 | 0.00978 | 0.00000 | 0.00000 | 0.00938 | 0.00000 | 0.06098 | 0.03285 | 0.02889 |
| **SLAMF8** | 0.00611 | 0.04723 | 0.01816 | 0.00606 | 0.02684 | 0.03777 | 0.02033 | 0.01039 | 0.01207 | 0.00527 | 0.01115 | 0.01506 |
| **SLAMF9** | 0.00374 | 0.03634 | 0.00390 | 0.00000 | 0.00183 | 0.08956 | 0.02476 | 0.03331 | 0.00000 | 0.05292 | 0.00901 | 0.00554 |
| **SLBP** | 4.63975 | 23.18935 | 13.58858 | 4.04885 | 15.85722 | 26.96358 | 6.42627 | 12.49919 | 24.24070 | 19.24168 | 27.80859 | 13.86017 |
| **SLC10A1** | 0.12665 | 0.00328 | 0.03450 | 0.04055 | 0.03747 | 0.00000 | 0.04612 | 0.10523 | 0.00000 | 0.07170 | 0.11613 | 0.08701 |
| **SLC10A3** | 0.65435 | 1.21880 | 3.48683 | 1.26822 | 3.09944 | 0.00000 | 2.82749 | 0.10898 | 3.80064 | 0.73887 | 4.52989 | 1.51753 |
| **SLC10A4** | 0.10779 | 0.15039 | 0.18398 | 0.00000 | 0.00000 | 0.27668 | 0.13948 | 0.45492 | 0.54009 | 0.38164 | 0.00019 | 0.00000 |
| **SLC10A5** | 0.01441 | 0.01787 | 0.03696 | 0.00818 | 0.01188 | 0.10525 | 0.01490 | 0.00711 | 0.00029 | 0.04089 | 0.04710 | 0.04828 |
| **SLC10A6** | 0.00316 | 0.01332 | 0.01101 | 0.01413 | 0.00914 | 0.02242 | 0.00921 | 0.02535 | 0.02131 | 0.04949 | 0.01323 | 0.02540 |
| **SLC10A7** | 0.91096 | 0.36374 | 4.05338 | 1.26935 | 0.00000 | 4.37292 | 0.00000 | 0.83430 | 4.28873 | 0.34004 | 1.47954 | 1.00320 |
| **SLC11A1** | 0.03793 | 0.19787 | 0.09695 | 0.07085 | 0.08268 | 0.15257 | 0.00000 | 0.00418 | 0.00000 | 0.27442 | 0.05259 | 0.11374 |
| **SLC11A2** | 2.11401 | 0.00000 | 10.86861 | 1.10965 | 12.35235 | 2.95742 | 9.04876 | 0.61908 | 26.19056 | 4.47488 | 23.51214 | 0.63731 |
| **SLC12A1** | 0.02904 | 0.01344 | 0.01470 | 0.01336 | 0.01820 | 0.00998 | 0.01929 | 0.05534 | 0.02216 | 0.00984 | 0.01523 | 0.04860 |
| **SLC12A2** | 5.28530 | 0.41056 | 4.43441 | 2.01052 | 0.00000 | 5.83039 | 2.02540 | 3.74076 | 5.19636 | 7.54218 | 0.00000 | 0.93795 |
| **SLC12A3** | 0.00000 | 17.77230 | 0.00000 | 0.00000 | 37.43512 | 5.47922 | 2.16301 | 0.00000 | 22.83885 | 13.00756 | 15.12398 | 0.00000 |
| **SLC12A4** | 0.14738 | 1.61135 | 3.28541 | 1.40201 | 4.01578 | 0.00000 | 0.00000 | 0.14944 | 2.91918 | 2.56058 | 0.96473 | 0.77149 |
| **SLC12A5** | 4.60800 | 0.13050 | 0.01829 | 0.05144 | 0.04889 | 0.00000 | 0.02077 | 0.08719 | 0.02894 | 0.01778 | 0.03079 | 0.10760 |
| **SLC12A6** | 1.51804 | 1.01429 | 0.26858 | 3.06576 | 0.00000 | 1.80957 | 0.00000 | 0.51550 | 1.86455 | 3.36902 | 0.56560 | 0.02226 |
| **SLC12A7** | 1.74193 | 7.70740 | 0.00000 | 2.48162 | 9.89845 | 0.00000 | 0.00000 | 0.36841 | 1.60105 | 5.98620 | 0.61564 | 3.34659 |
| **SLC12A8** | 3.00547 | 0.00122 | 0.33047 | 0.03028 | 0.00000 | 0.00000 | 0.00000 | 0.68589 | 0.37940 | 0.00720 | 1.00155 | 0.00000 |
| **SLC12A9** | 0.00132 | 0.48804 | 1.48040 | 0.19036 | 4.11352 | 0.00000 | 1.66462 | 0.01648 | 1.27867 | 8.89785 | 1.43066 | 0.19241 |
| **SLC13A1** | 0.00622 | 0.01840 | 0.01560 | 0.01179 | 0.01674 | 0.01503 | 0.01696 | 0.01045 | 0.01983 | 0.01648 | 0.01879 | 0.01692 |
| **SLC13A2** | 0.00856 | 0.01105 | 0.01403 | 0.01146 | 0.09657 | 0.00000 | 0.02556 | 0.01033 | 0.01159 | 0.03760 | 0.01013 | 0.03483 |
| **SLC13A3** | 0.73740 | 0.03739 | 0.50624 | 0.00219 | 0.20599 | 0.00000 | 0.00006 | 0.76486 | 0.43178 | 0.00263 | 0.00219 | 0.03183 |
| **SLC13A4** | 2.55529 | 0.01646 | 7.75222 | 0.03643 | 0.12642 | 0.00000 | 0.95793 | 0.68383 | 0.68169 | 0.00000 | 0.00421 | 0.21518 |
| **SLC13A5** | 0.00552 | 0.01494 | 0.05659 | 0.01798 | 0.01888 | 0.00568 | 0.01212 | 0.01180 | 0.01389 | 0.02976 | 0.02300 | 0.01926 |
| **SLC14A1** | 0.88797 | 0.02219 | 0.01045 | 0.00364 | 0.00780 | 0.04174 | 0.03759 | 0.01098 | 0.02492 | 0.03786 | 0.00801 | 0.00571 |
| **SLC14A2** | 0.55040 | 0.01470 | 0.02096 | 0.00170 | 0.05392 | 0.01447 | 0.00620 | 0.00434 | 0.06119 | 0.08327 | 0.02323 | 0.02060 |
| **SLC15A1** | 0.00781 | 0.00000 | 0.00000 | 0.01075 | 0.35581 | 0.05488 | 0.00000 | 0.04098 | 0.00000 | 0.13697 | 0.45034 | 0.01819 |
| **SLC15A2** | 0.00000 | 0.00000 | 0.00000 | 0.00000 | 4.72607 | 0.38222 | 0.39603 | 0.22932 | 7.81499 | 0.97627 | 12.20436 | 0.01915 |
| **SLC15A3** | 0.01087 | 0.55196 | 0.08199 | 0.01528 | 0.09550 | 0.00000 | 0.00548 | 0.24690 | 0.00000 | 0.02129 | 0.02035 | 2.57418 |
| **SLC15A4** | 1.02865 | 0.89579 | 2.23902 | 2.25054 | 4.88185 | 5.47331 | 0.09739 | 0.14851 | 1.88065 | 8.39967 | 1.74330 | 0.00000 |
| **SLC15A5** | 0.01298 | 0.00989 | 0.04837 | 0.01388 | 0.00000 | 0.03087 | 0.01307 | 0.04128 | 0.05834 | 0.05899 | 0.04777 | 0.01172 |
| **SLC16A1** | 3.93290 | 37.82204 | 25.26622 | 10.51094 | 14.63290 | 73.39324 | 0.00000 | 35.29401 | 75.69786 | 36.48085 | 31.77585 | 48.73881 |
| **SLC16A1-AS1** | 0.20960 | 0.43633 | 0.39466 | 0.20226 | 0.75923 | 0.00000 | 0.07565 | 0.54673 | 2.38543 | 0.00000 | 0.66509 | 2.12459 |
| **SLC16A10** | 0.69909 | 0.79873 | 0.91708 | 4.31315 | 0.89267 | 0.00009 | 0.23504 | 0.38512 | 1.73233 | 3.59200 | 0.00000 | 0.02544 |
| **SLC16A11** | 0.06272 | 0.01495 | 0.71881 | 0.00962 | 0.03331 | 0.00000 | 0.00000 | 0.01559 | 0.00000 | 0.03515 | 0.01615 | 0.01449 |

| | | | | | | | | | | | | |
|---|---|---|---|---|---|---|---|---|---|---|---|---|
| **SLC16A12** | 0.04564 | 0.11628 | 0.04966 | 0.10167 | 0.00386 | 0.00727 | 0.02578 | 0.16787 | 0.00378 | 0.10832 | 0.03619 | 0.05758 |
| **SLC16A13** | 0.29419 | 0.02895 | 1.33954 | 0.00571 | 0.01127 | 0.00000 | 0.11422 | 0.06661 | 0.81683 | 0.00000 | 0.07665 | 0.10678 |
| **SLC16A14** | 0.00136 | 0.78749 | 0.00000 | 0.23231 | 0.00000 | 0.37768 | 0.00000 | 0.38671 | 0.00000 | 0.04087 | 0.00176 | 0.00060 |
| **SLC16A2** | 0.00193 | 0.96703 | 0.05609 | 11.33254 | 5.48379 | 0.00000 | 0.00279 | 0.00169 | 0.00000 | 0.83025 | 0.00000 | 0.12347 |
| **SLC16A3** | 0.05690 | 0.00000 | 2.20448 | 0.00000 | 0.21247 | 0.00333 | 0.99458 | 0.01039 | 3.15801 | 0.87322 | 1.30610 | 0.00755 |
| **SLC16A4** | 0.04715 | 0.00000 | 0.11919 | 0.00000 | 0.43041 | 0.62281 | 0.38969 | 0.44471 | 0.13343 | 0.74461 | 0.51780 | 0.23017 |
| **SLC16A5** | 3.14545 | 0.05899 | 0.20722 | 0.56147 | 2.22571 | 0.00174 | 0.35990 | 2.62446 | 1.36752 | 0.88891 | 2.04138 | 0.02452 |
| **SLC16A6** | 0.03660 | 0.00000 | 1.15864 | 0.00000 | 0.60395 | 0.70400 | 0.00000 | 19.54155 | 0.00000 | 0.06578 | 0.24388 | 1.94425 |
| **SLC16A7** | 0.00945 | 0.01343 | 0.06842 | 0.01840 | 0.00000 | 0.03618 | 0.02374 | 0.01017 | 0.00000 | 0.04412 | 0.03039 | 0.01426 |
| **SLC16A8** | 0.22208 | 0.01099 | 0.03021 | 0.01076 | 0.00366 | 0.04691 | 0.01391 | 0.05497 | 0.01005 | 0.03634 | 0.01186 | 0.06143 |
| **SLC16A9** | 0.00120 | 0.87680 | 0.09391 | 11.86880 | 0.00000 | 0.52751 | 0.00000 | 2.71798 | 0.07895 | 0.17516 | 0.00000 | 11.78121 |
| **SLC17A2** | 0.00405 | 0.02575 | 0.01841 | 0.25805 | 0.06385 | 0.06525 | 0.00000 | 0.01641 | 0.00000 | 0.01492 | 0.06824 | 0.13012 |
| **SLC17A4** | 0.00495 | 0.00970 | 0.00840 | 0.00478 | 0.00000 | 0.03287 | 0.06692 | 0.00998 | 0.00000 | 0.08551 | 0.01362 | 0.01435 |
| **SLC17A5** | 0.29363 | 27.84977 | 0.00000 | 7.01140 | 22.71982 | 17.56020 | 0.00000 | 0.61459 | 4.25720 | 3.90287 | 7.78122 | 1.80878 |
| **SLC17A7** | 0.50924 | 0.01238 | 0.09738 | 0.03840 | 0.07362 | 0.00000 | 0.08000 | 0.91466 | 0.04983 | 0.00000 | 0.01331 | 0.76066 |
| **SLC17A8** | 0.83380 | 0.01813 | 0.01850 | 0.01885 | 0.01925 | 0.00000 | 0.04317 | 0.01111 | 0.01948 | 0.00651 | 0.01421 | 0.02538 |
| **SLC17A9** | 0.00459 | 0.02395 | 0.50917 | 0.03780 | 0.00000 | 0.00000 | 0.05790 | 0.01060 | 0.00000 | 0.09074 | 0.17775 | 0.01334 |
| **SLC18A1** | 0.00600 | 0.03062 | 0.01764 | 0.01526 | 0.01361 | 0.02345 | 0.01609 | 0.01040 | 0.01669 | 0.01557 | 0.01399 | 0.01698 |
| **SLC18A2** | 0.00000 | 0.11162 | 0.71537 | 0.16954 | 0.00000 | 2.22889 | 0.72741 | 1.10482 | 0.55283 | 1.56062 | 0.00000 | 0.60593 |
| **SLC18A3** | 0.00543 | 0.02623 | 0.00908 | 0.01164 | 0.00001 | 0.04655 | 0.00952 | 0.01025 | 0.00969 | 0.05020 | 0.01201 | 0.01071 |
| **SLC18B1** | 0.08801 | 0.02502 | 1.06431 | 0.00544 | 0.00000 | 0.02026 | 1.65993 | 0.23412 | 7.13238 | 0.00000 | 1.29898 | 0.24316 |
| **SLC19A1** | 1.24420 | 0.44509 | 4.56216 | 0.04042 | 0.70103 | 0.00000 | 0.29544 | 0.10303 | 1.07425 | 0.00000 | 1.53161 | 0.21347 |
| **SLC19A2** | 2.49346 | 0.00000 | 12.99023 | 0.12051 | 9.59247 | 0.00000 | 1.80185 | 2.22368 | 21.76471 | 4.43943 | 13.98940 | 0.00000 |
| **SLC19A3** | 0.00000 | 1.73813 | 1.02305 | 0.00000 | 1.72516 | 3.70298 | 0.00000 | 0.06707 | 7.66525 | 0.45569 | 10.75560 | 1.64342 |
| **SLC1A1** | 0.12739 | 0.38824 | 0.10338 | 0.02487 | 0.00000 | 0.01100 | 0.00388 | 0.13085 | 0.02340 | 0.01076 | 0.01652 | 0.58259 |
| **SLC1A2** | 0.00455 | 0.03609 | 0.21936 | 0.93074 | 0.15216 | 0.04245 | 0.00000 | 0.03164 | 0.00000 | 0.17818 | 0.00111 | 0.06587 |
| **SLC1A3** | 0.00000 | 0.00000 | 8.45418 | 6.26193 | 19.61579 | 3.21444 | 7.17718 | 0.00000 | 53.10218 | 1.62005 | 13.29513 | 0.00000 |
| **SLC1A4** | 0.42648 | 0.00711 | 0.28929 | 0.47062 | 0.00000 | 0.00000 | 0.25843 | 0.03554 | 0.10592 | 0.46781 | 0.01265 | 0.22839 |
| **SLC1A5** | 11.51161 | 3.69286 | 26.25342 | 6.54375 | 6.84543 | 1.50860 | 20.00740 | 0.75832 | 64.33255 | 8.90347 | 9.58968 | 17.52471 |
| **SLC1A6** | 1.72675 | 0.13187 | 1.22959 | 0.05170 | 0.41811 | 0.00000 | 0.00018 | 3.58362 | 0.00000 | 0.02043 | 0.01373 | 0.35202 |
| **SLC1A7** | 0.00399 | 0.19144 | 0.02239 | 0.00832 | 0.09603 | 0.09312 | 0.00223 | 0.01301 | 0.00004 | 0.02818 | 0.00836 | 0.12397 |
| **SLC20A1** | 1.14308 | 5.91012 | 14.00638 | 3.57136 | 3.12348 | 7.25412 | 0.97826 | 3.93462 | 19.68355 | 8.62926 | 8.23129 | 9.45302 |
| **SLC20A2** | 0.20553 | 3.76230 | 20.39401 | 2.55861 | 2.40651 | 2.06644 | 0.94273 | 0.42024 | 8.68151 | 0.00000 | 5.90152 | 0.80100 |
| **SLC22A1** | 0.00449 | 0.00688 | 0.17346 | 0.18045 | 0.19821 | 0.00000 | 0.11774 | 0.01006 | 0.00000 | 0.06993 | 0.09070 | 0.02088 |
| **SLC22A10** | 0.00597 | 0.04944 | 0.01502 | 0.01091 | 0.01508 | 0.00619 | 0.01894 | 0.00941 | 0.02236 | 0.01297 | 0.01590 | 0.04217 |
| **SLC22A11** | 0.00806 | 0.00000 | 4.60978 | 0.01678 | 0.04332 | 0.03003 | 0.12095 | 0.03040 | 0.72729 | 0.00000 | 0.27738 | 0.04662 |
| **SLC22A12** | 0.00556 | 0.01342 | 0.02558 | 0.00838 | 0.01416 | 0.02238 | 0.04174 | 0.01197 | 0.00001 | 0.02546 | 0.02145 | 0.01502 |
| **SLC22A13** | 0.06193 | 0.00000 | 0.06732 | 0.01410 | 0.07636 | 0.03337 | 0.00000 | 0.00565 | 0.03392 | 0.02547 | 0.20751 | 0.07999 |
| **SLC22A14** | 0.00535 | 0.00237 | 0.00669 | 0.00166 | 0.01157 | 0.06127 | 0.04672 | 0.04832 | 0.09498 | 0.02493 | 0.07668 | 0.04684 |
| **SLC22A15** | 0.06198 | 0.15616 | 0.01106 | 0.30889 | 0.00000 | 0.00460 | 0.07967 | 0.00420 | 0.31317 | 0.00000 | 0.00000 | 0.05151 |
| **SLC22A16** | 0.01754 | 0.02623 | 0.08854 | 0.01909 | 0.03394 | 0.00000 | 0.01518 | 0.09292 | 0.00866 | 0.00000 | 0.01853 | 0.05997 |
| **SLC22A17** | 0.09577 | 0.00040 | 0.23884 | 0.00141 | 0.00000 | 0.01129 | 0.00000 | 0.01378 | 0.14953 | 0.85797 | 0.00935 | 0.01645 |
| **SLC22A18** | 0.00310 | 0.45527 | 0.38181 | 1.12486 | 0.38832 | 0.53045 | 0.00000 | 0.00846 | 0.52874 | 0.13745 | 0.09284 | 0.08276 |
| **SLC22A18AS** | 0.00570 | 0.00422 | 0.08056 | 0.06772 | 0.00955 | 0.07124 | 0.00000 | 0.02194 | 0.01684 | 0.04011 | 0.01341 | 0.02826 |
| **SLC22A2** | 0.93202 | 0.00000 | 0.00000 | 0.00051 | 0.00000 | 0.07815 | 0.00116 | 0.09866 | 0.02211 | 0.20347 | 0.08868 | 0.12166 |
| **SLC22A20** | 0.00641 | 0.30660 | 0.45938 | 1.22459 | 0.03046 | 0.00000 | 0.20541 | 0.35548 | 0.10640 | 0.00000 | 0.03759 | 3.67795 |
| **SLC22A23** | 1.37880 | 0.15439 | 5.97049 | 0.82372 | 2.16381 | 2.59641 | 1.31824 | 0.36148 | 1.12543 | 4.91217 | 0.00000 | 0.00667 |
| **SLC22A24** | 0.01671 | 0.01378 | 0.02785 | 0.00618 | 0.03933 | 0.13428 | 0.02391 | 0.00902 | 0.00000 | 0.00000 | 0.02229 | 0.04366 |

| | | | | | | | | | | | | |
|---|---|---|---|---|---|---|---|---|---|---|---|---|
| **SLC22A25** | 0.00604 | 0.01744 | 0.03901 | 0.00749 | 0.01766 | 0.03041 | 0.02104 | 0.01036 | 0.01776 | 0.01243 | 0.01277 | 0.01878 |
| **SLC22A3** | 0.00800 | 0.01763 | 0.11665 | 0.01946 | 0.03035 | 0.00000 | 0.01036 | 0.01033 | 0.01325 | 0.00678 | 0.01746 | 0.01868 |
| **SLC22A31** | 0.00147 | 0.41419 | 0.28556 | 0.15768 | 0.38820 | 0.00000 | 0.00000 | 0.04463 | 0.00000 | 0.12616 | 0.03594 | 0.07671 |
| **SLC22A4** | 0.00445 | 0.19327 | 0.01272 | 0.14572 | 0.00000 | 0.00000 | 0.01811 | 0.01588 | 0.02013 | 0.15064 | 0.04030 | 0.04400 |
| **SLC22A5** | 0.11389 | 0.64832 | 0.25097 | 0.02998 | 1.75780 | 1.27070 | 0.00000 | 0.04995 | 1.99743 | 2.41610 | 1.85734 | 0.08269 |
| **SLC22A6** | 0.00629 | 0.01737 | 0.01687 | 0.01184 | 0.01603 | 0.01640 | 0.01610 | 0.01044 | 0.02021 | 0.01458 | 0.01860 | 0.01686 |
| **SLC22A7** | 0.00404 | 0.00051 | 0.00000 | 0.31604 | 0.00000 | 0.05999 | 0.02007 | 0.00932 | 0.00000 | 0.12837 | 0.02695 | 0.01294 |
| **SLC22A8** | 0.05141 | 0.02962 | 0.03014 | 0.01150 | 0.03109 | 0.00020 | 0.02329 | 0.00467 | 0.01640 | 0.01190 | 0.01744 | 0.03948 |
| **SLC22A9** | 0.01035 | 0.00175 | 0.00246 | 0.01098 | 0.00001 | 0.06356 | 0.00475 | 0.00777 | 0.00000 | 0.15381 | 0.00779 | 0.03149 |
| **SLC23A1** | 0.00477 | 0.03884 | 0.06360 | 0.03028 | 0.00000 | 0.16410 | 0.00000 | 0.20972 | 0.15993 | 0.38456 | 0.18992 | 0.02020 |
| **SLC23A2** | 0.00000 | 7.17051 | 0.59477 | 12.06544 | 18.18352 | 4.78325 | 0.00000 | 0.00000 | 4.32238 | 6.12553 | 5.96007 | 6.65414 |
| **SLC23A3** | 0.00533 | 0.03131 | 0.15448 | 0.02954 | 0.01113 | 0.00000 | 0.00000 | 0.00745 | 0.39067 | 0.08120 | 0.06543 | 0.07704 |
| **SLC24A1** | 0.13840 | 0.16274 | 0.30715 | 0.89831 | 0.75789 | 0.48394 | 0.04063 | 0.01378 | 0.00000 | 1.63342 | 0.29800 | 0.01772 |
| **SLC24A2** | 1.21377 | 0.17717 | 0.09437 | 0.62560 | 0.00000 | 0.00000 | 0.01128 | 0.11644 | 0.04628 | 0.19224 | 0.00027 | 0.26336 |
| **SLC24A3** | 0.00615 | 0.04755 | 0.08743 | 0.00601 | 0.01424 | 0.02495 | 0.00681 | 0.01033 | 0.01013 | 0.01646 | 0.01661 | 0.01441 |
| **SLC24A4** | 0.30706 | 0.07096 | 0.05588 | 0.00000 | 0.00000 | 0.04415 | 0.00828 | 0.00000 | 0.00000 | 0.20646 | 0.08486 | 0.14255 |
| **SLC24A5** | 0.17077 | 0.01518 | 0.01046 | 0.04873 | 0.03887 | 0.01955 | 0.04199 | 0.14442 | 0.00122 | 0.00000 | 0.01706 | 0.01208 |
| **SLC25A1** | 0.88878 | 4.71812 | 22.55624 | 14.91025 | 30.40764 | 0.00000 | 22.71603 | 0.34636 | 19.72475 | 7.23452 | 21.57866 | 1.40929 |
| **SLC25A10** | 0.19623 | 1.50286 | 2.78105 | 5.16217 | 3.76772 | 0.00000 | 2.84484 | 0.00000 | 4.97590 | 3.69159 | 10.19162 | 0.68910 |
| **SLC25A11** | 1.37680 | 25.64141 | 15.02541 | 18.52410 | 19.77374 | 4.76799 | 14.82720 | 1.92880 | 12.79026 | 0.00000 | 25.80534 | 9.39629 |
| **SLC25A12** | 0.82393 | 7.82065 | 0.28881 | 7.46958 | 0.00000 | 0.57622 | 0.00000 | 3.21315 | 0.00000 | 0.12350 | 0.00000 | 5.36005 |
| **SLC25A13** | 4.27290 | 6.21208 | 11.47574 | 6.10833 | 4.43159 | 9.04548 | 20.75904 | 0.13077 | 15.90507 | 0.55151 | 8.62796 | 0.00000 |
| **SLC25A14** | 3.62131 | 1.61436 | 2.47067 | 0.79212 | 0.73522 | 3.23429 | 2.05686 | 3.68981 | 2.85784 | 0.00399 | 2.03679 | 3.72557 |
| **SLC25A15** | 0.10224 | 2.29045 | 0.61521 | 0.57929 | 0.75567 | 1.49228 | 1.07623 | 0.00000 | 0.00000 | 2.36992 | 1.32049 | 0.07602 |
| **SLC25A16** | 0.74864 | 13.02008 | 4.73442 | 9.36818 | 12.90945 | 16.74299 | 0.00000 | 1.46712 | 0.00000 | 7.22569 | 4.82747 | 7.40361 |
| **SLC25A17** | 0.00000 | 5.40343 | 11.26535 | 3.98351 | 3.82391 | 15.19658 | 6.48594 | 4.24100 | 11.73694 | 3.48960 | 9.92203 | 9.10264 |
| **SLC25A18** | 0.28739 | 2.83412 | 0.65573 | 3.74447 | 0.00000 | 0.00000 | 0.00000 | 0.05634 | 0.16727 | 0.06491 | 0.02454 | 0.66206 |
| **SLC25A19** | 1.18633 | 8.71237 | 3.74543 | 2.11528 | 2.41893 | 5.44291 | 3.56252 | 0.01783 | 15.08221 | 0.00000 | 12.28493 | 10.68461 |
| **SLC25A2** | 0.00635 | 0.02299 | 0.01835 | 0.01505 | 0.02723 | 0.00979 | 0.01829 | 0.01067 | 0.02466 | 0.01036 | 0.01756 | 0.01647 |
| **SLC25A20** | 1.38922 | 2.04962 | 0.98688 | 0.16389 | 0.80613 | 1.30684 | 1.77611 | 0.57881 | 0.59078 | 1.40800 | 1.09906 | 3.06613 |
| **SLC25A21** | 0.36944 | 0.00000 | 0.05464 | 0.00141 | 0.00277 | 0.14776 | 0.11953 | 0.22614 | 0.13218 | 0.00000 | 0.09786 | 0.30545 |
| **SLC25A21-AS1** | 0.00814 | 0.04629 | 0.02182 | 0.01182 | 0.04050 | 0.01469 | 0.02485 | 0.00722 | 0.02102 | 0.00000 | 0.01309 | 0.09059 |
| **SLC25A22** | 1.10633 | 1.95153 | 2.23735 | 4.26499 | 5.42787 | 0.00000 | 0.00246 | 0.00160 | 7.34057 | 2.47709 | 2.87654 | 0.00000 |
| **SLC25A23** | 1.85165 | 1.38428 | 0.00000 | 3.36762 | 9.23828 | 0.00000 | 1.79661 | 0.00000 | 2.48913 | 6.64587 | 2.24463 | 1.91071 |
| **SLC25A24** | 0.00000 | 2.43621 | 3.21607 | 0.28179 | 0.00000 | 6.61726 | 3.07690 | 0.56193 | 0.59566 | 0.14139 | 1.00454 | 0.27954 |
| **SLC25A25** | 2.43191 | 1.40974 | 3.04103 | 2.10853 | 1.91021 | 0.00000 | 1.18097 | 3.96250 | 0.97129 | 2.75095 | 0.00000 | 3.10516 |
| **SLC25A26** | 3.18668 | 6.04592 | 10.38494 | 6.85896 | 4.69599 | 6.47775 | 13.91920 | 4.57830 | 10.39044 | 1.54469 | 6.54050 | 5.21221 |
| **SLC25A27** | 0.00646 | 0.03184 | 0.01870 | 0.02394 | 0.00000 | 0.12012 | 0.00066 | 0.00924 | 0.00568 | 0.02093 | 0.04366 | 0.03479 |
| **SLC25A28** | 0.00000 | 0.74234 | 0.15655 | 3.13443 | 1.18343 | 2.60292 | 1.14763 | 0.80346 | 0.00000 | 7.06658 | 0.09447 | 1.69168 |
| **SLC25A29** | 0.13536 | 6.02699 | 0.00000 | 9.95311 | 11.16372 | 0.00000 | 5.79583 | 0.00000 | 0.00000 | 16.11514 | 3.60753 | 0.19305 |
| **SLC25A3** | 10.20530 | 37.06765 | 34.71554 | 30.59352 | 39.22957 | 54.94213 | 48.24274 | 13.84132 | 52.95867 | 24.56071 | 43.17094 | 39.94128 |
| **SLC25A30** | 1.35349 | 0.39641 | 0.00000 | 0.15405 | 0.00000 | 0.57740 | 0.63383 | 0.26707 | 0.00000 | 1.69979 | 0.45841 | 0.19176 |
| **SLC25A30-AS1** | 0.01003 | 0.01760 | 0.01248 | 0.00988 | 0.02270 | 0.00000 | 0.01863 | 0.00993 | 0.06827 | 0.06605 | 0.01718 | 0.01502 |
| **SLC25A31** | 0.11513 | 0.02645 | 0.35558 | 0.01017 | 0.00000 | 0.02342 | 0.00000 | 0.53778 | 0.00000 | 0.06513 | 0.02269 | 0.87573 |
| **SLC25A32** | 1.05086 | 4.48703 | 8.12929 | 0.00000 | 0.00000 | 14.60286 | 9.70991 | 9.27481 | 12.63489 | 4.21593 | 7.97848 | 8.79858 |
| **SLC25A33** | 0.66298 | 0.37007 | 1.70958 | 1.03747 | 2.82626 | 4.12767 | 4.90042 | 1.11679 | 0.90185 | 6.18583 | 0.36632 | 1.31414 |
| **SLC25A34** | 0.00635 | 0.00000 | 0.05844 | 0.01075 | 0.12679 | 0.26171 | 0.01199 | 0.04792 | 0.19718 | 0.00000 | 0.09336 | 0.09781 |
| **SLC25A35** | 0.07051 | 0.89626 | 1.27498 | 1.72248 | 0.85085 | 0.00000 | 0.52254 | 0.07227 | 0.87699 | 0.00000 | 3.21936 | 0.39940 |

| | | | | | | | | | | | | |
|---|---|---|---|---|---|---|---|---|---|---|---|---|
| **SLC25A36** | 0.00000 | 3.71708 | 3.70800 | 3.04780 | 3.52347 | 23.94558 | 8.48748 | 13.34506 | 15.19887 | 15.10212 | 9.32458 | 9.07231 |
| **SLC25A37** | 0.79172 | 4.25601 | 4.39322 | 0.47073 | 3.77091 | 0.03930 | 6.02008 | 0.02683 | 5.04151 | 2.81210 | 7.60835 | 1.71241 |
| **SLC25A38** | 3.89314 | 17.38076 | 10.79718 | 18.78229 | 8.90049 | 9.00699 | 15.53241 | 4.93587 | 12.47000 | 0.00000 | 4.40518 | 14.37143 |
| **SLC25A39** | 7.16526 | 42.50107 | 17.06072 | 34.78220 | 47.60841 | 18.53629 | 22.91275 | 11.48692 | 37.91480 | 3.10921 | 29.50854 | 37.94070 |
| **SLC25A3P1** | 0.20563 | 0.00865 | 0.21099 | 0.06475 | 0.01917 | 0.01871 | 0.00000 | 0.15222 | 0.00000 | 0.01193 | 0.11617 | 0.03463 |
| **SLC25A4** | 0.39664 | 3.04633 | 7.86438 | 5.09967 | 2.21340 | 5.29824 | 2.31101 | 0.28239 | 0.83297 | 0.00000 | 1.70004 | 2.82944 |
| **SLC25A40** | 0.21499 | 1.65140 | 1.09600 | 0.32163 | 0.00000 | 3.68672 | 0.00569 | 0.00000 | 3.06142 | 2.30845 | 2.02943 | 1.04875 |
| **SLC25A41** | 0.00249 | 0.02424 | 0.01468 | 0.01848 | 0.06746 | 0.08624 | 0.03315 | 0.02192 | 0.00000 | 0.08758 | 0.02556 | 0.03382 |
| **SLC25A42** | 1.67053 | 0.43441 | 0.17298 | 0.73918 | 0.00000 | 0.00000 | 0.09462 | 0.05273 | 0.29442 | 2.19183 | 0.38544 | 0.20711 |
| **SLC25A43** | 0.00000 | 3.15455 | 0.54949 | 0.87674 | 5.44175 | 0.93586 | 1.37936 | 0.39742 | 1.58052 | 2.57926 | 4.88000 | 2.30801 |
| **SLC25A44** | 4.43038 | 18.40851 | 4.52434 | 22.23246 | 11.05147 | 21.93528 | 10.12920 | 24.55525 | 26.02259 | 6.15457 | 16.32370 | 15.08820 |
| **SLC25A45** | 0.01027 | 0.15973 | 0.06369 | 0.09337 | 0.24697 | 0.00000 | 0.03449 | 0.50174 | 0.12744 | 0.00000 | 0.38278 | 0.10172 |
| **SLC25A46** | 9.07327 | 3.01949 | 7.21059 | 1.51496 | 0.70755 | 12.62007 | 6.91285 | 8.88811 | 6.24860 | 0.34214 | 4.31609 | 2.88366 |
| **SLC25A47** | 0.01882 | 0.52007 | 0.00386 | 0.01656 | 0.00000 | 0.02839 | 0.05976 | 0.00951 | 0.00000 | 0.09251 | 0.00174 | 0.00461 |
| **SLC25A48** | 1.02154 | 0.32580 | 0.01153 | 0.00000 | 0.01842 | 0.03644 | 0.04000 | 0.06858 | 0.00785 | 0.00244 | 0.00532 | 0.00000 |
| **SLC25A5** | 10.51357 | 33.17391 | 26.96396 | 34.37938 | 33.15996 | 56.19084 | 49.81643 | 22.77744 | 37.10721 | 9.92504 | 50.23697 | 19.70184 |
| **SLC25A5-AS1** | 0.08921 | 0.32701 | 0.14767 | 0.03977 | 0.07243 | 0.06068 | 0.00000 | 0.11576 | 0.21977 | 0.00000 | 0.08810 | 0.15588 |
| **SLC25A51** | 0.84891 | 2.15009 | 0.75695 | 0.59267 | 0.00000 | 3.05463 | 0.68297 | 0.02036 | 1.16003 | 1.20286 | 1.51799 | 1.43681 |
| **SLC25A51P1** | 0.00593 | 0.01239 | 0.01581 | 0.01377 | 0.00785 | 0.04588 | 0.01307 | 0.00976 | 0.01334 | 0.01837 | 0.01847 | 0.02182 |
| **SLC25A52** | 0.08799 | 0.00262 | 0.06979 | 0.00000 | 0.03524 | 0.08732 | 0.00000 | 0.03117 | 0.00756 | 0.11427 | 0.01651 | 0.17518 |
| **SLC25A53** | 0.08155 | 0.18891 | 0.20423 | 0.01814 | 0.08774 | 0.00093 | 0.04419 | 0.75746 | 0.16252 | 0.00000 | 0.14908 | 0.35472 |
| **SLC26A1** | 0.02698 | 0.01287 | 0.04107 | 0.00955 | 0.01151 | 0.01833 | 0.02872 | 0.00961 | 0.01232 | 0.02984 | 0.01467 | 0.02062 |
| **SLC26A10** | 0.00565 | 0.04759 | 0.02456 | 0.04518 | 0.00000 | 0.00865 | 0.02108 | 0.00954 | 0.09837 | 0.06850 | 0.00288 | 0.04637 |
| **SLC26A11** | 0.17202 | 2.72563 | 0.70493 | 2.60001 | 1.59039 | 0.00000 | 0.10447 | 0.03623 | 7.99199 | 2.48428 | 0.35092 | 1.74081 |
| **SLC26A2** | 0.68338 | 0.02665 | 1.26882 | 0.00000 | 0.00000 | 0.00000 | 0.64977 | 0.00703 | 0.00000 | 0.68897 | 0.88498 | 0.20852 |
| **SLC26A3** | 0.00593 | 0.01227 | 0.00867 | 0.02354 | 0.01277 | 0.04242 | 0.02966 | 0.03375 | 0.01094 | 0.02241 | 0.01140 | 0.01325 |
| **SLC26A4** | 0.01596 | 0.31287 | 0.01063 | 0.00000 | 0.00000 | 0.01263 | 0.00000 | 0.01121 | 0.24875 | 0.05850 | 0.10270 | 0.11878 |
| **SLC26A4-AS1** | 0.00426 | 0.15600 | 0.05009 | 0.00000 | 0.00000 | 0.01842 | 0.04429 | 0.01742 | 0.01569 | 0.01925 | 0.08084 | 0.02929 |
| **SLC26A5** | 0.18955 | 0.01472 | 0.05098 | 0.01178 | 0.02497 | 0.00000 | 0.00849 | 0.00000 | 0.04624 | 0.02327 | 0.03225 | 0.18107 |
| **SLC26A6** | 4.97486 | 3.15424 | 3.52816 | 6.09001 | 6.08107 | 0.00000 | 0.39746 | 3.65685 | 16.89212 | 5.12244 | 3.76087 | 1.17682 |
| **SLC26A7** | 0.15604 | 0.00943 | 0.03979 | 0.00432 | 0.02919 | 0.02087 | 0.10241 | 0.10590 | 0.02514 | 0.03237 | 0.00723 | 0.24121 |
| **SLC26A8** | 0.50012 | 1.28279 | 0.00000 | 0.00000 | 1.50776 | 0.53311 | 0.00000 | 0.81079 | 0.02871 | 0.43811 | 0.15638 | 0.00000 |
| **SLC26A9** | 0.00596 | 0.01391 | 0.03184 | 0.01410 | 0.00652 | 0.01225 | 0.01633 | 0.01014 | 0.00000 | 0.03819 | 0.05902 | 0.01899 |
| **SLC27A1** | 0.40030 | 0.94465 | 0.14422 | 0.42441 | 0.00000 | 0.00000 | 0.24981 | 0.72884 | 0.00000 | 0.33678 | 0.27834 | 1.29046 |
| **SLC27A2** | 0.08598 | 4.80730 | 15.57724 | 0.79634 | 4.95659 | 0.00000 | 0.00000 | 0.17631 | 7.89646 | 0.00000 | 7.87429 | 0.00000 |
| **SLC27A3** | 0.00293 | 0.37243 | 0.51068 | 1.78315 | 1.09944 | 0.00000 | 0.38344 | 0.00000 | 0.61207 | 2.30494 | 0.29114 | 0.55035 |
| **SLC27A4** | 0.36936 | 17.69824 | 4.73240 | 4.78660 | 19.44488 | 0.00000 | 2.66076 | 0.00000 | 10.51368 | 23.23110 | 2.26050 | 9.36652 |
| **SLC27A5** | 0.18790 | 0.64058 | 0.09240 | 1.14496 | 0.00000 | 0.30517 | 1.28852 | 0.09425 | 1.74718 | 0.00000 | 0.91692 | 0.64142 |
| **SLC27A6** | 0.60270 | 0.15131 | 1.40870 | 0.00000 | 0.00000 | 0.09200 | 0.00000 | 4.14038 | 0.34291 | 0.04017 | 0.00000 | 0.00000 |
| **SLC28A1** | 0.00852 | 0.00857 | 0.07418 | 0.00678 | 0.29752 | 0.06495 | 0.00000 | 0.01089 | 0.00000 | 0.04501 | 0.01596 | 0.02927 |
| **SLC28A2** | 0.00242 | 0.01358 | 0.18316 | 0.02765 | 0.00000 | 0.02937 | 0.00753 | 0.03448 | 0.01826 | 0.03935 | 0.03494 | 0.16260 |
| **SLC28A3** | 1.27168 | 7.88433 | 0.00000 | 0.00000 | 21.06758 | 5.70619 | 4.14745 | 1.48517 | 20.31244 | 9.60788 | 20.78694 | 6.34926 |
| **SLC29A1** | 11.47721 | 28.96532 | 29.31693 | 6.84521 | 13.09097 | 6.37736 | 7.71387 | 4.15047 | 24.63500 | 0.00000 | 13.82832 | 7.96410 |
| **SLC29A2** | 0.15717 | 6.91073 | 4.64250 | 5.25051 | 13.21856 | 0.00000 | 1.56615 | 0.10092 | 16.30423 | 3.36008 | 8.74068 | 0.54877 |
| **SLC29A3** | 2.10134 | 2.49988 | 4.77735 | 1.18273 | 1.95858 | 0.00000 | 0.37984 | 0.29742 | 6.81140 | 0.00000 | 3.75596 | 2.51430 |
| **SLC29A4** | 3.11820 | 0.10405 | 0.01078 | 0.06662 | 0.00000 | 0.00000 | 0.15773 | 1.15802 | 0.11063 | 0.00000 | 0.05660 | 0.36070 |
| **SLC2A1** | 0.38830 | 45.63286 | 56.19080 | 17.82525 | 43.91962 | 29.57026 | 11.41848 | 0.22157 | 72.11373 | 7.95320 | 31.30467 | 6.00384 |
| **SLC2A1-AS1** | 0.00619 | 0.15466 | 0.53423 | 0.00235 | 0.13432 | 0.00416 | 0.00000 | 0.01112 | 0.03057 | 0.01019 | 0.07140 | 0.01320 |

| | | | | | | | | | | | | |
|---|---|---|---|---|---|---|---|---|---|---|---|---|
| **SLC2A10** | 0.08985 | 0.27103 | 0.03415 | 0.00000 | 0.00000 | 0.02705 | 0.00000 | 0.01550 | 0.00000 | 0.12695 | 0.03208 | 0.00871 |
| **SLC2A11** | 0.09833 | 3.81485 | 1.13111 | 2.88006 | 5.17075 | 3.74335 | 1.86224 | 0.00000 | 2.19784 | 1.00976 | 1.64381 | 0.62958 |
| **SLC2A12** | 0.03499 | 0.46566 | 0.00000 | 0.00000 | 0.03359 | 0.96662 | 0.58024 | 0.47111 | 0.33830 | 0.48867 | 0.32470 | 0.59812 |
| **SLC2A13** | 0.20249 | 0.00000 | 0.40012 | 0.51940 | 0.00000 | 0.94070 | 0.00000 | 0.22554 | 0.42961 | 1.26530 | 0.00042 | 0.00981 |
| **SLC2A14** | 0.35243 | 1.16647 | 1.55522 | 0.65233 | 0.00000 | 0.15297 | 0.00000 | 0.72910 | 0.72812 | 0.88563 | 1.30767 | 8.05694 |
| **SLC2A2** | 0.03704 | 0.02541 | 0.01584 | 0.14112 | 0.00000 | 0.04319 | 0.14097 | 0.00885 | 0.01219 | 0.01797 | 0.00761 | 0.01373 |
| **SLC2A3** | 0.00000 | 27.75383 | 32.75095 | 4.47709 | 21.14699 | 34.86219 | 17.06257 | 35.49215 | 67.24927 | 42.23355 | 30.54725 | 66.55774 |
| **SLC2A4** | 0.00000 | 0.74083 | 0.97010 | 0.75081 | 1.03899 | 2.15770 | 0.00083 | 0.73442 | 0.00000 | 1.54537 | 0.25042 | 1.14808 |
| **SLC2A4RG** | 0.80325 | 0.19747 | 0.54343 | 0.18146 | 0.49452 | 0.21286 | 0.00590 | 0.00000 | 0.24618 | 2.61572 | 0.00938 | 0.21295 |
| **SLC2A5** | 0.10276 | 0.15814 | 0.24726 | 0.07691 | 0.15706 | 0.00000 | 0.01722 | 0.50982 | 0.04614 | 0.00000 | 0.02674 | 2.03148 |
| **SLC2A6** | 0.00687 | 0.01262 | 0.07891 | 0.07183 | 0.06108 | 0.00000 | 0.02387 | 0.01958 | 0.03649 | 0.04913 | 0.03003 | 0.01627 |
| **SLC2A7** | 0.02203 | 0.01811 | 0.01986 | 0.02831 | 0.01793 | 0.00025 | 0.03291 | 0.02286 | 0.01278 | 0.01075 | 0.01682 | 0.04781 |
| **SLC2A8** | 1.57935 | 1.26277 | 2.17580 | 0.34766 | 2.98787 | 1.00459 | 1.66487 | 0.54572 | 6.27924 | 0.00886 | 0.00000 | 1.06130 |
| **SLC2A9** | 0.00585 | 0.22926 | 0.32134 | 0.11038 | 0.00131 | 0.12361 | 0.00000 | 0.01134 | 0.21423 | 0.11170 | 0.08576 | 0.05061 |
| **SLC30A1** | 2.04028 | 1.29466 | 6.65755 | 0.61774 | 2.91940 | 1.54501 | 0.00000 | 0.59552 | 0.00000 | 3.52839 | 0.00000 | 0.28871 |
| **SLC30A10** | 0.00218 | 0.00981 | 0.03002 | 0.02608 | 0.00212 | 0.04112 | 0.01004 | 0.11701 | 0.00799 | 0.03438 | 0.01829 | 0.00848 |
| **SLC30A2** | 0.00201 | 0.66763 | 0.07870 | 0.00000 | 9.35357 | 0.00008 | 0.00067 | 0.01427 | 0.00000 | 0.89157 | 1.53756 | 0.00020 |
| **SLC30A3** | 1.41129 | 0.08652 | 0.03535 | 0.00262 | 0.04303 | 0.10620 | 0.00000 | 0.32912 | 0.02466 | 0.00000 | 0.01810 | 0.05823 |
| **SLC30A4** | 0.88856 | 0.01536 | 0.01507 | 0.01007 | 0.11230 | 0.00648 | 0.02363 | 0.22157 | 0.01298 | 0.00289 | 0.02119 | 0.59445 |
| **SLC30A5** | 0.00000 | 5.50730 | 7.72802 | 1.77169 | 2.47566 | 11.85051 | 3.04543 | 0.67966 | 6.47471 | 10.11524 | 4.32428 | 2.85815 |
| **SLC30A6** | 1.72371 | 9.23770 | 5.08534 | 2.33489 | 1.80758 | 18.70521 | 0.00000 | 1.36783 | 8.18263 | 4.72152 | 3.31469 | 7.67323 |
| **SLC30A7** | 1.04408 | 4.86532 | 7.76801 | 1.47066 | 0.00155 | 21.86794 | 0.00000 | 9.98688 | 13.60395 | 7.14594 | 4.47820 | 13.85997 |
| **SLC30A8** | 1.14568 | 0.03603 | 0.03255 | 0.00000 | 0.04238 | 0.23801 | 0.00000 | 0.03618 | 0.00000 | 0.83542 | 0.09727 | 0.25455 |
| **SLC30A9** | 2.02467 | 4.85551 | 5.79653 | 4.26582 | 3.51615 | 14.87107 | 4.25364 | 1.32127 | 4.41759 | 0.00003 | 11.76292 | 2.64464 |
| **SLC31A1** | 1.03699 | 6.26701 | 12.53172 | 2.87747 | 10.40363 | 13.67368 | 2.22986 | 2.28802 | 22.18871 | 7.34031 | 13.01262 | 2.87449 |
| **SLC31A2** | 2.19055 | 1.93314 | 6.32263 | 0.64474 | 0.00000 | 2.32196 | 0.00004 | 1.44217 | 3.90539 | 0.00000 | 1.09801 | 2.29217 |
| **SLC32A1** | 0.00590 | 0.02103 | 0.01441 | 0.01384 | 0.01466 | 0.01577 | 0.01891 | 0.00873 | 0.01875 | 0.01197 | 0.01508 | 0.06402 |
| **SLC33A1** | 2.82853 | 3.15526 | 3.76315 | 2.59291 | 1.78495 | 9.85771 | 0.00000 | 0.72377 | 8.29838 | 0.75357 | 2.76228 | 6.47475 |
| **SLC34A1** | 0.00562 | 0.01434 | 0.06837 | 0.00843 | 0.01117 | 0.02277 | 0.00753 | 0.01211 | 0.00435 | 0.03696 | 0.01915 | 0.01451 |
| **SLC34A2** | 8.67030 | 12.83462 | 3.55752 | 0.00000 | 42.67944 | 0.00000 | 1.04080 | 43.87689 | 13.11802 | 0.91935 | 24.58193 | 6.53558 |
| **SLC34A3** | 0.00670 | 0.01776 | 0.01207 | 0.03633 | 0.01614 | 0.00034 | 0.02262 | 0.01032 | 0.06515 | 0.00000 | 0.01461 | 0.02368 |
| **SLC35A1** | 1.72539 | 3.97828 | 0.41999 | 5.27960 | 5.80130 | 10.51259 | 0.00000 | 1.59631 | 6.75868 | 0.00000 | 5.65377 | 0.47646 |
| **SLC35A2** | 0.26061 | 7.99056 | 5.67830 | 2.53737 | 5.42972 | 4.05236 | 0.84973 | 0.26166 | 20.23298 | 0.00000 | 5.82042 | 5.17341 |
| **SLC35A3** | 0.55783 | 1.44496 | 2.64436 | 0.00000 | 0.00000 | 7.81104 | 2.18842 | 0.10792 | 5.42927 | 4.85690 | 1.56140 | 0.00000 |
| **SLC35A4** | 3.92546 | 20.50099 | 21.50829 | 14.22203 | 9.88721 | 0.03354 | 21.34362 | 0.48087 | 24.30569 | 0.00000 | 19.69943 | 21.97971 |
| **SLC35A5** | 2.22746 | 9.53396 | 2.16687 | 8.79211 | 2.06112 | 12.46102 | 1.01234 | 7.46393 | 3.75053 | 1.50502 | 6.07241 | 15.44677 |
| **SLC35B1** | 4.65335 | 16.12889 | 12.51124 | 9.46819 | 11.76124 | 23.35458 | 0.00000 | 10.24352 | 22.42565 | 0.00000 | 10.54403 | 25.76240 |
| **SLC35B2** | 1.14031 | 10.86713 | 7.80047 | 5.39957 | 7.04492 | 0.00000 | 0.00000 | 0.00000 | 13.64435 | 0.00000 | 7.14381 | 5.58232 |
| **SLC35B3** | 0.00000 | 1.67689 | 2.92840 | 0.96037 | 0.63582 | 6.79707 | 1.19955 | 1.30179 | 3.17260 | 1.56604 | 4.03271 | 3.48522 |
| **SLC35B4** | 1.14025 | 7.59750 | 2.46777 | 4.38088 | 7.43758 | 15.00500 | 3.11319 | 0.23358 | 18.82400 | 6.09232 | 13.19457 | 2.32177 |
| **SLC35C1** | 5.53313 | 1.31191 | 3.07256 | 0.42507 | 3.51325 | 1.37501 | 5.56010 | 0.83873 | 16.15859 | 4.16960 | 6.84027 | 3.48555 |
| **SLC35C2** | 3.20106 | 10.78432 | 11.28901 | 3.47010 | 5.05450 | 0.00000 | 3.52717 | 0.23455 | 8.16535 | 0.00000 | 2.56917 | 10.41013 |
| **SLC35D1** | 8.86077 | 0.64401 | 3.00699 | 1.78923 | 0.00000 | 4.38844 | 3.99066 | 1.07818 | 4.85903 | 1.72471 | 2.31720 | 0.02582 |
| **SLC35D2** | 0.35244 | 0.51461 | 1.36828 | 1.93489 | 0.76497 | 2.33528 | 1.46047 | 0.00000 | 1.01138 | 2.63493 | 1.16758 | 0.04376 |
| **SLC35D3** | 0.04381 | 0.01173 | 0.01212 | 0.01791 | 0.00709 | 0.02177 | 0.01110 | 0.00939 | 0.01228 | 0.05310 | 0.01873 | 0.01785 |
| **SLC35E1** | 0.13662 | 1.49864 | 5.66236 | 0.95266 | 6.68057 | 2.88879 | 4.07356 | 2.01456 | 4.42561 | 16.97202 | 3.98771 | 3.76829 |
| **SLC35E2** | 0.07501 | 0.39632 | 0.46589 | 0.37553 | 0.13012 | 0.00000 | 0.55847 | 0.09810 | 0.58432 | 0.81724 | 0.23960 | 0.15773 |
| **SLC35E2B** | 0.46119 | 1.11007 | 1.03532 | 0.91137 | 0.69720 | 0.64039 | 0.05214 | 0.24317 | 1.96365 | 0.36392 | 1.25576 | 0.65678 |

| | | | | | | | | | | | | |
|---|---|---|---|---|---|---|---|---|---|---|---|---|
| SLC35E3 | 0.00000 | 2.08374 | 6.54722 | 3.24744 | 1.50066 | 4.22016 | 0.00000 | 0.13170 | 6.46471 | 1.37213 | 4.18632 | 1.80742 |
| SLC35E4 | 1.65787 | 15.72560 | 0.00000 | 11.89844 | 6.29470 | 0.00000 | 0.00000 | 5.05024 | 0.02479 | 5.54296 | 0.21001 | 22.58515 |
| SLC35F1 | 0.07358 | 0.50474 | 0.28848 | 0.00000 | 0.68124 | 0.00000 | 0.11747 | 0.10103 | 0.04327 | 0.05700 | 0.18278 | 0.02409 |
| SLC35F2 | 0.33927 | 18.30057 | 7.55160 | 7.12642 | 0.00000 | 18.42696 | 0.00000 | 21.56003 | 28.32200 | 11.33027 | 13.97351 | 39.67568 |
| SLC35F3 | 0.00208 | 0.05350 | 0.07558 | 0.00550 | 0.01102 | 0.01123 | 0.00824 | 0.80969 | 0.00841 | 0.03227 | 0.01588 | 0.52115 |
| SLC35F4 | 0.68651 | 0.03819 | 0.02698 | 0.00653 | 0.02640 | 0.00716 | 0.01033 | 0.18728 | 0.04053 | 0.00727 | 0.02030 | 0.02771 |
| SLC35F5 | 0.72051 | 5.87001 | 5.71537 | 1.38875 | 0.50682 | 11.04535 | 0.00000 | 3.82382 | 14.24966 | 0.75097 | 11.92769 | 3.73753 |
| SLC35F6 | 1.13060 | 23.53311 | 13.57078 | 27.77270 | 7.93491 | 13.39875 | 1.66299 | 0.05930 | 13.31080 | 4.25800 | 13.06217 | 15.52358 |
| SLC35G1 | 5.62238 | 2.30313 | 2.48801 | 0.53258 | 0.46184 | 2.82704 | 0.00000 | 0.56433 | 5.79544 | 0.00002 | 2.94160 | 3.64869 |
| SLC35G2 | 1.10074 | 0.06058 | 0.30922 | 0.00235 | 0.00000 | 0.06034 | 0.07195 | 0.00000 | 0.05773 | 0.02335 | 0.00000 | 0.48691 |
| SLC35G3 | 0.00641 | 0.19686 | 0.00957 | 0.57665 | 0.00000 | 0.00000 | 0.00497 | 0.00069 | 0.00000 | 0.27110 | 0.02026 | 1.08917 |
| SLC35G4P | 0.00206 | 0.01506 | 0.02427 | 0.01158 | 0.05052 | 0.00975 | 0.02249 | 0.10597 | 0.01225 | 0.00451 | 0.01367 | 0.06934 |
| SLC35G5 | 0.00303 | 0.01207 | 0.02817 | 0.07250 | 0.04470 | 0.00000 | 0.03159 | 0.10570 | 0.00775 | 0.01640 | 0.01641 | 0.26710 |
| SLC35G6 | 0.00906 | 0.01916 | 0.02477 | 0.02709 | 0.02228 | 0.01098 | 0.03754 | 0.01142 | 0.03144 | 0.00516 | 0.01374 | 0.03608 |
| SLC36A1 | 0.21005 | 0.33281 | 2.07520 | 2.42437 | 1.14995 | 2.14000 | 2.22561 | 1.15889 | 7.15851 | 6.29379 | 1.60948 | 1.21863 |
| SLC36A2 | 0.09472 | 0.00000 | 0.05754 | 0.00000 | 0.85644 | 0.00000 | 0.57609 | 1.66648 | 0.29890 | 2.27046 | 0.22733 | 0.35997 |
| SLC36A3 | 0.00505 | 0.00486 | 0.00853 | 0.01892 | 0.03624 | 0.04340 | 0.01307 | 0.00978 | 0.04614 | 0.05765 | 0.07435 | 0.01794 |
| SLC36A4 | 0.00000 | 0.45819 | 6.58386 | 0.00000 | 0.00000 | 0.60501 | 3.57149 | 0.00420 | 0.61191 | 1.21434 | 0.00000 | 0.47495 |
| SLC37A1 | 1.54570 | 3.21006 | 0.65918 | 0.15047 | 2.20898 | 0.73873 | 0.00000 | 0.13633 | 1.02376 | 0.00000 | 0.02734 | 0.00000 |
| SLC37A2 | 0.11260 | 0.01089 | 0.08002 | 0.01288 | 0.00887 | 0.02747 | 0.03608 | 0.32651 | 0.00179 | 0.03211 | 0.01893 | 0.23320 |
| SLC37A3 | 0.06165 | 17.47445 | 9.25955 | 10.34438 | 1.76882 | 20.37489 | 0.00000 | 2.95483 | 22.62411 | 6.12182 | 5.24414 | 10.37362 |
| SLC37A4 | 0.17466 | 18.75346 | 7.87536 | 19.62956 | 7.08275 | 5.40433 | 1.63392 | 0.00000 | 12.22343 | 6.26925 | 8.89507 | 8.07627 |
| SLC38A1 | 0.00144 | 0.00000 | 42.11224 | 0.00000 | 2.03828 | 2.93297 | 35.14016 | 0.00000 | 51.73296 | 0.04427 | 16.56695 | 0.00000 |
| SLC38A10 | 0.00000 | 2.95911 | 3.48293 | 1.15517 | 3.98703 | 0.00000 | 0.00000 | 0.15975 | 3.66666 | 7.56410 | 1.32356 | 0.56719 |
| SLC38A11 | 0.00576 | 0.01396 | 0.02175 | 0.01004 | 0.02261 | 0.02675 | 0.01414 | 0.01195 | 0.01609 | 0.01528 | 0.01360 | 0.01554 |
| SLC38A2 | 0.66963 | 0.00000 | 27.20996 | 13.79611 | 0.58222 | 15.16796 | 11.37782 | 1.65897 | 46.93801 | 15.66209 | 12.76375 | 3.97147 |
| SLC38A3 | 0.00318 | 0.53094 | 2.22707 | 0.00531 | 0.00926 | 0.00000 | 0.00000 | 0.13955 | 0.00000 | 0.00395 | 0.09929 | 1.83698 |
| SLC38A4 | 0.13741 | 0.00751 | 0.34208 | 0.03266 | 0.00000 | 0.04664 | 0.00000 | 0.00881 | 0.00000 | 0.03230 | 0.02295 | 0.01088 |
| SLC38A5 | 0.00476 | 0.08343 | 0.03956 | 0.09573 | 0.28818 | 0.00000 | 0.00000 | 0.01013 | 0.02319 | 0.27139 | 0.12186 | 0.01471 |
| SLC38A6 | 0.37691 | 0.34577 | 5.54556 | 0.30469 | 0.85316 | 1.42396 | 0.00000 | 1.54181 | 2.09606 | 0.02212 | 1.06240 | 1.47308 |
| SLC38A7 | 3.00684 | 8.87506 | 3.84022 | 4.30928 | 4.66234 | 0.00000 | 0.00000 | 0.07807 | 11.56461 | 7.92489 | 5.83270 | 2.65356 |
| SLC38A8 | 6.02973 | 0.03189 | 0.14362 | 0.06603 | 0.00868 | 0.00000 | 0.00000 | 2.41941 | 0.15340 | 0.00000 | 0.00000 | 0.15243 |
| SLC38A9 | 1.56187 | 4.62168 | 5.41677 | 4.22153 | 5.99721 | 12.86803 | 0.00000 | 0.01855 | 12.12591 | 0.85432 | 6.31165 | 0.62326 |
| SLC39A1 | 1.34916 | 28.68330 | 28.71691 | 18.82428 | 21.15242 | 7.95342 | 10.73467 | 2.77660 | 55.99797 | 0.00000 | 27.88354 | 25.73395 |
| SLC39A10 | 8.27676 | 3.60595 | 2.92776 | 20.74007 | 0.70176 | 14.89504 | 0.00000 | 7.90432 | 10.65190 | 9.43779 | 0.91635 | 9.18785 |
| SLC39A11 | 0.81571 | 2.75964 | 0.89164 | 1.91068 | 4.10453 | 1.01212 | 0.00000 | 0.10629 | 3.79938 | 0.22953 | 4.15291 | 1.62560 |
| SLC39A12 | 0.28596 | 0.00985 | 0.03176 | 0.00594 | 0.00712 | 0.01762 | 0.00000 | 0.04805 | 0.05727 | 0.09357 | 0.00579 | 0.01314 |
| SLC39A13 | 0.25497 | 0.77897 | 1.35050 | 1.78923 | 0.53220 | 0.00000 | 0.16472 | 0.00000 | 1.04140 | 0.92630 | 0.80494 | 1.33401 |
| SLC39A14 | 4.19709 | 12.96865 | 9.84433 | 2.12761 | 4.77491 | 6.31085 | 3.67486 | 0.78496 | 45.41362 | 6.16322 | 14.84655 | 14.34708 |
| SLC39A2 | 0.09889 | 0.02508 | 0.13320 | 0.00241 | 0.05048 | 0.34906 | 0.00000 | 0.02343 | 0.18200 | 0.00000 | 0.00182 | 0.01761 |
| SLC39A3 | 0.98099 | 18.65586 | 8.81485 | 3.57139 | 4.60120 | 0.00000 | 0.10141 | 1.82781 | 18.80096 | 0.37985 | 9.85687 | 16.51357 |
| SLC39A4 | 2.36500 | 5.48717 | 3.17671 | 7.72050 | 13.08780 | 0.88408 | 2.86263 | 0.87392 | 5.81136 | 1.48917 | 4.62952 | 2.40213 |
| SLC39A5 | 0.00547 | 0.00844 | 0.07781 | 0.03686 | 0.00000 | 0.02260 | 0.10308 | 0.00921 | 0.00000 | 0.05207 | 0.01535 | 0.02861 |
| SLC39A6 | 8.70571 | 3.97689 | 5.03064 | 1.46794 | 0.00000 | 5.94713 | 0.00000 | 5.16449 | 17.60717 | 4.35281 | 2.29367 | 12.29514 |
| SLC39A7 | 0.57669 | 8.14033 | 13.72876 | 7.84269 | 5.97318 | 0.00000 | 4.43584 | 0.68471 | 16.06778 | 2.03567 | 16.50434 | 12.93607 |
| SLC39A8 | 0.03203 | 0.01182 | 0.90126 | 0.19903 | 0.00239 | 0.00000 | 0.00000 | 0.00915 | 0.25332 | 0.00972 | 0.01436 | 0.03830 |
| SLC39A9 | 0.00000 | 6.21173 | 12.09667 | 3.62421 | 6.95788 | 20.61062 | 5.73908 | 8.59410 | 19.80034 | 7.02402 | 6.48591 | 8.75452 |
| SLC3A1 | 7.62813 | 0.00000 | 0.14162 | 0.61602 | 0.00000 | 0.00000 | 0.00000 | 2.55133 | 0.00000 | 0.15164 | 0.06903 | 0.35684 |

| | | | | | | | | | | | | |
|---|---|---|---|---|---|---|---|---|---|---|---|---|
| SLC3A2 | 1.66399 | 45.61429 | 48.02931 | 24.52454 | 48.27021 | 36.71326 | 18.33547 | 33.66764 | 93.06560 | 13.83141 | 44.62001 | 54.62469 |
| SLC40A1 | 0.20080 | 0.24466 | 30.08957 | 0.53014 | 4.16043 | 3.96075 | 1.75923 | 0.33415 | 0.00000 | 0.00000 | 9.40331 | 0.00000 |
| SLC41A1 | 2.66355 | 1.45211 | 1.44085 | 1.53345 | 1.00310 | 0.00000 | 1.16962 | 0.27703 | 0.00000 | 2.19528 | 0.68677 | 8.98337 |
| SLC41A2 | 3.47324 | 0.50213 | 0.35959 | 0.01699 | 0.02272 | 1.72584 | 0.00000 | 0.89848 | 0.48934 | 0.00000 | 0.24324 | 0.62129 |
| SLC41A3 | 1.27002 | 2.03573 | 5.64289 | 3.26278 | 1.42182 | 0.00078 | 3.12615 | 0.00000 | 7.32633 | 3.65896 | 1.04087 | 0.77312 |
| SLC43A1 | 0.00712 | 0.10820 | 0.16800 | 0.30376 | 0.00000 | 0.00000 | 0.00000 | 0.15369 | 0.34101 | 0.12693 | 0.00001 | 0.38361 |
| SLC43A2 | 1.19088 | 2.01471 | 9.97814 | 0.00000 | 4.14027 | 0.00000 | 0.00000 | 1.01732 | 2.38941 | 4.59194 | 0.34627 | 12.51207 |
| SLC43A3 | 5.86403 | 7.17094 | 0.24161 | 0.00000 | 0.32437 | 0.79139 | 0.00000 | 1.39027 | 0.31317 | 0.00000 | 0.29225 | 5.18648 |
| SLC44A1 | 0.01136 | 0.83977 | 1.68512 | 6.02406 | 0.78918 | 9.12124 | 0.81436 | 0.15552 | 0.00000 | 7.34988 | 3.00270 | 0.00000 |
| SLC44A2 | 0.03990 | 27.96511 | 22.00874 | 13.24277 | 35.83447 | 7.12124 | 3.54832 | 0.00000 | 30.32087 | 19.83302 | 12.02592 | 7.22515 |
| SLC44A3 | 0.11382 | 0.04388 | 0.00000 | 2.08634 | 0.11404 | 0.98284 | 0.00000 | 0.10168 | 0.62946 | 0.29404 | 1.50385 | 0.00000 |
| SLC44A4 | 0.03076 | 0.33247 | 0.10760 | 0.00000 | 0.00000 | 0.00129 | 0.00502 | 0.00251 | 0.09394 | 0.02310 | 0.02217 | 6.26063 |
| SLC44A5 | 0.12014 | 0.44603 | 0.12355 | 0.05787 | 0.13600 | 0.83978 | 0.00000 | 0.23225 | 0.29612 | 0.15398 | 0.21340 | 0.16859 |
| SLC45A1 | 0.26060 | 0.00774 | 0.04941 | 0.03563 | 0.01394 | 0.02968 | 0.00111 | 0.01434 | 0.05279 | 0.04431 | 0.01337 | 0.02383 |
| SLC45A2 | 0.00609 | 0.01484 | 0.02063 | 0.01172 | 0.01200 | 0.01480 | 0.03615 | 0.01057 | 0.00403 | 0.01772 | 0.03583 | 0.01630 |
| SLC45A3 | 2.20382 | 0.02132 | 0.05597 | 0.26218 | 0.00000 | 0.00000 | 0.19493 | 0.19496 | 0.00000 | 0.06263 | 0.07655 | 0.20145 |
| SLC45A4 | 0.78058 | 0.39065 | 3.36024 | 4.70729 | 1.87415 | 0.00000 | 1.37348 | 0.24559 | 0.07229 | 4.89709 | 0.10137 | 1.22623 |
| SLC46A1 | 0.00254 | 4.60640 | 0.42674 | 0.00000 | 3.86717 | 0.00000 | 0.51899 | 0.00000 | 0.50964 | 2.91819 | 1.74369 | 0.73649 |
| SLC46A2 | 0.00360 | 0.01764 | 0.00000 | 0.12138 | 0.00000 | 0.02073 | 0.00182 | 0.00830 | 0.12712 | 0.17646 | 0.05067 | 0.04312 |
| SLC46A3 | 0.00596 | 0.01731 | 0.15988 | 0.11020 | 0.01072 | 0.03729 | 0.00000 | 0.01070 | 0.02358 | 0.00300 | 0.05780 | 0.02365 |
| SLC47A1 | 0.00270 | 14.11019 | 0.00000 | 5.17050 | 0.00143 | 0.75136 | 0.00000 | 0.06257 | 0.00000 | 0.14311 | 0.00000 | 3.57184 |
| SLC47A2 | 0.00237 | 0.31987 | 0.01092 | 0.00953 | 0.03774 | 0.00000 | 0.03992 | 0.04097 | 0.01705 | 0.06189 | 0.00901 | 0.20132 |
| SLC48A1 | 1.39588 | 4.33931 | 0.00038 | 0.00000 | 5.64429 | 1.26410 | 0.87851 | 0.00000 | 0.45146 | 0.26610 | 3.34791 | 0.40994 |
| SLC4A1 | 0.11313 | 0.04573 | 0.02054 | 0.35795 | 0.00000 | 0.03387 | 0.01409 | 0.02626 | 0.00000 | 0.07331 | 0.02172 | 0.17917 |
| SLC4A10 | 0.00759 | 0.01140 | 0.05420 | 0.02564 | 0.00585 | 0.03025 | 0.00718 | 0.01057 | 0.00000 | 0.04656 | 0.03426 | 0.01416 |
| SLC4A11 | 2.58373 | 24.75017 | 0.00000 | 7.34455 | 33.68189 | 0.00000 | 0.00000 | 0.00000 | 8.88614 | 20.53282 | 7.45656 | 15.14089 |
| SLC4A1AP | 0.75314 | 12.58956 | 5.09989 | 5.26062 | 3.64926 | 13.60059 | 7.82112 | 8.51127 | 5.78350 | 3.98361 | 7.73684 | 12.35320 |
| SLC4A2 | 0.36329 | 10.20370 | 15.71356 | 5.46476 | 21.77623 | 0.00000 | 9.24452 | 0.11532 | 14.52645 | 21.28047 | 5.18853 | 2.95106 |
| SLC4A3 | 0.13327 | 0.22840 | 0.08922 | 0.14978 | 0.07673 | 0.00000 | 0.00000 | 0.00559 | 0.03898 | 0.45291 | 0.00000 | 0.02135 |
| SLC4A4 | 0.28155 | 0.01466 | 0.03719 | 0.00142 | 0.05244 | 0.00563 | 0.05161 | 0.00214 | 0.01342 | 0.27295 | 0.00120 | 0.09233 |
| SLC4A5 | 0.55771 | 0.51954 | 0.74700 | 0.76970 | 0.00000 | 4.00704 | 13.01416 | 0.19391 | 4.69400 | 4.60954 | 1.96695 | 0.00000 |
| SLC4A7 | 13.41249 | 0.00000 | 1.57144 | 0.00000 | 0.66345 | 3.75247 | 1.01293 | 2.00731 | 1.13314 | 5.08046 | 0.23415 | 0.54845 |
| SLC4A8 | 0.23534 | 2.62307 | 0.06723 | 13.47461 | 1.47023 | 1.28428 | 0.00000 | 0.00010 | 0.20950 | 0.50658 | 0.06142 | 2.74398 |
| SLC4A9 | 0.01086 | 0.01751 | 0.07688 | 0.02122 | 0.01348 | 0.00911 | 0.00886 | 0.01189 | 0.04180 | 0.02594 | 0.01449 | 0.01683 |
| SLC50A1 | 0.85628 | 2.97127 | 5.24069 | 1.13758 | 0.24078 | 5.64650 | 2.05812 | 3.16313 | 4.20458 | 1.31169 | 5.36858 | 4.18601 |
| SLC51A | 0.35417 | 0.00005 | 0.04695 | 0.00000 | 0.00000 | 0.06887 | 0.10591 | 0.38206 | 0.00000 | 0.39618 | 0.00000 | 0.05909 |
| SLC51B | 0.04701 | 0.01437 | 0.01372 | 0.01209 | 0.01357 | 0.02444 | 0.01733 | 0.00637 | 0.01837 | 0.01001 | 0.01428 | 0.02645 |
| SLC52A1 | 0.36576 | 3.64444 | 0.46497 | 4.88254 | 1.53751 | 0.00000 | 0.00000 | 0.58626 | 3.83094 | 3.82845 | 1.41910 | 5.05955 |
| SLC52A2 | 1.28717 | 14.37690 | 12.93767 | 6.33539 | 14.47692 | 0.00000 | 4.14276 | 0.16522 | 14.83635 | 15.64761 | 3.41972 | 10.75697 |
| SLC52A3 | 0.01825 | 3.07526 | 1.47026 | 0.64001 | 3.22827 | 0.00000 | 0.00000 | 0.04152 | 1.46578 | 0.00000 | 2.48032 | 3.10184 |
| SLC5A1 | 0.00298 | 0.02338 | 0.01627 | 0.00620 | 0.03151 | 0.04109 | 0.05365 | 0.34983 | 0.02466 | 0.00000 | 0.01298 | 0.11774 |
| SLC5A10 | 0.00360 | 0.31967 | 1.53439 | 0.07617 | 1.39514 | 0.00000 | 1.02503 | 0.03351 | 0.55000 | 0.63098 | 0.39064 | 0.00000 |
| SLC5A11 | 4.26435 | 0.18053 | 0.50110 | 0.08191 | 0.15681 | 0.00000 | 0.10788 | 1.57039 | 0.00000 | 0.19300 | 0.09294 | 0.03200 |
| SLC5A12 | 0.33276 | 0.01350 | 0.00000 | 0.00000 | 0.00444 | 0.22045 | 0.00000 | 0.01091 | 0.02174 | 0.10626 | 0.00233 | 0.02889 |
| SLC5A3 | 1.96867 | 0.48066 | 2.00544 | 0.00000 | 0.00000 | 8.73751 | 0.00000 | 0.22967 | 6.52142 | 13.25688 | 2.37704 | 0.84252 |
| SLC5A4 | 0.07202 | 0.01153 | 0.01414 | 0.01052 | 0.00947 | 0.02763 | 0.01309 | 0.03689 | 0.02924 | 0.02016 | 0.01214 | 0.02465 |
| SLC5A5 | 0.00000 | 13.41722 | 0.12881 | 5.24038 | 1.01415 | 1.95056 | 0.69884 | 6.27259 | 0.11576 | 2.22093 | 0.32055 | 10.25020 |
| SLC5A6 | 2.09037 | 17.40690 | 32.37814 | 16.66319 | 32.01118 | 3.34530 | 15.00505 | 2.86505 | 69.93371 | 7.87149 | 29.56788 | 18.84014 |

| | | | | | | | | | | | | |
|---|---|---|---|---|---|---|---|---|---|---|---|---|
| **SLC5A7** | 0.00553 | 0.01618 | 0.05173 | 0.02372 | 0.01283 | 0.00000 | 0.07288 | 0.01024 | 0.00000 | 0.04482 | 0.01543 | 0.02319 |
| **SLC5A9** | 0.27638 | 0.01687 | 0.04737 | 0.06510 | 0.00000 | 0.03307 | 0.00000 | 0.12121 | 0.04420 | 0.03544 | 0.02576 | 0.03139 |
| **SLC6A1** | 0.00560 | 0.01011 | 0.01398 | 0.02293 | 0.00466 | 0.01945 | 0.00674 | 0.01378 | 0.02085 | 0.07097 | 0.01415 | 0.02203 |
| **SLC6A10P** | 0.02084 | 0.01501 | 0.00973 | 0.02771 | 0.05062 | 0.00000 | 0.09038 | 0.00790 | 0.00000 | 0.10458 | 0.00682 | 0.04068 |
| **SLC6A11** | 0.47368 | 0.14728 | 0.01707 | 0.00836 | 0.03072 | 0.02110 | 0.06599 | 0.00113 | 0.03281 | 0.00000 | 0.01331 | 0.05305 |
| **SLC6A12** | 0.20436 | 0.01647 | 0.03679 | 0.00491 | 0.12103 | 0.02931 | 0.00671 | 0.00000 | 0.01633 | 0.04354 | 0.00880 | 0.09287 |
| **SLC6A13** | 0.51804 | 0.00743 | 0.22089 | 0.09051 | 0.00213 | 0.06605 | 0.08354 | 0.23061 | 0.00000 | 0.07873 | 0.02808 | 0.12082 |
| **SLC6A14** | 0.00569 | 0.01697 | 0.01274 | 0.01057 | 0.02721 | 0.00796 | 0.04035 | 0.00984 | 0.01882 | 0.03329 | 0.01126 | 0.02667 |
| **SLC6A15** | 0.00610 | 0.03657 | 1.60212 | 0.00232 | 0.00000 | 0.04800 | 0.00000 | 0.01133 | 0.00000 | 0.07358 | 0.02543 | 0.06493 |
| **SLC6A16** | 1.26579 | 0.25676 | 0.22920 | 0.01510 | 0.00000 | 0.00000 | 0.12622 | 0.11093 | 0.58053 | 0.16933 | 0.00000 | 0.80604 |
| **SLC6A17** | 1.33435 | 0.00891 | 0.10388 | 0.00810 | 0.01131 | 0.00827 | 0.02422 | 0.00055 | 0.00000 | 0.02964 | 0.05290 | 0.01837 |
| **SLC6A18** | 0.03298 | 0.01769 | 0.01687 | 0.01295 | 0.01203 | 0.01162 | 0.02544 | 0.00843 | 0.00623 | 0.02842 | 0.04786 | 0.01582 |
| **SLC6A19** | 0.00000 | 17.81298 | 10.11932 | 8.92389 | 27.41769 | 21.11808 | 0.00000 | 0.00000 | 22.91413 | 6.74417 | 0.43092 | 3.80169 |
| **SLC6A2** | 0.00618 | 0.00913 | 0.01298 | 0.03471 | 0.04444 | 0.00000 | 0.02269 | 0.00888 | 0.08580 | 0.00000 | 0.12951 | 0.05581 |
| **SLC6A20** | 1.19084 | 0.02814 | 0.01357 | 0.01585 | 0.00000 | 0.02430 | 0.00000 | 0.32122 | 0.03962 | 0.41461 | 0.00000 | 0.00000 |
| **SLC6A3** | 0.52902 | 0.25847 | 0.03805 | 0.37815 | 0.10971 | 0.36457 | 0.00000 | 2.62493 | 0.01493 | 0.12515 | 0.00000 | 0.08563 |
| **SLC6A4** | 0.00445 | 0.00448 | 0.37092 | 0.00000 | 0.51380 | 0.12081 | 0.00000 | 0.11031 | 0.00000 | 0.26738 | 0.27577 | 0.03628 |
| **SLC6A5** | 14.94778 | 0.11932 | 0.00000 | 0.01115 | 0.00240 | 0.04640 | 0.00000 | 2.95908 | 0.00208 | 0.02614 | 0.00576 | 2.07666 |
| **SLC6A6** | 3.97299 | 1.22398 | 10.17592 | 15.59719 | 4.83383 | 0.83739 | 1.37232 | 0.25504 | 16.51427 | 5.01124 | 3.20409 | 0.00000 |
| **SLC6A7** | 0.00468 | 0.00946 | 0.00379 | 0.02819 | 0.00975 | 0.01523 | 0.00357 | 0.01009 | 0.02730 | 0.13457 | 0.00659 | 0.01289 |
| **SLC6A8** | 0.74752 | 11.23192 | 0.77907 | 4.96044 | 10.97584 | 0.00000 | 2.07858 | 1.58795 | 0.00000 | 26.61536 | 1.87839 | 24.19526 |
| **SLC6A9** | 3.95596 | 0.19056 | 1.06713 | 0.00000 | 0.29200 | 0.00000 | 1.21220 | 0.03176 | 3.84723 | 1.63199 | 1.76434 | 0.14953 |
| **SLC7A1** | 0.07742 | 0.00000 | 0.00000 | 0.28362 | 0.02149 | 0.03298 | 3.12821 | 0.35199 | 0.55248 | 4.18937 | 0.00000 | 0.00000 |
| **SLC7A10** | 0.00598 | 0.01211 | 0.12091 | 0.00744 | 0.00891 | 0.03552 | 0.01370 | 0.01028 | 0.00685 | 0.02292 | 0.01461 | 0.01432 |
| **SLC7A11** | 2.72184 | 0.00000 | 13.70596 | 1.20644 | 0.00000 | 0.00814 | 19.31074 | 0.51133 | 14.30681 | 3.89939 | 0.00000 | 0.00000 |
| **SLC7A11-AS1** | 1.32403 | 0.00000 | 0.47371 | 1.86180 | 0.00596 | 0.20951 | 0.00000 | 0.03873 | 0.22907 | 0.27225 | 0.00000 | 0.15301 |
| **SLC7A14** | 0.11893 | 0.00736 | 0.08063 | 0.00588 | 0.04107 | 0.02955 | 0.01394 | 0.04354 | 0.00000 | 0.04043 | 0.01329 | 0.02615 |
| **SLC7A2** | 0.04908 | 0.00000 | 29.80890 | 0.00000 | 35.63642 | 18.32575 | 25.85286 | 1.22527 | 52.90112 | 24.57299 | 22.91805 | 0.83980 |
| **SLC7A3** | 0.08243 | 6.49984 | 0.00000 | 17.77528 | 12.46700 | 2.91600 | 0.00000 | 0.00000 | 0.00000 | 0.00000 | 5.65427 | 4.78122 |
| **SLC7A4** | 0.00934 | 0.93528 | 3.99745 | 1.00871 | 18.99599 | 0.00000 | 7.95535 | 0.00506 | 40.00146 | 0.00965 | 30.69990 | 0.00000 |
| **SLC7A5** | 0.59552 | 16.32421 | 26.60709 | 0.00000 | 45.62633 | 0.00000 | 27.51687 | 0.55593 | 53.44377 | 39.78968 | 21.94847 | 3.96530 |
| **SLC7A5P1** | 0.00206 | 0.17520 | 0.69324 | 0.01162 | 0.12410 | 0.00000 | 0.00000 | 0.10485 | 0.29665 | 0.00000 | 0.17558 | 0.01636 |
| **SLC7A5P2** | 0.00549 | 0.00000 | 0.66649 | 0.05298 | 1.06274 | 0.00000 | 0.05251 | 0.11567 | 0.00000 | 3.30659 | 0.33335 | 0.06897 |
| **SLC7A6** | 1.27312 | 3.82360 | 19.77491 | 7.49688 | 2.64060 | 5.82979 | 4.32470 | 6.41285 | 12.71089 | 8.42653 | 3.28246 | 4.08218 |
| **SLC7A6OS** | 0.52084 | 5.38469 | 2.66956 | 1.37964 | 2.18719 | 3.53295 | 1.74131 | 16.61620 | 5.88339 | 0.00000 | 2.23764 | 4.85520 |
| **SLC7A7** | 7.72863 | 12.22703 | 3.84123 | 11.93323 | 41.46252 | 24.45580 | 14.16194 | 3.52808 | 34.65646 | 14.31108 | 20.69233 | 5.05051 |
| **SLC7A8** | 17.71305 | 4.39040 | 49.97224 | 24.87822 | 51.70325 | 22.92538 | 33.57814 | 2.99567 | 66.96717 | 16.60279 | 51.63312 | 0.00000 |
| **SLC7A9** | 3.71152 | 0.00344 | 0.00315 | 0.00426 | 0.00000 | 0.00000 | 0.00278 | 1.78856 | 0.14452 | 0.23910 | 0.00000 | 0.02560 |
| **SLC8A1** | 5.97899 | 0.00000 | 0.00000 | 0.00000 | 0.72605 | 0.48674 | 0.00000 | 0.41052 | 0.47390 | 0.88049 | 0.28723 | 0.42585 |
| **SLC8A1-AS1** | 0.92194 | 0.01030 | 0.00038 | 0.00000 | 0.00227 | 0.07602 | 0.00000 | 0.04917 | 0.06547 | 0.20411 | 0.00305 | 0.00000 |
| **SLC8A2** | 4.23659 | 0.01441 | 0.07136 | 0.07558 | 0.07831 | 0.00151 | 0.03309 | 1.27438 | 0.00000 | 0.03892 | 0.01636 | 0.25555 |
| **SLC8A3** | 0.00697 | 0.01582 | 1.77121 | 0.00826 | 0.01948 | 0.00000 | 0.00000 | 0.00953 | 0.00000 | 0.01331 | 0.04779 | 0.02371 |
| **SLC8B1** | 0.00350 | 22.72076 | 2.84547 | 11.86528 | 19.82687 | 0.00000 | 0.03225 | 0.00000 | 2.60833 | 3.84704 | 7.58755 | 4.13134 |
| **SLC9A1** | 3.04916 | 6.40346 | 4.64128 | 5.77133 | 16.72938 | 0.00000 | 4.31949 | 2.57073 | 8.03841 | 22.77176 | 0.00001 | 3.42355 |
| **SLC9A2** | 0.01832 | 0.05117 | 0.11795 | 0.00247 | 0.00774 | 0.02924 | 0.00000 | 0.01426 | 0.85729 | 0.00290 | 0.00290 | 0.01650 |
| **SLC9A3** | 0.85610 | 0.42612 | 0.06828 | 0.00000 | 0.54512 | 0.00000 | 0.00000 | 0.14176 | 0.00000 | 0.74186 | 0.07662 | 0.07697 |
| **SLC9A3R1** | 3.35563 | 28.13613 | 16.31484 | 10.02608 | 23.86193 | 0.00000 | 26.64389 | 0.70641 | 42.63614 | 18.38200 | 33.49911 | 6.72677 |
| **SLC9A3R2** | 0.28546 | 2.66623 | 0.00000 | 0.61179 | 1.49353 | 0.00000 | 0.80155 | 0.25932 | 0.21644 | 1.61086 | 0.15714 | 3.04666 |

| | | | | | | | | | | | | |
|---|---|---|---|---|---|---|---|---|---|---|---|---|
| **SLC9A4** | 0.00575 | 0.01383 | 0.02342 | 0.05149 | 0.00281 | 0.00673 | 0.01098 | 0.01404 | 0.05622 | 0.02946 | 0.06159 | 0.01921 |
| **SLC9A5** | 0.00345 | 0.17287 | 0.13232 | 0.58506 | 0.00000 | 0.00000 | 0.00392 | 0.01013 | 0.40331 | 0.09154 | 0.00886 | 0.10737 |
| **SLC9A6** | 5.09713 | 1.89197 | 2.70276 | 0.00000 | 1.24323 | 2.91507 | 0.00000 | 0.89248 | 6.89427 | 2.48449 | 1.25726 | 2.15779 |
| **SLC9A7** | 0.96302 | 0.23028 | 0.21369 | 0.70070 | 0.27515 | 0.00000 | 0.27152 | 0.79824 | 0.41568 | 2.73684 | 0.15933 | 0.08106 |
| **SLC9A7P1** | 0.01641 | 0.01840 | 0.03974 | 0.02947 | 0.01243 | 0.02396 | 0.00957 | 0.01734 | 0.01388 | 0.06846 | 0.01531 | 0.02026 |
| **SLC9A8** | 0.61271 | 3.16229 | 1.87535 | 1.58543 | 2.77376 | 0.00000 | 0.00000 | 0.85148 | 2.32119 | 4.67058 | 0.03696 | 3.80434 |
| **SLC9A9** | 2.82687 | 0.01737 | 0.01752 | 0.00836 | 0.06368 | 0.00167 | 0.01818 | 0.25558 | 0.01408 | 0.00000 | 0.05848 | 0.00236 |
| **SLC9B1** | 0.05184 | 0.10380 | 0.43693 | 0.12127 | 0.13089 | 0.37672 | 0.30890 | 0.06903 | 0.48911 | 0.00000 | 0.28759 | 0.12922 |
| **SLC9B2** | 0.10699 | 0.13468 | 0.30487 | 0.07360 | 0.00000 | 0.00000 | 0.00000 | 0.05576 | 0.60742 | 0.17533 | 0.09034 | 0.06915 |
| **SLC9C1** | 0.02158 | 0.00000 | 0.01582 | 0.00000 | 0.05793 | 0.06762 | 0.00000 | 0.00779 | 0.11866 | 0.22559 | 0.00975 | 0.01505 |
| **SLC9C2** | 0.03885 | 0.02404 | 0.03211 | 0.05395 | 0.00000 | 0.03367 | 0.02397 | 0.01147 | 0.00131 | 0.03550 | 0.01658 | 0.01138 |
| **SLCO1A2** | 0.02467 | 0.08529 | 0.00000 | 3.43256 | 0.00000 | 1.04985 | 0.35523 | 0.00000 | 0.05863 | 0.74895 | 0.66650 | 1.04744 |
| **SLCO1B1** | 0.00778 | 0.11097 | 0.02947 | 0.08340 | 0.00000 | 0.11607 | 0.01987 | 0.08500 | 0.00000 | 0.26949 | 0.02288 | 0.08925 |
| **SLCO1B3** | 0.41064 | 0.01338 | 0.20899 | 0.05986 | 0.00196 | 0.14639 | 0.00000 | 0.12444 | 0.06959 | 0.00000 | 0.08909 | 0.08124 |
| **SLCO1B7** | 0.11387 | 0.00833 | 0.01956 | 0.01761 | 0.00688 | 0.02193 | 0.01454 | 0.00418 | 0.01272 | 0.04817 | 0.01350 | 0.07704 |
| **SLCO1C1** | 0.02574 | 0.00481 | 0.01343 | 0.13779 | 0.01153 | 0.00000 | 0.00900 | 0.01725 | 0.01559 | 0.26026 | 0.00746 | 0.02722 |
| **SLCO2A1** | 0.48962 | 0.18201 | 2.07981 | 8.35651 | 0.35887 | 0.00000 | 0.00000 | 0.06759 | 5.10035 | 0.00000 | 0.51145 | 0.00000 |
| **SLCO2B1** | 0.00997 | 0.98047 | 0.06679 | 0.00412 | 3.92938 | 0.00000 | 0.00867 | 0.00213 | 1.14736 | 0.00000 | 3.64392 | 0.43588 |
| **SLCO3A1** | 4.66574 | 0.07841 | 0.01037 | 0.00000 | 0.00000 | 0.01301 | 0.00000 | 0.91781 | 0.00000 | 0.18642 | 0.41363 | 0.74930 |
| **SLCO4A1** | 5.28902 | 0.05499 | 17.95332 | 0.00436 | 0.00000 | 0.00000 | 0.70030 | 0.01329 | 0.00000 | 0.01646 | 0.34147 | 0.17546 |
| **SLCO4A1-AS1** | 0.09378 | 0.01475 | 0.06306 | 0.00941 | 0.01582 | 0.01528 | 0.01526 | 0.00828 | 0.01931 | 0.01801 | 0.01324 | 0.01483 |
| **SLCO4C1** | 0.00000 | 5.54194 | 0.00000 | 0.00000 | 3.01064 | 7.89700 | 0.00000 | 0.25283 | 2.40574 | 6.72110 | 10.19889 | 7.49539 |
| **SLCO5A1** | 0.00557 | 0.00000 | 0.00000 | 1.43146 | 0.21337 | 0.03297 | 0.00418 | 0.00865 | 0.00000 | 0.08955 | 0.00978 | 0.01611 |
| **SLCO6A1** | 0.00523 | 0.00082 | 0.00823 | 0.01595 | 0.00530 | 0.12717 | 0.00000 | 0.00925 | 0.03988 | 0.03737 | 0.04397 | 0.00904 |
| **SLFN11** | 0.09739 | 0.01156 | 0.02215 | 0.00915 | 0.01210 | 0.02216 | 0.01565 | 0.00439 | 0.01135 | 0.02673 | 0.02114 | 0.02782 |
| **SLFN12** | 0.00642 | 0.05127 | 0.19396 | 0.00000 | 0.00000 | 0.30798 | 0.10835 | 0.00475 | 0.00000 | 0.15994 | 0.00000 | 0.06212 |
| **SLFN12L** | 0.00309 | 0.01016 | 0.01281 | 0.01070 | 0.01323 | 0.01961 | 0.01546 | 0.34070 | 0.01549 | 0.02397 | 0.01485 | 0.04098 |
| **SLFN13** | 0.59492 | 0.13611 | 0.00000 | 0.00000 | 0.00000 | 0.13284 | 0.00000 | 0.24885 | 0.00000 | 0.19171 | 0.01950 | 0.67388 |
| **SLFN14** | 0.02470 | 0.01621 | 0.01443 | 0.01312 | 0.01764 | 0.01067 | 0.02525 | 0.14049 | 0.01715 | 0.01069 | 0.01541 | 0.01248 |
| **SLFN5** | 0.00567 | 0.33921 | 0.23657 | 0.00753 | 0.00000 | 0.24756 | 0.00000 | 0.00431 | 0.00928 | 0.10880 | 0.04660 | 0.14173 |
| **SLFNL1** | 0.00483 | 0.04177 | 0.03785 | 0.01873 | 0.03445 | 0.04789 | 0.01678 | 0.00896 | 0.05316 | 0.13324 | 0.08275 | 0.03179 |
| **SLFNL1-AS1** | 0.01790 | 0.11496 | 0.00000 | 0.01702 | 0.02032 | 0.00000 | 0.00258 | 0.01512 | 0.24604 | 0.95331 | 0.17825 | 0.15258 |
| **SLIRP** | 1.73480 | 35.35927 | 29.06578 | 31.73646 | 41.74491 | 56.49038 | 59.50398 | 22.55001 | 30.74915 | 32.52537 | 40.34914 | 42.39068 |
| **SLIT1** | 0.05003 | 0.02361 | 0.09851 | 0.01076 | 0.10142 | 0.00000 | 0.00000 | 0.03531 | 0.02850 | 0.09149 | 0.03681 | 0.01577 |
| **SLIT2** | 0.00066 | 0.20488 | 0.00000 | 0.00000 | 0.16957 | 0.03079 | 0.00000 | 0.09474 | 0.52369 | 0.56074 | 0.03048 | 0.19041 |
| **SLIT3** | 6.49603 | 0.00000 | 0.00275 | 0.06889 | 0.00509 | 0.01762 | 0.00000 | 2.00606 | 0.02485 | 0.04111 | 0.00997 | 0.20492 |
| **SLITRK1** | 0.08313 | 0.00000 | 0.28233 | 0.00000 | 0.00000 | 0.65193 | 0.07861 | 0.09639 | 0.00038 | 1.34568 | 0.10297 | 0.11950 |
| **SLITRK2** | 0.01453 | 0.80260 | 0.00000 | 1.58859 | 0.00000 | 0.49424 | 0.14570 | 0.00376 | 0.00000 | 0.66241 | 0.03801 | 0.07450 |
| **SLITRK4** | 0.13831 | 0.04719 | 0.01184 | 0.03282 | 0.00000 | 0.28097 | 0.03072 | 0.04050 | 0.00000 | 0.28347 | 0.07938 | 0.05086 |
| **SLITRK5** | 0.00644 | 0.00056 | 0.09600 | 0.04362 | 0.10798 | 0.05408 | 0.00000 | 0.00979 | 0.00000 | 0.09759 | 0.00450 | 0.01044 |
| **SLITRK6** | 0.20093 | 0.01348 | 0.01323 | 0.01385 | 0.01339 | 0.01925 | 0.01590 | 0.04509 | 0.01951 | 0.01621 | 0.01425 | 0.01147 |
| **SLK** | 1.20342 | 0.00000 | 4.47824 | 5.66829 | 6.78401 | 14.56090 | 14.47470 | 0.00000 | 0.40261 | 26.09632 | 8.06439 | 0.61379 |
| **SLMAP** | 0.62591 | 3.82505 | 3.80484 | 3.48918 | 0.42010 | 14.20837 | 6.75120 | 2.55788 | 3.69033 | 14.80991 | 2.76216 | 4.09011 |
| **SLMO1** | 0.00275 | 0.00000 | 0.10882 | 0.08175 | 0.00000 | 0.05675 | 0.13431 | 0.00508 | 0.00000 | 0.32953 | 0.28412 | 0.23956 |
| **SLMO2** | 1.68318 | 10.17070 | 10.53949 | 5.95161 | 2.24862 | 36.27789 | 19.69739 | 0.08710 | 14.83785 | 8.79401 | 23.89726 | 0.53791 |
| **SLN** | 0.00843 | 0.27146 | 0.60726 | 0.00225 | 0.04578 | 0.01922 | 0.21336 | 0.00565 | 0.04086 | 0.00000 | 0.01329 | 0.26069 |
| **SLPI** | 0.00079 | 0.08994 | 0.19857 | 0.00000 | 4.31636 | 0.53501 | 1.90805 | 0.10450 | 0.03450 | 0.00000 | 1.46415 | 0.00000 |
| **SLTM** | 1.38668 | 4.40296 | 18.42452 | 9.59074 | 10.51823 | 24.18189 | 27.97736 | 5.45024 | 11.75454 | 32.44899 | 7.38260 | 15.09166 |

| | | | | | | | | | | | | |
|---|---|---|---|---|---|---|---|---|---|---|---|---|
| **SLU7** | 7.72873 | 9.13547 | 7.03198 | 4.27350 | 3.22416 | 23.24159 | 12.76203 | 14.93920 | 5.61562 | 8.58417 | 6.55841 | 21.29154 |
| **SLX4** | 0.87239 | 0.77519 | 1.00938 | 0.35301 | 1.73463 | 0.00000 | 0.36278 | 0.56212 | 1.17746 | 4.66994 | 0.54440 | 2.57093 |
| **SLX4IP** | 2.06678 | 0.65086 | 0.55395 | 1.17623 | 0.28723 | 0.84911 | 0.70588 | 0.12707 | 1.16110 | 2.04017 | 0.28544 | 2.21040 |
| **SMA5** | 0.00636 | 0.02080 | 0.01602 | 0.01343 | 0.01756 | 0.01024 | 0.01975 | 0.01053 | 0.02266 | 0.00909 | 0.01546 | 0.01848 |
| **SMAD1** | 4.24451 | 0.00000 | 0.46355 | 0.99024 | 0.87157 | 0.56400 | 4.33742 | 0.29916 | 2.03795 | 1.89801 | 4.73446 | 0.43491 |
| **SMAD1-AS2** | 0.00548 | 0.00869 | 0.02320 | 0.00502 | 0.00195 | 0.03415 | 0.00432 | 0.01027 | 0.01386 | 0.05831 | 0.07628 | 0.01312 |
| **SMAD2** | 6.41082 | 5.00174 | 6.95877 | 5.19331 | 3.41871 | 20.40678 | 8.02686 | 2.12878 | 6.02254 | 11.54409 | 6.17065 | 7.58552 |
| **SMAD3** | 1.97822 | 1.69989 | 0.86231 | 5.69421 | 0.00000 | 0.00000 | 0.00000 | 0.02239 | 0.00008 | 1.66930 | 0.00000 | 1.22304 |
| **SMAD4** | 0.58586 | 1.27094 | 2.21519 | 2.21958 | 3.10698 | 14.19353 | 11.45079 | 2.18520 | 0.75834 | 10.25903 | 2.72897 | 0.82825 |
| **SMAD5** | 0.13921 | 13.47989 | 7.02588 | 0.00000 | 13.38806 | 34.25003 | 15.64255 | 6.10114 | 8.29235 | 23.11577 | 17.43347 | 1.14143 |
| **SMAD5-AS1** | 0.00061 | 0.49160 | 0.32744 | 0.02626 | 1.07871 | 0.00000 | 0.12605 | 0.05267 | 0.11239 | 0.67979 | 0.55489 | 0.01728 |
| **SMAD6** | 0.10437 | 0.04670 | 0.53683 | 0.94980 | 0.29460 | 0.00000 | 0.00000 | 0.00674 | 0.00000 | 0.04362 | 0.01168 | 0.00866 |
| **SMAD7** | 0.00049 | 0.00000 | 4.93332 | 5.44648 | 5.55029 | 0.00000 | 3.19350 | 0.01573 | 0.34530 | 4.10915 | 0.58429 | 0.00000 |
| **SMAD9** | 0.47039 | 0.48979 | 1.95170 | 2.65375 | 1.18364 | 0.82029 | 0.00000 | 0.03338 | 0.00000 | 1.93364 | 1.29216 | 0.59648 |
| **SMAGP** | 0.02798 | 0.00000 | 22.10533 | 0.00000 | 15.89041 | 6.82599 | 15.74236 | 0.44973 | 21.97279 | 1.56456 | 14.46618 | 0.00000 |
| **SMAP1** | 1.53053 | 11.34661 | 6.44307 | 3.66302 | 11.81914 | 20.39469 | 12.17171 | 0.76375 | 5.93059 | 18.82682 | 8.74386 | 5.44443 |
| **SMAP2** | 0.56385 | 0.89988 | 1.33215 | 0.72303 | 0.75424 | 1.03653 | 3.42370 | 0.29729 | 0.00000 | 2.95863 | 1.03867 | 0.53667 |
| **SMARCA1** | 0.00207 | 0.00000 | 0.28641 | 0.90542 | 0.00000 | 0.56451 | 0.55855 | 0.17838 | 0.73320 | 0.00000 | 0.62917 | 0.00000 |
| **SMARCA2** | 9.01244 | 0.00000 | 19.03095 | 0.31510 | 0.00001 | 1.87013 | 13.69930 | 3.02168 | 2.52285 | 6.10084 | 0.97115 | 1.18188 |
| **SMARCA4** | 2.93009 | 8.61188 | 15.22911 | 12.18094 | 18.57919 | 0.00000 | 27.46255 | 2.97924 | 9.67814 | 34.75136 | 12.03512 | 8.80898 |
| **SMARCA5** | 2.82645 | 10.28488 | 15.91180 | 8.22820 | 6.55343 | 30.56433 | 19.88503 | 13.22738 | 14.86366 | 12.18299 | 17.12108 | 13.26609 |
| **SMARCA5-AS1** | 0.00437 | 0.01128 | 0.04135 | 0.01085 | 0.00032 | 0.01132 | 0.06871 | 0.08807 | 0.03073 | 0.06481 | 0.02540 | 0.04787 |
| **SMARCAD1** | 0.58139 | 6.18503 | 8.25369 | 6.24653 | 0.00000 | 20.87833 | 11.56874 | 3.15788 | 14.42330 | 14.79050 | 7.38921 | 10.14041 |
| **SMARCAL1** | 0.07533 | 3.38332 | 2.72513 | 2.55065 | 1.64648 | 1.92054 | 5.42108 | 0.02870 | 4.22489 | 0.86526 | 6.19291 | 2.41623 |
| **SMARCB1** | 9.87254 | 14.53349 | 10.47465 | 12.47605 | 11.88364 | 3.71185 | 16.54512 | 6.92944 | 20.42120 | 0.00000 | 12.69075 | 14.26280 |
| **SMARCC1** | 3.39299 | 4.02143 | 14.87674 | 7.95242 | 14.61477 | 19.50179 | 33.57878 | 1.22287 | 9.52321 | 44.04126 | 9.41420 | 6.62568 |
| **SMARCC2** | 8.24458 | 1.99201 | 3.36544 | 4.78309 | 2.87322 | 0.61165 | 3.04356 | 1.63504 | 4.27691 | 15.88170 | 2.34402 | 5.43839 |
| **SMARCD1** | 3.22307 | 2.07880 | 1.96671 | 1.95419 | 4.19990 | 0.00000 | 3.18833 | 0.71671 | 2.06842 | 10.08156 | 5.08915 | 2.46242 |
| **SMARCD2** | 0.03648 | 2.76042 | 6.88674 | 7.03525 | 14.69037 | 0.36552 | 19.05403 | 4.60000 | 4.06883 | 17.65954 | 5.91726 | 2.96902 |
| **SMARCD3** | 0.26215 | 0.05098 | 0.20801 | 0.46438 | 0.58624 | 0.00000 | 0.29414 | 0.00755 | 0.01643 | 0.48286 | 0.20392 | 0.08744 |
| **SMARCE1** | 1.72041 | 13.40301 | 6.77091 | 6.71908 | 13.64148 | 16.02436 | 18.85773 | 1.45835 | 9.49355 | 5.82698 | 10.45131 | 11.80744 |
| **SMC1A** | 6.32419 | 8.36499 | 9.55856 | 12.98276 | 10.06476 | 23.31136 | 8.56987 | 3.42049 | 14.57395 | 26.09811 | 15.72368 | 8.56380 |
| **SMC1B** | 1.40229 | 0.00000 | 0.00000 | 0.00000 | 0.00000 | 0.71032 | 0.00000 | 0.04790 | 0.05873 | 1.36716 | 0.43437 | 0.92103 |
| **SMC2** | 7.03294 | 6.50645 | 9.85928 | 6.12795 | 4.58337 | 30.96999 | 19.30063 | 8.69748 | 5.93547 | 19.84119 | 12.19856 | 14.66958 |
| **SMC3** | 10.06469 | 9.43606 | 13.10874 | 6.22955 | 6.69447 | 38.96909 | 19.72906 | 16.43780 | 13.53603 | 26.69483 | 14.82990 | 19.24824 |
| **SMC4** | 4.58661 | 12.94659 | 8.04027 | 6.53085 | 12.59514 | 48.38130 | 19.38237 | 16.75779 | 14.49306 | 40.94834 | 9.68061 | 19.64630 |
| **SMC5** | 2.71675 | 1.07169 | 6.62312 | 6.19553 | 0.00000 | 15.90039 | 7.33674 | 6.54261 | 7.04395 | 10.68972 | 5.33642 | 6.43809 |
| **SMC5-AS1** | 0.19725 | 0.04580 | 0.14436 | 0.00512 | 0.00000 | 0.00797 | 0.00000 | 0.21419 | 0.01279 | 0.04958 | 0.16169 | 0.03294 |
| **SMC6** | 2.08058 | 3.64182 | 5.58775 | 3.96538 | 3.30506 | 23.59739 | 7.69492 | 0.53929 | 5.47429 | 22.44983 | 6.36963 | 3.54940 |
| **SMCHD1** | 1.23072 | 0.35688 | 6.05040 | 1.78945 | 5.38823 | 5.90199 | 2.08221 | 2.07844 | 0.38937 | 17.13316 | 2.78588 | 2.55628 |
| **SMCO2** | 0.00680 | 0.01056 | 0.06768 | 0.01188 | 0.01947 | 0.00959 | 0.00809 | 0.00939 | 0.02618 | 0.01911 | 0.08237 | 0.03305 |
| **SMCO3** | 0.61617 | 0.07590 | 0.00000 | 0.04235 | 0.02770 | 0.16531 | 0.00000 | 0.12686 | 0.06611 | 0.00000 | 0.02578 | 0.05584 |
| **SMCO4** | 2.25385 | 3.35402 | 2.15547 | 3.82389 | 1.54850 | 2.00666 | 0.00000 | 0.01707 | 0.41817 | 0.00000 | 0.49279 | 0.61243 |
| **SMCR5** | 0.00649 | 0.01286 | 0.02501 | 0.01065 | 0.01242 | 0.01598 | 0.01588 | 0.01013 | 0.01395 | 0.03702 | 0.01515 | 0.02308 |
| **SMCR8** | 1.21942 | 0.08078 | 1.77174 | 1.14469 | 0.29456 | 0.00011 | 0.78011 | 0.00000 | 0.95125 | 3.44343 | 0.87340 | 2.96461 |
| **SMCR9** | 0.00864 | 0.01583 | 0.01939 | 0.01160 | 0.01430 | 0.02256 | 0.01863 | 0.01016 | 0.01975 | 0.01027 | 0.01873 | 0.01873 |
| **SMDT1** | 0.40220 | 0.31950 | 4.56466 | 1.99951 | 1.29803 | 0.32692 | 3.39361 | 0.50863 | 2.26703 | 0.82331 | 2.72173 | 1.05201 |
| **SMEK1** | 3.25329 | 5.33237 | 7.42753 | 0.00000 | 0.00000 | 25.38016 | 18.85860 | 16.46949 | 1.45134 | 33.14082 | 4.10856 | 13.53009 |

| | | | | | | | | | | | | |
|---|---|---|---|---|---|---|---|---|---|---|---|---|
| **SMEK2** | 2.60067 | 7.61692 | 7.90283 | 5.32415 | 2.29992 | 18.72910 | 7.15925 | 15.01886 | 6.89244 | 9.58991 | 4.61422 | 16.14334 |
| **SMEK3P** | 0.33476 | 0.33993 | 0.54978 | 0.00173 | 0.00000 | 0.44606 | 0.52176 | 0.11690 | 0.02835 | 0.00000 | 0.37123 | 0.16188 |
| **SMG1** | 4.55838 | 2.43766 | 5.26492 | 4.39985 | 5.30302 | 28.56763 | 7.37696 | 13.16108 | 13.10760 | 43.20726 | 4.40260 | 9.33625 |
| **SMG1P1** | 0.01603 | 0.01455 | 0.03233 | 0.03723 | 0.08703 | 0.03348 | 0.00219 | 0.05136 | 0.02763 | 0.04019 | 0.05205 | 0.04013 |
| **SMG1P2.uid-20** | 0.00144 | 0.02422 | 0.23166 | 0.09488 | 0.00037 | 0.34529 | 0.00000 | 0.16340 | 0.22194 | 0.98392 | 0.03917 | 0.06758 |
| **SMG1P3** | 0.00243 | 0.14308 | 0.17481 | 0.16555 | 0.16085 | 0.14136 | 0.00000 | 2.80749 | 0.23212 | 0.44887 | 0.00000 | 0.23909 |
| **SMG1P5** | 0.00213 | 0.05339 | 0.06507 | 0.04383 | 0.06377 | 0.00000 | 0.00000 | 0.36220 | 0.09133 | 0.19551 | 0.03871 | 0.02146 |
| **SMG1P7** | 0.00285 | 1.48918 | 0.11902 | 2.58973 | 0.00000 | 0.16605 | 0.00000 | 0.40692 | 0.05276 | 0.73519 | 0.00000 | 4.17589 |
| **SMG5** | 0.87275 | 12.55795 | 14.25769 | 7.61366 | 10.91288 | 0.00000 | 11.14368 | 3.97100 | 12.54333 | 16.76355 | 7.92104 | 7.38507 |
| **SMG6** | 6.62585 | 1.55840 | 2.83794 | 3.51090 | 5.68545 | 0.00000 | 3.94193 | 2.72487 | 3.44832 | 8.47872 | 1.05811 | 4.31425 |
| **SMG7** | 5.63218 | 3.76483 | 16.22601 | 6.25941 | 9.15046 | 6.95214 | 13.36756 | 3.95007 | 9.16107 | 16.17110 | 2.77325 | 6.52518 |
| **SMG7-AS1** | 0.00586 | 0.04373 | 0.22100 | 0.03738 | 0.00000 | 0.18878 | 0.17105 | 0.21874 | 0.12176 | 0.01866 | 0.03893 | 0.39147 |
| **SMG8** | 1.69363 | 8.86403 | 2.87495 | 4.98460 | 0.09367 | 16.13169 | 2.81061 | 2.24872 | 6.42466 | 0.38982 | 7.83738 | 11.61081 |
| **SMG9** | 0.82741 | 2.54112 | 7.28726 | 3.29832 | 3.88963 | 0.00000 | 8.09925 | 0.31355 | 1.96533 | 5.71180 | 3.44631 | 3.73872 |
| **SMIM1** | 0.04923 | 0.00000 | 0.09479 | 0.00000 | 0.24291 | 0.01091 | 0.12892 | 0.04433 | 0.49871 | 0.17252 | 6.45231 | 0.03773 |
| **SMIM10** | 0.07410 | 0.00000 | 0.00784 | 0.00000 | 0.20409 | 0.11873 | 0.00000 | 0.03931 | 0.00000 | 0.25008 | 0.66667 | 0.00187 |
| **SMIM11** | 0.13838 | 2.67203 | 2.27249 | 0.82630 | 0.00000 | 3.12284 | 4.71646 | 0.63867 | 1.65187 | 0.00001 | 4.04566 | 3.06723 |
| **SMIM12** | 0.00000 | 18.13899 | 7.73392 | 12.41669 | 18.88932 | 17.23552 | 7.86278 | 4.43864 | 18.92747 | 2.50178 | 13.26046 | 17.44481 |
| **SMIM13** | 0.74580 | 4.10958 | 7.43104 | 0.00000 | 0.00000 | 9.20885 | 8.08383 | 3.85465 | 0.52007 | 9.42515 | 1.14630 | 6.56438 |
| **SMIM14** | 2.40220 | 0.57453 | 2.31947 | 0.80825 | 0.56903 | 2.22200 | 0.62454 | 0.21185 | 0.00000 | 0.00000 | 1.10539 | 0.37992 |
| **SMIM15** | 2.21632 | 10.08107 | 12.92232 | 0.25848 | 2.05306 | 20.38158 | 12.73541 | 10.04132 | 11.04128 | 1.02797 | 9.41531 | 17.97104 |
| **SMIM17** | 0.04540 | 0.04134 | 0.01512 | 0.00637 | 0.06983 | 0.00000 | 0.02430 | 0.01223 | 0.06007 | 0.03877 | 0.02283 | 0.02263 |
| **SMIM18** | 0.00129 | 0.01251 | 0.03092 | 0.00000 | 0.00000 | 0.05124 | 0.02166 | 0.21731 | 0.03025 | 0.35080 | 0.20870 | 0.01782 |
| **SMIM19** | 0.87539 | 6.12855 | 2.33699 | 3.90705 | 5.51228 | 9.79068 | 0.00000 | 0.17695 | 1.78661 | 2.67997 | 3.92397 | 7.85882 |
| **SMIM2** | 0.00625 | 0.00505 | 0.07595 | 0.02071 | 0.00000 | 0.10340 | 0.04586 | 0.00911 | 0.02840 | 0.01372 | 0.00580 | 0.01565 |
| **SMIM2-AS1** | 0.15413 | 0.01407 | 0.09225 | 0.45878 | 0.00000 | 0.31444 | 0.00000 | 0.02056 | 0.08280 | 0.00000 | 0.01975 | 0.07105 |
| **SMIM20** | 0.61306 | 11.14270 | 7.08952 | 3.44491 | 9.43424 | 17.03753 | 0.00000 | 1.54231 | 10.45915 | 1.29240 | 3.81882 | 7.10840 |
| **SMIM22** | 0.00271 | 0.00000 | 0.48473 | 0.55102 | 0.49468 | 0.09378 | 0.56705 | 0.01390 | 1.67484 | 0.04761 | 1.68623 | 0.04740 |
| **SMIM24** | 0.00209 | 0.05827 | 0.02024 | 0.00000 | 0.04260 | 0.03485 | 0.01994 | 0.11906 | 0.01814 | 0.03026 | 0.01787 | 0.11019 |
| **SMIM3** | 0.00561 | 0.01044 | 0.10890 | 0.00000 | 0.00000 | 0.01196 | 0.42998 | 0.01028 | 0.82686 | 0.01970 | 0.00000 | 0.01350 |
| **SMIM4** | 0.00000 | 4.60655 | 5.30102 | 3.13941 | 3.51473 | 2.02220 | 11.16785 | 1.63170 | 4.67092 | 5.02476 | 9.04719 | 5.62813 |
| **SMIM5** | 0.00275 | 0.00000 | 0.20096 | 0.02435 | 0.39757 | 0.00000 | 0.00000 | 0.00573 | 0.00000 | 0.59396 | 0.06219 | 0.08275 |
| **SMIM6** | 0.22065 | 0.01894 | 0.04765 | 0.02687 | 0.02679 | 0.00028 | 0.01160 | 0.00236 | 0.04018 | 0.00000 | 0.01786 | 0.01547 |
| **SMIM7** | 0.65846 | 8.99433 | 9.87250 | 5.11004 | 7.47573 | 13.11463 | 4.62229 | 2.95143 | 14.25662 | 0.81571 | 5.28186 | 15.29086 |
| **SMIM8** | 0.48972 | 1.46680 | 1.10011 | 0.43589 | 0.00000 | 6.38539 | 0.00000 | 0.31564 | 0.99841 | 1.99922 | 2.09500 | 2.88048 |
| **SMIM9** | 0.00634 | 0.01321 | 0.22899 | 0.00659 | 0.01131 | 0.03165 | 0.02189 | 0.01023 | 0.00000 | 0.00862 | 0.02873 | 0.01553 |
| **SMKR1** | 4.24239 | 0.03688 | 0.05277 | 0.01668 | 0.03337 | 0.00000 | 0.01214 | 0.40747 | 0.04955 | 0.00000 | 0.01196 | 0.04274 |
| **SMLR1** | 0.03336 | 0.00208 | 0.20253 | 4.61667 | 0.40510 | 0.00000 | 0.00000 | 0.02947 | 0.00000 | 0.01951 | 0.02065 | 0.01450 |
| **SMNDC1** | 1.58757 | 4.44067 | 8.19709 | 2.44374 | 0.00000 | 19.34935 | 14.15219 | 5.07534 | 8.71209 | 13.11932 | 4.12486 | 7.63733 |
| **SMO** | 0.00151 | 0.09606 | 0.14514 | 1.16691 | 0.34580 | 0.00000 | 0.57011 | 0.02897 | 0.00000 | 0.50058 | 0.02282 | 0.00258 |
| **SMOC1** | 0.12057 | 0.00585 | 0.06786 | 0.10332 | 0.02742 | 0.06774 | 0.01162 | 0.07651 | 0.00000 | 0.01712 | 0.01388 | 0.01192 |
| **SMOC2** | 0.04723 | 0.17049 | 0.04869 | 0.00000 | 0.02889 | 0.00000 | 0.09628 | 0.04879 | 0.52311 | 0.05186 | 0.22929 | 0.03673 |
| **SMOX** | 1.18412 | 27.47603 | 6.78054 | 5.75710 | 17.80195 | 0.00000 | 14.33929 | 11.41353 | 16.54546 | 18.83183 | 11.32507 | 9.22100 |
| **SMPD1** | 0.02650 | 1.11231 | 2.22773 | 0.13102 | 13.97201 | 0.00000 | 0.39208 | 0.77898 | 1.89817 | 5.84522 | 1.71955 | 0.84492 |
| **SMPD2** | 0.33093 | 4.42500 | 4.88353 | 2.72514 | 4.37945 | 0.00000 | 2.05268 | 0.06158 | 5.63990 | 1.35234 | 3.00227 | 1.20106 |
| **SMPD3** | 5.72926 | 1.10923 | 0.86625 | 0.44031 | 1.94237 | 0.00000 | 0.05029 | 0.47114 | 0.15792 | 1.56615 | 0.48131 | 1.50617 |
| **SMPD4** | 1.38807 | 16.01145 | 11.16242 | 9.32995 | 6.12040 | 0.00000 | 5.74599 | 2.09737 | 21.47838 | 0.00000 | 9.06551 | 13.47973 |
| **SMPDL3A** | 0.00000 | 8.85977 | 1.95813 | 0.00000 | 11.94292 | 35.29887 | 5.81823 | 0.40400 | 32.74074 | 10.99062 | 12.62359 | 1.69155 |

| | | | | | | | | | | | | |
|---|---|---|---|---|---|---|---|---|---|---|---|---|
| **SMPDL3B** | 0.40706 | 0.02676 | 13.31546 | 0.00348 | 0.00000 | 0.34401 | 0.05916 | 0.17120 | 0.00000 | 0.00660 | 0.00422 | 0.05422 |
| **SMPX** | 0.10573 | 0.01970 | 0.02436 | 0.01059 | 0.01641 | 0.00784 | 0.01117 | 0.01655 | 0.07070 | 0.00508 | 0.00882 | 0.01929 |
| **SMS** | 5.24403 | 0.98825 | 4.32441 | 3.86034 | 2.54825 | 6.05940 | 10.93698 | 3.14514 | 6.17416 | 0.00001 | 6.64768 | 0.25359 |
| **SMTN** | 0.00000 | 2.33958 | 4.69392 | 3.81390 | 2.31606 | 0.00000 | 2.26780 | 0.10545 | 8.00546 | 2.63555 | 1.34164 | 0.74047 |
| **SMTNL1** | 0.00894 | 0.00745 | 0.07007 | 0.03656 | 0.00000 | 0.05270 | 0.00000 | 0.01052 | 0.01362 | 0.04382 | 0.01408 | 0.01209 |
| **SMTNL2** | 0.00250 | 0.00715 | 0.15363 | 0.29974 | 0.00000 | 0.15039 | 2.60404 | 0.04107 | 3.06140 | 0.00000 | 0.00000 | 0.00409 |
| **SMU1** | 0.78167 | 8.47808 | 12.18204 | 6.49849 | 0.00000 | 18.12519 | 17.31122 | 14.25656 | 8.11656 | 0.47489 | 12.74759 | 6.66582 |
| **SMUG1** | 0.00000 | 10.07376 | 9.37262 | 6.36636 | 8.84784 | 5.15746 | 6.41355 | 0.14619 | 2.51048 | 0.00000 | 11.08261 | 10.52443 |
| **SMURF1** | 7.75580 | 1.60884 | 3.42138 | 1.61807 | 2.83598 | 0.69739 | 2.92337 | 0.93580 | 3.55123 | 1.83812 | 0.00000 | 1.13905 |
| **SMURF2** | 5.41118 | 1.85051 | 1.64180 | 1.64281 | 1.13405 | 6.57809 | 0.92702 | 1.62051 | 0.25621 | 5.55653 | 0.17697 | 1.88899 |
| **SMYD1** | 0.24085 | 0.02620 | 0.03799 | 0.01362 | 0.03455 | 0.00080 | 0.03360 | 0.02308 | 0.01529 | 0.00000 | 0.02048 | 0.06529 |
| **SMYD2** | 0.35356 | 9.65594 | 10.17770 | 0.55141 | 2.81903 | 8.79288 | 9.16185 | 0.32169 | 10.11603 | 2.03374 | 12.71073 | 4.58846 |
| **SMYD3** | 2.49078 | 1.09912 | 2.05982 | 1.43078 | 1.34186 | 4.65478 | 2.12570 | 1.36102 | 0.01449 | 0.00002 | 3.54635 | 6.00551 |
| **SMYD4** | 1.31894 | 3.47935 | 1.94137 | 3.07465 | 2.85726 | 0.00040 | 0.65580 | 0.53049 | 2.85949 | 0.65701 | 2.64996 | 2.92625 |
| **SMYD5** | 0.89238 | 9.68325 | 8.42438 | 6.58186 | 7.70847 | 0.00000 | 9.08355 | 1.45963 | 15.03351 | 4.05843 | 2.86757 | 8.46203 |
| **SNAI1** | 0.57184 | 0.91301 | 8.39901 | 3.50109 | 2.87596 | 0.00000 | 2.45049 | 16.35679 | 2.51713 | 0.00000 | 0.00000 | 16.40494 |
| **SNAI2** | 0.00614 | 0.01280 | 0.05023 | 0.00756 | 0.00637 | 0.02830 | 0.03179 | 0.01038 | 0.02950 | 0.01114 | 0.03532 | 0.01587 |
| **SNAI3** | 2.48057 | 0.40231 | 0.15926 | 0.00684 | 0.31092 | 0.00000 | 0.17972 | 0.00000 | 0.00000 | 0.00000 | 0.05482 | 0.13423 |
| **SNAI3-AS1** | 0.13759 | 0.09156 | 0.18995 | 0.00770 | 0.61236 | 0.00000 | 0.05944 | 0.14872 | 0.28097 | 0.06274 | 0.21804 | 0.14021 |
| **SNAP23** | 1.38276 | 13.60615 | 14.91420 | 7.81832 | 9.79317 | 43.25920 | 16.32928 | 0.23316 | 17.83491 | 4.29922 | 18.07743 | 18.04581 |
| **SNAP25** | 0.00233 | 0.16495 | 0.38130 | 0.03738 | 0.01046 | 0.01439 | 0.03691 | 0.05076 | 0.06572 | 0.00000 | 0.00891 | 0.14423 |
| **SNAP25-AS1** | 0.00536 | 0.01146 | 0.01711 | 0.14875 | 0.01845 | 0.00000 | 0.02561 | 0.01571 | 0.02754 | 0.00396 | 0.01274 | 0.05265 |
| **SNAP29** | 1.31330 | 4.43805 | 6.25042 | 1.36488 | 8.73963 | 5.79976 | 3.28435 | 0.66845 | 4.19176 | 1.91726 | 9.63936 | 1.11189 |
| **SNAP47** | 2.47610 | 17.15584 | 7.13169 | 6.29361 | 5.91755 | 8.12240 | 13.33747 | 2.85755 | 14.61223 | 0.00000 | 9.56888 | 18.49639 |
| **SNAP91** | 0.71810 | 0.04395 | 0.14585 | 0.00000 | 0.00000 | 0.03109 | 0.04111 | 0.12033 | 0.04872 | 0.14423 | 0.00000 | 0.02043 |
| **SNAPC1** | 4.02728 | 1.54138 | 2.83630 | 0.64944 | 0.00000 | 6.84598 | 1.85956 | 11.58223 | 3.79368 | 2.13376 | 1.87325 | 10.04875 |
| **SNAPC2** | 2.98732 | 5.23992 | 0.71015 | 1.58671 | 2.81825 | 0.00000 | 0.00039 | 16.28321 | 5.35695 | 0.00000 | 0.88184 | 20.02119 |
| **SNAPC3** | 0.30584 | 6.95765 | 2.65660 | 11.03633 | 2.58737 | 10.94086 | 0.00000 | 6.72016 | 5.56969 | 0.97043 | 4.62061 | 6.15574 |
| **SNAPC4** | 1.04138 | 0.35354 | 2.04831 | 0.58619 | 3.40572 | 0.00000 | 3.23242 | 2.92603 | 4.16717 | 15.53674 | 1.31465 | 2.25505 |
| **SNAPC5** | 1.10861 | 5.69822 | 3.43094 | 1.82147 | 4.38475 | 5.78712 | 0.56441 | 2.60112 | 2.39376 | 0.00000 | 3.12433 | 1.53951 |
| **SNAPIN** | 3.56079 | 5.81494 | 1.55981 | 3.68927 | 2.77503 | 8.85086 | 4.50822 | 1.64598 | 4.15385 | 0.66638 | 4.92025 | 4.22529 |
| **SNAR-B1+SNAF** | 0.00498 | 0.01027 | 0.02283 | 0.01386 | 0.02520 | 0.03136 | 0.01051 | 0.01664 | 0.03451 | 0.02547 | 0.01830 | 0.01646 |
| **SNAR-C3** | 0.00434 | 0.01542 | 0.02017 | 0.02793 | 0.09835 | 0.00000 | 0.02626 | 0.01485 | 0.02923 | 0.08137 | 0.01296 | 0.03764 |
| **SNAR-D** | 0.00220 | 0.29464 | 0.04159 | 0.54630 | 1.23455 | 0.00000 | 0.04901 | 0.01424 | 0.00000 | 1.37184 | 0.00000 | 0.01846 |
| **SNAR-E** | 0.00000 | 1.70841 | 3.47096 | 5.05985 | 8.12781 | 33.74046 | 10.89514 | 3.30445 | 8.19396 | 38.31140 | 0.00000 | 16.99050 |
| **SNAR-F** | 0.00057 | 0.21583 | 0.28953 | 0.87342 | 0.27811 | 0.34395 | 0.79341 | 0.99263 | 1.12319 | 0.64836 | 0.17513 | 0.59457 |
| **SNAR-G1** | 0.00000 | 2.52815 | 0.07806 | 4.01412 | 10.75268 | 0.00000 | 0.16498 | 1.20899 | 0.68618 | 6.14636 | 0.00000 | 3.77563 |
| **SNAR-G2** | 0.00146 | 0.04997 | 0.06686 | 0.08796 | 0.13817 | 0.18515 | 0.00490 | 0.03872 | 0.11749 | 0.53715 | 0.04945 | 0.10670 |
| **SNAR-H** | 0.03916 | 1.66035 | 2.89937 | 3.84311 | 9.70130 | 0.00000 | 5.25098 | 0.00000 | 4.66955 | 7.34777 | 0.60909 | 0.77669 |
| **SNAR-I** | 0.00000 | 1.59458 | 1.24019 | 7.14974 | 4.82776 | 0.00000 | 2.64534 | 0.43646 | 0.58903 | 6.37083 | 0.30762 | 3.07610 |
| **SNCA** | 0.04169 | 0.00000 | 0.00000 | 0.19373 | 0.04570 | 0.09153 | 0.07476 | 0.02874 | 0.00000 | 0.07891 | 0.04797 | 0.06331 |
| **SNCAIP** | 0.03486 | 0.03120 | 0.01619 | 0.46880 | 0.29813 | 0.00000 | 0.03681 | 0.00365 | 0.00000 | 0.00000 | 0.32198 | 0.16546 |
| **SNCB** | 1.43601 | 0.01442 | 0.01171 | 0.01365 | 0.01203 | 0.02022 | 0.01616 | 0.00775 | 0.02094 | 0.01780 | 0.01628 | 0.05069 |
| **SNCG** | 0.00230 | 0.00979 | 0.09448 | 0.52856 | 0.00000 | 0.12384 | 0.11830 | 0.07457 | 0.01684 | 0.00000 | 0.00000 | 0.00742 |
| **SND1** | 2.40655 | 14.55503 | 20.47143 | 15.30212 | 15.18480 | 0.00000 | 25.22703 | 1.09182 | 28.16997 | 0.00000 | 18.02731 | 7.90760 |
| **SND1-IT1** | 0.06932 | 0.00000 | 1.00934 | 0.43505 | 1.72672 | 0.49828 | 0.34061 | 0.09218 | 1.90167 | 8.04176 | 0.74561 | 0.27987 |
| **SNED1** | 0.03329 | 0.00000 | 0.02408 | 0.09648 | 0.06855 | 0.13359 | 0.02650 | 0.11889 | 0.00000 | 0.45082 | 0.16491 | 0.17437 |
| **SNF8** | 0.00000 | 10.37745 | 11.06989 | 3.40743 | 14.45414 | 0.00000 | 11.94810 | 10.06453 | 6.54695 | 22.17579 | 5.40973 | 18.11868 |

| | | | | | | | | | | | | |
|---|---|---|---|---|---|---|---|---|---|---|---|---|
| **SNHG1** | 3.61091 | 10.41527 | 10.74047 | 14.82426 | 14.41538 | 26.46360 | 14.82286 | 11.82548 | 23.69664 | 39.17067 | 5.25469 | 19.89136 |
| **SNHG10** | 0.00000 | 2.52180 | 2.66729 | 0.45374 | 1.46143 | 3.55792 | 0.82070 | 5.16486 | 7.48349 | 2.41751 | 2.51072 | 8.51464 |
| **SNHG11** | 0.00208 | 4.87507 | 0.11976 | 5.58670 | 3.63195 | 2.99669 | 1.65454 | 7.10892 | 2.98414 | 0.00000 | 2.98965 | 4.21638 |
| **SNHG12** | 0.07163 | 6.08685 | 2.83205 | 5.74369 | 8.35598 | 12.54942 | 3.13762 | 3.38926 | 20.62569 | 26.02455 | 1.83005 | 6.81298 |
| **SNHG15** | 2.22891 | 27.28278 | 6.74851 | 15.54365 | 17.35253 | 15.63055 | 6.45050 | 26.80296 | 8.15414 | 13.63700 | 4.28391 | 43.19323 |
| **SNHG16** | 2.10745 | 21.90347 | 14.54019 | 10.67819 | 21.10906 | 27.50244 | 33.32504 | 11.87341 | 20.75082 | 21.88980 | 23.00498 | 29.03429 |
| **SNHG17** | 1.13709 | 14.87122 | 13.57942 | 11.87816 | 13.45334 | 0.00000 | 16.29042 | 14.26536 | 20.18048 | 4.68565 | 2.76899 | 13.23777 |
| **SNHG18** | 1.61675 | 0.00000 | 0.06859 | 0.00969 | 0.00000 | 0.00626 | 0.30702 | 0.00325 | 0.17736 | 0.00000 | 0.63476 | 0.00939 |
| **SNHG20** | 0.00270 | 0.37398 | 0.14758 | 0.59724 | 0.60183 | 0.73302 | 0.75522 | 1.32649 | 0.75236 | 0.00000 | 1.92434 | 2.37251 |
| **SNHG21** | 0.01392 | 0.68015 | 1.96077 | 0.19838 | 1.57998 | 2.67819 | 1.10757 | 5.18739 | 2.36047 | 0.00000 | 2.16691 | 3.30685 |
| **SNHG22** | 0.00506 | 0.01225 | 0.00448 | 0.08195 | 0.17193 | 0.00000 | 0.00252 | 0.01963 | 0.06166 | 0.06035 | 0.00467 | 0.01426 |
| **SNHG24** | 0.00319 | 0.01551 | 0.04175 | 0.11658 | 0.00355 | 0.00000 | 0.02654 | 0.03499 | 0.01140 | 0.01847 | 0.07548 | 0.01658 |
| **SNHG3** | 0.31730 | 7.13211 | 7.58281 | 9.50628 | 8.35153 | 18.42077 | 3.85776 | 10.39745 | 19.49916 | 22.26202 | 6.85253 | 12.87257 |
| **SNHG4.uid-7432** | 0.00157 | 0.14672 | 0.12280 | 0.02213 | 0.06361 | 0.00000 | 0.00000 | 0.05544 | 0.76533 | 0.50447 | 0.26894 | 0.29662 |
| **SNHG4.uid-7434** | 0.00000 | 0.06115 | 0.29335 | 0.21772 | 0.03139 | 0.27300 | 0.00548 | 0.07497 | 1.26067 | 1.49282 | 1.64403 | 0.78083 |
| **SNHG5** | 0.35905 | 0.00000 | 21.68030 | 6.47561 | 8.28174 | 34.47621 | 31.20379 | 7.17674 | 15.30703 | 32.36169 | 0.00000 | 15.02161 |
| **SNHG6** | 1.21771 | 6.93483 | 5.80542 | 4.68309 | 7.43212 | 15.28199 | 14.83055 | 5.20619 | 13.64768 | 21.47279 | 7.36953 | 13.54917 |
| **SNHG7** | 0.98776 | 0.95185 | 0.93199 | 1.89713 | 8.19527 | 0.00000 | 5.52213 | 1.55384 | 5.48515 | 19.13143 | 1.23283 | 4.50557 |
| **SNHG8** | 4.27261 | 8.38871 | 12.82708 | 11.34701 | 9.35493 | 17.97428 | 20.47514 | 8.59875 | 20.30409 | 17.81437 | 13.54419 | 16.94921 |
| **SNHG9** | 0.97602 | 1.78663 | 2.24931 | 0.51376 | 2.78117 | 0.00000 | 4.54290 | 4.07106 | 4.60200 | 7.43327 | 6.08324 | 7.21665 |
| **SNIP1** | 2.97882 | 3.94339 | 2.96693 | 1.65528 | 0.00774 | 6.38605 | 1.56988 | 13.34819 | 2.84720 | 5.87678 | 1.93775 | 4.13339 |
| **SNN** | 0.00859 | 0.65195 | 0.06919 | 0.41497 | 0.46675 | 0.00000 | 0.28845 | 0.46458 | 0.30946 | 1.50288 | 0.00000 | 0.83790 |
| **SNORA1** | 0.06262 | 0.03120 | 0.23482 | 0.02347 | 0.13866 | 0.05118 | 0.00000 | 0.06498 | 0.14421 | 0.12804 | 0.00000 | 0.08941 |
| **SNORA10** | 0.16013 | 0.08263 | 0.08558 | 0.22963 | 0.29270 | 0.00000 | 0.12499 | 0.09011 | 0.61836 | 2.04904 | 0.00006 | 0.13026 |
| **SNORA11B** | 0.00636 | 0.01100 | 0.03166 | 0.01779 | 0.02212 | 0.03036 | 0.01785 | 0.00983 | 0.01157 | 0.00702 | 0.01926 | 0.04312 |
| **SNORA11C** | 0.00436 | 0.01493 | 0.01408 | 0.01182 | 0.01578 | 0.02110 | 0.01812 | 0.04147 | 0.01782 | 0.01139 | 0.01489 | 0.01848 |
| **SNORA12** | 0.98338 | 0.00940 | 0.03457 | 0.00000 | 0.01772 | 0.13159 | 0.00000 | 0.33828 | 0.11397 | 0.22907 | 0.01824 | 0.48989 |
| **SNORA13** | 0.01638 | 0.01540 | 0.13912 | 0.11430 | 0.00000 | 0.03135 | 0.00000 | 0.08404 | 0.25940 | 0.59890 | 0.01216 | 0.08393 |
| **SNORA14A** | 0.03817 | 0.00790 | 0.05889 | 0.03042 | 0.04495 | 0.00855 | 0.05263 | 0.00990 | 0.00001 | 0.10036 | 0.00965 | 0.02382 |
| **SNORA14B** | 0.04708 | 0.09272 | 0.07525 | 0.00000 | 0.05035 | 0.05028 | 0.01738 | 0.02114 | 0.08338 | 0.19835 | 0.00000 | 0.15979 |
| **SNORA15** | 0.00598 | 0.01338 | 0.02325 | 0.02041 | 0.01941 | 0.02304 | 0.01299 | 0.01032 | 0.03250 | 0.02177 | 0.02330 | 0.01628 |
| **SNORA16A** | 0.00290 | 0.15636 | 0.06336 | 0.00000 | 0.27571 | 0.09693 | 0.03515 | 0.00724 | 0.38519 | 1.15316 | 0.00295 | 0.08076 |
| **SNORA16B** | 0.00522 | 0.01247 | 0.02623 | 0.00496 | 0.00974 | 0.04735 | 0.00000 | 0.00927 | 0.04627 | 0.05298 | 0.03751 | 0.02629 |
| **SNORA18** | 0.00000 | 0.08487 | 0.25712 | 0.06686 | 0.82515 | 1.02845 | 0.00000 | 0.09628 | 0.49937 | 2.11205 | 0.00000 | 0.30401 |
| **SNORA19** | 0.00316 | 0.03202 | 0.24267 | 0.00000 | 0.00021 | 0.27238 | 0.00000 | 0.47459 | 0.12424 | 1.00131 | 0.00000 | 0.19386 |
| **SNORA20** | 0.14096 | 0.04557 | 0.19566 | 0.10789 | 0.21156 | 0.00000 | 0.00000 | 0.04430 | 0.14383 | 0.32570 | 0.03222 | 0.15321 |
| **SNORA21** | 0.20348 | 0.06820 | 0.21325 | 0.15811 | 0.24149 | 0.00000 | 0.03480 | 0.26723 | 0.52701 | 0.70756 | 0.00950 | 0.33732 |
| **SNORA22** | 0.01300 | 0.01790 | 0.03000 | 0.02949 | 0.02900 | 0.00000 | 0.01168 | 0.01106 | 0.04514 | 0.03070 | 0.02282 | 0.02108 |
| **SNORA23** | 0.81819 | 0.00000 | 0.18207 | 0.05202 | 0.05081 | 0.00000 | 0.00000 | 0.46057 | 0.29859 | 0.13680 | 0.00000 | 0.25625 |
| **SNORA24** | 0.00000 | 0.16985 | 0.21520 | 0.26026 | 0.00000 | 0.80658 | 0.00000 | 0.13961 | 0.78773 | 1.16164 | 0.03745 | 0.42808 |
| **SNORA25** | 0.00065 | 0.00000 | 0.52712 | 0.17227 | 0.43953 | 0.43174 | 0.00000 | 0.13190 | 0.20880 | 2.78140 | 0.05192 | 0.15050 |
| **SNORA26** | 0.00241 | 0.13664 | 0.32902 | 0.00000 | 0.03318 | 0.22113 | 0.00000 | 0.09525 | 0.91948 | 0.58851 | 0.00659 | 0.26499 |
| **SNORA27** | 0.00286 | 0.23096 | 0.25913 | 0.18913 | 0.21181 | 0.35931 | 0.00000 | 0.37595 | 0.48318 | 0.48321 | 0.12162 | 0.33821 |
| **SNORA28** | 0.50482 | 0.02789 | 0.14435 | 0.00467 | 0.09210 | 0.00000 | 0.00172 | 0.07521 | 0.05056 | 0.44168 | 0.01016 | 0.09887 |
| **SNORA29** | 0.00154 | 0.00203 | 0.02439 | 0.11715 | 0.00000 | 0.10977 | 0.00000 | 0.05191 | 0.07603 | 0.20915 | 0.11627 | 0.03446 |
| **SNORA2A** | 0.15505 | 0.01726 | 0.05293 | 0.01586 | 0.02056 | 0.00000 | 0.00790 | 0.14700 | 0.05249 | 0.00000 | 0.01442 | 0.04872 |
| **SNORA2B** | 0.62554 | 0.00000 | 0.01980 | 0.00563 | 0.02482 | 0.15571 | 0.00000 | 0.37664 | 0.03862 | 0.12259 | 0.00001 | 0.09944 |
| **SNORA30** | 0.02882 | 0.02360 | 0.04207 | 0.00871 | 0.05204 | 0.00000 | 0.04647 | 0.01838 | 0.00158 | 0.03796 | 0.01519 | 0.02111 |

| | | | | | | | | | | | | |
|---|---|---|---|---|---|---|---|---|---|---|---|---|
| **SNORA31** | 0.00268 | 0.00000 | 0.09217 | 0.04671 | 0.12540 | 0.20502 | 0.00000 | 0.00638 | 0.04159 | 0.50252 | 0.00000 | 0.04365 |
| **SNORA32** | 0.01706 | 0.00000 | 0.17817 | 0.00000 | 0.19938 | 0.54583 | 0.00000 | 0.06127 | 0.11030 | 1.02617 | 0.00714 | 0.22907 |
| **SNORA33** | 0.14391 | 0.00000 | 0.22432 | 0.00254 | 0.32281 | 0.27078 | 0.10102 | 0.14396 | 0.49921 | 2.02139 | 0.00000 | 0.15265 |
| **SNORA34** | 0.00840 | 0.02104 | 0.01560 | 0.03276 | 0.02319 | 0.00000 | 0.02465 | 0.05138 | 0.06152 | 0.15520 | 0.01151 | 0.02627 |
| **SNORA36B** | 0.00807 | 0.01387 | 0.01559 | 0.01573 | 0.01434 | 0.02418 | 0.01764 | 0.01472 | 0.01600 | 0.01747 | 0.01512 | 0.01526 |
| **SNORA36C** | 0.00612 | 0.01458 | 0.01535 | 0.01115 | 0.01716 | 0.02553 | 0.01512 | 0.01043 | 0.01378 | 0.01657 | 0.02709 | 0.01542 |
| **SNORA37** | 0.00328 | 0.01062 | 0.06419 | 0.00000 | 0.04529 | 0.02115 | 0.04320 | 0.00881 | 0.00000 | 0.35382 | 0.03166 | 0.02035 |
| **SNORA38** | 0.00632 | 0.01600 | 0.01947 | 0.01069 | 0.01559 | 0.01992 | 0.01782 | 0.01040 | 0.02017 | 0.01796 | 0.01382 | 0.01590 |
| **SNORA38B** | 0.07473 | 0.01352 | 0.01605 | 0.01077 | 0.01251 | 0.01875 | 0.01653 | 0.01665 | 0.01511 | 0.02469 | 0.02671 | 0.01864 |
| **SNORA4** | 0.24281 | 0.16910 | 0.30830 | 0.17979 | 0.59726 | 0.43619 | 0.00000 | 0.29159 | 0.35565 | 0.28724 | 0.00000 | 0.37088 |
| **SNORA40** | 0.01697 | 0.07730 | 0.31962 | 0.07958 | 0.31855 | 0.46445 | 0.00000 | 0.07344 | 0.25980 | 1.06437 | 0.00000 | 0.30517 |
| **SNORA41** | 0.12269 | 0.09772 | 0.12513 | 0.21061 | 0.14058 | 0.00000 | 0.03228 | 0.05538 | 0.28390 | 0.05066 | 0.03845 | 0.11873 |
| **SNORA44** | 0.12913 | 0.33838 | 0.18624 | 0.03997 | 0.48278 | 0.00000 | 0.00000 | 0.01788 | 0.26170 | 1.17991 | 0.00000 | 0.15192 |
| **SNORA45A** | 0.10222 | 0.20671 | 0.39781 | 0.03266 | 0.26105 | 0.00000 | 0.00000 | 0.13902 | 0.37206 | 0.02730 | 0.19343 | 0.23329 |
| **SNORA45B** | 0.00298 | 0.11598 | 0.28785 | 0.21641 | 0.27175 | 0.00000 | 0.00000 | 0.09866 | 0.56348 | 1.19871 | 0.00356 | 0.18260 |
| **SNORA46** | 0.00281 | 0.00975 | 0.01037 | 0.00345 | 0.06140 | 0.00000 | 0.00310 | 0.04279 | 0.01753 | 0.28305 | 0.00452 | 0.01716 |
| **SNORA47** | 0.00563 | 0.01360 | 0.01230 | 0.01944 | 0.01260 | 0.02834 | 0.01603 | 0.01624 | 0.01678 | 0.01307 | 0.01619 | 0.01518 |
| **SNORA48** | 1.93177 | 0.22742 | 0.62837 | 0.02238 | 0.24094 | 0.00000 | 0.00000 | 1.05768 | 0.48696 | 0.90042 | 0.00000 | 0.38397 |
| **SNORA49** | 0.06602 | 0.01163 | 0.02152 | 0.05380 | 0.00899 | 0.04117 | 0.00400 | 0.01272 | 0.00956 | 0.05662 | 0.01103 | 0.06002 |
| **SNORA51** | 1.29411 | 0.06690 | 0.07806 | 0.00000 | 0.30217 | 0.00000 | 0.01673 | 1.03916 | 0.23532 | 0.30800 | 0.00000 | 0.29225 |
| **SNORA52** | 0.00290 | 0.08369 | 0.29442 | 0.08062 | 0.37740 | 0.00000 | 0.00333 | 0.14156 | 0.51207 | 0.15404 | 0.07402 | 0.16761 |
| **SNORA53** | 0.00430 | 0.02259 | 0.02603 | 0.00876 | 0.30108 | 0.00000 | 0.02681 | 0.06211 | 0.13124 | 0.45855 | 0.00000 | 0.03726 |
| **SNORA54** | 0.01901 | 0.01794 | 0.02359 | 0.00581 | 0.08043 | 0.00000 | 0.00380 | 0.02632 | 0.07506 | 0.03381 | 0.01107 | 0.02451 |
| **SNORA55** | 0.00230 | 0.02150 | 0.00143 | 0.06652 | 0.18918 | 0.00000 | 0.02906 | 0.02823 | 0.02967 | 0.59351 | 0.00966 | 0.02673 |
| **SNORA57** | 0.38013 | 0.25498 | 0.64382 | 0.94359 | 1.15787 | 0.00000 | 0.63804 | 0.93645 | 2.56275 | 3.02761 | 0.65091 | 0.55056 |
| **SNORA58** | 0.00275 | 0.05619 | 0.08170 | 0.05007 | 0.61275 | 0.00000 | 0.33100 | 0.16615 | 0.00001 | 1.64712 | 0.08873 | 0.47153 |
| **SNORA5A** | 0.17895 | 0.06183 | 0.08884 | 0.02606 | 0.06230 | 0.03653 | 0.00545 | 0.01992 | 0.07491 | 0.00000 | 0.02148 | 0.05779 |
| **SNORA5B** | 0.01613 | 0.02669 | 0.01366 | 0.01428 | 0.02026 | 0.01152 | 0.02004 | 0.01344 | 0.03647 | 0.00625 | 0.01499 | 0.02519 |
| **SNORA5C** | 0.04783 | 0.06882 | 0.06746 | 0.04452 | 0.02394 | 0.00000 | 0.00845 | 0.00797 | 0.10202 | 0.00000 | 0.01279 | 0.07360 |
| **SNORA6** | 0.34973 | 0.17391 | 0.07605 | 0.13671 | 0.17990 | 0.00000 | 0.00000 | 0.01386 | 0.55418 | 0.91400 | 0.00000 | 0.34995 |
| **SNORA60** | 0.01157 | 0.02606 | 0.01524 | 0.01610 | 0.03988 | 0.00783 | 0.01349 | 0.01856 | 0.06043 | 0.00000 | 0.01586 | 0.04053 |
| **SNORA61** | 0.03578 | 1.04286 | 0.96946 | 0.69379 | 1.36810 | 1.61301 | 0.00000 | 0.64288 | 3.16797 | 5.60631 | 0.00000 | 0.93753 |
| **SNORA62** | 0.15633 | 0.23332 | 0.12274 | 0.36811 | 0.48825 | 0.00000 | 0.07759 | 0.21168 | 0.37028 | 1.03366 | 0.16436 | 0.16416 |
| **SNORA63** | 0.93567 | 0.26851 | 0.41738 | 0.00600 | 0.36655 | 0.88391 | 0.00000 | 1.05443 | 0.12329 | 1.42805 | 0.00000 | 0.43215 |
| **SNORA64** | 0.10567 | 0.07028 | 0.36417 | 0.25730 | 0.40457 | 0.00000 | 0.00000 | 0.06048 | 0.54992 | 1.06456 | 0.00000 | 0.14942 |
| **SNORA65** | 0.12770 | 0.04552 | 0.10001 | 0.12276 | 0.13179 | 0.00000 | 0.00000 | 0.04801 | 0.18927 | 0.34711 | 0.01057 | 0.12034 |
| **SNORA66** | 0.02917 | 0.09306 | 0.03238 | 0.14198 | 0.14848 | 0.08446 | 0.06415 | 0.09478 | 0.20280 | 1.09706 | 0.00000 | 0.15307 |
| **SNORA67** | 1.39669 | 1.02894 | 1.12548 | 0.81667 | 1.48220 | 0.00000 | 0.25514 | 1.03129 | 2.32539 | 2.72404 | 0.15795 | 1.16444 |
| **SNORA68** | 0.05820 | 0.04402 | 0.12199 | 0.07749 | 0.20732 | 0.00000 | 0.00000 | 0.37834 | 0.18633 | 0.36896 | 0.00000 | 0.11761 |
| **SNORA69** | 0.00433 | 0.00000 | 0.06409 | 0.01028 | 0.01035 | 0.04630 | 0.03146 | 0.06499 | 0.00000 | 0.17096 | 0.04828 | 0.03683 |
| **SNORA70** | 0.00374 | 0.73606 | 0.04879 | 0.81205 | 0.87577 | 0.00000 | 0.00000 | 0.16975 | 1.45193 | 0.93268 | 0.39884 | 0.67707 |
| **SNORA70B** | 0.01041 | 0.01219 | 0.01218 | 0.01618 | 0.00000 | 0.05308 | 0.00942 | 0.01592 | 0.02537 | 0.03127 | 0.03848 | 0.01746 |
| **SNORA70D** | 0.00474 | 0.01117 | 0.01077 | 0.00947 | 0.00702 | 0.01477 | 0.01017 | 0.01031 | 0.02369 | 0.12140 | 0.00847 | 0.01226 |
| **SNORA70F** | 0.00324 | 0.00877 | 0.05307 | 0.00000 | 0.09499 | 0.08654 | 0.00000 | 0.00919 | 0.07805 | 0.20694 | 0.02633 | 0.00426 |
| **SNORA70G** | 0.01231 | 0.04150 | 0.12672 | 0.02321 | 0.20280 | 0.04354 | 0.00000 | 0.05210 | 0.14919 | 0.58788 | 0.03417 | 0.10322 |
| **SNORA71A** | 0.01808 | 0.03091 | 0.10916 | 0.08476 | 0.06114 | 0.00000 | 0.01870 | 0.01304 | 0.02911 | 0.08725 | 0.01393 | 0.05319 |
| **SNORA71B** | 0.01487 | 0.03608 | 0.03144 | 0.09157 | 0.01895 | 0.17685 | 0.00000 | 0.04451 | 0.10205 | 0.52710 | 0.02592 | 0.01706 |
| **SNORA71C** | 0.01422 | 0.08903 | 0.08924 | 0.01834 | 0.04415 | 0.00000 | 0.00000 | 0.00728 | 0.08251 | 0.07318 | 0.03885 | 0.09843 |

| | | | | | | | | | | | | |
|---|---|---|---|---|---|---|---|---|---|---|---|---|
| **SNORA71D** | 0.00930 | 0.00356 | 0.04183 | 0.01237 | 0.04102 | 0.00000 | 0.00120 | 0.01342 | 0.02670 | 0.23003 | 0.01839 | 0.05968 |
| **SNORA71E** | 0.00394 | 0.05642 | 0.00892 | 0.04058 | 0.00175 | 0.04609 | 0.00262 | 0.01190 | 0.03470 | 0.06177 | 0.03070 | 0.04179 |
| **SNORA72** | 0.00296 | 0.03853 | 0.08994 | 0.01402 | 0.01768 | 0.12088 | 0.00000 | 0.08250 | 0.29581 | 0.70976 | 0.00522 | 0.07802 |
| **SNORA74A** | 0.00546 | 0.01098 | 0.08522 | 0.04002 | 0.05785 | 0.00000 | 0.00000 | 0.01308 | 0.15943 | 0.02724 | 0.01637 | 0.06099 |
| **SNORA74B** | 0.08595 | 0.00559 | 0.05317 | 0.02166 | 0.02582 | 0.00000 | 0.00000 | 0.02463 | 0.04430 | 0.34157 | 0.00000 | 0.01561 |
| **SNORA75** | 0.11440 | 0.01410 | 0.07248 | 0.01485 | 0.15363 | 0.00500 | 0.00925 | 0.34055 | 0.03424 | 0.53603 | 0.01371 | 0.02762 |
| **SNORA76A** | 0.00523 | 0.01835 | 0.02764 | 0.02581 | 0.04366 | 0.00001 | 0.00240 | 0.00989 | 0.08643 | 0.07760 | 0.00368 | 0.02175 |
| **SNORA76C** | 0.06550 | 0.04579 | 0.14063 | 0.00998 | 0.25119 | 0.00000 | 0.37645 | 0.03378 | 0.41880 | 0.66517 | 0.00000 | 0.26211 |
| **SNORA77** | 0.04352 | 0.02338 | 0.07124 | 0.02935 | 0.12938 | 0.05353 | 0.00000 | 0.00779 | 0.04497 | 0.05067 | 0.00928 | 0.03005 |
| **SNORA78** | 0.08470 | 0.04837 | 0.08939 | 0.11899 | 0.29832 | 0.00000 | 0.00000 | 0.18241 | 0.37274 | 0.18686 | 0.05376 | 0.10492 |
| **SNORA79** | 0.18207 | 0.00792 | 0.04218 | 0.00268 | 0.01977 | 0.02265 | 0.00000 | 0.00020 | 0.05099 | 0.09393 | 0.00465 | 0.16681 |
| **SNORA7B** | 0.04587 | 0.00955 | 0.08801 | 0.05212 | 0.07551 | 0.00000 | 0.01637 | 0.16645 | 0.02221 | 0.26225 | 0.02091 | 0.05164 |
| **SNORA8** | 0.00508 | 0.00000 | 0.33062 | 0.25626 | 0.45592 | 0.37481 | 0.00000 | 0.19219 | 0.39316 | 2.68196 | 0.00000 | 0.17057 |
| **SNORA80A** | 0.44314 | 0.00677 | 0.01957 | 0.02686 | 0.10612 | 0.00000 | 0.05802 | 0.04614 | 0.08224 | 0.36345 | 0.02111 | 0.17947 |
| **SNORA80B** | 0.88951 | 0.07772 | 0.09648 | 0.00000 | 0.24559 | 0.00000 | 0.11687 | 0.14509 | 0.07841 | 0.57454 | 0.00000 | 0.21590 |
| **SNORA80E** | 0.21708 | 0.01957 | 0.22469 | 0.00000 | 0.34824 | 0.00000 | 0.06363 | 1.47139 | 0.17880 | 1.08339 | 0.06643 | 0.17564 |
| **SNORA81** | 0.05263 | 0.02944 | 0.07088 | 0.05721 | 0.17077 | 0.18960 | 0.00000 | 0.02288 | 0.11633 | 0.17495 | 0.00000 | 0.09177 |
| **SNORA84** | 0.01340 | 0.00336 | 0.09413 | 0.05044 | 0.03228 | 0.02355 | 0.00000 | 0.05727 | 0.12134 | 0.24646 | 0.14368 | 0.10352 |
| **SNORA9** | 0.14933 | 0.64181 | 0.11678 | 0.26656 | 0.34383 | 0.00000 | 0.00173 | 0.19106 | 0.68634 | 0.00000 | 0.12881 | 0.84260 |
| **SNORD10** | 0.04114 | 0.38749 | 0.18439 | 0.30247 | 0.17918 | 0.00000 | 0.00233 | 0.05762 | 1.16664 | 1.03415 | 0.87899 | 0.39084 |
| **SNORD100** | 0.04231 | 0.03287 | 0.08821 | 0.03056 | 0.09420 | 0.19150 | 0.00000 | 0.07132 | 0.10451 | 0.76998 | 0.02862 | 0.05877 |
| **SNORD101** | 0.01986 | 0.04702 | 0.03657 | 0.01234 | 0.07360 | 0.03982 | 0.00000 | 0.01472 | 0.10577 | 0.18239 | 0.02663 | 0.09397 |
| **SNORD102** | 0.02935 | 0.04373 | 0.11335 | 0.13441 | 0.09813 | 0.35862 | 0.08709 | 0.29069 | 0.34517 | 0.33874 | 0.11180 | 0.17486 |
| **SNORD104** | 0.00253 | 0.65331 | 0.21361 | 0.68052 | 1.06756 | 0.00000 | 0.00000 | 0.13799 | 1.24629 | 2.05441 | 0.32879 | 0.95849 |
| **SNORD105** | 0.01475 | 0.01659 | 0.02797 | 0.00970 | 0.02982 | 0.00000 | 0.00392 | 0.01986 | 0.04641 | 0.04015 | 0.01388 | 0.02441 |
| **SNORD105B** | 0.01212 | 0.01417 | 0.01403 | 0.01033 | 0.02790 | 0.00838 | 0.01275 | 0.01201 | 0.02712 | 0.03965 | 0.01270 | 0.01579 |
| **SNORD109A+SI** | 0.00540 | 0.02291 | 0.02362 | 0.11198 | 0.02211 | 0.03381 | 0.01315 | 0.00856 | 0.03591 | 0.05168 | 0.00904 | 0.04168 |
| **SNORD11** | 0.13346 | 0.04200 | 0.12880 | 0.03390 | 0.40658 | 0.00000 | 0.04002 | 0.38117 | 0.23288 | 0.87667 | 0.00000 | 0.11363 |
| **SNORD110** | 0.07432 | 0.06082 | 0.06990 | 0.13356 | 0.37743 | 0.00000 | 0.04594 | 0.06192 | 0.33344 | 0.83238 | 0.00000 | 0.15677 |
| **SNORD111** | 0.00555 | 0.01760 | 0.02428 | 0.01651 | 0.01354 | 0.03729 | 0.00610 | 0.00995 | 0.02459 | 0.04903 | 0.01222 | 0.01581 |
| **SNORD111B** | 0.00680 | 0.01189 | 0.01174 | 0.01826 | 0.00778 | 0.03185 | 0.00638 | 0.00995 | 0.02226 | 0.05565 | 0.01349 | 0.01410 |
| **SNORD114-1** | 0.00633 | 0.01532 | 0.02612 | 0.03293 | 0.01263 | 0.01806 | 0.01514 | 0.01048 | 0.01825 | 0.01068 | 0.01585 | 0.01646 |
| **SNORD114-13** | 0.00599 | 0.01274 | 0.01667 | 0.05150 | 0.00994 | 0.02083 | 0.01384 | 0.01044 | 0.01944 | 0.02311 | 0.01481 | 0.01565 |
| **SNORD114-2** | 0.00738 | 0.01531 | 0.01632 | 0.02128 | 0.01473 | 0.00607 | 0.01239 | 0.01037 | 0.02494 | 0.03401 | 0.02136 | 0.01626 |
| **SNORD114-20** | 0.00541 | 0.01213 | 0.03019 | 0.07907 | 0.00593 | 0.01816 | 0.00736 | 0.01037 | 0.01974 | 0.05686 | 0.01450 | 0.01436 |
| **SNORD114-3** | 0.00604 | 0.01324 | 0.01538 | 0.13249 | 0.01894 | 0.00796 | 0.01541 | 0.01046 | 0.02132 | 0.01794 | 0.01598 | 0.01666 |
| **SNORD114-6** | 0.00631 | 0.01399 | 0.01354 | 0.01786 | 0.01353 | 0.02432 | 0.01553 | 0.01046 | 0.01989 | 0.01441 | 0.01485 | 0.01589 |
| **SNORD116-1** | 0.00906 | 0.02386 | 0.01679 | 0.01299 | 0.01748 | 0.02015 | 0.01726 | 0.02041 | 0.01756 | 0.01217 | 0.01580 | 0.01650 |
| **SNORD116-11** | 0.01765 | 0.01453 | 0.01614 | 0.01215 | 0.01523 | 0.02110 | 0.01727 | 0.00821 | 0.01875 | 0.01375 | 0.01476 | 0.01585 |
| **SNORD116-12** | 0.00778 | 0.01570 | 0.01656 | 0.01157 | 0.01401 | 0.02464 | 0.01704 | 0.01111 | 0.01696 | 0.01438 | 0.01442 | 0.01589 |
| **SNORD116-13** | 0.01014 | 0.01842 | 0.02560 | 0.02383 | 0.02858 | 0.01015 | 0.03442 | 0.03122 | 0.01403 | 0.01660 | 0.01538 | 0.02006 |
| **SNORD116-14** | 0.00848 | 0.01344 | 0.02066 | 0.00907 | 0.03031 | 0.01731 | 0.02625 | 0.01357 | 0.00507 | 0.03473 | 0.01346 | 0.02066 |
| **SNORD116-15** | 0.01160 | 0.01651 | 0.01512 | 0.01379 | 0.02573 | 0.00705 | 0.02077 | 0.01105 | 0.01836 | 0.01375 | 0.01492 | 0.02016 |
| **SNORD116-17+** | 0.00646 | 0.01469 | 0.02767 | 0.01280 | 0.02253 | 0.01908 | 0.01743 | 0.01215 | 0.01221 | 0.02003 | 0.01664 | 0.01565 |
| **SNORD116-18** | 0.00771 | 0.02001 | 0.02951 | 0.01971 | 0.01967 | 0.01186 | 0.01701 | 0.01039 | 0.02192 | 0.01147 | 0.01572 | 0.01642 |
| **SNORD116-2** | 0.02873 | 0.01483 | 0.02383 | 0.01111 | 0.02048 | 0.02075 | 0.01503 | 0.01046 | 0.01506 | 0.03141 | 0.01599 | 0.01996 |
| **SNORD116-2+S** | 0.01025 | 0.02612 | 0.03081 | 0.04579 | 0.01307 | 0.03111 | 0.01696 | 0.01296 | 0.00598 | 0.04997 | 0.01405 | 0.02108 |
| **SNORD116-20** | 0.00990 | 0.01369 | 0.01572 | 0.05528 | 0.01565 | 0.02705 | 0.00792 | 0.01080 | 0.02434 | 0.02429 | 0.01331 | 0.02382 |

| | | | | | | | | | | | | |
|---|---|---|---|---|---|---|---|---|---|---|---|---|
| **SNORD116-21** | 0.00721 | 0.02269 | 0.01618 | 0.02004 | 0.01733 | 0.01800 | 0.01789 | 0.01692 | 0.02630 | 0.00746 | 0.01602 | 0.01635 |
| **SNORD116-22** | 0.01276 | 0.02045 | 0.01688 | 0.01577 | 0.01583 | 0.02898 | 0.02728 | 0.00978 | 0.01367 | 0.00728 | 0.01473 | 0.02474 |
| **SNORD116-23** | 0.02261 | 0.02228 | 0.02304 | 0.01400 | 0.03075 | 0.00293 | 0.02886 | 0.01062 | 0.02046 | 0.00000 | 0.01799 | 0.01923 |
| **SNORD116-24** | 0.04907 | 0.01825 | 0.02559 | 0.00964 | 0.02812 | 0.04568 | 0.00000 | 0.02766 | 0.04716 | 0.03840 | 0.00846 | 0.03373 |
| **SNORD116-25** | 0.02554 | 0.01732 | 0.03253 | 0.03617 | 0.03545 | 0.01551 | 0.01536 | 0.01001 | 0.02389 | 0.00460 | 0.03828 | 0.03401 |
| **SNORD116-26** | 0.04187 | 0.31236 | 0.12899 | 0.41888 | 0.15337 | 0.09610 | 0.00000 | 0.02317 | 0.02009 | 0.04454 | 0.05113 | 0.16940 |
| **SNORD116-27** | 0.02353 | 0.02182 | 0.03455 | 0.01332 | 0.01752 | 0.01622 | 0.01395 | 0.01003 | 0.01868 | 0.01550 | 0.01445 | 0.02706 |
| **SNORD116-28** | 0.00559 | 0.01604 | 0.02269 | 0.02793 | 0.01226 | 0.01225 | 0.01634 | 0.01711 | 0.01150 | 0.02187 | 0.02585 | 0.01777 |
| **SNORD116-29** | 0.00911 | 0.02026 | 0.03078 | 0.00948 | 0.02732 | 0.04905 | 0.00000 | 0.01250 | 0.00974 | 0.07949 | 0.01942 | 0.01703 |
| **SNORD116-4** | 0.00778 | 0.04619 | 0.01661 | 0.02532 | 0.01916 | 0.00361 | 0.03619 | 0.01015 | 0.01836 | 0.01319 | 0.01710 | 0.03016 |
| **SNORD116-5+S** | 0.00605 | 0.01570 | 0.01440 | 0.01158 | 0.01626 | 0.02141 | 0.01823 | 0.01148 | 0.01700 | 0.01310 | 0.01492 | 0.01726 |
| **SNORD116-8** | 0.01103 | 0.01562 | 0.01930 | 0.01164 | 0.02428 | 0.01614 | 0.01551 | 0.02229 | 0.01766 | 0.02381 | 0.01311 | 0.01635 |
| **SNORD117** | 0.01120 | 0.02396 | 0.02039 | 0.02622 | 0.00620 | 0.03987 | 0.00703 | 0.01657 | 0.02743 | 0.04633 | 0.01654 | 0.02738 |
| **SNORD119** | 0.10643 | 0.00678 | 0.04425 | 0.03765 | 0.02840 | 0.04222 | 0.00000 | 0.03345 | 0.05977 | 0.06427 | 0.01525 | 0.02001 |
| **SNORD11B** | 0.03127 | 0.02474 | 0.09781 | 0.08028 | 0.02539 | 0.07712 | 0.00000 | 0.09659 | 0.01348 | 0.23218 | 0.03198 | 0.04017 |
| **SNORD12** | 0.03692 | 0.02414 | 0.07196 | 0.07051 | 0.05597 | 0.00000 | 0.00000 | 0.04169 | 0.21558 | 0.39394 | 0.00906 | 0.11289 |
| **SNORD121A** | 0.01062 | 0.01402 | 0.00818 | 0.02496 | 0.00000 | 0.02268 | 0.00474 | 0.01918 | 0.04620 | 0.13946 | 0.01549 | 0.04157 |
| **SNORD124** | 0.00698 | 0.00851 | 0.02595 | 0.00618 | 0.03925 | 0.04428 | 0.01049 | 0.01330 | 0.00360 | 0.02811 | 0.04100 | 0.01671 |
| **SNORD126** | 0.00459 | 0.01118 | 0.02522 | 0.00591 | 0.01081 | 0.03876 | 0.02367 | 0.01453 | 0.01470 | 0.02857 | 0.01069 | 0.01930 |
| **SNORD127** | 0.00685 | 0.01373 | 0.02630 | 0.01187 | 0.01524 | 0.02360 | 0.01617 | 0.01314 | 0.02457 | 0.01474 | 0.01290 | 0.01786 |
| **SNORD12B** | 0.01915 | 0.00000 | 0.15787 | 0.02657 | 0.08749 | 0.04776 | 0.09458 | 0.03131 | 0.19876 | 0.46521 | 0.06086 | 0.10574 |
| **SNORD12C** | 0.00830 | 0.06574 | 0.21118 | 0.28701 | 0.30193 | 0.00000 | 0.00000 | 0.20783 | 1.06925 | 0.98022 | 0.39383 | 0.31736 |
| **SNORD15A** | 0.03081 | 0.23645 | 0.10744 | 0.14471 | 0.42459 | 0.00000 | 0.01666 | 0.03372 | 0.29351 | 0.12403 | 0.02427 | 0.30229 |
| **SNORD15B** | 0.10307 | 0.09211 | 0.17599 | 0.09307 | 0.37774 | 0.00000 | 0.00000 | 0.04938 | 0.19634 | 0.57205 | 0.00000 | 0.11148 |
| **SNORD16** | 0.23644 | 0.11263 | 0.13467 | 0.14536 | 0.15048 | 0.11753 | 0.00000 | 0.29503 | 0.42235 | 0.31012 | 0.00042 | 0.26763 |
| **SNORD17** | 0.09990 | 1.50182 | 1.68730 | 2.73101 | 6.29467 | 2.64785 | 1.78076 | 0.67359 | 3.98121 | 0.40959 | 0.00000 | 2.60844 |
| **SNORD18A** | 0.00444 | 0.02935 | 0.03594 | 0.02072 | 0.01899 | 0.04017 | 0.00786 | 0.04745 | 0.04725 | 0.03033 | 0.03206 | 0.02823 |
| **SNORD18B** | 0.02343 | 0.01831 | 0.02700 | 0.03420 | 0.03678 | 0.04931 | 0.00000 | 0.02345 | 0.03148 | 0.10852 | 0.01097 | 0.03669 |
| **SNORD18C** | 0.03176 | 0.02431 | 0.06060 | 0.04372 | 0.00000 | 0.16331 | 0.00000 | 0.03561 | 0.03065 | 0.11538 | 0.07216 | 0.09210 |
| **SNORD19** | 0.00615 | 0.01959 | 0.05076 | 0.08038 | 0.08821 | 0.00000 | 0.00000 | 0.00708 | 0.09848 | 0.26870 | 0.07249 | 0.09254 |
| **SNORD19B** | 0.00057 | 0.14427 | 0.07182 | 0.01793 | 0.06115 | 0.55302 | 0.00000 | 0.06241 | 0.37202 | 0.17746 | 0.00000 | 0.25253 |
| **SNORD1A** | 0.00832 | 0.01388 | 0.06739 | 0.01901 | 0.07611 | 0.00000 | 0.00020 | 0.01774 | 0.14983 | 0.29130 | 0.04260 | 0.05167 |
| **SNORD1B** | 0.03965 | 0.01942 | 0.01708 | 0.00556 | 0.05717 | 0.00000 | 0.00771 | 0.01520 | 0.07771 | 0.13634 | 0.03336 | 0.02419 |
| **SNORD1C** | 0.00717 | 0.03328 | 0.05805 | 0.01369 | 0.10190 | 0.01666 | 0.00000 | 0.01059 | 0.08466 | 0.12206 | 0.11233 | 0.05399 |
| **SNORD2** | 0.04044 | 0.22734 | 0.00466 | 0.00000 | 0.15350 | 0.25757 | 0.00000 | 0.13327 | 0.42491 | 0.24850 | 0.01543 | 0.15659 |
| **SNORD20** | 0.04329 | 0.01992 | 0.01075 | 0.00731 | 0.05950 | 0.00123 | 0.02832 | 0.01863 | 0.08906 | 0.28668 | 0.00000 | 0.05347 |
| **SNORD21** | 0.07540 | 0.02377 | 0.01077 | 0.02884 | 0.13357 | 0.00000 | 0.00056 | 0.01832 | 0.27140 | 0.30400 | 0.00828 | 0.07411 |
| **SNORD22** | 0.01218 | 0.43377 | 0.46821 | 0.63562 | 0.49464 | 0.00000 | 0.00000 | 0.08947 | 0.88425 | 1.74980 | 0.07441 | 0.43706 |
| **SNORD23** | 0.07421 | 0.19563 | 0.18215 | 0.04713 | 0.22042 | 0.10999 | 0.17284 | 0.01990 | 0.51749 | 0.21827 | 0.00000 | 0.23384 |
| **SNORD24** | 0.04599 | 0.04437 | 0.00604 | 0.01195 | 0.00296 | 0.08459 | 0.00000 | 0.06573 | 0.07232 | 0.13774 | 0.02114 | 0.05328 |
| **SNORD25** | 0.02027 | 0.17732 | 0.13652 | 0.22900 | 0.06795 | 0.15978 | 0.01170 | 0.06736 | 0.19145 | 0.00000 | 0.10177 | 0.21498 |
| **SNORD26** | 0.03063 | 0.09102 | 0.12090 | 0.12043 | 0.04777 | 0.09308 | 0.00000 | 0.02244 | 0.22758 | 0.27512 | 0.02795 | 0.12321 |
| **SNORD27** | 0.25429 | 0.11256 | 0.16163 | 0.14690 | 0.21604 | 0.29499 | 0.00000 | 0.02930 | 0.35866 | 0.68392 | 0.02327 | 0.18887 |
| **SNORD28** | 0.01327 | 0.14017 | 0.17930 | 0.21124 | 0.21407 | 0.42525 | 0.00000 | 0.01014 | 0.30726 | 0.50854 | 0.12261 | 0.18376 |
| **SNORD29** | 0.04586 | 0.22168 | 0.18617 | 0.15388 | 0.20565 | 0.00000 | 0.00000 | 0.06439 | 0.47693 | 0.45558 | 0.01833 | 0.23370 |
| **SNORD30** | 0.08879 | 0.09597 | 0.14050 | 0.15203 | 0.02084 | 0.13183 | 0.00000 | 0.07172 | 0.26453 | 0.35192 | 0.07443 | 0.19669 |
| **SNORD31** | 0.08069 | 0.14924 | 0.08206 | 0.09509 | 0.14856 | 0.00000 | 0.00000 | 0.04282 | 0.27019 | 0.93635 | 0.02668 | 0.13898 |
| **SNORD32A** | 0.04854 | 0.36858 | 0.66848 | 0.44301 | 0.52924 | 0.00000 | 0.00000 | 0.24255 | 0.84537 | 0.25831 | 0.07641 | 0.43085 |

| | | | | | | | | | | | | |
|---|---|---|---|---|---|---|---|---|---|---|---|---|
| **SNORD33** | 0.03268 | 0.10985 | 0.16088 | 0.09050 | 0.17189 | 0.00000 | 0.01444 | 0.08655 | 0.25548 | 0.00000 | 0.02450 | 0.14954 |
| **SNORD34** | 0.06381 | 0.05371 | 0.25849 | 0.11865 | 0.09255 | 0.00000 | 0.00000 | 0.12910 | 0.27393 | 0.37908 | 0.03776 | 0.16882 |
| **SNORD35A** | 0.45569 | 0.05696 | 0.34886 | 0.35326 | 0.14441 | 0.00000 | 0.00002 | 0.64144 | 0.13102 | 0.95600 | 0.02354 | 0.10492 |
| **SNORD35B** | 0.00467 | 0.23536 | 0.36139 | 0.28208 | 0.44626 | 0.00000 | 0.27186 | 0.32302 | 0.73839 | 0.79188 | 0.33577 | 0.36480 |
| **SNORD36A** | 0.11462 | 0.30959 | 0.12170 | 0.29272 | 0.34590 | 0.34639 | 0.00000 | 0.22160 | 0.84403 | 1.06666 | 0.07071 | 0.28255 |
| **SNORD36B** | 0.01966 | 0.09199 | 0.03229 | 0.08175 | 0.08661 | 0.00000 | 0.00000 | 0.01594 | 0.16189 | 0.42522 | 0.02082 | 0.06038 |
| **SNORD36C** | 0.02983 | 0.03356 | 0.04057 | 0.12943 | 0.05426 | 0.16278 | 0.00000 | 0.02572 | 0.15566 | 0.52260 | 0.01276 | 0.05442 |
| **SNORD37** | 0.01301 | 0.01563 | 0.01149 | 0.00657 | 0.02507 | 0.01216 | 0.00517 | 0.01584 | 0.04851 | 0.06205 | 0.01014 | 0.02652 |
| **SNORD38A** | 0.05089 | 0.05159 | 0.05174 | 0.08317 | 0.12110 | 0.00000 | 0.00005 | 0.03564 | 0.10307 | 1.52460 | 0.01540 | 0.14528 |
| **SNORD38B** | 0.00506 | 0.01697 | 0.02573 | 0.05406 | 0.06095 | 0.00000 | 0.00961 | 0.13066 | 0.18164 | 0.72733 | 0.00000 | 0.05316 |
| **SNORD41** | 0.03917 | 0.01386 | 0.01059 | 0.00860 | 0.03140 | 0.01460 | 0.00494 | 0.05511 | 0.03108 | 0.11747 | 0.00642 | 0.01379 |
| **SNORD42A** | 0.03313 | 0.00000 | 0.19258 | 0.00000 | 0.13283 | 0.21458 | 0.25815 | 0.11653 | 0.00000 | 2.24456 | 0.01987 | 0.11549 |
| **SNORD42B** | 0.01933 | 0.01451 | 0.21036 | 0.12870 | 0.12147 | 0.03033 | 0.00000 | 0.06986 | 0.20683 | 0.28951 | 0.09620 | 0.04586 |
| **SNORD43** | 0.01146 | 0.03792 | 0.03509 | 0.00366 | 0.03054 | 0.05388 | 0.16311 | 0.04595 | 0.04487 | 0.80087 | 0.00000 | 0.13931 |
| **SNORD44** | 0.01492 | 0.06942 | 0.11329 | 0.09033 | 0.06467 | 0.11489 | 0.00000 | 0.05689 | 0.23563 | 0.39550 | 0.03953 | 0.11069 |
| **SNORD45A** | 0.01287 | 0.14095 | 0.07005 | 0.00810 | 0.10723 | 0.15347 | 0.05520 | 0.01943 | 0.15126 | 0.00000 | 0.04579 | 0.17912 |
| **SNORD45B** | 0.00212 | 0.21632 | 0.08517 | 0.03164 | 0.07201 | 0.08074 | 0.00000 | 0.02653 | 0.27211 | 0.23814 | 0.12653 | 0.17574 |
| **SNORD45C** | 0.02116 | 0.09931 | 0.00964 | 0.00750 | 0.03477 | 0.53817 | 0.00000 | 0.00948 | 0.35716 | 0.00000 | 0.00000 | 0.20454 |
| **SNORD46** | 0.14245 | 0.12771 | 0.06027 | 0.03403 | 0.12549 | 0.00000 | 0.00000 | 0.58877 | 0.35551 | 0.84399 | 0.00000 | 0.07708 |
| **SNORD47** | 0.00000 | 0.00000 | 0.58341 | 0.36881 | 0.43803 | 2.91218 | 0.00000 | 0.06084 | 0.85609 | 5.79214 | 0.00000 | 0.45615 |
| **SNORD48** | 0.00704 | 0.09051 | 0.02111 | 0.22628 | 0.30089 | 0.11395 | 0.00000 | 0.04613 | 0.34518 | 0.24398 | 0.00869 | 0.37895 |
| **SNORD49A** | 0.00224 | 0.09121 | 0.11322 | 0.25346 | 0.46607 | 0.35050 | 0.00000 | 0.02028 | 1.21790 | 1.57663 | 0.16747 | 0.33150 |
| **SNORD49B** | 0.10707 | 0.26975 | 0.29175 | 0.41444 | 0.42582 | 0.68468 | 0.12017 | 0.01181 | 0.97079 | 0.07431 | 0.23534 | 0.39583 |
| **SNORD4A** | 0.00677 | 0.05225 | 0.11088 | 0.10111 | 0.13071 | 0.00000 | 0.00294 | 0.10691 | 0.25722 | 0.45230 | 0.14279 | 0.12223 |
| **SNORD4B** | 0.02029 | 0.00342 | 0.03458 | 0.02259 | 0.03022 | 0.02243 | 0.00000 | 0.04698 | 0.13234 | 0.18390 | 0.02538 | 0.01563 |
| **SNORD5** | 0.01481 | 0.02832 | 0.05753 | 0.03832 | 0.21584 | 0.07376 | 0.00000 | 0.05158 | 0.10942 | 0.34908 | 0.00000 | 0.06285 |
| **SNORD50A** | 0.00097 | 0.00000 | 0.70111 | 0.00000 | 0.00000 | 0.90938 | 1.61301 | 0.08702 | 0.00000 | 8.76633 | 0.13264 | 0.13906 |
| **SNORD50B** | 0.00081 | 0.00000 | 0.15368 | 0.00000 | 0.08061 | 0.60999 | 0.11482 | 0.04541 | 0.52800 | 1.28893 | 0.00000 | 0.12651 |
| **SNORD51** | 0.01604 | 0.05276 | 0.02477 | 0.04625 | 0.02853 | 0.00126 | 0.02796 | 0.02291 | 0.05281 | 0.05379 | 0.04263 | 0.01819 |
| **SNORD52** | 0.02396 | 0.21977 | 0.05776 | 0.21081 | 0.20385 | 0.53558 | 0.00000 | 0.06594 | 0.57917 | 0.36937 | 0.01929 | 0.33696 |
| **SNORD53** | 0.00449 | 0.02665 | 0.02761 | 0.01077 | 0.02899 | 0.04537 | 0.05507 | 0.03217 | 0.01630 | 0.00339 | 0.01030 | 0.02984 |
| **SNORD54** | 0.01071 | 0.01922 | 0.03949 | 0.04049 | 0.00000 | 0.18278 | 0.02922 | 0.06658 | 0.15474 | 0.19073 | 0.03057 | 0.09299 |
| **SNORD55** | 0.11687 | 0.15206 | 0.16246 | 0.38886 | 0.35481 | 0.00000 | 0.00000 | 0.60262 | 0.50403 | 0.43103 | 0.11459 | 0.32424 |
| **SNORD56** | 0.01852 | 0.01418 | 0.09511 | 0.31706 | 0.47569 | 0.00000 | 0.00000 | 0.08358 | 0.56830 | 0.95218 | 0.00000 | 0.08033 |
| **SNORD56B** | 0.01781 | 0.01486 | 0.01586 | 0.02365 | 0.01217 | 0.01942 | 0.01704 | 0.01033 | 0.02739 | 0.01752 | 0.01650 | 0.01543 |
| **SNORD57** | 0.00343 | 0.03474 | 0.09087 | 0.06184 | 0.54521 | 0.00000 | 0.11774 | 0.18514 | 0.24237 | 1.48361 | 0.00000 | 0.22925 |
| **SNORD58A** | 0.00341 | 0.00824 | 0.06171 | 0.03419 | 0.02860 | 0.10169 | 0.00000 | 0.01460 | 0.10674 | 0.13410 | 0.00350 | 0.02828 |
| **SNORD58C** | 0.00141 | 0.00000 | 0.13228 | 0.04441 | 0.00000 | 0.22081 | 0.00000 | 0.02772 | 0.29849 | 0.90162 | 0.00000 | 0.18830 |
| **SNORD59A** | 0.01113 | 0.07016 | 0.01361 | 0.02510 | 0.03589 | 0.04394 | 0.00000 | 0.00970 | 0.08428 | 0.24336 | 0.00000 | 0.01853 |
| **SNORD59B** | 0.01218 | 0.04164 | 0.03105 | 0.01464 | 0.02813 | 0.02169 | 0.00280 | 0.01545 | 0.08013 | 0.01688 | 0.03362 | 0.02623 |
| **SNORD6** | 0.00551 | 0.00951 | 0.05656 | 0.00618 | 0.04536 | 0.10562 | 0.00000 | 0.02019 | 0.03723 | 0.32607 | 0.00632 | 0.03683 |
| **SNORD60** | 0.01847 | 0.06819 | 0.04418 | 0.07188 | 0.11617 | 0.26976 | 0.00000 | 0.03398 | 0.28843 | 1.08962 | 0.13492 | 0.14671 |
| **SNORD61** | 0.00604 | 0.01323 | 0.01405 | 0.00908 | 0.02137 | 0.02750 | 0.01646 | 0.01108 | 0.01251 | 0.02113 | 0.02854 | 0.02028 |
| **SNORD62A+SN** | 0.00922 | 0.02176 | 0.01943 | 0.01594 | 0.01495 | 0.02214 | 0.01637 | 0.02178 | 0.03611 | 0.01308 | 0.01196 | 0.02612 |
| **SNORD62A+SN** | 0.02142 | 0.07653 | 0.03114 | 0.01690 | 0.07763 | 0.00000 | 0.00685 | 0.02838 | 0.09758 | 0.07186 | 0.02316 | 0.05101 |
| **SNORD63** | 0.00173 | 0.01001 | 0.02576 | 0.13798 | 0.15428 | 0.49102 | 0.00000 | 0.12040 | 0.00000 | 2.04955 | 0.10143 | 0.15466 |
| **SNORD65** | 0.00708 | 0.00000 | 0.00000 | 0.25401 | 0.69347 | 0.65413 | 0.00000 | 0.00641 | 0.40567 | 1.26008 | 0.00000 | 0.17469 |
| **SNORD66** | 0.07238 | 0.01374 | 0.02249 | 0.01301 | 0.01483 | 0.01281 | 0.01571 | 0.00772 | 0.02547 | 0.02056 | 0.01420 | 0.02266 |

| | | | | | | | | | | | | |
|---|---|---|---|---|---|---|---|---|---|---|---|---|
| **SNORD67** | 0.11738 | 0.05549 | 0.02216 | 0.01587 | 0.00765 | 0.07595 | 0.06747 | 0.05874 | 0.01735 | 0.05528 | 0.00877 | 0.04747 |
| **SNORD68** | 0.13685 | 0.16130 | 0.43126 | 0.18098 | 0.40519 | 0.00000 | 0.14144 | 0.20385 | 0.41536 | 0.47411 | 0.04073 | 0.23547 |
| **SNORD69** | 0.00324 | 0.28636 | 0.04395 | 0.26694 | 0.25746 | 0.20057 | 0.00000 | 0.17675 | 0.43856 | 0.02779 | 0.18226 | 0.20193 |
| **SNORD7** | 0.01125 | 0.02517 | 0.04415 | 0.01809 | 0.08203 | 0.00596 | 0.08762 | 0.04732 | 0.07175 | 0.02485 | 0.05786 | 0.04123 |
| **SNORD70** | 0.01719 | 0.01497 | 0.06254 | 0.04862 | 0.01204 | 0.11607 | 0.04453 | 0.01769 | 0.02491 | 0.04645 | 0.01513 | 0.08967 |
| **SNORD71** | 0.10541 | 0.02225 | 0.01778 | 0.01018 | 0.06188 | 0.00489 | 0.01018 | 0.10392 | 0.02255 | 0.05216 | 0.02384 | 0.02259 |
| **SNORD72** | 0.00470 | 0.01093 | 0.03685 | 0.00000 | 0.02793 | 0.11935 | 0.00000 | 0.03232 | 0.06522 | 0.08039 | 0.04616 | 0.03595 |
| **SNORD74** | 0.00856 | 0.07522 | 0.11356 | 0.12851 | 0.10954 | 0.00000 | 0.00351 | 0.12242 | 0.34896 | 0.14005 | 0.09360 | 0.15889 |
| **SNORD75** | 0.00524 | 0.03320 | 0.05280 | 0.03720 | 0.04623 | 0.01799 | 0.00710 | 0.01852 | 0.09371 | 0.00000 | 0.05709 | 0.03462 |
| **SNORD76** | 0.00323 | 0.17936 | 0.26348 | 0.22923 | 0.29413 | 0.00000 | 0.00000 | 0.15926 | 0.69083 | 0.37483 | 0.06624 | 0.26879 |
| **SNORD77** | 0.00515 | 0.09226 | 0.15869 | 0.10774 | 0.10570 | 0.13281 | 0.00000 | 0.08483 | 0.23053 | 0.06296 | 0.01640 | 0.14164 |
| **SNORD78** | 0.02841 | 0.03425 | 0.19459 | 0.09449 | 0.20330 | 0.14453 | 0.06879 | 0.11109 | 0.10231 | 0.13483 | 0.13402 | 0.07442 |
| **SNORD79** | 0.02837 | 0.00350 | 0.20499 | 0.04285 | 0.06957 | 1.40503 | 0.00000 | 0.12951 | 0.62520 | 1.44025 | 0.00000 | 0.12534 |
| **SNORD8** | 0.00722 | 0.00000 | 0.06385 | 0.02317 | 0.00250 | 0.05055 | 0.00000 | 0.02256 | 0.02709 | 0.49908 | 0.00911 | 0.09577 |
| **SNORD80** | 0.01908 | 0.30634 | 0.37796 | 0.74149 | 0.83061 | 1.79313 | 0.00000 | 0.34654 | 1.81198 | 1.25694 | 0.00000 | 0.36417 |
| **SNORD81** | 0.00186 | 0.06777 | 0.21216 | 0.04905 | 0.00962 | 0.94848 | 0.00000 | 0.50274 | 0.36642 | 2.74473 | 0.00000 | 0.25923 |
| **SNORD82** | 0.05766 | 0.02065 | 0.02585 | 0.00703 | 0.04605 | 0.01298 | 0.02092 | 0.01526 | 0.01929 | 0.04162 | 0.01213 | 0.03380 |
| **SNORD83A** | 0.01371 | 0.10251 | 0.01012 | 0.07353 | 0.19914 | 0.00000 | 0.00014 | 0.08413 | 0.29109 | 0.13298 | 0.00000 | 0.09407 |
| **SNORD83B** | 0.16983 | 0.08554 | 0.08744 | 0.10004 | 0.43283 | 0.00000 | 0.00000 | 0.09551 | 0.23530 | 0.78952 | 0.02180 | 0.20861 |
| **SNORD84** | 0.03277 | 0.03863 | 0.04757 | 0.02251 | 0.01229 | 0.07629 | 0.02528 | 0.02660 | 0.07754 | 0.02108 | 0.00245 | 0.04577 |
| **SNORD85** | 0.01444 | 0.01706 | 0.01155 | 0.00649 | 0.00534 | 0.02192 | 0.01167 | 0.01469 | 0.02360 | 0.06863 | 0.01191 | 0.01712 |
| **SNORD87** | 0.04169 | 0.01203 | 0.04723 | 0.05808 | 0.06799 | 0.58628 | 0.00000 | 0.24118 | 0.42614 | 1.16694 | 0.24279 | 0.10418 |
| **SNORD88A** | 0.01917 | 0.01742 | 0.03225 | 0.00787 | 0.10741 | 0.00000 | 0.00494 | 0.01920 | 0.03711 | 0.11489 | 0.00259 | 0.02810 |
| **SNORD88B** | 0.00170 | 0.68518 | 0.24114 | 0.48472 | 0.23510 | 0.00000 | 0.19960 | 0.10402 | 1.19409 | 0.00000 | 0.57304 | 1.16712 |
| **SNORD88C** | 0.00688 | 0.04569 | 0.08311 | 0.03959 | 0.34923 | 0.00000 | 0.00000 | 0.00914 | 0.08662 | 0.24198 | 0.01546 | 0.12078 |
| **SNORD89** | 0.00343 | 0.06734 | 0.07780 | 0.04519 | 0.12018 | 0.19140 | 0.00013 | 0.06398 | 0.05012 | 0.02429 | 0.01366 | 0.03467 |
| **SNORD9** | 0.00446 | 0.01468 | 0.02231 | 0.01045 | 0.01215 | 0.03027 | 0.01890 | 0.02228 | 0.02278 | 0.01336 | 0.01251 | 0.01593 |
| **SNORD90** | 0.03713 | 0.02060 | 0.02379 | 0.01639 | 0.02194 | 0.01656 | 0.01670 | 0.01695 | 0.01852 | 0.01402 | 0.01473 | 0.02175 |
| **SNORD91A** | 0.00196 | 0.00006 | 0.08715 | 0.08380 | 0.24620 | 0.09886 | 0.00000 | 0.02483 | 0.22220 | 0.41199 | 0.00000 | 0.09154 |
| **SNORD91B** | 0.00746 | 0.01063 | 0.01420 | 0.06723 | 0.04747 | 0.02858 | 0.00571 | 0.02450 | 0.03285 | 0.08982 | 0.01516 | 0.02585 |
| **SNORD92** | 0.01035 | 0.01387 | 0.04954 | 0.02645 | 0.01012 | 0.02434 | 0.02932 | 0.02820 | 0.00015 | 0.06140 | 0.02770 | 0.03286 |
| **SNORD93** | 0.00648 | 0.02311 | 0.05001 | 0.01720 | 0.05630 | 0.04517 | 0.00000 | 0.01117 | 0.04749 | 0.00563 | 0.02034 | 0.02497 |
| **SNORD94** | 0.28821 | 0.08763 | 0.03952 | 0.05173 | 0.07536 | 0.00000 | 0.00669 | 0.09081 | 0.16570 | 0.00827 | 0.04193 | 0.06409 |
| **SNORD95** | 0.03371 | 0.06040 | 0.04711 | 0.10848 | 0.06224 | 0.00000 | 0.03009 | 0.18662 | 0.06999 | 0.00000 | 0.00810 | 0.08089 |
| **SNORD96A** | 0.01169 | 0.24202 | 0.10772 | 0.31798 | 0.36766 | 0.00000 | 0.00392 | 0.12397 | 0.67582 | 0.00000 | 0.00507 | 0.34249 |
| **SNORD96B** | 0.00580 | 0.01216 | 0.01158 | 0.03190 | 0.00754 | 0.02102 | 0.00586 | 0.01043 | 0.03117 | 0.03930 | 0.01739 | 0.01464 |
| **SNORD97** | 0.08107 | 0.01382 | 0.09202 | 0.01656 | 0.00906 | 0.01876 | 0.02166 | 0.02715 | 0.04186 | 0.02914 | 0.00748 | 0.02349 |
| **SNORD98** | 0.01909 | 0.09310 | 0.11328 | 0.20499 | 0.12589 | 0.06397 | 0.02142 | 0.04734 | 0.13995 | 0.09702 | 0.08650 | 0.28185 |
| **SNORD99** | 0.00000 | 0.21474 | 0.41554 | 1.04714 | 1.07497 | 1.50577 | 0.00000 | 0.94032 | 1.68027 | 4.60830 | 1.75604 | 0.92972 |
| **SNPH** | 1.18086 | 0.01122 | 0.00611 | 0.48960 | 0.17590 | 0.00000 | 0.01231 | 0.12517 | 0.00000 | 0.07619 | 0.03106 | 0.02675 |
| **SNRK** | 1.80732 | 0.96998 | 1.19900 | 1.52403 | 0.33993 | 5.47703 | 2.57043 | 0.67478 | 0.28139 | 4.26069 | 1.80303 | 1.63352 |
| **SNRK-AS1** | 0.03086 | 0.01675 | 0.13271 | 0.00462 | 0.02200 | 0.07290 | 0.01425 | 0.01409 | 0.00623 | 0.04910 | 0.04309 | 0.02668 |
| **SNRNP200** | 6.02759 | 16.93922 | 15.35680 | 17.98359 | 18.29140 | 9.04551 | 27.03772 | 5.43043 | 21.97749 | 21.21697 | 16.85098 | 17.96047 |
| **SNRNP25** | 0.42332 | 14.97023 | 14.03083 | 12.38054 | 16.53118 | 1.78603 | 22.94418 | 13.90136 | 16.11101 | 1.15068 | 20.65684 | 14.26873 |
| **SNRNP27** | 2.08341 | 5.26273 | 7.10715 | 2.02782 | 3.73630 | 17.98420 | 10.54151 | 5.54998 | 6.50347 | 7.77292 | 6.69236 | 14.24857 |
| **SNRNP35** | 1.97769 | 8.58245 | 3.35725 | 2.46775 | 3.08581 | 10.17538 | 4.13457 | 10.25695 | 0.97100 | 1.38806 | 5.23182 | 12.29435 |
| **SNRNP40** | 2.76501 | 16.74227 | 16.53776 | 10.50599 | 7.38713 | 23.30632 | 19.94932 | 11.26001 | 21.64067 | 0.00000 | 18.70600 | 14.08877 |
| **SNRNP48** | 2.25633 | 0.00000 | 4.28940 | 0.94843 | 1.37939 | 7.07434 | 7.97713 | 5.47313 | 2.96155 | 8.27046 | 2.08796 | 1.87078 |

| | | | | | | | | | | | | |
|---|---|---|---|---|---|---|---|---|---|---|---|---|
| **SNRNP70** | 1.19306 | 1.32524 | 2.77003 | 2.44607 | 8.60802 | 0.00000 | 8.08679 | 1.85313 | 0.00000 | 23.76647 | 1.21658 | 1.70104 |
| **SNRPA** | 1.89722 | 16.95149 | 12.59665 | 13.81152 | 20.97780 | 0.01216 | 23.75440 | 2.11835 | 23.19531 | 13.23534 | 11.43525 | 20.49320 |
| **SNRPA1** | 0.31175 | 6.21171 | 6.63737 | 3.38092 | 4.86148 | 8.95558 | 8.80817 | 7.98120 | 9.78864 | 0.00000 | 4.48688 | 5.75434 |
| **SNRPB** | 7.08785 | 37.09114 | 39.10279 | 34.27938 | 42.81352 | 28.79762 | 53.18186 | 30.33216 | 53.94658 | 10.04423 | 35.63333 | 45.35205 |
| **SNRPB2** | 5.70720 | 13.82953 | 15.39811 | 4.76192 | 8.77487 | 30.60108 | 26.49019 | 18.62457 | 13.38423 | 22.13605 | 13.47149 | 18.77367 |
| **SNRPC** | 3.78117 | 30.86851 | 20.83853 | 18.99017 | 23.33003 | 34.47218 | 34.85097 | 28.40090 | 26.18820 | 10.04268 | 25.23779 | 41.51122 |
| **SNRPD1** | 8.07438 | 17.39653 | 20.63493 | 13.45255 | 14.53383 | 40.38090 | 35.37939 | 17.45194 | 21.84433 | 15.59125 | 23.64898 | 26.94355 |
| **SNRPD2** | 4.38757 | 32.72300 | 31.27384 | 27.39858 | 38.26949 | 29.47321 | 45.87141 | 18.71842 | 32.47027 | 17.16715 | 33.62348 | 44.44773 |
| **SNRPD2P2** | 0.00246 | 0.01906 | 0.01242 | 0.01594 | 0.01571 | 0.04032 | 0.01713 | 0.04433 | 0.03278 | 0.02542 | 0.03593 | 0.01819 |
| **SNRPD3** | 8.95382 | 21.55114 | 22.27575 | 19.38558 | 22.26288 | 30.08414 | 36.62500 | 19.62000 | 26.94641 | 18.44572 | 25.55982 | 27.77414 |
| **SNRPE** | 3.78438 | 11.19717 | 15.04057 | 7.11006 | 10.48646 | 29.97689 | 24.98813 | 6.89651 | 10.93702 | 19.17370 | 17.01232 | 16.62234 |
| **SNRPF** | 8.89937 | 18.20611 | 17.31220 | 13.90250 | 18.01681 | 28.98249 | 32.89580 | 12.20422 | 18.12330 | 18.08597 | 19.11471 | 22.85758 |
| **SNRPG** | 1.64093 | 12.82901 | 15.00010 | 11.47129 | 14.48316 | 18.07899 | 25.55314 | 9.20941 | 13.16343 | 7.23691 | 18.30660 | 17.05644 |
| **SNRPN.uid-189(** | 0.04898 | 0.06318 | 0.04437 | 0.01155 | 0.01901 | 0.00000 | 0.01940 | 0.02133 | 0.03785 | 0.00189 | 0.01950 | 0.02104 |
| **SNTA1** | 0.99150 | 0.44878 | 0.25253 | 0.34994 | 0.00000 | 0.00000 | 0.00310 | 0.30798 | 0.31329 | 0.00654 | 0.32293 | 0.60042 |
| **SNTB1** | 0.00327 | 0.00000 | 0.00000 | 0.01198 | 2.21867 | 0.15798 | 0.19387 | 0.06378 | 0.00000 | 0.00000 | 2.85806 | 0.03351 |
| **SNTB2** | 0.36481 | 0.62388 | 4.37066 | 2.43595 | 0.95410 | 3.17441 | 8.81553 | 2.27439 | 0.03314 | 12.42569 | 2.38232 | 3.50146 |
| **SNTG1** | 0.01026 | 0.01607 | 0.02772 | 0.01093 | 0.01402 | 0.01739 | 0.01399 | 0.01000 | 0.01887 | 0.02253 | 0.01435 | 0.01707 |
| **SNTG2** | 0.15072 | 0.01589 | 0.00949 | 0.00635 | 0.00000 | 0.03440 | 0.02121 | 0.00776 | 0.00000 | 0.07667 | 0.03628 | 0.02066 |
| **SNTN** | 0.01275 | 0.02967 | 0.01355 | 0.01023 | 0.00736 | 0.02599 | 0.03746 | 0.01057 | 0.00199 | 0.02182 | 0.02292 | 0.01525 |
| **SNUPN** | 2.21095 | 11.40908 | 4.02392 | 4.72159 | 1.75818 | 10.58683 | 2.80681 | 16.88295 | 0.29242 | 1.34391 | 4.54930 | 19.34555 |
| **SNURF.uid-1921** | 0.00689 | 0.17332 | 0.12450 | 0.15458 | 0.08186 | 0.15889 | 0.02502 | 0.00772 | 0.03594 | 0.00000 | 0.09105 | 0.14746 |
| **SNW1** | 8.14063 | 21.44492 | 13.59018 | 13.04151 | 16.53916 | 34.73273 | 18.65632 | 12.49999 | 12.74147 | 12.79590 | 20.01699 | 21.85233 |
| **SNX1** | 2.74565 | 4.37395 | 2.47248 | 3.10371 | 3.70651 | 3.15009 | 5.34103 | 4.13769 | 5.75349 | 0.45898 | 6.01743 | 6.88631 |
| **SNX10** | 0.57205 | 3.25896 | 2.04740 | 0.09409 | 0.00000 | 4.88490 | 1.05140 | 0.00000 | 0.02136 | 1.21331 | 1.49790 | 0.58072 |
| **SNX11** | 0.16634 | 5.22020 | 6.44000 | 4.62402 | 4.38560 | 0.00006 | 6.54974 | 0.70148 | 7.45108 | 1.91362 | 4.00679 | 4.86638 |
| **SNX12** | 1.55024 | 2.13744 | 0.00000 | 2.02873 | 2.80545 | 3.35755 | 4.64671 | 0.39244 | 0.00000 | 7.97858 | 5.88472 | 2.39112 |
| **SNX13** | 4.02206 | 0.51674 | 6.76871 | 2.54910 | 0.04191 | 8.54276 | 2.71659 | 3.22593 | 5.68802 | 2.44426 | 3.13440 | 3.17030 |
| **SNX14** | 4.61689 | 0.81233 | 6.36836 | 1.00878 | 0.00000 | 6.73692 | 0.00000 | 8.42409 | 10.25182 | 2.86079 | 3.67382 | 0.38527 |
| **SNX16** | 0.74886 | 0.03063 | 1.14625 | 0.00000 | 0.00000 | 4.38801 | 0.00000 | 1.52874 | 0.65668 | 4.86920 | 0.00000 | 5.52964 |
| **SNX17** | 0.77668 | 11.94804 | 10.56198 | 12.89642 | 11.12517 | 9.16422 | 11.90894 | 0.00000 | 8.38649 | 0.00000 | 23.46416 | 6.34989 |
| **SNX18** | 0.66584 | 0.00000 | 3.53980 | 0.00000 | 5.32166 | 4.95538 | 4.36290 | 0.60029 | 0.00000 | 11.08268 | 3.75216 | 2.08854 |
| **SNX19** | 2.72945 | 1.34651 | 0.80189 | 0.01107 | 0.80536 | 2.79986 | 0.00000 | 0.00000 | 0.89447 | 0.88783 | 1.43701 | 4.17599 |
| **SNX2** | 1.36007 | 24.57531 | 14.22415 | 10.08378 | 16.83339 | 54.69991 | 40.07618 | 5.44780 | 37.27646 | 9.89717 | 39.81969 | 19.07519 |
| **SNX20** | 0.04455 | 0.01149 | 0.01753 | 0.01230 | 0.05355 | 0.00868 | 0.03204 | 0.01614 | 0.00354 | 0.04027 | 0.01218 | 0.01881 |
| **SNX21** | 3.43286 | 5.21743 | 0.07366 | 3.41791 | 4.71666 | 0.00000 | 0.00000 | 0.69293 | 0.31379 | 1.78413 | 1.20444 | 1.04223 |
| **SNX22** | 0.07152 | 1.77350 | 3.62599 | 1.37418 | 1.99068 | 1.04856 | 0.56064 | 0.25146 | 4.88429 | 2.14049 | 1.59071 | 1.30571 |
| **SNX24** | 1.09449 | 11.23762 | 0.19928 | 3.73989 | 6.90295 | 15.80547 | 4.99344 | 1.36327 | 4.93246 | 9.84816 | 4.75477 | 10.48612 |
| **SNX25** | 2.55128 | 0.22984 | 0.43487 | 1.90321 | 0.41482 | 0.64185 | 0.00000 | 0.86686 | 0.00000 | 0.40744 | 0.00000 | 0.83099 |
| **SNX27** | 5.47729 | 8.26553 | 14.37272 | 15.79251 | 9.25001 | 14.76907 | 13.83954 | 3.41581 | 3.53659 | 5.37881 | 12.95765 | 4.65269 |
| **SNX29** | 1.42309 | 1.11662 | 2.49937 | 0.32401 | 11.24455 | 0.00000 | 1.08811 | 1.73327 | 5.13603 | 3.11834 | 4.92886 | 1.05781 |
| **SNX29P1** | 0.00641 | 0.01639 | 0.01434 | 0.02321 | 0.01681 | 0.01246 | 0.02229 | 0.01041 | 0.02211 | 0.00693 | 0.01713 | 0.01882 |
| **SNX29P2** | 0.00984 | 0.01831 | 0.01498 | 0.01267 | 0.01852 | 0.02358 | 0.02012 | 0.01127 | 0.01980 | 0.00982 | 0.01524 | 0.01816 |
| **SNX3** | 1.73511 | 11.01370 | 15.18934 | 7.37216 | 13.64822 | 29.04501 | 18.20419 | 3.91470 | 12.92392 | 17.14619 | 20.05049 | 6.58262 |
| **SNX30** | 2.83207 | 1.35219 | 1.70048 | 3.50749 | 0.22436 | 6.16005 | 2.25854 | 1.12883 | 1.56994 | 3.69274 | 4.74774 | 0.61339 |
| **SNX31** | 0.04799 | 0.00755 | 0.00945 | 0.00749 | 0.00000 | 0.04944 | 0.05301 | 0.00896 | 0.02990 | 0.08275 | 0.00175 | 0.01541 |
| **SNX32** | 0.00439 | 0.00795 | 0.01916 | 0.06194 | 0.18980 | 0.00000 | 0.00021 | 0.01012 | 0.12753 | 0.14235 | 0.00000 | 0.01212 |
| **SNX33** | 0.00209 | 0.26654 | 1.91155 | 0.41352 | 1.85060 | 0.00000 | 3.73076 | 0.00780 | 0.32909 | 0.44106 | 0.56284 | 0.00000 |

| | | | | | | | | | | | | |
|---|---|---|---|---|---|---|---|---|---|---|---|---|
| **SNX4** | 4.27489 | 9.74178 | 8.30253 | 2.59859 | 1.90618 | 19.30060 | 10.54738 | 0.29809 | 7.16932 | 1.65765 | 10.42352 | 8.91382 |
| **SNX5** | 5.93824 | 15.81733 | 11.79101 | 13.17124 | 13.65484 | 32.20589 | 23.34434 | 4.43389 | 15.44078 | 26.42159 | 20.30593 | 12.24916 |
| **SNX6** | 2.44729 | 7.61409 | 12.43596 | 12.68688 | 4.28463 | 53.98282 | 22.68712 | 5.97589 | 22.00544 | 11.45609 | 20.20589 | 15.01442 |
| **SNX7** | 0.00000 | 1.52802 | 0.03436 | 3.68749 | 0.38336 | 2.34355 | 2.26574 | 0.00000 | 0.00000 | 3.06116 | 0.38196 | 0.00000 |
| **SNX8** | 0.45599 | 21.80817 | 0.94630 | 11.54985 | 32.52494 | 11.33906 | 11.42859 | 6.78236 | 9.63731 | 3.47107 | 24.45326 | 13.85995 |
| **SNX9** | 2.99162 | 0.97081 | 3.87194 | 1.13890 | 1.55903 | 6.48242 | 8.11607 | 0.00000 | 1.15343 | 11.48477 | 2.86908 | 0.00000 |
| **SOAT1** | 2.60857 | 22.02827 | 9.74226 | 2.40102 | 23.65760 | 61.29312 | 0.00000 | 10.52518 | 42.25091 | 23.09980 | 21.25571 | 39.39179 |
| **SOAT2** | 0.00078 | 6.80119 | 0.00000 | 4.76595 | 1.04178 | 0.37493 | 0.00000 | 3.95077 | 0.00000 | 0.57335 | 0.20335 | 19.14252 |
| **SOBP** | 0.13714 | 0.24920 | 0.00000 | 0.10883 | 0.00000 | 0.04412 | 0.01409 | 0.26149 | 0.06208 | 0.01557 | 0.00745 | 0.88893 |
| **SOCS1** | 0.24849 | 0.49619 | 5.51357 | 0.42618 | 1.74397 | 0.00000 | 0.63098 | 0.11366 | 0.00000 | 2.83665 | 0.26587 | 0.00000 |
| **SOCS2** | 4.81454 | 0.76252 | 1.05736 | 6.03519 | 0.38926 | 2.32771 | 1.00365 | 1.21980 | 1.94754 | 0.85791 | 0.76461 | 2.17124 |
| **SOCS2-AS1** | 0.26017 | 0.03139 | 0.02015 | 0.46157 | 0.11419 | 0.02351 | 0.04276 | 0.04318 | 0.06760 | 0.00000 | 0.01358 | 1.00786 |
| **SOCS3** | 0.28578 | 1.04791 | 4.81025 | 3.95308 | 4.13394 | 1.62102 | 2.27417 | 0.19252 | 0.00006 | 1.38201 | 0.00000 | 0.00000 |
| **SOCS4** | 2.27300 | 1.92401 | 4.67335 | 6.54198 | 1.40392 | 20.06104 | 0.00000 | 9.23757 | 4.40510 | 9.64639 | 1.57246 | 12.23005 |
| **SOCS5** | 0.43249 | 0.19987 | 0.87115 | 1.00666 | 0.00000 | 2.95863 | 0.00000 | 1.83298 | 1.78730 | 2.61658 | 0.40940 | 0.93911 |
| **SOCS6** | 0.39178 | 1.99849 | 9.41488 | 1.67161 | 1.18327 | 7.59122 | 8.01148 | 2.23480 | 4.12740 | 0.65465 | 5.71547 | 0.00000 |
| **SOCS7** | 5.92568 | 1.15222 | 2.52915 | 2.06558 | 1.51185 | 1.27813 | 7.89921 | 2.66346 | 1.83892 | 11.65886 | 2.40717 | 4.89788 |
| **SOD1** | 6.82626 | 28.69973 | 29.66614 | 8.82814 | 29.82055 | 32.46426 | 47.25431 | 4.38239 | 34.38580 | 12.65051 | 29.55728 | 19.84885 |
| **SOD2** | 1.77787 | 10.20236 | 7.33052 | 10.07959 | 4.90319 | 14.03549 | 19.25382 | 3.49671 | 8.51973 | 1.63127 | 9.77801 | 5.39361 |
| **SOD3** | 0.00603 | 0.01420 | 0.14842 | 0.01069 | 0.00970 | 0.01652 | 0.02692 | 0.01047 | 0.00000 | 0.01814 | 0.04207 | 0.01593 |
| **SOGA1** | 0.20090 | 0.46356 | 0.00000 | 0.81363 | 1.63777 | 0.00000 | 0.98359 | 0.00000 | 0.57386 | 2.61294 | 0.10146 | 0.23888 |
| **SOGA3** | 0.13494 | 0.09142 | 0.01243 | 0.02332 | 0.02069 | 0.00000 | 0.03730 | 0.45840 | 0.01138 | 0.02829 | 0.01130 | 0.01875 |
| **SOHLH1** | 0.53731 | 0.02689 | 0.03036 | 0.00864 | 0.03244 | 0.00000 | 0.03057 | 0.05882 | 0.04140 | 0.00445 | 0.00692 | 0.11505 |
| **SOHLH2** | 2.27422 | 0.90561 | 0.53629 | 0.12315 | 0.00000 | 0.36999 | 1.45213 | 0.22991 | 1.03172 | 0.24922 | 0.27812 | 0.27601 |
| **SON** | 7.71446 | 16.61994 | 18.45563 | 15.05628 | 20.97031 | 34.46328 | 30.99039 | 24.34774 | 19.16623 | 47.56310 | 22.19082 | 24.78443 |
| **SORBS1** | 4.23606 | 10.74973 | 0.00000 | 4.71254 | 34.15786 | 12.21834 | 1.54223 | 5.87484 | 0.00000 | 2.59066 | 9.49077 | 12.58499 |
| **SORBS2** | 0.59680 | 0.05407 | 0.28217 | 0.04803 | 0.00330 | 0.05127 | 0.03584 | 0.07068 | 0.12736 | 0.00000 | 0.10121 | 0.02631 |
| **SORBS3** | 0.41039 | 1.17501 | 1.63638 | 0.70795 | 1.32656 | 0.00000 | 0.42128 | 0.00000 | 0.57452 | 1.07685 | 0.21105 | 0.50447 |
| **SORCS1** | 0.49643 | 0.03432 | 0.02176 | 0.03238 | 0.00000 | 0.04403 | 0.03424 | 0.08601 | 0.00000 | 0.04766 | 0.02683 | 0.00470 |
| **SORCS2** | 0.39070 | 0.06496 | 0.00149 | 0.13434 | 0.00000 | 0.00000 | 0.00473 | 0.07069 | 0.02151 | 0.22025 | 0.01458 | 0.00812 |
| **SORCS3** | 0.96340 | 0.04041 | 0.04875 | 0.01119 | 0.20179 | 0.00000 | 0.04110 | 0.29735 | 0.01297 | 0.00000 | 0.01288 | 0.01971 |
| **SORD** | 0.00228 | 1.16928 | 1.33881 | 0.00039 | 0.00000 | 0.00886 | 0.44029 | 0.06860 | 0.56846 | 0.00000 | 0.00000 | 0.97134 |
| **SORL1** | 0.41356 | 0.99073 | 0.00000 | 1.49285 | 1.14030 | 0.39990 | 0.00000 | 0.46283 | 0.00000 | 1.61628 | 0.33370 | 0.07641 |
| **SORT1** | 2.25204 | 4.66288 | 12.26089 | 5.90170 | 6.52347 | 3.82815 | 2.05037 | 1.57990 | 9.05522 | 8.33997 | 5.66196 | 0.00000 |
| **1.00000** | 0.00000 | 1.69956 | 3.36561 | 1.89333 | 0.84439 | 8.38670 | 2.75462 | 0.41712 | 4.11650 | 13.51440 | 1.50677 | 1.23844 |
| **2.00000** | 0.29382 | 0.69677 | 0.24635 | 0.95564 | 0.00000 | 1.64101 | 1.40004 | 0.55541 | 0.04548 | 3.89700 | 0.20445 | 1.39841 |
| **SOST** | 0.61553 | 0.01144 | 0.02196 | 0.01309 | 0.01127 | 0.02277 | 0.01448 | 0.02458 | 0.01614 | 0.01617 | 0.01342 | 0.01182 |
| **SOSTDC1** | 0.32625 | 0.01518 | 0.02284 | 0.08714 | 0.02374 | 0.00000 | 0.02080 | 0.00375 | 0.03524 | 0.00000 | 0.01284 | 0.04465 |
| **SOWAHA** | 0.09498 | 0.00000 | 0.00000 | 0.00000 | 2.35511 | 0.01154 | 0.28124 | 0.07060 | 0.00000 | 0.20253 | 0.09182 | 0.03459 |
| **SOWAHB** | 0.00623 | 0.05529 | 0.20996 | 0.33389 | 0.32251 | 0.19585 | 0.00000 | 0.00710 | 0.00000 | 0.03013 | 0.00000 | 0.01118 |
| **SOWAHC** | 0.00893 | 0.00000 | 2.22207 | 0.50122 | 1.28938 | 0.98801 | 5.02125 | 0.20584 | 0.00000 | 6.32163 | 0.27681 | 0.00000 |
| **SOWAHD** | 0.01859 | 0.00000 | 0.11149 | 0.16027 | 0.02742 | 0.01469 | 0.12562 | 0.07081 | 0.03286 | 0.20483 | 0.00000 | 0.02568 |
| **SOX1** | 0.38793 | 0.00686 | 0.03255 | 0.01810 | 0.00580 | 0.05553 | 0.00000 | 0.00121 | 0.01257 | 0.04960 | 0.01384 | 0.01091 |
| **SOX10** | 0.00608 | 0.01586 | 0.01179 | 0.03024 | 0.00580 | 0.03045 | 0.03027 | 0.01029 | 0.01643 | 0.01819 | 0.01434 | 0.01573 |
| **SOX11** | 1.32104 | 0.00000 | 0.00000 | 0.13095 | 0.01097 | 0.03763 | 0.00000 | 3.29177 | 0.02418 | 0.49590 | 0.00270 | 0.00000 |
| **SOX12** | 0.00367 | 0.00000 | 0.00000 | 0.44925 | 0.12977 | 0.00000 | 0.46981 | 0.07009 | 0.04665 | 2.39783 | 0.31685 | 2.09421 |
| **SOX13** | 10.23571 | 1.26905 | 13.36063 | 2.96317 | 2.84559 | 0.00000 | 6.38588 | 2.54346 | 0.69101 | 8.41628 | 0.20909 | 5.01113 |
| **SOX14** | 0.00613 | 0.00693 | 0.03774 | 0.12392 | 0.01338 | 0.08614 | 0.00796 | 0.01418 | 0.00000 | 0.00483 | 0.03796 | 0.01225 |

| | | | | | | | | | | | | |
|---|---|---|---|---|---|---|---|---|---|---|---|---|
| **SOX15** | 9.69873 | 26.03656 | 7.50740 | 2.35308 | 2.54458 | 0.00000 | 12.30508 | 30.92736 | 17.58482 | 6.78462 | 1.55056 | 43.05954 |
| **SOX17** | 0.00431 | 0.00309 | 0.75157 | 6.43879 | 0.08205 | 0.00000 | 0.00000 | 0.00839 | 0.00000 | 0.07540 | 0.02470 | 0.00974 |
| **SOX2** | 1.00906 | 3.36356 | 0.20603 | 7.19070 | 0.05639 | 0.00000 | 0.00000 | 0.65298 | 0.00000 | 2.25655 | 0.00000 | 9.77162 |
| **SOX2-OT** | 0.02535 | 0.19291 | 0.13380 | 0.33207 | 0.00000 | 0.03948 | 0.07355 | 0.04424 | 0.00000 | 0.06373 | 0.01579 | 0.39121 |
| **SOX21** | 0.05564 | 0.00952 | 0.08912 | 0.44852 | 0.00000 | 0.02475 | 0.00000 | 0.02898 | 0.01076 | 0.06496 | 0.01162 | 0.01764 |
| **SOX21-AS1** | 0.10421 | 0.00000 | 0.03481 | 0.21381 | 0.00000 | 0.09330 | 0.00000 | 0.14988 | 0.11430 | 0.16963 | 0.07361 | 0.04129 |
| **SOX30** | 2.14884 | 0.17042 | 0.00756 | 0.00000 | 0.00451 | 0.09765 | 0.06577 | 0.32002 | 0.00000 | 0.02166 | 0.02882 | 1.06767 |
| **SOX4** | 6.25076 | 13.82762 | 22.82080 | 16.87245 | 19.07609 | 28.97244 | 57.51387 | 5.38537 | 24.95657 | 44.98446 | 22.88007 | 9.01619 |
| **SOX5** | 0.82817 | 0.00000 | 0.00000 | 0.06730 | 0.00000 | 0.05264 | 0.00033 | 0.04785 | 0.02903 | 0.19973 | 0.00000 | 0.21938 |
| **SOX6** | 0.33909 | 0.00480 | 0.01180 | 0.03665 | 0.00000 | 0.10332 | 0.05835 | 0.12722 | 0.00000 | 0.09739 | 0.00333 | 0.01433 |
| **SOX7** | 0.34950 | 0.27607 | 0.20610 | 0.07633 | 0.21332 | 0.00000 | 0.24286 | 0.00006 | 0.00000 | 0.00000 | 0.04508 | 0.05139 |
| **SOX8** | 0.01851 | 0.01533 | 0.04308 | 0.10157 | 0.01001 | 0.00000 | 0.02543 | 0.01090 | 0.07015 | 0.01786 | 0.02734 | 0.01754 |
| **SOX9** | 0.01478 | 0.00642 | 0.56576 | 2.33771 | 0.00775 | 0.00000 | 0.15107 | 0.03952 | 0.00000 | 0.14248 | 0.00616 | 0.00754 |
| **SOX9-AS1** | 0.04068 | 0.01420 | 0.01505 | 0.01101 | 0.01296 | 0.02017 | 0.01845 | 0.00770 | 0.01043 | 0.01891 | 0.03241 | 0.01496 |
| **SP1** | 2.32455 | 5.58509 | 9.21078 | 4.44077 | 4.27136 | 15.12388 | 13.79286 | 12.12753 | 2.56148 | 23.15057 | 5.61622 | 15.80444 |
| **SP100** | 0.00632 | 0.08790 | 0.06232 | 0.01996 | 0.00000 | 0.19129 | 0.22305 | 0.02626 | 0.00000 | 0.18520 | 0.36951 | 0.23527 |
| **SP110** | 0.75885 | 0.13319 | 0.08991 | 0.04274 | 0.02583 | 0.11234 | 0.01481 | 0.27039 | 0.00967 | 0.00000 | 0.00302 | 0.26677 |
| **SP140** | 0.01263 | 0.05586 | 0.02432 | 0.21761 | 0.00393 | 0.00090 | 0.00878 | 0.04790 | 0.01312 | 0.06416 | 0.01813 | 0.01537 |
| **SP140L** | 0.05725 | 0.62750 | 0.48192 | 0.00000 | 0.00000 | 0.87135 | 0.84526 | 0.03418 | 0.00000 | 0.29014 | 0.67126 | 0.08598 |
| **SP2** | 4.02784 | 3.46776 | 4.66303 | 2.30611 | 3.57124 | 1.63730 | 3.21459 | 1.26481 | 7.27452 | 0.00000 | 1.85385 | 5.88131 |
| **SP3** | 2.74747 | 3.20823 | 9.03694 | 1.73404 | 1.90631 | 19.06660 | 7.05217 | 7.04827 | 4.32916 | 10.32429 | 3.82304 | 5.62169 |
| **SP4** | 0.03193 | 0.21542 | 2.17547 | 0.93447 | 0.00000 | 4.74952 | 3.34185 | 3.40971 | 2.04725 | 0.00000 | 1.12286 | 0.31213 |
| **SP5** | 0.07773 | 0.15694 | 0.60341 | 1.59043 | 0.21568 | 0.00000 | 0.00000 | 0.01102 | 0.00000 | 0.41515 | 0.23501 | 0.00275 |
| **SP6** | 0.00000 | 0.00000 | 36.51075 | 0.00000 | 3.77711 | 1.42127 | 22.18618 | 0.19734 | 4.72002 | 1.63205 | 0.00000 | 0.05199 |
| **SP7** | 0.05907 | 0.01072 | 1.03251 | 0.26719 | 0.00000 | 0.26071 | 0.62476 | 0.00256 | 0.22806 | 0.00000 | 0.00000 | 0.11365 |
| **SP8** | 0.03491 | 0.19458 | 0.75069 | 5.31151 | 0.30857 | 0.67216 | 0.00000 | 0.00000 | 0.00000 | 0.65065 | 0.00000 | 1.14607 |
| **SP9** | 0.02804 | 0.01391 | 0.01046 | 0.12661 | 0.03645 | 0.00000 | 0.01898 | 0.01010 | 0.02011 | 0.02061 | 0.01443 | 0.02458 |
| **SPA17** | 0.10274 | 1.93942 | 1.17913 | 1.49448 | 1.61126 | 5.78230 | 3.02721 | 0.00000 | 0.26838 | 0.29741 | 0.98238 | 1.15195 |
| **SPACA1** | 0.00193 | 0.00576 | 0.01292 | 0.01061 | 0.01097 | 0.01429 | 0.02129 | 0.14490 | 0.01893 | 0.03685 | 0.01688 | 0.26746 |
| **SPACA4** | 0.00537 | 0.01785 | 0.03136 | 0.01404 | 0.02449 | 0.00472 | 0.02601 | 0.05743 | 0.03741 | 0.00106 | 0.01262 | 0.01786 |
| **SPACA6P** | 0.06090 | 0.10409 | 0.00558 | 0.08666 | 0.13041 | 0.06747 | 0.00000 | 0.00961 | 0.09358 | 0.41915 | 0.02806 | 0.07910 |
| **SPACA6P-AS** | 0.00780 | 0.01280 | 0.02752 | 0.03046 | 0.04522 | 0.00035 | 0.04112 | 0.01343 | 0.00000 | 0.03251 | 0.01869 | 0.02936 |
| **SPAG1** | 0.25009 | 0.00000 | 0.28932 | 0.14761 | 0.07130 | 1.71948 | 0.25529 | 0.70606 | 0.25112 | 1.30303 | 0.12711 | 0.52463 |
| **SPAG16** | 2.11328 | 0.01777 | 0.79449 | 0.57161 | 0.27460 | 1.76966 | 0.81218 | 0.67468 | 0.00000 | 0.00000 | 1.16157 | 0.67786 |
| **SPAG17** | 0.00421 | 0.04347 | 0.03601 | 0.08051 | 0.00000 | 0.09133 | 0.10282 | 0.40457 | 0.00000 | 0.25167 | 0.01018 | 0.06942 |
| **SPAG4** | 0.43924 | 0.00389 | 0.18343 | 0.02000 | 0.05330 | 0.11478 | 0.00000 | 2.11964 | 0.03557 | 0.13105 | 0.01518 | 0.18450 |
| **SPAG5** | 0.92658 | 25.59147 | 7.82996 | 16.02749 | 11.68955 | 21.87920 | 12.79982 | 6.26627 | 17.24120 | 6.19440 | 10.64216 | 21.99999 |
| **SPAG5-AS1** | 0.02331 | 0.11762 | 0.15066 | 0.30721 | 0.00000 | 0.19516 | 0.02008 | 0.07795 | 0.34579 | 0.44005 | 0.12181 | 0.16938 |
| **SPAG6** | 1.15620 | 0.00000 | 0.21246 | 0.01229 | 0.09489 | 0.18495 | 0.00000 | 0.69159 | 0.52998 | 0.00000 | 0.02538 | 1.08404 |
| **SPAG7** | 2.14020 | 5.42057 | 10.99734 | 9.11789 | 5.65801 | 5.06735 | 12.91300 | 1.73308 | 2.28666 | 1.76545 | 11.32075 | 4.61023 |
| **SPAG8** | 0.00609 | 0.02001 | 0.01391 | 0.01196 | 0.01019 | 0.02378 | 0.00710 | 0.00917 | 0.03858 | 0.00273 | 0.08641 | 0.04908 |
| **SPAG9** | 1.15063 | 1.01800 | 9.17623 | 2.24554 | 5.33738 | 8.93458 | 11.21516 | 0.86667 | 3.77790 | 15.63582 | 3.80446 | 2.14376 |
| **SPAM1** | 0.06453 | 0.01049 | 0.16066 | 0.02973 | 0.01831 | 0.02603 | 0.02641 | 0.09251 | 0.15386 | 0.02336 | 0.00000 | 0.05915 |
| **SPANXA2-OT1** | 0.00543 | 0.00819 | 0.01819 | 0.00715 | 0.01314 | 0.03082 | 0.00000 | 0.01787 | 0.09136 | 0.14295 | 0.00160 | 0.05268 |
| **SPANXN3** | 0.21004 | 0.01114 | 0.01125 | 0.00741 | 0.05388 | 0.00599 | 0.03901 | 0.01905 | 0.00000 | 0.04424 | 0.01782 | 0.01266 |
| **SPANXN5** | 0.00194 | 0.01577 | 0.10440 | 0.01042 | 0.01720 | 0.01166 | 0.01067 | 0.06827 | 0.00000 | 0.01314 | 0.03531 | 0.01447 |
| **SPARC** | 0.00000 | 27.93619 | 12.65072 | 41.31277 | 26.91968 | 14.47334 | 0.00000 | 0.00000 | 4.12900 | 4.77352 | 10.99873 | 2.29249 |
| **SPARCL1** | 0.00873 | 0.03247 | 0.05646 | 0.32878 | 0.01079 | 0.06942 | 0.03084 | 0.01015 | 0.09221 | 0.01218 | 0.01095 | 0.05368 |

| | | | | | | | | | | | | |
|---|---|---|---|---|---|---|---|---|---|---|---|---|
| **SPAST** | 1.68914 | 1.71952 | 3.95272 | 1.75028 | 0.00444 | 9.53505 | 1.46343 | 0.49489 | 4.21868 | 6.24729 | 2.80975 | 1.34845 |
| **SPATA12** | 0.00532 | 0.01974 | 0.09720 | 0.07784 | 0.00000 | 0.05480 | 0.00000 | 0.00958 | 0.09783 | 0.02955 | 0.04557 | 0.03307 |
| **SPATA13** | 0.12969 | 0.84475 | 13.56886 | 1.27915 | 0.07708 | 2.59371 | 2.44161 | 0.11649 | 2.51048 | 2.46049 | 0.00000 | 0.85963 |
| **SPATA13-AS1** | 0.00653 | 0.01426 | 0.01547 | 0.01085 | 0.01357 | 0.02613 | 0.01502 | 0.01035 | 0.04007 | 0.00364 | 0.01517 | 0.01653 |
| **SPATA16** | 0.13918 | 0.01537 | 0.01689 | 0.01375 | 0.01801 | 0.01076 | 0.02516 | 0.01343 | 0.01616 | 0.01201 | 0.01505 | 0.01521 |
| **SPATA17** | 0.01780 | 0.40435 | 0.00000 | 0.00000 | 0.51033 | 1.13780 | 0.24758 | 0.19347 | 0.00000 | 0.01472 | 0.51022 | 2.47060 |
| **SPATA18** | 0.03663 | 2.32744 | 0.57152 | 0.38360 | 1.02824 | 0.82796 | 0.00000 | 0.00000 | 0.34931 | 0.41946 | 0.19412 | 0.69022 |
| **SPATA2** | 8.24967 | 7.44073 | 4.44848 | 4.13019 | 3.57765 | 0.00000 | 2.83027 | 8.31529 | 6.44362 | 3.05890 | 0.84115 | 11.90403 |
| **SPATA20** | 0.00505 | 0.15221 | 0.06783 | 0.00000 | 0.01944 | 0.05913 | 0.00005 | 0.01060 | 0.00000 | 0.03110 | 0.18335 | 0.02554 |
| **SPATA21** | 0.00252 | 0.04332 | 0.02501 | 0.10005 | 0.01958 | 0.01798 | 0.01527 | 0.05611 | 0.03210 | 0.00636 | 0.02748 | 0.03493 |
| **SPATA22** | 0.93788 | 0.08369 | 0.60732 | 0.17127 | 0.00000 | 0.88962 | 0.77979 | 0.33435 | 0.64862 | 0.19885 | 0.34487 | 0.83100 |
| **SPATA24** | 0.08636 | 0.07928 | 0.63704 | 0.68328 | 0.66238 | 0.12411 | 0.15756 | 0.02398 | 0.13762 | 0.00000 | 0.22198 | 0.37628 |
| **SPATA25** | 0.00495 | 0.04179 | 0.03114 | 0.07030 | 0.02402 | 0.00000 | 0.00954 | 0.01024 | 0.03886 | 0.08943 | 0.03200 | 0.01462 |
| **SPATA2L** | 0.14380 | 1.34379 | 1.16372 | 2.12138 | 1.59079 | 0.00000 | 1.70876 | 0.00000 | 2.49498 | 0.00000 | 0.41031 | 0.83456 |
| **SPATA3** | 0.01121 | 0.01468 | 0.02195 | 0.01222 | 0.01511 | 0.01885 | 0.01712 | 0.01048 | 0.01629 | 0.01466 | 0.01529 | 0.01652 |
| **SPATA3-AS1** | 0.00592 | 0.10215 | 0.19607 | 0.01118 | 0.00375 | 0.01037 | 0.02133 | 0.01154 | 0.05844 | 0.00640 | 0.01509 | 0.01363 |
| **SPATA31A6** | 0.00630 | 0.01433 | 0.01423 | 0.01266 | 0.02399 | 0.02443 | 0.01704 | 0.01042 | 0.01843 | 0.01014 | 0.01417 | 0.01602 |
| **SPATA31C1** | 0.08778 | 0.03654 | 0.02214 | 0.04980 | 0.00920 | 0.02448 | 0.00120 | 0.01564 | 0.03872 | 0.04182 | 0.00885 | 0.09932 |
| **SPATA31C2** | 0.04621 | 0.00217 | 0.01219 | 0.02564 | 0.02571 | 0.03051 | 0.04634 | 0.05644 | 0.00081 | 0.20415 | 0.01155 | 0.12058 |
| **SPATA31D1** | 0.00544 | 0.00606 | 0.00000 | 0.03945 | 0.00000 | 0.08425 | 0.09834 | 0.00746 | 0.04353 | 0.00000 | 0.02984 | 0.18192 |
| **SPATA31D5P** | 0.00839 | 0.04008 | 0.01140 | 0.00474 | 0.00626 | 0.04175 | 0.01321 | 0.01064 | 0.01193 | 0.02660 | 0.01330 | 0.01184 |
| **SPATA31E1** | 0.00482 | 0.00992 | 0.03575 | 0.00881 | 0.00342 | 0.01927 | 0.01036 | 0.00922 | 0.00574 | 0.11659 | 0.01781 | 0.02683 |
| **SPATA33** | 0.84374 | 1.67353 | 1.66492 | 0.52707 | 1.14413 | 0.00000 | 1.14316 | 6.80933 | 3.21974 | 2.26324 | 2.45005 | 4.62555 |
| **SPATA4** | 0.03983 | 0.05398 | 0.04240 | 0.02472 | 0.01101 | 0.00000 | 0.01345 | 0.06266 | 0.05734 | 0.00000 | 0.02251 | 0.04171 |
| **SPATA41** | 0.02632 | 0.01514 | 0.04255 | 0.01456 | 0.01624 | 0.00961 | 0.02674 | 0.00668 | 0.01231 | 0.01390 | 0.01806 | 0.02333 |
| **SPATA42** | 0.00960 | 0.01445 | 0.01639 | 0.01084 | 0.01354 | 0.02180 | 0.02692 | 0.01757 | 0.01460 | 0.01331 | 0.01599 | 0.01579 |
| **SPATA45** | 0.00459 | 0.00387 | 0.04686 | 0.02045 | 0.03697 | 0.06141 | 0.00000 | 0.00775 | 0.00000 | 0.08650 | 0.00798 | 0.06182 |
| **SPATA5** | 4.84163 | 1.50091 | 5.05468 | 1.33466 | 1.90341 | 7.91257 | 4.88913 | 4.96932 | 0.90365 | 4.71163 | 2.41424 | 4.66231 |
| **SPATA5L1** | 0.58566 | 3.29194 | 4.63883 | 0.00000 | 1.86361 | 2.10516 | 2.87883 | 1.35606 | 1.42815 | 7.42161 | 1.41997 | 2.40906 |
| **SPATA6** | 0.74117 | 1.52460 | 0.29627 | 2.99640 | 0.60705 | 1.04760 | 0.01078 | 0.00000 | 0.00000 | 0.71731 | 0.38881 | 1.28471 |
| **SPATA6L** | 6.02870 | 0.06940 | 0.36299 | 0.06185 | 0.03017 | 0.00000 | 0.00000 | 0.78362 | 0.04189 | 0.07278 | 0.08379 | 0.02640 |
| **SPATA7** | 1.50302 | 1.92720 | 0.38467 | 0.81528 | 0.00071 | 2.40173 | 0.00001 | 0.22025 | 0.00000 | 0.00126 | 0.41591 | 1.88917 |
| **SPATA8** | 0.00615 | 0.01485 | 0.03751 | 0.01070 | 0.01332 | 0.01637 | 0.01487 | 0.01044 | 0.03130 | 0.02204 | 0.01165 | 0.01579 |
| **SPATA9** | 0.00222 | 0.00601 | 0.00000 | 0.00415 | 0.05531 | 0.07335 | 0.00000 | 0.05428 | 0.13934 | 0.00450 | 0.03314 | 0.01112 |
| **SPATC1** | 0.00430 | 0.00968 | 0.00344 | 0.00782 | 0.00326 | 0.01714 | 0.00799 | 0.01020 | 0.02855 | 0.16391 | 0.00965 | 0.01108 |
| **SPATC1L** | 0.68870 | 0.23188 | 1.58162 | 0.00000 | 0.14394 | 0.00000 | 0.00000 | 0.07092 | 1.18631 | 0.10478 | 0.59328 | 0.10934 |
| **SPATS1** | 0.00829 | 0.01716 | 0.13181 | 0.01822 | 0.01175 | 0.03922 | 0.01346 | 0.01039 | 0.01124 | 0.00844 | 0.01440 | 0.01491 |
| **SPATS2** | 0.46374 | 2.25434 | 2.43791 | 7.33366 | 1.73668 | 7.10329 | 5.81476 | 0.39463 | 3.10001 | 6.68138 | 2.34789 | 2.19724 |
| **SPATS2L** | 0.41110 | 1.99500 | 1.04467 | 11.41740 | 0.95935 | 2.21393 | 0.28957 | 0.00000 | 0.00000 | 2.30056 | 0.32753 | 0.00000 |
| **SPC24** | 0.16515 | 1.00265 | 0.83860 | 0.86897 | 0.61866 | 0.73368 | 0.00000 | 0.00047 | 2.05935 | 0.00000 | 0.80725 | 0.98941 |
| **SPC25** | 0.00000 | 4.00948 | 6.12873 | 2.00083 | 1.62439 | 15.76886 | 7.43896 | 0.00000 | 3.10396 | 2.71571 | 5.61157 | 2.96938 |
| **SPCS1** | 5.17627 | 22.42679 | 21.87182 | 9.93645 | 10.85555 | 47.50135 | 7.55012 | 2.92850 | 39.84064 | 14.30503 | 16.62869 | 23.22800 |
| **SPCS2** | 1.30475 | 3.21962 | 4.38715 | 2.51310 | 2.92612 | 4.10605 | 2.12579 | 2.48442 | 6.78639 | 0.48118 | 3.29090 | 4.21886 |
| **SPCS3** | 0.00000 | 2.80494 | 20.94832 | 0.01531 | 1.40441 | 29.68585 | 0.01101 | 9.27382 | 22.07376 | 16.08364 | 10.73705 | 19.61906 |
| **SPDEF** | 0.04439 | 0.01755 | 0.06129 | 0.16259 | 0.00862 | 0.05243 | 0.03224 | 0.00638 | 0.01056 | 0.01140 | 0.03521 | 0.01169 |
| **SPDL1** | 12.32453 | 8.74574 | 6.89244 | 0.00002 | 1.65990 | 21.88028 | 13.34286 | 19.31044 | 6.25883 | 8.87860 | 8.18609 | 10.37912 |
| **SPDYA** | 0.01003 | 0.00112 | 0.39790 | 0.08099 | 0.00002 | 0.35488 | 0.00000 | 0.18093 | 0.14840 | 0.47984 | 0.22841 | 0.17617 |
| **SPDYC** | 11.95767 | 3.05170 | 1.14040 | 0.00000 | 0.00595 | 0.00000 | 0.00000 | 2.68294 | 0.23548 | 0.00000 | 0.42117 | 5.01882 |

| | | | | | | | | | | | | |
|---|---|---|---|---|---|---|---|---|---|---|---|---|
| **SPDYE1** | 0.05689 | 0.06873 | 0.01206 | 0.00883 | 0.11599 | 0.00000 | 0.04649 | 0.02301 | 0.13485 | 0.27734 | 0.13639 | 0.11468 |
| **SPDYE3** | 0.03308 | 0.10982 | 0.05027 | 0.10578 | 0.00738 | 0.00000 | 0.01453 | 0.00480 | 0.04584 | 0.04956 | 0.03686 | 0.07333 |
| **SPDYE5** | 0.00334 | 0.18752 | 0.08352 | 0.08398 | 0.00001 | 0.00000 | 0.00000 | 0.02551 | 0.23106 | 0.32236 | 0.27343 | 0.02840 |
| **SPDYE6** | 0.01693 | 0.02048 | 0.02570 | 0.01090 | 0.00717 | 0.03100 | 0.01865 | 0.02702 | 0.05278 | 0.02843 | 0.01939 | 0.03374 |
| **SPDYE7P** | 0.00592 | 0.01987 | 0.01406 | 0.01061 | 0.01363 | 0.01659 | 0.01533 | 0.01062 | 0.01647 | 0.03023 | 0.01483 | 0.02241 |
| **SPECC1** | 8.28393 | 4.10130 | 8.84723 | 3.89253 | 7.77771 | 8.31599 | 7.18204 | 4.69622 | 9.98760 | 11.75398 | 5.07741 | 6.70490 |
| **SPECC1L** | 0.65234 | 4.49118 | 1.39513 | 5.97977 | 3.89367 | 0.00000 | 3.84031 | 1.13352 | 0.88720 | 9.13059 | 2.14280 | 3.90924 |
| **SPECC1L-ADOI** | 0.00736 | 0.01446 | 0.01508 | 0.01444 | 0.01072 | 0.01835 | 0.01505 | 0.01057 | 0.01501 | 0.02106 | 0.03051 | 0.01789 |
| **SPEF1** | 0.08862 | 0.31344 | 0.08638 | 0.00000 | 0.00000 | 0.00000 | 0.00081 | 0.00816 | 0.00000 | 0.08090 | 0.13651 | 0.01682 |
| **SPEF2** | 0.00211 | 0.55054 | 0.00743 | 0.00000 | 0.10745 | 0.72536 | 0.00000 | 0.01769 | 0.00000 | 0.02518 | 0.04721 | 0.13745 |
| **SPEG** | 0.00730 | 0.01308 | 0.05639 | 0.01579 | 0.05951 | 0.00000 | 0.05474 | 0.01175 | 0.00000 | 0.03519 | 0.02218 | 0.01611 |
| **SPEM1** | 0.00450 | 0.00856 | 0.00064 | 0.00115 | 0.00000 | 0.02941 | 0.04265 | 0.01003 | 0.04453 | 0.12884 | 0.00000 | 0.01119 |
| **SPEN** | 0.00000 | 0.93193 | 4.97483 | 6.03410 | 11.38875 | 4.27897 | 11.35542 | 1.14243 | 8.97600 | 29.89994 | 4.00509 | 2.68402 |
| **SPERT** | 0.67869 | 0.00000 | 0.02455 | 0.00000 | 0.12867 | 0.06431 | 0.00000 | 0.01345 | 0.00000 | 0.05090 | 0.07493 | 0.07726 |
| **SPESP1** | 0.27584 | 5.19167 | 0.43648 | 0.00000 | 0.76165 | 3.33981 | 0.00000 | 3.21791 | 0.00000 | 0.00000 | 0.77461 | 3.56198 |
| **SPG11** | 1.27210 | 1.34458 | 2.08597 | 1.86289 | 2.84881 | 6.21536 | 0.00000 | 1.71607 | 0.00291 | 6.43622 | 4.38702 | 3.42409 |
| **SPG20** | 0.70040 | 7.40923 | 4.69889 | 0.99078 | 7.05009 | 18.87943 | 14.40846 | 0.48861 | 2.60704 | 3.28741 | 14.53188 | 5.19748 |
| **SPG20OS** | 0.00475 | 0.03886 | 0.03571 | 0.00145 | 0.07660 | 0.00000 | 0.07101 | 0.01247 | 0.00000 | 0.08877 | 0.08714 | 0.02021 |
| **SPG21** | 3.59949 | 4.52688 | 12.95373 | 2.81067 | 9.84104 | 4.91615 | 10.61112 | 0.39504 | 10.34707 | 2.62672 | 7.80889 | 2.31309 |
| **SPG7** | 0.88411 | 4.70476 | 4.04879 | 4.92193 | 6.17180 | 0.00000 | 0.27641 | 2.84457 | 8.86856 | 17.74953 | 3.50715 | 3.11640 |
| **SPHK1** | 0.39263 | 2.44957 | 1.39448 | 5.73760 | 4.42883 | 0.00000 | 1.42372 | 0.89313 | 0.83273 | 1.30588 | 0.62834 | 2.23207 |
| **SPHK2** | 0.36264 | 1.52275 | 8.65891 | 1.35270 | 0.00000 | 0.00000 | 8.47900 | 1.10543 | 9.42455 | 3.51310 | 0.94452 | 4.16920 |
| **SPHKAP** | 0.81894 | 0.00538 | 0.01014 | 0.00000 | 0.00984 | 0.07146 | 0.01957 | 0.13018 | 0.00026 | 0.06908 | 0.00776 | 0.06336 |
| **SPI1** | 0.00611 | 0.01421 | 0.01612 | 0.01093 | 0.01334 | 0.02420 | 0.02827 | 0.01040 | 0.01545 | 0.01667 | 0.01522 | 0.01656 |
| **SPIB** | 0.00850 | 0.55169 | 0.96488 | 0.43030 | 0.00000 | 0.00000 | 0.00000 | 0.01341 | 0.33856 | 1.32412 | 0.00000 | 1.11628 |
| **SPIC** | 0.02405 | 1.76314 | 0.00000 | 4.88344 | 0.00000 | 0.04136 | 0.00000 | 0.00000 | 0.00000 | 0.02565 | 0.00000 | 1.61102 |
| **SPICE1** | 1.08827 | 4.84371 | 2.19853 | 0.81687 | 0.00000 | 8.34060 | 0.00000 | 4.10846 | 3.47108 | 2.39198 | 1.87626 | 4.95647 |
| **SPIDR** | 1.18777 | 2.09822 | 3.54161 | 2.07275 | 1.44709 | 4.13626 | 2.08187 | 1.25191 | 5.41505 | 0.55063 | 5.20422 | 1.10177 |
| **SPIN1** | 5.94131 | 8.97434 | 6.50384 | 7.44780 | 0.70385 | 32.44333 | 16.81338 | 3.66716 | 5.85358 | 18.68021 | 11.70988 | 11.54557 |
| **SPIN2B** | 0.00107 | 0.05808 | 0.16367 | 0.00000 | 0.00000 | 0.10099 | 0.22888 | 0.31614 | 0.03265 | 0.29685 | 0.21182 | 0.14307 |
| **SPIN3** | 3.62880 | 0.34410 | 0.55333 | 0.37750 | 0.00000 | 0.00000 | 1.00519 | 1.69942 | 0.82197 | 0.00000 | 0.62277 | 0.83884 |
| **SPIN4** | 0.00000 | 3.60111 | 8.59715 | 0.00000 | 2.23323 | 11.29054 | 4.35193 | 0.00000 | 9.70203 | 5.48518 | 7.58640 | 0.99350 |
| **SPINK1** | 0.00180 | 0.97950 | 0.30668 | 1.38193 | 0.52561 | 0.00000 | 0.01911 | 0.00920 | 0.00000 | 0.74683 | 0.09785 | 1.05424 |
| **SPINK14** | 0.00535 | 0.01191 | 0.01464 | 0.00969 | 0.01051 | 0.03118 | 0.02392 | 0.01208 | 0.01724 | 0.01168 | 0.01828 | 0.02987 |
| **SPINK2** | 0.36640 | 0.79129 | 0.00000 | 0.56272 | 0.05851 | 0.06148 | 0.10813 | 0.00571 | 0.00000 | 0.05506 | 0.00002 | 0.69279 |
| **SPINK4** | 0.06812 | 0.00363 | 0.10264 | 0.08980 | 0.15674 | 0.00000 | 0.00106 | 0.06139 | 0.00000 | 0.12223 | 0.04806 | 0.02500 |
| **SPINK5** | 0.38285 | 1.74824 | 0.00000 | 1.58341 | 0.00000 | 0.16845 | 0.00000 | 0.09927 | 0.00000 | 1.78697 | 0.00000 | 2.47082 |
| **SPINK6** | 0.00627 | 0.01399 | 0.01455 | 0.01014 | 0.01107 | 0.02228 | 0.01718 | 0.01025 | 0.03963 | 0.01327 | 0.01655 | 0.01785 |
| **SPINK7** | 0.47281 | 0.27263 | 0.00329 | 0.00000 | 0.08852 | 0.01601 | 0.00819 | 1.47287 | 0.00000 | 0.07966 | 0.00224 | 0.11876 |
| **SPINK8** | 0.39036 | 0.01456 | 0.01794 | 0.01272 | 0.01734 | 0.01019 | 0.02001 | 0.00888 | 0.02260 | 0.00530 | 0.01656 | 0.02706 |
| **SPINT1** | 1.58161 | 26.37126 | 37.41300 | 8.71774 | 25.85890 | 0.00000 | 4.05604 | 1.53015 | 40.01620 | 18.43700 | 14.83813 | 10.15620 |
| **SPINT2** | 13.35103 | 40.96287 | 40.36247 | 25.06106 | 41.28002 | 36.66932 | 16.06271 | 14.87139 | 62.34633 | 15.83021 | 29.16597 | 26.21392 |
| **SPIRE1** | 1.43424 | 0.89638 | 0.00000 | 0.29919 | 3.75662 | 3.91906 | 0.67387 | 0.00000 | 0.35016 | 5.00845 | 0.61938 | 0.22851 |
| **SPIRE2** | 0.03406 | 0.00000 | 2.48627 | 0.24159 | 1.68957 | 0.00000 | 0.34176 | 0.13210 | 0.58050 | 0.99280 | 0.48283 | 0.00000 |
| **SPN** | 0.00941 | 0.01425 | 0.06639 | 0.01776 | 0.01639 | 0.00805 | 0.01235 | 0.01248 | 0.01698 | 0.05886 | 0.01767 | 0.01708 |
| **SPNS1** | 0.79540 | 9.59206 | 18.27802 | 6.82816 | 10.80599 | 0.00000 | 2.74035 | 0.09965 | 18.72901 | 19.87465 | 6.88718 | 2.09518 |
| **SPNS2** | 0.35143 | 0.06568 | 0.61505 | 0.03527 | 0.17421 | 0.00000 | 0.08765 | 0.00349 | 0.12554 | 0.15273 | 0.00000 | 0.20718 |
| **SPNS3** | 0.02389 | 0.02884 | 0.02562 | 0.01159 | 0.55650 | 0.21546 | 0.00000 | 0.04939 | 0.00000 | 0.00000 | 0.07594 | 0.50587 |

| | | | | | | | | | | | | |
|---|---|---|---|---|---|---|---|---|---|---|---|---|
| **SPO11** | 0.11652 | 0.00846 | 0.01273 | 0.01051 | 0.01201 | 0.02320 | 0.01769 | 0.06305 | 0.01579 | 0.02153 | 0.01412 | 0.05503 |
| **SPOCD1** | 0.00538 | 0.00894 | 0.01162 | 0.01726 | 0.00349 | 0.01576 | 0.01808 | 0.00829 | 0.03750 | 0.04446 | 0.03164 | 0.06075 |
| **SPOCK1** | 0.86360 | 0.00000 | 0.12474 | 0.42922 | 0.04736 | 0.24460 | 0.00000 | 0.27650 | 0.00000 | 0.01605 | 0.29735 | 2.21238 |
| **SPOCK2** | 0.20521 | 1.33716 | 0.00000 | 0.05454 | 1.43456 | 0.00000 | 0.30904 | 0.00000 | 0.08759 | 0.58001 | 0.00000 | 1.10049 |
| **SPOCK3** | 1.17338 | 0.21708 | 0.08436 | 0.00000 | 0.15501 | 0.00000 | 0.13562 | 0.20203 | 0.00000 | 0.09231 | 0.00466 | 0.00000 |
| **SPON1** | 1.85641 | 0.05943 | 0.12978 | 0.47532 | 0.00000 | 0.00000 | 0.09364 | 0.00571 | 0.33239 | 1.06809 | 0.08917 | 0.19125 |
| **SPON2** | 0.09725 | 0.00000 | 1.00592 | 1.50901 | 0.15108 | 0.00000 | 0.01608 | 0.30453 | 0.72965 | 0.01677 | 0.12642 | 0.31734 |
| **SPOP** | 1.56529 | 3.00875 | 7.81859 | 3.16092 | 2.22300 | 2.72157 | 8.63903 | 6.88342 | 4.93050 | 0.10795 | 6.85375 | 4.37042 |
| **SPOPL** | 0.77953 | 0.58664 | 6.06647 | 0.00000 | 2.12068 | 14.27674 | 7.11677 | 0.37737 | 4.49287 | 9.79282 | 2.84618 | 5.87655 |
| **SPP1** | 0.00446 | 0.00000 | 3.44068 | 9.59156 | 0.00000 | 0.03120 | 0.04747 | 0.00813 | 0.01484 | 0.01854 | 0.04508 | 0.00850 |
| **SPPL2A** | 0.95221 | 7.52740 | 24.22622 | 0.65958 | 4.97511 | 18.99198 | 0.00000 | 2.02488 | 28.65612 | 11.45663 | 12.80539 | 9.05280 |
| **SPPL2B** | 0.71160 | 2.18885 | 1.38837 | 0.78415 | 2.37637 | 0.00000 | 0.00000 | 0.03888 | 5.43798 | 6.26840 | 3.12043 | 1.40514 |
| **SPPL2C** | 0.00476 | 0.01647 | 0.02210 | 0.01692 | 0.01722 | 0.00651 | 0.04743 | 0.01576 | 0.00322 | 0.02012 | 0.01659 | 0.01676 |
| **SPPL3** | 5.19445 | 5.96824 | 3.62995 | 3.03861 | 3.04140 | 7.50718 | 4.41590 | 4.58452 | 0.64162 | 19.93290 | 0.00588 | 14.59876 |
| **SPR** | 1.87911 | 12.89767 | 3.73266 | 0.22338 | 1.55094 | 0.00008 | 5.01158 | 0.13785 | 3.74803 | 1.50696 | 4.05914 | 9.64522 |
| **SPRED1** | 0.21156 | 0.00000 | 0.25322 | 2.60171 | 0.00000 | 2.44947 | 0.70854 | 0.00000 | 0.00000 | 2.32068 | 0.00000 | 0.00000 |
| **SPRED2** | 1.01902 | 3.82476 | 1.49822 | 8.86807 | 0.80123 | 8.25177 | 4.05529 | 2.36562 | 1.64708 | 11.70330 | 0.67129 | 6.10177 |
| **SPRED3** | 0.03529 | 0.01166 | 0.26859 | 0.00084 | 0.11824 | 0.00000 | 0.02371 | 0.08468 | 0.34337 | 0.65878 | 0.00000 | 0.02458 |
| **SPRN** | 0.00027 | 0.62435 | 1.25430 | 1.03497 | 0.67818 | 0.00000 | 1.09416 | 0.08356 | 0.96990 | 3.59344 | 1.39893 | 0.65583 |
| **SPRNP1** | 0.00731 | 0.02500 | 0.02693 | 0.01166 | 0.03843 | 0.00000 | 0.01646 | 0.00996 | 0.03474 | 0.01358 | 0.01687 | 0.02312 |
| **SPRR2F** | 0.01682 | 0.01559 | 0.01373 | 0.01152 | 0.01419 | 0.01983 | 0.01728 | 0.12003 | 0.01729 | 0.01568 | 0.01482 | 0.01082 |
| **SPRR2G** | 0.00599 | 0.01409 | 0.01700 | 0.01380 | 0.01258 | 0.02008 | 0.02354 | 0.01036 | 0.01772 | 0.01766 | 0.01595 | 0.01991 |
| **SPRR3** | 0.00646 | 0.02264 | 0.01528 | 0.01711 | 0.04263 | 0.00000 | 0.03665 | 0.01066 | 0.01609 | 0.00036 | 0.01422 | 0.01974 |
| **SPRR4** | 0.00561 | 0.01222 | 0.01069 | 0.01052 | 0.01157 | 0.02700 | 0.03961 | 0.01015 | 0.00001 | 0.05746 | 0.01453 | 0.01426 |
| **SPRTN** | 0.75450 | 4.42859 | 5.53034 | 1.30145 | 1.18720 | 7.63253 | 4.39232 | 11.74579 | 7.92765 | 1.28025 | 3.85588 | 5.49754 |
| **SPRY1** | 0.18381 | 0.00000 | 0.63783 | 0.82186 | 0.00000 | 0.00000 | 1.02027 | 0.01187 | 0.63486 | 0.55965 | 0.52553 | 0.05249 |
| **SPRY2** | 0.00025 | 0.00000 | 0.97113 | 8.75272 | 0.00000 | 0.49973 | 0.00000 | 0.17123 | 0.00000 | 2.69626 | 0.00000 | 1.33171 |
| **SPRY4** | 1.23986 | 0.15454 | 1.13338 | 6.87919 | 2.51114 | 0.00000 | 0.51561 | 8.81640 | 0.75135 | 1.53760 | 0.00000 | 2.02041 |
| **SPRYD3** | 1.13928 | 3.37837 | 3.00991 | 1.69972 | 1.86569 | 0.00000 | 2.47331 | 1.02671 | 3.94609 | 0.00000 | 0.78474 | 5.57659 |
| **SPRYD4** | 0.01011 | 1.49004 | 3.65444 | 3.46191 | 0.65766 | 1.43075 | 3.38157 | 0.63890 | 1.53015 | 0.00000 | 4.00358 | 5.16732 |
| **SPRYD7** | 1.55276 | 4.96340 | 7.58082 | 1.85422 | 10.77576 | 14.52699 | 2.98944 | 0.56234 | 0.00000 | 2.26549 | 6.15284 | 1.34522 |
| **SPSB1** | 1.79143 | 0.78846 | 3.10413 | 0.16308 | 5.30038 | 0.48705 | 2.38611 | 0.10100 | 0.32607 | 2.72573 | 0.76576 | 0.20069 |
| **SPSB2** | 0.51218 | 6.96059 | 2.43703 | 10.13007 | 2.83912 | 0.00000 | 1.05768 | 0.56329 | 0.44962 | 0.21523 | 1.99710 | 7.53579 |
| **SPSB3** | 0.54707 | 16.72200 | 8.09272 | 5.36370 | 15.78170 | 0.00000 | 4.43023 | 3.14755 | 19.06786 | 2.14823 | 9.10715 | 17.76697 |
| **SPSB4** | 0.20871 | 0.00000 | 0.00000 | 0.38002 | 3.00780 | 0.00000 | 6.23865 | 0.14454 | 4.67790 | 4.67905 | 5.71026 | 0.52001 |
| **SPTA1** | 0.00372 | 0.00590 | 0.05306 | 0.03052 | 0.00045 | 0.04361 | 0.00001 | 0.03226 | 0.00000 | 0.05364 | 0.03408 | 0.03039 |
| **SPTAN1** | 7.01495 | 11.96675 | 4.10568 | 6.12101 | 24.15512 | 6.83134 | 2.39410 | 0.97354 | 20.50160 | 23.83518 | 10.92722 | 8.08265 |
| **SPTB** | 5.05612 | 2.36631 | 0.00000 | 2.04810 | 6.81663 | 0.00000 | 0.58833 | 0.30108 | 0.36744 | 2.11879 | 0.59068 | 2.27807 |
| **SPTBN1** | 9.62748 | 15.45491 | 0.00000 | 13.81118 | 33.65237 | 22.83310 | 3.64270 | 3.86803 | 19.06965 | 38.91852 | 8.38314 | 6.90635 |
| **SPTBN2** | 0.03449 | 0.47178 | 0.00000 | 0.00000 | 0.66201 | 0.00000 | 0.04940 | 0.02831 | 0.65227 | 4.10700 | 0.27754 | 1.66333 |
| **SPTBN4** | 0.31984 | 0.37621 | 0.04303 | 0.17705 | 0.16508 | 0.00000 | 0.03989 | 0.13558 | 0.00000 | 0.00000 | 0.08521 | 0.56450 |
| **SPTBN5** | 0.63991 | 0.10672 | 0.17832 | 0.00000 | 0.00000 | 0.00000 | 0.27425 | 0.13232 | 0.41256 | 1.28664 | 0.50199 | 0.18480 |
| **SPTLC1** | 2.45445 | 12.45483 | 12.79956 | 7.36879 | 4.50056 | 29.71730 | 0.00000 | 1.78595 | 33.73529 | 5.31036 | 7.28476 | 8.24973 |
| **SPTLC2** | 0.00000 | 22.04719 | 0.00000 | 13.18992 | 27.53984 | 19.13146 | 0.00000 | 0.46752 | 5.69689 | 13.27492 | 18.35579 | 6.46016 |
| **SPTLC3** | 0.03451 | 0.00000 | 1.30127 | 0.20893 | 0.94412 | 0.23819 | 0.00000 | 0.00360 | 0.29258 | 0.33190 | 0.53428 | 0.03743 |
| **SPTSSA** | 0.00000 | 6.04344 | 4.34136 | 7.98837 | 1.81899 | 22.92324 | 9.94332 | 1.20513 | 0.00000 | 19.02590 | 24.22032 | 9.80085 |
| **SPTSSB** | 0.10573 | 0.00709 | 0.05212 | 0.00845 | 0.01852 | 0.05842 | 0.00009 | 0.30335 | 0.00813 | 0.02214 | 0.01776 | 0.42384 |
| **SPTY2D1** | 4.20900 | 2.16232 | 7.17574 | 2.44548 | 2.29173 | 13.25530 | 8.78441 | 12.61587 | 3.62876 | 5.10522 | 5.99024 | 5.81756 |

| | | | | | | | | | | | | |
|---|---|---|---|---|---|---|---|---|---|---|---|---|
| SPTY2D1-AS1 | 0.01233 | 0.03953 | 0.15751 | 0.02384 | 0.13414 | 0.00000 | 0.03707 | 0.12650 | 0.00000 | 0.41333 | 0.39895 | 0.10367 |
| SPX | 0.02492 | 0.01374 | 0.01493 | 0.03856 | 0.04000 | 0.01583 | 0.02055 | 0.06494 | 0.01921 | 0.00000 | 0.01518 | 0.03192 |
| SPZ1 | 0.00332 | 0.16700 | 0.02594 | 0.18515 | 0.01044 | 0.06383 | 0.00807 | 0.23647 | 0.00191 | 0.03100 | 0.02522 | 0.52188 |
| SQLE | 1.16265 | 0.00000 | 22.43131 | 22.67413 | 43.03624 | 3.44275 | 15.03703 | 1.65158 | 42.20240 | 7.79539 | 31.15892 | 0.07189 |
| SQRDL | 10.36279 | 0.01028 | 0.01534 | 0.00626 | 0.09060 | 0.00000 | 0.00880 | 5.82563 | 0.02075 | 0.00000 | 0.00283 | 0.19978 |
| SQSTM1 | 5.03780 | 19.03254 | 19.46952 | 12.72888 | 17.19536 | 12.41752 | 21.47857 | 3.48079 | 15.08911 | 3.20157 | 2.92084 | 8.74569 |
| SRA1 | 0.18345 | 15.60120 | 15.18133 | 12.85569 | 16.49132 | 10.06919 | 23.70148 | 10.84190 | 15.20968 | 0.00000 | 15.58053 | 14.42065 |
| SRBD1 | 1.65534 | 4.28003 | 1.00726 | 1.25874 | 0.68415 | 8.36925 | 0.00000 | 1.97410 | 1.56783 | 2.72648 | 2.45141 | 3.51903 |
| SRC | 0.06115 | 0.46109 | 8.33729 | 0.35595 | 1.44391 | 0.00000 | 2.55700 | 0.00456 | 4.13508 | 2.55263 | 0.32755 | 0.20324 |
| SRCAP | 6.31816 | 8.15431 | 11.56416 | 9.46781 | 13.84355 | 0.00000 | 8.43703 | 2.09131 | 23.27299 | 38.82865 | 6.89855 | 13.22683 |
| SRCIN1 | 0.17651 | 0.01786 | 0.03276 | 0.01151 | 0.04953 | 0.04489 | 0.04763 | 0.00635 | 0.00480 | 0.01724 | 0.01224 | 0.01514 |
| SRCRB4D | 0.10544 | 0.12670 | 0.09720 | 0.00000 | 0.10009 | 0.00000 | 0.08032 | 0.05618 | 0.01369 | 0.09280 | 0.08562 | 0.13179 |
| SRD5A1 | 0.86494 | 1.07718 | 0.87031 | 1.15094 | 1.06262 | 1.71169 | 1.40609 | 0.14366 | 2.22147 | 0.36727 | 0.78759 | 0.19223 |
| SRD5A1P1 | 0.02751 | 0.04920 | 0.03540 | 0.02481 | 0.02178 | 0.11112 | 0.02761 | 0.02132 | 0.09901 | 0.00000 | 0.04005 | 0.02496 |
| SRD5A2 | 0.03222 | 0.00000 | 0.01201 | 0.01057 | 0.01350 | 0.01757 | 0.02666 | 0.00269 | 0.01451 | 0.01696 | 0.01460 | 0.58786 |
| SRD5A3 | 0.00706 | 10.12508 | 0.64497 | 3.64996 | 3.85407 | 0.00000 | 0.00000 | 0.06504 | 5.59662 | 8.28390 | 19.02556 | 2.28093 |
| SRD5A3-AS1 | 0.12602 | 0.03224 | 0.01769 | 0.01958 | 0.03212 | 0.01214 | 0.00924 | 0.05050 | 0.01899 | 0.03623 | 0.03538 | 0.03022 |
| SREBF1 | 0.46870 | 0.00000 | 8.40686 | 2.61419 | 4.34438 | 0.00000 | 0.34637 | 0.24989 | 13.94403 | 7.25372 | 1.06088 | 0.01363 |
| SREBF2 | 8.39167 | 0.00000 | 11.42685 | 9.03307 | 17.77669 | 0.00000 | 6.61148 | 0.52084 | 13.65401 | 13.35014 | 11.66711 | 0.31765 |
| SREK1 | 6.46840 | 6.05776 | 8.64885 | 4.10246 | 8.36702 | 18.61786 | 14.84818 | 6.77534 | 8.34005 | 30.10207 | 10.09408 | 3.15252 |
| SREK1IP1 | 2.11497 | 0.00000 | 6.87581 | 4.40036 | 0.00000 | 5.72954 | 13.56753 | 4.48370 | 6.45136 | 8.60515 | 3.18855 | 5.20273 |
| SRF | 0.51568 | 2.60251 | 4.03660 | 4.30840 | 9.05646 | 0.00000 | 8.24544 | 0.15126 | 0.00000 | 15.88931 | 0.40553 | 2.12557 |
| SRFBP1 | 0.32635 | 3.12279 | 3.49228 | 0.29215 | 1.12464 | 15.69036 | 5.01665 | 4.53558 | 6.54116 | 5.92271 | 2.71991 | 6.87581 |
| SRGAP1 | 2.58156 | 2.54709 | 0.69987 | 12.39427 | 1.04925 | 0.00000 | 0.69753 | 0.00010 | 0.00000 | 1.32292 | 0.26375 | 4.21783 |
| SRGAP2 | 1.07595 | 0.89967 | 2.77860 | 7.03528 | 1.85435 | 0.17945 | 1.71543 | 0.00000 | 2.77813 | 2.80108 | 0.00000 | 1.48120 |
| SRGAP2-AS1 | 0.00609 | 0.01491 | 0.02129 | 0.01097 | 0.01383 | 0.01960 | 0.01338 | 0.01026 | 0.02355 | 0.02012 | 0.01287 | 0.02022 |
| SRGAP2D.uid-4 | 0.07760 | 0.18553 | 0.23612 | 1.54388 | 0.00000 | 1.94015 | 0.54744 | 0.36813 | 0.00000 | 0.37055 | 0.03881 | 0.32166 |
| SRGAP3 | 0.86905 | 0.11993 | 0.08510 | 0.00362 | 0.00000 | 0.02912 | 0.01248 | 0.08548 | 0.01216 | 0.08474 | 0.06819 | 0.06786 |
| SRGN | 0.01104 | 0.03686 | 0.01970 | 0.00161 | 0.00324 | 0.09206 | 0.01539 | 0.01316 | 0.00760 | 0.01966 | 0.00814 | 0.14207 |
| SRI | 4.68100 | 10.43705 | 7.32254 | 4.76952 | 0.00000 | 9.92332 | 20.73337 | 1.39269 | 6.84247 | 2.02531 | 12.88278 | 2.53631 |
| SRL | 0.00584 | 0.01637 | 0.01692 | 0.01238 | 0.01123 | 0.02457 | 0.01903 | 0.01208 | 0.03358 | 0.01033 | 0.01275 | 0.01639 |
| SRM | 0.25581 | 31.30447 | 18.41442 | 8.32325 | 59.90874 | 13.22824 | 33.53094 | 6.64459 | 12.61536 | 51.61360 | 31.94181 | 24.19259 |
| SRMS | 0.00496 | 0.01195 | 0.01891 | 0.00372 | 0.03437 | 0.01467 | 0.01277 | 0.01047 | 0.02929 | 0.08595 | 0.01243 | 0.01297 |
| SRP14 | 4.38761 | 23.46757 | 29.54272 | 19.04245 | 23.51248 | 46.78742 | 41.16095 | 31.44717 | 28.66400 | 10.68626 | 29.90339 | 34.20505 |
| SRP14-AS1 | 0.11468 | 0.62576 | 2.49721 | 0.00529 | 1.08534 | 2.02453 | 0.86325 | 0.34505 | 0.68087 | 0.00000 | 2.56346 | 1.17176 |
| SRP19 | 3.58582 | 7.35010 | 20.40780 | 3.66065 | 3.90805 | 29.48035 | 24.05585 | 10.23702 | 11.94489 | 12.31728 | 13.24413 | 14.85002 |
| SRP54 | 1.43017 | 5.37968 | 14.30606 | 6.83827 | 0.27679 | 34.18488 | 14.85369 | 10.91837 | 14.31324 | 5.85668 | 13.80484 | 15.66237 |
| SRP68 | 5.67279 | 13.31188 | 11.40422 | 7.36182 | 12.61743 | 9.05422 | 10.60372 | 2.54168 | 6.31037 | 11.25380 | 11.72296 | 0.71913 |
| SRP72 | 3.47959 | 8.03226 | 15.99716 | 7.26243 | 10.00481 | 29.62305 | 22.92951 | 6.00730 | 12.51138 | 16.19233 | 15.16234 | 17.93133 |
| SRP9 | 0.33542 | 8.74184 | 15.35623 | 4.77009 | 0.00000 | 42.00329 | 21.63458 | 10.65415 | 13.35078 | 4.55105 | 11.82919 | 16.70572 |
| SRPK1 | 12.30299 | 13.03829 | 16.55981 | 6.29001 | 17.96003 | 34.26984 | 25.92054 | 10.49031 | 25.50011 | 19.14617 | 23.19309 | 16.75434 |
| SRPK2 | 3.83942 | 17.95058 | 4.16464 | 13.80209 | 11.20936 | 19.86128 | 7.12994 | 6.64402 | 4.82700 | 12.35883 | 6.36306 | 16.55462 |
| SRPK3 | 0.28711 | 0.01354 | 0.07061 | 0.01452 | 0.00238 | 0.06498 | 0.00000 | 0.00109 | 0.00000 | 0.04755 | 0.01639 | 0.01927 |
| SRPR | 4.97313 | 16.68280 | 16.81211 | 9.17928 | 11.83724 | 11.42905 | 8.47379 | 6.18946 | 25.36908 | 0.20529 | 12.29061 | 15.99319 |
| SRPRB | 1.41518 | 17.83064 | 17.61211 | 11.40526 | 9.60592 | 24.00945 | 4.68555 | 0.09048 | 28.79005 | 12.24279 | 12.44136 | 10.12607 |
| SRPX | 0.04649 | 0.13671 | 0.04285 | 0.16099 | 0.01207 | 0.00000 | 0.01955 | 0.03533 | 0.02075 | 0.00000 | 0.02004 | 0.41890 |
| SRPX2 | 0.00317 | 0.02273 | 0.01311 | 0.01737 | 0.07260 | 0.00000 | 0.02706 | 0.07717 | 0.02321 | 0.03716 | 0.01449 | 0.03115 |
| SRR | 0.01551 | 1.06879 | 0.63983 | 0.65875 | 0.30887 | 1.04359 | 0.85823 | 0.54726 | 0.91209 | 0.55861 | 0.98503 | 1.44594 |

| | | | | | | | | | | | | |
|---|---|---|---|---|---|---|---|---|---|---|---|---|
| **SRRD** | 1.23131 | 2.49698 | 3.54891 | 3.52757 | 4.53684 | 6.55988 | 8.91452 | 2.32308 | 1.39407 | 12.28847 | 2.41012 | 3.64470 |
| **SRRM1** | 3.79109 | 12.07018 | 13.34457 | 10.14197 | 18.94576 | 27.66527 | 20.07633 | 14.43977 | 16.34361 | 34.68316 | 19.65464 | 17.50230 |
| **SRRM2** | 9.90136 | 12.08139 | 14.17758 | 16.97034 | 31.76266 | 9.89062 | 17.43385 | 17.24290 | 30.82974 | 52.98292 | 8.40555 | 24.28473 |
| **SRRM2-AS1** | 0.19731 | 0.85302 | 0.12128 | 0.00000 | 0.02881 | 0.02254 | 0.01099 | 0.26641 | 0.00000 | 0.02860 | 0.14433 | 0.95598 |
| **SRRM3** | 0.13377 | 0.26350 | 0.63258 | 0.00000 | 0.17679 | 0.00000 | 0.45482 | 0.31835 | 0.00000 | 0.75472 | 0.00000 | 0.15158 |
| **SRRM4** | 0.21480 | 0.02808 | 0.63027 | 0.05425 | 0.17740 | 0.04928 | 0.07905 | 0.01386 | 0.47860 | 0.08444 | 0.00000 | 0.01709 |
| **SRRM5** | 0.08678 | 0.05092 | 0.03300 | 0.16671 | 0.04850 | 0.10049 | 0.04998 | 1.42627 | 0.30857 | 0.02571 | 0.01208 | 0.24515 |
| **SRRT** | 5.06130 | 17.56873 | 16.58272 | 16.73278 | 20.80228 | 0.00000 | 23.95058 | 16.28036 | 26.12705 | 18.41621 | 14.61348 | 13.51438 |
| **SRSF1** | 10.28194 | 23.61686 | 24.84106 | 17.10388 | 22.39326 | 60.14668 | 31.22055 | 18.87639 | 44.13087 | 15.80625 | 26.37940 | 36.59875 |
| **SRSF10.uid-100** | 1.81839 | 15.37468 | 14.56751 | 15.55518 | 9.00672 | 45.22718 | 20.74063 | 22.18652 | 20.78628 | 18.52408 | 18.47528 | 21.74678 |
| **SRSF11** | 6.67726 | 14.16010 | 16.45668 | 11.68104 | 14.18765 | 32.78534 | 29.83842 | 14.21728 | 21.20150 | 48.99137 | 16.71935 | 19.23475 |
| **SRSF12** | 0.38678 | 1.12990 | 0.00000 | 5.38460 | 0.14192 | 0.49248 | 0.49078 | 1.14769 | 0.79750 | 0.48626 | 0.49837 | 0.70328 |
| **SRSF2** | 6.37874 | 32.99514 | 28.40595 | 15.22884 | 19.91594 | 41.37544 | 18.66768 | 21.72209 | 41.04093 | 31.85366 | 28.07478 | 29.76193 |
| **SRSF3** | 4.86921 | 27.78739 | 24.41778 | 17.28747 | 18.46335 | 59.18656 | 41.67207 | 26.13297 | 34.01294 | 21.72099 | 31.09850 | 34.64444 |
| **SRSF4** | 1.68978 | 16.18657 | 16.09087 | 12.45467 | 11.05978 | 15.10280 | 18.63681 | 11.64597 | 15.41503 | 10.30822 | 13.32448 | 9.87130 |
| **SRSF5** | 5.97404 | 6.02743 | 16.62345 | 11.85666 | 6.62470 | 30.16529 | 15.42396 | 22.35091 | 18.13611 | 14.61414 | 17.96683 | 25.15636 |
| **SRSF6** | 0.61939 | 18.11705 | 12.17320 | 7.05827 | 3.31580 | 45.16257 | 25.88951 | 19.04486 | 24.36095 | 15.64004 | 23.15779 | 14.93398 |
| **SRSF7** | 7.65613 | 33.80040 | 27.55280 | 25.54554 | 20.46676 | 47.32735 | 31.95122 | 26.37436 | 42.66549 | 14.81547 | 34.95964 | 39.08658 |
| **SRSF8** | 2.54805 | 0.00000 | 3.55883 | 0.49500 | 0.42419 | 2.36766 | 5.15438 | 1.61699 | 4.01455 | 2.05612 | 0.70710 | 0.00000 |
| **SRSF9** | 0.23356 | 7.40476 | 8.09937 | 5.71192 | 8.90495 | 1.78823 | 8.51098 | 8.41623 | 8.33644 | 1.96894 | 2.92801 | 6.88371 |
| **SRXN1** | 0.48167 | 2.40138 | 13.88699 | 0.00000 | 8.28180 | 4.90087 | 21.31407 | 0.02053 | 5.83843 | 8.88989 | 1.97494 | 0.21230 |
| **SRY** | 0.01373 | 0.01450 | 0.01269 | 0.01110 | 0.01568 | 0.02058 | 0.01819 | 0.09987 | 0.02192 | 0.01333 | 0.01651 | 0.01185 |
| **SS18** | 2.13314 | 10.63453 | 11.71902 | 9.63142 | 5.63495 | 24.98459 | 13.82435 | 13.30814 | 12.12091 | 1.53844 | 10.65261 | 13.04476 |
| **SS18L1** | 0.02587 | 0.75269 | 2.36771 | 1.27581 | 1.33934 | 1.28148 | 1.52623 | 1.74480 | 5.78422 | 3.44017 | 2.05153 | 0.68800 |
| **SS18L2** | 2.44652 | 7.17226 | 15.67687 | 7.95349 | 5.94678 | 12.79360 | 19.48982 | 4.47065 | 14.16902 | 3.06007 | 13.64483 | 12.76947 |
| **SSB** | 4.21817 | 8.93403 | 13.31724 | 5.44582 | 3.49482 | 38.64211 | 26.62045 | 7.67491 | 17.35380 | 25.82707 | 14.87397 | 12.92750 |
| **SSBP1** | 0.92034 | 26.83010 | 25.47579 | 17.11741 | 19.21568 | 50.68027 | 39.23813 | 4.59938 | 26.44602 | 7.99877 | 27.93629 | 28.55262 |
| **SSBP2** | 1.63359 | 1.25140 | 0.39955 | 1.85857 | 0.13293 | 0.00000 | 0.00000 | 0.52699 | 0.35352 | 0.11503 | 0.13154 | 0.72781 |
| **SSBP3** | 3.97956 | 6.03994 | 1.35274 | 4.24151 | 14.10663 | 5.92427 | 14.21373 | 0.57308 | 0.00000 | 19.67696 | 1.97616 | 1.90557 |
| **SSBP3-AS1** | 0.00107 | 0.11225 | 0.08464 | 0.35730 | 1.55176 | 0.31443 | 0.00000 | 0.02584 | 0.48338 | 2.84499 | 0.28050 | 0.41047 |
| **SSBP4** | 0.01885 | 0.46174 | 0.16550 | 0.25894 | 0.74663 | 0.00029 | 0.71259 | 0.03921 | 0.28566 | 1.32474 | 0.27477 | 0.00000 |
| **SSC5D** | 0.06631 | 0.00680 | 0.04809 | 0.01336 | 0.02886 | 0.00125 | 0.01420 | 0.15073 | 0.01388 | 0.01955 | 0.01913 | 0.28988 |
| **SSFA2** | 4.03861 | 0.84884 | 1.38727 | 0.45211 | 0.00000 | 3.14520 | 0.00000 | 3.83198 | 1.07327 | 1.84271 | 0.00000 | 1.54580 |
| **SSH1** | 5.47803 | 1.73943 | 0.79160 | 1.61373 | 0.00000 | 3.37899 | 6.28200 | 0.42812 | 2.24972 | 6.81380 | 2.35783 | 1.02003 |
| **SSH2** | 4.67728 | 1.69041 | 1.48063 | 1.68654 | 2.08459 | 1.92970 | 4.39426 | 1.69841 | 4.20272 | 3.97080 | 0.61836 | 0.97162 |
| **SSH3** | 0.27791 | 2.20203 | 0.73189 | 0.00000 | 0.75564 | 0.00000 | 0.78521 | 0.00000 | 2.94105 | 1.73574 | 3.07613 | 0.51960 |
| **SSMEM1** | 0.54421 | 0.02020 | 0.01369 | 0.00618 | 0.10071 | 0.00000 | 0.02360 | 0.34167 | 0.00998 | 0.01654 | 0.00943 | 0.14443 |
| **SSNA1** | 3.38326 | 18.24413 | 10.86560 | 11.72004 | 13.02226 | 0.00001 | 16.62124 | 3.65104 | 9.82193 | 5.12382 | 14.34068 | 19.32524 |
| **SSPN** | 0.18555 | 0.01224 | 0.03887 | 0.25124 | 0.00000 | 0.02419 | 0.00000 | 0.01647 | 0.02796 | 0.04322 | 0.01112 | 0.01029 |
| **SSPO** | 0.03956 | 0.00285 | 0.01104 | 0.04414 | 0.00000 | 0.00591 | 0.00091 | 0.01213 | 0.01324 | 0.38799 | 0.02164 | 0.03695 |
| **SSR1** | 8.05851 | 12.17079 | 14.65807 | 6.13265 | 0.76719 | 37.29099 | 0.00000 | 14.79083 | 26.87863 | 25.48778 | 12.50331 | 14.83312 |
| **SSR2** | 3.34577 | 20.76761 | 32.22918 | 14.83356 | 17.67570 | 23.15098 | 7.66937 | 13.97821 | 50.47766 | 7.42234 | 16.90053 | 18.32265 |
| **SSR3** | 2.36841 | 15.88215 | 19.43455 | 6.27422 | 13.48985 | 35.05742 | 5.11853 | 3.71968 | 33.50206 | 21.34263 | 12.75976 | 19.36600 |
| **SSR4** | 6.29953 | 12.09191 | 14.18242 | 11.49004 | 21.03209 | 6.46683 | 10.31028 | 10.85036 | 25.41412 | 17.33148 | 13.05318 | 13.91784 |
| **SSR4P1** | 0.11489 | 0.10895 | 0.00871 | 0.00453 | 0.00851 | 0.00000 | 0.00400 | 0.00620 | 0.07278 | 0.04243 | 0.03192 | 0.00701 |
| **SSRP1** | 2.84314 | 8.52427 | 4.75555 | 10.25425 | 25.76158 | 11.70347 | 17.53589 | 1.45223 | 6.33154 | 36.33110 | 8.76609 | 13.82488 |
| **SSSCA1** | 0.01912 | 14.43891 | 13.83250 | 9.35468 | 16.24773 | 2.10237 | 15.53084 | 16.52021 | 12.41505 | 0.00000 | 10.97537 | 25.41557 |
| **SSSCA1-AS1** | 0.21862 | 0.28922 | 0.38476 | 0.08202 | 0.10597 | 0.00000 | 0.21197 | 0.50459 | 0.20889 | 0.00000 | 0.54352 | 0.95419 |

| | | | | | | | | | | | | |
|---|---|---|---|---|---|---|---|---|---|---|---|---|
| **SSTR1** | 0.00251 | 0.04040 | 0.03769 | 0.12353 | 0.00000 | 0.01992 | 0.10052 | 0.02076 | 0.00000 | 0.25127 | 0.00000 | 0.00497 |
| **SSTR2** | 0.00991 | 0.13479 | 0.27163 | 0.61065 | 0.00000 | 0.00000 | 0.03690 | 0.12668 | 0.00000 | 0.04609 | 0.11193 | 0.02987 |
| **SSTR3** | 1.18039 | 0.03616 | 0.01014 | 0.01338 | 0.01192 | 0.01600 | 0.02290 | 0.00589 | 0.02744 | 0.01284 | 0.00999 | 0.01173 |
| **SSTR5-AS1** | 15.83663 | 0.00000 | 0.00243 | 0.00000 | 0.13894 | 0.00000 | 0.00723 | 2.30998 | 0.02803 | 0.02499 | 0.00278 | 0.98567 |
| **SSU72** | 3.55773 | 25.44415 | 20.95779 | 20.61428 | 19.03321 | 20.67322 | 30.70661 | 10.01724 | 14.35465 | 23.25464 | 15.99883 | 25.80647 |
| **SSUH2** | 0.02259 | 0.00834 | 0.06149 | 0.31052 | 0.00000 | 0.02382 | 0.16822 | 0.02891 | 0.04504 | 0.52985 | 0.00000 | 0.01407 |
| **SSX1** | 0.35706 | 0.03268 | 0.16901 | 0.00094 | 0.27369 | 0.18742 | 0.07214 | 0.61084 | 0.00000 | 0.00000 | 0.00000 | 0.19209 |
| **SSX2IP** | 3.36312 | 3.75197 | 0.00000 | 0.00000 | 0.00000 | 7.55325 | 3.74777 | 2.42083 | 3.49832 | 6.91355 | 1.78754 | 12.59403 |
| **SSX3** | 0.00471 | 0.22994 | 0.04164 | 0.00116 | 0.02234 | 0.07933 | 0.05180 | 0.04907 | 0.08574 | 0.00000 | 0.00649 | 1.66564 |
| **SSX5** | 0.00933 | 0.07114 | 0.01502 | 0.00340 | 0.01013 | 0.08139 | 0.01134 | 0.00220 | 0.01993 | 0.01126 | 0.00919 | 0.77170 |
| **SSX6** | 0.00374 | 0.07675 | 0.02179 | 0.01897 | 0.01242 | 0.04393 | 0.01403 | 1.88504 | 0.01515 | 0.01347 | 0.00906 | 0.84583 |
| **SSX8** | 0.03017 | 0.01449 | 0.01483 | 0.01493 | 0.01584 | 0.01501 | 0.01590 | 0.01253 | 0.02149 | 0.02432 | 0.01414 | 0.03201 |
| **ST13** | 2.91232 | 5.73345 | 12.01378 | 5.39587 | 3.84484 | 21.97515 | 22.94298 | 1.87558 | 9.58372 | 13.12767 | 11.43009 | 6.14276 |
| **ST13P4** | 0.00811 | 0.00579 | 0.05493 | 0.01913 | 0.07463 | 0.07371 | 0.03267 | 0.00872 | 0.06819 | 0.03004 | 0.03958 | 0.06036 |
| **ST14** | 7.91340 | 1.61276 | 0.00000 | 1.30311 | 14.72649 | 0.00000 | 7.12357 | 2.67300 | 10.18595 | 8.81259 | 3.81055 | 0.00000 |
| **ST18** | 0.02777 | 0.07588 | 0.02300 | 0.00199 | 0.07956 | 0.04094 | 0.05569 | 0.02137 | 0.00000 | 0.00000 | 0.00599 | 0.21218 |
| **ST20** | 0.00878 | 0.55935 | 1.57908 | 0.57083 | 0.00298 | 1.84829 | 1.65371 | 0.00000 | 2.26390 | 1.43998 | 0.43823 | 1.26146 |
| **ST3GAL1** | 0.86652 | 12.18833 | 9.68246 | 22.72014 | 6.60681 | 0.00000 | 6.13477 | 2.62070 | 7.57470 | 9.15085 | 4.31565 | 6.09888 |
| **ST3GAL2** | 0.00427 | 3.25368 | 0.00000 | 1.33080 | 3.77324 | 0.00000 | 0.48296 | 1.33809 | 6.25488 | 7.69863 | 3.67926 | 7.93137 |
| **ST3GAL3** | 0.15569 | 0.09499 | 0.04275 | 0.13426 | 0.00000 | 0.10917 | 0.00000 | 0.06386 | 0.00000 | 0.10998 | 0.00671 | 0.13852 |
| **ST3GAL4** | 4.10015 | 0.60192 | 6.15131 | 2.68676 | 1.95335 | 0.00000 | 1.53736 | 1.44139 | 9.77782 | 0.00000 | 4.19264 | 0.92063 |
| **ST3GAL4-AS1** | 0.00100 | 0.37543 | 0.27581 | 0.15549 | 0.00000 | 0.33328 | 0.91623 | 0.07096 | 0.71445 | 0.00000 | 0.55163 | 0.49838 |
| **ST3GAL5** | 0.16503 | 0.02863 | 0.15429 | 0.02827 | 0.18764 | 0.00000 | 0.03359 | 0.00481 | 0.00000 | 0.00000 | 0.02744 | 0.02561 |
| **ST3GAL6** | 0.00000 | 4.81069 | 5.72818 | 0.43510 | 0.00000 | 3.75715 | 0.00000 | 0.11433 | 1.94659 | 2.16402 | 0.00000 | 3.57224 |
| **ST3GAL6-AS1** | 0.00540 | 0.20374 | 1.07731 | 0.00190 | 0.22455 | 0.00000 | 0.20743 | 0.00801 | 0.26665 | 0.00000 | 0.00000 | 0.36679 |
| **ST5** | 0.93049 | 7.43126 | 0.19684 | 5.78558 | 3.94261 | 0.75085 | 0.00003 | 0.00000 | 0.57878 | 3.40625 | 2.08241 | 1.55809 |
| **ST6GAL1** | 0.00000 | 21.72860 | 9.19463 | 14.43868 | 19.15820 | 22.73991 | 9.93279 | 0.38007 | 27.24231 | 8.05729 | 12.32186 | 12.60590 |
| **ST6GAL2** | 0.07149 | 0.01354 | 0.02117 | 0.04641 | 0.00023 | 0.06299 | 0.00685 | 0.00673 | 0.00763 | 0.03007 | 0.01198 | 0.09371 |
| **ST6GALNAC1** | 0.00056 | 6.37899 | 0.00000 | 6.36392 | 13.96653 | 0.00399 | 0.91470 | 0.00000 | 2.10146 | 2.00865 | 7.94806 | 0.29685 |
| **ST6GALNAC2** | 0.11149 | 1.04808 | 0.19554 | 0.00000 | 1.42348 | 0.00000 | 0.19460 | 0.05516 | 1.32040 | 2.19227 | 2.09140 | 0.60459 |
| **ST6GALNAC3** | 0.67433 | 0.24803 | 0.06115 | 2.98937 | 0.00146 | 0.12830 | 0.00000 | 0.05021 | 0.00000 | 0.00000 | 0.18760 | 0.00000 |
| **ST6GALNAC4** | 0.00255 | 0.00814 | 0.27117 | 0.00921 | 0.01175 | 0.03453 | 0.02030 | 0.04181 | 0.00000 | 0.01958 | 0.02976 | 0.06622 |
| **ST6GALNAC5** | 0.35016 | 0.06731 | 0.00944 | 0.20994 | 0.01651 | 0.07781 | 0.00000 | 0.05097 | 0.08660 | 0.00000 | 0.00432 | 0.03161 |
| **ST6GALNAC6** | 0.01322 | 0.16364 | 0.25559 | 0.42818 | 0.00000 | 0.00000 | 0.02869 | 0.38978 | 0.00000 | 0.76030 | 0.06004 | 0.65744 |
| **ST7** | 0.16555 | 1.13409 | 3.41380 | 2.21184 | 2.10155 | 2.38786 | 1.53268 | 0.44504 | 2.83528 | 0.45945 | 0.58121 | 2.42544 |
| **ST7-AS1** | 0.00602 | 0.02663 | 0.03229 | 0.01859 | 0.03338 | 0.01622 | 0.01919 | 0.01280 | 0.01766 | 0.00311 | 0.01625 | 0.02424 |
| **ST7-AS2.uid-10** | 0.10069 | 0.00868 | 0.03720 | 0.00957 | 0.01706 | 0.07182 | 0.00000 | 0.02099 | 0.02350 | 0.04741 | 0.01226 | 0.05592 |
| **ST7-AS2.uid-93** | 0.00521 | 0.00837 | 0.01850 | 0.03552 | 0.00000 | 0.08161 | 0.00000 | 0.00888 | 0.02347 | 0.05911 | 0.02343 | 0.02837 |
| **ST7-OT3** | 0.00722 | 0.03344 | 0.03323 | 0.01435 | 0.05358 | 0.00000 | 0.06272 | 0.03601 | 0.00287 | 0.00000 | 0.03366 | 0.04665 |
| **ST7-OT4** | 0.02821 | 0.05194 | 0.09326 | 0.08981 | 0.00000 | 0.05459 | 0.01328 | 0.07811 | 0.02050 | 0.04485 | 0.08877 | 0.21770 |
| **ST7L** | 0.13541 | 1.56846 | 0.54720 | 1.53118 | 1.11281 | 3.15271 | 0.00000 | 0.85303 | 2.33763 | 2.06385 | 1.87532 | 0.56756 |
| **ST8SIA1** | 0.11107 | 0.17214 | 0.46037 | 0.00000 | 0.00000 | 0.02691 | 0.18911 | 0.02265 | 0.00000 | 0.07509 | 0.21163 | 0.01862 |
| **ST8SIA2** | 0.10701 | 0.40402 | 0.01326 | 0.06252 | 0.31344 | 0.12039 | 0.00000 | 0.00446 | 0.00000 | 0.02219 | 0.00836 | 0.03826 |
| **ST8SIA3** | 0.00643 | 0.00609 | 0.01060 | 0.00379 | 0.00000 | 0.00509 | 0.00184 | 0.00932 | 0.00000 | 0.29858 | 0.17836 | 0.01036 |
| **ST8SIA4** | 0.04673 | 0.11359 | 10.58857 | 0.28530 | 0.00000 | 0.18328 | 0.00000 | 0.00603 | 0.35430 | 0.00000 | 0.17599 | 0.02792 |
| **ST8SIA5** | 0.48446 | 0.00000 | 0.00158 | 0.31532 | 0.00000 | 0.00000 | 0.08220 | 1.13752 | 0.00000 | 0.42567 | 0.06326 | 0.49399 |
| **ST8SIA6** | 0.00525 | 0.00979 | 0.07261 | 0.16133 | 0.01761 | 0.01498 | 0.01148 | 0.01667 | 0.01889 | 0.03110 | 0.03065 | 0.02053 |
| **ST8SIA6-AS1** | 0.28869 | 1.13416 | 2.58566 | 0.55764 | 0.47745 | 1.62264 | 0.31892 | 0.21664 | 1.09013 | 0.00000 | 1.41981 | 0.68767 |

| | | | | | | | | | | | | |
|---|---|---|---|---|---|---|---|---|---|---|---|---|
| **STAB1** | 0.18495 | 0.02021 | 0.02645 | 0.01356 | 0.00000 | 0.02826 | 0.02470 | 0.00404 | 0.04475 | 0.11381 | 0.05378 | 0.01428 |
| **STAB2** | 1.37540 | 0.00000 | 0.01733 | 0.10130 | 0.00000 | 0.00000 | 0.22714 | 0.27994 | 0.00000 | 0.74793 | 0.18067 | 0.17051 |
| **STAC** | 0.00433 | 0.44270 | 0.00000 | 0.00197 | 1.98703 | 0.65176 | 0.29580 | 1.29346 | 0.00000 | 0.00000 | 0.75121 | 1.34131 |
| **STAC2** | 0.00417 | 0.01949 | 0.09412 | 0.22496 | 0.00645 | 0.00785 | 0.02790 | 0.01072 | 0.00069 | 0.09766 | 0.01484 | 0.02304 |
| **STAC3** | 0.00799 | 0.03036 | 0.08002 | 0.02251 | 0.01695 | 0.00114 | 0.01767 | 0.01204 | 0.05076 | 0.15505 | 0.15677 | 0.02013 |
| **STAG1** | 3.53055 | 1.77708 | 5.86812 | 1.04708 | 1.99675 | 8.18667 | 7.99332 | 4.25834 | 9.56517 | 7.90621 | 4.36549 | 2.17405 |
| **STAG2** | 0.00000 | 2.21139 | 9.54073 | 7.76676 | 2.29761 | 22.31801 | 14.54039 | 6.28917 | 0.45157 | 19.94124 | 8.51929 | 17.49145 |
| **STAG3** | 9.47433 | 2.32260 | 0.33407 | 0.00000 | 2.76686 | 0.00001 | 1.84064 | 3.68923 | 3.09593 | 2.84931 | 4.60507 | 1.15623 |
| **STAG3L1.uid-97** | 0.01148 | 0.02786 | 0.02648 | 0.01870 | 0.06618 | 0.00000 | 0.04825 | 0.15547 | 0.02643 | 0.04390 | 0.01465 | 0.03753 |
| **STAG3L2** | 0.05412 | 0.51084 | 0.08987 | 0.15848 | 0.26190 | 0.24654 | 0.24314 | 2.80603 | 0.46132 | 0.58912 | 0.25339 | 0.88543 |
| **STAG3L3** | 0.02254 | 0.02550 | 0.02305 | 0.00687 | 0.04964 | 0.00785 | 0.01521 | 0.06510 | 0.02250 | 0.04202 | 0.02333 | 0.03079 |
| **STAG3L4** | 0.12626 | 4.30242 | 2.10882 | 2.66406 | 4.14389 | 4.18293 | 0.37176 | 8.50338 | 3.88455 | 2.69008 | 2.07768 | 3.51442 |
| **STAG3L5P** | 0.16291 | 0.22935 | 0.14678 | 0.00000 | 0.05233 | 0.15876 | 0.04520 | 0.65232 | 0.12891 | 0.00498 | 0.07520 | 0.25669 |
| **STAG3L5P-PVR** | 0.40789 | 0.62386 | 0.87729 | 0.99907 | 0.25474 | 0.00000 | 0.00000 | 0.71299 | 1.42915 | 0.00000 | 0.22756 | 0.41409 |
| **STAM** | 0.00000 | 5.97423 | 6.76951 | 6.11997 | 0.00000 | 17.75163 | 4.87786 | 15.17655 | 9.66405 | 1.78124 | 5.52538 | 7.40723 |
| **STAM-AS1** | 0.02536 | 0.27836 | 0.31071 | 0.25952 | 0.43124 | 0.90146 | 0.00000 | 0.83424 | 0.91878 | 2.49264 | 0.34967 | 0.66252 |
| **STAM2** | 1.96926 | 6.10578 | 4.51615 | 4.69575 | 0.81845 | 13.92616 | 6.20377 | 2.01920 | 2.07504 | 2.36722 | 5.04969 | 8.95133 |
| **STAMBP** | 1.64616 | 8.00427 | 5.45420 | 2.34342 | 2.32736 | 11.13919 | 8.05264 | 5.50012 | 7.33252 | 2.18594 | 6.04906 | 7.03445 |
| **STAMBPL1** | 0.00398 | 0.28258 | 0.11159 | 0.32118 | 0.08721 | 0.00006 | 0.42378 | 0.00701 | 0.41160 | 0.15102 | 0.00000 | 0.09576 |
| **STAP1** | 0.00498 | 0.01367 | 0.01196 | 0.00649 | 0.05497 | 0.02483 | 0.00930 | 0.01350 | 0.02188 | 0.03062 | 0.01641 | 0.01447 |
| **STAP2** | 0.03389 | 0.73769 | 1.03073 | 0.19048 | 0.00000 | 0.00000 | 0.00000 | 2.34448 | 0.00000 | 0.25693 | 0.00000 | 1.78722 |
| **STAR** | 0.01934 | 0.06707 | 0.22527 | 0.41483 | 0.00000 | 0.09568 | 0.00000 | 0.00847 | 0.04047 | 0.01468 | 0.01551 | 0.21043 |
| **STARD10** | 0.11784 | 0.98648 | 14.24169 | 1.13604 | 19.70338 | 0.00000 | 6.07083 | 0.01079 | 18.28211 | 4.01198 | 19.67194 | 0.08845 |
| **STARD13** | 0.00577 | 0.00000 | 0.22248 | 0.01328 | 0.04723 | 0.14161 | 0.29482 | 0.01234 | 0.13830 | 0.00000 | 0.03832 | 0.01039 |
| **STARD3** | 3.53740 | 8.24120 | 9.53250 | 5.44161 | 7.43841 | 0.00000 | 2.52947 | 1.00384 | 12.05610 | 0.00000 | 3.59711 | 9.54961 |
| **STARD3NL** | 3.06717 | 2.18905 | 6.22435 | 1.97316 | 0.00000 | 7.37623 | 4.56951 | 0.24803 | 5.41178 | 0.33188 | 4.54969 | 2.53746 |
| **STARD4** | 0.04079 | 0.00000 | 2.38087 | 2.22041 | 2.83094 | 1.06776 | 1.92516 | 0.05436 | 5.35825 | 0.00000 | 2.60580 | 0.00000 |
| **STARD4-AS1** | 0.84758 | 0.00840 | 0.13069 | 0.47565 | 0.07123 | 0.00014 | 0.09924 | 0.07812 | 0.23911 | 0.01488 | 0.15085 | 0.01796 |
| **STARD5** | 0.97127 | 0.56250 | 0.51768 | 0.02946 | 0.25991 | 0.95543 | 0.27902 | 0.21515 | 0.32182 | 0.00000 | 0.46839 | 0.13006 |
| **STARD6** | 0.00270 | 0.09384 | 0.05281 | 0.00153 | 0.00125 | 0.06513 | 0.01261 | 0.02512 | 0.00002 | 0.01577 | 0.01596 | 0.06520 |
| **STARD7** | 9.71220 | 17.04634 | 19.11718 | 15.98694 | 18.37736 | 26.58066 | 31.86404 | 5.37052 | 17.52147 | 24.26745 | 16.07881 | 14.53138 |
| **STARD7-AS1** | 0.24766 | 0.03362 | 0.25339 | 0.40044 | 0.00000 | 0.64367 | 0.19168 | 0.02528 | 0.11581 | 0.00000 | 0.53073 | 0.03762 |
| **STARD8** | 0.00526 | 0.01615 | 0.00421 | 0.03137 | 0.00723 | 0.00000 | 0.04765 | 0.00930 | 0.08569 | 0.07489 | 0.00655 | 0.03088 |
| **STARD9** | 1.27245 | 0.19616 | 0.17813 | 1.43091 | 0.15118 | 0.00000 | 0.00020 | 3.25496 | 0.00000 | 1.12178 | 0.00454 | 1.02466 |
| **STAT1** | 1.72681 | 3.65629 | 4.70955 | 1.32736 | 0.00000 | 7.67283 | 5.55173 | 1.00445 | 3.76255 | 1.86262 | 7.48637 | 2.17643 |
| **STAT2** | 0.00000 | 1.33287 | 4.11085 | 6.92643 | 0.00000 | 3.44094 | 0.00000 | 0.26069 | 0.52865 | 3.17565 | 2.76341 | 2.63765 |
| **STAT3** | 1.93739 | 11.96104 | 5.76525 | 19.37848 | 2.18482 | 4.23279 | 5.78159 | 1.72241 | 5.29228 | 2.33900 | 2.20771 | 25.70328 |
| **STAT4** | 0.00549 | 0.02698 | 0.22083 | 0.00176 | 0.00000 | 0.03769 | 0.00000 | 0.04913 | 0.00000 | 0.29978 | 0.00479 | 0.06056 |
| **STAT5A** | 1.54191 | 0.00000 | 0.02489 | 0.14192 | 0.00000 | 0.00000 | 0.01802 | 0.22053 | 0.00000 | 0.42322 | 0.05514 | 0.13807 |
| **STAT5B** | 0.92536 | 1.12342 | 1.10417 | 0.90327 | 1.50123 | 1.17245 | 0.58740 | 0.03918 | 0.00000 | 0.88770 | 0.41617 | 0.14335 |
| **STAT6** | 0.00329 | 0.00000 | 0.24742 | 0.47137 | 0.07524 | 0.03773 | 0.10688 | 0.00839 | 0.59089 | 0.26625 | 0.00000 | 0.03200 |
| **STAU1** | 14.61049 | 13.88272 | 20.77420 | 12.36924 | 15.77311 | 25.48621 | 30.42274 | 12.11494 | 26.34337 | 20.10381 | 16.93495 | 28.80146 |
| **STAU2** | 2.80332 | 3.93663 | 0.00000 | 1.38286 | 4.15099 | 8.97945 | 5.00250 | 0.58252 | 1.53732 | 7.96733 | 8.44729 | 2.55672 |
| **STAU2-AS1** | 0.00626 | 0.01338 | 0.02532 | 0.01226 | 0.00166 | 0.03982 | 0.02983 | 0.70054 | 0.04240 | 0.03703 | 0.01861 | 0.00000 |
| **STBD1** | 0.05533 | 0.18739 | 0.04267 | 0.03632 | 0.00000 | 0.02965 | 0.01214 | 0.00556 | 0.01654 | 0.02918 | 0.01199 | 0.03663 |
| **STC1** | 0.15368 | 0.01572 | 0.09871 | 0.00986 | 0.02661 | 0.01802 | 0.03482 | 0.11624 | 0.09552 | 0.00000 | 0.19145 | 0.02945 |
| **STC2** | 0.38159 | 0.00000 | 7.56938 | 2.26108 | 4.88177 | 0.00004 | 6.15498 | 0.29586 | 25.18381 | 6.27177 | 7.72392 | 0.00000 |
| **STEAP1** | 0.00206 | 9.76537 | 0.58430 | 1.76158 | 0.00000 | 1.38907 | 0.00000 | 0.00000 | 0.00001 | 0.32556 | 0.00000 | 6.39938 |

| | | | | | | | | | | | | |
|---|---|---|---|---|---|---|---|---|---|---|---|---|
| **STEAP1B** | 0.00226 | 0.36733 | 0.83219 | 0.00139 | 0.08714 | 0.76681 | 0.00000 | 0.06319 | 0.08471 | 0.00000 | 0.37127 | 0.86629 |
| **STEAP2** | 0.00485 | 0.03437 | 0.02529 | 0.00765 | 0.00316 | 0.01757 | 0.01610 | 0.00984 | 0.01484 | 0.08663 | 0.01427 | 0.01997 |
| **STEAP3** | 0.00412 | 1.42144 | 3.71226 | 0.46936 | 6.05587 | 0.00000 | 0.00000 | 0.02460 | 13.91239 | 0.00000 | 2.90705 | 0.02311 |
| **STEAP4** | 0.00254 | 0.16410 | 0.00000 | 0.00000 | 0.14790 | 0.32049 | 0.00000 | 0.01782 | 0.00000 | 0.05295 | 0.07876 | 0.07285 |
| **STIL** | 2.05876 | 6.06331 | 3.42543 | 2.48208 | 1.70924 | 16.94560 | 3.59346 | 14.59062 | 10.51894 | 3.96787 | 7.10065 | 10.69891 |
| **STIM1** | 3.23080 | 1.49426 | 4.12692 | 0.36612 | 3.64934 | 0.00000 | 1.59773 | 0.37760 | 2.85592 | 3.54182 | 1.72241 | 8.30187 |
| **STIM2** | 1.06805 | 1.23120 | 1.04245 | 0.12961 | 1.58612 | 3.08187 | 1.40330 | 0.83392 | 1.17454 | 7.36890 | 0.48132 | 3.48643 |
| **STIP1** | 8.34382 | 32.34525 | 31.02906 | 28.46934 | 38.38368 | 16.04877 | 44.91122 | 16.92997 | 34.99203 | 0.69981 | 34.99691 | 24.75041 |
| **STK10** | 0.00237 | 6.49814 | 4.01120 | 5.85555 | 17.35763 | 0.00000 | 2.49485 | 3.45852 | 7.70908 | 5.65217 | 12.10921 | 6.56340 |
| **STK11** | 0.96121 | 0.25491 | 1.26362 | 0.45692 | 1.78755 | 0.00000 | 1.17702 | 0.14778 | 0.00000 | 5.27942 | 0.00000 | 0.55807 |
| **STK11IP** | 0.35202 | 1.68356 | 1.98616 | 0.82868 | 1.39741 | 0.00000 | 1.62875 | 0.66471 | 2.05041 | 0.67552 | 0.84786 | 2.90160 |
| **STK16** | 0.65910 | 3.32077 | 1.59860 | 2.18235 | 1.64557 | 0.08606 | 1.22372 | 0.30900 | 1.29900 | 0.00000 | 4.22503 | 0.98421 |
| **STK17A** | 0.00000 | 5.73420 | 3.87718 | 0.69186 | 3.77896 | 6.17401 | 3.77860 | 0.55727 | 6.52765 | 8.21618 | 0.08481 | 3.91145 |
| **STK17B** | 0.38603 | 1.35398 | 3.67920 | 0.65610 | 0.00000 | 5.93871 | 4.11802 | 8.21800 | 7.40903 | 3.33720 | 4.66355 | 0.90389 |
| **STK19.uid-8457** | 0.54957 | 1.60882 | 1.32843 | 0.03703 | 1.47529 | 0.72495 | 1.01605 | 3.24888 | 1.31946 | 0.00006 | 0.60025 | 2.32960 |
| **STK24** | 14.95585 | 5.60615 | 4.40255 | 0.74191 | 5.11639 | 12.42009 | 15.02958 | 0.37218 | 1.10263 | 18.07798 | 0.99009 | 0.30832 |
| **STK25** | 3.20879 | 5.93033 | 8.01877 | 6.81727 | 9.68191 | 0.00000 | 14.37746 | 0.33573 | 10.24365 | 14.73260 | 7.20054 | 3.62659 |
| **STK3** | 0.48880 | 1.01628 | 7.59861 | 0.00000 | 3.02528 | 7.45124 | 5.40986 | 0.79304 | 4.61863 | 1.37272 | 1.99461 | 0.35879 |
| **STK31** | 1.77273 | 1.19347 | 0.00000 | 0.23705 | 0.00344 | 0.28557 | 0.00000 | 4.04641 | 0.00000 | 0.01319 | 0.00000 | 10.58848 |
| **STK32A** | 0.02666 | 0.01251 | 0.02136 | 0.01519 | 0.00004 | 0.00312 | 0.01798 | 0.00860 | 0.00000 | 0.07082 | 0.16669 | 0.04286 |
| **STK32B** | 8.68307 | 0.32598 | 0.00000 | 0.02202 | 0.05179 | 0.02420 | 0.00000 | 2.12739 | 0.00000 | 0.10425 | 0.00000 | 0.78010 |
| **STK32C** | 0.62611 | 0.17909 | 0.01293 | 0.00231 | 0.00000 | 0.00000 | 0.03911 | 0.00599 | 0.11776 | 0.03770 | 0.00000 | 0.03442 |
| **STK33** | 1.21421 | 1.08213 | 0.00000 | 0.22319 | 0.00000 | 0.11332 | 0.00000 | 0.18023 | 0.00000 | 0.23777 | 0.08790 | 0.38447 |
| **STK35** | 7.61617 | 4.02945 | 4.94137 | 7.44501 | 6.55745 | 3.42565 | 11.69722 | 5.31228 | 3.16616 | 9.98223 | 3.92917 | 6.24215 |
| **STK36** | 2.04325 | 3.07105 | 0.98420 | 4.09668 | 1.22166 | 0.00000 | 0.52176 | 0.17651 | 0.47745 | 0.57994 | 0.19607 | 2.13518 |
| **STK38** | 4.40339 | 3.01672 | 4.02900 | 1.19610 | 3.82826 | 6.09850 | 7.51419 | 0.31303 | 6.16031 | 3.47829 | 4.90653 | 3.23392 |
| **STK38L** | 2.44407 | 2.05880 | 4.57795 | 1.31237 | 0.00000 | 8.03447 | 5.61009 | 1.16138 | 4.68290 | 2.37296 | 2.14818 | 1.56773 |
| **STK39** | 1.84166 | 0.17720 | 0.10601 | 1.25564 | 0.10791 | 0.00000 | 0.00000 | 0.33141 | 0.00000 | 0.16748 | 0.19719 | 0.20932 |
| **STK4** | 7.16593 | 0.16329 | 8.35809 | 1.54676 | 1.81345 | 11.53515 | 8.05725 | 3.10319 | 6.39557 | 12.99346 | 5.79827 | 3.85931 |
| **STK4-AS1** | 0.15652 | 0.03023 | 0.03657 | 0.00775 | 0.01196 | 0.00531 | 0.02620 | 0.09548 | 0.03758 | 0.00000 | 0.06537 | 0.28252 |
| **STK40** | 1.92681 | 0.29188 | 4.93948 | 1.70316 | 2.23157 | 0.00000 | 2.78482 | 0.98447 | 1.64607 | 5.48055 | 0.20021 | 2.12342 |
| **STKLD1** | 0.02626 | 0.29739 | 0.06657 | 0.52310 | 0.00000 | 0.00000 | 0.00000 | 0.03249 | 0.15270 | 0.30733 | 0.02707 | 0.02806 |
| **STL** | 0.00609 | 0.01215 | 0.08041 | 0.02281 | 0.04874 | 0.02361 | 0.09942 | 0.01039 | 0.00128 | 0.01121 | 0.03297 | 0.01590 |
| **STMN1** | 0.87133 | 36.95115 | 18.70520 | 33.76802 | 27.86088 | 79.99933 | 37.11984 | 9.70536 | 32.43773 | 30.43802 | 45.66436 | 46.86433 |
| **STMN2** | 0.03050 | 0.02731 | 1.56066 | 0.01446 | 0.00000 | 0.06662 | 0.00000 | 0.05789 | 0.00000 | 0.05267 | 0.00727 | 0.01866 |
| **STMN3** | 0.16971 | 0.63140 | 0.13743 | 0.21931 | 0.09476 | 0.12090 | 0.00000 | 0.44409 | 0.00000 | 0.09013 | 0.05415 | 0.83420 |
| **STMN4** | 0.00579 | 0.01303 | 0.02422 | 0.03010 | 0.00710 | 0.05819 | 0.00638 | 0.00993 | 0.00624 | 0.02834 | 0.01361 | 0.01286 |
| **STOM** | 0.27298 | 8.60568 | 4.51559 | 14.67058 | 0.03187 | 11.00899 | 0.00000 | 13.54233 | 10.14917 | 0.00000 | 1.15850 | 14.32880 |
| **STOML1** | 0.95289 | 2.88494 | 0.58523 | 1.02709 | 1.36294 | 0.00000 | 0.21798 | 0.26879 | 0.00000 | 0.13139 | 1.57425 | 1.40569 |
| **STOML2** | 8.94126 | 34.11704 | 28.56393 | 28.12726 | 31.57682 | 16.86469 | 42.22061 | 20.53773 | 41.40219 | 0.00091 | 31.21763 | 37.10842 |
| **STOML3** | 0.03538 | 0.02535 | 0.11141 | 0.00864 | 0.01756 | 0.01118 | 0.00904 | 0.00891 | 0.01127 | 0.00618 | 0.01816 | 0.02763 |
| **STON1** | 0.00168 | 0.65519 | 0.00000 | 0.00000 | 0.89993 | 1.88119 | 3.26274 | 0.13385 | 1.08565 | 1.00812 | 1.50688 | 0.00000 |
| **STON1-GTF2A1** | 0.00550 | 0.01699 | 0.01382 | 0.02088 | 0.01760 | 0.00437 | 0.03701 | 0.01346 | 0.01994 | 0.01405 | 0.01713 | 0.01732 |
| **STON2** | 0.07171 | 7.23731 | 0.00000 | 6.05286 | 3.60673 | 13.91202 | 16.70715 | 0.00000 | 12.59374 | 9.49260 | 14.11362 | 0.00000 |
| **STOX1** | 0.61534 | 0.47950 | 0.00381 | 0.00000 | 0.06621 | 0.17142 | 0.00000 | 0.03659 | 0.00000 | 0.91005 | 0.06351 | 0.00000 |
| **STOX2** | 0.14410 | 0.09640 | 0.07622 | 0.70363 | 0.02113 | 0.00000 | 0.01262 | 0.17613 | 0.00000 | 0.30351 | 0.00148 | 0.04830 |
| **STPG1** | 4.02843 | 0.91932 | 1.53938 | 0.18558 | 0.93281 | 0.83134 | 0.49233 | 0.54515 | 1.54742 | 0.00000 | 2.30303 | 0.67176 |
| **STPG2** | 0.22295 | 0.06601 | 0.13218 | 0.00147 | 0.02269 | 0.02736 | 0.01040 | 0.02394 | 0.00002 | 0.11450 | 0.03100 | 0.05131 |

| | | | | | | | | | | | | |
|---|---|---|---|---|---|---|---|---|---|---|---|---|
| **STPG2-AS1** | 0.00316 | 0.02445 | 0.01510 | 0.02164 | 0.01226 | 0.01207 | 0.01790 | 0.03871 | 0.01727 | 0.02294 | 0.01512 | 0.02951 |
| **STRA13** | 0.95453 | 9.65588 | 21.71676 | 11.92485 | 17.20783 | 2.02038 | 23.59002 | 1.84551 | 16.29785 | 0.00000 | 18.51405 | 6.61077 |
| **STRA6** | 0.00609 | 0.00000 | 4.66847 | 0.00000 | 0.00000 | 0.00000 | 0.46028 | 0.09896 | 4.98276 | 0.34828 | 0.99698 | 0.00000 |
| **STRA8** | 0.02559 | 0.01104 | 0.08474 | 0.02958 | 0.00991 | 0.02488 | 0.07990 | 0.08722 | 0.05444 | 0.00000 | 0.02879 | 0.04122 |
| **STRADA** | 0.68383 | 3.42055 | 1.49090 | 2.72997 | 2.94641 | 1.12779 | 1.15958 | 0.23290 | 0.38881 | 0.01770 | 2.69798 | 2.44996 |
| **STRADB** | 0.67204 | 5.04638 | 0.35375 | 1.26488 | 0.44475 | 3.84697 | 0.58934 | 0.04513 | 0.00000 | 2.29969 | 2.94376 | 2.10224 |
| **STRAP** | 8.31654 | 24.88299 | 27.47825 | 13.27067 | 13.17191 | 54.00364 | 35.04779 | 18.05160 | 34.26740 | 6.52719 | 24.38651 | 26.96477 |
| **STRBP** | 4.22242 | 4.59587 | 3.97966 | 1.58024 | 8.58887 | 14.68940 | 12.09631 | 1.21567 | 0.59859 | 22.06648 | 5.13542 | 4.15493 |
| **STRC** | 0.14836 | 0.02486 | 0.01408 | 0.00522 | 0.00869 | 0.05141 | 0.00089 | 0.00409 | 0.04974 | 0.01601 | 0.01639 | 0.01916 |
| **STRIP1** | 2.50648 | 5.57614 | 5.11356 | 8.23471 | 2.90788 | 0.00000 | 1.34186 | 2.89747 | 6.17998 | 5.67890 | 5.53938 | 2.56208 |
| **STRIP2** | 0.38639 | 1.89993 | 2.64267 | 1.30993 | 1.06559 | 1.26828 | 0.77206 | 0.62593 | 2.67812 | 0.05261 | 0.88993 | 1.46625 |
| **STRN** | 0.53616 | 0.00000 | 2.81843 | 1.57864 | 2.33411 | 5.97668 | 4.66788 | 0.91539 | 0.00711 | 10.41714 | 0.10279 | 4.83178 |
| **STRN3** | 1.58557 | 2.91251 | 5.71122 | 3.22589 | 0.00000 | 37.63795 | 10.65787 | 6.86918 | 4.60814 | 13.15360 | 7.25184 | 6.80469 |
| **STRN4** | 0.86206 | 4.70645 | 5.82497 | 6.01087 | 8.77931 | 0.00000 | 6.23838 | 0.00000 | 5.15057 | 12.58287 | 2.59328 | 3.12348 |
| **STS** | 0.30501 | 0.00000 | 21.46216 | 0.52111 | 9.38801 | 2.19430 | 14.57071 | 0.00000 | 32.67311 | 0.00000 | 13.20774 | 0.00000 |
| **STT3A** | 10.55186 | 13.23289 | 16.11074 | 10.82842 | 9.61713 | 21.72102 | 2.06243 | 10.21903 | 32.28654 | 7.86894 | 19.87655 | 11.40389 |
| **STT3B** | 0.92050 | 2.77490 | 11.47140 | 3.12429 | 5.39635 | 32.02551 | 0.00000 | 1.79429 | 10.57280 | 29.18629 | 1.84579 | 3.47286 |
| **STUB1** | 0.00013 | 1.28111 | 2.75373 | 0.73703 | 3.11768 | 0.00000 | 1.04013 | 0.40559 | 0.99917 | 2.90626 | 1.42171 | 0.51541 |
| **STX10** | 0.24982 | 3.43421 | 9.73256 | 1.61725 | 2.25183 | 0.00000 | 4.93793 | 1.36644 | 9.38274 | 7.33406 | 3.26390 | 2.30617 |
| **STX11** | 0.00171 | 2.57103 | 0.86514 | 0.00000 | 0.71335 | 0.01215 | 0.09923 | 4.33598 | 0.01921 | 0.00000 | 0.48691 | 4.92778 |
| **STX12** | 4.88860 | 4.40130 | 9.14529 | 0.49296 | 0.99907 | 15.35221 | 5.19484 | 5.97835 | 8.25551 | 1.23199 | 5.74020 | 5.78862 |
| **STX16** | 2.90546 | 2.67417 | 1.57191 | 2.05866 | 1.14621 | 8.24910 | 0.41311 | 0.00000 | 6.53177 | 13.29701 | 2.22245 | 3.40111 |
| **STX16-NPEPL1** | 0.00437 | 0.01593 | 0.01407 | 0.03660 | 0.01809 | 0.02139 | 0.00000 | 0.00986 | 0.04478 | 0.12879 | 0.01889 | 0.01649 |
| **STX17** | 5.19642 | 1.84703 | 2.26318 | 0.71562 | 0.00000 | 3.27768 | 0.00000 | 0.76870 | 0.00000 | 2.52003 | 1.43049 | 2.47053 |
| **STX18** | 3.61750 | 8.01690 | 2.49905 | 3.69322 | 3.15108 | 9.91554 | 9.10818 | 0.89434 | 6.26584 | 0.00000 | 6.90776 | 6.05643 |
| **STX18-AS1** | 0.53411 | 1.29909 | 0.17413 | 0.70048 | 1.35350 | 0.13474 | 0.37782 | 0.10952 | 0.63021 | 0.00000 | 1.36630 | 0.14364 |
| **STX19** | 0.00499 | 0.01432 | 0.01303 | 0.01416 | 0.00649 | 0.06424 | 0.01620 | 0.01165 | 0.04028 | 0.00000 | 0.00905 | 0.09265 |
| **STX1A** | 0.39806 | 1.76472 | 0.85598 | 0.30994 | 1.21931 | 0.00000 | 0.19776 | 0.22842 | 3.58210 | 2.31323 | 0.00007 | 0.85950 |
| **STX1B** | 0.02671 | 0.01118 | 0.02202 | 0.05754 | 0.00558 | 0.00000 | 0.01072 | 0.31334 | 0.01028 | 0.06382 | 0.12091 | 0.26146 |
| **STX2** | 3.14232 | 0.81636 | 0.68474 | 3.08847 | 0.00000 | 0.75412 | 0.32975 | 1.64260 | 0.14701 | 1.52183 | 0.64432 | 0.95151 |
| **STX3** | 4.54457 | 9.43412 | 10.14016 | 7.08474 | 8.88141 | 19.23725 | 14.44662 | 4.04229 | 13.65193 | 14.91142 | 3.75914 | 11.03136 |
| **STX4** | 0.50899 | 0.82402 | 3.13184 | 2.93730 | 2.35073 | 0.00000 | 1.19107 | 0.77461 | 2.10774 | 2.83273 | 0.33330 | 1.35081 |
| **STX5** | 0.16870 | 0.87425 | 3.27889 | 1.44696 | 4.63929 | 0.00000 | 2.32563 | 4.70699 | 0.66774 | 5.96613 | 0.65970 | 1.50901 |
| **STX6** | 14.43198 | 1.34924 | 4.66649 | 0.48837 | 2.65198 | 3.58754 | 4.26809 | 1.44454 | 0.00000 | 9.94171 | 0.10455 | 1.10950 |
| **STX7** | 3.40568 | 5.84170 | 3.37273 | 1.89623 | 1.77256 | 20.10764 | 9.00171 | 0.81653 | 8.35503 | 3.23057 | 16.01820 | 5.01600 |
| **STX8** | 6.77561 | 6.00249 | 1.72281 | 2.13988 | 4.84612 | 3.58737 | 5.62665 | 6.55083 | 6.36427 | 2.78873 | 3.63109 | 3.23745 |
| **STXBP1** | 2.82372 | 0.34486 | 2.01252 | 0.00000 | 0.05980 | 0.83114 | 0.00000 | 1.13710 | 0.42984 | 0.63688 | 1.28813 | 1.74971 |
| **STXBP2** | 2.34424 | 3.83440 | 6.16541 | 3.96398 | 5.72064 | 0.00000 | 3.24256 | 0.33191 | 5.11322 | 1.05229 | 5.84322 | 1.12185 |
| **STXBP3** | 0.67444 | 5.60040 | 6.52951 | 1.80931 | 0.64654 | 13.83789 | 0.02968 | 8.01440 | 7.75551 | 0.67449 | 3.61616 | 6.68627 |
| **STXBP4** | 0.53912 | 0.80693 | 2.60849 | 0.10052 | 0.00101 | 5.46720 | 3.26841 | 0.55227 | 4.64386 | 2.37729 | 2.65233 | 1.22012 |
| **STXBP5** | 1.02346 | 0.20473 | 1.47400 | 2.16064 | 1.44083 | 4.33046 | 4.50035 | 0.00002 | 1.28234 | 5.62095 | 0.41436 | 0.09198 |
| **STXBP5-AS1** | 0.32995 | 0.01309 | 0.00969 | 0.01487 | 0.09900 | 0.00000 | 0.02000 | 0.01663 | 0.02864 | 0.03760 | 0.00699 | 0.02058 |
| **STXBP5L** | 0.00226 | 0.39033 | 0.04195 | 0.72176 | 0.11127 | 0.00000 | 0.00941 | 1.61102 | 0.00000 | 0.12917 | 0.12341 | 2.70662 |
| **STXBP6** | 1.41197 | 1.98091 | 0.00000 | 0.92850 | 6.35210 | 3.07147 | 0.63294 | 0.20675 | 5.44247 | 4.87165 | 9.76513 | 1.66452 |
| **STYK1** | 0.15127 | 0.99478 | 0.29110 | 0.00000 | 0.85614 | 0.70339 | 0.00000 | 0.00000 | 0.42600 | 0.00000 | 0.00000 | 0.47158 |
| **STYX** | 0.41655 | 0.53281 | 3.95515 | 0.77013 | 6.22985 | 20.11181 | 5.15626 | 9.68885 | 3.37537 | 9.80664 | 7.14010 | 7.30235 |
| **STYXL1** | 0.96150 | 2.37191 | 3.57373 | 2.01926 | 2.81190 | 3.86466 | 0.00000 | 0.08429 | 2.58965 | 0.00000 | 2.74376 | 0.90549 |
| **SUB1** | 3.30649 | 12.34673 | 19.39911 | 8.96064 | 7.99970 | 53.70631 | 23.73415 | 22.31682 | 13.76886 | 18.84524 | 17.36121 | 24.70480 |

| | | | | | | | | | | | | |
|---|---|---|---|---|---|---|---|---|---|---|---|---|
| **SUCLA2** | 0.19665 | 6.03613 | 7.61786 | 10.66047 | 0.06222 | 26.63842 | 7.86517 | 1.70056 | 9.93114 | 0.00000 | 10.23580 | 4.47015 |
| **SUCLG1** | 2.41467 | 15.48316 | 14.81554 | 16.10699 | 19.44943 | 21.11936 | 21.19324 | 0.59314 | 20.51248 | 2.72394 | 20.05900 | 5.97994 |
| **SUCLG2** | 0.15386 | 2.91844 | 0.67709 | 2.68998 | 1.05753 | 8.90172 | 0.00037 | 0.00000 | 0.00000 | 1.69197 | 5.00703 | 0.09426 |
| **SUCLG2-AS1** | 0.20452 | 0.03551 | 0.04583 | 0.07015 | 0.13318 | 0.09402 | 0.00137 | 0.02799 | 0.00000 | 0.16589 | 0.01513 | 0.04499 |
| **SUCNR1** | 0.00552 | 0.02310 | 0.07876 | 0.02100 | 0.02172 | 0.00000 | 0.00000 | 0.01036 | 0.13612 | 0.06105 | 0.01456 | 0.01630 |
| **SUCO** | 1.79698 | 0.33205 | 12.67068 | 5.72925 | 1.56912 | 9.98103 | 1.25423 | 0.95190 | 4.63938 | 16.14601 | 1.58179 | 2.15315 |
| **SUDS3** | 11.16400 | 7.01439 | 4.16164 | 7.72930 | 6.03448 | 12.06151 | 7.66709 | 6.21775 | 4.88373 | 10.57358 | 4.52262 | 6.74973 |
| **SUFU** | 0.81915 | 1.82532 | 2.10592 | 1.68299 | 3.79455 | 1.14410 | 4.27946 | 0.78312 | 2.50690 | 5.37327 | 1.39329 | 1.17241 |
| **SUGCT** | 5.66291 | 0.54849 | 0.03418 | 0.87680 | 0.26726 | 1.25185 | 0.00000 | 0.92853 | 1.56108 | 0.00000 | 0.77312 | 0.66860 |
| **SUGP1** | 0.86036 | 9.92198 | 6.16163 | 7.38587 | 9.83773 | 0.00000 | 10.50187 | 5.43133 | 8.89194 | 3.01477 | 8.02294 | 13.39206 |
| **SUGP2** | 2.47724 | 4.63192 | 2.89320 | 4.53895 | 3.24584 | 3.59797 | 1.69894 | 0.57839 | 2.77201 | 11.10435 | 4.60116 | 4.97686 |
| **SUGT1** | 5.29112 | 6.90537 | 8.97218 | 4.58916 | 4.58555 | 26.14583 | 15.14325 | 2.83840 | 11.14519 | 6.17990 | 10.19236 | 10.01753 |
| **SUGT1P1** | 0.00634 | 0.01514 | 0.03839 | 0.01544 | 0.01555 | 0.02410 | 0.01654 | 0.01155 | 0.01616 | 0.01092 | 0.01499 | 0.01826 |
| **SUGT1P3** | 0.01270 | 0.03964 | 0.03642 | 0.03355 | 0.01289 | 0.16082 | 0.01421 | 0.03234 | 0.00315 | 0.05286 | 0.01623 | 0.00785 |
| **SULF1** | 0.03105 | 0.00712 | 0.10101 | 0.05128 | 0.00000 | 0.04554 | 0.00000 | 0.09120 | 0.00000 | 0.05502 | 0.01513 | 0.00766 |
| **SULF2** | 2.57877 | 0.58587 | 0.00000 | 1.20623 | 5.01809 | 0.55788 | 1.06488 | 1.76104 | 0.81030 | 1.42650 | 0.80164 | 0.00000 |
| **SULT1A1** | 0.01337 | 0.81662 | 0.43523 | 1.08246 | 0.54547 | 0.00001 | 0.01451 | 0.00912 | 0.06926 | 0.00000 | 0.38314 | 0.00759 |
| **SULT1A2** | 0.00419 | 0.06204 | 0.03019 | 0.01517 | 0.17734 | 0.08076 | 0.01840 | 0.01164 | 0.00000 | 0.06458 | 0.07234 | 0.00920 |
| **SULT1B1** | 0.00381 | 0.05630 | 0.22494 | 0.19430 | 0.00000 | 0.01936 | 0.00002 | 0.06060 | 0.00000 | 0.05451 | 0.01883 | 0.04161 |
| **SULT1C2** | 0.00309 | 0.02214 | 0.01341 | 0.01007 | 0.01431 | 0.02289 | 0.01704 | 0.02999 | 0.01877 | 0.01447 | 0.01563 | 0.01405 |
| **SULT1E1** | 0.00361 | 0.00125 | 0.01368 | 0.00000 | 0.11355 | 0.01916 | 0.00080 | 0.01343 | 0.00000 | 0.31100 | 0.00000 | 0.00866 |
| **SULT2A1** | 0.13770 | 0.01979 | 1.10638 | 0.01421 | 0.17511 | 0.00000 | 0.20153 | 0.57287 | 0.00000 | 0.00000 | 0.02062 | 0.02242 |
| **SULT2B1** | 0.00534 | 0.53976 | 0.99367 | 0.04287 | 0.19064 | 0.00000 | 0.00000 | 1.88855 | 0.13009 | 0.05219 | 0.00403 | 0.82601 |
| **SULT4A1** | 0.69660 | 0.18534 | 0.01975 | 0.03686 | 0.00000 | 0.00000 | 0.02642 | 0.04100 | 0.00000 | 0.09255 | 0.03266 | 0.82692 |
| **SULT6B1** | 0.00539 | 0.00874 | 0.04657 | 0.02523 | 0.00000 | 0.04717 | 0.01144 | 0.02892 | 0.00362 | 0.03575 | 0.02345 | 0.01168 |
| **SUMF1** | 0.40883 | 4.25759 | 2.06739 | 1.48514 | 2.86585 | 5.24045 | 1.02330 | 0.00000 | 5.74398 | 1.68713 | 8.13548 | 0.69144 |
| **SUMF2** | 0.08600 | 13.15715 | 6.28123 | 7.08501 | 6.66691 | 3.89103 | 0.00000 | 0.32915 | 8.40276 | 0.00000 | 10.09539 | 4.08193 |
| **SUMO1** | 3.70122 | 8.89947 | 11.88157 | 11.66068 | 3.00527 | 30.11057 | 18.81013 | 11.18463 | 14.70314 | 0.00000 | 12.45206 | 11.73293 |
| **SUMO1P1** | 0.00548 | 0.52776 | 0.43247 | 0.01096 | 0.00057 | 0.31671 | 0.00000 | 0.02186 | 0.13079 | 0.00000 | 0.03410 | 0.11656 |
| **SUMO1P3** | 0.50114 | 0.94708 | 1.51008 | 1.27022 | 0.00000 | 5.72827 | 1.98198 | 1.45717 | 2.60700 | 0.00000 | 2.08821 | 1.68248 |
| **SUMO2** | 3.81400 | 9.06306 | 7.80735 | 5.81717 | 4.06070 | 26.28893 | 11.83475 | 11.02754 | 10.42586 | 8.05626 | 6.84193 | 17.00917 |
| **SUMO3** | 1.51295 | 14.79767 | 8.43288 | 7.97724 | 15.97877 | 18.32441 | 23.36936 | 1.90733 | 25.80416 | 2.37965 | 22.60037 | 7.84411 |
| **SUMO4** | 0.16729 | 0.00000 | 0.28683 | 0.10123 | 0.00000 | 0.07154 | 0.05619 | 0.14252 | 0.00000 | 0.37923 | 0.08214 | 0.08056 |
| **SUN1** | 8.85241 | 9.96517 | 2.88352 | 6.60226 | 3.03738 | 11.42249 | 0.07381 | 6.82149 | 11.22256 | 1.45950 | 3.32370 | 3.35694 |
| **SUN2** | 0.00574 | 1.27879 | 1.48021 | 0.14666 | 9.19462 | 0.00000 | 0.00000 | 0.18891 | 6.13933 | 7.52061 | 1.96740 | 0.11770 |
| **SUN3** | 0.00000 | 4.10498 | 3.84233 | 0.00000 | 10.31886 | 11.13110 | 16.19476 | 0.00000 | 17.96041 | 1.15268 | 12.49728 | 0.00000 |
| **SUOX** | 0.00053 | 2.53637 | 4.71105 | 1.09995 | 0.00000 | 0.00000 | 1.03278 | 0.16853 | 2.06882 | 0.00000 | 2.04450 | 0.39798 |
| **SUPT16H** | 1.33756 | 8.22954 | 16.29435 | 11.98530 | 12.57647 | 28.86009 | 28.50157 | 10.51247 | 17.81003 | 18.26373 | 19.60638 | 18.77427 |
| **SUPT20H** | 4.56010 | 5.86356 | 6.27448 | 2.51336 | 2.55619 | 12.63016 | 2.28374 | 1.41296 | 7.41286 | 0.00000 | 2.78900 | 7.10379 |
| **SUPT20HL1** | 0.00513 | 0.01770 | 0.02209 | 0.01495 | 0.01819 | 0.00795 | 0.01363 | 0.01523 | 0.03916 | 0.00999 | 0.01758 | 0.02132 |
| **SUPT20HL2** | 1.04141 | 0.02814 | 0.02327 | 0.01059 | 0.01311 | 0.00000 | 0.11467 | 0.03691 | 0.07107 | 0.00000 | 0.00332 | 0.14532 |
| **SUPT3H** | 4.46814 | 1.13103 | 0.93720 | 0.85324 | 1.56755 | 2.99964 | 0.41020 | 0.67160 | 1.87085 | 0.16385 | 1.86719 | 1.74707 |
| **SUPT4H1** | 0.85090 | 15.04624 | 15.88682 | 7.41298 | 9.57643 | 15.76457 | 25.59215 | 20.77455 | 26.90260 | 2.97555 | 18.10257 | 24.26352 |
| **SUPT5H** | 2.03015 | 14.78910 | 17.73401 | 11.89638 | 16.34840 | 0.00000 | 24.67838 | 6.28722 | 16.72471 | 26.54533 | 16.38086 | 18.37315 |
| **SUPT6H** | 1.60618 | 5.36852 | 11.95254 | 10.27800 | 12.09402 | 0.00000 | 12.01611 | 6.14217 | 10.45582 | 17.79373 | 2.67531 | 10.65993 |
| **SUPT7L** | 0.61161 | 17.95495 | 7.90918 | 7.50944 | 5.86986 | 16.31687 | 7.96897 | 12.95250 | 8.93798 | 4.76220 | 9.87796 | 18.43217 |
| **SUPV3L1** | 4.14778 | 14.07884 | 10.18877 | 7.43042 | 4.71106 | 14.98278 | 12.64440 | 7.08617 | 22.93547 | 0.00000 | 12.87204 | 13.47929 |
| **SURF1** | 0.18890 | 4.82675 | 5.97110 | 4.41421 | 7.38203 | 0.00000 | 4.33983 | 0.23631 | 0.00000 | 11.03341 | 6.64382 | 1.37425 |

| | | | | | | | | | | | | |
|---|---|---|---|---|---|---|---|---|---|---|---|---|
| **SURF2** | 1.01167 | 15.86012 | 13.91407 | 12.05242 | 18.35580 | 1.47827 | 24.63839 | 6.24679 | 15.48359 | 8.62042 | 15.44290 | 18.40537 |
| **SURF4** | 8.78330 | 28.58397 | 30.45217 | 25.46218 | 21.44163 | 12.59627 | 3.14041 | 7.72674 | 53.32713 | 7.36687 | 15.15897 | 16.21630 |
| **SURF6** | 0.99064 | 16.70183 | 12.01173 | 10.98561 | 11.52041 | 0.00000 | 14.11189 | 11.92762 | 14.79905 | 17.27731 | 11.57159 | 13.99526 |
| **SUSD1** | 0.79745 | 0.17179 | 1.05164 | 1.17849 | 0.25264 | 3.02048 | 0.76591 | 0.06221 | 2.60026 | 1.50106 | 0.16446 | 0.67041 |
| **SUSD2** | 0.22050 | 26.83053 | 6.59406 | 25.35328 | 25.81647 | 0.00000 | 0.00000 | 0.54463 | 7.07571 | 1.39609 | 4.48357 | 11.48521 |
| **SUSD3** | 3.53448 | 1.20272 | 0.41989 | 3.08026 | 0.26975 | 0.05153 | 0.00000 | 0.73659 | 0.00000 | 0.00000 | 0.02421 | 0.00000 |
| **SUSD4** | 0.18060 | 0.13842 | 0.05620 | 0.01292 | 0.00000 | 0.04320 | 0.00000 | 0.11622 | 0.00000 | 0.06280 | 0.01192 | 0.10830 |
| **SUSD5** | 0.12712 | 0.29670 | 0.02262 | 0.19935 | 0.00000 | 0.22686 | 0.00000 | 0.11221 | 0.00000 | 0.13770 | 0.00000 | 0.33016 |
| **SUV39H1** | 0.76920 | 2.91650 | 2.43794 | 1.18477 | 3.56890 | 1.20425 | 3.18854 | 1.47819 | 4.84667 | 0.00000 | 2.76086 | 3.69556 |
| **SUV39H2** | 4.63151 | 4.67869 | 4.39560 | 2.06398 | 0.22928 | 15.82793 | 4.56888 | 1.73290 | 8.03966 | 0.00000 | 6.94587 | 4.42273 |
| **SUV420H1** | 0.79683 | 3.44968 | 2.56998 | 0.00000 | 1.85634 | 6.70759 | 4.73946 | 6.85336 | 0.72612 | 9.19107 | 0.00000 | 3.37409 |
| **SUV420H2** | 0.33865 | 0.00004 | 0.71108 | 0.40352 | 0.53880 | 0.00000 | 0.81669 | 0.13848 | 0.20768 | 1.96738 | 0.05793 | 0.00648 |
| **SUZ12** | 0.00000 | 0.00000 | 6.22540 | 0.66951 | 4.60617 | 20.92206 | 17.39924 | 0.77338 | 0.00000 | 30.77889 | 4.22119 | 1.89073 |
| **SUZ12P1** | 0.00000 | 1.45887 | 0.10144 | 0.94975 | 0.03624 | 0.55974 | 0.00000 | 1.18188 | 0.00386 | 1.46476 | 0.09605 | 0.80240 |
| **SV2A** | 0.00231 | 0.00000 | 0.16726 | 0.16049 | 0.04322 | 0.02074 | 0.25199 | 0.02320 | 0.87707 | 0.07214 | 0.19802 | 0.06426 |
| **SV2B** | 3.95647 | 0.00990 | 0.02266 | 0.00636 | 0.00688 | 0.00200 | 0.01659 | 1.15704 | 0.00745 | 0.04286 | 0.03683 | 0.08725 |
| **SV2C** | 0.20517 | 0.44736 | 0.09834 | 0.15246 | 0.37224 | 0.52746 | 0.00000 | 0.04256 | 0.63330 | 0.21008 | 0.66288 | 0.13166 |
| **SVEP1** | 2.13604 | 0.00192 | 0.05249 | 0.49456 | 0.09245 | 0.00000 | 0.07461 | 0.31509 | 0.08510 | 0.10452 | 0.17050 | 0.35421 |
| **SVIL** | 0.11319 | 3.54626 | 0.00000 | 2.93407 | 2.00811 | 1.52855 | 0.00000 | 0.12483 | 0.30438 | 4.57318 | 1.06382 | 0.37320 |
| **SVILP1** | 0.00604 | 0.01999 | 0.01306 | 0.01790 | 0.01699 | 0.00747 | 0.01756 | 0.01056 | 0.02497 | 0.02752 | 0.01507 | 0.01603 |
| **SVIP** | 4.16989 | 1.83818 | 2.60676 | 0.00000 | 1.62579 | 4.73940 | 4.64076 | 0.25641 | 0.00000 | 1.20868 | 1.59880 | 0.58881 |
| **SVOP** | 0.00587 | 0.02127 | 0.16941 | 0.00440 | 0.08983 | 0.09737 | 0.03207 | 0.00593 | 0.12493 | 0.00000 | 0.58502 | 0.11817 |
| **SVOPL** | 11.11368 | 0.43049 | 0.03142 | 0.32173 | 0.49666 | 0.52213 | 0.00000 | 1.12408 | 0.38337 | 0.00000 | 0.38486 | 0.68251 |
| **SWAP70** | 2.46479 | 3.06714 | 6.34897 | 0.84819 | 7.84495 | 11.09756 | 8.77047 | 3.49298 | 3.05976 | 14.46306 | 6.44645 | 8.80972 |
| **SWI5** | 1.57892 | 8.90411 | 13.20085 | 4.54099 | 8.97895 | 2.00507 | 10.76283 | 0.12192 | 5.43528 | 0.00000 | 4.32512 | 4.75291 |
| **SWSAP1** | 0.09626 | 3.85801 | 2.91108 | 3.80870 | 2.49950 | 0.03110 | 2.14380 | 0.78339 | 2.87058 | 0.00000 | 2.06841 | 2.54751 |
| **SWT1** | 2.68637 | 0.75104 | 3.37127 | 1.34295 | 0.00000 | 4.02792 | 3.55761 | 5.04704 | 0.44511 | 1.82252 | 0.81123 | 2.19882 |
| **SYAP1** | 1.66772 | 8.20588 | 6.75363 | 7.67350 | 6.27520 | 15.20778 | 11.53172 | 7.72000 | 9.56814 | 3.20990 | 6.56314 | 10.78876 |
| **SYBU** | 0.07936 | 1.67095 | 0.29819 | 0.00000 | 0.00000 | 0.10482 | 3.64614 | 0.06747 | 7.98202 | 0.00000 | 7.34890 | 1.44118 |
| **SYCE1** | 0.29645 | 4.09480 | 0.06862 | 0.32167 | 0.73123 | 0.45619 | 0.00000 | 0.27664 | 0.00000 | 0.68778 | 0.09603 | 4.15693 |
| **SYCE1L** | 0.00575 | 0.14593 | 0.08703 | 0.13611 | 0.02113 | 0.00000 | 0.02366 | 0.09017 | 0.00001 | 0.18256 | 0.05306 | 0.03379 |
| **SYCE2** | 0.17288 | 0.20622 | 0.10632 | 0.18643 | 0.07626 | 0.21803 | 0.00000 | 0.03663 | 0.38234 | 0.16072 | 0.12334 | 0.55246 |
| **SYCE3** | 0.22062 | 0.01211 | 0.01857 | 0.02670 | 0.02616 | 0.05316 | 0.00646 | 0.18843 | 0.02139 | 0.00000 | 0.00746 | 0.28502 |
| **SYCP1** | 0.08520 | 0.00000 | 0.00000 | 0.04820 | 0.04439 | 0.00000 | 0.31346 | 0.45720 | 0.15105 | 0.16254 | 0.00000 | 1.12697 |
| **SYCP2** | 7.83085 | 0.00000 | 0.00000 | 0.43235 | 0.43997 | 0.06796 | 0.00000 | 1.88597 | 0.00000 | 0.47721 | 0.00794 | 1.35413 |
| **SYCP2L** | 2.46312 | 0.76065 | 0.00000 | 0.03254 | 0.38487 | 1.90310 | 0.00000 | 2.18399 | 1.59922 | 3.23290 | 0.51688 | 1.80712 |
| **SYCP3** | 5.66169 | 9.14020 | 5.21955 | 2.42363 | 7.31187 | 35.97959 | 12.60982 | 10.26700 | 3.17875 | 6.43985 | 6.53386 | 13.27745 |
| **SYDE1** | 0.01310 | 0.00000 | 0.16592 | 0.00000 | 0.01294 | 0.03094 | 0.02177 | 0.02516 | 0.00000 | 0.32855 | 0.10963 | 0.01380 |
| **SYDE2** | 0.16026 | 0.00000 | 2.63306 | 0.00000 | 2.97863 | 0.00291 | 4.44768 | 0.00123 | 3.81476 | 4.65751 | 1.68000 | 0.00000 |
| **SYF2** | 5.26172 | 10.72257 | 7.64460 | 6.32039 | 6.20437 | 25.64993 | 16.66925 | 11.47247 | 4.69959 | 17.38329 | 7.37307 | 17.81774 |
| **SYK** | 0.00000 | 3.84203 | 0.20326 | 1.50952 | 2.46745 | 1.20482 | 1.23003 | 0.06218 | 0.00000 | 3.37887 | 1.75826 | 0.98423 |
| **SYMPK** | 0.57779 | 1.98341 | 3.90157 | 1.87844 | 3.44153 | 0.00000 | 0.45205 | 1.44983 | 3.29662 | 6.61524 | 3.25677 | 2.72689 |
| **SYN1** | 0.03399 | 0.01330 | 0.03012 | 0.06159 | 0.00937 | 0.00853 | 0.02195 | 0.00984 | 0.03829 | 0.04421 | 0.01886 | 0.03222 |
| **SYN2** | 2.14074 | 0.00140 | 0.35679 | 0.44435 | 0.00000 | 0.20699 | 0.00000 | 0.29193 | 0.44667 | 0.19167 | 0.00000 | 0.17359 |
| **SYN3** | 0.64158 | 0.29311 | 0.00000 | 0.52270 | 0.54749 | 0.00000 | 0.00000 | 0.44403 | 0.12268 | 0.22288 | 0.14369 | 0.66744 |
| **SYNC** | 0.00202 | 0.19496 | 0.02031 | 0.12995 | 0.50437 | 0.04006 | 0.00000 | 0.18699 | 0.00000 | 0.02172 | 0.08867 | 0.35307 |
| **SYNCRIP** | 2.99719 | 12.18545 | 13.21183 | 7.97464 | 12.32140 | 38.46866 | 29.04362 | 5.03356 | 14.75820 | 37.65807 | 15.14502 | 14.45121 |
| **SYNDIG1** | 0.00941 | 0.01520 | 0.01475 | 0.01226 | 0.01224 | 0.02781 | 0.01847 | 0.00966 | 0.01747 | 0.01223 | 0.01389 | 0.01749 |

| | | | | | | | | | | | | |
|---|---|---|---|---|---|---|---|---|---|---|---|---|
| **SYNDIG1L** | 0.00662 | 0.02613 | 0.00000 | 0.02115 | 0.16316 | 0.31585 | 0.00203 | 0.00726 | 0.02142 | 0.00000 | 0.00762 | 0.00629 |
| **SYNE1** | 9.28866 | 3.26537 | 0.63455 | 0.37634 | 2.67290 | 0.78418 | 0.00000 | 4.30533 | 0.86736 | 2.75347 | 0.35346 | 2.09784 |
| **SYNE2** | 13.64097 | 2.13309 | 16.93453 | 5.72153 | 7.89962 | 27.83912 | 3.05941 | 8.72439 | 15.46348 | 35.69727 | 4.88270 | 15.84219 |
| **SYNE3** | 0.00440 | 0.09431 | 0.34455 | 0.00735 | 0.46083 | 0.00000 | 0.00000 | 0.02257 | 0.61625 | 0.00000 | 0.44804 | 0.05955 |
| **SYNE4** | 0.00372 | 0.00000 | 0.87812 | 0.12521 | 0.00000 | 0.00000 | 0.50516 | 0.00923 | 1.02856 | 0.44936 | 0.61428 | 0.01451 |
| **SYNGAP1** | 0.19489 | 0.00000 | 0.01431 | 0.16620 | 0.12833 | 0.00000 | 0.00233 | 0.00697 | 0.03201 | 0.35876 | 0.04385 | 0.06758 |
| **SYNGR1** | 0.00122 | 0.46591 | 3.15649 | 0.02308 | 0.06055 | 0.00000 | 0.31295 | 0.24670 | 0.00000 | 0.00000 | 0.50566 | 0.63644 |
| **SYNGR2** | 0.84513 | 30.31169 | 7.41076 | 9.19427 | 26.35219 | 4.72840 | 8.27759 | 1.46974 | 30.13680 | 3.93481 | 16.03352 | 17.46009 |
| **SYNGR3** | 1.80069 | 3.56103 | 0.00000 | 1.60733 | 6.68649 | 0.00000 | 1.32158 | 0.24529 | 6.25877 | 0.00000 | 6.26770 | 2.26020 |
| **SYNGR4** | 0.00135 | 0.10229 | 0.03789 | 0.00353 | 0.00000 | 0.00044 | 0.01005 | 0.51131 | 0.00000 | 0.08835 | 0.18411 | 1.94694 |
| **SYNJ1** | 4.86422 | 1.86913 | 4.29397 | 0.00000 | 2.94379 | 2.12389 | 4.15098 | 7.25158 | 0.58836 | 5.62507 | 2.80257 | 3.99262 |
| **SYNJ2** | 3.63681 | 1.25572 | 1.12884 | 1.24412 | 7.72513 | 4.65665 | 4.08368 | 0.12306 | 0.15057 | 4.76429 | 1.76916 | 0.18504 |
| **SYNJ2-IT1** | 0.00905 | 0.01455 | 0.04476 | 0.00886 | 0.06548 | 0.02342 | 0.01061 | 0.01411 | 0.02864 | 0.03319 | 0.01038 | 0.01449 |
| **SYNJ2BP** | 3.47189 | 0.00000 | 2.22220 | 5.21059 | 0.00000 | 11.51370 | 5.08910 | 0.39523 | 3.33681 | 9.29058 | 9.75403 | 1.99056 |
| **SYNM** | 2.13814 | 0.00000 | 0.00434 | 0.28928 | 0.28398 | 0.00000 | 0.03731 | 2.93290 | 0.00000 | 0.13258 | 0.14410 | 0.27343 |
| **SYNPO** | 0.01203 | 0.00000 | 4.36360 | 0.00000 | 0.39404 | 0.00000 | 3.82476 | 0.00767 | 0.04903 | 0.21388 | 1.57681 | 0.02334 |
| **SYNPO2** | 0.05547 | 0.00000 | 0.02082 | 0.02451 | 0.00000 | 0.02666 | 0.00000 | 0.00725 | 0.01744 | 0.37288 | 0.00349 | 0.09312 |
| **SYNPO2L** | 0.11135 | 0.13849 | 0.02979 | 0.00000 | 0.07255 | 0.00000 | 0.00033 | 0.00887 | 0.12088 | 0.43423 | 0.28633 | 0.11238 |
| **SYNPR** | 0.05908 | 0.01237 | 0.04001 | 0.02059 | 0.00867 | 0.01936 | 0.01027 | 0.01062 | 0.00761 | 0.04001 | 0.02238 | 0.01323 |
| **SYNPR-AS1** | 0.00381 | 0.00000 | 0.00000 | 0.03890 | 0.00000 | 0.11505 | 0.00000 | 0.01011 | 0.05314 | 0.07950 | 0.02245 | 0.03672 |
| **SYNRG** | 6.47142 | 4.50697 | 3.98907 | 5.99456 | 1.75901 | 5.49622 | 1.68135 | 2.51163 | 2.12113 | 9.15521 | 3.15746 | 2.96274 |
| **SYP** | 0.00430 | 0.00803 | 0.08824 | 0.04877 | 0.00000 | 0.09330 | 0.06482 | 0.33309 | 0.07800 | 0.00000 | 0.04196 | 0.26277 |
| **SYP-AS1** | 0.00528 | 0.02178 | 0.01653 | 0.01193 | 0.01441 | 0.01751 | 0.01838 | 0.01890 | 0.02024 | 0.01409 | 0.01505 | 0.02785 |
| **SYPL1** | 3.26034 | 10.18565 | 10.33203 | 7.56670 | 0.57486 | 23.24703 | 0.00000 | 6.29254 | 23.57937 | 0.00000 | 7.25715 | 9.87665 |
| **SYPL2** | 1.17287 | 0.10382 | 0.03353 | 0.26476 | 0.02785 | 0.08945 | 0.06270 | 0.00000 | 0.00000 | 0.24353 | 0.01883 | 0.00000 |
| **SYS1** | 0.80323 | 5.49529 | 9.74645 | 1.48179 | 2.49408 | 10.45445 | 2.73005 | 7.78074 | 19.96262 | 6.75926 | 4.15459 | 20.88825 |
| **SYT1** | 1.83983 | 0.29016 | 0.26906 | 0.00000 | 0.00000 | 1.02598 | 0.00000 | 0.52287 | 0.25445 | 0.85437 | 0.02219 | 0.44310 |
| **SYT10** | 0.00642 | 0.01419 | 0.03349 | 0.01032 | 0.01978 | 0.00000 | 0.03320 | 0.50431 | 0.00000 | 0.03651 | 0.03401 | 0.06377 |
| **SYT11** | 0.65099 | 0.02074 | 0.10739 | 0.01605 | 0.00148 | 0.02928 | 0.05805 | 0.56490 | 0.00000 | 0.00360 | 0.08031 | 0.07310 |
| **SYT12** | 0.00709 | 0.02744 | 0.07131 | 0.02540 | 0.00491 | 0.00825 | 0.00295 | 0.01092 | 0.11831 | 0.00684 | 0.07118 | 0.02343 |
| **SYT13** | 0.00458 | 0.03657 | 0.49941 | 1.35715 | 0.00000 | 0.00077 | 0.05024 | 0.00857 | 0.04495 | 0.06918 | 0.04484 | 0.02730 |
| **SYT14** | 0.75853 | 0.26367 | 0.03703 | 0.01425 | 0.12901 | 0.27254 | 0.00000 | 0.00000 | 0.00000 | 0.20522 | 0.00000 | 0.65433 |
| **SYT14L** | 0.01187 | 0.01518 | 0.01463 | 0.01098 | 0.01618 | 0.02172 | 0.01853 | 0.01015 | 0.01755 | 0.01315 | 0.01423 | 0.03015 |
| **SYT15** | 0.00712 | 0.00000 | 0.03458 | 0.01280 | 0.13224 | 0.08190 | 0.14136 | 0.05948 | 0.00000 | 0.09451 | 0.13142 | 0.10322 |
| **SYT16** | 3.99325 | 0.13552 | 0.18387 | 0.00280 | 0.01132 | 0.00000 | 0.07476 | 0.43926 | 0.00022 | 0.00000 | 0.01226 | 0.02667 |
| **SYT17** | 3.86899 | 0.57542 | 0.39949 | 0.00000 | 0.22968 | 0.25403 | 0.12998 | 0.08466 | 0.00000 | 0.06920 | 0.04815 | 0.91166 |
| **SYT2** | 13.03876 | 1.91942 | 0.00000 | 7.24887 | 3.61002 | 0.00000 | 0.00000 | 2.51879 | 0.41528 | 2.65575 | 4.91845 | 2.47020 |
| **SYT3** | 0.08214 | 0.05074 | 0.01834 | 0.07028 | 0.03125 | 0.00043 | 0.02754 | 0.00450 | 0.02489 | 0.00000 | 0.01514 | 0.13937 |
| **SYT5** | 0.17317 | 0.10550 | 0.02275 | 0.01360 | 0.00886 | 0.02693 | 0.02214 | 0.05899 | 0.00652 | 0.03999 | 0.01235 | 0.05224 |
| **SYT6** | 0.00185 | 0.12739 | 0.03811 | 0.07122 | 0.00863 | 0.13553 | 0.00000 | 0.54648 | 0.00000 | 0.00004 | 0.10440 | 0.01647 |
| **SYT7** | 0.08338 | 0.33585 | 1.07324 | 0.00000 | 0.84019 | 0.00000 | 0.99013 | 0.05900 | 0.97123 | 1.83380 | 0.00000 | 0.00000 |
| **SYT8** | 0.00637 | 0.01366 | 0.08505 | 0.01580 | 0.01383 | 0.02643 | 0.05731 | 0.01038 | 0.02725 | 0.00000 | 0.00976 | 0.01653 |
| **SYT9** | 1.13816 | 0.01754 | 0.03048 | 0.02761 | 0.00583 | 0.01208 | 0.01644 | 0.15868 | 0.00014 | 0.02111 | 0.06050 | 0.01465 |
| **SYTL1** | 0.23694 | 0.00742 | 0.01670 | 0.10433 | 0.00000 | 0.05410 | 0.02429 | 0.00219 | 0.00001 | 0.02924 | 0.06327 | 0.01199 |
| **SYTL2** | 4.14548 | 0.56283 | 0.04685 | 1.22987 | 0.00000 | 0.20072 | 0.00000 | 1.52559 | 0.00022 | 0.12931 | 0.00000 | 0.98310 |
| **SYTL3** | 11.08514 | 0.03307 | 0.09015 | 0.13626 | 0.66681 | 0.00000 | 0.00000 | 3.34399 | 0.49451 | 0.62533 | 0.41229 | 0.68069 |
| **SYTL4** | 0.68483 | 1.95737 | 0.00000 | 1.40451 | 0.00000 | 0.00000 | 0.00000 | 0.28584 | 0.00000 | 0.44594 | 0.29586 | 1.43724 |
| **SYTL5** | 0.47341 | 0.02268 | 0.21179 | 0.05525 | 0.00000 | 0.06099 | 0.00000 | 0.83089 | 0.00550 | 0.01909 | 0.00925 | 0.02470 |

| | | | | | | | | | | | | |
|---|---|---|---|---|---|---|---|---|---|---|---|---|
| **SYVN1** | 2.12164 | 1.89151 | 8.07766 | 2.16789 | 4.96523 | 0.00000 | 0.00000 | 0.00000 | 12.58774 | 6.31473 | 3.70550 | 3.15216 |
| **SZRD1** | 3.65169 | 19.80778 | 17.73554 | 14.92353 | 20.32333 | 11.69390 | 24.02542 | 10.73751 | 21.90745 | 5.27329 | 19.34073 | 22.53725 |
| **SZT2** | 1.30632 | 1.17256 | 1.79965 | 1.86056 | 2.05536 | 0.00000 | 0.03424 | 0.19499 | 3.04130 | 4.79403 | 1.87501 | 1.60780 |
| **T** | 0.02618 | 0.01211 | 0.15564 | 0.03355 | 0.00543 | 0.04830 | 0.02125 | 0.01459 | 0.00000 | 0.02241 | 0.01677 | 0.01982 |
| **TAAR5** | 0.08126 | 0.00000 | 0.11392 | 0.00000 | 0.00000 | 0.28807 | 2.26894 | 0.04715 | 0.00000 | 2.87974 | 0.00000 | 0.42993 |
| **TAAR8** | 0.00261 | 0.01532 | 0.01435 | 0.01227 | 0.01383 | 0.01564 | 0.01683 | 0.09664 | 0.01704 | 0.01427 | 0.02631 | 0.01302 |
| **TAB1** | 2.73611 | 2.52065 | 1.71029 | 1.77931 | 2.84718 | 0.00000 | 0.34846 | 0.00000 | 2.73617 | 1.10265 | 2.39830 | 1.02735 |
| **TAB2** | 4.77895 | 0.61743 | 3.54139 | 2.77357 | 0.65196 | 11.03104 | 5.90551 | 2.64002 | 0.00000 | 11.87373 | 2.03406 | 3.36763 |
| **TAB3** | 1.38850 | 0.00000 | 1.95208 | 0.30079 | 0.88219 | 2.83692 | 2.25101 | 0.14985 | 0.00000 | 3.45481 | 1.73488 | 0.34373 |
| **TAC1** | 1.09978 | 0.01395 | 0.01297 | 0.01290 | 0.01504 | 0.00066 | 0.01823 | 0.58415 | 0.03437 | 0.00289 | 0.02420 | 0.00000 |
| **TAC3** | 0.00644 | 0.00969 | 0.08372 | 0.00807 | 0.03767 | 0.04608 | 0.17446 | 0.00985 | 0.21949 | 0.00000 | 0.00698 | 0.01435 |
| **TAC4** | 0.00517 | 0.00263 | 0.63340 | 0.00000 | 0.00000 | 0.04394 | 0.08250 | 0.01451 | 0.20795 | 0.02925 | 0.10962 | 0.04019 |
| **TACC1** | 6.08841 | 3.44381 | 18.22589 | 0.00000 | 19.92670 | 12.85123 | 23.24696 | 2.41561 | 14.40213 | 36.35122 | 14.52415 | 7.77084 |
| **TACC2** | 0.57387 | 3.24332 | 2.97251 | 2.24059 | 3.11174 | 0.07725 | 5.23411 | 0.30963 | 5.49682 | 5.96216 | 0.60715 | 0.09301 |
| **TACC3** | 16.24111 | 29.91243 | 9.96479 | 17.32135 | 28.07497 | 0.14767 | 21.25011 | 2.57507 | 19.10037 | 8.65537 | 14.52147 | 19.95420 |
| **TACO1** | 0.20697 | 10.56687 | 1.45957 | 5.30633 | 6.18508 | 0.14835 | 2.42650 | 0.07638 | 1.70716 | 5.04402 | 4.77321 | 4.56474 |
| **TACR1** | 0.00158 | 0.13346 | 0.18142 | 0.09301 | 0.06414 | 0.00000 | 0.00000 | 0.12513 | 0.13595 | 0.15274 | 0.04230 | 0.24206 |
| **TACR2** | 0.00390 | 0.00155 | 0.24002 | 0.01661 | 0.00000 | 0.11858 | 0.00000 | 0.00920 | 0.00000 | 0.08235 | 0.03623 | 0.03224 |
| **TACR3** | 1.18744 | 0.00705 | 0.33024 | 0.05539 | 0.03614 | 0.03617 | 0.00000 | 0.06628 | 0.00002 | 0.00126 | 0.02222 | 0.00598 |
| **TACSTD2** | 0.00000 | 0.00000 | 14.73494 | 0.00000 | 37.93100 | 0.52225 | 8.34339 | 0.13921 | 55.46937 | 0.00190 | 19.77860 | 0.00000 |
| **TADA1** | 1.29877 | 7.95247 | 1.59250 | 1.46514 | 0.00000 | 10.16981 | 7.60046 | 2.40703 | 1.28597 | 0.00000 | 3.22739 | 13.49237 |
| **TADA2A** | 6.03667 | 2.89766 | 1.94714 | 0.31552 | 1.79468 | 7.76038 | 0.00000 | 0.17566 | 1.78300 | 1.45624 | 1.71713 | 3.65563 |
| **TADA2B** | 3.31928 | 3.44027 | 0.88614 | 0.93406 | 2.81638 | 3.01568 | 2.75272 | 0.26110 | 0.00000 | 6.14569 | 0.49963 | 2.43362 |
| **TADA3** | 3.93293 | 2.97009 | 7.33617 | 7.27295 | 2.59458 | 0.34817 | 5.00304 | 1.03708 | 6.44782 | 2.61615 | 4.60241 | 4.79891 |
| **TAF1** | 2.54117 | 0.82826 | 0.12986 | 1.65163 | 0.79092 | 6.09956 | 0.61500 | 2.13113 | 0.00000 | 5.68950 | 0.35595 | 1.72717 |
| **TAF10** | 1.72322 | 4.56635 | 4.82864 | 5.17774 | 11.41083 | 0.00000 | 4.07615 | 0.13992 | 1.87518 | 14.14439 | 2.49894 | 3.85166 |
| **TAF11** | 1.74552 | 9.76005 | 17.11407 | 3.30782 | 5.22161 | 17.66800 | 20.75981 | 13.68370 | 16.49056 | 8.33212 | 10.05431 | 14.14648 |
| **TAF12** | 6.26623 | 13.49448 | 14.72887 | 0.87250 | 7.83693 | 20.47491 | 5.85329 | 9.40892 | 8.47868 | 0.54724 | 13.11274 | 22.72024 |
| **TAF13** | 1.31268 | 4.58541 | 5.16112 | 4.01932 | 2.59984 | 12.88172 | 7.59171 | 16.79396 | 0.00000 | 3.50181 | 6.34702 | 11.16731 |
| **TAF15** | 0.43956 | 0.24130 | 1.57234 | 1.10363 | 2.36293 | 0.85308 | 3.46027 | 0.21337 | 0.45360 | 11.19952 | 4.69122 | 2.69622 |
| **TAF1A** | 1.31026 | 2.57713 | 3.70248 | 0.79439 | 0.00000 | 3.35002 | 0.02310 | 1.51731 | 5.23494 | 2.21722 | 0.98521 | 5.61483 |
| **TAF1A-AS1** | 0.97472 | 0.74361 | 0.21533 | 0.08348 | 0.07125 | 0.41054 | 0.00000 | 0.51204 | 0.41674 | 0.00000 | 0.11272 | 2.50853 |
| **TAF1B** | 0.83003 | 3.04410 | 10.15912 | 2.20946 | 2.01237 | 15.81238 | 3.71241 | 4.06542 | 13.31435 | 3.29891 | 6.06620 | 4.27752 |
| **TAF1C** | 0.39109 | 5.03141 | 4.27004 | 3.31368 | 5.67508 | 0.00000 | 0.00000 | 2.16485 | 6.91017 | 7.38877 | 4.72611 | 4.17052 |
| **TAF1D** | 0.86393 | 3.80173 | 9.31094 | 6.34279 | 13.99605 | 16.98696 | 6.69643 | 8.29336 | 16.14993 | 39.34812 | 4.35255 | 11.59709 |
| **TAF1L** | 0.70861 | 0.17286 | 0.17885 | 0.09720 | 0.00000 | 0.16247 | 0.46770 | 0.39046 | 0.00000 | 0.25988 | 0.08913 | 0.30984 |
| **TAF2** | 2.29046 | 1.94823 | 6.78558 | 5.11087 | 3.35336 | 9.96705 | 9.24335 | 2.53293 | 8.77773 | 8.08104 | 6.49811 | 4.30162 |
| **TAF3** | 10.18566 | 4.04674 | 7.61851 | 3.15303 | 4.91689 | 3.03215 | 5.86428 | 9.65355 | 4.01413 | 15.18882 | 3.73329 | 8.01668 |
| **TAF4** | 3.15623 | 0.00000 | 0.17242 | 0.00000 | 0.87930 | 2.14224 | 2.13369 | 0.34246 | 0.25688 | 7.58282 | 0.00000 | 1.10541 |
| **TAF4B** | 1.59779 | 3.51004 | 8.60807 | 0.00000 | 0.00000 | 6.52600 | 3.95935 | 2.56988 | 0.00000 | 9.86391 | 1.04028 | 5.32535 |
| **TAF5** | 4.69745 | 0.62151 | 6.66346 | 0.68124 | 1.26790 | 1.97144 | 5.88345 | 6.51099 | 0.34490 | 5.49585 | 2.22027 | 4.32895 |
| **TAF5L** | 4.43353 | 2.32452 | 2.65994 | 0.00000 | 3.27590 | 2.12858 | 3.37598 | 9.20804 | 0.00000 | 8.07681 | 5.68907 | 6.54781 |
| **TAF6** | 2.95234 | 11.02711 | 6.53611 | 5.09472 | 4.65700 | 0.00000 | 2.50728 | 0.45309 | 9.69700 | 0.00000 | 2.91134 | 7.08040 |
| **TAF6L** | 0.39858 | 0.24154 | 1.43868 | 0.68852 | 0.98774 | 0.00000 | 0.51856 | 0.15699 | 0.00000 | 4.01748 | 0.52611 | 0.51546 |
| **TAF7** | 4.75744 | 19.64448 | 13.72574 | 1.81697 | 0.00000 | 17.86173 | 8.01531 | 21.73003 | 6.95134 | 9.09239 | 9.27498 | 29.09274 |
| **TAF7L** | 0.11524 | 0.03694 | 0.12700 | 0.22326 | 0.16889 | 0.10136 | 0.06011 | 0.73261 | 0.00000 | 0.04761 | 0.07181 | 0.17337 |
| **TAF8** | 2.81370 | 2.51089 | 3.92140 | 2.83693 | 1.11185 | 2.44287 | 4.60333 | 2.38021 | 3.27442 | 0.00000 | 5.18830 | 5.42513 |
| **TAF9** | 1.33650 | 7.41918 | 16.37618 | 4.90723 | 4.28731 | 33.28427 | 26.55468 | 10.58900 | 19.74275 | 6.51240 | 20.28104 | 20.04635 |

| | | | | | | | | | | | | |
|---|---|---|---|---|---|---|---|---|---|---|---|---|
| **TAF9B** | 2.47187 | 1.02861 | 2.44485 | 1.55553 | 0.00000 | 4.09191 | 2.90941 | 0.37655 | 1.75407 | 2.14848 | 0.00000 | 1.89035 |
| **TAGAP** | 0.03414 | 0.12483 | 0.00000 | 0.10974 | 0.33050 | 0.33287 | 0.00000 | 0.00379 | 0.00000 | 0.19192 | 0.05892 | 0.00000 |
| **TAGLN** | 0.17374 | 0.13348 | 0.82918 | 0.27911 | 0.87971 | 0.00000 | 0.47319 | 0.31003 | 0.66399 | 6.56686 | 0.00000 | 0.83346 |
| **TAGLN2** | 5.01034 | 36.53652 | 16.51469 | 14.89543 | 57.88767 | 16.06800 | 42.05208 | 3.55758 | 63.67693 | 0.00000 | 38.91118 | 23.21711 |
| **TAGLN3** | 0.00585 | 0.02154 | 0.01918 | 0.03445 | 0.01672 | 0.00863 | 0.02406 | 0.01075 | 0.03757 | 0.02300 | 0.02276 | 0.01603 |
| **TAL1** | 0.00875 | 0.01311 | 0.00704 | 0.00935 | 0.00885 | 0.01802 | 0.05051 | 0.01128 | 0.01786 | 0.03301 | 0.03138 | 0.01482 |
| **TAL2** | 0.00521 | 0.01668 | 0.02326 | 0.10501 | 0.08658 | 0.00000 | 0.01885 | 0.01256 | 0.00311 | 0.04523 | 0.02143 | 0.03543 |
| **TALDO1** | 12.52324 | 21.17770 | 29.28258 | 31.20355 | 43.71891 | 31.98724 | 43.50656 | 12.40654 | 42.85415 | 7.62212 | 41.24658 | 9.79450 |
| **TAMM41** | 1.28950 | 5.83325 | 0.00000 | 1.94366 | 5.91063 | 5.98764 | 0.55810 | 2.84358 | 4.87856 | 0.00039 | 2.89244 | 4.00693 |
| **TANC1** | 3.26180 | 2.83897 | 2.41127 | 0.00000 | 1.85713 | 1.11720 | 0.65836 | 2.11890 | 0.00000 | 2.43112 | 0.60035 | 3.82322 |
| **TANC2** | 5.34404 | 0.59511 | 6.63660 | 0.15578 | 0.88545 | 0.01419 | 3.62816 | 0.36053 | 0.00911 | 2.37734 | 0.00000 | 1.17055 |
| **TANGO2** | 0.00016 | 4.25632 | 5.17654 | 1.13762 | 4.29938 | 1.73178 | 2.59575 | 0.11623 | 1.99965 | 0.00000 | 7.68183 | 2.26502 |
| **TANGO6** | 0.17371 | 2.97988 | 1.73055 | 0.21929 | 2.09124 | 0.62219 | 1.56691 | 0.36615 | 1.19114 | 0.00000 | 3.68385 | 4.96551 |
| **TANK** | 0.46860 | 1.85252 | 8.84092 | 2.37170 | 0.00000 | 11.30029 | 5.28054 | 4.86576 | 5.45170 | 0.00033 | 5.12822 | 2.43268 |
| **TAOK1** | 6.55911 | 2.58875 | 6.36134 | 1.64906 | 3.15707 | 18.15562 | 12.11521 | 3.95461 | 3.58773 | 31.37792 | 2.14521 | 7.18062 |
| **TAOK2** | 1.48093 | 1.81619 | 4.10939 | 3.12085 | 8.05958 | 0.00000 | 3.06840 | 0.18631 | 4.07635 | 17.80870 | 1.70331 | 2.86361 |
| **TAOK3** | 8.60280 | 5.20831 | 1.42553 | 9.15252 | 16.32807 | 9.52311 | 9.82333 | 11.49400 | 2.95098 | 16.25734 | 14.58515 | 11.34331 |
| **TAP1** | 0.04563 | 0.64467 | 0.27159 | 0.18815 | 0.27254 | 0.10614 | 0.00000 | 0.00007 | 0.32104 | 0.00000 | 0.00000 | 0.49098 |
| **TAP2** | 0.00349 | 0.00000 | 0.03060 | 0.07790 | 0.00000 | 0.19451 | 0.10991 | 0.13165 | 0.00000 | 0.16414 | 0.10125 | 0.79327 |
| **TAPBP** | 0.65859 | 0.33488 | 0.95678 | 0.52824 | 0.42596 | 0.00000 | 0.25485 | 0.38952 | 0.50431 | 0.79580 | 0.20297 | 1.46568 |
| **TAPBPL** | 0.01814 | 0.01453 | 0.01875 | 0.14026 | 0.13640 | 0.00002 | 0.05364 | 0.14351 | 0.00891 | 0.00001 | 0.01616 | 0.20053 |
| **TAPT1** | 1.67642 | 0.38156 | 0.83090 | 0.58276 | 0.00000 | 1.38305 | 0.00000 | 0.05920 | 1.08704 | 3.16521 | 0.00000 | 1.02371 |
| **TAPT1-AS1** | 2.58985 | 8.86169 | 7.04787 | 6.72566 | 12.78822 | 0.15362 | 14.09297 | 2.58976 | 11.31668 | 11.45328 | 4.80598 | 10.43747 |
| **TARBP1** | 0.66684 | 0.44508 | 0.00000 | 1.46882 | 2.08563 | 2.82424 | 0.75136 | 0.80055 | 0.78692 | 2.54389 | 0.83927 | 0.36696 |
| **TARBP2** | 0.56714 | 13.93805 | 6.40721 | 11.44413 | 11.73597 | 0.01139 | 14.29817 | 2.11270 | 14.63041 | 3.00481 | 10.81254 | 14.57335 |
| **TARDBP** | 4.84151 | 18.46582 | 19.15497 | 13.51293 | 16.13668 | 28.72330 | 27.18913 | 14.76482 | 24.60995 | 5.04901 | 20.22929 | 23.66877 |
| **TARS** | 4.31188 | 12.16061 | 19.24939 | 13.88905 | 8.34860 | 37.83442 | 34.95269 | 7.15364 | 32.21985 | 14.60093 | 29.85912 | 3.18757 |
| **TARS2** | 0.59749 | 8.40660 | 1.96193 | 4.87256 | 2.51220 | 5.17088 | 0.00000 | 1.08353 | 5.21586 | 1.64652 | 3.97887 | 5.81257 |
| **TARSL2** | 0.81152 | 1.40461 | 0.87466 | 0.60496 | 0.34211 | 0.63763 | 1.06799 | 1.32874 | 0.74870 | 1.10475 | 0.24126 | 1.19787 |
| **TAS1R1** | 0.00667 | 0.27316 | 0.03706 | 0.00000 | 0.00000 | 0.00000 | 0.09294 | 5.25209 | 0.07492 | 0.06960 | 0.00000 | 10.58165 |
| **TAS1R2** | 0.00493 | 0.01359 | 0.02127 | 0.01471 | 0.02162 | 0.00000 | 0.00170 | 0.01153 | 0.04677 | 0.09375 | 0.00541 | 0.01451 |
| **TAS1R3** | 0.01319 | 0.01591 | 0.02392 | 0.01163 | 0.00993 | 0.00000 | 0.14757 | 0.00944 | 0.00000 | 0.04308 | 0.02928 | 0.01601 |
| **TAS2R1** | 0.00653 | 0.01739 | 0.01129 | 0.01016 | 0.01702 | 0.00000 | 0.01050 | 0.01048 | 0.13820 | 0.02320 | 0.00213 | 0.01666 |
| **TAS2R10** | 0.00590 | 0.01170 | 0.14923 | 0.02137 | 0.02481 | 0.00997 | 0.00544 | 0.01037 | 0.00367 | 0.05962 | 0.01267 | 0.01489 |
| **TAS2R13** | 0.00567 | 0.01857 | 0.02772 | 0.02718 | 0.00000 | 0.05706 | 0.00000 | 0.00999 | 0.01693 | 0.04390 | 0.00968 | 0.01096 |
| **TAS2R14** | 0.03070 | 0.00000 | 0.06722 | 0.02403 | 0.14942 | 0.08703 | 0.01173 | 0.00497 | 0.06085 | 0.26749 | 0.04926 | 0.05087 |
| **TAS2R19** | 0.00744 | 0.01150 | 0.02939 | 0.06137 | 0.00000 | 0.01570 | 0.08617 | 0.01015 | 0.02798 | 0.07514 | 0.19364 | 0.01989 |
| **TAS2R20** | 0.00455 | 0.00000 | 0.27422 | 0.36313 | 0.00000 | 0.51495 | 0.36340 | 0.00153 | 0.34348 | 0.12337 | 0.36892 | 0.17065 |
| **TAS2R3** | 0.00556 | 0.01021 | 0.01137 | 0.00727 | 0.00294 | 0.02612 | 0.00679 | 0.00818 | 0.00493 | 0.07961 | 0.04375 | 0.04644 |
| **TAS2R30** | 0.00541 | 0.02405 | 0.03041 | 0.00567 | 0.03031 | 0.01964 | 0.00789 | 0.00926 | 0.06318 | 0.03658 | 0.00848 | 0.03764 |
| **TAS2R31** | 0.00507 | 0.01991 | 0.07598 | 0.00788 | 0.02067 | 0.01569 | 0.00416 | 0.01167 | 0.02145 | 0.06130 | 0.02437 | 0.02033 |
| **TAS2R39** | 0.00349 | 0.01606 | 0.02646 | 0.01117 | 0.01698 | 0.01474 | 0.01368 | 0.03534 | 0.04705 | 0.00508 | 0.00894 | 0.01610 |
| **TAS2R4** | 0.00441 | 0.03508 | 0.07262 | 0.05712 | 0.00000 | 0.10394 | 0.00000 | 0.00735 | 0.07350 | 0.07751 | 0.09353 | 0.04462 |
| **TAS2R43** | 0.00612 | 0.01548 | 0.01837 | 0.01597 | 0.01246 | 0.01987 | 0.01373 | 0.01037 | 0.05304 | 0.02105 | 0.01760 | 0.01688 |
| **TAS2R46** | 0.00539 | 0.00983 | 0.05984 | 0.10076 | 0.00778 | 0.03902 | 0.00000 | 0.01078 | 0.02971 | 0.03471 | 0.02503 | 0.02169 |
| **TAS2R5** | 0.00254 | 0.30638 | 0.06777 | 0.19428 | 0.00000 | 0.00000 | 0.00220 | 0.02892 | 0.42976 | 0.30263 | 0.20956 | 0.33548 |
| **TAS2R50** | 0.00622 | 0.01615 | 0.02699 | 0.01127 | 0.01517 | 0.01389 | 0.01573 | 0.01048 | 0.02624 | 0.01534 | 0.01399 | 0.01861 |
| **TAS2R60** | 0.03022 | 0.01506 | 0.01412 | 0.01128 | 0.02595 | 0.01847 | 0.01614 | 0.00799 | 0.02266 | 0.01183 | 0.02286 | 0.01567 |

| | | | | | | | | | | | | |
|---|---|---|---|---|---|---|---|---|---|---|---|---|
| **TASP1** | 1.40632 | 2.56482 | 2.00265 | 1.48276 | 1.55344 | 7.37779 | 0.00000 | 5.62706 | 6.83031 | 3.25151 | 2.47791 | 5.87620 |
| **TAT** | 0.13627 | 0.01585 | 0.00647 | 0.00012 | 0.00000 | 0.03219 | 0.00711 | 0.11997 | 0.00969 | 0.16369 | 0.02568 | 0.09914 |
| **TATDN1** | 0.17912 | 1.36685 | 3.64030 | 0.23385 | 0.00000 | 3.37383 | 1.76522 | 1.38356 | 1.72062 | 0.00000 | 0.88976 | 3.84288 |
| **TATDN2** | 0.93300 | 0.61274 | 2.40786 | 0.57238 | 4.30718 | 1.82517 | 7.14010 | 2.56946 | 0.00033 | 7.95380 | 1.80764 | 0.61894 |
| **TATDN3** | 0.31318 | 6.63319 | 1.39078 | 1.51375 | 0.96519 | 10.09616 | 2.95917 | 2.17032 | 3.28857 | 2.56897 | 3.15901 | 3.90760 |
| **TAX1BP1** | 14.24716 | 18.70497 | 15.49446 | 14.78957 | 13.81562 | 43.77110 | 25.71361 | 13.28351 | 9.98231 | 27.67357 | 16.09937 | 32.12532 |
| **TAX1BP3** | 0.05869 | 1.95728 | 0.67531 | 1.95660 | 1.46552 | 1.06448 | 0.00000 | 0.01238 | 2.41703 | 0.00000 | 2.76539 | 0.00771 |
| **TAZ** | 0.17211 | 1.72313 | 0.55369 | 0.64031 | 0.94131 | 0.00000 | 0.01376 | 0.01907 | 0.24489 | 1.74896 | 1.22633 | 1.14652 |
| **TBC1D1** | 10.59373 | 2.42248 | 0.00000 | 4.31279 | 19.43750 | 6.11834 | 11.55718 | 3.74834 | 6.31430 | 14.70547 | 12.81987 | 0.00000 |
| **TBC1D10A** | 1.83079 | 1.32749 | 9.97961 | 3.17713 | 0.46956 | 0.00000 | 1.18202 | 1.32735 | 1.70570 | 0.15638 | 1.89276 | 1.59219 |
| **TBC1D10B** | 5.17069 | 2.00798 | 3.48742 | 3.03672 | 8.09413 | 0.00000 | 5.56970 | 0.79903 | 0.00000 | 17.37205 | 2.71486 | 2.65157 |
| **TBC1D10C** | 0.00635 | 0.01712 | 0.01735 | 0.01396 | 0.01826 | 0.00987 | 0.01881 | 0.01045 | 0.02293 | 0.01105 | 0.01892 | 0.01877 |
| **TBC1D12** | 0.24531 | 0.08426 | 1.51979 | 1.20301 | 0.00063 | 2.68747 | 3.33596 | 2.79374 | 0.00000 | 3.28380 | 0.43192 | 0.30174 |
| **TBC1D13** | 2.51847 | 10.20506 | 2.58090 | 6.95040 | 4.67757 | 1.63862 | 2.72859 | 2.33781 | 4.46287 | 7.77011 | 1.66088 | 12.47087 |
| **TBC1D14** | 7.34617 | 5.90105 | 0.89777 | 4.81384 | 1.13261 | 0.00748 | 0.51510 | 1.10639 | 3.69577 | 0.00000 | 2.01896 | 1.29733 |
| **TBC1D15** | 1.84099 | 3.11139 | 12.71693 | 4.42995 | 0.00000 | 13.11280 | 9.29621 | 8.86239 | 6.63989 | 4.18710 | 5.46446 | 5.84069 |
| **TBC1D16** | 0.20440 | 0.47506 | 0.69450 | 3.58114 | 0.00000 | 0.01985 | 0.09348 | 0.00000 | 0.82753 | 1.01152 | 0.00000 | 0.90884 |
| **TBC1D17** | 0.91828 | 1.07950 | 4.45024 | 2.62555 | 2.31270 | 0.00000 | 1.40217 | 5.31364 | 4.93972 | 1.92334 | 1.36044 | 1.52941 |
| **TBC1D19** | 0.19033 | 0.00000 | 0.03296 | 0.67558 | 0.09451 | 0.59308 | 0.00000 | 0.12453 | 0.46759 | 0.23615 | 0.22406 | 0.97108 |
| **TBC1D2** | 1.58323 | 0.36209 | 3.49628 | 0.35599 | 4.94639 | 0.00000 | 1.86781 | 0.00000 | 1.84280 | 0.27495 | 2.84175 | 1.46013 |
| **TBC1D20** | 1.51140 | 1.56516 | 5.42400 | 2.76300 | 5.05339 | 1.78603 | 4.97643 | 0.68568 | 1.44551 | 8.15994 | 1.67904 | 3.15991 |
| **TBC1D22A** | 0.77668 | 2.93495 | 3.11947 | 2.16230 | 2.09365 | 0.00000 | 1.67807 | 0.34408 | 3.78561 | 2.06946 | 2.54622 | 0.86440 |
| **TBC1D22B** | 0.82657 | 2.52080 | 2.93853 | 1.71889 | 1.82608 | 0.00000 | 2.58537 | 0.80071 | 3.21638 | 1.07110 | 1.62645 | 0.73664 |
| **TBC1D23** | 7.20466 | 21.45486 | 4.94393 | 15.55154 | 0.00000 | 33.87643 | 0.00728 | 12.22422 | 3.05817 | 9.73743 | 4.79255 | 26.04844 |
| **TBC1D24** | 0.21287 | 1.97170 | 2.18800 | 1.39129 | 0.00000 | 0.00000 | 4.49317 | 0.02046 | 0.49055 | 2.11990 | 1.87941 | 1.84182 |
| **TBC1D25** | 1.35771 | 2.81561 | 2.05236 | 3.06048 | 3.36032 | 0.00000 | 1.61465 | 0.40563 | 2.50461 | 0.90630 | 1.20868 | 4.10247 |
| **TBC1D26** | 0.00606 | 0.01425 | 0.01838 | 0.01762 | 0.01098 | 0.01261 | 0.01619 | 0.00985 | 0.09646 | 0.01265 | 0.00604 | 0.03375 |
| **TBC1D29** | 0.00455 | 0.01421 | 0.01318 | 0.01169 | 0.01479 | 0.01980 | 0.01577 | 0.01684 | 0.02913 | 0.01413 | 0.01502 | 0.02768 |
| **TBC1D2B** | 1.42464 | 0.13860 | 2.25964 | 0.21266 | 0.70057 | 0.00000 | 1.15254 | 0.08212 | 1.07636 | 0.74766 | 0.66913 | 0.84834 |
| **TBC1D30** | 0.06477 | 0.00000 | 0.00000 | 0.16884 | 0.04655 | 0.17665 | 0.00000 | 0.00016 | 0.35695 | 0.75298 | 0.18373 | 1.52127 |
| **TBC1D31** | 2.52408 | 2.22682 | 0.47272 | 0.71195 | 0.00000 | 3.93437 | 0.00000 | 0.37048 | 2.91066 | 2.31573 | 0.24752 | 0.62554 |
| **TBC1D32** | 0.43513 | 0.10808 | 0.00000 | 0.05827 | 0.00000 | 2.72796 | 0.40690 | 0.29401 | 2.00359 | 3.00404 | 0.23345 | 0.45745 |
| **TBC1D3B** | 0.00660 | 0.01466 | 0.02184 | 0.01257 | 0.01448 | 0.02111 | 0.02693 | 0.01062 | 0.01575 | 0.00872 | 0.01515 | 0.01789 |
| **TBC1D3P1-DHX** | 0.00509 | 0.01338 | 0.01822 | 0.01232 | 0.00852 | 0.01976 | 0.02127 | 0.03518 | 0.00000 | 0.02074 | 0.05387 | 0.01609 |
| **TBC1D3P2** | 0.00769 | 0.01911 | 0.04520 | 0.02148 | 0.01445 | 0.01474 | 0.02807 | 0.01182 | 0.01845 | 0.00403 | 0.01648 | 0.01724 |
| **TBC1D3P5** | 0.00686 | 0.01498 | 0.03860 | 0.01259 | 0.01770 | 0.01356 | 0.01443 | 0.01061 | 0.01111 | 0.01631 | 0.02495 | 0.01649 |
| **TBC1D4** | 0.03037 | 0.00000 | 0.28251 | 0.00000 | 0.00000 | 0.26508 | 3.81992 | 0.00088 | 1.11734 | 0.63830 | 0.11781 | 0.14022 |
| **TBC1D5** | 0.82682 | 1.86973 | 3.08107 | 4.13967 | 3.93199 | 5.31693 | 3.06818 | 0.00000 | 2.71590 | 5.34763 | 1.45617 | 1.70649 |
| **TBC1D7** | 3.04354 | 6.52921 | 3.15038 | 4.46019 | 5.60125 | 11.62179 | 5.53344 | 8.26524 | 2.57992 | 0.00000 | 7.38967 | 8.73671 |
| **TBC1D8** | 1.29996 | 2.39413 | 3.47868 | 1.50547 | 0.80250 | 3.18798 | 0.80477 | 0.31759 | 3.07196 | 4.01028 | 1.57690 | 0.95992 |
| **TBC1D8B** | 0.10089 | 0.06835 | 0.07028 | 0.15824 | 0.00000 | 0.16153 | 0.11275 | 0.00803 | 0.05089 | 0.00002 | 0.08667 | 0.03315 |
| **TBC1D9** | 0.24934 | 0.05725 | 0.92331 | 0.00000 | 0.10166 | 0.29459 | 0.62565 | 2.12253 | 0.73291 | 0.06063 | 0.47460 | 3.67325 |
| **TBC1D9B** | 1.16498 | 8.12209 | 5.78674 | 6.58506 | 6.89374 | 3.79609 | 8.98738 | 0.40666 | 8.22766 | 3.48885 | 8.56285 | 4.70095 |
| **TBCA** | 3.33675 | 11.35999 | 10.64196 | 21.34713 | 16.16359 | 31.71488 | 15.12141 | 1.67259 | 12.05820 | 19.34427 | 10.33350 | 8.88622 |
| **TBCB** | 5.85414 | 7.58250 | 13.71841 | 20.32484 | 13.51105 | 0.03210 | 16.66667 | 12.72194 | 15.04477 | 0.00000 | 10.97135 | 9.14920 |
| **TBCC** | 6.37441 | 12.53450 | 3.39630 | 11.09700 | 7.81217 | 15.23561 | 4.69716 | 19.38300 | 5.61641 | 0.00000 | 8.49687 | 21.16345 |
| **TBCCD1** | 2.28076 | 4.13818 | 0.21434 | 1.65983 | 0.85115 | 8.93095 | 0.00001 | 2.16942 | 2.55065 | 0.66846 | 4.59291 | 5.73805 |
| **TBCD** | 1.93978 | 11.86188 | 4.45093 | 16.29540 | 22.60202 | 0.00000 | 3.23701 | 1.00258 | 3.83976 | 7.59433 | 4.56120 | 11.11879 |

| | | | | | | | | | | | | |
|---|---|---|---|---|---|---|---|---|---|---|---|---|
| TBCE | 1.66207 | 4.21282 | 6.43132 | 0.00000 | 1.53362 | 10.51765 | 5.47915 | 5.05827 | 3.42051 | 0.00000 | 4.99837 | 9.00594 |
| TBCEL | 2.50357 | 0.52744 | 0.16072 | 0.26898 | 0.00000 | 0.70722 | 1.03938 | 0.00000 | 0.00000 | 0.60192 | 0.19223 | 0.31123 |
| TBCK | 0.42143 | 2.69479 | 0.55409 | 1.78951 | 0.95686 | 3.93088 | 0.00001 | 0.02164 | 1.10479 | 2.96952 | 1.70732 | 0.38933 |
| TBK1 | 0.64230 | 4.56332 | 12.65008 | 1.10060 | 0.00022 | 21.60101 | 3.94136 | 3.41422 | 14.15442 | 3.13023 | 8.08024 | 1.18295 |
| TBKBP1 | 0.52725 | 0.74351 | 0.03387 | 0.95589 | 0.59236 | 0.00000 | 0.00000 | 0.02923 | 0.09225 | 1.20085 | 0.09486 | 0.76807 |
| TBL1X | 3.12196 | 0.50729 | 7.41489 | 0.00000 | 0.00000 | 2.32834 | 9.50925 | 0.49456 | 3.87334 | 8.33258 | 2.14723 | 0.46850 |
| TBL1XR1 | 0.75738 | 12.75326 | 10.17434 | 7.86806 | 8.81361 | 43.47041 | 30.83017 | 15.26723 | 6.63560 | 31.45077 | 17.01469 | 22.75719 |
| TBL1Y | 0.39621 | 0.00842 | 0.09633 | 0.32984 | 0.00031 | 0.11343 | 0.02448 | 0.31541 | 0.00000 | 0.00426 | 0.01143 | 0.02618 |
| TBL2 | 0.28637 | 9.09175 | 12.03047 | 2.66573 | 6.02754 | 0.00000 | 0.00000 | 0.04420 | 14.95062 | 0.87828 | 6.06894 | 1.40694 |
| TBL3 | 1.17659 | 16.94891 | 12.15732 | 7.61249 | 14.68077 | 0.00000 | 7.34089 | 5.70924 | 13.95784 | 0.00000 | 20.60838 | 12.98495 |
| TBP | 2.58427 | 9.03507 | 5.38948 | 2.21905 | 5.73311 | 9.59166 | 7.74675 | 17.02139 | 16.12776 | 0.22148 | 3.40850 | 12.82268 |
| TBPL1 | 12.94701 | 8.94943 | 1.84933 | 4.22341 | 4.96310 | 22.82564 | 11.84022 | 20.67492 | 10.83102 | 0.00002 | 7.54129 | 25.78982 |
| TBPL2 | 0.22723 | 0.91829 | 0.03624 | 0.35776 | 0.08507 | 0.08639 | 0.00000 | 0.21344 | 0.00000 | 0.06737 | 0.18615 | 0.79879 |
| TBR1 | 0.00314 | 0.01430 | 0.00982 | 0.01185 | 0.01898 | 0.01167 | 0.00963 | 0.03742 | 0.05278 | 0.02006 | 0.03400 | 0.01455 |
| TBRG1 | 2.92730 | 9.73509 | 8.62925 | 9.71926 | 6.16690 | 11.85984 | 10.92515 | 9.81229 | 7.79379 | 3.73454 | 5.76785 | 14.06855 |
| TBRG4 | 1.65828 | 32.34083 | 19.70923 | 20.15794 | 18.41307 | 7.44522 | 24.87276 | 7.11801 | 30.75434 | 0.00000 | 23.38253 | 18.28650 |
| TBX1 | 0.00197 | 0.01281 | 0.27128 | 0.05887 | 0.02170 | 0.00000 | 0.03987 | 0.06356 | 0.06350 | 0.06817 | 0.04179 | 0.01398 |
| TBX15 | 0.00617 | 0.00000 | 0.05056 | 0.01081 | 0.07537 | 0.11712 | 0.00000 | 0.19118 | 0.00000 | 0.04800 | 0.00718 | 0.00310 |
| TBX18 | 0.00387 | 0.00000 | 0.40203 | 0.11395 | 0.18674 | 0.04322 | 0.42226 | 0.00679 | 0.00000 | 0.19133 | 0.01036 | 0.08480 |
| TBX19 | 0.00059 | 0.38368 | 0.00000 | 0.59324 | 1.64245 | 0.00000 | 0.21425 | 0.26671 | 0.91114 | 0.41137 | 0.37886 | 0.04915 |
| TBX2 | 0.00157 | 0.00030 | 2.68135 | 0.00000 | 1.85163 | 0.00000 | 3.38365 | 0.00647 | 0.00000 | 2.17842 | 0.00000 | 0.00250 |
| TBX20 | 0.39928 | 0.02557 | 0.28094 | 0.13774 | 0.47338 | 0.00000 | 0.00000 | 0.13548 | 0.20870 | 0.32745 | 0.07770 | 0.08222 |
| TBX21 | 0.27517 | 0.01158 | 0.03643 | 0.01006 | 0.01622 | 0.00952 | 0.00464 | 0.00411 | 0.03299 | 0.02477 | 0.01649 | 0.04273 |
| TBX22 | 0.00607 | 0.01596 | 0.04913 | 0.04060 | 0.00887 | 0.00601 | 0.06145 | 0.01048 | 0.01950 | 0.02301 | 0.01616 | 0.01658 |
| TBX3 | 1.97378 | 13.72341 | 12.74612 | 10.34957 | 29.60141 | 8.76718 | 6.21203 | 0.00000 | 9.32656 | 20.86799 | 13.45303 | 0.10475 |
| TBX5 | 3.02005 | 0.03439 | 0.00149 | 0.03464 | 0.00701 | 0.06136 | 0.00137 | 0.39002 | 0.01761 | 0.05218 | 0.00472 | 0.01080 |
| TBX6 | 0.00575 | 0.01488 | 0.02108 | 0.06661 | 0.00622 | 0.01364 | 0.00923 | 0.00951 | 0.05474 | 0.02306 | 0.03529 | 0.03453 |
| TBXA2R | 0.09934 | 0.07871 | 0.10349 | 0.08463 | 0.05399 | 0.03375 | 0.01037 | 0.00493 | 0.10124 | 0.00000 | 0.80608 | 0.02806 |
| TBXAS1 | 0.01790 | 0.02228 | 0.00833 | 0.00930 | 0.00000 | 0.03889 | 0.04473 | 0.00753 | 0.00000 | 0.02902 | 0.03079 | 0.04987 |
| TC2N | 6.47976 | 1.19200 | 0.13790 | 0.00000 | 0.00000 | 3.39638 | 1.21568 | 5.23504 | 1.01731 | 0.43147 | 0.76899 | 0.79710 |
| TCAIM | 1.10647 | 2.68314 | 5.33242 | 0.24952 | 0.62474 | 5.14549 | 3.98803 | 0.30344 | 4.20461 | 0.00000 | 3.77516 | 2.75524 |
| TCAM1P | 0.27105 | 0.01354 | 0.01591 | 0.07286 | 0.00787 | 0.00944 | 0.02427 | 0.06109 | 0.04990 | 0.01371 | 0.01654 | 0.01197 |
| TCAP | 0.05776 | 0.04698 | 0.10377 | 0.01114 | 0.01699 | 0.00000 | 0.01039 | 0.03675 | 0.24559 | 0.01407 | 0.00196 | 0.07784 |
| TCEA1 | 0.26760 | 7.36826 | 2.86471 | 13.75863 | 1.60561 | 18.51641 | 3.64998 | 5.70951 | 0.00000 | 10.30339 | 3.86258 | 9.08673 |
| TCEA2 | 1.10837 | 0.56066 | 0.00000 | 0.45391 | 0.19631 | 0.00000 | 0.66789 | 0.05495 | 0.00043 | 1.43829 | 0.03040 | 0.00000 |
| TCEA3 | 0.00317 | 0.11077 | 0.81440 | 0.16487 | 0.00000 | 0.26717 | 0.12377 | 0.02478 | 0.12738 | 0.00000 | 0.04096 | 0.00548 |
| TCEAL1 | 0.00354 | 0.10803 | 0.10604 | 0.19004 | 0.00000 | 1.43794 | 0.00000 | 0.00000 | 0.00000 | 0.20456 | 0.28514 | 0.29859 |
| TCEAL2 | 0.00301 | 0.01113 | 0.00862 | 0.00383 | 0.01231 | 0.04374 | 0.02604 | 0.06055 | 0.00516 | 0.05209 | 0.01003 | 0.04959 |
| TCEAL3 | 0.03284 | 0.01160 | 0.06977 | 0.05181 | 0.00000 | 0.00567 | 0.00950 | 0.00000 | 0.00781 | 0.14066 | 0.02621 | 0.24059 |
| TCEAL4 | 0.19284 | 0.75335 | 0.51037 | 0.00397 | 0.16686 | 2.78146 | 0.48091 | 0.00000 | 0.00000 | 0.65761 | 0.27882 | 2.33119 |
| TCEAL5 | 0.01378 | 0.01763 | 0.01304 | 0.04638 | 0.01749 | 0.00204 | 0.02307 | 0.01053 | 0.01982 | 0.00663 | 0.01630 | 0.01881 |
| TCEAL7 | 0.00567 | 0.00520 | 0.35782 | 0.00000 | 0.00298 | 0.06227 | 0.39583 | 0.00980 | 0.00000 | 0.01208 | 0.03957 | 0.01238 |
| TCEAL8 | 0.11792 | 1.13228 | 4.46793 | 6.70080 | 2.48808 | 9.47930 | 4.91639 | 0.00000 | 7.16052 | 0.61488 | 5.25721 | 2.50035 |
| TCEANC | 0.65980 | 0.54628 | 0.17836 | 0.04071 | 0.00000 | 0.19377 | 0.00000 | 1.60263 | 0.08462 | 0.07245 | 0.43686 | 0.94357 |
| TCEANC2 | 1.55835 | 3.94101 | 1.68675 | 1.99650 | 1.70513 | 4.63824 | 1.77588 | 5.41609 | 4.39333 | 0.00000 | 1.70079 | 4.04225 |
| TCEB1 | 3.40157 | 13.70572 | 15.46316 | 10.19166 | 10.79625 | 22.42968 | 23.47247 | 26.44546 | 14.28551 | 10.51859 | 13.26499 | 29.79050 |
| TCEB2 | 0.00000 | 31.75425 | 14.95901 | 24.94762 | 34.76816 | 9.85741 | 33.06935 | 24.35278 | 13.15019 | 19.71885 | 19.00574 | 43.79621 |
| TCEB3 | 8.84736 | 22.71212 | 15.06952 | 12.93011 | 15.74165 | 23.89970 | 15.73001 | 17.83502 | 24.55517 | 8.40706 | 15.10548 | 21.44031 |

| | | | | | | | | | | | | |
|---|---|---|---|---|---|---|---|---|---|---|---|---|
| TCEB3-AS1 | 0.45208 | 0.98485 | 0.74136 | 0.47376 | 0.82697 | 0.00000 | 0.00000 | 0.40765 | 0.21196 | 0.20415 | 0.83653 | 1.90630 |
| TCEB3B | 0.13034 | 0.01467 | 0.01465 | 0.01241 | 0.01485 | 0.01566 | 0.01629 | 0.00685 | 0.02149 | 0.01448 | 0.01420 | 0.01409 |
| TCERG1 | 3.76352 | 9.18100 | 17.35990 | 7.69204 | 12.83335 | 12.23640 | 16.94773 | 10.70850 | 18.81133 | 27.16276 | 15.30632 | 14.18900 |
| TCERG1L | 0.09401 | 0.94304 | 1.05828 | 0.14306 | 2.65539 | 0.29034 | 0.76774 | 0.03017 | 0.48788 | 2.67239 | 0.37450 | 0.00000 |
| TCF12 | 0.76505 | 2.44729 | 3.98931 | 5.06385 | 1.76165 | 8.69204 | 3.93332 | 0.76332 | 5.80848 | 1.21554 | 3.55037 | 0.00000 |
| TCF15 | 0.54394 | 0.00000 | 0.07223 | 0.00000 | 0.18360 | 0.00000 | 0.12974 | 0.02084 | 0.18783 | 1.68952 | 0.02565 | 0.81259 |
| TCF19 | 0.94234 | 3.02175 | 2.04908 | 6.95692 | 2.68831 | 1.78773 | 2.37636 | 0.00000 | 0.00205 | 0.00000 | 3.42376 | 1.60943 |
| TCF20 | 8.15074 | 0.00000 | 0.45526 | 0.92566 | 1.69436 | 1.49959 | 5.61344 | 0.83444 | 0.00000 | 12.64906 | 0.37669 | 1.25953 |
| TCF21 | 0.00542 | 0.00465 | 0.00000 | 0.00239 | 0.00000 | 0.09477 | 0.12102 | 0.00972 | 0.00923 | 0.02427 | 0.00396 | 0.01634 |
| TCF24 | 0.00163 | 0.00000 | 0.00000 | 0.00000 | 0.51730 | 0.57231 | 0.65307 | 2.00294 | 0.00000 | 2.46442 | 0.08234 | 1.43334 |
| TCF25 | 1.69592 | 8.12449 | 7.63300 | 16.06075 | 19.51780 | 0.00000 | 12.77048 | 4.20629 | 12.07905 | 18.67051 | 10.59513 | 5.81089 |
| TCF3 | 5.09250 | 5.21744 | 3.56333 | 4.98472 | 9.45625 | 0.00000 | 7.29026 | 0.28449 | 7.69005 | 15.02943 | 1.66605 | 3.74639 |
| TCF4 | 0.00000 | 0.04373 | 0.00000 | 0.55127 | 0.00000 | 1.27241 | 1.80543 | 0.08785 | 0.07319 | 3.82055 | 0.31849 | 0.42294 |
| TCF7 | 3.18037 | 1.68503 | 0.45438 | 3.74136 | 3.33913 | 0.29545 | 2.23912 | 0.72356 | 0.00000 | 6.99577 | 0.00000 | 1.20563 |
| TCF7L1 | 2.77879 | 0.52083 | 0.00000 | 1.21191 | 20.21765 | 0.00000 | 10.00542 | 0.13243 | 0.00000 | 12.14230 | 6.51565 | 0.01250 |
| TCF7L2 | 0.24637 | 0.53361 | 20.54772 | 0.00000 | 6.33790 | 8.29502 | 21.55032 | 0.45781 | 11.97610 | 15.95376 | 7.90723 | 0.11531 |
| TCFL5 | 0.29367 | 0.02679 | 0.09357 | 0.30566 | 0.69079 | 0.73629 | 0.52780 | 0.18618 | 0.00000 | 1.70311 | 0.08594 | 0.01219 |
| TCHH | 0.00729 | 0.51251 | 0.16320 | 0.00000 | 0.00000 | 0.00000 | 0.00000 | 3.68882 | 0.05377 | 0.01679 | 0.31281 | 1.35002 |
| TCHP | 3.40960 | 5.43410 | 0.11483 | 1.59038 | 4.13110 | 3.00493 | 3.85666 | 2.14712 | 0.90143 | 1.61513 | 1.71839 | 5.59588 |
| TCIRG1 | 0.00000 | 2.01228 | 5.12863 | 0.68832 | 2.92187 | 0.00000 | 0.00000 | 0.14716 | 4.56989 | 4.71273 | 2.00839 | 0.56378 |
| TCL1A | 42.97432 | 2.04858 | 0.00000 | 0.22583 | 0.84057 | 0.00000 | 0.00000 | 23.72135 | 0.00000 | 0.01486 | 0.00000 | 21.01351 |
| TCL1B | 8.35793 | 4.02219 | 0.36414 | 3.61883 | 0.75685 | 0.00000 | 0.00000 | 2.86744 | 0.37284 | 0.89679 | 0.05944 | 14.95628 |
| TCL6 | 0.00246 | 0.01495 | 0.46845 | 0.01775 | 0.12217 | 0.01308 | 0.17334 | 0.04061 | 0.00000 | 0.01007 | 0.20738 | 0.04183 |
| TCN1 | 0.00442 | 0.01031 | 0.04583 | 0.00517 | 0.00576 | 0.02584 | 0.00432 | 0.01062 | 0.02220 | 0.10238 | 0.01727 | 0.01728 |
| TCN2 | 0.24309 | 0.00000 | 0.09304 | 2.41882 | 0.56116 | 0.00000 | 0.00328 | 1.66391 | 0.00000 | 0.50025 | 0.24235 | 2.05317 |
| TCOF1 | 10.13397 | 20.81920 | 15.48969 | 14.84949 | 20.68887 | 0.00000 | 28.38360 | 12.82236 | 18.54932 | 31.49046 | 18.81899 | 21.25009 |
| TCP1 | 6.69001 | 34.06286 | 26.81882 | 24.82436 | 16.72859 | 60.80406 | 37.48568 | 7.30867 | 40.29686 | 0.00000 | 29.58415 | 25.18016 |
| TCP10L | 0.00010 | 0.28040 | 0.18682 | 0.00000 | 0.15794 | 0.27545 | 0.22274 | 0.03148 | 0.50155 | 0.55647 | 0.00000 | 0.05375 |
| TCP11 | 1.84009 | 0.04033 | 0.00302 | 0.05812 | 0.00000 | 0.03290 | 0.01177 | 0.09384 | 0.03068 | 0.02626 | 0.01137 | 0.09396 |
| TCP11L1 | 0.58810 | 0.41259 | 0.49833 | 1.97585 | 0.50298 | 0.74524 | 0.00000 | 0.17191 | 1.03582 | 0.00000 | 0.71416 | 1.61357 |
| TCP11L2 | 1.06601 | 0.13916 | 0.62082 | 0.22946 | 0.11809 | 0.20727 | 0.00000 | 0.50555 | 0.23689 | 0.00000 | 0.00000 | 0.33369 |
| TCTA | 1.12985 | 3.77509 | 1.35387 | 2.86173 | 5.23780 | 0.40798 | 2.45533 | 0.60807 | 0.00000 | 0.00000 | 4.10555 | 4.55227 |
| TCTE1 | 0.08235 | 0.01073 | 0.01780 | 0.00918 | 0.01560 | 0.02373 | 0.01586 | 0.00203 | 0.01631 | 0.03158 | 0.01394 | 0.11122 |
| TCTE3 | 0.04815 | 0.07565 | 0.08428 | 0.03879 | 0.00317 | 0.04910 | 0.02460 | 0.06400 | 0.05633 | 0.00000 | 0.13252 | 0.13440 |
| TCTEX1D2 | 0.25097 | 0.23500 | 0.69106 | 1.05133 | 0.00000 | 0.45434 | 1.06249 | 0.21670 | 0.83921 | 0.00000 | 0.62663 | 0.04036 |
| TCTEX1D4 | 0.02776 | 0.01558 | 0.01621 | 0.02120 | 0.04076 | 0.00000 | 0.01947 | 0.00816 | 0.02391 | 0.01878 | 0.01946 | 0.01949 |
| TCTN1 | 2.27592 | 0.53513 | 0.09149 | 0.89378 | 0.02133 | 0.00000 | 0.00000 | 0.09817 | 0.09760 | 0.00000 | 0.38136 | 0.35215 |
| TCTN2 | 1.23946 | 1.20354 | 1.36985 | 0.77219 | 0.00000 | 0.33621 | 1.21062 | 3.75829 | 1.35914 | 1.85288 | 0.47577 | 1.82831 |
| TCTN3 | 1.37365 | 1.91645 | 4.62092 | 0.70198 | 3.66910 | 7.57761 | 0.00000 | 0.00000 | 14.71516 | 0.44839 | 4.03795 | 0.85186 |
| TDG | 0.01805 | 3.46785 | 10.28849 | 5.00119 | 9.55080 | 8.10917 | 18.01551 | 0.00000 | 9.40739 | 5.28036 | 9.58357 | 0.00000 |
| TDGF1 | 0.44964 | 0.62970 | 3.07197 | 17.80255 | 2.44190 | 0.52541 | 0.00000 | 0.10108 | 0.00000 | 0.00000 | 2.47893 | 0.00000 |
| TDGF1P3 | 0.22287 | 0.19543 | 0.41084 | 6.03322 | 0.70108 | 0.00000 | 0.00125 | 0.07375 | 0.00000 | 0.00000 | 0.69002 | 0.00257 |
| TDH | 0.06832 | 0.47987 | 0.80838 | 1.39147 | 0.09271 | 1.36622 | 0.00000 | 0.03233 | 0.00000 | 0.00355 | 0.00602 | 0.39254 |
| TDO2 | 0.08298 | 0.00000 | 1.26433 | 0.30256 | 0.00000 | 0.04088 | 0.60243 | 0.02093 | 0.18390 | 0.00000 | 0.06061 | 0.16840 |
| TDP1 | 8.45491 | 6.31078 | 6.14273 | 2.98271 | 2.56018 | 9.64234 | 3.54434 | 3.15302 | 6.62819 | 0.00000 | 7.10282 | 6.79656 |
| TDP2 | 1.78509 | 4.20950 | 9.07767 | 7.40020 | 0.00000 | 15.12084 | 12.92041 | 6.57676 | 2.37574 | 3.80245 | 5.57506 | 9.13025 |
| TDRD1 | 4.19554 | 0.02585 | 0.00000 | 0.00000 | 0.19799 | 0.10533 | 0.20856 | 1.32929 | 0.00000 | 0.06600 | 0.11559 | 0.44386 |
| TDRD10 | 0.24527 | 0.00497 | 0.00000 | 0.21966 | 0.00000 | 0.06021 | 0.00563 | 0.30000 | 0.01567 | 0.01103 | 0.00957 | 0.11219 |

| | | | | | | | | | | | | |
|---|---|---|---|---|---|---|---|---|---|---|---|---|
| **TDRD12** | 0.19623 | 0.00914 | 0.04410 | 0.03640 | 0.01738 | 0.13377 | 0.00000 | 0.00000 | 0.00000 | 0.09697 | 0.00788 | 0.15122 |
| **TDRD3** | 0.93493 | 1.04153 | 2.07195 | 1.42329 | 1.25641 | 2.47418 | 3.40934 | 0.31051 | 0.04199 | 5.42175 | 0.74845 | 0.83011 |
| **TDRD5** | 0.42445 | 0.00606 | 0.06095 | 0.03400 | 0.03362 | 0.00000 | 0.01567 | 0.01148 | 0.01608 | 0.03086 | 0.05574 | 0.23053 |
| **TDRD6** | 4.20618 | 0.00000 | 0.47655 | 0.16825 | 0.00000 | 1.34781 | 0.37329 | 1.72951 | 1.05256 | 0.59576 | 0.00000 | 1.60468 |
| **TDRD7** | 2.21577 | 0.92612 | 0.00000 | 1.41639 | 0.00000 | 1.01273 | 0.19573 | 0.00000 | 0.01550 | 0.00000 | 0.07950 | 2.58852 |
| **TDRD9** | 1.27864 | 0.01423 | 0.04504 | 0.00629 | 0.13844 | 0.00000 | 0.06095 | 0.33593 | 0.41566 | 0.01299 | 0.00000 | 0.47368 |
| **TDRG1** | 0.32048 | 0.00000 | 0.00000 | 0.03653 | 0.01724 | 0.02509 | 0.00000 | 1.81922 | 0.00909 | 0.17741 | 0.00245 | 0.24543 |
| **TDRKH** | 1.01183 | 1.36881 | 0.80282 | 4.67533 | 0.18587 | 3.71586 | 0.80254 | 0.32602 | 0.90893 | 0.00000 | 1.46148 | 1.70539 |
| **TDRP** | 1.81439 | 0.59273 | 18.81941 | 0.00000 | 10.62224 | 2.46295 | 18.00459 | 0.03528 | 1.13065 | 0.00000 | 3.57116 | 0.00000 |
| **TEAD1** | 1.49189 | 1.29391 | 20.18336 | 0.00000 | 16.91128 | 13.63554 | 34.09093 | 1.66836 | 13.55947 | 28.91906 | 9.38062 | 4.89355 |
| **TEAD2** | 0.38312 | 0.58871 | 0.67366 | 1.97042 | 1.33262 | 0.13260 | 3.03131 | 0.84316 | 1.27661 | 6.70604 | 0.00058 | 2.02821 |
| **TEAD3** | 1.53121 | 0.00000 | 23.14184 | 0.43542 | 7.86184 | 0.26410 | 13.94146 | 0.34279 | 4.69589 | 4.42363 | 0.55107 | 0.17858 |
| **TEAD4** | 1.28058 | 29.14300 | 12.61638 | 22.64575 | 26.26218 | 0.00000 | 23.56407 | 2.56058 | 26.27004 | 6.25045 | 12.41551 | 28.80011 |
| **TEC** | 0.00000 | 3.58029 | 0.20154 | 5.32528 | 0.00000 | 2.34982 | 0.00000 | 0.90011 | 0.11165 | 0.82611 | 0.37747 | 1.10676 |
| **TECPR1** | 0.28474 | 1.32563 | 3.02483 | 0.41508 | 2.00016 | 0.00000 | 2.31962 | 0.00000 | 0.88787 | 3.63299 | 1.96142 | 0.92753 |
| **TECPR2** | 8.10282 | 0.99517 | 1.17358 | 3.43331 | 2.03871 | 0.00000 | 0.98872 | 0.50488 | 0.40430 | 3.39271 | 0.78591 | 1.80269 |
| **TECR** | 0.62305 | 18.93130 | 10.04759 | 12.95829 | 13.72233 | 11.93420 | 1.78680 | 4.43621 | 31.30923 | 0.00000 | 13.68895 | 13.25985 |
| **TECRL** | 0.00757 | 0.01486 | 0.02244 | 0.01793 | 0.01233 | 0.01285 | 0.02152 | 0.01073 | 0.01377 | 0.02008 | 0.03435 | 0.01627 |
| **TECTA** | 9.92416 | 0.02411 | 0.02875 | 0.04372 | 0.35490 | 0.00000 | 0.00017 | 2.14035 | 0.00000 | 0.06820 | 0.00634 | 0.05114 |
| **TECTB** | 0.00619 | 0.01693 | 0.02185 | 0.01607 | 0.01590 | 0.00431 | 0.00868 | 0.01063 | 0.03116 | 0.03342 | 0.02852 | 0.01631 |
| **TEDDM1** | 0.00699 | 0.01447 | 0.02567 | 0.01086 | 0.01788 | 0.02110 | 0.01563 | 0.01545 | 0.01861 | 0.01141 | 0.01381 | 0.04343 |
| **TEF** | 0.21005 | 0.90276 | 0.38962 | 0.05874 | 0.35984 | 0.00000 | 1.49006 | 0.23797 | 0.76418 | 2.15649 | 0.46734 | 1.48433 |
| **TEFM** | 0.28107 | 7.46809 | 4.32910 | 2.36588 | 0.62931 | 16.04541 | 8.53592 | 11.42277 | 3.43518 | 0.00000 | 7.89831 | 11.73855 |
| **TEK** | 2.52520 | 0.16147 | 0.24570 | 0.00030 | 0.00750 | 0.31171 | 0.00000 | 0.02168 | 0.83980 | 0.00000 | 0.40060 | 0.00000 |
| **TEKT2** | 0.10231 | 0.00000 | 0.18998 | 0.00803 | 0.00751 | 0.00963 | 0.02735 | 0.00222 | 0.29516 | 0.00000 | 0.94563 | 0.58200 |
| **TEKT3** | 0.03229 | 0.00000 | 0.14493 | 0.40077 | 0.04560 | 0.14716 | 0.00000 | 0.11138 | 0.00000 | 0.10483 | 0.04314 | 0.04302 |
| **TEKT4** | 0.01525 | 0.03714 | 0.02252 | 0.02858 | 0.23880 | 0.00000 | 0.12804 | 0.00712 | 0.07382 | 0.16769 | 0.09863 | 0.12544 |
| **TEKT4P2** | 0.73029 | 3.65971 | 2.13478 | 0.96684 | 3.64451 | 0.00093 | 2.43429 | 0.83224 | 3.41117 | 1.83331 | 3.90737 | 4.60186 |
| **TEKT5** | 0.58529 | 0.00000 | 0.02424 | 0.00000 | 0.00000 | 0.00000 | 0.68148 | 0.07541 | 0.52864 | 0.58373 | 1.18946 | 0.04561 |
| **TELO2** | 0.00000 | 5.91774 | 4.58056 | 4.65667 | 6.29885 | 0.00000 | 3.97033 | 0.09347 | 8.38098 | 9.82930 | 3.11262 | 3.10073 |
| **TEN1-CDK3** | 0.00694 | 0.18149 | 0.02947 | 0.01267 | 0.02930 | 0.00000 | 0.01981 | 0.01001 | 0.05649 | 0.01939 | 0.03521 | 0.03681 |
| **TENC1** | 0.05656 | 0.05592 | 0.18640 | 0.22113 | 0.53939 | 0.00000 | 0.00000 | 0.05513 | 0.36499 | 0.05518 | 0.01375 | 0.14166 |
| **TENM1** | 0.17594 | 0.02353 | 0.00784 | 0.07214 | 0.00000 | 0.00000 | 0.04075 | 0.15554 | 0.05642 | 0.18810 | 0.03638 | 0.01714 |
| **TENM2** | 1.16828 | 0.14479 | 0.00000 | 0.00000 | 0.00000 | 0.06952 | 0.00000 | 0.29022 | 0.03252 | 0.40502 | 0.00000 | 0.30468 |
| **TENM3** | 0.00000 | 0.30391 | 19.24445 | 0.00000 | 0.77038 | 0.00000 | 6.00682 | 0.32907 | 14.15493 | 2.55668 | 1.74644 | 0.21276 |
| **TENM4** | 1.37614 | 0.07310 | 0.11622 | 0.34962 | 0.00000 | 0.00000 | 0.00000 | 0.13560 | 0.00000 | 0.16763 | 0.14269 | 0.09302 |
| **TEP1** | 0.71702 | 0.55129 | 0.62041 | 1.69569 | 0.62441 | 0.00000 | 0.00000 | 0.07012 | 0.00000 | 0.32723 | 0.41703 | 2.59454 |
| **TEPP** | 0.00185 | 0.01662 | 0.18343 | 0.25320 | 0.10174 | 0.00014 | 0.00000 | 0.19223 | 0.00000 | 0.06827 | 0.01599 | 0.10108 |
| **TERC** | 0.19973 | 0.09848 | 0.18998 | 0.01209 | 0.16246 | 0.00000 | 0.00000 | 0.10546 | 0.32210 | 0.59977 | 0.00000 | 0.12416 |
| **TERF1** | 0.01728 | 1.70052 | 4.83140 | 1.82989 | 1.05184 | 1.43371 | 2.55193 | 0.88098 | 0.83123 | 5.17884 | 1.12947 | 1.11696 |
| **TERF2** | 1.54840 | 1.31212 | 2.41215 | 0.58338 | 0.71060 | 5.33444 | 6.79594 | 2.18821 | 0.38756 | 6.94392 | 2.98613 | 2.22216 |
| **TERF2IP** | 0.37523 | 2.13913 | 2.17971 | 0.49492 | 1.47936 | 0.00000 | 1.32492 | 8.66703 | 0.00000 | 8.97297 | 1.46593 | 7.24732 |
| **TERT** | 0.02317 | 3.15534 | 0.06111 | 0.00000 | 0.46750 | 0.00000 | 1.41288 | 0.00000 | 2.94215 | 5.75363 | 2.91115 | 2.05457 |
| **TES** | 1.49131 | 5.47913 | 14.88837 | 3.35957 | 11.76251 | 19.45984 | 14.23711 | 2.62481 | 18.28766 | 5.15680 | 8.71801 | 0.08428 |
| **TESC** | 2.11238 | 0.16830 | 0.01063 | 0.11693 | 0.06924 | 0.06304 | 0.01806 | 0.30659 | 0.00643 | 0.00000 | 0.00427 | 0.22846 |
| **TESK1** | 4.01221 | 1.26573 | 0.60974 | 1.75264 | 1.47283 | 0.00000 | 3.23842 | 1.18781 | 0.75022 | 3.79366 | 0.00000 | 7.14455 |
| **TESK2** | 0.56400 | 1.39326 | 1.01891 | 3.07196 | 0.05699 | 1.58566 | 0.57437 | 7.01158 | 1.77986 | 0.00000 | 0.00063 | 4.16264 |
| **TESPA1** | 0.00611 | 0.01401 | 0.01443 | 0.01228 | 0.01358 | 0.02329 | 0.01767 | 0.03610 | 0.02202 | 0.01137 | 0.01438 | 0.01908 |

| | | | | | | | | | | | | |
|---|---|---|---|---|---|---|---|---|---|---|---|---|
| **TET1** | 0.00000 | 7.87363 | 6.11488 | 16.37968 | 42.28599 | 21.05515 | 13.61139 | 0.99494 | 17.93045 | 32.22305 | 15.11643 | 0.67431 |
| **TET2** | 4.93662 | 8.70933 | 13.94776 | 0.00000 | 21.93767 | 40.72936 | 10.67198 | 0.70976 | 15.86713 | 27.99403 | 25.02567 | 4.20597 |
| **TET3** | 10.12550 | 1.19674 | 4.95291 | 1.23862 | 1.45545 | 8.96339 | 6.14905 | 0.56298 | 6.16861 | 14.43697 | 1.14987 | 3.01331 |
| **TEX10** | 1.92140 | 12.69393 | 15.49582 | 4.57843 | 7.24875 | 22.14547 | 19.00480 | 5.13552 | 9.38821 | 4.27968 | 9.86940 | 12.82944 |
| **TEX101** | 0.10560 | 0.06713 | 0.01492 | 0.01817 | 0.00000 | 0.07461 | 0.00000 | 0.00322 | 0.03307 | 0.02540 | 0.01281 | 0.54431 |
| **TEX11** | 0.00141 | 5.39354 | 0.00000 | 3.81854 | 1.55797 | 3.20097 | 0.00000 | 0.59812 | 0.00000 | 0.01693 | 0.00000 | 4.78924 |
| **TEX12** | 0.00105 | 0.12591 | 0.03856 | 0.04526 | 0.00000 | 0.38683 | 0.00000 | 0.23888 | 0.00000 | 0.06948 | 0.04065 | 0.38534 |
| **TEX14** | 1.72473 | 0.36284 | 0.06302 | 0.19614 | 0.00000 | 0.02749 | 0.11095 | 2.07744 | 0.12739 | 0.34752 | 0.00000 | 1.18646 |
| **TEX15** | 3.50619 | 0.00982 | 0.00717 | 0.00044 | 0.01198 | 0.27572 | 0.00000 | 0.66824 | 0.00000 | 0.09512 | 0.00000 | 0.77684 |
| **TEX19** | 2.67925 | 0.00000 | 0.00000 | 0.32079 | 0.00000 | 0.18264 | 0.00000 | 2.24830 | 0.00000 | 0.01981 | 0.00123 | 7.90806 |
| **TEX2** | 9.73613 | 3.19968 | 5.50030 | 1.31882 | 2.70819 | 3.80199 | 0.00000 | 1.11609 | 4.75761 | 2.44616 | 1.88015 | 2.14636 |
| **TEX21P** | 0.04066 | 0.08722 | 0.02705 | 0.00394 | 0.00000 | 0.02741 | 0.09349 | 0.03437 | 0.02419 | 0.02360 | 0.02381 | 0.18970 |
| **TEX22** | 0.00835 | 0.01829 | 0.03010 | 0.01690 | 0.01791 | 0.00634 | 0.01646 | 0.00850 | 0.03024 | 0.01032 | 0.02120 | 0.05197 |
| **TEX26** | 0.01037 | 0.00762 | 0.02001 | 0.00737 | 0.01102 | 0.03519 | 0.00000 | 0.01140 | 0.03115 | 0.03341 | 0.12614 | 0.01384 |
| **TEX26-AS1** | 0.00921 | 0.25318 | 0.00209 | 0.00000 | 0.00000 | 0.01277 | 0.00156 | 0.01995 | 0.00866 | 0.26757 | 0.02053 | 0.13050 |
| **TEX261** | 0.96625 | 6.82826 | 5.09106 | 1.94525 | 5.83937 | 0.00000 | 1.21856 | 0.25733 | 8.02756 | 10.91956 | 4.12834 | 5.55432 |
| **TEX264** | 0.81240 | 16.44301 | 6.71371 | 9.87451 | 17.43706 | 0.00000 | 4.18190 | 0.08947 | 10.26821 | 0.00000 | 12.27005 | 13.03045 |
| **TEX29** | 0.00549 | 0.00000 | 0.02726 | 0.08124 | 0.00000 | 0.01848 | 0.03758 | 0.00518 | 0.05987 | 0.48165 | 0.00002 | 0.11298 |
| **TEX30** | 1.69615 | 3.80704 | 6.69367 | 0.35800 | 0.00000 | 9.33340 | 5.99504 | 3.42533 | 3.72060 | 0.30137 | 5.80406 | 16.37201 |
| **TEX35** | 0.06104 | 0.09902 | 0.00000 | 0.00101 | 0.13108 | 0.26905 | 0.02145 | 0.00542 | 0.00000 | 0.00000 | 0.29092 | 0.01655 |
| **TEX37** | 5.46543 | 0.41538 | 1.01063 | 0.00000 | 0.54904 | 0.04954 | 0.00000 | 5.76227 | 0.84269 | 0.00000 | 0.27517 | 1.23459 |
| **TEX38** | 0.00778 | 0.03157 | 0.13821 | 0.01039 | 0.17318 | 0.00000 | 0.00011 | 0.03966 | 0.00668 | 0.09272 | 0.01941 | 0.01125 |
| **TEX40** | 0.00959 | 0.69652 | 0.61810 | 0.44947 | 1.99337 | 0.00000 | 0.00768 | 0.17045 | 1.99773 | 0.00000 | 0.40618 | 0.63506 |
| **TEX41** | 1.85787 | 0.55994 | 0.03813 | 0.00000 | 0.13568 | 0.47040 | 0.00000 | 0.17183 | 0.00000 | 0.28591 | 0.03472 | 0.54305 |
| **TEX43** | 0.00680 | 0.01512 | 0.01352 | 0.01203 | 0.00000 | 0.03820 | 0.02275 | 0.06408 | 0.03776 | 0.04306 | 0.01399 | 0.03247 |
| **TEX9** | 1.85915 | 0.23175 | 0.05625 | 0.16289 | 0.11640 | 0.56926 | 0.12428 | 0.18569 | 0.27620 | 0.00000 | 0.03330 | 0.27118 |
| **TF** | 0.00525 | 0.60538 | 0.05951 | 0.66947 | 0.00000 | 0.00817 | 0.20636 | 0.01073 | 0.00000 | 0.12164 | 0.00000 | 0.00000 |
| **TFAM** | 1.02581 | 13.01385 | 12.01677 | 5.88389 | 6.06179 | 34.63615 | 28.99884 | 21.34051 | 16.30295 | 20.78824 | 16.18944 | 21.93037 |
| **TFAMP1** | 0.01387 | 0.03368 | 0.10073 | 0.01428 | 0.00000 | 1.09132 | 0.52323 | 0.50420 | 0.21348 | 1.12767 | 0.25027 | 0.60082 |
| **TFAP2A** | 0.19722 | 0.00000 | 10.24439 | 0.00000 | 3.75990 | 0.58411 | 16.07330 | 0.75706 | 3.46072 | 3.62918 | 2.50735 | 0.62415 |
| **TFAP2A-AS1** | 0.00551 | 0.00488 | 0.17505 | 0.01223 | 0.11670 | 0.02271 | 0.04792 | 0.01841 | 0.18930 | 0.08629 | 0.00000 | 0.01622 |
| **TFAP2B** | 3.14476 | 0.01519 | 0.04277 | 0.00000 | 0.00000 | 0.17304 | 0.01542 | 0.10124 | 0.00000 | 0.10570 | 0.00182 | 0.01957 |
| **TFAP2C** | 0.11571 | 9.90917 | 30.68226 | 0.00000 | 14.75204 | 29.67019 | 36.21533 | 34.02323 | 0.00080 | 29.60980 | 35.14952 | 21.74624 |
| **TFAP2D** | 1.21077 | 0.01339 | 0.01030 | 0.00569 | 0.00217 | 0.03855 | 0.01024 | 0.02149 | 0.00870 | 0.03266 | 0.02829 | 0.03445 |
| **TFAP2E** | 0.35134 | 0.01069 | 0.09663 | 0.04247 | 0.00000 | 0.02530 | 0.00000 | 0.00531 | 0.14224 | 0.04457 | 0.04120 | 0.03235 |
| **TFAP4** | 3.29298 | 0.10228 | 0.13234 | 0.01748 | 0.11705 | 0.00000 | 0.31562 | 0.33825 | 0.00000 | 1.39164 | 0.16561 | 0.01270 |
| **TFB1M** | 1.71750 | 8.03461 | 3.98600 | 5.63588 | 4.07217 | 12.77989 | 3.14617 | 7.11711 | 2.66813 | 2.36503 | 5.52485 | 8.86630 |
| **TFB2M** | 0.59642 | 13.80110 | 8.61137 | 1.68319 | 2.26840 | 20.74200 | 14.91766 | 27.04993 | 14.99919 | 0.09258 | 7.85951 | 24.13189 |
| **TFCP2** | 0.40292 | 4.95609 | 9.64650 | 1.98822 | 4.53800 | 8.07314 | 11.16435 | 3.00766 | 5.33237 | 5.26822 | 7.10752 | 5.35923 |
| **TFCP2L1** | 0.08404 | 18.46740 | 0.86495 | 14.80615 | 27.16858 | 26.26573 | 18.62117 | 0.00000 | 20.91995 | 39.66361 | 17.97815 | 5.31528 |
| **TFDP1** | 2.76282 | 7.38798 | 7.16045 | 2.14508 | 6.17640 | 16.18065 | 14.03975 | 0.15433 | 7.89183 | 9.26338 | 7.07517 | 4.74760 |
| **TFDP2** | 0.00000 | 10.40994 | 14.35058 | 3.93951 | 10.27280 | 15.65987 | 11.46683 | 7.10635 | 0.00000 | 9.41614 | 10.27827 | 25.82523 |
| **TFDP3** | 0.00411 | 0.31649 | 0.69566 | 0.05907 | 0.00000 | 0.00001 | 0.09417 | 0.01963 | 0.34350 | 0.00000 | 0.57611 | 0.00585 |
| **TFE3** | 4.63138 | 1.17842 | 2.49835 | 1.70298 | 10.36421 | 0.00002 | 5.52977 | 0.10585 | 7.52839 | 7.93840 | 5.42010 | 0.15701 |
| **TFEB** | 0.08630 | 5.93062 | 16.91537 | 0.00234 | 2.07533 | 0.00000 | 5.90533 | 0.63007 | 3.39140 | 0.00000 | 0.00000 | 1.24610 |
| **TFEC** | 0.04978 | 0.08105 | 0.69161 | 0.00530 | 0.00347 | 0.03811 | 0.00449 | 0.05944 | 0.00000 | 0.01275 | 0.01308 | 0.00687 |
| **TFF1** | 0.00596 | 0.01183 | 1.49010 | 0.00269 | 0.00000 | 0.03013 | 0.23704 | 0.01007 | 0.11809 | 0.00000 | 0.00000 | 0.01465 |
| **TFG** | 5.44628 | 13.62360 | 21.33681 | 13.61736 | 8.76224 | 36.44210 | 26.40923 | 10.89316 | 23.89600 | 0.00000 | 19.73101 | 12.35710 |

| | | | | | | | | | | | | |
|---|---|---|---|---|---|---|---|---|---|---|---|---|
| TFIP11 | 4.38292 | 14.75760 | 8.48016 | 8.75784 | 7.62676 | 7.99263 | 10.86488 | 22.85530 | 9.23776 | 0.00000 | 8.45539 | 15.97151 |
| TFPI | 0.10523 | 2.01460 | 3.04679 | 6.77500 | 0.09456 | 1.18321 | 0.00000 | 0.21247 | 0.82531 | 0.14396 | 0.03709 | 0.00000 |
| TFPI2 | 0.80737 | 0.00984 | 0.40784 | 1.34997 | 0.00000 | 0.06811 | 0.00000 | 0.09040 | 0.15438 | 0.03185 | 0.00000 | 0.27979 |
| TFPT | 1.84440 | 5.78941 | 6.06774 | 5.83930 | 8.80780 | 0.00000 | 7.62312 | 16.51111 | 2.35443 | 1.25935 | 3.73433 | 7.59277 |
| TFR2 | 1.85495 | 0.07487 | 0.14741 | 0.23015 | 0.00000 | 0.00000 | 0.00000 | 0.13222 | 0.20825 | 0.10382 | 0.00577 | 0.14448 |
| TFRC | 2.68845 | 18.00176 | 22.56898 | 0.00000 | 13.35471 | 46.53541 | 10.35654 | 1.39913 | 67.03015 | 33.88348 | 26.71624 | 6.90410 |
| TG | 0.20430 | 0.09930 | 0.02278 | 0.00000 | 0.05572 | 0.00000 | 0.04536 | 0.01758 | 0.00000 | 0.12962 | 0.01026 | 0.03199 |
| TGDS | 0.00000 | 4.01087 | 3.35363 | 1.99272 | 0.00000 | 15.43409 | 2.69229 | 0.23500 | 5.05362 | 3.58396 | 5.56843 | 3.62196 |
| TGFA | 2.03210 | 0.18352 | 0.01502 | 0.00000 | 0.00000 | 0.00848 | 0.02199 | 0.16342 | 0.02328 | 0.02218 | 0.00890 | 0.15741 |
| TGFB1 | 0.02944 | 0.13100 | 0.71378 | 0.04051 | 0.15068 | 0.14653 | 0.00000 | 0.02633 | 0.00000 | 0.74374 | 0.00000 | 0.80644 |
| TGFB1I1 | 0.10273 | 0.00618 | 0.06820 | 0.01980 | 0.02427 | 0.00000 | 0.01388 | 0.60381 | 0.02118 | 0.00000 | 0.01664 | 0.18185 |
| TGFB2 | 0.52340 | 0.00907 | 0.04055 | 0.02000 | 0.02986 | 0.03114 | 0.02377 | 0.03007 | 0.00000 | 0.09215 | 0.00590 | 0.09906 |
| TGFB2-AS1 | 0.00616 | 0.01346 | 0.00775 | 0.10144 | 0.02531 | 0.00000 | 0.02328 | 0.01028 | 0.03127 | 0.01257 | 0.01304 | 0.02141 |
| TGFB3 | 0.04850 | 2.86399 | 0.67227 | 2.02349 | 0.00000 | 0.11905 | 0.00000 | 0.03253 | 0.28632 | 0.42932 | 0.00000 | 1.27130 |
| TGFBI | 0.08149 | 6.90498 | 0.01356 | 0.00000 | 3.14069 | 0.00011 | 0.01327 | 0.01628 | 0.00000 | 0.22459 | 0.56041 | 0.04835 |
| TGFBR1 | 2.09598 | 3.17998 | 0.70999 | 0.00267 | 0.65664 | 6.28098 | 0.00000 | 3.42976 | 2.08485 | 7.94206 | 0.00000 | 2.87541 |
| TGFBR2 | 0.00350 | 0.02032 | 10.92523 | 0.14294 | 0.00000 | 0.01192 | 0.52500 | 0.00805 | 0.00000 | 0.11555 | 0.01603 | 0.03147 |
| TGFBR3 | 0.87111 | 5.06001 | 2.44241 | 0.00000 | 26.54156 | 15.71943 | 4.79034 | 0.02313 | 25.92260 | 28.49179 | 13.03255 | 3.14375 |
| TGFBR3L | 0.05364 | 0.98177 | 0.04275 | 0.03443 | 0.03379 | 0.00000 | 0.08277 | 0.43738 | 0.03696 | 0.01799 | 0.00000 | 0.26709 |
| TGFBRAP1 | 1.60165 | 1.62165 | 6.48193 | 1.67938 | 4.99203 | 2.19778 | 9.53355 | 0.00000 | 2.62240 | 5.76872 | 3.35982 | 0.03895 |
| TGIF1 | 0.00000 | 13.13867 | 16.90589 | 16.10751 | 3.74127 | 19.94119 | 20.97438 | 15.51893 | 14.08907 | 0.00000 | 5.43145 | 28.17076 |
| TGIF2 | 1.94977 | 3.66129 | 5.69874 | 7.09170 | 10.15377 | 0.00077 | 9.92259 | 3.51150 | 4.23965 | 16.52344 | 8.63102 | 7.37809 |
| TGIF2LX | 0.38084 | 0.03437 | 0.02736 | 0.01179 | 0.00903 | 0.00000 | 0.01838 | 0.12071 | 0.06994 | 0.00000 | 0.04150 | 0.08434 |
| TGIF2LY | 0.04622 | 0.01069 | 0.02010 | 0.01471 | 0.01229 | 0.02394 | 0.01778 | 0.04596 | 0.02536 | 0.00692 | 0.01908 | 0.07598 |
| TGM1 | 0.00179 | 0.14150 | 2.18189 | 0.02166 | 0.51909 | 0.00000 | 1.29810 | 0.04458 | 0.77117 | 0.00000 | 0.34414 | 0.13685 |
| TGM2 | 0.00388 | 0.00000 | 1.39298 | 0.62899 | 0.00000 | 0.00000 | 9.41960 | 0.00865 | 6.55417 | 0.03984 | 9.62773 | 0.00676 |
| TGM3 | 0.03318 | 0.01283 | 0.03461 | 0.01214 | 0.01373 | 0.02183 | 0.01290 | 0.02739 | 0.01420 | 0.02039 | 0.01534 | 0.02277 |
| TGM4 | 2.37155 | 0.00649 | 0.03123 | 0.00595 | 0.00044 | 0.02323 | 0.00000 | 0.12719 | 0.17017 | 0.03994 | 0.01173 | 0.00695 |
| TGM6 | 0.00323 | 0.02407 | 0.05187 | 0.03756 | 0.04676 | 0.00000 | 0.01443 | 0.04270 | 0.02068 | 0.03184 | 0.01210 | 0.01986 |
| TGM7 | 3.30178 | 0.01622 | 0.01865 | 0.01169 | 0.01810 | 0.00000 | 0.01477 | 0.65369 | 0.02531 | 0.00000 | 0.01328 | 0.16500 |
| TGOLN2 | 1.05136 | 8.24546 | 10.64665 | 4.20608 | 7.06462 | 9.54354 | 0.81744 | 17.75054 | 23.19589 | 20.58041 | 9.54039 | 9.72635 |
| TGS1 | 3.99244 | 4.64164 | 7.64133 | 10.68535 | 4.88767 | 15.90806 | 7.26742 | 12.26955 | 10.42176 | 12.52879 | 7.56384 | 27.58005 |
| TH | 0.07053 | 0.01323 | 0.01310 | 0.01283 | 0.01087 | 0.01825 | 0.01210 | 0.00898 | 0.01906 | 0.03787 | 0.01368 | 0.01379 |
| THADA | 1.66544 | 2.16816 | 5.80905 | 1.26595 | 3.97695 | 6.37416 | 4.31440 | 1.65260 | 4.11854 | 5.55382 | 2.99711 | 7.34448 |
| THAP1 | 3.18351 | 3.26657 | 6.69349 | 2.30284 | 0.00000 | 13.33982 | 8.14207 | 17.67196 | 4.11274 | 0.00000 | 6.12002 | 10.29247 |
| THAP10 | 0.47717 | 0.71947 | 0.02778 | 0.96089 | 0.00000 | 0.19550 | 0.33420 | 0.09076 | 0.14222 | 0.58912 | 0.52291 | 0.13272 |
| THAP11 | 1.77307 | 4.91221 | 6.85753 | 6.04473 | 3.85580 | 1.87986 | 10.03416 | 8.53551 | 8.80548 | 9.70695 | 1.09556 | 4.90433 |
| THAP2 | 0.15420 | 0.49596 | 0.93782 | 0.00000 | 0.10934 | 1.65130 | 0.62938 | 0.37645 | 1.03247 | 0.51585 | 0.55433 | 0.97933 |
| THAP3 | 0.12189 | 3.86855 | 5.53198 | 1.04280 | 1.69705 | 0.00302 | 5.13981 | 0.36066 | 2.21117 | 2.16817 | 1.31390 | 2.45968 |
| THAP4 | 1.11012 | 8.65357 | 5.63552 | 7.43919 | 10.64316 | 0.00000 | 8.44649 | 1.76827 | 5.84098 | 8.91155 | 7.41378 | 7.39889 |
| THAP5 | 2.01936 | 1.43045 | 4.92211 | 0.00003 | 0.00000 | 6.56656 | 0.00000 | 8.54689 | 1.58503 | 0.58904 | 3.58238 | 2.68403 |
| THAP6 | 1.76332 | 0.68532 | 1.66148 | 0.00000 | 0.00000 | 5.65718 | 0.00000 | 3.85670 | 0.95819 | 0.82122 | 1.25195 | 4.58441 |
| THAP7 | 2.62675 | 10.03057 | 5.64971 | 5.30800 | 8.13954 | 0.00000 | 8.06089 | 1.14558 | 7.38225 | 2.36365 | 9.26913 | 10.24454 |
| THAP7-AS1 | 0.16511 | 0.65956 | 0.07032 | 0.16822 | 0.52059 | 0.23401 | 0.00000 | 0.08250 | 0.56843 | 0.00000 | 0.11010 | 0.44297 |
| THAP8 | 1.23995 | 1.19336 | 1.05918 | 0.85941 | 0.10055 | 0.42303 | 0.15467 | 0.00000 | 0.65480 | 0.00000 | 0.37648 | 1.41823 |
| THAP9 | 0.78768 | 4.29996 | 1.99804 | 2.28778 | 0.00000 | 9.03142 | 0.00000 | 2.13975 | 2.10970 | 0.51030 | 1.13137 | 4.93749 |
| THAP9-AS1 | 0.00000 | 2.14344 | 5.02802 | 1.52715 | 0.00012 | 14.98004 | 2.19024 | 0.36740 | 4.46127 | 6.29345 | 1.65647 | 3.22722 |
| THBD | 0.00658 | 0.01686 | 0.32060 | 0.01757 | 0.02506 | 0.00000 | 0.00000 | 0.01045 | 0.00000 | 0.02092 | 0.04875 | 0.01659 |

| | | | | | | | | | | | | |
|---|---|---|---|---|---|---|---|---|---|---|---|---|
| THBS1 | 0.00109 | 0.00000 | 0.01017 | 0.00651 | 0.01740 | 0.16452 | 0.00379 | 0.46688 | 0.00000 | 0.14595 | 0.02129 | 0.00435 |
| THBS2 | 2.37627 | 0.00045 | 0.30189 | 0.09687 | 0.00000 | 0.09400 | 0.02565 | 1.93682 | 0.00000 | 0.01946 | 0.01418 | 0.03797 |
| THBS3 | 0.06565 | 0.14580 | 0.12778 | 0.03627 | 0.00000 | 0.00000 | 0.15096 | 0.02075 | 0.04982 | 0.31975 | 0.07750 | 0.06286 |
| THBS4 | 4.78930 | 0.36751 | 0.05958 | 0.18843 | 0.00000 | 0.00923 | 0.00000 | 2.20699 | 0.00000 | 0.02172 | 0.07120 | 0.00000 |
| THEG | 0.47401 | 0.06659 | 0.05407 | 0.00630 | 0.05736 | 0.00000 | 0.01134 | 2.29230 | 0.01542 | 0.00001 | 0.01054 | 1.06764 |
| THEGL | 0.23204 | 0.06754 | 0.01411 | 0.01100 | 0.00000 | 0.05787 | 0.04792 | 0.33951 | 0.00000 | 0.01700 | 0.00946 | 0.05335 |
| THEM4 | 0.13904 | 0.47871 | 2.69181 | 1.54082 | 0.00000 | 0.77892 | 3.78143 | 0.05127 | 4.14907 | 1.94223 | 6.79908 | 1.95627 |
| THEM5 | 0.00594 | 0.00762 | 0.24179 | 0.00621 | 0.05764 | 0.02576 | 0.00000 | 0.04950 | 0.05890 | 0.06733 | 0.04903 | 0.01166 |
| THEM6 | 2.30388 | 0.00000 | 0.09058 | 0.00000 | 0.00000 | 0.00000 | 0.00668 | 0.21812 | 1.68120 | 0.61194 | 3.42348 | 1.23440 |
| THEMIS | 1.11684 | 0.00000 | 0.00631 | 0.00668 | 0.02956 | 0.00000 | 0.00154 | 0.50082 | 0.01843 | 0.14674 | 0.01183 | 0.40570 |
| THEMIS2 | 0.08446 | 0.97915 | 0.34121 | 4.51174 | 0.00000 | 0.31522 | 0.79526 | 1.10726 | 0.03048 | 0.00000 | 0.00000 | 5.63617 |
| THG1L | 1.93605 | 4.37497 | 0.84280 | 1.39508 | 2.60528 | 5.62342 | 0.48027 | 2.84103 | 0.90527 | 0.00000 | 2.28989 | 9.52620 |
| THNSL1 | 1.73077 | 0.27135 | 0.64564 | 0.14535 | 0.05129 | 1.06788 | 0.49114 | 0.80577 | 0.55987 | 0.03850 | 0.33371 | 0.00000 |
| THNSL2 | 0.03730 | 0.38836 | 0.06617 | 0.00000 | 0.13317 | 0.71465 | 0.00000 | 0.00000 | 0.00000 | 0.15038 | 0.22917 | 0.40365 |
| THOC1 | 2.07902 | 7.35502 | 4.83918 | 2.07956 | 2.02393 | 15.90624 | 7.60924 | 2.97446 | 8.24666 | 5.63209 | 8.49706 | 7.56304 |
| THOC2 | 9.13877 | 7.31419 | 5.47959 | 8.35833 | 0.00000 | 30.01571 | 12.68672 | 11.92362 | 7.71968 | 36.33919 | 15.55525 | 12.91480 |
| THOC3 | 1.23629 | 2.75366 | 3.09965 | 2.29118 | 2.52880 | 2.66544 | 0.92489 | 3.19659 | 2.46784 | 0.00000 | 2.71645 | 3.27449 |
| THOC5 | 1.49520 | 18.27303 | 7.21047 | 16.05411 | 8.35006 | 6.26514 | 10.00310 | 10.20379 | 18.57374 | 0.00013 | 8.79737 | 20.97608 |
| THOC6 | 3.25017 | 10.35576 | 12.69265 | 10.95465 | 10.10837 | 0.00000 | 14.22361 | 1.55316 | 14.61585 | 0.00000 | 16.60408 | 6.18983 |
| THOC7 | 3.27720 | 6.01077 | 3.78135 | 2.27632 | 4.63317 | 21.17989 | 8.47042 | 0.81539 | 9.80590 | 8.34645 | 6.63159 | 6.53859 |
| THOC7-AS1 | 0.00566 | 0.00317 | 0.02133 | 0.00018 | 0.12789 | 0.00786 | 0.07830 | 0.00922 | 0.00000 | 0.15708 | 0.01141 | 0.02194 |
| THOP1 | 0.00000 | 6.23781 | 8.03608 | 2.22538 | 9.50484 | 0.00000 | 10.36411 | 0.03305 | 7.45225 | 12.31641 | 10.74196 | 1.85543 |
| THPO | 0.01557 | 0.00629 | 0.28478 | 0.36894 | 0.16322 | 0.02648 | 0.00023 | 0.02108 | 0.02359 | 0.00000 | 0.03860 | 0.02449 |
| THRA | 1.09372 | 0.41792 | 0.48406 | 0.21838 | 0.20990 | 0.00000 | 0.12907 | 0.00000 | 0.25781 | 1.77670 | 0.05328 | 0.29583 |
| THRAP3 | 3.49775 | 18.44245 | 18.07019 | 22.40114 | 23.30074 | 27.30821 | 31.58361 | 18.64279 | 19.91295 | 14.23314 | 30.93921 | 26.55100 |
| THRB | 5.15975 | 0.04377 | 0.06968 | 0.08177 | 0.00000 | 0.43050 | 0.22853 | 0.75577 | 0.00000 | 0.38228 | 0.04160 | 0.07067 |
| THRIL | 0.66430 | 0.04844 | 0.21247 | 0.00000 | 0.00000 | 0.99307 | 0.69918 | 0.00000 | 1.98070 | 1.79000 | 1.06479 | 0.10535 |
| THRSP | 0.09646 | 0.01968 | 0.13082 | 0.00464 | 0.00040 | 0.05754 | 0.01934 | 0.08555 | 0.26799 | 0.01689 | 0.07010 | 0.07694 |
| THSD1 | 0.00236 | 0.99864 | 1.87038 | 0.22555 | 0.00000 | 0.78885 | 1.19366 | 0.33661 | 1.74115 | 0.00000 | 1.37293 | 0.63324 |
| THSD4 | 3.11605 | 0.23970 | 0.06087 | 0.35274 | 0.04008 | 0.00000 | 0.00000 | 0.21888 | 0.01425 | 0.21145 | 0.04365 | 0.58145 |
| THSD7A | 6.39357 | 0.00291 | 0.05293 | 0.00000 | 0.23747 | 0.05913 | 0.00008 | 0.63171 | 0.01151 | 0.01484 | 0.00000 | 0.26649 |
| THSD7B | 0.75015 | 0.02415 | 0.07381 | 0.01276 | 0.01754 | 0.00275 | 0.00592 | 0.16325 | 0.01582 | 0.03978 | 0.01958 | 0.05454 |
| THTPA | 1.04567 | 1.67963 | 0.70129 | 1.68781 | 2.27625 | 0.00000 | 0.35527 | 0.37266 | 2.20715 | 0.00000 | 1.65606 | 2.65408 |
| THUMPD1 | 4.29211 | 13.73741 | 9.00566 | 7.35055 | 0.00000 | 29.00287 | 18.05677 | 17.83699 | 12.25296 | 14.16394 | 13.24708 | 15.85746 |
| THUMPD2 | 0.06275 | 2.66066 | 2.72306 | 0.18166 | 0.00000 | 5.86359 | 5.60312 | 0.72313 | 2.88294 | 3.65874 | 1.99464 | 2.22542 |
| THUMPD3 | 2.73673 | 11.55652 | 6.78709 | 2.45929 | 6.07668 | 16.53968 | 12.56526 | 2.84204 | 2.15004 | 7.17617 | 11.25598 | 12.12394 |
| THUMPD3-AS1 | 0.29394 | 1.18979 | 2.81187 | 0.59158 | 0.00000 | 9.55074 | 0.00000 | 4.02981 | 1.86500 | 9.53080 | 3.14180 | 6.43050 |
| THY1 | 0.00446 | 0.01668 | 0.80435 | 1.29439 | 0.01913 | 0.00000 | 0.00000 | 0.29165 | 0.15800 | 0.00000 | 0.03157 | 1.37601 |
| THYN1 | 10.83532 | 7.48821 | 6.51175 | 4.61378 | 2.39052 | 10.85365 | 12.54567 | 6.27358 | 3.53020 | 0.00000 | 11.03994 | 6.85549 |
| TIA1 | 0.00000 | 5.93097 | 4.37191 | 4.35875 | 0.77532 | 24.09129 | 9.35030 | 7.20653 | 10.18112 | 7.06999 | 5.59499 | 10.14332 |
| TIAF1 | 0.01481 | 0.01977 | 0.06049 | 0.17980 | 0.00000 | 0.01678 | 0.00000 | 0.07579 | 0.06787 | 0.02466 | 0.05342 | 0.01879 |
| TIAL1 | 1.09415 | 2.96446 | 7.77854 | 3.61074 | 3.04881 | 19.35290 | 18.40301 | 10.01472 | 2.53762 | 36.41628 | 3.33323 | 7.55752 |
| TIAM1 | 14.72391 | 2.84180 | 0.10514 | 4.02644 | 7.28389 | 3.93411 | 12.87631 | 2.52472 | 11.82773 | 11.14861 | 7.52172 | 2.46720 |
| TIAM2 | 2.36324 | 1.40613 | 0.03126 | 0.39582 | 0.00000 | 0.19463 | 0.00000 | 0.29692 | 0.07587 | 0.27543 | 0.08434 | 1.72156 |
| TICAM1 | 0.69936 | 0.28424 | 0.03035 | 0.00000 | 0.05129 | 0.00000 | 0.00000 | 0.17159 | 0.19782 | 0.04253 | 0.02442 | 0.75913 |
| TICAM2 | 0.00613 | 0.01756 | 0.01112 | 0.00662 | 0.00815 | 0.03582 | 0.01050 | 0.01596 | 0.01253 | 0.04251 | 0.01370 | 0.01651 |
| TICRR | 3.85953 | 4.18565 | 3.89193 | 3.37589 | 7.49807 | 0.62516 | 3.71809 | 0.29948 | 3.74711 | 5.75246 | 2.25377 | 2.43255 |
| TIE1 | 0.00388 | 0.00710 | 0.00629 | 0.00000 | 0.05525 | 0.02191 | 0.01659 | 0.01000 | 0.00000 | 0.21846 | 0.00471 | 0.00930 |

| | | | | | | | | | | | | |
|---|---|---|---|---|---|---|---|---|---|---|---|---|
| **TIFA** | 5.85746 | 0.02736 | 0.24488 | 0.01698 | 0.00000 | 0.02137 | 0.11857 | 0.12577 | 0.00000 | 0.01659 | 0.00497 | 0.11470 |
| **TIGD1** | 0.29810 | 1.00993 | 0.50825 | 0.18119 | 0.16489 | 1.63361 | 0.00000 | 0.08499 | 1.43088 | 0.00012 | 0.69538 | 3.18713 |
| **TIGD2** | 0.93259 | 0.59432 | 0.69012 | 0.18837 | 0.38889 | 1.25938 | 0.63417 | 0.45891 | 0.00256 | 0.96507 | 0.00000 | 2.21663 |
| **TIGD3** | 0.01111 | 0.16848 | 0.22036 | 0.10197 | 0.00000 | 0.00065 | 0.17497 | 0.06255 | 0.11193 | 0.22210 | 0.09357 | 0.05704 |
| **TIGD4** | 0.06635 | 0.08830 | 0.08060 | 0.16848 | 0.00213 | 0.08557 | 0.00000 | 0.15992 | 0.00000 | 0.00819 | 0.31113 | 0.10369 |
| **TIGD5** | 0.00163 | 0.26388 | 0.28189 | 1.06032 | 1.51529 | 0.00000 | 0.93342 | 0.00759 | 0.41415 | 2.34868 | 0.29760 | 0.00000 |
| **TIGD6** | 0.51654 | 0.69152 | 0.67761 | 0.00547 | 0.00385 | 1.36086 | 0.00000 | 1.34296 | 0.17318 | 0.00000 | 0.40317 | 1.13042 |
| **TIGD7** | 0.62699 | 0.94166 | 1.31979 | 0.00000 | 0.79360 | 2.41209 | 1.09100 | 1.43999 | 1.47773 | 0.62556 | 1.26436 | 3.91477 |
| **TIGIT** | 4.48005 | 0.00572 | 0.05531 | 0.03492 | 0.01913 | 0.00000 | 0.01831 | 0.66385 | 0.01989 | 0.00000 | 0.04146 | 0.10288 |
| **TIMD4** | 0.87583 | 0.43531 | 0.06220 | 0.58904 | 1.56479 | 0.00000 | 0.00000 | 0.97949 | 2.19821 | 1.56205 | 0.46911 | 0.50559 |
| **TIMELESS** | 2.66472 | 9.46495 | 5.41748 | 10.78596 | 2.21998 | 6.46499 | 6.38662 | 0.34907 | 6.04543 | 7.37514 | 10.98482 | 10.32452 |
| **TIMM10** | 0.38571 | 26.41119 | 15.43222 | 21.52443 | 33.85208 | 14.88481 | 37.61197 | 17.46455 | 16.60967 | 8.97012 | 24.47075 | 33.68027 |
| **TIMM10B** | 2.94708 | 10.40021 | 12.57602 | 11.57282 | 9.23110 | 13.24042 | 11.92022 | 27.37379 | 12.26060 | 0.00000 | 15.02321 | 17.13269 |
| **TIMM13** | 1.20736 | 35.59288 | 29.44142 | 30.12463 | 39.87996 | 11.17483 | 51.63911 | 13.70410 | 40.03018 | 12.93005 | 32.47681 | 37.67119 |
| **TIMM17A** | 2.28749 | 20.97957 | 19.37869 | 11.60016 | 13.73964 | 48.83651 | 27.02162 | 15.39714 | 30.07682 | 13.41079 | 28.46087 | 24.10464 |
| **TIMM17B** | 2.30397 | 10.20582 | 4.56269 | 9.78210 | 12.49621 | 1.62626 | 7.14985 | 0.98826 | 8.38342 | 0.00000 | 9.93224 | 8.91757 |
| **TIMM21** | 1.11320 | 13.55631 | 6.76576 | 4.43877 | 5.28306 | 26.93508 | 7.95090 | 0.00000 | 16.15404 | 1.15576 | 9.06230 | 10.88025 |
| **TIMM22** | 2.13038 | 22.07888 | 4.50445 | 10.45564 | 13.87001 | 10.91399 | 9.09584 | 12.71397 | 10.54728 | 0.00000 | 14.56340 | 16.59576 |
| **TIMM23** | 3.40187 | 13.86125 | 7.39771 | 9.43395 | 8.40106 | 28.62085 | 21.58028 | 8.90821 | 20.11079 | 10.63098 | 12.24139 | 20.83693 |
| **TIMM23B** | 0.00114 | 0.11868 | 0.28832 | 0.25074 | 0.35721 | 0.60685 | 0.00000 | 0.84384 | 0.36426 | 0.24187 | 0.38424 | 0.36193 |
| **TIMM44** | 8.67035 | 21.74411 | 12.70121 | 11.13450 | 22.12313 | 0.00000 | 22.82825 | 4.38115 | 28.01038 | 8.38435 | 20.78756 | 19.53187 |
| **TIMM50** | 1.40224 | 31.63308 | 18.20742 | 24.87332 | 25.90898 | 10.44650 | 22.01166 | 21.51818 | 27.30480 | 0.00000 | 25.24084 | 42.33544 |
| **TIMM8A** | 0.00000 | 6.70834 | 5.21703 | 6.25782 | 9.83859 | 12.47365 | 9.27122 | 14.35162 | 5.99521 | 5.70100 | 13.28246 | 14.81233 |
| **TIMM8B** | 0.00000 | 11.70294 | 14.03414 | 11.87269 | 17.77561 | 20.59740 | 21.28533 | 5.66958 | 10.54240 | 8.50329 | 18.57075 | 21.52961 |
| **TIMM9** | 0.82029 | 8.66731 | 2.14207 | 2.82917 | 11.36374 | 25.59416 | 6.54045 | 0.03509 | 6.40397 | 8.02550 | 8.71214 | 6.45749 |
| **TIMMDC1** | 2.16692 | 20.77397 | 15.48665 | 9.15669 | 10.13927 | 32.36385 | 12.70167 | 8.63759 | 19.85147 | 3.10780 | 23.68497 | 18.73035 |
| **TIMP1** | 0.01543 | 1.10353 | 0.00000 | 1.74523 | 0.16179 | 0.39080 | 0.06483 | 1.33810 | 0.00000 | 0.00000 | 0.07510 | 2.37348 |
| **TIMP2** | 3.68913 | 2.53917 | 0.00000 | 0.00000 | 0.00000 | 0.71561 | 0.68850 | 0.25884 | 0.00000 | 2.42625 | 0.00000 | 0.17429 |
| **TIMP3** | 0.18359 | 0.00000 | 3.30184 | 0.00000 | 0.56992 | 0.00000 | 0.00000 | 0.03009 | 0.00000 | 0.83230 | 0.68963 | 0.08969 |
| **TIMP4** | 0.00331 | 0.00000 | 12.99787 | 0.41342 | 0.29191 | 0.00000 | 0.06158 | 0.00726 | 0.89146 | 0.08937 | 0.00000 | 0.09145 |
| **TINAG** | 0.00521 | 0.00893 | 0.01278 | 0.01540 | 0.02486 | 0.04626 | 0.01324 | 0.01261 | 0.00952 | 0.01719 | 0.01453 | 0.01620 |
| **TINAGL1** | 0.00365 | 0.00000 | 12.23808 | 0.00000 | 0.00000 | 0.05105 | 2.03299 | 0.01139 | 1.06881 | 0.01901 | 0.00000 | 0.01263 |
| **TINCR** | 0.00000 | 0.64760 | 11.20010 | 0.00000 | 6.26702 | 0.70119 | 10.59734 | 0.13696 | 8.16950 | 4.22807 | 3.06547 | 5.54126 |
| **TINF2** | 2.64572 | 0.92769 | 7.74953 | 5.32381 | 2.89719 | 3.24908 | 2.54241 | 2.81204 | 2.85303 | 0.00000 | 2.86425 | 2.90591 |
| **TIPARP** | 9.99989 | 0.00000 | 3.69160 | 1.30470 | 0.47341 | 2.64299 | 0.74252 | 5.73378 | 0.00000 | 1.17490 | 1.97957 | 0.38648 |
| **TIPARP-AS1** | 0.29691 | 0.05536 | 0.11287 | 0.00000 | 0.09006 | 0.06535 | 0.09732 | 0.13991 | 0.08265 | 0.09653 | 0.11827 | 0.06075 |
| **TIPIN** | 1.01396 | 5.15926 | 19.33439 | 1.48095 | 10.22387 | 24.60698 | 16.92782 | 14.21337 | 13.90417 | 3.41671 | 17.43271 | 7.86790 |
| **TIPRL** | 0.34019 | 13.63325 | 4.95344 | 5.65977 | 6.71024 | 31.09890 | 8.28791 | 5.22390 | 7.15417 | 4.17169 | 11.55580 | 12.50607 |
| **TIRAP** | 0.75168 | 0.62220 | 1.06569 | 0.33968 | 0.00000 | 0.00000 | 1.15800 | 0.02688 | 1.24245 | 0.00000 | 1.16882 | 0.38696 |
| **TJAP1** | 7.19369 | 5.79280 | 2.19418 | 3.70756 | 2.41673 | 1.95790 | 2.44946 | 0.04542 | 3.40411 | 4.55910 | 3.06097 | 5.80846 |
| **TJP1** | 11.63743 | 9.03667 | 12.39671 | 1.20221 | 13.50124 | 28.65156 | 18.20596 | 3.46125 | 5.51483 | 35.02957 | 13.39565 | 4.32119 |
| **TJP2** | 1.84301 | 5.86527 | 10.37341 | 9.98875 | 18.12235 | 12.39618 | 20.59843 | 0.49713 | 5.72558 | 21.10628 | 9.35011 | 0.00000 |
| **TJP3** | 0.96641 | 0.35911 | 6.11085 | 0.24869 | 10.80017 | 0.00000 | 6.28106 | 0.33272 | 8.88622 | 0.00000 | 15.06443 | 0.00000 |
| **TK1** | 0.58002 | 23.16365 | 10.50501 | 15.72527 | 27.22534 | 2.74374 | 13.07259 | 6.39240 | 16.11127 | 1.92510 | 8.96855 | 17.06943 |
| **TK2** | 1.52183 | 1.66607 | 0.00000 | 0.79431 | 0.00000 | 0.29240 | 0.00000 | 0.30821 | 0.00000 | 0.53189 | 0.65146 | 0.70580 |
| **TKT** | 2.03573 | 17.16806 | 29.40618 | 24.47713 | 35.66004 | 8.02104 | 50.25434 | 4.14473 | 54.90278 | 13.78521 | 21.75541 | 7.43275 |
| **TKTL1** | 1.28141 | 18.84351 | 0.00000 | 8.73725 | 11.90103 | 8.60985 | 0.00000 | 2.77319 | 0.00000 | 0.00000 | 2.18103 | 5.37856 |
| **TKTL2** | 0.00671 | 0.04836 | 0.02285 | 0.00459 | 0.00000 | 0.09308 | 0.02305 | 0.00950 | 0.16710 | 0.01092 | 0.04506 | 0.02827 |

| | | | | | | | | | | | | |
|---|---|---|---|---|---|---|---|---|---|---|---|---|
| **TLCD1** | 1.71583 | 7.98508 | 6.84441 | 5.33987 | 17.84720 | 0.00000 | 0.00000 | 0.22611 | 5.34479 | 2.33706 | 10.78813 | 0.95111 |
| **TLCD2** | 0.00230 | 0.44295 | 0.01273 | 0.17247 | 0.61318 | 0.00000 | 0.05861 | 0.03602 | 0.00000 | 1.28013 | 0.08234 | 0.44229 |
| **TLDC1** | 0.88305 | 0.00000 | 2.98392 | 0.00000 | 0.41617 | 0.95484 | 3.72099 | 0.02601 | 0.23416 | 0.00000 | 3.24630 | 0.09235 |
| **TLDC2** | 0.44497 | 0.07255 | 0.00000 | 0.30629 | 0.00000 | 0.14916 | 0.00000 | 0.12676 | 0.00000 | 0.14145 | 0.01626 | 0.00769 |
| **TLE1** | 0.57959 | 6.99953 | 4.67851 | 0.00000 | 16.37854 | 3.30793 | 15.79669 | 0.05832 | 9.32340 | 23.32238 | 7.83881 | 0.45241 |
| **TLE2** | 3.19145 | 0.00000 | 0.45716 | 0.85887 | 0.17063 | 0.00094 | 0.00000 | 0.20448 | 0.12693 | 0.00000 | 0.00976 | 1.38863 |
| **TLE3** | 0.26911 | 15.99900 | 18.87541 | 14.62275 | 20.73934 | 7.89625 | 31.70937 | 0.00000 | 11.09653 | 16.87042 | 8.03012 | 0.01463 |
| **TLE4** | 6.72229 | 0.10245 | 4.81217 | 1.57965 | 0.76948 | 4.16093 | 4.93007 | 2.41749 | 1.30198 | 5.82084 | 0.00000 | 2.70712 |
| **TLE6** | 16.24971 | 1.26548 | 0.00000 | 0.00000 | 0.00000 | 0.00000 | 0.00000 | 9.84308 | 0.00370 | 0.17590 | 0.00000 | 1.70671 |
| **TLK1** | 3.75623 | 2.99323 | 14.06690 | 4.15260 | 0.00000 | 20.68538 | 8.15822 | 10.20272 | 0.18492 | 12.39224 | 7.32209 | 7.20752 |
| **TLK2** | 0.72722 | 0.20785 | 1.06565 | 1.79979 | 0.00000 | 3.84060 | 2.79186 | 2.65814 | 0.00234 | 9.68838 | 0.65882 | 0.55533 |
| **TLL1** | 1.09137 | 0.00551 | 0.02379 | 0.04161 | 0.00598 | 0.05999 | 0.04657 | 0.03788 | 0.00012 | 0.01126 | 0.02017 | 0.01900 |
| **TLL2** | 0.00104 | 0.50396 | 0.55393 | 1.58394 | 0.33173 | 0.00000 | 0.00000 | 0.35585 | 0.00000 | 0.05789 | 0.14364 | 0.29999 |
| **TLN1** | 1.60810 | 0.47233 | 4.43613 | 6.75921 | 3.60547 | 0.00000 | 6.77861 | 0.05444 | 3.37791 | 5.50619 | 0.22847 | 1.98863 |
| **TLN2** | 4.14374 | 2.73357 | 2.94537 | 7.74764 | 2.01131 | 0.00000 | 1.17697 | 0.00000 | 7.87741 | 6.47003 | 3.69285 | 0.49406 |
| **TLR1** | 0.00596 | 0.03071 | 0.03730 | 0.01471 | 0.01872 | 0.01693 | 0.01674 | 0.01046 | 0.01097 | 0.01331 | 0.01519 | 0.04145 |
| **TLR10** | 0.00858 | 0.01515 | 0.01466 | 0.01220 | 0.01709 | 0.01767 | 0.01901 | 0.01019 | 0.01845 | 0.01237 | 0.01490 | 0.01768 |
| **TLR2** | 0.16014 | 0.00894 | 0.04150 | 0.01161 | 0.00446 | 0.05172 | 0.03508 | 0.00348 | 0.00012 | 0.01468 | 0.02367 | 0.01186 |
| **TLR3** | 0.01577 | 0.01906 | 0.01769 | 0.01166 | 0.01426 | 0.01807 | 0.01674 | 0.09043 | 0.02136 | 0.01296 | 0.01394 | 0.01262 |
| **TLR4** | 0.05125 | 0.01235 | 0.00345 | 0.28119 | 0.00000 | 0.02980 | 0.02122 | 0.00628 | 0.01630 | 0.01579 | 0.01203 | 0.01368 |
| **TLR5** | 0.49260 | 0.00000 | 0.22104 | 0.02722 | 0.00000 | 0.00000 | 0.00000 | 0.08995 | 0.00000 | 0.22298 | 0.71047 | 0.08721 |
| **TLR6** | 0.07234 | 0.04130 | 0.01509 | 0.01658 | 0.01836 | 0.00000 | 0.02562 | 0.00622 | 0.01807 | 0.01699 | 0.01546 | 0.01468 |
| **TLR7** | 0.00939 | 0.00000 | 0.12138 | 0.18340 | 0.46728 | 0.23553 | 0.02248 | 0.01716 | 0.85917 | 0.00000 | 2.07355 | 0.00466 |
| **TLR8** | 0.14283 | 0.00372 | 0.00000 | 0.06671 | 0.05474 | 0.16373 | 0.00000 | 0.00139 | 0.00000 | 0.01969 | 0.01183 | 0.02979 |
| **TLR8-AS1** | 0.59191 | 0.00000 | 0.00000 | 0.61655 | 0.14828 | 0.20403 | 0.02072 | 0.04051 | 0.00000 | 0.01212 | 0.11976 | 0.04261 |
| **TLR9** | 0.00604 | 0.01975 | 0.01549 | 0.02550 | 0.04728 | 0.01408 | 0.03619 | 0.01069 | 0.01359 | 0.01124 | 0.01889 | 0.01573 |
| **TLX1** | 0.00598 | 0.04661 | 0.02421 | 0.01100 | 0.00687 | 0.00205 | 0.02007 | 0.00730 | 0.01636 | 0.05756 | 0.01676 | 0.09794 |
| **TLX2** | 0.14630 | 0.20687 | 0.01802 | 0.05898 | 0.29014 | 0.00000 | 0.05175 | 0.13146 | 0.00000 | 0.00000 | 0.01278 | 0.10675 |
| **TLX3** | 0.02721 | 0.02548 | 0.00974 | 0.02242 | 0.08737 | 0.00044 | 0.00706 | 0.01404 | 0.03754 | 0.04938 | 0.00542 | 0.06510 |
| **TM2D1** | 0.68916 | 4.26356 | 2.15322 | 0.63017 | 0.00000 | 12.44769 | 0.00000 | 3.46148 | 7.61744 | 7.72188 | 1.71759 | 6.00898 |
| **TM2D2** | 1.48182 | 7.80080 | 18.33518 | 6.84880 | 10.93262 | 18.29053 | 3.76263 | 2.75337 | 32.34391 | 2.17106 | 11.94117 | 6.31895 |
| **TM2D3** | 2.56178 | 7.18615 | 9.33923 | 1.87181 | 1.60428 | 8.92870 | 0.00000 | 3.60430 | 13.38180 | 0.00000 | 4.95698 | 10.19850 |
| **TM4SF1** | 0.00194 | 0.17106 | 0.10708 | 0.37369 | 0.04363 | 0.01305 | 0.00000 | 0.35655 | 0.00000 | 0.14379 | 0.00264 | 0.37408 |
| **TM4SF1-AS1** | 0.01013 | 0.11495 | 0.00601 | 0.20189 | 0.00731 | 0.02851 | 0.02327 | 0.34883 | 0.01010 | 0.00407 | 0.01673 | 0.26029 |
| **TM4SF18** | 0.00709 | 0.00887 | 0.02587 | 0.49942 | 0.10339 | 0.00000 | 0.00230 | 0.00974 | 0.04293 | 0.02871 | 0.00713 | 0.01558 |
| **TM4SF19** | 0.00736 | 0.02880 | 0.01948 | 0.00785 | 0.00810 | 0.04669 | 0.03310 | 0.01556 | 0.01626 | 0.00960 | 0.02338 | 0.01973 |
| **TM4SF19-AS1** | 0.00584 | 0.03940 | 0.01510 | 0.01212 | 0.00338 | 0.03225 | 0.00889 | 0.01040 | 0.03342 | 0.03299 | 0.01181 | 0.01181 |
| **TM4SF19-TCTE** | 0.00609 | 0.01445 | 0.01728 | 0.01130 | 0.01576 | 0.01787 | 0.01602 | 0.01050 | 0.02081 | 0.02251 | 0.01365 | 0.01576 |
| **TM4SF20** | 0.01829 | 0.01878 | 0.06814 | 0.01241 | 0.01238 | 0.00713 | 0.02646 | 0.01413 | 0.02419 | 0.04293 | 0.01642 | 0.01753 |
| **TM4SF4** | 0.10372 | 0.00833 | 0.02470 | 0.00722 | 0.00200 | 0.03504 | 0.04165 | 0.00124 | 0.00000 | 0.04866 | 0.01335 | 0.13094 |
| **TM4SF5** | 0.02671 | 4.95692 | 0.00000 | 0.00009 | 0.84471 | 0.26113 | 0.00000 | 0.00000 | 0.00000 | 0.00355 | 0.12047 | 4.02704 |
| **TM6SF1** | 0.36341 | 0.05850 | 0.07590 | 0.09484 | 0.28680 | 0.03363 | 0.04858 | 0.01961 | 0.00000 | 0.05700 | 0.00054 | 0.05350 |
| **TM6SF2** | 0.00591 | 0.00723 | 0.29255 | 0.22458 | 0.00000 | 0.05695 | 0.04329 | 0.00963 | 0.00000 | 0.01374 | 0.04240 | 0.01335 |
| **TM7SF2** | 0.00087 | 0.00000 | 4.67909 | 2.17340 | 4.52224 | 0.00000 | 0.32493 | 0.00000 | 7.95662 | 2.17399 | 7.31023 | 0.23564 |
| **TM7SF3** | 0.67918 | 4.77639 | 4.67163 | 0.63703 | 7.11797 | 3.17042 | 1.78506 | 0.09523 | 8.67173 | 7.07902 | 1.43534 | 0.00000 |
| **TM9SF1** | 1.48342 | 4.97059 | 10.33525 | 8.01433 | 2.41555 | 9.83158 | 0.00000 | 0.56588 | 22.12686 | 0.00000 | 7.33203 | 6.09625 |
| **TM9SF2** | 4.65010 | 13.65459 | 14.91497 | 6.99513 | 5.20819 | 41.38191 | 0.00000 | 4.87758 | 34.06574 | 8.62682 | 11.81665 | 10.00079 |
| **TM9SF3** | 3.26184 | 3.10892 | 12.60803 | 3.30968 | 2.89557 | 35.45020 | 4.90197 | 4.46063 | 15.98835 | 43.25442 | 5.15900 | 7.31058 |

| | | | | | | | | | | | | |
|---|---|---|---|---|---|---|---|---|---|---|---|---|
| TM9SF4 | 5.21632 | 12.35433 | 9.68764 | 7.32460 | 5.58898 | 0.00000 | 0.00000 | 2.37076 | 19.51936 | 2.93420 | 6.91298 | 9.56153 |
| TMA16 | 0.66225 | 6.87122 | 3.78932 | 5.71866 | 12.14110 | 15.75581 | 9.58668 | 12.26955 | 4.81025 | 9.31023 | 8.63290 | 13.54846 |
| TMA7 | 0.00000 | 6.54748 | 4.39626 | 3.41596 | 5.72515 | 10.09093 | 12.73383 | 3.29811 | 4.82662 | 7.25863 | 6.52798 | 11.12345 |
| TMBIM1 | 5.90118 | 34.19853 | 5.13629 | 30.53166 | 9.58994 | 11.29118 | 0.00000 | 0.00000 | 14.98376 | 1.95480 | 0.42609 | 8.06605 |
| TMBIM4 | 0.51427 | 10.49453 | 3.76257 | 7.20049 | 8.77040 | 23.90602 | 0.00000 | 0.45310 | 9.35089 | 9.02859 | 11.65319 | 6.68900 |
| TMBIM6 | 11.25792 | 29.81199 | 31.56430 | 27.80919 | 23.75580 | 56.25992 | 15.08627 | 34.17808 | 62.90762 | 30.57178 | 30.97845 | 35.97043 |
| TMC1 | 0.08717 | 0.14043 | 0.14815 | 0.03696 | 0.45234 | 0.24573 | 0.00000 | 0.05851 | 0.22281 | 2.41690 | 0.03543 | 0.32857 |
| TMC2 | 0.13189 | 0.01152 | 0.01560 | 0.01938 | 0.02160 | 0.00643 | 0.04397 | 0.00940 | 0.01622 | 0.01356 | 0.02514 | 0.05412 |
| TMC3 | 0.89934 | 0.00848 | 0.06594 | 0.00894 | 0.01911 | 0.00000 | 0.00890 | 0.00892 | 0.04354 | 0.00512 | 0.14089 | 0.22896 |
| TMC4 | 0.01294 | 0.11940 | 0.36619 | 0.16172 | 0.57197 | 0.00000 | 0.12052 | 0.01258 | 0.28991 | 0.00000 | 0.04710 | 0.15756 |
| TMC5 | 3.02181 | 0.08747 | 2.16651 | 0.00000 | 0.71485 | 2.43031 | 2.83878 | 1.49672 | 3.82403 | 4.70294 | 4.47959 | 0.62339 |
| TMC6 | 0.27958 | 0.43856 | 0.72834 | 0.51614 | 0.01561 | 0.00000 | 0.00000 | 0.09190 | 1.20705 | 1.80696 | 0.03255 | 0.00000 |
| TMC7 | 0.00000 | 10.85785 | 0.30096 | 5.91141 | 8.61217 | 2.22215 | 0.00000 | 0.64836 | 1.63083 | 5.56213 | 5.32329 | 6.88781 |
| TMC8 | 0.00272 | 0.04669 | 0.04986 | 0.00925 | 0.01411 | 0.00183 | 0.06125 | 0.31542 | 0.00000 | 0.12542 | 0.01975 | 0.11235 |
| TMCC1 | 6.50690 | 0.83096 | 1.13403 | 1.65035 | 1.35611 | 1.03673 | 0.93969 | 0.58583 | 0.00000 | 1.55942 | 0.71116 | 0.78639 |
| TMCC1-AS1 | 0.01496 | 0.17690 | 0.14457 | 0.04278 | 0.00000 | 0.00000 | 0.00000 | 0.00298 | 0.00000 | 0.18132 | 0.06291 | 0.16796 |
| TMCC2 | 2.83561 | 0.30034 | 0.31400 | 0.00123 | 0.09582 | 0.00000 | 0.00398 | 0.04708 | 0.13893 | 0.00000 | 0.02741 | 0.21628 |
| TMCC3 | 0.00322 | 0.48140 | 0.00000 | 0.00000 | 1.88090 | 0.00000 | 6.74425 | 0.60136 | 2.00588 | 0.95609 | 3.03542 | 0.20514 |
| TMCO1 | 1.07049 | 14.39981 | 19.77095 | 6.32251 | 9.51596 | 36.00284 | 5.27934 | 21.82571 | 27.66824 | 20.32726 | 12.67522 | 26.66677 |
| TMCO2 | 0.06508 | 0.01449 | 0.02751 | 0.01172 | 0.02042 | 0.01067 | 0.01695 | 0.00761 | 0.02056 | 0.00880 | 0.01493 | 0.03650 |
| TMCO3 | 0.83951 | 0.62317 | 2.70152 | 0.00269 | 0.00000 | 1.77074 | 6.29901 | 0.25465 | 3.99058 | 0.00000 | 4.77303 | 0.15067 |
| TMCO4 | 0.00275 | 3.97340 | 0.15065 | 1.35031 | 1.43548 | 0.19274 | 0.19052 | 0.00624 | 0.00000 | 0.17131 | 0.28438 | 1.40935 |
| TMCO6 | 0.09306 | 1.77253 | 1.21629 | 1.24178 | 0.60170 | 0.08419 | 1.48535 | 0.01793 | 1.52632 | 1.44108 | 3.11657 | 0.93267 |
| TMED1 | 0.85553 | 7.66445 | 6.13573 | 12.65314 | 7.73653 | 0.67936 | 2.51818 | 3.80978 | 9.00982 | 0.00000 | 7.24379 | 6.83931 |
| TMED10 | 6.04275 | 15.95499 | 18.71044 | 8.98703 | 10.82657 | 35.62739 | 0.00000 | 1.44338 | 38.59231 | 11.40287 | 15.05040 | 13.62138 |
| TMED10P1 | 0.22868 | 0.76906 | 0.67901 | 0.35530 | 0.47711 | 2.36518 | 0.00000 | 0.21363 | 1.59557 | 0.66575 | 0.88134 | 0.83083 |
| TMED11P | 0.05009 | 0.00559 | 0.01704 | 0.01740 | 0.02225 | 0.06031 | 0.00000 | 0.01275 | 0.06565 | 0.33525 | 0.08500 | 0.04119 |
| TMED2 | 7.17151 | 25.36553 | 29.68414 | 12.72144 | 6.66436 | 60.35138 | 17.00594 | 12.87234 | 48.26798 | 11.16059 | 29.00800 | 20.21275 |
| TMED3 | 1.83837 | 15.81611 | 17.50996 | 7.04229 | 22.30515 | 0.00000 | 11.14314 | 0.69735 | 33.26543 | 4.01159 | 13.67553 | 13.07702 |
| TMED4 | 1.47418 | 8.80030 | 7.18984 | 3.95015 | 5.87376 | 0.00000 | 0.00000 | 0.84652 | 9.51457 | 0.00000 | 8.37342 | 1.68357 |
| TMED5 | 0.00000 | 7.06337 | 9.16585 | 2.29059 | 0.00000 | 26.46497 | 0.00000 | 6.00551 | 24.85560 | 11.91924 | 4.45976 | 12.99317 |
| TMED6 | 0.06266 | 0.00000 | 0.00000 | 0.85566 | 0.14233 | 0.55891 | 0.09648 | 0.15442 | 0.00000 | 1.58237 | 0.36540 | 0.15133 |
| TMED7 | 0.00000 | 4.25951 | 11.10943 | 0.00357 | 0.00000 | 29.36434 | 0.00000 | 1.91152 | 17.91949 | 8.66587 | 6.84469 | 7.47614 |
| TMED8 | 0.88833 | 1.45622 | 0.82734 | 0.72822 | 0.66987 | 0.00000 | 0.00000 | 0.00000 | 0.00000 | 0.51461 | 1.13854 | 0.28141 |
| TMED9 | 1.32015 | 26.33563 | 29.36601 | 27.43491 | 27.68552 | 0.00000 | 12.62277 | 5.52154 | 46.96732 | 0.01569 | 26.91743 | 26.13508 |
| TMEFF1 | 0.00641 | 0.08271 | 0.03797 | 0.02762 | 0.01846 | 0.00000 | 0.02043 | 0.01075 | 0.02092 | 0.00671 | 0.01657 | 0.02700 |
| TMEFF2 | 1.18576 | 0.02167 | 0.18753 | 0.24019 | 0.00000 | 0.04061 | 0.00000 | 1.27043 | 0.00000 | 0.06167 | 0.19602 | 0.00286 |
| TMEM100 | 0.00275 | 0.02181 | 0.00496 | 0.01545 | 0.00871 | 0.00214 | 0.19143 | 0.03343 | 0.00000 | 0.03307 | 0.01205 | 0.01411 |
| TMEM101 | 3.26025 | 6.20227 | 8.50117 | 3.56203 | 1.83576 | 0.75699 | 0.88873 | 5.18396 | 4.92489 | 0.00000 | 3.94979 | 11.97441 |
| TMEM102 | 0.00283 | 0.07302 | 0.42564 | 0.49371 | 0.25340 | 0.00000 | 0.10208 | 0.02892 | 0.90804 | 0.00000 | 0.04985 | 0.18892 |
| TMEM104 | 0.91971 | 2.65509 | 2.79539 | 0.85920 | 4.76535 | 0.00000 | 0.03235 | 1.89416 | 9.25764 | 3.99432 | 2.71244 | 2.92594 |
| TMEM105 | 0.00381 | 0.12683 | 0.04512 | 0.00075 | 0.02742 | 0.19173 | 0.00000 | 0.06403 | 0.13469 | 0.04765 | 0.00000 | 0.00934 |
| TMEM106A | 0.56634 | 0.02673 | 0.19868 | 0.96881 | 0.00000 | 0.00000 | 0.05356 | 0.16410 | 0.00000 | 0.06775 | 0.04451 | 0.23362 |
| TMEM106B | 0.72739 | 2.48785 | 1.96080 | 0.28249 | 0.00004 | 11.89285 | 0.00000 | 0.29846 | 2.98500 | 1.89978 | 3.74596 | 1.36570 |
| TMEM106C | 3.79567 | 11.80296 | 12.97847 | 0.00000 | 5.58773 | 14.34990 | 14.19489 | 2.25499 | 18.56661 | 0.00000 | 19.53852 | 4.12701 |
| TMEM107 | 1.23848 | 0.55861 | 2.40426 | 1.45917 | 0.38447 | 1.24417 | 0.00000 | 1.17411 | 0.95812 | 0.82228 | 1.22093 | 0.54230 |
| TMEM108 | 3.34609 | 0.53456 | 0.27681 | 1.82076 | 0.24272 | 0.27450 | 0.08321 | 0.93543 | 0.12305 | 0.00000 | 0.06031 | 1.41894 |
| TMEM108-AS1 | 0.00544 | 0.00179 | 0.00000 | 0.09755 | 0.00239 | 0.02711 | 0.00091 | 0.00901 | 0.00723 | 0.44779 | 0.00000 | 0.04414 |

| | | | | | | | | | | | | |
|---|---|---|---|---|---|---|---|---|---|---|---|---|
| TMEM109 | 9.23796 | 14.38590 | 12.18511 | 17.73812 | 20.49871 | 0.00000 | 1.20430 | 0.31357 | 38.52677 | 2.54207 | 15.81150 | 7.38590 |
| TMEM11 | 6.05482 | 16.99789 | 17.68341 | 18.09720 | 13.98215 | 6.73713 | 21.51562 | 17.64109 | 18.25774 | 0.00000 | 16.17765 | 28.91564 |
| TMEM110 | 0.32055 | 1.80184 | 1.41927 | 1.61520 | 1.88901 | 0.00000 | 2.26476 | 0.03878 | 0.59524 | 2.71821 | 0.81861 | 0.88460 |
| TMEM114.uid-2 | 0.00619 | 0.01531 | 0.01404 | 0.01285 | 0.02123 | 0.01152 | 0.02215 | 0.01051 | 0.01863 | 0.01657 | 0.01540 | 0.01783 |
| TMEM115 | 1.93954 | 10.51237 | 8.58610 | 6.39333 | 6.89299 | 0.00000 | 1.28781 | 0.57772 | 16.84275 | 4.80086 | 4.14351 | 6.63708 |
| TMEM116 | 1.16436 | 1.33441 | 1.27514 | 0.92948 | 0.00000 | 0.76142 | 1.50793 | 0.00000 | 0.32619 | 0.00000 | 1.53526 | 1.14126 |
| TMEM117 | 0.04902 | 0.16006 | 0.05990 | 0.62026 | 0.00000 | 0.72399 | 0.00000 | 0.03618 | 0.31672 | 0.33986 | 0.10126 | 0.17185 |
| TMEM119 | 0.00643 | 0.00956 | 0.02644 | 0.00537 | 0.07968 | 0.00561 | 0.03055 | 0.01028 | 0.00000 | 0.07639 | 0.01028 | 0.01676 |
| TMEM120A | 0.00102 | 6.06417 | 5.63247 | 4.17880 | 7.16393 | 0.00000 | 1.67397 | 0.00000 | 12.58889 | 2.19231 | 6.53953 | 0.84701 |
| TMEM120B | 2.07618 | 3.24783 | 2.73585 | 2.14595 | 5.94174 | 0.00000 | 0.00058 | 0.89124 | 4.78862 | 6.23837 | 3.13210 | 1.94417 |
| TMEM121 | 0.00609 | 0.06202 | 0.03464 | 0.01170 | 0.05408 | 0.00000 | 0.01372 | 0.00935 | 0.02186 | 0.01342 | 0.01224 | 0.04299 |
| TMEM123 | 6.22913 | 0.00000 | 9.69864 | 14.80805 | 0.00435 | 10.02676 | 5.74591 | 0.78669 | 13.66754 | 6.31998 | 4.23712 | 1.86774 |
| TMEM125 | 0.67517 | 0.00000 | 0.45558 | 0.00111 | 0.00000 | 0.00422 | 0.96188 | 0.22620 | 0.74126 | 0.00000 | 1.29020 | 0.02619 |
| TMEM126A | 0.32533 | 8.50064 | 7.93736 | 0.66718 | 3.25894 | 21.89695 | 6.95568 | 7.43324 | 8.07618 | 5.88412 | 7.53624 | 13.76223 |
| TMEM126B | 0.16785 | 4.94458 | 10.38760 | 3.46162 | 0.00000 | 15.63510 | 13.65532 | 4.90309 | 11.89779 | 3.41249 | 9.90596 | 7.53255 |
| TMEM127 | 1.36813 | 4.43927 | 3.32485 | 3.17753 | 8.11039 | 0.09580 | 3.06730 | 0.22360 | 6.09638 | 7.23968 | 1.03324 | 3.77739 |
| TMEM128 | 5.36789 | 6.66107 | 0.76552 | 3.26796 | 2.14564 | 13.59999 | 0.00000 | 1.76662 | 3.09385 | 1.15974 | 5.00269 | 2.60499 |
| TMEM129 | 2.75504 | 2.74484 | 0.94168 | 2.47326 | 1.73318 | 0.00000 | 0.68302 | 0.00000 | 0.72964 | 3.51182 | 1.39721 | 0.60227 |
| TMEM130 | 0.12139 | 0.01273 | 0.01339 | 0.01446 | 0.01230 | 0.02700 | 0.02501 | 0.00717 | 0.01573 | 0.02144 | 0.01543 | 0.03848 |
| TMEM131 | 2.68265 | 0.00000 | 2.77028 | 2.55761 | 6.70409 | 7.80520 | 0.00000 | 0.40805 | 8.80851 | 22.28689 | 0.15637 | 0.67389 |
| TMEM132A | 0.00290 | 0.31336 | 0.25438 | 0.88644 | 0.44456 | 0.00000 | 0.00000 | 0.00574 | 0.64832 | 1.53761 | 0.00000 | 0.06904 |
| TMEM132B | 8.43768 | 0.00916 | 0.03327 | 0.06294 | 0.01428 | 0.00000 | 0.05721 | 0.31653 | 0.00000 | 0.00000 | 0.06871 | 0.00790 |
| TMEM132C | 0.00330 | 0.10747 | 0.01589 | 1.13927 | 0.00000 | 0.04991 | 0.00000 | 0.01017 | 0.00000 | 0.18819 | 0.00190 | 0.00044 |
| TMEM132D | 4.92666 | 0.00569 | 0.01833 | 0.00875 | 0.05630 | 0.00000 | 0.00717 | 0.00000 | 0.00973 | 0.08400 | 0.05809 | 0.09689 |
| TMEM132E | 0.00699 | 0.00134 | 0.00000 | 0.26756 | 0.01505 | 0.12940 | 0.00000 | 0.00894 | 0.00000 | 0.07037 | 0.05266 | 0.00637 |
| TMEM133 | 0.00287 | 0.14480 | 0.21432 | 1.87029 | 0.00000 | 1.60223 | 0.59924 | 0.00000 | 0.09928 | 1.30962 | 0.38078 | 0.06658 |
| TMEM134 | 1.48875 | 6.31760 | 0.81139 | 3.29276 | 3.59707 | 1.40891 | 0.00329 | 3.89320 | 1.03831 | 0.56825 | 2.00429 | 11.03877 |
| TMEM135 | 0.72558 | 1.54382 | 4.57133 | 0.35969 | 1.10980 | 8.53805 | 2.64867 | 0.16422 | 15.16742 | 3.12854 | 0.90491 | 2.24809 |
| TMEM136 | 0.70710 | 2.10850 | 0.00000 | 0.65701 | 0.60774 | 0.69530 | 0.00000 | 0.00000 | 0.00000 | 0.40888 | 0.31897 | 0.18390 |
| TMEM138 | 1.17964 | 7.12864 | 9.28663 | 0.29971 | 4.77260 | 12.46246 | 0.00000 | 10.92240 | 14.82454 | 0.00000 | 6.03712 | 15.91847 |
| TMEM139 | 0.05399 | 0.00000 | 1.59506 | 2.96872 | 0.00000 | 0.17621 | 0.67173 | 0.34677 | 0.52928 | 0.05558 | 2.08407 | 0.28843 |
| TMEM140 | 0.36724 | 1.61643 | 0.65608 | 0.33711 | 0.39487 | 0.00000 | 0.20357 | 0.00000 | 2.19811 | 0.00000 | 0.83713 | 0.53534 |
| TMEM141 | 0.50183 | 11.35786 | 4.35982 | 8.01202 | 8.82187 | 0.00000 | 0.72432 | 0.20078 | 0.00000 | 0.89080 | 7.46078 | 3.47021 |
| TMEM143 | 1.45327 | 3.01171 | 0.46646 | 0.96940 | 0.09601 | 0.00000 | 0.16615 | 0.67407 | 1.76756 | 0.00000 | 1.34915 | 1.77197 |
| TMEM144 | 0.04308 | 0.31174 | 0.58317 | 2.01111 | 0.00000 | 2.13587 | 0.36869 | 0.00000 | 1.04647 | 0.20552 | 1.02973 | 0.00000 |
| TMEM145 | 0.08179 | 0.02408 | 0.01536 | 0.01230 | 0.00075 | 0.01537 | 0.07107 | 0.01477 | 0.01576 | 0.05853 | 0.01349 | 0.03498 |
| TMEM147 | 3.54858 | 20.28087 | 19.89583 | 14.62625 | 27.81621 | 6.61178 | 10.72385 | 1.49113 | 31.35966 | 10.18393 | 22.43320 | 4.89635 |
| TMEM14A | 2.84267 | 8.12724 | 1.90883 | 1.75424 | 0.44476 | 19.18232 | 6.95880 | 0.80566 | 4.03169 | 2.08733 | 5.54188 | 6.06857 |
| TMEM14B | 4.55421 | 6.91437 | 7.68219 | 6.64422 | 6.47898 | 18.16972 | 8.31464 | 0.00000 | 7.54517 | 1.72877 | 9.59767 | 5.29404 |
| TMEM14C | 2.43908 | 7.32936 | 6.45028 | 12.09417 | 13.26383 | 16.24123 | 4.91522 | 0.19605 | 11.04882 | 1.86856 | 9.12330 | 1.45236 |
| TMEM14E | 0.00713 | 0.00733 | 0.13484 | 0.01300 | 0.09439 | 0.00000 | 0.00042 | 0.06004 | 0.02259 | 0.06944 | 0.02106 | 0.05012 |
| TMEM150A | 3.65267 | 2.09630 | 3.22335 | 2.20553 | 1.87673 | 0.00000 | 2.39066 | 0.09684 | 7.94440 | 2.70784 | 6.60073 | 0.63069 |
| TMEM150B | 1.79688 | 0.06637 | 0.03241 | 0.00511 | 0.06824 | 0.00065 | 0.01357 | 0.32100 | 0.02037 | 0.00000 | 0.00892 | 0.05908 |
| TMEM150C | 0.00432 | 2.75387 | 0.03050 | 0.59169 | 0.08491 | 2.13453 | 0.00000 | 0.00000 | 0.40410 | 0.00000 | 0.61739 | 0.94670 |
| TMEM151A | 0.00628 | 0.01581 | 0.06679 | 0.01125 | 0.05165 | 0.00000 | 0.01198 | 0.01017 | 0.01240 | 0.01494 | 0.02085 | 0.02295 |
| TMEM151B | 0.02623 | 0.00547 | 0.02477 | 0.12713 | 0.00755 | 0.00000 | 0.00693 | 0.06377 | 0.00271 | 0.19938 | 0.01331 | 0.05266 |
| TMEM154 | 3.33440 | 0.00000 | 15.26664 | 1.64839 | 0.12934 | 0.83922 | 0.94583 | 0.03509 | 2.10944 | 0.00000 | 0.00000 | 0.00000 |
| TMEM155 | 0.00419 | 0.00683 | 0.21196 | 0.07492 | 0.00000 | 0.02376 | 0.02298 | 0.01712 | 0.06168 | 0.04563 | 0.02310 | 0.04804 |

| | | | | | | | | | | | | |
|---|---|---|---|---|---|---|---|---|---|---|---|---|
| **TMEM156** | 0.00317 | 0.01517 | 0.01603 | 0.01227 | 0.01778 | 0.01598 | 0.01918 | 0.22117 | 0.01849 | 0.01310 | 0.01494 | 0.01023 |
| **TMEM158** | 0.00514 | 0.26986 | 0.10932 | 0.00000 | 0.42411 | 0.00000 | 0.11629 | 0.30640 | 0.00000 | 0.82767 | 0.01548 | 0.88233 |
| **TMEM159** | 0.00205 | 2.69763 | 0.44367 | 3.96907 | 0.52869 | 0.40122 | 0.42237 | 7.40859 | 1.04595 | 0.00000 | 0.00833 | 7.72409 |
| **TMEM160** | 0.54635 | 6.66860 | 1.53749 | 6.19431 | 9.98284 | 0.00000 | 1.49370 | 0.79892 | 3.67997 | 8.18307 | 2.77239 | 1.12082 |
| **TMEM161A** | 0.81928 | 8.98039 | 8.25109 | 4.01874 | 7.35866 | 0.00000 | 0.28703 | 0.62529 | 15.96269 | 2.56395 | 5.14331 | 7.42179 |
| **TMEM161B** | 0.00000 | 2.39862 | 3.64372 | 1.08299 | 1.69294 | 7.46810 | 0.00000 | 1.34512 | 2.84652 | 9.56292 | 3.86287 | 1.93478 |
| **TMEM161B-AS1** | 0.00000 | 2.29860 | 1.85540 | 0.00000 | 1.42348 | 3.56829 | 1.22763 | 0.39255 | 1.76581 | 2.25245 | 1.97418 | 2.64038 |
| **TMEM163** | 5.82307 | 0.01393 | 0.01884 | 0.00478 | 0.02078 | 0.00000 | 0.00000 | 0.67298 | 0.14320 | 0.03277 | 0.00000 | 0.04717 |
| **TMEM164** | 0.71424 | 0.59029 | 0.37342 | 4.45144 | 2.15529 | 0.00000 | 4.17699 | 1.16050 | 7.45022 | 5.08883 | 2.69236 | 1.31706 |
| **TMEM165** | 0.00000 | 7.60593 | 10.62187 | 1.66311 | 2.53217 | 13.71340 | 0.00000 | 0.00576 | 21.16884 | 16.32314 | 4.12932 | 0.72582 |
| **TMEM167A** | 2.88881 | 12.22768 | 20.23560 | 3.21206 | 1.65126 | 50.79066 | 23.98367 | 17.26653 | 20.16122 | 14.26613 | 14.57885 | 21.37387 |
| **TMEM167B** | 0.04682 | 3.00864 | 4.25214 | 3.27392 | 6.34156 | 7.28122 | 1.68274 | 10.14581 | 5.20181 | 4.44176 | 2.95030 | 9.42597 |
| **TMEM168** | 3.30471 | 7.71542 | 2.00294 | 0.00000 | 0.43492 | 15.02290 | 0.00000 | 4.23501 | 9.77673 | 4.46067 | 4.81897 | 8.89963 |
| **TMEM169** | 0.72805 | 0.00836 | 0.48221 | 0.00000 | 0.26969 | 0.06122 | 0.68188 | 0.23284 | 0.31434 | 0.07111 | 0.12639 | 0.00000 |
| **TMEM17** | 0.00673 | 0.25572 | 0.16354 | 0.00113 | 0.07608 | 0.43601 | 0.00000 | 0.00410 | 0.25280 | 0.00000 | 0.00027 | 0.20754 |
| **TMEM170A** | 0.55585 | 6.03698 | 8.33746 | 4.73432 | 2.89306 | 19.95114 | 3.84136 | 0.23322 | 11.27872 | 6.68606 | 8.24621 | 7.26863 |
| **TMEM170B** | 0.17283 | 0.00000 | 0.00000 | 0.00000 | 0.51378 | 0.01284 | 0.21971 | 0.00430 | 0.08387 | 0.24023 | 0.05809 | 0.00739 |
| **TMEM171** | 1.74225 | 1.69751 | 0.00000 | 0.00000 | 0.65900 | 0.73925 | 0.00000 | 0.00000 | 6.47614 | 0.48170 | 4.28206 | 0.02166 |
| **TMEM173** | 0.00999 | 0.00658 | 0.03653 | 0.04725 | 0.00232 | 0.04172 | 0.00922 | 0.16336 | 0.00532 | 0.02241 | 0.01757 | 0.05789 |
| **TMEM175** | 0.78932 | 1.05907 | 0.55129 | 0.17126 | 1.28766 | 0.00000 | 0.00000 | 0.36949 | 0.60571 | 0.21797 | 0.77769 | 0.72622 |
| **TMEM176A** | 0.23989 | 0.00537 | 0.00000 | 0.08828 | 0.03342 | 0.10836 | 0.00222 | 0.00000 | 0.00735 | 0.00698 | 0.00876 | 0.03458 |
| **TMEM176B** | 0.00633 | 0.11628 | 0.04116 | 0.05435 | 0.00861 | 0.00814 | 0.01469 | 0.01099 | 0.00967 | 0.01758 | 0.02127 | 0.00926 |
| **TMEM177** | 0.10223 | 5.78090 | 0.66017 | 4.69240 | 5.38537 | 1.72202 | 4.40024 | 0.31249 | 5.80693 | 0.00000 | 6.94801 | 6.82447 |
| **TMEM178A** | 0.10316 | 0.02033 | 0.01474 | 0.01338 | 0.00753 | 0.05454 | 0.00947 | 0.00396 | 0.01062 | 0.02905 | 0.01259 | 0.02600 |
| **TMEM178B** | 0.92778 | 0.00858 | 0.05505 | 0.00656 | 0.06994 | 0.01226 | 0.09666 | 0.01002 | 0.07581 | 0.00000 | 0.00000 | 0.02549 |
| **TMEM179** | 2.01749 | 0.70267 | 0.00820 | 0.02422 | 0.00000 | 0.04767 | 0.01308 | 0.04098 | 0.01657 | 0.02136 | 0.02427 | 0.19667 |
| **TMEM179B** | 3.00628 | 17.15274 | 13.02036 | 11.39934 | 16.21830 | 11.08433 | 0.00000 | 3.72720 | 16.94387 | 0.00000 | 13.28802 | 18.00813 |
| **TMEM18** | 1.77775 | 5.32299 | 5.76852 | 5.93950 | 6.68364 | 9.30942 | 5.30779 | 2.56764 | 9.90880 | 5.81590 | 4.39055 | 3.22107 |
| **TMEM180** | 0.00000 | 5.74935 | 6.67103 | 1.30107 | 3.44747 | 0.00000 | 0.41630 | 0.49762 | 8.39952 | 0.65755 | 5.87175 | 9.85412 |
| **TMEM181** | 2.55993 | 1.88095 | 3.11447 | 0.28169 | 2.97049 | 10.88138 | 2.74246 | 1.29816 | 8.73674 | 18.33499 | 3.74872 | 1.30470 |
| **TMEM182** | 0.01740 | 0.12279 | 0.54510 | 0.25628 | 0.41623 | 1.19149 | 0.34063 | 0.00000 | 0.17205 | 0.00000 | 0.31826 | 0.43173 |
| **TMEM183A** | 0.00962 | 0.16036 | 0.20734 | 0.04786 | 0.00000 | 0.00000 | 0.80311 | 0.02854 | 0.00000 | 1.14905 | 0.26457 | 0.03246 |
| **TMEM184A** | 0.00439 | 0.00522 | 0.03833 | 0.42188 | 0.06216 | 0.00000 | 0.03571 | 0.01115 | 0.00000 | 0.15242 | 0.02495 | 0.01310 |
| **TMEM184B** | 4.62223 | 7.99472 | 7.88129 | 6.06296 | 6.62696 | 0.00000 | 2.02324 | 0.00000 | 8.00374 | 4.36660 | 3.13114 | 10.25988 |
| **TMEM184C** | 2.38234 | 7.23071 | 6.10301 | 1.10389 | 5.03407 | 18.12243 | 0.00000 | 5.86974 | 15.88722 | 0.00000 | 9.56231 | 13.23839 |
| **TMEM185A** | 0.33155 | 2.56648 | 0.17307 | 1.12879 | 0.01656 | 0.00000 | 0.00000 | 3.43444 | 1.76904 | 0.80306 | 0.00000 | 2.14493 |
| **TMEM185B** | 1.93157 | 0.19067 | 0.24998 | 0.55986 | 0.42314 | 0.75549 | 0.21684 | 1.18945 | 1.13787 | 1.44048 | 0.00000 | 1.67326 |
| **TMEM186** | 1.34942 | 5.56975 | 3.99581 | 0.21951 | 3.71179 | 7.53798 | 4.00600 | 2.60599 | 6.72022 | 3.57926 | 6.28201 | 9.34390 |
| **TMEM187** | 0.20788 | 0.11431 | 0.21698 | 0.00367 | 0.00423 | 0.12868 | 0.00000 | 0.08205 | 0.00000 | 0.00007 | 0.48158 | 0.12761 |
| **TMEM189** | 4.23153 | 4.95117 | 13.72274 | 2.82298 | 9.40058 | 2.21331 | 4.30637 | 1.36694 | 18.49466 | 6.51241 | 4.27646 | 3.88121 |
| **TMEM19** | 0.01343 | 4.00196 | 3.80361 | 2.40446 | 0.50790 | 5.15557 | 0.00000 | 0.22929 | 8.23580 | 0.00000 | 2.58590 | 6.37533 |
| **TMEM190** | 0.02889 | 0.01505 | 0.02599 | 0.01128 | 0.01427 | 0.01454 | 0.01917 | 0.01024 | 0.01856 | 0.02295 | 0.01568 | 0.01519 |
| **TMEM191A** | 0.03167 | 0.43249 | 0.11040 | 0.56311 | 0.51741 | 0.00000 | 0.78029 | 0.27497 | 0.79071 | 0.00000 | 0.15866 | 1.37848 |
| **TMEM191B** | 0.01559 | 0.05722 | 0.14509 | 0.13264 | 0.15006 | 0.00000 | 0.03701 | 0.02048 | 0.15676 | 0.07650 | 0.16170 | 0.03833 |
| **TMEM191C** | 0.04735 | 0.14091 | 0.18478 | 0.17358 | 0.38735 | 0.00000 | 0.26034 | 0.03928 | 0.38013 | 0.41515 | 0.40825 | 0.02978 |
| **TMEM192** | 0.62450 | 1.93343 | 4.30329 | 0.66070 | 3.82405 | 4.90051 | 0.00000 | 0.05523 | 5.99938 | 3.85941 | 5.10018 | 1.76575 |
| **TMEM194A** | 4.86281 | 5.37406 | 0.25042 | 1.71706 | 0.21459 | 6.96574 | 0.00000 | 1.47610 | 5.36870 | 1.04762 | 1.90604 | 5.09799 |
| **TMEM194B** | 0.55537 | 0.14707 | 0.41647 | 0.00000 | 0.26500 | 0.53464 | 0.00000 | 0.04731 | 0.09164 | 0.00855 | 1.07495 | 0.71809 |

| | | | | | | | | | | | | |
|---|---|---|---|---|---|---|---|---|---|---|---|---|
| **TMEM198** | 0.16541 | 0.01389 | 0.45050 | 0.00736 | 0.06075 | 0.00000 | 0.01116 | 0.03155 | 0.02997 | 0.01276 | 0.01230 | 0.05278 |
| **TMEM198B** | 0.81921 | 0.59804 | 0.84126 | 0.00000 | 0.16227 | 0.00000 | 0.00000 | 0.00000 | 0.85366 | 0.50415 | 0.62023 | 1.20105 |
| **TMEM199** | 0.00124 | 11.21150 | 9.36701 | 6.10384 | 8.55279 | 12.39299 | 8.82966 | 7.14997 | 10.35384 | 0.00000 | 8.76443 | 12.93449 |
| **TMEM2** | 12.13150 | 0.26031 | 12.62132 | 1.57313 | 1.56649 | 9.24023 | 0.00001 | 11.48988 | 7.75505 | 7.48757 | 0.85989 | 1.33660 |
| **TMEM200A** | 2.91010 | 0.02157 | 1.70525 | 0.00779 | 0.00451 | 0.07532 | 0.00332 | 0.03777 | 0.08324 | 0.00199 | 0.00048 | 0.00308 |
| **TMEM200B** | 0.15759 | 0.58653 | 0.00574 | 0.05588 | 0.74487 | 0.00000 | 0.00598 | 0.05699 | 0.00000 | 0.18659 | 0.00000 | 0.59748 |
| **TMEM200C** | 0.00637 | 0.01006 | 0.01572 | 0.01798 | 0.02211 | 0.00000 | 0.01700 | 0.00620 | 0.03438 | 0.03645 | 0.01296 | 0.18242 |
| **TMEM201** | 3.94306 | 8.35132 | 4.16206 | 0.68392 | 7.66042 | 0.00000 | 0.40727 | 0.26617 | 6.34976 | 11.51070 | 1.88214 | 5.77541 |
| **TMEM202** | 0.00700 | 0.13727 | 0.00000 | 0.25976 | 0.01084 | 0.16472 | 0.02448 | 0.09114 | 0.02532 | 0.00000 | 0.00913 | 0.23138 |
| **TMEM203** | 0.66400 | 3.11950 | 1.71010 | 0.84408 | 4.68782 | 2.68234 | 3.18738 | 0.00000 | 2.42612 | 7.33190 | 1.60668 | 2.53118 |
| **TMEM204** | 0.00560 | 0.11351 | 0.01355 | 0.01287 | 0.00609 | 0.00000 | 0.00488 | 0.08188 | 0.04230 | 0.08840 | 0.09364 | 0.24662 |
| **TMEM205** | 0.43747 | 8.63796 | 5.14691 | 5.78079 | 12.08229 | 2.83982 | 3.62445 | 0.91723 | 14.27677 | 4.35494 | 6.49189 | 2.13212 |
| **TMEM206** | 1.53469 | 1.41671 | 5.52030 | 0.03201 | 2.72503 | 9.56865 | 0.00000 | 1.30765 | 11.91587 | 1.73031 | 3.52331 | 0.89262 |
| **TMEM207** | 0.02843 | 0.01070 | 0.00871 | 0.03412 | 0.07197 | 0.04072 | 0.01195 | 0.00690 | 0.00516 | 0.02923 | 0.01058 | 0.09112 |
| **TMEM208** | 1.29972 | 22.07684 | 20.08145 | 10.66030 | 15.16116 | 11.13033 | 12.33921 | 15.00613 | 23.28696 | 6.47842 | 14.49813 | 25.66780 |
| **TMEM209** | 2.99261 | 14.79948 | 1.84700 | 0.00000 | 1.58350 | 19.24110 | 0.00000 | 15.61320 | 12.87004 | 6.33982 | 7.86783 | 20.14016 |
| **TMEM210** | 0.01221 | 0.01449 | 0.02187 | 0.01022 | 0.01297 | 0.01323 | 0.02723 | 0.01213 | 0.01610 | 0.03972 | 0.01333 | 0.01665 |
| **TMEM211** | 0.00563 | 0.29083 | 0.00005 | 0.00000 | 0.30615 | 0.10251 | 0.00000 | 0.00960 | 0.00000 | 0.01921 | 0.00000 | 0.00141 |
| **TMEM212** | 0.01248 | 0.01488 | 0.01496 | 0.01205 | 0.01570 | 0.01877 | 0.01797 | 0.01094 | 0.01807 | 0.01463 | 0.01525 | 0.01609 |
| **TMEM213** | 0.00227 | 0.03478 | 0.00409 | 0.00000 | 0.00337 | 0.03091 | 0.00359 | 0.02385 | 0.02206 | 0.21268 | 0.00271 | 0.00536 |
| **TMEM214** | 1.50369 | 2.58811 | 6.79037 | 2.24255 | 6.31022 | 0.00000 | 0.33927 | 0.11543 | 8.19384 | 14.49651 | 1.06567 | 1.10614 |
| **TMEM215** | 0.10097 | 0.01007 | 0.01540 | 0.02603 | 0.00000 | 0.03580 | 0.06555 | 0.00486 | 0.00898 | 0.07895 | 0.02098 | 0.01735 |
| **TMEM216** | 2.70239 | 1.88683 | 1.71615 | 1.23322 | 2.22844 | 4.75102 | 0.93568 | 0.73920 | 4.60033 | 0.00000 | 1.57039 | 0.66791 |
| **TMEM217** | 0.01358 | 0.01545 | 0.24405 | 0.03752 | 0.00000 | 0.00974 | 0.21413 | 0.01950 | 0.21181 | 0.05419 | 0.03452 | 0.10889 |
| **TMEM218** | 1.49170 | 2.51521 | 2.77391 | 0.45826 | 0.76847 | 0.78851 | 2.42467 | 0.41155 | 0.19730 | 0.00000 | 0.75807 | 4.80984 |
| **TMEM219** | 1.75210 | 6.49383 | 7.79215 | 7.11476 | 6.62859 | 0.00000 | 5.17409 | 0.07037 | 7.77572 | 0.00000 | 4.43636 | 4.98906 |
| **TMEM220** | 0.00058 | 1.93608 | 0.28839 | 0.85905 | 0.29744 | 0.00000 | 0.00000 | 0.04362 | 0.44065 | 0.87338 | 0.37036 | 2.54218 |
| **TMEM220-AS1** | 0.00595 | 0.04618 | 0.01974 | 0.03333 | 0.03146 | 0.00147 | 0.01354 | 0.01006 | 0.02361 | 0.02265 | 0.02522 | 0.04546 |
| **TMEM221** | 0.10760 | 0.03666 | 0.14226 | 0.03122 | 0.01543 | 0.00000 | 0.00551 | 0.00766 | 0.02216 | 0.01961 | 0.02584 | 0.02366 |
| **TMEM222** | 0.56055 | 9.26015 | 11.69069 | 7.75334 | 8.88342 | 0.00000 | 10.30720 | 2.70554 | 16.25819 | 2.52599 | 8.70505 | 8.49530 |
| **TMEM223** | 0.12931 | 3.39607 | 8.07719 | 5.65524 | 5.32275 | 5.41703 | 6.86047 | 0.42537 | 4.19728 | 0.00000 | 9.60976 | 5.18045 |
| **TMEM225** | 0.00609 | 0.01146 | 0.01614 | 0.01243 | 0.02932 | 0.01589 | 0.01493 | 0.01028 | 0.01365 | 0.01453 | 0.05516 | 0.02108 |
| **TMEM229A** | 0.00456 | 0.02266 | 0.00767 | 0.01605 | 0.00000 | 0.01616 | 0.02386 | 0.01117 | 0.01830 | 0.10601 | 0.01689 | 0.01186 |
| **TMEM229B** | 6.07096 | 2.01944 | 0.00000 | 1.43157 | 0.00000 | 0.21462 | 0.00000 | 0.00000 | 0.00000 | 0.04292 | 0.06393 | 6.20302 |
| **TMEM230** | 2.35347 | 9.44651 | 7.40625 | 5.47846 | 3.25888 | 22.26935 | 5.96164 | 0.72339 | 7.79806 | 0.01008 | 10.69842 | 10.68326 |
| **TMEM231** | 1.16990 | 0.19126 | 0.00000 | 0.00000 | 0.22479 | 0.70339 | 0.00000 | 0.51744 | 0.00000 | 0.39981 | 0.01741 | 0.27468 |
| **TMEM232** | 0.98420 | 0.05908 | 0.22884 | 0.00000 | 0.03284 | 0.17514 | 0.00000 | 0.28602 | 0.00925 | 0.02109 | 0.00041 | 0.06115 |
| **TMEM234** | 0.00389 | 1.48398 | 2.69556 | 1.62925 | 3.14211 | 0.71251 | 0.04610 | 1.79909 | 1.53116 | 0.00000 | 1.05867 | 1.44047 |
| **TMEM235** | 0.01222 | 0.01603 | 0.01715 | 0.01250 | 0.01513 | 0.01338 | 0.01666 | 0.00933 | 0.01823 | 0.01362 | 0.02790 | 0.01681 |
| **TMEM236.uid-1** | 0.00548 | 0.01594 | 0.01586 | 0.00990 | 0.01131 | 0.03408 | 0.01172 | 0.01496 | 0.01552 | 0.03490 | 0.01362 | 0.01689 |
| **TMEM236.uid-1** | 0.01199 | 0.01255 | 0.01004 | 0.01113 | 0.01104 | 0.02426 | 0.04544 | 0.03136 | 0.04384 | 0.02029 | 0.01357 | 0.02222 |
| **TMEM237** | 0.89387 | 3.33225 | 2.11814 | 0.75176 | 0.16016 | 9.10750 | 0.00000 | 0.28146 | 7.14905 | 0.73293 | 0.73235 | 3.17035 |
| **TMEM238** | 0.00529 | 0.03457 | 0.04287 | 0.00613 | 0.00970 | 0.01851 | 0.03289 | 0.01890 | 0.00695 | 0.02840 | 0.02473 | 0.01698 |
| **TMEM240** | 0.00547 | 0.01370 | 0.01030 | 0.06200 | 0.00880 | 0.00728 | 0.01633 | 0.01149 | 0.01861 | 0.03998 | 0.01486 | 0.01567 |
| **TMEM241** | 0.53670 | 0.27190 | 2.35057 | 0.21718 | 0.88390 | 0.84084 | 1.03755 | 0.00000 | 2.23202 | 0.00004 | 2.29230 | 1.64399 |
| **TMEM242** | 1.58358 | 5.87516 | 1.32130 | 1.42748 | 2.90259 | 11.10381 | 2.23293 | 2.29874 | 4.75863 | 1.76389 | 3.11500 | 5.93590 |
| **TMEM243** | 0.75341 | 0.74672 | 6.53912 | 0.11040 | 0.00062 | 0.00602 | 0.31414 | 0.55401 | 2.36877 | 0.00000 | 1.02659 | 1.14620 |
| **TMEM245** | 2.88070 | 0.00000 | 8.45669 | 3.28148 | 3.66188 | 12.87169 | 4.24678 | 1.20189 | 14.01532 | 23.98435 | 7.32513 | 4.69179 |

| | | | | | | | | | | | | |
|---|---|---|---|---|---|---|---|---|---|---|---|---|
| **TMEM246** | 0.00651 | 0.54535 | 0.11010 | 0.32173 | 0.00000 | 0.03792 | 0.00000 | 0.01060 | 0.06438 | 0.00179 | 0.00723 | 0.00613 |
| **TMEM247** | 0.00611 | 0.17269 | 1.56072 | 0.00716 | 0.00083 | 0.00000 | 1.00981 | 0.18339 | 0.00000 | 0.00057 | 0.20073 | 0.13343 |
| **TMEM248** | 12.72694 | 7.57217 | 7.71906 | 7.68639 | 7.70609 | 8.18261 | 7.88527 | 3.81794 | 8.17598 | 20.28105 | 1.34350 | 3.81997 |
| **TMEM249** | 0.22810 | 0.02486 | 0.16697 | 0.03381 | 0.00000 | 0.05326 | 0.00000 | 0.00000 | 0.09858 | 0.15185 | 0.00386 | 0.07138 |
| **TMEM25** | 2.18097 | 0.62742 | 0.00000 | 0.57550 | 0.58258 | 1.24723 | 0.00000 | 0.45182 | 1.11324 | 0.00000 | 0.79216 | 0.37018 |
| **TMEM251** | 0.68761 | 3.99018 | 4.76232 | 2.02914 | 0.77534 | 10.47189 | 2.99168 | 3.36023 | 5.19216 | 0.49248 | 6.60969 | 10.40256 |
| **TMEM252** | 1.90821 | 0.03074 | 0.01208 | 0.00936 | 0.00423 | 0.00000 | 0.01935 | 1.35427 | 0.00000 | 0.02587 | 0.06810 | 0.01991 |
| **TMEM253** | 0.33869 | 0.01942 | 0.07409 | 0.14431 | 0.00000 | 0.00494 | 0.00675 | 0.05893 | 0.06678 | 0.03107 | 0.02988 | 0.02047 |
| **TMEM254** | 0.39878 | 4.30867 | 3.03933 | 0.59308 | 0.29088 | 5.61512 | 0.00000 | 0.00000 | 6.42151 | 0.73460 | 1.71071 | 4.88109 |
| **TMEM254-AS1** | 0.16289 | 0.69179 | 1.98733 | 0.00000 | 0.00000 | 0.20419 | 1.76355 | 11.15585 | 2.85722 | 0.00000 | 0.10086 | 1.33654 |
| **TMEM255A** | 0.00560 | 0.00013 | 0.00000 | 0.00103 | 0.05981 | 0.29604 | 0.00000 | 0.00873 | 0.00000 | 0.01131 | 0.00255 | 0.00967 |
| **TMEM255B** | 0.94454 | 0.03593 | 0.00005 | 0.03592 | 0.11938 | 0.07653 | 0.00000 | 0.09960 | 0.03584 | 0.13159 | 0.05308 | 0.04978 |
| **TMEM256** | 4.08651 | 3.32421 | 6.14501 | 6.08649 | 6.72135 | 0.12935 | 6.22867 | 2.71308 | 3.14582 | 3.85999 | 7.00801 | 2.10013 |
| **TMEM256-PLSC** | 0.00463 | 0.01889 | 0.01383 | 0.02511 | 0.02179 | 0.01349 | 0.01583 | 0.01759 | 0.02519 | 0.00708 | 0.02321 | 0.01978 |
| **TMEM257** | 0.01918 | 0.00627 | 0.01957 | 0.11474 | 0.00095 | 0.06689 | 0.00630 | 0.00806 | 0.00595 | 0.02388 | 0.01229 | 0.03199 |
| **TMEM258** | 3.09575 | 15.15145 | 18.22821 | 17.51600 | 23.55364 | 18.71861 | 25.98699 | 3.40791 | 16.45330 | 9.90257 | 19.70287 | 14.08981 |
| **TMEM259** | 0.00143 | 3.25952 | 3.93109 | 2.88266 | 4.79746 | 0.00000 | 0.00000 | 0.01206 | 6.28643 | 16.35420 | 0.63845 | 1.22236 |
| **TMEM26** | 0.00696 | 0.01375 | 0.01561 | 0.02350 | 0.00766 | 0.00871 | 0.02261 | 0.01027 | 0.00000 | 0.02709 | 0.06871 | 0.01641 |
| **TMEM260** | 0.03389 | 0.07825 | 0.87622 | 1.64056 | 2.58857 | 2.80642 | 0.00000 | 0.25016 | 1.46188 | 5.84044 | 0.90478 | 0.53319 |
| **TMEM261** | 0.06973 | 17.91800 | 5.48979 | 14.45491 | 20.47382 | 11.58456 | 14.20264 | 0.03839 | 12.34260 | 5.78195 | 19.05626 | 14.37927 |
| **TMEM262** | 0.00715 | 0.01100 | 0.02812 | 0.01804 | 0.04874 | 0.05850 | 0.01056 | 0.01002 | 0.05451 | 0.00000 | 0.01107 | 0.01863 |
| **TMEM263** | 0.72885 | 1.99827 | 0.13954 | 1.77310 | 0.00000 | 1.80534 | 1.50304 | 2.04618 | 0.00070 | 0.15828 | 1.02834 | 1.03372 |
| **TMEM27** | 0.17103 | 0.40036 | 0.52650 | 0.15719 | 0.00994 | 0.32899 | 0.00000 | 0.21501 | 0.17867 | 0.00000 | 0.12117 | 0.02589 |
| **TMEM30A** | 0.00000 | 8.16661 | 10.15233 | 5.71132 | 2.81259 | 29.81395 | 0.00000 | 13.01202 | 23.85526 | 5.92463 | 10.83756 | 12.85523 |
| **TMEM30B** | 0.32310 | 0.06165 | 4.57163 | 0.09684 | 0.00000 | 0.32038 | 0.15132 | 0.01081 | 0.74781 | 0.03285 | 0.09360 | 0.00000 |
| **TMEM30C** | 0.00591 | 0.01084 | 0.01369 | 0.02061 | 0.03449 | 0.02566 | 0.01222 | 0.01040 | 0.02392 | 0.02542 | 0.01422 | 0.01504 |
| **TMEM31** | 0.19744 | 0.00000 | 0.00000 | 0.00354 | 1.76849 | 1.66534 | 0.00000 | 0.95691 | 0.40338 | 0.00006 | 0.20948 | 0.05366 |
| **TMEM33** | 0.00000 | 17.04972 | 15.60358 | 0.69235 | 4.70267 | 48.04196 | 4.38441 | 8.57510 | 36.64625 | 21.78779 | 21.32807 | 24.66473 |
| **TMEM37** | 0.00038 | 15.59405 | 0.00000 | 8.25027 | 23.08538 | 2.54421 | 0.00000 | 0.64562 | 0.00000 | 0.01202 | 5.17997 | 10.57348 |
| **TMEM38A** | 0.08923 | 0.02550 | 0.03988 | 0.49071 | 0.02249 | 0.00000 | 0.00549 | 0.00412 | 0.11393 | 0.06977 | 0.11558 | 0.03449 |
| **TMEM38B** | 1.97234 | 9.73687 | 5.82397 | 2.91394 | 2.39427 | 23.24725 | 0.00000 | 0.98843 | 16.86823 | 2.32145 | 5.01779 | 4.96262 |
| **TMEM39A** | 0.44443 | 6.98729 | 10.84148 | 2.65029 | 3.17620 | 11.94211 | 0.05320 | 3.75817 | 10.22724 | 0.00000 | 4.52435 | 8.97991 |
| **TMEM39B** | 0.97641 | 4.39982 | 2.82325 | 4.84713 | 4.55783 | 0.00000 | 3.08783 | 0.96927 | 7.27195 | 0.00000 | 1.23074 | 2.54706 |
| **TMEM40** | 0.00793 | 0.28428 | 1.07074 | 0.00000 | 0.07990 | 0.00000 | 0.00000 | 0.06255 | 0.00000 | 0.03081 | 0.09506 | 0.08926 |
| **TMEM41A** | 4.43149 | 12.50937 | 9.40667 | 6.55143 | 13.69226 | 16.71060 | 0.00000 | 1.78277 | 14.81869 | 18.27344 | 9.15327 | 8.96264 |
| **TMEM41B** | 2.16447 | 5.70586 | 12.94905 | 9.59149 | 0.94737 | 38.80577 | 0.00000 | 20.28201 | 23.48589 | 21.44466 | 10.94292 | 17.73643 |
| **TMEM42** | 0.33842 | 0.18165 | 1.05205 | 1.01407 | 0.00000 | 0.00598 | 0.72490 | 0.23150 | 0.01978 | 0.08728 | 1.19551 | 0.98372 |
| **TMEM43** | 3.40272 | 8.94205 | 8.56102 | 4.44092 | 3.62125 | 6.50259 | 1.85041 | 0.18396 | 23.96904 | 3.57255 | 10.11069 | 5.82735 |
| **TMEM44** | 0.07574 | 0.41913 | 1.42227 | 0.11444 | 0.45165 | 0.10908 | 0.65720 | 0.27461 | 0.52214 | 0.17394 | 0.21031 | 0.13193 |
| **TMEM44-AS1** | 0.52746 | 0.22515 | 0.31250 | 0.00180 | 0.25035 | 0.88017 | 0.00000 | 0.09872 | 0.00000 | 0.29311 | 0.65869 | 0.90076 |
| **TMEM45A** | 0.08818 | 0.25028 | 0.24704 | 0.35449 | 0.00000 | 0.13132 | 0.00000 | 0.03360 | 0.23587 | 0.00000 | 0.00883 | 0.10797 |
| **TMEM45B** | 0.00544 | 10.67498 | 0.00000 | 0.00000 | 3.33086 | 2.40550 | 0.00000 | 0.00000 | 7.05911 | 0.00000 | 2.60247 | 4.11090 |
| **TMEM47** | 0.00306 | 0.06313 | 0.11287 | 0.00000 | 0.00000 | 0.04867 | 0.00000 | 0.01658 | 0.00000 | 0.18297 | 0.00820 | 0.10701 |
| **TMEM5** | 0.12253 | 3.50447 | 3.20919 | 0.53815 | 1.91200 | 6.69059 | 1.08425 | 0.00000 | 1.46501 | 7.29477 | 2.56404 | 3.36657 |
| **TMEM50A** | 3.19309 | 15.11556 | 16.49335 | 8.94387 | 7.86376 | 36.89701 | 0.00000 | 10.49020 | 27.97253 | 9.18257 | 9.44525 | 20.94537 |
| **TMEM50B** | 7.28354 | 5.01049 | 1.71226 | 1.73360 | 4.78080 | 7.47694 | 0.00000 | 6.83085 | 4.81922 | 0.70940 | 2.10592 | 6.39149 |
| **TMEM51** | 1.28898 | 6.86653 | 0.56530 | 5.57154 | 0.52893 | 0.00000 | 0.00000 | 0.00000 | 1.11788 | 3.97866 | 0.00000 | 1.77389 |
| **TMEM51-AS1** | 0.03927 | 0.13649 | 0.01870 | 0.25094 | 0.05068 | 0.00000 | 0.00201 | 0.00819 | 0.07185 | 0.12731 | 0.00000 | 0.02102 |

| | | | | | | | | | | | | |
|---|---|---|---|---|---|---|---|---|---|---|---|---|
| TMEM52 | 1.94701 | 0.56032 | 2.72755 | 0.23181 | 1.54480 | 0.00000 | 2.47557 | 2.13841 | 2.40318 | 0.00005 | 1.33326 | 0.16483 |
| TMEM52B | 0.00379 | 0.29057 | 3.82985 | 0.00000 | 1.16276 | 0.02156 | 3.98762 | 0.00000 | 5.73403 | 0.43729 | 1.33132 | 0.37136 |
| TMEM53 | 0.69272 | 0.77391 | 0.69639 | 1.60918 | 0.33531 | 1.61885 | 0.25318 | 0.24816 | 1.28297 | 0.06104 | 0.00000 | 0.33460 |
| TMEM54 | 0.54526 | 3.75909 | 9.04890 | 0.00000 | 22.85447 | 0.00000 | 4.56658 | 0.66144 | 0.00000 | 11.27493 | 1.36889 | 4.08805 |
| TMEM55A | 0.01953 | 0.00000 | 2.11914 | 0.27458 | 0.00000 | 0.80333 | 4.04933 | 0.31328 | 1.07573 | 0.00000 | 0.00000 | 0.11258 |
| TMEM55B | 0.35319 | 1.06155 | 3.29722 | 4.22034 | 4.67294 | 0.80613 | 4.54318 | 4.25410 | 1.00715 | 8.39317 | 0.45557 | 2.51470 |
| TMEM56 | 0.77028 | 11.56149 | 0.00000 | 0.00000 | 4.40001 | 4.57994 | 0.11190 | 0.00000 | 4.17826 | 8.92331 | 3.30841 | 7.12487 |
| TMEM56-RWDD | 0.00958 | 0.29730 | 0.00587 | 0.00000 | 0.07486 | 0.30569 | 0.00000 | 0.02929 | 0.03698 | 0.02208 | 0.04639 | 0.11256 |
| TMEM57 | 3.18992 | 0.00000 | 5.60348 | 0.72215 | 1.69169 | 1.05209 | 1.26583 | 0.18742 | 2.98875 | 7.27132 | 0.00000 | 0.17144 |
| TMEM59 | 7.06082 | 8.79053 | 16.60465 | 9.87337 | 5.93756 | 36.15420 | 0.00000 | 14.56129 | 29.31302 | 4.90592 | 12.95217 | 12.49416 |
| TMEM60 | 5.30310 | 1.07265 | 2.87466 | 1.14999 | 0.89123 | 5.39865 | 0.83450 | 1.35920 | 1.39411 | 2.50295 | 0.67707 | 0.44614 |
| TMEM61 | 0.59757 | 0.02176 | 0.00888 | 0.03039 | 0.00970 | 0.01795 | 0.00478 | 0.12091 | 0.03213 | 0.03845 | 0.01581 | 0.06348 |
| TMEM62 | 0.11099 | 3.04781 | 13.59808 | 1.68133 | 5.90050 | 3.09714 | 5.04039 | 0.41328 | 10.55056 | 4.97284 | 4.20918 | 1.03492 |
| TMEM63A | 1.15701 | 2.36982 | 3.48498 | 9.03176 | 2.25061 | 0.00000 | 0.00000 | 0.00000 | 4.05464 | 3.49159 | 0.18221 | 0.54578 |
| TMEM63B | 1.89997 | 0.22190 | 1.36438 | 0.82770 | 1.62087 | 0.00000 | 0.05949 | 0.24891 | 2.69656 | 4.64627 | 1.90725 | 0.67412 |
| TMEM63C | 1.01416 | 0.22116 | 0.00271 | 0.00000 | 0.00000 | 0.02994 | 0.01846 | 0.08693 | 0.00000 | 0.26467 | 0.00296 | 0.00729 |
| TMEM64 | 0.10523 | 0.00000 | 0.70563 | 0.00000 | 1.96734 | 12.11142 | 1.39763 | 0.21706 | 7.80389 | 5.41645 | 6.11923 | 1.57832 |
| TMEM65 | 0.03653 | 0.00000 | 0.00000 | 0.00000 | 0.31401 | 2.58334 | 1.25405 | 0.26366 | 0.00000 | 6.64132 | 2.00812 | 1.56003 |
| TMEM66 | 1.55189 | 17.05868 | 8.85854 | 6.86667 | 7.69394 | 34.00254 | 0.00000 | 8.40862 | 22.49328 | 10.76178 | 7.64553 | 22.17233 |
| TMEM67 | 0.44107 | 0.38851 | 0.44468 | 0.40235 | 0.00000 | 1.79085 | 0.00000 | 0.60085 | 1.15607 | 0.00000 | 0.58603 | 1.47808 |
| TMEM68 | 0.14863 | 6.41614 | 12.56222 | 1.07251 | 0.00000 | 16.60573 | 0.00000 | 6.81045 | 14.97120 | 2.02502 | 4.50793 | 16.39998 |
| TMEM69 | 4.55422 | 11.46024 | 9.24258 | 4.90228 | 5.06079 | 22.28061 | 5.13332 | 1.01777 | 17.57001 | 0.00000 | 12.90384 | 13.00283 |
| TMEM70 | 0.75341 | 14.81371 | 19.00957 | 13.24712 | 9.61537 | 31.72434 | 22.99546 | 7.88937 | 27.19315 | 3.25361 | 16.19721 | 19.15411 |
| TMEM71 | 0.03348 | 0.00636 | 0.06213 | 0.03080 | 0.00120 | 0.05025 | 0.02942 | 0.01897 | 0.04424 | 0.01996 | 0.00599 | 0.04715 |
| TMEM72-AS1 | 0.00593 | 0.01514 | 0.01677 | 0.01459 | 0.03547 | 0.00000 | 0.04305 | 0.01065 | 0.01498 | 0.02046 | 0.03866 | 0.02072 |
| TMEM74 | 1.36764 | 0.04489 | 0.00000 | 0.00972 | 0.00146 | 0.46181 | 0.00000 | 1.22601 | 0.70172 | 0.00000 | 0.08607 | 1.84438 |
| TMEM74B | 0.00296 | 0.22116 | 0.06225 | 0.41579 | 0.11367 | 0.00000 | 0.00322 | 0.07417 | 0.27391 | 0.00000 | 0.04897 | 2.85201 |
| TMEM75 | 0.00222 | 0.07019 | 0.01183 | 0.00000 | 0.15294 | 0.00000 | 0.01314 | 0.00816 | 0.35327 | 0.71682 | 0.00000 | 0.03223 |
| TMEM79 | 0.96654 | 1.48988 | 6.14425 | 0.00494 | 1.65464 | 0.08047 | 1.63250 | 1.21873 | 4.36529 | 0.00000 | 5.91243 | 2.93643 |
| TMEM80 | 1.17016 | 1.27911 | 2.43613 | 0.00000 | 0.00000 | 0.00000 | 0.87342 | 0.10826 | 2.49939 | 1.41236 | 4.52954 | 0.63642 |
| TMEM81 | 0.00300 | 0.01383 | 0.29354 | 0.17684 | 0.00000 | 0.19765 | 0.44455 | 0.67894 | 0.48340 | 0.93117 | 0.18164 | 0.23652 |
| TMEM82 | 0.00256 | 0.08859 | 0.02389 | 0.05275 | 0.00000 | 0.07381 | 0.00000 | 0.02675 | 0.00000 | 0.05438 | 0.03619 | 0.00745 |
| TMEM86A | 0.81914 | 0.61911 | 0.45931 | 5.62708 | 6.14357 | 0.00059 | 0.00000 | 0.06883 | 0.00000 | 0.00000 | 0.27745 | 0.45550 |
| TMEM86B | 0.02946 | 0.27362 | 0.35020 | 0.00094 | 0.00000 | 0.00000 | 0.37865 | 0.06079 | 0.91902 | 0.98889 | 0.00000 | 0.05816 |
| TMEM87A | 2.53483 | 8.37630 | 14.36162 | 5.21475 | 1.51004 | 20.84767 | 0.00000 | 8.84643 | 20.96400 | 15.24951 | 4.79377 | 15.79185 |
| TMEM87B | 0.05585 | 2.07542 | 0.32141 | 1.71049 | 1.71365 | 7.99916 | 0.00000 | 0.10225 | 3.05084 | 2.22480 | 0.00000 | 2.17911 |
| TMEM88 | 0.00675 | 0.00000 | 2.21701 | 1.30927 | 0.00000 | 0.00000 | 0.43548 | 0.35468 | 3.15388 | 0.00616 | 2.13825 | 0.07162 |
| TMEM89 | 0.00938 | 0.01455 | 0.06678 | 0.00519 | 0.01027 | 0.01322 | 0.04159 | 0.01362 | 0.04773 | 0.09883 | 0.01137 | 0.03336 |
| TMEM8A | 1.21920 | 2.00069 | 3.08480 | 6.81720 | 6.97045 | 0.00000 | 1.15728 | 0.00000 | 0.00000 | 6.94632 | 0.74792 | 1.46215 |
| TMEM8B | 0.77820 | 0.14418 | 0.08580 | 0.06160 | 0.01947 | 0.00000 | 0.00000 | 0.07023 | 0.06859 | 0.02272 | 0.03296 | 0.12082 |
| TMEM9 | 10.47669 | 13.07658 | 6.62257 | 6.40764 | 5.99179 | 9.80080 | 2.37943 | 6.81258 | 22.73765 | 0.00000 | 7.68100 | 9.59245 |
| TMEM91 | 0.00554 | 0.18661 | 0.09214 | 0.00000 | 0.07118 | 0.02161 | 0.19724 | 0.17610 | 0.00000 | 0.39626 | 0.32359 | 0.51787 |
| TMEM92 | 0.00315 | 2.62899 | 0.14013 | 2.36608 | 0.41491 | 0.00000 | 0.14694 | 15.01726 | 0.38119 | 0.00000 | 0.00000 | 20.68014 |
| TMEM95 | 0.00349 | 0.01436 | 0.05498 | 0.00846 | 0.01367 | 0.01925 | 0.01078 | 0.02665 | 0.01251 | 0.02754 | 0.01638 | 0.01456 |
| TMEM97 | 10.09397 | 0.00000 | 18.29895 | 7.13945 | 21.68898 | 3.94581 | 13.80874 | 0.56695 | 39.27354 | 4.11198 | 23.40622 | 1.77594 |
| TMEM98 | 0.02060 | 0.01205 | 0.25644 | 0.19619 | 0.00000 | 0.09951 | 0.11594 | 0.00759 | 0.10912 | 0.00000 | 0.23730 | 0.06884 |
| TMEM99 | 0.56815 | 9.36778 | 3.50391 | 0.00000 | 3.27259 | 4.77025 | 3.01632 | 2.59732 | 4.65166 | 2.71635 | 3.11372 | 20.94613 |
| TMEM9B | 4.83425 | 14.18127 | 7.97879 | 6.48699 | 6.99335 | 26.03714 | 0.00000 | 4.00926 | 9.92733 | 5.40444 | 10.07825 | 11.06912 |

| | | | | | | | | | | | | |
|---|---|---|---|---|---|---|---|---|---|---|---|---|
| **TMEM9B-AS1** | 0.21527 | 0.00135 | 0.08610 | 0.09403 | 0.16803 | 0.19942 | 0.00000 | 0.08226 | 0.08762 | 0.16008 | 0.12253 | 0.07423 |
| **TMF1** | 4.13674 | 2.82022 | 10.39674 | 2.97341 | 2.03072 | 16.76406 | 10.26878 | 9.40320 | 5.83063 | 9.23772 | 7.38742 | 8.67579 |
| **TMIE** | 0.00598 | 0.01414 | 0.02432 | 0.01202 | 0.01782 | 0.02194 | 0.01425 | 0.01118 | 0.02594 | 0.01525 | 0.01426 | 0.01591 |
| **TMIGD2** | 0.00607 | 0.01251 | 0.13926 | 0.00953 | 0.01737 | 0.02396 | 0.02112 | 0.00957 | 0.00000 | 0.01246 | 0.03233 | 0.02710 |
| **TMLHE** | 0.00226 | 0.00000 | 0.52582 | 0.39858 | 0.00000 | 0.36500 | 0.26290 | 0.04949 | 1.67188 | 0.00000 | 1.65382 | 0.24907 |
| **TMOD1** | 1.15263 | 4.36593 | 0.00000 | 1.08349 | 6.79604 | 1.98481 | 0.00000 | 0.23390 | 0.00000 | 0.32541 | 4.63700 | 1.32585 |
| **TMOD2** | 1.32593 | 0.03018 | 0.02398 | 0.02600 | 0.01141 | 0.00758 | 0.01620 | 0.30392 | 0.05478 | 0.01853 | 0.01234 | 0.00000 |
| **TMOD3** | 2.09611 | 5.12142 | 6.60129 | 3.60519 | 5.90209 | 26.82735 | 11.03984 | 7.33336 | 6.27029 | 11.92788 | 9.14790 | 4.24224 |
| **TMOD4** | 0.00199 | 0.10497 | 0.14331 | 0.04623 | 0.00978 | 0.00000 | 0.00000 | 0.07413 | 0.08252 | 0.08299 | 0.22566 | 0.04656 |
| **TMPO** | 3.56984 | 17.15661 | 7.09759 | 10.07007 | 5.61206 | 33.18130 | 25.29540 | 12.89144 | 12.64825 | 23.30170 | 19.85390 | 16.65373 |
| **TMPO-AS1** | 0.58554 | 0.19743 | 0.23660 | 0.35745 | 0.00000 | 0.08642 | 0.00000 | 0.77410 | 0.04544 | 0.84812 | 0.16256 | 0.53171 |
| **TMPPE** | 0.63395 | 1.51548 | 0.04629 | 0.88321 | 1.47332 | 0.00000 | 0.00029 | 0.29125 | 2.19935 | 1.29946 | 2.61945 | 1.79021 |
| **TMPRSS11A** | 0.30676 | 0.01420 | 0.01627 | 0.01301 | 0.01491 | 0.01027 | 0.01495 | 0.55991 | 0.02005 | 0.01505 | 0.01436 | 0.00102 |
| **TMPRSS11B** | 0.00524 | 0.01286 | 0.01737 | 0.01373 | 0.01089 | 0.00739 | 0.01667 | 0.01027 | 0.02491 | 0.07747 | 0.02192 | 0.01609 |
| **TMPRSS11E** | 0.00353 | 0.08585 | 0.39845 | 1.57377 | 0.28424 | 0.28933 | 0.00000 | 0.00581 | 0.04858 | 0.09092 | 0.07063 | 0.00000 |
| **TMPRSS11F** | 0.00931 | 0.00449 | 0.08573 | 0.00438 | 0.03609 | 0.07278 | 0.05204 | 0.00912 | 0.00000 | 0.00000 | 0.01895 | 0.07385 |
| **TMPRSS12** | 0.00865 | 0.00968 | 0.27923 | 0.00223 | 0.05825 | 0.00000 | 0.16383 | 0.01159 | 0.02356 | 0.09075 | 0.01436 | 0.01298 |
| **TMPRSS13** | 0.05197 | 0.00000 | 0.64643 | 0.00000 | 8.98386 | 0.00000 | 5.70130 | 0.00480 | 20.08553 | 0.14871 | 7.99991 | 0.00000 |
| **TMPRSS15** | 0.17842 | 0.02641 | 0.05214 | 0.06963 | 0.00000 | 0.00000 | 0.03041 | 0.05052 | 0.05505 | 0.09387 | 0.11538 | 0.17171 |
| **TMPRSS2** | 0.00532 | 0.00000 | 0.77058 | 0.00000 | 17.58243 | 0.01505 | 1.23966 | 0.01404 | 17.09310 | 0.05341 | 13.03115 | 0.01984 |
| **TMPRSS3** | 0.00744 | 0.01774 | 0.00809 | 0.00986 | 0.07435 | 0.00000 | 0.04863 | 0.01016 | 0.00000 | 0.11136 | 0.08379 | 0.01229 |
| **TMPRSS4** | 0.00215 | 0.05627 | 0.00000 | 0.01049 | 0.14283 | 0.79742 | 0.00000 | 0.03776 | 5.29604 | 0.00333 | 0.00000 | 0.00000 |
| **TMPRSS4-AS1** | 0.00616 | 0.02295 | 0.01959 | 0.01199 | 0.01441 | 0.00597 | 0.02474 | 0.01067 | 0.00000 | 0.01740 | 0.07154 | 0.01610 |
| **TMPRSS5** | 0.08807 | 0.24448 | 0.03653 | 0.06463 | 0.00000 | 0.00000 | 0.00021 | 0.00892 | 0.09468 | 0.04628 | 0.04763 | 0.06169 |
| **TMPRSS6** | 0.12694 | 0.01628 | 0.03891 | 0.03266 | 0.00889 | 0.01070 | 0.01977 | 0.00093 | 0.00320 | 0.03932 | 0.04512 | 0.07425 |
| **TMPRSS7** | 0.03461 | 0.02504 | 0.01115 | 0.00859 | 0.01181 | 0.04943 | 0.00583 | 0.11906 | 0.08997 | 0.00000 | 0.02431 | 0.03400 |
| **TMPRSS9** | 0.05242 | 0.03102 | 0.04137 | 0.02227 | 0.04572 | 0.00000 | 0.02086 | 0.14697 | 0.01917 | 0.49955 | 0.04804 | 0.06242 |
| **TMSB10** | 1.63173 | 21.81401 | 50.50281 | 40.55795 | 26.94880 | 27.86940 | 63.41595 | 8.95635 | 36.49549 | 21.95453 | 16.14980 | 10.32669 |
| **TMSB15A** | 0.02714 | 0.25772 | 0.04924 | 0.00092 | 0.01539 | 0.31098 | 0.06389 | 0.43432 | 0.00000 | 0.00000 | 0.00000 | 0.80960 |
| **TMSB15B** | 0.01110 | 0.28354 | 0.08344 | 0.12830 | 0.02351 | 0.06021 | 0.12959 | 0.00346 | 0.04792 | 0.00000 | 0.00000 | 0.48655 |
| **TMSB4X** | 21.75065 | 0.00000 | 16.63462 | 24.07576 | 13.02376 | 21.58878 | 39.39972 | 24.15535 | 25.06657 | 2.75910 | 28.54120 | 10.40003 |
| **TMSB4Y** | 0.05557 | 0.32770 | 2.05456 | 0.01990 | 0.00000 | 0.36777 | 0.71401 | 0.12133 | 0.10908 | 0.00000 | 1.03283 | 0.10175 |
| **TMTC1** | 0.70162 | 0.00000 | 0.23250 | 0.50949 | 0.14726 | 0.52875 | 0.00000 | 0.72105 | 0.48606 | 0.00000 | 0.00000 | 0.03515 |
| **TMTC2** | 0.71317 | 0.14610 | 1.04558 | 0.31256 | 0.57111 | 0.00000 | 0.00000 | 0.41380 | 0.70783 | 0.78671 | 0.29977 | 0.57790 |
| **TMTC3** | 0.05091 | 4.72837 | 10.19774 | 0.00000 | 0.00000 | 41.52985 | 8.00648 | 1.11916 | 36.98752 | 23.72371 | 20.34132 | 7.24795 |
| **TMTC4** | 0.18569 | 1.99734 | 1.29654 | 1.36656 | 2.72827 | 5.86037 | 0.00000 | 0.00000 | 4.15669 | 0.32185 | 4.83628 | 0.00000 |
| **TMUB1** | 0.95397 | 1.30005 | 7.24508 | 0.91640 | 7.12667 | 0.00000 | 3.22619 | 0.00000 | 8.85283 | 5.06874 | 1.86359 | 0.35355 |
| **TMUB2** | 1.52723 | 4.13215 | 10.51114 | 4.38011 | 5.97005 | 0.00000 | 2.44598 | 11.34057 | 11.07909 | 0.94265 | 3.15117 | 10.98461 |
| **TMX1** | 0.67156 | 7.95918 | 16.33566 | 9.27197 | 1.05622 | 55.87329 | 0.00000 | 9.54201 | 30.63271 | 10.99369 | 14.76254 | 20.02853 |
| **TMX2** | 2.34643 | 24.29179 | 18.61811 | 12.31604 | 11.33144 | 29.08180 | 3.07154 | 7.58459 | 37.26783 | 11.94981 | 16.58239 | 23.89133 |
| **TMX3** | 4.62606 | 2.20535 | 7.74689 | 0.39122 | 0.06847 | 11.83700 | 0.00000 | 2.95932 | 8.96840 | 7.38338 | 0.47741 | 3.39792 |
| **TMX4** | 1.91783 | 1.56799 | 0.51262 | 6.91852 | 0.61099 | 3.15530 | 0.00000 | 0.04610 | 0.29021 | 2.58046 | 0.59388 | 0.02463 |
| **TNC** | 0.00261 | 0.01085 | 0.03775 | 0.00000 | 0.05055 | 0.13853 | 0.01152 | 0.02401 | 0.00083 | 0.05120 | 0.00876 | 0.00850 |
| **TNF** | 0.00315 | 0.04896 | 0.00253 | 0.01999 | 0.00000 | 0.00663 | 0.00946 | 0.00766 | 0.00000 | 0.28868 | 0.10158 | 0.06732 |
| **TNFAIP1** | 2.28814 | 10.47789 | 8.59636 | 2.95900 | 4.94586 | 5.19827 | 3.21386 | 10.32599 | 6.43048 | 0.00000 | 3.00210 | 12.32481 |
| **TNFAIP2** | 0.55499 | 0.00618 | 6.07844 | 0.00000 | 0.00000 | 0.00758 | 1.59599 | 0.03223 | 0.40111 | 0.00000 | 0.00000 | 0.01667 |
| **TNFAIP3** | 0.01450 | 2.01984 | 11.14639 | 2.39151 | 0.00000 | 0.00000 | 0.00000 | 0.00000 | 0.35423 | 1.10381 | 0.43000 | 0.64797 |
| **TNFAIP6** | 0.00213 | 0.00693 | 0.06400 | 0.00368 | 0.05608 | 0.00001 | 0.00000 | 0.25316 | 0.02953 | 0.17790 | 0.00048 | 0.03601 |

| | | | | | | | | | | | | |
|---|---|---|---|---|---|---|---|---|---|---|---|---|
| **TNFAIP8** | 0.38855 | 0.27575 | 11.27872 | 0.86487 | 0.00000 | 3.84155 | 11.77499 | 0.29788 | 4.40369 | 0.00000 | 3.09533 | 0.10269 |
| **TNFAIP8L1** | 0.00000 | 0.71228 | 0.25839 | 0.21921 | 0.22867 | 0.00000 | 1.02592 | 1.01812 | 1.00139 | 1.41428 | 1.00640 | 0.45361 |
| **TNFAIP8L2** | 0.00192 | 0.06371 | 0.24361 | 0.00021 | 0.00015 | 0.12665 | 0.00000 | 0.11226 | 0.08088 | 0.05030 | 0.02388 | 0.06830 |
| **TNFAIP8L3** | 0.01767 | 0.10460 | 0.02163 | 0.00000 | 0.07342 | 0.00000 | 0.00000 | 0.00492 | 0.06624 | 0.31597 | 0.00933 | 0.10089 |
| **TNFRSF10A** | 0.00595 | 0.08267 | 0.10657 | 0.03501 | 0.06248 | 0.03406 | 0.00203 | 0.01153 | 0.03512 | 0.00000 | 0.03709 | 0.01391 |
| **TNFRSF10B** | 1.58109 | 1.14449 | 5.06671 | 1.04869 | 1.54212 | 1.37379 | 0.00000 | 0.19033 | 5.96247 | 0.30103 | 0.00000 | 0.39419 |
| **TNFRSF10C** | 0.00488 | 0.04540 | 0.26321 | 0.02225 | 0.31727 | 0.00000 | 0.00000 | 0.01012 | 0.26865 | 0.09361 | 0.00000 | 0.01109 |
| **TNFRSF10D** | 0.33179 | 0.54611 | 0.42068 | 0.53501 | 0.10912 | 0.00000 | 0.00000 | 0.86403 | 0.48805 | 0.00000 | 0.00000 | 0.12192 |
| **TNFRSF11A** | 0.23587 | 0.03331 | 0.00000 | 0.22598 | 0.00000 | 0.10060 | 0.00000 | 0.00000 | 0.00000 | 0.08911 | 0.00483 | 0.28512 |
| **TNFRSF11B** | 0.09115 | 0.01485 | 0.01216 | 0.01429 | 0.00983 | 0.02590 | 0.01442 | 0.01650 | 0.01658 | 0.02350 | 0.01369 | 0.01676 |
| **TNFRSF12A** | 1.88403 | 11.44681 | 11.15266 | 5.86641 | 19.08471 | 0.00000 | 3.78612 | 17.71748 | 19.48818 | 1.36918 | 4.11451 | 8.48948 |
| **TNFRSF13B** | 0.00538 | 0.01691 | 0.02990 | 0.03856 | 0.01682 | 0.00443 | 0.02083 | 0.01485 | 0.02162 | 0.00369 | 0.01903 | 0.01820 |
| **TNFRSF13C** | 0.06730 | 0.00635 | 0.01243 | 0.01406 | 0.03706 | 0.00506 | 0.01202 | 0.00606 | 0.01227 | 0.09900 | 0.01789 | 0.07155 |
| **TNFRSF14** | 0.00615 | 0.01534 | 0.01812 | 0.01352 | 0.01960 | 0.01293 | 0.03263 | 0.01057 | 0.00827 | 0.01762 | 0.01578 | 0.01639 |
| **TNFRSF18** | 0.00847 | 0.12579 | 0.24505 | 0.18092 | 0.01165 | 0.00000 | 0.02169 | 0.03797 | 0.00000 | 0.10021 | 0.01005 | 0.13910 |
| **TNFRSF19** | 0.00258 | 0.17381 | 0.97629 | 0.00000 | 0.00000 | 0.05001 | 0.02305 | 0.30647 | 0.00000 | 0.02266 | 0.04082 | 0.01467 |
| **TNFRSF1A** | 0.00203 | 1.90528 | 2.23949 | 10.48321 | 3.42412 | 0.00000 | 0.00494 | 0.05285 | 2.63398 | 0.02042 | 0.36023 | 0.00000 |
| **TNFRSF1B** | 0.71288 | 0.01444 | 2.52958 | 0.01135 | 0.00469 | 0.00000 | 0.12247 | 0.01248 | 0.00000 | 0.03987 | 0.01647 | 0.05161 |
| **TNFRSF21** | 0.80150 | 1.12125 | 15.13453 | 3.65080 | 0.00000 | 2.39282 | 4.72057 | 0.00000 | 13.60624 | 4.18413 | 0.53033 | 4.10710 |
| **TNFRSF25** | 0.15504 | 0.51808 | 0.03399 | 0.91916 | 0.03648 | 0.00000 | 0.00228 | 0.04792 | 0.09570 | 0.06711 | 0.01012 | 0.33786 |
| **TNFRSF4** | 0.00533 | 0.01103 | 0.01298 | 0.07180 | 0.00086 | 0.02305 | 0.01374 | 0.01009 | 0.01993 | 0.04936 | 0.01238 | 0.02037 |
| **TNFRSF6B** | 0.00613 | 0.01936 | 0.04446 | 0.03267 | 0.01410 | 0.00787 | 0.00981 | 0.01039 | 0.03268 | 0.01326 | 0.01422 | 0.02339 |
| **TNFRSF8** | 0.53770 | 1.61048 | 0.59537 | 2.31581 | 0.00000 | 0.00000 | 0.01283 | 0.00000 | 0.00000 | 0.01276 | 0.14081 | 1.73150 |
| **TNFRSF9** | 0.00743 | 0.01529 | 0.02927 | 0.01226 | 0.02174 | 0.01704 | 0.02151 | 0.01025 | 0.01510 | 0.01237 | 0.01748 | 0.01693 |
| **TNFSF10** | 0.00435 | 0.01124 | 0.00844 | 0.00366 | 0.15735 | 0.00000 | 0.02539 | 0.01082 | 0.01896 | 0.14682 | 0.00283 | 0.01517 |
| **TNFSF11** | 0.02919 | 0.01637 | 0.01042 | 0.02334 | 0.00000 | 0.03366 | 0.00406 | 0.02265 | 0.01487 | 0.08388 | 0.01112 | 0.00992 |
| **TNFSF12** | 0.09042 | 0.19914 | 0.64921 | 0.00000 | 0.88891 | 0.00000 | 1.34099 | 0.05203 | 1.64946 | 0.22404 | 0.14159 | 0.00000 |
| **TNFSF13** | 0.86212 | 0.02177 | 0.09755 | 0.00852 | 0.02392 | 0.01263 | 0.00773 | 0.05609 | 0.05609 | 0.00000 | 0.01262 | 0.08622 |
| **TNFSF13B** | 0.06320 | 0.37102 | 0.00000 | 0.12159 | 0.15826 | 4.37180 | 0.00000 | 0.00394 | 0.00000 | 2.97235 | 0.05663 | 0.00000 |
| **TNFSF14** | 0.00556 | 0.01654 | 0.04876 | 0.05370 | 0.02619 | 0.00000 | 0.09732 | 0.01177 | 0.00000 | 0.00821 | 0.01511 | 0.02591 |
| **TNFSF15** | 0.00548 | 0.00000 | 0.00000 | 0.17322 | 0.68146 | 0.23425 | 0.00000 | 0.00774 | 0.00000 | 0.00449 | 0.00000 | 0.00727 |
| **TNFSF4** | 0.01242 | 0.00196 | 0.12387 | 0.00071 | 0.05867 | 0.03232 | 0.01549 | 0.02189 | 0.00000 | 0.16036 | 0.01283 | 0.00904 |
| **TNFSF8** | 0.14370 | 0.01819 | 0.02383 | 0.01739 | 0.04797 | 0.00000 | 0.06971 | 0.00319 | 0.10273 | 0.00000 | 0.01584 | 0.01565 |
| **TNFSF9** | 0.22818 | 2.13334 | 0.00000 | 3.27892 | 1.12683 | 0.57656 | 0.00000 | 4.86235 | 0.00000 | 0.68031 | 0.00000 | 3.29122 |
| **TNIK** | 11.79699 | 0.00000 | 0.52728 | 0.59060 | 0.36750 | 0.00000 | 4.55966 | 1.17381 | 4.25083 | 2.10779 | 2.64931 | 0.05246 |
| **TNIP1** | 0.16826 | 0.93723 | 1.81543 | 1.63995 | 2.99869 | 0.11559 | 2.96333 | 0.00000 | 0.12690 | 3.68868 | 1.17380 | 0.84629 |
| **TNIP2** | 0.66563 | 4.23317 | 13.58986 | 2.86714 | 7.83063 | 0.00000 | 11.29921 | 1.68528 | 9.97566 | 9.48839 | 3.00284 | 1.51286 |
| **TNIP3** | 0.03508 | 0.01002 | 0.49943 | 1.34951 | 0.00000 | 0.00000 | 0.15346 | 0.00749 | 0.04230 | 0.04539 | 0.06760 | 0.04801 |
| **TNK1** | 0.12424 | 0.19621 | 0.54136 | 0.02767 | 0.39759 | 0.26428 | 0.00000 | 0.08443 | 0.85501 | 0.00000 | 0.11666 | 0.50965 |
| **TNK2** | 1.78692 | 0.71708 | 6.54622 | 4.32489 | 2.04727 | 0.00000 | 3.21969 | 0.25053 | 0.16832 | 11.94569 | 0.20496 | 0.78018 |
| **TNKS** | 0.43553 | 1.36118 | 6.31714 | 2.46781 | 2.07391 | 3.08149 | 3.45244 | 1.01681 | 1.99678 | 5.26827 | 1.60608 | 1.39476 |
| **TNKS1BP1** | 0.01269 | 1.73089 | 7.23516 | 2.37057 | 5.15722 | 0.00000 | 2.34240 | 1.07102 | 2.55149 | 9.27684 | 0.98288 | 0.96502 |
| **TNKS2** | 3.65586 | 0.70233 | 7.52166 | 0.62838 | 0.00884 | 12.34092 | 5.10405 | 0.99459 | 4.35545 | 10.35903 | 2.49925 | 2.92503 |
| **TNKS2-AS1** | 0.17016 | 0.16809 | 0.02907 | 0.01619 | 0.03182 | 0.20831 | 0.00000 | 0.09964 | 0.07552 | 0.00000 | 0.04491 | 0.37457 |
| **TNN** | 0.09607 | 0.02249 | 0.01296 | 0.00182 | 0.18463 | 0.00000 | 0.00941 | 0.01509 | 0.03879 | 0.19976 | 0.03699 | 0.02092 |
| **TNNC1** | 0.27739 | 0.22278 | 3.44459 | 0.45648 | 0.90125 | 0.00000 | 3.86280 | 0.20480 | 4.08230 | 0.27671 | 0.92496 | 0.00000 |
| **TNNC2** | 0.49396 | 0.33210 | 0.00000 | 1.07216 | 4.39663 | 0.00000 | 0.14777 | 1.03648 | 0.00000 | 0.30351 | 0.53212 | 0.15318 |
| **TNNI1** | 0.07776 | 0.00522 | 0.22787 | 0.01394 | 0.00000 | 0.07859 | 0.00000 | 0.01141 | 0.00000 | 0.05452 | 0.02864 | 0.00983 |

| | | | | | | | | | | | | |
|---|---|---|---|---|---|---|---|---|---|---|---|---|
| **TNNI2** | 0.00617 | 0.01520 | 0.14214 | 0.01427 | 0.01576 | 0.01408 | 0.01368 | 0.01053 | 0.01169 | 0.01765 | 0.01634 | 0.01614 |
| **TNNI3** | 0.02609 | 0.23611 | 0.00083 | 0.00075 | 0.00329 | 0.08311 | 0.00000 | 7.99408 | 0.05391 | 0.00642 | 0.01236 | 2.35262 |
| **TNNT1** | 0.99877 | 0.92924 | 0.00149 | 0.51194 | 1.37079 | 0.12446 | 0.00000 | 5.01628 | 0.00000 | 0.00852 | 0.00000 | 4.99223 |
| **TNNT2** | 0.05484 | 0.11666 | 0.04302 | 0.00000 | 0.41335 | 0.05102 | 0.43245 | 0.08936 | 0.44948 | 0.00000 | 0.10705 | 0.00548 |
| **TNNT3** | 0.00628 | 0.02421 | 0.01939 | 0.01261 | 0.01449 | 0.01527 | 0.01777 | 0.01054 | 0.02759 | 0.01358 | 0.01279 | 0.01564 |
| **TNPO1** | 8.54515 | 11.68543 | 16.89849 | 6.99761 | 8.01315 | 27.83276 | 31.14202 | 5.83026 | 17.30052 | 26.35082 | 17.36093 | 19.55220 |
| **TNPO2** | 3.26047 | 4.26181 | 5.34665 | 6.32106 | 11.80356 | 0.00000 | 8.05563 | 8.43686 | 5.30060 | 31.44359 | 9.52048 | 12.34277 |
| **TNPO3** | 13.05903 | 11.47697 | 14.75158 | 11.98926 | 12.36159 | 7.53557 | 19.08867 | 15.31696 | 15.01515 | 3.53168 | 14.18242 | 12.93010 |
| **TNR** | 0.00617 | 0.00617 | 0.15559 | 0.00698 | 0.12134 | 0.01879 | 0.00030 | 0.00991 | 0.00000 | 0.02534 | 0.15462 | 0.01985 |
| **TNRC18** | 0.07851 | 0.27460 | 1.21747 | 0.35068 | 2.06091 | 0.00000 | 1.70550 | 0.43463 | 0.51492 | 5.08472 | 0.00000 | 0.79046 |
| **TNRC18P1** | 0.00194 | 0.00346 | 0.09637 | 0.01335 | 0.02034 | 0.04354 | 0.03362 | 0.23590 | 0.00000 | 0.14102 | 0.04443 | 0.01975 |
| **TNRC6A** | 5.90750 | 2.46198 | 4.53884 | 2.62443 | 4.12097 | 4.65327 | 12.68706 | 4.16444 | 3.77234 | 25.77137 | 3.18485 | 7.03733 |
| **TNRC6B** | 6.80252 | 1.66753 | 2.41003 | 7.30890 | 4.02113 | 9.07014 | 6.79260 | 2.06110 | 5.03491 | 18.30101 | 7.29991 | 5.88955 |
| **TNRC6C** | 1.06510 | 0.70426 | 0.97792 | 5.24532 | 0.00000 | 0.31428 | 0.71170 | 0.71197 | 0.39441 | 4.71036 | 0.17770 | 1.07022 |
| **TNRC6C-AS1** | 0.00633 | 0.02070 | 0.01588 | 0.02296 | 0.03092 | 0.00000 | 0.03087 | 0.01075 | 0.03789 | 0.07216 | 0.00567 | 0.01395 |
| **TNS1** | 0.00382 | 0.64043 | 0.93724 | 0.00000 | 0.49125 | 0.00000 | 0.86914 | 0.00604 | 0.59058 | 1.18651 | 0.18070 | 0.03800 |
| **TNS3** | 0.24200 | 13.02742 | 9.59817 | 4.12354 | 14.14807 | 0.46288 | 8.43975 | 1.09146 | 7.07782 | 13.33468 | 7.71058 | 0.00000 |
| **TNS4** | 0.00610 | 0.01561 | 0.01969 | 0.02315 | 0.01895 | 0.01247 | 0.02458 | 0.01021 | 0.03361 | 0.01915 | 0.01173 | 0.02000 |
| **TNXB.uid-8455** | 0.00855 | 0.01448 | 0.02253 | 0.01034 | 0.01430 | 0.02320 | 0.03629 | 0.00980 | 0.02171 | 0.00435 | 0.01397 | 0.02193 |
| **TNXB.uid-8469** | 0.01337 | 0.55481 | 0.07614 | 0.01653 | 0.61337 | 0.00000 | 0.02223 | 0.01316 | 0.50246 | 0.00000 | 0.12023 | 0.19309 |
| **TOB1** | 4.15312 | 0.00687 | 0.95383 | 2.26010 | 0.00000 | 4.05165 | 4.06946 | 1.09958 | 4.03735 | 0.39777 | 2.58756 | 0.35922 |
| **TOB1-AS1** | 0.05555 | 0.01394 | 0.03505 | 0.03563 | 0.00508 | 0.04206 | 0.02118 | 0.00744 | 0.01554 | 0.02612 | 0.02066 | 0.01264 |
| **TOB2** | 2.51466 | 5.72368 | 6.45211 | 5.46503 | 10.92776 | 0.00000 | 15.24689 | 5.58608 | 13.41854 | 17.71314 | 7.10412 | 5.47016 |
| **TOB2P1** | 0.02465 | 0.02256 | 0.30021 | 0.00320 | 0.00000 | 0.01541 | 0.08991 | 0.02987 | 0.00000 | 0.16679 | 0.00813 | 0.07561 |
| **TOE1** | 0.00000 | 12.98845 | 6.29735 | 5.40214 | 4.12497 | 12.13454 | 8.07780 | 6.85278 | 17.54741 | 0.00000 | 14.19247 | 13.23073 |
| **TOLLIP** | 2.19128 | 9.83125 | 8.03735 | 8.04586 | 14.75429 | 0.00000 | 4.10701 | 3.39578 | 8.46906 | 0.00000 | 7.28168 | 8.26166 |
| **TOLLIP-AS1** | 0.36898 | 0.10048 | 0.09790 | 0.09771 | 0.14924 | 0.54252 | 0.00000 | 0.09607 | 0.10880 | 0.00000 | 0.10635 | 0.30740 |
| **TOM1** | 0.69756 | 5.08779 | 13.90784 | 6.88661 | 5.69034 | 0.00000 | 7.82606 | 0.00000 | 5.69746 | 1.12336 | 4.39105 | 1.98513 |
| **TOM1L1** | 3.52495 | 2.76078 | 0.77352 | 0.00433 | 0.36285 | 1.82829 | 0.50284 | 0.72738 | 0.68930 | 0.00000 | 1.91813 | 0.52969 |
| **TOM1L2** | 1.65461 | 1.53423 | 0.37806 | 1.17910 | 2.55588 | 0.80513 | 0.00000 | 0.02526 | 0.00000 | 0.00000 | 1.10769 | 0.27097 |
| **TOMM20** | 2.47008 | 11.96761 | 11.03653 | 9.43235 | 12.01697 | 24.18492 | 19.51975 | 6.97132 | 23.05163 | 17.12091 | 12.58395 | 13.19296 |
| **TOMM20L** | 0.03964 | 0.22909 | 0.00000 | 0.09507 | 0.12040 | 0.79835 | 0.00000 | 0.10998 | 0.32983 | 0.19206 | 0.13429 | 0.14733 |
| **TOMM22** | 0.06414 | 29.46918 | 22.80330 | 23.45316 | 25.01613 | 33.76101 | 39.52611 | 13.32004 | 29.92254 | 7.92115 | 27.56392 | 29.57092 |
| **TOMM34** | 10.27331 | 7.69487 | 16.13337 | 5.57737 | 8.55932 | 11.55538 | 14.81819 | 0.89845 | 20.96131 | 0.00000 | 13.05111 | 2.48132 |
| **TOMM40** | 0.09338 | 14.74255 | 8.91860 | 8.73082 | 23.55274 | 0.00000 | 15.86186 | 4.81461 | 8.93651 | 35.51031 | 6.70236 | 11.22989 |
| **TOMM40L** | 0.36687 | 0.72086 | 4.02674 | 2.31346 | 0.80134 | 0.52306 | 3.71496 | 0.12666 | 0.36808 | 1.53871 | 0.52413 | 0.09645 |
| **TOMM5** | 4.97640 | 21.68168 | 24.24198 | 11.57191 | 17.35827 | 40.16461 | 35.67011 | 30.03045 | 27.61186 | 14.87578 | 27.53187 | 38.91048 |
| **TOMM6** | 2.65097 | 30.30970 | 22.97036 | 17.11009 | 27.36463 | 44.61611 | 33.18835 | 17.43256 | 29.97716 | 17.87858 | 29.06805 | 40.83724 |
| **TOMM7** | 6.93165 | 28.12474 | 21.87100 | 24.59860 | 28.15132 | 27.84250 | 36.09754 | 4.62137 | 25.03339 | 19.34427 | 25.58332 | 34.57665 |
| **TOMM70A** | 4.81013 | 4.60136 | 6.98936 | 0.64563 | 7.09823 | 23.40524 | 19.04888 | 4.12701 | 5.78261 | 30.55027 | 5.93344 | 6.10630 |
| **TONSL** | 0.06980 | 8.59266 | 2.66279 | 4.34690 | 7.67757 | 0.00000 | 0.34788 | 0.00000 | 9.64164 | 8.58466 | 4.20732 | 3.91576 |
| **1.00000** | 7.50445 | 9.13407 | 22.65428 | 6.75876 | 15.73874 | 15.39142 | 30.70123 | 14.12137 | 20.55857 | 30.81468 | 8.46201 | 13.10089 |
| **TOP1MT** | 1.11388 | 17.89127 | 3.59296 | 7.76607 | 6.94003 | 0.00384 | 10.85271 | 3.54452 | 12.33694 | 1.66018 | 5.59533 | 17.69882 |
| **TOP1P1** | 0.31490 | 0.72549 | 1.08043 | 0.17194 | 0.30004 | 0.87857 | 1.47605 | 1.06987 | 0.87541 | 0.92605 | 0.17999 | 0.23239 |
| **TOP1P2** | 0.04858 | 0.00000 | 0.04782 | 0.00000 | 0.19552 | 0.03379 | 0.09667 | 0.02093 | 0.00000 | 0.28294 | 0.31857 | 0.00628 |
| **TOP2A** | 8.35671 | 17.13341 | 14.01169 | 7.52005 | 15.00345 | 38.27459 | 27.94553 | 10.02939 | 22.40712 | 23.99882 | 11.10585 | 21.92295 |
| **TOP2B** | 4.98639 | 1.43976 | 5.88829 | 3.64064 | 5.20204 | 15.46128 | 11.47020 | 4.17628 | 2.13215 | 6.91194 | 3.35480 | 0.00000 |
| **TOP3A** | 1.16370 | 3.52368 | 5.51686 | 6.54450 | 3.67122 | 0.00000 | 2.76690 | 7.66285 | 5.54107 | 0.00000 | 4.04810 | 10.58739 |

| | | | | | | | | | | | | |
|---|---|---|---|---|---|---|---|---|---|---|---|---|
| **TOP3B** | 0.12892 | 4.90465 | 1.16862 | 2.42462 | 1.01056 | 0.00687 | 0.00000 | 0.04453 | 5.61454 | 0.00000 | 2.83377 | 3.92022 |
| **TOPAZ1** | 0.74842 | 0.29398 | 0.18857 | 0.00000 | 0.16984 | 0.01270 | 0.08908 | 0.24416 | 0.10371 | 0.34608 | 0.00000 | 0.90165 |
| **TOPBP1** | 1.05359 | 2.27983 | 4.10471 | 3.50638 | 3.39964 | 18.34948 | 10.44290 | 3.50291 | 0.37214 | 17.49256 | 4.07785 | 5.24044 |
| **TOPORS** | 8.01252 | 5.05236 | 6.85052 | 5.57814 | 0.00000 | 14.32868 | 6.59263 | 33.45322 | 8.74781 | 7.11696 | 5.17307 | 16.98009 |
| **TOPORS-AS1** | 0.49777 | 0.99277 | 1.16957 | 0.33670 | 0.79702 | 0.05032 | 1.66417 | 5.37676 | 0.00004 | 0.00000 | 0.68649 | 3.04292 |
| **TOR1A** | 5.46349 | 11.43298 | 14.34593 | 7.07725 | 4.72483 | 15.34333 | 3.49574 | 15.47998 | 26.06757 | 0.00000 | 8.73792 | 19.98831 |
| **TOR1AIP1** | 0.01328 | 5.69821 | 10.52881 | 0.67902 | 1.62980 | 17.02937 | 0.00000 | 6.66526 | 17.11760 | 0.00005 | 7.93009 | 9.03378 |
| **TOR1AIP2** | 0.92701 | 10.54636 | 21.34156 | 6.42468 | 14.77051 | 23.61215 | 25.57841 | 24.86120 | 19.48156 | 19.96429 | 17.35740 | 17.87477 |
| **TOR1B** | 3.75408 | 12.85660 | 12.80219 | 0.00000 | 1.70568 | 8.01471 | 0.00000 | 22.95383 | 22.46342 | 6.21489 | 6.74093 | 25.67532 |
| **TOR2A** | 3.78216 | 3.70247 | 1.04288 | 1.95447 | 5.73290 | 1.50687 | 0.99199 | 1.93149 | 0.00000 | 0.00000 | 1.98322 | 3.43971 |
| **TOR3A** | 10.13238 | 10.75186 | 14.33496 | 14.83445 | 7.68706 | 2.47574 | 0.62205 | 4.31334 | 18.58862 | 1.43357 | 7.68787 | 0.57317 |
| **TOR4A** | 0.00559 | 0.00000 | 0.00000 | 0.03966 | 0.11544 | 0.00000 | 0.06594 | 0.00544 | 0.09162 | 0.53064 | 0.00000 | 0.17568 |
| **TOX** | 2.38389 | 0.00691 | 0.02590 | 0.54839 | 0.01419 | 0.00000 | 0.02168 | 0.00112 | 0.01554 | 0.03996 | 0.00885 | 0.01006 |
| **TOX2** | 8.04824 | 0.41373 | 0.02542 | 0.00000 | 0.19252 | 0.00000 | 0.03095 | 1.69375 | 0.00000 | 0.00000 | 0.06107 | 1.86872 |
| **TOX3** | 0.00846 | 0.00441 | 0.06046 | 0.10457 | 0.00000 | 0.06465 | 0.00000 | 0.01024 | 0.00000 | 0.05410 | 0.02506 | 0.01106 |
| **TOX4** | 7.17866 | 9.60063 | 12.42023 | 9.55738 | 11.28350 | 10.02426 | 11.39737 | 28.90606 | 5.31303 | 7.84821 | 10.63071 | 20.32100 |
| **TP53** | 0.98625 | 6.08124 | 4.74133 | 5.10539 | 2.51482 | 0.00000 | 8.24647 | 0.05233 | 4.66862 | 1.15171 | 6.86096 | 2.90442 |
| **TP53AIP1** | 0.00591 | 0.01679 | 0.24493 | 0.03116 | 0.00884 | 0.01354 | 0.12608 | 0.01078 | 0.00000 | 0.00776 | 0.01404 | 0.01759 |
| **TP53BP1** | 1.93261 | 2.66707 | 4.06316 | 2.22369 | 2.36073 | 0.66796 | 4.19696 | 0.50305 | 3.43558 | 5.02073 | 1.56474 | 3.74009 |
| **TP53BP2** | 0.68778 | 1.45989 | 3.12735 | 4.61450 | 2.35822 | 4.87718 | 7.55015 | 4.21348 | 7.00578 | 8.11235 | 3.40253 | 3.24585 |
| **TP53I11** | 0.16163 | 0.70006 | 0.05652 | 0.00052 | 0.00000 | 0.00761 | 0.00459 | 0.02849 | 0.03261 | 0.02328 | 0.01211 | 0.44685 |
| **TP53I13** | 0.05543 | 7.56801 | 1.35673 | 1.89297 | 1.94928 | 0.00000 | 0.00000 | 0.00418 | 3.33609 | 0.50491 | 7.77531 | 1.61842 |
| **TP53I3** | 0.75980 | 0.31418 | 0.52182 | 0.56909 | 0.24123 | 0.00000 | 0.12452 | 0.02027 | 1.28422 | 0.00112 | 0.10443 | 0.00000 |
| **TP53INP1** | 3.86989 | 0.00000 | 3.15751 | 2.68258 | 0.54953 | 6.49440 | 4.43527 | 0.30395 | 4.15757 | 3.75374 | 2.74878 | 0.00000 |
| **TP53INP2** | 2.88289 | 1.87353 | 0.63378 | 3.34521 | 2.18263 | 0.85024 | 0.97837 | 0.00000 | 0.00000 | 1.37438 | 0.33505 | 3.89514 |
| **TP53RK** | 0.46206 | 10.10967 | 15.95739 | 5.95975 | 6.60697 | 7.57116 | 15.07912 | 8.94839 | 9.85130 | 0.00000 | 12.46577 | 7.13158 |
| **TP53TG1** | 0.43691 | 0.01589 | 0.48076 | 0.00656 | 0.16677 | 0.80250 | 0.00000 | 0.00766 | 0.34861 | 0.00000 | 0.25900 | 0.13378 |
| **TP53TG3+TP53T** | 0.00612 | 0.01533 | 0.01782 | 0.01291 | 0.01578 | 0.01705 | 0.01822 | 0.01033 | 0.01750 | 0.01179 | 0.01540 | 0.02359 |
| **TP53TG3D** | 0.03435 | 0.04184 | 0.08394 | 0.00429 | 0.00610 | 0.01921 | 0.08289 | 1.04870 | 0.00000 | 0.04403 | 0.14187 | 0.35766 |
| **TP53TG5** | 0.00494 | 0.01615 | 0.02575 | 0.02610 | 0.02504 | 0.00000 | 0.01992 | 0.01499 | 0.04393 | 0.00670 | 0.04299 | 0.03388 |
| **TP63** | 3.89701 | 0.00000 | 3.38801 | 0.00000 | 0.00000 | 0.36660 | 7.96449 | 0.37091 | 4.38568 | 0.00000 | 0.00000 | 0.00000 |
| **TP73** | 0.19678 | 0.21010 | 2.92959 | 0.33353 | 0.07412 | 0.00000 | 0.00000 | 0.00000 | 0.49786 | 0.24253 | 0.00000 | 0.44692 |
| **TP73-AS1** | 4.92947 | 1.04911 | 1.19084 | 0.90822 | 1.49416 | 0.00000 | 0.00000 | 0.03926 | 2.09088 | 2.66908 | 0.44690 | 0.28137 |
| **TPBG** | 0.08926 | 0.72632 | 2.19184 | 0.05557 | 0.00850 | 1.47567 | 0.00004 | 1.43154 | 9.37488 | 6.74319 | 2.72391 | 0.95037 |
| **TPBGL** | 0.00929 | 0.01577 | 0.01385 | 0.02764 | 0.02447 | 0.03038 | 0.01728 | 0.01717 | 0.01484 | 0.00837 | 0.01435 | 0.03999 |
| **TPCN1** | 0.22448 | 1.21916 | 1.59790 | 0.99175 | 4.35086 | 0.00000 | 0.00000 | 0.00000 | 3.20121 | 4.05029 | 4.04602 | 0.57724 |
| **TPCN2** | 0.00204 | 3.79001 | 3.95510 | 2.93540 | 6.39888 | 0.00000 | 0.00000 | 0.00000 | 4.77895 | 3.81450 | 6.83579 | 0.30215 |
| **TPD52** | 9.02702 | 4.50460 | 8.46039 | 1.51430 | 0.45553 | 12.57033 | 7.35561 | 3.45678 | 13.23395 | 0.40472 | 9.51021 | 0.85519 |
| **TPD52L1** | 0.00402 | 1.44416 | 7.48521 | 1.23252 | 0.87257 | 2.53327 | 4.87783 | 0.00000 | 10.59447 | 0.00000 | 8.22161 | 0.00000 |
| **TPD52L2** | 4.72512 | 19.84306 | 14.40172 | 14.80948 | 16.95581 | 12.80230 | 23.42104 | 19.23686 | 25.48016 | 0.00000 | 21.45759 | 19.56831 |
| **TPGS1** | 0.07982 | 0.02292 | 0.49738 | 0.08197 | 0.14975 | 0.00000 | 0.06648 | 0.02031 | 0.00000 | 0.43880 | 0.34466 | 0.13656 |
| **TPGS2** | 1.60044 | 3.45423 | 3.46107 | 0.56620 | 2.12127 | 8.98853 | 5.55778 | 0.00000 | 1.13587 | 0.00000 | 6.79303 | 0.70226 |
| **TPH1** | 0.00516 | 0.03933 | 0.02457 | 0.15792 | 0.00000 | 0.03905 | 0.00000 | 0.01041 | 0.05425 | 0.04120 | 0.09844 | 0.04368 |
| **TPH2** | 0.76297 | 0.00895 | 0.00110 | 0.00135 | 0.00000 | 0.04405 | 0.01010 | 0.00991 | 0.06568 | 0.08174 | 0.00000 | 0.01120 |
| **TPI1** | 1.75748 | 36.96639 | 36.99618 | 43.04817 | 43.21362 | 34.81540 | 55.11414 | 7.91087 | 39.97959 | 12.26558 | 39.49418 | 12.63897 |
| **TPI1P2** | 0.51382 | 1.56541 | 1.94988 | 2.89457 | 1.49433 | 2.04434 | 2.18037 | 0.25193 | 1.78840 | 0.00000 | 1.90858 | 0.61519 |
| **TPI1P3** | 0.00674 | 0.09810 | 0.10818 | 0.39544 | 0.15777 | 0.09613 | 0.21805 | 0.03356 | 0.09839 | 0.00000 | 0.18632 | 0.04073 |
| **TPK1** | 2.43386 | 2.54713 | 1.31235 | 0.26919 | 0.00000 | 0.11618 | 0.97913 | 0.62221 | 4.34931 | 0.00000 | 4.87029 | 2.18923 |

| | | | | | | | | | | | | |
|---|---|---|---|---|---|---|---|---|---|---|---|---|
| **TPM1** | 7.49570 | 18.96631 | 20.35798 | 27.64385 | 27.24191 | 13.03512 | 32.88963 | 1.00631 | 15.46512 | 8.61437 | 16.82498 | 0.00000 |
| **TPM2** | 2.48682 | 9.68210 | 0.84036 | 9.99046 | 10.06803 | 1.73969 | 7.68485 | 4.73903 | 4.60749 | 1.15159 | 2.18074 | 26.65013 |
| **TPM3** | 4.89788 | 34.33863 | 22.42934 | 27.87259 | 30.96227 | 44.25798 | 35.18968 | 25.47098 | 32.47674 | 5.39351 | 33.57915 | 25.73036 |
| **TPM3P9** | 0.02818 | 1.28386 | 2.84938 | 1.40918 | 2.83436 | 2.47708 | 1.96619 | 3.78524 | 1.29686 | 1.38376 | 0.79308 | 1.22377 |
| **TPM4** | 0.00000 | 0.00000 | 27.22213 | 14.35635 | 33.44600 | 1.35836 | 48.20986 | 0.00000 | 31.33928 | 0.20542 | 35.71724 | 0.00000 |
| **TPMT** | 0.76520 | 4.83354 | 4.37197 | 2.79579 | 1.39173 | 21.11423 | 6.80816 | 0.60761 | 1.17212 | 0.00000 | 5.74648 | 5.22231 |
| **TPO** | 0.59544 | 0.05130 | 0.01280 | 0.00488 | 0.00000 | 0.05431 | 0.02064 | 0.04548 | 0.00081 | 0.02523 | 0.02996 | 0.13409 |
| **TPP1** | 0.40325 | 2.15518 | 3.06621 | 6.10171 | 5.10240 | 1.98351 | 0.34561 | 0.00000 | 12.03371 | 1.53204 | 8.10941 | 0.00000 |
| **TPP2** | 3.84869 | 2.22492 | 7.60084 | 4.94835 | 1.38884 | 11.13956 | 7.78335 | 2.33058 | 6.57830 | 13.71761 | 9.86322 | 1.54170 |
| **TPPP** | 1.12953 | 0.63139 | 0.12190 | 0.57786 | 0.00275 | 0.76381 | 0.00000 | 2.07142 | 0.00000 | 0.00000 | 0.26908 | 1.76749 |
| **TPPP2** | 0.01855 | 0.00867 | 0.01171 | 0.01654 | 0.02505 | 0.04856 | 0.01648 | 0.00954 | 0.03244 | 0.02696 | 0.01738 | 0.01557 |
| **TPPP3** | 0.18169 | 0.01196 | 1.97320 | 0.01322 | 0.06424 | 0.00000 | 0.32598 | 0.61834 | 0.00000 | 0.03201 | 0.01314 | 0.00308 |
| **TPR** | 6.93317 | 7.00276 | 20.20259 | 11.33118 | 15.96229 | 30.00716 | 13.82968 | 3.35570 | 16.00014 | 34.61518 | 13.45157 | 9.98718 |
| **TPRA1** | 0.73451 | 9.05330 | 5.59027 | 5.57403 | 14.26868 | 0.00000 | 6.34104 | 0.00000 | 4.87818 | 10.41205 | 1.36552 | 1.03755 |
| **TPRG1** | 1.69632 | 0.17845 | 0.00000 | 0.43594 | 0.08010 | 0.14915 | 0.00000 | 0.72385 | 0.00000 | 0.04633 | 0.16409 | 0.49922 |
| **TPRG1-AS1** | 0.00536 | 0.00878 | 0.08869 | 0.01175 | 0.04900 | 0.00000 | 0.01591 | 0.00996 | 0.01775 | 0.00000 | 0.02290 | 0.16922 |
| **TPRG1L** | 0.00000 | 3.14008 | 1.55327 | 1.36900 | 4.40838 | 0.57839 | 3.37616 | 4.10498 | 0.00000 | 8.83000 | 0.77862 | 4.06887 |
| **TPRKB** | 0.00000 | 8.58867 | 7.45904 | 1.16852 | 2.63195 | 32.32975 | 11.46441 | 9.51492 | 12.68456 | 4.84370 | 5.46890 | 21.68616 |
| **TPRN** | 0.00000 | 0.59441 | 1.25431 | 1.72140 | 3.98967 | 0.00000 | 6.52199 | 0.02077 | 2.84809 | 7.83759 | 2.00389 | 0.15721 |
| **TPRX1** | 1.94570 | 0.00000 | 0.00792 | 0.00465 | 0.01475 | 0.00000 | 0.00554 | 41.41165 | 0.01514 | 0.00728 | 0.00000 | 7.45842 |
| **TPRXL** | 27.97923 | 0.00000 | 0.03973 | 0.44736 | 0.95287 | 1.08362 | 0.30921 | 7.03254 | 0.00000 | 0.62349 | 0.75526 | 1.40260 |
| **TPSB2** | 0.03632 | 0.01525 | 0.01499 | 0.01327 | 0.01540 | 0.01638 | 0.01825 | 0.01711 | 0.01914 | 0.01076 | 0.01736 | 0.01532 |
| **TPSD1** | 0.00254 | 0.01552 | 0.01645 | 0.01280 | 0.01359 | 0.01177 | 0.02189 | 0.04286 | 0.00962 | 0.01909 | 0.03362 | 0.01755 |
| **TPSG1** | 0.03617 | 0.01747 | 0.01337 | 0.01516 | 0.01317 | 0.01913 | 0.01690 | 0.00912 | 0.01784 | 0.01759 | 0.01448 | 0.01623 |
| **TPST1** | 0.07291 | 3.46620 | 4.44474 | 0.68298 | 0.30640 | 1.79970 | 0.00000 | 8.30277 | 4.38912 | 0.00000 | 0.19222 | 2.81573 |
| **TPST2** | 0.00000 | 10.56449 | 10.52316 | 33.96450 | 4.39789 | 0.00000 | 0.00002 | 3.48869 | 14.87246 | 3.78945 | 5.04706 | 6.46547 |
| **TPT1** | 14.55901 | 28.25987 | 29.62716 | 23.08564 | 36.19850 | 54.47141 | 54.48246 | 18.70087 | 28.35492 | 64.59006 | 21.46168 | 40.85600 |
| **TPT1-AS1** | 0.03732 | 0.25291 | 1.31001 | 1.29114 | 0.79669 | 0.79266 | 0.00000 | 0.05611 | 2.23473 | 0.00000 | 0.66417 | 0.04643 |
| **TPTE** | 0.13879 | 1.31640 | 1.48063 | 0.12588 | 0.03528 | 4.09119 | 0.62578 | 7.26753 | 1.60611 | 1.34502 | 0.00000 | 2.56702 |
| **TPTE2** | 0.00159 | 0.88693 | 0.03709 | 0.00000 | 0.00000 | 0.09374 | 0.10186 | 0.20208 | 0.29905 | 0.20289 | 0.05644 | 0.30708 |
| **TPTE2P1** | 1.07468 | 0.22197 | 0.00000 | 0.90741 | 0.50695 | 0.08546 | 0.01766 | 0.00000 | 0.00000 | 0.07440 | 0.00000 | 0.48380 |
| **TPTE2P3** | 0.01265 | 0.05981 | 0.04389 | 0.00000 | 0.00000 | 0.16648 | 0.00000 | 0.03371 | 0.15582 | 0.61162 | 0.04359 | 0.15163 |
| **TPTE2P5** | 0.03064 | 0.18714 | 0.10591 | 0.11417 | 0.00000 | 0.39830 | 0.00000 | 0.00233 | 0.00000 | 0.07430 | 0.09748 | 0.09516 |
| **TPTE2P6** | 0.01829 | 0.00000 | 0.04023 | 0.47721 | 0.22267 | 0.00000 | 0.05133 | 0.00224 | 0.00000 | 0.41316 | 0.07240 | 1.02961 |
| **TPTEP1** | 0.20492 | 4.03369 | 2.51148 | 4.69644 | 5.36420 | 2.25723 | 1.81460 | 0.40141 | 7.77842 | 0.36866 | 3.86849 | 7.86679 |
| **TPX2** | 8.83523 | 24.99926 | 18.04839 | 10.59887 | 17.98712 | 29.04768 | 26.47835 | 20.66843 | 26.12554 | 5.76929 | 14.25074 | 27.33710 |
| **TRA2A** | 1.52135 | 8.72985 | 14.14046 | 10.86967 | 9.68456 | 35.18211 | 24.80385 | 26.61243 | 10.96879 | 14.67889 | 20.74665 | 21.52200 |
| **TRA2B** | 4.71621 | 15.60734 | 16.93788 | 15.42749 | 18.69433 | 31.60946 | 28.48006 | 23.88782 | 20.67216 | 21.12504 | 21.19228 | 23.65319 |
| **TRABD** | 0.46213 | 8.10609 | 9.54234 | 8.18568 | 10.54100 | 0.00000 | 7.29380 | 0.28919 | 17.76302 | 4.21637 | 10.06264 | 2.58402 |
| **TRABD2A** | 0.01959 | 0.02972 | 2.86870 | 0.00854 | 0.01137 | 0.00012 | 0.00000 | 0.01043 | 0.27349 | 0.00000 | 0.05256 | 0.02140 |
| **TRABD2B** | 0.00342 | 0.48670 | 0.02231 | 1.04456 | 0.00000 | 0.05381 | 0.00000 | 0.00652 | 0.00000 | 0.19273 | 0.00022 | 0.04082 |
| **TRADD** | 0.03175 | 2.20423 | 1.70280 | 0.68064 | 1.78221 | 0.04054 | 1.07921 | 0.00000 | 1.50227 | 0.00000 | 1.09418 | 1.37637 |
| **TRAF1** | 0.88123 | 0.07247 | 0.00839 | 0.00545 | 0.01117 | 0.01291 | 0.02857 | 0.00932 | 0.04750 | 0.00000 | 0.01401 | 0.10901 |
| **TRAF2** | 3.82220 | 10.04243 | 8.26194 | 6.58204 | 9.22669 | 0.00000 | 11.75348 | 2.41818 | 16.49929 | 0.00000 | 6.91338 | 8.93165 |
| **TRAF3** | 2.40948 | 0.00000 | 2.91471 | 0.07281 | 3.76030 | 0.00000 | 5.81592 | 0.07451 | 4.44698 | 4.85350 | 5.93436 | 0.49619 |
| **TRAF3IP1** | 4.37033 | 0.30177 | 0.00000 | 0.67258 | 0.16725 | 0.98070 | 1.01735 | 0.40426 | 0.24173 | 4.38281 | 0.90937 | 0.67117 |
| **TRAF3IP2** | 3.48194 | 0.95298 | 2.44115 | 3.32158 | 3.53419 | 1.31665 | 1.71823 | 0.30664 | 1.79241 | 0.37968 | 1.74016 | 0.41209 |
| **TRAF3IP2-AS1** | 0.00171 | 0.30382 | 1.22937 | 0.78920 | 0.00000 | 1.94428 | 0.00000 | 1.19552 | 1.89663 | 2.02112 | 0.37704 | 1.60085 |

| | | | | | | | | | | | | |
|---|---|---|---|---|---|---|---|---|---|---|---|---|
| **TRAF3IP3** | 0.20881 | 0.08013 | 0.22692 | 0.06718 | 0.00487 | 0.07395 | 0.01175 | 0.00119 | 0.00918 | 0.05017 | 0.06401 | 0.23977 |
| **TRAF4** | 0.82281 | 5.88829 | 12.70332 | 5.63095 | 13.25791 | 0.00000 | 14.42048 | 1.74713 | 16.95753 | 10.76886 | 5.98115 | 4.63836 |
| **TRAF5** | 1.14380 | 4.17178 | 0.00000 | 0.26186 | 0.00000 | 0.19379 | 0.00000 | 0.00000 | 0.47889 | 1.07863 | 0.25878 | 4.12711 |
| **TRAF6** | 0.73435 | 2.27365 | 0.00039 | 1.97394 | 1.49087 | 8.37761 | 1.16959 | 8.08427 | 2.63475 | 0.58929 | 1.54473 | 5.34813 |
| **TRAF7** | 4.99777 | 6.99569 | 10.79117 | 7.24732 | 10.12024 | 0.00000 | 7.24871 | 0.56125 | 17.57757 | 0.00000 | 8.68590 | 3.67795 |
| **TRAFD1** | 9.50212 | 7.22986 | 6.52812 | 5.03367 | 3.16259 | 4.50764 | 5.94644 | 6.18829 | 2.99796 | 0.00000 | 6.89562 | 5.55515 |
| **TRAIP** | 1.75887 | 4.57341 | 3.48200 | 2.47016 | 1.91986 | 5.46510 | 3.63810 | 6.14515 | 3.48184 | 0.00000 | 3.22608 | 7.89355 |
| **TRAK1** | 8.68083 | 5.74279 | 4.69371 | 5.88125 | 5.39944 | 7.68486 | 6.07090 | 1.37316 | 2.51698 | 18.27724 | 0.97581 | 6.73043 |
| **TRAK2** | 5.92520 | 5.28328 | 2.23957 | 4.59662 | 2.12356 | 5.11541 | 1.63151 | 5.24374 | 3.20653 | 4.38724 | 3.70046 | 5.19571 |
| **TRAM1** | 2.26377 | 12.77333 | 13.48902 | 4.16153 | 9.40361 | 36.28503 | 0.00000 | 4.48696 | 18.58862 | 13.98083 | 13.39216 | 6.90882 |
| **TRAM1L1** | 0.01433 | 0.38752 | 0.43658 | 0.07733 | 0.04551 | 0.64582 | 0.00000 | 0.21353 | 0.29227 | 0.23590 | 0.23845 | 0.52459 |
| **TRAM2** | 14.82499 | 5.23020 | 10.30870 | 7.19486 | 7.16869 | 0.00000 | 3.68641 | 2.00009 | 10.39946 | 11.13195 | 5.39394 | 13.51109 |
| **TRAM2-AS1** | 0.14570 | 0.11258 | 0.03089 | 0.21146 | 0.04045 | 0.14835 | 0.00002 | 0.00562 | 0.00000 | 0.17734 | 0.02414 | 0.00084 |
| **TRANK1** | 1.13266 | 0.01090 | 0.03167 | 0.04442 | 0.04606 | 0.03692 | 0.01956 | 0.00781 | 0.00234 | 0.03230 | 0.01061 | 0.01199 |
| **TRAP1** | 4.86302 | 23.31147 | 16.88198 | 16.84003 | 25.30348 | 2.41199 | 23.19399 | 2.17255 | 42.23696 | 0.00000 | 32.67243 | 9.58131 |
| **TRAPPC1** | 4.38803 | 4.47097 | 8.77531 | 9.45093 | 8.60808 | 1.28383 | 10.31541 | 2.47411 | 9.94666 | 0.00054 | 9.24021 | 8.79028 |
| **TRAPPC10** | 3.13748 | 2.90091 | 3.53442 | 3.39098 | 4.38742 | 3.51280 | 5.84576 | 0.73664 | 3.42433 | 11.33191 | 2.61056 | 1.92084 |
| **TRAPPC11** | 1.34923 | 3.84809 | 2.93340 | 3.24822 | 0.00000 | 5.91094 | 4.97619 | 0.88979 | 1.39044 | 1.47467 | 4.38947 | 3.90645 |
| **TRAPPC12** | 0.29636 | 0.81762 | 0.82969 | 2.00268 | 1.70043 | 0.82999 | 2.11057 | 0.00000 | 1.61673 | 2.35753 | 0.00000 | 0.22980 |
| **TRAPPC13** | 1.16798 | 3.83733 | 5.52188 | 1.35052 | 3.01465 | 17.29382 | 4.35746 | 0.19606 | 6.70517 | 0.88779 | 6.68961 | 3.47537 |
| **TRAPPC2** | 0.66156 | 0.39446 | 2.98134 | 1.33206 | 0.00000 | 3.48171 | 1.98126 | 1.38148 | 3.84346 | 1.41841 | 3.20265 | 1.11389 |
| **TRAPPC2L** | 0.76264 | 21.88537 | 19.35820 | 18.98579 | 28.09053 | 9.36797 | 32.61009 | 0.62589 | 15.10634 | 4.73559 | 27.42939 | 15.49387 |
| **TRAPPC2P1** | 0.80402 | 2.44979 | 2.56062 | 0.33265 | 0.26806 | 6.89108 | 4.01830 | 5.27094 | 1.85156 | 0.00000 | 1.05547 | 3.32678 |
| **TRAPPC3** | 2.94699 | 14.00346 | 17.63230 | 11.37770 | 13.12510 | 8.85119 | 25.75875 | 5.99324 | 16.03486 | 0.00000 | 18.51292 | 24.06826 |
| **TRAPPC3L** | 0.05032 | 0.01339 | 0.01589 | 0.01269 | 0.01503 | 0.01705 | 0.01728 | 0.06621 | 0.01738 | 0.00728 | 0.02696 | 0.02510 |
| **TRAPPC4** | 1.57019 | 22.10473 | 11.58846 | 9.72006 | 7.23328 | 20.41921 | 21.62086 | 15.47303 | 17.15876 | 0.00000 | 17.74838 | 27.07968 |
| **TRAPPC5** | 0.34126 | 8.27209 | 8.38431 | 6.32697 | 11.29877 | 0.00000 | 13.24811 | 1.27735 | 6.81059 | 5.77113 | 7.66606 | 13.63942 |
| **TRAPPC6A** | 0.11662 | 1.20401 | 1.31771 | 1.23847 | 1.17823 | 0.00000 | 0.14374 | 1.05316 | 1.53383 | 0.00000 | 2.12820 | 0.45816 |
| **TRAPPC6B** | 1.92247 | 1.11014 | 1.91007 | 3.12083 | 0.32502 | 16.94867 | 2.03131 | 8.92436 | 6.50843 | 3.21356 | 3.16489 | 8.94463 |
| **TRAPPC8** | 1.29209 | 4.57059 | 5.24864 | 3.46389 | 0.00000 | 24.78881 | 7.13883 | 5.37453 | 4.30929 | 23.78799 | 4.33744 | 7.72585 |
| **TRAPPC9** | 1.09208 | 0.89983 | 0.90260 | 1.06248 | 1.11474 | 0.34656 | 0.00000 | 0.05808 | 0.85782 | 0.26088 | 0.70531 | 0.32884 |
| **TRDMT1** | 0.00838 | 1.18387 | 0.17672 | 0.00000 | 0.00000 | 2.87024 | 0.20873 | 1.18633 | 0.24889 | 1.17002 | 1.93726 | 2.26882 |
| **TRDN** | 0.46806 | 0.00256 | 0.00000 | 0.05500 | 0.13055 | 0.59708 | 0.00000 | 0.00287 | 0.00729 | 0.00193 | 0.01184 | 0.05998 |
| **TREH** | 0.03379 | 0.02539 | 0.24401 | 0.00745 | 0.00159 | 0.00000 | 0.00000 | 0.01373 | 0.02167 | 0.07950 | 0.08564 | 0.01283 |
| **TREM1** | 0.00527 | 0.00954 | 7.16698 | 0.04301 | 0.00000 | 0.00563 | 0.34810 | 0.01055 | 0.78488 | 0.00023 | 0.00000 | 0.01656 |
| **TREM2** | 0.00682 | 0.01865 | 0.42170 | 0.01657 | 0.01877 | 0.00000 | 0.00477 | 0.01060 | 0.04621 | 0.00004 | 0.01723 | 0.01834 |
| **TREML1** | 0.00657 | 0.01676 | 0.04360 | 0.01139 | 0.01399 | 0.01088 | 0.02381 | 0.01050 | 0.04324 | 0.00000 | 0.03037 | 0.01893 |
| **TREML2** | 0.06902 | 0.01139 | 2.06965 | 0.00774 | 0.01928 | 0.00000 | 0.00000 | 0.12573 | 0.00000 | 0.03760 | 0.01291 | 0.07000 |
| **TREML3P** | 0.00614 | 0.01522 | 0.31555 | 0.00646 | 0.01121 | 0.01406 | 0.22141 | 0.01049 | 0.00634 | 0.00749 | 0.00773 | 0.01603 |
| **TREML4** | 0.00602 | 0.01489 | 0.03195 | 0.00837 | 0.00918 | 0.01418 | 0.00938 | 0.01044 | 0.05616 | 0.02952 | 0.02025 | 0.01544 |
| **TRERF1** | 0.44324 | 1.25649 | 5.11268 | 4.59020 | 3.59936 | 3.17643 | 6.95708 | 0.00000 | 9.31634 | 9.01582 | 3.57936 | 0.50180 |
| **TREX1** | 0.05951 | 0.76833 | 1.59602 | 0.05609 | 0.20318 | 0.73256 | 1.82122 | 0.00000 | 0.98472 | 0.38101 | 0.67244 | 0.00000 |
| **TREX2** | 0.00496 | 0.01279 | 0.01560 | 0.07566 | 0.00000 | 0.00869 | 0.02720 | 0.00907 | 0.06888 | 0.08267 | 0.04449 | 0.03192 |
| **TRG-AS1** | 0.04222 | 0.00000 | 0.00000 | 0.00000 | 0.12425 | 0.18361 | 0.09054 | 0.00468 | 0.00000 | 0.13063 | 0.49344 | 0.01921 |
| **TRH** | 0.00605 | 0.05931 | 0.02701 | 0.28525 | 0.00041 | 0.00014 | 0.00896 | 0.00859 | 0.05220 | 0.01382 | 0.01921 | 0.05707 |
| **TRHDE** | 0.65220 | 0.02775 | 0.10800 | 0.03279 | 0.00000 | 0.10651 | 0.00000 | 0.06426 | 0.00000 | 0.03956 | 0.00599 | 0.04703 |
| **TRHDE-AS1** | 0.14161 | 0.00443 | 0.11118 | 0.01246 | 0.00000 | 0.04343 | 0.00000 | 0.04364 | 0.03323 | 0.07196 | 0.00672 | 0.05518 |
| **TRHR** | 0.01023 | 0.01236 | 0.01411 | 0.01260 | 0.00792 | 0.02898 | 0.05206 | 0.05738 | 0.01706 | 0.02841 | 0.01342 | 0.01148 |

| | | | | | | | | | | | | |
|---|---|---|---|---|---|---|---|---|---|---|---|---|
| **TRIAP1** | 3.15314 | 23.58073 | 18.56433 | 8.68659 | 7.73901 | 42.16071 | 20.12845 | 11.87538 | 21.66422 | 8.15433 | 16.45573 | 23.54479 |
| **TRIB1** | 0.41300 | 0.53552 | 2.79534 | 0.65623 | 0.24456 | 2.51811 | 1.42286 | 1.34353 | 0.00000 | 6.32531 | 0.23260 | 2.77907 |
| **TRIB2** | 0.00610 | 0.05115 | 0.35271 | 0.51813 | 0.21841 | 0.00000 | 0.00664 | 0.01122 | 0.29560 | 0.00000 | 0.04243 | 0.03158 |
| **TRIB3** | 0.00000 | 2.00756 | 8.35373 | 1.00842 | 4.10980 | 4.83696 | 12.23680 | 0.03489 | 0.22425 | 0.84957 | 0.00000 | 0.88256 |
| **TRIL** | 0.42916 | 0.00432 | 0.03705 | 0.01170 | 0.01757 | 0.04862 | 0.02525 | 0.00000 | 0.00854 | 0.00037 | 0.01410 | 0.09954 |
| **TRIM10** | 0.00592 | 0.01454 | 0.05238 | 0.01412 | 0.01701 | 0.01325 | 0.01297 | 0.01049 | 0.00866 | 0.03277 | 0.01583 | 0.01542 |
| **TRIM11** | 2.89991 | 4.01110 | 9.27961 | 2.58420 | 11.67295 | 0.00000 | 9.33731 | 0.06154 | 6.05186 | 9.96083 | 3.38433 | 3.68154 |
| **TRIM13** | 0.54569 | 6.47420 | 10.14930 | 8.20323 | 3.86208 | 29.49473 | 8.42504 | 8.18385 | 11.72345 | 8.86951 | 9.50924 | 11.68543 |
| **TRIM14** | 0.16681 | 2.48523 | 0.34597 | 6.12950 | 1.41766 | 0.00000 | 0.13747 | 0.00000 | 0.52237 | 0.87716 | 0.31253 | 0.59388 |
| **TRIM15** | 0.00898 | 0.00982 | 0.01795 | 0.01084 | 0.05229 | 0.03716 | 0.00971 | 0.01010 | 0.01624 | 0.01222 | 0.01511 | 0.02162 |
| **TRIM16** | 0.05279 | 1.98832 | 0.35193 | 3.84900 | 4.25032 | 1.85398 | 0.92469 | 0.38130 | 0.00000 | 0.00000 | 3.71147 | 2.56104 |
| **TRIM16L** | 0.01070 | 0.00000 | 0.18457 | 0.05294 | 0.10788 | 0.03985 | 0.33525 | 0.03568 | 0.08831 | 0.49360 | 0.21676 | 0.74511 |
| **TRIM17** | 0.00423 | 0.00971 | 0.54327 | 0.80018 | 0.58283 | 0.14399 | 0.00000 | 0.00669 | 0.00000 | 0.05941 | 0.74838 | 0.01553 |
| **TRIM2** | 0.00000 | 0.05150 | 2.69289 | 23.56910 | 8.08763 | 8.48483 | 0.00000 | 0.00000 | 0.00000 | 6.19854 | 6.18020 | 0.57926 |
| **TRIM21** | 0.01155 | 1.19968 | 0.62357 | 0.85903 | 0.00000 | 0.65259 | 0.42271 | 0.98433 | 0.80469 | 0.00000 | 0.10837 | 1.40486 |
| **TRIM22** | 0.01228 | 0.17404 | 0.63296 | 1.00807 | 0.00000 | 1.04861 | 0.15878 | 0.00000 | 0.44954 | 0.01085 | 0.02949 | 0.24806 |
| **TRIM23** | 0.15870 | 0.75522 | 1.18103 | 0.43477 | 0.00029 | 5.12776 | 3.87267 | 1.26870 | 0.00000 | 4.21867 | 2.31138 | 3.82475 |
| **TRIM24** | 0.00000 | 1.84186 | 5.63237 | 4.28061 | 1.33957 | 6.50551 | 5.98580 | 0.44229 | 1.95359 | 9.41673 | 1.35278 | 0.15767 |
| **TRIM25** | 0.00163 | 0.00370 | 0.49247 | 0.06101 | 0.11561 | 0.00000 | 0.43992 | 0.02166 | 0.00091 | 0.52289 | 0.05745 | 0.00937 |
| **TRIM26** | 1.30261 | 2.51773 | 1.87562 | 11.49717 | 6.64794 | 0.03283 | 5.87495 | 0.96096 | 2.92976 | 1.75013 | 1.90380 | 2.56272 |
| **TRIM27** | 0.90211 | 17.03987 | 14.91100 | 11.21803 | 11.93565 | 4.92235 | 16.37885 | 6.54986 | 21.27408 | 0.00000 | 16.34455 | 15.83286 |
| **TRIM28** | 3.66846 | 25.22332 | 15.49353 | 25.92742 | 42.21392 | 1.14764 | 43.35517 | 11.68989 | 15.68086 | 59.90901 | 15.71871 | 32.44826 |
| **TRIM29** | 0.20438 | 0.00000 | 0.72372 | 0.06220 | 0.15953 | 0.20461 | 0.78806 | 0.01687 | 2.06080 | 0.00000 | 0.00065 | 0.13895 |
| **TRIM3** | 0.60976 | 1.13841 | 0.95361 | 1.60084 | 0.00000 | 0.00000 | 0.33422 | 0.28032 | 1.77977 | 0.92295 | 0.45950 | 0.80040 |
| **TRIM31** | 0.04594 | 0.01246 | 0.01097 | 0.01359 | 0.00916 | 0.04778 | 0.01144 | 0.00572 | 0.01409 | 0.01897 | 0.01350 | 0.03189 |
| **TRIM32** | 5.68797 | 3.02634 | 2.04516 | 1.79284 | 2.11863 | 10.35438 | 1.81997 | 1.04997 | 4.97159 | 0.00002 | 7.04545 | 5.51179 |
| **TRIM33** | 7.22533 | 1.65038 | 19.21119 | 6.03697 | 11.13848 | 25.13494 | 16.88440 | 2.96169 | 0.22617 | 31.52534 | 7.65947 | 6.68414 |
| **TRIM34.uid-151** | 0.00624 | 0.01469 | 0.02995 | 0.01122 | 0.01271 | 0.02067 | 0.01305 | 0.01047 | 0.02911 | 0.01736 | 0.01297 | 0.01580 |
| **TRIM34.uid-151** | 0.00647 | 0.01787 | 0.02913 | 0.01146 | 0.01558 | 0.02347 | 0.01301 | 0.01305 | 0.02242 | 0.01130 | 0.01744 | 0.01549 |
| **TRIM34.uid-151** | 0.00638 | 0.01754 | 0.04705 | 0.01306 | 0.01674 | 0.02506 | 0.01844 | 0.01077 | 0.01605 | 0.00592 | 0.01697 | 0.01760 |
| **TRIM35** | 1.80735 | 4.51228 | 4.11999 | 1.94011 | 1.02939 | 4.58308 | 4.35637 | 0.89506 | 5.75209 | 0.00000 | 4.12959 | 4.69325 |
| **TRIM36** | 3.26315 | 0.78933 | 0.19236 | 0.00000 | 0.00000 | 0.04691 | 0.10329 | 9.37076 | 0.00000 | 0.03907 | 0.00000 | 0.71970 |
| **TRIM37** | 2.83377 | 3.99799 | 1.66659 | 5.11985 | 3.05245 | 9.47129 | 6.70308 | 1.62735 | 7.78346 | 8.33371 | 6.02725 | 2.79480 |
| **TRIM38** | 0.82890 | 1.11630 | 0.00000 | 3.10507 | 0.99063 | 1.07638 | 0.00000 | 1.38069 | 0.00000 | 0.00171 | 0.09201 | 1.28747 |
| **TRIM39** | 2.39809 | 1.38268 | 1.79187 | 1.96921 | 0.00000 | 2.10063 | 2.22198 | 2.30520 | 0.27450 | 3.89806 | 0.99766 | 2.82560 |
| **TRIM4** | 0.03541 | 0.01558 | 3.21281 | 0.73059 | 0.00002 | 0.78336 | 0.73428 | 0.24572 | 2.99153 | 0.00000 | 0.90971 | 0.43415 |
| **TRIM40** | 0.00725 | 0.00070 | 0.00021 | 0.00338 | 0.08593 | 0.13603 | 0.00000 | 0.00978 | 0.00000 | 0.00637 | 0.09443 | 0.01070 |
| **TRIM41** | 0.78046 | 0.89545 | 1.48774 | 2.03573 | 2.67040 | 0.00000 | 4.40090 | 1.66835 | 0.00000 | 9.19186 | 0.99241 | 2.22865 |
| **TRIM42** | 0.89502 | 0.01160 | 0.03321 | 0.00840 | 0.01640 | 0.01857 | 0.01075 | 0.00174 | 0.02180 | 0.01080 | 0.01807 | 0.03146 |
| **TRIM43** | 3.17650 | 0.01324 | 0.01329 | 0.00982 | 0.02045 | 0.00323 | 0.01857 | 0.28025 | 0.02334 | 0.00000 | 0.01303 | 0.00152 |
| **TRIM43B** | 3.12929 | 0.01462 | 0.01523 | 0.01286 | 0.03203 | 0.00280 | 0.02890 | 0.30474 | 0.02290 | 0.00000 | 0.01379 | 0.00680 |
| **TRIM44** | 0.05931 | 2.60239 | 0.31856 | 2.00277 | 3.79704 | 2.60290 | 0.00000 | 0.83484 | 0.00000 | 8.13685 | 0.00000 | 1.95066 |
| **TRIM45** | 1.67658 | 0.00000 | 0.04896 | 0.15061 | 0.99413 | 1.10831 | 0.00000 | 0.24131 | 0.55751 | 1.55275 | 1.40774 | 0.00000 |
| **TRIM46** | 0.00436 | 2.03413 | 0.00000 | 0.37024 | 0.57483 | 0.00000 | 0.07488 | 0.00694 | 0.15687 | 0.09305 | 0.59349 | 0.05676 |
| **TRIM47** | 0.00298 | 0.08961 | 0.22703 | 0.11988 | 0.00000 | 0.00000 | 0.06047 | 0.02832 | 0.03127 | 0.01898 | 0.08932 | 0.01022 |
| **TRIM48** | 0.93290 | 0.02605 | 0.00971 | 0.01415 | 0.01192 | 0.00000 | 0.04570 | 0.85146 | 0.01465 | 0.01658 | 0.02050 | 0.00488 |
| **TRIM49** | 1.31634 | 0.01261 | 0.01414 | 0.01220 | 0.02247 | 0.01149 | 0.01928 | 0.05261 | 0.02162 | 0.00096 | 0.01303 | 0.00764 |
| **TRIM49B** | 0.89522 | 0.02158 | 0.01435 | 0.01069 | 0.01276 | 0.00684 | 0.02115 | 0.28159 | 0.01339 | 0.00761 | 0.03263 | 0.01323 |

| | | | | | | | | | | | | |
|---|---|---|---|---|---|---|---|---|---|---|---|---|
| **TRIM49C** | 1.15969 | 0.01120 | 0.01058 | 0.02711 | 0.02585 | 0.02825 | 0.01548 | 0.08126 | 0.01654 | 0.00405 | 0.01217 | 0.00510 |
| **TRIM5** | 0.24588 | 1.88864 | 5.04443 | 1.17392 | 2.02167 | 4.89921 | 2.58358 | 0.00000 | 6.33528 | 0.74861 | 0.94031 | 3.18121 |
| **TRIM50** | 1.39214 | 0.34169 | 0.00892 | 0.00000 | 0.01165 | 0.00001 | 0.01346 | 0.00000 | 0.01892 | 0.02594 | 0.02195 | 0.67344 |
| **TRIM51** | 1.51976 | 0.02168 | 0.01685 | 0.01068 | 0.01674 | 0.00000 | 0.01660 | 0.87264 | 0.02200 | 0.00774 | 0.01308 | 0.00000 |
| **TRIM51HP** | 0.46953 | 0.01602 | 0.01351 | 0.01167 | 0.01603 | 0.00987 | 0.01687 | 0.23009 | 0.02433 | 0.01013 | 0.01415 | 0.00541 |
| **TRIM52** | 0.66038 | 0.55945 | 0.42132 | 1.10766 | 0.00000 | 0.63552 | 0.19284 | 1.49069 | 0.33449 | 0.00000 | 0.62447 | 1.52205 |
| **TRIM52-AS1** | 0.00109 | 3.42881 | 0.26050 | 2.29015 | 2.81441 | 0.00000 | 0.20144 | 0.41560 | 0.25564 | 0.00186 | 1.46955 | 3.95842 |
| **TRIM53AP.uid-1** | 0.88911 | 0.26530 | 0.00000 | 0.08795 | 0.00000 | 0.12493 | 0.00000 | 0.26184 | 0.00000 | 0.02671 | 0.00059 | 0.27732 |
| **TRIM53AP.uid-1** | 1.61883 | 0.41697 | 0.00000 | 0.10340 | 0.00000 | 0.07323 | 0.00000 | 0.70368 | 0.00077 | 0.02833 | 0.00292 | 0.41089 |
| **TRIM54** | 0.00286 | 0.11013 | 0.01897 | 0.27705 | 0.27372 | 0.00000 | 0.02909 | 2.89521 | 0.00000 | 0.03014 | 0.02514 | 2.14260 |
| **TRIM55** | 0.00495 | 0.00023 | 0.11373 | 0.01328 | 0.00000 | 0.21204 | 0.00001 | 0.00826 | 0.06162 | 0.06506 | 0.00564 | 0.00504 |
| **TRIM56** | 0.14910 | 0.94553 | 0.13860 | 2.75465 | 0.93102 | 0.00000 | 0.12114 | 0.00000 | 0.00000 | 2.69687 | 0.00000 | 0.85023 |
| **TRIM58** | 0.02780 | 0.34260 | 0.55322 | 0.48967 | 0.63248 | 0.99443 | 0.20122 | 0.08252 | 0.00000 | 1.08304 | 0.65867 | 0.25962 |
| **TRIM59** | 10.03103 | 11.81819 | 7.25271 | 1.66724 | 0.81511 | 30.95627 | 14.27170 | 1.83860 | 16.41722 | 3.04632 | 13.37824 | 17.69830 |
| **TRIM6** | 3.87920 | 3.69386 | 3.11727 | 4.06255 | 0.00033 | 14.16966 | 5.81631 | 0.81075 | 4.12404 | 6.47177 | 3.26788 | 0.33476 |
| **TRIM60** | 0.98947 | 14.19095 | 3.66099 | 8.76328 | 1.41243 | 23.73029 | 0.47548 | 15.76676 | 7.48350 | 0.72730 | 6.15446 | 28.36335 |
| **TRIM61** | 0.73962 | 7.31620 | 0.00000 | 3.35813 | 1.89654 | 4.85154 | 0.00000 | 7.25363 | 1.08824 | 0.47564 | 1.91051 | 15.47200 |
| **TRIM62** | 2.97810 | 0.30414 | 0.78972 | 0.12209 | 0.10351 | 0.00000 | 0.51469 | 0.00509 | 1.12502 | 0.39192 | 0.00125 | 0.00640 |
| **TRIM63** | 0.00477 | 0.20736 | 0.00000 | 0.16786 | 2.40557 | 0.00000 | 0.00049 | 0.05638 | 0.00000 | 0.46907 | 1.43830 | 0.10496 |
| **TRIM64B** | 0.27341 | 0.01640 | 0.01146 | 0.00997 | 0.00955 | 0.01084 | 0.01377 | 0.08286 | 0.02124 | 0.07274 | 0.01074 | 0.00704 |
| **TRIM64C** | 0.61921 | 0.01490 | 0.01441 | 0.01250 | 0.01708 | 0.00945 | 0.01917 | 0.05705 | 0.02241 | 0.00502 | 0.01398 | 0.00863 |
| **TRIM65** | 0.00195 | 0.16909 | 0.57209 | 1.10599 | 0.30281 | 0.00000 | 1.82131 | 0.00000 | 0.37618 | 2.06144 | 1.71876 | 0.33079 |
| **TRIM66** | 0.02897 | 0.59373 | 0.56535 | 0.13542 | 0.00000 | 0.32772 | 0.00000 | 0.00000 | 0.14293 | 0.85081 | 1.01604 | 0.61806 |
| **TRIM67** | 0.00516 | 0.00631 | 0.01615 | 0.01016 | 0.00514 | 0.01721 | 0.01675 | 0.00895 | 0.00000 | 0.04746 | 0.06913 | 0.08853 |
| **TRIM68** | 0.67087 | 2.49059 | 1.42148 | 0.73241 | 1.01403 | 0.30515 | 1.12445 | 0.00000 | 1.52113 | 1.21542 | 0.99776 | 2.99370 |
| **TRIM7** | 0.02268 | 0.20341 | 0.04295 | 0.03744 | 0.00000 | 0.15770 | 0.07026 | 1.07672 | 0.00000 | 0.05419 | 0.05329 | 0.29567 |
| **TRIM71** | 2.97182 | 0.00000 | 8.42642 | 12.53064 | 11.96900 | 1.02967 | 9.72059 | 1.29056 | 3.98549 | 16.64788 | 2.48322 | 9.33158 |
| **TRIM72** | 0.00199 | 0.01525 | 0.02164 | 0.02559 | 0.08663 | 0.00000 | 0.02651 | 0.07433 | 0.00948 | 0.04372 | 0.00963 | 0.01375 |
| **TRIM77** | 11.20352 | 0.00476 | 0.00559 | 0.00225 | 0.01144 | 0.04754 | 0.00956 | 0.64880 | 0.00886 | 0.00432 | 0.01591 | 0.07722 |
| **TRIM8** | 2.23207 | 5.99104 | 0.00000 | 3.54293 | 6.45795 | 1.83050 | 0.45096 | 0.00000 | 0.00000 | 5.25657 | 0.43180 | 0.25362 |
| **TRIM9** | 0.70791 | 0.00000 | 0.13282 | 0.13839 | 0.01498 | 0.08476 | 0.00000 | 0.06570 | 0.00000 | 0.09282 | 0.00639 | 0.12475 |
| **TRIML1** | 0.01925 | 0.00000 | 6.87682 | 0.00000 | 2.10513 | 0.02670 | 8.08715 | 0.00045 | 11.23378 | 0.00000 | 3.03565 | 0.60604 |
| **TRIML2** | 0.00000 | 5.72485 | 10.11011 | 13.01906 | 9.48418 | 10.50762 | 9.23157 | 6.25384 | 11.74757 | 9.09093 | 8.17936 | 13.42051 |
| **TRIO** | 9.67899 | 0.07267 | 5.45111 | 0.32308 | 1.37166 | 0.00000 | 1.46667 | 2.40192 | 1.39120 | 4.94426 | 0.00089 | 0.52503 |
| **TRIOBP** | 1.79012 | 0.65553 | 1.32539 | 4.08751 | 1.44367 | 0.00000 | 1.71266 | 0.29301 | 0.86007 | 4.87577 | 1.10967 | 0.40727 |
| **TRIP10** | 1.12598 | 14.42608 | 13.21519 | 14.25245 | 25.32685 | 0.00000 | 14.62017 | 4.14199 | 11.79307 | 7.88108 | 9.19488 | 5.01704 |
| **TRIP11** | 5.44644 | 3.89481 | 2.39532 | 1.89384 | 3.55663 | 16.45888 | 2.12403 | 2.40713 | 3.28654 | 14.53755 | 5.52769 | 7.75585 |
| **TRIP12** | 14.25097 | 15.25778 | 10.79232 | 6.18992 | 8.27585 | 32.54576 | 17.03842 | 16.15234 | 7.76017 | 28.18115 | 6.42818 | 26.06926 |
| **TRIP13** | 2.32771 | 14.39512 | 12.76401 | 2.76332 | 5.95980 | 15.35214 | 10.04557 | 1.39886 | 12.84447 | 0.00004 | 16.16288 | 7.37879 |
| **TRIP4** | 0.95231 | 8.68532 | 9.90720 | 0.95566 | 4.82813 | 18.12470 | 9.42413 | 1.93298 | 9.94746 | 0.00000 | 8.78912 | 9.30846 |
| **TRIP6** | 0.19459 | 0.00000 | 9.82396 | 0.00000 | 1.76961 | 0.00000 | 8.04295 | 0.01884 | 4.56246 | 2.60304 | 2.69915 | 0.02579 |
| **TRIQK** | 0.00000 | 0.17227 | 0.52020 | 1.66984 | 0.00000 | 1.71150 | 0.12817 | 0.19096 | 0.21862 | 0.13607 | 0.00026 | 0.14748 |
| **TRIT1** | 1.70116 | 5.58192 | 3.13819 | 5.01235 | 0.98456 | 10.54961 | 5.51714 | 2.89001 | 4.74516 | 3.31762 | 5.13855 | 6.66362 |
| **TRMT1** | 0.00000 | 13.33792 | 8.65514 | 10.21805 | 8.23407 | 0.00000 | 9.62688 | 3.03807 | 20.90388 | 5.78202 | 8.05268 | 18.47690 |
| **TRMT10A** | 0.00000 | 3.21076 | 5.58764 | 0.00000 | 0.00000 | 11.90588 | 6.38810 | 6.05748 | 3.32660 | 3.46526 | 4.49112 | 8.04942 |
| **TRMT10B** | 0.01886 | 0.39841 | 0.32679 | 0.89731 | 0.81067 | 1.14790 | 1.17537 | 0.24513 | 0.47402 | 1.05238 | 1.30618 | 0.48927 |
| **TRMT10C** | 1.49786 | 10.25851 | 15.27324 | 7.31421 | 9.44042 | 42.04937 | 22.66035 | 14.62591 | 18.49562 | 18.80892 | 18.57533 | 23.98252 |
| **TRMT11** | 2.26997 | 0.98481 | 8.17970 | 3.26301 | 0.00000 | 9.74860 | 3.25355 | 0.97961 | 10.63138 | 1.13810 | 7.28392 | 0.97662 |

| | | | | | | | | | | | | |
|---|---|---|---|---|---|---|---|---|---|---|---|---|
| **TRMT112** | 14.97863 | 21.22647 | 17.77211 | 13.97066 | 18.10099 | 19.12317 | 29.14474 | 23.25396 | 17.56385 | 5.56688 | 20.27394 | 33.69246 |
| **TRMT12** | 0.00101 | 3.07924 | 4.07141 | 0.97715 | 1.37879 | 5.43196 | 3.90338 | 2.16599 | 3.54868 | 0.00000 | 1.85039 | 4.49958 |
| **TRMT13** | 2.33334 | 2.16536 | 4.27967 | 2.19277 | 1.24513 | 9.35615 | 3.50109 | 2.85863 | 3.14823 | 3.21866 | 0.91613 | 3.28944 |
| **TRMT1L** | 1.47306 | 6.04663 | 2.98041 | 2.83415 | 0.59505 | 13.83008 | 0.00000 | 0.74268 | 1.89452 | 5.54671 | 4.31998 | 4.80878 |
| **TRMT2A** | 0.17200 | 9.84331 | 12.39170 | 4.41848 | 6.86391 | 0.00000 | 8.00137 | 0.56111 | 17.21939 | 3.12727 | 13.84806 | 5.20995 |
| **TRMT2B** | 0.79882 | 1.24956 | 1.87000 | 1.34753 | 1.59627 | 2.76113 | 2.37706 | 0.05553 | 3.17104 | 1.40472 | 3.80191 | 0.02721 |
| **TRMT44** | 7.71017 | 0.54076 | 1.49194 | 0.25913 | 2.11822 | 0.00000 | 1.21237 | 3.44437 | 1.39500 | 2.81244 | 0.80057 | 1.53745 |
| **TRMT5** | 0.96642 | 4.78311 | 3.68518 | 0.93881 | 0.18563 | 12.49793 | 2.12045 | 4.10776 | 3.69119 | 1.54670 | 4.67608 | 11.57808 |
| **TRMT6** | 2.29186 | 11.13575 | 14.94664 | 2.72115 | 6.10457 | 16.01828 | 14.21641 | 11.39576 | 10.22749 | 10.51878 | 13.64954 | 14.29900 |
| **TRMT61A** | 0.25019 | 8.24201 | 8.52453 | 3.53444 | 9.75701 | 0.00000 | 10.60002 | 4.20786 | 12.75242 | 3.23682 | 5.21698 | 14.97199 |
| **TRMT61B** | 0.51624 | 5.19641 | 5.31491 | 2.75858 | 0.00000 | 6.98154 | 3.20320 | 2.38786 | 4.89830 | 0.00000 | 4.12255 | 6.74330 |
| **TRMU** | 0.30059 | 7.92407 | 12.35549 | 7.18203 | 7.59084 | 0.00920 | 24.82336 | 3.75626 | 19.20047 | 0.00000 | 12.00229 | 6.30915 |
| **TRNAU1AP** | 1.48790 | 9.26840 | 8.91425 | 5.66668 | 9.95267 | 4.65741 | 8.26315 | 5.81027 | 7.94134 | 0.82442 | 9.95095 | 6.57631 |
| **TRNP1** | 0.13449 | 0.10040 | 0.12870 | 0.60338 | 0.75492 | 0.00000 | 0.43106 | 0.04460 | 0.00000 | 0.84316 | 0.04537 | 0.04142 |
| **TRNT1** | 0.79046 | 3.88777 | 3.03034 | 1.37038 | 0.16888 | 21.14744 | 6.56253 | 6.31286 | 2.95377 | 0.00000 | 9.63420 | 7.58452 |
| **TRO** | 0.00176 | 1.09010 | 0.14502 | 2.39379 | 0.00000 | 0.00000 | 0.02721 | 0.15502 | 0.76987 | 0.35635 | 0.17069 | 2.22066 |
| **TROAP** | 6.13504 | 16.17520 | 5.80047 | 12.76455 | 11.76049 | 0.00005 | 11.00963 | 6.06030 | 18.12628 | 0.00000 | 9.50664 | 16.00595 |
| **TROVE2** | 1.74965 | 4.66697 | 4.60213 | 4.05477 | 0.00000 | 12.99050 | 9.30888 | 6.11903 | 9.48046 | 5.60117 | 3.90039 | 6.31658 |
| **TRPA1** | 4.17733 | 0.03739 | 0.00551 | 0.00267 | 0.01367 | 0.05306 | 0.04285 | 0.60985 | 0.00202 | 0.00000 | 0.00876 | 0.04658 |
| **TRPC1** | 0.09474 | 0.18161 | 0.06138 | 0.34927 | 0.12486 | 0.25849 | 0.37807 | 0.04378 | 0.00000 | 0.80881 | 0.02004 | 0.17234 |
| **TRPC3** | 1.58643 | 0.00668 | 0.01115 | 0.01670 | 0.01654 | 0.01338 | 0.04916 | 0.04655 | 0.00163 | 0.02020 | 0.02736 | 0.04925 |
| **TRPC4** | 0.03072 | 0.04782 | 0.01120 | 0.00662 | 0.00725 | 0.02482 | 0.01395 | 0.03536 | 0.02398 | 0.02468 | 0.02122 | 0.00925 |
| **TRPC4AP** | 8.80155 | 9.24629 | 10.94235 | 9.75296 | 8.48706 | 0.00000 | 5.24829 | 1.90088 | 12.33439 | 5.84684 | 6.29682 | 4.43770 |
| **TRPC5** | 0.00653 | 0.01885 | 0.05530 | 0.02324 | 0.03877 | 0.01098 | 0.01602 | 0.00987 | 0.01410 | 0.01643 | 0.01253 | 0.03053 |
| **TRPC5OS** | 0.03658 | 0.01116 | 0.01861 | 0.02702 | 0.03054 | 0.01352 | 0.02910 | 0.00885 | 0.00269 | 0.03671 | 0.01385 | 0.02949 |
| **TRPC6** | 0.03828 | 0.06011 | 0.00928 | 0.07844 | 0.03479 | 0.11206 | 0.00000 | 0.01878 | 0.00861 | 0.01141 | 0.00798 | 0.00580 |
| **TRPC7** | 1.50294 | 0.06604 | 0.06599 | 0.68532 | 0.22644 | 0.00000 | 0.00000 | 0.43698 | 0.00000 | 0.22471 | 0.22138 | 2.46814 |
| **TRPM1** | 0.00887 | 0.24360 | 0.80681 | 0.00000 | 0.00000 | 0.31716 | 0.22459 | 0.00358 | 0.00000 | 0.07694 | 0.86194 | 0.18363 |
| **TRPM2** | 0.01719 | 0.01041 | 0.08566 | 0.02385 | 0.00114 | 0.19386 | 0.00000 | 0.05895 | 0.05571 | 0.00000 | 0.01503 | 0.05274 |
| **TRPM3** | 0.00526 | 0.02516 | 0.02513 | 0.01139 | 0.02880 | 0.01664 | 0.03907 | 0.15450 | 0.01849 | 0.00000 | 0.01482 | 0.02765 |
| **TRPM4** | 0.00049 | 1.08972 | 2.03883 | 2.48074 | 6.32173 | 0.00000 | 1.51965 | 1.13521 | 6.43181 | 0.03117 | 0.79907 | 2.36775 |
| **TRPM5** | 2.96441 | 0.02241 | 0.19507 | 0.00770 | 0.04654 | 0.00000 | 0.02075 | 0.56403 | 0.00000 | 0.03196 | 0.06151 | 0.01429 |
| **TRPM6** | 0.00000 | 3.37795 | 0.00000 | 0.00000 | 3.45038 | 6.56599 | 1.47087 | 2.74236 | 3.23717 | 7.62729 | 6.30785 | 7.66773 |
| **TRPM7** | 2.64548 | 1.17271 | 10.84484 | 0.39924 | 1.36174 | 17.38912 | 3.66151 | 6.54176 | 11.04005 | 16.22116 | 3.84259 | 5.08561 |
| **TRPM8** | 0.02743 | 0.01459 | 0.01598 | 0.00851 | 0.03419 | 0.01929 | 0.01707 | 0.05180 | 0.01429 | 0.02469 | 0.01223 | 0.01204 |
| **TRPS1** | 0.12327 | 0.05596 | 0.52065 | 0.87352 | 0.00000 | 0.29554 | 0.88382 | 0.00000 | 0.02372 | 0.36512 | 0.13134 | 0.57932 |
| **TRPT1** | 0.01440 | 5.74724 | 4.97725 | 5.86588 | 5.17863 | 1.07746 | 1.45423 | 0.12016 | 2.47690 | 0.00000 | 5.10766 | 2.73495 |
| **TRPV1** | 2.38789 | 3.49327 | 0.00000 | 3.47107 | 0.34719 | 0.75637 | 0.04369 | 0.00000 | 1.03731 | 3.07815 | 0.57624 | 3.26870 |
| **TRPV2** | 0.00580 | 0.09429 | 13.89515 | 0.00000 | 0.00000 | 0.00000 | 0.00000 | 0.03346 | 0.00000 | 0.62831 | 0.76820 | 0.15033 |
| **TRPV3** | 0.37728 | 5.12375 | 4.45385 | 2.17967 | 1.99360 | 3.55652 | 2.01945 | 0.25117 | 7.97981 | 5.53308 | 1.53949 | 0.00000 |
| **TRPV4** | 3.69952 | 0.14932 | 0.15442 | 0.33815 | 1.01296 | 0.00000 | 0.17082 | 0.48305 | 0.96621 | 0.56063 | 0.42176 | 0.00224 |
| **TRPV5** | 0.00556 | 0.01512 | 0.01371 | 0.01133 | 0.01122 | 0.00000 | 0.04439 | 0.01043 | 0.00000 | 0.05616 | 0.07943 | 0.01709 |
| **TRPV6** | 1.70996 | 0.01017 | 0.15905 | 0.00682 | 0.00000 | 0.01750 | 0.13223 | 0.26859 | 0.16551 | 0.00000 | 0.00000 | 0.01537 |
| **TRRAP** | 0.75533 | 5.29729 | 2.33153 | 6.91443 | 7.17750 | 2.54812 | 3.62068 | 10.23319 | 4.18305 | 10.26039 | 1.90702 | 8.53752 |
| **TRUB1** | 0.67576 | 4.64314 | 0.45940 | 0.63260 | 0.00000 | 14.18301 | 2.28133 | 0.94907 | 2.87100 | 3.86164 | 5.20474 | 2.03430 |
| **TRUB2** | 0.34794 | 18.34765 | 10.91622 | 11.80770 | 10.39160 | 11.39852 | 17.86995 | 6.17991 | 17.27786 | 0.00000 | 16.06014 | 19.45302 |
| **TRY2P** | 0.00232 | 0.00000 | 0.00000 | 0.00209 | 0.00284 | 0.01386 | 0.00088 | 0.67899 | 0.01057 | 0.60012 | 0.00000 | 0.23442 |
| **TSACC** | 0.03669 | 0.93402 | 0.68689 | 0.35359 | 0.85757 | 1.28661 | 0.00000 | 0.35651 | 1.74918 | 1.58578 | 1.70572 | 1.00936 |

| | | | | | | | | | | | | |
|---|---|---|---|---|---|---|---|---|---|---|---|---|
| **TSC1** | 7.58946 | 5.66840 | 5.23985 | 2.97795 | 4.86176 | 6.46176 | 5.35736 | 8.76749 | 5.31480 | 4.04619 | 5.24739 | 6.36983 |
| **TSC2** | 1.37649 | 3.80732 | 3.08773 | 5.39747 | 3.63134 | 0.00000 | 2.04560 | 0.59054 | 7.08702 | 5.64861 | 1.30040 | 3.24604 |
| **TSC22D1** | 4.46522 | 0.00000 | 13.85760 | 6.07741 | 4.21682 | 9.15555 | 16.53167 | 2.04614 | 19.82899 | 0.87048 | 0.72303 | 2.43409 |
| **TSC22D1-AS1** | 0.26889 | 0.01124 | 0.10582 | 0.01109 | 0.04824 | 0.00000 | 0.00964 | 0.02810 | 0.00000 | 0.05610 | 0.18865 | 0.08139 |
| **TSC22D2** | 3.48222 | 0.32333 | 7.53982 | 2.05163 | 3.12439 | 5.08584 | 4.65754 | 3.82881 | 6.97198 | 7.68793 | 3.91768 | 2.37024 |
| **TSC22D3** | 0.45761 | 0.00000 | 0.14536 | 0.15581 | 0.00000 | 0.03168 | 0.28223 | 0.12734 | 0.01501 | 0.00000 | 0.01261 | 1.12929 |
| **TSC22D4** | 0.00579 | 0.69641 | 0.69907 | 1.03799 | 0.98664 | 0.00000 | 1.50646 | 0.08791 | 0.15272 | 2.00541 | 0.34699 | 0.00000 |
| **TSEN15** | 0.00000 | 4.92532 | 0.00000 | 0.00000 | 2.72865 | 18.23575 | 4.59179 | 0.41192 | 2.79198 | 0.72588 | 6.20394 | 4.35094 |
| **TSEN2** | 0.84977 | 5.10707 | 2.51229 | 1.08885 | 0.88325 | 7.18418 | 0.00000 | 0.05296 | 6.11154 | 0.00000 | 6.39750 | 5.87904 |
| **TSEN34** | 0.66458 | 3.83534 | 8.79890 | 2.22667 | 5.24576 | 0.00000 | 4.18337 | 3.27871 | 4.08124 | 6.19221 | 4.16836 | 2.30639 |
| **TSEN54** | 0.00000 | 15.72772 | 0.82335 | 6.24585 | 9.65533 | 0.00000 | 6.88131 | 0.20334 | 17.43958 | 10.49595 | 14.23929 | 9.39243 |
| **TSFM** | 7.00874 | 15.84856 | 6.97122 | 9.34742 | 10.90116 | 21.11076 | 17.34834 | 18.46435 | 20.71096 | 3.68764 | 16.60075 | 18.74092 |
| **TSG1** | 0.15690 | 0.01496 | 0.03149 | 0.01021 | 0.01610 | 0.00916 | 0.01596 | 0.07704 | 0.02057 | 0.02185 | 0.01341 | 0.01635 |
| **TSG101** | 7.23736 | 12.54237 | 12.57425 | 9.57350 | 2.79295 | 23.24141 | 6.17193 | 2.32025 | 11.22687 | 0.00000 | 9.90291 | 12.97244 |
| **TSGA10** | 2.38693 | 1.27213 | 0.00002 | 0.17246 | 1.20063 | 1.76108 | 0.00000 | 0.53405 | 0.00000 | 2.82851 | 0.81269 | 1.84455 |
| **TSGA10IP** | 0.00558 | 0.00985 | 0.01574 | 0.11898 | 0.02854 | 0.00000 | 0.01209 | 0.01035 | 0.01177 | 0.04021 | 0.08002 | 0.01809 |
| **TSHR** | 0.15533 | 0.03386 | 0.03459 | 0.00387 | 0.00982 | 0.02939 | 0.00013 | 0.09984 | 0.01894 | 0.06215 | 0.01094 | 0.13391 |
| **TSHZ1** | 1.04247 | 0.17651 | 0.00000 | 0.00000 | 0.50184 | 0.33082 | 0.00000 | 0.08787 | 0.00000 | 0.14049 | 0.00000 | 0.37798 |
| **TSHZ2** | 0.44844 | 0.00817 | 0.07779 | 0.01289 | 0.00941 | 0.03307 | 0.01217 | 0.00000 | 0.00000 | 0.06131 | 0.03428 | 0.10635 |
| **TSHZ3** | 0.38613 | 0.04847 | 0.01572 | 0.33962 | 0.00000 | 0.00000 | 0.05546 | 0.11749 | 0.04239 | 0.02500 | 0.02834 | 0.00825 |
| **TSIX** | 0.02889 | 0.00000 | 0.10194 | 0.04223 | 0.00000 | 0.00000 | 0.01291 | 0.25142 | 0.00000 | 0.19548 | 0.36055 | 1.12771 |
| **TSKS** | 0.15802 | 0.02885 | 0.09162 | 0.21691 | 0.09861 | 0.00000 | 0.22518 | 1.24581 | 0.45363 | 0.21826 | 0.49108 | 0.65892 |
| **TSKU** | 3.46806 | 6.00056 | 4.29394 | 5.61504 | 0.19494 | 2.86034 | 2.39040 | 0.06130 | 6.14409 | 0.84137 | 0.00000 | 0.18162 |
| **TSLP** | 0.00585 | 0.01379 | 0.01329 | 0.01166 | 0.01511 | 0.02295 | 0.02440 | 0.01142 | 0.02049 | 0.00719 | 0.01355 | 0.05375 |
| **TSN** | 5.49121 | 15.94610 | 9.85865 | 5.02640 | 3.04558 | 25.33309 | 15.22397 | 6.85804 | 16.16978 | 2.88657 | 16.85197 | 21.16259 |
| **TSNARE1** | 0.00263 | 0.96354 | 0.19023 | 0.42293 | 0.50530 | 0.00000 | 0.30751 | 0.09750 | 0.00063 | 1.22512 | 0.21232 | 0.44048 |
| **TSNAX** | 0.98428 | 3.39377 | 3.83832 | 0.00000 | 0.00000 | 18.00999 | 7.32529 | 2.70557 | 1.98382 | 5.84769 | 6.06764 | 4.01516 |
| **TSNAX-DISC1.u** | 0.00792 | 0.02208 | 0.03746 | 0.01371 | 0.02277 | 0.00067 | 0.03086 | 0.02067 | 0.01419 | 0.00000 | 0.01835 | 0.03101 |
| **TSNAXIP1** | 0.20887 | 0.06561 | 0.01600 | 0.02676 | 0.00000 | 0.00000 | 0.00235 | 0.15895 | 0.00835 | 0.21666 | 0.03550 | 0.11747 |
| **TSPAN1** | 0.00134 | 0.98048 | 0.19391 | 0.00000 | 0.72875 | 0.00625 | 0.02104 | 0.04673 | 1.55496 | 0.02125 | 3.37453 | 0.00069 |
| **TSPAN10** | 0.02513 | 0.50621 | 0.00000 | 0.00000 | 0.01776 | 0.07130 | 0.00000 | 0.00851 | 0.00000 | 0.10052 | 0.01259 | 0.65727 |
| **TSPAN11** | 0.00344 | 0.00000 | 0.06031 | 0.00000 | 0.00000 | 0.05233 | 0.05585 | 0.00948 | 0.00000 | 0.27299 | 0.00000 | 0.00671 |
| **TSPAN12** | 3.05635 | 0.15586 | 0.82571 | 0.25120 | 0.00000 | 0.69379 | 0.31476 | 0.69465 | 0.62202 | 0.00000 | 0.00000 | 0.57260 |
| **TSPAN13** | 0.03080 | 6.67864 | 5.28756 | 22.41421 | 6.06694 | 27.76121 | 0.00000 | 3.62419 | 14.04134 | 7.36090 | 7.74459 | 1.70285 |
| **TSPAN14** | 5.09259 | 22.69166 | 14.82715 | 8.95646 | 25.44599 | 14.20669 | 2.57312 | 2.33292 | 31.89169 | 20.79211 | 8.33317 | 15.03546 |
| **TSPAN15** | 1.55096 | 0.00000 | 11.27991 | 2.99029 | 8.15645 | 0.00000 | 6.07784 | 0.39352 | 30.64569 | 0.01446 | 13.87074 | 0.00000 |
| **TSPAN16** | 0.06465 | 0.09837 | 0.00498 | 0.06327 | 0.00000 | 0.11416 | 0.00000 | 0.00000 | 0.00000 | 0.06131 | 0.00802 | 0.24035 |
| **TSPAN17** | 0.05538 | 3.26914 | 0.83824 | 0.57404 | 0.31102 | 0.55164 | 0.00000 | 0.26851 | 0.00000 | 0.90970 | 0.00401 | 2.68351 |
| **TSPAN18** | 1.31116 | 1.13892 | 1.65711 | 7.58278 | 6.33028 | 0.00000 | 0.01845 | 0.20441 | 0.00000 | 1.40960 | 0.39003 | 0.33145 |
| **TSPAN2** | 0.00296 | 0.00341 | 0.00060 | 0.00097 | 0.00227 | 0.02203 | 0.00472 | 0.01079 | 0.02180 | 0.35438 | 0.00164 | 0.01902 |
| **TSPAN3** | 0.00000 | 7.89771 | 12.09541 | 19.89002 | 5.71264 | 17.36031 | 0.00000 | 0.48975 | 22.85983 | 3.29446 | 9.15728 | 0.64907 |
| **TSPAN31** | 0.00044 | 0.41560 | 0.05388 | 0.77745 | 0.37308 | 0.73844 | 0.14794 | 0.10789 | 0.11799 | 0.80774 | 0.52300 | 0.41130 |
| **TSPAN32** | 0.00509 | 0.01198 | 0.01094 | 0.08157 | 0.00000 | 0.00159 | 0.01893 | 0.01045 | 0.00000 | 0.08212 | 0.06572 | 0.01469 |
| **TSPAN33** | 0.16354 | 0.00000 | 0.13816 | 0.72865 | 0.39826 | 0.19424 | 0.79851 | 0.02648 | 0.00000 | 0.00000 | 0.00000 | 0.00348 |
| **TSPAN4** | 0.79709 | 3.50826 | 2.65390 | 1.81389 | 5.08482 | 0.00000 | 0.00000 | 0.18962 | 5.77199 | 2.47594 | 9.06166 | 0.00000 |
| **TSPAN5** | 4.34168 | 0.19725 | 1.90543 | 0.07402 | 0.20753 | 0.00000 | 1.04371 | 0.19038 | 0.65961 | 2.57986 | 0.00000 | 0.83082 |
| **TSPAN6** | 0.00000 | 11.73993 | 6.35511 | 3.15522 | 6.96738 | 30.00269 | 0.00000 | 0.05524 | 21.48884 | 4.30013 | 15.85453 | 7.11736 |
| **TSPAN7** | 2.14485 | 0.17654 | 0.01831 | 0.00000 | 0.01083 | 0.08975 | 0.00000 | 0.44528 | 0.00000 | 0.05474 | 0.00676 | 0.14085 |

| | | | | | | | | | | | | |
|---|---|---|---|---|---|---|---|---|---|---|---|---|
| **TSPAN8** | 0.00385 | 0.00179 | 0.18134 | 0.00000 | 0.05298 | 0.07718 | 0.12983 | 0.01579 | 0.00000 | 0.03050 | 0.03059 | 0.01105 |
| **TSPAN9** | 1.03391 | 3.11627 | 10.08753 | 5.49876 | 2.90585 | 0.00000 | 0.14318 | 1.72452 | 1.03771 | 0.13547 | 0.02871 | 4.30798 |
| **TSPEAR** | 1.26317 | 0.03598 | 0.05234 | 0.00828 | 0.01149 | 0.00421 | 0.02827 | 0.03682 | 0.04751 | 0.00000 | 0.01990 | 0.01433 |
| **TSPEAR-AS1** | 0.30241 | 0.01614 | 0.02528 | 0.01111 | 0.03216 | 0.00739 | 0.02053 | 0.09417 | 0.02711 | 0.00000 | 0.02030 | 0.01783 |
| **TSPO** | 0.61746 | 0.79593 | 7.62036 | 0.13424 | 0.00000 | 0.00000 | 1.09774 | 0.07092 | 0.90932 | 0.29513 | 0.83817 | 0.66846 |
| **TSPO2** | 0.00696 | 0.01067 | 0.01454 | 0.05134 | 0.00000 | 0.05559 | 0.06048 | 0.01056 | 0.01061 | 0.06610 | 0.01583 | 0.01039 |
| **TSPY1** | 0.01093 | 0.03725 | 0.01405 | 0.01006 | 0.01376 | 0.04383 | 0.02844 | 0.01100 | 0.01638 | 0.00946 | 0.01256 | 0.01935 |
| **TSPY2** | 0.00345 | 0.32178 | 0.50700 | 0.00000 | 0.38175 | 0.00000 | 0.14075 | 0.01962 | 0.00000 | 0.40021 | 0.02380 | 0.09613 |
| **TSPY8** | 0.00606 | 0.01326 | 0.01671 | 0.00879 | 0.02431 | 0.03249 | 0.01851 | 0.01038 | 0.01064 | 0.01659 | 0.01699 | 0.01497 |
| **TSPYL1** | 5.72332 | 5.59369 | 9.24060 | 6.75733 | 0.00000 | 26.20299 | 17.43079 | 11.48294 | 1.04331 | 2.84086 | 11.92356 | 5.57769 |
| **TSPYL2** | 0.33940 | 1.76674 | 1.39549 | 2.21235 | 0.87851 | 0.29005 | 0.97381 | 2.05629 | 2.90462 | 2.33505 | 0.00000 | 3.50846 |
| **TSPYL4** | 2.23970 | 1.63066 | 5.15549 | 0.64964 | 0.25988 | 0.51006 | 2.77374 | 0.79368 | 1.98462 | 2.43763 | 3.01505 | 1.70631 |
| **TSPYL5** | 1.55716 | 0.24033 | 0.49743 | 0.00000 | 0.98816 | 0.90089 | 2.01497 | 0.06957 | 0.74413 | 1.33759 | 0.54702 | 1.86541 |
| **TSPYL6** | 0.00156 | 1.57078 | 2.02930 | 0.69488 | 0.08870 | 1.05482 | 0.54982 | 2.01413 | 1.45193 | 0.00000 | 1.21099 | 4.29779 |
| **TSR1** | 0.00399 | 22.40988 | 16.57107 | 9.70124 | 12.63007 | 20.60906 | 17.09492 | 14.85129 | 27.99336 | 0.00000 | 24.55580 | 23.98825 |
| **TSR2** | 2.35283 | 11.18615 | 7.08289 | 9.04477 | 7.48148 | 0.73187 | 14.69576 | 4.70372 | 9.64646 | 0.00000 | 11.44212 | 11.61665 |
| **TSR3** | 0.00084 | 2.74574 | 1.61965 | 1.53646 | 5.50713 | 0.00000 | 6.03151 | 0.00000 | 0.83942 | 9.01282 | 0.83338 | 1.24376 |
| **TSSC1** | 4.11220 | 8.72643 | 4.22341 | 7.64815 | 7.39203 | 5.18301 | 5.21371 | 0.64288 | 5.37275 | 0.00000 | 9.90856 | 2.26590 |
| **TSSC4** | 1.60767 | 13.23545 | 12.33804 | 5.07021 | 9.22298 | 0.00000 | 8.52895 | 2.19010 | 14.76754 | 0.09218 | 10.46486 | 10.15941 |
| **TSSK1B** | 0.00812 | 0.01375 | 0.04013 | 0.02593 | 0.00269 | 0.03985 | 0.01217 | 0.01044 | 0.00408 | 0.03627 | 0.01769 | 0.01399 |
| **TSSK3** | 0.00453 | 0.04724 | 0.04250 | 0.00013 | 0.00000 | 0.05988 | 0.00000 | 0.00881 | 0.05772 | 0.09603 | 0.03864 | 0.04171 |
| **TSSK4** | 0.06873 | 0.01683 | 0.06112 | 0.01705 | 0.11304 | 0.00000 | 0.10655 | 0.00174 | 0.15373 | 0.18832 | 0.09125 | 0.06174 |
| **TSSK6** | 0.00878 | 0.12397 | 0.04764 | 0.09654 | 0.00000 | 0.00000 | 0.05958 | 0.61492 | 0.14986 | 0.42179 | 0.01010 | 0.27266 |
| **TST** | 0.03646 | 0.77827 | 1.31293 | 0.62711 | 1.50634 | 0.09609 | 2.95585 | 0.03684 | 3.02486 | 0.00101 | 1.93383 | 0.10807 |
| **TSTA3** | 2.54959 | 9.53467 | 12.04195 | 13.94810 | 16.51214 | 0.01105 | 14.55722 | 0.48863 | 23.25076 | 0.00000 | 17.79157 | 5.80686 |
| **TSTD1** | 0.56273 | 6.42203 | 1.13246 | 5.08071 | 3.79062 | 0.46718 | 3.14938 | 4.25714 | 1.17913 | 2.54324 | 3.48465 | 3.09879 |
| **TSTD2** | 5.36370 | 3.00645 | 2.25831 | 1.64750 | 0.87415 | 2.84992 | 2.13929 | 3.75246 | 2.44770 | 0.00000 | 1.47076 | 3.55354 |
| **TSTD3** | 0.00239 | 0.29658 | 0.65005 | 0.14205 | 0.25659 | 0.99065 | 0.00000 | 0.18070 | 0.70956 | 1.09093 | 1.17354 | 0.21247 |
| **TTBK1** | 0.00509 | 0.01196 | 0.06880 | 0.01801 | 0.02022 | 0.02114 | 0.02743 | 0.01141 | 0.00595 | 0.02112 | 0.02441 | 0.04144 |
| **TTBK2** | 2.63670 | 0.22027 | 0.00000 | 0.65737 | 0.51464 | 0.97510 | 0.12536 | 0.22098 | 0.32826 | 1.67388 | 0.25536 | 0.71129 |
| **TTC1** | 3.67018 | 14.31485 | 19.07258 | 6.92293 | 9.11036 | 20.32956 | 32.95797 | 12.26415 | 12.96149 | 10.50604 | 14.26406 | 18.47715 |
| **TTC12** | 3.08115 | 1.65475 | 0.00000 | 0.87682 | 0.32117 | 0.42548 | 0.39968 | 0.62730 | 0.92470 | 0.00000 | 0.31026 | 1.24020 |
| **TTC13** | 0.02057 | 0.38429 | 0.88797 | 0.29811 | 0.41288 | 0.00000 | 0.00000 | 0.25654 | 1.60074 | 0.32855 | 0.78098 | 0.00000 |
| **TTC14** | 0.00000 | 1.27452 | 4.13949 | 0.56694 | 0.00000 | 10.46649 | 4.07944 | 5.97471 | 10.49347 | 12.35496 | 4.00372 | 7.08973 |
| **TTC16** | 0.07615 | 0.00695 | 0.01654 | 0.01305 | 0.02114 | 0.00907 | 0.01821 | 0.28481 | 0.02109 | 0.00701 | 0.01456 | 0.16374 |
| **TTC17** | 2.91192 | 5.07425 | 3.18801 | 4.66488 | 2.48401 | 5.12599 | 1.04430 | 1.90511 | 12.53725 | 6.09894 | 1.49206 | 6.39642 |
| **TTC18** | 0.33066 | 0.50094 | 0.18562 | 0.02869 | 0.10084 | 0.35143 | 0.00000 | 1.13804 | 0.35772 | 0.28535 | 0.40994 | 1.10714 |
| **TTC19** | 0.02231 | 2.70122 | 0.07238 | 6.13691 | 3.23073 | 10.87966 | 1.73750 | 1.25396 | 1.51759 | 14.51466 | 3.63889 | 1.77851 |
| **TTC21A** | 0.01015 | 0.07391 | 0.00000 | 0.19290 | 0.16550 | 0.23572 | 0.69590 | 0.06286 | 0.26980 | 1.01325 | 0.00000 | 0.11742 |
| **TTC21B** | 1.18203 | 1.19309 | 0.92201 | 0.93611 | 0.00003 | 2.34284 | 0.00000 | 0.47427 | 1.17803 | 0.71023 | 0.56282 | 1.37863 |
| **TTC21B-AS1** | 0.00641 | 0.04727 | 0.03094 | 0.01291 | 0.03193 | 0.00000 | 0.01951 | 0.01050 | 0.03763 | 0.00000 | 0.02078 | 0.02294 |
| **TTC22** | 0.00381 | 0.00948 | 0.01779 | 0.12495 | 0.00000 | 0.07942 | 0.00000 | 0.01115 | 0.00000 | 0.13106 | 0.01429 | 0.01538 |
| **TTC23** | 0.00283 | 1.77449 | 0.09042 | 0.59376 | 0.48525 | 0.00000 | 0.13107 | 0.31190 | 0.23103 | 0.00000 | 0.43280 | 0.75888 |
| **TTC23L** | 0.44235 | 0.04973 | 0.16048 | 0.00000 | 0.05361 | 0.08439 | 0.16431 | 0.91212 | 0.22068 | 0.00477 | 0.00000 | 0.03062 |
| **TTC24** | 0.06648 | 0.00842 | 0.01370 | 0.00857 | 0.00919 | 0.03450 | 0.02147 | 0.00618 | 0.01600 | 0.02502 | 0.01415 | 0.04638 |
| **TTC25** | 0.09033 | 0.28246 | 0.07034 | 0.00000 | 0.00000 | 0.25925 | 0.04702 | 0.27194 | 0.21241 | 0.00421 | 0.02546 | 0.16490 |
| **TTC26** | 0.48265 | 3.76785 | 0.34276 | 0.53900 | 0.78030 | 0.05452 | 0.28224 | 0.63187 | 0.48338 | 0.00000 | 0.67482 | 3.27871 |
| **TTC27** | 0.00000 | 9.06949 | 2.43083 | 1.24829 | 2.91162 | 12.70512 | 5.99938 | 4.30658 | 8.61812 | 3.11260 | 7.52418 | 13.53768 |

| | | | | | | | | | | | | |
|---|---|---|---|---|---|---|---|---|---|---|---|---|
| **TTC28** | 2.39580 | 0.85968 | 0.62720 | 0.46407 | 7.39187 | 1.44534 | 0.00000 | 0.28623 | 2.16502 | 1.59750 | 3.30301 | 0.02614 |
| **TTC28-AS1** | 1.65639 | 0.93607 | 3.48441 | 2.20695 | 1.66195 | 3.35738 | 1.26848 | 5.85045 | 1.00615 | 7.89516 | 0.99547 | 4.05004 |
| **TTC29** | 0.52536 | 0.01727 | 0.00934 | 0.00346 | 0.03102 | 0.02197 | 0.01945 | 0.02291 | 0.01306 | 0.05328 | 0.01345 | 0.00981 |
| **TTC3** | 5.91930 | 0.71180 | 3.73645 | 0.82881 | 1.82761 | 7.14162 | 8.75311 | 0.74569 | 3.21716 | 21.28142 | 5.90970 | 0.00000 |
| **TTC30A** | 0.36422 | 0.23317 | 0.42119 | 0.00000 | 0.32273 | 0.00000 | 0.00000 | 0.51988 | 0.12837 | 0.30172 | 0.01276 | 0.41901 |
| **TTC30B** | 0.18745 | 0.15986 | 0.05215 | 0.00000 | 0.00604 | 0.52757 | 0.00000 | 0.05971 | 0.08831 | 0.76719 | 0.24411 | 0.32065 |
| **TTC31** | 1.23725 | 5.89208 | 1.48972 | 5.54853 | 2.99783 | 1.03292 | 3.59619 | 0.31245 | 4.25581 | 0.00000 | 3.22505 | 5.67517 |
| **TTC32** | 0.65532 | 1.47211 | 2.71052 | 1.32673 | 0.00000 | 1.67743 | 4.49959 | 0.99032 | 0.40393 | 2.69176 | 1.42400 | 2.19867 |
| **TTC33** | 0.59295 | 2.30771 | 0.43374 | 0.23383 | 0.00000 | 5.31097 | 2.51703 | 3.69505 | 1.24222 | 0.30607 | 1.76446 | 3.20038 |
| **TTC36** | 0.00638 | 0.02540 | 0.01580 | 0.01255 | 0.03584 | 0.00000 | 0.01465 | 0.01069 | 0.03803 | 0.01368 | 0.01814 | 0.01650 |
| **TTC37** | 0.56548 | 1.46514 | 3.45102 | 2.67143 | 2.35005 | 10.69466 | 6.49816 | 0.33480 | 8.06558 | 6.21962 | 7.84602 | 0.62306 |
| **TTC38** | 0.84023 | 0.00000 | 2.49158 | 1.78318 | 0.73086 | 0.33954 | 2.09188 | 0.05939 | 4.94483 | 0.00000 | 2.42269 | 0.26744 |
| **TTC39A** | 0.82636 | 0.08417 | 3.42527 | 0.00285 | 0.11684 | 0.40098 | 1.22520 | 0.24688 | 0.01553 | 0.00000 | 0.98036 | 0.24268 |
| **TTC39B** | 0.17973 | 0.40734 | 0.00000 | 7.27011 | 0.22218 | 1.86188 | 0.00000 | 0.61278 | 0.00000 | 0.42464 | 0.00000 | 1.14052 |
| **TTC39C** | 1.76140 | 0.36979 | 0.40007 | 0.00000 | 0.00000 | 0.30626 | 0.26447 | 2.94357 | 0.00001 | 0.00142 | 0.03650 | 2.12914 |
| **TTC3P1** | 0.34973 | 0.00000 | 0.16345 | 0.00000 | 0.16471 | 0.21788 | 0.73022 | 0.04342 | 0.00000 | 1.47132 | 0.09346 | 0.00000 |
| **TTC4** | 0.34101 | 1.99287 | 0.43524 | 0.15064 | 1.01096 | 0.71878 | 0.73092 | 1.36156 | 0.06802 | 0.00000 | 0.52265 | 2.97178 |
| **TTC40** | 0.07684 | 0.30679 | 0.00000 | 1.24057 | 0.00000 | 0.14529 | 0.00000 | 0.02040 | 0.15573 | 0.04788 | 0.51419 | 0.24065 |
| **TTC5** | 5.71877 | 3.37276 | 0.70286 | 4.63852 | 4.16074 | 2.24645 | 0.72345 | 1.04397 | 2.44507 | 0.00000 | 3.52901 | 5.35641 |
| **TTC7A** | 0.10837 | 2.16519 | 2.41714 | 2.06475 | 3.68479 | 0.00000 | 2.81997 | 0.11363 | 4.01768 | 5.36951 | 1.23652 | 1.24027 |
| **TTC7B** | 0.65430 | 0.72213 | 3.96420 | 0.00000 | 3.09639 | 2.09340 | 7.05022 | 0.00000 | 2.62116 | 7.91776 | 3.00783 | 1.43151 |
| **TTC8** | 0.29700 | 1.73875 | 0.09789 | 0.04782 | 0.00660 | 1.33889 | 0.57657 | 0.06772 | 1.61843 | 0.00000 | 1.06076 | 1.85920 |
| **TTC9** | 0.59631 | 0.00000 | 0.76668 | 0.09699 | 0.71577 | 0.00743 | 1.34165 | 0.00709 | 0.82173 | 0.48925 | 0.13732 | 0.00703 |
| **TTC9B** | 0.14650 | 0.06157 | 0.01034 | 0.00828 | 0.03550 | 0.00000 | 0.00773 | 0.48690 | 0.07505 | 0.03035 | 0.00857 | 0.40510 |
| **TTC9C** | 5.80679 | 5.31241 | 6.01839 | 4.41211 | 7.53152 | 10.41719 | 7.72902 | 17.06110 | 8.55370 | 0.00000 | 7.63886 | 12.55480 |
| **TTF1** | 2.18865 | 6.22212 | 2.21366 | 2.86217 | 4.43189 | 14.96189 | 5.73687 | 7.81213 | 4.45762 | 11.14662 | 1.78383 | 11.97945 |
| **TTF2** | 3.94897 | 1.56526 | 3.22007 | 4.59143 | 7.88526 | 3.37314 | 1.72087 | 2.03255 | 6.90361 | 8.28898 | 7.82794 | 3.82390 |
| **TTI1** | 2.35586 | 9.05748 | 7.96749 | 4.46602 | 6.37713 | 6.15724 | 3.85837 | 7.54649 | 12.36013 | 0.00000 | 12.48224 | 11.08410 |
| **TTI2** | 0.00000 | 5.60780 | 2.78780 | 2.41688 | 2.22679 | 6.51131 | 4.28922 | 5.88739 | 4.24832 | 0.00000 | 6.04749 | 9.38274 |
| **TTK** | 5.63105 | 11.09057 | 5.08616 | 1.37566 | 1.26277 | 27.33329 | 10.88489 | 8.67411 | 7.35769 | 4.45049 | 4.95780 | 10.97662 |
| **TTL** | 0.54162 | 2.56038 | 0.00000 | 0.39445 | 4.94843 | 6.12648 | 4.43280 | 1.55837 | 0.07080 | 7.03978 | 4.00425 | 3.55122 |
| **TTLL1** | 1.38054 | 0.57830 | 0.00000 | 0.59284 | 0.10661 | 0.35977 | 0.22250 | 1.37378 | 0.00000 | 0.10571 | 0.11057 | 0.45677 |
| **TTLL10** | 0.06109 | 0.01566 | 0.09269 | 0.09687 | 0.01218 | 0.00000 | 0.00721 | 0.00613 | 0.01202 | 0.02653 | 0.01744 | 0.01608 |
| **TTLL11** | 0.08165 | 0.00000 | 0.45133 | 0.22313 | 0.14433 | 0.03561 | 0.13839 | 0.06813 | 0.17749 | 0.35446 | 0.08861 | 0.01114 |
| **TTLL12** | 2.16662 | 8.53363 | 8.06977 | 5.05294 | 16.15595 | 0.00000 | 12.52828 | 0.00000 | 3.22274 | 18.62298 | 5.97687 | 3.02022 |
| **TTLL13** | 0.00690 | 0.28331 | 0.06012 | 0.00839 | 0.06346 | 0.00000 | 0.00439 | 0.01346 | 0.20746 | 0.00000 | 0.08202 | 0.03898 |
| **TTLL2** | 0.16030 | 0.02926 | 0.04019 | 0.00674 | 0.01230 | 0.03133 | 0.01448 | 0.00255 | 0.01523 | 0.00981 | 0.01311 | 0.01182 |
| **TTLL3** | 0.00127 | 0.16220 | 0.35282 | 0.37558 | 0.06258 | 0.00000 | 0.00000 | 0.01498 | 0.40237 | 1.11413 | 0.27655 | 0.16378 |
| **TTLL4** | 8.69403 | 10.28657 | 8.56535 | 12.08137 | 11.16306 | 4.30132 | 10.55929 | 3.01629 | 16.66045 | 7.58585 | 10.70694 | 6.97879 |
| **TTLL5** | 2.43658 | 4.60371 | 5.75988 | 3.02737 | 3.18712 | 4.98806 | 5.99932 | 7.55245 | 7.74891 | 4.24902 | 4.26217 | 7.44902 |
| **TTLL6** | 0.00631 | 0.01601 | 0.02618 | 0.03385 | 0.01831 | 0.01818 | 0.01674 | 0.01055 | 0.02116 | 0.00819 | 0.01491 | 0.01661 |
| **TTLL7** | 0.00241 | 0.01479 | 1.18347 | 0.01122 | 0.00000 | 0.11481 | 0.25750 | 0.38196 | 0.47428 | 0.00000 | 0.01929 | 0.14459 |
| **TTLL9** | 0.25741 | 0.54897 | 0.05879 | 0.00000 | 0.21819 | 0.37612 | 0.00000 | 0.05833 | 0.00000 | 0.21093 | 0.00000 | 0.05310 |
| **TTN** | 2.01474 | 0.24656 | 4.65482 | 4.18379 | 4.33825 | 3.52926 | 0.19829 | 1.51604 | 6.69331 | 23.31800 | 5.98265 | 3.08958 |
| **TTN-AS1** | 0.62146 | 0.80541 | 0.69480 | 1.49564 | 0.38652 | 0.00000 | 0.00000 | 1.41505 | 1.72896 | 0.28952 | 0.45221 | 1.31293 |
| **TTPA** | 0.05252 | 0.02843 | 0.07691 | 0.01119 | 0.01605 | 0.00000 | 0.01142 | 0.00668 | 0.02269 | 0.01675 | 0.02908 | 0.10127 |
| **TTPAL** | 0.77366 | 2.63435 | 1.35573 | 1.62612 | 0.47307 | 4.36464 | 2.72329 | 2.53935 | 2.97973 | 3.81987 | 3.99063 | 4.26632 |
| **TTR** | 0.26210 | 0.01318 | 0.36720 | 0.01229 | 0.02872 | 0.00011 | 0.00080 | 0.15869 | 0.03042 | 0.00000 | 0.01495 | 0.23319 |

| | | | | | | | | | | | | |
|---|---|---|---|---|---|---|---|---|---|---|---|---|
| **TTTY14** | 0.00590 | 0.03696 | 0.47464 | 0.14512 | 0.21931 | 0.00031 | 0.00368 | 0.01044 | 0.47323 | 0.00000 | 0.07531 | 0.01416 |
| **TTTY15** | 0.00000 | 4.01808 | 1.08003 | 0.36583 | 0.86151 | 7.59812 | 3.65684 | 1.73340 | 3.14229 | 2.51326 | 1.99954 | 11.79291 |
| **TTYH1** | 0.00453 | 0.00000 | 0.14547 | 0.95601 | 0.20609 | 0.00000 | 0.02054 | 0.00955 | 0.49124 | 0.12302 | 1.12369 | 0.02540 |
| **TTYH2** | 3.19516 | 0.11033 | 0.11087 | 0.00000 | 0.00000 | 0.00000 | 0.00000 | 0.67547 | 0.17592 | 0.23224 | 0.08964 | 0.26566 |
| **TTYH3** | 0.18030 | 0.12890 | 1.47385 | 1.84169 | 0.07812 | 0.16355 | 0.11897 | 0.01475 | 0.44520 | 0.80517 | 0.00000 | 0.00000 |
| **TUB** | 0.80176 | 0.12658 | 0.24048 | 0.00000 | 0.00000 | 0.80550 | 0.21359 | 0.02680 | 0.14194 | 0.77198 | 0.05565 | 0.28055 |
| **TUBA1A** | 0.17067 | 0.83466 | 1.75985 | 4.92350 | 1.00219 | 1.19021 | 2.42146 | 0.08703 | 0.00000 | 0.00000 | 1.04996 | 1.61540 |
| **TUBA1B** | 1.65221 | 32.59813 | 32.47365 | 24.45691 | 34.03583 | 38.83759 | 62.75781 | 13.98622 | 47.76042 | 23.51784 | 42.28065 | 21.21332 |
| **TUBA1C** | 23.72167 | 52.06431 | 26.42604 | 33.59963 | 55.97539 | 46.41861 | 52.31255 | 14.09922 | 50.85507 | 30.33136 | 42.30451 | 35.56692 |
| **TUBA3C** | 0.00713 | 3.85811 | 0.15852 | 0.00000 | 0.00000 | 0.02524 | 0.00000 | 3.45251 | 0.00000 | 0.21258 | 0.00000 | 5.62974 |
| **TUBA3D** | 0.53772 | 3.49459 | 1.06940 | 1.42457 | 1.34907 | 0.08616 | 0.76990 | 4.76351 | 0.00000 | 0.00000 | 1.26623 | 5.70749 |
| **TUBA3E** | 0.00166 | 0.19102 | 0.20050 | 0.70079 | 0.00000 | 0.66353 | 0.00000 | 6.23824 | 0.00000 | 0.02125 | 0.00311 | 0.83022 |
| **TUBA3FP** | 0.00609 | 0.01513 | 0.09671 | 0.05377 | 0.00206 | 0.06462 | 0.02881 | 0.14077 | 0.12707 | 0.01440 | 0.03356 | 0.01706 |
| **TUBA4A** | 7.29340 | 0.00000 | 2.81024 | 0.00000 | 0.00000 | 0.92490 | 0.10010 | 0.35824 | 0.00000 | 0.14874 | 0.12367 | 0.11901 |
| **TUBA4B** | 9.72377 | 0.02463 | 0.12348 | 0.02550 | 0.02058 | 0.09208 | 0.00975 | 2.26892 | 0.01385 | 0.00000 | 0.06230 | 0.40158 |
| **TUBA8** | 0.07840 | 0.32398 | 0.40031 | 0.40857 | 0.57424 | 0.00000 | 0.42807 | 0.03821 | 0.32448 | 0.64210 | 0.33403 | 0.02893 |
| **TUBAL3** | 3.50048 | 0.01496 | 0.11638 | 0.00475 | 0.01200 | 0.02761 | 0.09166 | 0.96611 | 0.00000 | 0.00000 | 0.01162 | 0.00683 |
| **TUBB** | 0.00000 | 33.28755 | 31.72643 | 39.76019 | 37.86032 | 30.69732 | 55.90858 | 4.99251 | 54.79971 | 14.17274 | 48.75312 | 26.85897 |
| **TUBB1** | 0.12482 | 0.15282 | 0.10330 | 0.00134 | 0.18761 | 0.31943 | 0.00000 | 0.83156 | 0.50632 | 1.19201 | 0.13776 | 0.79681 |
| **TUBB2A** | 0.13934 | 1.66765 | 1.08440 | 13.91421 | 3.99639 | 4.81562 | 0.35229 | 0.24869 | 0.00000 | 3.53448 | 1.39787 | 0.51529 |
| **TUBB2B** | 0.00241 | 0.00000 | 1.23304 | 15.79052 | 6.37600 | 0.02733 | 1.19746 | 0.00896 | 0.73451 | 0.16189 | 2.35468 | 0.00000 |
| **TUBB3** | 3.03763 | 6.27623 | 7.22408 | 13.61196 | 5.36527 | 0.00000 | 2.69381 | 0.76076 | 6.22595 | 1.84539 | 0.57713 | 0.66251 |
| **TUBB4A** | 0.00000 | 57.29792 | 6.99962 | 55.74018 | 78.69824 | 11.83266 | 5.98612 | 0.00000 | 33.96367 | 30.93212 | 59.95863 | 28.09853 |
| **TUBB4B** | 11.33334 | 51.01881 | 35.32450 | 32.94612 | 37.53243 | 25.42496 | 42.90233 | 20.38490 | 55.03885 | 0.00000 | 37.00793 | 42.93696 |
| **TUBB6** | 0.00000 | 11.10094 | 11.35824 | 6.93664 | 6.60317 | 13.31143 | 19.40348 | 6.26377 | 29.75224 | 0.00000 | 22.29349 | 5.61879 |
| **TUBB8** | 29.30241 | 0.37501 | 0.00000 | 0.00000 | 0.00468 | 0.00000 | 0.00324 | 7.11220 | 0.00384 | 0.01995 | 0.00818 | 2.45160 |
| **TUBBP5** | 16.13672 | 0.97291 | 0.00000 | 0.42657 | 0.00000 | 0.02763 | 0.20905 | 5.59227 | 0.03213 | 0.06547 | 0.01994 | 2.21417 |
| **TUBD1** | 0.47405 | 2.63913 | 3.68792 | 2.50748 | 0.00000 | 10.28018 | 4.75244 | 1.01253 | 7.68323 | 0.43911 | 5.04954 | 8.76711 |
| **TUBE1** | 0.99303 | 3.11222 | 0.00000 | 2.27617 | 0.33657 | 8.33233 | 0.00000 | 0.00000 | 2.67413 | 1.59260 | 3.66476 | 4.37598 |
| **TUBG1** | 1.82033 | 15.37155 | 14.61038 | 11.71148 | 8.63359 | 6.07992 | 15.80426 | 1.88800 | 12.06802 | 0.00000 | 14.63581 | 6.18511 |
| **TUBG2** | 4.30769 | 0.87453 | 0.74710 | 2.02675 | 0.00000 | 0.18880 | 0.22713 | 0.28916 | 1.41318 | 0.92022 | 0.94060 | 0.77937 |
| **TUBGCP2** | 0.52156 | 10.40317 | 4.99585 | 8.21480 | 8.57388 | 0.06437 | 8.67419 | 0.92426 | 5.12221 | 2.34285 | 8.16967 | 6.78240 |
| **TUBGCP3** | 2.40393 | 6.45990 | 3.80917 | 1.21792 | 2.64460 | 7.92834 | 0.99265 | 1.12832 | 7.23986 | 3.98681 | 6.06746 | 3.91924 |
| **TUBGCP4** | 3.78852 | 4.13023 | 3.54995 | 2.33235 | 4.92289 | 3.06448 | 5.77995 | 0.87256 | 9.92275 | 0.00000 | 4.71739 | 3.77979 |
| **TUBGCP5** | 1.16892 | 3.10911 | 2.57062 | 2.21119 | 2.03665 | 3.87484 | 1.98518 | 0.09202 | 2.94510 | 0.96154 | 3.94232 | 2.59649 |
| **TUBGCP6** | 0.30707 | 2.37726 | 1.59849 | 1.13885 | 1.47765 | 0.00000 | 0.00000 | 0.42921 | 5.32932 | 7.64657 | 0.59137 | 2.58562 |
| **TUFM** | 0.65417 | 41.96422 | 33.12470 | 28.82712 | 37.67541 | 16.04214 | 37.36452 | 8.00803 | 51.97303 | 12.67883 | 33.47454 | 32.85044 |
| **TUFT1** | 0.46867 | 6.73584 | 13.62023 | 0.00000 | 12.78999 | 4.95227 | 4.68207 | 1.09953 | 7.20037 | 2.23569 | 4.94998 | 4.50078 |
| **TUG1** | 2.01911 | 3.67881 | 9.67368 | 3.23290 | 2.31216 | 18.60617 | 12.70418 | 9.98129 | 5.21746 | 29.16643 | 8.19155 | 15.89305 |
| **TULP1** | 0.00234 | 0.00892 | 0.86196 | 0.01054 | 0.03619 | 0.00000 | 0.00000 | 0.07179 | 0.16943 | 0.03535 | 0.00503 | 0.10486 |
| **TULP2** | 0.15837 | 1.35552 | 0.41237 | 2.03600 | 1.18903 | 0.00000 | 0.04699 | 0.53714 | 0.30296 | 0.19823 | 0.12598 | 1.18378 |
| **TULP3** | 15.64512 | 6.24647 | 4.05715 | 2.74048 | 4.18259 | 5.84632 | 6.04662 | 9.06903 | 3.03672 | 0.00000 | 4.73840 | 13.75171 |
| **TULP4** | 1.38952 | 12.40575 | 5.41264 | 8.15592 | 5.42750 | 23.05999 | 11.55343 | 4.24455 | 8.04656 | 14.16527 | 12.57660 | 6.29249 |
| **TUNAR** | 0.44103 | 0.01739 | 0.01875 | 0.00960 | 0.04912 | 0.01970 | 0.03511 | 0.00322 | 0.00353 | 0.00010 | 0.01571 | 0.01262 |
| **TUSC1** | 0.22381 | 0.26153 | 2.21534 | 1.06400 | 0.27344 | 0.65365 | 1.64152 | 0.16042 | 0.00000 | 3.83224 | 0.62754 | 0.48594 |
| **TUSC2** | 2.23543 | 10.62844 | 13.24778 | 6.91226 | 13.74211 | 0.17619 | 17.16121 | 7.50819 | 10.63365 | 3.70180 | 11.44921 | 16.28426 |
| **TUSC3** | 1.56157 | 1.24796 | 12.06345 | 2.39069 | 0.00136 | 7.63873 | 0.00061 | 0.13653 | 0.93900 | 6.00889 | 0.82326 | 0.75422 |
| **TUSC5** | 1.99726 | 0.01144 | 0.05110 | 0.00188 | 0.00453 | 0.02372 | 0.00000 | 0.27253 | 0.01563 | 0.07188 | 0.01058 | 0.01808 |

| | | | | | | | | | | | | |
|---|---|---|---|---|---|---|---|---|---|---|---|---|
| **TUSC8** | 0.34382 | 0.01713 | 0.01666 | 0.07180 | 0.01797 | 0.00000 | 0.02340 | 0.04117 | 0.01923 | 0.00000 | 0.01643 | 0.02886 |
| **TUT1** | 0.84113 | 8.34972 | 4.14858 | 3.22676 | 6.47151 | 0.00000 | 4.29530 | 5.30881 | 3.20400 | 0.00000 | 6.26872 | 6.33726 |
| **TVP23A** | 0.07562 | 0.07658 | 0.13637 | 0.09832 | 0.00019 | 0.24416 | 0.39989 | 0.03251 | 0.12249 | 0.12968 | 0.13384 | 0.22839 |
| **TVP23B** | 0.73990 | 1.96611 | 2.96350 | 0.46299 | 0.00000 | 4.43276 | 0.00000 | 4.24049 | 3.84031 | 0.00000 | 2.33738 | 4.82815 |
| **TVP23C.uid-209** | 0.09879 | 0.28775 | 0.16818 | 0.04181 | 0.19789 | 0.49423 | 0.00000 | 0.40724 | 0.11077 | 0.28406 | 0.06512 | 0.62384 |
| **TVP23C.uid-209** | 0.00086 | 1.39720 | 0.14620 | 0.37126 | 1.40429 | 0.00000 | 0.13567 | 0.44041 | 0.51886 | 1.29057 | 0.23077 | 0.42442 |
| **TWF1** | 1.20477 | 3.82421 | 2.92178 | 0.32256 | 0.52163 | 16.34526 | 5.18563 | 1.23333 | 5.72307 | 0.00209 | 3.28085 | 4.00497 |
| **TWF2** | 0.17520 | 17.51924 | 8.06255 | 22.10442 | 19.29410 | 0.17530 | 16.38629 | 1.87472 | 11.24831 | 0.00000 | 13.71121 | 6.44536 |
| **TWIST1** | 0.06948 | 0.00000 | 0.43572 | 2.17920 | 0.00000 | 0.14921 | 0.16350 | 0.06638 | 0.00000 | 0.29948 | 0.00000 | 0.07657 |
| **TWIST2** | 0.00510 | 0.01844 | 0.10054 | 0.05080 | 0.00740 | 0.00662 | 0.03595 | 0.01077 | 0.00000 | 0.05494 | 0.01875 | 0.01800 |
| **TWISTNB** | 6.51848 | 9.63412 | 13.06651 | 6.80837 | 8.64293 | 19.51923 | 23.54392 | 5.79139 | 6.06316 | 10.96603 | 12.89158 | 11.90305 |
| **TWSG1** | 0.03677 | 0.15071 | 4.62224 | 0.03210 | 0.00006 | 2.64698 | 0.00000 | 4.65510 | 2.59405 | 0.34413 | 0.00000 | 1.52757 |
| **TXK** | 0.10216 | 0.33232 | 0.00000 | 1.59069 | 0.00000 | 0.90160 | 0.10114 | 0.00000 | 0.00000 | 0.24514 | 0.00000 | 0.60296 |
| **TXLNA** | 0.00000 | 11.72973 | 11.97684 | 14.11349 | 9.37067 | 3.82391 | 17.07021 | 5.21873 | 17.30154 | 2.68346 | 18.05008 | 13.25461 |
| **TXLNB** | 0.08094 | 0.00569 | 0.03918 | 0.00170 | 0.07520 | 0.12196 | 0.01700 | 0.10171 | 0.00000 | 0.00590 | 0.03718 | 0.04991 |
| **TXLNG** | 4.98363 | 4.83945 | 6.44457 | 3.27814 | 5.78695 | 10.58405 | 12.96885 | 3.79667 | 6.66598 | 12.13495 | 9.65252 | 3.82296 |
| **TXLNGY** | 0.00002 | 0.00001 | 1.73195 | 1.71446 | 0.00000 | 2.48775 | 1.05918 | 0.00000 | 0.00000 | 3.10354 | 0.57978 | 0.02410 |
| **TXN** | 0.41734 | 25.33662 | 24.79781 | 28.74805 | 28.04817 | 53.81178 | 33.02617 | 14.29235 | 30.17647 | 17.27883 | 31.13040 | 41.58661 |
| **TXN2** | 6.06528 | 23.57986 | 14.36035 | 16.71308 | 25.06168 | 7.20975 | 26.79120 | 13.83582 | 22.40816 | 3.35577 | 25.15587 | 25.59829 |
| **TXNDC11** | 2.09347 | 2.98789 | 2.27405 | 3.34134 | 3.84088 | 0.00000 | 0.00000 | 1.61692 | 3.00971 | 4.02678 | 1.44646 | 1.55143 |
| **TXNDC12** | 0.77392 | 16.06370 | 14.43605 | 17.47813 | 8.98867 | 40.80462 | 0.00000 | 1.39129 | 31.85615 | 11.96753 | 12.41961 | 13.83543 |
| **TXNDC15** | 0.00000 | 6.94729 | 3.37303 | 5.80895 | 2.72339 | 6.94060 | 0.00000 | 0.60244 | 7.43980 | 2.62028 | 1.02894 | 6.48477 |
| **TXNDC16** | 0.64710 | 0.00000 | 0.17234 | 0.18920 | 1.26889 | 1.13018 | 0.29142 | 0.68470 | 0.00000 | 1.48388 | 0.13792 | 0.56575 |
| **TXNDC17** | 11.45913 | 13.13345 | 14.96125 | 14.37827 | 15.70258 | 21.31612 | 29.35001 | 14.14020 | 21.53042 | 7.40012 | 20.63188 | 8.45918 |
| **TXNDC2** | 0.00347 | 0.00000 | 0.00462 | 0.00000 | 0.02361 | 0.05945 | 0.09183 | 0.00946 | 0.02822 | 0.22779 | 0.01826 | 0.00579 |
| **TXNDC5.uid-81** | 0.00390 | 0.16913 | 0.07392 | 0.00775 | 0.26092 | 0.00000 | 0.00186 | 0.02987 | 0.19694 | 0.00000 | 0.05872 | 0.01081 |
| **TXNDC8** | 0.00392 | 0.01587 | 0.01393 | 0.03890 | 0.02271 | 0.00051 | 0.02357 | 0.01863 | 0.02211 | 0.00289 | 0.02275 | 0.07515 |
| **TXNDC9** | 2.41607 | 6.73284 | 11.52480 | 1.74652 | 0.00000 | 36.44988 | 13.69556 | 8.45321 | 9.67124 | 6.52253 | 12.14949 | 8.21584 |
| **TXNIP** | 0.00000 | 6.32990 | 0.00000 | 8.03673 | 1.28954 | 20.73871 | 5.59952 | 0.00000 | 7.09926 | 1.79286 | 0.00000 | 5.02120 |
| **TXNL1** | 1.05656 | 13.12470 | 24.58022 | 6.91027 | 8.71998 | 33.71025 | 29.70894 | 12.62029 | 20.04247 | 0.00000 | 14.31562 | 11.42634 |
| **TXNL4A** | 0.21669 | 6.42102 | 7.40490 | 7.44591 | 10.88567 | 5.89145 | 20.17500 | 0.20124 | 6.73475 | 21.89012 | 7.11749 | 3.52145 |
| **TXNL4B** | 2.87425 | 5.65122 | 8.40685 | 0.83304 | 3.03694 | 5.87197 | 9.64453 | 15.58537 | 4.39236 | 3.47726 | 3.11052 | 9.06853 |
| **TXNRD1** | 2.50474 | 31.87228 | 18.35299 | 10.18660 | 20.73797 | 60.13254 | 34.16863 | 0.51449 | 32.79178 | 21.48649 | 24.05662 | 29.99246 |
| **TXNRD2** | 0.00236 | 5.16073 | 0.62848 | 1.01520 | 4.60024 | 0.00000 | 0.86781 | 0.00000 | 0.00000 | 3.36574 | 3.15197 | 0.23781 |
| **TXNRD3** | 0.00956 | 0.04159 | 0.01463 | 0.03724 | 0.05658 | 0.11045 | 0.00000 | 0.00902 | 0.01806 | 0.12582 | 0.00084 | 0.00336 |
| **TYK2** | 0.14014 | 3.00321 | 1.76040 | 5.68634 | 5.60943 | 0.00000 | 1.19355 | 0.67250 | 4.11094 | 4.98927 | 1.94388 | 2.98281 |
| **TYMP** | 0.00796 | 0.08747 | 0.04014 | 0.07788 | 0.00387 | 0.00000 | 0.00123 | 0.01847 | 0.39714 | 0.35968 | 0.13784 | 0.01634 |
| **TYMS** | 0.33581 | 13.29061 | 4.84104 | 4.24830 | 1.77521 | 8.22524 | 0.00000 | 1.27409 | 9.04720 | 0.00000 | 5.00436 | 6.38292 |
| **TYMSOS** | 0.00000 | 0.58119 | 0.57744 | 1.18560 | 0.57176 | 1.08298 | 0.00000 | 0.35179 | 0.00000 | 0.79689 | 0.28285 | 1.86282 |
| **TYR** | 0.01575 | 0.00800 | 0.00594 | 0.00498 | 0.00420 | 0.03410 | 0.04539 | 0.01147 | 0.00000 | 0.10980 | 0.02612 | 0.03323 |
| **TYRO3** | 0.00000 | 10.17162 | 17.81199 | 5.72935 | 11.72432 | 0.00000 | 5.99566 | 0.14437 | 18.44235 | 10.71080 | 2.54389 | 8.55949 |
| **TYRO3P** | 0.02215 | 0.05544 | 0.08630 | 0.07403 | 0.09505 | 0.00000 | 0.12058 | 0.01901 | 0.17540 | 0.12817 | 0.00852 | 0.06212 |
| **TYROBP** | 0.00225 | 0.09994 | 0.07685 | 0.00585 | 0.00033 | 0.02810 | 0.00134 | 0.03694 | 0.01950 | 0.03991 | 0.01228 | 0.04570 |
| **TYRP1** | 0.48762 | 0.07487 | 0.00000 | 0.00217 | 0.03524 | 0.20208 | 0.00000 | 0.21240 | 0.00000 | 0.00963 | 0.01394 | 0.06749 |
| **TYSND1** | 1.10494 | 1.36355 | 2.60607 | 2.71055 | 0.63033 | 0.00000 | 6.44027 | 1.27835 | 0.31313 | 3.97416 | 2.10267 | 1.72600 |
| **TYW1** | 1.36770 | 4.18415 | 2.12096 | 2.45955 | 2.90370 | 4.34940 | 0.00000 | 3.65800 | 7.35285 | 2.20074 | 0.78317 | 3.68951 |
| **TYW1B** | 0.40485 | 0.72187 | 0.48275 | 0.22668 | 1.11461 | 1.03393 | 0.16351 | 5.23157 | 0.97750 | 1.40570 | 0.00000 | 0.63311 |
| **TYW3** | 0.85445 | 5.62266 | 3.74507 | 4.93433 | 3.62785 | 14.62378 | 4.19793 | 1.50458 | 5.91566 | 11.87547 | 2.29286 | 6.82926 |

| | | | | | | | | | | | | |
|---|---|---|---|---|---|---|---|---|---|---|---|---|
| **TYW5** | 0.39004 | 3.21119 | 2.55275 | 0.83579 | 0.00368 | 12.76297 | 0.00000 | 1.04242 | 3.52785 | 3.19514 | 2.58079 | 3.37902 |
| **U2AF1** | 2.28192 | 28.18353 | 26.61693 | 24.14310 | 30.45966 | 15.18768 | 46.26512 | 21.88771 | 29.68700 | 7.94440 | 27.95951 | 29.39071 |
| **U2AF1L4** | 0.07195 | 2.32968 | 1.59383 | 4.29017 | 1.48575 | 1.02072 | 3.25268 | 2.98419 | 2.70072 | 0.00000 | 2.08843 | 3.85266 |
| **U2AF2** | 4.89465 | 4.10658 | 7.53106 | 1.80911 | 10.69979 | 0.27909 | 12.47716 | 0.89318 | 8.92474 | 22.15894 | 0.83089 | 10.86393 |
| **U2SURP** | 3.63640 | 11.20740 | 10.07571 | 6.99490 | 6.52363 | 42.31851 | 18.62490 | 13.65465 | 7.46548 | 34.53752 | 15.52694 | 21.62001 |
| **UACA** | 5.11234 | 17.54721 | 5.84131 | 55.45983 | 11.57536 | 15.21279 | 0.00000 | 2.63852 | 0.00000 | 14.90335 | 5.74587 | 5.10262 |
| **UAP1** | 2.13233 | 15.45965 | 4.60895 | 5.78695 | 3.59975 | 18.01775 | 4.33281 | 7.04237 | 4.54640 | 0.90090 | 7.10242 | 7.91489 |
| **UAP1L1** | 0.00000 | 9.89432 | 0.27906 | 4.80058 | 15.52202 | 0.00000 | 1.55821 | 0.17285 | 3.62090 | 2.86448 | 1.92004 | 0.00000 |
| **UBA1** | 3.92724 | 20.17172 | 16.05838 | 12.97449 | 18.43768 | 1.34530 | 23.37632 | 4.38404 | 15.95749 | 17.09966 | 16.67163 | 22.34521 |
| **UBA2** | 0.00000 | 3.33624 | 9.83791 | 2.58151 | 6.20665 | 19.45786 | 14.97611 | 1.72546 | 6.54008 | 19.57120 | 3.26984 | 6.21794 |
| **UBA3** | 0.82426 | 5.23469 | 4.59645 | 3.84157 | 0.56088 | 22.33485 | 8.63167 | 2.47696 | 7.92481 | 3.03988 | 14.33632 | 8.23823 |
| **UBA5** | 2.59004 | 6.86076 | 5.52703 | 1.54022 | 1.58569 | 11.40436 | 2.84116 | 0.44849 | 5.30647 | 4.39304 | 4.27308 | 3.00592 |
| **UBA52** | 9.55099 | 38.52760 | 31.00582 | 29.38820 | 39.92923 | 29.98149 | 53.91819 | 17.63925 | 44.08299 | 35.86350 | 30.30008 | 44.41390 |
| **UBA6** | 3.32776 | 5.84816 | 5.51939 | 4.12151 | 0.00000 | 19.12355 | 5.14710 | 3.42156 | 4.11321 | 9.04953 | 5.05247 | 4.89596 |
| **UBA6-AS1** | 0.15019 | 0.90548 | 1.46694 | 0.60175 | 0.00000 | 1.09247 | 1.45747 | 0.01097 | 0.50986 | 0.00000 | 1.03723 | 0.22923 |
| **UBA7** | 0.07408 | 0.01355 | 0.19269 | 0.01680 | 0.13595 | 0.00000 | 0.04643 | 0.01206 | 0.01655 | 0.00000 | 0.01405 | 0.04175 |
| **UBAC1** | 6.82059 | 9.93240 | 9.31298 | 6.49758 | 9.65326 | 4.58412 | 16.70230 | 4.07603 | 6.40974 | 10.71868 | 10.64208 | 3.67615 |
| **UBAC2** | 2.30322 | 19.17819 | 16.82545 | 11.81803 | 9.41489 | 12.49201 | 1.80698 | 0.22463 | 24.69874 | 2.53243 | 5.17339 | 8.56413 |
| **UBAC2-AS1** | 0.05136 | 0.73779 | 0.20251 | 0.44233 | 0.26777 | 1.13393 | 0.08138 | 0.08576 | 0.00000 | 0.00000 | 0.71709 | 0.93207 |
| **UBALD1** | 1.33254 | 0.52521 | 3.70713 | 0.27994 | 2.25521 | 0.00000 | 0.72103 | 0.00000 | 0.54912 | 5.84688 | 0.23552 | 0.66358 |
| **UBALD2** | 0.18694 | 0.00000 | 4.91924 | 1.42968 | 4.74840 | 0.00000 | 5.55977 | 0.34364 | 0.00000 | 11.22738 | 0.00000 | 0.67059 |
| **UBAP1** | 2.25778 | 11.92326 | 14.51847 | 10.02328 | 6.98706 | 4.80844 | 9.49007 | 10.89231 | 11.55617 | 0.00000 | 8.35413 | 11.26572 |
| **UBAP1L** | 0.23692 | 0.00753 | 0.03072 | 0.07257 | 0.00000 | 0.00000 | 0.30216 | 0.28554 | 0.06059 | 0.86772 | 0.09579 | 0.08686 |
| **UBAP2** | 4.36052 | 19.65329 | 11.26387 | 19.97463 | 12.13962 | 14.09404 | 12.21572 | 6.12635 | 19.54545 | 21.37694 | 10.74535 | 25.61179 |
| **UBAP2L** | 3.03277 | 15.35563 | 16.28039 | 18.15765 | 17.30111 | 13.36299 | 22.43188 | 9.00545 | 22.30628 | 7.46810 | 19.68616 | 16.41339 |
| **UBASH3A** | 0.00370 | 0.00677 | 0.00001 | 0.00265 | 0.00206 | 0.02399 | 0.00821 | 0.00986 | 0.02066 | 0.26615 | 0.00127 | 0.00880 |
| **UBASH3B** | 0.00169 | 20.10484 | 2.62318 | 18.76257 | 5.33034 | 5.73999 | 0.00000 | 0.48416 | 0.00000 | 0.00250 | 3.45340 | 5.79698 |
| **UBB** | 18.05576 | 26.55675 | 38.39645 | 36.13483 | 33.58535 | 43.16777 | 50.73427 | 27.14548 | 49.73031 | 2.86730 | 40.78998 | 39.00966 |
| **UBC** | 7.07486 | 13.85847 | 22.53960 | 14.11256 | 19.10807 | 27.58645 | 32.13250 | 8.90728 | 24.79949 | 6.89126 | 16.20619 | 21.94962 |
| **UBD** | 0.04649 | 0.00807 | 0.07142 | 0.02093 | 0.00000 | 0.05252 | 0.00913 | 0.01759 | 0.00000 | 0.03209 | 0.01811 | 0.01337 |
| **UBE2A** | 6.58834 | 7.65872 | 16.63638 | 9.42526 | 8.59563 | 19.21034 | 13.34543 | 0.36850 | 15.79454 | 0.65141 | 12.76950 | 1.06985 |
| **UBE2B** | 0.24492 | 5.94825 | 10.83396 | 3.47531 | 3.63474 | 29.43247 | 10.60958 | 22.90307 | 1.99619 | 8.31857 | 8.28727 | 14.02353 |
| **UBE2C** | 6.77084 | 40.97041 | 22.13432 | 18.66503 | 34.53123 | 40.73449 | 27.41936 | 41.26959 | 25.42347 | 0.84986 | 17.72785 | 51.14135 |
| **UBE2D1** | 1.06782 | 5.15430 | 5.07859 | 2.07149 | 0.00000 | 11.34733 | 2.43857 | 0.00000 | 4.09924 | 4.34369 | 6.78641 | 3.59050 |
| **UBE2D2** | 4.95437 | 6.95900 | 15.54509 | 5.31967 | 9.32754 | 8.89204 | 13.89716 | 0.62528 | 11.70457 | 17.18383 | 9.40939 | 3.39708 |
| **UBE2D3** | 15.52910 | 18.27084 | 26.46045 | 16.78215 | 15.07172 | 38.59543 | 33.81295 | 20.01329 | 31.74314 | 6.21416 | 23.58827 | 25.07271 |
| **UBE2D4** | 0.27094 | 4.36205 | 0.15021 | 0.00000 | 0.60829 | 1.38693 | 1.52053 | 0.27576 | 0.00000 | 1.37115 | 3.44729 | 0.87690 |
| **UBE2DNL** | 0.00252 | 0.01212 | 0.03040 | 0.04274 | 0.00301 | 0.05233 | 0.03071 | 0.06698 | 0.10872 | 0.00000 | 0.00376 | 0.03227 |
| **UBE2E1** | 0.14713 | 4.49404 | 3.79492 | 2.20084 | 3.22867 | 4.63264 | 10.52489 | 0.47474 | 0.00000 | 13.67725 | 11.89454 | 4.24868 |
| **UBE2E1-AS1** | 0.00524 | 0.13071 | 0.01815 | 0.00000 | 0.01443 | 0.08366 | 0.00000 | 0.00778 | 0.02812 | 0.02456 | 0.02171 | 0.05958 |
| **UBE2E2** | 1.69691 | 4.54161 | 5.70548 | 0.28006 | 4.29417 | 6.94544 | 2.34860 | 1.16842 | 2.31747 | 0.06610 | 3.60352 | 5.16022 |
| **UBE2E3** | 2.13966 | 3.11846 | 10.11203 | 2.10443 | 4.26108 | 9.05478 | 9.98969 | 0.00000 | 3.44961 | 6.25619 | 5.66319 | 3.59507 |
| **UBE2E4P** | 0.48018 | 0.49173 | 1.18164 | 0.37217 | 0.11500 | 1.33052 | 0.62714 | 0.04632 | 0.86344 | 0.00000 | 0.72430 | 0.35859 |
| **UBE2F** | 1.01278 | 7.27606 | 6.57973 | 3.76884 | 7.96211 | 15.88697 | 10.23301 | 1.34010 | 9.45719 | 9.47439 | 8.00282 | 4.52940 |
| **UBE2F-SCLY** | 0.01252 | 0.01605 | 0.03141 | 0.01244 | 0.01461 | 0.01729 | 0.01152 | 0.01784 | 0.02311 | 0.01787 | 0.01529 | 0.01520 |
| **UBE2G1** | 6.20222 | 14.89223 | 18.47464 | 7.91036 | 3.67804 | 25.45736 | 18.19406 | 20.23291 | 15.19703 | 14.98066 | 14.65961 | 33.31445 |
| **UBE2G2** | 0.00000 | 11.93361 | 15.24289 | 2.88440 | 3.03859 | 3.94198 | 21.41118 | 0.98109 | 19.27106 | 5.90561 | 17.60280 | 3.42283 |
| **UBE2H** | 9.16991 | 3.00082 | 14.08582 | 4.25499 | 1.75248 | 5.83632 | 15.11673 | 1.01614 | 0.00000 | 7.98719 | 0.30201 | 2.74745 |

| | | | | | | | | | | | | |
|---|---|---|---|---|---|---|---|---|---|---|---|---|
| **UBE2I** | 16.03206 | 26.75345 | 22.41639 | 19.64194 | 27.06029 | 18.08019 | 39.33774 | 5.37577 | 30.47462 | 8.48752 | 25.47811 | 15.76916 |
| **UBE2J1** | 0.00000 | 0.00000 | 10.95013 | 0.00000 | 0.00000 | 0.51787 | 20.67134 | 0.07680 | 11.90384 | 1.03729 | 9.16870 | 0.00000 |
| **UBE2J2** | 4.37211 | 17.38675 | 11.91571 | 8.40540 | 17.00678 | 1.46027 | 16.83404 | 1.43014 | 16.63141 | 8.15140 | 10.73320 | 12.69375 |
| **UBE2K** | 4.12472 | 12.06770 | 19.91363 | 9.64220 | 13.11453 | 32.92070 | 33.06979 | 9.92376 | 11.75646 | 16.61663 | 13.23712 | 16.49871 |
| **UBE2L3** | 2.91207 | 13.49219 | 16.63690 | 12.19290 | 11.01918 | 16.55486 | 27.11201 | 8.13022 | 17.59579 | 0.17217 | 14.94401 | 14.63226 |
| **UBE2L6** | 0.04094 | 0.00637 | 0.57366 | 0.00422 | 0.00965 | 0.00513 | 0.00937 | 0.76144 | 0.49390 | 0.00000 | 0.00000 | 0.24746 |
| **UBE2M** | 0.15427 | 1.41539 | 1.20520 | 3.07515 | 7.38874 | 0.00000 | 13.14334 | 1.95892 | 0.05977 | 20.84905 | 1.58131 | 1.58076 |
| **UBE2MP1** | 0.08179 | 0.30941 | 0.33190 | 0.43309 | 0.78850 | 0.00027 | 0.80652 | 0.32762 | 0.38517 | 0.89830 | 0.16282 | 0.21870 |
| **UBE2N** | 5.80445 | 15.50373 | 19.68596 | 11.26258 | 8.15089 | 38.30373 | 26.13696 | 8.96667 | 20.03097 | 4.52084 | 21.43356 | 14.67163 |
| **UBE2NL** | 0.06259 | 0.22173 | 0.73598 | 0.26514 | 0.00000 | 1.31837 | 0.07025 | 0.23960 | 0.85500 | 0.00000 | 0.56788 | 0.26617 |
| **UBE2O** | 2.47824 | 5.73557 | 4.90340 | 2.91480 | 7.61724 | 0.00014 | 8.37632 | 1.67852 | 5.98712 | 19.58628 | 1.12185 | 5.21599 |
| **UBE2Q1** | 1.21888 | 0.83578 | 4.06951 | 2.46256 | 3.68612 | 3.65491 | 9.01367 | 0.98283 | 0.00000 | 14.30890 | 0.53400 | 5.29553 |
| **UBE2Q2** | 18.55623 | 7.88927 | 5.83811 | 2.84243 | 3.27611 | 18.15261 | 4.82922 | 3.69670 | 1.87655 | 19.61755 | 0.33881 | 0.18773 |
| **UBE2Q2L** | 0.04524 | 0.01445 | 0.01412 | 0.01196 | 0.01517 | 0.01823 | 0.01778 | 0.00857 | 0.01856 | 0.01394 | 0.01469 | 0.01676 |
| **UBE2Q2P1** | 0.37111 | 0.12639 | 0.00080 | 0.14383 | 0.09262 | 0.26281 | 0.00000 | 0.03143 | 0.00000 | 0.04586 | 0.00612 | 0.03766 |
| **UBE2QL1** | 0.00684 | 0.01789 | 0.01742 | 0.01489 | 0.02510 | 0.00948 | 0.03248 | 0.01105 | 0.01512 | 0.00681 | 0.01515 | 0.01931 |
| **UBE2R2** | 5.27006 | 2.32542 | 5.52171 | 4.37724 | 7.68748 | 10.09497 | 15.49778 | 1.87950 | 0.00000 | 27.11950 | 2.39605 | 4.36719 |
| **UBE2S** | 0.68385 | 1.82891 | 1.51318 | 0.58667 | 2.42270 | 0.00000 | 1.38333 | 0.57288 | 1.81597 | 3.61710 | 0.00000 | 1.15182 |
| **UBE2T** | 1.82883 | 18.95847 | 12.31130 | 14.88469 | 15.07306 | 30.16290 | 10.83849 | 8.87850 | 18.11718 | 0.00000 | 12.93372 | 17.61775 |
| **UBE2U** | 1.80718 | 0.08710 | 0.00000 | 0.75035 | 0.28588 | 0.16935 | 0.00000 | 0.50708 | 0.00000 | 0.06065 | 0.00000 | 0.30158 |
| **UBE2V1** | 3.53983 | 1.29019 | 1.88084 | 1.01643 | 0.73800 | 5.16091 | 0.80007 | 0.98238 | 1.70877 | 0.00000 | 1.37618 | 3.29104 |
| **UBE2V2** | 0.31369 | 8.02316 | 9.73989 | 3.35121 | 0.00000 | 42.17427 | 16.71098 | 5.30064 | 13.44536 | 6.99344 | 12.68853 | 11.99448 |
| **UBE2W** | 7.51892 | 2.06444 | 4.82913 | 2.97099 | 0.00000 | 12.53003 | 2.85515 | 3.19297 | 7.59661 | 2.06719 | 3.90783 | 10.21578 |
| **UBE2Z** | 0.97050 | 5.48758 | 4.52125 | 7.88550 | 11.43988 | 8.64922 | 10.89217 | 2.79926 | 3.05532 | 21.93441 | 4.43697 | 6.92952 |
| **UBE3A** | 5.33572 | 4.25088 | 8.60767 | 3.70355 | 3.57031 | 13.06829 | 13.79143 | 4.48558 | 0.00000 | 23.33882 | 6.60163 | 4.03146 |
| **UBE3B** | 2.31201 | 9.04339 | 5.30770 | 3.31479 | 0.00000 | 10.07369 | 0.00000 | 0.00000 | 6.38705 | 8.73587 | 3.21982 | 4.83781 |
| **UBE3C** | 0.75989 | 5.88374 | 7.52079 | 5.83843 | 6.87486 | 9.31183 | 15.62413 | 6.94444 | 3.80308 | 11.98583 | 5.34956 | 6.43548 |
| **UBE3D** | 0.24461 | 1.58091 | 2.09671 | 0.17288 | 0.00000 | 3.87718 | 2.08915 | 0.77943 | 5.17054 | 0.00000 | 1.88162 | 0.95694 |
| **UBE4A** | 0.45650 | 6.03261 | 8.22416 | 4.25103 | 4.18429 | 18.02361 | 0.00000 | 9.51884 | 15.72204 | 9.25997 | 8.64747 | 14.84355 |
| **UBE4B** | 0.71109 | 11.99035 | 11.72554 | 5.29437 | 3.95063 | 6.91696 | 15.08365 | 7.34687 | 9.05655 | 7.87226 | 3.19323 | 7.62326 |
| **UBFD1** | 5.32809 | 5.17795 | 7.58946 | 7.93733 | 8.15166 | 13.02102 | 20.32352 | 2.76921 | 7.23999 | 16.49051 | 3.33014 | 4.23738 |
| **UBIAD1** | 2.62758 | 14.25070 | 10.59990 | 6.40280 | 7.17793 | 10.23520 | 5.37099 | 6.77079 | 16.42420 | 0.00000 | 8.05222 | 9.96927 |
| **UBL3** | 0.32188 | 0.19584 | 0.00000 | 0.00000 | 4.11994 | 6.15584 | 11.28481 | 1.38561 | 1.12232 | 11.47900 | 4.72748 | 0.09504 |
| **UBL4A** | 0.00317 | 8.89456 | 9.34188 | 8.23576 | 7.87019 | 0.00000 | 17.95803 | 0.00091 | 15.88595 | 0.00000 | 17.72558 | 4.36676 |
| **UBL4B** | 0.00418 | 0.01528 | 0.01428 | 0.01546 | 0.01098 | 0.01930 | 0.03809 | 0.07469 | 0.01790 | 0.01919 | 0.01198 | 0.01284 |
| **UBL5** | 2.77443 | 27.13319 | 14.17866 | 14.08143 | 26.13787 | 26.02225 | 29.36398 | 22.79964 | 24.08805 | 17.99417 | 19.47706 | 37.81178 |
| **UBL7** | 2.75594 | 12.00845 | 14.19645 | 9.19870 | 13.84486 | 8.71018 | 15.08305 | 13.68390 | 7.48612 | 0.00000 | 14.71300 | 16.48972 |
| **UBL7-AS1** | 1.74042 | 0.70001 | 3.53428 | 0.74035 | 0.89009 | 2.41018 | 0.00000 | 0.39832 | 1.61711 | 0.48357 | 0.35626 | 1.56583 |
| **UBLCP1** | 1.66700 | 5.02252 | 5.85627 | 0.69256 | 0.00000 | 13.95166 | 4.50352 | 0.27181 | 3.16738 | 4.20688 | 5.07766 | 5.05877 |
| **UBN1** | 3.14913 | 3.09378 | 5.62140 | 5.87345 | 6.78104 | 0.00180 | 10.44294 | 3.58329 | 4.45052 | 20.64691 | 3.03668 | 4.10115 |
| **UBN2** | 5.36552 | 2.82621 | 2.59692 | 2.89840 | 0.00000 | 7.35785 | 5.83973 | 7.85114 | 0.77989 | 17.59756 | 0.75549 | 7.17254 |
| **UBOX5** | 0.85869 | 2.53681 | 1.74283 | 2.18687 | 1.15992 | 0.00000 | 2.70554 | 1.56342 | 0.00000 | 0.00000 | 2.33050 | 1.89744 |
| **UBOX5-AS1** | 0.00783 | 0.03880 | 0.01997 | 0.13517 | 0.01938 | 0.00000 | 0.05949 | 0.07430 | 0.00000 | 0.07228 | 0.13480 | 0.07451 |
| **UBP1** | 0.89280 | 1.09221 | 2.67921 | 1.42737 | 0.00000 | 11.13729 | 8.48365 | 5.72507 | 0.00000 | 12.78307 | 2.58382 | 3.03910 |
| **UBQLN1** | 13.24101 | 19.21967 | 19.06118 | 14.16932 | 20.69960 | 33.73070 | 17.50521 | 16.79534 | 24.58421 | 6.66375 | 15.56699 | 23.27759 |
| **UBQLN2** | 0.00000 | 4.03791 | 5.18524 | 0.00000 | 0.01488 | 7.25949 | 3.70938 | 1.34890 | 6.21364 | 4.05834 | 4.56653 | 5.57468 |
| **UBQLN4** | 0.02590 | 2.47497 | 0.58276 | 0.00000 | 2.11488 | 2.49236 | 1.55369 | 0.00000 | 1.69051 | 6.77244 | 0.41317 | 1.95027 |
| **UBQLNL** | 0.02611 | 0.04883 | 0.13488 | 0.08502 | 0.00000 | 0.10330 | 0.00000 | 0.00745 | 0.16549 | 0.01597 | 0.03856 | 0.01171 |

| | | | | | | | | | | | | |
|---|---|---|---|---|---|---|---|---|---|---|---|---|
| **UBR1** | 2.45830 | 1.82297 | 9.42306 | 5.90287 | 1.93480 | 17.20078 | 2.54650 | 2.61215 | 6.24824 | 19.21807 | 2.77599 | 5.93609 |
| **UBR2** | 4.30208 | 6.12354 | 5.09824 | 9.08276 | 2.00666 | 10.10529 | 7.48041 | 1.85264 | 4.39699 | 12.56767 | 7.74760 | 8.24377 |
| **UBR3** | 0.48363 | 0.00000 | 4.19109 | 3.25631 | 0.00000 | 7.93465 | 6.90843 | 0.76510 | 1.94124 | 10.85252 | 1.10488 | 0.26228 |
| **UBR4** | 2.89343 | 5.55948 | 16.69568 | 15.61176 | 19.52399 | 2.02970 | 19.08809 | 1.36244 | 10.64777 | 29.03708 | 6.41132 | 9.07253 |
| **UBR5** | 6.69217 | 1.88165 | 11.67480 | 4.26581 | 2.97653 | 21.44741 | 4.92312 | 6.21910 | 2.49683 | 26.66287 | 0.32016 | 8.79241 |
| **UBR7** | 5.21218 | 9.34985 | 6.91042 | 4.04120 | 7.12258 | 13.12314 | 3.22490 | 16.32881 | 6.02812 | 0.00000 | 9.91917 | 7.46567 |
| **UBTD1** | 0.00000 | 2.73335 | 1.32842 | 5.12918 | 0.59960 | 0.00000 | 0.98473 | 5.55143 | 0.00000 | 2.43047 | 0.00000 | 7.40150 |
| **UBTD2** | 2.27403 | 1.11638 | 2.24867 | 0.51024 | 3.39554 | 9.14770 | 6.55363 | 3.36014 | 3.05732 | 10.85218 | 1.64310 | 3.40683 |
| **UBTF** | 4.26890 | 10.00548 | 3.90485 | 9.94159 | 9.50834 | 0.00002 | 12.23337 | 3.37383 | 5.33102 | 17.08145 | 5.64274 | 8.82096 |
| **UBTFL1** | 0.59112 | 0.01394 | 0.01396 | 0.01243 | 0.01879 | 0.00000 | 0.01761 | 1.05107 | 0.02118 | 0.00928 | 0.01436 | 0.00000 |
| **UBXN1** | 4.08703 | 24.20672 | 19.70988 | 17.51173 | 25.82700 | 5.48205 | 26.53240 | 27.14622 | 24.14674 | 4.70704 | 13.06022 | 31.70190 |
| **UBXN10** | 1.06075 | 0.02759 | 0.01030 | 0.01823 | 0.01604 | 0.02970 | 0.01421 | 0.22783 | 0.02125 | 0.00000 | 0.01118 | 0.08801 |
| **UBXN11** | 14.91574 | 4.17958 | 2.13691 | 2.40891 | 5.73499 | 0.00000 | 0.29331 | 13.16384 | 2.56577 | 0.31670 | 2.52458 | 4.32575 |
| **UBXN2A** | 0.00000 | 2.72524 | 4.62241 | 1.41823 | 1.78284 | 8.61889 | 3.83044 | 10.19192 | 4.52318 | 2.02651 | 3.60751 | 6.35059 |
| **UBXN2B** | 0.83156 | 3.73780 | 3.03116 | 3.31602 | 0.00000 | 10.69851 | 9.73226 | 0.29588 | 5.45639 | 0.90613 | 5.91741 | 1.79787 |
| **UBXN4** | 9.77367 | 8.65410 | 15.08852 | 10.75521 | 10.41356 | 21.76663 | 15.97417 | 7.04105 | 15.54744 | 21.51479 | 13.02095 | 8.50951 |
| **UBXN6** | 2.38638 | 1.64271 | 0.76815 | 2.23459 | 4.61905 | 0.44313 | 2.50086 | 2.09860 | 0.42717 | 8.90587 | 0.75006 | 5.32153 |
| **UBXN7** | 1.46682 | 2.77892 | 4.64340 | 2.55090 | 0.00000 | 12.07941 | 8.24211 | 19.25188 | 5.64860 | 6.63134 | 5.60220 | 15.40941 |
| **UBXN8** | 1.14467 | 10.95534 | 7.68493 | 0.76503 | 0.00041 | 13.36161 | 1.38097 | 6.36734 | 8.47471 | 4.96916 | 4.32204 | 14.73384 |
| **UCA1** | 0.00473 | 0.00000 | 1.94171 | 0.00000 | 0.71379 | 0.09166 | 2.80569 | 0.01534 | 3.35493 | 0.00000 | 3.62065 | 0.00670 |
| **UCHL1** | 16.92670 | 0.21002 | 7.34247 | 0.16778 | 0.00000 | 0.00000 | 13.33069 | 5.25647 | 3.08908 | 0.00000 | 0.00000 | 1.38677 |
| **UCHL3** | 1.00455 | 5.54645 | 14.16275 | 3.50953 | 5.88157 | 26.34579 | 18.03536 | 2.12317 | 8.86706 | 5.37638 | 15.03389 | 4.37144 |
| **UCHL5** | 1.05065 | 8.48770 | 6.89753 | 4.87157 | 4.52112 | 22.96785 | 11.12414 | 0.02425 | 8.66825 | 1.37435 | 12.97423 | 1.51575 |
| **UCK1** | 1.38331 | 8.47683 | 9.48032 | 7.68025 | 4.04536 | 0.00000 | 6.52439 | 4.51457 | 5.77918 | 5.74544 | 6.32809 | 5.64220 |
| **UCK2** | 0.80582 | 9.61309 | 12.25162 | 4.31595 | 8.95147 | 0.34706 | 15.69312 | 1.24655 | 11.22075 | 10.33818 | 6.35650 | 8.64946 |
| **UCKL1** | 0.01008 | 5.76350 | 2.87201 | 8.83486 | 6.19800 | 2.90961 | 7.58794 | 3.06672 | 11.87592 | 5.87602 | 5.72788 | 2.37891 |
| **UCKL1-AS1** | 0.00092 | 0.05193 | 0.32313 | 0.43088 | 0.69681 | 0.00000 | 0.00000 | 0.05485 | 0.16844 | 1.08894 | 0.00000 | 0.03087 |
| **UCMA** | 0.98367 | 0.04021 | 0.02023 | 0.00589 | 0.01512 | 0.01035 | 0.01493 | 0.21283 | 0.01798 | 0.01949 | 0.01134 | 0.75293 |
| **UCN** | 0.00982 | 0.02460 | 0.01534 | 0.00132 | 0.04243 | 0.00000 | 0.03531 | 0.02838 | 0.00000 | 0.20758 | 0.04805 | 0.01030 |
| **UCN2** | 0.00553 | 0.00955 | 0.05573 | 0.09065 | 0.00000 | 0.04202 | 0.00000 | 0.00989 | 0.10870 | 0.04941 | 0.00655 | 0.01317 |
| **UCP1** | 0.00168 | 0.07744 | 0.00961 | 0.00827 | 0.01960 | 0.02761 | 0.03591 | 0.12555 | 0.09101 | 0.00000 | 0.03543 | 0.10071 |
| **UCP2** | 8.60332 | 6.59570 | 2.53554 | 11.32126 | 7.50326 | 9.22310 | 2.08906 | 6.16836 | 0.00000 | 0.00000 | 7.65395 | 4.75677 |
| **UCP3** | 0.00358 | 0.00000 | 0.23941 | 0.08879 | 0.00000 | 0.00000 | 0.00723 | 0.06525 | 0.00000 | 0.45935 | 0.09377 | 0.29208 |
| **UEVLD** | 0.10160 | 1.48368 | 1.46565 | 0.00000 | 0.89091 | 4.14381 | 0.00000 | 0.64289 | 2.02749 | 0.00000 | 0.78707 | 1.95708 |
| **UFC1** | 2.75166 | 14.52540 | 31.56240 | 10.83383 | 14.97434 | 17.41856 | 45.20205 | 3.94786 | 27.72537 | 2.86147 | 25.27636 | 19.04921 |
| **UFD1L** | 0.00000 | 9.84978 | 19.63927 | 8.11292 | 10.19895 | 15.49715 | 23.78928 | 3.82853 | 17.78173 | 1.22964 | 12.49761 | 11.57022 |
| **UFL1** | 2.68937 | 1.46072 | 4.97233 | 2.19599 | 2.56421 | 13.29531 | 0.00000 | 0.67614 | 4.50180 | 9.74692 | 4.02732 | 1.56301 |
| **UFM1** | 1.64015 | 8.57450 | 9.78617 | 7.25913 | 2.70530 | 30.07763 | 10.19670 | 3.92506 | 5.87310 | 4.08232 | 7.05635 | 18.92375 |
| **UFSP1** | 0.21140 | 0.20673 | 0.34656 | 0.23602 | 0.00000 | 0.00000 | 0.46185 | 0.13959 | 0.52178 | 0.00000 | 0.95298 | 0.15295 |
| **UFSP2** | 2.06791 | 3.65427 | 3.86182 | 1.56843 | 1.20827 | 10.29967 | 5.26568 | 1.36734 | 2.89715 | 0.87134 | 4.76612 | 4.94867 |
| **UG0898H09** | 0.00600 | 0.02305 | 0.01472 | 0.01233 | 0.01435 | 0.01599 | 0.02015 | 0.01095 | 0.02564 | 0.00741 | 0.02311 | 0.02135 |
| **UGCG** | 0.81339 | 9.46073 | 0.00000 | 5.42289 | 28.24480 | 11.89602 | 0.00000 | 0.14223 | 3.09612 | 13.92070 | 16.00773 | 0.00000 |
| **UGDH** | 2.64567 | 0.84196 | 6.97093 | 8.06740 | 0.00000 | 7.34312 | 10.40289 | 7.37037 | 6.66333 | 0.00000 | 8.05931 | 3.64092 |
| **UGDH-AS1** | 0.02240 | 0.00000 | 0.10123 | 0.07774 | 0.09090 | 0.00000 | 0.00083 | 0.01390 | 0.04482 | 0.29704 | 0.07692 | 0.06653 |
| **UGGT1** | 0.81188 | 3.91487 | 7.33272 | 3.35774 | 6.37723 | 11.76341 | 0.00000 | 5.23341 | 17.93127 | 19.41994 | 7.33733 | 6.22139 |
| **UGGT2** | 1.22210 | 0.46222 | 1.88993 | 1.42813 | 0.00000 | 4.76890 | 0.00000 | 0.74056 | 5.69547 | 5.12320 | 1.48337 | 1.09476 |
| **UGP2** | 6.89716 | 11.69784 | 15.48853 | 18.58598 | 8.15206 | 43.02273 | 20.44244 | 4.24637 | 10.36926 | 11.55017 | 13.62385 | 7.09954 |
| **UGT2A1** | 0.06746 | 0.01471 | 0.02694 | 0.01266 | 0.01846 | 0.00724 | 0.01889 | 0.02104 | 0.01815 | 0.01108 | 0.01555 | 0.10163 |

| | | | | | | | | | | | | |
|---|---|---|---|---|---|---|---|---|---|---|---|---|
| **UGT2B11** | 0.00617 | 0.01285 | 0.02555 | 0.00901 | 0.01003 | 0.03421 | 0.03988 | 0.01034 | 0.01232 | 0.01042 | 0.01279 | 0.01542 |
| **UGT2B15** | 0.00754 | 0.01475 | 0.01544 | 0.01813 | 0.01254 | 0.02303 | 0.01835 | 0.01068 | 0.01799 | 0.01426 | 0.01510 | 0.01592 |
| **UGT2B28** | 0.00723 | 0.01394 | 0.04572 | 0.00847 | 0.01367 | 0.02357 | 0.08617 | 0.01072 | 0.00000 | 0.01699 | 0.01342 | 0.01534 |
| **UGT2B7** | 0.00375 | 0.00551 | 0.07449 | 0.00000 | 0.00000 | 0.03284 | 0.02659 | 0.00976 | 0.00000 | 0.24968 | 0.00483 | 0.00805 |
| **UGT3A1** | 0.00231 | 0.01689 | 0.00000 | 0.00583 | 0.01316 | 0.16009 | 0.00000 | 0.09324 | 0.44774 | 0.01853 | 0.09438 | 0.00260 |
| **UGT3A2** | 0.00626 | 0.25265 | 0.13209 | 0.00364 | 0.04103 | 0.00928 | 0.03405 | 0.01116 | 0.05525 | 0.00000 | 0.04654 | 0.00648 |
| **UGT8** | 2.09514 | 1.03915 | 0.00000 | 1.73709 | 0.20023 | 0.82289 | 0.00000 | 0.69308 | 0.47441 | 0.63271 | 0.06124 | 1.62857 |
| **UHMK1** | 3.41042 | 1.69326 | 4.04507 | 9.00970 | 0.48243 | 23.15107 | 0.00089 | 13.36374 | 8.33188 | 13.98006 | 3.61904 | 9.44847 |
| **UHRF1** | 13.79515 | 2.54911 | 1.90645 | 3.25182 | 2.62261 | 0.00504 | 3.06538 | 2.63951 | 8.31237 | 0.00000 | 5.75739 | 0.52164 |
| **UHRF1BP1** | 4.75043 | 6.36067 | 1.77355 | 3.66875 | 2.78774 | 8.28405 | 5.52523 | 0.99393 | 0.73373 | 12.08739 | 4.76975 | 2.66691 |
| **UHRF1BP1L** | 5.52337 | 1.80154 | 2.80066 | 6.12510 | 0.40643 | 4.02344 | 3.74663 | 3.50284 | 0.92296 | 0.00029 | 0.83711 | 2.46307 |
| **UHRF2** | 5.32325 | 1.19659 | 2.15138 | 1.10359 | 1.54476 | 5.60902 | 3.32182 | 2.78975 | 2.99657 | 10.62686 | 2.39949 | 2.16775 |
| **UIMC1** | 3.12899 | 4.02706 | 2.27292 | 4.99565 | 1.16674 | 7.49174 | 2.95111 | 9.62600 | 10.00602 | 0.00000 | 3.72455 | 6.26211 |
| **ULBP1** | 0.00283 | 0.01303 | 0.29195 | 0.04479 | 0.15612 | 0.00000 | 0.00000 | 0.01447 | 0.12280 | 0.28565 | 0.00000 | 0.01139 |
| **ULBP2** | 0.02527 | 0.21892 | 0.11472 | 0.11561 | 0.00000 | 0.01235 | 0.00000 | 0.00427 | 0.02645 | 0.01748 | 0.01128 | 0.36542 |
| **ULBP3** | 0.00158 | 0.01811 | 0.02138 | 0.01021 | 0.01807 | 0.04204 | 0.01779 | 0.14182 | 0.02960 | 0.00000 | 0.01333 | 0.04068 |
| **ULK1** | 4.17194 | 7.14790 | 7.27675 | 7.05369 | 19.12662 | 2.88393 | 11.12839 | 0.95683 | 4.84265 | 30.13546 | 0.00000 | 6.70836 |
| **ULK2** | 2.78753 | 8.65967 | 0.00000 | 0.42420 | 0.00000 | 0.75955 | 0.00000 | 13.22539 | 0.08304 | 1.34163 | 0.00000 | 22.25576 |
| **ULK3** | 0.70750 | 4.21386 | 4.11191 | 2.21922 | 2.79386 | 0.00000 | 0.00000 | 0.10669 | 3.79007 | 3.36974 | 2.74418 | 1.05558 |
| **ULK4** | 4.29573 | 1.65036 | 0.83992 | 3.03472 | 2.92567 | 2.98246 | 1.10042 | 3.99997 | 1.37904 | 2.66158 | 0.54725 | 5.79587 |
| **UMOD** | 0.00622 | 0.05399 | 0.03342 | 0.01991 | 0.01758 | 0.00656 | 0.01842 | 0.01078 | 0.02130 | 0.00885 | 0.01628 | 0.01616 |
| **UMODL1** | 0.48905 | 0.00447 | 0.00001 | 0.05444 | 0.00013 | 0.06278 | 0.12173 | 0.01968 | 0.02261 | 0.00000 | 0.01501 | 0.04158 |
| **UMPS** | 2.26423 | 22.87093 | 14.10986 | 12.61617 | 11.89773 | 37.76608 | 16.45624 | 3.37975 | 19.14620 | 0.00000 | 21.27641 | 20.49379 |
| **UNC119** | 0.00000 | 3.58037 | 2.63704 | 4.82672 | 0.65731 | 0.00000 | 1.90831 | 1.97577 | 0.52550 | 3.48748 | 0.92312 | 0.26601 |
| **UNC119B** | 0.00000 | 2.27740 | 2.35087 | 1.60652 | 2.02715 | 1.36832 | 4.08988 | 0.11618 | 0.00154 | 4.92087 | 2.28389 | 2.23394 |
| **UNC13A** | 0.00856 | 0.00039 | 0.02113 | 0.01448 | 0.01111 | 0.00000 | 0.08377 | 0.17843 | 0.00000 | 0.17381 | 0.13430 | 0.15359 |
| **UNC13B** | 1.15254 | 2.30383 | 1.51764 | 7.02342 | 3.51896 | 1.50154 | 3.21430 | 0.49267 | 0.00000 | 4.36387 | 7.81835 | 2.05730 |
| **UNC13C** | 21.47942 | 0.02971 | 0.00108 | 0.00000 | 0.00153 | 0.00000 | 0.00000 | 5.98081 | 0.04045 | 0.23665 | 0.00000 | 0.33310 |
| **UNC13D** | 1.05887 | 1.02411 | 0.94106 | 1.74639 | 2.22181 | 0.00000 | 0.17228 | 0.94958 | 0.86194 | 0.00000 | 0.73128 | 0.11144 |
| **UNC45A** | 0.10085 | 3.47887 | 11.45238 | 8.82339 | 2.00680 | 0.00000 | 10.72513 | 0.02894 | 12.39457 | 2.21384 | 8.51798 | 1.91087 |
| **UNC45B** | 0.10462 | 0.00219 | 0.00428 | 0.03914 | 0.00000 | 0.04076 | 0.00000 | 0.01217 | 0.00000 | 0.10840 | 0.09691 | 0.12781 |
| **UNC50** | 0.37738 | 9.12597 | 10.83003 | 1.17435 | 0.03610 | 21.66516 | 0.00000 | 1.09821 | 17.57914 | 6.43431 | 5.78734 | 5.77343 |
| **UNC5A** | 0.83012 | 0.11919 | 0.03889 | 0.00003 | 0.24071 | 0.00000 | 0.01887 | 0.02327 | 0.00302 | 0.00781 | 0.00907 | 0.08960 |
| **UNC5B** | 0.00487 | 0.35477 | 0.37152 | 10.04202 | 0.74196 | 0.00000 | 0.07513 | 0.00505 | 0.00000 | 1.29070 | 0.00000 | 0.01861 |
| **UNC5B-AS1** | 0.36712 | 0.07715 | 0.00000 | 0.86052 | 0.00066 | 0.07455 | 0.00000 | 0.08740 | 0.00864 | 0.01193 | 0.01098 | 0.00000 |
| **UNC5C** | 7.63355 | 0.06146 | 0.40430 | 0.00479 | 0.00000 | 0.00000 | 0.44454 | 1.67147 | 0.00031 | 0.03659 | 0.05560 | 0.00000 |
| **UNC5CL** | 0.00339 | 0.13500 | 0.22120 | 0.14757 | 0.01479 | 0.08788 | 0.00000 | 0.00211 | 0.26173 | 0.35005 | 0.45007 | 0.43804 |
| **UNC5D** | 0.85665 | 0.07181 | 0.03988 | 0.18607 | 0.02109 | 0.04708 | 0.01226 | 0.04539 | 0.04579 | 0.06170 | 0.00349 | 0.06862 |
| **UNC79** | 0.53642 | 0.00741 | 0.08560 | 0.00399 | 0.11119 | 0.00000 | 0.02620 | 0.12908 | 0.06027 | 0.10525 | 0.01336 | 0.09466 |
| **UNC80** | 0.19852 | 0.00892 | 0.03350 | 0.01309 | 0.05274 | 0.00000 | 0.04289 | 0.19049 | 0.00442 | 0.01164 | 0.01748 | 0.08349 |
| **UNC93A** | 1.41595 | 0.10011 | 0.00000 | 1.79682 | 0.00000 | 0.28063 | 0.00000 | 1.64535 | 0.13719 | 0.00000 | 0.00000 | 0.00000 |
| **UNC93B1** | 0.00324 | 0.00799 | 0.29141 | 0.00000 | 0.08905 | 0.00000 | 0.00094 | 0.00935 | 0.18038 | 0.41782 | 0.01117 | 0.00825 |
| **UNG** | 10.98653 | 23.97475 | 10.78661 | 13.92887 | 5.68974 | 12.63092 | 11.35349 | 9.88194 | 22.01699 | 5.01065 | 17.27583 | 21.97910 |
| **UNK** | 2.78885 | 1.99297 | 5.29406 | 1.29390 | 1.83465 | 7.71458 | 4.53171 | 1.62927 | 1.63376 | 0.72653 | 1.99178 | 6.45753 |
| **UNKL** | 8.04498 | 3.25906 | 7.49407 | 1.02219 | 6.32338 | 0.00000 | 6.12035 | 4.01455 | 6.74109 | 4.09094 | 2.52358 | 3.71910 |
| **UNQ6494** | 0.01308 | 0.01805 | 0.12309 | 0.10200 | 0.04795 | 0.00000 | 0.00382 | 0.00874 | 0.05724 | 0.00000 | 0.01382 | 0.03343 |
| **UOX** | 0.02309 | 0.00000 | 0.01667 | 0.00678 | 0.08199 | 0.05054 | 0.01283 | 0.07981 | 0.01905 | 0.02302 | 0.05464 | 0.10705 |
| **UPB1** | 2.76081 | 0.07344 | 0.00778 | 0.04758 | 0.05650 | 0.00632 | 0.01756 | 0.24356 | 0.02287 | 0.00000 | 0.01144 | 0.04652 |

| | | | | | | | | | | | | |
|---|---|---|---|---|---|---|---|---|---|---|---|---|
| **UPF1** | 1.24968 | 4.09495 | 7.17787 | 6.91449 | 9.98078 | 0.00000 | 15.80779 | 2.91925 | 2.51500 | 23.20370 | 3.27841 | 9.04704 |
| **UPF2** | 1.98401 | 1.60040 | 8.34885 | 3.30611 | 6.53994 | 9.22169 | 15.79802 | 10.25811 | 5.08153 | 27.24166 | 9.55084 | 7.95080 |
| **UPF3A** | 1.20162 | 1.75066 | 1.40658 | 1.38488 | 1.11308 | 2.38018 | 3.67352 | 0.93226 | 2.10986 | 4.17603 | 1.65782 | 0.70314 |
| **UPF3B** | 1.83980 | 2.61347 | 7.28830 | 3.56157 | 4.77158 | 6.30713 | 14.37768 | 5.38419 | 3.31995 | 12.84053 | 2.82702 | 4.60441 |
| **UPK1A** | 0.00275 | 0.08512 | 0.10217 | 0.08342 | 0.00000 | 0.00000 | 0.07473 | 0.05032 | 0.00000 | 0.06895 | 0.15649 | 0.09228 |
| **UPK1A-AS1** | 0.00220 | 0.01151 | 0.01680 | 0.01455 | 0.01168 | 0.01259 | 0.01876 | 0.05923 | 0.00782 | 0.03105 | 0.03778 | 0.03340 |
| **UPK1B** | 0.00787 | 0.02374 | 0.01709 | 0.04708 | 0.01486 | 0.00029 | 0.02071 | 0.00852 | 0.02665 | 0.01268 | 0.01631 | 0.06158 |
| **UPK2** | 0.04089 | 0.00000 | 0.54497 | 0.04694 | 0.34968 | 0.11558 | 0.00000 | 0.01178 | 0.90967 | 0.00000 | 0.26479 | 0.00947 |
| **UPK3B** | 0.00786 | 0.01396 | 0.01588 | 0.01208 | 0.01307 | 0.02177 | 0.02804 | 0.01030 | 0.01568 | 0.01890 | 0.01472 | 0.01640 |
| **UPP1** | 0.27691 | 35.13158 | 10.25607 | 33.18150 | 31.83736 | 11.12736 | 23.40163 | 0.98682 | 20.25815 | 5.83383 | 12.79756 | 23.10471 |
| **UPP2** | 0.06812 | 0.00707 | 0.05014 | 0.00635 | 0.01084 | 0.02629 | 0.02757 | 0.03728 | 0.04751 | 0.01906 | 0.02252 | 0.06844 |
| **UPRT** | 0.03908 | 0.13230 | 0.59163 | 0.77355 | 0.51732 | 3.28167 | 3.37195 | 0.40987 | 3.01136 | 0.00000 | 1.69089 | 0.10515 |
| **UQCC1** | 0.42752 | 13.90337 | 11.52731 | 5.97545 | 3.97227 | 9.45703 | 14.64205 | 0.32996 | 12.76330 | 0.00000 | 9.94653 | 7.53204 |
| **UQCC2** | 1.54454 | 6.28187 | 3.86346 | 5.26140 | 8.81864 | 4.80475 | 4.90963 | 0.39837 | 0.00000 | 2.54452 | 6.61842 | 6.19481 |
| **UQCR10** | 0.75975 | 22.96066 | 21.80816 | 29.38303 | 30.31630 | 25.21117 | 41.29338 | 11.95352 | 23.91043 | 19.23699 | 35.42529 | 27.68729 |
| **UQCR11** | 0.61286 | 23.99039 | 27.64490 | 28.49342 | 33.44671 | 19.79052 | 42.88821 | 1.04376 | 34.96555 | 13.01791 | 36.52442 | 14.41375 |
| **UQCRB** | 1.48667 | 10.27734 | 11.90333 | 12.33628 | 8.11970 | 47.19830 | 24.96546 | 1.12173 | 12.54062 | 24.37210 | 19.76792 | 17.54289 |
| **UQCRBP1** | 0.04797 | 0.78614 | 1.32522 | 1.86643 | 1.04801 | 4.09427 | 2.59083 | 0.09056 | 0.10182 | 1.57860 | 2.29602 | 1.33822 |
| **UQCRC1** | 2.03250 | 37.01387 | 30.65886 | 36.45136 | 43.75111 | 19.66723 | 45.23053 | 6.49467 | 46.46429 | 0.00000 | 42.16519 | 23.68519 |
| **UQCRC2** | 3.75204 | 23.31227 | 14.39277 | 14.43523 | 15.87678 | 27.38297 | 25.64323 | 9.29973 | 33.11052 | 4.49469 | 28.85731 | 11.28173 |
| **UQCRFS1** | 2.43269 | 20.76704 | 16.86495 | 16.97945 | 16.63162 | 27.31271 | 24.89603 | 4.85544 | 26.06459 | 1.47517 | 20.89959 | 16.29899 |
| **UQCRH** | 0.33400 | 26.75205 | 24.36280 | 35.41975 | 27.81773 | 56.15677 | 42.76966 | 13.78328 | 31.83930 | 28.31179 | 29.43352 | 28.15264 |
| **UQCRHL** | 0.00996 | 12.94359 | 10.47901 | 22.09693 | 14.43360 | 30.02250 | 18.28686 | 5.28504 | 16.59850 | 0.00468 | 14.13739 | 12.94570 |
| **UQCRQ** | 3.27519 | 21.23445 | 25.14741 | 17.28973 | 30.27332 | 34.62048 | 46.48236 | 12.79785 | 23.33863 | 29.37597 | 36.20847 | 20.86553 |
| **URAD** | 0.01357 | 0.00401 | 0.06318 | 0.00000 | 0.87548 | 0.00000 | 0.17878 | 0.01291 | 0.47325 | 0.07242 | 1.23825 | 0.01934 |
| **URAHP** | 0.02787 | 0.03700 | 0.04116 | 0.00795 | 0.01063 | 0.02261 | 0.00836 | 0.03159 | 0.04979 | 0.03379 | 0.03721 | 0.03081 |
| **URB1** | 0.09670 | 6.13953 | 3.22011 | 1.18402 | 3.06746 | 2.60883 | 4.06142 | 4.76396 | 4.87773 | 7.25975 | 3.83493 | 6.09384 |
| **URB2** | 0.00000 | 6.96624 | 7.96654 | 4.15989 | 4.30844 | 0.93194 | 10.59005 | 3.90303 | 8.36022 | 0.00000 | 8.15171 | 6.71123 |
| **URGCP** | 3.79854 | 6.90981 | 0.71789 | 1.64794 | 2.18488 | 0.00000 | 0.83757 | 0.10805 | 0.00000 | 0.00000 | 1.99093 | 9.01372 |
| **URI1** | 0.41938 | 0.77042 | 3.59962 | 0.78871 | 0.80780 | 13.06299 | 9.66354 | 1.01403 | 1.02087 | 17.14872 | 1.63222 | 4.47446 |
| **URM1** | 0.76633 | 11.71560 | 11.20332 | 6.57126 | 11.06625 | 0.00000 | 14.16081 | 1.93972 | 17.60681 | 3.99245 | 9.27266 | 14.42134 |
| **UROC1** | 0.00652 | 0.01376 | 0.02957 | 0.01089 | 0.01369 | 0.02360 | 0.01472 | 0.01155 | 0.01213 | 0.01937 | 0.01556 | 0.01718 |
| **UROD** | 2.13495 | 19.14033 | 12.16246 | 14.84925 | 6.82914 | 5.34605 | 19.29548 | 8.21411 | 12.93622 | 0.00000 | 19.76770 | 14.94523 |
| **UROS** | 0.35356 | 1.73447 | 3.00464 | 0.77990 | 5.54022 | 0.00000 | 0.08202 | 0.00000 | 2.63312 | 2.22422 | 4.25134 | 1.10275 |
| **USB1** | 9.18006 | 7.67287 | 9.31514 | 2.37078 | 2.87624 | 5.80528 | 2.36563 | 0.77227 | 8.47465 | 0.00000 | 1.64199 | 4.79323 |
| **USE1** | 0.00000 | 3.51419 | 2.71505 | 1.56928 | 3.22240 | 0.32429 | 3.61692 | 1.56333 | 3.30104 | 0.27492 | 3.03398 | 4.22438 |
| **USF1** | 0.76088 | 9.02264 | 6.51317 | 6.57556 | 3.62153 | 4.13348 | 5.45716 | 5.51039 | 11.19830 | 0.00000 | 0.18103 | 7.61478 |
| **USF2** | 2.69639 | 4.43869 | 4.28089 | 3.78436 | 9.72180 | 0.00000 | 7.46964 | 0.00000 | 2.92143 | 17.09760 | 2.44464 | 3.00818 |
| **USH1C** | 0.10963 | 0.00397 | 0.03814 | 0.10569 | 0.00000 | 0.01035 | 0.19839 | 0.09921 | 0.53795 | 0.00000 | 0.00000 | 0.30613 |
| **USH1G** | 0.02576 | 0.01661 | 0.02173 | 0.02596 | 0.07528 | 0.00000 | 0.03132 | 0.01737 | 0.00000 | 0.01453 | 0.06889 | 0.05170 |
| **USH2A** | 1.94205 | 0.18393 | 0.00000 | 0.03604 | 0.45592 | 0.10917 | 0.00000 | 0.73820 | 0.04004 | 0.41287 | 0.08299 | 0.14724 |
| **USHBP1** | 0.02214 | 0.01099 | 0.46820 | 0.07651 | 0.10982 | 0.01795 | 0.25020 | 0.00790 | 0.07437 | 0.00000 | 0.21090 | 0.04741 |
| **USMG5** | 0.13988 | 17.95758 | 19.19808 | 13.11075 | 16.55827 | 43.15347 | 36.98300 | 8.75491 | 22.89987 | 25.88796 | 28.08668 | 21.74375 |
| **USO1** | 2.12545 | 4.68227 | 7.46617 | 3.86284 | 1.63994 | 18.37636 | 5.65708 | 1.27982 | 7.68427 | 7.31529 | 7.08666 | 4.09732 |
| **USP1** | 6.51059 | 7.84199 | 5.78403 | 5.60506 | 5.69236 | 31.19457 | 12.64542 | 16.29438 | 10.95512 | 21.07411 | 8.30953 | 18.36992 |
| **USP10** | 5.26655 | 20.17256 | 20.84513 | 11.92480 | 11.29719 | 24.33957 | 22.21597 | 9.67527 | 18.60290 | 4.42137 | 19.44986 | 18.70691 |
| **USP11** | 5.68751 | 3.92158 | 3.04332 | 6.61621 | 2.35086 | 0.00049 | 4.21969 | 3.54192 | 2.14227 | 2.92081 | 1.70460 | 7.45573 |
| **USP12** | 0.03978 | 1.63031 | 0.08258 | 0.50183 | 0.00139 | 2.24802 | 0.00000 | 0.95465 | 0.23689 | 3.11284 | 0.10215 | 3.41844 |

| | | | | | | | | | | | | |
|---|---|---|---|---|---|---|---|---|---|---|---|---|
| **USP12-AS1** | 0.01395 | 0.02098 | 0.00903 | 0.00803 | 0.00709 | 0.03501 | 0.01152 | 0.01393 | 0.01599 | 0.07629 | 0.02004 | 0.02743 |
| **USP12-AS2** | 0.00486 | 0.01881 | 0.02308 | 0.01207 | 0.01941 | 0.01290 | 0.02321 | 0.03801 | 0.01348 | 0.01206 | 0.01655 | 0.01491 |
| **USP13** | 9.22272 | 1.40794 | 0.85689 | 1.52390 | 0.00000 | 0.25046 | 0.76184 | 4.26144 | 0.01042 | 2.71166 | 0.43801 | 1.03546 |
| **USP14** | 5.03430 | 13.20513 | 20.12112 | 6.96413 | 5.91209 | 35.65663 | 27.82546 | 2.62827 | 19.25441 | 8.59928 | 17.06941 | 11.19353 |
| **USP15** | 4.06347 | 4.35626 | 8.45504 | 0.97539 | 5.58773 | 25.64673 | 7.80068 | 4.33876 | 10.09177 | 5.29185 | 7.86904 | 8.16363 |
| **USP16** | 4.51909 | 13.29797 | 7.94689 | 5.31430 | 2.72240 | 26.69866 | 14.18444 | 9.60378 | 7.52638 | 17.95358 | 7.08753 | 18.24105 |
| **USP18** | 0.17918 | 0.32639 | 0.00764 | 0.38379 | 0.20899 | 0.10054 | 0.02810 | 0.05580 | 0.27994 | 0.33055 | 0.14934 | 0.26442 |
| **USP19** | 0.92438 | 6.29890 | 3.37754 | 4.70176 | 6.32403 | 0.00000 | 2.74131 | 1.15172 | 7.41031 | 10.24783 | 1.18237 | 4.25761 |
| **USP2** | 21.57373 | 2.80646 | 0.26630 | 0.35164 | 3.06853 | 1.20030 | 0.42421 | 7.06101 | 0.18333 | 3.03194 | 0.78165 | 1.66793 |
| **USP2-AS1** | 0.55239 | 0.30227 | 0.40686 | 0.00000 | 0.36977 | 0.04049 | 0.55746 | 1.72966 | 0.52423 | 0.22153 | 0.07565 | 3.07053 |
| **USP20** | 4.19399 | 1.84484 | 0.89954 | 0.00000 | 2.10571 | 0.00000 | 0.73404 | 0.00000 | 1.05546 | 0.00000 | 0.65538 | 0.77845 |
| **USP21** | 2.57432 | 2.22134 | 1.43492 | 1.51792 | 0.44339 | 1.39368 | 0.70826 | 0.19896 | 1.61403 | 0.50630 | 1.82085 | 1.09011 |
| **USP22** | 3.58857 | 18.16516 | 16.27302 | 22.00482 | 18.83410 | 12.30512 | 35.86121 | 1.94741 | 12.98240 | 23.86651 | 17.54097 | 15.22394 |
| **USP24** | 4.19570 | 1.82295 | 3.30548 | 1.30618 | 3.35376 | 11.23894 | 8.76672 | 2.63478 | 4.24416 | 16.58725 | 4.20929 | 2.12695 |
| **USP25** | 2.03685 | 0.69929 | 6.94790 | 0.52754 | 1.84227 | 9.57105 | 6.50694 | 0.41469 | 0.00449 | 9.08572 | 1.89615 | 0.09750 |
| **USP26** | 0.00388 | 0.01364 | 0.01643 | 0.00993 | 0.02284 | 0.01596 | 0.03406 | 0.02079 | 0.02260 | 0.01747 | 0.02231 | 0.01858 |
| **USP27X** | 0.03769 | 0.00000 | 1.58042 | 0.12717 | 0.00000 | 0.20190 | 0.45044 | 0.01134 | 0.00000 | 0.13489 | 0.00000 | 0.05530 |
| **USP27X-AS1** | 0.11472 | 0.04746 | 0.49884 | 0.03626 | 0.00540 | 0.19489 | 0.00000 | 0.11408 | 0.03171 | 0.00000 | 0.00697 | 0.12894 |
| **USP28** | 0.37430 | 9.73902 | 1.03101 | 27.58285 | 2.12960 | 14.47039 | 10.10705 | 0.87930 | 9.71998 | 8.31276 | 13.07556 | 5.55153 |
| **USP29** | 2.45287 | 0.03560 | 0.01388 | 0.01336 | 0.02038 | 0.00000 | 0.01507 | 0.81426 | 0.03811 | 0.00000 | 0.01010 | 0.02792 |
| **USP3** | 4.64090 | 0.78595 | 5.17385 | 1.26073 | 2.07519 | 5.77442 | 10.76654 | 2.50042 | 12.92026 | 3.41921 | 4.96843 | 0.94008 |
| **USP3-AS1** | 2.45438 | 0.00000 | 0.17307 | 0.00604 | 0.00000 | 0.50636 | 0.03422 | 0.32390 | 0.68067 | 0.10014 | 0.08138 | 0.14998 |
| **USP30** | 1.98920 | 4.00032 | 1.59067 | 1.31915 | 0.77161 | 4.07339 | 1.13868 | 2.16404 | 2.39803 | 0.00000 | 1.89733 | 3.65002 |
| **USP30-AS1** | 0.00486 | 0.04413 | 0.03154 | 0.00699 | 0.05223 | 0.00000 | 0.00270 | 0.01077 | 0.04759 | 0.06434 | 0.05255 | 0.02661 |
| **USP31** | 2.20191 | 2.76290 | 0.00000 | 0.15341 | 1.08361 | 3.12946 | 1.58522 | 0.90954 | 2.90335 | 6.28038 | 1.42926 | 2.04901 |
| **USP32** | 1.27152 | 0.79637 | 1.07522 | 2.81205 | 1.48679 | 3.62777 | 3.49252 | 1.16341 | 3.79119 | 6.34693 | 1.30641 | 1.92264 |
| **USP32P1** | 0.13034 | 0.03132 | 0.06144 | 0.00149 | 0.00000 | 0.36575 | 0.00000 | 0.15808 | 0.04942 | 0.08644 | 0.00000 | 0.52246 |
| **USP32P2** | 0.00219 | 0.00000 | 0.02995 | 0.07323 | 0.00647 | 0.01378 | 0.01420 | 0.03144 | 0.06162 | 0.08581 | 0.06356 | 0.14830 |
| **USP33** | 0.00000 | 6.67286 | 8.07498 | 5.21527 | 5.48412 | 10.39882 | 8.61640 | 3.52276 | 9.71170 | 3.06270 | 5.92421 | 3.76480 |
| **USP34** | 20.37973 | 0.84135 | 7.14490 | 1.76701 | 2.90608 | 18.48825 | 4.28421 | 8.42785 | 8.01337 | 29.14429 | 2.42075 | 5.20112 |
| **USP35** | 2.59980 | 0.23553 | 0.51977 | 0.24904 | 0.05236 | 0.00000 | 0.00683 | 0.14314 | 0.82454 | 0.78482 | 0.07559 | 0.11728 |
| **USP36** | 7.32335 | 6.08603 | 7.49777 | 2.87468 | 11.55405 | 0.00000 | 13.09776 | 5.10876 | 9.01665 | 20.89064 | 10.64334 | 9.70120 |
| **USP37** | 1.79266 | 1.80520 | 3.68135 | 2.65503 | 0.00419 | 13.84792 | 1.41599 | 5.32703 | 4.41663 | 11.68454 | 3.28854 | 6.30879 |
| **USP38** | 1.98881 | 4.77826 | 5.47450 | 2.06035 | 0.00000 | 8.20427 | 6.46055 | 6.99836 | 3.22898 | 4.56183 | 5.02399 | 6.20796 |
| **USP39** | 1.70873 | 16.22796 | 11.51152 | 12.24083 | 11.27112 | 19.50630 | 19.77200 | 5.88422 | 20.68229 | 0.00020 | 13.97933 | 12.71437 |
| **USP4** | 1.08985 | 6.67787 | 1.46653 | 1.67024 | 6.16180 | 1.90162 | 0.84798 | 1.85296 | 5.90508 | 3.57102 | 5.04341 | 5.90444 |
| **USP40** | 3.15798 | 2.28598 | 1.75267 | 0.98957 | 2.62082 | 5.07904 | 0.98902 | 0.61610 | 1.12320 | 2.64673 | 3.76965 | 3.85711 |
| **USP42** | 0.25498 | 0.32252 | 4.31578 | 0.93527 | 0.80948 | 1.46077 | 4.22016 | 4.62831 | 1.73589 | 9.92551 | 1.39361 | 1.27670 |
| **USP43** | 1.14506 | 3.99336 | 0.55761 | 0.00000 | 4.92671 | 2.66587 | 1.00110 | 0.86646 | 1.21330 | 6.09541 | 1.91266 | 5.65500 |
| **USP44** | 1.93432 | 0.07173 | 0.00000 | 0.60364 | 0.35570 | 0.57801 | 0.32749 | 0.21717 | 0.00000 | 0.00071 | 0.14670 | 0.41228 |
| **USP45** | 2.76593 | 1.35845 | 1.35404 | 0.61320 | 0.00000 | 5.74887 | 0.00000 | 0.44471 | 2.66334 | 2.14942 | 1.70713 | 1.79390 |
| **USP46** | 0.66968 | 2.41793 | 1.37463 | 2.44513 | 0.00000 | 6.56533 | 3.88845 | 0.44272 | 1.16975 | 1.81752 | 3.21420 | 4.57774 |
| **USP47** | 3.16741 | 3.34824 | 12.52149 | 0.46773 | 4.06407 | 11.94581 | 15.86845 | 8.06986 | 3.51238 | 11.49856 | 8.76110 | 5.54253 |
| **USP48** | 2.37329 | 6.41842 | 4.19667 | 5.69555 | 6.41584 | 8.81095 | 5.24839 | 0.71419 | 4.70289 | 20.47756 | 6.54616 | 5.70675 |
| **USP49** | 0.76575 | 0.66666 | 3.07638 | 0.90914 | 0.52240 | 2.86410 | 0.47677 | 0.83042 | 1.99064 | 3.90553 | 0.62917 | 1.97943 |
| **USP5** | 2.05540 | 10.45826 | 13.54708 | 9.98408 | 7.14044 | 0.00000 | 13.06695 | 3.28155 | 7.13598 | 4.65868 | 10.86907 | 4.92003 |
| **USP50** | 0.56949 | 0.00617 | 0.01778 | 0.01319 | 0.01927 | 0.01741 | 0.00071 | 0.04211 | 0.04142 | 0.13865 | 0.00798 | 0.01413 |
| **USP51** | 0.01197 | 0.01872 | 0.07899 | 0.03548 | 0.00892 | 0.00000 | 0.00355 | 0.01082 | 0.02436 | 0.06090 | 0.03023 | 0.20278 |

| | | | | | | | | | | | | |
|---|---|---|---|---|---|---|---|---|---|---|---|---|
| USP53 | 2.69663 | 0.21303 | 6.67313 | 2.06435 | 4.93047 | 5.49965 | 7.92837 | 0.28478 | 7.10460 | 6.43051 | 2.62888 | 0.36024 |
| USP54 | 3.47680 | 1.84241 | 1.59197 | 1.97306 | 1.72987 | 0.60539 | 2.32979 | 4.25832 | 3.97015 | 0.85220 | 1.85406 | 6.92571 |
| USP6 | 0.00086 | 0.06869 | 0.05806 | 0.64209 | 0.00000 | 0.05762 | 0.00000 | 0.14519 | 0.10826 | 0.48459 | 0.21973 | 0.32111 |
| USP6NL | 2.52154 | 1.99488 | 1.79600 | 0.00000 | 0.00000 | 1.72098 | 2.62410 | 0.61370 | 0.62146 | 4.13751 | 2.15751 | 1.12231 |
| USP7 | 4.65007 | 22.32482 | 6.10029 | 7.83883 | 15.02567 | 24.09606 | 22.39212 | 29.86476 | 1.94390 | 42.91582 | 5.57291 | 43.26783 |
| USP8 | 1.55249 | 1.98221 | 8.22580 | 3.37084 | 3.33358 | 25.85208 | 8.11457 | 4.52443 | 2.44365 | 20.35672 | 9.49281 | 5.74431 |
| USP9X | 6.02666 | 4.42619 | 4.57931 | 5.79263 | 4.57014 | 18.90254 | 6.71812 | 6.50262 | 3.50314 | 20.35355 | 7.20425 | 5.32713 |
| USP9Y | 0.00000 | 0.00000 | 0.18923 | 1.03598 | 0.49513 | 5.44705 | 0.00017 | 0.16065 | 1.01957 | 4.41740 | 0.00000 | 1.88245 |
| USPL1 | 0.25389 | 2.85326 | 3.85290 | 0.17050 | 0.60333 | 9.39516 | 2.70602 | 11.50109 | 5.77833 | 2.32087 | 3.70546 | 5.61830 |
| UST | 0.80377 | 0.00000 | 0.64401 | 0.89852 | 5.37727 | 1.56438 | 1.55916 | 0.08775 | 0.35267 | 0.54227 | 0.51468 | 0.00000 |
| UTF1 | 0.02432 | 0.00000 | 0.57746 | 3.60988 | 0.12494 | 0.00000 | 0.05400 | 0.00938 | 0.00000 | 0.24450 | 0.00000 | 0.01001 |
| UTP11L | 1.88006 | 12.84510 | 15.21559 | 5.84133 | 5.75221 | 35.20233 | 16.73291 | 20.52010 | 13.40404 | 3.21304 | 10.80101 | 14.23305 |
| UTP14A | 3.97328 | 10.99861 | 5.21192 | 3.82691 | 6.93814 | 9.18909 | 11.63491 | 9.55405 | 14.51039 | 6.10066 | 10.80665 | 13.95142 |
| UTP14C | 0.05346 | 0.97455 | 3.07640 | 0.00064 | 0.00000 | 5.78430 | 1.46151 | 0.34985 | 4.60305 | 7.66780 | 0.61196 | 1.56212 |
| UTP15 | 0.00000 | 10.16071 | 7.15703 | 0.82550 | 0.00000 | 21.32834 | 9.76572 | 5.90710 | 10.45058 | 0.00000 | 7.91680 | 16.35322 |
| UTP18 | 0.00000 | 14.22568 | 13.25307 | 6.80791 | 6.65912 | 22.08206 | 16.88438 | 8.42213 | 15.30996 | 3.61133 | 10.60377 | 17.75734 |
| UTP20 | 0.00000 | 3.72129 | 8.93423 | 2.44676 | 2.14208 | 11.16247 | 15.66674 | 7.16758 | 7.30486 | 19.90812 | 6.13884 | 9.89469 |
| UTP23 | 2.05398 | 4.54500 | 9.45550 | 3.79524 | 3.54824 | 10.00714 | 9.20151 | 8.57625 | 7.26415 | 3.33834 | 5.13479 | 10.49241 |
| UTP3 | 0.06563 | 2.07364 | 3.79158 | 3.29685 | 0.00000 | 8.87582 | 7.12770 | 7.20447 | 0.00000 | 9.52958 | 2.35677 | 11.11005 |
| UTP6 | 4.08766 | 14.44575 | 16.20675 | 7.49949 | 10.65660 | 26.43721 | 23.57587 | 20.00825 | 22.06978 | 0.00001 | 19.50589 | 15.69723 |
| UTRN | 0.15389 | 0.34788 | 8.48685 | 3.62083 | 12.37417 | 11.47923 | 0.00000 | 0.15776 | 10.77075 | 31.02646 | 5.21523 | 3.76393 |
| UTS2 | 0.00243 | 0.05680 | 0.02230 | 0.00000 | 0.00588 | 0.05077 | 0.01515 | 0.25861 | 0.00000 | 0.03448 | 0.01584 | 0.33272 |
| UTS2B | 2.42349 | 0.00433 | 0.00940 | 0.00610 | 0.00130 | 0.03565 | 0.00426 | 0.26300 | 0.01269 | 0.06017 | 0.01003 | 0.01512 |
| UTY | 0.32148 | 1.26790 | 2.72872 | 1.16876 | 1.04433 | 3.07925 | 0.00000 | 0.81120 | 2.96436 | 3.45953 | 0.69067 | 2.83280 |
| UVRAG | 1.00655 | 4.24419 | 1.62863 | 3.08911 | 2.41091 | 0.87388 | 0.75066 | 0.23741 | 2.10069 | 3.51041 | 0.53643 | 2.49942 |
| UVSSA | 0.21249 | 0.22606 | 0.02723 | 0.52290 | 0.53635 | 0.38483 | 0.50074 | 0.43534 | 0.12233 | 1.70575 | 0.36067 | 0.30142 |
| UXS1 | 5.30963 | 7.21476 | 7.92419 | 2.69999 | 2.44364 | 12.30592 | 0.00000 | 5.34498 | 14.93878 | 7.50717 | 7.41979 | 3.27853 |
| UXT | 3.91113 | 9.56457 | 8.99352 | 9.30396 | 13.64095 | 11.68177 | 14.18430 | 11.61412 | 10.85816 | 0.49616 | 10.49124 | 17.25958 |
| UXT-AS1 | 0.01510 | 0.01525 | 0.01723 | 0.01346 | 0.01370 | 0.01511 | 0.02149 | 0.01215 | 0.01052 | 0.01475 | 0.03232 | 0.01854 |
| VAC14 | 0.09250 | 2.07781 | 3.23903 | 2.66414 | 9.05888 | 0.00000 | 5.74439 | 0.36601 | 1.97638 | 11.87272 | 2.55483 | 1.63715 |
| VAC14-AS1 | 0.06444 | 0.16461 | 0.17094 | 0.01216 | 0.00000 | 0.15929 | 0.02874 | 0.07379 | 0.00000 | 0.93647 | 0.12065 | 0.08708 |
| VAMP1 | 0.99230 | 3.33298 | 0.20756 | 2.39184 | 1.52083 | 0.00000 | 0.20275 | 8.28879 | 1.04221 | 0.00000 | 1.23939 | 5.24313 |
| VAMP2 | 2.11945 | 0.99761 | 3.69759 | 1.80327 | 0.00000 | 0.00000 | 3.01036 | 4.45236 | 1.83932 | 4.12760 | 2.27652 | 3.65091 |
| VAMP3 | 5.48357 | 14.02832 | 9.86125 | 4.09147 | 4.06720 | 25.89771 | 7.27638 | 3.75088 | 9.71167 | 7.70389 | 5.69339 | 13.51156 |
| VAMP4 | 1.11112 | 1.07234 | 0.19964 | 0.33888 | 0.27489 | 2.56980 | 0.00075 | 0.39528 | 0.26478 | 0.00000 | 0.97001 | 1.75424 |
| VAMP5 | 0.00071 | 0.31823 | 0.50691 | 0.20723 | 1.32106 | 0.00000 | 0.58998 | 0.09918 | 1.69448 | 0.99218 | 0.00000 | 0.37101 |
| VAMP8 | 0.81723 | 17.73309 | 7.74121 | 1.20243 | 28.02669 | 28.28360 | 31.52337 | 0.34815 | 20.43464 | 13.74422 | 17.16593 | 11.86329 |
| VANGL1 | 0.49263 | 4.42241 | 0.59742 | 3.72838 | 2.92640 | 0.01459 | 0.27815 | 0.29045 | 6.39407 | 2.15910 | 1.08032 | 1.41708 |
| VANGL2 | 0.00325 | 0.01156 | 0.07838 | 2.70248 | 0.00000 | 0.02798 | 0.21012 | 0.00755 | 0.70238 | 0.39991 | 0.00000 | 0.12043 |
| VAPA | 2.41205 | 22.81792 | 18.46082 | 14.47786 | 16.63064 | 51.36946 | 28.93975 | 17.65311 | 24.05788 | 6.94333 | 28.06006 | 22.27537 |
| VAPB | 2.93103 | 10.98607 | 7.96555 | 6.84366 | 8.45750 | 4.79985 | 12.56602 | 0.70410 | 5.75507 | 6.15960 | 6.89294 | 11.04845 |
| VARS | 0.00088 | 9.55434 | 12.87783 | 9.70053 | 20.01170 | 0.00000 | 16.02958 | 0.00000 | 11.38301 | 19.25822 | 6.46007 | 7.41844 |
| VARS2 | 0.14351 | 1.28329 | 0.34700 | 1.61024 | 1.96044 | 0.00000 | 0.62053 | 0.07281 | 1.44959 | 8.72465 | 0.00000 | 1.49952 |
| VASH1 | 0.03921 | 0.95776 | 0.00000 | 2.41143 | 0.96417 | 0.00000 | 0.00195 | 0.28766 | 0.00000 | 0.48701 | 0.02440 | 1.04596 |
| VASH2 | 0.00220 | 0.00000 | 1.32758 | 2.77665 | 0.07816 | 0.00000 | 0.21339 | 0.60880 | 0.00000 | 0.47815 | 0.28107 | 0.02824 |
| VASN | 0.07440 | 0.00000 | 0.70939 | 0.00000 | 0.06425 | 0.00000 | 0.00000 | 0.05110 | 0.59862 | 0.40483 | 0.01863 | 0.11584 |
| VASP | 0.63813 | 5.36833 | 7.44925 | 1.58328 | 9.83774 | 0.00000 | 4.93949 | 1.34926 | 2.89385 | 17.57339 | 0.97587 | 7.54812 |
| VAT1 | 9.98450 | 7.79913 | 21.56130 | 23.75130 | 26.93219 | 0.39822 | 30.59296 | 3.21968 | 30.38131 | 1.61104 | 9.54034 | 17.76471 |

| | | | | | | | | | | | | |
|---|---|---|---|---|---|---|---|---|---|---|---|---|
| **VAT1L** | 0.01646 | 0.04743 | 0.07181 | 0.00000 | 0.39527 | 0.01545 | 0.00000 | 0.00601 | 0.00000 | 0.01581 | 0.42394 | 0.07245 |
| **VAV1** | 0.00533 | 0.09258 | 0.85599 | 0.00032 | 0.29709 | 0.51263 | 0.00000 | 0.00373 | 0.54402 | 0.00000 | 0.21290 | 0.07953 |
| **VAV2** | 0.07534 | 11.40360 | 1.56903 | 7.40314 | 10.13253 | 0.00000 | 2.24020 | 0.11124 | 6.38455 | 7.69734 | 1.77499 | 2.77428 |
| **VAV3** | 9.73070 | 0.10011 | 0.42586 | 0.00000 | 0.00000 | 0.06371 | 1.30357 | 0.40656 | 0.34572 | 0.11649 | 0.00000 | 0.44228 |
| **VAV3-AS1** | 0.50395 | 0.01903 | 0.01348 | 0.00996 | 0.01133 | 0.01572 | 0.03359 | 0.00456 | 0.01160 | 0.02871 | 0.01768 | 0.01354 |
| **VAX1** | 0.12619 | 0.00963 | 0.01893 | 0.01165 | 0.01382 | 0.01304 | 0.01563 | 0.10838 | 0.01890 | 0.02039 | 0.02013 | 0.07637 |
| **VAX2** | 0.00588 | 0.01889 | 0.00899 | 0.06671 | 0.11172 | 0.05228 | 0.01451 | 0.03610 | 0.00483 | 0.00048 | 0.01607 | 0.01223 |
| **VBP1** | 0.08551 | 9.86528 | 11.86381 | 7.95248 | 0.42380 | 39.26172 | 18.33554 | 5.84840 | 7.15804 | 4.19788 | 15.80952 | 14.92528 |
| **VCAM1** | 0.00629 | 0.01717 | 0.01430 | 0.04399 | 0.01561 | 0.00583 | 0.02104 | 0.01059 | 0.01562 | 0.01449 | 0.02165 | 0.01725 |
| **VCAN** | 0.10879 | 0.68525 | 3.65095 | 21.55894 | 0.00000 | 0.00000 | 0.00000 | 0.05502 | 0.00000 | 0.70528 | 0.15310 | 5.03671 |
| **VCL** | 2.06460 | 6.29988 | 8.09271 | 13.04480 | 6.57089 | 11.53713 | 0.69520 | 0.11043 | 12.32368 | 9.36751 | 5.08211 | 1.89441 |
| **VCP** | 1.59473 | 35.91527 | 42.33280 | 33.81692 | 35.19458 | 28.55293 | 53.91399 | 23.91012 | 45.38306 | 5.30935 | 35.88163 | 24.85142 |
| **VCPIP1** | 5.65386 | 2.99473 | 3.47579 | 0.00000 | 0.49320 | 10.00575 | 5.68623 | 6.18728 | 3.12696 | 4.82645 | 3.76919 | 4.31176 |
| **VCPKMT** | 0.08960 | 1.28299 | 3.54637 | 1.80902 | 0.00337 | 6.73601 | 1.74577 | 3.97179 | 4.84279 | 1.47937 | 1.45908 | 4.14399 |
| **VCX** | 0.00168 | 0.01840 | 0.00000 | 0.95431 | 0.11715 | 0.00000 | 0.02120 | 0.27078 | 0.13631 | 0.00000 | 0.00388 | 0.00910 |
| **VCX2** | 0.00591 | 0.01082 | 0.01891 | 0.01661 | 0.01365 | 0.02577 | 0.01649 | 0.05194 | 0.01751 | 0.01344 | 0.02474 | 0.03189 |
| **VCX3A** | 0.00628 | 0.00000 | 0.10117 | 0.09029 | 0.47305 | 0.00000 | 0.21430 | 0.16904 | 0.28746 | 0.24689 | 0.03716 | 0.00245 |
| **VCX3B** | 0.00126 | 0.00041 | 0.02809 | 0.00000 | 0.26852 | 0.18068 | 0.10384 | 0.06961 | 0.00000 | 0.15017 | 0.22672 | 0.00479 |
| **VDAC1** | 6.94191 | 17.09722 | 16.18242 | 12.35864 | 18.18500 | 34.00987 | 26.17078 | 0.75963 | 24.23031 | 11.18641 | 18.08618 | 7.35103 |
| **VDAC2** | 0.00000 | 15.47047 | 9.82921 | 8.48609 | 10.50756 | 28.06776 | 17.41400 | 4.55882 | 20.30019 | 8.92053 | 15.57912 | 13.54892 |
| **VDAC3** | 5.67941 | 22.19056 | 21.28662 | 16.62140 | 13.67110 | 42.26855 | 30.72727 | 1.59490 | 23.01497 | 2.87680 | 31.69783 | 12.86243 |
| **VDR** | 0.03950 | 0.00338 | 0.00431 | 0.01564 | 0.00000 | 0.08286 | 0.00000 | 0.04407 | 0.03980 | 0.36118 | 0.00000 | 0.12561 |
| **VEGFA** | 0.01065 | 0.00000 | 4.72340 | 0.00000 | 0.45626 | 0.01357 | 5.52256 | 0.02029 | 0.11227 | 4.55754 | 0.58650 | 0.00000 |
| **VEGFB** | 0.04208 | 0.07586 | 0.66533 | 0.56804 | 0.28254 | 0.00000 | 1.81006 | 0.02942 | 0.00000 | 2.03547 | 0.23784 | 0.03765 |
| **VEGFC** | 0.09233 | 0.08619 | 0.11626 | 0.04861 | 0.00000 | 0.00000 | 0.03174 | 0.32161 | 0.01888 | 0.02147 | 0.03137 | 0.25213 |
| **VENTX** | 0.21561 | 0.00692 | 0.37108 | 13.58398 | 0.05810 | 0.00000 | 0.07172 | 0.00662 | 0.00000 | 0.00000 | 0.01913 | 0.01562 |
| **VENTXP1** | 0.01564 | 0.01444 | 0.04099 | 0.02157 | 0.01442 | 0.01169 | 0.01686 | 0.01100 | 0.01134 | 0.03140 | 0.01634 | 0.01548 |
| **VENTXP7** | 0.00460 | 0.00966 | 0.00317 | 0.12930 | 0.00000 | 0.08545 | 0.09134 | 0.02764 | 0.00000 | 0.05963 | 0.00485 | 0.04389 |
| **VEPH1** | 2.36288 | 0.29799 | 0.00000 | 0.35114 | 0.90713 | 0.41348 | 0.00000 | 0.16573 | 0.49629 | 0.11949 | 1.45915 | 0.00000 |
| **VEZF1** | 3.08993 | 1.77914 | 3.80397 | 0.54719 | 2.04604 | 6.95490 | 2.20400 | 0.52635 | 3.93273 | 6.52465 | 3.17859 | 2.16556 |
| **VEZT** | 0.00875 | 3.03620 | 9.20345 | 5.10392 | 0.83923 | 11.32789 | 2.88960 | 1.81442 | 8.28926 | 3.51077 | 1.11213 | 2.83756 |
| **VGF** | 0.11945 | 0.12579 | 0.10001 | 0.18863 | 0.02992 | 0.00000 | 0.03371 | 0.05276 | 0.13289 | 1.46605 | 0.01284 | 0.12509 |
| **VGLL1** | 0.00645 | 0.00000 | 31.02846 | 0.00000 | 4.04461 | 0.12776 | 16.92171 | 0.00473 | 13.63598 | 0.00000 | 5.91662 | 0.00250 |
| **VGLL2** | 0.07192 | 0.01160 | 0.01478 | 0.01456 | 0.02473 | 0.00000 | 0.01172 | 0.01835 | 0.04112 | 0.01629 | 0.01404 | 0.06972 |
| **VGLL3** | 0.00418 | 0.00269 | 0.03523 | 0.05116 | 0.05101 | 0.03062 | 0.04030 | 0.04785 | 0.00000 | 0.15011 | 0.00990 | 0.02157 |
| **VGLL4** | 3.03124 | 8.72849 | 6.67778 | 0.85768 | 8.73998 | 0.00000 | 6.40434 | 0.00000 | 11.16065 | 1.16413 | 8.27843 | 7.38316 |
| **VHL** | 1.77113 | 1.98114 | 6.76881 | 2.50042 | 2.27004 | 3.90134 | 13.21427 | 2.38999 | 0.00000 | 19.82424 | 3.62184 | 0.70094 |
| **VHLL** | 0.00626 | 0.01408 | 0.01613 | 0.01326 | 0.01589 | 0.02181 | 0.01839 | 0.01052 | 0.02828 | 0.01179 | 0.01429 | 0.01631 |
| **VIL1** | 1.35374 | 0.02929 | 0.09140 | 3.02157 | 0.02346 | 0.00000 | 0.00577 | 0.21121 | 0.02610 | 0.00000 | 0.09275 | 0.02619 |
| **VILL** | 0.00595 | 0.28986 | 0.18053 | 0.27576 | 0.01988 | 0.00000 | 0.05397 | 0.00353 | 0.32228 | 0.35256 | 0.17520 | 0.27564 |
| **VIM** | 0.20382 | 0.01880 | 0.25750 | 1.08037 | 0.00000 | 0.10818 | 0.00000 | 0.08354 | 0.00000 | 0.09427 | 0.01723 | 0.20835 |
| **VIM-AS1** | 0.25356 | 0.01173 | 0.06341 | 0.01439 | 0.01316 | 0.01624 | 0.00890 | 0.00095 | 0.01582 | 0.01891 | 0.01455 | 0.04008 |
| **VIMP** | 1.40456 | 14.24898 | 16.65003 | 6.21288 | 12.86768 | 15.27448 | 0.00000 | 13.10993 | 22.40665 | 11.92549 | 7.52488 | 9.77871 |
| **VIP** | 0.00714 | 0.01558 | 0.00000 | 0.00467 | 0.00000 | 0.08426 | 0.08433 | 0.01022 | 0.02528 | 0.01641 | 0.05793 | 0.01367 |
| **VIPAS39** | 1.00131 | 5.97895 | 2.52329 | 0.40340 | 2.21007 | 9.91769 | 1.26886 | 0.69844 | 3.38106 | 0.00000 | 1.12909 | 3.22124 |
| **VIPR1** | 0.00528 | 0.01286 | 0.00616 | 0.06585 | 0.00536 | 0.01624 | 0.01646 | 0.01005 | 0.01388 | 0.05653 | 0.01346 | 0.01898 |
| **VIPR2** | 0.38358 | 0.06689 | 0.05602 | 0.00639 | 0.01199 | 0.00000 | 0.01113 | 0.00282 | 0.01013 | 0.03985 | 0.01921 | 0.11959 |
| **VIT** | 0.00267 | 0.00000 | 0.05234 | 0.00000 | 0.19196 | 0.03672 | 0.02732 | 0.29323 | 0.00000 | 0.17270 | 0.01063 | 0.00567 |

| | | | | | | | | | | | | |
|---|---|---|---|---|---|---|---|---|---|---|---|---|
| **VKORC1** | 4.34117 | 20.73000 | 11.93616 | 8.65821 | 8.48348 | 0.31580 | 20.24467 | 14.38442 | 29.00180 | 4.58308 | 19.87941 | 30.01956 |
| **VKORC1L1** | 0.91745 | 2.15618 | 5.72697 | 0.00000 | 2.49565 | 8.45650 | 4.44762 | 0.91651 | 4.16048 | 17.74013 | 1.60182 | 4.96910 |
| **VLDLR** | 2.41005 | 0.05896 | 0.25176 | 0.90398 | 0.00000 | 0.05124 | 0.98995 | 1.08748 | 0.61461 | 0.00000 | 0.00000 | 0.17335 |
| **VLDLR-AS1** | 1.22878 | 0.00956 | 0.03239 | 0.02299 | 0.00000 | 0.03396 | 0.11356 | 0.43663 | 0.04683 | 0.01559 | 0.00000 | 0.05839 |
| **VMA21** | 3.02014 | 9.86503 | 6.38360 | 1.16571 | 4.50634 | 15.82965 | 11.86454 | 2.03647 | 12.66050 | 7.61628 | 9.03409 | 13.36605 |
| **VMAC** | 0.99391 | 0.39789 | 0.25803 | 0.21162 | 0.55760 | 0.00000 | 0.18787 | 0.11371 | 0.03294 | 0.00000 | 0.24788 | 0.15581 |
| **VMO1** | 0.00383 | 1.71612 | 0.00000 | 0.00000 | 2.36637 | 0.15717 | 0.00014 | 3.89441 | 0.00000 | 0.12286 | 0.28023 | 10.25163 |
| **VMP1** | 0.33754 | 11.73910 | 18.56681 | 9.46541 | 3.31824 | 32.50337 | 0.00000 | 16.11500 | 35.56931 | 5.97816 | 10.70075 | 17.36768 |
| **VN1R1** | 0.11588 | 0.09142 | 0.02177 | 0.12498 | 0.00000 | 0.00012 | 0.33826 | 0.01622 | 0.01067 | 0.44648 | 0.08357 | 0.27838 |
| **VN1R10P** | 0.00761 | 0.01538 | 0.01439 | 0.01666 | 0.01512 | 0.01517 | 0.01993 | 0.01080 | 0.02165 | 0.01337 | 0.01460 | 0.01645 |
| **VN1R2** | 0.00386 | 0.01092 | 0.02531 | 0.00120 | 0.03948 | 0.05552 | 0.03013 | 0.01732 | 0.00000 | 0.03105 | 0.00918 | 0.01699 |
| **VN1R4** | 0.00774 | 0.01499 | 0.01496 | 0.01159 | 0.02435 | 0.01563 | 0.01780 | 0.04005 | 0.01778 | 0.01663 | 0.01417 | 0.03148 |
| **VN1R5** | 0.00307 | 0.01381 | 0.00850 | 0.00734 | 0.00228 | 0.02897 | 0.01348 | 0.08075 | 0.02656 | 0.05373 | 0.02255 | 0.00971 |
| **1.00000** | 0.02199 | 0.00340 | 0.01206 | 0.07119 | 0.03940 | 0.08827 | 0.00674 | 0.21466 | 0.00000 | 0.00763 | 0.02712 | 0.00617 |
| **2.00000** | 0.00599 | 0.01150 | 0.01653 | 0.01115 | 0.00579 | 0.02331 | 0.02014 | 0.01038 | 0.00514 | 0.02193 | 0.08216 | 0.01658 |
| **3.00000** | 0.00993 | 0.00989 | 0.01450 | 0.00969 | 0.00876 | 0.01980 | 0.01796 | 0.56903 | 0.00000 | 0.02578 | 0.05739 | 0.02092 |
| **VOPP1** | 0.13584 | 1.50410 | 0.11151 | 0.32925 | 1.95971 | 0.00127 | 2.34970 | 0.04758 | 0.00000 | 4.49646 | 1.04880 | 0.21550 |
| **VPRBP** | 8.79562 | 5.17348 | 7.39803 | 4.46864 | 3.80036 | 13.45575 | 9.67410 | 0.68752 | 8.26250 | 5.61962 | 7.67329 | 4.59765 |
| **VPREB1** | 0.01733 | 0.00876 | 0.01309 | 0.00971 | 0.01075 | 0.03588 | 0.01605 | 0.03590 | 0.02040 | 0.01167 | 0.01430 | 0.04546 |
| **VPREB3** | 0.00226 | 0.12711 | 0.09566 | 0.00000 | 0.04816 | 0.01889 | 0.13710 | 0.03303 | 0.00000 | 0.06081 | 0.02120 | 0.17630 |
| **VPS11** | 0.00000 | 8.12277 | 7.75052 | 5.10156 | 6.70483 | 4.87136 | 2.60159 | 2.57758 | 6.51534 | 1.50680 | 6.82358 | 5.71921 |
| **VPS13A** | 2.06722 | 0.00000 | 2.05140 | 1.27952 | 1.08914 | 7.61892 | 2.41588 | 1.67542 | 4.94774 | 12.02378 | 0.08864 | 0.25368 |
| **VPS13A-AS1** | 0.00680 | 0.01924 | 0.03107 | 0.02430 | 0.01677 | 0.00000 | 0.00833 | 0.01145 | 0.06275 | 0.01487 | 0.01973 | 0.02083 |
| **VPS13B** | 9.29508 | 0.87967 | 1.73476 | 3.56102 | 1.80111 | 4.16682 | 0.00000 | 1.56001 | 3.19384 | 4.84974 | 0.38717 | 1.84079 |
| **VPS13C** | 2.43435 | 0.31251 | 0.08717 | 1.07642 | 0.74219 | 3.04867 | 0.00000 | 0.00000 | 0.00000 | 3.53582 | 0.54731 | 0.12162 |
| **VPS13D** | 3.94859 | 2.60342 | 0.00000 | 2.09047 | 4.60350 | 5.74738 | 1.25774 | 0.39960 | 2.04214 | 8.72689 | 1.37829 | 4.17791 |
| **VPS16** | 1.83938 | 4.77862 | 2.80939 | 3.46158 | 2.77113 | 1.60427 | 3.22304 | 0.29153 | 5.61504 | 1.36772 | 4.49807 | 3.07700 |
| **VPS18** | 0.97196 | 9.43414 | 10.11080 | 6.02502 | 7.58214 | 0.00000 | 8.80622 | 0.00000 | 10.53011 | 3.68412 | 7.62573 | 5.73686 |
| **VPS25** | 2.43797 | 19.19230 | 12.34507 | 12.34180 | 18.56274 | 25.79501 | 12.11507 | 1.37127 | 14.39782 | 0.00000 | 14.58475 | 18.14094 |
| **VPS26A** | 0.57175 | 8.87298 | 14.98008 | 4.55865 | 1.84507 | 37.94933 | 22.25536 | 13.76367 | 11.89627 | 2.71326 | 14.63724 | 12.19794 |
| **VPS26B** | 0.89411 | 1.88340 | 3.39980 | 4.86562 | 4.80991 | 0.00002 | 4.03964 | 0.00000 | 0.02065 | 6.15313 | 2.84393 | 1.48561 |
| **VPS28** | 1.97576 | 10.39698 | 6.80402 | 14.79075 | 13.21445 | 0.00001 | 8.34310 | 0.45600 | 12.13380 | 8.38311 | 10.87748 | 5.87152 |
| **VPS29** | 2.82659 | 10.74901 | 17.43485 | 10.36477 | 8.24406 | 38.98096 | 24.67069 | 1.40117 | 14.01770 | 1.82852 | 24.85410 | 9.26124 |
| **VPS33A** | 0.63391 | 15.44378 | 11.00011 | 7.22921 | 4.36919 | 22.06698 | 11.16390 | 1.31086 | 12.51627 | 0.00672 | 15.97337 | 11.13128 |
| **VPS33B** | 0.58047 | 2.71764 | 2.72344 | 2.85008 | 3.19022 | 2.26730 | 2.06757 | 0.59398 | 2.28603 | 2.17000 | 2.77933 | 2.56550 |
| **VPS35** | 0.88834 | 14.88944 | 12.06877 | 11.67992 | 3.46696 | 31.93646 | 17.93914 | 5.50219 | 14.37039 | 12.20042 | 17.02005 | 6.53625 |
| **VPS36** | 1.26624 | 3.11884 | 5.51714 | 2.16607 | 1.49499 | 13.84277 | 9.51204 | 1.31359 | 9.22430 | 5.50602 | 6.27723 | 2.83681 |
| **VPS37A** | 3.33795 | 5.19930 | 12.15133 | 2.79756 | 2.82509 | 18.25167 | 8.89694 | 2.55499 | 8.79330 | 0.00037 | 6.34782 | 5.18475 |
| **VPS37B** | 5.11177 | 2.35430 | 6.46745 | 3.77770 | 4.62905 | 0.00000 | 7.68698 | 1.14415 | 2.81367 | 9.17013 | 0.32151 | 2.38036 |
| **VPS37C** | 2.74615 | 2.35617 | 3.63501 | 3.34839 | 8.36669 | 0.00000 | 5.83220 | 1.57948 | 3.56284 | 11.68404 | 2.36753 | 4.49417 |
| **VPS37D** | 0.03655 | 0.00000 | 0.00000 | 0.00000 | 0.00000 | 0.03849 | 0.08015 | 0.03696 | 0.00676 | 0.36076 | 0.00000 | 0.03700 |
| **VPS39** | 0.56978 | 6.96910 | 4.15518 | 4.72151 | 3.65536 | 2.02951 | 2.13312 | 0.45023 | 4.96022 | 2.51094 | 5.49933 | 4.84946 |
| **VPS41** | 0.33658 | 4.53357 | 5.27280 | 1.40752 | 2.86347 | 10.79814 | 4.15919 | 0.11318 | 7.14410 | 0.00000 | 4.49651 | 2.30258 |
| **VPS45** | 0.72941 | 4.92524 | 2.69567 | 0.26735 | 0.46670 | 8.99712 | 4.16779 | 1.58357 | 4.17813 | 0.00000 | 4.82528 | 5.33681 |
| **VPS4A** | 2.06242 | 15.04323 | 5.55702 | 8.01480 | 11.66264 | 0.00000 | 2.22849 | 3.30373 | 5.62907 | 7.92187 | 7.05183 | 5.56916 |
| **VPS4B** | 4.74459 | 8.62253 | 10.36457 | 3.69558 | 6.99151 | 21.14293 | 17.96921 | 18.37724 | 7.21642 | 12.16655 | 9.78783 | 12.50335 |
| **VPS51** | 2.66769 | 4.62956 | 4.62189 | 6.83268 | 8.77846 | 0.00000 | 2.61936 | 0.19238 | 3.25008 | 4.21424 | 2.99086 | 2.39443 |
| **VPS52** | 0.19830 | 7.14666 | 6.58914 | 5.93068 | 6.65051 | 1.79996 | 2.99398 | 3.80672 | 4.39915 | 0.96012 | 8.06342 | 6.62665 |

| | | | | | | | | | | | | |
|---|---|---|---|---|---|---|---|---|---|---|---|---|
| **VPS53** | 0.41808 | 4.27205 | 4.42823 | 5.12265 | 4.08966 | 0.00007 | 5.34449 | 10.25457 | 2.16259 | 8.13560 | 8.18653 | 2.41601 |
| **VPS54** | 0.28565 | 0.33068 | 0.00000 | 1.49270 | 1.20105 | 7.11136 | 0.97802 | 7.41413 | 0.00000 | 6.89151 | 0.42755 | 2.30933 |
| **VPS72** | 8.56851 | 10.81672 | 10.83839 | 7.84554 | 11.24140 | 7.26122 | 14.72214 | 1.45860 | 12.60019 | 0.00000 | 15.51746 | 9.66613 |
| **VPS8** | 0.69927 | 1.34465 | 0.84624 | 0.97595 | 0.00002 | 1.67806 | 1.82185 | 1.69393 | 3.73895 | 4.23611 | 0.85827 | 1.07624 |
| **VPS9D1** | 0.28130 | 0.26288 | 0.12636 | 0.23903 | 0.20023 | 0.00000 | 0.22480 | 0.12825 | 0.18952 | 0.47349 | 0.02936 | 0.14112 |
| **VPS9D1-AS1** | 0.49014 | 3.07373 | 1.23911 | 0.17245 | 3.58761 | 0.00000 | 0.00000 | 0.00000 | 1.82069 | 3.55830 | 0.26044 | 1.26235 |
| **VRK1** | 4.47965 | 8.87953 | 7.89974 | 6.63542 | 5.40576 | 39.40505 | 11.32775 | 2.09984 | 7.80111 | 7.33903 | 15.72336 | 11.52706 |
| **VRK2** | 0.58326 | 5.75359 | 0.89131 | 3.36185 | 3.63133 | 22.21722 | 0.22229 | 0.92918 | 3.96824 | 1.87992 | 7.01259 | 3.14051 |
| **VRK3** | 2.70012 | 11.56656 | 6.55448 | 3.77011 | 8.94372 | 6.31802 | 8.97824 | 5.94546 | 7.48493 | 0.00000 | 6.30072 | 9.89017 |
| **VRTN** | 0.42063 | 0.66838 | 0.58076 | 0.13595 | 0.00000 | 0.00000 | 0.00130 | 1.35651 | 0.00000 | 0.18084 | 0.01229 | 14.84973 |
| **VSIG1** | 0.00728 | 0.01560 | 0.00428 | 0.01157 | 0.01386 | 0.00603 | 0.08894 | 0.04228 | 0.09231 | 0.00000 | 0.00000 | 0.02844 |
| **VSIG10** | 0.00000 | 17.37800 | 0.48493 | 24.18961 | 15.10345 | 22.89463 | 0.00000 | 11.89832 | 0.00000 | 6.80477 | 12.85260 | 13.14850 |
| **VSIG10L** | 0.44867 | 0.03056 | 0.03647 | 0.11773 | 0.02664 | 0.06604 | 0.06452 | 0.06478 | 0.00000 | 0.16077 | 0.00001 | 0.08780 |
| **VSIG2** | 0.03209 | 0.00493 | 0.05354 | 0.79653 | 0.03973 | 0.02100 | 0.05478 | 0.01915 | 0.00000 | 0.05439 | 0.01663 | 0.01780 |
| **VSNL1** | 0.00559 | 0.01523 | 1.88542 | 1.98976 | 0.01105 | 0.00000 | 0.00000 | 0.02099 | 0.05179 | 0.00000 | 0.01695 | 0.01658 |
| **VSTM2A** | 0.00454 | 0.09527 | 0.00585 | 0.14436 | 0.00491 | 0.02083 | 0.01521 | 0.03037 | 0.01926 | 0.00934 | 0.01546 | 0.00890 |
| **VSTM2A-OT1** | 0.00580 | 0.03071 | 0.00836 | 0.00963 | 0.00695 | 0.01745 | 0.08850 | 0.01047 | 0.00000 | 0.03181 | 0.01376 | 0.01380 |
| **VSTM2L** | 0.00256 | 0.02103 | 0.00000 | 0.00000 | 0.00819 | 0.04412 | 0.19224 | 0.00885 | 0.07958 | 0.49193 | 0.03479 | 0.00455 |
| **VSTM4** | 0.00352 | 0.00771 | 0.02783 | 0.01020 | 0.01503 | 0.01196 | 0.14849 | 0.01377 | 0.00000 | 0.10815 | 0.00582 | 0.04477 |
| **VSTM5** | 0.00737 | 0.01247 | 0.01767 | 0.01289 | 0.00632 | 0.02318 | 0.01011 | 0.01142 | 0.04043 | 0.02739 | 0.02193 | 0.01985 |
| **VSX1** | 0.00645 | 0.00897 | 0.02902 | 0.03654 | 0.00000 | 0.03163 | 0.01035 | 0.01026 | 0.00000 | 0.04170 | 0.06260 | 0.01990 |
| **VSX2** | 0.20585 | 0.00318 | 0.03694 | 0.00971 | 0.01178 | 0.00975 | 0.00677 | 0.00000 | 0.01386 | 0.05227 | 0.02007 | 0.19835 |
| **VTA1** | 1.79410 | 14.80834 | 15.16367 | 4.94719 | 2.89831 | 39.02390 | 20.10507 | 8.62405 | 15.36354 | 2.05487 | 19.39586 | 16.57385 |
| **VTCN1** | 3.37527 | 0.00000 | 0.02535 | 0.00000 | 0.28416 | 0.00000 | 0.21329 | 0.50273 | 3.85978 | 0.05686 | 1.57377 | 0.00000 |
| **VTI1A** | 0.50368 | 2.83341 | 4.82825 | 2.04958 | 2.73577 | 7.45562 | 1.73561 | 3.11018 | 5.34381 | 3.22069 | 4.79657 | 3.70099 |
| **VTI1B** | 0.47336 | 4.11584 | 2.59351 | 9.33359 | 11.45661 | 13.66674 | 6.14032 | 1.13074 | 5.17964 | 0.00003 | 7.60890 | 11.13758 |
| **VTN** | 0.09204 | 0.02817 | 0.13999 | 0.10711 | 0.01502 | 0.02620 | 0.00000 | 0.00784 | 0.06934 | 0.01858 | 0.01018 | 0.02014 |
| **VTRNA1-1** | 0.01778 | 0.03083 | 0.03130 | 0.03647 | 0.07180 | 0.00992 | 0.05932 | 0.02375 | 0.02743 | 0.00620 | 0.02228 | 0.03164 |
| **VTRNA1-2** | 0.00128 | 0.18674 | 0.32656 | 0.08754 | 0.12586 | 0.00000 | 0.25726 | 0.07072 | 0.00000 | 0.29423 | 0.06870 | 0.11151 |
| **VTRNA1-3** | 0.00333 | 0.00540 | 3.79628 | 0.29387 | 0.43471 | 0.00000 | 0.00000 | 0.02475 | 0.00006 | 0.89056 | 0.00621 | 0.00416 |
| **VTRNA2-1** | 0.00508 | 0.01134 | 0.02693 | 0.01369 | 0.05650 | 0.01039 | 0.06388 | 0.01235 | 0.00000 | 0.05420 | 0.00953 | 0.01429 |
| **VWA1** | 0.00195 | 0.00000 | 5.02741 | 1.67886 | 6.05296 | 0.00000 | 3.54198 | 0.00063 | 2.99187 | 12.84782 | 0.52355 | 0.10019 |
| **VWA2** | 0.12440 | 0.00898 | 0.02025 | 0.00447 | 0.01151 | 0.05470 | 0.01420 | 0.08790 | 0.00776 | 0.04008 | 0.01312 | 0.04022 |
| **VWA3A** | 0.55351 | 0.00000 | 0.00000 | 0.00000 | 0.00000 | 0.05520 | 0.00000 | 0.47133 | 0.02934 | 0.23245 | 0.03430 | 0.00000 |
| **VWA3B** | 0.02412 | 0.01079 | 0.06328 | 0.16033 | 0.02909 | 0.00712 | 0.00973 | 0.07713 | 0.06742 | 0.00000 | 0.03259 | 0.07312 |
| **VWA5A** | 4.41057 | 0.29562 | 0.00000 | 5.98860 | 0.64093 | 0.29622 | 0.00000 | 1.65783 | 0.00000 | 0.00920 | 0.00000 | 1.62026 |
| **VWA5B1** | 0.00531 | 0.01665 | 0.04429 | 0.01563 | 0.00898 | 0.00260 | 0.05593 | 0.01074 | 0.00940 | 0.05932 | 0.02209 | 0.01482 |
| **VWA5B2** | 0.02574 | 0.04516 | 0.04610 | 0.30975 | 0.01232 | 0.00000 | 0.00113 | 0.02052 | 0.01648 | 0.08555 | 0.02289 | 0.01293 |
| **VWA7** | 0.14867 | 0.03285 | 0.07221 | 0.00647 | 0.04268 | 0.00000 | 0.00795 | 0.23554 | 0.29637 | 0.05944 | 0.00000 | 0.40775 |
| **VWA8** | 0.98174 | 2.07344 | 0.30532 | 1.55736 | 1.27825 | 6.19984 | 0.00000 | 0.00000 | 0.81358 | 5.23952 | 2.26534 | 0.88658 |
| **VWA8-AS1** | 0.01718 | 0.35659 | 0.01066 | 0.00000 | 0.00000 | 0.00995 | 0.02073 | 0.00767 | 0.01750 | 0.03529 | 0.01105 | 0.04612 |
| **VWA9** | 1.87811 | 6.99061 | 12.99005 | 4.47649 | 1.21363 | 13.93027 | 12.25331 | 1.90807 | 11.09263 | 0.00000 | 10.18362 | 6.61235 |
| **VWC2** | 0.00368 | 0.00165 | 0.01456 | 0.05766 | 0.02317 | 0.00889 | 0.02083 | 0.02556 | 0.03930 | 0.00000 | 0.00948 | 0.82246 |
| **VWCE** | 0.17015 | 0.49132 | 0.53911 | 0.79518 | 0.00000 | 0.00000 | 0.00000 | 0.22857 | 0.60538 | 0.66031 | 0.04039 | 0.61532 |
| **VWDE** | 0.00462 | 0.08896 | 0.22303 | 1.02328 | 0.00000 | 0.00000 | 0.00000 | 0.00858 | 0.14172 | 0.15666 | 0.11125 | 0.02451 |
| **VWF** | 0.37284 | 0.05430 | 0.03380 | 0.03277 | 0.00000 | 0.00087 | 0.10721 | 0.01518 | 0.00000 | 0.13538 | 0.00993 | 0.03148 |
| **WAC** | 8.64368 | 5.99927 | 9.75769 | 7.11981 | 9.99994 | 26.69595 | 19.14153 | 7.51232 | 10.99000 | 34.26280 | 6.04837 | 15.00046 |
| **WAC-AS1** | 0.22628 | 0.30262 | 0.70950 | 0.16255 | 1.02354 | 0.46067 | 0.34325 | 0.18297 | 0.00000 | 2.30338 | 1.71597 | 1.08819 |

| | | | | | | | | | | | | |
|---|---|---|---|---|---|---|---|---|---|---|---|---|
| **WAPAL** | 6.93852 | 2.44935 | 11.85696 | 5.76688 | 6.32846 | 21.58417 | 18.14714 | 9.67564 | 3.63326 | 30.76110 | 9.36495 | 12.62706 |
| **WARS** | 1.48831 | 40.69988 | 8.47408 | 35.25649 | 14.47726 | 23.23205 | 21.92576 | 7.85821 | 20.26914 | 4.19822 | 29.44057 | 23.41499 |
| **WARS2** | 0.40476 | 2.13286 | 2.71767 | 0.91664 | 0.24052 | 6.12186 | 3.28848 | 0.69573 | 4.27614 | 0.46027 | 2.97682 | 2.63729 |
| **WAS** | 0.00282 | 0.00918 | 0.00581 | 0.03583 | 0.00000 | 0.05764 | 0.00000 | 0.02788 | 0.04890 | 0.08057 | 0.01465 | 0.02000 |
| **WASF1** | 3.64924 | 3.12077 | 2.79823 | 0.24909 | 0.12165 | 4.35297 | 3.84717 | 7.76425 | 7.01955 | 3.89642 | 4.50271 | 3.48709 |
| **WASF2** | 5.18472 | 4.41304 | 9.13987 | 4.93671 | 16.50623 | 4.06407 | 20.04566 | 0.22088 | 7.91548 | 13.55755 | 10.20292 | 2.06584 |
| **WASF3** | 5.44091 | 0.82652 | 0.39360 | 1.02796 | 0.06466 | 1.03628 | 0.00000 | 0.00000 | 0.00000 | 1.82432 | 0.00006 | 6.47372 |
| **WASH1** | 0.03109 | 0.13266 | 0.00465 | 0.10108 | 0.27773 | 0.00000 | 0.18807 | 0.10550 | 0.00000 | 1.24287 | 0.03656 | 0.34110 |
| **WASH2P** | 0.05598 | 0.28952 | 0.04068 | 0.10171 | 0.35335 | 0.00000 | 0.04328 | 0.02724 | 0.13336 | 0.27031 | 0.08006 | 0.38467 |
| **WASH3P** | 0.02465 | 0.03006 | 0.03008 | 0.03231 | 0.13268 | 0.02118 | 0.00413 | 0.03890 | 0.11853 | 0.10104 | 0.00794 | 0.18359 |
| **WASH5P** | 0.00634 | 0.01709 | 0.01439 | 0.01355 | 0.02324 | 0.01100 | 0.01977 | 0.01052 | 0.02405 | 0.00904 | 0.01482 | 0.01931 |
| **WASH7P** | 0.00705 | 0.01446 | 0.01895 | 0.01349 | 0.01865 | 0.01251 | 0.01611 | 0.01020 | 0.02237 | 0.01726 | 0.01544 | 0.03098 |
| **WASL** | 3.75042 | 13.30690 | 11.49969 | 12.54500 | 11.08322 | 19.51151 | 18.17142 | 3.69389 | 4.71505 | 15.15084 | 10.95337 | 6.55529 |
| **WBP1** | 0.01804 | 0.15307 | 0.06314 | 0.07491 | 0.01252 | 0.08399 | 0.00000 | 0.01469 | 0.11425 | 0.04363 | 0.04978 | 0.09814 |
| **WBP11** | 6.45503 | 9.13328 | 14.02905 | 7.57230 | 11.75856 | 12.70977 | 18.24234 | 16.68070 | 9.38390 | 9.46311 | 11.75604 | 14.55564 |
| **WBP11P1** | 0.58337 | 0.21217 | 0.73064 | 0.20983 | 0.26677 | 1.60558 | 0.72798 | 1.40931 | 0.21341 | 0.85311 | 0.77590 | 0.92347 |
| **WBP1L** | 0.11744 | 4.91200 | 6.20549 | 4.70465 | 4.99145 | 3.96343 | 15.22243 | 3.98357 | 7.01232 | 4.48064 | 6.83654 | 5.01895 |
| **WBP2** | 3.80529 | 5.86303 | 13.08141 | 8.03354 | 9.23831 | 0.04512 | 9.40704 | 0.55255 | 14.67852 | 0.00000 | 6.26059 | 7.61102 |
| **WBP2NL** | 0.03022 | 0.01077 | 0.01816 | 0.01249 | 0.00678 | 0.04224 | 0.01322 | 0.08553 | 0.01015 | 0.01794 | 0.01366 | 0.29449 |
| **WBP4** | 2.05523 | 5.39360 | 6.68220 | 3.11053 | 3.37356 | 14.71669 | 9.75411 | 7.30108 | 0.46013 | 12.47576 | 4.91705 | 10.56862 |
| **WBP5** | 0.00000 | 0.00000 | 17.17797 | 0.01073 | 0.96060 | 21.44666 | 29.08545 | 9.87968 | 11.41550 | 12.72405 | 15.37957 | 13.31375 |
| **WBSCR16** | 0.18202 | 0.72136 | 1.02026 | 0.55589 | 1.40529 | 0.22589 | 1.17439 | 0.08548 | 0.54999 | 0.77635 | 0.51482 | 0.13097 |
| **WBSCR17** | 0.05815 | 0.00000 | 0.01846 | 0.08538 | 0.48098 | 0.00351 | 0.00128 | 0.08010 | 0.00000 | 0.16941 | 0.07129 | 0.03549 |
| **WBSCR22** | 6.41276 | 22.85627 | 19.03848 | 8.78416 | 17.96837 | 16.41131 | 26.28933 | 19.21438 | 24.42433 | 1.21742 | 14.85732 | 19.76244 |
| **WBSCR27** | 0.00664 | 1.05764 | 0.07572 | 0.43507 | 0.00000 | 0.37668 | 0.00049 | 0.00186 | 0.24608 | 0.00000 | 0.00905 | 0.13077 |
| **WBSCR28** | 0.00525 | 0.01351 | 0.02272 | 0.01247 | 0.03434 | 0.02360 | 0.00688 | 0.01169 | 0.01749 | 0.04117 | 0.01081 | 0.03363 |
| **WDFY1** | 21.78676 | 0.49103 | 4.19117 | 2.22620 | 2.59236 | 9.46040 | 2.38333 | 3.08188 | 4.64522 | 0.19568 | 1.77318 | 0.04821 |
| **WDFY2** | 2.29372 | 1.52958 | 3.67253 | 2.27289 | 2.73132 | 3.47526 | 1.42876 | 9.05706 | 3.72581 | 1.35669 | 3.42452 | 2.21139 |
| **WDFY3** | 5.24239 | 0.36974 | 3.69616 | 4.08971 | 0.96677 | 7.06057 | 2.04346 | 1.24729 | 0.86001 | 12.61652 | 0.00000 | 1.73409 |
| **WDFY3-AS2** | 0.20898 | 0.14020 | 0.10222 | 0.04454 | 0.00000 | 0.34183 | 0.03161 | 0.04468 | 0.00000 | 0.02467 | 0.06230 | 0.09789 |
| **WDFY4** | 0.03325 | 0.01011 | 0.04033 | 0.01765 | 0.00512 | 0.01505 | 0.06464 | 0.03162 | 0.00000 | 0.03591 | 0.06006 | 0.02608 |
| **WDHD1** | 1.86412 | 6.44063 | 5.04142 | 6.09597 | 1.73026 | 26.02631 | 0.00024 | 4.30347 | 4.38571 | 12.57229 | 7.08414 | 12.03954 |
| **WDPCP** | 0.69824 | 0.98886 | 0.97094 | 0.18065 | 0.09860 | 1.04391 | 0.00000 | 0.06404 | 0.55101 | 0.00000 | 0.32884 | 0.52911 |
| **WDR1** | 12.44603 | 30.21782 | 28.73080 | 26.56585 | 27.54933 | 12.54766 | 26.12695 | 5.04941 | 28.70617 | 2.44363 | 18.24002 | 17.95298 |
| **WDR11** | 4.16271 | 9.84886 | 0.00000 | 8.10901 | 1.39075 | 19.91624 | 1.88276 | 2.25368 | 0.67170 | 5.36752 | 5.95617 | 5.71126 |
| **WDR11-AS1** | 0.01071 | 3.81882 | 0.03010 | 2.40633 | 0.59838 | 0.44444 | 0.00000 | 0.16289 | 0.00000 | 0.02651 | 0.00073 | 1.53319 |
| **WDR12** | 2.25639 | 12.71379 | 5.39704 | 6.27158 | 4.58996 | 29.51083 | 10.35168 | 6.74212 | 14.34600 | 2.71687 | 12.71619 | 15.09380 |
| **WDR13** | 0.17045 | 4.35150 | 1.32142 | 4.83346 | 3.48858 | 0.00000 | 3.30449 | 0.68003 | 1.72812 | 0.00000 | 5.55856 | 2.68290 |
| **WDR16** | 0.00642 | 0.17245 | 0.00000 | 0.08747 | 0.02437 | 0.11911 | 0.04558 | 0.13567 | 0.00000 | 0.21665 | 0.03513 | 0.25908 |
| **WDR17** | 0.06108 | 0.00819 | 0.47906 | 0.00095 | 0.00000 | 0.05558 | 0.08041 | 0.01917 | 0.14746 | 0.00784 | 0.00000 | 0.03201 |
| **WDR18** | 1.19572 | 22.73359 | 21.38575 | 17.47839 | 23.18778 | 0.00000 | 17.92593 | 0.68202 | 31.17107 | 0.00000 | 24.93676 | 18.51347 |
| **WDR19** | 1.17951 | 0.73901 | 0.65558 | 0.25273 | 0.14923 | 1.17574 | 0.56823 | 1.28727 | 1.07499 | 1.53939 | 0.23669 | 0.91830 |
| **WDR20** | 6.99065 | 9.58522 | 5.93800 | 6.64522 | 2.14100 | 13.90792 | 4.92611 | 18.79079 | 8.52509 | 0.00000 | 5.30925 | 8.16754 |
| **WDR24** | 0.93224 | 8.33437 | 1.14731 | 3.17508 | 4.59955 | 0.00000 | 2.85935 | 0.00000 | 5.28061 | 0.00000 | 2.87937 | 4.03899 |
| **WDR25** | 4.53691 | 13.28101 | 1.02427 | 2.65980 | 3.34950 | 2.12661 | 0.52512 | 1.36294 | 1.31103 | 0.00000 | 3.43734 | 7.04047 |
| **WDR26** | 4.98004 | 1.77419 | 6.73928 | 3.81381 | 2.14802 | 15.41862 | 8.57821 | 3.81071 | 4.30328 | 22.46748 | 1.94374 | 7.53010 |
| **WDR27** | 0.27853 | 0.47786 | 0.28965 | 0.28815 | 0.23535 | 0.00000 | 0.09240 | 0.44285 | 0.81565 | 0.57785 | 0.36869 | 0.89432 |
| **WDR3** | 0.00000 | 10.02146 | 12.23706 | 2.19050 | 4.21233 | 18.72609 | 11.25919 | 14.79671 | 17.47959 | 0.00000 | 13.50306 | 14.75551 |

| | | | | | | | | | | | | |
|---|---|---|---|---|---|---|---|---|---|---|---|---|
| **WDR31** | 0.54755 | 0.03747 | 0.00928 | 0.14053 | 0.00517 | 0.00000 | 0.20024 | 0.16049 | 0.00000 | 0.07741 | 0.06166 | 0.08472 |
| **WDR33** | 4.96873 | 8.69194 | 12.69173 | 11.08235 | 12.00612 | 17.28250 | 15.26868 | 1.49266 | 9.15884 | 11.59173 | 13.09840 | 10.91748 |
| **WDR34** | 4.80982 | 30.96306 | 18.80943 | 11.85898 | 24.94479 | 0.00000 | 11.48294 | 0.00000 | 16.21162 | 6.21002 | 23.64993 | 13.26909 |
| **WDR35** | 1.44927 | 0.09375 | 0.01773 | 0.00000 | 0.00000 | 0.60273 | 0.46525 | 1.55990 | 0.47639 | 0.15198 | 0.52163 | 0.82991 |
| **WDR36** | 0.00000 | 6.86942 | 8.74557 | 1.70488 | 0.00000 | 17.98376 | 12.14311 | 8.05754 | 11.22974 | 11.56691 | 10.36246 | 9.15236 |
| **WDR37** | 3.00689 | 1.78479 | 2.24191 | 1.47346 | 1.11004 | 0.00000 | 2.07624 | 5.17675 | 3.00475 | 1.24067 | 1.07344 | 2.12340 |
| **WDR38** | 1.51213 | 0.01696 | 0.02608 | 0.01794 | 0.02353 | 0.00000 | 0.02276 | 0.53869 | 0.06484 | 0.08668 | 0.02834 | 0.08867 |
| **WDR4** | 0.58487 | 13.06978 | 7.12323 | 6.99811 | 13.40113 | 0.00000 | 8.12535 | 4.40344 | 6.96294 | 4.00740 | 9.17549 | 15.06127 |
| **WDR41** | 6.45541 | 7.99314 | 2.42327 | 4.16874 | 2.33738 | 10.84355 | 3.63463 | 0.88334 | 3.25595 | 0.46187 | 2.84907 | 5.94534 |
| **WDR43** | 1.31817 | 17.93057 | 21.26574 | 13.31061 | 16.48381 | 31.31228 | 33.89733 | 24.40072 | 18.21969 | 25.04808 | 26.66443 | 27.38411 |
| **WDR44** | 2.45789 | 1.05930 | 0.76858 | 1.17507 | 0.00000 | 4.36759 | 0.00002 | 2.04036 | 2.82452 | 1.21867 | 0.82561 | 3.36469 |
| **WDR45** | 0.65677 | 11.69064 | 4.72236 | 2.59326 | 3.23032 | 3.60963 | 7.04175 | 11.42633 | 9.79631 | 0.00000 | 6.83117 | 33.56982 |
| **WDR45B** | 14.58619 | 18.19737 | 10.57991 | 12.55852 | 12.70057 | 13.24675 | 16.42888 | 21.73740 | 18.23933 | 6.30193 | 14.26157 | 17.93521 |
| **WDR46** | 2.03031 | 24.02900 | 17.78695 | 17.12585 | 21.74097 | 0.00000 | 29.25013 | 12.52085 | 24.94483 | 0.00000 | 19.07287 | 27.88249 |
| **WDR47** | 5.63852 | 7.24458 | 4.72171 | 4.41340 | 4.03501 | 12.50532 | 0.00000 | 32.74101 | 0.22545 | 3.14208 | 1.90050 | 20.47662 |
| **WDR48** | 4.41796 | 6.24828 | 4.78515 | 4.18583 | 0.75140 | 13.52239 | 8.28157 | 7.82411 | 12.68392 | 0.17311 | 5.25165 | 9.49084 |
| **WDR49** | 0.33198 | 0.01042 | 0.03790 | 0.00629 | 0.05532 | 0.00027 | 0.02532 | 0.06700 | 0.00612 | 0.06276 | 0.01447 | 0.03033 |
| **WDR5** | 5.82925 | 22.06882 | 8.99945 | 14.48273 | 24.57998 | 13.09596 | 10.18926 | 6.16248 | 17.98029 | 22.43191 | 13.46120 | 18.47304 |
| **WDR52** | 0.01334 | 0.00000 | 0.35812 | 0.00000 | 0.02141 | 0.47027 | 0.09553 | 0.32192 | 0.71456 | 4.47262 | 0.09089 | 0.68502 |
| **WDR52-AS1** | 0.00526 | 0.01225 | 0.01184 | 0.01584 | 0.00814 | 0.04180 | 0.01600 | 0.01430 | 0.01967 | 0.02476 | 0.01364 | 0.01351 |
| **WDR53** | 1.40848 | 7.19865 | 2.01560 | 3.36958 | 1.28785 | 7.06277 | 2.50103 | 0.47746 | 2.26987 | 0.55640 | 1.78260 | 6.07629 |
| **WDR54** | 0.55610 | 2.85846 | 1.88230 | 1.88778 | 0.66022 | 0.00000 | 1.12467 | 0.23063 | 0.55325 | 0.00000 | 1.29044 | 0.38783 |
| **WDR55** | 0.00000 | 10.50782 | 9.42482 | 9.83070 | 6.88859 | 8.77876 | 10.17676 | 11.27845 | 16.55300 | 0.00000 | 17.72741 | 12.62120 |
| **WDR59** | 2.06285 | 0.98679 | 3.08980 | 1.47819 | 5.86937 | 1.44886 | 5.40961 | 0.83843 | 2.94315 | 13.50118 | 0.96941 | 1.90276 |
| **WDR5B** | 0.07503 | 0.98116 | 0.72299 | 0.00000 | 0.00000 | 2.02838 | 0.00000 | 0.15971 | 0.49700 | 0.04895 | 0.93753 | 1.16922 |
| **WDR6** | 0.33248 | 2.43685 | 14.60166 | 10.84337 | 1.08099 | 0.00000 | 7.46669 | 0.25778 | 15.22444 | 7.83495 | 5.36548 | 2.46185 |
| **WDR60** | 3.94293 | 0.63107 | 0.46506 | 2.05946 | 1.20468 | 0.00000 | 1.26056 | 1.02570 | 0.51081 | 5.31145 | 0.18708 | 1.11687 |
| **WDR61** | 2.19165 | 8.64206 | 8.48623 | 2.56514 | 5.07026 | 21.26785 | 5.70589 | 4.15623 | 9.47802 | 0.31981 | 15.11146 | 9.35584 |
| **WDR62** | 1.16412 | 4.61252 | 3.57029 | 8.22035 | 2.66623 | 0.00000 | 1.71916 | 0.16102 | 6.50966 | 1.32755 | 0.00000 | 3.78650 |
| **WDR63** | 0.49405 | 0.01378 | 0.01760 | 0.06178 | 0.04580 | 0.00000 | 0.07223 | 0.03260 | 0.00001 | 0.00000 | 0.05264 | 0.01401 |
| **WDR64** | 0.07324 | 0.00006 | 0.43764 | 0.00311 | 0.07704 | 0.19647 | 0.06858 | 0.00454 | 0.00000 | 0.00000 | 0.02413 | 0.01207 |
| **WDR65** | 0.01083 | 0.09496 | 0.13949 | 0.01262 | 0.00000 | 0.00973 | 0.16530 | 1.11966 | 0.00000 | 0.11572 | 0.04568 | 0.13925 |
| **WDR66** | 1.07032 | 9.34072 | 0.00000 | 9.83537 | 8.08155 | 10.62724 | 2.77250 | 7.98815 | 0.00000 | 4.52279 | 2.42000 | 12.60268 |
| **WDR7** | 5.42404 | 1.26376 | 0.00000 | 1.76373 | 2.28953 | 3.06104 | 0.30229 | 0.39481 | 0.82634 | 1.78256 | 1.11389 | 1.92375 |
| **WDR70** | 4.71465 | 7.98829 | 2.33404 | 4.53898 | 6.10702 | 3.73724 | 6.20749 | 4.16146 | 4.39319 | 5.84369 | 6.63502 | 7.47501 |
| **WDR72** | 1.69733 | 2.83007 | 0.24870 | 1.87044 | 0.66624 | 5.16495 | 0.00000 | 0.10865 | 1.66533 | 1.35385 | 0.00000 | 0.71123 |
| **WDR73** | 0.00000 | 9.19697 | 9.47043 | 6.00983 | 6.96056 | 6.20556 | 8.00779 | 5.05915 | 11.81426 | 1.15981 | 10.32459 | 14.84987 |
| **WDR74** | 1.50547 | 24.85266 | 21.98331 | 13.49256 | 21.22680 | 7.81011 | 28.80763 | 12.45194 | 26.91730 | 5.46052 | 20.76929 | 29.02436 |
| **WDR75** | 2.75427 | 11.74736 | 4.84721 | 4.69543 | 2.33008 | 27.03386 | 11.93653 | 13.23824 | 11.02644 | 2.69209 | 11.93780 | 15.87692 |
| **WDR76** | 6.84544 | 3.98073 | 3.95582 | 0.00000 | 2.34877 | 5.79489 | 1.09205 | 6.26294 | 0.00000 | 3.42974 | 4.94339 | 5.47321 |
| **WDR77** | 0.00000 | 21.65118 | 24.17855 | 17.34784 | 16.46978 | 21.89965 | 26.16538 | 8.22123 | 31.24314 | 0.00000 | 23.08552 | 22.91305 |
| **WDR78** | 0.19203 | 0.21927 | 0.31015 | 0.00786 | 0.02631 | 0.05446 | 0.00000 | 0.08786 | 0.32538 | 0.09572 | 0.22287 | 0.19902 |
| **WDR81** | 0.00180 | 0.31119 | 1.74827 | 1.18180 | 0.00000 | 0.00000 | 2.36979 | 0.02922 | 0.00000 | 2.14283 | 1.10197 | 1.33620 |
| **WDR82** | 1.85639 | 4.27858 | 4.31307 | 6.59058 | 7.54542 | 4.20651 | 16.16360 | 4.29423 | 1.53329 | 21.42341 | 11.85420 | 8.88244 |
| **WDR83** | 2.20674 | 7.88447 | 4.91775 | 5.14349 | 5.60661 | 2.10201 | 5.63118 | 13.26314 | 4.66223 | 0.00000 | 7.54380 | 8.82346 |
| **WDR83OS** | 0.96929 | 18.94841 | 18.07017 | 17.42772 | 26.90992 | 11.51773 | 21.42314 | 5.70466 | 23.21585 | 6.26680 | 16.38755 | 18.86124 |
| **WDR86** | 0.04284 | 0.01443 | 0.01994 | 0.01203 | 0.01328 | 0.02108 | 0.01851 | 0.00737 | 0.01752 | 0.01579 | 0.01443 | 0.01750 |
| **WDR86-AS1** | 0.06967 | 0.01478 | 0.01552 | 0.01520 | 0.01415 | 0.01687 | 0.01736 | 0.11552 | 0.01768 | 0.01526 | 0.01471 | 0.01023 |

| | | | | | | | | | | | | |
|---|---|---|---|---|---|---|---|---|---|---|---|---|
| **WDR87** | 0.15838 | 0.00000 | 0.08126 | 0.00000 | 0.36651 | 0.00000 | 0.28872 | 0.10865 | 0.29879 | 1.04762 | 0.05945 | 0.04118 |
| **WDR88** | 0.03724 | 0.01340 | 0.07593 | 0.00585 | 0.01876 | 0.01828 | 0.00831 | 0.01052 | 0.03793 | 0.02905 | 0.02381 | 0.03198 |
| **WDR89** | 1.53304 | 1.50912 | 9.44673 | 0.56177 | 1.40670 | 13.52608 | 6.97789 | 7.55233 | 10.63333 | 0.40246 | 6.58652 | 7.11100 |
| **WDR90** | 0.02478 | 0.76200 | 0.34531 | 1.08257 | 0.97565 | 0.00000 | 0.48708 | 0.43892 | 1.60365 | 3.36947 | 0.16661 | 0.99060 |
| **WDR91** | 0.20763 | 1.47820 | 1.50032 | 0.79791 | 0.00000 | 1.53633 | 0.68123 | 0.34977 | 0.48061 | 0.00000 | 0.09388 | 1.07967 |
| **WDR92** | 1.49606 | 0.84856 | 0.87574 | 2.41097 | 0.00213 | 4.78919 | 1.00312 | 0.37362 | 3.54426 | 2.34904 | 2.44847 | 7.06339 |
| **WDR93** | 0.00136 | 0.32998 | 0.00586 | 0.04137 | 0.00000 | 0.00000 | 0.02743 | 0.77876 | 0.04598 | 0.35930 | 0.08743 | 0.67174 |
| **WDR96** | 0.00408 | 0.52740 | 0.03380 | 0.00000 | 0.03049 | 0.16777 | 0.00000 | 0.16893 | 0.00000 | 0.04681 | 0.00320 | 0.56059 |
| **WDSUB1** | 1.70068 | 2.48495 | 0.97544 | 2.59550 | 0.00000 | 7.85145 | 4.45096 | 0.00000 | 6.12769 | 1.69917 | 6.41525 | 0.68013 |
| **WDTC1** | 3.40244 | 8.58491 | 0.53282 | 2.90869 | 5.71144 | 0.00006 | 1.52080 | 2.90087 | 0.74860 | 4.30954 | 0.24352 | 9.17767 |
| **WDYHV1** | 1.25960 | 2.78868 | 1.98786 | 0.46446 | 2.79725 | 8.44925 | 0.00000 | 0.25445 | 3.24776 | 0.00000 | 2.06100 | 2.16659 |
| **WEE1** | 0.00000 | 0.34082 | 6.37343 | 1.50921 | 0.00000 | 12.45227 | 12.84782 | 1.03461 | 0.00000 | 13.32361 | 0.00000 | 1.91792 |
| **WEE2** | 44.61830 | 0.20443 | 0.00000 | 0.00000 | 0.10741 | 0.00000 | 0.00069 | 20.42073 | 0.00000 | 0.03816 | 0.00035 | 6.21487 |
| **WEE2-AS1** | 3.46999 | 0.07768 | 0.42245 | 0.01794 | 0.05757 | 0.05798 | 0.08614 | 0.71514 | 0.00842 | 0.00000 | 0.09040 | 0.03379 |
| **WFDC10A** | 0.16796 | 0.00789 | 0.01530 | 0.01313 | 0.02019 | 0.00247 | 0.01629 | 0.05848 | 0.02191 | 0.01945 | 0.01500 | 0.22525 |
| **WFDC10B** | 0.01466 | 0.02907 | 0.01855 | 0.01086 | 0.01134 | 0.01761 | 0.01826 | 0.01030 | 0.01547 | 0.01680 | 0.01603 | 0.05323 |
| **WFDC13** | 0.00659 | 0.00889 | 0.01372 | 0.01130 | 0.01298 | 0.02664 | 0.01727 | 0.02140 | 0.01702 | 0.00738 | 0.01802 | 0.12450 |
| **WFDC2** | 11.23268 | 0.75127 | 0.76044 | 0.00000 | 0.57616 | 0.00056 | 1.17421 | 2.91998 | 0.74510 | 1.18328 | 0.29790 | 5.60813 |
| **WFDC21P** | 0.02844 | 0.01420 | 0.04939 | 0.01860 | 0.02133 | 0.02357 | 0.01407 | 0.00796 | 0.01330 | 0.01323 | 0.01466 | 0.01535 |
| **WFDC3** | 0.03839 | 0.04280 | 0.03055 | 0.02840 | 0.08460 | 0.03312 | 0.01533 | 0.03814 | 0.00000 | 0.00000 | 0.04540 | 0.49769 |
| **WFDC6** | 0.00790 | 0.02756 | 0.03523 | 0.01186 | 0.00000 | 0.00000 | 0.01151 | 0.00696 | 0.00000 | 0.07556 | 0.18117 | 0.08086 |
| **WFDC8** | 0.04935 | 0.01529 | 0.01727 | 0.01298 | 0.01439 | 0.01555 | 0.01647 | 0.01831 | 0.04146 | 0.01237 | 0.01553 | 0.05365 |
| **WFDC9** | 0.00620 | 0.01460 | 0.01425 | 0.01211 | 0.01518 | 0.02074 | 0.01864 | 0.01297 | 0.01842 | 0.00914 | 0.01501 | 0.02990 |
| **WFIKKN1** | 0.02198 | 0.03902 | 0.06201 | 0.00498 | 0.06786 | 0.00188 | 0.19978 | 0.00824 | 0.11680 | 0.00070 | 0.00000 | 0.02126 |
| **WFIKKN2** | 0.01590 | 0.00953 | 0.00000 | 0.00000 | 0.00000 | 0.00426 | 0.00137 | 0.02233 | 0.15422 | 0.39412 | 0.01225 | 0.00709 |
| **WFS1** | 9.32540 | 5.97784 | 2.81030 | 3.31211 | 6.07118 | 0.00000 | 0.01780 | 1.29800 | 6.34416 | 4.28057 | 2.15881 | 2.33412 |
| **WHAMM** | 0.93343 | 0.29359 | 1.46825 | 0.38694 | 1.34380 | 0.78196 | 1.13697 | 1.85734 | 0.33404 | 5.89667 | 0.00009 | 0.33601 |
| **WHAMMP1** | 0.01901 | 0.01541 | 0.01109 | 0.04955 | 0.02435 | 0.02743 | 0.04874 | 0.03127 | 0.00825 | 0.09386 | 0.01562 | 0.02373 |
| **WHAMMP2** | 0.02328 | 0.01759 | 0.01309 | 0.01183 | 0.01458 | 0.02276 | 0.01639 | 0.01757 | 0.01534 | 0.02647 | 0.01439 | 0.01717 |
| **WHAMMP3** | 0.01258 | 0.01994 | 0.02687 | 0.01915 | 0.00000 | 0.01180 | 0.01131 | 0.01063 | 0.00747 | 0.10919 | 0.04777 | 0.01330 |
| **WHSC1** | 14.28909 | 4.93423 | 2.58482 | 0.00020 | 7.12157 | 8.47892 | 5.47668 | 0.95294 | 1.95210 | 25.18941 | 2.54288 | 2.00836 |
| **WHSC1L1** | 1.81609 | 1.42125 | 7.52532 | 2.13673 | 4.43438 | 3.34863 | 8.09803 | 0.08412 | 4.24148 | 12.89278 | 1.25119 | 0.00000 |
| **WIBG** | 0.59605 | 14.12548 | 7.88849 | 11.34487 | 8.62634 | 19.81251 | 13.46046 | 3.83835 | 13.61179 | 0.00000 | 16.73058 | 10.80334 |
| **WIF1** | 0.24027 | 0.01668 | 0.01569 | 0.02070 | 0.03603 | 0.00633 | 0.02207 | 0.00364 | 0.02174 | 0.00000 | 0.01389 | 0.01989 |
| **WIPF1** | 0.00129 | 0.00159 | 0.55989 | 0.05416 | 0.32173 | 0.10468 | 0.96388 | 0.07977 | 0.47254 | 0.12788 | 0.15284 | 0.06475 |
| **WIPF2** | 9.50068 | 2.06726 | 4.81966 | 3.49046 | 7.55228 | 2.80426 | 8.74826 | 3.54780 | 2.53854 | 17.43470 | 2.93550 | 4.93657 |
| **WIPF3** | 0.05084 | 0.93092 | 0.00000 | 1.52548 | 2.46751 | 0.98545 | 0.00000 | 0.00000 | 0.00000 | 0.87873 | 2.84707 | 0.18476 |
| **WIPI1** | 0.30841 | 0.18190 | 7.95960 | 1.48780 | 2.25232 | 0.00000 | 0.00000 | 0.07536 | 0.00000 | 0.02805 | 0.16692 | 0.40868 |
| **WIPI2** | 5.51793 | 9.55730 | 6.20282 | 7.37196 | 12.65218 | 0.43468 | 15.91961 | 11.25345 | 6.63118 | 15.48302 | 2.61184 | 16.70414 |
| **WISP3** | 0.00529 | 0.02969 | 0.01632 | 0.00073 | 0.01600 | 0.08094 | 0.00060 | 0.00907 | 0.01731 | 0.04156 | 0.01258 | 0.01857 |
| **WIZ** | 1.96913 | 0.88090 | 1.25350 | 1.23345 | 1.99728 | 0.00000 | 3.49774 | 0.36909 | 0.00009 | 6.13614 | 2.14665 | 1.88236 |
| **WLS** | 7.67995 | 0.00000 | 16.88446 | 0.00000 | 11.25728 | 0.66070 | 3.52795 | 3.03396 | 11.22547 | 0.00000 | 17.11077 | 2.44025 |
| **WNK1** | 4.89609 | 5.50682 | 7.64420 | 1.92200 | 5.60670 | 10.75858 | 14.32321 | 2.59106 | 3.79251 | 29.05544 | 7.89412 | 12.35576 |
| **WNK2** | 0.73029 | 0.14505 | 1.25780 | 0.00000 | 0.61415 | 0.00000 | 0.01295 | 0.00086 | 0.98358 | 2.11893 | 0.00000 | 0.37601 |
| **WNK3** | 1.62496 | 0.09471 | 0.00000 | 2.11158 | 0.00000 | 0.22733 | 0.00000 | 4.77892 | 0.49368 | 1.16394 | 0.12371 | 6.04034 |
| **WNK4** | 0.00178 | 0.11973 | 0.20962 | 0.00000 | 0.00000 | 0.04193 | 0.27949 | 0.09625 | 0.00000 | 0.31085 | 0.18619 | 0.26285 |
| **WNT1** | 0.00463 | 0.01087 | 0.04337 | 0.01460 | 0.00000 | 0.03344 | 0.00000 | 0.01727 | 0.09017 | 0.07766 | 0.01018 | 0.01227 |
| **WNT10A** | 0.00623 | 0.03291 | 0.03249 | 0.00896 | 0.01409 | 0.02186 | 0.01207 | 0.01055 | 0.02405 | 0.01160 | 0.01690 | 0.01460 |

| | | | | | | | | | | | | |
|---|---|---|---|---|---|---|---|---|---|---|---|---|
| **WNT10B** | 0.00104 | 0.57465 | 2.50272 | 0.00137 | 0.04039 | 0.59597 | 0.04964 | 0.05918 | 0.00000 | 0.00000 | 0.00000 | 0.75289 |
| **WNT11** | 0.00000 | 6.86323 | 0.14923 | 4.63386 | 2.52416 | 0.00000 | 1.73484 | 1.49827 | 3.61143 | 2.56085 | 2.80285 | 2.41152 |
| **WNT16** | 0.03473 | 0.16455 | 0.00000 | 0.41334 | 2.34858 | 0.63052 | 0.00000 | 0.01075 | 0.00094 | 0.00520 | 1.75701 | 0.09875 |
| **WNT2** | 0.00465 | 0.00000 | 0.00000 | 0.00000 | 0.04147 | 0.02270 | 0.00020 | 0.00817 | 0.26469 | 0.08482 | 1.00672 | 0.02921 |
| **WNT2B** | 0.22587 | 0.08309 | 0.13562 | 0.19508 | 0.05191 | 0.00000 | 0.02078 | 0.07519 | 0.15893 | 0.22737 | 0.00000 | 0.27294 |
| **WNT3** | 0.00303 | 0.00000 | 0.80339 | 3.20331 | 0.00000 | 0.00001 | 0.34821 | 1.18426 | 0.08386 | 0.00000 | 0.13647 | 2.03181 |
| **WNT3A** | 0.00109 | 0.04987 | 0.01731 | 0.07340 | 0.09525 | 0.15095 | 0.08430 | 0.13387 | 0.06300 | 0.04930 | 0.17171 | 0.00234 |
| **WNT4** | 0.00372 | 0.01475 | 0.00952 | 0.00322 | 0.08595 | 0.00000 | 0.07475 | 0.07499 | 0.07873 | 0.09396 | 0.00676 | 0.04796 |
| **WNT5A** | 0.38771 | 0.00000 | 0.00000 | 0.04960 | 0.00000 | 0.00519 | 0.48384 | 0.08791 | 0.39573 | 0.52701 | 0.00000 | 0.00072 |
| **WNT5B** | 0.32232 | 6.41796 | 0.97787 | 1.98208 | 2.10557 | 0.00000 | 2.17193 | 0.00000 | 0.58600 | 1.32524 | 3.80296 | 1.65103 |
| **WNT6** | 0.00111 | 0.00000 | 0.86857 | 0.00000 | 1.07798 | 0.00000 | 3.98327 | 0.00541 | 2.83393 | 2.79363 | 0.25988 | 0.00000 |
| **WNT7A** | 0.00255 | 0.00000 | 0.00000 | 0.00000 | 3.22451 | 0.00000 | 1.15091 | 0.00805 | 4.72683 | 0.17426 | 1.94843 | 0.00554 |
| **WNT7B** | 0.00157 | 0.00000 | 0.47718 | 0.00000 | 2.49444 | 0.00000 | 1.90632 | 0.00637 | 1.78132 | 2.64514 | 3.02700 | 0.00000 |
| **WNT8A** | 0.00256 | 0.00290 | 0.01138 | 0.21319 | 0.00000 | 0.05976 | 0.00001 | 0.20456 | 0.01740 | 0.01734 | 0.01202 | 0.10346 |
| **WNT8B** | 0.35487 | 0.01286 | 0.00065 | 0.11457 | 0.00000 | 0.00972 | 0.05126 | 0.00312 | 0.00038 | 0.17689 | 0.00685 | 0.00758 |
| **WNT9A** | 0.76578 | 0.81189 | 0.00053 | 0.00290 | 0.24700 | 0.08900 | 0.00206 | 0.15897 | 0.00000 | 0.00000 | 0.47833 | 0.33547 |
| **WRAP53** | 0.70099 | 8.48201 | 8.83124 | 5.58091 | 9.05265 | 0.00000 | 11.41939 | 11.17883 | 17.29968 | 0.00000 | 9.99956 | 12.70725 |
| **WRAP73** | 0.99169 | 10.24778 | 8.52642 | 2.72207 | 4.18475 | 1.19302 | 7.21716 | 0.13456 | 11.26431 | 0.00000 | 9.38321 | 5.41959 |
| **WRB** | 4.04794 | 4.96747 | 5.18183 | 1.73195 | 0.00000 | 7.48619 | 3.33560 | 0.75292 | 13.36616 | 0.00000 | 7.48280 | 3.90426 |
| **WRN** | 2.88890 | 1.69749 | 2.34342 | 1.67601 | 0.00000 | 4.42623 | 2.49352 | 0.90198 | 4.92625 | 4.75109 | 0.48739 | 2.25037 |
| **WRNIP1** | 1.20652 | 0.12136 | 3.84727 | 2.20868 | 2.08607 | 5.25315 | 8.48875 | 0.30595 | 0.33653 | 10.41454 | 1.96621 | 0.75334 |
| **WSB1** | 1.62366 | 12.29966 | 27.77255 | 16.40584 | 13.38519 | 37.80759 | 23.42870 | 14.72864 | 22.20468 | 11.21835 | 12.22124 | 17.90017 |
| **WSB2** | 1.20161 | 0.77025 | 6.05187 | 2.67439 | 0.00000 | 6.97117 | 5.51025 | 0.35675 | 1.25466 | 4.68366 | 2.95922 | 1.12157 |
| **WSCD1** | 1.24152 | 0.04868 | 0.00200 | 0.42887 | 0.02418 | 0.00000 | 0.02454 | 0.02519 | 0.01430 | 0.06078 | 0.08810 | 0.00311 |
| **WSCD2** | 0.14096 | 0.02417 | 0.05129 | 0.10065 | 0.01006 | 0.01235 | 0.00690 | 0.02395 | 0.01167 | 0.03781 | 0.01929 | 0.02644 |
| **WT1** | 0.01330 | 0.01117 | 0.00947 | 0.00949 | 0.00515 | 0.02159 | 0.01688 | 0.00852 | 0.01199 | 0.08493 | 0.02861 | 0.01251 |
| **WT1-AS** | 0.00545 | 0.01024 | 0.01075 | 0.00465 | 0.02173 | 0.01954 | 0.00455 | 0.01028 | 0.03579 | 0.06976 | 0.02646 | 0.01340 |
| **WTAP** | 3.71322 | 12.16538 | 11.69661 | 7.84891 | 3.20389 | 33.16167 | 25.78437 | 18.03544 | 8.14237 | 20.58639 | 16.00577 | 14.56004 |
| **WTAPP1** | 0.00621 | 0.01517 | 0.01743 | 0.01151 | 0.01385 | 0.02020 | 0.01853 | 0.01046 | 0.01085 | 0.01348 | 0.03425 | 0.01642 |
| **WTH3DI** | 0.31247 | 0.82951 | 0.50384 | 0.34928 | 0.00000 | 1.15144 | 0.25101 | 1.44557 | 0.54254 | 0.00000 | 0.50054 | 0.99452 |
| **WTIP** | 0.10259 | 0.58471 | 0.00000 | 0.02690 | 3.08699 | 0.00000 | 0.78910 | 0.02539 | 0.93386 | 2.77278 | 0.08561 | 0.04859 |
| **WWC1** | 0.09049 | 2.96367 | 2.91954 | 0.00000 | 12.74501 | 1.53041 | 6.72086 | 0.05511 | 2.23635 | 11.33660 | 2.65292 | 3.87429 |
| **WWC2** | 3.11946 | 2.22656 | 5.91358 | 2.75297 | 12.19340 | 4.34382 | 14.32990 | 0.23021 | 9.38517 | 17.37014 | 11.00206 | 1.38133 |
| **WWC2-AS2** | 0.01364 | 0.01028 | 0.23007 | 0.28229 | 0.05869 | 0.00000 | 0.36530 | 0.00907 | 0.50231 | 0.00000 | 0.16788 | 0.03537 |
| **WWC3** | 5.10259 | 0.00000 | 0.68045 | 0.00000 | 4.04718 | 1.53898 | 5.27716 | 0.07999 | 4.16157 | 3.29558 | 3.48482 | 0.17596 |
| **WWOX** | 5.43092 | 1.07403 | 1.71090 | 0.07422 | 3.12423 | 1.35571 | 2.83546 | 0.34119 | 1.30402 | 0.36716 | 3.26380 | 0.66883 |
| **WWP1** | 0.46297 | 1.32308 | 3.18507 | 0.00000 | 0.50750 | 12.50081 | 7.46280 | 0.58855 | 2.17154 | 10.66174 | 2.63593 | 0.04181 |
| **WWP2** | 4.38240 | 3.61599 | 5.32787 | 3.48701 | 5.22036 | 0.00000 | 5.24697 | 4.27094 | 7.34098 | 7.88582 | 4.77323 | 1.54545 |
| **WWTR1** | 0.00136 | 18.19145 | 22.09283 | 14.33869 | 8.33433 | 13.10176 | 19.43090 | 1.28736 | 14.75324 | 1.04252 | 11.39691 | 23.57782 |
| **WWTR1-AS1** | 0.00515 | 0.03209 | 0.18066 | 0.01210 | 0.00000 | 0.00000 | 0.18703 | 0.05144 | 0.01517 | 0.17035 | 0.02530 | 0.22978 |
| **XAB2** | 1.89231 | 12.15091 | 6.70073 | 3.86759 | 8.92136 | 0.00000 | 9.98908 | 2.56519 | 5.36711 | 5.04090 | 9.97943 | 10.21245 |
| **XAF1** | 0.00517 | 0.01271 | 0.02533 | 0.01045 | 0.00000 | 0.00435 | 0.02291 | 0.00975 | 0.00000 | 0.05797 | 0.16055 | 0.03129 |
| **XAGE2.uid-1286** | 0.00604 | 0.01318 | 0.03283 | 0.00894 | 0.02454 | 0.02585 | 0.03643 | 0.01070 | 0.03947 | 0.00739 | 0.04319 | 0.01692 |
| **XAGE3** | 0.00293 | 2.70491 | 0.49701 | 0.00000 | 2.36525 | 0.76446 | 0.83447 | 0.01818 | 1.72919 | 0.00000 | 0.65149 | 0.92370 |
| **XAGE5** | 0.02661 | 0.63040 | 0.25518 | 0.01435 | 0.00000 | 0.04900 | 0.38312 | 0.01913 | 0.14058 | 0.00000 | 0.00000 | 0.72126 |
| **XBP1** | 0.00000 | 7.95131 | 18.71806 | 3.15434 | 6.75789 | 10.92766 | 8.59064 | 0.19585 | 19.83957 | 27.23550 | 6.69150 | 3.51300 |
| **XCL1** | 0.08204 | 0.02185 | 0.05793 | 0.09738 | 0.11997 | 0.05039 | 0.01935 | 0.23623 | 0.08140 | 0.08899 | 0.03314 | 0.00893 |
| **XCL2** | 0.02059 | 0.02109 | 0.01562 | 0.01849 | 0.03865 | 0.00000 | 0.00911 | 0.01881 | 0.04407 | 0.06132 | 0.02668 | 0.06526 |

| | | | | | | | | | | | | |
|---|---|---|---|---|---|---|---|---|---|---|---|---|
| **XCR1** | 1.28366 | 0.01124 | 0.06278 | 0.01180 | 0.01551 | 0.01570 | 0.02336 | 0.05113 | 0.01663 | 0.00000 | 0.01483 | 0.02254 |
| **XDH** | 0.04418 | 0.15202 | 0.00000 | 0.00000 | 0.05804 | 0.00000 | 0.00319 | 0.00506 | 0.14769 | 0.25177 | 0.00000 | 0.18410 |
| **XG** | 0.09198 | 0.02927 | 0.01925 | 0.00000 | 0.13646 | 0.01692 | 0.11197 | 0.03346 | 0.00000 | 0.09873 | 0.14347 | 0.01961 |
| **XGY2.uid-13475** | 0.00636 | 0.01513 | 0.01545 | 0.01132 | 0.01437 | 0.02053 | 0.01695 | 0.01040 | 0.02093 | 0.01454 | 0.01413 | 0.01639 |
| **XIAP** | 0.60935 | 0.75374 | 6.05507 | 1.79050 | 0.00000 | 7.89207 | 5.71947 | 0.00000 | 0.00000 | 17.92589 | 3.01222 | 4.64793 |
| **XIRP1** | 0.09952 | 0.01234 | 0.04877 | 0.00742 | 0.02059 | 0.01588 | 0.01374 | 0.00514 | 0.00000 | 0.03275 | 0.03699 | 0.01374 |
| **XIRP2** | 0.23426 | 0.02370 | 0.26317 | 0.01160 | 0.01324 | 0.02644 | 0.09928 | 0.06308 | 0.05970 | 0.00000 | 0.04299 | 0.03126 |
| **XIST** | 0.00000 | 3.03487 | 5.43083 | 0.28829 | 0.00000 | 0.42359 | 0.87940 | 0.07313 | 0.00000 | 14.04624 | 8.60028 | 2.78523 |
| **XK** | 0.04136 | 0.04808 | 0.02788 | 0.00000 | 0.05004 | 0.08035 | 0.00000 | 0.01875 | 0.00806 | 0.01573 | 0.00616 | 0.02501 |
| **XKR4** | 0.00544 | 0.02409 | 0.01671 | 0.01558 | 0.01334 | 0.02309 | 0.02626 | 0.02514 | 0.00702 | 0.04712 | 0.01803 | 0.06040 |
| **XKR5** | 0.00680 | 0.01323 | 0.06994 | 0.01326 | 0.02507 | 0.02797 | 0.02661 | 0.01110 | 0.00000 | 0.01751 | 0.01587 | 0.01499 |
| **XKR6** | 0.08155 | 0.05910 | 0.04213 | 0.19105 | 0.00000 | 0.09752 | 0.09160 | 0.00297 | 0.02086 | 0.09736 | 0.32821 | 0.01732 |
| **XKR7** | 0.00639 | 0.01854 | 0.01646 | 0.01811 | 0.02812 | 0.00287 | 0.02157 | 0.01059 | 0.02212 | 0.00856 | 0.01464 | 0.01930 |
| **XKR8** | 2.05278 | 3.08358 | 8.13020 | 3.60705 | 1.21783 | 0.00000 | 0.00000 | 0.27783 | 7.83132 | 0.19447 | 0.34952 | 0.33567 |
| **XKR9** | 0.01001 | 0.07973 | 0.12106 | 0.06036 | 0.03078 | 0.13654 | 0.00000 | 0.06850 | 0.00000 | 0.00000 | 0.04219 | 0.01029 |
| **XKRX** | 0.18383 | 0.00000 | 0.00743 | 0.17739 | 0.02994 | 0.01790 | 0.02509 | 0.00000 | 0.02885 | 0.00000 | 0.01729 | 0.62005 |
| **XPA** | 0.07544 | 1.38343 | 1.88504 | 0.00000 | 1.25629 | 3.43909 | 3.22135 | 0.03684 | 2.05907 | 3.37128 | 3.50024 | 1.03799 |
| **XPC** | 2.23864 | 5.73631 | 5.75470 | 4.55970 | 3.23304 | 8.25519 | 3.92307 | 1.84836 | 4.05382 | 6.17571 | 6.17801 | 4.45994 |
| **XPNPEP1** | 0.44280 | 9.69093 | 6.53927 | 0.00000 | 7.49160 | 9.88865 | 6.66305 | 0.00000 | 9.66523 | 1.43766 | 12.77392 | 0.24209 |
| **XPNPEP2** | 0.00569 | 0.00703 | 0.01356 | 0.00375 | 0.09593 | 0.04501 | 0.06108 | 0.01145 | 0.00440 | 0.00132 | 0.04359 | 0.01566 |
| **XPNPEP3** | 0.43494 | 5.74731 | 3.37924 | 2.72699 | 3.11649 | 8.36076 | 5.71051 | 1.52417 | 2.55030 | 7.49713 | 6.72995 | 7.56667 |
| **XPO1** | 2.53693 | 13.31904 | 16.59746 | 6.81364 | 7.41248 | 42.51071 | 30.48489 | 10.37324 | 14.32464 | 37.02941 | 9.36736 | 21.05838 |
| **XPO4** | 3.35110 | 3.06308 | 6.56923 | 2.69209 | 2.64527 | 21.18277 | 12.17840 | 2.04148 | 5.14130 | 16.47911 | 5.82056 | 4.40807 |
| **XPO5** | 3.90902 | 16.65476 | 10.02773 | 10.01851 | 10.70298 | 12.62094 | 14.25627 | 4.68607 | 19.43670 | 6.99316 | 12.96327 | 13.43961 |
| **XPO6** | 3.60239 | 6.37131 | 7.79654 | 8.91556 | 16.42623 | 1.10945 | 15.14737 | 0.99116 | 12.00391 | 18.67151 | 5.31622 | 6.77291 |
| **XPO7** | 5.40300 | 6.72202 | 3.78111 | 6.23840 | 7.44544 | 11.79220 | 9.97206 | 1.19967 | 9.62533 | 12.07356 | 5.31794 | 11.81143 |
| **XPOT** | 5.47816 | 8.19873 | 14.35073 | 3.95976 | 0.00000 | 23.74360 | 18.53637 | 4.38261 | 15.05145 | 5.83325 | 12.28692 | 6.72967 |
| **XPR1** | 6.43187 | 2.64433 | 4.30084 | 3.59310 | 0.92458 | 4.15947 | 0.48552 | 1.99839 | 2.55235 | 5.30327 | 7.43955 | 3.96911 |
| **XRCC1** | 6.26661 | 6.08940 | 4.10311 | 3.14403 | 2.12837 | 0.00000 | 1.60162 | 2.51582 | 3.06044 | 0.00202 | 4.96755 | 1.73995 |
| **XRCC2** | 0.48131 | 4.33578 | 3.32002 | 1.82242 | 3.03739 | 5.04637 | 0.00000 | 1.05382 | 3.23505 | 2.62205 | 5.21022 | 5.74053 |
| **XRCC3** | 0.36561 | 1.28332 | 2.47402 | 1.11234 | 3.92219 | 0.00000 | 0.99802 | 0.93955 | 4.72452 | 1.91071 | 0.89874 | 3.88582 |
| **XRCC4** | 1.71883 | 3.82885 | 4.86117 | 0.46785 | 2.32094 | 18.72765 | 6.06149 | 3.15387 | 2.83327 | 4.43770 | 4.78306 | 5.71331 |
| **XRCC5** | 0.87334 | 27.24375 | 28.70964 | 19.56464 | 18.97995 | 49.39052 | 43.17759 | 12.79707 | 37.07908 | 14.39854 | 31.98977 | 34.19847 |
| **XRCC6** | 6.53557 | 32.10981 | 26.70058 | 22.99825 | 22.39605 | 30.45436 | 37.50436 | 14.22979 | 39.72953 | 0.00000 | 32.59681 | 32.24291 |
| **XRCC6BP1** | 0.58022 | 3.90529 | 0.26260 | 1.33409 | 0.73297 | 2.89808 | 0.14894 | 0.98616 | 0.00000 | 0.79155 | 3.00409 | 3.61147 |
| **XRCC6P5** | 0.19096 | 1.12093 | 0.84505 | 0.58983 | 0.06399 | 1.89761 | 0.95865 | 0.35132 | 1.65578 | 0.00000 | 1.14725 | 1.12725 |
| **XRN1** | 7.93584 | 0.98814 | 0.12199 | 1.47302 | 1.58319 | 8.83104 | 0.74779 | 0.73343 | 0.00000 | 7.08056 | 0.32384 | 1.59429 |
| **XRN2** | 8.07376 | 11.42318 | 13.04192 | 10.66828 | 3.64792 | 32.94498 | 19.84724 | 3.01339 | 17.82898 | 14.35586 | 14.46105 | 15.54349 |
| **XRRA1** | 1.91041 | 0.85869 | 0.52522 | 0.96170 | 2.27204 | 0.00000 | 0.74902 | 0.74000 | 1.69267 | 2.96816 | 0.80784 | 1.49584 |
| **XXYLT1** | 0.93399 | 1.95504 | 0.68773 | 2.11730 | 3.34833 | 0.02032 | 0.62364 | 0.30131 | 3.42534 | 2.52256 | 0.71438 | 0.36453 |
| **XXYLT1-AS2** | 0.00784 | 0.01001 | 0.00588 | 0.00412 | 0.00638 | 0.05922 | 0.01314 | 0.00935 | 0.01857 | 0.05161 | 0.01893 | 0.01322 |
| **XYLB** | 0.01086 | 2.42327 | 2.91873 | 1.10904 | 2.82668 | 0.98271 | 0.00000 | 0.05808 | 1.77701 | 0.05758 | 2.58398 | 0.87536 |
| **XYLT1** | 2.48328 | 0.14401 | 0.39599 | 0.14360 | 0.14597 | 0.00000 | 0.00000 | 0.47733 | 0.00000 | 0.31244 | 0.05007 | 0.01277 |
| **XYLT2** | 0.69848 | 2.43160 | 1.70508 | 3.39292 | 2.14574 | 0.00000 | 0.00000 | 2.43969 | 1.04166 | 2.83585 | 0.66336 | 9.34015 |
| **YAE1D1** | 0.00000 | 3.82851 | 7.95516 | 1.19792 | 0.00000 | 5.90440 | 10.50809 | 1.30254 | 3.05484 | 0.00000 | 6.06349 | 5.44399 |
| **YAF2** | 1.27261 | 4.31236 | 2.51670 | 5.54500 | 3.64114 | 6.42417 | 5.51956 | 8.32690 | 1.14020 | 4.30524 | 0.26163 | 1.93697 |
| **YAP1** | 4.80789 | 0.00000 | 7.92122 | 2.45887 | 2.67690 | 6.09028 | 25.40405 | 1.30164 | 0.01827 | 20.10520 | 4.60467 | 1.12101 |
| **YARS** | 0.00000 | 20.62821 | 16.27170 | 21.92334 | 25.21026 | 16.37123 | 23.19480 | 1.15393 | 35.33060 | 8.49471 | 27.68951 | 15.96875 |

| | | | | | | | | | | | | |
|---|---|---|---|---|---|---|---|---|---|---|---|---|
| YARS2 | 4.57578 | 9.95343 | 6.79878 | 4.83754 | 4.46380 | 16.17656 | 8.75539 | 9.67078 | 15.85382 | 0.00000 | 7.65920 | 15.38769 |
| YBEY | 0.25009 | 8.87169 | 2.36177 | 1.55893 | 2.75612 | 1.18802 | 1.06244 | 0.47422 | 4.17102 | 4.64127 | 2.46544 | 7.98676 |
| YBX1 | 0.50459 | 1.81615 | 7.00076 | 5.05078 | 12.96417 | 13.50718 | 20.67736 | 7.23596 | 0.01582 | 32.32343 | 7.51982 | 9.37591 |
| YBX2 | 6.16387 | 0.20450 | 0.00000 | 0.16652 | 0.67333 | 0.00000 | 0.06139 | 0.30731 | 0.00000 | 0.85027 | 0.00000 | 0.63350 |
| YBX3 | 1.35683 | 4.15178 | 3.03758 | 5.93198 | 8.23882 | 5.79821 | 0.64512 | 0.54932 | 0.00000 | 17.93684 | 0.00000 | 4.62002 |
| YBX3P1 | 0.14605 | 0.37326 | 0.35618 | 0.79822 | 0.60779 | 0.31698 | 0.00000 | 0.10694 | 0.00000 | 0.76311 | 0.00000 | 0.38109 |
| YDJC | 0.17004 | 11.71060 | 10.31392 | 9.72179 | 12.82743 | 0.00000 | 10.53868 | 0.00000 | 27.85611 | 3.26929 | 13.04461 | 2.94020 |
| YEATS2 | 4.03021 | 6.78206 | 4.44085 | 2.01080 | 1.84816 | 0.00000 | 5.30701 | 6.65712 | 8.78730 | 6.93478 | 5.08322 | 9.45580 |
| YEATS4 | 6.48141 | 9.75267 | 3.67110 | 4.92858 | 4.93544 | 28.60859 | 5.69826 | 0.08487 | 7.60314 | 5.40157 | 10.22366 | 8.72598 |
| YES1 | 7.72181 | 3.76102 | 8.30707 | 0.64763 | 2.96947 | 20.50525 | 12.75420 | 0.66173 | 11.48745 | 10.20871 | 13.88971 | 2.59563 |
| YIF1A | 0.77981 | 15.18158 | 12.73328 | 13.91085 | 22.03837 | 2.89476 | 10.40538 | 0.73443 | 18.67867 | 0.00000 | 17.07861 | 11.82128 |
| YIF1B | 0.65743 | 22.96741 | 3.89327 | 15.77785 | 19.60006 | 2.23095 | 5.53227 | 6.90151 | 21.19653 | 0.00000 | 11.64206 | 21.07701 |
| YIPF1 | 2.74390 | 6.13020 | 4.18244 | 1.87231 | 8.23883 | 9.98561 | 0.86594 | 0.24843 | 11.50467 | 0.00000 | 7.48570 | 4.80264 |
| YIPF2 | 1.49142 | 5.32280 | 7.03684 | 4.38493 | 7.42910 | 0.00000 | 5.98806 | 2.09979 | 4.20334 | 8.27292 | 7.63568 | 3.72497 |
| YIPF3 | 0.34362 | 13.41049 | 15.10922 | 9.23392 | 8.39608 | 7.85011 | 4.65427 | 4.05762 | 24.61967 | 1.33817 | 5.93522 | 10.25775 |
| YIPF4 | 0.23800 | 22.06146 | 4.71870 | 14.95019 | 1.05605 | 24.73328 | 0.00000 | 3.95806 | 8.16534 | 4.39845 | 4.90676 | 29.41199 |
| YIPF5 | 2.08003 | 8.20183 | 9.78149 | 2.77512 | 1.02517 | 19.53842 | 0.00000 | 6.11457 | 13.82874 | 9.78540 | 3.48207 | 12.28170 |
| YIPF6 | 4.08312 | 2.46013 | 6.71482 | 3.02818 | 4.04632 | 17.33631 | 3.51027 | 1.33409 | 15.16791 | 3.95151 | 8.60602 | 4.23316 |
| YIPF7 | 0.00623 | 0.01413 | 0.02070 | 0.00994 | 0.01146 | 0.02691 | 0.04339 | 0.01048 | 0.01606 | 0.00865 | 0.01408 | 0.01598 |
| YJEFN3 | 0.12809 | 0.64869 | 1.09115 | 0.00000 | 0.00000 | 0.00000 | 1.56355 | 0.45340 | 2.13623 | 1.02812 | 0.72683 | 0.62845 |
| YKT6 | 2.97165 | 8.99340 | 12.03357 | 4.70883 | 12.42682 | 0.00000 | 17.94787 | 10.92195 | 1.85296 | 19.99690 | 6.53356 | 6.66068 |
| YLPM1 | 6.62483 | 3.15163 | 4.25920 | 2.23242 | 8.59436 | 11.79959 | 10.96188 | 6.24145 | 5.65636 | 20.66611 | 3.37635 | 6.88831 |
| YME1L1 | 1.88628 | 10.98244 | 13.48484 | 7.82463 | 9.48629 | 30.32093 | 17.07300 | 14.85127 | 20.84962 | 20.34361 | 11.47776 | 19.97237 |
| YOD1 | 0.00000 | 0.73211 | 3.58539 | 0.00000 | 0.59862 | 6.10554 | 5.90173 | 4.85854 | 2.94449 | 4.32806 | 2.56221 | 2.62643 |
| YPEL1 | 3.34297 | 0.07085 | 0.00000 | 0.21749 | 0.40175 | 0.36081 | 0.14421 | 0.07216 | 0.00000 | 0.74192 | 0.12939 | 0.16317 |
| YPEL2 | 6.58474 | 22.27540 | 0.00000 | 13.25028 | 19.00171 | 8.87872 | 1.88616 | 22.26623 | 3.78565 | 20.40544 | 5.61017 | 35.61493 |
| YPEL3 | 0.75655 | 0.90680 | 0.89695 | 2.28209 | 3.36926 | 0.00000 | 1.70886 | 0.39039 | 1.00246 | 6.39980 | 0.21989 | 0.81490 |
| YPEL4 | 0.00460 | 0.00003 | 0.09830 | 0.00000 | 0.17590 | 0.08684 | 0.00000 | 0.01009 | 0.06949 | 0.05521 | 0.02176 | 0.02058 |
| YPEL5 | 16.79840 | 10.08595 | 17.77050 | 5.51468 | 4.00584 | 17.15282 | 14.12772 | 36.97540 | 5.87177 | 3.89896 | 1.76403 | 31.72782 |
| YRDC | 0.00000 | 4.47775 | 8.97673 | 2.42472 | 4.82240 | 6.31288 | 16.45690 | 14.93429 | 4.55823 | 10.10214 | 3.56183 | 12.83367 |
| YTHDC1 | 4.08221 | 3.65249 | 12.95219 | 10.47481 | 7.75897 | 12.44721 | 20.61534 | 7.88482 | 2.64431 | 34.48504 | 7.93567 | 8.34586 |
| YTHDC2 | 1.14999 | 2.39170 | 2.64112 | 0.25254 | 0.00000 | 9.64568 | 2.12797 | 0.20397 | 0.99274 | 10.70091 | 3.12367 | 0.39033 |
| YTHDF1 | 6.60453 | 9.92742 | 4.59531 | 6.42368 | 15.27467 | 16.97031 | 21.37413 | 5.57769 | 0.00937 | 39.32243 | 6.14571 | 11.65192 |
| YTHDF2 | 2.25033 | 14.31919 | 21.57887 | 14.04119 | 16.09290 | 24.99534 | 27.81043 | 25.40866 | 23.64744 | 6.51419 | 19.43516 | 21.71444 |
| YTHDF3 | 3.58658 | 17.70086 | 11.62641 | 8.57646 | 2.81954 | 21.82117 | 16.32862 | 24.74843 | 12.03524 | 0.71050 | 14.07047 | 18.87032 |
| YTHDF3-AS1 | 0.36625 | 0.37115 | 0.04226 | 0.25310 | 0.03833 | 0.54079 | 0.00000 | 1.43088 | 0.00000 | 0.34463 | 0.18923 | 0.81813 |
| YWHAB | 11.59928 | 21.05302 | 24.82130 | 18.00038 | 27.32228 | 41.81113 | 45.04056 | 19.46898 | 21.87148 | 28.49059 | 25.92400 | 24.16624 |
| YWHAE | 8.82509 | 27.14460 | 32.09612 | 25.98019 | 23.94783 | 42.09141 | 48.91168 | 30.00761 | 25.95945 | 25.31840 | 37.28244 | 38.29219 |
| YWHAEP1 | 0.11057 | 0.06647 | 0.11260 | 0.01162 | 0.00000 | 0.28532 | 0.04891 | 0.45238 | 0.00000 | 0.31565 | 0.06415 | 0.32154 |
| YWHAEP7 | 0.04181 | 0.00866 | 0.22737 | 0.03097 | 0.00000 | 0.09828 | 0.00000 | 0.02109 | 0.00000 | 0.03550 | 0.01110 | 0.01074 |
| YWHAG | 8.14488 | 2.60594 | 7.20514 | 3.15566 | 2.50866 | 8.30468 | 20.96458 | 5.94236 | 0.00001 | 25.88664 | 6.63574 | 5.17339 |
| YWHAH | 2.51011 | 6.79283 | 13.19664 | 9.49504 | 10.41357 | 19.34796 | 23.47587 | 3.18197 | 4.82590 | 35.27685 | 10.36338 | 4.03843 |
| YWHAQ | 10.31924 | 23.47148 | 22.51514 | 13.85790 | 14.00323 | 49.06802 | 30.73533 | 18.06229 | 30.97180 | 0.00000 | 28.38818 | 24.61450 |
| YWHAZ | 2.70874 | 20.64233 | 28.05487 | 14.91454 | 13.43301 | 62.51868 | 33.13306 | 11.51019 | 27.92988 | 12.70594 | 24.96496 | 18.02147 |
| YY1 | 1.27095 | 1.66081 | 5.81347 | 5.78329 | 5.69973 | 16.58823 | 19.73816 | 14.88899 | 0.00000 | 42.07389 | 6.08044 | 13.43302 |
| YY1AP1 | 3.90313 | 14.11671 | 10.80690 | 10.40855 | 10.09717 | 8.08096 | 8.43536 | 8.37711 | 5.15705 | 10.43778 | 6.96503 | 19.76495 |
| YY1P2 | 0.00932 | 0.01240 | 0.02074 | 0.00769 | 0.00696 | 0.02572 | 0.02165 | 0.01617 | 0.02072 | 0.05929 | 0.01297 | 0.02261 |
| YY2 | 0.29980 | 0.07784 | 0.12727 | 0.00928 | 0.03014 | 0.05031 | 0.08076 | 0.99601 | 0.40751 | 0.00000 | 0.05961 | 0.38382 |

| | | | | | | | | | | | | |
|---|---|---|---|---|---|---|---|---|---|---|---|---|
| **ZACN** | 0.01931 | 0.00724 | 0.00177 | 0.00000 | 0.00280 | 0.03605 | 0.00073 | 0.00834 | 0.10096 | 0.14782 | 0.00000 | 0.00912 |
| **ZADH2** | 0.07470 | 0.45311 | 0.09512 | 0.48345 | 0.10700 | 0.18306 | 0.00000 | 0.04814 | 0.00000 | 0.41038 | 0.15151 | 0.21772 |
| **ZAK** | 2.64481 | 1.42140 | 3.04154 | 0.00000 | 0.00000 | 1.96993 | 0.00000 | 0.39115 | 0.74875 | 1.33944 | 0.63641 | 1.67838 |
| **ZAN** | 0.28971 | 0.02183 | 0.04405 | 0.07423 | 0.00000 | 0.00000 | 0.10932 | 0.08813 | 0.00000 | 0.43731 | 0.05973 | 0.17995 |
| **ZAP70** | 0.00678 | 0.06315 | 0.16152 | 0.10879 | 0.00102 | 0.02356 | 0.00911 | 0.01030 | 0.00000 | 0.05309 | 0.06948 | 0.00819 |
| **1.00000** | 1.80529 | 0.01020 | 0.02515 | 0.02278 | 0.00756 | 0.00000 | 0.08843 | 0.15289 | 0.01982 | 0.05008 | 0.00912 | 0.13819 |
| **ZAR1L** | 11.62620 | 0.03253 | 0.00000 | 0.00033 | 0.00229 | 0.11895 | 0.00000 | 0.87449 | 0.08227 | 0.00000 | 0.00122 | 0.52849 |
| **ZBBX** | 0.37553 | 0.03158 | 0.01406 | 0.00144 | 0.04765 | 0.04383 | 0.00562 | 0.00802 | 0.01155 | 0.03665 | 0.01475 | 0.08165 |
| **ZBED3** | 0.17022 | 0.00000 | 0.00325 | 0.88175 | 0.06437 | 0.00000 | 0.03907 | 0.00000 | 0.00000 | 0.51402 | 0.00140 | 0.04919 |
| **ZBED3-AS1** | 0.00123 | 0.04348 | 0.00461 | 0.09240 | 0.00000 | 0.10906 | 0.11010 | 0.19900 | 0.00000 | 0.11450 | 0.08935 | 0.02962 |
| **ZBED4** | 1.81295 | 1.55521 | 2.69734 | 1.37532 | 0.78087 | 3.66786 | 3.90727 | 0.89186 | 4.86482 | 4.85149 | 1.33507 | 0.85949 |
| **ZBED5** | 0.46250 | 1.26841 | 3.49281 | 1.91286 | 0.00000 | 4.23267 | 3.47953 | 4.67546 | 2.90528 | 5.59096 | 1.57445 | 4.29328 |
| **ZBED5-AS1** | 0.00735 | 0.61951 | 0.23356 | 0.46398 | 0.00000 | 0.31648 | 0.08697 | 3.09390 | 0.05586 | 0.22505 | 0.25301 | 2.58797 |
| **ZBED6** | 0.00000 | 0.92076 | 1.85051 | 1.42707 | 0.00007 | 3.98219 | 0.00000 | 1.11654 | 2.47355 | 4.30089 | 0.74887 | 0.05089 |
| **ZBED6CL** | 0.01021 | 1.44378 | 3.20139 | 1.18438 | 0.97384 | 0.00000 | 2.35401 | 0.00000 | 0.00000 | 0.54333 | 2.06669 | 0.68389 |
| **ZBED8** | 0.00224 | 0.16514 | 0.41726 | 0.03590 | 0.00000 | 0.02046 | 0.00000 | 0.04206 | 0.00000 | 0.02211 | 0.16774 | 0.20443 |
| **ZBED9** | 0.00000 | 0.85844 | 0.01116 | 0.67452 | 0.00000 | 2.01125 | 0.00000 | 0.11858 | 0.27294 | 0.90604 | 0.34082 | 1.34234 |
| **ZBP1** | 0.00606 | 0.01357 | 0.02269 | 0.01284 | 0.01271 | 0.01875 | 0.01771 | 0.01008 | 0.01246 | 0.02546 | 0.01764 | 0.01979 |
| **ZBTB1** | 0.00075 | 1.53816 | 3.30356 | 2.47849 | 0.00576 | 9.85004 | 1.74316 | 7.10745 | 7.22315 | 0.00000 | 4.21125 | 4.70446 |
| **ZBTB10** | 3.27640 | 1.34535 | 7.86227 | 0.24407 | 0.00000 | 17.60823 | 7.08510 | 8.38986 | 0.00289 | 23.62701 | 0.00006 | 6.66431 |
| **ZBTB11** | 2.35938 | 4.20917 | 2.93827 | 0.96638 | 1.86723 | 9.42379 | 3.23648 | 12.83018 | 2.87479 | 7.82533 | 2.82032 | 7.24982 |
| **ZBTB11-AS1** | 0.35724 | 2.28828 | 0.32788 | 0.86684 | 2.15112 | 1.70010 | 0.08985 | 0.79501 | 0.54007 | 1.77454 | 0.99043 | 3.10936 |
| **ZBTB12** | 0.00543 | 0.01362 | 0.01590 | 0.01432 | 0.01323 | 0.01792 | 0.02474 | 0.00981 | 0.01962 | 0.04708 | 0.01609 | 0.03269 |
| **ZBTB14** | 0.79331 | 2.42540 | 2.39085 | 2.20345 | 0.80831 | 5.73005 | 1.59897 | 2.70677 | 2.22728 | 1.76204 | 2.97215 | 4.56006 |
| **ZBTB16** | 9.26707 | 0.03532 | 0.00113 | 0.16020 | 0.00000 | 0.00000 | 0.00991 | 3.17299 | 0.01727 | 0.01400 | 0.02963 | 0.47788 |
| **ZBTB17** | 2.89188 | 5.46455 | 4.14424 | 3.07531 | 3.09916 | 0.00000 | 4.76751 | 3.22255 | 3.33112 | 3.98204 | 2.57535 | 6.67951 |
| **ZBTB18** | 0.66270 | 0.00000 | 0.00000 | 0.25982 | 0.58593 | 0.54920 | 0.51451 | 0.42184 | 0.00000 | 2.64219 | 0.00000 | 0.20830 |
| **ZBTB2** | 3.34528 | 5.37539 | 3.15863 | 4.50829 | 2.15365 | 11.37171 | 1.51942 | 4.38954 | 3.35969 | 6.62231 | 2.77400 | 4.44777 |
| **ZBTB20** | 0.16984 | 0.00000 | 0.31806 | 0.32259 | 0.00000 | 1.67003 | 0.17308 | 0.04482 | 0.00000 | 2.56215 | 0.00000 | 0.18330 |
| **ZBTB20-AS1** | 0.00624 | 0.01805 | 0.01478 | 0.01134 | 0.01570 | 0.02170 | 0.01686 | 0.01048 | 0.01820 | 0.01342 | 0.01744 | 0.01570 |
| **ZBTB20-AS4** | 0.00611 | 0.01350 | 0.04367 | 0.01719 | 0.01330 | 0.02619 | 0.02384 | 0.01059 | 0.01053 | 0.01244 | 0.01409 | 0.01585 |
| **ZBTB21** | 3.19902 | 5.87045 | 1.09145 | 0.19004 | 1.37637 | 5.69429 | 0.00000 | 7.10920 | 4.24500 | 0.50432 | 2.00595 | 7.11064 |
| **ZBTB22** | 0.38812 | 0.33842 | 1.05391 | 0.00000 | 1.11328 | 0.00000 | 0.22297 | 0.00000 | 0.37132 | 0.48956 | 0.02699 | 0.38994 |
| **ZBTB24** | 4.58184 | 1.58788 | 1.03782 | 2.08779 | 1.06220 | 6.67189 | 1.46546 | 4.14192 | 0.00168 | 5.69214 | 0.84178 | 2.12347 |
| **ZBTB25** | 0.11129 | 0.58372 | 0.56747 | 0.95244 | 0.41234 | 2.35605 | 0.00000 | 0.00000 | 0.51131 | 0.00000 | 0.96883 | 0.75624 |
| **ZBTB26** | 0.54800 | 0.03102 | 0.63312 | 0.01444 | 0.00229 | 1.86033 | 0.00000 | 0.00000 | 0.61990 | 0.00000 | 0.41727 | 0.60827 |
| **ZBTB3** | 0.71277 | 1.95743 | 2.73928 | 1.92382 | 1.09697 | 0.00000 | 0.00000 | 1.04408 | 0.83981 | 1.02302 | 1.22995 | 2.41655 |
| **ZBTB32** | 0.00555 | 0.00554 | 0.10765 | 0.03122 | 0.00000 | 0.04528 | 0.05294 | 0.00931 | 0.00000 | 0.11670 | 0.00857 | 0.01993 |
| **ZBTB33** | 3.08700 | 2.85583 | 1.96108 | 0.00000 | 0.00000 | 3.84745 | 4.67346 | 3.91523 | 2.35839 | 4.66442 | 3.40287 | 2.66216 |
| **ZBTB34** | 0.11311 | 0.14397 | 1.66705 | 0.83565 | 0.00000 | 3.33652 | 3.90226 | 1.37731 | 1.55150 | 4.65721 | 0.26973 | 1.47132 |
| **ZBTB37** | 2.24590 | 0.50033 | 0.39607 | 0.16737 | 0.00000 | 1.07199 | 0.00000 | 0.70921 | 0.15878 | 0.00687 | 0.08415 | 1.71986 |
| **ZBTB38** | 2.72608 | 0.28610 | 1.53001 | 0.00000 | 0.25954 | 1.29071 | 1.10888 | 0.38858 | 0.51707 | 0.69263 | 0.42297 | 0.21497 |
| **ZBTB39** | 5.71030 | 2.73859 | 0.08410 | 1.44936 | 1.07751 | 0.56561 | 1.52726 | 0.13010 | 0.00000 | 1.59616 | 0.85146 | 2.27708 |
| **ZBTB4** | 1.02042 | 0.07719 | 0.96155 | 0.00000 | 0.33344 | 0.72952 | 1.16045 | 0.01992 | 1.01412 | 3.08523 | 0.65970 | 0.39776 |
| **ZBTB40** | 2.80568 | 4.91913 | 9.03797 | 3.41008 | 7.78836 | 1.81804 | 8.23995 | 2.84747 | 7.50406 | 13.71470 | 4.55095 | 6.64329 |
| **ZBTB41** | 0.81539 | 0.00450 | 0.62882 | 0.00117 | 0.00000 | 4.17327 | 0.82146 | 0.36763 | 0.36457 | 1.28873 | 0.28245 | 1.62434 |
| **ZBTB42** | 0.00597 | 0.10119 | 0.19355 | 0.04810 | 0.11085 | 0.00000 | 0.08153 | 0.01095 | 0.00000 | 0.00000 | 0.08650 | 0.01106 |
| **ZBTB43** | 4.87828 | 3.08029 | 4.81883 | 2.55846 | 2.54940 | 7.33776 | 5.40178 | 4.44664 | 6.12892 | 7.06967 | 3.75879 | 5.23116 |

| | | | | | | | | | | | | |
|---|---|---|---|---|---|---|---|---|---|---|---|---|
| **ZBTB44** | 3.56928 | 0.45625 | 2.56436 | 2.51089 | 0.00000 | 15.24597 | 11.57933 | 0.38374 | 0.00000 | 21.53327 | 3.92205 | 0.68661 |
| **ZBTB45** | 0.71048 | 1.12517 | 0.53337 | 1.40470 | 4.40477 | 0.00000 | 1.88665 | 0.00000 | 2.52043 | 3.99349 | 2.51037 | 2.24966 |
| **ZBTB46** | 0.33179 | 0.20982 | 0.40239 | 0.77041 | 0.16062 | 0.00000 | 0.76842 | 0.00000 | 0.19617 | 1.35195 | 0.06487 | 0.30982 |
| **ZBTB46-AS1** | 0.01238 | 0.70373 | 0.02000 | 0.00000 | 0.06699 | 0.10946 | 0.15896 | 0.00295 | 0.00000 | 0.03999 | 0.04123 | 0.27053 |
| **ZBTB47** | 0.11936 | 0.02080 | 0.03346 | 0.06326 | 0.02166 | 0.00000 | 0.03620 | 0.04463 | 0.01845 | 0.02665 | 0.02844 | 0.05563 |
| **ZBTB48** | 0.87173 | 4.14819 | 1.61317 | 2.74212 | 1.17880 | 0.00000 | 2.81946 | 0.19863 | 1.94867 | 0.00000 | 0.86556 | 4.80389 |
| **ZBTB49** | 3.66611 | 1.44281 | 1.34705 | 0.31919 | 0.26696 | 0.37351 | 0.36659 | 1.55272 | 1.79216 | 0.62602 | 0.33597 | 2.03447 |
| **ZBTB5** | 4.67301 | 1.01609 | 1.45650 | 0.20051 | 0.19096 | 3.34656 | 0.00000 | 0.69507 | 2.21670 | 3.99238 | 0.28625 | 1.20349 |
| **ZBTB6** | 0.64714 | 0.62378 | 1.84980 | 0.18173 | 0.04659 | 2.85323 | 3.90216 | 2.16842 | 0.95604 | 0.04372 | 0.72896 | 2.61462 |
| **ZBTB7A** | 0.00081 | 0.00000 | 0.61272 | 0.18877 | 1.72707 | 0.00000 | 2.02875 | 0.09094 | 0.00000 | 8.88559 | 0.11620 | 0.01350 |
| **ZBTB7B** | 0.37763 | 0.00000 | 2.56616 | 0.21560 | 1.81827 | 0.00000 | 1.19854 | 0.08626 | 1.93779 | 3.62899 | 0.32069 | 0.00278 |
| **ZBTB7C** | 0.43199 | 0.00000 | 0.96142 | 0.00000 | 0.00000 | 0.00000 | 1.60981 | 0.04555 | 2.48691 | 0.67369 | 1.58403 | 0.00000 |
| **ZBTB8A** | 0.31603 | 2.86701 | 0.00000 | 3.01401 | 0.00000 | 2.87400 | 1.94822 | 1.15732 | 1.18560 | 1.26148 | 0.08507 | 10.52152 |
| **ZBTB8B** | 1.55042 | 0.00000 | 0.03630 | 0.26736 | 0.29888 | 0.00528 | 0.52042 | 0.20688 | 0.02626 | 0.00000 | 0.26225 | 0.65022 |
| **ZBTB8OS** | 0.48023 | 4.34398 | 5.28194 | 7.80506 | 1.79957 | 13.67126 | 6.63984 | 1.77495 | 1.05506 | 6.60808 | 4.88593 | 9.68652 |
| **ZBTB9** | 0.32901 | 4.04674 | 1.06217 | 3.68952 | 0.00569 | 2.59881 | 3.81227 | 1.03237 | 6.06486 | 0.00000 | 3.88101 | 7.58835 |
| **ZC2HC1A** | 0.13469 | 0.75501 | 4.92766 | 0.00000 | 0.00000 | 1.99688 | 1.69237 | 2.55038 | 3.29450 | 0.72644 | 0.05092 | 2.50543 |
| **ZC2HC1B** | 7.16831 | 0.00012 | 0.00095 | 0.00915 | 0.03139 | 0.00000 | 0.14507 | 0.98969 | 0.00000 | 0.09220 | 0.02301 | 0.11510 |
| **ZC2HC1C** | 1.23405 | 0.31252 | 4.68408 | 0.21660 | 0.00000 | 1.49089 | 0.00000 | 0.05157 | 0.36520 | 0.00000 | 0.71640 | 0.02606 |
| **ZC3H10** | 1.84930 | 1.92498 | 1.29218 | 1.79183 | 0.75198 | 1.54598 | 0.77099 | 3.22036 | 0.00000 | 0.00000 | 1.09016 | 3.06213 |
| **ZC3H11A** | 0.60705 | 2.00840 | 7.87533 | 3.15577 | 1.89706 | 7.85984 | 5.09659 | 6.86532 | 4.50166 | 8.48814 | 2.71250 | 4.53174 |
| **ZC3H12A** | 1.64968 | 2.01615 | 3.31934 | 1.54564 | 3.58646 | 0.00000 | 1.02760 | 0.00000 | 4.88647 | 0.73051 | 0.00001 | 0.90497 |
| **ZC3H12B** | 0.00000 | 0.57304 | 0.00000 | 0.00000 | 1.20520 | 0.45318 | 0.00000 | 0.07597 | 1.31091 | 4.52717 | 0.76397 | 0.07582 |
| **ZC3H12C** | 0.00544 | 0.13463 | 0.07769 | 0.61399 | 0.11323 | 1.24832 | 0.00000 | 0.07325 | 0.00000 | 0.47662 | 0.14213 | 0.62601 |
| **ZC3H12D** | 0.01978 | 0.01441 | 0.01600 | 0.01206 | 0.04820 | 0.01841 | 0.01879 | 0.08969 | 0.03144 | 0.03234 | 0.02489 | 0.02957 |
| **ZC3H13** | 7.05054 | 3.93527 | 10.70882 | 5.98077 | 11.11118 | 15.97967 | 15.70554 | 15.25414 | 7.69686 | 35.21762 | 9.51271 | 14.47149 |
| **ZC3H14** | 3.47559 | 8.52884 | 8.37385 | 4.48841 | 6.96550 | 16.89955 | 10.64140 | 4.86560 | 13.99183 | 5.58307 | 15.59543 | 8.51837 |
| **ZC3H15** | 6.42194 | 17.86785 | 15.75797 | 12.60750 | 8.33098 | 44.73200 | 29.24431 | 11.90961 | 19.01085 | 25.35703 | 16.31296 | 24.76936 |
| **ZC3H18** | 11.78506 | 4.12700 | 4.61593 | 3.75842 | 6.20487 | 0.74357 | 7.44174 | 5.99602 | 1.64576 | 13.80184 | 0.86440 | 6.32034 |
| **ZC3H3** | 3.79166 | 7.00728 | 2.84256 | 2.03137 | 6.35933 | 0.00000 | 2.51682 | 0.00000 | 6.88737 | 1.22145 | 1.70065 | 5.93898 |
| **ZC3H4** | 1.15295 | 3.26350 | 4.19076 | 4.22030 | 2.30915 | 10.32826 | 12.69300 | 11.36220 | 1.23394 | 19.33888 | 7.09766 | 7.58804 |
| **ZC3H6** | 1.28870 | 0.07482 | 0.00000 | 0.00000 | 0.01542 | 0.15145 | 0.00000 | 0.11708 | 0.00772 | 0.17176 | 0.00008 | 0.05432 |
| **ZC3H7A** | 4.39226 | 6.00301 | 3.79942 | 3.87404 | 1.33669 | 8.13041 | 2.01368 | 0.43745 | 2.91502 | 8.32073 | 6.69295 | 0.72856 |
| **ZC3H7B** | 0.22641 | 0.63042 | 0.79007 | 1.03364 | 2.01266 | 0.00000 | 0.24389 | 0.21886 | 1.68940 | 5.05569 | 1.99669 | 0.00000 |
| **ZC3H8** | 2.37232 | 1.50958 | 4.75068 | 0.00000 | 0.00000 | 5.61213 | 6.75184 | 0.47971 | 3.13342 | 5.32788 | 3.95713 | 3.74082 |
| **ZC3HAV1** | 0.01629 | 10.24348 | 3.86508 | 36.43013 | 3.41110 | 2.51360 | 1.95498 | 0.25012 | 0.00000 | 2.96163 | 1.08830 | 0.17267 |
| **ZC3HAV1L** | 0.00715 | 0.02717 | 0.06316 | 0.06656 | 0.00000 | 0.00734 | 0.11622 | 0.01183 | 0.00000 | 0.01768 | 0.06754 | 0.03311 |
| **ZC3HC1** | 3.52111 | 11.89157 | 10.10483 | 5.66938 | 2.92020 | 8.25309 | 12.80826 | 4.42531 | 7.70687 | 0.00000 | 14.38231 | 12.51313 |
| **ZC4H2** | 0.11211 | 1.25449 | 0.00000 | 1.47552 | 0.19846 | 1.39212 | 2.30111 | 0.03147 | 1.06691 | 0.00000 | 3.02728 | 0.95477 |
| **ZCCHC10** | 1.54127 | 3.55372 | 9.98208 | 3.76398 | 0.00000 | 15.54989 | 9.07818 | 12.88891 | 2.49066 | 7.05720 | 5.87977 | 7.78343 |
| **ZCCHC11** | 0.67011 | 0.94291 | 1.12872 | 1.92655 | 0.00000 | 5.00145 | 3.71769 | 0.72543 | 1.50489 | 10.19350 | 0.10735 | 2.26612 |
| **ZCCHC12** | 1.10109 | 0.09727 | 0.00670 | 0.00000 | 0.00188 | 0.01839 | 0.01916 | 0.00388 | 0.01913 | 0.13370 | 0.00793 | 0.00335 |
| **ZCCHC14** | 6.76827 | 0.70136 | 0.00000 | 0.75218 | 2.79017 | 2.39089 | 0.87502 | 1.79312 | 2.29874 | 4.52305 | 0.00000 | 0.71115 |
| **ZCCHC17** | 8.38003 | 14.13625 | 12.49535 | 9.87286 | 15.30920 | 14.10141 | 24.27645 | 11.56840 | 10.97926 | 5.04179 | 10.56420 | 12.69897 |
| **ZCCHC18** | 0.00239 | 0.01643 | 0.08978 | 0.01268 | 0.03006 | 0.06447 | 0.00000 | 0.08449 | 0.00000 | 0.04320 | 0.01426 | 0.00703 |
| **ZCCHC2** | 8.46032 | 0.00000 | 0.23817 | 0.58788 | 2.73796 | 1.44700 | 2.35378 | 1.88719 | 0.00000 | 7.20972 | 0.95743 | 0.37622 |
| **ZCCHC24** | 0.01284 | 0.00000 | 0.22738 | 0.95135 | 0.00000 | 0.18310 | 0.06864 | 0.00695 | 0.00000 | 0.37481 | 0.04894 | 0.02813 |
| **ZCCHC3** | 0.90898 | 0.25681 | 1.11647 | 0.00000 | 1.35145 | 2.17062 | 6.94627 | 0.39407 | 0.00000 | 4.92919 | 1.98867 | 4.48194 |

| | | | | | | | | | | | | |
|---|---|---|---|---|---|---|---|---|---|---|---|---|
| **ZCCHC4** | 1.99597 | 3.44026 | 1.90391 | 1.11110 | 0.90020 | 7.21636 | 0.00000 | 0.10036 | 1.31444 | 0.00006 | 1.93028 | 6.26893 |
| **ZCCHC6** | 3.32241 | 1.52900 | 5.87833 | 3.28334 | 2.41722 | 7.93668 | 4.77297 | 13.08962 | 5.11404 | 7.34282 | 1.17120 | 4.82586 |
| **ZCCHC7** | 7.10305 | 8.26604 | 5.04678 | 2.70524 | 3.78088 | 17.15120 | 6.10266 | 5.31091 | 3.22232 | 6.27365 | 6.50878 | 14.13319 |
| **ZCCHC8** | 3.04621 | 10.01194 | 17.24944 | 5.53481 | 0.54259 | 19.57266 | 6.83984 | 9.11377 | 8.49513 | 1.82754 | 9.75923 | 10.60699 |
| **ZCCHC9** | 1.31437 | 4.89423 | 6.64919 | 3.13008 | 1.93313 | 10.92032 | 6.19862 | 5.37163 | 2.77520 | 6.66167 | 8.06441 | 9.22263 |
| **ZCRB1** | 4.94877 | 9.68859 | 7.66902 | 7.62479 | 13.30143 | 26.69316 | 11.63645 | 9.04723 | 6.91795 | 11.08272 | 10.98843 | 13.85846 |
| **ZCWPW1** | 5.28727 | 0.42985 | 0.24028 | 0.40288 | 0.71242 | 0.00000 | 0.31413 | 0.69569 | 0.00000 | 0.00000 | 0.12751 | 0.61190 |
| **ZCWPW2** | 0.19336 | 0.14734 | 0.02850 | 0.00031 | 0.00000 | 0.15094 | 0.01291 | 0.04990 | 0.00382 | 0.00912 | 0.06733 | 0.32271 |
| **ZDBF2** | 0.97607 | 1.83909 | 0.75494 | 1.94692 | 0.00000 | 6.66983 | 1.08877 | 0.69922 | 4.32632 | 6.97375 | 1.91194 | 0.65515 |
| **ZDHHC1** | 0.03648 | 0.01120 | 0.11110 | 0.01372 | 0.03316 | 0.01761 | 0.02221 | 0.00671 | 0.00000 | 0.02443 | 0.03846 | 0.02024 |
| **ZDHHC11** | 0.00782 | 0.77957 | 1.74020 | 0.00000 | 0.00000 | 0.00000 | 1.69407 | 0.06498 | 5.78473 | 4.69273 | 0.53922 | 0.61764 |
| **ZDHHC12** | 0.49419 | 9.83889 | 8.73213 | 10.77666 | 9.34565 | 1.53720 | 4.50249 | 0.41600 | 9.91198 | 0.00000 | 10.41052 | 4.03499 |
| **ZDHHC13** | 0.46139 | 0.92553 | 4.59533 | 1.17360 | 0.34789 | 6.39191 | 0.00000 | 0.27983 | 2.16139 | 0.00000 | 1.19020 | 0.95086 |
| **ZDHHC14** | 0.24305 | 0.00000 | 1.44777 | 0.00000 | 0.55064 | 0.04401 | 3.52619 | 0.10190 | 0.65425 | 1.12734 | 0.47344 | 0.00000 |
| **ZDHHC15** | 0.08186 | 0.04192 | 0.02466 | 0.01783 | 0.00000 | 0.06156 | 0.01394 | 0.00456 | 0.00797 | 0.03735 | 0.01002 | 0.03320 |
| **ZDHHC16** | 0.40188 | 7.14149 | 9.64780 | 4.53735 | 3.81698 | 0.00000 | 0.75087 | 1.47209 | 5.67399 | 3.79282 | 3.59835 | 3.72847 |
| **ZDHHC17** | 0.00000 | 0.49866 | 1.68019 | 0.82294 | 0.00000 | 8.82935 | 1.55650 | 4.40880 | 4.40539 | 5.43332 | 5.03405 | 6.66414 |
| **ZDHHC18** | 2.69556 | 3.47452 | 3.30225 | 2.71432 | 5.51116 | 0.00000 | 6.88500 | 0.00000 | 2.88097 | 13.84354 | 2.20554 | 5.46650 |
| **ZDHHC19** | 0.00807 | 0.21780 | 0.15027 | 0.04425 | 0.00368 | 0.00000 | 0.00503 | 0.01407 | 0.00866 | 0.02960 | 0.01669 | 0.12926 |
| **ZDHHC2** | 0.15337 | 0.30953 | 0.46214 | 0.36215 | 0.00000 | 1.62109 | 0.89714 | 0.37183 | 0.00000 | 0.66289 | 0.10169 | 0.33758 |
| **ZDHHC20** | 0.00000 | 1.55386 | 6.94017 | 3.45682 | 2.78923 | 23.27782 | 7.04246 | 1.04942 | 2.97855 | 14.80034 | 3.92278 | 0.30930 |
| **ZDHHC21** | 0.35866 | 0.00000 | 1.36928 | 0.00000 | 0.00000 | 3.93541 | 0.13812 | 0.13362 | 2.91686 | 3.00784 | 1.13504 | 0.52314 |
| **ZDHHC22** | 0.00532 | 0.00765 | 0.01294 | 0.01177 | 0.00702 | 0.03036 | 0.00700 | 0.00901 | 0.01387 | 0.05822 | 0.02209 | 0.07424 |
| **ZDHHC23** | 0.08344 | 0.79306 | 0.00000 | 0.90445 | 0.85200 | 0.41135 | 0.00000 | 0.00553 | 0.25067 | 2.02378 | 0.43776 | 0.30360 |
| **ZDHHC24** | 0.38690 | 1.91721 | 1.37913 | 1.92573 | 3.05085 | 0.00000 | 0.00000 | 0.03793 | 2.09724 | 0.00000 | 1.92185 | 0.81118 |
| **ZDHHC3** | 3.35038 | 10.30107 | 13.14250 | 7.35531 | 8.46700 | 3.46348 | 9.80605 | 1.89626 | 10.74253 | 8.05736 | 8.95534 | 4.59567 |
| **ZDHHC4** | 3.13613 | 2.75261 | 4.18204 | 5.94829 | 5.30571 | 8.38488 | 2.63991 | 0.43919 | 8.54587 | 0.00000 | 8.20833 | 3.85975 |
| **ZDHHC5** | 4.92393 | 11.20862 | 13.43550 | 8.38395 | 9.97225 | 3.36735 | 11.84800 | 5.76955 | 3.43567 | 9.72150 | 6.87448 | 9.06891 |
| **ZDHHC6** | 0.29174 | 6.86548 | 11.16464 | 3.30675 | 0.40661 | 19.49946 | 13.22614 | 2.11669 | 17.82238 | 0.00000 | 10.82867 | 6.99130 |
| **ZDHHC7** | 4.85961 | 4.80983 | 5.72901 | 4.54187 | 5.89160 | 2.59695 | 7.40585 | 1.81163 | 7.05752 | 12.74804 | 4.66343 | 4.03297 |
| **ZDHHC8** | 2.29446 | 0.00000 | 0.18609 | 0.70148 | 0.06190 | 0.05253 | 0.15702 | 0.00000 | 0.75693 | 1.33258 | 0.10703 | 0.06043 |
| **ZDHHC8P1** | 0.00304 | 0.07907 | 0.01702 | 0.16075 | 0.00000 | 0.00000 | 0.15590 | 0.01019 | 0.22539 | 0.47637 | 0.00272 | 0.03791 |
| **ZDHHC9** | 0.10008 | 12.14336 | 2.56517 | 9.61872 | 8.71273 | 0.00000 | 0.01354 | 1.15590 | 11.05850 | 8.90465 | 3.04265 | 6.63208 |
| **ZEB1** | 1.24180 | 0.17031 | 0.00000 | 0.01148 | 0.00000 | 0.12314 | 0.00000 | 0.55944 | 0.00041 | 0.13541 | 0.00061 | 0.62301 |
| **ZEB1-AS1** | 0.42509 | 0.81024 | 0.00000 | 0.00000 | 0.00000 | 0.42744 | 0.00000 | 0.04941 | 0.00000 | 0.11495 | 0.05818 | 1.87144 |
| **ZEB2** | 0.00332 | 0.21244 | 0.00000 | 0.00000 | 0.14199 | 0.00000 | 0.01891 | 0.13101 | 0.00000 | 0.56737 | 0.13321 | 0.10067 |
| **ZER1** | 0.51505 | 4.53723 | 2.40007 | 1.96439 | 3.22766 | 0.00000 | 1.97153 | 0.39707 | 1.22701 | 1.61394 | 1.29658 | 3.07192 |
| **ZFAND1** | 0.40003 | 4.03476 | 3.28602 | 2.06617 | 0.44015 | 14.79896 | 2.90292 | 0.86737 | 5.93170 | 2.04783 | 4.91680 | 2.18412 |
| **ZFAND2A** | 13.32997 | 5.87129 | 15.81383 | 1.02559 | 2.74700 | 12.26370 | 13.51617 | 19.67110 | 0.76178 | 1.48067 | 3.27369 | 8.41867 |
| **ZFAND2B** | 0.00000 | 4.63646 | 2.28457 | 5.19973 | 2.54496 | 0.00000 | 2.26740 | 1.10400 | 0.62762 | 1.10039 | 2.52232 | 3.20023 |
| **ZFAND3** | 3.02039 | 0.64341 | 2.70763 | 2.09412 | 3.36903 | 0.00000 | 3.78389 | 1.27941 | 0.76563 | 8.34975 | 0.46432 | 1.87177 |
| **ZFAND4** | 0.51131 | 0.52067 | 0.07357 | 0.06037 | 0.70427 | 2.75699 | 0.00000 | 1.58129 | 0.62200 | 0.00000 | 0.12232 | 1.67059 |
| **ZFAND5** | 7.76815 | 26.09999 | 16.14984 | 0.00360 | 1.67103 | 22.05860 | 15.05537 | 23.23679 | 4.16175 | 29.60736 | 4.73240 | 29.66282 |
| **ZFAND6** | 5.48532 | 7.40486 | 13.90724 | 6.39532 | 4.87261 | 32.72234 | 9.46197 | 11.51207 | 14.30883 | 2.92635 | 8.48109 | 10.96206 |
| **ZFAS1** | 4.34755 | 9.92880 | 17.77819 | 14.92690 | 19.11033 | 23.33555 | 24.01067 | 10.53986 | 28.59583 | 33.41886 | 6.10836 | 21.00229 |
| **ZFAT** | 1.64521 | 1.36101 | 8.71717 | 1.73954 | 0.58483 | 0.13159 | 2.39420 | 0.11230 | 1.01124 | 0.00000 | 1.46678 | 1.42214 |
| **ZFAT-AS1** | 0.00848 | 0.02126 | 0.08071 | 0.02790 | 0.01754 | 0.00756 | 0.01995 | 0.01904 | 0.03652 | 0.00606 | 0.01747 | 0.02555 |
| **ZFC3H1** | 3.56930 | 0.00000 | 6.74801 | 1.48095 | 2.13488 | 9.37475 | 5.16978 | 4.07398 | 8.35870 | 18.61765 | 3.77682 | 4.15741 |

| | | | | | | | | | | | | |
|---|---|---|---|---|---|---|---|---|---|---|---|---|
| ZFHX2 | 0.27804 | 0.07458 | 0.26027 | 0.61387 | 0.00000 | 0.13980 | 0.00000 | 0.00170 | 0.28337 | 1.51422 | 0.00000 | 0.00772 |
| ZFHX3 | 4.41202 | 1.43705 | 19.08153 | 0.00000 | 10.60973 | 5.44094 | 21.60026 | 0.82992 | 3.98485 | 27.13831 | 8.33594 | 0.00000 |
| ZFHX4 | 0.00274 | 0.00000 | 0.03595 | 0.30552 | 0.16449 | 0.02402 | 0.02463 | 0.89302 | 0.11428 | 0.02266 | 0.01486 | 0.35575 |
| ZFP1 | 0.45966 | 0.60539 | 1.90399 | 0.27427 | 0.00000 | 2.90587 | 0.00002 | 0.44277 | 1.63413 | 1.48778 | 1.31773 | 1.42475 |
| ZFP14 | 0.71774 | 0.00542 | 0.25591 | 0.30567 | 0.14968 | 0.00000 | 0.13561 | 0.00000 | 0.25513 | 1.56037 | 0.58121 | 0.35621 |
| ZFP2 | 0.03799 | 0.20236 | 0.22923 | 0.00000 | 0.06095 | 0.00117 | 0.00000 | 0.14224 | 0.00000 | 0.27557 | 0.00569 | 0.06293 |
| ZFP28 | 3.67203 | 0.40058 | 0.12579 | 0.46534 | 0.26143 | 0.00110 | 0.00000 | 4.88196 | 0.72477 | 0.87541 | 0.06988 | 1.12842 |
| ZFP3 | 1.57763 | 0.54589 | 0.00000 | 0.08272 | 0.05362 | 1.87745 | 0.00000 | 1.76785 | 0.00000 | 0.09880 | 0.19787 | 1.97780 |
| ZFP30 | 0.06188 | 0.05311 | 0.46322 | 0.79979 | 0.00000 | 0.86393 | 0.00000 | 0.00000 | 0.00000 | 0.86562 | 1.17460 | 1.03682 |
| ZFP36 | 1.14769 | 0.16135 | 5.28431 | 1.09495 | 0.93357 | 0.00000 | 0.37958 | 0.00000 | 2.36012 | 0.68836 | 0.16420 | 0.48746 |
| ZFP36L1 | 0.77926 | 1.04201 | 20.11478 | 6.22127 | 6.66001 | 4.77165 | 13.74210 | 0.00000 | 3.52918 | 11.17538 | 7.15160 | 7.42455 |
| ZFP36L2 | 0.67173 | 3.95294 | 6.36999 | 7.01480 | 13.21613 | 12.01303 | 9.11223 | 1.45048 | 0.00000 | 29.21327 | 7.33376 | 2.45164 |
| ZFP37 | 0.00000 | 1.04577 | 1.33412 | 0.22292 | 0.00000 | 1.08987 | 1.23086 | 6.52865 | 0.00000 | 0.68444 | 0.27147 | 1.68622 |
| ZFP41 | 0.99186 | 0.28778 | 0.00000 | 0.04889 | 0.00000 | 0.00000 | 0.36992 | 0.11556 | 0.39841 | 1.00632 | 0.17613 | 1.26479 |
| ZFP42 | 0.00000 | 26.97819 | 14.47550 | 23.58627 | 18.00511 | 44.13336 | 28.62664 | 13.22898 | 37.38546 | 8.04078 | 29.47856 | 25.54700 |
| ZFP57 | 0.00173 | 14.64726 | 1.66539 | 11.13295 | 2.71545 | 2.02691 | 5.84475 | 0.00000 | 7.60147 | 0.00000 | 8.08126 | 6.93071 |
| ZFP62 | 2.82567 | 2.72675 | 3.15972 | 0.38484 | 0.00000 | 0.33446 | 2.51811 | 3.56017 | 5.22872 | 1.93248 | 3.05598 | 7.39645 |
| ZFP64 | 1.16322 | 9.19101 | 3.29042 | 6.57330 | 4.76229 | 1.56085 | 8.98041 | 0.00000 | 12.53516 | 0.00000 | 9.80513 | 2.15025 |
| ZFP69 | 0.29543 | 0.29428 | 0.10538 | 0.00000 | 0.00000 | 1.32970 | 0.39751 | 0.00000 | 0.32975 | 0.58665 | 0.09245 | 0.27320 |
| ZFP69B | 0.16235 | 1.28258 | 0.22634 | 0.00000 | 0.00000 | 0.74358 | 0.95983 | 0.40120 | 0.16360 | 0.54504 | 0.65428 | 2.88940 |
| ZFP82 | 0.00130 | 3.41849 | 0.35353 | 1.50536 | 0.00000 | 5.43811 | 0.45604 | 6.71284 | 6.26900 | 0.00000 | 6.58256 | 4.65254 |
| ZFP90 | 0.84065 | 0.85562 | 0.00000 | 0.00000 | 0.20249 | 1.21954 | 0.37024 | 1.04432 | 0.00000 | 0.46164 | 0.26001 | 1.02252 |
| ZFP91 | 0.68572 | 0.00000 | 3.70616 | 0.42365 | 2.84267 | 4.85777 | 7.52212 | 2.84294 | 2.20020 | 14.60920 | 1.24087 | 3.09684 |
| ZFPL1 | 1.32025 | 7.52520 | 5.98728 | 4.74330 | 9.54054 | 0.62416 | 7.75897 | 1.01284 | 5.08750 | 0.00000 | 6.60526 | 6.06415 |
| ZFPM1 | 0.11392 | 0.09971 | 0.00765 | 0.09863 | 0.35808 | 0.00000 | 0.02829 | 0.00554 | 0.00000 | 0.48567 | 0.00769 | 0.03422 |
| ZFPM2 | 6.68945 | 0.00341 | 0.00980 | 0.00372 | 0.00168 | 0.02972 | 0.02170 | 0.79106 | 0.09758 | 0.01440 | 0.01629 | 0.08996 |
| ZFR | 4.61178 | 10.64174 | 12.19048 | 8.71532 | 6.44680 | 25.70694 | 18.42233 | 8.96385 | 13.03469 | 11.60887 | 11.52528 | 15.21187 |
| ZFR2 | 0.87331 | 0.28497 | 0.13687 | 0.05809 | 0.06756 | 0.05650 | 0.00000 | 1.56492 | 0.01458 | 0.04671 | 0.00000 | 7.52333 |
| ZFX | 0.55732 | 0.64671 | 2.59625 | 1.67545 | 0.00000 | 6.93091 | 2.82443 | 1.77886 | 4.17930 | 3.32486 | 3.89999 | 2.07273 |
| ZFX-AS1 | 0.01420 | 0.01577 | 0.02293 | 0.01546 | 0.01268 | 0.01809 | 0.01680 | 0.01637 | 0.01432 | 0.02300 | 0.01515 | 0.01880 |
| ZFY | 0.09757 | 0.31508 | 0.81978 | 0.81330 | 0.56750 | 4.69468 | 3.00385 | 0.06069 | 1.28001 | 3.44994 | 1.76217 | 0.58574 |
| ZFYVE1 | 1.00383 | 16.54871 | 1.31201 | 5.74773 | 0.97741 | 2.96339 | 2.07849 | 17.19009 | 3.08605 | 0.33646 | 2.33308 | 25.57156 |
| ZFYVE16 | 0.72838 | 0.17765 | 2.82876 | 0.16996 | 0.73187 | 9.37620 | 2.21090 | 1.63729 | 2.33881 | 4.03139 | 1.93519 | 1.94866 |
| ZFYVE19 | 1.15446 | 12.66577 | 9.17323 | 1.50179 | 1.69807 | 0.00000 | 2.86916 | 10.83375 | 4.62660 | 7.90822 | 6.65000 | 18.58956 |
| ZFYVE20 | 3.66112 | 5.63191 | 6.01593 | 4.72950 | 3.46313 | 4.23536 | 6.89095 | 0.42539 | 4.00660 | 2.19481 | 5.45326 | 3.03993 |
| ZFYVE21 | 7.54425 | 13.09164 | 6.86230 | 9.68011 | 15.86956 | 16.96871 | 13.85070 | 2.12539 | 13.80582 | 2.52217 | 19.81354 | 2.22083 |
| ZFYVE26 | 0.48473 | 0.21274 | 1.96828 | 1.56368 | 1.80069 | 3.39480 | 0.00000 | 0.00000 | 2.11255 | 3.17549 | 1.66427 | 0.61554 |
| ZFYVE27 | 0.00107 | 5.31210 | 4.46727 | 2.69859 | 1.14935 | 0.00000 | 2.01659 | 3.60676 | 9.40172 | 0.00000 | 4.69554 | 6.95369 |
| ZFYVE28 | 0.03656 | 0.02375 | 0.06874 | 0.15316 | 0.00748 | 0.01523 | 0.02375 | 0.01679 | 0.00000 | 0.05952 | 0.11166 | 0.01463 |
| ZFYVE9 | 2.83228 | 1.18847 | 0.00000 | 1.88007 | 0.00000 | 2.34117 | 0.89941 | 0.00000 | 0.82728 | 5.59211 | 1.54144 | 1.49601 |
| ZG16 | 0.00378 | 0.01293 | 0.09735 | 0.00000 | 0.34121 | 0.00000 | 0.17786 | 0.00676 | 0.00000 | 0.19268 | 0.66196 | 0.06824 |
| ZG16B | 0.00615 | 0.13278 | 0.02201 | 0.00498 | 0.02044 | 0.00000 | 0.01648 | 0.01110 | 0.01769 | 0.02741 | 0.01434 | 0.00859 |
| ZGLP1 | 1.54334 | 0.19741 | 0.26796 | 0.00000 | 0.00000 | 0.10725 | 0.35959 | 0.34272 | 0.98403 | 0.50022 | 0.05228 | 0.48840 |
| ZGPAT | 0.54010 | 9.13052 | 5.97811 | 8.09477 | 7.65778 | 0.00000 | 5.33216 | 6.81302 | 6.15922 | 3.14557 | 6.21206 | 13.19579 |
| ZGRF1 | 0.34017 | 1.33945 | 0.10803 | 1.93587 | 1.14098 | 7.99768 | 0.00000 | 2.78651 | 0.60248 | 10.66914 | 1.54147 | 5.36567 |
| ZHX1 | 5.91612 | 1.85879 | 0.23434 | 5.15955 | 1.12558 | 5.07408 | 0.00000 | 2.39017 | 0.40328 | 2.87483 | 0.23540 | 2.60204 |
| ZHX1-C8orf76 | 0.37163 | 0.34269 | 0.37839 | 0.32054 | 0.16777 | 0.32079 | 0.39829 | 0.47727 | 0.43342 | 1.01661 | 0.06392 | 0.93988 |
| ZHX2 | 1.98331 | 0.21459 | 0.21817 | 0.34733 | 0.04655 | 0.17841 | 0.02633 | 0.46787 | 0.10507 | 0.64571 | 0.16731 | 0.17309 |

| | | | | | | | | | | | | |
|---|---|---|---|---|---|---|---|---|---|---|---|---|
| ZHX3 | 15.48434 | 0.98751 | 0.15856 | 0.49060 | 1.40198 | 1.50665 | 0.00000 | 2.97827 | 0.54005 | 2.36141 | 0.11369 | 0.00000 |
| ZIC1 | 0.06627 | 0.00581 | 0.02040 | 0.03348 | 0.00000 | 0.03402 | 0.00000 | 0.00798 | 0.00797 | 0.15894 | 0.01402 | 0.01461 |
| ZIC2 | 0.00370 | 0.04901 | 0.52801 | 1.04422 | 0.00000 | 0.09305 | 0.00000 | 0.48836 | 0.00000 | 0.46815 | 0.00000 | 0.04627 |
| ZIC3 | 0.00446 | 0.00369 | 0.33391 | 2.66252 | 0.00000 | 0.02277 | 0.01622 | 0.05462 | 0.00000 | 0.02908 | 0.03216 | 0.04265 |
| ZIC4 | 0.00619 | 0.01389 | 0.01429 | 0.02290 | 0.01815 | 0.03061 | 0.01506 | 0.01035 | 0.01694 | 0.01325 | 0.01405 | 0.01529 |
| ZIC5 | 0.00569 | 0.15089 | 0.00000 | 0.89768 | 0.13514 | 0.87046 | 0.00000 | 0.18873 | 0.00000 | 0.69550 | 0.00000 | 0.07049 |
| ZIK1 | 0.85557 | 2.97397 | 1.53053 | 1.38035 | 1.11208 | 0.85772 | 3.03429 | 15.73550 | 0.07139 | 0.14542 | 1.50343 | 11.97867 |
| ZIM2 | 2.33550 | 0.00000 | 0.08689 | 0.00000 | 0.24055 | 0.00000 | 0.05900 | 1.05290 | 0.02659 | 0.09036 | 0.02536 | 1.91987 |
| ZIM3 | 0.03851 | 0.05553 | 0.00292 | 0.00000 | 0.00000 | 0.01065 | 0.00110 | 30.75961 | 0.00976 | 0.03522 | 0.00518 | 10.89582 |
| ZKSCAN1 | 0.51049 | 0.49466 | 2.03906 | 1.86292 | 0.65558 | 3.81438 | 1.29892 | 2.64278 | 4.93943 | 5.28883 | 3.48344 | 1.69235 |
| ZKSCAN2 | 2.34483 | 0.67693 | 1.62111 | 0.63455 | 0.50198 | 1.22208 | 1.19175 | 0.96671 | 2.58521 | 2.59374 | 0.36296 | 1.28975 |
| ZKSCAN3 | 0.08300 | 2.69863 | 0.21275 | 0.00000 | 0.02279 | 1.05992 | 0.00000 | 0.00000 | 0.63869 | 1.48876 | 0.23604 | 0.02391 |
| ZKSCAN4 | 0.43577 | 3.97145 | 0.16632 | 0.11950 | 0.33955 | 0.00000 | 0.00000 | 1.73944 | 0.32895 | 0.12318 | 0.30970 | 4.37019 |
| ZKSCAN5 | 1.14447 | 3.20033 | 1.35051 | 2.31942 | 2.88585 | 0.93855 | 2.11033 | 8.64980 | 5.43048 | 2.00482 | 2.29413 | 5.70409 |
| ZKSCAN7 | 0.20328 | 0.00492 | 0.01980 | 0.10333 | 0.01778 | 0.00186 | 0.02323 | 0.00000 | 0.00000 | 0.00015 | 0.16520 | 0.21268 |
| ZKSCAN8 | 0.86294 | 1.43886 | 0.07808 | 0.10055 | 1.60708 | 3.23792 | 0.03726 | 2.51123 | 2.17435 | 2.92142 | 2.60454 | 4.11532 |
| ZMAT1 | 0.00453 | 0.01296 | 0.00783 | 0.02970 | 0.00000 | 0.05742 | 0.00000 | 0.00877 | 0.01012 | 0.09872 | 0.00710 | 0.03186 |
| ZMAT2 | 5.23135 | 4.46165 | 11.14116 | 5.44905 | 7.69873 | 11.27848 | 21.02636 | 6.57181 | 3.03464 | 19.00345 | 8.98418 | 13.45211 |
| ZMAT3 | 0.14495 | 1.22110 | 2.37417 | 3.72850 | 4.10374 | 10.38600 | 2.94447 | 1.06476 | 0.00000 | 14.29936 | 0.34419 | 2.53239 |
| ZMAT4 | 0.00372 | 0.02068 | 0.01590 | 0.02340 | 0.03137 | 0.00076 | 0.06154 | 0.04104 | 0.01411 | 0.00092 | 0.01588 | 0.02556 |
| ZMAT5 | 0.59337 | 6.65506 | 4.79288 | 0.90939 | 4.41868 | 1.08487 | 5.79361 | 18.15330 | 2.68915 | 0.00000 | 4.10482 | 10.70072 |
| ZMIZ1 | 0.00000 | 5.83093 | 0.09815 | 9.05332 | 13.09278 | 7.19261 | 10.51573 | 0.00000 | 8.99442 | 15.29962 | 3.03000 | 0.75019 |
| ZMIZ1-AS1 | 0.00622 | 0.01338 | 0.01096 | 0.12303 | 0.00741 | 0.01690 | 0.01582 | 0.01044 | 0.01898 | 0.01084 | 0.01680 | 0.01655 |
| ZMIZ2 | 1.21798 | 1.11154 | 0.40353 | 2.29871 | 5.11875 | 0.00000 | 0.52856 | 0.06180 | 1.87256 | 6.17809 | 0.94028 | 0.09358 |
| ZMPSTE24 | 1.01668 | 10.41482 | 8.49150 | 2.94886 | 2.10562 | 41.19893 | 0.00000 | 7.46135 | 30.04599 | 15.83703 | 10.44807 | 8.79933 |
| ZMYM1 | 0.10090 | 8.76441 | 0.36320 | 0.68390 | 0.00008 | 17.14220 | 0.00000 | 3.04047 | 4.81812 | 6.99458 | 4.72694 | 13.95003 |
| ZMYM2 | 6.93614 | 5.04928 | 2.94647 | 3.48675 | 0.37412 | 23.61552 | 0.00008 | 6.68835 | 7.24919 | 15.91352 | 1.59904 | 8.09760 |
| ZMYM3 | 1.08790 | 1.95794 | 0.27655 | 4.13339 | 3.59306 | 0.63049 | 1.49364 | 0.00000 | 2.71652 | 5.86018 | 3.24472 | 2.43915 |
| ZMYM4 | 3.72588 | 1.50765 | 1.70100 | 1.34209 | 3.29845 | 9.79146 | 7.60496 | 0.46245 | 1.62963 | 13.64702 | 0.98175 | 1.67458 |
| ZMYM5 | 3.98360 | 3.54365 | 2.27461 | 1.58723 | 0.09751 | 8.78978 | 2.46631 | 3.79957 | 1.48834 | 2.22669 | 3.87281 | 5.58634 |
| ZMYM6 | 0.67293 | 0.99984 | 0.08673 | 1.66574 | 0.76938 | 3.19845 | 1.82459 | 5.13010 | 1.95388 | 0.00000 | 1.68026 | 2.97011 |
| ZMYM6NB | 0.88605 | 4.79331 | 0.52215 | 3.61631 | 0.69596 | 2.10925 | 0.00000 | 0.00000 | 0.00000 | 0.00000 | 0.51436 | 7.41178 |
| ZMYND10 | 0.00314 | 0.21345 | 0.20890 | 0.00000 | 0.00000 | 0.00152 | 0.03171 | 0.00510 | 0.46268 | 0.45473 | 0.00000 | 0.14293 |
| ZMYND11 | 1.97595 | 2.19934 | 1.99571 | 0.00000 | 1.72653 | 3.99873 | 8.70931 | 0.38796 | 5.15528 | 8.86449 | 7.12852 | 0.41347 |
| ZMYND12 | 0.55653 | 0.14824 | 0.12206 | 0.10239 | 0.20602 | 0.15143 | 0.00000 | 0.15289 | 0.00000 | 0.01300 | 0.41251 | 0.00000 |
| ZMYND15 | 0.10128 | 0.00143 | 0.08350 | 0.02882 | 0.00000 | 0.05241 | 0.00004 | 0.01663 | 0.00000 | 0.05511 | 0.02610 | 0.08155 |
| ZMYND19 | 3.93917 | 2.24370 | 2.32427 | 2.03155 | 3.99552 | 0.00000 | 6.11466 | 2.60103 | 0.00000 | 11.50330 | 1.61745 | 3.21293 |
| ZMYND8 | 7.98607 | 5.53534 | 11.73909 | 14.11682 | 5.92035 | 5.31583 | 7.86360 | 13.51275 | 10.03615 | 20.34321 | 7.77691 | 7.24151 |
| ZNF10 | 2.61467 | 1.12708 | 0.97276 | 0.22151 | 0.00000 | 2.25187 | 0.74045 | 1.38664 | 0.30924 | 0.00005 | 0.24136 | 2.05748 |
| ZNF100 | 0.00000 | 7.90368 | 2.24184 | 3.21963 | 0.00000 | 11.77345 | 0.00000 | 8.21899 | 2.79860 | 7.27707 | 2.01221 | 8.50733 |
| ZNF101 | 0.99069 | 5.32899 | 2.06534 | 2.36045 | 1.60334 | 6.67630 | 0.00000 | 3.97519 | 0.00000 | 2.15929 | 3.34924 | 8.57520 |
| ZNF106 | 7.22748 | 15.89370 | 12.25133 | 8.87050 | 16.60392 | 28.91195 | 19.21314 | 4.47907 | 17.02388 | 33.04229 | 18.52269 | 31.03345 |
| ZNF107 | 10.51363 | 4.80846 | 2.49777 | 1.45589 | 0.00000 | 7.12527 | 0.00000 | 2.34637 | 4.71944 | 3.98487 | 1.01961 | 5.29594 |
| ZNF112 | 0.20073 | 0.57288 | 0.73328 | 0.03228 | 0.00000 | 1.54372 | 0.26952 | 0.00000 | 0.28988 | 0.00000 | 0.07965 | 0.86849 |
| ZNF114 | 0.00000 | 3.39321 | 7.47203 | 2.68180 | 11.85179 | 7.59760 | 7.53796 | 0.59066 | 4.72439 | 0.00000 | 4.92868 | 5.23462 |
| ZNF117 | 0.04351 | 0.00000 | 1.81312 | 0.07506 | 0.00000 | 1.09107 | 0.90751 | 0.19799 | 0.96451 | 0.85456 | 0.00000 | 0.20078 |
| ZNF12 | 0.00000 | 0.32187 | 0.55584 | 0.63864 | 0.00000 | 4.96261 | 0.37942 | 1.52373 | 1.79977 | 5.73924 | 1.29938 | 1.93901 |
| ZNF121 | 0.00000 | 1.34363 | 0.74171 | 0.52737 | 0.35472 | 4.79249 | 0.00000 | 1.90994 | 2.73284 | 2.30717 | 1.46872 | 1.19567 |

| | | | | | | | | | | | | |
|---|---|---|---|---|---|---|---|---|---|---|---|---|
| **ZNF124** | 0.24347 | 1.69686 | 1.92235 | 1.65255 | 0.00000 | 6.86361 | 1.30301 | 4.53153 | 5.65452 | 2.67630 | 3.80577 | 4.50976 |
| **ZNF131** | 1.93076 | 3.11353 | 1.83503 | 2.50664 | 2.28807 | 10.77935 | 2.44829 | 11.10919 | 5.16120 | 13.24318 | 4.58351 | 5.49501 |
| **ZNF132** | 0.57291 | 0.52283 | 0.10606 | 0.04337 | 0.00000 | 0.32338 | 0.14466 | 1.22650 | 0.65669 | 0.00000 | 0.37831 | 2.16465 |
| **ZNF133** | 0.22871 | 1.98271 | 2.19880 | 1.29636 | 2.27017 | 0.00000 | 0.04792 | 0.72446 | 3.89980 | 1.24272 | 2.92734 | 2.58130 |
| **ZNF134** | 2.38019 | 12.21853 | 2.72828 | 3.88835 | 2.55826 | 2.81961 | 3.15692 | 8.59709 | 2.97214 | 0.00000 | 3.78206 | 17.55660 |
| **ZNF135** | 0.74744 | 0.66484 | 0.00000 | 0.00000 | 0.05398 | 0.16245 | 0.09742 | 0.04957 | 0.02906 | 0.01259 | 0.00000 | 0.06822 |
| **ZNF136** | 3.48592 | 2.77416 | 1.47406 | 0.73123 | 0.00000 | 8.20288 | 0.00000 | 9.04330 | 3.14884 | 4.48561 | 2.21220 | 3.52243 |
| **ZNF137P** | 0.00922 | 0.05555 | 0.19116 | 0.48675 | 0.00000 | 0.48718 | 1.22412 | 0.56299 | 0.00000 | 1.54538 | 0.57142 | 0.23692 |
| **ZNF138** | 0.24075 | 9.04172 | 2.71232 | 3.41760 | 0.00000 | 10.37778 | 1.95259 | 4.72358 | 3.53914 | 3.03848 | 2.75415 | 6.80876 |
| **ZNF14** | 0.18845 | 2.31502 | 0.04123 | 0.25212 | 0.31332 | 4.49371 | 0.02861 | 0.40698 | 1.05817 | 0.00000 | 1.72100 | 1.33287 |
| **ZNF140** | 0.90740 | 2.77216 | 0.48181 | 2.50347 | 0.00000 | 3.14056 | 2.25912 | 13.02597 | 2.45303 | 0.00000 | 0.00000 | 3.39486 |
| **ZNF141** | 0.62650 | 1.65091 | 0.17588 | 0.82768 | 0.33946 | 4.08967 | 0.00000 | 0.79648 | 0.83544 | 0.00000 | 1.31749 | 2.16868 |
| **ZNF142** | 5.82369 | 4.39343 | 0.65114 | 3.96014 | 9.04528 | 0.00000 | 0.11944 | 4.00884 | 5.75566 | 5.86981 | 1.24272 | 8.39511 |
| **ZNF143** | 3.42585 | 4.07747 | 18.05869 | 2.53388 | 0.00000 | 13.02679 | 10.47856 | 8.65747 | 15.23005 | 1.76307 | 8.22583 | 6.96733 |
| **ZNF146** | 4.35114 | 11.13243 | 7.89576 | 9.96114 | 0.00000 | 42.67536 | 22.18120 | 3.11464 | 25.26668 | 10.13229 | 15.23068 | 11.72102 |
| **ZNF148** | 4.52623 | 0.19500 | 0.42509 | 1.26444 | 0.00000 | 10.01329 | 6.73229 | 8.04958 | 2.64848 | 16.77456 | 4.33258 | 4.55788 |
| **ZNF154** | 0.00036 | 0.00000 | 0.29351 | 0.67214 | 0.28815 | 0.14900 | 0.00255 | 0.14771 | 0.47612 | 5.40586 | 0.04149 | 0.26738 |
| **ZNF155** | 5.79517 | 0.91321 | 0.00000 | 0.00000 | 0.30526 | 1.24081 | 0.17588 | 1.51460 | 0.38679 | 0.00000 | 0.00124 | 1.17573 |
| **ZNF157** | 0.01716 | 0.00628 | 0.06264 | 0.06232 | 0.00000 | 0.03744 | 0.00000 | 4.14815 | 0.04331 | 0.03068 | 0.04239 | 0.67473 |
| **ZNF16** | 2.64214 | 2.64059 | 0.30190 | 0.95234 | 0.00000 | 0.09466 | 0.92894 | 0.86787 | 0.94230 | 0.00000 | 0.68181 | 5.67168 |
| **ZNF160** | 1.49195 | 1.64972 | 1.97480 | 1.26085 | 1.44779 | 3.63718 | 1.70835 | 1.95183 | 0.91362 | 2.66776 | 1.64208 | 1.27530 |
| **ZNF165** | 0.00000 | 6.52139 | 15.81906 | 0.00000 | 1.47866 | 8.84052 | 14.15703 | 0.41844 | 14.88681 | 0.00000 | 11.42953 | 2.41550 |
| **ZNF169** | 0.16172 | 0.59668 | 0.06915 | 0.09762 | 0.10895 | 0.00000 | 0.18952 | 1.12094 | 0.97122 | 0.86195 | 1.11341 | 1.38597 |
| **ZNF17** | 1.17567 | 2.91776 | 0.00000 | 0.88224 | 0.23494 | 4.25593 | 0.00000 | 1.82528 | 1.46420 | 0.00000 | 0.93707 | 3.53554 |
| **ZNF174** | 0.00086 | 5.71437 | 1.34563 | 1.62630 | 0.79352 | 0.00000 | 2.55333 | 6.10664 | 3.45022 | 0.00000 | 2.86891 | 7.62550 |
| **ZNF175** | 0.36341 | 0.38084 | 0.44617 | 0.30915 | 0.00000 | 0.96732 | 0.22554 | 1.36745 | 0.47267 | 0.77968 | 0.07815 | 1.28088 |
| **ZNF177.uid-232** | 0.00574 | 0.03542 | 0.01365 | 0.01555 | 0.01566 | 0.01206 | 0.01816 | 0.01866 | 0.02143 | 0.02030 | 0.01654 | 0.02156 |
| **ZNF18** | 1.38237 | 2.72522 | 0.62004 | 3.63470 | 0.00000 | 1.97584 | 0.00000 | 2.30575 | 0.00000 | 0.73098 | 0.45216 | 3.52397 |
| **ZNF180** | 0.26153 | 2.37920 | 0.66916 | 1.32223 | 0.00000 | 7.11749 | 1.77956 | 3.12153 | 4.41348 | 0.00000 | 1.88847 | 3.97290 |
| **ZNF181** | 0.33675 | 0.22924 | 0.00000 | 0.27085 | 0.00000 | 1.17785 | 0.00762 | 0.00000 | 0.00000 | 0.07262 | 0.35654 | 0.01530 |
| **ZNF182** | 0.84965 | 0.63078 | 0.00000 | 0.00000 | 0.00000 | 1.86000 | 1.60624 | 4.31117 | 0.06997 | 1.15887 | 1.44846 | 0.90985 |
| **ZNF184** | 0.00347 | 1.31025 | 1.15705 | 0.03575 | 0.40134 | 1.28286 | 0.60730 | 6.45619 | 0.78668 | 0.00000 | 0.88624 | 3.71419 |
| **ZNF185** | 0.87105 | 0.24505 | 0.30644 | 0.00094 | 0.27602 | 0.00000 | 0.00967 | 0.13203 | 0.16985 | 0.00059 | 0.14328 | 0.00000 |
| **ZNF189** | 0.43001 | 0.50718 | 1.14691 | 0.33871 | 0.00000 | 1.02859 | 0.38159 | 2.13450 | 0.56701 | 1.92715 | 0.15701 | 1.33984 |
| **ZNF19** | 1.16820 | 1.63106 | 0.00000 | 0.00000 | 0.58569 | 0.40501 | 0.00000 | 0.28044 | 0.00000 | 0.03020 | 0.11415 | 3.95163 |
| **ZNF192P1** | 0.30548 | 0.17905 | 0.00156 | 0.00114 | 0.00000 | 0.45297 | 0.00001 | 0.12906 | 0.00000 | 0.01369 | 0.00404 | 0.15227 |
| **ZNF195** | 1.82776 | 9.77177 | 8.87471 | 5.76488 | 0.00000 | 19.59735 | 13.12077 | 3.71984 | 13.57766 | 4.53749 | 7.08071 | 9.27749 |
| **ZNF197** | 0.23318 | 0.86068 | 0.22553 | 0.45585 | 0.45081 | 0.23011 | 0.00000 | 0.13254 | 0.17069 | 0.07795 | 0.16708 | 1.19351 |
| **ZNF197-AS1** | 0.01737 | 0.13187 | 0.02103 | 0.00509 | 0.00000 | 0.02796 | 0.01779 | 0.00839 | 0.01239 | 0.01663 | 0.01710 | 0.00749 |
| **ZNF2** | 0.43391 | 1.48436 | 1.00940 | 0.35011 | 0.08487 | 0.00000 | 0.00844 | 2.44500 | 0.86069 | 0.70384 | 1.64510 | 2.18073 |
| **ZNF20** | 0.01083 | 0.02173 | 0.01605 | 0.02414 | 0.00000 | 0.03732 | 0.00955 | 0.01393 | 0.01049 | 0.05942 | 0.02937 | 0.02859 |
| **ZNF200** | 4.45827 | 9.87622 | 4.32204 | 5.41909 | 0.00000 | 7.47091 | 8.07032 | 7.08712 | 7.13971 | 1.34735 | 4.33677 | 18.81075 |
| **ZNF202** | 2.97448 | 2.32206 | 1.28288 | 1.77710 | 2.22568 | 0.52773 | 2.39719 | 4.10919 | 1.96415 | 2.70429 | 1.59758 | 4.12800 |
| **ZNF204P** | 0.01186 | 0.00000 | 0.97731 | 0.43002 | 0.00000 | 0.00397 | 0.36004 | 0.01045 | 0.74616 | 0.15555 | 0.29766 | 0.00792 |
| **ZNF205** | 0.01341 | 0.36736 | 0.17313 | 0.23023 | 0.51963 | 0.00000 | 0.17029 | 0.01382 | 0.11772 | 1.00147 | 0.11485 | 0.04488 |
| **ZNF205-AS1** | 0.01965 | 0.01786 | 0.03785 | 0.01454 | 0.04283 | 0.01364 | 0.02105 | 0.01047 | 0.01739 | 0.00000 | 0.03838 | 0.07462 |
| **ZNF207** | 2.72552 | 20.25973 | 17.86647 | 14.33992 | 18.38730 | 38.15794 | 28.37697 | 19.77155 | 26.18436 | 3.40138 | 16.50857 | 27.95142 |
| **ZNF208** | 0.48314 | 1.46687 | 2.15246 | 2.76217 | 0.00010 | 8.42868 | 0.00000 | 2.26496 | 5.78812 | 22.96077 | 4.29385 | 2.60596 |

| | | | | | | | | | | | | |
|---|---|---|---|---|---|---|---|---|---|---|---|---|
| **ZNF211** | 1.51567 | 5.14165 | 3.75192 | 3.21184 | 2.68386 | 0.15046 | 1.65671 | 8.24667 | 4.61010 | 0.00000 | 1.88646 | 4.25315 |
| **ZNF212** | 0.19862 | 7.91537 | 1.49084 | 1.66945 | 3.11210 | 0.61817 | 4.49725 | 4.50187 | 2.78098 | 1.21338 | 2.43470 | 11.21822 |
| **ZNF213** | 0.09896 | 0.10671 | 0.13926 | 0.16265 | 0.00000 | 0.19831 | 0.05892 | 0.02341 | 0.19472 | 0.36014 | 0.18401 | 0.31740 |
| **ZNF213-AS1** | 0.02548 | 0.49091 | 0.25557 | 0.13133 | 0.14088 | 0.00000 | 0.29299 | 0.24708 | 0.85880 | 0.00069 | 0.92244 | 1.21822 |
| **ZNF214** | 3.42891 | 0.28474 | 0.39169 | 0.02562 | 0.00000 | 0.56364 | 0.00000 | 1.61695 | 0.76592 | 0.18291 | 0.00000 | 0.92871 |
| **ZNF215** | 5.99070 | 1.72212 | 2.90348 | 1.00012 | 0.00000 | 6.12124 | 0.00354 | 3.79077 | 3.94177 | 1.91319 | 1.95064 | 4.82760 |
| **ZNF217** | 3.94261 | 2.77182 | 4.93896 | 6.04680 | 0.00025 | 12.01129 | 6.74623 | 2.06854 | 0.00000 | 12.80400 | 4.25386 | 11.40837 |
| **ZNF219** | 0.02355 | 0.02429 | 0.77515 | 0.47396 | 0.06206 | 0.06016 | 0.00000 | 0.00623 | 0.00001 | 1.23155 | 0.42828 | 0.03759 |
| **ZNF22** | 2.21686 | 6.49970 | 0.00000 | 4.67596 | 5.60108 | 9.34613 | 0.00000 | 3.68311 | 0.00000 | 0.74542 | 1.71047 | 13.35868 |
| **ZNF221** | 0.05875 | 0.47284 | 0.35712 | 0.03078 | 0.00000 | 0.00000 | 0.00000 | 1.53451 | 0.37847 | 0.28587 | 0.11795 | 0.71197 |
| **ZNF222** | 2.10559 | 0.65391 | 1.99634 | 0.26738 | 0.00000 | 0.83338 | 0.00000 | 0.10747 | 2.44290 | 0.07652 | 0.00000 | 1.04538 |
| **ZNF223** | 0.58969 | 0.50718 | 0.06296 | 0.16535 | 0.00001 | 1.07479 | 0.00000 | 0.27494 | 0.57331 | 0.00000 | 0.42280 | 0.12568 |
| **ZNF224** | 0.00193 | 0.41214 | 0.20465 | 0.04649 | 0.13154 | 0.07189 | 0.00000 | 2.21741 | 0.13643 | 0.00000 | 0.27931 | 0.25836 |
| **ZNF225** | 0.00386 | 0.98331 | 0.08522 | 0.53081 | 0.00000 | 1.61016 | 0.00000 | 0.63303 | 0.60317 | 0.00000 | 0.68002 | 1.88383 |
| **ZNF226** | 0.46062 | 5.40937 | 0.96771 | 1.50812 | 0.15577 | 1.71604 | 0.00000 | 3.65465 | 3.68100 | 0.00000 | 0.14867 | 7.60772 |
| **ZNF227** | 3.29087 | 5.17604 | 3.11210 | 0.21734 | 0.00000 | 1.83392 | 3.29615 | 0.84444 | 7.79997 | 0.00000 | 2.61930 | 6.56064 |
| **ZNF229** | 3.24247 | 3.32369 | 2.00590 | 0.94078 | 0.00000 | 1.43521 | 1.87813 | 3.32004 | 1.52275 | 0.14943 | 0.00000 | 4.30835 |
| **ZNF23** | 0.26393 | 4.09078 | 0.16332 | 0.76440 | 1.24776 | 2.26459 | 0.00000 | 5.71631 | 1.62709 | 1.77383 | 0.79299 | 7.25481 |
| **ZNF230** | 0.00394 | 1.72412 | 0.32936 | 0.02306 | 0.00000 | 2.02339 | 0.63191 | 3.29274 | 0.54169 | 0.00000 | 0.00076 | 4.18236 |
| **ZNF232** | 1.43551 | 7.87071 | 5.89964 | 6.92119 | 3.52794 | 5.42844 | 9.05365 | 0.39967 | 8.52333 | 0.00000 | 10.29653 | 17.10887 |
| **ZNF233** | 0.28879 | 0.15049 | 0.55480 | 0.00000 | 0.99813 | 0.00000 | 0.21157 | 0.00000 | 0.15760 | 0.24424 | 0.12364 | 0.56888 |
| **ZNF234** | 0.00057 | 1.42634 | 1.02509 | 0.00001 | 0.00000 | 3.95388 | 2.02151 | 0.76108 | 4.84064 | 0.00000 | 0.30158 | 2.29275 |
| **ZNF235** | 0.11129 | 1.56927 | 1.75979 | 1.14402 | 0.04897 | 0.40538 | 0.33460 | 0.50661 | 3.28961 | 0.00000 | 1.02012 | 4.12350 |
| **ZNF236** | 1.10638 | 0.25924 | 2.41400 | 2.87137 | 2.66479 | 2.41499 | 4.70954 | 0.24763 | 2.92164 | 7.46088 | 1.55924 | 3.85083 |
| **ZNF239** | 0.64758 | 0.22523 | 0.11903 | 0.94112 | 0.00000 | 0.58999 | 1.03144 | 0.76637 | 0.51776 | 0.00000 | 0.76298 | 1.70327 |
| **ZNF24** | 1.21973 | 5.04997 | 4.88374 | 3.60872 | 3.59644 | 19.15189 | 5.39536 | 8.76929 | 9.02875 | 13.26485 | 2.80631 | 12.19971 |
| **ZNF248** | 0.04666 | 0.80471 | 1.76160 | 0.07031 | 0.00000 | 0.39534 | 0.06769 | 0.12743 | 2.32555 | 0.02138 | 1.64254 | 0.37218 |
| **ZNF25** | 1.12527 | 0.02679 | 0.09281 | 0.02134 | 0.00375 | 0.00000 | 0.00221 | 0.11626 | 0.05733 | 0.04430 | 0.03250 | 0.00775 |
| **ZNF250** | 0.39208 | 0.93436 | 0.00000 | 0.64543 | 0.00231 | 2.11043 | 0.39101 | 1.34690 | 0.17790 | 0.00000 | 1.16656 | 1.47615 |
| **ZNF251** | 2.34027 | 2.64552 | 1.57212 | 1.45377 | 1.24934 | 0.49846 | 1.97204 | 1.63680 | 1.07800 | 0.39484 | 1.85590 | 2.01366 |
| **ZNF252P** | 2.29203 | 1.28511 | 0.21182 | 0.80150 | 1.00796 | 0.27229 | 0.00000 | 2.83978 | 0.49196 | 0.00000 | 1.03738 | 3.12721 |
| **ZNF252P-AS1** | 0.25154 | 0.02225 | 0.00000 | 0.00596 | 0.04298 | 0.13359 | 0.00000 | 0.08533 | 0.00000 | 0.05551 | 0.02757 | 0.13018 |
| **ZNF253** | 0.37079 | 0.85391 | 0.73720 | 0.59131 | 0.00000 | 3.47000 | 0.00000 | 1.34107 | 4.85567 | 0.00000 | 2.05923 | 2.20273 |
| **ZNF254** | 0.00000 | 3.85231 | 5.84415 | 1.83372 | 0.00000 | 12.65056 | 2.42046 | 6.30060 | 8.06883 | 2.68341 | 2.57662 | 5.36594 |
| **ZNF256** | 1.72693 | 7.59194 | 0.82543 | 0.41500 | 0.46437 | 2.02704 | 0.77450 | 12.37919 | 4.93702 | 0.00000 | 0.00002 | 11.98473 |
| **ZNF257** | 0.00000 | 5.06730 | 1.96251 | 2.22641 | 0.00000 | 12.38173 | 0.00000 | 1.15941 | 5.85406 | 7.74269 | 3.11237 | 5.28327 |
| **ZNF26.uid-1752** | 0.00000 | 3.10192 | 1.43974 | 1.13015 | 0.04950 | 1.87435 | 0.68829 | 3.40771 | 1.49389 | 1.91166 | 0.53617 | 2.23390 |
| **ZNF260** | 1.15208 | 5.59463 | 1.91437 | 2.98800 | 0.00000 | 13.52367 | 0.00000 | 3.26187 | 7.64257 | 3.53499 | 1.13760 | 11.76648 |
| **ZNF263** | 0.00000 | 3.93410 | 1.64168 | 4.69209 | 3.14771 | 0.13199 | 5.29706 | 4.54941 | 4.61660 | 11.04882 | 0.14692 | 5.95770 |
| **ZNF264** | 0.07680 | 0.00000 | 0.94960 | 0.25940 | 0.27249 | 2.33730 | 2.10589 | 1.30915 | 0.52468 | 6.16094 | 0.41901 | 2.37146 |
| **ZNF266** | 1.80535 | 5.95240 | 1.27891 | 1.66315 | 0.61774 | 4.04275 | 1.36608 | 7.01255 | 3.66209 | 0.76000 | 1.06761 | 7.29995 |
| **ZNF267** | 1.89347 | 7.45490 | 2.68326 | 2.06057 | 0.32199 | 21.14375 | 2.57566 | 10.44726 | 5.95547 | 4.30327 | 3.02688 | 9.84837 |
| **ZNF268** | 0.91434 | 9.71649 | 2.19114 | 1.43331 | 0.00000 | 5.90895 | 2.98676 | 10.57637 | 2.44421 | 2.48970 | 1.18908 | 15.96823 |
| **ZNF271** | 0.99652 | 4.85685 | 2.39050 | 0.94107 | 0.98261 | 12.40214 | 6.24829 | 3.41983 | 5.43658 | 5.85921 | 4.60343 | 8.31944 |
| **ZNF273** | 0.59801 | 4.15335 | 1.58049 | 2.35395 | 0.00000 | 6.46387 | 0.82475 | 3.95747 | 3.25667 | 1.28008 | 1.29771 | 6.38171 |
| **ZNF274** | 1.50300 | 6.48714 | 1.06229 | 0.90565 | 0.00000 | 6.31588 | 1.25292 | 1.75454 | 8.26419 | 0.89097 | 0.57963 | 5.64754 |
| **ZNF275** | 0.08913 | 0.05292 | 0.95968 | 0.26116 | 0.09094 | 0.04780 | 0.66505 | 0.00000 | 0.00000 | 0.06316 | 0.07907 | 0.52768 |
| **ZNF276** | 0.04872 | 0.89854 | 0.03445 | 2.10141 | 0.44806 | 0.19953 | 1.23661 | 0.47807 | 2.59519 | 3.67504 | 0.28049 | 0.65598 |

| | | | | | | | | | | | | |
|---|---|---|---|---|---|---|---|---|---|---|---|---|
| ZNF277 | 1.91981 | 1.86704 | 15.47212 | 0.59720 | 3.80703 | 12.85967 | 2.19930 | 0.32545 | 6.47478 | 3.08768 | 1.82682 | 2.11565 |
| ZNF28 | 0.00000 | 8.86392 | 4.53848 | 4.13169 | 0.00000 | 16.96615 | 3.83351 | 6.65116 | 7.54120 | 5.48258 | 4.51745 | 11.00244 |
| ZNF280A | 1.48960 | 4.48362 | 1.27916 | 1.51648 | 0.00000 | 0.72622 | 0.00000 | 42.53613 | 0.40628 | 0.00000 | 0.21768 | 28.70137 |
| ZNF280B | 12.74706 | 2.90125 | 0.96077 | 4.24692 | 0.00000 | 0.53747 | 0.23459 | 23.49022 | 0.00000 | 0.00000 | 0.36748 | 4.89941 |
| ZNF280C | 4.41138 | 0.13005 | 0.23809 | 0.00000 | 0.00000 | 2.06447 | 0.00000 | 1.67575 | 0.92964 | 0.54870 | 0.45188 | 0.57727 |
| ZNF280D | 1.38660 | 0.74883 | 0.03430 | 0.79817 | 0.76633 | 0.29166 | 0.00652 | 0.18158 | 0.10143 | 0.79710 | 0.28526 | 0.37611 |
| ZNF281 | 5.88382 | 6.56599 | 7.47222 | 9.63507 | 3.81206 | 21.53145 | 12.15858 | 12.88956 | 6.80008 | 23.20240 | 11.14006 | 15.01852 |
| ZNF282 | 0.85582 | 0.85381 | 1.09152 | 1.52511 | 1.59169 | 0.00000 | 1.25257 | 0.00000 | 1.26017 | 3.88111 | 0.71372 | 0.15238 |
| ZNF283 | 0.26260 | 0.99306 | 0.61489 | 0.00411 | 0.00000 | 1.37920 | 1.74824 | 0.38093 | 1.33529 | 0.00000 | 0.33197 | 1.10095 |
| ZNF284 | 0.19986 | 0.23742 | 0.05727 | 0.00101 | 0.00000 | 0.18694 | 0.19525 | 1.22044 | 0.45239 | 0.00000 | 0.21459 | 0.74581 |
| ZNF285 | 1.41842 | 1.76558 | 0.18146 | 0.00000 | 0.00000 | 0.39403 | 0.00000 | 0.58790 | 0.41767 | 0.25976 | 0.21369 | 1.14999 |
| ZNF286A | 0.32261 | 1.31439 | 3.87982 | 1.84408 | 0.00000 | 5.16797 | 4.28122 | 3.38399 | 4.46764 | 2.09997 | 3.34183 | 2.67496 |
| ZNF286B | 0.00170 | 0.04190 | 0.62768 | 0.23590 | 0.00000 | 0.23208 | 1.67802 | 1.29416 | 0.53038 | 0.99768 | 0.81691 | 0.28895 |
| ZNF287 | 0.00324 | 0.19206 | 0.31385 | 0.09930 | 0.00646 | 0.11787 | 0.00000 | 0.31114 | 0.00010 | 0.00000 | 0.06737 | 1.11528 |
| ZNF292 | 0.44180 | 0.86900 | 18.11504 | 4.92608 | 0.00000 | 8.26464 | 9.74369 | 4.16200 | 4.88326 | 15.85068 | 1.82538 | 5.06007 |
| ZNF295-AS1 | 0.10042 | 0.04847 | 0.03620 | 0.00350 | 0.00140 | 0.04004 | 0.01360 | 0.06549 | 0.05711 | 0.02907 | 0.01161 | 0.05493 |
| ZNF296 | 10.15885 | 19.77704 | 2.49144 | 12.99424 | 19.89273 | 0.00000 | 4.52799 | 17.36132 | 8.32413 | 1.52542 | 9.07697 | 26.79732 |
| ZNF3 | 1.25579 | 7.38244 | 3.33486 | 5.61475 | 5.19541 | 6.94417 | 6.17129 | 0.88832 | 5.94493 | 7.06768 | 3.96806 | 4.38392 |
| ZNF30 | 0.27490 | 2.69017 | 0.08466 | 0.54966 | 0.00000 | 0.81254 | 0.00000 | 1.29583 | 0.00000 | 0.30635 | 0.13053 | 2.63532 |
| ZNF300 | 0.39239 | 1.23686 | 2.50107 | 0.00000 | 0.00000 | 1.08392 | 1.13923 | 4.17653 | 1.52905 | 4.30840 | 0.34456 | 1.54422 |
| ZNF300P1 | 0.00283 | 0.15945 | 0.84952 | 0.00000 | 0.00000 | 1.63235 | 0.89233 | 0.40253 | 1.57473 | 0.05068 | 0.44873 | 0.56831 |
| ZNF302 | 0.22741 | 3.48948 | 0.11278 | 0.28670 | 0.00009 | 2.90526 | 0.00000 | 0.98240 | 2.47009 | 1.85506 | 0.93234 | 3.16334 |
| ZNF304 | 2.96103 | 5.43030 | 0.84966 | 1.15021 | 2.00261 | 1.61873 | 3.23244 | 6.62502 | 1.82438 | 1.00445 | 1.59509 | 7.33108 |
| ZNF311 | 0.00245 | 1.39964 | 0.00000 | 0.75305 | 0.11393 | 0.02410 | 0.00000 | 1.52145 | 0.00000 | 0.24678 | 0.00778 | 1.50713 |
| ZNF316 | 0.17692 | 0.00000 | 0.38747 | 0.27236 | 0.32906 | 0.00000 | 1.18929 | 0.03588 | 0.54945 | 2.28959 | 0.03284 | 0.16184 |
| ZNF317 | 0.00000 | 4.44088 | 4.04324 | 4.05701 | 3.15132 | 0.51112 | 6.68388 | 0.84595 | 9.19956 | 6.13587 | 2.75352 | 2.62376 |
| ZNF318 | 3.56927 | 2.04983 | 0.27985 | 5.91345 | 2.79464 | 5.59364 | 1.35876 | 1.81036 | 2.30163 | 10.55825 | 0.09479 | 6.59651 |
| ZNF319 | 0.19039 | 0.00000 | 0.11976 | 0.26016 | 0.04547 | 0.05564 | 0.11468 | 0.00865 | 0.00000 | 0.11610 | 0.00000 | 0.02757 |
| ZNF32 | 0.05699 | 1.21883 | 0.39056 | 0.83397 | 0.00000 | 1.94885 | 0.28757 | 0.00000 | 1.34400 | 0.00000 | 1.57420 | 0.74391 |
| ZNF32-AS1 | 0.00600 | 0.01706 | 0.01300 | 0.01365 | 0.00972 | 0.01979 | 0.02149 | 0.01048 | 0.02388 | 0.02046 | 0.04677 | 0.01524 |
| ZNF32-AS2 | 0.01682 | 0.07793 | 0.00764 | 0.04589 | 0.00046 | 0.05163 | 0.01581 | 0.00701 | 0.15366 | 0.01543 | 0.07553 | 0.08628 |
| ZNF32-AS3 | 0.00597 | 0.02817 | 0.01573 | 0.00894 | 0.01168 | 0.03515 | 0.01443 | 0.01021 | 0.01389 | 0.01772 | 0.01359 | 0.01639 |
| ZNF320 | 0.00000 | 3.64450 | 3.71911 | 2.48275 | 2.03371 | 6.67730 | 0.93565 | 1.39446 | 0.00610 | 9.27164 | 6.28528 | 2.44391 |
| ZNF321P | 0.00415 | 0.05066 | 0.07929 | 0.06253 | 0.07398 | 0.00000 | 0.01435 | 0.06766 | 0.13680 | 0.05174 | 0.02492 | 0.07055 |
| ZNF322 | 0.00203 | 0.30659 | 0.31350 | 0.11277 | 0.00000 | 0.84650 | 0.19982 | 0.51809 | 0.33950 | 0.04763 | 0.23619 | 0.35717 |
| ZNF324 | 2.19484 | 5.13360 | 0.70366 | 3.12067 | 3.21334 | 0.00000 | 1.83135 | 2.29157 | 4.51254 | 4.72750 | 1.55306 | 3.48556 |
| ZNF324B | 0.21327 | 0.53379 | 0.17463 | 0.18477 | 0.20591 | 0.00000 | 0.22280 | 0.01796 | 0.21193 | 0.45823 | 0.31037 | 0.68165 |
| ZNF326 | 1.38568 | 1.22341 | 14.09000 | 2.70753 | 6.64985 | 14.99390 | 12.15734 | 0.84163 | 6.25915 | 10.84202 | 6.74085 | 15.67136 |
| ZNF329 | 2.03507 | 1.27808 | 0.31370 | 0.27439 | 0.00000 | 0.00000 | 0.04936 | 0.34062 | 0.53583 | 0.01620 | 0.00000 | 2.68170 |
| ZNF330 | 4.90172 | 9.51781 | 6.55690 | 2.64922 | 8.12212 | 14.70547 | 7.56427 | 10.67398 | 4.79986 | 2.80604 | 6.55572 | 11.98136 |
| ZNF331 | 0.00000 | 4.46108 | 11.90032 | 3.72549 | 3.25298 | 5.65865 | 2.00278 | 0.88223 | 16.09959 | 10.66776 | 1.28096 | 4.89111 |
| ZNF333 | 0.15167 | 0.00000 | 0.45273 | 0.84464 | 0.00356 | 0.24885 | 0.00000 | 0.08583 | 0.71460 | 1.19735 | 0.69829 | 0.37558 |
| ZNF334 | 0.00000 | 1.11743 | 0.27641 | 0.40657 | 0.00000 | 0.99677 | 0.00000 | 0.01022 | 0.01367 | 0.93732 | 0.02688 | 1.23276 |
| ZNF335 | 2.08715 | 8.17080 | 10.45877 | 10.75240 | 5.77019 | 0.00000 | 4.55201 | 4.11252 | 12.89470 | 13.99525 | 3.16386 | 7.77867 |
| ZNF337 | 1.49564 | 3.72094 | 0.50538 | 0.83271 | 0.00000 | 3.94699 | 1.34847 | 0.62308 | 2.16767 | 0.39040 | 1.80960 | 4.24460 |
| ZNF33A | 0.17892 | 1.02819 | 1.07206 | 1.38779 | 0.18425 | 4.41652 | 2.09096 | 0.21365 | 2.77528 | 5.52356 | 2.53995 | 2.98354 |
| ZNF33B | 0.00000 | 0.96899 | 0.77252 | 0.00000 | 0.00000 | 1.70025 | 1.01181 | 0.87252 | 1.27767 | 1.14452 | 1.27204 | 0.72298 |
| ZNF33BP1 | 0.00770 | 0.01450 | 0.01715 | 0.01264 | 0.01562 | 0.00377 | 0.02441 | 0.01033 | 0.00201 | 0.02628 | 0.08508 | 0.01755 |

| | | | | | | | | | | | | |
|---|---|---|---|---|---|---|---|---|---|---|---|---|
| **ZNF34** | 0.55925 | 0.53360 | 0.57443 | 3.02773 | 1.72523 | 0.00000 | 0.00000 | 1.18632 | 0.67938 | 1.29681 | 0.00000 | 1.81107 |
| **ZNF341** | 0.15085 | 0.74091 | 2.40186 | 0.99765 | 0.67130 | 0.05436 | 4.36868 | 0.22085 | 4.75961 | 0.25981 | 1.27250 | 1.25483 |
| **ZNF341-AS1** | 0.00627 | 0.01862 | 0.03966 | 0.01962 | 0.01483 | 0.00000 | 0.02593 | 0.01068 | 0.05694 | 0.01467 | 0.03091 | 0.01726 |
| **ZNF343** | 3.02856 | 7.58744 | 0.85057 | 2.07099 | 0.00000 | 3.52393 | 0.00000 | 7.33439 | 2.64454 | 1.10016 | 3.37709 | 6.21347 |
| **ZNF345** | 0.60074 | 0.71517 | 1.44909 | 0.13414 | 0.19502 | 1.42305 | 0.26483 | 1.22905 | 1.99101 | 0.61940 | 0.99796 | 2.08223 |
| **ZNF346** | 3.67840 | 4.54441 | 3.39757 | 2.60537 | 3.52111 | 0.17637 | 4.32334 | 1.24324 | 4.26753 | 3.33052 | 4.24987 | 1.96662 |
| **ZNF347** | 0.00845 | 1.94526 | 1.05440 | 0.02290 | 0.43846 | 2.11436 | 0.00000 | 1.10100 | 0.54604 | 1.35435 | 0.74007 | 1.81415 |
| **ZNF35** | 0.17713 | 0.00000 | 0.21579 | 0.24114 | 0.00799 | 0.74010 | 0.00000 | 0.38777 | 0.09353 | 0.07713 | 0.25190 | 0.24053 |
| **ZNF350** | 0.59755 | 9.77204 | 0.78420 | 1.10328 | 0.00000 | 4.53543 | 0.00006 | 15.22015 | 1.27554 | 2.18691 | 1.44273 | 11.85468 |
| **ZNF350-AS1** | 0.03174 | 0.18034 | 0.21038 | 0.13114 | 0.00000 | 0.00000 | 0.24079 | 0.47574 | 0.00070 | 0.15045 | 0.12419 | 0.30957 |
| **ZNF354A** | 1.47459 | 1.06098 | 0.00000 | 0.59289 | 0.39326 | 3.88602 | 0.65614 | 0.67713 | 0.46360 | 1.81041 | 0.80802 | 3.56610 |
| **ZNF354B** | 0.27465 | 0.08228 | 0.03304 | 0.18858 | 0.00000 | 0.26698 | 0.22670 | 0.20875 | 0.00000 | 1.07292 | 0.03265 | 0.54535 |
| **ZNF354C** | 0.00425 | 0.01051 | 0.00306 | 0.04711 | 0.03291 | 0.00000 | 0.01017 | 0.22825 | 0.29161 | 0.00000 | 0.03279 | 0.15852 |
| **ZNF358** | 0.07295 | 0.06921 | 0.00851 | 0.23095 | 0.40593 | 0.00001 | 0.64505 | 0.00467 | 0.00084 | 0.03660 | 0.11383 | 0.17024 |
| **ZNF362** | 1.97131 | 0.12745 | 0.54344 | 0.00000 | 0.80512 | 0.06673 | 3.24881 | 0.05988 | 0.84723 | 0.25347 | 0.69779 | 0.03489 |
| **ZNF365** | 0.17123 | 0.04974 | 0.09345 | 0.02697 | 0.01099 | 0.00000 | 0.00732 | 0.00000 | 0.03826 | 0.00563 | 0.03013 | 0.34878 |
| **ZNF366** | 0.00577 | 0.01001 | 0.02283 | 0.00703 | 0.08653 | 0.01552 | 0.01355 | 0.01036 | 0.01287 | 0.03437 | 0.01364 | 0.01613 |
| **ZNF367** | 0.07346 | 0.24910 | 0.60762 | 0.53289 | 0.80679 | 1.04463 | 0.43111 | 0.00000 | 0.00000 | 0.73510 | 0.67568 | 0.00000 |
| **ZNF37A** | 0.00000 | 0.91267 | 2.90502 | 1.60059 | 0.50879 | 5.05229 | 4.35988 | 0.74503 | 6.32075 | 8.00322 | 3.17084 | 2.43870 |
| **ZNF37BP** | 0.00001 | 0.46609 | 2.09237 | 0.50771 | 0.00000 | 0.69100 | 0.32728 | 0.22112 | 3.40075 | 2.76980 | 2.08029 | 1.87587 |
| **ZNF382** | 0.12596 | 1.40479 | 0.13435 | 0.00000 | 0.06183 | 0.16265 | 0.30392 | 0.05284 | 0.42414 | 0.05278 | 0.82675 | 1.49404 |
| **ZNF383** | 0.11369 | 1.20473 | 0.10950 | 0.64887 | 0.00000 | 1.44265 | 0.00000 | 0.00000 | 2.68452 | 0.82281 | 0.90779 | 2.00192 |
| **ZNF384** | 1.64679 | 7.57317 | 6.74414 | 8.90977 | 8.07762 | 1.89156 | 6.32769 | 4.52380 | 10.38572 | 2.01478 | 5.20898 | 10.61267 |
| **ZNF385A** | 0.04634 | 0.00000 | 3.01475 | 0.00000 | 2.08937 | 0.00000 | 4.28202 | 0.00000 | 5.79483 | 1.77629 | 0.35314 | 0.44387 |
| **ZNF385B** | 1.05437 | 0.29387 | 0.06670 | 0.14984 | 0.18060 | 0.08456 | 0.02433 | 0.15692 | 0.01013 | 0.00000 | 0.04182 | 0.20278 |
| **ZNF385C** | 0.07503 | 0.06194 | 0.04067 | 0.00835 | 0.00000 | 0.00000 | 0.08141 | 0.01811 | 0.00000 | 0.10299 | 0.16927 | 0.02618 |
| **ZNF385D** | 0.06998 | 0.02921 | 0.03108 | 0.01619 | 0.00000 | 0.02128 | 0.04051 | 0.03913 | 0.01928 | 0.08474 | 0.01876 | 0.01013 |
| **ZNF385D-AS1** | 1.57988 | 0.39012 | 0.32640 | 1.02993 | 0.00000 | 1.29394 | 0.00000 | 0.09386 | 0.84286 | 0.08746 | 0.34059 | 0.52691 |
| **ZNF391** | 0.75173 | 0.53935 | 0.42828 | 0.63192 | 0.08715 | 0.13039 | 0.16985 | 0.14176 | 0.63463 | 0.00000 | 0.42649 | 1.10820 |
| **ZNF394** | 2.14241 | 10.97765 | 4.64492 | 4.75573 | 3.84451 | 6.20149 | 3.66501 | 27.63103 | 4.91817 | 0.00000 | 3.20574 | 14.77503 |
| **ZNF395** | 3.25375 | 2.19882 | 0.31149 | 1.01823 | 2.74402 | 0.00180 | 2.21691 | 0.57437 | 0.00000 | 1.82707 | 1.33274 | 0.45532 |
| **ZNF396** | 0.01180 | 0.05859 | 0.05554 | 0.03108 | 0.05548 | 0.06264 | 0.00233 | 0.03158 | 0.02033 | 0.00000 | 0.04982 | 0.05540 |
| **ZNF397** | 2.92616 | 1.69902 | 1.11042 | 0.95253 | 2.11968 | 2.84975 | 0.15146 | 1.63835 | 1.42115 | 2.22602 | 1.52937 | 4.88642 |
| **ZNF398** | 1.21715 | 4.24154 | 0.41546 | 4.67616 | 4.31479 | 4.98389 | 0.82941 | 1.32468 | 0.00000 | 4.73627 | 0.71746 | 7.56352 |
| **ZNF404** | 0.51956 | 0.65012 | 0.45639 | 0.00658 | 0.02531 | 0.29603 | 0.15428 | 0.14066 | 0.26159 | 0.00000 | 0.15137 | 0.05744 |
| **ZNF407** | 1.44864 | 0.88831 | 3.10082 | 0.30349 | 0.83922 | 1.21452 | 0.00000 | 3.27766 | 3.53214 | 6.01900 | 1.18319 | 3.70172 |
| **ZNF408** | 1.07393 | 9.47969 | 5.11468 | 5.29722 | 7.24617 | 0.00000 | 2.08914 | 14.23124 | 6.52785 | 1.67273 | 3.91292 | 13.35479 |
| **ZNF41** | 0.16318 | 0.70861 | 0.35624 | 0.00611 | 0.00000 | 0.93608 | 0.69272 | 0.04067 | 1.55068 | 0.00003 | 1.80798 | 1.11464 |
| **ZNF410** | 2.69725 | 7.04365 | 11.25665 | 2.18062 | 6.55123 | 21.34973 | 12.55233 | 19.36745 | 17.13114 | 0.00000 | 9.97377 | 12.77957 |
| **ZNF414** | 0.42926 | 0.61967 | 0.33946 | 0.00000 | 0.67594 | 0.00000 | 0.03874 | 0.07158 | 0.18219 | 1.04564 | 0.36719 | 0.31448 |
| **ZNF415** | 1.27803 | 3.05339 | 2.53067 | 0.10847 | 1.50967 | 4.13309 | 0.00000 | 0.80407 | 3.82968 | 0.00000 | 1.15767 | 3.97995 |
| **ZNF416** | 2.15279 | 6.72772 | 0.32755 | 1.27513 | 0.00000 | 2.88937 | 1.76088 | 10.38342 | 1.27307 | 0.00000 | 1.49120 | 12.91067 |
| **ZNF417** | 0.00000 | 0.58937 | 0.46094 | 0.39330 | 0.26880 | 1.42646 | 0.55982 | 0.46857 | 0.40018 | 1.19403 | 0.45689 | 1.27594 |
| **ZNF418** | 0.29183 | 0.48993 | 0.00000 | 0.29567 | 0.42216 | 0.55887 | 0.00000 | 0.56417 | 1.00963 | 0.71100 | 1.15435 | 0.61288 |
| **ZNF419** | 1.53891 | 4.39775 | 1.33045 | 0.73454 | 0.50798 | 2.41565 | 0.99283 | 3.52477 | 2.08902 | 2.94641 | 2.02930 | 4.61446 |
| **ZNF420** | 1.04012 | 1.68515 | 0.71260 | 0.82832 | 0.00000 | 5.31399 | 0.00000 | 0.24191 | 4.63028 | 0.56164 | 2.72927 | 1.35518 |
| **ZNF423** | 0.01615 | 0.04199 | 0.04140 | 0.45786 | 0.00000 | 0.05445 | 0.13886 | 0.00846 | 0.00000 | 0.06762 | 0.00192 | 0.00720 |
| **ZNF425** | 1.15330 | 0.54373 | 0.00000 | 0.37056 | 0.32461 | 0.09928 | 0.23367 | 0.00000 | 0.32518 | 0.00000 | 0.13494 | 0.61240 |

| | | | | | | | | | | | | |
|---|---|---|---|---|---|---|---|---|---|---|---|---|
| **ZNF426** | 4.69090 | 4.12450 | 1.49173 | 1.83404 | 0.58954 | 6.22623 | 2.57961 | 1.05221 | 3.31811 | 0.43320 | 1.24888 | 7.54830 |
| **ZNF428** | 0.38175 | 2.38374 | 1.61849 | 2.50638 | 0.97941 | 0.00000 | 1.09693 | 0.31606 | 0.47157 | 2.11909 | 1.19697 | 2.66788 |
| **ZNF429** | 0.14516 | 2.20739 | 0.03804 | 0.78657 | 0.00000 | 1.93330 | 0.00000 | 0.63688 | 0.60786 | 1.06292 | 0.75980 | 1.49691 |
| **ZNF43** | 0.65354 | 5.99721 | 3.09042 | 3.53745 | 0.46224 | 15.08829 | 1.40779 | 3.14504 | 7.67733 | 9.40517 | 4.58132 | 5.51439 |
| **ZNF430** | 0.76133 | 2.26361 | 2.56820 | 3.83544 | 0.89074 | 4.82999 | 4.69824 | 2.52104 | 5.00780 | 6.74916 | 3.16650 | 1.54571 |
| **ZNF431** | 1.18644 | 1.27979 | 2.60281 | 1.06738 | 0.09675 | 6.23190 | 1.29355 | 0.48900 | 1.62061 | 1.27754 | 1.14593 | 2.21181 |
| **ZNF432** | 0.00073 | 1.47458 | 0.70952 | 1.89542 | 0.00000 | 1.63603 | 0.93555 | 2.90431 | 1.22854 | 0.16037 | 0.27734 | 0.25141 |
| **ZNF433** | 1.54416 | 0.17850 | 1.88930 | 0.01076 | 0.00476 | 3.94572 | 0.00000 | 0.77580 | 1.25583 | 0.00000 | 0.97706 | 0.86695 |
| **ZNF436** | 1.09828 | 0.71128 | 6.45754 | 0.14433 | 0.90677 | 1.44126 | 0.55008 | 6.19289 | 1.62878 | 0.00000 | 1.32138 | 1.41789 |
| **ZNF438** | 1.19610 | 1.16460 | 0.14138 | 0.29844 | 0.00400 | 0.66601 | 0.00000 | 0.91372 | 0.00000 | 0.42314 | 0.35778 | 0.64559 |
| **ZNF439** | 0.04744 | 0.00000 | 2.56454 | 0.00000 | 0.00000 | 0.29717 | 0.77494 | 0.24729 | 0.83135 | 0.00000 | 0.84918 | 0.15679 |
| **ZNF44** | 0.74199 | 4.42395 | 2.81512 | 2.99237 | 2.13505 | 6.30205 | 1.35851 | 1.76350 | 2.68658 | 7.84977 | 1.85537 | 1.94620 |
| **ZNF440** | 0.03316 | 0.04737 | 2.73459 | 0.13082 | 3.34640 | 2.26884 | 2.33470 | 0.00000 | 2.59740 | 0.00152 | 2.24138 | 0.00000 |
| **ZNF441** | 0.20752 | 0.13671 | 0.18636 | 0.23917 | 0.00816 | 0.58581 | 0.00000 | 0.34308 | 0.32400 | 0.38786 | 0.48793 | 1.50202 |
| **ZNF442** | 0.18157 | 0.00000 | 0.07626 | 0.05457 | 0.04639 | 0.00000 | 0.00219 | 0.00000 | 0.00000 | 0.14303 | 0.03577 | 0.16321 |
| **ZNF443** | 1.45394 | 1.97756 | 1.17458 | 0.80222 | 0.00000 | 1.66087 | 0.49246 | 1.23596 | 0.22199 | 0.03851 | 0.64253 | 1.85292 |
| **ZNF444** | 0.16968 | 2.25814 | 2.85205 | 1.35031 | 2.14651 | 0.00000 | 1.79941 | 0.05044 | 3.02919 | 4.41000 | 0.75364 | 0.79722 |
| **ZNF445** | 3.26736 | 1.59426 | 0.51870 | 1.92568 | 1.89123 | 4.01495 | 0.02701 | 0.20890 | 1.93175 | 2.30760 | 6.19190 | 2.90216 |
| **ZNF446** | 0.70200 | 0.90515 | 0.49546 | 1.31871 | 0.51257 | 0.00000 | 1.41625 | 0.05753 | 1.31110 | 0.00000 | 0.97536 | 0.68178 |
| **ZNF449** | 1.96528 | 1.33457 | 0.48401 | 0.59074 | 0.00000 | 1.98159 | 0.00000 | 0.81624 | 0.95557 | 0.00000 | 0.29489 | 1.45188 |
| **ZNF45** | 0.18612 | 3.09615 | 1.42475 | 1.40850 | 0.00000 | 0.71305 | 3.45473 | 2.26403 | 2.07172 | 0.00000 | 0.75733 | 2.60596 |
| **ZNF451** | 3.70123 | 3.11451 | 4.05255 | 6.57751 | 0.00000 | 6.24240 | 4.85036 | 7.37046 | 6.47143 | 7.38482 | 2.29667 | 14.24039 |
| **ZNF454** | 0.00793 | 0.05478 | 0.03270 | 0.09319 | 0.00000 | 0.09230 | 0.10790 | 0.16329 | 0.02800 | 0.05886 | 0.10998 | 0.12224 |
| **ZNF460** | 1.19311 | 2.30395 | 1.72077 | 0.17345 | 0.00000 | 3.53029 | 1.48363 | 1.45437 | 3.95919 | 1.88222 | 4.06948 | 1.01020 |
| **ZNF461** | 1.73896 | 1.96239 | 1.98398 | 0.36355 | 0.00000 | 3.60251 | 0.00000 | 1.05539 | 0.88281 | 0.47318 | 1.56389 | 2.84707 |
| **ZNF462** | 2.92307 | 0.00000 | 0.26317 | 4.05975 | 0.31887 | 0.11919 | 0.00000 | 0.87372 | 0.00000 | 0.28770 | 0.00000 | 1.38872 |
| **ZNF467** | 0.03301 | 0.95451 | 0.00000 | 0.67020 | 0.68382 | 0.17511 | 0.00000 | 0.00525 | 0.00000 | 0.69076 | 0.10489 | 0.00000 |
| **ZNF468** | 0.00000 | 10.29077 | 9.53551 | 2.70360 | 4.28981 | 25.29168 | 17.42003 | 8.51160 | 9.41564 | 0.82764 | 16.94236 | 11.39063 |
| **ZNF469** | 0.00624 | 0.01249 | 0.02488 | 0.01280 | 0.00929 | 0.01500 | 0.02328 | 0.01123 | 0.00000 | 0.02219 | 0.07626 | 0.01785 |
| **ZNF470** | 0.34157 | 1.09255 | 1.17564 | 0.77762 | 0.00000 | 1.28285 | 0.51993 | 0.94261 | 0.93788 | 1.88144 | 0.23215 | 0.32152 |
| **ZNF471** | 0.14900 | 0.42385 | 0.44155 | 0.00000 | 0.00000 | 0.05405 | 1.22489 | 0.00057 | 0.00000 | 0.19597 | 0.07279 | 0.34341 |
| **ZNF473** | 0.82914 | 7.70126 | 2.66293 | 3.50583 | 3.52150 | 3.80468 | 4.48562 | 22.45497 | 7.86965 | 0.00000 | 4.77072 | 11.98400 |
| **ZNF474** | 0.02285 | 0.01587 | 0.02317 | 0.01448 | 0.01774 | 0.00851 | 0.02173 | 0.04069 | 0.02756 | 0.00755 | 0.01633 | 0.02237 |
| **ZNF479** | 0.17309 | 0.01476 | 0.01663 | 0.01357 | 0.01811 | 0.01349 | 0.01684 | 0.43888 | 0.01286 | 0.01628 | 0.02790 | 0.01148 |
| **ZNF48** | 0.00092 | 1.24656 | 0.00000 | 2.48208 | 1.35809 | 0.00000 | 0.53885 | 0.59501 | 0.00000 | 2.87636 | 0.32111 | 0.58321 |
| **ZNF480** | 0.00000 | 6.88435 | 4.42676 | 4.39106 | 3.95688 | 24.39652 | 0.00000 | 0.74609 | 10.68450 | 9.96856 | 4.46512 | 7.10217 |
| **ZNF483** | 0.77296 | 0.01446 | 1.22556 | 0.62601 | 0.44421 | 0.00000 | 0.29503 | 0.21247 | 0.25218 | 0.65916 | 0.71324 | 0.02246 |
| **ZNF484** | 0.06199 | 1.03556 | 0.52946 | 0.93795 | 0.00000 | 1.99481 | 0.35623 | 0.43972 | 0.64657 | 0.07942 | 0.54445 | 2.01950 |
| **ZNF485** | 0.00384 | 1.68610 | 0.20854 | 0.29491 | 0.00535 | 1.46983 | 0.56098 | 1.37919 | 0.59723 | 0.00000 | 2.15651 | 3.37762 |
| **ZNF486** | 1.05996 | 6.64104 | 0.67507 | 5.55879 | 0.00000 | 17.13006 | 0.00000 | 4.99423 | 2.49923 | 9.37909 | 2.23823 | 4.01572 |
| **ZNF487** | 0.00362 | 2.40672 | 0.43757 | 0.00418 | 0.00169 | 1.33874 | 0.00000 | 4.24901 | 0.36349 | 0.00000 | 0.11156 | 3.54045 |
| **ZNF488** | 0.00439 | 0.39762 | 1.51049 | 0.00701 | 0.07279 | 0.18894 | 0.39393 | 0.01026 | 0.02422 | 0.00000 | 0.00000 | 0.05317 |
| **ZNF490** | 1.26149 | 4.58092 | 1.18340 | 5.59116 | 0.00000 | 1.45685 | 1.42842 | 5.72079 | 0.51282 | 1.53757 | 0.60272 | 3.92307 |
| **ZNF491** | 0.06515 | 0.00000 | 0.15968 | 0.00000 | 0.00000 | 0.08260 | 0.09397 | 0.07328 | 0.00000 | 0.13461 | 0.04295 | 0.05638 |
| **ZNF492** | 0.02875 | 0.79534 | 0.79921 | 0.52165 | 0.00000 | 0.41204 | 1.01156 | 0.40536 | 0.91087 | 0.76645 | 0.86592 | 0.59903 |
| **ZNF493** | 0.49358 | 0.98750 | 0.49892 | 0.51669 | 0.00000 | 4.60241 | 0.00000 | 0.79641 | 4.48086 | 2.72950 | 1.16772 | 2.07634 |
| **ZNF496** | 3.29954 | 2.73140 | 0.33173 | 1.94685 | 6.24077 | 0.00000 | 0.00000 | 0.55698 | 4.69729 | 1.37292 | 0.81012 | 0.58122 |
| **ZNF497** | 0.05829 | 0.03671 | 0.04840 | 0.00000 | 0.11939 | 0.14416 | 0.05903 | 0.08016 | 0.01163 | 0.31275 | 0.04595 | 0.40455 |

| | | | | | | | | | | | | |
|---|---|---|---|---|---|---|---|---|---|---|---|---|
| **ZNF500** | 0.43508 | 1.80293 | 0.36551 | 2.41179 | 1.16422 | 0.00000 | 0.36261 | 0.05597 | 5.66480 | 3.72146 | 1.90190 | 1.07350 |
| **ZNF501** | 2.80391 | 1.07086 | 0.16900 | 0.05364 | 0.00150 | 0.75118 | 0.00000 | 2.74452 | 0.09086 | 0.00000 | 0.15255 | 1.55003 |
| **ZNF502** | 2.59804 | 2.11755 | 0.25034 | 0.81058 | 0.00000 | 0.24465 | 0.67478 | 3.45161 | 0.18547 | 0.00000 | 0.72328 | 2.95018 |
| **ZNF503** | 0.98438 | 0.00000 | 0.04535 | 0.38963 | 0.04761 | 0.00000 | 0.01719 | 0.00833 | 0.06903 | 0.18718 | 0.00019 | 0.11927 |
| **ZNF503-AS1** | 0.18850 | 0.03420 | 0.01496 | 0.01064 | 0.01076 | 0.00986 | 0.02168 | 0.06319 | 0.02072 | 0.02292 | 0.01363 | 0.10025 |
| **ZNF503-AS2** | 0.57641 | 0.03918 | 0.10512 | 0.59522 | 0.14920 | 0.09481 | 0.00000 | 0.05663 | 0.08973 | 0.02054 | 0.00000 | 0.00003 |
| **ZNF506** | 4.77941 | 1.46490 | 1.45453 | 1.08340 | 0.00000 | 6.07785 | 0.00000 | 8.67081 | 5.20765 | 3.06715 | 0.60777 | 5.62044 |
| **ZNF507** | 1.09345 | 3.46004 | 1.78618 | 1.55326 | 1.75775 | 6.27465 | 2.43382 | 0.49457 | 1.89451 | 5.38984 | 1.58442 | 5.22201 |
| **ZNF510** | 0.59522 | 1.74672 | 0.76338 | 1.34382 | 0.00000 | 3.28344 | 0.63161 | 0.46783 | 0.46031 | 1.54165 | 1.15979 | 1.92849 |
| **ZNF511** | 0.00425 | 9.74749 | 6.39492 | 6.46640 | 8.82915 | 0.00000 | 9.23341 | 1.06470 | 10.98109 | 1.96702 | 6.51629 | 4.16928 |
| **ZNF512** | 1.20009 | 1.79642 | 1.32959 | 1.63811 | 0.00000 | 3.14727 | 2.37058 | 1.99093 | 2.42773 | 1.08689 | 3.28436 | 1.32329 |
| **ZNF512B** | 1.81038 | 0.22056 | 0.61591 | 0.69598 | 0.40865 | 0.00000 | 0.00454 | 0.00000 | 0.00000 | 2.40523 | 0.60031 | 0.47459 |
| **ZNF513** | 0.72361 | 0.33012 | 1.44611 | 1.12589 | 1.42160 | 0.00000 | 0.00597 | 0.30246 | 1.97037 | 2.50942 | 0.05134 | 1.02178 |
| **ZNF514** | 0.09465 | 0.03376 | 0.23610 | 0.10371 | 0.05801 | 0.71721 | 0.08926 | 0.14164 | 0.68836 | 0.68546 | 0.11493 | 0.07278 |
| **ZNF516** | 0.04415 | 0.61415 | 0.14992 | 1.80907 | 0.54611 | 1.69413 | 0.40571 | 0.12240 | 0.00001 | 0.00000 | 0.00000 | 3.13110 |
| **ZNF517** | 0.16746 | 0.10838 | 0.02769 | 0.24078 | 0.00000 | 0.00000 | 0.12146 | 0.06860 | 0.00000 | 0.46142 | 0.13450 | 0.19601 |
| **ZNF518A** | 1.23344 | 3.14595 | 5.27634 | 3.30212 | 0.08604 | 27.00372 | 5.34563 | 8.95159 | 5.36990 | 22.60611 | 5.25913 | 11.89743 |
| **ZNF518B** | 4.61604 | 2.92953 | 7.42137 | 1.43234 | 0.00000 | 8.89862 | 8.71186 | 1.75080 | 7.45133 | 1.95657 | 4.03376 | 5.88271 |
| **ZNF519** | 0.16462 | 0.67894 | 1.27961 | 0.54889 | 0.00000 | 3.31343 | 1.04510 | 2.01670 | 4.17253 | 3.25368 | 0.57179 | 1.11894 |
| **ZNF521** | 0.03919 | 0.03879 | 0.03025 | 0.09215 | 0.00000 | 0.00007 | 0.05493 | 0.00950 | 0.00000 | 0.08014 | 0.15186 | 0.03508 |
| **ZNF524** | 0.48393 | 1.87556 | 0.04549 | 0.38004 | 0.49570 | 0.20418 | 0.51954 | 0.00000 | 1.32649 | 0.00000 | 0.40155 | 0.85325 |
| **ZNF525** | 2.11085 | 2.18362 | 1.02921 | 0.46974 | 0.44898 | 8.05911 | 0.35244 | 1.01099 | 3.47833 | 3.88827 | 0.00325 | 3.74867 |
| **ZNF526** | 2.35866 | 7.21431 | 3.14418 | 1.40538 | 5.05945 | 0.00000 | 2.03196 | 0.00000 | 3.29258 | 3.07270 | 4.15494 | 10.56363 |
| **ZNF527** | 0.48627 | 0.55054 | 0.05107 | 1.06608 | 0.00000 | 0.93712 | 0.68952 | 0.26820 | 0.00000 | 0.49333 | 0.91945 | 0.29644 |
| **ZNF528** | 1.84885 | 4.65981 | 0.28128 | 0.00344 | 0.00000 | 2.50284 | 0.18984 | 1.84218 | 0.05275 | 1.66452 | 0.00000 | 9.82394 |
| **ZNF529** | 0.65009 | 1.17307 | 3.24541 | 2.19025 | 0.93737 | 3.43701 | 2.03791 | 0.54277 | 2.48292 | 2.59340 | 2.54576 | 3.62761 |
| **ZNF530** | 2.21350 | 3.11387 | 1.64879 | 0.50370 | 0.77931 | 0.98354 | 1.84973 | 9.68970 | 1.26478 | 0.00000 | 1.50898 | 3.91540 |
| **ZNF532** | 2.38920 | 1.55758 | 3.07367 | 1.69975 | 2.27083 | 6.11617 | 2.45792 | 0.38977 | 5.74716 | 7.78548 | 2.15925 | 1.80351 |
| **ZNF534** | 0.54606 | 6.16345 | 1.18490 | 1.43847 | 1.73201 | 3.52094 | 1.08569 | 5.08660 | 0.97377 | 0.99128 | 2.34766 | 15.44098 |
| **ZNF536** | 0.00895 | 0.01018 | 0.09802 | 0.03712 | 0.00338 | 0.03485 | 0.00000 | 0.01412 | 0.00000 | 0.04458 | 0.01907 | 0.01479 |
| **ZNF540** | 0.41619 | 0.58565 | 0.11538 | 0.00272 | 0.00000 | 0.72758 | 0.21726 | 0.00000 | 0.11361 | 0.00000 | 0.07067 | 0.64782 |
| **ZNF541** | 0.92844 | 0.01112 | 0.05463 | 0.02944 | 0.01564 | 0.00000 | 0.00962 | 0.00000 | 0.01171 | 0.02394 | 0.03667 | 1.07388 |
| **ZNF542P** | 2.50107 | 3.55056 | 1.30793 | 1.10731 | 0.15601 | 2.07736 | 1.86913 | 2.43328 | 5.06049 | 1.76663 | 0.00000 | 6.08929 |
| **ZNF543** | 0.11446 | 1.35453 | 0.48457 | 0.68027 | 0.00000 | 3.36047 | 0.00000 | 1.03896 | 0.40371 | 0.00000 | 1.14329 | 2.32056 |
| **ZNF544** | 1.43121 | 3.48129 | 3.31080 | 2.24322 | 0.00000 | 1.67818 | 6.67880 | 6.87238 | 11.03524 | 0.00000 | 4.08800 | 4.35979 |
| **ZNF546** | 0.50166 | 0.17394 | 0.12467 | 0.11585 | 0.00000 | 0.00000 | 0.00000 | 0.06987 | 0.07695 | 0.30028 | 0.48674 | 0.60255 |
| **ZNF547** | 0.62563 | 2.10032 | 0.17058 | 0.56387 | 0.66586 | 0.47681 | 1.18842 | 2.37406 | 0.09750 | 0.00000 | 0.50892 | 1.61975 |
| **ZNF548** | 1.48768 | 1.99849 | 0.62577 | 0.12368 | 0.00000 | 3.00609 | 0.00000 | 3.20946 | 3.00403 | 1.84760 | 0.23134 | 3.35517 |
| **ZNF549** | 4.49498 | 3.87008 | 1.09969 | 1.29117 | 0.00000 | 1.81105 | 3.14479 | 3.50103 | 1.66010 | 0.00000 | 1.99502 | 7.34323 |
| **ZNF550** | 1.61675 | 2.74212 | 0.67170 | 1.16163 | 0.00028 | 2.24135 | 0.66410 | 2.03405 | 2.55086 | 1.88570 | 2.47029 | 3.07100 |
| **ZNF551** | 0.46376 | 4.08307 | 2.50595 | 2.31209 | 3.91214 | 2.57294 | 3.41555 | 2.20700 | 5.79387 | 0.00000 | 3.53158 | 4.58325 |
| **ZNF552** | 1.89688 | 5.80428 | 1.88929 | 3.27957 | 3.26382 | 7.66873 | 1.16593 | 0.05598 | 2.54836 | 1.76858 | 3.97112 | 8.80229 |
| **ZNF554** | 1.68342 | 1.26089 | 0.08165 | 0.00000 | 0.19981 | 0.00000 | 0.04484 | 2.01376 | 0.00000 | 1.01447 | 1.11643 | 0.22876 |
| **ZNF555** | 2.37213 | 2.15118 | 0.00000 | 0.00696 | 0.00938 | 1.61047 | 0.62716 | 4.99689 | 0.01788 | 0.00000 | 1.41453 | 1.61358 |
| **ZNF556** | 0.24221 | 0.01072 | 0.00998 | 0.02883 | 0.00505 | 0.01017 | 0.79110 | 0.08084 | 0.00000 | 0.00260 | 0.12881 | 0.01276 |
| **ZNF557** | 0.19655 | 2.57461 | 2.21706 | 2.40496 | 1.25516 | 4.70255 | 2.35999 | 8.48439 | 2.31938 | 4.37661 | 3.83309 | 7.26467 |
| **ZNF558** | 0.80306 | 0.75356 | 0.37105 | 0.62271 | 0.00000 | 1.09592 | 0.61374 | 0.77907 | 0.97463 | 0.89524 | 0.21064 | 0.53998 |
| **ZNF559** | 0.55791 | 2.41430 | 0.07447 | 0.07988 | 0.00000 | 5.14429 | 1.86191 | 3.12719 | 0.77071 | 0.00000 | 0.00247 | 5.18006 |

| | | | | | | | | | | | | |
|---|---|---|---|---|---|---|---|---|---|---|---|---|
| ZNF559-ZNF177 | 0.01039 | 0.03551 | 0.01475 | 0.03458 | 0.00642 | 0.01762 | 0.02753 | 0.03015 | 0.01985 | 0.03258 | 0.01576 | 0.02715 |
| ZNF559-ZNF177 | 0.00468 | 0.01526 | 0.00382 | 0.00713 | 0.00000 | 0.03083 | 0.00478 | 0.03036 | 0.02371 | 0.14476 | 0.01060 | 0.03590 |
| ZNF560 | 0.68175 | 1.24603 | 1.91451 | 0.64495 | 0.00000 | 3.18280 | 1.29311 | 2.66457 | 2.72446 | 0.00000 | 0.30482 | 4.56108 |
| ZNF561 | 1.25735 | 4.23605 | 0.67142 | 0.31348 | 0.00000 | 3.13312 | 1.42860 | 3.53085 | 2.03807 | 0.84148 | 1.74599 | 5.90676 |
| ZNF562 | 2.27531 | 5.31322 | 4.35439 | 1.75340 | 0.00000 | 3.02272 | 5.26644 | 0.89808 | 4.92758 | 1.47189 | 6.08096 | 5.60036 |
| ZNF563 | 0.05778 | 0.01141 | 0.04889 | 0.01512 | 0.01306 | 0.03039 | 0.01797 | 0.10805 | 0.00820 | 0.01377 | 0.02102 | 0.02903 |
| ZNF564 | 0.82206 | 4.74602 | 0.95997 | 2.89087 | 0.94122 | 5.75825 | 0.00000 | 1.85436 | 4.13291 | 0.00000 | 1.83841 | 7.19038 |
| ZNF565 | 0.16323 | 3.27777 | 1.44045 | 0.36123 | 0.29194 | 3.06332 | 0.88649 | 1.27729 | 1.01011 | 0.91808 | 1.19721 | 4.66166 |
| ZNF566 | 0.00000 | 6.49235 | 3.11821 | 1.29565 | 0.00000 | 8.48827 | 4.80795 | 3.29733 | 3.25111 | 3.31286 | 6.67691 | 12.65085 |
| ZNF567 | 0.00000 | 3.32549 | 3.83263 | 0.72161 | 0.00000 | 8.10350 | 4.99333 | 0.97606 | 6.79683 | 3.29677 | 2.18449 | 2.63953 |
| ZNF568 | 0.55202 | 0.95957 | 0.70281 | 0.63524 | 0.00000 | 3.41599 | 2.30786 | 5.77790 | 0.31001 | 0.35605 | 1.17472 | 2.11141 |
| ZNF569 | 0.95200 | 4.17393 | 0.00000 | 0.77870 | 0.00000 | 2.67621 | 1.40891 | 2.37215 | 0.68163 | 0.00000 | 1.38613 | 4.36998 |
| ZNF57 | 0.21200 | 23.56732 | 1.23673 | 2.20099 | 5.46521 | 19.35748 | 0.00000 | 3.43469 | 4.17814 | 1.39142 | 5.17486 | 5.95352 |
| ZNF570 | 1.68839 | 0.95833 | 1.75802 | 0.43282 | 0.46388 | 0.00000 | 0.54886 | 0.02759 | 0.24907 | 0.00000 | 0.30954 | 1.03048 |
| ZNF571 | 2.14883 | 1.11087 | 0.38811 | 0.79464 | 0.00000 | 2.27266 | 0.70977 | 0.13031 | 0.63571 | 0.06609 | 0.72709 | 1.35367 |
| ZNF571-AS1 | 0.64218 | 0.18533 | 0.27472 | 0.51845 | 0.00000 | 1.15089 | 0.40399 | 0.00000 | 0.06446 | 0.19169 | 0.18495 | 0.33217 |
| ZNF572 | 0.00056 | 0.16604 | 0.00000 | 0.00000 | 0.00000 | 0.62494 | 0.09324 | 0.11413 | 0.00000 | 0.11467 | 0.01123 | 0.31554 |
| ZNF573 | 1.90853 | 0.46879 | 0.20823 | 1.31834 | 0.00000 | 2.56881 | 0.00000 | 0.16835 | 2.33247 | 1.02418 | 0.97727 | 1.74752 |
| ZNF574 | 8.08512 | 17.42194 | 4.91710 | 7.37691 | 11.78255 | 0.00000 | 2.71553 | 8.07979 | 11.45939 | 0.00000 | 9.99698 | 23.31741 |
| ZNF575 | 0.08538 | 0.01428 | 0.04595 | 0.07736 | 0.01148 | 0.01569 | 0.01064 | 0.01699 | 0.01083 | 0.01925 | 0.02926 | 0.12240 |
| ZNF576 | 0.99437 | 15.69034 | 5.00132 | 7.27693 | 14.03693 | 0.92547 | 11.63853 | 8.79820 | 11.78757 | 0.00000 | 7.14432 | 17.04962 |
| ZNF577 | 0.00000 | 1.99316 | 0.21014 | 1.17717 | 0.00000 | 2.28876 | 0.00000 | 1.45554 | 1.42247 | 2.74368 | 0.58273 | 3.64510 |
| ZNF578 | 0.00000 | 3.96035 | 1.21346 | 1.67180 | 0.00000 | 0.00000 | 0.89361 | 12.53564 | 0.92188 | 2.28772 | 2.86446 | 10.13842 |
| ZNF579 | 0.00283 | 0.05079 | 0.00711 | 0.00132 | 0.00000 | 0.04936 | 0.00445 | 0.02723 | 0.00000 | 0.06762 | 0.06178 | 0.17771 |
| ZNF580 | 0.70841 | 0.18386 | 0.12236 | 0.01924 | 0.20120 | 0.00000 | 0.58821 | 0.00000 | 0.66339 | 2.45798 | 0.13693 | 0.24061 |
| ZNF581 | 2.39527 | 23.15414 | 0.34505 | 9.08837 | 14.67409 | 2.06848 | 11.82137 | 4.35476 | 10.22201 | 6.75818 | 5.77113 | 26.18220 |
| ZNF582 | 0.34076 | 0.80067 | 0.33966 | 0.30982 | 0.00000 | 1.09261 | 0.00000 | 0.69528 | 0.52913 | 0.00000 | 0.15162 | 1.07907 |
| ZNF582-AS1 | 0.01025 | 0.96297 | 0.27406 | 1.47225 | 0.00000 | 0.00000 | 1.02064 | 0.15620 | 0.31710 | 0.00000 | 0.28579 | 0.88849 |
| ZNF583 | 0.64839 | 1.49840 | 0.86304 | 0.08971 | 0.00000 | 0.50491 | 0.00000 | 0.09028 | 0.60989 | 0.00000 | 0.13333 | 1.12942 |
| ZNF584 | 1.84467 | 5.99294 | 0.36776 | 1.32719 | 1.15649 | 0.35129 | 0.00000 | 5.55125 | 2.21773 | 0.00000 | 1.36234 | 8.77305 |
| ZNF585A | 2.14364 | 3.75157 | 1.70908 | 2.02174 | 0.87795 | 4.59850 | 4.96788 | 0.68485 | 2.33640 | 1.78992 | 2.39892 | 2.98591 |
| ZNF585B | 0.02394 | 1.19524 | 0.31015 | 0.49916 | 0.25765 | 2.92118 | 0.75541 | 1.32651 | 1.05017 | 0.13288 | 0.95683 | 2.21230 |
| ZNF586 | 3.43286 | 7.81199 | 2.37861 | 3.23273 | 3.29064 | 2.75912 | 2.58591 | 8.16580 | 4.06184 | 0.00000 | 5.78498 | 12.11032 |
| ZNF587 | 1.28671 | 1.30810 | 2.14418 | 4.48234 | 1.66936 | 1.69069 | 2.23710 | 1.17276 | 1.80185 | 12.30059 | 1.86781 | 4.21987 |
| ZNF587B | 1.34115 | 1.71228 | 2.52529 | 2.22594 | 0.67057 | 3.68507 | 3.49714 | 0.51392 | 0.55144 | 4.08579 | 4.33026 | 6.05681 |
| ZNF589 | 5.80158 | 12.42463 | 0.49143 | 7.35115 | 2.78182 | 7.00637 | 5.59331 | 2.37930 | 13.47255 | 8.89794 | 8.56166 | 11.92619 |
| ZNF592 | 1.42890 | 4.80173 | 5.28887 | 3.64547 | 7.11657 | 0.00059 | 6.59562 | 0.23736 | 6.28969 | 5.40220 | 3.97695 | 8.85658 |
| ZNF593 | 3.12826 | 21.05527 | 21.86464 | 12.51086 | 22.91771 | 0.00003 | 28.41289 | 8.50122 | 16.71144 | 11.40556 | 12.21851 | 27.31702 |
| ZNF594 | 0.01014 | 0.00000 | 0.74179 | 0.19641 | 0.00000 | 0.11608 | 0.25895 | 0.00546 | 0.30255 | 0.04121 | 0.77729 | 0.13743 |
| ZNF595 | 0.01595 | 3.20207 | 1.12335 | 1.01478 | 0.00000 | 5.16753 | 2.46780 | 7.00603 | 0.81905 | 0.00061 | 2.93703 | 7.41146 |
| ZNF596 | 0.01272 | 1.67916 | 0.10729 | 0.00000 | 0.00000 | 1.34986 | 0.13440 | 4.79730 | 0.03809 | 0.20592 | 0.10045 | 1.05370 |
| ZNF597 | 4.49624 | 0.55624 | 1.07997 | 0.84436 | 0.00000 | 0.80054 | 2.02554 | 15.86433 | 1.00417 | 0.50008 | 0.42800 | 8.21509 |
| ZNF598 | 1.02732 | 0.73746 | 2.03579 | 0.78082 | 7.05477 | 0.00000 | 5.08415 | 0.45466 | 1.30658 | 15.53881 | 0.22433 | 1.27688 |
| ZNF599 | 0.15005 | 0.91679 | 0.22779 | 0.01427 | 0.00000 | 0.00690 | 0.20512 | 0.96627 | 0.05400 | 0.00000 | 0.69095 | 1.21046 |
| ZNF600 | 1.24605 | 8.62109 | 9.22550 | 5.99013 | 1.59026 | 15.53969 | 0.00005 | 1.96671 | 4.95295 | 6.10085 | 2.66536 | 5.66986 |
| ZNF605 | 1.03553 | 0.38968 | 0.00000 | 0.25196 | 0.00000 | 1.47737 | 0.00000 | 4.25823 | 1.90847 | 1.47653 | 0.00000 | 1.96797 |
| ZNF606 | 1.30708 | 1.70558 | 0.17516 | 0.58028 | 0.00000 | 1.97023 | 0.00000 | 2.93504 | 0.38818 | 0.00000 | 1.03734 | 2.44798 |
| ZNF607 | 0.51069 | 3.71783 | 2.24808 | 1.27276 | 0.01807 | 1.84897 | 0.00000 | 0.24835 | 2.64177 | 3.61382 | 1.47361 | 2.54109 |

| | | | | | | | | | | | | |
|---|---|---|---|---|---|---|---|---|---|---|---|---|
| **ZNF608** | 0.01775 | 0.00000 | 0.02448 | 1.17225 | 0.00000 | 0.00000 | 0.13730 | 1.06409 | 0.10291 | 1.33500 | 0.06739 | 1.24850 |
| **ZNF609** | 2.31095 | 0.80500 | 2.92820 | 3.84100 | 2.82649 | 0.57975 | 5.42212 | 0.04958 | 3.20767 | 11.69890 | 0.56668 | 1.53619 |
| **ZNF610** | 2.07628 | 1.02653 | 0.43071 | 0.23336 | 0.03528 | 0.31974 | 0.00000 | 0.43316 | 0.26151 | 0.00000 | 0.35238 | 2.87334 |
| **ZNF611** | 0.42600 | 0.74785 | 4.03880 | 3.28989 | 2.11633 | 2.99430 | 2.28633 | 4.50330 | 0.70973 | 10.31845 | 4.17829 | 4.40517 |
| **ZNF613** | 0.00086 | 1.48638 | 0.38663 | 2.02804 | 0.31175 | 2.03256 | 0.44209 | 2.96330 | 1.02522 | 0.00000 | 0.37395 | 3.42245 |
| **ZNF614** | 0.06170 | 3.78019 | 1.40868 | 4.91555 | 0.00174 | 6.32594 | 0.17148 | 2.57200 | 3.25774 | 1.52084 | 1.01971 | 8.75970 |
| **ZNF615** | 0.05288 | 2.59111 | 0.95137 | 0.76048 | 0.00000 | 1.63718 | 0.65302 | 2.98738 | 1.34751 | 0.86773 | 0.66080 | 2.57405 |
| **ZNF616** | 0.38132 | 0.97320 | 0.77246 | 0.29446 | 0.00000 | 1.46295 | 0.00000 | 6.02916 | 1.74931 | 0.31979 | 1.68611 | 3.35007 |
| **ZNF618** | 10.51591 | 0.78937 | 0.00000 | 0.26207 | 0.37513 | 0.00000 | 0.00000 | 1.13544 | 0.00000 | 0.18957 | 0.00000 | 0.90400 |
| **ZNF619** | 0.07957 | 1.63364 | 0.00000 | 0.46687 | 1.71037 | 0.00000 | 0.00000 | 0.43175 | 0.00000 | 0.41078 | 0.18394 | 0.95585 |
| **ZNF620** | 0.58353 | 2.11972 | 0.87364 | 0.54864 | 0.84318 | 2.37281 | 0.00000 | 0.27209 | 0.00000 | 0.75186 | 2.41252 | 2.22351 |
| **ZNF621** | 0.00000 | 0.80056 | 0.52447 | 0.00000 | 3.59861 | 3.92420 | 0.69661 | 2.23021 | 0.00000 | 4.64822 | 1.48193 | 0.80975 |
| **ZNF622** | 0.15023 | 16.92835 | 13.49643 | 8.74052 | 10.36738 | 15.63839 | 15.49407 | 28.19540 | 12.21864 | 0.00000 | 12.45348 | 14.05639 |
| **ZNF623** | 2.27505 | 2.32094 | 0.63876 | 1.17746 | 2.49184 | 1.48342 | 0.69523 | 1.46259 | 0.47764 | 2.30619 | 0.53028 | 3.86155 |
| **ZNF624** | 0.25542 | 0.26960 | 0.71436 | 0.24978 | 0.00000 | 1.14262 | 0.15407 | 0.66098 | 0.38396 | 0.00000 | 0.21953 | 0.88638 |
| **ZNF625** | 0.00152 | 1.43047 | 0.48434 | 0.05309 | 0.00275 | 3.17986 | 0.00000 | 1.65605 | 0.02268 | 0.00000 | 1.67147 | 0.90650 |
| **ZNF625-ZNF20** | 0.00139 | 0.46378 | 0.14354 | 0.45548 | 0.28012 | 1.43954 | 0.16866 | 0.30011 | 0.35296 | 0.54651 | 0.79491 | 0.66144 |
| **ZNF626** | 1.23290 | 2.36945 | 4.08920 | 0.39182 | 0.15609 | 8.53714 | 3.31045 | 1.71863 | 8.55717 | 2.41740 | 4.23735 | 7.83464 |
| **ZNF627** | 0.00283 | 0.97433 | 0.82589 | 0.76441 | 0.00000 | 1.36056 | 1.79860 | 0.26473 | 2.28415 | 0.07738 | 0.67115 | 0.09783 |
| **ZNF628** | 0.76206 | 0.05519 | 0.27673 | 0.32791 | 1.79357 | 0.00000 | 1.69819 | 0.20324 | 0.00000 | 3.94881 | 0.00000 | 0.95253 |
| **ZNF629** | 0.04820 | 0.03263 | 0.00000 | 0.53264 | 0.38436 | 0.00000 | 0.01734 | 0.07421 | 0.17162 | 0.04502 | 0.00000 | 0.07966 |
| **ZNF630** | 0.00221 | 0.03276 | 0.13042 | 0.06156 | 0.07855 | 0.16786 | 0.00003 | 0.80331 | 0.05262 | 0.00989 | 0.00000 | 0.07792 |
| **ZNF630-AS1** | 0.00572 | 0.01426 | 0.01527 | 0.01278 | 0.01301 | 0.02380 | 0.02289 | 0.05089 | 0.01532 | 0.01359 | 0.01471 | 0.01348 |
| **ZNF638** | 4.61696 | 3.82800 | 6.54457 | 4.91596 | 3.45626 | 28.54898 | 8.34535 | 10.03257 | 8.39846 | 19.28438 | 8.26627 | 12.97139 |
| **ZNF639** | 1.91726 | 4.53507 | 0.36681 | 1.52731 | 1.72092 | 11.16314 | 4.35254 | 14.20712 | 0.00000 | 16.12901 | 2.10226 | 7.32876 |
| **ZNF641** | 1.52154 | 0.96349 | 0.20979 | 0.96778 | 1.18887 | 1.62911 | 1.25244 | 0.11778 | 0.36619 | 2.93001 | 0.53808 | 1.76729 |
| **ZNF644** | 0.09379 | 0.55793 | 6.23821 | 3.03569 | 0.00000 | 4.33533 | 9.55122 | 0.90699 | 4.80364 | 5.12195 | 3.50859 | 4.38493 |
| **ZNF646** | 2.33915 | 1.60409 | 0.95205 | 0.00000 | 0.95048 | 0.00708 | 0.51232 | 0.00000 | 0.62563 | 2.42654 | 0.43144 | 1.44958 |
| **ZNF648** | 0.00571 | 0.14096 | 0.04824 | 0.05108 | 0.00717 | 0.01038 | 0.02689 | 0.00934 | 0.00765 | 0.00289 | 0.04955 | 0.04958 |
| **ZNF649** | 0.77699 | 3.32508 | 0.00000 | 3.52146 | 0.44351 | 5.99528 | 1.02222 | 0.28239 | 1.38124 | 1.19221 | 1.14743 | 3.57784 |
| **ZNF652** | 11.08811 | 0.96607 | 1.11897 | 2.87459 | 0.00000 | 9.68277 | 7.32525 | 4.85652 | 2.15815 | 9.39762 | 9.68278 | 4.98083 |
| **ZNF653** | 0.04518 | 0.00249 | 0.06744 | 0.00633 | 0.02526 | 0.01300 | 0.05612 | 0.04250 | 0.12925 | 0.21651 | 0.00000 | 0.09380 |
| **ZNF654** | 4.01881 | 0.84339 | 1.29895 | 1.10292 | 0.95447 | 3.27866 | 2.20944 | 6.82480 | 0.62746 | 3.78739 | 0.00000 | 2.05132 |
| **ZNF655** | 1.43843 | 0.66795 | 9.78392 | 3.96812 | 2.24601 | 0.76914 | 5.07695 | 9.05440 | 7.13931 | 3.76062 | 4.09084 | 6.72411 |
| **ZNF658** | 0.02479 | 0.01390 | 0.05921 | 0.06602 | 0.00000 | 0.12559 | 0.00000 | 0.01390 | 0.00000 | 0.12441 | 0.01377 | 0.00471 |
| **ZNF660** | 0.00570 | 0.01363 | 0.03643 | 0.00061 | 0.02148 | 0.05910 | 0.00004 | 0.01094 | 0.01448 | 0.05751 | 0.00769 | 0.02244 |
| **ZNF662** | 1.86071 | 1.03786 | 0.00000 | 1.34062 | 0.11181 | 1.75693 | 0.00000 | 0.44231 | 0.00000 | 0.04754 | 0.26231 | 0.90721 |
| **ZNF663P** | 0.00684 | 0.01197 | 0.01284 | 0.01092 | 0.01290 | 0.00644 | 0.00328 | 0.00889 | 0.11611 | 0.01098 | 0.03490 | 0.06802 |
| **ZNF664** | 3.62228 | 3.41492 | 4.05344 | 0.85954 | 0.00000 | 6.29512 | 6.94043 | 0.30239 | 1.46811 | 9.53996 | 9.40031 | 4.44755 |
| **ZNF664-FAM10** | 0.09441 | 0.06371 | 0.00592 | 0.00192 | 0.00187 | 0.03668 | 0.00101 | 0.02000 | 0.04253 | 0.05566 | 0.00865 | 0.04514 |
| **ZNF665** | 0.80915 | 1.64724 | 0.67105 | 0.00006 | 0.51513 | 0.04202 | 0.23986 | 0.36920 | 0.65381 | 0.00000 | 0.75250 | 1.43047 |
| **ZNF667** | 0.60658 | 1.67970 | 0.58806 | 0.66127 | 0.00000 | 0.44407 | 1.80791 | 1.85367 | 0.73290 | 0.32994 | 0.53661 | 1.43918 |
| **ZNF667-AS1** | 1.43511 | 3.37526 | 1.66954 | 0.00000 | 1.26071 | 0.45237 | 1.71352 | 2.05332 | 2.60140 | 0.00000 | 0.27861 | 9.67162 |
| **ZNF668** | 2.10988 | 4.78272 | 5.21391 | 2.01425 | 3.50915 | 0.00000 | 4.11838 | 6.59713 | 7.88745 | 0.00000 | 3.45453 | 11.75965 |
| **ZNF669** | 0.01015 | 1.46030 | 0.96803 | 0.41740 | 0.72246 | 0.01509 | 1.47624 | 10.94079 | 1.78865 | 0.16970 | 0.68885 | 5.44404 |
| **ZNF670** | 0.10877 | 4.34950 | 4.07489 | 1.07947 | 0.00000 | 8.37955 | 4.89422 | 8.00361 | 4.92060 | 0.08053 | 1.73923 | 9.30196 |
| **ZNF670-ZNF695** | 0.00325 | 0.00839 | 0.01812 | 0.04413 | 0.00000 | 0.09308 | 0.00000 | 0.01045 | 0.06514 | 0.19030 | 0.11853 | 0.01074 |
| **ZNF671** | 0.00388 | 0.49476 | 0.01684 | 0.58989 | 0.14714 | 0.34004 | 0.21649 | 0.08630 | 1.76969 | 0.00000 | 1.54935 | 2.71936 |

| | | | | | | | | | | | | |
|---|---|---|---|---|---|---|---|---|---|---|---|---|
| **ZNF672** | 0.47275 | 7.18325 | 4.10482 | 2.54544 | 2.43957 | 1.05004 | 3.70418 | 0.06011 | 5.69526 | 4.97000 | 3.81023 | 2.42159 |
| **ZNF674** | 0.42035 | 0.29796 | 0.22022 | 0.62404 | 0.00000 | 0.97602 | 0.00000 | 0.51397 | 0.00000 | 0.46469 | 0.26555 | 0.39179 |
| **ZNF674-AS1** | 0.16452 | 1.62616 | 0.36419 | 0.19945 | 0.09821 | 0.70053 | 0.00000 | 1.08291 | 0.98948 | 0.00000 | 0.84054 | 3.06114 |
| **ZNF675** | 2.20239 | 7.64947 | 1.46325 | 2.90375 | 0.00000 | 13.52964 | 0.00000 | 6.04516 | 0.00000 | 1.51322 | 1.68874 | 17.75555 |
| **ZNF676** | 0.00000 | 2.49654 | 1.69728 | 0.61526 | 0.07979 | 5.06367 | 0.00000 | 2.50546 | 3.41959 | 2.11347 | 2.17751 | 2.83538 |
| **ZNF677** | 4.84153 | 5.11963 | 1.57192 | 0.00000 | 0.56366 | 2.74541 | 1.63109 | 2.13048 | 1.11065 | 0.33021 | 1.07353 | 8.03709 |
| **ZNF678** | 0.11752 | 0.11502 | 0.85155 | 0.52983 | 0.17361 | 1.88128 | 1.12745 | 0.28230 | 0.10172 | 1.92647 | 0.96056 | 0.80476 |
| **ZNF679** | 2.41348 | 0.00000 | 0.00173 | 0.01189 | 0.00859 | 0.00000 | 0.00000 | 26.59024 | 0.01399 | 0.09645 | 0.01537 | 0.73182 |
| **ZNF680** | 0.00000 | 6.88681 | 1.54310 | 2.13359 | 0.00000 | 14.72078 | 0.18523 | 2.92769 | 4.52879 | 2.65622 | 2.42668 | 6.71916 |
| **ZNF681** | 0.18150 | 3.25286 | 1.86493 | 0.00000 | 0.00000 | 8.06262 | 0.00000 | 0.65512 | 0.00000 | 3.50526 | 2.12608 | 2.76906 |
| **ZNF682** | 1.15057 | 0.81779 | 2.75049 | 3.68537 | 0.65365 | 10.83461 | 3.27635 | 6.94671 | 3.75292 | 1.53465 | 0.21351 | 2.21402 |
| **ZNF683** | 0.05390 | 0.00340 | 0.00000 | 0.00097 | 0.00000 | 0.02982 | 0.00000 | 0.00663 | 0.02006 | 0.27823 | 0.00447 | 0.01667 |
| **ZNF684** | 0.25847 | 5.31792 | 0.03918 | 3.03202 | 0.00000 | 3.97818 | 0.00000 | 5.99275 | 0.95056 | 0.68096 | 0.85607 | 8.10063 |
| **ZNF687** | 1.08218 | 2.90107 | 2.59920 | 1.89197 | 0.71356 | 0.00000 | 1.01608 | 0.54682 | 1.85830 | 0.00000 | 3.30356 | 1.57816 |
| **ZNF688** | 0.00451 | 0.21512 | 0.00000 | 0.48854 | 0.06057 | 0.00000 | 0.18338 | 0.00106 | 0.36981 | 0.60955 | 0.01100 | 0.53533 |
| **ZNF689** | 7.81661 | 4.51351 | 1.15422 | 6.85916 | 3.44489 | 0.00000 | 3.70939 | 10.63739 | 5.00123 | 0.70740 | 3.74904 | 10.83338 |
| **ZNF69** | 0.00245 | 0.00000 | 1.27574 | 0.23954 | 0.47674 | 0.00000 | 0.90564 | 0.02417 | 0.38267 | 1.04976 | 0.41620 | 0.03612 |
| **ZNF691** | 0.00439 | 4.86741 | 1.69198 | 1.91876 | 1.16482 | 5.56083 | 1.50438 | 4.95826 | 3.78066 | 0.00000 | 4.30965 | 6.98568 |
| **ZNF692** | 0.24780 | 5.01693 | 5.30990 | 0.00000 | 2.39670 | 0.00000 | 4.74830 | 0.46417 | 8.88877 | 5.70366 | 5.77027 | 3.73982 |
| **ZNF695** | 0.00000 | 1.28260 | 1.51684 | 0.67347 | 1.26744 | 6.85332 | 0.00000 | 0.07985 | 2.32638 | 1.62400 | 1.07698 | 0.27610 |
| **ZNF696** | 0.00000 | 2.83579 | 0.00000 | 1.69184 | 0.49083 | 0.00000 | 0.46415 | 0.03989 | 0.95863 | 3.77447 | 1.67416 | 1.73598 |
| **ZNF697** | 0.02690 | 0.18165 | 3.47761 | 0.76676 | 0.56441 | 2.21591 | 2.56273 | 0.02239 | 0.20092 | 1.31363 | 0.92265 | 0.00153 |
| **ZNF699** | 0.00216 | 0.04567 | 0.07912 | 0.14255 | 0.05370 | 0.00000 | 0.03774 | 0.09678 | 0.00000 | 0.00000 | 0.19828 | 0.08360 |
| **ZNF7** | 1.22980 | 8.95738 | 0.15043 | 4.06745 | 0.97887 | 7.74294 | 0.00000 | 4.07883 | 4.24787 | 2.38266 | 2.54356 | 10.13384 |
| **ZNF70** | 0.88438 | 0.27134 | 0.14243 | 0.16806 | 0.84060 | 0.14217 | 0.00000 | 0.04925 | 0.00000 | 0.04274 | 0.27485 | 0.59090 |
| **ZNF700** | 0.00000 | 1.38687 | 1.12782 | 0.25379 | 0.00000 | 5.84763 | 0.00000 | 1.06212 | 1.81134 | 2.34401 | 1.85006 | 2.49263 |
| **ZNF701** | 0.01138 | 3.04887 | 2.10405 | 1.75585 | 1.46341 | 3.25587 | 0.91499 | 9.12237 | 1.06231 | 1.83152 | 1.10214 | 8.13066 |
| **ZNF702P** | 0.09478 | 0.48809 | 3.52128 | 0.00059 | 0.22396 | 2.63253 | 0.72051 | 0.18830 | 2.79045 | 0.00000 | 1.58440 | 0.45062 |
| **ZNF703** | 0.04607 | 2.80718 | 0.22662 | 0.00000 | 10.90823 | 0.00000 | 1.40854 | 0.74520 | 0.00000 | 4.46132 | 0.00000 | 0.00000 |
| **ZNF704** | 0.44068 | 0.00000 | 2.59890 | 0.00000 | 4.82992 | 0.00793 | 4.69056 | 0.00353 | 1.07249 | 4.29826 | 5.98408 | 0.00000 |
| **ZNF705A** | 2.56645 | 0.01044 | 0.01192 | 0.01005 | 0.01947 | 0.00000 | 0.01664 | 1.82316 | 0.02182 | 0.00001 | 0.01343 | 0.00000 |
| **ZNF705B** | 0.01163 | 0.01470 | 0.01390 | 0.01176 | 0.01456 | 0.01958 | 0.01757 | 0.01762 | 0.01761 | 0.01622 | 0.01487 | 0.01537 |
| **ZNF705D** | 0.02141 | 0.03262 | 0.00938 | 0.00651 | 0.00084 | 0.05583 | 0.01308 | 0.11667 | 0.01648 | 0.02058 | 0.01416 | 0.07311 |
| **ZNF705E** | 2.56749 | 0.10065 | 0.02197 | 0.00121 | 0.02531 | 0.00000 | 0.02252 | 2.40573 | 0.01243 | 0.00000 | 0.01792 | 0.00000 |
| **ZNF705G** | 0.63495 | 0.01440 | 0.01406 | 0.01193 | 0.01677 | 0.00819 | 0.01822 | 0.31430 | 0.02138 | 0.00757 | 0.01398 | 0.00295 |
| **ZNF706** | 1.59403 | 10.53986 | 13.27578 | 5.99599 | 12.37127 | 19.14236 | 10.73360 | 1.59094 | 8.36117 | 0.15888 | 9.61785 | 8.40476 |
| **ZNF707** | 2.36473 | 1.77852 | 0.85871 | 0.62596 | 1.33345 | 0.00000 | 0.04528 | 7.62738 | 2.96268 | 0.00000 | 0.40080 | 2.85831 |
| **ZNF708** | 0.00000 | 0.00000 | 2.03530 | 0.93896 | 0.00000 | 2.69512 | 0.84924 | 0.13826 | 3.08122 | 0.81473 | 0.63660 | 0.59432 |
| **ZNF709** | 0.00509 | 1.28215 | 1.02959 | 0.50981 | 0.00000 | 2.48861 | 1.04428 | 2.56363 | 0.83971 | 0.11160 | 0.76273 | 2.21132 |
| **ZNF71** | 0.29583 | 1.68188 | 1.68460 | 1.25176 | 0.73699 | 0.00000 | 1.45209 | 0.01171 | 2.07751 | 0.01704 | 0.86424 | 0.10191 |
| **ZNF710** | 0.75729 | 0.45293 | 0.13794 | 2.58958 | 0.13491 | 0.00000 | 0.24184 | 0.03725 | 0.05766 | 0.25675 | 0.12313 | 0.75267 |
| **ZNF711** | 0.00140 | 0.00128 | 0.04828 | 0.00579 | 0.01791 | 0.40882 | 0.28721 | 0.47167 | 0.01027 | 0.00000 | 0.00000 | 0.37540 |
| **ZNF713** | 0.00109 | 0.00000 | 0.32361 | 0.03452 | 0.07582 | 0.09272 | 0.28445 | 0.04281 | 0.00000 | 0.96574 | 0.13981 | 0.04999 |
| **ZNF714** | 0.16198 | 0.00000 | 2.52729 | 2.59285 | 0.00000 | 1.28381 | 5.17307 | 0.28935 | 4.42712 | 1.89722 | 2.36690 | 0.03851 |
| **ZNF716** | 0.03976 | 0.79357 | 0.10874 | 0.00000 | 0.32452 | 0.00000 | 0.00000 | 12.77325 | 0.00000 | 0.28171 | 0.48058 | 5.04851 |
| **ZNF717** | 0.71369 | 0.80954 | 0.66676 | 0.77402 | 0.00000 | 0.83112 | 0.98031 | 0.40456 | 0.23935 | 0.92355 | 1.29565 | 1.46886 |
| **ZNF718** | 0.03499 | 3.28346 | 1.22000 | 0.83590 | 0.00000 | 2.81998 | 1.66889 | 3.65587 | 2.20470 | 2.69923 | 3.04761 | 4.45375 |
| **ZNF720** | 0.62085 | 0.19958 | 0.88888 | 0.06997 | 0.28978 | 1.00983 | 1.04767 | 0.07315 | 0.42147 | 0.00000 | 0.55464 | 0.58162 |

| | | | | | | | | | | | | |
|---|---|---|---|---|---|---|---|---|---|---|---|---|
| **ZNF721** | 2.98660 | 3.79233 | 2.10207 | 0.98815 | 0.23059 | 12.24504 | 0.00000 | 2.62187 | 7.60216 | 10.46112 | 3.62609 | 4.92348 |
| **ZNF724P** | 0.41814 | 0.23252 | 1.05551 | 0.53863 | 0.19803 | 1.74034 | 0.00000 | 1.39888 | 1.23747 | 1.83537 | 0.95805 | 2.76761 |
| **ZNF726** | 0.04381 | 0.51328 | 1.38407 | 0.57604 | 0.28412 | 0.81361 | 0.00000 | 0.01504 | 0.12578 | 0.00000 | 0.88148 | 0.09374 |
| **ZNF727P** | 0.60448 | 0.44679 | 0.09031 | 0.00000 | 0.00000 | 0.23434 | 0.00000 | 4.35158 | 0.36488 | 0.18356 | 0.01627 | 0.21790 |
| **ZNF728** | 0.01767 | 4.63503 | 0.91175 | 0.88980 | 0.00000 | 5.13314 | 0.00000 | 14.03872 | 2.46732 | 0.00097 | 2.90664 | 4.81065 |
| **ZNF729** | 0.00000 | 2.64907 | 0.00000 | 3.50117 | 0.91484 | 7.25237 | 0.00000 | 1.94124 | 0.00028 | 0.39087 | 2.01134 | 4.04915 |
| **ZNF730** | 0.18628 | 0.54864 | 0.38968 | 1.72300 | 0.00027 | 3.75131 | 0.00000 | 0.66904 | 2.27801 | 3.71733 | 1.15975 | 0.90045 |
| **ZNF732** | 0.45443 | 0.40370 | 0.07805 | 0.35802 | 0.07593 | 2.48290 | 0.00000 | 0.84637 | 1.44783 | 1.28637 | 1.11131 | 2.17128 |
| **ZNF733P** | 0.03846 | 0.01169 | 0.01282 | 0.00841 | 0.01232 | 0.03536 | 0.01966 | 0.51404 | 0.01632 | 0.00686 | 0.01273 | 0.06448 |
| **ZNF735P** | 0.91610 | 0.00000 | 0.00841 | 0.02625 | 0.02182 | 0.00000 | 0.01404 | 23.63519 | 0.01381 | 0.00581 | 0.00699 | 2.14536 |
| **ZNF736** | 0.20910 | 0.46349 | 0.36967 | 0.00234 | 0.06415 | 0.90194 | 0.00000 | 8.62183 | 0.09540 | 0.00000 | 0.19022 | 0.70114 |
| **ZNF737** | 0.55084 | 1.28166 | 0.79343 | 1.71470 | 0.00000 | 3.48206 | 1.78080 | 1.30656 | 1.41480 | 0.46050 | 0.95025 | 2.69724 |
| **ZNF738** | 4.97602 | 10.22804 | 5.31142 | 1.25330 | 0.00000 | 13.17727 | 6.53658 | 6.59119 | 6.60749 | 7.73629 | 9.54250 | 4.60920 |
| **ZNF74** | 0.59839 | 2.57142 | 0.81374 | 1.94917 | 0.00000 | 0.00000 | 2.66783 | 0.00000 | 3.63345 | 2.34286 | 3.26784 | 0.31152 |
| **ZNF740** | 0.24937 | 0.43503 | 0.29303 | 0.90962 | 2.47264 | 0.16402 | 2.32156 | 0.00000 | 0.71861 | 4.69849 | 3.89746 | 0.39637 |
| **ZNF746** | 2.29200 | 1.09270 | 1.09548 | 0.96495 | 1.67490 | 1.21656 | 1.89031 | 0.00000 | 0.00000 | 5.49254 | 0.09254 | 1.81736 |
| **ZNF747** | 0.35237 | 0.27885 | 0.42949 | 1.18823 | 0.39241 | 0.00000 | 0.02657 | 0.00000 | 0.00000 | 0.00000 | 0.02650 | 0.25120 |
| **ZNF749** | 0.65019 | 3.04190 | 0.84972 | 1.27002 | 1.81217 | 1.00271 | 0.22229 | 1.70644 | 2.93435 | 0.00000 | 1.85679 | 3.28794 |
| **ZNF750** | 0.00514 | 0.00000 | 15.61379 | 0.00000 | 7.23430 | 0.11662 | 3.31322 | 0.00000 | 5.90659 | 0.13436 | 0.58635 | 0.30369 |
| **ZNF75A** | 0.41391 | 2.09485 | 1.27824 | 0.44051 | 0.68197 | 1.97673 | 1.09263 | 0.71003 | 1.97130 | 2.44677 | 1.99118 | 3.87024 |
| **ZNF75D** | 0.00576 | 0.20809 | 0.49455 | 2.41661 | 1.49909 | 4.66421 | 0.00000 | 1.47022 | 2.91079 | 0.25015 | 0.46075 | 0.17753 |
| **ZNF76** | 4.88769 | 5.71241 | 4.18957 | 5.69833 | 4.60444 | 0.00000 | 4.93990 | 0.05039 | 5.85298 | 0.00000 | 5.50169 | 2.69786 |
| **ZNF761** | 1.72267 | 4.69210 | 5.28222 | 3.38332 | 0.00026 | 9.94448 | 4.90194 | 3.49852 | 8.00189 | 5.35217 | 2.10701 | 4.35010 |
| **ZNF763** | 0.00689 | 0.00002 | 0.15553 | 0.00525 | 0.08598 | 0.14756 | 0.00958 | 0.07519 | 0.05921 | 0.03533 | 0.00338 | 0.26054 |
| **ZNF764** | 1.32683 | 4.48484 | 1.04620 | 2.28287 | 0.63359 | 1.83695 | 0.09023 | 0.00000 | 3.32976 | 1.35732 | 1.24693 | 4.63401 |
| **ZNF765** | 0.46010 | 4.04371 | 1.27931 | 1.70728 | 0.86205 | 8.82467 | 0.00075 | 4.84654 | 6.23279 | 2.22480 | 1.10632 | 7.15575 |
| **ZNF766** | 0.18534 | 8.06496 | 1.21394 | 2.64481 | 1.17920 | 8.36108 | 0.00000 | 9.31210 | 2.50087 | 2.39932 | 1.65016 | 13.87592 |
| **ZNF767P** | 0.00266 | 0.53236 | 0.31510 | 0.42179 | 0.70999 | 0.00000 | 0.00647 | 0.24118 | 0.00000 | 2.67280 | 0.65928 | 0.27248 |
| **ZNF768** | 0.59625 | 0.66619 | 0.00000 | 1.05623 | 1.52510 | 0.00000 | 0.15050 | 0.00000 | 0.00000 | 1.27767 | 0.04362 | 0.32101 |
| **ZNF77** | 0.87521 | 1.04891 | 0.00000 | 0.77054 | 1.18014 | 3.48141 | 0.83205 | 2.67463 | 0.63365 | 0.00000 | 1.62936 | 1.62472 |
| **ZNF770** | 5.59401 | 0.42227 | 3.32102 | 1.96318 | 1.85830 | 19.85569 | 11.60718 | 0.92067 | 3.09333 | 15.09096 | 5.03874 | 11.10878 |
| **ZNF771** | 0.03580 | 0.98212 | 0.84007 | 1.15773 | 0.18692 | 0.00000 | 1.89313 | 0.01976 | 0.06346 | 0.11815 | 0.54727 | 4.02805 |
| **ZNF772** | 2.96673 | 1.25635 | 0.00000 | 0.01216 | 0.25321 | 2.87842 | 0.63692 | 0.24811 | 0.54675 | 0.74500 | 0.88072 | 1.46094 |
| **ZNF773** | 1.19928 | 2.32208 | 0.86686 | 1.37310 | 1.67106 | 0.36372 | 2.58513 | 2.66492 | 1.10685 | 0.00000 | 2.26014 | 3.96370 |
| **ZNF774** | 0.05887 | 0.53125 | 0.35209 | 0.00000 | 0.19073 | 0.00000 | 0.00000 | 0.06757 | 0.06625 | 0.01991 | 0.26226 | 0.03867 |
| **ZNF775** | 0.01119 | 0.12837 | 0.00456 | 0.03919 | 0.13267 | 0.00000 | 0.46983 | 0.00646 | 0.00000 | 0.47230 | 0.00000 | 0.03094 |
| **ZNF776** | 0.60106 | 2.10468 | 0.59391 | 1.21761 | 0.00000 | 5.12181 | 1.67666 | 4.43779 | 1.85008 | 1.67806 | 2.91219 | 3.72282 |
| **ZNF777** | 1.60482 | 0.16156 | 0.10332 | 0.07034 | 1.39705 | 0.00000 | 0.47381 | 0.03003 | 0.00000 | 3.95365 | 0.00000 | 0.92063 |
| **ZNF778** | 0.89252 | 2.04282 | 0.61715 | 1.82587 | 0.58415 | 0.00000 | 0.00000 | 6.20124 | 2.84737 | 0.00000 | 1.04039 | 3.00870 |
| **ZNF780A** | 0.00000 | 1.36521 | 1.96325 | 0.89507 | 0.19871 | 2.28890 | 1.50728 | 0.00000 | 1.97360 | 1.67086 | 1.29240 | 0.79973 |
| **ZNF780B** | 0.00115 | 0.23347 | 0.51277 | 0.88936 | 0.25175 | 0.00002 | 0.11749 | 0.04225 | 0.42470 | 0.89379 | 0.49329 | 0.51706 |
| **ZNF781** | 0.28504 | 0.08402 | 0.89423 | 0.15944 | 0.00000 | 0.20158 | 0.16652 | 0.00000 | 0.52674 | 0.12433 | 0.24052 | 1.53411 |
| **ZNF782** | 0.35489 | 0.05863 | 0.00000 | 0.00000 | 0.00000 | 1.52753 | 0.00000 | 0.01941 | 0.25829 | 1.04086 | 0.45299 | 0.27540 |
| **ZNF783** | 0.07944 | 1.58060 | 0.00000 | 1.70408 | 2.43818 | 1.43670 | 0.48324 | 0.00000 | 0.00000 | 1.42347 | 2.10119 | 2.37775 |
| **ZNF784** | 0.19049 | 0.04013 | 0.33202 | 0.08184 | 0.21360 | 0.00000 | 0.13871 | 0.00228 | 0.00000 | 0.53425 | 0.00000 | 0.12116 |
| **ZNF785** | 0.04262 | 0.18953 | 0.04476 | 0.39862 | 0.01130 | 0.04951 | 0.02754 | 0.02693 | 0.01420 | 0.00000 | 0.19770 | 0.11675 |
| **ZNF786** | 0.18171 | 0.63626 | 1.28183 | 0.36093 | 0.62913 | 0.00010 | 0.73256 | 0.23638 | 0.56367 | 0.23284 | 2.10120 | 1.11670 |
| **ZNF787** | 0.34458 | 0.44949 | 0.09364 | 0.38610 | 1.54231 | 0.34609 | 2.00815 | 0.40776 | 0.00000 | 3.52415 | 0.00000 | 0.06763 |

| | | | | | | | | | | | | |
|---|---|---|---|---|---|---|---|---|---|---|---|---|
| **ZNF788** | 6.29077 | 1.32419 | 2.21493 | 0.29066 | 1.23813 | 1.75130 | 2.46482 | 1.71434 | 2.35333 | 0.00000 | 2.65352 | 1.14557 |
| **ZNF789** | 0.50532 | 3.25184 | 1.75008 | 0.65461 | 4.70084 | 2.88957 | 2.84810 | 3.20102 | 3.06410 | 0.67020 | 2.33820 | 2.47466 |
| **ZNF79** | 0.91602 | 3.38511 | 2.66174 | 1.25439 | 1.23948 | 0.00027 | 0.00000 | 7.48608 | 2.83266 | 0.69224 | 1.87243 | 2.87709 |
| **ZNF790** | 0.15726 | 0.18584 | 0.94730 | 0.12042 | 0.00629 | 1.98541 | 0.00105 | 0.49479 | 1.27272 | 1.09805 | 0.56264 | 0.45621 |
| **ZNF790-AS1** | 0.00000 | 0.12631 | 1.60753 | 1.08032 | 0.00143 | 1.31686 | 1.00684 | 8.16072 | 0.98971 | 0.90894 | 0.19658 | 1.28449 |
| **ZNF791** | 0.00954 | 6.00972 | 3.75145 | 5.00340 | 3.20924 | 6.79538 | 0.00000 | 8.73813 | 6.70341 | 0.00000 | 2.68420 | 6.87807 |
| **ZNF792** | 0.67819 | 0.48010 | 0.04129 | 0.19042 | 0.00000 | 0.00000 | 0.02972 | 0.19310 | 0.03649 | 0.19880 | 0.08196 | 0.05919 |
| **ZNF793** | 0.10163 | 1.54430 | 0.53247 | 0.80263 | 0.00000 | 0.92219 | 0.53329 | 0.74316 | 0.57271 | 0.36761 | 0.21920 | 1.85612 |
| **ZNF799** | 1.62996 | 0.99633 | 0.76559 | 0.25224 | 0.04942 | 2.31963 | 0.00000 | 0.85454 | 0.00000 | 0.93731 | 0.06657 | 0.46582 |
| **ZNF8** | 0.33976 | 3.78582 | 1.88316 | 0.05453 | 2.10406 | 1.73326 | 0.00000 | 1.74831 | 5.74682 | 0.01064 | 0.36352 | 4.90119 |
| **ZNF80** | 0.00486 | 0.01936 | 0.13043 | 0.01843 | 0.01610 | 0.00000 | 0.02724 | 0.01560 | 0.01440 | 0.03213 | 0.01750 | 0.01671 |
| **ZNF800** | 1.11535 | 2.50778 | 5.45960 | 0.00000 | 4.05401 | 3.87084 | 0.00000 | 0.23481 | 6.08610 | 10.58599 | 2.11236 | 1.25094 |
| **ZNF804A** | 0.00735 | 0.03548 | 0.01570 | 0.01126 | 0.01842 | 0.00446 | 0.02127 | 0.02066 | 0.01802 | 0.02067 | 0.01824 | 0.01371 |
| **ZNF804B** | 0.08257 | 0.01414 | 0.03619 | 0.00963 | 0.04416 | 0.00643 | 0.01411 | 0.09773 | 0.01027 | 0.02386 | 0.01853 | 0.01058 |
| **ZNF805** | 2.66627 | 2.62854 | 1.31704 | 1.21775 | 0.11870 | 3.59819 | 2.43332 | 2.49211 | 2.87009 | 0.89635 | 3.43457 | 4.45338 |
| **ZNF808** | 0.24174 | 0.67781 | 1.02087 | 0.62493 | 0.00000 | 1.52085 | 0.00000 | 2.41804 | 1.54169 | 0.86696 | 1.60005 | 1.89840 |
| **ZNF81** | 0.00000 | 0.15724 | 0.29051 | 0.83457 | 0.00000 | 0.84136 | 0.42730 | 0.08777 | 0.62785 | 0.23936 | 0.43600 | 0.24386 |
| **ZNF812** | 0.09737 | 0.00000 | 0.19256 | 0.04289 | 0.07783 | 0.00000 | 0.06158 | 0.13589 | 0.00000 | 0.06731 | 0.06988 | 0.18875 |
| **ZNF813** | 0.90144 | 1.46626 | 0.41417 | 1.19337 | 0.00000 | 4.32718 | 0.96932 | 0.62346 | 0.69852 | 0.00000 | 1.06048 | 1.93366 |
| **ZNF814** | 2.01893 | 9.91302 | 3.70894 | 5.37352 | 4.19700 | 5.71737 | 9.29127 | 8.30826 | 5.02969 | 7.24361 | 9.61388 | 17.48103 |
| **ZNF815P** | 0.56640 | 0.04487 | 0.01906 | 0.00000 | 0.05198 | 0.15521 | 0.11538 | 1.08458 | 0.05281 | 1.14245 | 0.00950 | 0.98117 |
| **ZNF816** | 0.39595 | 2.91220 | 2.03817 | 0.21193 | 0.87757 | 7.33425 | 1.12733 | 3.75109 | 2.22165 | 2.24033 | 3.62281 | 8.66318 |
| **ZNF818P** | 0.01636 | 0.52815 | 0.46207 | 0.00000 | 0.59220 | 0.00000 | 0.22303 | 0.42634 | 0.00000 | 0.84641 | 0.29548 | 1.86887 |
| **ZNF821** | 1.60671 | 1.15429 | 1.30685 | 0.53254 | 0.22628 | 0.00000 | 0.80755 | 1.19291 | 0.00000 | 1.65893 | 0.06439 | 1.56316 |
| **ZNF823** | 1.10142 | 4.45299 | 3.78938 | 2.17829 | 4.32294 | 13.43348 | 0.00000 | 9.66120 | 5.00502 | 3.42555 | 4.83256 | 3.21740 |
| **ZNF826P** | 0.17060 | 2.34679 | 0.16737 | 1.40584 | 0.50633 | 1.21165 | 0.00000 | 0.59130 | 1.01242 | 0.00000 | 0.34953 | 1.87854 |
| **ZNF827** | 0.81693 | 0.00000 | 0.10642 | 1.27498 | 0.01339 | 0.00000 | 0.10595 | 0.10498 | 0.00000 | 0.06649 | 0.06679 | 0.16900 |
| **ZNF829** | 0.00000 | 1.79861 | 0.47846 | 1.21603 | 0.00000 | 3.85477 | 1.70354 | 7.99443 | 0.66553 | 3.66392 | 2.29785 | 5.69907 |
| **ZNF83** | 1.43362 | 4.02624 | 3.42186 | 0.57521 | 0.18838 | 9.30846 | 1.45652 | 0.53793 | 2.58899 | 0.88626 | 1.43491 | 8.52422 |
| **ZNF830** | 2.82705 | 7.06029 | 4.37870 | 1.62227 | 0.00000 | 10.48522 | 0.00000 | 13.18502 | 10.31733 | 4.26321 | 3.38081 | 12.87695 |
| **ZNF831** | 0.05613 | 0.01514 | 0.05990 | 0.01179 | 0.02033 | 0.01189 | 0.01196 | 0.00737 | 0.01222 | 0.01750 | 0.02146 | 0.01525 |
| **ZNF833P** | 1.22818 | 0.00000 | 0.53886 | 0.00000 | 0.14700 | 0.41826 | 0.30262 | 0.82461 | 0.19199 | 0.00000 | 0.31047 | 0.01803 |
| **ZNF835** | 0.79504 | 0.59880 | 0.29209 | 0.00120 | 0.10330 | 0.00000 | 0.02108 | 0.00000 | 0.89088 | 0.00000 | 0.98772 | 0.04859 |
| **ZNF836** | 0.22468 | 0.43096 | 1.15817 | 0.21220 | 0.00000 | 0.21044 | 0.85583 | 0.61738 | 1.07606 | 0.00000 | 0.95681 | 1.02901 |
| **ZNF837** | 0.00578 | 0.16875 | 0.01981 | 0.17221 | 0.26882 | 0.00000 | 0.01186 | 0.00442 | 0.01498 | 0.15935 | 0.00000 | 0.21949 |
| **ZNF839** | 0.33831 | 2.22972 | 0.02610 | 0.00000 | 1.45524 | 0.76278 | 0.32721 | 0.29024 | 0.05180 | 2.48706 | 0.81411 | 1.73807 |
| **ZNF84.uid-1752** | 0.54577 | 5.02455 | 2.53534 | 0.97251 | 0.00000 | 3.12237 | 1.38665 | 5.81152 | 3.15314 | 2.58293 | 0.99928 | 5.45584 |
| **ZNF841** | 0.00246 | 1.21883 | 2.03967 | 0.34187 | 0.61166 | 0.00050 | 0.00689 | 4.52627 | 0.38144 | 1.20991 | 1.95644 | 1.85958 |
| **ZNF843** | 0.00548 | 0.16679 | 0.01147 | 0.00000 | 0.07896 | 0.00000 | 0.02734 | 0.88661 | 0.00511 | 0.04991 | 0.00937 | 0.36173 |
| **ZNF844** | 2.10618 | 0.26000 | 0.96451 | 0.10244 | 0.07847 | 3.17546 | 0.00000 | 2.51456 | 0.62118 | 1.56884 | 0.00000 | 1.12992 |
| **ZNF845** | 0.72666 | 4.35290 | 2.49990 | 1.27833 | 1.19689 | 1.55551 | 2.05982 | 5.00222 | 3.06351 | 0.00000 | 0.69788 | 10.24593 |
| **ZNF846** | 0.08618 | 0.18022 | 0.04769 | 0.07209 | 0.00000 | 0.02186 | 0.02943 | 0.21656 | 0.00335 | 0.00954 | 0.06114 | 0.42066 |
| **ZNF847P** | 0.00795 | 0.01644 | 0.11429 | 0.03779 | 0.00000 | 0.03568 | 0.00000 | 0.00995 | 0.00000 | 0.10770 | 0.01351 | 0.01996 |
| **ZNF85** | 0.14955 | 2.15305 | 0.46246 | 1.69049 | 0.00000 | 4.39739 | 0.00000 | 1.67319 | 2.51047 | 0.69503 | 0.77571 | 1.81593 |
| **ZNF850** | 2.38129 | 6.07798 | 3.39556 | 3.29072 | 1.65648 | 9.83438 | 6.92132 | 3.03110 | 5.87027 | 7.80635 | 1.40965 | 6.85443 |
| **ZNF852** | 0.00000 | 0.12294 | 0.14662 | 0.14819 | 0.00000 | 1.62783 | 0.00000 | 2.27270 | 0.66008 | 1.25705 | 0.86021 | 1.33443 |
| **ZNF853** | 0.04825 | 0.01626 | 0.06014 | 0.03943 | 0.01982 | 0.00000 | 0.01430 | 0.02418 | 0.01826 | 0.00950 | 0.01614 | 0.02250 |
| **ZNF860** | 0.12946 | 0.09881 | 0.73532 | 0.43431 | 0.00000 | 0.09104 | 0.61286 | 0.13045 | 0.59106 | 0.06036 | 0.47298 | 0.05824 |

| | | | | | | | | | | | | |
|---|---|---|---|---|---|---|---|---|---|---|---|---|
| **ZNF862** | 0.25864 | 0.07890 | 0.06914 | 0.24982 | 0.03947 | 0.20244 | 0.00000 | 0.00097 | 0.00074 | 0.01216 | 0.01944 | 0.13428 |
| **ZNF865** | 0.96275 | 0.07145 | 0.01204 | 0.02156 | 0.25250 | 0.00000 | 0.10199 | 0.16336 | 0.13970 | 0.40826 | 0.00000 | 0.27016 |
| **ZNF876P** | 2.20978 | 3.54501 | 1.59334 | 0.00000 | 0.00000 | 3.49034 | 0.59221 | 0.47040 | 2.80970 | 1.98665 | 2.36806 | 2.59677 |
| **ZNF878** | 0.00984 | 0.00000 | 1.58217 | 1.19670 | 0.00000 | 0.02400 | 0.31414 | 5.76517 | 1.40072 | 0.00000 | 1.01001 | 0.98360 |
| **ZNF879** | 0.53361 | 0.17749 | 0.00000 | 0.30729 | 0.00000 | 1.32266 | 0.00107 | 0.00000 | 0.00000 | 0.94667 | 0.56518 | 0.39836 |
| **ZNF880** | 0.20272 | 7.84390 | 1.18599 | 0.21283 | 0.00000 | 0.59339 | 1.91490 | 5.44014 | 1.46698 | 3.30521 | 3.35741 | 7.75710 |
| **ZNF883** | 0.36129 | 0.23305 | 0.21937 | 0.01696 | 0.00131 | 0.46580 | 0.00000 | 0.13261 | 0.06064 | 0.00000 | 0.00290 | 0.07007 |
| **ZNF890P** | 0.01528 | 0.01669 | 0.01869 | 0.01034 | 0.01196 | 0.02084 | 0.01459 | 0.01851 | 0.01708 | 0.03783 | 0.01406 | 0.04594 |
| **ZNF891** | 1.65191 | 0.83318 | 0.81340 | 0.47550 | 0.00000 | 0.65821 | 0.50152 | 2.87565 | 0.94325 | 0.17627 | 0.12610 | 2.99443 |
| **ZNF90** | 0.64186 | 11.28997 | 1.07476 | 5.38067 | 0.00000 | 27.14577 | 0.00000 | 5.50728 | 5.97932 | 13.90146 | 1.69214 | 13.85396 |
| **ZNF91** | 1.05291 | 0.37174 | 4.67069 | 1.80713 | 0.20591 | 5.41621 | 0.00000 | 1.49232 | 2.98353 | 6.18127 | 3.20624 | 2.42147 |
| **ZNF92** | 2.07194 | 9.44095 | 0.62379 | 0.38841 | 0.00000 | 20.18668 | 0.00000 | 10.03837 | 7.29043 | 7.60150 | 3.57576 | 13.39798 |
| **ZNF93** | 3.30825 | 5.85575 | 3.95061 | 3.13752 | 0.69707 | 12.35127 | 0.00455 | 8.73350 | 8.60697 | 2.68602 | 3.58023 | 8.66242 |
| **ZNF98** | 0.29847 | 0.00000 | 0.31084 | 0.61845 | 0.11607 | 0.45709 | 0.00000 | 0.41941 | 0.69073 | 1.01957 | 1.65166 | 1.16260 |
| **ZNF99** | 0.01306 | 1.66550 | 0.20323 | 0.00000 | 0.00000 | 2.47242 | 0.00000 | 0.74540 | 0.43016 | 1.17828 | 1.82336 | 0.60530 |
| **ZNFX1** | 1.86913 | 4.12115 | 4.50740 | 3.57408 | 5.06078 | 0.50380 | 7.05017 | 12.27572 | 10.77509 | 8.59033 | 3.48312 | 9.83230 |
| **ZNHIT1** | 0.51848 | 19.41908 | 16.54491 | 19.49778 | 20.44970 | 2.79806 | 28.35676 | 18.42982 | 15.04780 | 0.00000 | 14.40948 | 23.50967 |
| **ZNHIT2** | 1.28641 | 3.42010 | 4.11468 | 2.90174 | 3.68406 | 0.00000 | 3.20312 | 3.43959 | 3.35787 | 0.00000 | 3.32490 | 9.92863 |
| **ZNHIT3** | 8.89295 | 11.21032 | 11.12966 | 8.73062 | 7.85636 | 26.56678 | 17.90408 | 20.26543 | 13.11431 | 1.81817 | 7.99091 | 16.61708 |
| **ZNHIT6** | 4.43990 | 2.82948 | 2.89053 | 1.74610 | 3.21223 | 6.69119 | 8.86874 | 9.13812 | 8.07689 | 14.74283 | 5.47555 | 9.30555 |
| **ZNRD1** | 4.41329 | 10.19525 | 4.27497 | 5.12117 | 11.44816 | 6.76550 | 6.54010 | 3.67533 | 6.24201 | 1.12546 | 5.53215 | 11.93241 |
| **ZNRD1-AS1** | 0.32266 | 0.17128 | 0.10386 | 0.00000 | 0.33764 | 0.27916 | 0.07753 | 0.11607 | 0.00000 | 0.22193 | 0.10174 | 0.76625 |
| **ZNRF1** | 0.16213 | 0.68305 | 5.64189 | 1.46277 | 4.77230 | 0.01015 | 13.30532 | 0.42702 | 0.00000 | 20.05721 | 0.62787 | 3.02664 |
| **ZNRF2** | 0.38024 | 0.00000 | 1.44258 | 0.00000 | 0.00000 | 0.88205 | 2.69877 | 0.02588 | 0.25280 | 1.69482 | 0.00000 | 0.10355 |
| **ZNRF2P1** | 0.01460 | 0.03255 | 0.09349 | 0.03050 | 0.05661 | 0.00001 | 0.03380 | 0.01089 | 0.06731 | 0.04663 | 0.02938 | 0.01845 |
| **ZNRF3** | 0.16068 | 2.16689 | 4.23853 | 6.39700 | 1.62668 | 0.05186 | 1.23119 | 0.04359 | 8.19421 | 14.04795 | 0.07285 | 1.76885 |
| **ZNRF3-AS1** | 0.00378 | 0.00000 | 0.06569 | 0.13345 | 0.03684 | 0.06952 | 0.00000 | 0.05269 | 0.24053 | 0.44812 | 0.00000 | 0.01578 |
| **ZNRF4** | 0.00831 | 0.01290 | 0.01774 | 0.01360 | 0.01786 | 0.00922 | 0.01836 | 0.01363 | 0.01696 | 0.02044 | 0.01592 | 0.03857 |
| **ZP1** | 5.28309 | 1.18358 | 0.00000 | 0.12276 | 0.12737 | 0.00000 | 0.04105 | 10.99240 | 0.00000 | 0.00000 | 0.01832 | 1.65805 |
| **ZP2** | 15.85655 | 4.29225 | 0.00000 | 0.30815 | 0.56394 | 0.00000 | 0.01766 | 22.51361 | 0.00000 | 0.04884 | 0.05160 | 4.97206 |
| **ZP3** | 18.06028 | 5.67209 | 0.00000 | 0.19495 | 0.62766 | 0.00000 | 0.03835 | 28.58947 | 0.00000 | 0.00000 | 0.57136 | 10.53200 |
| **ZP4** | 6.26033 | 1.71739 | 0.00000 | 0.00000 | 0.00000 | 0.00000 | 0.00000 | 23.56533 | 0.00000 | 0.47960 | 0.00000 | 1.25459 |
| **ZPBP** | 0.03989 | 0.02974 | 0.00981 | 0.01025 | 0.00412 | 0.02427 | 0.03838 | 0.02421 | 0.01949 | 0.00000 | 0.09106 | 0.14682 |
| **ZPBP2** | 0.00281 | 0.00000 | 0.01807 | 0.00795 | 0.02086 | 0.02882 | 0.01527 | 0.44573 | 0.00978 | 0.01472 | 0.01615 | 0.63458 |
| **ZPLD1** | 0.92521 | 0.02261 | 0.01466 | 0.01268 | 0.04042 | 0.00359 | 0.02065 | 0.31224 | 0.02157 | 0.00000 | 0.01102 | 0.00549 |
| **ZPR1** | 0.09593 | 11.51169 | 14.93145 | 4.94745 | 13.70343 | 0.81443 | 8.74040 | 0.67490 | 12.05680 | 7.65623 | 12.92512 | 4.97316 |
| **ZRANB1** | 3.76096 | 0.00000 | 0.29737 | 2.15316 | 3.41694 | 8.62587 | 6.39178 | 5.00015 | 0.25031 | 13.26736 | 1.07902 | 3.20424 |
| **ZRANB2** | 0.14648 | 9.92632 | 13.58880 | 2.26764 | 3.39149 | 32.86580 | 18.25257 | 15.32505 | 8.49768 | 21.44997 | 12.83151 | 21.26144 |
| **ZRANB2-AS1** | 0.00231 | 0.02159 | 0.06452 | 0.00000 | 0.01200 | 0.37406 | 0.00000 | 0.03579 | 0.08822 | 0.09243 | 0.05069 | 0.04612 |
| **ZRANB2-AS2** | 0.00444 | 0.09794 | 0.07772 | 0.00201 | 0.00904 | 0.32030 | 0.00000 | 0.17627 | 0.00000 | 0.04189 | 0.03178 | 0.48386 |
| **ZRANB3** | 0.00000 | 2.61514 | 1.67279 | 1.68738 | 0.46085 | 3.50059 | 1.56007 | 1.07451 | 0.36942 | 0.97658 | 2.55867 | 2.11253 |
| **ZRSR2** | 1.06192 | 0.10033 | 0.78854 | 0.07093 | 0.17576 | 0.54489 | 0.90117 | 1.00840 | 0.00000 | 1.73015 | 0.27565 | 0.94219 |
| **ZSCAN1** | 0.01551 | 0.13418 | 0.01552 | 0.00000 | 0.00882 | 0.00000 | 0.00073 | 0.01803 | 0.10872 | 0.18986 | 0.00000 | 0.26206 |
| **ZSCAN10** | 0.43178 | 0.00000 | 1.22688 | 5.33672 | 0.00000 | 0.00000 | 0.48774 | 0.04354 | 0.59375 | 0.00262 | 1.65176 | 0.01597 |
| **ZSCAN12** | 0.00000 | 0.38980 | 0.00000 | 0.00004 | 0.88503 | 2.17894 | 0.31689 | 0.40466 | 0.00000 | 0.34353 | 0.30164 | 0.21197 |
| **ZSCAN12P1** | 0.03578 | 0.00000 | 1.87531 | 0.19726 | 0.00000 | 0.17865 | 0.87022 | 0.00000 | 0.93976 | 0.00000 | 1.00015 | 0.42625 |
| **ZSCAN16** | 0.63123 | 4.15021 | 0.04478 | 0.37625 | 1.00880 | 1.44930 | 0.38815 | 0.37483 | 0.14030 | 0.00000 | 1.61474 | 0.22094 |
| **ZSCAN16-AS1** | 0.01111 | 0.79633 | 0.00000 | 0.23013 | 1.05006 | 1.10377 | 0.17049 | 0.00352 | 0.00000 | 0.44992 | 0.71319 | 1.96505 |

| | | | | | | | | | | | | |
|---|---|---|---|---|---|---|---|---|---|---|---|---|
| **ZSCAN18** | 0.07260 | 0.78032 | 0.22378 | 0.00000 | 0.19079 | 0.00000 | 0.38885 | 2.11933 | 0.00000 | 1.04461 | 0.07529 | 1.75970 |
| **ZSCAN2** | 3.45662 | 1.67171 | 1.91150 | 3.91815 | 0.00000 | 0.89053 | 1.47412 | 1.05964 | 0.91369 | 0.00000 | 1.47307 | 3.95958 |
| **ZSCAN20** | 0.00509 | 0.20929 | 0.31072 | 0.25702 | 0.27920 | 0.17258 | 0.24978 | 0.34974 | 0.38442 | 0.00000 | 0.17596 | 0.50244 |
| **ZSCAN21** | 1.84608 | 5.50009 | 1.13591 | 0.66837 | 1.83393 | 2.50675 | 0.65485 | 0.60538 | 0.50259 | 0.17117 | 1.17130 | 6.43382 |
| **ZSCAN22** | 0.48785 | 1.29900 | 1.02394 | 1.30032 | 0.99988 | 0.02813 | 1.11483 | 0.61402 | 1.63837 | 0.00000 | 1.30957 | 1.43422 |
| **ZSCAN23** | 0.04944 | 0.11267 | 0.01416 | 0.02446 | 0.03353 | 0.01276 | 0.05602 | 0.05525 | 0.24713 | 0.00000 | 0.02207 | 0.10432 |
| **ZSCAN25** | 1.20615 | 5.39083 | 0.42437 | 0.49966 | 0.00000 | 4.11477 | 1.28923 | 1.71666 | 4.15644 | 1.71320 | 3.74727 | 5.48573 |
| **ZSCAN26** | 1.70357 | 1.50069 | 0.07215 | 0.40597 | 0.00999 | 0.52235 | 0.00000 | 0.85629 | 0.00000 | 0.34649 | 0.30282 | 2.03295 |
| **ZSCAN29** | 4.20978 | 1.45767 | 2.01221 | 1.51882 | 1.94443 | 5.15721 | 4.36674 | 0.76262 | 0.99564 | 5.37148 | 1.09405 | 3.02858 |
| **ZSCAN30** | 0.01036 | 0.81180 | 0.16106 | 0.30916 | 0.79248 | 0.79926 | 0.00000 | 0.40653 | 0.00000 | 0.00000 | 0.19848 | 0.76865 |
| **ZSCAN31** | 0.00420 | 0.31063 | 0.47357 | 0.00239 | 0.00000 | 0.47920 | 0.76633 | 0.24599 | 1.15756 | 0.00000 | 0.01162 | 0.19291 |
| **ZSCAN32** | 0.16477 | 3.42322 | 1.90109 | 2.46147 | 1.35306 | 3.01926 | 2.40363 | 7.05216 | 2.36331 | 0.00000 | 1.32635 | 5.69656 |
| **ZSCAN4** | 2.08321 | 0.00000 | 0.09539 | 0.00718 | 0.03241 | 0.00000 | 0.00000 | 47.67049 | 0.01879 | 0.00326 | 0.00000 | 5.65278 |
| **ZSCAN5A** | 3.78497 | 3.58816 | 0.81426 | 0.22252 | 0.86926 | 2.46850 | 2.49977 | 6.09579 | 2.04882 | 0.00000 | 0.00000 | 4.62021 |
| **ZSCAN5B** | 3.62106 | 0.04661 | 0.00652 | 0.00166 | 0.02804 | 0.00000 | 0.00745 | 30.99460 | 0.07257 | 0.00000 | 0.00000 | 9.99724 |
| **ZSCAN9** | 1.76071 | 1.94627 | 0.21007 | 0.45606 | 0.00000 | 3.76332 | 0.00000 | 0.64913 | 0.84345 | 1.04554 | 1.04184 | 3.80774 |
| **ZSWIM1** | 0.00283 | 1.51633 | 2.04267 | 0.31887 | 2.02965 | 1.87308 | 2.41090 | 0.07772 | 1.51092 | 0.33193 | 2.60442 | 3.08736 |
| **ZSWIM3** | 4.79704 | 3.50271 | 1.11061 | 5.39930 | 0.00530 | 2.24673 | 0.00000 | 0.00000 | 0.83777 | 0.22625 | 0.81868 | 4.01995 |
| **ZSWIM4** | 0.87959 | 0.00000 | 1.03164 | 0.11730 | 0.00000 | 0.07183 | 0.31277 | 0.00000 | 0.00857 | 0.91992 | 0.04879 | 0.26109 |
| **ZSWIM5** | 0.00006 | 0.45764 | 0.00000 | 0.77008 | 0.31581 | 0.16830 | 0.03602 | 0.17154 | 0.00000 | 0.51719 | 0.00000 | 0.31478 |
| **ZSWIM6** | 0.00783 | 0.00518 | 0.22821 | 0.93303 | 0.05994 | 0.28957 | 0.17915 | 0.23082 | 0.10453 | 0.43205 | 0.00000 | 0.00000 |
| **ZSWIM7** | 2.59615 | 8.99964 | 4.15622 | 10.66903 | 7.48071 | 12.51852 | 5.86631 | 2.87418 | 5.61275 | 5.85392 | 9.98389 | 5.14307 |
| **ZSWIM8** | 0.72114 | 0.00000 | 5.55972 | 0.00000 | 2.65207 | 0.00000 | 2.35936 | 0.50833 | 4.24430 | 7.33482 | 0.62167 | 1.60725 |
| **ZSWIM8-AS1** | 0.11818 | 0.10011 | 0.06766 | 0.00000 | 0.12579 | 0.04969 | 0.36832 | 0.02822 | 0.10471 | 0.25572 | 0.00000 | 0.07106 |
| **ZUFSP** | 0.00245 | 3.62075 | 2.56049 | 1.45525 | 0.89082 | 11.54993 | 0.00000 | 3.32646 | 6.66971 | 0.00000 | 3.57455 | 5.83373 |
| **ZW10** | 2.03684 | 6.13594 | 5.71480 | 1.61332 | 0.00000 | 13.28938 | 6.60831 | 4.03525 | 5.06977 | 0.00000 | 8.07827 | 4.73550 |
| **ZWILCH** | 1.03079 | 8.20387 | 9.66097 | 6.13023 | 4.44431 | 29.09443 | 13.12815 | 2.71428 | 7.52894 | 7.63651 | 10.43417 | 11.25323 |
| **ZWINT** | 7.17687 | 19.77833 | 20.96832 | 13.46959 | 14.99245 | 38.80425 | 28.90393 | 17.43841 | 28.22927 | 0.00000 | 22.15215 | 27.86087 |
| **ZXDA** | 0.00573 | 0.01254 | 0.00000 | 0.00302 | 0.17036 | 0.13970 | 0.04538 | 0.00897 | 0.13656 | 0.00000 | 0.00000 | 0.01138 |
| **ZXDB** | 0.16486 | 0.08049 | 0.14924 | 0.17931 | 0.39040 | 0.32103 | 0.18948 | 0.00000 | 0.00000 | 1.56853 | 0.20733 | 0.00000 |
| **ZXDC** | 1.23245 | 1.31595 | 7.24319 | 2.82795 | 3.42272 | 3.34717 | 5.69380 | 0.00000 | 0.08778 | 7.39690 | 1.89764 | 1.11217 |
| **ZYG11A** | 3.00003 | 7.29986 | 0.93019 | 13.07160 | 3.24470 | 4.45252 | 1.33233 | 2.11948 | 2.75458 | 4.43310 | 1.12012 | 5.65189 |
| **ZYG11B** | 0.00000 | 0.87216 | 3.07219 | 1.15787 | 2.58658 | 5.97110 | 1.59145 | 0.28908 | 0.00000 | 5.55608 | 2.50733 | 0.17661 |
| **ZYX** | 4.79902 | 13.99515 | 9.35087 | 9.55776 | 18.27421 | 0.00000 | 8.77596 | 1.39245 | 7.90539 | 14.97017 | 1.69559 | 5.18088 |
| **ZZEF1** | 2.57975 | 2.07202 | 2.60997 | 4.14785 | 4.03651 | 0.11987 | 2.83621 | 0.89814 | 2.19311 | 8.60374 | 3.10045 | 3.05848 |
| **ZZZ3** | 0.48354 | 4.97527 | 2.85180 | 2.49075 | 0.87771 | 16.98532 | 9.09559 | 13.26565 | 8.91609 | 6.00504 | 6.47639 | 5.56532 |

### Supplementary Table S11

#### Human female NMF coefficient matrix

| Gene | Cluster 1 | Cluster 2 | Cluster 3 | Cluster 4 | Cluster 5 | Cluster 6 | Cluster 7 |
|---|---|---|---|---|---|---|---|
| A1BG | 0.00000 | 0.09312 | 1.17750 | 0.63035 | 8.57770 | 0.45704 | 0.05919 |
| A1BG-AS1 | 0.04762 | 0.00194 | 0.10790 | 0.10843 | 0.23029 | 0.03922 | 0.00163 |
| A1CF | 0.00000 | 0.04317 | 0.00000 | 0.11007 | 1.07607 | 0.04331 | 0.06853 |
| A2M | 0.00789 | 0.05163 | 0.03856 | 0.11234 | 0.01345 | 0.45918 | 0.00557 |
| A2M-AS1 | 0.01255 | 0.04448 | 0.03819 | 0.07086 | 0.00554 | 0.01267 | 0.06451 |
| A2ML1 | 0.61338 | 0.00000 | 0.06982 | 1.58774 | 0.51199 | 0.02054 | 0.31262 |
| A3GALT2 | 0.01313 | 0.01548 | 0.21323 | 0.01841 | 0.18932 | 0.01044 | 0.02287 |
| A4GALT | 7.14825 | 27.94214 | 8.10132 | 7.24734 | 0.64582 | 12.65962 | 33.23882 |
| A4GNT | 0.03763 | 0.01922 | 0.04635 | 0.02130 | 0.01164 | 0.11247 | 0.00709 |
| AAAS | 16.48837 | 17.05797 | 9.17568 | 0.88818 | 2.45853 | 7.72716 | 7.77320 |
| AACS | 7.46654 | 1.04718 | 0.23184 | 1.97483 | 1.29474 | 2.05144 | 6.19759 |
| AACSP1 | 0.02075 | 0.00000 | 0.20507 | 0.05391 | 0.51689 | 0.00267 | 0.03535 |
| AADAC | 0.06540 | 0.00388 | 0.01389 | 0.00013 | 0.00945 | 0.01767 | 0.09489 |
| AADACL2 | 0.05053 | 0.05969 | 0.01458 | 0.02180 | 0.00741 | 0.02976 | 0.09736 |
| AADACL2-AS1 | 0.24596 | 0.24514 | 0.11867 | 0.28334 | 0.06970 | 0.10345 | 0.33783 |
| AADACL3 | 0.96165 | 0.36503 | 0.00029 | 0.00043 | 0.00582 | 0.56743 | 0.00000 |
| AADACP1 | 0.04205 | 0.02615 | 0.01129 | 0.02465 | 0.00717 | 0.00399 | 0.07475 |
| AADAT | 5.21977 | 0.35412 | 0.50267 | 2.03911 | 0.12063 | 1.58069 | 0.00000 |
| AAED1 | 0.01803 | 0.00000 | 1.81232 | 1.38971 | 0.07667 | 0.26628 | 1.88757 |
| AAGAB | 19.31638 | 6.80503 | 6.63983 | 14.98670 | 1.83630 | 5.63051 | 0.00000 |
| AAK1 | 11.97999 | 1.68271 | 2.71164 | 14.85838 | 2.95298 | 1.62833 | 26.02898 |
| AAMDC | 1.51317 | 10.19067 | 8.11042 | 0.39220 | 4.19293 | 5.13781 | 1.44172 |
| AAMP | 44.04338 | 40.62208 | 17.34833 | 42.27853 | 8.63516 | 19.09067 | 0.00000 |
| AANAT | 0.01303 | 0.01176 | 0.01719 | 0.02037 | 0.01142 | 0.00762 | 0.21530 |
| AAR2 | 19.71431 | 12.44382 | 9.73011 | 12.88251 | 3.84606 | 5.88579 | 0.00000 |
| AARD | 0.10237 | 0.11552 | 0.03070 | 0.04317 | 0.03000 | 0.02759 | 0.04842 |
| AARS | 28.89499 | 5.70521 | 4.46021 | 19.52587 | 0.71651 | 8.87704 | 11.24959 |
| AARS2 | 2.02302 | 10.97623 | 6.59466 | 2.11685 | 0.56685 | 15.40287 | 5.26510 |
| AARSD1 | 0.67198 | 0.33360 | 0.18654 | 0.20573 | 0.12438 | 0.17219 | 0.00000 |
| AASDH | 4.71553 | 0.67385 | 4.81116 | 0.91948 | 1.18814 | 2.84440 | 3.12777 |
| AASDHPPT | 20.91809 | 3.74631 | 9.78016 | 10.06558 | 0.35425 | 5.59841 | 0.98943 |
| AASS | 3.78756 | 0.28344 | 1.47890 | 8.52331 | 0.59020 | 3.69165 | 2.04394 |
| AATF | 26.66974 | 25.26784 | 11.63847 | 21.39178 | 6.29970 | 9.54388 | 0.00000 |

| | | | | | | | |
|---|---|---|---|---|---|---|---|
| **AATK** | 0.01952 | 0.01820 | 0.00861 | 0.02492 | 0.04812 | 0.02249 | 0.02739 |
| **AATK-AS1** | 0.02762 | 0.01753 | 0.01262 | 0.01900 | 0.00777 | 0.01174 | 0.02637 |
| **ABAT** | 0.00000 | 0.71810 | 1.09896 | 0.04321 | 0.74989 | 0.87430 | 0.02157 |
| **ABCA1** | 7.01516 | 0.00000 | 2.79501 | 1.47660 | 1.70554 | 16.32350 | 4.10411 |
| **ABCA10** | 0.00000 | 0.02380 | 0.90432 | 0.00994 | 0.09890 | 0.02670 | 0.04642 |
| **ABCA11P** | 2.20429 | 0.49333 | 1.15434 | 0.59494 | 1.23353 | 0.44946 | 0.20166 |
| **ABCA12** | 0.16563 | 0.04103 | 0.14118 | 0.00000 | 0.24677 | 0.11334 | 0.12566 |
| **ABCA13** | 1.44563 | 0.67459 | 0.95989 | 0.00000 | 0.14721 | 0.23375 | 0.60329 |
| **ABCA17P** | 0.00000 | 0.53962 | 8.53954 | 0.00250 | 1.32689 | 0.02261 | 0.00000 |
| **ABCA2** | 0.03643 | 0.12782 | 0.26921 | 0.07169 | 0.34323 | 0.19950 | 0.53837 |
| **ABCA3** | 0.20094 | 0.00000 | 1.08096 | 0.00000 | 0.82520 | 0.10099 | 0.22907 |
| **ABCA4** | 1.06762 | 0.00000 | 0.00000 | 0.95669 | 0.19531 | 0.00000 | 0.92923 |
| **ABCA5** | 0.63439 | 0.00000 | 0.50346 | 0.17055 | 1.22627 | 0.39609 | 0.50622 |
| **ABCA6** | 0.04158 | 0.00223 | 0.04358 | 0.02286 | 0.09688 | 0.01105 | 0.01907 |
| **ABCA7** | 0.42886 | 0.58758 | 0.17598 | 1.15590 | 0.12184 | 1.09407 | 3.16937 |
| **ABCA8** | 0.02441 | 0.01732 | 0.01016 | 0.02641 | 0.04108 | 0.01481 | 0.01374 |
| **ABCA9** | 0.02962 | 0.00000 | 0.08189 | 0.02240 | 0.06798 | 0.02461 | 0.01810 |
| **ABCB1** | 0.04804 | 0.13645 | 0.46561 | 0.00077 | 0.46914 | 0.00000 | 0.00878 |
| **ABCB10** | 3.86379 | 1.54699 | 1.72045 | 0.00000 | 0.00000 | 1.46341 | 9.13207 |
| **ABCB11** | 0.01804 | 0.01676 | 0.00663 | 0.02286 | 0.08943 | 0.01182 | 0.03395 |
| **ABCB4** | 0.02290 | 0.07981 | 0.00000 | 0.21951 | 2.44906 | 0.00271 | 0.00000 |
| **ABCB5** | 0.02729 | 0.00000 | 0.09766 | 0.01486 | 0.00822 | 0.30699 | 0.04662 |
| **ABCB6** | 12.91748 | 7.88293 | 2.23003 | 17.47515 | 0.59209 | 2.66636 | 5.69782 |
| **ABCB7** | 11.02239 | 5.01400 | 3.76757 | 0.00000 | 3.36699 | 5.11582 | 5.10429 |
| **ABCB8** | 2.91213 | 19.28588 | 7.09322 | 6.71801 | 0.00000 | 9.75982 | 5.45010 |
| **ABCB9** | 0.01401 | 0.07321 | 0.25058 | 0.55932 | 2.62618 | 0.29359 | 0.53501 |
| **ABCC1** | 0.30126 | 0.62037 | 0.62860 | 1.04882 | 1.43103 | 2.10049 | 1.28890 |
| **ABCC10** | 0.41174 | 1.79549 | 0.29336 | 2.71181 | 0.02334 | 4.23941 | 2.97924 |
| **ABCC11** | 0.01458 | 0.02604 | 0.07235 | 0.02848 | 0.04227 | 0.05292 | 0.02780 |
| **ABCC12** | 0.02740 | 0.00119 | 0.04151 | 0.03955 | 0.10797 | 0.05727 | 0.01340 |
| **ABCC13** | 0.20727 | 1.18436 | 0.27381 | 0.00000 | 0.00000 | 1.19244 | 0.00000 |
| **ABCC2** | 1.28988 | 0.84879 | 0.26476 | 0.18154 | 0.13132 | 4.27509 | 0.00000 |
| **ABCC3** | 0.02237 | 0.09589 | 0.00000 | 0.01230 | 0.59600 | 0.01256 | 0.02119 |
| **ABCC4** | 11.76562 | 5.95945 | 0.02154 | 0.00000 | 2.37069 | 0.00000 | 8.73973 |
| **ABCC5** | 13.74492 | 4.30127 | 9.05442 | 19.48828 | 3.99204 | 5.90548 | 7.59270 |

| | | | | | | | |
|---|---|---|---|---|---|---|---|
| **ABCC5-AS1** | 0.02272 | 0.00000 | 0.04529 | 0.00953 | 0.00841 | 0.07059 | 0.16568 |
| **ABCC6** | 0.04013 | 0.04561 | 0.14878 | 0.00632 | 5.16983 | 0.00763 | 0.11157 |
| **ABCC6P1** | 0.00003 | 0.00008 | 0.56744 | 0.00937 | 12.73765 | 0.00000 | 0.03122 |
| **ABCC8** | 0.02565 | 0.00000 | 0.03386 | 0.03148 | 0.13240 | 0.01661 | 0.05287 |
| **ABCC9** | 0.12571 | 0.00408 | 0.01672 | 0.00000 | 0.00915 | 0.04072 | 0.28285 |
| **ABCD1** | 0.00001 | 0.98303 | 0.00000 | 1.81754 | 0.06699 | 0.00000 | 3.26053 |
| **ABCD3** | 16.74289 | 0.68488 | 7.10301 | 8.12416 | 1.31386 | 3.88692 | 8.47022 |
| **ABCD4** | 6.72023 | 2.23678 | 2.34166 | 1.95984 | 4.58514 | 3.91633 | 2.18296 |
| **ABCE1** | 33.56994 | 0.62594 | 14.21681 | 11.01074 | 2.34311 | 9.27210 | 10.98937 |
| **ABCF1** | 23.54766 | 16.77576 | 25.17365 | 28.52169 | 11.06141 | 13.26749 | 33.27166 |
| **ABCF2** | 37.57622 | 23.92666 | 25.11330 | 25.27502 | 15.85134 | 11.88537 | 5.51131 |
| **ABCF3** | 17.92810 | 20.27027 | 7.03217 | 18.07244 | 3.18766 | 9.84615 | 0.00000 |
| **ABCG1** | 0.02190 | 0.00253 | 0.02822 | 0.15656 | 0.05614 | 0.01171 | 0.08516 |
| **ABCG2** | 38.36430 | 0.00000 | 0.00000 | 31.36520 | 0.00000 | 0.00000 | 17.70148 |
| **ABCG4** | 0.00000 | 0.02947 | 0.13461 | 0.06234 | 0.21322 | 0.00000 | 0.18504 |
| **ABCG5** | 0.03089 | 0.05935 | 0.03056 | 0.01970 | 0.12803 | 0.00996 | 0.01530 |
| **ABCG8** | 0.05071 | 0.09595 | 0.08202 | 0.00281 | 1.34616 | 0.00522 | 0.02101 |
| **ABHD1** | 0.02656 | 0.02431 | 0.01679 | 0.04517 | 0.06365 | 0.04706 | 0.11838 |
| **ABHD10** | 11.72512 | 5.62062 | 1.96411 | 3.28406 | 0.16279 | 2.59333 | 0.00000 |
| **ABHD11** | 18.39003 | 13.44404 | 1.94344 | 18.29304 | 1.60256 | 4.77072 | 0.00000 |
| **ABHD12** | 1.78643 | 10.28018 | 1.25244 | 6.64849 | 1.09032 | 8.12305 | 24.97441 |
| **ABHD12B** | 4.36140 | 4.94860 | 0.81163 | 0.42132 | 0.66238 | 9.34251 | 0.00002 |
| **ABHD13** | 5.84037 | 0.00000 | 4.00560 | 1.37116 | 2.28582 | 1.63182 | 3.88869 |
| **ABHD14A** | 0.91758 | 0.19439 | 0.13929 | 1.08689 | 0.00207 | 0.09399 | 0.00000 |
| **ABHD14B** | 5.61219 | 0.35830 | 8.95128 | 2.71835 | 0.00000 | 0.32437 | 2.31662 |
| **ABHD15** | 1.27629 | 1.91693 | 2.05269 | 0.00000 | 0.00000 | 1.94402 | 1.42817 |
| **ABHD16A** | 4.38777 | 3.49139 | 2.57022 | 7.76669 | 1.99089 | 5.16110 | 0.73707 |
| **ABHD16B** | 0.00000 | 0.15921 | 0.07271 | 0.23851 | 0.06834 | 0.00000 | 0.38471 |
| **ABHD17A** | 0.00000 | 0.14230 | 0.02985 | 0.01041 | 0.01384 | 0.14718 | 1.09455 |
| **ABHD17B** | 1.84482 | 0.86966 | 1.36019 | 1.40494 | 2.00610 | 3.58950 | 7.37531 |
| **ABHD17C** | 3.30420 | 2.45387 | 4.22562 | 2.34107 | 0.00000 | 2.51785 | 14.57213 |
| **ABHD2** | 11.08995 | 4.91570 | 4.15953 | 5.47365 | 3.48494 | 4.87355 | 1.00130 |
| **ABHD3** | 7.92649 | 6.09473 | 8.89246 | 6.15931 | 8.04915 | 4.12976 | 0.00000 |
| **ABHD4** | 7.12030 | 0.02815 | 0.00000 | 15.65560 | 0.56278 | 1.19366 | 0.00000 |
| **ABHD5** | 12.78724 | 1.67789 | 3.53838 | 15.88531 | 1.14807 | 5.15691 | 6.70882 |

| | | | | | | | |
|---|---|---|---|---|---|---|---|
| **ABHD6** | 3.98120 | 2.17913 | 1.25150 | 3.05851 | 0.08645 | 3.73986 | 3.22524 |
| **ABHD8** | 1.74356 | 0.09271 | 0.11351 | 2.62025 | 0.33824 | 0.00000 | 0.18435 |
| **ABI1** | 19.27660 | 5.47896 | 5.87130 | 15.38762 | 3.90571 | 9.74039 | 0.00754 |
| **ABI2** | 2.93276 | 0.88167 | 1.94403 | 5.23986 | 1.09888 | 4.97067 | 1.02500 |
| **ABI3** | 0.00000 | 0.20996 | 0.06620 | 0.01896 | 0.09887 | 0.52849 | 0.04457 |
| **ABI3BP** | 0.12503 | 0.00000 | 0.01196 | 0.01797 | 1.37464 | 0.11016 | 0.06887 |
| **ABL1** | 0.99895 | 3.22627 | 2.32053 | 3.62711 | 1.63121 | 4.06231 | 12.18747 |
| **ABL2** | 4.52231 | 5.43757 | 1.69655 | 2.02060 | 2.15935 | 6.40514 | 9.51425 |
| **ABLIM1** | 0.96482 | 0.88043 | 2.41173 | 0.23161 | 0.46200 | 0.59421 | 0.25859 |
| **ABLIM2** | 0.04976 | 0.00170 | 0.00081 | 0.17206 | 0.44380 | 0.21119 | 0.06208 |
| **ABLIM3** | 0.22033 | 0.03859 | 0.07034 | 1.49343 | 0.01293 | 0.61070 | 0.00000 |
| **ABO** | 0.02871 | 0.01284 | 0.21076 | 0.08243 | 0.07069 | 0.04253 | 0.00000 |
| **ABR** | 4.90317 | 3.05879 | 0.87376 | 6.30632 | 1.77835 | 10.48694 | 20.62057 |
| **ABRA** | 0.03046 | 0.01242 | 0.01380 | 0.03365 | 0.00825 | 0.05012 | 0.00421 |
| **ABRACL** | 14.23124 | 12.96598 | 3.88715 | 6.24950 | 0.00000 | 8.55015 | 2.24495 |
| **ABT1** | 34.81035 | 26.80513 | 20.79096 | 27.11453 | 11.31234 | 13.86045 | 0.00000 |
| **ABTB1** | 0.00000 | 0.00000 | 1.47644 | 1.77741 | 0.09084 | 1.12931 | 0.08955 |
| **ABTB2** | 0.00000 | 5.45679 | 4.34834 | 3.65691 | 3.39099 | 4.49690 | 5.35776 |
| **AC093375.1** | 0.18792 | 0.19226 | 0.09030 | 0.01061 | 0.00462 | 0.05884 | 0.00000 |
| **ACAA1** | 9.99932 | 10.79497 | 0.28105 | 13.61866 | 2.42875 | 10.81434 | 14.45978 |
| **ACAA2** | 32.85573 | 16.94169 | 9.06331 | 13.80869 | 1.26454 | 11.94710 | 1.72127 |
| **ACACA** | 19.56918 | 2.38352 | 3.53256 | 13.11856 | 1.71812 | 5.10167 | 23.75348 |
| **ACACB** | 0.27097 | 0.23213 | 0.44688 | 0.00048 | 1.05158 | 0.49591 | 0.13666 |
| **ACAD10** | 2.95864 | 4.43008 | 2.42333 | 0.92534 | 1.19006 | 2.45867 | 0.31130 |
| **ACAD11** | 0.16931 | 0.10413 | 0.10352 | 0.06692 | 0.46074 | 0.18580 | 0.00000 |
| **ACAD8** | 7.79481 | 6.24959 | 2.24825 | 7.23152 | 0.99152 | 5.12789 | 0.00000 |
| **ACAD9** | 12.94190 | 7.60009 | 4.42405 | 7.74718 | 1.65540 | 5.95424 | 2.80638 |
| **ACADL** | 0.11088 | 0.11086 | 0.00532 | 0.00247 | 0.03042 | 0.03458 | 0.01804 |
| **ACADM** | 19.68028 | 0.14931 | 0.50220 | 8.79012 | 0.00000 | 10.52560 | 4.47384 |
| **ACADS** | 0.00000 | 6.57962 | 0.64807 | 0.00000 | 0.09945 | 7.39260 | 2.30440 |
| **ACADSB** | 11.64553 | 1.71669 | 1.65946 | 1.90849 | 3.70431 | 2.31934 | 0.00000 |
| **ACADVL** | 2.63096 | 5.98062 | 0.00000 | 13.12924 | 0.39077 | 6.09064 | 16.09419 |
| **ACAN** | 0.02149 | 0.03259 | 0.08470 | 0.02368 | 0.06629 | 0.20982 | 0.07875 |
| **ACAP1** | 0.00000 | 0.67343 | 0.42250 | 0.10831 | 0.20286 | 0.09932 | 0.27048 |
| **ACAP2** | 8.40580 | 3.17042 | 3.01059 | 6.45063 | 2.16776 | 4.52407 | 10.77742 |

| | | | | | | | |
|---|---|---|---|---|---|---|---|
| **ACAP3** | 0.00000 | 0.37269 | 0.00377 | 0.51619 | 0.01474 | 0.20277 | 4.13148 |
| **ACAT1** | 23.51115 | 6.08578 | 2.55007 | 15.66741 | 1.44729 | 3.38879 | 0.00000 |
| **ACAT2** | 61.98151 | 12.74233 | 0.51117 | 20.01131 | 5.77606 | 13.19434 | 0.00000 |
| **ACBD3** | 6.39534 | 0.46992 | 5.55427 | 4.83407 | 6.17045 | 5.79129 | 9.65532 |
| **ACBD4** | 0.07478 | 0.07207 | 0.00853 | 0.21363 | 0.09285 | 0.04498 | 0.00000 |
| **ACBD5** | 6.11908 | 0.00000 | 4.39360 | 2.44774 | 2.64079 | 2.27762 | 0.38329 |
| **ACBD6** | 3.05268 | 4.08290 | 2.92600 | 6.88576 | 0.00000 | 2.74236 | 5.03087 |
| **ACBD7** | 0.00000 | 0.28477 | 0.16433 | 0.39201 | 0.25602 | 0.28814 | 0.05791 |
| **ACCS** | 0.05781 | 0.15308 | 0.02990 | 0.04601 | 0.00549 | 0.13697 | 0.03012 |
| **ACCSL** | 0.00587 | 0.18623 | 0.07040 | 0.00291 | 11.03990 | 0.00000 | 0.01415 |
| **ACD** | 9.95623 | 13.35591 | 3.05984 | 12.22337 | 2.56432 | 6.80333 | 0.00000 |
| **ACE** | 0.00000 | 7.26854 | 8.39867 | 0.00000 | 0.00000 | 6.49493 | 0.00000 |
| **ACE2** | 17.82440 | 8.61709 | 0.00000 | 0.00000 | 0.55607 | 0.00000 | 1.85460 |
| **ACER1** | 0.02397 | 0.00000 | 0.21770 | 0.03209 | 0.04851 | 0.00748 | 0.01547 |
| **ACER2** | 0.20211 | 0.65764 | 2.06001 | 1.41534 | 0.67462 | 1.09483 | 0.00000 |
| **ACER3** | 7.20370 | 0.00000 | 5.57225 | 9.36722 | 0.41580 | 0.34862 | 1.90161 |
| **ACHE** | 0.00000 | 0.48271 | 0.23455 | 0.39970 | 0.09710 | 0.86043 | 0.12736 |
| **ACIN1** | 11.72820 | 13.82357 | 6.40631 | 16.64049 | 3.76260 | 7.40771 | 33.24398 |
| **ACKR2** | 5.79336 | 0.00000 | 0.00000 | 8.28225 | 0.00000 | 0.00000 | 3.73002 |
| **ACKR3** | 2.85836 | 0.03491 | 0.01564 | 0.00000 | 0.00186 | 1.22495 | 0.18579 |
| **ACKR4** | 0.03972 | 0.02281 | 0.03205 | 0.04150 | 0.06055 | 0.05287 | 0.02678 |
| **ACLY** | 36.49714 | 17.77791 | 5.37534 | 27.50027 | 4.82703 | 10.11866 | 25.30032 |
| **ACMSD** | 0.09627 | 0.00061 | 0.02713 | 0.04118 | 0.01359 | 0.00628 | 0.03905 |
| **ACN9** | 18.39782 | 5.67655 | 16.86431 | 8.75971 | 3.17694 | 5.60445 | 0.44848 |
| **ACO1** | 27.55477 | 6.12206 | 1.04069 | 5.44331 | 1.31641 | 4.96042 | 0.00000 |
| **ACO2** | 19.15830 | 15.79313 | 2.52705 | 14.17769 | 1.17430 | 8.77062 | 0.00000 |
| **ACOT1** | 0.09244 | 0.00000 | 0.12090 | 0.31864 | 0.12008 | 0.06156 | 0.00000 |
| **ACOT11** | 0.11346 | 0.48237 | 2.78672 | 0.08449 | 2.09603 | 5.45604 | 0.00000 |
| **ACOT12** | 0.02698 | 0.06313 | 0.02760 | 0.03424 | 0.10238 | 0.02932 | 0.00000 |
| **ACOT13** | 6.24198 | 6.92607 | 5.07567 | 7.47364 | 2.41997 | 5.55107 | 3.66922 |
| **ACOT2** | 0.88488 | 0.36335 | 2.76749 | 2.23556 | 1.31365 | 0.00000 | 0.00000 |
| **ACOT4** | 2.35198 | 1.36913 | 0.99261 | 1.35568 | 1.40374 | 0.00000 | 0.35833 |
| **ACOT6** | 0.03717 | 0.03297 | 0.01525 | 0.03670 | 0.01830 | 0.05539 | 0.04951 |
| **ACOT7** | 17.10885 | 17.83232 | 1.36813 | 19.79079 | 0.90329 | 7.91022 | 3.66587 |
| **ACOT8** | 3.19427 | 8.82478 | 2.69264 | 1.65360 | 0.09329 | 7.15683 | 0.00000 |

| | | | | | | | |
|---|---|---|---|---|---|---|---|
| **ACOT9** | 1.22519 | 3.54447 | 11.95485 | 0.33640 | 2.53078 | 0.21824 | 0.66368 |
| **ACOX1** | 19.44507 | 6.93470 | 1.09389 | 15.77707 | 0.60824 | 7.09084 | 7.10719 |
| **ACOX2** | 0.01286 | 0.01384 | 0.04796 | 0.04468 | 0.03903 | 0.28844 | 0.00712 |
| **ACOX3** | 3.05461 | 2.43785 | 1.57562 | 2.76409 | 1.78531 | 0.66867 | 0.00000 |
| **ACOXL** | 5.13312 | 1.31191 | 0.23310 | 7.06592 | 0.12475 | 4.77687 | 1.44982 |
| **ACP1** | 34.40909 | 26.21651 | 23.29467 | 26.35099 | 7.21255 | 14.89253 | 6.86058 |
| **ACP2** | 6.96934 | 3.09186 | 2.47872 | 1.50178 | 0.15755 | 2.91258 | 0.00000 |
| **ACP5** | 7.24428 | 21.75379 | 2.21017 | 6.45475 | 0.00237 | 5.25146 | 0.01047 |
| **ACP6** | 0.76588 | 0.67678 | 0.18944 | 4.29567 | 0.24561 | 0.86420 | 0.62672 |
| **ACPP** | 1.07403 | 0.30948 | 0.03879 | 2.11084 | 0.02683 | 4.59809 | 0.00000 |
| **ACPT** | 0.00000 | 0.14101 | 0.04024 | 0.03751 | 0.01805 | 0.14124 | 0.57765 |
| **ACR** | 0.03019 | 0.00977 | 0.02328 | 0.01600 | 0.01605 | 0.03100 | 0.02355 |
| **ACRBP** | 0.01756 | 0.05421 | 0.16919 | 0.04304 | 0.00973 | 0.00000 | 0.04525 |
| **ACRC** | 2.20341 | 0.52281 | 1.14115 | 0.00000 | 3.92768 | 0.65203 | 1.81114 |
| **ACRV1** | 0.19448 | 0.00000 | 0.10082 | 0.13698 | 0.05350 | 0.15734 | 0.12936 |
| **ACSBG1** | 0.51189 | 0.04771 | 0.06751 | 0.20478 | 0.36520 | 0.02362 | 0.81660 |
| **ACSBG2** | 0.06988 | 0.01163 | 0.14943 | 0.03838 | 0.01368 | 0.01884 | 0.01553 |
| **ACSF2** | 0.59890 | 0.30381 | 0.00000 | 0.29369 | 0.04795 | 1.75574 | 0.35789 |
| **ACSF3** | 3.23896 | 3.61232 | 1.44293 | 0.00000 | 0.00000 | 0.77545 | 4.28805 |
| **ACSL1** | 0.08381 | 0.18241 | 0.83569 | 1.78227 | 1.35813 | 1.04596 | 0.44057 |
| **ACSL3** | 35.96450 | 19.07095 | 21.63492 | 9.38655 | 2.91907 | 16.27704 | 20.17754 |
| **ACSL4** | 15.39750 | 0.00000 | 11.19636 | 0.00000 | 0.33669 | 5.07581 | 5.40177 |
| **ACSL5** | 0.08412 | 0.20944 | 0.02925 | 0.01829 | 0.05976 | 0.04861 | 0.05099 |
| **ACSL6** | 8.79922 | 0.00000 | 0.00141 | 0.00000 | 0.00128 | 0.00000 | 0.59278 |
| **ACSM1** | 0.06250 | 0.00000 | 0.02296 | 0.08664 | 0.00982 | 0.00000 | 0.20545 |
| **ACSM2B** | 0.02337 | 0.02169 | 0.01623 | 0.01886 | 0.00762 | 0.01809 | 0.01817 |
| **ACSM3** | 0.16084 | 0.00000 | 0.02761 | 0.09454 | 0.11353 | 0.73213 | 0.00000 |
| **ACSM4** | 0.02747 | 0.01646 | 0.01324 | 0.02211 | 0.01116 | 0.01298 | 0.01652 |
| **ACSM5** | 0.00000 | 0.01756 | 0.14481 | 0.04860 | 0.00624 | 0.00304 | 0.06184 |
| **ACSM6** | 0.08271 | 0.00539 | 0.05279 | 0.01704 | 0.00484 | 0.06243 | 0.08081 |
| **ACSS1** | 0.01290 | 0.00000 | 0.05876 | 0.02694 | 0.00533 | 0.00781 | 0.08924 |
| **ACSS2** | 10.31567 | 2.48647 | 5.29608 | 1.54078 | 1.82762 | 0.81534 | 10.48761 |
| **ACSS3** | 0.02295 | 0.09543 | 0.02890 | 0.05397 | 0.27091 | 0.00000 | 0.08640 |
| **ACTA1** | 0.06491 | 0.06779 | 0.13706 | 0.21759 | 0.18406 | 0.00550 | 0.31008 |
| **ACTA2** | 0.02627 | 0.03822 | 0.05192 | 0.05134 | 0.05496 | 0.03362 | 0.13554 |

| | | | | | | | |
|---|---|---|---|---|---|---|---|
| **ACTB** | 117.56101 | 102.95042 | 30.48177 | 88.85083 | 12.48825 | 51.16032 | 59.36004 |
| **ACTBL2** | 0.01255 | 0.03518 | 0.01163 | 0.03642 | 0.01715 | 0.05308 | 0.11281 |
| **ACTC1** | 0.51090 | 0.09517 | 0.01099 | 6.78327 | 0.01000 | 0.00667 | 0.00000 |
| **ACTG1** | 63.37823 | 83.50464 | 32.60361 | 64.16492 | 18.68734 | 50.01316 | 25.08791 |
| **ACTG1P4** | 0.06030 | 0.06658 | 0.02410 | 0.08170 | 0.12163 | 0.05255 | 0.12944 |
| **ACTG2** | 3.35626 | 5.51673 | 0.00000 | 0.62125 | 1.92674 | 0.00000 | 0.10851 |
| **ACTL10** | 0.01163 | 0.00194 | 0.03002 | 0.05873 | 0.03983 | 0.04591 | 0.10871 |
| **ACTL6A** | 32.59354 | 5.43875 | 12.95916 | 18.03815 | 1.94031 | 9.72895 | 0.00000 |
| **ACTL6B** | 0.01421 | 0.01593 | 0.01255 | 0.03049 | 0.00854 | 0.03162 | 0.03004 |
| **ACTL7B** | 0.02456 | 0.01201 | 0.00988 | 0.02690 | 0.14172 | 0.01305 | 0.02064 |
| **ACTL8** | 0.00000 | 0.50796 | 5.37369 | 0.00000 | 38.62298 | 0.00000 | 0.00000 |
| **ACTL9** | 0.02818 | 0.01706 | 0.01309 | 0.02035 | 0.00897 | 0.01692 | 0.01589 |
| **ACTN1** | 19.33634 | 29.85244 | 0.00000 | 33.11843 | 3.67366 | 4.83543 | 27.27988 |
| **ACTN1-AS1** | 0.03510 | 0.05152 | 0.00744 | 0.01890 | 0.05802 | 0.00821 | 0.01270 |
| **ACTN2** | 1.43797 | 0.00000 | 0.20248 | 0.35776 | 0.24292 | 0.00000 | 0.58382 |
| **ACTN3** | 0.04928 | 0.01587 | 0.15441 | 0.00638 | 0.17459 | 0.59775 | 0.01660 |
| **ACTN4** | 12.60427 | 10.97778 | 4.95414 | 15.42204 | 1.74823 | 17.93745 | 37.92603 |
| **ACTR10** | 16.35959 | 4.66876 | 10.01237 | 5.32357 | 0.28125 | 4.93036 | 1.51046 |
| **ACTR1A** | 24.62851 | 22.62732 | 6.56719 | 27.95352 | 2.54362 | 12.28624 | 4.29535 |
| **ACTR1B** | 2.01173 | 2.46361 | 0.29349 | 0.58237 | 0.30025 | 2.78351 | 6.36892 |
| **ACTR2** | 10.56569 | 3.01360 | 5.87589 | 7.31321 | 3.12746 | 6.36208 | 14.85111 |
| **ACTR3** | 29.16009 | 9.60619 | 11.57025 | 14.87385 | 2.11983 | 12.17215 | 10.74054 |
| **ACTR3B** | 0.31647 | 1.08116 | 0.39792 | 1.15609 | 0.89853 | 1.01314 | 2.66850 |
| **ACTR3C** | 0.03235 | 0.02665 | 0.01344 | 0.06116 | 0.00800 | 0.07777 | 0.00000 |
| **ACTR5** | 1.98538 | 4.96163 | 4.06670 | 3.97750 | 2.53314 | 3.07171 | 4.45766 |
| **ACTR6** | 19.06211 | 0.00000 | 4.84850 | 15.68458 | 2.67101 | 4.52226 | 2.38980 |
| **ACTR8** | 10.47356 | 6.37905 | 7.52547 | 7.36931 | 0.69290 | 5.39168 | 0.00000 |
| **ACTRT3** | 1.97795 | 0.00000 | 0.67605 | 1.30170 | 0.28180 | 0.60888 | 0.00000 |
| **ACVR1** | 4.13412 | 0.40286 | 0.68557 | 1.58942 | 0.32347 | 2.62180 | 5.10168 |
| **ACVR1B** | 0.20174 | 1.74835 | 1.46844 | 1.66406 | 2.36568 | 3.07828 | 3.65344 |
| **ACVR1C** | 0.10870 | 0.00000 | 0.86811 | 0.02741 | 0.12921 | 0.44500 | 0.18300 |
| **ACVR2A** | 5.59340 | 1.81726 | 1.67571 | 1.79797 | 3.48028 | 2.36805 | 2.28956 |
| **ACVR2B** | 1.69360 | 3.14065 | 4.65868 | 10.48461 | 2.81083 | 7.88093 | 14.19509 |
| **ACVR2B-AS1** | 0.01263 | 0.20588 | 0.13160 | 0.16277 | 0.03665 | 0.02191 | 0.05699 |
| **ACVRL1** | 0.00000 | 0.07040 | 0.00067 | 0.77768 | 0.07370 | 0.95829 | 0.00000 |

| | | | | | | | |
|---|---|---|---|---|---|---|---|
| **ACY1** | 1.17280 | 0.24075 | 0.16185 | 1.33305 | 0.00074 | 0.31043 | 0.00000 |
| **ACY3** | 0.01079 | 0.01338 | 0.01298 | 0.03292 | 0.00804 | 0.03580 | 0.03399 |
| **ACYP1** | 4.22630 | 0.78595 | 0.48318 | 10.06846 | 0.60304 | 0.83983 | 1.01906 |
| **ACYP2** | 0.19250 | 0.14240 | 0.19746 | 0.35965 | 0.10198 | 0.40009 | 0.00000 |
| **ADA** | 1.69507 | 19.32008 | 3.39102 | 6.38766 | 2.09446 | 2.93813 | 3.18556 |
| **ADAD1** | 0.26444 | 0.00000 | 2.66055 | 0.03400 | 4.31249 | 0.01609 | 0.01474 |
| **ADAD2** | 0.04205 | 0.03233 | 0.24246 | 0.05971 | 0.51777 | 0.17120 | 0.05783 |
| **ADAL** | 3.27614 | 1.29591 | 4.43155 | 2.19816 | 2.16238 | 1.45922 | 4.15503 |
| **ADAM10** | 25.30256 | 3.18875 | 10.33208 | 11.27644 | 6.29264 | 7.85374 | 6.15422 |
| **ADAM11** | 0.11838 | 0.00000 | 0.07819 | 0.25962 | 0.03119 | 0.05880 | 0.12598 |
| **ADAM12** | 0.00000 | 0.00564 | 0.20607 | 0.24146 | 0.49551 | 0.00000 | 0.32269 |
| **ADAM15** | 11.60664 | 1.55596 | 0.00000 | 15.93282 | 0.37581 | 0.00000 | 22.52310 |
| **ADAM17** | 14.27861 | 5.95041 | 4.33797 | 11.88814 | 8.86685 | 8.03843 | 4.86610 |
| **ADAM18** | 1.46359 | 0.15194 | 0.00291 | 0.76045 | 0.00058 | 3.17422 | 0.47015 |
| **ADAM19** | 1.55058 | 2.20962 | 3.27839 | 2.28053 | 3.57728 | 3.27302 | 1.17172 |
| **ADAM1A** | 0.75920 | 0.15134 | 0.61100 | 0.27303 | 0.00001 | 0.22410 | 1.05838 |
| **ADAM2** | 0.01640 | 0.01729 | 0.01272 | 0.03056 | 0.00783 | 0.01290 | 0.02927 |
| **ADAM20** | 0.04736 | 0.00000 | 0.08793 | 0.06951 | 0.04064 | 0.01535 | 0.19067 |
| **ADAM20P1** | 0.11315 | 0.07444 | 0.03443 | 0.00313 | 0.06050 | 0.03854 | 0.09805 |
| **ADAM21** | 0.00000 | 0.00013 | 0.57059 | 0.17233 | 0.10648 | 0.00000 | 0.13423 |
| **ADAM21P1** | 0.02608 | 0.01016 | 0.05893 | 0.02134 | 0.01881 | 0.01472 | 0.02899 |
| **ADAM22** | 0.14539 | 0.00000 | 0.12441 | 0.02483 | 0.12135 | 0.00899 | 0.38645 |
| **ADAM23** | 0.00000 | 1.29033 | 0.89515 | 0.00000 | 0.07201 | 2.35861 | 1.13768 |
| **ADAM28** | 0.10271 | 0.00119 | 0.01208 | 0.06328 | 0.11878 | 0.07011 | 0.00000 |
| **ADAM32** | 0.16379 | 0.00000 | 0.05389 | 0.00000 | 0.05141 | 0.05228 | 0.28645 |
| **ADAM33** | 0.02595 | 0.02317 | 0.02673 | 0.02141 | 0.00808 | 0.01258 | 0.01598 |
| **ADAM3A** | 0.02549 | 0.01791 | 0.01263 | 0.02307 | 0.00797 | 0.01249 | 0.01866 |
| **ADAM5** | 0.02527 | 0.00000 | 0.13200 | 0.01648 | 0.00903 | 0.01160 | 0.02351 |
| **ADAM7** | 0.02816 | 0.01850 | 0.00975 | 0.01801 | 0.03314 | 0.01335 | 0.01678 |
| **ADAM8** | 0.00000 | 0.25419 | 0.44515 | 1.60588 | 0.04825 | 0.31376 | 1.40192 |
| **ADAM9** | 30.96834 | 2.37023 | 4.70683 | 10.69182 | 1.76367 | 0.59869 | 15.26758 |
| **ADAMTS1** | 1.71373 | 0.00000 | 0.35030 | 7.04284 | 0.27125 | 0.39375 | 1.30574 |
| **ADAMTS10** | 0.01913 | 0.01478 | 0.01072 | 0.00840 | 0.06060 | 0.00804 | 0.08842 |
| **ADAMTS12** | 0.40984 | 0.00000 | 0.00634 | 0.00000 | 0.10657 | 2.01865 | 0.03813 |
| **ADAMTS13** | 0.04897 | 0.15572 | 0.11797 | 0.13885 | 0.08128 | 0.01807 | 0.41136 |

| | | | | | | | |
|---|---|---|---|---|---|---|---|
| **ADAMTS14** | 0.00650 | 0.02616 | 0.34763 | 0.01506 | 0.50736 | 0.10489 | 0.02608 |
| **ADAMTS15** | 0.00960 | 0.24257 | 0.07857 | 0.02189 | 0.04876 | 0.10861 | 0.00251 |
| **ADAMTS16** | 0.03029 | 0.00614 | 0.02230 | 0.02001 | 0.00757 | 0.04795 | 0.02461 |
| **ADAMTS17** | 0.07070 | 0.48223 | 0.24738 | 0.00000 | 1.27929 | 0.00000 | 0.19798 |
| **ADAMTS18** | 0.05093 | 0.07294 | 0.00896 | 0.02202 | 0.25230 | 0.02644 | 0.00811 |
| **ADAMTS19** | 0.04263 | 0.06793 | 0.15301 | 0.04475 | 0.04204 | 0.00618 | 0.00000 |
| **ADAMTS2** | 0.11547 | 2.26225 | 0.25516 | 0.00000 | 0.05605 | 0.38763 | 0.06429 |
| **ADAMTS20** | 0.05123 | 0.00000 | 0.34277 | 2.14181 | 0.04618 | 0.04230 | 0.00000 |
| **ADAMTS3** | 0.40325 | 0.00000 | 0.24768 | 0.19787 | 0.11910 | 0.53886 | 0.65235 |
| **ADAMTS4** | 0.00000 | 0.01192 | 0.06919 | 0.28687 | 0.03410 | 1.71017 | 0.04865 |
| **ADAMTS5** | 0.25789 | 0.06804 | 0.01355 | 0.00000 | 0.11249 | 0.23034 | 0.26327 |
| **ADAMTS6** | 1.32532 | 0.00000 | 0.52064 | 0.12551 | 0.06689 | 0.09508 | 0.38768 |
| **ADAMTS7** | 0.02034 | 0.04866 | 0.00827 | 0.09203 | 0.08168 | 0.01025 | 0.01352 |
| **ADAMTS8** | 0.02849 | 0.00000 | 0.10253 | 0.00679 | 0.00418 | 0.07182 | 0.10581 |
| **ADAMTS9** | 0.29916 | 0.03598 | 0.25302 | 0.00000 | 4.54001 | 0.96822 | 0.98367 |
| **ADAMTS9-AS2** | 0.06724 | 0.01210 | 0.01168 | 0.02690 | 0.01336 | 0.04169 | 0.04181 |
| **ADAMTSL1** | 0.00948 | 0.06780 | 0.06967 | 0.02796 | 1.32193 | 0.00654 | 0.02448 |
| **ADAMTSL2** | 0.00000 | 0.23637 | 0.00034 | 0.03281 | 0.02990 | 0.51980 | 0.01177 |
| **ADAMTSL3** | 0.06403 | 0.00000 | 0.53720 | 0.06782 | 1.76151 | 0.91809 | 0.99247 |
| **ADAMTSL4** | 1.73133 | 0.00000 | 0.00000 | 3.14055 | 0.00650 | 0.00000 | 5.81793 |
| **ADAMTSL4-AS1** | 0.00000 | 0.03605 | 0.13193 | 0.12691 | 0.01824 | 0.15070 | 0.09630 |
| **ADAMTSL5** | 0.00000 | 2.96790 | 3.11986 | 0.49814 | 0.05526 | 1.64440 | 1.35692 |
| **ADAP1** | 0.00000 | 0.34162 | 0.37482 | 0.05240 | 2.18629 | 0.18537 | 0.00620 |
| **ADAP2** | 1.19430 | 0.65078 | 0.73023 | 10.02033 | 0.37912 | 0.08523 | 0.62137 |
| **ADAR** | 18.19453 | 11.10374 | 4.45842 | 15.50917 | 7.51754 | 10.38983 | 21.44747 |
| **ADARB1** | 3.30707 | 0.77657 | 0.09603 | 2.18949 | 0.03967 | 0.46911 | 2.83419 |
| **ADARB2** | 0.02855 | 0.01322 | 0.00903 | 0.01520 | 0.10523 | 0.01195 | 0.02735 |
| **ADAT1** | 4.10350 | 2.35038 | 3.16272 | 2.05427 | 3.19875 | 1.90661 | 1.22602 |
| **ADAT2** | 6.16395 | 0.00000 | 3.95498 | 2.46955 | 0.22305 | 0.14734 | 1.62060 |
| **ADAT3** | 0.00000 | 1.90633 | 0.26745 | 1.95797 | 0.65168 | 1.06203 | 8.32160 |
| **ADCK1** | 2.18702 | 14.21361 | 3.44339 | 0.00000 | 0.00000 | 2.01032 | 4.55599 |
| **ADCK2** | 0.00000 | 14.00119 | 7.53247 | 3.27513 | 0.28207 | 0.95270 | 6.46353 |
| **ADCK3** | 7.51245 | 3.95496 | 9.93579 | 10.53467 | 0.38132 | 3.35114 | 2.07981 |
| **ADCK4** | 2.03314 | 1.65022 | 1.57287 | 2.39818 | 0.00000 | 1.87178 | 1.20294 |
| **ADCK5** | 2.86396 | 5.24802 | 0.47804 | 0.68533 | 0.40853 | 1.21120 | 0.58987 |

| | | | | | | | |
|---|---|---|---|---|---|---|---|
| **ADCY1** | 0.00000 | 0.03560 | 0.00000 | 0.23042 | 0.06933 | 0.35315 | 0.27052 |
| **ADCY10** | 0.10685 | 0.36262 | 0.35745 | 0.97367 | 0.39577 | 0.82689 | 2.72154 |
| **ADCY10P1** | 0.06442 | 0.14255 | 0.07242 | 0.35914 | 0.45772 | 0.33996 | 0.79202 |
| **ADCY2** | 0.01372 | 0.03146 | 0.01604 | 0.01934 | 0.57829 | 0.00965 | 0.06913 |
| **ADCY3** | 0.83540 | 0.52352 | 0.32462 | 1.21227 | 0.61607 | 0.74205 | 0.11116 |
| **ADCY4** | 0.02327 | 0.01731 | 0.01530 | 0.02135 | 0.01343 | 0.01454 | 0.02054 |
| **ADCY5** | 0.00000 | 0.00000 | 0.36534 | 1.15567 | 5.24168 | 0.00000 | 0.51093 |
| **ADCY6** | 0.00000 | 0.03167 | 0.14934 | 0.26263 | 0.04131 | 0.38398 | 0.23651 |
| **ADCY7** | 0.38861 | 0.17981 | 0.12943 | 0.42328 | 1.27773 | 0.23691 | 0.87514 |
| **ADCY8** | 0.02362 | 0.02308 | 0.01224 | 0.07145 | 0.00879 | 0.03929 | 0.00035 |
| **ADCY9** | 0.43433 | 1.99586 | 0.13609 | 0.26346 | 3.67145 | 0.00000 | 2.14267 |
| **ADCYAP1** | 0.02436 | 0.01847 | 0.01056 | 0.02392 | 0.02983 | 0.01478 | 0.01923 |
| **ADCYAP1R1** | 0.02219 | 0.00515 | 0.01343 | 0.03081 | 0.08511 | 0.08409 | 0.01924 |
| **ADD1** | 7.59118 | 10.73290 | 3.32055 | 9.27702 | 0.66744 | 8.87431 | 20.18522 |
| **ADD2** | 0.00000 | 0.31039 | 0.00000 | 0.00000 | 1.01241 | 1.99563 | 0.03413 |
| **ADD3** | 12.51466 | 4.45280 | 2.98289 | 1.63160 | 6.57865 | 7.29919 | 4.37849 |
| **ADD3-AS1** | 0.04832 | 0.01877 | 0.01232 | 0.01028 | 0.00778 | 0.10077 | 0.01307 |
| **ADGB** | 0.02682 | 0.01646 | 0.01039 | 0.02184 | 0.06660 | 0.01303 | 0.01528 |
| **ADH1B** | 0.02670 | 0.01314 | 0.01314 | 0.01606 | 0.00778 | 0.01245 | 0.03389 |
| **ADH4** | 0.01724 | 0.01473 | 0.01861 | 0.03697 | 0.01386 | 0.01068 | 0.03820 |
| **ADH5** | 17.30573 | 8.91692 | 6.15734 | 11.31853 | 1.63345 | 9.43150 | 6.09638 |
| **ADH6** | 0.00948 | 0.00000 | 0.02715 | 0.06699 | 0.00690 | 0.02134 | 0.12753 |
| **ADHFE1** | 0.03484 | 0.06995 | 0.51676 | 0.13742 | 0.00469 | 0.00492 | 0.00000 |
| **ADI1** | 34.89634 | 41.01341 | 8.15443 | 13.62991 | 0.05843 | 5.82354 | 10.43905 |
| **ADIPOQ** | 0.02269 | 0.01073 | 0.01460 | 0.02559 | 0.02472 | 0.04493 | 0.03271 |
| **ADIPOQ-AS1** | 0.00807 | 0.21494 | 0.09279 | 0.00973 | 0.01219 | 0.23547 | 0.01823 |
| **ADIPOR1** | 20.48720 | 15.79046 | 10.92820 | 26.06643 | 7.08850 | 13.83226 | 0.00073 |
| **ADIPOR2** | 8.10077 | 4.96485 | 5.92741 | 8.96257 | 3.78011 | 3.63457 | 21.02710 |
| **ADIRF** | 0.03734 | 0.03851 | 0.00804 | 0.05166 | 0.04895 | 0.00940 | 0.01536 |
| **ADK** | 27.72238 | 1.65958 | 0.27796 | 15.88549 | 0.57500 | 0.00000 | 6.02609 |
| **ADM** | 18.29623 | 5.22402 | 15.18495 | 32.53260 | 0.35706 | 2.67428 | 6.12973 |
| **ADM2** | 0.00000 | 0.13222 | 0.00000 | 1.29548 | 0.07518 | 0.00000 | 1.48611 |
| **ADM5** | 0.80430 | 0.12216 | 0.21532 | 1.48049 | 0.00000 | 0.79548 | 3.03546 |
| **ADNP** | 22.08420 | 6.67978 | 10.71498 | 8.59843 | 4.02619 | 11.70418 | 11.03816 |
| **ADNP2** | 4.13151 | 2.44876 | 4.82191 | 6.48501 | 4.37315 | 5.74417 | 11.11385 |

| | | | | | | | |
|---|---|---|---|---|---|---|---|
| **ADO** | 0.67032 | 0.25360 | 1.74015 | 0.82366 | 0.18151 | 2.51776 | 6.18223 |
| **ADORA1** | 0.01575 | 0.02422 | 0.01082 | 0.01258 | 0.13567 | 0.06080 | 0.04990 |
| **ADORA2A-AS1** | 0.00896 | 0.13894 | 0.06330 | 0.04095 | 0.09915 | 0.05721 | 0.09417 |
| **ADORA2A.uid-2** | 0.02409 | 0.05473 | 0.01624 | 0.02377 | 0.01959 | 0.01361 | 0.00683 |
| **ADORA2B** | 2.21300 | 2.72321 | 3.76391 | 0.29125 | 0.00000 | 5.60444 | 0.00000 |
| **ADORA3** | 0.00000 | 1.23109 | 0.14130 | 0.02966 | 0.01547 | 0.11998 | 0.00148 |
| **ADPGK** | 23.15554 | 9.35994 | 8.02705 | 12.08773 | 8.26486 | 7.14435 | 3.54355 |
| **ADPGK-AS1** | 0.00000 | 0.00000 | 0.05363 | 0.09268 | 0.00709 | 0.09824 | 1.19610 |
| **ADPRH** | 0.00000 | 0.01882 | 0.01514 | 0.42975 | 0.00747 | 0.10903 | 0.01354 |
| **ADPRHL1** | 0.22306 | 0.59631 | 0.12302 | 0.37287 | 0.20083 | 0.47717 | 0.78685 |
| **ADPRHL2** | 16.59910 | 14.16143 | 8.99313 | 11.23778 | 0.73090 | 3.98133 | 0.00000 |
| **ADPRM** | 2.75431 | 0.00000 | 2.37293 | 1.58912 | 0.52476 | 0.46227 | 0.60164 |
| **ADRA1A** | 0.01872 | 0.01995 | 0.01128 | 0.01722 | 0.01028 | 0.01124 | 0.04040 |
| **ADRA1B** | 0.02233 | 0.01795 | 0.00718 | 0.08771 | 0.08992 | 0.01312 | 0.00078 |
| **ADRA1D** | 0.00000 | 0.03181 | 0.01225 | 0.08541 | 0.00912 | 0.01167 | 0.11293 |
| **ADRA2A** | 0.02250 | 0.01678 | 0.01205 | 0.01900 | 0.01450 | 0.01635 | 0.02117 |
| **ADRA2B** | 0.04493 | 0.01192 | 0.00257 | 0.01547 | 0.28061 | 0.01168 | 0.02990 |
| **ADRA2C** | 0.00392 | 0.04063 | 0.03918 | 0.01666 | 0.00484 | 0.48093 | 0.05827 |
| **ADRB1** | 0.02059 | 0.02317 | 0.01459 | 0.02131 | 0.08358 | 0.01611 | 0.03201 |
| **ADRB2** | 0.02425 | 0.00951 | 0.00990 | 0.05231 | 0.40148 | 0.02257 | 0.00483 |
| **ADRB3** | 0.04237 | 0.00000 | 0.04707 | 0.01876 | 0.01011 | 0.05288 | 0.03142 |
| **ADRBK1** | 0.00000 | 7.38216 | 2.84564 | 2.51319 | 0.07378 | 2.42549 | 12.08206 |
| **ADRBK2** | 2.43119 | 1.72113 | 0.36184 | 5.29905 | 0.16452 | 3.16071 | 6.36652 |
| **ADRM1** | 39.30791 | 52.42661 | 15.29789 | 39.53772 | 9.60291 | 20.02752 | 8.04153 |
| **ADSL** | 39.99284 | 19.46989 | 12.24020 | 24.35133 | 2.83067 | 12.18085 | 7.02395 |
| **ADSS** | 13.56578 | 2.11560 | 4.30227 | 13.31617 | 2.43107 | 5.78685 | 3.30625 |
| **ADSSL1** | 0.01092 | 0.00000 | 0.17183 | 0.36394 | 0.02957 | 0.00000 | 0.07656 |
| **ADTRP** | 18.95009 | 3.93874 | 1.15906 | 4.70509 | 0.50346 | 1.26014 | 0.00000 |
| **AEBP1** | 0.03623 | 0.02451 | 0.14236 | 0.10189 | 0.02921 | 0.54759 | 0.08203 |
| **AEBP2** | 17.88454 | 0.00000 | 8.54781 | 12.77463 | 6.65206 | 6.74519 | 5.93800 |
| **AEN** | 23.95660 | 31.35900 | 16.10130 | 16.86430 | 5.65558 | 14.38482 | 6.81148 |
| **AES** | 0.00000 | 0.68676 | 11.31027 | 3.22358 | 1.49889 | 0.38340 | 0.86733 |
| **AFAP1** | 3.77644 | 1.74946 | 1.09105 | 2.70097 | 2.18399 | 2.06406 | 7.27737 |
| **AFAP1-AS1** | 1.21017 | 1.12160 | 1.15721 | 0.00000 | 3.22252 | 0.00000 | 0.55241 |
| **AFAP1L1** | 0.00000 | 1.20416 | 0.91117 | 3.72805 | 0.00174 | 1.44318 | 1.33084 |

| | | | | | | | |
|---|---|---|---|---|---|---|---|
| **AFAP1L2** | 0.00000 | 0.34268 | 0.00000 | 1.66988 | 0.91920 | 0.16010 | 0.00000 |
| **AFF1** | 6.28562 | 3.74934 | 8.99940 | 5.48448 | 8.15624 | 8.28529 | 9.12843 |
| **AFF2** | 0.07386 | 0.06113 | 0.08414 | 0.10212 | 0.23288 | 0.91595 | 0.01917 |
| **AFF3** | 0.00000 | 0.00173 | 0.03189 | 0.03255 | 0.59230 | 0.37074 | 0.06690 |
| **AFF4** | 10.55406 | 1.11645 | 4.97545 | 15.17255 | 3.50138 | 6.94592 | 25.40898 |
| **AFG3L1P** | 3.81668 | 3.27757 | 2.46865 | 0.76970 | 0.42686 | 1.67044 | 6.19026 |
| **AFG3L2** | 15.48636 | 17.01374 | 2.81305 | 6.80906 | 0.49898 | 16.19154 | 23.90165 |
| **AFMID** | 6.74281 | 2.39144 | 0.65229 | 5.56325 | 11.82336 | 1.78664 | 0.72534 |
| **AFP** | 0.01086 | 0.04531 | 0.01097 | 0.02713 | 0.00759 | 0.04923 | 0.02283 |
| **AFTPH** | 5.20418 | 1.27011 | 11.60161 | 5.40094 | 4.61956 | 7.64621 | 9.12273 |
| **AGA** | 2.12798 | 1.31829 | 0.25299 | 0.67567 | 0.26768 | 2.05287 | 0.00006 |
| **AGAP1** | 1.22233 | 3.62602 | 0.34378 | 4.47012 | 0.27763 | 2.87277 | 10.55477 |
| **AGAP11** | 0.18097 | 0.00000 | 0.01023 | 0.00000 | 0.08824 | 0.00083 | 0.15289 |
| **AGAP2** | 0.00000 | 0.01044 | 0.01794 | 0.06222 | 0.00724 | 0.04565 | 0.10701 |
| **AGAP2-AS1** | 0.01086 | 0.07111 | 0.05369 | 0.24954 | 0.00540 | 0.02669 | 0.00000 |
| **AGAP3** | 2.43938 | 5.39138 | 0.26285 | 6.63472 | 0.05580 | 0.73008 | 14.85375 |
| **AGAP4** | 0.00578 | 0.09882 | 0.06559 | 0.03903 | 0.05174 | 0.11060 | 0.26930 |
| **AGAP5** | 0.06248 | 0.15430 | 0.05943 | 0.05872 | 0.04242 | 0.00778 | 0.07784 |
| **AGAP6** | 0.90637 | 0.11431 | 0.67609 | 0.43943 | 0.13180 | 0.51799 | 1.30510 |
| **AGAP7P** | 0.12820 | 0.23446 | 0.02514 | 0.05669 | 0.00688 | 0.02732 | 0.13376 |
| **AGAP9.uid-1418** | 0.02377 | 0.01965 | 0.01328 | 0.01920 | 0.00787 | 0.01391 | 0.01679 |
| **AGBL1** | 0.02372 | 0.02558 | 0.00861 | 0.01949 | 0.06116 | 0.04233 | 0.01911 |
| **AGBL2** | 0.00957 | 0.01813 | 0.09003 | 0.04306 | 0.32053 | 0.01103 | 0.07344 |
| **AGBL3** | 0.28121 | 0.00000 | 0.09384 | 0.08226 | 0.38950 | 0.11032 | 0.24980 |
| **AGBL4** | 0.04221 | 0.01706 | 0.00813 | 0.04770 | 0.05795 | 0.18450 | 0.01411 |
| **AGBL5** | 0.28253 | 0.35477 | 1.34203 | 2.55137 | 0.46259 | 0.90337 | 1.94471 |
| **AGBL5-AS1** | 0.02437 | 0.01965 | 0.01266 | 0.02340 | 0.00791 | 0.01254 | 0.01583 |
| **AGER** | 7.24587 | 9.40678 | 0.38010 | 2.73609 | 0.14327 | 0.00000 | 0.00000 |
| **AGFG1** | 5.18797 | 2.47316 | 4.51898 | 4.93373 | 0.74338 | 7.89669 | 22.15271 |
| **AGFG2** | 0.15486 | 0.41755 | 0.45893 | 0.42300 | 1.00855 | 0.63788 | 0.16556 |
| **AGGF1** | 5.73107 | 1.98707 | 5.58222 | 2.47076 | 1.97115 | 7.04116 | 3.66073 |
| **AGK** | 9.04080 | 4.20564 | 3.76470 | 0.58703 | 3.04728 | 1.80875 | 0.00000 |
| **AGL** | 41.00862 | 7.61106 | 25.14552 | 7.46670 | 1.55476 | 34.36726 | 30.85365 |
| **AGMAT** | 0.17404 | 0.16513 | 0.04719 | 0.11746 | 0.32282 | 0.08836 | 0.17696 |
| **AGMO** | 0.06441 | 0.01589 | 0.02039 | 0.06534 | 0.97533 | 0.05005 | 0.00000 |

| | | | | | | | |
|---|---|---|---|---|---|---|---|
| **AGO1** | 6.29229 | 0.89768 | 4.31144 | 5.42699 | 2.21705 | 6.01656 | 2.13243 |
| **AGO2** | 3.97813 | 5.63437 | 4.04844 | 3.61488 | 2.95334 | 3.35263 | 18.90551 |
| **AGO3** | 3.04074 | 0.42469 | 2.30303 | 5.47639 | 3.41745 | 1.49626 | 0.57715 |
| **AGO4** | 0.99071 | 0.00000 | 0.15085 | 0.99353 | 0.18070 | 0.45995 | 0.63432 |
| **AGPAT1** | 4.08981 | 2.05530 | 4.33600 | 10.73432 | 0.27356 | 3.33228 | 5.35489 |
| **AGPAT2** | 13.09752 | 34.43950 | 0.00000 | 1.35737 | 1.85563 | 14.19784 | 4.08266 |
| **AGPAT3** | 0.00000 | 1.89656 | 0.00000 | 0.17089 | 0.62681 | 1.42411 | 1.01487 |
| **AGPAT4** | 0.50039 | 0.00000 | 0.32617 | 1.52754 | 0.78577 | 1.10144 | 1.48291 |
| **AGPAT4-IT1** | 0.11200 | 0.04871 | 0.03013 | 0.06207 | 0.01190 | 0.36356 | 0.70668 |
| **AGPAT5** | 18.40059 | 0.02775 | 1.87210 | 13.33107 | 0.29725 | 0.42580 | 10.40810 |
| **AGPAT6** | 16.35354 | 19.72266 | 4.94959 | 14.36420 | 3.11144 | 10.41715 | 7.71941 |
| **AGPAT9** | 2.42237 | 10.87926 | 6.72432 | 0.00000 | 2.01052 | 0.17156 | 0.00000 |
| **AGPS** | 10.02532 | 2.06743 | 2.45731 | 7.58383 | 0.76667 | 2.83926 | 9.46751 |
| **AGR2** | 0.03077 | 0.01335 | 0.01113 | 0.03014 | 0.01091 | 0.02817 | 0.05718 |
| **AGR3** | 0.01351 | 0.01508 | 0.01202 | 0.02727 | 0.00976 | 0.01222 | 0.06286 |
| **AGRN** | 0.15610 | 0.80159 | 0.00000 | 0.53661 | 0.02314 | 0.00000 | 6.87642 |
| **AGRP** | 0.01340 | 0.02608 | 0.01906 | 0.06047 | 0.00726 | 0.01097 | 0.01711 |
| **AGT** | 0.01446 | 0.00618 | 0.12156 | 0.01791 | 0.00897 | 0.03441 | 0.02686 |
| **AGTPBP1** | 3.33645 | 0.70517 | 2.41715 | 2.10775 | 0.99151 | 2.44096 | 4.40446 |
| **AGTR1** | 0.04096 | 0.00144 | 0.04470 | 0.00912 | 0.11719 | 0.03056 | 0.06874 |
| **AGTRAP** | 5.40457 | 7.99103 | 0.00000 | 2.79927 | 1.48640 | 7.06298 | 1.15132 |
| **AGXT2** | 0.02248 | 0.05678 | 0.00517 | 0.08277 | 0.01892 | 0.01084 | 0.01898 |
| **AHCTF1** | 4.48872 | 0.36857 | 3.90227 | 1.54552 | 3.89587 | 2.13871 | 11.52465 |
| **AHCTF1P1** | 1.23030 | 0.10080 | 0.95354 | 0.49310 | 1.49928 | 0.43618 | 1.77256 |
| **AHCY** | 35.31421 | 22.56158 | 13.91673 | 32.18445 | 3.66942 | 17.35508 | 19.05096 |
| **AHCYL1** | 11.15008 | 2.53197 | 1.39374 | 16.64208 | 0.00000 | 8.92530 | 26.58455 |
| **AHCYL2** | 1.78511 | 0.07441 | 0.91049 | 0.66245 | 0.85042 | 4.74217 | 1.25810 |
| **AHDC1** | 0.00000 | 0.08888 | 0.11275 | 0.90794 | 0.17596 | 0.40657 | 4.77852 |
| **AHI1** | 3.30763 | 0.00448 | 1.37097 | 1.17384 | 1.86398 | 0.20149 | 3.38840 |
| **AHNAK** | 21.23317 | 17.28429 | 0.22709 | 7.54702 | 1.19153 | 2.13700 | 44.08081 |
| **AHNAK2** | 0.02118 | 0.00000 | 0.02063 | 0.00693 | 0.04903 | 0.00341 | 0.27959 |
| **AHR** | 0.31056 | 0.00000 | 0.91460 | 0.63020 | 0.70689 | 1.13873 | 0.68541 |
| **AHRR** | 0.15150 | 1.20202 | 0.16147 | 0.07760 | 0.47010 | 0.69167 | 0.34067 |
| **AHSA1** | 45.61452 | 33.18409 | 23.15739 | 42.48276 | 12.90367 | 19.07315 | 7.74000 |
| **AHSA2** | 0.17072 | 0.06180 | 0.23647 | 1.25522 | 0.14225 | 0.85064 | 1.58596 |

| | | | | | | | |
|---|---|---|---|---|---|---|---|
| **AHSG** | 0.03139 | 0.01780 | 0.03844 | 0.02096 | 0.00671 | 0.01635 | 0.01153 |
| **AHSP** | 0.02699 | 0.10117 | 0.33469 | 0.00922 | 0.00225 | 0.00347 | 0.01256 |
| **AICDA** | 0.08329 | 0.73027 | 0.02742 | 0.08243 | 0.02812 | 2.67190 | 0.00000 |
| **AIDA** | 0.50260 | 0.00000 | 0.61518 | 0.17678 | 0.21035 | 0.55241 | 1.59653 |
| **AIF1** | 0.04030 | 0.06219 | 0.00865 | 0.01126 | 0.01523 | 0.13162 | 0.00643 |
| **AIF1L** | 37.32635 | 44.24990 | 8.78671 | 35.47901 | 0.79700 | 13.08248 | 16.79883 |
| **AIFM1** | 27.07080 | 21.67911 | 8.11249 | 8.68147 | 3.95445 | 11.40696 | 8.60591 |
| **AIFM2** | 1.41190 | 3.92315 | 0.02070 | 2.49828 | 0.07522 | 5.64165 | 0.00000 |
| **AIFM3** | 0.04396 | 0.05351 | 0.02420 | 0.04718 | 0.04621 | 0.08478 | 0.02491 |
| **AIG1** | 0.12045 | 0.00000 | 0.25041 | 0.81020 | 0.17083 | 0.05110 | 0.00000 |
| **AIM1** | 0.00000 | 0.17502 | 0.00000 | 6.20304 | 3.95452 | 2.30801 | 0.67764 |
| **AIM1L** | 7.23667 | 1.59564 | 0.00000 | 8.92220 | 0.26078 | 1.18171 | 18.44371 |
| **AIM2** | 0.00000 | 0.26158 | 0.00636 | 0.01490 | 0.00758 | 0.04417 | 0.01416 |
| **AIMP1** | 21.21606 | 12.87450 | 7.60864 | 5.46763 | 0.00000 | 11.19439 | 22.35636 |
| **AIMP2** | 18.81068 | 17.65237 | 3.88183 | 7.90005 | 2.14818 | 6.48051 | 0.00000 |
| **AIP** | 1.82689 | 1.83892 | 6.62503 | 5.43898 | 5.59247 | 4.17819 | 2.03254 |
| **AIPL1** | 0.03296 | 0.02020 | 0.01366 | 0.03042 | 0.00940 | 0.02268 | 0.00454 |
| **AIRE** | 0.00000 | 4.06363 | 0.00071 | 0.00065 | 0.00500 | 1.63048 | 0.72360 |
| **AIRN** | 0.00000 | 0.00902 | 0.09595 | 0.10583 | 0.06415 | 0.03773 | 0.11343 |
| **AJAP1** | 0.04782 | 0.01499 | 0.00432 | 0.06301 | 0.04136 | 0.00000 | 1.20535 |
| **AJUBA** | 2.30929 | 1.53606 | 1.17664 | 9.36002 | 3.35659 | 0.00000 | 5.30600 |
| **AK1** | 4.24841 | 7.04421 | 0.19009 | 1.17090 | 1.64635 | 1.14659 | 0.00000 |
| **AK2** | 34.42106 | 21.45786 | 10.92425 | 13.71597 | 4.98280 | 13.90364 | 3.77836 |
| **AK3** | 0.76527 | 0.00000 | 0.05188 | 1.97777 | 0.20412 | 0.68879 | 1.82229 |
| **AK4** | 29.82692 | 28.31865 | 14.49972 | 0.00000 | 0.90241 | 29.20475 | 7.33720 |
| **AK5** | 0.00000 | 0.34405 | 0.65449 | 0.51214 | 1.41377 | 1.61640 | 0.88279 |
| **AK6** | 25.55467 | 0.00000 | 14.63820 | 8.88019 | 1.26826 | 7.12292 | 5.54860 |
| **AK7** | 0.56516 | 0.08318 | 1.84844 | 2.65226 | 0.62521 | 0.00000 | 0.00000 |
| **AK8** | 0.03367 | 0.37551 | 0.28902 | 0.04673 | 0.21896 | 0.77729 | 0.00000 |
| **AK9** | 0.33891 | 0.06514 | 0.09289 | 0.83781 | 0.89651 | 0.06971 | 0.10396 |
| **AKAP1** | 13.59300 | 18.61715 | 6.58675 | 6.97287 | 3.19110 | 20.06928 | 20.55727 |
| **AKAP10** | 5.93155 | 2.94175 | 5.32020 | 3.99163 | 2.08968 | 5.71452 | 4.43335 |
| **AKAP11** | 1.64980 | 0.00000 | 1.45985 | 1.41953 | 3.00881 | 2.33026 | 6.26521 |
| **AKAP12** | 16.70907 | 29.81569 | 16.07323 | 13.44636 | 0.44210 | 29.53644 | 33.39725 |
| **AKAP13** | 13.34704 | 0.00000 | 9.33198 | 18.64474 | 6.78279 | 2.36751 | 18.49296 |

| | | | | | | | |
|---|---|---|---|---|---|---|---|
| **AKAP14** | 0.00000 | 0.00000 | 0.58351 | 0.01925 | 0.15495 | 0.01152 | 0.12450 |
| **AKAP2.uid-1139** | 0.08069 | 0.37725 | 0.11074 | 0.06479 | 0.06037 | 0.03685 | 0.36750 |
| **AKAP2.uid-1192** | 0.12098 | 0.28056 | 0.00269 | 0.13397 | 0.01125 | 0.00247 | 0.00000 |
| **AKAP3** | 0.07670 | 0.69112 | 0.64372 | 0.12470 | 0.02037 | 0.18393 | 0.00000 |
| **AKAP4** | 0.02396 | 0.03151 | 0.02375 | 0.01919 | 0.00685 | 0.01187 | 0.01455 |
| **AKAP5** | 0.07719 | 0.00000 | 0.79260 | 0.21573 | 0.03259 | 0.14558 | 0.63670 |
| **AKAP6** | 0.06554 | 0.33353 | 0.06306 | 0.00720 | 0.23558 | 0.07503 | 0.00000 |
| **AKAP7** | 0.12421 | 0.23420 | 0.59448 | 0.03295 | 0.15887 | 1.41970 | 0.21316 |
| **AKAP8** | 14.43114 | 17.97848 | 11.80468 | 16.60736 | 12.92414 | 8.83347 | 9.49026 |
| **AKAP8L** | 0.12475 | 2.16267 | 0.86447 | 1.95629 | 1.15113 | 1.41113 | 4.67298 |
| **AKAP9** | 9.14478 | 5.17864 | 7.13217 | 10.52680 | 0.71728 | 8.58317 | 24.92686 |
| **AKIP1** | 10.46140 | 10.18505 | 15.17368 | 6.16257 | 2.69685 | 3.31381 | 0.34845 |
| **AKIRIN1** | 7.41576 | 8.99990 | 12.70065 | 10.22654 | 2.67275 | 14.86766 | 24.09284 |
| **AKIRIN2** | 3.95037 | 8.27932 | 15.70199 | 10.53380 | 11.91599 | 9.49627 | 13.68913 |
| **AKNA** | 0.00000 | 0.04736 | 0.00000 | 0.65117 | 0.10195 | 4.14423 | 1.25067 |
| **AKNAD1** | 0.01327 | 0.00909 | 0.06073 | 0.05596 | 0.03120 | 0.01377 | 0.03256 |
| **AKR1A1** | 40.22035 | 32.20451 | 8.75429 | 24.75350 | 3.24174 | 12.28323 | 4.93502 |
| **AKR1B1** | 1.70975 | 2.33918 | 0.32436 | 3.09726 | 1.15377 | 3.69173 | 0.00000 |
| **AKR1B10** | 0.02243 | 0.01732 | 0.00215 | 0.02137 | 0.20657 | 0.01373 | 0.02211 |
| **AKR1B15** | 0.03503 | 0.01708 | 0.00940 | 0.02640 | 0.05294 | 0.01313 | 0.00835 |
| **AKR1C1** | 0.02790 | 0.02695 | 0.01257 | 0.01856 | 0.01108 | 0.01966 | 0.01276 |
| **AKR1C2** | 0.02913 | 0.01810 | 0.01246 | 0.01863 | 0.01124 | 0.01298 | 0.01694 |
| **AKR1C3** | 0.01691 | 0.00000 | 0.34218 | 0.01014 | 1.86947 | 0.00380 | 0.03335 |
| **AKR1CL1** | 0.03228 | 0.01478 | 0.00897 | 0.01505 | 0.05905 | 0.01812 | 0.01883 |
| **AKR1D1** | 0.05231 | 0.00000 | 0.18473 | 0.02859 | 0.01027 | 0.01128 | 0.00374 |
| **AKR1E2** | 0.10681 | 0.10524 | 0.30451 | 0.01017 | 0.08473 | 0.02553 | 0.00000 |
| **AKR7A2** | 5.94866 | 1.85775 | 1.74370 | 13.83634 | 0.39091 | 2.58424 | 6.74727 |
| **AKR7A2P1** | 0.27076 | 0.14792 | 0.12925 | 0.64849 | 0.04364 | 0.15150 | 0.27593 |
| **AKR7A3** | 0.04019 | 0.03580 | 0.70300 | 0.13109 | 0.00159 | 0.09081 | 0.17748 |
| **AKR7L** | 0.21321 | 0.00000 | 0.20788 | 0.28045 | 0.00357 | 0.00000 | 0.25880 |
| **AKT1** | 3.01345 | 5.84767 | 0.00000 | 8.52585 | 1.22326 | 2.67445 | 10.56967 |
| **AKT1S1** | 0.00000 | 6.30130 | 5.56460 | 9.07862 | 6.98902 | 1.98035 | 19.68900 |
| **AKT2** | 0.75532 | 2.57647 | 0.93488 | 6.87647 | 2.42026 | 4.73810 | 22.69369 |
| **AKT3** | 0.16461 | 0.65483 | 1.03600 | 0.77758 | 1.66237 | 0.71645 | 1.74401 |
| **AKTIP** | 7.36647 | 2.62982 | 1.77831 | 4.55124 | 2.19085 | 0.02941 | 1.07774 |

| | | | | | | | |
|---|---|---|---|---|---|---|---|
| **ALAD** | 28.14741 | 33.73446 | 11.37504 | 7.47160 | 2.25675 | 1.04117 | 0.00000 |
| **ALAS1** | 25.81357 | 34.31876 | 10.55175 | 32.87828 | 6.22203 | 1.28064 | 0.00000 |
| **ALAS2** | 0.04198 | 0.01723 | 0.02660 | 0.03531 | 0.00992 | 0.00975 | 0.01884 |
| **ALB** | 0.00548 | 0.00000 | 0.08361 | 0.04483 | 0.00649 | 0.11197 | 0.09488 |
| **ALCAM** | 0.34822 | 0.00000 | 0.10368 | 0.00000 | 0.05496 | 0.69239 | 0.37332 |
| **ALDH16A1** | 0.00000 | 2.56545 | 0.31766 | 3.95696 | 0.66013 | 2.85896 | 8.13733 |
| **ALDH18A1** | 13.18933 | 3.74174 | 1.08554 | 12.98130 | 2.19387 | 1.85472 | 0.00000 |
| **ALDH1A1** | 7.82951 | 0.00000 | 0.01502 | 0.09535 | 0.00195 | 2.48697 | 0.00000 |
| **ALDH1A2** | 0.23865 | 0.03321 | 0.00000 | 0.00000 | 2.56600 | 0.00000 | 0.18951 |
| **ALDH1A3** | 0.16394 | 0.00000 | 0.04370 | 0.00580 | 0.02730 | 0.62333 | 0.05729 |
| **ALDH1B1** | 3.40052 | 3.29891 | 0.00000 | 5.17519 | 0.10490 | 2.24963 | 0.00000 |
| **ALDH1L1** | 0.04319 | 0.00123 | 0.02602 | 0.06240 | 0.05282 | 0.01075 | 0.02856 |
| **ALDH1L2** | 0.74405 | 0.00000 | 0.05206 | 0.22301 | 0.01516 | 0.40180 | 0.85510 |
| **ALDH2** | 19.48067 | 2.61817 | 0.00000 | 5.60905 | 0.00000 | 27.40190 | 0.24144 |
| **ALDH3A1** | 0.00946 | 0.17010 | 0.00395 | 0.02421 | 0.01055 | 0.02408 | 0.02967 |
| **ALDH3A2** | 10.87796 | 16.62364 | 2.96426 | 2.78384 | 0.92472 | 12.60677 | 0.00000 |
| **ALDH3B1** | 0.02664 | 0.15468 | 0.00597 | 0.12514 | 0.00941 | 0.01130 | 0.00000 |
| **ALDH3B2** | 0.04289 | 0.01125 | 0.01034 | 0.90919 | 0.00662 | 0.00055 | 0.07516 |
| **ALDH4A1** | 5.70806 | 2.12939 | 0.00000 | 11.35877 | 0.09954 | 1.27127 | 11.23285 |
| **ALDH5A1** | 0.64297 | 0.42601 | 0.04212 | 0.61296 | 0.57233 | 0.17751 | 1.96407 |
| **ALDH6A1** | 13.19692 | 9.38377 | 2.50761 | 3.52375 | 1.13711 | 2.76715 | 0.00000 |
| **ALDH7A1** | 10.39664 | 3.85976 | 0.31505 | 8.95560 | 4.95698 | 5.74983 | 0.77656 |
| **ALDH8A1** | 0.07718 | 0.02255 | 0.00000 | 0.04468 | 0.60000 | 0.01936 | 0.00000 |
| **ALDH9A1** | 17.04222 | 1.46795 | 11.06213 | 10.45111 | 3.26385 | 2.59234 | 2.65413 |
| **ALDOA** | 71.55528 | 63.37383 | 35.63981 | 65.10476 | 17.11144 | 41.64143 | 47.59503 |
| **ALDOB** | 0.03609 | 0.01828 | 0.01101 | 0.01917 | 0.01741 | 0.01047 | 0.02672 |
| **ALDOC** | 4.53037 | 0.00000 | 0.38090 | 2.80587 | 3.60861 | 3.30439 | 0.44559 |
| **ALG1** | 10.00728 | 6.34320 | 1.55816 | 6.98612 | 0.08804 | 1.78763 | 0.00000 |
| **ALG10** | 0.47998 | 0.14531 | 0.44199 | 0.15978 | 0.04772 | 0.11051 | 0.00000 |
| **ALG10B** | 0.61577 | 0.00000 | 1.01141 | 0.16365 | 0.02887 | 0.27149 | 0.13115 |
| **ALG11** | 4.59278 | 2.51765 | 3.75066 | 3.13090 | 0.48959 | 1.66950 | 0.29494 |
| **ALG12** | 4.56728 | 6.71125 | 1.74653 | 7.44731 | 0.79897 | 2.12377 | 1.11618 |
| **ALG13** | 18.34267 | 2.10027 | 10.01981 | 10.03888 | 3.74962 | 3.80742 | 8.13361 |
| **ALG14** | 10.68957 | 3.80980 | 2.20861 | 6.15711 | 0.00000 | 0.68951 | 2.75241 |
| **ALG1L** | 0.07206 | 0.68038 | 0.56381 | 0.06914 | 0.04763 | 0.49272 | 0.00521 |

| | | | | | | | |
|---|---|---|---|---|---|---|---|
| **ALG1L2** | 0.01551 | 0.01552 | 0.01546 | 0.02182 | 0.01291 | 0.03370 | 0.03949 |
| **ALG1L9P** | 0.04021 | 0.01538 | 0.04895 | 0.03620 | 0.05955 | 0.06564 | 0.00000 |
| **ALG2** | 22.71254 | 13.42571 | 13.22508 | 16.18732 | 7.10953 | 11.72011 | 1.10635 |
| **ALG3** | 16.69201 | 16.03656 | 14.48402 | 16.45099 | 1.15115 | 9.21285 | 4.90050 |
| **ALG5** | 20.31869 | 5.74918 | 9.69501 | 6.87185 | 2.53183 | 4.85244 | 5.40597 |
| **ALG6** | 10.82954 | 0.15888 | 8.04612 | 2.03352 | 0.72808 | 3.03903 | 2.67390 |
| **ALG8** | 17.12918 | 3.56453 | 8.36749 | 11.78895 | 0.20491 | 7.77379 | 0.49982 |
| **ALG9** | 3.94333 | 1.97617 | 0.70741 | 3.94272 | 0.51314 | 3.45653 | 6.44557 |
| **ALK** | 0.01745 | 0.05394 | 0.04076 | 0.03690 | 0.13801 | 0.03172 | 0.01093 |
| **ALKBH1** | 10.49734 | 11.98158 | 13.96347 | 7.51178 | 3.89460 | 6.26754 | 0.00000 |
| **ALKBH2** | 4.36838 | 3.88625 | 7.43945 | 4.02339 | 3.49756 | 1.99165 | 1.27893 |
| **ALKBH3** | 3.08388 | 1.33735 | 1.96859 | 1.44756 | 2.11347 | 0.70928 | 0.37981 |
| **ALKBH3-AS1** | 0.03127 | 0.00737 | 0.02900 | 0.03034 | 0.00845 | 0.01518 | 0.04764 |
| **ALKBH4** | 5.17502 | 4.36664 | 2.66623 | 5.31718 | 0.92174 | 3.54160 | 0.00024 |
| **ALKBH5** | 7.98632 | 11.19755 | 5.56108 | 6.96121 | 5.74359 | 8.74627 | 19.11804 |
| **ALKBH6** | 1.37936 | 1.20926 | 0.57948 | 4.90906 | 0.05672 | 1.78577 | 3.03617 |
| **ALKBH7** | 6.94797 | 22.51985 | 9.19867 | 10.81490 | 7.89219 | 17.89346 | 0.72668 |
| **ALKBH8** | 5.49914 | 0.38385 | 3.07518 | 1.40853 | 3.06974 | 1.84656 | 1.86853 |
| **ALLC** | 0.02469 | 0.03047 | 0.00568 | 0.01902 | 0.06302 | 0.01289 | 0.01613 |
| **ALMS1** | 2.09882 | 0.82326 | 3.40059 | 2.27850 | 7.16401 | 2.08981 | 6.50498 |
| **ALMS1-IT1** | 0.11553 | 0.17618 | 0.09307 | 0.05219 | 0.65972 | 0.14083 | 0.25681 |
| **ALMS1P** | 0.00184 | 0.02468 | 0.01347 | 0.11089 | 0.00787 | 0.06926 | 0.07869 |
| **ALOX12** | 2.00049 | 0.00000 | 0.25760 | 3.08880 | 0.10681 | 0.34119 | 0.00000 |
| **ALOX12B** | 0.01922 | 0.01827 | 0.01293 | 0.01536 | 0.01571 | 0.00308 | 0.14362 |
| **ALOX12P2** | 0.03972 | 0.01320 | 0.03856 | 0.06363 | 0.07249 | 0.00662 | 0.03010 |
| **ALOX15** | 0.03504 | 0.07774 | 0.21101 | 0.03363 | 5.81449 | 0.00000 | 0.14824 |
| **ALOX15B** | 0.00543 | 0.00000 | 0.38572 | 0.01176 | 0.01604 | 0.00295 | 0.07698 |
| **ALOX15P1** | 0.02211 | 0.03293 | 0.01633 | 0.02503 | 0.01351 | 0.01193 | 0.01204 |
| **ALOX5** | 0.37493 | 0.98271 | 0.40449 | 0.00000 | 0.00000 | 2.36276 | 0.29154 |
| **ALOX5AP** | 0.03045 | 0.01496 | 0.01159 | 0.01265 | 0.03172 | 0.07202 | 0.02039 |
| **ALOXE3** | 0.00000 | 0.19042 | 0.00000 | 0.34008 | 1.77758 | 0.00296 | 0.00000 |
| **ALPI** | 0.03954 | 0.03910 | 0.02357 | 0.05272 | 0.09670 | 0.02797 | 0.06557 |
| **ALPK1** | 0.02048 | 0.02023 | 0.00834 | 0.02646 | 0.06290 | 0.01272 | 0.02321 |
| **ALPK2** | 0.05323 | 0.05392 | 0.00882 | 0.02721 | 0.10364 | 0.01286 | 0.03009 |
| **ALPK3** | 0.17323 | 0.20033 | 0.26409 | 1.85619 | 0.02435 | 0.11947 | 2.59792 |

| | | | | | | | |
|---|---|---|---|---|---|---|---|
| **ALPL** | 15.77708 | 54.30803 | 19.37549 | 0.00000 | 0.00000 | 24.50082 | 5.28500 |
| **ALPP** | 31.65726 | 23.19310 | 0.95113 | 5.68223 | 1.80106 | 0.00000 | 15.48048 |
| **ALPPL2** | 45.57414 | 64.77467 | 29.74677 | 11.58843 | 16.74099 | 16.98907 | 43.61730 |
| **ALS2** | 4.77808 | 1.33733 | 4.96825 | 4.02566 | 5.89435 | 3.48376 | 1.96981 |
| **ALS2CL** | 0.03519 | 0.31320 | 0.13759 | 1.78189 | 0.00312 | 0.08753 | 0.00000 |
| **ALS2CR11** | 0.16772 | 0.00000 | 2.34242 | 0.21863 | 0.59026 | 0.39588 | 0.00000 |
| **ALS2CR12** | 0.20211 | 0.00000 | 0.40253 | 0.03147 | 0.11088 | 0.05743 | 0.01568 |
| **ALX1** | 0.56691 | 0.56176 | 2.56836 | 0.31286 | 0.00739 | 0.32315 | 0.00000 |
| **ALX3** | 0.06439 | 0.27574 | 0.33623 | 0.52159 | 0.00532 | 0.23787 | 0.13295 |
| **ALX4** | 0.05128 | 0.00901 | 0.02739 | 0.05874 | 0.03377 | 0.03147 | 0.00000 |
| **ALYREF** | 1.23856 | 8.46707 | 3.22014 | 5.82073 | 6.92500 | 5.23638 | 27.40132 |
| **AMACR** | 0.39459 | 0.67373 | 0.12819 | 0.03362 | 0.33884 | 0.15487 | 0.00000 |
| **AMBN** | 0.49709 | 0.00000 | 0.01136 | 0.04137 | 0.00726 | 1.41351 | 0.00000 |
| **AMBP** | 0.03233 | 0.00000 | 0.02217 | 0.01997 | 0.65203 | 0.03867 | 0.02803 |
| **AMBRA1** | 3.45023 | 5.12912 | 7.67841 | 2.59929 | 4.14707 | 6.02963 | 8.89999 |
| **1.00000** | 47.01595 | 7.36533 | 28.99091 | 18.97450 | 17.96690 | 12.07395 | 21.28279 |
| **AMDHD1** | 0.85747 | 0.00000 | 1.18439 | 0.06190 | 0.58411 | 4.56304 | 0.00000 |
| **AMDHD2** | 6.78565 | 7.01077 | 4.13105 | 5.57458 | 2.24817 | 1.75180 | 4.75626 |
| **AMER1** | 0.22474 | 0.14153 | 0.25453 | 0.02952 | 0.07082 | 0.51704 | 0.99295 |
| **AMER2** | 0.82710 | 0.00000 | 0.05154 | 0.86891 | 0.00373 | 0.00000 | 0.63322 |
| **AMER3** | 0.02299 | 0.12341 | 0.00668 | 0.03839 | 0.00897 | 0.00580 | 0.00000 |
| **AMFR** | 3.87779 | 9.83674 | 6.13687 | 1.61613 | 1.77421 | 4.84119 | 15.99444 |
| **AMH** | 0.00000 | 0.10221 | 0.05816 | 0.09021 | 0.00449 | 0.09818 | 0.58662 |
| **AMHR2** | 0.00000 | 0.03506 | 0.36424 | 2.91128 | 0.21146 | 1.69836 | 0.00000 |
| **AMICA1** | 0.27344 | 0.15170 | 0.33252 | 0.24918 | 0.01736 | 0.97598 | 0.15600 |
| **AMIGO1** | 0.14776 | 0.04421 | 0.03579 | 0.05089 | 0.27876 | 0.03225 | 0.01487 |
| **AMIGO2** | 0.03495 | 0.00000 | 0.38872 | 0.01566 | 0.18952 | 0.17368 | 0.01877 |
| **AMIGO3** | 0.75877 | 0.57490 | 0.34422 | 1.02363 | 0.00000 | 0.84334 | 1.48472 |
| **AMMECR1** | 10.58173 | 0.00000 | 1.46644 | 3.66612 | 0.08671 | 0.57818 | 8.11711 |
| **AMMECR1L** | 3.30455 | 2.68699 | 3.56538 | 5.13981 | 5.85767 | 2.60287 | 7.93768 |
| **AMN** | 0.02779 | 0.03917 | 0.01249 | 0.05728 | 0.01013 | 0.05787 | 0.22260 |
| **AMN1** | 2.30286 | 0.00000 | 1.02056 | 2.69626 | 1.67710 | 0.51797 | 0.00000 |
| **AMOT** | 9.72710 | 12.09211 | 9.35265 | 0.57649 | 2.05986 | 1.65421 | 16.53967 |
| **AMOTL1** | 0.62316 | 0.00000 | 0.59148 | 2.02323 | 2.32331 | 3.41169 | 3.43245 |
| **AMOTL2** | 4.58776 | 18.41241 | 1.61403 | 27.43742 | 0.00000 | 0.00000 | 14.19663 |

| | | | | | | | |
|---|---|---|---|---|---|---|---|
| **AMPD1** | 0.02502 | 0.00000 | 0.12208 | 0.02109 | 0.12610 | 0.01002 | 0.01878 |
| **AMPD2** | 2.38271 | 1.10148 | 3.22025 | 7.41452 | 2.58410 | 1.12437 | 3.08739 |
| **AMPD3** | 0.24275 | 0.00000 | 0.17207 | 0.31879 | 1.34644 | 0.58205 | 0.00000 |
| **AMPH** | 0.00000 | 0.59777 | 1.83768 | 0.00000 | 0.39708 | 0.00000 | 0.00356 |
| **AMT** | 7.29654 | 8.22544 | 0.00000 | 0.58394 | 0.02650 | 7.77724 | 0.67206 |
| **AMY2B** | 0.13675 | 0.00000 | 0.14262 | 0.01780 | 0.08078 | 0.10668 | 0.68274 |
| **AMZ1** | 0.00000 | 0.00000 | 0.14153 | 0.20134 | 0.18631 | 0.04177 | 0.56408 |
| **AMZ2** | 5.47218 | 4.36551 | 5.93529 | 4.21094 | 3.18905 | 4.70913 | 2.44278 |
| **AMZ2P1** | 0.48861 | 0.37556 | 0.96470 | 0.50949 | 0.79464 | 0.98569 | 0.10433 |
| **ANAPC1** | 4.25234 | 2.39343 | 1.86483 | 2.20398 | 1.35165 | 2.18962 | 1.33612 |
| **ANAPC10** | 9.44692 | 0.00000 | 7.94833 | 4.13205 | 1.20909 | 3.23805 | 2.28782 |
| **ANAPC11** | 34.14766 | 29.89521 | 5.31421 | 32.14757 | 11.77331 | 20.23077 | 19.79747 |
| **ANAPC13** | 22.36605 | 0.71900 | 15.23718 | 17.39471 | 4.21118 | 7.67627 | 1.36835 |
| **ANAPC15** | 3.99429 | 4.81356 | 2.76278 | 4.22952 | 3.59033 | 1.32445 | 0.00000 |
| **ANAPC16** | 13.68163 | 8.34858 | 9.79924 | 10.78700 | 0.47908 | 9.05930 | 5.35591 |
| **ANAPC1P1** | 0.02574 | 0.01106 | 0.02489 | 0.02377 | 0.12469 | 0.01314 | 0.01273 |
| **ANAPC2** | 0.00000 | 2.50761 | 1.68484 | 1.55104 | 0.59276 | 0.55995 | 2.51461 |
| **ANAPC4** | 14.24075 | 2.63477 | 1.76006 | 6.47605 | 1.98003 | 6.37630 | 7.52170 |
| **ANAPC5** | 38.52549 | 25.17801 | 12.99964 | 18.66823 | 2.41528 | 15.22748 | 8.81373 |
| **ANAPC7** | 21.63264 | 5.71761 | 12.22837 | 13.99957 | 2.63011 | 5.69685 | 8.49239 |
| **ANG** | 0.16725 | 0.25956 | 0.67858 | 0.33200 | 0.07057 | 0.00000 | 0.00000 |
| **ANGEL1** | 2.15074 | 4.54327 | 2.14918 | 2.69973 | 0.64434 | 3.83876 | 0.01239 |
| **ANGEL2** | 8.85226 | 0.65958 | 4.92815 | 5.21656 | 1.65777 | 5.13640 | 0.00000 |
| **ANGPT1** | 6.94006 | 0.00000 | 1.60667 | 2.03132 | 0.05403 | 0.91461 | 0.30715 |
| **ANGPT2** | 0.35980 | 0.09621 | 0.38311 | 0.00000 | 0.48789 | 0.12473 | 1.65088 |
| **ANGPT4** | 0.00000 | 0.02243 | 0.01258 | 0.93756 | 0.00908 | 0.01180 | 0.00364 |
| **ANGPTL1** | 0.02588 | 0.00000 | 0.11736 | 0.00000 | 0.00426 | 0.14900 | 0.18356 |
| **ANGPTL2** | 0.00985 | 0.01246 | 0.01404 | 0.17872 | 0.00633 | 0.00707 | 0.16771 |
| **ANGPTL3** | 0.04228 | 0.00408 | 0.01365 | 0.01049 | 0.00709 | 0.00735 | 0.09108 |
| **ANGPTL4** | 0.01421 | 0.08177 | 0.04754 | 0.07251 | 0.32614 | 0.13979 | 0.11005 |
| **ANGPTL5** | 0.04537 | 0.00000 | 0.07807 | 0.29956 | 0.11622 | 0.00000 | 0.38841 |
| **ANGPTL6** | 0.09258 | 0.00000 | 0.11298 | 0.00000 | 0.21184 | 0.03584 | 0.06556 |
| **ANGPTL7** | 0.02449 | 0.00588 | 0.07145 | 0.02731 | 0.00583 | 0.04409 | 0.06454 |
| **ANHX** | 0.02662 | 0.02550 | 0.00822 | 0.01796 | 0.03553 | 0.03094 | 0.01138 |
| **ANK1** | 0.14140 | 2.47732 | 0.37521 | 0.67579 | 0.02410 | 0.36897 | 0.05578 |

| | | | | | | | |
|---|---|---|---|---|---|---|---|
| **ANK2** | 0.16197 | 0.00000 | 0.01059 | 0.17564 | 0.27579 | 1.74614 | 0.41249 |
| **ANK3** | 1.67244 | 0.00000 | 0.00000 | 0.58955 | 2.54354 | 2.04598 | 3.28476 |
| **ANKAR** | 0.24350 | 0.00000 | 0.24391 | 0.33850 | 0.22391 | 0.14328 | 0.38678 |
| **ANKDD1A** | 0.14823 | 0.00000 | 0.12469 | 0.31950 | 0.04125 | 0.07852 | 0.64182 |
| **ANKDD1B** | 0.30056 | 0.62016 | 0.92936 | 0.03337 | 0.20280 | 0.79899 | 0.09356 |
| **ANKEF1** | 1.36558 | 0.06248 | 1.06397 | 1.68492 | 0.10256 | 0.39736 | 0.00000 |
| **ANKFN1** | 0.00000 | 0.01799 | 0.02260 | 0.77125 | 0.02883 | 0.09236 | 0.01417 |
| **ANKFY1** | 11.35336 | 2.87572 | 4.45436 | 6.31439 | 1.75471 | 4.31915 | 8.04756 |
| **ANKH** | 0.75045 | 0.76404 | 0.13648 | 1.32632 | 2.50834 | 0.16442 | 2.93274 |
| **ANKHD1-EIF4E** | 0.09538 | 0.19815 | 0.04737 | 0.13422 | 0.02835 | 0.02516 | 0.10264 |
| **ANKHD1.uid-68** | 0.49697 | 0.00000 | 0.31402 | 1.26910 | 0.04151 | 0.81991 | 3.14977 |
| **ANKHD1.uid-74** | 0.00000 | 0.61950 | 0.71979 | 0.36942 | 0.05117 | 1.14440 | 3.38750 |
| **ANKIB1** | 6.29908 | 2.25683 | 3.99677 | 3.39047 | 1.10507 | 5.50804 | 12.60265 |
| **ANKK1** | 0.00469 | 0.00000 | 0.41295 | 0.02503 | 0.00153 | 0.77529 | 0.04263 |
| **ANKLE1** | 1.09800 | 0.00000 | 0.17011 | 3.89554 | 0.08914 | 0.96009 | 0.42843 |
| **ANKLE2** | 5.85864 | 4.80566 | 1.92505 | 4.69426 | 0.59593 | 6.98363 | 17.59844 |
| **ANKMY1** | 0.95590 | 0.36825 | 0.09797 | 0.74712 | 0.32327 | 0.06235 | 0.07172 |
| **ANKMY2** | 7.21720 | 0.00000 | 0.88150 | 13.54423 | 2.84324 | 1.93920 | 0.00000 |
| **ANKRA2** | 3.08416 | 1.90608 | 2.33000 | 3.46594 | 0.78672 | 3.05703 | 0.00000 |
| **ANKRD1** | 0.00000 | 0.51332 | 0.00476 | 0.48476 | 0.03942 | 1.03046 | 1.00969 |
| **ANKRD10** | 28.90985 | 5.88574 | 5.62473 | 18.98662 | 6.75000 | 15.40286 | 6.89580 |
| **ANKRD11** | 11.55317 | 14.64823 | 3.75894 | 9.96860 | 1.80740 | 10.29708 | 49.69025 |
| **ANKRD12** | 7.56243 | 0.00000 | 4.76155 | 9.03705 | 6.04695 | 6.68283 | 11.20180 |
| **ANKRD13A** | 4.62811 | 3.44331 | 1.16892 | 2.71860 | 2.70483 | 2.35906 | 8.54524 |
| **ANKRD13B** | 0.00000 | 0.53085 | 0.73617 | 0.65737 | 0.45154 | 0.51009 | 0.62813 |
| **ANKRD13C** | 17.25892 | 2.04921 | 5.07004 | 9.78275 | 6.56674 | 4.50411 | 2.27723 |
| **ANKRD13D** | 1.04353 | 0.64291 | 0.16810 | 1.81052 | 0.00000 | 1.62404 | 5.67686 |
| **ANKRD16** | 0.56106 | 0.42912 | 0.07532 | 0.27518 | 0.00000 | 0.64716 | 3.49276 |
| **ANKRD17** | 18.42385 | 5.32883 | 10.45765 | 11.80954 | 4.05344 | 11.93818 | 25.87235 |
| **ANKRD18A** | 0.01827 | 0.02079 | 0.01891 | 0.02327 | 0.05957 | 0.01919 | 0.02698 |
| **ANKRD18B** | 0.00000 | 0.01143 | 0.02046 | 0.04739 | 0.01355 | 1.16757 | 0.07362 |
| **ANKRD18DP** | 0.02141 | 0.03500 | 0.07238 | 0.01945 | 0.03037 | 0.00847 | 0.01098 |
| **ANKRD19P** | 2.72438 | 2.60244 | 0.94290 | 3.23313 | 1.85572 | 1.71625 | 9.01817 |
| **ANKRD2** | 0.60613 | 0.00000 | 0.00950 | 0.37323 | 0.00434 | 0.29805 | 0.13538 |
| **ANKRD20A11P** | 0.01536 | 0.06215 | 0.05440 | 0.18655 | 0.00733 | 0.29109 | 0.00466 |

| | | | | | | | |
|---|---|---|---|---|---|---|---|
| **ANKRD20A12P** | 0.06832 | 0.00683 | 0.01994 | 0.02126 | 0.04875 | 0.05164 | 0.03835 |
| **ANKRD20A19P** | 0.28526 | 0.00000 | 0.12221 | 2.54196 | 0.15075 | 0.04678 | 0.00000 |
| **ANKRD20A4** | 0.02079 | 0.01257 | 0.05230 | 0.02100 | 0.04086 | 0.01029 | 0.03092 |
| **ANKRD20A5P** | 0.09030 | 0.01463 | 0.19591 | 0.00466 | 0.06825 | 0.57740 | 0.08523 |
| **ANKRD20A8P** | 0.02540 | 0.04930 | 0.01771 | 0.01919 | 0.04324 | 0.01402 | 0.02895 |
| **ANKRD20A9P** | 0.02641 | 0.02342 | 0.04676 | 0.05040 | 0.10908 | 0.10577 | 0.15392 |
| **ANKRD22** | 0.21098 | 0.00000 | 0.02324 | 0.07991 | 0.23501 | 0.01934 | 0.00000 |
| **ANKRD23** | 0.02032 | 0.11876 | 0.27857 | 1.83241 | 0.02962 | 0.18795 | 1.16150 |
| **ANKRD24** | 0.00000 | 0.00000 | 0.08841 | 0.04117 | 0.04040 | 0.00052 | 0.92260 |
| **ANKRD26** | 1.23752 | 0.76190 | 3.90822 | 0.27822 | 1.46493 | 2.84929 | 9.10866 |
| **ANKRD26P1** | 0.29744 | 0.12203 | 0.59520 | 0.38079 | 0.66837 | 0.53378 | 0.88764 |
| **ANKRD26P3** | 0.05818 | 0.00000 | 0.14042 | 0.00629 | 0.00493 | 0.03492 | 0.03317 |
| **ANKRD27** | 7.90960 | 1.50642 | 1.76402 | 6.17082 | 1.28806 | 2.69791 | 3.16160 |
| **ANKRD28** | 4.66596 | 3.07337 | 4.70389 | 0.39163 | 0.37064 | 5.62970 | 6.81648 |
| **ANKRD29** | 0.02253 | 0.05495 | 0.07491 | 0.03160 | 0.00619 | 0.00667 | 0.00864 |
| **ANKRD30A** | 0.05640 | 0.06879 | 0.07788 | 0.00000 | 0.06696 | 0.00000 | 0.21120 |
| **ANKRD30BL** | 0.00000 | 0.15504 | 0.19232 | 0.32301 | 0.00186 | 1.92936 | 9.76411 |
| **ANKRD30BP2** | 0.02859 | 0.01438 | 0.01559 | 0.02128 | 0.01061 | 0.01215 | 0.02042 |
| **ANKRD31** | 0.04763 | 0.00711 | 0.02841 | 0.04545 | 0.12495 | 0.02285 | 0.00312 |
| **ANKRD32** | 7.48245 | 0.51521 | 4.60373 | 5.01393 | 4.35399 | 4.58071 | 6.04844 |
| **ANKRD33** | 0.03683 | 0.01703 | 0.01130 | 0.05535 | 0.00762 | 0.13692 | 0.02100 |
| **ANKRD33B** | 7.40409 | 6.39660 | 1.30223 | 1.79634 | 0.31171 | 0.00000 | 13.01698 |
| **ANKRD34A** | 0.02800 | 0.00000 | 0.04059 | 0.17294 | 0.03218 | 1.29049 | 0.01978 |
| **ANKRD34B** | 0.06183 | 0.00079 | 0.01899 | 0.00560 | 0.01225 | 0.01119 | 0.07091 |
| **ANKRD34C** | 0.04137 | 0.01147 | 0.01154 | 0.01775 | 0.02964 | 0.01394 | 0.02313 |
| **ANKRD35** | 0.04290 | 0.00000 | 0.10710 | 0.01755 | 7.48033 | 0.22771 | 0.00000 |
| **ANKRD36** | 1.03088 | 0.00000 | 1.96959 | 0.65537 | 0.04954 | 0.03806 | 1.94386 |
| **ANKRD36B** | 1.05068 | 0.00000 | 2.15783 | 1.14046 | 0.14843 | 0.21925 | 1.36162 |
| **ANKRD36BP1** | 1.37643 | 1.02393 | 0.38657 | 0.08995 | 0.34411 | 0.10661 | 0.53881 |
| **ANKRD36BP2** | 0.00000 | 0.07316 | 0.30047 | 0.00000 | 0.46064 | 0.09669 | 0.36160 |
| **ANKRD37** | 0.31511 | 0.00528 | 1.37575 | 1.09603 | 5.45357 | 0.00000 | 0.77592 |
| **ANKRD39** | 6.33322 | 5.67629 | 1.94846 | 10.99901 | 1.22507 | 3.62812 | 1.24376 |
| **ANKRD40** | 22.44237 | 12.02343 | 9.73526 | 5.82856 | 5.16373 | 8.39680 | 4.08847 |
| **ANKRD42** | 0.90738 | 0.87002 | 1.28733 | 1.13941 | 2.36470 | 0.42817 | 0.00000 |
| **ANKRD44** | 0.09604 | 0.00000 | 0.04723 | 0.16289 | 0.37732 | 0.12399 | 0.07132 |

| | | | | | | | |
|---|---|---|---|---|---|---|---|
| **ANKRD44-IT1** | 0.03023 | 0.01303 | 0.01293 | 0.01646 | 0.00764 | 0.01343 | 0.03405 |
| **ANKRD45** | 0.17139 | 0.00000 | 0.00980 | 0.00000 | 0.14896 | 3.22860 | 0.04269 |
| **ANKRD46** | 3.78726 | 0.15069 | 2.25201 | 0.00000 | 0.77512 | 0.99894 | 1.59615 |
| **ANKRD49** | 8.22477 | 0.45645 | 4.57654 | 3.33339 | 1.11320 | 2.34227 | 2.66275 |
| **ANKRD50** | 2.37995 | 0.27919 | 2.82285 | 2.91285 | 2.51797 | 1.58335 | 2.64413 |
| **ANKRD52** | 0.00000 | 0.00000 | 0.81037 | 2.44282 | 1.54719 | 1.94306 | 5.89997 |
| **ANKRD53** | 0.04451 | 0.00466 | 0.01286 | 0.13866 | 0.14512 | 0.08050 | 0.01850 |
| **ANKRD54** | 2.79665 | 5.44894 | 0.73196 | 5.02602 | 1.35187 | 3.42733 | 6.47925 |
| **ANKRD55** | 0.00000 | 1.14623 | 2.09623 | 0.03075 | 0.00000 | 4.65264 | 0.00000 |
| **ANKRD6** | 5.92017 | 0.00000 | 0.00000 | 8.90969 | 1.11420 | 0.00000 | 4.56620 |
| **ANKRD61** | 0.00000 | 0.02889 | 0.12606 | 0.15335 | 0.00768 | 0.16817 | 0.18503 |
| **ANKRD62** | 0.04226 | 0.00000 | 0.34192 | 0.01631 | 0.13042 | 0.05512 | 0.03361 |
| **ANKRD62P1-PA** | 0.01414 | 0.00000 | 0.05214 | 0.05340 | 0.01175 | 0.06677 | 0.43610 |
| **ANKRD65** | 0.00000 | 0.11069 | 0.11521 | 1.41972 | 0.00677 | 0.05293 | 0.03928 |
| **ANKRD7** | 0.07434 | 0.02878 | 0.07751 | 0.00818 | 0.41697 | 0.01383 | 0.00554 |
| **ANKRD9** | 0.00000 | 1.82385 | 0.01749 | 0.01668 | 0.00239 | 0.65404 | 2.72659 |
| **ANKS1A** | 10.26184 | 8.18653 | 1.45094 | 12.67611 | 1.03530 | 2.92364 | 11.23137 |
| **ANKS1B** | 0.00000 | 0.10910 | 0.11307 | 0.02090 | 0.18397 | 0.01612 | 0.07509 |
| **ANKS3** | 0.26425 | 1.75905 | 0.99898 | 3.48675 | 0.11193 | 0.38678 | 5.03995 |
| **ANKS4B** | 0.11316 | 0.00000 | 0.06697 | 0.00000 | 1.26055 | 0.63627 | 0.01424 |
| **ANKS6** | 0.84906 | 1.88477 | 0.65444 | 0.10867 | 0.15876 | 0.81211 | 2.07182 |
| **ANKUB1** | 0.06828 | 0.00160 | 0.01387 | 0.00552 | 0.00717 | 0.00391 | 0.07961 |
| **ANKZF1** | 11.73401 | 5.90400 | 10.63063 | 11.64430 | 4.66000 | 3.87108 | 0.00101 |
| **ANLN** | 30.22855 | 6.71208 | 5.43778 | 13.07767 | 2.81310 | 5.40291 | 8.28431 |
| **ANO1** | 1.51609 | 0.95901 | 0.00000 | 1.72690 | 0.11350 | 0.00000 | 1.25492 |
| **ANO10** | 1.94551 | 2.22307 | 3.51431 | 6.83024 | 2.91616 | 3.56442 | 2.80887 |
| **ANO2** | 0.07125 | 0.00000 | 0.02557 | 0.00717 | 0.00565 | 0.07423 | 0.14104 |
| **ANO3** | 0.02538 | 0.00580 | 0.01663 | 0.41764 | 0.23293 | 0.01564 | 0.00000 |
| **ANO4** | 0.00000 | 0.16224 | 0.31151 | 0.01238 | 0.04456 | 0.25459 | 0.07152 |
| **ANO5** | 0.08898 | 0.00000 | 0.12603 | 0.09416 | 0.36826 | 0.30359 | 0.00000 |
| **ANO6** | 15.32951 | 2.38931 | 2.61402 | 14.48122 | 9.71975 | 2.89615 | 20.86509 |
| **ANO7** | 1.50437 | 0.27742 | 0.00941 | 1.25475 | 0.19757 | 0.00000 | 4.30985 |
| **ANO8** | 0.00065 | 0.02421 | 0.01812 | 0.13013 | 0.01520 | 0.06775 | 0.40431 |
| **ANO9** | 0.31625 | 0.00000 | 0.00096 | 0.06434 | 0.00542 | 0.00000 | 0.99313 |
| **ANP32A** | 5.35174 | 12.01289 | 3.48661 | 12.41581 | 0.20540 | 7.12974 | 18.47979 |

| | | | | | | | |
|---|---|---|---|---|---|---|---|
| **ANP32A-IT1** | 0.00000 | 0.34746 | 0.19295 | 0.33418 | 0.00720 | 0.47560 | 1.58966 |
| **ANP32AP1** | 0.84944 | 1.51978 | 0.47561 | 1.67935 | 0.02311 | 0.98915 | 1.33194 |
| **ANP32B** | 29.57633 | 26.84184 | 19.91294 | 25.78413 | 4.72968 | 11.49810 | 42.14303 |
| **ANP32C** | 0.11430 | 0.07812 | 0.10842 | 0.17057 | 0.00406 | 0.15278 | 0.28304 |
| **ANP32D** | 0.02763 | 0.03131 | 0.02301 | 0.02799 | 0.00966 | 0.01035 | 0.03242 |
| **ANP32E** | 39.46367 | 0.00000 | 2.06103 | 32.79576 | 0.60258 | 14.82455 | 11.62822 |
| **ANPEP** | 24.72630 | 61.74052 | 0.00000 | 4.56577 | 0.12451 | 16.44199 | 30.98399 |
| **ANTXR1** | 0.00000 | 0.05057 | 0.01594 | 0.24120 | 0.02173 | 0.02353 | 0.23592 |
| **ANTXR2** | 0.00000 | 0.35452 | 0.10213 | 0.74112 | 0.22240 | 1.53849 | 0.70657 |
| **ANTXRL** | 0.04402 | 0.01879 | 0.01223 | 0.08786 | 0.00790 | 0.00766 | 0.01270 |
| **ANTXRLP1** | 0.02468 | 0.01869 | 0.01280 | 0.03357 | 0.00800 | 0.01451 | 0.01219 |
| **ANXA1** | 0.85808 | 0.00000 | 0.00917 | 24.70940 | 0.01542 | 0.03827 | 0.00000 |
| **ANXA10** | 0.02682 | 0.01461 | 0.01866 | 0.01786 | 0.02907 | 0.01326 | 0.01748 |
| **ANXA11** | 11.61040 | 13.23056 | 1.11124 | 6.86819 | 0.12606 | 3.60960 | 32.00640 |
| **ANXA13** | 0.02153 | 0.01607 | 0.01446 | 0.01859 | 0.01664 | 0.01186 | 0.02730 |
| **ANXA2** | 63.06545 | 46.69928 | 0.00000 | 43.26509 | 2.18996 | 28.46211 | 24.10487 |
| **ANXA2P1** | 8.64542 | 2.63780 | 0.00000 | 3.76181 | 0.04803 | 2.59590 | 0.00000 |
| **ANXA2P2** | 54.03197 | 29.96485 | 0.00000 | 30.84987 | 0.82167 | 19.30180 | 0.00025 |
| **ANXA2P3** | 7.58454 | 2.59493 | 0.00000 | 3.75795 | 0.03697 | 2.62655 | 0.40123 |
| **ANXA2R** | 0.00000 | 0.00000 | 0.93181 | 0.06076 | 0.29581 | 0.14093 | 0.12654 |
| **ANXA3** | 42.91197 | 13.54131 | 8.08247 | 16.03572 | 0.00000 | 6.24107 | 5.35599 |
| **ANXA4** | 1.40254 | 0.15471 | 1.01861 | 0.00000 | 0.02454 | 3.98519 | 0.00000 |
| **ANXA5** | 48.28759 | 11.17587 | 19.57653 | 40.26725 | 4.61379 | 9.16746 | 11.51953 |
| **ANXA6** | 49.05931 | 7.46823 | 0.00000 | 21.58443 | 0.10286 | 3.62133 | 3.75116 |
| **ANXA7** | 36.11826 | 13.86482 | 9.75647 | 19.27274 | 4.45639 | 10.83710 | 1.71530 |
| **ANXA8.uid-1418** | 0.36612 | 0.06719 | 0.00442 | 0.50815 | 0.00739 | 0.00063 | 0.00000 |
| **ANXA8L1** | 0.67280 | 0.31083 | 0.00054 | 0.93623 | 0.00633 | 0.00036 | 0.00000 |
| **ANXA9** | 1.13524 | 0.00000 | 0.01998 | 1.73213 | 0.00282 | 0.00000 | 0.07913 |
| **AOAH** | 0.02917 | 0.00918 | 0.02318 | 0.04825 | 1.64769 | 0.01207 | 0.01199 |
| **AOC2** | 0.69431 | 1.51747 | 0.50131 | 3.06886 | 0.45768 | 0.79807 | 2.60614 |
| **AOC3** | 0.49567 | 0.00000 | 1.07236 | 1.53296 | 0.10988 | 0.34185 | 3.16925 |
| **AOC4P** | 0.00656 | 0.01360 | 0.00634 | 0.02961 | 0.08069 | 0.50041 | 0.03424 |
| **AOX1** | 0.00000 | 0.38467 | 5.27306 | 0.00000 | 0.59284 | 0.52543 | 0.00249 |
| **AOX2P** | 0.03087 | 0.03318 | 0.01469 | 0.01580 | 0.04894 | 0.01795 | 0.01268 |
| **AP1AR** | 10.50059 | 2.74632 | 3.09851 | 2.62757 | 0.96592 | 3.51380 | 0.31761 |

| | | | | | | | |
|---|---|---|---|---|---|---|---|
| **AP1B1** | 7.39142 | 13.08908 | 2.05882 | 9.71349 | 0.73350 | 9.57195 | 20.39544 |
| **AP1B1P1** | 0.02360 | 0.01799 | 0.01073 | 0.01932 | 0.03066 | 0.01315 | 0.01645 |
| **AP1G1** | 19.24720 | 4.28403 | 11.38844 | 11.48047 | 7.73992 | 7.67442 | 10.61421 |
| **AP1G2** | 4.95592 | 2.76187 | 0.03756 | 8.20151 | 0.00000 | 5.20136 | 2.77364 |
| **AP1M1** | 8.05604 | 6.82321 | 5.10743 | 11.98628 | 6.02902 | 2.81520 | 2.16344 |
| **AP1M2** | 22.53364 | 15.65427 | 7.20750 | 15.10341 | 4.43555 | 14.59183 | 5.27978 |
| **AP1S1** | 4.79369 | 7.58249 | 1.14886 | 23.10660 | 0.71571 | 11.45954 | 2.95107 |
| **AP1S2** | 1.26732 | 0.00000 | 1.04641 | 0.99002 | 1.78407 | 1.26691 | 0.00000 |
| **AP1S3** | 2.03292 | 0.00000 | 3.20941 | 0.59309 | 0.18132 | 0.08570 | 0.21355 |
| **AP2A1** | 3.03142 | 6.58273 | 0.86420 | 6.02259 | 0.15769 | 2.05136 | 24.83286 |
| **AP2A2** | 0.00000 | 1.99608 | 1.13554 | 2.89211 | 0.37593 | 1.37569 | 11.28856 |
| **AP2B1** | 27.68903 | 12.24936 | 16.41079 | 20.75566 | 3.27524 | 17.41817 | 7.92889 |
| **AP2M1** | 50.12108 | 55.82931 | 16.86095 | 36.15417 | 4.90480 | 32.39872 | 17.40884 |
| **AP2S1** | 32.57053 | 37.76255 | 19.92968 | 34.68352 | 5.48741 | 21.00468 | 28.21297 |
| **AP3B1** | 16.81708 | 1.22218 | 7.47464 | 11.08278 | 2.06510 | 5.66875 | 12.24173 |
| **AP3B2** | 0.00000 | 0.05361 | 0.52407 | 0.87139 | 0.68954 | 0.03120 | 0.00000 |
| **AP3D1** | 9.51004 | 20.08971 | 3.69054 | 12.49979 | 0.31326 | 11.88527 | 42.67199 |
| **AP3M1** | 14.85588 | 1.75243 | 6.32689 | 9.87628 | 0.70316 | 2.04576 | 0.00000 |
| **AP3M2** | 6.18524 | 2.58428 | 4.55475 | 4.54134 | 2.76452 | 0.71985 | 1.45387 |
| **AP3S1** | 15.93732 | 5.40810 | 11.34841 | 7.61515 | 2.87561 | 8.59989 | 5.82279 |
| **AP3S2** | 0.65275 | 0.66310 | 0.49987 | 0.54393 | 0.16787 | 0.12449 | 0.00000 |
| **AP4B1** | 4.63164 | 0.00014 | 4.27843 | 2.61306 | 4.37947 | 1.29117 | 0.00000 |
| **AP4B1-AS1** | 0.16848 | 0.00755 | 0.05052 | 0.05004 | 0.13649 | 0.04508 | 0.01122 |
| **AP4E1** | 5.13289 | 1.23579 | 2.42810 | 4.50182 | 1.14658 | 1.05442 | 5.11763 |
| **AP4M1** | 5.00277 | 5.51864 | 4.26036 | 6.50957 | 5.66771 | 2.87645 | 0.29097 |
| **AP4S1** | 0.67097 | 0.26304 | 0.31637 | 1.42639 | 0.03886 | 0.74582 | 1.21747 |
| **AP5B1** | 0.00000 | 2.21102 | 0.05062 | 7.01984 | 0.02787 | 2.07609 | 3.33015 |
| **AP5M1** | 17.31458 | 4.09704 | 7.97424 | 6.96987 | 3.33382 | 7.78397 | 0.00000 |
| **AP5S1** | 2.63996 | 2.25482 | 3.88515 | 2.07238 | 0.73166 | 0.88334 | 1.55831 |
| **AP5Z1** | 1.44453 | 3.68889 | 1.62088 | 6.79119 | 0.42295 | 2.38529 | 9.62813 |
| **APAF1** | 0.00000 | 0.55083 | 1.05178 | 0.53076 | 0.69605 | 1.41137 | 0.76873 |
| **APBA1** | 0.00000 | 0.07964 | 0.27873 | 0.02006 | 1.36462 | 0.04331 | 0.14219 |
| **APBA2** | 0.00000 | 0.05634 | 0.31637 | 0.04087 | 1.07527 | 0.22656 | 0.02087 |
| **APBA3** | 0.00000 | 3.39262 | 2.02235 | 2.88292 | 0.00000 | 1.40074 | 4.74170 |
| **APBB1** | 3.34671 | 2.69487 | 0.09939 | 1.35950 | 0.14804 | 0.36992 | 2.22495 |

| | | | | | | | |
|---|---|---|---|---|---|---|---|
| **APBB1IP** | 0.84066 | 0.00000 | 0.04288 | 0.42883 | 0.03733 | 0.00000 | 1.16744 |
| **APBB2** | 4.29647 | 2.14138 | 9.43068 | 5.85290 | 4.91667 | 3.59329 | 6.54961 |
| **APBB3** | 0.94757 | 0.18940 | 0.65185 | 2.88760 | 0.23858 | 0.10024 | 1.26496 |
| **APC** | 6.29712 | 0.00000 | 3.85223 | 2.32485 | 4.38990 | 2.76991 | 8.09212 |
| **APC2** | 0.00000 | 0.36158 | 0.07922 | 0.11365 | 0.56852 | 0.10186 | 0.22522 |
| **APCDD1** | 0.05005 | 0.05878 | 0.03218 | 0.11879 | 0.00865 | 0.02558 | 0.01848 |
| **APCDD1L** | 0.05749 | 0.03501 | 0.04349 | 0.01672 | 0.00681 | 0.01174 | 0.00779 |
| **APCDD1L-AS1** | 0.02854 | 0.10194 | 0.01761 | 0.03363 | 0.08284 | 0.01571 | 0.01746 |
| **APEH** | 20.93206 | 20.88752 | 6.91427 | 13.58183 | 0.26116 | 14.40997 | 5.82253 |
| **APELA** | 4.01940 | 2.03675 | 0.00002 | 0.00000 | 0.00530 | 5.05791 | 0.00000 |
| **APEX1** | 41.93626 | 19.99351 | 28.23954 | 26.81432 | 10.64412 | 26.18465 | 13.50467 |
| **APEX2** | 13.18900 | 15.53263 | 13.24387 | 10.69342 | 6.61652 | 6.31843 | 1.20061 |
| **APH1A** | 7.35530 | 3.78960 | 5.51072 | 11.19590 | 2.43207 | 8.21339 | 7.06322 |
| **APH1B** | 0.54840 | 0.00809 | 0.12791 | 0.17869 | 0.54088 | 0.00000 | 0.52211 |
| **API5** | 28.75850 | 3.77069 | 12.90605 | 15.81541 | 8.99870 | 6.92671 | 0.00000 |
| **APIP** | 8.61714 | 1.24393 | 2.23411 | 1.76610 | 0.02678 | 3.93891 | 0.86570 |
| **APITD1** | 1.71287 | 0.16533 | 2.37849 | 0.23615 | 0.81984 | 1.69548 | 1.01789 |
| **APLF** | 0.18543 | 0.00000 | 0.39156 | 0.03760 | 0.34148 | 0.30936 | 0.20080 |
| **APLN** | 0.03714 | 0.01527 | 0.01329 | 0.01500 | 0.01180 | 0.02204 | 0.01576 |
| **APLNR** | 0.02861 | 0.00809 | 0.01012 | 0.04936 | 0.00727 | 0.59045 | 0.04200 |
| **APLP1** | 0.00000 | 0.00000 | 0.16438 | 0.05213 | 0.31568 | 0.77838 | 0.11987 |
| **APLP2** | 51.55987 | 50.37464 | 20.47696 | 39.60697 | 10.30277 | 34.41122 | 36.13059 |
| **APMAP** | 18.20879 | 17.38130 | 7.96443 | 21.15367 | 1.84234 | 9.57191 | 0.00000 |
| **APOA1** | 4.39679 | 0.00000 | 0.09507 | 9.40981 | 0.00000 | 11.86266 | 0.00000 |
| **APOA1BP** | 20.62314 | 17.09631 | 0.86199 | 23.12283 | 0.47752 | 2.95974 | 1.39038 |
| **APOA2** | 1.07520 | 0.00261 | 0.00000 | 6.27320 | 1.66507 | 7.53997 | 0.00000 |
| **APOA4** | 2.24162 | 0.93019 | 0.00028 | 0.00000 | 0.00315 | 0.42803 | 0.00154 |
| **APOA5** | 0.04777 | 0.00488 | 0.05221 | 0.61094 | 0.00633 | 0.03487 | 0.00000 |
| **APOB** | 0.01978 | 0.00555 | 0.01074 | 0.19861 | 0.02348 | 0.12086 | 0.04512 |
| **APOBEC1** | 0.06281 | 0.01748 | 0.01329 | 0.01229 | 0.02633 | 0.01163 | 0.01221 |
| **APOBEC2** | 0.57394 | 0.19195 | 0.21902 | 3.04250 | 0.05206 | 0.02182 | 1.95523 |
| **APOBEC3A** | 0.03148 | 0.01907 | 0.02634 | 0.01657 | 0.00707 | 0.01619 | 0.01889 |
| **APOBEC3B** | 0.02528 | 0.00742 | 0.01954 | 0.02417 | 0.00937 | 0.20840 | 0.00628 |
| **APOBEC3B-AS1** | 0.01749 | 0.00000 | 0.04188 | 0.03666 | 0.01043 | 0.42132 | 0.01075 |
| **APOBEC3C** | 0.06703 | 0.00664 | 0.00829 | 0.09737 | 0.00757 | 4.49909 | 0.00000 |

| | | | | | | | |
|---|---|---|---|---|---|---|---|
| **APOBEC3D** | 0.03657 | 0.07527 | 0.00000 | 0.03742 | 0.05422 | 3.18029 | 0.02460 |
| **APOBEC3F** | 0.00000 | 0.34006 | 0.99122 | 0.01954 | 0.03620 | 2.96568 | 0.01447 |
| **APOBEC3G** | 0.05673 | 0.00000 | 0.02460 | 0.11964 | 0.00747 | 2.01388 | 0.00000 |
| **APOBEC3H** | 0.02002 | 0.03304 | 0.01012 | 0.01948 | 0.00898 | 0.04847 | 0.01199 |
| **APOBEC4** | 0.03240 | 0.03321 | 0.00744 | 0.00267 | 0.00768 | 0.00387 | 0.10890 |
| **APOBR** | 0.00000 | 1.02200 | 2.89790 | 0.00000 | 0.04336 | 0.72281 | 0.28163 |
| **APOC1** | 27.79624 | 44.55365 | 15.99973 | 13.24584 | 0.00000 | 36.08754 | 28.24238 |
| **APOC1P1** | 0.00000 | 5.71603 | 3.87032 | 0.45523 | 0.00000 | 3.28724 | 0.87882 |
| **APOC3** | 3.74716 | 0.67175 | 0.03363 | 5.35969 | 0.00064 | 0.02722 | 0.02711 |
| **APOC4-APOC2** | 0.04261 | 0.03781 | 0.01548 | 0.03751 | 0.00787 | 0.03132 | 0.00000 |
| **APOD** | 5.37339 | 0.21743 | 0.00000 | 0.96537 | 0.44953 | 0.52739 | 0.00000 |
| **APOE** | 0.00000 | 3.80670 | 0.00000 | 3.35365 | 0.00299 | 19.28469 | 17.73838 |
| **APOF** | 0.02740 | 0.01532 | 0.04410 | 0.01921 | 0.03810 | 0.01351 | 0.03389 |
| **APOH** | 0.07860 | 0.17348 | 0.47787 | 0.02777 | 0.00383 | 0.00004 | 0.00000 |
| **APOL1** | 0.05587 | 0.00196 | 0.00637 | 0.03768 | 0.01611 | 0.91043 | 0.13101 |
| **APOL2** | 1.95497 | 0.00000 | 0.00379 | 0.87415 | 0.03753 | 1.15173 | 0.00005 |
| **APOL3** | 0.04390 | 0.01378 | 0.01180 | 0.03751 | 0.02068 | 0.01407 | 0.00984 |
| **APOL4** | 0.61829 | 0.00000 | 0.01058 | 1.31812 | 0.00612 | 0.05911 | 0.00000 |
| **APOL5** | 0.02388 | 0.00000 | 0.21736 | 0.02473 | 0.01550 | 0.00863 | 0.02203 |
| **APOL6** | 0.04493 | 0.01929 | 0.05042 | 0.07546 | 0.02245 | 0.02967 | 0.08481 |
| **APOLD1** | 0.10608 | 0.00850 | 0.01800 | 0.02589 | 0.00830 | 0.02526 | 0.17215 |
| **APOM** | 0.10272 | 0.02893 | 0.03755 | 0.12320 | 0.00807 | 1.11640 | 0.00000 |
| **APOO** | 4.85596 | 5.20589 | 8.85134 | 0.19615 | 2.55830 | 5.63108 | 0.12935 |
| **APOOL** | 13.74855 | 6.83815 | 2.53169 | 5.50149 | 0.06206 | 4.25180 | 7.20951 |
| **APOOP5** | 0.32826 | 0.29081 | 0.77524 | 0.02364 | 0.20104 | 0.49090 | 0.24631 |
| **APOPT1** | 14.01776 | 9.43017 | 5.42136 | 8.96872 | 1.65709 | 5.64769 | 5.69546 |
| **APP** | 2.64850 | 5.17553 | 0.87569 | 4.04841 | 1.98197 | 1.55236 | 12.11463 |
| **APPBP2** | 0.23334 | 0.59315 | 1.95548 | 1.93898 | 1.57040 | 1.95107 | 4.42516 |
| **APPL1** | 8.55261 | 3.76830 | 2.84445 | 3.40623 | 0.24321 | 9.26517 | 15.62873 |
| **APPL2** | 0.48709 | 1.23239 | 0.43854 | 1.75842 | 0.95155 | 0.91312 | 1.33021 |
| **APRT** | 31.01101 | 28.56722 | 2.74382 | 22.90424 | 0.00000 | 16.79254 | 14.42215 |
| **APTR** | 3.55116 | 0.74530 | 1.69595 | 7.18787 | 8.55284 | 1.28061 | 0.00000 |
| **APTX** | 23.36928 | 19.55530 | 16.68851 | 15.61330 | 5.34858 | 6.77984 | 0.00000 |
| **AQP1** | 0.03222 | 0.00000 | 0.01554 | 0.00000 | 0.00757 | 0.06178 | 0.36682 |
| **AQP10** | 0.02105 | 0.03124 | 0.05345 | 0.02606 | 0.00966 | 0.02245 | 0.01421 |

| | | | | | | | |
|---|---|---|---|---|---|---|---|
| **AQP11** | 1.85523 | 0.00000 | 1.17774 | 0.47952 | 0.71917 | 0.26587 | 0.92103 |
| **AQP2** | 0.02327 | 0.01861 | 0.01352 | 0.01951 | 0.00960 | 0.01271 | 0.02019 |
| **AQP3** | 13.23671 | 59.39377 | 8.12606 | 26.06505 | 0.03572 | 8.02271 | 13.11425 |
| **AQP4** | 0.04085 | 0.02056 | 0.01687 | 0.05336 | 0.01248 | 0.11721 | 0.00000 |
| **AQP4-AS1** | 0.02370 | 0.01527 | 0.01844 | 0.03555 | 0.01164 | 0.08756 | 0.03548 |
| **AQP5** | 0.02423 | 0.01657 | 0.01336 | 0.02207 | 0.04082 | 0.01542 | 0.01904 |
| **AQP6** | 0.20236 | 0.11866 | 0.18676 | 0.24841 | 0.90302 | 0.04692 | 1.13218 |
| **AQP7** | 0.09214 | 0.09072 | 0.29620 | 0.07672 | 0.00345 | 0.11922 | 0.00000 |
| **AQP7P1** | 0.01896 | 0.02513 | 0.01746 | 0.02820 | 0.00808 | 0.01731 | 0.01738 |
| **AQP8** | 0.00799 | 0.11182 | 0.01096 | 0.06389 | 0.01204 | 0.01510 | 0.21973 |
| **AQP9** | 0.06311 | 0.01111 | 0.01284 | 0.00839 | 0.00757 | 0.01162 | 0.02916 |
| **AQPEP** | 1.46753 | 0.97793 | 0.10937 | 0.00000 | 0.00000 | 1.22784 | 2.11318 |
| **AQR** | 21.40408 | 3.70440 | 12.60117 | 9.57752 | 4.94671 | 7.63598 | 6.07946 |
| **AR** | 0.03366 | 0.01087 | 0.05016 | 0.02162 | 0.01041 | 0.01304 | 0.01594 |
| **ARAF** | 14.17372 | 11.59372 | 1.59513 | 9.67339 | 0.28150 | 7.68639 | 3.40437 |
| **ARAP1** | 1.07521 | 8.45939 | 2.70581 | 3.33411 | 0.00427 | 3.21263 | 12.14737 |
| **ARAP2** | 0.08845 | 0.07975 | 0.10509 | 0.01894 | 2.72105 | 0.00000 | 0.17625 |
| **ARAP3** | 0.01557 | 0.04451 | 0.01317 | 0.01176 | 0.00843 | 0.21395 | 0.05598 |
| **ARC** | 0.00000 | 0.09376 | 0.00654 | 0.04790 | 0.08221 | 0.09297 | 0.04014 |
| **ARCN1** | 31.11146 | 12.92377 | 13.76489 | 16.19860 | 8.03483 | 8.31064 | 2.09041 |
| **AREL1** | 5.33896 | 3.42193 | 3.86352 | 4.89293 | 3.90848 | 4.05364 | 5.51872 |
| **ARF1** | 61.72159 | 48.10540 | 25.64450 | 46.54409 | 16.87657 | 29.87312 | 24.88153 |
| **ARF3** | 3.67785 | 7.32295 | 6.27104 | 7.37651 | 10.39967 | 17.38196 | 30.25919 |
| **ARF4** | 49.76202 | 15.22982 | 16.55636 | 29.62660 | 8.14176 | 15.34121 | 11.30358 |
| **ARF5** | 15.50990 | 22.68353 | 7.29328 | 21.93256 | 2.43447 | 15.95797 | 18.29184 |
| **ARF6** | 20.40755 | 13.66142 | 9.10165 | 11.59190 | 9.08397 | 9.09607 | 21.80216 |
| **ARFGAP1** | 3.15153 | 12.63655 | 4.98506 | 14.68603 | 0.22875 | 6.35093 | 17.25831 |
| **ARFGAP2** | 15.51173 | 17.58482 | 9.96598 | 14.30601 | 2.91366 | 7.57834 | 6.27701 |
| **ARFGAP3** | 2.33419 | 1.34027 | 5.36873 | 2.78809 | 3.22081 | 7.40896 | 1.59817 |
| **ARFGEF1** | 7.81544 | 10.35268 | 10.64432 | 0.81919 | 2.54873 | 18.81389 | 14.10071 |
| **ARFGEF2** | 9.60910 | 3.47665 | 4.79521 | 3.34171 | 3.19805 | 11.99573 | 13.64700 |
| **ARFIP1** | 13.74047 | 5.46848 | 5.02740 | 3.78554 | 2.81014 | 8.11664 | 3.17885 |
| **ARFIP2** | 19.26214 | 7.93252 | 3.60583 | 18.79256 | 2.73191 | 6.79229 | 7.82872 |
| **ARFRP1** | 5.92893 | 16.04469 | 1.34712 | 7.43198 | 0.68471 | 5.01882 | 1.96114 |
| **ARG1** | 0.10512 | 0.00000 | 0.08695 | 0.02929 | 0.00769 | 0.15538 | 0.07720 |

| | | | | | | | |
|---|---|---|---|---|---|---|---|
| **ARG2** | 1.79230 | 0.00000 | 3.38247 | 8.70433 | 5.45152 | 3.42515 | 0.00000 |
| **ARGFX** | 0.00000 | 1.21049 | 31.16744 | 0.00000 | 13.01481 | 13.87978 | 0.00000 |
| **ARGFXP2** | 0.11415 | 0.02712 | 2.09047 | 0.00000 | 0.99085 | 0.69359 | 0.08516 |
| **ARGLU1** | 20.99405 | 10.00880 | 12.71454 | 14.27035 | 9.38079 | 10.49355 | 17.63853 |
| **ARHGAP1** | 12.22952 | 22.35763 | 6.76137 | 6.16616 | 3.86401 | 13.75465 | 5.28871 |
| **ARHGAP10** | 7.28585 | 0.31136 | 0.10455 | 9.66290 | 4.80728 | 0.48030 | 16.17344 |
| **ARHGAP11A** | 8.12896 | 0.00000 | 9.66346 | 3.36258 | 3.48888 | 3.27909 | 14.94357 |
| **ARHGAP11B** | 0.22110 | 0.00000 | 0.36520 | 0.16850 | 0.06764 | 0.25009 | 2.28249 |
| **ARHGAP12** | 10.35092 | 1.28166 | 3.23803 | 1.94977 | 2.36921 | 8.11193 | 4.15447 |
| **ARHGAP15** | 0.09761 | 0.00000 | 0.01798 | 0.01087 | 0.72345 | 0.01136 | 0.00782 |
| **ARHGAP17** | 1.41435 | 5.72523 | 1.68556 | 9.35657 | 2.95939 | 1.74797 | 11.07398 |
| **ARHGAP18** | 38.96203 | 17.06437 | 0.00000 | 13.90963 | 9.34958 | 0.00000 | 15.82974 |
| **ARHGAP19** | 4.46430 | 4.02580 | 2.81537 | 2.73390 | 0.59884 | 3.34203 | 3.25788 |
| **ARHGAP20** | 0.03032 | 0.00000 | 0.10639 | 0.04593 | 0.39540 | 0.02445 | 0.00787 |
| **ARHGAP21** | 2.51351 | 3.02256 | 5.56154 | 2.28590 | 0.71033 | 3.61203 | 10.67565 |
| **ARHGAP22** | 0.00000 | 1.06804 | 0.01244 | 0.11148 | 0.03167 | 0.29587 | 0.00189 |
| **ARHGAP23** | 0.30001 | 1.50912 | 0.00000 | 1.75477 | 0.02035 | 0.00000 | 8.60099 |
| **ARHGAP24** | 1.05648 | 0.00000 | 0.02463 | 7.55843 | 0.15767 | 0.00000 | 1.21182 |
| **ARHGAP25** | 0.04827 | 0.00999 | 0.06262 | 0.07188 | 0.02149 | 0.11969 | 0.00000 |
| **ARHGAP26** | 0.65612 | 0.24485 | 0.20209 | 0.28433 | 1.84571 | 1.35139 | 2.29434 |
| **ARHGAP26-AS1** | 0.02021 | 0.05357 | 0.01400 | 0.01981 | 0.21066 | 0.02480 | 0.02478 |
| **ARHGAP27** | 1.80718 | 0.71887 | 1.34281 | 2.57183 | 0.00030 | 0.00000 | 4.71815 |
| **ARHGAP28** | 0.31455 | 0.14475 | 0.06467 | 1.74992 | 0.02680 | 0.33098 | 0.90515 |
| **ARHGAP29** | 8.15554 | 0.05574 | 1.04963 | 7.36466 | 2.21843 | 0.49582 | 16.83793 |
| **ARHGAP30** | 0.00000 | 0.00000 | 0.20281 | 0.72424 | 0.16231 | 1.38858 | 1.52840 |
| **ARHGAP31** | 0.09012 | 0.10848 | 0.11739 | 0.37050 | 0.09595 | 0.65890 | 0.20378 |
| **ARHGAP31-AS1** | 0.02184 | 0.01664 | 0.00000 | 0.02334 | 0.43136 | 0.02397 | 0.01097 |
| **ARHGAP32** | 0.42948 | 4.03422 | 0.63625 | 8.52242 | 2.93731 | 2.99271 | 5.43787 |
| **ARHGAP33** | 0.03786 | 0.00000 | 0.03571 | 0.07496 | 0.63476 | 1.23437 | 0.26995 |
| **ARHGAP35** | 0.28816 | 0.51391 | 2.45905 | 2.55699 | 1.16040 | 2.12411 | 12.76027 |
| **ARHGAP36** | 0.05313 | 0.06495 | 0.00141 | 0.01497 | 0.08595 | 0.00818 | 0.00379 |
| **ARHGAP39** | 0.00000 | 0.41360 | 0.62659 | 0.02095 | 0.41026 | 0.57642 | 2.35028 |
| **ARHGAP4** | 0.01304 | 0.00074 | 0.13059 | 0.11003 | 0.01267 | 0.20784 | 0.04996 |
| **ARHGAP40** | 0.07528 | 0.01038 | 0.00756 | 0.07127 | 0.06914 | 0.06269 | 0.00000 |
| **ARHGAP42** | 0.69597 | 0.44254 | 0.89940 | 0.46866 | 0.00000 | 5.70049 | 4.16438 |

| | | | | | | | |
|---|---|---|---|---|---|---|---|
| **ARHGAP44** | 0.91712 | 2.02280 | 0.00214 | 0.00000 | 0.47442 | 0.60610 | 2.24269 |
| **ARHGAP5** | 7.58769 | 0.00000 | 11.01183 | 4.26718 | 2.98900 | 8.11968 | 21.69589 |
| **ARHGAP5-AS1** | 0.22861 | 1.05725 | 0.92831 | 0.59124 | 0.17676 | 0.23227 | 0.25547 |
| **ARHGAP6** | 0.01906 | 0.04219 | 0.01813 | 0.02375 | 0.29954 | 0.02170 | 0.08232 |
| **ARHGAP8** | 0.34706 | 0.71345 | 0.02933 | 0.50057 | 0.01550 | 0.15491 | 0.00000 |
| **ARHGAP9** | 0.00000 | 3.07807 | 0.60508 | 0.05806 | 0.07299 | 0.08374 | 0.00325 |
| **ARHGDIA** | 7.69761 | 7.28707 | 0.98344 | 25.76826 | 2.88405 | 9.23634 | 36.15648 |
| **ARHGDIB** | 1.26851 | 0.34780 | 0.00000 | 2.89867 | 0.00041 | 0.00000 | 0.90964 |
| **ARHGDIG** | 0.00000 | 0.08133 | 0.01664 | 0.01528 | 0.12291 | 0.23371 | 0.05960 |
| **ARHGEF1** | 1.87092 | 2.81487 | 1.29410 | 2.35293 | 0.04244 | 2.23886 | 7.59130 |
| **ARHGEF10** | 0.78856 | 0.09021 | 0.21731 | 0.09346 | 0.00963 | 1.88170 | 0.35356 |
| **ARHGEF10L** | 0.00005 | 0.00348 | 0.01531 | 0.61468 | 0.04237 | 0.13345 | 0.13451 |
| **ARHGEF11** | 0.23417 | 3.80408 | 2.79748 | 1.36844 | 0.98149 | 1.71744 | 8.89157 |
| **ARHGEF12** | 2.11518 | 0.91627 | 1.21019 | 2.49929 | 2.27849 | 2.03124 | 11.64444 |
| **ARHGEF15** | 0.02181 | 0.00000 | 0.05802 | 0.07436 | 0.01122 | 1.19880 | 0.03156 |
| **ARHGEF16** | 1.83055 | 3.37734 | 0.00000 | 1.42727 | 0.50781 | 6.92498 | 7.11340 |
| **ARHGEF17** | 0.50626 | 1.51659 | 0.00000 | 2.75601 | 0.01503 | 0.00000 | 1.62888 |
| **ARHGEF18** | 1.43453 | 9.39808 | 3.68589 | 3.75669 | 2.01762 | 2.48423 | 10.76416 |
| **ARHGEF19** | 0.55804 | 0.46754 | 0.07901 | 0.84472 | 0.16593 | 0.55560 | 0.00000 |
| **ARHGEF2** | 1.08376 | 0.40869 | 0.00000 | 2.50521 | 0.01552 | 0.41361 | 4.17275 |
| **ARHGEF25** | 0.07043 | 0.00000 | 0.05507 | 0.01608 | 0.01656 | 0.25812 | 0.00340 |
| **ARHGEF26** | 9.55036 | 3.41221 | 1.94182 | 0.00000 | 1.54731 | 0.00000 | 2.20921 |
| **ARHGEF26-AS1** | 0.03689 | 0.11555 | 0.02162 | 0.01498 | 0.16382 | 0.03283 | 0.02464 |
| **ARHGEF28** | 0.02968 | 0.01787 | 0.02099 | 0.01947 | 0.73024 | 0.12020 | 0.01040 |
| **ARHGEF3** | 1.33326 | 0.32620 | 1.52374 | 1.36798 | 0.78690 | 1.96698 | 0.03394 |
| **ARHGEF33** | 0.00000 | 0.07318 | 3.44177 | 0.00000 | 0.87995 | 0.01677 | 0.32805 |
| **ARHGEF34P** | 0.01112 | 0.03166 | 0.01341 | 0.03370 | 0.01040 | 0.02821 | 0.07291 |
| **ARHGEF35** | 0.01743 | 0.01379 | 0.01353 | 0.14235 | 0.00632 | 0.07102 | 0.14020 |
| **ARHGEF37** | 0.00000 | 0.26221 | 0.33519 | 1.31263 | 0.08333 | 0.00000 | 0.00000 |
| **ARHGEF38** | 0.59837 | 0.35922 | 0.06437 | 0.06055 | 0.14330 | 0.12335 | 0.06900 |
| **ARHGEF38-IT1** | 0.09649 | 0.19174 | 0.00005 | 0.00000 | 0.06253 | 0.01328 | 0.02495 |
| **ARHGEF39** | 4.98656 | 8.14858 | 3.63039 | 3.67831 | 0.51678 | 2.79458 | 0.00000 |
| **ARHGEF4** | 0.00000 | 0.01466 | 0.14729 | 1.37405 | 0.00672 | 0.06547 | 0.00822 |
| **ARHGEF40** | 0.03746 | 0.02004 | 0.02944 | 0.10610 | 0.00718 | 0.15154 | 0.17000 |
| **ARHGEF5** | 0.00000 | 0.04308 | 0.12920 | 0.47504 | 0.05770 | 0.30597 | 0.65865 |

| | | | | | | | |
|---|---|---|---|---|---|---|---|
| **ARHGEF6** | 0.04145 | 0.00000 | 0.37177 | 0.01740 | 0.32774 | 0.00486 | 0.01978 |
| **ARHGEF7** | 9.91784 | 2.53887 | 7.61858 | 8.17759 | 5.57244 | 4.95437 | 10.69806 |
| **ARHGEF9** | 0.71249 | 0.39065 | 0.35140 | 0.39879 | 0.54464 | 0.30966 | 1.23596 |
| **ARHGEF9-IT1** | 0.02466 | 0.01878 | 0.01247 | 0.02889 | 0.00813 | 0.01335 | 0.01556 |
| **ARID1A** | 13.98002 | 15.56339 | 9.93385 | 12.20051 | 3.44907 | 9.19136 | 40.41726 |
| **ARID1B** | 3.11217 | 3.95236 | 2.98245 | 5.05031 | 0.47507 | 5.03570 | 9.45998 |
| **ARID2** | 10.36548 | 0.99175 | 6.08741 | 6.98972 | 2.83606 | 3.23807 | 11.90754 |
| **ARID3A** | 0.66816 | 1.51242 | 1.43322 | 18.22581 | 0.18524 | 2.03030 | 33.25726 |
| **ARID3B** | 2.45499 | 0.74579 | 3.21167 | 9.07576 | 2.03723 | 3.49815 | 6.23466 |
| **ARID3C** | 0.02259 | 0.01386 | 0.01618 | 0.02039 | 0.04632 | 0.01790 | 0.01751 |
| **ARID4A** | 5.00144 | 1.15722 | 3.39019 | 4.74651 | 10.08459 | 3.26294 | 7.10749 |
| **ARID4B** | 4.62924 | 3.77662 | 3.65472 | 10.80071 | 6.65134 | 6.41571 | 13.74242 |
| **ARID5A** | 1.08710 | 2.78746 | 0.57311 | 1.35829 | 0.13274 | 1.28884 | 0.13315 |
| **ARID5B** | 1.54683 | 0.69115 | 0.00000 | 4.75106 | 0.16267 | 1.22342 | 0.00007 |
| **ARIH1** | 6.29070 | 2.21440 | 4.37756 | 4.46202 | 2.84992 | 5.01208 | 18.37969 |
| **ARIH2** | 32.14848 | 16.32888 | 15.97174 | 23.10507 | 5.04968 | 12.23248 | 9.91884 |
| **ARIH2OS** | 0.47067 | 0.33366 | 0.22597 | 0.64175 | 0.55790 | 0.35381 | 0.02654 |
| **1.00000** | 17.63451 | 3.53495 | 8.59349 | 5.29503 | 0.89936 | 10.45521 | 6.29518 |
| **10.00000** | 0.07163 | 0.00568 | 0.00935 | 0.04177 | 0.01095 | 0.10223 | 0.09656 |
| **11.00000** | 1.63479 | 0.00000 | 0.18699 | 0.99013 | 0.00317 | 0.49446 | 0.00000 |
| **ARL13A** | 0.00143 | 0.00000 | 0.10670 | 0.01032 | 0.07782 | 0.10410 | 0.17114 |
| **ARL13B** | 3.46842 | 0.56627 | 0.24852 | 1.81342 | 3.44183 | 3.78497 | 2.07834 |
| **14.00000** | 0.01804 | 0.02718 | 0.01411 | 0.01370 | 0.00689 | 0.01519 | 0.06057 |
| **ARL14EP** | 5.67903 | 3.50884 | 7.75548 | 1.24396 | 2.97511 | 1.08075 | 0.57806 |
| **ARL14EPL** | 0.02411 | 0.01051 | 0.01325 | 0.02059 | 0.00769 | 0.08219 | 0.02640 |
| **15.00000** | 10.19169 | 1.26358 | 13.23179 | 5.73059 | 0.41419 | 1.46154 | 0.93541 |
| **16.00000** | 10.22389 | 12.09816 | 3.54282 | 9.29806 | 2.25355 | 5.03824 | 0.00000 |
| **ARL17A.uid-210** | 0.01224 | 0.01093 | 0.11210 | 0.04894 | 0.20155 | 0.02832 | 0.06078 |
| **ARL17A.uid-219** | 0.09451 | 0.00000 | 0.08703 | 0.07771 | 0.12611 | 0.08636 | 0.03392 |
| **ARL17A.uid-219** | 0.14133 | 0.20016 | 0.08021 | 0.01215 | 0.06697 | 0.06501 | 0.06607 |
| **2.00000** | 12.59790 | 21.93031 | 3.80327 | 12.24729 | 0.15198 | 9.67393 | 7.31354 |
| **ARL2BP** | 9.23346 | 10.40514 | 6.52188 | 4.20679 | 1.36615 | 5.57898 | 0.00000 |
| **3.00000** | 5.53856 | 2.25738 | 4.99034 | 11.40835 | 2.99829 | 5.64860 | 0.27323 |
| **ARL4A** | 3.00745 | 2.07160 | 2.31534 | 0.13360 | 1.39920 | 2.91242 | 0.00000 |
| **ARL4C** | 0.00000 | 1.74639 | 0.62297 | 4.86251 | 0.82420 | 1.06528 | 3.58371 |

| | | | | | | | |
|---|---|---|---|---|---|---|---|
| **ARL4D** | 19.90109 | 36.13460 | 14.39550 | 7.31392 | 23.33612 | 8.12397 | 0.01304 |
| **ARL5A** | 5.29947 | 0.00000 | 3.96987 | 8.57867 | 1.34542 | 8.52721 | 25.84387 |
| **ARL5B** | 3.19230 | 0.00000 | 6.70392 | 6.22017 | 6.14050 | 1.20632 | 5.77303 |
| **ARL5C** | 0.00822 | 0.03899 | 0.01051 | 0.05727 | 0.01187 | 0.04165 | 0.03033 |
| **6.00000** | 0.82024 | 0.10979 | 0.90256 | 0.92673 | 0.06762 | 0.52823 | 0.30266 |
| **ARL6IP1** | 41.46281 | 23.69981 | 29.31534 | 30.93000 | 18.48413 | 17.94344 | 15.80189 |
| **ARL6IP4** | 6.97453 | 10.54301 | 1.43919 | 7.09428 | 0.12540 | 9.99400 | 12.87737 |
| **ARL6IP5** | 14.60426 | 1.63143 | 11.44090 | 4.84627 | 0.30188 | 1.11164 | 8.44157 |
| **ARL6IP6** | 12.18610 | 1.81285 | 10.77198 | 11.58001 | 2.70357 | 8.97163 | 0.00000 |
| **ARL8A** | 0.00000 | 1.31102 | 1.15621 | 0.26523 | 0.03600 | 1.93551 | 3.63417 |
| **ARL8B** | 20.36659 | 7.43350 | 28.67473 | 15.68122 | 10.27922 | 16.23410 | 9.85910 |
| **9.00000** | 0.03952 | 0.00000 | 0.02627 | 0.04062 | 0.75422 | 0.00066 | 0.18150 |
| **ARMC1** | 22.05434 | 1.28837 | 12.47568 | 5.49664 | 9.00297 | 9.70561 | 5.23465 |
| **ARMC10** | 0.62098 | 0.48797 | 0.89859 | 1.17695 | 0.04599 | 0.88952 | 3.33024 |
| **ARMC12** | 0.01313 | 0.03906 | 0.00551 | 0.00853 | 0.05034 | 0.16966 | 0.07775 |
| **ARMC2** | 0.46614 | 0.09282 | 0.96539 | 0.14158 | 6.27624 | 1.07964 | 0.00000 |
| **ARMC2-AS1** | 0.01669 | 0.01424 | 0.01410 | 0.02203 | 0.00770 | 0.04743 | 0.02501 |
| **ARMC3** | 0.07624 | 0.08301 | 0.01207 | 0.03261 | 0.01941 | 0.03180 | 0.00000 |
| **ARMC4** | 0.07694 | 0.00450 | 0.01278 | 0.00679 | 0.02747 | 0.07477 | 0.01214 |
| **ARMC5** | 0.10243 | 0.50806 | 0.68428 | 0.49649 | 0.50195 | 0.66291 | 0.89885 |
| **ARMC6** | 12.01587 | 9.60461 | 9.69065 | 10.11302 | 0.36928 | 6.08747 | 7.17008 |
| **ARMC7** | 11.07591 | 6.57733 | 5.60405 | 4.14071 | 1.06255 | 1.70723 | 3.95842 |
| **ARMC8** | 20.14540 | 5.69145 | 10.71898 | 9.22073 | 5.61622 | 8.67296 | 0.00000 |
| **ARMC9** | 2.66869 | 1.17645 | 0.82783 | 1.04144 | 2.87426 | 2.32502 | 0.00000 |
| **ARMCX1** | 0.52526 | 0.00000 | 0.07001 | 0.07332 | 0.01131 | 0.01639 | 0.00000 |
| **ARMCX2** | 0.05767 | 0.00000 | 0.23174 | 0.00229 | 0.00771 | 1.44674 | 0.02616 |
| **ARMCX3** | 0.85666 | 0.00000 | 0.06296 | 0.21087 | 0.05477 | 0.63382 | 0.00000 |
| **ARMCX4** | 0.93028 | 0.33374 | 0.12165 | 0.00000 | 0.01093 | 0.00000 | 0.57204 |
| **ARMCX5** | 0.88945 | 0.37510 | 1.10603 | 0.06264 | 0.00238 | 0.00000 | 0.00000 |
| **ARMCX5-GPRA** | 0.06721 | 0.06022 | 0.03540 | 0.00804 | 0.00728 | 0.02696 | 0.02187 |
| **ARMCX6** | 0.19284 | 0.02033 | 0.01444 | 0.01756 | 0.01490 | 0.05186 | 0.00000 |
| **ARMS2** | 0.03181 | 0.17037 | 0.00464 | 0.05147 | 0.01132 | 0.00451 | 0.00000 |
| **ARNT** | 8.39162 | 5.68971 | 3.68691 | 6.39847 | 0.78430 | 5.10603 | 0.00000 |
| **ARNT2** | 0.09497 | 0.00994 | 0.10904 | 0.01406 | 0.29567 | 0.00906 | 0.01244 |
| **ARNTL** | 1.27131 | 1.10784 | 0.17502 | 1.24308 | 1.39964 | 0.81606 | 0.00000 |

| | | | | | | | |
|---|---|---|---|---|---|---|---|
| **ARNTL2** | 0.79006 | 0.85470 | 0.55375 | 0.20183 | 0.09867 | 0.60095 | 2.54434 |
| **ARNTL2-AS1** | 0.02778 | 0.01597 | 0.00971 | 0.02122 | 0.05957 | 0.01538 | 0.02525 |
| **ARPC1A** | 28.01280 | 16.68233 | 8.32141 | 19.50977 | 0.49851 | 12.87915 | 11.07290 |
| **ARPC1B** | 45.02469 | 40.17614 | 10.92165 | 40.06574 | 1.64016 | 16.46917 | 7.70506 |
| **ARPC2** | 55.55204 | 40.42456 | 24.76328 | 41.81902 | 8.41360 | 27.40243 | 21.19300 |
| **ARPC3** | 22.64669 | 7.11478 | 15.42837 | 18.79402 | 0.19357 | 12.18154 | 9.28978 |
| **ARPC4** | 26.74654 | 32.98580 | 8.58686 | 27.48619 | 5.21494 | 17.05134 | 11.31327 |
| **ARPC5** | 43.57700 | 17.73013 | 13.93042 | 29.51897 | 0.45029 | 12.62622 | 3.12790 |
| **ARPC5L** | 5.90980 | 15.99348 | 9.68115 | 14.16761 | 6.55575 | 11.04873 | 34.55701 |
| **ARPIN** | 0.07358 | 0.13515 | 0.04953 | 0.07273 | 0.05832 | 0.09113 | 0.38749 |
| **ARPP19** | 22.32758 | 1.85578 | 17.33602 | 10.97604 | 8.12463 | 10.58082 | 21.88825 |
| **ARPP21** | 0.01818 | 0.02264 | 0.01082 | 0.02551 | 0.03286 | 0.01657 | 0.02197 |
| **ARR3** | 0.10815 | 0.03915 | 0.25149 | 0.00230 | 0.92426 | 0.00688 | 0.07532 |
| **ARRB1** | 2.22784 | 9.65250 | 4.02037 | 10.29253 | 1.08002 | 10.97584 | 5.88984 |
| **ARRB2** | 0.68505 | 0.40290 | 0.31599 | 4.34784 | 0.00263 | 1.35773 | 1.66565 |
| **ARRDC1** | 2.94111 | 3.23988 | 0.31587 | 7.59895 | 0.05058 | 1.72635 | 5.29806 |
| **ARRDC2** | 0.00000 | 3.10036 | 3.62817 | 0.00000 | 4.99398 | 4.69023 | 0.67671 |
| **ARRDC3** | 23.15280 | 5.93604 | 2.16319 | 3.74341 | 2.46501 | 2.39607 | 0.00000 |
| **ARRDC3-AS1** | 0.02331 | 0.01603 | 0.06497 | 0.08972 | 0.02703 | 0.15139 | 0.00578 |
| **ARRDC4** | 1.39190 | 2.54046 | 0.94694 | 1.70638 | 0.00000 | 0.81860 | 1.42998 |
| **ARRDC5** | 0.02191 | 0.03452 | 0.00868 | 0.01615 | 0.00844 | 0.05645 | 0.01689 |
| **ARSA** | 0.45021 | 0.60217 | 0.42394 | 1.74607 | 0.31539 | 0.78424 | 0.61338 |
| **ARSB** | 0.89443 | 0.16174 | 0.18365 | 2.08218 | 0.00954 | 0.62401 | 1.98583 |
| **ARSD** | 0.02699 | 0.12633 | 0.94653 | 0.27403 | 0.41811 | 0.15217 | 0.52320 |
| **ARSE** | 0.11639 | 0.00000 | 0.03814 | 0.01842 | 0.00485 | 3.08548 | 0.00000 |
| **ARSF** | 0.03797 | 0.01507 | 0.01351 | 0.01827 | 0.01173 | 0.01309 | 0.00906 |
| **ARSG** | 0.15132 | 0.30390 | 0.36136 | 0.01994 | 0.40933 | 0.16009 | 0.06245 |
| **ARSI** | 0.00000 | 0.31037 | 1.22159 | 0.00000 | 0.07254 | 0.30706 | 0.13186 |
| **ARSJ** | 0.02655 | 0.00510 | 0.03486 | 0.02896 | 0.10990 | 0.01301 | 0.01277 |
| **ARSK** | 5.30943 | 0.23081 | 0.74635 | 2.19199 | 1.75381 | 0.68998 | 1.08451 |
| **ART1** | 0.02151 | 0.00000 | 0.06955 | 0.00841 | 0.01076 | 0.00760 | 0.08597 |
| **ART3** | 0.01618 | 0.00000 | 0.16842 | 0.03941 | 0.85689 | 0.00740 | 0.03113 |
| **ART4** | 0.15050 | 0.00000 | 0.04550 | 0.29870 | 0.06324 | 0.03901 | 0.00000 |
| **ART5** | 0.00000 | 0.00000 | 0.47510 | 0.15517 | 0.84481 | 0.41550 | 0.09475 |
| **ARTN** | 0.02855 | 0.01151 | 0.01365 | 0.02209 | 0.00781 | 0.04389 | 0.02772 |

| | | | | | | | |
|---|---|---|---|---|---|---|---|
| **ARV1** | 5.84457 | 2.73985 | 2.56835 | 4.93886 | 0.22637 | 0.96249 | 0.00000 |
| **ARVCF** | 0.04520 | 0.20369 | 0.00574 | 0.04796 | 0.00612 | 0.00000 | 0.64231 |
| **ARX** | 0.03811 | 0.03778 | 0.22861 | 0.00000 | 0.10975 | 1.48515 | 0.16067 |
| **AS3MT** | 0.02226 | 0.01688 | 0.01599 | 0.02252 | 0.00896 | 0.01219 | 0.02145 |
| **ASAH1** | 38.68970 | 3.60786 | 0.97772 | 19.07757 | 1.59978 | 1.48286 | 2.12154 |
| **ASAH2** | 0.08104 | 0.00356 | 0.02089 | 0.02411 | 0.61314 | 0.03691 | 0.00000 |
| **ASAH2B** | 0.29412 | 0.26552 | 0.34452 | 0.15666 | 0.07785 | 0.01671 | 0.00000 |
| **ASAP1** | 7.74106 | 4.27137 | 2.44749 | 17.31902 | 1.61910 | 6.38985 | 30.81603 |
| **ASAP1-IT1** | 0.00000 | 0.07091 | 0.02683 | 0.18067 | 0.00832 | 0.11868 | 1.42789 |
| **ASAP1-IT2** | 0.17345 | 0.21091 | 0.15636 | 0.16375 | 0.05074 | 0.28169 | 1.25818 |
| **ASAP2** | 3.50162 | 0.87900 | 0.56820 | 0.00000 | 1.87525 | 4.53840 | 14.70532 |
| **ASAP3** | 0.00000 | 2.72254 | 2.35433 | 3.51762 | 1.76611 | 1.16097 | 0.08422 |
| **ASB1** | 0.00000 | 5.34894 | 3.02895 | 3.84782 | 0.79563 | 5.86074 | 1.58997 |
| **ASB11** | 0.02683 | 0.02179 | 0.01072 | 0.02178 | 0.01865 | 0.01164 | 0.02418 |
| **ASB12** | 0.06297 | 0.11311 | 0.10409 | 0.10841 | 0.02641 | 0.16314 | 0.00000 |
| **ASB13** | 0.40264 | 0.00000 | 0.00117 | 0.30930 | 0.00600 | 0.00000 | 0.41882 |
| **ASB14** | 0.04010 | 0.07883 | 0.06735 | 0.10841 | 0.02076 | 0.44858 | 0.83120 |
| **ASB15** | 0.48887 | 0.15468 | 0.05435 | 0.33467 | 0.09549 | 0.00030 | 0.00000 |
| **ASB16** | 0.02783 | 0.07050 | 0.12952 | 0.12570 | 0.02110 | 0.02264 | 0.00000 |
| **ASB16-AS1** | 0.91213 | 0.83419 | 1.36699 | 2.32711 | 0.25680 | 0.12741 | 0.07459 |
| **ASB18** | 0.01979 | 0.04145 | 0.01880 | 0.01904 | 0.00864 | 0.02131 | 0.02598 |
| **ASB2** | 0.00000 | 0.07668 | 0.05661 | 0.11863 | 0.57811 | 0.12380 | 0.08067 |
| **ASB3** | 1.63122 | 0.00000 | 1.61241 | 1.09842 | 0.52424 | 0.84096 | 1.33257 |
| **ASB4** | 0.05425 | 0.01704 | 0.01221 | 0.01726 | 0.00920 | 0.02288 | 0.00555 |
| **ASB5** | 0.01104 | 0.00893 | 0.01136 | 0.17793 | 0.02117 | 0.00557 | 0.09175 |
| **ASB6** | 4.79138 | 18.28078 | 4.41201 | 9.63300 | 3.73144 | 4.01917 | 7.90499 |
| **ASB7** | 3.25424 | 2.38255 | 2.28877 | 3.23643 | 5.82503 | 2.00121 | 0.82850 |
| **ASB8** | 5.94051 | 2.31104 | 3.46160 | 3.28685 | 2.88849 | 1.44719 | 3.46298 |
| **ASB9** | 0.10588 | 0.00154 | 0.01331 | 0.02499 | 0.04711 | 0.00842 | 0.02440 |
| **ASB9P1** | 0.02047 | 0.10721 | 0.00212 | 0.02624 | 0.16065 | 0.02155 | 0.06934 |
| **ASCC1** | 11.35488 | 0.76220 | 12.38407 | 6.99993 | 1.10353 | 4.26159 | 5.08021 |
| **ASCC2** | 15.87243 | 17.27519 | 6.28846 | 11.80847 | 3.82133 | 8.27202 | 0.31134 |
| **ASCC3** | 6.80302 | 3.83553 | 5.44106 | 8.67590 | 4.29051 | 7.42120 | 12.67217 |
| **ASCL1** | 0.02643 | 0.01542 | 0.02059 | 0.02298 | 0.00751 | 0.02831 | 0.01690 |
| **ASCL2** | 0.00520 | 0.09309 | 0.24201 | 0.00000 | 0.00313 | 0.00000 | 0.21365 |



1532

| | | | | | | | |
|---|---|---|---|---|---|---|---|
| **ASCL3** | 0.02675 | 0.02838 | 0.01918 | 0.02232 | 0.03495 | 0.01084 | 0.01136 |
| **ASF1A** | 15.81607 | 5.11631 | 16.97492 | 9.57922 | 4.56466 | 3.24446 | 0.33553 |
| **ASF1B** | 10.58100 | 8.15708 | 4.36169 | 16.96204 | 3.35204 | 2.23872 | 0.00000 |
| **ASGR1** | 0.18277 | 0.07535 | 0.00000 | 0.23670 | 0.25261 | 0.00000 | 0.43145 |
| **ASGR2** | 0.14045 | 0.85110 | 0.13872 | 0.00000 | 0.04512 | 0.00000 | 0.11622 |
| **ASH1L** | 6.97705 | 1.88800 | 6.00927 | 6.74564 | 3.00130 | 5.65535 | 19.15006 |
| **ASH1L-AS1** | 0.48410 | 1.41340 | 0.73643 | 1.34267 | 3.77565 | 0.23126 | 0.49000 |
| **ASH2L** | 10.73413 | 11.77547 | 16.46691 | 8.61999 | 0.73091 | 23.23826 | 0.00000 |
| **ASIC1** | 0.03742 | 0.10440 | 0.02966 | 0.24793 | 0.20175 | 0.00000 | 0.43700 |
| **ASIC2** | 0.03647 | 0.02412 | 0.06223 | 0.02245 | 0.03812 | 0.21156 | 0.07495 |
| **ASIC3** | 0.00000 | 1.15165 | 0.91724 | 1.24433 | 0.00059 | 0.13512 | 0.20052 |
| **ASIC4** | 0.01929 | 0.02930 | 0.01105 | 0.03951 | 0.00967 | 0.03514 | 0.02143 |
| **ASIC5** | 0.02506 | 0.02039 | 0.01372 | 0.02062 | 0.00788 | 0.01266 | 0.01664 |
| **ASIP** | 0.03834 | 0.02705 | 0.05156 | 0.00000 | 0.00469 | 0.01697 | 0.17990 |
| **ASL** | 4.12709 | 3.73841 | 3.29612 | 4.11101 | 0.85504 | 0.73067 | 1.31061 |
| **ASNA1** | 21.38936 | 27.33568 | 18.19485 | 24.36844 | 9.68076 | 8.49018 | 2.45416 |
| **ASNS** | 15.20651 | 0.00000 | 0.00000 | 10.15202 | 0.19620 | 3.70545 | 15.36045 |
| **ASNSD1** | 23.33041 | 9.36485 | 13.51987 | 18.59125 | 4.40531 | 8.06293 | 0.00000 |
| **ASPA** | 0.10815 | 0.29839 | 0.10291 | 0.00000 | 0.01288 | 0.63171 | 0.02458 |
| **ASPDH** | 0.01582 | 0.01852 | 0.01150 | 0.02513 | 0.00750 | 0.00926 | 0.05870 |
| **ASPG** | 0.04150 | 0.01302 | 0.02353 | 0.01120 | 0.00693 | 0.00975 | 0.03223 |
| **ASPH** | 9.76367 | 2.72269 | 4.51260 | 1.17334 | 2.50964 | 5.12246 | 0.77864 |
| **ASPHD1** | 0.00000 | 0.35473 | 0.43704 | 0.18183 | 0.02984 | 0.00909 | 0.17688 |
| **ASPHD2** | 0.01141 | 0.21542 | 0.30503 | 0.33766 | 2.02619 | 1.52894 | 0.65083 |
| **ASPM** | 10.39601 | 0.00000 | 13.51966 | 3.61939 | 3.08952 | 5.12892 | 25.22210 |
| **ASPN** | 0.02171 | 0.00000 | 0.06621 | 0.00000 | 0.00656 | 0.24278 | 1.19441 |
| **ASPRV1** | 0.29794 | 0.00000 | 0.09050 | 2.32424 | 0.02892 | 0.29459 | 0.01192 |
| **ASPSCR1** | 0.00002 | 0.93902 | 0.49821 | 1.89857 | 1.38728 | 1.11674 | 4.11358 |
| **ASRGL1** | 26.43647 | 45.40393 | 16.41549 | 7.12518 | 1.41904 | 36.79136 | 0.57153 |
| **ASS1** | 12.62428 | 0.00000 | 0.00000 | 11.08897 | 0.00574 | 0.25049 | 5.37756 |
| **ASTE1** | 4.23022 | 1.21329 | 2.39994 | 1.65239 | 1.68416 | 0.91627 | 0.00000 |
| **ASTL** | 0.02828 | 0.00404 | 0.42264 | 0.01109 | 2.46345 | 0.02010 | 0.06193 |
| **ASTN1** | 0.01643 | 0.01701 | 0.00831 | 0.03563 | 0.04282 | 0.01014 | 0.05710 |
| **ASTN2** | 4.09154 | 9.17625 | 0.90986 | 0.00000 | 1.72471 | 1.99095 | 0.00000 |
| **ASUN** | 15.45093 | 4.86644 | 7.96517 | 6.48452 | 0.55762 | 4.63809 | 1.53366 |

| | | | | | | | |
|---|---|---|---|---|---|---|---|
| **ASXL1** | 5.32169 | 5.88229 | 3.88698 | 3.89503 | 3.52588 | 1.56110 | 19.79877 |
| **ASXL2** | 2.96975 | 1.72157 | 6.09301 | 4.37511 | 2.67733 | 4.20370 | 16.23695 |
| **ASXL3** | 0.10307 | 0.00000 | 0.16188 | 0.36817 | 1.02681 | 0.03888 | 0.00000 |
| **ASZ1** | 0.01573 | 0.02188 | 0.44233 | 0.09292 | 0.08125 | 0.00415 | 0.02727 |
| **ATAD1** | 17.65821 | 0.20236 | 11.93511 | 10.24585 | 2.96494 | 5.78630 | 3.34611 |
| **ATAD2** | 15.36631 | 6.20367 | 11.07250 | 8.69868 | 10.33197 | 3.73779 | 8.90895 |
| **ATAD2B** | 6.93963 | 1.05267 | 2.42096 | 3.23908 | 2.20944 | 1.53540 | 8.14349 |
| **ATAD3A** | 4.16845 | 11.11174 | 0.44697 | 4.06528 | 0.10744 | 4.18898 | 16.94840 |
| **ATAD3B** | 11.25301 | 36.29199 | 13.41991 | 12.46053 | 0.27711 | 18.22165 | 37.90049 |
| **ATAD3C** | 0.00000 | 0.33277 | 0.00000 | 0.26061 | 3.72293 | 0.49153 | 0.83076 |
| **ATAD5** | 9.64193 | 3.26175 | 5.49821 | 3.21911 | 0.61428 | 5.24803 | 15.08488 |
| **ATAT1** | 3.89152 | 0.90738 | 1.03416 | 5.33725 | 1.72907 | 0.94484 | 1.87686 |
| **ATCAY** | 0.05546 | 0.00000 | 0.06269 | 0.01636 | 0.01815 | 0.17251 | 0.01441 |
| **ATE1** | 12.47986 | 3.32530 | 4.28073 | 5.76918 | 0.54766 | 5.74554 | 3.21843 |
| **ATE1-AS1** | 0.01880 | 0.04786 | 0.02441 | 0.04768 | 0.00878 | 0.02316 | 0.03672 |
| **ATF1** | 17.12225 | 0.00000 | 5.55277 | 14.60775 | 4.79666 | 6.75126 | 7.63606 |
| **ATF2** | 8.91762 | 0.36472 | 2.76892 | 3.41251 | 8.56720 | 3.39078 | 1.70441 |
| **ATF3** | 2.19842 | 5.30627 | 9.39434 | 9.43127 | 2.33441 | 1.92017 | 8.78615 |
| **ATF4** | 27.92351 | 13.14856 | 9.15004 | 29.17958 | 8.30921 | 19.00859 | 44.75625 |
| **ATF5** | 3.01257 | 14.12817 | 1.54321 | 1.50756 | 0.34798 | 1.46859 | 18.64485 |
| **ATF6** | 15.03875 | 6.72836 | 5.23312 | 9.39805 | 1.26712 | 5.73612 | 5.81869 |
| **ATF6B** | 2.36336 | 2.15670 | 2.11206 | 5.72003 | 0.65942 | 2.05475 | 1.65596 |
| **ATF7** | 5.76074 | 1.31259 | 1.23680 | 2.74328 | 2.42532 | 2.15922 | 3.03856 |
| **ATF7IP** | 22.96769 | 5.01450 | 16.57859 | 12.15986 | 7.22338 | 11.63208 | 12.44330 |
| **ATF7IP2** | 2.85564 | 0.00000 | 0.60195 | 4.74886 | 0.48995 | 0.08570 | 3.91600 |
| **ATG10** | 10.00485 | 1.02356 | 8.28817 | 1.40951 | 0.02793 | 5.11568 | 1.32655 |
| **ATG101** | 11.21470 | 20.78334 | 14.94356 | 17.81696 | 11.41777 | 9.95542 | 7.70260 |
| **ATG12** | 14.62014 | 2.22747 | 12.74043 | 14.65681 | 11.52265 | 10.02157 | 7.52882 |
| **ATG13** | 13.24220 | 9.38260 | 17.89957 | 17.60538 | 10.74019 | 10.02070 | 7.16129 |
| **ATG14** | 8.71155 | 11.12049 | 13.96568 | 4.26431 | 13.57007 | 1.59757 | 9.47434 |
| **ATG16L1** | 17.57329 | 16.44230 | 13.63104 | 10.61283 | 5.06044 | 7.79053 | 0.01779 |
| **ATG16L2** | 0.00000 | 0.12570 | 0.00903 | 0.53790 | 0.06915 | 0.24202 | 0.10879 |
| **ATG2A** | 0.00000 | 19.67068 | 22.47752 | 4.43458 | 0.98031 | 1.28227 | 8.65435 |
| **ATG2B** | 2.49016 | 0.09728 | 0.70777 | 3.42801 | 2.07582 | 1.85188 | 4.06278 |
| **ATG3** | 30.73648 | 36.00248 | 42.26913 | 14.90112 | 14.41753 | 37.73581 | 0.00000 |

| | | | | | | | |
|---|---|---|---|---|---|---|---|
| **ATG4A** | 9.23934 | 1.69190 | 4.58572 | 3.28630 | 3.75002 | 1.89771 | 0.00000 |
| **ATG4B** | 11.31500 | 13.98772 | 5.62096 | 17.50645 | 1.60994 | 8.16023 | 11.94132 |
| **ATG4C** | 7.88875 | 0.85162 | 4.15333 | 2.76265 | 0.16049 | 1.70098 | 0.00000 |
| **ATG4D** | 0.00000 | 2.33821 | 0.53242 | 1.02951 | 0.46972 | 1.89436 | 1.74630 |
| **ATG5** | 30.61587 | 9.80499 | 29.79749 | 19.09470 | 9.85129 | 24.53600 | 1.52863 |
| **ATG7** | 9.69561 | 7.00350 | 6.03322 | 5.66615 | 3.10177 | 4.44716 | 0.00000 |
| **ATG9A** | 3.95284 | 9.37025 | 20.86052 | 13.93578 | 5.98170 | 7.27202 | 15.65478 |
| **ATG9B** | 0.00000 | 16.07603 | 15.50716 | 4.59386 | 0.23485 | 8.66193 | 2.73533 |
| **ATHL1** | 0.11244 | 0.07728 | 0.06142 | 0.11954 | 0.00242 | 0.49585 | 0.39338 |
| **ATIC** | 37.15457 | 15.46234 | 17.08813 | 31.00429 | 0.95953 | 12.56364 | 16.48580 |
| **ATL1** | 0.24773 | 0.00000 | 0.15764 | 0.10876 | 0.20878 | 0.15758 | 0.16967 |
| **ATL2** | 17.70793 | 5.66109 | 12.37101 | 7.74962 | 7.27007 | 14.44071 | 0.00000 |
| **ATL3** | 10.22037 | 8.66881 | 8.47788 | 0.67835 | 2.71995 | 11.06137 | 0.61028 |
| **ATM** | 2.73436 | 0.00000 | 1.73024 | 1.02917 | 1.89999 | 1.93974 | 4.94289 |
| **ATMIN** | 5.73417 | 4.64197 | 6.21905 | 5.93443 | 0.77323 | 4.63439 | 21.67471 |
| **ATN1** | 0.00000 | 0.47971 | 0.78522 | 6.74306 | 0.29647 | 3.89098 | 21.33191 |
| **ATOH7** | 0.01607 | 0.00695 | 0.24086 | 0.01788 | 0.07358 | 0.00609 | 0.02263 |
| **ATOH8** | 0.00994 | 0.00345 | 0.02956 | 0.07024 | 0.01922 | 0.09052 | 0.05265 |
| **ATOX1** | 14.40598 | 13.58280 | 7.27847 | 16.73419 | 2.66375 | 10.54967 | 12.76188 |
| **ATP10A** | 0.00000 | 0.00000 | 0.09615 | 0.02112 | 1.11348 | 0.43171 | 0.04406 |
| **ATP10B** | 0.01050 | 0.01192 | 0.00000 | 0.78920 | 0.32668 | 0.00000 | 0.37885 |
| **ATP10D** | 0.00000 | 0.01297 | 0.00000 | 1.68530 | 3.20202 | 0.01027 | 0.00000 |
| **ATP11A** | 16.54709 | 16.12845 | 3.96314 | 3.76337 | 0.17705 | 9.28958 | 36.22421 |
| **ATP11A-AS1** | 0.15574 | 1.09924 | 0.35847 | 0.00000 | 0.00000 | 0.31781 | 3.34030 |
| **ATP11AUN** | 0.10020 | 1.20969 | 0.36685 | 0.02060 | 0.13035 | 0.27137 | 0.00000 |
| **ATP11B** | 11.44390 | 0.26811 | 7.01205 | 0.00000 | 1.34622 | 0.31252 | 13.33170 |
| **ATP11C** | 18.36125 | 0.09582 | 5.49778 | 2.42293 | 0.37770 | 14.24139 | 27.34866 |
| **ATP12A** | 0.29892 | 0.10605 | 0.03294 | 0.04458 | 0.00350 | 3.47668 | 0.14912 |
| **ATP13A1** | 1.92368 | 8.94438 | 4.91899 | 7.00788 | 0.73561 | 8.86003 | 19.42412 |
| **ATP13A2** | 6.94991 | 20.43539 | 2.61658 | 1.23757 | 0.00000 | 6.75778 | 23.57928 |
| **ATP13A3** | 13.87421 | 2.95699 | 8.00073 | 7.00807 | 2.36906 | 3.78716 | 24.85420 |
| **ATP13A5** | 0.02119 | 0.01787 | 0.00497 | 0.02050 | 0.12362 | 0.01447 | 0.01912 |
| **ATP1A1** | 22.17819 | 49.52662 | 16.13987 | 19.37581 | 4.87194 | 15.58711 | 40.00136 |
| **ATP1A1-AS1** | 0.52202 | 2.09823 | 0.35103 | 0.54695 | 0.04528 | 0.15992 | 1.23755 |
| **ATP1A2** | 0.01460 | 0.07463 | 0.04069 | 0.02118 | 0.01446 | 0.00732 | 0.05017 |

| | | | | | | | |
|---|---|---|---|---|---|---|---|
| **ATP1A3** | 0.23206 | 0.25947 | 0.00000 | 0.29633 | 0.02248 | 0.76661 | 0.07067 |
| **ATP1A4** | 0.03111 | 0.00879 | 0.02447 | 1.62648 | 0.00746 | 0.01381 | 0.00000 |
| **ATP1B1** | 16.75175 | 24.21446 | 4.66072 | 0.00000 | 3.47705 | 20.63075 | 12.69914 |
| **ATP1B2** | 0.02464 | 0.47370 | 3.96409 | 0.00000 | 1.95722 | 0.74452 | 0.00053 |
| **ATP1B3** | 41.90185 | 23.49605 | 24.05215 | 34.74251 | 4.34473 | 11.98092 | 20.23572 |
| **ATP2A1** | 0.10088 | 1.44175 | 0.26189 | 0.16369 | 0.32451 | 0.69324 | 0.57962 |
| **ATP2A2** | 4.80764 | 5.67965 | 8.44397 | 6.24658 | 5.41448 | 6.23328 | 28.40017 |
| **ATP2A3** | 0.00000 | 0.00000 | 0.69684 | 0.29976 | 0.78097 | 0.23344 | 0.22986 |
| **ATP2B1** | 48.20494 | 5.49781 | 10.06077 | 17.33710 | 5.59729 | 14.20749 | 41.84425 |
| **ATP2B2** | 0.02874 | 0.01791 | 0.00739 | 0.02591 | 0.02846 | 0.00516 | 0.16405 |
| **ATP2B3** | 0.06980 | 0.01001 | 0.06329 | 0.01105 | 0.05106 | 0.01085 | 0.02618 |
| **ATP2B4** | 5.00683 | 1.99733 | 0.00000 | 8.04826 | 2.54136 | 6.21265 | 10.39536 |
| **ATP2C1** | 9.70058 | 5.98278 | 6.84034 | 9.72088 | 1.45858 | 8.34484 | 8.97129 |
| **ATP2C2** | 4.73622 | 0.00000 | 0.00000 | 5.34816 | 1.21423 | 0.00000 | 5.15892 |
| **ATP4A** | 0.06051 | 0.00666 | 0.04726 | 0.04680 | 0.08242 | 0.06481 | 0.02402 |
| **ATP4B** | 0.02590 | 0.04054 | 0.01991 | 0.00204 | 0.00597 | 0.03779 | 0.17049 |
| **ATP5A1** | 86.07454 | 52.99160 | 20.22322 | 49.57729 | 9.02708 | 37.75397 | 15.94497 |
| **ATP5B** | 102.15997 | 59.59623 | 18.99723 | 69.89746 | 6.93090 | 49.13859 | 35.89407 |
| **ATP5C1** | 60.15974 | 26.84603 | 13.06939 | 33.74481 | 5.56987 | 28.02747 | 17.20112 |
| **ATP5D** | 22.44089 | 24.82878 | 3.32584 | 25.76006 | 1.82599 | 17.57813 | 40.21578 |
| **ATP5E** | 9.70990 | 15.26481 | 4.29939 | 9.46804 | 0.33170 | 5.78615 | 13.29374 |
| **ATP5EP2** | 18.97031 | 13.47914 | 8.25318 | 14.63877 | 0.24591 | 10.92984 | 21.83017 |
| **ATP5F1** | 51.50878 | 22.79890 | 18.84348 | 30.33824 | 1.36610 | 20.60972 | 9.93043 |
| **ATP5G1** | 27.83783 | 16.99624 | 3.26901 | 28.00385 | 0.69606 | 16.38490 | 9.57136 |
| **ATP5G2** | 21.55674 | 10.76168 | 10.77222 | 9.75041 | 1.64433 | 16.49489 | 23.93986 |
| **ATP5G3** | 75.78323 | 44.17626 | 23.53070 | 48.49670 | 8.50434 | 41.36299 | 49.79706 |
| **ATP5H** | 16.93146 | 9.79823 | 10.56769 | 14.10292 | 0.30464 | 13.06016 | 17.82028 |
| **ATP5I** | 41.79878 | 41.11059 | 14.49324 | 25.10323 | 1.18630 | 28.62067 | 42.10352 |
| **ATP5J** | 56.51428 | 29.33722 | 16.17861 | 32.01922 | 4.30873 | 23.89113 | 18.79482 |
| **ATP5J2** | 31.16004 | 28.91371 | 8.57985 | 28.40912 | 1.61093 | 18.94867 | 23.95686 |
| **ATP5L** | 44.36256 | 31.88516 | 18.32394 | 31.42427 | 4.55019 | 26.86287 | 30.98322 |
| **ATP5L2** | 0.04413 | 0.02441 | 0.05052 | 0.03949 | 0.00490 | 0.03173 | 0.63377 |
| **ATP5O** | 64.50179 | 36.36403 | 16.81812 | 35.32007 | 2.26933 | 31.29698 | 21.65087 |
| **ATP5S** | 3.50070 | 2.03659 | 2.79987 | 2.73009 | 0.62216 | 1.38456 | 0.00000 |
| **ATP5SL** | 18.52972 | 11.02160 | 8.42251 | 16.77534 | 1.10691 | 7.27313 | 1.18698 |

| | | | | | | | |
|---|---|---|---|---|---|---|---|
| **ATP6AP1** | 44.26707 | 53.21089 | 13.98010 | 22.72046 | 1.57915 | 14.62349 | 10.49145 |
| **ATP6AP1L** | 0.13893 | 0.21137 | 0.01871 | 0.02623 | 0.68955 | 0.10443 | 0.08638 |
| **ATP6AP2** | 29.85489 | 16.09865 | 17.10370 | 4.99512 | 3.36552 | 17.97166 | 4.56236 |
| **ATP6V0A1** | 2.98863 | 4.23386 | 2.89228 | 7.59271 | 0.96229 | 6.78079 | 5.19728 |
| **ATP6V0A2** | 1.63491 | 2.30303 | 1.07714 | 1.62132 | 0.28068 | 4.03878 | 4.85880 |
| **ATP6V0A4** | 33.69112 | 9.24655 | 0.00000 | 3.01320 | 0.18330 | 0.00000 | 10.66361 |
| **ATP6V0B** | 40.57218 | 43.45469 | 14.32614 | 38.83167 | 10.86493 | 20.96164 | 20.67729 |
| **ATP6V0C** | 1.26135 | 19.78218 | 6.06897 | 16.45745 | 1.15711 | 7.89191 | 50.47091 |
| **ATP6V0CP3** | 0.01936 | 0.02181 | 0.01143 | 0.02265 | 0.01067 | 0.01310 | 0.04353 |
| **ATP6V0D1** | 27.36226 | 18.52521 | 7.23952 | 26.08575 | 9.55647 | 11.31833 | 4.78877 |
| **ATP6V0D2** | 9.73825 | 17.69748 | 0.00000 | 0.00000 | 0.00000 | 0.00000 | 0.28955 |
| **ATP6V0E1** | 38.41048 | 23.20412 | 16.93453 | 31.84889 | 10.52478 | 17.91257 | 14.30146 |
| **ATP6V0E2** | 3.26855 | 0.57826 | 0.00000 | 1.44233 | 0.64020 | 0.53437 | 2.98614 |
| **ATP6V0E2-AS1** | 0.02466 | 0.01529 | 0.02291 | 0.04808 | 0.00741 | 0.02295 | 0.01023 |
| **ATP6V1A** | 36.65943 | 12.30358 | 10.65873 | 15.87485 | 0.40071 | 5.62896 | 11.19156 |
| **ATP6V1B1** | 27.84840 | 17.43127 | 0.00000 | 1.95535 | 0.00000 | 0.00000 | 5.99171 |
| **ATP6V1B2** | 26.80927 | 16.08623 | 7.84434 | 23.39817 | 7.47125 | 6.70500 | 0.00000 |
| **ATP6V1C1** | 25.82865 | 2.39101 | 13.88961 | 13.92110 | 3.26154 | 7.07105 | 3.28385 |
| **ATP6V1C2** | 7.21545 | 5.50439 | 0.00000 | 9.72691 | 0.09715 | 0.00000 | 22.74251 |
| **ATP6V1D** | 23.95470 | 13.58152 | 14.94925 | 18.66997 | 10.00924 | 11.06135 | 9.97598 |
| **ATP6V1E1** | 21.65992 | 19.63078 | 16.59590 | 16.21452 | 6.67689 | 11.68578 | 10.08836 |
| **ATP6V1E2** | 0.01323 | 0.04790 | 0.05655 | 0.02331 | 0.00502 | 0.03352 | 0.07566 |
| **ATP6V1F** | 28.10973 | 24.42669 | 10.78068 | 25.58342 | 2.99671 | 16.29195 | 11.49345 |
| **ATP6V1G1** | 44.99412 | 22.16866 | 15.77888 | 30.31571 | 7.67577 | 23.52988 | 15.02459 |
| **ATP6V1G2** | 0.02980 | 0.13185 | 0.02757 | 0.02642 | 0.33562 | 0.11676 | 0.00797 |
| **ATP6V1H** | 21.39452 | 8.60369 | 3.71268 | 9.50502 | 0.74268 | 6.41745 | 4.51884 |
| **ATP7A** | 0.66794 | 0.54968 | 1.71994 | 0.00000 | 0.68677 | 1.87219 | 2.17927 |
| **ATP7B** | 2.98788 | 2.02599 | 0.00000 | 9.92560 | 0.00243 | 0.81376 | 6.85299 |
| **ATP8A1** | 0.11502 | 0.01041 | 0.06913 | 0.02933 | 1.72372 | 0.42407 | 0.11868 |
| **ATP8A2** | 2.34041 | 0.21311 | 0.32094 | 0.55650 | 0.59170 | 0.78003 | 1.20219 |
| **ATP8B1** | 9.47877 | 9.09338 | 5.75048 | 0.00000 | 4.03051 | 0.00000 | 19.14718 |
| **ATP8B2** | 0.00000 | 0.47552 | 0.27639 | 0.69328 | 0.00209 | 7.84724 | 0.34154 |
| **ATP8B3** | 0.14102 | 0.00000 | 0.14583 | 0.30231 | 0.16693 | 0.50677 | 0.01182 |
| **ATP8B4** | 0.10268 | 0.00000 | 0.14856 | 0.05150 | 0.13023 | 0.04393 | 0.00000 |
| **ATP8B5P** | 0.99574 | 0.11648 | 0.44458 | 0.69057 | 0.48973 | 0.54096 | 0.09037 |

| | | | | | | | |
|---|---|---|---|---|---|---|---|
| **ATP9A** | 0.55951 | 2.85536 | 0.00000 | 0.31891 | 1.64748 | 0.41714 | 2.86132 |
| **ATP9B** | 1.60535 | 3.90359 | 3.61802 | 1.60649 | 1.18603 | 3.93362 | 1.95823 |
| **ATPAF1** | 1.22145 | 3.94799 | 1.86283 | 0.00000 | 0.16104 | 4.62554 | 10.26439 |
| **ATPAF2** | 11.95267 | 10.07414 | 0.95753 | 11.73413 | 2.38609 | 5.22883 | 0.00000 |
| **ATPIF1** | 20.91246 | 13.29491 | 16.56544 | 31.28942 | 18.21664 | 16.09833 | 14.00976 |
| **ATR** | 1.93062 | 0.62239 | 1.67808 | 0.34473 | 0.46096 | 1.14079 | 3.79659 |
| **ATRAID** | 12.88828 | 16.22709 | 5.89823 | 5.33580 | 0.86801 | 13.90026 | 6.95822 |
| **ATRIP** | 1.80141 | 6.49690 | 1.06327 | 0.97491 | 0.78564 | 1.58008 | 6.26652 |
| **ATRN** | 5.18130 | 1.42505 | 2.01391 | 3.32686 | 1.23901 | 3.40949 | 7.93450 |
| **ATRNL1** | 0.00027 | 0.00000 | 0.06702 | 0.01826 | 0.61954 | 0.11269 | 0.10195 |
| **ATRX** | 17.08248 | 0.00000 | 9.55112 | 5.09543 | 4.30414 | 12.72879 | 27.83866 |
| **ATXN1** | 0.00000 | 0.07850 | 0.15495 | 0.07320 | 0.81851 | 0.45519 | 0.30990 |
| **ATXN10** | 12.75089 | 11.78178 | 4.52855 | 8.06383 | 0.43657 | 10.58279 | 0.00000 |
| **ATXN1L** | 4.40075 | 2.50972 | 4.55534 | 5.19283 | 5.42022 | 3.23944 | 10.95299 |
| **ATXN2** | 2.50123 | 5.51856 | 8.33649 | 7.39025 | 1.96245 | 8.63138 | 17.94870 |
| **ATXN2L** | 2.61737 | 4.87811 | 6.24340 | 15.21006 | 1.78081 | 9.47983 | 36.49966 |
| **ATXN3** | 8.42002 | 3.07601 | 5.74527 | 5.61058 | 2.10261 | 7.28369 | 2.36746 |
| **ATXN3L** | 0.04647 | 0.01114 | 0.01295 | 0.01993 | 0.01646 | 0.00848 | 0.04603 |
| **ATXN7** | 2.25627 | 0.97779 | 1.71453 | 2.38499 | 0.85561 | 2.37972 | 9.52167 |
| **ATXN7L1** | 1.35862 | 0.00000 | 2.01295 | 1.04502 | 0.54179 | 1.61523 | 1.39905 |
| **ATXN7L2** | 0.84677 | 0.97705 | 1.27000 | 3.74921 | 1.28693 | 0.53600 | 1.89854 |
| **ATXN7L3** | 0.00000 | 1.14222 | 1.59140 | 3.96867 | 0.43440 | 5.43100 | 15.30649 |
| **ATXN7L3B** | 14.67388 | 7.47750 | 8.42914 | 12.39763 | 3.64107 | 2.14309 | 1.86108 |
| **ATXN8OS** | 0.02112 | 0.02054 | 0.02055 | 0.03884 | 0.00700 | 0.01131 | 0.04226 |
| **AUH** | 0.83182 | 1.28596 | 2.28842 | 1.66524 | 0.33529 | 3.90389 | 6.20782 |
| **AUNIP** | 11.39696 | 6.45710 | 8.92779 | 7.15131 | 4.26356 | 3.73599 | 3.42541 |
| **AUP1** | 33.93405 | 30.18522 | 7.53358 | 30.00597 | 3.03619 | 18.76361 | 13.70661 |
| **AURKA** | 37.55212 | 25.19198 | 22.00103 | 24.22433 | 8.94188 | 5.65760 | 5.94821 |
| **AURKAIP1** | 25.38431 | 46.08828 | 15.27906 | 32.41085 | 4.81663 | 19.75607 | 26.99343 |
| **AURKAPS1** | 2.60405 | 1.18714 | 1.97901 | 1.59082 | 1.09893 | 0.70198 | 0.43908 |
| **AURKB** | 34.75693 | 20.51820 | 15.95755 | 26.79484 | 8.04757 | 11.10997 | 9.40495 |
| **AURKC** | 0.54735 | 2.20666 | 7.97722 | 1.38058 | 11.16788 | 0.75949 | 0.07710 |
| **AUTS2** | 0.28808 | 0.90422 | 1.34541 | 0.15093 | 1.06897 | 1.99739 | 3.87687 |
| **AVEN** | 0.37453 | 0.23963 | 0.07496 | 1.07918 | 1.06443 | 0.32400 | 3.89012 |
| **AVIL** | 0.25335 | 0.50044 | 0.48609 | 0.02986 | 0.74956 | 0.18401 | 0.08683 |

| | | | | | | | |
|---|---|---|---|---|---|---|---|
| **AVL9** | 5.53679 | 0.78293 | 1.49782 | 5.76065 | 3.10197 | 5.03707 | 0.00000 |
| **AVP** | 0.01784 | 0.02660 | 0.01021 | 0.02928 | 0.01406 | 0.04129 | 0.01161 |
| **AVPI1** | 3.45540 | 6.48865 | 1.86717 | 14.84928 | 4.46813 | 1.29633 | 0.00000 |
| **AVPR1A** | 0.03650 | 0.02476 | 0.01125 | 0.04124 | 0.01399 | 0.09233 | 0.02606 |
| **AVPR2** | 0.02151 | 0.01596 | 0.02933 | 0.02304 | 0.04216 | 0.01216 | 0.01561 |
| **AWAT1** | 0.01963 | 0.02583 | 0.01093 | 0.02298 | 0.00805 | 0.02694 | 0.02333 |
| **AWAT2** | 0.02455 | 0.01081 | 0.03173 | 0.04567 | 0.04721 | 0.01645 | 0.00589 |
| **AXDND1** | 0.00000 | 0.09741 | 0.45007 | 0.00000 | 0.22169 | 0.44328 | 0.20290 |
| **AXIN1** | 1.60060 | 5.71880 | 1.42849 | 5.44539 | 1.52181 | 2.93987 | 14.07305 |
| **AXIN2** | 0.63575 | 0.81937 | 0.65838 | 5.90834 | 0.49971 | 7.15564 | 4.49669 |
| **AXL** | 0.01047 | 0.11910 | 0.37310 | 0.00000 | 0.07380 | 0.79798 | 3.82072 |
| **AZGP1** | 0.00983 | 0.02401 | 0.00775 | 0.08388 | 0.04971 | 0.01319 | 0.01266 |
| **AZI2** | 8.81858 | 0.42776 | 6.68125 | 8.69519 | 5.43663 | 5.77304 | 4.84191 |
| **AZIN1** | 46.35960 | 6.08974 | 25.88770 | 24.63950 | 13.11414 | 11.02504 | 6.28106 |
| **AZIN2** | 0.39898 | 0.00000 | 0.78731 | 0.59456 | 3.08177 | 0.92708 | 0.30908 |
| **AZU1** | 0.02737 | 0.01987 | 0.01177 | 0.01996 | 0.01136 | 0.01218 | 0.02233 |
| **B2M** | 0.45678 | 0.00003 | 0.71691 | 7.45496 | 2.02312 | 9.72877 | 0.00000 |
| **B3GALNT1** | 0.24790 | 0.20252 | 0.23641 | 0.38000 | 0.76888 | 0.07977 | 0.00000 |
| **B3GALNT2** | 12.34466 | 4.35229 | 2.62679 | 9.04554 | 3.11824 | 4.31096 | 3.41328 |
| **B3GALT1** | 0.03967 | 0.02400 | 0.01107 | 0.02530 | 0.13750 | 0.01387 | 0.00494 |
| **B3GALT2** | 0.00000 | 0.00000 | 0.32757 | 0.27536 | 0.04095 | 0.10651 | 0.60505 |
| **B3GALT4** | 0.18175 | 0.01444 | 0.04566 | 0.42280 | 0.07617 | 0.13607 | 0.20708 |
| **B3GALT6** | 0.00000 | 0.08749 | 0.20882 | 1.61656 | 0.05294 | 1.03767 | 4.18084 |
| **B3GALTL** | 1.04425 | 1.11093 | 0.45548 | 0.05020 | 0.27034 | 0.94707 | 1.49406 |
| **B3GAT1** | 0.01037 | 0.07553 | 0.04965 | 0.02669 | 0.03220 | 0.00937 | 0.02560 |
| **B3GAT2** | 0.01476 | 0.09623 | 0.04656 | 0.02020 | 0.22570 | 0.01634 | 0.00325 |
| **B3GAT3** | 1.14749 | 2.68156 | 1.07157 | 5.81531 | 0.49219 | 1.10073 | 2.15130 |
| **B3GNT1** | 7.88061 | 3.01681 | 0.61841 | 4.97195 | 0.32101 | 1.61691 | 2.42279 |
| **B3GNT2** | 18.45362 | 0.00000 | 0.44367 | 7.78648 | 20.89670 | 1.09677 | 19.00404 |
| **B3GNT3** | 0.01162 | 0.02415 | 0.00209 | 0.00382 | 0.12542 | 0.00457 | 0.19191 |
| **B3GNT4** | 0.36446 | 0.00000 | 0.20527 | 0.21702 | 1.34399 | 0.26249 | 0.75619 |
| **B3GNT5** | 0.47804 | 0.00000 | 0.00000 | 11.02411 | 0.33336 | 0.50377 | 1.14733 |
| **B3GNT6** | 0.01781 | 0.02231 | 0.01062 | 0.01999 | 0.01434 | 0.01800 | 0.08618 |
| **B3GNT7** | 12.64434 | 24.81811 | 0.00000 | 0.00000 | 1.35958 | 24.68101 | 0.00000 |
| **B3GNT8** | 0.09594 | 0.03776 | 0.02161 | 0.21886 | 0.00659 | 0.00000 | 0.14094 |

| | | | | | | | |
|---|---|---|---|---|---|---|---|
| **B3GNT9** | 0.00974 | 0.08868 | 0.01656 | 0.03851 | 0.15375 | 0.26933 | 0.00046 |
| **B3GNTL1** | 1.12453 | 1.13067 | 0.07424 | 0.74869 | 0.19945 | 0.24259 | 0.26243 |
| **B4GALNT1** | 0.00000 | 0.21230 | 0.48730 | 0.33307 | 0.09819 | 0.00107 | 0.05290 |
| **B4GALNT2** | 0.10825 | 0.00000 | 0.89368 | 0.03804 | 0.07033 | 0.53195 | 0.00000 |
| **B4GALNT3** | 0.90755 | 0.55910 | 0.08535 | 0.77730 | 0.00000 | 0.78191 | 1.09566 |
| **B4GALNT4** | 0.03932 | 0.00418 | 0.01740 | 0.01721 | 0.00826 | 0.01455 | 0.08054 |
| **B4GALT1** | 34.71896 | 54.48692 | 1.16930 | 32.97211 | 11.20438 | 7.70960 | 21.33589 |
| **B4GALT1-AS1** | 0.39792 | 0.29681 | 0.14222 | 0.06952 | 0.02869 | 0.06627 | 0.16084 |
| **B4GALT2** | 0.08638 | 0.95406 | 0.08677 | 2.45013 | 0.16103 | 0.84319 | 3.77338 |
| **B4GALT3** | 9.99246 | 4.69099 | 7.98035 | 17.89785 | 4.16205 | 4.07046 | 14.08237 |
| **B4GALT4** | 7.55959 | 5.80916 | 3.14746 | 6.54591 | 2.26162 | 11.02828 | 0.00000 |
| **B4GALT4-AS1** | 0.02369 | 0.03140 | 0.06279 | 0.04223 | 0.00702 | 0.02513 | 0.06415 |
| **B4GALT5** | 9.54757 | 13.30786 | 6.01388 | 3.93191 | 0.68092 | 4.67009 | 32.11629 |
| **B4GALT6** | 1.55499 | 0.00000 | 0.05942 | 0.73802 | 0.58926 | 0.27354 | 1.09314 |
| **B4GALT7** | 2.39754 | 6.69887 | 1.45722 | 1.02576 | 0.00000 | 3.06443 | 1.51385 |
| **B9D1** | 1.87715 | 2.35089 | 0.33673 | 1.12212 | 1.17267 | 2.38808 | 0.00000 |
| **B9D2** | 1.57156 | 2.31339 | 3.02718 | 3.21014 | 4.20759 | 1.60424 | 0.00000 |
| **BAALC** | 0.01197 | 0.02730 | 0.03124 | 0.03729 | 1.30858 | 0.02033 | 0.02881 |
| **BAAT** | 0.20356 | 0.00000 | 0.18620 | 0.17310 | 0.13828 | 0.01278 | 0.00000 |
| **BABAM1** | 25.58559 | 15.70825 | 6.13961 | 31.28547 | 4.00064 | 11.25207 | 2.61223 |
| **BACE1** | 0.21668 | 2.22447 | 3.16298 | 0.12419 | 0.30955 | 4.57887 | 0.65702 |
| **BACE1-AS** | 0.10661 | 0.09978 | 0.20045 | 0.00000 | 0.01004 | 0.00838 | 0.22754 |
| **BACE2** | 0.01592 | 0.11133 | 0.01170 | 0.10548 | 0.10544 | 0.39558 | 0.01434 |
| **BACH1** | 7.34408 | 0.86505 | 2.30197 | 4.19284 | 4.31185 | 2.72499 | 1.84081 |
| **BACH1-IT2** | 0.04824 | 0.00912 | 0.01289 | 0.03937 | 0.01519 | 0.01024 | 0.03746 |
| **BACH2** | 0.03287 | 0.02930 | 0.09718 | 0.00537 | 0.55320 | 0.53235 | 0.43823 |
| **BAD** | 1.64815 | 4.28220 | 2.68828 | 7.54687 | 1.44101 | 5.47979 | 2.23135 |
| **BAG1** | 4.02107 | 10.46157 | 1.51884 | 6.44486 | 1.31672 | 5.31273 | 18.05721 |
| **BAG2** | 1.09683 | 0.00000 | 0.84720 | 3.44902 | 0.32096 | 0.38299 | 0.36341 |
| **BAG3** | 2.15894 | 2.84460 | 0.86771 | 14.11283 | 0.36015 | 4.68658 | 10.23241 |
| **BAG4** | 15.01109 | 0.00000 | 18.78765 | 5.78950 | 14.41591 | 9.75478 | 1.26631 |
| **BAG5** | 13.31365 | 7.21960 | 12.07198 | 5.82985 | 5.54790 | 5.85042 | 10.37488 |
| **BAG6** | 2.04018 | 13.90942 | 0.80628 | 10.58233 | 0.32690 | 9.73069 | 28.17794 |
| **BAGE** | 0.04573 | 0.00000 | 0.37736 | 0.05253 | 0.11591 | 0.01755 | 0.01696 |
| **BAHCC1** | 0.02819 | 0.02373 | 0.04450 | 0.02313 | 0.07924 | 0.07851 | 0.13288 |

| | | | | | | | |
|---|---|---|---|---|---|---|---|
| **BAHD1** | 1.35670 | 1.56068 | 0.24552 | 3.17278 | 0.43033 | 2.43689 | 14.14989 |
| **BAI1** | 0.01120 | 0.04073 | 0.04524 | 0.02250 | 0.00532 | 0.02349 | 0.03829 |
| **BAI2** | 0.00714 | 0.03530 | 0.00129 | 0.02331 | 0.15750 | 0.07007 | 0.03450 |
| **BAI3** | 0.02644 | 0.00000 | 0.07486 | 0.02077 | 0.08156 | 0.01242 | 0.01841 |
| **BAIAP2** | 11.89244 | 3.23260 | 4.66118 | 17.24436 | 8.57612 | 0.00000 | 19.68956 |
| **BAIAP2-AS1** | 0.00000 | 0.05248 | 0.48880 | 0.20211 | 0.26167 | 0.12970 | 0.62656 |
| **BAIAP2L1** | 8.83081 | 16.50711 | 4.48177 | 12.11057 | 5.54697 | 0.00000 | 33.66534 |
| **BAIAP2L2** | 0.00000 | 4.70174 | 10.11716 | 0.00000 | 1.55174 | 0.07583 | 0.00000 |
| **BAIAP3** | 0.00000 | 0.61825 | 0.26136 | 0.18453 | 0.24951 | 0.09313 | 0.59634 |
| **BAK1** | 1.83542 | 13.23727 | 23.70074 | 3.55498 | 0.00000 | 12.51591 | 2.03397 |
| **BAMBI** | 0.00000 | 0.00000 | 8.11476 | 1.00843 | 1.15454 | 21.89710 | 2.01542 |
| **BANCR** | 0.02375 | 0.04000 | 0.01368 | 0.14614 | 0.00997 | 0.49196 | 0.00180 |
| **BANF1** | 23.08623 | 25.85021 | 16.42464 | 19.91105 | 7.45680 | 16.21038 | 9.66839 |
| **BANF2** | 0.03180 | 0.02827 | 0.03030 | 0.02506 | 0.00733 | 0.00979 | 0.00700 |
| **BANK1** | 0.06531 | 0.00279 | 0.02012 | 0.01419 | 0.29411 | 0.01630 | 0.00909 |
| **BANP** | 3.74091 | 9.40044 | 5.27534 | 5.89633 | 1.03637 | 1.89762 | 0.86487 |
| **BAP1** | 5.14762 | 9.53313 | 5.07417 | 9.55865 | 2.36226 | 9.08496 | 15.27387 |
| **BARD1** | 16.04026 | 18.46035 | 13.14940 | 9.31716 | 4.69728 | 17.20154 | 0.01018 |
| **BARHL2** | 0.01858 | 0.01205 | 0.01742 | 0.01730 | 0.11103 | 0.04411 | 0.02815 |
| **BARX1** | 0.00000 | 0.57259 | 0.79581 | 0.00000 | 0.32350 | 0.05778 | 0.00596 |
| **BARX2** | 0.05155 | 0.02735 | 0.08925 | 0.00447 | 1.00756 | 0.12475 | 0.07802 |
| **BASP1** | 22.71953 | 29.49454 | 10.84089 | 14.53040 | 5.72255 | 0.00000 | 62.94132 |
| **BASP1P1** | 0.02886 | 0.02773 | 0.03419 | 0.02186 | 0.00590 | 0.02048 | 0.02432 |
| **BATF** | 0.73649 | 0.13095 | 0.06097 | 3.23602 | 0.02687 | 0.00000 | 0.05020 |
| **BATF2** | 0.00000 | 0.00550 | 0.50482 | 0.05446 | 0.61992 | 0.01203 | 0.03541 |
| **BATF3** | 2.37810 | 9.89085 | 11.32117 | 0.00000 | 0.16749 | 0.85782 | 0.99886 |
| **BAX** | 0.00000 | 9.72878 | 1.56780 | 4.36988 | 0.04395 | 5.79168 | 8.71297 |
| **BAZ1A** | 9.37506 | 3.70408 | 8.54816 | 5.84371 | 11.69496 | 2.72485 | 24.53119 |
| **BAZ1B** | 13.54603 | 6.26438 | 10.28350 | 6.44276 | 6.11755 | 8.75048 | 38.06979 |
| **BAZ2A** | 4.16021 | 4.91196 | 12.37068 | 9.62590 | 8.20692 | 9.18923 | 30.10745 |
| **BAZ2B.uid-2643** | 0.16166 | 0.06630 | 0.02689 | 0.39588 | 0.19116 | 0.14052 | 0.47613 |
| **BAZ2B.uid-2983** | 0.68795 | 0.70587 | 0.92645 | 0.00000 | 0.56114 | 0.34942 | 0.53282 |
| **BBC3** | 0.00000 | 2.40588 | 0.15373 | 0.35948 | 0.09087 | 4.51533 | 1.91771 |
| **BBIP1** | 1.52192 | 0.17563 | 3.82302 | 2.36156 | 3.42035 | 1.21565 | 1.34414 |
| **BBOX1** | 0.05955 | 0.03984 | 0.02579 | 0.13936 | 0.00847 | 0.03955 | 0.08040 |

| | | | | | | | |
|---|---|---|---|---|---|---|---|
| **BBS1** | 1.77667 | 0.58202 | 1.56941 | 1.12009 | 0.66768 | 1.29970 | 0.20122 |
| **BBS10** | 1.04815 | 0.00000 | 1.23120 | 0.81183 | 0.30882 | 0.00515 | 0.00000 |
| **BBS12** | 0.61925 | 0.11856 | 0.57972 | 0.98400 | 0.03602 | 0.58647 | 0.00000 |
| **BBS2** | 3.25886 | 1.26937 | 1.69179 | 2.27038 | 0.04277 | 2.12838 | 0.00000 |
| **BBS4** | 3.45336 | 2.77838 | 4.01947 | 1.38891 | 2.04281 | 1.33670 | 0.91928 |
| **BBS5** | 0.32899 | 0.02537 | 0.73657 | 0.40609 | 1.62479 | 0.00000 | 0.29368 |
| **BBS7** | 2.03754 | 2.39363 | 3.21676 | 2.19462 | 2.40231 | 2.22637 | 0.16941 |
| **BBS9** | 1.83071 | 2.86606 | 0.00000 | 4.61508 | 5.56138 | 2.42159 | 0.00000 |
| **BBX** | 5.57842 | 1.02791 | 5.73927 | 5.10734 | 4.01957 | 5.21270 | 4.48701 |
| **BCAM** | 0.00000 | 0.16284 | 0.04216 | 4.01940 | 0.00215 | 0.00000 | 1.66083 |
| **BCAN** | 0.00805 | 0.01746 | 0.08912 | 0.04664 | 0.05776 | 0.07365 | 0.02033 |
| **BCAP29** | 11.76622 | 0.52587 | 8.77476 | 2.53790 | 0.50231 | 2.44573 | 4.78538 |
| **BCAP31** | 29.96065 | 41.10507 | 21.28064 | 25.57787 | 9.16611 | 20.15032 | 14.15151 |
| **BCAR1** | 0.66846 | 5.12200 | 0.10610 | 7.41149 | 0.06315 | 2.63310 | 12.98796 |
| **BCAR3** | 2.35649 | 0.00000 | 0.10738 | 4.09072 | 0.00000 | 1.45081 | 3.93580 |
| **BCAR4** | 0.80285 | 0.00000 | 0.04782 | 11.73355 | 9.84587 | 0.00000 | 0.08736 |
| **BCAS1** | 0.07865 | 0.00430 | 0.01169 | 0.23885 | 0.79630 | 0.05759 | 0.02018 |
| **BCAS2** | 28.34810 | 9.73368 | 12.56351 | 21.66600 | 2.24918 | 9.20522 | 2.17667 |
| **BCAS3** | 1.02521 | 1.48680 | 2.28902 | 1.88610 | 1.76502 | 1.06059 | 0.52551 |
| **BCAS4** | 0.00000 | 0.87028 | 1.01322 | 0.20913 | 0.09542 | 0.64612 | 0.09192 |
| **BCAT1** | 49.59607 | 8.35900 | 30.70533 | 29.72126 | 3.04889 | 25.17367 | 30.54057 |
| **BCAT2** | 9.59113 | 6.28032 | 1.64964 | 10.30014 | 1.85399 | 0.00000 | 3.92071 |
| **BCCIP** | 38.16235 | 17.19735 | 20.99219 | 28.63527 | 7.70698 | 13.14219 | 9.38044 |
| **BCDIN3D** | 1.46451 | 0.44250 | 2.04522 | 1.13732 | 0.18912 | 0.30630 | 0.16045 |
| **BCDIN3D-AS1** | 0.12224 | 0.08807 | 0.14239 | 0.09409 | 0.10672 | 0.06186 | 0.33271 |
| **BCKDHA** | 9.76838 | 7.81069 | 6.48546 | 13.31650 | 0.31079 | 5.40517 | 0.00000 |
| **BCKDHB** | 1.33566 | 0.77711 | 0.07206 | 1.08182 | 0.46382 | 0.78297 | 0.00000 |
| **BCKDK** | 8.58892 | 6.19154 | 3.39461 | 14.79644 | 0.19745 | 11.72678 | 0.66051 |
| **BCL10** | 16.58797 | 7.84864 | 5.03020 | 8.45102 | 10.06904 | 6.48645 | 0.23368 |
| **BCL11A** | 0.22181 | 7.53694 | 8.24661 | 0.00000 | 0.47033 | 2.86096 | 0.08624 |
| **BCL11B** | 0.01486 | 0.01559 | 0.02710 | 0.02450 | 0.34834 | 0.64099 | 0.04699 |
| **BCL2** | 0.00000 | 0.10513 | 0.00627 | 0.04423 | 0.00770 | 0.07007 | 0.10669 |
| **BCL2A1** | 0.04435 | 0.00740 | 0.01558 | 0.00000 | 0.04865 | 0.01963 | 0.09079 |
| **BCL2L1** | 4.31055 | 12.70762 | 1.71700 | 14.98101 | 1.90080 | 5.44027 | 12.43874 |
| **BCL2L10** | 0.00000 | 0.09661 | 0.09395 | 0.01082 | 5.57114 | 0.01360 | 0.02122 |

| | | | | | | | |
|---|---|---|---|---|---|---|---|
| **BCL2L11** | 6.34573 | 0.61001 | 2.71761 | 6.09693 | 6.44973 | 1.54475 | 6.18383 |
| **BCL2L12** | 8.48231 | 17.19798 | 4.15807 | 21.26155 | 3.10295 | 5.67293 | 16.97464 |
| **BCL2L13** | 8.24795 | 11.59431 | 10.96157 | 11.59072 | 4.81030 | 4.94814 | 0.00000 |
| **BCL2L14** | 0.09201 | 0.07004 | 0.14349 | 0.00821 | 0.10400 | 0.15242 | 0.00000 |
| **BCL2L15** | 0.08917 | 0.03126 | 0.04998 | 0.05501 | 0.15381 | 0.65446 | 0.00000 |
| **BCL2L2** | 0.92236 | 0.00000 | 0.00096 | 1.69791 | 0.07134 | 0.00000 | 1.40134 |
| **BCL3** | 0.00000 | 0.96164 | 0.02876 | 0.27424 | 0.11199 | 6.13312 | 0.00842 |
| **BCL6** | 0.11393 | 0.20657 | 0.53353 | 2.32649 | 0.01804 | 0.57230 | 0.56928 |
| **BCL6B** | 0.39557 | 0.00000 | 0.07461 | 0.00000 | 0.24038 | 0.00014 | 0.07992 |
| **BCL7A** | 0.98687 | 3.23506 | 0.00000 | 1.67090 | 0.30011 | 3.20569 | 5.95618 |
| **BCL7B** | 3.25544 | 7.60218 | 2.78473 | 5.75212 | 0.52672 | 4.93760 | 8.62954 |
| **BCL7C** | 0.03466 | 1.73048 | 0.14470 | 0.27632 | 0.00284 | 0.28275 | 3.42826 |
| **BCL9** | 4.35684 | 0.00000 | 0.76998 | 1.96512 | 1.91657 | 4.64257 | 7.93762 |
| **BCL9L** | 3.20415 | 4.80240 | 0.12810 | 8.51495 | 0.10871 | 1.56309 | 23.71431 |
| **BCLAF1** | 37.84493 | 9.35450 | 17.75833 | 13.48302 | 7.91353 | 10.88477 | 25.47351 |
| **BCMO1** | 0.02084 | 0.35467 | 0.00090 | 0.05561 | 0.24567 | 0.52605 | 0.00000 |
| **BCO2** | 0.17649 | 0.01631 | 0.07717 | 0.35178 | 0.22399 | 0.16334 | 0.25768 |
| **BCOR** | 4.12804 | 6.07079 | 6.19221 | 0.00000 | 4.73193 | 18.88059 | 6.77197 |
| **BCORL1** | 2.06734 | 5.03237 | 2.90116 | 1.11355 | 2.34330 | 0.00000 | 9.72715 |
| **BCR** | 4.44546 | 7.97391 | 1.24942 | 8.07783 | 0.56325 | 2.02578 | 15.06616 |
| **BCRP3** | 0.02908 | 0.00484 | 0.02084 | 0.03401 | 0.00736 | 1.06411 | 0.00180 |
| **BCS1L** | 17.25771 | 16.62968 | 7.69543 | 2.92265 | 5.08745 | 4.69609 | 6.16164 |
| **BCYRN1** | 0.02501 | 0.02090 | 0.01293 | 0.01846 | 0.00816 | 0.02007 | 0.02271 |
| **BDH1** | 0.29156 | 0.68726 | 0.00000 | 0.32674 | 1.14109 | 0.21224 | 0.07530 |
| **BDH2** | 0.95643 | 0.14279 | 0.03021 | 0.35633 | 0.70218 | 0.41369 | 0.26761 |
| **BDKRB1** | 0.02853 | 0.01456 | 0.02247 | 0.08169 | 0.06548 | 0.01417 | 0.00417 |
| **BDKRB2** | 0.03273 | 0.01371 | 0.01288 | 0.01577 | 0.00805 | 0.05910 | 0.02074 |
| **BDNF** | 0.16013 | 0.19214 | 0.50219 | 0.19432 | 0.32513 | 0.73923 | 0.00000 |
| **BDNF-AS** | 0.12267 | 0.25981 | 0.28687 | 0.03426 | 0.87289 | 0.09430 | 0.00000 |
| **BDP1** | 12.01592 | 5.28759 | 10.30379 | 7.39588 | 6.60857 | 12.88922 | 29.63633 |
| **BEAN1** | 0.00000 | 0.04716 | 0.03165 | 0.19859 | 0.37850 | 0.06211 | 0.01427 |
| **BECN1** | 18.95130 | 11.16746 | 6.10355 | 10.51118 | 3.80533 | 3.40617 | 0.00000 |
| **BECN1P1** | 0.00000 | 0.28999 | 17.42837 | 0.00000 | 14.74417 | 0.00000 | 0.00000 |
| **BEGAIN** | 0.01674 | 0.01655 | 0.02045 | 0.03054 | 0.03850 | 0.01457 | 0.03096 |
| **BEND2** | 0.01995 | 0.00000 | 0.05753 | 0.06391 | 0.21003 | 0.00823 | 0.10657 |

| | | | | | | | |
|---|---|---|---|---|---|---|---|
| **BEND3** | 4.49554 | 4.65915 | 6.88943 | 2.50737 | 1.64796 | 8.07406 | 12.66563 |
| **BEND3P3** | 0.27350 | 0.00000 | 0.09974 | 0.24671 | 0.05873 | 0.02147 | 0.16710 |
| **BEND4** | 0.00000 | 0.22686 | 0.00000 | 0.05593 | 0.25006 | 3.52886 | 0.06878 |
| **BEND5** | 0.02868 | 0.02622 | 0.02247 | 0.03455 | 0.01288 | 0.03973 | 0.00336 |
| **BEND6** | 0.07056 | 0.00000 | 0.08318 | 0.04687 | 0.18122 | 0.01472 | 0.00000 |
| **BEND7** | 0.00000 | 0.12331 | 0.07341 | 0.10088 | 0.13404 | 0.15823 | 1.03010 |
| **BEST1** | 0.00000 | 0.01322 | 0.01312 | 0.14387 | 0.00740 | 0.03261 | 0.07547 |
| **BEST2** | 0.02435 | 0.01961 | 0.05447 | 0.02923 | 0.03041 | 0.00853 | 0.05338 |
| **BEST3** | 0.04481 | 0.03771 | 0.01522 | 0.01403 | 0.05450 | 0.01091 | 0.00524 |
| **BEST4** | 0.00000 | 0.15172 | 0.03621 | 0.10656 | 0.01007 | 0.00819 | 0.07094 |
| **BET1** | 9.20085 | 1.81570 | 8.11276 | 8.72425 | 1.66631 | 3.75051 | 0.58426 |
| **BET1L** | 8.72500 | 10.51938 | 7.89864 | 7.35872 | 2.05451 | 4.93228 | 6.13039 |
| **BEX1** | 0.01829 | 0.01489 | 0.15124 | 0.16069 | 0.03894 | 0.18730 | 0.02193 |
| **BEX2** | 1.05466 | 0.03569 | 2.16473 | 7.45636 | 0.99572 | 0.02702 | 2.24824 |
| **BEX4** | 14.07005 | 0.00000 | 7.29568 | 8.88855 | 0.77931 | 1.40564 | 6.04991 |
| **BEX5** | 0.05023 | 0.06375 | 0.04489 | 0.02188 | 0.03769 | 0.01407 | 0.00588 |
| **BFAR** | 12.03169 | 7.22044 | 15.02097 | 8.98689 | 2.49100 | 9.23885 | 8.19434 |
| **BFSP1** | 0.03801 | 0.16804 | 0.34804 | 0.04409 | 0.08094 | 0.67081 | 0.05947 |
| **BFSP2** | 0.03261 | 0.00003 | 0.38467 | 0.02644 | 0.04829 | 0.20190 | 0.04444 |
| **BGLAP** | 0.03184 | 0.04448 | 0.00980 | 0.02204 | 0.00807 | 0.02228 | 0.01603 |
| **BGN** | 0.02171 | 0.01204 | 0.02716 | 0.09025 | 0.05106 | 0.01366 | 0.04161 |
| **BHLHA15** | 0.01650 | 0.01564 | 0.01257 | 0.02539 | 0.00754 | 0.00992 | 0.04633 |
| **BHLHB9** | 0.19635 | 0.06109 | 0.00000 | 0.05809 | 2.39064 | 0.08611 | 0.08623 |
| **BHLHE22** | 0.00000 | 0.32179 | 0.41086 | 0.00000 | 0.21005 | 0.46892 | 0.05196 |
| **BHLHE40** | 0.00000 | 1.18981 | 0.00000 | 14.58093 | 2.05566 | 8.12536 | 2.35953 |
| **BHLHE40-AS1** | 0.05378 | 0.01258 | 0.01165 | 0.06481 | 0.00746 | 0.03280 | 0.09426 |
| **BHLHE41** | 0.01808 | 0.01526 | 0.01554 | 0.04458 | 0.01872 | 0.01677 | 0.02255 |
| **BHMT** | 0.00000 | 3.34899 | 1.78303 | 0.00000 | 0.37798 | 2.56261 | 0.00000 |
| **BHMT2** | 0.01491 | 0.00711 | 0.02518 | 1.36454 | 0.02884 | 0.10527 | 0.00000 |
| **BICC1** | 0.01802 | 0.06230 | 0.03003 | 0.01453 | 0.74558 | 0.00491 | 0.02539 |
| **BICD1** | 0.75138 | 4.57189 | 0.62299 | 2.20522 | 0.29426 | 8.24707 | 2.23411 |
| **BICD2** | 3.40224 | 5.44122 | 4.05669 | 5.33679 | 1.45372 | 5.83225 | 18.78526 |
| **BID** | 0.63373 | 0.19238 | 0.05102 | 0.49494 | 0.07226 | 0.29924 | 0.36761 |
| **BIK** | 6.49990 | 1.60950 | 43.49124 | 9.64246 | 17.89067 | 0.00000 | 0.00000 |
| **BIN1** | 8.96888 | 14.38695 | 0.00000 | 7.58443 | 0.00000 | 5.36072 | 20.41844 |

| | | | | | | | |
|---|---|---|---|---|---|---|---|
| **BIN2** | 4.38128 | 0.00000 | 0.42272 | 8.39013 | 0.19205 | 0.13491 | 5.59419 |
| **BIN3** | 6.82763 | 4.98017 | 3.28005 | 9.19440 | 2.36563 | 2.07827 | 0.72540 |
| **BIN3-IT1** | 0.00000 | 0.08280 | 0.08628 | 0.31622 | 0.03598 | 0.04785 | 0.77956 |
| **BIRC2** | 9.93153 | 0.00000 | 9.04411 | 6.31586 | 6.60365 | 3.68947 | 5.40335 |
| **BIRC3** | 0.01133 | 0.00000 | 0.10180 | 0.00000 | 0.28045 | 0.11156 | 0.67607 |
| **BIRC5** | 32.25296 | 35.44683 | 14.36242 | 26.88168 | 10.10021 | 16.29835 | 3.39360 |
| **BIRC6** | 13.82076 | 5.67458 | 7.71085 | 9.84178 | 4.60575 | 17.25045 | 29.18784 |
| **BIRC7** | 0.04529 | 0.00219 | 0.01615 | 0.03358 | 0.00719 | 0.01632 | 0.06144 |
| **BIRC8** | 0.01443 | 0.07113 | 0.02768 | 0.01934 | 0.01146 | 0.00862 | 0.01447 |
| **BIVM** | 0.39289 | 0.04214 | 0.15297 | 0.39283 | 0.02111 | 0.24428 | 0.36371 |
| **BLACAT1** | 0.05472 | 0.01232 | 0.00885 | 0.08822 | 0.00750 | 0.00152 | 0.17998 |
| **BLACE** | 0.00000 | 0.67233 | 1.24522 | 0.00000 | 0.00033 | 0.57701 | 0.00912 |
| **BLCAP** | 11.94673 | 11.79365 | 9.68352 | 19.05238 | 10.00042 | 4.22256 | 0.04500 |
| **BLK** | 0.02397 | 0.00417 | 0.02028 | 0.01797 | 0.44008 | 0.01318 | 0.01973 |
| **BLM** | 10.17540 | 2.69696 | 5.45244 | 5.70352 | 2.51767 | 1.32347 | 4.51208 |
| **BLMH** | 27.93143 | 16.29084 | 7.95185 | 25.19017 | 2.47068 | 8.91451 | 11.57759 |
| **BLNK** | 0.07691 | 0.00000 | 0.15081 | 0.06630 | 0.14701 | 0.05221 | 0.00178 |
| **BLOC1S1** | 3.09629 | 6.65938 | 1.95659 | 5.12586 | 0.43246 | 4.18848 | 3.74915 |
| **BLOC1S2** | 19.89897 | 8.85327 | 9.52580 | 15.98051 | 1.94294 | 10.06047 | 0.79001 |
| **BLOC1S3** | 6.54907 | 9.10199 | 2.35482 | 7.14876 | 10.11750 | 2.74679 | 0.00000 |
| **BLOC1S4** | 0.61976 | 1.14472 | 0.92411 | 1.76679 | 0.18732 | 2.28665 | 7.35315 |
| **BLOC1S5.uid-8** | 0.03778 | 0.01776 | 0.01839 | 0.04394 | 0.00738 | 0.01789 | 0.00839 |
| **BLOC1S6** | 7.83890 | 0.39991 | 6.97896 | 7.64032 | 2.62798 | 3.88912 | 8.61824 |
| **BLVRA** | 0.25826 | 0.37209 | 0.54780 | 0.05196 | 0.03881 | 0.00000 | 0.08176 |
| **BLVRB** | 2.01812 | 8.79936 | 13.29351 | 15.52775 | 0.80096 | 1.64249 | 0.00000 |
| **BLZF1** | 6.21323 | 0.72225 | 5.82128 | 4.27992 | 3.88903 | 1.21243 | 0.00000 |
| **BMF** | 0.54574 | 0.00000 | 0.00000 | 2.81094 | 0.09687 | 0.24475 | 1.43041 |
| **BMI1** | 0.12637 | 0.04479 | 0.05608 | 0.07007 | 0.01902 | 0.04921 | 0.02471 |
| **BMP1** | 0.09691 | 0.23166 | 0.01355 | 1.15177 | 0.00529 | 0.00000 | 1.63383 |
| **BMP10** | 0.02672 | 0.02139 | 0.01744 | 0.01651 | 0.00740 | 0.06469 | 0.01219 |
| **BMP15** | 0.00000 | 0.11784 | 0.00001 | 0.00358 | 15.07098 | 0.01389 | 0.01940 |
| **BMP2** | 0.00000 | 0.65021 | 0.00144 | 0.03408 | 0.00649 | 13.87247 | 0.02044 |
| **BMP2K** | 3.16338 | 0.96793 | 0.99002 | 0.00000 | 0.61404 | 3.27912 | 3.74228 |
| **BMP3** | 0.03145 | 0.00658 | 0.02683 | 0.01605 | 0.00705 | 0.02221 | 0.02598 |
| **BMP4** | 2.25390 | 0.62642 | 0.10830 | 3.11423 | 4.49730 | 1.55115 | 0.00000 |

| | | | | | | | |
|---|---|---|---|---|---|---|---|
| **BMP5** | 0.01120 | 0.01975 | 0.01019 | 0.01275 | 0.00768 | 0.06625 | 0.05943 |
| **BMP6** | 0.32922 | 0.20967 | 0.00000 | 0.38262 | 6.20332 | 4.03559 | 0.00000 |
| **BMP7** | 0.00000 | 0.00000 | 0.70510 | 1.22810 | 0.25316 | 0.15727 | 0.49270 |
| **BMP8A** | 0.00001 | 0.98798 | 0.20097 | 0.72869 | 0.05530 | 0.43050 | 1.01821 |
| **BMP8B** | 0.03813 | 0.01218 | 0.01378 | 0.07923 | 0.00676 | 0.02093 | 0.08474 |
| **BMPER** | 0.03291 | 0.06116 | 0.01129 | 0.02278 | 0.12575 | 0.07249 | 0.01086 |
| **BMPR1A** | 4.50109 | 0.00000 | 1.65337 | 0.00000 | 0.32381 | 2.29466 | 2.49449 |
| **BMPR1B** | 0.07310 | 0.06118 | 0.26538 | 0.00785 | 0.59978 | 0.01698 | 0.15522 |
| **BMPR2** | 2.46455 | 0.59649 | 0.80613 | 3.74782 | 0.44282 | 3.21384 | 8.47024 |
| **BMS1** | 19.56164 | 15.29193 | 14.47349 | 10.99613 | 4.84668 | 7.06714 | 14.74716 |
| **BMS1P20** | 2.11357 | 3.22020 | 2.52236 | 1.40808 | 1.87853 | 0.48717 | 1.36465 |
| **BMS1P4** | 0.66837 | 0.71259 | 1.01906 | 0.25424 | 0.15968 | 0.42773 | 0.75632 |
| **BMS1P5.uid-14** | 0.01519 | 0.01148 | 0.02083 | 0.01489 | 0.00662 | 0.02285 | 0.07711 |
| **BMS1P5.uid-14** | 0.04500 | 0.01743 | 0.02482 | 0.01521 | 0.00615 | 0.03359 | 0.08478 |
| **BMX** | 0.03035 | 0.07117 | 0.00712 | 0.01278 | 0.00751 | 0.01353 | 0.04575 |
| **BNC1** | 0.00000 | 0.04623 | 0.00000 | 0.52608 | 0.46941 | 0.06803 | 0.10127 |
| **BNC2** | 0.00000 | 0.00000 | 0.58443 | 0.11679 | 0.23982 | 0.66921 | 0.62123 |
| **BNIP1** | 5.31163 | 4.14741 | 2.67302 | 1.40243 | 1.21727 | 2.27704 | 0.00001 |
| **BNIP2** | 16.54236 | 1.35481 | 16.39624 | 13.26777 | 12.51672 | 8.25040 | 2.36157 |
| **BNIP3** | 28.42926 | 2.86157 | 9.80979 | 27.04001 | 5.94793 | 10.14189 | 6.06354 |
| **BNIP3L** | 3.16932 | 1.66211 | 6.82510 | 10.38796 | 9.77276 | 1.37795 | 0.16365 |
| **BNIPL** | 0.19851 | 0.00779 | 0.08607 | 0.08372 | 0.01975 | 0.09922 | 0.05151 |
| **BOC** | 0.03416 | 0.09921 | 0.01603 | 0.05542 | 0.00926 | 0.09350 | 0.00000 |
| **BOD1** | 0.67245 | 1.71972 | 0.20988 | 2.61051 | 5.99455 | 3.17556 | 12.69498 |
| **BOD1L1** | 5.18025 | 0.44511 | 7.31754 | 2.17425 | 4.40077 | 7.86086 | 22.08419 |
| **BOD1L2** | 0.00744 | 0.09759 | 0.02489 | 0.03115 | 0.02887 | 0.03278 | 0.03115 |
| **BOK** | 0.00000 | 0.83409 | 0.00888 | 1.02268 | 0.05594 | 0.60545 | 1.29485 |
| **BOK-AS1** | 0.85146 | 0.00000 | 0.00718 | 0.44652 | 0.09775 | 0.00000 | 0.45772 |
| **BOLA1** | 4.46431 | 3.56199 | 3.38076 | 5.78146 | 1.88980 | 0.30504 | 0.74112 |
| **BOLA2+BOLA2I** | 0.03111 | 0.02102 | 0.03414 | 0.01972 | 0.01353 | 0.02972 | 0.05181 |
| **BOLA2+BOLA2I** | 0.06140 | 0.04425 | 0.22199 | 0.04761 | 0.00294 | 0.21400 | 0.22292 |
| **BOLA3** | 20.47016 | 21.07523 | 3.65527 | 0.00000 | 0.00000 | 9.71321 | 7.08206 |
| **BOLA3-AS1** | 1.17888 | 0.33960 | 0.20951 | 1.63527 | 0.09879 | 0.44982 | 0.74495 |
| **BOLL** | 0.01699 | 0.17591 | 0.82418 | 0.00000 | 0.06473 | 0.00000 | 0.12992 |
| **1.00000** | 0.00000 | 1.47764 | 0.02528 | 0.37240 | 0.09956 | 0.39873 | 1.07372 |

| | | | | | | | |
|---|---|---|---|---|---|---|---|
| **BORA** | 10.17580 | 1.15225 | 9.29500 | 3.47021 | 3.54426 | 1.72558 | 0.00000 |
| **BPESC1** | 0.00877 | 0.00002 | 0.04311 | 0.00000 | 0.22024 | 0.09130 | 0.34291 |
| **BPGM** | 4.90217 | 0.00000 | 3.01591 | 12.46780 | 10.66675 | 0.94622 | 0.00000 |
| **BPHL** | 1.54988 | 1.18129 | 1.23442 | 0.30664 | 0.16941 | 1.08516 | 0.00000 |
| **BPI** | 0.02100 | 0.01869 | 0.01242 | 0.02053 | 0.00766 | 0.01316 | 0.03203 |
| **BPIFA2** | 0.09080 | 0.45609 | 0.19161 | 0.08298 | 0.09897 | 0.30251 | 0.00910 |
| **BPIFA3** | 0.01333 | 0.02204 | 0.01107 | 0.01434 | 0.00777 | 0.02965 | 0.05778 |
| **BPIFB1** | 0.00972 | 0.03511 | 0.01226 | 0.02466 | 0.00756 | 0.02436 | 0.03639 |
| **BPIFB2** | 0.02198 | 0.01866 | 0.01209 | 0.01824 | 0.00784 | 0.01875 | 0.02576 |
| **BPIFB4** | 0.04252 | 0.00689 | 0.22087 | 0.01259 | 0.00868 | 0.04393 | 0.02133 |
| **BPIFC** | 0.02287 | 0.00000 | 0.08752 | 0.02384 | 0.17784 | 0.01227 | 0.01785 |
| **BPNT1** | 4.20947 | 0.00000 | 2.39128 | 4.84676 | 0.46380 | 1.18845 | 2.06833 |
| **BPTF** | 11.08059 | 6.17869 | 8.01975 | 6.25506 | 4.22860 | 7.43141 | 37.11646 |
| **BRAF** | 0.61544 | 1.42117 | 2.23392 | 2.17001 | 0.34939 | 1.55441 | 2.23334 |
| **BRAP** | 15.36169 | 17.02224 | 12.96786 | 12.18584 | 9.24524 | 8.70826 | 0.00000 |
| **BRAT1** | 0.00000 | 5.56328 | 4.69435 | 3.82851 | 0.82222 | 5.03975 | 10.67247 |
| **BRCA1** | 9.04790 | 2.34119 | 7.47319 | 3.82186 | 2.97347 | 5.76654 | 9.95070 |
| **BRCA2** | 6.80077 | 0.00000 | 12.91302 | 0.90269 | 6.42807 | 2.56259 | 11.47878 |
| **BRCC3** | 21.36967 | 4.71609 | 3.27378 | 9.27950 | 0.18882 | 5.07861 | 3.06591 |
| **BRD1** | 1.94490 | 2.88808 | 2.02836 | 3.74616 | 0.89878 | 2.50382 | 7.55125 |
| **BRD2** | 22.30189 | 17.30025 | 13.43750 | 28.54411 | 14.29672 | 19.21643 | 33.31052 |
| **BRD3** | 11.77789 | 37.99696 | 5.27952 | 11.03532 | 1.20917 | 18.98641 | 11.01012 |
| **BRD4** | 1.48136 | 10.54856 | 4.64570 | 13.11901 | 6.56111 | 9.51709 | 40.02266 |
| **BRD7** | 2.06224 | 4.07716 | 1.99403 | 2.00076 | 1.61991 | 2.62991 | 9.09226 |
| **BRD7P3** | 0.04734 | 0.00703 | 0.18432 | 0.03384 | 0.02429 | 0.05262 | 0.03733 |
| **BRD8** | 20.12430 | 12.03613 | 12.95976 | 17.35339 | 5.35760 | 8.34954 | 0.07476 |
| **BRD9** | 4.25827 | 9.69654 | 2.00768 | 4.41739 | 1.22669 | 5.67017 | 14.69347 |
| **BRDT** | 13.08320 | 12.48348 | 24.41123 | 11.22289 | 21.12136 | 19.79511 | 0.67724 |
| **BRE** | 9.02925 | 4.67012 | 2.22478 | 2.62428 | 0.14067 | 3.02044 | 1.46602 |
| **BRE-AS1** | 0.18833 | 0.17035 | 0.10027 | 0.08292 | 0.02942 | 0.05674 | 0.00000 |
| **BREA2** | 0.01790 | 0.00789 | 0.04740 | 0.08867 | 0.05590 | 0.00000 | 0.23157 |
| **BRF1** | 0.00000 | 2.29459 | 1.01311 | 1.70767 | 0.70042 | 1.41052 | 6.11645 |
| **BRF2** | 6.54284 | 5.88852 | 6.06111 | 8.73477 | 2.77644 | 2.48485 | 0.00000 |
| **BRI3** | 0.00000 | 2.10150 | 0.15126 | 3.24290 | 0.42401 | 1.13316 | 13.21065 |
| **BRI3BP** | 0.61397 | 0.42143 | 0.71394 | 2.28045 | 0.10694 | 0.13574 | 4.71835 |

| | | | | | | | |
|---|---|---|---|---|---|---|---|
| **BRICD5** | 0.16979 | 0.86285 | 0.00558 | 0.81452 | 0.01460 | 0.25303 | 0.99594 |
| **BRINP1** | 0.14273 | 0.04589 | 0.00000 | 0.05963 | 2.65050 | 1.13023 | 0.00000 |
| **BRINP2** | 0.00840 | 0.02576 | 0.01013 | 0.05441 | 0.03012 | 0.01263 | 0.02894 |
| **BRIP1** | 2.84481 | 0.00000 | 1.50363 | 0.64914 | 0.06850 | 1.00491 | 1.22314 |
| **BRIX1** | 31.62041 | 9.11361 | 24.30768 | 14.51101 | 10.47489 | 8.17771 | 8.16076 |
| **BRK1** | 23.49609 | 14.55732 | 20.00317 | 24.19531 | 1.58675 | 15.79140 | 9.15776 |
| **BRMS1** | 26.26395 | 32.01184 | 16.19596 | 22.75234 | 5.29773 | 14.56985 | 4.23461 |
| **BRMS1L** | 7.93737 | 3.54759 | 7.64099 | 7.55095 | 1.31941 | 10.03205 | 0.00000 |
| **BROX** | 23.94368 | 1.60349 | 10.48892 | 9.73593 | 5.84715 | 5.97876 | 0.00000 |
| **BRPF1** | 2.06710 | 4.90680 | 3.97067 | 4.47553 | 3.47320 | 5.08419 | 9.61605 |
| **BRPF3** | 7.54298 | 4.13499 | 5.59925 | 2.87190 | 0.15427 | 6.78004 | 13.92887 |
| **BRS3** | 0.08290 | 0.00000 | 0.01457 | 0.00000 | 0.02018 | 0.01008 | 0.05696 |
| **BRSK1** | 0.00000 | 0.00000 | 0.06681 | 0.01296 | 0.21093 | 0.46243 | 1.19170 |
| **BRSK2** | 0.00000 | 0.00000 | 0.20040 | 0.14879 | 0.08398 | 0.09920 | 0.22364 |
| **BRWD1** | 8.07744 | 2.40304 | 9.94965 | 4.91072 | 8.14765 | 6.20467 | 19.25852 |
| **BRWD1-AS1** | 0.02902 | 0.01687 | 0.02618 | 0.02015 | 0.01281 | 0.01019 | 0.03960 |
| **BRWD1-IT2** | 0.01214 | 0.03248 | 0.07609 | 0.01804 | 0.08416 | 0.02694 | 0.10896 |
| **BRWD3** | 2.40223 | 0.00125 | 18.75932 | 0.00000 | 1.48752 | 14.82383 | 14.86315 |
| **BSCL2.uid-1559** | 0.03432 | 0.00000 | 0.24180 | 0.24906 | 0.09099 | 0.13252 | 0.15620 |
| **BSCL2.uid-1560** | 0.07225 | 0.00000 | 0.05072 | 0.12994 | 0.03160 | 0.02119 | 0.30789 |
| **BSCL2.uid-1560** | 0.28701 | 0.68795 | 0.15460 | 0.48331 | 0.22707 | 0.12941 | 0.00000 |
| **BSDC1** | 6.65854 | 3.91304 | 9.91884 | 15.86815 | 3.07826 | 6.03860 | 5.11751 |
| **BSG** | 44.33786 | 58.06860 | 45.02218 | 43.96636 | 18.08420 | 30.85422 | 31.10695 |
| **BSN** | 0.00000 | 0.27942 | 0.30919 | 0.33473 | 0.42971 | 0.08669 | 0.35360 |
| **BSN-AS2** | 0.01813 | 0.04888 | 0.05202 | 0.04952 | 0.00762 | 0.01136 | 0.01053 |
| **BSND** | 0.03110 | 0.02465 | 0.04494 | 0.01643 | 0.00556 | 0.21725 | 0.00732 |
| **BSPRY** | 0.41891 | 8.82204 | 1.27196 | 0.48722 | 0.00000 | 0.00000 | 8.81441 |
| **BST1** | 0.06663 | 0.16021 | 0.01077 | 0.08118 | 0.03300 | 0.52444 | 0.00000 |
| **BST2** | 0.00000 | 0.10110 | 0.00000 | 0.02933 | 0.20083 | 0.90743 | 0.07117 |
| **BTAF1** | 9.99059 | 0.00000 | 7.51594 | 7.11004 | 6.68558 | 5.18549 | 10.74744 |
| **BTBD1** | 4.24067 | 0.54680 | 1.45227 | 0.88818 | 0.07486 | 2.24918 | 6.10310 |
| **BTBD10** | 12.95626 | 2.62549 | 4.96722 | 9.34115 | 4.05418 | 3.27006 | 1.15690 |
| **BTBD11** | 0.15727 | 0.05362 | 0.00000 | 0.00000 | 0.46602 | 0.17281 | 0.02928 |
| **BTBD16** | 0.03506 | 0.06846 | 0.00890 | 0.02287 | 0.01148 | 0.06348 | 0.00574 |
| **BTBD17** | 0.02875 | 0.02510 | 0.01007 | 0.01162 | 0.00771 | 0.00819 | 0.04194 |

| | | | | | | | |
|---|---|---|---|---|---|---|---|
| **BTBD18** | 0.07737 | 0.22190 | 0.98021 | 0.01443 | 1.35466 | 0.16168 | 0.81113 |
| **BTBD19** | 0.00000 | 0.10552 | 0.04271 | 0.06605 | 0.01738 | 0.06735 | 0.26692 |
| **BTBD2** | 0.00000 | 0.49774 | 1.04611 | 2.84462 | 1.51203 | 0.87780 | 4.92716 |
| **BTBD3** | 9.59962 | 1.53295 | 0.68806 | 3.77744 | 0.60956 | 0.77586 | 7.61515 |
| **BTBD6** | 0.26702 | 2.41345 | 2.11069 | 0.82353 | 0.42448 | 0.00000 | 4.83402 |
| **BTBD7** | 2.77784 | 2.93223 | 2.17228 | 4.59251 | 1.04461 | 3.01221 | 10.34617 |
| **BTBD8** | 0.11422 | 0.00000 | 0.49728 | 0.00875 | 0.06555 | 0.25324 | 0.26935 |
| **BTBD9** | 0.49621 | 0.86361 | 0.77723 | 1.02983 | 0.61341 | 0.49149 | 2.83610 |
| **BTC** | 0.02600 | 0.01731 | 0.01288 | 0.01907 | 0.01673 | 0.01406 | 0.01545 |
| **BTD** | 3.55512 | 0.46721 | 1.36095 | 4.17942 | 0.23033 | 0.59425 | 0.30840 |
| **BTF3** | 46.38198 | 18.57210 | 17.38577 | 40.78807 | 10.01575 | 23.06424 | 34.09746 |
| **BTF3L4** | 25.17406 | 0.17344 | 11.55796 | 10.81383 | 0.74639 | 9.12932 | 7.54667 |
| **BTF3P11** | 0.05744 | 0.00000 | 0.01848 | 0.02946 | 0.00746 | 0.02663 | 0.13662 |
| **BTG1** | 6.24378 | 1.54710 | 1.07879 | 16.71031 | 4.52738 | 2.49508 | 32.91365 |
| **BTG2** | 1.36895 | 7.36499 | 0.01953 | 12.10514 | 0.00106 | 8.36376 | 0.00000 |
| **BTG3** | 12.43751 | 6.04522 | 14.95194 | 8.58164 | 3.19698 | 5.47884 | 2.65628 |
| **BTG4** | 0.03122 | 0.10638 | 0.09917 | 0.03338 | 2.50163 | 0.03569 | 0.02561 |
| **BTK** | 0.88699 | 1.10888 | 1.40747 | 0.00000 | 2.37376 | 2.10778 | 0.00000 |
| **BTLA** | 0.02344 | 0.09334 | 2.64062 | 0.00000 | 0.91706 | 4.31497 | 0.00533 |
| **BTN1A1** | 0.00000 | 0.00969 | 12.26909 | 0.00500 | 0.13075 | 0.00000 | 0.02498 |
| **BTN2A1** | 1.74161 | 0.00000 | 0.09219 | 2.16185 | 0.00062 | 3.35055 | 0.00000 |
| **BTN2A2** | 0.01871 | 0.02271 | 0.01118 | 0.07318 | 0.00770 | 0.01077 | 0.02859 |
| **BTN2A3P** | 0.02628 | 0.02051 | 0.01272 | 0.01922 | 0.00786 | 0.01724 | 0.02362 |
| **BTN3A1** | 0.03413 | 0.01785 | 0.01354 | 0.02497 | 0.00811 | 0.01636 | 0.01317 |
| **BTNL2** | 0.02387 | 0.01764 | 0.00954 | 0.01926 | 0.04495 | 0.01303 | 0.01747 |
| **BTNL3** | 0.00575 | 0.00000 | 0.02259 | 0.01192 | 0.33403 | 0.00683 | 0.17274 |
| **BTNL8** | 0.02021 | 0.06439 | 0.02231 | 0.04457 | 0.07751 | 0.07369 | 0.00000 |
| **BTNL9** | 0.03045 | 0.09101 | 0.60748 | 0.02354 | 0.21007 | 0.03478 | 0.00000 |
| **BTRC** | 5.33999 | 9.16453 | 19.86830 | 4.79944 | 8.01817 | 4.36600 | 1.53071 |
| **BUB1** | 32.49696 | 17.54318 | 14.08180 | 11.05954 | 6.77322 | 10.21023 | 9.10074 |
| **BUB1B** | 25.96366 | 1.96620 | 7.12832 | 5.27911 | 3.25406 | 4.44256 | 14.58689 |
| **BUB3** | 47.31478 | 33.03609 | 23.64056 | 34.05796 | 12.29494 | 17.91363 | 7.39810 |
| **BUD13** | 7.50273 | 3.49339 | 5.83401 | 4.79468 | 2.35954 | 2.32755 | 1.95674 |
| **BUD31** | 24.97385 | 13.36098 | 8.20095 | 20.09246 | 14.73136 | 12.98693 | 1.64806 |
| **BVES** | 0.04805 | 0.15974 | 0.10543 | 0.03571 | 0.77519 | 0.15906 | 0.00000 |

| | | | | | | | |
|---|---|---|---|---|---|---|---|
| **BVES-AS1** | 0.02513 | 0.00663 | 0.04694 | 0.01895 | 0.00944 | 0.01255 | 0.01778 |
| **BYSL** | 28.29732 | 19.58525 | 20.95771 | 25.13770 | 5.30674 | 8.77727 | 6.70837 |
| **BZRAP1** | 0.05389 | 0.02137 | 0.01115 | 0.17467 | 0.00874 | 0.00986 | 0.08072 |
| **BZRAP1-AS1** | 0.26812 | 0.20375 | 0.33588 | 0.77743 | 0.23969 | 0.01299 | 0.00000 |
| **BZW1** | 24.43705 | 7.42270 | 12.78788 | 14.70037 | 13.42427 | 8.44061 | 0.00000 |
| **BZW2** | 35.00088 | 2.73084 | 16.11828 | 35.69752 | 9.97679 | 11.44786 | 21.35708 |
| **C10orf10** | 9.36053 | 11.24482 | 0.00020 | 6.09821 | 0.00266 | 2.55228 | 0.00000 |
| **C10orf105** | 0.02426 | 0.01901 | 0.01312 | 0.02129 | 0.00806 | 0.01437 | 0.01764 |
| **C10orf11** | 0.04894 | 0.00407 | 0.01231 | 0.02111 | 0.21208 | 0.02732 | 0.03913 |
| **C10orf111** | 0.15458 | 0.26946 | 0.23092 | 0.13942 | 0.13648 | 0.09388 | 0.11954 |
| **C10orf113** | 0.02245 | 0.06015 | 0.00823 | 0.01504 | 0.00776 | 0.04947 | 0.03195 |
| **C10orf12** | 2.18201 | 1.29097 | 3.01688 | 2.03618 | 1.24789 | 1.97456 | 2.40646 |
| **C10orf120** | 0.02499 | 0.00982 | 0.01996 | 0.01801 | 0.22687 | 0.01318 | 0.01679 |
| **C10orf126** | 0.03536 | 0.00622 | 0.01084 | 0.03670 | 0.03344 | 0.14035 | 0.03820 |
| **C10orf131** | 0.25999 | 0.04699 | 0.12997 | 0.08611 | 0.00335 | 0.04038 | 0.33923 |
| **C10orf2** | 7.77003 | 5.21842 | 7.04629 | 5.78825 | 3.03911 | 2.68676 | 6.51320 |
| **C10orf25** | 0.59692 | 0.06725 | 0.62855 | 0.05573 | 0.04940 | 0.00033 | 0.00000 |
| **C10orf32** | 2.76771 | 0.00000 | 2.52070 | 4.35067 | 0.00000 | 1.42306 | 2.03696 |
| **C10orf35** | 1.77438 | 1.07129 | 0.00795 | 9.67176 | 0.33756 | 0.03325 | 0.00000 |
| **C10orf40** | 0.02548 | 0.01717 | 0.00977 | 0.02190 | 0.03896 | 0.02385 | 0.01697 |
| **C10orf53** | 0.05773 | 0.01271 | 0.01347 | 0.02114 | 0.00802 | 0.01642 | 0.00431 |
| **C10orf54** | 0.00000 | 1.03782 | 0.02152 | 2.57791 | 0.00619 | 0.00080 | 0.00099 |
| **C10orf55** | 0.02882 | 0.04310 | 0.03074 | 0.05113 | 0.00936 | 0.09035 | 0.02178 |
| **C10orf67** | 0.02343 | 0.03776 | 0.03106 | 0.01674 | 0.00661 | 0.08951 | 0.01824 |
| **C10orf71** | 0.02669 | 0.01966 | 0.01214 | 0.06388 | 0.01110 | 0.04121 | 0.00378 |
| **C10orf71-AS1** | 0.02406 | 0.01778 | 0.01311 | 0.03086 | 0.00808 | 0.01352 | 0.01318 |
| **C10orf76** | 2.86033 | 1.88850 | 5.76998 | 3.54576 | 1.54567 | 4.27085 | 5.97727 |
| **C10orf82** | 0.55363 | 0.05800 | 0.01923 | 0.14537 | 1.82218 | 0.07369 | 0.00000 |
| **C10orf88** | 5.93008 | 0.32789 | 5.07984 | 4.14579 | 1.56521 | 0.62510 | 0.00000 |
| **C10orf90** | 0.00000 | 0.24280 | 0.00000 | 0.00523 | 0.05027 | 0.98413 | 0.01097 |
| **C10orf91** | 0.00664 | 0.06020 | 0.05309 | 0.03251 | 0.01872 | 0.01344 | 0.07121 |
| **C10orf95** | 0.00000 | 1.08135 | 0.32906 | 0.41842 | 0.08985 | 0.51457 | 0.38868 |
| **C10orf99** | 0.05485 | 0.06474 | 0.03050 | 0.04671 | 0.00809 | 0.00911 | 0.00000 |
| **C11orf1** | 1.67593 | 0.41543 | 1.76253 | 1.17940 | 0.16215 | 0.63906 | 0.10226 |
| **C11orf16** | 0.12503 | 0.24140 | 0.18726 | 0.05830 | 0.44223 | 0.07022 | 0.00000 |

| | | | | | | | |
|---|---|---|---|---|---|---|---|
| **C11orf24** | 7.34146 | 9.09064 | 5.71647 | 10.25372 | 1.22187 | 3.07810 | 1.15118 |
| **C11orf30** | 2.10367 | 1.20985 | 1.95282 | 4.68579 | 1.66756 | 1.35178 | 5.29160 |
| **C11orf31** | 15.96896 | 30.64923 | 14.55549 | 18.55592 | 15.06173 | 13.16081 | 13.50958 |
| **C11orf40** | 0.00000 | 0.36579 | 0.37261 | 0.00279 | 5.50191 | 0.00000 | 0.00594 |
| **C11orf42** | 0.03288 | 0.03282 | 0.02700 | 0.10001 | 0.00878 | 0.01646 | 0.06331 |
| **C11orf44** | 0.02130 | 0.02336 | 0.01365 | 0.02853 | 0.01548 | 0.01440 | 0.03425 |
| **C11orf45** | 0.28810 | 0.08510 | 0.16668 | 0.07134 | 0.08285 | 0.08123 | 0.14288 |
| **C11orf48** | 0.85507 | 0.17812 | 0.78192 | 1.12948 | 0.95912 | 0.17517 | 0.18244 |
| **C11orf49** | 4.48478 | 1.02023 | 0.85383 | 2.55347 | 0.12780 | 1.51578 | 1.37395 |
| **C11orf53** | 0.02444 | 0.04211 | 0.05271 | 0.01762 | 0.02736 | 0.00449 | 0.03502 |
| **C11orf54** | 7.92853 | 1.01901 | 2.41480 | 8.72610 | 0.98521 | 3.52362 | 0.00000 |
| **C11orf57** | 4.93733 | 0.00000 | 7.46957 | 8.54759 | 2.32796 | 3.16502 | 9.79280 |
| **C11orf58** | 34.22230 | 14.79014 | 21.68912 | 20.71884 | 3.28594 | 16.77787 | 18.47445 |
| **C11orf63** | 0.04602 | 0.00893 | 0.03681 | 0.04169 | 0.03525 | 0.01895 | 0.00000 |
| **C11orf65** | 0.10452 | 0.05441 | 1.53384 | 0.00086 | 0.23009 | 0.04338 | 0.01455 |
| **C11orf68** | 0.00000 | 0.81084 | 0.37504 | 0.69255 | 0.16459 | 1.24405 | 2.29578 |
| **C11orf70** | 0.28747 | 0.00000 | 0.09340 | 2.32664 | 0.03287 | 0.00402 | 0.00000 |
| **C11orf71** | 2.77183 | 3.99971 | 5.17024 | 4.36481 | 4.97666 | 0.91281 | 0.00000 |
| **C11orf73** | 14.52342 | 5.37271 | 3.87832 | 10.25135 | 0.04372 | 4.43923 | 1.60382 |
| **C11orf74** | 2.82748 | 2.31247 | 2.00176 | 0.67089 | 1.76522 | 1.43303 | 1.34061 |
| **C11orf80** | 0.00000 | 0.05440 | 0.36974 | 0.26104 | 0.15290 | 0.31714 | 0.97466 |
| **C11orf83** | 7.09235 | 28.96051 | 9.17162 | 0.00000 | 11.67859 | 6.45562 | 9.91277 |
| **C11orf84** | 0.00027 | 1.39184 | 0.62016 | 0.77048 | 0.86475 | 0.43097 | 3.27292 |
| **C11orf85** | 0.02665 | 0.00000 | 0.28829 | 0.00037 | 0.94176 | 0.02201 | 0.07171 |
| **C11orf86** | 0.04218 | 0.00787 | 0.15842 | 0.01976 | 0.52967 | 0.10558 | 0.00000 |
| **C11orf87** | 0.02790 | 0.02332 | 0.00893 | 0.01895 | 0.02638 | 0.03356 | 0.01632 |
| **C11orf88** | 0.51777 | 0.00025 | 0.00266 | 0.36434 | 0.07434 | 0.00000 | 0.14809 |
| **C11orf91** | 0.07208 | 0.00387 | 0.04477 | 0.06840 | 0.01567 | 0.00391 | 0.04749 |
| **C11orf94** | 0.01526 | 0.05648 | 0.01485 | 0.06039 | 0.00748 | 0.03922 | 0.01708 |
| **C11orf95** | 2.14390 | 2.65702 | 1.24834 | 0.39541 | 0.25804 | 2.76667 | 10.55103 |
| **C11orf96** | 0.00000 | 0.01678 | 0.00466 | 0.33657 | 0.00701 | 0.46459 | 0.46820 |
| **C12orf10** | 8.85445 | 8.58042 | 7.16656 | 13.73781 | 4.82298 | 5.17796 | 4.88000 |
| **C12orf29** | 14.32803 | 3.93771 | 7.77519 | 9.73040 | 0.75069 | 1.64592 | 0.00000 |
| **C12orf36** | 0.04654 | 0.14122 | 0.86777 | 0.00149 | 0.05249 | 0.92948 | 0.00000 |
| **C12orf4** | 8.48858 | 1.12656 | 6.51078 | 5.85604 | 1.29043 | 1.80019 | 0.00000 |

| | | | | | | | |
|---|---|---|---|---|---|---|---|
| **C12orf40** | 0.22128 | 0.00000 | 0.62320 | 0.04441 | 0.52353 | 0.02369 | 0.32173 |
| **C12orf42** | 0.00000 | 0.08829 | 0.01480 | 0.01359 | 0.09418 | 0.07436 | 0.05977 |
| **C12orf43** | 11.25993 | 7.57977 | 12.03506 | 9.40219 | 6.53999 | 3.35882 | 0.23494 |
| **C12orf45** | 19.26094 | 16.39153 | 12.75964 | 14.85022 | 5.94383 | 5.67107 | 1.22363 |
| **C12orf49** | 0.76855 | 2.14730 | 2.38281 | 3.75488 | 1.55265 | 1.09614 | 6.79869 |
| **C12orf5** | 49.82523 | 13.20583 | 24.10322 | 27.50070 | 4.65718 | 5.65022 | 12.88088 |
| **C12orf50** | 0.02480 | 0.01695 | 0.01092 | 0.04099 | 0.03623 | 0.01255 | 0.01099 |
| **C12orf54** | 0.01132 | 0.04968 | 0.03945 | 0.02985 | 0.00586 | 0.04775 | 0.02908 |
| **C12orf56** | 0.38616 | 0.85738 | 1.26938 | 0.03086 | 0.01480 | 0.42788 | 0.24645 |
| **C12orf57** | 4.56615 | 9.54247 | 7.14518 | 12.10692 | 6.03468 | 11.01627 | 6.58318 |
| **C12orf60** | 0.98214 | 0.00000 | 2.31158 | 0.45915 | 5.07668 | 0.21221 | 0.00000 |
| **C12orf61** | 0.00000 | 0.11269 | 0.10255 | 0.07019 | 0.00700 | 0.01636 | 0.09226 |
| **C12orf65** | 5.04985 | 6.98660 | 0.72758 | 3.21210 | 1.83325 | 3.96526 | 4.59656 |
| **C12orf66** | 1.60031 | 1.30260 | 1.48118 | 0.75931 | 0.69016 | 0.49953 | 0.00000 |
| **C12orf71** | 0.00103 | 0.04129 | 0.03836 | 0.05084 | 0.01001 | 0.00405 | 0.08047 |
| **C12orf73** | 4.31978 | 1.78878 | 1.85250 | 4.40644 | 0.00000 | 1.83024 | 3.20797 |
| **C12orf74** | 0.02190 | 0.01166 | 0.02482 | 0.01658 | 0.03134 | 0.06266 | 0.01922 |
| **C12orf75** | 0.53191 | 0.23087 | 0.00053 | 2.28546 | 0.27435 | 2.69870 | 1.09047 |
| **C12orf76** | 4.92942 | 0.00000 | 1.89822 | 0.62279 | 1.19853 | 0.06855 | 1.42989 |
| **C12orf77** | 0.00779 | 0.08603 | 0.04113 | 0.02319 | 0.91082 | 0.00556 | 0.01382 |
| **C12orf79** | 0.00000 | 0.00220 | 0.02263 | 0.17605 | 0.09610 | 0.02018 | 0.11176 |
| **C14orf1** | 30.84082 | 13.87544 | 2.41922 | 17.63529 | 2.88889 | 6.60998 | 6.35395 |
| **C14orf105** | 0.02792 | 0.00813 | 0.01446 | 0.01281 | 0.00770 | 0.01034 | 0.05671 |
| **C14orf119** | 36.49362 | 24.29814 | 23.20375 | 28.72537 | 12.84833 | 15.01801 | 2.73353 |
| **C14orf132** | 0.03176 | 0.01580 | 0.01583 | 0.04330 | 0.04415 | 0.02660 | 0.01777 |
| **C14orf142** | 10.60348 | 1.28348 | 6.59249 | 8.33005 | 3.15720 | 3.77568 | 0.00000 |
| **C14orf159** | 0.75347 | 3.16688 | 1.07671 | 0.41535 | 0.15731 | 2.99140 | 1.54845 |
| **C14orf166** | 31.14598 | 20.41158 | 19.48836 | 28.73077 | 14.85019 | 14.26732 | 19.71185 |
| **C14orf169** | 1.89697 | 4.00006 | 3.21545 | 4.18693 | 2.72375 | 4.89825 | 5.12698 |
| **C14orf178** | 0.06760 | 0.01003 | 0.15921 | 0.05913 | 0.08574 | 0.23022 | 0.00000 |
| **C14orf180** | 0.02054 | 0.02217 | 0.01158 | 0.03373 | 0.00750 | 0.02607 | 0.04494 |
| **C14orf182** | 0.14285 | 0.09762 | 0.12687 | 0.47762 | 0.03242 | 0.00000 | 0.57466 |
| **C14orf183** | 0.03269 | 0.01897 | 0.03855 | 0.06377 | 0.01966 | 0.03068 | 0.03367 |
| **C14orf2** | 50.37206 | 25.95509 | 27.29082 | 35.34279 | 13.68471 | 24.06632 | 32.22218 |
| **C14orf23** | 0.01913 | 0.00560 | 0.03158 | 0.02380 | 0.04411 | 0.01181 | 0.03348 |

| | | | | | | | |
|---|---|---|---|---|---|---|---|
| **C14orf28** | 0.15933 | 0.00000 | 0.26478 | 0.39480 | 0.17490 | 0.23161 | 0.21979 |
| **C14orf37** | 0.00000 | 2.69440 | 1.32522 | 0.00000 | 2.02605 | 0.17660 | 0.05158 |
| **C14orf39** | 0.02626 | 0.02705 | 0.00915 | 0.01837 | 0.03388 | 0.01071 | 0.02231 |
| **C14orf64** | 0.01770 | 0.01768 | 0.01308 | 0.01699 | 0.00760 | 0.01940 | 0.03723 |
| **C14orf79** | 0.06578 | 0.70570 | 0.68773 | 0.08046 | 0.29293 | 0.26882 | 0.11412 |
| **C14orf80** | 0.00000 | 3.94537 | 0.17426 | 0.27953 | 0.01020 | 1.04468 | 5.89941 |
| **C14orf93** | 3.15817 | 0.83117 | 0.04821 | 3.02742 | 0.15662 | 0.44930 | 0.24804 |
| **C15orf26** | 0.02365 | 0.04701 | 0.11691 | 0.01634 | 0.07274 | 0.02988 | 0.00738 |
| **C15orf27** | 0.08106 | 0.00215 | 0.02153 | 0.03705 | 0.11972 | 0.01152 | 0.00000 |
| **C15orf37** | 0.23339 | 1.24584 | 0.40033 | 0.49949 | 0.03261 | 0.12103 | 0.24570 |
| **C15orf39** | 6.86185 | 7.76364 | 6.86198 | 2.22864 | 0.16365 | 4.14067 | 18.78583 |
| **C15orf40** | 4.51900 | 6.16959 | 4.54622 | 2.58745 | 0.48967 | 1.99673 | 0.00000 |
| **C15orf41** | 7.10755 | 0.11014 | 8.89292 | 3.19728 | 2.53129 | 1.23513 | 0.00000 |
| **C15orf43** | 0.02811 | 0.06088 | 0.08005 | 0.05248 | 0.01126 | 0.01042 | 0.07092 |
| **C15orf48** | 0.00907 | 0.04620 | 0.06507 | 0.09720 | 0.12608 | 0.07141 | 0.08895 |
| **C15orf52** | 0.04276 | 0.06070 | 0.00651 | 0.03886 | 0.02240 | 0.01574 | 0.00376 |
| **C15orf53** | 0.03988 | 0.01446 | 0.01251 | 0.01365 | 0.01429 | 0.01291 | 0.01831 |
| **C15orf56** | 0.04016 | 0.02740 | 0.01100 | 0.06990 | 0.00810 | 0.01142 | 0.00442 |
| **C15orf57** | 5.68290 | 7.21214 | 0.97383 | 9.52072 | 1.15190 | 0.65371 | 0.05091 |
| **C15orf61** | 4.82594 | 5.28648 | 2.55653 | 8.49717 | 0.55029 | 0.80669 | 7.21193 |
| **C15orf62** | 0.00358 | 0.04163 | 0.04214 | 0.14658 | 0.01518 | 0.00327 | 0.12859 |
| **C15orf65** | 0.01465 | 0.00000 | 2.81138 | 0.04945 | 1.01699 | 0.02632 | 0.16230 |
| **C16orf13** | 13.36884 | 11.41923 | 1.03861 | 13.93341 | 0.23162 | 6.17267 | 7.59944 |
| **C16orf3** | 0.04235 | 0.01856 | 0.01460 | 0.01714 | 0.01279 | 0.01079 | 0.01608 |
| **C16orf45** | 0.06690 | 0.04602 | 0.23355 | 0.03138 | 0.03915 | 0.00916 | 0.00000 |
| **C16orf46** | 0.89667 | 0.00000 | 1.75972 | 1.79102 | 2.79314 | 0.30673 | 0.98830 |
| **C16orf47** | 0.27938 | 0.00000 | 0.01381 | 1.04213 | 0.00997 | 0.01178 | 0.01352 |
| **C16orf52** | 0.20472 | 0.00000 | 0.67275 | 0.64768 | 0.08502 | 0.53793 | 2.84680 |
| **C16orf54** | 0.00000 | 1.44263 | 1.17733 | 0.23306 | 0.14754 | 0.00000 | 0.00000 |
| **C16orf58** | 0.00000 | 2.65352 | 0.81150 | 0.29431 | 0.28660 | 0.82309 | 2.28847 |
| **C16orf59** | 4.59310 | 0.00000 | 0.95199 | 4.01165 | 0.28099 | 1.63389 | 1.10735 |
| **C16orf62** | 7.43846 | 5.58779 | 1.94713 | 2.74268 | 1.83464 | 4.59059 | 0.00000 |
| **C16orf70** | 15.47491 | 6.22030 | 7.18330 | 11.77596 | 3.97688 | 4.51133 | 0.00000 |
| **C16orf71** | 0.08066 | 0.43956 | 0.34080 | 0.32428 | 0.39493 | 0.05863 | 0.32959 |
| **C16orf72** | 4.91098 | 0.00000 | 12.60066 | 3.44750 | 5.78403 | 7.44088 | 18.41769 |

| | | | | | | | |
|---|---|---|---|---|---|---|---|
| **C16orf74** | 0.19097 | 1.97442 | 0.00000 | 1.65840 | 3.49448 | 0.00000 | 4.23574 |
| **C16orf80** | 21.93028 | 14.88803 | 12.09617 | 13.54297 | 13.79716 | 8.40998 | 0.00000 |
| **C16orf86** | 0.05856 | 0.05744 | 0.02485 | 0.10620 | 0.01269 | 0.17757 | 0.00000 |
| **C16orf87** | 8.58841 | 0.00000 | 8.43415 | 4.22209 | 1.09124 | 3.20897 | 1.97090 |
| **C16orf89** | 0.00330 | 0.10584 | 0.00491 | 0.00599 | 0.00712 | 0.00453 | 0.14365 |
| **C16orf90** | 0.02457 | 0.01842 | 0.01254 | 0.02630 | 0.00800 | 0.01404 | 0.01763 |
| **C16orf91** | 14.91030 | 17.27779 | 13.11236 | 11.83414 | 13.69578 | 2.48529 | 0.00000 |
| **C16orf93** | 0.03619 | 0.06212 | 0.03288 | 0.11573 | 0.16198 | 0.02460 | 0.01285 |
| **C16orf95** | 0.57096 | 0.86881 | 0.56282 | 0.27588 | 0.27624 | 1.05636 | 1.67861 |
| **C16orf96** | 0.02183 | 0.00430 | 0.03918 | 0.02454 | 0.00683 | 0.01097 | 0.03567 |
| **C17orf100** | 0.35263 | 0.00000 | 0.23188 | 0.16790 | 1.82579 | 0.02050 | 0.03128 |
| **C17orf104** | 1.18056 | 0.00000 | 0.20348 | 0.43624 | 0.60596 | 0.28183 | 1.65839 |
| **C17orf105** | 0.02412 | 0.01684 | 0.01121 | 0.02117 | 0.02483 | 0.01288 | 0.01774 |
| **C17orf107** | 0.00002 | 0.01564 | 0.20570 | 0.13728 | 0.43843 | 0.32402 | 0.28683 |
| **C17orf112** | 0.02165 | 0.02220 | 0.00662 | 0.02194 | 0.14967 | 0.01278 | 0.01468 |
| **C17orf47** | 0.02354 | 0.02964 | 0.04161 | 0.03330 | 0.00640 | 0.01086 | 0.00897 |
| **C17orf51** | 6.86305 | 3.03838 | 5.97564 | 6.44592 | 4.13439 | 4.54348 | 5.25109 |
| **C17orf53** | 6.57130 | 5.96037 | 5.18987 | 8.19829 | 4.41191 | 4.43841 | 0.00000 |
| **C17orf58** | 5.91464 | 1.97019 | 1.04193 | 4.15877 | 0.22941 | 0.70023 | 3.03460 |
| **C17orf59** | 0.00000 | 3.05726 | 0.00000 | 3.71958 | 2.68422 | 2.18476 | 6.40862 |
| **C17orf62** | 4.87999 | 7.62395 | 4.06587 | 8.10885 | 0.24794 | 4.42193 | 5.40108 |
| **C17orf64** | 0.02907 | 0.01419 | 0.02768 | 0.01364 | 0.02995 | 0.02984 | 0.02754 |
| **C17orf67** | 0.02568 | 0.01485 | 0.27790 | 0.02816 | 0.38869 | 0.01325 | 0.00237 |
| **C17orf70** | 1.42949 | 8.09211 | 0.85509 | 2.42965 | 0.12379 | 3.65907 | 6.24645 |
| **C17orf74** | 0.03055 | 0.01937 | 0.01636 | 0.01498 | 0.00836 | 0.01067 | 0.02683 |
| **C17orf75** | 7.25276 | 7.48502 | 0.62813 | 0.00000 | 0.88490 | 1.82978 | 1.64325 |
| **C17orf77** | 0.03415 | 0.01645 | 0.01306 | 0.01727 | 0.00799 | 0.01262 | 0.01602 |
| **C17orf78** | 0.02171 | 0.02106 | 0.01207 | 0.04375 | 0.00809 | 0.01347 | 0.01280 |
| **C17orf80** | 11.97392 | 4.78337 | 12.59681 | 4.79335 | 2.36435 | 4.30615 | 1.07720 |
| **C17orf85** | 4.05984 | 6.78663 | 3.99229 | 8.39254 | 3.26578 | 7.90079 | 17.53045 |
| **C17orf89** | 0.12414 | 12.33512 | 5.99371 | 4.41576 | 0.60375 | 2.63253 | 17.54535 |
| **C17orf96** | 0.00000 | 1.64996 | 0.23864 | 1.66100 | 0.14836 | 1.07479 | 3.42769 |
| **C17orf97** | 0.00000 | 0.07156 | 0.09425 | 0.03436 | 0.01361 | 0.01940 | 0.18998 |
| **C17orf98** | 0.01397 | 0.05694 | 0.07505 | 0.24678 | 0.23419 | 0.03406 | 0.16347 |
| **C17orf99** | 0.01400 | 0.09320 | 0.06111 | 0.01170 | 1.00269 | 0.00532 | 0.04331 |

| | | | | | | | |
|---|---|---|---|---|---|---|---|
| **C18orf21** | 8.99928 | 5.52882 | 10.78981 | 4.69996 | 6.35041 | 3.75764 | 2.28033 |
| **C18orf25** | 11.81381 | 7.88930 | 5.31765 | 6.39946 | 7.17255 | 6.82331 | 3.44516 |
| **C18orf32** | 0.88596 | 1.83339 | 0.19721 | 0.84658 | 0.14152 | 0.26153 | 0.43962 |
| **C18orf42** | 0.02571 | 0.01246 | 0.00529 | 0.01709 | 0.12061 | 0.06178 | 0.01797 |
| **C18orf54** | 5.61747 | 0.74076 | 4.98851 | 3.46781 | 0.36712 | 2.48184 | 2.54001 |
| **C18orf61** | 0.04359 | 0.00710 | 0.04686 | 0.02673 | 0.01510 | 0.04003 | 0.02321 |
| **C18orf63** | 0.02451 | 0.00793 | 0.01801 | 0.02258 | 0.24624 | 0.01472 | 0.01305 |
| **C18orf64+LINC(** | 0.03508 | 0.01446 | 0.03860 | 0.05838 | 0.01263 | 0.01269 | 0.00000 |
| **C18orf8** | 10.58581 | 7.60515 | 2.50991 | 8.68115 | 2.89522 | 2.04179 | 2.75156 |
| **C19orf10** | 33.48563 | 31.95684 | 10.71449 | 27.61640 | 10.05020 | 14.67476 | 0.00000 |
| **C19orf12** | 3.75130 | 3.15202 | 1.68965 | 5.34666 | 0.26633 | 0.92228 | 9.63730 |
| **C19orf18** | 0.06492 | 0.37218 | 0.12237 | 0.11634 | 0.42688 | 0.00000 | 0.03047 |
| **C19orf24** | 14.24697 | 15.87552 | 8.46171 | 16.80588 | 2.05321 | 7.70775 | 16.85929 |
| **C19orf25** | 5.87111 | 10.49076 | 3.78396 | 10.66534 | 0.86087 | 4.43312 | 3.03306 |
| **C19orf26** | 0.02374 | 0.02294 | 0.01637 | 0.01925 | 0.00878 | 0.01265 | 0.01767 |
| **C19orf35** | 0.01277 | 0.00904 | 0.01714 | 0.01208 | 0.00735 | 0.01003 | 0.17021 |
| **C19orf38** | 0.00000 | 0.04554 | 0.80216 | 0.30486 | 0.55839 | 0.00000 | 0.24426 |
| **C19orf40** | 2.49137 | 2.82949 | 0.89435 | 1.89001 | 0.93587 | 0.91621 | 0.00000 |
| **C19orf43** | 0.00000 | 6.84142 | 10.74200 | 5.79229 | 3.19085 | 4.66712 | 35.04883 |
| **C19orf44** | 0.27765 | 0.81509 | 0.40049 | 2.08357 | 0.45245 | 0.15934 | 1.30067 |
| **C19orf45** | 0.02471 | 0.00924 | 0.00597 | 0.01916 | 0.28308 | 0.01659 | 0.04229 |
| **C19orf47** | 0.86882 | 4.06843 | 5.76569 | 9.00092 | 5.24066 | 2.10344 | 7.34622 |
| **C19orf48** | 49.42291 | 49.11972 | 32.39716 | 37.78505 | 16.01893 | 20.42826 | 6.93153 |
| **C19orf52** | 0.00945 | 3.13391 | 0.76028 | 1.45432 | 0.87092 | 0.82117 | 4.64831 |
| **C19orf53** | 27.60556 | 35.43620 | 26.40094 | 29.78909 | 18.36005 | 19.10263 | 20.54185 |
| **C19orf54** | 0.72220 | 6.23175 | 5.35646 | 3.47691 | 0.49596 | 2.58830 | 2.04810 |
| **C19orf57** | 0.17320 | 1.01042 | 0.49303 | 0.00000 | 0.07477 | 1.40825 | 0.30313 |
| **C19orf60** | 3.63313 | 12.62056 | 0.29887 | 3.91098 | 0.27650 | 7.38038 | 8.45145 |
| **C19orf66** | 0.02152 | 0.07310 | 0.00756 | 0.02391 | 0.17600 | 0.02831 | 0.00532 |
| **C19orf67** | 0.01369 | 0.04533 | 0.03218 | 0.03752 | 1.24820 | 0.00000 | 0.09510 |
| **C19orf68** | 0.00000 | 0.01057 | 0.08658 | 0.38115 | 0.00628 | 0.03213 | 0.49389 |
| **C19orf70** | 14.54452 | 16.20669 | 1.92556 | 7.31253 | 4.59360 | 8.73946 | 7.50752 |
| **C19orf71** | 0.01262 | 0.03557 | 0.04364 | 0.08099 | 0.00632 | 0.01204 | 0.06580 |
| **C19orf73** | 0.19428 | 0.91745 | 0.48668 | 0.20126 | 0.80273 | 0.12748 | 0.00000 |
| **C19orf80** | 0.00000 | 0.00559 | 0.03597 | 0.00287 | 0.00524 | 0.20397 | 0.27476 |

| | | | | | | | |
|---|---|---|---|---|---|---|---|
| **C19orf81** | 0.00000 | 0.02336 | 0.11464 | 0.04278 | 0.03351 | 0.05605 | 0.11301 |
| **C19orf82** | 0.06612 | 0.21099 | 0.71446 | 0.10735 | 0.00253 | 0.04015 | 0.04073 |
| **C19orf83** | 0.05326 | 0.31194 | 0.61223 | 0.00512 | 0.82088 | 0.01674 | 0.00000 |
| **C19orf84** | 0.00484 | 0.02712 | 0.04544 | 0.03628 | 0.10185 | 0.11895 | 0.04867 |
| **C1D** | 6.78104 | 0.00000 | 5.11555 | 1.77963 | 2.38411 | 1.70889 | 2.13782 |
| **C1GALT1** | 11.50377 | 0.00000 | 10.93032 | 4.53602 | 5.16110 | 2.29871 | 4.43144 |
| **C1GALT1C1** | 15.80178 | 0.02064 | 2.27207 | 4.79231 | 0.66965 | 4.54757 | 0.93941 |
| **C1QA** | 0.00648 | 0.24645 | 0.12216 | 0.01785 | 0.00417 | 0.01802 | 0.00456 |
| **C1QBP** | 69.00180 | 48.29250 | 49.69012 | 51.76813 | 10.14837 | 28.88403 | 15.39736 |
| **C1QL1** | 0.00104 | 0.00000 | 0.05406 | 0.04403 | 0.07559 | 0.04466 | 0.08094 |
| **C1QL3** | 0.11852 | 0.17198 | 0.00763 | 0.00000 | 0.01341 | 0.00000 | 0.11539 |
| **C1QL4** | 0.01453 | 0.03736 | 0.02767 | 0.02626 | 0.01165 | 0.00965 | 0.03044 |
| **C1QTNF1** | 0.00206 | 0.01999 | 0.00704 | 0.21868 | 0.24143 | 0.01100 | 0.03592 |
| **C1QTNF1-AS1** | 0.03949 | 0.04383 | 0.01284 | 0.01636 | 0.00903 | 0.01266 | 0.00288 |
| **C1QTNF2** | 0.01647 | 0.00512 | 0.31281 | 0.01875 | 0.01094 | 0.00796 | 0.02425 |
| **C1QTNF3** | 0.04506 | 0.01687 | 0.01411 | 0.01747 | 0.00745 | 0.00598 | 0.11204 |
| **C1QTNF3-AMA(** | 0.03000 | 0.01661 | 0.01331 | 0.01452 | 0.00766 | 0.01159 | 0.02726 |
| **C1QTNF4** | 0.00000 | 0.55631 | 0.00000 | 0.57700 | 0.03305 | 0.28219 | 0.52393 |
| **C1QTNF5+MFR** | 0.04739 | 0.00923 | 0.01584 | 0.01751 | 0.01910 | 0.00955 | 0.02594 |
| **C1QTNF6** | 6.95868 | 3.81934 | 1.36006 | 3.49202 | 2.07837 | 0.00000 | 5.38048 |
| **C1QTNF7** | 0.03594 | 0.01690 | 0.01169 | 0.00922 | 0.00827 | 0.01054 | 0.12886 |
| **C1QTNF8** | 0.02297 | 0.01780 | 0.01224 | 0.01996 | 0.01987 | 0.01270 | 0.01900 |
| **C1QTNF9** | 0.00000 | 0.65454 | 0.00000 | 0.02105 | 0.01238 | 0.30248 | 0.03095 |
| **C1QTNF9B** | 0.05127 | 0.02573 | 0.03076 | 0.08466 | 0.01948 | 0.03638 | 0.06563 |
| **C1QTNF9B-AS1** | 0.18248 | 0.24415 | 0.07134 | 0.11679 | 0.04791 | 0.29308 | 0.00000 |
| **C1R** | 0.00000 | 0.33787 | 0.19062 | 0.01039 | 0.00592 | 1.53123 | 0.01592 |
| **C1RL** | 0.03111 | 0.07673 | 0.01143 | 0.02539 | 0.01037 | 0.19058 | 0.01315 |
| **C1RL-AS1** | 0.00013 | 0.04701 | 0.00836 | 0.00641 | 0.00705 | 0.02426 | 0.18957 |
| **C1S** | 0.53595 | 3.29526 | 1.07445 | 0.06234 | 0.33223 | 1.63335 | 0.64184 |
| **C1orf100** | 0.02255 | 0.01471 | 0.01971 | 0.02923 | 0.07798 | 0.01367 | 0.02040 |
| **C1orf101** | 0.01681 | 0.00000 | 0.06481 | 0.08901 | 0.06700 | 0.04986 | 0.08072 |
| **C1orf105** | 0.53135 | 0.00000 | 0.03618 | 1.75572 | 0.00648 | 0.00334 | 0.00213 |
| **C1orf106** | 1.32943 | 5.03695 | 0.00000 | 4.95791 | 0.00000 | 1.08824 | 3.81828 |
| **C1orf109** | 19.68514 | 11.75883 | 7.32589 | 13.03632 | 0.61951 | 7.72485 | 0.00000 |
| **C1orf110** | 0.06686 | 0.00022 | 0.03451 | 0.00653 | 1.11656 | 0.23601 | 0.00507 |

| | | | | | | | |
|---|---|---|---|---|---|---|---|
| **C1orf111** | 0.04213 | 0.05429 | 0.01186 | 0.01552 | 0.00732 | 0.01738 | 0.02854 |
| **C1orf112** | 9.17505 | 1.57273 | 4.35009 | 2.51629 | 0.22631 | 1.79261 | 0.00000 |
| **C1orf115** | 1.57987 | 1.08990 | 0.00000 | 2.56849 | 0.27869 | 0.00000 | 2.68857 |
| **C1orf116** | 0.25554 | 1.09744 | 0.00000 | 0.31299 | 0.08542 | 0.00000 | 0.23132 |
| **C1orf122** | 0.00000 | 4.10553 | 0.79202 | 1.82645 | 0.43283 | 4.85022 | 20.18726 |
| **C1orf123** | 9.65746 | 7.00874 | 4.69776 | 6.79408 | 1.12841 | 2.83307 | 0.59731 |
| **C1orf127** | 0.02043 | 0.00139 | 0.03013 | 0.02412 | 0.17177 | 0.01531 | 0.03432 |
| **C1orf131** | 16.88236 | 11.96706 | 14.24445 | 7.94880 | 2.47265 | 4.75674 | 3.05820 |
| **C1orf140** | 0.03210 | 0.01690 | 0.01279 | 0.02355 | 0.00897 | 0.02520 | 0.00763 |
| **C1orf141** | 0.09707 | 0.00619 | 0.01363 | 0.03241 | 0.01907 | 0.02210 | 0.00000 |
| **C1orf145** | 0.01788 | 0.02089 | 0.01090 | 0.01513 | 0.07086 | 0.01417 | 0.07078 |
| **C1orf146** | 0.05557 | 0.17552 | 0.08502 | 0.24296 | 0.01233 | 0.06203 | 0.09936 |
| **C1orf158** | 0.07419 | 0.00799 | 0.01446 | 0.01937 | 0.00773 | 0.02446 | 0.01151 |
| **C1orf159** | 0.04255 | 2.98064 | 3.35144 | 3.40523 | 0.00000 | 0.91315 | 4.36903 |
| **C1orf162** | 0.01753 | 0.00000 | 2.46585 | 0.01685 | 0.51797 | 0.28494 | 0.26129 |
| **C1orf167** | 0.01695 | 0.05025 | 0.00976 | 0.04599 | 0.00872 | 0.01194 | 0.00759 |
| **C1orf168** | 0.70106 | 0.00000 | 0.00000 | 0.72206 | 0.30120 | 0.00000 | 1.68587 |
| **C1orf174** | 22.55525 | 13.55919 | 7.20393 | 11.11486 | 1.60441 | 7.51851 | 0.00000 |
| **C1orf177** | 0.01818 | 0.00655 | 0.01547 | 0.02427 | 0.00707 | 0.03254 | 0.12278 |
| **C1orf180** | 0.02563 | 0.01561 | 0.01758 | 0.01948 | 0.01964 | 0.01265 | 0.01877 |
| **C1orf186** | 0.00718 | 0.01657 | 0.01323 | 0.06591 | 0.01264 | 0.50106 | 0.02300 |
| **C1orf189** | 0.02821 | 0.02830 | 0.03646 | 0.04910 | 0.00694 | 0.00928 | 0.00837 |
| **C1orf192** | 0.06464 | 0.03995 | 0.10692 | 0.14369 | 0.01228 | 0.08138 | 0.16456 |
| **C1orf194** | 0.01729 | 0.00000 | 0.34497 | 0.01955 | 0.03867 | 0.01966 | 0.02824 |
| **C1orf195** | 0.01278 | 0.03698 | 0.03529 | 0.01544 | 0.00585 | 0.03724 | 0.07093 |
| **C1orf198** | 0.09082 | 1.55911 | 2.90176 | 6.30748 | 0.72067 | 0.00000 | 0.00000 |
| **C1orf200** | 0.02255 | 0.00600 | 0.05273 | 0.01892 | 0.01053 | 0.01202 | 0.02280 |
| **C1orf21** | 2.93479 | 0.00000 | 0.00000 | 9.11981 | 0.38116 | 0.00000 | 3.97174 |
| **C1orf210** | 1.97278 | 0.86590 | 0.00000 | 1.15307 | 2.39150 | 1.31641 | 0.00000 |
| **C1orf213** | 0.21013 | 0.04398 | 0.49369 | 1.48659 | 0.16196 | 0.01742 | 0.00000 |
| **C1orf216** | 1.19755 | 0.34868 | 1.94667 | 1.64868 | 2.28401 | 1.65278 | 2.00471 |
| **C1orf220** | 0.05290 | 0.00000 | 0.08023 | 0.03101 | 0.09073 | 0.00952 | 0.05311 |
| **C1orf226** | 4.76481 | 1.57042 | 0.62136 | 0.69202 | 1.10293 | 0.84860 | 2.12889 |
| **C1orf228** | 0.00000 | 0.19436 | 0.04011 | 0.01429 | 0.03742 | 0.10329 | 0.07814 |
| **C1orf229** | 0.02831 | 0.00600 | 0.03252 | 0.02211 | 0.09256 | 0.01294 | 0.02696 |

| | | | | | | | |
|---|---|---|---|---|---|---|---|
| **C1orf233** | 0.00000 | 2.92916 | 0.43865 | 0.00000 | 0.12075 | 3.11711 | 2.64793 |
| **C1orf234** | 0.02810 | 0.01605 | 0.01706 | 0.05468 | 0.02669 | 0.01560 | 0.00294 |
| **C1orf27** | 13.50321 | 2.96911 | 1.04800 | 13.73353 | 0.12482 | 3.80753 | 5.79678 |
| **C1orf35** | 0.00000 | 15.02527 | 2.37859 | 8.91084 | 0.62962 | 3.01114 | 13.86401 |
| **C1orf43** | 24.85463 | 9.70257 | 15.09529 | 19.39495 | 4.29734 | 11.70301 | 4.03148 |
| **C1orf50** | 1.02411 | 3.03983 | 2.99764 | 1.32839 | 0.43909 | 0.95892 | 2.29661 |
| **C1orf52** | 21.24035 | 8.85173 | 12.88584 | 13.39972 | 13.85568 | 5.88664 | 4.22003 |
| **C1orf53** | 0.04741 | 0.32816 | 0.09881 | 0.17767 | 0.07551 | 0.39467 | 0.17767 |
| **C1orf54** | 0.04303 | 0.07676 | 0.86498 | 0.10554 | 0.09760 | 0.02753 | 0.03038 |
| **C1orf56** | 3.61592 | 0.46829 | 0.03059 | 2.22346 | 1.31378 | 0.42331 | 0.20872 |
| **C1orf61** | 3.22948 | 0.19703 | 0.08963 | 4.40054 | 0.07994 | 0.00000 | 6.74815 |
| **C1orf64** | 0.00000 | 0.90153 | 0.76915 | 0.00000 | 0.44852 | 2.50105 | 0.00739 |
| **C1orf74** | 0.88084 | 1.02501 | 2.56488 | 0.81206 | 0.17176 | 0.12616 | 0.00000 |
| **C1orf85** | 7.90353 | 7.76903 | 2.66531 | 1.86297 | 0.07399 | 4.63287 | 1.42002 |
| **C1orf86** | 0.00000 | 0.76011 | 0.27092 | 3.71699 | 0.25139 | 1.09952 | 4.34413 |
| **C1orf87** | 0.02809 | 0.01747 | 0.01285 | 0.02044 | 0.00873 | 0.01296 | 0.01526 |
| **C1orf95** | 0.02468 | 0.01872 | 0.01184 | 0.01977 | 0.07969 | 0.02815 | 0.01468 |
| **C2** | 0.00000 | 0.99349 | 0.94949 | 0.38842 | 0.03933 | 1.05673 | 0.00000 |
| **C20orf144** | 0.01759 | 0.01545 | 0.01560 | 0.02126 | 0.01840 | 0.01186 | 0.04632 |
| **C20orf166-AS1** | 0.02349 | 0.01661 | 0.01270 | 0.02118 | 0.01983 | 0.01439 | 0.01901 |
| **C20orf173** | 0.02435 | 0.02766 | 0.01301 | 0.03599 | 0.00817 | 0.06119 | 0.00702 |
| **C20orf194** | 1.65548 | 0.83607 | 2.47203 | 0.94607 | 3.47685 | 1.71602 | 0.87377 |
| **C20orf195** | 0.03226 | 0.01098 | 0.01468 | 0.01919 | 0.00763 | 0.02764 | 0.02764 |
| **C20orf196** | 1.90728 | 5.34123 | 3.58171 | 0.85277 | 0.65672 | 1.95720 | 0.82862 |
| **C20orf202** | 0.01523 | 0.04262 | 0.07038 | 0.03918 | 0.00446 | 0.00100 | 0.07479 |
| **C20orf203** | 0.01369 | 0.09936 | 0.00390 | 0.10649 | 0.06083 | 0.00280 | 0.11341 |
| **C20orf24** | 0.00000 | 4.87947 | 0.66769 | 2.55077 | 0.77348 | 3.15952 | 4.43586 |
| **C20orf26** | 0.00896 | 0.17801 | 0.51013 | 0.07186 | 0.23950 | 0.00000 | 0.01599 |
| **C20orf27** | 20.22976 | 15.60182 | 8.27920 | 25.63135 | 0.71260 | 8.98246 | 10.05003 |
| **C20orf96** | 0.65551 | 0.00000 | 1.06423 | 0.61325 | 2.12048 | 0.00000 | 1.74043 |
| **C21orf119** | 0.34988 | 0.71601 | 2.36298 | 0.71696 | 1.29315 | 0.08681 | 0.10950 |
| **C21orf140** | 0.02504 | 0.02935 | 0.01834 | 0.06005 | 0.02234 | 0.00465 | 0.03956 |
| **C21orf15** | 0.01581 | 0.05827 | 0.04408 | 0.03915 | 0.00723 | 0.16613 | 0.14510 |
| **C21orf2** | 0.37503 | 3.10969 | 0.61419 | 1.30206 | 0.00000 | 1.12266 | 4.20113 |
| **C21orf33** | 24.48517 | 24.45643 | 3.31818 | 10.02759 | 0.75844 | 12.02924 | 1.74626 |

| | | | | | | | |
|---|---|---|---|---|---|---|---|
| **C21orf37** | 0.02768 | 0.06817 | 0.01528 | 0.02206 | 0.00753 | 0.00881 | 0.00816 |
| **C21orf49** | 0.09278 | 0.02939 | 0.01153 | 0.04438 | 0.41290 | 0.00971 | 0.00000 |
| **C21orf58** | 0.24982 | 0.95534 | 0.63566 | 2.11708 | 0.17559 | 0.45884 | 3.44837 |
| **C21orf59** | 34.52814 | 19.76059 | 17.31827 | 25.45926 | 3.11655 | 10.01607 | 1.67547 |
| **C21orf62** | 0.04084 | 0.04826 | 0.00790 | 0.00683 | 0.01277 | 0.00521 | 0.05209 |
| **C21orf67** | 0.30762 | 1.05486 | 1.67703 | 0.06762 | 0.01858 | 0.04985 | 0.01053 |
| **C21orf90** | 0.04056 | 0.00192 | 0.03617 | 0.02702 | 0.12483 | 0.02847 | 0.01113 |
| **C21orf91** | 6.44191 | 0.00000 | 9.59201 | 6.54034 | 11.86846 | 2.95038 | 0.84759 |
| **C21orf91-OT1** | 0.05344 | 0.01222 | 0.02473 | 0.01676 | 0.02566 | 0.02632 | 0.03875 |
| **C22orf15** | 0.03838 | 0.06717 | 0.00275 | 0.02539 | 0.05138 | 0.01716 | 0.07107 |
| **C22orf23** | 0.43592 | 0.65913 | 0.40425 | 0.44181 | 0.48446 | 0.02821 | 0.00000 |
| **C22orf24** | 0.01212 | 0.05140 | 0.10151 | 0.04004 | 0.00687 | 0.02494 | 0.03638 |
| **C22orf29** | 8.43735 | 9.24712 | 1.03291 | 6.70568 | 1.36567 | 0.49505 | 0.36790 |
| **C22orf31** | 0.04759 | 0.01112 | 0.01313 | 0.02592 | 0.00769 | 0.01104 | 0.02991 |
| **C22orf34** | 0.13069 | 0.11411 | 0.59900 | 0.14674 | 0.39520 | 0.06368 | 0.26728 |
| **C22orf39** | 6.31983 | 3.55953 | 0.91697 | 7.73319 | 0.44884 | 1.64939 | 1.04135 |
| **C22orf42** | 0.02395 | 0.01725 | 0.01074 | 0.01830 | 0.03615 | 0.01309 | 0.01953 |
| **C22orf46** | 0.00000 | 1.61351 | 3.11674 | 0.80074 | 0.16807 | 0.78655 | 1.56261 |
| **C2CD2** | 3.86673 | 0.63321 | 0.09279 | 0.76519 | 0.00056 | 0.00000 | 6.06501 |
| **C2CD2L** | 0.25243 | 0.39159 | 0.61786 | 1.69843 | 0.70320 | 0.57233 | 2.96248 |
| **C2CD3** | 1.91626 | 2.19745 | 1.35561 | 3.94057 | 2.95571 | 1.78848 | 2.75262 |
| **C2CD4A** | 0.01927 | 0.01487 | 0.01296 | 0.02335 | 0.00858 | 0.09682 | 0.01603 |
| **C2CD4B** | 0.00607 | 0.01977 | 0.01314 | 0.01954 | 0.01169 | 0.18498 | 0.03897 |
| **C2CD4C** | 0.00000 | 0.13942 | 0.09674 | 0.00000 | 0.00548 | 1.07301 | 0.05227 |
| **C2CD4D** | 0.01072 | 0.00774 | 0.01220 | 0.02006 | 0.01154 | 0.00908 | 0.12081 |
| **C2CD5** | 4.36341 | 0.80686 | 0.84618 | 0.07590 | 3.27593 | 4.53531 | 0.01632 |
| **C2orf15** | 1.18186 | 0.00000 | 0.12320 | 0.67871 | 1.55068 | 0.19936 | 0.54492 |
| **C2orf16** | 0.03703 | 0.14996 | 0.12008 | 0.11857 | 2.03645 | 0.17330 | 0.00000 |
| **C2orf27A** | 0.02963 | 0.00000 | 0.06946 | 0.02683 | 0.00700 | 0.01945 | 0.09611 |
| **C2orf40** | 0.00000 | 0.04590 | 0.02184 | 0.06534 | 0.02122 | 0.00533 | 0.07312 |
| **C2orf42** | 2.64024 | 2.24367 | 1.72210 | 2.19503 | 1.55023 | 0.51274 | 0.00000 |
| **C2orf43** | 6.10487 | 0.90258 | 3.28488 | 3.17476 | 0.49280 | 1.33072 | 1.86790 |
| **C2orf44** | 5.84981 | 0.19357 | 5.36952 | 5.17739 | 3.18459 | 1.38391 | 0.61929 |
| **C2orf47** | 18.31403 | 10.84450 | 5.69373 | 8.46361 | 0.85834 | 6.42135 | 0.76542 |
| **C2orf48** | 0.81915 | 0.59211 | 0.09718 | 0.46476 | 0.26643 | 0.30824 | 0.51533 |

| | | | | | | | |
|---|---|---|---|---|---|---|---|
| **C2orf49** | 12.66355 | 7.43045 | 12.08116 | 9.17213 | 7.36592 | 5.79141 | 6.48725 |
| **C2orf50** | 0.00676 | 0.12993 | 0.01470 | 0.03657 | 0.01010 | 0.10950 | 0.00297 |
| **C2orf54** | 0.01092 | 0.01106 | 0.01385 | 0.03395 | 0.00733 | 0.01238 | 0.06253 |
| **C2orf57** | 0.02590 | 0.01737 | 0.00884 | 0.02014 | 0.04457 | 0.02405 | 0.01255 |
| **C2orf61** | 0.06272 | 0.01798 | 0.00936 | 0.02287 | 0.08979 | 0.01204 | 0.02424 |
| **C2orf66** | 0.16964 | 0.03887 | 0.04113 | 0.12420 | 0.08528 | 0.03356 | 0.00000 |
| **C2orf68** | 9.14445 | 16.32146 | 11.46055 | 10.75498 | 2.08851 | 4.76693 | 4.79635 |
| **C2orf69** | 14.47676 | 0.73295 | 13.07533 | 6.00048 | 10.49903 | 4.45203 | 8.90095 |
| **C2orf71** | 0.00872 | 0.03030 | 0.00810 | 0.14403 | 0.08643 | 0.01257 | 0.01036 |
| **C2orf72** | 0.00000 | 0.16944 | 0.00024 | 3.72379 | 0.00846 | 0.45422 | 0.21017 |
| **C2orf74** | 3.85889 | 1.49096 | 4.65722 | 1.45196 | 0.02158 | 1.29852 | 0.00000 |
| **C2orf76** | 2.28549 | 1.14725 | 4.66607 | 2.51236 | 1.76267 | 0.63186 | 0.00000 |
| **C2orf80** | 0.03662 | 0.21428 | 0.55814 | 0.00372 | 0.12046 | 0.24524 | 0.00065 |
| **C2orf81** | 0.06054 | 0.06970 | 0.01410 | 0.13382 | 0.06206 | 0.09307 | 0.00000 |
| **C2orf82** | 0.00773 | 0.01501 | 0.01683 | 0.01643 | 0.00652 | 0.02644 | 0.11302 |
| **C2orf83** | 0.01812 | 0.00718 | 0.00595 | 0.03220 | 0.12157 | 0.06944 | 0.04498 |
| **C2orf88** | 0.02071 | 0.00000 | 0.26489 | 0.49698 | 0.98609 | 0.26539 | 0.00000 |
| **C2orf91** | 0.02432 | 0.04234 | 0.20928 | 0.02497 | 0.01408 | 0.10011 | 0.00632 |
| **C3** | 0.00000 | 0.00000 | 0.58535 | 0.12018 | 2.89342 | 0.14120 | 0.02409 |
| **C3AR1** | 0.00000 | 0.47093 | 1.70942 | 0.05482 | 0.00112 | 0.00000 | 0.04155 |
| **C3P1** | 0.00478 | 0.01059 | 0.01886 | 0.01281 | 0.02704 | 0.02711 | 0.16511 |
| **C3orf14** | 6.69524 | 3.05884 | 6.88656 | 9.69972 | 0.69509 | 3.58449 | 0.00000 |
| **C3orf17** | 10.32552 | 1.32154 | 16.50846 | 5.11036 | 5.80063 | 6.98066 | 5.97522 |
| **C3orf18** | 0.24035 | 0.38482 | 0.23203 | 0.34546 | 0.01572 | 0.26509 | 0.00000 |
| **C3orf20** | 0.06221 | 0.00000 | 0.06738 | 0.22053 | 0.09777 | 0.07873 | 0.04295 |
| **C3orf27** | 0.00134 | 0.11125 | 0.00344 | 0.00665 | 0.00757 | 0.05419 | 0.16134 |
| **C3orf33** | 0.58180 | 0.00000 | 1.65309 | 0.15195 | 0.54308 | 0.18448 | 0.00000 |
| **C3orf35** | 0.04992 | 0.15599 | 0.03452 | 0.14195 | 0.00557 | 0.10591 | 0.33704 |
| **C3orf36** | 0.01378 | 0.03324 | 0.01085 | 0.02125 | 0.00739 | 0.02665 | 0.05188 |
| **C3orf38** | 8.62981 | 0.75273 | 8.33615 | 6.91208 | 1.78561 | 4.01674 | 0.00000 |
| **C3orf43** | 0.02732 | 0.00000 | 0.01151 | 0.01960 | 2.71617 | 0.20410 | 0.01884 |
| **C3orf49** | 0.09638 | 0.00000 | 0.02569 | 0.01548 | 0.05975 | 0.01219 | 0.07170 |
| **C3orf52** | 9.56640 | 3.78039 | 0.45557 | 4.35127 | 0.53441 | 1.48175 | 1.06496 |
| **C3orf55** | 0.12831 | 0.02972 | 0.21823 | 0.01049 | 0.00580 | 0.03221 | 0.00000 |
| **C3orf56** | 0.00000 | 0.83896 | 0.00000 | 0.07589 | 10.40420 | 0.00448 | 0.00000 |

| | | | | | | | |
|---|---|---|---|---|---|---|---|
| **C3orf58** | 4.11276 | 0.00000 | 1.22023 | 0.93017 | 2.96360 | 1.25175 | 3.78522 |
| **C3orf62** | 1.18118 | 0.34023 | 0.35594 | 0.61836 | 0.01904 | 0.00000 | 1.19065 |
| **C3orf65** | 0.03909 | 0.74130 | 0.06337 | 0.22351 | 0.16552 | 0.37159 | 0.00000 |
| **C3orf67** | 0.44442 | 0.54134 | 1.20013 | 0.42762 | 0.35189 | 0.40964 | 0.29243 |
| **C3orf70** | 0.22061 | 0.00000 | 0.57325 | 0.21780 | 0.02800 | 0.38056 | 0.11730 |
| **C3orf80** | 0.12294 | 0.53495 | 0.00000 | 0.14030 | 0.02421 | 0.07765 | 0.48707 |
| **C4A+C4B+C4B-** | 0.01972 | 0.07752 | 0.00919 | 0.23619 | 0.00784 | 0.02289 | 0.00233 |
| **C4A+C4B+C4B-** | 0.00739 | 0.04022 | 0.01378 | 0.30647 | 0.00847 | 0.02861 | 0.00115 |
| **C4BPA** | 0.05784 | 0.00905 | 0.01607 | 0.04494 | 0.00811 | 0.02551 | 0.00000 |
| **C4BPB** | 12.98930 | 0.15593 | 0.00000 | 1.44465 | 0.01019 | 0.00000 | 0.46385 |
| **C4orf19** | 4.35033 | 1.50683 | 0.04199 | 6.52095 | 0.26423 | 0.31551 | 5.72200 |
| **C4orf22** | 0.01300 | 0.06411 | 0.02010 | 0.01951 | 0.14713 | 0.00757 | 0.02218 |
| **C4orf26** | 0.12088 | 0.00492 | 0.01301 | 0.14411 | 0.24099 | 0.01216 | 0.00000 |
| **C4orf27** | 7.39296 | 2.70234 | 6.82112 | 2.34983 | 4.95113 | 3.69213 | 4.13883 |
| **C4orf29** | 4.37092 | 1.13546 | 1.17350 | 2.61891 | 0.64298 | 1.95965 | 1.12160 |
| **C4orf3** | 21.51218 | 19.11012 | 16.73736 | 18.90554 | 0.66539 | 10.99255 | 11.65033 |
| **C4orf32** | 0.00000 | 0.21690 | 1.76566 | 0.08247 | 0.09124 | 0.11066 | 0.36979 |
| **C4orf33** | 3.06443 | 0.00000 | 0.66112 | 0.47365 | 1.34854 | 0.48862 | 0.69560 |
| **C4orf36** | 0.02398 | 0.05361 | 0.11513 | 0.05074 | 0.02970 | 0.01667 | 0.02247 |
| **C4orf45** | 0.05391 | 0.02194 | 0.04116 | 0.05385 | 0.16112 | 0.01433 | 0.10122 |
| **C4orf46** | 12.21257 | 0.74182 | 5.48747 | 6.78838 | 5.01857 | 3.72983 | 1.05864 |
| **C4orf47** | 0.11733 | 0.00000 | 0.20438 | 0.24337 | 1.71733 | 0.22797 | 0.00000 |
| **C4orf48** | 0.00000 | 5.83552 | 0.07481 | 2.98861 | 0.04854 | 1.72857 | 11.70173 |
| **C4orf51** | 0.25752 | 0.06610 | 0.10635 | 0.04688 | 0.13636 | 0.37166 | 0.24037 |
| **C5** | 0.03124 | 0.00000 | 0.17806 | 0.03402 | 0.33630 | 0.01973 | 0.01435 |
| **C5AR1** | 0.30058 | 0.00000 | 0.08711 | 0.24823 | 0.28144 | 0.00000 | 1.08329 |
| **C5orf15** | 19.37212 | 1.94325 | 8.97980 | 8.66874 | 2.51470 | 7.36840 | 0.00000 |
| **C5orf20** | 0.02855 | 0.01882 | 0.01277 | 0.01976 | 0.00801 | 0.01315 | 0.01280 |
| **C5orf22** | 13.93527 | 2.84726 | 5.12091 | 6.01194 | 3.27913 | 5.05193 | 0.00000 |
| **C5orf24** | 4.36540 | 0.00000 | 4.49820 | 4.55238 | 0.93114 | 0.28208 | 8.90327 |
| **C5orf27** | 0.03565 | 0.01425 | 0.02173 | 0.04588 | 0.08714 | 0.01452 | 0.01977 |
| **C5orf28** | 3.15811 | 0.00000 | 2.97007 | 0.94843 | 1.42742 | 1.44626 | 1.16561 |
| **C5orf30** | 0.93217 | 0.00000 | 0.77194 | 9.46667 | 0.36276 | 0.17789 | 0.00000 |
| **C5orf34** | 3.26621 | 2.16739 | 4.38590 | 1.04185 | 0.87436 | 3.42636 | 0.00000 |
| **C5orf38** | 0.00000 | 0.01133 | 0.01191 | 0.16239 | 0.00706 | 0.01973 | 0.11164 |

| | | | | | | | |
|---|---|---|---|---|---|---|---|
| **C5orf42** | 0.00000 | 0.34905 | 0.84436 | 0.21710 | 0.32057 | 2.26503 | 0.42098 |
| **C5orf45** | 0.37069 | 1.02233 | 0.53965 | 0.53738 | 1.15755 | 0.30253 | 0.00000 |
| **C5orf47** | 0.01986 | 0.18394 | 0.08043 | 0.01179 | 0.25850 | 0.16067 | 0.00006 |
| **C5orf49** | 0.00000 | 0.06019 | 0.02604 | 0.10028 | 0.11592 | 0.14278 | 0.10403 |
| **C5orf51** | 13.00739 | 0.00000 | 3.48883 | 11.77176 | 0.44916 | 3.96770 | 7.83561 |
| **C5orf52** | 0.01417 | 0.01419 | 0.14379 | 0.01469 | 0.00346 | 0.02654 | 0.04406 |
| **C5orf55** | 0.00000 | 0.23724 | 1.78042 | 0.06342 | 0.17203 | 0.00656 | 0.06458 |
| **C5orf56** | 0.00732 | 0.01728 | 0.01519 | 0.03624 | 0.00721 | 0.02882 | 0.04925 |
| **C5orf58** | 0.81991 | 0.15111 | 0.38689 | 0.00000 | 0.19092 | 0.22444 | 0.43100 |
| **C5orf60** | 0.01815 | 0.10430 | 0.06956 | 0.02511 | 0.05517 | 0.09972 | 0.00537 |
| **C5orf63** | 0.77354 | 0.14878 | 1.75239 | 1.24207 | 0.13060 | 0.47493 | 0.74966 |
| **C5orf64** | 0.02692 | 0.01390 | 0.02245 | 0.01855 | 0.00723 | 0.01270 | 0.01651 |
| **C5orf66** | 0.12949 | 0.28357 | 0.09909 | 0.24643 | 0.08624 | 0.27408 | 1.73100 |
| **C5orf66-AS1** | 0.00000 | 0.16820 | 0.24264 | 0.13909 | 0.01987 | 0.19116 | 0.03314 |
| **C5orf66-AS2** | 0.02382 | 0.01263 | 0.00911 | 0.02666 | 0.07267 | 0.04556 | 0.00943 |
| **C6orf1** | 0.96288 | 1.63912 | 2.26214 | 4.25743 | 1.18580 | 3.19343 | 0.00000 |
| **C6orf10** | 0.04158 | 0.09714 | 0.44392 | 0.01141 | 0.16186 | 0.34744 | 0.00000 |
| **C6orf100** | 0.03498 | 0.01523 | 0.01025 | 0.06702 | 0.03398 | 0.02860 | 0.04328 |
| **C6orf106** | 9.39988 | 12.57144 | 10.83122 | 13.30946 | 4.08684 | 16.10490 | 24.96780 |
| **C6orf120** | 0.45289 | 0.00001 | 3.73989 | 1.64600 | 1.10594 | 0.91545 | 6.50108 |
| **C6orf123** | 0.01762 | 0.09790 | 0.01838 | 0.04004 | 0.00891 | 0.01409 | 0.00160 |
| **C6orf132** | 4.00327 | 18.89393 | 0.49402 | 6.55793 | 0.00000 | 0.32307 | 20.39097 |
| **C6orf136** | 2.52470 | 7.53660 | 2.23622 | 4.08131 | 1.23036 | 3.77067 | 5.90403 |
| **C6orf141** | 0.00000 | 0.00000 | 0.31371 | 0.07738 | 0.11926 | 0.01682 | 0.36011 |
| **C6orf147** | 2.86597 | 0.69827 | 1.56502 | 4.77727 | 1.34965 | 0.23379 | 0.68502 |
| **C6orf15** | 0.02346 | 0.01769 | 0.00608 | 0.01929 | 0.11289 | 0.01341 | 0.01518 |
| **C6orf163** | 0.35018 | 0.19172 | 0.20222 | 0.04989 | 0.04816 | 0.16752 | 0.46035 |
| **C6orf164** | 0.04848 | 0.00000 | 0.12114 | 0.08991 | 0.00370 | 0.04774 | 0.08325 |
| **C6orf165** | 0.34280 | 0.00000 | 0.72246 | 0.09716 | 0.21240 | 0.00241 | 0.00000 |
| **C6orf201** | 0.43835 | 0.00000 | 0.03036 | 0.01213 | 0.03341 | 0.02012 | 0.14455 |
| **C6orf203** | 3.08572 | 2.63753 | 10.10776 | 0.72331 | 2.31420 | 3.80228 | 1.09812 |
| **C6orf211** | 48.46835 | 9.52369 | 28.41490 | 8.20865 | 2.43561 | 24.85370 | 1.15680 |
| **C6orf222** | 0.03333 | 0.00116 | 0.04185 | 0.02425 | 0.03059 | 0.04978 | 0.00868 |
| **C6orf223** | 0.45221 | 0.00000 | 0.04756 | 0.20803 | 0.00358 | 0.23669 | 0.51723 |
| **C6orf226** | 0.30744 | 1.62141 | 0.67682 | 1.68042 | 1.78239 | 0.44194 | 0.62296 |

| | | | | | | | |
|---|---|---|---|---|---|---|---|
| **C6orf229** | 0.01322 | 0.01252 | 0.02228 | 0.04744 | 0.04415 | 0.01045 | 0.04417 |
| **C6orf25** | 0.00000 | 0.02163 | 0.02227 | 0.17759 | 0.06779 | 0.00698 | 0.27845 |
| **C6orf47** | 2.05101 | 1.92750 | 0.11392 | 3.12719 | 0.12731 | 1.62491 | 1.99320 |
| **C6orf48** | 19.20533 | 16.94095 | 22.45733 | 13.93888 | 9.63299 | 15.26356 | 15.65494 |
| **C6orf52** | 0.21573 | 0.44257 | 0.95578 | 0.53081 | 9.76520 | 0.11032 | 0.24764 |
| **C6orf57** | 0.37361 | 0.47033 | 0.62479 | 0.00000 | 0.12051 | 0.45064 | 0.38235 |
| **C6orf58** | 0.02409 | 0.02448 | 0.00878 | 0.02046 | 0.03488 | 0.02831 | 0.01256 |
| **C6orf62** | 60.57657 | 22.34875 | 17.14877 | 28.59744 | 18.58400 | 20.28369 | 0.00000 |
| **C6orf89** | 6.14161 | 4.10604 | 6.23873 | 4.68977 | 0.87245 | 2.90999 | 2.48119 |
| **C6orf99** | 2.79239 | 0.00000 | 0.56536 | 3.92055 | 0.16359 | 0.52862 | 0.87541 |
| **C7** | 0.04350 | 0.05050 | 0.00889 | 0.01627 | 0.04467 | 0.01657 | 0.02901 |
| **C7orf13** | 0.02504 | 0.00910 | 2.61464 | 0.58210 | 0.43818 | 0.00000 | 0.00000 |
| **C7orf25** | 2.63406 | 0.19570 | 3.93785 | 1.62664 | 0.91280 | 0.56491 | 0.00000 |
| **C7orf26** | 0.00000 | 2.46100 | 1.01314 | 1.93226 | 0.45367 | 1.30404 | 8.59105 |
| **C7orf31** | 0.21791 | 0.00000 | 1.00932 | 0.07070 | 0.25762 | 0.91707 | 0.91260 |
| **C7orf33** | 0.04460 | 0.00000 | 0.12545 | 0.04563 | 0.02544 | 0.05470 | 0.00000 |
| **C7orf43** | 0.12451 | 0.72187 | 0.11588 | 2.74163 | 0.34934 | 1.18946 | 2.97376 |
| **C7orf49** | 6.87815 | 12.83884 | 6.72856 | 5.29792 | 0.97846 | 2.96735 | 0.00000 |
| **C7orf50** | 10.58137 | 17.69432 | 5.33677 | 13.11594 | 0.23469 | 7.39585 | 8.89590 |
| **C7orf55** | 2.49419 | 3.47269 | 1.56653 | 2.33475 | 0.52157 | 1.41166 | 0.38616 |
| **C7orf57** | 0.13115 | 0.21950 | 0.06449 | 0.07314 | 0.00390 | 0.48339 | 0.00066 |
| **C7orf60** | 0.10721 | 0.00000 | 0.45535 | 0.91276 | 1.92425 | 0.46024 | 1.30927 |
| **C7orf61** | 0.03963 | 0.15040 | 0.00572 | 0.01684 | 0.00779 | 0.10793 | 0.00000 |
| **C7orf63** | 0.21837 | 0.00000 | 0.08659 | 0.20318 | 0.08284 | 0.25486 | 0.06810 |
| **C7orf65** | 0.02329 | 0.01935 | 0.01238 | 0.03061 | 0.00782 | 0.02122 | 0.02126 |
| **C7orf66** | 0.02383 | 0.02711 | 0.01075 | 0.01885 | 0.02835 | 0.01611 | 0.01496 |
| **C7orf69** | 0.02945 | 0.01650 | 0.02254 | 0.01799 | 0.04135 | 0.01255 | 0.01369 |
| **C7orf71** | 0.02893 | 0.01451 | 0.01137 | 0.06169 | 0.03188 | 0.02146 | 0.04623 |
| **C7orf73** | 14.05732 | 5.86220 | 10.59556 | 12.24061 | 0.46598 | 6.36143 | 7.20567 |
| **C7orf76** | 0.02254 | 0.01748 | 0.00919 | 0.02246 | 0.05054 | 0.01341 | 0.01757 |
| **C8A** | 0.01672 | 0.01336 | 0.01100 | 0.01110 | 0.00730 | 0.03871 | 0.09261 |
| **C8B** | 0.02517 | 0.00353 | 0.03077 | 0.01854 | 0.00648 | 0.00942 | 0.05324 |
| **C8G** | 0.03063 | 0.25365 | 0.17298 | 0.03836 | 0.00858 | 0.04365 | 0.16553 |
| **C8orf12** | 0.02938 | 0.05496 | 0.00000 | 0.01532 | 2.46965 | 0.04467 | 0.05290 |
| **C8orf22** | 0.02672 | 0.00000 | 0.09037 | 0.01640 | 0.00515 | 0.01127 | 0.02580 |

| | | | | | | | |
|---|---|---|---|---|---|---|---|
| **C8orf31** | 0.93691 | 0.08924 | 0.01358 | 0.71449 | 0.03477 | 0.00121 | 0.00000 |
| **C8orf33** | 23.09906 | 26.11876 | 16.35983 | 12.75416 | 14.37476 | 8.25061 | 0.00000 |
| **C8orf34** | 0.03799 | 0.00000 | 0.13264 | 0.01458 | 0.04905 | 0.02088 | 0.01231 |
| **C8orf37** | 0.95956 | 0.14078 | 0.97893 | 0.62607 | 0.22634 | 0.00000 | 0.00000 |
| **C8orf37-AS1** | 0.00000 | 0.28300 | 2.39303 | 0.01759 | 0.65890 | 0.00000 | 0.02543 |
| **C8orf4** | 0.03501 | 0.01221 | 0.01338 | 0.06267 | 0.00810 | 0.03055 | 0.02909 |
| **C8orf44** | 0.88576 | 0.36587 | 0.77285 | 0.33947 | 0.20968 | 0.05766 | 0.00000 |
| **C8orf44-SGK3** | 0.02487 | 0.01773 | 0.01345 | 0.02315 | 0.00793 | 0.01276 | 0.01517 |
| **C8orf46** | 0.03103 | 0.01567 | 0.00862 | 0.06310 | 0.09691 | 0.01211 | 0.01591 |
| **C8orf48** | 0.07695 | 0.00000 | 0.08300 | 0.09967 | 0.04489 | 0.00829 | 0.00000 |
| **C8orf49** | 0.01944 | 0.01272 | 0.01200 | 0.02963 | 0.00800 | 0.14913 | 0.01800 |
| **C8orf58** | 0.00000 | 0.00413 | 0.00000 | 1.27973 | 0.65679 | 0.34807 | 0.77105 |
| **C8orf59** | 22.76201 | 0.11631 | 11.67448 | 12.82575 | 0.92227 | 7.52046 | 16.41312 |
| **C8orf76** | 3.87909 | 1.49790 | 3.21601 | 2.34151 | 0.66992 | 3.50898 | 3.25110 |
| **C8orf82** | 0.00000 | 0.32625 | 0.08425 | 0.30938 | 0.51449 | 0.50671 | 3.57271 |
| **C8orf88** | 0.21149 | 0.00000 | 0.39431 | 0.28526 | 0.22875 | 0.14835 | 0.00000 |
| **C9** | 0.07541 | 0.02488 | 0.03266 | 0.07453 | 0.01657 | 0.01445 | 0.04883 |
| **C9orf114** | 10.58535 | 13.61313 | 7.00897 | 9.27144 | 3.87773 | 5.73726 | 4.77921 |
| **C9orf116** | 0.35446 | 1.68606 | 0.79599 | 0.00000 | 0.00200 | 0.05536 | 0.03590 |
| **C9orf117** | 0.06436 | 0.12425 | 0.14523 | 0.16480 | 0.02792 | 0.08138 | 0.19204 |
| **C9orf129** | 0.02829 | 0.02815 | 0.03921 | 0.02850 | 0.05134 | 0.06276 | 0.02362 |
| **C9orf131** | 0.02238 | 0.09514 | 0.04715 | 0.09451 | 0.02242 | 0.02698 | 0.26806 |
| **C9orf135** | 0.27801 | 0.36622 | 0.00000 | 0.00000 | 0.09418 | 4.51405 | 0.50280 |
| **C9orf135-AS1** | 0.06081 | 0.10068 | 0.02231 | 0.00640 | 0.00793 | 0.00480 | 0.00768 |
| **C9orf139** | 0.02675 | 0.01825 | 0.01264 | 0.01824 | 0.00809 | 0.01319 | 0.01794 |
| **C9orf142** | 12.24701 | 6.27373 | 3.12310 | 13.66207 | 0.06435 | 1.09878 | 19.54166 |
| **C9orf152** | 0.00000 | 0.00000 | 0.07418 | 0.40140 | 0.00962 | 0.02819 | 0.23394 |
| **C9orf153** | 0.09993 | 0.12962 | 0.11390 | 0.10001 | 0.18974 | 0.10203 | 0.00000 |
| **C9orf156** | 8.17801 | 5.11822 | 4.27907 | 5.13134 | 0.98103 | 1.45027 | 0.00000 |
| **C9orf16** | 3.34757 | 9.29867 | 4.25978 | 10.35147 | 1.74486 | 3.21548 | 6.51460 |
| **C9orf163** | 0.01381 | 0.05776 | 0.02668 | 0.14556 | 0.00624 | 0.00934 | 0.01063 |
| **C9orf170** | 0.02734 | 0.01871 | 0.01230 | 0.02512 | 0.00790 | 0.01162 | 0.01875 |
| **C9orf171** | 0.04700 | 0.02083 | 0.05539 | 0.01937 | 0.28233 | 0.10061 | 0.02886 |
| **C9orf172** | 0.00079 | 0.03237 | 0.01111 | 0.02191 | 0.01598 | 0.04614 | 0.12053 |
| **C9orf173** | 0.00000 | 0.12048 | 0.03493 | 0.02988 | 0.00791 | 0.06980 | 0.01955 |

| | | | | | | | |
|---|---|---|---|---|---|---|---|
| **C9orf24** | 0.04063 | 0.00771 | 0.01271 | 0.10213 | 0.03054 | 0.01975 | 0.02882 |
| **C9orf3** | 7.35551 | 12.17234 | 2.80614 | 9.66893 | 2.06496 | 2.38553 | 2.42351 |
| **C9orf37** | 2.90871 | 5.83409 | 4.33462 | 5.93703 | 1.80480 | 0.80329 | 0.00000 |
| **C9orf40** | 0.99028 | 0.43379 | 2.78527 | 0.81209 | 0.12895 | 1.20201 | 9.18734 |
| **C9orf41** | 1.96428 | 0.27932 | 0.68586 | 0.70036 | 0.18927 | 1.00799 | 2.15865 |
| **C9orf43** | 0.17740 | 0.20551 | 0.32764 | 0.18569 | 0.30824 | 0.01896 | 0.45408 |
| **C9orf47** | 0.00000 | 0.05714 | 0.34225 | 0.05255 | 0.41586 | 0.27358 | 0.09916 |
| **C9orf53** | 0.02029 | 0.01326 | 0.01866 | 0.03823 | 0.02050 | 0.01398 | 0.01624 |
| **C9orf64** | 10.44888 | 5.52915 | 12.76367 | 3.93318 | 2.86500 | 3.23437 | 0.00000 |
| **C9orf66** | 0.04655 | 0.00000 | 0.05367 | 0.08526 | 0.06699 | 0.06613 | 0.11949 |
| **C9orf69** | 0.18278 | 4.47565 | 0.17653 | 1.43614 | 0.01913 | 2.20908 | 11.92001 |
| **C9orf72** | 2.15826 | 0.11586 | 2.61102 | 3.54404 | 1.52792 | 2.32207 | 0.00000 |
| **C9orf78** | 31.27505 | 22.01326 | 14.85539 | 21.64185 | 8.13622 | 12.32242 | 6.53155 |
| **C9orf84** | 0.24224 | 0.00000 | 1.20092 | 0.00000 | 0.10530 | 0.04134 | 0.48923 |
| **C9orf85** | 4.95494 | 1.81676 | 3.47236 | 3.74233 | 5.89437 | 2.13041 | 0.70088 |
| **C9orf89** | 0.52028 | 2.11887 | 0.32671 | 2.84482 | 0.12616 | 3.30370 | 1.46695 |
| **C9orf9** | 0.02610 | 0.70087 | 0.65180 | 0.26827 | 0.06010 | 0.09901 | 0.28501 |
| **C9orf91** | 0.83633 | 0.00000 | 0.07131 | 0.10283 | 1.06955 | 0.28025 | 1.45413 |
| **C9orf92** | 0.02106 | 0.00544 | 0.05126 | 0.02291 | 0.07224 | 0.01200 | 0.03792 |
| **CA1** | 0.02640 | 0.00000 | 0.06455 | 0.02535 | 0.04216 | 0.02590 | 0.01599 |
| **CA10** | 0.02524 | 0.01951 | 0.00223 | 0.08026 | 0.29541 | 0.01302 | 0.00006 |
| **CA11** | 0.09861 | 0.06991 | 0.06187 | 0.02699 | 0.10545 | 0.12883 | 0.28369 |
| **CA12** | 21.99291 | 0.00000 | 0.00000 | 16.41060 | 0.38543 | 0.00000 | 9.85005 |
| **CA13** | 0.07809 | 0.01528 | 0.11387 | 0.14640 | 0.09657 | 0.98314 | 0.02240 |
| **CA14** | 0.08308 | 0.00000 | 0.36800 | 0.09452 | 0.06418 | 0.02183 | 0.11308 |
| **CA2** | 45.17930 | 28.83875 | 14.27613 | 22.92033 | 0.00000 | 15.72291 | 0.00000 |
| **CA3** | 0.02729 | 0.01753 | 0.01301 | 0.01576 | 0.00769 | 0.02235 | 0.02498 |
| **CA4** | 0.02058 | 0.00668 | 0.00770 | 0.02258 | 0.00705 | 3.78125 | 0.00893 |
| **CA5A** | 0.02134 | 0.01617 | 0.01077 | 0.02114 | 0.04068 | 0.01747 | 0.02109 |
| **CA5B** | 3.27101 | 0.00000 | 1.66533 | 1.45305 | 0.65919 | 0.56791 | 0.10577 |
| **CA5BP1** | 4.87767 | 3.16385 | 3.67050 | 4.96185 | 1.18194 | 1.67873 | 0.00000 |
| **CA6** | 0.01434 | 0.01979 | 0.00000 | 0.02082 | 0.98144 | 0.06718 | 0.01259 |
| **CA7** | 0.01389 | 0.06016 | 0.00916 | 0.01956 | 0.00773 | 0.01539 | 0.03258 |
| **CA8** | 2.60902 | 0.24639 | 0.50464 | 1.06325 | 0.09100 | 0.29440 | 0.00000 |
| **CA9** | 0.02813 | 0.30989 | 0.00581 | 0.04316 | 0.03628 | 0.01700 | 0.00000 |

| | | | | | | | |
|---|---|---|---|---|---|---|---|
| **CAAP1** | 14.55551 | 1.20867 | 8.32728 | 4.40009 | 3.28529 | 6.89814 | 1.76444 |
| **CAB39** | 4.72482 | 1.24233 | 1.10660 | 2.72236 | 0.76903 | 8.96371 | 13.44037 |
| **CAB39L** | 3.51250 | 0.00000 | 6.11278 | 4.24743 | 5.15553 | 0.45254 | 2.07154 |
| **CABIN1** | 3.62339 | 3.10018 | 2.47026 | 6.87251 | 1.96556 | 6.25947 | 14.03909 |
| **CABLES1** | 0.13188 | 0.72810 | 0.28812 | 0.19545 | 0.06942 | 1.34153 | 0.94188 |
| **CABLES2** | 0.00000 | 0.08117 | 0.05485 | 0.49139 | 0.33438 | 0.16933 | 1.04199 |
| **CABP1** | 0.00000 | 0.03024 | 0.26943 | 0.01722 | 0.04299 | 0.08343 | 0.06544 |
| **CABP4** | 0.05936 | 0.11825 | 0.08617 | 0.02973 | 0.05229 | 0.05055 | 0.04861 |
| **CABP5** | 0.01961 | 0.00919 | 0.01455 | 0.02037 | 0.08173 | 0.01265 | 0.03837 |
| **CABP7** | 0.12846 | 0.26059 | 0.48815 | 0.24019 | 1.51975 | 0.01059 | 0.00000 |
| **CABYR** | 1.35819 | 3.21912 | 4.82634 | 2.23990 | 5.96541 | 0.41035 | 0.00000 |
| **CACFD1** | 0.00000 | 1.26473 | 1.42364 | 1.22307 | 0.39632 | 2.26042 | 0.10753 |
| **CACHD1** | 4.84331 | 3.01692 | 3.83299 | 0.00000 | 0.00000 | 4.49911 | 3.40941 |
| **CACNA1A** | 0.00939 | 0.00000 | 0.21321 | 0.07786 | 0.10219 | 0.10057 | 0.71117 |
| **CACNA1B** | 0.00000 | 2.02037 | 1.49549 | 0.00000 | 0.00000 | 1.46606 | 1.18187 |
| **CACNA1C** | 0.02354 | 0.07755 | 0.00000 | 0.02760 | 1.14643 | 0.02374 | 0.01457 |
| **CACNA1C-AS1** | 0.02285 | 0.01689 | 0.00880 | 0.01857 | 0.05917 | 0.01322 | 0.01989 |
| **CACNA1C-AS2** | 0.02563 | 0.01305 | 0.01730 | 0.03638 | 0.06121 | 0.01225 | 0.01395 |
| **CACNA1D** | 0.02298 | 0.00000 | 0.33899 | 0.03041 | 1.45415 | 0.04909 | 0.03115 |
| **CACNA1E** | 0.00000 | 0.01180 | 0.00754 | 0.03927 | 0.59424 | 0.24055 | 0.07354 |
| **CACNA1F** | 0.05282 | 0.01227 | 0.02792 | 0.02756 | 0.02623 | 0.01163 | 0.06515 |
| **CACNA1G** | 0.02237 | 0.01963 | 0.00946 | 0.02192 | 0.10870 | 0.02070 | 0.02563 |
| **CACNA1H** | 0.00000 | 0.28994 | 0.08835 | 0.00000 | 0.83057 | 0.15723 | 0.22331 |
| **CACNA1I** | 0.00000 | 0.04521 | 0.00754 | 0.06077 | 0.06258 | 0.06412 | 0.06948 |
| **CACNA1S** | 0.02664 | 0.02051 | 0.04615 | 0.01737 | 0.01630 | 0.05913 | 0.04491 |
| **CACNA2D1** | 0.00000 | 0.11781 | 0.69295 | 0.00000 | 2.27791 | 1.07593 | 0.02517 |
| **CACNA2D2** | 0.00000 | 1.39067 | 0.18870 | 0.00000 | 0.77593 | 6.63925 | 0.70100 |
| **CACNA2D3** | 1.87626 | 1.48669 | 0.00000 | 0.86102 | 0.28425 | 1.43820 | 0.66007 |
| **CACNA2D3-AS1** | 0.01590 | 0.03618 | 0.00909 | 0.01715 | 0.00796 | 0.04240 | 0.05106 |
| **CACNA2D4** | 0.02165 | 0.05386 | 0.01111 | 0.02056 | 0.01165 | 0.01608 | 0.07200 |
| **CACNB1** | 0.00000 | 0.07143 | 0.20616 | 0.30986 | 0.38496 | 0.79098 | 0.01255 |
| **CACNB2** | 8.71329 | 0.67613 | 0.81298 | 0.00000 | 0.41111 | 0.00000 | 1.94622 |
| **CACNB3** | 0.00000 | 2.42601 | 9.02320 | 0.71119 | 0.25615 | 6.53570 | 0.43319 |
| **CACNB4** | 0.00000 | 0.03173 | 0.10685 | 0.07287 | 0.09025 | 0.17285 | 0.02628 |
| **CACNG1** | 0.02341 | 0.02002 | 0.00800 | 0.02323 | 0.05791 | 0.01351 | 0.01336 |

| | | | | | | | |
|---|---|---|---|---|---|---|---|
| **CACNG2** | 0.05750 | 0.03609 | 0.03652 | 0.00000 | 0.02109 | 0.03349 | 0.04690 |
| **CACNG3** | 0.03546 | 0.01518 | 0.01118 | 0.01655 | 0.03333 | 0.03550 | 0.00618 |
| **CACNG4** | 0.04759 | 0.11084 | 0.01349 | 0.01858 | 0.00802 | 0.00210 | 0.05128 |
| **CACNG5** | 0.02448 | 0.01451 | 0.00996 | 0.01800 | 0.09435 | 0.01358 | 0.02061 |
| **CACNG6** | 0.02597 | 0.01967 | 0.01357 | 0.01837 | 0.00789 | 0.01782 | 0.01470 |
| **CACNG7** | 0.01563 | 0.01866 | 0.02053 | 0.01747 | 0.26364 | 0.02914 | 0.03382 |
| **CACNG8** | 0.00415 | 0.44008 | 0.04876 | 0.00000 | 0.58423 | 0.00634 | 0.30470 |
| **CACTIN** | 0.19796 | 3.01969 | 0.79752 | 0.92767 | 2.96223 | 1.75692 | 7.34984 |
| **CACTIN-AS1** | 0.01586 | 0.03855 | 0.01841 | 0.02671 | 0.01621 | 0.02315 | 0.02600 |
| **CACUL1** | 24.67618 | 16.60535 | 10.48494 | 14.46916 | 11.35051 | 12.53285 | 2.86437 |
| **CACYBP** | 43.83745 | 18.02487 | 11.38211 | 31.72034 | 9.36553 | 15.96481 | 5.93730 |
| **CAD** | 13.17642 | 5.37299 | 7.70408 | 12.12390 | 0.53360 | 7.31349 | 27.16121 |
| **CADM1** | 0.79375 | 11.03403 | 2.83048 | 11.89121 | 4.75599 | 13.77048 | 0.00000 |
| **CADM2** | 0.00000 | 0.33192 | 0.23002 | 0.06254 | 0.10300 | 0.48948 | 0.01648 |
| **CADM2-AS2** | 0.00583 | 0.01877 | 0.01127 | 0.03445 | 0.01038 | 0.01157 | 0.05685 |
| **CADM3** | 0.00521 | 0.02634 | 0.00904 | 0.17085 | 0.00780 | 0.07346 | 0.03557 |
| **CADM4** | 0.05053 | 0.02028 | 0.11063 | 0.01297 | 0.05248 | 0.11125 | 0.15527 |
| **CADPS** | 0.02052 | 0.07193 | 0.00000 | 0.06254 | 0.30929 | 0.03460 | 0.02170 |
| **CADPS2** | 0.26642 | 0.02035 | 0.01241 | 0.02216 | 0.97374 | 0.27963 | 0.43486 |
| **CAGE1** | 0.08405 | 0.09795 | 0.01400 | 0.07329 | 0.12209 | 0.00320 | 0.00000 |
| **CAHM** | 0.03253 | 0.03581 | 0.03918 | 0.03264 | 0.05226 | 0.02204 | 0.12687 |
| **CALB1** | 2.51032 | 0.00000 | 0.02408 | 0.00140 | 0.18765 | 6.32118 | 0.03008 |
| **CALB2** | 0.26956 | 0.00000 | 0.00166 | 5.89212 | 0.05530 | 0.00000 | 0.15650 |
| **CALCA** | 0.03787 | 0.00658 | 0.00969 | 0.01187 | 0.03995 | 0.14586 | 0.01679 |
| **CALCB** | 0.06482 | 0.00516 | 0.00000 | 0.36806 | 0.41315 | 0.00832 | 0.07529 |
| **CALCOCO1** | 2.07049 | 2.04176 | 2.34657 | 8.89497 | 2.29289 | 0.76429 | 0.00000 |
| **CALCOCO2** | 17.40294 | 0.31482 | 8.60192 | 8.61993 | 7.55235 | 8.97284 | 1.66409 |
| **CALCR** | 0.03554 | 0.02605 | 0.00350 | 0.02116 | 0.10800 | 0.43850 | 0.00267 |
| **CALCRL** | 0.04297 | 0.00532 | 0.02272 | 0.01750 | 0.00948 | 0.01090 | 0.05478 |
| **CALD1** | 22.43668 | 1.32405 | 0.00000 | 13.59969 | 0.00000 | 0.60016 | 11.93563 |
| **CALHM1** | 0.00000 | 0.00000 | 0.05770 | 0.03868 | 0.01423 | 0.04660 | 0.14592 |
| **CALHM2** | 0.00000 | 0.35596 | 0.13751 | 1.48306 | 0.01555 | 0.72501 | 0.00000 |
| **CALHM3** | 0.00000 | 0.09129 | 0.00000 | 0.06574 | 0.08697 | 0.29353 | 0.00301 |
| **CALM1** | 53.70988 | 42.55006 | 13.65514 | 43.14923 | 17.07751 | 26.19006 | 38.26619 |
| **CALM2** | 98.91617 | 42.02032 | 29.77044 | 41.07984 | 14.71935 | 33.20475 | 31.80628 |

| | | | | | | | |
|---|---|---|---|---|---|---|---|
| **CALM3** | 16.70841 | 37.06610 | 14.08991 | 12.81420 | 10.68395 | 20.59694 | 22.02534 |
| **CALML3** | 0.00000 | 0.01656 | 0.01355 | 0.29075 | 0.02030 | 0.02413 | 0.18255 |
| **CALML3-AS1** | 0.00198 | 0.01396 | 0.01294 | 0.12009 | 0.01123 | 0.02487 | 0.03939 |
| **CALML4** | 1.90715 | 1.02119 | 0.51443 | 0.67163 | 2.80181 | 1.13477 | 6.66833 |
| **CALML5** | 0.00080 | 0.81513 | 0.00295 | 0.55905 | 0.01021 | 0.00213 | 0.00010 |
| **CALML6** | 0.02208 | 0.01125 | 0.01774 | 0.01129 | 0.01798 | 0.00724 | 0.11759 |
| **CALN1** | 0.28658 | 0.37274 | 0.86961 | 0.00000 | 1.82040 | 0.00000 | 0.00000 |
| **CALR** | 42.18681 | 40.78801 | 9.46935 | 44.96725 | 17.58353 | 24.52857 | 26.73685 |
| **CALR3** | 0.03144 | 0.10027 | 0.32162 | 0.06065 | 0.05630 | 0.00084 | 0.00000 |
| **CALU** | 39.49004 | 6.72206 | 16.08701 | 26.32726 | 12.53258 | 17.05235 | 9.05375 |
| **CALY** | 0.01414 | 0.03848 | 0.00852 | 0.02190 | 0.00699 | 0.03065 | 0.12877 |
| **CAMK1** | 0.03499 | 0.12467 | 0.04750 | 0.11693 | 0.05111 | 0.14550 | 0.62298 |
| **CAMK1D** | 0.01363 | 0.10766 | 0.14547 | 0.00582 | 0.50370 | 0.11712 | 0.16425 |
| **CAMK2A** | 0.03885 | 0.00000 | 0.03176 | 0.01569 | 2.15212 | 0.05616 | 0.01515 |
| **CAMK2B** | 0.01815 | 0.00349 | 0.02742 | 0.03341 | 0.03254 | 0.09686 | 0.27746 |
| **CAMK2D** | 6.89599 | 0.00000 | 0.00000 | 0.36412 | 0.00135 | 1.82222 | 2.40581 |
| **CAMK2G** | 14.90867 | 6.34660 | 0.62333 | 13.16395 | 4.61314 | 1.27647 | 7.92767 |
| **CAMK2N1** | 0.60381 | 0.98502 | 0.00000 | 0.00000 | 0.06554 | 0.00000 | 1.34646 |
| **CAMK2N2** | 0.00000 | 0.31359 | 0.36491 | 0.47841 | 0.13655 | 0.00000 | 0.32188 |
| **CAMK4** | 0.02509 | 0.00000 | 0.10472 | 0.01501 | 0.26712 | 0.05078 | 0.02466 |
| **CAMKK1** | 0.60232 | 0.00368 | 0.13749 | 0.17062 | 0.00152 | 0.27969 | 0.19613 |
| **CAMKK2** | 3.27134 | 7.41155 | 1.60187 | 3.70209 | 0.14043 | 0.00000 | 5.66175 |
| **CAMKMT** | 2.42584 | 1.70017 | 1.55238 | 1.44011 | 1.08147 | 0.84599 | 1.38217 |
| **CAMKV** | 0.13279 | 0.13502 | 0.19735 | 0.01996 | 0.03033 | 3.93052 | 0.00000 |
| **CAMLG** | 10.94335 | 1.69784 | 6.80141 | 11.99478 | 3.58009 | 11.69571 | 7.91222 |
| **CAMP** | 0.56255 | 1.78292 | 0.46035 | 0.23946 | 0.01103 | 1.91545 | 3.12932 |
| **CAMSAP1** | 4.32862 | 4.65852 | 2.59990 | 2.70742 | 3.02202 | 3.97533 | 7.26644 |
| **CAMSAP2** | 4.83359 | 0.00000 | 4.35243 | 6.68636 | 0.24234 | 1.73495 | 15.80520 |
| **CAMSAP3** | 0.21532 | 3.51089 | 0.00000 | 0.00000 | 0.04469 | 0.43441 | 5.45428 |
| **CAMTA1** | 6.92473 | 5.53023 | 7.10104 | 8.04410 | 6.97157 | 8.13140 | 3.72025 |
| **CAMTA2** | 1.30901 | 4.24709 | 1.20788 | 4.22041 | 6.03301 | 1.92384 | 10.34522 |
| **CAND1** | 17.90090 | 1.92900 | 9.46517 | 10.66534 | 0.34404 | 9.16107 | 17.20703 |
| **CAND1.11** | 0.05118 | 0.01463 | 0.12743 | 0.05420 | 0.03307 | 0.01613 | 0.02168 |
| **CAND2** | 0.00000 | 0.00000 | 2.38015 | 0.24004 | 0.15686 | 0.06066 | 0.83321 |
| **CANT1** | 1.81922 | 4.23058 | 2.57455 | 6.06201 | 1.24054 | 1.61807 | 9.58277 |

| | | | | | | | |
|---|---|---|---|---|---|---|---|
| **CANX** | 67.92490 | 15.42384 | 18.24182 | 39.07824 | 9.29795 | 29.15984 | 28.13642 |
| **CAP1** | 38.47656 | 26.61543 | 14.31696 | 35.03087 | 5.47506 | 22.09183 | 12.51684 |
| **CAP2** | 0.80324 | 0.28785 | 0.00000 | 0.63472 | 0.75955 | 0.06139 | 0.55876 |
| **CAPG** | 11.09672 | 3.58017 | 0.00000 | 2.67619 | 0.01492 | 20.24202 | 1.24045 |
| **CAPN1** | 5.86885 | 11.62786 | 0.41865 | 17.40223 | 0.00000 | 9.32779 | 17.81924 |
| **CAPN10** | 0.07041 | 0.53986 | 0.02694 | 3.45213 | 0.23059 | 1.35760 | 4.97678 |
| **CAPN10-AS1** | 0.66063 | 0.73212 | 0.54956 | 1.63003 | 0.15282 | 0.00000 | 1.49140 |
| **CAPN11** | 0.02614 | 0.01846 | 0.01336 | 0.02372 | 0.00781 | 0.01537 | 0.01858 |
| **CAPN12** | 0.42544 | 0.18925 | 0.25308 | 0.53100 | 1.86421 | 0.79250 | 1.54420 |
| **CAPN13** | 0.00000 | 0.00000 | 0.51591 | 0.04366 | 8.37455 | 0.39955 | 0.03724 |
| **CAPN14** | 0.02504 | 0.01886 | 0.01530 | 0.01858 | 0.01381 | 0.01667 | 0.01642 |
| **CAPN15** | 0.00065 | 4.06991 | 0.34027 | 3.25740 | 0.00000 | 1.37463 | 18.15588 |
| **CAPN2** | 13.97350 | 3.07277 | 0.00000 | 7.90910 | 0.19608 | 3.88250 | 14.92034 |
| **CAPN3** | 0.31369 | 0.00000 | 0.82456 | 0.16246 | 1.38210 | 0.17301 | 0.24693 |
| **CAPN5** | 0.00000 | 3.98986 | 0.04632 | 0.21090 | 0.12645 | 11.92780 | 1.58576 |
| **CAPN6** | 0.00000 | 0.02037 | 0.01405 | 1.64850 | 0.00947 | 0.01576 | 0.00182 |
| **CAPN7** | 4.99026 | 1.72864 | 0.22458 | 4.02403 | 0.72182 | 1.48565 | 2.58960 |
| **CAPN8** | 0.02076 | 0.02137 | 0.01029 | 0.02120 | 0.02826 | 0.01030 | 0.03465 |
| **CAPN9** | 0.01533 | 0.02210 | 0.01125 | 0.02493 | 0.00786 | 0.02673 | 0.02844 |
| **CAPNS1** | 1.93515 | 8.92579 | 4.31132 | 3.24350 | 0.35696 | 10.39033 | 21.39763 |
| **CAPNS2** | 0.02765 | 0.03275 | 0.01258 | 0.02999 | 0.00751 | 0.00987 | 0.01864 |
| **CAPRIN1** | 38.24169 | 10.92238 | 12.61339 | 26.88790 | 6.06828 | 14.08745 | 12.26521 |
| **CAPRIN2** | 0.41924 | 0.44775 | 1.29183 | 0.17792 | 1.92623 | 1.95135 | 1.28122 |
| **CAPS** | 0.04972 | 0.15688 | 0.07162 | 0.32723 | 0.01855 | 0.07019 | 0.35966 |
| **CAPS2** | 0.05639 | 0.01736 | 0.01614 | 0.13598 | 0.00794 | 0.02275 | 0.03544 |
| **CAPZA1** | 43.16988 | 11.34494 | 13.64216 | 25.41971 | 4.64797 | 15.15648 | 4.50401 |
| **CAPZA2** | 29.98482 | 2.92858 | 6.18635 | 11.76255 | 0.94490 | 9.32609 | 0.00000 |
| **CAPZB** | 17.46852 | 17.08222 | 8.16326 | 21.52168 | 2.02217 | 16.44135 | 19.51018 |
| **CARD10** | 0.00000 | 4.84147 | 0.00181 | 1.84499 | 0.00468 | 2.24976 | 4.59439 |
| **CARD11** | 0.00000 | 0.11106 | 0.08210 | 0.03410 | 0.00781 | 0.76430 | 0.15960 |
| **CARD14** | 0.07419 | 0.29493 | 0.06110 | 0.34082 | 0.00545 | 0.03698 | 0.35466 |
| **CARD16** | 0.01572 | 0.02900 | 0.01105 | 0.03990 | 0.00819 | 0.04618 | 0.01889 |
| **CARD17** | 0.02625 | 0.01566 | 0.01316 | 0.02394 | 0.00786 | 0.01278 | 0.01901 |
| **CARD18** | 0.14525 | 0.01652 | 0.05817 | 0.06418 | 0.00553 | 0.01611 | 0.08928 |
| **CARD6** | 0.02632 | 0.01802 | 0.00002 | 0.02978 | 0.24514 | 0.02556 | 0.01964 |

| | | | | | | | |
|---|---|---|---|---|---|---|---|
| **CARD8** | 2.80701 | 0.00000 | 0.63062 | 1.71799 | 0.02131 | 0.32965 | 0.85513 |
| **CARD9** | 0.01580 | 0.00000 | 0.45452 | 0.27638 | 0.00257 | 0.00000 | 1.07488 |
| **CARF** | 0.07452 | 0.00000 | 0.60724 | 0.27577 | 0.34727 | 0.09159 | 0.12707 |
| **CARHSP1** | 12.77036 | 10.88919 | 1.53683 | 22.82336 | 6.21852 | 18.55519 | 6.89029 |
| **CARKD** | 12.21852 | 6.32645 | 4.39080 | 9.55460 | 4.74246 | 10.91951 | 2.01869 |
| **CARM1** | 0.02023 | 8.17027 | 2.40776 | 6.67029 | 1.27835 | 4.04881 | 16.91942 |
| **CARNS1** | 0.14352 | 0.01009 | 0.01657 | 0.00939 | 0.07192 | 0.02504 | 0.01580 |
| **CARS** | 23.93987 | 7.02197 | 4.40100 | 18.99965 | 0.86368 | 7.35480 | 8.15661 |
| **CARS2** | 1.63988 | 1.13632 | 0.10774 | 1.50183 | 0.24596 | 0.50373 | 8.26484 |
| **CASC1** | 0.00000 | 0.00000 | 0.30059 | 0.04896 | 1.44209 | 0.05705 | 0.15048 |
| **CASC10** | 0.00000 | 0.15447 | 0.00981 | 0.88309 | 0.05285 | 0.05910 | 0.60157 |
| **CASC11** | 0.06283 | 0.00000 | 0.16831 | 0.05395 | 2.88662 | 0.01564 | 0.00149 |
| **CASC14** | 0.01160 | 0.01622 | 0.01680 | 0.03723 | 0.01629 | 0.00977 | 0.08458 |
| **CASC15** | 0.36887 | 0.00000 | 0.24643 | 1.05488 | 1.08456 | 0.98877 | 0.03482 |
| **CASC18** | 0.02582 | 0.00368 | 0.03317 | 0.01781 | 0.20675 | 0.01281 | 0.01831 |
| **CASC2** | 0.20556 | 0.07987 | 0.53044 | 0.09491 | 0.05001 | 0.04452 | 0.00000 |
| **CASC20** | 0.04313 | 0.01375 | 0.02111 | 0.02970 | 0.04732 | 0.03880 | 0.01089 |
| **CASC21** | 0.01900 | 0.02398 | 0.01562 | 0.01984 | 0.01762 | 0.05036 | 0.01752 |
| **CASC3** | 11.89949 | 5.57792 | 5.19525 | 13.08187 | 3.43829 | 6.57280 | 13.29729 |
| **CASC4** | 1.34599 | 0.00000 | 0.21936 | 5.12160 | 0.47352 | 2.23151 | 2.24276 |
| **CASC5** | 16.78518 | 0.00000 | 13.16227 | 0.00000 | 7.87725 | 6.22725 | 32.04855 |
| **CASC8** | 0.06268 | 0.00563 | 0.06071 | 0.00000 | 2.01135 | 0.18072 | 0.06548 |
| **CASC9** | 0.13333 | 0.00000 | 0.83987 | 0.07651 | 0.00739 | 0.73914 | 0.36862 |
| **CASD1** | 4.96850 | 0.00000 | 0.98829 | 4.06667 | 0.01815 | 1.44372 | 2.00337 |
| **CASK** | 2.64324 | 1.63784 | 0.98813 | 0.28106 | 0.05995 | 4.22077 | 3.55833 |
| **CASKIN1** | 0.00000 | 0.70502 | 0.19734 | 0.00000 | 0.33809 | 0.57644 | 0.19656 |
| **CASKIN2** | 0.67147 | 1.19033 | 0.86197 | 1.03859 | 0.17743 | 0.77809 | 4.10297 |
| **CASP1** | 0.04301 | 0.01962 | 0.01254 | 0.03498 | 0.00841 | 0.01254 | 0.00000 |
| **CASP10** | 0.49819 | 0.69027 | 1.04275 | 0.04682 | 0.11633 | 3.10203 | 0.00000 |
| **CASP12** | 0.02507 | 0.01410 | 0.01207 | 0.01834 | 0.08645 | 0.01360 | 0.01737 |
| **CASP14** | 0.02628 | 0.01872 | 0.01259 | 0.02237 | 0.00807 | 0.01327 | 0.01516 |
| **CASP2** | 24.16175 | 5.48334 | 0.95708 | 10.66475 | 1.36862 | 4.48006 | 0.00000 |
| **CASP3** | 24.82289 | 7.38602 | 16.68573 | 11.15473 | 4.38664 | 6.28331 | 0.00000 |
| **CASP4** | 0.02917 | 0.00189 | 0.03422 | 0.11834 | 0.00864 | 0.01083 | 0.03190 |
| **CASP5** | 0.02059 | 0.01748 | 0.01335 | 0.03001 | 0.00794 | 0.01331 | 0.01750 |

| | | | | | | | |
|---|---|---|---|---|---|---|---|
| **CASP6** | 18.64451 | 3.26752 | 18.58851 | 6.73589 | 0.30962 | 1.48198 | 3.61358 |
| **CASP7** | 1.03865 | 2.41539 | 2.13535 | 0.13009 | 0.02611 | 0.77936 | 0.00000 |
| **CASP8** | 0.26441 | 0.60219 | 2.25241 | 0.06188 | 0.48198 | 3.87998 | 0.13505 |
| **CASP8AP2** | 12.26349 | 0.00000 | 7.38506 | 7.87093 | 6.70853 | 4.16612 | 8.55852 |
| **CASP9** | 2.07739 | 4.59076 | 2.84315 | 2.67365 | 0.26128 | 1.99732 | 0.14468 |
| **CASQ1** | 0.31012 | 0.00000 | 2.39355 | 1.73589 | 0.00044 | 2.21893 | 0.15546 |
| **CASQ2** | 0.14072 | 0.16996 | 0.00658 | 0.61370 | 0.00683 | 0.02117 | 0.00000 |
| **CASR** | 0.01805 | 0.01628 | 0.02445 | 0.03848 | 0.02108 | 0.01958 | 0.01969 |
| **CASS4** | 0.03940 | 0.08717 | 0.11923 | 0.09536 | 0.00417 | 0.02623 | 0.00000 |
| **CAST** | 21.78875 | 0.64908 | 0.76008 | 8.32148 | 2.57421 | 0.00000 | 6.53935 |
| **CASZ1** | 0.03386 | 0.00000 | 0.55850 | 0.24245 | 0.11592 | 0.00000 | 0.14150 |
| **CAT** | 4.60375 | 0.50828 | 0.48883 | 1.28780 | 0.00000 | 3.51730 | 0.00000 |
| **CATIP** | 0.01473 | 0.12647 | 0.12404 | 0.02041 | 0.27467 | 0.02707 | 0.02756 |
| **CATIP-AS1** | 0.03817 | 0.08179 | 0.41380 | 0.02572 | 0.37670 | 0.02997 | 0.02093 |
| **CATSPER1** | 0.03804 | 0.00119 | 0.09546 | 0.02466 | 0.07174 | 0.00924 | 0.01773 |
| **CATSPER2** | 0.06493 | 0.00000 | 0.10923 | 0.08984 | 0.15326 | 0.00000 | 0.19621 |
| **CATSPER2P1** | 0.10648 | 0.14650 | 0.07537 | 0.10695 | 0.07412 | 0.00156 | 0.00000 |
| **CATSPER3** | 0.21501 | 0.24235 | 0.30997 | 0.00000 | 0.01966 | 0.67823 | 0.26478 |
| **CATSPER4** | 0.02355 | 0.01309 | 0.02388 | 0.03155 | 0.00731 | 0.01226 | 0.01564 |
| **CATSPERB** | 0.08899 | 0.00000 | 0.05341 | 0.01625 | 0.23255 | 0.00695 | 0.02141 |
| **CATSPERD** | 0.04033 | 0.00000 | 0.01896 | 0.05632 | 0.77511 | 0.02761 | 0.06598 |
| **CATSPERG** | 0.30552 | 0.73622 | 0.20296 | 0.96503 | 0.22819 | 0.27739 | 0.34348 |
| **CAV1** | 0.02957 | 0.05965 | 0.08386 | 3.68419 | 0.01223 | 1.01858 | 0.00000 |
| **CAV2** | 0.05294 | 0.00114 | 0.01533 | 0.35439 | 0.00871 | 0.29684 | 0.00000 |
| **CAV3** | 0.02549 | 0.01698 | 0.01237 | 0.02205 | 0.00777 | 0.01271 | 0.02625 |
| **CBFA2T2** | 2.73571 | 6.40329 | 3.10401 | 1.12665 | 2.13343 | 21.41837 | 8.39309 |
| **CBFA2T3** | 0.04377 | 0.01646 | 0.00000 | 0.13666 | 0.90651 | 0.06479 | 0.01629 |
| **CBFB** | 7.79783 | 0.00000 | 5.48646 | 4.64756 | 3.07749 | 2.87357 | 12.13153 |
| **CBL** | 6.92697 | 6.43892 | 14.30348 | 14.07073 | 2.17696 | 4.26426 | 26.85472 |
| **CBLB** | 9.15465 | 0.00000 | 0.31601 | 36.00701 | 0.00000 | 3.08645 | 5.66926 |
| **CBLC** | 0.61326 | 0.00000 | 0.08345 | 1.46589 | 0.10328 | 0.68302 | 1.73014 |
| **CBLL1** | 19.35137 | 3.60993 | 11.19818 | 8.99475 | 9.03212 | 5.08702 | 0.98803 |
| **CBLN1** | 0.00000 | 0.00000 | 2.73558 | 0.02026 | 2.73839 | 0.00000 | 0.20828 |
| **CBLN2** | 0.17602 | 0.00000 | 0.00476 | 0.34015 | 0.00657 | 0.00000 | 0.29097 |
| **CBLN3** | 0.05801 | 0.03578 | 0.02892 | 0.00399 | 0.00612 | 0.01685 | 0.03475 |

| | | | | | | | |
|---|---|---|---|---|---|---|---|
| **CBLN4** | 0.02066 | 0.05432 | 0.00000 | 0.01889 | 0.46782 | 0.00770 | 0.02068 |
| **CBR1** | 10.17115 | 1.23637 | 3.94612 | 22.93072 | 1.79853 | 0.39833 | 0.31851 |
| **CBR3** | 0.04563 | 1.13535 | 2.40090 | 0.05107 | 0.10304 | 4.27975 | 0.00000 |
| **CBR3-AS1** | 0.78622 | 0.26535 | 0.27949 | 0.66121 | 0.47411 | 0.00000 | 0.36743 |
| **CBR4** | 10.56825 | 3.03959 | 3.18014 | 1.36424 | 1.94823 | 4.56282 | 2.67000 |
| **CBS** | 9.88218 | 0.00000 | 0.00000 | 15.46487 | 0.00000 | 6.43517 | 13.51455 |
| **CBWD1** | 6.03395 | 0.00000 | 1.61600 | 5.99819 | 0.36990 | 0.79039 | 1.42916 |
| **CBWD2** | 3.87413 | 0.19699 | 1.00816 | 4.74608 | 0.68453 | 1.06791 | 0.00000 |
| **CBWD3** | 0.02667 | 0.01866 | 0.01496 | 0.01966 | 0.00867 | 0.01309 | 0.01474 |
| **CBWD5.uid-117** | 1.32027 | 1.68007 | 0.44790 | 2.54087 | 0.51560 | 0.17487 | 0.00000 |
| **CBWD6** | 0.55422 | 0.13495 | 0.16403 | 0.77422 | 0.11894 | 0.14282 | 0.16081 |
| **CBX1** | 15.55693 | 7.53008 | 6.91937 | 11.61434 | 4.44928 | 9.94095 | 14.95665 |
| **CBX2** | 2.57530 | 0.74699 | 1.30202 | 8.44716 | 0.29497 | 0.39717 | 8.22766 |
| **CBX3** | 28.41712 | 16.66668 | 19.13283 | 17.72074 | 8.18556 | 16.87174 | 32.52805 |
| **CBX3P2** | 0.04647 | 0.02530 | 0.01440 | 0.01409 | 0.01599 | 0.09086 | 0.00833 |
| **CBX4** | 0.00012 | 0.00000 | 0.16341 | 0.04045 | 0.24086 | 0.74413 | 3.02412 |
| **CBX5** | 9.47179 | 0.00000 | 16.21778 | 5.63480 | 7.94620 | 4.74938 | 8.04817 |
| **CBX6** | 0.00000 | 0.82109 | 0.96648 | 0.14231 | 0.45536 | 0.95887 | 1.18685 |
| **CBX7** | 0.04357 | 0.05308 | 0.08311 | 0.16940 | 0.00944 | 0.38182 | 0.22982 |
| **CBX8** | 1.21208 | 0.27977 | 2.46001 | 0.72215 | 0.58811 | 0.52097 | 0.45965 |
| **CBY1** | 5.08780 | 5.41936 | 0.73356 | 2.18453 | 1.38596 | 2.61818 | 0.59824 |
| **CBY3** | 0.06251 | 0.14590 | 0.12718 | 0.00522 | 0.00752 | 0.10277 | 0.00000 |
| **CC2D1A** | 0.98824 | 6.03943 | 0.18509 | 3.18390 | 0.14161 | 3.21645 | 5.73001 |
| **CC2D1B** | 3.29315 | 6.04041 | 0.89706 | 5.88297 | 0.34957 | 3.60122 | 7.42755 |
| **CC2D2A** | 0.76406 | 0.30742 | 0.26579 | 0.27943 | 0.57008 | 0.77070 | 0.58628 |
| **CC2D2B** | 0.10453 | 0.04314 | 0.05467 | 0.00000 | 0.00635 | 0.01323 | 0.11033 |
| **CCAR1** | 35.79824 | 8.46485 | 17.54600 | 23.30079 | 3.31808 | 13.90836 | 26.09914 |
| **CCAR2** | 9.30785 | 9.55620 | 8.69983 | 11.87908 | 2.15286 | 5.81487 | 8.33276 |
| **CCAT1** | 0.01965 | 0.00847 | 0.01108 | 0.02815 | 0.00883 | 0.42861 | 0.01006 |
| **CCAT2** | 0.02162 | 0.01943 | 0.01373 | 0.01524 | 0.00832 | 0.02324 | 0.03615 |
| **CCBE1** | 0.56749 | 0.00000 | 0.00000 | 3.04183 | 0.58963 | 0.00000 | 1.57017 |
| **CCBL1** | 4.11979 | 1.07144 | 1.69222 | 3.08756 | 0.55567 | 0.82487 | 0.51653 |
| **CCBL2** | 6.05231 | 0.23523 | 5.50953 | 7.40068 | 0.09437 | 2.16751 | 0.85301 |
| **CCDC101** | 6.25834 | 8.74899 | 4.34011 | 4.67386 | 2.07678 | 3.86461 | 0.61812 |
| **CCDC102A** | 0.78874 | 0.31399 | 0.02708 | 2.01198 | 0.00481 | 1.12684 | 0.00000 |

| | | | | | | | |
|---|---|---|---|---|---|---|---|
| **CCDC102B** | 0.21687 | 0.03138 | 0.07774 | 0.05382 | 0.08119 | 0.05665 | 0.10484 |
| **CCDC103** | 0.70479 | 0.15017 | 1.00700 | 0.27171 | 0.20536 | 0.57908 | 0.10588 |
| **CCDC104** | 2.61045 | 1.77325 | 2.96792 | 3.55508 | 0.34233 | 2.27665 | 1.00059 |
| **CCDC106** | 0.00000 | 0.71755 | 0.01111 | 0.24808 | 0.07587 | 0.34650 | 1.59583 |
| **CCDC107** | 0.53941 | 0.60207 | 0.20531 | 1.38599 | 0.28046 | 0.39501 | 0.23011 |
| **CCDC108** | 0.11133 | 0.00000 | 0.40610 | 0.15381 | 0.22924 | 0.05046 | 0.00441 |
| **CCDC109B** | 4.85939 | 8.06567 | 0.90430 | 3.82017 | 0.18493 | 3.47827 | 11.10666 |
| **CCDC11** | 0.01574 | 0.05536 | 0.28810 | 0.48375 | 0.46774 | 0.12366 | 0.12208 |
| **CCDC110** | 0.01914 | 0.00000 | 0.11415 | 0.03486 | 1.27805 | 0.01790 | 0.01413 |
| **CCDC112** | 0.63122 | 0.00000 | 1.03852 | 0.43276 | 0.00009 | 0.64460 | 1.06873 |
| **CCDC113** | 0.21361 | 0.40049 | 2.43377 | 0.63589 | 0.29604 | 0.42569 | 0.00000 |
| **CCDC114** | 0.00034 | 0.00000 | 0.05774 | 0.05550 | 0.24403 | 0.04408 | 0.08004 |
| **CCDC115** | 5.67930 | 2.81211 | 4.09016 | 4.64649 | 3.84514 | 5.84703 | 0.00000 |
| **CCDC116** | 2.76590 | 0.00000 | 0.08639 | 2.23970 | 0.23983 | 0.20017 | 0.00000 |
| **CCDC117** | 10.69227 | 1.53240 | 12.42892 | 6.63869 | 3.35947 | 6.28107 | 9.54107 |
| **CCDC12** | 17.91800 | 23.00252 | 13.77151 | 25.03966 | 13.82423 | 11.83594 | 5.12496 |
| **CCDC120** | 0.27692 | 2.71861 | 0.62798 | 0.94938 | 0.24084 | 1.81887 | 5.15888 |
| **CCDC121** | 0.39304 | 0.00000 | 0.55966 | 0.36615 | 1.41287 | 0.14695 | 0.00000 |
| **CCDC122** | 0.05296 | 0.04516 | 0.86265 | 0.13057 | 0.08504 | 0.00000 | 0.34061 |
| **CCDC124** | 20.87093 | 41.26042 | 17.75524 | 22.19711 | 1.20617 | 13.98804 | 11.28069 |
| **CCDC125** | 2.99660 | 0.17109 | 0.56528 | 2.33065 | 0.00000 | 0.00000 | 1.94122 |
| **CCDC126** | 6.83984 | 0.77335 | 4.98474 | 4.46313 | 4.02810 | 4.46398 | 0.00000 |
| **CCDC127** | 4.56498 | 4.36081 | 5.27084 | 2.17357 | 1.85122 | 2.05587 | 0.20181 |
| **CCDC129** | 0.04423 | 0.00000 | 0.02963 | 0.02201 | 0.11778 | 0.03423 | 0.02612 |
| **CCDC13** | 0.03707 | 0.13906 | 0.31110 | 0.39830 | 0.69050 | 0.06286 | 0.04080 |
| **CCDC130** | 4.93348 | 6.32337 | 4.48761 | 6.77215 | 0.60207 | 6.39339 | 1.16283 |
| **CCDC132** | 5.36999 | 0.18041 | 1.94717 | 3.07159 | 0.62264 | 2.16885 | 1.82704 |
| **CCDC134** | 2.48028 | 1.51259 | 0.38875 | 1.10741 | 0.18700 | 0.52092 | 0.74847 |
| **CCDC135** | 0.08697 | 0.00000 | 0.37358 | 0.06503 | 0.41272 | 0.00874 | 0.10718 |
| **CCDC136** | 0.50645 | 0.00000 | 0.26924 | 0.56014 | 1.54438 | 0.85183 | 0.10256 |
| **CCDC137** | 10.66394 | 23.15299 | 5.20234 | 11.51460 | 5.44975 | 6.41538 | 25.23241 |
| **CCDC138** | 8.76462 | 0.00000 | 3.51627 | 5.34287 | 1.21006 | 1.38549 | 0.00000 |
| **CCDC14** | 8.98901 | 0.00000 | 6.02089 | 4.86199 | 1.77921 | 2.68256 | 8.40517 |
| **CCDC141** | 1.47327 | 0.00000 | 0.30705 | 1.41239 | 0.86027 | 0.00000 | 2.34769 |
| **CCDC142** | 3.06534 | 1.54674 | 1.03472 | 7.41698 | 0.49254 | 1.57620 | 1.67826 |

| | | | | | | | |
|---|---|---|---|---|---|---|---|
| **CCDC144A** | 0.03230 | 0.17289 | 0.59276 | 0.05728 | 0.06759 | 0.01225 | 0.00243 |
| **CCDC144B** | 0.13292 | 0.03913 | 1.02428 | 0.13126 | 0.21414 | 0.11812 | 0.45727 |
| **CCDC144CP** | 0.17763 | 0.37699 | 1.07204 | 0.67387 | 0.13139 | 0.33239 | 1.03837 |
| **CCDC144NL** | 0.06942 | 0.13550 | 0.21621 | 0.35540 | 0.20391 | 0.00000 | 0.02357 |
| **CCDC146** | 0.05475 | 0.26427 | 0.21815 | 0.01590 | 2.88656 | 0.00000 | 0.53924 |
| **CCDC147** | 0.06661 | 0.00000 | 0.39857 | 0.00070 | 8.98385 | 0.00057 | 0.01864 |
| **CCDC147-AS1** | 0.04460 | 0.00000 | 0.11721 | 0.03659 | 0.13872 | 0.01373 | 0.00501 |
| **CCDC148** | 0.00000 | 0.10187 | 0.35688 | 0.03287 | 1.20520 | 0.19365 | 0.18993 |
| **CCDC149** | 0.05211 | 0.05919 | 0.00390 | 0.01437 | 0.33991 | 0.06517 | 0.00000 |
| **CCDC15** | 0.23121 | 0.49414 | 1.24035 | 0.24809 | 5.41702 | 0.43502 | 0.52448 |
| **CCDC150** | 2.51335 | 0.00000 | 1.01606 | 1.45681 | 0.85954 | 0.61440 | 5.40907 |
| **CCDC151** | 0.11563 | 0.13701 | 0.13389 | 0.03881 | 0.07797 | 0.02783 | 0.00000 |
| **CCDC152** | 0.48756 | 0.10346 | 0.21058 | 0.00000 | 0.00193 | 0.39338 | 0.36737 |
| **CCDC153** | 0.21786 | 0.83643 | 0.48211 | 0.24106 | 0.04085 | 0.00000 | 0.06870 |
| **CCDC154** | 0.12023 | 0.09070 | 0.06643 | 0.08596 | 0.00350 | 0.00000 | 0.09370 |
| **CCDC155** | 0.00465 | 0.00000 | 0.15638 | 0.02007 | 0.16906 | 0.04445 | 0.20769 |
| **CCDC157** | 0.10985 | 1.03288 | 0.41864 | 0.47194 | 0.71360 | 0.14457 | 0.00000 |
| **CCDC158** | 0.03040 | 0.00000 | 0.15612 | 0.01753 | 0.00769 | 0.01030 | 0.01595 |
| **CCDC159** | 0.00209 | 0.42085 | 0.00826 | 0.36315 | 0.01844 | 0.35767 | 0.00000 |
| **CCDC160** | 0.00131 | 0.00318 | 0.01846 | 0.12753 | 0.04768 | 0.11138 | 0.07933 |
| **CCDC162P** | 0.03637 | 0.00943 | 0.04949 | 0.02944 | 0.08213 | 0.01248 | 0.01291 |
| **CCDC163P** | 1.91442 | 6.82827 | 8.46244 | 1.10823 | 4.54594 | 5.20008 | 0.16369 |
| **CCDC166** | 0.02622 | 0.00900 | 0.01556 | 0.01500 | 0.01544 | 0.03680 | 0.04240 |
| **CCDC167** | 9.78276 | 11.82726 | 3.18347 | 18.61693 | 0.36383 | 11.65896 | 4.80204 |
| **CCDC168** | 0.00055 | 0.00000 | 0.14633 | 0.10986 | 0.08460 | 0.00000 | 0.63365 |
| **CCDC169** | 0.31943 | 0.00000 | 0.02216 | 0.12232 | 0.00531 | 0.07326 | 0.27960 |
| **CCDC17** | 0.23710 | 0.00000 | 1.00445 | 0.62080 | 0.14966 | 0.18881 | 1.03640 |
| **CCDC170** | 0.02114 | 0.00000 | 0.20709 | 0.01420 | 0.53733 | 0.05382 | 0.08829 |
| **CCDC171** | 0.11902 | 0.03749 | 0.45442 | 0.15530 | 0.89013 | 0.64734 | 0.65755 |
| **CCDC172** | 0.02077 | 0.00000 | 0.27123 | 0.01711 | 0.07487 | 0.01569 | 0.02444 |
| **CCDC173** | 0.56333 | 0.00000 | 0.64817 | 0.08448 | 0.84253 | 0.00000 | 0.96496 |
| **CCDC174** | 7.97226 | 1.67272 | 5.16899 | 4.82942 | 4.12082 | 2.84837 | 14.01746 |
| **CCDC175** | 0.01573 | 0.00080 | 0.06910 | 0.02065 | 0.22931 | 0.02235 | 0.11684 |
| **CCDC176** | 0.34660 | 0.31755 | 0.81295 | 0.41966 | 0.67852 | 0.00000 | 0.05685 |
| **CCDC177** | 0.02330 | 0.01715 | 0.00786 | 0.04503 | 0.14433 | 0.02264 | 0.01943 |

| | | | | | | | |
|---|---|---|---|---|---|---|---|
| **CCDC178** | 0.02549 | 0.07684 | 0.27658 | 0.00000 | 0.92317 | 0.07488 | 0.41536 |
| **CCDC18** | 1.80173 | 0.46127 | 2.72335 | 0.95829 | 1.02733 | 1.67657 | 3.72838 |
| **CCDC181** | 0.23785 | 0.00000 | 0.35535 | 0.18904 | 0.96580 | 0.03297 | 0.02285 |
| **CCDC183** | 0.00804 | 0.14658 | 0.01262 | 0.07677 | 0.02695 | 0.00000 | 0.29019 |
| **CCDC183-AS1** | 0.00000 | 0.42159 | 0.03128 | 0.15686 | 0.01129 | 0.04641 | 0.92047 |
| **CCDC184** | 0.02163 | 0.00000 | 0.24237 | 0.01815 | 0.13793 | 0.00710 | 0.02298 |
| **CCDC186** | 5.28297 | 0.00000 | 4.44246 | 5.45195 | 4.93701 | 2.16716 | 9.50786 |
| **CCDC19** | 0.48811 | 0.20401 | 0.20734 | 0.90508 | 0.15160 | 0.00000 | 0.14847 |
| **CCDC22** | 4.58171 | 2.93516 | 0.00000 | 0.39064 | 0.00000 | 2.52265 | 3.06558 |
| **CCDC23** | 7.84630 | 2.69037 | 2.61025 | 10.33895 | 1.97969 | 1.97998 | 2.12019 |
| **CCDC24** | 0.51285 | 0.84300 | 0.25214 | 0.44720 | 0.04401 | 0.54397 | 0.42479 |
| **CCDC25** | 5.24795 | 3.18736 | 4.61784 | 3.18892 | 12.41169 | 3.96311 | 2.16279 |
| **CCDC27** | 0.00966 | 0.06308 | 0.00000 | 0.01485 | 5.74259 | 0.02226 | 0.02784 |
| **CCDC28A** | 7.59494 | 3.91700 | 5.89456 | 10.97556 | 4.00517 | 1.79722 | 0.00000 |
| **CCDC28B** | 0.33879 | 0.00000 | 0.05626 | 0.17370 | 0.24749 | 1.08648 | 0.29778 |
| **CCDC3** | 0.02684 | 0.45189 | 0.26901 | 0.02510 | 0.44261 | 0.18680 | 0.03398 |
| **CCDC30** | 0.09078 | 0.02723 | 0.04430 | 0.10294 | 0.82632 | 0.04804 | 0.00000 |
| **CCDC33** | 0.00639 | 0.01823 | 0.01429 | 0.06548 | 0.02868 | 0.02321 | 0.05174 |
| **CCDC34** | 8.74329 | 18.81339 | 11.70328 | 6.75306 | 4.33773 | 1.68973 | 8.60697 |
| **CCDC36** | 0.01702 | 0.00000 | 0.06006 | 0.03744 | 0.19638 | 0.05841 | 0.04169 |
| **CCDC37** | 0.01094 | 0.00476 | 0.03614 | 0.05986 | 0.00556 | 0.12805 | 0.13872 |
| **CCDC37-AS1** | 0.02988 | 0.01701 | 0.01233 | 0.02004 | 0.00806 | 0.03097 | 0.01170 |
| **CCDC38** | 0.05754 | 0.13874 | 0.14929 | 0.01981 | 0.14855 | 0.08364 | 0.18423 |
| **CCDC39** | 0.24024 | 0.00000 | 0.16503 | 0.02996 | 0.84224 | 0.07386 | 0.29291 |
| **CCDC40** | 0.21144 | 0.10131 | 0.48080 | 0.08431 | 0.15011 | 0.00000 | 0.03325 |
| **CCDC42** | 0.09206 | 0.20662 | 0.00137 | 0.38229 | 0.02898 | 0.01976 | 0.00000 |
| **CCDC42B** | 0.00877 | 0.02047 | 0.02281 | 0.03632 | 0.01812 | 0.01730 | 0.13696 |
| **CCDC43** | 20.43642 | 1.98270 | 8.23802 | 11.85346 | 2.79875 | 5.55757 | 7.51895 |
| **CCDC47** | 39.76778 | 13.28905 | 13.56534 | 28.81573 | 9.14048 | 13.21324 | 16.25949 |
| **CCDC50** | 2.05374 | 2.68056 | 0.00000 | 3.22469 | 0.80124 | 4.11143 | 9.01080 |
| **CCDC51** | 29.18436 | 24.09283 | 13.82627 | 17.58297 | 1.85527 | 3.23262 | 0.00000 |
| **CCDC53** | 4.69134 | 2.69075 | 4.83633 | 1.77437 | 0.53923 | 5.34118 | 1.24006 |
| **CCDC57** | 0.00000 | 4.46889 | 0.12999 | 0.34075 | 0.00000 | 1.13437 | 2.53831 |
| **CCDC58** | 10.49468 | 8.80148 | 16.32077 | 8.72524 | 0.03533 | 4.34203 | 9.21061 |
| **CCDC59** | 14.74981 | 4.95509 | 13.00721 | 12.62992 | 6.80089 | 6.01681 | 11.24865 |

| | | | | | | | |
|---|---|---|---|---|---|---|---|
| **CCDC6** | 11.89099 | 3.39096 | 5.95449 | 3.70020 | 3.36578 | 4.73405 | 17.05263 |
| **CCDC60** | 0.02960 | 0.04488 | 0.00077 | 0.04712 | 0.12666 | 0.21484 | 0.00751 |
| **CCDC61** | 0.25055 | 0.31861 | 0.19681 | 0.39395 | 0.16961 | 0.13760 | 1.36550 |
| **CCDC62** | 0.29702 | 0.17045 | 0.54024 | 0.14866 | 1.77029 | 0.25386 | 0.08239 |
| **CCDC63** | 0.01596 | 0.19845 | 0.04316 | 0.03663 | 0.14665 | 0.10760 | 0.00000 |
| **CCDC64** | 0.70115 | 1.08115 | 1.60034 | 2.12998 | 1.67442 | 1.22925 | 4.78105 |
| **CCDC64B** | 0.03155 | 0.04368 | 0.00000 | 1.25033 | 0.14113 | 0.14073 | 1.02356 |
| **CCDC65** | 0.00000 | 0.27301 | 0.97995 | 0.07252 | 0.57512 | 0.35308 | 0.25894 |
| **CCDC66** | 5.96027 | 0.23923 | 0.44216 | 2.77797 | 0.11492 | 2.22700 | 2.98278 |
| **CCDC67** | 0.02420 | 0.01571 | 0.01651 | 0.02141 | 0.01118 | 0.01280 | 0.01709 |
| **CCDC68** | 0.00977 | 0.18228 | 0.00714 | 0.03403 | 0.00701 | 0.19717 | 0.02275 |
| **CCDC69** | 0.60377 | 0.07080 | 0.33315 | 0.01467 | 0.00239 | 0.32021 | 0.45692 |
| **CCDC7** | 0.82456 | 0.09697 | 0.74633 | 0.48987 | 0.48446 | 0.32933 | 0.42546 |
| **CCDC70** | 0.02688 | 0.10819 | 0.01613 | 0.02752 | 0.00609 | 0.00381 | 0.07064 |
| **CCDC71** | 0.29123 | 0.00000 | 0.42219 | 0.32585 | 0.03546 | 0.07056 | 0.06038 |
| **CCDC71L** | 0.01328 | 0.00000 | 0.48372 | 0.45778 | 0.00166 | 0.22220 | 0.84212 |
| **CCDC73** | 0.73014 | 0.80515 | 4.50521 | 0.66466 | 2.76880 | 0.65349 | 1.78354 |
| **CCDC74A** | 0.01996 | 0.00593 | 0.06980 | 0.06466 | 0.04366 | 0.05940 | 0.00788 |
| **CCDC74B** | 0.02870 | 0.01147 | 0.04967 | 0.04567 | 0.01177 | 0.01436 | 0.01161 |
| **CCDC77** | 10.35657 | 0.00000 | 5.19800 | 9.26469 | 4.55751 | 2.40857 | 3.38527 |
| **CCDC78** | 0.02025 | 0.06916 | 0.02087 | 0.08285 | 0.00543 | 0.13493 | 0.11993 |
| **CCDC79** | 0.08009 | 0.05326 | 0.08865 | 0.00288 | 0.63490 | 0.03807 | 0.01932 |
| **CCDC8** | 0.54175 | 0.00000 | 0.00000 | 0.00004 | 0.14658 | 0.65956 | 1.20397 |
| **CCDC80** | 1.03857 | 1.72447 | 0.31547 | 0.18986 | 0.45303 | 8.08084 | 0.00000 |
| **CCDC81** | 0.57712 | 0.00000 | 0.14693 | 0.15267 | 0.01400 | 0.12432 | 0.01693 |
| **CCDC82** | 4.26381 | 0.00000 | 4.23019 | 2.58575 | 7.57025 | 2.17027 | 0.00000 |
| **CCDC83** | 0.06106 | 0.00000 | 0.07892 | 0.00921 | 0.05228 | 0.01496 | 0.05091 |
| **CCDC84** | 2.65940 | 4.27879 | 2.53200 | 5.98144 | 0.83795 | 3.96202 | 5.46200 |
| **CCDC85A** | 0.12730 | 0.04903 | 0.14809 | 0.00000 | 0.17907 | 0.01714 | 0.05108 |
| **CCDC85B** | 0.00000 | 3.11999 | 0.86300 | 0.34947 | 0.73105 | 1.57682 | 10.40064 |
| **CCDC85C** | 5.05014 | 6.34713 | 0.51703 | 4.88571 | 0.02855 | 2.21866 | 20.31470 |
| **CCDC86** | 26.02865 | 22.35913 | 18.58216 | 27.03423 | 4.53511 | 7.24708 | 1.22348 |
| **CCDC87** | 0.04539 | 0.41859 | 0.39138 | 0.35422 | 0.17233 | 0.11325 | 0.05143 |
| **CCDC88A** | 0.88013 | 0.00000 | 6.07070 | 0.00000 | 4.76523 | 4.39857 | 5.10827 |
| **CCDC88B** | 0.06040 | 0.04393 | 0.06184 | 0.54224 | 0.11907 | 0.00000 | 0.80783 |

| | | | | | | | |
|---|---|---|---|---|---|---|---|
| **CCDC88C** | 0.70950 | 0.00000 | 0.92762 | 4.46119 | 2.72510 | 0.00000 | 10.82759 |
| **CCDC89** | 0.03565 | 0.01646 | 0.01393 | 0.04792 | 0.00829 | 0.01248 | 0.00000 |
| **CCDC9** | 2.30507 | 1.11897 | 0.38718 | 3.08636 | 0.89122 | 3.78855 | 6.61759 |
| **CCDC90B** | 7.91849 | 5.45267 | 6.47836 | 3.56258 | 0.82094 | 6.53258 | 0.42991 |
| **CCDC91** | 2.30918 | 0.35912 | 2.20207 | 2.99836 | 0.78776 | 1.60839 | 4.79316 |
| **CCDC92** | 0.02870 | 0.00000 | 0.15527 | 0.25080 | 0.29871 | 0.05802 | 0.10808 |
| **CCDC93** | 10.39441 | 4.13587 | 4.35467 | 7.32663 | 2.17493 | 4.01234 | 5.95108 |
| **CCDC94** | 11.25437 | 11.44259 | 7.24616 | 7.28066 | 6.16041 | 4.08649 | 0.00000 |
| **CCDC96** | 0.06586 | 0.05136 | 0.01528 | 0.01585 | 0.31880 | 0.17881 | 0.00000 |
| **CCDC97** | 2.67048 | 5.81890 | 5.91149 | 14.12792 | 2.52377 | 0.94285 | 12.65769 |
| **CCER1** | 0.02354 | 0.02011 | 0.02117 | 0.03992 | 0.00864 | 0.00382 | 0.07602 |
| **CCER2** | 0.02608 | 0.00000 | 0.06629 | 0.04555 | 0.09996 | 0.01741 | 0.11863 |
| **CCHCR1** | 1.98598 | 6.98533 | 3.40508 | 2.76494 | 0.18908 | 4.80858 | 0.25194 |
| **CCIN** | 1.52187 | 1.39118 | 3.65818 | 0.57723 | 0.00000 | 0.00000 | 0.02841 |
| **CCK** | 0.01133 | 0.00014 | 0.36769 | 0.01371 | 3.37241 | 0.00694 | 0.02904 |
| **CCKBR** | 11.86623 | 0.00000 | 0.17391 | 72.14993 | 0.00000 | 0.00000 | 8.76290 |
| **CCL13** | 0.04177 | 0.01888 | 0.01119 | 0.03254 | 0.01844 | 0.01164 | 0.00644 |
| **CCL15** | 0.01804 | 0.00000 | 0.34606 | 0.01560 | 0.00798 | 0.00753 | 0.02495 |
| **CCL16** | 0.03442 | 0.00308 | 0.07492 | 0.01876 | 0.00616 | 0.01240 | 0.01495 |
| **CCL17** | 0.02100 | 0.01748 | 0.01583 | 0.02125 | 0.00764 | 0.02562 | 0.02100 |
| **CCL19** | 0.01800 | 0.02491 | 0.01462 | 0.02440 | 0.02082 | 0.02665 | 0.02224 |
| **CCL2** | 0.29870 | 0.00000 | 0.03124 | 0.22692 | 0.27045 | 0.34937 | 0.00000 |
| **CCL20** | 0.01414 | 0.01847 | 0.00261 | 0.02899 | 0.17642 | 0.03054 | 0.02071 |
| **CCL22** | 0.00000 | 0.10416 | 0.02213 | 0.05488 | 0.04481 | 0.00641 | 0.25418 |
| **CCL25** | 0.02442 | 0.00000 | 0.09998 | 0.01816 | 0.22864 | 0.00774 | 0.04401 |
| **CCL26** | 0.65964 | 0.00000 | 0.12457 | 0.00034 | 0.18882 | 1.00267 | 0.00964 |
| **CCL27** | 0.04496 | 0.01901 | 0.03739 | 0.14808 | 0.10489 | 0.01776 | 0.05937 |
| **CCL28** | 0.59818 | 0.19570 | 0.94192 | 0.49613 | 0.02700 | 0.04544 | 0.45673 |
| **CCL5** | 0.14161 | 0.00000 | 0.02743 | 0.02308 | 0.04954 | 0.00000 | 0.26850 |
| **CCL8** | 0.10827 | 0.00000 | 0.00706 | 0.00000 | 0.00714 | 0.00052 | 0.38052 |
| **CCM2** | 0.00000 | 13.73886 | 1.89058 | 0.86500 | 0.00047 | 3.83913 | 5.43922 |
| **CCM2L** | 0.00000 | 0.02157 | 0.00000 | 0.04235 | 0.34168 | 0.44034 | 0.03860 |
| **CCNA1** | 1.07927 | 7.89576 | 5.00274 | 0.00000 | 22.22260 | 0.66216 | 0.72968 |
| **CCNA2** | 38.76797 | 20.29221 | 26.09512 | 17.38184 | 10.89211 | 15.31899 | 7.39029 |
| **CCNB1** | 61.98970 | 37.10355 | 42.78161 | 48.95620 | 27.78919 | 22.29404 | 5.34955 |

| | | | | | | | |
|---|---|---|---|---|---|---|---|
| **CCNB1IP1** | 10.34282 | 6.07985 | 18.16932 | 9.49155 | 6.11545 | 9.07651 | 10.25801 |
| **CCNB2** | 29.07413 | 12.42665 | 17.90053 | 18.75583 | 4.05864 | 8.56056 | 3.37339 |
| **CCNB3** | 0.11705 | 0.46108 | 0.18412 | 0.00000 | 4.40122 | 0.00000 | 0.00000 |
| **CCNC** | 43.70922 | 6.64910 | 18.54557 | 23.00059 | 4.58907 | 14.06132 | 9.15001 |
| **CCND1** | 0.62323 | 3.96796 | 5.29699 | 13.42122 | 1.54958 | 1.95438 | 6.94707 |
| **CCND2** | 0.04791 | 0.23546 | 0.40019 | 0.02937 | 4.95070 | 0.12315 | 0.03422 |
| **CCND3** | 20.49805 | 19.43401 | 8.26449 | 25.83611 | 0.73078 | 14.28842 | 26.94707 |
| **CCNDBP1** | 17.69334 | 22.50470 | 20.31805 | 11.51503 | 2.61126 | 12.16250 | 0.00000 |
| **CCNE1** | 0.80421 | 11.12525 | 7.73899 | 24.86141 | 12.08749 | 9.02616 | 26.00428 |
| **CCNE2** | 8.77302 | 0.00000 | 2.33066 | 5.76623 | 7.59982 | 0.25535 | 0.00000 |
| **CCNF** | 15.07720 | 24.36736 | 12.14493 | 21.66410 | 7.44422 | 12.04301 | 12.82458 |
| **CCNG1** | 24.35292 | 0.31817 | 4.80021 | 14.98556 | 3.99392 | 9.50637 | 9.64314 |
| **CCNG2** | 1.12329 | 0.00000 | 0.36502 | 1.75261 | 0.11995 | 0.95228 | 2.71708 |
| **CCNH** | 11.98823 | 3.59979 | 6.87672 | 6.64330 | 4.11452 | 3.33477 | 0.00000 |
| **CCNI** | 4.12722 | 5.90050 | 4.32159 | 4.31944 | 9.79575 | 9.99410 | 34.95519 |
| **CCNI2** | 0.61129 | 1.14149 | 0.16902 | 0.00000 | 0.82278 | 0.08784 | 0.56208 |
| **CCNJ** | 5.33932 | 0.05442 | 0.91185 | 4.48516 | 2.86106 | 1.82038 | 5.04594 |
| **CCNJL** | 0.02189 | 0.01329 | 0.02650 | 0.02385 | 0.07857 | 0.02393 | 0.01441 |
| **CCNK** | 8.06188 | 0.00000 | 5.72374 | 9.22124 | 5.98865 | 2.75413 | 14.01989 |
| **CCNL1** | 14.61038 | 3.36266 | 11.70863 | 12.72863 | 6.48836 | 13.19294 | 11.88629 |
| **CCNL2** | 7.73829 | 13.80585 | 7.10639 | 9.83696 | 7.03171 | 11.39158 | 17.50311 |
| **CCNO** | 0.42957 | 0.00000 | 0.66547 | 0.20523 | 3.47803 | 0.84356 | 0.00000 |
| **CCNT1** | 5.47377 | 0.00000 | 5.90128 | 4.27750 | 7.98456 | 4.26133 | 12.90403 |
| **CCNT2** | 9.91631 | 0.99681 | 6.55111 | 11.20330 | 4.72980 | 3.81723 | 8.40716 |
| **CCNT2-AS1** | 0.11350 | 0.07025 | 0.05435 | 0.76586 | 0.16707 | 0.14674 | 0.10133 |
| **CCNY** | 2.05006 | 8.44860 | 2.53123 | 3.84238 | 0.03821 | 5.69456 | 16.73359 |
| **CCNYL1** | 1.53671 | 0.00000 | 6.66643 | 3.49441 | 4.19404 | 2.24526 | 5.43491 |
| **CCP110** | 8.67577 | 1.33372 | 13.71608 | 5.37623 | 5.26149 | 1.83809 | 3.85213 |
| **CCPG1** | 1.04641 | 0.00000 | 16.25619 | 0.00000 | 1.43143 | 3.46861 | 2.16297 |
| **CCR1** | 0.29236 | 0.00000 | 0.01085 | 1.29672 | 0.00546 | 0.00880 | 0.06074 |
| **CCR10** | 0.03495 | 0.02111 | 0.00000 | 0.04580 | 0.13185 | 0.00728 | 0.10273 |
| **CCR2** | 0.01912 | 0.01846 | 0.01037 | 0.08468 | 0.02318 | 0.01053 | 0.02169 |
| **CCR3** | 0.05170 | 0.04928 | 0.00000 | 0.05775 | 0.82574 | 0.06016 | 0.00000 |
| **CCR4** | 0.00417 | 0.09552 | 0.01025 | 0.10106 | 0.01552 | 0.00816 | 0.00258 |
| **CCR6** | 0.08735 | 0.20578 | 0.00000 | 0.10789 | 0.02013 | 0.05095 | 0.11887 |

| | | | | | | | |
|---|---|---|---|---|---|---|---|
| **CCR7** | 4.84110 | 0.00000 | 0.00905 | 31.76999 | 0.00418 | 0.00764 | 0.00000 |
| **CCR8** | 0.11596 | 0.03064 | 0.00000 | 0.00000 | 0.44906 | 6.61976 | 0.00090 |
| **CCR9** | 0.02877 | 0.01471 | 0.01349 | 0.02646 | 0.00886 | 0.02608 | 0.00794 |
| **CCRN4L** | 8.18637 | 13.46618 | 7.52434 | 6.36739 | 11.68843 | 6.45837 | 0.64245 |
| **CCS** | 6.79230 | 20.78331 | 5.54014 | 6.89285 | 1.15032 | 8.14151 | 3.94997 |
| **CCSAP** | 4.25335 | 0.00000 | 2.27316 | 2.48921 | 8.24693 | 1.46275 | 7.29235 |
| **CCSER1** | 0.11977 | 0.00000 | 0.01491 | 0.40775 | 0.21799 | 0.00669 | 0.02402 |
| **CCSER2** | 8.66346 | 3.74223 | 6.60628 | 9.16278 | 2.93540 | 4.03354 | 15.56217 |
| **CCT2** | 68.27069 | 34.51764 | 30.60730 | 45.12961 | 13.97472 | 25.96693 | 24.17530 |
| **CCT3** | 68.60638 | 43.76001 | 27.13940 | 50.45733 | 10.51087 | 31.82366 | 23.23585 |
| **CCT4** | 61.54096 | 31.96696 | 20.44566 | 39.49687 | 1.02633 | 24.52911 | 16.31574 |
| **CCT5** | 66.72830 | 36.54203 | 26.85212 | 48.92569 | 7.85350 | 24.10189 | 22.56138 |
| **CCT6A** | 65.90683 | 28.23111 | 26.42448 | 44.35717 | 2.83190 | 25.56062 | 24.52175 |
| **CCT6B** | 0.08676 | 0.30790 | 0.13287 | 0.02311 | 0.25815 | 0.09603 | 0.07798 |
| **CCT6P1** | 0.83554 | 0.00000 | 0.40691 | 0.89820 | 0.39173 | 0.87941 | 0.47732 |
| **CCT6P3** | 0.43491 | 0.08023 | 0.31218 | 0.31457 | 0.14164 | 0.19854 | 0.00000 |
| **CCT7** | 56.84317 | 42.25401 | 24.53677 | 28.38665 | 2.35701 | 25.01828 | 22.86806 |
| **CCT8** | 59.64561 | 16.26295 | 25.82982 | 33.27841 | 6.96555 | 16.59894 | 13.50268 |
| **CCZ1** | 0.15291 | 0.17503 | 0.16398 | 0.45829 | 0.14642 | 1.07195 | 2.98251 |
| **CCZ1B** | 0.23357 | 0.17309 | 0.18059 | 0.09629 | 0.09626 | 0.42709 | 1.32188 |
| **CD101** | 0.00000 | 0.34470 | 0.02350 | 0.03583 | 0.01649 | 0.28692 | 0.34376 |
| **CD109** | 0.19655 | 0.00000 | 0.26877 | 0.03024 | 0.93727 | 1.17903 | 0.00000 |
| **CD14** | 0.12418 | 0.00000 | 1.08780 | 0.30276 | 0.32758 | 0.20544 | 0.00000 |
| **CD151** | 1.38181 | 12.88037 | 0.47582 | 8.73715 | 0.00000 | 5.64602 | 18.99756 |
| **CD160** | 0.07562 | 0.00000 | 0.07545 | 0.00940 | 0.01728 | 0.15684 | 0.27569 |
| **CD163** | 0.00000 | 0.08982 | 0.05934 | 0.05802 | 0.00590 | 0.36167 | 0.02464 |
| **CD163L1** | 0.43025 | 0.02401 | 0.00620 | 0.19864 | 0.29000 | 0.11015 | 0.00000 |
| **CD164** | 47.82524 | 14.46947 | 23.01397 | 27.99412 | 12.14428 | 12.52011 | 6.00894 |
| **CD164L2** | 0.02481 | 0.00000 | 0.05963 | 0.02218 | 0.07034 | 0.01494 | 0.01816 |
| **CD177** | 0.03397 | 0.02310 | 0.01488 | 0.02472 | 0.00789 | 0.01456 | 0.00654 |
| **CD180** | 0.02290 | 0.01872 | 0.00730 | 0.03151 | 0.07400 | 0.04575 | 0.00458 |
| **CD19** | 0.72739 | 0.13448 | 0.11634 | 0.24349 | 0.02612 | 0.71479 | 0.10550 |
| **CD1D** | 0.00000 | 0.07485 | 0.01065 | 0.02942 | 0.00736 | 0.12984 | 0.04770 |
| **CD200** | 0.00021 | 0.14508 | 0.82925 | 0.18922 | 0.00000 | 4.59052 | 0.00000 |
| **CD200R1** | 0.39043 | 0.20896 | 0.27745 | 0.00000 | 0.31073 | 0.31025 | 0.28156 |

| | | | | | | | |
|---|---|---|---|---|---|---|---|
| **CD200R1L** | 0.04430 | 0.00469 | 0.04048 | 0.00000 | 0.03541 | 0.02714 | 0.26012 |
| **CD209** | 0.03761 | 0.06111 | 0.06795 | 0.05991 | 0.04293 | 0.02159 | 0.14817 |
| **CD22** | 0.00000 | 3.04475 | 0.00970 | 0.10031 | 0.00679 | 0.00338 | 0.00508 |
| **CD226** | 0.03839 | 0.01092 | 0.02066 | 0.01854 | 0.01459 | 0.01062 | 0.02724 |
| **CD24** | 33.97006 | 0.00000 | 0.00000 | 46.59462 | 0.15880 | 0.00000 | 15.04461 |
| **CD244** | 0.03559 | 0.04214 | 0.03855 | 0.03200 | 0.00673 | 0.03597 | 0.00584 |
| **CD247** | 0.06046 | 0.00000 | 0.07377 | 0.05719 | 0.05945 | 0.00218 | 0.06409 |
| **CD248** | 0.02258 | 0.01307 | 0.01126 | 0.01594 | 0.00760 | 0.28251 | 0.02764 |
| **CD27** | 0.02738 | 0.01623 | 0.01447 | 0.01961 | 0.01044 | 0.01525 | 0.01715 |
| **CD27-AS1** | 0.25292 | 0.32665 | 0.05511 | 0.00536 | 0.22901 | 0.07690 | 0.00514 |
| **CD274** | 3.14956 | 0.00000 | 0.01212 | 9.64299 | 0.00966 | 0.00000 | 0.21636 |
| **CD276** | 6.57504 | 1.22486 | 0.07154 | 14.74050 | 0.00000 | 0.77012 | 12.19742 |
| **CD28** | 0.01947 | 0.02840 | 0.00671 | 0.01773 | 0.07415 | 0.01531 | 0.03120 |
| **CD2AP** | 10.49831 | 0.00000 | 11.13109 | 2.76377 | 1.18477 | 13.36481 | 17.93387 |
| **CD2BP2** | 15.36868 | 15.86212 | 10.81782 | 20.00899 | 7.42696 | 11.91466 | 1.60899 |
| **CD300A** | 0.02960 | 0.01492 | 0.01863 | 0.01832 | 0.00754 | 0.01494 | 0.01425 |
| **CD300E** | 0.02986 | 0.00208 | 0.04237 | 0.01154 | 0.30248 | 0.01612 | 0.03711 |
| **CD300LB** | 0.01482 | 0.01239 | 0.01423 | 0.01665 | 0.00751 | 0.04867 | 0.03678 |
| **CD300LF** | 0.86239 | 2.06228 | 0.03140 | 0.00045 | 0.11264 | 0.00056 | 0.00000 |
| **CD300LG** | 0.01920 | 0.01774 | 0.01305 | 0.03034 | 0.00886 | 0.01207 | 0.02423 |
| **CD302** | 0.02573 | 0.02607 | 0.03572 | 0.03413 | 0.02648 | 0.01141 | 0.01922 |
| **CD320** | 21.10928 | 45.17612 | 9.42048 | 24.83898 | 0.32465 | 21.65517 | 5.24446 |
| **CD33** | 0.01828 | 0.10900 | 0.02099 | 0.01534 | 0.00599 | 0.01431 | 0.05103 |
| **CD34** | 0.03243 | 0.02292 | 0.01017 | 0.00225 | 0.01186 | 0.03294 | 0.09351 |
| **CD36** | 0.40545 | 0.00000 | 0.00323 | 0.00666 | 4.08189 | 0.01407 | 0.03692 |
| **CD37** | 0.09569 | 0.00000 | 4.67209 | 0.00000 | 2.06632 | 0.00000 | 0.36002 |
| **CD38** | 0.04702 | 0.76726 | 0.42132 | 0.00000 | 0.44735 | 0.90893 | 0.02411 |
| **CD3D** | 0.01723 | 0.01989 | 0.01889 | 0.02760 | 0.01546 | 0.15834 | 0.01006 |
| **CD3E** | 0.01571 | 0.02182 | 0.04055 | 0.02070 | 0.00657 | 0.02829 | 0.02451 |
| **CD3EAP** | 12.03315 | 4.95703 | 9.15462 | 2.99750 | 2.22070 | 2.78118 | 4.25701 |
| **CD3G** | 0.01374 | 0.02571 | 0.03803 | 0.01571 | 0.00560 | 0.21083 | 0.01949 |
| **CD4** | 0.04644 | 0.01114 | 0.01946 | 0.03026 | 0.01052 | 0.01707 | 0.01077 |
| **CD40** | 0.02115 | 0.02197 | 0.45054 | 0.05146 | 0.11094 | 0.00539 | 0.00000 |
| **CD40LG** | 0.01292 | 0.01342 | 0.04276 | 0.03512 | 0.00759 | 0.01378 | 0.03503 |
| **CD44** | 0.00000 | 0.08952 | 0.39297 | 0.03350 | 0.00084 | 3.49063 | 0.04350 |

| | | | | | | | |
|---|---|---|---|---|---|---|---|
| **CD46** | 26.02789 | 4.36629 | 7.90144 | 6.71783 | 1.56358 | 6.36953 | 6.81405 |
| **CD47** | 25.38288 | 0.00000 | 4.50195 | 11.24832 | 0.19919 | 2.58075 | 8.19387 |
| **CD48** | 0.13378 | 0.18841 | 0.22098 | 0.00000 | 0.00027 | 1.44175 | 0.00003 |
| **CD5** | 0.00000 | 0.12896 | 0.03712 | 0.03043 | 0.11339 | 0.04032 | 0.08824 |
| **CD52** | 1.26013 | 0.51470 | 0.22742 | 0.81219 | 0.19863 | 0.44379 | 0.87634 |
| **CD53** | 64.41073 | 17.90601 | 2.67049 | 0.92548 | 0.11489 | 7.78245 | 0.00000 |
| **CD55** | 32.83425 | 11.67220 | 1.62310 | 26.05243 | 1.32667 | 1.48043 | 21.51199 |
| **CD58** | 1.29735 | 0.00000 | 0.83031 | 0.23431 | 0.00066 | 0.50285 | 0.00000 |
| **CD59** | 16.89605 | 6.96098 | 3.47608 | 15.51762 | 8.17397 | 3.77620 | 2.89630 |
| **CD6** | 0.00086 | 0.20102 | 0.01668 | 0.01431 | 0.05516 | 0.00495 | 0.05484 |
| **CD63** | 20.70374 | 26.85042 | 13.35816 | 42.12202 | 3.47565 | 27.84290 | 22.77419 |
| **CD68** | 0.76102 | 0.89620 | 1.13619 | 2.83147 | 1.04005 | 0.45368 | 0.00000 |
| **CD69** | 0.01723 | 0.01994 | 0.01298 | 0.08959 | 0.00805 | 0.02312 | 0.01306 |
| **CD7** | 0.01182 | 0.05576 | 0.02087 | 0.02160 | 0.00643 | 0.22867 | 0.01406 |
| **CD70** | 0.02427 | 0.03527 | 0.14717 | 0.07992 | 0.00649 | 2.44941 | 0.01122 |
| **CD72** | 0.00021 | 0.00000 | 0.54827 | 0.02909 | 0.21455 | 0.03601 | 0.14422 |
| **CD74** | 0.03579 | 0.10248 | 0.02673 | 1.58872 | 0.04960 | 0.06656 | 0.04220 |
| **CD79A** | 0.04304 | 0.00000 | 0.10448 | 0.05907 | 0.16149 | 0.08059 | 0.07219 |
| **CD79B** | 0.02275 | 0.00000 | 0.07676 | 0.06188 | 0.10831 | 0.10203 | 0.07059 |
| **CD80** | 0.02165 | 0.03032 | 0.04280 | 0.00342 | 0.07685 | 0.13000 | 0.09061 |
| **CD81** | 7.60745 | 22.28504 | 2.29257 | 5.41245 | 0.00000 | 22.36244 | 48.56079 |
| **CD81-AS1** | 0.01719 | 0.03263 | 0.02523 | 0.01541 | 0.00682 | 0.06924 | 0.03765 |
| **CD82** | 0.00000 | 0.19631 | 9.46892 | 0.00000 | 3.62280 | 0.04584 | 0.00623 |
| **CD83** | 1.08715 | 0.18436 | 1.77578 | 0.00000 | 2.25595 | 0.20893 | 0.04115 |
| **CD84** | 0.04247 | 0.01673 | 0.01300 | 0.02152 | 0.01827 | 0.01488 | 0.01092 |
| **CD86** | 0.01508 | 0.00000 | 0.63720 | 0.02261 | 0.00310 | 0.00303 | 0.02275 |
| **CD8A** | 0.02195 | 0.02004 | 0.02199 | 0.02270 | 0.00717 | 0.01289 | 0.01723 |
| **CD9** | 15.37344 | 30.31959 | 26.14082 | 8.84911 | 6.96329 | 33.19457 | 1.11778 |
| **CD96** | 0.03811 | 0.03756 | 0.06035 | 0.05282 | 0.01596 | 0.06054 | 0.01490 |
| **CD97** | 5.81827 | 5.99652 | 0.19942 | 1.14489 | 0.06723 | 0.88664 | 9.15241 |
| **CD99L2** | 0.36306 | 0.12881 | 0.02077 | 0.00000 | 0.26333 | 0.03600 | 0.31623 |
| **CDA** | 0.00000 | 4.06124 | 3.55350 | 0.00000 | 1.59977 | 7.20365 | 0.00000 |
| **CDADC1** | 3.66818 | 1.13871 | 2.35414 | 1.07667 | 0.30920 | 1.04457 | 1.81009 |
| **CDAN1** | 0.06055 | 3.88058 | 1.80913 | 1.92667 | 2.02795 | 1.32681 | 2.71273 |
| **CDC123** | 46.35565 | 21.56044 | 21.79645 | 39.72035 | 4.54763 | 16.52002 | 0.00000 |

| | | | | | | | |
|---|---|---|---|---|---|---|---|
| **CDC14A** | 1.02276 | 0.73689 | 0.67232 | 1.30405 | 0.25912 | 0.81886 | 0.14630 |
| **CDC14B** | 0.06435 | 0.75695 | 0.81204 | 1.26418 | 0.25749 | 1.51121 | 0.58210 |
| **CDC14C** | 0.01297 | 0.02948 | 0.01514 | 0.02684 | 0.01248 | 0.04404 | 0.03477 |
| **CDC16** | 5.66486 | 4.09260 | 3.30312 | 0.00000 | 0.79029 | 5.20714 | 21.42195 |
| **CDC20** | 31.59556 | 44.22548 | 22.16153 | 34.62180 | 15.27339 | 12.30037 | 7.74035 |
| **CDC20B** | 0.00842 | 0.02303 | 0.13993 | 0.01263 | 0.78649 | 0.00928 | 0.06612 |
| **CDC23** | 16.76215 | 1.53219 | 12.20919 | 14.99750 | 2.90843 | 3.86204 | 0.00000 |
| **CDC25A** | 5.78784 | 11.75374 | 11.20136 | 3.93634 | 4.65476 | 6.08425 | 25.26722 |
| **CDC25B** | 4.22385 | 7.78553 | 15.29849 | 10.15137 | 5.85612 | 5.36834 | 13.16897 |
| **CDC25C** | 4.16151 | 0.93097 | 3.97182 | 4.06247 | 3.01366 | 2.75483 | 0.37924 |
| **CDC26** | 2.99995 | 4.28067 | 4.32405 | 3.10234 | 5.53212 | 0.71936 | 0.00000 |
| **CDC27** | 17.32462 | 3.51381 | 8.31102 | 8.35650 | 3.68049 | 7.64475 | 8.88973 |
| **CDC34** | 0.00000 | 3.67727 | 1.67897 | 1.19550 | 1.40171 | 3.17082 | 16.10085 |
| **CDC37** | 25.96097 | 34.82198 | 14.44094 | 35.16022 | 14.16370 | 13.64948 | 14.45743 |
| **CDC37L1** | 9.18241 | 5.90065 | 4.84707 | 4.75022 | 2.32208 | 4.48001 | 0.08872 |
| **CDC37L1-AS1** | 0.04690 | 0.03397 | 0.01294 | 0.11500 | 0.00690 | 0.02008 | 0.05133 |
| **CDC40** | 7.49775 | 0.25543 | 4.22752 | 6.53042 | 4.01432 | 2.82432 | 5.11481 |
| **CDC42** | 32.58214 | 12.52198 | 12.48882 | 20.65922 | 8.20048 | 15.07752 | 13.82468 |
| **CDC42BPA** | 2.49671 | 2.77296 | 4.13685 | 1.45678 | 3.33609 | 6.27072 | 12.29469 |
| **CDC42BPB** | 0.00000 | 0.72420 | 0.78026 | 1.56675 | 3.49313 | 2.53226 | 15.73238 |
| **CDC42BPG** | 0.20832 | 1.19367 | 0.95872 | 2.53643 | 0.07552 | 0.49413 | 5.92744 |
| **CDC42EP1** | 0.00000 | 5.88112 | 2.60158 | 18.56328 | 0.00000 | 0.00000 | 14.95146 |
| **CDC42EP2** | 3.09290 | 2.28482 | 1.79871 | 7.26007 | 0.71517 | 1.32017 | 1.31217 |
| **CDC42EP3** | 0.69326 | 0.00001 | 0.45924 | 6.92317 | 0.38332 | 1.38378 | 2.42736 |
| **CDC42EP4** | 0.00000 | 16.03880 | 4.15669 | 9.47392 | 0.00000 | 23.23389 | 3.25300 |
| **CDC42EP5** | 0.03660 | 0.00658 | 0.01804 | 0.02058 | 0.03282 | 0.01424 | 0.04898 |
| **CDC42P3** | 0.11191 | 0.02042 | 0.02367 | 0.05440 | 0.01639 | 0.04321 | 0.03931 |
| **CDC42SE1** | 13.28230 | 6.53078 | 0.00000 | 17.10704 | 1.67710 | 4.09868 | 4.11657 |
| **CDC42SE2** | 26.81110 | 3.68060 | 6.06281 | 11.74738 | 1.53169 | 3.86797 | 8.83359 |
| **CDC45** | 13.09666 | 9.03212 | 6.16376 | 12.69656 | 1.84141 | 4.93480 | 1.95050 |
| **CDC5L** | 23.54996 | 9.23110 | 13.30920 | 13.27690 | 9.66887 | 10.31058 | 9.32879 |
| **CDC6** | 35.01820 | 14.20549 | 13.83088 | 18.74026 | 11.65958 | 6.16635 | 0.00000 |
| **CDC7** | 11.70366 | 2.81698 | 7.16514 | 3.58717 | 2.81346 | 2.94786 | 0.00000 |
| **CDC73** | 6.66372 | 0.00000 | 6.10370 | 5.97451 | 0.70812 | 3.70366 | 16.86262 |
| **CDCA2** | 13.73704 | 8.93699 | 9.88461 | 7.86407 | 4.78843 | 4.83910 | 7.24820 |

| | | | | | | | |
|---|---|---|---|---|---|---|---|
| **CDCA3** | 23.32804 | 20.99080 | 8.69788 | 22.07155 | 2.60243 | 9.73336 | 0.00000 |
| **CDCA4** | 30.82547 | 22.96458 | 13.73378 | 15.33567 | 6.62002 | 11.88155 | 0.43870 |
| **CDCA5** | 25.03859 | 15.42028 | 11.52799 | 26.59961 | 6.26620 | 9.28513 | 15.50848 |
| **CDCA7** | 20.90415 | 1.10570 | 8.87621 | 5.95858 | 0.00000 | 4.02410 | 4.75001 |
| **CDCA7L** | 0.06818 | 0.01370 | 0.37034 | 0.04675 | 5.96230 | 4.75177 | 0.00000 |
| **CDCA8** | 26.85773 | 19.80806 | 17.15770 | 19.17948 | 10.45921 | 8.84240 | 0.88552 |
| **CDCP1** | 1.52107 | 2.11229 | 0.75861 | 0.33964 | 0.01624 | 1.58805 | 0.80031 |
| **CDCP2** | 0.03180 | 0.11713 | 0.00889 | 0.04605 | 0.00806 | 0.00561 | 0.00000 |
| **CDH1** | 50.58667 | 43.01822 | 7.08827 | 36.96182 | 7.15238 | 11.70665 | 18.00683 |
| **CDH10** | 0.09787 | 0.00000 | 0.08627 | 0.01059 | 0.78805 | 0.05606 | 0.00000 |
| **CDH11** | 0.03246 | 0.00116 | 0.00877 | 0.13106 | 0.00855 | 4.01998 | 0.00000 |
| **CDH12** | 0.01756 | 0.00000 | 0.02862 | 0.05157 | 1.61045 | 0.01687 | 0.01741 |
| **CDH13** | 0.01611 | 0.00857 | 0.00750 | 0.01698 | 0.11755 | 0.06608 | 0.05721 |
| **CDH15** | 0.02434 | 0.00407 | 0.03708 | 0.04716 | 0.08586 | 0.01115 | 0.06233 |
| **CDH16** | 0.00925 | 0.01729 | 0.00885 | 0.08969 | 0.01544 | 0.32209 | 0.20432 |
| **CDH17** | 0.03414 | 0.01483 | 0.10969 | 0.00730 | 2.10250 | 0.09736 | 0.01502 |
| **CDH18** | 0.05002 | 0.05540 | 0.05029 | 0.19137 | 0.22955 | 0.00547 | 0.00254 |
| **CDH19** | 0.06259 | 0.00927 | 0.02359 | 0.02509 | 0.00758 | 0.01678 | 0.00282 |
| **CDH2** | 0.16165 | 0.00000 | 2.42219 | 0.13542 | 1.56752 | 2.46474 | 0.04088 |
| **CDH20** | 0.03207 | 0.01368 | 0.01540 | 0.03387 | 0.04731 | 0.01512 | 0.01192 |
| **CDH22** | 0.01130 | 0.03670 | 0.00917 | 0.02415 | 0.03608 | 0.02816 | 0.03478 |
| **CDH23** | 0.00725 | 1.49324 | 0.67802 | 0.09485 | 0.06891 | 0.00000 | 0.32442 |
| **CDH24** | 0.00000 | 0.00000 | 0.27057 | 0.32174 | 0.51113 | 0.00000 | 2.21957 |
| **CDH26** | 0.07878 | 0.07073 | 0.01854 | 0.04701 | 0.00641 | 0.24399 | 0.00000 |
| **CDH3** | 4.24966 | 6.38455 | 12.49241 | 3.32999 | 12.24437 | 6.07573 | 0.11112 |
| **CDH4** | 0.24372 | 0.00534 | 0.02213 | 0.13750 | 0.20753 | 0.07850 | 0.82453 |
| **CDH5** | 0.00000 | 0.11457 | 0.00000 | 0.46330 | 0.21067 | 1.27132 | 0.06566 |
| **CDH6** | 0.05298 | 0.00203 | 0.01818 | 0.00845 | 0.05234 | 0.02360 | 0.14367 |
| **CDH7** | 0.08825 | 0.00733 | 0.02839 | 0.01119 | 0.01427 | 0.08508 | 0.00000 |
| **CDH8** | 0.14123 | 0.91888 | 0.39340 | 0.00439 | 0.38870 | 2.86685 | 0.00000 |
| **CDHR1** | 0.04690 | 0.28034 | 0.04525 | 0.02716 | 0.00225 | 13.35430 | 0.00000 |
| **CDHR2** | 0.04357 | 0.00223 | 0.01953 | 0.07293 | 0.05288 | 0.17535 | 0.00000 |
| **CDHR3** | 0.02548 | 0.01915 | 0.02045 | 0.03836 | 0.01420 | 0.07069 | 0.00777 |
| **CDHR4** | 0.02511 | 0.01903 | 0.01033 | 0.04378 | 0.03006 | 0.01295 | 0.00752 |
| **CDHR5** | 0.01024 | 0.01356 | 0.01398 | 0.02075 | 0.01202 | 0.02326 | 0.13357 |

| | | | | | | | |
|---|---|---|---|---|---|---|---|
| **CDIP1** | 0.00000 | 3.25331 | 2.42671 | 0.66902 | 3.47102 | 3.83442 | 0.68856 |
| **CDIPT** | 0.95737 | 11.68131 | 6.19114 | 4.95684 | 2.64376 | 8.34615 | 26.18208 |
| **CDIPT-AS1** | 0.17645 | 0.17104 | 0.59546 | 0.22356 | 0.48841 | 0.00000 | 0.01518 |
| **CDK1** | 46.16364 | 3.76559 | 31.30132 | 12.93851 | 8.16953 | 8.97384 | 14.13099 |
| **CDK10** | 4.28184 | 2.98447 | 2.91095 | 4.77998 | 0.74807 | 4.63628 | 3.32306 |
| **CDK11A** | 0.23959 | 0.95354 | 0.27950 | 1.23591 | 0.52512 | 0.70818 | 1.32632 |
| **CDK11B** | 0.83343 | 1.56487 | 1.32203 | 1.97031 | 0.96031 | 0.81791 | 1.37858 |
| **CDK12** | 10.30660 | 11.64632 | 9.20116 | 8.68497 | 9.95308 | 8.55572 | 24.99077 |
| **CDK13** | 4.35078 | 1.57602 | 2.91695 | 3.10641 | 1.01420 | 6.80696 | 11.17157 |
| **CDK14** | 0.50506 | 0.00000 | 0.11505 | 0.19737 | 0.05844 | 0.19971 | 0.00000 |
| **CDK15** | 0.02139 | 0.03655 | 0.00000 | 0.01770 | 0.38252 | 0.01098 | 0.01976 |
| **CDK16** | 9.40043 | 21.59133 | 5.46477 | 7.87555 | 3.43702 | 13.28444 | 24.39968 |
| **CDK17** | 1.40457 | 0.00000 | 3.33571 | 2.39372 | 2.83603 | 1.50183 | 4.05731 |
| **CDK18** | 0.01378 | 0.10887 | 0.00000 | 0.01602 | 0.12519 | 0.15850 | 0.01435 |
| **CDK19** | 0.32683 | 0.16434 | 0.36430 | 1.61946 | 0.33996 | 0.75622 | 1.09535 |
| **CDK2** | 11.39798 | 5.39914 | 1.94283 | 14.31854 | 3.09712 | 2.74356 | 0.00000 |
| **CDK20** | 1.66852 | 0.00000 | 1.69672 | 0.77054 | 1.27962 | 1.02546 | 0.00056 |
| **CDK2AP1** | 8.18249 | 2.54746 | 12.41333 | 4.69405 | 2.61129 | 4.94976 | 10.93932 |
| **CDK2AP2** | 15.62155 | 18.26811 | 13.18973 | 22.56317 | 18.42048 | 15.06523 | 1.84276 |
| **CDK4** | 35.21036 | 21.83013 | 15.41400 | 30.47397 | 7.32586 | 16.03123 | 4.81362 |
| **CDK5** | 6.52556 | 6.95810 | 1.99825 | 5.04195 | 1.10051 | 6.03485 | 2.75547 |
| **CDK5R1** | 0.85518 | 0.00000 | 0.51615 | 0.92627 | 0.30605 | 0.16797 | 1.85474 |
| **CDK5R2** | 0.00000 | 0.02721 | 0.21414 | 0.01836 | 0.30500 | 0.04146 | 0.11298 |
| **CDK5RAP1** | 22.63617 | 11.62182 | 10.15208 | 12.30517 | 2.85292 | 5.68337 | 0.00000 |
| **CDK5RAP2** | 5.35988 | 15.81162 | 15.44056 | 2.70930 | 0.59269 | 3.70424 | 12.27157 |
| **CDK5RAP3** | 8.45769 | 8.15612 | 3.33593 | 3.68161 | 0.74685 | 5.82444 | 0.00002 |
| **CDK6** | 22.95228 | 0.00000 | 0.17402 | 3.16130 | 0.49266 | 0.00000 | 27.78369 |
| **CDK7** | 19.67821 | 1.50475 | 11.95607 | 18.00683 | 7.97788 | 6.48520 | 7.81767 |
| **CDK8** | 1.74017 | 1.81407 | 1.02077 | 1.16891 | 0.64654 | 0.74431 | 2.67845 |
| **CDK9** | 2.78422 | 6.36563 | 3.11989 | 3.95069 | 4.12134 | 3.79928 | 6.48545 |
| **CDKAL1** | 7.36909 | 1.17921 | 4.14499 | 5.55170 | 0.64779 | 3.21146 | 1.81351 |
| **CDKL1** | 0.16854 | 0.25049 | 0.42600 | 0.09236 | 0.07893 | 0.01490 | 0.00116 |
| **CDKL2** | 0.20330 | 0.00000 | 0.08738 | 0.00862 | 0.27364 | 0.00148 | 0.07346 |
| **CDKL3** | 0.28768 | 0.02784 | 0.47936 | 0.27098 | 0.54195 | 0.05728 | 0.13640 |
| **CDKL4** | 0.03188 | 0.00343 | 0.19374 | 0.02211 | 0.00797 | 0.03475 | 0.04368 |

| | | | | | | | |
|---|---|---|---|---|---|---|---|
| **CDKL5** | 0.26422 | 0.35104 | 0.90232 | 0.36264 | 1.20225 | 0.45908 | 1.27192 |
| **CDKN1A** | 0.21383 | 4.76627 | 0.00000 | 8.94626 | 2.72508 | 4.50341 | 1.46148 |
| **CDKN1B** | 1.24588 | 0.00000 | 0.18140 | 0.41348 | 0.00490 | 1.26990 | 1.94382 |
| **CDKN1C** | 0.00000 | 0.21481 | 0.00000 | 20.50505 | 0.79113 | 1.86584 | 2.98009 |
| **CDKN2A** | 0.02009 | 0.06909 | 0.01194 | 0.06075 | 0.02317 | 0.02713 | 0.00000 |
| **CDKN2AIP** | 6.64702 | 0.60436 | 4.06409 | 4.27355 | 14.69740 | 3.61479 | 2.34162 |
| **CDKN2AIPNL** | 8.72375 | 6.90750 | 3.37114 | 0.39787 | 0.43133 | 3.05751 | 1.58343 |
| **CDKN2B** | 0.06275 | 0.22832 | 0.06981 | 0.09379 | 0.16673 | 0.00194 | 0.00000 |
| **CDKN2B-AS1** | 0.04318 | 0.00468 | 0.03941 | 0.02804 | 0.01116 | 0.01252 | 0.01159 |
| **CDKN2C** | 0.00000 | 0.18099 | 0.10529 | 0.16039 | 0.31436 | 0.45195 | 0.03542 |
| **CDKN2D** | 3.68761 | 1.86609 | 2.44893 | 6.63193 | 2.32653 | 2.10458 | 3.91275 |
| **CDKN3** | 11.39623 | 4.77950 | 4.18874 | 3.29026 | 0.54017 | 5.89068 | 0.00000 |
| **CDNF** | 0.18905 | 0.24101 | 0.15427 | 0.13265 | 0.11891 | 0.07467 | 0.48995 |
| **CDO1** | 1.11297 | 0.00000 | 0.63463 | 6.93356 | 0.07776 | 1.01705 | 0.00000 |
| **CDON** | 0.36240 | 0.28972 | 0.21471 | 0.39644 | 0.88126 | 0.84879 | 0.46222 |
| **CDPF1** | 2.34585 | 1.15797 | 2.38865 | 1.64054 | 0.05151 | 0.26465 | 0.66845 |
| **CDR1** | 1.56647 | 0.00000 | 7.74950 | 0.00000 | 4.06945 | 2.82095 | 8.57078 |
| **CDR2** | 0.76663 | 0.00000 | 6.84036 | 2.30791 | 1.22131 | 4.97714 | 3.67854 |
| **CDR2L** | 8.50416 | 9.54586 | 0.45807 | 9.48526 | 0.46787 | 2.45783 | 0.04537 |
| **CDRT1** | 0.12406 | 0.16221 | 0.03992 | 0.14379 | 0.03772 | 0.06191 | 0.14798 |
| **CDRT15** | 0.03148 | 0.01776 | 0.01180 | 0.02631 | 0.00771 | 0.01263 | 0.03791 |
| **CDRT15P1** | 0.04885 | 0.02238 | 0.00982 | 0.06856 | 0.01772 | 0.01781 | 0.05621 |
| **CDRT4** | 0.01226 | 0.11031 | 0.04805 | 0.01565 | 0.01230 | 0.00482 | 0.08081 |
| **CDS1** | 3.39859 | 2.37393 | 1.05647 | 2.46343 | 1.00838 | 1.69356 | 14.43025 |
| **CDS2** | 9.83172 | 0.01185 | 12.72225 | 10.98495 | 2.94551 | 2.88516 | 2.90716 |
| **CDSN** | 0.00002 | 0.02857 | 0.03195 | 0.00000 | 0.08664 | 0.10080 | 0.24110 |
| **CDT1** | 2.00825 | 16.35194 | 9.18548 | 6.71497 | 1.28575 | 9.19799 | 26.84096 |
| **CDV3** | 40.31121 | 15.88176 | 20.77694 | 15.91794 | 15.17159 | 17.57036 | 14.62926 |
| **CDX1** | 6.71876 | 0.00000 | 0.00000 | 3.17848 | 0.00000 | 0.97883 | 10.91697 |
| **CDX2** | 21.72420 | 4.07000 | 0.00000 | 0.00000 | 0.03846 | 0.40401 | 14.50372 |
| **CDX4** | 0.02509 | 0.00687 | 0.02188 | 0.01752 | 0.00724 | 0.09074 | 0.01694 |
| **CDYL** | 7.39369 | 10.84123 | 3.83212 | 7.76912 | 0.35355 | 3.26123 | 13.85726 |
| **CDYL2** | 0.06354 | 0.37809 | 0.00000 | 0.05743 | 0.13231 | 0.00000 | 0.36510 |
| **CEACAM1** | 0.64851 | 2.48050 | 0.00054 | 2.62246 | 0.01332 | 10.30216 | 0.00000 |
| **CEACAM16** | 0.01698 | 0.01732 | 0.01134 | 0.02194 | 0.00783 | 0.00845 | 0.06477 |

| | | | | | | | |
|---|---|---|---|---|---|---|---|
| **CEACAM19** | 0.15477 | 0.08322 | 0.07874 | 0.08803 | 0.01203 | 0.16020 | 0.11135 |
| **CEACAM20** | 0.02760 | 0.02951 | 0.00960 | 0.18175 | 0.02492 | 0.07214 | 0.00000 |
| **CEACAM21** | 0.00000 | 0.00000 | 0.18761 | 0.12502 | 0.14553 | 0.10844 | 0.09856 |
| **CEACAM22P** | 0.30392 | 0.06282 | 0.00784 | 0.16757 | 0.25461 | 0.02944 | 0.00000 |
| **CEACAM3** | 0.01519 | 0.01899 | 0.00854 | 0.02037 | 0.03591 | 0.03809 | 0.02748 |
| **CEACAM4** | 0.02530 | 0.01860 | 0.01206 | 0.01761 | 0.00933 | 0.01206 | 0.02217 |
| **CEACAM5** | 0.01347 | 0.02199 | 0.01417 | 0.02157 | 0.00992 | 0.20094 | 0.04278 |
| **CEACAM6** | 0.00564 | 1.42374 | 0.03382 | 0.01135 | 0.02072 | 11.16033 | 0.00000 |
| **CEACAM7** | 0.01306 | 0.03335 | 0.02580 | 0.01308 | 0.02237 | 0.18882 | 0.02640 |
| **CEACAM8** | 0.02357 | 0.06057 | 0.01264 | 0.01442 | 0.01423 | 0.01500 | 0.02232 |
| **CEBPA** | 6.78300 | 7.38335 | 0.00000 | 18.25966 | 0.00000 | 0.00000 | 30.62530 |
| **CEBPA-AS1** | 0.21310 | 0.06819 | 0.03573 | 0.20745 | 0.00406 | 0.00000 | 0.30505 |
| **CEBPB** | 0.00000 | 0.08793 | 0.06711 | 2.99203 | 0.06334 | 0.18431 | 5.75120 |
| **CEBPD** | 0.00000 | 0.43201 | 0.00028 | 2.96320 | 0.00541 | 0.72671 | 3.59154 |
| **CEBPE** | 0.00000 | 0.01223 | 0.01879 | 0.16853 | 0.00611 | 0.10768 | 0.15365 |
| **CEBPG** | 10.24372 | 8.70309 | 9.26959 | 16.70771 | 2.43256 | 3.58427 | 22.73834 |
| **CEBPZ** | 27.08322 | 12.25028 | 21.65837 | 14.71190 | 8.17171 | 24.05262 | 18.58152 |
| **CEBPZOS** | 4.05874 | 1.08440 | 7.18882 | 2.52301 | 1.33406 | 4.53952 | 6.21369 |
| **CECR1** | 0.06470 | 0.74611 | 0.30874 | 0.00000 | 0.04651 | 4.64024 | 0.00154 |
| **CECR2** | 0.14166 | 1.38428 | 0.54335 | 1.23447 | 1.32494 | 3.94813 | 1.26588 |
| **CECR5** | 4.13835 | 9.70351 | 6.85905 | 3.28007 | 0.54683 | 8.66899 | 3.70376 |
| **CECR5-AS1** | 0.03729 | 0.04641 | 0.07002 | 0.02353 | 0.01990 | 0.02151 | 0.00051 |
| **CECR6** | 0.00000 | 0.00989 | 0.12930 | 0.01224 | 0.00488 | 0.07020 | 0.18498 |
| **CECR7** | 0.02057 | 0.00000 | 0.16604 | 0.03124 | 0.25573 | 0.01181 | 0.01717 |
| **CEL** | 0.00624 | 0.00590 | 0.03196 | 0.02990 | 0.00869 | 0.01304 | 0.17525 |
| **CELA1** | 1.79938 | 1.29243 | 0.61237 | 0.89280 | 0.15994 | 0.00000 | 0.75946 |
| **CELA2A** | 0.02634 | 0.01458 | 0.01979 | 0.02224 | 0.04902 | 0.01273 | 0.01542 |
| **CELA2B** | 0.00891 | 0.03562 | 0.00654 | 0.03685 | 0.03136 | 0.02776 | 0.03868 |
| **CELA3A** | 0.00855 | 0.10958 | 0.96265 | 0.06200 | 0.39292 | 1.07411 | 0.00000 |
| **CELA3B** | 0.00043 | 0.41098 | 0.78700 | 0.03133 | 0.41357 | 0.74583 | 0.00576 |
| **CELF1** | 13.82878 | 8.79241 | 9.83347 | 14.88314 | 4.18441 | 11.47682 | 31.41078 |
| **CELF2** | 0.00000 | 1.00035 | 0.00000 | 0.58917 | 2.22421 | 1.28333 | 0.81652 |
| **CELF2-AS2** | 0.00858 | 0.04389 | 0.02028 | 0.02877 | 0.01025 | 0.06805 | 0.04420 |
| **CELF3** | 0.00000 | 0.12481 | 0.02660 | 0.06526 | 0.06609 | 0.03045 | 0.15633 |
| **CELF4** | 0.11075 | 0.18360 | 0.03554 | 0.01018 | 0.01337 | 0.30337 | 0.25530 |

| | | | | | | | |
|---|---|---|---|---|---|---|---|
| **CELF5** | 0.00737 | 0.00994 | 0.09549 | 0.01418 | 0.17786 | 0.02220 | 0.16741 |
| **CELF6** | 0.17587 | 0.00000 | 0.06988 | 0.48455 | 0.01994 | 0.06139 | 0.55875 |
| **CELP** | 0.02182 | 0.01624 | 0.01371 | 0.02097 | 0.02560 | 0.01630 | 0.01909 |
| **CELSR1** | 0.00000 | 0.50663 | 0.26910 | 0.01643 | 1.70844 | 0.52127 | 1.61128 |
| **CELSR2** | 0.00000 | 0.85290 | 0.74048 | 2.63268 | 0.00000 | 2.82026 | 4.00327 |
| **CELSR3** | 0.25943 | 0.00000 | 0.58062 | 1.23369 | 0.00101 | 1.28999 | 1.43874 |
| **CELSR3-AS1** | 0.01816 | 0.07904 | 0.05995 | 0.04501 | 0.02749 | 0.01864 | 0.06191 |
| **CEMIP** | 0.30985 | 0.00000 | 0.02829 | 0.03733 | 0.08203 | 0.00000 | 0.61236 |
| **CEMP1** | 0.37069 | 0.84744 | 0.52392 | 2.29877 | 0.20397 | 0.30774 | 7.75997 |
| **CEND1** | 0.03078 | 0.01897 | 0.00265 | 0.01166 | 0.12283 | 0.32622 | 0.01106 |
| **CENPA** | 0.00000 | 4.03938 | 8.83809 | 0.96203 | 3.78845 | 9.66044 | 6.01261 |
| **CENPB** | 0.00000 | 1.68719 | 0.41034 | 1.71008 | 0.00000 | 1.48879 | 8.68041 |
| **CENPBD1** | 1.56620 | 0.41307 | 1.81409 | 2.06600 | 0.47115 | 1.25674 | 0.59335 |
| **CENPBD1P1** | 1.84264 | 3.74101 | 1.43809 | 4.83366 | 0.19875 | 3.07870 | 4.29226 |
| **CENPC** | 4.47373 | 0.00000 | 2.47190 | 1.78762 | 2.52061 | 1.41814 | 4.80954 |
| **CENPE** | 17.83290 | 0.48126 | 12.54536 | 9.83224 | 6.80948 | 6.82499 | 25.71717 |
| **CENPF** | 21.31693 | 7.67098 | 25.22164 | 23.28096 | 6.93053 | 13.17353 | 37.78680 |
| **CENPH** | 1.79426 | 1.72057 | 5.42009 | 0.87194 | 0.80212 | 3.60339 | 6.49670 |
| **CENPI** | 3.67577 | 1.04470 | 1.74586 | 0.37236 | 1.10473 | 0.12290 | 1.18660 |
| **CENPJ** | 7.62825 | 2.01446 | 5.75677 | 1.87147 | 2.11032 | 8.23109 | 12.17538 |
| **CENPK** | 13.70857 | 0.00000 | 5.78226 | 5.86840 | 1.80857 | 4.23879 | 4.10657 |
| **CENPL** | 10.18010 | 0.92175 | 5.16247 | 5.67269 | 4.01991 | 2.25139 | 0.00000 |
| **CENPM** | 6.82580 | 9.96395 | 1.94506 | 1.80395 | 0.53016 | 6.10420 | 0.89322 |
| **CENPN** | 20.61441 | 15.03786 | 11.58780 | 20.17626 | 7.52376 | 12.52010 | 7.00215 |
| **CENPO** | 8.24476 | 5.29720 | 4.70173 | 9.88694 | 4.40338 | 2.88026 | 4.76315 |
| **CENPP** | 4.27630 | 2.83826 | 1.26261 | 1.38814 | 3.52549 | 1.56075 | 0.00000 |
| **CENPQ** | 6.97609 | 0.00000 | 7.65776 | 7.70722 | 2.58888 | 2.07239 | 0.63377 |
| **CENPT** | 10.67048 | 12.40796 | 3.19187 | 18.63020 | 7.17491 | 7.35732 | 5.48813 |
| **CENPU** | 14.99769 | 3.55200 | 8.71801 | 12.34828 | 4.52220 | 8.42930 | 8.90460 |
| **CENPV** | 0.00000 | 4.29533 | 2.59172 | 2.47394 | 0.00000 | 6.66999 | 13.27404 |
| **CENPW** | 20.21191 | 16.81945 | 16.65794 | 18.66650 | 8.96238 | 10.54163 | 2.77462 |
| **CEP104** | 2.35594 | 6.19323 | 1.31282 | 1.73528 | 1.26275 | 3.00728 | 8.82322 |
| **CEP112** | 0.08901 | 0.46811 | 0.32266 | 0.00000 | 0.43694 | 0.64892 | 0.50232 |
| **CEP120** | 1.60295 | 0.21677 | 0.33913 | 0.00000 | 1.09235 | 1.28557 | 2.81052 |
| **CEP128** | 2.06968 | 0.76466 | 1.45511 | 0.81284 | 1.16894 | 1.29305 | 2.74246 |

| | | | | | | | |
|---|---|---|---|---|---|---|---|
| **CEP131** | 7.95993 | 7.50390 | 0.05261 | 2.20087 | 0.33212 | 1.58351 | 5.73430 |
| **CEP135** | 4.41990 | 0.00000 | 3.68583 | 1.55028 | 0.38887 | 3.38603 | 3.81180 |
| **CEP152** | 5.11127 | 0.65719 | 3.67248 | 2.45627 | 3.23734 | 2.77889 | 8.40526 |
| **CEP162** | 2.91506 | 0.00000 | 1.50383 | 2.28687 | 2.39415 | 1.08128 | 2.85423 |
| **CEP164** | 0.14064 | 3.82167 | 10.14647 | 0.53155 | 0.28059 | 8.10208 | 5.99225 |
| **CEP170** | 1.81808 | 0.00000 | 0.90851 | 0.96929 | 2.43296 | 1.59022 | 1.07374 |
| **CEP170B** | 0.00000 | 1.56726 | 0.00000 | 2.94268 | 0.34977 | 0.08481 | 6.45744 |
| **CEP170P1** | 0.06719 | 0.01209 | 0.02469 | 0.01949 | 0.02629 | 0.09446 | 0.00000 |
| **CEP19** | 0.24957 | 0.00000 | 1.61930 | 0.34184 | 0.23111 | 0.67901 | 0.00000 |
| **CEP192** | 4.04367 | 3.49185 | 4.22802 | 0.96798 | 1.49919 | 3.90965 | 6.07474 |
| **CEP250** | 1.85020 | 3.28682 | 1.27833 | 4.30230 | 1.19205 | 1.48803 | 2.97535 |
| **CEP290** | 3.68082 | 1.07657 | 1.41882 | 2.10909 | 0.00000 | 0.19512 | 13.39246 |
| **CEP350** | 8.23640 | 0.65995 | 7.47446 | 4.72000 | 2.14117 | 6.95912 | 19.24420 |
| **CEP41** | 5.40112 | 2.75431 | 3.27327 | 3.61831 | 0.72116 | 1.02790 | 0.00000 |
| **CEP44** | 4.58675 | 0.00000 | 5.11654 | 1.79442 | 2.35936 | 4.47297 | 2.54621 |
| **CEP55** | 1.57667 | 0.00000 | 3.93173 | 3.91239 | 1.10066 | 0.40921 | 6.57180 |
| **CEP57** | 15.86217 | 3.63704 | 9.93966 | 7.86543 | 4.95280 | 7.77258 | 7.59557 |
| **CEP57L1** | 6.00988 | 0.00000 | 3.67489 | 0.61373 | 0.75739 | 0.83389 | 0.92461 |
| **CEP63** | 8.00610 | 2.33423 | 6.96421 | 2.86833 | 1.72023 | 3.57486 | 8.05751 |
| **CEP68** | 6.71028 | 3.76809 | 1.75570 | 6.31944 | 0.58731 | 0.68455 | 0.82219 |
| **CEP70** | 1.61788 | 0.00000 | 1.45394 | 1.86560 | 2.03634 | 0.87997 | 0.00000 |
| **CEP72** | 3.58612 | 3.06710 | 0.31508 | 3.16089 | 0.08795 | 0.66355 | 2.00077 |
| **CEP76** | 6.07659 | 0.80221 | 5.17136 | 5.13993 | 0.85190 | 2.74396 | 3.55392 |
| **CEP78** | 4.86205 | 3.58720 | 6.18728 | 4.14713 | 2.41447 | 2.55853 | 4.64749 |
| **CEP83** | 4.20463 | 0.00000 | 1.76616 | 2.32646 | 1.40072 | 1.21433 | 2.32593 |
| **CEP83-AS1** | 0.08149 | 0.02001 | 0.13378 | 0.02716 | 0.20878 | 0.00349 | 0.00378 |
| **CEP85** | 12.58275 | 4.97053 | 4.51495 | 24.36311 | 6.38645 | 3.18405 | 8.53329 |
| **CEP85L** | 1.47438 | 0.00000 | 7.92907 | 1.98670 | 9.66080 | 7.27704 | 3.56194 |
| **CEP89** | 1.21831 | 0.67624 | 0.25225 | 2.16301 | 1.03420 | 1.25143 | 0.69557 |
| **CEP95** | 9.86645 | 1.14616 | 4.57533 | 2.98837 | 4.88762 | 5.95318 | 6.34583 |
| **CEP97** | 2.16278 | 0.00000 | 4.52619 | 3.21447 | 5.54756 | 1.12545 | 1.22408 |
| **CEPT1** | 16.65635 | 0.00002 | 8.68980 | 9.85930 | 0.24005 | 6.03957 | 3.01416 |
| **CERCAM** | 1.50449 | 0.00000 | 0.62533 | 1.45466 | 0.27418 | 0.18078 | 1.91493 |
| **CERK** | 0.02408 | 0.15036 | 0.00710 | 0.03320 | 0.01689 | 0.01042 | 0.07690 |
| **CERKL** | 0.04712 | 0.00000 | 0.22895 | 0.00128 | 0.40883 | 0.11219 | 0.04500 |

| | | | | | | | |
|---|---|---|---|---|---|---|---|
| **CERS1.uid-2288** | 0.00000 | 0.59741 | 0.11932 | 0.09524 | 0.06757 | 0.07368 | 0.18119 |
| **CERS2** | 8.44395 | 5.97924 | 12.39258 | 9.45493 | 5.12319 | 5.05890 | 15.82005 |
| **CERS3** | 0.02422 | 0.01764 | 0.00545 | 0.05233 | 0.19773 | 0.01483 | 0.00286 |
| **CERS4** | 10.11723 | 6.00861 | 2.02853 | 1.12631 | 0.06447 | 0.67093 | 7.29786 |
| **CERS5** | 27.85605 | 20.64200 | 24.19236 | 27.09332 | 7.62015 | 1.55132 | 8.00331 |
| **CERS6** | 0.07504 | 0.00000 | 0.58863 | 0.07374 | 0.80879 | 1.29391 | 0.01015 |
| **CES1** | 0.04265 | 0.06718 | 0.00261 | 0.37430 | 0.09444 | 0.02580 | 0.00000 |
| **CES1P1** | 0.01978 | 0.01841 | 0.01339 | 0.03501 | 0.00775 | 0.01164 | 0.02209 |
| **CES1P2** | 0.02416 | 0.01744 | 0.01325 | 0.02609 | 0.00777 | 0.01257 | 0.02721 |
| **CES2** | 2.38117 | 5.62995 | 2.57080 | 6.00611 | 0.00000 | 3.70659 | 4.42821 |
| **CES3** | 0.10805 | 0.88656 | 0.13168 | 0.00777 | 0.04939 | 0.01886 | 0.00000 |
| **CES4A** | 0.18100 | 0.24461 | 0.43153 | 0.15059 | 0.02539 | 0.06507 | 0.00000 |
| **CES5A** | 0.02720 | 0.01249 | 0.01296 | 0.03350 | 0.05803 | 0.01176 | 0.02788 |
| **CES5AP1** | 0.02801 | 0.01908 | 0.01235 | 0.04819 | 0.01172 | 0.02026 | 0.00120 |
| **CETN1** | 0.01570 | 0.42495 | 0.28181 | 0.05854 | 0.01438 | 0.00580 | 0.00149 |
| **CETN2** | 15.20721 | 8.00994 | 11.30699 | 5.14033 | 8.89052 | 4.85079 | 6.59647 |
| **CETN3** | 10.16779 | 0.00000 | 6.04797 | 3.82149 | 0.64191 | 4.81917 | 5.10296 |
| **CETN4P** | 0.07327 | 0.02886 | 0.04649 | 0.03593 | 0.02179 | 0.02309 | 0.00000 |
| **CETP** | 0.03054 | 0.02690 | 0.01135 | 0.10128 | 0.00846 | 0.01929 | 0.00321 |
| **CFB** | 0.03644 | 0.00000 | 0.23268 | 0.18629 | 0.21883 | 0.45861 | 0.00000 |
| **CFD** | 2.70162 | 1.69678 | 8.81387 | 0.00000 | 0.86757 | 0.00000 | 2.55101 |
| **CFDP1** | 12.57539 | 2.95814 | 10.13157 | 13.18389 | 5.37850 | 8.11614 | 12.86048 |
| **CFH** | 0.02399 | 0.04411 | 0.00850 | 0.02733 | 0.00845 | 0.01301 | 0.01817 |
| **CFHR1** | 0.02533 | 0.01251 | 0.02270 | 0.02161 | 0.00989 | 0.01298 | 0.01503 |
| **CFHR2** | 0.02105 | 0.01413 | 0.01311 | 0.02137 | 0.00795 | 0.02569 | 0.02399 |
| **CFHR3** | 0.02721 | 0.04794 | 0.03196 | 0.01529 | 0.02324 | 0.00667 | 0.02396 |
| **CFHR4** | 0.02386 | 0.02820 | 0.02598 | 0.02417 | 0.00693 | 0.01136 | 0.01102 |
| **CFI** | 0.02217 | 0.01634 | 0.01244 | 0.01911 | 0.00928 | 0.02544 | 0.01757 |
| **CFL1** | 77.45456 | 64.42893 | 32.91823 | 67.33154 | 22.81603 | 45.55604 | 37.38469 |
| **CFL1P1** | 0.06585 | 0.00000 | 0.12730 | 0.23221 | 0.19720 | 0.01891 | 0.19087 |
| **CFL2** | 21.60109 | 1.69168 | 14.51818 | 6.65971 | 3.69614 | 2.90157 | 6.37565 |
| **CFLAR** | 0.38949 | 4.68547 | 14.79231 | 0.23983 | 0.96409 | 23.95114 | 0.41124 |
| **CFLAR-AS1** | 0.01358 | 0.00000 | 0.11623 | 0.02227 | 0.07774 | 0.17111 | 0.04457 |
| **CFP** | 0.01901 | 0.00000 | 0.08889 | 0.10364 | 0.02275 | 0.18403 | 0.46130 |
| **CFTR** | 0.10429 | 0.00000 | 0.07619 | 0.19392 | 0.18760 | 0.00982 | 0.01878 |

| | | | | | | | |
|---|---|---|---|---|---|---|---|
| **CGA** | 0.00000 | 0.00000 | 0.06536 | 72.37302 | 0.02934 | 0.14185 | 0.00854 |
| **CGB** | 0.00000 | 0.00963 | 0.14535 | 4.54047 | 0.06066 | 0.00431 | 0.00000 |
| **CGB1** | 0.00098 | 0.01799 | 0.01147 | 0.10336 | 0.01410 | 0.01191 | 0.05289 |
| **CGB2** | 0.01337 | 0.02033 | 0.00865 | 0.04956 | 0.04822 | 0.01912 | 0.01726 |
| **CGB5** | 0.00000 | 0.01130 | 0.04000 | 5.41354 | 0.01163 | 0.01278 | 0.00000 |
| **CGB7** | 0.00000 | 0.00662 | 0.07895 | 2.16989 | 0.17154 | 0.00577 | 0.00000 |
| **CGB8** | 0.00000 | 0.00530 | 0.13546 | 3.54376 | 0.02896 | 0.00337 | 0.00076 |
| **CGGBP1** | 7.49545 | 9.31741 | 4.79819 | 9.26476 | 3.21376 | 6.03323 | 6.53832 |
| **CGN** | 3.03691 | 13.62170 | 0.43287 | 1.26607 | 0.95878 | 7.39762 | 3.95287 |
| **CGNL1** | 0.00000 | 0.09889 | 4.40108 | 0.05927 | 0.21075 | 4.24493 | 0.36496 |
| **CGREF1** | 0.79355 | 0.77379 | 0.00000 | 0.39339 | 1.62709 | 0.09554 | 0.82215 |
| **CGRRF1** | 5.30715 | 2.49261 | 3.40868 | 2.17328 | 0.67490 | 2.50540 | 1.40695 |
| **CHAC1** | 11.87855 | 0.64690 | 0.00000 | 6.08539 | 2.04720 | 0.00000 | 12.34560 |
| **CHAC2** | 4.93204 | 0.90247 | 0.93735 | 0.71669 | 0.03798 | 1.34176 | 0.00000 |
| **CHAD** | 0.09908 | 0.05390 | 0.05946 | 0.00000 | 0.00458 | 0.08097 | 0.01973 |
| **CHADL** | 0.01919 | 0.12001 | 0.01859 | 0.07842 | 0.02376 | 0.03020 | 0.13720 |
| **CHAF1A** | 6.57711 | 14.25925 | 4.78081 | 5.30635 | 8.07431 | 5.06435 | 19.15926 |
| **CHAF1B** | 13.47456 | 5.94922 | 3.55520 | 4.05787 | 2.91728 | 2.83637 | 0.95738 |
| **CHAMP1** | 1.03608 | 0.00000 | 1.88607 | 0.07768 | 2.12281 | 0.36990 | 4.53832 |
| **CHAT** | 0.03154 | 0.00768 | 0.03024 | 0.01996 | 0.04375 | 0.01036 | 0.02893 |
| **CHCHD1** | 23.06444 | 9.46783 | 12.80862 | 22.13719 | 0.12099 | 8.47957 | 17.07695 |
| **CHCHD10** | 0.55381 | 6.10687 | 0.09417 | 1.61513 | 0.06792 | 3.75089 | 30.32201 |
| **CHCHD2** | 44.04859 | 27.75140 | 21.70782 | 29.43246 | 11.95297 | 21.51629 | 21.55682 |
| **CHCHD3** | 26.82802 | 14.33336 | 3.23732 | 14.53515 | 0.74613 | 16.75836 | 0.00000 |
| **CHCHD4** | 16.58452 | 18.71845 | 10.64242 | 6.03200 | 0.00000 | 7.92896 | 0.00000 |
| **CHCHD5** | 6.53841 | 13.69551 | 6.23998 | 6.32095 | 0.53944 | 4.41053 | 3.37516 |
| **CHCHD6** | 3.71102 | 2.37578 | 0.51805 | 3.33473 | 2.21512 | 1.49425 | 0.67888 |
| **CHCHD7** | 29.14221 | 10.66457 | 12.44406 | 8.67286 | 8.12837 | 8.18372 | 12.81135 |
| **CHD1** | 11.89332 | 1.82991 | 8.05606 | 6.88215 | 4.64298 | 7.48793 | 26.51528 |
| **CHD1L** | 7.80760 | 6.05078 | 2.60557 | 3.56768 | 0.12565 | 9.30778 | 3.89270 |
| **CHD2** | 8.08454 | 0.72166 | 2.18850 | 8.79969 | 4.55815 | 5.39149 | 18.69329 |
| **CHD3** | 0.87802 | 0.00000 | 0.83722 | 3.22714 | 0.32888 | 1.63442 | 8.51604 |
| **CHD4** | 18.04469 | 18.68991 | 10.93194 | 12.36538 | 3.96962 | 14.72229 | 38.96748 |
| **CHD5** | 0.00000 | 0.00000 | 0.47394 | 0.01272 | 0.02964 | 0.50790 | 0.46290 |
| **CHD6** | 2.61846 | 3.65053 | 6.36658 | 3.32145 | 2.49678 | 7.97164 | 10.16662 |

| | | | | | | | |
|---|---|---|---|---|---|---|---|
| **CHD7** | 18.38900 | 11.20377 | 12.60480 | 14.12585 | 4.57289 | 0.00000 | 29.01773 |
| **CHD8** | 7.49367 | 5.76796 | 6.79649 | 10.38447 | 7.11033 | 5.09992 | 25.09500 |
| **CHD9** | 1.67255 | 0.26327 | 1.27679 | 0.14832 | 1.13975 | 1.50335 | 1.66617 |
| **CHDC2** | 0.03257 | 0.00000 | 0.08153 | 0.05437 | 0.04445 | 0.08314 | 0.13173 |
| **CHDH** | 0.62322 | 1.79836 | 0.01581 | 0.28401 | 0.00455 | 2.46200 | 0.94365 |
| **CHEK1** | 16.13071 | 3.75441 | 12.15484 | 6.55274 | 4.59707 | 9.26356 | 10.74876 |
| **CHEK2** | 8.58418 | 3.05767 | 1.79288 | 13.40788 | 0.67498 | 5.79057 | 0.00000 |
| **CHEK2P2** | 0.00699 | 0.24198 | 0.00585 | 0.01830 | 0.23755 | 0.00469 | 0.01501 |
| **CHERP** | 5.47270 | 8.20765 | 4.97706 | 8.58438 | 2.29835 | 4.85932 | 12.73990 |
| **CHFR** | 8.16669 | 5.09594 | 1.96961 | 4.70478 | 0.98747 | 6.04130 | 6.00744 |
| **CHGA** | 0.00000 | 6.81123 | 2.25253 | 0.00000 | 0.15268 | 5.74439 | 0.22917 |
| **CHI3L1** | 0.71135 | 0.15640 | 0.00000 | 0.00510 | 0.00538 | 0.00000 | 0.10687 |
| **CHI3L2** | 15.94670 | 9.24430 | 6.14236 | 0.96873 | 0.00000 | 4.01849 | 0.00000 |
| **CHIA** | 0.09115 | 0.11832 | 0.03439 | 0.02747 | 0.01040 | 0.32305 | 0.00000 |
| **CHIAP2** | 0.08469 | 0.01099 | 0.01402 | 0.03009 | 0.00789 | 0.01516 | 0.00000 |
| **CHIC1** | 0.07182 | 0.00000 | 0.87650 | 0.00000 | 0.65380 | 0.13014 | 0.04293 |
| **CHIC2** | 2.55057 | 3.07492 | 5.17020 | 3.82664 | 4.51728 | 3.21995 | 3.22735 |
| **CHID1** | 21.54469 | 22.88808 | 7.45924 | 11.66913 | 1.16668 | 7.33781 | 3.51456 |
| **CHIT1** | 0.08222 | 0.00473 | 0.01618 | 0.01341 | 0.00787 | 0.08249 | 0.00039 |
| **CHKA** | 0.98423 | 0.12926 | 0.07851 | 5.21346 | 0.39592 | 0.85861 | 3.14304 |
| **CHKB** | 0.24168 | 0.57113 | 0.01033 | 0.18381 | 0.01465 | 0.22608 | 1.62055 |
| **CHKB-AS1** | 0.00000 | 0.46584 | 0.08410 | 0.08509 | 0.05273 | 0.00000 | 0.49534 |
| **CHL1** | 0.04216 | 0.01991 | 0.02223 | 0.01280 | 0.03562 | 0.01262 | 0.02816 |
| **CHM** | 4.69084 | 0.00000 | 4.81927 | 3.63489 | 0.28927 | 1.96348 | 2.78465 |
| **CHML** | 2.36555 | 0.00000 | 2.59329 | 2.27635 | 0.75073 | 1.02486 | 9.21206 |
| **CHMP1A** | 19.28815 | 16.74944 | 14.02636 | 29.55866 | 7.07206 | 14.21030 | 13.12243 |
| **CHMP1B** | 25.55656 | 8.42997 | 17.89104 | 30.97867 | 10.83779 | 8.05938 | 3.74672 |
| **CHMP1B2P** | 0.49415 | 0.33447 | 0.21202 | 0.42376 | 0.05082 | 0.30399 | 0.76557 |
| **CHMP2A** | 11.30785 | 6.57320 | 7.96685 | 8.04469 | 4.18297 | 6.31510 | 3.55889 |
| **CHMP2B** | 15.60867 | 2.82857 | 8.90100 | 15.60739 | 2.93831 | 7.93999 | 13.50194 |
| **CHMP3** | 0.73013 | 1.30946 | 0.42238 | 0.66013 | 0.27237 | 0.44633 | 1.02964 |
| **CHMP4A** | 16.33140 | 10.74344 | 1.02683 | 12.49734 | 0.00000 | 7.06079 | 2.08529 |
| **CHMP4B** | 0.29587 | 0.19819 | 1.03952 | 0.00000 | 0.02089 | 3.19605 | 15.26567 |
| **CHMP4C** | 2.57093 | 0.22068 | 0.00000 | 3.20716 | 0.00334 | 1.95845 | 0.12262 |
| **CHMP5** | 18.69124 | 2.97119 | 19.75022 | 22.12054 | 6.36307 | 7.49677 | 2.24989 |

| | | | | | | | |
|---|---|---|---|---|---|---|---|
| **CHMP6** | 3.00346 | 17.59927 | 4.17742 | 1.96694 | 1.05529 | 2.04446 | 0.00000 |
| **CHMP7** | 6.10324 | 6.15682 | 4.73202 | 12.19338 | 1.08729 | 1.30497 | 4.15785 |
| **CHN1** | 0.12047 | 0.58181 | 0.08033 | 0.18271 | 0.23251 | 0.94370 | 0.30731 |
| **CHN2** | 0.28455 | 1.15201 | 0.30408 | 1.01522 | 1.15517 | 0.40034 | 0.53800 |
| **CHODL** | 0.16977 | 0.22893 | 0.01419 | 0.00300 | 0.07332 | 0.65535 | 0.00000 |
| **CHORDC1** | 32.17271 | 3.61302 | 14.14909 | 18.52823 | 5.26282 | 8.08623 | 0.00000 |
| **CHP1** | 52.94283 | 36.42972 | 20.50348 | 22.60709 | 16.74618 | 32.49012 | 0.05786 |
| **CHP2** | 0.04476 | 0.01207 | 0.03949 | 0.00039 | 0.08205 | 0.00431 | 0.10878 |
| **CHPF** | 0.00000 | 1.67343 | 0.10923 | 4.80627 | 0.33993 | 2.18116 | 4.45164 |
| **CHPF2** | 0.00000 | 0.79340 | 0.30809 | 1.13458 | 0.19899 | 1.24047 | 2.35509 |
| **CHPT1** | 3.86378 | 0.00000 | 1.57948 | 2.58013 | 0.21112 | 1.73791 | 11.75284 |
| **CHRAC1** | 3.66265 | 2.46840 | 9.94534 | 4.46428 | 0.78083 | 3.70212 | 7.97727 |
| **CHRD** | 0.04664 | 0.25991 | 0.52806 | 0.04845 | 0.05018 | 0.34254 | 0.12057 |
| **CHRDL1** | 0.02114 | 0.02918 | 0.00847 | 0.01845 | 0.02519 | 0.01114 | 0.02274 |
| **CHRDL2** | 0.01299 | 0.03581 | 0.04498 | 0.02196 | 0.09998 | 0.02468 | 0.02885 |
| **CHRFAM7A** | 0.03268 | 0.01724 | 0.01657 | 0.01959 | 0.01388 | 0.01959 | 0.01454 |
| **CHRM1** | 0.00000 | 0.00000 | 0.19753 | 0.19491 | 0.03083 | 0.00003 | 0.10987 |
| **CHRM2** | 0.09929 | 0.00000 | 0.03378 | 0.01486 | 0.00665 | 0.03420 | 0.04046 |
| **CHRM3** | 0.28190 | 0.01900 | 0.17750 | 0.17611 | 0.00665 | 0.24875 | 0.00000 |
| **CHRM3-AS2** | 0.02414 | 0.02015 | 0.01347 | 0.02086 | 0.00779 | 0.01405 | 0.01829 |
| **CHRM4** | 0.01535 | 0.01807 | 0.02450 | 0.04910 | 0.00694 | 0.03389 | 0.03060 |
| **CHRM5** | 0.13726 | 0.00000 | 0.58644 | 0.55624 | 0.01758 | 0.04688 | 0.09050 |
| **CHRNA1** | 0.04189 | 0.00373 | 0.01729 | 0.01905 | 0.02109 | 0.00553 | 0.16691 |
| **CHRNA10** | 0.24520 | 0.00000 | 3.07997 | 0.39402 | 0.40691 | 0.03355 | 2.21670 |
| **CHRNA3** | 0.00000 | 0.22321 | 0.56954 | 0.26513 | 0.42363 | 0.63618 | 0.00000 |
| **CHRNA4** | 0.02273 | 0.01286 | 0.01224 | 0.02012 | 0.01560 | 0.04773 | 0.02994 |
| **CHRNA5** | 0.07461 | 0.26682 | 2.14236 | 1.31438 | 0.54783 | 1.01874 | 0.89653 |
| **CHRNA6** | 0.01374 | 0.05617 | 0.03274 | 0.01707 | 0.13032 | 0.05042 | 0.02362 |
| **CHRNA7** | 0.06387 | 0.16048 | 0.19706 | 0.18424 | 0.75926 | 0.24081 | 0.00000 |
| **CHRNA9** | 0.01613 | 0.04551 | 0.01488 | 0.09285 | 0.02115 | 0.01028 | 0.02178 |
| **CHRNB1** | 1.96324 | 0.00000 | 2.14640 | 4.20375 | 4.37308 | 0.50735 | 0.59437 |
| **CHRNB2** | 0.04777 | 0.03796 | 0.09836 | 0.06770 | 0.00592 | 0.01418 | 0.00000 |
| **CHRNB3** | 0.01760 | 0.05994 | 0.09580 | 0.01633 | 0.02532 | 0.06282 | 0.03115 |
| **CHRNB4** | 0.00719 | 0.05762 | 0.20092 | 0.02482 | 0.24240 | 0.17387 | 0.00670 |
| **CHRND** | 0.02015 | 0.01498 | 0.01187 | 0.01742 | 0.01294 | 0.04639 | 0.02189 |

| | | | | | | | |
|---|---|---|---|---|---|---|---|
| **CHRNE** | 0.13142 | 0.21766 | 0.40496 | 0.12989 | 0.64089 | 0.15522 | 0.48487 |
| **CHRNG** | 0.02707 | 0.00564 | 0.01422 | 0.03195 | 0.00744 | 0.02139 | 0.06567 |
| **CHST1** | 0.00000 | 0.32842 | 0.17770 | 0.00000 | 0.00210 | 0.10830 | 0.11301 |
| **CHST10** | 0.11004 | 0.70143 | 0.14545 | 0.18720 | 0.17048 | 0.03584 | 1.18873 |
| **CHST11** | 0.85674 | 0.91479 | 0.32100 | 0.18927 | 0.26480 | 0.97504 | 0.99039 |
| **CHST12** | 0.00000 | 21.85731 | 2.33386 | 4.00539 | 3.55600 | 18.30235 | 5.65169 |
| **CHST13** | 0.00851 | 0.01591 | 0.01683 | 0.06871 | 0.24096 | 0.07454 | 0.06843 |
| **CHST14** | 0.86927 | 0.52029 | 0.00000 | 1.40126 | 0.00162 | 0.52179 | 4.37855 |
| **CHST15** | 2.25276 | 1.16154 | 5.87817 | 1.89367 | 1.12503 | 2.72888 | 0.64019 |
| **CHST2** | 0.00000 | 14.76701 | 1.07135 | 0.61223 | 0.48775 | 18.03506 | 5.43614 |
| **CHST3** | 0.00980 | 0.03843 | 0.01429 | 0.14531 | 0.01809 | 0.40849 | 0.05667 |
| **CHST4** | 0.00000 | 0.68217 | 2.05061 | 0.00000 | 0.12662 | 0.47907 | 0.00028 |
| **CHST5** | 0.00000 | 0.17517 | 0.30471 | 0.01905 | 0.00465 | 0.00238 | 0.02940 |
| **CHST6** | 0.00000 | 1.47065 | 1.62786 | 0.03836 | 0.39120 | 0.49072 | 0.05189 |
| **CHST7** | 0.05414 | 0.73128 | 1.18442 | 0.06702 | 0.25041 | 2.35519 | 0.17187 |
| **CHST8** | 0.00000 | 4.12106 | 0.66483 | 0.00000 | 0.02482 | 1.74905 | 0.80281 |
| **CHST9** | 0.07112 | 0.00145 | 0.01153 | 0.00943 | 0.00780 | 0.73222 | 0.00000 |
| **CHSY1** | 5.38625 | 1.55220 | 3.73654 | 3.98643 | 0.58220 | 1.50568 | 5.45090 |
| **CHSY3** | 0.02819 | 0.01720 | 0.01115 | 0.02145 | 0.03214 | 0.01248 | 0.01626 |
| **CHTF18** | 0.00000 | 3.54049 | 0.64970 | 2.87401 | 0.15345 | 3.77662 | 9.84920 |
| **CHTF8** | 10.49994 | 5.48611 | 8.80947 | 14.32466 | 1.68720 | 5.77436 | 8.90924 |
| **CHTOP** | 43.73986 | 29.08770 | 21.81822 | 29.13506 | 16.22776 | 16.59977 | 5.73553 |
| **CHUK** | 6.41202 | 0.84547 | 3.07295 | 2.86899 | 0.06042 | 3.82512 | 12.38434 |
| **CHURC1** | 10.44807 | 3.00177 | 9.82700 | 4.09664 | 0.40191 | 7.40189 | 8.46771 |
| **CIAO1** | 30.63899 | 25.67969 | 10.69298 | 24.36125 | 1.67650 | 12.83036 | 6.30057 |
| **CIAPIN1** | 28.38101 | 17.03496 | 9.65929 | 21.35948 | 3.47121 | 9.14805 | 0.00000 |
| **CIART** | 0.04197 | 0.00000 | 3.99857 | 0.04585 | 3.64788 | 0.00000 | 0.16839 |
| **CIB1** | 13.98044 | 30.72353 | 13.89499 | 12.59182 | 3.42236 | 11.42697 | 8.08580 |
| **CIB2** | 0.00000 | 3.53274 | 0.50409 | 0.17067 | 0.03040 | 1.27949 | 2.27804 |
| **CIB3** | 0.07917 | 0.04599 | 0.03635 | 0.07996 | 0.00549 | 0.00167 | 0.03952 |
| **CIB4** | 0.03626 | 0.01490 | 0.01415 | 0.01757 | 0.00915 | 0.04202 | 0.00909 |
| **CIC** | 0.00000 | 1.07412 | 0.50782 | 1.68660 | 0.00000 | 3.39606 | 14.55308 |
| **CIDEC** | 7.93489 | 2.50062 | 0.00000 | 0.46949 | 0.01008 | 0.38798 | 0.39014 |
| **CIDECP** | 2.88162 | 4.59870 | 4.89384 | 3.78105 | 6.62130 | 0.97152 | 0.00000 |
| **CIITA** | 0.02555 | 0.00595 | 0.05025 | 0.02340 | 0.00792 | 0.02622 | 0.03157 |

| | | | | | | | |
|---|---|---|---|---|---|---|---|
| **CILP** | 0.02779 | 0.01773 | 0.50710 | 0.04019 | 0.24888 | 0.02177 | 0.00000 |
| **CILP2** | 0.00387 | 0.29178 | 0.62334 | 0.09325 | 0.02482 | 0.29234 | 1.06323 |
| **CINP** | 13.74309 | 13.16057 | 12.30458 | 8.40623 | 0.91871 | 5.92694 | 4.46062 |
| **CIPC** | 7.49346 | 2.15858 | 2.39542 | 5.05271 | 3.29236 | 1.03779 | 0.49106 |
| **CIR1** | 5.76224 | 2.12418 | 9.48820 | 7.02442 | 5.35566 | 4.79672 | 4.96476 |
| **CIRBP** | 7.41548 | 12.27717 | 8.60362 | 10.56351 | 2.43263 | 7.06366 | 11.91723 |
| **CIRBP-AS1** | 0.00000 | 0.13893 | 0.71244 | 0.36663 | 0.00051 | 0.28758 | 0.86207 |
| **CIRH1A** | 36.87481 | 23.15422 | 15.82722 | 26.13094 | 7.53592 | 12.61871 | 0.00000 |
| **CISD1** | 21.01850 | 6.65381 | 12.05483 | 15.32361 | 4.96017 | 14.77073 | 15.35844 |
| **CISD2** | 8.47627 | 1.90067 | 1.07221 | 6.89881 | 0.50728 | 3.11608 | 0.04346 |
| **CISD3** | 4.99394 | 10.84358 | 0.99188 | 5.08529 | 0.00000 | 1.50659 | 2.54004 |
| **CISH** | 1.05293 | 1.72619 | 3.19561 | 1.98283 | 0.27748 | 1.84325 | 1.61680 |
| **CIT** | 4.59590 | 10.65738 | 3.54181 | 3.07814 | 2.26070 | 5.28354 | 2.84874 |
| **CITED1** | 0.04396 | 0.02528 | 0.05124 | 0.01361 | 0.32011 | 0.03649 | 0.06905 |
| **CITED2** | 0.00000 | 1.93384 | 0.00000 | 8.04369 | 0.13910 | 0.00000 | 12.02946 |
| **CITED4** | 0.00000 | 8.19704 | 0.17147 | 0.22296 | 0.00510 | 0.00000 | 17.37964 |
| **CIZ1** | 1.77664 | 5.94915 | 6.64725 | 3.27860 | 1.62852 | 1.36276 | 11.18856 |
| **CKAP2** | 27.85313 | 7.55916 | 17.95457 | 10.55104 | 10.38099 | 8.18875 | 6.94769 |
| **CKAP2L** | 5.54455 | 1.42694 | 7.49829 | 3.89837 | 3.69970 | 3.79796 | 0.00000 |
| **CKAP4** | 0.00000 | 1.73043 | 1.81831 | 0.32055 | 0.07639 | 1.48645 | 0.84731 |
| **CKAP5** | 27.93794 | 16.61575 | 16.35473 | 21.23466 | 8.04563 | 13.31989 | 21.94532 |
| **CKB** | 19.55331 | 47.89057 | 4.03890 | 31.79563 | 1.56312 | 26.14766 | 26.17222 |
| **CKLF** | 0.47482 | 0.20594 | 0.60459 | 0.29506 | 0.12783 | 0.71694 | 0.30762 |
| **CKM** | 0.00000 | 0.09660 | 0.00000 | 0.20461 | 0.68544 | 0.00125 | 0.15058 |
| **CKMT1A** | 0.84930 | 0.00061 | 0.14503 | 1.33284 | 0.03963 | 0.65795 | 0.00000 |
| **CKMT1B** | 0.02111 | 0.04905 | 0.05912 | 0.78130 | 0.04606 | 0.73324 | 0.07648 |
| **CKMT2** | 0.39658 | 0.07413 | 0.05305 | 0.05942 | 1.57787 | 0.68248 | 0.53236 |
| **CKMT2-AS1** | 2.29258 | 0.53870 | 2.05261 | 1.22500 | 4.62110 | 0.70164 | 0.00000 |
| **CKS1B** | 10.94808 | 11.53168 | 6.57972 | 13.88913 | 3.25653 | 7.52060 | 3.71490 |
| **CKS2** | 34.55443 | 21.15663 | 19.79391 | 29.51035 | 16.67263 | 14.86150 | 11.57335 |
| **CLASP1** | 3.74465 | 1.96779 | 3.22120 | 3.12550 | 2.08232 | 3.25801 | 6.50042 |
| **CLASP2** | 6.19425 | 2.46577 | 3.87291 | 4.07385 | 0.34541 | 2.73679 | 8.24896 |
| **CLASRP** | 0.00000 | 1.90695 | 0.58051 | 2.61739 | 0.98497 | 1.13846 | 5.53053 |
| **CLC** | 0.03209 | 0.00000 | 0.40138 | 0.04069 | 1.31982 | 0.09400 | 0.01066 |
| **CLCA1** | 0.01386 | 0.01887 | 0.02811 | 0.11481 | 0.01206 | 0.12611 | 0.00000 |

| | | | | | | | |
|---|---|---|---|---|---|---|---|
| **CLCA2** | 0.13418 | 0.00000 | 0.02060 | 0.13868 | 0.00966 | 0.07347 | 0.00501 |
| **CLCA3P** | 0.03838 | 0.01948 | 0.01157 | 0.01393 | 0.01253 | 0.02215 | 0.02575 |
| **CLCA4** | 0.01952 | 0.01881 | 0.01201 | 0.01519 | 0.00744 | 0.02035 | 0.04993 |
| **CLCC1** | 9.54861 | 2.96200 | 0.45055 | 6.78159 | 0.36895 | 0.42552 | 0.00000 |
| **CLCF1** | 0.00151 | 0.00000 | 0.38306 | 0.02264 | 0.19436 | 0.02628 | 0.20398 |
| **CLCN1** | 0.16180 | 0.00000 | 0.11701 | 0.00000 | 0.00589 | 0.01303 | 0.07734 |
| **CLCN2** | 0.00000 | 0.00000 | 0.24662 | 0.67445 | 0.19179 | 0.65040 | 0.77254 |
| **CLCN3** | 5.85435 | 2.57492 | 4.57188 | 7.60258 | 2.40487 | 5.44515 | 7.92160 |
| **CLCN4** | 0.00000 | 0.00000 | 0.76237 | 0.01408 | 2.33179 | 0.00000 | 0.07835 |
| **CLCN5** | 5.48178 | 6.26326 | 2.27022 | 0.00000 | 3.11379 | 3.39159 | 6.71843 |
| **CLCN6** | 5.60669 | 0.09686 | 0.00000 | 5.87830 | 0.01515 | 0.93841 | 6.64290 |
| **CLCN7** | 0.00000 | 1.41018 | 1.45078 | 0.30605 | 0.00000 | 1.48954 | 8.27195 |
| **CLCNKA** | 0.06569 | 0.01658 | 0.00660 | 0.00000 | 0.00688 | 0.01748 | 0.28448 |
| **CLCNKB** | 0.05363 | 0.01610 | 0.01080 | 0.02568 | 0.00778 | 0.01570 | 0.10249 |
| **CLDN1** | 0.58701 | 0.00000 | 0.00000 | 0.26730 | 0.00424 | 0.00000 | 0.66232 |
| **CLDN10** | 33.44512 | 55.79252 | 0.00000 | 0.00000 | 1.27476 | 0.00000 | 0.29207 |
| **CLDN10-AS1** | 0.05974 | 0.29730 | 0.00638 | 0.00000 | 0.00662 | 0.00000 | 0.15946 |
| **CLDN11** | 0.25415 | 0.00000 | 2.53056 | 0.09747 | 0.17508 | 0.49309 | 2.94523 |
| **CLDN12** | 11.72132 | 0.00000 | 5.04838 | 4.60226 | 3.73241 | 2.56486 | 0.00000 |
| **CLDN14** | 0.02553 | 0.02694 | 0.00860 | 0.04615 | 0.02840 | 0.00958 | 0.02345 |
| **CLDN15** | 0.00000 | 0.39856 | 0.66168 | 1.66622 | 0.12914 | 0.35649 | 1.86012 |
| **CLDN16** | 0.02952 | 0.00062 | 0.06125 | 0.01974 | 0.01599 | 0.01973 | 0.01567 |
| **CLDN18** | 0.00857 | 0.01437 | 0.08308 | 0.01541 | 1.12761 | 0.06243 | 0.02982 |
| **CLDN19** | 0.00000 | 9.42370 | 0.43136 | 0.00000 | 0.00242 | 10.98136 | 0.27642 |
| **CLDN2** | 0.04263 | 0.06546 | 0.00408 | 0.00451 | 0.00879 | 0.00000 | 0.15913 |
| **CLDN20** | 0.08266 | 0.02480 | 0.18378 | 0.00000 | 0.02447 | 0.07990 | 0.09632 |
| **CLDN23** | 0.00000 | 0.61753 | 0.09063 | 0.20339 | 0.00190 | 0.13663 | 0.45035 |
| **CLDN24** | 0.02387 | 0.01719 | 0.03071 | 0.02101 | 0.00662 | 0.01186 | 0.01659 |
| **CLDN3** | 1.46762 | 0.00000 | 0.00000 | 13.27563 | 0.00000 | 0.00000 | 9.90880 |
| **CLDN4** | 63.27986 | 31.64732 | 0.00000 | 57.57270 | 0.00000 | 0.00000 | 28.91073 |
| **CLDN5** | 0.06195 | 0.00504 | 0.03136 | 0.04862 | 0.00685 | 0.01299 | 0.00609 |
| **CLDN6** | 2.37110 | 15.33290 | 15.81270 | 14.11038 | 2.25626 | 9.64857 | 38.39034 |
| **CLDN7** | 30.35897 | 42.09280 | 11.11547 | 39.01463 | 0.89889 | 13.97771 | 15.45257 |
| **CLDN8** | 0.03083 | 0.01386 | 0.00884 | 0.06643 | 0.01038 | 0.03827 | 0.14425 |
| **CLDN9** | 0.10597 | 0.00899 | 0.01670 | 0.37779 | 0.01465 | 0.26270 | 0.01229 |

| | | | | | | | |
|---|---|---|---|---|---|---|---|
| **CLDND1** | 20.34894 | 1.90020 | 14.05845 | 20.92278 | 3.73222 | 6.50139 | 0.00000 |
| **CLDND2** | 0.11120 | 0.05190 | 0.00389 | 0.06623 | 0.08427 | 0.08283 | 0.14173 |
| **CLEC10A** | 0.01456 | 0.02948 | 0.00000 | 0.01014 | 2.82012 | 0.00652 | 0.03218 |
| **CLEC11A** | 0.18430 | 2.15025 | 0.95578 | 2.04564 | 0.57462 | 0.37381 | 0.00323 |
| **CLEC12A** | 0.40832 | 0.00000 | 0.09454 | 0.41898 | 0.00477 | 0.43081 | 0.00992 |
| **CLEC12B** | 0.02654 | 0.01437 | 0.04110 | 0.03039 | 0.02211 | 0.04178 | 0.01329 |
| **CLEC16A** | 6.69700 | 8.80817 | 2.60926 | 3.71392 | 2.51923 | 6.85064 | 4.64549 |
| **CLEC17A** | 0.04339 | 0.02962 | 0.01616 | 0.02398 | 0.66725 | 0.00859 | 0.00000 |
| **CLEC18A** | 0.03359 | 0.01161 | 0.01858 | 0.01863 | 0.00683 | 0.00901 | 0.06443 |
| **CLEC18B** | 0.02651 | 0.05643 | 0.02193 | 0.04556 | 0.00714 | 0.01210 | 0.03891 |
| **CLEC18C** | 0.01844 | 0.04592 | 0.01800 | 0.02504 | 0.00820 | 0.01396 | 0.01286 |
| **CLEC19A** | 0.01817 | 0.01859 | 0.01246 | 0.02979 | 0.00783 | 0.01108 | 0.03455 |
| **CLEC1A** | 0.01764 | 0.05523 | 0.34648 | 6.27202 | 0.05330 | 0.02663 | 0.00000 |
| **CLEC1B** | 0.01256 | 0.02125 | 0.12738 | 0.25775 | 0.02843 | 0.03642 | 0.00000 |
| **CLEC2A** | 0.02647 | 0.01226 | 0.01299 | 0.02135 | 0.00825 | 0.07334 | 0.01415 |
| **CLEC2B** | 0.03690 | 0.01309 | 0.01302 | 0.03161 | 0.01431 | 0.01526 | 0.01390 |
| **CLEC2D** | 1.39055 | 0.00000 | 0.53501 | 0.27680 | 0.08197 | 0.31913 | 0.11976 |
| **CLEC2L** | 0.12875 | 0.78526 | 2.75837 | 1.03753 | 0.53159 | 0.00000 | 0.29738 |
| **CLEC3B** | 0.03634 | 0.01551 | 0.01389 | 0.01251 | 0.00751 | 0.01458 | 0.03051 |
| **CLEC4A** | 0.04169 | 0.00000 | 0.20717 | 0.03998 | 0.54260 | 0.06272 | 0.10356 |
| **CLEC4C** | 0.06513 | 0.07680 | 0.11590 | 0.02323 | 1.55801 | 0.04083 | 0.00001 |
| **CLEC4D** | 0.29191 | 0.12539 | 1.25346 | 0.00000 | 0.00000 | 1.69051 | 0.00385 |
| **CLEC4E** | 0.02833 | 0.00478 | 0.00915 | 0.01961 | 0.00732 | 1.90757 | 0.00873 |
| **CLEC4F** | 0.02062 | 0.01626 | 0.01312 | 0.04155 | 0.00793 | 0.01283 | 0.01916 |
| **CLEC4G** | 0.03592 | 0.02595 | 0.01848 | 0.01789 | 0.03039 | 0.05870 | 0.00337 |
| **CLEC4GP1** | 0.05328 | 0.00000 | 0.00397 | 2.39029 | 0.01165 | 0.00000 | 0.40947 |
| **CLEC4M** | 0.08099 | 0.01305 | 0.01335 | 0.03108 | 0.00796 | 0.01001 | 0.00000 |
| **CLEC5A** | 0.03222 | 0.02669 | 0.01230 | 0.02617 | 0.00697 | 0.03448 | 0.05767 |
| **CLEC6A** | 0.52463 | 0.76174 | 1.56751 | 0.00000 | 0.00000 | 1.03251 | 0.00000 |
| **CLEC7A** | 0.42251 | 0.00000 | 0.07685 | 0.28039 | 0.01769 | 0.90135 | 0.05669 |
| **CLEC9A** | 0.04178 | 0.00397 | 0.38718 | 0.08097 | 0.00649 | 0.01128 | 0.00058 |
| **CLECL1** | 0.02750 | 0.01168 | 0.02275 | 0.01671 | 0.00762 | 0.02018 | 0.01934 |
| **CLGN** | 0.14702 | 0.16954 | 0.53632 | 0.08055 | 0.31290 | 0.00167 | 0.00000 |
| **CLHC1** | 0.65153 | 0.00000 | 1.55473 | 0.55568 | 0.49373 | 0.22736 | 0.92550 |
| **CLIC1** | 49.45901 | 14.43218 | 0.00000 | 39.89715 | 0.00000 | 26.15750 | 6.50841 |

| | | | | | | | |
|---|---|---|---|---|---|---|---|
| **CLIC2** | 0.25003 | 0.00000 | 0.58755 | 0.00000 | 0.00120 | 0.25774 | 0.06837 |
| **CLIC3** | 1.38559 | 0.00000 | 0.00000 | 6.70357 | 0.00035 | 0.00000 | 2.46585 |
| **CLIC4** | 41.09739 | 18.47021 | 9.71105 | 20.17884 | 2.74650 | 7.38301 | 1.41937 |
| **CLIC5** | 0.15004 | 0.42348 | 0.00000 | 0.00000 | 0.01936 | 0.12834 | 0.00702 |
| **CLIC6** | 0.04791 | 0.00857 | 0.02174 | 0.02534 | 0.02192 | 0.02572 | 0.01342 |
| **CLINT1** | 19.02960 | 8.76720 | 9.64856 | 15.90070 | 4.72556 | 13.72978 | 11.57896 |
| **CLIP1** | 13.05892 | 16.20540 | 13.58430 | 11.11668 | 4.00430 | 8.38839 | 31.58552 |
| **CLIP1-AS1** | 0.04951 | 0.13911 | 0.03442 | 0.16891 | 0.00741 | 0.01057 | 0.00433 |
| **CLIP2** | 0.01273 | 0.00021 | 0.81696 | 0.00987 | 0.25865 | 0.33588 | 0.02989 |
| **CLIP3** | 0.00000 | 0.16961 | 0.00000 | 0.16951 | 1.11606 | 0.23683 | 0.19946 |
| **CLIP4** | 0.07272 | 0.00000 | 0.06890 | 0.01337 | 1.63141 | 0.01059 | 0.14999 |
| **CLK1** | 12.54942 | 0.00000 | 19.07571 | 10.50364 | 10.24312 | 9.68129 | 10.44659 |
| **CLK2** | 2.49713 | 1.13403 | 1.58699 | 2.06832 | 0.40752 | 2.44294 | 5.83149 |
| **CLK2P** | 0.33676 | 0.15304 | 0.13314 | 0.20525 | 0.04805 | 0.18060 | 0.20850 |
| **CLK3** | 12.85101 | 10.28213 | 6.35541 | 13.98820 | 8.78486 | 5.25069 | 0.00000 |
| **CLK4** | 4.76406 | 0.00000 | 1.55806 | 2.15844 | 5.25353 | 2.02811 | 0.00000 |
| **CLMN** | 0.54513 | 0.35629 | 0.03206 | 0.00000 | 0.46861 | 0.50299 | 0.35789 |
| **CLMP** | 0.00000 | 0.07081 | 0.06388 | 2.87202 | 0.03185 | 0.10188 | 0.15245 |
| **CLN3** | 15.27802 | 11.24892 | 4.80697 | 22.78477 | 0.07251 | 6.33831 | 3.63627 |
| **CLN5** | 11.85146 | 1.87599 | 1.25334 | 5.75062 | 0.96670 | 1.86610 | 2.31540 |
| **CLN6** | 2.34498 | 9.06129 | 6.36510 | 5.28341 | 0.31211 | 7.54389 | 19.07198 |
| **CLN8** | 12.47734 | 8.12812 | 1.96485 | 9.11151 | 1.87827 | 4.41232 | 6.28966 |
| **CLNK** | 0.02974 | 0.01520 | 0.01447 | 0.01919 | 0.00795 | 0.03549 | 0.01503 |
| **CLNS1A** | 29.38115 | 8.83473 | 15.39841 | 13.75309 | 0.84690 | 12.34372 | 9.49029 |
| **CLOCK** | 4.60925 | 0.00000 | 3.80110 | 0.54306 | 8.34129 | 2.39997 | 6.26572 |
| **1.00000** | 20.01556 | 5.84189 | 13.28594 | 16.25622 | 11.17985 | 8.65750 | 0.00000 |
| **CLPB** | 14.77976 | 15.15581 | 5.92447 | 12.04988 | 4.74582 | 4.59043 | 0.00000 |
| **CLPP** | 13.75763 | 23.04134 | 16.45194 | 20.01168 | 9.46100 | 12.85598 | 11.05318 |
| **CLPS** | 1.40806 | 0.00000 | 0.23369 | 0.00000 | 0.45706 | 0.03956 | 0.67055 |
| **CLPSL1** | 1.35915 | 0.09878 | 0.00000 | 0.00000 | 0.00905 | 0.19053 | 0.21669 |
| **CLPSL2** | 0.20759 | 0.02147 | 0.01239 | 0.01776 | 0.00549 | 0.07355 | 0.15680 |
| **CLPTM1** | 0.69289 | 12.31681 | 4.82154 | 7.67126 | 1.17575 | 11.36570 | 28.21382 |
| **CLPTM1L** | 7.73800 | 18.52852 | 10.03053 | 7.48873 | 3.42026 | 5.92751 | 20.32155 |
| **CLPX** | 15.88403 | 8.18048 | 16.15572 | 8.31646 | 3.74173 | 12.18210 | 2.86182 |
| **CLRN1** | 0.05837 | 0.01068 | 0.01074 | 0.00000 | 0.00737 | 0.00524 | 0.08517 |

| | | | | | | | |
|---|---|---|---|---|---|---|---|
| **CLRN1-AS1** | 0.04671 | 0.00889 | 0.03372 | 0.07271 | 0.00870 | 0.01985 | 0.01804 |
| **CLSPN** | 20.06656 | 9.87222 | 10.47632 | 15.70622 | 14.45892 | 10.95858 | 12.49577 |
| **CLSTN1** | 0.82702 | 10.42908 | 5.39269 | 3.72893 | 5.70466 | 8.79861 | 12.72528 |
| **CLSTN2** | 0.01489 | 0.01652 | 0.01445 | 0.00476 | 1.51406 | 0.00575 | 0.08987 |
| **CLSTN2-AS1** | 0.02734 | 0.01617 | 0.01854 | 0.02084 | 0.02017 | 0.01971 | 0.01422 |
| **CLSTN3** | 0.35822 | 0.00000 | 0.65946 | 0.00000 | 1.01570 | 0.56303 | 0.02943 |
| **CLTA** | 23.78320 | 29.90810 | 20.36713 | 31.82588 | 2.91226 | 13.83942 | 50.03733 |
| **CLTB** | 0.42860 | 23.30134 | 1.67335 | 0.40202 | 1.57590 | 5.70479 | 13.88273 |
| **CLTC** | 49.33526 | 16.95527 | 10.59754 | 23.48199 | 4.76763 | 16.96392 | 35.77554 |
| **CLTCL1** | 0.51310 | 1.71992 | 0.58142 | 1.20144 | 0.33517 | 0.34425 | 0.62302 |
| **CLU** | 1.51446 | 0.60544 | 0.01812 | 2.60116 | 0.00000 | 0.03730 | 0.97135 |
| **CLUAP1** | 1.91421 | 1.42139 | 3.02238 | 0.10654 | 1.05027 | 1.68184 | 1.43907 |
| **CLUH** | 1.55595 | 18.38237 | 7.21013 | 6.25392 | 0.20377 | 4.79529 | 44.57961 |
| **CLUHP3** | 1.04392 | 0.00000 | 0.00979 | 1.20040 | 0.00545 | 0.23497 | 0.12514 |
| **CLUL1** | 0.04930 | 0.11041 | 0.01366 | 0.03780 | 0.31091 | 0.02475 | 0.05198 |
| **CLVS1** | 0.08099 | 0.00058 | 0.03967 | 0.18189 | 0.00752 | 0.01571 | 0.00146 |
| **CLVS2** | 0.16779 | 0.14420 | 0.02402 | 0.21791 | 0.00568 | 0.67193 | 0.00000 |
| **CLYBL** | 0.54548 | 0.04257 | 0.01914 | 0.00000 | 0.07154 | 0.28805 | 0.01140 |
| **CMA1** | 0.02204 | 0.02964 | 0.01024 | 0.02288 | 0.00830 | 0.01003 | 0.03171 |
| **CMAHP** | 0.05202 | 0.00000 | 0.08693 | 0.38905 | 0.08816 | 0.00955 | 0.01925 |
| **CMAS** | 7.09686 | 4.35447 | 7.10048 | 15.21133 | 0.51380 | 7.71959 | 2.24689 |
| **CMBL** | 0.43915 | 1.93731 | 0.12404 | 0.49577 | 0.07441 | 4.88856 | 0.00000 |
| **CMC1** | 5.96212 | 3.88860 | 3.44527 | 2.57358 | 0.00000 | 3.37445 | 2.01916 |
| **CMC2** | 19.16171 | 8.68088 | 14.45301 | 13.67707 | 11.61191 | 10.47128 | 9.48861 |
| **CMC4** | 0.80464 | 0.65698 | 0.36830 | 0.76169 | 0.43651 | 1.21143 | 0.80661 |
| **CMIP** | 0.00000 | 0.41520 | 0.02428 | 0.56481 | 0.19483 | 1.20617 | 6.17034 |
| **CMKLR1** | 0.04744 | 0.00000 | 0.01247 | 0.03024 | 1.43372 | 0.01678 | 0.05097 |
| **CMPK1** | 22.05701 | 1.47891 | 8.92032 | 14.83954 | 4.38732 | 7.21940 | 8.16523 |
| **CMPK2** | 0.02982 | 0.03488 | 0.05404 | 0.01047 | 0.00458 | 0.00000 | 0.10406 |
| **CMSS1** | 14.80395 | 4.95893 | 8.78280 | 4.93488 | 0.29425 | 6.04166 | 14.14688 |
| **CMTM1** | 0.02603 | 0.06790 | 0.00000 | 0.01358 | 0.12549 | 0.01004 | 0.06314 |
| **CMTM2** | 0.04691 | 0.06160 | 0.01771 | 0.04199 | 0.00778 | 0.01307 | 0.01126 |
| **CMTM3** | 0.00000 | 0.66706 | 0.04691 | 1.09584 | 0.00357 | 0.84608 | 1.19677 |
| **CMTM4** | 4.46525 | 3.79006 | 0.69019 | 0.76383 | 4.18224 | 4.14641 | 11.84192 |
| **CMTM6** | 32.19078 | 7.99301 | 15.92931 | 23.51291 | 9.45154 | 11.53208 | 14.52284 |

| | | | | | | | |
|---|---|---|---|---|---|---|---|
| **CMTM7** | 8.50611 | 6.52342 | 0.00000 | 10.38318 | 0.00000 | 3.33407 | 9.00911 |
| **CMTM8** | 0.21610 | 0.00000 | 0.11510 | 0.83079 | 0.09341 | 0.27320 | 2.90526 |
| **CMTR1** | 2.19802 | 3.78751 | 1.26304 | 3.00950 | 1.52921 | 2.38471 | 2.47141 |
| **CMTR2** | 4.61482 | 0.00000 | 2.22947 | 3.49645 | 0.21938 | 0.51288 | 0.01498 |
| **CMYA5** | 0.00000 | 0.19375 | 0.23546 | 0.26021 | 1.04805 | 0.35116 | 1.20398 |
| **CNBD1** | 0.01747 | 0.00000 | 0.07678 | 0.01831 | 0.86245 | 0.01124 | 0.02956 |
| **CNBD2** | 0.22040 | 0.00731 | 0.31746 | 0.27373 | 0.12449 | 0.00801 | 0.00000 |
| **CNBP** | 71.49496 | 33.98852 | 47.50487 | 44.07288 | 19.28235 | 31.43657 | 7.40156 |
| **CNDP1** | 0.35847 | 0.03771 | 0.04798 | 0.00000 | 0.14429 | 0.07577 | 0.26342 |
| **CNDP2** | 33.81992 | 13.07248 | 4.57676 | 31.81828 | 0.00464 | 7.13166 | 8.26938 |
| **CNEP1R1** | 9.48663 | 0.60380 | 12.50321 | 2.37451 | 4.44300 | 3.64685 | 0.00000 |
| **CNFN** | 0.00000 | 1.09366 | 1.96785 | 0.09778 | 1.34471 | 0.25425 | 0.00000 |
| **CNGA1** | 0.25681 | 0.02029 | 0.63103 | 0.00000 | 0.00195 | 0.49191 | 0.03695 |
| **CNGA2** | 0.02021 | 0.00814 | 0.03599 | 0.01867 | 0.08086 | 0.02761 | 0.02983 |
| **CNGA3** | 0.05753 | 0.00015 | 0.03813 | 0.02037 | 0.00949 | 0.04090 | 0.01949 |
| **CNGA4** | 0.00000 | 0.01532 | 0.02834 | 0.64032 | 0.02965 | 0.00696 | 0.06166 |
| **CNGB1** | 0.00763 | 0.03049 | 0.01961 | 0.05191 | 0.06035 | 0.03816 | 0.06451 |
| **CNGB3** | 0.01644 | 0.02877 | 0.01141 | 0.02193 | 0.29553 | 0.03939 | 0.01786 |
| **CNIH1** | 38.00065 | 3.78889 | 23.20475 | 14.89051 | 5.90019 | 12.34427 | 11.60969 |
| **CNIH2** | 0.00000 | 0.14027 | 0.03425 | 0.34945 | 0.06080 | 0.00477 | 0.48283 |
| **CNIH3** | 0.03933 | 0.08637 | 0.08883 | 0.23173 | 0.02679 | 4.32429 | 0.10754 |
| **CNIH4** | 26.77573 | 19.75863 | 18.80090 | 12.83612 | 1.84068 | 29.87159 | 6.70437 |
| **CNKSR1** | 0.22394 | 1.57387 | 3.92382 | 6.96213 | 1.44842 | 0.49167 | 0.00446 |
| **CNKSR2** | 0.05061 | 0.05183 | 0.13133 | 0.01216 | 0.96094 | 0.04994 | 0.10144 |
| **CNKSR3** | 0.00000 | 0.18751 | 1.62930 | 0.01552 | 1.30954 | 1.11606 | 0.03352 |
| **CNN1** | 0.15841 | 0.05941 | 0.00000 | 0.00000 | 1.12573 | 0.04657 | 0.35806 |
| **CNN2** | 4.00173 | 8.35901 | 3.10581 | 22.51768 | 0.59391 | 8.11435 | 0.00000 |
| **CNN3** | 54.37378 | 39.23274 | 10.10595 | 19.43539 | 6.47555 | 15.88879 | 0.00000 |
| **CNNM1** | 0.00000 | 0.34591 | 0.35195 | 0.02943 | 0.25619 | 0.62149 | 0.00300 |
| **CNNM2** | 0.00000 | 1.42914 | 1.65321 | 3.72058 | 14.06922 | 0.21343 | 3.75975 |
| **CNNM3** | 4.06218 | 10.63172 | 5.09075 | 3.63721 | 0.05950 | 6.63632 | 20.74838 |
| **CNNM4** | 6.50546 | 9.16326 | 1.63941 | 4.30715 | 0.87067 | 5.51826 | 8.53483 |
| **CNOT1** | 25.50035 | 16.41674 | 7.99150 | 15.77687 | 5.83000 | 13.46270 | 19.41189 |
| **CNOT10** | 26.36673 | 8.66375 | 12.12551 | 7.80450 | 3.28626 | 7.90386 | 0.00000 |
| **CNOT11** | 2.93767 | 1.20321 | 2.52776 | 4.65215 | 1.22438 | 5.23989 | 16.15865 |

| | | | | | | | |
|---|---|---|---|---|---|---|---|
| **CNOT2** | 18.93422 | 2.65321 | 13.31324 | 9.34107 | 4.16337 | 7.57161 | 11.59348 |
| **CNOT3** | 0.00000 | 2.61514 | 2.46387 | 1.14449 | 0.65242 | 1.24198 | 12.16791 |
| **CNOT4** | 5.99342 | 2.98336 | 8.66732 | 4.29255 | 4.74433 | 1.71550 | 5.70866 |
| **CNOT6** | 12.07011 | 1.42208 | 6.31238 | 9.09400 | 7.10351 | 6.38011 | 13.17612 |
| **CNOT6L** | 1.05489 | 0.77539 | 1.24993 | 0.16947 | 3.87296 | 2.53688 | 4.90735 |
| **CNOT7** | 40.71547 | 14.52773 | 17.44329 | 22.52694 | 6.02311 | 14.92216 | 4.81711 |
| **CNOT8** | 18.08190 | 3.22216 | 9.73312 | 6.93441 | 6.06275 | 7.92317 | 0.00004 |
| **CNP** | 1.84259 | 8.37792 | 5.92611 | 7.62461 | 3.62663 | 6.30542 | 0.00000 |
| **CNPPD1** | 2.56526 | 6.85415 | 2.73874 | 11.87333 | 1.24039 | 2.94369 | 4.77035 |
| **CNPY1** | 0.03615 | 0.00936 | 0.03787 | 0.02405 | 0.08428 | 0.01682 | 0.00518 |
| **CNPY2** | 37.40423 | 25.46229 | 10.09494 | 32.74463 | 1.01812 | 18.62226 | 11.83609 |
| **CNPY3** | 13.80033 | 8.19219 | 7.00394 | 12.68279 | 0.94494 | 9.57416 | 1.30973 |
| **CNPY4** | 2.88079 | 0.00000 | 2.40412 | 1.40814 | 5.35139 | 2.17899 | 0.61711 |
| **CNR1** | 0.16282 | 0.03140 | 0.02386 | 0.07740 | 0.08562 | 0.66671 | 0.00000 |
| **CNR2** | 0.00835 | 1.94925 | 0.05031 | 0.02968 | 0.00269 | 3.64182 | 0.00000 |
| **CNRIP1** | 0.05871 | 0.01425 | 0.30398 | 0.00936 | 2.19997 | 0.00271 | 0.00390 |
| **CNST** | 6.86696 | 2.29818 | 5.74311 | 8.85491 | 10.68355 | 2.33883 | 1.05974 |
| **CNTD1** | 0.24131 | 0.17428 | 0.25634 | 0.33655 | 0.28077 | 0.29272 | 0.89493 |
| **CNTD2** | 0.00000 | 0.17975 | 0.02327 | 0.13139 | 1.50102 | 0.07767 | 0.01265 |
| **CNTF** | 0.02772 | 0.01594 | 0.01926 | 0.01957 | 0.01109 | 0.01495 | 0.01856 |
| **CNTFR** | 0.00000 | 0.52073 | 0.06708 | 0.02636 | 0.01125 | 0.51998 | 0.02339 |
| **CNTLN** | 0.39558 | 0.00000 | 2.72755 | 0.02694 | 0.69910 | 1.72952 | 0.70580 |
| **CNTN1** | 0.02713 | 0.00858 | 0.01520 | 0.01918 | 0.24214 | 0.01394 | 0.01434 |
| **CNTN2** | 0.03977 | 0.24222 | 0.00938 | 0.06534 | 0.02346 | 0.08505 | 0.08526 |
| **CNTN3** | 0.02527 | 0.03531 | 0.04340 | 0.01977 | 0.00641 | 0.01240 | 0.01787 |
| **CNTN4** | 0.29161 | 0.00000 | 0.00000 | 0.00000 | 1.52983 | 0.04914 | 0.62049 |
| **CNTN4-AS1** | 0.02867 | 0.01280 | 0.01539 | 0.01752 | 0.01594 | 0.03409 | 0.01539 |
| **CNTN4-AS2** | 0.01495 | 0.02762 | 0.00902 | 0.01254 | 0.00768 | 0.04885 | 0.05619 |
| **CNTN5** | 0.03039 | 0.02912 | 0.05728 | 0.02289 | 0.40190 | 0.01067 | 0.02184 |
| **CNTN6** | 1.74739 | 1.83027 | 1.19757 | 0.00000 | 0.44469 | 1.10465 | 0.00000 |
| **CNTNAP1** | 0.06621 | 0.00000 | 0.29738 | 0.00855 | 0.00305 | 0.05870 | 0.34578 |
| **CNTNAP2** | 0.42778 | 0.00000 | 0.00335 | 2.13689 | 1.58019 | 0.87468 | 0.00000 |
| **CNTNAP3** | 0.01814 | 0.05200 | 0.11300 | 0.01816 | 0.00602 | 0.00888 | 0.01753 |
| **CNTNAP4** | 0.27318 | 0.00000 | 0.04801 | 0.06795 | 0.13323 | 0.02844 | 0.00001 |
| **CNTNAP5** | 0.08095 | 0.23649 | 0.27242 | 0.00132 | 0.06323 | 0.37932 | 0.00000 |

| | | | | | | | |
|---|---|---|---|---|---|---|---|
| **CNTRL** | 1.09864 | 0.00000 | 1.43093 | 0.42899 | 4.40157 | 0.33034 | 1.20789 |
| **CNTROB** | 1.40730 | 1.55501 | 2.28167 | 2.34466 | 1.58831 | 1.79408 | 3.66445 |
| **COA1** | 18.94027 | 7.51573 | 11.13743 | 5.80453 | 4.85423 | 5.69650 | 5.42198 |
| **COA3** | 23.97257 | 28.28935 | 5.42754 | 20.72948 | 0.93579 | 13.55575 | 2.84789 |
| **COA4** | 24.86209 | 12.85728 | 10.02370 | 25.16272 | 2.54528 | 13.20141 | 13.59600 |
| **COA5** | 6.67158 | 4.20370 | 6.90901 | 6.45279 | 1.11105 | 3.36753 | 3.21841 |
| **COA6** | 23.47726 | 10.49576 | 17.51879 | 19.93343 | 5.52147 | 9.06964 | 7.52323 |
| **COA7** | 17.88896 | 13.08148 | 16.02627 | 15.15375 | 0.66605 | 5.19061 | 0.00000 |
| **COASY** | 23.06605 | 23.20422 | 10.16847 | 24.96260 | 2.30823 | 14.18174 | 7.05477 |
| **COBL** | 9.62309 | 9.39210 | 0.30893 | 2.43242 | 0.00000 | 3.75454 | 16.26444 |
| **COBLL1** | 6.32641 | 3.20771 | 0.00000 | 6.85556 | 1.15107 | 0.00000 | 7.63971 |
| **COCH** | 2.44882 | 0.00000 | 0.15576 | 5.10841 | 0.65499 | 0.08871 | 0.80281 |
| **COG1** | 6.01995 | 3.55560 | 1.90802 | 4.81655 | 0.41104 | 1.90645 | 1.26802 |
| **COG2** | 8.20216 | 5.17646 | 3.77570 | 2.78339 | 0.57953 | 2.93698 | 1.24934 |
| **COG3** | 18.11510 | 4.38537 | 1.95717 | 7.38506 | 2.29366 | 1.85700 | 1.80478 |
| **COG4** | 9.63130 | 11.63628 | 3.89517 | 7.10837 | 0.71689 | 6.92461 | 1.83217 |
| **COG5** | 5.81346 | 1.18036 | 2.55765 | 3.74007 | 3.20690 | 2.23682 | 0.42972 |
| **COG6** | 5.08934 | 1.12526 | 1.87268 | 2.36075 | 0.54970 | 1.62731 | 0.84318 |
| **COG7** | 6.95337 | 6.06919 | 2.54756 | 7.00473 | 1.61932 | 4.24417 | 0.00000 |
| **COG8** | 0.90479 | 2.54992 | 1.45606 | 1.88967 | 2.14998 | 1.61716 | 1.45380 |
| **COIL** | 14.73964 | 6.82818 | 5.18354 | 4.57981 | 2.55906 | 1.61856 | 4.03681 |
| **COL10A1** | 0.02058 | 0.00000 | 0.05321 | 0.01384 | 0.00495 | 0.04694 | 0.14664 |
| **COL11A1** | 0.02960 | 0.01921 | 0.01719 | 0.02791 | 0.06808 | 0.02498 | 0.08719 |
| **COL11A2** | 0.00000 | 0.00000 | 0.23695 | 0.69189 | 0.10788 | 0.36700 | 0.52341 |
| **COL12A1** | 0.24046 | 0.00000 | 0.00000 | 0.39828 | 1.11000 | 0.00281 | 0.13330 |
| **COL13A1** | 0.01354 | 0.01230 | 0.03745 | 0.03768 | 0.05140 | 0.09260 | 0.01696 |
| **COL14A1** | 0.07857 | 0.00000 | 0.02050 | 0.06108 | 0.15470 | 0.20923 | 0.04987 |
| **COL15A1** | 0.00000 | 0.42302 | 0.07559 | 0.05012 | 0.11349 | 0.17431 | 0.45523 |
| **COL16A1** | 0.02955 | 0.01621 | 0.04201 | 0.30422 | 0.02186 | 0.01288 | 0.00000 |
| **COL17A1** | 0.00809 | 0.08458 | 0.11032 | 0.02950 | 0.00521 | 0.70085 | 0.00692 |
| **COL18A1** | 0.00000 | 0.02078 | 0.00000 | 0.66477 | 0.34430 | 2.27547 | 0.24991 |
| **COL19A1** | 0.72890 | 0.03571 | 0.12427 | 0.15879 | 0.04183 | 0.29978 | 0.00000 |
| **COL1A1** | 0.21773 | 0.01826 | 0.04567 | 2.40348 | 0.00544 | 0.22083 | 1.39640 |
| **COL1A2** | 0.04398 | 0.04567 | 0.00889 | 0.75118 | 0.00720 | 0.45361 | 0.00000 |
| **COL20A1** | 0.02705 | 0.06543 | 0.16025 | 0.05874 | 0.00305 | 0.01001 | 0.02667 |

| | | | | | | | |
|---|---|---|---|---|---|---|---|
| **COL21A1** | 2.48655 | 0.00000 | 0.00000 | 8.08687 | 0.01277 | 0.00000 | 2.51735 |
| **COL22A1** | 0.01802 | 0.07936 | 0.01051 | 0.02078 | 0.99260 | 0.01749 | 0.01090 |
| **COL23A1** | 0.08134 | 0.00000 | 0.01682 | 0.00000 | 0.18524 | 1.18115 | 0.07562 |
| **COL24A1** | 0.08471 | 0.03400 | 0.21878 | 0.01858 | 0.08917 | 0.02493 | 0.00000 |
| **COL25A1** | 0.06612 | 0.04399 | 0.15193 | 0.00114 | 0.42861 | 0.00246 | 0.02148 |
| **COL26A1** | 0.06972 | 1.09848 | 0.04843 | 0.18986 | 0.19996 | 2.56249 | 0.07441 |
| **COL27A1** | 0.00000 | 1.65519 | 0.97191 | 0.01729 | 0.03718 | 2.63227 | 0.44004 |
| **COL28A1** | 0.06158 | 0.17034 | 0.17053 | 0.03636 | 2.06691 | 0.02404 | 0.11237 |
| **COL2A1** | 0.00000 | 0.10225 | 0.18863 | 0.13538 | 0.00548 | 0.11573 | 0.06013 |
| **COL3A1** | 0.02009 | 0.01066 | 0.03176 | 0.02880 | 0.00733 | 0.01254 | 0.04532 |
| **COL4A1** | 0.00000 | 0.00000 | 0.00912 | 0.96205 | 0.02529 | 9.12877 | 0.46140 |
| **COL4A2** | 0.00000 | 0.43188 | 0.00000 | 1.55497 | 0.11986 | 6.72056 | 1.24867 |
| **COL4A2-AS1** | 0.01857 | 0.01734 | 0.01226 | 0.01879 | 0.00779 | 0.02280 | 0.02965 |
| **COL4A3** | 0.01871 | 0.01539 | 0.03376 | 0.03170 | 0.00952 | 0.05988 | 0.12020 |
| **COL4A3BP** | 9.66825 | 7.74104 | 4.92015 | 7.35845 | 2.49061 | 11.66520 | 6.63597 |
| **COL4A4** | 0.02106 | 0.00000 | 0.06624 | 0.01893 | 0.14186 | 0.13013 | 0.04398 |
| **COL4A5** | 0.17114 | 0.00000 | 0.02407 | 0.15397 | 0.01137 | 0.53438 | 0.00000 |
| **COL4A6** | 0.02980 | 0.01286 | 0.00846 | 0.01495 | 0.05012 | 0.02552 | 0.03196 |
| **COL5A1** | 0.00000 | 0.01327 | 0.00566 | 0.06177 | 0.39707 | 0.96535 | 0.16233 |
| **COL5A2** | 0.73523 | 0.00000 | 0.87888 | 0.24821 | 0.00028 | 0.05198 | 0.56640 |
| **COL5A3** | 0.01953 | 0.02054 | 0.01369 | 0.03525 | 0.00834 | 0.01444 | 0.03435 |
| **COL6A1** | 0.00000 | 0.32351 | 5.23567 | 0.27690 | 0.83183 | 2.11804 | 4.00265 |
| **COL6A2** | 0.00000 | 0.66709 | 0.05994 | 0.03628 | 0.32844 | 1.04189 | 0.53780 |
| **COL6A3** | 0.00000 | 0.17317 | 0.01017 | 0.03720 | 0.41023 | 0.01220 | 0.01338 |
| **COL6A4P1** | 0.03032 | 0.00634 | 0.05214 | 0.03234 | 0.08391 | 0.01224 | 0.01328 |
| **COL6A4P2** | 0.01986 | 0.03866 | 0.27486 | 0.04679 | 0.03796 | 0.03716 | 0.01078 |
| **COL6A5** | 0.00900 | 0.00000 | 0.03655 | 0.15276 | 0.94298 | 0.03532 | 0.01452 |
| **COL6A6** | 0.00000 | 0.00000 | 0.28632 | 0.06058 | 1.58005 | 0.13055 | 0.17535 |
| **COL7A1** | 0.00000 | 0.94841 | 0.09983 | 1.77880 | 0.12129 | 1.14041 | 6.02249 |
| **COL8A1** | 0.01859 | 0.03370 | 0.04858 | 0.01663 | 0.00620 | 0.01973 | 0.02686 |
| **COL8A2** | 0.01160 | 0.02023 | 0.01334 | 0.03227 | 0.00895 | 0.01067 | 0.03842 |
| **COL9A1** | 0.02690 | 0.06283 | 0.00580 | 0.03740 | 0.01368 | 0.06742 | 0.00561 |
| **COL9A2** | 0.00246 | 0.01285 | 0.01805 | 0.59485 | 0.03489 | 0.00253 | 0.39216 |
| **COL9A3** | 0.00000 | 3.37885 | 3.12582 | 0.61830 | 0.00000 | 1.60937 | 3.98856 |
| **COLCA1** | 0.06114 | 0.01397 | 0.01430 | 0.02153 | 0.01258 | 0.01455 | 0.00000 |

| | | | | | | | |
|---|---|---|---|---|---|---|---|
| **COLEC11** | 0.00303 | 0.06853 | 0.01104 | 0.04676 | 0.35900 | 0.14709 | 0.13868 |
| **COLEC12** | 3.89833 | 0.00000 | 0.37395 | 2.09131 | 0.05062 | 0.00000 | 17.29222 |
| **COLGALT1** | 25.61333 | 25.27543 | 5.71552 | 15.21175 | 11.14632 | 7.94254 | 1.00974 |
| **COLGALT2** | 0.05071 | 0.01533 | 0.00938 | 0.02192 | 0.02110 | 0.01232 | 0.04832 |
| **COLQ** | 0.67915 | 0.00000 | 0.20634 | 0.86991 | 0.21522 | 0.11932 | 0.00000 |
| **COMMD1** | 9.73207 | 9.85527 | 1.18166 | 1.73231 | 3.67119 | 3.42373 | 3.04427 |
| **COMMD10** | 11.26650 | 0.00000 | 4.44433 | 5.90335 | 0.54962 | 5.06012 | 0.36035 |
| **COMMD2** | 24.48332 | 13.45607 | 9.56311 | 12.75771 | 0.75617 | 7.06489 | 1.58004 |
| **COMMD3** | 7.59421 | 1.30999 | 2.24721 | 1.72595 | 0.00000 | 0.79841 | 4.91000 |
| **COMMD4** | 12.28959 | 12.27524 | 3.09183 | 7.13580 | 1.54733 | 9.82239 | 8.52888 |
| **COMMD5** | 6.59080 | 7.94332 | 5.93653 | 8.72134 | 4.30384 | 4.30615 | 0.00000 |
| **COMMD6** | 13.84851 | 4.44306 | 5.16924 | 11.23141 | 8.28982 | 7.62657 | 12.27971 |
| **COMMD7** | 7.43073 | 9.50164 | 0.57640 | 6.36110 | 0.60955 | 4.39442 | 19.91403 |
| **COMMD8** | 11.94510 | 0.89879 | 4.71134 | 5.54012 | 1.08559 | 2.30318 | 1.88804 |
| **COMMD9** | 24.66636 | 11.72120 | 6.77538 | 16.29415 | 3.79363 | 5.87035 | 0.00000 |
| **COMP** | 0.01838 | 0.04833 | 0.08747 | 0.01971 | 0.56056 | 0.12216 | 0.03308 |
| **COMT** | 9.54779 | 5.75641 | 0.78127 | 0.98545 | 0.00000 | 0.00000 | 0.52959 |
| **COMTD1** | 0.60446 | 2.42248 | 0.00000 | 2.93664 | 0.01428 | 3.04138 | 11.25297 |
| **COPA** | 23.13760 | 8.91735 | 7.73595 | 18.08908 | 1.56701 | 11.76517 | 7.21160 |
| **COPB1** | 31.48655 | 5.14249 | 16.07367 | 15.98276 | 3.23549 | 12.17976 | 5.32153 |
| **COPB2** | 36.03735 | 12.53382 | 9.50356 | 23.82651 | 9.82366 | 13.49428 | 2.68109 |
| **COPE** | 30.61055 | 34.92584 | 14.19887 | 33.24162 | 7.98918 | 18.17310 | 6.84613 |
| **COPG1** | 24.75685 | 11.55205 | 4.79664 | 15.00370 | 2.93316 | 8.98863 | 0.00000 |
| **COPG2** | 1.13734 | 0.15720 | 0.66919 | 0.42121 | 0.41618 | 0.45135 | 0.86678 |
| **COPRS** | 0.65962 | 3.53293 | 0.51975 | 4.22227 | 3.36617 | 1.40032 | 16.83212 |
| **COPS2** | 21.88463 | 0.00000 | 13.30595 | 13.03968 | 6.34777 | 8.58463 | 11.09397 |
| **COPS3** | 35.31164 | 13.34484 | 11.82943 | 22.52300 | 8.80021 | 14.76877 | 3.27464 |
| **COPS4** | 21.81995 | 5.55683 | 8.89617 | 4.63681 | 6.44188 | 8.81552 | 1.82961 |
| **COPS5** | 28.48928 | 6.16400 | 14.22080 | 16.31122 | 5.15515 | 9.83806 | 3.47861 |
| **COPS6** | 29.75468 | 22.15365 | 11.48456 | 29.95067 | 5.79413 | 15.00428 | 0.12368 |
| **COPS7A** | 16.58632 | 4.82336 | 9.11147 | 10.23355 | 4.63865 | 3.86239 | 0.00000 |
| **COPS7B** | 5.40872 | 10.02782 | 6.91453 | 5.92807 | 6.01042 | 5.87705 | 5.39843 |
| **COPS8** | 19.47305 | 11.28095 | 2.85841 | 8.93724 | 5.64842 | 8.09873 | 0.92455 |
| **COPZ1** | 22.88202 | 11.09798 | 12.26272 | 12.84998 | 0.21189 | 11.53996 | 6.96394 |
| **COPZ2** | 0.02539 | 0.01449 | 0.01956 | 0.02805 | 0.00757 | 0.01110 | 0.02612 |

| | | | | | | | |
|---|---|---|---|---|---|---|---|
| **COQ10A** | 0.11711 | 0.00000 | 0.38077 | 0.90784 | 0.76510 | 0.68158 | 1.48797 |
| **COQ10B** | 16.55757 | 4.10319 | 8.82641 | 11.00988 | 9.10049 | 6.50491 | 0.00000 |
| **COQ2** | 2.14985 | 0.68733 | 2.73945 | 0.62303 | 0.03484 | 3.19000 | 7.08380 |
| **COQ3** | 5.55401 | 4.22693 | 3.81366 | 0.00000 | 0.48745 | 2.22131 | 0.69087 |
| **COQ4** | 9.93702 | 26.05820 | 16.82072 | 8.51433 | 0.98235 | 5.53715 | 1.28987 |
| **COQ5** | 8.12053 | 8.70621 | 7.10085 | 0.45035 | 0.02539 | 6.20144 | 2.64250 |
| **COQ6** | 6.03274 | 5.72210 | 2.24077 | 3.53834 | 1.56682 | 2.72501 | 0.00000 |
| **COQ7** | 10.61005 | 10.19935 | 10.13128 | 4.15406 | 3.28790 | 2.36654 | 0.00000 |
| **COQ9** | 9.63757 | 9.71430 | 3.93844 | 5.70063 | 0.35403 | 3.36175 | 0.43277 |
| **CORIN** | 0.13008 | 0.07111 | 0.05933 | 0.13016 | 0.04229 | 0.00259 | 0.00000 |
| **CORO1A** | 0.76724 | 7.11033 | 11.92346 | 0.00000 | 0.43174 | 6.14945 | 0.02716 |
| **CORO1B** | 1.88718 | 11.11718 | 5.73118 | 9.08199 | 0.00000 | 4.00507 | 7.89895 |
| **CORO1C** | 52.63881 | 35.86496 | 21.91760 | 24.26688 | 6.35916 | 19.36597 | 10.94872 |
| **CORO2A** | 20.66208 | 8.72575 | 0.00000 | 11.64110 | 0.37488 | 5.63062 | 3.14324 |
| **CORO2B** | 0.03947 | 0.00000 | 0.07009 | 0.01287 | 0.04151 | 0.01947 | 0.02766 |
| **CORO6** | 0.00000 | 0.25581 | 0.14660 | 0.25528 | 0.06772 | 0.33278 | 0.12670 |
| **CORO7** | 0.82817 | 0.93083 | 0.01657 | 1.59637 | 0.00205 | 0.86534 | 2.17727 |
| **CORT** | 0.03877 | 0.16187 | 0.12134 | 0.05262 | 0.18578 | 0.09663 | 0.20457 |
| **COTL1** | 2.24460 | 0.09628 | 0.35405 | 12.99606 | 0.90619 | 1.56816 | 10.51362 |
| **COX10** | 18.24130 | 9.09787 | 6.12780 | 11.45025 | 3.94681 | 4.49509 | 0.95412 |
| **COX10-AS1** | 1.61023 | 0.72144 | 0.18486 | 1.33083 | 0.38686 | 0.04113 | 0.00000 |
| **COX11** | 26.84748 | 7.37333 | 5.95264 | 8.98943 | 0.29016 | 10.13159 | 0.00000 |
| **COX14** | 19.05646 | 18.21047 | 8.19002 | 19.78634 | 3.34720 | 11.64837 | 8.97003 |
| **COX15** | 16.07098 | 10.21125 | 7.90350 | 9.65503 | 0.93548 | 6.27478 | 0.00000 |
| **COX16** | 10.37167 | 14.25886 | 8.13846 | 8.58459 | 5.86359 | 6.06286 | 6.21252 |
| **COX17** | 7.16596 | 6.75580 | 4.51607 | 5.03115 | 1.08785 | 5.15363 | 6.35751 |
| **COX18** | 2.73467 | 0.47967 | 0.54751 | 0.34178 | 0.07656 | 1.49212 | 3.05106 |
| **COX19** | 7.26106 | 9.05777 | 5.11788 | 8.80477 | 3.09323 | 3.14541 | 1.67254 |
| **COX20** | 2.66896 | 3.70099 | 0.13301 | 2.59501 | 0.00000 | 0.71875 | 0.64342 |
| **COX4I1** | 63.14662 | 61.70681 | 28.38502 | 48.33413 | 8.59440 | 39.10467 | 44.67527 |
| **COX4I2** | 0.00000 | 0.00426 | 0.13191 | 0.18356 | 4.88650 | 0.00125 | 0.01454 |
| **COX5A** | 54.12365 | 29.14265 | 18.26957 | 40.00090 | 1.98607 | 27.94737 | 34.93323 |
| **COX5B** | 53.44194 | 49.43710 | 24.56847 | 40.21162 | 16.97980 | 34.74062 | 33.62565 |
| **COX6A1** | 30.52695 | 25.98980 | 18.77953 | 23.33288 | 10.46372 | 20.91192 | 21.78933 |
| **COX6B1** | 49.44930 | 39.63116 | 18.81832 | 42.37307 | 7.75539 | 33.39019 | 41.15728 |

| | | | | | | | |
|---|---|---|---|---|---|---|---|
| **COX6B2** | 0.05038 | 0.16913 | 0.59754 | 0.00000 | 1.18625 | 0.02533 | 0.01629 |
| **COX6C** | 46.50477 | 25.28147 | 22.82418 | 28.90104 | 12.14330 | 29.46507 | 37.29075 |
| **COX7A1** | 0.05497 | 0.27017 | 0.02891 | 0.00000 | 0.02414 | 0.32209 | 0.02060 |
| **COX7A2** | 47.60413 | 26.29009 | 21.57916 | 32.61069 | 8.48295 | 26.52515 | 31.94080 |
| **COX7A2L** | 42.84942 | 24.77278 | 19.66700 | 25.88885 | 11.77779 | 17.72160 | 12.58662 |
| **COX7B** | 58.84707 | 26.50505 | 25.06753 | 30.57413 | 13.69480 | 31.15846 | 33.51044 |
| **COX7B2** | 17.72804 | 3.14413 | 10.22659 | 9.96907 | 0.88212 | 4.14419 | 13.27322 |
| **COX7C** | 42.14674 | 13.40173 | 10.18236 | 32.78971 | 3.40088 | 28.00245 | 42.92202 |
| **COX8A** | 36.29921 | 35.15405 | 13.11444 | 37.72013 | 6.87834 | 28.99804 | 33.67376 |
| **COX8C** | 0.73520 | 0.02080 | 1.42770 | 1.78372 | 0.32735 | 0.00000 | 0.61803 |
| **CP** | 0.28808 | 0.29911 | 0.00205 | 0.01476 | 0.00976 | 0.02923 | 0.00000 |
| **CPA1** | 0.00000 | 0.02273 | 0.05959 | 0.10306 | 0.72843 | 0.05225 | 0.26249 |
| **CPA2** | 0.24805 | 0.04485 | 0.05629 | 0.04686 | 0.49871 | 0.13433 | 0.00000 |
| **CPA4** | 0.07177 | 0.00000 | 0.00940 | 0.04178 | 0.07771 | 0.04039 | 0.16168 |
| **CPA5** | 0.02323 | 0.06946 | 0.00800 | 0.12906 | 0.05878 | 0.01591 | 0.00002 |
| **CPA6** | 0.02303 | 0.00000 | 0.03273 | 0.02280 | 0.04054 | 0.04217 | 0.04213 |
| **CPAMD8** | 0.04422 | 0.01009 | 0.01599 | 0.14897 | 0.00781 | 0.00518 | 0.09431 |
| **CPB1** | 0.05642 | 0.01066 | 0.03142 | 0.01392 | 0.01644 | 0.01146 | 0.01536 |
| **CPB2** | 0.04526 | 0.00000 | 0.08409 | 0.04263 | 0.05350 | 0.02884 | 0.14741 |
| **CPB2-AS1** | 0.06425 | 0.00000 | 0.17308 | 0.16208 | 0.27796 | 0.00463 | 0.16327 |
| **CPD** | 2.36972 | 1.95265 | 5.49768 | 1.30011 | 3.45564 | 5.48487 | 3.94831 |
| **CPE** | 0.03090 | 0.00713 | 0.03088 | 0.00989 | 0.01791 | 0.09597 | 0.04410 |
| **CPEB1** | 0.00000 | 0.39115 | 1.43559 | 0.00000 | 1.85496 | 0.00941 | 0.04372 |
| **CPEB2** | 0.60793 | 0.00000 | 2.29703 | 0.30005 | 1.57727 | 1.01187 | 1.93600 |
| **CPEB2-AS1** | 0.02260 | 0.00559 | 0.05920 | 0.02880 | 0.01320 | 0.01546 | 0.01943 |
| **CPEB3** | 0.77254 | 0.00000 | 1.03720 | 0.27197 | 0.49340 | 0.12912 | 2.30509 |
| **CPEB4** | 0.10845 | 0.88176 | 1.92933 | 4.63842 | 5.22531 | 1.75728 | 1.98819 |
| **CPED1** | 0.27661 | 0.00000 | 2.61970 | 0.00000 | 2.30100 | 0.08048 | 0.34968 |
| **CPLX1** | 0.00000 | 0.26968 | 0.21508 | 0.08003 | 0.11600 | 0.08552 | 0.09040 |
| **CPLX2** | 0.00000 | 0.22705 | 0.03247 | 0.19037 | 0.00614 | 0.08922 | 0.90745 |
| **CPLX3** | 0.00000 | 0.28045 | 0.13235 | 0.01292 | 0.00321 | 0.00663 | 0.02510 |
| **CPLX4** | 0.01591 | 0.05110 | 0.00000 | 0.01624 | 1.00715 | 0.00812 | 0.01694 |
| **CPM** | 0.00000 | 0.16095 | 0.54297 | 4.09856 | 1.30038 | 0.00000 | 0.00000 |
| **CPN1** | 0.04728 | 0.05204 | 0.13300 | 0.12024 | 0.07190 | 1.69543 | 0.00743 |
| **CPN2** | 0.00000 | 0.15271 | 0.01721 | 0.01098 | 0.00641 | 0.25165 | 0.02555 |

| | | | | | | | |
|---|---|---|---|---|---|---|---|
| **CPNE1** | 31.83642 | 22.37633 | 12.28583 | 22.81030 | 6.75457 | 8.98535 | 6.03930 |
| **CPNE2** | 0.00000 | 1.10739 | 0.46687 | 0.00000 | 0.19022 | 0.89939 | 0.12563 |
| **CPNE3** | 17.34211 | 0.97123 | 6.92327 | 6.08972 | 0.79059 | 6.56425 | 5.80780 |
| **CPNE4** | 0.06448 | 0.19815 | 0.00000 | 0.01106 | 0.50848 | 0.02280 | 0.00000 |
| **CPNE5** | 0.03621 | 0.02138 | 0.01288 | 0.11333 | 0.01848 | 0.01813 | 0.00054 |
| **CPNE6** | 0.02576 | 0.02057 | 0.01258 | 0.03475 | 0.00790 | 0.01433 | 0.01683 |
| **CPNE7** | 0.03114 | 0.14270 | 0.04405 | 0.05456 | 0.00638 | 0.01207 | 0.01715 |
| **CPNE8** | 2.25418 | 0.00000 | 0.09950 | 1.36441 | 0.01264 | 0.37439 | 0.09976 |
| **CPNE9** | 0.10448 | 0.45495 | 0.43151 | 0.28085 | 2.31981 | 0.29867 | 0.00000 |
| **CPO** | 0.02730 | 0.00801 | 0.02522 | 0.02696 | 0.01107 | 0.17664 | 0.02947 |
| **CPOX** | 1.97100 | 0.00000 | 3.06046 | 1.22368 | 0.00000 | 2.51019 | 9.04035 |
| **CPPED1** | 10.43636 | 6.58718 | 9.20622 | 0.51845 | 0.43307 | 7.27707 | 0.00000 |
| **CPQ** | 0.01361 | 0.14176 | 0.00000 | 0.01214 | 0.46135 | 0.02305 | 0.02144 |
| **CPS1** | 0.07296 | 0.21215 | 0.04614 | 0.00000 | 1.86684 | 0.00000 | 0.03331 |
| **CPSF1** | 4.38824 | 10.17830 | 3.25141 | 7.93396 | 0.57459 | 7.93424 | 20.02406 |
| **CPSF2** | 11.77995 | 1.49242 | 7.73875 | 10.61253 | 2.21312 | 7.16345 | 11.03143 |
| **CPSF3** | 20.96392 | 11.42877 | 8.58566 | 7.49830 | 0.12130 | 7.77370 | 3.81866 |
| **CPSF3L** | 21.55514 | 15.29259 | 8.65144 | 16.75266 | 2.96891 | 9.98045 | 2.79751 |
| **CPSF4** | 2.72721 | 6.41168 | 3.61956 | 12.14685 | 2.27428 | 8.06611 | 12.38817 |
| **CPSF4L** | 0.00325 | 0.40811 | 0.27803 | 0.02897 | 0.42405 | 0.04396 | 0.00000 |
| **CPSF6** | 32.94200 | 10.52087 | 15.54816 | 17.30643 | 3.58438 | 12.39712 | 12.36531 |
| **CPSF7** | 9.01827 | 3.92682 | 6.95071 | 8.90008 | 1.69350 | 6.05371 | 15.09725 |
| **CPT1A** | 0.23503 | 0.55841 | 1.05960 | 0.80100 | 0.98791 | 5.28806 | 0.00000 |
| **CPT1B** | 0.03539 | 0.02783 | 0.03611 | 0.03169 | 0.00672 | 0.01901 | 0.11134 |
| **CPT1C** | 0.35293 | 0.00000 | 0.42012 | 0.29627 | 6.09196 | 0.26289 | 0.29516 |
| **CPT2** | 1.83471 | 1.14438 | 0.60731 | 2.85461 | 0.14830 | 1.65243 | 3.06013 |
| **CPTP** | 0.03143 | 4.76142 | 0.03965 | 1.43179 | 0.00000 | 0.98085 | 5.58963 |
| **CPVL** | 27.20798 | 4.67803 | 0.16261 | 7.29654 | 0.23949 | 5.80005 | 3.67518 |
| **CPXM1** | 0.03544 | 0.01974 | 0.01538 | 0.11492 | 0.00802 | 0.01163 | 0.05384 |
| **CPXM2** | 0.00000 | 0.10301 | 0.04592 | 0.02809 | 0.17980 | 0.38879 | 0.01332 |
| **CPZ** | 0.07198 | 0.00000 | 0.00943 | 6.29457 | 0.00960 | 0.00000 | 1.34519 |
| **CR1** | 0.04357 | 0.15173 | 0.18928 | 0.00000 | 0.02385 | 0.13570 | 0.01512 |
| **CR1L** | 0.03998 | 0.04868 | 0.03751 | 0.00000 | 0.01833 | 1.20728 | 0.02219 |
| **CR2** | 0.08321 | 0.00180 | 0.00769 | 0.00888 | 3.12329 | 0.05564 | 0.00662 |
| **CRABP1** | 0.00000 | 0.00000 | 0.01630 | 0.04766 | 1.61695 | 0.53149 | 0.05376 |

| | | | | | | | |
|---|---|---|---|---|---|---|---|
| **CRABP2** | 0.10594 | 0.00000 | 0.32125 | 0.56528 | 0.27424 | 0.35709 | 0.60814 |
| **CRADD** | 6.12999 | 5.12266 | 1.58304 | 2.22249 | 0.08753 | 2.13759 | 0.00000 |
| **CRAMP1L** | 1.41963 | 4.28694 | 3.50459 | 3.49743 | 1.37292 | 4.24207 | 10.95675 |
| **CRAT** | 0.00000 | 0.00005 | 0.16399 | 2.49594 | 0.27748 | 0.24335 | 0.48133 |
| **CRB1** | 0.03237 | 0.02290 | 0.01615 | 0.01593 | 0.00752 | 0.02966 | 0.02310 |
| **CRB2** | 0.05034 | 0.01631 | 0.01465 | 0.05241 | 0.00861 | 0.02943 | 0.02204 |
| **CRB3** | 5.63614 | 23.15037 | 6.27034 | 11.10749 | 0.51796 | 6.05893 | 6.88014 |
| **CRBN** | 12.49381 | 5.13007 | 5.32364 | 6.29348 | 0.84465 | 5.36271 | 0.00000 |
| **CRCP** | 8.05343 | 3.65750 | 8.70711 | 6.23832 | 5.82225 | 2.30941 | 3.44319 |
| **CRCT1** | 0.01937 | 0.03261 | 0.01781 | 0.04019 | 0.00705 | 0.02079 | 0.01546 |
| **CREB1** | 11.11794 | 1.30400 | 4.64165 | 3.76762 | 5.07157 | 4.72982 | 5.53102 |
| **CREB3** | 8.22629 | 17.65306 | 7.26552 | 5.70310 | 3.94834 | 7.03209 | 0.00000 |
| **CREB3L1** | 0.06453 | 0.10415 | 0.01742 | 0.41190 | 0.02386 | 2.44277 | 0.72308 |
| **CREB3L2** | 1.23481 | 0.00000 | 0.24316 | 6.36653 | 2.46889 | 1.74148 | 0.51071 |
| **CREB3L3** | 0.19908 | 0.77801 | 0.00022 | 0.07386 | 0.33570 | 0.66922 | 0.16955 |
| **CREB3L4** | 1.29044 | 0.87269 | 0.82563 | 2.54693 | 0.00000 | 1.61231 | 0.41790 |
| **CREB5** | 0.06205 | 0.00000 | 0.07989 | 0.07863 | 1.81245 | 0.07707 | 0.11509 |
| **CREBBP** | 2.34921 | 12.98694 | 10.36564 | 4.59066 | 2.99494 | 7.50210 | 33.88140 |
| **CREBL2** | 1.07969 | 1.42183 | 2.09826 | 0.16858 | 0.24155 | 9.72372 | 3.78149 |
| **CREBRF** | 0.34339 | 0.11782 | 1.52257 | 1.50611 | 1.71259 | 1.80087 | 1.94825 |
| **CREBZF** | 6.35937 | 0.63906 | 1.49081 | 3.95072 | 1.32738 | 3.93688 | 16.16513 |
| **CREG1** | 0.43529 | 0.00000 | 6.68716 | 2.99175 | 0.27454 | 2.12155 | 4.12894 |
| **CREG2** | 0.06663 | 0.06155 | 0.00557 | 0.01321 | 0.00790 | 0.00000 | 0.06695 |
| **CRELD1** | 1.97624 | 5.49658 | 10.18620 | 3.57685 | 0.39381 | 3.46132 | 2.49083 |
| **CRELD2** | 2.63495 | 8.19204 | 0.77393 | 7.59896 | 3.73245 | 2.11245 | 4.95142 |
| **CREM** | 13.59192 | 3.09451 | 11.28228 | 9.32853 | 5.94225 | 7.54630 | 5.33434 |
| **CRH** | 0.00629 | 0.02841 | 0.01390 | 0.42262 | 0.00735 | 0.02440 | 0.00943 |
| **CRHBP** | 0.00891 | 0.00000 | 0.27908 | 0.04197 | 0.50691 | 0.83804 | 0.05766 |
| **CRHR1** | 0.03251 | 0.02987 | 0.02040 | 0.01173 | 0.00688 | 0.07800 | 0.01696 |
| **CRHR1-IT1** | 0.32342 | 0.10128 | 0.28533 | 0.03646 | 0.06579 | 0.11810 | 0.55739 |
| **CRHR2** | 0.01344 | 0.11083 | 0.05319 | 0.17442 | 0.00820 | 0.00000 | 0.22440 |
| **CRIM1** | 2.73225 | 0.34318 | 1.73880 | 5.95242 | 0.37701 | 1.05155 | 10.54888 |
| **CRIP1** | 29.33636 | 34.23982 | 0.00000 | 20.56705 | 0.00000 | 22.07371 | 39.37177 |
| **CRIP2** | 0.00000 | 0.43749 | 0.28756 | 1.05810 | 0.00390 | 1.10144 | 0.03079 |
| **CRIP3** | 0.27336 | 0.00000 | 0.20244 | 0.22202 | 0.01457 | 0.71429 | 0.00000 |

| | | | | | | | |
|---|---|---|---|---|---|---|---|
| **CRIPAK** | 0.04034 | 1.01408 | 0.07640 | 0.34032 | 0.06125 | 0.34360 | 2.16032 |
| **CRIPT** | 9.40825 | 0.33894 | 12.70796 | 8.31332 | 8.08452 | 3.84174 | 3.03157 |
| **CRISP1** | 0.06654 | 0.02035 | 0.01140 | 0.02204 | 0.00906 | 0.00883 | 0.01364 |
| **CRISP2** | 0.01110 | 0.00000 | 0.14525 | 0.08040 | 0.08806 | 0.00762 | 0.02652 |
| **CRISPLD1** | 0.08923 | 0.00000 | 1.03712 | 0.20568 | 1.21817 | 0.08280 | 0.00733 |
| **CRISPLD2** | 0.08960 | 0.00000 | 0.09313 | 0.72724 | 2.89903 | 1.55008 | 0.00000 |
| **CRK** | 13.17786 | 7.87821 | 6.59027 | 12.56187 | 4.02726 | 7.69676 | 18.89034 |
| **CRKL** | 31.23470 | 22.60759 | 11.82946 | 17.01263 | 17.06857 | 13.16710 | 4.31235 |
| **CRLF1** | 0.00838 | 0.10074 | 0.14345 | 0.03862 | 0.09817 | 0.01896 | 0.19128 |
| **CRLF3** | 11.71318 | 1.55995 | 3.44458 | 9.01643 | 2.93558 | 2.67020 | 0.65346 |
| **CRLS1** | 2.09302 | 1.54449 | 1.75992 | 1.11533 | 3.41851 | 4.66040 | 14.15958 |
| **CRMP1** | 0.03500 | 0.07949 | 0.00000 | 0.00000 | 4.99573 | 0.19738 | 0.02988 |
| **CRNDE** | 0.45964 | 0.00000 | 1.33108 | 0.00000 | 0.72240 | 0.15628 | 0.81447 |
| **CRNKL1** | 12.13919 | 6.86693 | 9.37878 | 11.06782 | 6.44685 | 4.44852 | 0.95537 |
| **CROCC** | 0.00871 | 0.06095 | 0.15259 | 0.17599 | 0.15177 | 0.07378 | 0.61854 |
| **CROCCP2** | 1.23415 | 1.11007 | 1.17602 | 2.43001 | 0.28133 | 0.67525 | 3.09767 |
| **CROCCP3** | 0.24358 | 0.11570 | 0.12073 | 0.08861 | 0.07519 | 0.07048 | 0.20378 |
| **CROT** | 0.28974 | 0.46654 | 0.08192 | 0.61625 | 0.35796 | 0.13785 | 0.00000 |
| **CRP** | 0.00248 | 0.00000 | 0.48454 | 0.01371 | 0.30903 | 0.02916 | 0.06709 |
| **CRSP8P** | 2.73712 | 1.81366 | 1.51394 | 2.35701 | 0.90003 | 0.91846 | 0.00000 |
| **CRTAC1** | 0.03757 | 0.03102 | 0.03277 | 0.02211 | 0.43637 | 0.09058 | 0.02582 |
| **CRTAM** | 0.02710 | 0.00768 | 0.05194 | 0.02052 | 0.01627 | 0.01408 | 0.01636 |
| **CRTAP** | 12.01367 | 2.30607 | 1.03086 | 3.62414 | 0.01193 | 8.79540 | 9.74272 |
| **CRTC1** | 0.00000 | 1.45130 | 0.40501 | 0.98175 | 0.53445 | 0.71855 | 3.69204 |
| **CRTC2** | 0.00000 | 3.25416 | 3.33373 | 10.20300 | 1.09678 | 4.61450 | 11.41086 |
| **CRTC3** | 0.21128 | 1.01348 | 1.20595 | 7.43673 | 0.37341 | 0.11487 | 7.85392 |
| **CRX** | 0.02157 | 0.05129 | 0.08720 | 0.01447 | 0.14738 | 0.01129 | 0.08932 |
| **CRY1** | 7.47520 | 3.93051 | 2.24020 | 8.35691 | 2.36247 | 3.91292 | 0.00000 |
| **CRY2** | 9.57746 | 6.26107 | 4.73957 | 5.93617 | 2.96373 | 2.14640 | 1.39792 |
| **CRYAB** | 0.00000 | 0.00773 | 0.00724 | 0.19140 | 0.11245 | 0.18112 | 0.06391 |
| **CRYBA1** | 0.08449 | 0.00000 | 0.01393 | 0.76873 | 0.13507 | 0.01793 | 0.01936 |
| **CRYBA2** | 0.00000 | 1.57972 | 0.19699 | 3.15372 | 0.00000 | 0.00000 | 1.50968 |
| **CRYBA4** | 0.07530 | 0.13296 | 0.02026 | 0.02533 | 0.08457 | 1.33289 | 0.00853 |
| **CRYBB1** | 0.02280 | 0.04558 | 0.01809 | 0.09319 | 0.10241 | 2.09246 | 0.00000 |
| **CRYBB2** | 0.08375 | 0.06982 | 0.01296 | 0.07670 | 0.01516 | 0.17591 | 0.02960 |

| | | | | | | | |
|---|---|---|---|---|---|---|---|
| **CRYBB2P1** | 3.37290 | 3.68170 | 0.61614 | 5.40243 | 1.10095 | 2.08087 | 0.00000 |
| **CRYBB3** | 0.00000 | 0.33018 | 0.02462 | 0.05301 | 0.01365 | 0.05764 | 0.06993 |
| **CRYBG3** | 0.02852 | 0.00000 | 0.99678 | 0.05442 | 1.10955 | 0.20122 | 0.00000 |
| **CRYGB** | 0.07912 | 0.01472 | 0.10006 | 0.00105 | 0.05647 | 1.05256 | 0.09975 |
| **CRYGC** | 0.46458 | 0.19335 | 0.74131 | 0.08059 | 0.52257 | 1.10865 | 0.00000 |
| **CRYGD** | 0.44032 | 0.01039 | 0.13484 | 0.12298 | 0.04192 | 1.56743 | 0.00000 |
| **CRYGS** | 0.10116 | 0.11576 | 0.07126 | 0.07284 | 0.14574 | 0.01622 | 0.23329 |
| **CRYL1** | 0.02188 | 0.01171 | 0.02094 | 0.07201 | 0.05058 | 0.03404 | 0.02675 |
| **CRYM** | 0.10827 | 0.00000 | 0.20344 | 0.04457 | 0.86743 | 2.52560 | 0.00000 |
| **CRYM-AS1** | 0.09619 | 0.03882 | 0.03955 | 0.06030 | 0.00586 | 0.03855 | 0.00000 |
| **CRYZ** | 13.59267 | 0.00000 | 4.30308 | 3.92888 | 2.32943 | 3.97886 | 0.00000 |
| **CRYZL1** | 7.03236 | 1.66629 | 1.98447 | 0.74803 | 0.68259 | 2.78395 | 0.00000 |
| **CS** | 35.61328 | 14.19276 | 15.30490 | 12.18436 | 7.45456 | 17.80991 | 15.88160 |
| **CSAD** | 3.98537 | 0.54258 | 1.00700 | 0.29086 | 0.00000 | 0.00000 | 6.44029 |
| **CSAG1** | 0.01778 | 0.00000 | 3.40245 | 0.48971 | 4.92485 | 0.02595 | 0.00000 |
| **CSAG3.uid-1334** | 0.02187 | 0.01434 | 0.03487 | 0.01995 | 0.01046 | 0.01170 | 0.02399 |
| **CSAG4** | 0.04474 | 0.00786 | 0.03259 | 0.03180 | 0.18198 | 0.01812 | 0.00000 |
| **CSDC2** | 0.01533 | 0.01775 | 0.01225 | 0.03332 | 0.00759 | 0.01027 | 0.04082 |
| **CSDE1** | 50.60165 | 23.09811 | 26.05793 | 34.52947 | 15.71510 | 23.35502 | 19.67745 |
| **CSE1L** | 57.20492 | 17.88618 | 21.14933 | 27.37648 | 2.80766 | 19.33330 | 13.10102 |
| **CSF1** | 0.11126 | 0.00259 | 0.06995 | 0.33120 | 0.02944 | 0.10070 | 0.02981 |
| **CSF1R** | 0.18174 | 0.00000 | 0.74767 | 0.79571 | 6.63206 | 0.00000 | 0.14284 |
| **CSF2RB** | 0.01426 | 0.01078 | 0.01491 | 0.16406 | 0.00860 | 0.05316 | 0.02428 |
| **CSF3R** | 0.00000 | 0.57690 | 0.09185 | 10.21866 | 0.00645 | 0.12291 | 0.00000 |
| **CSGALNACT1** | 0.75012 | 0.35023 | 0.00000 | 1.44295 | 0.49136 | 0.00000 | 0.68919 |
| **CSGALNACT2** | 0.68579 | 0.00000 | 1.02740 | 0.50443 | 1.89381 | 0.00000 | 1.73119 |
| **CSH1** | 0.01490 | 0.01160 | 0.02570 | 0.01904 | 0.00766 | 0.07861 | 0.02733 |
| **CSH2** | 0.01714 | 0.01480 | 0.01688 | 0.02245 | 0.02562 | 0.03074 | 0.03073 |
| **CSK** | 0.00000 | 5.54166 | 3.58417 | 8.17536 | 0.63718 | 5.48712 | 12.90855 |
| **CSMD1** | 0.48925 | 0.00000 | 0.11973 | 0.02858 | 0.70683 | 0.00000 | 0.54030 |
| **CSMD2** | 0.01159 | 0.01382 | 0.02293 | 0.09118 | 0.03702 | 0.06912 | 0.12615 |
| **CSMD3** | 0.08259 | 0.00000 | 0.11838 | 0.14252 | 0.06192 | 0.11960 | 0.15171 |
| **CSN3** | 0.04369 | 0.01476 | 0.01285 | 0.07443 | 0.01446 | 0.01650 | 0.00000 |
| **CSNK1A1** | 37.32255 | 16.18425 | 19.73320 | 27.39691 | 7.84868 | 14.06125 | 11.00553 |
| **CSNK1A1L** | 1.76921 | 0.26906 | 1.06759 | 1.22328 | 0.29774 | 0.62055 | 0.00000 |

| | | | | | | | |
|---|---|---|---|---|---|---|---|
| **CSNK1A1P1** | 2.32160 | 1.52499 | 1.53987 | 1.05486 | 0.35363 | 0.73645 | 0.00000 |
| **CSNK1D** | 5.70601 | 10.10291 | 5.94504 | 11.84703 | 2.48257 | 10.09539 | 24.58412 |
| **CSNK1E** | 0.00000 | 14.46899 | 4.68243 | 8.81495 | 3.39832 | 1.53030 | 15.25321 |
| **CSNK1G1** | 5.91250 | 1.01571 | 4.93313 | 5.81751 | 6.49374 | 0.00000 | 8.91830 |
| **CSNK1G2** | 0.43744 | 1.61295 | 0.78680 | 2.45885 | 1.04164 | 1.25337 | 19.81662 |
| **CSNK1G2-AS1** | 0.03922 | 0.00136 | 0.02699 | 0.00616 | 0.02624 | 0.04106 | 0.30173 |
| **CSNK1G3** | 6.76110 | 0.00000 | 5.52345 | 6.29682 | 3.44923 | 3.22081 | 6.50580 |
| **CSNK2A1** | 20.61669 | 11.04043 | 16.13647 | 9.52024 | 4.03847 | 9.22596 | 11.93226 |
| **CSNK2A2** | 0.00000 | 3.91642 | 6.10396 | 2.65865 | 3.08740 | 2.15954 | 10.99341 |
| **CSNK2A3** | 8.00336 | 2.52259 | 6.50396 | 3.24724 | 0.97817 | 2.56149 | 0.00000 |
| **CSNK2B** | 30.13900 | 25.16074 | 20.96708 | 29.79179 | 13.03756 | 20.03720 | 21.96736 |
| **CSPG4** | 0.00977 | 0.06151 | 0.00618 | 0.02502 | 0.12422 | 0.00805 | 0.24557 |
| **CSPG5** | 0.00000 | 0.02043 | 0.75544 | 0.09159 | 0.15944 | 0.04814 | 0.12928 |
| **CSPP1** | 9.07954 | 0.43594 | 5.48731 | 1.61572 | 5.37410 | 5.12764 | 13.56304 |
| **CSRNP1** | 0.18706 | 7.69524 | 3.56224 | 0.65793 | 2.65175 | 4.18661 | 4.96909 |
| **CSRNP2** | 2.29092 | 1.35889 | 2.58910 | 6.13330 | 2.34333 | 2.36569 | 8.34773 |
| **CSRNP3** | 0.20232 | 0.05668 | 0.20108 | 0.19585 | 0.00662 | 0.01419 | 0.00000 |
| **CSRP1** | 0.89938 | 3.42023 | 0.49735 | 1.68499 | 4.01503 | 0.18806 | 1.70499 |
| **CSRP2** | 2.07991 | 3.54681 | 1.32058 | 2.57864 | 3.19908 | 0.81180 | 0.00000 |
| **CSRP2BP** | 14.31939 | 15.84444 | 5.71122 | 15.30050 | 0.56841 | 1.55850 | 7.37629 |
| **CSRP3** | 0.02512 | 0.00745 | 0.03572 | 0.01970 | 0.05561 | 0.01275 | 0.01683 |
| **CST1** | 0.05619 | 0.00890 | 0.03999 | 0.35018 | 0.03601 | 0.42669 | 0.00000 |
| **CST11** | 0.03159 | 0.06083 | 0.01129 | 0.01619 | 0.07780 | 0.00870 | 0.00494 |
| **CST13P** | 0.02289 | 0.02451 | 0.02934 | 0.01872 | 0.00653 | 0.01101 | 0.02174 |
| **CST2** | 0.02111 | 0.02200 | 0.01130 | 0.03086 | 0.00831 | 0.16394 | 0.00719 |
| **CST3** | 0.00000 | 3.17632 | 0.00037 | 6.37908 | 0.00377 | 8.40571 | 8.05748 |
| **CST4** | 0.04719 | 0.00565 | 0.01331 | 0.06698 | 0.00831 | 0.14258 | 0.01263 |
| **CST5** | 0.03535 | 0.01654 | 0.01183 | 0.01663 | 0.01939 | 0.01547 | 0.01201 |
| **CST6** | 0.12492 | 0.01450 | 0.00511 | 0.45044 | 0.01507 | 0.00000 | 0.12191 |
| **CST7** | 0.02207 | 0.00706 | 0.02945 | 0.04312 | 0.01048 | 0.02487 | 0.03685 |
| **CST8** | 0.02454 | 0.01776 | 0.01026 | 0.01879 | 0.06491 | 0.01324 | 0.01608 |
| **CSTA** | 0.41296 | 0.00000 | 0.04753 | 0.18130 | 0.06907 | 0.21981 | 0.48356 |
| **CSTB** | 34.33943 | 14.61428 | 4.18631 | 24.73494 | 1.47999 | 13.34640 | 30.59922 |
| **CSTF1** | 18.70518 | 11.50782 | 14.12418 | 9.26614 | 3.58647 | 4.76292 | 0.00000 |
| **CSTF2** | 37.95292 | 28.20227 | 15.87093 | 17.24883 | 8.41406 | 7.51570 | 0.00000 |

| | | | | | | | |
|---|---|---|---|---|---|---|---|
| **CSTF2T** | 9.27142 | 0.00000 | 9.11435 | 6.75923 | 1.69233 | 2.92273 | 0.00000 |
| **CSTF3** | 27.63146 | 3.51159 | 13.24714 | 14.57240 | 5.59878 | 5.22068 | 2.85765 |
| **CSTF3-AS1** | 0.02616 | 0.00935 | 0.03608 | 0.03120 | 0.42155 | 0.03993 | 0.01097 |
| **CSTL1** | 0.02338 | 0.02197 | 0.05124 | 0.01883 | 0.00785 | 0.00917 | 0.02157 |
| **CT45A1** | 0.02459 | 0.02531 | 0.01490 | 0.01908 | 0.01160 | 0.01198 | 0.01378 |
| **CT47A6.uid-131** | 0.02072 | 0.01887 | 0.02572 | 0.02799 | 0.02072 | 0.01792 | 0.03763 |
| **CT47B1** | 0.01502 | 0.01544 | 0.01374 | 0.01320 | 0.81789 | 0.01954 | 0.04488 |
| **CT55** | 0.15689 | 1.50625 | 1.79888 | 0.00087 | 0.18256 | 0.00519 | 0.00000 |
| **CT62** | 0.04294 | 0.01276 | 0.01919 | 0.01879 | 0.00701 | 0.01256 | 0.04061 |
| **CT83** | 0.06064 | 0.00000 | 0.15248 | 0.09546 | 0.46771 | 0.00215 | 0.02922 |
| **CTAG2** | 0.00000 | 0.62717 | 0.59480 | 0.21141 | 0.04274 | 0.15593 | 1.70603 |
| **CTAGE1** | 0.09281 | 0.00800 | 0.03255 | 0.07460 | 0.03151 | 0.04830 | 0.01425 |
| **CTAGE10P** | 0.03672 | 0.01814 | 0.01331 | 0.02150 | 0.00792 | 0.02009 | 0.02934 |
| **CTAGE11P** | 0.05192 | 0.01717 | 0.01300 | 0.02428 | 0.01873 | 0.02380 | 0.00848 |
| **CTAGE15** | 0.02424 | 0.01812 | 0.01553 | 0.02217 | 0.00773 | 0.01260 | 0.01617 |
| **CTAGE4.uid-101** | 0.02309 | 0.01890 | 0.01325 | 0.01934 | 0.00777 | 0.01231 | 0.02296 |
| **CTAGE5** | 3.80705 | 0.41807 | 0.78740 | 2.67552 | 1.74058 | 2.51272 | 2.13653 |
| **CTAGE7P** | 0.05865 | 0.01506 | 0.01925 | 0.03222 | 0.04013 | 0.04369 | 0.02529 |
| **CTAGE8** | 0.02455 | 0.01867 | 0.01325 | 0.02007 | 0.00785 | 0.01297 | 0.01673 |
| **CTAGE9** | 0.02594 | 0.01640 | 0.01293 | 0.02324 | 0.00795 | 0.02013 | 0.01917 |
| **CTB-113P19.1** | 0.17940 | 0.34055 | 0.03577 | 0.03861 | 0.00847 | 0.51252 | 0.00474 |
| **CTB-118N6.3** | 0.25421 | 0.00000 | 0.02545 | 0.03500 | 0.00456 | 0.05257 | 0.64636 |
| **CTB-12O2.1** | 0.03809 | 0.01100 | 0.04226 | 0.02227 | 0.00833 | 0.15971 | 0.03905 |
| **CTB-178M22.2** | 0.04304 | 0.06984 | 0.54323 | 0.01645 | 0.23164 | 0.00208 | 0.00003 |
| **CTB-7E3.1** | 0.02694 | 0.01676 | 0.01154 | 0.01950 | 0.04158 | 0.01406 | 0.01282 |
| **CTBP1** | 5.61736 | 15.02670 | 1.67712 | 1.91972 | 0.34925 | 3.61005 | 12.49437 |
| **CTBP1-AS** | 0.02380 | 0.08423 | 0.04430 | 0.07274 | 0.00613 | 0.05907 | 0.26301 |
| **CTBP1-AS2** | 0.00000 | 0.78333 | 0.70171 | 0.28745 | 0.00180 | 0.71636 | 2.16982 |
| **CTBP2** | 4.19557 | 4.03701 | 1.32162 | 3.19512 | 0.40365 | 6.80297 | 16.18555 |
| **CTBS** | 0.89618 | 0.00000 | 2.19135 | 0.19262 | 0.78891 | 0.13292 | 0.00000 |
| **CTC-338M12.4** | 0.98704 | 0.89113 | 1.29768 | 2.18447 | 0.26576 | 1.36882 | 1.38985 |
| **CTC-360P9.3** | 0.02040 | 0.01705 | 0.02450 | 0.02942 | 0.05203 | 0.01195 | 0.01580 |
| **CTC-436P18.1** | 0.03209 | 0.00166 | 0.06648 | 0.02143 | 0.04354 | 0.01821 | 0.02523 |
| **CTC1** | 2.70580 | 1.14740 | 0.35891 | 1.57076 | 0.14667 | 1.31557 | 5.05844 |
| **CTCF** | 9.60527 | 6.32111 | 5.80202 | 4.66164 | 9.91795 | 7.30493 | 20.00235 |

| | | | | | | | |
|---|---|---|---|---|---|---|---|
| **CTCFL** | 0.32046 | 0.11697 | 0.93280 | 0.00000 | 2.20430 | 0.10241 | 0.16422 |
| **CTD-2151A2.1** | 0.05545 | 0.01757 | 0.05418 | 0.04191 | 0.02275 | 0.02088 | 0.00633 |
| **CTD-2194D22.4** | 0.03454 | 0.01664 | 0.01228 | 0.11918 | 0.00786 | 0.01003 | 0.01853 |
| **CTD-2201I18.1** | 0.05213 | 0.07071 | 0.00000 | 0.09468 | 1.77804 | 0.03136 | 0.00000 |
| **CTD-2258A20.5** | 0.02324 | 0.01931 | 0.01298 | 0.04875 | 0.00791 | 0.01381 | 0.01444 |
| **CTD-2350J17.1** | 0.02120 | 0.03964 | 0.01287 | 0.02986 | 0.01564 | 0.01039 | 0.01830 |
| **CTD-3080P12.3** | 0.06860 | 0.00590 | 0.01627 | 0.06948 | 0.01422 | 0.04280 | 0.00000 |
| **CTDNEP1** | 0.53676 | 5.07025 | 2.74351 | 4.95536 | 1.25877 | 4.77829 | 23.05315 |
| **CTDP1** | 0.00000 | 3.25058 | 1.81616 | 1.08539 | 1.11197 | 3.02041 | 9.75074 |
| **CTDSP1** | 1.74718 | 10.69072 | 3.97752 | 2.83585 | 0.60063 | 8.02982 | 14.10536 |
| **CTDSP2** | 3.73543 | 3.48769 | 1.53847 | 4.21146 | 4.33351 | 3.89082 | 12.94692 |
| **CTDSPL** | 3.93023 | 0.24575 | 0.00000 | 8.30759 | 0.86470 | 1.20181 | 10.57446 |
| **CTDSPL2** | 10.79742 | 0.00000 | 10.04296 | 6.97296 | 6.49881 | 3.35092 | 5.99988 |
| **CTF1** | 0.01953 | 0.01383 | 0.02447 | 0.08820 | 0.00946 | 0.02151 | 0.06355 |
| **CTGF** | 0.00000 | 3.53727 | 2.75395 | 0.69193 | 0.00000 | 3.01780 | 2.37007 |
| **CTGLF12P** | 0.03403 | 0.00574 | 0.02658 | 0.01928 | 0.01348 | 0.01363 | 0.07827 |
| **CTH** | 19.83347 | 2.35478 | 1.93839 | 16.30222 | 0.00000 | 7.47175 | 3.21945 |
| **CTHRC1** | 0.05928 | 0.00000 | 0.27165 | 0.66931 | 0.00346 | 0.28273 | 0.11394 |
| **CTIF** | 0.60194 | 1.08221 | 0.00000 | 0.87525 | 0.41196 | 0.42091 | 6.17731 |
| **CTLA4** | 0.02820 | 0.01405 | 0.01985 | 0.04054 | 0.00706 | 0.01148 | 0.04719 |
| **CTNNA1** | 43.48652 | 25.87738 | 13.92232 | 32.45003 | 10.30835 | 16.05815 | 12.56284 |
| **CTNNA2** | 0.02601 | 0.02519 | 0.00826 | 0.01247 | 1.29587 | 0.13876 | 0.00886 |
| **CTNNA3** | 0.05152 | 0.02651 | 0.00000 | 0.04640 | 1.59192 | 0.08894 | 0.01484 |
| **CTNNAL1** | 20.52972 | 9.49146 | 3.70076 | 20.78922 | 1.91054 | 3.93267 | 9.98246 |
| **CTNNB1** | 43.10584 | 18.17551 | 17.00633 | 40.62002 | 8.80923 | 11.82630 | 22.63317 |
| **CTNNBIP1** | 0.00000 | 0.21505 | 0.00000 | 0.22035 | 0.38349 | 0.49014 | 1.33424 |
| **CTNNBL1** | 18.15902 | 15.60098 | 12.25523 | 22.45034 | 11.11769 | 10.20127 | 0.00000 |
| **CTNND1** | 0.70898 | 0.00000 | 1.68796 | 2.37282 | 0.27465 | 0.62375 | 0.97372 |
| **CTNND2** | 0.01153 | 0.16631 | 0.00000 | 0.01256 | 0.38410 | 0.08980 | 0.02075 |
| **CTNS** | 5.54731 | 10.46854 | 4.24351 | 6.96874 | 3.12835 | 7.46031 | 2.07509 |
| **CTPS1** | 36.86386 | 17.90121 | 15.59048 | 20.07425 | 3.08161 | 9.69763 | 8.81819 |
| **CTPS2** | 0.06785 | 0.00000 | 0.11906 | 0.04288 | 0.02418 | 0.09573 | 0.15607 |
| **CTR9** | 17.81749 | 7.25690 | 9.06375 | 10.80039 | 5.07821 | 8.11875 | 4.92955 |
| **CTRB1** | 0.01786 | 0.03009 | 0.00000 | 0.01616 | 0.45144 | 0.02278 | 0.02372 |
| **CTRB2** | 0.00000 | 0.19732 | 0.68909 | 0.01466 | 3.11775 | 0.00441 | 0.03326 |

| | | | | | | | |
|---|---|---|---|---|---|---|---|
| **CTRC** | 0.03131 | 0.01709 | 0.01073 | 0.01225 | 0.02485 | 0.01252 | 0.03261 |
| **CTRL** | 1.84551 | 0.80040 | 0.02469 | 3.00685 | 0.00229 | 0.73311 | 0.00000 |
| **CTSA** | 20.98079 | 32.63569 | 7.34266 | 9.66015 | 2.12198 | 12.52636 | 7.57957 |
| **CTSB** | 69.50463 | 70.69122 | 25.08383 | 20.95572 | 10.50413 | 19.88764 | 13.83451 |
| **CTSC** | 51.42691 | 13.87591 | 19.62294 | 31.29793 | 1.80558 | 32.96103 | 2.22656 |
| **CTSD** | 15.30496 | 25.73836 | 0.53352 | 31.41965 | 0.41085 | 15.86994 | 27.95953 |
| **CTSE** | 0.87741 | 0.60380 | 0.00045 | 1.01985 | 0.00635 | 9.19933 | 0.00000 |
| **CTSF** | 0.00000 | 5.77599 | 5.84027 | 0.00000 | 0.28876 | 5.60932 | 1.47260 |
| **CTSH** | 15.02286 | 1.05142 | 1.10713 | 6.57889 | 0.40946 | 0.00000 | 2.96883 |
| **CTSK** | 0.38753 | 0.00000 | 0.38038 | 0.39364 | 0.06197 | 0.32243 | 0.55098 |
| **CTSL** | 52.27224 | 43.05798 | 17.02079 | 51.18088 | 2.38993 | 19.85954 | 19.70891 |
| **CTSL3P** | 0.20400 | 0.00000 | 0.44026 | 0.56452 | 0.00225 | 0.09155 | 0.00000 |
| **CTSLP8** | 0.06916 | 0.02862 | 0.02548 | 0.02963 | 0.00768 | 0.01588 | 0.04213 |
| **CTSO** | 0.04072 | 0.03234 | 0.00970 | 0.01593 | 0.00790 | 0.02319 | 0.01709 |
| **CTSS** | 0.00000 | 0.58977 | 0.24677 | 0.03499 | 0.62676 | 0.01662 | 0.00089 |
| **CTSV** | 50.56843 | 48.62305 | 34.55947 | 76.76299 | 11.75343 | 12.69258 | 4.44159 |
| **CTSW** | 0.00000 | 0.24828 | 0.53625 | 0.00000 | 0.01387 | 0.03213 | 0.03070 |
| **CTSZ** | 0.00000 | 1.16212 | 0.00000 | 0.26656 | 0.12961 | 0.10323 | 1.45779 |
| **CTTN** | 48.21779 | 48.18030 | 6.92023 | 32.84242 | 7.47923 | 29.99164 | 0.00000 |
| **CTTNBP2** | 4.72696 | 0.00000 | 0.17538 | 3.01130 | 1.25443 | 0.00000 | 6.93743 |
| **CTTNBP2NL** | 4.57242 | 3.74275 | 1.78727 | 4.36298 | 7.10069 | 1.71005 | 0.95923 |
| **CTU1** | 0.00000 | 0.35959 | 0.13109 | 0.00000 | 0.06332 | 0.10933 | 1.01361 |
| **CTU2** | 7.45829 | 9.67304 | 4.13919 | 8.65850 | 1.61196 | 5.44646 | 3.89669 |
| **CTXN1** | 0.00000 | 0.74367 | 0.43047 | 0.00000 | 0.00268 | 0.14126 | 0.70945 |
| **CTXN2** | 0.02360 | 0.01748 | 0.01460 | 0.02631 | 0.00800 | 0.01364 | 0.01431 |
| **CUBN** | 0.00801 | 0.02826 | 0.01950 | 0.04706 | 0.19426 | 0.39709 | 0.01670 |
| **CUEDC1** | 0.00000 | 0.02436 | 0.08082 | 0.04161 | 0.27898 | 0.08660 | 0.25874 |
| **CUEDC2** | 5.09450 | 2.17110 | 1.00229 | 1.49028 | 0.07813 | 2.65110 | 1.14277 |
| **CUL1** | 15.90174 | 12.59641 | 8.26491 | 13.85600 | 6.81306 | 10.30901 | 10.06451 |
| **CUL2** | 20.91059 | 0.00000 | 21.24203 | 11.05029 | 17.09501 | 6.37248 | 0.00000 |
| **CUL3** | 8.65955 | 2.22211 | 5.54749 | 4.85283 | 2.79106 | 2.73191 | 22.49156 |
| **CUL4A** | 11.81607 | 6.21385 | 7.27994 | 6.65678 | 1.71507 | 8.15864 | 17.58980 |
| **CUL4B** | 8.20446 | 0.00000 | 7.85607 | 3.85174 | 3.81572 | 4.28939 | 8.02204 |
| **CUL5** | 15.02112 | 1.15008 | 6.69765 | 8.31989 | 16.63277 | 3.61104 | 8.88777 |
| **CUL7** | 0.00000 | 0.13252 | 0.47426 | 1.09947 | 0.08327 | 1.21093 | 1.25126 |

| | | | | | | | |
|---|---|---|---|---|---|---|---|
| **CUL9** | 0.00000 | 0.00000 | 0.70297 | 0.64203 | 0.03153 | 0.66895 | 2.45969 |
| **CUTA** | 5.79808 | 4.82036 | 6.05571 | 8.62170 | 3.03449 | 3.04813 | 2.95700 |
| **CUTC** | 14.33256 | 3.48085 | 10.79517 | 6.97385 | 0.37113 | 5.41282 | 0.00000 |
| **CUX1** | 2.05226 | 0.47421 | 0.73784 | 1.43791 | 2.19171 | 2.43762 | 7.28134 |
| **CUX2** | 0.00000 | 0.28395 | 0.21329 | 0.00458 | 0.30993 | 0.28228 | 0.04100 |
| **CWC15** | 28.71255 | 15.74685 | 20.74122 | 17.66383 | 12.17095 | 10.62126 | 9.84093 |
| **CWC22** | 15.03611 | 0.00000 | 8.87004 | 9.15162 | 3.33790 | 7.42796 | 2.30825 |
| **CWC25** | 10.42374 | 9.63150 | 5.33225 | 9.68726 | 13.14962 | 4.13535 | 6.22891 |
| **CWC27** | 10.12129 | 3.07208 | 5.26372 | 6.06601 | 1.38186 | 5.27682 | 5.37566 |
| **CWF19L1** | 14.60702 | 9.74563 | 7.83786 | 9.19292 | 2.98604 | 6.21301 | 2.32929 |
| **CWF19L2** | 5.76771 | 2.64514 | 5.47527 | 5.43769 | 4.24836 | 3.93375 | 9.51940 |
| **CX3CL1** | 0.02652 | 0.01600 | 0.01349 | 0.07402 | 0.00912 | 0.05240 | 0.00718 |
| **CX3CR1** | 0.09796 | 0.00000 | 0.18826 | 0.01260 | 0.01200 | 0.29862 | 0.07890 |
| **CXADR** | 41.06174 | 17.37273 | 30.56194 | 0.00001 | 14.36522 | 15.64339 | 0.00000 |
| **CXADRP2** | 0.02758 | 0.01748 | 0.02236 | 0.02052 | 0.01520 | 0.01288 | 0.01757 |
| **CXADRP3** | 0.29377 | 0.11524 | 0.49741 | 0.00355 | 0.28560 | 0.05549 | 0.00000 |
| **CXCL12** | 0.01806 | 0.02484 | 0.12023 | 0.00243 | 0.01115 | 6.08269 | 0.02878 |
| **CXCL14** | 0.30287 | 0.00000 | 0.18441 | 0.04656 | 1.18799 | 0.27674 | 0.00000 |
| **CXCL16** | 0.00000 | 0.21739 | 0.99553 | 0.13324 | 0.49033 | 0.17266 | 0.08085 |
| **CXCL17** | 0.05892 | 0.05521 | 0.01222 | 0.04134 | 0.00800 | 0.22838 | 0.00000 |
| **CXCL3** | 0.02277 | 0.00000 | 0.08502 | 0.02478 | 0.03157 | 0.01162 | 0.01975 |
| **CXCL5** | 0.03400 | 0.01369 | 0.01350 | 0.02080 | 0.04027 | 0.01231 | 0.03642 |
| **CXCL8** | 0.06180 | 0.00738 | 0.01085 | 0.01021 | 0.00702 | 0.01282 | 0.09120 |
| **CXCL9** | 0.02649 | 0.00442 | 0.06290 | 0.01882 | 0.00727 | 0.01243 | 0.01711 |
| **CXCR1** | 0.01416 | 0.00000 | 0.31902 | 0.02222 | 0.06677 | 0.03399 | 0.02193 |
| **CXCR2** | 0.03483 | 0.01358 | 0.02631 | 0.02197 | 0.00722 | 0.03831 | 0.02411 |
| **CXCR2P1** | 0.02638 | 0.01849 | 0.01256 | 0.01911 | 0.00793 | 0.01265 | 0.01688 |
| **CXCR4** | 0.25238 | 0.05848 | 0.17083 | 0.00000 | 0.51909 | 1.11684 | 0.01314 |
| **CXCR5** | 0.02025 | 0.00772 | 0.01446 | 0.09400 | 0.00695 | 0.02234 | 0.09313 |
| **CXCR6** | 0.02681 | 0.03650 | 0.00770 | 2.17642 | 0.00585 | 0.00000 | 0.11499 |
| **CXXC1** | 9.57068 | 12.56197 | 7.08212 | 19.37427 | 1.20587 | 8.29521 | 13.68658 |
| **CXXC4** | 0.03762 | 0.01806 | 0.01317 | 0.04554 | 0.48480 | 0.42150 | 0.00000 |
| **CXXC5** | 0.24929 | 1.79589 | 0.00000 | 1.52868 | 0.22719 | 1.63395 | 0.58338 |
| **CXorf22** | 0.24350 | 0.23817 | 0.47944 | 0.06165 | 0.02210 | 0.18203 | 0.00001 |
| **CXorf23** | 1.82080 | 0.00000 | 0.53659 | 0.93789 | 0.88298 | 1.10611 | 3.47738 |

| | | | | | | | |
|---|---|---|---|---|---|---|---|
| **CXorf30** | 0.00737 | 0.18468 | 0.05618 | 0.03681 | 0.05474 | 0.01691 | 0.13340 |
| **CXorf31** | 0.05561 | 0.00066 | 0.07431 | 0.02119 | 0.20609 | 0.01418 | 0.01935 |
| **CXorf36** | 0.02161 | 0.01832 | 0.01134 | 0.02777 | 0.02544 | 0.01295 | 0.01578 |
| **CXorf38** | 5.78680 | 3.71813 | 3.42269 | 0.99265 | 1.75012 | 2.50419 | 0.18816 |
| **CXorf40A** | 1.00366 | 1.60107 | 2.40788 | 0.51619 | 2.06346 | 0.15240 | 0.00000 |
| **CXorf40B** | 7.34084 | 10.95154 | 7.56367 | 4.35710 | 11.90844 | 2.92317 | 0.00000 |
| **CXorf49+CXorf4** | 0.02391 | 0.02179 | 0.01857 | 0.01973 | 0.01113 | 0.01354 | 0.01655 |
| **CXorf49+CXorf4** | 0.02490 | 0.01929 | 0.01449 | 0.01902 | 0.00942 | 0.01444 | 0.01537 |
| **CXorf56** | 15.85178 | 1.17164 | 10.57364 | 6.58769 | 9.97469 | 4.65285 | 3.94943 |
| **CXorf57** | 0.24371 | 0.00000 | 2.00111 | 0.60238 | 0.19390 | 0.21302 | 0.00000 |
| **CXorf58** | 0.07294 | 0.00000 | 0.15383 | 0.01265 | 0.18153 | 0.00492 | 0.04687 |
| **CXorf65** | 1.73483 | 6.55572 | 1.35392 | 0.00000 | 0.00000 | 0.00000 | 2.04233 |
| **CXorf67** | 0.13453 | 0.29074 | 4.06360 | 0.41284 | 10.37331 | 0.00000 | 0.00000 |
| **CYB561** | 0.00000 | 7.20558 | 9.92483 | 1.97570 | 0.28587 | 5.92382 | 0.74705 |
| **CYB561A3** | 3.43306 | 2.54168 | 3.62911 | 7.84130 | 0.28504 | 1.51795 | 0.02784 |
| **CYB561D1** | 0.71856 | 1.68791 | 1.42986 | 1.36683 | 1.21322 | 0.28825 | 2.03018 |
| **CYB561D2** | 17.22617 | 19.05168 | 5.36139 | 20.65354 | 1.40643 | 4.06639 | 0.00000 |
| **CYB5A** | 16.88298 | 3.17456 | 0.00000 | 10.11247 | 0.57169 | 3.74846 | 4.72504 |
| **CYB5B** | 51.42353 | 21.80858 | 15.60745 | 25.27538 | 13.48462 | 9.90134 | 0.00005 |
| **CYB5D1** | 8.91269 | 4.13183 | 1.82761 | 8.29967 | 1.78536 | 1.72648 | 2.52022 |
| **CYB5D2** | 0.83859 | 2.40191 | 2.99091 | 1.73123 | 1.25934 | 0.40237 | 0.91456 |
| **CYB5R1** | 6.50656 | 4.86630 | 1.48112 | 21.35888 | 1.99704 | 0.99646 | 0.90276 |
| **CYB5R2** | 1.66949 | 1.06810 | 0.29626 | 2.15763 | 0.25926 | 0.75609 | 0.46550 |
| **CYB5R3** | 13.07482 | 19.01133 | 1.66423 | 20.05524 | 2.30209 | 10.39579 | 11.25381 |
| **CYB5R4** | 10.32350 | 0.00000 | 4.64968 | 7.39618 | 0.62561 | 1.97517 | 0.00000 |
| **CYB5RL** | 1.49275 | 1.93113 | 2.07938 | 1.02076 | 0.31071 | 0.00000 | 0.75509 |
| **CYBA** | 0.02526 | 0.06402 | 0.00000 | 1.76777 | 0.00486 | 1.14783 | 1.99389 |
| **CYBB** | 0.01866 | 0.00000 | 0.20393 | 0.02488 | 0.00476 | 0.07250 | 0.01659 |
| **CYBRD1** | 0.51170 | 0.00000 | 0.83895 | 1.46036 | 0.23305 | 0.10470 | 0.00000 |
| **CYC1** | 28.22031 | 26.55977 | 1.25145 | 24.76010 | 0.18170 | 19.39929 | 34.80146 |
| **CYCS** | 59.80533 | 21.32466 | 34.54442 | 27.56451 | 17.30309 | 24.02958 | 22.90722 |
| **CYCSP52** | 0.07306 | 0.05850 | 0.07676 | 0.00477 | 0.00634 | 0.09361 | 0.21135 |
| **CYFIP1** | 14.27545 | 9.53776 | 3.65572 | 14.08940 | 2.11960 | 6.16233 | 3.11941 |
| **CYFIP2** | 0.00000 | 3.61165 | 0.05044 | 1.11062 | 0.76394 | 1.82739 | 0.44547 |
| **CYGB** | 0.00000 | 0.23634 | 1.75464 | 0.06368 | 0.06279 | 0.00000 | 0.13264 |

| | | | | | | | |
|---|---|---|---|---|---|---|---|
| **CYHR1** | 1.31670 | 5.93371 | 2.13934 | 1.49000 | 4.10775 | 1.09673 | 3.40131 |
| **CYLC2** | 0.03025 | 0.01259 | 0.01435 | 0.02026 | 0.00897 | 0.01580 | 0.02800 |
| **CYLD** | 1.14560 | 1.00632 | 1.82563 | 0.30358 | 3.18013 | 0.56028 | 0.48295 |
| **CYMP** | 0.02773 | 0.02217 | 0.01133 | 0.02445 | 0.01256 | 0.02939 | 0.00891 |
| **CYP11A1** | 13.80801 | 0.54563 | 0.52153 | 35.39052 | 0.00000 | 0.00000 | 3.49736 |
| **CYP17A1** | 0.04968 | 0.03320 | 0.01722 | 0.06393 | 0.00728 | 0.00868 | 0.01176 |
| **CYP19A1** | 0.48869 | 0.00000 | 0.03639 | 29.71803 | 0.01632 | 0.04049 | 0.00000 |
| **CYP1A1** | 0.00000 | 0.65735 | 0.00000 | 24.02236 | 0.10747 | 6.45696 | 0.00000 |
| **CYP1A2** | 0.02484 | 0.02956 | 0.01192 | 0.30414 | 0.00981 | 0.01770 | 0.00000 |
| **CYP1B1** | 0.00000 | 19.67987 | 2.51525 | 8.05650 | 0.00000 | 12.46248 | 0.00000 |
| **CYP1B1-AS1** | 0.01896 | 0.05345 | 0.10781 | 0.01817 | 0.01688 | 0.02429 | 0.00671 |
| **CYP20A1** | 6.21823 | 7.83248 | 1.19490 | 3.00986 | 0.15401 | 7.19708 | 5.77826 |
| **CYP21A1P** | 0.02482 | 0.01549 | 0.01663 | 0.02171 | 0.00775 | 0.01663 | 0.01405 |
| **CYP21A2.uid-84** | 0.00000 | 0.02185 | 0.01413 | 0.09106 | 0.00744 | 0.02582 | 0.04215 |
| **CYP24A1** | 0.01361 | 0.08992 | 0.00000 | 0.01637 | 0.42717 | 0.00642 | 0.02307 |
| **CYP26A1** | 0.00000 | 24.65022 | 0.29687 | 0.04111 | 0.03344 | 0.00000 | 0.00077 |
| **CYP26B1** | 0.02374 | 0.08685 | 0.23375 | 0.00694 | 0.03181 | 0.01208 | 0.14229 |
| **CYP26C1** | 0.02871 | 0.05744 | 0.01665 | 0.02570 | 0.00897 | 0.01046 | 0.00520 |
| **CYP27A1** | 0.15725 | 0.03299 | 0.00417 | 0.01945 | 0.02272 | 0.39200 | 0.05863 |
| **CYP27B1** | 0.55320 | 0.34053 | 0.94597 | 0.70084 | 0.00000 | 0.33759 | 1.52503 |
| **CYP27C1** | 0.00000 | 0.00000 | 1.58472 | 0.07209 | 0.05005 | 0.37887 | 0.09487 |
| **CYP2A13** | 0.02668 | 0.01403 | 0.01352 | 0.01829 | 0.00786 | 0.02947 | 0.01932 |
| **CYP2A6** | 0.02134 | 0.01961 | 0.01215 | 0.03600 | 0.00825 | 0.01569 | 0.01462 |
| **CYP2B6** | 0.04128 | 0.00668 | 0.01511 | 0.14619 | 0.00925 | 0.44469 | 0.00000 |
| **CYP2B7P** | 0.10736 | 0.00138 | 0.01803 | 0.05399 | 0.00766 | 0.05055 | 0.00000 |
| **CYP2C18** | 0.84730 | 0.00000 | 0.09022 | 3.03902 | 1.20016 | 0.00000 | 0.12943 |
| **CYP2C19** | 0.07397 | 0.07602 | 0.01245 | 0.00000 | 0.00728 | 0.04104 | 0.18087 |
| **CYP2C8** | 0.24610 | 0.00000 | 0.03696 | 0.15897 | 0.00566 | 0.25693 | 0.00000 |
| **CYP2C9** | 0.04985 | 0.00000 | 0.04283 | 0.32996 | 0.06904 | 0.01400 | 0.00028 |
| **CYP2D7P** | 0.13373 | 0.09096 | 0.10309 | 0.14841 | 0.04414 | 0.11905 | 0.30802 |
| **20.00000** | 0.15353 | 2.61902 | 3.22909 | 0.00000 | 0.00402 | 4.96239 | 0.32039 |
| **CYP2F1** | 0.01758 | 0.50064 | 0.07889 | 0.23820 | 0.07122 | 0.99392 | 0.00000 |
| **CYP2J2** | 0.01678 | 0.06356 | 0.09846 | 0.01921 | 0.03604 | 0.01615 | 0.05324 |
| **CYP2R1** | 0.05527 | 0.00000 | 0.35649 | 0.01474 | 1.77147 | 0.32892 | 0.01347 |
| **CYP2S1** | 22.74727 | 48.07015 | 5.69912 | 2.14942 | 0.45138 | 16.79213 | 16.72136 |

| | | | | | | | |
|---|---|---|---|---|---|---|---|
| **CYP2U1** | 0.00000 | 0.08091 | 0.01061 | 0.04017 | 0.03958 | 0.15402 | 0.08284 |
| **CYP2W1** | 0.01454 | 0.02962 | 0.09169 | 0.01860 | 0.00442 | 0.00828 | 0.02796 |
| **CYP39A1** | 0.02460 | 0.01398 | 0.02623 | 0.04839 | 0.01479 | 0.01080 | 0.02492 |
| **CYP3A4** | 0.05250 | 0.01033 | 0.01652 | 0.01381 | 0.00737 | 0.01024 | 0.03161 |
| **CYP3A43** | 0.00403 | 0.05237 | 0.01158 | 0.03648 | 0.00759 | 0.04900 | 0.02590 |
| **CYP3A5** | 0.52466 | 0.00000 | 0.14996 | 0.18996 | 0.07584 | 0.37386 | 0.58760 |
| **CYP3A7** | 0.16826 | 0.01384 | 0.01128 | 0.14559 | 0.00758 | 0.02524 | 0.00000 |
| **CYP3A7-CYP3A** | 0.02849 | 0.01272 | 0.01533 | 0.02238 | 0.00795 | 0.04686 | 0.01760 |
| **CYP46A1** | 0.01625 | 0.07439 | 0.01027 | 0.02666 | 0.18900 | 0.00542 | 0.03569 |
| **CYP4A22** | 0.03136 | 0.01604 | 0.01284 | 0.01830 | 0.00794 | 0.01197 | 0.02018 |
| **CYP4B1** | 0.02307 | 0.03083 | 0.04502 | 0.03468 | 0.38790 | 0.01180 | 0.01096 |
| **CYP4F11** | 0.02154 | 0.01707 | 0.00923 | 0.02530 | 0.06304 | 0.01370 | 0.01712 |
| **CYP4F12** | 0.02234 | 0.03102 | 0.02321 | 0.02483 | 0.03526 | 0.01055 | 0.01451 |
| **CYP4F2** | 0.01462 | 0.05733 | 0.00474 | 0.02138 | 0.05197 | 0.06203 | 0.01809 |
| **CYP4F22** | 0.00741 | 0.13834 | 0.15433 | 0.01469 | 0.03788 | 0.02876 | 0.01092 |
| **CYP4F24P** | 0.01663 | 0.02944 | 0.02364 | 0.03847 | 0.01992 | 0.00922 | 0.02553 |
| **CYP4F3** | 0.01092 | 0.02626 | 0.01717 | 0.01472 | 0.01824 | 0.00832 | 0.07322 |
| **CYP4F35P** | 0.02467 | 0.00242 | 0.03474 | 0.01424 | 0.01340 | 0.79395 | 0.01373 |
| **CYP4F62P** | 0.02359 | 0.01742 | 0.01478 | 0.02030 | 0.00783 | 0.01333 | 0.01678 |
| **CYP4F8** | 0.04128 | 0.02731 | 0.00993 | 0.01119 | 0.00786 | 0.00835 | 0.02663 |
| **CYP4V2** | 0.06702 | 0.04968 | 0.16031 | 0.00665 | 0.28832 | 0.22469 | 0.14088 |
| **CYP4X1** | 0.02001 | 0.01439 | 0.01347 | 0.04573 | 0.27758 | 0.24002 | 0.00000 |
| **CYP4Z1** | 0.02379 | 0.01411 | 0.01281 | 0.01688 | 0.00807 | 0.03228 | 0.02250 |
| **CYP51A1** | 43.32027 | 4.87911 | 0.87165 | 8.85332 | 0.75277 | 8.73708 | 6.52902 |
| **CYP51A1-AS1** | 0.48242 | 0.01300 | 0.12295 | 0.79937 | 0.12383 | 0.02097 | 0.12819 |
| **CYP7B1** | 0.13744 | 0.04072 | 0.02976 | 0.08895 | 0.04324 | 0.03502 | 0.14425 |
| **CYP8B1** | 0.04941 | 0.00644 | 0.00590 | 0.22677 | 0.00635 | 0.01535 | 0.52815 |
| **CYR61** | 0.00000 | 1.18666 | 0.00000 | 7.40327 | 0.01393 | 1.67605 | 3.43990 |
| **CYS1** | 0.02135 | 0.02003 | 0.04224 | 0.02815 | 0.00608 | 0.02452 | 0.01314 |
| **CYSRT1** | 0.03266 | 0.06586 | 0.03887 | 0.07083 | 0.13643 | 0.01208 | 0.00000 |
| **CYSTM1** | 19.18264 | 17.45883 | 3.78797 | 6.09874 | 7.09062 | 4.09207 | 4.93973 |
| **CYTH1** | 3.01077 | 0.96702 | 4.21859 | 1.73911 | 3.60431 | 0.77284 | 3.67892 |
| **CYTH2** | 5.43729 | 14.56341 | 2.87481 | 15.36312 | 5.52438 | 5.61794 | 5.70132 |
| **CYTH3** | 1.05779 | 6.19580 | 5.87642 | 2.67262 | 2.88931 | 1.13880 | 9.60633 |
| **CYTH4** | 0.02104 | 0.01765 | 0.01228 | 0.03090 | 0.01118 | 0.01704 | 0.01824 |

| | | | | | | | |
|---|---|---|---|---|---|---|---|
| **CYTIP** | 0.04939 | 0.00000 | 0.02617 | 0.01105 | 0.08080 | 0.01224 | 0.04825 |
| **CYTL1** | 0.02728 | 0.01801 | 0.01257 | 0.01855 | 0.00868 | 0.01292 | 0.01643 |
| **CYYR1** | 0.02721 | 0.01712 | 0.01318 | 0.02357 | 0.00800 | 0.01397 | 0.01267 |
| **D2HGDH** | 0.00000 | 0.19243 | 0.11994 | 0.04549 | 0.38034 | 0.07043 | 0.59951 |
| **DAAM1** | 9.99583 | 1.91753 | 4.51440 | 4.96140 | 7.93456 | 4.45213 | 10.64004 |
| **DAAM2** | 0.25458 | 0.00000 | 0.91021 | 0.31926 | 2.16361 | 0.49144 | 3.45962 |
| **DAB1** | 0.03752 | 0.00000 | 0.02661 | 0.05980 | 0.71631 | 0.33898 | 0.00000 |
| **DAB2** | 60.02107 | 16.03316 | 12.02824 | 32.44881 | 0.97677 | 0.00000 | 25.77112 |
| **DAB2IP** | 0.03621 | 3.23890 | 0.01167 | 0.46157 | 0.51861 | 1.65434 | 5.94755 |
| **DACH1** | 0.11505 | 0.09117 | 0.04145 | 0.02457 | 0.17474 | 1.02954 | 0.00000 |
| **DACH2** | 0.02441 | 0.00979 | 0.05026 | 0.08840 | 0.15441 | 0.00980 | 0.01035 |
| **DACT1** | 0.28281 | 0.32130 | 0.00000 | 0.81458 | 0.00598 | 2.81121 | 2.56019 |
| **DACT2** | 0.13302 | 0.16214 | 0.00000 | 0.00000 | 0.00498 | 0.52890 | 3.36534 |
| **DACT3** | 0.00859 | 0.02257 | 0.02157 | 0.02050 | 0.03761 | 0.03618 | 0.07062 |
| **DACT3-AS1** | 0.02929 | 0.00000 | 0.06705 | 0.03071 | 0.03722 | 0.06370 | 0.01615 |
| **DAD1** | 39.44410 | 25.40553 | 22.52067 | 30.20271 | 1.85876 | 16.90388 | 15.94366 |
| **DAG1** | 23.36730 | 27.40191 | 4.33566 | 28.20288 | 2.85762 | 7.30009 | 43.70147 |
| **DAGLA** | 0.02853 | 0.43608 | 0.19983 | 0.00000 | 0.11721 | 1.66444 | 0.46108 |
| **DAGLB** | 3.67701 | 6.96984 | 7.15247 | 7.68002 | 7.76099 | 6.91849 | 1.03377 |
| **DAK** | 1.57081 | 1.73628 | 1.88878 | 1.32995 | 0.02702 | 1.41138 | 1.89546 |
| **DALRD3** | 6.02719 | 4.37813 | 1.36344 | 5.86056 | 0.37091 | 3.26929 | 0.26663 |
| **DANCR** | 43.16820 | 28.27202 | 27.37451 | 33.62827 | 6.01264 | 11.00228 | 33.28137 |
| **DAND5** | 0.02196 | 0.10233 | 0.00000 | 0.03164 | 0.82304 | 0.00696 | 0.00000 |
| **DAO** | 0.03700 | 0.01968 | 0.01203 | 0.18283 | 0.00790 | 0.00883 | 0.01038 |
| **DAP** | 4.26221 | 5.36852 | 2.27669 | 8.16335 | 1.10768 | 2.55837 | 7.19982 |
| **DAP3** | 28.56394 | 16.06961 | 19.16799 | 28.31878 | 6.70349 | 13.30165 | 3.72048 |
| **DAPK1** | 3.50255 | 7.17901 | 3.41585 | 5.51319 | 0.00000 | 6.26655 | 4.19501 |
| **DAPK2** | 0.03015 | 0.00365 | 0.03325 | 0.02455 | 0.03042 | 0.53271 | 0.00570 |
| **DAPK3** | 0.00000 | 8.59116 | 0.97076 | 6.72283 | 1.19900 | 3.61709 | 8.46183 |
| **DAPP1** | 0.29876 | 0.56297 | 0.00790 | 0.00758 | 0.18495 | 0.00451 | 0.00000 |
| **DARS** | 39.46265 | 14.33427 | 10.94352 | 17.35453 | 2.65304 | 14.62770 | 15.38043 |
| **DARS2** | 15.65829 | 4.41242 | 4.33708 | 9.34635 | 0.50799 | 5.42395 | 2.41922 |
| **DAW1** | 0.01432 | 0.11333 | 0.00000 | 0.02558 | 1.76886 | 0.01716 | 0.05907 |
| **DAXX** | 22.34932 | 19.10328 | 19.15679 | 30.50839 | 5.49265 | 14.03790 | 3.13412 |
| **DAZAP1** | 0.15384 | 7.03667 | 2.10808 | 5.73301 | 0.00000 | 6.72460 | 23.65111 |

| | | | | | | | |
|---|---|---|---|---|---|---|---|
| **DAZAP2** | 13.46713 | 5.80979 | 9.79145 | 7.56142 | 13.04774 | 3.95142 | 0.00000 |
| **DAZL** | 0.00000 | 0.00356 | 2.75654 | 0.87394 | 2.13436 | 0.00708 | 0.00000 |
| **DBET** | 0.02277 | 0.04645 | 0.02295 | 0.04936 | 0.04352 | 0.04099 | 0.11579 |
| **DBF4** | 11.35099 | 1.80585 | 13.55554 | 6.75377 | 2.37707 | 4.43409 | 9.96249 |
| **DBF4B** | 2.13900 | 1.19938 | 3.43763 | 3.84207 | 4.21525 | 1.50563 | 6.76196 |
| **DBH** | 0.01677 | 0.01444 | 0.00525 | 0.01969 | 0.23247 | 0.03367 | 0.03505 |
| **DBH-AS1** | 0.02789 | 0.01714 | 0.01212 | 0.12323 | 0.01565 | 0.01095 | 0.00766 |
| **DBI** | 55.42968 | 19.62758 | 6.37180 | 26.24979 | 9.91318 | 19.40958 | 27.16735 |
| **DBIL5P** | 0.03244 | 0.00000 | 0.20578 | 0.23299 | 0.15554 | 0.02962 | 0.07386 |
| **DBIL5P2** | 0.00951 | 0.02562 | 0.04419 | 0.03971 | 0.00634 | 0.06568 | 0.03875 |
| **DBN1** | 6.97302 | 9.98933 | 0.78683 | 4.41567 | 0.02910 | 11.37884 | 14.95472 |
| **DBNDD1** | 0.04083 | 0.06486 | 0.02341 | 0.38504 | 0.00937 | 0.37322 | 0.62240 |
| **DBNDD2** | 0.62079 | 0.69169 | 0.02638 | 0.72076 | 0.07911 | 0.00000 | 0.33034 |
| **DBNL** | 12.61504 | 19.68221 | 4.65557 | 9.87361 | 0.20416 | 5.33823 | 0.00000 |
| **DBP** | 0.08944 | 0.69252 | 5.52101 | 0.00000 | 0.80899 | 1.17964 | 0.96002 |
| **DBR1** | 16.63342 | 3.19425 | 6.57430 | 9.71483 | 18.31994 | 5.05479 | 0.00000 |
| **DBT** | 8.41410 | 0.31731 | 4.20185 | 5.90686 | 1.24594 | 3.42741 | 2.67754 |
| **DBX1** | 0.02855 | 0.04514 | 0.02257 | 0.02327 | 0.06884 | 0.00951 | 0.00286 |
| **DBX2** | 0.02288 | 0.00500 | 0.04573 | 0.02367 | 0.01435 | 0.01389 | 0.01952 |
| **DCAF10** | 0.87874 | 0.62943 | 0.19955 | 0.50423 | 0.15518 | 1.24572 | 2.69034 |
| **DCAF11** | 8.28337 | 3.97691 | 1.39621 | 12.65383 | 0.51174 | 4.96848 | 0.00000 |
| **DCAF12** | 15.32843 | 8.81101 | 7.86158 | 18.79097 | 4.59810 | 5.99275 | 0.54616 |
| **DCAF12L1** | 0.04322 | 0.00000 | 0.09259 | 0.02822 | 0.13860 | 0.25976 | 0.00473 |
| **DCAF12L2** | 0.02588 | 0.01899 | 0.01176 | 0.02568 | 0.01986 | 0.02478 | 0.00955 |
| **DCAF13** | 35.36561 | 3.51404 | 19.99469 | 22.41410 | 12.29250 | 11.00885 | 0.00000 |
| **DCAF13P3** | 3.02858 | 0.35968 | 1.82183 | 2.29278 | 1.70236 | 0.65316 | 0.00000 |
| **DCAF15** | 0.00000 | 0.63703 | 0.37327 | 0.85003 | 0.51786 | 1.00865 | 3.84080 |
| **DCAF16** | 7.78987 | 3.10624 | 4.41809 | 4.81517 | 4.27236 | 0.92656 | 8.01066 |
| **DCAF17** | 5.66576 | 1.16562 | 3.97470 | 3.21490 | 1.65821 | 2.15535 | 4.22238 |
| **DCAF4** | 23.99259 | 4.60243 | 7.55447 | 21.99123 | 5.91613 | 9.08991 | 5.20428 |
| **DCAF4L1** | 2.86936 | 0.00000 | 7.23119 | 1.27518 | 10.93185 | 0.39518 | 0.87052 |
| **DCAF4L2** | 2.29232 | 0.83897 | 5.02973 | 2.38707 | 1.71421 | 0.22675 | 0.00000 |
| **DCAF5** | 2.52780 | 4.77664 | 0.69121 | 2.80149 | 5.59319 | 2.46007 | 7.27963 |
| **DCAF6** | 6.93895 | 4.05173 | 5.46733 | 12.83060 | 3.25258 | 3.20374 | 3.57381 |
| **DCAF7** | 8.87025 | 12.40645 | 12.22242 | 13.83497 | 7.90024 | 9.88411 | 6.42988 |

| | | | | | | | |
|---|---|---|---|---|---|---|---|
| **DCAF8** | 12.88763 | 8.99284 | 5.09982 | 13.45126 | 4.69285 | 4.90160 | 6.07771 |
| **DCAF8L1** | 0.25649 | 0.05312 | 0.05246 | 0.12806 | 0.54111 | 0.00071 | 0.04479 |
| **DCAF8L2** | 0.08371 | 0.00000 | 0.07824 | 0.08726 | 0.18454 | 0.06278 | 0.00000 |
| **DCAKD** | 6.46087 | 4.64487 | 1.85030 | 7.03830 | 2.21130 | 1.14826 | 3.74780 |
| **DCBLD1** | 1.53390 | 0.00000 | 0.11219 | 0.63173 | 0.19214 | 0.02711 | 1.37595 |
| **DCBLD2** | 1.51018 | 0.75166 | 0.72690 | 1.79545 | 0.41302 | 2.44011 | 3.66823 |
| **DCC** | 0.02615 | 0.02081 | 0.08882 | 0.02193 | 1.18033 | 0.04921 | 0.05893 |
| **DCD** | 0.03296 | 0.00995 | 0.01606 | 0.00963 | 0.00823 | 0.01234 | 0.08475 |
| **DCDC1** | 0.03917 | 0.00000 | 0.14431 | 0.04499 | 0.02890 | 0.05077 | 0.02132 |
| **DCDC2** | 1.72031 | 0.47774 | 0.26186 | 0.16257 | 9.16594 | 0.28509 | 0.00000 |
| **DCDC2B** | 0.12790 | 0.03300 | 0.07227 | 0.12319 | 0.10886 | 0.03193 | 0.15327 |
| **DCDC2C** | 0.00865 | 0.02682 | 0.00667 | 0.01404 | 0.30420 | 0.02709 | 0.07136 |
| **DCDC5** | 0.02764 | 0.06299 | 0.00770 | 0.01134 | 0.00826 | 0.04079 | 0.02591 |
| **DCHS1** | 0.00000 | 0.22231 | 0.24421 | 0.25053 | 0.02851 | 0.78728 | 1.14800 |
| **DCHS2** | 0.00000 | 0.05407 | 0.00000 | 0.02790 | 1.17403 | 0.08093 | 0.04683 |
| **DCK** | 7.38225 | 0.00000 | 3.38076 | 3.18454 | 1.32834 | 1.43399 | 0.90685 |
| **DCLK1** | 0.00000 | 0.00809 | 0.00880 | 0.09857 | 0.01197 | 0.72919 | 0.05991 |
| **DCLK2** | 0.01582 | 0.05471 | 0.00000 | 0.00345 | 2.83811 | 0.51135 | 0.04958 |
| **DCLK3** | 0.02613 | 0.04433 | 0.02700 | 0.03670 | 0.00711 | 0.01007 | 0.00636 |
| **DCLRE1A** | 5.33365 | 1.10491 | 1.27086 | 0.37492 | 2.92799 | 1.12648 | 0.50339 |
| **DCLRE1B** | 7.52966 | 2.01672 | 6.42307 | 5.09495 | 11.62807 | 3.07518 | 1.53039 |
| **DCLRE1C** | 11.72156 | 3.21071 | 2.50540 | 6.47067 | 1.40400 | 1.92678 | 1.21034 |
| **DCN** | 0.04694 | 0.01615 | 0.01393 | 0.01955 | 0.00828 | 0.01140 | 0.01494 |
| **DCP1A** | 17.09326 | 9.12296 | 11.44835 | 18.85347 | 18.55233 | 7.62466 | 9.34674 |
| **DCP1B** | 1.16062 | 3.08031 | 2.02728 | 0.23162 | 0.02521 | 1.32749 | 0.00000 |
| **DCP2** | 21.85077 | 5.45044 | 10.72398 | 10.67922 | 13.83394 | 8.21248 | 6.23171 |
| **DCPS** | 11.37586 | 4.08845 | 2.07269 | 10.20596 | 1.00813 | 5.19879 | 1.11562 |
| **DCST1** | 0.04749 | 0.01653 | 0.02235 | 0.04710 | 0.01435 | 0.00000 | 0.10522 |
| **DCST2** | 0.05032 | 0.05971 | 0.06965 | 0.11059 | 0.00549 | 0.00000 | 0.28790 |
| **DCT** | 0.09558 | 0.00000 | 0.14274 | 0.00361 | 2.24052 | 0.00206 | 0.02712 |
| **DCTD** | 5.56031 | 2.79879 | 2.48870 | 14.08016 | 0.12032 | 0.63681 | 5.13348 |
| **DCTN1** | 7.73084 | 15.39779 | 7.08796 | 9.32944 | 0.75902 | 14.16675 | 18.03241 |
| **DCTN1-AS1** | 0.00172 | 0.05795 | 0.03882 | 0.02321 | 0.01127 | 0.06087 | 0.07412 |
| **DCTN2** | 14.64609 | 12.58848 | 7.86708 | 21.12784 | 0.10382 | 16.67748 | 13.20089 |
| **DCTN3** | 10.08329 | 12.18802 | 8.78860 | 9.48163 | 7.30585 | 4.58831 | 3.67569 |

| | | | | | | | |
|---|---|---|---|---|---|---|---|
| **DCTN4** | 11.72392 | 1.60562 | 4.79037 | 8.56998 | 1.14593 | 2.46835 | 5.77159 |
| **DCTN5** | 26.53187 | 22.15271 | 23.45652 | 21.66869 | 6.60825 | 22.06790 | 1.04490 |
| **DCTN6** | 15.00263 | 1.49851 | 6.06199 | 8.99536 | 0.00000 | 4.70955 | 2.14076 |
| **DCTPP1** | 31.86780 | 14.91674 | 10.40542 | 25.04548 | 0.95192 | 17.97188 | 6.81285 |
| **DCUN1D1** | 9.39758 | 0.00000 | 5.20223 | 5.02385 | 0.62939 | 2.93395 | 1.58957 |
| **DCUN1D2** | 2.95559 | 0.98320 | 0.33055 | 1.51287 | 0.29965 | 3.25068 | 5.34881 |
| **DCUN1D3** | 0.51152 | 1.55097 | 0.56222 | 1.40169 | 3.09182 | 3.17733 | 0.89924 |
| **DCUN1D4** | 13.66950 | 2.72636 | 5.43190 | 1.79067 | 2.75490 | 5.73853 | 1.65710 |
| **DCUN1D5** | 32.91747 | 21.57552 | 9.52436 | 20.28282 | 3.74873 | 12.09190 | 6.09903 |
| **DCX** | 0.05062 | 0.00860 | 0.00698 | 0.00859 | 0.21941 | 0.01665 | 0.06021 |
| **DCXR** | 19.37077 | 6.85030 | 0.00000 | 17.96882 | 1.45414 | 9.97046 | 19.42165 |
| **DDA1** | 4.11792 | 10.34013 | 9.09982 | 17.85815 | 10.96924 | 5.09114 | 7.30975 |
| **DDAH1** | 0.54657 | 0.00000 | 0.10868 | 0.16551 | 1.05624 | 0.30692 | 0.39360 |
| **DDAH2** | 0.27518 | 0.29672 | 1.40386 | 5.75133 | 4.64329 | 3.35876 | 2.22550 |
| **DDB1** | 29.75801 | 24.46190 | 12.51321 | 46.77442 | 4.25114 | 17.82825 | 21.34102 |
| **DDB2** | 1.53400 | 6.35781 | 5.15698 | 2.77325 | 1.49238 | 4.95074 | 0.00066 |
| **DDC** | 0.14139 | 0.00000 | 0.12877 | 0.16962 | 0.02299 | 0.50758 | 0.00000 |
| **DDC-AS1** | 0.03449 | 0.01253 | 0.01846 | 0.01992 | 0.00825 | 0.02206 | 0.00986 |
| **DDHD1** | 6.79802 | 2.21671 | 12.13044 | 4.80273 | 0.33230 | 8.78078 | 14.28861 |
| **DDHD2** | 3.04942 | 0.00000 | 0.37307 | 2.01925 | 0.65329 | 0.05155 | 5.07621 |
| **DDI2** | 0.36450 | 0.62526 | 0.75298 | 0.50141 | 0.97248 | 0.31369 | 1.95703 |
| **DDIAS** | 8.99533 | 0.00000 | 10.01581 | 7.87915 | 11.72351 | 3.15137 | 0.29106 |
| **DDIT3** | 0.82870 | 6.63283 | 5.25861 | 9.79902 | 17.06746 | 8.07271 | 1.12741 |
| **DDIT4** | 33.86270 | 2.05134 | 0.00000 | 33.41509 | 1.04305 | 4.21648 | 10.19150 |
| **DDIT4L** | 12.67396 | 4.47187 | 0.00000 | 1.18804 | 0.03460 | 1.36246 | 0.51527 |
| **DDN** | 0.11410 | 0.02399 | 0.03248 | 0.33805 | 0.04790 | 0.01500 | 0.12921 |
| **DDOST** | 50.89252 | 32.97651 | 9.76657 | 39.03202 | 6.80387 | 21.70600 | 0.67874 |
| **DDR1** | 0.47763 | 0.00000 | 2.82420 | 6.34627 | 0.55639 | 3.51662 | 10.04972 |
| **DDR2** | 0.03647 | 0.03327 | 0.03630 | 0.31722 | 0.10935 | 0.04053 | 0.12700 |
| **DDRGK1** | 0.00000 | 5.70958 | 0.62481 | 6.91681 | 0.43163 | 5.61851 | 17.59454 |
| **DDT** | 0.02641 | 0.02219 | 0.01326 | 0.02565 | 0.00807 | 0.01279 | 0.01619 |
| **DDTL** | 0.57982 | 1.26095 | 0.21919 | 0.61904 | 0.57362 | 0.42041 | 0.90575 |
| **DDX1** | 45.49578 | 23.64583 | 17.41655 | 19.29601 | 1.44618 | 18.88431 | 12.40335 |
| **DDX10** | 15.86144 | 3.68138 | 10.35676 | 10.03434 | 12.68456 | 5.45807 | 14.38122 |
| **DDX11** | 1.48473 | 2.08681 | 0.48122 | 2.96240 | 0.14901 | 1.76175 | 6.03073 |

| | | | | | | | |
|---|---|---|---|---|---|---|---|
| **DDX11-AS1** | 0.06550 | 0.08109 | 0.03083 | 0.08664 | 0.03418 | 0.12593 | 0.00000 |
| **DDX11L10** | 0.02481 | 0.01638 | 0.01453 | 0.01889 | 0.01026 | 0.01369 | 0.01714 |
| **DDX11L2** | 0.03732 | 0.00000 | 0.21766 | 0.09982 | 0.35705 | 0.09002 | 0.00921 |
| **DDX11L5** | 0.02277 | 0.01497 | 0.03239 | 0.01894 | 0.01291 | 0.01386 | 0.02325 |
| **DDX12P** | 0.15593 | 0.24591 | 0.30971 | 0.63581 | 0.01449 | 0.22502 | 1.26075 |
| **DDX17** | 37.03700 | 4.71212 | 15.34787 | 26.89720 | 4.53800 | 16.62462 | 37.01960 |
| **DDX18** | 33.19337 | 18.12889 | 19.58631 | 19.56148 | 6.33178 | 8.53590 | 22.51086 |
| **DDX19A** | 18.99151 | 7.95172 | 6.18428 | 11.33930 | 1.31617 | 7.07834 | 4.97039 |
| **DDX19B** | 5.76276 | 2.70254 | 3.86888 | 4.99507 | 0.17211 | 2.44482 | 0.00000 |
| **DDX20** | 14.45660 | 7.52608 | 13.54724 | 9.01523 | 5.51797 | 6.94922 | 0.01652 |
| **DDX21** | 52.45778 | 16.84327 | 30.18799 | 27.84216 | 13.87663 | 21.94913 | 46.02687 |
| **DDX23** | 16.32868 | 16.21739 | 13.35397 | 20.28424 | 1.25168 | 12.78864 | 9.39210 |
| **DDX24** | 27.52788 | 22.10995 | 15.62951 | 21.71913 | 13.78885 | 8.84647 | 10.05033 |
| **DDX25** | 0.27071 | 0.13324 | 3.86501 | 0.00000 | 0.22660 | 0.23627 | 0.00280 |
| **DDX26B** | 0.63834 | 0.00000 | 6.11424 | 0.00000 | 5.67764 | 1.57242 | 0.33545 |
| **DDX26B-AS1** | 0.02165 | 0.00751 | 0.03439 | 0.01793 | 0.01422 | 0.03059 | 0.02224 |
| **DDX27** | 25.45609 | 24.97667 | 14.10845 | 26.04957 | 5.63495 | 9.26370 | 15.59234 |
| **DDX28** | 12.77896 | 12.32983 | 9.52061 | 11.76649 | 1.98795 | 6.37168 | 0.00000 |
| **DDX31** | 7.84498 | 4.44488 | 9.86899 | 5.74066 | 0.40736 | 5.87713 | 5.48367 |
| **DDX39A** | 49.10377 | 44.51401 | 26.11311 | 32.41879 | 20.83998 | 17.87469 | 5.27536 |
| **DDX39B** | 0.13584 | 0.09501 | 0.09266 | 0.24409 | 0.12436 | 0.03108 | 0.02589 |
| **DDX3X** | 58.03274 | 24.20898 | 26.91176 | 29.45213 | 23.24737 | 22.18951 | 20.95705 |
| **DDX3Y** | 0.00000 | 0.00000 | 0.31913 | 0.23767 | 0.01163 | 0.08145 | 0.09631 |
| **DDX4** | 0.00000 | 0.00000 | 2.67335 | 0.08754 | 3.53379 | 0.17381 | 0.10768 |
| **DDX41** | 24.07488 | 14.40828 | 7.06671 | 9.19984 | 5.35052 | 12.61583 | 7.11888 |
| **DDX42** | 9.86217 | 8.81295 | 7.68162 | 4.70272 | 5.00497 | 6.22460 | 17.01947 |
| **DDX43** | 4.16752 | 2.16507 | 8.19011 | 2.63243 | 3.69137 | 5.50204 | 0.00000 |
| **DDX46** | 22.80895 | 11.73812 | 15.88813 | 19.36853 | 4.96542 | 10.88469 | 33.96131 |
| **DDX47** | 30.42713 | 12.82226 | 14.73637 | 17.97456 | 8.44815 | 9.42821 | 0.00000 |
| **DDX49** | 19.99177 | 21.18799 | 11.95167 | 17.17179 | 7.43047 | 7.27278 | 5.62496 |
| **DDX5** | 70.23805 | 32.14680 | 38.98331 | 43.13810 | 27.80220 | 30.71855 | 4.16971 |
| **DDX50** | 7.24392 | 2.82582 | 3.06843 | 2.79101 | 4.59373 | 2.28104 | 4.62671 |
| **DDX51** | 1.90976 | 6.37458 | 1.49258 | 2.83114 | 1.21270 | 3.39699 | 13.00594 |
| **DDX52** | 19.11583 | 6.42581 | 13.74208 | 15.63020 | 3.57361 | 7.86693 | 7.70188 |
| **DDX53** | 0.04563 | 0.09063 | 0.27081 | 0.39313 | 0.10797 | 0.02908 | 0.00000 |

| | | | | | | | |
|---|---|---|---|---|---|---|---|
| **DDX54** | 3.32980 | 12.43664 | 5.31435 | 11.41295 | 4.37789 | 5.24244 | 22.86242 |
| **DDX55** | 16.55256 | 14.90345 | 9.04336 | 9.85236 | 3.64541 | 6.75383 | 5.37244 |
| **DDX56** | 37.98756 | 28.70494 | 16.79900 | 29.55036 | 5.16212 | 13.81474 | 3.87403 |
| **DDX58** | 1.57544 | 0.12596 | 1.39203 | 0.09030 | 2.24743 | 0.75045 | 0.70780 |
| **DDX59** | 4.34648 | 0.00000 | 6.28710 | 6.69383 | 3.72036 | 2.74472 | 1.47160 |
| **DDX6** | 24.55821 | 2.18574 | 16.27713 | 12.41640 | 6.84341 | 8.31090 | 14.59178 |
| **DDX60** | 0.02278 | 0.01982 | 0.02685 | 0.02319 | 0.01412 | 0.01196 | 0.02798 |
| **DDX60L** | 0.15311 | 0.01325 | 0.35984 | 0.15773 | 0.36482 | 0.00000 | 0.00979 |
| **DEAF1** | 0.72200 | 0.86215 | 1.46632 | 2.13538 | 2.25414 | 0.64166 | 5.65837 |
| **DEAR** | 0.03547 | 0.02005 | 0.01947 | 0.02180 | 0.00979 | 0.01454 | 0.02114 |
| **DEC1** | 0.02718 | 0.01683 | 0.00893 | 0.02751 | 0.09472 | 0.01302 | 0.01057 |
| **DECR1** | 2.59421 | 0.00000 | 0.00000 | 1.74167 | 0.14523 | 1.56251 | 0.26292 |
| **DECR2** | 13.44364 | 19.22353 | 9.43812 | 7.69281 | 0.19408 | 2.80460 | 10.07280 |
| **DEDD** | 12.49038 | 6.57731 | 15.01345 | 11.62777 | 7.81792 | 5.86694 | 1.84172 |
| **DEDD2** | 0.78769 | 6.49550 | 1.34528 | 16.33312 | 0.80389 | 3.10289 | 10.77605 |
| **DEF6** | 0.23951 | 0.00000 | 0.31762 | 0.90413 | 0.67566 | 0.24562 | 0.76593 |
| **DEF8** | 16.24028 | 25.54297 | 2.50860 | 6.00824 | 3.35968 | 10.93555 | 7.16265 |
| **DEFA6** | 0.03535 | 0.01062 | 0.01339 | 0.11625 | 0.00823 | 0.06363 | 0.00000 |
| **DEFB1** | 0.07022 | 0.00536 | 0.01009 | 0.01034 | 0.00836 | 0.65202 | 0.00000 |
| **DEFB118** | 0.02721 | 0.01153 | 0.02026 | 0.00562 | 0.01347 | 0.03716 | 0.10422 |
| **DEFB119** | 0.02136 | 0.21248 | 0.01839 | 0.01156 | 0.01254 | 0.01165 | 0.00452 |
| **DEFB121** | 0.02801 | 0.02472 | 0.00921 | 0.02049 | 0.06976 | 0.01156 | 0.01483 |
| **DEFB122** | 0.00000 | 4.06676 | 1.94178 | 0.00000 | 0.03400 | 1.11466 | 0.00046 |
| **DEFB123** | 0.04195 | 0.05005 | 0.02745 | 0.00296 | 0.02832 | 0.53520 | 0.01739 |
| **DEFB124** | 0.00000 | 0.78337 | 0.57957 | 0.00000 | 0.00997 | 1.05553 | 0.00543 |
| **DEFB126** | 0.03393 | 0.00322 | 0.06224 | 0.01274 | 0.01183 | 0.00991 | 0.03803 |
| **DEGS1** | 34.53533 | 16.27083 | 9.54035 | 16.62786 | 5.49089 | 11.30209 | 0.00000 |
| **DEGS2** | 0.00000 | 1.03667 | 0.20376 | 0.03365 | 5.58570 | 0.08809 | 0.03038 |
| **DEK** | 26.68498 | 16.49513 | 19.33999 | 20.55374 | 4.28879 | 16.11263 | 23.12630 |
| **DENND1A** | 1.46885 | 3.24765 | 0.00000 | 2.96250 | 0.76114 | 1.18334 | 4.15981 |
| **DENND1B** | 0.42708 | 0.18562 | 0.55897 | 0.31612 | 0.00778 | 2.79811 | 1.22526 |
| **DENND1C** | 2.31690 | 6.93511 | 2.77716 | 0.08989 | 0.26983 | 0.01078 | 0.16005 |
| **DENND2A** | 0.14722 | 0.41450 | 1.15028 | 0.16158 | 0.54893 | 2.42854 | 0.00000 |
| **DENND2C** | 0.16113 | 0.00000 | 0.59417 | 0.10279 | 0.13956 | 2.14149 | 0.00000 |
| **DENND2D** | 1.19602 | 2.36589 | 0.66628 | 5.63638 | 0.00589 | 0.01082 | 0.00000 |

| | | | | | | | |
|---|---|---|---|---|---|---|---|
| **DENND3** | 0.00857 | 0.00000 | 0.46408 | 0.53973 | 0.13199 | 0.48904 | 0.57003 |
| **DENND4A** | 1.53876 | 0.23876 | 1.44464 | 1.92179 | 0.92153 | 0.97094 | 3.68220 |
| **DENND4B** | 1.64266 | 2.66246 | 0.04531 | 0.00000 | 0.06649 | 4.47474 | 5.85679 |
| **DENND4C** | 12.04343 | 4.99047 | 6.49052 | 11.67782 | 0.92423 | 5.29572 | 15.48232 |
| **DENND5A** | 7.28365 | 2.74445 | 4.45482 | 10.06790 | 2.69619 | 6.00259 | 15.46583 |
| **DENND5B** | 0.63006 | 0.14054 | 1.68050 | 0.06764 | 1.69355 | 0.73456 | 0.88229 |
| **DENND5B-AS1** | 0.15968 | 0.09768 | 0.08195 | 0.16988 | 0.01481 | 0.00000 | 0.02509 |
| **DENND6A** | 0.85700 | 0.08510 | 0.33562 | 0.56380 | 0.36219 | 0.43334 | 3.56121 |
| **DENND6B** | 0.19894 | 0.00000 | 0.00629 | 0.00418 | 0.01690 | 0.00000 | 0.31449 |
| **DENR** | 39.29854 | 18.30724 | 21.84556 | 17.42678 | 8.26640 | 22.12980 | 10.92163 |
| **DEPDC1** | 8.64382 | 2.38575 | 1.03421 | 4.93071 | 0.00000 | 0.65984 | 5.73123 |
| **DEPDC1-AS1** | 0.17136 | 0.01233 | 0.07396 | 0.04308 | 0.00570 | 0.05786 | 0.00000 |
| **DEPDC1B** | 7.70361 | 0.64553 | 5.98173 | 9.56851 | 2.69161 | 2.54358 | 0.00000 |
| **DEPDC4** | 0.30156 | 0.05885 | 0.66206 | 0.27493 | 0.48972 | 0.04377 | 0.07719 |
| **DEPDC5** | 4.40509 | 2.90804 | 2.05086 | 3.03806 | 2.25447 | 2.95317 | 4.90735 |
| **DEPDC7** | 0.76840 | 0.00000 | 3.79922 | 0.02800 | 6.24219 | 3.72527 | 0.00000 |
| **DEPTOR** | 2.23297 | 7.35482 | 13.08330 | 0.00000 | 0.18206 | 13.56565 | 0.00000 |
| **DERA** | 9.54530 | 10.18290 | 6.31508 | 1.01465 | 1.25132 | 7.53077 | 0.00000 |
| **DERL1** | 17.97934 | 8.43097 | 6.74229 | 15.71200 | 2.00768 | 9.93884 | 6.35193 |
| **DERL2** | 15.48326 | 8.39540 | 11.43241 | 13.16158 | 0.74619 | 7.78955 | 0.00000 |
| **DERL3** | 1.63892 | 0.69066 | 0.32972 | 0.99804 | 6.69897 | 0.43766 | 0.00000 |
| **DES** | 0.03783 | 0.00000 | 0.00340 | 0.02110 | 0.70998 | 0.64120 | 0.00751 |
| **DESI1** | 9.59756 | 10.60348 | 1.20072 | 2.98830 | 0.26222 | 3.89977 | 12.32188 |
| **DESI2** | 9.82863 | 2.14765 | 5.07894 | 6.85300 | 2.01105 | 8.78991 | 28.46506 |
| **DET1** | 2.41694 | 1.92152 | 2.20002 | 0.64658 | 0.95069 | 3.71484 | 0.00000 |
| **DEXI** | 0.04814 | 1.08880 | 0.18872 | 0.68680 | 0.05689 | 0.54724 | 2.13965 |
| **DFFA** | 35.73172 | 21.05348 | 9.19520 | 23.79836 | 3.77664 | 16.64539 | 0.00001 |
| **DFFB** | 8.80864 | 4.04863 | 0.48889 | 3.43182 | 0.29148 | 0.66994 | 0.00000 |
| **DFNA5** | 0.00000 | 0.25098 | 0.06290 | 1.79972 | 1.68181 | 0.32400 | 0.29885 |
| **DFNB31** | 0.00000 | 0.26701 | 0.00000 | 0.32865 | 2.66292 | 1.07998 | 0.20759 |
| **DFNB59** | 0.27025 | 0.00000 | 0.18359 | 0.05814 | 0.00856 | 0.10658 | 0.30095 |
| **DGAT1** | 0.89629 | 1.43119 | 0.03232 | 2.78017 | 0.27002 | 3.32676 | 7.27243 |
| **DGAT2** | 0.00000 | 0.00000 | 2.66913 | 0.06400 | 0.10137 | 2.55653 | 0.06214 |
| **DGCR10** | 0.05132 | 0.00414 | 0.03576 | 0.00364 | 0.00579 | 0.00178 | 0.11318 |
| **DGCR11** | 0.48556 | 0.11589 | 0.20725 | 0.25838 | 0.05391 | 0.73323 | 0.18199 |

| | | | | | | | |
|---|---|---|---|---|---|---|---|
| **DGCR14** | 8.12601 | 9.35848 | 7.39588 | 6.34607 | 6.80529 | 4.43460 | 0.74436 |
| **DGCR2** | 4.27765 | 9.78144 | 2.50603 | 3.22652 | 0.24653 | 5.36907 | 11.75230 |
| **DGCR5** | 0.02592 | 0.14947 | 0.06822 | 0.01995 | 0.01427 | 0.00650 | 0.01502 |
| **DGCR6** | 4.29605 | 6.74612 | 0.23875 | 0.00000 | 0.35575 | 2.37833 | 0.02241 |
| **DGCR6L** | 3.45798 | 8.56576 | 2.23265 | 3.32425 | 0.24978 | 2.51894 | 0.67946 |
| **DGCR8** | 7.55677 | 6.42672 | 0.00000 | 7.95264 | 0.61225 | 3.05140 | 8.72325 |
| **DGCR9** | 0.18796 | 0.03703 | 0.03208 | 0.09906 | 0.00484 | 0.00000 | 0.19408 |
| **DGKA** | 0.06147 | 0.00000 | 0.05582 | 0.69184 | 0.04047 | 0.71694 | 0.00000 |
| **DGKB** | 0.06617 | 0.00000 | 0.02118 | 0.02657 | 0.80339 | 0.00000 | 0.25710 |
| **DGKD** | 0.48767 | 3.58651 | 4.15231 | 7.47027 | 0.70995 | 2.18388 | 7.67356 |
| **DGKE** | 0.04651 | 0.04087 | 0.90717 | 0.00000 | 1.67515 | 0.06183 | 0.14738 |
| **DGKG** | 0.01657 | 0.00998 | 0.01514 | 0.02947 | 0.10521 | 0.05169 | 0.02846 |
| **DGKH** | 0.13543 | 0.24025 | 0.27078 | 0.02875 | 0.17857 | 0.08952 | 0.05282 |
| **DGKI** | 0.05740 | 0.02298 | 0.00000 | 0.00000 | 0.70740 | 0.00348 | 0.08695 |
| **DGKK** | 0.63928 | 0.36621 | 0.02207 | 0.00026 | 0.03704 | 0.11744 | 0.00000 |
| **DGKQ** | 0.15318 | 1.09862 | 0.20405 | 0.30202 | 0.03349 | 0.53120 | 1.96261 |
| **DGKZ** | 0.00000 | 2.25500 | 0.24033 | 2.76856 | 0.00000 | 2.22371 | 9.47021 |
| **DGUOK** | 12.83393 | 12.29177 | 9.51285 | 13.33473 | 2.34656 | 8.90019 | 2.78815 |
| **DGUOK-AS1** | 0.69488 | 0.33615 | 0.16698 | 0.46586 | 0.06031 | 0.82098 | 0.00000 |
| **DHCR24** | 58.50731 | 25.84760 | 4.88765 | 27.18979 | 1.93938 | 27.85263 | 26.45655 |
| **DHCR7** | 38.99224 | 6.63705 | 0.00000 | 29.46488 | 1.47097 | 11.07974 | 15.16540 |
| **DHDDS** | 12.41083 | 11.01329 | 8.23582 | 9.50417 | 4.67470 | 2.12015 | 0.00000 |
| **DHDH** | 0.00000 | 0.64200 | 0.37290 | 0.44126 | 0.22287 | 0.17151 | 0.41079 |
| **DHFR** | 0.39054 | 1.24162 | 0.74336 | 0.22733 | 0.09280 | 0.55642 | 1.40595 |
| **DHFRL1** | 0.70429 | 0.00000 | 0.43783 | 0.16866 | 0.23259 | 0.12074 | 0.00000 |
| **DHH** | 0.00423 | 0.15297 | 0.08306 | 0.02571 | 0.22227 | 0.10226 | 0.06708 |
| **DHODH** | 8.58312 | 5.45610 | 4.13655 | 2.59084 | 4.02411 | 2.70486 | 0.00000 |
| **DHPS** | 6.57923 | 2.81169 | 3.93044 | 9.21670 | 1.13044 | 4.18910 | 2.01603 |
| **DHRS1** | 2.10778 | 0.70875 | 0.70487 | 3.47802 | 0.60581 | 0.83222 | 0.00000 |
| **DHRS11** | 0.41615 | 0.31454 | 0.84155 | 1.32051 | 0.41678 | 0.88964 | 3.18430 |
| **DHRS12** | 0.10366 | 1.01645 | 0.09783 | 0.17334 | 0.06444 | 0.33356 | 0.88240 |
| **DHRS13** | 1.43502 | 4.66584 | 3.09890 | 4.42587 | 0.00000 | 4.37536 | 7.03460 |
| **DHRS2** | 0.00515 | 0.16278 | 0.01482 | 0.15543 | 0.01792 | 0.00463 | 0.00000 |
| **DHRS3** | 1.19132 | 26.58615 | 1.01667 | 0.26320 | 0.07683 | 4.78904 | 0.00000 |
| **DHRS4** | 0.05105 | 0.20437 | 0.05542 | 0.04443 | 0.00714 | 0.06901 | 0.00000 |

| | | | | | | | |
|---|---|---|---|---|---|---|---|
| **DHRS4-AS1** | 0.09941 | 0.53721 | 0.65743 | 0.41464 | 0.05090 | 0.03316 | 0.01097 |
| **DHRS4L1** | 0.04197 | 0.02617 | 0.01868 | 0.05186 | 0.00731 | 0.01295 | 0.00589 |
| **DHRS4L2** | 0.05949 | 0.04504 | 0.03931 | 0.12152 | 0.00602 | 0.06054 | 0.00000 |
| **DHRS7** | 3.23070 | 1.29662 | 3.39865 | 2.65304 | 0.30790 | 1.01522 | 0.82509 |
| **DHRS7B** | 3.30681 | 7.70170 | 8.64034 | 5.73019 | 1.03741 | 1.62543 | 0.00000 |
| **DHRS9** | 0.02645 | 0.01656 | 0.01177 | 0.02515 | 0.01626 | 0.01309 | 0.02096 |
| **DHTKD1** | 10.86861 | 5.96422 | 4.70957 | 12.34027 | 0.46467 | 1.80006 | 12.23322 |
| **DHX15** | 47.19835 | 10.98998 | 21.21682 | 22.35021 | 12.09625 | 17.68763 | 19.32768 |
| **DHX16** | 12.63737 | 16.17240 | 7.15433 | 12.46213 | 6.29006 | 6.91978 | 2.99028 |
| **DHX29** | 15.76554 | 3.97498 | 9.74852 | 8.28258 | 4.53042 | 8.42184 | 14.08098 |
| **DHX30** | 18.35522 | 20.24488 | 8.60452 | 16.34776 | 0.65891 | 7.80681 | 13.80451 |
| **DHX32** | 2.06622 | 1.51455 | 0.52133 | 2.29597 | 0.19856 | 1.39692 | 4.51680 |
| **DHX33** | 11.87474 | 7.11142 | 6.60668 | 8.91006 | 2.10138 | 10.05214 | 21.68020 |
| **DHX34** | 0.30325 | 3.62118 | 3.51319 | 3.30183 | 3.04374 | 2.53240 | 10.60105 |
| **DHX35** | 5.71625 | 1.20907 | 2.12258 | 1.79761 | 0.22192 | 2.50856 | 0.73482 |
| **DHX36** | 8.59754 | 0.00000 | 6.46721 | 5.71272 | 1.10939 | 5.61033 | 24.03218 |
| **DHX37** | 3.14784 | 9.00081 | 4.72319 | 11.83907 | 0.94293 | 5.66591 | 19.03421 |
| **DHX38** | 10.22260 | 10.90466 | 4.90712 | 15.22222 | 4.28358 | 6.74350 | 7.56578 |
| **DHX40** | 6.25126 | 0.59518 | 5.06922 | 5.15475 | 3.93128 | 4.16140 | 3.93650 |
| **DHX57** | 5.62408 | 0.61286 | 2.54200 | 5.13099 | 0.53059 | 2.11282 | 5.38074 |
| **DHX58** | 0.48988 | 0.00000 | 4.07799 | 1.49103 | 2.10238 | 0.00056 | 0.00000 |
| **DHX8** | 14.62916 | 12.69767 | 7.67135 | 11.53778 | 8.73726 | 8.27650 | 2.31594 |
| **DHX9** | 36.98472 | 14.41690 | 17.83229 | 21.87547 | 10.33328 | 16.73725 | 18.55684 |
| **DIABLO** | 29.03694 | 20.67185 | 24.03148 | 21.16398 | 12.16258 | 14.68938 | 4.37636 |
| **DIAPH1** | 9.46485 | 11.42831 | 6.84238 | 8.46468 | 1.37172 | 9.47782 | 23.31883 |
| **DIAPH2** | 0.52501 | 0.00000 | 2.61307 | 0.40526 | 2.02963 | 4.15205 | 1.30707 |
| **DIAPH3** | 15.18078 | 4.78014 | 5.46801 | 6.09605 | 1.65600 | 2.82710 | 16.82191 |
| **DIAPH3-AS1** | 0.03227 | 0.11118 | 0.09460 | 0.09836 | 0.01143 | 0.03172 | 0.33215 |
| **DIAPH3-AS2** | 0.05449 | 0.05444 | 0.03493 | 0.01757 | 0.00701 | 0.01186 | 0.09791 |
| **DICER1** | 7.38662 | 3.65152 | 2.05975 | 1.56763 | 1.45116 | 2.96577 | 15.84519 |
| **DICER1-AS1** | 0.17652 | 0.14823 | 0.01248 | 0.01349 | 0.01263 | 0.01207 | 0.01956 |
| **DIDO1** | 4.67239 | 2.90287 | 4.05336 | 3.74419 | 5.70513 | 2.94020 | 13.33251 |
| **DIEXF** | 19.96854 | 2.60475 | 10.25005 | 8.31978 | 5.67909 | 3.41146 | 0.00000 |
| **DIMT1** | 30.94157 | 7.28058 | 12.18799 | 19.36771 | 5.06839 | 9.18288 | 0.00000 |
| **DIO1** | 0.19198 | 0.01661 | 0.01848 | 0.01155 | 0.01104 | 0.02293 | 0.00000 |

| | | | | | | | |
|---|---|---|---|---|---|---|---|
| **DIO2** | 0.19804 | 0.00000 | 0.01460 | 0.21390 | 0.15424 | 0.27822 | 0.00000 |
| **DIO2-AS1** | 0.02423 | 0.01347 | 0.02068 | 0.01955 | 0.00725 | 0.03614 | 0.02307 |
| **DIO3** | 0.00270 | 0.01245 | 0.00000 | 0.14660 | 0.36986 | 0.00853 | 0.21653 |
| **DIO3OS** | 0.02879 | 0.01792 | 0.01245 | 0.01874 | 0.00792 | 0.01559 | 0.01708 |
| **DIP2A** | 3.14427 | 1.92016 | 1.14522 | 4.69132 | 1.42212 | 2.62665 | 7.13020 |
| **DIP2A-IT1** | 0.05923 | 0.00000 | 0.12964 | 0.04470 | 0.24467 | 0.08083 | 0.15155 |
| **DIP2B** | 8.37715 | 0.00000 | 5.22961 | 17.56772 | 2.36592 | 1.24431 | 7.50901 |
| **DIP2C** | 0.54832 | 0.19544 | 1.57883 | 1.36005 | 2.51386 | 0.38123 | 1.94292 |
| **DIRAS1** | 0.00000 | 0.28157 | 0.05887 | 0.01391 | 0.22722 | 0.12431 | 0.03198 |
| **DIRAS2** | 0.02497 | 0.00577 | 0.02857 | 0.01837 | 0.17246 | 0.01286 | 0.01840 |
| **DIRAS3** | 0.13659 | 0.00000 | 0.05662 | 0.03670 | 2.32306 | 0.68255 | 0.03258 |
| **DIRC2** | 2.96980 | 0.92208 | 2.36070 | 7.11776 | 0.09135 | 4.21774 | 14.81435 |
| **DIRC3** | 0.04918 | 0.10634 | 0.02448 | 0.09483 | 0.07517 | 0.00283 | 0.00002 |
| **DIS3** | 15.09940 | 0.71368 | 10.23352 | 7.57445 | 7.80152 | 5.60979 | 13.42027 |
| **DIS3L** | 8.97736 | 1.01484 | 14.92559 | 4.06672 | 1.71952 | 0.20756 | 1.66158 |
| **DIS3L2** | 2.96668 | 0.51426 | 4.82154 | 3.00819 | 1.60783 | 1.37611 | 4.21810 |
| **DISC1** | 0.02478 | 0.00070 | 0.02316 | 0.01174 | 0.02359 | 0.00828 | 0.12037 |
| **DISC1FP1** | 0.02068 | 0.02095 | 0.01681 | 0.03791 | 0.09626 | 0.03819 | 0.07373 |
| **DISC2** | 0.02448 | 0.01971 | 0.01631 | 0.02088 | 0.00749 | 0.01200 | 0.01768 |
| **DISP1** | 0.41474 | 0.30635 | 0.53101 | 0.72346 | 0.26065 | 0.12034 | 0.12119 |
| **DISP2** | 0.00333 | 0.00000 | 0.04757 | 0.21622 | 0.08133 | 0.01128 | 0.11763 |
| **DIXDC1** | 0.30065 | 0.00000 | 0.92626 | 0.27085 | 0.30273 | 1.00910 | 0.01402 |
| **DKC1** | 51.70706 | 29.72369 | 26.84672 | 34.88247 | 13.09914 | 19.67383 | 21.09163 |
| **DKFZP434A062** | 0.01563 | 0.00925 | 0.01697 | 0.05205 | 0.00856 | 0.00975 | 0.05497 |
| **DKFZP434H168** | 0.01400 | 0.01215 | 0.01663 | 0.01813 | 0.01232 | 0.07703 | 0.02865 |
| **DKFZP434I0714** | 0.72786 | 0.17547 | 0.98254 | 0.42929 | 0.58911 | 0.18356 | 0.15590 |
| **DKFZP586I1420** | 1.31719 | 0.68250 | 0.37067 | 2.04437 | 0.10295 | 0.56872 | 3.16442 |
| **DKFZp434J022** | 0.00556 | 0.09438 | 0.03465 | 0.07558 | 0.01135 | 0.04496 | 0.04141 |
| **DKFZp434L192** | 0.02532 | 0.02190 | 0.01176 | 0.01856 | 0.01860 | 0.01271 | 0.02047 |
| **DKFZp686K168** | 0.02349 | 0.01616 | 0.01320 | 0.01673 | 0.00861 | 0.01416 | 0.02717 |
| **1.00000** | 0.03016 | 0.00122 | 0.01264 | 0.31203 | 0.00764 | 0.05498 | 0.12639 |
| **3.00000** | 0.26963 | 0.31568 | 0.00000 | 0.00000 | 0.07964 | 0.64413 | 0.13851 |
| **DKKL1** | 0.10731 | 0.09401 | 0.59769 | 0.07602 | 0.03931 | 0.13992 | 0.19706 |
| **DLAT** | 34.92139 | 15.41577 | 14.08665 | 17.28007 | 0.00000 | 15.93392 | 9.90273 |
| **DLC1** | 19.76444 | 23.27126 | 5.64452 | 9.59456 | 2.94133 | 0.61517 | 11.82156 |

| | | | | | | | |
|---|---|---|---|---|---|---|---|
| **DLD** | 39.93738 | 8.78511 | 13.93768 | 18.60286 | 4.67988 | 13.91021 | 0.00000 |
| **DLEC1** | 0.00000 | 0.00000 | 1.29029 | 0.06969 | 1.19103 | 0.32244 | 0.42568 |
| **DLEU1** | 17.45236 | 4.39917 | 4.16348 | 3.96517 | 1.13462 | 2.28132 | 5.42042 |
| **DLEU2** | 1.36783 | 0.00000 | 1.21550 | 0.00000 | 0.73927 | 0.56053 | 0.70229 |
| **DLEU2L** | 0.47393 | 0.27981 | 0.48044 | 0.00000 | 0.21495 | 0.20659 | 0.56173 |
| **DLEU7** | 0.03405 | 0.02053 | 0.00000 | 0.01787 | 2.30345 | 0.02189 | 0.00913 |
| **DLEU7-AS1** | 0.02100 | 0.01708 | 0.01775 | 0.02380 | 0.01311 | 0.01125 | 0.04064 |
| **DLG1** | 7.58474 | 0.48711 | 2.24238 | 2.08892 | 1.55421 | 3.23382 | 9.64637 |
| **DLG1-AS1** | 0.05179 | 0.07868 | 0.03214 | 0.03684 | 0.13238 | 0.02062 | 0.01178 |
| **DLG2** | 2.80780 | 0.00000 | 0.00000 | 3.63320 | 0.77953 | 0.00000 | 1.95806 |
| **DLG3** | 8.89422 | 22.54100 | 0.96507 | 0.00000 | 1.70973 | 11.85832 | 5.20839 |
| **DLG3-AS1** | 0.03895 | 0.17271 | 0.01029 | 0.02219 | 0.00699 | 0.10690 | 0.03286 |
| **DLG4** | 0.05514 | 0.00000 | 0.04970 | 0.22240 | 0.01348 | 0.06644 | 0.58691 |
| **DLG5** | 16.37552 | 4.72880 | 2.75077 | 38.30782 | 0.45567 | 3.33514 | 20.87415 |
| **DLG5-AS1** | 0.00000 | 0.01034 | 0.11163 | 0.77546 | 0.04596 | 0.00000 | 0.44861 |
| **DLGAP1** | 0.18003 | 0.77175 | 0.16643 | 0.06119 | 0.21245 | 0.24469 | 0.18965 |
| **DLGAP1-AS1** | 0.14545 | 0.13934 | 3.34342 | 0.36573 | 0.07126 | 0.48527 | 0.18731 |
| **DLGAP1-AS2** | 0.45831 | 0.97978 | 2.43562 | 0.00000 | 0.00000 | 1.30357 | 0.17820 |
| **DLGAP1-AS3** | 0.02492 | 0.01954 | 0.01160 | 0.01254 | 0.00841 | 0.01768 | 0.05006 |
| **DLGAP1-AS4** | 0.01990 | 0.02093 | 0.01424 | 0.02059 | 0.00721 | 0.00831 | 0.06494 |
| **DLGAP1-AS5** | 0.05335 | 0.01857 | 0.02721 | 0.01053 | 0.00785 | 0.05701 | 0.03575 |
| **DLGAP2** | 0.05386 | 0.00000 | 0.15306 | 0.00000 | 0.20091 | 0.00153 | 0.13475 |
| **DLGAP3** | 0.00000 | 0.92895 | 0.12444 | 0.00000 | 0.00535 | 3.23782 | 0.21297 |
| **DLGAP4** | 9.83430 | 15.23865 | 2.42539 | 17.87344 | 1.23437 | 3.18759 | 10.03520 |
| **DLGAP5** | 24.12421 | 7.18526 | 12.36279 | 13.70619 | 12.55505 | 9.12476 | 5.89302 |
| **DLK1** | 0.00589 | 0.01463 | 0.05295 | 0.03547 | 0.01782 | 0.00000 | 0.21029 |
| **DLK2** | 0.00000 | 0.71312 | 2.97081 | 0.02264 | 0.98077 | 0.01468 | 0.15914 |
| **DLL1** | 0.01724 | 0.01191 | 0.00886 | 0.03864 | 0.08723 | 0.07092 | 0.01432 |
| **DLL3** | 0.65609 | 0.00000 | 5.98781 | 0.47414 | 1.29318 | 2.16127 | 0.46222 |
| **DLL4** | 0.14044 | 0.11882 | 0.00000 | 0.07045 | 0.28851 | 0.26613 | 0.02586 |
| **DLST** | 0.75966 | 0.83390 | 0.39543 | 1.54207 | 2.86617 | 0.65850 | 1.60466 |
| **DLX1** | 0.04527 | 0.02326 | 0.00000 | 0.00000 | 1.31742 | 0.05581 | 0.07687 |
| **DLX2** | 0.00372 | 0.45270 | 0.00000 | 0.00000 | 0.29965 | 0.00218 | 0.70257 |
| **DLX3** | 21.40069 | 0.00000 | 0.00000 | 35.83141 | 0.00000 | 0.30676 | 15.66710 |
| **DLX4** | 2.28030 | 0.77317 | 0.00000 | 6.11992 | 0.00000 | 0.80382 | 5.41416 |

| | | | | | | | |
|---|---|---|---|---|---|---|---|
| **DLX5** | 2.55668 | 0.10314 | 0.02035 | 13.80989 | 0.07893 | 0.00000 | 2.94788 |
| **DLX6** | 2.55951 | 3.06533 | 0.97670 | 5.46040 | 1.61596 | 0.00000 | 0.22544 |
| **DLX6-AS1** | 0.01747 | 0.07146 | 0.08137 | 0.24053 | 0.00478 | 0.00000 | 0.53513 |
| **DMAP1** | 9.23311 | 14.82766 | 6.72476 | 8.48754 | 2.06663 | 6.25247 | 1.25652 |
| **DMBT1** | 0.02249 | 0.00000 | 0.06845 | 0.03879 | 0.07620 | 0.01665 | 0.02583 |
| **DMBT1P1** | 0.02720 | 0.01697 | 0.02023 | 0.02512 | 0.00737 | 0.01749 | 0.01808 |
| **DMBX1** | 0.01312 | 0.03195 | 0.13109 | 0.02529 | 0.04774 | 0.63375 | 0.01811 |
| **DMC1** | 0.38668 | 1.06571 | 0.04017 | 0.01503 | 0.00710 | 0.57580 | 0.10660 |
| **DMD** | 5.65147 | 0.14070 | 6.69215 | 3.54186 | 2.86219 | 0.67716 | 5.34306 |
| **DMGDH** | 0.00962 | 0.03581 | 0.00000 | 0.04808 | 0.49377 | 0.01055 | 0.02281 |
| **DMKN** | 0.56406 | 0.39963 | 0.00000 | 0.83092 | 1.35875 | 0.59625 | 0.06489 |
| **DMP1** | 0.01264 | 0.05997 | 0.01539 | 0.04368 | 0.04197 | 0.04374 | 0.01607 |
| **DMPK** | 0.00000 | 0.50422 | 0.29670 | 1.41831 | 0.09654 | 0.29163 | 6.13268 |
| **DMRT1** | 0.00000 | 0.00000 | 0.17452 | 0.03219 | 0.30946 | 0.11255 | 0.09647 |
| **DMRT2** | 0.00502 | 0.00823 | 0.01608 | 0.03650 | 0.02973 | 0.11924 | 0.04220 |
| **DMRTA1** | 0.01383 | 0.08681 | 0.01969 | 0.07891 | 0.13010 | 0.07044 | 0.00000 |
| **DMRTA2** | 0.00000 | 0.01028 | 0.00867 | 0.01828 | 0.00733 | 0.52605 | 0.14514 |
| **DMRTB1** | 0.00000 | 0.62560 | 5.53700 | 0.00000 | 1.09987 | 0.22692 | 0.57141 |
| **DMRTC2** | 0.02649 | 0.00000 | 0.33693 | 0.00959 | 0.60912 | 0.00251 | 0.03346 |
| **DMTF1** | 9.28385 | 0.00000 | 3.73839 | 3.49585 | 2.53395 | 2.82992 | 2.36796 |
| **DMTN** | 0.00000 | 0.01294 | 0.06937 | 0.08805 | 0.52203 | 0.53079 | 0.03792 |
| **DMWD** | 0.00000 | 1.89346 | 1.37715 | 2.57461 | 0.48860 | 1.55338 | 11.93710 |
| **DMXL1** | 5.05589 | 1.20901 | 3.12806 | 2.46630 | 2.41218 | 8.33436 | 9.10568 |
| **DMXL2** | 0.43358 | 0.00000 | 1.36621 | 1.48569 | 0.32671 | 1.11512 | 2.48744 |
| **DNA2** | 10.82133 | 0.00000 | 4.59524 | 6.89375 | 1.80764 | 4.83346 | 2.87581 |
| **DNAAF1** | 0.01872 | 0.01036 | 0.02081 | 0.02531 | 0.08594 | 0.02574 | 0.04645 |
| **DNAAF2** | 2.82463 | 2.27337 | 2.92850 | 7.92211 | 0.68432 | 3.99605 | 12.87648 |
| **DNAAF3** | 0.04565 | 0.00000 | 0.23375 | 0.04551 | 0.63924 | 0.04175 | 0.00000 |
| **DNAH1** | 0.15297 | 0.14679 | 0.26170 | 0.25841 | 0.05255 | 0.04459 | 0.47606 |
| **DNAH10** | 0.10212 | 0.02655 | 0.18971 | 0.03770 | 3.51316 | 0.06116 | 0.00000 |
| **DNAH11** | 0.00000 | 0.23086 | 0.15119 | 0.00000 | 3.11568 | 0.19305 | 0.20244 |
| **DNAH12** | 0.00000 | 0.00000 | 0.66472 | 0.00538 | 1.25053 | 0.22232 | 0.28120 |
| **DNAH14** | 2.32304 | 0.00000 | 1.27059 | 2.31155 | 0.45885 | 0.62293 | 3.23262 |
| **DNAH17** | 0.12305 | 1.19520 | 0.04637 | 0.40513 | 1.72554 | 0.28432 | 0.80174 |
| **DNAH17-AS1** | 0.01091 | 0.43922 | 0.00358 | 0.01954 | 0.00789 | 0.13642 | 0.00850 |

| | | | | | | | |
|---|---|---|---|---|---|---|---|
| **DNAH2** | 0.05845 | 0.00000 | 0.18760 | 0.08255 | 0.11607 | 0.11882 | 0.02769 |
| **DNAH3** | 3.80017 | 4.13825 | 0.00673 | 0.02255 | 1.48727 | 1.86629 | 0.48968 |
| **DNAH5** | 0.01144 | 0.01780 | 0.07196 | 0.01850 | 0.14280 | 0.10480 | 0.04484 |
| **DNAH6** | 0.06032 | 0.00000 | 0.09380 | 0.07220 | 0.01537 | 0.00720 | 0.06669 |
| **DNAH7** | 0.01154 | 0.00000 | 0.23785 | 0.01496 | 0.32204 | 0.22323 | 0.28961 |
| **DNAH8** | 0.10296 | 0.00000 | 0.02627 | 0.10301 | 0.25248 | 0.02281 | 0.00000 |
| **DNAH9** | 0.05694 | 0.08538 | 0.07861 | 0.06410 | 0.82532 | 0.54160 | 0.00000 |
| **DNAI1** | 0.28912 | 0.00000 | 0.05570 | 0.26031 | 0.34434 | 0.39808 | 0.09416 |
| **DNAI2** | 0.01796 | 0.00421 | 0.05742 | 0.02764 | 0.00986 | 0.01395 | 0.02697 |
| **DNAJA1** | 46.04670 | 21.18896 | 22.91286 | 40.20732 | 19.88479 | 20.37692 | 15.74521 |
| **DNAJA1P5** | 0.06230 | 0.02606 | 0.02685 | 0.04789 | 0.02226 | 0.02361 | 0.00488 |
| **DNAJA2** | 36.25135 | 15.74497 | 19.51642 | 21.98612 | 10.59217 | 20.43690 | 11.58049 |
| **DNAJA3** | 20.97923 | 12.34470 | 7.79395 | 17.62230 | 1.58911 | 8.98218 | 4.41679 |
| **DNAJA4** | 1.90907 | 1.37231 | 0.38110 | 1.05441 | 1.32369 | 1.22190 | 1.02463 |
| **DNAJB1** | 28.60407 | 28.53160 | 20.04589 | 32.74033 | 30.26037 | 13.03582 | 3.81713 |
| **DNAJB11** | 11.92608 | 12.10750 | 10.62079 | 11.72249 | 6.75588 | 10.31811 | 20.13487 |
| **DNAJB12** | 11.73355 | 7.32260 | 8.13403 | 12.53641 | 2.80127 | 4.45377 | 3.98272 |
| **DNAJB13** | 0.04992 | 0.21120 | 0.07314 | 0.01370 | 0.29362 | 0.07779 | 0.00000 |
| **DNAJB14** | 4.43183 | 0.00000 | 5.34335 | 2.31778 | 2.35530 | 2.34592 | 3.47787 |
| **DNAJB2** | 0.00000 | 2.43989 | 0.73327 | 1.08429 | 0.51492 | 1.39328 | 6.19282 |
| **DNAJB3** | 0.02079 | 0.01695 | 0.00953 | 0.05638 | 0.05159 | 0.03757 | 0.00763 |
| **DNAJB4** | 4.87058 | 0.00000 | 7.18763 | 2.30421 | 1.75045 | 3.10373 | 0.00000 |
| **DNAJB5** | 0.00000 | 0.21159 | 0.07629 | 0.06686 | 0.01679 | 0.36274 | 0.19598 |
| **DNAJB5-AS1** | 0.00553 | 0.17971 | 0.08764 | 0.06774 | 0.13456 | 0.00000 | 0.05208 |
| **DNAJB6** | 22.76082 | 10.19545 | 7.09879 | 22.35221 | 6.77926 | 15.96878 | 1.11356 |
| **DNAJB7** | 0.42120 | 0.10397 | 0.25079 | 0.09596 | 0.00680 | 0.12926 | 0.30847 |
| **DNAJB8** | 0.03701 | 0.01887 | 0.01210 | 0.03300 | 0.00819 | 0.03876 | 0.00000 |
| **DNAJB8-AS1** | 0.01959 | 0.02735 | 0.01044 | 0.03161 | 0.01160 | 0.01711 | 0.01711 |
| **DNAJB9** | 9.62953 | 0.00000 | 7.00809 | 13.81342 | 6.03082 | 2.88009 | 0.00000 |
| **DNAJC1** | 10.79505 | 0.90526 | 1.39139 | 7.35677 | 2.93068 | 1.86264 | 14.72121 |
| **DNAJC10** | 16.35804 | 1.01529 | 8.05164 | 8.37225 | 1.77552 | 5.75512 | 7.46676 |
| **DNAJC11** | 22.10289 | 16.26203 | 7.65662 | 8.38749 | 1.12049 | 6.94460 | 0.00000 |
| **DNAJC12** | 0.04191 | 0.01548 | 0.07550 | 0.05560 | 0.02883 | 0.10113 | 0.00000 |
| **DNAJC13** | 4.11204 | 1.21785 | 2.70014 | 1.13304 | 4.27697 | 3.74072 | 7.17277 |
| **DNAJC14** | 9.14876 | 5.00420 | 5.50341 | 9.07434 | 2.51092 | 4.42059 | 0.00000 |

| | | | | | | | |
|---|---|---|---|---|---|---|---|
| **DNAJC15** | 78.68698 | 77.27995 | 45.34898 | 35.01280 | 14.95445 | 31.32239 | 44.35222 |
| **DNAJC16** | 4.35966 | 3.84850 | 4.28228 | 0.23079 | 1.03350 | 3.57345 | 3.44054 |
| **DNAJC17** | 3.84742 | 2.55240 | 4.45664 | 4.61182 | 3.03446 | 1.16187 | 0.00016 |
| **DNAJC18** | 0.52312 | 0.01571 | 0.13072 | 0.64809 | 0.61455 | 0.41273 | 0.00000 |
| **DNAJC19** | 11.79046 | 9.95858 | 7.87163 | 4.80573 | 0.02660 | 4.06925 | 4.80949 |
| **DNAJC2** | 23.83850 | 9.80720 | 12.54182 | 17.52563 | 3.68742 | 10.34509 | 19.79860 |
| **DNAJC21** | 3.35947 | 2.06024 | 3.50100 | 1.50321 | 1.12090 | 3.93994 | 18.49506 |
| **DNAJC22** | 0.01038 | 0.51481 | 0.01909 | 0.09688 | 0.04219 | 2.13953 | 0.00000 |
| **DNAJC24** | 2.99792 | 0.26290 | 2.50520 | 3.19357 | 0.72809 | 2.10104 | 3.90327 |
| **DNAJC25** | 0.97504 | 1.52597 | 3.95481 | 0.60115 | 0.76966 | 6.33267 | 6.63425 |
| **DNAJC27** | 1.79006 | 0.00000 | 0.37058 | 0.42083 | 2.77047 | 1.21079 | 1.29525 |
| **DNAJC27-AS1** | 0.00000 | 0.06822 | 0.20948 | 0.03578 | 0.02546 | 0.12230 | 0.07130 |
| **DNAJC28** | 1.25233 | 0.35477 | 1.00553 | 0.37498 | 0.14087 | 0.61096 | 0.12609 |
| **DNAJC3** | 29.15400 | 15.15125 | 10.61262 | 15.99167 | 7.41043 | 17.53882 | 0.00001 |
| **DNAJC30** | 4.17649 | 5.81088 | 6.27816 | 4.60377 | 2.16675 | 1.91258 | 0.00000 |
| **DNAJC4** | 0.79698 | 0.33937 | 0.29512 | 1.33613 | 0.38646 | 1.07788 | 0.15399 |
| **DNAJC5** | 4.73201 | 7.14884 | 1.84751 | 6.74380 | 0.89142 | 1.68209 | 19.92729 |
| **DNAJC5B** | 0.01838 | 0.01477 | 0.00656 | 0.05948 | 0.07696 | 0.00928 | 0.05547 |
| **DNAJC5G** | 0.05114 | 0.11247 | 0.00000 | 0.00386 | 0.31981 | 0.12191 | 0.06391 |
| **DNAJC6** | 10.12916 | 0.18056 | 0.01060 | 0.00000 | 0.07258 | 0.00000 | 4.12889 |
| **DNAJC7** | 27.56993 | 6.37326 | 12.24034 | 16.98293 | 6.36225 | 9.35113 | 6.16136 |
| **DNAJC8** | 26.10470 | 12.84828 | 17.77685 | 23.37613 | 12.04565 | 11.74618 | 9.63010 |
| **DNAJC9** | 22.22821 | 7.81076 | 9.92381 | 17.26016 | 8.29555 | 7.17288 | 2.50098 |
| **DNAJC9-AS1** | 0.03252 | 0.07860 | 0.17558 | 0.13101 | 0.04387 | 0.02809 | 0.33674 |
| **DNAL1** | 0.89902 | 0.28414 | 3.63845 | 1.80486 | 7.04981 | 1.36534 | 1.58996 |
| **DNAL4** | 0.48612 | 2.55751 | 3.22203 | 1.39106 | 2.19532 | 0.45562 | 1.05588 |
| **DNALI1** | 0.34983 | 0.01238 | 0.07272 | 1.82738 | 0.06445 | 0.27431 | 0.00000 |
| **DNASE1** | 0.48845 | 0.45884 | 0.98223 | 1.29242 | 0.47598 | 0.16456 | 0.85715 |
| **DNASE1L1** | 0.18369 | 0.42686 | 0.37671 | 0.16247 | 0.00556 | 0.30716 | 0.32560 |
| **DNASE1L2** | 0.00000 | 0.58808 | 0.28974 | 0.07487 | 0.00520 | 0.15947 | 0.19318 |
| **DNASE1L3** | 0.23367 | 0.00000 | 0.20510 | 0.55294 | 0.02319 | 0.05152 | 0.00000 |
| **DNASE2** | 15.33842 | 18.10656 | 6.86192 | 11.47224 | 4.70100 | 3.15376 | 0.00000 |
| **DNASE2B** | 0.01996 | 0.00000 | 0.29312 | 0.01770 | 0.08729 | 0.00708 | 0.01739 |
| **DND1** | 1.76716 | 0.73417 | 0.30374 | 2.18429 | 0.86637 | 12.59458 | 0.00000 |
| **DNER** | 0.25847 | 0.44331 | 0.05452 | 0.00000 | 0.09247 | 0.12230 | 0.20848 |

| | | | | | | | |
|---|---|---|---|---|---|---|---|
| **DNHD1** | 1.78103 | 0.00000 | 1.42121 | 3.54589 | 0.48952 | 1.24776 | 2.72844 |
| **DNLZ** | 0.00000 | 1.65112 | 0.08956 | 0.37612 | 0.10558 | 1.27177 | 3.33531 |
| **DNM1** | 0.00000 | 0.01536 | 0.06276 | 0.16936 | 0.11769 | 0.58989 | 0.24382 |
| **DNM1L** | 24.45448 | 8.33868 | 5.09931 | 18.96554 | 1.69233 | 8.26974 | 11.26485 |
| **DNM1P35** | 0.01998 | 0.01268 | 0.01348 | 0.02351 | 0.02551 | 0.01010 | 0.05271 |
| **DNM1P41** | 0.02469 | 0.01610 | 0.01198 | 0.01836 | 0.01458 | 0.01192 | 0.02694 |
| **DNM1P46** | 0.02618 | 0.01833 | 0.02131 | 0.01698 | 0.03557 | 0.01354 | 0.03559 |
| **DNM2** | 0.00000 | 10.09309 | 6.06834 | 11.98267 | 0.18920 | 7.66489 | 23.78283 |
| **DNM3** | 0.21342 | 0.04725 | 0.00000 | 0.07273 | 2.28612 | 0.05231 | 0.00815 |
| **DNM3OS** | 0.10333 | 0.00000 | 0.03818 | 0.00000 | 0.05578 | 0.01578 | 0.21646 |
| **DNMBP** | 3.96091 | 1.92862 | 1.75219 | 0.22577 | 1.46371 | 3.59507 | 2.74385 |
| **DNMBP-AS1** | 0.07054 | 0.05873 | 0.04830 | 0.04211 | 0.04787 | 0.00706 | 0.09772 |
| **DNMT1** | 6.34024 | 6.29868 | 16.91960 | 29.52558 | 34.97215 | 6.43519 | 22.49196 |
| **DNMT3A** | 24.59013 | 30.01494 | 8.05644 | 3.54163 | 2.20174 | 10.20032 | 12.31088 |
| **DNMT3B** | 37.88145 | 28.28076 | 0.00000 | 23.01988 | 0.59571 | 9.23032 | 42.26054 |
| **DNMT3L** | 86.85049 | 108.46675 | 13.93425 | 46.90074 | 0.00000 | 41.11677 | 19.02179 |
| **DNPEP** | 1.83505 | 3.22854 | 1.61265 | 5.00917 | 3.31828 | 5.44184 | 3.93055 |
| **DNPH1** | 7.86314 | 8.89777 | 0.57911 | 2.52420 | 0.00000 | 3.06131 | 2.96794 |
| **DNTT** | 0.00832 | 0.11511 | 0.02770 | 0.00000 | 0.00465 | 0.46526 | 0.14417 |
| **DNTTIP1** | 0.00000 | 8.15647 | 2.78176 | 1.97741 | 0.23150 | 3.10301 | 9.15823 |
| **DNTTIP2** | 44.33455 | 13.73661 | 27.35631 | 28.25663 | 13.34645 | 13.14540 | 11.01860 |
| **DOC2A** | 0.00000 | 0.09676 | 0.06926 | 0.15259 | 0.04962 | 0.30799 | 1.00473 |
| **DOC2B** | 0.01763 | 0.01580 | 0.01446 | 0.01755 | 0.25082 | 0.00902 | 0.05031 |
| **DOCK1** | 1.81487 | 2.72270 | 2.18323 | 2.52865 | 1.15118 | 3.29946 | 2.91744 |
| **DOCK10** | 0.02620 | 0.00704 | 0.01656 | 0.01725 | 0.31409 | 0.01364 | 0.02487 |
| **DOCK11** | 0.08722 | 0.00000 | 0.87468 | 0.02117 | 0.24641 | 0.11749 | 0.00160 |
| **DOCK2** | 0.05076 | 0.05544 | 0.07880 | 0.04939 | 1.27639 | 0.14284 | 0.03236 |
| **DOCK3** | 0.00000 | 2.62955 | 1.00255 | 0.27107 | 0.08930 | 2.67127 | 2.59713 |
| **DOCK4** | 0.99653 | 0.00000 | 0.35457 | 4.05940 | 0.59277 | 3.09913 | 3.97737 |
| **DOCK4-AS1** | 0.01783 | 0.00264 | 0.01178 | 0.02774 | 0.01452 | 0.06817 | 0.10778 |
| **DOCK5** | 0.18573 | 0.89203 | 0.27733 | 0.33042 | 0.48168 | 0.29795 | 0.01112 |
| **DOCK6** | 0.28763 | 6.17731 | 1.32869 | 1.17415 | 0.09704 | 11.24104 | 6.71618 |
| **DOCK7** | 4.02358 | 0.00000 | 1.05200 | 3.53791 | 0.51817 | 2.03686 | 4.55635 |
| **DOCK8** | 0.27877 | 0.12083 | 0.03423 | 0.96044 | 0.40434 | 0.05702 | 1.16054 |
| **DOCK9** | 10.12635 | 0.00000 | 0.00000 | 16.58668 | 1.66229 | 0.62274 | 11.14110 |

| | | | | | | | |
|---|---|---|---|---|---|---|---|
| **DOCK9-AS1** | 0.01724 | 0.01612 | 0.01146 | 0.02127 | 0.00738 | 0.00698 | 0.07932 |
| **DOCK9-AS2** | 0.00000 | 0.04471 | 0.01921 | 0.22183 | 0.02273 | 0.00567 | 0.14157 |
| **DOHH** | 0.00000 | 3.81772 | 0.49760 | 6.64740 | 0.72940 | 1.68704 | 7.14653 |
| **DOK1** | 0.35741 | 0.19893 | 0.38224 | 1.87458 | 0.72602 | 0.96006 | 0.01583 |
| **DOK2** | 1.12283 | 0.20827 | 0.14597 | 0.05945 | 0.31056 | 0.00000 | 0.07497 |
| **DOK3** | 1.27265 | 0.25134 | 2.31587 | 1.47668 | 0.07809 | 1.05683 | 3.83021 |
| **DOK4** | 1.49528 | 0.72766 | 0.22003 | 6.07662 | 3.00403 | 1.59171 | 1.63397 |
| **DOK5** | 0.00000 | 0.07429 | 0.07874 | 0.03674 | 0.95842 | 0.06868 | 0.01714 |
| **DOK6** | 0.03517 | 0.04966 | 0.01162 | 0.01050 | 0.13710 | 0.01138 | 0.08485 |
| **DOK7** | 0.00000 | 0.02673 | 0.01783 | 0.16325 | 0.00972 | 0.06454 | 0.13898 |
| **DOLK** | 5.00686 | 13.90335 | 4.64716 | 8.63634 | 0.80126 | 4.22941 | 0.00000 |
| **DOLPP1** | 9.93674 | 13.00957 | 1.00615 | 16.45354 | 0.73387 | 4.85297 | 0.00000 |
| **DONSON** | 3.29997 | 0.54930 | 2.45051 | 4.11229 | 0.07040 | 2.54137 | 7.76291 |
| **DOPEY1** | 0.66429 | 0.29449 | 0.15911 | 0.47305 | 4.40681 | 0.58440 | 1.07922 |
| **DOPEY2** | 1.95539 | 0.00000 | 0.43031 | 2.26220 | 1.58192 | 0.06776 | 2.34890 |
| **DOT1L** | 1.98491 | 8.55598 | 5.39851 | 4.77298 | 3.41271 | 0.01305 | 33.35451 |
| **DPAGT1** | 19.32385 | 8.31692 | 8.48690 | 17.18683 | 0.35110 | 8.80313 | 0.00000 |
| **DPCD** | 4.85436 | 7.67873 | 7.17645 | 5.23737 | 2.57246 | 3.19978 | 0.00000 |
| **DPCR1** | 0.01359 | 0.02685 | 0.15332 | 0.01378 | 0.05146 | 0.09592 | 0.02908 |
| **DPEP1** | 0.00000 | 0.61415 | 1.61002 | 0.06035 | 1.76402 | 0.00000 | 0.10950 |
| **DPEP2** | 0.01058 | 0.03790 | 0.00685 | 0.06756 | 0.00718 | 0.11194 | 0.17539 |
| **DPEP3** | 0.04300 | 0.10202 | 0.21784 | 0.07752 | 0.02737 | 0.01559 | 0.01090 |
| **DPF1** | 0.00000 | 2.64697 | 0.00000 | 0.00000 | 0.86862 | 5.73530 | 0.41883 |
| **DPF2** | 21.20823 | 16.35694 | 11.84783 | 19.54800 | 3.67420 | 8.58995 | 0.00004 |
| **DPF3** | 2.78097 | 0.00000 | 0.07094 | 9.24714 | 0.26316 | 0.00000 | 7.03883 |
| **DPH1** | 0.69441 | 4.30360 | 1.80060 | 1.84812 | 0.13271 | 1.64527 | 2.87615 |
| **DPH2** | 33.28192 | 18.79379 | 17.22071 | 27.82499 | 5.88331 | 12.94302 | 0.00000 |
| **DPH3** | 21.08744 | 10.97962 | 18.90556 | 19.30058 | 9.61268 | 9.83663 | 8.61345 |
| **DPH3P1** | 0.79870 | 0.69709 | 1.12132 | 0.78016 | 0.41367 | 0.52238 | 1.12541 |
| **DPH5** | 13.11769 | 5.21154 | 6.43890 | 4.80277 | 0.00000 | 4.62039 | 3.45757 |
| **DPH6** | 4.27175 | 2.61635 | 5.77593 | 0.00000 | 0.00000 | 2.18309 | 5.21996 |
| **DPH6-AS1** | 0.03425 | 0.03244 | 0.01019 | 0.02467 | 0.06138 | 0.01490 | 0.00966 |
| **DPH7** | 9.27293 | 9.96018 | 2.66368 | 10.06999 | 0.99300 | 6.53317 | 6.48427 |
| **DPM1** | 28.98387 | 0.33577 | 20.46213 | 15.88431 | 4.91806 | 10.56913 | 4.82806 |
| **DPM2** | 12.84093 | 18.81594 | 10.57825 | 16.84463 | 3.64698 | 6.15881 | 0.08762 |

| | | | | | | | |
|---|---|---|---|---|---|---|---|
| **DPM3** | 2.47465 | 4.03245 | 1.85798 | 5.40570 | 1.38988 | 1.46547 | 3.93163 |
| **DPP10** | 0.00352 | 0.05565 | 0.00000 | 0.01223 | 0.65150 | 0.06950 | 0.04417 |
| **DPP10-AS1** | 0.03553 | 0.03649 | 0.00806 | 0.04094 | 0.02007 | 0.01761 | 0.01314 |
| **DPP3** | 11.17707 | 5.06017 | 1.66934 | 14.28858 | 2.59803 | 7.81055 | 0.00000 |
| **DPP4** | 1.52970 | 0.00000 | 3.91717 | 1.89912 | 0.00015 | 8.24737 | 0.00000 |
| **DPP6** | 0.01655 | 0.00265 | 0.00672 | 0.01394 | 1.05762 | 0.11430 | 0.20688 |
| **DPP7** | 0.00006 | 0.10352 | 0.15146 | 0.24016 | 0.07112 | 0.08872 | 0.33533 |
| **DPP8** | 9.31084 | 1.55736 | 6.26868 | 4.48925 | 2.37028 | 4.13936 | 3.01711 |
| **DPP9** | 7.00924 | 9.23452 | 1.85584 | 11.68929 | 0.17499 | 5.35438 | 13.75312 |
| **DPP9-AS1** | 0.02700 | 0.22022 | 0.04281 | 0.17176 | 0.00603 | 0.10656 | 0.23566 |
| **DPPA2** | 5.17587 | 0.85958 | 8.10802 | 1.61837 | 2.80556 | 15.96005 | 0.00000 |
| **DPPA2P3** | 0.08671 | 0.00000 | 0.09777 | 0.02557 | 0.01678 | 0.22268 | 0.02187 |
| **DPPA3** | 54.77783 | 12.30296 | 37.06339 | 10.66148 | 20.94044 | 16.37677 | 17.29529 |
| **DPPA4** | 22.61710 | 5.46477 | 19.07780 | 14.59410 | 0.46340 | 22.01375 | 0.17038 |
| **DPPA5** | 0.00000 | 18.04373 | 35.07119 | 0.00000 | 36.94478 | 45.44711 | 0.00000 |
| **DPRX** | 0.00000 | 0.08021 | 18.01195 | 0.61409 | 3.54026 | 0.04609 | 0.00000 |
| **DPRXP4** | 0.00000 | 0.08497 | 0.97368 | 0.30515 | 0.05690 | 0.02927 | 0.11373 |
| **DPY19L1** | 0.19676 | 0.57460 | 0.37687 | 0.66749 | 0.01445 | 4.72469 | 3.08945 |
| **DPY19L1P1** | 0.00000 | 0.08786 | 0.01836 | 0.18381 | 0.12030 | 0.15842 | 0.35710 |
| **DPY19L2** | 0.08506 | 0.00494 | 0.01885 | 0.02386 | 0.00829 | 0.07564 | 0.00000 |
| **DPY19L2P1** | 0.02866 | 0.00830 | 0.12250 | 0.03564 | 0.02286 | 0.08380 | 0.01311 |
| **DPY19L2P2** | 0.01486 | 0.00000 | 0.11685 | 0.07391 | 0.02563 | 0.04771 | 0.05584 |
| **DPY19L2P3** | 0.03333 | 0.00999 | 0.02821 | 0.07712 | 0.00704 | 0.02616 | 0.02711 |
| **DPY19L2P4** | 0.02462 | 0.01388 | 0.01783 | 0.01968 | 0.03591 | 0.01444 | 0.01529 |
| **DPY19L3** | 3.19261 | 0.00000 | 0.88412 | 1.04306 | 1.45656 | 3.58960 | 3.74983 |
| **DPY19L4** | 3.15029 | 0.00005 | 0.58666 | 0.00000 | 0.31439 | 1.91548 | 4.19398 |
| **DPY30** | 21.24993 | 12.88586 | 14.09847 | 19.78526 | 1.72150 | 11.75189 | 12.86945 |
| **DPYD** | 0.03225 | 0.01412 | 0.01099 | 0.05121 | 0.10190 | 0.04841 | 0.01807 |
| **DPYS** | 0.03431 | 0.00718 | 0.01488 | 0.04513 | 0.14340 | 0.02558 | 0.00962 |
| **DPYSL2** | 4.55196 | 1.69938 | 6.67531 | 3.79049 | 2.97475 | 5.93102 | 0.00000 |
| **DPYSL3** | 0.33836 | 0.00000 | 0.03894 | 0.75818 | 0.00585 | 1.52401 | 0.00000 |
| **DPYSL4** | 0.01507 | 0.00000 | 0.10274 | 0.03371 | 0.34764 | 0.01148 | 0.02198 |
| **DPYSL5** | 0.03465 | 0.78631 | 0.00000 | 0.58862 | 0.24326 | 0.18826 | 0.95956 |
| **DQX1** | 0.09896 | 0.06651 | 0.27092 | 0.47399 | 0.03154 | 0.28967 | 0.10894 |
| **DR1** | 12.52736 | 5.23523 | 14.93527 | 12.85606 | 7.12798 | 6.18714 | 9.60903 |

| | | | | | | | |
|---|---|---|---|---|---|---|---|
| **DRAM1** | 2.01759 | 1.41550 | 11.96359 | 7.85461 | 2.49916 | 1.08092 | 7.14899 |
| **DRAM2** | 17.57316 | 2.97785 | 14.12029 | 7.09885 | 0.28167 | 6.44954 | 4.94766 |
| **DRAP1** | 1.40594 | 11.52592 | 2.98785 | 2.96761 | 2.66877 | 8.04471 | 8.00955 |
| **DRAXIN** | 0.00423 | 0.00906 | 0.01489 | 0.01797 | 0.01676 | 0.14318 | 0.09278 |
| **DRC1** | 0.08131 | 0.08613 | 0.00405 | 0.06181 | 0.09418 | 0.00884 | 0.00432 |
| **DRD1** | 0.08895 | 0.02205 | 0.00692 | 0.07903 | 0.02728 | 0.00245 | 0.03003 |
| **DRD2** | 0.00000 | 0.06947 | 0.01664 | 0.02933 | 0.00615 | 0.36969 | 0.05687 |
| **DRD3** | 0.03402 | 0.01440 | 0.01520 | 0.01500 | 0.00771 | 0.03042 | 0.01400 |
| **DRD4** | 0.00000 | 0.07130 | 0.04613 | 0.09650 | 0.00398 | 0.01008 | 0.45041 |
| **DRD5P2+LOC1(** | 0.02373 | 0.01000 | 0.01301 | 0.01305 | 0.00753 | 0.01875 | 0.05533 |
| **DRG1** | 41.30116 | 23.41897 | 17.06037 | 29.25703 | 8.38006 | 15.60128 | 4.73277 |
| **DRG2** | 17.01775 | 12.25557 | 5.45919 | 15.52760 | 1.44847 | 6.41165 | 3.18307 |
| **DRGX** | 0.02526 | 0.01640 | 0.01041 | 0.01943 | 0.03994 | 0.02676 | 0.01254 |
| **DRICH1** | 0.11497 | 0.01853 | 0.16098 | 0.04567 | 0.00737 | 0.00000 | 0.44580 |
| **DROSHA** | 9.23925 | 5.19587 | 1.99907 | 9.33885 | 0.63298 | 5.84212 | 16.91697 |
| **DRP2** | 0.06506 | 0.00000 | 0.08080 | 0.08849 | 0.12057 | 0.00000 | 0.12373 |
| **DSC1** | 0.38691 | 0.00000 | 0.00888 | 0.15177 | 0.11715 | 0.05030 | 0.00000 |
| **DSC2** | 16.54130 | 9.11827 | 0.00000 | 24.10554 | 0.15455 | 8.61503 | 11.82367 |
| **DSC3** | 0.12205 | 0.00000 | 0.02089 | 1.88686 | 0.17816 | 0.00000 | 0.80872 |
| **DSCAM** | 0.04224 | 0.02293 | 0.00850 | 0.04950 | 0.34315 | 0.01021 | 0.01082 |
| **DSCAM-AS1** | 0.02582 | 0.01741 | 0.01383 | 0.01935 | 0.00780 | 0.01299 | 0.01884 |
| **DSCAML1** | 0.09974 | 1.48828 | 0.02682 | 0.00000 | 0.18410 | 0.43050 | 0.00476 |
| **DSCC1** | 12.59342 | 0.30833 | 6.92565 | 6.38637 | 1.53550 | 4.65032 | 1.49336 |
| **DSCR10** | 0.02579 | 0.01206 | 0.01027 | 0.05494 | 0.05593 | 0.01232 | 0.01870 |
| **DSCR3** | 17.37547 | 3.60212 | 1.51320 | 9.83141 | 0.99485 | 4.47579 | 0.00000 |
| **DSCR4** | 2.69724 | 0.58997 | 0.15619 | 1.22237 | 0.13884 | 0.58963 | 0.47836 |
| **DSCR8** | 6.79135 | 3.34255 | 0.49645 | 2.00716 | 0.88227 | 4.50152 | 2.16131 |
| **DSCR9** | 0.31939 | 0.17400 | 0.25994 | 0.24234 | 0.61155 | 0.00000 | 0.03496 |
| **DSE** | 1.17133 | 0.19586 | 0.13459 | 0.82256 | 1.02885 | 0.49399 | 0.51928 |
| **DSEL** | 0.05393 | 0.00000 | 1.56979 | 0.79705 | 2.65055 | 1.07147 | 0.19165 |
| **DSG1** | 0.00102 | 0.00949 | 0.12287 | 0.05782 | 0.06549 | 0.01849 | 0.07732 |
| **DSG2** | 33.03544 | 3.52229 | 4.76010 | 8.01466 | 0.21848 | 6.08878 | 30.42061 |
| **DSG3** | 1.55843 | 0.00000 | 0.05889 | 0.83649 | 0.03272 | 0.00818 | 0.00956 |
| **DSG4** | 0.02938 | 0.02074 | 0.01223 | 0.02999 | 0.00825 | 0.01640 | 0.00639 |
| **DSN1** | 12.71796 | 10.90047 | 5.37570 | 4.18314 | 2.63727 | 5.38300 | 0.00000 |

| | | | | | | | |
|---|---|---|---|---|---|---|---|
| **DSP** | 25.59885 | 21.49180 | 6.90323 | 30.92265 | 0.00000 | 16.23363 | 31.32424 |
| **DSPP** | 0.02629 | 0.00355 | 0.03593 | 0.02257 | 0.00816 | 0.05753 | 0.04481 |
| **DST** | 2.07988 | 8.80025 | 2.99260 | 1.28621 | 0.74729 | 16.44629 | 14.76993 |
| **DSTN** | 30.89482 | 6.33755 | 6.96266 | 22.71749 | 4.20070 | 15.28358 | 17.97277 |
| **DSTNP2** | 1.70358 | 2.49841 | 2.49645 | 0.79359 | 0.37980 | 5.68423 | 0.00223 |
| **DSTYK** | 3.09304 | 2.70438 | 1.43656 | 0.00000 | 0.93919 | 1.09155 | 3.93382 |
| **DTD1** | 10.01830 | 6.92899 | 2.49428 | 2.91319 | 0.70695 | 4.53597 | 2.87916 |
| **DTD2** | 1.08380 | 0.78183 | 0.45762 | 0.07953 | 0.07474 | 0.31336 | 1.16433 |
| **DTL** | 22.41762 | 9.14183 | 16.98270 | 11.08376 | 9.72550 | 12.93990 | 0.00000 |
| **DTNA** | 0.01592 | 0.00000 | 0.01568 | 0.14241 | 0.01092 | 0.31236 | 0.11227 |
| **DTNB** | 6.23689 | 7.59904 | 2.47339 | 0.00019 | 0.87738 | 4.57240 | 0.34608 |
| **DTNBP1** | 0.32386 | 1.72231 | 0.96800 | 1.20062 | 0.84381 | 2.26952 | 2.64587 |
| **DTWD1** | 5.61995 | 0.13743 | 2.56676 | 2.70196 | 0.00000 | 1.49442 | 2.52466 |
| **DTWD2** | 0.48080 | 0.00000 | 0.93102 | 0.07282 | 0.39621 | 1.24445 | 1.23487 |
| **DTX1** | 0.00889 | 0.26187 | 0.00147 | 0.02932 | 0.05162 | 0.02205 | 0.00216 |
| **DTX2** | 0.00000 | 3.77761 | 2.00957 | 3.76417 | 1.97670 | 0.99432 | 4.13256 |
| **DTX2P1-UPK3B** | 0.13828 | 0.14949 | 0.15938 | 0.17927 | 0.06594 | 0.07062 | 0.01650 |
| **DTX3** | 0.00696 | 0.02104 | 0.00878 | 0.04599 | 0.00738 | 0.03872 | 0.14542 |
| **DTX3L** | 0.01277 | 0.30619 | 0.04589 | 0.01974 | 0.00589 | 1.42875 | 0.00000 |
| **DTX4** | 5.21931 | 3.27559 | 0.02294 | 7.37967 | 0.00000 | 0.43131 | 9.18655 |
| **DTYMK** | 26.57870 | 29.50327 | 14.29751 | 22.09276 | 1.85401 | 11.47973 | 6.20675 |
| **DUOX1** | 0.09536 | 0.09397 | 0.03023 | 0.34841 | 0.02222 | 0.08550 | 0.11417 |
| **DUOX2** | 0.00000 | 0.02027 | 0.00548 | 0.08588 | 0.06176 | 0.00361 | 0.14123 |
| **DUOXA1** | 0.02063 | 0.01315 | 0.01411 | 0.68152 | 0.00868 | 0.01385 | 0.02549 |
| **DUOXA2** | 0.02226 | 0.01746 | 0.01247 | 0.01908 | 0.00786 | 0.01569 | 0.02476 |
| **DUPD1** | 0.03908 | 0.01958 | 0.01721 | 0.01049 | 0.03083 | 0.00973 | 0.07457 |
| **DUS1L** | 4.49414 | 10.81057 | 2.15380 | 2.65999 | 0.03937 | 3.57205 | 22.83489 |
| **DUS2** | 21.27790 | 10.46848 | 6.44995 | 14.07898 | 0.51968 | 6.10392 | 0.00000 |
| **DUS3L** | 9.73467 | 22.66383 | 8.18842 | 13.49355 | 2.61392 | 5.60448 | 7.64288 |
| **DUS4L** | 1.83435 | 0.99670 | 4.78706 | 1.39166 | 0.60543 | 1.23203 | 1.53594 |
| **DUSP1** | 0.00000 | 11.34377 | 0.99069 | 3.11559 | 0.25277 | 1.02806 | 5.79076 |
| **DUSP10** | 0.22080 | 0.17236 | 0.00000 | 0.04007 | 2.18288 | 1.17031 | 0.00000 |
| **DUSP11** | 15.84863 | 4.10355 | 10.51937 | 9.37914 | 4.50280 | 5.80537 | 0.00000 |
| **DUSP12** | 9.14912 | 8.68849 | 13.40949 | 3.22449 | 11.28640 | 4.64766 | 0.41169 |
| **DUSP13** | 0.48170 | 0.08011 | 0.01899 | 2.71963 | 0.02475 | 0.01941 | 0.00000 |

| | | | | | | | |
|---|---|---|---|---|---|---|---|
| **DUSP14** | 20.77267 | 22.21644 | 18.27674 | 17.94076 | 6.13143 | 5.13521 | 0.35382 |
| **DUSP15** | 0.02422 | 0.01251 | 0.02943 | 0.02007 | 0.01911 | 0.05847 | 0.01568 |
| **DUSP16** | 0.86891 | 5.25982 | 3.90286 | 0.00000 | 0.37155 | 9.35453 | 5.66903 |
| **DUSP18** | 0.58854 | 2.35568 | 7.60563 | 1.77109 | 6.62668 | 2.12441 | 1.38604 |
| **DUSP19** | 0.51226 | 0.00000 | 0.91864 | 0.36288 | 2.07002 | 0.04134 | 0.00000 |
| **DUSP2** | 0.00000 | 0.47584 | 0.00071 | 0.02119 | 0.00849 | 0.03639 | 0.06159 |
| **DUSP22** | 0.62136 | 2.22880 | 1.35661 | 0.94517 | 2.18970 | 1.63266 | 0.98871 |
| **DUSP23** | 3.65958 | 7.81479 | 2.20419 | 5.38677 | 0.78748 | 3.02790 | 0.00001 |
| **DUSP26** | 0.02000 | 0.01553 | 0.03629 | 0.03768 | 0.02891 | 0.05073 | 0.01060 |
| **DUSP27** | 0.25522 | 1.23451 | 7.20482 | 0.00000 | 0.00000 | 0.00000 | 0.99503 |
| **DUSP28** | 0.12621 | 0.48406 | 0.24786 | 0.40018 | 0.05419 | 0.52198 | 1.20022 |
| **DUSP3** | 5.64915 | 6.01233 | 7.23050 | 8.66298 | 1.87490 | 3.27185 | 0.87960 |
| **DUSP4** | 0.00000 | 10.66215 | 4.63452 | 1.38346 | 0.19127 | 6.59253 | 0.00000 |
| **DUSP5** | 0.00000 | 4.97486 | 8.05781 | 10.73741 | 1.79521 | 1.96842 | 11.53432 |
| **DUSP5P1** | 0.12392 | 0.03312 | 0.02579 | 0.17154 | 0.00749 | 0.00000 | 0.20547 |
| **DUSP6** | 33.29410 | 0.77496 | 0.00000 | 6.98827 | 0.02467 | 2.12069 | 5.90555 |
| **DUSP7** | 3.13753 | 7.18695 | 1.87361 | 0.08440 | 3.41608 | 6.29785 | 3.43973 |
| **DUSP8** | 0.00000 | 0.11604 | 0.09001 | 0.19391 | 0.10423 | 0.00000 | 1.09182 |
| **DUSP9** | 0.00000 | 6.57398 | 0.00539 | 0.00000 | 0.05257 | 0.22899 | 0.33641 |
| **DUT** | 25.03615 | 9.13263 | 18.21382 | 16.06492 | 3.60839 | 10.89206 | 21.05105 |
| **DUXA** | 0.00000 | 0.00395 | 12.19280 | 0.00206 | 26.46736 | 0.00000 | 0.03012 |
| **DUXAP8** | 0.02856 | 0.03496 | 0.08695 | 0.06210 | 0.01248 | 0.00549 | 0.01625 |
| **DVL1** | 1.26052 | 1.56281 | 0.00000 | 2.40595 | 0.07011 | 1.68183 | 21.73377 |
| **DVL2** | 0.00000 | 3.53458 | 1.04252 | 2.80812 | 0.56647 | 7.71209 | 9.87068 |
| **DVL3** | 1.55879 | 5.66164 | 3.81314 | 8.38008 | 0.05212 | 3.45926 | 34.62965 |
| **DXO** | 4.74103 | 1.17440 | 1.45657 | 8.25626 | 0.63436 | 1.93130 | 0.18393 |
| **DYDC2** | 0.09127 | 0.10234 | 0.00850 | 0.00415 | 0.12024 | 0.00503 | 0.02253 |
| **DYM** | 10.00358 | 1.57349 | 2.32829 | 5.82872 | 1.52022 | 5.13937 | 5.69598 |
| **DYNAP** | 0.02327 | 0.02885 | 0.02750 | 0.01520 | 0.05628 | 0.06579 | 0.01415 |
| **DYNC1H1** | 16.07329 | 15.67783 | 1.92104 | 16.92711 | 2.94691 | 12.02683 | 20.50726 |
| **DYNC1I1** | 0.24502 | 0.16346 | 0.28803 | 0.06175 | 0.30034 | 0.39697 | 0.00000 |
| **DYNC1I2** | 8.71929 | 8.77006 | 1.85345 | 8.15673 | 5.17098 | 5.50329 | 9.81052 |
| **DYNC1LI1** | 14.72879 | 5.72702 | 24.20285 | 10.91220 | 2.65265 | 5.69888 | 5.68182 |
| **DYNC1LI2** | 21.52301 | 12.19696 | 11.85284 | 17.50253 | 2.82463 | 9.79970 | 15.51930 |
| **DYNC2H1** | 1.66729 | 0.00000 | 0.42726 | 0.53794 | 3.60667 | 0.92287 | 1.66923 |

| | | | | | | | |
|---|---|---|---|---|---|---|---|
| **DYNC2LI1** | 2.79064 | 0.00008 | 1.68812 | 0.72330 | 0.53158 | 0.96738 | 1.43241 |
| **DYNLL1** | 17.85897 | 8.82555 | 14.52476 | 22.42833 | 11.61229 | 9.08462 | 13.30361 |
| **DYNLL1-AS1** | 1.31387 | 0.00130 | 3.13324 | 2.89324 | 8.06422 | 0.00000 | 0.00000 |
| **DYNLL2** | 0.06424 | 5.27046 | 0.43811 | 0.84127 | 4.97783 | 1.09513 | 9.70323 |
| **DYNLRB1** | 12.67616 | 11.24356 | 4.84790 | 14.60321 | 1.35550 | 6.98529 | 14.80377 |
| **DYNLRB2** | 0.03002 | 0.06414 | 0.06998 | 0.02457 | 0.07103 | 0.00000 | 0.27466 |
| **DYNLT1** | 17.90440 | 9.15672 | 15.85394 | 12.95405 | 1.19262 | 7.15448 | 4.94772 |
| **DYNLT3** | 0.63009 | 0.31304 | 4.58344 | 0.59120 | 1.15354 | 2.07156 | 0.31094 |
| **DYRK1A** | 3.54874 | 0.00000 | 3.73279 | 1.73409 | 0.53824 | 2.83632 | 12.08454 |
| **DYRK1B** | 0.00000 | 0.86713 | 0.24268 | 0.83148 | 0.13600 | 0.19793 | 1.75639 |
| **DYRK2** | 1.89588 | 0.53395 | 2.89508 | 2.41308 | 1.27452 | 1.54709 | 6.69541 |
| **DYRK3** | 0.00478 | 0.02852 | 0.01379 | 0.06469 | 0.16623 | 0.34142 | 0.02635 |
| **DYRK4** | 1.62470 | 2.12831 | 2.79854 | 0.01749 | 0.35569 | 1.19709 | 0.60403 |
| **DYSF** | 13.73935 | 0.00000 | 0.00000 | 7.24074 | 1.40014 | 3.26121 | 19.42342 |
| **DYX1C1** | 0.08110 | 0.00000 | 0.48664 | 0.05419 | 0.03602 | 0.04280 | 0.05182 |
| **DZANK1** | 2.11099 | 0.65693 | 0.37405 | 0.63709 | 0.98682 | 2.24859 | 0.00000 |
| **DZIP1** | 0.02408 | 0.23917 | 0.21875 | 0.00000 | 1.42787 | 0.56124 | 0.60344 |
| **DZIP1L** | 0.00000 | 0.10184 | 0.59176 | 0.05816 | 0.02524 | 1.30518 | 0.03920 |
| **DZIP3** | 3.84419 | 2.08005 | 2.60529 | 1.73967 | 1.48194 | 2.37096 | 2.13138 |
| **E2F1** | 0.39512 | 5.21882 | 2.03472 | 3.10320 | 0.35948 | 4.65346 | 9.44637 |
| **E2F2** | 0.56587 | 4.27497 | 0.66833 | 2.34494 | 0.00000 | 1.50507 | 4.15779 |
| **E2F3** | 5.06992 | 0.48995 | 1.30648 | 3.09223 | 0.39417 | 1.69886 | 9.95806 |
| **E2F4** | 6.21183 | 14.13844 | 6.25961 | 11.96142 | 0.00000 | 12.05126 | 20.27154 |
| **E2F5** | 0.11627 | 0.02831 | 0.06358 | 0.22679 | 0.06362 | 0.11641 | 0.86024 |
| **E2F6** | 1.15881 | 0.96825 | 1.20826 | 1.05682 | 0.95943 | 0.67792 | 2.33966 |
| **E2F7** | 2.12006 | 1.35585 | 0.52836 | 3.36251 | 0.29983 | 0.93565 | 1.39083 |
| **E2F8** | 8.58506 | 2.12473 | 1.94029 | 11.36222 | 0.22471 | 1.03971 | 0.90250 |
| **E4F1** | 1.02082 | 14.46373 | 8.47677 | 8.40283 | 3.05305 | 7.85219 | 11.86359 |
| **EAF1** | 20.34134 | 4.71057 | 6.98172 | 19.84323 | 7.82639 | 5.02984 | 0.00000 |
| **EAF2** | 5.11818 | 0.06451 | 1.02625 | 3.53452 | 0.07291 | 0.05329 | 0.00000 |
| **EAPP** | 13.79125 | 3.89358 | 12.17070 | 11.95780 | 3.58133 | 7.77349 | 2.08469 |
| **EARS2** | 14.34696 | 13.56126 | 5.23097 | 7.88102 | 1.31743 | 8.00914 | 0.00000 |
| **EBAG9** | 9.51135 | 5.02116 | 6.50830 | 3.46840 | 1.85862 | 2.18248 | 3.40946 |
| **EBF1** | 0.00899 | 0.05703 | 0.02928 | 0.01255 | 3.00103 | 0.02494 | 0.06227 |
| **EBF2** | 0.01080 | 0.16555 | 0.00000 | 0.01371 | 2.79800 | 0.03418 | 0.00156 |

| | | | | | | | |
|---|---|---|---|---|---|---|---|
| **EBF3** | 0.02745 | 0.06712 | 0.06794 | 0.03168 | 0.13992 | 0.13171 | 0.00000 |
| **EBF4** | 0.00000 | 0.15158 | 0.01279 | 0.03963 | 0.00884 | 0.03971 | 0.11610 |
| **EBI3** | 0.00000 | 0.28649 | 0.06254 | 0.80514 | 0.02834 | 0.01122 | 0.00000 |
| **EBLN1** | 0.09210 | 0.02624 | 0.03225 | 0.03572 | 2.12263 | 0.00000 | 0.06212 |
| **EBLN2** | 0.77413 | 0.49564 | 0.50115 | 0.35495 | 0.26741 | 0.42118 | 2.47840 |
| **EBLN3** | 7.68076 | 0.00000 | 12.61228 | 6.05478 | 6.28851 | 3.59308 | 5.05149 |
| **EBNA1BP2** | 49.32175 | 39.83022 | 24.65025 | 43.45181 | 15.52558 | 15.94616 | 13.95062 |
| **EBP** | 72.15880 | 53.74217 | 10.10965 | 26.42197 | 12.98989 | 26.07290 | 16.96119 |
| **EBPL** | 1.76928 | 0.70532 | 1.77125 | 2.64166 | 0.53939 | 2.90699 | 10.23236 |
| **ECD** | 15.84724 | 3.25346 | 10.22963 | 9.23582 | 0.94285 | 3.06451 | 0.00000 |
| **ECE1** | 0.38720 | 4.01306 | 0.45793 | 5.95681 | 0.32170 | 6.22979 | 7.08779 |
| **ECE2** | 17.30487 | 14.04630 | 11.83358 | 22.42435 | 1.51907 | 6.46828 | 0.00024 |
| **ECEL1** | 0.02458 | 0.03165 | 0.00651 | 0.00199 | 0.00789 | 0.06073 | 0.15517 |
| **ECEL1P2** | 0.00000 | 0.11447 | 0.05817 | 2.40166 | 0.01619 | 0.00944 | 2.21486 |
| **ECH1** | 24.88320 | 14.03987 | 7.82507 | 17.47758 | 3.22197 | 9.32267 | 1.03297 |
| **ECHDC1** | 6.88784 | 0.00000 | 8.11874 | 9.81714 | 1.29145 | 1.28088 | 0.00106 |
| **ECHDC2** | 3.48877 | 2.40109 | 1.60630 | 7.21328 | 0.10600 | 0.00000 | 5.05736 |
| **ECHDC3** | 0.25872 | 0.33081 | 0.05264 | 0.68271 | 0.04759 | 0.00000 | 1.37584 |
| **ECHS1** | 50.32278 | 34.16341 | 0.38472 | 33.20975 | 1.82332 | 12.43655 | 22.83779 |
| **ECI1** | 0.64507 | 6.40294 | 1.40541 | 6.45975 | 0.12410 | 5.14556 | 23.22119 |
| **ECI2** | 14.48014 | 0.00000 | 1.10328 | 2.25456 | 0.82530 | 1.21246 | 0.61417 |
| **ECM1** | 2.65281 | 0.00000 | 0.02754 | 2.08149 | 0.12055 | 0.09134 | 0.90416 |
| **ECM2** | 0.35089 | 0.00000 | 0.28121 | 0.06347 | 0.05399 | 0.29997 | 0.59163 |
| **ECSCR** | 0.03647 | 0.01393 | 0.00714 | 0.01761 | 0.12776 | 0.01577 | 0.01031 |
| **ECSIT** | 16.83681 | 16.15284 | 5.10867 | 14.56821 | 2.20019 | 8.65916 | 4.92153 |
| **ECT2** | 18.49984 | 0.00000 | 14.62446 | 8.58709 | 7.72637 | 11.13557 | 8.69482 |
| **ECT2L** | 0.09570 | 0.00000 | 0.08672 | 0.04822 | 0.09511 | 0.06244 | 0.17496 |
| **EDA** | 0.18360 | 0.12463 | 2.75777 | 0.00000 | 1.29403 | 0.76610 | 0.34351 |
| **EDA2R** | 1.92053 | 3.88517 | 0.00000 | 2.68434 | 0.16949 | 2.33275 | 0.12063 |
| **EDAR** | 0.03872 | 0.06505 | 0.00000 | 0.00000 | 0.54775 | 0.00576 | 0.25326 |
| **EDARADD** | 46.75241 | 31.00610 | 17.84352 | 35.35595 | 4.83086 | 20.99580 | 1.78428 |
| **EDC3** | 10.57695 | 9.57070 | 9.73877 | 17.08440 | 4.69089 | 4.20080 | 4.27262 |
| **EDC4** | 3.43926 | 13.80815 | 1.43538 | 9.79825 | 0.46811 | 6.64165 | 18.03554 |
| **EDDM3A** | 0.09559 | 0.01107 | 0.01404 | 0.02910 | 0.00812 | 0.01723 | 0.00000 |
| **EDDM3B** | 0.12192 | 0.00627 | 0.01228 | 0.03165 | 0.00716 | 0.00593 | 0.03811 |

| | | | | | | | |
|---|---|---|---|---|---|---|---|
| **EDEM1** | 9.04639 | 3.76976 | 0.91552 | 5.92183 | 2.42044 | 5.27562 | 0.83618 |
| **EDEM2** | 13.27936 | 10.15900 | 5.67732 | 10.97903 | 3.37222 | 7.24939 | 0.00000 |
| **EDEM3** | 8.85233 | 0.00000 | 2.81857 | 3.29061 | 0.80021 | 2.88673 | 3.06086 |
| **EDF1** | 27.73742 | 26.47359 | 9.02461 | 36.36700 | 3.35837 | 15.38271 | 30.44809 |
| **EDIL3** | 0.24521 | 0.00000 | 0.01582 | 0.52196 | 0.22981 | 0.04082 | 0.00000 |
| **EDN1** | 0.00000 | 0.17593 | 0.35777 | 0.08587 | 0.03209 | 0.29789 | 0.22455 |
| **EDN3** | 0.00485 | 0.10237 | 0.00624 | 0.01441 | 0.03351 | 0.20029 | 0.01488 |
| **EDNRA** | 0.03116 | 0.01160 | 0.00953 | 0.03855 | 0.03754 | 0.10043 | 0.02404 |
| **EDNRB** | 1.94166 | 0.02439 | 0.03697 | 7.97762 | 0.12179 | 0.02401 | 0.00013 |
| **EDNRB-AS1** | 0.01949 | 0.00000 | 0.05431 | 0.02682 | 0.47198 | 0.01724 | 0.08556 |
| **EDRF1** | 8.78702 | 1.02300 | 2.97696 | 5.72442 | 3.80496 | 1.59561 | 3.37541 |
| **EDRF1-AS1** | 0.08847 | 0.02117 | 0.06003 | 0.04817 | 0.01664 | 0.02845 | 0.06741 |
| **EEA1** | 13.04530 | 0.00000 | 7.62524 | 5.30535 | 1.33469 | 8.51152 | 23.72636 |
| **EED** | 10.29072 | 2.75526 | 7.17865 | 9.76150 | 1.24823 | 2.84217 | 3.21060 |
| **EEF1A1** | 29.13911 | 11.35861 | 16.45285 | 21.91632 | 3.75323 | 15.33668 | 13.70175 |
| **EEF1A2** | 0.00000 | 0.15490 | 0.06396 | 0.05345 | 0.26815 | 0.02817 | 0.42583 |
| **EEF1B2** | 29.65188 | 26.53937 | 12.35558 | 23.93705 | 7.88581 | 16.37613 | 15.66098 |
| **EEF1D** | 33.57547 | 34.83658 | 12.15155 | 29.59251 | 4.01544 | 18.92148 | 42.90712 |
| **EEF1DP3** | 0.01990 | 0.10194 | 0.03057 | 0.01351 | 0.06722 | 0.04413 | 0.01918 |
| **EEF1E1.uid-815** | 0.04567 | 0.05294 | 0.00850 | 0.06042 | 0.01910 | 0.00203 | 0.05839 |
| **EEF1E1.uid-816** | 17.45195 | 0.00000 | 5.88850 | 10.06484 | 0.70996 | 3.62257 | 8.19843 |
| **EEF1G** | 30.43988 | 28.40918 | 12.79247 | 35.15584 | 7.76387 | 18.73242 | 29.35845 |
| **EEF2** | 40.87202 | 44.59487 | 30.56527 | 59.36248 | 15.14491 | 32.95112 | 64.94592 |
| **EEF2K** | 4.32955 | 0.58215 | 1.43107 | 2.23990 | 2.49796 | 1.45450 | 7.86521 |
| **EEFSEC** | 5.43074 | 4.64092 | 0.36814 | 1.77508 | 0.08106 | 3.03746 | 2.79177 |
| **EEPD1** | 0.11635 | 0.00000 | 0.93189 | 0.69634 | 1.68211 | 0.29614 | 0.00000 |
| **EFCAB1** | 0.02508 | 0.00517 | 0.03957 | 0.01937 | 0.03394 | 0.01262 | 0.01984 |
| **EFCAB10** | 0.15574 | 0.00000 | 0.25156 | 0.04303 | 0.77474 | 0.15345 | 0.07641 |
| **EFCAB11** | 6.10591 | 1.54672 | 4.40022 | 3.73932 | 2.28299 | 3.74590 | 1.93471 |
| **EFCAB12** | 0.00342 | 0.08301 | 0.01379 | 0.21131 | 0.35390 | 0.01279 | 0.00000 |
| **EFCAB13** | 0.43831 | 0.00000 | 0.94322 | 0.05834 | 0.35140 | 0.45730 | 0.74018 |
| **EFCAB14** | 10.94534 | 3.85071 | 9.87264 | 7.34945 | 7.04077 | 7.66510 | 13.40388 |
| **EFCAB14-AS1** | 0.05247 | 0.02281 | 0.02504 | 0.02913 | 0.01965 | 0.04605 | 0.01650 |
| **EFCAB2** | 2.77151 | 0.11718 | 0.87413 | 2.56538 | 0.30301 | 1.60961 | 2.01100 |
| **EFCAB3** | 0.03028 | 0.01644 | 0.01274 | 0.01636 | 0.00796 | 0.01270 | 0.02064 |

| | | | | | | | |
|---|---|---|---|---|---|---|---|
| **EFCAB4A** | 1.45211 | 0.48809 | 0.00000 | 0.00000 | 0.14994 | 0.69565 | 0.50745 |
| **EFCAB4B** | 0.07636 | 0.09745 | 0.00000 | 0.03804 | 0.09588 | 0.02335 | 0.00000 |
| **EFCAB5** | 0.11341 | 0.05908 | 0.01906 | 0.02914 | 0.10909 | 0.05689 | 0.04077 |
| **EFCAB6** | 0.03362 | 0.01786 | 0.21255 | 0.06547 | 0.90477 | 0.04196 | 0.00000 |
| **EFCAB7** | 1.45631 | 0.00000 | 0.63892 | 1.33964 | 0.79094 | 0.09888 | 0.00000 |
| **EFCAB9** | 0.02751 | 0.01148 | 0.01355 | 0.01807 | 0.01113 | 0.09869 | 0.04280 |
| **EFCC1** | 0.01836 | 0.00953 | 0.02465 | 0.02655 | 0.05310 | 0.11676 | 0.07314 |
| **EFEMP1** | 0.66059 | 0.32251 | 0.02636 | 0.08364 | 0.00368 | 1.42417 | 0.00000 |
| **EFEMP2** | 0.00000 | 0.00000 | 0.39743 | 0.05724 | 0.00908 | 0.15022 | 0.25025 |
| **EFHB** | 0.02727 | 0.01018 | 0.03724 | 0.03848 | 0.09056 | 0.01073 | 0.01995 |
| **EFHC1** | 1.79357 | 0.11645 | 1.53568 | 2.12657 | 1.74744 | 1.56187 | 0.12071 |
| **EFHC2** | 0.04069 | 0.00000 | 0.21340 | 0.03662 | 1.05696 | 0.03189 | 0.02883 |
| **EFHD1** | 5.34494 | 16.16597 | 0.00000 | 0.83322 | 0.71836 | 0.67477 | 9.84296 |
| **EFHD2** | 0.24342 | 7.95439 | 0.01518 | 2.06380 | 0.36006 | 2.98674 | 16.21447 |
| **EFNA1** | 38.20000 | 18.32913 | 0.00000 | 27.60862 | 1.03683 | 0.00000 | 3.46804 |
| **EFNA2** | 0.10357 | 0.15794 | 1.04414 | 0.01287 | 0.03352 | 0.18512 | 0.32037 |
| **EFNA3** | 0.35504 | 1.34008 | 0.38709 | 0.44074 | 0.52197 | 0.34057 | 0.80818 |
| **EFNA4** | 7.02153 | 9.84288 | 12.20858 | 2.62813 | 0.80590 | 2.48153 | 2.41412 |
| **EFNA5** | 0.16527 | 0.08816 | 0.19762 | 0.01687 | 0.55245 | 0.49245 | 0.24520 |
| **EFNB1** | 0.00000 | 1.27845 | 0.31588 | 0.00000 | 0.59033 | 0.52776 | 0.94990 |
| **EFNB2** | 2.86045 | 1.85703 | 0.00000 | 0.00000 | 3.00537 | 1.76661 | 7.79412 |
| **EFNB3** | 0.00000 | 0.00000 | 2.31699 | 0.01231 | 1.32786 | 0.18830 | 0.36584 |
| **EFR3A** | 10.71719 | 1.15889 | 2.52678 | 1.36583 | 0.75161 | 2.83166 | 8.36577 |
| **EFR3B** | 7.27644 | 20.21098 | 6.84348 | 0.00000 | 1.22053 | 11.40171 | 0.00000 |
| **EFS** | 0.49736 | 0.00000 | 0.18427 | 1.36191 | 0.10574 | 0.00000 | 5.88183 |
| **EFTUD1** | 4.53563 | 1.10524 | 1.11271 | 2.37023 | 1.90237 | 2.82060 | 1.90544 |
| **EFTUD1P1** | 0.02761 | 0.01846 | 0.01808 | 0.01784 | 0.01865 | 0.01552 | 0.01959 |
| **EFTUD2** | 35.33044 | 21.79619 | 15.91667 | 24.81313 | 3.22047 | 13.50308 | 8.92851 |
| **EGF** | 0.02415 | 0.06545 | 0.00000 | 0.01789 | 0.97901 | 0.01296 | 0.01790 |
| **EGFEM1P** | 0.02176 | 0.03291 | 0.01208 | 0.01667 | 0.16080 | 0.02003 | 0.02431 |
| **EGFL6** | 0.00051 | 0.67246 | 0.33436 | 0.00000 | 0.05501 | 0.06084 | 0.03125 |
| **EGFL7** | 3.47397 | 8.47185 | 0.32207 | 6.14135 | 0.02849 | 5.22753 | 1.05482 |
| **EGFL8.uid-8475** | 0.02521 | 0.03280 | 0.02753 | 0.02115 | 0.00714 | 0.01207 | 0.01742 |
| **EGFLAM** | 0.97139 | 5.14617 | 2.79090 | 0.00000 | 0.00000 | 8.42217 | 2.32191 |
| **EGFLAM-AS2** | 0.01896 | 0.02024 | 0.01862 | 0.01799 | 0.00710 | 0.02073 | 0.02916 |

| | | | | | | | |
|---|---|---|---|---|---|---|---|
| **EGFLAM-AS4** | 0.06482 | 0.01055 | 0.11543 | 0.02582 | 0.01079 | 0.04209 | 0.10079 |
| **EGFR** | 0.22455 | 0.00000 | 0.00000 | 0.61096 | 0.27462 | 0.00000 | 0.98477 |
| **EGFR-AS1** | 0.02344 | 0.01318 | 0.01222 | 0.02051 | 0.00800 | 0.04801 | 0.02982 |
| **EGLN1** | 2.06221 | 0.09018 | 2.99613 | 0.08840 | 0.27537 | 0.72100 | 7.51076 |
| **EGLN2.uid-2379** | 0.16573 | 0.64150 | 0.27672 | 0.56421 | 0.10261 | 0.26342 | 0.00000 |
| **EGLN2.uid-2379** | 0.04575 | 0.07376 | 0.02316 | 0.04679 | 0.00629 | 0.03233 | 0.01292 |
| **EGLN3** | 16.57731 | 29.87397 | 4.56309 | 18.48073 | 1.10245 | 35.92302 | 0.00000 |
| **EGR1** | 0.00000 | 0.12856 | 0.16477 | 2.05512 | 5.94029 | 4.23118 | 1.36553 |
| **EGR2** | 0.00924 | 0.00009 | 0.00487 | 0.87840 | 1.08663 | 0.51992 | 0.00510 |
| **EGR3** | 0.04241 | 0.00000 | 0.01696 | 0.10161 | 0.01893 | 0.04142 | 0.08565 |
| **EGR4** | 0.01673 | 0.02144 | 0.00974 | 0.02086 | 0.01972 | 0.04664 | 0.02390 |
| **EHBP1** | 4.37454 | 1.09900 | 3.88540 | 2.90182 | 2.05624 | 3.20053 | 5.91267 |
| **EHBP1L1** | 0.00000 | 0.01116 | 0.00517 | 0.20402 | 0.06489 | 0.43855 | 0.42720 |
| **EHD1** | 4.64453 | 15.45702 | 2.87606 | 19.19064 | 5.27691 | 0.71699 | 10.48023 |
| **EHD2** | 0.00000 | 0.38715 | 0.42091 | 0.00000 | 0.03301 | 0.01223 | 0.19146 |
| **EHD3** | 0.02133 | 0.67710 | 0.07214 | 1.09248 | 0.17336 | 0.99804 | 0.00000 |
| **EHD4** | 32.19777 | 21.64600 | 1.02173 | 18.16648 | 0.40143 | 5.40169 | 8.74051 |
| **EHF** | 0.01393 | 0.10919 | 0.00303 | 0.01320 | 0.01257 | 0.03063 | 0.03222 |
| **EHHADH** | 0.29645 | 0.53862 | 1.03376 | 0.04493 | 0.02451 | 1.52572 | 0.00000 |
| **EHMT1** | 3.79449 | 5.02684 | 3.63702 | 6.35325 | 3.18471 | 5.35503 | 15.21758 |
| **EHMT1-IT1** | 0.00000 | 0.03940 | 0.01528 | 0.22903 | 0.11209 | 0.09302 | 0.18650 |
| **EHMT2** | 2.69312 | 10.28111 | 12.92899 | 4.47872 | 1.71329 | 7.41547 | 21.39983 |
| **EI24** | 12.69139 | 14.63063 | 9.16223 | 9.40927 | 0.65469 | 12.27538 | 27.83427 |
| **EID1** | 21.89734 | 15.59309 | 14.06731 | 13.73296 | 2.44909 | 12.28230 | 10.68978 |
| **EID2** | 0.67719 | 0.48164 | 1.30725 | 0.10139 | 0.01416 | 0.45382 | 4.96666 |
| **EID2B** | 0.04757 | 0.00000 | 0.50074 | 0.03987 | 0.31926 | 0.22128 | 0.02006 |
| **EID3** | 0.15717 | 0.00000 | 9.33868 | 0.15344 | 5.17728 | 0.46500 | 1.57721 |
| **EIF1** | 45.95518 | 31.56668 | 38.53776 | 51.42442 | 23.70384 | 29.69319 | 46.06043 |
| **EIF1AD** | 25.13460 | 18.06789 | 13.34908 | 15.62870 | 19.04501 | 7.63309 | 0.00000 |
| **EIF1AX** | 19.70490 | 5.00762 | 10.92192 | 5.67140 | 4.23773 | 7.44551 | 23.15993 |
| **EIF1AX-AS1** | 0.05542 | 0.00000 | 0.05560 | 0.01621 | 0.03368 | 0.12232 | 0.06163 |
| **EIF1AY** | 0.04872 | 0.00224 | 0.04581 | 0.04344 | 0.00653 | 0.02533 | 0.04987 |
| **EIF1B** | 26.73016 | 7.14973 | 19.45723 | 21.35647 | 14.50419 | 10.99975 | 15.52493 |
| **EIF1B-AS1** | 0.07307 | 0.13569 | 0.00707 | 0.01259 | 0.31236 | 0.05721 | 0.00000 |
| **EIF2A** | 30.49512 | 3.46018 | 14.30799 | 19.54362 | 6.65895 | 8.37587 | 4.13743 |

| | | | | | | | |
|---|---|---|---|---|---|---|---|
| **EIF2AK1** | 5.00323 | 6.24768 | 2.32989 | 16.05866 | 0.22594 | 6.44572 | 22.36571 |
| **EIF2AK2** | 1.97723 | 0.00000 | 0.52184 | 3.69336 | 1.48425 | 0.23226 | 0.87380 |
| **EIF2AK3** | 1.70167 | 0.48744 | 0.04666 | 0.70093 | 0.41772 | 1.27178 | 2.61774 |
| **EIF2AK4** | 4.65280 | 7.33808 | 0.01882 | 5.88804 | 0.94177 | 8.72298 | 9.43986 |
| **EIF2B1** | 31.67557 | 15.16726 | 14.90916 | 21.49208 | 5.70906 | 8.38814 | 0.00000 |
| **EIF2B2** | 21.16455 | 12.62831 | 10.70205 | 26.54383 | 4.15954 | 6.00387 | 0.00000 |
| **EIF2B3** | 24.70509 | 15.91346 | 11.25210 | 9.69751 | 1.87271 | 5.72102 | 1.60478 |
| **EIF2B4** | 24.47468 | 24.96057 | 5.63073 | 18.86638 | 1.73503 | 12.82506 | 0.00000 |
| **EIF2B5** | 2.42640 | 6.88937 | 1.81879 | 2.67729 | 0.31674 | 7.81331 | 13.69380 |
| **EIF2D** | 5.46709 | 3.11357 | 1.89965 | 0.00000 | 0.46049 | 1.86162 | 0.96677 |
| **EIF2S1** | 59.90751 | 27.50177 | 25.65137 | 33.29866 | 12.50434 | 18.91283 | 0.00004 |
| **EIF2S2** | 20.67488 | 17.91552 | 13.16730 | 18.82114 | 4.43483 | 10.82570 | 25.04120 |
| **EIF2S3** | 56.48563 | 28.62682 | 26.26488 | 17.60323 | 10.75962 | 23.14633 | 24.43871 |
| **EIF3A** | 37.91356 | 22.25745 | 22.87983 | 27.99348 | 8.85450 | 19.01421 | 40.21257 |
| **EIF3B** | 19.09800 | 27.96822 | 11.28027 | 19.93251 | 4.08102 | 16.64746 | 42.02080 |
| **EIF3D** | 60.34535 | 39.33230 | 26.07867 | 42.86382 | 6.15444 | 27.29989 | 22.56876 |
| **EIF3E** | 46.48861 | 6.14049 | 21.03562 | 23.99652 | 2.85078 | 15.65043 | 12.39282 |
| **EIF3F** | 15.81351 | 10.99727 | 10.17245 | 19.42779 | 7.40916 | 9.16882 | 12.36604 |
| **EIF3G** | 31.21710 | 30.04184 | 21.51164 | 33.70969 | 11.75109 | 15.69939 | 11.21399 |
| **EIF3H** | 40.87218 | 19.18709 | 23.92598 | 27.12524 | 10.24943 | 21.57077 | 22.61646 |
| **EIF3I** | 53.53870 | 36.17353 | 26.91035 | 43.18420 | 14.91259 | 20.64477 | 5.82041 |
| **EIF3IP1** | 4.85438 | 3.02520 | 2.48394 | 3.86603 | 1.47958 | 1.48446 | 0.00000 |
| **EIF3J** | 9.71837 | 5.35679 | 11.76738 | 14.87275 | 7.83437 | 6.07326 | 44.73737 |
| **EIF3J-AS1** | 2.13537 | 0.38118 | 0.37440 | 3.75704 | 2.78031 | 0.10714 | 0.37228 |
| **EIF3K** | 43.52649 | 47.91770 | 28.27636 | 36.08379 | 10.77857 | 35.95916 | 24.89130 |
| **EIF3L** | 46.53634 | 21.24755 | 19.80894 | 35.39054 | 11.27173 | 21.42365 | 22.66528 |
| **EIF3M** | 49.27538 | 22.78143 | 23.76228 | 34.92132 | 7.88837 | 17.02296 | 17.08817 |
| **EIF4A2** | 32.29834 | 15.06120 | 20.87439 | 18.01924 | 15.24485 | 16.87098 | 24.17716 |
| **EIF4A3** | 40.32744 | 33.62807 | 23.38234 | 25.96818 | 27.67374 | 19.34476 | 13.76788 |
| **EIF4B** | 21.89270 | 5.88072 | 9.54352 | 13.51528 | 4.62780 | 7.28087 | 7.45926 |
| **EIF4E** | 23.35595 | 0.00000 | 11.07328 | 5.78642 | 3.41039 | 7.47858 | 7.50805 |
| **EIF4E1B** | 0.02249 | 0.06157 | 0.19703 | 0.08756 | 6.69313 | 0.06307 | 0.00000 |
| **EIF4E2** | 32.70762 | 26.33297 | 19.79104 | 25.54122 | 6.25056 | 13.55544 | 10.04973 |
| **EIF4E3** | 0.81758 | 0.39046 | 0.00000 | 0.25002 | 0.19291 | 1.48150 | 0.90316 |
| **EIF4EBP1** | 27.27655 | 38.26480 | 8.55040 | 21.05886 | 0.02820 | 20.78322 | 14.82150 |

| | | | | | | | |
|---|---|---|---|---|---|---|---|
| **EIF4EBP2** | 13.23222 | 7.54937 | 4.75573 | 12.63487 | 0.00000 | 17.52188 | 20.55894 |
| **EIF4EBP3** | 0.78840 | 1.36147 | 1.34736 | 1.09948 | 0.07802 | 0.40676 | 0.17556 |
| **EIF4ENIF1** | 3.65662 | 3.11784 | 2.99011 | 1.34125 | 5.00328 | 3.63587 | 8.04107 |
| **EIF4G1** | 33.02749 | 32.36414 | 27.56310 | 40.97352 | 8.32722 | 23.21088 | 47.24623 |
| **EIF4G2** | 78.20089 | 41.27526 | 38.66205 | 64.97640 | 22.03264 | 30.96233 | 27.02153 |
| **EIF4G3** | 7.64887 | 4.68820 | 5.88826 | 5.50741 | 1.10554 | 7.00267 | 21.49934 |
| **EIF4H** | 26.12969 | 31.21361 | 11.83281 | 21.06088 | 4.11011 | 13.25636 | 5.45144 |
| **EIF5** | 56.04874 | 49.21354 | 34.27151 | 46.82726 | 27.26555 | 22.88294 | 19.33406 |
| **EIF5A** | 35.86976 | 54.04727 | 20.03208 | 41.38146 | 15.88720 | 23.39039 | 38.98270 |
| **EIF5A2** | 5.82021 | 0.00000 | 10.35496 | 3.65304 | 6.67827 | 3.90467 | 0.00000 |
| **EIF5AL1** | 3.96417 | 2.37380 | 2.31617 | 2.43505 | 1.09227 | 2.43312 | 5.27622 |
| **EIF5B** | 31.37132 | 21.19147 | 22.41304 | 28.89645 | 15.97233 | 14.42563 | 45.63034 |
| **EIF6** | 41.32460 | 49.46250 | 20.11454 | 32.23919 | 14.58178 | 25.69508 | 16.18253 |
| **ELAC1** | 0.58162 | 1.32599 | 0.76970 | 0.00000 | 0.00000 | 0.10531 | 0.28956 |
| **ELAC2** | 33.93127 | 27.07269 | 15.43976 | 22.75781 | 8.43638 | 14.00808 | 6.61679 |
| **ELANE** | 0.01205 | 0.00000 | 0.37354 | 0.01486 | 1.45342 | 0.00291 | 0.02787 |
| **ELAVL1** | 3.68479 | 9.85201 | 2.67607 | 7.34361 | 0.77570 | 7.20153 | 28.13466 |
| **ELAVL2** | 0.00000 | 0.18189 | 0.38789 | 0.00000 | 0.76946 | 0.43776 | 0.11755 |
| **ELAVL3** | 0.00000 | 0.03448 | 0.11643 | 0.11045 | 0.55899 | 0.09252 | 0.07128 |
| **ELAVL4** | 0.02459 | 0.01799 | 0.01016 | 0.03066 | 0.03448 | 0.02178 | 0.01675 |
| **ELF1** | 8.79518 | 0.34708 | 4.63889 | 4.47577 | 5.31632 | 3.24709 | 8.43754 |
| **ELF2** | 3.78198 | 1.91801 | 3.29151 | 2.88060 | 1.66897 | 3.62322 | 8.61235 |
| **ELF3** | 28.48492 | 36.15361 | 2.25536 | 11.18615 | 0.00000 | 18.30536 | 27.18687 |
| **ELF4** | 2.74962 | 5.67966 | 0.31707 | 3.84589 | 0.02837 | 0.00000 | 6.92080 |
| **ELF5** | 0.16136 | 0.06759 | 0.01004 | 0.09163 | 0.04206 | 0.01106 | 0.00000 |
| **ELFN1** | 0.00000 | 0.17034 | 0.00000 | 0.00546 | 0.19629 | 0.28352 | 0.06251 |
| **ELFN1-AS1** | 0.02337 | 0.06393 | 0.00990 | 0.02116 | 0.00811 | 0.01124 | 0.00432 |
| **ELFN2** | 0.00000 | 0.04011 | 0.00000 | 0.11027 | 0.25003 | 0.02258 | 0.06105 |
| **ELK1** | 2.54650 | 2.00803 | 3.72828 | 3.96568 | 1.49472 | 3.08095 | 4.69143 |
| **ELK2AP** | 0.02362 | 0.01635 | 0.01484 | 0.02173 | 0.02058 | 0.01307 | 0.01582 |
| **ELK3** | 0.26176 | 0.48527 | 1.87641 | 0.15827 | 1.53233 | 1.85250 | 0.84316 |
| **ELK4** | 3.86533 | 0.30709 | 1.87533 | 0.85377 | 2.07362 | 1.48756 | 6.37790 |
| **ELL** | 2.79687 | 8.92177 | 3.31104 | 8.09042 | 2.49975 | 1.75783 | 4.67603 |
| **ELL2** | 1.17569 | 0.66224 | 1.02876 | 0.09104 | 2.62821 | 2.02516 | 2.59175 |
| **ELL3** | 0.69933 | 0.00000 | 9.75197 | 0.94225 | 3.81160 | 0.00000 | 0.50443 |

| | | | | | | | |
|---|---|---|---|---|---|---|---|
| **ELMO1** | 3.77901 | 0.82467 | 0.38658 | 8.87801 | 4.17897 | 2.66040 | 0.25554 |
| **ELMO1-AS1** | 0.02604 | 0.00189 | 0.01358 | 0.05185 | 0.00693 | 0.03328 | 0.12056 |
| **ELMO2** | 3.03203 | 4.09629 | 3.80381 | 2.46642 | 5.96589 | 4.65111 | 0.00000 |
| **ELMO3** | 0.31523 | 1.24154 | 0.10081 | 0.86012 | 0.01013 | 0.73505 | 1.72945 |
| **ELMOD1** | 0.08993 | 1.57051 | 0.79338 | 0.00000 | 0.35685 | 2.27029 | 0.79329 |
| **ELMOD2** | 8.91146 | 0.39466 | 5.42729 | 0.59350 | 0.27145 | 1.96306 | 1.70601 |
| **ELMOD3** | 1.85403 | 3.17707 | 1.29342 | 3.23963 | 1.12008 | 1.69181 | 0.00000 |
| **ELMSAN1** | 18.97590 | 10.16535 | 7.24129 | 15.13600 | 10.52029 | 5.18230 | 2.36959 |
| **ELN** | 0.00810 | 0.02666 | 0.01299 | 0.04331 | 0.00793 | 0.26112 | 0.02632 |
| **ELOF1** | 27.81869 | 28.20782 | 23.89392 | 25.68664 | 16.24939 | 12.47291 | 8.28757 |
| **ELOVL1** | 25.56956 | 14.77888 | 6.13460 | 30.98128 | 1.77947 | 3.63731 | 6.37422 |
| **ELOVL2** | 0.07898 | 0.01666 | 0.09806 | 0.10925 | 0.53677 | 0.03163 | 0.18543 |
| **ELOVL2-AS1** | 0.01271 | 0.07314 | 0.01286 | 0.01940 | 0.07104 | 0.01219 | 0.02200 |
| **ELOVL3** | 0.51450 | 0.40518 | 0.96219 | 0.55104 | 3.59644 | 0.31727 | 0.00000 |
| **ELOVL4** | 0.79936 | 0.30302 | 0.84533 | 1.26596 | 7.28723 | 2.81653 | 0.00000 |
| **ELOVL5** | 19.96052 | 0.00000 | 5.25815 | 16.88300 | 6.49605 | 5.86549 | 10.41951 |
| **ELOVL6** | 28.47033 | 0.00000 | 5.22283 | 10.57696 | 2.02366 | 3.37264 | 14.57173 |
| **ELOVL7** | 0.03955 | 0.10089 | 3.74065 | 0.03285 | 0.20687 | 0.34346 | 0.00227 |
| **ELP2** | 20.31911 | 7.88198 | 8.58134 | 12.72798 | 0.67647 | 4.46961 | 1.46665 |
| **ELP3** | 15.88608 | 5.33074 | 8.26656 | 5.91365 | 5.55616 | 2.84808 | 2.19735 |
| **ELP4** | 6.73413 | 0.85242 | 3.16688 | 2.62006 | 2.48025 | 2.21832 | 0.00000 |
| **ELP5** | 18.90988 | 16.01826 | 8.94938 | 13.53902 | 4.33486 | 8.64108 | 10.74565 |
| **ELP6** | 9.47167 | 8.58390 | 7.52513 | 8.53112 | 4.61707 | 3.15935 | 0.75399 |
| **ELSPBP1** | 0.03073 | 0.07069 | 0.04104 | 0.02413 | 0.18257 | 0.00499 | 0.00570 |
| **ELTD1** | 0.02830 | 0.01394 | 0.01182 | 0.12641 | 0.07005 | 0.01319 | 0.00000 |
| **EMB** | 0.03197 | 0.26282 | 0.04120 | 0.00979 | 0.15808 | 0.21301 | 0.05775 |
| **EMBP1** | 0.03445 | 0.00427 | 0.03923 | 0.00885 | 0.01257 | 0.02141 | 0.08285 |
| **EMC1** | 15.73127 | 9.47905 | 4.97425 | 14.65387 | 1.04123 | 8.20745 | 9.40237 |
| **EMC10** | 0.01612 | 2.47113 | 1.56917 | 0.69113 | 0.00000 | 3.71566 | 15.52020 |
| **EMC2** | 9.65601 | 1.11209 | 11.50223 | 4.30612 | 0.95366 | 4.99075 | 2.68921 |
| **EMC3** | 24.53882 | 20.82832 | 21.55053 | 16.90206 | 9.02396 | 7.67558 | 0.42408 |
| **EMC3-AS1** | 1.74404 | 2.51302 | 2.36164 | 4.12678 | 3.53568 | 0.18978 | 5.60011 |
| **EMC4** | 34.91507 | 20.98735 | 21.42155 | 29.09311 | 13.19778 | 13.42296 | 1.81077 |
| **EMC6** | 6.15094 | 9.98102 | 3.47517 | 6.58032 | 0.68146 | 2.73422 | 0.00002 |
| **EMC7** | 24.79124 | 14.11570 | 14.69539 | 23.82405 | 4.74864 | 9.52470 | 0.00004 |

| | | | | | | | |
|---|---|---|---|---|---|---|---|
| **EMC8** | 1.97766 | 12.38403 | 8.75455 | 9.99358 | 3.68148 | 11.29593 | 27.71764 |
| **EMC9** | 1.58258 | 3.78651 | 3.82458 | 2.73669 | 0.90788 | 3.61609 | 3.66806 |
| **EMCN** | 0.06765 | 0.00714 | 0.01416 | 0.00081 | 0.00842 | 0.01072 | 0.04487 |
| **EMD** | 6.38015 | 11.44466 | 3.23139 | 5.71846 | 0.85464 | 4.73208 | 11.62553 |
| **EME1** | 3.85336 | 6.71719 | 5.84568 | 1.21329 | 0.06810 | 0.52421 | 0.00000 |
| **EME2** | 0.00000 | 0.19095 | 0.03400 | 0.18055 | 0.00442 | 0.00578 | 0.23877 |
| **EMG1** | 27.49897 | 18.52517 | 19.60261 | 28.84667 | 4.70499 | 14.51470 | 3.93021 |
| **EMID1** | 0.00000 | 0.17925 | 0.01168 | 0.02727 | 0.00613 | 0.42963 | 0.03870 |
| **EMILIN1** | 0.00000 | 0.15206 | 0.05622 | 0.03379 | 0.01168 | 0.00875 | 0.07787 |
| **EMILIN2** | 3.80544 | 3.51495 | 0.27843 | 5.91122 | 3.79461 | 0.00000 | 12.40782 |
| **EMILIN3** | 0.12413 | 0.02864 | 0.16630 | 0.01424 | 0.02848 | 0.00000 | 0.02213 |
| **EML1** | 0.09899 | 0.13276 | 0.00904 | 0.01962 | 0.56963 | 0.00000 | 0.11514 |
| **EML2** | 0.78376 | 3.61597 | 1.35740 | 0.51977 | 0.32141 | 1.96409 | 0.35719 |
| **EML3** | 2.15577 | 1.89372 | 0.00000 | 1.23258 | 0.89561 | 1.12965 | 9.07801 |
| **EML4** | 8.42344 | 3.91287 | 6.97659 | 6.42235 | 5.17894 | 6.85975 | 9.96616 |
| **EML5** | 0.40147 | 0.00000 | 0.03240 | 0.20604 | 1.67823 | 0.29245 | 1.47504 |
| **EML6** | 0.34229 | 0.16599 | 1.78163 | 0.00000 | 0.40669 | 1.62707 | 0.42140 |
| **EMP1** | 0.03952 | 0.01238 | 0.01734 | 0.02093 | 0.01426 | 0.01265 | 0.02715 |
| **EMP2** | 25.65831 | 3.77899 | 1.16843 | 36.96953 | 0.42272 | 0.00000 | 17.07418 |
| **EMP3** | 0.00000 | 0.09024 | 1.40704 | 0.06384 | 0.02919 | 0.92943 | 0.05310 |
| **EMR1** | 0.32028 | 0.00000 | 0.23450 | 0.11354 | 1.82753 | 0.12360 | 0.00000 |
| **EMR2** | 0.96401 | 0.32289 | 0.09398 | 2.05668 | 0.05916 | 0.00000 | 0.46017 |
| **EMR3** | 0.02240 | 0.01499 | 0.01566 | 0.02448 | 0.02919 | 0.01340 | 0.02198 |
| **EMR4P** | 0.02231 | 0.00964 | 0.04948 | 0.02077 | 0.01871 | 0.01518 | 0.02331 |
| **EMX1** | 0.02152 | 0.19636 | 0.00000 | 0.15694 | 3.35534 | 0.00000 | 0.06008 |
| **EMX2** | 0.03213 | 0.00759 | 0.01270 | 0.00747 | 0.03676 | 0.01234 | 0.10329 |
| **EMX2OS** | 0.04759 | 0.05210 | 0.01025 | 0.03021 | 0.01075 | 0.04991 | 0.02790 |
| **EN1** | 0.03775 | 0.00000 | 1.12704 | 0.00000 | 1.49366 | 0.07064 | 0.84239 |
| **EN2** | 0.01562 | 0.11898 | 0.02976 | 0.01981 | 0.09461 | 0.17535 | 0.01480 |
| **ENAH** | 23.24927 | 2.00330 | 17.65879 | 13.37682 | 3.46507 | 12.25800 | 24.36437 |
| **ENAM** | 0.02267 | 0.01392 | 0.01435 | 0.02385 | 0.08434 | 0.01259 | 0.01891 |
| **ENC1** | 0.00000 | 0.00000 | 0.05041 | 0.67700 | 0.12512 | 0.99375 | 0.11530 |
| **ENDOD1** | 1.38130 | 1.81806 | 3.92159 | 1.06015 | 0.24750 | 0.00000 | 0.48897 |
| **ENDOG** | 0.24424 | 1.84168 | 0.00000 | 0.76221 | 0.09905 | 1.07277 | 1.23653 |
| **ENDOU** | 0.50667 | 0.00647 | 0.01014 | 3.10434 | 0.00587 | 0.01116 | 0.00000 |

| | | | | | | | |
|---|---|---|---|---|---|---|---|
| **ENDOV** | 1.51986 | 9.11279 | 6.48204 | 3.86598 | 0.31422 | 1.07871 | 0.37209 |
| **ENG** | 0.00000 | 0.17538 | 0.00781 | 1.21860 | 5.22623 | 0.00580 | 0.00000 |
| **ENGASE** | 0.22260 | 0.85218 | 0.11898 | 0.97956 | 0.12374 | 0.11378 | 1.82571 |
| **ENHO** | 0.00000 | 0.07884 | 0.21106 | 0.09549 | 0.05436 | 0.62358 | 0.17158 |
| **ENKD1** | 0.00000 | 1.48236 | 0.91250 | 0.79289 | 0.32280 | 0.54786 | 2.26161 |
| **ENKUR** | 0.04537 | 0.02365 | 0.03439 | 0.05566 | 0.00602 | 0.00895 | 0.03233 |
| **ENO1** | 98.54321 | 79.05375 | 44.69211 | 83.37231 | 16.68757 | 52.42676 | 50.12113 |
| **ENO1-AS1** | 0.01426 | 0.08102 | 0.10944 | 0.04311 | 0.00915 | 0.00000 | 0.11798 |
| **ENO2** | 6.34446 | 4.16258 | 18.91935 | 0.00000 | 9.27364 | 24.85389 | 0.23644 |
| **ENO3** | 3.33732 | 0.00000 | 1.32817 | 0.11925 | 0.97304 | 1.36568 | 2.68459 |
| **ENO4** | 0.46457 | 0.00000 | 0.29334 | 0.36463 | 0.03981 | 0.00000 | 0.58868 |
| **ENOPH1** | 30.14606 | 5.50619 | 9.28686 | 11.79997 | 1.86657 | 10.65470 | 8.79706 |
| **ENOSF1** | 10.15161 | 5.61443 | 4.46554 | 3.81910 | 1.03802 | 3.71314 | 1.64581 |
| **ENOX1** | 0.02780 | 0.00000 | 0.24995 | 0.01418 | 0.55589 | 0.01228 | 0.03933 |
| **ENOX2** | 9.90184 | 6.90401 | 1.28453 | 2.70851 | 0.27293 | 2.80116 | 1.70758 |
| **ENPEP** | 25.06111 | 0.67861 | 0.00000 | 3.96963 | 0.00000 | 0.00000 | 7.11351 |
| **ENPP1** | 6.17751 | 0.00000 | 0.04613 | 0.00000 | 0.66876 | 5.07564 | 2.52174 |
| **ENPP2** | 0.03040 | 0.00141 | 0.00000 | 0.04792 | 3.98116 | 1.08595 | 0.00000 |
| **ENPP3** | 0.02493 | 0.01394 | 0.01155 | 0.01788 | 0.02221 | 0.02566 | 0.02227 |
| **ENPP4** | 0.79320 | 0.00000 | 0.39660 | 0.24427 | 0.16835 | 0.00000 | 0.54354 |
| **ENPP5** | 0.34778 | 0.00000 | 0.05362 | 0.19676 | 0.00436 | 0.20560 | 0.00347 |
| **ENPP6** | 0.02363 | 0.02043 | 0.01309 | 0.02134 | 0.00896 | 0.01683 | 0.01682 |
| **ENPP7** | 0.02198 | 0.01512 | 0.01308 | 0.01886 | 0.00790 | 0.02778 | 0.02008 |
| **ENSA** | 16.89605 | 13.32398 | 12.92043 | 22.61133 | 6.87933 | 14.11962 | 9.84648 |
| **ENTHD1** | 0.03883 | 0.01005 | 0.01866 | 0.01229 | 0.02098 | 0.01064 | 0.02851 |
| **ENTHD2** | 0.00000 | 3.98599 | 1.15750 | 1.39280 | 0.18959 | 0.84738 | 5.98083 |
| **ENTPD1** | 19.23407 | 0.00000 | 4.73158 | 12.30289 | 0.09677 | 0.00004 | 11.68453 |
| **ENTPD1-AS1** | 0.22072 | 0.21698 | 0.27157 | 0.08362 | 0.29961 | 0.05654 | 0.00000 |
| **ENTPD2** | 0.01606 | 0.09887 | 0.03796 | 0.07343 | 0.06186 | 0.00642 | 0.05201 |
| **ENTPD3** | 0.02305 | 0.02009 | 0.01630 | 0.02191 | 0.00893 | 0.01637 | 0.02511 |
| **ENTPD3-AS1** | 0.36174 | 0.14393 | 0.42512 | 0.09224 | 0.05922 | 0.15768 | 0.15602 |
| **ENTPD4** | 3.17556 | 4.85567 | 5.30299 | 3.66894 | 0.41786 | 0.59879 | 2.07160 |
| **ENTPD5** | 0.36525 | 0.71164 | 0.40345 | 0.20742 | 0.32702 | 0.33274 | 0.25809 |
| **ENTPD6** | 3.68578 | 19.13855 | 0.38218 | 5.00042 | 0.00000 | 15.56241 | 10.11549 |
| **ENTPD7** | 4.50576 | 1.16552 | 0.47338 | 3.26607 | 0.25458 | 1.06651 | 0.39983 |

| | | | | | | | |
|---|---|---|---|---|---|---|---|
| **ENTPD8** | 0.40325 | 0.59398 | 0.00284 | 0.72600 | 0.00643 | 0.00038 | 0.00000 |
| **ENY2** | 29.78391 | 12.59280 | 19.36531 | 18.67130 | 13.43700 | 15.47338 | 16.49807 |
| **EOGT** | 3.42731 | 0.30949 | 2.57323 | 2.04174 | 2.52502 | 0.55525 | 1.70133 |
| **EOMES** | 0.09636 | 0.01941 | 0.14066 | 0.07033 | 0.27000 | 0.21475 | 0.00000 |
| **EP300** | 12.56062 | 10.78589 | 11.91740 | 15.25382 | 5.79522 | 11.35623 | 28.82104 |
| **EP300-AS1** | 0.05870 | 0.16627 | 0.23932 | 0.28461 | 0.21066 | 0.11157 | 0.08579 |
| **EP400** | 4.60487 | 4.69284 | 3.57470 | 5.20102 | 2.43317 | 4.32738 | 22.66858 |
| **EP400NL** | 0.35972 | 0.18400 | 0.05982 | 0.28064 | 0.08951 | 0.21387 | 0.78205 |
| **EPAS1** | 0.29869 | 2.20501 | 3.22634 | 8.88040 | 1.77376 | 0.00000 | 3.03967 |
| **EPB41** | 1.17655 | 1.19989 | 0.97537 | 1.30550 | 1.39122 | 1.30179 | 2.09220 |
| **EPB41L1** | 5.21429 | 3.01184 | 0.40879 | 4.27861 | 0.29683 | 5.65032 | 10.96537 |
| **EPB41L2** | 5.58927 | 3.38508 | 7.88250 | 0.87285 | 7.97884 | 11.23626 | 1.92141 |
| **EPB41L3** | 9.07093 | 0.00000 | 0.00000 | 5.88264 | 0.40389 | 0.00000 | 11.01456 |
| **EPB41L4A** | 3.67961 | 2.46168 | 2.22067 | 0.60795 | 2.33364 | 2.36523 | 3.33081 |
| **EPB41L4A-AS1** | 5.97491 | 1.00561 | 3.87871 | 9.78501 | 2.24013 | 6.52422 | 13.92298 |
| **EPB41L4A-AS2** | 0.00675 | 0.05256 | 0.05269 | 0.00199 | 0.02310 | 0.06096 | 0.18013 |
| **EPB41L4B** | 0.64698 | 0.51417 | 1.16037 | 3.30307 | 0.30491 | 1.23454 | 2.54005 |
| **EPB41L5** | 36.52249 | 15.83741 | 11.39952 | 3.41145 | 5.20299 | 9.41365 | 0.00000 |
| **EPB42** | 0.02245 | 0.02142 | 0.01205 | 0.03539 | 0.00807 | 0.01672 | 0.01070 |
| **EPC1** | 3.71265 | 2.98831 | 4.57513 | 2.08699 | 3.65164 | 2.63856 | 5.26531 |
| **EPC2** | 1.82464 | 0.87543 | 2.01154 | 1.91576 | 0.80199 | 1.27227 | 5.81112 |
| **EPCAM** | 56.35204 | 12.14214 | 0.00000 | 31.79578 | 2.92116 | 21.57463 | 21.22519 |
| **EPDR1** | 0.01890 | 0.01329 | 0.01617 | 0.02509 | 0.07172 | 0.01115 | 0.03258 |
| **EPG5** | 2.28138 | 1.12020 | 1.82231 | 2.38625 | 0.66550 | 1.92833 | 4.24145 |
| **EPGN** | 0.11070 | 0.01119 | 0.00855 | 0.06777 | 0.00840 | 0.00000 | 0.09777 |
| **EPHA1** | 3.03015 | 0.00000 | 0.37734 | 0.89959 | 1.19695 | 4.22238 | 1.08705 |
| **EPHA1-AS1** | 0.15549 | 0.03724 | 0.07050 | 0.02283 | 0.30807 | 0.08705 | 0.23215 |
| **EPHA10** | 0.00000 | 0.00000 | 1.01372 | 0.00802 | 0.25374 | 0.62749 | 0.17180 |
| **EPHA2** | 0.00000 | 1.10804 | 1.27296 | 2.64043 | 0.09981 | 0.00000 | 1.12821 |
| **EPHA3** | 0.01636 | 0.01153 | 0.01382 | 0.03571 | 0.01085 | 0.06963 | 0.02753 |
| **EPHA4** | 0.00000 | 0.10182 | 0.37123 | 0.20884 | 2.04512 | 10.18847 | 0.00000 |
| **EPHA5** | 0.02747 | 0.01914 | 0.01271 | 0.01896 | 0.01079 | 0.01732 | 0.01735 |
| **EPHA6** | 0.03157 | 0.01968 | 0.02461 | 0.03280 | 0.01006 | 0.03254 | 0.00927 |
| **EPHA7** | 0.00000 | 0.00000 | 0.22759 | 0.00840 | 0.36470 | 2.48591 | 0.04549 |
| **EPHA8** | 0.00000 | 0.18750 | 0.02459 | 0.04875 | 0.39687 | 0.03133 | 0.20781 |

| | | | | | | | |
|---|---|---|---|---|---|---|---|
| **EPHB1** | 0.00000 | 0.73014 | 1.43131 | 0.06520 | 3.36829 | 2.01091 | 0.00000 |
| **EPHB2** | 0.84816 | 1.56578 | 0.00000 | 0.00000 | 0.04638 | 3.95557 | 0.85688 |
| **EPHB3** | 0.00000 | 0.16108 | 0.00588 | 1.36767 | 0.18824 | 0.25952 | 0.81643 |
| **EPHB4** | 0.00000 | 4.90305 | 3.13633 | 14.88908 | 0.77475 | 12.68862 | 8.77872 |
| **EPHB6** | 0.00000 | 0.33617 | 0.16490 | 0.00000 | 0.02569 | 0.26055 | 0.05341 |
| **EPHX1** | 0.00000 | 1.27647 | 2.38078 | 0.32559 | 0.72370 | 0.86099 | 0.36740 |
| **EPHX2** | 0.10205 | 0.27891 | 0.01109 | 0.03392 | 0.23540 | 0.49291 | 0.00000 |
| **EPHX3** | 1.49970 | 0.95013 | 1.90384 | 1.51861 | 0.67119 | 4.27310 | 0.42485 |
| **EPHX4** | 0.27261 | 0.00000 | 0.17706 | 0.13694 | 1.17976 | 0.31003 | 0.34579 |
| **EPM2A** | 0.19284 | 0.03252 | 0.03698 | 0.09142 | 0.03871 | 0.07491 | 0.00000 |
| **EPM2AIP1** | 6.00983 | 0.00000 | 2.94021 | 2.20200 | 4.92754 | 0.83703 | 4.36497 |
| **EPN1** | 0.00000 | 3.56701 | 0.85427 | 0.62731 | 0.11201 | 3.29678 | 11.98920 |
| **EPN2** | 2.29225 | 1.37200 | 5.11699 | 2.71877 | 3.61907 | 3.35619 | 3.83251 |
| **EPN2-AS1** | 0.02108 | 0.01807 | 0.01422 | 0.02713 | 0.04319 | 0.01430 | 0.02715 |
| **EPN2-IT1** | 0.14235 | 0.00000 | 0.49528 | 0.09643 | 3.46191 | 0.09041 | 0.00000 |
| **EPN3** | 0.11336 | 0.02897 | 0.02565 | 0.35604 | 0.01063 | 0.00000 | 0.08420 |
| **EPO** | 0.00000 | 0.00000 | 0.06628 | 0.92791 | 0.03326 | 0.35011 | 0.14628 |
| **EPOR** | 0.07597 | 0.10195 | 0.78286 | 0.96346 | 2.38921 | 1.45447 | 0.00001 |
| **EPPIN** | 0.07286 | 0.00652 | 0.01105 | 0.07843 | 0.00793 | 0.00943 | 0.09935 |
| **EPPIN-WFDC6** | 0.03890 | 0.01464 | 0.01374 | 0.02532 | 0.03210 | 0.01371 | 0.01840 |
| **EPPK1** | 0.00000 | 25.93182 | 0.27231 | 5.71462 | 0.01957 | 10.94024 | 16.89989 |
| **EPRS** | 38.41984 | 10.42961 | 12.87013 | 20.26542 | 2.99011 | 11.68169 | 23.78022 |
| **EPS15** | 12.35484 | 0.00000 | 5.26127 | 7.74999 | 4.19680 | 4.96819 | 7.54330 |
| **EPS15L1** | 9.41158 | 9.41713 | 1.43937 | 9.08159 | 2.96018 | 1.56042 | 7.03679 |
| **EPS8** | 4.01815 | 0.00000 | 0.28173 | 1.50563 | 0.25248 | 1.51957 | 0.52798 |
| **EPS8L1** | 0.00000 | 0.00734 | 0.00000 | 0.79058 | 6.96730 | 0.51806 | 0.36427 |
| **EPS8L2** | 4.79143 | 7.76666 | 0.00000 | 1.38298 | 0.00000 | 0.29060 | 10.63523 |
| **EPS8L3** | 0.02809 | 0.01919 | 0.01233 | 0.10262 | 0.00827 | 0.03714 | 0.00000 |
| **EPSTI1** | 6.16041 | 1.74285 | 0.00000 | 0.00000 | 3.00458 | 1.66265 | 1.33911 |
| **EPT1** | 23.65118 | 3.74574 | 9.92373 | 11.92016 | 2.88023 | 7.34325 | 13.44128 |
| **EPX** | 0.05213 | 0.05783 | 0.00788 | 0.01319 | 0.01388 | 0.01667 | 0.00000 |
| **EQTN** | 0.08798 | 0.00592 | 0.01420 | 0.07397 | 0.01047 | 0.03151 | 0.07158 |
| **ERAL1** | 22.96504 | 26.33780 | 14.43299 | 16.39036 | 3.20117 | 10.78673 | 4.02979 |
| **ERAP1** | 0.15139 | 0.03102 | 0.00251 | 0.00000 | 0.07778 | 0.06019 | 0.19426 |
| **ERAP2** | 0.02396 | 0.01958 | 0.01316 | 0.02235 | 0.00931 | 0.01299 | 0.01620 |

| | | | | | | | |
|---|---|---|---|---|---|---|---|
| **ERAS** | 0.04563 | 0.00448 | 0.02871 | 0.02592 | 0.09839 | 0.01314 | 0.00165 |
| **ERBB2** | 3.15977 | 2.72134 | 0.18939 | 2.56191 | 0.38251 | 0.74976 | 7.57514 |
| **ERBB2IP** | 7.42490 | 7.08571 | 7.36443 | 5.97486 | 1.58603 | 11.93771 | 19.54105 |
| **ERBB3** | 3.06623 | 10.59527 | 8.67026 | 13.49577 | 2.09980 | 3.91331 | 8.19794 |
| **ERBB4** | 0.07185 | 0.00000 | 0.69343 | 0.00379 | 5.72365 | 2.10620 | 0.69174 |
| **ERC1** | 6.27726 | 3.24207 | 3.53235 | 3.91054 | 3.18117 | 2.72144 | 21.77351 |
| **ERC2** | 0.12435 | 0.00000 | 0.54100 | 0.23749 | 1.00019 | 0.52006 | 0.77048 |
| **ERC2-IT1** | 0.02072 | 0.01451 | 0.01045 | 0.03077 | 0.00747 | 0.00397 | 0.12578 |
| **ERCC1** | 2.52658 | 7.75171 | 3.98570 | 1.37558 | 2.25038 | 3.06823 | 2.25366 |
| **ERCC2** | 4.87195 | 3.23383 | 1.70387 | 2.45932 | 2.15913 | 1.23412 | 1.33633 |
| **ERCC3** | 20.25732 | 12.85027 | 7.96879 | 17.38696 | 0.59335 | 10.70793 | 0.00000 |
| **ERCC4** | 3.52946 | 1.68486 | 3.16102 | 1.88547 | 1.20150 | 3.47222 | 4.49822 |
| **ERCC5** | 0.65241 | 0.50162 | 0.33137 | 0.31593 | 0.38706 | 0.00150 | 0.00000 |
| **ERCC6** | 2.85131 | 0.37740 | 2.10168 | 1.44001 | 1.07255 | 0.85107 | 2.88076 |
| **ERCC6L** | 17.69840 | 1.77669 | 7.41030 | 3.61539 | 6.91349 | 5.17731 | 5.08709 |
| **ERCC6L2** | 4.60571 | 0.00000 | 2.02046 | 2.47265 | 0.74513 | 1.48561 | 2.84830 |
| **ERCC8** | 4.79453 | 0.00000 | 3.14340 | 3.66724 | 0.71671 | 1.60847 | 0.00000 |
| **EREG** | 0.04292 | 0.00961 | 0.01571 | 0.00928 | 0.00702 | 0.00710 | 0.05813 |
| **ERF** | 0.00000 | 3.97240 | 2.97340 | 0.90090 | 0.12173 | 3.80158 | 18.33642 |
| **ERG** | 0.00873 | 0.00680 | 0.14332 | 0.02214 | 0.47830 | 0.00697 | 0.05574 |
| **ERGIC1** | 3.22480 | 1.53774 | 0.45923 | 0.58138 | 0.09176 | 0.67792 | 2.41347 |
| **ERGIC2** | 20.55911 | 1.07460 | 14.18574 | 13.63554 | 2.20989 | 7.19926 | 2.01684 |
| **ERGIC3** | 22.91997 | 22.86880 | 13.21074 | 24.28958 | 3.41605 | 13.03656 | 13.27887 |
| **ERH** | 39.66732 | 15.65985 | 23.51458 | 30.65295 | 7.57874 | 18.59937 | 19.89030 |
| **ERI1** | 20.81168 | 0.00000 | 10.07075 | 8.56899 | 8.12153 | 5.81420 | 4.57629 |
| **ERI2** | 5.27327 | 1.27561 | 4.38775 | 1.77493 | 0.78154 | 1.55057 | 0.00000 |
| **ERI3** | 10.94686 | 9.47390 | 3.93443 | 12.28519 | 0.05980 | 8.16857 | 5.67857 |
| **ERI3-IT1** | 0.01752 | 0.05299 | 0.06949 | 0.05961 | 0.00825 | 0.05180 | 0.11305 |
| **ERICH1** | 7.38145 | 13.33561 | 2.11811 | 7.35109 | 2.57502 | 4.22537 | 13.48292 |
| **ERICH1-AS1** | 0.02948 | 0.03142 | 0.09493 | 0.09867 | 0.13279 | 0.00000 | 0.26803 |
| **ERICH2** | 0.31471 | 1.15469 | 0.97386 | 0.25829 | 0.04820 | 0.28138 | 0.18076 |
| **ERICH3** | 0.14265 | 0.10384 | 0.01993 | 0.02789 | 0.00588 | 0.11051 | 0.00510 |
| **ERICH3-AS1** | 0.04970 | 0.01643 | 0.01458 | 0.01883 | 0.00783 | 0.02012 | 0.00527 |
| **ERICH4** | 0.02817 | 0.02040 | 0.01209 | 0.03052 | 0.00869 | 0.01203 | 0.01548 |
| **ERICH5** | 1.90621 | 0.10713 | 0.42386 | 1.01208 | 0.75480 | 2.33716 | 0.00000 |

| | | | | | | | |
|---|---|---|---|---|---|---|---|
| **ERICH6** | 0.02527 | 0.02273 | 0.02210 | 0.02383 | 0.00811 | 0.02479 | 0.02787 |
| **ERICH6-AS1** | 0.06863 | 0.02138 | 0.12976 | 0.04170 | 0.01066 | 0.15872 | 0.00000 |
| **ERICH6B** | 0.01954 | 0.01902 | 0.01131 | 0.02659 | 0.02108 | 0.03062 | 0.02014 |
| **ERLEC1** | 11.46141 | 4.80545 | 7.84532 | 11.84096 | 2.64107 | 5.95883 | 5.42994 |
| **ERLIN1** | 23.19755 | 7.39965 | 12.94017 | 12.77418 | 3.94158 | 4.77686 | 5.48615 |
| **ERLIN2** | 2.90327 | 0.51393 | 5.12080 | 0.21537 | 0.13401 | 2.04884 | 8.23208 |
| **ERMAP** | 1.33534 | 0.00000 | 1.07379 | 0.47184 | 0.18860 | 0.00787 | 0.20622 |
| **ERMARD** | 3.97175 | 1.34541 | 2.94286 | 0.72786 | 1.02996 | 0.77286 | 1.27775 |
| **ERMN** | 0.24838 | 0.00000 | 0.01520 | 0.78153 | 0.00644 | 0.01382 | 0.01817 |
| **ERMP1** | 4.99570 | 0.00000 | 1.29151 | 0.72528 | 0.35890 | 0.85126 | 6.06192 |
| **1.00000** | 0.00000 | 0.51922 | 0.46093 | 1.30656 | 2.00186 | 0.06839 | 0.76773 |
| **2.00000** | 0.03294 | 0.13565 | 0.04953 | 0.03578 | 0.01574 | 0.00000 | 0.08176 |
| **ERO1L** | 2.99958 | 0.00000 | 0.55106 | 1.59823 | 1.22975 | 0.26747 | 6.22177 |
| **ERO1LB** | 0.18784 | 0.00000 | 0.23196 | 0.51559 | 2.65963 | 0.40144 | 0.80164 |
| **ERP27** | 0.74456 | 0.00000 | 0.01280 | 1.50603 | 0.01020 | 0.00535 | 0.00000 |
| **ERP29** | 40.27876 | 19.18196 | 12.76660 | 33.71924 | 2.51720 | 22.50326 | 29.96673 |
| **ERP44** | 20.46116 | 11.00877 | 11.12176 | 13.92569 | 5.50624 | 13.73436 | 1.12259 |
| **ERRFI1** | 7.61697 | 0.00000 | 2.13086 | 26.18010 | 3.88383 | 6.21094 | 22.46227 |
| **ERV3-1** | 0.72113 | 0.00000 | 0.19691 | 1.75189 | 0.26939 | 0.03370 | 1.30460 |
| **ERVFRD-1** | 0.00000 | 0.47014 | 0.08513 | 13.77601 | 0.16267 | 0.00000 | 0.17790 |
| **ERVK13-1** | 0.41312 | 0.31867 | 0.55615 | 0.48976 | 2.99201 | 0.26814 | 0.21167 |
| **ERVMER34-1** | 3.38983 | 0.44333 | 0.45835 | 12.92387 | 0.00000 | 0.00000 | 0.24474 |
| **ERVV-1** | 0.08513 | 0.08991 | 0.01235 | 8.22768 | 0.00610 | 0.00000 | 0.02394 |
| **ERVV-2** | 1.01803 | 0.02857 | 0.01620 | 10.75439 | 0.02239 | 0.06206 | 0.00000 |
| **ESAM** | 0.00000 | 0.37635 | 0.13792 | 0.29467 | 0.01098 | 3.16258 | 0.00000 |
| **ESCO1** | 11.11887 | 0.20711 | 5.94857 | 4.82904 | 0.85938 | 11.98171 | 7.74110 |
| **ESCO2** | 9.03364 | 1.99322 | 6.89073 | 3.38458 | 9.36096 | 1.15793 | 0.64453 |
| **ESD** | 13.06256 | 0.16165 | 2.01655 | 7.00668 | 0.00000 | 4.01110 | 3.77233 |
| **ESF1** | 19.39929 | 0.62146 | 18.05112 | 9.30167 | 4.85564 | 9.82896 | 21.11603 |
| **ESPL1** | 6.85486 | 6.13069 | 10.27751 | 7.73053 | 1.30551 | 1.52920 | 16.72206 |
| **ESPN** | 0.01822 | 0.11022 | 0.19274 | 0.03617 | 0.06071 | 0.07799 | 0.10036 |
| **ESPNL** | 0.03098 | 0.01430 | 0.01225 | 0.04218 | 0.00792 | 0.03446 | 0.02700 |
| **ESPNP** | 0.02416 | 0.02007 | 0.01205 | 0.02261 | 0.00893 | 0.01227 | 0.01877 |
| **ESR1** | 0.09785 | 0.00000 | 0.19997 | 0.01860 | 0.06282 | 0.01819 | 0.10556 |
| **ESR2** | 0.39317 | 0.19391 | 0.26918 | 0.46163 | 0.38080 | 0.10909 | 0.67320 |

| | | | | | | | |
|---|---|---|---|---|---|---|---|
| **ESRG** | 6.96697 | 1.73150 | 0.65546 | 2.34077 | 0.00000 | 19.76802 | 2.96773 |
| **ESRP1** | 20.66527 | 13.06368 | 15.05676 | 10.94103 | 5.36597 | 12.52812 | 15.82200 |
| **ESRP2** | 5.76608 | 8.67050 | 2.96049 | 7.30181 | 0.05561 | 2.58985 | 14.01918 |
| **ESRRA** | 2.57482 | 10.02180 | 0.21945 | 7.38208 | 0.00000 | 4.85310 | 14.25493 |
| **ESRRB** | 1.66438 | 18.25900 | 4.35479 | 0.00000 | 0.00000 | 0.37353 | 1.54684 |
| **ESRRG** | 2.52432 | 1.09781 | 0.00000 | 6.48915 | 0.56800 | 0.01802 | 0.00000 |
| **ESX1** | 0.02457 | 0.00926 | 0.02712 | 0.01916 | 0.02899 | 0.01297 | 0.01853 |
| **ESYT1** | 3.75509 | 0.00000 | 13.71514 | 14.26046 | 1.40266 | 5.90202 | 3.35058 |
| **ESYT2** | 7.79243 | 3.50470 | 3.76370 | 6.82476 | 1.97021 | 5.22889 | 15.15719 |
| **ESYT3** | 0.05955 | 0.00000 | 0.07503 | 0.21536 | 1.23332 | 0.79015 | 0.00000 |
| **ETAA1** | 3.77686 | 0.38753 | 1.81860 | 1.62284 | 1.30506 | 0.85855 | 0.00000 |
| **ETF1** | 34.42579 | 14.94367 | 16.94586 | 23.75984 | 11.90267 | 13.35113 | 23.77124 |
| **ETFA** | 33.51624 | 9.86548 | 5.66449 | 23.47895 | 9.44495 | 8.96499 | 9.05860 |
| **ETFB** | 10.06299 | 18.90999 | 0.98589 | 9.45395 | 0.74472 | 8.93293 | 2.18987 |
| **ETFDH** | 7.44400 | 0.87618 | 1.56933 | 5.14711 | 3.43505 | 2.17712 | 0.00000 |
| **ETHE1** | 4.64122 | 18.71214 | 0.96204 | 0.73544 | 2.20507 | 11.75019 | 4.71858 |
| **ETNK1** | 18.47045 | 1.52159 | 4.28500 | 8.20855 | 6.73497 | 2.97791 | 0.00000 |
| **ETNK2** | 0.17037 | 0.37144 | 0.49930 | 2.16854 | 0.04800 | 5.69135 | 2.41354 |
| **ETNPPL** | 0.00000 | 3.40645 | 22.13133 | 0.00000 | 0.05549 | 0.00000 | 0.00000 |
| **ETS1** | 0.89057 | 0.00000 | 0.33391 | 0.12262 | 0.27207 | 0.43551 | 0.09914 |
| **ETS2** | 1.46348 | 0.52383 | 0.51818 | 3.95608 | 0.12317 | 0.27076 | 3.38324 |
| **ETV1** | 0.10071 | 0.00000 | 0.00104 | 0.07515 | 0.26392 | 5.17689 | 0.00000 |
| **ETV2** | 0.02624 | 0.28539 | 0.01561 | 0.16508 | 0.13772 | 0.08483 | 0.00406 |
| **ETV3** | 0.84502 | 1.06929 | 1.46706 | 0.00000 | 2.27341 | 1.19290 | 5.44755 |
| **ETV3L** | 0.01927 | 0.07296 | 0.00746 | 0.01574 | 0.03923 | 0.00561 | 0.04448 |
| **ETV4** | 0.00000 | 0.46486 | 2.58031 | 3.13736 | 1.65046 | 23.39208 | 0.00000 |
| **ETV5** | 0.59754 | 0.47421 | 1.23119 | 1.00659 | 6.15715 | 6.39097 | 0.43958 |
| **ETV6** | 0.72420 | 1.31865 | 1.26030 | 2.27757 | 0.21380 | 3.10662 | 3.97271 |
| **ETV7** | 0.03268 | 0.00000 | 0.02462 | 0.07895 | 0.06041 | 0.04151 | 0.12313 |
| **EVA1A** | 0.03395 | 0.00549 | 0.04095 | 0.03100 | 0.04257 | 0.02559 | 0.01109 |
| **EVA1B** | 0.00000 | 0.30334 | 0.05447 | 0.32490 | 0.07417 | 0.16339 | 1.62066 |
| **EVA1C** | 0.03362 | 0.01914 | 0.01342 | 0.01851 | 0.02807 | 0.01241 | 0.01617 |
| **EVC** | 0.06552 | 0.06089 | 0.24971 | 0.03562 | 0.05284 | 0.12206 | 0.10035 |
| **EVC2** | 0.01101 | 0.00000 | 0.11908 | 0.07067 | 0.47822 | 0.65664 | 0.08953 |
| **EVI2A** | 0.08950 | 0.00000 | 0.21523 | 0.16791 | 0.00645 | 0.08283 | 0.59982 |

| | | | | | | | |
|---|---|---|---|---|---|---|---|
| **EVI2B** | 0.02883 | 0.00000 | 0.06258 | 0.01208 | 0.00445 | 0.03370 | 0.25102 |
| **EVI5** | 5.61152 | 0.00000 | 3.31538 | 6.66529 | 0.74508 | 1.07641 | 10.99885 |
| **EVI5L** | 0.00000 | 0.02301 | 0.03548 | 0.16376 | 0.20918 | 0.00000 | 0.25553 |
| **EVL** | 0.36341 | 0.51845 | 0.00973 | 0.52039 | 0.33267 | 0.18336 | 2.84065 |
| **EVPL** | 0.00000 | 0.04360 | 0.00177 | 0.29337 | 0.00847 | 0.06233 | 0.92752 |
| **EVPLL** | 0.00387 | 0.08845 | 0.02082 | 0.02385 | 0.00583 | 0.01395 | 0.12185 |
| **EVX1-AS** | 0.02562 | 0.01609 | 0.01283 | 0.02124 | 0.00911 | 0.02654 | 0.01302 |
| **EVX2** | 0.01692 | 0.01084 | 0.05005 | 0.02876 | 0.02447 | 0.01322 | 0.02746 |
| **EWSR1** | 47.58065 | 47.03617 | 17.02743 | 40.59242 | 10.25772 | 22.10494 | 0.23872 |
| **EXD1** | 0.04514 | 0.00000 | 0.25839 | 0.01832 | 0.61145 | 0.09721 | 0.03095 |
| **EXD2** | 8.03530 | 0.08271 | 5.24054 | 8.76907 | 1.15919 | 1.00280 | 1.46659 |
| **EXD3** | 0.19540 | 0.94091 | 0.46209 | 1.18410 | 1.78028 | 0.26004 | 0.49190 |
| **EXO1** | 13.53997 | 8.82752 | 9.05166 | 15.68539 | 5.74229 | 4.40902 | 0.00000 |
| **EXO5** | 3.17298 | 1.30398 | 4.11222 | 2.43021 | 4.15868 | 0.00000 | 0.00000 |
| **EXOC1** | 7.56249 | 1.64772 | 3.21783 | 2.27594 | 2.54953 | 3.56038 | 3.41825 |
| **EXOC2** | 5.56255 | 3.50602 | 1.10534 | 2.88824 | 1.08517 | 3.53992 | 0.12342 |
| **EXOC3** | 4.61562 | 7.57996 | 1.75115 | 3.81587 | 1.49492 | 1.23003 | 3.51710 |
| **EXOC3L1** | 0.14642 | 0.00994 | 0.02985 | 0.04117 | 0.00957 | 1.46229 | 0.00553 |
| **EXOC3L2** | 0.00000 | 0.15349 | 0.47596 | 0.03459 | 4.28593 | 0.00000 | 0.00837 |
| **EXOC3L4** | 0.01838 | 0.01279 | 0.01288 | 0.03730 | 0.00770 | 0.03162 | 0.03759 |
| **EXOC4** | 3.05408 | 2.59639 | 1.24000 | 4.17619 | 1.97618 | 3.16571 | 3.37077 |
| **EXOC5** | 15.31364 | 0.73160 | 5.58667 | 9.76024 | 3.39781 | 8.26250 | 9.94021 |
| **EXOC6** | 4.12728 | 0.00000 | 1.82997 | 1.06897 | 0.22168 | 0.04100 | 3.91596 |
| **EXOC6B** | 1.04120 | 0.69184 | 1.32769 | 2.90512 | 0.70115 | 0.00000 | 0.56336 |
| **EXOC7** | 6.41476 | 8.23838 | 0.30869 | 10.19779 | 0.30155 | 5.34407 | 4.58119 |
| **EXOC8** | 1.61690 | 0.00000 | 1.91275 | 2.43842 | 1.59591 | 1.53926 | 4.36837 |
| **EXOG** | 1.88806 | 0.80559 | 0.44261 | 2.40491 | 0.24514 | 1.68718 | 0.00000 |
| **EXOSC1** | 16.32995 | 10.37237 | 13.15331 | 17.25713 | 2.99599 | 7.75039 | 3.21757 |
| **EXOSC10** | 32.82695 | 17.60483 | 14.59455 | 21.77848 | 7.29825 | 10.51672 | 0.70722 |
| **EXOSC2** | 25.37441 | 7.73778 | 11.89515 | 9.70351 | 2.29619 | 7.19366 | 4.51681 |
| **EXOSC3** | 28.20635 | 16.73391 | 19.83867 | 20.21132 | 11.75336 | 6.29594 | 0.00000 |
| **EXOSC4** | 25.61637 | 29.83387 | 15.51813 | 21.83848 | 3.10918 | 9.01389 | 7.14695 |
| **EXOSC5** | 22.68951 | 19.30741 | 17.36634 | 25.72012 | 1.08247 | 8.27516 | 0.00000 |
| **EXOSC6** | 1.47584 | 13.03377 | 7.75938 | 2.98729 | 2.80149 | 5.06801 | 5.42351 |
| **EXOSC7** | 22.22510 | 16.61362 | 10.54116 | 17.13653 | 5.24369 | 11.82548 | 2.63091 |

| | | | | | | | |
|---|---|---|---|---|---|---|---|
| **EXOSC8** | 37.84553 | 20.33005 | 11.87333 | 14.00075 | 1.43127 | 15.16374 | 5.92118 |
| **EXOSC9** | 23.81104 | 8.64810 | 14.47534 | 12.93023 | 2.86537 | 8.31629 | 5.18649 |
| **EXPH5** | 17.10026 | 0.00000 | 3.50745 | 6.79301 | 0.00000 | 0.92030 | 3.21500 |
| **EXT1** | 1.46984 | 1.40023 | 1.41251 | 1.04189 | 2.37772 | 6.95608 | 4.22017 |
| **EXT2** | 11.96490 | 11.83986 | 1.32632 | 8.63675 | 3.40234 | 5.54870 | 1.33282 |
| **EXTL1** | 0.00000 | 0.14333 | 0.06014 | 0.16310 | 0.12064 | 0.04464 | 0.13404 |
| **EXTL2** | 5.27497 | 0.00000 | 3.86690 | 5.95152 | 0.89601 | 1.71648 | 1.76978 |
| **EXTL3** | 4.22581 | 4.92626 | 9.72783 | 1.43291 | 4.59147 | 4.23827 | 4.57994 |
| **EYA1** | 0.09869 | 0.00000 | 0.05590 | 0.00000 | 1.52547 | 0.09903 | 0.14409 |
| **EYA2** | 0.02173 | 0.01186 | 0.09348 | 0.02683 | 1.70486 | 0.03249 | 0.00946 |
| **EYA3** | 21.02130 | 8.34596 | 8.27523 | 18.67852 | 10.23411 | 6.11290 | 0.00000 |
| **EYA4** | 0.06982 | 0.00000 | 0.14952 | 0.05762 | 0.03493 | 0.10958 | 0.00814 |
| **EYA4-AS1** | 0.02260 | 0.02343 | 0.00000 | 0.02152 | 0.33445 | 0.01255 | 0.01392 |
| **EYS** | 0.13152 | 0.00000 | 0.08247 | 0.00000 | 0.36595 | 0.06182 | 0.75157 |
| **EZH1** | 1.18067 | 0.00000 | 1.06522 | 2.92748 | 0.07125 | 0.15081 | 1.92818 |
| **EZH2** | 13.66944 | 8.92468 | 11.02936 | 10.22687 | 2.52721 | 4.22464 | 13.35184 |
| **EZR** | 82.59021 | 55.81214 | 14.82709 | 55.33717 | 9.70625 | 26.09379 | 36.58930 |
| **EZR-AS1** | 0.08790 | 0.10109 | 0.01629 | 0.05817 | 0.01144 | 0.01389 | 0.05324 |
| **F10** | 0.02297 | 0.01397 | 0.01566 | 0.01965 | 0.07253 | 0.01414 | 0.01978 |
| **F11** | 0.02452 | 0.01149 | 0.01285 | 0.01719 | 0.04495 | 0.01316 | 0.03343 |
| **F11-AS1** | 0.02532 | 0.05416 | 0.05374 | 0.04212 | 0.04736 | 0.00000 | 0.11023 |
| **F11R** | 42.28173 | 35.84583 | 35.13062 | 45.43746 | 14.96170 | 29.82310 | 17.52524 |
| **F12** | 0.04672 | 0.04246 | 0.14759 | 0.10683 | 0.09297 | 0.19828 | 0.03649 |
| **F13A1** | 0.01441 | 0.01364 | 0.00000 | 0.02270 | 0.69238 | 0.02131 | 0.02890 |
| **F13B** | 0.04063 | 0.02692 | 0.02500 | 0.01182 | 0.07503 | 0.03723 | 0.01537 |
| **F2** | 0.05499 | 0.00725 | 0.07548 | 0.07482 | 0.60809 | 0.58044 | 0.00000 |
| **F2R** | 0.43767 | 0.50362 | 0.63810 | 1.29312 | 0.01063 | 1.05123 | 0.00000 |
| **F2RL1** | 29.69787 | 9.45809 | 20.55598 | 12.46537 | 1.96883 | 24.42529 | 0.00000 |
| **F2RL2** | 0.35242 | 0.27325 | 0.08479 | 0.04560 | 0.02257 | 0.05899 | 0.55509 |
| **F2RL3** | 0.02857 | 0.01960 | 0.01471 | 0.02167 | 0.00760 | 0.01831 | 0.02982 |
| **F3** | 14.37171 | 3.99764 | 1.27508 | 17.50605 | 0.00000 | 8.99263 | 0.00000 |
| **F5** | 0.36507 | 0.00000 | 0.01588 | 0.08732 | 0.00722 | 0.00710 | 0.00000 |
| **F7** | 0.02542 | 0.01828 | 0.01378 | 0.01979 | 0.00788 | 0.01272 | 0.01588 |
| **F8** | 0.38894 | 0.14312 | 0.91813 | 0.05380 | 0.02980 | 0.19216 | 0.04798 |
| **F8A1.uid-13426** | 0.00733 | 0.06534 | 0.08103 | 0.03546 | 0.00930 | 0.04982 | 0.08201 |

| | | | | | | | |
|---|---|---|---|---|---|---|---|
| **F9** | 0.02342 | 0.01238 | 0.01787 | 0.01635 | 0.03874 | 0.01245 | 0.02896 |
| **FA2H** | 0.00000 | 5.10376 | 4.85522 | 0.00000 | 0.04243 | 1.63635 | 0.35281 |
| **FAAH** | 0.15418 | 1.13646 | 1.41043 | 0.35181 | 0.25428 | 2.43820 | 1.31619 |
| **FAAH2** | 0.05021 | 0.17082 | 0.00190 | 0.02549 | 0.00739 | 0.44617 | 0.08931 |
| **FAAHP1** | 0.17328 | 0.08190 | 0.03109 | 0.12300 | 0.04956 | 0.15847 | 0.00000 |
| **FABP2** | 0.04929 | 0.00000 | 0.07643 | 0.01512 | 0.00591 | 0.00927 | 0.02073 |
| **FABP3** | 75.41548 | 0.27380 | 0.00000 | 15.84312 | 0.00000 | 0.00000 | 29.60782 |
| **FABP4** | 0.04038 | 0.00949 | 0.02223 | 0.16505 | 0.01058 | 0.01700 | 0.00000 |
| **FABP5** | 14.74983 | 5.48468 | 0.00000 | 10.35468 | 0.02281 | 8.49451 | 4.87485 |
| **FABP5P3** | 0.40361 | 0.00000 | 0.04177 | 0.15513 | 0.00636 | 0.15804 | 0.21766 |
| **FABP6** | 0.18659 | 0.00000 | 0.08625 | 0.09601 | 0.15542 | 0.53401 | 0.00000 |
| **FABP7** | 10.45061 | 0.00000 | 0.00000 | 4.84989 | 0.01284 | 0.00005 | 2.33281 |
| **FADD** | 5.03806 | 5.55497 | 5.12416 | 7.75220 | 0.13463 | 3.50076 | 5.28833 |
| **FADS1** | 6.87225 | 0.00000 | 0.31400 | 29.68307 | 1.83455 | 0.00000 | 4.47470 |
| **FADS2** | 0.05273 | 0.00000 | 0.00000 | 10.23433 | 0.00564 | 0.00000 | 9.85209 |
| **FADS3** | 0.00000 | 1.24184 | 0.26409 | 7.73575 | 1.48402 | 2.46741 | 9.42548 |
| **FADS6** | 0.03824 | 0.00797 | 0.01444 | 0.01543 | 0.01671 | 0.06402 | 0.01419 |
| **FAF1** | 2.16673 | 8.02390 | 5.36102 | 4.79034 | 0.00000 | 10.51654 | 17.31196 |
| **FAF2** | 17.44208 | 6.81890 | 15.93369 | 13.86577 | 5.15538 | 6.72788 | 6.99071 |
| **FAH** | 3.87619 | 1.48919 | 0.18908 | 3.03685 | 0.00115 | 0.00000 | 0.22415 |
| **FAHD1** | 6.41506 | 8.59968 | 0.41451 | 3.52430 | 0.30748 | 4.04099 | 2.61860 |
| **FAHD2A** | 2.94394 | 2.62485 | 0.49528 | 1.53866 | 1.18101 | 1.14573 | 0.00000 |
| **FAHD2B** | 0.27437 | 0.34165 | 0.13324 | 0.28153 | 0.20064 | 0.35710 | 0.00000 |
| **FAHD2CP** | 0.02875 | 0.22177 | 0.09622 | 0.05532 | 0.00831 | 0.02374 | 0.00264 |
| **FAIM** | 5.99963 | 1.91501 | 0.04718 | 4.18316 | 0.08830 | 0.82525 | 1.33849 |
| **FAIM2** | 0.01209 | 0.01281 | 0.03393 | 0.03450 | 0.57738 | 0.02227 | 0.03264 |
| **FAIM3** | 0.01634 | 0.05385 | 0.03131 | 0.02388 | 0.07704 | 0.00913 | 0.01207 |
| **FAM101B** | 14.60826 | 11.05487 | 0.00000 | 2.37158 | 0.00000 | 0.00000 | 10.47418 |
| **FAM102A** | 0.12459 | 4.24746 | 1.03178 | 3.60584 | 1.88914 | 3.19304 | 11.08743 |
| **FAM102B** | 1.20969 | 2.10060 | 7.94423 | 1.46410 | 0.00003 | 9.76044 | 4.17131 |
| **FAM103A1** | 4.64567 | 5.03285 | 4.55578 | 2.66042 | 2.64961 | 3.21136 | 0.00000 |
| **FAM104A** | 18.06087 | 13.62457 | 14.57122 | 19.97570 | 15.79928 | 6.00304 | 0.01210 |
| **FAM104B** | 6.24947 | 4.90176 | 12.81336 | 3.16714 | 1.10911 | 0.93647 | 3.08539 |
| **FAM105A** | 0.05704 | 0.84114 | 1.03654 | 0.30161 | 1.93825 | 0.36750 | 0.00000 |
| **FAM106A** | 0.02289 | 0.01359 | 0.03510 | 0.02863 | 0.00657 | 0.02619 | 0.05986 |

| | | | | | | | |
|---|---|---|---|---|---|---|---|
| **FAM106CP** | 0.02630 | 0.01240 | 0.05810 | 0.01351 | 0.00714 | 0.00942 | 0.03836 |
| **FAM107A** | 0.02554 | 0.02714 | 0.01541 | 0.00753 | 0.05206 | 0.07706 | 0.08222 |
| **FAM107B** | 9.11602 | 1.08639 | 2.38641 | 0.00000 | 0.64022 | 5.66878 | 11.15259 |
| **FAM109A** | 0.00000 | 1.67665 | 1.25424 | 0.70147 | 0.06542 | 2.88758 | 4.19495 |
| **FAM109B** | 0.36162 | 1.90921 | 1.73221 | 0.02083 | 0.12090 | 1.48262 | 0.00000 |
| **FAM110A** | 0.67643 | 2.30004 | 0.00000 | 13.05850 | 0.38256 | 0.00000 | 11.96343 |
| **FAM110B** | 7.67431 | 1.81899 | 0.12457 | 4.22254 | 0.25244 | 0.34554 | 1.57379 |
| **FAM110D** | 0.01671 | 0.00000 | 0.20029 | 0.01873 | 0.36513 | 0.00749 | 0.02187 |
| **FAM111A** | 0.00272 | 0.00000 | 0.11414 | 0.01052 | 0.03881 | 0.12791 | 0.08176 |
| **FAM111B** | 0.79038 | 0.12625 | 0.07100 | 0.03631 | 0.76114 | 1.12748 | 0.59462 |
| **FAM114A1** | 0.02464 | 0.12192 | 0.18402 | 0.01446 | 0.13448 | 0.05247 | 0.00260 |
| **FAM114A2** | 8.30301 | 1.60891 | 4.89372 | 2.28405 | 1.69727 | 0.83906 | 0.00000 |
| **FAM115A** | 0.51546 | 0.00671 | 0.00422 | 2.65711 | 0.11748 | 0.00000 | 0.95762 |
| **FAM115C** | 0.02999 | 0.00989 | 0.01429 | 0.17974 | 0.01840 | 0.01168 | 0.02391 |
| **FAM117A** | 0.84709 | 0.00000 | 0.71471 | 0.70583 | 0.97550 | 1.11927 | 3.29034 |
| **FAM117B** | 4.56512 | 1.54633 | 8.09319 | 0.13728 | 6.89881 | 5.29785 | 3.30583 |
| **FAM118A** | 15.58401 | 10.41794 | 3.59091 | 14.47359 | 2.49799 | 0.00000 | 10.09261 |
| **FAM118B** | 18.60058 | 5.42457 | 8.17646 | 13.40073 | 7.38378 | 4.33031 | 0.00000 |
| **FAM120A** | 9.62064 | 6.11548 | 2.83347 | 6.38397 | 0.00000 | 4.82467 | 30.01692 |
| **FAM120AOS** | 5.42914 | 3.65500 | 4.41819 | 6.11763 | 0.65826 | 1.80539 | 6.01446 |
| **FAM120B** | 2.70425 | 0.00000 | 2.93181 | 1.18546 | 1.14457 | 0.08432 | 0.24125 |
| **FAM120C** | 1.68497 | 1.10783 | 1.13593 | 3.86655 | 1.16666 | 2.53683 | 5.50371 |
| **FAM122A** | 0.48201 | 1.60556 | 1.68904 | 2.46128 | 2.33988 | 2.14818 | 6.60344 |
| **FAM122B** | 9.54214 | 0.91377 | 1.67964 | 1.40752 | 2.09208 | 4.22015 | 5.62173 |
| **FAM122C** | 0.85804 | 5.18108 | 11.04511 | 0.00000 | 6.26090 | 3.68187 | 0.00000 |
| **FAM124A** | 0.00000 | 0.36529 | 1.59740 | 0.00000 | 0.23830 | 5.99259 | 0.39284 |
| **FAM124B** | 0.00000 | 0.22015 | 0.46733 | 0.00000 | 0.00068 | 0.96462 | 0.01329 |
| **FAM126A** | 10.86429 | 7.17597 | 9.04032 | 3.98383 | 2.68368 | 7.63422 | 3.79157 |
| **FAM126B** | 10.91978 | 2.14356 | 4.83635 | 3.56072 | 7.50637 | 9.40947 | 3.49007 |
| **FAM127A** | 7.96208 | 9.83414 | 0.80489 | 9.95162 | 2.36832 | 11.19953 | 5.13215 |
| **FAM127B** | 4.49532 | 6.99649 | 4.52767 | 4.44461 | 3.54300 | 4.50989 | 0.88140 |
| **FAM127C** | 0.57891 | 0.06999 | 1.07239 | 0.00000 | 0.79067 | 2.09449 | 0.05187 |
| **FAM129A** | 0.02572 | 0.02433 | 0.00388 | 0.10337 | 0.13931 | 0.08797 | 0.00601 |
| **FAM129B** | 0.00000 | 0.00000 | 0.31069 | 2.90144 | 0.23253 | 0.01851 | 2.34204 |
| **FAM129C** | 0.03639 | 0.06371 | 0.04384 | 0.03648 | 0.00657 | 0.00976 | 0.00000 |

| | | | | | | | |
|---|---|---|---|---|---|---|---|
| **FAM131A** | 0.06519 | 0.39108 | 0.07730 | 3.70124 | 0.15706 | 0.40583 | 0.00000 |
| **FAM131B** | 0.01672 | 0.00621 | 0.01249 | 0.09952 | 0.01018 | 0.03292 | 0.08495 |
| **FAM132A** | 0.00000 | 0.64008 | 0.02350 | 0.00000 | 0.04967 | 2.61740 | 0.37717 |
| **FAM132B** | 0.09954 | 0.05673 | 0.27783 | 0.00000 | 0.00513 | 0.05169 | 0.02513 |
| **FAM133A** | 2.85985 | 1.05541 | 2.90818 | 0.00000 | 0.14093 | 1.07974 | 1.37346 |
| **FAM133B** | 0.20955 | 0.00000 | 0.22030 | 0.14002 | 0.24276 | 0.20358 | 0.57173 |
| **FAM133CP** | 0.21245 | 0.00000 | 0.21791 | 0.06179 | 0.22137 | 0.14136 | 0.47759 |
| **FAM134A** | 1.50900 | 5.06202 | 3.71358 | 5.71039 | 1.06052 | 7.14143 | 17.57487 |
| **FAM134B** | 0.23812 | 0.00000 | 0.00251 | 0.00230 | 0.00917 | 0.00000 | 1.33195 |
| **FAM134C** | 4.29851 | 4.39455 | 4.19974 | 9.43068 | 1.28169 | 5.87557 | 4.78098 |
| **FAM135A** | 8.99425 | 0.70758 | 3.43884 | 4.21388 | 2.71219 | 2.62341 | 5.26425 |
| **FAM135B** | 0.02273 | 0.01770 | 0.00000 | 0.01674 | 0.25819 | 0.02097 | 0.03471 |
| **FAM136A** | 43.18473 | 32.65656 | 16.81848 | 32.41324 | 4.95831 | 15.75752 | 5.09752 |
| **FAM13A** | 0.12868 | 0.27260 | 0.83224 | 0.47355 | 7.61843 | 0.00000 | 0.04636 |
| **FAM13A-AS1** | 0.29315 | 0.00000 | 0.02807 | 0.00000 | 0.00579 | 0.00143 | 0.46011 |
| **FAM13B** | 2.53298 | 0.00000 | 1.74235 | 0.87419 | 3.31226 | 2.49746 | 5.20312 |
| **FAM13C** | 0.00783 | 0.02227 | 0.01070 | 0.01642 | 0.02064 | 0.10772 | 0.03971 |
| **FAM149A** | 0.01287 | 0.03406 | 0.06156 | 0.01400 | 0.69869 | 0.01002 | 0.03469 |
| **FAM149B1** | 0.98953 | 0.02967 | 0.88199 | 0.29071 | 0.17623 | 1.20406 | 2.10785 |
| **FAM150A** | 0.02239 | 0.03118 | 0.04467 | 0.02349 | 0.09192 | 0.01212 | 0.00774 |
| **FAM150B** | 0.01988 | 0.00000 | 0.15202 | 0.08410 | 0.03570 | 0.01132 | 0.06460 |
| **FAM151A** | 0.00000 | 45.09926 | 40.98173 | 0.00000 | 14.25127 | 24.34748 | 0.00000 |
| **FAM151B** | 0.69282 | 0.00000 | 0.01448 | 0.68370 | 0.21135 | 0.07345 | 0.36173 |
| **FAM153A** | 0.03511 | 0.02266 | 0.01394 | 0.03900 | 0.00895 | 0.01607 | 0.00027 |
| **FAM153C** | 0.03089 | 0.01710 | 0.01336 | 0.02054 | 0.00796 | 0.01973 | 0.01117 |
| **FAM154A** | 0.00000 | 0.27366 | 0.10908 | 0.01722 | 3.93599 | 0.03376 | 0.01102 |
| **FAM154B** | 0.00000 | 0.21439 | 0.05120 | 0.13820 | 1.17582 | 0.06586 | 0.22667 |
| **FAM155A** | 0.02598 | 0.03727 | 0.02553 | 0.00799 | 0.01690 | 0.68328 | 0.04179 |
| **FAM155A-IT1** | 0.02006 | 0.00000 | 0.08354 | 0.00555 | 0.00438 | 0.26255 | 0.07273 |
| **FAM155B** | 0.02634 | 0.65196 | 0.86820 | 0.23942 | 0.00000 | 0.54366 | 0.59796 |
| **FAM156A.uid-12** | 0.07017 | 0.02508 | 0.01712 | 0.03293 | 0.01417 | 0.00877 | 0.03384 |
| **FAM157A** | 0.05886 | 0.01401 | 0.06603 | 0.07333 | 0.04222 | 0.10490 | 0.00004 |
| **FAM157B** | 0.02487 | 0.01743 | 0.01221 | 0.01912 | 0.01102 | 0.01372 | 0.01900 |
| **FAM159A** | 0.01236 | 0.01732 | 0.02452 | 0.03772 | 0.03899 | 0.03260 | 0.02636 |
| **FAM159B** | 0.00000 | 0.11716 | 0.00499 | 0.24348 | 0.01071 | 0.19228 | 0.01989 |

| | | | | | | | |
|---|---|---|---|---|---|---|---|
| **FAM160A1** | 0.85877 | 1.62649 | 1.57956 | 0.69033 | 1.06690 | 2.21789 | 0.00004 |
| **FAM160A2** | 0.00000 | 1.41864 | 1.97821 | 16.02592 | 1.54759 | 0.65927 | 7.20749 |
| **FAM160B1** | 4.70108 | 1.64694 | 3.70656 | 5.11176 | 0.26290 | 1.90996 | 9.85169 |
| **FAM160B2** | 0.00000 | 1.82553 | 2.48537 | 1.19353 | 0.11422 | 2.12646 | 4.92204 |
| **FAM161A** | 1.10043 | 0.00000 | 1.44449 | 2.12896 | 1.91315 | 1.25122 | 2.12920 |
| **FAM161B** | 0.20935 | 0.36656 | 0.40036 | 0.45419 | 1.18812 | 0.14803 | 0.48170 |
| **FAM162A** | 26.91442 | 6.63233 | 8.77421 | 13.36463 | 0.88014 | 13.73768 | 8.11675 |
| **FAM162B** | 0.00000 | 0.43140 | 12.04168 | 0.00000 | 0.56867 | 3.33094 | 0.00002 |
| **FAM163A** | 0.01807 | 0.02071 | 0.01188 | 0.02276 | 0.01846 | 0.01569 | 0.03104 |
| **FAM163B** | 0.00000 | 0.34926 | 0.29814 | 0.00890 | 0.00234 | 0.04849 | 0.01570 |
| **FAM166A** | 0.00259 | 0.02352 | 0.02695 | 0.04438 | 0.00626 | 0.01965 | 0.08652 |
| **FAM166B** | 0.01199 | 0.00000 | 0.09903 | 0.04692 | 0.11184 | 0.01571 | 0.03139 |
| **FAM167A** | 0.02621 | 0.10565 | 0.00000 | 0.03598 | 1.11852 | 0.00387 | 0.00059 |
| **FAM167B** | 0.02908 | 0.01277 | 0.02516 | 0.03379 | 0.00719 | 0.01307 | 0.01237 |
| **FAM168A** | 1.09061 | 4.09785 | 0.85138 | 2.35646 | 1.89665 | 2.69450 | 4.69924 |
| **FAM168B** | 15.95263 | 9.45384 | 5.53806 | 9.70708 | 2.15240 | 5.78393 | 14.92459 |
| **FAM169A** | 2.24705 | 3.19420 | 3.30867 | 4.54644 | 1.90713 | 5.83523 | 2.99632 |
| **FAM169B** | 0.02415 | 0.00375 | 0.03024 | 0.01895 | 0.26649 | 0.01313 | 0.02051 |
| **FAM171A1** | 0.00000 | 2.59757 | 2.43775 | 0.20030 | 0.19638 | 1.56744 | 0.76282 |
| **FAM171A2** | 0.02592 | 0.00934 | 0.04815 | 0.02262 | 0.00684 | 0.01338 | 0.01766 |
| **FAM171B** | 0.02507 | 0.06491 | 0.01220 | 0.03499 | 0.40581 | 0.02272 | 0.00000 |
| **FAM172A** | 3.99917 | 0.00000 | 3.64313 | 3.21118 | 1.06432 | 0.96909 | 3.29356 |
| **FAM172BP** | 0.03142 | 0.02923 | 0.09645 | 0.00529 | 0.71413 | 0.00000 | 0.05701 |
| **FAM173A** | 3.15279 | 5.07337 | 0.03998 | 3.02300 | 0.02096 | 1.57123 | 7.45096 |
| **FAM173B** | 2.11466 | 0.62370 | 3.25708 | 0.55527 | 0.22889 | 1.41830 | 0.88604 |
| **FAM174A** | 2.46225 | 2.17929 | 0.78600 | 2.50056 | 1.06230 | 1.24898 | 0.82314 |
| **FAM174B** | 0.00000 | 0.72238 | 0.36813 | 0.00000 | 1.23470 | 1.39900 | 0.12673 |
| **FAM175A** | 2.58901 | 0.00000 | 3.21677 | 0.48225 | 0.36301 | 1.09053 | 0.24739 |
| **FAM175B** | 9.51701 | 3.50979 | 9.58283 | 4.15218 | 4.00382 | 4.31851 | 3.65333 |
| **FAM177A1** | 6.48223 | 5.17962 | 2.79955 | 9.37963 | 6.47708 | 6.45226 | 9.92068 |
| **FAM178A** | 9.40094 | 3.04202 | 4.28088 | 3.13866 | 1.73406 | 3.46134 | 3.44348 |
| **FAM178B** | 0.22026 | 0.89058 | 2.09637 | 0.65338 | 0.13307 | 0.28241 | 1.12668 |
| **FAM179A** | 0.01111 | 0.08853 | 0.00442 | 0.11210 | 0.44067 | 0.00864 | 0.03004 |
| **FAM179B** | 1.56878 | 0.17239 | 2.82660 | 1.57694 | 0.00000 | 0.94503 | 4.21519 |
| **FAM180A** | 0.00289 | 0.01706 | 0.00776 | 0.43495 | 0.04483 | 0.00370 | 0.08171 |

| | | | | | | | |
|---|---|---|---|---|---|---|---|
| **FAM180B** | 0.02752 | 0.02045 | 0.00416 | 0.04036 | 0.11323 | 0.01628 | 0.03217 |
| **FAM181A-AS1** | 0.01855 | 0.01542 | 0.01569 | 0.02349 | 0.00903 | 0.02546 | 0.03342 |
| **FAM181B** | 0.01338 | 0.03011 | 0.01884 | 0.02807 | 0.02153 | 0.02247 | 0.02151 |
| **FAM182A** | 0.03168 | 0.03150 | 0.01345 | 0.03465 | 0.00763 | 0.06605 | 0.00458 |
| **FAM182B** | 0.02134 | 0.02870 | 0.02610 | 0.07552 | 0.00654 | 0.07158 | 0.01962 |
| **FAM183A** | 0.04182 | 0.02456 | 0.02134 | 0.03663 | 0.01125 | 0.03737 | 0.05846 |
| **FAM183B** | 0.05433 | 0.00317 | 0.04667 | 0.03688 | 0.00710 | 0.00741 | 0.02301 |
| **FAM183CP** | 0.04625 | 0.01274 | 0.01338 | 0.01728 | 0.01281 | 0.01269 | 0.01075 |
| **FAM184A** | 1.58765 | 0.00000 | 0.37086 | 2.91133 | 0.01722 | 0.00000 | 5.78516 |
| **FAM184B** | 0.60105 | 0.00000 | 0.22833 | 0.53628 | 0.95767 | 0.30137 | 1.32657 |
| **FAM185A** | 1.32882 | 0.66909 | 0.25244 | 1.17030 | 0.11868 | 0.50224 | 0.19544 |
| **FAM186A** | 0.01779 | 0.10011 | 0.02028 | 0.02438 | 0.02441 | 0.00682 | 0.01812 |
| **FAM186B** | 0.16768 | 0.00000 | 0.13827 | 0.17179 | 0.12248 | 0.00000 | 0.37978 |
| **FAM188A** | 7.09255 | 1.56952 | 8.69379 | 3.75657 | 4.43997 | 0.37803 | 0.00000 |
| **FAM188B** | 0.05420 | 0.03910 | 0.09811 | 0.14017 | 0.01425 | 0.00815 | 0.12656 |
| **FAM189A1** | 0.03225 | 0.01838 | 0.05241 | 0.03107 | 0.02609 | 0.04368 | 0.02000 |
| **FAM189A2** | 0.27804 | 0.00000 | 0.00927 | 0.46626 | 2.00332 | 0.00000 | 0.05460 |
| **FAM189B** | 0.21261 | 0.00000 | 0.01972 | 2.46490 | 0.05253 | 1.13977 | 6.21032 |
| **FAM192A** | 13.56096 | 15.38312 | 11.46796 | 9.94520 | 2.17456 | 6.38349 | 4.05081 |
| **FAM193A** | 3.67297 | 8.65840 | 2.54647 | 1.26689 | 1.89621 | 3.83115 | 12.38015 |
| **FAM193B** | 0.56154 | 1.64660 | 1.87164 | 3.74485 | 0.06217 | 8.46070 | 10.79005 |
| **FAM195A** | 6.59281 | 21.41160 | 8.92398 | 1.46207 | 0.00000 | 4.19754 | 9.03893 |
| **FAM195B** | 2.61219 | 0.00000 | 0.66891 | 16.43005 | 0.69598 | 3.77424 | 5.93596 |
| **FAM196A** | 0.12216 | 0.00000 | 0.29532 | 0.27036 | 0.06308 | 0.34512 | 1.46594 |
| **FAM196B** | 0.02889 | 0.01890 | 0.01303 | 0.03541 | 0.00788 | 0.01344 | 0.01593 |
| **FAM198A** | 0.06094 | 0.16904 | 0.08150 | 0.16286 | 0.11031 | 0.03664 | 0.00000 |
| **FAM198B** | 0.15588 | 0.16309 | 0.05948 | 0.31448 | 0.14912 | 0.90853 | 0.00000 |
| **FAM199X** | 6.28839 | 2.56191 | 1.29231 | 0.43531 | 0.62678 | 5.38229 | 11.21915 |
| **FAM19A1** | 0.02555 | 0.01699 | 0.01213 | 0.01831 | 0.01753 | 0.01292 | 0.01766 |
| **FAM19A2** | 0.00440 | 0.01300 | 0.01567 | 0.15433 | 0.42085 | 0.01919 | 0.03011 |
| **FAM19A3** | 0.00000 | 0.09504 | 4.90274 | 0.00136 | 0.90990 | 0.00000 | 0.01893 |
| **FAM19A4** | 0.06289 | 0.00000 | 0.67132 | 0.11443 | 2.60386 | 0.04132 | 0.17102 |
| **FAM19A5** | 0.00000 | 0.25697 | 0.00000 | 0.00000 | 0.45041 | 0.35669 | 0.26314 |
| **FAM200A** | 1.98312 | 0.00000 | 4.34401 | 0.81332 | 3.38531 | 0.62671 | 0.24343 |
| **FAM200B** | 8.01581 | 4.73196 | 5.58302 | 3.40930 | 3.13655 | 2.91396 | 3.86914 |

| | | | | | | | |
|---|---|---|---|---|---|---|---|
| **FAM201A** | 0.03807 | 0.00851 | 0.05079 | 0.03020 | 0.01122 | 0.01007 | 0.01789 |
| **FAM204A** | 12.31924 | 2.99033 | 7.11374 | 6.79029 | 2.05833 | 4.04857 | 2.38771 |
| **FAM205B** | 0.04034 | 0.01043 | 0.01465 | 0.01737 | 0.00731 | 0.00968 | 0.04142 |
| **FAM206A** | 8.63515 | 2.26899 | 8.05473 | 7.99264 | 0.49704 | 3.01934 | 0.00000 |
| **FAM207A** | 0.11151 | 6.12217 | 0.96660 | 0.00000 | 0.15977 | 1.67370 | 7.05272 |
| **FAM208A** | 4.98415 | 0.31796 | 7.25762 | 0.24800 | 3.00976 | 5.40368 | 6.63029 |
| **FAM208B** | 11.19096 | 2.70367 | 2.82699 | 2.32914 | 5.25294 | 4.65659 | 14.11310 |
| **FAM209A** | 0.02845 | 0.06517 | 0.01889 | 0.03898 | 0.01758 | 0.01491 | 0.02449 |
| **FAM209B** | 0.10346 | 0.17522 | 0.23863 | 0.07089 | 0.02701 | 0.06456 | 0.00000 |
| **FAM20A** | 0.04416 | 0.09730 | 0.00775 | 0.03193 | 0.01446 | 0.51925 | 0.15663 |
| **FAM20B** | 4.50360 | 1.41289 | 1.17421 | 2.11380 | 0.51458 | 5.30356 | 13.94573 |
| **FAM20C** | 0.02467 | 0.01821 | 0.01291 | 0.01826 | 0.00792 | 0.01805 | 0.01942 |
| **FAM210A** | 23.07441 | 11.87663 | 4.32717 | 7.66968 | 5.44569 | 6.42248 | 3.81995 |
| **FAM210B** | 0.00000 | 0.84905 | 0.28578 | 0.62176 | 0.09460 | 0.57950 | 1.05269 |
| **FAM212A** | 0.05283 | 0.09565 | 0.16163 | 0.08474 | 0.15910 | 0.00749 | 0.00000 |
| **FAM212B** | 0.00000 | 1.05458 | 1.31888 | 0.00000 | 0.00071 | 0.21089 | 0.01142 |
| **FAM212B-AS1** | 0.00000 | 0.13630 | 1.21995 | 0.00000 | 0.18291 | 0.00000 | 0.05938 |
| **FAM213A** | 31.17130 | 2.13914 | 4.43410 | 13.41032 | 0.06489 | 5.59964 | 12.80894 |
| **FAM213B** | 0.01034 | 0.00000 | 0.41436 | 0.40683 | 0.12182 | 0.35774 | 0.09361 |
| **FAM214A** | 0.46284 | 0.00000 | 8.23489 | 2.49226 | 5.11070 | 0.97763 | 1.08024 |
| **FAM214B** | 0.00000 | 0.33774 | 3.61061 | 0.48468 | 1.77530 | 3.92931 | 1.72247 |
| **FAM215A** | 0.03016 | 0.02246 | 0.01022 | 0.01844 | 0.08755 | 0.02300 | 0.00763 |
| **FAM216A** | 2.35738 | 0.00000 | 1.71012 | 3.99258 | 0.24624 | 1.30219 | 0.00000 |
| **FAM217A** | 0.02368 | 0.02446 | 0.00993 | 0.02657 | 0.02278 | 0.01237 | 0.01243 |
| **FAM217B** | 5.33517 | 0.58018 | 0.84567 | 2.10774 | 0.14740 | 1.72631 | 4.55852 |
| **FAM218A** | 0.31148 | 0.52369 | 0.86693 | 0.18236 | 3.70309 | 0.45480 | 0.00921 |
| **FAM219A** | 0.00000 | 1.11023 | 0.02218 | 4.49302 | 0.15033 | 0.70983 | 5.28602 |
| **FAM219B** | 2.25959 | 2.98318 | 1.18810 | 4.24819 | 0.37405 | 2.70662 | 8.05059 |
| **FAM21A** | 0.58794 | 0.47084 | 0.24178 | 0.73352 | 0.31488 | 0.28856 | 0.00000 |
| **FAM21C** | 1.23023 | 0.41897 | 0.37984 | 0.69771 | 0.20376 | 0.53562 | 0.00000 |
| **FAM21EP** | 0.05624 | 0.10289 | 0.01061 | 0.02351 | 0.00897 | 0.03377 | 0.01628 |
| **FAM220A** | 1.77672 | 0.59060 | 1.52326 | 0.32694 | 7.13361 | 1.56350 | 3.27249 |
| **FAM221A** | 0.94139 | 0.24498 | 0.23239 | 0.12138 | 0.15336 | 3.75860 | 0.00000 |
| **FAM221B** | 0.00022 | 0.01733 | 0.01171 | 0.05175 | 0.02628 | 0.00915 | 0.10355 |
| **FAM222A** | 0.18478 | 1.34084 | 0.38437 | 0.78668 | 1.02440 | 0.81752 | 4.78053 |

| | | | | | | | |
|---|---|---|---|---|---|---|---|
| **FAM222A-AS1** | 0.06564 | 0.03916 | 0.01522 | 0.04025 | 0.46508 | 0.09019 | 0.10784 |
| **FAM222B** | 12.85360 | 10.34749 | 7.03767 | 11.09431 | 6.66722 | 11.33204 | 17.00111 |
| **FAM223A+FAM2** | 0.02172 | 0.05430 | 0.01071 | 0.02273 | 0.02540 | 0.01453 | 0.00997 |
| **FAM223A+FAM2** | 0.02220 | 0.01727 | 0.02458 | 0.01981 | 0.00859 | 0.01873 | 0.02120 |
| **FAM225A** | 0.02439 | 0.00000 | 0.03214 | 0.05097 | 0.00680 | 0.02223 | 0.09888 |
| **FAM225B** | 0.02308 | 0.01448 | 0.02886 | 0.02209 | 0.01202 | 0.01243 | 0.01616 |
| **FAM226A+FAM2** | 0.03062 | 0.00671 | 0.03679 | 0.01704 | 0.00652 | 0.02005 | 0.02454 |
| **FAM227A** | 0.34696 | 0.00000 | 0.00592 | 0.21703 | 0.41518 | 0.32396 | 0.00000 |
| **FAM227B** | 0.27815 | 0.09896 | 0.21000 | 0.05268 | 0.14379 | 0.29264 | 0.00000 |
| **FAM228A** | 0.00000 | 0.10922 | 0.67983 | 0.10907 | 2.03922 | 0.23067 | 0.01221 |
| **FAM228B** | 0.13981 | 0.03788 | 0.04217 | 0.07873 | 0.03575 | 0.02181 | 0.00000 |
| **FAM229A** | 0.03014 | 0.11976 | 0.04885 | 0.14744 | 0.01137 | 0.00000 | 0.19933 |
| **FAM229B** | 0.89505 | 0.50971 | 1.02572 | 0.04920 | 0.06970 | 0.26784 | 0.80420 |
| **FAM230B** | 0.02336 | 0.03004 | 0.01728 | 0.01811 | 0.00731 | 0.01235 | 0.01897 |
| **FAM24A** | 0.03237 | 0.01954 | 0.01361 | 0.01502 | 0.00769 | 0.00991 | 0.02650 |
| **FAM24B** | 1.10046 | 0.43900 | 0.32399 | 1.46643 | 0.95739 | 0.66284 | 0.68480 |
| **FAM24B-CUZD1** | 0.02550 | 0.01675 | 0.01278 | 0.02357 | 0.00798 | 0.02031 | 0.01531 |
| **FAM25A** | 0.01721 | 0.02246 | 0.01059 | 0.01583 | 0.01980 | 0.01127 | 0.04040 |
| **FAM25BP** | 0.02428 | 0.01780 | 0.01265 | 0.02042 | 0.00929 | 0.01302 | 0.01584 |
| **FAM26D** | 0.18280 | 0.10595 | 0.01212 | 0.00000 | 0.16459 | 0.00000 | 0.12020 |
| **FAM26E** | 0.38279 | 0.00000 | 0.00633 | 0.00000 | 0.00561 | 0.00039 | 0.24626 |
| **FAM26F** | 0.16200 | 0.03052 | 0.19747 | 0.15754 | 0.00343 | 0.01781 | 0.00000 |
| **FAM27L** | 0.02384 | 0.01832 | 0.01271 | 0.02038 | 0.00799 | 0.01303 | 0.01709 |
| **FAM32A** | 36.39051 | 24.53660 | 23.42986 | 30.60167 | 20.47905 | 15.47539 | 1.32510 |
| **FAM35A** | 2.70493 | 0.00000 | 2.15886 | 2.17754 | 0.54628 | 2.55935 | 2.83938 |
| **FAM35BP** | 0.02450 | 0.01716 | 0.01288 | 0.02127 | 0.01024 | 0.01590 | 0.01846 |
| **FAM35DP** | 0.03409 | 0.01825 | 0.01767 | 0.01952 | 0.01692 | 0.01469 | 0.01339 |
| **FAM3A** | 1.28789 | 4.74922 | 1.07693 | 1.87385 | 0.00000 | 3.61446 | 2.29259 |
| **FAM3B** | 0.11030 | 0.29235 | 0.19021 | 0.00889 | 0.14790 | 0.00350 | 0.05766 |
| **FAM3C** | 2.54604 | 0.00000 | 0.18620 | 0.73572 | 0.03244 | 0.61897 | 6.11082 |
| **FAM3D** | 0.02726 | 0.01102 | 0.02594 | 0.01900 | 0.03418 | 0.05860 | 0.01145 |
| **FAM41C** | 0.00289 | 0.02086 | 0.01380 | 0.02623 | 0.12782 | 0.02843 | 0.06454 |
| **FAM43A** | 0.00000 | 0.01189 | 0.00000 | 0.26476 | 1.16820 | 0.29926 | 0.06025 |
| **FAM43B** | 0.00000 | 0.16659 | 0.15020 | 0.01386 | 0.11595 | 0.06485 | 0.01656 |
| **FAM45A** | 0.06652 | 0.30321 | 0.08297 | 0.06785 | 0.03050 | 0.05290 | 0.00000 |

| | | | | | | | |
|---|---|---|---|---|---|---|---|
| **FAM45B.uid-124** | 3.04955 | 2.69197 | 1.50993 | 0.76498 | 0.07643 | 2.15634 | 0.00000 |
| **FAM45B.uid-145** | 0.02616 | 0.02183 | 0.02434 | 0.01736 | 0.00825 | 0.02589 | 0.03600 |
| **FAM46A** | 10.59713 | 3.42224 | 1.56587 | 10.52064 | 2.64920 | 0.00000 | 6.03682 |
| **FAM46B** | 0.00000 | 24.59536 | 0.00000 | 1.33156 | 2.30200 | 18.34936 | 8.97884 |
| **FAM46C** | 0.06561 | 1.02659 | 2.94647 | 0.23027 | 16.67106 | 2.17890 | 0.00000 |
| **FAM46D** | 0.06598 | 0.00000 | 0.25385 | 0.01806 | 0.07566 | 0.00693 | 0.01979 |
| **FAM47C** | 0.02371 | 0.01440 | 0.01591 | 0.02290 | 0.02622 | 0.01303 | 0.01666 |
| **FAM47E** | 0.02995 | 0.02915 | 0.01890 | 0.01265 | 0.06088 | 0.01983 | 0.06956 |
| **FAM49A** | 0.18017 | 0.08290 | 0.31268 | 8.47348 | 0.10665 | 0.10637 | 0.00000 |
| **FAM49B** | 17.62818 | 1.14272 | 12.52480 | 17.53823 | 1.17400 | 9.32781 | 12.05816 |
| **FAM50A** | 16.96966 | 26.76573 | 5.55294 | 17.91651 | 0.00000 | 8.85837 | 17.54618 |
| **FAM50B** | 0.21074 | 0.03573 | 0.29389 | 0.30238 | 0.12764 | 0.02968 | 0.04395 |
| **FAM53A** | 0.00000 | 0.20789 | 0.18464 | 0.02014 | 0.04782 | 0.08988 | 0.18629 |
| **FAM53B** | 0.81897 | 5.33145 | 1.07432 | 1.17065 | 4.31836 | 0.87378 | 9.77592 |
| **FAM53B-AS1** | 0.05019 | 0.00000 | 0.03370 | 0.03929 | 0.28816 | 0.09041 | 0.23418 |
| **FAM53C** | 7.54725 | 1.06898 | 7.95036 | 9.51211 | 8.92071 | 6.03016 | 15.55573 |
| **FAM57A** | 1.10959 | 1.13284 | 0.21183 | 1.23693 | 0.12443 | 0.00000 | 3.65680 |
| **FAM57B** | 0.02404 | 0.01405 | 0.02016 | 0.01881 | 0.01090 | 0.01238 | 0.02101 |
| **FAM58A** | 0.01270 | 4.35538 | 5.96408 | 0.60670 | 6.09341 | 1.69837 | 11.47079 |
| **FAM60A** | 22.27829 | 5.48279 | 8.93483 | 13.81249 | 3.65878 | 9.02143 | 3.82981 |
| **FAM63A** | 0.39364 | 0.00000 | 0.16327 | 0.25907 | 0.28035 | 0.00446 | 0.43095 |
| **FAM63B** | 4.05506 | 7.13550 | 5.76438 | 8.86748 | 2.11209 | 10.61695 | 15.87969 |
| **FAM64A** | 0.46483 | 0.00000 | 0.06823 | 0.37270 | 2.59832 | 7.32679 | 0.00000 |
| **FAM65A** | 0.00000 | 0.00000 | 0.64631 | 3.33658 | 0.00047 | 0.42085 | 0.77924 |
| **FAM65B** | 0.41096 | 1.53837 | 0.10819 | 7.66951 | 0.85023 | 0.00000 | 0.23542 |
| **FAM65C** | 0.11337 | 0.11479 | 0.23080 | 0.03915 | 0.53471 | 0.04794 | 0.12484 |
| **FAM66A** | 0.02605 | 0.01345 | 0.03286 | 0.01814 | 0.00846 | 0.01197 | 0.01831 |
| **FAM66C** | 0.01655 | 0.03258 | 0.11897 | 0.01616 | 0.07617 | 0.15692 | 0.01312 |
| **FAM66D** | 0.01030 | 0.06880 | 0.04490 | 0.02040 | 0.05683 | 0.03058 | 0.02030 |
| **FAM69A** | 0.77883 | 0.49701 | 0.55662 | 0.51407 | 0.44574 | 0.89789 | 0.05969 |
| **FAM69B** | 0.00000 | 4.37724 | 0.02577 | 1.53497 | 0.71278 | 6.44117 | 6.86936 |
| **FAM69C** | 0.04944 | 0.01631 | 0.00944 | 0.01594 | 0.05200 | 0.01321 | 0.00522 |
| **FAM71A** | 0.02123 | 0.05142 | 0.02007 | 0.02175 | 0.01412 | 0.00900 | 0.01183 |
| **FAM71C** | 0.02587 | 0.00527 | 0.03632 | 0.01731 | 0.04132 | 0.01427 | 0.02411 |
| **FAM71D** | 0.01491 | 0.02440 | 0.01086 | 0.03936 | 0.00849 | 0.00761 | 0.04821 |

| | | | | | | | |
|---|---|---|---|---|---|---|---|
| **FAM71E1** | 3.08573 | 4.42670 | 1.25297 | 3.22207 | 2.08364 | 2.72514 | 1.50281 |
| **FAM71E2** | 0.02621 | 0.01241 | 0.01221 | 0.02439 | 0.03470 | 0.01925 | 0.02270 |
| **FAM71F1** | 0.06168 | 0.06100 | 0.02427 | 0.05778 | 0.02184 | 0.26730 | 0.02254 |
| **FAM71F2** | 0.33843 | 0.08734 | 0.25741 | 0.07038 | 0.00261 | 0.00000 | 0.56986 |
| **FAM72B** | 0.48651 | 0.54132 | 0.96753 | 0.36510 | 0.06128 | 0.15942 | 0.09375 |
| **FAM73A** | 6.10410 | 0.00000 | 3.79842 | 6.68323 | 0.95794 | 1.57890 | 10.17067 |
| **FAM73B** | 0.75367 | 2.67381 | 1.68117 | 2.70897 | 0.06302 | 2.68102 | 6.94118 |
| **FAM76A** | 22.44221 | 0.43061 | 10.82230 | 11.35066 | 7.65372 | 2.11728 | 5.74512 |
| **FAM76B** | 5.07430 | 0.00000 | 3.65216 | 1.26721 | 1.55384 | 2.79016 | 5.56704 |
| **FAM78A** | 0.32489 | 0.09972 | 0.09197 | 2.14900 | 0.12523 | 0.00000 | 2.27312 |
| **FAM78B** | 0.00000 | 0.02516 | 0.25902 | 0.00491 | 0.14332 | 0.52731 | 0.08534 |
| **FAM81A** | 0.30370 | 0.00563 | 0.00880 | 0.05869 | 0.39334 | 0.11461 | 0.00000 |
| **FAM81B** | 0.02116 | 0.00064 | 0.02951 | 0.02673 | 0.56941 | 0.03428 | 0.01452 |
| **FAM83A** | 0.09426 | 0.00000 | 0.21815 | 0.14263 | 0.07474 | 0.09235 | 0.11008 |
| **FAM83A-AS1** | 0.03678 | 0.00000 | 0.06936 | 0.02867 | 0.02385 | 0.01985 | 0.07550 |
| **FAM83B** | 4.05176 | 0.00000 | 0.56112 | 4.59446 | 0.29987 | 0.00000 | 3.35474 |
| **FAM83C** | 0.03303 | 0.01481 | 0.02254 | 0.01551 | 0.00613 | 0.04492 | 0.06489 |
| **FAM83C-AS1** | 0.02876 | 0.01327 | 0.01433 | 0.06297 | 0.00699 | 0.00508 | 0.07108 |
| **FAM83D** | 20.39633 | 16.34629 | 5.31243 | 8.91665 | 3.72016 | 7.84297 | 12.92806 |
| **FAM83E** | 0.00000 | 0.00000 | 0.15366 | 0.42661 | 1.25327 | 0.07008 | 0.16775 |
| **FAM83F** | 0.00000 | 1.24990 | 0.00000 | 0.64862 | 0.05350 | 1.94393 | 3.25731 |
| **FAM83G** | 0.00000 | 2.03890 | 0.00000 | 2.63740 | 0.07005 | 0.36987 | 2.15215 |
| **FAM83H** | 0.05004 | 4.94914 | 0.07617 | 3.52059 | 0.00000 | 1.03327 | 17.78364 |
| **FAM83H-AS1** | 1.17426 | 1.08619 | 0.00072 | 4.36230 | 0.00000 | 0.00000 | 6.04081 |
| **FAM84A** | 0.12514 | 0.00000 | 0.02254 | 0.01103 | 0.01324 | 0.00420 | 0.09396 |
| **FAM84B** | 7.63201 | 17.59755 | 3.06619 | 0.93761 | 1.27516 | 9.75199 | 10.66487 |
| **FAM86A** | 2.17135 | 3.41243 | 1.81608 | 1.13099 | 0.07720 | 0.53055 | 0.00000 |
| **FAM86B1** | 0.02309 | 0.07398 | 0.06457 | 0.03378 | 0.04465 | 0.00497 | 0.02033 |
| **FAM86B2** | 0.02205 | 0.02634 | 0.03268 | 0.02069 | 0.05430 | 0.01383 | 0.01382 |
| **FAM86B3P** | 0.33809 | 0.13094 | 0.02066 | 0.08229 | 0.19932 | 0.08854 | 0.04274 |
| **FAM86C1** | 0.93816 | 2.55287 | 1.02775 | 0.42584 | 0.21378 | 0.26729 | 0.00000 |
| **FAM86C2P** | 0.11982 | 0.18719 | 0.36456 | 0.20044 | 0.05580 | 0.04345 | 0.05596 |
| **FAM86DP** | 0.00000 | 0.69428 | 0.40883 | 0.30629 | 0.09145 | 0.18725 | 0.54727 |
| **FAM86EP** | 0.12036 | 0.79424 | 1.06943 | 0.00000 | 0.12106 | 1.64933 | 0.40298 |
| **FAM86FP** | 0.04081 | 0.05809 | 0.08738 | 0.07582 | 0.09118 | 0.04685 | 0.00000 |

| | | | | | | | |
|---|---|---|---|---|---|---|---|
| **FAM86HP** | 0.04456 | 0.02049 | 0.06375 | 0.06312 | 0.01319 | 0.11841 | 0.06519 |
| **FAM86JP** | 0.04918 | 0.31557 | 0.50393 | 0.07505 | 0.11669 | 0.12826 | 0.03147 |
| **FAM87A** | 0.03904 | 0.02589 | 0.01115 | 0.04122 | 0.03094 | 0.01218 | 0.01456 |
| **FAM87B** | 0.00730 | 0.04706 | 0.02565 | 0.06666 | 0.04191 | 0.02074 | 0.05439 |
| **FAM89A** | 0.00000 | 0.00000 | 0.38969 | 4.96338 | 1.22059 | 0.47942 | 0.94500 |
| **FAM89B** | 0.00000 | 0.97127 | 0.77738 | 2.50527 | 2.88956 | 1.20035 | 5.29455 |
| **FAM8A1** | 3.44866 | 0.00000 | 0.81974 | 1.63200 | 0.64361 | 0.81109 | 4.27645 |
| **FAM90A1** | 0.02468 | 0.00722 | 0.01437 | 0.01745 | 0.41171 | 0.02901 | 0.02596 |
| **FAM90A25P** | 0.02406 | 0.01767 | 0.01202 | 0.01867 | 0.02329 | 0.01274 | 0.01884 |
| **FAM90A27P** | 0.02022 | 0.01729 | 0.00000 | 0.03138 | 0.66081 | 0.01345 | 0.01404 |
| **FAM91A1** | 7.12075 | 0.00000 | 4.24225 | 2.90735 | 0.35723 | 1.55431 | 8.21286 |
| **FAM92A1** | 1.44754 | 0.15744 | 0.56758 | 0.05915 | 0.14687 | 2.16439 | 0.82014 |
| **FAM92A1P2** | 0.01967 | 0.01141 | 0.02670 | 0.01177 | 0.01069 | 0.02735 | 0.06025 |
| **FAM92B** | 0.01077 | 0.02022 | 0.01199 | 0.03240 | 0.00780 | 0.02786 | 0.03658 |
| **FAM95A** | 0.01194 | 0.21032 | 0.01063 | 0.01954 | 0.00880 | 0.11174 | 0.00000 |
| **FAM95C** | 0.03484 | 0.01238 | 0.08688 | 0.05475 | 0.01515 | 0.16810 | 0.00072 |
| **FAM96A** | 31.30356 | 15.41560 | 12.45663 | 15.11683 | 1.66238 | 14.13778 | 4.09444 |
| **FAM96B** | 30.65336 | 40.48305 | 18.69228 | 27.36931 | 10.75883 | 19.95811 | 14.65640 |
| **FAM98A** | 22.21574 | 13.45709 | 9.05613 | 9.66609 | 1.86901 | 5.89907 | 5.31098 |
| **FAM98B** | 1.25216 | 0.00000 | 1.23770 | 1.81040 | 0.10496 | 1.41015 | 9.78895 |
| **FAM98C** | 0.00000 | 1.40840 | 0.32098 | 0.40659 | 0.15138 | 0.42779 | 0.71574 |
| **FAM9A** | 0.01887 | 0.00000 | 0.18029 | 0.06424 | 1.78562 | 0.00536 | 0.08979 |
| **FAM9B** | 0.27788 | 0.15612 | 0.34005 | 0.01375 | 1.26560 | 0.03990 | 0.00000 |
| **FAM9C** | 1.51543 | 0.52497 | 0.22748 | 2.93435 | 0.18180 | 0.28709 | 0.00000 |
| **FAN1** | 2.46552 | 1.50066 | 2.12530 | 0.04789 | 0.08252 | 1.11168 | 1.20599 |
| **FANCA** | 2.54213 | 7.45130 | 2.16558 | 2.61222 | 1.53299 | 9.11574 | 1.89505 |
| **FANCB** | 6.54148 | 0.00000 | 3.93343 | 4.70133 | 3.03760 | 2.27980 | 2.84706 |
| **FANCC** | 2.00352 | 1.22286 | 2.29524 | 1.76007 | 0.32398 | 0.17776 | 0.96472 |
| **FANCD2** | 14.76431 | 7.32374 | 7.91819 | 5.53965 | 2.96155 | 10.80344 | 9.07576 |
| **FANCD2OS** | 0.04860 | 0.03774 | 0.05239 | 0.05761 | 0.01651 | 0.08454 | 0.00170 |
| **FANCE** | 2.86298 | 3.15555 | 10.59876 | 7.00137 | 0.57275 | 2.07333 | 7.43527 |
| **FANCF** | 3.28493 | 0.52801 | 4.00432 | 1.27598 | 0.60913 | 1.01960 | 0.00000 |
| **FANCG** | 7.04955 | 3.81844 | 1.39117 | 9.98650 | 0.20861 | 6.03938 | 0.24954 |
| **FANCI** | 14.63127 | 4.72215 | 6.70447 | 12.26376 | 6.73789 | 4.13987 | 3.25784 |
| **FANCL** | 4.23078 | 0.00000 | 2.58851 | 4.55376 | 0.97094 | 1.29742 | 0.00000 |

| | | | | | | | |
|---|---|---|---|---|---|---|---|
| **FANCM** | 4.73956 | 0.76150 | 3.88125 | 2.21135 | 7.42623 | 2.28590 | 4.52569 |
| **FANK1** | 0.45430 | 0.39155 | 0.63186 | 0.66145 | 0.14179 | 0.29991 | 0.11128 |
| **FAP** | 0.02389 | 0.01701 | 0.01141 | 0.02498 | 0.02166 | 0.01352 | 0.01570 |
| **FAR1** | 23.12240 | 0.00000 | 4.45569 | 8.49892 | 4.76235 | 2.72302 | 1.62174 |
| **FAR2** | 0.12195 | 0.12321 | 0.06029 | 1.17542 | 0.33034 | 0.00762 | 0.00000 |
| **FAR2P1** | 0.00000 | 0.40532 | 0.19663 | 0.44727 | 0.07811 | 0.07908 | 0.02787 |
| **FAR2P2** | 0.02574 | 0.01914 | 0.02969 | 0.02539 | 0.01339 | 0.02285 | 0.02095 |
| **FARP1** | 4.61290 | 5.96793 | 0.00000 | 0.00000 | 1.06225 | 2.13443 | 12.56279 |
| **FARP2** | 2.30496 | 2.07738 | 0.39512 | 1.84003 | 2.86499 | 0.56795 | 0.00000 |
| **FARS2** | 5.70222 | 3.03205 | 4.00837 | 3.68053 | 0.82517 | 2.97941 | 0.00000 |
| **FARSA** | 39.52737 | 44.63573 | 9.75624 | 32.08371 | 0.63713 | 14.75758 | 2.64012 |
| **FARSB** | 30.62932 | 13.42270 | 14.52784 | 11.17216 | 1.70545 | 8.08117 | 4.41362 |
| **FAS** | 0.03886 | 0.01114 | 0.21301 | 0.05663 | 0.12585 | 0.05478 | 0.00006 |
| **FAS-AS1** | 0.02230 | 0.01826 | 0.01059 | 0.02050 | 0.04253 | 0.01310 | 0.01789 |
| **FASLG** | 0.00438 | 0.01428 | 0.01445 | 0.54793 | 0.00781 | 0.02425 | 0.01301 |
| **FASN** | 0.73939 | 7.86812 | 0.00000 | 17.05593 | 0.01199 | 3.29369 | 73.46486 |
| **FASTK** | 3.67618 | 6.57974 | 4.44575 | 6.73273 | 0.00000 | 1.72993 | 8.11539 |
| **FASTKD1** | 22.84240 | 10.52716 | 13.37503 | 3.98539 | 1.67606 | 8.56644 | 9.48458 |
| **FASTKD2** | 26.12689 | 6.11970 | 12.88863 | 8.99170 | 2.96450 | 7.86617 | 0.00000 |
| **FASTKD3** | 13.78578 | 0.16629 | 6.85160 | 3.52238 | 0.90763 | 2.54490 | 0.00000 |
| **FASTKD5** | 20.99217 | 8.41770 | 11.60083 | 10.67038 | 10.39464 | 8.30957 | 0.00662 |
| **FAT1** | 6.35380 | 3.62488 | 1.46675 | 0.00000 | 4.83913 | 0.00000 | 12.15396 |
| **FAT2** | 0.43553 | 0.02551 | 0.00000 | 1.04112 | 1.88992 | 0.00102 | 0.69257 |
| **FAT3** | 0.05135 | 0.00318 | 0.00000 | 0.02929 | 0.34572 | 0.10112 | 0.12300 |
| **FAT4** | 0.02010 | 0.01280 | 0.01682 | 0.02389 | 0.03010 | 0.16401 | 0.02112 |
| **FATE1** | 0.02393 | 0.02923 | 0.01815 | 0.02353 | 0.02189 | 0.01044 | 0.01514 |
| **FAU** | 54.68708 | 52.75692 | 33.13388 | 54.10195 | 22.24499 | 29.77366 | 39.93612 |
| **FAXC** | 0.00794 | 0.13673 | 1.30757 | 0.36442 | 0.02554 | 0.30616 | 0.20056 |
| **FAXDC2** | 0.13472 | 0.13446 | 0.00000 | 0.07107 | 0.22386 | 0.08933 | 0.55328 |
| **FBF1** | 0.07245 | 0.79870 | 0.16683 | 0.77265 | 0.00000 | 0.43159 | 1.15998 |
| **FBL** | 15.67218 | 26.36243 | 19.11027 | 29.74449 | 11.07771 | 12.37475 | 48.15863 |
| **FBLIM1** | 0.00000 | 0.19854 | 0.00000 | 2.88631 | 0.19055 | 1.12551 | 0.49887 |
| **FBLL1** | 0.01677 | 0.02824 | 0.04809 | 0.13339 | 0.07173 | 0.09621 | 0.24110 |
| **FBLN1** | 3.31987 | 0.83224 | 0.00000 | 19.37874 | 0.19100 | 8.04578 | 14.77093 |
| **FBLN2** | 0.30814 | 0.67891 | 0.00000 | 1.75244 | 0.74070 | 0.00000 | 1.69722 |

| | | | | | | | |
|---|---|---|---|---|---|---|---|
| **FBLN5** | 0.00000 | 0.31466 | 0.15960 | 0.19661 | 1.39518 | 0.15381 | 0.00000 |
| **FBLN7** | 0.01268 | 0.03021 | 0.01381 | 0.04868 | 3.03814 | 0.00697 | 0.00948 |
| **FBN1** | 0.53204 | 0.00000 | 0.12023 | 0.04659 | 0.30682 | 0.22744 | 0.05614 |
| **FBN2** | 0.27561 | 0.00000 | 0.14161 | 0.90820 | 0.12973 | 0.01416 | 0.00000 |
| **FBN3** | 0.02294 | 0.01882 | 0.01258 | 0.03576 | 0.06168 | 0.01639 | 0.02691 |
| **FBP1** | 0.34876 | 0.00000 | 0.11944 | 0.12843 | 0.30065 | 14.09811 | 0.00000 |
| **FBP2** | 0.01848 | 0.03348 | 0.18987 | 0.22059 | 0.27895 | 0.00000 | 0.04702 |
| **FBRS** | 0.00041 | 1.82648 | 2.76978 | 9.29434 | 0.61345 | 1.15118 | 13.29672 |
| **FBRSL1** | 0.18907 | 0.60055 | 0.01819 | 0.00000 | 0.05839 | 0.10086 | 8.46281 |
| **FBXL12** | 12.10023 | 14.78730 | 8.33739 | 7.92012 | 9.15976 | 6.35689 | 0.00000 |
| **FBXL13** | 0.09775 | 0.18417 | 0.07658 | 0.04463 | 0.22059 | 0.05946 | 0.01093 |
| **FBXL14** | 0.12558 | 0.00000 | 0.06690 | 0.05123 | 0.02980 | 0.81257 | 0.82432 |
| **FBXL15** | 8.15412 | 17.02474 | 0.73830 | 12.54320 | 0.86672 | 5.72716 | 1.55697 |
| **FBXL16** | 0.00003 | 0.20332 | 0.07984 | 0.05452 | 0.02765 | 0.09370 | 0.58940 |
| **FBXL17** | 0.05153 | 0.00000 | 0.06082 | 0.19349 | 0.01622 | 0.21773 | 0.57710 |
| **FBXL18** | 3.65687 | 7.05955 | 0.63967 | 10.55587 | 1.52288 | 0.00000 | 15.01529 |
| **FBXL19** | 0.00000 | 0.17108 | 0.48303 | 0.00001 | 0.01068 | 0.25363 | 3.07111 |
| **FBXL19-AS1** | 0.00000 | 0.55429 | 0.30534 | 1.62229 | 3.61475 | 0.47172 | 1.74319 |
| **FBXL2** | 0.20091 | 0.00000 | 0.79347 | 0.38653 | 0.73077 | 0.08184 | 0.10298 |
| **FBXL20** | 4.43184 | 22.37391 | 21.61287 | 1.89185 | 7.08642 | 9.84319 | 1.47640 |
| **FBXL21** | 1.36645 | 0.00000 | 0.03799 | 2.18604 | 0.12769 | 0.00000 | 0.20102 |
| **FBXL22** | 0.01779 | 0.02114 | 0.06373 | 0.01822 | 0.64219 | 0.02030 | 0.01868 |
| **FBXL3** | 2.53431 | 0.00000 | 3.13135 | 0.05807 | 2.58622 | 1.20365 | 5.15002 |
| **FBXL4** | 2.49721 | 0.32367 | 2.43241 | 1.34238 | 0.45581 | 0.73087 | 1.18497 |
| **FBXL5** | 10.70632 | 2.25473 | 4.43128 | 2.75088 | 4.61567 | 4.66014 | 0.00000 |
| **FBXL6** | 0.69694 | 6.47837 | 0.87559 | 0.89732 | 0.76135 | 2.20501 | 5.99551 |
| **FBXL7** | 0.02558 | 0.03371 | 0.01405 | 0.04490 | 0.56755 | 0.13910 | 0.00262 |
| **FBXL8** | 0.00000 | 0.24371 | 0.02401 | 0.13650 | 0.00791 | 0.06322 | 0.05788 |
| **FBXO10** | 0.57056 | 0.63969 | 1.62116 | 1.51179 | 1.33339 | 0.53768 | 0.82620 |
| **FBXO11** | 8.96827 | 0.00000 | 3.31345 | 5.86842 | 1.30831 | 2.55021 | 17.89011 |
| **FBXO15** | 0.05237 | 0.00000 | 0.12079 | 0.06668 | 0.64908 | 0.10716 | 0.01866 |
| **FBXO16** | 0.26245 | 0.34204 | 0.37400 | 0.00086 | 0.44811 | 0.00000 | 0.00000 |
| **FBXO17** | 0.30650 | 0.91013 | 1.40246 | 0.51699 | 0.00000 | 1.71267 | 0.00000 |
| **FBXO18** | 1.10464 | 6.65563 | 2.07398 | 5.15290 | 1.26762 | 2.61973 | 12.46904 |
| **FBXO2** | 0.03120 | 0.16494 | 0.19350 | 0.00000 | 0.27799 | 0.39232 | 0.03565 |

| | | | | | | | |
|---|---|---|---|---|---|---|---|
| **FBXO21** | 20.78439 | 7.07036 | 4.82679 | 14.28189 | 1.87024 | 6.93424 | 0.00000 |
| **FBXO22** | 14.57293 | 11.95608 | 6.80438 | 11.64827 | 0.77438 | 5.14521 | 0.00000 |
| **FBXO22-AS1** | 0.02685 | 0.01855 | 0.01262 | 0.02210 | 0.01246 | 0.01821 | 0.01794 |
| **FBXO24** | 0.04360 | 0.40880 | 0.04031 | 0.16441 | 0.07447 | 0.12628 | 0.00000 |
| **FBXO25** | 4.13233 | 5.76571 | 3.32030 | 3.52368 | 0.87449 | 1.58044 | 6.04999 |
| **FBXO27** | 0.03624 | 0.00000 | 0.56364 | 0.07929 | 1.55790 | 0.49465 | 0.26563 |
| **FBXO28** | 19.32033 | 3.10625 | 10.67317 | 6.59554 | 5.42104 | 6.70764 | 3.20759 |
| **FBXO3** | 7.50053 | 0.64280 | 2.13110 | 4.01097 | 1.14349 | 0.88765 | 0.00000 |
| **FBXO30** | 3.70526 | 0.00000 | 1.22115 | 5.36387 | 1.57272 | 1.53615 | 4.10641 |
| **FBXO31** | 0.00000 | 1.59903 | 2.97459 | 1.11045 | 2.93056 | 2.84582 | 8.86028 |
| **FBXO32** | 0.00000 | 2.99399 | 2.43296 | 2.57178 | 0.08470 | 0.00000 | 1.38209 |
| **FBXO33** | 3.05651 | 0.21323 | 0.99646 | 2.74696 | 5.54644 | 1.18165 | 5.49776 |
| **FBXO34** | 6.65753 | 3.35902 | 2.95888 | 2.93840 | 1.05158 | 1.81484 | 3.41729 |
| **FBXO36** | 4.82209 | 6.55764 | 8.56415 | 0.47507 | 0.00000 | 4.71619 | 0.00000 |
| **FBXO38** | 5.21706 | 0.79771 | 2.71928 | 4.74169 | 1.33916 | 2.42353 | 0.29310 |
| **FBXO39** | 0.00889 | 0.03130 | 0.00677 | 0.00885 | 0.02006 | 0.00956 | 0.10316 |
| **FBXO4** | 0.99489 | 0.16968 | 0.16981 | 0.18019 | 0.04043 | 0.00105 | 0.15893 |
| **FBXO40** | 0.05104 | 0.00000 | 0.07892 | 0.03580 | 0.20210 | 0.02187 | 0.00565 |
| **FBXO41** | 0.00000 | 0.39122 | 0.04761 | 0.39317 | 0.05314 | 0.52022 | 0.13042 |
| **FBXO42** | 5.20720 | 4.41938 | 1.75318 | 4.37286 | 1.19214 | 2.36957 | 0.00000 |
| **FBXO43** | 0.01049 | 0.00000 | 0.10432 | 0.06726 | 1.16872 | 0.01729 | 0.06935 |
| **FBXO44** | 0.00000 | 0.29255 | 0.16984 | 0.17027 | 0.43413 | 1.75118 | 0.00000 |
| **FBXO45** | 12.04905 | 1.15426 | 4.56453 | 4.15048 | 6.68893 | 7.24300 | 11.28585 |
| **FBXO46** | 0.69761 | 2.75895 | 1.17200 | 3.98158 | 1.90765 | 1.03508 | 6.00142 |
| **FBXO47** | 0.02672 | 0.00802 | 0.01926 | 0.02210 | 0.22301 | 0.01400 | 0.01338 |
| **FBXO48** | 1.06558 | 1.05119 | 0.97685 | 0.27482 | 0.21958 | 0.83477 | 0.00000 |
| **FBXO5** | 19.76278 | 0.00000 | 19.15616 | 7.48324 | 17.41647 | 3.78612 | 0.05244 |
| **FBXO6** | 1.41596 | 7.93166 | 1.79568 | 0.00000 | 3.01992 | 1.14270 | 0.00000 |
| **FBXO7** | 25.05394 | 19.18852 | 11.34036 | 16.97834 | 3.79315 | 15.74839 | 8.58803 |
| **FBXO8** | 5.08228 | 1.65394 | 3.22545 | 1.56804 | 2.44205 | 4.00494 | 0.36613 |
| **FBXO9** | 6.46874 | 4.08085 | 3.31758 | 12.84000 | 0.32571 | 5.59542 | 8.31162 |
| **FBXW10** | 0.05686 | 0.00789 | 0.07559 | 0.02173 | 0.03597 | 0.00687 | 0.04119 |
| **FBXW11** | 6.52130 | 2.29133 | 6.41601 | 3.53165 | 1.78789 | 4.30076 | 7.59048 |
| **FBXW12** | 0.01658 | 0.10149 | 0.00000 | 0.03228 | 4.63666 | 0.04638 | 0.00000 |
| **FBXW2** | 14.92992 | 8.69101 | 7.65213 | 9.57225 | 4.19003 | 6.48569 | 8.18977 |

| | | | | | | | |
|---|---|---|---|---|---|---|---|
| **FBXW4** | 1.01814 | 2.36376 | 0.60744 | 1.12143 | 0.00849 | 1.59446 | 3.29640 |
| **FBXW4P1** | 0.00000 | 0.08081 | 0.55703 | 0.55629 | 0.31991 | 0.60474 | 1.84380 |
| **FBXW5** | 7.40745 | 17.73778 | 1.83674 | 11.87220 | 0.00000 | 9.74158 | 18.46736 |
| **FBXW7** | 5.38985 | 3.09505 | 3.39366 | 3.65330 | 8.44841 | 5.65749 | 8.70502 |
| **FBXW8** | 0.64245 | 0.25372 | 0.56527 | 0.93565 | 0.24762 | 0.16247 | 0.31396 |
| **FBXW9** | 2.47142 | 2.57209 | 1.47442 | 4.15753 | 0.59331 | 0.87434 | 3.51688 |
| **FCAMR** | 0.05488 | 0.00733 | 0.01674 | 0.02078 | 0.06349 | 0.02417 | 0.00000 |
| **FCAR** | 0.02646 | 0.01702 | 0.05883 | 0.02005 | 0.00589 | 0.01395 | 0.03107 |
| **FCER1G** | 0.63184 | 0.00000 | 0.02192 | 0.47872 | 0.03358 | 1.74097 | 0.00000 |
| **FCER2** | 0.37504 | 0.63840 | 0.00063 | 0.00000 | 0.00909 | 0.05249 | 0.00559 |
| **FCF1** | 7.44407 | 2.22547 | 6.38997 | 4.87340 | 1.21965 | 2.71751 | 6.15077 |
| **FCGBP** | 0.17316 | 0.47443 | 0.13132 | 1.21957 | 0.10621 | 6.09082 | 0.00000 |
| **FCGR1A** | 0.02727 | 0.00541 | 0.03952 | 0.03145 | 0.03432 | 0.01269 | 0.03425 |
| **FCGR1B** | 0.02715 | 0.00399 | 0.03408 | 0.02621 | 0.02973 | 0.02206 | 0.01888 |
| **FCGR1C** | 0.03409 | 0.01448 | 0.01790 | 0.01754 | 0.00990 | 0.01596 | 0.01172 |
| **FCGR2A** | 0.02331 | 0.01349 | 0.02791 | 0.02162 | 0.03820 | 0.01183 | 0.02371 |
| **FCGR2B** | 0.02074 | 0.01111 | 0.03691 | 0.03832 | 0.00655 | 0.11552 | 0.00809 |
| **FCGR3B** | 0.03183 | 0.01603 | 0.01306 | 0.01789 | 0.00792 | 0.01469 | 0.01521 |
| **FCGRT** | 0.00000 | 0.31017 | 0.32208 | 0.22619 | 0.31425 | 1.00813 | 0.01556 |
| **FCHO1** | 0.01928 | 1.02240 | 0.22507 | 0.39450 | 1.59878 | 0.33682 | 0.00000 |
| **FCHO2** | 2.39812 | 0.00000 | 1.40627 | 2.60272 | 0.70131 | 0.91572 | 1.40814 |
| **FCHSD1** | 0.01977 | 0.06905 | 0.03973 | 0.87004 | 0.55434 | 0.50372 | 0.70614 |
| **FCHSD2** | 1.68645 | 0.99818 | 0.77687 | 0.94874 | 0.00769 | 0.97551 | 2.07935 |
| **FCRL3** | 0.02091 | 0.01609 | 0.01276 | 0.02299 | 0.00944 | 0.01386 | 0.02388 |
| **FCRL4** | 0.01844 | 0.01891 | 0.01262 | 0.02666 | 0.00783 | 0.01592 | 0.02253 |
| **FCRL5** | 0.05829 | 0.01153 | 0.01425 | 0.02283 | 0.00796 | 0.02233 | 0.00589 |
| **FCRL6** | 0.03049 | 0.00851 | 0.01350 | 0.02502 | 0.00724 | 0.00706 | 0.06119 |
| **FCRLA** | 0.02019 | 0.01437 | 0.01037 | 0.01490 | 0.02683 | 0.01182 | 0.04709 |
| **FCRLB** | 0.17505 | 0.02569 | 0.02177 | 0.01980 | 0.08583 | 0.01459 | 0.08252 |
| **FDCSP** | 0.02063 | 0.01680 | 0.01283 | 0.02865 | 0.00893 | 0.01417 | 0.02079 |
| **FDFT1** | 52.19036 | 7.84537 | 0.31394 | 13.67094 | 6.53695 | 12.13841 | 21.87852 |
| **FDPS** | 51.56999 | 15.59179 | 2.39493 | 26.08140 | 0.00138 | 10.49192 | 3.98175 |
| **FDPSP2** | 0.00005 | 0.10129 | 0.05707 | 0.09364 | 0.03227 | 0.08525 | 0.27605 |
| **FDX1** | 2.40948 | 2.73258 | 0.82394 | 4.47608 | 0.05725 | 4.00101 | 10.49541 |
| **FDX1L** | 0.74408 | 9.04170 | 5.20844 | 5.94615 | 2.42790 | 3.96220 | 6.43190 |

| | | | | | | | |
|---|---|---|---|---|---|---|---|
| **FDXACB1** | 3.49916 | 0.00000 | 1.76139 | 1.23213 | 1.32155 | 0.31128 | 0.00000 |
| **FDXR** | 6.33106 | 26.11365 | 0.84629 | 13.23234 | 0.28736 | 5.07582 | 0.29397 |
| **FECH** | 7.15383 | 0.13212 | 0.71068 | 4.97228 | 0.17542 | 0.16935 | 1.55147 |
| **FEM1A** | 7.83280 | 9.02909 | 4.88973 | 8.41004 | 0.65487 | 6.47088 | 8.41333 |
| **FEM1B** | 4.30746 | 0.00000 | 4.01086 | 5.19096 | 1.77881 | 6.70645 | 11.32729 |
| **FEM1C** | 4.03308 | 0.00005 | 3.52771 | 1.41710 | 1.20407 | 1.08983 | 3.50078 |
| **FEN1** | 35.04516 | 23.65441 | 12.37603 | 28.74868 | 2.95969 | 12.74782 | 0.00000 |
| **FENDRR** | 0.02485 | 0.01398 | 0.01964 | 0.01917 | 0.00816 | 0.01275 | 0.01836 |
| **FER** | 3.12142 | 1.48031 | 2.16735 | 1.50802 | 1.83215 | 1.28870 | 3.65458 |
| **FER1L4** | 0.00000 | 0.09878 | 0.16032 | 0.35370 | 0.17037 | 0.06530 | 0.06476 |
| **FER1L5** | 0.00000 | 0.09736 | 0.13474 | 0.14114 | 0.01245 | 0.09633 | 0.08383 |
| **FER1L6** | 0.00747 | 0.02908 | 0.00000 | 0.02286 | 0.82098 | 0.02825 | 0.12949 |
| **FER1L6-AS2** | 0.07824 | 0.05168 | 0.03626 | 0.09231 | 0.17689 | 0.01471 | 0.00000 |
| **FERMT1** | 0.12593 | 0.00000 | 0.01846 | 0.49428 | 0.05221 | 0.12123 | 0.46777 |
| **FERMT2** | 10.49987 | 14.50499 | 12.99479 | 2.48459 | 3.37207 | 13.15281 | 8.94130 |
| **FERMT3** | 0.05011 | 1.15968 | 0.32129 | 0.24580 | 0.00489 | 1.22656 | 3.07082 |
| **FES** | 1.10368 | 5.23700 | 0.09375 | 0.97764 | 0.00779 | 3.30103 | 1.03719 |
| **FETUB** | 0.01276 | 0.11657 | 0.01530 | 0.03995 | 0.02185 | 0.04834 | 0.02275 |
| **FEV** | 0.00593 | 0.01838 | 0.01140 | 0.02830 | 0.02597 | 0.02219 | 0.09153 |
| **FEZ1** | 0.03404 | 0.56780 | 1.04561 | 0.15676 | 0.19603 | 0.16180 | 0.00000 |
| **FEZ2** | 24.76892 | 10.39526 | 12.05734 | 0.00000 | 6.90664 | 7.43856 | 27.58423 |
| **FEZF1** | 0.02425 | 0.00092 | 0.06872 | 0.01937 | 0.01265 | 0.01200 | 0.02046 |
| **FEZF1-AS1** | 0.02612 | 0.00035 | 0.05637 | 0.02170 | 0.13254 | 0.01270 | 0.01614 |
| **FEZF2** | 0.01904 | 0.18434 | 0.00000 | 0.00000 | 0.87563 | 0.02794 | 0.16708 |
| **FFAR2** | 0.02266 | 0.02264 | 0.01248 | 0.02798 | 0.14266 | 0.04352 | 0.01823 |
| **FFAR4** | 0.02387 | 0.09859 | 0.03430 | 0.13433 | 0.01037 | 0.01589 | 0.00000 |
| **FGA** | 0.14655 | 0.00000 | 0.00114 | 0.02516 | 0.20448 | 1.47549 | 0.00000 |
| **FGB** | 0.05414 | 0.00883 | 0.01202 | 0.07269 | 0.00936 | 0.35156 | 0.00000 |
| **FGD1** | 0.00000 | 1.48789 | 5.18525 | 0.10666 | 0.13849 | 2.20897 | 0.91031 |
| **FGD2** | 0.03549 | 0.07544 | 0.01152 | 0.01451 | 0.02115 | 0.17597 | 0.00966 |
| **FGD3** | 0.61259 | 0.00000 | 0.56353 | 0.79273 | 0.27054 | 0.16536 | 0.51938 |
| **FGD4** | 1.89879 | 1.29010 | 3.28670 | 0.00000 | 1.84004 | 5.54605 | 5.33366 |
| **FGD5** | 0.01957 | 0.00184 | 0.05013 | 0.08861 | 0.40410 | 0.02862 | 0.03848 |
| **FGD5-AS1** | 8.52809 | 3.70685 | 9.21303 | 7.33684 | 3.73281 | 6.48870 | 10.88095 |
| **FGD5P1** | 0.04704 | 0.00228 | 0.01237 | 0.04008 | 0.14197 | 0.08738 | 0.00000 |

| | | | | | | | |
|---|---|---|---|---|---|---|---|
| **FGD6** | 5.43254 | 0.09055 | 3.37680 | 7.58727 | 4.27437 | 5.90481 | 4.67975 |
| **FGF1** | 0.00799 | 0.02167 | 0.00017 | 0.01300 | 0.09582 | 1.59662 | 0.01488 |
| **FGF10** | 0.05586 | 0.01200 | 0.01435 | 0.03429 | 0.00793 | 0.01186 | 0.00425 |
| **FGF10-AS1** | 0.02732 | 0.01859 | 0.01261 | 0.02480 | 0.00804 | 0.01287 | 0.01407 |
| **FGF11** | 0.00000 | 0.00000 | 0.17682 | 1.08941 | 0.00706 | 0.36347 | 0.31628 |
| **FGF12** | 0.09194 | 0.39934 | 0.05200 | 0.06475 | 0.04971 | 0.64748 | 0.27887 |
| **FGF13** | 0.96806 | 5.65568 | 5.41282 | 0.00000 | 0.73265 | 2.71487 | 0.39073 |
| **FGF13-AS1** | 0.08072 | 0.26934 | 0.10924 | 0.00299 | 0.00361 | 0.08149 | 0.00000 |
| **FGF14** | 0.00000 | 0.06777 | 0.03555 | 0.01183 | 4.86965 | 0.07807 | 0.03612 |
| **FGF14-AS2** | 0.15154 | 0.49274 | 0.37208 | 0.19194 | 0.27090 | 0.03579 | 0.00000 |
| **FGF14-IT1** | 0.01427 | 0.06824 | 0.01911 | 0.02551 | 0.02422 | 0.00571 | 0.04081 |
| **FGF16** | 0.01972 | 0.03527 | 0.00815 | 0.01595 | 0.03204 | 0.02231 | 0.02506 |
| **FGF17** | 0.00000 | 0.01547 | 0.01231 | 0.03435 | 0.03103 | 0.05327 | 0.10559 |
| **FGF18** | 0.28446 | 1.95051 | 0.00000 | 0.36267 | 0.75382 | 1.37169 | 0.56105 |
| **FGF19** | 0.02174 | 0.01525 | 0.01293 | 0.02647 | 0.01466 | 0.02134 | 0.02085 |
| **FGF2** | 0.03485 | 0.00000 | 0.13431 | 0.02814 | 0.00652 | 0.60476 | 0.03900 |
| **FGF20** | 0.01381 | 0.00185 | 0.06559 | 0.04673 | 0.00828 | 0.01374 | 0.02409 |
| **FGF21** | 0.21090 | 0.05414 | 0.00054 | 0.01433 | 0.00643 | 0.00000 | 0.38605 |
| **FGF23** | 0.01662 | 0.00000 | 0.12662 | 0.00000 | 0.07215 | 0.16473 | 0.63709 |
| **FGF3** | 0.01154 | 0.01536 | 0.01306 | 0.03177 | 0.00754 | 0.01545 | 0.04253 |
| **FGF4** | 0.00000 | 0.04726 | 0.00602 | 0.01793 | 0.00866 | 1.63824 | 0.01776 |
| **FGF5** | 0.02193 | 0.01738 | 0.01394 | 0.01890 | 0.00840 | 0.02707 | 0.02441 |
| **FGF7** | 0.03135 | 0.01768 | 0.01315 | 0.01778 | 0.00906 | 0.01430 | 0.01289 |
| **FGF8** | 0.00000 | 0.39653 | 0.43017 | 0.12768 | 0.07107 | 0.92384 | 0.08320 |
| **FGF9** | 0.08868 | 0.22257 | 0.01346 | 0.05602 | 0.85834 | 0.00000 | 0.03106 |
| **FGFBP1** | 1.04039 | 0.00000 | 0.00082 | 0.54065 | 0.00414 | 0.00000 | 0.28750 |
| **FGFBP2** | 1.23764 | 1.19377 | 0.00000 | 0.00000 | 0.04837 | 0.00000 | 0.04804 |
| **FGFBP3** | 0.04340 | 0.00000 | 0.17382 | 0.00786 | 0.13949 | 0.15430 | 0.15474 |
| **FGFR1** | 0.00000 | 7.05576 | 2.99708 | 2.88856 | 0.91747 | 23.99478 | 9.35412 |
| **FGFR1OP** | 20.78568 | 12.67468 | 7.84717 | 11.03701 | 5.57137 | 7.03970 | 0.00000 |
| **FGFR1OP2** | 9.56350 | 1.09663 | 3.80635 | 6.38421 | 8.75086 | 4.80516 | 3.57486 |
| **FGFR2** | 0.00000 | 11.54800 | 3.38223 | 4.42142 | 0.66718 | 13.32926 | 4.65161 |
| **FGFR3** | 0.00000 | 11.98930 | 3.06209 | 0.00000 | 0.00073 | 4.05067 | 25.69559 |
| **FGFR4** | 15.45813 | 2.26471 | 0.00000 | 15.32390 | 0.12204 | 6.30111 | 25.99869 |
| **FGFRL1** | 0.42663 | 0.91507 | 0.34268 | 0.00000 | 1.11003 | 0.00000 | 2.43987 |

| | | | | | | | |
|---|---|---|---|---|---|---|---|
| **FGG** | 0.23331 | 0.00000 | 0.01065 | 0.04183 | 0.00804 | 0.83467 | 0.00000 |
| **FGGY** | 0.27966 | 0.96288 | 0.31229 | 0.24777 | 1.47413 | 0.12971 | 0.04356 |
| **FGL1** | 0.02965 | 0.00057 | 0.04401 | 0.04031 | 0.02329 | 0.05489 | 0.00569 |
| **FGL2** | 0.02554 | 0.01872 | 0.01396 | 0.02574 | 0.00775 | 0.01437 | 0.01733 |
| **FGR** | 0.01643 | 0.03083 | 0.03588 | 0.02043 | 0.01204 | 0.03450 | 0.01738 |
| **FH** | 28.74556 | 10.88450 | 2.88382 | 10.44274 | 5.27990 | 10.67125 | 5.11512 |
| **FHAD1** | 0.07308 | 0.00875 | 0.02849 | 0.01191 | 0.25990 | 0.15542 | 0.02072 |
| **FHDC1** | 0.00000 | 0.50285 | 0.01345 | 12.38218 | 0.00677 | 2.92258 | 0.85256 |
| **FHIT** | 0.45994 | 0.20613 | 0.00000 | 0.23316 | 0.87371 | 0.24843 | 0.60348 |
| **FHL1** | 0.03294 | 0.00000 | 0.30582 | 0.12317 | 1.58127 | 0.02115 | 0.00087 |
| **FHL2** | 13.06897 | 7.84048 | 0.77357 | 15.79523 | 0.00000 | 0.00000 | 4.18280 |
| **FHL3** | 1.08840 | 3.36498 | 1.32421 | 2.52147 | 0.00000 | 3.16901 | 5.50539 |
| **FHL5** | 0.03498 | 0.01864 | 0.01146 | 0.02346 | 0.03630 | 0.02616 | 0.00880 |
| **FHOD1** | 0.59612 | 1.85582 | 3.43937 | 2.81292 | 0.30690 | 1.38376 | 7.34600 |
| **FHOD3** | 0.02988 | 0.06527 | 0.03699 | 0.02122 | 2.36719 | 0.20113 | 0.09937 |
| **FIBCD1** | 0.14306 | 0.28205 | 0.00000 | 0.46983 | 0.41521 | 0.20100 | 0.73583 |
| **FIBP** | 14.30416 | 11.22456 | 4.11640 | 12.09098 | 2.91041 | 9.60302 | 1.08965 |
| **FICD** | 0.72688 | 0.27439 | 0.26350 | 0.60183 | 0.06574 | 0.47798 | 0.00000 |
| **FIG4** | 1.65502 | 0.16092 | 0.56600 | 1.62361 | 0.32822 | 0.68317 | 0.00000 |
| **FIGF** | 0.03633 | 0.01515 | 0.01317 | 0.01613 | 0.00781 | 0.01277 | 0.01820 |
| **FIGLA** | 0.01428 | 0.00000 | 0.14877 | 0.01219 | 2.53524 | 0.02190 | 0.03027 |
| **FIGN** | 1.72628 | 0.61136 | 0.75626 | 7.90005 | 0.51932 | 0.00000 | 7.32349 |
| **FIGNL1** | 10.73634 | 0.15684 | 7.56596 | 2.69625 | 6.44983 | 2.15288 | 1.93806 |
| **FIGNL2** | 0.00000 | 0.00400 | 0.02243 | 2.36484 | 0.00267 | 0.14391 | 5.05607 |
| **FILIP1** | 5.33073 | 3.81547 | 2.75350 | 1.02147 | 0.15543 | 11.53196 | 4.51363 |
| **FILIP1L** | 0.00000 | 0.00000 | 0.23754 | 0.18043 | 0.01162 | 0.16404 | 1.05472 |
| **FIP1L1** | 23.16852 | 21.09330 | 15.20626 | 23.41541 | 3.31589 | 11.44785 | 9.83006 |
| **FIRRE** | 0.28853 | 0.46499 | 0.14993 | 0.05703 | 0.07374 | 0.11880 | 0.00000 |
| **FIS1** | 9.78630 | 12.40816 | 5.03630 | 11.11165 | 7.88141 | 11.04958 | 9.84173 |
| **FITM1** | 0.00000 | 0.01007 | 0.01391 | 0.08110 | 0.00638 | 0.12809 | 0.16655 |
| **FITM2** | 0.92934 | 2.08112 | 2.00392 | 1.42650 | 0.62605 | 0.17759 | 0.00000 |
| **FIZ1** | 0.00000 | 0.28231 | 0.39818 | 0.25210 | 0.06654 | 0.38396 | 1.03747 |
| **FJX1** | 0.18289 | 0.10604 | 0.05244 | 0.22365 | 0.00351 | 0.12253 | 0.08248 |
| **FKBP10** | 0.03115 | 0.06401 | 0.02366 | 0.08500 | 0.01387 | 0.06120 | 0.15606 |
| **FKBP11** | 3.20036 | 9.39706 | 5.73477 | 4.68333 | 0.00000 | 7.51091 | 1.18409 |

| | | | | | | | |
|---|---|---|---|---|---|---|---|
| **FKBP14** | 4.81133 | 0.00000 | 1.43841 | 3.88344 | 0.07364 | 1.17267 | 2.20448 |
| **FKBP15** | 7.47350 | 5.44598 | 2.68012 | 5.43699 | 2.04576 | 2.87780 | 2.89273 |
| **FKBP1A** | 21.89413 | 14.08172 | 9.90139 | 27.76514 | 2.06661 | 19.25632 | 12.36991 |
| **FKBP1AP1** | 2.59485 | 0.62745 | 0.88727 | 2.99808 | 0.17742 | 2.03145 | 0.78637 |
| **FKBP1B** | 0.00000 | 0.35114 | 2.78455 | 1.47072 | 0.21212 | 1.43227 | 2.33906 |
| **FKBP2** | 11.07636 | 20.12465 | 6.88720 | 16.79389 | 3.60975 | 10.80715 | 12.11105 |
| **FKBP3** | 32.73552 | 4.62751 | 6.47617 | 20.76971 | 0.60428 | 10.32707 | 11.01747 |
| **FKBP4** | 22.18625 | 26.38972 | 13.77894 | 20.57131 | 7.73854 | 20.81083 | 25.65389 |
| **FKBP5** | 8.12438 | 2.66203 | 1.77775 | 2.63689 | 0.95681 | 2.06904 | 0.54890 |
| **FKBP6** | 0.00000 | 0.00809 | 3.65431 | 0.00947 | 1.72111 | 0.00000 | 0.07767 |
| **FKBP7** | 0.28103 | 0.00000 | 0.33402 | 0.47170 | 0.01801 | 0.08906 | 0.00000 |
| **FKBP8** | 0.00000 | 2.62022 | 0.44190 | 1.84578 | 0.03084 | 3.63535 | 5.66643 |
| **FKBP9** | 1.41692 | 0.84731 | 0.14458 | 0.84232 | 0.07008 | 0.60193 | 0.47453 |
| **FKBP9P1** | 0.02537 | 0.01957 | 0.01366 | 0.02748 | 0.00760 | 0.00988 | 0.04136 |
| **FKBPL** | 2.05215 | 10.16743 | 12.74424 | 6.84258 | 5.71560 | 1.93499 | 0.00404 |
| **FKRP** | 1.05713 | 1.85610 | 0.83048 | 1.25446 | 0.00135 | 0.95247 | 3.29329 |
| **FKTN** | 1.89225 | 0.54696 | 2.68225 | 0.28404 | 0.01658 | 0.81277 | 1.46864 |
| **FLAD1** | 26.99170 | 23.05363 | 15.28461 | 15.25617 | 2.85157 | 12.95983 | 2.21548 |
| **FLCN** | 2.23483 | 2.31746 | 1.73190 | 5.64918 | 2.21396 | 3.05883 | 2.67943 |
| **FLG** | 0.07865 | 0.31424 | 0.11780 | 0.07873 | 0.01262 | 0.04790 | 0.01500 |
| **FLG-AS1** | 0.12728 | 0.68578 | 0.17763 | 0.06771 | 0.06523 | 0.11202 | 0.14235 |
| **FLG2** | 0.05708 | 0.08545 | 0.01941 | 0.00009 | 0.01182 | 0.00013 | 0.08285 |
| **FLI1** | 0.02459 | 0.00641 | 0.07127 | 0.03085 | 0.18320 | 0.01756 | 0.03863 |
| **FLII** | 7.97306 | 17.52095 | 1.50793 | 12.22670 | 1.76115 | 9.54949 | 4.86725 |
| **FLJ10038** | 0.00354 | 0.07922 | 0.51554 | 0.21677 | 0.13307 | 0.14296 | 2.17437 |
| **FLJ12825** | 0.08409 | 0.12962 | 0.08464 | 0.06059 | 0.00930 | 0.03920 | 0.00074 |
| **FLJ13224** | 0.00000 | 0.10007 | 0.02358 | 0.07983 | 0.10583 | 0.08376 | 0.36577 |
| **FLJ16171** | 0.03607 | 0.01874 | 0.01291 | 0.01063 | 0.00763 | 0.03469 | 0.02924 |
| **FLJ16779** | 0.05834 | 0.00937 | 0.01536 | 0.11845 | 0.00629 | 0.00000 | 0.13869 |
| **FLJ20021** | 0.03888 | 0.10459 | 0.06216 | 0.05060 | 0.19016 | 0.00925 | 0.00000 |
| **FLJ20518.uid-5** | 0.02137 | 0.03202 | 0.00531 | 0.01768 | 0.05914 | 0.01667 | 0.01867 |
| **FLJ21408** | 0.02879 | 0.01603 | 0.10926 | 0.02492 | 0.01007 | 0.01081 | 0.00886 |
| **FLJ22184** | 0.00257 | 0.01714 | 0.00000 | 0.01692 | 0.37181 | 0.01222 | 0.25176 |
| **FLJ22447** | 0.03912 | 0.00605 | 0.01515 | 0.09086 | 0.01062 | 0.04355 | 0.01560 |
| **FLJ22763** | 0.03445 | 0.01410 | 0.05594 | 0.01603 | 0.02090 | 0.19428 | 0.00802 |

| | | | | | | | |
|---|---|---|---|---|---|---|---|
| **FLJ23867** | 0.00436 | 0.03226 | 0.02027 | 0.19885 | 0.06317 | 0.09918 | 0.07389 |
| **FLJ25758** | 0.01793 | 0.00000 | 1.59246 | 0.07445 | 3.41781 | 0.00247 | 0.00000 |
| **FLJ26850** | 0.02010 | 0.03807 | 0.06084 | 0.03936 | 0.27869 | 0.09369 | 0.00844 |
| **FLJ27354** | 0.02263 | 0.00804 | 0.05356 | 0.02432 | 0.03588 | 0.02110 | 0.01742 |
| **FLJ30403** | 0.11270 | 0.06942 | 0.12852 | 0.14425 | 0.17433 | 0.10418 | 0.00000 |
| **FLJ30679** | 0.00002 | 0.00000 | 0.09724 | 0.03044 | 0.11465 | 0.00689 | 0.21042 |
| **FLJ31104** | 0.00000 | 0.10501 | 0.01280 | 0.16158 | 0.02877 | 0.03708 | 0.02236 |
| **FLJ31306** | 6.69578 | 0.63719 | 4.58178 | 3.95604 | 3.39714 | 3.19583 | 4.52994 |
| **FLJ31356** | 0.03060 | 0.02218 | 0.01157 | 0.02361 | 0.00894 | 0.01185 | 0.02273 |
| **FLJ32255** | 0.00000 | 0.15041 | 0.22697 | 0.01234 | 0.42853 | 0.11431 | 0.09854 |
| **FLJ33360** | 0.05708 | 0.00896 | 0.14990 | 0.12901 | 0.00683 | 0.06259 | 0.00000 |
| **FLJ33534** | 0.02546 | 0.04523 | 0.05370 | 0.02859 | 0.14699 | 0.02203 | 0.01401 |
| **FLJ33581** | 0.03289 | 0.01672 | 0.01354 | 0.01646 | 0.00774 | 0.01806 | 0.02331 |
| **FLJ34503** | 0.01624 | 0.02172 | 0.01375 | 0.04220 | 0.00964 | 0.01181 | 0.02935 |
| **FLJ35934** | 0.01804 | 0.01965 | 0.02000 | 0.01285 | 0.00996 | 0.04285 | 0.04714 |
| **FLJ36000** | 0.01904 | 0.05119 | 0.01421 | 0.03385 | 0.11353 | 0.03396 | 0.02675 |
| **FLJ36777** | 0.03303 | 0.02046 | 0.01485 | 0.01641 | 0.01231 | 0.04416 | 0.01422 |
| **FLJ37035** | 0.01738 | 0.07663 | 0.17996 | 0.02750 | 0.00486 | 0.00565 | 0.03483 |
| **FLJ37201** | 0.07582 | 0.00930 | 0.09817 | 0.00840 | 0.24947 | 0.03285 | 0.03436 |
| **FLJ37453** | 0.11341 | 1.22776 | 1.00185 | 0.35964 | 0.36314 | 0.65945 | 1.62996 |
| **FLJ38576** | 0.02303 | 0.01361 | 0.02628 | 0.01971 | 0.00759 | 0.02540 | 0.01493 |
| **FLJ39080** | 0.05636 | 0.03979 | 0.08005 | 0.03274 | 0.00619 | 0.06743 | 0.01369 |
| **FLJ40194** | 0.00569 | 0.01631 | 0.06837 | 0.01217 | 0.10392 | 0.02590 | 0.10164 |
| **FLJ41200** | 0.02660 | 0.01723 | 0.01276 | 0.01893 | 0.00793 | 0.01249 | 0.01896 |
| **FLJ41278** | 0.04655 | 0.04641 | 0.02983 | 0.01255 | 0.01518 | 0.03478 | 0.03041 |
| **FLJ41941** | 0.00641 | 0.02681 | 0.01172 | 0.16345 | 0.00829 | 0.01317 | 0.00373 |
| **FLJ42102** | 0.01841 | 0.02266 | 0.01158 | 0.02329 | 0.00810 | 0.01294 | 0.02556 |
| **FLJ42351** | 0.02309 | 0.02474 | 0.01305 | 0.01969 | 0.00772 | 0.01435 | 0.02138 |
| **FLJ42393** | 0.15391 | 0.00000 | 0.47513 | 0.00000 | 0.05097 | 0.45138 | 1.38095 |
| **FLJ42627** | 0.00000 | 0.12447 | 0.01015 | 0.17587 | 0.88980 | 0.01981 | 0.12020 |
| **FLJ42969** | 0.01680 | 0.06076 | 0.00141 | 0.69829 | 0.05462 | 0.00126 | 0.08983 |
| **FLJ43681** | 0.03179 | 0.01327 | 0.02446 | 0.01829 | 0.01505 | 0.01275 | 0.02198 |
| **FLJ43879** | 0.04132 | 0.01085 | 0.01157 | 0.00996 | 0.02360 | 0.01045 | 0.03041 |
| **FLJ44313** | 0.03224 | 0.00000 | 0.18360 | 0.01304 | 0.10284 | 0.01278 | 0.02554 |
| **FLJ44511** | 0.04588 | 0.01497 | 0.06841 | 0.07580 | 0.04345 | 0.00000 | 0.20601 |

| | | | | | | | |
|---|---|---|---|---|---|---|---|
| **FLJ44635** | 0.97792 | 0.00000 | 3.09812 | 0.77048 | 0.73553 | 2.17371 | 11.09925 |
| **FLJ45079** | 0.02409 | 0.02012 | 0.01235 | 0.02900 | 0.00840 | 0.01349 | 0.01037 |
| **FLJ45513** | 0.36994 | 0.00000 | 0.43840 | 0.32972 | 0.14157 | 0.10749 | 0.38564 |
| **FLJ46066** | 0.02134 | 0.01976 | 0.03917 | 0.02834 | 0.01876 | 0.00815 | 0.03924 |
| **FLJ46284** | 0.03021 | 0.01604 | 0.01408 | 0.04642 | 0.00810 | 0.01401 | 0.00743 |
| **FLJ46906** | 0.89595 | 0.60598 | 0.12062 | 1.07227 | 0.05834 | 0.46220 | 0.38293 |
| **FLNA** | 5.59938 | 16.30957 | 0.00000 | 8.70186 | 0.46164 | 7.74155 | 24.42750 |
| **FLNB** | 29.28345 | 34.59294 | 0.76737 | 25.31027 | 1.54328 | 11.38060 | 34.05185 |
| **FLNC** | 0.00000 | 2.24410 | 0.00000 | 1.86070 | 0.32287 | 1.16677 | 2.45485 |
| **FLOT1** | 1.71336 | 5.37111 | 2.64879 | 4.09687 | 7.72519 | 8.32272 | 4.72774 |
| **FLOT2** | 7.39016 | 7.24485 | 0.38721 | 10.26812 | 0.25549 | 7.42920 | 12.08028 |
| **FLRT1** | 0.01379 | 0.52285 | 0.03666 | 0.12001 | 0.02110 | 0.21797 | 0.00000 |
| **FLRT2** | 0.01029 | 0.00782 | 0.00506 | 0.05498 | 0.13245 | 0.45331 | 0.02409 |
| **FLRT3** | 0.20196 | 0.00000 | 0.09696 | 0.12436 | 0.00553 | 3.51422 | 0.13004 |
| **FLT1** | 0.00000 | 0.10402 | 0.20931 | 0.07592 | 0.41019 | 0.03003 | 0.16628 |
| **FLT3** | 0.14193 | 0.07243 | 0.15867 | 0.00000 | 0.36836 | 0.00533 | 0.01597 |
| **FLT3LG** | 0.39525 | 0.00000 | 0.86025 | 0.18399 | 1.00801 | 0.05190 | 0.86869 |
| **FLT4** | 0.00000 | 1.06040 | 0.36071 | 0.06816 | 0.00091 | 0.24254 | 0.05195 |
| **FLVCR1** | 6.43860 | 1.29363 | 0.43775 | 2.72999 | 0.05045 | 9.43293 | 7.40642 |
| **FLVCR1-AS1** | 0.95862 | 0.74249 | 0.00518 | 0.59759 | 0.40448 | 1.65003 | 0.00000 |
| **FLVCR2** | 4.81804 | 0.07172 | 2.05197 | 7.75560 | 2.09722 | 0.02675 | 2.64548 |
| **FLYWCH1** | 0.11436 | 2.17195 | 1.25529 | 1.92243 | 0.43895 | 2.42998 | 3.57092 |
| **FLYWCH2** | 2.13981 | 8.95640 | 3.39312 | 5.03663 | 0.82276 | 3.08518 | 0.80581 |
| **FMN1** | 0.06196 | 0.00000 | 1.32846 | 0.37361 | 3.30621 | 1.18034 | 0.44150 |
| **FMN2** | 0.00000 | 0.15469 | 0.14001 | 0.00239 | 0.82049 | 0.30831 | 0.11307 |
| **FMNL1** | 0.00000 | 0.09349 | 0.11831 | 0.91439 | 0.35776 | 0.00000 | 0.22087 |
| **FMNL2** | 1.15008 | 0.99123 | 2.02713 | 6.66211 | 0.24189 | 3.85901 | 8.33329 |
| **FMNL3** | 0.35325 | 0.28790 | 0.23804 | 0.08038 | 0.37854 | 0.38772 | 0.21660 |
| **FMO1** | 0.03810 | 0.01612 | 0.01209 | 0.02031 | 0.01624 | 0.01412 | 0.01285 |
| **FMO2** | 0.01494 | 0.00000 | 0.46408 | 0.01813 | 0.91600 | 0.00696 | 0.03679 |
| **FMO3** | 0.02770 | 0.00922 | 0.04410 | 0.01846 | 0.04763 | 0.01253 | 0.01595 |
| **FMO4** | 0.09352 | 0.17218 | 0.06220 | 0.18893 | 0.16947 | 0.00102 | 0.00000 |
| **FMO5** | 0.00000 | 0.00000 | 0.39629 | 0.12229 | 0.01423 | 2.34827 | 0.07836 |
| **FMO6P** | 0.01824 | 0.02916 | 0.01375 | 0.01851 | 0.02185 | 0.02673 | 0.02095 |
| **FMO9P** | 0.01156 | 0.01781 | 0.01305 | 0.06706 | 0.00794 | 0.01521 | 0.01887 |

| | | | | | | | |
|---|---|---|---|---|---|---|---|
| **FMOD** | 0.01001 | 0.09277 | 0.01345 | 0.04046 | 0.00714 | 0.02797 | 0.00801 |
| **FMR1** | 4.62005 | 0.48567 | 5.25933 | 3.85944 | 2.67227 | 2.53194 | 8.89220 |
| **FMR1-AS1** | 0.02255 | 0.05304 | 0.15985 | 0.03277 | 0.17060 | 0.19761 | 0.09969 |
| **FMR1NB** | 12.24188 | 19.44639 | 17.56699 | 4.83360 | 2.93704 | 3.15883 | 2.23232 |
| **FN1** | 0.65036 | 0.14785 | 1.31276 | 1.72906 | 13.80916 | 6.42972 | 0.52808 |
| **FN3K** | 0.50582 | 1.96646 | 0.00000 | 0.10966 | 0.36657 | 0.66399 | 1.09912 |
| **FN3KRP** | 6.83097 | 8.40963 | 4.36809 | 4.46103 | 1.69986 | 6.28690 | 1.48297 |
| **FNBP1** | 2.03183 | 6.59163 | 12.04421 | 2.11950 | 3.63920 | 1.95227 | 11.21540 |
| **FNBP1L** | 29.00748 | 7.76811 | 9.66986 | 10.93863 | 1.77818 | 4.28202 | 13.65962 |
| **FNBP4** | 5.01293 | 2.86384 | 6.78131 | 10.06589 | 4.01977 | 10.65810 | 31.26184 |
| **FNDC1** | 0.04420 | 0.01668 | 0.01314 | 0.01018 | 0.00773 | 0.01326 | 0.02489 |
| **FNDC3A** | 6.21945 | 0.00000 | 2.41292 | 3.98865 | 0.05916 | 2.90217 | 5.56027 |
| **FNDC3B** | 9.68598 | 3.41101 | 0.78594 | 5.46438 | 3.97683 | 4.04722 | 8.52218 |
| **FNDC4** | 0.00000 | 0.07671 | 0.00000 | 0.04844 | 0.43555 | 4.62601 | 0.01785 |
| **FNDC5** | 0.10930 | 0.06458 | 0.04611 | 0.03850 | 0.05163 | 0.00931 | 0.18194 |
| **FNDC7** | 0.03596 | 0.00614 | 0.00528 | 0.00892 | 0.11953 | 0.00696 | 0.12601 |
| **FNDC8** | 0.09419 | 0.00000 | 0.07025 | 0.13753 | 0.00358 | 0.35780 | 0.43116 |
| **FNIP1** | 9.18239 | 0.00000 | 1.50084 | 3.59549 | 1.44198 | 2.22301 | 0.57856 |
| **FNIP2** | 3.72203 | 8.62106 | 4.29722 | 0.03772 | 2.69633 | 3.72848 | 7.15556 |
| **FNTA** | 4.50133 | 4.25440 | 2.84237 | 5.27295 | 0.66521 | 7.71264 | 11.65156 |
| **FNTB** | 1.26670 | 0.77780 | 0.39181 | 0.30694 | 0.05979 | 0.62024 | 0.00000 |
| **FOCAD** | 4.44672 | 3.21206 | 2.37049 | 4.05418 | 1.50071 | 1.78683 | 4.02441 |
| **FOCAD-AS1** | 0.00669 | 0.03198 | 0.00921 | 0.04242 | 0.15192 | 0.01685 | 0.07906 |
| **FOLH1** | 0.05689 | 0.03311 | 0.12630 | 0.01287 | 0.23604 | 0.02703 | 0.00000 |
| **FOLH1B** | 0.03183 | 0.01590 | 0.01854 | 0.01757 | 0.03941 | 0.01858 | 0.01372 |
| **FOLR1** | 31.77486 | 16.90485 | 0.00000 | 26.84175 | 0.88562 | 0.00000 | 16.14363 |
| **FOLR2** | 0.06623 | 0.02412 | 0.01690 | 0.02039 | 0.01384 | 0.03346 | 0.08882 |
| **FOLR3** | 0.15923 | 0.77761 | 0.00000 | 0.00000 | 0.09822 | 0.76401 | 0.01375 |
| **FOLR4** | 0.01452 | 0.08076 | 0.00000 | 0.02213 | 2.50187 | 0.01031 | 0.00941 |
| **FOPNL** | 14.73449 | 2.24933 | 9.94566 | 8.20432 | 1.29342 | 4.79347 | 0.00000 |
| **FOS** | 2.00600 | 8.17982 | 0.59510 | 8.30356 | 6.47949 | 5.92483 | 9.35251 |
| **FOSB** | 0.00000 | 0.00000 | 0.04803 | 4.55279 | 0.69726 | 0.02941 | 1.21999 |
| **FOSL1** | 0.00000 | 0.04349 | 0.00512 | 3.91370 | 0.46867 | 0.58445 | 0.51744 |
| **FOSL2** | 0.30100 | 4.20989 | 0.71158 | 0.90904 | 0.00854 | 0.29636 | 5.46964 |
| **FOXA1** | 0.01521 | 0.02544 | 0.01735 | 0.02381 | 0.00653 | 0.04983 | 0.12637 |

| | | | | | | | |
|---|---|---|---|---|---|---|---|
| **FOXA2** | 0.00000 | 0.00122 | 0.00968 | 0.03346 | 0.03008 | 2.96971 | 0.09543 |
| **FOXA3** | 0.00000 | 0.13562 | 0.06749 | 0.43117 | 0.00779 | 0.31143 | 0.00124 |
| **FOXB1** | 0.01781 | 0.02422 | 0.01097 | 0.07989 | 0.00825 | 0.05250 | 0.00292 |
| **FOXB2** | 0.01110 | 0.01527 | 0.01100 | 0.01725 | 0.02007 | 0.13995 | 0.02889 |
| **FOXC1** | 0.00019 | 0.34085 | 0.02689 | 0.11881 | 0.12597 | 0.81303 | 0.50410 |
| **FOXC2** | 0.02149 | 0.01841 | 0.01642 | 0.01903 | 0.01120 | 0.02002 | 0.01885 |
| **FOXD1** | 0.00000 | 0.13534 | 0.00514 | 0.25410 | 0.17235 | 0.00000 | 0.45899 |
| **FOXD2** | 0.12629 | 0.16805 | 0.25316 | 0.08146 | 0.28668 | 0.03735 | 0.12966 |
| **FOXD2-AS1** | 0.00000 | 0.24512 | 1.68399 | 0.63080 | 0.71732 | 0.09082 | 0.57248 |
| **FOXD3** | 0.00000 | 0.04405 | 0.39656 | 0.00301 | 0.01070 | 0.81284 | 0.09113 |
| **FOXD3-AS1** | 0.03449 | 0.47058 | 0.40879 | 0.05217 | 0.42196 | 3.36647 | 0.00000 |
| **FOXD4** | 0.04537 | 0.02584 | 0.01843 | 0.02114 | 0.01144 | 0.00187 | 0.10402 |
| **FOXD4L1** | 0.03532 | 0.01576 | 0.01224 | 0.02339 | 0.01588 | 0.01619 | 0.00878 |
| **FOXE1** | 0.06733 | 1.70009 | 0.84405 | 0.00000 | 0.00025 | 0.00000 | 1.15019 |
| **FOXE3** | 0.00009 | 0.46269 | 0.02650 | 0.05902 | 0.00526 | 0.01006 | 0.02757 |
| **FOXF1** | 0.03184 | 0.02273 | 0.01043 | 0.00701 | 0.00728 | 0.01469 | 0.07151 |
| **FOXF2** | 0.00000 | 0.10814 | 0.01970 | 0.02658 | 0.02830 | 0.01561 | 0.11985 |
| **FOXG1** | 0.28750 | 0.00000 | 0.20452 | 0.10038 | 0.36911 | 0.06143 | 0.38413 |
| **FOXH1** | 3.59727 | 18.11865 | 1.84470 | 0.00000 | 0.00000 | 14.02230 | 1.12772 |
| **FOXI2** | 0.02638 | 0.02313 | 0.01869 | 0.01956 | 0.00729 | 0.01530 | 0.01360 |
| **FOXI3** | 0.00000 | 2.63618 | 1.55283 | 1.22658 | 0.16118 | 0.00000 | 0.12298 |
| **FOXJ1** | 0.05133 | 0.00858 | 0.01071 | 0.06568 | 0.05271 | 0.02239 | 0.02472 |
| **FOXJ2** | 2.60452 | 0.00000 | 1.70185 | 1.19601 | 0.29698 | 0.72371 | 2.89847 |
| **FOXJ3** | 4.59205 | 0.85417 | 4.19657 | 5.14703 | 5.06734 | 2.98357 | 9.02202 |
| **FOXK1** | 10.67695 | 13.86288 | 2.58816 | 7.80117 | 0.25598 | 3.15171 | 8.46892 |
| **FOXK2** | 7.63012 | 13.75784 | 2.79820 | 6.64218 | 3.16704 | 5.71024 | 20.34157 |
| **FOXL2** | 0.03552 | 0.01274 | 0.01246 | 0.04109 | 0.00701 | 0.00396 | 0.09637 |
| **FOXL2NB** | 0.02281 | 0.03269 | 0.02235 | 0.03011 | 0.00607 | 0.01569 | 0.06078 |
| **FOXM1** | 6.21454 | 11.99260 | 6.54142 | 9.04928 | 0.25690 | 8.92225 | 21.99934 |
| **FOXN1** | 0.01254 | 0.01579 | 0.02087 | 0.03767 | 0.00909 | 0.01287 | 0.04729 |
| **FOXN2** | 8.09787 | 0.00000 | 4.18737 | 3.21968 | 11.12127 | 4.57412 | 7.42273 |
| **FOXN3** | 3.44635 | 8.65554 | 0.00000 | 2.64958 | 0.21012 | 8.98449 | 21.25723 |
| **FOXN3-AS1** | 0.00000 | 0.22298 | 0.03486 | 0.01364 | 0.00450 | 0.04774 | 0.12816 |
| **FOXN3-AS2** | 0.02759 | 0.01869 | 0.01240 | 0.01913 | 0.00787 | 0.01224 | 0.02444 |
| **FOXN4** | 0.17050 | 0.55680 | 0.00000 | 0.02899 | 0.06782 | 0.00000 | 0.09844 |

| | | | | | | | |
|---|---|---|---|---|---|---|---|
| **FOXO1** | 0.21632 | 6.25074 | 6.45601 | 1.15494 | 1.59087 | 3.67272 | 5.32283 |
| **FOXO3** | 0.92984 | 0.53800 | 2.80327 | 0.73782 | 2.03999 | 1.35977 | 4.65156 |
| **FOXO3B** | 0.05519 | 0.00000 | 0.13922 | 0.70253 | 0.63795 | 0.41772 | 0.96517 |
| **FOXO4** | 0.37075 | 1.95804 | 0.06595 | 0.95705 | 0.00655 | 0.00000 | 4.50140 |
| **FOXO6** | 0.00000 | 0.61195 | 0.01129 | 1.89389 | 0.00487 | 0.00000 | 4.23014 |
| **FOXP1** | 0.86196 | 1.37846 | 0.00000 | 0.25884 | 0.35739 | 5.81828 | 2.27996 |
| **FOXP2** | 0.05856 | 0.01830 | 0.03007 | 0.03118 | 0.01832 | 0.18393 | 0.02587 |
| **FOXP3** | 0.02117 | 0.16840 | 0.03629 | 0.02182 | 0.00811 | 0.00113 | 0.08764 |
| **FOXP4** | 0.00000 | 2.05566 | 0.85919 | 0.58855 | 0.42842 | 1.64998 | 8.38716 |
| **FOXQ1** | 0.01516 | 0.05507 | 0.00340 | 0.02002 | 0.15682 | 0.05770 | 0.01561 |
| **FOXR1** | 2.73048 | 36.63494 | 28.96566 | 0.00000 | 23.90429 | 7.70437 | 0.00000 |
| **FOXR2** | 0.09711 | 0.00000 | 0.22717 | 0.05698 | 0.25319 | 0.16574 | 0.00000 |
| **FOXRED1** | 17.59641 | 18.09707 | 3.65793 | 17.03858 | 0.05901 | 12.18604 | 4.08842 |
| **FOXRED2** | 4.48602 | 0.73963 | 4.26738 | 2.77741 | 1.21251 | 0.93145 | 3.90543 |
| **FOXS1** | 0.01469 | 0.03991 | 0.07869 | 0.02168 | 0.00455 | 0.00558 | 0.03202 |
| **FPGS** | 2.50710 | 7.36509 | 4.48262 | 9.25479 | 1.07469 | 3.68894 | 6.87722 |
| **FPGT** | 6.41327 | 0.00000 | 3.46905 | 1.69130 | 0.41774 | 0.92809 | 0.00000 |
| **FPR1** | 0.02526 | 0.01498 | 0.02082 | 0.02369 | 0.00776 | 0.01660 | 0.01474 |
| **FPR3** | 0.01428 | 0.01255 | 0.10911 | 0.01436 | 0.04966 | 0.03300 | 0.04708 |
| **FRA10AC1** | 2.01835 | 1.95962 | 3.70042 | 1.56093 | 0.18140 | 1.50526 | 5.21392 |
| **FRAS1** | 1.41479 | 0.00000 | 0.04860 | 2.69623 | 0.21536 | 4.94325 | 2.54956 |
| **FRAT1** | 0.00000 | 0.07954 | 0.64997 | 0.62832 | 0.22274 | 0.91658 | 0.52307 |
| **FRAT2** | 0.00000 | 11.68206 | 22.77188 | 3.15743 | 5.44018 | 17.03384 | 18.74750 |
| **FREM1** | 0.03337 | 0.01570 | 0.00862 | 0.02746 | 0.05895 | 0.02445 | 0.01191 |
| **FREM2** | 0.47034 | 0.00000 | 0.12647 | 1.10787 | 0.34044 | 0.00000 | 1.09520 |
| **FREM3** | 0.01967 | 0.01894 | 0.01479 | 0.04134 | 0.00991 | 0.03488 | 0.02068 |
| **FRG1** | 9.38044 | 16.94055 | 9.35977 | 7.85078 | 10.25794 | 6.53199 | 7.75938 |
| **FRG1B** | 2.69493 | 4.28845 | 2.37649 | 2.18743 | 2.20763 | 3.49037 | 5.14402 |
| **FRG2.uid-6569** | 0.02945 | 0.01467 | 0.00549 | 0.01378 | 0.42800 | 0.01298 | 0.02512 |
| **FRG2B** | 0.01581 | 0.01985 | 0.00000 | 0.01561 | 0.39866 | 0.01652 | 0.04104 |
| **FRK** | 0.50705 | 0.00000 | 0.11760 | 0.08084 | 0.00539 | 0.04673 | 0.06115 |
| **FRMD1** | 0.02045 | 0.01567 | 0.01356 | 0.01553 | 0.00682 | 0.00917 | 0.10251 |
| **FRMD3** | 0.09607 | 0.25001 | 0.35215 | 0.23649 | 4.63511 | 0.57629 | 0.03309 |
| **FRMD4A** | 3.75755 | 0.31841 | 0.00000 | 8.98453 | 0.90890 | 0.00000 | 9.99801 |
| **FRMD4B** | 17.61384 | 6.12780 | 7.63648 | 13.89714 | 2.76696 | 0.00000 | 10.34552 |

| | | | | | | | |
|---|---|---|---|---|---|---|---|
| **FRMD5** | 0.96998 | 0.00000 | 1.51433 | 0.41705 | 0.58386 | 0.06977 | 0.90677 |
| **FRMD6** | 12.47348 | 1.43191 | 2.26037 | 8.03852 | 1.77895 | 9.29532 | 0.72303 |
| **FRMD6-AS1** | 0.33341 | 0.08718 | 0.10996 | 0.84304 | 0.03677 | 0.60480 | 0.71447 |
| **FRMD6-AS2** | 0.05646 | 0.00939 | 0.01242 | 0.04295 | 0.00719 | 0.02386 | 0.05921 |
| **FRMD8** | 2.66693 | 4.43378 | 4.08922 | 6.89743 | 1.10569 | 2.17499 | 11.72286 |
| **FRMD8P1** | 0.01201 | 0.24340 | 0.08473 | 0.16272 | 0.02115 | 0.08006 | 0.21323 |
| **FRMPD1** | 0.01960 | 0.00641 | 0.02214 | 0.05827 | 0.01968 | 0.01856 | 0.05125 |
| **FRMPD2.uid-13** | 0.02100 | 0.03265 | 0.01617 | 0.03127 | 0.00734 | 0.01175 | 0.01429 |
| **FRMPD3** | 0.00000 | 0.45193 | 0.98276 | 0.00000 | 0.00115 | 0.54710 | 0.02658 |
| **FRMPD4** | 0.02558 | 0.01239 | 0.00782 | 0.01776 | 0.05162 | 0.11209 | 0.01718 |
| **FRRS1** | 1.49960 | 0.00028 | 0.35570 | 1.07162 | 0.09945 | 0.00000 | 0.47702 |
| **FRRS1L** | 0.02274 | 0.01682 | 0.00833 | 0.01993 | 0.06380 | 0.01272 | 0.02285 |
| **FRS2** | 8.53853 | 0.00000 | 5.01210 | 10.94653 | 10.36774 | 2.06711 | 6.71658 |
| **FRS3** | 0.03721 | 0.12540 | 0.19470 | 0.10443 | 0.37775 | 0.46308 | 0.34381 |
| **FRY** | 0.00000 | 0.09447 | 0.07217 | 0.06933 | 0.08340 | 0.84049 | 0.13282 |
| **FRY-AS1** | 0.05793 | 0.12106 | 0.05658 | 0.01025 | 0.04671 | 0.24231 | 0.00001 |
| **FRYL** | 4.83869 | 0.00000 | 2.92579 | 3.69265 | 0.88093 | 4.27428 | 9.30532 |
| **FRZB** | 0.01396 | 0.00021 | 0.00734 | 0.12466 | 0.05342 | 2.90396 | 0.00074 |
| **FSCN1** | 2.70728 | 7.12015 | 1.47510 | 25.54632 | 0.38765 | 9.31702 | 42.65003 |
| **FSCN2** | 0.00000 | 0.00799 | 0.01565 | 0.02513 | 0.00660 | 0.13737 | 0.15881 |
| **FSCN3** | 0.04428 | 0.02259 | 0.01087 | 0.07950 | 0.00787 | 0.02083 | 0.01121 |
| **FSD1** | 0.00000 | 0.02883 | 0.02416 | 1.65296 | 0.10201 | 0.58306 | 0.41118 |
| **FSD1L** | 0.86779 | 0.00000 | 0.29277 | 0.13975 | 0.44987 | 0.05844 | 0.48369 |
| **FSD2** | 2.26620 | 0.00000 | 0.93621 | 0.45470 | 0.60034 | 0.37492 | 0.00000 |
| **FSHR** | 0.07530 | 0.04284 | 0.01014 | 0.01937 | 0.01104 | 0.01117 | 0.00000 |
| **FSIP1** | 0.30303 | 0.00000 | 0.20953 | 0.09619 | 0.34073 | 0.03864 | 0.00000 |
| **FSIP2** | 0.05943 | 0.00000 | 0.24349 | 0.02142 | 0.49923 | 0.17595 | 0.33243 |
| **FST** | 0.00000 | 0.00069 | 0.00414 | 6.43793 | 0.28529 | 1.08025 | 0.31691 |
| **FSTL1** | 7.59671 | 0.00000 | 0.00000 | 6.38934 | 0.00285 | 0.33662 | 1.37339 |
| **FSTL3** | 0.00000 | 0.89503 | 1.40174 | 0.64035 | 0.08236 | 0.04104 | 0.68484 |
| **FSTL4** | 0.00000 | 0.02526 | 0.12401 | 0.04245 | 2.12523 | 0.08356 | 0.03945 |
| **FSTL5** | 0.05707 | 0.04093 | 0.00000 | 0.02844 | 0.23049 | 0.02714 | 0.03022 |
| **FTCD** | 0.01730 | 0.02705 | 0.03214 | 0.02416 | 2.22598 | 0.03759 | 0.01322 |
| **FTCDNL1** | 0.00528 | 0.17950 | 0.00224 | 0.03087 | 0.04583 | 0.02289 | 0.08272 |
| **FTH1** | 38.03001 | 46.50670 | 17.89550 | 40.30178 | 6.24236 | 27.71380 | 9.94040 |

| | | | | | | | |
|---|---|---|---|---|---|---|---|
| **FTH1P3** | 0.05361 | 0.41343 | 0.22946 | 0.16319 | 1.25352 | 0.42792 | 0.67280 |
| **FTHL17** | 0.01017 | 0.19956 | 0.28330 | 0.43927 | 0.52861 | 0.13251 | 0.00000 |
| **FTL** | 60.73286 | 36.46738 | 17.31353 | 78.17073 | 25.11378 | 23.46123 | 26.45750 |
| **FTLP10** | 0.02429 | 0.02251 | 0.01265 | 0.02146 | 0.00807 | 0.01203 | 0.01665 |
| **FTMT** | 0.02128 | 0.01778 | 0.01383 | 0.02089 | 0.00768 | 0.01242 | 0.02459 |
| **FTO** | 5.76339 | 0.79932 | 1.78238 | 4.12199 | 2.67726 | 3.84403 | 0.00000 |
| **FTO-IT1** | 0.13442 | 0.00000 | 0.28665 | 0.18906 | 0.31216 | 0.14245 | 0.39406 |
| **FTSJ1** | 23.63323 | 9.85871 | 7.74940 | 13.75907 | 0.78747 | 11.87391 | 0.00000 |
| **FTSJ2** | 27.19601 | 17.85805 | 14.39853 | 17.06193 | 4.08679 | 10.12251 | 0.00000 |
| **FTSJ3** | 31.71701 | 25.64208 | 14.74122 | 30.26097 | 8.33563 | 10.28089 | 5.48117 |
| **FTX** | 1.29180 | 0.07097 | 0.65497 | 0.68659 | 0.16676 | 0.51740 | 0.92214 |
| **FUBP1** | 26.23146 | 13.71102 | 24.33629 | 19.10911 | 8.14530 | 13.43374 | 15.12066 |
| **FUBP3** | 8.98508 | 3.71769 | 3.83999 | 4.13253 | 2.00036 | 4.81035 | 10.92393 |
| **FUCA1** | 9.89788 | 18.55364 | 9.69606 | 2.46357 | 2.84049 | 11.12220 | 0.00000 |
| **FUCA2** | 26.94477 | 4.40400 | 0.00000 | 17.79499 | 0.16206 | 1.19382 | 0.00000 |
| **FUK** | 0.00000 | 0.63909 | 0.34852 | 0.79874 | 0.00000 | 0.42038 | 2.37368 |
| **FUNDC1** | 17.56946 | 14.32377 | 14.54876 | 1.19806 | 6.07646 | 12.88888 | 4.12278 |
| **FUNDC2** | 15.91256 | 13.88952 | 9.02890 | 11.29475 | 2.21963 | 7.72103 | 7.60930 |
| **FUNDC2P2** | 0.56799 | 0.46791 | 0.23900 | 0.31142 | 0.31649 | 0.19445 | 0.00000 |
| **FUOM** | 10.14087 | 9.33437 | 0.00000 | 0.54198 | 0.00000 | 3.40202 | 3.12425 |
| **FURIN** | 3.51549 | 16.64375 | 5.73089 | 12.53217 | 1.88638 | 6.11383 | 39.62746 |
| **FUS** | 32.75870 | 37.77677 | 21.89545 | 34.16507 | 14.87818 | 25.27068 | 25.92487 |
| **FUT1** | 4.82700 | 0.02209 | 0.00000 | 0.00000 | 0.00000 | 0.00000 | 4.51328 |
| **FUT10** | 4.27856 | 0.16960 | 1.96208 | 1.36129 | 1.56345 | 0.86102 | 0.00000 |
| **FUT11** | 0.79826 | 0.13567 | 0.04662 | 2.36148 | 0.09903 | 0.18406 | 1.15584 |
| **FUT2** | 0.01084 | 0.14508 | 0.11995 | 0.04662 | 0.26109 | 0.37360 | 0.00524 |
| **FUT3** | 0.25690 | 0.00000 | 9.05342 | 0.02192 | 0.27762 | 0.35300 | 0.00000 |
| **FUT4** | 11.89552 | 0.00000 | 7.05339 | 3.21778 | 2.80430 | 0.00000 | 13.16185 |
| **FUT5** | 0.00177 | 0.02108 | 0.00441 | 0.01861 | 0.69632 | 0.19443 | 0.04855 |
| **FUT6** | 0.04670 | 0.08566 | 0.19574 | 0.02213 | 0.07974 | 0.39086 | 0.02668 |
| **FUT8** | 12.15135 | 2.04057 | 1.18069 | 18.60571 | 1.74538 | 3.45771 | 9.13136 |
| **FUT8-AS1** | 0.04859 | 0.10536 | 0.07188 | 0.14997 | 0.09552 | 0.04863 | 0.00000 |
| **FUT9** | 0.13090 | 0.09609 | 0.00000 | 0.01125 | 0.48280 | 0.01508 | 0.00000 |
| **FUZ** | 1.02556 | 1.31994 | 0.71027 | 3.92411 | 0.02689 | 1.61816 | 0.42660 |
| **FXN** | 16.23847 | 21.79381 | 7.61328 | 8.08254 | 0.00000 | 12.29972 | 4.85420 |

| | | | | | | | |
|---|---|---|---|---|---|---|---|
| **FXR1** | 20.72751 | 5.78247 | 14.11872 | 9.73535 | 4.59317 | 10.91641 | 13.19955 |
| **FXR2** | 0.71540 | 2.50312 | 4.37237 | 6.46086 | 0.80361 | 1.27129 | 13.92532 |
| **FXYD1** | 0.01636 | 0.01871 | 0.02500 | 0.03080 | 0.04809 | 0.03460 | 0.04641 |
| **FXYD3** | 0.51040 | 0.43685 | 0.03887 | 0.31626 | 0.05083 | 0.62075 | 0.32585 |
| **FXYD4** | 8.99168 | 19.42597 | 6.74385 | 0.05468 | 0.25914 | 1.18661 | 14.01845 |
| **FXYD5** | 1.70674 | 0.22145 | 0.15469 | 13.96840 | 0.00111 | 6.07157 | 3.60333 |
| **FXYD6** | 0.00000 | 0.04921 | 0.00537 | 0.02397 | 0.00781 | 0.46737 | 0.03410 |
| **FXYD7** | 0.00000 | 0.00389 | 0.03146 | 0.09680 | 0.10326 | 0.07731 | 0.17671 |
| **FYB** | 2.64730 | 0.00000 | 0.00000 | 25.53687 | 0.00000 | 0.00000 | 0.49972 |
| **FYCO1** | 2.33720 | 0.00000 | 0.50706 | 1.40118 | 0.00246 | 0.10068 | 2.07337 |
| **FYN** | 0.00000 | 0.85345 | 1.08899 | 0.83513 | 1.68705 | 2.38729 | 1.62394 |
| **FYTTD1** | 6.21337 | 3.00659 | 10.18623 | 0.15830 | 5.09282 | 1.11000 | 4.78849 |
| **FZD1** | 0.03660 | 0.22493 | 0.14741 | 0.22879 | 0.31650 | 0.30584 | 0.15076 |
| **FZD10** | 0.00000 | 0.04418 | 0.00248 | 1.19789 | 0.07199 | 0.33382 | 0.02779 |
| **FZD10-AS1** | 0.00000 | 0.05005 | 0.00903 | 0.07425 | 0.03816 | 0.08406 | 0.06561 |
| **FZD2** | 0.01131 | 0.05828 | 0.03228 | 0.04460 | 0.00524 | 0.00285 | 0.10431 |
| **FZD3** | 7.23494 | 0.00000 | 5.09548 | 2.35493 | 2.56077 | 0.75119 | 9.58352 |
| **FZD4** | 4.94659 | 5.13144 | 0.54542 | 0.20710 | 2.19392 | 0.75409 | 0.60498 |
| **FZD5** | 13.48988 | 11.22413 | 0.46559 | 19.79793 | 0.73185 | 26.84579 | 12.51500 |
| **FZD6** | 1.77442 | 0.01347 | 0.00000 | 3.13690 | 0.43510 | 0.63091 | 1.31833 |
| **FZD7** | 0.10525 | 0.98801 | 0.15832 | 0.00000 | 0.78286 | 1.03625 | 3.83920 |
| **FZD8** | 0.08741 | 0.13975 | 0.01256 | 0.00000 | 0.00501 | 0.74781 | 0.14137 |
| **FZD9** | 0.00642 | 0.01762 | 0.03662 | 0.03614 | 0.06607 | 0.25016 | 0.05128 |
| **FZR1** | 0.00000 | 4.54598 | 1.41580 | 6.22135 | 0.41886 | 1.31316 | 11.91544 |
| **G0S2** | 0.05934 | 0.08518 | 0.00553 | 0.00994 | 0.01390 | 0.01190 | 0.00000 |
| **G2E3** | 17.11990 | 0.00000 | 10.45192 | 9.90582 | 3.65265 | 5.40069 | 9.63521 |
| **G3BP1** | 71.52739 | 31.78454 | 17.54940 | 29.64929 | 3.86320 | 24.48868 | 2.74989 |
| **G3BP2** | 36.37720 | 9.32745 | 28.95654 | 8.89516 | 7.55324 | 17.19992 | 11.91431 |
| **G6PC** | 0.05958 | 0.00288 | 0.06042 | 0.01257 | 0.00813 | 0.02609 | 0.00648 |
| **G6PC2** | 0.02042 | 0.01944 | 0.01264 | 0.02314 | 0.00776 | 0.01809 | 0.02429 |
| **G6PC3** | 4.57696 | 7.62633 | 1.71910 | 9.60938 | 0.01305 | 11.52640 | 4.53967 |
| **G6PD** | 10.68080 | 10.79252 | 2.87277 | 9.91697 | 0.76157 | 5.00454 | 11.21173 |
| **GAA** | 0.00000 | 0.62751 | 0.42351 | 1.96653 | 0.27456 | 0.54052 | 2.77666 |
| **GAB1** | 1.88681 | 7.49594 | 3.94869 | 3.30694 | 7.77185 | 4.76213 | 7.21048 |
| **GAB2** | 8.05599 | 1.08654 | 0.18255 | 19.21127 | 2.23991 | 0.00000 | 24.84928 |

| | | | | | | | |
|---|---|---|---|---|---|---|---|
| **GAB3** | 0.04239 | 0.00000 | 0.34114 | 0.07168 | 0.01140 | 0.06455 | 0.09394 |
| **GABARAP** | 19.93165 | 4.50518 | 1.07663 | 25.55524 | 5.97805 | 13.15428 | 16.00711 |
| **GABARAPL1** | 4.68619 | 15.26207 | 41.60940 | 22.76413 | 1.55496 | 26.96060 | 2.81981 |
| **GABARAPL2** | 14.85585 | 8.15596 | 16.64701 | 13.56321 | 14.88085 | 11.87026 | 9.84492 |
| **GABARAPL3** | 0.04023 | 0.03620 | 0.11703 | 0.05372 | 0.02667 | 0.04205 | 0.00000 |
| **GABBR1** | 0.25047 | 0.00000 | 0.35413 | 0.30989 | 0.07643 | 3.26231 | 1.13478 |
| **GABBR2** | 7.88397 | 0.00000 | 0.00000 | 0.00000 | 2.75834 | 0.00000 | 7.03962 |
| **GABPA** | 8.77379 | 0.00000 | 5.58362 | 3.25404 | 6.34334 | 3.08633 | 1.65176 |
| **GABPB1** | 19.80986 | 7.42185 | 9.10284 | 9.25750 | 3.81536 | 3.98633 | 1.42521 |
| **GABPB1-AS1** | 0.52362 | 0.64809 | 2.23948 | 2.81944 | 0.67677 | 0.64312 | 6.35982 |
| **GABPB2** | 0.75512 | 1.17025 | 1.49388 | 0.63054 | 1.01468 | 0.40385 | 0.12239 |
| **GABRA1** | 0.02373 | 0.00537 | 0.03614 | 0.01986 | 0.18692 | 0.01485 | 0.01658 |
| **GABRA2** | 0.01684 | 0.00460 | 0.02338 | 0.03253 | 0.04108 | 0.12469 | 0.14081 |
| **GABRA3** | 0.13450 | 0.02464 | 0.05074 | 0.00000 | 0.48870 | 0.07422 | 0.03509 |
| **GABRA4** | 0.02606 | 0.00654 | 0.03617 | 0.02909 | 0.00708 | 0.01527 | 0.02166 |
| **GABRA5** | 0.55651 | 0.37266 | 0.00381 | 0.04455 | 0.07590 | 0.22830 | 0.00000 |
| **GABRA6** | 0.02968 | 0.06450 | 0.01140 | 0.01836 | 0.04413 | 0.00970 | 0.00005 |
| **GABRB1** | 0.03642 | 0.02083 | 0.00672 | 0.03878 | 0.05978 | 0.02722 | 0.03839 |
| **GABRB2** | 0.36723 | 0.00000 | 0.00672 | 0.00000 | 0.00800 | 0.05327 | 0.42471 |
| **GABRB3** | 2.95869 | 2.40910 | 1.83346 | 3.50399 | 0.91604 | 4.96375 | 1.98288 |
| **GABRD** | 0.00000 | 0.06307 | 0.49886 | 0.00427 | 0.00412 | 0.28132 | 0.04985 |
| **GABRE** | 0.05183 | 0.03117 | 0.00000 | 1.52119 | 0.15680 | 0.15948 | 0.31000 |
| **GABRG1** | 0.18084 | 0.00000 | 0.01182 | 0.13639 | 0.00650 | 0.00000 | 0.06627 |
| **GABRG2** | 0.30163 | 0.00000 | 0.04016 | 0.01324 | 0.00588 | 0.19467 | 0.00829 |
| **GABRG3** | 0.03927 | 0.11944 | 0.02940 | 0.02849 | 0.10831 | 0.00805 | 0.00000 |
| **GABRP** | 0.03551 | 0.00000 | 0.05801 | 0.00479 | 0.34654 | 0.38163 | 0.02704 |
| **GABRQ** | 0.00954 | 0.03706 | 0.32154 | 0.08203 | 1.27708 | 0.62573 | 0.00000 |
| **GABRR1** | 0.16751 | 0.09628 | 0.32792 | 0.03020 | 0.01953 | 0.05819 | 0.00000 |
| **GABRR2** | 0.05428 | 0.01905 | 0.01119 | 0.09118 | 0.00809 | 0.03312 | 0.00000 |
| **GABRR3** | 0.01920 | 0.01201 | 0.00000 | 0.01576 | 2.82101 | 0.01071 | 0.01487 |
| **GAD1** | 0.13404 | 0.19414 | 0.73064 | 0.00000 | 0.53077 | 0.52821 | 0.00002 |
| **GAD2** | 0.03853 | 0.01009 | 0.02532 | 0.01653 | 0.00869 | 0.25974 | 0.01225 |
| **GADD45A** | 0.39872 | 3.08193 | 0.25446 | 6.65051 | 3.71262 | 3.95020 | 9.85722 |
| **GADD45B** | 1.75269 | 2.57829 | 0.00000 | 2.13383 | 9.16901 | 0.85044 | 0.24992 |
| **GADD45G** | 0.00000 | 2.18277 | 0.00000 | 4.29340 | 0.00426 | 1.01472 | 1.71348 |

| | | | | | | | |
|---|---|---|---|---|---|---|---|
| **GADD45GIP1** | 0.00000 | 18.33800 | 5.23972 | 6.55072 | 0.47410 | 11.73885 | 30.29953 |
| **GADL1** | 0.00597 | 0.01175 | 0.03811 | 0.04955 | 0.09154 | 0.07393 | 0.04394 |
| **GAGE1** | 1.45498 | 4.24991 | 1.45235 | 0.00000 | 0.13038 | 0.00000 | 1.00052 |
| **GAGE10** | 0.04971 | 0.08383 | 0.01731 | 0.01792 | 0.01230 | 0.01604 | 0.01288 |
| **GAGE12J** | 0.12077 | 5.21510 | 0.53677 | 0.07593 | 0.05770 | 0.09925 | 0.00000 |
| **GAGE2A.uid-12** | 0.00000 | 9.58562 | 0.37152 | 0.00975 | 0.12151 | 0.03805 | 0.00579 |
| **GAGE2B** | 0.00000 | 0.86294 | 0.05566 | 0.01123 | 0.02203 | 0.16931 | 0.01376 |
| **GAK** | 5.35772 | 12.82013 | 2.33411 | 5.12479 | 0.72669 | 6.45572 | 24.73965 |
| **GAL** | 0.00000 | 5.75410 | 1.12023 | 0.00000 | 0.08679 | 2.56428 | 6.56311 |
| **GAL3ST1** | 0.00000 | 0.01153 | 0.03530 | 0.07435 | 0.00968 | 0.07693 | 0.04302 |
| **GAL3ST2** | 0.02286 | 0.01672 | 0.01297 | 0.01956 | 0.00793 | 0.01607 | 0.01944 |
| **GAL3ST3** | 0.02522 | 0.01488 | 0.27466 | 0.00591 | 0.00399 | 0.00000 | 0.14024 |
| **GAL3ST4** | 0.54515 | 0.18402 | 0.00286 | 1.49914 | 0.17757 | 0.16448 | 0.91721 |
| **GALC** | 0.89178 | 0.00271 | 0.78818 | 1.30860 | 2.33180 | 0.58013 | 0.00000 |
| **GALE** | 11.72208 | 14.33100 | 1.04171 | 3.80257 | 0.00000 | 5.37949 | 1.27828 |
| **GALK1** | 8.03473 | 37.68688 | 8.90842 | 2.06255 | 0.03206 | 2.95331 | 0.00000 |
| **GALK2** | 6.59312 | 1.27248 | 1.95129 | 3.84467 | 0.44825 | 3.79177 | 0.00000 |
| **GALM** | 13.98143 | 9.72112 | 5.73121 | 0.93235 | 3.15570 | 6.12391 | 0.00000 |
| **GALNS** | 1.18537 | 0.51254 | 0.32869 | 5.10185 | 0.11274 | 0.62996 | 1.68889 |
| **GALNT1** | 4.46444 | 1.51664 | 2.65408 | 0.86702 | 0.52945 | 1.44651 | 12.89897 |
| **GALNT10** | 18.87143 | 1.90209 | 0.00000 | 0.08571 | 0.13662 | 0.00000 | 11.82414 |
| **GALNT11** | 18.55825 | 0.00000 | 1.70329 | 9.52026 | 1.64502 | 0.13294 | 6.32124 |
| **GALNT12** | 0.17090 | 0.71924 | 0.11202 | 0.00000 | 0.00470 | 0.13640 | 0.36349 |
| **GALNT13** | 0.03434 | 0.00000 | 0.02615 | 0.07283 | 0.91853 | 0.03032 | 0.21071 |
| **GALNT14** | 0.21800 | 0.51654 | 0.04757 | 0.68397 | 0.27360 | 0.11367 | 1.13479 |
| **GALNT15** | 0.02760 | 0.01057 | 0.01106 | 0.01873 | 0.08647 | 0.01563 | 0.03263 |
| **GALNT16** | 0.06531 | 0.04871 | 0.00761 | 0.02183 | 0.03836 | 0.03531 | 0.15724 |
| **GALNT18** | 0.44659 | 0.11500 | 0.03668 | 0.05719 | 0.23311 | 2.77681 | 0.35106 |
| **GALNT2** | 0.00000 | 5.88779 | 1.63792 | 8.76418 | 0.77935 | 2.53841 | 5.35296 |
| **GALNT3** | 1.70108 | 0.02168 | 0.42560 | 1.48854 | 0.40286 | 0.09278 | 0.53327 |
| **GALNT4** | 0.15472 | 0.08503 | 0.00582 | 0.03533 | 0.00791 | 0.04605 | 0.00005 |
| **GALNT5** | 0.08485 | 0.00000 | 0.18858 | 0.08648 | 0.33377 | 0.01300 | 0.03128 |
| **GALNT6** | 17.60009 | 14.85192 | 5.08574 | 0.75935 | 0.00000 | 6.47596 | 9.33381 |
| **GALNT7** | 3.49023 | 0.18770 | 0.36815 | 1.22271 | 0.85395 | 2.52680 | 1.62439 |
| **GALNT8** | 0.47184 | 0.36702 | 0.64684 | 0.52241 | 0.11563 | 1.28919 | 0.00000 |

| | | | | | | | |
|---|---|---|---|---|---|---|---|
| **GALNT9** | 0.02217 | 0.01520 | 0.01740 | 0.02087 | 0.04708 | 0.01221 | 0.02168 |
| **GALNTL5** | 0.02337 | 0.02313 | 0.01835 | 0.01666 | 0.00925 | 0.01545 | 0.02409 |
| **GALNTL6** | 0.02307 | 0.01350 | 0.01537 | 0.01902 | 0.34859 | 0.01434 | 0.01444 |
| **GALP** | 0.02258 | 0.01766 | 0.01140 | 0.01943 | 0.04672 | 0.01303 | 0.01871 |
| **GALR1** | 0.04660 | 0.06311 | 0.00959 | 0.01635 | 0.00695 | 0.06151 | 0.04086 |
| **GALR2** | 0.19260 | 0.15250 | 0.00574 | 0.91970 | 0.00860 | 0.03189 | 0.00000 |
| **GALR3** | 0.02203 | 0.02836 | 0.01448 | 0.02768 | 0.02733 | 0.01148 | 0.01406 |
| **GALT** | 8.59550 | 3.49114 | 0.76067 | 6.51431 | 0.85734 | 3.41073 | 0.97844 |
| **GAMT** | 0.74700 | 1.97721 | 0.12203 | 0.26688 | 0.02860 | 1.14776 | 0.60008 |
| **GAN** | 2.01069 | 1.64852 | 1.75087 | 3.64507 | 0.80775 | 0.00000 | 5.18748 |
| **GANAB** | 23.22862 | 6.29427 | 11.22453 | 30.80657 | 0.38842 | 15.56295 | 22.69021 |
| **GANC** | 1.13787 | 0.03756 | 1.65984 | 0.55322 | 1.04111 | 0.71481 | 0.40094 |
| **GAP43** | 0.10401 | 0.00000 | 0.09598 | 0.00000 | 0.32049 | 0.36375 | 0.07420 |
| **GAPDH** | 86.47831 | 68.60128 | 22.33004 | 77.97769 | 5.46899 | 44.38331 | 68.30662 |
| **GAPDHS** | 0.15247 | 0.23001 | 0.02556 | 0.34157 | 0.00771 | 0.01530 | 0.13230 |
| **GAPVD1** | 22.40894 | 11.87794 | 8.26823 | 15.07062 | 7.69477 | 8.84033 | 8.47819 |
| **GAR1** | 13.41445 | 9.69134 | 11.06442 | 13.72729 | 4.86104 | 3.31514 | 10.74206 |
| **GAREM** | 0.24440 | 0.00000 | 0.88129 | 0.87132 | 0.43460 | 2.82147 | 0.78118 |
| **GAREML** | 0.07959 | 0.18701 | 0.81969 | 0.16577 | 0.67625 | 0.09626 | 0.39980 |
| **GARNL3** | 0.21342 | 0.00000 | 0.04373 | 0.02586 | 0.32054 | 0.13147 | 0.00000 |
| **GARS** | 37.58783 | 12.45972 | 6.33352 | 24.95640 | 4.03982 | 20.38581 | 25.29874 |
| **GART** | 55.31789 | 17.84106 | 15.38610 | 25.94673 | 2.74681 | 10.59358 | 2.58147 |
| **GAS1** | 0.00053 | 0.07393 | 0.01450 | 0.01408 | 0.00604 | 0.13476 | 0.53556 |
| **GAS2** | 0.08134 | 0.00000 | 0.00884 | 0.00000 | 0.11626 | 0.02970 | 0.16078 |
| **GAS2L1** | 0.30477 | 2.24582 | 0.72684 | 0.08668 | 0.13509 | 1.70881 | 6.40855 |
| **GAS2L1P2** | 0.00543 | 0.02379 | 0.00881 | 0.04939 | 0.00759 | 0.06934 | 0.07874 |
| **GAS2L2** | 0.02469 | 0.01298 | 0.01339 | 0.01991 | 0.00764 | 0.02913 | 0.03024 |
| **GAS2L3** | 1.60780 | 0.26221 | 0.07805 | 4.43643 | 0.12021 | 0.02677 | 0.30051 |
| **GAS5** | 12.53172 | 6.60009 | 10.29526 | 14.34321 | 8.18734 | 14.64245 | 23.60013 |
| **GAS5-AS1** | 0.00151 | 0.08439 | 0.04019 | 0.09527 | 0.02598 | 0.01748 | 0.23652 |
| **GAS6** | 1.40869 | 16.08598 | 0.24724 | 0.00000 | 0.00809 | 10.48620 | 15.97949 |
| **GAS6-AS1** | 0.01114 | 0.07444 | 0.04096 | 0.01212 | 0.04544 | 0.02447 | 0.10036 |
| **GAS6-AS2** | 0.33349 | 1.59807 | 0.34221 | 0.28726 | 0.00234 | 0.03211 | 0.00000 |
| **GAS7** | 4.99493 | 8.71114 | 1.00728 | 0.88484 | 2.05109 | 9.97397 | 11.58596 |
| **GAS8** | 0.34432 | 0.30958 | 0.59174 | 0.96572 | 0.48316 | 0.30575 | 0.41290 |

| | | | | | | | |
|---|---|---|---|---|---|---|---|
| **GAST** | 0.00000 | 0.01220 | 0.00213 | 13.01690 | 0.34487 | 0.02142 | 0.00000 |
| **GATA2** | 33.68866 | 6.17472 | 0.00000 | 43.05912 | 0.44032 | 0.00000 | 61.65101 |
| **GATA3** | 27.75376 | 26.36403 | 0.25006 | 32.09330 | 0.14423 | 0.00000 | 54.78636 |
| **GATA3-AS1** | 0.58264 | 0.86146 | 0.00000 | 2.51457 | 3.19604 | 0.00000 | 1.43798 |
| **GATA4** | 0.01777 | 0.00000 | 0.00000 | 0.19478 | 0.24175 | 6.14081 | 0.06943 |
| **GATA5** | 0.13800 | 0.01736 | 0.01087 | 0.03481 | 0.01188 | 0.12519 | 0.00473 |
| **GATA6** | 12.67780 | 15.49916 | 26.03651 | 2.72015 | 1.70483 | 5.07559 | 16.81289 |
| **GATA6-AS1** | 1.23475 | 5.56653 | 4.42741 | 2.47922 | 0.01342 | 0.62464 | 1.03245 |
| **GATAD1** | 2.51204 | 0.95959 | 0.00000 | 2.52676 | 0.00000 | 2.61517 | 8.20546 |
| **GATAD2A** | 9.60728 | 10.28858 | 6.75263 | 14.94529 | 3.51641 | 9.17788 | 30.36837 |
| **GATAD2B** | 3.18435 | 1.44682 | 2.03807 | 2.89862 | 3.02133 | 2.13259 | 4.23888 |
| **GATC** | 11.09562 | 7.33698 | 6.09397 | 14.14943 | 6.23437 | 4.92323 | 3.17563 |
| **GATM** | 0.50013 | 0.00000 | 0.18552 | 0.18427 | 0.32776 | 0.00000 | 0.12344 |
| **GATS** | 0.33449 | 0.69096 | 0.15531 | 0.99052 | 1.14847 | 1.37418 | 1.81311 |
| **GATSL2** | 0.02932 | 0.02167 | 0.01410 | 0.02172 | 0.00799 | 0.01471 | 0.01279 |
| **GATSL3** | 0.00000 | 1.52789 | 0.58532 | 5.15273 | 0.26341 | 1.12398 | 0.00000 |
| **GBA** | 5.35338 | 2.33932 | 1.35456 | 4.96501 | 0.00377 | 2.26545 | 0.00000 |
| **GBA2** | 3.76879 | 3.75591 | 1.03642 | 6.53856 | 0.19836 | 5.52187 | 7.81222 |
| **GBA3** | 0.01127 | 0.03892 | 0.04856 | 0.05762 | 0.05259 | 1.03602 | 0.00714 |
| **GBAP1** | 0.13089 | 0.08391 | 0.02929 | 0.29508 | 0.04144 | 0.13189 | 0.05572 |
| **GBAS** | 5.60045 | 1.09020 | 0.47694 | 4.05210 | 0.52648 | 4.40596 | 3.70517 |
| **GBE1** | 2.57585 | 1.06726 | 1.79392 | 1.68933 | 0.63882 | 1.99494 | 0.43767 |
| **GBF1** | 5.47644 | 6.82192 | 4.03902 | 7.66181 | 2.40628 | 5.17216 | 11.33593 |
| **GBGT1** | 2.62030 | 0.00000 | 0.00000 | 0.88561 | 0.00000 | 0.00000 | 2.47353 |
| **GBP1** | 0.00578 | 0.07852 | 0.15443 | 0.27888 | 0.00381 | 0.02749 | 0.00000 |
| **GBP1P1** | 0.10625 | 0.02742 | 0.31018 | 0.01510 | 0.09280 | 0.01124 | 0.00000 |
| **GBP2** | 0.03480 | 0.00613 | 0.29292 | 0.17826 | 0.01424 | 0.02667 | 0.00000 |
| **GBP3** | 0.02162 | 0.02021 | 0.02041 | 0.02332 | 0.00953 | 0.02101 | 0.01835 |
| **GBP4** | 0.01093 | 0.01700 | 0.01583 | 0.15144 | 0.01103 | 0.01764 | 0.01330 |
| **GBP5** | 0.06449 | 0.03570 | 0.01586 | 0.12946 | 0.00733 | 0.02791 | 0.00000 |
| **GBP6** | 0.02547 | 0.01624 | 0.01250 | 0.01975 | 0.03847 | 0.01451 | 0.02046 |
| **GBP7** | 0.01947 | 0.01788 | 0.01305 | 0.03005 | 0.00778 | 0.01239 | 0.02736 |
| **GBX1** | 0.01907 | 0.02311 | 0.01089 | 0.01646 | 0.25172 | 0.01948 | 0.02516 |
| **GBX2** | 0.04721 | 0.11417 | 1.53005 | 0.06760 | 1.30133 | 1.63855 | 0.00000 |
| **GC** | 0.03276 | 0.02663 | 0.00968 | 0.01918 | 0.01290 | 0.04543 | 0.00262 |

| | | | | | | | |
|---|---|---|---|---|---|---|---|
| **GCA** | 4.87079 | 0.29963 | 0.91843 | 2.20060 | 0.55224 | 3.47588 | 0.00000 |
| **GCAT** | 5.27127 | 7.15568 | 3.61453 | 6.06920 | 3.18260 | 2.29448 | 0.00000 |
| **GCC1** | 7.88356 | 5.26725 | 4.38720 | 7.84936 | 2.48208 | 4.65517 | 1.88046 |
| **GCC2** | 11.10985 | 1.88765 | 7.24589 | 8.53302 | 5.36566 | 7.37481 | 12.02988 |
| **GCDH** | 15.16642 | 16.12702 | 5.30268 | 11.24435 | 0.41738 | 5.18216 | 0.00000 |
| **GCFC2** | 3.44423 | 1.41763 | 4.84715 | 0.80463 | 1.51355 | 2.42815 | 6.44567 |
| **GCG** | 0.02033 | 0.00820 | 0.04019 | 0.01867 | 0.01141 | 0.01954 | 0.03189 |
| **GCGR** | 0.00000 | 0.29996 | 0.11425 | 0.00853 | 0.04733 | 0.34801 | 0.01331 |
| **GCH1** | 21.77707 | 0.00000 | 0.49962 | 2.30522 | 0.57261 | 0.00000 | 18.70923 |
| **GCHFR** | 0.90924 | 9.59827 | 14.74072 | 0.00000 | 0.65308 | 4.84950 | 0.00000 |
| **GCK** | 0.47480 | 0.06968 | 1.12438 | 0.00026 | 0.75765 | 0.00000 | 0.00000 |
| **GCKR** | 0.02831 | 0.02915 | 0.02248 | 0.05008 | 0.15604 | 0.65443 | 0.00000 |
| **GCLC** | 4.33878 | 0.23462 | 0.93935 | 3.74234 | 0.31892 | 1.19961 | 8.20133 |
| **GCLM** | 3.20989 | 1.63394 | 0.12022 | 4.87871 | 0.02557 | 1.85492 | 15.60731 |
| **GCM1** | 6.74448 | 0.44708 | 0.00000 | 49.54171 | 0.25635 | 9.94456 | 0.00000 |
| **GCM2** | 0.33229 | 0.03290 | 0.24717 | 0.00174 | 0.00230 | 0.71093 | 0.00000 |
| **GCN1L1** | 16.97316 | 12.17338 | 9.46693 | 15.36083 | 0.99072 | 9.05098 | 30.68844 |
| **GCNT1** | 8.82694 | 0.02016 | 0.21028 | 1.89750 | 4.41719 | 3.58715 | 0.00000 |
| **GCNT2** | 7.25950 | 0.02345 | 0.39055 | 4.60524 | 0.00000 | 3.55033 | 2.06765 |
| **GCNT3** | 0.29783 | 0.23326 | 2.06765 | 0.00713 | 6.15701 | 2.73950 | 0.00000 |
| **GCNT4** | 0.01432 | 0.01124 | 0.02405 | 0.07555 | 0.14736 | 0.01446 | 0.02266 |
| **GCNT7** | 0.17961 | 0.00000 | 0.19008 | 0.00545 | 0.10225 | 0.20402 | 0.25111 |
| **GCOM1.uid-190** | 0.02218 | 0.01698 | 0.00898 | 0.02163 | 0.04648 | 0.01335 | 0.02107 |
| **GCSAM** | 0.50999 | 0.29092 | 0.00299 | 0.00554 | 0.00677 | 0.17073 | 0.00000 |
| **GCSAML** | 0.02673 | 0.00458 | 0.14429 | 0.01564 | 0.11526 | 0.01642 | 0.03446 |
| **GCSAML-AS1** | 0.02384 | 0.00714 | 0.04358 | 0.01988 | 0.02107 | 0.03240 | 0.02155 |
| **GCSH** | 4.04410 | 3.98426 | 0.92954 | 2.25753 | 0.07625 | 0.78621 | 1.09357 |
| **GCSHP3** | 0.84818 | 0.21092 | 0.37980 | 0.31970 | 0.04309 | 0.77921 | 1.74816 |
| **GDA** | 0.77714 | 0.00000 | 0.09345 | 0.77238 | 0.32300 | 0.00000 | 0.70880 |
| **GDAP1** | 0.70920 | 0.30884 | 1.04710 | 2.18830 | 9.55792 | 0.02210 | 0.00000 |
| **GDAP1L1** | 0.07710 | 0.00005 | 0.02699 | 0.01015 | 0.00837 | 0.00768 | 0.05303 |
| **GDAP2** | 11.17799 | 0.08202 | 4.95184 | 6.41928 | 4.11459 | 1.86357 | 0.00000 |
| **GDE1** | 16.96411 | 23.25768 | 22.59369 | 12.30849 | 0.00000 | 5.73366 | 13.39386 |
| **GDF10** | 0.02487 | 0.01704 | 0.01331 | 0.03883 | 0.00787 | 0.01143 | 0.02022 |
| **GDF11** | 0.17629 | 1.43422 | 1.05127 | 2.09137 | 1.08475 | 2.15102 | 5.83093 |

| | | | | | | | |
|---|---|---|---|---|---|---|---|
| **GDF15** | 0.00000 | 10.05490 | 0.00000 | 12.05727 | 0.66890 | 7.06327 | 2.63703 |
| **GDF3** | 1.56229 | 0.00000 | 0.32179 | 1.21423 | 0.26099 | 34.91590 | 0.00000 |
| **GDF5** | 0.01017 | 0.04145 | 0.00705 | 0.07972 | 0.02075 | 0.03556 | 0.01154 |
| **GDF6** | 0.01811 | 0.02002 | 0.01618 | 0.04382 | 0.00739 | 0.06223 | 0.01696 |
| **GDF7** | 0.02111 | 0.01429 | 0.01300 | 0.02337 | 0.00827 | 0.01360 | 0.04295 |
| **GDF9** | 2.33607 | 4.11551 | 1.49393 | 1.79743 | 34.28913 | 0.75395 | 0.64867 |
| **GDI1** | 0.11754 | 4.03911 | 6.45465 | 6.37099 | 2.36974 | 5.35323 | 10.89814 |
| **GDI2** | 48.75276 | 19.93618 | 20.65920 | 26.94572 | 14.02987 | 18.74064 | 23.06022 |
| **GDNF** | 0.02915 | 0.01822 | 0.01072 | 0.01901 | 0.03171 | 0.01345 | 0.01551 |
| **GDPD1** | 2.00391 | 0.00000 | 11.68579 | 0.56504 | 0.42069 | 3.67056 | 0.43053 |
| **GDPD2** | 1.38293 | 20.31094 | 0.17973 | 0.00000 | 0.00000 | 14.97084 | 0.00000 |
| **GDPD3** | 7.58923 | 0.30600 | 0.04995 | 1.52017 | 0.00091 | 0.00000 | 2.60413 |
| **GDPD4** | 0.01520 | 0.16363 | 0.10541 | 0.02234 | 0.07263 | 0.00191 | 0.01883 |
| **GDPD5** | 0.33823 | 5.95912 | 0.02250 | 0.99653 | 0.17342 | 1.98442 | 7.28300 |
| **GDPGP1** | 0.68388 | 0.21964 | 0.39260 | 0.14504 | 0.01381 | 0.03698 | 0.00000 |
| **GEM** | 0.00384 | 0.00929 | 3.75881 | 0.14489 | 0.69473 | 1.17752 | 0.00000 |
| **GEMIN2** | 8.41622 | 6.52793 | 5.77653 | 4.14564 | 1.36511 | 8.00588 | 0.52426 |
| **GEMIN4** | 17.30755 | 10.79564 | 9.96006 | 13.89435 | 0.68045 | 5.31107 | 3.76737 |
| **GEMIN5** | 13.65048 | 3.92700 | 3.91420 | 5.20396 | 9.53935 | 4.87666 | 6.60954 |
| **GEMIN6** | 18.16039 | 3.88456 | 15.35267 | 10.09546 | 4.58523 | 6.31297 | 3.46409 |
| **GEMIN7** | 10.37556 | 17.50891 | 9.91564 | 6.66757 | 1.86512 | 6.28323 | 3.46071 |
| **GEMIN8** | 6.70819 | 2.36827 | 2.86920 | 5.09350 | 1.04572 | 3.68618 | 1.61116 |
| **GEMIN8P4** | 1.09849 | 0.31650 | 2.65044 | 4.34619 | 2.09644 | 0.38405 | 0.10158 |
| **GEN1** | 4.98229 | 0.23350 | 3.06873 | 2.05656 | 0.14226 | 0.92657 | 4.87294 |
| **GET4** | 2.48956 | 18.78746 | 1.69216 | 11.84726 | 4.10201 | 7.65790 | 28.08900 |
| **GFAP** | 0.03303 | 0.06085 | 0.01095 | 0.05780 | 0.01117 | 0.01110 | 0.03705 |
| **GFER** | 0.22529 | 8.80059 | 2.14594 | 2.16131 | 0.01112 | 3.35245 | 14.73772 |
| **GFI1** | 0.01588 | 0.45565 | 0.08339 | 0.02037 | 0.09400 | 0.14085 | 0.01015 |
| **GFI1B** | 0.02685 | 0.01853 | 0.02227 | 0.00000 | 0.00548 | 0.18720 | 0.13552 |
| **GFM1** | 26.99262 | 5.51275 | 11.87011 | 8.35528 | 4.12492 | 8.40543 | 2.10857 |
| **GFM2** | 12.95672 | 1.56838 | 2.70376 | 4.34915 | 5.64620 | 5.73882 | 0.00000 |
| **GFOD1** | 0.00000 | 0.04166 | 0.23594 | 0.01352 | 0.33182 | 0.60558 | 0.09248 |
| **GFOD2** | 5.76850 | 2.27647 | 4.26775 | 10.35420 | 3.84134 | 3.53195 | 1.46005 |
| **GFPT1** | 17.12111 | 12.22652 | 3.27907 | 13.52638 | 6.07380 | 10.32540 | 0.00000 |
| **GFPT2** | 1.69692 | 0.00000 | 0.63832 | 1.65291 | 1.39557 | 1.26282 | 1.06491 |

| | | | | | | | |
|---|---|---|---|---|---|---|---|
| **GFRA1** | 0.01628 | 0.02551 | 0.07703 | 0.02229 | 0.37646 | 0.00454 | 0.04248 |
| **GFRA2** | 0.00000 | 0.00000 | 0.04867 | 0.05277 | 2.87874 | 0.64090 | 0.04621 |
| **GFRA3** | 0.01339 | 0.02853 | 0.17285 | 0.18093 | 0.08598 | 0.03899 | 0.57420 |
| **GFRAL** | 0.01619 | 0.01519 | 0.01359 | 0.01922 | 0.01198 | 0.01985 | 0.04818 |
| **GFY** | 0.00000 | 0.24007 | 0.00020 | 0.01346 | 0.65652 | 0.08909 | 0.01405 |
| **GGA1** | 2.82175 | 3.67262 | 0.44063 | 7.87010 | 1.10140 | 3.81118 | 18.12480 |
| **GGA2** | 12.46732 | 16.70416 | 4.23341 | 4.23822 | 2.41366 | 8.47647 | 18.48226 |
| **GGA3** | 3.89706 | 3.95990 | 6.25723 | 4.76245 | 1.16557 | 3.16032 | 12.20038 |
| **GGACT** | 0.04419 | 0.87469 | 0.00000 | 0.00000 | 3.84038 | 0.00000 | 0.48531 |
| **GGCT** | 19.17114 | 1.45124 | 1.56384 | 20.42872 | 0.93062 | 4.26789 | 7.06212 |
| **GGCX** | 7.36034 | 5.55888 | 0.20897 | 12.02251 | 1.14931 | 2.70606 | 1.69877 |
| **GGH** | 12.22107 | 2.31352 | 7.52718 | 7.10015 | 0.47824 | 2.08690 | 5.70836 |
| **GGN** | 0.00478 | 0.06282 | 0.04447 | 0.08413 | 0.05424 | 0.06964 | 0.05827 |
| **GGNBP1** | 0.02420 | 0.00608 | 0.06146 | 0.01973 | 0.00550 | 0.01299 | 0.02515 |
| **GGNBP2** | 9.36288 | 14.21559 | 7.93693 | 10.67682 | 12.90739 | 9.17503 | 26.78298 |
| **GGPS1** | 8.03997 | 0.00000 | 7.37044 | 10.65057 | 3.58284 | 3.65329 | 0.00000 |
| **GGT1** | 0.00000 | 0.22276 | 0.16242 | 0.41274 | 0.07962 | 2.52640 | 0.02785 |
| **GGT3P** | 0.02459 | 0.02060 | 0.01259 | 0.02232 | 0.00807 | 0.03647 | 0.01059 |
| **GGT5** | 0.23569 | 0.00000 | 0.16830 | 0.80002 | 0.33179 | 0.37081 | 0.05763 |
| **GGT6** | 0.00192 | 0.19628 | 0.00000 | 0.00942 | 0.31353 | 0.00162 | 0.08545 |
| **GGT7** | 1.06424 | 0.36130 | 0.64684 | 1.58617 | 0.69335 | 0.24394 | 0.24108 |
| **GGT8P** | 0.01969 | 0.01665 | 0.01283 | 0.03225 | 0.00826 | 0.02521 | 0.02139 |
| **GGTA1P** | 0.07942 | 0.02233 | 0.01613 | 0.01057 | 0.17972 | 0.02148 | 0.00449 |
| **GGTLC1** | 0.01262 | 0.01016 | 0.30555 | 0.01999 | 0.00234 | 0.01180 | 0.02986 |
| **GGTLC2** | 0.02245 | 0.02091 | 0.01802 | 0.02259 | 0.00776 | 0.01318 | 0.01496 |
| **GH2** | 0.01364 | 0.01452 | 0.01244 | 0.02881 | 0.01943 | 0.05693 | 0.02766 |
| **GHDC** | 0.13281 | 0.17788 | 0.01663 | 0.21767 | 0.00601 | 0.22398 | 1.47659 |
| **GHITM** | 73.27700 | 36.57650 | 24.64275 | 46.57693 | 9.60042 | 37.58951 | 7.46600 |
| **GHR** | 0.15059 | 0.00000 | 0.00723 | 0.28786 | 0.01111 | 0.04786 | 0.13962 |
| **GHRHR** | 0.00922 | 0.02606 | 0.06054 | 0.01597 | 0.33422 | 0.00787 | 0.04222 |
| **GHRL** | 0.03720 | 0.15940 | 0.05809 | 0.10905 | 0.01958 | 0.04703 | 0.02485 |
| **GHRLOS** | 0.75363 | 1.53719 | 0.37511 | 0.68874 | 0.02962 | 0.46084 | 2.39490 |
| **GHSR** | 0.02203 | 0.00883 | 0.01435 | 0.03462 | 0.00764 | 0.04087 | 0.04595 |
| **GID4** | 0.73850 | 2.80830 | 2.40798 | 2.89016 | 0.34616 | 3.43123 | 4.61295 |
| **GID8** | 12.41147 | 8.70144 | 6.48308 | 10.04415 | 3.48843 | 9.04421 | 19.67595 |

| | | | | | | | |
|---|---|---|---|---|---|---|---|
| **GIF** | 0.06477 | 0.01369 | 0.01654 | 0.03492 | 0.01617 | 0.02058 | 0.00000 |
| **GIGYF1** | 4.45747 | 7.44868 | 1.68441 | 11.84794 | 1.22206 | 10.08053 | 11.88402 |
| **GIGYF2** | 16.06292 | 8.94721 | 9.02754 | 11.44076 | 4.03520 | 5.42156 | 16.49290 |
| **GIMAP1** | 0.02561 | 0.00420 | 0.04024 | 0.03794 | 0.01463 | 0.02738 | 0.02070 |
| **GIMAP6** | 0.02657 | 0.00120 | 0.06838 | 0.02002 | 0.00860 | 0.01222 | 0.01779 |
| **GIMAP7** | 0.03333 | 0.02230 | 0.07064 | 0.01278 | 0.07347 | 0.01556 | 0.02014 |
| **GIMAP8** | 0.06810 | 0.00000 | 0.11014 | 0.03158 | 0.00669 | 0.00974 | 0.00000 |
| **GIN1** | 2.64103 | 0.00000 | 2.25755 | 0.74979 | 1.24749 | 0.11873 | 0.18363 |
| **GINM1** | 5.01794 | 1.42051 | 3.76790 | 3.26285 | 0.44647 | 1.56027 | 1.29582 |
| **GINS1** | 32.22094 | 9.54730 | 8.30243 | 12.49572 | 1.53262 | 10.60959 | 3.03538 |
| **GINS2** | 27.38583 | 32.62872 | 9.90709 | 13.36178 | 7.34110 | 8.45801 | 0.00037 |
| **GINS3** | 14.76564 | 9.12019 | 23.03978 | 2.36437 | 10.48611 | 10.35170 | 1.48111 |
| **GINS4** | 9.74690 | 6.13623 | 6.79598 | 7.53526 | 4.81611 | 8.87945 | 2.85547 |
| **GIP** | 0.00000 | 2.07786 | 6.93961 | 1.68043 | 7.74319 | 0.00000 | 0.15272 |
| **GIPC1** | 1.38498 | 2.68443 | 0.04377 | 2.48125 | 0.07426 | 2.91023 | 9.29033 |
| **GIPC2** | 1.66932 | 1.95941 | 2.31290 | 0.00000 | 0.51733 | 1.14414 | 1.43719 |
| **GIPC3** | 0.02528 | 0.01634 | 0.02542 | 0.02331 | 0.02333 | 0.01071 | 0.02132 |
| **GIPR** | 0.01779 | 0.11463 | 0.16927 | 0.04991 | 0.14897 | 0.06331 | 0.09951 |
| **GIT1** | 0.58509 | 9.21044 | 2.02156 | 3.89191 | 0.51019 | 6.03020 | 8.98548 |
| **GIT2** | 9.72596 | 5.99111 | 4.99815 | 3.53390 | 13.51678 | 2.23987 | 0.30271 |
| **GJA1** | 30.53956 | 5.26868 | 28.24783 | 19.94003 | 1.35626 | 20.85730 | 41.13979 |
| **GJA10** | 0.19037 | 0.12974 | 0.10049 | 0.05442 | 0.00301 | 0.35388 | 0.00000 |
| **GJA3** | 0.02163 | 0.01347 | 0.02670 | 0.01983 | 0.06797 | 0.01299 | 0.02453 |
| **GJA4** | 0.10432 | 0.00000 | 0.19903 | 0.00557 | 0.00300 | 0.04686 | 0.01608 |
| **GJA5** | 1.56323 | 0.38667 | 0.00080 | 16.16738 | 0.00302 | 1.07523 | 0.00000 |
| **GJA8** | 0.00937 | 0.01989 | 0.00887 | 0.02317 | 0.02449 | 0.05164 | 0.03653 |
| **GJA9-MYCBP.ui** | 0.02348 | 0.00239 | 0.06171 | 0.05971 | 0.13296 | 0.02559 | 0.04306 |
| **GJB1** | 0.04212 | 0.00174 | 0.00982 | 0.02695 | 0.01736 | 0.70035 | 0.00357 |
| **GJB2** | 0.01458 | 0.01231 | 0.05776 | 0.02048 | 0.02273 | 0.01692 | 0.03349 |
| **GJB3** | 0.17246 | 0.04078 | 0.07334 | 0.14680 | 0.00816 | 0.07494 | 0.00000 |
| **GJB4** | 0.00916 | 0.29273 | 0.01106 | 0.06909 | 0.00677 | 0.26751 | 0.00000 |
| **GJB5** | 0.19217 | 0.46383 | 0.08436 | 0.12539 | 0.00814 | 0.13317 | 0.00000 |
| **GJB7** | 0.90312 | 0.00000 | 0.51349 | 0.70332 | 0.10468 | 0.30367 | 0.00000 |
| **GJC1** | 5.59915 | 0.00000 | 1.41063 | 3.17679 | 4.18505 | 0.00000 | 14.71046 |
| **GJC2** | 0.02924 | 0.00000 | 0.01039 | 0.04067 | 0.00667 | 0.00346 | 0.27419 |

| | | | | | | | |
|---|---|---|---|---|---|---|---|
| **GJD3** | 0.01578 | 0.02105 | 0.00641 | 0.08815 | 0.02912 | 0.11009 | 0.21019 |
| **GJD4** | 0.04449 | 0.07535 | 0.06260 | 0.09480 | 0.01474 | 0.02199 | 0.00461 |
| **GK** | 0.00000 | 0.65176 | 0.53396 | 0.00653 | 0.02028 | 2.90941 | 0.04047 |
| **GK2** | 0.02299 | 0.01879 | 0.01558 | 0.01915 | 0.00769 | 0.01288 | 0.01962 |
| **GK3P** | 0.03141 | 0.04654 | 0.07709 | 0.02287 | 0.00647 | 0.28036 | 0.00478 |
| **GK5** | 5.21806 | 1.92412 | 2.63929 | 1.16562 | 0.00000 | 0.51558 | 5.62652 |
| **GKAP1** | 0.00000 | 1.64929 | 1.59217 | 0.00000 | 0.71759 | 2.35949 | 2.49044 |
| **GLA** | 37.60547 | 3.31388 | 8.50878 | 21.87051 | 11.85507 | 5.01528 | 11.83058 |
| **GLB1** | 29.42239 | 3.33236 | 2.63854 | 15.31737 | 5.70011 | 9.03155 | 7.20564 |
| **GLB1L** | 0.36383 | 0.00000 | 1.14778 | 0.32862 | 1.89797 | 0.14941 | 0.00000 |
| **GLB1L2** | 0.18520 | 0.04145 | 0.04659 | 0.18239 | 0.27394 | 0.00832 | 0.08608 |
| **GLB1L3** | 1.38367 | 0.12285 | 0.54355 | 0.73549 | 0.17553 | 0.12336 | 1.10071 |
| **GLCCI1** | 1.25336 | 0.00000 | 0.36636 | 0.36400 | 2.07040 | 0.68972 | 2.15669 |
| **GLCE** | 1.28434 | 0.10060 | 0.70527 | 0.98866 | 2.38367 | 0.08387 | 0.00120 |
| **GLDC** | 7.96492 | 1.61344 | 0.14248 | 10.74516 | 0.06939 | 13.08818 | 14.12014 |
| **GLDN** | 0.29025 | 0.00000 | 0.02108 | 2.41893 | 0.32129 | 0.00000 | 0.29549 |
| **GLE1** | 10.04575 | 5.99392 | 6.09959 | 9.13709 | 3.01680 | 2.14294 | 0.66732 |
| **GLG1** | 15.03901 | 18.23376 | 11.38589 | 8.70248 | 6.64696 | 12.66139 | 14.31855 |
| **GLI1** | 0.10164 | 0.17687 | 0.02386 | 0.01707 | 0.00491 | 0.00000 | 0.20465 |
| **GLI2** | 0.00365 | 0.00382 | 0.01147 | 0.01930 | 0.00849 | 0.77801 | 0.08341 |
| **GLI3** | 0.20469 | 0.02785 | 0.18179 | 0.32395 | 0.35134 | 0.31278 | 0.06142 |
| **GLI4** | 0.02762 | 0.53956 | 0.37265 | 0.04732 | 0.02717 | 0.11147 | 0.61278 |
| **GLIDR** | 0.01847 | 0.10919 | 0.07870 | 0.14360 | 0.03317 | 0.10970 | 0.03418 |
| **GLIPR1** | 0.23203 | 0.03911 | 0.25520 | 2.69246 | 0.98312 | 0.17935 | 0.04111 |
| **GLIPR1L1** | 0.03369 | 0.02051 | 0.01258 | 0.05564 | 0.00820 | 0.01413 | 0.00139 |
| **GLIPR1L2** | 0.05887 | 0.00445 | 0.02931 | 0.02174 | 0.00960 | 0.01545 | 0.00416 |
| **GLIPR2** | 27.48352 | 46.14112 | 9.77676 | 3.12233 | 0.00000 | 0.50409 | 6.93767 |
| **GLIS1** | 0.01830 | 0.00000 | 0.00962 | 0.14772 | 0.03358 | 0.14405 | 0.17872 |
| **GLIS2** | 0.00000 | 0.37868 | 0.24188 | 0.72876 | 0.06022 | 3.43027 | 2.02682 |
| **GLIS3** | 0.51030 | 0.00000 | 0.02735 | 0.62767 | 0.07211 | 0.00000 | 1.10216 |
| **GLIS3-AS1** | 0.00830 | 0.02098 | 0.00963 | 0.03050 | 0.01159 | 0.00647 | 0.09683 |
| **GLMN** | 12.44361 | 0.00000 | 6.60240 | 6.90557 | 2.29302 | 1.88171 | 0.40979 |
| **GLO1** | 44.30467 | 2.72989 | 24.23839 | 31.00910 | 4.21934 | 15.90335 | 9.88462 |
| **GLOD4** | 35.82951 | 15.20122 | 13.93510 | 19.77120 | 1.98570 | 11.07580 | 6.00284 |
| **GLOD5** | 0.01846 | 0.11279 | 0.00718 | 0.01738 | 0.00870 | 0.00718 | 0.00840 |

| | | | | | | | |
|---|---|---|---|---|---|---|---|
| **GLP1R** | 0.02628 | 0.03742 | 0.01237 | 0.02659 | 0.00871 | 0.01153 | 0.00745 |
| **GLP2R** | 0.02362 | 0.01674 | 0.00984 | 0.01976 | 0.06161 | 0.01350 | 0.01613 |
| **GLRA1** | 0.05515 | 0.00353 | 0.01128 | 0.00983 | 0.06819 | 0.02958 | 0.06787 |
| **GLRA2** | 0.05617 | 0.00000 | 0.03689 | 0.03730 | 0.23432 | 0.02008 | 0.18845 |
| **GLRA3** | 0.04946 | 0.04891 | 0.00362 | 0.01394 | 0.03974 | 0.00453 | 0.04797 |
| **GLRA4** | 0.10848 | 0.08302 | 0.00170 | 0.12607 | 0.11058 | 0.03237 | 0.00000 |
| **GLRB** | 0.03052 | 0.00247 | 0.11465 | 0.01938 | 0.00749 | 0.01077 | 0.01326 |
| **GLRX** | 10.46878 | 0.00000 | 0.00000 | 0.03900 | 0.88509 | 2.42254 | 1.90160 |
| **GLRX2** | 6.89479 | 1.90489 | 3.90881 | 6.74601 | 0.34930 | 2.55235 | 3.00094 |
| **GLRX3** | 35.32874 | 10.71237 | 14.27162 | 24.64122 | 1.98022 | 10.32612 | 1.96932 |
| **GLRX5** | 4.45893 | 6.30381 | 3.41150 | 11.54431 | 2.31462 | 6.21763 | 31.94377 |
| **GLS** | 3.45402 | 0.00000 | 1.83376 | 7.99396 | 3.66140 | 3.06756 | 8.34338 |
| **GLS2** | 0.16094 | 0.00000 | 0.15969 | 0.21862 | 0.90916 | 1.77481 | 0.00000 |
| **GLT1D1** | 0.94620 | 0.29237 | 0.00000 | 0.00000 | 0.03661 | 1.52461 | 1.39129 |
| **GLT6D1** | 0.02410 | 0.01409 | 0.02475 | 0.04869 | 0.01777 | 0.01212 | 0.00769 |
| **GLT8D1** | 3.37210 | 1.24615 | 2.15911 | 3.62258 | 0.99264 | 2.25800 | 4.37779 |
| **GLT8D2** | 0.14173 | 0.01908 | 0.04923 | 0.10749 | 0.03539 | 0.00000 | 0.10956 |
| **GLTP** | 8.41437 | 1.61524 | 0.24785 | 3.70652 | 1.12947 | 0.25201 | 1.67979 |
| **GLTPD2** | 0.00000 | 1.75879 | 0.24597 | 0.00000 | 0.06750 | 0.51128 | 0.39464 |
| **GLTSCR1** | 0.00000 | 0.56321 | 0.64586 | 0.76908 | 0.61567 | 0.46745 | 3.91026 |
| **GLTSCR1L** | 1.20861 | 0.00000 | 0.65111 | 0.94950 | 0.35400 | 1.27629 | 2.09683 |
| **GLTSCR2** | 0.00000 | 8.04343 | 1.94734 | 2.04535 | 3.36144 | 4.33158 | 23.39566 |
| **GLUD1** | 4.29472 | 13.93243 | 16.94825 | 1.97231 | 1.52923 | 13.58693 | 17.31204 |
| **GLUD1P3** | 0.00000 | 0.11624 | 0.31570 | 0.55881 | 0.19529 | 0.05627 | 1.63018 |
| **GLUD2** | 0.91397 | 4.31013 | 3.09698 | 0.24722 | 0.50196 | 2.30270 | 0.76959 |
| **GLUL** | 43.34647 | 37.60748 | 34.10017 | 38.26015 | 21.65592 | 24.33019 | 1.62296 |
| **GLYATL1** | 0.02203 | 0.01153 | 0.13897 | 0.01078 | 3.86478 | 0.00197 | 0.02436 |
| **GLYCTK** | 1.53031 | 1.72016 | 1.40612 | 1.43088 | 0.05348 | 0.83638 | 0.01263 |
| **GLYCTK-AS1** | 0.01871 | 0.01240 | 0.03069 | 0.03600 | 0.00872 | 0.01415 | 0.04729 |
| **GLYR1** | 5.12648 | 4.79766 | 3.96224 | 8.72172 | 1.49660 | 4.74743 | 19.81124 |
| **GM2A** | 22.11238 | 7.75784 | 0.00000 | 7.97048 | 1.77437 | 0.48164 | 18.99012 |
| **GMCL1** | 2.57737 | 0.95677 | 2.01638 | 3.28460 | 1.10843 | 1.55160 | 1.07123 |
| **GMCL1P1** | 0.23453 | 0.02451 | 0.17535 | 0.23191 | 0.07599 | 0.13835 | 0.16314 |
| **GMDS** | 1.89516 | 2.60009 | 0.49547 | 4.26936 | 0.99736 | 1.68164 | 3.37658 |
| **GMDS-AS1** | 0.04171 | 0.05646 | 0.02388 | 0.02070 | 0.06661 | 0.03918 | 0.00388 |

| | | | | | | | |
|---|---|---|---|---|---|---|---|
| GMEB1 | 7.60863 | 1.59347 | 3.87624 | 8.99165 | 1.69001 | 4.58078 | 1.32595 |
| GMEB2 | 0.00000 | 2.69276 | 2.71534 | 2.98591 | 0.34967 | 1.56502 | 5.62793 |
| GMFB | 11.60222 | 5.09103 | 11.49161 | 0.99749 | 8.77237 | 11.85913 | 3.52809 |
| GMFG | 0.08938 | 0.00486 | 0.03781 | 0.08915 | 0.03617 | 0.08434 | 0.01915 |
| GMIP | 0.00000 | 1.05016 | 0.70651 | 0.59656 | 0.04033 | 0.72932 | 1.20533 |
| GML | 0.02145 | 0.01994 | 0.00000 | 0.01885 | 0.25826 | 0.01321 | 0.01660 |
| GMNC | 0.01315 | 0.01647 | 0.01338 | 0.04070 | 0.00929 | 0.01468 | 0.02501 |
| GMNN | 41.31390 | 14.34353 | 20.27386 | 20.27636 | 15.40278 | 14.07728 | 2.42136 |
| GMPPA | 9.89912 | 13.14571 | 3.51051 | 9.02043 | 1.08953 | 4.85499 | 0.77889 |
| GMPPB | 1.37337 | 5.31522 | 0.73821 | 5.42978 | 0.28629 | 3.45747 | 5.81208 |
| GMPR | 0.05737 | 0.00000 | 0.19409 | 0.02244 | 0.24060 | 0.58578 | 0.02421 |
| GMPR2 | 12.67268 | 4.79799 | 5.42013 | 15.36911 | 5.50785 | 5.23845 | 0.00600 |
| GMPS | 29.91602 | 5.96796 | 23.66325 | 12.31907 | 3.01679 | 6.16104 | 0.00000 |
| GNA11 | 0.00000 | 13.49422 | 6.41108 | 1.78549 | 1.98734 | 5.44634 | 19.39972 |
| GNA12 | 4.16515 | 0.23442 | 2.64126 | 29.39085 | 1.96620 | 2.71338 | 6.79144 |
| GNA13 | 8.73525 | 4.00583 | 10.63987 | 6.17171 | 6.70905 | 12.32057 | 13.04137 |
| GNA14 | 0.00000 | 0.05059 | 0.00000 | 0.02259 | 1.99887 | 0.49663 | 0.04724 |
| GNA14-AS1 | 0.03181 | 0.01734 | 0.01169 | 0.01338 | 0.00842 | 0.02809 | 0.02421 |
| GNA15 | 0.03133 | 0.01443 | 0.01279 | 0.01474 | 0.00762 | 0.00991 | 0.03682 |
| GNAI1 | 0.13836 | 0.54374 | 1.44171 | 5.25778 | 4.88404 | 0.71177 | 0.00000 |
| GNAI2 | 0.17468 | 7.35529 | 0.97166 | 2.15778 | 2.78845 | 10.28305 | 13.62812 |
| GNAI3 | 36.80405 | 10.30026 | 17.29466 | 24.84573 | 13.73930 | 12.44173 | 0.00000 |
| GNAL | 0.17418 | 1.48579 | 2.22653 | 3.04371 | 1.67924 | 1.66220 | 1.57310 |
| GNAO1 | 0.00000 | 0.22678 | 0.11918 | 0.00000 | 0.52170 | 0.16053 | 0.24098 |
| GNAQ | 1.20844 | 2.12602 | 2.36633 | 0.90574 | 0.37343 | 4.77980 | 8.74134 |
| GNAS | 15.39169 | 13.55373 | 1.55496 | 20.48759 | 1.58038 | 18.95694 | 46.43609 |
| GNAS-AS1 | 0.29611 | 0.00000 | 0.23979 | 0.13179 | 0.11365 | 0.03739 | 0.24789 |
| GNAT1 | 0.00221 | 0.03073 | 0.03358 | 0.02293 | 0.01094 | 0.03288 | 0.09941 |
| GNAT2 | 0.13587 | 0.10413 | 0.03578 | 0.35833 | 1.36785 | 0.09749 | 0.51918 |
| GNAT3 | 0.03296 | 0.01529 | 0.01277 | 0.01620 | 0.01371 | 0.01105 | 0.02454 |
| GNAZ | 0.01815 | 0.04421 | 0.02011 | 0.03004 | 0.22304 | 0.22023 | 0.04663 |
| GNB1 | 11.84638 | 12.78295 | 11.66652 | 12.17405 | 1.20061 | 5.56164 | 33.71061 |
| GNB1L | 7.89452 | 10.40951 | 1.14268 | 12.29078 | 0.21593 | 1.88831 | 3.02645 |
| GNB2 | 0.00000 | 10.46314 | 3.93065 | 7.55021 | 0.97434 | 9.32784 | 29.74106 |
| GNB2L1 | 66.47069 | 51.96552 | 42.31578 | 63.03678 | 15.76519 | 39.57391 | 54.20483 |

| | | | | | | | |
|---|---|---|---|---|---|---|---|
| **GNB3** | 0.29048 | 0.14958 | 0.19291 | 0.46683 | 0.07777 | 0.08671 | 0.10608 |
| **GNB4** | 1.95902 | 0.00000 | 4.97832 | 4.40032 | 2.68347 | 0.00000 | 2.71688 |
| **GNB5** | 2.27362 | 0.48460 | 1.87779 | 1.69038 | 0.94863 | 2.10997 | 0.00000 |
| **GNE** | 0.00000 | 0.00717 | 0.61553 | 5.71227 | 0.28145 | 0.01043 | 0.49667 |
| **GNG10** | 1.62600 | 4.36648 | 0.24133 | 0.76538 | 0.00873 | 0.00278 | 0.00000 |
| **GNG11** | 0.02954 | 0.00573 | 0.03457 | 0.02289 | 0.26339 | 0.01431 | 0.01495 |
| **GNG12** | 1.91403 | 0.00000 | 3.01610 | 8.92944 | 3.89221 | 0.41228 | 0.98010 |
| **GNG12-AS1** | 0.03235 | 0.00187 | 0.04041 | 0.06571 | 0.12221 | 0.00724 | 0.04526 |
| **GNG13** | 0.02514 | 0.02002 | 0.01275 | 0.05081 | 0.00796 | 0.01220 | 0.00980 |
| **GNG2** | 0.00000 | 0.00653 | 0.02165 | 0.40293 | 2.41886 | 0.41946 | 0.01789 |
| **GNG3** | 0.18578 | 0.04058 | 0.38566 | 0.15354 | 2.61990 | 0.00000 | 0.30197 |
| **GNG4** | 0.10789 | 1.32288 | 0.10911 | 0.29702 | 0.10797 | 0.00477 | 0.00000 |
| **GNG5** | 23.64039 | 19.58195 | 22.24684 | 22.24640 | 9.06209 | 11.06554 | 18.09812 |
| **GNG7** | 0.03051 | 0.38662 | 0.66855 | 0.00000 | 0.22529 | 0.25247 | 0.08887 |
| **GNG8** | 0.02320 | 0.01855 | 0.00777 | 0.01925 | 0.06218 | 0.01321 | 0.01644 |
| **GNGT1** | 0.04942 | 0.01242 | 0.01024 | 0.02953 | 0.10399 | 0.01171 | 0.09753 |
| **GNGT2** | 0.07412 | 0.05171 | 0.00000 | 0.03373 | 0.76731 | 0.09620 | 0.00539 |
| **GNL1** | 20.05636 | 21.69225 | 4.91432 | 23.53267 | 9.45038 | 9.32421 | 0.00000 |
| **GNL2** | 39.84784 | 17.13476 | 18.07237 | 23.12543 | 9.10695 | 11.65959 | 0.93992 |
| **GNL3** | 56.55647 | 19.56975 | 25.56087 | 32.60024 | 7.92755 | 17.16321 | 10.57035 |
| **GNL3L** | 15.43199 | 5.65839 | 10.16632 | 9.33257 | 2.83725 | 4.40191 | 23.08059 |
| **GNLY** | 0.00000 | 0.29650 | 0.36400 | 0.00432 | 0.00156 | 0.00000 | 0.07419 |
| **GNMT** | 0.04116 | 0.12453 | 0.00193 | 0.09874 | 0.00720 | 0.06621 | 0.18935 |
| **GNN** | 0.01697 | 0.23310 | 0.12435 | 0.15442 | 0.20067 | 0.03380 | 0.00000 |
| **GNPAT** | 10.30767 | 0.93797 | 2.63581 | 7.70598 | 3.09958 | 2.43415 | 1.63544 |
| **GNPDA1** | 40.45008 | 16.94072 | 8.56529 | 12.43033 | 0.18151 | 6.81657 | 8.59623 |
| **GNPDA2** | 2.36440 | 0.00000 | 1.60355 | 2.64445 | 2.88919 | 0.01374 | 0.00000 |
| **GNPNAT1** | 22.09219 | 9.64957 | 11.32727 | 5.98839 | 4.72488 | 11.22311 | 8.62693 |
| **GNPTAB** | 4.70937 | 3.52100 | 6.92115 | 1.30047 | 3.50737 | 7.31208 | 8.79644 |
| **GNPTG** | 7.87201 | 6.31822 | 0.79979 | 4.62891 | 0.64347 | 3.44407 | 3.59454 |
| **GNRH1** | 0.32583 | 0.00000 | 0.20958 | 0.25928 | 0.16257 | 0.19062 | 0.36659 |
| **GNRH2** | 0.00000 | 0.37785 | 0.01508 | 0.06272 | 0.08356 | 2.06012 | 0.00658 |
| **GNRHR** | 0.03054 | 0.00268 | 0.01196 | 0.01381 | 0.00617 | 0.03688 | 0.30232 |
| **GNRHR2** | 0.02897 | 0.04392 | 0.02237 | 0.02830 | 0.01404 | 0.01557 | 0.01382 |
| **GNS** | 12.76603 | 3.49493 | 4.43249 | 13.18865 | 13.71188 | 1.57074 | 3.08894 |

| | | | | | | | |
|---|---|---|---|---|---|---|---|
| **GOLGA1** | 4.88406 | 4.61024 | 1.43098 | 2.82683 | 0.41857 | 1.07561 | 3.74532 |
| **GOLGA2** | 3.27388 | 5.69358 | 1.98721 | 9.11738 | 2.95774 | 2.88562 | 8.51347 |
| **GOLGA2P5** | 0.15371 | 0.00000 | 0.01763 | 0.63095 | 0.88319 | 0.19761 | 0.51317 |
| **GOLGA2P6** | 0.01517 | 0.02099 | 0.01175 | 0.02227 | 0.01254 | 0.01162 | 0.06030 |
| **GOLGA2P7** | 0.00000 | 0.04023 | 0.01664 | 0.05977 | 0.03126 | 0.07150 | 0.32615 |
| **GOLGA2P9** | 0.03471 | 0.01032 | 0.01214 | 0.02353 | 0.29908 | 0.01483 | 0.01922 |
| **GOLGA3** | 17.88221 | 14.86246 | 4.93361 | 11.59204 | 10.00965 | 6.50421 | 5.43956 |
| **GOLGA4** | 21.41996 | 3.00531 | 10.94483 | 16.39648 | 4.97569 | 14.37077 | 44.93546 |
| **GOLGA5** | 11.26428 | 4.47051 | 8.44120 | 10.00384 | 2.34531 | 5.14264 | 0.01383 |
| **GOLGA6L10** | 0.02850 | 0.01752 | 0.01299 | 0.02131 | 0.00786 | 0.01438 | 0.01725 |
| **GOLGA6L22.uid** | 0.01811 | 0.01312 | 0.01050 | 0.01792 | 0.03911 | 0.03994 | 0.02576 |
| **GOLGA6L5P** | 0.03058 | 0.04663 | 0.01491 | 0.02719 | 0.00814 | 0.01528 | 0.00854 |
| **GOLGA6L7P** | 0.01728 | 0.03235 | 0.01659 | 0.01932 | 0.01654 | 0.02324 | 0.01755 |
| **GOLGA6L9.uid-** | 0.02655 | 0.01643 | 0.01295 | 0.01922 | 0.00787 | 0.01234 | 0.01941 |
| **GOLGA7** | 18.27603 | 8.55777 | 19.04216 | 26.04760 | 4.13699 | 14.30483 | 0.00000 |
| **GOLGA7B** | 0.00000 | 0.75010 | 3.03898 | 0.18514 | 0.00523 | 0.67935 | 0.00000 |
| **GOLGA8A** | 1.28659 | 0.00000 | 1.33603 | 2.48831 | 0.39631 | 1.35943 | 6.72507 |
| **GOLGA8B** | 0.53983 | 0.00000 | 1.11108 | 0.94322 | 0.09718 | 0.90197 | 2.09440 |
| **GOLGA8EP** | 0.02513 | 0.01703 | 0.01234 | 0.01879 | 0.01291 | 0.01244 | 0.01981 |
| **GOLGA8H** | 0.02634 | 0.04032 | 0.00677 | 0.01674 | 0.04726 | 0.00959 | 0.02083 |
| **GOLGA8I** | 0.02503 | 0.02226 | 0.00808 | 0.02051 | 0.06351 | 0.01277 | 0.01250 |
| **GOLGA8M** | 0.02360 | 0.01746 | 0.01242 | 0.01964 | 0.01679 | 0.01314 | 0.01668 |
| **GOLGA8N** | 0.02598 | 0.01695 | 0.01669 | 0.01594 | 0.01894 | 0.01876 | 0.02552 |
| **GOLGA8S** | 0.02544 | 0.02921 | 0.00943 | 0.01870 | 0.01166 | 0.01148 | 0.03044 |
| **GOLGB1** | 11.63506 | 4.48416 | 6.79095 | 10.93628 | 5.53434 | 10.80429 | 20.14480 |
| **GOLIM4** | 1.32735 | 0.11530 | 0.38933 | 1.04997 | 0.31838 | 1.64682 | 0.43420 |
| **GOLM1** | 0.98763 | 2.32632 | 1.15586 | 1.40550 | 10.39429 | 2.32069 | 4.49321 |
| **GOLPH3** | 8.62340 | 1.63922 | 8.56356 | 8.84021 | 1.88519 | 11.70601 | 27.79932 |
| **GOLPH3L** | 6.15999 | 0.30799 | 1.97656 | 2.33049 | 0.85219 | 4.01616 | 3.07454 |
| **GOLT1A** | 0.13996 | 0.03302 | 0.11880 | 0.27223 | 0.14700 | 0.00103 | 0.00000 |
| **GOLT1B** | 15.34906 | 0.00000 | 13.11587 | 11.42692 | 5.63477 | 5.60993 | 0.00000 |
| **GON4L** | 6.26340 | 1.32973 | 8.54486 | 7.06934 | 4.88122 | 8.38955 | 11.44185 |
| **GOPC** | 3.34838 | 0.45484 | 3.14289 | 4.71099 | 1.90402 | 3.94508 | 12.93376 |
| **GORAB** | 2.94416 | 0.18717 | 1.86203 | 1.11461 | 0.21831 | 1.45057 | 0.00000 |
| **GORASP1** | 8.12572 | 8.83698 | 2.54251 | 3.50209 | 0.78411 | 3.60366 | 2.36981 |

| | | | | | | | |
|---|---|---|---|---|---|---|---|
| **GORASP2** | 29.17205 | 15.68820 | 14.12532 | 22.31192 | 6.70016 | 9.47922 | 4.30062 |
| **GOSR1** | 16.90360 | 12.39904 | 16.05423 | 8.26720 | 1.22552 | 8.12397 | 7.32953 |
| **GOSR2** | 17.38155 | 14.92981 | 14.29464 | 11.58194 | 7.18331 | 6.71558 | 0.00000 |
| **GOT1** | 29.62085 | 16.62611 | 4.41292 | 26.10754 | 1.59227 | 9.78433 | 1.85078 |
| **GOT1L1** | 0.03027 | 0.01847 | 0.01351 | 0.01909 | 0.00959 | 0.01343 | 0.01126 |
| **GOT2** | 38.72051 | 19.84513 | 5.51528 | 21.24465 | 4.02386 | 16.36763 | 12.78754 |
| **GP1BA** | 0.19013 | 0.00000 | 0.11488 | 0.02692 | 0.29781 | 0.03945 | 0.36933 |
| **GP2** | 0.01487 | 0.08156 | 0.04663 | 0.00969 | 0.00792 | 0.24668 | 0.02255 |
| **GP5** | 0.01929 | 0.07352 | 0.02652 | 0.00009 | 0.00533 | 0.10836 | 0.10648 |
| **GP6** | 0.07332 | 0.01565 | 0.18816 | 0.10122 | 0.00226 | 0.29059 | 0.42887 |
| **GPA33** | 0.01202 | 0.32699 | 0.01037 | 0.08388 | 0.03482 | 0.07311 | 0.00848 |
| **GPAA1** | 12.83050 | 21.24566 | 4.01557 | 10.46102 | 0.00000 | 12.21677 | 13.05263 |
| **GPALPP1** | 2.80270 | 0.55622 | 1.68551 | 1.86218 | 1.28661 | 0.51598 | 0.76615 |
| **GPAM** | 12.39951 | 4.25188 | 4.67710 | 4.42706 | 1.17332 | 0.69935 | 4.76914 |
| **GPANK1** | 11.59983 | 17.20359 | 13.97894 | 18.17994 | 7.06939 | 7.98692 | 2.01834 |
| **GPAT2** | 0.06405 | 0.10168 | 0.04755 | 0.01913 | 0.03186 | 0.08730 | 0.18981 |
| **GPATCH1** | 7.84418 | 7.22118 | 3.39245 | 2.90245 | 1.05880 | 2.11662 | 1.95971 |
| **GPATCH11** | 3.62836 | 0.66501 | 4.14697 | 2.34068 | 5.80837 | 0.66792 | 2.01419 |
| **GPATCH2** | 5.60464 | 2.69513 | 7.65099 | 2.21003 | 3.75269 | 2.91939 | 5.87976 |
| **GPATCH2L** | 3.92495 | 1.59851 | 2.65294 | 5.24890 | 6.21555 | 1.72336 | 3.78176 |
| **GPATCH3** | 11.27779 | 6.42552 | 13.15270 | 16.00720 | 14.91805 | 6.73599 | 0.05557 |
| **GPATCH4** | 25.42800 | 22.33097 | 17.75510 | 12.25800 | 4.84654 | 11.60383 | 18.54290 |
| **GPATCH8** | 15.10293 | 6.94355 | 6.41570 | 8.95498 | 2.81839 | 9.47218 | 14.80815 |
| **GPBAR1** | 0.00862 | 0.00000 | 0.04293 | 0.11125 | 0.20162 | 0.10727 | 0.32766 |
| **GPBP1** | 24.91393 | 9.33847 | 21.01580 | 15.24106 | 11.49402 | 7.93284 | 7.65895 |
| **GPBP1L1** | 16.36592 | 6.65865 | 9.49204 | 12.60345 | 10.09563 | 11.64414 | 12.26309 |
| **GPC1** | 0.20056 | 1.15372 | 0.00000 | 2.12339 | 0.41424 | 0.13294 | 10.31155 |
| **GPC2** | 0.12401 | 0.00000 | 0.00445 | 0.20308 | 0.01196 | 0.07455 | 0.44489 |
| **GPC3** | 0.93256 | 0.71228 | 0.84589 | 0.00340 | 3.31149 | 6.65895 | 0.00000 |
| **GPC4** | 0.02075 | 0.00000 | 0.18565 | 0.12248 | 0.87022 | 3.83404 | 0.00000 |
| **GPC5** | 0.02196 | 0.01971 | 0.05897 | 0.09635 | 0.45371 | 0.01293 | 0.03155 |
| **GPC5-AS1** | 0.02417 | 0.01524 | 0.06499 | 0.01920 | 0.14321 | 0.00993 | 0.01331 |
| **GPC6** | 0.08874 | 0.01956 | 0.00000 | 0.01394 | 1.04835 | 0.07629 | 0.00000 |
| **GPC6-AS1** | 0.02315 | 0.01047 | 0.01965 | 0.01902 | 0.09434 | 0.01308 | 0.02011 |
| **GPC6-AS2** | 0.02553 | 0.01615 | 0.01315 | 0.01915 | 0.03547 | 0.01327 | 0.01503 |

| | | | | | | | |
|---|---|---|---|---|---|---|---|
| **GPCPD1** | 4.55346 | 16.89348 | 15.38636 | 0.00000 | 2.26290 | 8.97453 | 6.64442 |
| **GPD1** | 0.04663 | 0.00000 | 0.18722 | 0.08498 | 0.53035 | 0.00000 | 0.17190 |
| **GPD1L** | 5.46334 | 4.27429 | 1.15293 | 3.76150 | 1.19220 | 0.17491 | 0.00000 |
| **GPD2** | 12.45236 | 21.83739 | 3.76913 | 0.00000 | 3.70212 | 8.65217 | 7.90156 |
| **GPER1** | 0.00000 | 0.13233 | 0.11579 | 0.09802 | 0.01919 | 0.01268 | 0.10794 |
| **GPHA2** | 0.07429 | 1.46221 | 0.38649 | 0.00000 | 2.33427 | 0.37047 | 0.00000 |
| **GPHB5** | 0.00000 | 0.14478 | 0.07616 | 0.05097 | 0.03507 | 0.09749 | 0.04125 |
| **GPHN** | 2.96807 | 2.59680 | 0.59237 | 2.01149 | 0.65326 | 1.96507 | 2.91906 |
| **GPI** | 44.96249 | 26.03216 | 6.55246 | 35.51672 | 0.60844 | 23.80350 | 31.00131 |
| **GPKOW** | 7.55184 | 16.46078 | 13.59162 | 0.24444 | 3.82237 | 4.24287 | 2.03863 |
| **GPLD1** | 0.96143 | 0.55399 | 0.06491 | 0.92958 | 2.92795 | 0.00000 | 1.28032 |
| **GPM6A** | 0.00000 | 0.00403 | 0.03802 | 0.00000 | 0.05282 | 0.83704 | 1.35382 |
| **GPM6B** | 0.90187 | 0.39192 | 0.69809 | 0.27205 | 1.83017 | 2.94515 | 1.40739 |
| **GPN1** | 26.22876 | 9.56744 | 13.61546 | 10.29618 | 5.17252 | 9.00081 | 0.00000 |
| **GPN2** | 7.03165 | 10.75274 | 5.62563 | 7.48537 | 2.92896 | 4.59209 | 1.15423 |
| **GPN3** | 18.57353 | 4.87127 | 12.79717 | 15.31403 | 7.52015 | 7.55187 | 0.50230 |
| **GPNMB** | 0.97603 | 0.00000 | 1.77555 | 0.04836 | 1.02859 | 1.30538 | 0.00000 |
| **GPR1** | 0.15399 | 0.16877 | 0.23183 | 0.10733 | 0.43615 | 0.36339 | 0.00000 |
| **GPR1-AS** | 1.06910 | 0.52602 | 0.97124 | 1.13614 | 0.34510 | 1.36842 | 1.30787 |
| **GPR107** | 4.94271 | 4.01902 | 5.36851 | 7.28870 | 1.00721 | 5.36632 | 10.73464 |
| **GPR108** | 9.81004 | 11.51060 | 1.54182 | 10.04478 | 1.39853 | 11.19112 | 3.85665 |
| **GPR110** | 0.01414 | 0.02233 | 0.01501 | 0.03554 | 0.00896 | 0.01872 | 0.04853 |
| **GPR111** | 0.08947 | 0.00000 | 0.02792 | 0.01921 | 0.00732 | 0.21909 | 0.00000 |
| **GPR112** | 0.01397 | 0.05967 | 0.01439 | 0.01921 | 0.01629 | 0.01121 | 0.02934 |
| **GPR113** | 0.00000 | 0.15098 | 0.09909 | 0.20248 | 0.75839 | 0.22755 | 0.72647 |
| **GPR114** | 0.28455 | 0.10482 | 0.00317 | 1.66798 | 0.00414 | 0.17027 | 0.33787 |
| **GPR115** | 0.22392 | 0.00000 | 0.01046 | 0.29190 | 0.00709 | 0.07296 | 0.05351 |
| **GPR116** | 0.00399 | 0.00667 | 0.01042 | 0.20447 | 0.00885 | 0.00434 | 0.22582 |
| **GPR119** | 0.02192 | 0.00809 | 0.03495 | 0.02666 | 0.01492 | 0.01142 | 0.02606 |
| **GPR12** | 0.01801 | 0.13426 | 0.20777 | 0.00000 | 0.00694 | 0.07940 | 0.07375 |
| **GPR123** | 0.03003 | 0.01162 | 0.01290 | 0.02143 | 0.01254 | 0.01202 | 0.03574 |
| **GPR124** | 0.05375 | 0.03318 | 0.05436 | 0.11614 | 0.03076 | 0.28430 | 0.01294 |
| **GPR125** | 0.64627 | 1.80097 | 0.82041 | 0.95666 | 1.09382 | 3.93340 | 4.43442 |
| **GPR126** | 4.44250 | 0.00000 | 0.00000 | 0.00000 | 0.61710 | 0.00000 | 2.20936 |
| **GPR128** | 0.03821 | 0.03866 | 0.04389 | 0.02836 | 0.01374 | 0.01877 | 0.00845 |

| | | | | | | | |
|---|---|---|---|---|---|---|---|
| **GPR132** | 0.00531 | 0.15295 | 0.00631 | 0.03050 | 0.01099 | 0.02603 | 0.01618 |
| **GPR133** | 0.53677 | 1.25141 | 0.92235 | 0.00000 | 0.43415 | 1.15498 | 0.00000 |
| **GPR135** | 0.00000 | 0.04435 | 0.01953 | 0.10842 | 0.00662 | 0.01698 | 0.13708 |
| **GPR137** | 0.00000 | 1.02283 | 1.40909 | 2.32118 | 0.32636 | 2.14414 | 2.58283 |
| **GPR137B** | 0.00000 | 0.10084 | 0.07535 | 0.00805 | 0.53418 | 0.13299 | 0.11907 |
| **GPR137C** | 0.05586 | 0.00000 | 0.17967 | 0.01907 | 0.17528 | 0.06648 | 0.63721 |
| **GPR141** | 0.04177 | 0.01568 | 0.01607 | 0.02394 | 0.00801 | 0.01462 | 0.00417 |
| **GPR142** | 0.02460 | 0.02401 | 0.01098 | 0.01866 | 0.00786 | 0.01099 | 0.02478 |
| **GPR143** | 0.00000 | 0.09887 | 1.12849 | 0.02698 | 8.58220 | 0.00000 | 0.30506 |
| **GPR146** | 0.00000 | 0.07008 | 0.14844 | 0.19115 | 0.04646 | 0.19347 | 0.33998 |
| **GPR151** | 0.02258 | 0.01969 | 0.01859 | 0.03267 | 0.00747 | 0.03433 | 0.01750 |
| **GPR152** | 0.02471 | 0.01853 | 0.01303 | 0.02248 | 0.00791 | 0.01301 | 0.01534 |
| **GPR153** | 0.00000 | 1.39872 | 0.16408 | 0.00000 | 0.11784 | 2.58095 | 0.53278 |
| **GPR155** | 3.53320 | 0.00000 | 1.94342 | 0.26349 | 8.33409 | 0.00000 | 0.28404 |
| **GPR156** | 0.03268 | 0.02179 | 0.02550 | 0.11046 | 0.23260 | 0.01505 | 0.01452 |
| **GPR157** | 0.10205 | 0.12479 | 0.05552 | 0.38788 | 0.16909 | 0.00054 | 0.00000 |
| **GPR158** | 0.03885 | 0.03424 | 0.08117 | 0.02715 | 1.18605 | 0.00270 | 0.01716 |
| **GPR160** | 3.01136 | 4.43579 | 8.56647 | 0.00000 | 0.59245 | 19.22499 | 0.93673 |
| **GPR161** | 0.00000 | 1.02506 | 2.58010 | 0.85595 | 0.55586 | 1.91264 | 1.07519 |
| **GPR162** | 0.02069 | 0.01927 | 0.01359 | 0.01906 | 0.00770 | 0.01540 | 0.02419 |
| **GPR17** | 0.01831 | 0.00798 | 0.02320 | 0.01825 | 0.01082 | 0.01056 | 0.04794 |
| **GPR171** | 0.05774 | 0.00000 | 0.39482 | 0.76127 | 0.02811 | 0.03483 | 0.89221 |
| **GPR173** | 0.01592 | 0.02604 | 0.01181 | 0.02207 | 0.00762 | 0.02644 | 0.03407 |
| **GPR176** | 0.00000 | 3.12181 | 1.07707 | 0.00000 | 0.04190 | 9.68050 | 0.00935 |
| **GPR179** | 0.07228 | 0.00000 | 0.07028 | 0.14260 | 0.00347 | 0.00000 | 0.40544 |
| **GPR18** | 0.00000 | 0.67789 | 0.49327 | 0.15947 | 0.00000 | 0.34817 | 1.80597 |
| **GPR180** | 3.54085 | 0.00000 | 1.09639 | 0.31563 | 0.26514 | 0.76072 | 1.17613 |
| **GPR183** | 0.24603 | 0.41480 | 0.23741 | 0.10602 | 0.00139 | 0.12105 | 0.00611 |
| **GPR19** | 0.82235 | 0.00000 | 0.49496 | 0.43877 | 0.18069 | 1.48928 | 0.00000 |
| **GPR20** | 0.02208 | 0.02051 | 0.01218 | 0.02607 | 0.00791 | 0.01291 | 0.01836 |
| **GPR21** | 0.02892 | 0.00522 | 0.02222 | 0.02442 | 0.01017 | 0.01093 | 0.05660 |
| **GPR22** | 0.53046 | 0.00000 | 0.50762 | 0.00000 | 0.04185 | 0.59135 | 1.11810 |
| **GPR25** | 0.02640 | 0.01890 | 0.01087 | 0.02523 | 0.00747 | 0.00827 | 0.05222 |
| **GPR26** | 0.02685 | 0.01711 | 0.01332 | 0.02700 | 0.00869 | 0.01594 | 0.01941 |
| **GPR27** | 0.00000 | 0.18388 | 0.02446 | 0.02348 | 0.00592 | 0.04653 | 0.10003 |

| | | | | | | | |
|---|---|---|---|---|---|---|---|
| **GPR3** | 0.00000 | 0.00000 | 0.13994 | 0.33042 | 0.67973 | 0.63247 | 0.13609 |
| **GPR32** | 0.00000 | 0.01932 | 0.01254 | 2.15726 | 0.00860 | 0.01033 | 0.00500 |
| **GPR34** | 0.00000 | 0.28091 | 0.05669 | 0.00989 | 0.00339 | 0.17069 | 0.20453 |
| **GPR35** | 0.00000 | 0.03635 | 0.05317 | 0.12168 | 0.08204 | 0.00621 | 0.20515 |
| **GPR37** | 1.75671 | 0.00000 | 0.00000 | 0.62328 | 1.77033 | 0.00000 | 5.28459 |
| **GPR37L1** | 0.59488 | 2.24651 | 1.84248 | 0.43447 | 0.45389 | 0.00000 | 0.00000 |
| **GPR39** | 0.04866 | 0.00262 | 0.02868 | 0.01424 | 1.95965 | 0.05478 | 0.00000 |
| **GPR4** | 0.04791 | 0.00000 | 0.60717 | 0.06748 | 0.16054 | 0.04565 | 0.31083 |
| **GPR45** | 0.05381 | 0.01220 | 0.01757 | 0.08718 | 0.00762 | 0.11394 | 0.00252 |
| **GPR50** | 0.03848 | 0.00491 | 0.00000 | 0.01691 | 2.06264 | 0.49749 | 0.00000 |
| **GPR52** | 0.02896 | 0.01834 | 0.01227 | 0.03021 | 0.00797 | 0.03360 | 0.01126 |
| **GPR55** | 0.03117 | 0.01735 | 0.01603 | 0.02753 | 0.00779 | 0.01940 | 0.01089 |
| **GPR56** | 0.51090 | 0.00000 | 0.00000 | 5.12115 | 0.20474 | 0.00000 | 0.60160 |
| **GPR61** | 0.02009 | 0.01833 | 0.01378 | 0.01843 | 0.00769 | 0.01576 | 0.02659 |
| **GPR62** | 0.02289 | 0.01823 | 0.00989 | 0.01784 | 0.04128 | 0.01429 | 0.02173 |
| **GPR63** | 0.77694 | 0.00000 | 0.62386 | 0.37111 | 0.29322 | 0.30897 | 0.00000 |
| **GPR64** | 0.00000 | 0.00000 | 0.37775 | 0.11662 | 0.50827 | 0.20108 | 0.05475 |
| **GPR65** | 0.02625 | 0.01787 | 0.01265 | 0.01999 | 0.00806 | 0.01372 | 0.01631 |
| **GPR68** | 0.00108 | 0.04420 | 0.21813 | 0.03186 | 0.05789 | 0.02760 | 0.10764 |
| **GPR75** | 0.05489 | 0.00000 | 0.15245 | 0.65965 | 0.00106 | 0.67586 | 1.13567 |
| **GPR78** | 0.03930 | 0.00000 | 0.00000 | 4.43237 | 0.22135 | 0.00000 | 0.22348 |
| **GPR82** | 0.11254 | 0.04589 | 0.04027 | 0.00725 | 0.04966 | 0.03775 | 0.07274 |
| **GPR83** | 0.36074 | 0.19304 | 0.27319 | 0.43379 | 0.35192 | 0.06968 | 0.00000 |
| **GPR84** | 0.01689 | 0.01606 | 0.01378 | 0.05919 | 0.00781 | 0.01095 | 0.02424 |
| **GPR85** | 0.01203 | 0.02207 | 0.01841 | 0.03360 | 0.00668 | 0.03945 | 0.09198 |
| **GPR87** | 0.14138 | 0.00000 | 0.03294 | 0.48259 | 0.00468 | 0.00000 | 0.18038 |
| **GPR89A** | 2.27469 | 0.49072 | 0.76827 | 2.24585 | 0.69713 | 0.70005 | 0.77376 |
| **GPR89B** | 0.46149 | 0.23550 | 0.09690 | 0.72225 | 0.21040 | 0.40363 | 0.10662 |
| **GPR97** | 0.01848 | 0.02180 | 0.01162 | 0.03768 | 0.00783 | 0.03362 | 0.02224 |
| **GPR98** | 0.60514 | 0.53742 | 0.26059 | 0.46203 | 1.02262 | 1.21620 | 0.04687 |
| **GPRASP1** | 0.02706 | 0.00818 | 0.01936 | 0.01535 | 0.05062 | 0.02270 | 0.03922 |
| **GPRC5A** | 31.38167 | 26.46254 | 0.00000 | 0.00000 | 0.17108 | 0.00000 | 5.11259 |
| **GPRC5B** | 0.00000 | 0.85071 | 1.87389 | 0.31412 | 0.41406 | 21.62660 | 0.00000 |
| **GPRC5C** | 1.40597 | 3.82047 | 3.33160 | 2.75138 | 0.00000 | 16.29960 | 0.57192 |
| **GPRC5D** | 0.58756 | 0.22288 | 0.31861 | 0.36158 | 0.08034 | 0.00062 | 0.00000 |

| | | | | | | | |
|---|---|---|---|---|---|---|---|
| **GPRC6A** | 0.02446 | 0.00000 | 0.13943 | 0.02359 | 0.07252 | 0.03649 | 0.01214 |
| **GPRIN1** | 0.03094 | 0.00000 | 0.12035 | 0.51911 | 0.00309 | 0.00000 | 1.23855 |
| **GPRIN2** | 0.02679 | 0.00756 | 0.01686 | 0.00575 | 0.00658 | 0.00336 | 0.14658 |
| **GPRIN3** | 0.02433 | 0.01632 | 0.01252 | 0.01764 | 0.00807 | 0.02138 | 0.03115 |
| **GPS1** | 25.11195 | 27.93334 | 9.65014 | 25.16319 | 1.81007 | 11.81553 | 13.39940 |
| **GPS2** | 11.66459 | 19.47246 | 7.22492 | 15.13751 | 4.88814 | 8.29410 | 11.25879 |
| **GPSM1** | 0.00000 | 0.10699 | 0.01791 | 0.10700 | 0.00933 | 0.02279 | 0.10053 |
| **GPSM2** | 0.40819 | 0.00000 | 0.38486 | 1.62152 | 0.45271 | 0.10265 | 0.80927 |
| **GPSM3** | 0.00000 | 1.20515 | 2.34912 | 0.08275 | 0.28553 | 1.56404 | 1.14936 |
| **GPT** | 0.02789 | 0.01850 | 0.01245 | 0.06158 | 0.00823 | 0.03194 | 0.04803 |
| **GPT2** | 3.53826 | 1.56001 | 1.69320 | 5.27554 | 0.62192 | 7.46649 | 13.78407 |
| **GPX1** | 14.46642 | 33.29077 | 16.28634 | 12.73413 | 8.93413 | 21.43569 | 16.77028 |
| **GPX2** | 1.01777 | 0.00000 | 0.06440 | 2.90411 | 0.03030 | 47.17120 | 0.00000 |
| **GPX3** | 0.34473 | 0.08316 | 0.03975 | 2.55608 | 0.40647 | 2.52951 | 0.00000 |
| **GPX4** | 26.17341 | 34.49846 | 16.90201 | 33.69815 | 4.47016 | 14.71292 | 23.59295 |
| **GPX5** | 0.05204 | 0.01516 | 0.01322 | 0.01169 | 0.00787 | 0.01344 | 0.01283 |
| **GPX6** | 0.03329 | 0.12912 | 0.08560 | 0.05147 | 0.00649 | 0.05780 | 0.00000 |
| **GPX7** | 0.02124 | 0.16556 | 0.12868 | 0.03810 | 0.12496 | 0.21036 | 0.00000 |
| **GPX8** | 1.24888 | 0.00000 | 0.22750 | 0.92567 | 0.00105 | 0.00000 | 0.78286 |
| **GRAMD1A** | 0.00000 | 0.48249 | 0.00000 | 1.73850 | 0.83426 | 7.42447 | 1.15580 |
| **GRAMD1B** | 0.02795 | 0.00000 | 1.40641 | 0.06924 | 1.63951 | 0.10402 | 0.01413 |
| **GRAMD1C** | 0.35441 | 0.00000 | 0.35872 | 0.19969 | 0.38965 | 0.89353 | 0.00000 |
| **GRAMD2** | 2.69960 | 1.72942 | 0.01861 | 19.85623 | 0.00000 | 0.12602 | 3.04848 |
| **GRAMD3** | 4.81364 | 0.00000 | 0.04176 | 11.69944 | 0.11285 | 0.00000 | 0.73380 |
| **GRAMD4** | 0.00000 | 0.05678 | 0.10044 | 0.25041 | 0.10001 | 0.14045 | 2.61737 |
| **GRAP** | 0.19585 | 0.56071 | 0.00816 | 0.01320 | 0.06223 | 0.00193 | 0.00000 |
| **GRAP2** | 0.01845 | 0.01115 | 0.02922 | 0.05961 | 0.35054 | 0.01376 | 0.01114 |
| **GRASP** | 0.00000 | 0.03724 | 0.02460 | 0.13890 | 0.06176 | 0.31968 | 0.51378 |
| **GRB10** | 7.97832 | 7.24831 | 5.10622 | 3.51069 | 0.78030 | 3.77567 | 21.40678 |
| **GRB14** | 0.57918 | 0.03725 | 0.42194 | 0.09558 | 0.07114 | 0.00000 | 0.51607 |
| **GRB2** | 36.00377 | 29.54619 | 10.97065 | 35.31143 | 19.02200 | 16.97071 | 11.55666 |
| **GRB7** | 1.07198 | 9.58114 | 0.12610 | 2.50000 | 1.62382 | 11.87071 | 2.74882 |
| **GREB1** | 0.46759 | 0.00000 | 0.03181 | 0.26586 | 0.67635 | 0.11599 | 0.50286 |
| **GREB1L** | 7.07606 | 0.30329 | 0.34397 | 8.20295 | 0.00000 | 0.00000 | 10.12155 |
| **GREM1** | 0.02477 | 0.02223 | 0.01133 | 0.02104 | 0.01060 | 0.02916 | 0.02395 |

| | | | | | | | |
|---|---|---|---|---|---|---|---|
| **GREM2** | 3.22645 | 0.14705 | 0.00000 | 25.43880 | 0.03584 | 1.19018 | 0.01642 |
| **GRHL1** | 21.86018 | 1.13773 | 0.00000 | 23.91479 | 0.85346 | 0.00000 | 8.88115 |
| **GRHL2** | 14.36282 | 13.53877 | 0.00000 | 6.17888 | 1.85461 | 0.00000 | 6.88276 |
| **GRHL3** | 5.64600 | 7.42766 | 0.00000 | 18.66113 | 0.88624 | 0.00000 | 0.00303 |
| **GRHPR** | 21.71980 | 4.16277 | 0.10761 | 15.24199 | 1.99629 | 1.97924 | 4.99356 |
| **GRIA1** | 0.00786 | 0.06332 | 0.00000 | 0.02182 | 0.63662 | 0.01393 | 0.03294 |
| **GRIA2** | 0.02094 | 0.00000 | 0.08573 | 0.03551 | 0.09865 | 0.04477 | 0.02148 |
| **GRIA3** | 0.01806 | 0.07409 | 0.00752 | 0.00000 | 0.01787 | 0.06076 | 0.15831 |
| **GRIA4** | 0.00784 | 0.01707 | 0.06461 | 0.03628 | 0.02058 | 0.05235 | 0.05069 |
| **GRID1** | 0.02396 | 0.02597 | 0.03928 | 0.08104 | 0.04798 | 0.04340 | 0.00827 |
| **GRID2** | 0.02170 | 0.07914 | 0.00000 | 0.00630 | 0.19931 | 0.14366 | 0.07381 |
| **GRID2IP** | 0.02954 | 0.02944 | 0.01198 | 0.06517 | 0.04535 | 0.00459 | 0.05840 |
| **GRIK1** | 0.49448 | 0.00000 | 0.01990 | 0.12473 | 0.01003 | 0.03425 | 0.08665 |
| **GRIK1-AS2** | 0.05751 | 0.00960 | 0.01078 | 0.14664 | 0.03792 | 0.03168 | 0.04981 |
| **GRIK2** | 0.02952 | 0.02438 | 0.00000 | 0.01282 | 0.68809 | 0.01104 | 0.09062 |
| **GRIK3** | 0.00171 | 0.02937 | 0.00913 | 0.00475 | 0.00809 | 0.26195 | 0.14209 |
| **GRIK4** | 0.05081 | 0.02093 | 0.00000 | 0.06601 | 0.84486 | 0.02730 | 0.06130 |
| **GRIK5** | 0.01473 | 0.03299 | 0.02222 | 0.02315 | 0.02910 | 0.02200 | 0.07754 |
| **GRIN1** | 0.01515 | 0.02502 | 0.01209 | 0.02095 | 0.01501 | 0.01691 | 0.04674 |
| **GRIN2A** | 0.14199 | 0.17145 | 0.00000 | 0.00000 | 0.98777 | 0.85564 | 0.25426 |
| **GRIN2B** | 0.02063 | 0.05976 | 0.00000 | 0.07115 | 1.34816 | 0.03081 | 0.01736 |
| **GRIN2C** | 0.10289 | 0.12989 | 0.03339 | 0.07491 | 0.02709 | 0.08617 | 0.25834 |
| **GRIN2D** | 0.11238 | 0.27498 | 0.40193 | 0.61097 | 0.36432 | 0.87972 | 4.06072 |
| **GRIN3A** | 0.02414 | 0.01137 | 0.03784 | 0.01547 | 0.03560 | 0.07490 | 0.16022 |
| **GRIN3B** | 0.03368 | 0.00782 | 0.01776 | 0.03456 | 0.00988 | 0.01924 | 0.03908 |
| **GRINA** | 0.51865 | 0.64114 | 0.85163 | 12.65146 | 0.42981 | 4.96697 | 8.86266 |
| **GRIP1** | 0.36666 | 0.00000 | 0.12037 | 0.11829 | 0.47473 | 0.45654 | 0.52875 |
| **GRIP2** | 0.00000 | 0.10481 | 0.01440 | 0.13995 | 0.00916 | 1.69806 | 0.32135 |
| **GRIPAP1** | 8.27589 | 7.78048 | 3.49272 | 2.41125 | 1.10199 | 4.22480 | 6.71214 |
| **GRK1** | 0.02273 | 0.01202 | 0.01171 | 0.01866 | 0.02642 | 0.06514 | 0.01417 |
| **GRK4** | 0.04330 | 0.11429 | 0.05566 | 0.08523 | 0.07814 | 0.20615 | 0.11086 |
| **GRK5** | 0.00000 | 1.18353 | 1.01155 | 0.16317 | 1.15777 | 3.80505 | 0.86169 |
| **GRK6** | 1.05398 | 3.29334 | 0.00000 | 4.18545 | 0.12184 | 4.05498 | 10.20053 |
| **GRK7** | 0.01815 | 0.00056 | 0.05381 | 0.07256 | 0.35391 | 0.08916 | 0.00720 |
| **GRM1** | 0.02299 | 0.01911 | 0.01218 | 0.09413 | 0.03551 | 0.01229 | 0.00253 |

| | | | | | | | |
|---|---|---|---|---|---|---|---|
| **GRM2** | 0.05008 | 0.00000 | 0.30686 | 0.21919 | 0.00149 | 0.30876 | 0.16422 |
| **GRM3** | 0.01971 | 0.01695 | 0.02438 | 0.01021 | 0.00620 | 0.01913 | 0.07868 |
| **GRM4** | 0.38423 | 1.72070 | 0.00000 | 0.00000 | 0.96409 | 1.70929 | 0.08862 |
| **GRM5** | 0.02775 | 0.01251 | 0.01194 | 0.01889 | 0.08999 | 0.01287 | 0.02271 |
| **GRM6** | 0.03235 | 0.02906 | 0.03904 | 0.04764 | 0.00981 | 0.02084 | 0.01834 |
| **GRM7** | 0.02732 | 0.02435 | 0.02008 | 0.01113 | 1.15742 | 0.00798 | 0.02868 |
| **GRM8** | 0.02000 | 0.04330 | 0.00287 | 0.02311 | 0.37799 | 0.01373 | 0.02918 |
| **GRN** | 8.91667 | 25.03086 | 24.53824 | 10.41491 | 6.92319 | 18.47227 | 4.21519 |
| **GRP** | 0.01345 | 0.10841 | 0.29668 | 0.01196 | 0.03899 | 0.00000 | 0.02247 |
| **GRPEL1** | 46.67714 | 30.65036 | 20.99319 | 23.78498 | 12.16992 | 18.19705 | 8.05682 |
| **GRPEL2** | 22.04959 | 1.03873 | 10.25928 | 10.67620 | 7.81356 | 5.65425 | 5.12656 |
| **GRPR** | 0.03473 | 0.00000 | 0.28480 | 0.04147 | 0.53439 | 0.31361 | 0.01103 |
| **GRSF1** | 33.46308 | 10.87752 | 27.60233 | 16.83053 | 4.01050 | 20.61107 | 26.81951 |
| **GRTP1** | 5.14308 | 4.87184 | 0.51215 | 0.82611 | 2.33387 | 1.82972 | 5.83116 |
| **GRTP1-AS1** | 0.00880 | 0.11961 | 0.02189 | 0.05109 | 0.33431 | 0.04976 | 0.06329 |
| **GRWD1** | 33.87991 | 27.05613 | 17.90384 | 28.70903 | 9.79444 | 11.51028 | 2.49857 |
| **GS1-259H13.2** | 0.02410 | 0.00479 | 0.08407 | 0.14348 | 0.06370 | 0.04289 | 0.01279 |
| **GS1-279B7.1** | 0.02403 | 0.02789 | 0.01021 | 0.02070 | 0.01029 | 0.01198 | 0.02225 |
| **GS1-600G8.3** | 0.00836 | 0.00000 | 0.05784 | 0.08500 | 0.02963 | 0.02774 | 0.24616 |
| **GSAP** | 0.05487 | 0.00000 | 0.20585 | 0.03542 | 0.65435 | 0.06938 | 0.04323 |
| **GSC** | 0.00000 | 1.62103 | 0.27564 | 0.00000 | 1.03523 | 4.61587 | 0.00586 |
| **GSDMA** | 0.04113 | 0.20170 | 0.15699 | 0.00640 | 0.00629 | 0.00000 | 0.02467 |
| **GSDMB** | 0.39901 | 0.49467 | 0.32332 | 0.72578 | 0.43882 | 0.56568 | 2.45722 |
| **GSDMC** | 0.02179 | 0.01437 | 0.01226 | 0.01875 | 0.00800 | 0.03686 | 0.04109 |
| **GSDMD** | 0.18736 | 0.00000 | 0.00824 | 0.20822 | 0.00587 | 1.07942 | 0.13205 |
| **GSE1** | 8.93439 | 13.24813 | 1.25105 | 24.22063 | 2.84118 | 1.05139 | 27.07945 |
| **GSG1** | 0.03798 | 0.00000 | 0.35031 | 0.03663 | 0.13090 | 0.07847 | 0.00073 |
| **GSG1L** | 0.01723 | 0.02231 | 0.02456 | 0.04254 | 0.01169 | 0.02240 | 0.09524 |
| **GSG2** | 2.74967 | 2.83821 | 3.78078 | 5.30180 | 0.62944 | 2.98086 | 2.49090 |
| **GSK3A** | 0.00000 | 1.97309 | 2.33231 | 1.73260 | 0.49128 | 2.04076 | 14.49048 |
| **GSK3B** | 1.71679 | 3.45526 | 2.82186 | 14.79083 | 0.77833 | 3.56947 | 15.57677 |
| **GSKIP** | 26.83211 | 3.97015 | 7.92866 | 18.03084 | 14.20741 | 6.13367 | 0.00000 |
| **GSN** | 0.76288 | 3.71718 | 0.71765 | 0.61972 | 0.76674 | 8.22912 | 0.97463 |
| **GSN-AS1** | 0.02239 | 0.03964 | 0.11922 | 0.00000 | 0.23382 | 0.09307 | 0.43521 |
| **GSPT1** | 21.21418 | 10.52604 | 8.24807 | 15.66743 | 7.15067 | 12.54958 | 36.70483 |

| | | | | | | | |
|---|---|---|---|---|---|---|---|
| **GSPT2** | 0.67347 | 0.34931 | 1.04024 | 0.45765 | 0.70911 | 0.67286 | 0.36309 |
| **GSR** | 3.18659 | 2.39682 | 0.70901 | 4.65657 | 0.32203 | 2.79467 | 6.61320 |
| **GSS** | 4.82689 | 4.46401 | 2.14899 | 5.53218 | 0.02508 | 3.98312 | 0.29194 |
| **GSTA1** | 0.03027 | 0.01452 | 0.01267 | 0.01548 | 0.00801 | 0.06544 | 0.01378 |
| **GSTA2** | 0.05643 | 0.05378 | 0.00688 | 0.00599 | 0.00760 | 0.86236 | 0.00000 |
| **GSTA3** | 3.86859 | 0.00000 | 0.00868 | 0.07226 | 0.00154 | 0.02138 | 0.86097 |
| **GSTA4** | 5.31767 | 0.00000 | 0.17366 | 2.65024 | 0.09817 | 0.90219 | 0.00141 |
| **GSTA5** | 0.02547 | 0.01794 | 0.01280 | 0.01899 | 0.00798 | 0.01310 | 0.01643 |
| **GSTCD** | 7.55204 | 1.02276 | 3.71647 | 2.44654 | 1.69824 | 1.83844 | 1.62960 |
| **GSTK1** | 17.12863 | 5.17717 | 3.16237 | 14.97357 | 0.30117 | 3.23601 | 5.41619 |
| **GSTM1** | 0.16952 | 0.00000 | 0.02501 | 0.28961 | 0.02071 | 0.05667 | 0.00000 |
| **GSTM2** | 0.03407 | 0.01163 | 0.04283 | 0.10168 | 0.05266 | 0.03618 | 0.00000 |
| **GSTM2P1** | 0.14169 | 0.03223 | 0.01518 | 0.15034 | 0.04678 | 0.17576 | 0.00000 |
| **GSTM3** | 1.47304 | 0.00000 | 0.94180 | 10.35556 | 6.20374 | 0.00193 | 0.00000 |
| **GSTM4** | 1.81691 | 0.52936 | 0.02044 | 2.37541 | 0.19819 | 3.91283 | 0.00000 |
| **GSTM5** | 0.04476 | 0.01787 | 0.01369 | 0.02358 | 0.00788 | 0.01149 | 0.01056 |
| **GSTO1** | 11.01501 | 14.48597 | 0.62162 | 0.39809 | 0.00000 | 16.21436 | 1.12329 |
| **GSTO2** | 0.58497 | 1.07228 | 6.14175 | 0.00727 | 0.00000 | 1.03695 | 0.00280 |
| **GSTP1** | 74.11029 | 79.64354 | 11.29892 | 37.30886 | 0.69858 | 45.98497 | 41.28480 |
| **GSTT1** | 1.96994 | 0.00000 | 0.00000 | 16.86711 | 0.00385 | 0.26142 | 0.23432 |
| **GSTTP1** | 0.02874 | 0.01423 | 0.01394 | 0.01698 | 0.00772 | 0.01172 | 0.02411 |
| **GSTTP2** | 0.02447 | 0.01761 | 0.01314 | 0.04470 | 0.00803 | 0.01415 | 0.01056 |
| **GSTZ1** | 5.66768 | 15.09366 | 2.87721 | 3.88199 | 0.07749 | 5.45236 | 4.28729 |
| **GTDC1** | 8.41323 | 0.49814 | 4.45655 | 2.97378 | 1.72035 | 2.83629 | 2.03625 |
| **GTF2A1** | 3.49928 | 0.00000 | 6.26821 | 2.60968 | 2.46192 | 2.20436 | 8.49542 |
| **GTF2A1L** | 0.03521 | 0.02895 | 0.00281 | 0.01749 | 0.31124 | 0.01011 | 0.01342 |
| **GTF2A2** | 52.39960 | 13.28355 | 29.69400 | 26.34078 | 14.05806 | 13.25602 | 22.15898 |
| **GTF2B** | 17.68073 | 1.98360 | 15.55628 | 12.77887 | 6.90609 | 7.71150 | 5.34386 |
| **GTF2E1** | 17.42440 | 1.74983 | 9.90929 | 6.08833 | 5.70829 | 4.49851 | 0.00000 |
| **GTF2E2** | 2.68133 | 4.24699 | 2.90212 | 2.98701 | 3.01021 | 4.13028 | 9.04247 |
| **GTF2F1** | 12.01432 | 20.24054 | 7.94408 | 16.05541 | 9.29177 | 10.57830 | 14.39617 |
| **GTF2F2** | 18.60193 | 3.19001 | 13.45460 | 8.02176 | 12.08310 | 8.86367 | 11.75723 |
| **GTF2H1** | 22.91095 | 2.69243 | 10.03034 | 14.80927 | 9.90728 | 6.80938 | 0.00000 |
| **GTF2H2** | 1.03959 | 0.03454 | 0.39620 | 0.50087 | 0.00991 | 0.30285 | 0.00002 |
| **GTF2H2B** | 0.11792 | 0.00757 | 0.02756 | 0.06535 | 0.00597 | 0.07391 | 0.06082 |

| | | | | | | | |
|---|---|---|---|---|---|---|---|
| **GTF2H2C** | 0.02002 | 0.02851 | 0.01223 | 0.03397 | 0.00777 | 0.02491 | 0.04374 |
| **GTF2H2C-2** | 0.30025 | 0.27194 | 0.10522 | 0.11343 | 0.00757 | 0.06984 | 0.04615 |
| **GTF2H3** | 9.49734 | 0.45764 | 3.27649 | 3.09827 | 0.17863 | 1.20154 | 3.86848 |
| **GTF2H4** | 3.53855 | 0.89652 | 1.66280 | 1.20197 | 0.00000 | 1.75323 | 2.27543 |
| **GTF2H5** | 16.87812 | 11.49309 | 14.12529 | 8.44197 | 0.24142 | 9.03258 | 10.15506 |
| **GTF2I** | 0.17118 | 0.39656 | 0.41487 | 0.91030 | 0.32113 | 2.25166 | 3.22649 |
| **GTF2IP1.uid-92(** | 0.02251 | 0.02265 | 0.01174 | 0.02201 | 0.00954 | 0.02018 | 0.01678 |
| **GTF2IRD1** | 0.10729 | 0.44977 | 1.58524 | 0.65450 | 0.17840 | 1.15861 | 0.56421 |
| **GTF2IRD1P1** | 0.03149 | 0.00548 | 0.07574 | 0.05872 | 0.00506 | 0.02429 | 0.04337 |
| **GTF2IRD2** | 0.04031 | 0.04028 | 0.01365 | 0.03022 | 0.15302 | 0.02374 | 0.00000 |
| **GTF2IRD2+GTF** | 0.02558 | 0.02055 | 0.01400 | 0.02058 | 0.00906 | 0.01518 | 0.01314 |
| **GTF3A** | 39.75753 | 19.02009 | 18.70987 | 15.25322 | 0.05166 | 13.69392 | 17.33296 |
| **GTF3C1** | 5.40920 | 11.44025 | 8.41891 | 10.66349 | 2.84571 | 16.70725 | 12.15088 |
| **GTF3C2** | 18.79583 | 11.66796 | 15.12712 | 17.64931 | 4.47504 | 10.26280 | 13.51837 |
| **GTF3C2-AS1** | 0.11429 | 0.10314 | 0.08892 | 0.13437 | 0.07394 | 0.10903 | 0.06143 |
| **GTF3C3** | 16.19636 | 4.71177 | 9.79786 | 5.13229 | 1.95012 | 3.97338 | 5.02794 |
| **GTF3C4** | 4.55130 | 5.54007 | 6.65600 | 4.70386 | 1.76690 | 4.32368 | 8.99158 |
| **GTF3C5** | 20.00812 | 26.54495 | 13.68989 | 25.63431 | 2.97149 | 7.74074 | 1.60683 |
| **GTF3C6** | 22.33930 | 7.35414 | 21.86242 | 17.36123 | 2.78609 | 8.05759 | 5.39704 |
| **GTPBP1** | 0.71819 | 4.46574 | 0.63691 | 4.73010 | 1.49495 | 4.18849 | 7.16505 |
| **GTPBP10** | 13.28203 | 2.34046 | 6.97592 | 5.78808 | 2.03639 | 2.22803 | 2.90087 |
| **GTPBP2** | 1.46060 | 1.83948 | 0.93526 | 2.45163 | 0.38639 | 1.81317 | 2.99712 |
| **GTPBP3** | 8.67254 | 0.00000 | 5.31530 | 3.40962 | 2.76584 | 3.78970 | 6.91154 |
| **GTPBP4** | 46.68747 | 17.65409 | 21.43036 | 25.42561 | 10.47340 | 12.67249 | 9.97765 |
| **GTPBP8** | 11.62087 | 3.88881 | 8.86557 | 2.40181 | 1.55413 | 4.98331 | 1.39928 |
| **GTSE1** | 11.23829 | 12.08382 | 4.73553 | 13.91930 | 2.61328 | 3.49162 | 17.42205 |
| **GTSE1-AS1** | 0.00000 | 0.72278 | 0.53863 | 0.00000 | 0.00294 | 0.04838 | 0.02632 |
| **GTSF1** | 35.70757 | 17.69853 | 9.17043 | 21.91445 | 5.10109 | 9.14363 | 5.32245 |
| **GTSF1L** | 0.01934 | 0.02610 | 0.00659 | 0.02794 | 0.06762 | 0.01870 | 0.02111 |
| **GUCA1A** | 0.64979 | 0.00000 | 0.66305 | 0.62435 | 0.22804 | 4.59816 | 0.00000 |
| **GUCA1B** | 0.07493 | 0.20375 | 0.05045 | 0.33448 | 0.12448 | 0.19554 | 0.10135 |
| **GUCA2A** | 0.00000 | 0.12235 | 0.00311 | 0.27332 | 0.00779 | 0.67698 | 0.00194 |
| **GUCD1** | 6.39132 | 23.61474 | 2.90804 | 9.94748 | 10.41587 | 7.13250 | 4.01165 |
| **GUCY1A2** | 0.02921 | 0.00771 | 0.03036 | 0.12448 | 1.10411 | 0.08912 | 0.03853 |
| **GUCY1A3** | 0.04147 | 0.00590 | 0.01851 | 0.02862 | 0.16916 | 0.01311 | 0.02972 |

| | | | | | | | |
|---|---|---|---|---|---|---|---|
| **GUCY1B2** | 0.03361 | 0.00000 | 0.12155 | 0.03697 | 0.01375 | 0.03858 | 0.03512 |
| **GUCY1B3** | 0.03373 | 0.01083 | 0.01337 | 0.06169 | 0.01922 | 0.09407 | 0.00418 |
| **GUCY2C** | 0.12575 | 0.03777 | 0.17794 | 0.03674 | 0.02429 | 0.29468 | 0.00000 |
| **GUCY2D** | 0.08150 | 0.00678 | 0.05975 | 0.03362 | 0.06149 | 0.00313 | 0.07045 |
| **GUCY2EP** | 0.03432 | 0.01317 | 0.01514 | 0.02520 | 0.00804 | 0.01892 | 0.01034 |
| **GUCY2F** | 0.02297 | 0.01722 | 0.01257 | 0.01821 | 0.01055 | 0.01643 | 0.02179 |
| **GUCY2GP** | 0.03399 | 0.01809 | 0.01581 | 0.02001 | 0.02604 | 0.01258 | 0.01595 |
| **GUF1** | 22.06999 | 8.98327 | 12.22237 | 6.09184 | 3.53582 | 4.67254 | 8.00762 |
| **GUK1** | 11.37849 | 15.95804 | 1.83558 | 17.54069 | 0.83012 | 9.00956 | 30.46439 |
| **GULP1** | 7.07623 | 0.00000 | 3.13262 | 6.21908 | 0.00000 | 1.16522 | 2.98557 |
| **GUSB** | 4.26376 | 9.16384 | 6.52972 | 5.81592 | 1.37740 | 5.14456 | 7.23471 |
| **GUSBP1** | 3.32270 | 2.43668 | 3.21564 | 4.02021 | 6.70364 | 0.22497 | 0.00000 |
| **GUSBP10** | 0.02921 | 0.03015 | 0.07521 | 0.03371 | 0.13769 | 0.02715 | 0.00000 |
| **GUSBP11** | 2.73426 | 1.14745 | 0.99727 | 3.37330 | 0.49965 | 1.35665 | 2.42642 |
| **GUSBP2** | 0.03740 | 0.01571 | 0.01600 | 0.02493 | 0.00720 | 0.00633 | 0.06727 |
| **GUSBP3.uid-67** | 0.02402 | 0.01782 | 0.01355 | 0.01951 | 0.00787 | 0.01315 | 0.01671 |
| **GUSBP4** | 0.32025 | 0.15706 | 0.35617 | 0.29126 | 1.72243 | 0.16735 | 0.28036 |
| **GUSBP5** | 0.05743 | 0.04831 | 0.40908 | 0.03939 | 0.06776 | 0.00000 | 0.27880 |
| **GVINP1** | 0.79767 | 0.00000 | 0.54408 | 0.23230 | 0.01768 | 0.07716 | 0.48923 |
| **GXYLT1** | 2.64772 | 2.39549 | 2.57889 | 2.04495 | 0.20553 | 3.63797 | 13.15899 |
| **GXYLT2** | 0.01477 | 0.00863 | 0.01439 | 0.03632 | 0.01922 | 0.00359 | 0.18282 |
| **GYG1** | 9.46824 | 6.15128 | 4.32341 | 5.39464 | 6.67743 | 9.56368 | 7.74793 |
| **GYG2** | 0.96536 | 0.39590 | 1.38192 | 1.70615 | 0.04763 | 3.60292 | 0.59266 |
| **GYLTL1B** | 14.26085 | 18.49115 | 0.00000 | 11.64878 | 0.00000 | 0.00000 | 32.38972 |
| **GYPC** | 0.00000 | 0.02515 | 0.00527 | 0.01305 | 0.00625 | 8.84868 | 0.04489 |
| **GYPE** | 0.02430 | 0.01762 | 0.01068 | 0.01924 | 0.03242 | 0.01307 | 0.01724 |
| **GYS1** | 10.95047 | 16.66166 | 1.97415 | 19.63025 | 2.04897 | 1.07761 | 17.17515 |
| **GYS2** | 0.07121 | 0.00000 | 0.02617 | 0.07829 | 0.00634 | 0.06613 | 0.05485 |
| **GZF1** | 5.29375 | 0.93602 | 1.17591 | 1.19966 | 0.56677 | 1.02872 | 3.17336 |
| **GZMM** | 0.07528 | 0.10089 | 0.06682 | 0.07805 | 0.00426 | 0.07787 | 0.02132 |
| **H19** | 0.00000 | 0.02728 | 0.00038 | 17.99747 | 0.00332 | 0.72392 | 4.56468 |
| **H1F0** | 16.04100 | 39.45317 | 5.59117 | 28.56177 | 7.77472 | 18.07933 | 0.00000 |
| **H1FNT** | 0.01836 | 0.00000 | 0.08449 | 0.02431 | 0.84879 | 0.00741 | 0.03514 |
| **H1FOO** | 0.00000 | 0.02785 | 1.11781 | 0.01172 | 23.06653 | 0.00000 | 0.02556 |
| **H1FX** | 0.00000 | 3.20702 | 0.78996 | 0.23723 | 0.00378 | 2.72955 | 9.03948 |

| | | | | | | | |
|---|---|---|---|---|---|---|---|
| **H1FX-AS1** | 0.07252 | 0.00000 | 0.09701 | 0.36656 | 0.00292 | 0.01282 | 0.49223 |
| **H2AFJ** | 4.57715 | 5.43058 | 1.49363 | 21.66719 | 1.29963 | 3.81222 | 5.83695 |
| **H2AFV** | 5.39956 | 13.61523 | 12.41316 | 6.65290 | 7.70393 | 10.56081 | 23.33857 |
| **H2AFX** | 0.00000 | 13.26345 | 4.66718 | 7.15717 | 1.98099 | 5.69615 | 29.48505 |
| **H2AFY** | 13.00650 | 11.85661 | 3.60876 | 8.61810 | 0.00000 | 7.13692 | 27.42487 |
| **H2AFY2** | 0.77876 | 1.59036 | 0.00000 | 0.00000 | 0.46945 | 4.47460 | 0.84787 |
| **H2AFZ** | 76.25785 | 41.00710 | 39.27384 | 48.39002 | 30.42125 | 32.74720 | 29.20416 |
| **H2BFM** | 0.01888 | 0.01312 | 0.07139 | 0.01917 | 0.00683 | 0.01099 | 0.02665 |
| **H2BFWT** | 0.06987 | 0.01683 | 0.09916 | 0.00274 | 0.00392 | 0.02141 | 0.06679 |
| **H3F3A** | 0.63981 | 0.89825 | 1.22011 | 1.77378 | 0.77120 | 1.00618 | 1.30698 |
| **H3F3B** | 73.96772 | 57.71973 | 38.11631 | 59.96434 | 37.17118 | 31.49726 | 20.57641 |
| **H3F3C** | 10.77376 | 20.85964 | 5.33759 | 13.58285 | 15.87557 | 3.49803 | 0.00000 |
| **H6PD** | 0.16740 | 0.07258 | 0.06294 | 0.03903 | 0.18855 | 0.07548 | 0.27293 |
| **HAAO** | 0.05234 | 0.00000 | 0.16409 | 0.00422 | 0.00820 | 0.54917 | 0.04375 |
| **HABP2** | 0.00797 | 0.03442 | 0.00000 | 0.16475 | 8.02564 | 1.47196 | 0.00000 |
| **HABP4** | 0.00000 | 0.47241 | 0.25504 | 0.22860 | 0.43161 | 0.37060 | 0.81363 |
| **HACE1** | 0.87049 | 0.23644 | 0.61984 | 1.65139 | 1.45244 | 1.03622 | 0.59364 |
| **HACL1** | 12.61406 | 2.57814 | 7.40251 | 14.26247 | 0.96985 | 4.37243 | 0.00000 |
| **HADH** | 12.14422 | 3.00829 | 1.06907 | 6.60874 | 0.97254 | 5.46421 | 5.16101 |
| **HADHA** | 21.43782 | 13.59634 | 6.10206 | 15.90708 | 1.47745 | 10.56087 | 6.16400 |
| **HADHB** | 17.70671 | 8.03220 | 5.51081 | 8.48754 | 0.65324 | 9.18615 | 1.60679 |
| **HAGH** | 2.17186 | 3.07993 | 0.00000 | 4.54954 | 4.30242 | 2.68953 | 0.82779 |
| **HAGHL** | 0.30499 | 0.03518 | 0.00750 | 0.59466 | 0.00631 | 0.00000 | 0.94399 |
| **HAGLR** | 0.00000 | 0.06230 | 0.98865 | 0.02701 | 0.21465 | 0.46066 | 0.02687 |
| **HAGLROS** | 0.00000 | 0.01460 | 0.05549 | 0.02540 | 0.00925 | 0.25845 | 0.03952 |
| **HAL** | 0.14131 | 0.00000 | 0.00670 | 0.09887 | 2.22996 | 0.00000 | 0.10023 |
| **HAMP** | 0.00472 | 0.23923 | 0.01644 | 0.16213 | 0.20074 | 0.13155 | 0.00000 |
| **HAND1** | 12.61730 | 0.00000 | 0.00000 | 14.44041 | 0.18348 | 0.00000 | 14.29928 |
| **HAND2** | 0.02549 | 0.01553 | 0.01397 | 0.02035 | 0.03493 | 0.01338 | 0.01493 |
| **HAO2** | 0.03161 | 0.00943 | 0.01403 | 0.01913 | 0.00755 | 0.01500 | 0.03641 |
| **HAP1** | 0.00000 | 0.10817 | 1.37425 | 4.45404 | 5.18333 | 0.12026 | 0.24253 |
| **HAPLN1** | 15.62414 | 0.00000 | 0.00000 | 21.67987 | 0.00186 | 0.00000 | 0.13167 |
| **HAPLN2** | 0.01781 | 0.01681 | 0.01465 | 0.03450 | 0.00869 | 0.01241 | 0.01967 |
| **HAPLN3** | 0.00000 | 0.50441 | 0.30185 | 0.00000 | 0.73163 | 0.68930 | 0.01303 |
| **HAPLN4** | 0.00666 | 0.01814 | 0.01387 | 0.06001 | 0.00739 | 0.02250 | 0.04329 |

| | | | | | | | |
|---|---|---|---|---|---|---|---|
| **HAR1A** | 0.01758 | 0.01427 | 0.01328 | 0.02409 | 0.00827 | 0.02038 | 0.03183 |
| **HARBI1** | 2.76314 | 2.97934 | 2.84403 | 1.60773 | 1.51173 | 1.68581 | 0.00000 |
| **HARS** | 39.98854 | 26.82035 | 10.47122 | 23.40795 | 5.86733 | 10.65348 | 0.00000 |
| **HARS2** | 11.59290 | 11.52783 | 2.14709 | 5.95286 | 0.73008 | 5.07289 | 1.64445 |
| **HAS1** | 0.02341 | 0.02738 | 0.03748 | 0.02504 | 0.01454 | 0.01203 | 0.01986 |
| **HAS2** | 1.61378 | 0.00000 | 0.00000 | 0.00302 | 0.38618 | 4.81962 | 0.00000 |
| **HAS2-AS1** | 0.07727 | 0.01031 | 0.01428 | 0.01553 | 0.01161 | 0.18394 | 0.00000 |
| **HAS3** | 0.00000 | 0.20130 | 1.80861 | 2.93693 | 0.51188 | 0.00000 | 0.86004 |
| **HAT1** | 23.23912 | 2.61115 | 26.25619 | 14.78056 | 8.46977 | 19.78158 | 0.79967 |
| **HAUS1** | 17.85369 | 4.03110 | 2.30225 | 11.05702 | 0.09461 | 9.03706 | 2.29928 |
| **HAUS2** | 11.39359 | 1.78542 | 13.33513 | 10.64897 | 2.75665 | 4.75185 | 8.52924 |
| **HAUS3** | 7.32369 | 0.00481 | 5.68296 | 2.15733 | 4.99782 | 3.53421 | 2.81491 |
| **HAUS4** | 10.78872 | 6.39681 | 0.70410 | 7.01604 | 5.18222 | 4.49048 | 4.38158 |
| **HAUS5** | 5.30482 | 6.87658 | 3.59818 | 6.09160 | 0.70530 | 4.48426 | 0.27738 |
| **HAUS6** | 9.70982 | 0.00000 | 6.44037 | 4.54618 | 0.62786 | 2.94443 | 5.70603 |
| **HAUS7** | 16.40517 | 19.61996 | 5.80639 | 12.21197 | 1.03715 | 3.70674 | 0.00000 |
| **HAUS8** | 6.54933 | 5.54232 | 4.02337 | 8.78171 | 2.29849 | 1.28422 | 3.63435 |
| **HAVCR1** | 41.88130 | 28.84571 | 8.62624 | 10.43919 | 0.68199 | 3.58006 | 0.00000 |
| **HAVCR1P1** | 5.24508 | 3.35481 | 0.91628 | 0.86943 | 0.03327 | 0.13814 | 0.00000 |
| **HAVCR2** | 4.14614 | 0.85342 | 0.37833 | 0.00000 | 0.02053 | 0.00000 | 0.11271 |
| **HAX1** | 21.95237 | 14.80874 | 14.27342 | 10.59023 | 0.65222 | 11.91663 | 11.24083 |
| **HBA1** | 0.00000 | 2.16396 | 0.44880 | 0.24702 | 0.03068 | 0.01063 | 0.07768 |
| **HBA2** | 0.00000 | 2.42981 | 0.20823 | 0.05002 | 0.19809 | 0.01785 | 0.03675 |
| **HBB** | 0.04154 | 0.00005 | 0.00971 | 0.00838 | 0.03160 | 0.03947 | 0.30513 |
| **HBBP1** | 0.02017 | 0.01726 | 0.01243 | 0.01847 | 0.00781 | 0.01629 | 0.02703 |
| **HBD** | 0.02371 | 0.00000 | 0.01674 | 0.02861 | 0.00624 | 0.05237 | 0.35601 |
| **HBE1** | 0.50104 | 0.08375 | 0.00453 | 0.21846 | 0.00344 | 0.00000 | 1.75627 |
| **HBEGF** | 0.00000 | 0.06289 | 0.87366 | 2.69899 | 1.96042 | 0.39935 | 0.24368 |
| **HBG1** | 0.02309 | 0.03045 | 0.01003 | 0.01812 | 0.02652 | 0.01171 | 0.01781 |
| **HBG2** | 0.02895 | 0.04000 | 0.01124 | 0.01860 | 0.00798 | 0.03727 | 0.00715 |
| **HBM** | 0.07828 | 0.35862 | 0.00000 | 0.14558 | 0.43497 | 0.00112 | 0.00000 |
| **HBP1** | 5.50671 | 3.97989 | 8.28981 | 12.87279 | 6.65878 | 4.02618 | 0.00000 |
| **HBQ1** | 0.05237 | 0.00000 | 0.00329 | 0.33482 | 0.31587 | 0.01202 | 0.15622 |
| **HBS1L** | 21.28993 | 3.72191 | 9.77645 | 12.79259 | 7.54218 | 6.54800 | 0.00000 |
| **HBZ** | 0.69029 | 0.00068 | 0.08992 | 0.24529 | 0.12144 | 0.14777 | 0.00000 |

| | | | | | | | |
|---|---|---|---|---|---|---|---|
| **HCAR1** | 0.05173 | 0.04771 | 0.00763 | 0.00774 | 0.01034 | 0.68007 | 0.08424 |
| **HCAR2** | 0.80239 | 1.50495 | 0.07126 | 0.00000 | 0.21932 | 0.62605 | 0.00664 |
| **HCAR3** | 0.25354 | 0.29162 | 0.01411 | 0.17443 | 0.04458 | 0.14229 | 0.00000 |
| **HCCS** | 32.13129 | 27.00693 | 14.53688 | 17.38090 | 9.01948 | 11.32446 | 0.00000 |
| **HCFC1** | 9.40776 | 12.00362 | 16.15318 | 12.18903 | 1.12015 | 12.82499 | 41.99883 |
| **HCFC1-AS1** | 0.00812 | 0.04264 | 0.01769 | 0.01384 | 0.01146 | 0.02980 | 0.05886 |
| **HCFC1R1** | 4.34211 | 7.64416 | 1.49333 | 7.03176 | 0.32821 | 5.29935 | 6.81734 |
| **HCFC2** | 1.50047 | 0.00000 | 1.53942 | 1.80998 | 2.74838 | 0.44763 | 0.00000 |
| **HCG11** | 0.55457 | 0.35899 | 0.84689 | 0.58282 | 0.79713 | 0.00000 | 0.88290 |
| **HCG14** | 0.02020 | 0.01486 | 0.00939 | 0.04979 | 0.22340 | 0.01067 | 0.12960 |
| **HCG17** | 0.12151 | 0.18884 | 0.18953 | 0.08224 | 0.82814 | 0.10524 | 0.00000 |
| **HCG18** | 3.94674 | 4.20096 | 5.91039 | 5.41068 | 6.56469 | 3.13410 | 3.72777 |
| **HCG2040054** | 0.06098 | 0.01553 | 0.02923 | 0.04248 | 0.00696 | 0.00830 | 0.00865 |
| **HCG22** | 0.01989 | 0.01832 | 0.01004 | 0.02312 | 0.02695 | 0.01308 | 0.04321 |
| **HCG23** | 0.02302 | 0.05782 | 0.02324 | 0.02477 | 0.00737 | 0.00841 | 0.00738 |
| **HCG25** | 0.08514 | 0.08396 | 0.06671 | 0.11694 | 0.02282 | 0.09794 | 0.10810 |
| **HCG26** | 0.01594 | 0.00946 | 0.04900 | 0.03076 | 0.00645 | 0.01805 | 0.02333 |
| **HCG27** | 0.02808 | 0.00080 | 0.09169 | 0.01785 | 0.00903 | 0.01212 | 0.02017 |
| **HCG4B** | 0.01857 | 0.01093 | 0.05161 | 0.03551 | 0.01835 | 0.03867 | 0.01456 |
| **HCG8** | 0.08794 | 0.28815 | 0.16918 | 0.00000 | 0.04119 | 0.05463 | 0.14111 |
| **HCG9** | 0.01360 | 0.01849 | 0.01283 | 0.02075 | 0.00713 | 0.00837 | 0.08718 |
| **HCK** | 0.02744 | 0.05174 | 0.02871 | 0.03711 | 0.03743 | 0.00634 | 0.01998 |
| **HCLS1** | 3.30740 | 6.02363 | 0.00420 | 2.96333 | 0.11801 | 3.57197 | 0.00000 |
| **HCN1** | 0.13073 | 0.93092 | 0.16003 | 0.00000 | 0.12268 | 0.04974 | 0.16590 |
| **HCN2** | 0.00000 | 0.05295 | 0.04667 | 0.01943 | 0.22330 | 0.09894 | 0.18204 |
| **HCN3** | 0.18870 | 0.02074 | 0.20975 | 0.49670 | 0.09157 | 0.17071 | 0.00000 |
| **HCN4** | 0.02532 | 0.03710 | 0.00000 | 0.00262 | 0.88200 | 0.19693 | 0.06011 |
| **HCP5** | 0.08118 | 0.00000 | 0.38672 | 0.01408 | 0.00281 | 0.01610 | 0.00000 |
| **HCRTR1** | 0.00346 | 0.01929 | 0.01188 | 0.07796 | 0.00920 | 0.01097 | 0.03997 |
| **HCRTR2** | 0.04749 | 0.01972 | 0.01189 | 0.02488 | 0.00797 | 0.01432 | 0.00828 |
| **HCST** | 0.03987 | 0.01700 | 0.06478 | 0.01883 | 0.08277 | 0.01444 | 0.04058 |
| **HDAC1** | 42.51658 | 14.63559 | 17.15142 | 30.70360 | 7.23569 | 16.96985 | 0.00000 |
| **HDAC10** | 0.00000 | 0.04029 | 0.00294 | 0.23842 | 0.01355 | 0.43339 | 1.13789 |
| **HDAC11** | 0.44175 | 0.24829 | 0.11947 | 0.25329 | 0.70646 | 0.02990 | 0.17259 |
| **HDAC11-AS1** | 0.00573 | 0.06644 | 0.01719 | 0.06226 | 0.10321 | 0.01081 | 0.04677 |

| | | | | | | | |
|---|---|---|---|---|---|---|---|
| **HDAC2** | 36.21923 | 9.85500 | 23.83005 | 28.36376 | 12.83782 | 18.89271 | 19.39617 |
| **HDAC3** | 8.53265 | 6.14857 | 5.35553 | 12.05656 | 2.24980 | 7.42490 | 6.08326 |
| **HDAC4** | 0.00000 | 0.58373 | 0.09606 | 0.73715 | 0.18838 | 0.43890 | 1.06564 |
| **HDAC5** | 0.00000 | 3.31494 | 2.37186 | 12.19823 | 0.83296 | 3.29801 | 4.85495 |
| **HDAC6** | 4.27808 | 4.48474 | 1.66979 | 1.97840 | 1.43434 | 5.17822 | 1.92483 |
| **HDAC7** | 0.00000 | 1.36103 | 0.20457 | 0.51724 | 0.00000 | 4.20250 | 1.85588 |
| **HDAC8** | 8.09193 | 5.90966 | 14.68633 | 1.07402 | 0.71706 | 5.55977 | 5.04057 |
| **HDAC9** | 0.14072 | 0.00000 | 0.08563 | 0.00000 | 3.44021 | 0.20083 | 0.02086 |
| **HDC** | 0.01131 | 0.00175 | 0.00430 | 0.01229 | 7.77581 | 0.21375 | 0.03047 |
| **HDDC2** | 25.77547 | 20.40447 | 15.38655 | 18.57842 | 0.00198 | 12.15202 | 10.16736 |
| **HDDC3** | 10.26523 | 9.12303 | 1.21972 | 0.00000 | 0.00000 | 3.04415 | 1.38239 |
| **HDGF** | 7.62897 | 7.09600 | 4.97631 | 13.29168 | 0.09033 | 5.97425 | 40.67170 |
| **HDGFL1** | 0.00000 | 0.05783 | 0.39842 | 0.03474 | 0.12108 | 0.02857 | 0.14006 |
| **HDGFRP2** | 0.00000 | 1.48304 | 0.62559 | 0.07363 | 0.98417 | 1.25953 | 4.22703 |
| **HDGFRP3** | 0.61561 | 1.21279 | 0.74028 | 0.16062 | 0.41805 | 2.67634 | 1.24567 |
| **HDHD1** | 14.32497 | 5.82006 | 4.59559 | 9.68646 | 0.00000 | 5.96776 | 8.36618 |
| **HDHD2** | 6.39521 | 2.61684 | 5.67497 | 2.95807 | 1.83918 | 1.95132 | 0.00000 |
| **HDHD3** | 13.59462 | 14.73492 | 6.46578 | 11.79581 | 5.16141 | 2.47438 | 1.14615 |
| **HDLBP** | 34.18463 | 34.49708 | 13.99479 | 31.30128 | 5.43377 | 17.14570 | 31.64694 |
| **HDX** | 0.17455 | 0.09693 | 0.04159 | 0.04075 | 0.15993 | 0.04705 | 0.00000 |
| **HEATR1** | 17.56928 | 5.16394 | 11.33036 | 10.83214 | 8.01165 | 4.21275 | 4.38712 |
| **HEATR2** | 9.24400 | 5.16968 | 2.36618 | 3.70293 | 0.62907 | 1.73436 | 13.95599 |
| **HEATR3** | 1.19456 | 1.14545 | 2.43345 | 0.99860 | 0.44400 | 0.64075 | 2.78908 |
| **HEATR4** | 0.06475 | 0.03028 | 0.07601 | 0.12030 | 0.54542 | 0.05422 | 0.07254 |
| **HEATR5A** | 18.50239 | 7.36669 | 3.68723 | 4.98974 | 1.20771 | 5.94768 | 5.45673 |
| **HEATR5B** | 7.00312 | 0.44746 | 4.00655 | 3.96583 | 1.11312 | 3.69131 | 2.91680 |
| **HEATR6** | 5.81935 | 3.03719 | 1.95819 | 3.96699 | 0.09579 | 1.97742 | 0.33988 |
| **HEATR9** | 0.01054 | 0.00000 | 0.25019 | 0.01537 | 0.03204 | 0.02771 | 0.07721 |
| **HEBP1** | 5.73227 | 0.00000 | 0.75268 | 0.41577 | 0.08022 | 0.33738 | 2.43973 |
| **HEBP2** | 2.12600 | 1.32426 | 1.00886 | 5.93091 | 0.24669 | 1.13605 | 15.17229 |
| **HECA** | 2.15456 | 0.00000 | 1.83308 | 2.78959 | 1.72788 | 0.22273 | 4.09176 |
| **HECTD1** | 13.39288 | 4.09532 | 8.15698 | 9.95941 | 1.37143 | 9.35449 | 23.77742 |
| **HECTD2** | 0.00000 | 0.71999 | 0.61502 | 0.87964 | 0.92222 | 0.14026 | 0.17922 |
| **HECTD3** | 4.49532 | 14.46631 | 4.89971 | 7.34097 | 2.36721 | 7.19443 | 13.98321 |
| **HECTD4** | 1.56067 | 1.76702 | 0.38041 | 2.92950 | 0.44211 | 2.54200 | 9.25012 |

| | | | | | | | |
|---|---|---|---|---|---|---|---|
| **HECW1** | 0.01889 | 0.02119 | 0.06289 | 0.02711 | 0.08770 | 0.01458 | 0.02016 |
| **HECW2** | 0.55745 | 0.00000 | 0.06067 | 0.49066 | 0.21029 | 0.00000 | 1.03711 |
| **HEG1** | 0.80550 | 0.51992 | 0.19373 | 5.05956 | 0.00000 | 0.00000 | 4.02868 |
| **HEIH** | 0.00000 | 0.00000 | 0.45410 | 1.25236 | 0.38608 | 0.00000 | 0.50943 |
| **HELB** | 1.76849 | 0.00000 | 3.18151 | 2.13011 | 2.26283 | 1.18727 | 0.00000 |
| **HELLS** | 13.66800 | 2.09141 | 5.75474 | 9.30510 | 2.84366 | 7.20391 | 17.36312 |
| **HELQ** | 4.50907 | 4.00468 | 5.17090 | 3.27472 | 2.66914 | 2.03225 | 0.40357 |
| **HELT** | 0.02377 | 0.02476 | 0.02644 | 0.02026 | 0.00703 | 0.01187 | 0.01584 |
| **HELZ** | 12.72040 | 1.93151 | 7.81851 | 9.66364 | 6.18353 | 7.59078 | 17.92134 |
| **HELZ2** | 0.00000 | 0.14746 | 0.03413 | 0.02514 | 0.00459 | 0.24066 | 0.15253 |
| **HEMGN** | 0.02390 | 0.01672 | 0.01403 | 0.02177 | 0.00817 | 0.01282 | 0.01890 |
| **HEMK1** | 4.02534 | 3.89326 | 0.13273 | 0.34701 | 0.01312 | 1.91238 | 1.01126 |
| **HENMT1** | 8.87332 | 7.76236 | 7.28331 | 2.80792 | 13.52370 | 8.91232 | 0.00000 |
| **HEPACAM** | 0.01876 | 0.01533 | 0.01458 | 0.02291 | 0.00775 | 0.02586 | 0.02597 |
| **HEPACAM2** | 0.01240 | 0.01226 | 0.02075 | 0.07762 | 0.00780 | 0.01246 | 0.02702 |
| **HEPH** | 0.47979 | 0.13424 | 0.34339 | 0.13882 | 0.03147 | 0.06436 | 0.00000 |
| **HEPHL1** | 0.03800 | 0.00000 | 0.03639 | 0.02521 | 0.26087 | 0.06300 | 0.02711 |
| **HERC1** | 4.68773 | 2.17361 | 0.00000 | 3.33626 | 1.88375 | 4.12980 | 8.80621 |
| **HERC2** | 3.60240 | 6.28241 | 1.58665 | 0.11162 | 1.11242 | 4.01294 | 7.92252 |
| **HERC2P10** | 0.05133 | 0.00000 | 0.06187 | 0.00992 | 0.06618 | 0.02630 | 0.13162 |
| **HERC2P2** | 0.42665 | 0.04913 | 0.30851 | 0.00835 | 0.05949 | 0.61652 | 1.32718 |
| **HERC2P3** | 0.07621 | 0.12116 | 0.02644 | 0.07479 | 0.02254 | 0.02928 | 0.09341 |
| **HERC2P4** | 0.08463 | 0.03426 | 0.06637 | 0.04044 | 0.06335 | 0.02977 | 0.00000 |
| **HERC2P9** | 0.32555 | 0.25155 | 0.12034 | 0.01738 | 0.05506 | 0.17714 | 0.16438 |
| **HERC3** | 4.95959 | 0.58682 | 2.38459 | 3.50981 | 2.86710 | 0.53288 | 3.72140 |
| **HERC4** | 10.90924 | 0.00000 | 4.55488 | 4.62693 | 1.35283 | 2.35912 | 6.35252 |
| **HERC5** | 3.11680 | 0.34322 | 5.61861 | 5.72767 | 2.96237 | 3.54615 | 3.14417 |
| **HERC6** | 0.13834 | 0.35093 | 0.60745 | 0.00000 | 8.48474 | 0.09344 | 0.12782 |
| **HERPUD1** | 38.78366 | 2.00302 | 1.56100 | 31.84937 | 2.23185 | 3.74120 | 4.20805 |
| **HERPUD2** | 1.68414 | 0.31453 | 0.76974 | 1.70434 | 0.58758 | 0.60455 | 1.98713 |
| **HES1** | 1.71522 | 1.23660 | 0.00000 | 2.37849 | 0.67143 | 0.00000 | 4.59544 |
| **HES2** | 0.00000 | 1.88081 | 3.79180 | 1.22243 | 0.35272 | 0.39995 | 0.01802 |
| **HES3** | 0.02761 | 0.02625 | 0.01308 | 0.01693 | 0.00765 | 0.02459 | 0.01711 |
| **HES4** | 0.00000 | 0.24310 | 0.05470 | 0.04348 | 0.00268 | 0.00364 | 0.68578 |
| **HES5** | 0.01504 | 0.01581 | 0.03127 | 0.05120 | 0.09127 | 0.01127 | 0.01404 |

| | | | | | | | |
|---|---|---|---|---|---|---|---|
| **HES6** | 0.62268 | 0.59412 | 0.00000 | 1.92514 | 0.82440 | 1.83490 | 8.04284 |
| **HES7** | 0.02488 | 0.02751 | 0.06536 | 0.04531 | 0.06530 | 0.01203 | 0.08449 |
| **HESX1** | 0.14206 | 0.00000 | 0.53350 | 0.08628 | 0.97622 | 4.40995 | 0.00000 |
| **HEXA** | 14.53742 | 6.24620 | 2.29888 | 10.59482 | 2.85792 | 6.16618 | 2.26057 |
| **HEXA-AS1** | 0.05477 | 0.08508 | 0.05740 | 0.15053 | 0.20820 | 0.00654 | 0.14980 |
| **HEXB** | 20.62269 | 4.49237 | 1.50712 | 17.70472 | 2.60908 | 5.25372 | 5.69632 |
| **HEXDC** | 0.03509 | 0.28256 | 0.05014 | 0.27418 | 0.05088 | 0.42586 | 1.34909 |
| **HEXIM1** | 9.87789 | 4.25938 | 3.23375 | 21.33136 | 15.41886 | 3.05984 | 5.87771 |
| **HEXIM2** | 3.28690 | 6.65640 | 6.40395 | 2.29476 | 1.63161 | 2.33992 | 0.00000 |
| **HEY1** | 0.00000 | 0.00000 | 0.89139 | 0.45034 | 1.58999 | 0.42872 | 0.00544 |
| **HEY2** | 0.00000 | 0.62729 | 3.18486 | 0.00000 | 0.47424 | 2.78188 | 0.28729 |
| **HEYL** | 0.07881 | 0.18963 | 0.22980 | 0.16894 | 0.05077 | 0.00129 | 0.03009 |
| **HFE** | 0.02162 | 0.11013 | 0.00085 | 0.04478 | 0.06909 | 0.00670 | 0.00918 |
| **HFE2** | 0.05812 | 0.00000 | 0.76600 | 0.09759 | 0.03190 | 0.08909 | 0.00000 |
| **HFM1** | 0.06649 | 0.12028 | 0.09870 | 0.03826 | 0.00922 | 0.03678 | 0.00000 |
| **HGD** | 0.07318 | 0.01014 | 0.00918 | 0.02790 | 0.02016 | 0.42038 | 0.00000 |
| **HGFAC** | 0.03896 | 0.02292 | 0.01130 | 0.01716 | 0.00785 | 0.02840 | 0.01541 |
| **HGH1.uid-11150** | 0.01111 | 0.14508 | 0.01747 | 0.03024 | 0.00791 | 0.02644 | 0.04919 |
| **HGH1.uid-11154** | 0.02522 | 0.02113 | 0.01263 | 0.01853 | 0.00778 | 0.01654 | 0.02236 |
| **HGS** | 6.24442 | 3.58997 | 4.86192 | 23.41729 | 5.37578 | 9.43148 | 11.11290 |
| **HGSNAT** | 0.70856 | 0.76981 | 0.39795 | 0.09108 | 0.39650 | 1.17890 | 2.19519 |
| **HHAT** | 0.09211 | 0.10740 | 0.22719 | 0.31545 | 3.06478 | 0.29792 | 0.06948 |
| **HHATL** | 0.01197 | 0.02095 | 0.00968 | 0.05934 | 0.01389 | 0.06244 | 0.03270 |
| **HHEX** | 0.00000 | 0.18247 | 0.21374 | 0.12598 | 0.58323 | 0.80415 | 0.15231 |
| **HHIP** | 0.07290 | 0.02827 | 0.01041 | 0.01469 | 0.07086 | 0.00784 | 0.02919 |
| **HHIP-AS1** | 0.03327 | 0.00397 | 0.04595 | 0.02353 | 0.00536 | 0.00478 | 0.07375 |
| **HHIPL1** | 0.10766 | 0.00000 | 0.08905 | 0.00491 | 0.00769 | 0.00000 | 0.16968 |
| **HHIPL2** | 0.29183 | 0.00000 | 0.07602 | 0.03982 | 0.05310 | 0.00000 | 0.06970 |
| **HHLA1** | 0.05234 | 0.00000 | 0.00592 | 0.09518 | 0.16382 | 1.74917 | 0.00000 |
| **HHLA2** | 0.00000 | 0.00970 | 0.96858 | 0.01319 | 8.86270 | 0.00000 | 0.06576 |
| **HHLA3** | 0.90131 | 1.00848 | 1.06131 | 2.30829 | 0.51426 | 1.21963 | 0.74529 |
| **HIAT1** | 21.81169 | 3.88537 | 5.32683 | 10.21829 | 7.79067 | 8.96259 | 0.00000 |
| **HIATL1** | 9.15053 | 1.46892 | 7.83195 | 2.23203 | 4.90157 | 6.48673 | 14.63393 |
| **HIATL2** | 0.00000 | 0.12382 | 0.06170 | 0.15889 | 0.33372 | 0.17273 | 0.32486 |
| **HIBADH** | 5.83912 | 1.33716 | 2.17034 | 2.25638 | 0.24474 | 3.05537 | 3.42653 |

| | | | | | | | |
|---|---|---|---|---|---|---|---|
| **HIBCH** | 5.18656 | 0.07223 | 3.06256 | 7.01349 | 2.71948 | 4.87682 | 0.00000 |
| **HIC1** | 0.01501 | 0.02919 | 0.02624 | 0.04111 | 0.10248 | 0.01852 | 0.04689 |
| **HIC2** | 22.36543 | 35.36854 | 5.79399 | 26.79314 | 2.77627 | 7.25218 | 64.14511 |
| **HID1** | 0.68662 | 0.08141 | 0.09160 | 0.14944 | 0.28827 | 0.11225 | 1.00194 |
| **HIF1A** | 16.72203 | 2.40084 | 3.73073 | 16.05589 | 3.58121 | 4.02660 | 0.00000 |
| **HIF1A-AS2** | 0.02814 | 0.01591 | 0.01246 | 0.03014 | 0.01428 | 0.01464 | 0.01758 |
| **HIF1AN** | 15.43193 | 9.85229 | 10.06822 | 7.22380 | 1.76012 | 6.52407 | 5.58713 |
| **HIF3A** | 0.00000 | 0.00000 | 0.15464 | 0.11177 | 0.29056 | 0.02497 | 0.33670 |
| **HIGD1A** | 41.69876 | 10.52879 | 27.39292 | 19.74206 | 2.67474 | 17.77551 | 11.29357 |
| **HIGD1B** | 0.04631 | 0.03925 | 0.09585 | 0.12306 | 0.02148 | 0.01353 | 0.06059 |
| **HIGD1C** | 0.05141 | 0.00000 | 0.00703 | 0.13218 | 0.07912 | 0.00766 | 0.11763 |
| **HIGD2A** | 20.70688 | 16.41421 | 9.92365 | 20.17301 | 8.02535 | 15.17187 | 15.22398 |
| **HIGD2B** | 0.02648 | 0.01758 | 0.03236 | 0.01461 | 0.08472 | 0.01485 | 0.20617 |
| **HILPDA** | 12.71200 | 13.27764 | 21.84580 | 17.58552 | 3.15142 | 4.74535 | 5.79895 |
| **HINFP** | 7.01742 | 3.50432 | 5.50960 | 9.41278 | 4.05546 | 4.74144 | 0.00000 |
| **HINT1** | 42.33998 | 0.00000 | 1.77777 | 47.06801 | 20.09026 | 16.46055 | 24.41876 |
| **HINT2** | 1.24264 | 2.52992 | 2.44858 | 0.79915 | 0.48920 | 1.33654 | 4.56467 |
| **HINT3** | 8.18051 | 4.29910 | 4.69711 | 6.55132 | 0.11155 | 1.92291 | 0.00000 |
| **HIP1** | 16.97403 | 4.41638 | 0.51298 | 2.16640 | 1.25426 | 0.00000 | 17.46778 |
| **HIP1R** | 2.16917 | 6.29379 | 1.64993 | 7.27434 | 2.78646 | 4.74185 | 5.71720 |
| **HIPK1** | 11.07547 | 0.01957 | 7.49007 | 5.02694 | 3.80714 | 7.10740 | 15.61435 |
| **HIPK1-AS1** | 0.00000 | 0.09209 | 0.00598 | 0.06917 | 0.04783 | 0.07092 | 0.05060 |
| **HIPK2** | 0.00000 | 2.14417 | 0.91016 | 0.00000 | 2.07737 | 1.93106 | 0.18686 |
| **HIPK3** | 9.96104 | 0.00000 | 6.99938 | 2.71449 | 21.10946 | 3.65066 | 7.60661 |
| **HIPK4** | 0.00000 | 0.00000 | 0.62243 | 0.05941 | 0.35094 | 0.00328 | 0.04574 |
| **HIRA** | 1.89291 | 0.91908 | 0.76793 | 1.91030 | 0.41060 | 2.26458 | 8.54694 |
| **HIRIP3** | 3.40051 | 4.60045 | 2.90389 | 3.26893 | 6.25392 | 3.27037 | 0.96249 |
| **HIST1H1A** | 5.64320 | 5.10708 | 2.78974 | 4.24218 | 11.70702 | 1.35526 | 9.52826 |
| **HIST1H1B** | 1.18229 | 0.41398 | 0.77790 | 1.29043 | 1.15030 | 1.47664 | 6.35237 |
| **HIST1H1C** | 0.04682 | 2.59941 | 1.76310 | 8.48577 | 2.86646 | 2.01050 | 6.88890 |
| **HIST1H1D** | 3.66723 | 0.61703 | 1.91475 | 3.81956 | 0.00000 | 0.72677 | 9.03552 |
| **HIST1H1E** | 0.67281 | 1.02778 | 0.82582 | 2.03423 | 2.21763 | 1.17491 | 7.97652 |
| **HIST1H1T** | 0.00000 | 0.04765 | 0.00000 | 0.08629 | 0.17429 | 0.01568 | 0.28898 |
| **HIST1H2AA** | 0.39198 | 0.29718 | 0.86764 | 4.47220 | 1.55529 | 0.64308 | 0.96338 |
| **HIST1H2AB** | 0.03607 | 0.02100 | 0.03514 | 0.19729 | 0.28936 | 0.11648 | 0.08352 |

| | | | | | | | |
|---|---|---|---|---|---|---|---|
| **HIST1H2AC** | 0.23497 | 1.08455 | 0.12654 | 4.69353 | 9.76696 | 2.05566 | 0.00000 |
| **HIST1H2AD** | 0.01183 | 0.03820 | 0.01320 | 0.06580 | 0.03810 | 0.05041 | 0.05083 |
| **HIST1H2AE** | 0.00000 | 0.30839 | 0.21226 | 0.86985 | 0.98452 | 0.27306 | 1.01659 |
| **HIST1H2AG** | 0.01076 | 0.18593 | 0.00000 | 0.40880 | 1.72988 | 0.45718 | 0.86487 |
| **HIST1H2AH** | 0.00004 | 1.65586 | 0.07694 | 1.77211 | 2.13548 | 0.96328 | 6.57531 |
| **HIST1H2AI** | 0.00000 | 0.07620 | 0.00000 | 0.17847 | 0.77438 | 0.19205 | 0.31495 |
| **HIST1H2AJ** | 0.61923 | 0.15609 | 0.54165 | 0.96141 | 0.22368 | 0.36122 | 3.29613 |
| **HIST1H2AK** | 0.03702 | 0.10399 | 0.02356 | 0.00000 | 0.00800 | 0.20925 | 0.24345 |
| **HIST1H2AL** | 0.00000 | 0.38629 | 0.02932 | 0.63350 | 1.36623 | 0.25398 | 0.39184 |
| **HIST1H2AM** | 0.61627 | 1.09660 | 0.40240 | 2.64710 | 9.18368 | 0.66910 | 4.59167 |
| **HIST1H2APS1** | 0.02126 | 0.06253 | 0.00506 | 0.02600 | 0.05488 | 0.00997 | 0.00503 |
| **HIST1H2BA** | 0.00000 | 0.01531 | 0.03996 | 0.04694 | 0.01202 | 0.05854 | 0.21647 |
| **HIST1H2BB** | 0.00000 | 0.00973 | 0.05013 | 0.36673 | 1.13263 | 0.27747 | 0.51694 |
| **HIST1H2BC** | 0.00000 | 0.89294 | 0.45557 | 0.82163 | 9.14139 | 0.38156 | 0.87025 |
| **HIST1H2BD** | 0.07943 | 0.00000 | 1.02456 | 0.70842 | 3.86183 | 0.67298 | 0.54159 |
| **HIST1H2BE** | 0.00000 | 0.11406 | 0.06907 | 0.17468 | 0.78800 | 0.08077 | 0.65644 |
| **HIST1H2BF** | 0.02081 | 0.09171 | 0.06432 | 0.26047 | 0.04322 | 0.18017 | 0.61841 |
| **HIST1H2BG** | 0.00000 | 0.64310 | 0.00000 | 1.60927 | 5.85546 | 0.65915 | 0.79932 |
| **HIST1H2BH** | 0.00000 | 1.67954 | 0.53189 | 2.43568 | 7.41539 | 0.51313 | 0.28760 |
| **HIST1H2BI** | 0.00000 | 0.06765 | 0.03737 | 0.11637 | 0.21864 | 0.05281 | 0.31592 |
| **HIST1H2BJ** | 0.32141 | 0.32557 | 0.01974 | 1.66532 | 3.68573 | 0.81475 | 0.00000 |
| **HIST1H2BK** | 2.25783 | 0.00000 | 3.84339 | 9.47946 | 7.52298 | 2.60865 | 0.55914 |
| **HIST1H2BL** | 0.00020 | 0.05695 | 0.04758 | 0.11444 | 0.34057 | 0.05921 | 0.20598 |
| **HIST1H2BM** | 0.00000 | 0.32640 | 0.03236 | 0.17958 | 0.02838 | 0.19098 | 0.39788 |
| **HIST1H2BN** | 0.00000 | 0.10435 | 0.00000 | 0.26924 | 0.53553 | 0.21348 | 0.19295 |
| **HIST1H2BO** | 0.09626 | 0.04266 | 0.05523 | 0.38767 | 1.63169 | 0.10858 | 0.12995 |
| **HIST1H3A** | 0.00000 | 0.21359 | 0.00000 | 0.35989 | 6.49234 | 0.16051 | 0.31063 |
| **HIST1H3B** | 0.00000 | 0.11502 | 0.02807 | 0.15079 | 0.36559 | 0.18545 | 2.09329 |
| **HIST1H3C** | 0.00000 | 0.65562 | 0.08289 | 2.10566 | 4.28131 | 0.89440 | 1.85859 |
| **HIST1H3D** | 0.00000 | 0.77168 | 0.00000 | 1.59152 | 1.42316 | 0.34257 | 0.83456 |
| **HIST1H3E** | 0.04202 | 0.05087 | 0.01252 | 0.10475 | 0.39770 | 0.09392 | 0.14816 |
| **HIST1H3F** | 0.00000 | 0.01497 | 0.01857 | 0.12599 | 0.24407 | 0.39278 | 0.21485 |
| **HIST1H3G** | 0.14599 | 1.41141 | 0.00000 | 1.57311 | 5.69728 | 0.70382 | 0.62001 |
| **HIST1H3H** | 0.02943 | 0.40411 | 0.00000 | 0.78859 | 1.02688 | 0.46831 | 0.58773 |
| **HIST1H3I** | 0.09950 | 0.00000 | 0.36644 | 0.96132 | 0.29535 | 0.41318 | 0.65515 |

| | | | | | | | |
|---|---|---|---|---|---|---|---|
| **HIST1H3J** | 0.00000 | 0.81430 | 0.10350 | 1.61869 | 0.69050 | 0.35316 | 0.99651 |
| **HIST1H4A** | 0.00000 | 0.03053 | 0.00000 | 0.05142 | 0.12721 | 0.10955 | 0.29608 |
| **HIST1H4B** | 0.00000 | 0.06824 | 0.00000 | 0.08000 | 0.29662 | 0.28748 | 0.51067 |
| **HIST1H4C** | 74.13849 | 54.20595 | 21.83402 | 54.75204 | 6.39522 | 25.59397 | 27.25754 |
| **HIST1H4D** | 0.04132 | 0.02870 | 0.04587 | 0.06364 | 0.14591 | 0.23200 | 0.72308 |
| **HIST1H4E** | 0.19919 | 0.61652 | 0.15524 | 0.24451 | 0.33415 | 0.37812 | 1.05087 |
| **HIST1H4F** | 0.00000 | 0.18072 | 0.00000 | 0.03775 | 0.16668 | 0.20305 | 0.83224 |
| **HIST1H4G** | 0.02461 | 0.01789 | 0.01216 | 0.02050 | 0.02986 | 0.01423 | 0.01556 |
| **HIST1H4H** | 0.07014 | 0.38656 | 0.00000 | 0.76162 | 4.08791 | 0.13365 | 0.25056 |
| **HIST1H4I** | 0.28253 | 0.00000 | 0.05702 | 0.23770 | 0.03371 | 0.28156 | 0.41411 |
| **HIST1H4J** | 0.00000 | 0.10062 | 0.04096 | 0.23700 | 0.24341 | 0.06322 | 0.07925 |
| **HIST1H4K** | 0.01088 | 0.04626 | 0.00174 | 0.02906 | 0.15888 | 0.03527 | 0.05040 |
| **HIST1H4L** | 0.00000 | 0.04163 | 0.02453 | 0.12722 | 0.06797 | 0.10386 | 0.12031 |
| **HIST2H2AB** | 0.03069 | 0.06609 | 0.04189 | 0.00952 | 0.14374 | 0.04271 | 0.16271 |
| **HIST2H2AC** | 0.00000 | 0.28529 | 0.13329 | 1.04131 | 0.16182 | 0.20782 | 2.18463 |
| **HIST2H2BA** | 0.00679 | 0.02130 | 0.08012 | 0.02202 | 0.28941 | 0.12230 | 0.06505 |
| **HIST2H2BC** | 0.05223 | 0.01912 | 0.02208 | 0.12120 | 0.07552 | 0.02547 | 0.06354 |
| **HIST2H2BE** | 0.16257 | 0.00000 | 0.05469 | 0.44152 | 9.98355 | 0.53234 | 0.73118 |
| **HIST2H2BF** | 0.00132 | 0.04892 | 0.01000 | 0.03249 | 0.33418 | 0.06994 | 0.24616 |
| **HIST2H3D** | 0.00741 | 0.02921 | 0.01115 | 0.04593 | 0.02597 | 0.06604 | 0.09888 |
| **HIST3H2A** | 0.51125 | 0.06946 | 0.00000 | 3.52253 | 0.00604 | 0.50182 | 1.23193 |
| **HIST3H2BB** | 0.00417 | 0.08624 | 0.00276 | 0.16080 | 0.02685 | 0.00736 | 0.13663 |
| **HIST3H3** | 0.00000 | 0.02096 | 0.00635 | 0.38754 | 0.04519 | 0.07932 | 0.10469 |
| **HIST4H4** | 0.17877 | 0.00112 | 0.07520 | 0.18711 | 0.32453 | 0.08255 | 0.13037 |
| **HIVEP1** | 4.92332 | 0.00000 | 2.37180 | 3.78186 | 1.47122 | 0.00000 | 11.98276 |
| **HIVEP2** | 1.05961 | 0.19815 | 0.00000 | 0.47889 | 0.12721 | 0.00000 | 1.12510 |
| **HIVEP3** | 0.08379 | 0.03748 | 0.16353 | 0.09643 | 0.03011 | 0.21634 | 0.21568 |
| **HJURP** | 16.35958 | 16.11959 | 22.68670 | 11.53008 | 11.00334 | 3.14664 | 0.00001 |
| **HK1** | 7.49693 | 21.20148 | 12.83589 | 15.29374 | 0.91396 | 14.88841 | 7.33098 |
| **HK2** | 26.64236 | 27.73716 | 23.80037 | 2.75088 | 8.78148 | 9.48733 | 14.33054 |
| **HK3** | 0.03400 | 0.00988 | 0.01587 | 0.00725 | 0.00698 | 0.00900 | 0.06917 |
| **HKDC1** | 0.00000 | 0.87572 | 2.94265 | 0.16744 | 0.03477 | 0.00000 | 0.00000 |
| **HKR1** | 5.05655 | 3.08009 | 11.35852 | 4.59308 | 3.30126 | 4.95096 | 0.00000 |
| **HLA-A** | 0.02404 | 0.00000 | 0.09473 | 0.12749 | 0.21440 | 0.58374 | 0.00236 |
| **HLA-B** | 0.00000 | 0.33008 | 0.66077 | 0.04267 | 0.23658 | 0.99237 | 0.18940 |

| | | | | | | | |
|---|---|---|---|---|---|---|---|
| **HLA-C** | 0.17463 | 1.00625 | 0.37690 | 5.19039 | 0.13129 | 3.35635 | 0.00000 |
| **HLA-DMA** | 0.00000 | 0.39919 | 0.26785 | 0.72232 | 0.08596 | 0.07042 | 0.00000 |
| **HLA-DOA** | 0.02415 | 0.01951 | 0.01082 | 0.01932 | 0.03594 | 0.01349 | 0.01841 |
| **HLA-DOB** | 0.03381 | 0.00729 | 0.02450 | 0.03219 | 0.00816 | 0.01170 | 0.02785 |
| **HLA-DPA1** | 0.00267 | 0.04270 | 0.01690 | 0.12064 | 0.00676 | 0.18996 | 0.02554 |
| **HLA-DPB1** | 0.04229 | 0.00000 | 0.15518 | 0.14473 | 0.02344 | 0.28338 | 0.00000 |
| **HLA-DPB2** | 0.05221 | 0.01843 | 0.01172 | 0.01062 | 0.02845 | 0.36910 | 0.00177 |
| **HLA-DQA1** | 0.01843 | 0.01324 | 0.01216 | 0.01831 | 0.01758 | 0.03364 | 0.02766 |
| **HLA-DQA2** | 0.03639 | 0.02460 | 0.01348 | 0.02603 | 0.01046 | 0.01321 | 0.00785 |
| **HLA-DQB1** | 0.00000 | 0.01306 | 0.97106 | 0.00000 | 0.00168 | 0.15351 | 0.19521 |
| **HLA-DQB2** | 0.05693 | 0.00204 | 0.14501 | 0.01653 | 0.00450 | 0.00886 | 0.00717 |
| **HLA-DRB1** | 0.00067 | 0.01360 | 0.18189 | 3.09784 | 0.27134 | 0.34249 | 0.01175 |
| **HLA-DRB5** | 0.02059 | 0.03377 | 0.05872 | 0.28152 | 0.25417 | 0.04204 | 0.00000 |
| **HLA-DRB6** | 0.14261 | 0.01141 | 0.02876 | 0.59733 | 0.05532 | 0.00418 | 0.10846 |
| **HLA-E** | 7.75959 | 15.55883 | 1.42694 | 13.25654 | 1.27964 | 16.02786 | 3.28093 |
| **HLA-F** | 2.25985 | 17.70676 | 3.97403 | 0.00000 | 0.59277 | 12.15710 | 0.00000 |
| **HLA-F-AS1** | 0.09403 | 0.36050 | 0.24454 | 0.00000 | 0.18464 | 0.54946 | 0.01096 |
| **HLA-G** | 0.00000 | 0.09391 | 0.01941 | 0.29316 | 0.01586 | 0.28413 | 0.41671 |
| **HLA-H** | 0.01964 | 0.02177 | 0.04198 | 0.03280 | 0.00838 | 0.04380 | 0.01881 |
| **HLA-J** | 0.04567 | 0.08582 | 0.11984 | 0.00675 | 0.03042 | 0.12759 | 0.04826 |
| **HLA-L** | 0.08787 | 0.06658 | 0.07033 | 0.09930 | 0.04481 | 0.19025 | 0.14030 |
| **HLCS** | 4.54329 | 0.68797 | 0.85452 | 2.56120 | 0.45903 | 0.54814 | 0.42580 |
| **HLF** | 0.05422 | 0.00978 | 0.00977 | 0.02660 | 0.10061 | 0.03176 | 0.00569 |
| **HLTF** | 4.17249 | 1.02021 | 4.03455 | 0.00000 | 1.68569 | 7.31057 | 1.32067 |
| **HLX** | 0.07377 | 0.00000 | 0.04494 | 0.21998 | 0.09641 | 0.04422 | 0.01471 |
| **HLX-AS1** | 0.01981 | 0.01619 | 0.01232 | 0.01777 | 0.00772 | 0.02692 | 0.03214 |
| **HM13** | 24.59774 | 26.28047 | 12.03512 | 26.33320 | 4.03714 | 16.02165 | 13.70554 |
| **HM13-AS1** | 0.02208 | 0.08065 | 0.05541 | 0.04231 | 0.07851 | 0.02843 | 0.06918 |
| **HMBOX1** | 2.21728 | 0.00000 | 1.78959 | 4.49631 | 1.46701 | 3.41112 | 4.36080 |
| **HMBS** | 16.31988 | 17.42021 | 2.32281 | 9.67065 | 0.00000 | 6.52370 | 0.23128 |
| **HMCES** | 15.41746 | 17.81429 | 6.00248 | 22.49460 | 2.74395 | 6.60725 | 0.00000 |
| **HMCN1** | 0.04677 | 0.00598 | 0.76205 | 0.00880 | 2.22643 | 0.49683 | 0.03151 |
| **HMCN2** | 0.13021 | 0.71548 | 0.06538 | 0.08792 | 0.02240 | 0.06972 | 0.27943 |
| **HMG20A** | 9.90834 | 3.92273 | 4.04844 | 5.29044 | 1.29705 | 2.38590 | 0.00000 |
| **HMG20B** | 1.44052 | 6.28983 | 1.41036 | 8.44772 | 3.06748 | 3.02563 | 14.25022 |

| | | | | | | | |
|---|---|---|---|---|---|---|---|
| **HMGA1** | 24.25080 | 47.99042 | 24.25177 | 55.74926 | 3.86732 | 29.34205 | 63.98470 |
| **HMGA1P7** | 0.01985 | 0.02153 | 0.01482 | 0.03645 | 0.00762 | 0.01201 | 0.02023 |
| **HMGA2** | 0.00000 | 0.00521 | 0.00946 | 0.09573 | 0.01789 | 0.76306 | 0.06372 |
| **HMGB1** | 24.98559 | 10.52776 | 17.81828 | 9.01727 | 2.49818 | 8.73795 | 17.83602 |
| **HMGB2** | 37.57746 | 21.12896 | 28.12528 | 26.17259 | 11.28515 | 13.84460 | 25.50135 |
| **HMGB3** | 37.72798 | 46.47240 | 19.40837 | 8.72367 | 15.86091 | 27.11628 | 11.67589 |
| **HMGB3P1** | 0.01692 | 0.01890 | 0.01864 | 0.04289 | 0.01395 | 0.02244 | 0.05608 |
| **HMGCL** | 1.32828 | 3.75438 | 1.62017 | 1.18945 | 0.69141 | 1.99027 | 0.00000 |
| **HMGCLL1** | 0.01745 | 0.03205 | 0.01238 | 0.01838 | 0.15645 | 0.01127 | 0.02141 |
| **HMGCR** | 47.18422 | 10.79914 | 3.23586 | 15.58488 | 6.02708 | 24.27805 | 8.96929 |
| **HMGCS1** | 62.83670 | 5.00889 | 0.85584 | 10.27465 | 3.27445 | 7.60455 | 11.87897 |
| **HMGCS2** | 0.03344 | 0.01257 | 0.01224 | 0.03531 | 0.00836 | 0.10846 | 0.00000 |
| **HMGN1** | 28.72336 | 10.43976 | 13.30235 | 14.34165 | 0.12961 | 11.83465 | 12.24743 |
| **HMGN2** | 16.27689 | 26.84308 | 7.31609 | 7.06951 | 6.29429 | 4.93069 | 0.00000 |
| **HMGN2P46** | 0.03308 | 0.18101 | 0.09881 | 0.03974 | 0.25683 | 0.00871 | 0.02280 |
| **HMGN3** | 5.80598 | 0.00000 | 0.35504 | 4.31614 | 0.04837 | 6.15282 | 1.23371 |
| **HMGN3-AS1** | 0.21754 | 0.20912 | 0.00743 | 0.28569 | 0.01908 | 0.26454 | 0.00000 |
| **HMGN4** | 15.47849 | 2.63702 | 19.68699 | 21.81905 | 8.33388 | 7.61650 | 0.00000 |
| **HMGN5** | 3.19275 | 0.83291 | 0.81344 | 0.00000 | 0.00000 | 2.96766 | 3.80737 |
| **HMGXB3** | 7.20675 | 5.58957 | 7.42676 | 5.96208 | 3.68527 | 3.90432 | 8.83926 |
| **HMGXB4** | 8.86507 | 6.32155 | 5.93082 | 4.40197 | 4.41336 | 6.91700 | 10.14978 |
| **HMHA1** | 0.69730 | 7.87001 | 0.00000 | 9.04961 | 3.60421 | 3.18393 | 6.34457 |
| **HMMR** | 18.65052 | 5.62624 | 18.70263 | 6.34978 | 2.90599 | 5.54406 | 10.38206 |
| **HMMR-AS1** | 0.40483 | 0.05452 | 0.59747 | 0.16987 | 0.08404 | 0.14365 | 0.47991 |
| **HMOX1** | 18.26869 | 18.38951 | 2.67011 | 43.68778 | 0.00000 | 4.28470 | 32.78648 |
| **HMOX2** | 38.81767 | 25.24559 | 8.26850 | 24.19793 | 7.80369 | 10.26690 | 0.00000 |
| **HMP19** | 0.01463 | 0.02297 | 0.01109 | 0.01998 | 0.01453 | 0.01917 | 0.04040 |
| **HMX1** | 0.01591 | 0.07178 | 0.01052 | 0.00905 | 0.00687 | 0.04387 | 0.05970 |
| **HMX2** | 0.03488 | 0.00960 | 0.04971 | 0.01675 | 0.00620 | 0.01066 | 0.02515 |
| **HMX3** | 0.02537 | 0.01724 | 0.01366 | 0.02149 | 0.03131 | 0.01374 | 0.01607 |
| **HN1** | 49.22021 | 55.22856 | 37.24114 | 37.70809 | 5.87991 | 31.25611 | 34.97186 |
| **HN1L** | 32.12083 | 12.53925 | 7.77154 | 32.22777 | 0.43589 | 10.48657 | 10.53820 |
| **HNF1A** | 0.26490 | 0.70037 | 0.39351 | 0.57694 | 2.88227 | 0.53611 | 0.00000 |
| **HNF1A-AS1** | 0.02236 | 0.00852 | 0.00679 | 0.03741 | 0.05613 | 0.66360 | 0.01825 |
| **HNF1B** | 0.30811 | 0.71698 | 0.00004 | 0.08375 | 0.01409 | 6.17417 | 0.00000 |

| | | | | | | | |
|---|---|---|---|---|---|---|---|
| **HNF4A** | 2.63823 | 9.37545 | 18.60991 | 0.00000 | 0.51710 | 9.92618 | 0.00448 |
| **HNF4A-AS1** | 0.01471 | 0.06297 | 0.13055 | 0.01237 | 0.02248 | 0.01381 | 0.06247 |
| **HNF4G** | 0.02975 | 0.01365 | 0.05184 | 0.01687 | 0.00478 | 0.35711 | 0.04417 |
| **HNMT** | 0.03908 | 0.20307 | 0.26296 | 0.00693 | 0.23834 | 0.26565 | 0.00000 |
| **HNRNPA0** | 4.42863 | 4.03335 | 5.23110 | 12.49475 | 2.05553 | 11.46197 | 35.76676 |
| **HNRNPA1** | 12.24997 | 0.00000 | 10.88221 | 8.48924 | 2.42186 | 6.64015 | 16.40149 |
| **HNRNPA1L2** | 1.57225 | 1.51900 | 1.89753 | 0.00000 | 1.83099 | 1.01535 | 0.00001 |
| **HNRNPA1P33** | 0.70114 | 0.46441 | 0.02543 | 0.17410 | 0.01833 | 0.00000 | 1.92043 |
| **HNRNPA2B1** | 49.78974 | 23.00439 | 33.71964 | 42.57934 | 17.67385 | 36.00661 | 52.62377 |
| **HNRNPA3** | 7.87916 | 0.43705 | 5.05709 | 2.67674 | 1.13583 | 4.03800 | 20.52029 |
| **HNRNPA3P1** | 0.03705 | 0.02611 | 0.01864 | 0.02759 | 0.01238 | 0.02216 | 0.03601 |
| **HNRNPAB** | 12.96047 | 15.85398 | 10.21720 | 12.86325 | 9.72898 | 13.94654 | 54.23076 |
| **HNRNPC** | 57.46838 | 28.34457 | 29.48214 | 39.78891 | 16.79066 | 25.00916 | 16.15232 |
| **HNRNPCL1** | 0.01120 | 0.03632 | 0.00000 | 0.01451 | 4.79460 | 0.00721 | 0.01250 |
| **HNRNPCL2** | 0.02720 | 0.05060 | 0.06141 | 0.02513 | 16.11270 | 0.01563 | 0.03410 |
| **HNRNPD** | 23.93352 | 4.07612 | 19.29484 | 15.08762 | 1.75343 | 15.80960 | 50.46199 |
| **HNRNPDL** | 42.77130 | 10.91597 | 31.35386 | 21.28253 | 12.94998 | 22.03045 | 33.45217 |
| **HNRNPF** | 69.31294 | 40.24435 | 36.39248 | 42.99776 | 22.45918 | 28.54912 | 19.59533 |
| **HNRNPH1** | 59.59960 | 15.01363 | 29.39333 | 39.07244 | 19.61368 | 31.08786 | 21.23146 |
| **HNRNPH2** | 3.80967 | 1.19147 | 1.24658 | 0.94652 | 1.06213 | 0.56320 | 0.00000 |
| **HNRNPH3** | 31.31537 | 18.27540 | 23.41227 | 25.19710 | 4.48197 | 11.43536 | 13.52929 |
| **HNRNPK** | 54.90404 | 25.53362 | 20.99969 | 33.59789 | 12.42997 | 24.48859 | 7.77225 |
| **HNRNPKP3** | 0.17122 | 0.09446 | 0.09713 | 0.15960 | 0.11286 | 0.08905 | 0.33358 |
| **HNRNPL** | 8.81770 | 7.83075 | 12.99254 | 13.98052 | 7.46791 | 11.21378 | 28.73947 |
| **HNRNPLL** | 2.44528 | 0.47180 | 2.93573 | 1.59048 | 0.68003 | 4.12768 | 8.50807 |
| **HNRNPM** | 47.67954 | 47.37547 | 22.30776 | 43.42750 | 15.04997 | 27.16801 | 23.49352 |
| **HNRNPR** | 44.52350 | 18.54159 | 22.51603 | 32.33717 | 6.45254 | 17.22776 | 16.31739 |
| **HNRNPU** | 30.29922 | 15.84060 | 20.64665 | 20.33542 | 10.77854 | 18.81306 | 53.03422 |
| **HNRNPU-AS1** | 3.68506 | 0.00000 | 1.33953 | 0.88204 | 0.63749 | 1.73456 | 9.06177 |
| **HNRNPUL1** | 5.28700 | 12.40335 | 11.20796 | 12.43953 | 2.19991 | 10.60999 | 25.94821 |
| **HNRNPUL2** | 0.00000 | 0.82544 | 0.82179 | 0.73100 | 0.21918 | 1.13796 | 7.11490 |
| **HOGA1** | 0.06475 | 0.06278 | 0.02420 | 0.01483 | 0.01643 | 0.00516 | 0.02840 |
| **HOMER1** | 0.95457 | 0.13648 | 0.74793 | 0.92794 | 0.47950 | 1.99536 | 6.62500 |
| **HOMER2** | 0.00000 | 2.91439 | 1.30803 | 0.13296 | 2.66698 | 5.66178 | 0.01413 |
| **HOMER3** | 0.00000 | 0.12522 | 0.58794 | 0.11595 | 0.06074 | 0.96752 | 0.32993 |

| | | | | | | | |
|---|---|---|---|---|---|---|---|
| **HOMEZ** | 0.18223 | 0.61910 | 0.01373 | 1.91817 | 0.05556 | 0.10121 | 0.37126 |
| **HOOK1** | 4.33010 | 0.00000 | 0.41088 | 1.87061 | 0.84272 | 0.00000 | 5.65641 |
| **HOOK2** | 4.17008 | 2.09045 | 0.74247 | 6.30197 | 0.29532 | 1.94633 | 3.59723 |
| **HOOK3** | 4.77936 | 0.89866 | 3.98673 | 1.53866 | 2.14464 | 6.83124 | 19.68858 |
| **HOPX** | 0.00000 | 0.00561 | 0.03164 | 8.92794 | 0.01434 | 0.35824 | 0.00184 |
| **HORMAD1** | 0.96840 | 1.85584 | 4.50143 | 1.05758 | 0.14694 | 2.98388 | 0.25816 |
| **HORMAD2** | 0.01552 | 0.10728 | 0.21834 | 0.02778 | 0.04756 | 0.07046 | 0.00058 |
| **HORMAD2-AS1** | 1.25497 | 6.72066 | 0.42849 | 0.00000 | 0.00000 | 8.99794 | 0.00000 |
| **HOTAIR** | 0.02793 | 0.00002 | 0.02904 | 0.00145 | 0.00883 | 0.00890 | 0.20062 |
| **HOTAIRM1** | 0.00000 | 0.88137 | 1.49297 | 0.48789 | 0.53794 | 0.56608 | 0.00009 |
| **HOTTIP** | 0.02642 | 0.01497 | 0.01927 | 0.01878 | 0.00951 | 0.01312 | 0.02184 |
| **HOXA-AS2** | 0.02465 | 0.01611 | 0.02035 | 0.02846 | 0.00837 | 0.03201 | 0.00931 |
| **HOXA-AS3** | 0.02139 | 0.01919 | 0.01608 | 0.01859 | 0.01411 | 0.01474 | 0.02415 |
| **HOXA1** | 0.14580 | 0.09596 | 1.97143 | 0.33385 | 0.54573 | 0.50451 | 0.47802 |
| **HOXA10** | 0.09943 | 0.02312 | 0.14710 | 0.01142 | 0.02582 | 0.00616 | 0.02726 |
| **HOXA10-AS** | 0.04495 | 0.01665 | 0.01465 | 0.01779 | 0.00790 | 0.01433 | 0.00688 |
| **HOXA11** | 0.05300 | 0.01006 | 0.09791 | 0.00773 | 0.04451 | 0.03961 | 0.02262 |
| **HOXA11-AS** | 0.03923 | 0.01392 | 0.05591 | 0.01929 | 0.01564 | 0.03108 | 0.00345 |
| **HOXA13** | 0.02531 | 0.00976 | 0.02950 | 0.01776 | 0.00761 | 0.05905 | 0.01718 |
| **HOXA2** | 0.02326 | 0.00000 | 0.06855 | 0.02093 | 0.02728 | 0.01327 | 0.02221 |
| **HOXA3** | 0.00113 | 0.00819 | 0.33887 | 0.01147 | 0.12358 | 0.13113 | 0.06623 |
| **HOXA4** | 0.01276 | 0.10083 | 0.02902 | 0.02160 | 0.15894 | 0.03400 | 0.11814 |
| **HOXA5** | 0.00635 | 0.04302 | 0.05191 | 0.03299 | 0.02650 | 0.02666 | 0.03354 |
| **HOXA7** | 0.06522 | 0.46258 | 0.12339 | 0.00000 | 1.66719 | 0.00562 | 0.03754 |
| **HOXA9** | 0.02003 | 0.03070 | 0.01267 | 0.02022 | 0.17747 | 0.02217 | 0.01491 |
| **HOXB-AS1** | 0.01784 | 0.02660 | 0.01510 | 0.11441 | 0.02102 | 0.10166 | 0.08009 |
| **HOXB-AS3** | 0.04505 | 0.00000 | 0.04560 | 0.37066 | 0.04180 | 0.00000 | 0.13604 |
| **HOXB1** | 0.04474 | 0.01580 | 0.01112 | 0.04120 | 0.00807 | 0.14612 | 0.00000 |
| **HOXB13** | 0.35282 | 0.00000 | 0.46334 | 0.06049 | 0.92587 | 0.29660 | 0.13619 |
| **HOXB2** | 0.00000 | 0.12348 | 0.10597 | 0.59575 | 0.09206 | 0.41333 | 0.98026 |
| **HOXB3** | 0.15016 | 0.00000 | 0.02746 | 0.16708 | 0.01074 | 0.75489 | 1.19912 |
| **HOXB4** | 0.45980 | 0.00760 | 0.34498 | 0.76701 | 0.11954 | 0.00000 | 2.63590 |
| **HOXB5** | 0.02764 | 0.00000 | 0.16685 | 0.18072 | 0.00730 | 0.00000 | 0.12638 |
| **HOXB6** | 0.06151 | 0.03181 | 0.02893 | 0.04336 | 0.32621 | 0.10680 | 0.02061 |
| **HOXB7** | 1.62707 | 3.01030 | 0.82489 | 0.04710 | 0.29568 | 1.21083 | 0.78670 |

| | | | | | | | |
|---|---|---|---|---|---|---|---|
| **HOXB8** | 0.01404 | 0.01843 | 0.02370 | 0.01883 | 0.03039 | 0.12237 | 0.02001 |
| **HOXB9** | 0.19272 | 0.71324 | 0.74473 | 0.00000 | 0.00931 | 0.00000 | 0.17313 |
| **HOXC-AS1** | 0.03390 | 0.07685 | 0.02501 | 0.05112 | 0.00555 | 0.02230 | 0.12878 |
| **HOXC-AS3** | 0.04751 | 0.01421 | 0.01958 | 0.02491 | 0.01637 | 0.01324 | 0.03428 |
| **HOXC10** | 0.18236 | 0.33719 | 0.35126 | 0.00000 | 1.07858 | 0.00000 | 0.52518 |
| **HOXC11** | 0.02170 | 0.00000 | 0.03260 | 0.00882 | 0.07434 | 0.03600 | 0.08656 |
| **HOXC12** | 0.03929 | 0.03093 | 0.01818 | 0.03601 | 0.00659 | 0.00397 | 0.05006 |
| **HOXC13** | 1.06700 | 0.56020 | 0.06904 | 1.02097 | 0.44161 | 0.00000 | 4.29729 |
| **HOXC13-AS** | 0.25882 | 0.04514 | 0.00163 | 0.54238 | 0.03819 | 0.00000 | 0.79867 |
| **HOXC4** | 0.03896 | 0.09893 | 0.00377 | 0.01747 | 0.02447 | 0.01022 | 0.00661 |
| **HOXC5** | 0.07306 | 0.07949 | 0.04213 | 0.00618 | 0.00472 | 0.10404 | 0.06177 |
| **HOXC6** | 0.02035 | 0.00579 | 0.03706 | 0.01864 | 0.00760 | 0.01547 | 0.06060 |
| **HOXC8** | 0.02616 | 0.03620 | 0.00667 | 0.08876 | 0.01649 | 0.00000 | 0.12693 |
| **HOXC9** | 0.45673 | 1.78612 | 1.52780 | 1.52819 | 0.48235 | 0.00000 | 1.60639 |
| **HOXD1** | 0.00000 | 0.27432 | 0.07866 | 0.28736 | 0.46921 | 0.97993 | 0.00130 |
| **HOXD11** | 0.02585 | 0.01522 | 0.01138 | 0.01720 | 0.02635 | 0.01133 | 0.02960 |
| **HOXD13** | 0.12982 | 0.07233 | 0.08044 | 0.02569 | 0.04181 | 0.78290 | 0.72978 |
| **HOXD3** | 0.02078 | 0.02515 | 0.01165 | 0.02258 | 0.00791 | 0.01771 | 0.03102 |
| **HOXD4** | 0.02101 | 0.01238 | 0.05650 | 0.01745 | 0.05648 | 0.11051 | 0.01503 |
| **HOXD8** | 0.00000 | 0.27704 | 0.14780 | 0.00000 | 0.20148 | 0.04465 | 0.07080 |
| **HOXD9** | 0.00000 | 0.62936 | 0.05496 | 0.00000 | 0.02768 | 0.08424 | 0.16453 |
| **HP** | 0.03497 | 0.01474 | 0.01193 | 0.02492 | 0.02366 | 0.02322 | 0.00739 |
| **HP08942** | 0.03750 | 0.06840 | 0.00845 | 0.04265 | 0.01177 | 0.01386 | 0.00880 |
| **HP09025** | 0.01496 | 0.02917 | 0.01447 | 0.01391 | 0.00821 | 0.00723 | 0.07048 |
| **HP09053** | 0.19746 | 0.19288 | 0.49647 | 0.02946 | 0.18030 | 0.13574 | 0.22488 |
| **HP1BP3** | 11.04051 | 1.96884 | 9.14600 | 9.91917 | 1.27086 | 6.53009 | 17.54382 |
| **HPCA** | 0.02126 | 0.03543 | 0.13622 | 0.02435 | 0.02538 | 0.02389 | 0.04272 |
| **HPCAL1** | 0.58957 | 5.04727 | 0.55744 | 3.54738 | 0.02952 | 1.02001 | 10.14282 |
| **HPCAL4** | 0.03082 | 0.00040 | 0.05136 | 0.04965 | 0.65666 | 0.02016 | 0.00000 |
| **HPD** | 0.01042 | 0.02266 | 0.16644 | 3.83471 | 0.06961 | 0.18835 | 0.00000 |
| **HPDL** | 11.62793 | 20.17157 | 12.49084 | 15.45339 | 0.00000 | 7.22535 | 1.08929 |
| **HPGD** | 23.29065 | 11.99652 | 0.72617 | 4.68151 | 0.00000 | 14.98385 | 4.19754 |
| **HPGDS** | 0.03169 | 0.00256 | 0.03874 | 0.01514 | 0.00880 | 0.01091 | 0.03466 |
| **HPN** | 0.02933 | 0.00693 | 0.02766 | 0.03506 | 0.09725 | 0.16403 | 0.00000 |
| **HPN-AS1** | 0.02652 | 0.00126 | 0.04343 | 0.02802 | 0.11738 | 0.02840 | 0.08258 |

| | | | | | | | |
|---|---|---|---|---|---|---|---|
| **HPR** | 0.08290 | 0.00000 | 0.05019 | 0.01178 | 0.07459 | 0.03278 | 0.09867 |
| **HPRT1** | 15.29510 | 10.08434 | 18.00682 | 0.00000 | 4.56273 | 0.87191 | 5.43638 |
| **HPS1** | 8.38162 | 11.46249 | 2.93685 | 8.14981 | 0.34542 | 4.70077 | 2.89165 |
| **HPS3** | 3.81913 | 1.88936 | 0.21805 | 2.76904 | 1.32762 | 0.88457 | 0.00000 |
| **HPS4** | 4.26477 | 5.70044 | 2.44514 | 3.73296 | 3.38565 | 2.34862 | 1.71946 |
| **HPS5** | 6.03319 | 2.45431 | 3.99372 | 3.16268 | 3.73510 | 1.62367 | 2.31072 |
| **HPS6** | 0.69205 | 4.84986 | 2.31395 | 1.62008 | 0.02717 | 2.13118 | 3.68886 |
| **HPSE** | 0.86526 | 0.00000 | 0.31350 | 0.00000 | 0.35558 | 2.22485 | 0.06741 |
| **HPSE2** | 0.00727 | 0.03925 | 0.03222 | 0.02400 | 1.54886 | 0.07101 | 0.02540 |
| **HPX** | 0.40662 | 0.33645 | 0.08757 | 0.00000 | 0.01706 | 0.32093 | 0.27406 |
| **HR** | 0.03814 | 0.02053 | 0.01630 | 0.06167 | 0.03308 | 0.00000 | 0.22655 |
| **HRAS** | 0.51723 | 2.48919 | 0.00000 | 7.00798 | 0.97163 | 2.47553 | 9.20055 |
| **HRASLS** | 0.02133 | 0.00000 | 0.09859 | 0.02924 | 0.13877 | 0.02846 | 0.09761 |
| **HRASLS2** | 0.07647 | 0.02565 | 0.04098 | 0.05235 | 0.02813 | 0.02130 | 0.00000 |
| **HRASLS5** | 5.00523 | 4.37879 | 20.09314 | 0.00000 | 0.18809 | 0.95473 | 0.49620 |
| **HRC** | 0.09470 | 0.01436 | 0.01316 | 0.03852 | 0.10333 | 0.01942 | 0.00000 |
| **HRCT1** | 0.02245 | 0.01762 | 0.01149 | 0.01893 | 0.01784 | 0.01466 | 0.02067 |
| **HRG** | 0.13548 | 0.31969 | 0.03667 | 0.01050 | 0.58949 | 0.30773 | 0.00000 |
| **HRH1** | 3.05265 | 0.00310 | 0.12405 | 1.19134 | 0.80520 | 0.00109 | 0.00000 |
| **HRH2** | 0.04261 | 0.68707 | 0.00000 | 0.27960 | 3.28636 | 0.08276 | 0.00000 |
| **HRH3** | 0.01695 | 0.05765 | 0.00762 | 0.01540 | 0.01006 | 0.01033 | 0.04592 |
| **HRH4** | 0.02465 | 0.02315 | 0.01537 | 0.01943 | 0.07906 | 0.01131 | 0.02794 |
| **HRK** | 0.00000 | 0.20819 | 0.05601 | 0.10526 | 0.00337 | 0.54548 | 0.23559 |
| **HRNR** | 0.09279 | 0.33183 | 0.67168 | 0.11386 | 0.00048 | 0.06376 | 0.14416 |
| **HRSP12** | 23.29887 | 1.53643 | 3.67651 | 7.04820 | 0.01106 | 7.39682 | 2.48844 |
| **HS1BP3** | 1.19157 | 2.33581 | 1.13045 | 0.82837 | 0.05185 | 0.00000 | 1.33875 |
| **HS1BP3-IT1** | 0.02677 | 0.01591 | 0.01049 | 0.04132 | 0.03708 | 0.01116 | 0.01766 |
| **HS2ST1** | 2.72232 | 0.30930 | 2.29566 | 1.31275 | 0.97549 | 4.05004 | 6.36250 |
| **HS3ST1** | 0.05892 | 0.09098 | 0.05929 | 0.00000 | 0.00429 | 0.80842 | 0.12994 |
| **HS3ST2** | 0.01754 | 0.00000 | 0.07769 | 0.01668 | 0.00823 | 0.22833 | 0.02140 |
| **HS3ST3A1** | 0.30425 | 0.06279 | 0.00651 | 0.41851 | 0.00382 | 1.45027 | 0.50525 |
| **HS3ST3B1** | 0.00000 | 0.03753 | 0.01948 | 0.85133 | 0.03603 | 0.07670 | 0.02068 |
| **HS3ST4** | 0.01081 | 0.00757 | 0.00905 | 0.00557 | 0.00735 | 0.58721 | 0.07939 |
| **HS3ST5** | 0.16456 | 0.19291 | 0.01176 | 0.00289 | 0.01125 | 0.00989 | 0.00557 |
| **HS3ST6** | 0.02227 | 0.00000 | 0.09279 | 0.01759 | 0.17673 | 0.01153 | 0.02344 |

| | | | | | | | |
|---|---|---|---|---|---|---|---|
| **HS6ST1** | 0.00000 | 15.07673 | 1.33096 | 0.00000 | 0.00000 | 6.91052 | 4.25803 |
| **HS6ST2** | 1.07230 | 0.07274 | 0.00000 | 1.22211 | 0.00210 | 0.00000 | 1.36227 |
| **HS6ST3** | 0.00000 | 0.06565 | 0.00571 | 0.02654 | 0.06720 | 0.12010 | 0.08629 |
| **HSBP1** | 39.08499 | 23.22726 | 14.77331 | 30.42915 | 7.35556 | 19.33302 | 13.16210 |
| **HSBP1L1** | 0.35826 | 0.24452 | 0.86056 | 0.00000 | 0.00972 | 0.94567 | 3.35447 |
| **HSCB** | 5.12741 | 2.84881 | 2.60783 | 6.39752 | 1.00831 | 3.45127 | 2.47914 |
| **HSD11B1L** | 0.23078 | 0.00000 | 0.26164 | 0.65561 | 0.47934 | 0.18712 | 1.30036 |
| **HSD11B2** | 0.59173 | 1.02315 | 0.00000 | 2.09482 | 0.04109 | 0.64944 | 0.00000 |
| **HSD17B1** | 3.01774 | 3.90512 | 0.05124 | 7.75694 | 0.00000 | 0.00000 | 20.25239 |
| **HSD17B10** | 39.71614 | 31.73449 | 13.19884 | 40.33820 | 4.78249 | 19.59646 | 5.16030 |
| **HSD17B11** | 12.43353 | 0.00000 | 2.16145 | 13.16431 | 0.00000 | 3.52716 | 0.00000 |
| **HSD17B12** | 23.39831 | 0.00000 | 0.64659 | 13.26077 | 2.26569 | 9.71679 | 8.81001 |
| **HSD17B13** | 0.02577 | 0.01954 | 0.01338 | 0.07079 | 0.00768 | 0.00948 | 0.02105 |
| **HSD17B14** | 0.01418 | 0.12097 | 0.25180 | 0.30731 | 0.74652 | 0.43828 | 0.22781 |
| **HSD17B2** | 0.03877 | 0.00204 | 0.01689 | 0.02049 | 0.00850 | 0.97173 | 0.01333 |
| **HSD17B3** | 0.20519 | 0.73011 | 0.16867 | 0.00000 | 0.01748 | 0.76989 | 0.00400 |
| **HSD17B4** | 19.50062 | 0.31122 | 0.00000 | 13.16261 | 0.89100 | 8.89536 | 4.27764 |
| **HSD17B6** | 0.27955 | 0.15643 | 0.81108 | 0.53278 | 0.04702 | 0.62823 | 0.22009 |
| **HSD17B7** | 7.08375 | 0.59173 | 0.00000 | 2.94994 | 0.14989 | 0.95382 | 0.00000 |
| **HSD17B7P2** | 0.53707 | 0.01847 | 0.00000 | 0.10248 | 0.03997 | 0.03316 | 0.22875 |
| **HSD17B8** | 0.24424 | 0.13816 | 0.24458 | 0.56959 | 0.03844 | 0.30279 | 0.73944 |
| **HSD3B1** | 0.17703 | 0.00000 | 0.00962 | 20.12406 | 0.00536 | 0.00000 | 0.02881 |
| **HSD3B2** | 0.02506 | 0.01342 | 0.01464 | 0.03785 | 0.00799 | 0.03068 | 0.01153 |
| **HSD3B7** | 0.20364 | 0.70763 | 0.41861 | 0.40943 | 0.00000 | 0.01295 | 0.63281 |
| **HSD3BP4** | 0.09004 | 0.03829 | 0.00826 | 0.12701 | 0.00801 | 0.04401 | 0.00000 |
| **HSDL1** | 1.47375 | 0.00000 | 2.50262 | 1.71876 | 1.68662 | 1.38937 | 0.37259 |
| **HSDL2** | 9.26948 | 0.00000 | 0.61064 | 8.66469 | 0.15151 | 4.00158 | 6.20709 |
| **HSF1** | 1.37167 | 9.51016 | 2.93895 | 11.08959 | 0.96340 | 8.02216 | 14.97503 |
| **HSF2** | 7.24189 | 0.00000 | 4.36476 | 4.85731 | 0.98695 | 0.99227 | 0.72662 |
| **HSF2BP** | 0.62640 | 2.80411 | 2.39324 | 0.64569 | 9.25057 | 0.00000 | 0.00000 |
| **HSF4** | 0.02393 | 0.07316 | 0.07870 | 0.13723 | 0.04181 | 0.02712 | 0.30163 |
| **HSF5** | 0.01864 | 0.00000 | 0.32452 | 0.02251 | 0.82181 | 0.13883 | 0.06010 |
| **HSH2D** | 0.94673 | 0.78915 | 0.21172 | 1.96487 | 0.07095 | 1.41989 | 0.80913 |
| **HSP90AA1** | 77.85982 | 32.78452 | 23.52058 | 56.54712 | 17.07306 | 41.45984 | 83.78470 |
| **HSP90AB1** | 88.53044 | 59.49990 | 24.84160 | 65.23424 | 13.21842 | 40.77417 | 50.93437 |

| | | | | | | | |
|---|---|---|---|---|---|---|---|
| **HSP90AB4P** | 0.03530 | 0.13733 | 0.10167 | 0.27429 | 0.49743 | 0.37023 | 1.17212 |
| **HSP90B1** | 48.64810 | 11.38479 | 4.92467 | 25.55340 | 3.11988 | 31.92939 | 31.15820 |
| **HSP90B2P** | 10.90975 | 0.07835 | 0.77211 | 3.74644 | 0.54331 | 7.71921 | 6.53312 |
| **HSPA12A** | 0.07146 | 0.00621 | 0.04441 | 0.01214 | 0.43096 | 0.21185 | 0.30397 |
| **HSPA12B** | 0.00000 | 0.10636 | 0.00000 | 0.13189 | 0.30696 | 0.59190 | 0.27215 |
| **HSPA13** | 10.42275 | 0.00000 | 4.67627 | 9.05834 | 2.72703 | 4.13331 | 0.00000 |
| **HSPA14** | 35.75348 | 12.65371 | 16.82552 | 19.76387 | 9.33571 | 14.96260 | 5.58322 |
| **HSPA1A** | 0.21918 | 1.50702 | 0.00000 | 4.24594 | 2.26256 | 2.83148 | 2.61841 |
| **HSPA1B** | 1.57317 | 0.74858 | 0.11624 | 8.69677 | 1.28047 | 3.54150 | 7.12072 |
| **HSPA1L** | 0.03601 | 0.00000 | 0.18643 | 0.12974 | 0.51643 | 0.13345 | 0.43724 |
| **HSPA2** | 2.94679 | 2.18069 | 2.73867 | 13.50237 | 10.53519 | 1.35712 | 0.00000 |
| **HSPA4** | 46.02429 | 19.76935 | 15.57742 | 27.80929 | 3.11413 | 17.92531 | 13.57447 |
| **HSPA4L** | 4.35811 | 2.61674 | 1.76182 | 0.87832 | 3.65521 | 0.38912 | 0.39817 |
| **HSPA5** | 65.79637 | 40.15069 | 19.11926 | 49.44073 | 28.60484 | 29.58393 | 0.00000 |
| **HSPA6** | 0.00000 | 0.00315 | 0.03444 | 0.85145 | 0.01357 | 0.34138 | 0.01279 |
| **HSPA7** | 0.00000 | 0.01026 | 0.01961 | 0.10076 | 0.01575 | 0.07148 | 0.02600 |
| **HSPA8** | 97.00971 | 67.81649 | 44.24363 | 63.31491 | 26.81323 | 36.89050 | 41.25762 |
| **HSPA9** | 63.25344 | 23.56589 | 24.02411 | 37.99989 | 8.51777 | 28.74955 | 31.31241 |
| **HSPB1** | 0.00000 | 8.41583 | 6.38337 | 4.20620 | 14.34362 | 10.31428 | 0.00000 |
| **HSPB11** | 15.17269 | 2.85305 | 4.79192 | 2.16698 | 4.91286 | 0.00000 | 3.46780 |
| **HSPB2** | 0.02783 | 0.00783 | 0.02928 | 0.02520 | 0.00857 | 0.01822 | 0.02145 |
| **HSPB2-C11orf5** | 0.04219 | 0.02783 | 0.01155 | 0.02125 | 0.00793 | 0.01031 | 0.01360 |
| **HSPB3** | 0.95001 | 0.17986 | 0.00163 | 0.00193 | 0.00583 | 0.15332 | 0.00000 |
| **HSPB6** | 0.00482 | 0.05500 | 0.03104 | 0.02503 | 0.40568 | 0.01065 | 0.05663 |
| **HSPB7** | 0.19412 | 0.06553 | 0.00120 | 0.00000 | 0.00799 | 0.00000 | 0.37455 |
| **HSPB8** | 9.06545 | 0.00000 | 0.00000 | 19.46703 | 0.00086 | 0.00000 | 0.12486 |
| **HSPB9** | 0.19974 | 0.00000 | 0.05846 | 0.27235 | 0.13951 | 0.20674 | 0.19058 |
| **HSPBAP1** | 2.92393 | 0.00000 | 3.20830 | 6.07930 | 0.81707 | 1.61025 | 1.56382 |
| **HSPBP1** | 2.05440 | 20.29239 | 6.83989 | 17.30977 | 3.20835 | 6.19251 | 21.64600 |
| **HSPD1** | 52.17966 | 37.20002 | 6.97875 | 35.97269 | 0.95000 | 27.13208 | 34.65376 |
| **HSPE1** | 23.42182 | 16.63929 | 4.79449 | 22.66689 | 0.61703 | 15.90354 | 30.47656 |
| **HSPG2** | 0.00000 | 0.00000 | 0.02314 | 1.83677 | 0.00766 | 0.19637 | 1.50838 |
| **HSPH1** | 11.54256 | 5.79978 | 4.77924 | 8.81197 | 6.67718 | 12.05139 | 36.14520 |
| **HTATIP2** | 19.01691 | 0.03678 | 0.08316 | 18.75925 | 0.07585 | 1.67920 | 8.41678 |
| **HTATSF1** | 8.22991 | 6.71882 | 6.84757 | 7.91728 | 5.49111 | 6.35058 | 17.08492 |

| | | | | | | | |
|---|---|---|---|---|---|---|---|
| **HTATSF1P2** | 0.19102 | 0.03403 | 0.12320 | 0.22432 | 0.06073 | 0.41425 | 0.52370 |
| **HTR1D** | 0.00000 | 0.02541 | 0.00000 | 0.45487 | 0.16111 | 0.58229 | 0.00092 |
| **HTR2A** | 0.01705 | 0.02712 | 0.00000 | 0.02492 | 0.39647 | 0.01123 | 0.02448 |
| **HTR2A-AS1** | 0.02087 | 0.02405 | 0.00521 | 0.01897 | 0.17968 | 0.01253 | 0.01655 |
| **HTR2B** | 0.16655 | 0.05630 | 0.04560 | 0.25597 | 0.29192 | 0.24280 | 0.50823 |
| **HTR2C** | 0.02609 | 0.29167 | 0.13676 | 0.01635 | 0.08233 | 0.20498 | 0.02062 |
| **HTR3A** | 0.03415 | 0.00700 | 0.02601 | 0.01982 | 0.01297 | 0.06241 | 0.00835 |
| **HTR3B** | 0.18001 | 0.00000 | 0.02088 | 0.00173 | 0.01005 | 0.31542 | 0.00488 |
| **HTR3C** | 0.01815 | 0.08395 | 0.27615 | 0.00156 | 0.27707 | 0.01629 | 0.05848 |
| **HTR3D** | 0.00667 | 0.00000 | 0.24546 | 0.01460 | 0.01024 | 0.01522 | 0.07660 |
| **HTR3E** | 0.03172 | 0.00000 | 0.28469 | 0.01624 | 0.07052 | 0.00651 | 0.01362 |
| **HTR4** | 0.05096 | 0.00841 | 0.01525 | 0.02365 | 0.00818 | 0.05039 | 0.00000 |
| **HTR5A** | 0.00000 | 0.15397 | 0.00916 | 0.01628 | 0.07477 | 0.07494 | 0.01787 |
| **HTR5AOS** | 0.00965 | 0.01665 | 0.04680 | 0.01534 | 0.00609 | 0.19655 | 0.02746 |
| **HTR7** | 0.00243 | 0.00000 | 0.11524 | 0.02891 | 0.05720 | 0.02340 | 0.19121 |
| **HTR7P1** | 0.13082 | 0.00000 | 0.08560 | 0.15402 | 0.17914 | 0.00019 | 0.03723 |
| **HTRA2** | 7.03255 | 10.60245 | 2.06039 | 10.82577 | 1.40188 | 4.18755 | 3.11022 |
| **HTRA3** | 0.01714 | 0.03716 | 0.00669 | 0.03318 | 0.01921 | 0.06200 | 0.07575 |
| **HTRA4** | 0.00000 | 0.01787 | 0.01293 | 0.74767 | 0.01598 | 0.01038 | 0.01117 |
| **HTT** | 3.28968 | 5.39991 | 3.08751 | 1.92310 | 2.59080 | 11.68407 | 13.46824 |
| **HULC** | 0.05531 | 0.01425 | 0.01509 | 0.01683 | 0.00766 | 0.00865 | 0.02561 |
| **HUNK** | 3.13004 | 3.55072 | 1.03322 | 9.09155 | 0.02987 | 0.00000 | 6.46635 |
| **HUS1** | 18.29563 | 9.66096 | 11.16292 | 11.65566 | 1.29389 | 18.92591 | 0.81131 |
| **HUS1B** | 0.11269 | 0.00603 | 0.46146 | 0.10427 | 0.06774 | 0.22233 | 0.09548 |
| **HUWE1** | 36.10203 | 32.38070 | 24.72189 | 24.39422 | 3.46806 | 30.54137 | 58.04097 |
| **HVCN1** | 0.32046 | 0.23059 | 4.11628 | 0.06831 | 1.67205 | 0.36227 | 0.00000 |
| **HYAL1** | 0.19093 | 1.01331 | 1.03328 | 1.57540 | 0.18585 | 1.93418 | 0.09766 |
| **HYAL2** | 1.73119 | 9.78360 | 4.22602 | 3.71740 | 0.58498 | 4.84541 | 9.06752 |
| **HYAL3** | 0.85044 | 0.44965 | 0.97580 | 3.27039 | 0.18496 | 1.23718 | 0.00000 |
| **HYAL4** | 4.44361 | 1.58095 | 0.73468 | 0.34519 | 0.14620 | 3.54893 | 0.00000 |
| **HYALP1** | 0.06354 | 0.11741 | 0.06929 | 0.03703 | 0.03865 | 0.18493 | 0.00000 |
| **HYDIN** | 0.07293 | 0.01950 | 0.50191 | 0.04741 | 0.44227 | 0.24351 | 0.00000 |
| **HYKK** | 0.10271 | 0.81691 | 0.56525 | 0.09217 | 0.18510 | 0.65729 | 0.71383 |
| **HYLS1** | 14.14862 | 4.13398 | 26.22101 | 1.99354 | 6.27533 | 4.24113 | 0.00000 |
| **HYMAI** | 0.02037 | 0.02132 | 0.01812 | 0.01738 | 0.00718 | 0.03493 | 0.02704 |

| | | | | | | | |
|---|---|---|---|---|---|---|---|
| **HYOU1** | 6.66398 | 24.84437 | 12.48644 | 20.57740 | 3.80259 | 24.30973 | 27.37229 |
| **HYPK** | 0.03074 | 0.04160 | 0.04499 | 0.02236 | 0.02094 | 0.02150 | 0.05649 |
| **HYPM** | 0.05258 | 0.04133 | 0.04529 | 0.02164 | 0.10034 | 0.00771 | 0.00000 |
| **IAH1** | 4.60596 | 2.29988 | 0.09295 | 0.56085 | 3.76678 | 1.45240 | 2.67499 |
| **IAPP** | 0.03063 | 0.00738 | 0.03901 | 0.01490 | 0.00685 | 0.02234 | 0.02504 |
| **IARS** | 46.67244 | 10.85244 | 11.42508 | 22.95876 | 1.42949 | 17.44605 | 9.72010 |
| **IARS2** | 24.08972 | 7.64792 | 9.61862 | 14.00008 | 2.16202 | 11.56145 | 14.84755 |
| **IBA57** | 1.54806 | 3.63760 | 0.94957 | 1.59898 | 0.00000 | 0.78418 | 5.38630 |
| **IBA57-AS1** | 0.00000 | 0.01604 | 0.01277 | 0.03659 | 0.00693 | 0.00601 | 0.28066 |
| **IBSP** | 0.02974 | 0.01226 | 0.02674 | 0.01702 | 0.00723 | 0.01289 | 0.01753 |
| **IBTK** | 15.80719 | 2.12550 | 5.01464 | 0.00000 | 0.00000 | 3.59255 | 8.90094 |
| **ICA1** | 0.61018 | 0.47207 | 0.03076 | 0.22205 | 1.50567 | 0.08748 | 0.31543 |
| **ICA1L** | 3.71917 | 1.09874 | 1.34638 | 1.70283 | 1.27657 | 0.13803 | 1.92978 |
| **ICAM1** | 0.05784 | 0.00000 | 0.07042 | 0.27471 | 2.18990 | 0.10929 | 0.29980 |
| **ICAM2** | 0.03803 | 0.05756 | 0.02131 | 0.02559 | 0.09941 | 0.00977 | 0.00002 |
| **ICAM3** | 2.80105 | 0.91757 | 0.77680 | 3.44233 | 1.16089 | 0.33592 | 1.44890 |
| **ICAM4** | 0.11910 | 0.00000 | 0.01677 | 0.05326 | 0.00608 | 0.08532 | 0.12192 |
| **ICAM5** | 0.00804 | 0.00355 | 0.00000 | 0.01386 | 0.81054 | 0.10841 | 0.24428 |
| **ICE1** | 8.59406 | 3.10888 | 6.46440 | 2.48150 | 2.06360 | 5.10057 | 17.69643 |
| **ICE2** | 10.14055 | 2.40068 | 5.78446 | 4.04003 | 4.05835 | 4.38849 | 2.43761 |
| **ICK** | 7.04708 | 2.56792 | 1.97243 | 5.19468 | 0.24351 | 1.58463 | 11.29025 |
| **ICMT** | 21.63820 | 20.14618 | 14.22428 | 13.83890 | 1.22192 | 14.06634 | 21.66089 |
| **ICOSLG** | 0.00000 | 6.11127 | 4.69118 | 1.08834 | 0.00885 | 3.41406 | 2.43642 |
| **ICT1** | 21.00626 | 13.61965 | 12.52217 | 4.18178 | 0.76869 | 10.33498 | 7.37726 |
| **ID1** | 5.15368 | 9.47079 | 8.69457 | 7.87763 | 1.76479 | 21.96444 | 3.01222 |
| **ID2** | 4.89920 | 0.72423 | 0.00000 | 9.96415 | 5.26076 | 5.79177 | 2.59767 |
| **ID3** | 7.01920 | 3.67246 | 0.00000 | 12.47375 | 0.87304 | 14.76624 | 0.00000 |
| **ID4** | 0.05372 | 0.85711 | 0.66887 | 0.39323 | 2.26516 | 1.77639 | 0.72657 |
| **IDE** | 24.86653 | 4.39272 | 5.47443 | 10.17056 | 3.11391 | 4.95646 | 1.74301 |
| **IDH1** | 72.48245 | 45.96841 | 5.76252 | 12.53156 | 2.85562 | 34.25389 | 0.00000 |
| **IDH1-AS1** | 0.17160 | 0.17664 | 0.05797 | 0.05252 | 0.04167 | 0.07991 | 0.02187 |
| **IDH2** | 6.68558 | 1.58199 | 0.32956 | 8.84130 | 6.34196 | 0.31743 | 0.00000 |
| **IDH3A** | 34.30828 | 19.86516 | 6.43220 | 11.21658 | 1.80625 | 9.69984 | 0.00000 |
| **IDH3B** | 31.69207 | 23.91440 | 14.67712 | 27.03335 | 2.07798 | 21.37924 | 12.30533 |
| **IDH3G** | 24.73686 | 18.95414 | 3.33192 | 10.40905 | 3.18545 | 12.36745 | 1.02419 |

| | | | | | | | |
|---|---|---|---|---|---|---|---|
| **IDI1** | 49.08601 | 6.92665 | 6.32023 | 18.70519 | 8.26804 | 10.15654 | 9.29494 |
| **IDI2** | 0.29240 | 0.40501 | 0.29612 | 0.90245 | 0.04573 | 0.00000 | 1.77492 |
| **IDI2-AS1** | 1.45117 | 0.10443 | 0.15230 | 0.73724 | 0.48053 | 0.33020 | 0.00000 |
| **IDNK** | 0.07110 | 0.41849 | 0.27070 | 0.02869 | 0.05914 | 0.00000 | 1.16762 |
| **IDO1** | 0.04407 | 0.01297 | 0.95944 | 0.00247 | 0.01750 | 0.34635 | 0.01638 |
| **IDS** | 0.10586 | 1.08449 | 2.03050 | 0.60482 | 2.66252 | 0.31050 | 0.37868 |
| **IDUA** | 0.02265 | 0.01784 | 0.01228 | 0.02537 | 0.00783 | 0.01515 | 0.02968 |
| **IER2** | 1.23661 | 28.78017 | 6.19159 | 19.66245 | 1.18248 | 18.89352 | 27.31303 |
| **IER3** | 0.00000 | 0.52133 | 10.87640 | 6.12029 | 9.48980 | 4.91781 | 4.76370 |
| **IER3IP1** | 19.42872 | 4.00103 | 14.93241 | 17.49420 | 7.42609 | 7.58679 | 9.26012 |
| **IER5** | 0.00000 | 0.00000 | 8.19241 | 0.87758 | 2.82500 | 4.46300 | 1.03818 |
| **IER5L** | 0.00000 | 0.03313 | 0.00931 | 0.04955 | 0.10371 | 0.04969 | 0.89138 |
| **IFFO1** | 0.19516 | 0.11360 | 0.46730 | 0.18009 | 0.46685 | 0.72353 | 0.20134 |
| **IFFO2** | 0.00000 | 0.43621 | 0.15662 | 0.49941 | 0.30816 | 0.05742 | 0.01421 |
| **IFI16** | 0.00000 | 2.00090 | 0.17329 | 0.06229 | 0.05443 | 22.16554 | 0.03241 |
| **IFI27** | 0.08390 | 0.44201 | 0.00000 | 0.21863 | 0.12134 | 0.52960 | 0.21544 |
| **IFI27L1** | 1.27239 | 1.19728 | 0.24742 | 2.51916 | 1.51634 | 0.82914 | 1.23693 |
| **IFI27L2** | 0.01291 | 0.02881 | 0.03493 | 0.20027 | 0.00692 | 0.02911 | 0.01759 |
| **IFI30** | 20.04313 | 32.44846 | 16.86360 | 11.53696 | 4.51764 | 5.16409 | 0.00000 |
| **IFI35** | 0.15439 | 0.03502 | 0.00831 | 0.16862 | 0.11142 | 0.17359 | 0.00000 |
| **IFI44L** | 0.02070 | 0.01542 | 0.01463 | 0.03380 | 0.00781 | 0.01650 | 0.02103 |
| **IFI6** | 0.28459 | 0.14404 | 0.04180 | 6.70227 | 0.11641 | 0.41901 | 0.43250 |
| **IFIH1** | 0.02009 | 0.02839 | 0.03356 | 0.02120 | 0.00630 | 0.01269 | 0.01574 |
| **IFIT1** | 0.03859 | 0.01942 | 0.01255 | 0.01032 | 0.00784 | 0.00902 | 0.03244 |
| **IFIT3** | 0.02637 | 0.01742 | 0.01319 | 0.02996 | 0.00831 | 0.01251 | 0.02079 |
| **IFIT5** | 0.02581 | 0.00000 | 0.44269 | 0.15226 | 0.15854 | 0.03349 | 0.00000 |
| **IFITM1** | 10.93654 | 1.99907 | 0.00000 | 19.51987 | 0.05532 | 52.44634 | 0.00018 |
| **IFITM10** | 0.00000 | 0.20397 | 0.02289 | 0.11266 | 0.00396 | 0.10063 | 0.60189 |
| **IFITM2** | 7.51767 | 16.99865 | 0.00000 | 13.67576 | 0.00000 | 6.42209 | 7.02485 |
| **IFITM3** | 21.63589 | 7.10552 | 0.00000 | 19.64043 | 0.00000 | 34.16685 | 12.61543 |
| **IFITM4P** | 0.02534 | 0.01873 | 0.01251 | 0.03350 | 0.00809 | 0.02454 | 0.01185 |
| **IFITM5** | 0.01892 | 0.01323 | 0.01255 | 0.03197 | 0.00806 | 0.11529 | 0.01443 |
| **IFNAR1** | 11.35821 | 1.85140 | 4.47803 | 0.84124 | 0.21590 | 0.99389 | 0.93008 |
| **IFNAR2** | 4.29757 | 0.94983 | 0.69821 | 0.73089 | 2.61939 | 0.55589 | 1.28194 |
| **IFNB1** | 0.02818 | 0.00952 | 0.01085 | 0.01740 | 0.03487 | 0.01005 | 0.04787 |

| | | | | | | | |
|---|---|---|---|---|---|---|---|
| **IFNGR1** | 12.94597 | 0.60533 | 7.41396 | 12.26655 | 2.69730 | 4.29642 | 0.00000 |
| **IFNGR2** | 0.33013 | 0.89109 | 0.48785 | 0.17699 | 1.69077 | 0.76907 | 0.96579 |
| **IFNK** | 0.01572 | 0.01570 | 0.02072 | 0.02231 | 0.00652 | 0.01393 | 0.05996 |
| **IFNL1** | 0.03480 | 0.01488 | 0.01723 | 0.06924 | 0.00791 | 0.01146 | 0.00593 |
| **IFNL2** | 0.02335 | 0.01776 | 0.01293 | 0.04078 | 0.00796 | 0.01225 | 0.01260 |
| **IFNL3** | 0.02291 | 0.02251 | 0.01223 | 0.05803 | 0.00809 | 0.01265 | 0.00598 |
| **IFNL4** | 0.03251 | 0.01961 | 0.01205 | 0.02526 | 0.00788 | 0.01149 | 0.01654 |
| **IFNLR1** | 0.09294 | 0.24211 | 0.95032 | 0.06138 | 0.00972 | 0.38038 | 0.70428 |
| **IFNW1** | 0.02230 | 0.01611 | 0.01284 | 0.01914 | 0.01903 | 0.01292 | 0.02163 |
| **IFRD1** | 16.36731 | 5.06232 | 11.17506 | 26.60058 | 7.98314 | 8.32693 | 3.50919 |
| **IFRD2** | 27.90100 | 19.34303 | 8.08769 | 23.61407 | 1.86201 | 7.75965 | 0.00000 |
| **IFT122** | 1.80417 | 0.55642 | 2.38688 | 1.60773 | 0.83621 | 2.41573 | 0.54685 |
| **IFT140** | 0.19872 | 2.71185 | 0.66658 | 0.70350 | 0.42051 | 1.55126 | 0.80261 |
| **IFT172** | 1.92647 | 2.13941 | 3.54421 | 1.32196 | 1.90959 | 7.93855 | 0.01954 |
| **IFT20** | 4.93879 | 3.59460 | 5.72040 | 6.76876 | 3.27790 | 2.57594 | 0.00000 |
| **IFT22** | 4.69668 | 7.38054 | 12.98581 | 3.36810 | 5.36416 | 5.61415 | 0.57699 |
| **IFT27** | 0.78036 | 1.06141 | 0.00050 | 1.28739 | 0.86009 | 1.49031 | 0.00000 |
| **IFT43** | 0.75021 | 4.05673 | 0.76041 | 2.19716 | 0.00000 | 2.90049 | 0.27164 |
| **IFT46** | 2.32733 | 1.87967 | 4.99082 | 0.00000 | 3.01336 | 1.01979 | 0.00000 |
| **IFT52** | 4.56291 | 2.08361 | 3.46617 | 3.76649 | 1.52616 | 4.35253 | 0.86097 |
| **IFT57** | 3.24226 | 1.46807 | 0.85216 | 3.45238 | 2.35629 | 3.70139 | 0.00000 |
| **IFT74** | 1.59084 | 0.00000 | 0.37567 | 2.40048 | 0.37492 | 0.55802 | 2.04275 |
| **IFT74-AS1** | 0.03122 | 0.01594 | 0.01178 | 0.01927 | 0.03409 | 0.01969 | 0.01051 |
| **IFT80** | 2.92256 | 0.00000 | 1.70049 | 1.38587 | 1.62511 | 1.39369 | 0.00000 |
| **IFT81** | 0.55379 | 0.00000 | 1.13430 | 0.25752 | 0.87297 | 0.67802 | 0.92140 |
| **IFT88** | 0.97425 | 0.00000 | 2.07376 | 0.82809 | 0.35520 | 0.88980 | 2.46637 |
| **IGBP1** | 15.49245 | 9.65330 | 11.15822 | 12.24645 | 2.72439 | 11.23269 | 9.97101 |
| **IGBP1P1** | 0.74392 | 0.23613 | 0.52881 | 0.56701 | 0.21855 | 0.41336 | 0.23244 |
| **IGDCC3** | 1.74876 | 0.00000 | 0.01761 | 0.54474 | 0.97524 | 0.00000 | 5.76219 |
| **IGDCC4** | 0.26345 | 0.05876 | 0.01057 | 0.03040 | 0.07987 | 0.00751 | 0.01307 |
| **IGF1** | 0.22260 | 0.00000 | 0.09713 | 0.21493 | 0.00316 | 3.15233 | 0.42962 |
| **IGF1R** | 1.65868 | 6.77548 | 4.08759 | 1.34443 | 5.13829 | 2.92702 | 1.34478 |
| **IGF2.uid-15072** | 0.01779 | 0.03651 | 0.01323 | 0.02451 | 0.04081 | 0.01246 | 0.01946 |
| **IGF2.uid-15081** | 0.02242 | 0.01853 | 0.01216 | 0.02605 | 0.01343 | 0.01280 | 0.01606 |
| **IGF2BP1** | 41.12502 | 31.45553 | 21.44990 | 33.21888 | 3.09681 | 15.71800 | 31.98714 |

| | | | | | | | |
|---|---|---|---|---|---|---|---|
| **IGF2BP2** | 0.08620 | 5.17153 | 2.16019 | 4.78791 | 0.89869 | 5.04524 | 12.50805 |
| **IGF2BP3** | 7.04006 | 0.00000 | 1.19243 | 11.42730 | 1.20518 | 2.70624 | 12.63982 |
| **IGF2R** | 0.79520 | 1.53260 | 2.52132 | 1.98071 | 1.44059 | 4.34569 | 14.96848 |
| **IGFALS** | 0.02371 | 0.01665 | 0.01330 | 0.02580 | 0.05893 | 0.01259 | 0.02043 |
| **IGFBP1** | 0.01533 | 0.02524 | 0.00000 | 0.02280 | 0.96785 | 0.01721 | 0.01384 |
| **IGFBP2** | 0.08252 | 0.12175 | 0.09730 | 2.13009 | 1.45206 | 5.30650 | 2.88553 |
| **IGFBP3** | 0.03020 | 0.04028 | 0.09184 | 0.00000 | 4.80605 | 0.58799 | 0.10008 |
| **IGFBP4** | 0.00631 | 0.01061 | 0.01565 | 0.17481 | 0.00718 | 0.02336 | 0.05509 |
| **IGFBP5** | 0.02808 | 0.17580 | 0.01066 | 0.00197 | 0.08090 | 0.03443 | 0.08438 |
| **IGFBP6** | 0.20564 | 0.00000 | 0.03732 | 0.23512 | 1.83710 | 0.23276 | 1.44384 |
| **IGFBP7** | 0.04359 | 0.00000 | 0.02858 | 0.02448 | 0.02058 | 0.02810 | 0.16570 |
| **IGFBP7-AS1** | 0.02241 | 0.03842 | 0.33234 | 0.01014 | 0.01006 | 0.35095 | 0.01144 |
| **IGFBPL1** | 0.00000 | 1.03738 | 0.11517 | 0.12728 | 2.68755 | 0.71887 | 0.00000 |
| **IGFL2** | 0.00984 | 0.01656 | 0.10625 | 0.04489 | 0.14108 | 0.01276 | 0.01827 |
| **IGFL3** | 0.02557 | 0.01251 | 0.02041 | 0.01875 | 0.01542 | 0.01346 | 0.02014 |
| **IGFL4** | 0.03130 | 0.00000 | 0.07213 | 0.03458 | 0.00707 | 0.01132 | 0.02417 |
| **IGFLR1** | 0.50645 | 0.63053 | 0.19005 | 1.12111 | 0.22760 | 1.85392 | 0.00000 |
| **IGFN1** | 0.01880 | 0.01772 | 0.01090 | 0.02489 | 0.01125 | 0.07022 | 0.01819 |
| **IGHMBP2** | 2.23446 | 7.59813 | 5.65285 | 7.17476 | 2.19673 | 4.35023 | 5.32751 |
| **IGIP** | 0.23427 | 0.04906 | 0.05147 | 0.08415 | 0.00570 | 0.00000 | 0.61174 |
| **IGJ** | 0.02479 | 0.02063 | 0.01065 | 0.01959 | 0.11959 | 0.01223 | 0.01788 |
| **IGLL1** | 0.08758 | 0.01726 | 0.02075 | 0.16781 | 0.00741 | 0.01852 | 0.00225 |
| **IGLL3P** | 0.00000 | 0.20434 | 0.00000 | 0.08748 | 0.91208 | 0.14748 | 0.11659 |
| **IGLON5** | 0.01299 | 0.01322 | 0.01590 | 0.02217 | 0.01073 | 0.00890 | 0.06671 |
| **IGSF1** | 0.11925 | 0.00000 | 0.33018 | 0.00000 | 0.16088 | 6.38529 | 0.08764 |
| **IGSF10** | 0.16416 | 0.06317 | 0.10199 | 0.16986 | 3.09720 | 0.08810 | 0.21256 |
| **IGSF11** | 0.10772 | 0.00000 | 0.11327 | 0.03534 | 0.24907 | 0.13940 | 0.00000 |
| **IGSF22** | 0.04842 | 0.00000 | 0.14907 | 0.03730 | 0.01850 | 0.07420 | 0.01473 |
| **IGSF23** | 0.05101 | 0.00402 | 0.05200 | 0.02451 | 0.02415 | 0.06038 | 0.00000 |
| **IGSF3** | 4.51455 | 1.62852 | 0.00000 | 4.90887 | 0.07256 | 9.00167 | 7.46839 |
| **IGSF5** | 0.02267 | 0.01378 | 0.03047 | 0.03139 | 0.11789 | 0.02167 | 0.02788 |
| **IGSF6** | 0.01772 | 0.02068 | 0.04902 | 0.03749 | 0.02187 | 0.03001 | 0.02929 |
| **IGSF8** | 0.00000 | 0.75372 | 0.52340 | 3.83724 | 0.05160 | 2.37376 | 1.31465 |
| **IGSF9** | 0.28167 | 0.92131 | 1.72650 | 2.80072 | 0.85197 | 0.00000 | 4.18610 |
| **IGSF9B** | 0.00000 | 0.13061 | 0.05822 | 0.10591 | 0.02430 | 0.17537 | 0.32423 |

| | | | | | | | |
|---|---|---|---|---|---|---|---|
| **IHH** | 0.00000 | 0.02200 | 0.00734 | 0.03682 | 0.00719 | 0.14634 | 0.18649 |
| **IK** | 24.10274 | 13.98401 | 7.38425 | 14.31720 | 5.83612 | 10.46119 | 4.92628 |
| **IKBIP** | 0.13424 | 0.00000 | 0.71571 | 0.52850 | 0.55166 | 1.37908 | 0.80489 |
| **IKBKAP** | 14.50974 | 4.82854 | 5.22179 | 12.83595 | 0.91519 | 7.44479 | 4.33235 |
| **IKBKB** | 1.60627 | 4.05176 | 1.97233 | 3.77663 | 0.84350 | 2.19719 | 4.63311 |
| **IKBKE** | 0.02601 | 0.08006 | 0.02706 | 0.02918 | 0.20327 | 0.02009 | 0.00756 |
| **IKBKG** | 0.63206 | 1.91690 | 0.46493 | 1.04417 | 0.30164 | 0.68882 | 0.00000 |
| **IKZF1** | 0.02458 | 0.01576 | 0.01164 | 0.01200 | 0.00975 | 0.01476 | 0.04504 |
| **IKZF2** | 0.15125 | 0.10644 | 0.00440 | 0.00000 | 0.00778 | 0.00000 | 0.13350 |
| **IKZF3** | 0.01270 | 0.00000 | 0.31391 | 0.03164 | 0.12674 | 0.23062 | 0.08334 |
| **IKZF4** | 0.48435 | 0.11946 | 0.68883 | 1.41562 | 0.45924 | 1.40909 | 1.84127 |
| **IKZF5** | 10.18173 | 5.44747 | 4.43818 | 9.10589 | 6.75112 | 2.70782 | 0.00000 |
| **IL10** | 0.01823 | 0.02201 | 0.01375 | 0.01743 | 0.00756 | 0.01702 | 0.03025 |
| **IL10RA** | 0.01641 | 0.04392 | 0.00951 | 0.05790 | 0.00872 | 0.01004 | 0.01810 |
| **IL10RB** | 6.58321 | 2.40372 | 3.11118 | 1.80382 | 0.16871 | 0.50707 | 0.00000 |
| **IL10RB-AS1** | 0.09202 | 0.00002 | 0.04104 | 0.04731 | 0.13541 | 0.00000 | 0.07273 |
| **IL11** | 0.02546 | 0.01463 | 0.00898 | 0.01938 | 0.07284 | 0.01196 | 0.04419 |
| **IL11RA** | 0.33881 | 0.51592 | 0.01071 | 3.92179 | 0.07043 | 0.94726 | 2.99510 |
| **IL12A** | 0.09929 | 0.02851 | 0.02658 | 0.18334 | 0.02583 | 0.02246 | 0.01117 |
| **IL12A-AS1** | 0.01844 | 0.02690 | 0.01073 | 0.03136 | 0.00979 | 0.02369 | 0.01828 |
| **IL12B** | 0.02468 | 0.01631 | 0.00827 | 0.02235 | 0.07503 | 0.01612 | 0.01856 |
| **IL12RB1** | 0.02060 | 0.01079 | 0.02996 | 0.03587 | 0.00752 | 0.01237 | 0.02591 |
| **IL12RB2** | 0.06241 | 0.00000 | 0.05135 | 0.03422 | 0.04290 | 0.01742 | 0.21971 |
| **IL13RA1** | 0.18566 | 0.00000 | 0.60289 | 0.04711 | 1.85506 | 0.83494 | 0.56472 |
| **IL13RA2** | 0.00000 | 0.00000 | 2.51285 | 0.00641 | 0.36661 | 0.23208 | 0.03203 |
| **IL15** | 0.01998 | 0.24267 | 0.04045 | 0.03460 | 0.01241 | 2.23984 | 0.00000 |
| **IL15RA** | 0.01595 | 0.01292 | 0.01438 | 0.03863 | 0.02971 | 0.01572 | 0.02903 |
| **IL16** | 0.23673 | 0.05591 | 0.07729 | 0.03838 | 1.49933 | 0.15262 | 0.21159 |
| **IL17B** | 0.06057 | 0.26819 | 3.72917 | 0.00000 | 9.53475 | 0.00000 | 0.00000 |
| **IL17C** | 0.01184 | 0.01111 | 0.00740 | 0.02744 | 0.07457 | 0.11792 | 0.02416 |
| **IL17D** | 0.06505 | 0.13616 | 0.07032 | 0.62410 | 0.02077 | 0.00000 | 0.38557 |
| **IL17F** | 0.02225 | 0.01809 | 0.05239 | 0.06580 | 0.04536 | 0.02737 | 0.00572 |
| **IL17RA** | 0.00010 | 1.75369 | 0.86151 | 0.88019 | 0.84111 | 0.97647 | 3.63443 |
| **IL17RB** | 0.61221 | 0.54414 | 0.16656 | 0.67238 | 0.05902 | 2.39638 | 0.43659 |
| **IL17RC** | 0.00000 | 0.22559 | 0.47227 | 0.13929 | 0.06721 | 0.51190 | 0.32155 |

| | | | | | | | |
|---|---|---|---|---|---|---|---|
| **IL17RD** | 0.06225 | 0.00000 | 0.59489 | 0.42540 | 1.00468 | 2.77656 | 0.21435 |
| **IL17RE** | 0.30196 | 0.00000 | 0.01421 | 0.88315 | 0.02294 | 0.00000 | 0.46249 |
| **IL17REL** | 0.02069 | 0.02497 | 0.01232 | 0.02414 | 0.00809 | 0.01920 | 0.01553 |
| **IL18** | 0.14227 | 0.01909 | 0.04368 | 0.01657 | 0.05334 | 0.06020 | 0.11182 |
| **IL18BP** | 0.06526 | 0.00000 | 0.45123 | 1.41986 | 0.17779 | 0.85349 | 2.49487 |
| **IL18R1** | 0.06257 | 0.02645 | 0.01652 | 0.02141 | 0.00746 | 0.01954 | 0.00008 |
| **IL18RAP** | 0.07804 | 0.01184 | 0.01305 | 0.01428 | 0.00774 | 0.01131 | 0.01198 |
| **IL19** | 0.02233 | 0.01762 | 0.01271 | 0.01970 | 0.00795 | 0.02979 | 0.01861 |
| **IL1A** | 0.35411 | 0.00000 | 0.42705 | 0.01833 | 0.02944 | 0.03498 | 0.00000 |
| **IL1B** | 0.02894 | 0.02023 | 0.01585 | 0.02051 | 0.09405 | 0.03499 | 0.01138 |
| **IL1R1** | 0.00000 | 0.00000 | 0.05867 | 5.80304 | 0.05085 | 0.01228 | 0.12656 |
| **IL1R2** | 0.01066 | 0.01824 | 0.01202 | 0.16555 | 0.00816 | 0.09205 | 0.00263 |
| **IL1RAP** | 0.00000 | 0.00000 | 1.01634 | 0.44105 | 0.76407 | 1.68339 | 0.03365 |
| **IL1RAPL1** | 0.04972 | 0.01788 | 0.00429 | 0.02343 | 0.59160 | 0.02316 | 0.02323 |
| **IL1RAPL2** | 0.02608 | 0.02112 | 0.03240 | 0.01707 | 0.20700 | 0.01542 | 0.01469 |
| **IL1RL1** | 0.13295 | 0.00698 | 0.01264 | 0.00158 | 0.02945 | 0.08424 | 0.00684 |
| **IL1RL2** | 0.02099 | 0.00000 | 0.04262 | 0.01929 | 0.02201 | 0.00876 | 0.07221 |
| **IL1RN** | 0.00000 | 0.15398 | 0.11588 | 0.05453 | 0.38346 | 0.00272 | 0.00895 |
| **IL20RA** | 0.01711 | 0.01799 | 0.01322 | 0.01534 | 0.06974 | 0.01356 | 0.04443 |
| **IL20RB** | 0.67490 | 0.22559 | 0.37169 | 2.12756 | 0.35717 | 0.32063 | 0.00000 |
| **IL21-AS1** | 0.02968 | 0.00000 | 0.06185 | 0.04262 | 0.06281 | 0.07489 | 0.00399 |
| **IL21R** | 0.06963 | 0.00000 | 0.02504 | 0.19345 | 0.00571 | 0.00000 | 0.20354 |
| **IL21R-AS1** | 0.04698 | 0.01436 | 0.02787 | 0.01457 | 0.00660 | 0.00847 | 0.03557 |
| **IL22RA1** | 0.00000 | 0.22488 | 0.58281 | 0.00459 | 0.00094 | 0.43191 | 0.00992 |
| **IL23A** | 0.07979 | 0.00000 | 0.04739 | 0.02276 | 0.57076 | 1.71856 | 0.00000 |
| **IL23R** | 0.05077 | 0.00000 | 0.26150 | 0.01585 | 1.92532 | 0.00908 | 0.00950 |
| **IL24** | 0.02985 | 0.01423 | 0.01742 | 0.01733 | 0.00752 | 0.01336 | 0.01828 |
| **IL27** | 0.01768 | 0.03083 | 0.05078 | 0.10802 | 0.00982 | 0.01256 | 0.05825 |
| **IL27RA** | 0.13873 | 0.00000 | 0.06264 | 0.02847 | 0.01181 | 0.08957 | 0.00000 |
| **IL2RA** | 0.02685 | 0.01470 | 0.01863 | 0.01374 | 0.42825 | 0.01324 | 0.03523 |
| **IL2RB** | 0.52547 | 0.08499 | 0.00493 | 1.38267 | 0.03099 | 0.00000 | 2.00845 |
| **IL2RG** | 0.08080 | 0.70602 | 0.86737 | 0.00000 | 0.32163 | 0.14234 | 0.00000 |
| **IL3** | 0.03866 | 0.03200 | 0.02982 | 0.01441 | 0.00637 | 0.01013 | 0.01384 |
| **IL31RA** | 0.03069 | 0.16230 | 0.01287 | 0.03492 | 0.15754 | 0.04752 | 0.00000 |
| **IL32** | 4.48563 | 0.00000 | 0.00442 | 1.70074 | 0.00243 | 3.27731 | 0.00000 |

| | | | | | | | |
|---|---|---|---|---|---|---|---|
| **IL34** | 4.56100 | 8.48806 | 1.94608 | 1.71720 | 0.07524 | 1.67124 | 0.60748 |
| **IL36B** | 0.06561 | 0.01138 | 0.01339 | 0.02404 | 0.00774 | 0.01156 | 0.01329 |
| **IL36G** | 0.02909 | 0.06643 | 0.00543 | 0.02227 | 0.02639 | 0.01593 | 0.00868 |
| **IL36RN** | 0.00574 | 0.11653 | 0.00497 | 1.31175 | 0.00802 | 0.00396 | 0.00000 |
| **IL37** | 0.05879 | 0.02599 | 0.02818 | 0.00623 | 0.01228 | 0.02593 | 0.05525 |
| **IL4** | 0.02560 | 0.01627 | 0.00775 | 0.02046 | 0.11780 | 0.01454 | 0.01396 |
| **IL4I1** | 0.07204 | 0.07839 | 0.24466 | 0.12785 | 0.28093 | 0.07813 | 0.00000 |
| **IL4R** | 0.00000 | 0.58028 | 0.27412 | 0.21311 | 0.03759 | 4.65499 | 0.70741 |
| **IL5** | 0.02167 | 0.01570 | 0.03424 | 0.01793 | 0.01179 | 0.02633 | 0.02005 |
| **IL5RA** | 0.01877 | 0.04528 | 0.00244 | 0.01743 | 0.17121 | 0.01242 | 0.03362 |
| **IL6** | 0.41713 | 3.15308 | 0.65599 | 5.92303 | 0.00000 | 0.41410 | 0.73083 |
| **IL6R** | 0.00000 | 2.38108 | 4.00676 | 0.18167 | 1.33080 | 34.42829 | 0.00000 |
| **IL6ST** | 5.45353 | 2.33860 | 2.62317 | 6.40202 | 0.70440 | 15.14598 | 8.33833 |
| **IL7** | 0.03350 | 0.01469 | 0.01557 | 0.01705 | 0.01786 | 0.01266 | 0.01401 |
| **IL7R** | 0.01740 | 0.01517 | 0.01299 | 0.02726 | 0.00779 | 0.04768 | 0.02403 |
| **ILDR1** | 0.02657 | 0.01701 | 0.01839 | 0.01351 | 0.00699 | 0.01604 | 0.06392 |
| **ILDR2** | 0.00000 | 0.02250 | 0.02979 | 0.00761 | 1.39390 | 0.26615 | 0.10659 |
| **ILF2** | 66.79754 | 31.17065 | 28.76804 | 45.33678 | 21.46231 | 25.45114 | 10.20602 |
| **ILF3** | 34.22612 | 33.24507 | 25.68557 | 36.89524 | 9.14019 | 21.49160 | 46.35521 |
| **ILF3-AS1** | 0.15320 | 2.77512 | 2.93221 | 2.48936 | 0.93299 | 0.78082 | 2.83309 |
| **ILK** | 14.47699 | 12.47765 | 2.85524 | 10.81319 | 3.16370 | 11.34056 | 5.07000 |
| **ILKAP** | 10.64050 | 10.38296 | 6.41483 | 9.64633 | 2.29629 | 4.45650 | 9.00046 |
| **ILVBL** | 8.01985 | 4.73536 | 1.69031 | 10.34932 | 0.00000 | 2.63158 | 4.55922 |
| **IMMP1L** | 3.92228 | 1.35481 | 3.38752 | 4.14810 | 1.19074 | 1.00730 | 1.29942 |
| **IMMP2L** | 1.85322 | 0.65398 | 0.95116 | 0.95531 | 0.15456 | 0.49452 | 1.04919 |
| **IMMT** | 35.50850 | 16.72998 | 14.01198 | 22.93213 | 8.17640 | 15.22753 | 12.49494 |
| **IMP3** | 38.00178 | 22.63225 | 23.97393 | 28.31439 | 18.34506 | 15.59708 | 18.10436 |
| **IMP4** | 38.70459 | 37.96898 | 19.53358 | 29.07163 | 13.03774 | 15.97201 | 5.39376 |
| **IMPA1** | 7.32462 | 0.00000 | 5.18091 | 2.77769 | 3.47466 | 3.65974 | 2.16959 |
| **IMPA2** | 0.83250 | 5.09359 | 1.60039 | 14.82525 | 0.37742 | 4.67696 | 18.33257 |
| **IMPACT** | 7.53740 | 1.98580 | 4.93668 | 1.46991 | 0.80044 | 1.79003 | 3.55593 |
| **IMPAD1** | 10.02677 | 1.63518 | 5.96485 | 3.54492 | 1.07551 | 13.39637 | 21.74087 |
| **IMPDH1** | 13.88013 | 33.51940 | 22.51268 | 13.83966 | 5.13008 | 1.13579 | 10.37306 |
| **IMPDH2** | 61.14878 | 32.83966 | 16.47468 | 47.73935 | 8.81880 | 20.18934 | 25.61019 |
| **IMPG1** | 0.11738 | 0.00000 | 0.01719 | 0.04313 | 0.19060 | 0.01657 | 0.02558 |

| | | | | | | | |
|---|---|---|---|---|---|---|---|
| **IMPG2** | 0.02804 | 0.01487 | 0.03161 | 0.02075 | 0.05469 | 0.01262 | 0.04618 |
| **INA** | 10.87831 | 0.00000 | 0.00000 | 5.01321 | 0.21919 | 0.00000 | 16.14975 |
| **INADL** | 8.00915 | 5.32165 | 7.36549 | 6.09960 | 0.36671 | 2.33154 | 8.93371 |
| **INCA1** | 0.22562 | 0.08582 | 0.10196 | 0.33139 | 0.14847 | 0.13823 | 0.15133 |
| **INCENP** | 0.04793 | 0.16343 | 1.86274 | 3.10118 | 1.84934 | 1.03888 | 10.53498 |
| **INE1** | 0.30988 | 0.23028 | 0.35400 | 0.09479 | 0.03339 | 0.24926 | 0.10795 |
| **INF2** | 2.52804 | 7.03160 | 0.00000 | 10.52622 | 0.00696 | 0.45576 | 13.54622 |
| **ING1** | 3.76857 | 4.00950 | 0.87552 | 4.03508 | 5.26171 | 2.92739 | 4.64059 |
| **ING2** | 1.43835 | 1.27281 | 7.25989 | 1.33001 | 2.58291 | 2.88521 | 8.38478 |
| **ING3** | 10.52316 | 6.53076 | 7.61964 | 5.02527 | 10.47354 | 4.58937 | 0.00000 |
| **ING4** | 1.33382 | 0.00000 | 2.25704 | 2.01231 | 0.25185 | 0.67568 | 0.20660 |
| **ING5** | 3.04432 | 2.39599 | 1.13373 | 4.02868 | 0.33153 | 1.54769 | 6.89650 |
| **INGX** | 0.01328 | 0.04255 | 0.04948 | 0.02431 | 0.05510 | 0.01551 | 0.02661 |
| **INHA** | 0.01546 | 0.02155 | 0.01202 | 0.17299 | 0.00734 | 0.01485 | 0.04653 |
| **INHBA** | 0.02201 | 0.01414 | 0.01372 | 0.01962 | 0.00754 | 0.01073 | 0.03729 |
| **INHBA-AS1** | 0.02252 | 0.01510 | 0.01607 | 0.02117 | 0.00804 | 0.01762 | 0.01840 |
| **INHBB** | 0.03164 | 0.02229 | 0.00986 | 0.01926 | 0.02025 | 0.02206 | 0.01140 |
| **INHBC** | 0.02349 | 0.01778 | 0.02681 | 0.03021 | 0.02929 | 0.01218 | 0.01459 |
| **INHBE** | 0.07984 | 0.00000 | 0.03182 | 0.05194 | 0.00664 | 0.02297 | 0.02661 |
| **INIP** | 9.64747 | 2.85447 | 5.79571 | 5.60465 | 3.23146 | 2.53385 | 0.32283 |
| **INMT** | 0.13254 | 0.06790 | 0.05911 | 0.22731 | 0.01477 | 0.00000 | 0.10833 |
| **INO80** | 5.91634 | 5.01630 | 4.26499 | 7.08756 | 3.18849 | 4.20332 | 11.19614 |
| **INO80B** | 0.00000 | 0.02887 | 0.25294 | 0.11200 | 0.03835 | 0.04643 | 0.69103 |
| **INO80C** | 7.51620 | 4.95667 | 12.25687 | 3.98632 | 5.22716 | 1.72807 | 2.12901 |
| **INO80D** | 5.55408 | 0.32205 | 7.16305 | 3.59701 | 2.93459 | 6.75139 | 10.25685 |
| **INO80E** | 0.03536 | 2.04154 | 0.72435 | 2.52265 | 0.05369 | 1.78734 | 4.13242 |
| **INPP1** | 0.00000 | 0.03848 | 0.04540 | 0.08649 | 0.27275 | 0.04917 | 0.04810 |
| **INPP4A** | 2.06973 | 1.63220 | 0.44513 | 2.21269 | 0.36672 | 2.47654 | 2.76802 |
| **INPP4B** | 0.00000 | 0.00000 | 0.06500 | 0.05815 | 0.27857 | 0.07432 | 0.09443 |
| **INPP5A** | 2.19435 | 4.26513 | 0.68064 | 0.00000 | 0.31744 | 1.11306 | 3.30939 |
| **INPP5B** | 3.80707 | 3.10841 | 3.97028 | 3.04332 | 0.62795 | 3.17057 | 2.04628 |
| **INPP5D** | 2.37564 | 13.42355 | 2.05594 | 0.00000 | 0.02566 | 8.81311 | 2.70636 |
| **INPP5E** | 0.19839 | 1.38172 | 0.49670 | 0.25938 | 0.07915 | 2.38057 | 6.06299 |
| **INPP5F** | 5.60256 | 6.75721 | 14.00821 | 5.75881 | 8.39756 | 11.20474 | 20.64603 |
| **INPP5J** | 0.38808 | 0.00000 | 1.36535 | 0.38086 | 2.50940 | 0.39912 | 0.46962 |

| | | | | | | | |
|---|---|---|---|---|---|---|---|
| **INPP5K** | 7.05015 | 10.68738 | 6.55669 | 10.45595 | 5.03894 | 4.87323 | 0.24639 |
| **INPPL1** | 0.10609 | 3.65019 | 0.08673 | 6.54006 | 0.00000 | 3.96312 | 19.99366 |
| **INSC** | 0.12579 | 0.00013 | 0.00465 | 0.12401 | 0.35967 | 0.00000 | 0.09759 |
| **INSIG1** | 18.14125 | 6.52182 | 2.56203 | 8.39450 | 4.62031 | 6.62494 | 15.01735 |
| **INSIG2** | 4.65234 | 0.06261 | 2.61176 | 2.92679 | 0.64579 | 1.30512 | 0.30453 |
| **INSL3** | 0.03033 | 0.00458 | 0.03385 | 0.02114 | 0.00656 | 0.01624 | 0.03926 |
| **INSL4** | 1.09384 | 0.00000 | 0.06767 | 0.94904 | 0.00344 | 0.00298 | 0.00000 |
| **INSL6** | 0.05157 | 0.02250 | 0.42885 | 0.18085 | 0.03660 | 0.00288 | 0.00000 |
| **INSM1** | 0.00000 | 0.54307 | 0.57427 | 0.00000 | 0.08110 | 0.50521 | 0.04582 |
| **INSM2** | 0.02832 | 0.01039 | 0.06762 | 0.01524 | 0.07398 | 0.00551 | 0.07821 |
| **INSR** | 4.77492 | 8.00010 | 2.73758 | 4.13834 | 3.81842 | 13.14121 | 3.25540 |
| **INSRR** | 0.00952 | 0.01783 | 0.02638 | 0.01666 | 0.00894 | 0.01704 | 0.11035 |
| **INTS1** | 0.00000 | 7.48967 | 2.76021 | 5.34613 | 1.36152 | 6.78094 | 20.21183 |
| **INTS10** | 15.66528 | 7.99751 | 5.61251 | 10.73104 | 1.49973 | 7.56212 | 1.54157 |
| **INTS12** | 9.87997 | 4.01812 | 8.90631 | 4.72512 | 2.44847 | 1.38786 | 0.94468 |
| **INTS2** | 7.37467 | 1.79613 | 2.45844 | 4.71312 | 2.73374 | 3.21757 | 0.00000 |
| **INTS3** | 4.17379 | 2.57388 | 2.82682 | 6.56011 | 0.90023 | 3.48556 | 7.85246 |
| **INTS4** | 6.94466 | 2.70718 | 5.71240 | 9.36377 | 3.34016 | 3.03162 | 0.71365 |
| **INTS4L2** | 0.03003 | 0.03997 | 0.01770 | 0.01739 | 0.01082 | 0.01462 | 0.00883 |
| **INTS5** | 1.10062 | 2.05275 | 2.50953 | 3.50788 | 0.30273 | 2.33056 | 7.25971 |
| **INTS6** | 2.93078 | 0.00000 | 2.73737 | 0.99431 | 3.14042 | 2.46218 | 7.97874 |
| **INTS6-AS1** | 0.01194 | 0.10159 | 0.06126 | 0.10979 | 0.33663 | 0.03438 | 0.28962 |
| **INTS7** | 20.56285 | 12.42572 | 13.25216 | 10.16451 | 1.74625 | 9.74027 | 0.00000 |
| **INTS8** | 6.25890 | 3.08718 | 5.72692 | 1.67293 | 3.16306 | 6.17945 | 9.72570 |
| **INTS9** | 9.47167 | 3.27811 | 5.50761 | 4.49323 | 3.14750 | 3.03621 | 0.00000 |
| **INTU** | 0.89698 | 0.04891 | 0.94543 | 1.19882 | 0.30410 | 0.92169 | 1.39842 |
| **INVS** | 1.39301 | 2.36541 | 0.44172 | 1.59617 | 0.34854 | 1.18373 | 0.44720 |
| **IP6K1** | 2.37535 | 8.93837 | 2.05632 | 8.00521 | 0.34018 | 4.71744 | 13.60077 |
| **IP6K2** | 25.51708 | 17.85036 | 12.41618 | 24.39630 | 3.88308 | 12.63628 | 1.52515 |
| **IP6K3** | 0.03807 | 0.00590 | 0.06049 | 0.04657 | 0.01307 | 0.01813 | 0.00714 |
| **IPCEF1** | 0.00000 | 0.12484 | 0.00000 | 0.01970 | 0.32462 | 0.15299 | 0.08712 |
| **IPMK** | 1.94728 | 0.00000 | 3.82263 | 0.12766 | 0.00000 | 4.57283 | 4.98884 |
| **IPO11** | 13.61905 | 0.73042 | 6.43806 | 4.97543 | 2.71049 | 2.80049 | 0.00000 |
| **IPO13** | 1.71255 | 3.66967 | 1.08049 | 3.03062 | 0.86726 | 3.49546 | 8.45908 |
| **IPO4** | 16.43019 | 11.93162 | 9.27162 | 15.99310 | 1.29695 | 9.66473 | 10.88796 |

| | | | | | | | |
|---|---|---|---|---|---|---|---|
| **IPO5** | 40.89813 | 20.25204 | 17.92830 | 17.06307 | 2.62325 | 15.22336 | 29.27160 |
| **IPO5P1** | 1.86375 | 2.34280 | 4.12699 | 4.30294 | 6.72510 | 1.83403 | 1.82581 |
| **IPO7** | 9.10636 | 0.00000 | 7.19019 | 8.36981 | 1.79539 | 6.27905 | 32.61505 |
| **IPO8** | 3.95531 | 1.01320 | 1.99723 | 2.18467 | 7.81699 | 2.88800 | 8.96203 |
| **IPO9** | 15.88146 | 7.11622 | 7.14827 | 13.79808 | 1.51167 | 9.62894 | 15.19876 |
| **IPO9-AS1** | 0.01143 | 0.15014 | 0.06397 | 0.03535 | 0.01519 | 0.05431 | 0.03114 |
| **IPP** | 1.79266 | 2.85656 | 3.44570 | 0.87747 | 1.24222 | 3.42931 | 0.17279 |
| **IPPK** | 5.09685 | 4.25394 | 2.26285 | 7.97415 | 3.26038 | 3.85852 | 0.13443 |
| **IPW** | 1.89063 | 0.00000 | 1.82986 | 0.29976 | 0.00018 | 3.43713 | 1.97861 |
| **IQCA1** | 1.12636 | 0.55336 | 0.00000 | 0.19196 | 4.62419 | 0.27766 | 0.44129 |
| **IQCB1** | 11.81986 | 2.25657 | 4.29969 | 6.56176 | 0.90927 | 2.83060 | 2.35503 |
| **IQCC** | 2.42293 | 2.16470 | 1.40531 | 1.53854 | 0.47942 | 0.77300 | 0.00000 |
| **IQCD** | 0.47121 | 0.60796 | 0.32246 | 1.72032 | 2.10927 | 0.14305 | 0.00000 |
| **IQCE** | 0.07163 | 1.51841 | 1.89368 | 0.61817 | 0.61091 | 1.27732 | 1.27352 |
| **IQCF1** | 0.08166 | 0.22047 | 0.01117 | 0.02634 | 0.00764 | 0.08958 | 0.00000 |
| **IQCF2** | 0.01111 | 0.04334 | 0.15700 | 0.01859 | 0.00963 | 0.01241 | 0.02361 |
| **IQCF4** | 0.02157 | 0.01152 | 0.02006 | 0.02292 | 0.04272 | 0.01368 | 0.02022 |
| **IQCF5** | 0.02173 | 0.01964 | 0.01239 | 0.01653 | 0.00772 | 0.02367 | 0.02779 |
| **IQCG** | 6.18866 | 0.10030 | 2.24353 | 4.26657 | 1.44468 | 0.58092 | 0.00000 |
| **IQCH** | 0.28581 | 0.00000 | 0.37194 | 0.10568 | 0.16731 | 0.14652 | 0.00000 |
| **IQCH-AS1** | 2.25337 | 1.84618 | 0.21156 | 1.33088 | 0.05161 | 0.02020 | 1.19802 |
| **IQCJ-SCHIP1-A** | 0.01064 | 0.05768 | 0.08805 | 0.01852 | 0.00793 | 0.03399 | 0.03433 |
| **IQCJ.uid-4645** | 0.01465 | 0.05749 | 0.01158 | 0.01793 | 0.13905 | 0.01046 | 0.01994 |
| **IQCK** | 1.72416 | 2.39235 | 3.12298 | 1.80699 | 1.67367 | 3.07691 | 0.98895 |
| **IQGAP1** | 23.54424 | 12.73791 | 7.22198 | 20.80521 | 9.24302 | 6.51635 | 26.10030 |
| **IQGAP2** | 20.18324 | 8.09864 | 0.26630 | 0.52689 | 6.79470 | 3.88751 | 36.64171 |
| **IQGAP3** | 5.91519 | 12.36235 | 5.96189 | 1.09314 | 0.88062 | 11.02852 | 11.95059 |
| **IQSEC1** | 2.59751 | 5.42693 | 0.15153 | 2.65151 | 2.40392 | 0.55559 | 12.47975 |
| **IQSEC2** | 0.03516 | 0.00000 | 0.04490 | 0.01649 | 0.01963 | 0.56496 | 0.17727 |
| **IQSEC3** | 0.02140 | 0.00000 | 0.11298 | 0.02325 | 0.00456 | 0.00849 | 0.03738 |
| **IQUB** | 0.05116 | 0.00000 | 0.08736 | 0.08261 | 3.09176 | 0.06629 | 0.07657 |
| **IRAK1** | 0.00000 | 7.06076 | 2.02088 | 6.60901 | 0.00000 | 1.64260 | 6.71979 |
| **IRAK1BP1** | 0.47130 | 0.46220 | 0.51952 | 0.13525 | 0.50790 | 0.44795 | 0.41896 |
| **IRAK2** | 0.26161 | 0.00000 | 0.00956 | 3.12753 | 0.27341 | 0.01640 | 0.00913 |
| **IRAK3** | 0.04486 | 0.00000 | 0.63791 | 0.01537 | 0.00330 | 0.00289 | 0.01919 |

| | | | | | | | |
|---|---|---|---|---|---|---|---|
| **IRAK4** | 1.89344 | 0.35534 | 0.59362 | 1.05121 | 0.25606 | 1.20297 | 1.10102 |
| **IREB2** | 16.37983 | 0.00000 | 8.41045 | 5.41647 | 1.07067 | 5.92486 | 17.90588 |
| **IRF1** | 0.74377 | 3.08723 | 1.19030 | 0.58930 | 1.82303 | 2.13328 | 2.14088 |
| **IRF2** | 0.00885 | 0.01071 | 0.05286 | 0.25452 | 0.01270 | 0.40667 | 0.12510 |
| **IRF2BP1** | 0.00000 | 0.66440 | 1.00260 | 0.65360 | 0.20958 | 1.40397 | 4.66900 |
| **IRF2BP2** | 3.62546 | 2.50641 | 1.24697 | 3.64851 | 0.23918 | 8.12940 | 11.99469 |
| **IRF2BPL** | 1.88923 | 2.79530 | 4.05143 | 3.74004 | 1.27535 | 5.52370 | 13.71273 |
| **IRF3** | 16.26874 | 18.93441 | 3.00480 | 22.90097 | 0.00000 | 6.95876 | 3.82330 |
| **IRF4** | 0.01609 | 0.00854 | 0.00000 | 0.02506 | 6.12110 | 0.53185 | 0.00478 |
| **IRF5** | 0.00875 | 0.00755 | 0.05773 | 0.03045 | 0.00486 | 0.01884 | 0.06908 |
| **IRF6** | 1.94366 | 0.44040 | 0.38868 | 6.98027 | 1.70616 | 0.77895 | 0.53409 |
| **IRF7** | 0.08199 | 0.73211 | 0.15053 | 1.80763 | 0.16483 | 1.81193 | 0.88685 |
| **IRF8** | 0.05357 | 0.00000 | 1.38389 | 0.62390 | 0.54578 | 0.28579 | 0.00000 |
| **IRF9** | 1.27799 | 0.93018 | 0.42832 | 1.47112 | 0.38402 | 2.63279 | 2.99951 |
| **IRG1** | 0.02000 | 0.02177 | 0.01278 | 0.01781 | 0.00780 | 0.01743 | 0.03063 |
| **IRGC** | 0.05937 | 0.01367 | 0.01437 | 0.03506 | 0.00790 | 0.02294 | 0.00000 |
| **IRGM** | 0.01886 | 0.01916 | 0.00000 | 0.01783 | 0.27352 | 0.01733 | 0.02982 |
| **IRGQ** | 2.59921 | 12.20527 | 18.04123 | 4.09324 | 4.21112 | 2.23430 | 5.91693 |
| **IRS1** | 0.08863 | 0.00000 | 0.22995 | 3.61707 | 0.03505 | 0.15564 | 2.37967 |
| **IRS2** | 0.16353 | 0.00000 | 0.64248 | 0.00000 | 0.10342 | 1.79631 | 1.19533 |
| **IRS4** | 0.01196 | 0.02291 | 0.01194 | 0.04373 | 0.00761 | 0.01269 | 0.03456 |
| **IRX1** | 0.00000 | 0.02061 | 0.01027 | 0.01427 | 0.01403 | 0.13995 | 0.08300 |
| **IRX2** | 0.53143 | 0.00000 | 0.00000 | 6.58894 | 0.15039 | 0.00000 | 13.97652 |
| **IRX3** | 0.00000 | 0.00000 | 0.99391 | 0.20743 | 0.19692 | 1.24275 | 0.96371 |
| **IRX4** | 0.00000 | 0.00645 | 0.00445 | 1.67864 | 0.02296 | 0.00119 | 3.98335 |
| **IRX5** | 0.00000 | 0.00000 | 0.65930 | 0.00836 | 0.48578 | 0.02952 | 0.06554 |
| **IRX6** | 0.02362 | 0.02846 | 0.01043 | 0.02228 | 0.00844 | 0.08569 | 0.00596 |
| **ISCA1** | 15.05869 | 6.41859 | 8.45939 | 7.69582 | 2.42877 | 5.08202 | 0.00000 |
| **ISCA2** | 14.67202 | 14.16374 | 6.60666 | 7.93325 | 1.47724 | 6.22401 | 0.00000 |
| **ISCU** | 27.05869 | 21.38939 | 17.76403 | 18.32376 | 5.45070 | 13.34983 | 10.53820 |
| **ISG15** | 0.48807 | 1.39788 | 0.00000 | 1.06801 | 0.08766 | 0.94355 | 0.00331 |
| **ISG20** | 0.09681 | 0.20426 | 0.06124 | 0.65919 | 0.11888 | 0.42870 | 0.30075 |
| **ISG20L2** | 15.04539 | 7.03041 | 10.30917 | 15.45832 | 4.05293 | 4.62578 | 3.21404 |
| **ISL1** | 0.21786 | 0.00000 | 0.27218 | 0.21429 | 0.64492 | 1.10945 | 0.21893 |
| **ISL2** | 0.02295 | 0.00000 | 0.88904 | 0.03390 | 0.71322 | 0.00000 | 0.45952 |

| | | | | | | | |
|---|---|---|---|---|---|---|---|
| **ISLR** | 0.01322 | 0.02595 | 0.01046 | 0.01636 | 0.00763 | 0.01000 | 0.05287 |
| **ISLR2** | 0.03123 | 0.02585 | 0.00000 | 0.01012 | 0.21873 | 0.02443 | 0.03702 |
| **ISM1** | 0.00000 | 0.14549 | 0.12787 | 0.02928 | 0.08545 | 0.14253 | 0.13969 |
| **ISM1-AS1** | 0.02174 | 0.01345 | 0.02217 | 0.01397 | 0.00624 | 0.05271 | 0.10179 |
| **ISM2** | 0.00000 | 0.85147 | 0.24972 | 1.85638 | 0.00031 | 0.63811 | 0.62394 |
| **ISOC1** | 19.46461 | 7.07077 | 12.68988 | 15.33049 | 1.24626 | 2.06570 | 0.00000 |
| **ISOC2** | 17.11571 | 7.68739 | 1.21878 | 2.52087 | 0.08109 | 14.87759 | 4.52304 |
| **ISPD** | 0.72568 | 0.00000 | 0.23675 | 0.01141 | 1.81571 | 0.00000 | 1.63413 |
| **ISPD-AS1** | 0.06975 | 0.00000 | 0.02166 | 0.01140 | 0.09273 | 0.01876 | 0.06790 |
| **IST1** | 31.98803 | 16.18979 | 16.18792 | 24.74454 | 13.31004 | 13.18505 | 3.98146 |
| **ISX** | 0.09780 | 0.00570 | 0.01517 | 0.02834 | 0.00791 | 0.02076 | 0.00000 |
| **ISY1** | 8.42598 | 3.63327 | 4.75040 | 8.43571 | 6.43257 | 3.56079 | 5.17405 |
| **ISYNA1** | 8.08704 | 4.63369 | 0.00000 | 16.99844 | 0.02811 | 3.46072 | 5.69658 |
| **ITCH** | 11.82578 | 4.72688 | 11.36133 | 8.87403 | 2.20010 | 7.08740 | 5.57453 |
| **ITFG1** | 6.58409 | 3.50308 | 5.83049 | 2.93533 | 0.30486 | 4.36018 | 4.37791 |
| **ITFG2** | 3.93195 | 4.25191 | 1.90476 | 2.77407 | 1.04466 | 2.33937 | 1.28772 |
| **ITFG3** | 0.04804 | 12.75788 | 19.59546 | 1.90252 | 4.13241 | 13.39255 | 3.56756 |
| **ITGA1** | 0.00436 | 0.00864 | 0.01012 | 1.18371 | 0.01110 | 0.69458 | 0.00059 |
| **ITGA10** | 0.04749 | 0.12097 | 0.46201 | 0.19019 | 0.30530 | 0.05554 | 0.14161 |
| **ITGA11** | 0.01360 | 0.00110 | 0.04587 | 0.02660 | 0.63135 | 0.13980 | 0.03547 |
| **ITGA2** | 0.15555 | 0.00000 | 0.00495 | 0.00000 | 0.01495 | 2.69274 | 0.05720 |
| **ITGA2B** | 0.82837 | 0.00000 | 0.59337 | 0.50359 | 0.03119 | 0.04490 | 0.63325 |
| **ITGA3** | 0.00000 | 0.00000 | 0.20735 | 0.10394 | 0.01917 | 0.30249 | 0.71124 |
| **ITGA4** | 0.01270 | 0.00000 | 0.01380 | 0.01269 | 1.83343 | 0.11749 | 0.04048 |
| **ITGA5** | 2.89694 | 0.00000 | 8.98722 | 23.70688 | 0.20500 | 0.00000 | 2.17372 |
| **ITGA6** | 3.93468 | 3.98966 | 2.44191 | 1.38209 | 1.29483 | 13.95299 | 15.02734 |
| **ITGA7** | 0.00000 | 0.15913 | 0.03678 | 0.21323 | 0.00600 | 0.37624 | 0.98178 |
| **ITGA8** | 0.02032 | 0.02938 | 0.01008 | 0.03440 | 0.06878 | 0.02725 | 0.01893 |
| **ITGA9** | 0.00679 | 0.06797 | 0.02235 | 0.10795 | 0.66198 | 0.34640 | 0.04877 |
| **ITGA9-AS1** | 0.03377 | 0.01176 | 0.02032 | 0.08065 | 0.04223 | 0.01254 | 0.02106 |
| **ITGAD** | 0.00917 | 0.03115 | 0.01709 | 0.10038 | 0.05629 | 0.03784 | 0.01328 |
| **ITGAE** | 8.08657 | 4.21223 | 4.68878 | 10.93657 | 1.76915 | 6.18683 | 5.17772 |
| **ITGAL** | 0.01203 | 0.03060 | 0.11431 | 0.07755 | 0.73911 | 0.51503 | 0.00000 |
| **ITGAM** | 0.00000 | 2.24589 | 1.16933 | 0.00000 | 0.08976 | 8.53084 | 0.00344 |
| **ITGAV** | 0.44426 | 0.00000 | 0.02283 | 0.60764 | 1.90602 | 1.97706 | 1.36139 |

| | | | | | | | |
|---|---|---|---|---|---|---|---|
| **ITGAX** | 0.00000 | 0.47568 | 0.40496 | 0.00000 | 0.16354 | 0.61687 | 0.01060 |
| **ITGB1** | 13.91080 | 2.64366 | 3.34782 | 8.94184 | 2.48855 | 5.13683 | 2.97066 |
| **ITGB1BP1** | 20.41468 | 11.75281 | 6.84486 | 17.02113 | 0.45877 | 6.68964 | 3.82073 |
| **ITGB1BP2** | 0.46628 | 0.00000 | 1.46635 | 0.03862 | 0.49020 | 0.40677 | 0.48522 |
| **ITGB2** | 0.06887 | 0.00897 | 0.01144 | 0.02084 | 0.00793 | 0.13306 | 0.03183 |
| **ITGB2-AS1** | 0.02110 | 0.01766 | 0.01172 | 0.02312 | 0.01467 | 0.01310 | 0.02183 |
| **ITGB3** | 0.00000 | 0.05430 | 0.67193 | 0.01817 | 0.00529 | 0.14071 | 0.03099 |
| **ITGB3BP** | 9.31986 | 0.42135 | 6.60894 | 5.79716 | 1.69385 | 4.09479 | 5.38282 |
| **ITGB4** | 0.00000 | 0.00101 | 0.01663 | 9.29360 | 0.02389 | 0.00000 | 1.51829 |
| **ITGB5** | 1.72565 | 7.37243 | 2.36856 | 8.95048 | 0.22211 | 9.04441 | 13.35963 |
| **ITGB6** | 0.02822 | 0.01433 | 0.01505 | 0.01523 | 0.00770 | 0.06294 | 0.02634 |
| **ITGB7** | 0.16066 | 0.00000 | 1.07540 | 0.04102 | 0.00069 | 0.06560 | 0.07308 |
| **ITGB8** | 0.00000 | 0.00000 | 0.21743 | 0.05891 | 0.11860 | 0.39704 | 0.06520 |
| **ITGBL1** | 0.10515 | 0.00000 | 0.02837 | 0.07206 | 0.74745 | 0.01931 | 0.03757 |
| **ITIH1** | 0.02669 | 0.02279 | 0.01268 | 0.01615 | 0.00788 | 0.01859 | 0.01999 |
| **ITIH2** | 0.05683 | 0.00000 | 0.03487 | 0.02872 | 0.16381 | 0.02022 | 0.00344 |
| **ITIH3** | 0.03372 | 0.01640 | 0.02215 | 0.03590 | 0.00730 | 0.01413 | 0.01472 |
| **ITIH4** | 0.07512 | 0.17383 | 0.74197 | 0.22953 | 0.00922 | 0.18273 | 0.21871 |
| **ITIH4-AS1** | 0.02804 | 0.01611 | 0.01848 | 0.01920 | 0.00898 | 0.01365 | 0.01519 |
| **ITIH5** | 1.02381 | 0.87505 | 0.00000 | 0.23396 | 1.44428 | 2.58442 | 0.00000 |
| **ITIH6** | 0.01269 | 0.01212 | 0.21778 | 0.00812 | 0.00284 | 0.03968 | 0.07837 |
| **ITK** | 0.04819 | 0.00206 | 0.01541 | 0.01254 | 0.00749 | 0.11012 | 0.02596 |
| **ITLN1** | 0.04918 | 0.24916 | 0.15586 | 0.05166 | 0.14360 | 0.10590 | 0.00000 |
| **ITLN2** | 0.00000 | 2.07472 | 2.06156 | 0.34360 | 0.29288 | 2.53173 | 0.00000 |
| **ITM2A** | 0.73089 | 0.81504 | 0.00000 | 0.01152 | 0.00341 | 0.00000 | 0.12236 |
| **ITM2B** | 15.93533 | 4.02189 | 13.86589 | 14.42462 | 1.03651 | 14.62279 | 38.92387 |
| **ITM2C** | 0.00000 | 26.22306 | 10.87399 | 2.74543 | 4.55410 | 18.98783 | 8.37312 |
| **ITPA** | 22.27893 | 17.57837 | 12.72256 | 19.52915 | 0.53992 | 9.66797 | 5.23421 |
| **ITPK1** | 0.76762 | 14.38112 | 2.84600 | 5.55422 | 1.01570 | 5.80766 | 14.08222 |
| **ITPK1-AS1** | 0.00000 | 0.86029 | 0.17524 | 0.14363 | 0.00143 | 0.28106 | 0.20803 |
| **ITPKA** | 0.00000 | 0.31650 | 0.37669 | 0.06946 | 1.02672 | 0.18774 | 0.15610 |
| **ITPKB** | 0.02275 | 0.01388 | 0.01240 | 0.04071 | 0.02636 | 0.05835 | 0.01889 |
| **ITPKC** | 0.45829 | 5.17076 | 2.53741 | 5.06441 | 0.02574 | 1.99515 | 7.34207 |
| **ITPR1** | 1.14926 | 0.57915 | 0.06890 | 0.56252 | 2.14345 | 1.16873 | 1.96212 |
| **ITPR1-AS1** | 0.04079 | 0.05674 | 0.03304 | 0.01230 | 0.00619 | 0.11901 | 0.08929 |

| | | | | | | | |
|---|---|---|---|---|---|---|---|
| **ITPR2** | 0.32974 | 0.40771 | 0.00000 | 1.04314 | 0.06502 | 3.35156 | 1.98637 |
| **ITPR3** | 6.61632 | 4.70354 | 0.11105 | 0.43759 | 0.79425 | 7.71770 | 11.09897 |
| **ITPRIP** | 0.69988 | 0.45330 | 1.08069 | 0.73056 | 1.51022 | 1.27927 | 2.06078 |
| **ITPRIPL1** | 1.12401 | 0.15902 | 0.46119 | 0.57249 | 0.04136 | 0.02753 | 0.00000 |
| **ITPRIPL2** | 0.03533 | 0.39021 | 0.21465 | 1.88533 | 1.02964 | 0.00000 | 3.27926 |
| **ITSN1** | 29.98125 | 10.73979 | 7.14690 | 21.59509 | 10.51945 | 7.29462 | 27.55169 |
| **ITSN2** | 6.17485 | 2.09004 | 3.73940 | 6.58968 | 1.25243 | 3.94443 | 16.87441 |
| **IVD** | 6.26997 | 1.08472 | 1.47693 | 2.63390 | 1.56367 | 4.16015 | 1.33358 |
| **IVL** | 0.03304 | 0.01846 | 0.01409 | 0.03058 | 0.00796 | 0.01148 | 0.01223 |
| **IVNS1ABP** | 17.30622 | 0.24156 | 3.44796 | 18.25930 | 3.47662 | 2.66277 | 1.30726 |
| **IWS1** | 17.75919 | 14.68853 | 12.10230 | 15.57855 | 11.11608 | 8.70101 | 13.67624 |
| **IYD** | 0.02112 | 0.01624 | 0.01122 | 0.09558 | 0.04056 | 0.01335 | 0.00604 |
| **IZUMO1** | 0.05825 | 0.00000 | 0.00742 | 0.03077 | 0.03184 | 0.00000 | 0.22001 |
| **IZUMO2** | 0.01768 | 0.09106 | 0.02775 | 0.00741 | 0.01857 | 0.00099 | 0.08992 |
| **IZUMO4** | 0.03991 | 0.00000 | 0.17577 | 0.15614 | 0.09763 | 0.00000 | 0.14490 |
| **JADE1** | 15.89317 | 2.73696 | 3.16781 | 14.12042 | 0.22333 | 4.98448 | 11.67359 |
| **JADE2** | 2.78755 | 0.91690 | 0.00000 | 0.00000 | 0.06309 | 8.00651 | 7.23736 |
| **JADE3** | 4.32437 | 3.67911 | 2.66359 | 7.28163 | 1.07965 | 0.00000 | 7.11088 |
| **JAG1** | 0.00000 | 0.13771 | 2.79440 | 1.21984 | 6.70463 | 5.12784 | 0.86217 |
| **JAG2** | 0.00000 | 8.14533 | 1.30341 | 0.00000 | 0.41115 | 3.24031 | 4.20013 |
| **JAGN1** | 24.56892 | 13.16855 | 19.63817 | 17.38192 | 1.23032 | 7.28799 | 8.06520 |
| **JAK1** | 1.98338 | 3.63118 | 2.71535 | 2.88085 | 2.08768 | 8.90260 | 8.73993 |
| **JAK2** | 0.73452 | 0.00000 | 0.69692 | 0.27260 | 0.76410 | 0.63213 | 1.19953 |
| **JAK3** | 0.08295 | 0.00000 | 0.17231 | 0.03281 | 0.13678 | 0.49437 | 0.01464 |
| **JAKMIP1** | 0.05552 | 0.00000 | 0.04232 | 0.00000 | 0.12252 | 0.13117 | 0.35171 |
| **JAKMIP2** | 0.10450 | 0.01899 | 0.01107 | 0.01535 | 0.02772 | 0.03189 | 0.01006 |
| **JAKMIP2-AS1** | 0.05525 | 0.01594 | 0.01277 | 0.02206 | 0.00864 | 0.01091 | 0.00804 |
| **JAKMIP3** | 0.01876 | 0.11283 | 0.00000 | 0.02457 | 0.19790 | 0.00553 | 0.01478 |
| **JAM2** | 0.13681 | 0.00000 | 0.09662 | 0.12884 | 0.57748 | 0.07103 | 0.09495 |
| **JAM3** | 3.95108 | 4.46056 | 4.94620 | 8.80623 | 4.35527 | 8.40712 | 2.14439 |
| **JARID2** | 5.42315 | 9.85754 | 7.28871 | 14.71368 | 7.52407 | 18.05963 | 23.30358 |
| **JARID2-AS1** | 0.04348 | 0.02908 | 0.04695 | 0.03442 | 0.01555 | 0.01615 | 0.00000 |
| **JAZF1** | 0.23163 | 1.85677 | 0.81909 | 0.00000 | 3.14497 | 1.60299 | 0.46725 |
| **JAZF1-AS1** | 0.01741 | 0.05152 | 0.01941 | 0.01648 | 0.01599 | 0.02802 | 0.03395 |
| **JDP2** | 7.33349 | 2.99585 | 0.00000 | 2.67742 | 0.14385 | 0.00000 | 15.98447 |

| | | | | | | | |
|---|---|---|---|---|---|---|---|
| **JHDM1D-AS1** | 0.00000 | 0.37512 | 2.49717 | 0.18659 | 0.28728 | 0.43113 | 0.36959 |
| **JKAMP** | 23.40939 | 0.33780 | 14.54795 | 12.53548 | 4.22532 | 5.75956 | 0.00000 |
| **JMJD1C** | 4.53072 | 1.11470 | 3.62677 | 2.63873 | 0.73878 | 6.87662 | 13.16153 |
| **JMJD1C-AS1** | 0.01782 | 0.02125 | 0.02966 | 0.01660 | 0.06109 | 0.00913 | 0.11148 |
| **JMJD4** | 1.45117 | 7.92019 | 4.00195 | 2.78881 | 0.20354 | 1.19609 | 1.91268 |
| **JMJD6** | 25.60067 | 22.94134 | 6.82796 | 22.27949 | 6.98670 | 7.84156 | 0.04183 |
| **JMJD7** | 0.61145 | 0.08912 | 0.20543 | 0.63028 | 0.29575 | 0.16695 | 0.75701 |
| **JMJD8** | 0.88949 | 3.43908 | 0.73711 | 3.69916 | 1.42077 | 2.81388 | 8.54111 |
| **JMY** | 0.21978 | 0.00000 | 19.58354 | 1.45652 | 3.17089 | 7.02997 | 16.56534 |
| **JOSD1** | 1.74016 | 2.65628 | 1.33851 | 4.11153 | 0.82401 | 1.59370 | 6.11603 |
| **JOSD2** | 0.00000 | 0.80250 | 0.51884 | 5.27369 | 0.05283 | 0.79717 | 4.27446 |
| **JPH1** | 0.00000 | 0.00000 | 0.72251 | 0.06452 | 0.02528 | 0.20397 | 0.30083 |
| **JPH2** | 0.00014 | 0.01218 | 0.00000 | 0.37705 | 0.41384 | 0.00198 | 0.17239 |
| **JPH3** | 0.00000 | 1.04673 | 0.09751 | 0.00000 | 0.80336 | 0.52350 | 0.96382 |
| **JPH4** | 0.00849 | 0.05171 | 0.04364 | 0.02148 | 0.00798 | 0.03082 | 0.03896 |
| **JPX** | 5.06515 | 3.58779 | 2.93528 | 0.57961 | 0.12669 | 2.59050 | 2.50689 |
| **JRK** | 0.76966 | 1.21069 | 1.40775 | 1.33332 | 0.24433 | 0.44629 | 1.81088 |
| **JRKL** | 1.16139 | 0.00000 | 2.68918 | 0.79007 | 1.03809 | 0.00000 | 0.44751 |
| **JSRP1** | 0.03207 | 0.02996 | 0.00882 | 0.05581 | 0.00769 | 0.00000 | 0.08969 |
| **JTB** | 4.37651 | 14.67076 | 8.54653 | 16.38809 | 0.30113 | 7.21865 | 44.96051 |
| **JUN** | 0.00000 | 0.00000 | 0.29406 | 0.93156 | 0.14516 | 1.97800 | 3.21184 |
| **JUNB** | 0.00000 | 6.41485 | 2.14217 | 2.19025 | 0.72409 | 5.20465 | 7.03859 |
| **JUND** | 0.00000 | 3.85700 | 1.40478 | 4.28135 | 0.48911 | 1.52348 | 21.47193 |
| **JUP** | 27.24621 | 32.92446 | 0.00000 | 41.01296 | 0.00000 | 5.29867 | 44.70598 |
| **KAAG1** | 0.10221 | 0.11617 | 0.03154 | 0.02682 | 0.03951 | 0.02150 | 0.00000 |
| **KAL1** | 0.01302 | 0.00000 | 0.14697 | 0.00000 | 0.36280 | 0.21606 | 0.28913 |
| **KALRN** | 2.81143 | 1.37025 | 0.00000 | 7.44863 | 4.62078 | 1.38597 | 1.12338 |
| **KANK1** | 9.29575 | 9.33542 | 3.90902 | 8.86100 | 0.47657 | 6.71744 | 1.39025 |
| **KANK2** | 0.70128 | 3.79188 | 2.02267 | 1.36720 | 0.63851 | 3.39397 | 1.86100 |
| **KANK3** | 0.04294 | 0.66455 | 0.40054 | 0.44341 | 0.22156 | 0.44296 | 0.82445 |
| **KANK4** | 7.01255 | 0.12534 | 0.55434 | 7.33322 | 0.00000 | 0.00000 | 10.73084 |
| **KANSL1** | 1.95298 | 0.00000 | 4.06643 | 2.09146 | 1.25041 | 6.02833 | 12.46877 |
| **KANSL1-AS1** | 0.39194 | 0.08310 | 0.04127 | 0.12092 | 0.06035 | 0.43511 | 0.51604 |
| **KANSL1L** | 0.91156 | 0.00000 | 0.47555 | 0.45006 | 0.18114 | 1.05905 | 1.34485 |
| **KANSL2** | 22.81835 | 10.70824 | 12.56524 | 12.34693 | 11.09775 | 11.56429 | 0.00000 |

| | | | | | | | |
|---|---|---|---|---|---|---|---|
| **KANSL3** | 8.03484 | 2.57326 | 5.97083 | 5.66996 | 3.51399 | 5.35578 | 7.48042 |
| **KANTR** | 0.35873 | 0.19468 | 0.27634 | 0.27309 | 0.23712 | 0.06421 | 0.42048 |
| **KARS** | 48.61554 | 20.88430 | 30.09311 | 33.23768 | 4.51144 | 16.47386 | 19.92397 |
| **KAT2A** | 3.09689 | 7.59202 | 2.46662 | 3.12966 | 0.00000 | 3.92445 | 14.78829 |
| **KAT2B** | 1.24631 | 0.00000 | 3.45301 | 3.09444 | 0.00177 | 0.00000 | 1.88649 |
| **KAT5** | 2.95440 | 2.93558 | 1.93556 | 3.36820 | 4.52119 | 1.24282 | 0.00000 |
| **KAT6A** | 2.53293 | 1.63019 | 4.21571 | 3.16180 | 1.53637 | 3.72322 | 13.16190 |
| **KAT6B** | 5.16628 | 2.84158 | 8.02817 | 2.60553 | 1.43422 | 8.66443 | 12.36076 |
| **KAT7** | 10.88013 | 1.40683 | 7.15015 | 6.24310 | 5.68425 | 12.16115 | 0.00000 |
| **KAT8** | 0.00000 | 2.99905 | 1.49528 | 2.57716 | 0.46657 | 2.10175 | 7.47734 |
| **KATNA1** | 17.24918 | 8.57554 | 5.94325 | 13.78463 | 1.04479 | 3.41881 | 0.00000 |
| **KATNAL1** | 1.62867 | 3.00088 | 2.87483 | 0.00000 | 5.98248 | 2.06501 | 0.76208 |
| **KATNAL2** | 0.09515 | 0.12301 | 0.01204 | 0.11320 | 0.02918 | 0.03866 | 0.02531 |
| **KATNB1** | 6.34605 | 8.83020 | 0.71242 | 1.16387 | 3.30326 | 4.21868 | 3.46645 |
| **KATNBL1** | 3.40812 | 0.91625 | 0.79939 | 10.56983 | 0.24584 | 0.36353 | 1.66380 |
| **KATNBL1P6** | 1.51301 | 0.11008 | 0.37816 | 4.49473 | 0.08971 | 0.08733 | 0.00689 |
| **KAZALD1** | 0.02376 | 0.01537 | 0.00947 | 0.10069 | 0.04798 | 0.03727 | 0.01685 |
| **KAZN** | 0.00000 | 5.04768 | 0.20292 | 1.34315 | 0.00000 | 9.49764 | 3.48268 |
| **KBTBD11** | 0.00516 | 0.22512 | 0.00468 | 0.05836 | 0.14984 | 0.00001 | 0.23386 |
| **KBTBD12** | 0.02176 | 0.01913 | 0.01951 | 0.01764 | 0.00715 | 0.04072 | 0.01959 |
| **KBTBD13** | 0.03678 | 0.02009 | 0.01207 | 0.03030 | 0.00852 | 0.01443 | 0.01157 |
| **KBTBD2** | 0.39423 | 0.00000 | 2.98161 | 6.08338 | 2.99201 | 8.15508 | 2.26840 |
| **KBTBD3** | 0.49590 | 0.00000 | 0.32068 | 0.41001 | 0.26387 | 0.25531 | 0.01056 |
| **KBTBD4** | 3.70885 | 0.21774 | 1.41484 | 7.34028 | 1.97498 | 0.86392 | 0.00000 |
| **KBTBD6** | 9.42056 | 0.10937 | 3.27269 | 3.31479 | 7.11173 | 1.59996 | 0.00000 |
| **KBTBD7** | 3.13306 | 0.16300 | 1.91112 | 1.91292 | 2.20547 | 1.40036 | 0.00000 |
| **KBTBD8** | 4.96884 | 0.65572 | 0.75234 | 1.75962 | 4.84700 | 1.91578 | 0.00000 |
| **KC6** | 0.09787 | 0.07217 | 0.01546 | 0.01681 | 0.01237 | 0.07992 | 0.01188 |
| **KCMF1** | 2.85770 | 4.97058 | 0.26884 | 4.80946 | 0.43577 | 6.41728 | 26.00484 |
| **KCNA1** | 0.02271 | 0.08301 | 0.00000 | 0.01761 | 0.47429 | 0.00850 | 0.00025 |
| **KCNA2** | 0.02082 | 0.00872 | 0.01602 | 0.02324 | 0.20478 | 0.01362 | 0.02401 |
| **KCNA3** | 0.02316 | 0.02102 | 0.00989 | 0.02280 | 0.03427 | 0.01226 | 0.02491 |
| **KCNA4** | 0.02568 | 0.06723 | 0.00819 | 0.04809 | 0.05492 | 0.01011 | 0.00000 |
| **KCNA6** | 0.09786 | 0.24179 | 0.05499 | 0.26558 | 0.01299 | 0.46334 | 0.00000 |
| **KCNA7** | 0.00000 | 0.01061 | 0.04601 | 0.19063 | 0.49621 | 0.01260 | 0.06789 |

| | | | | | | | |
|---|---|---|---|---|---|---|---|
| **KCNAB1** | 0.37057 | 0.32063 | 0.18160 | 0.02386 | 0.20573 | 0.06003 | 1.22466 |
| **KCNAB1-AS2** | 0.02105 | 0.02106 | 0.01120 | 0.01873 | 0.01295 | 0.01873 | 0.02136 |
| **KCNAB2** | 0.00000 | 0.14029 | 0.11256 | 0.19521 | 0.04026 | 0.14970 | 0.00462 |
| **KCNAB3** | 0.00000 | 0.00000 | 0.16280 | 0.50861 | 0.15373 | 0.25945 | 0.33552 |
| **KCNB1** | 0.02421 | 0.00442 | 0.04445 | 0.02105 | 0.13686 | 0.01448 | 0.01899 |
| **KCNC1** | 0.01011 | 0.00899 | 0.01377 | 0.02980 | 0.00779 | 0.02665 | 0.08191 |
| **KCNC2** | 0.02851 | 0.01661 | 0.01361 | 0.01754 | 0.01515 | 0.01478 | 0.01580 |
| **KCNC3** | 0.00000 | 0.21718 | 0.21888 | 0.95844 | 0.15078 | 0.27347 | 4.10118 |
| **KCNC4** | 0.00000 | 2.26205 | 2.59508 | 0.17767 | 1.25289 | 2.37908 | 0.49469 |
| **KCNC4-AS1** | 0.00853 | 0.08569 | 0.31340 | 0.04438 | 0.24219 | 0.01385 | 0.01726 |
| **KCND1** | 0.34416 | 0.10097 | 0.26160 | 0.69761 | 0.01109 | 0.25722 | 1.79085 |
| **KCND2** | 0.02333 | 0.00000 | 0.04463 | 0.01885 | 0.07190 | 0.07977 | 0.01586 |
| **KCND3** | 0.01818 | 0.27046 | 0.06358 | 0.01058 | 0.00937 | 0.02193 | 0.03697 |
| **KCNE1** | 0.03942 | 0.00000 | 0.04972 | 0.01898 | 0.04329 | 0.01203 | 0.02155 |
| **KCNE1L** | 0.06784 | 0.18911 | 0.18622 | 0.00405 | 0.06651 | 0.03497 | 0.04946 |
| **KCNE2** | 0.02806 | 0.14506 | 0.32181 | 0.03082 | 0.14564 | 0.00000 | 0.12223 |
| **KCNE3** | 0.33055 | 0.02607 | 0.02668 | 3.94494 | 0.08926 | 1.22698 | 0.01028 |
| **KCNE4** | 0.05010 | 0.00000 | 0.05847 | 0.06336 | 0.03587 | 0.01807 | 0.00182 |
| **KCNF1** | 0.02579 | 0.00000 | 0.30523 | 0.06407 | 0.00258 | 0.06616 | 0.01439 |
| **KCNG1** | 0.00000 | 0.46395 | 0.05724 | 0.01186 | 0.00568 | 0.04217 | 0.05684 |
| **KCNG2** | 0.03717 | 0.02874 | 0.00692 | 0.01008 | 0.21346 | 0.02540 | 0.08025 |
| **KCNG3** | 0.00000 | 0.14977 | 0.05860 | 0.18504 | 0.11705 | 0.93179 | 0.00992 |
| **KCNG4** | 0.02250 | 0.01729 | 0.00916 | 0.01897 | 0.04513 | 0.01301 | 0.01996 |
| **KCNH1** | 0.06983 | 0.09546 | 0.03879 | 0.15887 | 0.36615 | 0.00000 | 0.24426 |
| **KCNH2** | 0.12110 | 0.00000 | 0.01126 | 0.03963 | 0.01573 | 1.82179 | 0.58498 |
| **KCNH3** | 0.00526 | 0.03147 | 0.01880 | 0.03325 | 0.09040 | 0.05277 | 0.08906 |
| **KCNH4** | 0.02652 | 0.14176 | 0.09412 | 0.02856 | 0.00439 | 0.03322 | 0.02773 |
| **KCNH5** | 0.81882 | 0.10035 | 2.29943 | 0.28298 | 0.80556 | 0.00583 | 0.04190 |
| **KCNH6** | 0.01619 | 0.08056 | 0.01299 | 0.03444 | 0.02424 | 0.11914 | 0.00002 |
| **KCNH7** | 0.02299 | 0.03545 | 0.03528 | 0.02201 | 0.13441 | 0.00920 | 0.03064 |
| **KCNH8** | 0.12330 | 0.07850 | 0.19288 | 0.00652 | 0.41919 | 0.08703 | 0.01214 |
| **KCNIP2** | 0.15518 | 0.10623 | 0.08387 | 0.05839 | 0.47840 | 0.14859 | 0.57424 |
| **KCNIP2-AS1** | 0.13329 | 0.00000 | 0.02315 | 0.15015 | 0.97609 | 0.00000 | 0.10400 |
| **KCNIP3** | 0.01212 | 0.00674 | 0.02210 | 0.01931 | 0.00675 | 0.21934 | 0.04406 |
| **KCNIP4** | 0.11251 | 0.05476 | 0.04489 | 0.05578 | 0.27655 | 0.17058 | 0.14252 |

| | | | | | | | |
|---|---|---|---|---|---|---|---|
| **KCNIP4-IT1** | 0.02340 | 0.01223 | 0.01256 | 0.01453 | 0.00738 | 0.01043 | 0.07464 |
| **KCNJ1** | 0.03685 | 0.02898 | 0.00905 | 0.01439 | 0.00821 | 0.10790 | 0.00268 |
| **KCNJ10** | 0.01586 | 0.03821 | 0.06546 | 0.02245 | 0.00516 | 0.00821 | 0.01764 |
| **KCNJ11** | 0.09058 | 0.00000 | 0.01658 | 0.07286 | 0.00634 | 0.00000 | 0.11105 |
| **KCNJ12** | 0.01813 | 0.01900 | 0.01159 | 0.02808 | 0.03923 | 0.01316 | 0.02777 |
| **KCNJ13** | 2.14166 | 0.35282 | 0.15626 | 0.93342 | 0.12429 | 1.27367 | 0.69492 |
| **KCNJ14** | 0.10188 | 0.00000 | 0.16585 | 0.44376 | 0.05303 | 0.06579 | 0.94383 |
| **KCNJ15** | 0.06453 | 0.00882 | 0.01204 | 0.09036 | 0.05051 | 0.02367 | 0.00000 |
| **KCNJ16** | 0.05231 | 0.00000 | 0.00585 | 0.00000 | 0.04498 | 0.89451 | 0.14139 |
| **KCNJ2** | 0.01508 | 0.00000 | 0.04064 | 0.01942 | 0.03174 | 0.08255 | 0.18092 |
| **KCNJ3** | 0.12776 | 0.00000 | 0.23224 | 0.00000 | 0.22305 | 0.17596 | 0.86632 |
| **KCNJ4** | 0.02421 | 0.01694 | 0.00660 | 0.01787 | 0.08840 | 0.01268 | 0.02078 |
| **KCNJ6** | 0.11868 | 0.00000 | 0.03905 | 0.03663 | 0.24666 | 0.10413 | 0.00000 |
| **KCNJ8** | 0.09042 | 0.00000 | 0.01758 | 0.08624 | 0.08587 | 0.63555 | 0.10521 |
| **KCNJ9** | 0.03291 | 0.00805 | 0.01200 | 0.00917 | 0.02135 | 0.00889 | 0.06586 |
| **KCNK1** | 9.04261 | 2.55156 | 3.27911 | 2.37121 | 0.57697 | 0.81098 | 10.43691 |
| **KCNK10** | 0.01346 | 0.01315 | 0.02603 | 0.04502 | 0.03627 | 0.01349 | 0.03272 |
| **KCNK12** | 0.00000 | 0.06792 | 0.05393 | 0.03364 | 0.09923 | 0.94526 | 0.03469 |
| **KCNK13** | 0.00000 | 0.04799 | 0.05677 | 0.00297 | 0.01890 | 0.84219 | 0.17736 |
| **KCNK15** | 0.02389 | 0.03774 | 0.03378 | 0.01824 | 0.10017 | 0.01061 | 0.01036 |
| **KCNK16** | 0.04455 | 0.00000 | 0.05170 | 0.00769 | 0.29754 | 0.00833 | 0.03670 |
| **KCNK17** | 0.02187 | 0.02326 | 0.00985 | 0.01830 | 0.01982 | 0.01334 | 0.02080 |
| **KCNK18** | 0.02244 | 0.01801 | 0.00117 | 0.01802 | 0.24169 | 0.01470 | 0.01813 |
| **KCNK2** | 0.16104 | 0.00000 | 0.03616 | 0.13381 | 0.00571 | 0.04321 | 0.07334 |
| **KCNK3** | 0.00250 | 0.02538 | 0.01738 | 0.02374 | 0.00697 | 0.14560 | 0.04562 |
| **KCNK4** | 0.01299 | 0.05923 | 0.19622 | 0.03716 | 0.00464 | 0.04828 | 0.01514 |
| **KCNK5** | 0.72409 | 0.00000 | 3.48842 | 0.67050 | 2.89556 | 2.11858 | 1.10820 |
| **KCNK6** | 0.00000 | 2.59752 | 1.42185 | 1.03224 | 0.20413 | 5.11699 | 2.45902 |
| **KCNK7** | 0.02360 | 0.07296 | 0.00815 | 0.15874 | 0.06381 | 0.01863 | 0.05203 |
| **KCNK9** | 0.00753 | 0.39764 | 0.00000 | 0.04010 | 4.14116 | 0.00404 | 0.00000 |
| **KCNMA1** | 0.02768 | 0.02160 | 0.00664 | 0.03137 | 0.05550 | 0.20993 | 0.03452 |
| **KCNMB2** | 0.09932 | 0.03643 | 0.06584 | 0.00000 | 0.01293 | 0.02941 | 0.12958 |
| **KCNMB3** | 0.04575 | 0.00000 | 0.30820 | 0.16429 | 0.01940 | 0.08550 | 1.07474 |
| **KCNMB4** | 1.69820 | 2.08104 | 1.59257 | 0.16520 | 0.33525 | 2.49889 | 1.22187 |
| **KCNN1** | 0.00000 | 1.78444 | 0.00121 | 0.00000 | 0.02174 | 1.53895 | 0.16955 |

| | | | | | | | |
|---|---|---|---|---|---|---|---|
| **KCNN2** | 0.07769 | 0.27805 | 0.38121 | 0.08677 | 0.63044 | 0.47283 | 0.00000 |
| **KCNN3** | 0.02074 | 0.06495 | 0.00000 | 0.02051 | 0.28908 | 0.29261 | 0.10195 |
| **KCNN4** | 2.91885 | 0.00000 | 0.00000 | 11.12637 | 0.01602 | 0.00000 | 1.05300 |
| **KCNQ1** | 0.02624 | 0.02569 | 0.00405 | 0.02387 | 0.16919 | 0.01405 | 0.00978 |
| **KCNQ1OT1** | 1.34690 | 0.72635 | 0.09906 | 4.07003 | 1.69850 | 4.14632 | 7.61147 |
| **KCNQ2** | 0.00000 | 0.90430 | 1.09137 | 0.01673 | 0.00255 | 0.10712 | 0.20778 |
| **KCNQ3** | 0.03232 | 0.02166 | 0.01724 | 0.01620 | 0.15662 | 0.02059 | 0.06070 |
| **KCNQ4** | 0.77968 | 1.32641 | 0.50445 | 0.09900 | 0.00000 | 0.00000 | 4.35300 |
| **KCNQ5** | 0.19629 | 0.00000 | 0.15655 | 0.36175 | 0.04661 | 0.11637 | 0.29357 |
| **KCNQ5-AS1** | 0.02333 | 0.01636 | 0.01030 | 0.02680 | 0.02648 | 0.01278 | 0.02634 |
| **KCNQ5-IT1** | 0.07199 | 0.00000 | 0.15431 | 0.08936 | 0.00462 | 0.01478 | 0.51919 |
| **KCNRG** | 0.40333 | 0.00000 | 0.48499 | 0.40354 | 0.35054 | 0.22261 | 0.83687 |
| **KCNS1** | 0.00016 | 0.00000 | 0.18046 | 0.09556 | 0.00739 | 0.00303 | 0.26455 |
| **KCNS2** | 0.06050 | 0.00375 | 0.02711 | 0.01761 | 0.00934 | 0.00953 | 0.04663 |
| **KCNS3** | 0.00000 | 0.26915 | 0.04353 | 0.98346 | 0.15441 | 3.21862 | 0.19956 |
| **KCNT1** | 0.01697 | 0.01050 | 0.01530 | 0.02425 | 0.05511 | 0.01441 | 0.05096 |
| **KCNT2** | 0.02410 | 0.06233 | 0.09049 | 0.01214 | 0.03971 | 0.01445 | 0.03801 |
| **KCNU1** | 0.01798 | 0.02170 | 0.01204 | 0.03893 | 0.00810 | 0.01243 | 0.01561 |
| **KCNV1** | 3.01474 | 0.00000 | 0.00000 | 0.07641 | 0.06005 | 0.00000 | 0.89162 |
| **KCNV2** | 0.24606 | 0.14483 | 1.15872 | 0.80395 | 0.14354 | 2.46047 | 0.30126 |
| **KCP** | 0.00000 | 0.59160 | 0.11070 | 0.01435 | 0.12822 | 0.10839 | 0.14507 |
| **KCTD1** | 1.19220 | 2.04021 | 0.62004 | 0.16686 | 0.15187 | 1.07745 | 5.96303 |
| **KCTD10** | 16.56463 | 8.38943 | 9.47328 | 21.45310 | 5.97112 | 5.86020 | 2.27028 |
| **KCTD11** | 0.00000 | 0.44653 | 0.38880 | 1.32485 | 0.00245 | 0.00000 | 0.18657 |
| **KCTD12** | 2.04492 | 0.00000 | 0.39114 | 1.96665 | 0.00352 | 0.00000 | 4.75532 |
| **KCTD13** | 0.00000 | 2.25948 | 1.22324 | 5.00922 | 2.24085 | 0.10923 | 5.00088 |
| **KCTD14.uid-159** | 0.03996 | 0.02417 | 0.01733 | 0.01882 | 0.01050 | 0.01323 | 0.02656 |
| **KCTD15** | 4.21777 | 7.02467 | 2.80332 | 1.14193 | 0.00000 | 3.23994 | 8.78855 |
| **KCTD16** | 0.03598 | 0.04170 | 0.02775 | 0.01474 | 0.01178 | 0.01622 | 0.03344 |
| **KCTD17** | 0.00000 | 0.79240 | 0.03688 | 0.01053 | 0.08695 | 0.39811 | 0.21598 |
| **KCTD18** | 0.24319 | 0.80550 | 0.51104 | 0.30972 | 0.25621 | 0.20208 | 0.45176 |
| **KCTD19** | 0.01910 | 0.01149 | 0.12433 | 0.02351 | 0.04767 | 0.00923 | 0.02844 |
| **KCTD2** | 2.91141 | 5.27984 | 2.46810 | 3.34021 | 1.46836 | 4.71487 | 10.65378 |
| **KCTD20** | 9.50044 | 2.37437 | 0.66415 | 10.18620 | 2.79479 | 0.11910 | 2.33987 |
| **KCTD21** | 0.44861 | 1.06219 | 0.15194 | 0.97427 | 0.79730 | 0.52214 | 0.05652 |

| | | | | | | | |
|---|---|---|---|---|---|---|---|
| **KCTD21-AS1** | 0.17483 | 0.00000 | 0.22018 | 0.15728 | 0.01662 | 0.11892 | 0.00000 |
| **KCTD3** | 6.53924 | 0.31928 | 4.61845 | 0.00000 | 0.46158 | 3.20388 | 12.11884 |
| **KCTD4** | 0.05841 | 0.03133 | 0.06396 | 0.00913 | 0.11268 | 0.13081 | 0.15142 |
| **KCTD5** | 2.30853 | 7.42908 | 2.95571 | 4.94988 | 4.12040 | 3.60635 | 9.19716 |
| **KCTD6** | 2.26265 | 0.38463 | 5.17043 | 0.00000 | 1.10653 | 4.38209 | 1.49301 |
| **KCTD7** | 0.00000 | 1.08482 | 1.17159 | 0.55901 | 0.19313 | 2.64080 | 0.12590 |
| **KCTD8** | 0.00000 | 0.14289 | 0.01435 | 0.02352 | 0.02995 | 0.05881 | 0.02511 |
| **KCTD9** | 3.77152 | 0.00000 | 1.73062 | 2.48232 | 2.35294 | 1.33383 | 1.96906 |
| **KDELC1** | 3.14583 | 0.00000 | 0.38633 | 1.80325 | 0.69984 | 1.72974 | 0.00000 |
| **KDELC2** | 2.54965 | 0.00000 | 0.27573 | 0.57460 | 0.05293 | 0.33239 | 2.90190 |
| **KDELR1** | 5.62099 | 7.41621 | 2.75548 | 12.73501 | 0.91567 | 11.47859 | 21.78966 |
| **KDELR2** | 33.91162 | 10.61769 | 14.17967 | 24.87102 | 4.68600 | 14.41418 | 15.91501 |
| **KDELR3** | 0.30362 | 0.00266 | 0.14184 | 0.09744 | 0.11605 | 0.08054 | 0.03304 |
| **KDF1** | 5.02789 | 7.30919 | 3.13884 | 6.81360 | 0.00001 | 2.58940 | 1.52652 |
| **KDM1A** | 8.51524 | 7.67554 | 5.54584 | 15.86366 | 4.89777 | 8.35883 | 26.19214 |
| **KDM1B** | 0.21152 | 0.43510 | 2.02657 | 0.96410 | 3.36754 | 0.64760 | 1.90507 |
| **KDM2A** | 3.94598 | 7.15597 | 3.10927 | 10.21412 | 2.17715 | 7.46028 | 22.12510 |
| **KDM2B** | 3.94178 | 2.77353 | 3.84962 | 5.49073 | 0.79603 | 4.82187 | 14.07427 |
| **KDM3A** | 3.85182 | 2.62654 | 6.38984 | 4.28820 | 6.98763 | 2.79733 | 4.76457 |
| **KDM3B** | 8.25789 | 5.17434 | 1.64829 | 3.20031 | 1.05882 | 5.80880 | 10.31415 |
| **KDM4A** | 7.10948 | 1.88496 | 5.72481 | 6.83429 | 7.37308 | 7.94825 | 2.48605 |
| **KDM4A-AS1** | 0.33224 | 0.10895 | 0.17065 | 0.55940 | 0.21031 | 0.27439 | 0.00000 |
| **KDM4B** | 0.00000 | 1.06141 | 0.01290 | 1.05388 | 0.00274 | 0.60483 | 6.45223 |
| **KDM4C** | 3.29075 | 4.16564 | 5.13396 | 3.78145 | 3.32003 | 8.68251 | 5.89899 |
| **KDM4D** | 0.56271 | 0.04162 | 2.23578 | 0.41284 | 1.98084 | 0.12569 | 0.00000 |
| **KDM4E** | 0.00957 | 0.16237 | 3.42826 | 0.04631 | 2.90160 | 0.51167 | 0.00000 |
| **KDM5A** | 12.39786 | 10.51743 | 6.71766 | 8.81468 | 11.78607 | 9.00715 | 15.60305 |
| **KDM5B** | 27.98342 | 30.47777 | 18.92146 | 19.11833 | 19.31248 | 12.05215 | 25.94468 |
| **KDM5C** | 0.48301 | 0.93037 | 4.07585 | 6.22526 | 2.04067 | 3.67975 | 15.79205 |
| **KDM5D** | 0.00513 | 0.09353 | 0.00530 | 0.16083 | 0.00811 | 0.01004 | 0.01672 |
| **KDM6A** | 8.15375 | 3.80742 | 3.74208 | 7.48646 | 1.60606 | 6.54613 | 5.51653 |
| **KDM6B** | 0.21255 | 3.92854 | 0.00000 | 6.19244 | 1.49025 | 4.21328 | 14.27427 |
| **KDM7A** | 0.76843 | 0.00000 | 6.95161 | 2.19283 | 2.56111 | 7.73338 | 3.91655 |
| **KDM8** | 1.44732 | 1.15883 | 1.25280 | 0.31723 | 0.07352 | 0.48107 | 0.07291 |
| **KDR** | 0.06729 | 0.05753 | 0.00630 | 0.02530 | 0.02900 | 0.39649 | 0.00000 |

| | | | | | | | |
|---|---|---|---|---|---|---|---|
| **KDSR** | 8.98179 | 1.64227 | 6.49365 | 8.55997 | 0.10460 | 4.98208 | 7.94037 |
| **KEAP1** | 15.11260 | 12.87603 | 2.36082 | 20.55994 | 1.86466 | 7.11824 | 1.29342 |
| **KEL** | 0.03088 | 0.00794 | 0.02641 | 0.03548 | 0.00726 | 0.02216 | 0.02081 |
| **KGFLP2** | 0.02803 | 0.02487 | 0.02583 | 0.02979 | 0.00756 | 0.01240 | 0.01184 |
| **KHDC1** | 3.17844 | 2.62285 | 2.79557 | 6.09172 | 12.10165 | 0.82001 | 0.00000 |
| **KHDC1L** | 81.03058 | 82.59712 | 52.66629 | 15.18036 | 38.87787 | 61.76843 | 31.22456 |
| **KHDC3L** | 0.00000 | 0.03378 | 1.81430 | 0.83759 | 25.66506 | 12.64433 | 0.00000 |
| **KHDRBS1** | 15.33469 | 4.93190 | 7.77642 | 13.79284 | 1.93707 | 9.28424 | 36.54013 |
| **KHDRBS2** | 0.03198 | 0.01105 | 0.01351 | 0.01475 | 0.01567 | 0.01618 | 0.03040 |
| **KHDRBS3** | 10.88884 | 3.71374 | 0.53252 | 1.85185 | 0.44220 | 0.65105 | 26.19360 |
| **KHK** | 3.26262 | 0.27700 | 0.00000 | 1.50840 | 1.00858 | 0.28770 | 0.00000 |
| **KHNYN** | 1.53771 | 1.37092 | 0.17355 | 2.41011 | 0.08069 | 2.27045 | 2.79806 |
| **KHSRP** | 1.73320 | 7.91015 | 5.62671 | 5.85753 | 1.66529 | 9.89039 | 44.01945 |
| **KIAA0020** | 37.57891 | 10.10883 | 21.40729 | 19.93311 | 7.91789 | 11.74919 | 17.68550 |
| **KIAA0040** | 4.26079 | 1.15344 | 4.11718 | 1.03260 | 0.92319 | 1.00130 | 0.19894 |
| **KIAA0087** | 0.02189 | 0.01805 | 0.01055 | 0.01882 | 0.02760 | 0.01486 | 0.02092 |
| **KIAA0100** | 8.04337 | 12.73959 | 4.46834 | 11.60193 | 2.69889 | 9.36129 | 18.91992 |
| **KIAA0101** | 16.43959 | 0.00000 | 0.07239 | 17.68334 | 0.25848 | 0.00000 | 8.67274 |
| **KIAA0125** | 0.02784 | 0.01656 | 0.01530 | 0.02427 | 0.00987 | 0.01573 | 0.01833 |
| **KIAA0141** | 4.77067 | 0.95320 | 0.27335 | 5.87169 | 0.32556 | 2.80777 | 2.61304 |
| **KIAA0195** | 0.74783 | 0.13641 | 1.64851 | 5.37215 | 0.33259 | 3.78694 | 12.02191 |
| **KIAA0196** | 19.15467 | 2.22606 | 4.80748 | 3.98294 | 2.13778 | 5.14250 | 2.50927 |
| **KIAA0226** | 3.59143 | 5.55039 | 1.74253 | 5.17195 | 4.01274 | 3.35731 | 1.25145 |
| **KIAA0226L** | 0.11379 | 1.23684 | 0.67744 | 0.12860 | 0.00192 | 0.12040 | 0.00000 |
| **KIAA0232** | 4.25036 | 3.91577 | 1.92003 | 0.81641 | 1.09755 | 4.45100 | 16.42499 |
| **KIAA0247** | 1.14291 | 4.95822 | 1.62975 | 1.30433 | 6.11251 | 2.12425 | 2.11721 |
| **KIAA0319** | 0.00000 | 0.00000 | 0.39470 | 0.24769 | 1.70912 | 0.13395 | 0.13298 |
| **KIAA0319L** | 4.02038 | 5.64474 | 0.40957 | 4.60284 | 0.63727 | 3.46152 | 4.18337 |
| **KIAA0355** | 0.33279 | 0.35884 | 2.32612 | 3.96224 | 1.34452 | 1.88121 | 3.41221 |
| **KIAA0368** | 13.43836 | 4.79024 | 2.18696 | 9.79761 | 0.25127 | 9.38997 | 15.08963 |
| **KIAA0391** | 15.30458 | 4.75438 | 12.68657 | 6.63713 | 1.06836 | 5.66252 | 1.01426 |
| **KIAA0430** | 2.38006 | 0.77566 | 3.91561 | 4.60013 | 2.67912 | 2.12346 | 7.67253 |
| **KIAA0513** | 0.00000 | 0.10952 | 0.56669 | 0.59251 | 0.99314 | 1.64236 | 0.54310 |
| **KIAA0556** | 1.14111 | 0.78517 | 2.34020 | 2.30099 | 0.83399 | 0.98799 | 1.89521 |
| **KIAA0586** | 4.45344 | 0.48447 | 2.29439 | 0.85387 | 1.41486 | 1.34300 | 5.08408 |

| | | | | | | | |
|---|---|---|---|---|---|---|---|
| **KIAA0753** | 2.36702 | 1.09979 | 2.18466 | 1.92207 | 1.05390 | 0.65785 | 1.33219 |
| **KIAA0754** | 0.00000 | 0.17552 | 0.17146 | 0.10366 | 0.15461 | 0.59426 | 1.52468 |
| **KIAA0825** | 0.29223 | 0.00000 | 0.11362 | 0.08941 | 0.62951 | 0.19153 | 0.59536 |
| **KIAA0895** | 0.30289 | 0.10496 | 0.00341 | 0.70869 | 0.08974 | 0.05352 | 0.13650 |
| **KIAA0895L** | 0.23342 | 1.30333 | 0.41604 | 1.13935 | 0.03751 | 0.67096 | 1.39452 |
| **KIAA0907** | 7.83454 | 1.55645 | 3.53047 | 6.31397 | 4.76432 | 3.19005 | 9.13810 |
| **KIAA0922** | 6.71631 | 4.19685 | 1.12301 | 0.64688 | 5.26313 | 4.39833 | 1.37946 |
| **KIAA0930** | 0.42486 | 6.07374 | 0.00000 | 1.70427 | 0.84313 | 2.46138 | 11.52557 |
| **KIAA1024** | 0.10623 | 0.00000 | 0.14014 | 0.21919 | 0.33134 | 0.10880 | 0.53894 |
| **KIAA1024L** | 0.02456 | 0.01787 | 0.01359 | 0.06857 | 0.00820 | 0.01364 | 0.00239 |
| **KIAA1033** | 15.73123 | 0.84532 | 4.13960 | 5.27197 | 4.03705 | 6.30737 | 7.01092 |
| **KIAA1045** | 0.08459 | 0.40685 | 0.31962 | 0.05098 | 0.00157 | 0.12389 | 0.26538 |
| **KIAA1107** | 0.12241 | 0.04526 | 0.94159 | 0.15061 | 0.31220 | 0.35734 | 0.33703 |
| **KIAA1109** | 2.14452 | 0.00000 | 1.87192 | 1.16976 | 2.75282 | 1.83789 | 7.14488 |
| **KIAA1143** | 7.09415 | 0.00000 | 6.46838 | 7.91623 | 1.04561 | 1.82363 | 4.07161 |
| **KIAA1147** | 0.87197 | 0.00000 | 0.49818 | 0.12179 | 3.57841 | 0.05139 | 0.83128 |
| **KIAA1161** | 3.70218 | 3.01763 | 0.06961 | 1.51879 | 0.00000 | 0.00000 | 5.85242 |
| **KIAA1191** | 9.72411 | 0.47938 | 12.73735 | 9.95777 | 18.09608 | 7.99021 | 5.30487 |
| **KIAA1210** | 0.02142 | 0.01120 | 0.00868 | 0.06838 | 0.35526 | 0.01215 | 0.03786 |
| **KIAA1211** | 0.54127 | 0.00000 | 0.48630 | 0.54771 | 0.06897 | 0.18273 | 1.16024 |
| **KIAA1211L** | 0.00238 | 0.06459 | 0.03138 | 0.08636 | 0.02593 | 0.00345 | 0.12277 |
| **KIAA1217** | 1.46892 | 0.00000 | 0.00000 | 21.09262 | 0.00198 | 10.41336 | 3.43829 |
| **KIAA1244** | 0.23397 | 0.00000 | 0.19909 | 0.26221 | 1.56819 | 0.21015 | 0.05417 |
| **KIAA1257** | 0.14707 | 0.29594 | 1.37332 | 0.00000 | 2.09552 | 0.32640 | 0.05662 |
| **KIAA1279** | 5.10369 | 7.04651 | 2.01023 | 0.88394 | 0.66347 | 5.80385 | 3.83804 |
| **KIAA1324** | 0.02863 | 0.07123 | 0.15659 | 0.94012 | 0.13272 | 0.00000 | 0.10204 |
| **KIAA1324L** | 0.39815 | 0.25124 | 0.05261 | 0.15960 | 1.75444 | 0.21957 | 0.52167 |
| **KIAA1328** | 0.83613 | 0.47411 | 0.52755 | 0.78949 | 0.22464 | 0.13737 | 0.06700 |
| **KIAA1377** | 0.25035 | 0.00000 | 0.49326 | 1.11761 | 0.06527 | 0.01416 | 0.41555 |
| **KIAA1407** | 0.20355 | 0.15697 | 0.34734 | 0.05813 | 0.78172 | 0.03141 | 0.00000 |
| **KIAA1429** | 14.56971 | 1.85102 | 7.60934 | 5.94484 | 3.55898 | 4.07456 | 8.59053 |
| **KIAA1430** | 0.49937 | 0.00000 | 9.35305 | 0.00000 | 8.81972 | 0.98546 | 1.23266 |
| **KIAA1432** | 3.12322 | 0.67421 | 1.41840 | 4.45717 | 0.53565 | 0.66667 | 5.36492 |
| **KIAA1456** | 0.05020 | 0.00000 | 0.01998 | 0.04006 | 0.11131 | 0.05365 | 0.06010 |
| **KIAA1462** | 4.77814 | 0.00000 | 0.14503 | 3.73988 | 0.36036 | 0.86810 | 6.76844 |

| | | | | | | | |
|---|---|---|---|---|---|---|---|
| **KIAA1467** | 5.43323 | 0.32108 | 4.01969 | 2.51134 | 2.07404 | 0.62402 | 0.00000 |
| **KIAA1468** | 4.05350 | 1.61856 | 1.01694 | 2.11370 | 1.00513 | 2.38315 | 4.04222 |
| **KIAA1522** | 0.00000 | 0.52001 | 3.58610 | 3.80276 | 0.19088 | 1.54322 | 7.58920 |
| **KIAA1524** | 15.98990 | 1.30480 | 6.88989 | 6.70149 | 4.38242 | 3.97407 | 5.38239 |
| **KIAA1549** | 0.35483 | 0.24662 | 0.03349 | 2.43510 | 0.17223 | 1.56469 | 3.71171 |
| **KIAA1549L** | 0.00000 | 0.00000 | 0.12701 | 0.02910 | 2.01472 | 1.19352 | 0.15807 |
| **KIAA1551** | 21.63144 | 2.09334 | 25.73561 | 3.80565 | 5.32176 | 19.10507 | 26.87171 |
| **KIAA1586** | 0.99800 | 0.00000 | 2.95135 | 1.88084 | 0.72374 | 1.69524 | 0.00348 |
| **KIAA1598** | 16.80645 | 0.00000 | 13.29327 | 12.28420 | 1.93160 | 2.63357 | 14.75285 |
| **KIAA1614** | 0.02241 | 0.05731 | 0.00960 | 0.13424 | 0.02816 | 0.13243 | 0.09804 |
| **KIAA1644** | 0.03471 | 0.00435 | 0.13389 | 0.00604 | 0.05760 | 0.00192 | 0.14621 |
| **KIAA1656** | 0.04046 | 0.21688 | 0.01293 | 0.14837 | 0.07187 | 0.22972 | 0.45900 |
| **KIAA1671** | 1.74079 | 4.00356 | 1.15019 | 0.84543 | 0.71721 | 6.13618 | 3.50077 |
| **KIAA1683** | 0.04607 | 0.02163 | 0.01153 | 0.07729 | 0.00800 | 0.00941 | 0.00747 |
| **KIAA1715** | 4.09356 | 0.43931 | 2.83369 | 2.03327 | 3.19242 | 0.65729 | 2.01751 |
| **KIAA1731** | 4.64758 | 0.45699 | 5.09947 | 2.71049 | 2.35202 | 3.81577 | 17.55226 |
| **KIAA1751** | 0.00000 | 0.01853 | 0.33675 | 0.06401 | 0.24792 | 0.00000 | 0.07821 |
| **KIAA1755** | 0.05829 | 0.00443 | 0.01670 | 0.00787 | 0.01562 | 0.02318 | 0.04631 |
| **KIAA1804** | 0.79266 | 0.00000 | 1.43097 | 0.00000 | 0.26140 | 1.24827 | 1.33327 |
| **KIAA1841** | 0.34999 | 0.44159 | 0.27478 | 0.94754 | 0.60190 | 1.11457 | 0.30173 |
| **KIAA1875** | 0.00465 | 0.00000 | 0.07596 | 0.11847 | 0.00565 | 0.01578 | 0.41533 |
| **KIAA1919** | 2.27582 | 0.28646 | 4.52907 | 1.89866 | 1.85904 | 2.84083 | 7.71017 |
| **KIAA1958** | 1.10234 | 1.82821 | 0.67363 | 1.90862 | 0.14431 | 3.66546 | 5.80351 |
| **KIAA2013** | 1.59219 | 9.69902 | 3.51082 | 3.36213 | 0.29101 | 6.00998 | 17.00528 |
| **KIAA2018** | 7.56771 | 1.17702 | 1.37124 | 4.60490 | 1.77731 | 6.52151 | 3.34889 |
| **KIAA2022** | 0.05786 | 0.01428 | 0.01014 | 0.00491 | 0.02553 | 0.00370 | 0.15501 |
| **KIAA2026** | 1.80842 | 0.00000 | 2.69188 | 0.34399 | 1.61643 | 3.31075 | 7.49731 |
| **KIDINS220** | 5.07962 | 2.96061 | 2.90747 | 3.42517 | 6.72418 | 1.48887 | 1.32422 |
| **KIF11** | 15.32986 | 4.38441 | 8.57804 | 8.32821 | 2.09234 | 5.84060 | 7.13649 |
| **KIF12** | 0.00000 | 0.68999 | 0.88844 | 0.00000 | 0.22976 | 0.91781 | 0.01170 |
| **KIF13A** | 0.82377 | 0.36377 | 0.74739 | 2.20477 | 0.25618 | 0.82258 | 5.04132 |
| **KIF13B** | 0.60479 | 1.39676 | 0.00001 | 2.07522 | 2.48324 | 0.49593 | 5.55373 |
| **KIF14** | 4.88340 | 0.00000 | 4.73637 | 10.05714 | 1.23594 | 0.25032 | 5.79543 |
| **KIF15** | 9.81633 | 1.39959 | 5.22087 | 5.54254 | 3.00728 | 3.04832 | 5.24827 |
| **KIF16B** | 3.76729 | 2.34882 | 4.87208 | 6.04694 | 0.68792 | 3.19422 | 2.18444 |

| | | | | | | | |
|---|---|---|---|---|---|---|---|
| **KIF17** | 1.53710 | 3.57214 | 0.21975 | 0.00000 | 0.03437 | 1.00105 | 1.19419 |
| **KIF18A** | 6.74315 | 0.00000 | 6.19554 | 4.53119 | 4.94750 | 1.58977 | 0.22839 |
| **KIF18B** | 4.22788 | 1.97546 | 3.34578 | 6.62496 | 0.49804 | 2.08746 | 11.47008 |
| **KIF19** | 0.02401 | 0.01080 | 0.01395 | 0.01883 | 0.54610 | 0.01324 | 0.01757 |
| **KIF1A** | 0.04922 | 0.38904 | 0.00000 | 0.00000 | 1.70112 | 0.00000 | 0.38331 |
| **KIF1B** | 3.53595 | 6.15149 | 4.04794 | 4.51306 | 4.77641 | 6.28396 | 13.89528 |
| **KIF1C** | 0.00000 | 5.76104 | 0.41290 | 4.79289 | 0.04277 | 9.22946 | 14.04488 |
| **KIF20A** | 28.66883 | 18.88230 | 15.83515 | 22.27175 | 2.14043 | 5.63776 | 0.00000 |
| **KIF20B** | 17.90657 | 0.00000 | 11.73092 | 10.48451 | 10.88017 | 6.62893 | 16.32582 |
| **KIF21A** | 9.79436 | 0.51670 | 0.00000 | 10.53739 | 0.05806 | 6.55080 | 9.06382 |
| **KIF21B** | 0.04089 | 0.00000 | 0.35703 | 0.12504 | 0.96709 | 0.02159 | 0.28790 |
| **KIF22** | 31.06090 | 24.90771 | 14.35119 | 30.89304 | 3.86207 | 12.77675 | 4.66322 |
| **KIF23** | 26.88147 | 6.60600 | 21.74461 | 13.05180 | 12.49780 | 6.76627 | 5.07902 |
| **KIF24** | 1.86845 | 1.10302 | 0.99794 | 1.71924 | 1.46629 | 0.36466 | 0.50708 |
| **KIF25** | 0.01122 | 0.01573 | 0.01417 | 0.03926 | 0.00855 | 0.01121 | 0.03961 |
| **KIF25-AS1** | 0.02367 | 0.01659 | 0.01393 | 0.02020 | 0.06677 | 0.01303 | 0.01523 |
| **KIF26A** | 0.00000 | 4.66462 | 0.78735 | 0.00000 | 0.00000 | 0.48153 | 4.46842 |
| **KIF26B** | 0.03849 | 0.01109 | 0.14871 | 0.00821 | 1.70651 | 0.13448 | 0.10196 |
| **KIF27** | 0.84566 | 0.52513 | 0.70774 | 0.72306 | 2.37456 | 0.68778 | 0.05940 |
| **KIF2A** | 2.96286 | 0.00000 | 2.45228 | 4.68148 | 1.52466 | 3.55621 | 5.63521 |
| **KIF2C** | 25.01154 | 12.37321 | 13.99311 | 28.68423 | 19.03040 | 10.00746 | 0.00000 |
| **KIF3A** | 6.41678 | 0.51417 | 2.58482 | 5.76100 | 6.48738 | 3.91017 | 1.94890 |
| **KIF3B** | 7.13661 | 5.95095 | 5.09250 | 6.38246 | 2.96893 | 2.96497 | 6.44078 |
| **KIF3C** | 0.75505 | 1.44035 | 0.86136 | 1.64137 | 2.90973 | 2.61279 | 1.23168 |
| **KIF4A** | 7.34655 | 9.85851 | 12.07891 | 2.70956 | 5.93754 | 4.17889 | 11.43574 |
| **KIF4B** | 1.79506 | 1.60121 | 2.60172 | 0.40633 | 1.40928 | 0.79564 | 0.94456 |
| **KIF5A** | 0.01964 | 0.03838 | 0.08889 | 0.05228 | 0.07621 | 1.14129 | 0.03195 |
| **KIF5B** | 35.13328 | 22.55876 | 19.32557 | 23.12932 | 6.46360 | 15.26656 | 24.31431 |
| **KIF5C** | 0.05427 | 0.35714 | 0.00000 | 0.18707 | 1.72220 | 1.63064 | 0.10311 |
| **KIF6** | 0.01928 | 0.15499 | 0.07983 | 0.08627 | 0.21142 | 0.03099 | 0.00000 |
| **KIF7** | 0.25080 | 0.07835 | 0.00000 | 0.25396 | 0.11961 | 0.57454 | 0.87526 |
| **KIF9** | 2.36753 | 8.76862 | 7.13102 | 1.92473 | 0.94290 | 5.63834 | 0.55891 |
| **KIF9-AS1** | 0.06719 | 0.52122 | 1.30881 | 0.13131 | 0.28210 | 0.18987 | 0.27368 |
| **KIFAP3** | 7.20244 | 1.45209 | 3.09074 | 2.39486 | 1.86127 | 6.33777 | 2.22718 |
| **KIFC1** | 10.49200 | 7.76134 | 8.99836 | 12.80743 | 2.10518 | 5.40972 | 3.21350 |

| | | | | | | | |
|---|---|---|---|---|---|---|---|
| **KIFC2** | 0.04074 | 0.18314 | 0.01815 | 0.10119 | 0.15335 | 0.21851 | 1.73226 |
| **KIFC3** | 0.00000 | 0.09218 | 1.76534 | 0.57449 | 0.63116 | 1.54638 | 1.30486 |
| **KIN** | 9.46832 | 4.21825 | 7.35426 | 7.71655 | 4.88925 | 4.12101 | 4.87360 |
| **KIR2DL1** | 0.06685 | 0.00632 | 0.03441 | 0.01670 | 0.00768 | 0.02070 | 0.00000 |
| **KIR2DL3** | 0.05040 | 0.01043 | 0.02602 | 0.01265 | 0.00842 | 0.02145 | 0.01115 |
| **KIR2DL4.uid-24** | 0.02260 | 0.02230 | 0.01175 | 0.02067 | 0.00795 | 0.01844 | 0.02410 |
| **KIR3DL1** | 0.03801 | 0.00510 | 0.04065 | 0.02118 | 0.00659 | 0.02661 | 0.01464 |
| **KIR3DL2** | 0.03307 | 0.00741 | 0.05603 | 0.07214 | 0.00830 | 0.04456 | 0.00161 |
| **KIR3DX1** | 0.04035 | 0.02411 | 0.01332 | 0.01515 | 0.00774 | 0.01134 | 0.01502 |
| **KIRREL** | 0.05407 | 0.00325 | 0.01091 | 0.10319 | 0.02533 | 0.50647 | 0.00000 |
| **KIRREL2** | 0.00000 | 0.05785 | 0.01395 | 0.09768 | 0.14273 | 0.20727 | 0.10200 |
| **KIRREL3** | 0.06157 | 0.01644 | 0.01951 | 0.07636 | 0.03091 | 0.32289 | 0.00000 |
| **KIRREL3-AS3** | 0.02230 | 0.01744 | 0.00913 | 0.01938 | 0.04325 | 0.01297 | 0.02026 |
| **KISS1R** | 0.02601 | 0.01576 | 0.01270 | 0.01755 | 0.00942 | 0.01676 | 0.02294 |
| **KIT** | 0.00000 | 3.98876 | 0.49999 | 0.00000 | 3.95333 | 5.61378 | 0.11839 |
| **KITLG** | 2.71630 | 0.00000 | 7.61272 | 0.00000 | 3.29137 | 0.00000 | 0.04927 |
| **KIZ** | 2.74503 | 0.32887 | 3.63728 | 3.63875 | 1.53783 | 2.80422 | 0.84370 |
| **KL** | 0.01662 | 0.00956 | 0.04851 | 0.01371 | 0.00665 | 0.01084 | 0.05357 |
| **KLB** | 0.48150 | 0.40311 | 0.15451 | 0.84456 | 0.27112 | 0.84795 | 0.34743 |
| **KLC1** | 0.00000 | 3.92851 | 1.52750 | 4.02598 | 2.32093 | 3.18682 | 12.79055 |
| **KLC2** | 0.00000 | 0.41837 | 1.77492 | 1.56171 | 0.91405 | 0.94124 | 6.22954 |
| **KLC3** | 0.28084 | 0.21848 | 0.10214 | 0.50799 | 0.13860 | 0.51450 | 0.28428 |
| **KLC4** | 3.15955 | 1.83137 | 0.35448 | 6.33731 | 1.62497 | 3.69820 | 0.00000 |
| **KLF1** | 0.11866 | 0.02285 | 0.06567 | 0.09345 | 0.02954 | 0.14689 | 0.16962 |
| **KLF10** | 15.75184 | 8.55460 | 2.16888 | 10.81389 | 8.34184 | 2.37260 | 0.00000 |
| **KLF11** | 0.97420 | 4.71272 | 4.93416 | 0.00000 | 4.79597 | 4.70653 | 3.10898 |
| **KLF12** | 0.03695 | 0.00000 | 0.80020 | 0.03866 | 1.30075 | 0.19578 | 0.04516 |
| **KLF13** | 0.00000 | 2.87833 | 0.74714 | 0.50707 | 0.12818 | 1.77100 | 9.60728 |
| **KLF14** | 0.00000 | 0.12481 | 0.04313 | 0.01514 | 0.04768 | 0.08121 | 0.09794 |
| **KLF15** | 0.00952 | 0.09423 | 0.02183 | 0.29896 | 0.08373 | 0.11034 | 0.59318 |
| **KLF16** | 0.00000 | 5.84702 | 3.46317 | 1.40202 | 1.78989 | 4.22019 | 15.26374 |
| **KLF17** | 0.15098 | 0.00000 | 13.17702 | 0.00000 | 25.36785 | 10.22114 | 0.00796 |
| **KLF2** | 0.00000 | 0.01447 | 0.00000 | 0.02654 | 0.50860 | 0.40121 | 0.27607 |
| **KLF3** | 3.71617 | 6.42146 | 10.01622 | 0.13419 | 0.87266 | 5.24927 | 10.25505 |
| **KLF3-AS1** | 0.02416 | 0.07069 | 0.02014 | 0.01645 | 0.01393 | 0.01529 | 0.01655 |

| | | | | | | | |
|---|---|---|---|---|---|---|---|
| **KLF4** | 0.00000 | 27.40158 | 17.15910 | 5.58971 | 1.76033 | 32.94114 | 0.00000 |
| **KLF5** | 14.40135 | 18.03987 | 13.36103 | 0.50756 | 4.83688 | 6.23767 | 43.10881 |
| **KLF6** | 14.87905 | 6.20251 | 0.00000 | 13.25175 | 0.00000 | 13.28797 | 11.23434 |
| **KLF7** | 0.09680 | 0.00000 | 0.12375 | 0.05091 | 0.22157 | 2.86926 | 0.80378 |
| **KLF8** | 3.40220 | 9.97715 | 5.93360 | 0.00000 | 0.00000 | 2.95599 | 2.63610 |
| **KLF9** | 0.44070 | 0.00000 | 4.37352 | 0.23900 | 1.68994 | 2.24090 | 2.12346 |
| **KLHDC1** | 0.02768 | 0.01597 | 0.01220 | 0.01762 | 0.03445 | 0.01560 | 0.01925 |
| **KLHDC10** | 1.57041 | 0.06145 | 0.67447 | 1.95004 | 0.31562 | 1.10909 | 6.13438 |
| **KLHDC2** | 0.27498 | 2.44541 | 5.84608 | 1.92306 | 0.38253 | 4.47459 | 7.39416 |
| **KLHDC3** | 18.98983 | 14.73824 | 12.66410 | 25.40936 | 6.82255 | 15.05221 | 9.38492 |
| **KLHDC4** | 7.22505 | 5.66698 | 1.31401 | 4.39447 | 0.99847 | 3.07959 | 2.80787 |
| **KLHDC7A** | 0.04978 | 0.00358 | 0.01338 | 0.02469 | 0.09509 | 0.04981 | 0.01752 |
| **KLHDC7B** | 0.02648 | 0.01856 | 0.01256 | 0.02014 | 0.00793 | 0.01622 | 0.01586 |
| **KLHDC8A** | 0.02290 | 0.01796 | 0.01580 | 0.03275 | 0.00866 | 0.01394 | 0.01365 |
| **KLHDC8B** | 0.98006 | 0.49125 | 0.31938 | 2.39016 | 0.43124 | 1.18926 | 0.31312 |
| **KLHDC9** | 0.00000 | 0.85433 | 0.34669 | 0.01576 | 0.04533 | 0.52755 | 0.09328 |
| **KLHL1** | 0.05190 | 0.01107 | 0.02578 | 0.02503 | 0.02734 | 0.01128 | 0.00554 |
| **KLHL10** | 0.03693 | 0.00385 | 0.03132 | 0.02574 | 0.00729 | 0.01608 | 0.03337 |
| **KLHL11** | 0.00000 | 3.58326 | 3.81630 | 5.29395 | 9.09750 | 0.74348 | 3.91997 |
| **KLHL12** | 6.23234 | 2.31026 | 4.07806 | 3.81854 | 2.40442 | 3.43402 | 0.94114 |
| **KLHL13** | 0.08621 | 0.00000 | 0.28323 | 0.07998 | 0.02679 | 0.02493 | 0.06083 |
| **KLHL14** | 0.26907 | 0.23762 | 0.00798 | 0.09403 | 0.02342 | 0.00654 | 0.00000 |
| **KLHL15** | 4.39433 | 0.00000 | 7.85664 | 0.23707 | 11.88746 | 1.46070 | 3.97598 |
| **KLHL17** | 0.50805 | 1.30633 | 0.97692 | 1.26994 | 0.60840 | 1.28417 | 4.90335 |
| **KLHL18** | 7.53234 | 18.25349 | 15.86671 | 7.77993 | 6.53535 | 20.36427 | 6.06298 |
| **KLHL2** | 2.41543 | 0.00000 | 1.55098 | 0.68145 | 0.66879 | 0.14698 | 3.89410 |
| **KLHL20** | 4.33329 | 0.13004 | 3.93609 | 3.16984 | 3.44950 | 2.58286 | 2.21174 |
| **KLHL21** | 8.93061 | 7.32212 | 2.86607 | 9.50633 | 3.62832 | 0.67333 | 16.48283 |
| **KLHL22** | 0.00000 | 1.84152 | 0.57786 | 1.50391 | 0.36915 | 0.34087 | 0.98457 |
| **KLHL23** | 0.28953 | 0.04375 | 0.00984 | 0.97969 | 0.03407 | 0.00318 | 0.43451 |
| **KLHL24** | 2.81250 | 2.09487 | 6.47658 | 8.05463 | 5.24386 | 5.44878 | 3.25609 |
| **KLHL25** | 0.51038 | 1.31577 | 0.55936 | 2.43459 | 0.33616 | 0.79638 | 2.46358 |
| **KLHL26** | 0.22795 | 0.62614 | 0.52125 | 0.91789 | 1.11208 | 0.00000 | 0.79404 |
| **KLHL28** | 4.41605 | 0.00000 | 2.72544 | 3.00103 | 3.58637 | 1.50541 | 0.87728 |
| **KLHL29** | 0.05841 | 0.00962 | 0.03199 | 0.48311 | 0.07233 | 0.00000 | 0.69094 |

| | | | | | | | |
|---|---|---|---|---|---|---|---|
| **KLHL3** | 0.11079 | 0.26930 | 0.00798 | 0.02151 | 0.07442 | 3.65371 | 0.00901 |
| **KLHL30** | 0.02121 | 0.02826 | 0.01775 | 0.02630 | 0.00668 | 0.01683 | 0.04588 |
| **KLHL31** | 0.12034 | 0.46827 | 0.00236 | 0.01409 | 0.00846 | 0.00000 | 0.02266 |
| **KLHL32** | 0.01995 | 0.00184 | 0.04440 | 0.01903 | 1.13340 | 0.06750 | 0.03156 |
| **KLHL34** | 0.02476 | 0.03876 | 0.00523 | 0.01106 | 0.04084 | 0.00750 | 0.04522 |
| **KLHL35** | 0.00000 | 1.24051 | 2.50300 | 0.00000 | 0.32750 | 0.00000 | 0.07364 |
| **KLHL36** | 0.00000 | 1.46900 | 0.00000 | 0.89689 | 0.50979 | 0.68032 | 1.66905 |
| **KLHL4** | 0.01058 | 0.00897 | 0.02533 | 0.04569 | 0.00696 | 0.09228 | 0.03418 |
| **KLHL40** | 0.02165 | 0.01779 | 0.01297 | 0.02198 | 0.00798 | 0.01392 | 0.01896 |
| **KLHL41** | 0.15141 | 0.00003 | 0.09076 | 0.04575 | 0.27023 | 0.01415 | 0.00000 |
| **KLHL42** | 1.43152 | 2.20029 | 1.50578 | 2.26596 | 0.04544 | 4.21581 | 9.07687 |
| **KLHL5** | 0.73405 | 0.88481 | 0.00000 | 0.00000 | 0.00801 | 0.27767 | 2.28361 |
| **KLHL6** | 0.01903 | 0.02466 | 0.01139 | 0.04070 | 0.01585 | 0.02798 | 0.03284 |
| **KLHL7** | 12.70956 | 8.13446 | 9.96465 | 11.14746 | 7.63816 | 3.72853 | 0.00000 |
| **KLHL7-AS1** | 0.44919 | 0.00000 | 0.28412 | 0.08916 | 0.00164 | 0.11038 | 0.07818 |
| **KLHL8** | 2.95076 | 0.06832 | 0.74093 | 2.38460 | 2.05728 | 1.28474 | 2.87566 |
| **KLHL9** | 2.73020 | 3.63297 | 6.09761 | 2.44806 | 0.70424 | 0.78919 | 3.44412 |
| **KLK1** | 0.00000 | 0.21250 | 0.11248 | 0.00884 | 0.03379 | 0.12351 | 0.01802 |
| **KLK10** | 0.00000 | 0.05162 | 1.12667 | 0.11983 | 3.58507 | 0.00000 | 0.00229 |
| **KLK11** | 0.00000 | 0.00000 | 2.82817 | 0.00315 | 11.67798 | 0.00000 | 0.03694 |
| **KLK12** | 0.02228 | 0.01224 | 0.02415 | 0.02399 | 0.01790 | 0.01491 | 0.01716 |
| **KLK13** | 0.04301 | 0.22072 | 0.14699 | 0.00000 | 0.10318 | 3.86083 | 0.00000 |
| **KLK14** | 0.02493 | 0.04526 | 0.05050 | 0.03395 | 0.01984 | 0.01656 | 0.01143 |
| **KLK15** | 0.03523 | 0.00000 | 0.17179 | 0.01547 | 0.01914 | 0.01001 | 0.01551 |
| **KLK2** | 0.03361 | 0.13308 | 0.31186 | 0.07837 | 0.09198 | 0.00000 | 0.03713 |
| **KLK4** | 0.02455 | 0.00000 | 0.04936 | 0.01946 | 0.41304 | 0.01237 | 0.01564 |
| **KLK5** | 0.03291 | 0.22923 | 0.03032 | 0.00425 | 0.00523 | 0.00052 | 0.03110 |
| **KLK6** | 0.00000 | 0.06729 | 0.01183 | 0.05079 | 0.00992 | 2.09171 | 0.01763 |
| **KLK7** | 1.51692 | 0.73902 | 0.07029 | 0.46132 | 0.39294 | 0.35427 | 0.28842 |
| **KLK8** | 4.16120 | 5.47653 | 2.66020 | 1.39805 | 1.59481 | 3.06816 | 2.37713 |
| **KLK9** | 0.02142 | 0.00880 | 0.02134 | 0.03563 | 0.09763 | 0.01644 | 0.06481 |
| **KLKB1** | 0.00868 | 0.01434 | 0.01021 | 0.04482 | 0.03987 | 0.02226 | 0.03878 |
| **KLKP1** | 0.02478 | 0.00000 | 0.06881 | 0.01761 | 0.23347 | 0.01180 | 0.01935 |
| **KLLN** | 0.00000 | 0.30199 | 0.16559 | 0.08523 | 0.65729 | 0.18158 | 0.04553 |
| **KLRAP1** | 0.34212 | 0.00000 | 0.29656 | 0.12881 | 0.06722 | 0.02325 | 0.24010 |

| | | | | | | | |
|---|---|---|---|---|---|---|---|
| **KLRB1** | 0.01827 | 0.01202 | 0.15604 | 0.01515 | 0.11353 | 0.03977 | 0.05570 |
| **KLRC1** | 0.05418 | 0.01237 | 0.01990 | 0.04339 | 0.01089 | 0.02717 | 0.01004 |
| **KLRC2** | 0.02979 | 0.01692 | 0.01366 | 0.02159 | 0.00828 | 0.01331 | 0.01886 |
| **KLRC3** | 0.03794 | 0.01530 | 0.01274 | 0.01648 | 0.00781 | 0.01127 | 0.01989 |
| **KLRD1** | 0.04679 | 0.02294 | 0.00246 | 0.09246 | 0.11245 | 0.00171 | 0.07834 |
| **KLRF1** | 0.02477 | 0.02002 | 0.00540 | 0.01879 | 0.10331 | 0.02072 | 0.01491 |
| **KLRF2** | 0.02535 | 0.01829 | 0.01116 | 0.01995 | 0.02753 | 0.01347 | 0.01540 |
| **KLRG1** | 0.12428 | 0.00000 | 0.53496 | 0.45444 | 0.89296 | 0.10886 | 0.00000 |
| **KLRG2** | 0.00000 | 14.00420 | 1.27368 | 0.39366 | 0.02971 | 7.12203 | 11.27374 |
| **KMO** | 0.01950 | 0.02852 | 0.02757 | 0.19533 | 0.01050 | 0.18291 | 0.02302 |
| **KMT2A** | 3.38783 | 0.73181 | 0.44067 | 2.24398 | 1.36665 | 3.05153 | 11.00455 |
| **KMT2B** | 0.07237 | 3.59757 | 2.20883 | 3.01436 | 0.99013 | 4.79466 | 20.43129 |
| **KMT2C** | 10.13985 | 7.64314 | 15.00606 | 8.06215 | 15.20300 | 14.01809 | 34.05391 |
| **KMT2D** | 0.00000 | 0.66265 | 4.08452 | 7.24702 | 1.13818 | 5.64969 | 33.36470 |
| **KMT2E** | 7.26665 | 4.95711 | 3.69766 | 2.96109 | 2.89720 | 6.51942 | 14.28993 |
| **KMT2E-AS1** | 2.76946 | 3.30447 | 0.94214 | 1.75376 | 0.30699 | 0.80801 | 0.97762 |
| **KNCN** | 0.00960 | 0.04705 | 0.00594 | 0.05489 | 0.22566 | 0.01151 | 0.01037 |
| **KNDC1** | 0.02873 | 0.00925 | 0.01197 | 0.13005 | 0.01131 | 0.23653 | 0.00000 |
| **KNG1** | 0.15366 | 0.24307 | 0.66091 | 0.01427 | 0.01142 | 0.47471 | 0.54072 |
| **KNOP1** | 20.56694 | 17.38007 | 16.15810 | 11.44101 | 5.02541 | 8.43548 | 16.15856 |
| **KNSTRN** | 32.63324 | 16.45266 | 28.30957 | 26.49893 | 6.89362 | 4.16947 | 0.00000 |
| **KNTC1** | 7.60768 | 2.22719 | 3.07925 | 2.33452 | 3.30041 | 3.89413 | 6.06092 |
| **KPNA1** | 23.87055 | 5.29367 | 13.48161 | 11.82807 | 4.83881 | 8.59953 | 3.05386 |
| **KPNA2** | 68.25633 | 32.02140 | 33.24136 | 39.85272 | 29.62406 | 22.32809 | 6.73640 |
| **KPNA3** | 22.12124 | 6.97960 | 19.37419 | 8.25132 | 4.04849 | 13.85361 | 21.31583 |
| **KPNA4** | 10.72158 | 3.47362 | 3.42795 | 5.40394 | 1.55912 | 6.01343 | 16.23615 |
| **KPNA5** | 1.54678 | 0.00000 | 0.88431 | 0.79783 | 0.00284 | 0.29101 | 1.67532 |
| **KPNA6** | 35.56522 | 13.86969 | 21.92598 | 29.45525 | 16.94446 | 16.86832 | 15.87879 |
| **KPNA7** | 0.00000 | 0.17226 | 0.14367 | 0.00000 | 27.35690 | 0.04660 | 0.01907 |
| **KPNB1** | 36.01752 | 25.41638 | 14.18806 | 27.91654 | 7.03490 | 21.66467 | 44.59252 |
| **KPTN** | 4.46521 | 6.50231 | 4.97857 | 7.91794 | 7.76475 | 1.73573 | 2.47128 |
| **KRAS** | 18.05996 | 3.44402 | 21.12688 | 10.45028 | 5.15813 | 11.20406 | 23.93172 |
| **KRBA1** | 0.00001 | 0.57300 | 0.13543 | 1.84116 | 0.00093 | 0.50906 | 3.30796 |
| **KRBA2** | 0.02239 | 0.55817 | 0.46111 | 0.01374 | 0.42037 | 0.24944 | 0.10062 |
| **KRBOX1** | 1.31366 | 6.11745 | 6.99812 | 0.00000 | 2.02803 | 0.70134 | 0.87212 |

| | | | | | | | |
|---|---|---|---|---|---|---|---|
| **KRBOX1-AS1** | 0.01847 | 0.01663 | 0.11034 | 0.03548 | 0.03476 | 0.00721 | 0.02577 |
| **KRBOX4** | 2.08566 | 1.37656 | 5.10448 | 0.05810 | 2.39790 | 0.66683 | 0.59256 |
| **KRCC1** | 0.16136 | 0.01885 | 0.55148 | 0.21446 | 0.10836 | 0.01036 | 0.23460 |
| **KREMEN1** | 0.35538 | 0.59454 | 0.00000 | 3.14726 | 0.03232 | 3.93046 | 3.23501 |
| **KREMEN2** | 0.11126 | 0.29651 | 0.00000 | 0.00000 | 0.01328 | 1.55629 | 1.64414 |
| **KRI1** | 13.55109 | 12.56062 | 4.98764 | 12.90059 | 2.87688 | 8.49061 | 8.12352 |
| **KRIT1** | 8.56582 | 1.09578 | 7.61394 | 5.06057 | 0.93106 | 3.43854 | 8.09381 |
| **KRR1** | 12.75917 | 0.23860 | 9.95651 | 12.85745 | 4.33641 | 4.78919 | 5.65087 |
| **KRT1** | 0.03507 | 0.03321 | 0.01170 | 0.03599 | 0.00791 | 0.01044 | 0.00905 |
| **KRT10** | 1.28834 | 0.30322 | 5.86649 | 4.39442 | 1.80874 | 0.91081 | 12.79152 |
| **KRT12** | 0.06302 | 0.01234 | 0.01385 | 0.08236 | 0.06995 | 0.03552 | 0.00000 |
| **KRT13** | 0.02369 | 0.01038 | 0.01179 | 0.04184 | 0.01759 | 0.02137 | 0.05425 |
| **KRT14** | 0.01125 | 0.03027 | 0.00861 | 0.21965 | 0.04542 | 0.01193 | 0.00006 |
| **KRT15** | 0.13203 | 0.00038 | 0.02644 | 0.51536 | 0.03999 | 0.00000 | 0.17472 |
| **KRT16** | 0.00942 | 0.03180 | 0.01012 | 0.40235 | 0.00951 | 0.01867 | 0.00000 |
| **KRT16P1** | 0.01285 | 0.00948 | 0.01119 | 0.02185 | 0.00779 | 0.29957 | 0.02854 |
| **KRT16P2** | 0.04659 | 0.01358 | 0.01167 | 0.01264 | 0.00944 | 0.06078 | 0.03916 |
| **KRT16P3** | 0.03295 | 0.01136 | 0.00954 | 0.00996 | 0.00767 | 0.41042 | 0.02844 |
| **KRT17** | 0.00237 | 0.02158 | 0.00201 | 1.50496 | 0.25874 | 0.01375 | 0.00000 |
| **KRT17P5** | 0.02362 | 0.01815 | 0.01276 | 0.02098 | 0.00802 | 0.01383 | 0.01804 |
| **KRT18** | 95.01073 | 95.08193 | 0.00000 | 73.33522 | 0.00000 | 14.72555 | 32.10374 |
| **KRT18P55** | 0.28618 | 0.51548 | 0.04868 | 0.27535 | 0.00565 | 0.00000 | 0.35061 |
| **KRT19** | 62.08695 | 0.00000 | 0.00000 | 75.98727 | 0.00000 | 0.00000 | 25.34689 |
| **KRT19P2** | 0.05878 | 0.00879 | 0.01766 | 0.04307 | 0.00750 | 0.01049 | 0.02677 |
| **KRT2** | 0.02481 | 0.01882 | 0.01261 | 0.02079 | 0.00794 | 0.01302 | 0.01649 |
| **KRT222** | 0.02262 | 0.00000 | 0.12752 | 0.01916 | 0.45708 | 0.00921 | 0.01510 |
| **KRT23** | 0.00000 | 0.23659 | 0.00034 | 18.00525 | 0.02153 | 2.13766 | 0.00000 |
| **KRT24** | 0.02198 | 0.01731 | 0.01095 | 0.02170 | 0.03873 | 0.01319 | 0.01968 |
| **KRT26** | 0.03689 | 0.01644 | 0.01311 | 0.01834 | 0.00815 | 0.01346 | 0.00948 |
| **KRT27** | 0.05446 | 0.02171 | 0.15662 | 0.02017 | 0.01614 | 0.00370 | 0.02487 |
| **KRT28** | 0.02663 | 0.01790 | 0.01319 | 0.01874 | 0.00852 | 0.01459 | 0.01687 |
| **KRT3** | 0.02933 | 0.01924 | 0.01203 | 0.02709 | 0.00776 | 0.01193 | 0.02879 |
| **KRT32** | 0.02242 | 0.01398 | 0.01488 | 0.02373 | 0.00769 | 0.02545 | 0.02029 |
| **KRT33B** | 0.02261 | 0.01851 | 0.01270 | 0.02912 | 0.00794 | 0.01243 | 0.01597 |
| **KRT34** | 0.03123 | 0.02297 | 0.01238 | 0.01880 | 0.00812 | 0.01248 | 0.01220 |

| | | | | | | | |
|---|---|---|---|---|---|---|---|
| **KRT35** | 0.02732 | 0.01779 | 0.01291 | 0.02523 | 0.00800 | 0.01328 | 0.01221 |
| **KRT36** | 0.02788 | 0.02101 | 0.00500 | 0.03271 | 0.18382 | 0.01143 | 0.01155 |
| **KRT38** | 0.02496 | 0.01349 | 0.02230 | 0.02077 | 0.00766 | 0.01278 | 0.01651 |
| **KRT4** | 0.01466 | 0.02120 | 0.01114 | 0.04551 | 0.01018 | 0.01457 | 0.05358 |
| **KRT42P** | 0.02199 | 0.03794 | 0.00933 | 0.01958 | 0.00878 | 0.08450 | 0.01036 |
| **KRT5** | 0.01565 | 0.01846 | 0.01268 | 0.02946 | 0.00783 | 0.01473 | 0.02601 |
| **KRT6A** | 0.03156 | 0.01542 | 0.01317 | 0.02349 | 0.01373 | 0.01105 | 0.02560 |
| **KRT6B** | 0.02790 | 0.01814 | 0.01373 | 0.01952 | 0.00839 | 0.01195 | 0.02791 |
| **KRT7** | 1.28584 | 4.52277 | 0.00000 | 25.73828 | 0.14362 | 4.59908 | 2.89234 |
| **KRT71** | 0.00462 | 0.02464 | 0.01211 | 0.15954 | 0.00783 | 0.01450 | 0.02012 |
| **KRT72** | 0.05119 | 0.02989 | 0.01247 | 0.01164 | 0.00760 | 0.00998 | 0.02008 |
| **KRT73** | 0.01208 | 0.03620 | 0.06699 | 0.02220 | 0.00452 | 0.00856 | 0.03444 |
| **KRT74** | 0.01504 | 0.02262 | 0.01177 | 0.04361 | 0.00801 | 0.01290 | 0.01891 |
| **KRT75** | 0.04548 | 0.03713 | 0.01599 | 0.02688 | 0.00713 | 0.01407 | 0.02842 |
| **KRT76** | 0.03620 | 0.01383 | 0.01352 | 0.01540 | 0.00782 | 0.01191 | 0.01800 |
| **KRT77** | 0.02812 | 0.01109 | 0.01244 | 0.01845 | 0.02147 | 0.12071 | 0.01171 |
| **KRT78** | 0.01262 | 0.02199 | 0.01140 | 0.02969 | 0.00758 | 0.01054 | 0.04413 |
| **KRT79** | 0.02083 | 0.01618 | 0.01300 | 0.03246 | 0.00998 | 0.06462 | 0.01656 |
| **KRT8** | 93.21020 | 99.86882 | 0.52437 | 80.17029 | 0.00000 | 17.51700 | 34.20392 |
| **KRT80** | 0.00000 | 0.06257 | 0.00040 | 4.07880 | 0.13253 | 0.00930 | 0.00000 |
| **KRT81** | 0.01137 | 0.02139 | 0.01359 | 0.24333 | 0.00778 | 0.00876 | 0.01201 |
| **KRT83** | 0.01693 | 0.05119 | 0.02178 | 0.01229 | 0.00637 | 0.01328 | 0.04552 |
| **KRT84** | 0.02611 | 0.02362 | 0.01190 | 0.02593 | 0.00780 | 0.00875 | 0.03725 |
| **KRT85** | 0.01241 | 0.01855 | 0.01209 | 0.02704 | 0.00764 | 0.01155 | 0.04361 |
| **KRT86** | 0.00298 | 0.02114 | 0.01210 | 0.39848 | 0.00767 | 0.00770 | 0.02238 |
| **KRT8P41** | 0.31918 | 0.28917 | 0.04746 | 0.26492 | 0.00474 | 0.03195 | 0.08639 |
| **KRT9** | 0.04702 | 0.00075 | 0.02289 | 0.05707 | 0.00914 | 0.00354 | 0.12934 |
| **KRTAP10-2** | 0.02215 | 0.01874 | 0.01240 | 0.01990 | 0.01235 | 0.01289 | 0.01948 |
| **KRTAP10-6** | 0.02245 | 0.01507 | 0.03601 | 0.01886 | 0.02262 | 0.01204 | 0.01891 |
| **KRTAP11-1** | 0.02168 | 0.03265 | 0.01068 | 0.02166 | 0.05774 | 0.01204 | 0.01253 |
| **KRTAP19-8** | 0.04132 | 0.03629 | 0.00415 | 0.01690 | 0.08366 | 0.00682 | 0.03376 |
| **KRTAP4-1** | 0.04017 | 0.02101 | 0.02556 | 0.01419 | 0.00725 | 0.03094 | 0.00869 |
| **KRTAP5-1** | 0.02042 | 0.01723 | 0.01252 | 0.02675 | 0.00787 | 0.01207 | 0.02767 |
| **KRTAP5-10** | 0.03278 | 0.01552 | 0.02383 | 0.01964 | 0.00693 | 0.02249 | 0.01632 |
| **KRTAP5-7** | 0.01829 | 0.01738 | 0.01406 | 0.02337 | 0.00766 | 0.01317 | 0.03074 |

| | | | | | | | |
|---|---|---|---|---|---|---|---|
| **KRTAP5-9** | 0.02333 | 0.00436 | 0.10731 | 0.02289 | 0.00505 | 0.02720 | 0.01962 |
| **KRTAP5-AS1** | 0.02719 | 0.03473 | 0.01724 | 0.04166 | 0.00772 | 0.01361 | 0.03695 |
| **KRTAP7-1** | 0.04988 | 0.00891 | 0.01275 | 0.01644 | 0.01850 | 0.01016 | 0.02693 |
| **KRTCAP2** | 11.96759 | 14.91222 | 8.27040 | 12.80217 | 1.88527 | 7.75133 | 16.39055 |
| **KRTCAP3** | 3.94229 | 6.42813 | 2.27360 | 2.77386 | 0.43959 | 9.26854 | 0.00000 |
| **KRTDAP** | 0.04076 | 0.01497 | 0.03038 | 0.03471 | 0.07633 | 0.01952 | 0.01155 |
| **KSR1** | 0.02902 | 0.48990 | 0.00000 | 2.96054 | 0.68799 | 0.40045 | 3.64858 |
| **KSR2** | 0.00000 | 0.08126 | 0.06067 | 0.13967 | 2.69975 | 0.06639 | 0.16508 |
| **KTN1** | 19.38465 | 5.90724 | 8.92701 | 11.03406 | 4.67112 | 15.88676 | 48.59897 |
| **KTN1-AS1** | 0.47453 | 0.98520 | 0.43534 | 0.42487 | 0.21499 | 0.41201 | 0.74722 |
| **KU-MEL-3** | 0.03188 | 0.01963 | 0.01203 | 0.03069 | 0.00804 | 0.01099 | 0.01614 |
| **KXD1** | 26.66323 | 26.81508 | 11.22947 | 23.64660 | 0.26596 | 18.29380 | 12.66581 |
| **KY** | 0.01996 | 0.00798 | 0.03239 | 0.02715 | 0.03766 | 0.01759 | 0.02830 |
| **KYNU** | 0.05046 | 0.00000 | 0.12434 | 1.91841 | 0.18587 | 0.18539 | 0.00000 |
| **L1CAM** | 0.16755 | 0.24192 | 0.10901 | 0.11686 | 0.15896 | 0.00000 | 2.08569 |
| **L1TD1** | 26.63482 | 6.25744 | 25.51445 | 26.03670 | 2.74642 | 38.09489 | 18.94669 |
| **L2HGDH** | 5.32003 | 1.90200 | 2.18999 | 5.66816 | 0.28492 | 2.97760 | 2.01728 |
| **L3HYPDH** | 10.27580 | 0.00383 | 0.98737 | 5.67052 | 0.71396 | 0.66837 | 0.00000 |
| **L3MBTL1** | 0.26726 | 0.00000 | 0.22433 | 0.51253 | 0.02678 | 0.00000 | 0.31201 |
| **L3MBTL2** | 12.88516 | 8.75324 | 3.19597 | 11.42918 | 1.33243 | 3.98401 | 3.38067 |
| **L3MBTL3** | 0.94104 | 0.15990 | 0.22037 | 0.00000 | 0.15851 | 0.72398 | 2.95242 |
| **LACC1** | 0.30926 | 0.32876 | 0.15841 | 0.56960 | 3.01252 | 0.00000 | 0.35251 |
| **LACE1** | 3.32043 | 1.14831 | 1.06704 | 0.80790 | 0.21796 | 1.14151 | 0.00000 |
| **LACTB** | 1.21764 | 0.61583 | 0.07840 | 0.50101 | 0.09855 | 5.03215 | 2.35076 |
| **LACTB2** | 3.70279 | 2.95389 | 0.07795 | 1.21273 | 0.10397 | 1.85492 | 0.17401 |
| **LAD1** | 11.17067 | 5.33711 | 0.00000 | 9.54738 | 7.90987 | 0.00000 | 12.84430 |
| **LAG3** | 0.03315 | 0.00110 | 0.03136 | 0.04948 | 0.07699 | 0.04090 | 0.03328 |
| **LAGE3** | 1.92599 | 6.61982 | 0.95830 | 6.11147 | 1.55558 | 7.74213 | 20.16139 |
| **LAIR1** | 0.01855 | 0.03184 | 0.03561 | 0.03610 | 0.00665 | 0.00575 | 0.04888 |
| **LAIR2** | 0.02062 | 0.01297 | 0.01042 | 0.10327 | 0.00746 | 0.00377 | 0.12109 |
| **LALBA** | 0.00000 | 0.55562 | 0.26982 | 0.00000 | 0.00216 | 0.00000 | 0.06769 |
| **LAMA1** | 47.10178 | 30.18554 | 7.48215 | 28.51167 | 0.65113 | 6.74948 | 48.28037 |
| **LAMA2** | 0.35827 | 0.14023 | 0.00000 | 0.00000 | 0.74631 | 0.17576 | 0.39211 |
| **LAMA3** | 0.19050 | 0.46176 | 0.43000 | 0.00000 | 0.03839 | 0.14612 | 0.05789 |
| **LAMA4** | 0.00000 | 1.90271 | 1.16906 | 0.00000 | 0.96061 | 7.13263 | 0.00032 |

| | | | | | | | |
|---|---|---|---|---|---|---|---|
| **LAMA5** | 0.30142 | 6.35443 | 0.08496 | 7.82134 | 0.25697 | 6.32227 | 43.75510 |
| **LAMA5-AS1** | 0.02566 | 0.02375 | 0.01139 | 0.01549 | 0.00785 | 0.01916 | 0.03059 |
| **LAMB1** | 42.80708 | 12.43113 | 1.43199 | 20.73874 | 5.45634 | 8.26416 | 32.45727 |
| **LAMB2** | 3.01919 | 0.94373 | 0.07959 | 2.02193 | 0.07209 | 1.05097 | 2.79534 |
| **LAMB2P1** | 0.00837 | 0.00501 | 0.02077 | 0.03736 | 0.00714 | 0.08834 | 0.04974 |
| **LAMB3** | 1.34302 | 0.37383 | 0.00000 | 3.07745 | 0.00026 | 0.00000 | 1.22757 |
| **LAMB4** | 0.18603 | 0.02672 | 0.01492 | 0.05176 | 0.17022 | 0.08021 | 0.00000 |
| **LAMC1** | 38.19095 | 18.01379 | 3.68824 | 15.78326 | 2.67933 | 12.19125 | 47.71456 |
| **LAMC2** | 1.08576 | 1.43102 | 0.08219 | 3.28119 | 0.07945 | 2.87446 | 0.41672 |
| **LAMC3** | 0.00000 | 3.44592 | 0.08965 | 0.15227 | 0.00859 | 0.14531 | 1.73244 |
| **LAMP1** | 0.53181 | 3.31967 | 2.59812 | 3.32206 | 1.01918 | 3.92416 | 33.40444 |
| **LAMP2** | 69.97150 | 14.93600 | 4.90109 | 18.89826 | 4.49740 | 4.83906 | 16.04865 |
| **LAMP3** | 0.06168 | 0.10615 | 0.15979 | 0.02834 | 0.39081 | 0.01869 | 0.00000 |
| **LAMP5** | 0.02472 | 0.00843 | 0.03264 | 0.02222 | 0.01763 | 0.01268 | 0.01904 |
| **LAMP5-AS1** | 0.02269 | 0.02565 | 0.01104 | 0.01964 | 0.00811 | 0.01291 | 0.02087 |
| **LAMTOR1** | 28.98142 | 19.98240 | 6.98935 | 25.74360 | 0.96919 | 14.60380 | 16.13218 |
| **LAMTOR2** | 23.24869 | 34.63863 | 7.79615 | 18.81581 | 2.87867 | 16.04147 | 3.99035 |
| **LAMTOR3** | 23.97850 | 0.52590 | 14.45992 | 9.01022 | 8.81509 | 5.71108 | 1.13369 |
| **LAMTOR4** | 34.82539 | 25.85130 | 6.04366 | 22.82698 | 7.96264 | 12.64230 | 15.42583 |
| **LAMTOR5** | 37.70331 | 21.63800 | 23.41671 | 21.91090 | 12.08267 | 14.72433 | 16.31499 |
| **LAMTOR5-AS1** | 0.14173 | 0.26470 | 0.49122 | 0.03326 | 0.05962 | 0.14954 | 0.11564 |
| **LANCL1** | 1.63790 | 0.17259 | 0.67625 | 0.53592 | 0.07312 | 1.02635 | 0.00000 |
| **LANCL1-AS1** | 0.02904 | 0.02236 | 0.01639 | 0.03124 | 0.00940 | 0.01121 | 0.01575 |
| **LANCL2** | 4.62234 | 9.57265 | 2.74152 | 2.93228 | 0.59180 | 1.76095 | 1.01931 |
| **LANCL3** | 0.00951 | 0.00314 | 0.04548 | 0.03824 | 0.01386 | 0.35851 | 0.02403 |
| **LAP3** | 19.27471 | 5.31016 | 6.21440 | 8.92272 | 3.72119 | 8.79259 | 2.16073 |
| **LAPTM4A** | 36.82139 | 9.89371 | 23.23007 | 19.73011 | 13.30025 | 22.94460 | 2.57579 |
| **LAPTM4B** | 3.99498 | 6.23985 | 15.81011 | 1.11026 | 3.20567 | 23.41610 | 18.15050 |
| **LAPTM5** | 0.00000 | 4.14640 | 0.38682 | 0.50507 | 0.03729 | 1.17902 | 0.33715 |
| **LARGE** | 0.68368 | 0.91481 | 0.01857 | 0.54830 | 0.37581 | 1.65534 | 3.88001 |
| **LARGE-AS1** | 0.06810 | 0.24911 | 0.16375 | 0.02758 | 0.03326 | 0.13509 | 0.56926 |
| **LARP1** | 39.61726 | 26.95912 | 3.78974 | 21.72954 | 4.25918 | 18.27792 | 55.83188 |
| **LARP1B** | 5.21126 | 5.69320 | 24.61666 | 5.05053 | 4.78353 | 7.73501 | 14.13003 |
| **LARP4** | 27.34069 | 7.82184 | 15.63340 | 18.04006 | 11.60411 | 8.59651 | 13.50658 |
| **LARP4B** | 21.54782 | 20.77484 | 5.37394 | 15.22219 | 10.15782 | 10.21334 | 4.07904 |

| | | | | | | | |
|---|---|---|---|---|---|---|---|
| **LARP6** | 0.00000 | 4.98300 | 0.23289 | 0.00000 | 0.00659 | 10.03865 | 1.49751 |
| **LARP7** | 24.85963 | 20.54038 | 18.27470 | 3.59796 | 3.57416 | 25.28389 | 20.53839 |
| **LARS** | 36.38412 | 6.29153 | 4.81252 | 22.79735 | 2.18857 | 16.19030 | 26.07621 |
| **LARS2** | 11.31550 | 4.36788 | 3.52785 | 5.23944 | 2.50276 | 6.79657 | 3.65509 |
| **LARS2-AS1** | 0.21418 | 0.00000 | 0.03031 | 0.11133 | 0.07876 | 0.17895 | 0.56110 |
| **LAS1L** | 37.62935 | 38.49430 | 17.14397 | 20.98664 | 5.15402 | 12.89110 | 0.00000 |
| **LASP1** | 6.41910 | 36.78761 | 12.14518 | 11.52075 | 2.96696 | 16.07369 | 26.73862 |
| **LAT** | 0.41552 | 0.60062 | 0.39418 | 0.78804 | 0.00000 | 0.18199 | 1.63168 |
| **LAT2** | 0.33410 | 0.28202 | 0.53166 | 0.25419 | 0.49309 | 0.15821 | 0.67138 |
| **LATS1** | 4.30772 | 0.00000 | 5.44232 | 3.40062 | 4.89980 | 3.11179 | 2.85603 |
| **LATS2** | 5.02661 | 3.24584 | 5.92033 | 2.82036 | 0.75557 | 0.78290 | 12.54437 |
| **LAX1** | 0.05885 | 0.00000 | 0.34010 | 0.11830 | 0.05075 | 0.10217 | 0.00000 |
| **LAYN** | 2.65825 | 6.70005 | 0.36258 | 1.41983 | 0.16169 | 0.12617 | 0.31124 |
| **LBH** | 0.00000 | 2.92594 | 2.22410 | 0.31101 | 4.15121 | 18.79582 | 0.00000 |
| **LBP** | 0.00000 | 0.03666 | 0.33852 | 0.05323 | 0.00231 | 0.19146 | 0.09384 |
| **LBR** | 29.79103 | 2.55646 | 10.99489 | 10.76103 | 5.92472 | 8.55766 | 8.78168 |
| **LBX1-AS1** | 0.01848 | 0.00603 | 0.01253 | 0.02083 | 0.02141 | 0.06380 | 0.06064 |
| **LBX2** | 0.00000 | 0.26538 | 0.03911 | 0.09712 | 0.00475 | 0.25029 | 0.09613 |
| **LBX2-AS1** | 0.00000 | 1.00088 | 0.02200 | 1.17169 | 0.07578 | 0.22881 | 0.30503 |
| **LCA5** | 0.34022 | 0.00000 | 0.12072 | 1.08958 | 0.99150 | 0.22239 | 0.00112 |
| **LCA5L** | 0.09452 | 0.00000 | 0.96791 | 0.09839 | 0.24535 | 0.08216 | 0.08135 |
| **LCAT** | 0.00000 | 1.25408 | 0.00698 | 1.43506 | 0.02381 | 0.52620 | 1.29944 |
| **LCE2B** | 0.01869 | 0.01182 | 0.01358 | 0.01933 | 0.00775 | 0.04241 | 0.02919 |
| **LCE3D** | 0.02566 | 0.01344 | 0.01465 | 0.01801 | 0.00774 | 0.02451 | 0.02387 |
| **LCK** | 0.03590 | 0.20172 | 0.01322 | 0.16259 | 0.04056 | 0.23582 | 0.06637 |
| **LCLAT1** | 7.95795 | 1.21416 | 5.92472 | 2.12563 | 0.93634 | 5.55918 | 1.26954 |
| **LCMT1** | 13.24990 | 8.33084 | 4.22439 | 14.22031 | 6.86273 | 7.41384 | 6.36023 |
| **LCMT2** | 3.15037 | 1.22715 | 4.54348 | 3.51296 | 0.71990 | 0.17488 | 0.00000 |
| **LCN10** | 0.02986 | 0.01149 | 0.01354 | 0.01671 | 0.00856 | 0.01016 | 0.03779 |
| **LCN12** | 0.02152 | 0.02773 | 0.02239 | 0.01593 | 0.01245 | 0.01021 | 0.03153 |
| **LCN15** | 0.03783 | 0.05836 | 0.00684 | 0.01731 | 0.00823 | 0.00997 | 0.00864 |
| **LCN2** | 0.01610 | 0.01409 | 0.01441 | 0.02370 | 0.00759 | 0.02245 | 0.03482 |
| **LCNL1** | 0.03995 | 0.01776 | 0.01406 | 0.02460 | 0.00906 | 0.01296 | 0.01179 |
| **LCOR** | 3.91403 | 0.00000 | 3.59785 | 2.78255 | 6.83442 | 3.07802 | 4.18058 |
| **LCORL** | 3.34745 | 0.45001 | 1.81209 | 1.26296 | 2.94523 | 1.72121 | 2.84716 |

| | | | | | | | |
|---|---|---|---|---|---|---|---|
| **LCP1** | 26.77931 | 0.00000 | 0.36992 | 25.60902 | 0.22856 | 17.58061 | 0.01268 |
| **LCP2** | 0.02164 | 0.03427 | 0.01921 | 0.01911 | 0.10300 | 0.01392 | 0.01825 |
| **LCT** | 0.06976 | 0.02100 | 0.08423 | 0.03680 | 0.01333 | 0.01432 | 0.02663 |
| **LCTL** | 0.60857 | 0.00000 | 0.27341 | 0.08694 | 0.10573 | 0.10631 | 0.13400 |
| **LDB1** | 1.75503 | 7.61184 | 1.55341 | 2.71852 | 0.94959 | 4.52728 | 17.43694 |
| **LDB2** | 0.01536 | 0.11538 | 0.01478 | 0.02122 | 0.04250 | 0.03645 | 0.01932 |
| **LDB3** | 0.13273 | 0.02992 | 0.13464 | 0.00000 | 0.06489 | 0.03754 | 0.07572 |
| **LDHA** | 78.91403 | 32.83524 | 60.38383 | 55.36565 | 11.52864 | 32.23811 | 28.05090 |
| **LDHAL6A** | 0.15541 | 0.00000 | 0.05982 | 0.00000 | 3.32347 | 0.06198 | 0.08320 |
| **LDHAL6B** | 0.00825 | 0.02491 | 0.03494 | 0.05892 | 0.02169 | 0.01332 | 0.05369 |
| **LDHB** | 85.28799 | 41.01300 | 39.65305 | 54.11992 | 2.81928 | 41.72235 | 38.78896 |
| **LDHC** | 0.00000 | 0.24377 | 0.73280 | 0.08319 | 1.15119 | 0.00000 | 0.06584 |
| **LDHD** | 0.00000 | 1.92265 | 0.61842 | 0.53036 | 0.15028 | 1.13251 | 0.00000 |
| **LDLR** | 20.25798 | 0.00000 | 0.00000 | 12.92264 | 0.58177 | 14.47559 | 17.57607 |
| **LDLRAD1** | 0.01856 | 0.02513 | 0.00899 | 0.02810 | 0.02246 | 0.01281 | 0.06927 |
| **LDLRAD2** | 0.02062 | 0.02692 | 0.01818 | 0.04444 | 0.00697 | 0.10265 | 0.01936 |
| **LDLRAD3** | 1.00885 | 0.94999 | 0.55413 | 2.60662 | 0.75417 | 1.11756 | 0.20694 |
| **LDLRAD4** | 0.00182 | 0.00000 | 0.63571 | 0.04378 | 0.00974 | 0.00000 | 0.15160 |
| **LDLRAD4-AS1** | 0.03125 | 0.01537 | 0.01162 | 0.01569 | 0.02379 | 0.01528 | 0.01935 |
| **LDLRAP1** | 0.00000 | 8.61217 | 4.87141 | 1.02950 | 1.16207 | 8.18292 | 2.95691 |
| **LDOC1** | 9.45641 | 1.52173 | 0.96776 | 11.65472 | 1.19104 | 1.51365 | 1.96985 |
| **LDOC1L** | 0.29143 | 0.00000 | 0.02284 | 1.36350 | 0.05078 | 0.00000 | 0.79025 |
| **LEAP2** | 0.54624 | 0.25744 | 0.29140 | 0.74397 | 0.82796 | 0.44094 | 0.56964 |
| **LECT1** | 0.01783 | 0.07316 | 0.03313 | 0.07098 | 0.03897 | 0.22391 | 0.02461 |
| **LECT2** | 0.03759 | 0.00000 | 0.00932 | 0.22493 | 0.00950 | 0.00279 | 0.26400 |
| **LEF1** | 0.02081 | 0.08893 | 0.00000 | 0.00061 | 2.56058 | 0.00650 | 0.05703 |
| **LEF1-AS1** | 0.02343 | 0.01185 | 0.02913 | 0.01849 | 0.05176 | 0.01241 | 0.01979 |
| **LEFTY1** | 0.04995 | 0.00000 | 0.02092 | 0.90222 | 0.00428 | 15.40784 | 0.00000 |
| **LEFTY2** | 0.01114 | 0.00241 | 0.00688 | 1.78626 | 0.00825 | 12.24597 | 0.00000 |
| **LEKR1** | 0.04353 | 0.01831 | 0.02339 | 0.03004 | 0.22162 | 0.05796 | 0.00000 |
| **LEMD1** | 0.02576 | 0.01194 | 0.01459 | 0.32614 | 0.01507 | 0.06808 | 0.00000 |
| **LEMD1-AS1** | 0.02898 | 0.05957 | 0.01221 | 0.01543 | 0.00717 | 0.01113 | 0.02298 |
| **LEMD2** | 0.00000 | 2.77575 | 0.61493 | 1.10413 | 0.04520 | 2.96824 | 8.28528 |
| **LEMD3** | 2.06285 | 0.01670 | 1.84700 | 0.07378 | 0.94507 | 4.34656 | 3.90398 |
| **LENEP** | 0.00546 | 0.00649 | 0.02842 | 0.05917 | 0.02294 | 0.00991 | 0.06209 |

| | | | | | | | |
|---|---|---|---|---|---|---|---|
| **LENG1** | 3.87657 | 5.88033 | 0.97761 | 5.65701 | 3.23805 | 3.06508 | 0.00000 |
| **LENG8** | 0.00000 | 2.28892 | 2.19291 | 3.02930 | 2.52914 | 1.49726 | 8.24290 |
| **LENG9** | 0.00000 | 0.00501 | 0.01016 | 0.00876 | 0.00676 | 0.14372 | 0.24417 |
| **LEO1** | 21.90164 | 3.07753 | 13.07527 | 11.71780 | 3.22311 | 6.31086 | 0.87084 |
| **LEP** | 3.49980 | 0.00000 | 0.00250 | 4.05211 | 0.00804 | 0.00000 | 0.09069 |
| **LEPR** | 0.75283 | 0.00000 | 0.25029 | 2.09550 | 0.37006 | 0.05697 | 0.31863 |
| **LEPRE1** | 8.11957 | 7.45729 | 4.20025 | 4.42002 | 0.14025 | 8.60881 | 2.33556 |
| **LEPREL1** | 0.07578 | 0.00000 | 0.00698 | 0.33030 | 0.11914 | 1.43540 | 0.00000 |
| **LEPREL2** | 0.01040 | 0.01540 | 0.01274 | 0.02687 | 0.00760 | 0.10607 | 0.03543 |
| **LEPREL4** | 0.10358 | 0.40613 | 0.02461 | 0.30141 | 0.01197 | 0.06773 | 0.46705 |
| **LEPROT** | 9.11155 | 2.95633 | 5.76282 | 7.64703 | 1.42358 | 1.96279 | 3.43590 |
| **LEPROTL1** | 17.78754 | 3.98832 | 8.85411 | 11.59477 | 1.11507 | 5.57109 | 9.20661 |
| **LETM1** | 18.83160 | 42.02992 | 10.56960 | 10.16723 | 0.00000 | 6.44847 | 20.11697 |
| **LETM2** | 0.00000 | 0.00000 | 0.31252 | 0.28820 | 1.06358 | 0.32683 | 0.16650 |
| **LETMD1** | 9.51207 | 0.65459 | 3.70910 | 3.86405 | 0.76601 | 2.17648 | 5.69967 |
| **LEUTX** | 0.00928 | 0.01171 | 0.00000 | 0.01002 | 16.64676 | 0.00972 | 0.01852 |
| **LFNG** | 0.00000 | 0.06317 | 0.68240 | 0.03604 | 0.58450 | 1.03136 | 0.16989 |
| **LGALS1** | 10.05318 | 0.17206 | 0.00000 | 8.82968 | 13.07198 | 13.25701 | 2.09192 |
| **LGALS12** | 0.43781 | 0.97083 | 0.47885 | 0.35554 | 0.33763 | 3.91129 | 0.22694 |
| **LGALS13** | 0.01350 | 0.01560 | 0.01305 | 1.03542 | 0.00786 | 0.01956 | 0.00000 |
| **LGALS16** | 0.00000 | 0.00501 | 0.00903 | 14.30293 | 0.00483 | 0.20861 | 0.11496 |
| **LGALS17A** | 0.02166 | 0.01755 | 0.01254 | 0.04792 | 0.01011 | 0.01286 | 0.01710 |
| **LGALS2** | 0.02387 | 0.00000 | 0.02496 | 0.29906 | 0.04602 | 7.35571 | 0.00000 |
| **LGALS3** | 0.55163 | 0.00000 | 0.12172 | 31.09778 | 0.02478 | 0.60138 | 0.13368 |
| **LGALS3BP** | 0.29003 | 0.00000 | 0.04417 | 0.38012 | 0.00451 | 1.05674 | 0.00000 |
| **LGALS4** | 0.00000 | 1.54245 | 5.97665 | 0.00000 | 0.42604 | 0.62469 | 0.00000 |
| **LGALS7** | 0.02368 | 0.03915 | 0.03238 | 0.02529 | 0.02775 | 0.01121 | 0.00931 |
| **LGALS7B** | 0.01670 | 0.03932 | 0.02225 | 0.03748 | 0.02243 | 0.03332 | 0.03509 |
| **LGALS8** | 0.53256 | 0.10118 | 0.56904 | 0.41769 | 2.90588 | 0.09958 | 1.25122 |
| **LGALS8-AS1** | 0.02102 | 0.06381 | 0.01540 | 0.02489 | 0.01648 | 0.01502 | 0.00558 |
| **LGALS9** | 0.00000 | 0.01186 | 0.01335 | 0.77288 | 0.00936 | 0.06999 | 1.15831 |
| **LGALS9C** | 0.02290 | 0.01632 | 0.01345 | 0.02297 | 0.01229 | 0.01676 | 0.02013 |
| **LGALSL** | 0.90089 | 0.33484 | 0.28188 | 1.32090 | 0.07024 | 1.05038 | 0.00000 |
| **LGI1** | 0.03233 | 0.01580 | 0.00929 | 0.01898 | 0.05090 | 0.01697 | 0.01698 |
| **LGI2** | 0.01701 | 0.00000 | 0.03904 | 0.01683 | 0.08926 | 0.30248 | 0.04073 |

| | | | | | | | |
|---|---|---|---|---|---|---|---|
| **LGI3** | 0.03930 | 0.01286 | 0.00948 | 0.02370 | 0.03527 | 0.02333 | 0.03040 |
| **LGI4** | 0.00912 | 0.01644 | 0.02087 | 0.04977 | 0.03051 | 0.01878 | 0.04875 |
| **LGMN** | 16.86927 | 3.70899 | 20.10647 | 24.54270 | 14.87285 | 4.68938 | 0.00000 |
| **LGR4** | 10.52128 | 0.00000 | 2.05700 | 5.94287 | 0.23958 | 5.49326 | 13.74500 |
| **LGR5** | 0.04326 | 0.00978 | 0.00597 | 0.05465 | 0.22745 | 0.04724 | 0.00000 |
| **LGR6** | 0.00000 | 0.02324 | 0.01464 | 0.02211 | 0.01298 | 0.92634 | 0.03210 |
| **LGSN** | 0.81995 | 0.14186 | 0.07194 | 0.09597 | 0.00388 | 0.00000 | 0.09291 |
| **LHB** | 0.00000 | 0.00549 | 0.60758 | 0.28075 | 0.15286 | 0.02504 | 0.02888 |
| **LHCGR** | 0.04630 | 0.00643 | 0.01806 | 0.01244 | 0.01061 | 0.12083 | 0.00433 |
| **LHFP** | 0.95772 | 0.00000 | 0.71070 | 2.36486 | 0.09625 | 1.12244 | 0.36404 |
| **LHFPL1** | 0.02801 | 0.07624 | 0.06932 | 0.02147 | 0.11003 | 0.03767 | 0.00434 |
| **LHFPL2** | 3.99197 | 0.00000 | 0.47975 | 2.91174 | 0.09641 | 0.00000 | 5.76919 |
| **LHFPL3** | 0.51732 | 0.00000 | 1.16639 | 0.13746 | 0.57519 | 0.54050 | 0.23984 |
| **LHFPL3-AS1** | 0.06169 | 0.00000 | 0.12814 | 0.00378 | 0.05577 | 0.03763 | 0.04120 |
| **LHFPL3-AS2** | 1.62724 | 0.71572 | 3.24026 | 1.77668 | 3.77520 | 1.51371 | 0.71272 |
| **LHFPL4** | 0.00000 | 0.10470 | 2.50459 | 0.09946 | 0.00003 | 1.71743 | 0.17050 |
| **LHFPL5** | 0.08415 | 0.08100 | 0.00486 | 0.03316 | 0.00668 | 0.12880 | 0.14856 |
| **LHPP** | 1.81987 | 2.49352 | 0.00155 | 0.60428 | 0.26293 | 1.18904 | 0.19508 |
| **LHX1** | 0.03699 | 0.01359 | 0.01186 | 0.02491 | 0.02908 | 0.01404 | 0.01381 |
| **LHX2** | 0.00684 | 0.00000 | 1.09452 | 0.05268 | 1.64992 | 0.00482 | 0.14118 |
| **LHX3** | 0.00395 | 0.01918 | 0.01016 | 0.03436 | 0.01056 | 0.02347 | 0.07372 |
| **LHX4** | 0.56613 | 0.62092 | 0.00000 | 0.30013 | 0.29040 | 0.00000 | 0.46197 |
| **LHX4-AS1** | 0.16446 | 0.09325 | 0.17377 | 0.34128 | 0.00105 | 0.05062 | 0.53161 |
| **LHX5** | 0.02344 | 0.01681 | 0.01302 | 0.02109 | 0.01083 | 0.01621 | 0.01802 |
| **LHX6** | 0.05033 | 0.04632 | 1.02043 | 0.00000 | 0.38180 | 0.20984 | 0.37184 |
| **LHX8** | 0.02058 | 0.02197 | 0.08790 | 0.03145 | 0.52209 | 0.21418 | 0.00467 |
| **LHX9** | 0.03578 | 0.01188 | 0.01191 | 0.08324 | 0.03292 | 0.02219 | 0.00000 |
| **LIAS** | 10.86699 | 2.48881 | 9.77923 | 2.27860 | 1.16737 | 2.52038 | 2.00139 |
| **LIF** | 0.00000 | 1.39996 | 0.25260 | 0.21488 | 0.00558 | 0.80496 | 0.09352 |
| **LIFR** | 2.27343 | 0.00000 | 4.20674 | 1.02387 | 1.71204 | 0.63869 | 6.40750 |
| **LIFR-AS1** | 0.03846 | 0.05836 | 0.03004 | 0.03591 | 0.05318 | 0.00000 | 0.20497 |
| **LIG1** | 10.68420 | 9.18858 | 3.50421 | 8.00412 | 2.08355 | 6.06344 | 4.73970 |
| **LIG3** | 17.94252 | 13.06083 | 4.45806 | 11.41978 | 1.56816 | 6.01965 | 2.02639 |
| **LIG4** | 4.21230 | 0.00000 | 1.54924 | 1.06514 | 0.41130 | 0.36436 | 1.06313 |
| **LILRA2** | 0.02504 | 0.03724 | 0.01728 | 0.01809 | 0.00713 | 0.01031 | 0.01629 |

| | | | | | | | |
|---|---|---|---|---|---|---|---|
| **LILRA3** | 0.04774 | 0.16567 | 0.20513 | 0.01542 | 0.00294 | 0.00000 | 0.00977 |
| **LILRA4** | 0.01265 | 0.02149 | 0.00815 | 0.03387 | 0.13611 | 0.02633 | 0.02336 |
| **LILRA5** | 0.04814 | 0.00636 | 0.01451 | 0.01056 | 0.01218 | 0.07137 | 0.01571 |
| **LILRA6** | 0.01603 | 0.04837 | 0.00710 | 0.04094 | 0.02879 | 0.02430 | 0.08835 |
| **LILRB1** | 0.00261 | 0.03545 | 0.04114 | 0.03113 | 0.00689 | 0.03101 | 0.04957 |
| **LILRB2** | 0.02469 | 0.01269 | 0.01442 | 0.01980 | 0.00787 | 0.08629 | 0.00892 |
| **LILRB3** | 0.00000 | 0.50172 | 0.00000 | 0.12448 | 0.25472 | 0.45740 | 0.40819 |
| **LILRB4** | 0.02766 | 0.00965 | 0.04152 | 0.02788 | 0.00736 | 0.01548 | 0.01165 |
| **LILRB5** | 0.01563 | 0.01266 | 0.02061 | 0.03308 | 0.04951 | 0.02111 | 0.06898 |
| **LILRP2** | 0.03471 | 0.07524 | 0.00335 | 0.01615 | 0.05612 | 0.01071 | 0.00000 |
| **LIM2** | 0.00000 | 0.18675 | 0.70260 | 0.00702 | 0.30416 | 2.89297 | 0.00656 |
| **LIMA1** | 30.62573 | 0.00000 | 7.82525 | 17.71061 | 15.07016 | 2.16315 | 21.72069 |
| **LIMCH1** | 0.07770 | 0.00000 | 0.00000 | 0.01171 | 0.53855 | 8.03798 | 0.13491 |
| **LIMD1** | 2.30720 | 3.08684 | 0.34125 | 1.94807 | 0.01388 | 2.69504 | 6.44732 |
| **LIMD1-AS1** | 0.05418 | 0.01576 | 0.01878 | 0.04574 | 0.08702 | 0.00995 | 0.00831 |
| **LIMD2** | 0.00000 | 0.02793 | 0.58667 | 0.37702 | 0.61060 | 1.00339 | 0.11621 |
| **LIME1** | 0.00000 | 1.03189 | 0.22748 | 0.59683 | 0.04297 | 0.37588 | 2.65720 |
| **LIMK1** | 0.18816 | 2.55333 | 1.28397 | 0.60601 | 1.36790 | 1.03733 | 3.11320 |
| **LIMK2** | 17.54900 | 16.24054 | 2.86585 | 17.52738 | 0.37599 | 2.91099 | 13.01650 |
| **LIMS1** | 4.23750 | 2.44437 | 1.68331 | 4.93815 | 0.33456 | 4.12270 | 11.43556 |
| **LIMS2** | 0.00000 | 0.65400 | 0.19163 | 0.03481 | 0.89516 | 0.19170 | 0.05090 |
| **LIN28A** | 33.60598 | 13.89876 | 26.62791 | 19.10435 | 2.28638 | 18.64410 | 36.03024 |
| **LIN28B** | 11.77550 | 0.00000 | 7.78188 | 13.97143 | 1.01316 | 2.39360 | 5.21315 |
| **LIN37** | 4.35699 | 5.58046 | 0.78934 | 6.98005 | 0.83001 | 2.48185 | 0.29823 |
| **LIN52** | 3.68241 | 1.51202 | 5.06166 | 3.18058 | 2.87619 | 1.22571 | 0.00000 |
| **LIN54** | 4.51656 | 0.10592 | 2.17752 | 3.96034 | 3.65195 | 1.58996 | 2.88276 |
| **LIN7A** | 0.03329 | 0.00808 | 0.02297 | 0.01881 | 0.05339 | 0.01244 | 0.01780 |
| **LIN7B** | 0.62393 | 0.09353 | 0.47151 | 0.47161 | 0.18847 | 0.03759 | 0.15777 |
| **LIN7C** | 26.72450 | 5.08708 | 7.93849 | 7.44644 | 3.98169 | 5.28397 | 0.00000 |
| **LIN9** | 8.03837 | 0.00000 | 6.48872 | 2.92447 | 2.55659 | 1.78544 | 0.40835 |
| **LINC-PINT.uid-1** | 0.06088 | 0.00564 | 0.02974 | 0.04956 | 0.00684 | 0.01504 | 0.01291 |
| **LINC-PINT.uid-9** | 0.01972 | 0.19521 | 0.36930 | 0.44569 | 0.35978 | 0.54612 | 0.83659 |
| **LINC-ROR** | 0.08998 | 0.00000 | 0.01765 | 0.00592 | 0.05357 | 2.59250 | 0.00535 |
| **LINC00028** | 0.02787 | 0.04164 | 0.05597 | 0.01629 | 0.00565 | 0.00904 | 0.01277 |
| **LINC00029** | 0.02341 | 0.00000 | 0.08708 | 0.02067 | 0.04087 | 0.01157 | 0.02475 |

| | | | | | | | |
|---|---|---|---|---|---|---|---|
| **LINC00032** | 0.01631 | 0.00000 | 0.07731 | 0.03308 | 0.24063 | 0.02971 | 0.13859 |
| **LINC00035** | 0.01969 | 0.01127 | 0.01279 | 0.05476 | 0.00739 | 0.01202 | 0.08178 |
| **LINC00052** | 0.14754 | 0.02724 | 0.12936 | 0.04654 | 0.00421 | 0.01850 | 0.00000 |
| **LINC00086** | 0.00000 | 0.57738 | 0.64742 | 0.00000 | 0.01361 | 1.40187 | 0.02180 |
| **LINC00087** | 0.07916 | 0.09595 | 0.06792 | 0.00000 | 0.40481 | 0.00000 | 0.17267 |
| **LINC00092** | 0.02059 | 0.02175 | 0.01125 | 0.02113 | 0.02807 | 0.01156 | 0.02413 |
| **LINC00094** | 0.43842 | 0.89583 | 1.61716 | 1.66004 | 1.39417 | 2.25425 | 0.22015 |
| **LINC00113** | 0.04268 | 0.01980 | 0.01404 | 0.02349 | 0.00799 | 0.01261 | 0.00271 |
| **LINC00115** | 0.29767 | 0.28108 | 0.18466 | 0.26868 | 0.48840 | 0.07164 | 0.07959 |
| **LINC00116** | 4.22383 | 16.28948 | 3.48586 | 2.31274 | 0.82472 | 2.58331 | 7.21172 |
| **LINC00152** | 1.15595 | 1.16924 | 0.00000 | 4.40165 | 0.00102 | 0.45765 | 1.33100 |
| **LINC00158** | 0.03573 | 0.01023 | 0.02531 | 0.01159 | 0.06356 | 0.03424 | 0.03278 |
| **LINC00159** | 0.03335 | 0.01673 | 0.01304 | 0.01886 | 0.01961 | 0.01282 | 0.01179 |
| **LINC00162** | 0.02990 | 0.02333 | 0.01518 | 0.01741 | 0.05645 | 0.01283 | 0.01467 |
| **LINC00163** | 0.01264 | 0.01238 | 0.03743 | 0.02788 | 0.00631 | 0.04337 | 0.02968 |
| **LINC00167** | 0.02048 | 0.12961 | 0.01452 | 0.05214 | 0.13543 | 0.07983 | 0.09560 |
| **LINC00173** | 0.05120 | 0.04275 | 0.04882 | 0.00000 | 0.04012 | 0.06866 | 0.17749 |
| **LINC00174** | 0.06545 | 0.25630 | 0.83787 | 0.26406 | 0.04778 | 0.54149 | 1.47980 |
| **LINC00176** | 0.00000 | 0.09328 | 0.08149 | 0.04582 | 0.23175 | 0.09920 | 0.23450 |
| **LINC00184** | 0.01632 | 0.02228 | 0.01150 | 0.01935 | 0.00780 | 0.02528 | 0.03035 |
| **LINC00189** | 0.47874 | 0.03712 | 0.48809 | 0.00000 | 0.24232 | 0.60671 | 0.05516 |
| **LINC00200** | 0.15472 | 0.93336 | 1.73981 | 1.60203 | 0.00349 | 0.01896 | 0.00000 |
| **LINC00202-1** | 0.03893 | 0.00629 | 0.01269 | 0.08170 | 0.00876 | 0.04765 | 0.17292 |
| **LINC00202-2** | 0.07331 | 0.06148 | 0.05031 | 0.34088 | 0.00475 | 0.00000 | 0.14991 |
| **LINC00211** | 0.02493 | 0.01786 | 0.01265 | 0.02104 | 0.00816 | 0.01432 | 0.01784 |
| **LINC00221** | 4.18069 | 2.08802 | 5.05795 | 1.61416 | 2.85622 | 0.77028 | 3.32336 |
| **LINC00222** | 0.03573 | 0.01213 | 0.01448 | 0.01724 | 0.00788 | 0.05686 | 0.00999 |
| **LINC00226** | 0.02677 | 0.00000 | 0.09615 | 0.01684 | 0.23322 | 0.01191 | 0.01896 |
| **LINC00235** | 0.02086 | 0.03721 | 0.01696 | 0.02493 | 0.00724 | 0.02964 | 0.00989 |
| **LINC00238** | 0.02124 | 0.01690 | 0.01681 | 0.02188 | 0.01605 | 0.03102 | 0.02479 |
| **LINC00240** | 0.03901 | 0.01820 | 0.02457 | 0.12927 | 0.02547 | 0.01424 | 0.00440 |
| **LINC00242** | 0.03228 | 0.02113 | 0.01780 | 0.01869 | 0.00770 | 0.06696 | 0.01435 |
| **LINC00260** | 0.11465 | 0.35381 | 0.01716 | 0.32055 | 0.03657 | 0.16733 | 1.22254 |
| **LINC00261** | 0.04170 | 0.00000 | 0.01025 | 0.24945 | 0.00687 | 4.07493 | 0.00000 |
| **LINC00263** | 2.05130 | 0.00000 | 1.27728 | 3.68967 | 0.23636 | 2.03086 | 0.13558 |

| | | | | | | | |
|---|---|---|---|---|---|---|---|
| **LINC00264** | 0.01779 | 0.01902 | 0.01724 | 0.02650 | 0.01955 | 0.01109 | 0.03224 |
| **LINC00265** | 0.06061 | 0.00000 | 0.06223 | 0.21839 | 0.06846 | 0.20297 | 0.00981 |
| **LINC00266-3.ui** | 0.02371 | 0.01438 | 0.01462 | 0.01734 | 0.03001 | 0.01329 | 0.02689 |
| **LINC00269** | 0.01198 | 0.01070 | 0.01108 | 0.01450 | 0.01077 | 0.00909 | 0.12270 |
| **LINC00271** | 0.02411 | 0.01582 | 0.01501 | 0.02245 | 0.00873 | 0.01255 | 0.03284 |
| **LINC00272** | 0.02599 | 0.01859 | 0.01552 | 0.01785 | 0.01414 | 0.01271 | 0.01793 |
| **LINC00273** | 0.00000 | 0.00004 | 0.04653 | 0.17819 | 0.00388 | 0.25970 | 1.01958 |
| **LINC00276** | 0.02431 | 0.00733 | 0.03741 | 0.01421 | 0.16803 | 0.01188 | 0.04454 |
| **LINC00277** | 0.01933 | 0.00874 | 0.02886 | 0.02453 | 0.02161 | 0.01514 | 0.02745 |
| **LINC00282** | 0.04832 | 0.06084 | 0.00536 | 0.12273 | 0.04323 | 0.02754 | 0.28955 |
| **LINC00284** | 0.02569 | 0.00985 | 0.05718 | 0.04220 | 0.00726 | 0.01413 | 0.01182 |
| **LINC00293** | 0.02534 | 0.01792 | 0.01541 | 0.02008 | 0.00970 | 0.01257 | 0.01737 |
| **LINC00294** | 0.47549 | 0.11848 | 0.20018 | 1.17667 | 0.02586 | 0.37499 | 0.12151 |
| **LINC00298** | 0.02502 | 0.00781 | 0.02185 | 0.01897 | 0.14980 | 0.01337 | 0.01585 |
| **LINC00299** | 0.03915 | 0.01133 | 0.01110 | 0.01153 | 0.08874 | 0.04892 | 0.01921 |
| **LINC00301** | 0.02395 | 0.01886 | 0.02142 | 0.01818 | 0.02785 | 0.01337 | 0.01901 |
| **LINC00303** | 0.02014 | 0.00655 | 0.02271 | 0.07100 | 0.02883 | 0.13608 | 0.04301 |
| **LINC00304** | 0.02555 | 0.01855 | 0.01268 | 0.01969 | 0.00791 | 0.01300 | 0.01762 |
| **LINC00308** | 0.04449 | 0.01924 | 0.01544 | 0.03163 | 0.00858 | 0.01388 | 0.00555 |
| **LINC00309** | 0.03030 | 0.02473 | 0.02233 | 0.01915 | 0.01104 | 0.04669 | 0.01367 |
| **LINC00310** | 0.07826 | 0.02499 | 0.08014 | 0.01234 | 0.14533 | 0.00244 | 0.03162 |
| **LINC00311** | 0.00000 | 0.20321 | 0.00360 | 0.00263 | 0.01243 | 0.03078 | 0.30726 |
| **LINC00312** | 0.01269 | 0.00918 | 0.05326 | 0.01653 | 0.07025 | 0.18115 | 0.05739 |
| **LINC00313** | 0.01755 | 0.06691 | 0.03152 | 0.02968 | 0.02154 | 0.03802 | 0.03217 |
| **LINC00319** | 0.02520 | 0.02772 | 0.02491 | 0.02090 | 0.00703 | 0.01095 | 0.01811 |
| **LINC00320** | 0.06140 | 0.01119 | 0.02314 | 0.01888 | 0.00784 | 0.12561 | 0.00000 |
| **LINC00324** | 0.09102 | 0.03423 | 0.18937 | 0.05516 | 7.23319 | 0.00000 | 0.06820 |
| **LINC00327** | 0.02235 | 0.01907 | 0.01686 | 0.04732 | 0.00823 | 0.01651 | 0.04924 |
| **LINC00330** | 0.01862 | 0.61034 | 1.00566 | 0.00000 | 0.01931 | 0.00000 | 0.00254 |
| **LINC00333** | 0.02114 | 0.01781 | 0.01385 | 0.03922 | 0.00942 | 0.01367 | 0.02086 |
| **LINC00336** | 0.03476 | 0.06414 | 0.07296 | 0.02240 | 0.00848 | 0.00951 | 0.00000 |
| **LINC00337** | 0.23052 | 4.53172 | 4.32495 | 0.82602 | 1.09475 | 1.26773 | 1.67352 |
| **LINC00339** | 0.49253 | 0.72572 | 3.37862 | 0.04409 | 6.02314 | 1.74566 | 0.00000 |
| **LINC00341** | 0.10688 | 0.00000 | 0.00356 | 0.58440 | 0.00999 | 0.00000 | 0.81064 |
| **LINC00342** | 0.00589 | 0.01458 | 0.06475 | 0.01367 | 0.00500 | 0.00622 | 0.29604 |

| | | | | | | | |
|---|---|---|---|---|---|---|---|
| **LINC00346** | 0.11362 | 0.00391 | 0.00674 | 0.00456 | 0.04187 | 0.00445 | 0.13660 |
| **LINC00347** | 0.00224 | 0.00000 | 0.02703 | 0.04459 | 0.13034 | 0.00724 | 0.11609 |
| **LINC00348** | 0.03968 | 0.01587 | 0.01205 | 0.03711 | 0.01930 | 0.01234 | 0.00718 |
| **LINC00351** | 0.02335 | 0.01838 | 0.00845 | 0.02190 | 0.06191 | 0.01315 | 0.01487 |
| **LINC00354** | 0.02325 | 0.00887 | 0.02778 | 0.01914 | 0.03084 | 0.01223 | 0.02852 |
| **LINC00366** | 0.02487 | 0.02453 | 0.00482 | 0.04600 | 0.14870 | 0.02937 | 0.00213 |
| **LINC00367** | 0.03887 | 0.02490 | 0.01969 | 0.17991 | 0.01047 | 0.01453 | 0.00000 |
| **LINC00368** | 0.01968 | 0.00545 | 0.00978 | 0.00000 | 0.02315 | 0.00622 | 0.29634 |
| **LINC00371** | 0.39232 | 0.64795 | 0.16059 | 0.00000 | 0.12797 | 0.60206 | 0.00000 |
| **LINC00378** | 0.02231 | 0.01429 | 0.01966 | 0.02077 | 0.05223 | 0.01866 | 0.01911 |
| **LINC00379** | 0.30258 | 0.00000 | 0.01010 | 1.48533 | 0.05989 | 0.00761 | 0.00000 |
| **LINC00381** | 0.02421 | 0.01701 | 0.02210 | 0.01943 | 0.00732 | 0.01866 | 0.01559 |
| **LINC00392** | 0.00000 | 0.09293 | 0.15336 | 0.11349 | 0.01311 | 0.09013 | 0.03833 |
| **LINC00400** | 0.02524 | 0.05663 | 0.01256 | 0.01769 | 0.06709 | 0.00947 | 0.10576 |
| **LINC00408** | 0.02474 | 0.01067 | 0.02295 | 0.04497 | 0.03592 | 0.01213 | 0.01162 |
| **LINC00412** | 0.04929 | 0.03972 | 0.02643 | 0.02224 | 0.00732 | 0.03472 | 0.00129 |
| **LINC00417** | 0.02827 | 0.01760 | 0.01359 | 0.02081 | 0.00795 | 0.02717 | 0.01083 |
| **LINC00421** | 0.02430 | 0.00917 | 0.03421 | 0.01845 | 0.00671 | 0.01261 | 0.02051 |
| **LINC00423** | 0.02668 | 0.01787 | 0.01081 | 0.01854 | 0.02564 | 0.01449 | 0.01530 |
| **LINC00426** | 0.01115 | 0.03706 | 0.28814 | 0.00135 | 0.27306 | 0.17583 | 0.05471 |
| **LINC00441** | 0.02120 | 0.05694 | 0.01824 | 0.04499 | 0.00746 | 0.00681 | 0.03444 |
| **LINC00446** | 0.02547 | 0.01530 | 0.01274 | 0.01891 | 0.06621 | 0.01345 | 0.01503 |
| **LINC00448** | 0.02725 | 0.01815 | 0.01112 | 0.01844 | 0.07352 | 0.01367 | 0.01544 |
| **LINC00452** | 0.02063 | 0.03938 | 0.02258 | 0.01793 | 0.01241 | 0.02328 | 0.02458 |
| **LINC00457** | 0.05826 | 0.02549 | 0.02188 | 0.01357 | 0.00848 | 0.03158 | 0.00331 |
| **LINC00458** | 0.02000 | 0.00000 | 0.02270 | 1.37203 | 0.09448 | 0.64784 | 0.00000 |
| **LINC00461** | 0.01951 | 0.09159 | 0.00151 | 0.00000 | 0.01820 | 0.57261 | 0.15640 |
| **LINC00463** | 0.02979 | 0.01720 | 0.01838 | 0.01762 | 0.00991 | 0.01278 | 0.01894 |
| **LINC00466** | 0.03227 | 0.01176 | 0.01323 | 0.02067 | 0.09356 | 0.09020 | 0.02664 |
| **LINC00467** | 0.07822 | 0.12705 | 1.81832 | 0.13970 | 0.52672 | 1.20115 | 0.02254 |
| **LINC00469** | 0.02136 | 0.01797 | 0.01248 | 0.02038 | 0.01164 | 0.02030 | 0.01919 |
| **LINC00470** | 1.01884 | 0.00000 | 0.28146 | 1.46091 | 0.02329 | 0.24329 | 1.27645 |
| **LINC00471** | 0.13462 | 0.32067 | 0.59852 | 0.08497 | 0.82674 | 0.16633 | 0.15188 |
| **LINC00472** | 0.04478 | 0.04336 | 0.07284 | 0.04641 | 0.15575 | 0.00944 | 0.00214 |
| **LINC00475** | 0.00000 | 0.14379 | 0.00784 | 0.01451 | 0.01017 | 0.27261 | 0.02015 |

| | | | | | | | |
|---|---|---|---|---|---|---|---|
| **LINC00476.uid-** | 0.35090 | 0.82472 | 0.39431 | 0.59117 | 0.32405 | 0.00000 | 0.00000 |
| **LINC00476.uid-** | 0.07777 | 0.00000 | 0.07559 | 0.13841 | 0.03840 | 0.03458 | 0.15199 |
| **LINC00477** | 0.07063 | 0.05444 | 0.03227 | 0.02070 | 0.06545 | 0.01776 | 0.02519 |
| **LINC00478** | 0.01801 | 0.05286 | 0.01160 | 0.00980 | 0.07810 | 0.18960 | 0.08664 |
| **LINC00479** | 0.65178 | 2.52291 | 0.00000 | 1.74962 | 0.09850 | 0.16362 | 0.00000 |
| **LINC00482** | 0.01683 | 0.01783 | 0.03236 | 0.23206 | 0.00743 | 0.00000 | 0.16954 |
| **LINC00483** | 0.00983 | 0.12924 | 0.00000 | 0.01624 | 0.45119 | 0.01653 | 0.01132 |
| **LINC00485** | 0.03270 | 0.01794 | 0.01481 | 0.02009 | 0.00803 | 0.02773 | 0.01828 |
| **LINC00486** | 0.05902 | 0.01905 | 0.02556 | 0.00363 | 0.04750 | 0.00847 | 0.04189 |
| **LINC00488** | 0.02635 | 0.01727 | 0.01412 | 0.02184 | 0.05074 | 0.01647 | 0.01786 |
| **LINC00491** | 0.36648 | 0.10328 | 0.19780 | 0.06616 | 0.16170 | 0.17835 | 0.31345 |
| **LINC00492** | 0.02822 | 0.02567 | 0.01613 | 0.01867 | 0.00734 | 0.01342 | 0.01875 |
| **LINC00493** | 18.33093 | 10.46295 | 12.54393 | 20.35412 | 1.28130 | 9.84223 | 11.99180 |
| **LINC00501** | 0.04328 | 1.18343 | 1.54472 | 0.03425 | 0.19949 | 0.37694 | 0.10151 |
| **LINC00506** | 0.02505 | 0.01847 | 0.01393 | 0.02415 | 0.00799 | 0.01473 | 0.01442 |
| **LINC00511** | 0.06881 | 0.06983 | 0.24827 | 0.00000 | 0.00399 | 0.19320 | 0.02367 |
| **LINC00514** | 0.01324 | 0.04692 | 0.01381 | 0.03401 | 0.01021 | 0.01192 | 0.04166 |
| **LINC00515** | 0.03641 | 0.01404 | 0.01325 | 0.02294 | 0.00791 | 0.04608 | 0.01374 |
| **LINC00518** | 0.03381 | 0.54632 | 0.19369 | 0.18329 | 0.00150 | 0.00000 | 0.08957 |
| **LINC00520** | 0.02489 | 0.01963 | 0.00910 | 0.01846 | 0.03714 | 0.01428 | 0.01659 |
| **LINC00521** | 0.00820 | 0.03421 | 0.00907 | 0.01742 | 0.00787 | 0.08611 | 0.03131 |
| **LINC00525** | 0.03630 | 0.01203 | 0.03867 | 0.02167 | 0.00641 | 0.02824 | 0.01032 |
| **LINC00526** | 0.15748 | 0.62203 | 0.33934 | 0.38849 | 0.51405 | 0.19926 | 0.32928 |
| **LINC00528** | 0.02373 | 0.01702 | 0.02519 | 0.01904 | 0.00732 | 0.01261 | 0.01936 |
| **LINC00534** | 0.03383 | 0.01740 | 0.01189 | 0.01411 | 0.01999 | 0.01124 | 0.03636 |
| **LINC00535** | 0.02727 | 0.12044 | 0.00873 | 0.01225 | 0.08071 | 0.03798 | 0.03375 |
| **LINC00536** | 0.03062 | 0.01627 | 0.00940 | 0.02261 | 0.05161 | 0.01688 | 0.01063 |
| **LINC00539** | 0.03863 | 0.18084 | 0.02700 | 0.03558 | 0.12474 | 0.00610 | 0.00000 |
| **LINC00540** | 2.00782 | 0.98344 | 3.07046 | 2.68517 | 0.50163 | 0.45913 | 0.00000 |
| **LINC00545** | 0.01777 | 0.03537 | 0.02520 | 0.01858 | 0.00649 | 0.03818 | 0.01683 |
| **LINC00548** | 0.03048 | 0.01749 | 0.01271 | 0.02805 | 0.00800 | 0.01309 | 0.01200 |
| **LINC00550** | 0.01994 | 0.01657 | 0.01330 | 0.01690 | 0.00797 | 0.05498 | 0.02794 |
| **LINC00551** | 0.04458 | 0.03598 | 0.00905 | 0.01341 | 0.00824 | 0.01595 | 0.00825 |
| **LINC00552** | 0.02714 | 0.00859 | 0.02382 | 0.02682 | 0.09185 | 0.01283 | 0.01833 |
| **LINC00554** | 0.02301 | 0.01761 | 0.01545 | 0.01693 | 0.00746 | 0.02480 | 0.02629 |

| | | | | | | | |
|---|---|---|---|---|---|---|---|
| **LINC00558** | 0.02959 | 0.01866 | 0.00888 | 0.01456 | 0.21165 | 0.01288 | 0.02052 |
| **LINC00559** | 0.03103 | 0.01714 | 0.01325 | 0.02131 | 0.00825 | 0.01375 | 0.01741 |
| **LINC00563** | 0.02805 | 0.03314 | 0.01002 | 0.10398 | 0.00821 | 0.01333 | 0.00093 |
| **LINC00568** | 0.03010 | 0.07252 | 0.09680 | 0.02073 | 0.02115 | 0.01392 | 0.23546 |
| **LINC00570** | 0.04093 | 0.00000 | 0.04429 | 0.03154 | 0.08779 | 0.01163 | 0.01711 |
| **LINC00571** | 0.02176 | 0.02670 | 0.01660 | 0.02035 | 0.00747 | 0.01222 | 0.01582 |
| **LINC00574** | 0.04891 | 0.00034 | 0.04608 | 0.03979 | 0.06143 | 0.01913 | 0.05250 |
| **LINC00577** | 0.02129 | 0.00778 | 0.10364 | 0.02023 | 0.06601 | 0.01070 | 0.01824 |
| **LINC00578** | 0.02929 | 0.02554 | 0.12199 | 0.02955 | 0.03540 | 0.03297 | 0.05923 |
| **LINC00581** | 0.03066 | 0.01781 | 0.01295 | 0.02914 | 0.00777 | 0.01077 | 0.02304 |
| **LINC00592** | 0.02726 | 0.00744 | 0.01576 | 0.06533 | 0.00697 | 0.00605 | 0.07209 |
| **LINC00597** | 0.12192 | 0.00000 | 0.07044 | 0.25822 | 0.00467 | 0.00000 | 1.17399 |
| **LINC00598** | 0.04118 | 0.02859 | 0.01545 | 0.01407 | 0.04548 | 0.08614 | 0.02758 |
| **LINC00599** | 0.05174 | 0.01729 | 0.01154 | 0.01146 | 0.11406 | 0.04338 | 0.00982 |
| **LINC00601** | 0.02339 | 0.01808 | 0.00409 | 0.01902 | 0.15650 | 0.01347 | 0.01530 |
| **LINC00605** | 0.05465 | 0.08634 | 0.03921 | 0.00000 | 0.01648 | 0.17691 | 0.73395 |
| **LINC00607** | 0.11872 | 0.03594 | 0.01331 | 0.01687 | 0.05641 | 0.15960 | 0.00000 |
| **LINC00612** | 0.02651 | 0.01909 | 0.01538 | 0.02104 | 0.01171 | 0.00551 | 0.09657 |
| **LINC00615** | 0.02447 | 0.02032 | 0.01305 | 0.01930 | 0.04361 | 0.01308 | 0.01392 |
| **LINC00616** | 0.03208 | 0.01682 | 0.00993 | 0.01749 | 0.03840 | 0.01415 | 0.01291 |
| **LINC00618** | 0.00000 | 0.10407 | 0.05039 | 0.05210 | 0.00680 | 0.21304 | 0.56255 |
| **LINC00620** | 0.03747 | 0.01408 | 0.01268 | 0.03562 | 0.01111 | 0.07840 | 0.00147 |
| **LINC00622** | 0.02673 | 0.02527 | 0.00941 | 0.28342 | 0.00754 | 0.07225 | 0.02781 |
| **LINC00623.uid-** | 0.02856 | 0.01572 | 0.01293 | 0.01746 | 0.00787 | 0.01437 | 0.01951 |
| **LINC00624** | 3.92905 | 2.34894 | 1.22960 | 2.09754 | 1.76690 | 2.01728 | 0.87917 |
| **LINC00628** | 0.01217 | 0.02175 | 0.01542 | 0.02789 | 0.02269 | 0.08216 | 0.02700 |
| **LINC00629** | 0.04208 | 0.04537 | 0.01799 | 0.02260 | 0.17163 | 0.00742 | 0.00557 |
| **LINC00630** | 1.13286 | 0.55481 | 1.08343 | 0.67408 | 1.33947 | 0.45240 | 0.21570 |
| **LINC00632** | 0.07355 | 0.02346 | 0.01521 | 0.00499 | 0.12915 | 0.04919 | 0.03977 |
| **LINC00634** | 0.02082 | 0.04906 | 0.03258 | 0.02904 | 0.02917 | 0.03836 | 0.00637 |
| **LINC00635** | 0.03019 | 0.01485 | 0.01295 | 0.02497 | 0.02681 | 0.01271 | 0.01402 |
| **LINC00636** | 0.02387 | 0.01990 | 0.01326 | 0.02015 | 0.00805 | 0.01297 | 0.01663 |
| **LINC00637** | 0.00389 | 0.01056 | 0.01583 | 0.17347 | 0.02294 | 0.00712 | 0.09392 |
| **LINC00638** | 0.01761 | 0.06898 | 0.01365 | 0.01220 | 0.01099 | 0.00256 | 0.17776 |
| **LINC00639** | 0.00000 | 0.01791 | 0.00544 | 0.06999 | 0.12893 | 0.05032 | 0.04654 |

| | | | | | | | |
|---|---|---|---|---|---|---|---|
| **LINC00640** | 0.04468 | 0.02708 | 0.00952 | 0.05789 | 0.09856 | 0.14148 | 0.00987 |
| **LINC00641** | 2.38462 | 0.00000 | 0.56008 | 1.38990 | 0.26105 | 1.24027 | 5.86167 |
| **LINC00642** | 0.02979 | 0.01866 | 0.00997 | 0.02636 | 0.03319 | 0.01279 | 0.01053 |
| **LINC00643** | 0.11869 | 0.00000 | 0.02070 | 5.05179 | 0.04085 | 0.00000 | 0.17133 |
| **LINC00645** | 0.18620 | 0.00000 | 0.04914 | 0.06384 | 0.00559 | 0.01034 | 0.35417 |
| **LINC00648** | 0.28471 | 0.12070 | 0.09555 | 0.51884 | 0.07974 | 0.00000 | 0.15124 |
| **LINC00649** | 0.18571 | 0.03595 | 0.33351 | 0.14364 | 0.02124 | 0.27296 | 0.28596 |
| **LINC00652** | 0.06235 | 0.05209 | 0.01336 | 0.05532 | 0.02714 | 0.01201 | 0.00000 |
| **LINC00654** | 0.05329 | 0.04402 | 0.01760 | 0.07121 | 0.03819 | 0.09689 | 0.00000 |
| **LINC00657** | 1.01677 | 1.50642 | 4.76547 | 2.85696 | 1.83068 | 3.02097 | 13.16235 |
| **LINC00661** | 0.02448 | 0.01638 | 0.01299 | 0.01938 | 0.00800 | 0.02548 | 0.01791 |
| **LINC00662** | 1.35981 | 1.29965 | 1.00349 | 0.86207 | 1.02386 | 0.49308 | 0.43022 |
| **LINC00663** | 0.02892 | 0.01788 | 0.09559 | 0.06936 | 0.17447 | 0.01124 | 0.07424 |
| **LINC00664** | 0.04077 | 0.16304 | 0.96479 | 0.01820 | 0.25660 | 0.56061 | 0.23935 |
| **LINC00665** | 2.34374 | 0.41708 | 1.05456 | 6.42149 | 1.63095 | 2.11582 | 0.00000 |
| **LINC00667** | 2.65693 | 4.66502 | 4.40390 | 3.99702 | 0.56461 | 0.52223 | 0.00000 |
| **LINC00668** | 10.36777 | 3.64645 | 1.89425 | 11.42880 | 0.52154 | 0.00000 | 3.87424 |
| **LINC00669** | 0.37963 | 0.01897 | 0.09140 | 0.27035 | 0.02795 | 0.08253 | 0.18063 |
| **LINC00670** | 0.04058 | 0.02175 | 0.01438 | 0.01581 | 0.06375 | 0.03038 | 0.02497 |
| **LINC00671** | 0.04136 | 0.03128 | 0.02265 | 0.01557 | 0.00693 | 0.00950 | 0.04466 |
| **LINC00672** | 0.04767 | 0.17071 | 0.13549 | 0.05771 | 0.04367 | 0.42741 | 0.26979 |
| **LINC00673** | 0.14067 | 1.16061 | 0.00966 | 0.00000 | 0.01016 | 2.79040 | 0.00000 |
| **LINC00674** | 0.58571 | 0.07206 | 0.07459 | 0.33112 | 0.46851 | 0.99789 | 0.00827 |
| **LINC00676** | 0.02070 | 0.01493 | 0.01085 | 0.01698 | 0.02130 | 0.04409 | 0.02264 |
| **LINC00678** | 0.13558 | 1.80962 | 0.47263 | 0.81977 | 0.02086 | 1.25316 | 0.00000 |
| **LINC00682** | 0.01595 | 0.01541 | 0.00000 | 0.01266 | 0.60597 | 0.03410 | 0.04223 |
| **LINC00687** | 0.00371 | 0.01825 | 0.01413 | 0.11495 | 0.00820 | 0.01084 | 0.02677 |
| **LINC00689** | 0.00970 | 0.00000 | 0.86463 | 0.12114 | 0.13398 | 0.03306 | 0.14243 |
| **LINC00690** | 0.03106 | 0.01751 | 0.01226 | 0.01827 | 0.01044 | 0.01394 | 0.01694 |
| **LINC00691** | 0.02995 | 0.01646 | 0.09733 | 0.04203 | 0.01715 | 0.07359 | 0.02884 |
| **LINC00692** | 0.01742 | 0.00802 | 0.02063 | 0.03161 | 0.02250 | 0.02044 | 0.05487 |
| **LINC00693** | 0.00321 | 0.01123 | 0.01499 | 0.02828 | 0.02630 | 0.08839 | 0.08453 |
| **LINC00696** | 0.04820 | 0.01723 | 0.04246 | 0.01344 | 0.01005 | 0.00922 | 0.02181 |
| **LINC00698** | 0.05661 | 0.00300 | 0.02223 | 0.03852 | 0.02696 | 0.18651 | 0.00000 |
| **LINC00701** | 0.04826 | 0.00655 | 0.31034 | 0.05065 | 0.01799 | 0.00631 | 0.00342 |

| | | | | | | | |
|---|---|---|---|---|---|---|---|
| **LINC00707** | 0.00000 | 0.68835 | 0.77673 | 0.00000 | 0.64540 | 0.63022 | 0.02101 |
| **LINC00836** | 0.05679 | 0.02574 | 0.01053 | 0.19226 | 0.00788 | 0.01142 | 0.01094 |
| **LINC00837** | 0.02724 | 0.00304 | 0.03651 | 0.01959 | 0.04009 | 0.03305 | 0.01656 |
| **LINC00839** | 0.15574 | 0.00000 | 0.05048 | 0.14120 | 1.33035 | 0.19529 | 0.00000 |
| **LINC00842** | 1.15262 | 0.84320 | 0.00000 | 1.35365 | 0.00000 | 0.00000 | 1.62822 |
| **LINC00844** | 0.05386 | 0.02666 | 0.04607 | 0.01160 | 0.01127 | 0.01290 | 0.00460 |
| **LINC00847** | 0.00000 | 0.00688 | 0.33699 | 0.37961 | 0.13098 | 0.00620 | 0.08045 |
| **LINC00852** | 0.49858 | 0.47810 | 0.17504 | 0.70220 | 0.07385 | 0.53592 | 0.54904 |
| **LINC00853** | 0.02424 | 0.01303 | 0.01831 | 0.01828 | 0.01010 | 0.01808 | 0.02124 |
| **LINC00854** | 0.02134 | 0.07544 | 0.04079 | 0.01256 | 0.00590 | 0.04841 | 0.04862 |
| **LINC00857** | 1.11072 | 0.42165 | 0.00007 | 0.23308 | 0.00985 | 0.00000 | 0.02483 |
| **LINC00858** | 0.07341 | 0.30595 | 0.39020 | 0.07201 | 0.00167 | 0.10753 | 0.00000 |
| **LINC00861** | 0.02362 | 0.01835 | 0.01288 | 0.01999 | 0.00791 | 0.01288 | 0.01723 |
| **LINC00862** | 0.07177 | 0.05415 | 0.00664 | 0.06462 | 0.05586 | 0.21149 | 0.03783 |
| **LINC00865** | 0.02724 | 0.03772 | 0.01093 | 0.02075 | 0.08558 | 0.00997 | 0.01414 |
| **LINC00867** | 0.00512 | 0.05861 | 0.05219 | 0.02986 | 0.00804 | 0.08015 | 0.02293 |
| **LINC00869.uid-1** | 0.02571 | 0.01597 | 0.01352 | 0.02302 | 0.00806 | 0.01908 | 0.01889 |
| **LINC00870** | 0.02805 | 0.00962 | 0.01936 | 0.01746 | 0.01064 | 0.02589 | 0.02452 |
| **LINC00871** | 0.03166 | 0.00000 | 0.10054 | 0.01767 | 0.11425 | 0.02350 | 0.01647 |
| **LINC00877** | 0.02166 | 0.01915 | 0.01213 | 0.03482 | 0.01201 | 0.01008 | 0.03104 |
| **LINC00879** | 0.03039 | 0.01993 | 0.01455 | 0.03805 | 0.00776 | 0.01602 | 0.01169 |
| **LINC00882** | 0.01607 | 0.08618 | 0.03894 | 0.02618 | 0.11374 | 0.00780 | 0.00872 |
| **LINC00883** | 0.12734 | 0.00000 | 0.18369 | 0.32655 | 0.48527 | 0.28438 | 0.00000 |
| **LINC00884** | 0.00000 | 0.66411 | 1.08308 | 0.23348 | 0.32150 | 0.00000 | 0.00000 |
| **LINC00885** | 0.01544 | 0.15851 | 0.01256 | 0.07371 | 0.18219 | 0.00851 | 0.10597 |
| **LINC00886** | 0.03588 | 0.00081 | 0.09890 | 0.02084 | 0.03979 | 0.01640 | 0.01517 |
| **LINC00887** | 0.02523 | 0.00000 | 0.06199 | 0.01424 | 1.48084 | 0.00997 | 0.02647 |
| **LINC00888** | 1.86803 | 0.81846 | 1.17075 | 2.26311 | 0.09480 | 3.26024 | 2.06850 |
| **LINC00889** | 0.03173 | 0.09703 | 0.01474 | 0.02855 | 0.00714 | 0.01740 | 0.00000 |
| **LINC00890** | 0.02619 | 0.04573 | 0.01290 | 0.01912 | 0.00979 | 0.06516 | 0.00725 |
| **LINC00892** | 0.00652 | 0.01812 | 0.01280 | 0.08008 | 0.00818 | 0.01512 | 0.02282 |
| **LINC00893** | 0.03553 | 0.01995 | 0.01396 | 0.02996 | 0.19547 | 0.01493 | 0.02184 |
| **LINC00894** | 0.09470 | 0.11453 | 0.07862 | 0.07264 | 0.27862 | 0.14397 | 0.09061 |
| **LINC00896** | 0.02681 | 0.01861 | 0.04178 | 0.02035 | 0.00613 | 0.01513 | 0.01181 |
| **LINC00898** | 0.02416 | 0.01894 | 0.01362 | 0.01933 | 0.00774 | 0.01401 | 0.02022 |

| | | | | | | | |
|---|---|---|---|---|---|---|---|
| **LINC00899** | 0.02881 | 0.01673 | 0.01257 | 0.01776 | 0.00787 | 0.01207 | 0.02417 |
| **LINC00907** | 0.02549 | 0.08737 | 0.10834 | 0.01396 | 0.47584 | 0.10653 | 0.03883 |
| **LINC00908** | 0.02499 | 0.01741 | 0.01260 | 0.02050 | 0.00796 | 0.01747 | 0.01789 |
| **LINC00909** | 1.47181 | 0.54023 | 2.15637 | 1.58144 | 1.04236 | 0.18877 | 0.00000 |
| **LINC00910** | 0.08087 | 0.00000 | 0.67623 | 0.19942 | 0.07214 | 0.04459 | 0.47017 |
| **LINC00917** | 0.04344 | 0.00653 | 0.07991 | 0.01965 | 0.00577 | 0.01189 | 0.01165 |
| **LINC00921** | 0.04212 | 0.00000 | 0.20326 | 0.04333 | 0.11139 | 0.04461 | 0.03680 |
| **LINC00922** | 0.01640 | 0.02810 | 0.03563 | 0.01334 | 0.22549 | 0.14595 | 0.02942 |
| **LINC00923** | 0.04301 | 0.07087 | 0.01928 | 0.01686 | 0.11111 | 0.00815 | 0.00249 |
| **LINC00924** | 0.02425 | 0.01812 | 0.01296 | 0.01969 | 0.00791 | 0.01299 | 0.01691 |
| **LINC00925** | 0.61358 | 0.00611 | 2.48256 | 0.58023 | 0.89396 | 0.00000 | 0.00000 |
| **LINC00926** | 0.00000 | 0.02604 | 0.00000 | 0.09465 | 0.54157 | 0.06123 | 0.11371 |
| **LINC00933** | 0.02000 | 0.02362 | 0.01711 | 0.01415 | 0.03147 | 0.01581 | 0.05064 |
| **LINC00934** | 0.02417 | 0.01644 | 0.01283 | 0.01827 | 0.00795 | 0.01724 | 0.01897 |
| **LINC00935** | 0.00673 | 0.04260 | 0.62287 | 0.00840 | 0.44761 | 0.13104 | 0.02056 |
| **LINC00936** | 1.40375 | 1.04546 | 0.45446 | 1.10653 | 0.36845 | 0.17090 | 1.03884 |
| **LINC00937** | 0.00287 | 0.03554 | 0.04519 | 0.02493 | 0.02284 | 0.25248 | 0.04790 |
| **LINC00938** | 2.54255 | 1.89530 | 0.90967 | 1.84142 | 0.21682 | 0.51065 | 0.21988 |
| **LINC00939** | 0.02173 | 0.01644 | 0.01098 | 0.02197 | 0.01645 | 0.02932 | 0.02200 |
| **LINC00940** | 0.02218 | 0.02124 | 0.01274 | 0.02392 | 0.00804 | 0.01700 | 0.01964 |
| **LINC00941** | 0.01715 | 0.02303 | 0.06308 | 0.02852 | 0.01112 | 0.22696 | 0.01308 |
| **LINC00942** | 0.01213 | 0.02277 | 0.01495 | 0.08366 | 0.00738 | 0.03458 | 0.12890 |
| **LINC00943** | 0.02127 | 0.02219 | 0.00000 | 0.01852 | 0.67181 | 0.01360 | 0.01621 |
| **LINC00948** | 0.01883 | 0.00000 | 3.07519 | 0.78667 | 2.42138 | 0.16375 | 0.61507 |
| **LINC00950** | 0.02274 | 0.00552 | 0.00851 | 0.00648 | 0.03534 | 0.01287 | 0.16667 |
| **LINC00951** | 0.01243 | 0.01645 | 0.01267 | 0.04992 | 0.00770 | 0.01447 | 0.03362 |
| **LINC00954** | 0.01109 | 0.01426 | 0.00629 | 0.03756 | 0.18842 | 0.01023 | 0.04395 |
| **LINC00955** | 0.01119 | 0.02186 | 0.01333 | 0.01628 | 0.00733 | 0.03720 | 0.04795 |
| **LINC00957** | 0.02680 | 0.08508 | 0.03742 | 0.12777 | 0.03107 | 0.04411 | 0.00000 |
| **LINC00958** | 0.80707 | 0.00000 | 0.90746 | 1.34101 | 0.76833 | 0.00000 | 2.44631 |
| **LINC00959** | 0.19903 | 0.00000 | 0.25293 | 0.02984 | 0.30392 | 0.17718 | 0.00000 |
| **LINC00960.uid-!** | 0.12909 | 0.34623 | 0.08203 | 0.17387 | 0.01159 | 0.16994 | 0.04372 |
| **LINC00961** | 0.04378 | 0.01189 | 0.01687 | 0.01554 | 0.02872 | 0.01225 | 0.01087 |
| **LINC00963** | 0.67616 | 0.08696 | 0.00407 | 0.56177 | 0.00894 | 0.00680 | 0.03300 |
| **LINC00964** | 0.02092 | 0.01750 | 0.01318 | 0.02172 | 0.00780 | 0.02042 | 0.02112 |

| | | | | | | | |
|---|---|---|---|---|---|---|---|
| **LINC00967** | 0.05509 | 0.06179 | 0.08661 | 0.15733 | 0.04152 | 0.04231 | 0.00000 |
| **LINC00969** | 0.27515 | 0.17661 | 0.16327 | 0.14194 | 0.02788 | 0.19574 | 0.37495 |
| **LINC00971** | 0.03134 | 0.02830 | 0.01815 | 0.08361 | 0.02274 | 0.03923 | 0.00000 |
| **LINC00974** | 1.19163 | 0.00000 | 0.02798 | 0.93015 | 0.00308 | 0.00000 | 0.03861 |
| **LINC00977** | 0.01834 | 0.02420 | 0.00000 | 0.01784 | 0.50649 | 0.01178 | 0.01944 |
| **LINC00982** | 0.03097 | 0.00000 | 0.05121 | 0.01776 | 0.00603 | 0.01153 | 0.03944 |
| **LINC00987** | 0.08881 | 0.10951 | 0.03672 | 0.11162 | 0.02883 | 0.06459 | 0.15794 |
| **LINC00989** | 0.02033 | 0.04909 | 0.15242 | 0.01036 | 0.05227 | 0.37851 | 0.00807 |
| **LINC00992** | 0.05562 | 0.03865 | 0.02837 | 0.01181 | 0.01364 | 0.03032 | 0.04315 |
| **LINC00997** | 0.07742 | 0.00608 | 0.04821 | 0.79232 | 0.03115 | 0.30797 | 0.54923 |
| **LINC00998** | 6.18696 | 0.13351 | 3.95993 | 3.42200 | 1.30212 | 1.61981 | 0.00000 |
| **LINC00999** | 0.02069 | 0.16534 | 0.03209 | 0.03050 | 0.02059 | 0.20638 | 0.19267 |
| **LINC01000** | 0.00000 | 0.92186 | 0.73874 | 1.12931 | 0.01789 | 1.34252 | 5.94983 |
| **LINC01001** | 0.02231 | 0.01481 | 0.02001 | 0.01951 | 0.01230 | 0.01724 | 0.02095 |
| **LINC01003** | 0.00000 | 0.17621 | 0.22588 | 0.03618 | 0.00184 | 0.13650 | 0.70474 |
| **LINC01004** | 1.53823 | 0.00000 | 0.70557 | 0.46822 | 0.00000 | 0.04531 | 2.38946 |
| **LINC01005** | 0.02621 | 0.01854 | 0.01233 | 0.01835 | 0.04265 | 0.01285 | 0.01637 |
| **LINC01006** | 0.02008 | 0.00319 | 0.04279 | 0.03247 | 0.09597 | 0.03654 | 0.01659 |
| **LINC01007** | 0.04014 | 0.01339 | 0.03761 | 0.04193 | 0.05738 | 0.01015 | 0.00252 |
| **LINC01010** | 0.01182 | 0.14925 | 0.29813 | 0.01478 | 0.04245 | 0.00123 | 0.01009 |
| **LINC01011** | 0.07882 | 0.79778 | 0.51016 | 0.27357 | 0.20035 | 0.34740 | 0.00000 |
| **LINC01012** | 0.67837 | 0.05534 | 0.00579 | 0.41566 | 0.05768 | 0.26576 | 0.20296 |
| **LINC01015** | 0.01673 | 0.02410 | 0.01143 | 0.02945 | 0.00783 | 0.01118 | 0.02479 |
| **LINC01017** | 0.02606 | 0.02296 | 0.01693 | 0.02257 | 0.00746 | 0.01246 | 0.01411 |
| **LINC01019** | 0.01115 | 0.06661 | 0.01434 | 0.26071 | 0.88123 | 0.08972 | 0.34698 |
| **LINC01021** | 0.60875 | 1.92365 | 0.00000 | 2.65655 | 1.07436 | 0.56287 | 0.00000 |
| **LINC01023** | 0.01810 | 0.63123 | 0.02643 | 0.13874 | 0.11374 | 0.06347 | 0.54319 |
| **LINC01024** | 0.22606 | 0.00000 | 0.67783 | 0.31607 | 0.27582 | 0.25014 | 0.55394 |
| **LINC01028** | 0.02138 | 0.01858 | 0.01306 | 0.03350 | 0.00942 | 0.02278 | 0.01319 |
| **LINC01029** | 0.05667 | 0.00000 | 0.10170 | 0.58294 | 0.10172 | 0.00000 | 0.16748 |
| **LINC01030** | 0.60923 | 0.06774 | 0.23883 | 0.29226 | 2.58724 | 0.45459 | 0.04692 |
| **LINC01049** | 0.04258 | 0.01466 | 0.01442 | 0.03117 | 0.00839 | 0.01545 | 0.00575 |
| **LINC01056** | 0.02991 | 0.01334 | 0.08018 | 0.03687 | 0.28148 | 0.00849 | 0.00611 |
| **LINC01057** | 0.07219 | 0.05060 | 0.01362 | 0.08371 | 0.00896 | 0.24039 | 0.04574 |
| **LINC01060** | 0.08587 | 0.02414 | 0.03911 | 0.02981 | 0.00595 | 0.12709 | 0.00000 |

| | | | | | | | |
|---|---|---|---|---|---|---|---|
| **LINC01061** | 0.06762 | 0.13657 | 0.22092 | 0.13392 | 0.00162 | 0.13680 | 0.52596 |
| **LINC01081** | 0.02363 | 0.01148 | 0.01114 | 0.01954 | 0.21601 | 0.01407 | 0.01631 |
| **LINC01082** | 0.02185 | 0.01719 | 0.01117 | 0.02635 | 0.02816 | 0.01272 | 0.01767 |
| **LINC01087** | 0.10311 | 0.02134 | 0.15276 | 0.06024 | 0.02265 | 0.00000 | 0.03327 |
| **LINC01088** | 0.06954 | 0.00000 | 0.07391 | 0.08222 | 0.05587 | 0.03767 | 0.05616 |
| **LINC01089** | 0.58613 | 1.09852 | 1.19354 | 2.94169 | 0.38329 | 1.21648 | 3.49292 |
| **LINC01091** | 0.14396 | 0.01713 | 0.07030 | 0.43851 | 0.07011 | 0.08855 | 0.73872 |
| **LINC01094** | 0.19965 | 0.40060 | 0.01554 | 0.00000 | 0.03768 | 0.01747 | 0.04939 |
| **LINC01096** | 0.04227 | 0.00871 | 0.01233 | 0.03073 | 0.19863 | 0.01278 | 0.00520 |
| **LINC01097** | 0.01600 | 0.01545 | 0.01204 | 0.02418 | 0.00820 | 0.12414 | 0.01609 |
| **LINC01098** | 0.03165 | 0.05727 | 0.08493 | 0.09254 | 0.21232 | 0.01873 | 0.00000 |
| **LINC01099** | 0.03250 | 0.07900 | 0.09835 | 0.01819 | 0.21593 | 0.03391 | 0.00087 |
| **LINC01104** | 0.02420 | 0.01812 | 0.01264 | 0.03007 | 0.00865 | 0.01278 | 0.01841 |
| **LINC01105** | 0.04892 | 0.00503 | 0.03908 | 0.01554 | 0.00713 | 0.01638 | 0.01514 |
| **LINC01108** | 0.39466 | 3.09478 | 0.65914 | 0.00000 | 0.00000 | 9.49616 | 0.00000 |
| **LINC01111** | 0.03023 | 0.00664 | 0.01984 | 0.01163 | 0.01505 | 0.01415 | 0.08131 |
| **LINC01115** | 0.02497 | 0.01470 | 0.01338 | 0.02288 | 0.07991 | 0.01255 | 0.01849 |
| **LINC01116** | 0.02422 | 0.01022 | 0.04431 | 0.01708 | 0.01009 | 0.01084 | 0.02716 |
| **LINC01118** | 0.00000 | 0.08432 | 0.00000 | 0.01415 | 5.56451 | 0.03474 | 0.01501 |
| **LINC01120** | 0.02319 | 0.01499 | 0.01412 | 0.02201 | 0.03385 | 0.01326 | 0.01741 |
| **LINC01121** | 0.02762 | 0.01809 | 0.01429 | 0.01994 | 0.02661 | 0.01561 | 0.01466 |
| **LINC01122** | 0.02885 | 0.01670 | 0.02204 | 0.01803 | 0.15146 | 0.01281 | 0.01747 |
| **LINC01123.uid-** | 0.11299 | 0.00000 | 0.07812 | 0.03081 | 0.00928 | 0.00768 | 0.03324 |
| **LINC01123.uid-** | 0.05713 | 0.01502 | 0.05771 | 0.03020 | 0.00521 | 0.00957 | 0.02464 |
| **LINC01124** | 0.01914 | 0.05153 | 0.02211 | 0.02238 | 0.00681 | 0.01229 | 0.01111 |
| **LINC01125** | 0.06424 | 0.12536 | 0.06210 | 0.09380 | 0.02698 | 0.00685 | 0.00196 |
| **LINC01126** | 0.00182 | 0.01429 | 0.06570 | 0.01749 | 0.00411 | 0.15847 | 0.14646 |
| **LINC01128** | 3.83971 | 2.15785 | 2.02272 | 3.21187 | 3.45778 | 0.33578 | 2.32653 |
| **LINC01133** | 0.23869 | 0.23825 | 0.00862 | 0.00571 | 0.01766 | 0.05451 | 0.00000 |
| **LINC01134** | 0.00000 | 0.20686 | 0.24580 | 0.20238 | 0.00850 | 0.00000 | 0.03074 |
| **LINC01135** | 0.02020 | 0.01760 | 0.01223 | 0.02702 | 0.00797 | 0.01279 | 0.02962 |
| **LINC01136** | 0.04195 | 0.03124 | 0.01355 | 0.02684 | 0.01183 | 0.01109 | 0.01556 |
| **LINC01137** | 0.18424 | 0.28274 | 0.03996 | 0.05020 | 0.03513 | 0.10016 | 0.00000 |
| **LINC01138** | 0.02409 | 0.02013 | 0.01472 | 0.02677 | 0.00906 | 0.01500 | 0.01444 |
| **LINC01140** | 0.31515 | 0.00000 | 0.09969 | 0.29142 | 0.07035 | 0.17043 | 0.19008 |

| | | | | | | | |
|---|---|---|---|---|---|---|---|
| **LINC01141** | 0.02054 | 0.01778 | 0.00830 | 0.03132 | 0.04801 | 0.05920 | 0.01151 |
| **LINC01142** | 0.02500 | 0.01802 | 0.01252 | 0.01684 | 0.00795 | 0.01796 | 0.02177 |
| **LINC01144** | 0.00000 | 0.11487 | 0.25621 | 0.09177 | 0.14709 | 0.01590 | 0.07298 |
| **LINC01150** | 0.00000 | 0.09218 | 0.04648 | 0.02326 | 0.00624 | 0.12774 | 0.02554 |
| **LINC01152** | 0.01931 | 0.01476 | 0.13109 | 0.01072 | 3.44844 | 0.00274 | 0.02511 |
| **LINC01159** | 0.03393 | 0.02174 | 0.03296 | 0.01716 | 0.02192 | 0.22677 | 0.00000 |
| **LINC01160** | 0.03688 | 9.03061 | 1.26198 | 0.55274 | 0.00000 | 7.56062 | 0.00000 |
| **LINC01164** | 0.03797 | 0.01823 | 0.01286 | 0.02071 | 0.03664 | 0.00577 | 0.05723 |
| **LINC01166** | 0.01071 | 0.02397 | 0.40568 | 0.01270 | 0.00641 | 0.03954 | 0.02720 |
| **LINC01168** | 0.03258 | 0.00000 | 0.15101 | 0.02079 | 0.00529 | 0.02435 | 0.03403 |
| **LINC01169** | 0.03870 | 0.01546 | 0.00770 | 0.02670 | 0.02899 | 0.01118 | 0.17211 |
| **LINC01176** | 0.01983 | 0.04562 | 0.03233 | 0.28112 | 0.00401 | 0.11418 | 0.66000 |
| **LINC01179** | 0.02586 | 0.01226 | 0.02470 | 0.02021 | 0.00869 | 0.01272 | 0.01607 |
| **LINC01180** | 0.02715 | 0.01644 | 0.01190 | 0.02383 | 0.01438 | 0.00961 | 0.04216 |
| **LINC01182** | 0.03152 | 0.01620 | 0.01315 | 0.01350 | 0.03712 | 0.04169 | 0.02331 |
| **LINC01184** | 0.04807 | 0.01156 | 0.02623 | 0.01533 | 0.00752 | 0.07929 | 0.02008 |
| **LINC01185** | 0.02366 | 0.03214 | 0.02514 | 0.02120 | 0.00702 | 0.01472 | 0.00945 |
| **LINC01186** | 0.02245 | 0.29280 | 0.69973 | 0.00000 | 0.00175 | 0.00000 | 0.01648 |
| **LINC01187** | 0.02462 | 0.01937 | 0.00983 | 0.01888 | 0.03080 | 0.01325 | 0.01614 |
| **LINC01191** | 0.02618 | 0.00000 | 0.04594 | 0.07361 | 0.07344 | 0.04523 | 0.05822 |
| **LINC01192** | 0.02797 | 0.01715 | 0.01368 | 0.02048 | 0.00861 | 0.01462 | 0.01442 |
| **LINC01193** | 0.03221 | 0.01725 | 0.01780 | 0.01830 | 0.22773 | 0.01334 | 0.00796 |
| **LINC01194** | 0.23895 | 0.13421 | 0.20904 | 0.41619 | 0.01278 | 0.07337 | 0.31771 |
| **LINC01203** | 0.08351 | 0.00226 | 0.08151 | 0.25871 | 0.03734 | 0.00573 | 0.09528 |
| **LINC01204** | 0.02138 | 0.01084 | 0.01252 | 0.03324 | 0.07230 | 0.04311 | 0.02822 |
| **LINC01205** | 1.30720 | 1.05081 | 0.16550 | 0.36598 | 0.00362 | 0.90557 | 0.49986 |
| **LINC01206** | 0.07289 | 0.11139 | 0.08800 | 0.00913 | 0.27910 | 0.95956 | 0.00000 |
| **LINC01207** | 0.04017 | 0.01723 | 0.01308 | 0.03278 | 0.00969 | 0.01510 | 0.01440 |
| **LINC01208** | 0.02147 | 0.00533 | 0.03573 | 0.01969 | 0.29065 | 0.01211 | 0.02214 |
| **LINC01209** | 0.02231 | 0.01092 | 0.02548 | 0.02017 | 0.02036 | 0.01412 | 0.02204 |
| **LINC01212** | 0.02635 | 0.01618 | 0.01362 | 0.01918 | 0.07444 | 0.01572 | 0.01330 |
| **LINC01214** | 0.05283 | 0.01144 | 0.01309 | 0.03855 | 0.01502 | 0.01263 | 0.04442 |
| **LINC01220** | 0.01976 | 0.03411 | 0.02957 | 0.08004 | 0.01069 | 0.02712 | 0.00907 |
| **LINC01225** | 0.02498 | 0.01785 | 0.01271 | 0.02625 | 0.00793 | 0.01264 | 0.01586 |
| **LINC01226** | 0.02432 | 0.01836 | 0.00483 | 0.02709 | 0.14945 | 0.01449 | 0.01196 |

| | | | | | | | |
|---|---|---|---|---|---|---|---|
| **LINC01230** | 0.02545 | 0.02443 | 0.02755 | 0.01153 | 0.01793 | 0.01337 | 0.04542 |
| **LINC01231** | 0.01865 | 0.02117 | 0.00929 | 0.01871 | 0.05088 | 0.02290 | 0.02235 |
| **LINC01232** | 0.07064 | 0.06073 | 0.02219 | 0.08223 | 0.06105 | 0.01634 | 0.00000 |
| **LINC01233** | 0.01949 | 0.03744 | 0.02304 | 0.03240 | 0.03995 | 0.00926 | 0.01560 |
| **LINC01234** | 5.00451 | 0.58552 | 0.54447 | 0.87633 | 0.00000 | 0.79670 | 0.00000 |
| **LINC01239** | 0.03753 | 0.00000 | 0.08296 | 0.00148 | 0.07265 | 0.04795 | 0.07350 |
| **LINC01241** | 0.02984 | 0.01853 | 0.01476 | 0.02900 | 0.02215 | 0.01647 | 0.00993 |
| **LINC01242** | 0.05189 | 0.02171 | 0.04471 | 0.01830 | 0.00651 | 0.01191 | 0.01590 |
| **LINC01248** | 0.02545 | 0.01940 | 0.01242 | 0.01861 | 0.00895 | 0.01275 | 0.01695 |
| **LINC01249** | 0.02572 | 0.12129 | 0.05685 | 0.01980 | 0.09560 | 0.00824 | 0.00006 |
| **LINC01250** | 0.04715 | 0.05284 | 0.05886 | 0.00902 | 0.00987 | 0.15742 | 0.03616 |
| **LINC01251** | 0.02913 | 0.00369 | 0.04576 | 0.02580 | 0.01810 | 0.01531 | 0.01315 |
| **LINC01252** | 0.01698 | 0.01894 | 0.02406 | 0.01628 | 1.19533 | 0.00688 | 0.05815 |
| **LINC01254** | 0.02529 | 0.01615 | 0.00933 | 0.02070 | 0.08680 | 0.01333 | 0.01391 |
| **LINC01257** | 0.02427 | 0.00502 | 0.02835 | 0.01846 | 0.22226 | 0.01353 | 0.01787 |
| **LINC01260** | 0.57837 | 0.35691 | 0.65229 | 0.14210 | 0.03216 | 0.00000 | 0.00000 |
| **LINC01263** | 0.03361 | 0.05455 | 0.03948 | 0.00626 | 0.07255 | 0.25510 | 0.04823 |
| **LINC01266** | 0.01958 | 0.01481 | 0.01427 | 0.02570 | 0.01861 | 0.02101 | 0.02186 |
| **LINC01268** | 0.01943 | 0.02606 | 0.00978 | 0.01663 | 0.05860 | 0.01413 | 0.03030 |
| **LINC01270** | 0.04587 | 0.01801 | 0.01449 | 0.05415 | 0.00793 | 0.02750 | 0.00000 |
| **LINC01272** | 0.01690 | 0.01825 | 0.01299 | 0.04181 | 0.00987 | 0.01867 | 0.01676 |
| **LINC01273** | 0.00875 | 0.00000 | 0.04982 | 0.31093 | 0.01604 | 0.00873 | 0.14126 |
| **LINC01277** | 0.02558 | 0.01685 | 0.00941 | 0.02095 | 0.05882 | 0.01331 | 0.01482 |
| **LINC01278** | 4.54252 | 2.64853 | 0.59149 | 4.14000 | 0.38101 | 0.00000 | 5.55386 |
| **LINC01279** | 0.07008 | 0.04558 | 0.10596 | 0.06928 | 0.12090 | 0.18475 | 0.03033 |
| **LINC01280** | 0.02770 | 0.00637 | 0.03205 | 0.02329 | 0.03716 | 0.01373 | 0.01887 |
| **LINC01285** | 0.02513 | 0.01318 | 0.01801 | 0.01920 | 0.03473 | 0.01324 | 0.01571 |
| **LINC01289** | 0.01748 | 0.01139 | 0.01492 | 0.16552 | 0.00745 | 0.00873 | 0.04243 |
| **LINC01296.uid-** | 0.01364 | 0.02464 | 0.02633 | 0.02706 | 0.02540 | 0.00908 | 0.04141 |
| **LINC01299** | 0.02559 | 0.01598 | 0.01355 | 0.01893 | 0.00776 | 0.01245 | 0.02801 |
| **LINC01300** | 0.23467 | 0.10930 | 0.00276 | 0.11669 | 0.00765 | 0.20450 | 0.00000 |
| **LINC01301** | 0.12825 | 0.00000 | 0.12362 | 0.07271 | 0.01679 | 0.01823 | 0.06473 |
| **LINC01304** | 0.02156 | 0.01575 | 0.01317 | 0.02410 | 0.00798 | 0.01267 | 0.02540 |
| **LINC01305** | 0.00703 | 0.00643 | 0.01671 | 0.02971 | 0.01860 | 0.18628 | 0.04081 |
| **LINC01310** | 0.02317 | 0.01567 | 0.01239 | 0.02050 | 0.01161 | 0.01852 | 0.02051 |

| | | | | | | | |
|---|---|---|---|---|---|---|---|
| **LINC01311** | 0.47367 | 0.96971 | 0.24287 | 0.76461 | 0.00000 | 0.00000 | 1.68595 |
| **LINC01312** | 0.02525 | 0.02318 | 0.00407 | 0.01874 | 0.14068 | 0.01340 | 0.01331 |
| **LINC01314** | 0.03938 | 0.00987 | 0.01096 | 0.02674 | 0.02377 | 0.01064 | 0.05180 |
| **LINC01315** | 0.29117 | 0.22989 | 0.01902 | 0.70651 | 0.03409 | 0.04195 | 0.09899 |
| **LINC01330** | 0.03927 | 0.01620 | 0.01347 | 0.02235 | 0.00805 | 0.01492 | 0.00518 |
| **LINC01333** | 0.02762 | 0.01718 | 0.02184 | 0.01939 | 0.00736 | 0.01915 | 0.01342 |
| **LINC01338** | 0.03631 | 0.01906 | 0.01802 | 0.02566 | 0.02434 | 0.01057 | 0.01521 |
| **LINC01340** | 0.04069 | 0.02654 | 0.01132 | 0.01359 | 0.01839 | 0.01327 | 0.01585 |
| **LINC01341** | 0.05827 | 0.00806 | 0.01753 | 0.06542 | 0.00768 | 0.02005 | 0.00933 |
| **LINC01344** | 0.00782 | 0.13279 | 0.05343 | 0.01049 | 0.10883 | 0.00836 | 0.03998 |
| **LINC01347** | 0.01113 | 0.00704 | 0.05904 | 0.12385 | 0.02387 | 0.01737 | 0.13179 |
| **LINC01352** | 0.02383 | 0.01022 | 0.01299 | 0.05796 | 0.00733 | 0.00612 | 0.06863 |
| **LINC01354** | 0.02462 | 0.00517 | 0.03449 | 0.03191 | 0.07685 | 0.01413 | 0.02064 |
| **LINC01355** | 1.45847 | 0.00000 | 0.23628 | 0.35582 | 0.03464 | 0.00000 | 1.42480 |
| **LINC01356** | 2.39251 | 0.09214 | 0.24956 | 10.71496 | 0.05565 | 15.09353 | 0.17533 |
| **LINC01358** | 0.02606 | 0.01815 | 0.01229 | 0.01853 | 0.01022 | 0.01326 | 0.01755 |
| **LINC01359** | 0.02395 | 0.00456 | 0.07001 | 0.02374 | 0.04504 | 0.01525 | 0.01698 |
| **LINC01360** | 0.02666 | 0.02142 | 0.02556 | 0.03904 | 0.00696 | 0.01463 | 0.00984 |
| **LINC01361** | 0.02713 | 0.01789 | 0.01247 | 0.01784 | 0.00798 | 0.01450 | 0.01864 |
| **LINC01363** | 0.15350 | 0.17591 | 0.03413 | 0.04673 | 0.14017 | 0.00164 | 0.00000 |
| **LINC01364** | 0.04595 | 0.01715 | 0.01332 | 0.04865 | 0.00962 | 0.01482 | 0.00000 |
| **LINC01372** | 0.05091 | 0.00979 | 0.00985 | 0.04227 | 0.05666 | 0.01373 | 0.03823 |
| **LINC01391** | 0.03749 | 0.02147 | 0.01373 | 0.02121 | 0.00767 | 0.00951 | 0.02164 |
| **LINC01393** | 0.03662 | 0.03696 | 0.03151 | 0.01444 | 0.00705 | 0.01884 | 0.01008 |
| **LINC01395** | 0.06632 | 0.03890 | 0.04014 | 0.02458 | 0.00607 | 0.01940 | 0.00435 |
| **LINC01405** | 0.21119 | 0.00006 | 0.00123 | 0.00000 | 0.01847 | 0.00187 | 0.88806 |
| **LINC01410** | 0.09488 | 0.04230 | 0.01581 | 0.03567 | 0.01589 | 0.03705 | 0.02416 |
| **LINC01413** | 0.08155 | 0.04247 | 0.00797 | 0.15953 | 0.04784 | 0.07006 | 0.00000 |
| **LINC01419** | 0.16128 | 0.04954 | 0.10440 | 0.31610 | 0.02096 | 0.00728 | 0.06559 |
| **LINC01420** | 0.56231 | 0.60407 | 1.96318 | 0.69143 | 0.00378 | 0.71362 | 0.00731 |
| **LINC01422** | 0.04294 | 0.06157 | 0.06602 | 0.01165 | 0.04354 | 0.13112 | 0.00000 |
| **LINC01424** | 0.08302 | 0.01816 | 0.07519 | 0.03455 | 0.00980 | 0.01342 | 0.00964 |
| **LINC01425** | 0.02538 | 0.01565 | 0.00804 | 0.01664 | 0.10536 | 0.02072 | 0.02362 |
| **LINC01426** | 0.24626 | 0.13603 | 0.00583 | 0.05569 | 0.00667 | 0.47953 | 0.00000 |
| **LINC01428** | 0.02581 | 0.01315 | 0.01063 | 0.02392 | 0.03499 | 0.03353 | 0.01612 |

| | | | | | | | |
|---|---|---|---|---|---|---|---|
| **LINC01429** | 0.01896 | 0.02617 | 0.01162 | 0.02213 | 0.00721 | 0.00972 | 0.06454 |
| **LINC01431** | 0.03206 | 0.05012 | 0.16045 | 0.05282 | 0.19143 | 0.00000 | 0.03808 |
| **LINC01433** | 0.04264 | 0.10546 | 0.00509 | 0.04461 | 0.04122 | 0.19423 | 0.60574 |
| **LINC01436** | 0.04324 | 0.01192 | 0.01226 | 0.01737 | 0.01619 | 0.01583 | 0.02130 |
| **LINC01446** | 0.59623 | 0.14421 | 0.06719 | 0.14238 | 0.05505 | 0.22906 | 0.00000 |
| **LINC01447** | 0.02867 | 0.31181 | 0.14809 | 0.96463 | 0.06446 | 0.12701 | 0.00000 |
| **LINC01448** | 0.03215 | 0.00941 | 0.07509 | 0.01785 | 0.00933 | 0.01280 | 0.02245 |
| **LINC01452** | 0.03313 | 0.03580 | 0.01093 | 0.01641 | 0.00858 | 0.01241 | 0.01118 |
| **LINCR-0001** | 0.03086 | 0.00741 | 0.01543 | 0.05536 | 0.00892 | 0.06445 | 0.03733 |
| **LINCR-0002** | 0.02874 | 0.01746 | 0.01428 | 0.02982 | 0.01531 | 0.01317 | 0.01058 |
| **LINCR-0003** | 0.02223 | 0.01812 | 0.00000 | 0.01857 | 0.36855 | 0.01326 | 0.01631 |
| **LINGO1** | 0.00000 | 0.01713 | 0.01000 | 0.00000 | 0.05680 | 0.46650 | 1.25408 |
| **LINGO2** | 0.00919 | 0.01961 | 0.15302 | 0.02218 | 3.57029 | 0.01036 | 0.04457 |
| **LINGO3** | 0.02169 | 0.02015 | 0.01559 | 0.02261 | 0.00774 | 0.01141 | 0.02350 |
| **LINGO4** | 0.02397 | 0.03666 | 0.01114 | 0.02258 | 0.00973 | 0.00976 | 0.02206 |
| **LINS** | 3.84140 | 0.51391 | 3.76876 | 2.60373 | 2.65331 | 0.92846 | 0.85467 |
| **LIPA** | 0.12515 | 0.00000 | 0.79555 | 0.55334 | 0.58062 | 0.14830 | 0.11010 |
| **LIPC** | 0.50031 | 0.33963 | 0.00000 | 0.37924 | 0.00456 | 0.26998 | 0.19640 |
| **LIPE** | 0.02744 | 0.30436 | 0.02493 | 0.04389 | 0.41440 | 0.04371 | 0.02022 |
| **LIPE-AS1** | 0.05558 | 0.00000 | 0.29259 | 0.02693 | 0.65558 | 0.47639 | 0.02262 |
| **LIPF** | 0.04155 | 0.01347 | 0.01144 | 0.01694 | 0.02377 | 0.01306 | 0.02420 |
| **LIPG** | 1.32572 | 0.00000 | 0.08491 | 5.51109 | 2.46641 | 0.68316 | 0.00000 |
| **LIPH** | 4.98376 | 3.75054 | 0.00000 | 0.15674 | 0.10699 | 0.00000 | 0.66656 |
| **LIPI** | 0.03368 | 0.00975 | 0.02730 | 0.03734 | 0.00991 | 0.01069 | 0.01832 |
| **LIPJ** | 0.01992 | 0.00730 | 0.02539 | 0.01892 | 0.01317 | 0.02846 | 0.03505 |
| **LIPK** | 0.02491 | 0.01229 | 0.02300 | 0.01876 | 0.01776 | 0.01345 | 0.01779 |
| **LIPM** | 0.03114 | 0.01766 | 0.04100 | 0.02053 | 0.22347 | 0.01165 | 0.00533 |
| **LIPT1** | 4.81045 | 0.93301 | 6.21925 | 0.52318 | 1.40262 | 1.12816 | 0.86313 |
| **LIPT2** | 0.05200 | 1.03665 | 0.52992 | 0.51434 | 0.55493 | 0.39376 | 1.09318 |
| **LITAF** | 0.54463 | 0.00000 | 0.88192 | 1.32553 | 0.13492 | 1.26836 | 0.00000 |
| **LIX1** | 0.06619 | 0.34021 | 0.03448 | 0.21230 | 0.24528 | 0.07717 | 0.05067 |
| **LIX1L** | 0.08313 | 0.00000 | 0.21329 | 0.03316 | 0.05017 | 1.70494 | 0.00000 |
| **LL22NC01-81G** | 0.02015 | 0.02108 | 0.01240 | 0.01836 | 0.00780 | 0.02224 | 0.02316 |
| **LLGL1** | 0.24747 | 0.15650 | 0.64745 | 1.54143 | 0.00000 | 1.43075 | 2.16643 |
| **LLGL2** | 11.58412 | 14.41851 | 2.85274 | 9.94136 | 0.93510 | 5.31386 | 28.83190 |

| | | | | | | | |
|---|---|---|---|---|---|---|---|
| **LLPH** | 17.33290 | 3.95666 | 12.01380 | 9.97165 | 4.07567 | 6.15944 | 3.99007 |
| **LMAN1** | 23.53025 | 6.37894 | 13.82730 | 16.28965 | 4.73349 | 8.58760 | 15.06565 |
| **LMAN1L** | 0.03606 | 0.01050 | 0.01430 | 0.01351 | 0.00757 | 0.01719 | 0.04346 |
| **LMAN2** | 31.11631 | 30.34132 | 12.33464 | 29.20135 | 11.83278 | 22.74147 | 15.38136 |
| **LMAN2L** | 7.54200 | 7.36239 | 4.87121 | 9.01955 | 0.50219 | 5.92819 | 0.00000 |
| **LMBR1** | 3.81416 | 0.12269 | 2.39738 | 1.31025 | 1.58445 | 3.41345 | 8.05250 |
| **LMBR1L** | 12.96196 | 10.25863 | 18.50409 | 4.92774 | 8.50395 | 10.86693 | 5.98800 |
| **LMBRD1** | 12.21469 | 0.00000 | 6.78783 | 7.72504 | 3.23177 | 8.71511 | 3.42751 |
| **LMBRD2** | 2.46312 | 0.24811 | 0.76956 | 0.59842 | 0.41471 | 0.98659 | 1.82076 |
| **LMCD1** | 0.06967 | 0.00000 | 1.32785 | 0.01361 | 0.62675 | 0.31434 | 0.03813 |
| **LMCD1-AS1** | 0.01756 | 0.00984 | 0.05779 | 0.17858 | 0.04649 | 0.03093 | 0.00259 |
| **LMF1** | 0.96919 | 1.40673 | 0.17723 | 0.30310 | 0.02968 | 0.69995 | 0.24660 |
| **LMF1-AS1** | 0.02314 | 0.01177 | 0.02088 | 0.03028 | 0.02235 | 0.01857 | 0.03101 |
| **LMF2** | 2.26249 | 2.14057 | 0.73994 | 5.15857 | 0.41220 | 2.72952 | 8.16116 |
| **LMLN** | 0.27699 | 0.90353 | 1.52239 | 0.48344 | 0.88303 | 1.14445 | 0.37888 |
| **LMLN-AS1** | 0.02649 | 0.01847 | 0.01767 | 0.02219 | 0.00925 | 0.01470 | 0.01378 |
| **LMNA** | 39.37133 | 33.69579 | 3.50249 | 39.63895 | 6.06037 | 8.28870 | 20.01522 |
| **LMNB1** | 11.53370 | 3.08934 | 6.98702 | 8.40833 | 2.02514 | 6.19825 | 3.12521 |
| **LMNB2** | 6.69334 | 18.28348 | 2.92670 | 15.45693 | 0.13772 | 10.38830 | 40.75961 |
| **LMNTD1** | 0.01100 | 0.00000 | 0.87703 | 0.01332 | 0.00151 | 0.00413 | 0.03982 |
| **LMNTD2** | 0.03453 | 0.00000 | 0.01623 | 0.04166 | 0.04158 | 0.02727 | 0.13840 |
| **LMO1** | 0.09911 | 0.05286 | 1.00683 | 0.26091 | 0.60781 | 0.04312 | 0.00000 |
| **LMO2** | 0.00000 | 0.00000 | 1.09765 | 1.69266 | 0.00156 | 0.20443 | 0.00728 |
| **LMO3** | 0.09982 | 0.00000 | 0.01484 | 0.01087 | 0.18245 | 0.07610 | 0.03588 |
| **LMO4** | 2.22595 | 9.68373 | 4.35517 | 12.91233 | 4.21913 | 11.86475 | 23.84559 |
| **LMO7** | 0.25777 | 0.07220 | 2.10315 | 0.35057 | 3.15061 | 0.32408 | 0.59163 |
| **LMO7-AS1** | 0.81666 | 0.23682 | 0.20286 | 0.40124 | 0.21842 | 0.33433 | 0.33426 |
| **LMO7DN** | 0.02438 | 0.01818 | 0.01314 | 0.01891 | 0.00791 | 0.01410 | 0.01728 |
| **LMOD1** | 0.06663 | 0.06875 | 0.06342 | 0.40079 | 0.00635 | 0.05956 | 0.00475 |
| **LMOD2** | 3.17549 | 0.30082 | 0.03077 | 0.53423 | 0.07927 | 0.00000 | 0.38394 |
| **LMOD3** | 0.00000 | 0.02574 | 0.44891 | 0.00000 | 2.21204 | 2.44406 | 3.15289 |
| **LMTK2** | 2.43178 | 3.70092 | 2.52509 | 4.43787 | 1.40484 | 1.92722 | 10.95235 |
| **LMTK3** | 0.01091 | 0.00000 | 0.03227 | 0.07495 | 0.12135 | 0.09184 | 0.46004 |
| **LMX1A** | 0.02291 | 0.01923 | 0.00665 | 0.03968 | 0.09075 | 0.01173 | 0.01525 |
| **LMX1B** | 0.15194 | 0.38455 | 0.06184 | 0.55151 | 0.00272 | 0.00000 | 2.78494 |

| | | | | | | | |
|---|---|---|---|---|---|---|---|
| **LNP1** | 0.12126 | 0.00000 | 0.18979 | 0.38993 | 0.79427 | 0.29451 | 0.01667 |
| **LNPEP** | 0.27503 | 0.47803 | 0.82071 | 0.49267 | 1.96296 | 0.03206 | 0.00000 |
| **LNX1** | 1.82420 | 0.79645 | 0.16114 | 1.43483 | 0.16826 | 1.15193 | 0.50635 |
| **LNX1-AS1** | 0.02018 | 0.01625 | 0.01682 | 0.03256 | 0.00737 | 0.01034 | 0.05058 |
| **LNX1-AS2** | 0.03235 | 0.01443 | 0.01580 | 0.02406 | 0.01045 | 0.01208 | 0.01565 |
| **LNX2** | 1.71247 | 0.00000 | 1.63013 | 0.48498 | 1.15227 | 2.12139 | 2.93499 |
| **LOC100126784** | 0.01023 | 0.12115 | 0.00449 | 0.01502 | 0.12814 | 0.01299 | 0.02477 |
| **LOC100128006** | 0.02145 | 0.01781 | 0.01272 | 0.02470 | 0.01527 | 0.01164 | 0.02565 |
| **LOC100128076** | 0.01979 | 0.01045 | 0.03016 | 0.03747 | 0.01400 | 0.01428 | 0.02275 |
| **LOC100128127** | 0.02176 | 0.01913 | 0.01237 | 0.02413 | 0.00798 | 0.03392 | 0.01638 |
| **LOC100128164** | 0.01711 | 0.25604 | 0.47725 | 0.03833 | 0.25892 | 0.06864 | 0.00000 |
| **LOC100128176** | 0.02841 | 0.00908 | 0.01402 | 0.01804 | 0.00778 | 0.01566 | 0.05006 |
| **LOC100128233** | 0.02725 | 0.03110 | 0.01960 | 0.01394 | 0.01073 | 0.04306 | 0.03404 |
| **LOC100128239** | 0.03940 | 0.00297 | 0.02965 | 0.01819 | 0.20170 | 0.01329 | 0.01028 |
| **LOC100128288** | 0.00000 | 0.78657 | 0.59277 | 0.00000 | 0.08262 | 0.90140 | 0.25285 |
| **LOC100128361** | 0.06781 | 0.07109 | 0.09403 | 0.09216 | 0.15656 | 0.22881 | 0.61884 |
| **LOC100128398** | 0.08984 | 0.08490 | 0.08675 | 0.16979 | 0.07933 | 0.00068 | 0.31552 |
| **LOC100128494** | 0.06894 | 0.00000 | 0.02352 | 0.02879 | 0.00531 | 0.12546 | 0.31875 |
| **LOC100128505** | 0.02514 | 0.08405 | 0.01399 | 0.01003 | 0.01474 | 0.02629 | 0.10120 |
| **LOC100128531** | 0.03239 | 0.02795 | 0.01712 | 0.01274 | 0.00784 | 0.01426 | 0.04677 |
| **LOC100128554** | 0.02429 | 0.01788 | 0.01015 | 0.01971 | 0.03324 | 0.01322 | 0.01523 |
| **LOC100128568** | 0.06148 | 0.08591 | 0.11134 | 0.13598 | 0.01326 | 0.01818 | 0.02261 |
| **LOC100128573** | 0.01491 | 0.01058 | 0.01592 | 0.02404 | 0.00719 | 0.00856 | 0.08178 |
| **LOC100128593** | 0.02284 | 0.01992 | 0.01272 | 0.01933 | 0.00869 | 0.01369 | 0.01828 |
| **LOC100128682** | 0.02044 | 0.01363 | 0.03163 | 0.03057 | 0.25657 | 0.01104 | 0.02032 |
| **LOC100128714** | 0.02099 | 0.01524 | 0.01349 | 0.03419 | 0.00786 | 0.01364 | 0.02429 |
| **LOC100128770** | 0.00449 | 0.04754 | 0.00500 | 0.02345 | 0.00691 | 0.02083 | 0.32015 |
| **LOC100128885** | 13.95771 | 2.09776 | 3.41978 | 4.53757 | 0.65468 | 6.13924 | 10.18340 |
| **LOC100129027** | 0.05323 | 0.03068 | 0.04015 | 0.00500 | 0.00653 | 0.04532 | 0.04885 |
| **LOC100129034** | 2.94693 | 3.67770 | 1.60714 | 3.26370 | 0.35307 | 1.49492 | 2.68046 |
| **LOC100129046** | 0.04702 | 0.01352 | 0.01194 | 0.12988 | 0.00748 | 0.05448 | 0.04306 |
| **LOC100129055** | 0.02852 | 0.01114 | 0.03671 | 0.01757 | 0.00729 | 0.01657 | 0.01688 |
| **LOC100129083** | 0.02012 | 0.02190 | 0.02177 | 0.01933 | 0.01206 | 0.02313 | 0.01770 |
| **LOC100129138** | 0.09201 | 0.21582 | 0.06552 | 0.13877 | 0.01569 | 0.03657 | 0.00206 |
| **LOC100129148** | 0.13579 | 0.08136 | 0.07739 | 0.19944 | 0.16387 | 0.21990 | 0.02685 |

| | | | | | | | |
|---|---|---|---|---|---|---|---|
| **LOC100129203** | 0.03924 | 0.00000 | 0.42190 | 0.01517 | 1.01905 | 0.14651 | 0.01020 |
| **LOC100129213** | 0.02954 | 0.02200 | 0.01338 | 0.05971 | 0.00800 | 0.01920 | 0.00016 |
| **LOC100129216** | 0.02118 | 0.01637 | 0.01701 | 0.01678 | 0.01415 | 0.03155 | 0.02900 |
| **LOC100129316** | 0.01801 | 0.01848 | 0.01315 | 0.03855 | 0.01474 | 0.01181 | 0.02695 |
| **LOC100129345** | 0.02955 | 0.01637 | 0.01287 | 0.01749 | 0.00795 | 0.01277 | 0.01843 |
| **LOC100129361** | 13.11991 | 5.52575 | 11.64398 | 11.47029 | 0.52771 | 2.53497 | 2.01880 |
| **LOC100129518** | 0.10389 | 0.00665 | 0.00870 | 0.30721 | 0.04624 | 0.06853 | 0.11313 |
| **LOC100129520** | 0.55516 | 0.63821 | 0.37040 | 0.93576 | 0.00112 | 0.06417 | 0.00000 |
| **LOC100129534** | 0.16870 | 0.77813 | 0.18851 | 0.16960 | 0.03002 | 0.27564 | 0.00000 |
| **LOC100129550** | 0.82257 | 0.08428 | 0.27728 | 1.28133 | 0.00988 | 0.25742 | 1.96713 |
| **LOC100129617** | 0.00000 | 0.00620 | 0.01434 | 0.06363 | 0.00723 | 0.17133 | 0.30729 |
| **LOC100129636** | 0.03874 | 0.01750 | 0.01285 | 0.02391 | 0.00883 | 0.01082 | 0.01880 |
| **LOC100129722** | 0.01505 | 0.03976 | 0.05231 | 0.02188 | 0.00744 | 0.02735 | 0.03648 |
| **LOC100129917** | 0.10306 | 0.32299 | 0.35478 | 0.08628 | 0.15239 | 0.22778 | 0.32093 |
| **LOC100129924** | 0.81101 | 0.00000 | 0.01409 | 0.18104 | 0.00383 | 0.00000 | 0.18310 |
| **LOC100129931** | 0.02315 | 0.00000 | 0.02018 | 0.00882 | 0.00747 | 0.03475 | 0.24693 |
| **LOC100129935** | 0.00000 | 0.01086 | 0.01166 | 9.21608 | 0.00621 | 0.00674 | 0.01901 |
| **LOC100129940.** | 0.11372 | 0.00394 | 0.02024 | 0.04022 | 0.00857 | 0.02656 | 0.00000 |
| **LOC100129940.** | 0.01883 | 0.01645 | 0.01220 | 0.03807 | 0.00804 | 0.03281 | 0.02198 |
| **LOC100129973** | 0.07143 | 0.01536 | 0.05815 | 0.02908 | 0.02773 | 0.01485 | 0.00000 |
| **LOC100130075** | 0.00000 | 0.20196 | 0.06147 | 0.70733 | 0.23216 | 0.03422 | 0.01808 |
| **LOC100130093** | 0.27523 | 1.96104 | 0.36640 | 0.49719 | 0.48266 | 0.87730 | 0.01631 |
| **LOC100130264** | 0.02681 | 0.01436 | 0.01669 | 0.01893 | 0.00898 | 0.03345 | 0.01078 |
| **LOC100130298** | 0.02168 | 0.01953 | 0.01696 | 0.01824 | 0.12071 | 0.01165 | 0.02086 |
| **LOC100130301** | 0.02782 | 0.00634 | 0.01820 | 0.01779 | 0.19366 | 0.01637 | 0.01576 |
| **LOC100130331** | 0.32012 | 0.46933 | 0.25081 | 0.32697 | 0.69485 | 0.35882 | 0.00000 |
| **LOC100130357** | 0.40315 | 0.00000 | 0.57079 | 0.29015 | 0.01471 | 0.14304 | 0.00863 |
| **LOC100130370** | 0.00000 | 0.02394 | 0.01090 | 0.00664 | 0.00671 | 0.09465 | 0.16913 |
| **LOC100130452** | 0.06583 | 0.00694 | 0.02163 | 0.01909 | 0.00724 | 0.01880 | 0.05195 |
| **LOC100130476** | 0.00000 | 0.09179 | 0.02759 | 0.43505 | 0.00643 | 0.24767 | 0.00053 |
| **LOC100130587** | 0.02129 | 0.01448 | 0.01261 | 0.01665 | 0.00755 | 0.01025 | 0.04576 |
| **LOC100130673** | 0.09511 | 0.00000 | 0.13180 | 0.20918 | 0.13050 | 0.21297 | 0.00000 |
| **LOC100130691** | 0.01079 | 0.04857 | 0.01238 | 0.04362 | 0.01636 | 0.12445 | 0.04878 |
| **LOC100130700** | 0.02750 | 0.01739 | 0.15621 | 0.09116 | 0.11881 | 0.01588 | 0.00000 |
| **LOC100130705** | 0.02126 | 0.02026 | 0.03980 | 0.05427 | 0.01287 | 0.19256 | 0.09994 |

| | | | | | | | |
|---|---|---|---|---|---|---|---|
| **LOC100130744** | 0.16350 | 0.00000 | 0.45042 | 0.35543 | 0.13349 | 0.00000 | 0.50262 |
| **LOC100130849** | 0.02382 | 0.01666 | 0.01530 | 0.02953 | 0.02225 | 0.01595 | 0.01209 |
| **LOC100130872** | 0.04330 | 0.29622 | 0.11407 | 0.00253 | 0.01341 | 0.00000 | 0.04258 |
| **LOC100130880** | 0.02626 | 0.01135 | 0.01320 | 0.01730 | 0.00788 | 0.03127 | 0.02971 |
| **LOC100130899** | 1.18022 | 0.91673 | 0.00000 | 1.25983 | 0.00187 | 1.41166 | 1.01176 |
| **LOC100130950** | 0.11039 | 0.00000 | 0.23904 | 0.12533 | 0.14325 | 0.00458 | 0.21120 |
| **LOC100130964** | 0.02978 | 0.03297 | 0.02581 | 0.01721 | 0.00662 | 0.03166 | 0.01338 |
| **LOC100130987** | 0.21117 | 0.28425 | 0.02151 | 0.24013 | 0.08346 | 0.02391 | 0.00000 |
| **LOC100130992** | 0.03368 | 0.04047 | 0.01015 | 0.01988 | 0.00989 | 0.01670 | 0.00185 |
| **LOC100131047** | 0.02809 | 0.01312 | 0.01733 | 0.02055 | 0.00782 | 0.02790 | 0.01586 |
| **LOC100131257** | 0.02131 | 0.01766 | 0.01806 | 0.03309 | 0.02030 | 0.10635 | 0.10138 |
| **LOC100131289** | 0.12674 | 0.00697 | 0.09434 | 0.39533 | 0.04649 | 0.04483 | 0.00000 |
| **LOC100131303** | 0.00000 | 0.00454 | 0.04215 | 0.40220 | 0.10904 | 0.02224 | 0.01593 |
| **LOC100131347** | 0.00000 | 0.12442 | 0.02432 | 0.02880 | 0.02992 | 0.00515 | 0.12657 |
| **LOC100131496** | 0.03806 | 0.27036 | 0.10343 | 0.05495 | 0.02289 | 0.94008 | 0.12957 |
| **LOC100131564** | 0.53154 | 0.00000 | 1.65681 | 0.24947 | 0.00054 | 0.00000 | 1.11380 |
| **LOC100131626** | 0.03761 | 0.02824 | 0.02491 | 0.02553 | 0.01941 | 0.01862 | 0.03231 |
| **LOC100131655** | 0.00691 | 0.07872 | 0.04809 | 0.10025 | 0.00832 | 0.03077 | 0.06358 |
| **LOC100131691** | 0.00000 | 0.57649 | 0.45676 | 0.32845 | 0.35396 | 0.11656 | 0.26746 |
| **LOC100132057.** | 0.02361 | 0.01497 | 0.01752 | 0.01984 | 0.00760 | 0.01454 | 0.01762 |
| **LOC100132062+** | 0.02898 | 0.02492 | 0.01364 | 0.02292 | 0.00863 | 0.00998 | 0.03484 |
| **LOC100132077** | 0.07107 | 0.20443 | 0.07563 | 0.00358 | 0.01282 | 0.06761 | 0.00138 |
| **LOC100132078** | 0.02540 | 0.01607 | 0.01265 | 0.01919 | 0.00867 | 0.02177 | 0.02423 |
| **LOC100132111** | 0.03611 | 0.01879 | 0.01419 | 0.05122 | 0.01748 | 0.03329 | 0.00000 |
| **LOC100132146** | 0.00523 | 0.05680 | 0.00500 | 0.00499 | 0.01067 | 0.03494 | 0.12514 |
| **LOC100132215** | 0.13097 | 0.09959 | 0.20950 | 0.27482 | 0.06096 | 0.14168 | 0.40869 |
| **LOC100132352** | 0.35358 | 0.00000 | 0.30596 | 0.09234 | 0.20150 | 0.13594 | 0.13652 |
| **LOC100132356** | 0.19582 | 0.26105 | 0.29372 | 0.12920 | 0.61692 | 0.18120 | 0.14509 |
| **LOC100132529** | 0.00000 | 0.01694 | 0.10852 | 0.06656 | 0.00394 | 0.01231 | 0.11563 |
| **LOC100132735** | 1.34208 | 0.00000 | 0.00073 | 0.99048 | 0.00285 | 0.00000 | 0.29703 |
| **LOC100132781** | 0.04542 | 0.00000 | 0.02655 | 0.05266 | 0.04372 | 0.21200 | 0.21051 |
| **LOC100132831** | 0.08433 | 0.16823 | 0.05934 | 0.29365 | 0.01769 | 0.09896 | 0.26890 |
| **LOC100132891** | 0.02513 | 0.01165 | 0.02067 | 0.02032 | 0.04138 | 0.01895 | 0.01524 |
| **LOC100133077** | 0.00000 | 0.19162 | 0.71191 | 0.00000 | 0.04804 | 0.07163 | 0.13780 |
| **LOC100133091** | 0.00307 | 0.07112 | 0.07404 | 0.02685 | 0.01291 | 0.07958 | 0.39359 |

| | | | | | | | |
|---|---|---|---|---|---|---|---|
| **LOC100133286** | 0.02757 | 0.04925 | 0.04841 | 0.09998 | 0.15144 | 0.00591 | 0.00650 |
| **LOC100133315** | 0.40120 | 0.00000 | 0.40572 | 0.38199 | 0.87290 | 0.76041 | 0.00000 |
| **LOC100133331** | 0.03672 | 0.03344 | 0.07931 | 0.01898 | 0.01180 | 0.04647 | 0.11615 |
| **LOC100133669** | 0.07553 | 0.01394 | 0.01288 | 0.06559 | 0.00803 | 0.02987 | 0.00000 |
| **LOC100133985** | 0.08314 | 0.04671 | 0.01462 | 0.13012 | 0.00946 | 0.06990 | 0.00000 |
| **LOC100134317** | 0.07965 | 0.01497 | 0.03696 | 0.03815 | 0.00438 | 0.20697 | 0.21140 |
| **LOC100134368** | 0.04559 | 0.02464 | 0.01108 | 0.01279 | 0.01429 | 0.02727 | 0.01388 |
| **LOC100134868** | 0.14861 | 0.01435 | 0.01276 | 0.26031 | 0.00676 | 0.05303 | 0.02377 |
| **LOC100144595** | 0.02550 | 0.01624 | 0.01560 | 0.01963 | 0.01996 | 0.01389 | 0.01646 |
| **LOC100188947** | 0.02130 | 0.01596 | 0.01400 | 0.01910 | 0.03379 | 0.01912 | 0.02547 |
| **LOC100190940** | 0.02379 | 0.01794 | 0.01278 | 0.02104 | 0.01006 | 0.01300 | 0.01609 |
| **LOC100190986** | 0.02317 | 0.01638 | 0.01304 | 0.01927 | 0.00778 | 0.01329 | 0.02587 |
| **LOC100240734** | 0.02291 | 0.01616 | 0.00946 | 0.01895 | 0.07763 | 0.01318 | 0.01827 |
| **LOC100240735** | 0.03222 | 0.01947 | 0.01553 | 0.01877 | 0.01880 | 0.03619 | 0.00678 |
| **LOC100268168** | 0.05846 | 0.26226 | 0.14261 | 0.12080 | 0.35723 | 0.00000 | 0.05147 |
| **LOC100270746** | 0.17164 | 0.03085 | 0.07657 | 0.20549 | 0.05384 | 0.00000 | 0.08934 |
| **LOC100270804** | 0.15599 | 0.10681 | 0.06334 | 0.12685 | 0.05677 | 0.00000 | 0.25753 |
| **LOC100271832** | 0.01197 | 0.08387 | 0.00464 | 0.01113 | 0.00829 | 0.00442 | 0.08631 |
| **LOC100272216** | 0.07969 | 0.05130 | 0.01916 | 0.03061 | 0.00772 | 0.01232 | 0.08311 |
| **LOC100272217** | 0.02316 | 0.31436 | 0.24835 | 0.38580 | 0.16031 | 0.01153 | 0.00000 |
| **LOC100286922** | 0.06919 | 0.00635 | 0.01308 | 0.00634 | 0.00892 | 0.02086 | 0.04445 |
| **LOC100287015** | 0.51899 | 3.54314 | 2.83938 | 2.05653 | 1.63631 | 0.10214 | 0.00000 |
| **LOC100287036** | 0.00779 | 0.25450 | 0.00835 | 0.00386 | 0.00896 | 0.00000 | 0.17500 |
| **LOC100287042** | 0.27200 | 0.00000 | 0.07871 | 0.16564 | 0.03203 | 0.06021 | 0.23315 |
| **LOC100287072** | 0.08507 | 0.13018 | 0.10603 | 0.01860 | 0.04401 | 0.30257 | 0.11160 |
| **LOC100287098** | 0.00137 | 0.02145 | 0.02666 | 0.02071 | 0.01019 | 0.02390 | 0.17520 |
| **LOC100287225** | 0.05743 | 0.04117 | 0.00000 | 0.02593 | 0.84440 | 0.15129 | 0.34164 |
| **LOC100287632** | 0.02230 | 0.01879 | 0.01694 | 0.03919 | 0.00948 | 0.01293 | 0.02483 |
| **LOC100287704+** | 0.02389 | 0.02180 | 0.01440 | 0.02114 | 0.00768 | 0.01277 | 0.01620 |
| **LOC100287728** | 0.02836 | 0.01930 | 0.01224 | 0.02686 | 0.00814 | 0.01948 | 0.01035 |
| **LOC100287792** | 0.10914 | 0.00000 | 0.11583 | 0.05036 | 0.05671 | 0.18963 | 1.13275 |
| **LOC100287846** | 0.02750 | 0.06648 | 0.04921 | 0.02520 | 0.00609 | 0.16339 | 0.00571 |
| **LOC100287944** | 0.03413 | 0.01642 | 0.01443 | 0.01884 | 0.00772 | 0.01893 | 0.01742 |
| **LOC100288069** | 0.04999 | 0.01213 | 0.09228 | 0.03059 | 0.00811 | 0.02662 | 0.04707 |
| **LOC100288123** | 0.00856 | 0.00833 | 0.01267 | 0.04342 | 0.00891 | 0.01945 | 0.13449 |

| | | | | | | | |
|---|---|---|---|---|---|---|---|
| **LOC100288162.** | 0.02908 | 0.01889 | 0.01230 | 0.01737 | 0.00815 | 0.01161 | 0.02162 |
| **LOC100288162.** | 0.02298 | 0.01529 | 0.01562 | 0.02273 | 0.00829 | 0.01377 | 0.03280 |
| **LOC100288181** | 0.04426 | 0.01629 | 0.03699 | 0.00815 | 0.03780 | 0.01189 | 0.03508 |
| **LOC100288637** | 2.22432 | 0.00000 | 0.62414 | 0.86146 | 0.28424 | 0.36176 | 2.41881 |
| **LOC100288748** | 0.33990 | 0.03361 | 0.42249 | 0.30980 | 0.46091 | 0.06301 | 0.20788 |
| **LOC100288778** | 0.00000 | 0.30983 | 0.11320 | 0.01527 | 0.04050 | 0.04330 | 0.34697 |
| **LOC100288814** | 0.02685 | 0.01422 | 0.01267 | 0.02247 | 0.00757 | 0.01309 | 0.03793 |
| **LOC100288842** | 0.36653 | 0.18005 | 0.66917 | 0.17752 | 0.13010 | 0.00433 | 0.01287 |
| **LOC100288846** | 0.11513 | 0.15956 | 0.00000 | 0.07177 | 0.35278 | 0.09720 | 0.00000 |
| **LOC100288866** | 0.02488 | 0.01893 | 0.01072 | 0.02235 | 0.01929 | 0.01226 | 0.01917 |
| **LOC100288911** | 0.00000 | 0.10109 | 0.01330 | 0.22293 | 0.03756 | 0.32075 | 0.13794 |
| **LOC100288974** | 0.15626 | 0.00000 | 0.06874 | 0.00428 | 0.02027 | 0.00656 | 0.05538 |
| **LOC100289019** | 0.84692 | 1.34379 | 0.87440 | 0.29094 | 0.01945 | 0.73239 | 1.73413 |
| **LOC100289092** | 0.14004 | 0.08083 | 0.64605 | 0.33897 | 0.18916 | 0.23939 | 0.05751 |
| **LOC100289230** | 0.10144 | 1.04076 | 0.35742 | 0.18012 | 0.23572 | 0.09320 | 0.32955 |
| **LOC100289333** | 0.06802 | 0.05082 | 0.10345 | 0.12191 | 0.80777 | 0.02352 | 0.00000 |
| **LOC100289361** | 0.17475 | 0.63407 | 0.51646 | 0.02334 | 1.03904 | 0.24073 | 0.22585 |
| **LOC100289473** | 0.03263 | 0.01748 | 0.01780 | 0.02325 | 0.00755 | 0.02822 | 0.00837 |
| **LOC100289495** | 0.00537 | 0.12109 | 0.02877 | 0.06192 | 0.21719 | 0.03124 | 0.07132 |
| **LOC100289511** | 0.08486 | 0.15997 | 0.03486 | 0.18521 | 0.14017 | 0.07002 | 0.04416 |
| **LOC100289561** | 0.00431 | 0.06397 | 0.03131 | 0.01207 | 0.00555 | 0.02566 | 0.29589 |
| **LOC100289580** | 0.04535 | 0.03163 | 0.02247 | 0.01509 | 0.03771 | 0.01865 | 0.44032 |
| **LOC100289650** | 0.01463 | 0.01624 | 0.01179 | 0.02040 | 0.02505 | 0.03536 | 0.03243 |
| **LOC100289656** | 0.01500 | 0.04367 | 0.01467 | 0.01643 | 0.01007 | 0.01717 | 0.04073 |
| **LOC100289673** | 0.02652 | 0.01508 | 0.01163 | 0.01886 | 0.06904 | 0.01290 | 0.01687 |
| **LOC100294145** | 2.96382 | 1.72327 | 3.02027 | 2.53387 | 3.00366 | 2.41088 | 0.00000 |
| **LOC100294362** | 0.05158 | 0.20849 | 0.07132 | 0.18646 | 0.44197 | 0.19055 | 0.00000 |
| **LOC100303749** | 0.00000 | 0.00000 | 0.15995 | 0.05516 | 0.00921 | 0.25676 | 0.33877 |
| **LOC100335030** | 0.22742 | 0.02320 | 0.48329 | 0.11081 | 0.32962 | 0.38350 | 0.63663 |
| **LOC100379224** | 0.19789 | 0.12189 | 0.92670 | 0.77138 | 0.10221 | 0.20601 | 0.00000 |
| **LOC100420587** | 0.16451 | 0.03564 | 0.03474 | 0.29913 | 0.03668 | 0.49109 | 0.00000 |
| **LOC100422556** | 0.03121 | 0.09272 | 0.06356 | 0.04760 | 0.15425 | 0.00670 | 0.01816 |
| **LOC100422737** | 0.02521 | 0.01057 | 0.01406 | 0.04232 | 0.01371 | 0.02686 | 0.02046 |
| **LOC100499194** | 0.02776 | 0.02063 | 0.00488 | 0.02596 | 0.12025 | 0.01307 | 0.00836 |
| **LOC100499484** | 0.03303 | 0.02040 | 0.08918 | 0.02672 | 0.09884 | 0.10307 | 0.00029 |

| | | | | | | | |
|---|---|---|---|---|---|---|---|
| **LOC100499484-** | 0.02387 | 0.01896 | 0.01173 | 0.01906 | 0.01145 | 0.01300 | 0.02004 |
| **LOC100499489** | 0.02716 | 0.01546 | 0.00939 | 0.01758 | 0.07562 | 0.01267 | 0.01934 |
| **LOC100500773** | 0.04011 | 0.00058 | 0.01619 | 0.00000 | 0.00651 | 0.00119 | 0.47301 |
| **LOC100505474** | 0.02476 | 0.02056 | 0.01192 | 0.02096 | 0.00803 | 0.01677 | 0.01616 |
| **LOC100505549** | 1.63374 | 0.06717 | 1.31343 | 2.99053 | 0.16709 | 2.28670 | 2.31647 |
| **LOC100505592** | 0.00233 | 0.00000 | 0.37487 | 0.05335 | 0.00494 | 0.02834 | 0.08747 |
| **LOC100505658** | 0.03953 | 0.10053 | 0.13412 | 0.19473 | 0.01666 | 0.12278 | 0.38547 |
| **LOC100505666** | 0.17653 | 0.07909 | 0.14586 | 0.57092 | 0.11586 | 0.00209 | 0.00000 |
| **LOC100505715** | 0.84808 | 1.11749 | 0.31501 | 1.88028 | 0.99606 | 0.73656 | 0.00044 |
| **LOC100505716** | 0.05727 | 0.00000 | 0.08979 | 0.05326 | 0.00390 | 0.03587 | 0.16856 |
| **LOC100505718** | 0.02737 | 0.01654 | 0.01461 | 0.01833 | 0.00855 | 0.01434 | 0.01519 |
| **LOC100505736** | 0.04516 | 0.01055 | 0.14809 | 0.05242 | 0.01120 | 0.11096 | 0.20171 |
| **LOC100505795** | 0.00457 | 0.00640 | 0.01264 | 0.01218 | 0.00696 | 0.23584 | 0.11276 |
| **LOC100505811** | 0.02526 | 0.02661 | 0.01578 | 0.02058 | 0.01504 | 0.01626 | 0.01110 |
| **LOC100505812** | 0.11322 | 0.00000 | 0.12726 | 0.13816 | 0.00501 | 0.02984 | 0.03836 |
| **LOC100505817** | 2.43245 | 1.11861 | 0.14282 | 0.00000 | 1.01118 | 5.69405 | 0.00000 |
| **LOC100505835** | 0.02416 | 0.01739 | 0.01363 | 0.01956 | 0.00810 | 0.01286 | 0.01809 |
| **LOC100505841** | 0.02762 | 0.01654 | 0.00642 | 0.01952 | 0.10863 | 0.01322 | 0.01504 |
| **LOC100505878** | 0.03349 | 0.03120 | 0.00611 | 0.01534 | 0.15285 | 0.02027 | 0.00821 |
| **LOC100505887** | 0.02050 | 0.03275 | 0.01308 | 0.01852 | 0.06927 | 0.01549 | 0.01618 |
| **LOC100505912** | 0.00671 | 0.00416 | 0.01641 | 0.05992 | 0.01767 | 0.24203 | 0.16164 |
| **LOC100505918** | 0.09904 | 0.12227 | 0.04543 | 0.00000 | 0.00800 | 0.02272 | 0.11997 |
| **LOC100505921** | 0.03360 | 0.01140 | 0.01504 | 0.04642 | 0.00796 | 0.01817 | 0.02492 |
| **LOC100505942** | 0.03842 | 0.00000 | 0.06339 | 0.24813 | 0.03031 | 0.00547 | 0.03328 |
| **LOC100505978** | 0.07825 | 0.01645 | 0.19416 | 0.04104 | 0.00957 | 0.02566 | 0.00000 |
| **LOC100505984** | 0.04955 | 0.01851 | 0.01995 | 0.02969 | 0.00683 | 0.10517 | 0.01229 |
| **LOC100506022** | 0.02441 | 0.00986 | 0.05018 | 0.02605 | 0.01821 | 0.02450 | 0.01251 |
| **LOC100506023** | 0.05478 | 0.00000 | 0.15805 | 0.30793 | 0.25519 | 0.24568 | 0.10245 |
| **LOC100506071** | 0.17907 | 0.00174 | 0.01761 | 0.23797 | 0.03434 | 0.02633 | 0.05929 |
| **LOC100506082** | 0.13111 | 0.22055 | 0.50810 | 0.00000 | 0.09757 | 0.14331 | 0.01900 |
| **LOC100506083** | 0.23986 | 0.64944 | 0.72730 | 0.14734 | 0.02958 | 0.08382 | 0.00000 |
| **LOC100506085** | 0.00359 | 0.02454 | 0.02378 | 0.00000 | 0.01924 | 0.27469 | 0.30713 |
| **LOC100506100** | 2.48846 | 1.32403 | 0.89438 | 4.38873 | 0.57300 | 3.26201 | 0.56509 |
| **LOC100506119** | 0.02486 | 0.02001 | 0.01139 | 0.07626 | 0.00836 | 0.08528 | 0.00000 |
| **LOC100506123** | 0.05300 | 0.00369 | 0.05086 | 0.02471 | 0.00816 | 0.01629 | 0.11227 |

| | | | | | | | |
|---|---|---|---|---|---|---|---|
| **LOC100506124** | 0.04482 | 0.00000 | 0.02296 | 0.00000 | 0.00548 | 0.00363 | 0.32634 |
| **LOC100506125** | 0.01349 | 0.06407 | 0.02096 | 0.02043 | 0.06385 | 0.01144 | 0.01488 |
| **LOC100506127** | 0.10627 | 0.15053 | 0.18205 | 0.13720 | 0.15715 | 0.03490 | 0.03049 |
| **LOC100506136** | 0.16500 | 0.00000 | 0.18155 | 0.32148 | 0.46265 | 0.03999 | 0.00384 |
| **LOC100506142** | 0.03426 | 0.00351 | 0.03978 | 0.04439 | 0.05090 | 0.02124 | 0.05002 |
| **LOC100506172** | 0.02734 | 0.01342 | 0.01484 | 0.01736 | 0.02907 | 0.01287 | 0.02118 |
| **LOC100506175** | 0.02669 | 0.00819 | 0.02412 | 0.01761 | 0.01537 | 0.01932 | 0.02907 |
| **LOC100506178** | 2.59677 | 1.12340 | 0.19152 | 4.50152 | 0.00000 | 0.36989 | 0.05985 |
| **LOC100506207** | 0.11099 | 0.00000 | 0.09629 | 0.07266 | 0.07367 | 0.04472 | 0.00000 |
| **LOC100506258** | 0.02141 | 0.00595 | 0.04753 | 0.02483 | 0.08257 | 0.02712 | 0.03349 |
| **LOC100506271** | 0.01436 | 0.01576 | 0.01127 | 0.05086 | 0.02025 | 0.02088 | 0.02372 |
| **LOC100506272** | 0.00000 | 0.52410 | 0.16499 | 0.00715 | 0.00800 | 0.05829 | 0.04062 |
| **LOC100506281** | 0.00801 | 0.05053 | 0.00697 | 0.01658 | 0.00779 | 0.00620 | 0.13476 |
| **LOC100506299** | 0.00557 | 0.10114 | 0.03025 | 0.09437 | 0.05957 | 0.00928 | 0.02211 |
| **LOC100506314** | 2.32964 | 0.46383 | 1.17793 | 1.02068 | 0.03516 | 0.06167 | 1.46328 |
| **LOC100506321** | 0.12242 | 0.61954 | 0.17226 | 0.13960 | 0.10322 | 0.21993 | 0.24314 |
| **LOC100506325** | 0.04677 | 0.06758 | 0.16697 | 0.08396 | 0.03266 | 0.00000 | 0.10958 |
| **LOC100506368** | 0.09840 | 0.00000 | 0.10032 | 0.01404 | 0.01239 | 0.02628 | 0.01522 |
| **LOC100506403** | 0.03229 | 0.00936 | 0.02678 | 0.01787 | 0.01299 | 0.03337 | 0.01113 |
| **LOC100506422** | 0.02914 | 0.01468 | 0.01511 | 0.01656 | 0.00778 | 0.01580 | 0.02330 |
| **LOC100506457** | 0.02540 | 0.02287 | 0.01581 | 0.01250 | 0.21984 | 0.07199 | 0.02161 |
| **LOC100506469** | 0.77755 | 0.97819 | 0.00550 | 1.44674 | 0.00287 | 0.34040 | 0.85788 |
| **LOC100506472** | 0.12150 | 0.00000 | 0.14231 | 0.08452 | 0.00238 | 0.18565 | 0.82356 |
| **LOC100506476** | 0.01961 | 0.02474 | 0.08824 | 0.02489 | 0.02903 | 0.00224 | 0.07469 |
| **LOC100506497** | 0.10718 | 0.00969 | 0.01718 | 0.00647 | 0.02781 | 0.00736 | 0.03177 |
| **LOC100506499** | 0.34688 | 0.12783 | 0.00806 | 0.76756 | 0.02472 | 0.00000 | 0.40718 |
| **LOC100506548** | 2.95944 | 0.00000 | 2.93533 | 4.93491 | 4.49741 | 2.40793 | 9.95027 |
| **LOC100506551** | 0.17443 | 0.00000 | 0.18887 | 0.06702 | 0.00698 | 0.00000 | 0.15163 |
| **LOC100506585** | 0.02143 | 0.01570 | 0.01320 | 0.02355 | 0.00770 | 0.01283 | 0.02849 |
| **LOC100506603** | 0.10194 | 0.34828 | 0.23181 | 0.13693 | 0.06025 | 0.58750 | 0.59994 |
| **LOC100506606** | 0.03481 | 0.12020 | 0.02062 | 0.04076 | 0.04149 | 0.00236 | 0.02603 |
| **LOC100506639** | 0.35237 | 0.00000 | 0.35562 | 0.15857 | 0.07679 | 0.11259 | 0.08026 |
| **LOC100506655** | 3.39497 | 0.20511 | 0.03521 | 20.35081 | 0.01374 | 0.00000 | 2.37990 |
| **LOC100506674** | 0.14059 | 0.09283 | 0.08498 | 0.17463 | 0.36362 | 0.03555 | 0.00000 |
| **LOC100506675** | 0.15059 | 0.00000 | 0.30143 | 0.00000 | 0.10606 | 0.35364 | 2.12298 |

| | | | | | | | |
|---|---|---|---|---|---|---|---|
| **LOC100506682** | 0.02225 | 0.02802 | 0.00692 | 0.02006 | 0.05268 | 0.02763 | 0.01050 |
| **LOC100506688** | 1.31417 | 0.34392 | 0.25043 | 0.11251 | 0.00000 | 0.00000 | 0.13531 |
| **LOC100506700** | 0.15304 | 0.07474 | 0.06583 | 0.19136 | 0.00673 | 0.04509 | 0.14745 |
| **LOC100506713** | 0.39692 | 0.20234 | 0.16804 | 0.34099 | 0.06919 | 0.10600 | 0.14068 |
| **LOC100506730** | 0.12789 | 0.00000 | 0.22135 | 0.78620 | 0.00125 | 0.00000 | 0.74241 |
| **LOC100506746** | 0.00000 | 0.45280 | 0.80223 | 0.23420 | 0.41393 | 0.04152 | 0.20910 |
| **LOC100506747** | 0.16031 | 0.04370 | 0.02987 | 0.01543 | 0.12291 | 0.09727 | 0.07018 |
| **LOC100506787** | 0.02942 | 0.00721 | 0.03407 | 0.02873 | 0.01120 | 0.01519 | 0.02435 |
| **LOC100506790** | 0.00000 | 0.61541 | 5.68485 | 0.00000 | 8.20419 | 3.98500 | 0.00000 |
| **LOC100506801** | 0.05817 | 0.03563 | 0.00000 | 0.06064 | 0.09503 | 0.20271 | 0.35829 |
| **LOC100506804** | 0.21126 | 0.17523 | 0.09637 | 0.07148 | 0.18309 | 0.02113 | 0.00000 |
| **LOC100506834** | 0.02832 | 0.01972 | 0.01102 | 0.02261 | 0.01935 | 0.01342 | 0.01975 |
| **LOC100506844** | 0.00000 | 0.25083 | 0.49752 | 0.01973 | 1.42416 | 0.15566 | 0.01020 |
| **LOC100506860** | 0.04780 | 0.05809 | 0.01270 | 0.03874 | 0.00746 | 0.01719 | 0.12254 |
| **LOC100506895** | 0.04619 | 0.00000 | 0.06338 | 0.00461 | 0.31711 | 0.00611 | 0.05646 |
| **LOC100506985** | 0.27167 | 0.11076 | 0.04733 | 0.04125 | 0.05420 | 0.03300 | 0.01929 |
| **LOC100506990** | 0.06763 | 0.00000 | 1.60638 | 0.39938 | 0.80851 | 0.45429 | 0.12528 |
| **LOC100507002** | 0.00000 | 2.27629 | 0.12910 | 0.14659 | 0.06745 | 3.32766 | 0.00018 |
| **LOC100507006** | 0.04401 | 0.02102 | 0.01130 | 0.00872 | 0.04547 | 0.22979 | 0.01804 |
| **LOC100507012** | 0.01121 | 0.05109 | 0.01985 | 0.02049 | 0.07857 | 0.00931 | 0.03067 |
| **LOC100507053** | 0.23493 | 0.00001 | 0.27533 | 0.16330 | 1.35287 | 0.02085 | 0.00000 |
| **LOC100507065** | 0.03073 | 0.01595 | 0.01302 | 0.01735 | 0.00788 | 0.01217 | 0.01775 |
| **LOC100507073** | 0.02520 | 0.01380 | 0.01406 | 0.01730 | 0.04954 | 0.01223 | 0.02357 |
| **LOC100507091** | 0.04645 | 0.01039 | 0.01433 | 0.04336 | 0.00792 | 0.05478 | 0.01077 |
| **LOC100507140** | 0.02728 | 0.01652 | 0.00862 | 0.01546 | 0.04593 | 0.01298 | 0.02832 |
| **LOC100507156** | 0.04486 | 0.05805 | 0.07989 | 0.01929 | 0.02250 | 0.01063 | 0.02805 |
| **LOC100507175** | 0.02843 | 0.00887 | 0.01351 | 0.01521 | 0.21124 | 0.01483 | 0.02828 |
| **LOC100507194** | 0.06137 | 0.04102 | 0.00805 | 0.00000 | 0.01576 | 0.01808 | 0.08313 |
| **LOC100507195** | 0.02861 | 0.03527 | 0.10028 | 0.03429 | 0.47472 | 0.15419 | 0.00000 |
| **LOC100507205** | 0.03278 | 0.01618 | 0.01325 | 0.01705 | 0.01571 | 0.01546 | 0.01562 |
| **LOC100507217** | 2.43995 | 0.00000 | 0.95362 | 2.56180 | 0.51983 | 2.33584 | 13.96255 |
| **LOC100507250** | 0.03802 | 0.26507 | 0.51348 | 0.03478 | 0.08140 | 0.08581 | 0.00000 |
| **LOC100507283** | 0.03567 | 0.03249 | 0.01714 | 0.02408 | 0.02575 | 0.02542 | 0.01168 |
| **LOC100507291** | 0.31242 | 0.35254 | 0.39633 | 0.11268 | 0.15530 | 0.04774 | 0.21185 |
| **LOC100507316** | 0.03331 | 0.00000 | 1.02610 | 0.00166 | 0.24333 | 0.00000 | 0.18400 |

| | | | | | | | |
|---|---|---|---|---|---|---|---|
| **LOC100507334** | 0.01969 | 0.01951 | 0.02768 | 0.02494 | 0.01319 | 0.01184 | 0.02066 |
| **LOC100507346** | 0.02348 | 0.00616 | 0.02382 | 0.03601 | 0.01797 | 0.07801 | 0.03334 |
| **LOC100507351** | 0.02319 | 0.00366 | 0.07197 | 0.01926 | 0.03198 | 0.09886 | 0.01510 |
| **LOC100507373** | 0.06409 | 0.05968 | 0.46777 | 0.10364 | 0.11658 | 0.09142 | 0.54983 |
| **LOC100507377** | 0.17328 | 0.17338 | 0.21361 | 0.17116 | 0.68438 | 0.19941 | 0.00000 |
| **LOC100507387** | 0.02339 | 0.01897 | 0.01145 | 0.01955 | 0.01795 | 0.01314 | 0.01758 |
| **LOC100507388** | 0.02768 | 0.00942 | 0.05014 | 0.03632 | 0.04109 | 0.01289 | 0.01580 |
| **LOC100507389** | 0.01301 | 0.01611 | 0.01082 | 0.02909 | 0.02233 | 0.01893 | 0.04115 |
| **LOC100507391** | 0.02092 | 0.01193 | 0.00599 | 0.01999 | 0.10744 | 0.01487 | 0.04694 |
| **LOC100507420.** | 0.05338 | 0.26753 | 0.00009 | 0.00284 | 0.00949 | 0.14309 | 0.04309 |
| **LOC100507420.** | 0.43461 | 1.01950 | 0.00000 | 0.00000 | 0.03343 | 0.58623 | 0.00262 |
| **LOC100507424** | 0.23212 | 0.00000 | 0.10223 | 0.17084 | 0.00593 | 0.07861 | 0.59815 |
| **LOC100507443** | 0.04876 | 0.01801 | 0.05223 | 0.02323 | 0.00620 | 0.03207 | 0.00000 |
| **LOC100507459** | 0.41603 | 0.51011 | 0.27182 | 0.07080 | 0.00668 | 0.00000 | 0.39743 |
| **LOC100507462** | 0.43243 | 0.00025 | 0.14650 | 0.44622 | 0.10658 | 0.09512 | 0.19136 |
| **LOC100507477** | 0.02281 | 0.00994 | 0.02869 | 0.09018 | 0.00812 | 0.01818 | 0.02628 |
| **LOC100507501** | 0.09820 | 0.33935 | 0.13959 | 0.15781 | 0.15284 | 0.01310 | 0.00000 |
| **LOC100507506** | 0.04866 | 0.01430 | 0.00936 | 0.01558 | 0.06638 | 0.01349 | 0.00799 |
| **LOC100507534** | 0.03012 | 0.05159 | 0.00715 | 0.02168 | 0.00834 | 0.03244 | 0.01089 |
| **LOC100507537** | 0.03828 | 0.01474 | 0.01472 | 0.01353 | 0.00753 | 0.01737 | 0.03431 |
| **LOC100507540** | 0.06362 | 0.20570 | 0.11400 | 0.14820 | 0.17665 | 0.07209 | 0.00000 |
| **LOC100507547** | 0.00000 | 0.50528 | 0.36401 | 0.13622 | 0.14617 | 0.38129 | 0.16523 |
| **LOC100507557** | 0.30948 | 0.00000 | 0.09746 | 0.02043 | 0.01085 | 0.05818 | 0.37452 |
| **LOC100507564** | 0.00955 | 0.01583 | 0.44109 | 0.06612 | 0.10899 | 0.02069 | 0.16497 |
| **LOC100507577** | 0.22732 | 0.12576 | 0.15195 | 0.09340 | 0.44571 | 0.14280 | 0.33382 |
| **LOC100507600** | 0.02148 | 0.01531 | 0.02851 | 0.01943 | 0.00691 | 0.03312 | 0.01910 |
| **LOC100507634** | 0.06693 | 0.15200 | 0.20825 | 0.38708 | 0.27893 | 0.01877 | 0.09780 |
| **LOC100507639** | 0.15653 | 0.18250 | 0.00000 | 0.37411 | 0.17151 | 0.00807 | 0.02011 |
| **LOC100507642** | 0.02396 | 0.02310 | 0.01116 | 0.04386 | 0.00825 | 0.04926 | 0.00431 |
| **LOC100507651** | 0.04796 | 0.00982 | 0.04040 | 0.03746 | 0.04542 | 0.04848 | 0.00000 |
| **LOC100507661** | 0.02724 | 0.01288 | 0.01603 | 0.02175 | 0.00854 | 0.08185 | 0.00789 |
| **LOC100508046** | 0.02567 | 0.01920 | 0.01186 | 0.02299 | 0.01063 | 0.01185 | 0.02018 |
| **LOC100630918** | 0.00000 | 0.39189 | 0.16629 | 0.28540 | 0.67709 | 0.07514 | 0.25070 |
| **LOC100631378** | 0.08580 | 0.01150 | 0.42581 | 0.01761 | 0.14314 | 0.12518 | 0.00630 |
| **LOC100652758** | 0.00000 | 0.31970 | 0.14048 | 0.03773 | 0.62246 | 0.13489 | 0.18938 |

| | | | | | | | |
|---|---|---|---|---|---|---|---|
| **LOC100652768** | 0.03885 | 0.05596 | 0.07762 | 0.08304 | 0.03658 | 0.00968 | 0.25036 |
| **LOC100652770** | 0.19449 | 0.22396 | 0.00000 | 0.04405 | 1.83595 | 0.11667 | 0.08546 |
| **LOC100652824** | 0.01243 | 0.08672 | 0.10758 | 0.03832 | 0.65500 | 0.06916 | 0.01577 |
| **LOC100652999** | 0.03060 | 0.01455 | 0.01299 | 0.01723 | 0.00773 | 0.01168 | 0.02641 |
| **LOC100653515** | 0.02087 | 0.03305 | 0.01262 | 0.03696 | 0.03358 | 0.01282 | 0.03321 |
| **LOC100862671** | 2.20807 | 1.42482 | 0.66753 | 9.19555 | 0.14061 | 0.77808 | 1.80099 |
| **LOC100996249** | 0.00000 | 0.13218 | 1.96675 | 0.00000 | 0.00281 | 0.13271 | 0.03419 |
| **LOC100996251** | 0.20304 | 0.05374 | 0.13893 | 0.08420 | 0.12717 | 0.06953 | 0.00000 |
| **LOC100996255** | 0.04165 | 0.03707 | 0.01668 | 0.03173 | 0.01919 | 0.01171 | 0.00412 |
| **LOC100996266** | 0.02171 | 0.02727 | 0.01054 | 0.01908 | 0.00808 | 0.01349 | 0.02191 |
| **LOC100996286** | 0.02336 | 0.01267 | 0.02047 | 0.01977 | 0.01153 | 0.02027 | 0.02145 |
| **LOC100996291** | 0.02653 | 0.02425 | 0.01418 | 0.01658 | 0.00752 | 0.03121 | 0.01931 |
| **LOC100996324** | 0.05831 | 0.05506 | 0.55423 | 0.10288 | 2.43887 | 0.59183 | 0.00000 |
| **LOC100996342** | 0.03570 | 0.02854 | 0.02181 | 0.01883 | 0.00723 | 0.01978 | 0.00270 |
| **LOC100996348** | 0.12330 | 0.03620 | 0.13361 | 0.06890 | 0.00360 | 0.02250 | 0.05891 |
| **LOC100996349** | 0.03825 | 0.02449 | 0.01004 | 0.03113 | 0.02837 | 0.01264 | 0.00761 |
| **LOC100996351** | 0.02286 | 0.01686 | 0.01254 | 0.02086 | 0.00991 | 0.01405 | 0.02032 |
| **LOC100996385** | 0.02400 | 0.01487 | 0.01771 | 0.02106 | 0.00972 | 0.01285 | 0.01724 |
| **LOC100996437** | 0.20603 | 0.34409 | 0.20607 | 0.12757 | 0.04124 | 0.14817 | 0.02116 |
| **LOC100996447** | 2.72438 | 1.02188 | 0.71616 | 0.28570 | 0.03953 | 1.27810 | 1.86453 |
| **LOC100996455** | 0.22572 | 0.07611 | 0.08532 | 0.53379 | 0.00245 | 0.00000 | 0.07621 |
| **LOC100996579** | 0.18694 | 0.01794 | 0.01289 | 0.00000 | 0.02059 | 0.13220 | 0.04773 |
| **LOC100996590** | 0.02434 | 0.01700 | 0.01629 | 0.01921 | 0.02403 | 0.01306 | 0.01586 |
| **LOC100996634** | 0.11889 | 0.14038 | 0.21365 | 0.06024 | 0.13544 | 0.06065 | 0.00000 |
| **LOC100996654** | 0.01273 | 0.04108 | 0.01995 | 0.05764 | 0.03300 | 0.02379 | 0.02544 |
| **LOC100996669** | 0.01158 | 0.01812 | 0.01389 | 0.03310 | 0.00934 | 0.01364 | 0.03999 |
| **LOC100996693** | 0.01279 | 0.05799 | 0.01120 | 0.08835 | 0.06919 | 0.01749 | 0.04736 |
| **LOC100996694** | 0.05777 | 0.00000 | 0.07193 | 0.00729 | 0.00696 | 0.02055 | 0.02849 |
| **LOC101054525** | 0.02345 | 0.01765 | 0.01313 | 0.02362 | 0.00972 | 0.01257 | 0.01717 |
| **LOC101059948** | 0.01177 | 0.00127 | 0.03897 | 0.02267 | 0.05298 | 0.00787 | 0.09472 |
| **LOC101060019** | 0.02270 | 0.01363 | 0.01065 | 0.02089 | 0.21339 | 0.01302 | 0.01742 |
| **LOC101060091** | 0.02496 | 0.02325 | 0.01270 | 0.08121 | 0.00886 | 0.01150 | 0.00490 |
| **LOC101060385** | 0.02546 | 0.01291 | 0.03406 | 0.01803 | 0.00925 | 0.01179 | 0.01964 |
| **LOC101060498** | 0.06446 | 0.14657 | 0.16474 | 0.00745 | 0.02465 | 0.00367 | 0.09152 |
| **LOC101060542** | 0.03243 | 0.02178 | 0.01297 | 0.02025 | 0.01104 | 0.02432 | 0.01257 |

| | | | | | | | |
|---|---|---|---|---|---|---|---|
| **LOC101060553** | 0.04176 | 0.01551 | 0.01150 | 0.02816 | 0.03918 | 0.01357 | 0.00800 |
| **LOC101060691** | 1.62768 | 0.00000 | 1.81118 | 1.25629 | 0.37178 | 0.42503 | 3.24887 |
| **LOC101241902** | 0.03362 | 0.01484 | 0.01017 | 0.01856 | 0.04267 | 0.01220 | 0.01981 |
| **LOC101243545** | 0.06229 | 0.00000 | 0.05580 | 0.02733 | 0.01673 | 0.00960 | 0.03294 |
| **LOC101409256** | 0.07029 | 0.00000 | 0.05711 | 0.02343 | 0.00563 | 0.02337 | 0.03100 |
| **LOC101448202** | 0.02923 | 0.01812 | 0.01591 | 0.06903 | 0.01816 | 0.01269 | 0.02076 |
| **LOC101559451** | 0.07115 | 0.21169 | 0.41148 | 0.03704 | 0.10723 | 0.00000 | 0.09163 |
| **LOC101593348** | 0.03054 | 0.08795 | 0.03042 | 0.07761 | 0.32420 | 0.02467 | 0.11781 |
| **LOC101805491** | 0.16428 | 0.00862 | 0.17450 | 1.00544 | 0.72979 | 0.19420 | 0.25219 |
| **LOC101926889** | 0.03377 | 0.05754 | 0.00918 | 0.05462 | 0.00850 | 0.04323 | 0.00000 |
| **LOC101926892** | 0.01331 | 0.01942 | 0.02001 | 0.99819 | 0.00720 | 0.07484 | 0.00000 |
| **LOC101926895** | 0.11185 | 0.02183 | 0.10878 | 0.08053 | 0.05544 | 0.00000 | 0.16116 |
| **LOC101926896** | 0.00547 | 0.04390 | 0.00834 | 0.02218 | 0.00836 | 0.04898 | 0.09726 |
| **LOC101926905** | 0.01780 | 0.01359 | 0.50695 | 0.02639 | 0.08784 | 0.00655 | 0.01638 |
| **LOC101926911** | 0.12564 | 0.03060 | 0.04818 | 0.11217 | 0.04488 | 0.00000 | 0.11407 |
| **LOC101926913** | 0.08543 | 0.04905 | 0.04058 | 0.02558 | 0.00846 | 0.01163 | 0.01098 |
| **LOC101926928** | 0.02631 | 0.01794 | 0.01163 | 0.02004 | 0.02574 | 0.01329 | 0.01607 |
| **LOC101926933** | 0.05259 | 0.00000 | 0.09601 | 0.07352 | 0.03243 | 0.00489 | 0.10452 |
| **LOC101926935** | 0.00864 | 0.04063 | 0.08277 | 0.08967 | 0.00784 | 0.21183 | 0.04421 |
| **LOC101926940** | 0.10320 | 0.00068 | 0.02274 | 0.02529 | 0.00703 | 0.01917 | 0.00898 |
| **LOC101926941** | 0.01644 | 0.02850 | 0.01891 | 0.02693 | 0.04765 | 0.02280 | 0.02855 |
| **LOC101926943** | 0.02738 | 0.04558 | 0.05238 | 0.04958 | 0.01858 | 0.14286 | 0.23465 |
| **LOC101926950** | 0.01864 | 0.04174 | 0.00784 | 0.01906 | 0.02135 | 0.01241 | 0.01721 |
| **LOC101926955** | 0.01707 | 0.02216 | 0.01421 | 0.02523 | 0.00776 | 0.01921 | 0.04248 |
| **LOC101926960** | 0.03131 | 0.01931 | 0.01421 | 0.01913 | 0.00968 | 0.01645 | 0.01166 |
| **LOC101926962** | 0.02908 | 0.01833 | 0.01281 | 0.01936 | 0.00797 | 0.01252 | 0.01491 |
| **LOC101926963** | 0.03135 | 0.01452 | 0.01288 | 0.02437 | 0.00777 | 0.01089 | 0.02805 |
| **LOC101926964** | 0.02388 | 0.01970 | 0.01438 | 0.02135 | 0.00962 | 0.01725 | 0.01687 |
| **LOC101926966** | 0.52343 | 0.06087 | 0.50798 | 0.18631 | 0.02545 | 0.00000 | 0.00000 |
| **LOC101926975** | 0.02272 | 0.01620 | 0.01008 | 0.01911 | 0.09551 | 0.01316 | 0.01772 |
| **LOC101926987** | 0.00000 | 0.00000 | 0.26713 | 0.09011 | 0.25158 | 0.39099 | 0.09198 |
| **LOC101926996** | 0.00827 | 0.02850 | 0.03736 | 0.04189 | 0.06098 | 0.15525 | 0.06861 |
| **LOC101927018** | 0.05517 | 0.00000 | 0.04015 | 0.00429 | 0.00840 | 0.00149 | 0.17857 |
| **LOC101927021** | 0.00000 | 0.22257 | 0.17612 | 0.16506 | 0.07497 | 0.24573 | 0.31801 |
| **LOC101927023** | 0.02439 | 0.01225 | 0.03049 | 0.01189 | 0.11697 | 0.01618 | 0.04264 |

| | | | | | | | |
|---|---|---|---|---|---|---|---|
| **LOC101927027** | 1.07824 | 0.00000 | 1.00250 | 0.04650 | 0.00050 | 0.84544 | 0.93753 |
| **LOC101927040** | 0.02593 | 0.01629 | 0.00862 | 0.03226 | 0.06011 | 0.01101 | 0.02411 |
| **LOC101927043** | 0.17244 | 0.00000 | 0.02565 | 0.07564 | 0.04770 | 0.00000 | 0.16981 |
| **LOC101927045** | 0.51964 | 0.27333 | 0.49622 | 0.29625 | 0.07767 | 0.06869 | 0.00000 |
| **LOC101927049** | 0.00000 | 0.03477 | 0.01444 | 0.00680 | 0.00632 | 0.07275 | 0.14744 |
| **LOC101927051** | 0.00547 | 0.04701 | 0.01142 | 0.04754 | 0.02766 | 0.09495 | 0.06433 |
| **LOC101927053** | 0.00754 | 0.02637 | 0.05359 | 0.06803 | 0.00688 | 0.00755 | 0.06134 |
| **LOC101927055** | 0.02704 | 0.01636 | 0.01317 | 0.02775 | 0.00786 | 0.01206 | 0.02048 |
| **LOC101927069** | 0.07254 | 0.16425 | 0.45550 | 0.01415 | 0.00126 | 0.00000 | 1.20582 |
| **LOC101927081** | 0.09136 | 0.00000 | 0.14725 | 0.05486 | 0.01717 | 0.00923 | 0.26405 |
| **LOC101927087** | 0.02306 | 0.01797 | 0.00625 | 0.01831 | 0.09421 | 0.02131 | 0.02333 |
| **LOC101927100** | 0.08300 | 0.00000 | 0.21832 | 0.08675 | 0.16935 | 0.13653 | 0.13958 |
| **LOC101927102** | 0.03579 | 0.01528 | 0.01534 | 0.01734 | 0.00786 | 0.01934 | 0.01438 |
| **LOC101927112** | 0.03202 | 0.01787 | 0.01636 | 0.01693 | 0.00878 | 0.01479 | 0.01374 |
| **LOC101927123** | 0.02428 | 0.02076 | 0.01233 | 0.01690 | 0.00748 | 0.03139 | 0.04004 |
| **LOC101927124** | 0.25687 | 0.14158 | 0.02614 | 0.09869 | 0.03803 | 0.00214 | 0.03413 |
| **LOC101927129** | 0.09147 | 0.09420 | 0.05200 | 0.00334 | 0.04775 | 0.08351 | 0.00000 |
| **LOC101927131** | 0.04203 | 0.03189 | 0.01053 | 0.03055 | 0.01027 | 0.02720 | 0.00000 |
| **LOC101927132** | 0.04584 | 0.00000 | 0.10159 | 0.06125 | 0.00843 | 0.00522 | 0.03759 |
| **LOC101927134** | 0.03695 | 0.01584 | 0.04047 | 0.01742 | 0.03013 | 0.01086 | 0.01031 |
| **LOC101927136** | 0.02940 | 0.01994 | 0.01327 | 0.01754 | 0.00782 | 0.01291 | 0.01741 |
| **LOC101927142** | 0.02522 | 0.01601 | 0.01580 | 0.02687 | 0.01432 | 0.01231 | 0.01823 |
| **LOC101927143** | 0.00000 | 0.48109 | 0.05154 | 0.04857 | 0.02273 | 0.08409 | 0.01368 |
| **LOC101927151** | 1.64995 | 2.65604 | 3.22175 | 3.32492 | 1.80630 | 0.00000 | 0.08237 |
| **LOC101927153** | 0.03926 | 0.01657 | 0.03283 | 0.01755 | 0.00731 | 0.01217 | 0.00968 |
| **LOC101927164** | 0.11840 | 0.04112 | 0.05652 | 0.07459 | 0.03493 | 0.00676 | 0.00000 |
| **LOC101927166** | 0.04444 | 0.01452 | 0.01051 | 0.01398 | 0.10707 | 0.00851 | 0.02983 |
| **LOC101927178** | 0.06761 | 0.00000 | 0.04628 | 0.06527 | 0.03074 | 0.08139 | 0.10845 |
| **LOC101927181** | 0.00000 | 0.68107 | 0.22600 | 0.12310 | 0.06332 | 0.25812 | 0.05448 |
| **LOC101927190** | 0.02664 | 0.00000 | 0.06610 | 0.00608 | 0.00436 | 0.03928 | 0.18754 |
| **LOC101927196** | 0.01258 | 0.01652 | 0.01527 | 0.02599 | 0.01291 | 0.04573 | 0.05838 |
| **LOC101927207** | 0.04282 | 0.01144 | 0.02588 | 0.01948 | 0.01504 | 0.01405 | 0.00489 |
| **LOC101927211** | 0.05032 | 0.01077 | 0.04397 | 0.04545 | 0.05085 | 0.03824 | 0.02367 |
| **LOC101927217** | 0.02928 | 0.01640 | 0.01822 | 0.01858 | 0.12606 | 0.03566 | 0.02399 |
| **LOC101927230** | 0.09681 | 0.26964 | 0.30029 | 0.00000 | 0.00220 | 0.19655 | 0.00718 |

| | | | | | | | |
|---|---|---|---|---|---|---|---|
| **LOC101927237** | 0.10660 | 0.00000 | 0.01888 | 0.02278 | 0.11907 | 0.02812 | 0.00000 |
| **LOC101927239** | 0.11443 | 0.64902 | 0.21583 | 0.00708 | 0.16188 | 0.13741 | 0.01558 |
| **LOC101927243** | 0.02369 | 0.01578 | 0.01210 | 0.02363 | 0.06124 | 0.01380 | 0.01454 |
| **LOC101927248** | 0.03958 | 0.03973 | 0.00561 | 0.01167 | 0.41295 | 0.02494 | 0.00874 |
| **LOC101927257** | 0.00000 | 0.06157 | 0.02459 | 0.09938 | 0.23069 | 0.14633 | 0.47677 |
| **LOC101927258** | 0.09066 | 0.00000 | 0.06251 | 0.02424 | 0.77398 | 0.03025 | 0.00002 |
| **LOC101927267** | 0.18800 | 0.08877 | 0.40696 | 0.06700 | 0.45432 | 0.08343 | 0.22157 |
| **LOC101927273** | 0.41328 | 0.05428 | 0.12576 | 0.00028 | 0.00378 | 0.05616 | 0.11951 |
| **LOC101927274** | 0.02341 | 0.01935 | 0.03244 | 0.02230 | 0.01010 | 0.01196 | 0.02053 |
| **LOC101927275** | 0.03770 | 0.05581 | 0.02789 | 0.27114 | 0.10992 | 0.03277 | 0.15095 |
| **LOC101927280** | 0.01816 | 0.00723 | 0.00707 | 0.01489 | 1.78108 | 0.01524 | 0.02373 |
| **LOC101927282** | 0.02967 | 0.00881 | 0.03104 | 0.02725 | 0.06221 | 0.01965 | 0.01712 |
| **LOC101927284** | 0.11082 | 0.00000 | 0.05117 | 0.03643 | 0.16754 | 0.12300 | 0.97717 |
| **LOC101927285** | 0.06241 | 0.01978 | 0.01159 | 0.04579 | 0.04608 | 0.00766 | 0.19473 |
| **LOC101927286** | 0.03887 | 0.01290 | 0.01215 | 0.03293 | 0.19212 | 0.04896 | 0.02581 |
| **LOC101927292** | 0.02781 | 0.06538 | 0.01482 | 0.01864 | 0.00783 | 0.01351 | 0.00918 |
| **LOC101927296** | 0.02868 | 0.02934 | 0.01611 | 0.02206 | 0.00785 | 0.01582 | 0.00605 |
| **LOC101927318** | 0.03464 | 0.01773 | 0.01098 | 0.01946 | 0.02170 | 0.01203 | 0.01266 |
| **LOC101927322** | 0.04141 | 0.00676 | 0.23443 | 0.06419 | 0.22180 | 0.02519 | 0.00705 |
| **LOC101927323** | 0.02175 | 0.01990 | 0.01214 | 0.02477 | 0.00792 | 0.01266 | 0.02074 |
| **LOC101927324** | 0.03297 | 0.04483 | 0.02682 | 0.01601 | 0.01111 | 0.01276 | 0.01353 |
| **LOC101927332** | 0.02532 | 0.01771 | 0.00457 | 0.02388 | 0.16588 | 0.01297 | 0.01315 |
| **LOC101927347** | 0.04455 | 0.00469 | 0.01700 | 0.00876 | 0.00693 | 0.00406 | 0.09783 |
| **LOC101927354** | 0.40618 | 0.04735 | 0.03816 | 0.26652 | 0.04765 | 0.06599 | 0.00000 |
| **LOC101927356** | 0.02152 | 0.03789 | 0.01994 | 0.01955 | 0.00697 | 0.01835 | 0.01118 |
| **LOC101927374** | 0.02279 | 0.01657 | 0.01237 | 0.01998 | 0.01028 | 0.01301 | 0.02444 |
| **LOC101927378** | 0.01621 | 0.03153 | 0.00894 | 0.02516 | 0.00768 | 0.01514 | 0.05888 |
| **LOC101927379** | 1.13164 | 0.33655 | 0.00000 | 0.05904 | 0.00648 | 0.00000 | 0.15180 |
| **LOC101927380** | 0.01534 | 0.00000 | 0.05564 | 0.06191 | 0.00578 | 0.24406 | 0.47660 |
| **LOC101927391** | 0.54435 | 0.01408 | 0.02061 | 0.00000 | 0.07377 | 0.00000 | 0.38947 |
| **LOC101927394** | 0.02251 | 0.01701 | 0.00746 | 0.01889 | 0.08611 | 0.01321 | 0.01872 |
| **LOC101927396** | 0.02811 | 0.01518 | 0.02408 | 0.02716 | 0.00729 | 0.01655 | 0.01640 |
| **LOC101927415** | 0.20418 | 0.01125 | 0.21828 | 0.31634 | 0.16864 | 0.12360 | 0.00979 |
| **LOC101927416** | 0.04891 | 0.01823 | 0.01523 | 0.00897 | 0.00887 | 0.01848 | 0.01898 |
| **LOC101927418** | 0.15454 | 0.09392 | 0.25606 | 0.03198 | 1.11012 | 0.03421 | 0.00000 |

| | | | | | | | |
|---|---|---|---|---|---|---|---|
| **LOC101927419** | 0.02214 | 0.01880 | 0.01214 | 0.02271 | 0.00992 | 0.03082 | 0.01973 |
| **LOC101927421** | 0.02425 | 0.01214 | 0.01190 | 0.01812 | 0.46557 | 0.01856 | 0.01402 |
| **LOC101927437** | 0.02218 | 0.01603 | 0.00986 | 0.02075 | 0.04941 | 0.02093 | 0.01667 |
| **LOC101927438** | 0.08735 | 0.07061 | 0.00463 | 0.02821 | 0.02173 | 0.00425 | 0.00000 |
| **LOC101927446** | 0.03172 | 0.01719 | 0.01474 | 0.03582 | 0.00868 | 0.01258 | 0.00738 |
| **LOC101927457** | 0.02991 | 0.01352 | 0.00918 | 0.01668 | 0.01780 | 0.00393 | 0.09266 |
| **LOC101927460** | 0.03834 | 0.01499 | 0.01167 | 0.01302 | 0.01381 | 0.03088 | 0.01578 |
| **LOC101927472** | 0.01626 | 0.02602 | 0.04294 | 0.03049 | 0.12765 | 0.01274 | 0.01750 |
| **LOC101927476** | 0.02398 | 0.10221 | 0.03468 | 0.02579 | 0.02353 | 0.01433 | 0.00864 |
| **LOC101927477** | 0.01044 | 0.04423 | 0.02817 | 0.02795 | 0.02661 | 0.00764 | 0.03502 |
| **LOC101927478** | 0.00992 | 0.02192 | 0.02477 | 0.05188 | 0.06724 | 0.00799 | 0.06158 |
| **LOC101927482** | 4.69965 | 0.08769 | 0.06006 | 0.00000 | 0.00029 | 0.00000 | 0.12956 |
| **LOC101927492** | 1.44858 | 1.15910 | 0.29034 | 0.94785 | 0.06651 | 0.28962 | 0.00000 |
| **LOC101927495** | 0.02091 | 0.00982 | 0.02758 | 0.02811 | 0.00728 | 0.01931 | 0.02164 |
| **LOC101927497** | 0.18275 | 0.00460 | 0.01673 | 0.04290 | 0.00525 | 0.00000 | 0.18153 |
| **LOC101927501** | 0.04854 | 1.34833 | 0.08783 | 4.22748 | 0.05041 | 0.00112 | 0.00000 |
| **LOC101927502** | 0.06127 | 0.01106 | 0.00848 | 0.02728 | 0.12884 | 0.01469 | 0.00000 |
| **LOC101927523** | 0.02331 | 0.01694 | 0.00789 | 0.02009 | 0.09248 | 0.01354 | 0.01533 |
| **LOC101927539** | 0.02362 | 0.01561 | 0.01142 | 0.01822 | 0.02377 | 0.01206 | 0.02569 |
| **LOC101927550** | 0.06577 | 0.81008 | 0.32770 | 0.31233 | 0.17269 | 0.44678 | 0.47914 |
| **LOC101927557** | 0.01277 | 0.02205 | 0.01078 | 0.02780 | 0.00726 | 0.01756 | 0.07205 |
| **LOC101927560** | 0.02678 | 0.03456 | 0.01611 | 0.02029 | 0.07313 | 0.01172 | 0.00637 |
| **LOC101927571** | 0.16822 | 0.04524 | 0.12287 | 0.02616 | 0.19913 | 0.04998 | 0.00000 |
| **LOC101927572** | 0.27094 | 0.02936 | 0.08099 | 0.29153 | 0.01134 | 0.05172 | 0.17365 |
| **LOC101927575** | 0.00909 | 0.04089 | 0.04103 | 0.01305 | 0.28863 | 0.01014 | 0.07620 |
| **LOC101927577** | 0.02267 | 0.02245 | 0.01340 | 0.01792 | 0.01057 | 0.01227 | 0.02157 |
| **LOC101927580** | 0.23239 | 0.00000 | 0.29477 | 0.00000 | 0.04935 | 0.08775 | 0.14180 |
| **LOC101927583** | 0.02272 | 0.01766 | 0.01197 | 0.02412 | 0.01615 | 0.01572 | 0.01801 |
| **LOC101927587** | 0.02569 | 0.01878 | 0.01420 | 0.03491 | 0.01910 | 0.01297 | 0.00871 |
| **LOC101927592** | 0.04053 | 0.07177 | 0.01734 | 0.02182 | 0.17866 | 0.08441 | 0.00896 |
| **LOC101927599** | 1.11460 | 1.08447 | 1.59493 | 1.46177 | 0.68230 | 0.10303 | 0.47546 |
| **LOC101927616** | 0.02439 | 0.01757 | 0.01303 | 0.02606 | 0.00799 | 0.01306 | 0.01351 |
| **LOC101927619** | 0.03938 | 0.01148 | 0.01396 | 0.02426 | 0.00785 | 0.01388 | 0.01901 |
| **LOC101927620** | 0.01745 | 0.12894 | 0.07060 | 0.05353 | 0.00511 | 0.07641 | 0.01222 |
| **LOC101927621** | 0.73712 | 0.15307 | 0.13470 | 0.92983 | 0.05760 | 0.04098 | 0.00000 |

| | | | | | | | |
|---|---|---|---|---|---|---|---|
| **LOC101927623** | 0.02672 | 0.02987 | 0.05333 | 0.01695 | 0.00594 | 0.03017 | 0.01969 |
| **LOC101927630** | 0.02274 | 0.01466 | 0.01157 | 0.02378 | 0.02822 | 0.01308 | 0.02454 |
| **LOC101927631** | 0.06057 | 0.24601 | 0.07732 | 0.11695 | 0.01981 | 0.07603 | 0.16710 |
| **LOC101927641** | 0.03220 | 0.00000 | 0.04298 | 0.01824 | 0.33647 | 0.01266 | 0.01287 |
| **LOC101927650** | 0.02884 | 0.01489 | 0.01309 | 0.02161 | 0.00809 | 0.02370 | 0.01368 |
| **LOC101927653** | 0.02737 | 0.02305 | 0.01656 | 0.01706 | 0.00748 | 0.01539 | 0.02215 |
| **LOC101927661** | 0.02558 | 0.00574 | 0.05338 | 0.01818 | 0.00724 | 0.01285 | 0.01926 |
| **LOC101927666** | 0.05841 | 0.12189 | 0.00589 | 0.00354 | 0.01761 | 0.01312 | 0.01824 |
| **LOC101927667** | 0.05900 | 0.00105 | 0.12294 | 0.15885 | 0.13413 | 0.04278 | 0.06608 |
| **LOC101927668** | 0.03430 | 0.01429 | 0.01120 | 0.03606 | 0.21374 | 0.35875 | 0.00000 |
| **LOC101927688** | 0.03112 | 0.02438 | 0.01836 | 0.02236 | 0.00756 | 0.01688 | 0.01012 |
| **LOC101927692** | 0.03226 | 0.00573 | 0.01434 | 0.00762 | 0.01201 | 0.00699 | 0.08012 |
| **LOC101927697** | 0.02198 | 0.01286 | 0.01962 | 0.02155 | 0.00992 | 0.01427 | 0.02312 |
| **LOC101927698** | 0.07951 | 0.01252 | 0.01508 | 0.02723 | 0.01116 | 0.01289 | 0.00000 |
| **LOC101927700** | 0.28226 | 0.00000 | 0.05946 | 0.03425 | 0.00475 | 0.00529 | 0.11726 |
| **LOC101927701** | 0.02433 | 0.01844 | 0.00263 | 0.01807 | 0.18663 | 0.01358 | 0.01640 |
| **LOC101927709** | 0.02637 | 0.01754 | 0.01259 | 0.01814 | 0.00785 | 0.01240 | 0.02137 |
| **LOC101927718** | 0.08044 | 0.07633 | 0.04748 | 0.04183 | 0.00489 | 0.05392 | 0.04245 |
| **LOC101927720** | 0.71516 | 1.31285 | 2.06374 | 0.97883 | 0.72179 | 1.00104 | 0.00000 |
| **LOC101927722** | 0.01378 | 0.11221 | 0.07657 | 0.01254 | 1.34011 | 0.00000 | 0.02470 |
| **LOC101927728** | 0.00003 | 0.04380 | 0.21625 | 0.04561 | 0.03858 | 0.07434 | 0.12708 |
| **LOC101927730** | 0.00949 | 0.02564 | 0.01803 | 0.14114 | 0.00753 | 0.06381 | 0.00494 |
| **LOC101927740** | 0.02327 | 0.01852 | 0.01316 | 0.01838 | 0.01274 | 0.01254 | 0.02122 |
| **LOC101927746** | 6.88072 | 7.18051 | 1.62616 | 2.96082 | 1.75140 | 7.75228 | 0.00000 |
| **LOC101927750** | 0.03411 | 0.08936 | 0.11789 | 0.12360 | 0.00314 | 0.00000 | 0.15962 |
| **LOC101927755** | 0.62362 | 0.01869 | 1.57731 | 0.21134 | 0.44687 | 0.00000 | 0.00000 |
| **LOC101927762** | 0.01999 | 0.02885 | 0.01452 | 0.02368 | 0.01152 | 0.01542 | 0.01637 |
| **LOC101927769** | 0.02519 | 0.01024 | 0.03281 | 0.02591 | 0.00768 | 0.01759 | 0.01172 |
| **LOC101927770** | 0.00000 | 0.07954 | 0.06918 | 0.04783 | 0.53005 | 0.06503 | 0.15240 |
| **LOC101927780** | 0.00364 | 0.01335 | 0.01099 | 0.04533 | 0.01916 | 0.01969 | 0.08221 |
| **LOC101927795** | 0.01917 | 0.03355 | 0.01191 | 0.02334 | 0.04362 | 0.01334 | 0.01743 |
| **LOC101927797** | 0.93595 | 0.22603 | 0.03205 | 0.33147 | 0.01039 | 0.01293 | 0.34197 |
| **LOC101927811** | 0.01649 | 0.06866 | 0.08171 | 0.04029 | 0.07579 | 0.01101 | 0.00977 |
| **LOC101927814** | 0.01185 | 0.04838 | 0.00610 | 0.02331 | 0.01442 | 0.01032 | 0.06136 |
| **LOC101927817** | 0.03836 | 0.00000 | 0.07741 | 0.02551 | 0.02439 | 0.24118 | 0.58425 |

| | | | | | | | |
|---|---|---|---|---|---|---|---|
| **LOC101927829** | 0.04012 | 0.01694 | 0.01080 | 0.01977 | 0.02981 | 0.01409 | 0.01093 |
| **LOC101927830-** | 0.02493 | 0.01632 | 0.01510 | 0.02148 | 0.00884 | 0.01420 | 0.03002 |
| **LOC101927839** | 0.02213 | 0.01964 | 0.01254 | 0.02212 | 0.01006 | 0.01441 | 0.01952 |
| **LOC101927843** | 0.02539 | 0.06542 | 0.00676 | 0.02495 | 0.04577 | 0.00936 | 0.01346 |
| **LOC101927844** | 0.02278 | 0.01218 | 0.01265 | 0.02011 | 0.00948 | 0.01366 | 0.04496 |
| **LOC101927854** | 0.00982 | 0.00328 | 0.05172 | 0.33129 | 0.06640 | 0.11190 | 0.04660 |
| **LOC101927865** | 0.02072 | 0.03727 | 0.03566 | 0.01557 | 0.08950 | 0.05400 | 0.02312 |
| **LOC101927870** | 0.03754 | 0.00630 | 0.01194 | 0.01706 | 0.31787 | 0.01523 | 0.01083 |
| **LOC101927873** | 0.12283 | 0.76074 | 0.65729 | 0.00000 | 0.87303 | 0.73908 | 0.00000 |
| **LOC101927881** | 0.01892 | 0.07636 | 0.02973 | 0.02020 | 0.00704 | 0.01584 | 0.00624 |
| **LOC101927901** | 0.00984 | 0.00373 | 0.01230 | 0.02014 | 0.07131 | 0.00543 | 0.19583 |
| **LOC101927905** | 0.02799 | 0.01957 | 0.03371 | 0.02541 | 0.01644 | 0.02040 | 0.00706 |
| **LOC101927907** | 0.02787 | 0.02096 | 0.01386 | 0.02602 | 0.02818 | 0.03741 | 0.03430 |
| **LOC101927911** | 0.07111 | 0.03737 | 0.02957 | 0.02073 | 0.00702 | 0.01726 | 0.00304 |
| **LOC101927914** | 0.03780 | 0.01440 | 0.01315 | 0.02752 | 0.03047 | 0.01821 | 0.01038 |
| **LOC101927919** | 0.39288 | 0.23058 | 0.36722 | 0.57646 | 0.13142 | 0.07260 | 0.00000 |
| **LOC101927921** | 0.02610 | 0.01481 | 0.01231 | 0.01622 | 1.23459 | 0.01089 | 0.01128 |
| **LOC101927926** | 0.02807 | 0.01543 | 0.01289 | 0.02557 | 0.01081 | 0.01673 | 0.01509 |
| **LOC101927931** | 0.03225 | 0.01930 | 0.00182 | 0.02015 | 0.26887 | 0.01955 | 0.00892 |
| **LOC101927934** | 0.04508 | 0.50106 | 0.02646 | 0.00000 | 0.01974 | 0.00000 | 0.14799 |
| **LOC101927948** | 0.12885 | 0.01956 | 0.02610 | 0.04234 | 0.00524 | 0.01107 | 0.16858 |
| **LOC101927950** | 0.02681 | 0.01180 | 0.02002 | 0.02622 | 0.00727 | 0.02260 | 0.03308 |
| **LOC101927954** | 0.02643 | 0.01553 | 0.02255 | 0.04783 | 0.05797 | 0.01423 | 0.02252 |
| **LOC101927964** | 0.03740 | 0.01930 | 0.01264 | 0.01413 | 0.00829 | 0.01205 | 0.02313 |
| **LOC101927967** | 0.02369 | 0.00304 | 0.02094 | 0.02172 | 0.00709 | 0.05359 | 0.07865 |
| **LOC101927990** | 0.02359 | 0.01140 | 0.02492 | 0.01735 | 0.00972 | 0.01210 | 0.02746 |
| **LOC101928001** | 0.02349 | 0.01746 | 0.03090 | 0.00000 | 0.01001 | 0.02329 | 0.19740 |
| **LOC101928002** | 0.01459 | 0.00674 | 0.01909 | 0.01296 | 0.00799 | 0.00696 | 0.11192 |
| **LOC101928008** | 0.03470 | 0.00729 | 0.03169 | 0.02818 | 0.00685 | 0.06422 | 0.08516 |
| **LOC101928012** | 0.64614 | 0.24141 | 0.84026 | 0.00000 | 0.01957 | 0.52517 | 0.01854 |
| **LOC101928020** | 0.15932 | 0.00000 | 0.08597 | 0.07569 | 0.23433 | 0.06786 | 0.37515 |
| **LOC101928035** | 0.02911 | 0.04692 | 0.02188 | 0.02190 | 0.01306 | 0.02327 | 0.01016 |
| **LOC101928036** | 0.02513 | 0.02383 | 0.01139 | 0.02742 | 0.00786 | 0.01175 | 0.02872 |
| **LOC101928043** | 0.01472 | 0.04377 | 0.26847 | 0.02002 | 0.35732 | 0.00471 | 0.09936 |
| **LOC101928053** | 0.12006 | 0.00000 | 0.28522 | 0.27666 | 0.92861 | 0.02195 | 0.00000 |

| | | | | | | | |
|---|---|---|---|---|---|---|---|
| **LOC101928062** | 0.94144 | 0.99188 | 1.65541 | 0.89509 | 1.77073 | 0.31530 | 0.03674 |
| **LOC101928063** | 0.02264 | 0.05484 | 0.02910 | 0.08848 | 0.02616 | 0.00681 | 0.00862 |
| **LOC101928068** | 0.07356 | 0.00000 | 0.07731 | 0.32264 | 0.22269 | 0.10077 | 0.00000 |
| **LOC101928069** | 0.03858 | 0.05065 | 0.04969 | 0.08217 | 0.00723 | 0.03809 | 0.06300 |
| **LOC101928075** | 0.01632 | 0.01782 | 0.01477 | 0.03954 | 0.00819 | 0.01432 | 0.02332 |
| **LOC101928093** | 0.01785 | 0.02810 | 0.01239 | 0.06835 | 0.01036 | 0.01321 | 0.01029 |
| **LOC101928099** | 0.07509 | 0.26116 | 2.27193 | 1.28411 | 1.75705 | 0.14148 | 0.04923 |
| **LOC101928100** | 0.45566 | 0.08078 | 0.13419 | 0.00000 | 0.19007 | 0.00000 | 0.44400 |
| **LOC101928103** | 0.00000 | 0.52868 | 0.17821 | 0.08325 | 0.09252 | 0.43856 | 0.00079 |
| **LOC101928104** | 0.02945 | 0.02228 | 0.01707 | 0.01765 | 0.00870 | 0.01600 | 0.01822 |
| **LOC101928105** | 0.08144 | 0.00632 | 0.01401 | 0.03496 | 0.05011 | 0.00887 | 0.01178 |
| **LOC101928107** | 0.00936 | 0.01145 | 0.01228 | 0.03055 | 0.01286 | 0.00927 | 0.08281 |
| **LOC101928122** | 0.10483 | 0.00000 | 0.25962 | 0.00378 | 0.33253 | 0.07885 | 0.03198 |
| **LOC101928128** | 0.02635 | 0.01196 | 0.02790 | 0.01861 | 0.00716 | 0.01251 | 0.01673 |
| **LOC101928131** | 0.04702 | 0.01068 | 0.00294 | 0.02122 | 0.21711 | 0.00832 | 0.03489 |
| **LOC101928135** | 0.00843 | 0.01325 | 0.00613 | 0.00976 | 0.06118 | 0.06753 | 0.09520 |
| **LOC101928136** | 0.02134 | 0.01635 | 0.01335 | 0.02466 | 0.00845 | 0.02112 | 0.01629 |
| **LOC101928137** | 0.03024 | 0.01467 | 0.02051 | 0.02030 | 0.01557 | 0.01613 | 0.03160 |
| **LOC101928150** | 0.04039 | 0.00748 | 0.03558 | 0.02256 | 0.03374 | 0.03934 | 0.03114 |
| **LOC101928163** | 0.04475 | 0.00000 | 0.04046 | 0.02770 | 0.04092 | 0.00976 | 0.08970 |
| **LOC101928167** | 0.03476 | 0.02148 | 0.01558 | 0.02107 | 0.00892 | 0.01325 | 0.00865 |
| **LOC101928168** | 0.09516 | 0.02481 | 0.14593 | 0.00000 | 0.03053 | 0.28533 | 1.07451 |
| **LOC101928174** | 0.03337 | 0.01338 | 0.01761 | 0.03922 | 0.01477 | 0.01479 | 0.00784 |
| **LOC101928203** | 0.00729 | 0.00000 | 0.16837 | 0.04206 | 0.18753 | 0.13801 | 0.11662 |
| **LOC101928222** | 0.02401 | 0.03126 | 0.02418 | 0.02168 | 0.04092 | 0.00987 | 0.02236 |
| **LOC101928243** | 0.63365 | 0.53747 | 0.12549 | 0.47022 | 0.03736 | 0.21507 | 0.00000 |
| **LOC101928244** | 0.03416 | 0.02030 | 0.01253 | 0.01737 | 0.00891 | 0.01237 | 0.01925 |
| **LOC101928254** | 0.01744 | 0.31808 | 0.04450 | 0.00000 | 0.04936 | 1.71045 | 0.00986 |
| **LOC101928259** | 0.01780 | 0.10453 | 0.56897 | 0.00000 | 0.13084 | 0.25361 | 0.47333 |
| **LOC101928266** | 0.03446 | 0.00840 | 0.01155 | 0.02623 | 0.02182 | 0.01488 | 0.04106 |
| **LOC101928269** | 0.02128 | 0.01616 | 0.01222 | 0.03088 | 0.07492 | 0.01222 | 0.02351 |
| **LOC101928272** | 0.00000 | 0.11586 | 0.23068 | 0.01057 | 0.04252 | 0.13607 | 0.02264 |
| **LOC101928283** | 0.06316 | 0.01005 | 0.06166 | 0.02729 | 0.00526 | 0.11456 | 0.00811 |
| **LOC101928290** | 0.04110 | 0.05190 | 0.10172 | 0.06124 | 0.06535 | 0.03231 | 0.05440 |
| **LOC101928295** | 0.10130 | 0.00000 | 0.35593 | 0.01731 | 0.21001 | 0.01565 | 0.16780 |

| | | | | | | | |
|---|---|---|---|---|---|---|---|
| **LOC101928303** | 0.00903 | 0.23073 | 0.01811 | 0.50240 | 0.05078 | 0.06869 | 0.00000 |
| **LOC101928335** | 0.02328 | 0.01460 | 0.01678 | 0.01911 | 0.01767 | 0.01349 | 0.02024 |
| **LOC101928336** | 0.00004 | 0.00419 | 0.02512 | 0.03618 | 0.01413 | 0.04911 | 0.64798 |
| **LOC101928358** | 0.02468 | 0.00000 | 0.06020 | 0.03218 | 0.01143 | 0.01160 | 0.03102 |
| **LOC101928363** | 0.14971 | 0.09852 | 0.00000 | 0.00000 | 0.02743 | 0.05514 | 0.75826 |
| **LOC101928371** | 0.07985 | 0.27081 | 0.00000 | 0.15209 | 0.25471 | 0.04321 | 0.28402 |
| **LOC101928372** | 0.00000 | 0.26763 | 1.44523 | 0.06861 | 0.24213 | 0.52267 | 0.19031 |
| **LOC101928378** | 0.00000 | 0.55348 | 0.05120 | 0.74706 | 0.08275 | 0.35257 | 2.87787 |
| **LOC101928386** | 0.07301 | 0.00459 | 0.02985 | 0.03108 | 0.00733 | 0.00819 | 0.05307 |
| **LOC101928388** | 0.17379 | 0.26261 | 0.01455 | 0.03995 | 0.00594 | 0.00337 | 0.00912 |
| **LOC101928401** | 0.08228 | 0.03075 | 0.23050 | 0.06219 | 0.00566 | 0.00000 | 0.04559 |
| **LOC101928404** | 0.12092 | 0.00000 | 0.17550 | 0.00158 | 0.00762 | 0.00296 | 0.04087 |
| **LOC101928414** | 0.02878 | 0.00382 | 0.05194 | 0.02139 | 0.03940 | 0.01438 | 0.01818 |
| **LOC101928418** | 0.03133 | 0.02004 | 0.01435 | 0.01949 | 0.01267 | 0.01031 | 0.02545 |
| **LOC101928420** | 0.02367 | 0.01577 | 0.02976 | 0.01717 | 0.00704 | 0.01196 | 0.02382 |
| **LOC101928437** | 0.03927 | 0.01128 | 0.02173 | 0.01473 | 0.11420 | 0.05181 | 0.01347 |
| **LOC101928438** | 0.02810 | 0.03260 | 0.00969 | 0.01852 | 0.01420 | 0.01964 | 0.01848 |
| **LOC101928440** | 0.02545 | 0.01749 | 0.01206 | 0.01837 | 0.01641 | 0.01333 | 0.01747 |
| **LOC101928443** | 0.01384 | 0.08045 | 0.00662 | 0.03918 | 0.04419 | 0.00983 | 0.05610 |
| **LOC101928445** | 0.10388 | 0.15170 | 0.14593 | 0.07396 | 0.03150 | 0.19811 | 0.09396 |
| **LOC101928446** | 0.02037 | 0.01423 | 0.00847 | 0.01905 | 0.04964 | 0.03781 | 0.03386 |
| **LOC101928448** | 0.02670 | 0.01759 | 0.00795 | 0.01807 | 0.08623 | 0.01358 | 0.01508 |
| **LOC101928453** | 0.01971 | 0.02438 | 0.00944 | 0.07247 | 0.02519 | 0.01223 | 0.00496 |
| **LOC101928460** | 0.06669 | 0.02366 | 0.03590 | 0.08881 | 0.10917 | 0.02058 | 0.00507 |
| **LOC101928470** | 0.03138 | 0.01729 | 0.01287 | 0.01826 | 0.00815 | 0.01772 | 0.01246 |
| **LOC101928489** | 0.02885 | 0.01369 | 0.01076 | 0.01970 | 0.05411 | 0.00731 | 0.05606 |
| **LOC101928491** | 0.00396 | 0.03573 | 0.00694 | 0.05046 | 0.20762 | 0.00582 | 0.07610 |
| **LOC101928495** | 0.02020 | 0.11760 | 0.02696 | 0.02415 | 0.00891 | 0.06911 | 0.02935 |
| **LOC101928496** | 0.02249 | 0.01926 | 0.01780 | 0.02295 | 0.01366 | 0.01786 | 0.02632 |
| **LOC101928505** | 0.01094 | 0.11837 | 0.01638 | 0.02458 | 0.00846 | 0.04177 | 0.00825 |
| **LOC101928514** | 0.01871 | 0.04517 | 0.03395 | 0.01799 | 0.11279 | 0.00461 | 0.07596 |
| **LOC101928517** | 0.03976 | 0.00500 | 0.02794 | 0.01635 | 0.52042 | 0.01448 | 0.00963 |
| **LOC101928519** | 0.02760 | 0.01618 | 0.01395 | 0.01696 | 0.12619 | 0.01273 | 0.01718 |
| **LOC101928523** | 0.00947 | 0.03917 | 0.58557 | 0.03647 | 0.64918 | 0.00000 | 0.01494 |
| **LOC101928525** | 0.58505 | 0.63658 | 0.37987 | 0.07456 | 0.00389 | 0.29034 | 0.08900 |

| | | | | | | | |
|---|---|---|---|---|---|---|---|
| **LOC101928530** | 0.03779 | 0.17256 | 0.24655 | 0.02661 | 0.21082 | 0.00000 | 0.03773 |
| **LOC101928535** | 0.04867 | 0.00000 | 0.06347 | 0.04601 | 0.08662 | 0.04891 | 0.00640 |
| **LOC101928537** | 0.00969 | 0.19603 | 0.69823 | 0.00330 | 0.66963 | 0.00401 | 0.00679 |
| **LOC101928539** | 0.03556 | 0.01714 | 0.02095 | 0.02265 | 0.00759 | 0.01180 | 0.01719 |
| **LOC101928545** | 0.05865 | 0.02156 | 0.07350 | 0.05515 | 0.27423 | 0.02278 | 0.00000 |
| **LOC101928550** | 0.02928 | 0.01894 | 0.01350 | 0.02888 | 0.00762 | 0.02231 | 0.02526 |
| **LOC101928559** | 0.02038 | 0.01248 | 0.00000 | 0.00328 | 3.96918 | 0.01465 | 0.06347 |
| **LOC101928564** | 0.02398 | 0.01474 | 0.02712 | 0.02170 | 0.00926 | 0.01437 | 0.01912 |
| **LOC101928567** | 0.17784 | 0.12947 | 0.07901 | 0.16015 | 0.03294 | 0.02959 | 0.04083 |
| **LOC101928571** | 0.02837 | 0.04346 | 0.09684 | 0.03353 | 0.00699 | 0.03505 | 0.00000 |
| **LOC101928580** | 0.02530 | 0.00979 | 0.04937 | 0.02002 | 0.11566 | 0.01089 | 0.02128 |
| **LOC101928597.** | 0.03586 | 0.02057 | 0.01216 | 0.01342 | 0.04493 | 0.01013 | 0.03498 |
| **LOC101928600** | 0.02603 | 0.02113 | 0.01290 | 0.02114 | 0.00813 | 0.01270 | 0.01318 |
| **LOC101928663** | 0.18307 | 0.10674 | 0.15790 | 0.00000 | 0.03312 | 0.11455 | 0.23165 |
| **LOC101928674** | 0.00000 | 0.21447 | 0.00000 | 0.27905 | 0.03124 | 0.15903 | 0.24554 |
| **LOC101928682** | 0.03033 | 0.04468 | 0.01334 | 0.02460 | 0.00760 | 0.00982 | 0.01029 |
| **LOC101928694** | 0.01453 | 0.05250 | 0.00000 | 0.02907 | 0.69488 | 0.02832 | 0.01860 |
| **LOC101928696** | 0.02662 | 0.01272 | 0.01550 | 0.01547 | 0.00727 | 0.00912 | 0.05069 |
| **LOC101928700** | 0.04037 | 0.00659 | 0.02467 | 0.01583 | 0.07048 | 0.01505 | 0.01362 |
| **LOC101928710** | 0.04384 | 0.00000 | 0.05654 | 0.06004 | 0.04173 | 0.02413 | 0.11050 |
| **LOC101928720** | 0.03750 | 0.02653 | 0.01727 | 0.04732 | 0.01847 | 0.01618 | 0.00965 |
| **LOC101928731** | 0.07647 | 0.03740 | 0.02503 | 0.00001 | 0.00535 | 0.01085 | 0.13511 |
| **LOC101928736** | 0.15369 | 0.00000 | 3.04829 | 0.08222 | 4.22433 | 0.00000 | 0.28163 |
| **LOC101928737** | 0.03209 | 0.07470 | 0.01694 | 0.01932 | 0.01729 | 0.02542 | 0.01736 |
| **LOC101928738** | 0.02413 | 0.01941 | 0.01398 | 0.02053 | 0.00800 | 0.01681 | 0.01535 |
| **LOC101928766** | 0.03741 | 0.02233 | 0.00966 | 0.00533 | 0.04761 | 0.04387 | 0.04847 |
| **LOC101928767** | 0.02386 | 0.01914 | 0.01828 | 0.01561 | 0.07486 | 0.10387 | 0.02453 |
| **LOC101928769** | 0.01821 | 0.01860 | 0.02349 | 0.01698 | 1.57077 | 0.01011 | 0.01259 |
| **LOC101928775** | 0.00432 | 0.21471 | 0.43250 | 0.10852 | 0.50175 | 0.02826 | 0.00000 |
| **LOC101928778** | 0.02310 | 0.01741 | 0.01027 | 0.01908 | 0.03504 | 0.01316 | 0.01810 |
| **LOC101928782** | 0.02447 | 0.01795 | 0.02215 | 0.02191 | 0.04727 | 0.01306 | 0.01305 |
| **LOC101928784** | 0.26717 | 0.00000 | 0.44472 | 0.05294 | 0.55605 | 0.10422 | 0.31130 |
| **LOC101928786** | 0.00000 | 0.25824 | 0.13957 | 0.01416 | 0.00617 | 0.06693 | 0.05583 |
| **LOC101928794** | 0.09509 | 0.00000 | 0.01777 | 0.32329 | 0.01247 | 0.11542 | 0.01893 |
| **LOC101928796** | 0.03214 | 0.10130 | 0.13096 | 0.04231 | 0.06848 | 0.00105 | 0.01074 |

| | | | | | | | |
|---|---|---|---|---|---|---|---|
| **LOC101928812** | 0.05346 | 0.02835 | 0.09951 | 0.08680 | 0.00337 | 0.05006 | 0.05354 |
| **LOC101928823** | 0.01565 | 0.05287 | 0.05561 | 0.01609 | 0.49166 | 0.09301 | 0.01879 |
| **LOC101928837** | 0.00164 | 0.02136 | 0.01245 | 0.06818 | 0.00805 | 0.07165 | 0.07077 |
| **LOC101928844** | 0.01805 | 0.00000 | 2.06061 | 0.01415 | 1.00905 | 0.00895 | 0.01075 |
| **LOC101928847** | 0.02616 | 0.01918 | 0.01285 | 0.02383 | 0.00790 | 0.01238 | 0.01508 |
| **LOC101928855** | 0.00000 | 0.04258 | 0.36111 | 0.00010 | 0.00257 | 0.23090 | 1.39942 |
| **LOC101928861** | 0.02357 | 0.01663 | 0.01807 | 0.05834 | 0.00696 | 0.01178 | 0.05412 |
| **LOC101928865** | 0.11724 | 0.00000 | 0.01833 | 0.06821 | 0.06094 | 0.00000 | 0.17057 |
| **LOC101928869** | 0.02317 | 0.02747 | 0.01755 | 0.02169 | 0.00765 | 0.01605 | 0.01643 |
| **LOC101928882** | 0.02405 | 0.01662 | 0.01078 | 0.01794 | 0.02953 | 0.01720 | 0.02013 |
| **LOC101928886** | 0.02829 | 0.02088 | 0.01360 | 0.03487 | 0.00790 | 0.01269 | 0.00762 |
| **LOC101928891** | 0.02138 | 0.01864 | 0.01287 | 0.01057 | 0.00727 | 0.01661 | 0.06294 |
| **LOC101928894** | 1.73071 | 0.21448 | 0.00087 | 0.33815 | 0.02883 | 0.00000 | 0.10546 |
| **LOC101928896** | 0.10014 | 0.03895 | 0.05436 | 0.03172 | 0.04379 | 0.11426 | 0.04990 |
| **LOC101928909** | 0.04332 | 0.00000 | 0.12860 | 0.13464 | 0.00329 | 0.00000 | 0.90810 |
| **LOC101928911** | 0.01688 | 0.05866 | 0.01512 | 0.01774 | 0.01779 | 0.01731 | 0.01669 |
| **LOC101928913** | 0.06395 | 0.02019 | 0.18602 | 0.04949 | 0.02296 | 0.16328 | 0.00000 |
| **LOC101928936** | 0.02605 | 0.01557 | 0.01330 | 0.01339 | 0.00760 | 0.01316 | 0.04690 |
| **LOC101928940** | 0.02405 | 0.05670 | 0.02129 | 0.03276 | 0.05694 | 0.01183 | 0.00030 |
| **LOC101928943** | 0.01915 | 0.01279 | 0.01843 | 0.03148 | 0.00722 | 0.01049 | 0.03715 |
| **LOC101928961** | 0.02419 | 0.01560 | 0.01306 | 0.02491 | 0.00961 | 0.01226 | 0.02434 |
| **LOC101928989** | 0.03050 | 0.02706 | 0.05970 | 0.02057 | 0.15603 | 0.01672 | 0.02601 |
| **LOC101928992** | 0.02139 | 0.02206 | 0.01571 | 0.03393 | 0.00920 | 0.02068 | 0.01250 |
| **LOC101928994** | 0.00306 | 0.02377 | 0.01246 | 0.17487 | 0.04593 | 0.01785 | 0.01781 |
| **LOC101929010** | 0.01205 | 0.00831 | 0.01223 | 0.03921 | 0.00856 | 0.09041 | 0.06067 |
| **LOC101929025** | 0.00963 | 0.02757 | 0.00861 | 0.00721 | 0.00718 | 0.00952 | 0.13404 |
| **LOC101929057** | 0.05021 | 0.04619 | 0.03609 | 0.20183 | 0.04854 | 0.07966 | 0.28195 |
| **LOC101929059** | 0.21511 | 0.14365 | 0.62792 | 0.14194 | 0.05824 | 0.09885 | 0.00000 |
| **LOC101929076** | 0.04150 | 0.02128 | 0.05182 | 0.08262 | 0.96285 | 0.03133 | 0.00000 |
| **LOC101929089** | 0.06156 | 0.01375 | 0.03377 | 0.02001 | 0.00913 | 0.01900 | 0.02041 |
| **LOC101929099** | 0.04947 | 0.02501 | 0.03389 | 0.04656 | 0.00993 | 0.08985 | 0.00000 |
| **LOC101929106** | 0.01743 | 0.06297 | 0.04432 | 0.01349 | 0.00984 | 0.13926 | 0.02064 |
| **LOC101929116** | 0.02141 | 0.01937 | 0.01184 | 0.02362 | 0.00893 | 0.01198 | 0.03283 |
| **LOC101929122** | 0.01967 | 0.01995 | 0.00162 | 0.03644 | 0.20448 | 0.02909 | 0.01223 |
| **LOC101929124** | 0.02454 | 0.07199 | 0.02913 | 0.01663 | 0.16235 | 0.02421 | 0.01309 |

| | | | | | | | |
|---|---|---|---|---|---|---|---|
| **LOC101929125** | 0.02331 | 0.00000 | 0.10868 | 0.19533 | 0.02081 | 0.00789 | 0.05456 |
| **LOC101929140** | 0.04830 | 0.06164 | 0.08193 | 0.18478 | 0.00541 | 0.11972 | 0.32691 |
| **LOC101929144** | 0.01651 | 0.01841 | 0.01317 | 0.04570 | 0.00881 | 0.01800 | 0.01568 |
| **LOC101929153** | 0.21771 | 0.00000 | 0.03866 | 0.05233 | 0.00498 | 0.04309 | 0.07870 |
| **LOC101929162** | 0.04629 | 0.04785 | 0.04108 | 0.05742 | 0.00613 | 0.16463 | 0.50272 |
| **LOC101929173** | 0.01299 | 0.01693 | 0.00880 | 0.01412 | 0.03198 | 0.02127 | 0.07306 |
| **LOC101929181** | 0.00000 | 0.01278 | 0.01627 | 0.14706 | 0.01015 | 0.01230 | 0.07513 |
| **LOC101929196** | 0.01677 | 0.04930 | 0.07665 | 0.01661 | 0.00548 | 0.04355 | 0.01782 |
| **LOC101929207** | 0.02140 | 0.01674 | 0.01292 | 0.02395 | 0.00784 | 0.01448 | 0.02226 |
| **LOC101929218** | 0.02696 | 0.01123 | 0.02188 | 0.02070 | 0.01292 | 0.02231 | 0.01537 |
| **LOC101929221** | 0.02954 | 0.01873 | 0.00952 | 0.02706 | 0.04053 | 0.01345 | 0.00911 |
| **LOC101929224** | 0.05836 | 0.04424 | 0.03749 | 0.02351 | 0.03299 | 0.01180 | 0.01359 |
| **LOC101929225** | 0.16607 | 0.03165 | 0.15146 | 0.08604 | 0.01386 | 0.10123 | 0.56343 |
| **LOC101929231** | 0.10049 | 0.09994 | 0.05316 | 0.18378 | 0.22734 | 0.10115 | 0.00000 |
| **LOC101929234** | 0.05602 | 0.01937 | 0.00533 | 0.02723 | 0.12945 | 0.04266 | 0.03182 |
| **LOC101929259** | 0.02933 | 0.01007 | 0.02756 | 0.01656 | 0.00729 | 0.01857 | 0.06376 |
| **LOC101929260** | 0.02582 | 0.01560 | 0.01729 | 0.02419 | 0.00774 | 0.01284 | 0.01640 |
| **LOC101929279** | 0.02226 | 0.01885 | 0.01401 | 0.01770 | 0.00798 | 0.01356 | 0.02978 |
| **LOC101929295** | 0.01560 | 0.05094 | 0.02140 | 0.04934 | 0.00965 | 0.00890 | 0.01162 |
| **LOC101929302** | 0.02972 | 0.01759 | 0.01302 | 0.02122 | 0.01605 | 0.01193 | 0.01619 |
| **LOC101929319** | 0.02423 | 0.01584 | 0.01152 | 0.02157 | 0.08102 | 0.01342 | 0.01461 |
| **LOC101929331** | 0.02803 | 0.01933 | 0.01247 | 0.02651 | 0.00809 | 0.01598 | 0.00820 |
| **LOC101929335** | 0.03610 | 0.05218 | 0.01033 | 0.01746 | 0.02851 | 0.01076 | 0.03031 |
| **LOC101929340** | 0.12844 | 0.06041 | 0.13382 | 0.15184 | 0.00556 | 0.21495 | 0.05020 |
| **LOC101929344** | 0.00001 | 0.02797 | 0.07440 | 0.12656 | 0.00389 | 2.12862 | 0.01275 |
| **LOC101929372** | 0.08362 | 0.23031 | 0.06399 | 0.07283 | 0.04202 | 0.02430 | 0.00000 |
| **LOC101929374** | 0.01111 | 0.36971 | 0.18312 | 0.02091 | 0.00294 | 0.01556 | 0.00057 |
| **LOC101929380** | 0.01277 | 0.00858 | 0.01401 | 0.05556 | 0.14291 | 0.03457 | 0.04273 |
| **LOC101929395** | 0.02466 | 0.00590 | 0.05222 | 0.03521 | 0.03120 | 0.01199 | 0.01206 |
| **LOC101929397** | 0.13855 | 0.00000 | 1.31482 | 0.80947 | 1.57248 | 0.00000 | 0.06520 |
| **LOC101929412** | 0.04541 | 0.00909 | 0.00246 | 0.04705 | 0.58821 | 0.01625 | 0.00555 |
| **LOC101929413** | 0.03703 | 0.02160 | 0.01341 | 0.03005 | 0.00825 | 0.04850 | 0.00000 |
| **LOC101929420** | 0.01014 | 0.05961 | 0.02747 | 0.03896 | 0.00701 | 0.01240 | 0.08608 |
| **LOC101929431** | 0.01379 | 0.02335 | 0.09124 | 0.05043 | 0.00699 | 0.00826 | 0.04248 |
| **LOC101929439** | 0.01479 | 0.00856 | 0.01350 | 0.01871 | 0.00717 | 0.01341 | 0.08204 |

| | | | | | | | |
|---|---|---|---|---|---|---|---|
| **LOC101929454** | 0.02180 | 0.00998 | 0.01274 | 0.01818 | 0.26388 | 0.01304 | 0.02098 |
| **LOC101929486** | 0.01738 | 0.04015 | 0.03235 | 0.02498 | 0.03363 | 0.05954 | 0.01450 |
| **LOC101929497** | 0.03492 | 0.04161 | 0.01150 | 0.02595 | 0.00762 | 0.00883 | 0.02021 |
| **LOC101929505** | 0.03054 | 0.00929 | 0.01907 | 0.02411 | 0.04448 | 0.01065 | 0.03238 |
| **LOC101929512** | 0.02662 | 0.01312 | 0.02002 | 0.01815 | 0.01909 | 0.01222 | 0.02091 |
| **LOC101929524** | 0.00840 | 0.05047 | 0.06077 | 0.00874 | 2.08592 | 0.00734 | 0.05049 |
| **LOC101929526** | 0.01899 | 0.01474 | 0.01640 | 0.02781 | 0.00750 | 0.01519 | 0.03081 |
| **LOC101929549** | 0.33463 | 0.94177 | 1.36553 | 0.00000 | 0.04988 | 1.64119 | 0.13775 |
| **LOC101929555** | 0.02385 | 0.02598 | 0.01080 | 0.02152 | 0.01447 | 0.01581 | 0.02272 |
| **LOC101929563** | 0.01508 | 0.02454 | 0.05448 | 0.05303 | 0.05246 | 0.05416 | 0.01803 |
| **LOC101929567** | 0.03576 | 0.01789 | 0.01224 | 0.01976 | 0.00927 | 0.01884 | 0.02096 |
| **LOC101929570** | 0.02733 | 0.01176 | 0.01327 | 0.02520 | 0.00962 | 0.01011 | 0.03707 |
| **LOC101929574** | 0.02614 | 0.03432 | 0.08759 | 0.03771 | 0.00427 | 0.03266 | 0.02521 |
| **LOC101929584** | 0.02556 | 0.01589 | 0.01805 | 0.01696 | 0.00776 | 0.01573 | 0.02743 |
| **LOC101929586** | 0.02328 | 0.01806 | 0.01333 | 0.02282 | 0.00806 | 0.01293 | 0.01614 |
| **LOC101929591** | 0.03446 | 0.00001 | 0.02171 | 0.02204 | 0.01778 | 0.08508 | 0.08810 |
| **LOC101929596** | 0.05858 | 0.00758 | 0.01235 | 0.06505 | 0.07424 | 0.08251 | 0.00313 |
| **LOC101929607** | 0.02388 | 0.01861 | 0.01155 | 0.01993 | 0.01679 | 0.01304 | 0.01626 |
| **LOC101929613** | 0.01169 | 0.06445 | 0.01608 | 0.08163 | 0.02213 | 0.00882 | 0.00913 |
| **LOC101929631** | 0.02809 | 0.01586 | 0.01769 | 0.01720 | 0.01229 | 0.01245 | 0.01812 |
| **LOC101929633** | 0.00000 | 0.14902 | 0.01013 | 0.35085 | 0.00768 | 0.07461 | 0.00588 |
| **LOC101929645** | 0.03505 | 0.01172 | 0.01255 | 0.02591 | 0.00862 | 0.02338 | 0.03353 |
| **LOC101929653** | 0.02004 | 0.02525 | 0.00257 | 0.02713 | 0.21775 | 0.01595 | 0.01993 |
| **LOC101929655** | 0.04481 | 0.02360 | 0.07313 | 0.09757 | 0.32296 | 0.00638 | 0.00000 |
| **LOC101929657** | 0.02795 | 0.02459 | 0.03681 | 0.01927 | 0.00593 | 0.45211 | 0.03172 |
| **LOC101929679** | 0.03623 | 0.01985 | 0.01706 | 0.02077 | 0.00700 | 0.01723 | 0.05869 |
| **LOC101929684** | 0.02609 | 0.01734 | 0.01613 | 0.03365 | 0.00774 | 0.01266 | 0.00993 |
| **LOC101929690** | 0.02539 | 0.01666 | 0.01344 | 0.01948 | 0.00894 | 0.01462 | 0.01676 |
| **LOC101929694** | 0.04464 | 0.26036 | 0.12877 | 0.02965 | 0.00766 | 0.10138 | 0.02258 |
| **LOC101929698** | 0.02548 | 0.02135 | 0.01184 | 0.01597 | 0.01944 | 0.01353 | 0.02872 |
| **LOC101929715** | 0.02869 | 0.11093 | 0.11661 | 0.03193 | 0.99018 | 0.00062 | 0.00669 |
| **LOC101929717** | 0.03113 | 0.03146 | 0.01672 | 0.01951 | 0.00748 | 0.02788 | 0.00974 |
| **LOC101929718** | 0.58058 | 0.14336 | 0.15624 | 0.23192 | 0.09399 | 0.42999 | 0.10588 |
| **LOC101929723** | 0.01865 | 0.02561 | 0.00714 | 0.01005 | 0.05499 | 0.03719 | 0.05987 |
| **LOC101929733** | 0.02247 | 0.00541 | 0.06025 | 0.02702 | 0.09625 | 0.06614 | 0.00969 |

| | | | | | | | |
|---|---|---|---|---|---|---|---|
| **LOC101929743** | 0.00000 | 0.58783 | 0.50134 | 0.00000 | 0.00068 | 0.41361 | 0.05986 |
| **LOC101929760** | 0.03345 | 0.01683 | 0.01425 | 0.02294 | 0.01050 | 0.01735 | 0.01262 |
| **LOC101929765** | 0.00000 | 0.01375 | 0.02394 | 0.06994 | 0.03225 | 0.01095 | 0.07793 |
| **LOC101929767** | 0.38786 | 0.13704 | 0.37991 | 0.84922 | 0.00000 | 2.57313 | 0.16048 |
| **LOC101930452** | 0.05267 | 0.00000 | 0.12270 | 0.22982 | 0.12224 | 0.06768 | 0.63098 |
| **LOC102060414** | 0.01185 | 0.04152 | 0.02628 | 0.10444 | 0.03692 | 0.08205 | 0.01058 |
| **LOC102288414** | 5.73187 | 6.18875 | 3.22137 | 5.24567 | 2.40532 | 2.84729 | 0.00017 |
| **LOC102467073** | 0.01787 | 0.07959 | 0.32505 | 0.11939 | 0.15314 | 0.01630 | 0.01562 |
| **LOC102467212** | 0.01809 | 0.01071 | 0.03052 | 0.02788 | 0.00662 | 0.01168 | 0.03127 |
| **LOC102467213** | 0.00818 | 0.17309 | 0.11588 | 0.01507 | 0.01457 | 0.28727 | 0.00852 |
| **LOC102467223** | 0.02975 | 0.00705 | 0.01627 | 0.02856 | 0.24070 | 0.01404 | 0.00880 |
| **LOC102467224** | 0.02719 | 0.01645 | 0.01073 | 0.01790 | 0.04425 | 0.01332 | 0.01663 |
| **LOC102467225** | 0.02601 | 0.01864 | 0.01002 | 0.01751 | 0.03116 | 0.01237 | 0.02035 |
| **LOC102467655** | 0.02437 | 0.01682 | 0.01309 | 0.02025 | 0.00926 | 0.01352 | 0.01818 |
| **LOC102503427** | 0.02422 | 0.01696 | 0.01270 | 0.01968 | 0.02545 | 0.01317 | 0.01568 |
| **LOC102546226** | 0.04140 | 0.00365 | 0.06557 | 0.02104 | 0.01944 | 0.05632 | 0.03863 |
| **LOC102546229** | 0.72075 | 0.00000 | 0.24172 | 0.00000 | 0.11244 | 0.24993 | 0.41643 |
| **LOC102546294** | 0.01406 | 0.02439 | 0.03989 | 0.13683 | 0.38768 | 0.00241 | 0.04289 |
| **LOC102546298** | 0.06178 | 0.02294 | 0.02661 | 0.06954 | 0.04319 | 0.06928 | 0.08484 |
| **LOC102577426** | 0.03932 | 0.03747 | 0.05837 | 0.02220 | 0.02455 | 0.00741 | 0.01018 |
| **LOC102606465** | 0.57049 | 0.14470 | 0.22842 | 0.13193 | 0.15946 | 0.01275 | 0.24999 |
| **LOC102659288** | 0.02063 | 0.01580 | 0.00000 | 0.02058 | 0.70201 | 0.01365 | 0.01420 |
| **LOC102723344** | 0.04463 | 0.01321 | 0.01292 | 0.01738 | 0.04194 | 0.01227 | 0.01075 |
| **LOC102723354** | 0.00000 | 0.92598 | 1.36222 | 0.03393 | 0.03839 | 0.03007 | 0.02241 |
| **LOC102723373** | 0.06350 | 0.00000 | 0.02560 | 0.09284 | 0.03033 | 0.00517 | 0.04332 |
| **LOC102723385** | 0.04510 | 0.20230 | 0.14236 | 0.02293 | 0.02511 | 0.16362 | 0.11674 |
| **LOC102723439** | 0.13947 | 0.00000 | 0.00905 | 0.05042 | 0.03625 | 0.00000 | 0.40069 |
| **LOC102723483** | 0.03334 | 0.03180 | 0.04408 | 0.03777 | 0.02520 | 0.01360 | 0.00696 |
| **LOC102723487** | 0.03217 | 0.01872 | 0.01391 | 0.01540 | 0.01893 | 0.01079 | 0.02567 |
| **LOC102723505** | 0.03276 | 0.01379 | 0.01646 | 0.01637 | 0.04028 | 0.01977 | 0.01322 |
| **LOC102723510** | 0.17129 | 0.14486 | 0.18411 | 0.40505 | 0.03772 | 0.01341 | 0.00000 |
| **LOC102723513** | 0.02413 | 0.02403 | 0.01984 | 0.01953 | 0.01282 | 0.01174 | 0.01402 |
| **LOC102723517** | 0.02327 | 0.01229 | 0.01444 | 0.01837 | 0.41084 | 0.01268 | 0.01575 |
| **LOC102723562** | 0.02534 | 0.01728 | 0.01278 | 0.01983 | 0.00796 | 0.01311 | 0.01968 |
| **LOC102723631** | 0.02256 | 0.00816 | 0.13654 | 0.02040 | 0.05293 | 0.00849 | 0.01559 |

| | | | | | | | |
|---|---|---|---|---|---|---|---|
| **LOC102723641** | 0.02535 | 0.01815 | 0.01884 | 0.02076 | 0.00759 | 0.01362 | 0.01670 |
| **LOC102723644** | 0.00531 | 0.03279 | 0.00828 | 0.01146 | 0.01190 | 0.01594 | 0.09063 |
| **LOC102723703** | 0.05926 | 0.39151 | 0.46520 | 0.09600 | 0.09259 | 0.29050 | 0.47074 |
| **LOC102723742** | 0.02430 | 0.01765 | 0.01042 | 0.01877 | 0.02499 | 0.01767 | 0.01733 |
| **LOC102723769** | 0.05058 | 0.03450 | 0.03445 | 0.01892 | 0.01199 | 0.03793 | 0.11834 |
| **LOC102723809** | 0.02731 | 0.01806 | 0.01824 | 0.02701 | 0.01426 | 0.00975 | 0.02954 |
| **LOC102723824** | 0.02997 | 0.01611 | 0.01280 | 0.01515 | 0.01625 | 0.01325 | 0.03373 |
| **LOC102723854** | 0.26521 | 0.12724 | 0.00000 | 0.84962 | 0.22571 | 0.05591 | 0.01561 |
| **LOC102723885** | 0.41667 | 0.43341 | 0.03547 | 0.71659 | 0.06155 | 0.36824 | 0.00589 |
| **LOC102723895** | 0.02242 | 0.01637 | 0.01331 | 0.02045 | 0.00794 | 0.03257 | 0.01599 |
| **LOC102723927** | 0.00828 | 0.02673 | 0.00950 | 0.01837 | 0.00783 | 0.09147 | 0.03710 |
| **LOC102724009** | 0.01147 | 0.20286 | 0.01473 | 0.04617 | 0.04160 | 0.03628 | 0.05541 |
| **LOC102724020** | 0.03568 | 0.01904 | 0.04025 | 0.02225 | 0.04172 | 0.01430 | 0.00267 |
| **LOC102724034.** | 0.02432 | 0.01782 | 0.01197 | 0.01902 | 0.01461 | 0.01318 | 0.01684 |
| **LOC102724050** | 0.00505 | 0.00000 | 0.44505 | 0.12136 | 0.06456 | 0.00000 | 0.57989 |
| **LOC102724058** | 0.01783 | 0.03047 | 0.02158 | 0.02099 | 0.00997 | 0.01121 | 0.02055 |
| **LOC102724084** | 0.03335 | 0.01215 | 0.01560 | 0.01711 | 0.03339 | 0.01513 | 0.01529 |
| **LOC102724094** | 0.02106 | 0.00000 | 0.03704 | 0.03885 | 0.37265 | 0.04473 | 0.01470 |
| **LOC102724153** | 0.00000 | 0.08584 | 2.03512 | 0.00000 | 0.24507 | 0.00000 | 0.31921 |
| **LOC102724163** | 0.02414 | 0.01849 | 0.01268 | 0.01929 | 0.00909 | 0.01277 | 0.01869 |
| **LOC102724190** | 0.01448 | 0.04827 | 0.01333 | 0.02064 | 0.01222 | 0.02474 | 0.01559 |
| **LOC102724246** | 0.01720 | 0.03395 | 0.01123 | 0.03015 | 0.00782 | 0.01111 | 0.02065 |
| **LOC102724301** | 0.02937 | 0.00027 | 0.05153 | 0.65234 | 0.17548 | 0.00750 | 0.00000 |
| **LOC102724316** | 6.23077 | 4.20891 | 0.72197 | 4.41660 | 0.25791 | 1.67884 | 0.98856 |
| **LOC102724321** | 0.02355 | 0.01877 | 0.01194 | 0.01915 | 0.02738 | 0.01286 | 0.01683 |
| **LOC102724323** | 0.02014 | 0.05279 | 0.02475 | 0.01877 | 0.01567 | 0.03541 | 0.00911 |
| **LOC102724467** | 0.06433 | 0.01495 | 0.01417 | 0.02183 | 0.04320 | 0.01854 | 0.07636 |
| **LOC102724484** | 0.01944 | 0.01181 | 0.02663 | 0.05967 | 0.00687 | 0.01824 | 0.01611 |
| **LOC102724508** | 0.03238 | 0.04137 | 0.03058 | 0.02008 | 0.14902 | 0.02810 | 0.02024 |
| **LOC102724550** | 0.01956 | 0.01868 | 0.00674 | 0.05185 | 0.07382 | 0.01315 | 0.02722 |
| **LOC102724589** | 0.02776 | 0.01799 | 0.01173 | 0.01732 | 0.01406 | 0.01943 | 0.01971 |
| **LOC102724596** | 0.02698 | 0.04262 | 0.02975 | 0.02056 | 0.01462 | 0.00730 | 0.02720 |
| **LOC102724601** | 0.01034 | 0.01098 | 0.05371 | 0.05394 | 0.03500 | 0.03610 | 0.05295 |
| **LOC102724691** | 0.04507 | 0.03791 | 0.01972 | 0.07788 | 0.01496 | 0.02186 | 0.00000 |
| **LOC102724719** | 0.02318 | 0.02084 | 0.01420 | 0.01979 | 0.03416 | 0.01558 | 0.01797 |

| | | | | | | | |
|---|---|---|---|---|---|---|---|
| **LOC102724784** | 0.00250 | 0.14004 | 0.07848 | 0.00454 | 0.00485 | 0.01720 | 0.39660 |
| **LOC102724814** | 0.26871 | 0.23115 | 0.12846 | 0.80396 | 0.00025 | 0.40227 | 0.45449 |
| **LOC102724890** | 0.02343 | 0.01709 | 0.01127 | 0.02053 | 0.04359 | 0.01322 | 0.01750 |
| **LOC102724927** | 0.01517 | 0.05222 | 0.00568 | 0.03041 | 0.06602 | 0.00274 | 0.08238 |
| **LOC102724933** | 0.02329 | 0.02450 | 0.02382 | 0.01821 | 0.00905 | 0.01030 | 0.02538 |
| **LOC102724957** | 0.02645 | 0.02531 | 0.00898 | 0.02761 | 0.05508 | 0.01131 | 0.01416 |
| **LOC102800447** | 0.02590 | 0.00653 | 0.04447 | 0.01848 | 0.00765 | 0.01266 | 0.02420 |
| **LOC103091866** | 0.02355 | 0.01499 | 0.01313 | 0.01672 | 0.02825 | 0.01599 | 0.03277 |
| **LOC103191607** | 0.03101 | 0.01464 | 0.02903 | 0.02864 | 0.00679 | 0.01300 | 0.01115 |
| **LOC103344931** | 0.09756 | 0.01726 | 0.03784 | 0.22035 | 0.00719 | 0.28798 | 0.50511 |
| **LOC113230** | 0.00000 | 0.05070 | 0.13359 | 0.15368 | 0.24865 | 0.02281 | 0.20809 |
| **LOC143666** | 0.01051 | 0.04974 | 0.01809 | 0.10545 | 0.01703 | 0.09690 | 0.10085 |
| **LOC145474** | 0.42556 | 0.00000 | 0.12962 | 0.10392 | 0.19626 | 0.21307 | 1.72592 |
| **LOC145783** | 0.06631 | 0.46596 | 0.09760 | 0.53703 | 0.25345 | 0.42651 | 0.00000 |
| **LOC145837** | 0.03594 | 0.02580 | 0.11154 | 0.10621 | 0.60299 | 0.07101 | 0.00000 |
| **LOC145845** | 0.02379 | 0.01545 | 0.01200 | 0.01476 | 0.00970 | 0.01134 | 0.04330 |
| **LOC146513** | 0.02736 | 0.01404 | 0.01770 | 0.02067 | 0.00834 | 0.02994 | 0.06293 |
| **LOC146880** | 0.49722 | 0.39575 | 0.39908 | 0.49179 | 0.10027 | 0.36368 | 1.83838 |
| **LOC148413** | 4.96889 | 8.52976 | 6.67433 | 8.63056 | 4.49333 | 2.32356 | 0.00000 |
| **LOC148696** | 0.01853 | 0.01021 | 0.00000 | 0.01189 | 0.78992 | 0.00000 | 0.19056 |
| **LOC148709** | 1.34640 | 0.80429 | 0.00000 | 0.79526 | 0.71220 | 0.14278 | 0.73064 |
| **LOC149950** | 0.02372 | 0.01770 | 0.01079 | 0.01966 | 0.02412 | 0.01294 | 0.01877 |
| **LOC150776** | 1.32083 | 1.10534 | 1.68588 | 2.03523 | 0.54858 | 2.31536 | 4.07943 |
| **LOC151121** | 0.03010 | 0.01711 | 0.00621 | 0.03691 | 0.03005 | 0.13228 | 0.15027 |
| **LOC151174** | 0.25616 | 0.13753 | 0.00883 | 0.11444 | 0.04785 | 0.21991 | 0.03862 |
| **LOC151475** | 0.01231 | 0.02543 | 0.00953 | 0.21466 | 0.01641 | 0.01120 | 0.01667 |
| **LOC151484** | 0.02141 | 0.01980 | 0.01347 | 0.02313 | 0.00783 | 0.01336 | 0.01713 |
| **LOC152225** | 0.03663 | 0.01164 | 0.02179 | 0.03242 | 0.05319 | 0.01495 | 0.00926 |
| **LOC152578** | 0.05792 | 0.01389 | 0.03082 | 0.01482 | 0.00773 | 0.02075 | 0.04704 |
| **LOC153684** | 0.01759 | 0.00000 | 0.19547 | 0.01989 | 0.04126 | 0.02558 | 0.02813 |
| **LOC154761** | 0.02549 | 0.01664 | 0.01287 | 0.02517 | 0.00914 | 0.01235 | 0.02008 |
| **LOC155060** | 0.13787 | 0.16944 | 0.07929 | 0.00000 | 0.01141 | 0.02693 | 0.59913 |
| **LOC157273** | 0.08130 | 0.01559 | 0.04357 | 0.02816 | 0.00635 | 0.02093 | 0.00000 |
| **LOC158434** | 0.05057 | 0.01097 | 0.02077 | 0.01164 | 0.01437 | 0.01190 | 0.01628 |
| **LOC158435** | 0.04362 | 0.00490 | 0.03537 | 0.01650 | 0.00669 | 0.03889 | 0.00606 |

| | | | | | | | |
|---|---|---|---|---|---|---|---|
| **LOC158960** | 1.28360 | 2.94013 | 0.34570 | 0.26197 | 0.12172 | 0.03832 | 0.65360 |
| **LOC1720** | 0.02655 | 0.02267 | 0.01253 | 0.01912 | 0.00785 | 0.01218 | 0.01594 |
| **LOC200726** | 0.03261 | 0.01852 | 0.01296 | 0.01716 | 0.00786 | 0.01490 | 0.02033 |
| **LOC202181** | 0.53041 | 0.43057 | 0.24463 | 0.17550 | 0.09055 | 0.41183 | 0.39745 |
| **LOC220729** | 2.32221 | 1.58871 | 2.04519 | 1.00246 | 1.14613 | 1.33696 | 2.06614 |
| **LOC253573** | 0.06932 | 0.00119 | 0.02300 | 0.00730 | 0.00672 | 0.01063 | 0.04902 |
| **LOC254028** | 0.04346 | 0.00943 | 0.03472 | 0.07675 | 0.02035 | 0.06068 | 0.00000 |
| **LOC254896** | 0.03090 | 0.01393 | 0.02162 | 0.01162 | 0.00893 | 0.00922 | 0.05734 |
| **LOC255187** | 0.02948 | 0.00000 | 0.05601 | 0.08667 | 0.00587 | 0.02016 | 0.03174 |
| **LOC256880** | 0.30523 | 0.00000 | 0.15043 | 0.06980 | 3.49682 | 0.17432 | 0.04117 |
| **LOC257396** | 0.20786 | 0.14051 | 0.19730 | 0.09152 | 0.39142 | 0.00737 | 0.13825 |
| **LOC283038** | 0.00840 | 0.04223 | 0.24226 | 0.01886 | 0.00415 | 0.04533 | 0.14794 |
| **LOC283045** | 0.02592 | 0.01770 | 0.01290 | 0.02628 | 0.00795 | 0.01268 | 0.01487 |
| **LOC283070** | 0.02181 | 0.03123 | 0.11318 | 0.01806 | 0.02278 | 0.12394 | 0.03960 |
| **LOC283177** | 0.02290 | 0.02193 | 0.02172 | 0.01842 | 0.00964 | 0.01140 | 0.02219 |
| **LOC283194** | 0.02105 | 0.02026 | 0.01253 | 0.03307 | 0.00829 | 0.02152 | 0.01064 |
| **LOC283299** | 0.04873 | 0.02386 | 0.27531 | 0.29676 | 6.77123 | 0.00758 | 0.00000 |
| **LOC283335** | 0.23640 | 0.27075 | 0.01398 | 0.37183 | 0.03430 | 0.00438 | 0.00000 |
| **LOC283352** | 0.05567 | 0.00000 | 0.09394 | 0.01212 | 0.01605 | 0.03969 | 0.04154 |
| **LOC283440** | 0.02826 | 0.01330 | 0.01987 | 0.02070 | 0.00852 | 0.01499 | 0.01439 |
| **LOC283575** | 0.02977 | 0.02192 | 0.01642 | 0.02753 | 0.00800 | 0.02140 | 0.01187 |
| **LOC283585** | 0.02406 | 0.01693 | 0.01624 | 0.02119 | 0.01228 | 0.01305 | 0.01672 |
| **LOC283683** | 0.03695 | 0.09979 | 0.02019 | 0.09679 | 0.03616 | 0.29888 | 0.00272 |
| **LOC283692** | 0.09882 | 0.56738 | 0.20285 | 0.28880 | 0.22907 | 0.25226 | 0.44721 |
| **LOC283693** | 0.03694 | 0.13819 | 0.03676 | 0.05947 | 0.04460 | 0.01685 | 0.02809 |
| **LOC283731** | 0.02309 | 0.01670 | 0.01317 | 0.01968 | 0.00782 | 0.01872 | 0.02671 |
| **LOC283856** | 0.02806 | 0.02478 | 0.00882 | 0.02480 | 0.03402 | 0.01211 | 0.02527 |
| **LOC283914** | 0.00704 | 0.01410 | 0.01312 | 0.06164 | 0.00780 | 0.01050 | 0.05384 |
| **LOC283922** | 0.11139 | 0.00000 | 0.28251 | 0.19785 | 0.04642 | 0.07875 | 0.00000 |
| **LOC284009** | 0.05532 | 0.19147 | 0.22583 | 0.12038 | 0.00610 | 0.00626 | 0.20938 |
| **LOC284023** | 0.53599 | 0.19730 | 0.93769 | 0.09768 | 1.92087 | 0.03649 | 0.17533 |
| **LOC284294** | 0.01975 | 0.03694 | 0.00803 | 0.01315 | 0.10640 | 0.01181 | 0.04527 |
| **LOC284344** | 0.01774 | 0.01747 | 0.01310 | 0.03451 | 0.00784 | 0.01179 | 0.02659 |
| **LOC284379** | 0.01231 | 0.01762 | 0.01436 | 2.68431 | 0.00942 | 0.00953 | 0.00402 |
| **LOC284412** | 0.19188 | 0.01604 | 0.07338 | 0.07123 | 0.01582 | 0.20468 | 0.18191 |

| | | | | | | | |
|---|---|---|---|---|---|---|---|
| **LOC284454** | 0.55517 | 1.54985 | 0.00000 | 4.29776 | 0.05572 | 0.60279 | 1.91340 |
| **LOC284578** | 0.02561 | 0.00000 | 0.11099 | 0.02454 | 0.10087 | 0.02491 | 0.01417 |
| **LOC284581** | 0.08959 | 0.00000 | 0.63565 | 0.15892 | 0.28148 | 0.14071 | 0.16369 |
| **LOC284632** | 0.02459 | 0.01667 | 0.01257 | 0.02396 | 0.00914 | 0.03048 | 0.01403 |
| **LOC284648** | 0.03774 | 0.01270 | 0.01460 | 0.07221 | 0.00979 | 0.01670 | 0.00000 |
| **LOC284798** | 0.02816 | 0.01445 | 0.01382 | 0.01939 | 0.08274 | 0.01320 | 0.01309 |
| **LOC284837** | 0.00000 | 0.00000 | 0.29302 | 0.24389 | 0.00507 | 0.40266 | 1.15765 |
| **LOC284865** | 0.02303 | 0.00764 | 0.04657 | 0.02211 | 0.00640 | 0.01854 | 0.01938 |
| **LOC284889** | 11.53258 | 31.65063 | 4.24829 | 16.53358 | 1.89456 | 18.26945 | 8.63638 |
| **LOC284930** | 0.23759 | 0.33450 | 0.28014 | 0.07115 | 0.19386 | 0.18731 | 0.00719 |
| **LOC284950** | 0.02164 | 0.01755 | 0.00715 | 0.02034 | 0.07909 | 0.01418 | 0.01873 |
| **LOC285000** | 0.02823 | 0.02004 | 0.01184 | 0.02649 | 0.00801 | 0.01882 | 0.01402 |
| **LOC285043** | 0.01143 | 0.01927 | 0.03977 | 0.02210 | 0.00594 | 0.07908 | 0.03285 |
| **LOC285074** | 0.04880 | 0.01040 | 0.04909 | 0.08194 | 0.13915 | 0.18532 | 0.22133 |
| **LOC285484** | 0.01410 | 0.06018 | 0.05165 | 0.01086 | 0.00442 | 0.00000 | 0.14015 |
| **LOC285626** | 0.06096 | 0.01498 | 0.01319 | 0.03098 | 0.00829 | 0.01365 | 0.00351 |
| **LOC285629** | 0.35798 | 0.00000 | 0.01749 | 0.05847 | 0.03233 | 0.01818 | 0.00000 |
| **LOC285692** | 0.02007 | 0.02032 | 0.01152 | 0.03866 | 0.01783 | 0.01371 | 0.01286 |
| **LOC285696** | 0.07218 | 0.07353 | 0.05956 | 0.16342 | 0.00492 | 0.00000 | 0.06463 |
| **LOC285740** | 0.04381 | 0.00000 | 0.03868 | 0.02032 | 0.03558 | 0.15656 | 0.02011 |
| **LOC285762** | 0.02015 | 0.02362 | 0.01024 | 0.02111 | 0.00770 | 0.00872 | 0.05152 |
| **LOC285819** | 0.04534 | 0.00824 | 0.02713 | 0.01705 | 0.00788 | 0.01385 | 0.01618 |
| **LOC285847** | 0.03191 | 0.02442 | 0.01066 | 0.01791 | 0.01020 | 0.03433 | 0.00861 |
| **LOC285889** | 0.01760 | 0.01411 | 0.00000 | 0.01695 | 1.93115 | 0.01179 | 0.01690 |
| **LOC286059** | 0.01989 | 0.02176 | 0.00927 | 0.02509 | 0.06290 | 0.01183 | 0.03071 |
| **LOC286083** | 0.03183 | 0.10268 | 0.00631 | 0.02162 | 0.02261 | 0.03637 | 0.00360 |
| **LOC286189** | 0.01984 | 0.03091 | 0.02625 | 0.02040 | 0.13821 | 0.01126 | 0.01409 |
| **LOC286190** | 0.02019 | 0.17449 | 0.06953 | 0.02093 | 0.00473 | 0.05368 | 0.04129 |
| **LOC286238** | 0.01867 | 0.01224 | 0.09948 | 0.01877 | 0.01781 | 0.01897 | 0.02147 |
| **LOC286359** | 0.02193 | 0.01484 | 0.02687 | 0.01817 | 0.03872 | 0.01422 | 0.02246 |
| **LOC286367** | 0.11085 | 0.00060 | 0.02453 | 0.18633 | 0.00690 | 0.01498 | 0.03729 |
| **LOC286437** | 0.26866 | 0.14641 | 0.00000 | 0.42025 | 0.77663 | 0.00000 | 0.55225 |
| **LOC338963** | 0.02566 | 0.02170 | 0.01532 | 0.02513 | 0.02399 | 0.01277 | 0.01491 |
| **LOC339059** | 0.02234 | 0.06170 | 0.00641 | 0.00954 | 0.00733 | 0.00609 | 0.08667 |
| **LOC339166** | 0.00467 | 0.44923 | 0.00000 | 0.11048 | 0.57502 | 0.34663 | 0.00000 |

| | | | | | | | |
|---|---|---|---|---|---|---|---|
| **LOC339298** | 0.02576 | 0.01815 | 0.01280 | 0.01896 | 0.00802 | 0.01389 | 0.01576 |
| **LOC339529** | 0.03554 | 0.01929 | 0.01299 | 0.03152 | 0.00803 | 0.01613 | 0.01017 |
| **LOC339539** | 0.13364 | 0.02598 | 0.01006 | 0.05969 | 0.00719 | 0.03752 | 0.09025 |
| **LOC339622** | 0.02535 | 0.01804 | 0.01279 | 0.02546 | 0.00798 | 0.01384 | 0.01331 |
| **LOC339666** | 0.01439 | 0.06078 | 0.02047 | 0.04897 | 0.00583 | 0.00000 | 0.13399 |
| **LOC339803** | 2.59759 | 2.61853 | 2.67002 | 3.01143 | 0.26706 | 2.40235 | 4.10388 |
| **LOC339807** | 0.06252 | 0.01276 | 0.01388 | 0.02706 | 0.00809 | 0.01056 | 0.00417 |
| **LOC339862** | 0.09204 | 0.03470 | 0.17546 | 0.08840 | 0.02343 | 0.07294 | 0.04352 |
| **LOC339874** | 0.08943 | 0.02979 | 0.01870 | 0.00935 | 0.12075 | 0.03263 | 0.02763 |
| **LOC339975** | 0.03527 | 0.16191 | 0.07628 | 0.11337 | 0.00747 | 0.00000 | 0.15909 |
| **LOC340113** | 0.02741 | 0.01776 | 0.01169 | 0.02060 | 0.01516 | 0.01246 | 0.01632 |
| **LOC340357** | 0.02151 | 0.01496 | 0.01002 | 0.01800 | 0.07447 | 0.02790 | 0.02063 |
| **LOC340515** | 0.02408 | 0.02068 | 0.01072 | 0.03175 | 0.01995 | 0.02485 | 0.00570 |
| **LOC341056** | 1.45605 | 0.29066 | 0.42181 | 1.11574 | 0.32230 | 0.52375 | 0.00000 |
| **LOC344887** | 0.02516 | 0.01711 | 0.00611 | 0.01627 | 0.08732 | 0.01422 | 0.02984 |
| **LOC344967** | 1.43710 | 1.58722 | 0.07088 | 1.79949 | 0.11783 | 0.63311 | 0.00000 |
| **LOC349160** | 0.06062 | 0.03194 | 0.03769 | 0.04990 | 0.00888 | 0.01149 | 0.06480 |
| **LOC374443** | 0.58969 | 0.39903 | 0.76492 | 0.84244 | 0.47557 | 0.47429 | 0.28514 |
| **LOC375196** | 0.06219 | 0.03420 | 0.90659 | 0.07256 | 0.55838 | 0.00788 | 0.00000 |
| **LOC387810** | 0.03723 | 0.01518 | 0.01165 | 0.01675 | 0.02392 | 0.01618 | 0.01121 |
| **LOC388282** | 0.00953 | 0.01575 | 0.01234 | 0.06616 | 0.00770 | 0.00864 | 0.05399 |
| **LOC388436+LO** | 0.01966 | 0.02652 | 0.04737 | 0.02094 | 0.03466 | 0.02029 | 0.01735 |
| **LOC388553** | 0.47784 | 1.85769 | 0.71273 | 0.24643 | 0.06354 | 0.03227 | 0.07284 |
| **LOC388692** | 0.03264 | 0.01394 | 0.01604 | 0.01694 | 0.00951 | 0.01167 | 0.02226 |
| **LOC388780** | 0.01051 | 0.13227 | 0.94179 | 0.03870 | 0.33695 | 0.01268 | 0.00851 |
| **LOC388813** | 0.02546 | 0.01763 | 0.00232 | 0.01904 | 0.22442 | 0.01457 | 0.01370 |
| **LOC388882** | 0.02553 | 0.09127 | 0.05677 | 0.01848 | 0.00883 | 0.01553 | 0.00720 |
| **LOC388942** | 0.02313 | 0.01749 | 0.01197 | 0.02012 | 0.01997 | 0.01275 | 0.01917 |
| **LOC389033** | 0.02406 | 0.09376 | 0.00827 | 0.02347 | 0.01518 | 0.01222 | 0.01394 |
| **LOC389247** | 0.00207 | 0.08438 | 0.22805 | 0.11162 | 0.10910 | 0.07247 | 0.29544 |
| **LOC389332** | 0.11974 | 0.28579 | 0.00000 | 0.01508 | 0.01178 | 0.00002 | 0.14594 |
| **LOC389641** | 0.05328 | 0.02239 | 0.03797 | 0.03374 | 0.01016 | 0.00816 | 0.00993 |
| **LOC389705** | 0.01333 | 0.05458 | 0.06655 | 0.01445 | 0.34218 | 0.00479 | 0.04521 |
| **LOC389765** | 0.01340 | 0.07821 | 0.15433 | 0.08928 | 0.00829 | 0.11596 | 0.30367 |
| **LOC389895** | 0.01298 | 0.00428 | 0.13132 | 0.02506 | 0.08349 | 0.04227 | 0.02251 |

| | | | | | | | |
|---|---|---|---|---|---|---|---|
| **LOC389906** | 0.14926 | 1.09223 | 0.33908 | 0.35474 | 0.16213 | 0.56188 | 3.56995 |
| **LOC391003.uid-** | 0.00000 | 1.26933 | 3.04346 | 0.00000 | 3.18844 | 0.00000 | 0.00002 |
| **LOC391322** | 0.26825 | 0.04973 | 0.09161 | 0.36690 | 0.06409 | 0.15923 | 0.00000 |
| **LOC392232** | 0.02214 | 0.02908 | 0.02567 | 0.02227 | 0.00972 | 0.01206 | 0.01582 |
| **LOC392364** | 0.05241 | 0.01679 | 0.02300 | 0.02442 | 0.00732 | 0.01395 | 0.00812 |
| **LOC392452** | 0.02479 | 0.01682 | 0.01563 | 0.01908 | 0.00786 | 0.01276 | 0.01762 |
| **LOC399715** | 0.00767 | 0.16061 | 0.33278 | 0.01239 | 1.52921 | 0.43883 | 0.00229 |
| **LOC399815** | 2.91162 | 0.21372 | 0.62682 | 0.87656 | 0.02703 | 0.00001 | 0.59951 |
| **LOC399886** | 0.02451 | 0.01467 | 0.01993 | 0.02044 | 0.01064 | 0.01303 | 0.01585 |
| **LOC400043** | 0.06682 | 0.51430 | 0.98230 | 0.17816 | 0.00000 | 2.44141 | 0.58309 |
| **LOC400511** | 0.03691 | 0.00965 | 0.01409 | 0.01095 | 0.00901 | 0.03778 | 0.05416 |
| **LOC400548** | 0.02486 | 0.01813 | 0.01412 | 0.01906 | 0.00868 | 0.01330 | 0.01756 |
| **LOC400558** | 0.00771 | 0.02389 | 0.01066 | 0.02752 | 0.00765 | 0.01585 | 0.04887 |
| **LOC400620** | 0.02258 | 0.02128 | 0.01562 | 0.01892 | 0.08072 | 0.01282 | 0.01622 |
| **LOC400655** | 4.01185 | 1.03859 | 0.68397 | 4.72204 | 0.20158 | 1.20032 | 0.00000 |
| **LOC400661** | 0.05764 | 0.00785 | 0.02863 | 0.00710 | 0.08725 | 0.00999 | 0.02087 |
| **LOC400684** | 0.01679 | 0.16780 | 0.12504 | 0.12646 | 0.03437 | 0.01041 | 0.00000 |
| **LOC400685** | 0.03191 | 0.00867 | 0.03790 | 0.02175 | 0.11306 | 0.02057 | 0.00747 |
| **LOC400736+PR** | 0.02342 | 0.01774 | 0.01262 | 0.01966 | 0.02522 | 0.01295 | 0.01718 |
| **LOC400736+PR** | 0.02251 | 0.06094 | 0.00000 | 0.01399 | 0.72295 | 0.01230 | 0.01163 |
| **LOC400794** | 0.02722 | 0.01444 | 0.01039 | 0.04403 | 0.02904 | 0.13781 | 0.01471 |
| **LOC400863** | 0.04714 | 0.01501 | 0.06111 | 0.02264 | 0.00604 | 0.00775 | 0.04327 |
| **LOC400867** | 0.02170 | 0.02314 | 0.00213 | 0.02116 | 0.25977 | 0.01233 | 0.01660 |
| **LOC400891** | 0.00630 | 0.03444 | 0.02599 | 0.11782 | 0.00778 | 0.02504 | 0.02119 |
| **LOC400927** | 0.04728 | 0.13447 | 0.11475 | 0.08110 | 0.05237 | 0.06644 | 0.00000 |
| **LOC400940** | 0.04079 | 0.01934 | 0.01237 | 0.02029 | 0.00820 | 0.01562 | 0.00568 |
| **LOC400958** | 0.00000 | 0.21064 | 0.14815 | 0.03160 | 0.00654 | 0.01678 | 0.00907 |
| **LOC400997** | 0.31637 | 0.05185 | 0.01819 | 0.09183 | 0.00501 | 0.00000 | 0.27036 |
| **LOC401010** | 1.43708 | 2.36524 | 1.28150 | 2.38032 | 0.77476 | 0.80061 | 0.34482 |
| **LOC401052** | 0.08850 | 0.03622 | 0.04993 | 0.03493 | 0.00620 | 0.00453 | 0.16000 |
| **LOC401127** | 0.77898 | 1.75126 | 1.31821 | 0.40341 | 0.20716 | 0.88308 | 1.63480 |
| **LOC401177** | 0.02986 | 0.07292 | 0.00616 | 0.02178 | 0.03599 | 0.00557 | 0.02609 |
| **LOC401242** | 0.03770 | 0.05278 | 0.03879 | 0.05214 | 0.00961 | 0.04272 | 0.11385 |
| **LOC401286** | 0.05073 | 0.00606 | 0.01468 | 0.00729 | 0.00743 | 0.00865 | 0.04500 |
| **LOC401312** | 0.48285 | 0.62958 | 0.53731 | 0.41824 | 0.00000 | 0.00000 | 0.93567 |

| | | | | | | | |
|---|---|---|---|---|---|---|---|
| **LOC401320** | 0.18475 | 0.00000 | 0.69136 | 0.28242 | 0.01268 | 0.21152 | 0.26634 |
| **LOC401463** | 0.06538 | 0.03117 | 0.08784 | 0.00961 | 0.00490 | 0.02228 | 0.00000 |
| **LOC402160** | 0.02650 | 0.01475 | 0.01419 | 0.02718 | 0.03754 | 0.01175 | 0.01874 |
| **LOC403323** | 0.16421 | 0.09969 | 0.00467 | 0.05136 | 0.01967 | 0.03945 | 0.00000 |
| **LOC407835** | 0.28898 | 1.34310 | 0.15100 | 0.78545 | 0.16803 | 0.54535 | 1.82325 |
| **LOC414300** | 0.03898 | 0.00847 | 0.01740 | 0.04551 | 0.04337 | 0.05083 | 0.00496 |
| **LOC439933** | 0.00984 | 0.01797 | 0.01549 | 0.03743 | 0.01496 | 0.01263 | 0.04583 |
| **LOC439994** | 0.05267 | 0.31014 | 0.06373 | 0.10684 | 0.33709 | 0.03675 | 0.03547 |
| **LOC440028** | 0.00000 | 0.60476 | 1.50196 | 0.00957 | 0.20650 | 0.00000 | 0.05607 |
| **LOC440040** | 0.02904 | 0.06226 | 0.01606 | 0.01947 | 0.01199 | 0.01471 | 0.00518 |
| **LOC440117** | 0.02547 | 0.01756 | 0.01298 | 0.01852 | 0.00842 | 0.01310 | 0.01848 |
| **LOC440173** | 0.06643 | 0.00370 | 0.13420 | 0.04013 | 0.01952 | 0.01865 | 0.20062 |
| **LOC440300** | 0.01964 | 0.03190 | 0.00940 | 0.01750 | 0.00790 | 0.02478 | 0.03109 |
| **LOC440311** | 0.00000 | 1.01262 | 0.22268 | 0.04053 | 0.27841 | 0.67583 | 2.94757 |
| **LOC440416** | 1.15592 | 0.00000 | 1.39525 | 2.34791 | 0.36744 | 0.00000 | 1.09645 |
| **LOC440434** | 1.24476 | 0.00000 | 1.14483 | 1.14039 | 0.35180 | 1.55828 | 2.36896 |
| **LOC440446** | 0.02288 | 0.01669 | 0.01044 | 0.02207 | 0.04831 | 0.01329 | 0.01719 |
| **LOC440461** | 0.01441 | 0.02869 | 0.01141 | 0.02278 | 0.00775 | 0.02675 | 0.02954 |
| **LOC440600** | 0.42970 | 0.28167 | 0.50592 | 0.29635 | 0.09179 | 0.11743 | 0.04785 |
| **LOC440602** | 0.02616 | 0.01443 | 0.01299 | 0.02180 | 0.01166 | 0.03366 | 0.01134 |
| **LOC440700** | 0.03230 | 0.01410 | 0.01411 | 0.03619 | 0.09991 | 0.01565 | 0.01045 |
| **LOC440896** | 0.01780 | 0.01609 | 0.01325 | 0.02850 | 0.00802 | 0.01186 | 0.02754 |
| **LOC440910** | 0.04902 | 0.00698 | 0.06589 | 0.02427 | 0.04526 | 0.00025 | 0.11237 |
| **LOC441081** | 0.02389 | 0.01847 | 0.01264 | 0.02394 | 0.01071 | 0.01485 | 0.01580 |
| **LOC441155** | 0.02386 | 0.01634 | 0.01360 | 0.02328 | 0.01356 | 0.01312 | 0.01727 |
| **LOC441178** | 0.04371 | 0.01390 | 0.01464 | 0.02009 | 0.00916 | 0.01529 | 0.00462 |
| **LOC441204** | 0.02770 | 0.01758 | 0.00658 | 0.01725 | 0.09589 | 0.01391 | 0.01623 |
| **LOC441242** | 0.13709 | 0.21075 | 0.06572 | 0.24287 | 0.02339 | 0.25059 | 0.10048 |
| **LOC441454** | 0.09609 | 0.08011 | 0.03345 | 0.08207 | 0.01335 | 0.04141 | 0.07662 |
| **LOC441455** | 0.05578 | 0.00000 | 2.17881 | 0.01059 | 0.49715 | 0.44042 | 0.22615 |
| **LOC441461** | 0.04332 | 0.00629 | 0.04892 | 0.12868 | 0.01336 | 0.03302 | 0.00000 |
| **LOC441601** | 3.78294 | 4.55641 | 1.65909 | 5.46567 | 1.64700 | 1.81574 | 0.00000 |
| **LOC441666** | 0.68743 | 0.00000 | 0.09329 | 0.07813 | 0.01966 | 0.16009 | 0.36103 |
| **LOC442028** | 0.02111 | 0.06040 | 0.05098 | 0.05173 | 0.02121 | 0.00876 | 0.00905 |
| **LOC442132** | 0.01814 | 0.02970 | 0.01677 | 0.01851 | 0.03202 | 0.01125 | 0.02618 |

| | | | | | | | |
|---|---|---|---|---|---|---|---|
| **LOC493754** | 2.81108 | 4.18790 | 5.26900 | 3.85212 | 9.01926 | 6.39556 | 4.10995 |
| **LOC494127** | 0.64523 | 0.67785 | 0.45759 | 0.41229 | 0.12496 | 0.29061 | 0.04547 |
| **LOC494141** | 0.02821 | 0.06705 | 0.05572 | 0.02713 | 0.00590 | 0.00979 | 0.00365 |
| **LOC54944** | 0.00000 | 0.07063 | 0.17268 | 0.06797 | 0.02181 | 0.15871 | 0.18113 |
| **LOC553103** | 0.09402 | 0.16630 | 0.02262 | 0.01078 | 0.00641 | 0.00625 | 0.00004 |
| **LOC554206** | 0.00717 | 0.62247 | 0.02691 | 0.32397 | 0.11440 | 0.05274 | 0.00000 |
| **LOC613037** | 0.02085 | 0.01877 | 0.01934 | 0.02602 | 0.00852 | 0.02680 | 0.09803 |
| **LOC63930** | 0.03557 | 0.00000 | 0.09812 | 0.04419 | 0.01332 | 0.01915 | 0.00268 |
| **LOC641367** | 0.12692 | 0.22922 | 0.42438 | 0.01674 | 0.15505 | 0.05556 | 0.61281 |
| **LOC641515** | 0.30496 | 0.05358 | 0.09921 | 0.11356 | 0.00444 | 0.00445 | 0.00000 |
| **LOC641746** | 0.24535 | 0.45537 | 0.73289 | 0.08284 | 0.27439 | 0.29599 | 0.42575 |
| **LOC642236** | 0.33335 | 0.00000 | 0.31265 | 0.10207 | 0.15462 | 0.22647 | 0.48501 |
| **LOC642361** | 0.00000 | 0.26460 | 0.03952 | 0.10676 | 0.86670 | 0.00000 | 0.17320 |
| **LOC642366** | 0.06717 | 0.01246 | 0.03390 | 0.02647 | 0.03746 | 0.12288 | 0.00196 |
| **LOC642757** | 0.01629 | 0.02232 | 0.10613 | 0.06638 | 0.00454 | 0.01154 | 0.17612 |
| **LOC642846** | 0.19410 | 0.34130 | 0.05787 | 0.43733 | 0.01422 | 0.23418 | 0.79342 |
| **LOC642852** | 0.57607 | 0.33835 | 1.36296 | 1.06341 | 0.00795 | 0.38475 | 1.14636 |
| **LOC642943** | 0.03009 | 0.01776 | 0.01943 | 0.03212 | 0.03984 | 0.01278 | 0.01364 |
| **LOC643072** | 0.03395 | 0.00000 | 0.06400 | 0.07435 | 0.00871 | 0.02454 | 0.12276 |
| **LOC643201** | 1.12888 | 0.00000 | 0.45264 | 0.25234 | 0.21056 | 0.21964 | 0.00000 |
| **LOC643339** | 0.02587 | 0.21830 | 0.12080 | 0.11359 | 0.12156 | 0.10086 | 0.00000 |
| **LOC643355** | 0.02896 | 0.00000 | 0.02658 | 0.01193 | 0.15192 | 0.08779 | 0.21140 |
| **LOC643387** | 0.31002 | 0.15688 | 0.12985 | 0.26241 | 0.18659 | 0.08172 | 0.00000 |
| **LOC643406** | 0.02271 | 0.12185 | 0.02133 | 0.01663 | 0.03984 | 0.01746 | 0.00173 |
| **LOC643441** | 1.54846 | 0.65149 | 0.10404 | 0.99819 | 0.00329 | 0.02332 | 0.11580 |
| **LOC643486** | 0.05382 | 0.00000 | 0.10434 | 0.03733 | 0.21173 | 0.11979 | 0.07262 |
| **LOC643623** | 0.00000 | 2.46222 | 1.39456 | 0.00000 | 0.61352 | 0.63757 | 0.00010 |
| **LOC643711** | 0.02876 | 0.00103 | 0.03111 | 0.01871 | 0.13088 | 0.01701 | 0.02387 |
| **LOC643770** | 0.02702 | 0.26939 | 0.19835 | 0.02325 | 0.00347 | 0.08108 | 0.00000 |
| **LOC643802** | 0.02538 | 0.01685 | 0.01304 | 0.01904 | 0.00985 | 0.01461 | 0.01684 |
| **LOC643923** | 0.06581 | 0.43366 | 0.44731 | 0.01451 | 0.11929 | 0.08502 | 0.02299 |
| **LOC644050** | 0.04785 | 0.02629 | 0.01061 | 0.03338 | 0.00805 | 0.08440 | 0.00000 |
| **LOC644172.uid-** | 0.04517 | 0.04255 | 0.00459 | 0.01040 | 0.01310 | 0.08223 | 0.42126 |
| **LOC644172.uid-** | 0.01859 | 0.08788 | 0.01898 | 0.01880 | 0.01195 | 0.11612 | 0.16471 |
| **LOC644189** | 0.08131 | 0.04021 | 0.03151 | 0.18520 | 0.04793 | 0.03181 | 0.01356 |

| | | | | | | | |
|---|---|---|---|---|---|---|---|
| **LOC644554** | 0.02828 | 0.00000 | 0.05967 | 0.24108 | 0.19965 | 0.07753 | 0.04404 |
| **LOC644656** | 0.82677 | 0.85424 | 0.39242 | 1.52139 | 0.55935 | 0.29998 | 0.00000 |
| **LOC644762** | 1.24931 | 0.20502 | 0.33890 | 0.87625 | 0.10653 | 0.49083 | 0.68147 |
| **LOC644838** | 0.02583 | 0.00835 | 0.03822 | 0.01973 | 0.00700 | 0.01278 | 0.01672 |
| **LOC644919** | 0.16354 | 0.18278 | 0.14043 | 0.09298 | 0.03012 | 0.28453 | 0.38835 |
| **LOC644936** | 29.51005 | 21.02296 | 0.93831 | 16.84743 | 0.44076 | 7.30385 | 2.06470 |
| **LOC644961** | 6.09126 | 13.75929 | 2.81048 | 7.30597 | 2.92961 | 6.78704 | 0.05213 |
| **LOC645166** | 0.28097 | 0.84819 | 0.39959 | 0.51007 | 0.34219 | 0.00000 | 0.16504 |
| **LOC645354+LO** | 0.02207 | 0.02431 | 0.00000 | 0.01476 | 0.63404 | 0.01544 | 0.02106 |
| **LOC645354+LO** | 0.02507 | 0.06439 | 0.00000 | 0.00473 | 1.08328 | 0.01562 | 0.03455 |
| **LOC645355** | 0.02251 | 0.01794 | 0.01298 | 0.02160 | 0.00796 | 0.02085 | 0.01911 |
| **LOC645434** | 0.03983 | 0.01511 | 0.01323 | 0.01694 | 0.01096 | 0.01327 | 0.01164 |
| **LOC645513** | 1.31058 | 0.32586 | 0.97187 | 0.26818 | 0.75956 | 1.08151 | 0.00000 |
| **LOC645752** | 0.03849 | 0.01448 | 0.00937 | 0.01841 | 0.08857 | 0.02433 | 0.01688 |
| **LOC645949** | 0.02372 | 0.01783 | 0.00473 | 0.01965 | 0.13309 | 0.01316 | 0.01722 |
| **LOC646214** | 1.12286 | 0.06299 | 0.67814 | 1.41346 | 1.79440 | 1.15954 | 2.04276 |
| **LOC646471** | 0.30349 | 0.42407 | 0.06327 | 0.25972 | 0.13038 | 0.34278 | 0.19770 |
| **LOC646522** | 0.02327 | 0.02180 | 0.01289 | 0.02682 | 0.03212 | 0.01211 | 0.01535 |
| **LOC646626** | 0.10404 | 0.04710 | 0.07878 | 0.20083 | 0.07292 | 0.08653 | 0.05498 |
| **LOC646719** | 0.18122 | 0.31417 | 0.71989 | 0.94955 | 1.63755 | 0.21341 | 1.31013 |
| **LOC646736** | 0.02374 | 0.01719 | 0.01205 | 0.03442 | 0.01236 | 0.01160 | 0.01965 |
| **LOC646762** | 2.95597 | 1.23849 | 0.12822 | 4.98087 | 0.53176 | 0.06945 | 1.29853 |
| **LOC646813** | 0.02480 | 0.01634 | 0.01378 | 0.02094 | 0.00907 | 0.01470 | 0.02103 |
| **LOC646862** | 0.02556 | 0.01557 | 0.01310 | 0.01736 | 0.00774 | 0.01551 | 0.02591 |
| **LOC646903** | 0.10490 | 0.00000 | 0.04704 | 0.02838 | 0.04787 | 0.02604 | 0.08587 |
| **LOC647859** | 0.05335 | 0.01047 | 0.01368 | 0.00877 | 0.00742 | 0.01736 | 0.03835 |
| **LOC648691** | 0.02109 | 0.00000 | 0.06216 | 0.01693 | 0.16998 | 0.04591 | 0.02260 |
| **LOC648987** | 0.14856 | 0.22280 | 0.88124 | 0.32850 | 0.26602 | 0.15618 | 0.30718 |
| **LOC649133** | 0.02179 | 0.01770 | 0.01290 | 0.04015 | 0.00795 | 0.01209 | 0.01619 |
| **LOC649330** | 0.02169 | 0.01044 | 0.00405 | 0.01898 | 1.33449 | 0.01286 | 0.01487 |
| **LOC649352** | 0.02540 | 0.01883 | 0.01328 | 0.02059 | 0.02037 | 0.01330 | 0.01679 |
| **LOC650226** | 2.47078 | 0.55130 | 3.20705 | 5.14316 | 1.67382 | 3.01717 | 13.76188 |
| **LOC650368** | 0.01371 | 0.21619 | 0.13029 | 0.00030 | 0.12292 | 1.10668 | 0.43386 |
| **LOC652276** | 0.34803 | 1.61389 | 0.56191 | 2.80836 | 0.08393 | 1.31308 | 1.04344 |
| **LOC653160** | 0.01415 | 0.00312 | 0.01438 | 0.03309 | 0.01372 | 0.01513 | 0.20612 |

| | | | | | | | |
|---|---|---|---|---|---|---|---|
| **LOC653513** | 0.02415 | 0.03758 | 0.04952 | 0.05816 | 0.52162 | 0.34647 | 0.01310 |
| **LOC653653** | 0.15580 | 0.03053 | 0.18047 | 0.09632 | 0.18199 | 0.25902 | 0.00000 |
| **LOC653712** | 0.21912 | 0.15608 | 0.17164 | 0.03202 | 0.04874 | 0.26763 | 0.27365 |
| **LOC653786** | 0.00000 | 0.48935 | 0.07810 | 0.01014 | 0.06396 | 0.35200 | 0.01259 |
| **LOC654342** | 1.34594 | 3.59560 | 1.67981 | 2.17569 | 0.41812 | 0.05101 | 0.14093 |
| **LOC654841** | 0.09332 | 0.12510 | 0.07426 | 0.02845 | 0.04834 | 0.08581 | 0.03019 |
| **LOC727896** | 6.24282 | 1.23038 | 2.65200 | 3.91651 | 0.84452 | 0.84125 | 0.00000 |
| **LOC727924** | 0.02390 | 0.01690 | 0.01071 | 0.01986 | 0.05080 | 0.01326 | 0.01553 |
| **LOC728024** | 2.56582 | 0.37840 | 1.82740 | 0.99725 | 2.24550 | 0.32856 | 0.00000 |
| **LOC728084** | 0.20286 | 0.00000 | 0.00661 | 0.27379 | 0.00589 | 0.00000 | 0.20611 |
| **LOC728175** | 0.03638 | 0.01680 | 0.01203 | 0.01667 | 0.01469 | 0.02046 | 0.02847 |
| **LOC728323** | 0.34969 | 0.13028 | 0.09663 | 0.39001 | 0.36014 | 0.15397 | 1.97783 |
| **LOC728392** | 0.00000 | 0.20178 | 0.11803 | 0.19312 | 0.03083 | 1.23256 | 0.16521 |
| **LOC728554** | 3.41207 | 2.59203 | 1.96129 | 2.02206 | 1.24665 | 2.53720 | 0.00000 |
| **LOC728613** | 0.55447 | 0.75484 | 0.76392 | 0.41942 | 1.20195 | 0.08694 | 0.23259 |
| **LOC728637** | 2.63504 | 0.00000 | 3.22098 | 0.00000 | 2.66276 | 0.00000 | 0.33732 |
| **LOC728730** | 0.14148 | 0.30212 | 0.15363 | 0.08716 | 0.56006 | 0.04217 | 0.21372 |
| **LOC728739** | 0.18842 | 0.08143 | 0.07083 | 0.23989 | 0.18226 | 0.08849 | 0.09835 |
| **LOC728743** | 0.00000 | 2.75224 | 0.50320 | 0.06247 | 0.00000 | 1.09659 | 0.63063 |
| **LOC728752** | 0.00000 | 0.29354 | 0.63070 | 0.40070 | 1.94669 | 0.18419 | 0.04503 |
| **LOC728819** | 0.08613 | 0.00000 | 1.08923 | 0.00000 | 0.11150 | 0.25547 | 0.01343 |
| **LOC728989** | 0.03943 | 0.02233 | 0.01752 | 0.03769 | 0.00769 | 0.01618 | 0.01035 |
| **LOC729080** | 0.09147 | 0.04139 | 0.07613 | 0.03635 | 0.00892 | 0.02197 | 0.01990 |
| **LOC729083** | 0.01056 | 0.00000 | 0.32277 | 0.04053 | 0.46423 | 0.00288 | 0.06443 |
| **LOC729218** | 0.52075 | 0.89678 | 0.45904 | 0.59605 | 0.25649 | 0.32965 | 0.87302 |
| **LOC729296** | 0.02413 | 0.01422 | 0.02882 | 0.01844 | 0.06067 | 0.01232 | 0.01792 |
| **LOC729506** | 0.02674 | 0.01731 | 0.02355 | 0.04557 | 0.00835 | 0.01311 | 0.00531 |
| **LOC729603** | 0.27655 | 0.30237 | 0.61436 | 0.45739 | 0.06491 | 0.30418 | 0.73948 |
| **LOC729609** | 0.01584 | 0.02699 | 0.02006 | 0.03051 | 0.00680 | 0.02616 | 0.17951 |
| **LOC729683** | 0.02395 | 0.01885 | 0.03334 | 0.02948 | 0.05193 | 0.04457 | 0.01598 |
| **LOC729732** | 0.00000 | 0.05996 | 0.02994 | 0.06401 | 0.04928 | 0.05244 | 0.05876 |
| **LOC729737** | 0.03482 | 0.02950 | 0.06955 | 0.05907 | 0.02864 | 0.07022 | 0.13360 |
| **LOC729739** | 0.42234 | 0.13190 | 0.36358 | 1.26135 | 0.07316 | 0.00000 | 0.36241 |
| **LOC729930** | 0.18363 | 0.10071 | 0.10046 | 0.04971 | 0.13520 | 0.09991 | 0.00451 |
| **LOC729966** | 0.03386 | 0.01409 | 0.02459 | 0.01913 | 0.01613 | 0.02270 | 0.00750 |

| | | | | | | | |
|---|---|---|---|---|---|---|---|
| **LOC729970** | 0.10891 | 0.08721 | 0.02837 | 0.03090 | 0.01438 | 0.01074 | 0.07797 |
| **LOC729987** | 0.02414 | 0.01799 | 0.01327 | 0.01969 | 0.00789 | 0.01295 | 0.01642 |
| **LOC730101** | 4.75048 | 2.07027 | 0.13817 | 3.01497 | 0.17241 | 0.58292 | 0.00000 |
| **LOC730102** | 0.37659 | 0.00000 | 0.20226 | 0.02125 | 0.03122 | 0.05088 | 0.05149 |
| **LOC730159** | 0.02529 | 0.01555 | 0.01362 | 0.01695 | 0.00779 | 0.03967 | 0.01692 |
| **LOC730183** | 0.10193 | 0.94287 | 0.31388 | 0.30016 | 0.10517 | 0.00000 | 0.30188 |
| **LOC730668** | 0.01571 | 0.00649 | 0.01317 | 0.01150 | 0.00755 | 0.20608 | 0.04083 |
| **LOC731157** | 0.05604 | 0.06546 | 0.06825 | 0.09301 | 0.02121 | 0.04897 | 0.08946 |
| **LOC731424** | 0.11998 | 0.09842 | 0.00358 | 0.01272 | 0.07778 | 0.00000 | 0.17472 |
| **LOC80154.uid-1** | 0.01990 | 0.02863 | 0.00975 | 0.01795 | 0.00792 | 0.02824 | 0.02834 |
| **LOC81691** | 1.83549 | 1.35706 | 1.70187 | 0.43196 | 1.26850 | 0.39241 | 0.00000 |
| **LOC90246** | 0.05155 | 0.02087 | 0.00970 | 0.23069 | 0.00996 | 0.00364 | 0.05344 |
| **LOC90784** | 1.03920 | 1.53505 | 1.62745 | 2.97304 | 0.46399 | 0.30516 | 6.15691 |
| **LOC90834** | 0.00000 | 0.29900 | 0.02719 | 0.28774 | 0.12756 | 0.28957 | 0.46287 |
| **LOC91450** | 0.03035 | 0.01313 | 0.02109 | 0.01905 | 0.00705 | 0.01185 | 0.02511 |
| **LOC93432** | 0.02431 | 0.00000 | 0.06880 | 0.03091 | 0.40875 | 0.06052 | 0.05315 |
| **LOC93622** | 1.17633 | 0.67532 | 0.52549 | 0.33369 | 0.04021 | 0.85513 | 4.17509 |
| **LOH12CR1** | 2.54073 | 3.48051 | 2.91148 | 1.26603 | 5.37387 | 1.86689 | 0.00000 |
| **LOH12CR2** | 0.06215 | 0.00000 | 0.32587 | 0.03486 | 0.12150 | 0.02257 | 0.00000 |
| **LONP1** | 10.60555 | 16.60874 | 2.33432 | 14.48161 | 1.44494 | 6.69780 | 22.27340 |
| **LONP2** | 10.91893 | 7.69156 | 3.29189 | 1.50714 | 3.92751 | 3.90644 | 3.21517 |
| **LONRF1** | 1.39638 | 0.00000 | 0.01631 | 0.00000 | 0.13078 | 2.78761 | 4.64541 |
| **LONRF2** | 0.00897 | 0.00000 | 0.01550 | 0.23002 | 0.08047 | 0.00326 | 0.32349 |
| **LONRF3** | 0.04720 | 0.04294 | 0.20357 | 0.01071 | 0.35190 | 0.05115 | 0.01595 |
| **LOR** | 0.01884 | 0.03540 | 0.01353 | 0.02045 | 0.03384 | 0.02209 | 0.02096 |
| **LOX** | 0.08587 | 0.04189 | 0.11793 | 0.03712 | 0.06517 | 0.01943 | 0.61878 |
| **LOXHD1** | 0.02960 | 0.01273 | 0.00473 | 0.02890 | 0.98072 | 0.09892 | 0.00277 |
| **LOXL1** | 0.01122 | 0.01450 | 0.14536 | 0.03491 | 0.24256 | 0.18087 | 0.01400 |
| **LOXL1-AS1** | 0.01899 | 0.00000 | 0.27122 | 0.01781 | 0.10538 | 0.18948 | 0.05419 |
| **LOXL2** | 0.00000 | 0.78846 | 2.82746 | 2.28704 | 0.29521 | 1.03163 | 0.00000 |
| **LOXL3** | 0.33078 | 0.40506 | 0.12335 | 0.76841 | 0.22473 | 0.60454 | 0.37563 |
| **LOXL4** | 0.02018 | 0.01482 | 0.02203 | 0.04159 | 0.01280 | 0.09814 | 0.03535 |
| **LPA** | 0.01391 | 0.03304 | 0.00888 | 0.01667 | 0.29252 | 0.00994 | 0.03146 |
| **LPAL2** | 0.02675 | 0.02139 | 0.01905 | 0.02818 | 0.09701 | 0.01851 | 0.00488 |
| **LPAR1** | 0.01444 | 0.28867 | 3.71337 | 0.00000 | 0.10482 | 1.53833 | 0.00000 |

| | | | | | | | |
|---|---|---|---|---|---|---|---|
| **LPAR2** | 4.88168 | 8.89579 | 10.22365 | 5.37522 | 0.47093 | 7.62475 | 2.91592 |
| **LPAR3** | 0.00890 | 0.18867 | 0.12654 | 0.01384 | 0.03026 | 0.06735 | 0.02074 |
| **LPAR4** | 0.14624 | 0.00000 | 0.11153 | 0.00000 | 0.08808 | 0.34421 | 0.02846 |
| **LPAR5** | 0.53017 | 0.22360 | 0.07928 | 0.00000 | 0.00491 | 0.00000 | 0.44046 |
| **LPAR6** | 0.08998 | 0.01111 | 0.03341 | 0.02508 | 0.01910 | 0.06457 | 0.06620 |
| **LPCAT1** | 0.00000 | 6.16999 | 0.45643 | 1.14630 | 1.60646 | 5.94929 | 13.28611 |
| **LPCAT2** | 0.29117 | 0.00000 | 1.62815 | 0.86818 | 1.04898 | 0.43761 | 0.74293 |
| **LPCAT3** | 29.91342 | 34.88937 | 17.56764 | 5.65758 | 0.00417 | 10.48651 | 2.96824 |
| **LPCAT4** | 2.42629 | 8.16351 | 3.24793 | 0.00000 | 5.70734 | 7.31328 | 11.06145 |
| **LPGAT1** | 4.04037 | 3.55936 | 4.14410 | 0.00000 | 0.50227 | 1.58306 | 6.01072 |
| **LPHN1** | 0.00000 | 1.68107 | 3.94211 | 0.33778 | 0.76935 | 3.78549 | 8.92826 |
| **LPHN2** | 11.28094 | 0.00000 | 0.08336 | 4.97842 | 0.25589 | 0.16584 | 1.56560 |
| **LPHN3** | 0.71867 | 0.00000 | 1.77313 | 2.19835 | 0.82696 | 0.00000 | 0.58975 |
| **LPHN3-AS1** | 0.00966 | 0.01630 | 0.01635 | 0.03360 | 0.01255 | 0.02252 | 0.05542 |
| **LPIN1** | 10.95057 | 17.83431 | 3.25176 | 8.47123 | 0.51714 | 17.14873 | 12.41256 |
| **LPIN2** | 9.21830 | 11.23660 | 7.93660 | 0.36528 | 2.71173 | 6.76256 | 11.20503 |
| **LPIN3** | 1.54504 | 6.18542 | 0.29870 | 1.69007 | 1.77025 | 2.71309 | 1.08571 |
| **LPL** | 0.03988 | 0.02981 | 0.01669 | 0.02274 | 0.00810 | 0.01390 | 0.01122 |
| **LPO** | 0.02231 | 0.01616 | 0.01525 | 0.02171 | 0.00767 | 0.01210 | 0.02326 |
| **LPP** | 6.74652 | 4.01365 | 8.21078 | 7.67754 | 0.10858 | 10.98246 | 8.35782 |
| **LPP-AS1** | 0.01857 | 0.02245 | 0.01909 | 0.02457 | 0.00692 | 0.00834 | 0.05681 |
| **LPP-AS2** | 0.11809 | 0.51186 | 1.09929 | 0.22706 | 0.12570 | 0.02361 | 0.09087 |
| **LPPR1** | 0.03321 | 0.05570 | 0.03073 | 0.02811 | 0.58040 | 0.07944 | 0.00000 |
| **LPPR2** | 0.00000 | 1.22619 | 0.03185 | 1.25737 | 0.74106 | 1.66574 | 0.29702 |
| **LPPR3** | 0.00000 | 0.03141 | 0.04759 | 0.10712 | 0.00722 | 0.01769 | 0.22132 |
| **LPPR4** | 0.02835 | 0.01094 | 0.04063 | 0.02391 | 0.00814 | 0.01416 | 0.01188 |
| **LPXN** | 0.23120 | 0.52216 | 0.08631 | 0.28786 | 0.07891 | 0.05331 | 0.00000 |
| **LRAT** | 0.05665 | 0.08973 | 0.19729 | 0.03702 | 3.21784 | 0.21434 | 0.01537 |
| **LRBA** | 5.26230 | 1.19355 | 1.27968 | 1.70226 | 0.12010 | 4.05318 | 8.96543 |
| **LRCH1** | 3.65702 | 1.11495 | 4.13690 | 1.47308 | 0.61585 | 3.51809 | 8.87169 |
| **LRCH2** | 0.04616 | 0.00413 | 0.02572 | 0.01051 | 0.05487 | 0.01274 | 0.03430 |
| **LRCH3** | 3.74282 | 2.27219 | 1.70217 | 2.66536 | 1.77572 | 1.43618 | 0.00000 |
| **LRCH4** | 0.00000 | 0.96777 | 0.34682 | 0.37055 | 0.74964 | 1.05487 | 8.03581 |
| **LRFN1** | 0.00893 | 0.06271 | 0.00456 | 0.47031 | 0.02938 | 0.02387 | 2.04161 |
| **LRFN2** | 0.02076 | 0.02077 | 0.01221 | 0.01920 | 0.00793 | 0.02436 | 0.01769 |

| | | | | | | | |
|---|---|---|---|---|---|---|---|
| **LRFN3** | 0.00000 | 0.00000 | 0.30687 | 0.03633 | 0.00840 | 0.61128 | 0.26853 |
| **LRFN4** | 0.00000 | 2.32004 | 0.95837 | 0.00000 | 0.00000 | 0.00000 | 1.84848 |
| **LRFN5** | 0.01606 | 0.01361 | 0.01132 | 0.36898 | 0.00791 | 0.24605 | 0.00000 |
| **LRG1** | 0.75077 | 1.96587 | 0.00000 | 0.00000 | 0.12195 | 1.45044 | 0.00732 |
| **LRGUK** | 0.01770 | 0.01908 | 0.01636 | 0.09969 | 0.15327 | 0.16596 | 0.11949 |
| **LRIF1** | 6.40486 | 0.80676 | 6.45860 | 5.16698 | 3.58962 | 2.90285 | 4.09256 |
| **LRIG1** | 0.41786 | 0.00000 | 0.33637 | 0.17078 | 4.29689 | 3.51757 | 1.27088 |
| **LRIG2** | 2.19449 | 5.01270 | 4.46539 | 3.62397 | 5.52452 | 3.85829 | 2.19709 |
| **LRIG3** | 11.44511 | 12.71266 | 8.76837 | 0.94318 | 0.67442 | 0.00000 | 5.23976 |
| **LRIT1** | 0.03658 | 0.01596 | 0.02288 | 0.01917 | 0.00810 | 0.01199 | 0.00998 |
| **LRIT2** | 0.03921 | 0.00378 | 0.02040 | 0.01368 | 0.00729 | 0.18774 | 0.01737 |
| **LRIT3** | 0.12960 | 0.00000 | 0.36079 | 0.10772 | 0.15553 | 0.00413 | 0.00000 |
| **LRMP** | 0.05651 | 0.05707 | 0.04138 | 0.00000 | 1.44971 | 0.01548 | 0.18449 |
| **LRP1** | 1.30032 | 10.91873 | 0.65634 | 6.40769 | 0.35866 | 1.32893 | 17.17674 |
| **LRP10** | 0.00000 | 2.66920 | 1.18625 | 5.54642 | 0.91391 | 4.74816 | 9.80419 |
| **LRP11** | 0.10450 | 0.01816 | 0.14699 | 0.00000 | 0.04849 | 0.13963 | 0.28170 |
| **LRP12** | 1.08691 | 0.12700 | 1.48923 | 0.14099 | 1.71284 | 0.16090 | 1.93955 |
| **LRP1B** | 0.05958 | 0.10806 | 0.08100 | 0.04932 | 1.12497 | 0.21475 | 0.04269 |
| **LRP2** | 62.79992 | 0.01045 | 0.00000 | 18.75588 | 0.00000 | 0.00000 | 49.27935 |
| **LRP2BP** | 0.06587 | 0.01616 | 0.38786 | 0.09868 | 0.83026 | 0.32053 | 0.52855 |
| **LRP3** | 0.00000 | 0.14017 | 0.16612 | 0.10514 | 0.26728 | 0.12751 | 0.22693 |
| **LRP4** | 0.36078 | 0.48824 | 3.45219 | 0.48545 | 1.93073 | 15.88117 | 0.39955 |
| **LRP4-AS1** | 0.00476 | 0.06926 | 0.08317 | 0.05192 | 0.36071 | 0.18709 | 0.30051 |
| **LRP5** | 0.00000 | 0.36290 | 0.26753 | 0.31106 | 2.41961 | 0.00000 | 1.49811 |
| **LRP5L** | 0.03785 | 0.18319 | 0.00107 | 0.72673 | 0.00606 | 0.92233 | 0.64665 |
| **LRP6** | 4.38172 | 0.00000 | 3.43971 | 5.03676 | 7.16885 | 3.71945 | 14.37975 |
| **LRP8** | 0.58280 | 0.00000 | 0.02327 | 2.17636 | 0.01256 | 0.74073 | 2.23622 |
| **LRPAP1** | 49.48755 | 59.92225 | 10.62974 | 22.78122 | 5.61885 | 19.65982 | 16.18841 |
| **LRPPRC** | 46.13658 | 20.42026 | 25.35584 | 23.34719 | 2.60275 | 17.54673 | 32.72339 |
| **LRR1** | 23.15958 | 7.27847 | 15.79610 | 12.04781 | 3.15140 | 8.59967 | 2.77195 |
| **LRRC1** | 4.52883 | 4.53867 | 0.18343 | 2.32768 | 0.69953 | 1.22302 | 8.27243 |
| **LRRC10** | 0.02737 | 0.02014 | 0.01269 | 0.01859 | 0.00787 | 0.01415 | 0.01588 |
| **LRRC10B** | 0.01238 | 0.03024 | 0.01861 | 0.02580 | 0.04928 | 0.04184 | 0.03880 |
| **LRRC14** | 4.51631 | 4.59776 | 3.06508 | 5.32933 | 0.51398 | 5.78754 | 7.15674 |
| **LRRC15** | 0.02002 | 0.01848 | 0.00982 | 0.08684 | 0.03786 | 0.01282 | 0.00524 |

| | | | | | | | |
|---|---|---|---|---|---|---|---|
| **LRRC16A** | 13.89160 | 5.46568 | 5.50002 | 2.47686 | 11.05172 | 6.86396 | 1.14883 |
| **LRRC16B** | 0.01668 | 0.01476 | 0.01258 | 0.03093 | 0.00762 | 0.04644 | 0.03888 |
| **LRRC17** | 0.02339 | 0.04802 | 0.01334 | 0.02990 | 3.02409 | 0.06795 | 0.00603 |
| **LRRC19** | 0.03820 | 0.00000 | 0.04055 | 0.06201 | 0.01628 | 0.02339 | 0.27224 |
| **LRRC2** | 0.23758 | 1.54961 | 0.18852 | 0.00048 | 0.00394 | 0.81302 | 0.00000 |
| **LRRC2-AS1** | 0.04548 | 0.02511 | 0.01691 | 0.01006 | 0.00971 | 0.02678 | 0.01949 |
| **LRRC20** | 0.00000 | 0.02106 | 0.30203 | 0.12628 | 0.12803 | 0.37472 | 0.20997 |
| **LRRC23** | 0.07612 | 0.44599 | 1.18896 | 0.53764 | 0.34903 | 0.64622 | 0.13824 |
| **LRRC25** | 0.00000 | 0.01046 | 0.01139 | 0.02987 | 0.00763 | 0.19935 | 0.08866 |
| **LRRC26** | 0.01943 | 0.02769 | 0.01349 | 0.03347 | 0.00786 | 0.01237 | 0.03870 |
| **LRRC27** | 0.23544 | 0.33764 | 0.08885 | 0.06631 | 0.67236 | 0.00000 | 0.79574 |
| **LRRC28** | 2.34399 | 1.34906 | 1.40631 | 2.16518 | 1.03128 | 1.34624 | 3.60180 |
| **LRRC29** | 0.03221 | 0.00171 | 0.05839 | 0.03692 | 0.00939 | 0.01731 | 0.01801 |
| **LRRC3** | 0.15986 | 0.08929 | 0.55683 | 0.00000 | 0.05198 | 0.38833 | 0.30328 |
| **LRRC3-AS1** | 0.02728 | 0.00301 | 0.02220 | 0.01886 | 0.16528 | 0.06833 | 0.03331 |
| **LRRC31** | 0.03031 | 0.01870 | 0.01536 | 0.02004 | 0.01822 | 0.04029 | 0.00890 |
| **LRRC32** | 0.00000 | 1.60526 | 0.06910 | 0.01144 | 0.00312 | 4.49771 | 0.01042 |
| **LRRC34** | 0.43928 | 0.00000 | 0.56803 | 0.02841 | 0.76555 | 0.66551 | 0.19178 |
| **LRRC36** | 0.01952 | 0.04040 | 0.22331 | 0.01611 | 0.04744 | 0.02740 | 0.00904 |
| **LRRC37A** | 0.02289 | 0.03038 | 0.01492 | 0.02707 | 0.01046 | 0.01480 | 0.02062 |
| **LRRC37A11P** | 0.26530 | 2.56776 | 0.38679 | 0.84594 | 0.01512 | 0.19070 | 0.00000 |
| **LRRC37A2** | 0.10136 | 0.10596 | 0.08427 | 0.09241 | 0.06274 | 0.19280 | 0.14688 |
| **LRRC37A3** | 0.18309 | 0.38701 | 0.12346 | 0.03807 | 0.06574 | 0.18739 | 0.20903 |
| **LRRC37A4P** | 0.01421 | 0.01946 | 0.30184 | 0.01874 | 0.04493 | 0.00270 | 0.05158 |
| **LRRC37A6P** | 0.13320 | 0.04670 | 0.07752 | 0.15117 | 0.03488 | 0.17694 | 0.00000 |
| **LRRC37B** | 0.14531 | 0.16245 | 0.07976 | 0.42659 | 0.02718 | 0.24215 | 0.58765 |
| **LRRC37BP1** | 0.98232 | 1.30876 | 1.80890 | 0.22284 | 2.01476 | 3.17670 | 0.30019 |
| **LRRC38** | 0.00980 | 0.00547 | 0.32854 | 0.02337 | 1.39442 | 0.12340 | 0.02163 |
| **LRRC39** | 0.11003 | 0.00000 | 0.06115 | 0.03469 | 0.05893 | 0.09880 | 0.19789 |
| **LRRC3B** | 0.01677 | 0.01250 | 0.00685 | 0.03214 | 0.11436 | 0.01450 | 0.04364 |
| **LRRC4** | 0.61123 | 0.25055 | 1.95109 | 0.00000 | 0.90303 | 0.00000 | 2.55124 |
| **LRRC40** | 10.20253 | 0.52442 | 1.39550 | 1.89177 | 1.46096 | 3.24812 | 0.00000 |
| **LRRC41** | 7.08844 | 6.29235 | 1.23125 | 13.00745 | 2.48987 | 11.55715 | 3.93660 |
| **LRRC42** | 7.12558 | 6.84222 | 6.12251 | 5.26909 | 2.53743 | 1.41152 | 4.65309 |
| **LRRC43** | 0.02561 | 0.06527 | 0.03259 | 0.08091 | 0.10093 | 0.04526 | 0.01851 |

| | | | | | | | |
|---|---|---|---|---|---|---|---|
| **LRRC45** | 0.33823 | 0.74561 | 0.00000 | 0.91998 | 0.22521 | 0.52769 | 4.27259 |
| **LRRC46** | 0.11424 | 0.22086 | 0.51033 | 0.14631 | 0.31322 | 0.08123 | 0.16320 |
| **LRRC47** | 8.30045 | 20.63990 | 4.60512 | 4.46074 | 1.22113 | 13.55984 | 12.21794 |
| **LRRC48** | 0.25559 | 0.35922 | 0.26161 | 0.39359 | 1.00922 | 0.34614 | 0.00000 |
| **LRRC49** | 0.37888 | 0.18027 | 0.09592 | 0.00000 | 0.91040 | 1.18825 | 0.03281 |
| **LRRC4B** | 0.00000 | 0.15776 | 0.08860 | 0.00000 | 0.11968 | 0.14649 | 0.17432 |
| **LRRC4C** | 0.02529 | 0.00402 | 0.04072 | 0.01826 | 0.01116 | 0.01851 | 0.02992 |
| **LRRC55** | 0.03144 | 0.01711 | 0.01238 | 0.01644 | 0.02385 | 0.01406 | 0.01549 |
| **LRRC56** | 0.00000 | 0.14695 | 0.02441 | 0.00716 | 0.13775 | 0.31220 | 1.09153 |
| **LRRC57** | 6.36462 | 2.77692 | 6.31147 | 5.55531 | 3.54300 | 2.11333 | 0.00000 |
| **LRRC58** | 7.78994 | 0.00000 | 0.37815 | 18.13145 | 0.00000 | 3.67438 | 8.63314 |
| **LRRC59** | 24.35634 | 27.76385 | 10.12763 | 18.89560 | 6.41517 | 15.12495 | 7.95584 |
| **LRRC6** | 0.58195 | 0.31050 | 0.34030 | 0.54784 | 0.50933 | 0.18358 | 0.05565 |
| **LRRC61** | 0.00000 | 8.75151 | 0.59330 | 9.72516 | 0.02601 | 5.29068 | 8.48706 |
| **LRRC63** | 0.03946 | 0.08383 | 0.08347 | 0.08789 | 0.03208 | 0.06522 | 0.00000 |
| **LRRC66** | 0.01185 | 0.01585 | 0.14361 | 0.02216 | 0.00389 | 0.13478 | 0.08853 |
| **LRRC69** | 0.61420 | 0.13526 | 0.73070 | 0.16652 | 0.06255 | 0.35048 | 0.48673 |
| **LRRC7** | 0.02737 | 0.00028 | 0.04032 | 0.01929 | 1.13147 | 0.01702 | 0.01708 |
| **LRRC70** | 0.04149 | 0.14271 | 0.11328 | 0.01667 | 0.39628 | 0.23777 | 0.18929 |
| **LRRC71** | 0.03593 | 0.03040 | 0.02224 | 0.04664 | 0.00676 | 0.00782 | 0.04108 |
| **LRRC72** | 0.02175 | 0.01415 | 0.01455 | 0.02884 | 0.02641 | 0.01211 | 0.02339 |
| **LRRC73** | 0.02799 | 0.00698 | 0.04946 | 0.04359 | 0.03461 | 0.03643 | 0.00000 |
| **LRRC74** | 0.02374 | 0.00152 | 0.06244 | 0.02106 | 0.03462 | 0.01190 | 0.02127 |
| **LRRC75A** | 0.18859 | 0.48163 | 0.13373 | 0.11482 | 0.09861 | 0.11700 | 0.65603 |
| **LRRC75A-AS1** | 57.00853 | 42.63447 | 33.78900 | 48.68779 | 14.77251 | 34.75227 | 46.70866 |
| **LRRC75B** | 0.01889 | 0.01537 | 0.01942 | 0.03299 | 0.00655 | 0.20642 | 0.04115 |
| **LRRC8A** | 6.13035 | 13.20655 | 3.49661 | 13.70570 | 6.03518 | 6.32934 | 8.84135 |
| **LRRC8B** | 1.46126 | 0.00000 | 0.02406 | 4.00149 | 0.83622 | 0.00000 | 3.69328 |
| **LRRC8C** | 0.13697 | 0.05186 | 1.03899 | 0.40182 | 0.34419 | 0.00000 | 0.03572 |
| **LRRC8D** | 5.86426 | 1.01888 | 1.10702 | 6.87309 | 1.09004 | 2.63064 | 5.56871 |
| **LRRC8E** | 0.50605 | 0.16338 | 0.10397 | 0.84601 | 0.99608 | 0.46398 | 0.53133 |
| **LRRC9** | 0.09623 | 0.07837 | 0.13344 | 0.09005 | 0.08630 | 0.04814 | 0.00000 |
| **LRRCC1** | 2.74622 | 1.08490 | 4.80578 | 0.00000 | 0.19979 | 1.87245 | 3.79778 |
| **LRRD1** | 0.04680 | 0.00545 | 0.02515 | 0.05055 | 0.05531 | 0.07326 | 0.00000 |
| **LRRFIP1** | 24.35895 | 8.28730 | 0.73828 | 22.64912 | 1.56795 | 3.19763 | 37.53727 |

| | | | | | | | |
|---|---|---|---|---|---|---|---|
| **LRRFIP2** | 5.72046 | 6.12476 | 6.55600 | 6.19966 | 3.98175 | 3.90448 | 6.31867 |
| **LRRIQ1** | 0.04520 | 0.00000 | 0.14475 | 0.00266 | 0.05753 | 0.04735 | 0.18839 |
| **LRRIQ3** | 0.09862 | 0.17740 | 1.25971 | 0.10555 | 0.23345 | 0.12874 | 0.08454 |
| **LRRIQ4** | 0.09188 | 0.00808 | 0.05543 | 0.01569 | 0.08872 | 0.01336 | 0.00750 |
| **LRRK1** | 0.00000 | 0.00000 | 0.06844 | 0.14857 | 1.24810 | 0.26462 | 0.39652 |
| **LRRK2** | 0.10163 | 0.00819 | 0.03583 | 0.00648 | 0.01378 | 0.61551 | 0.01036 |
| **LRRN1** | 0.08158 | 0.00000 | 0.00749 | 0.08889 | 0.13935 | 0.79632 | 0.02915 |
| **LRRN2** | 0.02290 | 0.04364 | 0.01868 | 0.04251 | 0.00644 | 0.00614 | 0.04751 |
| **LRRN3** | 0.02239 | 0.03980 | 0.02206 | 0.02972 | 0.00475 | 0.00000 | 0.93017 |
| **LRRN4** | 0.02589 | 0.00000 | 0.06659 | 0.10394 | 0.55243 | 0.02306 | 0.04583 |
| **LRRN4CL** | 0.02891 | 0.00000 | 0.03207 | 0.05807 | 0.01017 | 0.02509 | 0.18655 |
| **LRRTM1** | 0.00458 | 0.00000 | 0.02138 | 0.20404 | 0.00659 | 0.58234 | 0.10827 |
| **LRRTM2** | 0.10758 | 0.00000 | 0.28745 | 0.24952 | 0.00105 | 0.19617 | 2.25695 |
| **LRRTM3** | 0.07014 | 0.01204 | 0.01128 | 0.01459 | 0.03244 | 0.02304 | 0.00823 |
| **LRRTM4** | 0.00000 | 0.89001 | 0.04531 | 0.04439 | 8.80390 | 0.35455 | 0.00177 |
| **LRSAM1** | 1.51738 | 1.65546 | 0.92221 | 3.33609 | 0.33762 | 0.93369 | 2.74394 |
| **LRTM1** | 0.09773 | 0.14229 | 0.01204 | 0.03803 | 0.00662 | 0.22721 | 0.00000 |
| **LRTM2** | 0.00000 | 0.23211 | 0.25770 | 0.00000 | 0.00207 | 0.02844 | 0.09713 |
| **LRTOMT** | 0.44517 | 0.53492 | 2.57959 | 1.57752 | 3.75781 | 0.57187 | 0.92398 |
| **LRWD1** | 6.95777 | 9.01663 | 0.45547 | 13.40602 | 2.04183 | 4.56614 | 0.00000 |
| **LSAMP** | 0.00000 | 0.11434 | 0.01219 | 0.02196 | 0.02873 | 0.49788 | 0.03157 |
| **LSAMP-AS1** | 0.03494 | 0.29017 | 0.11451 | 0.01279 | 0.00483 | 0.13907 | 0.00000 |
| **LSG1** | 18.80563 | 14.19804 | 12.65743 | 15.06112 | 8.25643 | 7.17159 | 7.70124 |
| **1.00000** | 24.21130 | 12.57512 | 20.93088 | 15.31998 | 6.61040 | 11.55898 | 2.33429 |
| **10.00000** | 19.95880 | 14.83411 | 11.64902 | 15.20603 | 8.85366 | 7.53674 | 0.00000 |
| **11.00000** | 0.10828 | 0.51457 | 0.44402 | 0.14750 | 0.47221 | 0.04852 | 1.11272 |
| **12.00000** | 5.66520 | 1.34203 | 2.08728 | 2.32310 | 0.98934 | 1.21836 | 0.00000 |
| **LSM14A** | 26.08256 | 11.70487 | 16.24402 | 19.14859 | 9.26031 | 11.14409 | 16.46255 |
| **LSM14B** | 0.00001 | 0.39823 | 0.38394 | 0.51363 | 0.18482 | 1.25002 | 3.92232 |
| **2.00000** | 25.97590 | 13.45309 | 14.35461 | 25.91665 | 3.89229 | 11.56479 | 11.78782 |
| **3.00000** | 32.44781 | 7.91120 | 17.13357 | 21.79640 | 1.72163 | 15.43442 | 17.79664 |
| **4.00000** | 22.89304 | 41.56392 | 10.93491 | 29.09004 | 0.28297 | 14.78239 | 25.68182 |
| **5.00000** | 28.18073 | 8.00839 | 11.69312 | 13.12896 | 0.93297 | 13.41700 | 17.59436 |
| **6.00000** | 16.59052 | 4.96878 | 7.79022 | 3.38081 | 0.00000 | 8.39299 | 10.87877 |
| **7.00000** | 20.20299 | 24.98825 | 5.08653 | 25.47759 | 10.01498 | 10.70663 | 14.16088 |

| 8.00000 | 18.57045 | 1.24213 | 12.48311 | 13.43347 | 1.16531 | 5.82599 | 17.25972 |
|---|---|---|---|---|---|---|---|
| LSMEM1 | 0.13545 | 0.04639 | 0.07042 | 0.44055 | 0.87104 | 0.43827 | 0.26088 |
| LSMEM2 | 0.03248 | 0.07543 | 0.03245 | 0.06202 | 0.02578 | 0.01435 | 0.01184 |
| LSP1 | 0.02440 | 0.01225 | 0.01479 | 0.04367 | 0.00778 | 0.03649 | 0.02220 |
| LSP1P3 | 0.09555 | 0.23560 | 0.16142 | 0.10343 | 0.05487 | 0.01863 | 0.00807 |
| LSR | 0.00000 | 5.26767 | 3.44706 | 7.39832 | 1.19258 | 5.76295 | 14.57669 |
| LSS | 23.22474 | 1.77722 | 1.89749 | 8.36770 | 1.90068 | 3.35968 | 13.79456 |
| LTA | 0.02356 | 0.60422 | 0.01041 | 0.06603 | 0.01753 | 0.00109 | 0.01211 |
| LTA4H | 24.73591 | 5.50686 | 0.00000 | 14.19197 | 4.91705 | 7.82570 | 2.01319 |
| LTB | 0.03266 | 0.02118 | 0.01400 | 0.01991 | 0.00804 | 0.03146 | 0.00968 |
| LTB4R | 0.11635 | 0.32770 | 0.97064 | 1.18536 | 0.15105 | 0.03789 | 5.31625 |
| LTB4R2 | 0.00911 | 0.05815 | 0.02782 | 0.10550 | 0.06899 | 0.06054 | 0.33943 |
| LTBP1 | 0.17838 | 0.28636 | 0.03312 | 0.17380 | 2.95273 | 2.23401 | 1.54973 |
| LTBP2 | 0.25967 | 0.27740 | 0.29223 | 0.14429 | 0.96563 | 0.18894 | 0.08262 |
| LTBP3 | 1.09928 | 0.39652 | 0.40740 | 2.18009 | 1.27602 | 0.54501 | 4.52258 |
| LTBP4 | 0.31353 | 4.04042 | 4.17469 | 1.56831 | 3.76029 | 4.09709 | 2.70310 |
| LTBR | 1.47552 | 0.25172 | 0.00000 | 2.39198 | 0.04156 | 1.96425 | 4.26005 |
| LTC4S | 0.04114 | 0.01293 | 0.01440 | 0.02183 | 0.00758 | 0.01346 | 0.02408 |
| LTF | 0.04654 | 0.04789 | 0.01250 | 0.07837 | 0.00758 | 0.00699 | 0.00000 |
| LTK | 0.03561 | 0.00000 | 0.04369 | 0.16381 | 0.17330 | 0.08292 | 0.09335 |
| LTN1 | 13.14617 | 0.00000 | 6.96382 | 5.19747 | 2.49376 | 5.61498 | 9.09449 |
| LTV1 | 33.27535 | 9.42827 | 17.42915 | 16.16644 | 8.35477 | 8.04013 | 4.11157 |
| LUC7L | 17.55160 | 11.51464 | 8.01302 | 12.49861 | 7.22871 | 8.25193 | 10.24892 |
| LUC7L2.uid-939 | 0.05489 | 0.03699 | 0.05091 | 0.03944 | 0.07098 | 0.01358 | 0.00328 |
| LUC7L2.uid-939 | 1.70183 | 5.47470 | 0.87361 | 2.88735 | 2.67778 | 0.55758 | 0.00000 |
| LUC7L3 | 25.09649 | 8.11919 | 17.78511 | 18.85871 | 9.30026 | 12.87515 | 32.75616 |
| LUCAT1 | 0.41599 | 0.86225 | 0.20010 | 0.18411 | 0.00000 | 0.19550 | 0.32851 |
| LUM | 0.03930 | 0.01689 | 0.01289 | 0.01696 | 0.00812 | 0.01299 | 0.00954 |
| LURAP1 | 0.86979 | 0.91025 | 0.36345 | 0.03034 | 0.11066 | 0.30113 | 0.65240 |
| LURAP1L | 0.00000 | 0.02857 | 0.01506 | 2.00800 | 0.02408 | 0.00995 | 0.00000 |
| LUZP1 | 19.72237 | 19.05218 | 11.04610 | 10.04390 | 13.25723 | 8.31798 | 21.95889 |
| LUZP2 | 0.01602 | 0.02026 | 0.02378 | 0.03086 | 0.12845 | 0.04669 | 0.02264 |
| LUZP4 | 0.09150 | 0.05783 | 5.12998 | 0.00000 | 0.07907 | 0.52467 | 0.00608 |
| LUZP6+MTPN | 8.98012 | 4.58763 | 12.29386 | 13.56839 | 5.51635 | 6.49402 | 23.98487 |
| LXN | 0.45114 | 0.00000 | 0.05329 | 1.81954 | 0.00974 | 0.00981 | 0.00000 |

| | | | | | | | |
|---|---|---|---|---|---|---|---|
| **LY6D** | 0.02955 | 0.01759 | 0.01201 | 0.03403 | 0.01085 | 0.02162 | 0.00827 |
| **LY6E** | 25.51876 | 14.58846 | 0.20685 | 9.12322 | 0.11449 | 15.77168 | 6.40252 |
| **LY6G5B** | 1.73064 | 0.33614 | 0.94601 | 1.19083 | 1.55922 | 1.16867 | 1.71799 |
| **LY6G5C** | 0.29558 | 0.00000 | 0.08318 | 0.27715 | 0.09986 | 0.15831 | 0.00000 |
| **LY6G6C** | 0.75616 | 0.19629 | 0.00098 | 4.51198 | 0.00509 | 0.00000 | 0.02456 |
| **LY6G6D** | 0.00000 | 0.00984 | 0.01289 | 0.12581 | 0.00694 | 0.00534 | 0.11434 |
| **LY6G6E** | 0.03173 | 0.01957 | 0.01256 | 0.05338 | 0.00809 | 0.01188 | 0.00519 |
| **LY6G6F** | 0.02344 | 0.00765 | 0.03455 | 0.01743 | 0.00818 | 0.01191 | 0.02970 |
| **LY6H** | 0.01986 | 0.02420 | 0.00952 | 0.02221 | 0.02217 | 0.01747 | 0.02545 |
| **LY6K** | 0.02117 | 0.01374 | 0.06012 | 0.01771 | 0.09314 | 0.03298 | 0.03738 |
| **LY75** | 0.02506 | 0.01410 | 0.00977 | 0.02359 | 0.03587 | 0.06437 | 0.01032 |
| **LY86-AS1** | 0.03076 | 0.01614 | 0.01459 | 0.03202 | 0.00991 | 0.01859 | 0.01854 |
| **LY9** | 0.86673 | 0.10982 | 0.16435 | 0.41203 | 0.04016 | 0.00079 | 0.00000 |
| **LY96** | 0.01014 | 0.00208 | 0.04726 | 0.03460 | 0.00577 | 0.23602 | 0.03102 |
| **LYAR** | 27.97379 | 11.82882 | 16.93913 | 13.47624 | 7.12051 | 6.38872 | 17.42982 |
| **LYG1** | 0.06052 | 0.00000 | 0.34958 | 1.24502 | 0.73445 | 0.54069 | 0.56313 |
| **LYG2** | 0.01834 | 0.32930 | 1.88190 | 0.07612 | 0.00139 | 0.20118 | 0.18740 |
| **LYL1** | 0.01634 | 0.01211 | 0.01478 | 0.03069 | 0.00716 | 0.01279 | 0.06668 |
| **LYN** | 18.01977 | 7.64738 | 0.00000 | 23.22798 | 0.00000 | 2.38646 | 40.41819 |
| **LYNX1** | 0.06142 | 0.01701 | 0.02882 | 0.02107 | 0.15298 | 0.04926 | 0.02976 |
| **LYPD1** | 0.01252 | 0.00000 | 0.10093 | 0.07434 | 1.43925 | 0.27723 | 0.00691 |
| **LYPD2** | 0.18315 | 0.01798 | 0.00201 | 0.05866 | 0.00651 | 0.00000 | 0.50244 |
| **LYPD3** | 15.61392 | 0.31339 | 0.00000 | 12.14013 | 0.27602 | 0.79893 | 4.59001 |
| **LYPD4** | 0.01286 | 0.07067 | 0.19698 | 0.02197 | 1.85349 | 0.00445 | 0.02436 |
| **LYPD5** | 0.00000 | 0.07022 | 0.27005 | 5.12461 | 0.12189 | 0.04659 | 0.00000 |
| **LYPD6** | 0.24364 | 0.01882 | 0.00000 | 0.06393 | 0.83836 | 0.19567 | 0.00000 |
| **LYPD6B** | 3.21916 | 3.10082 | 0.07274 | 0.70956 | 0.00143 | 0.61758 | 0.01966 |
| **LYPLA1** | 26.42164 | 0.00000 | 14.64245 | 6.19954 | 1.87650 | 16.32351 | 12.64641 |
| **LYPLA2** | 15.76049 | 9.71277 | 1.51415 | 21.15575 | 0.11446 | 7.26999 | 1.91651 |
| **LYPLA2P2** | 0.92110 | 0.44093 | 0.06931 | 1.61374 | 0.02496 | 0.38768 | 0.00000 |
| **LYPLAL1** | 6.63970 | 3.47966 | 3.70537 | 4.80149 | 0.00000 | 1.04094 | 2.86787 |
| **LYPLAL1-AS1** | 0.02888 | 0.01746 | 0.01264 | 0.02205 | 0.00933 | 0.01264 | 0.01418 |
| **LYRM1** | 5.34915 | 2.43609 | 7.26299 | 3.29739 | 1.28847 | 4.10037 | 1.63047 |
| **LYRM2** | 8.00302 | 4.93516 | 6.24579 | 8.48244 | 1.10010 | 2.47514 | 1.58551 |
| **LYRM4** | 15.65054 | 8.18370 | 9.59679 | 17.23279 | 0.82250 | 5.57979 | 6.02376 |

| | | | | | | | |
|---|---|---|---|---|---|---|---|
| **LYRM5** | 2.79749 | 0.48857 | 0.65622 | 0.87887 | 0.47443 | 0.86472 | 0.33018 |
| **LYRM7** | 12.39095 | 0.00000 | 3.35829 | 4.07933 | 0.33200 | 3.18849 | 3.19586 |
| **LYRM9** | 0.04153 | 0.11874 | 0.01525 | 0.08599 | 0.26600 | 0.00788 | 0.00000 |
| **LYSMD1** | 2.43084 | 2.43175 | 2.44723 | 4.32212 | 8.76834 | 1.98858 | 2.40745 |
| **LYSMD2** | 0.01521 | 0.05696 | 0.07965 | 0.05342 | 0.00589 | 0.00734 | 0.01792 |
| **LYSMD3** | 9.40817 | 0.14943 | 3.17046 | 3.60570 | 4.07728 | 2.18229 | 0.01273 |
| **LYSMD4** | 1.82853 | 3.70614 | 1.32932 | 2.19127 | 0.83541 | 0.86929 | 0.00000 |
| **LYST** | 0.68954 | 0.00000 | 0.85153 | 0.42944 | 1.04723 | 1.23067 | 0.96771 |
| **LYVE1** | 0.02655 | 0.01659 | 0.01254 | 0.02021 | 0.00803 | 0.01902 | 0.01962 |
| **LYZ** | 0.04168 | 0.00000 | 0.00000 | 0.00000 | 0.62641 | 3.06603 | 0.13371 |
| **LYZL6** | 0.03204 | 0.00981 | 0.03424 | 0.01767 | 0.00738 | 0.01363 | 0.01431 |
| **LZIC** | 11.99147 | 3.91127 | 10.91586 | 7.26206 | 2.78084 | 3.79474 | 4.59642 |
| **LZTFL1** | 1.04594 | 0.00000 | 2.93480 | 0.84177 | 1.11439 | 0.77877 | 0.00000 |
| **LZTR1** | 1.52529 | 0.73854 | 0.03872 | 3.60075 | 0.00000 | 1.40608 | 5.64169 |
| **LZTS1** | 0.00000 | 0.06402 | 0.00000 | 0.25676 | 0.12465 | 5.04251 | 0.00003 |
| **LZTS1-AS1** | 0.01468 | 0.02288 | 0.01080 | 0.03050 | 0.00792 | 0.15331 | 0.01974 |
| **LZTS2** | 0.00000 | 0.70443 | 0.09487 | 3.96267 | 0.37866 | 1.05090 | 2.96235 |
| **LZTS3** | 0.00000 | 0.04052 | 1.20020 | 0.16984 | 0.25474 | 1.12286 | 0.53562 |
| **M1AP** | 2.83621 | 0.00000 | 5.12541 | 0.59605 | 0.39751 | 0.06123 | 0.00000 |
| **M6PR** | 18.27159 | 3.33601 | 7.31809 | 17.22052 | 1.86147 | 5.29377 | 0.00000 |
| **MAATS1** | 0.03592 | 0.02428 | 0.02102 | 0.14492 | 0.26411 | 0.37667 | 0.00000 |
| **MAB21L1** | 0.02607 | 0.00677 | 0.07318 | 0.01632 | 0.02978 | 0.00881 | 0.03148 |
| **MAB21L2** | 0.05013 | 0.02924 | 0.02280 | 0.00555 | 0.07531 | 0.00848 | 0.04771 |
| **MAB21L3** | 0.28615 | 0.00000 | 0.01120 | 0.93511 | 0.01305 | 0.05236 | 0.15130 |
| **MACC1** | 0.00450 | 0.02372 | 0.00000 | 0.00000 | 0.88199 | 0.20438 | 0.26520 |
| **MACF1** | 6.60477 | 5.36128 | 7.60740 | 5.88161 | 0.44787 | 11.22301 | 14.41658 |
| **MACROD1** | 0.03981 | 0.53764 | 0.00954 | 0.02252 | 0.03623 | 0.36313 | 0.58800 |
| **MACROD2** | 0.08078 | 0.18292 | 0.05068 | 0.04722 | 0.12281 | 0.07410 | 0.00000 |
| **MACROD2-AS1** | 0.01349 | 0.00000 | 0.09297 | 0.02240 | 0.00543 | 0.01493 | 0.05474 |
| **MACROD2-IT1** | 0.02228 | 0.02050 | 0.00938 | 0.02224 | 0.06236 | 0.01392 | 0.01465 |
| **MAD1L1** | 12.09814 | 15.32064 | 2.66755 | 9.24537 | 5.55658 | 5.94770 | 0.57575 |
| **MAD2L1** | 38.67058 | 1.76477 | 19.92279 | 15.19788 | 9.68777 | 10.09733 | 3.50414 |
| **MAD2L1BP** | 25.02214 | 17.59208 | 18.23759 | 11.79933 | 23.10145 | 10.44646 | 0.00000 |
| **MAD2L2** | 5.49943 | 11.60309 | 6.41504 | 4.90756 | 5.03997 | 12.29392 | 0.74277 |
| **MADCAM1** | 0.01983 | 0.02086 | 0.01594 | 0.02521 | 0.03748 | 0.02881 | 0.01780 |

| | | | | | | | |
|---|---|---|---|---|---|---|---|
| **MADD** | 2.58078 | 1.31741 | 1.03118 | 5.72751 | 2.23671 | 0.92011 | 4.56336 |
| **MAEA** | 26.42027 | 20.88482 | 10.19244 | 14.07424 | 3.25268 | 7.28684 | 9.90352 |
| **MAEL** | 1.02464 | 5.94137 | 16.15933 | 0.08900 | 2.48654 | 5.99371 | 1.41087 |
| **MAF** | 0.00000 | 0.06834 | 0.01488 | 0.11982 | 0.18141 | 0.28450 | 0.29840 |
| **1.00000** | 0.00000 | 16.59073 | 2.84843 | 12.36211 | 0.22073 | 10.44969 | 26.02149 |
| **MAFA** | 0.00000 | 1.06151 | 0.28133 | 0.00000 | 0.02017 | 0.01892 | 1.73016 |
| **MAFB** | 0.02045 | 0.08497 | 0.00000 | 0.00000 | 0.08293 | 0.02107 | 0.74354 |
| **MAFF** | 0.00000 | 1.72394 | 0.34789 | 4.61946 | 0.65441 | 0.91499 | 0.94217 |
| **MAFG** | 0.25582 | 4.77802 | 1.08373 | 11.04766 | 0.34419 | 6.50924 | 22.76730 |
| **MAFG-AS1** | 0.00000 | 0.17342 | 0.04067 | 0.41199 | 0.20776 | 0.19451 | 0.83685 |
| **MAFK** | 0.00000 | 0.34030 | 0.04397 | 0.78083 | 0.12206 | 0.08280 | 1.37191 |
| **MAG** | 0.00000 | 8.85763 | 0.60873 | 0.31482 | 0.00123 | 0.00000 | 0.01576 |
| **MAGEA1** | 0.68796 | 0.14321 | 1.71603 | 0.08448 | 2.19228 | 0.00000 | 0.00000 |
| **MAGEA10** | 0.55157 | 0.29853 | 3.30829 | 1.10314 | 1.14556 | 0.00000 | 0.00000 |
| **MAGEA10-MAG** | 0.02647 | 0.01088 | 0.04174 | 0.02104 | 0.02318 | 0.01210 | 0.01394 |
| **MAGEA11** | 0.02876 | 0.05043 | 0.00000 | 0.41540 | 0.65304 | 0.00547 | 0.00000 |
| **MAGEA12** | 0.09367 | 0.00000 | 1.32165 | 0.36738 | 10.88418 | 0.02558 | 0.00000 |
| **MAGEA2.uid-13** | 0.02430 | 0.01987 | 0.01693 | 0.02375 | 0.03022 | 0.01337 | 0.01363 |
| **MAGEA3** | 0.06746 | 0.00000 | 0.53887 | 0.15105 | 5.11068 | 0.00281 | 0.00000 |
| **MAGEA4** | 15.90585 | 8.03492 | 33.77544 | 0.00000 | 9.05473 | 0.00000 | 0.00000 |
| **MAGEA6** | 0.07440 | 0.00000 | 0.84972 | 0.08811 | 4.61903 | 0.00178 | 0.00000 |
| **MAGEA8** | 0.12634 | 0.00000 | 0.35873 | 0.22186 | 0.07761 | 0.02099 | 0.00000 |
| **MAGEA8-AS1** | 0.02143 | 0.01888 | 0.03703 | 0.04056 | 0.01693 | 0.01189 | 0.02125 |
| **MAGEB1** | 1.50299 | 0.97803 | 8.05350 | 1.13277 | 0.44057 | 0.03727 | 0.00000 |
| **MAGEB10** | 0.00818 | 0.00000 | 1.89296 | 0.01018 | 0.21405 | 0.00000 | 0.03907 |
| **MAGEB16** | 0.24285 | 0.08194 | 0.02743 | 0.36405 | 0.32751 | 0.00000 | 0.12359 |
| **MAGEB17** | 0.04677 | 0.00000 | 0.10106 | 0.17288 | 0.46316 | 0.15621 | 0.00000 |
| **MAGEB2** | 7.12386 | 10.89358 | 23.93796 | 5.76911 | 2.47276 | 2.76313 | 0.00000 |
| **MAGEB3** | 0.00792 | 0.00000 | 1.84531 | 0.00933 | 0.74177 | 0.00000 | 0.05550 |
| **MAGEB4** | 0.02550 | 0.01994 | 0.02322 | 0.01653 | 0.00694 | 0.01209 | 0.02371 |
| **MAGEB6** | 0.08513 | 0.00000 | 0.23127 | 0.26713 | 0.00970 | 0.16352 | 0.07541 |
| **MAGEC1** | 0.48898 | 2.05906 | 0.00000 | 0.75877 | 0.04633 | 0.00137 | 0.00000 |
| **MAGEC2** | 0.00000 | 5.37694 | 3.11642 | 3.87336 | 1.16735 | 0.00000 | 0.00000 |
| **MAGEC3** | 0.06969 | 0.01203 | 0.01425 | 0.09142 | 0.00824 | 0.01276 | 0.00000 |
| **MAGED1** | 7.43111 | 6.22825 | 3.75163 | 14.79132 | 1.74103 | 0.63549 | 0.00000 |

| | | | | | | | |
|---|---|---|---|---|---|---|---|
| **MAGED2** | 1.94330 | 4.18613 | 2.43268 | 0.02853 | 0.16014 | 4.84003 | 0.00000 |
| **MAGEE1** | 0.02134 | 0.02425 | 0.01805 | 0.01935 | 0.01599 | 0.01503 | 0.01631 |
| **MAGEF1** | 0.00000 | 2.30135 | 0.37968 | 2.81319 | 0.00000 | 0.91983 | 6.72383 |
| **MAGEH1** | 0.34493 | 0.15072 | 1.28300 | 0.12135 | 0.17656 | 0.00000 | 0.00000 |
| **MAGI1** | 0.00000 | 6.89581 | 3.59459 | 4.10496 | 0.34576 | 2.33095 | 9.54665 |
| **MAGI1-AS1** | 0.00000 | 0.27994 | 0.10048 | 0.10153 | 0.00862 | 0.06603 | 0.35723 |
| **MAGI2** | 0.00000 | 0.00000 | 0.07927 | 0.03896 | 0.25427 | 0.20030 | 0.13595 |
| **MAGI2-AS3** | 0.01720 | 0.00000 | 0.32329 | 0.02043 | 1.90030 | 0.00603 | 0.02562 |
| **MAGI3** | 4.22462 | 0.28500 | 0.45834 | 5.29242 | 0.10285 | 0.56813 | 14.45636 |
| **MAGIX** | 0.34258 | 0.34406 | 0.35286 | 0.12822 | 0.01468 | 0.18117 | 0.06457 |
| **MAGOH** | 14.72492 | 6.20983 | 12.92643 | 14.90190 | 5.64008 | 7.96991 | 7.60568 |
| **MAGOH2** | 2.05657 | 0.20924 | 1.76700 | 1.57873 | 0.50584 | 0.78870 | 0.54926 |
| **MAGOHB** | 15.04072 | 3.47867 | 11.59960 | 7.11937 | 0.01709 | 5.05225 | 6.23195 |
| **MAGT1** | 12.30553 | 4.06768 | 9.40424 | 1.44071 | 5.54633 | 3.64752 | 7.21964 |
| **MAK** | 0.68603 | 0.30188 | 1.40308 | 0.26462 | 8.01158 | 0.82623 | 1.72467 |
| **MAK16** | 24.39948 | 1.97059 | 10.20388 | 10.65204 | 3.54030 | 4.78681 | 7.02245 |
| **MAL** | 0.00000 | 0.64295 | 0.09160 | 0.00000 | 0.00383 | 1.15118 | 0.03028 |
| **MAL2** | 0.51406 | 0.00000 | 0.51237 | 0.90689 | 0.25784 | 0.00000 | 1.47555 |
| **MALAT1** | 70.96193 | 24.47094 | 42.63008 | 48.98209 | 10.65186 | 37.97520 | 88.28931 |
| **MALL** | 0.02246 | 0.01066 | 0.01921 | 0.02321 | 0.01411 | 0.03298 | 0.01668 |
| **MALRD1** | 0.06848 | 0.00000 | 0.16499 | 0.02432 | 0.72431 | 0.01704 | 0.01659 |
| **MALSU1** | 19.90614 | 15.60314 | 15.99677 | 10.46811 | 6.55994 | 8.62023 | 12.09180 |
| **MALT1** | 1.33706 | 0.00000 | 0.69014 | 0.75436 | 0.65779 | 1.77917 | 3.70344 |
| **MAMDC2** | 0.15484 | 0.00622 | 0.10953 | 0.03385 | 0.10060 | 0.00607 | 0.00000 |
| **MAMDC4** | 0.00000 | 0.88546 | 0.06248 | 0.30162 | 0.00254 | 0.36755 | 1.97205 |
| **MAML1** | 6.98704 | 0.00000 | 2.56108 | 4.42896 | 0.63560 | 4.80871 | 11.23075 |
| **MAML2** | 0.00000 | 0.03351 | 0.05651 | 0.13115 | 0.09772 | 0.04479 | 0.09917 |
| **MAML3** | 0.00000 | 0.37877 | 0.07492 | 1.23383 | 0.07094 | 0.67935 | 1.19455 |
| **MAMLD1** | 0.02832 | 0.00974 | 0.03268 | 0.02109 | 0.09498 | 0.01340 | 0.01608 |
| **MAMSTR** | 0.04155 | 0.00000 | 0.07819 | 0.11295 | 0.27970 | 0.23381 | 0.26011 |
| **MAN1A1** | 1.82508 | 0.00000 | 0.28432 | 0.00000 | 0.64148 | 0.40436 | 2.85610 |
| **MAN1A2** | 5.53008 | 2.67183 | 2.96789 | 11.99139 | 1.46672 | 6.20975 | 13.05756 |
| **MAN1B1** | 6.41038 | 8.65297 | 1.99231 | 5.22443 | 0.58263 | 5.02962 | 2.76953 |
| **MAN1B1-AS1** | 0.00000 | 0.09371 | 0.03402 | 0.09936 | 0.00531 | 0.16171 | 0.07777 |
| **MAN1C1** | 0.00000 | 2.54554 | 0.65342 | 0.07594 | 0.21688 | 16.11459 | 0.00000 |

| | | | | | | | |
|---|---|---|---|---|---|---|---|
| **MAN2A1** | 9.06552 | 0.00000 | 4.27108 | 4.36159 | 0.94516 | 5.48961 | 17.36678 |
| **MAN2A2** | 1.82583 | 1.48586 | 0.56048 | 3.26027 | 0.14050 | 1.68717 | 9.35131 |
| **MAN2B1** | 0.56736 | 2.23699 | 0.53999 | 3.35134 | 0.00000 | 1.66578 | 6.68657 |
| **MAN2B2** | 2.00247 | 1.35577 | 0.41895 | 1.85352 | 0.17945 | 0.64191 | 4.73182 |
| **MAN2C1** | 2.02746 | 1.80354 | 1.09735 | 1.09156 | 0.21354 | 2.45948 | 4.31546 |
| **MANBA** | 11.80052 | 0.00000 | 0.00000 | 1.56748 | 0.22750 | 2.25237 | 3.73254 |
| **MANBAL** | 9.45618 | 7.51249 | 10.35536 | 15.31825 | 1.40273 | 5.05577 | 3.23254 |
| **MANEA** | 6.58959 | 0.00000 | 7.12490 | 0.88434 | 1.24184 | 0.91543 | 0.65855 |
| **MANEA-AS1** | 0.17623 | 0.00000 | 0.25525 | 0.08192 | 0.06499 | 0.27172 | 0.35912 |
| **MANEAL** | 0.00000 | 0.38601 | 1.79469 | 1.14228 | 1.89819 | 0.90059 | 2.05629 |
| **MANF** | 2.20908 | 2.32186 | 1.71982 | 8.22933 | 4.03182 | 6.02260 | 13.88152 |
| **MANSC1** | 2.33887 | 0.00000 | 0.20823 | 0.95507 | 0.47871 | 0.21774 | 0.60269 |
| **MANSC4** | 0.02140 | 0.00000 | 0.09483 | 0.01811 | 0.16943 | 0.00999 | 0.02819 |
| **MAOA** | 1.12740 | 0.79258 | 2.16076 | 0.71562 | 0.19039 | 0.00000 | 0.99285 |
| **MAP10** | 0.13390 | 0.00000 | 0.27144 | 0.20610 | 0.04998 | 0.02007 | 0.00000 |
| **MAP1A** | 0.15633 | 0.02699 | 0.13454 | 0.60129 | 1.90542 | 0.94763 | 1.10503 |
| **MAP1B** | 1.08657 | 0.00000 | 4.18700 | 0.60924 | 4.47282 | 6.60622 | 0.29298 |
| **MAP1LC3A** | 0.00000 | 0.55363 | 0.78753 | 0.82792 | 0.73988 | 1.26501 | 0.79267 |
| **MAP1LC3B** | 19.38949 | 2.09884 | 24.43407 | 19.86807 | 17.91901 | 9.52824 | 4.27139 |
| **MAP1LC3B2** | 4.49576 | 0.91584 | 8.20265 | 5.12713 | 6.60618 | 2.11180 | 0.85085 |
| **MAP1LC3C** | 0.00626 | 1.81064 | 0.33558 | 0.01922 | 0.13958 | 0.81224 | 0.00000 |
| **MAP1S** | 0.00000 | 8.60105 | 7.92432 | 4.08481 | 1.20223 | 4.86222 | 10.83717 |
| **MAP2** | 0.02121 | 0.00794 | 0.01142 | 0.01988 | 0.22788 | 0.00817 | 0.08265 |
| **MAP2K1** | 3.27017 | 1.39080 | 0.71962 | 4.63269 | 2.15951 | 1.61856 | 16.33663 |
| **MAP2K2** | 1.46493 | 11.49841 | 1.58911 | 6.63148 | 0.11768 | 3.91767 | 19.82895 |
| **MAP2K3** | 5.80234 | 0.37004 | 0.21789 | 6.46320 | 1.26888 | 6.23248 | 3.07314 |
| **MAP2K4** | 2.18622 | 1.84522 | 2.86606 | 3.38946 | 0.74074 | 4.20208 | 10.11976 |
| **MAP2K4P1** | 0.04587 | 0.00000 | 0.23462 | 0.11184 | 0.03916 | 0.19358 | 0.49682 |
| **MAP2K5** | 0.97571 | 1.49265 | 0.45183 | 2.75064 | 0.17690 | 0.85267 | 3.10462 |
| **MAP2K6** | 2.74839 | 0.00000 | 0.74597 | 0.00000 | 0.50684 | 2.75590 | 0.22429 |
| **MAP2K7** | 0.00000 | 4.54075 | 0.87427 | 2.99116 | 0.59798 | 2.07184 | 10.36477 |
| **MAP3K1** | 2.04645 | 1.13687 | 2.85445 | 0.63433 | 0.51450 | 3.19229 | 2.26494 |
| **MAP3K10** | 0.00000 | 0.06198 | 0.01848 | 0.11522 | 0.13458 | 0.14214 | 0.75536 |
| **MAP3K11** | 0.00000 | 1.61425 | 0.91041 | 1.68159 | 0.20742 | 2.00091 | 7.77469 |
| **MAP3K12** | 0.00000 | 0.28594 | 0.28491 | 0.38488 | 0.45118 | 0.00001 | 0.96446 |

| | | | | | | | |
|---|---|---|---|---|---|---|---|
| **MAP3K13** | 5.65653 | 0.74502 | 12.15992 | 3.71825 | 2.50493 | 1.81177 | 5.00458 |
| **MAP3K14** | 0.07457 | 1.70443 | 0.78974 | 1.04638 | 0.22115 | 0.00000 | 0.83747 |
| **MAP3K14-AS1** | 0.00350 | 0.11768 | 0.00883 | 0.03156 | 0.01043 | 0.00572 | 0.25485 |
| **MAP3K15** | 0.29816 | 0.00000 | 0.19123 | 0.00000 | 0.36055 | 0.25128 | 0.43577 |
| **MAP3K19** | 0.01286 | 0.10439 | 0.09513 | 0.01757 | 0.00951 | 0.31942 | 0.03101 |
| **MAP3K2** | 5.59037 | 1.13199 | 3.70565 | 1.21044 | 0.13789 | 7.43235 | 13.05619 |
| **MAP3K3** | 0.00000 | 2.07761 | 1.29514 | 0.60829 | 1.74553 | 2.61500 | 0.00000 |
| **MAP3K4** | 3.49617 | 7.16638 | 4.62645 | 7.30891 | 2.24398 | 6.79300 | 9.60383 |
| **MAP3K5** | 0.30150 | 1.64612 | 1.63580 | 0.00000 | 0.22903 | 1.50523 | 1.04363 |
| **MAP3K6** | 0.00000 | 1.12344 | 0.31417 | 0.35953 | 0.38327 | 1.34810 | 0.56742 |
| **MAP3K7** | 17.59663 | 3.31628 | 5.59350 | 11.52032 | 4.83958 | 6.72954 | 0.00000 |
| **MAP3K7CL** | 0.00000 | 0.10284 | 0.00000 | 0.01860 | 6.45093 | 0.00873 | 0.06524 |
| **MAP3K8** | 3.66389 | 0.57452 | 2.95999 | 7.91203 | 0.62895 | 0.53564 | 0.00000 |
| **MAP3K9** | 1.75797 | 0.25466 | 0.60884 | 2.99830 | 1.00667 | 4.35196 | 4.77548 |
| **MAP4** | 22.60326 | 31.45815 | 12.28191 | 12.15468 | 1.70994 | 17.01821 | 36.79657 |
| **MAP4K1** | 0.16673 | 0.66228 | 0.29743 | 0.15728 | 0.46736 | 5.89606 | 0.00000 |
| **MAP4K2** | 0.41094 | 1.33660 | 6.10244 | 1.37133 | 0.50164 | 1.40752 | 3.98236 |
| **MAP4K3** | 3.32874 | 0.00000 | 1.61499 | 0.59687 | 0.72692 | 4.29607 | 8.87252 |
| **MAP4K4** | 1.20403 | 15.63876 | 3.62659 | 6.83958 | 0.85894 | 5.56004 | 16.68726 |
| **MAP4K5** | 3.97819 | 0.00000 | 2.43635 | 3.03723 | 0.17650 | 1.03221 | 8.39340 |
| **MAP6** | 0.01612 | 0.01144 | 0.08377 | 0.10650 | 0.00537 | 0.28489 | 0.05406 |
| **MAP6D1** | 0.43478 | 0.27040 | 0.16615 | 0.54550 | 0.23175 | 0.00000 | 1.16027 |
| **MAP7** | 7.79997 | 2.97007 | 3.40202 | 5.44353 | 1.39658 | 0.31419 | 8.03696 |
| **MAP7D1** | 0.00000 | 0.56228 | 0.89181 | 4.29705 | 3.61110 | 1.58880 | 3.64581 |
| **MAP7D2** | 1.89379 | 3.84248 | 3.58967 | 0.66109 | 4.21179 | 0.63444 | 2.50485 |
| **MAP7D3** | 2.57690 | 5.91483 | 11.60577 | 1.95538 | 1.26623 | 6.06367 | 2.73601 |
| **MAP9** | 1.43364 | 0.00000 | 1.14711 | 0.47280 | 2.23034 | 3.09013 | 1.63095 |
| **MAPK1** | 11.17090 | 3.50955 | 13.96190 | 2.17520 | 3.51476 | 9.02221 | 13.47667 |
| **MAPK10** | 0.00000 | 0.22220 | 0.01616 | 0.06303 | 0.30971 | 0.02784 | 0.00995 |
| **MAPK11** | 0.02420 | 0.00574 | 0.02415 | 0.01823 | 0.00681 | 0.08566 | 0.03501 |
| **MAPK12** | 0.07508 | 1.28466 | 0.02695 | 0.09563 | 0.39645 | 0.70132 | 0.37327 |
| **MAPK13** | 0.95395 | 1.84168 | 4.40958 | 0.08953 | 0.28560 | 0.60182 | 1.33170 |
| **MAPK14** | 12.56801 | 6.22962 | 3.08766 | 4.09826 | 1.04339 | 2.09945 | 19.08826 |
| **MAPK15** | 0.07584 | 0.00000 | 0.02370 | 0.13242 | 0.02984 | 0.02818 | 0.32467 |
| **MAPK1IP1L** | 43.53115 | 15.68274 | 18.20607 | 23.61639 | 8.37609 | 17.43154 | 4.01445 |

| | | | | | | | |
|---|---|---|---|---|---|---|---|
| **MAPK3** | 6.55605 | 20.91210 | 2.03642 | 1.00136 | 0.00000 | 3.78940 | 9.10900 |
| **MAPK4** | 0.00000 | 0.24901 | 0.01027 | 0.12161 | 0.50508 | 0.58342 | 0.05552 |
| **MAPK6** | 10.89654 | 0.00000 | 12.52853 | 8.78045 | 8.52465 | 3.66925 | 7.09763 |
| **MAPK7** | 0.00000 | 1.83499 | 0.40133 | 1.23058 | 0.76872 | 1.99347 | 2.30550 |
| **MAPK8** | 4.81592 | 0.00000 | 4.43976 | 4.44717 | 6.64464 | 1.96506 | 6.48229 |
| **MAPK8IP1** | 0.01868 | 0.84886 | 0.09398 | 0.69799 | 0.23800 | 0.28345 | 1.39894 |
| **MAPK8IP2** | 0.00000 | 0.51082 | 0.12921 | 0.15242 | 0.19154 | 0.19629 | 0.00000 |
| **MAPK8IP3** | 0.00000 | 1.27699 | 0.08412 | 1.92949 | 0.16280 | 1.49994 | 4.07275 |
| **MAPK9** | 4.49297 | 1.87610 | 0.49451 | 1.61331 | 0.54844 | 2.40839 | 13.89083 |
| **MAPKAP1** | 2.33392 | 13.06041 | 2.01600 | 5.38074 | 1.56303 | 9.61139 | 13.86388 |
| **MAPKAPK2** | 0.00000 | 2.09514 | 0.19523 | 0.45189 | 1.96327 | 2.01016 | 6.85811 |
| **MAPKAPK3** | 0.43252 | 0.16518 | 0.08768 | 3.39009 | 0.03161 | 1.93937 | 3.22052 |
| **MAPKAPK5** | 3.10430 | 3.09446 | 3.36169 | 2.01286 | 0.84820 | 4.99696 | 11.90212 |
| **MAPKAPK5-AS** | 8.74437 | 0.00000 | 4.25999 | 6.38128 | 5.13774 | 2.33796 | 3.90181 |
| **MAPKBP1** | 0.03844 | 0.00000 | 2.58134 | 4.08718 | 2.43318 | 0.41058 | 4.26355 |
| **MAPRE1** | 42.09693 | 16.07118 | 17.33359 | 23.59393 | 9.81693 | 13.28268 | 10.52873 |
| **MAPRE2** | 1.13486 | 8.31999 | 3.14709 | 2.16512 | 4.20065 | 0.38367 | 0.00000 |
| **MAPRE3** | 0.00000 | 0.22046 | 0.00000 | 0.29714 | 0.56944 | 0.21059 | 0.35588 |
| **MAPT** | 0.00000 | 0.66564 | 0.64249 | 0.74644 | 0.15261 | 0.00000 | 0.15925 |
| **MAPT-IT1** | 0.01930 | 0.02144 | 0.01735 | 0.02217 | 0.01832 | 0.01498 | 0.02422 |
| **MARC1** | 0.44513 | 0.07580 | 0.04193 | 2.08307 | 0.04979 | 0.09351 | 2.63560 |
| **MARC2** | 0.00000 | 0.02407 | 0.00273 | 1.67464 | 0.51706 | 0.01018 | 0.02438 |
| **MARCH1** | 0.80566 | 0.00000 | 0.55441 | 0.00000 | 0.64936 | 0.00000 | 1.18937 |
| **MARCH10** | 0.03837 | 0.02306 | 0.06490 | 0.07334 | 0.04004 | 0.04596 | 0.00000 |
| **MARCH11** | 0.00000 | 0.01349 | 0.33142 | 0.03504 | 0.16634 | 0.12747 | 0.05046 |
| **MARCH2** | 0.00000 | 0.61287 | 1.60384 | 0.38365 | 0.16371 | 0.49300 | 0.46207 |
| **MARCH3** | 1.22959 | 2.52574 | 0.16716 | 0.60235 | 1.58560 | 2.22183 | 0.00000 |
| **MARCH4** | 0.02094 | 0.00505 | 0.07104 | 0.01935 | 0.61367 | 0.00839 | 0.06725 |
| **MARCH5** | 9.39802 | 3.64710 | 4.14563 | 6.90280 | 3.82100 | 3.51260 | 3.32837 |
| **MARCH6** | 26.41268 | 4.71387 | 4.71314 | 14.43563 | 1.97330 | 7.25447 | 8.74803 |
| **MARCH7** | 23.52601 | 1.14343 | 14.07345 | 12.18517 | 8.88115 | 8.73495 | 6.74998 |
| **MARCH8** | 7.63343 | 7.78875 | 12.85814 | 1.01362 | 6.22627 | 7.95457 | 9.63675 |
| **MARCH9** | 0.51704 | 1.95081 | 0.32190 | 0.31091 | 0.25936 | 1.31896 | 0.18060 |
| **MARCKS** | 13.34505 | 13.22232 | 4.42355 | 0.00000 | 0.22245 | 9.01791 | 12.36498 |
| **MARCKSL1** | 13.79192 | 11.24811 | 14.38571 | 25.11534 | 0.93942 | 13.18241 | 16.62654 |

| | | | | | | | |
|---|---|---|---|---|---|---|---|
| **MARCO** | 0.01354 | 0.00777 | 0.01790 | 0.05917 | 0.00721 | 0.04774 | 0.05738 |
| **MARK1** | 1.56007 | 0.93260 | 2.56752 | 0.00000 | 0.17664 | 2.86922 | 3.95110 |
| **MARK2** | 0.00000 | 4.32538 | 1.63091 | 2.57014 | 0.20089 | 3.14197 | 18.00217 |
| **MARK2P9** | 0.08351 | 0.02714 | 0.09672 | 0.13273 | 0.24764 | 0.05493 | 0.54881 |
| **MARK3** | 3.56380 | 4.25832 | 2.57567 | 5.29704 | 4.10530 | 6.00190 | 19.19457 |
| **MARK4** | 0.00000 | 0.68348 | 0.27481 | 1.23800 | 1.57282 | 0.32157 | 3.22025 |
| **MARS** | 46.45893 | 35.45170 | 19.59009 | 29.58510 | 9.68131 | 25.41597 | 5.03336 |
| **MARS2** | 10.89482 | 4.99274 | 3.39253 | 4.05673 | 3.01814 | 3.34950 | 0.00000 |
| **MARVELD1** | 0.00000 | 0.81774 | 1.87003 | 0.05258 | 0.28017 | 1.45316 | 0.36801 |
| **MARVELD2** | 7.12390 | 1.58198 | 1.57687 | 2.19690 | 0.00000 | 2.46249 | 3.56037 |
| **MARVELD3** | 0.40913 | 3.43685 | 0.85405 | 2.88478 | 0.00000 | 0.25531 | 6.53698 |
| **MAS1** | 0.05241 | 0.00786 | 0.01088 | 0.01569 | 0.01974 | 0.20395 | 0.00000 |
| **MASP1** | 0.05220 | 0.03365 | 0.03887 | 0.00000 | 0.01527 | 0.08015 | 0.04877 |
| **MASP2** | 2.53779 | 1.91128 | 0.76272 | 2.08435 | 0.23720 | 0.00000 | 0.60455 |
| **MAST1** | 0.00000 | 0.00000 | 0.31907 | 0.21330 | 0.00219 | 0.37133 | 0.45016 |
| **MAST2** | 0.00000 | 12.71416 | 10.26150 | 4.23143 | 1.91311 | 11.09520 | 12.77976 |
| **MAST3** | 1.42583 | 0.29445 | 4.88713 | 0.02152 | 6.80399 | 0.68972 | 1.26076 |
| **MAST4** | 0.43308 | 0.83982 | 0.25680 | 1.60918 | 3.03862 | 0.87871 | 1.80619 |
| **MASTL** | 16.31313 | 3.71352 | 11.06273 | 8.32198 | 12.83433 | 6.78073 | 4.56294 |
| **MAT1A** | 0.03515 | 0.04406 | 0.02235 | 0.00260 | 0.72642 | 0.01242 | 0.07043 |
| **MAT2A** | 35.43471 | 8.62400 | 14.65835 | 19.76616 | 14.03445 | 7.92897 | 0.00000 |
| **MAT2B** | 19.04656 | 3.92281 | 8.40557 | 12.97094 | 3.33015 | 8.01772 | 0.00000 |
| **MATK** | 0.04124 | 0.20183 | 0.00000 | 0.00000 | 0.22260 | 0.16485 | 0.51958 |
| **MATN1** | 0.02918 | 0.02269 | 0.05519 | 0.06934 | 0.11250 | 0.02019 | 0.00018 |
| **MATN1-AS1** | 0.03909 | 0.01682 | 0.01375 | 0.04695 | 0.00844 | 0.03146 | 0.01731 |
| **MATN2** | 0.81861 | 0.72263 | 0.60647 | 0.00000 | 0.61806 | 0.19011 | 0.00000 |
| **MATN3** | 0.27677 | 0.23238 | 0.00000 | 0.00000 | 0.17099 | 0.00000 | 0.33678 |
| **MATN4** | 0.00000 | 0.16256 | 0.46652 | 0.00321 | 0.00144 | 0.00645 | 0.10566 |
| **MATR3** | 41.13959 | 4.39828 | 18.54040 | 20.24471 | 13.91229 | 16.00764 | 16.30665 |
| **MAU2** | 4.26111 | 4.51308 | 4.01568 | 9.04976 | 0.01204 | 2.81327 | 9.43740 |
| **MAVS** | 1.56372 | 10.44718 | 7.12661 | 7.48430 | 0.02405 | 4.94014 | 11.34205 |
| **MAX** | 30.21938 | 26.80454 | 17.96464 | 16.89343 | 8.93238 | 10.25647 | 0.00000 |
| **MAZ** | 7.69555 | 19.91070 | 6.72099 | 18.11996 | 0.35933 | 13.50454 | 44.83119 |
| **MB** | 0.03243 | 0.79167 | 0.01948 | 0.00000 | 0.00546 | 0.08975 | 0.06055 |
| **MB21D1** | 0.01561 | 0.03121 | 0.00906 | 0.04954 | 0.19262 | 0.11683 | 0.03411 |

| | | | | | | | |
|---|---|---|---|---|---|---|---|
| **MB21D2** | 1.50178 | 0.49060 | 0.35317 | 1.58367 | 0.82499 | 0.98640 | 0.77144 |
| **MBD1** | 5.58984 | 5.95766 | 4.36628 | 11.01896 | 2.03320 | 3.97405 | 8.62981 |
| **MBD2** | 0.71925 | 1.82574 | 1.53120 | 2.58229 | 0.29769 | 1.76057 | 8.13499 |
| **MBD3** | 0.89501 | 23.24533 | 3.91822 | 13.89870 | 0.03843 | 8.89450 | 37.22927 |
| **MBD3L2** | 0.00768 | 0.02993 | 0.00000 | 0.00641 | 8.24838 | 0.01558 | 0.04120 |
| **MBD3L3** | 0.01019 | 0.01004 | 0.00000 | 0.00990 | 8.93333 | 0.00791 | 0.02650 |
| **MBD3L4** | 0.02388 | 0.01841 | 0.01102 | 0.01921 | 0.02205 | 0.01311 | 0.01648 |
| **MBD3L5** | 0.01757 | 0.01429 | 0.00000 | 0.01405 | 3.09894 | 0.01207 | 0.01700 |
| **MBD4** | 15.23253 | 5.12278 | 9.81947 | 15.96043 | 9.10483 | 9.50822 | 5.21812 |
| **MBD5** | 0.53170 | 0.00000 | 0.38705 | 0.73653 | 0.59851 | 0.09759 | 0.77739 |
| **MBD6** | 0.00000 | 0.57647 | 0.63591 | 2.24607 | 0.29210 | 1.64581 | 6.74711 |
| **MBIP** | 7.42669 | 0.00000 | 10.57131 | 3.32229 | 5.99718 | 4.10127 | 0.00000 |
| **MBL1P** | 0.09268 | 0.21313 | 0.01104 | 0.01300 | 0.01135 | 0.00000 | 0.02278 |
| **MBL2** | 0.02356 | 0.01917 | 0.01312 | 0.01949 | 0.00788 | 0.01298 | 0.01655 |
| **MBLAC1** | 0.00000 | 0.31355 | 0.33484 | 0.20778 | 0.41589 | 0.11508 | 0.39100 |
| **MBLAC2** | 0.31118 | 0.38944 | 0.21170 | 0.02472 | 0.01496 | 0.18297 | 0.87851 |
| **MBNL1** | 4.66038 | 0.00000 | 4.72242 | 2.22662 | 3.07813 | 1.36227 | 4.51843 |
| **MBNL1-AS1** | 0.03385 | 0.01714 | 0.01427 | 0.03161 | 0.01501 | 0.01114 | 0.03500 |
| **MBNL2** | 0.05555 | 0.00000 | 0.00325 | 4.75922 | 0.06586 | 0.00000 | 0.33152 |
| **MBNL3** | 5.98009 | 5.62055 | 2.33743 | 8.41769 | 0.25416 | 0.00000 | 5.17364 |
| **MBOAT1** | 6.32182 | 9.70280 | 0.85007 | 1.79395 | 0.25029 | 4.08203 | 0.00000 |
| **MBOAT2** | 0.91892 | 0.98832 | 1.10161 | 1.17686 | 0.94154 | 3.33555 | 6.01864 |
| **MBOAT4** | 0.10407 | 0.06824 | 0.06601 | 0.25947 | 0.03604 | 0.02938 | 0.00000 |
| **MBOAT7** | 5.29695 | 4.29034 | 1.32237 | 19.44428 | 0.00000 | 4.98910 | 23.30348 |
| **MBP** | 0.75750 | 0.28132 | 0.00000 | 0.25516 | 0.58873 | 0.69157 | 1.10630 |
| **MBTD1** | 0.43453 | 0.00000 | 2.95985 | 1.38377 | 0.77784 | 1.09708 | 5.49142 |
| **MBTPS1** | 11.95712 | 6.61277 | 1.30756 | 10.78141 | 2.72354 | 4.25867 | 3.38899 |
| **MBTPS2** | 5.52986 | 2.69814 | 0.01785 | 0.00000 | 0.15240 | 1.47432 | 0.63559 |
| **MC1R** | 0.14495 | 0.79104 | 0.56099 | 1.53014 | 0.00000 | 1.10327 | 3.21161 |
| **MC2R** | 0.02626 | 0.02054 | 0.12250 | 0.01699 | 0.00399 | 0.02127 | 0.01352 |
| **MC3R** | 0.02470 | 0.01696 | 0.01410 | 0.01939 | 0.01314 | 0.01317 | 0.01849 |
| **MC5R** | 0.01463 | 0.07669 | 0.10714 | 0.01369 | 0.03293 | 0.00146 | 0.03456 |
| **MCAM** | 0.00000 | 3.94968 | 4.56778 | 1.65364 | 1.14061 | 6.27164 | 1.01369 |
| **MCAT** | 1.33553 | 5.39796 | 3.03065 | 5.59711 | 1.06532 | 1.62544 | 12.96783 |
| **MCC** | 2.32964 | 0.43292 | 1.89103 | 0.44019 | 1.91363 | 0.23189 | 0.76679 |

| | | | | | | | |
|---|---|---|---|---|---|---|---|
| **MCCC1** | 3.33902 | 0.00000 | 0.01047 | 3.60248 | 0.49176 | 0.00000 | 0.37964 |
| **MCCC2** | 24.41953 | 3.03730 | 2.02827 | 16.33210 | 0.93644 | 4.88497 | 8.17823 |
| **MCEE** | 6.93868 | 0.37533 | 0.85173 | 4.31678 | 0.14934 | 0.58810 | 1.88610 |
| **MCEMP1** | 0.03116 | 0.01660 | 0.01348 | 0.02420 | 0.00801 | 0.01413 | 0.01094 |
| **2.00000** | 0.27469 | 0.00000 | 0.00706 | 0.00000 | 0.08306 | 0.22513 | 0.19139 |
| **MCF2L** | 0.07527 | 0.02210 | 0.03665 | 0.00880 | 0.02872 | 0.34924 | 0.05556 |
| **MCF2L-AS1** | 0.41226 | 1.22737 | 0.14741 | 0.05766 | 0.00425 | 0.19790 | 0.00000 |
| **MCF2L2** | 0.01975 | 0.01993 | 0.01223 | 0.04513 | 0.06965 | 0.02837 | 0.03947 |
| **MCFD2** | 25.26264 | 1.44426 | 4.05455 | 23.39936 | 0.67231 | 2.81468 | 10.97615 |
| **MCHR1** | 0.00927 | 0.02121 | 0.05034 | 0.03369 | 0.02759 | 0.20430 | 0.01703 |
| **MCHR2** | 0.02991 | 0.02073 | 0.01116 | 0.01812 | 0.01737 | 0.01202 | 0.01553 |
| **MCHR2-AS1** | 0.02713 | 0.01888 | 0.01321 | 0.02792 | 0.01205 | 0.01298 | 0.01610 |
| **MCIDAS** | 0.02998 | 0.00000 | 0.42200 | 0.09348 | 0.03892 | 0.01171 | 0.07958 |
| **MCL1** | 9.08890 | 8.74187 | 23.41208 | 14.34364 | 8.53652 | 28.21811 | 23.41319 |
| **MCM10** | 30.74841 | 16.58852 | 14.62190 | 13.24813 | 5.07572 | 11.08901 | 0.00000 |
| **MCM2** | 20.09433 | 26.34108 | 14.02098 | 23.10458 | 3.65246 | 12.46223 | 21.83477 |
| **MCM3** | 39.72384 | 25.12412 | 16.68851 | 24.56215 | 10.02467 | 22.48812 | 13.51455 |
| **MCM3AP** | 13.02824 | 3.08486 | 6.92051 | 8.34994 | 1.04082 | 6.17781 | 18.11285 |
| **MCM3AP-AS1** | 1.23007 | 0.00000 | 1.15035 | 0.84885 | 0.01849 | 0.47468 | 1.79704 |
| **MCM4** | 51.18405 | 32.08545 | 16.55928 | 21.85641 | 7.81580 | 20.11192 | 15.80455 |
| **MCM5** | 27.81303 | 38.93351 | 17.03597 | 22.76895 | 5.53904 | 17.61282 | 15.97490 |
| **MCM6** | 36.22250 | 19.48426 | 16.71616 | 11.60126 | 3.66032 | 14.75237 | 4.01176 |
| **MCM7** | 45.30970 | 37.23785 | 17.59187 | 41.20519 | 11.98414 | 26.25943 | 17.17142 |
| **MCM8** | 3.33014 | 1.38019 | 2.83027 | 2.66508 | 0.82985 | 2.29692 | 12.35119 |
| **MCM9** | 3.86326 | 0.00000 | 1.25163 | 3.10125 | 1.95818 | 1.07875 | 0.94627 |
| **MCMBP** | 7.20255 | 5.81383 | 6.53589 | 5.55400 | 1.82937 | 8.54328 | 16.31086 |
| **MCMDC2** | 0.05801 | 0.00000 | 0.40853 | 0.08189 | 0.42700 | 0.02151 | 0.29845 |
| **MCOLN1** | 17.76971 | 22.14105 | 5.94697 | 11.20853 | 1.84443 | 2.23292 | 7.14122 |
| **MCOLN2** | 0.24177 | 0.13697 | 0.00584 | 0.20139 | 0.40365 | 0.01680 | 0.00000 |
| **MCOLN3** | 0.01423 | 0.03925 | 0.02613 | 0.03886 | 0.02565 | 0.05339 | 0.00831 |
| **MCPH1** | 10.00274 | 3.91702 | 8.36214 | 5.23216 | 4.53556 | 2.81925 | 3.62490 |
| **MCRS1** | 19.94833 | 26.38720 | 12.58294 | 19.38478 | 5.59326 | 11.70075 | 7.33769 |
| **MCTP1** | 0.20336 | 0.00000 | 0.12196 | 0.00000 | 1.71425 | 0.17150 | 0.51655 |
| **MCTP2** | 0.00000 | 0.02032 | 0.00000 | 0.16403 | 1.28085 | 0.12354 | 0.01320 |
| **MCTS1** | 27.36086 | 8.36398 | 15.49093 | 9.51524 | 2.86843 | 13.71774 | 24.51899 |

| | | | | | | | |
|---|---|---|---|---|---|---|---|
| **MCTS2P** | 3.69956 | 0.42685 | 1.48807 | 1.61467 | 0.11364 | 1.62175 | 1.49262 |
| **MCU** | 2.25683 | 1.59916 | 1.24891 | 2.34340 | 1.08113 | 3.88172 | 4.35091 |
| **MCUR1** | 6.41673 | 9.56945 | 1.04785 | 8.80371 | 0.19887 | 9.09492 | 22.71998 |
| **1.00000** | 5.99348 | 3.95556 | 5.19588 | 4.48059 | 0.82633 | 1.28635 | 9.32916 |
| **MDFI** | 0.00000 | 0.41377 | 0.00057 | 2.92746 | 0.02444 | 6.34081 | 0.00000 |
| **MDFIC** | 0.01986 | 0.00760 | 0.01294 | 0.13138 | 0.04117 | 0.01997 | 0.11091 |
| **MDGA1** | 0.01327 | 0.01946 | 0.02074 | 0.06207 | 0.06371 | 0.13749 | 0.01925 |
| **MDGA2** | 0.10356 | 0.00000 | 0.04785 | 0.08438 | 0.70231 | 1.18752 | 0.00000 |
| **MDH1** | 52.30608 | 28.66195 | 15.17307 | 24.90810 | 1.25940 | 22.33993 | 10.05945 |
| **MDH1B** | 0.11110 | 0.10106 | 0.27752 | 0.11852 | 0.33327 | 0.17645 | 0.04255 |
| **MDH2** | 61.76658 | 41.83340 | 1.72240 | 46.25281 | 4.56108 | 31.51196 | 22.13698 |
| **MDK** | 21.73794 | 28.13638 | 0.61172 | 24.51236 | 0.88909 | 29.97829 | 11.56633 |
| **MDM1** | 2.62553 | 0.00000 | 2.09978 | 1.70424 | 2.12176 | 0.65900 | 1.07376 |
| **MDM2** | 21.30500 | 12.51706 | 12.53568 | 29.25580 | 11.71523 | 12.83456 | 5.20496 |
| **MDM4** | 24.64445 | 4.13098 | 14.43610 | 9.63249 | 8.38276 | 9.32715 | 23.84265 |
| **MDN1** | 15.79594 | 6.77922 | 11.54564 | 9.11599 | 3.53083 | 13.02194 | 28.88991 |
| **MDP1** | 0.14565 | 0.53326 | 0.28323 | 0.13917 | 0.01426 | 1.15140 | 0.01009 |
| **MDS2** | 0.02010 | 0.02017 | 0.01997 | 0.02118 | 0.02082 | 0.01243 | 0.02313 |
| **ME1** | 0.22240 | 0.00118 | 0.00000 | 0.88144 | 0.30224 | 0.07850 | 0.11387 |
| **ME2** | 4.02841 | 0.00000 | 0.07864 | 4.91902 | 0.01392 | 1.73393 | 8.98676 |
| **ME3** | 0.04453 | 0.00000 | 0.06192 | 0.04194 | 0.74376 | 0.01028 | 0.18502 |
| **MEA1** | 26.44646 | 20.06808 | 10.91042 | 24.62346 | 7.35782 | 12.85654 | 0.00000 |
| **MEAF6** | 16.47107 | 7.38306 | 10.80303 | 8.12488 | 2.82190 | 9.56498 | 0.66007 |
| **MECOM** | 0.00012 | 0.00000 | 0.46563 | 0.07978 | 0.51860 | 0.25370 | 2.00497 |
| **MECP2** | 0.39538 | 3.64609 | 2.08184 | 1.17025 | 2.37526 | 2.94536 | 7.66001 |
| **MECR** | 16.23264 | 11.80515 | 3.02204 | 8.38150 | 0.56503 | 6.34840 | 0.77947 |
| **MED1** | 11.37457 | 0.52197 | 8.37412 | 10.21634 | 2.46935 | 6.85406 | 16.83386 |
| **MED10** | 12.15989 | 18.28359 | 12.69945 | 22.26799 | 8.65228 | 7.03602 | 0.00000 |
| **MED11** | 6.80616 | 4.75917 | 7.77772 | 6.96545 | 1.57848 | 2.86142 | 0.14789 |
| **MED12** | 6.42955 | 9.10750 | 7.61979 | 2.42226 | 0.11326 | 0.00000 | 13.92717 |
| **MED12L** | 6.07209 | 1.18455 | 3.75534 | 18.89952 | 3.15174 | 0.08919 | 7.01654 |
| **MED13** | 12.40993 | 1.70031 | 11.81472 | 13.58287 | 5.39123 | 11.83636 | 24.18215 |
| **MED13L** | 12.96409 | 5.63610 | 5.80391 | 3.90800 | 2.34096 | 10.68925 | 23.04901 |
| **MED14** | 5.37303 | 1.08015 | 7.60292 | 4.61189 | 5.15807 | 4.36538 | 4.32276 |
| **MED14OS** | 0.03855 | 0.04872 | 0.12389 | 0.09190 | 0.07338 | 0.04325 | 0.15007 |

| | | | | | | | |
|---|---|---|---|---|---|---|---|
| **MED15** | 2.08896 | 20.49228 | 13.03636 | 11.44848 | 5.81292 | 13.45322 | 16.30014 |
| **MED15P9** | 0.02415 | 0.01234 | 0.04688 | 0.03009 | 0.26237 | 0.01174 | 0.01157 |
| **MED16** | 4.76137 | 12.20675 | 3.04332 | 12.72628 | 5.05014 | 4.26781 | 6.65014 |
| **MED17** | 7.09778 | 0.00000 | 5.67089 | 6.49752 | 2.98232 | 2.96949 | 9.02341 |
| **MED18** | 7.95255 | 2.92458 | 8.83916 | 6.20720 | 0.91734 | 2.63060 | 0.00000 |
| **MED19** | 11.27964 | 16.36973 | 14.99785 | 14.97110 | 15.73342 | 6.87688 | 5.77460 |
| **MED20** | 12.78450 | 6.20037 | 9.07880 | 5.96881 | 3.68906 | 6.42340 | 0.00000 |
| **MED21** | 17.89453 | 2.84048 | 8.44257 | 13.77078 | 9.16628 | 6.52199 | 0.00000 |
| **MED22** | 1.30101 | 5.99477 | 4.00053 | 7.32866 | 0.57818 | 4.04279 | 13.57808 |
| **MED23** | 4.98327 | 0.39327 | 4.01807 | 2.80801 | 1.58498 | 2.45573 | 1.52871 |
| **MED24** | 8.73005 | 5.72595 | 3.57509 | 13.31018 | 2.10405 | 4.67112 | 9.30510 |
| **MED25** | 0.00000 | 2.01811 | 1.58510 | 3.10773 | 0.71525 | 1.60299 | 6.82323 |
| **MED26** | 0.00000 | 1.57478 | 0.78755 | 0.86289 | 2.20480 | 1.50594 | 7.98358 |
| **MED27** | 6.46324 | 8.14805 | 3.54898 | 7.17556 | 1.93319 | 2.86703 | 0.00000 |
| **MED28** | 24.59609 | 13.53123 | 12.75975 | 13.85989 | 6.73991 | 9.18996 | 0.95722 |
| **MED29** | 10.21762 | 15.95550 | 9.71832 | 13.86133 | 3.89514 | 6.68752 | 1.41309 |
| **MED30** | 7.38741 | 6.40614 | 3.68876 | 6.36938 | 1.99139 | 3.20497 | 0.00000 |
| **MED31** | 3.59445 | 0.25078 | 2.99652 | 4.82335 | 7.54824 | 0.50547 | 1.69252 |
| **MED4** | 15.08286 | 1.46472 | 14.01140 | 8.30810 | 8.37846 | 7.31075 | 4.70048 |
| **MED4-AS1** | 0.00138 | 0.02989 | 0.02522 | 0.10755 | 0.05575 | 0.06584 | 0.21541 |
| **MED6** | 17.43107 | 0.00000 | 18.62201 | 14.59161 | 9.41117 | 5.05567 | 0.88414 |
| **MED7** | 10.26259 | 1.43889 | 8.58076 | 3.96746 | 6.51389 | 2.66214 | 2.85223 |
| **MED8** | 15.62613 | 10.76564 | 11.66253 | 16.40205 | 4.03822 | 4.23329 | 0.00000 |
| **MED9** | 5.92160 | 8.16786 | 7.68628 | 5.73019 | 0.81314 | 3.66122 | 0.32370 |
| **MEDAG** | 0.05945 | 0.01218 | 0.01337 | 0.01367 | 0.00772 | 0.01261 | 0.02277 |
| **MEF2A** | 3.26841 | 1.94053 | 2.91797 | 1.16647 | 0.85262 | 2.50189 | 4.78092 |
| **MEF2B** | 0.02622 | 0.01793 | 0.01287 | 0.02043 | 0.00791 | 0.01200 | 0.01973 |
| **MEF2BNB.uid-2** | 0.84241 | 1.08063 | 1.25521 | 1.38831 | 0.80840 | 0.54339 | 0.73299 |
| **MEF2BNB.uid-2** | 1.65080 | 3.83857 | 2.28900 | 0.98190 | 0.00000 | 0.43281 | 1.36323 |
| **MEF2C** | 0.02503 | 0.00699 | 0.03028 | 0.02122 | 0.04141 | 0.01690 | 0.02496 |
| **MEF2C-AS1** | 0.02895 | 0.01332 | 0.03969 | 0.01624 | 0.00697 | 0.01119 | 0.02296 |
| **MEF2D** | 0.00000 | 2.99749 | 1.95230 | 1.12145 | 1.05671 | 3.21675 | 4.82236 |
| **MEFV** | 0.03770 | 0.03144 | 0.01269 | 0.02785 | 0.00828 | 0.05532 | 0.00000 |
| **MEG3** | 0.00000 | 1.07325 | 0.00000 | 0.06917 | 0.22860 | 20.24343 | 0.00837 |
| **MEG8** | 0.00000 | 0.07859 | 0.00382 | 0.01739 | 0.00907 | 0.45584 | 0.03615 |

| | | | | | | | |
|---|---|---|---|---|---|---|---|
| **MEG9** | 0.00434 | 0.01867 | 0.00766 | 0.01614 | 0.01555 | 0.93902 | 0.05549 |
| **MEGF10** | 0.02264 | 0.01328 | 0.00838 | 0.02381 | 0.52149 | 0.03563 | 0.01359 |
| **MEGF11** | 0.27365 | 0.11761 | 0.01600 | 0.17507 | 1.17074 | 0.21741 | 0.56933 |
| **MEGF6** | 0.00000 | 0.02102 | 0.08302 | 0.18229 | 0.00816 | 0.03045 | 0.01583 |
| **MEGF8** | 0.36988 | 2.11666 | 0.19889 | 0.00000 | 0.17937 | 1.42777 | 6.14705 |
| **MEGF9** | 0.16584 | 0.21587 | 0.17908 | 0.35984 | 0.08386 | 0.70705 | 0.39350 |
| **MEI1** | 0.82285 | 4.73629 | 2.73979 | 0.00000 | 0.69692 | 2.36472 | 1.82512 |
| **MEI4** | 0.26773 | 0.08810 | 0.21067 | 0.00000 | 0.28553 | 0.13129 | 0.09356 |
| **MEIG1** | 0.05774 | 0.03459 | 0.06338 | 0.04277 | 0.02678 | 0.06419 | 0.00773 |
| **MEIOB** | 0.05855 | 0.00000 | 0.51470 | 0.00000 | 0.16666 | 0.09693 | 0.03263 |
| **MEIS1** | 4.24311 | 0.03553 | 0.83013 | 1.06775 | 0.23360 | 1.58986 | 0.00000 |
| **MEIS1-AS3** | 0.02027 | 0.01567 | 0.01206 | 0.05031 | 0.00775 | 0.02280 | 0.03375 |
| **MEIS2** | 0.18280 | 0.16898 | 0.00000 | 0.89899 | 0.02707 | 2.39434 | 0.17703 |
| **MEIS3** | 0.00000 | 0.23442 | 0.00000 | 0.36112 | 0.31669 | 0.14242 | 0.39561 |
| **MEIS3P1** | 0.05084 | 0.00366 | 0.01969 | 0.07752 | 0.02676 | 0.15737 | 0.14216 |
| **MELK** | 26.50292 | 11.94712 | 14.06019 | 18.62019 | 6.47114 | 13.59904 | 0.00000 |
| **MEMO1** | 2.94627 | 1.18208 | 2.60354 | 2.31227 | 0.13004 | 1.22721 | 1.49753 |
| **MEN1** | 0.37979 | 1.31327 | 2.08515 | 2.07640 | 0.10797 | 1.68714 | 11.78084 |
| **MEOX2** | 0.02304 | 0.01957 | 0.01251 | 0.02100 | 0.00796 | 0.01276 | 0.01697 |
| **MEP1B** | 0.02332 | 0.02105 | 0.00700 | 0.02018 | 0.10640 | 0.07867 | 0.01466 |
| **MEPCE** | 3.28916 | 7.47955 | 5.10510 | 4.14459 | 2.55963 | 4.62159 | 15.62810 |
| **MEPE** | 0.02506 | 0.00650 | 0.05127 | 0.01788 | 0.00664 | 0.07255 | 0.02208 |
| **MERTK** | 9.49951 | 7.03478 | 6.01683 | 4.19607 | 0.25372 | 10.52777 | 7.72145 |
| **MESDC1** | 0.00000 | 0.08110 | 1.08273 | 0.63251 | 0.80089 | 1.94391 | 1.80740 |
| **MESDC2** | 38.67372 | 19.07539 | 12.34135 | 15.16505 | 8.32532 | 11.36949 | 14.18360 |
| **MESP1** | 0.28608 | 0.31396 | 0.00000 | 0.72857 | 0.14914 | 0.00000 | 0.50979 |
| **MESP2** | 0.00000 | 0.42826 | 0.06861 | 0.05676 | 0.00252 | 0.47015 | 0.27223 |
| **MEST** | 24.56628 | 10.53617 | 14.81904 | 1.23335 | 5.16990 | 11.27894 | 1.60064 |
| **MESTIT1** | 0.05296 | 0.00000 | 0.05472 | 0.00000 | 0.02930 | 0.06299 | 0.07563 |
| **MET** | 1.97188 | 0.00000 | 2.29299 | 2.64928 | 0.00000 | 0.00000 | 2.77880 |
| **METAP1** | 28.87324 | 8.84722 | 7.39661 | 12.12168 | 1.49155 | 9.62769 | 9.57510 |
| **METAP1D** | 2.15467 | 1.06476 | 1.23869 | 0.29375 | 0.00000 | 0.26162 | 0.78056 |
| **METAP2** | 27.22718 | 9.16449 | 21.22246 | 17.28648 | 11.26783 | 14.64127 | 29.91986 |
| **METRN** | 0.00000 | 0.53159 | 0.01548 | 0.00000 | 0.07661 | 0.66908 | 0.15913 |
| **METRNL** | 0.03344 | 4.73760 | 0.00000 | 1.83981 | 0.09946 | 0.00000 | 8.45038 |

| | | | | | | | |
|---|---|---|---|---|---|---|---|
| **METTL1** | 14.89164 | 12.48786 | 14.35421 | 7.31493 | 1.93456 | 3.88844 | 2.38191 |
| **METTL10** | 7.63941 | 1.79468 | 3.64014 | 0.00000 | 6.48786 | 3.83330 | 2.50752 |
| **METTL11B** | 0.04509 | 0.00000 | 0.04548 | 0.02041 | 0.06322 | 0.09911 | 0.00524 |
| **METTL12** | 6.46987 | 4.43260 | 1.14538 | 10.06284 | 2.39736 | 1.12100 | 3.47327 |
| **METTL13** | 24.16945 | 16.40260 | 11.57105 | 17.06533 | 2.20089 | 7.29695 | 0.00000 |
| **METTL14** | 9.99130 | 3.71172 | 5.80975 | 5.08342 | 2.31995 | 2.75120 | 0.71989 |
| **METTL15** | 2.66598 | 1.35651 | 1.14419 | 0.30957 | 0.12957 | 0.74502 | 1.50991 |
| **METTL16** | 11.38001 | 4.47523 | 9.99493 | 11.09074 | 2.94468 | 10.25269 | 8.50361 |
| **METTL17** | 9.95220 | 4.45540 | 5.08402 | 8.50577 | 1.07372 | 6.07147 | 0.00001 |
| **METTL18** | 3.98060 | 1.01162 | 5.80771 | 1.65424 | 0.60915 | 0.60243 | 0.00000 |
| **METTL20** | 0.36027 | 0.00000 | 1.45737 | 0.19209 | 1.41170 | 0.20779 | 0.00000 |
| **METTL21A** | 4.41839 | 0.00000 | 4.86497 | 3.35957 | 0.99891 | 0.86091 | 3.19294 |
| **METTL21B** | 0.49715 | 0.00000 | 0.38939 | 0.65482 | 0.42643 | 0.10118 | 0.27627 |
| **METTL21C** | 0.10136 | 0.00467 | 0.01318 | 0.04428 | 0.00708 | 0.00965 | 0.04600 |
| **METTL21EP** | 0.01569 | 0.02444 | 0.01082 | 0.02991 | 0.00801 | 0.02631 | 0.01840 |
| **METTL22** | 2.88295 | 2.87087 | 5.61952 | 4.42821 | 1.65174 | 2.21355 | 2.69777 |
| **METTL23** | 1.79333 | 4.62535 | 2.13416 | 3.38259 | 0.11657 | 4.17627 | 1.09727 |
| **METTL25** | 2.43590 | 0.00000 | 0.26152 | 1.35045 | 0.06675 | 1.05794 | 0.89492 |
| **METTL2A** | 10.02887 | 4.44845 | 2.97213 | 5.64954 | 0.50596 | 2.36242 | 4.86834 |
| **METTL2B** | 8.24453 | 3.28648 | 2.85541 | 4.24346 | 0.68207 | 2.75240 | 2.81970 |
| **METTL3** | 11.39028 | 4.88495 | 2.66184 | 15.36071 | 0.44866 | 1.91385 | 0.00000 |
| **METTL4** | 7.74346 | 0.00000 | 6.20996 | 8.33664 | 2.21033 | 1.86800 | 2.10919 |
| **METTL5** | 26.68374 | 13.95218 | 8.07023 | 11.36518 | 0.06343 | 8.72209 | 2.31929 |
| **METTL6** | 4.46181 | 1.43542 | 4.83397 | 3.91459 | 0.94696 | 1.48118 | 1.32420 |
| **METTL7A** | 0.13026 | 0.00000 | 0.02822 | 0.00611 | 0.17445 | 0.61485 | 0.40100 |
| **METTL7B** | 0.08065 | 0.25184 | 0.00897 | 0.00394 | 0.00779 | 0.07302 | 0.00000 |
| **METTL8** | 10.23391 | 2.51000 | 3.45975 | 7.56360 | 0.07931 | 2.48769 | 5.32081 |
| **METTL9** | 2.08406 | 4.61764 | 7.33637 | 5.84438 | 3.59071 | 10.46665 | 14.76681 |
| **MEX3A** | 0.00000 | 0.33372 | 0.23168 | 2.57680 | 0.47954 | 0.60701 | 5.91748 |
| **MEX3B** | 0.19586 | 0.00000 | 0.14993 | 0.04306 | 1.21123 | 0.17298 | 0.41081 |
| **MEX3C** | 8.66180 | 0.00000 | 1.09373 | 4.77496 | 3.41256 | 3.07256 | 11.15010 |
| **MEX3D** | 0.00001 | 0.00000 | 0.42579 | 0.10863 | 0.05942 | 0.40908 | 4.04296 |
| **MFAP1** | 23.86029 | 7.79000 | 18.13365 | 15.09303 | 19.39472 | 4.96122 | 5.85018 |
| **MFAP2** | 0.69363 | 1.63727 | 0.00000 | 1.94050 | 0.36783 | 0.42558 | 0.28697 |
| **MFAP3** | 9.98049 | 0.00000 | 9.17619 | 4.41314 | 3.08861 | 1.06652 | 1.79096 |

| | | | | | | | |
|---|---|---|---|---|---|---|---|
| **MFAP3L** | 5.45732 | 2.57226 | 2.14490 | 0.00000 | 2.39444 | 4.95977 | 2.66051 |
| **MFAP4** | 0.05158 | 0.01573 | 0.04164 | 0.02232 | 0.14072 | 0.01325 | 0.02370 |
| **MFAP5** | 0.92216 | 1.10055 | 1.23046 | 0.42796 | 0.00081 | 0.11327 | 0.00000 |
| **MFF** | 11.80830 | 3.65926 | 2.88362 | 8.27465 | 1.21260 | 5.39568 | 4.23313 |
| **MFGE8** | 0.15492 | 1.15699 | 0.49162 | 0.80344 | 0.20094 | 4.23076 | 0.00000 |
| **MFHAS1** | 2.71878 | 1.48544 | 0.26644 | 0.60600 | 0.92039 | 6.08335 | 9.05660 |
| **MFI2** | 0.00000 | 0.75103 | 0.14461 | 0.00000 | 0.03027 | 0.94549 | 0.30795 |
| **MFI2-AS1** | 0.00229 | 0.04539 | 0.01799 | 0.04611 | 0.06861 | 0.02444 | 0.12560 |
| **MFN1** | 17.36725 | 13.51101 | 32.85414 | 4.58126 | 13.51429 | 2.76061 | 1.50831 |
| **MFN2** | 18.00681 | 36.80905 | 12.36227 | 14.33369 | 1.75038 | 6.17258 | 20.78642 |
| **MFNG** | 0.11870 | 0.08355 | 0.02984 | 0.08071 | 0.03807 | 0.08279 | 0.11476 |
| **MFSD1** | 18.00744 | 3.43758 | 4.10117 | 11.19242 | 0.62989 | 3.58445 | 0.00000 |
| **MFSD10** | 4.73472 | 1.40486 | 0.67159 | 12.65655 | 6.01734 | 3.60150 | 10.01066 |
| **MFSD11** | 11.13973 | 3.20374 | 5.50549 | 5.28373 | 5.07373 | 3.13627 | 3.30378 |
| **MFSD12** | 0.59480 | 2.03358 | 0.28324 | 4.71566 | 0.26587 | 0.30965 | 6.98697 |
| **MFSD2A** | 0.00634 | 0.00000 | 15.07893 | 0.11569 | 19.22032 | 1.22480 | 0.00467 |
| **MFSD2B** | 0.03364 | 0.03801 | 0.02246 | 0.02950 | 0.02812 | 0.01706 | 0.01163 |
| **MFSD3** | 0.23259 | 0.36300 | 0.04734 | 0.38365 | 0.05079 | 0.91020 | 3.48790 |
| **MFSD4** | 0.04493 | 0.00000 | 0.45334 | 0.03891 | 0.83914 | 0.32373 | 0.01734 |
| **MFSD5** | 1.36091 | 8.63076 | 4.86145 | 9.84785 | 2.57248 | 3.15971 | 11.94367 |
| **MFSD6** | 0.00000 | 0.00000 | 0.17367 | 0.76147 | 0.04540 | 0.12320 | 0.11058 |
| **MFSD6L** | 0.03470 | 0.01739 | 0.00549 | 0.07637 | 0.10210 | 0.01707 | 0.01304 |
| **MFSD7** | 0.02643 | 0.18879 | 0.00346 | 0.03653 | 0.00897 | 0.35933 | 0.67311 |
| **MFSD8** | 9.75926 | 1.65281 | 4.30204 | 4.14867 | 0.66593 | 4.13348 | 2.30662 |
| **MFSD9** | 4.59892 | 6.01960 | 1.57608 | 4.37970 | 0.06360 | 2.38464 | 0.00000 |
| **MGA** | 11.40627 | 3.12017 | 13.74263 | 8.24683 | 7.37567 | 6.74358 | 20.31040 |
| **MGAM** | 0.05734 | 0.00000 | 0.04178 | 0.02072 | 0.10557 | 0.99433 | 0.04880 |
| **MGARP** | 0.03902 | 0.02204 | 0.00079 | 0.07878 | 0.10646 | 0.11459 | 0.10407 |
| **MGAT1** | 9.60267 | 13.55757 | 6.63250 | 32.55428 | 2.91178 | 9.42481 | 14.29810 |
| **MGAT2** | 4.84942 | 0.52282 | 5.51500 | 3.36537 | 2.81952 | 2.76033 | 2.82731 |
| **MGAT3** | 0.00000 | 0.44160 | 0.00000 | 1.07934 | 0.02849 | 0.26874 | 1.28458 |
| **MGAT4A** | 1.01467 | 0.00000 | 0.05054 | 5.55716 | 2.20513 | 1.89682 | 0.56643 |
| **MGAT4B** | 0.00000 | 0.83339 | 0.00000 | 0.10472 | 0.32221 | 2.40479 | 5.08707 |
| **MGAT4C** | 0.03367 | 0.02267 | 0.02865 | 0.02962 | 0.03290 | 0.01039 | 0.01439 |
| **MGAT5** | 0.98741 | 1.50785 | 0.33664 | 2.35991 | 3.42589 | 1.20459 | 6.23156 |

| | | | | | | | |
|---|---|---|---|---|---|---|---|
| **MGAT5B** | 0.00000 | 0.07187 | 0.24168 | 0.08757 | 1.59811 | 0.07489 | 0.12119 |
| **MGC12916** | 0.00208 | 0.13010 | 0.01013 | 0.73095 | 0.23853 | 0.13629 | 0.26963 |
| **MGC15885** | 0.08812 | 0.02281 | 0.01383 | 0.02056 | 0.00762 | 0.03993 | 0.01569 |
| **MGC16025** | 0.04368 | 0.06188 | 0.04543 | 0.01883 | 0.00582 | 0.02749 | 0.00000 |
| **MGC16142** | 0.15845 | 0.00000 | 0.14447 | 0.12200 | 0.21754 | 0.00907 | 0.01801 |
| **MGC16275** | 0.32414 | 0.53316 | 0.24656 | 0.71418 | 0.23660 | 0.05452 | 0.07968 |
| **MGC27345** | 5.59265 | 2.65477 | 0.88072 | 2.01304 | 0.07503 | 0.92130 | 2.58700 |
| **MGC27382** | 0.02317 | 0.01881 | 0.01241 | 0.01944 | 0.05324 | 0.01317 | 0.01564 |
| **MGC2889** | 0.02735 | 0.01419 | 0.02164 | 0.01820 | 0.00829 | 0.01329 | 0.01605 |
| **MGC32805** | 0.06021 | 0.11738 | 0.07740 | 0.00000 | 0.01233 | 0.38983 | 0.03528 |
| **MGC34796** | 0.03935 | 0.04856 | 0.01925 | 0.04273 | 0.03951 | 0.04380 | 0.05317 |
| **MGC45800** | 0.00000 | 0.01433 | 0.02626 | 1.55368 | 0.00973 | 0.00356 | 0.04921 |
| **MGC45922** | 0.04845 | 0.00000 | 0.08259 | 0.05918 | 0.05006 | 0.00450 | 0.03392 |
| **MGC57346** | 1.58343 | 0.38684 | 0.67070 | 0.76037 | 0.44999 | 1.06501 | 0.33140 |
| **MGC72080** | 0.29162 | 0.81518 | 0.04220 | 1.36590 | 0.46786 | 0.38030 | 1.83598 |
| **MGEA5** | 9.68144 | 3.87721 | 12.14992 | 8.04687 | 3.78548 | 10.58914 | 21.43658 |
| **MGLL** | 5.02147 | 0.82976 | 0.00000 | 2.88481 | 0.00000 | 4.98653 | 5.28239 |
| **MGME1** | 9.76780 | 8.76024 | 10.20865 | 7.53289 | 2.58393 | 6.19152 | 0.00000 |
| **MGMT** | 1.49620 | 2.48012 | 0.24161 | 1.32506 | 1.50978 | 1.54411 | 0.19590 |
| **MGP** | 0.02925 | 0.02425 | 0.01679 | 0.02892 | 0.11509 | 0.00912 | 0.07616 |
| **MGRN1** | 0.57385 | 1.49433 | 0.28590 | 4.19135 | 3.90999 | 2.37598 | 8.04407 |
| **MGST1** | 29.55561 | 0.00000 | 0.14727 | 9.16494 | 0.00000 | 0.00000 | 6.25118 |
| **MGST2** | 6.95597 | 3.69627 | 6.22801 | 0.00000 | 3.26556 | 5.60349 | 2.87095 |
| **MGST3** | 30.06671 | 5.38410 | 6.06411 | 39.51586 | 4.79567 | 3.11974 | 16.19861 |
| **MIA** | 0.03159 | 0.01934 | 0.01799 | 0.02593 | 0.02262 | 0.01123 | 0.01687 |
| **MIA3** | 9.15813 | 3.48700 | 5.04244 | 6.22583 | 5.24916 | 5.21670 | 3.77366 |
| **MIAT** | 0.00000 | 0.08784 | 0.10215 | 0.01686 | 0.19378 | 5.51487 | 0.18867 |
| **MIATNB** | 0.12902 | 0.02551 | 0.33136 | 0.04349 | 0.12591 | 0.14969 | 0.00000 |
| **MIB1** | 2.86598 | 0.00000 | 2.23757 | 0.65041 | 1.29048 | 1.55881 | 8.45532 |
| **MIB2** | 0.00000 | 2.63694 | 0.17583 | 0.12903 | 1.54656 | 1.93736 | 3.59776 |
| **MICA** | 0.00686 | 0.20846 | 1.55898 | 0.30242 | 0.04712 | 1.19317 | 0.00000 |
| **MICAL1** | 0.00000 | 1.91933 | 1.62194 | 1.92315 | 0.32509 | 2.10988 | 2.89655 |
| **MICAL2** | 3.02763 | 2.81239 | 0.00000 | 6.00290 | 1.33395 | 0.28304 | 9.34642 |
| **MICAL3** | 0.66646 | 2.74434 | 2.22022 | 3.62934 | 2.95475 | 2.42997 | 5.68938 |
| **MICALCL** | 0.11223 | 0.27455 | 0.00000 | 0.26681 | 2.05997 | 0.00000 | 1.37885 |

| | | | | | | | |
|---|---|---|---|---|---|---|---|
| **MICALL1** | 0.75532 | 5.61557 | 0.19576 | 7.96359 | 1.41368 | 1.50587 | 13.70829 |
| **MICALL2** | 0.15151 | 0.46343 | 0.06842 | 1.28172 | 0.38993 | 0.22212 | 2.09044 |
| **MICB** | 0.43477 | 0.41726 | 2.69508 | 1.31223 | 0.00005 | 3.68931 | 0.00000 |
| **MICU1** | 4.86584 | 4.31334 | 2.72845 | 3.53474 | 6.43288 | 4.27832 | 2.64040 |
| **MICU2** | 11.41805 | 0.00000 | 6.95868 | 6.04041 | 1.07701 | 5.48848 | 9.05002 |
| **MICU3** | 0.35836 | 0.00000 | 0.00843 | 0.31064 | 0.00445 | 0.11319 | 0.53192 |
| **MID1** | 4.33518 | 1.40257 | 6.83648 | 19.43707 | 1.45360 | 3.49036 | 2.39975 |
| **MID1IP1** | 0.68938 | 0.12413 | 0.01639 | 0.73989 | 0.00075 | 1.49874 | 10.00781 |
| **MID1IP1-AS1** | 0.04110 | 0.02431 | 0.01062 | 0.02268 | 0.00800 | 0.01309 | 0.01491 |
| **MID2** | 0.06727 | 0.07746 | 0.02282 | 0.01921 | 0.05758 | 0.03193 | 0.00091 |
| **MIDN** | 0.00000 | 22.64184 | 2.18905 | 8.78773 | 0.78903 | 6.59822 | 32.91206 |
| **MIEF1** | 25.20794 | 17.91566 | 10.07027 | 21.46711 | 4.15956 | 9.21893 | 13.94510 |
| **MIEF2** | 1.12123 | 2.81304 | 0.53055 | 1.91035 | 0.00000 | 0.91739 | 0.00035 |
| **MIEN1** | 16.59028 | 17.83347 | 7.06172 | 13.79051 | 2.67988 | 5.75384 | 2.28098 |
| **MIER1** | 16.90141 | 0.53503 | 6.03102 | 10.44863 | 3.73301 | 5.21801 | 5.96893 |
| **MIER2** | 0.00000 | 1.14406 | 1.65569 | 1.52049 | 0.25753 | 2.73444 | 6.74106 |
| **MIER3** | 4.40585 | 0.00000 | 4.33745 | 7.98934 | 1.66132 | 2.96787 | 3.68028 |
| **MIF4GD** | 1.30779 | 1.99846 | 0.06828 | 0.91473 | 1.05895 | 1.27209 | 0.57826 |
| **MIIP** | 25.41708 | 46.28746 | 11.05622 | 20.14678 | 1.53622 | 14.38553 | 14.64876 |
| **MILR1** | 0.02390 | 0.08384 | 0.00956 | 0.01963 | 0.03669 | 0.12785 | 0.00153 |
| **MIMT1** | 0.03728 | 0.01297 | 0.03318 | 0.02301 | 0.01553 | 0.01424 | 0.01842 |
| **MINA** | 5.31955 | 0.89211 | 4.99038 | 3.84083 | 2.27344 | 0.08545 | 0.00000 |
| **MINK1** | 0.00000 | 2.35490 | 1.30758 | 3.38539 | 0.18539 | 0.42025 | 8.06333 |
| **MINOS1** | 22.57411 | 9.76801 | 5.33542 | 19.17409 | 0.96998 | 12.20119 | 13.34272 |
| **MINOS1P1** | 0.36717 | 0.25435 | 0.41499 | 0.48058 | 0.09217 | 0.37136 | 1.07771 |
| **MINPP1** | 7.15505 | 1.29438 | 4.00200 | 3.97566 | 1.85162 | 3.13540 | 0.00000 |
| **MIOS** | 5.99710 | 2.35050 | 2.29608 | 1.81384 | 1.00023 | 0.20844 | 7.74750 |
| **MIOX** | 0.01869 | 0.06148 | 0.00448 | 0.12519 | 0.08338 | 0.00769 | 0.00000 |
| **MIP** | 0.08634 | 0.00000 | 0.61586 | 0.32028 | 0.23175 | 0.22212 | 0.00000 |
| **MIPEP** | 11.03983 | 6.88251 | 3.06632 | 0.62988 | 1.29305 | 3.77582 | 1.45194 |
| **MIPEPP3** | 0.93923 | 1.33470 | 1.32560 | 0.32589 | 1.39534 | 0.07125 | 0.00000 |
| **MIPOL1** | 0.48115 | 0.00000 | 0.58349 | 0.51125 | 0.03749 | 0.00000 | 0.59252 |
| **MIR100HG** | 0.02680 | 0.01159 | 0.00711 | 0.02099 | 0.08419 | 0.03105 | 0.04301 |
| **MIR103A2** | 0.17113 | 0.02026 | 0.07993 | 0.08247 | 0.04886 | 0.11967 | 0.14906 |
| **MIR106B** | 0.03950 | 0.03367 | 0.02400 | 0.05484 | 0.01225 | 0.01514 | 0.00000 |

| | | | | | | | |
|---|---|---|---|---|---|---|---|
| **MIR1178** | 0.02156 | 0.01697 | 0.01522 | 0.01884 | 0.00764 | 0.01699 | 0.02124 |
| **MIR1204** | 0.06748 | 0.25612 | 0.01488 | 0.19175 | 0.05114 | 0.05604 | 0.10275 |
| **MIR1206** | 0.03750 | 0.00819 | 0.01655 | 0.01007 | 0.00754 | 0.01036 | 0.05892 |
| **MIR1207** | 0.02602 | 0.01693 | 0.01364 | 0.02310 | 0.00785 | 0.01426 | 0.01681 |
| **MIR1226** | 0.22133 | 0.31232 | 0.11634 | 0.21829 | 0.00340 | 0.12535 | 0.37224 |
| **MIR1227** | 0.02395 | 0.06160 | 0.01847 | 0.05301 | 0.01017 | 0.01336 | 0.05944 |
| **MIR1228** | 0.02579 | 0.02730 | 0.01156 | 0.02412 | 0.00773 | 0.01161 | 0.02573 |
| **MIR1229** | 0.01093 | 0.03444 | 0.01264 | 0.02566 | 0.01005 | 0.04115 | 0.03473 |
| **MIR1231** | 0.02791 | 0.01945 | 0.01274 | 0.02289 | 0.00797 | 0.01971 | 0.01174 |
| **MIR1234** | 0.01694 | 0.02614 | 0.01470 | 0.02586 | 0.00735 | 0.02207 | 0.05522 |
| **MIR1236** | 0.02766 | 0.03959 | 0.01517 | 0.03382 | 0.03466 | 0.01668 | 0.01255 |
| **MIR1237** | 0.02264 | 0.02037 | 0.01191 | 0.02342 | 0.00749 | 0.01861 | 0.04050 |
| **MIR1238** | 0.01809 | 0.06899 | 0.01586 | 0.02634 | 0.01717 | 0.04621 | 0.03231 |
| **MIR124-2HG** | 0.02660 | 0.01530 | 0.00918 | 0.01872 | 0.09410 | 0.01746 | 0.01609 |
| **MIR1244-1+MIR** | 0.02235 | 0.02004 | 0.01346 | 0.02065 | 0.00781 | 0.01269 | 0.01746 |
| **MIR1244-1+MIR** | 0.02505 | 0.01822 | 0.01243 | 0.02105 | 0.01027 | 0.01243 | 0.01769 |
| **MIR126** | 0.02697 | 0.01792 | 0.01258 | 0.01998 | 0.00789 | 0.01216 | 0.01841 |
| **MIR1272** | 0.02240 | 0.02590 | 0.03497 | 0.01917 | 0.00665 | 0.01431 | 0.02539 |
| **MIR1277** | 0.02607 | 0.01789 | 0.01327 | 0.01946 | 0.00800 | 0.01347 | 0.01512 |
| **MIR128-1** | 0.02060 | 0.01814 | 0.01709 | 0.02138 | 0.00880 | 0.01260 | 0.02705 |
| **MIR1281** | 0.01345 | 0.01428 | 0.01226 | 0.01714 | 0.00756 | 0.02653 | 0.05156 |
| **MIR1283-1** | 0.02118 | 0.01587 | 0.01534 | 0.02438 | 0.00749 | 0.01688 | 0.02704 |
| **MIR1283-2** | 0.01735 | 0.01773 | 0.01376 | 0.04382 | 0.00726 | 0.01213 | 0.05650 |
| **MIR1291** | 0.02927 | 0.02078 | 0.01176 | 0.02138 | 0.00797 | 0.01933 | 0.01616 |
| **MIR1292** | 0.38348 | 0.57518 | 0.33332 | 0.43290 | 0.15614 | 0.23080 | 0.06689 |
| **MIR1296** | 0.03051 | 0.02767 | 0.02093 | 0.01964 | 0.00694 | 0.01225 | 0.02512 |
| **MIR1304** | 0.14028 | 0.21474 | 0.04340 | 0.09938 | 0.00832 | 0.10308 | 0.28320 |
| **MIR130A** | 0.02123 | 0.03130 | 0.02237 | 0.02868 | 0.00726 | 0.02466 | 0.01824 |
| **MIR130B** | 0.02193 | 0.01646 | 0.01467 | 0.06021 | 0.00798 | 0.03149 | 0.00897 |
| **MIR1322** | 0.02512 | 0.01448 | 0.01781 | 0.02222 | 0.00761 | 0.01632 | 0.01709 |
| **MIR1323** | 0.00742 | 0.02135 | 0.02529 | 0.04117 | 0.00622 | 0.03261 | 0.09990 |
| **MIR133A1HG** | 0.09614 | 0.00000 | 0.06015 | 0.06598 | 0.05905 | 0.06379 | 0.19695 |
| **MIR1343** | 0.01682 | 0.03870 | 0.01130 | 0.02154 | 0.00772 | 0.02815 | 0.02531 |
| **MIR137HG** | 0.02740 | 0.02082 | 0.01171 | 0.01779 | 0.01000 | 0.01201 | 0.01972 |
| **MIR138-1** | 0.01907 | 0.02025 | 0.01168 | 0.01823 | 0.00781 | 0.01174 | 0.03423 |

| | | | | | | | |
|---|---|---|---|---|---|---|---|
| **MIR141** | 0.02363 | 0.01845 | 0.01409 | 0.01743 | 0.00759 | 0.01503 | 0.03086 |
| **MIR142** | 0.03011 | 0.01812 | 0.01277 | 0.02915 | 0.00796 | 0.01196 | 0.01221 |
| **MIR143HG** | 0.02176 | 0.02947 | 0.01537 | 0.02614 | 0.00958 | 0.01177 | 0.01107 |
| **MIR144** | 0.01953 | 0.01810 | 0.01174 | 0.01311 | 0.01069 | 0.00997 | 0.05459 |
| **MIR1469** | 0.01901 | 0.01706 | 0.01324 | 0.03265 | 0.00987 | 0.01250 | 0.02418 |
| **MIR146B** | 0.01552 | 0.02914 | 0.01540 | 0.01652 | 0.00714 | 0.03685 | 0.03373 |
| **MIR148B** | 0.02521 | 0.01899 | 0.01716 | 0.02397 | 0.00771 | 0.01440 | 0.01214 |
| **MIR1539** | 0.02364 | 0.01735 | 0.01479 | 0.01991 | 0.00772 | 0.01369 | 0.01847 |
| **MIR155HG** | 0.05962 | 0.00558 | 0.05039 | 0.01250 | 0.00768 | 0.01243 | 0.02638 |
| **MIR15A** | 0.02527 | 0.01769 | 0.01272 | 0.01940 | 0.00792 | 0.01278 | 0.01783 |
| **MIR15B** | 0.02655 | 0.01834 | 0.01427 | 0.01916 | 0.00930 | 0.01301 | 0.01566 |
| **MIR16-2** | 0.04206 | 0.00000 | 0.05952 | 0.02860 | 0.01614 | 0.06557 | 0.16682 |
| **MIR17HG** | 9.17127 | 0.00000 | 1.78579 | 3.92052 | 0.12289 | 3.75165 | 19.01142 |
| **MIR181A1** | 0.03154 | 0.02077 | 0.01197 | 0.02167 | 0.00791 | 0.01164 | 0.01662 |
| **MIR181A1HG** | 0.02450 | 0.02649 | 0.01061 | 0.02211 | 0.01138 | 0.02121 | 0.00939 |
| **MIR181A2** | 0.02285 | 0.01820 | 0.01853 | 0.03108 | 0.01541 | 0.01924 | 0.02997 |
| **MIR181A2HG** | 0.00773 | 0.01137 | 0.04432 | 0.15134 | 0.01764 | 0.04311 | 0.20684 |
| **MIR181B2** | 0.02157 | 0.01560 | 0.01729 | 0.02464 | 0.00771 | 0.01180 | 0.02672 |
| **MIR186** | 0.02559 | 0.01153 | 0.01728 | 0.02380 | 0.00905 | 0.02069 | 0.06281 |
| **MIR188** | 0.02295 | 0.02199 | 0.01738 | 0.02054 | 0.00964 | 0.01914 | 0.02618 |
| **MIR1909** | 0.01078 | 0.02007 | 0.01473 | 0.03397 | 0.00709 | 0.03539 | 0.11177 |
| **MIR191** | 0.02187 | 0.02122 | 0.01073 | 0.03499 | 0.01254 | 0.01233 | 0.02807 |
| **MIR1910** | 0.02621 | 0.02243 | 0.01295 | 0.02839 | 0.00776 | 0.01103 | 0.01717 |
| **MIR1914** | 0.03712 | 0.06498 | 0.01328 | 0.04259 | 0.10196 | 0.02457 | 0.05551 |
| **MIR192** | 0.01222 | 0.17473 | 0.42250 | 0.06846 | 0.00308 | 0.00298 | 0.00491 |
| **MIR193A** | 0.02545 | 0.01653 | 0.01281 | 0.02067 | 0.00790 | 0.01406 | 0.02082 |
| **MIR193B** | 0.02350 | 0.02029 | 0.01275 | 0.01849 | 0.00798 | 0.01501 | 0.02007 |
| **MIR194-2** | 0.02397 | 0.01578 | 0.01822 | 0.01967 | 0.00756 | 0.01297 | 0.01655 |
| **MIR197** | 0.02684 | 0.01751 | 0.01670 | 0.02208 | 0.00943 | 0.01005 | 0.03303 |
| **MIR1972-1+MIR** | 0.02646 | 0.01809 | 0.01261 | 0.01864 | 0.00847 | 0.01324 | 0.01945 |
| **MIR1973** | 0.06152 | 0.03593 | 0.09594 | 0.06631 | 0.02674 | 0.10472 | 0.93115 |
| **MIR202HG** | 0.03145 | 0.01316 | 0.01178 | 0.03459 | 0.02104 | 0.02443 | 0.01672 |
| **MIR205** | 0.03144 | 0.01979 | 0.01112 | 0.07120 | 0.00776 | 0.02356 | 0.02881 |
| **MIR2052** | 0.02406 | 0.01786 | 0.01284 | 0.02135 | 0.00795 | 0.01311 | 0.01583 |
| **MIR205HG** | 1.57111 | 0.00000 | 0.00000 | 5.41742 | 0.00000 | 0.48465 | 3.83301 |

| | | | | | | | |
|---|---|---|---|---|---|---|---|
| **MIR21** | 0.01966 | 0.01689 | 0.01127 | 0.01500 | 0.01054 | 0.01429 | 0.05790 |
| **MIR210** | 0.02445 | 0.01976 | 0.01218 | 0.01861 | 0.00792 | 0.01216 | 0.01978 |
| **MIR210HG** | 0.03379 | 0.00633 | 0.00969 | 0.62958 | 0.00937 | 0.08967 | 0.34772 |
| **MIR2116** | 0.02301 | 0.02020 | 0.01258 | 0.02201 | 0.00785 | 0.01146 | 0.02328 |
| **MIR2117** | 0.02441 | 0.02868 | 0.01632 | 0.02510 | 0.00821 | 0.01043 | 0.02150 |
| **MIR221** | 0.01815 | 0.01532 | 0.01309 | 0.02830 | 0.00778 | 0.01551 | 0.03656 |
| **MIR2278** | 0.02419 | 0.02412 | 0.01524 | 0.02114 | 0.00756 | 0.01538 | 0.01478 |
| **MIR22HG** | 5.14172 | 3.98892 | 1.21829 | 9.19853 | 4.01303 | 3.25555 | 4.99865 |
| **MIR23B** | 0.02310 | 0.02033 | 0.01201 | 0.02243 | 0.00777 | 0.01147 | 0.02685 |
| **MIR23C** | 0.02202 | 0.01333 | 0.02463 | 0.02126 | 0.01078 | 0.01208 | 0.02458 |
| **MIR24-2** | 0.01846 | 0.01640 | 0.01285 | 0.02721 | 0.00788 | 0.01167 | 0.02862 |
| **MIR25** | 0.17550 | 0.30040 | 0.11296 | 0.31766 | 0.04816 | 0.22545 | 0.10015 |
| **MIR26A1** | 0.02804 | 0.01536 | 0.01311 | 0.03119 | 0.00782 | 0.01463 | 0.02187 |
| **MIR26A2** | 0.02784 | 0.01585 | 0.01516 | 0.02339 | 0.02123 | 0.01353 | 0.01460 |
| **MIR26B** | 0.02189 | 0.02655 | 0.01396 | 0.01966 | 0.00771 | 0.02331 | 0.01942 |
| **MIR27B** | 0.01809 | 0.01113 | 0.01435 | 0.02273 | 0.00674 | 0.00791 | 0.11567 |
| **MIR2861** | 0.02348 | 0.01848 | 0.01246 | 0.01889 | 0.00794 | 0.01480 | 0.01841 |
| **MIR2909** | 0.02448 | 0.02130 | 0.01303 | 0.02154 | 0.00780 | 0.01162 | 0.01981 |
| **MIR296** | 0.03175 | 0.01630 | 0.01312 | 0.02070 | 0.00783 | 0.01573 | 0.01721 |
| **MIR29A** | 0.02257 | 0.01708 | 0.01601 | 0.02547 | 0.00848 | 0.01229 | 0.01814 |
| **MIR29B1** | 0.02329 | 0.01887 | 0.01211 | 0.01827 | 0.00788 | 0.01166 | 0.02675 |
| **MIR29C** | 0.02979 | 0.01679 | 0.01351 | 0.01840 | 0.01261 | 0.01381 | 0.01244 |
| **MIR300** | 0.01977 | 0.01511 | 0.01224 | 0.02012 | 0.01120 | 0.03920 | 0.02037 |
| **MIR301A** | 0.02691 | 0.01716 | 0.01676 | 0.01822 | 0.00776 | 0.01632 | 0.01488 |
| **MIR302A** | 0.08780 | 0.06659 | 0.06289 | 0.00700 | 0.00591 | 0.09057 | 0.00000 |
| **MIR302B** | 0.08033 | 0.03775 | 0.07655 | 0.00950 | 0.00534 | 0.09210 | 0.00000 |
| **MIR302C** | 0.06041 | 0.13423 | 0.05075 | 0.01305 | 0.00765 | 0.09954 | 0.00000 |
| **MIR302D** | 0.08106 | 0.07318 | 0.04725 | 0.00918 | 0.00755 | 0.08587 | 0.00000 |
| **MIR3064** | 0.33770 | 0.30898 | 0.31321 | 0.18279 | 0.76869 | 0.20865 | 0.00000 |
| **MIR3074** | 0.02640 | 0.03528 | 0.01336 | 0.02292 | 0.00919 | 0.01169 | 0.01440 |
| **MIR30A** | 0.03782 | 0.03353 | 0.00782 | 0.00000 | 0.00728 | 0.01103 | 0.14190 |
| **MIR30C1** | 0.02489 | 0.01867 | 0.01361 | 0.02040 | 0.00790 | 0.01493 | 0.01631 |
| **MIR3115** | 0.02446 | 0.01949 | 0.01288 | 0.01994 | 0.01059 | 0.01435 | 0.01490 |
| **MIR3124** | 0.02295 | 0.02511 | 0.02123 | 0.02277 | 0.01293 | 0.01314 | 0.01262 |
| **MIR3127** | 0.02157 | 0.03286 | 0.01096 | 0.02115 | 0.00790 | 0.02308 | 0.01593 |

| | | | | | | | |
|---|---|---|---|---|---|---|---|
| **MIR3128** | 0.02715 | 0.01627 | 0.01523 | 0.01910 | 0.00755 | 0.01263 | 0.02543 |
| **MIR3129** | 0.02554 | 0.01746 | 0.01385 | 0.01931 | 0.00786 | 0.01298 | 0.01579 |
| **MIR3140** | 0.02322 | 0.01633 | 0.01330 | 0.02334 | 0.00769 | 0.01333 | 0.02793 |
| **MIR3143** | 0.01960 | 0.01728 | 0.01309 | 0.02530 | 0.01306 | 0.02200 | 0.02331 |
| **MIR3145** | 0.02491 | 0.01861 | 0.01281 | 0.01962 | 0.00789 | 0.01616 | 0.01717 |
| **MIR3153** | 0.02418 | 0.01783 | 0.01828 | 0.02135 | 0.00755 | 0.01393 | 0.01778 |
| **MIR3162** | 0.02389 | 0.01900 | 0.01423 | 0.01865 | 0.00789 | 0.01456 | 0.01782 |
| **MIR3173** | 0.02386 | 0.01430 | 0.01985 | 0.01898 | 0.00738 | 0.01454 | 0.01960 |
| **MIR3175** | 0.02350 | 0.01095 | 0.01662 | 0.02663 | 0.02302 | 0.01734 | 0.03290 |
| **MIR3176** | 0.02902 | 0.00454 | 0.01934 | 0.01535 | 0.00685 | 0.01789 | 0.05634 |
| **MIR3178** | 0.01986 | 0.02054 | 0.01078 | 0.01556 | 0.02171 | 0.01007 | 0.04364 |
| **MIR3188** | 0.00608 | 0.01490 | 0.01176 | 0.05796 | 0.00745 | 0.01573 | 0.08501 |
| **MIR3189** | 0.01062 | 0.01488 | 0.01355 | 0.04367 | 0.00775 | 0.01615 | 0.04223 |
| **MIR3192** | 0.01819 | 0.01328 | 0.01878 | 0.01789 | 0.00735 | 0.03608 | 0.02659 |
| **MIR31HG** | 0.03183 | 0.00686 | 0.01467 | 0.02777 | 0.00845 | 0.10200 | 0.03837 |
| **MIR32** | 0.02801 | 0.01790 | 0.01270 | 0.01891 | 0.00791 | 0.01289 | 0.01723 |
| **MIR320A** | 0.01892 | 0.02004 | 0.01535 | 0.02188 | 0.02769 | 0.01395 | 0.03298 |
| **MIR320B2** | 0.02374 | 0.01897 | 0.01951 | 0.02035 | 0.00806 | 0.01871 | 0.02954 |
| **MIR320C1** | 0.02443 | 0.01853 | 0.01525 | 0.01940 | 0.01085 | 0.01326 | 0.01641 |
| **MIR320D1** | 0.02402 | 0.01549 | 0.01328 | 0.01875 | 0.00999 | 0.01752 | 0.02691 |
| **MIR320D2** | 0.03026 | 0.01587 | 0.01690 | 0.01840 | 0.00746 | 0.01144 | 0.02480 |
| **MIR320E** | 0.02434 | 0.02023 | 0.01830 | 0.02610 | 0.01275 | 0.03554 | 0.02661 |
| **MIR324** | 0.02525 | 0.01274 | 0.01318 | 0.03136 | 0.00735 | 0.03672 | 0.05067 |
| **MIR326** | 0.02440 | 0.01430 | 0.01326 | 0.01923 | 0.00782 | 0.01307 | 0.04257 |
| **MIR328** | 0.02473 | 0.01980 | 0.01383 | 0.01775 | 0.00776 | 0.01750 | 0.01933 |
| **MIR330** | 0.03650 | 0.05357 | 0.02104 | 0.01447 | 0.00901 | 0.02112 | 0.01190 |
| **MIR331** | 0.02596 | 0.01787 | 0.01556 | 0.02709 | 0.00854 | 0.02334 | 0.01567 |
| **MIR33B** | 0.02385 | 0.01891 | 0.01252 | 0.01986 | 0.00793 | 0.01375 | 0.01740 |
| **MIR345** | 0.01567 | 0.03454 | 0.01858 | 0.02703 | 0.00732 | 0.03031 | 0.02099 |
| **MIR34A** | 0.02601 | 0.02025 | 0.01505 | 0.04330 | 0.00775 | 0.01176 | 0.01501 |
| **MIR34C** | 0.02391 | 0.01927 | 0.01234 | 0.03315 | 0.00875 | 0.01221 | 0.01405 |
| **MIR3605** | 0.02504 | 0.01740 | 0.01645 | 0.03217 | 0.00803 | 0.01530 | 0.01113 |
| **MIR3607** | 0.07471 | 0.00000 | 0.03058 | 0.02027 | 0.00773 | 0.03524 | 0.17081 |
| **MIR3609** | 0.01391 | 0.03499 | 0.01457 | 0.02091 | 0.01723 | 0.03620 | 0.02601 |
| **MIR3620** | 1.07562 | 0.66586 | 0.31320 | 0.69385 | 0.24678 | 0.48495 | 0.00000 |

| | | | | | | | |
|---|---|---|---|---|---|---|---|
| **MIR3646** | 0.02184 | 0.01064 | 0.02484 | 0.02153 | 0.00726 | 0.01785 | 0.02149 |
| **MIR3648** | 0.00088 | 0.00753 | 0.02627 | 0.00370 | 0.01783 | 0.01136 | 0.79881 |
| **MIR3651** | 0.01974 | 0.03247 | 0.01790 | 0.03447 | 0.00772 | 0.01984 | 0.03453 |
| **MIR3653** | 0.02456 | 0.01777 | 0.01281 | 0.01919 | 0.00902 | 0.01283 | 0.01726 |
| **MIR3656** | 0.02404 | 0.01965 | 0.01371 | 0.02065 | 0.00835 | 0.01291 | 0.01843 |
| **MIR3660** | 0.02875 | 0.01213 | 0.01809 | 0.02081 | 0.00769 | 0.02216 | 0.01515 |
| **MIR3661** | 0.11826 | 0.27618 | 0.13388 | 0.31488 | 0.22059 | 0.10831 | 0.22201 |
| **MIR3662** | 0.02616 | 0.02269 | 0.01264 | 0.01818 | 0.01256 | 0.02021 | 0.02803 |
| **MIR3668** | 0.02544 | 0.01751 | 0.01263 | 0.01963 | 0.01128 | 0.01295 | 0.01598 |
| **MIR367** | 0.11774 | 0.06140 | 0.10655 | 0.00548 | 0.00500 | 0.09830 | 0.00000 |
| **MIR3671** | 0.03634 | 0.01903 | 0.03986 | 0.01372 | 0.00985 | 0.05817 | 0.06010 |
| **MIR3675.uid-91** | 0.02228 | 0.01654 | 0.01326 | 0.03330 | 0.00783 | 0.01196 | 0.02243 |
| **MIR3675.uid-91** | 0.01400 | 0.01696 | 0.01247 | 0.03881 | 0.00770 | 0.01044 | 0.04215 |
| **MIR3679** | 0.02486 | 0.04080 | 0.01589 | 0.03373 | 0.00676 | 0.01765 | 0.05700 |
| **MIR3682** | 0.03428 | 0.00000 | 0.04311 | 0.01928 | 0.01065 | 0.06980 | 0.29498 |
| **MIR3685** | 0.06004 | 0.00472 | 0.02056 | 0.01697 | 0.00971 | 0.02584 | 0.01875 |
| **MIR3687** | 0.00183 | 0.01562 | 0.01527 | 0.02012 | 0.00996 | 0.03011 | 0.14626 |
| **MIR371A** | 0.01858 | 0.03743 | 0.01401 | 0.02057 | 0.00769 | 0.02334 | 0.03180 |
| **MIR371B** | 0.02649 | 0.02447 | 0.01366 | 0.02359 | 0.00870 | 0.01492 | 0.01718 |
| **MIR372** | 0.43245 | 0.99579 | 0.17038 | 0.12557 | 0.07281 | 0.48695 | 0.40572 |
| **MIR373** | 0.31221 | 0.36692 | 0.16945 | 0.10476 | 0.02566 | 0.13665 | 0.30890 |
| **MIR378J** | 0.03085 | 0.02050 | 0.02253 | 0.02362 | 0.01019 | 0.02279 | 0.01102 |
| **MIR3911** | 0.02039 | 0.08566 | 0.01588 | 0.03651 | 0.02075 | 0.00453 | 0.05685 |
| **MIR3916** | 0.11314 | 0.02415 | 0.07281 | 0.10896 | 0.15349 | 0.12490 | 0.03970 |
| **MIR3917** | 3.25051 | 3.29233 | 2.19832 | 0.88965 | 0.07791 | 3.37183 | 0.00000 |
| **MIR3921** | 0.02566 | 0.01732 | 0.01611 | 0.01860 | 0.00756 | 0.01220 | 0.01993 |
| **MIR3928** | 0.02465 | 0.01793 | 0.01491 | 0.01983 | 0.00757 | 0.01129 | 0.02550 |
| **MIR3939** | 0.09493 | 0.01578 | 0.01879 | 0.00000 | 0.00660 | 0.01381 | 0.07260 |
| **MIR3940** | 0.02770 | 0.02548 | 0.02502 | 0.02476 | 0.00970 | 0.02290 | 0.08667 |
| **MIR3941** | 0.01732 | 0.03091 | 0.01618 | 0.02831 | 0.00765 | 0.01155 | 0.02401 |
| **MIR3942** | 0.03119 | 0.01466 | 0.02297 | 0.01882 | 0.00931 | 0.01243 | 0.01241 |
| **MIR3944** | 0.08933 | 0.02980 | 0.01154 | 0.01218 | 0.00759 | 0.01218 | 0.00994 |
| **MIR3975** | 0.02122 | 0.02208 | 0.01324 | 0.01746 | 0.00934 | 0.01323 | 0.03570 |
| **MIR421** | 0.04542 | 0.02483 | 0.02483 | 0.02932 | 0.00645 | 0.02218 | 0.03968 |
| **MIR423** | 0.02443 | 0.01983 | 0.02040 | 0.01970 | 0.00782 | 0.01551 | 0.03341 |

| | | | | | | | |
|---|---|---|---|---|---|---|---|
| **MIR425** | 0.01220 | 0.02521 | 0.01605 | 0.06074 | 0.00720 | 0.00856 | 0.04373 |
| **MIR4256** | 0.02377 | 0.01920 | 0.01245 | 0.02038 | 0.00795 | 0.01362 | 0.01683 |
| **MIR4257** | 0.03484 | 0.01198 | 0.01318 | 0.02662 | 0.00753 | 0.00980 | 0.03661 |
| **MIR4258** | 0.05632 | 0.00110 | 0.02326 | 0.03749 | 0.01252 | 0.06489 | 0.10661 |
| **MIR4263** | 0.05594 | 0.03737 | 0.01161 | 0.02198 | 0.00732 | 0.02734 | 0.02564 |
| **MIR4269** | 0.02129 | 0.02441 | 0.01502 | 0.02026 | 0.00891 | 0.01226 | 0.01994 |
| **MIR4276** | 0.02669 | 0.01776 | 0.01300 | 0.01826 | 0.00785 | 0.01237 | 0.01840 |
| **MIR4278** | 0.02728 | 0.01578 | 0.01346 | 0.01955 | 0.00797 | 0.02543 | 0.01222 |
| **MIR4279** | 0.03243 | 0.02283 | 0.01441 | 0.03236 | 0.00729 | 0.01064 | 0.03882 |
| **MIR4284** | 0.01433 | 0.02247 | 0.01248 | 0.02566 | 0.00695 | 0.00911 | 0.09800 |
| **MIR4292** | 0.03172 | 0.01630 | 0.01572 | 0.00658 | 0.00698 | 0.01812 | 0.08590 |
| **MIR4295** | 0.02777 | 0.01562 | 0.01372 | 0.02037 | 0.00782 | 0.01230 | 0.01851 |
| **MIR4296** | 0.02902 | 0.01358 | 0.01300 | 0.02145 | 0.00796 | 0.02013 | 0.02966 |
| **MIR4307** | 0.03688 | 0.01388 | 0.01723 | 0.01791 | 0.00942 | 0.01376 | 0.02230 |
| **MIR4308** | 0.02908 | 0.01829 | 0.01280 | 0.02100 | 0.00789 | 0.01212 | 0.01701 |
| **MIR4312** | 0.02215 | 0.01677 | 0.01295 | 0.01622 | 0.00733 | 0.00830 | 0.05513 |
| **MIR4324** | 0.02044 | 0.01501 | 0.02382 | 0.03508 | 0.02032 | 0.01832 | 0.04016 |
| **MIR4326** | 0.02350 | 0.02017 | 0.01264 | 0.01972 | 0.00868 | 0.01290 | 0.01815 |
| **MIR4329** | 0.02370 | 0.01981 | 0.01510 | 0.01977 | 0.00851 | 0.01248 | 0.01657 |
| **MIR4427** | 0.01412 | 0.01000 | 0.01199 | 0.01036 | 0.00728 | 0.00718 | 0.11716 |
| **MIR4434** | 0.01909 | 0.03816 | 0.02980 | 0.02012 | 0.01384 | 0.01101 | 0.03160 |
| **MIR4435-1HG** | 1.27790 | 1.44955 | 0.00000 | 3.32955 | 0.00000 | 1.02863 | 1.23164 |
| **MIR4442** | 0.02256 | 0.02260 | 0.01190 | 0.02314 | 0.00781 | 0.02229 | 0.02317 |
| **MIR4444-1+MIR** | 0.02542 | 0.01716 | 0.01457 | 0.01950 | 0.00780 | 0.01452 | 0.01611 |
| **MIR4449** | 0.01526 | 0.04283 | 0.02758 | 0.04061 | 0.01140 | 0.02270 | 0.07645 |
| **MIR4450** | 0.03860 | 0.03643 | 0.01416 | 0.01917 | 0.00745 | 0.02859 | 0.01683 |
| **MIR4451** | 0.02122 | 0.01872 | 0.01249 | 0.02207 | 0.00788 | 0.01244 | 0.02112 |
| **MIR4458HG** | 1.17638 | 0.76384 | 0.88750 | 1.09564 | 0.00000 | 0.00000 | 0.21884 |
| **MIR4461** | 0.04962 | 0.02633 | 0.13962 | 0.00000 | 0.20409 | 0.10763 | 0.44099 |
| **MIR4465** | 0.02426 | 0.01994 | 0.01280 | 0.02328 | 0.00796 | 0.01249 | 0.01591 |
| **MIR4467** | 0.01937 | 0.01612 | 0.01317 | 0.02536 | 0.00764 | 0.01221 | 0.03294 |
| **MIR4469** | 0.01965 | 0.02267 | 0.01374 | 0.01829 | 0.00743 | 0.01381 | 0.03696 |
| **MIR4472-2** | 0.01748 | 0.03158 | 0.01519 | 0.02526 | 0.00791 | 0.03646 | 0.03063 |
| **MIR4482** | 0.03245 | 0.01786 | 0.01791 | 0.02933 | 0.01217 | 0.01992 | 0.02859 |
| **MIR4497** | 0.02509 | 0.02010 | 0.01223 | 0.01835 | 0.00788 | 0.01252 | 0.01983 |

| | | | | | | | |
|---|---|---|---|---|---|---|---|
| **MIR4500HG** | 0.02346 | 0.00000 | 0.04570 | 0.01670 | 0.57655 | 0.01179 | 0.02241 |
| **MIR4517** | 0.06226 | 0.08335 | 0.01421 | 0.03505 | 0.03325 | 0.02051 | 0.05375 |
| **MIR4519** | 0.03313 | 0.03533 | 0.01924 | 0.01742 | 0.02218 | 0.00929 | 0.01368 |
| **MIR4521** | 0.02418 | 0.01424 | 0.02018 | 0.01933 | 0.01047 | 0.01298 | 0.01719 |
| **MIR454** | 0.02115 | 0.00975 | 0.01235 | 0.01523 | 0.01226 | 0.00906 | 0.07218 |
| **MIR4632** | 0.02323 | 0.02029 | 0.00884 | 0.02351 | 0.04427 | 0.01338 | 0.01331 |
| **MIR4639** | 0.05732 | 0.01720 | 0.01580 | 0.02229 | 0.01756 | 0.01806 | 0.07114 |
| **MIR4640** | 0.01623 | 0.01963 | 0.01585 | 0.03871 | 0.00782 | 0.02224 | 0.04089 |
| **MIR4641** | 0.02166 | 0.02112 | 0.02015 | 0.02317 | 0.00968 | 0.01516 | 0.02248 |
| **MIR4646** | 0.02558 | 0.01916 | 0.01272 | 0.02009 | 0.00794 | 0.01419 | 0.01513 |
| **MIR4648** | 0.02512 | 0.01839 | 0.01375 | 0.01909 | 0.00984 | 0.01571 | 0.01778 |
| **MIR4651** | 0.02695 | 0.01662 | 0.01362 | 0.02154 | 0.01528 | 0.01238 | 0.01660 |
| **MIR4652** | 0.02441 | 0.01490 | 0.01977 | 0.01874 | 0.00690 | 0.02098 | 0.04496 |
| **MIR4653** | 0.01486 | 0.04639 | 0.01443 | 0.03276 | 0.00872 | 0.02398 | 0.02145 |
| **MIR4655** | 0.02355 | 0.02110 | 0.01447 | 0.01969 | 0.00768 | 0.01246 | 0.01658 |
| **MIR4656** | 0.02550 | 0.02117 | 0.01345 | 0.01831 | 0.00798 | 0.01664 | 0.02310 |
| **MIR4660** | 0.02675 | 0.03019 | 0.01094 | 0.01897 | 0.00799 | 0.01158 | 0.01220 |
| **MIR4664** | 0.02821 | 0.01289 | 0.01342 | 0.02237 | 0.00778 | 0.01746 | 0.02533 |
| **MIR4667** | 0.01878 | 0.02589 | 0.03098 | 0.02384 | 0.01344 | 0.01894 | 0.03359 |
| **MIR4668** | 0.15893 | 0.25641 | 0.00731 | 0.00000 | 0.01274 | 0.11719 | 0.27863 |
| **MIR4670** | 0.02132 | 0.02155 | 0.01263 | 0.01638 | 0.00756 | 0.01792 | 0.03698 |
| **MIR4676** | 0.01967 | 0.01935 | 0.02167 | 0.01784 | 0.00857 | 0.02083 | 0.03079 |
| **MIR4677** | 0.02335 | 0.01861 | 0.01283 | 0.02084 | 0.00791 | 0.01657 | 0.01703 |
| **MIR4685** | 0.03172 | 0.05160 | 0.02031 | 0.01298 | 0.00616 | 0.00575 | 0.08686 |
| **MIR4690** | 0.02187 | 0.05301 | 0.01456 | 0.02565 | 0.00674 | 0.00702 | 0.04850 |
| **MIR4691** | 0.06315 | 0.11264 | 0.03992 | 0.06415 | 0.00653 | 0.05106 | 0.11029 |
| **MIR4697HG** | 0.00000 | 0.08150 | 0.11687 | 0.12954 | 0.00462 | 0.28608 | 0.28582 |
| **MIR4712** | 0.03341 | 0.02220 | 0.02715 | 0.03720 | 0.00747 | 0.01750 | 0.03750 |
| **MIR4720** | 0.03300 | 0.03182 | 0.01470 | 0.03082 | 0.00837 | 0.01223 | 0.02548 |
| **MIR4721** | 0.60725 | 0.40380 | 0.18296 | 0.53261 | 0.03580 | 0.25502 | 0.12407 |
| **MIR4722** | 0.02769 | 0.03394 | 0.01737 | 0.01946 | 0.00679 | 0.01131 | 0.05575 |
| **MIR4723** | 0.23791 | 0.10131 | 0.14667 | 0.13662 | 0.11021 | 0.10359 | 0.00000 |
| **MIR4727** | 0.02851 | 0.02121 | 0.01164 | 0.02191 | 0.01511 | 0.00978 | 0.02907 |
| **MIR4728** | 0.02599 | 0.01863 | 0.01254 | 0.02363 | 0.00838 | 0.01272 | 0.01480 |
| **MIR4729** | 0.02291 | 0.01718 | 0.01750 | 0.01639 | 0.00744 | 0.02225 | 0.02559 |

| | | | | | | | |
|---|---|---|---|---|---|---|---|
| **MIR4730** | 0.02456 | 0.01837 | 0.01348 | 0.02034 | 0.00787 | 0.01360 | 0.01667 |
| **MIR4737** | 0.04862 | 0.05186 | 0.02579 | 0.03893 | 0.00685 | 0.03375 | 0.00000 |
| **MIR4742** | 0.02446 | 0.02015 | 0.01214 | 0.02080 | 0.00795 | 0.01377 | 0.01774 |
| **MIR4743** | 0.02450 | 0.02121 | 0.01024 | 0.01609 | 0.01201 | 0.01390 | 0.04863 |
| **MIR4745** | 0.02582 | 0.01717 | 0.01524 | 0.01973 | 0.00766 | 0.01284 | 0.01725 |
| **MIR4746** | 0.00000 | 0.18685 | 0.05041 | 0.03204 | 0.03415 | 0.07577 | 0.15657 |
| **MIR4747** | 0.02592 | 0.01716 | 0.01333 | 0.02136 | 0.00791 | 0.01677 | 0.01487 |
| **MIR4748** | 0.01081 | 0.06336 | 0.00692 | 0.01991 | 0.01671 | 0.02266 | 0.02897 |
| **MIR4749** | 0.01976 | 0.04463 | 0.01183 | 0.04266 | 0.01169 | 0.05096 | 0.08225 |
| **MIR4750** | 0.01108 | 0.03398 | 0.02304 | 0.04266 | 0.01417 | 0.02222 | 0.05510 |
| **MIR4754** | 0.04651 | 0.02909 | 0.02282 | 0.04167 | 0.00914 | 0.02611 | 0.01188 |
| **MIR4755** | 0.02646 | 0.02099 | 0.02151 | 0.02055 | 0.02108 | 0.01756 | 0.02389 |
| **MIR4762** | 0.03482 | 0.01207 | 0.01353 | 0.01563 | 0.00778 | 0.01409 | 0.02311 |
| **MIR4766** | 0.03405 | 0.01919 | 0.01455 | 0.02429 | 0.00730 | 0.01560 | 0.03347 |
| **MIR4767** | 0.01950 | 0.04608 | 0.00751 | 0.02456 | 0.01277 | 0.05225 | 0.01064 |
| **MIR4781** | 0.02388 | 0.01727 | 0.01315 | 0.02071 | 0.01295 | 0.01300 | 0.01636 |
| **MIR4784** | 0.02418 | 0.03048 | 0.01245 | 0.02939 | 0.00784 | 0.02128 | 0.01250 |
| **MIR4785** | 0.02177 | 0.02323 | 0.01158 | 0.02157 | 0.00796 | 0.01411 | 0.02164 |
| **MIR4786** | 0.03034 | 0.06786 | 0.02231 | 0.01926 | 0.00737 | 0.01201 | 0.02688 |
| **MIR4787** | 0.01830 | 0.03122 | 0.01119 | 0.01875 | 0.00784 | 0.01990 | 0.02151 |
| **MIR4793** | 0.02329 | 0.01728 | 0.01378 | 0.02415 | 0.00778 | 0.01366 | 0.01955 |
| **MIR4794** | 0.02284 | 0.01923 | 0.01789 | 0.01858 | 0.00733 | 0.01961 | 0.01828 |
| **MIR4802** | 0.02372 | 0.01873 | 0.01580 | 0.02990 | 0.00788 | 0.01433 | 0.01329 |
| **MIR491** | 0.02616 | 0.03527 | 0.01894 | 0.02283 | 0.00817 | 0.00892 | 0.01952 |
| **MIR492** | 2.33685 | 0.01280 | 0.00601 | 4.67352 | 0.00221 | 0.00000 | 0.07795 |
| **MIR493** | 0.02212 | 0.01699 | 0.01260 | 0.01952 | 0.00797 | 0.02338 | 0.01723 |
| **MIR497HG** | 0.08103 | 0.12512 | 0.10056 | 0.21331 | 0.12336 | 0.13546 | 0.19756 |
| **MIR498** | 0.02923 | 0.01403 | 0.01736 | 0.07384 | 0.00827 | 0.04160 | 0.09745 |
| **MIR5000** | 0.02631 | 0.02977 | 0.03525 | 0.02424 | 0.00660 | 0.01658 | 0.01099 |
| **MIR5001** | 0.03345 | 0.05168 | 0.02770 | 0.04486 | 0.04641 | 0.01726 | 0.02272 |
| **MIR5004** | 0.02406 | 0.01614 | 0.01296 | 0.01774 | 0.01182 | 0.01274 | 0.02441 |
| **MIR5006** | 0.02312 | 0.01976 | 0.01277 | 0.02039 | 0.00975 | 0.01399 | 0.01707 |
| **MIR5010** | 0.02453 | 0.02068 | 0.01328 | 0.02479 | 0.00854 | 0.01902 | 0.10807 |
| **MIR503** | 0.02349 | 0.01773 | 0.01519 | 0.02038 | 0.00759 | 0.01425 | 0.02463 |
| **MIR503HG** | 0.03131 | 0.00717 | 0.08563 | 0.00000 | 0.02721 | 0.00000 | 0.48950 |

| | | | | | | | |
|---|---|---|---|---|---|---|---|
| **MIR5047** | 0.33773 | 0.61450 | 0.78091 | 0.42302 | 0.79249 | 0.21060 | 0.60085 |
| **MIR5087** | 0.01074 | 0.01272 | 0.01624 | 0.01132 | 0.00660 | 0.03642 | 0.10217 |
| **MIR5088** | 0.50469 | 0.32650 | 0.17106 | 0.47026 | 0.09693 | 0.15307 | 0.00011 |
| **MIR5090** | 0.02278 | 0.03613 | 0.02615 | 0.03905 | 0.00714 | 0.01974 | 0.00863 |
| **MIR5091** | 0.03196 | 0.03512 | 0.03210 | 0.01678 | 0.01024 | 0.00607 | 0.04962 |
| **MIR5094** | 0.02863 | 0.01673 | 0.01410 | 0.01190 | 0.00715 | 0.01196 | 0.05491 |
| **MIR512-2.uid-24** | 0.02433 | 0.01783 | 0.01341 | 0.01925 | 0.00784 | 0.01290 | 0.01790 |
| **MIR516A1** | 0.03393 | 0.01768 | 0.01617 | 0.02560 | 0.00826 | 0.01508 | 0.01629 |
| **MIR516A2** | 0.02997 | 0.01703 | 0.01297 | 0.01966 | 0.00787 | 0.01235 | 0.01710 |
| **MIR516B1** | 0.02552 | 0.01692 | 0.01550 | 0.02279 | 0.00777 | 0.01353 | 0.01562 |
| **MIR516B2** | 0.03751 | 0.01067 | 0.01521 | 0.02040 | 0.00752 | 0.01404 | 0.02856 |
| **MIR517A** | 0.03365 | 0.01300 | 0.02204 | 0.04506 | 0.00753 | 0.01892 | 0.02641 |
| **MIR517B** | 0.02723 | 0.01348 | 0.01685 | 0.02983 | 0.00719 | 0.02731 | 0.04147 |
| **MIR517C** | 0.02955 | 0.01246 | 0.01585 | 0.02524 | 0.00740 | 0.01388 | 0.05720 |
| **MIR5187** | 0.02407 | 0.02229 | 0.01238 | 0.02536 | 0.00797 | 0.01890 | 0.01255 |
| **MIR5188** | 0.01962 | 0.01530 | 0.01300 | 0.03107 | 0.00813 | 0.02346 | 0.02046 |
| **MIR518A1** | 0.01687 | 0.02559 | 0.01930 | 0.03710 | 0.00711 | 0.02294 | 0.03649 |
| **MIR518A2** | 0.02298 | 0.02607 | 0.01360 | 0.02814 | 0.00781 | 0.01519 | 0.01228 |
| **MIR518C** | 0.02814 | 0.01639 | 0.01309 | 0.01723 | 0.00793 | 0.01540 | 0.01859 |
| **MIR518D** | 0.03649 | 0.01777 | 0.01443 | 0.01844 | 0.00760 | 0.01450 | 0.02318 |
| **MIR518E** | 0.02343 | 0.01764 | 0.01265 | 0.02299 | 0.00785 | 0.01223 | 0.02249 |
| **MIR518F** | 0.02614 | 0.01482 | 0.02017 | 0.02075 | 0.00741 | 0.01207 | 0.01888 |
| **MIR5192** | 0.02741 | 0.01713 | 0.01273 | 0.01934 | 0.00788 | 0.01206 | 0.01930 |
| **MIR5196** | 0.02290 | 0.03446 | 0.01080 | 0.01901 | 0.00789 | 0.01274 | 0.01907 |
| **MIR519A1** | 0.02392 | 0.01738 | 0.01292 | 0.02111 | 0.00792 | 0.01282 | 0.01728 |
| **MIR519A2** | 0.04831 | 0.00000 | 0.02514 | 0.04469 | 0.00769 | 0.03566 | 0.04411 |
| **MIR519B** | 0.02191 | 0.01725 | 0.01635 | 0.03701 | 0.00751 | 0.01285 | 0.03227 |
| **MIR519C** | 0.02476 | 0.01789 | 0.01285 | 0.02413 | 0.00796 | 0.01314 | 0.01487 |
| **MIR519D** | 0.02271 | 0.01750 | 0.01323 | 0.02669 | 0.00789 | 0.01282 | 0.02133 |
| **MIR519E** | 0.02308 | 0.01806 | 0.01277 | 0.02084 | 0.00790 | 0.01291 | 0.01899 |
| **MIR520A** | 0.02756 | 0.00833 | 0.01441 | 0.03206 | 0.00730 | 0.01374 | 0.05187 |
| **MIR520C** | 0.01501 | 0.03348 | 0.01271 | 0.03600 | 0.00715 | 0.02229 | 0.04907 |
| **MIR520D** | 0.08646 | 0.00808 | 0.05871 | 0.10685 | 0.01159 | 0.06290 | 0.12490 |
| **MIR520E** | 0.02330 | 0.01231 | 0.02162 | 0.04550 | 0.00680 | 0.01972 | 0.04884 |
| **MIR520F** | 0.03418 | 0.00000 | 0.05238 | 0.13200 | 0.00726 | 0.05395 | 0.12901 |

| | | | | | | | |
|---|---|---|---|---|---|---|---|
| **MIR520G** | 0.02483 | 0.01675 | 0.01439 | 0.03506 | 0.00735 | 0.01467 | 0.06938 |
| **MIR520H** | 0.03153 | 0.00608 | 0.01884 | 0.02744 | 0.00674 | 0.02694 | 0.07121 |
| **MIR521-1** | 0.02385 | 0.01714 | 0.01263 | 0.01947 | 0.00784 | 0.01209 | 0.02549 |
| **MIR521-2** | 0.01956 | 0.01771 | 0.01336 | 0.04120 | 0.00888 | 0.01222 | 0.03388 |
| **MIR522** | 0.02603 | 0.01414 | 0.01848 | 0.02363 | 0.00736 | 0.01967 | 0.02382 |
| **MIR523** | 0.02290 | 0.01692 | 0.01453 | 0.03153 | 0.00856 | 0.01234 | 0.01944 |
| **MIR525** | 0.02043 | 0.01896 | 0.01240 | 0.02413 | 0.00792 | 0.01637 | 0.01955 |
| **MIR526A1** | 0.02344 | 0.00746 | 0.02310 | 0.05165 | 0.00669 | 0.02483 | 0.07550 |
| **MIR526A2** | 0.02606 | 0.01071 | 0.01798 | 0.03324 | 0.00724 | 0.01966 | 0.03422 |
| **MIR526B** | 0.02916 | 0.01496 | 0.01453 | 0.01902 | 0.00781 | 0.01528 | 0.01942 |
| **MIR527** | 0.02278 | 0.01921 | 0.01460 | 0.02423 | 0.00791 | 0.01411 | 0.01805 |
| **MIR542** | 0.02234 | 0.02092 | 0.01498 | 0.01932 | 0.01256 | 0.01338 | 0.01722 |
| **MIR545** | 0.02707 | 0.01622 | 0.01370 | 0.02441 | 0.00778 | 0.01186 | 0.02105 |
| **MIR548AC** | 0.02802 | 0.01750 | 0.01386 | 0.01955 | 0.00829 | 0.01327 | 0.01406 |
| **MIR548AE2** | 0.02410 | 0.01605 | 0.01497 | 0.01952 | 0.01778 | 0.01305 | 0.01656 |
| **MIR548AZ** | 0.02372 | 0.01611 | 0.01277 | 0.02351 | 0.00796 | 0.01743 | 0.02329 |
| **MIR548C+MIR5** | 0.02319 | 0.01916 | 0.01208 | 0.01853 | 0.00783 | 0.01186 | 0.02971 |
| **MIR548F2** | 0.03462 | 0.01326 | 0.01373 | 0.01598 | 0.00816 | 0.01203 | 0.02128 |
| **MIR548G** | 0.02615 | 0.01602 | 0.01546 | 0.01848 | 0.00772 | 0.01392 | 0.01761 |
| **MIR548H2** | 0.02943 | 0.00961 | 0.01326 | 0.01044 | 0.00774 | 0.00894 | 0.05621 |
| **MIR548I2** | 0.01990 | 0.02822 | 0.01461 | 0.01883 | 0.00748 | 0.02056 | 0.01880 |
| **MIR548J** | 0.01996 | 0.01456 | 0.01367 | 0.01973 | 0.00757 | 0.02469 | 0.03562 |
| **MIR548K** | 0.02551 | 0.01625 | 0.01707 | 0.01969 | 0.00736 | 0.01562 | 0.02234 |
| **MIR548N** | 0.02397 | 0.01672 | 0.01341 | 0.01930 | 0.00873 | 0.01300 | 0.02130 |
| **MIR548Y** | 0.02435 | 0.01515 | 0.01326 | 0.01890 | 0.00755 | 0.01040 | 0.03761 |
| **MIR553** | 0.02628 | 0.01738 | 0.01307 | 0.01895 | 0.00789 | 0.01564 | 0.01678 |
| **MIR554** | 0.03010 | 0.00628 | 0.05790 | 0.02414 | 0.15943 | 0.01355 | 0.01060 |
| **MIR555** | 0.04126 | 0.02573 | 0.01620 | 0.02299 | 0.00904 | 0.01870 | 0.02956 |
| **MIR557** | 0.03194 | 0.01623 | 0.01376 | 0.01996 | 0.00777 | 0.01330 | 0.01840 |
| **MIR558** | 0.02469 | 0.01622 | 0.01852 | 0.02089 | 0.01231 | 0.02039 | 0.03062 |
| **MIR5581** | 0.03845 | 0.01932 | 0.01311 | 0.02088 | 0.01005 | 0.01227 | 0.01667 |
| **MIR5586** | 0.02450 | 0.01836 | 0.01349 | 0.01843 | 0.01044 | 0.01375 | 0.01910 |
| **MIR561** | 0.01885 | 0.01862 | 0.01563 | 0.02794 | 0.00934 | 0.00969 | 0.04089 |
| **MIR564** | 0.02799 | 0.02554 | 0.02007 | 0.02810 | 0.00942 | 0.01613 | 0.00665 |
| **MIR567** | 0.01631 | 0.01425 | 0.01286 | 0.01907 | 0.00741 | 0.01996 | 0.05816 |

| | | | | | | | |
|---|---|---|---|---|---|---|---|
| **MIR5687** | 0.04712 | 0.00627 | 0.02472 | 0.00781 | 0.00665 | 0.12470 | 0.03568 |
| **MIR5691** | 0.02466 | 0.03088 | 0.01173 | 0.01864 | 0.00801 | 0.01304 | 0.01377 |
| **MIR5695** | 0.02126 | 0.01718 | 0.01318 | 0.01879 | 0.00776 | 0.02113 | 0.02456 |
| **MIR570** | 0.10196 | 0.02365 | 0.07190 | 0.06487 | 0.00549 | 0.03566 | 0.13449 |
| **MIR5703** | 0.02262 | 0.01731 | 0.01362 | 0.02274 | 0.00785 | 0.01309 | 0.02267 |
| **MIR572** | 0.02314 | 0.08064 | 0.01656 | 0.02277 | 0.00873 | 0.00801 | 0.00771 |
| **MIR573** | 0.05886 | 0.01594 | 0.02586 | 0.05015 | 0.01273 | 0.02312 | 0.02131 |
| **MIR580** | 0.02738 | 0.01751 | 0.01291 | 0.01798 | 0.00792 | 0.01812 | 0.01849 |
| **MIR581** | 0.02905 | 0.01342 | 0.01892 | 0.01900 | 0.00766 | 0.01251 | 0.01690 |
| **MIR589** | 0.03233 | 0.04613 | 0.02351 | 0.09292 | 0.01148 | 0.06012 | 0.04351 |
| **MIR590** | 0.02712 | 0.01939 | 0.02077 | 0.02248 | 0.01285 | 0.02380 | 0.01735 |
| **MIR593** | 0.01582 | 0.01719 | 0.01396 | 0.02716 | 0.00831 | 0.01520 | 0.03416 |
| **MIR600HG** | 1.05792 | 1.07214 | 0.12754 | 1.76954 | 0.00000 | 0.35554 | 2.57276 |
| **MIR603** | 0.02763 | 0.01103 | 0.01379 | 0.03308 | 0.00753 | 0.01660 | 0.03852 |
| **MIR604** | 0.02422 | 0.02072 | 0.01661 | 0.02044 | 0.00753 | 0.01394 | 0.01489 |
| **MIR610** | 0.02090 | 0.01672 | 0.01386 | 0.02089 | 0.00946 | 0.01357 | 0.04062 |
| **MIR611** | 0.41703 | 0.90147 | 0.17188 | 0.49393 | 0.09413 | 0.47248 | 0.39410 |
| **MIR612** | 0.02229 | 0.01837 | 0.01171 | 0.02023 | 0.01155 | 0.02141 | 0.02175 |
| **MIR6124** | 0.02764 | 0.02965 | 0.01150 | 0.01324 | 0.00739 | 0.01377 | 0.04335 |
| **MIR6126** | 0.02438 | 0.04306 | 0.02948 | 0.10928 | 0.01662 | 0.06770 | 0.05131 |
| **MIR6132** | 0.02140 | 0.01590 | 0.01747 | 0.02997 | 0.00748 | 0.01874 | 0.01935 |
| **MIR614** | 1.08577 | 1.62264 | 0.00000 | 0.00000 | 0.00465 | 0.00000 | 0.01697 |
| **MIR6165.uid-77** | 0.02662 | 0.01642 | 0.01318 | 0.01884 | 0.00797 | 0.01219 | 0.02034 |
| **MIR622** | 0.05361 | 0.06004 | 0.00997 | 0.04840 | 0.00778 | 0.00691 | 0.02487 |
| **MIR623** | 0.02487 | 0.01214 | 0.01792 | 0.02261 | 0.00746 | 0.01500 | 0.02864 |
| **MIR624** | 0.03614 | 0.01991 | 0.02230 | 0.02780 | 0.01409 | 0.02341 | 0.01363 |
| **MIR626** | 0.02857 | 0.01375 | 0.01380 | 0.01921 | 0.00780 | 0.01833 | 0.02201 |
| **MIR629** | 0.02229 | 0.02021 | 0.01234 | 0.02198 | 0.00794 | 0.01258 | 0.01714 |
| **MIR630** | 0.03394 | 0.01926 | 0.01832 | 0.01709 | 0.00866 | 0.01077 | 0.02492 |
| **MIR631** | 0.02892 | 0.01391 | 0.01856 | 0.02790 | 0.00758 | 0.06106 | 0.02446 |
| **MIR632** | 0.02665 | 0.03685 | 0.03098 | 0.02250 | 0.02264 | 0.01229 | 0.01121 |
| **MIR635** | 0.02600 | 0.02193 | 0.01224 | 0.02444 | 0.00798 | 0.01259 | 0.01289 |
| **MIR636** | 0.01946 | 0.03197 | 0.01323 | 0.01971 | 0.01453 | 0.01581 | 0.02006 |
| **MIR637** | 0.01819 | 0.03937 | 0.01310 | 0.03194 | 0.00714 | 0.01801 | 0.04622 |
| **MIR639** | 0.02856 | 0.02753 | 0.01783 | 0.02426 | 0.00751 | 0.01231 | 0.01135 |

| | | | | | | | |
|---|---|---|---|---|---|---|---|
| **MIR644A** | 0.01160 | 0.00000 | 0.03700 | 0.02503 | 0.00700 | 0.03691 | 0.10026 |
| **MIR646HG** | 0.17425 | 0.39033 | 0.24034 | 0.14563 | 0.63159 | 0.04141 | 0.00069 |
| **MIR647** | 0.00281 | 0.06937 | 0.04236 | 0.02954 | 0.00428 | 0.01861 | 0.31375 |
| **MIR6499** | 0.02670 | 0.01551 | 0.01545 | 0.02143 | 0.00765 | 0.01181 | 0.02078 |
| **MIR6505** | 0.06379 | 0.02762 | 0.02190 | 0.03625 | 0.00944 | 0.01555 | 0.00000 |
| **MIR6514** | 0.02340 | 0.03825 | 0.01385 | 0.03552 | 0.01398 | 0.02281 | 0.03584 |
| **MIR6515** | 0.51929 | 0.33301 | 0.07114 | 0.74935 | 0.07777 | 0.40495 | 0.15995 |
| **MIR6516** | 0.02258 | 0.01749 | 0.01267 | 0.02153 | 0.00892 | 0.01281 | 0.01937 |
| **MIR658** | 0.01772 | 0.01727 | 0.01239 | 0.02114 | 0.00764 | 0.02311 | 0.03622 |
| **MIR659** | 0.02606 | 0.05130 | 0.01067 | 0.01810 | 0.01116 | 0.01746 | 0.02949 |
| **MIR663A** | 0.00009 | 0.00543 | 0.01837 | 0.00567 | 0.00534 | 0.01741 | 1.58877 |
| **MIR663AHG** | 0.00000 | 0.02820 | 0.04796 | 0.00597 | 0.02496 | 0.53596 | 2.73052 |
| **MIR663B** | 0.01933 | 0.01696 | 0.01380 | 0.01957 | 0.00756 | 0.01085 | 0.03843 |
| **MIR6716** | 0.03819 | 0.02085 | 0.01717 | 0.02606 | 0.00906 | 0.01244 | 0.01930 |
| **MIR6717** | 0.02338 | 0.01947 | 0.01272 | 0.02017 | 0.00787 | 0.01266 | 0.01857 |
| **MIR6719** | 0.05102 | 0.03297 | 0.02113 | 0.03009 | 0.00711 | 0.01983 | 0.01846 |
| **MIR6721** | 0.02562 | 0.02161 | 0.01382 | 0.02364 | 0.01175 | 0.01226 | 0.01296 |
| **MIR6723** | 0.07822 | 0.05155 | 0.09296 | 0.02092 | 0.07707 | 0.07726 | 0.28050 |
| **MIR6727** | 0.02815 | 0.02087 | 0.01470 | 0.02801 | 0.01100 | 0.02011 | 0.02242 |
| **MIR6728** | 0.05450 | 0.03843 | 0.01570 | 0.02366 | 0.01259 | 0.04352 | 0.02565 |
| **MIR6729** | 0.05932 | 0.08526 | 0.03171 | 0.07003 | 0.01119 | 0.07123 | 0.00000 |
| **MIR6730** | 0.02400 | 0.02868 | 0.01485 | 0.02190 | 0.00760 | 0.02181 | 0.01105 |
| **MIR6732** | 0.03121 | 0.03130 | 0.01185 | 0.04975 | 0.01229 | 0.02046 | 0.02308 |
| **MIR6733** | 0.04596 | 0.06639 | 0.01757 | 0.06295 | 0.05601 | 0.10358 | 0.09764 |
| **MIR6734** | 0.06793 | 0.02967 | 0.01418 | 0.03326 | 0.00754 | 0.01517 | 0.00927 |
| **MIR6735** | 0.02906 | 0.02133 | 0.01660 | 0.02468 | 0.00926 | 0.01317 | 0.01755 |
| **MIR6736** | 0.04559 | 0.04846 | 0.02190 | 0.04491 | 0.02079 | 0.04881 | 0.00415 |
| **MIR6737** | 0.02329 | 0.01824 | 0.01318 | 0.02959 | 0.00775 | 0.02093 | 0.02904 |
| **MIR6739** | 0.08890 | 0.06888 | 0.03834 | 0.07690 | 0.01093 | 0.02971 | 0.00000 |
| **MIR6740** | 0.05834 | 0.04798 | 0.02417 | 0.01941 | 0.01028 | 0.03802 | 0.14307 |
| **MIR6741** | 0.03241 | 0.01614 | 0.01451 | 0.02274 | 0.00775 | 0.01218 | 0.01672 |
| **MIR6742** | 0.02391 | 0.02471 | 0.01263 | 0.03159 | 0.00761 | 0.02020 | 0.02196 |
| **MIR6743** | 0.07676 | 0.14110 | 0.02382 | 0.08732 | 0.13137 | 0.06801 | 0.00000 |
| **MIR6745** | 0.02375 | 0.01798 | 0.01339 | 0.02085 | 0.01141 | 0.01259 | 0.01924 |
| **MIR6746** | 0.02094 | 0.01485 | 0.01337 | 0.03180 | 0.00793 | 0.03817 | 0.01905 |

| | | | | | | | |
|---|---|---|---|---|---|---|---|
| **MIR6747** | 0.02544 | 0.03024 | 0.01563 | 0.03156 | 0.00698 | 0.02231 | 0.04275 |
| **MIR6748** | 0.24228 | 0.19904 | 0.14789 | 0.03264 | 0.10585 | 0.08601 | 0.11233 |
| **MIR6749** | 0.02033 | 0.02640 | 0.02416 | 0.02009 | 0.00760 | 0.01494 | 0.02581 |
| **MIR6750** | 0.01492 | 0.06564 | 0.03511 | 0.02105 | 0.01776 | 0.00902 | 0.01722 |
| **MIR6751** | 0.02364 | 0.01791 | 0.01290 | 0.02143 | 0.00800 | 0.01631 | 0.01621 |
| **MIR6752** | 0.05573 | 0.03631 | 0.10895 | 0.07932 | 0.07057 | 0.07105 | 0.00533 |
| **MIR6753** | 0.02718 | 0.02090 | 0.01381 | 0.03218 | 0.00891 | 0.01562 | 0.02479 |
| **MIR6754** | 0.04378 | 0.05267 | 0.01134 | 0.05200 | 0.00932 | 0.02634 | 0.00952 |
| **MIR6758** | 1.01261 | 0.59247 | 0.43821 | 0.34802 | 0.09214 | 0.69339 | 0.00000 |
| **MIR6759** | 0.02655 | 0.03059 | 0.01370 | 0.03263 | 0.01753 | 0.02495 | 0.03300 |
| **MIR6761** | 0.16414 | 0.05636 | 0.00524 | 0.05041 | 0.00757 | 0.45601 | 0.02991 |
| **MIR6762** | 0.02294 | 0.02679 | 0.01445 | 0.02842 | 0.00781 | 0.01605 | 0.02374 |
| **MIR6763** | 0.01882 | 0.02067 | 0.01281 | 0.01548 | 0.00716 | 0.00826 | 0.06730 |
| **MIR6764** | 0.00000 | 0.15740 | 0.08844 | 0.04253 | 0.10021 | 0.29252 | 0.61376 |
| **MIR6765** | 0.02212 | 0.02041 | 0.01647 | 0.02004 | 0.00822 | 0.01499 | 0.02148 |
| **MIR6766** | 0.02390 | 0.01993 | 0.01248 | 0.01905 | 0.00777 | 0.01159 | 0.02463 |
| **MIR6767** | 0.09202 | 0.07040 | 0.06012 | 0.11396 | 0.01368 | 0.07983 | 0.00000 |
| **MIR6768** | 0.07648 | 0.02461 | 0.01823 | 0.07870 | 0.01383 | 0.02065 | 0.00373 |
| **MIR6773** | 0.03226 | 0.01892 | 0.01527 | 0.01837 | 0.00773 | 0.01669 | 0.01500 |
| **MIR6774** | 0.02386 | 0.02119 | 0.02382 | 0.02386 | 0.00735 | 0.01623 | 0.01105 |
| **MIR6775** | 0.00622 | 0.09280 | 0.00453 | 0.03954 | 0.00856 | 0.01195 | 0.08642 |
| **MIR6776** | 0.02536 | 0.03149 | 0.02229 | 0.02029 | 0.00663 | 0.01319 | 0.03667 |
| **MIR6777** | 0.02063 | 0.00986 | 0.01251 | 0.05070 | 0.00732 | 0.02978 | 0.07280 |
| **MIR6778** | 0.02439 | 0.03232 | 0.00984 | 0.02709 | 0.01566 | 0.01678 | 0.01887 |
| **MIR6779** | 0.01833 | 0.13775 | 0.04179 | 0.04700 | 0.02258 | 0.02902 | 0.02169 |
| **MIR6780B** | 0.01827 | 0.01823 | 0.02818 | 0.02912 | 0.00835 | 0.05271 | 0.04373 |
| **MIR6781** | 0.05685 | 0.10439 | 0.01258 | 0.06532 | 0.05458 | 0.06417 | 0.00000 |
| **MIR6783** | 0.04253 | 0.01439 | 0.01682 | 0.02797 | 0.00753 | 0.01867 | 0.01331 |
| **MIR6784** | 0.02064 | 0.04227 | 0.01818 | 0.03764 | 0.00712 | 0.01385 | 0.01635 |
| **MIR6785** | 0.04258 | 0.03290 | 0.01814 | 0.06443 | 0.00874 | 0.03457 | 0.03049 |
| **MIR6786** | 0.05343 | 0.03051 | 0.03891 | 0.13737 | 0.00982 | 0.04619 | 0.00354 |
| **MIR6787** | 0.03442 | 0.02946 | 0.01199 | 0.03325 | 0.00759 | 0.01405 | 0.01952 |
| **MIR6789** | 0.01907 | 0.07862 | 0.01279 | 0.04341 | 0.00737 | 0.03251 | 0.02106 |
| **MIR6790** | 0.11257 | 0.41404 | 0.13625 | 0.25759 | 0.11222 | 0.14105 | 0.53802 |
| **MIR6791** | 0.01941 | 0.01931 | 0.01823 | 0.02964 | 0.00757 | 0.03939 | 0.01821 |

| | | | | | | | |
|---|---|---|---|---|---|---|---|
| **MIR6792** | 0.02691 | 0.01721 | 0.01260 | 0.01791 | 0.00787 | 0.01309 | 0.02176 |
| **MIR6793** | 0.02299 | 0.02423 | 0.01857 | 0.01984 | 0.01380 | 0.01611 | 0.01501 |
| **MIR6794** | 0.02639 | 0.01036 | 0.02269 | 0.02105 | 0.00750 | 0.01754 | 0.02147 |
| **MIR6796** | 0.10182 | 0.18648 | 0.21199 | 0.46181 | 0.03092 | 0.02837 | 0.00345 |
| **MIR6797** | 2.21062 | 9.05496 | 4.68636 | 9.84332 | 2.49070 | 4.84807 | 12.86732 |
| **MIR6798** | 0.23207 | 0.34301 | 0.04922 | 0.18121 | 0.02859 | 0.06615 | 0.00000 |
| **MIR6799** | 0.03693 | 0.07568 | 0.01205 | 0.08829 | 0.00898 | 0.00300 | 0.28285 |
| **MIR6800** | 0.02387 | 0.02046 | 0.01337 | 0.02267 | 0.00783 | 0.01462 | 0.01517 |
| **MIR6801** | 0.03072 | 0.04303 | 0.01438 | 0.02479 | 0.00756 | 0.01884 | 0.02997 |
| **MIR6802** | 0.02209 | 0.01753 | 0.01252 | 0.02631 | 0.00797 | 0.01868 | 0.01911 |
| **MIR6803** | 0.00994 | 0.02470 | 0.01353 | 0.02932 | 0.00853 | 0.01656 | 0.06977 |
| **MIR6806** | 0.02620 | 0.02330 | 0.01393 | 0.02829 | 0.01205 | 0.01225 | 0.01075 |
| **MIR6807** | 0.15205 | 1.73274 | 0.87034 | 0.32785 | 0.22036 | 1.51957 | 4.83731 |
| **MIR6810** | 0.02763 | 0.02200 | 0.03825 | 0.03243 | 0.00930 | 0.01582 | 0.02439 |
| **MIR6812** | 0.03062 | 0.02152 | 0.01420 | 0.02943 | 0.01039 | 0.01777 | 0.02449 |
| **MIR6816** | 0.02481 | 0.01931 | 0.01241 | 0.02669 | 0.00795 | 0.01260 | 0.01657 |
| **MIR6817** | 0.10052 | 0.12581 | 0.03070 | 0.12909 | 0.03332 | 0.05838 | 0.00000 |
| **MIR6818** | 0.02577 | 0.03532 | 0.01440 | 0.02076 | 0.02623 | 0.01400 | 0.01624 |
| **MIR6820** | 0.60980 | 1.83501 | 0.80560 | 1.34224 | 0.79106 | 0.42295 | 0.62179 |
| **MIR6821** | 0.00000 | 0.16101 | 0.01927 | 0.11277 | 0.05461 | 0.02800 | 0.25169 |
| **MIR6822** | 0.02493 | 0.01862 | 0.01488 | 0.01970 | 0.00808 | 0.01231 | 0.01984 |
| **MIR6825** | 0.02272 | 0.02395 | 0.01299 | 0.02409 | 0.00774 | 0.02280 | 0.01844 |
| **MIR6826** | 0.22573 | 0.10721 | 0.03342 | 0.10736 | 0.02003 | 0.09464 | 0.00000 |
| **MIR6828** | 0.02481 | 0.01395 | 0.02362 | 0.01935 | 0.01120 | 0.01276 | 0.01599 |
| **MIR6831** | 0.02482 | 0.02083 | 0.01688 | 0.03277 | 0.00724 | 0.01158 | 0.02692 |
| **MIR6832** | 0.01636 | 0.02545 | 0.03053 | 0.03545 | 0.00876 | 0.02476 | 0.07688 |
| **MIR6833** | 0.04003 | 0.01163 | 0.04943 | 0.04885 | 0.00607 | 0.04319 | 0.02386 |
| **MIR6834** | 0.11687 | 0.04344 | 0.09981 | 0.17115 | 0.03446 | 0.10359 | 0.00432 |
| **MIR6835** | 0.03195 | 0.08808 | 0.03262 | 0.04739 | 0.00694 | 0.03100 | 0.03470 |
| **MIR6836** | 0.02333 | 0.02067 | 0.01648 | 0.02300 | 0.00753 | 0.01997 | 0.01698 |
| **MIR6837** | 0.13857 | 0.27170 | 0.04916 | 0.09576 | 0.00889 | 0.07187 | 0.00000 |
| **MIR6839** | 0.04029 | 0.01711 | 0.03614 | 0.03526 | 0.07513 | 0.01862 | 0.04298 |
| **MIR6840** | 0.01491 | 0.07857 | 0.02689 | 0.04275 | 0.03191 | 0.02233 | 0.05691 |
| **MIR6842** | 0.02121 | 0.01429 | 0.01447 | 0.02021 | 0.01572 | 0.01529 | 0.04560 |
| **MIR6845** | 0.01652 | 0.03572 | 0.01729 | 0.02071 | 0.00849 | 0.03486 | 0.01783 |

| | | | | | | | |
|---|---|---|---|---|---|---|---|
| **MIR6847** | 0.17835 | 0.63703 | 0.21838 | 0.18671 | 0.01240 | 0.07188 | 0.00419 |
| **MIR6848** | 0.01935 | 0.01856 | 0.01388 | 0.03861 | 0.00748 | 0.01094 | 0.03396 |
| **MIR6849** | 0.02420 | 0.01567 | 0.01281 | 0.02132 | 0.00746 | 0.02342 | 0.05514 |
| **MIR6850** | 6.58997 | 8.59619 | 2.15675 | 6.52972 | 1.98026 | 5.32674 | 0.36005 |
| **MIR6851** | 0.02342 | 0.01910 | 0.01634 | 0.02304 | 0.00767 | 0.01309 | 0.01950 |
| **MIR6853** | 0.10177 | 0.25532 | 0.07867 | 0.06586 | 0.04484 | 0.08798 | 0.00000 |
| **MIR6855** | 0.02340 | 0.01833 | 0.01308 | 0.02575 | 0.00995 | 0.01886 | 0.01926 |
| **MIR6856** | 0.05577 | 0.10660 | 0.01891 | 0.03839 | 0.03282 | 0.02278 | 0.00000 |
| **MIR6857** | 0.19569 | 0.14263 | 0.06079 | 0.07229 | 0.03606 | 0.17149 | 0.02728 |
| **MIR6858** | 0.02900 | 0.03425 | 0.02158 | 0.02699 | 0.00690 | 0.02115 | 0.05371 |
| **MIR6861** | 0.01861 | 0.01546 | 0.01257 | 0.01987 | 0.00770 | 0.01452 | 0.04674 |
| **MIR6863** | 0.02784 | 0.14717 | 0.02211 | 0.00692 | 0.00792 | 0.00230 | 0.07380 |
| **MIR6864** | 0.03132 | 0.01975 | 0.01526 | 0.01613 | 0.00742 | 0.01083 | 0.02902 |
| **MIR6865** | 0.02716 | 0.02393 | 0.01443 | 0.01605 | 0.04098 | 0.01153 | 0.02093 |
| **MIR6866** | 0.05420 | 0.02629 | 0.03025 | 0.07351 | 0.01019 | 0.01810 | 0.00953 |
| **MIR6869** | 0.30042 | 0.63030 | 0.17055 | 0.84842 | 0.03991 | 0.38188 | 0.00000 |
| **MIR6871** | 0.02864 | 0.01622 | 0.01284 | 0.02775 | 0.01219 | 0.01935 | 0.02846 |
| **MIR6873** | 0.04650 | 0.01581 | 0.01287 | 0.04351 | 0.01039 | 0.03980 | 0.02918 |
| **MIR6875** | 0.03297 | 0.01552 | 0.01194 | 0.04415 | 0.00769 | 0.01509 | 0.03527 |
| **MIR6877** | 0.10100 | 0.16323 | 0.13992 | 0.21838 | 0.01249 | 0.09538 | 0.00000 |
| **MIR6878** | 0.18920 | 0.06865 | 0.05500 | 0.07496 | 0.04839 | 0.08316 | 0.00000 |
| **MIR6879** | 0.37818 | 0.48083 | 0.09061 | 0.31839 | 0.01332 | 0.26054 | 0.00000 |
| **MIR6880** | 0.02496 | 0.01998 | 0.01341 | 0.01928 | 0.00771 | 0.01201 | 0.02189 |
| **MIR6883** | 0.02473 | 0.01725 | 0.01281 | 0.02251 | 0.00788 | 0.01247 | 0.01957 |
| **MIR6884** | 0.02512 | 0.01722 | 0.01293 | 0.02033 | 0.00790 | 0.01340 | 0.01754 |
| **MIR6886** | 0.13247 | 0.06258 | 0.00918 | 0.11183 | 0.01382 | 0.07287 | 0.00000 |
| **MIR6887** | 0.03935 | 0.03115 | 0.01277 | 0.03103 | 0.01018 | 0.04684 | 0.04721 |
| **MIR6888** | 0.02506 | 0.02123 | 0.01414 | 0.01879 | 0.00781 | 0.01262 | 0.01636 |
| **MIR6889** | 0.02734 | 0.01665 | 0.01402 | 0.01849 | 0.00774 | 0.01547 | 0.02107 |
| **MIR6890** | 0.05038 | 0.04591 | 0.01180 | 0.01996 | 0.00777 | 0.07044 | 0.02902 |
| **MIR6892** | 0.02116 | 0.08260 | 0.01857 | 0.05153 | 0.01105 | 0.05765 | 0.05859 |
| **MIR6893** | 0.02008 | 0.02122 | 0.01248 | 0.01730 | 0.00759 | 0.01861 | 0.03453 |
| **MIR6894** | 0.02024 | 0.01859 | 0.01504 | 0.02790 | 0.00793 | 0.01801 | 0.01628 |
| **MIR7-1** | 0.06698 | 0.01845 | 0.02387 | 0.03363 | 0.00710 | 0.02472 | 0.02791 |
| **MIR7-3HG** | 0.03748 | 0.04313 | 0.00000 | 0.04053 | 0.40669 | 0.00929 | 0.00000 |

| | | | | | | | |
|---|---|---|---|---|---|---|---|
| **MIR7107** | 0.02459 | 0.01559 | 0.01440 | 0.02094 | 0.00780 | 0.01691 | 0.01731 |
| **MIR7108** | 0.02254 | 0.09339 | 0.01770 | 0.03854 | 0.01103 | 0.04030 | 0.02859 |
| **MIR7109** | 0.04250 | 0.04507 | 0.01257 | 0.04710 | 0.00915 | 0.03404 | 0.01457 |
| **MIR7110** | 0.03505 | 0.02158 | 0.01027 | 0.04970 | 0.00655 | 0.05194 | 0.25714 |
| **MIR7111** | 0.03989 | 0.05383 | 0.04330 | 0.07907 | 0.01132 | 0.31455 | 0.32850 |
| **MIR7113** | 0.23392 | 0.20384 | 0.04424 | 0.24364 | 0.02247 | 0.08616 | 0.00000 |
| **MIR7114** | 0.02429 | 0.01557 | 0.01611 | 0.01936 | 0.01025 | 0.01607 | 0.01573 |
| **MIR7150** | 0.02554 | 0.01755 | 0.01306 | 0.01858 | 0.01877 | 0.01132 | 0.02865 |
| **MIR7161** | 0.02505 | 0.01925 | 0.01250 | 0.02500 | 0.00766 | 0.01163 | 0.03381 |
| **MIR744** | 0.02325 | 0.01411 | 0.02039 | 0.02094 | 0.00726 | 0.02251 | 0.02875 |
| **MIR761** | 0.02314 | 0.02592 | 0.01309 | 0.02413 | 0.00812 | 0.02509 | 0.01123 |
| **MIR765** | 0.02260 | 0.02330 | 0.01159 | 0.01739 | 0.00812 | 0.01758 | 0.02423 |
| **MIR7703** | 0.01589 | 0.02567 | 0.00940 | 0.03924 | 0.03250 | 0.17811 | 0.02967 |
| **MIR7846** | 0.02156 | 0.01793 | 0.01275 | 0.02363 | 0.00790 | 0.01279 | 0.01942 |
| **MIR7851.uid-20** | 0.02610 | 0.01970 | 0.01521 | 0.02476 | 0.00771 | 0.01237 | 0.01409 |
| **MIR7851.uid-46** | 0.02756 | 0.01441 | 0.01368 | 0.01582 | 0.00800 | 0.01168 | 0.02649 |
| **MIR7854** | 0.02571 | 0.02096 | 0.01563 | 0.02314 | 0.00736 | 0.01529 | 0.03061 |
| **MIR8055** | 0.02558 | 0.02130 | 0.01783 | 0.02134 | 0.00834 | 0.01451 | 0.02831 |
| **MIR8069.uid-25** | 0.01957 | 0.03392 | 0.01966 | 0.02036 | 0.00938 | 0.01111 | 0.01595 |
| **MIR8072** | 0.37941 | 0.34753 | 0.16662 | 0.42707 | 0.21697 | 0.06989 | 0.04983 |
| **MIR8076** | 0.02738 | 0.01492 | 0.01865 | 0.02445 | 0.00875 | 0.01391 | 0.01337 |
| **MIR8079** | 0.01429 | 0.01330 | 0.01199 | 0.05588 | 0.00739 | 0.01443 | 0.06098 |
| **MIR8085** | 0.01846 | 0.02363 | 0.01206 | 0.02571 | 0.00795 | 0.03201 | 0.01355 |
| **MIR8087** | 0.02843 | 0.01344 | 0.01738 | 0.01704 | 0.00866 | 0.01245 | 0.02036 |
| **MIR877** | 0.20527 | 0.11934 | 0.16007 | 0.13681 | 0.09894 | 0.07815 | 0.00000 |
| **MIR9-1** | 0.02338 | 0.01728 | 0.01366 | 0.02144 | 0.00798 | 0.01273 | 0.01810 |
| **MIR921** | 0.02285 | 0.02180 | 0.01248 | 0.01724 | 0.00772 | 0.01320 | 0.02463 |
| **MIR93** | 0.02896 | 0.02609 | 0.02677 | 0.05088 | 0.01277 | 0.01107 | 0.03819 |
| **MIR934** | 0.02268 | 0.01609 | 0.01302 | 0.02112 | 0.00786 | 0.01237 | 0.02400 |
| **MIR937** | 0.02132 | 0.01788 | 0.01320 | 0.01750 | 0.00771 | 0.01858 | 0.02784 |
| **MIR939** | 0.00000 | 0.08754 | 0.01332 | 0.04457 | 0.00701 | 0.10914 | 0.09607 |
| **MIR940** | 0.03014 | 0.01365 | 0.03980 | 0.02979 | 0.00658 | 0.01451 | 0.01587 |
| **MIR943** | 0.02384 | 0.03513 | 0.00937 | 0.01637 | 0.00813 | 0.01231 | 0.02723 |
| **MIR944** | 0.02444 | 0.01793 | 0.01272 | 0.02090 | 0.00792 | 0.01264 | 0.01801 |
| **MIR98** | 0.03866 | 0.05225 | 0.05582 | 0.03329 | 0.00458 | 0.08466 | 0.08304 |

| | | | | | | | |
|---|---|---|---|---|---|---|---|
| **MIRLET7A1** | 0.03159 | 0.01477 | 0.01684 | 0.03076 | 0.00770 | 0.01693 | 0.01546 |
| **MIRLET7BHG** | 0.01783 | 0.06301 | 0.02093 | 0.05253 | 0.00912 | 0.00641 | 0.02284 |
| **MIRLET7D** | 0.09125 | 0.04934 | 0.04996 | 0.00889 | 0.00485 | 0.09193 | 0.07131 |
| **MIRLET7DHG** | 0.53740 | 0.07460 | 0.22656 | 0.18667 | 0.16667 | 0.44657 | 0.60199 |
| **MIRLET7F1** | 0.03430 | 0.02393 | 0.03286 | 0.04011 | 0.00864 | 0.03243 | 0.01117 |
| **MIRLET7F2** | 0.02749 | 0.01998 | 0.01452 | 0.02146 | 0.00780 | 0.01359 | 0.01329 |
| **MIRLET7G** | 0.02448 | 0.02074 | 0.01184 | 0.02967 | 0.00818 | 0.02657 | 0.00781 |
| **MIRLET7I** | 0.00915 | 0.03826 | 0.01308 | 0.03856 | 0.00932 | 0.00814 | 0.05109 |
| **MIS12** | 10.80400 | 0.03290 | 6.07164 | 7.89135 | 2.16847 | 3.79899 | 0.00000 |
| **MIS18A** | 18.40835 | 18.27587 | 14.48128 | 11.17418 | 5.00016 | 8.86056 | 8.35080 |
| **MIS18BP1** | 13.40436 | 0.00000 | 9.01541 | 8.45270 | 4.99672 | 6.11610 | 13.33499 |
| **MISP** | 5.11511 | 6.52512 | 1.17930 | 1.92951 | 0.77440 | 0.00000 | 4.63960 |
| **MITD1** | 8.76932 | 1.53650 | 2.49115 | 5.05151 | 0.41227 | 2.99070 | 3.57607 |
| **MITF** | 5.10015 | 0.17560 | 1.30607 | 1.69396 | 3.42188 | 0.16514 | 2.01833 |
| **MIXL1** | 0.00000 | 0.00000 | 0.13247 | 0.00757 | 3.33689 | 1.46416 | 0.40481 |
| **MKI67** | 15.87602 | 14.89643 | 16.84674 | 17.60389 | 5.47270 | 9.71065 | 40.64003 |
| **MKKS** | 27.02210 | 18.30216 | 19.90492 | 5.18374 | 0.00000 | 9.80633 | 10.74331 |
| **MKL1** | 0.51816 | 3.25966 | 0.77036 | 5.49993 | 2.85357 | 3.09952 | 9.83040 |
| **MKL2** | 5.97952 | 0.08451 | 1.61084 | 3.02857 | 1.16908 | 0.06261 | 4.50271 |
| **MKLN1** | 17.27851 | 4.05701 | 5.93991 | 5.85436 | 2.29138 | 5.54462 | 5.96588 |
| **MKNK1** | 7.74390 | 9.97929 | 6.77592 | 6.72040 | 1.50201 | 4.76086 | 0.00000 |
| **MKNK1-AS1** | 0.07379 | 0.03577 | 0.06001 | 0.03199 | 0.02498 | 0.03089 | 0.00000 |
| **MKNK2** | 2.70587 | 1.80512 | 0.76869 | 8.32349 | 2.04356 | 3.76228 | 31.10706 |
| **MKRN1** | 5.60416 | 12.82254 | 19.34883 | 4.84367 | 11.66338 | 22.33145 | 21.34651 |
| **MKRN2** | 13.40060 | 7.31982 | 6.20051 | 9.98455 | 9.37344 | 2.67844 | 0.00000 |
| **MKRN2OS** | 0.18267 | 0.00000 | 0.02823 | 0.04435 | 0.00671 | 0.06496 | 0.02900 |
| **MKRN3** | 5.23252 | 1.77880 | 2.62391 | 4.85675 | 0.62139 | 0.12167 | 0.00000 |
| **MKRN7P** | 0.02141 | 0.02317 | 0.02134 | 0.02900 | 0.00915 | 0.02365 | 0.01207 |
| **MKRN9P** | 0.16999 | 0.44328 | 0.61871 | 0.30074 | 0.59251 | 0.53837 | 0.21693 |
| **MKS1** | 5.49221 | 5.54863 | 5.16207 | 2.56665 | 1.19405 | 2.39437 | 1.25377 |
| **MKX** | 0.01944 | 0.01681 | 0.13682 | 0.00465 | 0.00402 | 0.09635 | 0.13449 |
| **MLANA** | 0.03823 | 0.02718 | 0.01337 | 0.01865 | 0.02095 | 0.01884 | 0.01549 |
| **MLC1** | 0.00944 | 0.04088 | 0.07419 | 0.01910 | 0.02943 | 0.00717 | 0.04295 |
| **MLEC** | 8.80072 | 13.26146 | 5.41888 | 6.15029 | 2.10665 | 16.51243 | 25.90057 |
| **1.00000** | 2.34235 | 0.11320 | 8.90427 | 1.22597 | 12.33465 | 0.45161 | 0.00000 |

| | | | | | | | |
|---|---|---|---|---|---|---|---|
| **2.00000** | 33.19852 | 24.69871 | 13.60765 | 37.05029 | 12.83970 | 20.96059 | 4.17787 |
| **MLH1** | 9.74164 | 1.08915 | 6.51879 | 3.94713 | 1.74896 | 3.20664 | 0.00000 |
| **MLH3** | 3.28958 | 0.14970 | 2.03626 | 4.25312 | 0.11206 | 1.12195 | 3.93665 |
| **MLIP** | 0.01660 | 0.02435 | 0.01086 | 0.01744 | 0.03852 | 0.01189 | 0.03562 |
| **MLIP-IT1** | 0.02356 | 0.01187 | 0.04261 | 0.01947 | 0.01099 | 0.01196 | 0.01765 |
| **MLK7-AS1** | 0.21015 | 0.08755 | 0.13373 | 1.43869 | 1.39625 | 0.26911 | 0.72860 |
| **MLKL** | 0.05437 | 0.00000 | 0.04058 | 0.04201 | 0.34173 | 0.30387 | 0.00000 |
| **MLLT1** | 0.00000 | 2.49537 | 2.47473 | 9.06529 | 0.11532 | 1.97272 | 16.57957 |
| **MLLT10** | 4.18325 | 2.06145 | 6.07946 | 7.82479 | 1.92015 | 6.34532 | 20.10619 |
| **MLLT10P1** | 0.17073 | 0.00000 | 0.28836 | 0.02598 | 0.00494 | 0.19487 | 0.33232 |
| **MLLT11** | 0.77315 | 0.00000 | 0.99423 | 13.47539 | 1.51214 | 0.16527 | 0.00000 |
| **MLLT3** | 0.13935 | 0.07188 | 0.27376 | 0.00000 | 0.66206 | 3.92565 | 2.52388 |
| **MLLT4** | 4.89636 | 2.28891 | 3.90177 | 4.65690 | 1.42028 | 4.31462 | 27.77247 |
| **MLLT4-AS1** | 0.36054 | 0.04714 | 0.09773 | 0.37580 | 0.24015 | 0.02749 | 0.05512 |
| **MLLT6** | 0.00075 | 1.64549 | 0.72351 | 0.50698 | 0.09934 | 1.94909 | 3.11403 |
| **MLN** | 0.02262 | 0.00777 | 0.00902 | 0.00847 | 0.00745 | 1.25790 | 0.01721 |
| **MLNR** | 0.01667 | 0.01489 | 0.57697 | 0.01059 | 0.06064 | 0.05548 | 0.03140 |
| **MLPH** | 0.00000 | 0.00000 | 0.04643 | 0.13149 | 0.11796 | 0.22298 | 0.03388 |
| **MLST8** | 13.70034 | 13.55672 | 2.91438 | 12.76171 | 0.15070 | 9.39860 | 9.76851 |
| **MLX** | 14.83572 | 5.78970 | 10.15091 | 14.77717 | 4.10286 | 8.64654 | 0.00000 |
| **MLXIP** | 3.32748 | 14.26404 | 6.05844 | 1.51411 | 1.42422 | 10.57026 | 13.91494 |
| **MLXIPL** | 0.46494 | 0.00344 | 0.00000 | 0.00000 | 1.26200 | 0.00000 | 0.40445 |
| **MLYCD** | 0.37538 | 0.97518 | 0.31006 | 0.30880 | 0.39183 | 0.38713 | 1.44635 |
| **MMAA** | 0.72965 | 0.58338 | 3.55209 | 0.10763 | 0.66322 | 0.06921 | 0.44788 |
| **MMAB** | 15.58547 | 3.42499 | 0.00100 | 7.87286 | 0.13211 | 3.05180 | 3.55488 |
| **MMACHC** | 4.84130 | 0.00000 | 6.79222 | 3.74923 | 3.29011 | 1.02573 | 0.73887 |
| **MMADHC** | 38.74175 | 7.20196 | 21.54494 | 24.50829 | 10.58366 | 12.55640 | 6.29665 |
| **MMD** | 1.77720 | 0.00000 | 0.41301 | 1.85924 | 0.38719 | 2.96037 | 1.65955 |
| **MMD2** | 0.02028 | 0.04187 | 0.03961 | 0.01833 | 0.00790 | 0.00921 | 0.01950 |
| **MME** | 0.14215 | 0.05246 | 0.00061 | 1.35681 | 0.16957 | 0.06612 | 0.00000 |
| **MMEL1** | 0.02121 | 0.02057 | 0.01956 | 0.02520 | 0.00966 | 0.01784 | 0.01851 |
| **MMGT1** | 15.70727 | 3.21681 | 13.79391 | 3.31817 | 3.82136 | 2.14985 | 8.80248 |
| **MMP11** | 0.00000 | 0.09457 | 0.02698 | 0.15930 | 0.06812 | 0.04637 | 0.17107 |
| **MMP12** | 0.02827 | 0.01725 | 0.01285 | 0.02118 | 0.00808 | 0.01373 | 0.01565 |
| **MMP13** | 0.02193 | 0.01819 | 0.00233 | 0.02130 | 0.23180 | 0.01431 | 0.01410 |

| | | | | | | | |
|---|---|---|---|---|---|---|---|
| **MMP14** | 0.00000 | 0.08510 | 0.15221 | 2.48796 | 0.05815 | 0.01709 | 1.14594 |
| **MMP15** | 0.00000 | 3.39156 | 0.49604 | 0.95574 | 0.06913 | 3.61694 | 0.00000 |
| **MMP16** | 0.37395 | 0.00000 | 0.00722 | 0.00000 | 0.00586 | 0.00520 | 0.15764 |
| **MMP17** | 0.00000 | 0.47110 | 0.01964 | 0.18860 | 0.00794 | 0.43057 | 2.82292 |
| **MMP19** | 0.32868 | 0.06252 | 0.11088 | 3.85991 | 0.07232 | 0.00000 | 0.03099 |
| **MMP2** | 0.00000 | 0.55607 | 0.01553 | 0.35825 | 0.11970 | 10.04187 | 0.00000 |
| **MMP20** | 0.03700 | 0.02003 | 0.01262 | 0.04386 | 0.00813 | 0.01272 | 0.00167 |
| **MMP21** | 0.02176 | 0.01126 | 0.01094 | 0.01450 | 0.04404 | 0.06768 | 0.03518 |
| **MMP24** | 0.09693 | 1.12926 | 1.66410 | 0.03815 | 0.06417 | 1.94285 | 0.00200 |
| **MMP24-AS1** | 0.00000 | 1.85259 | 0.73164 | 0.10200 | 0.20455 | 1.47638 | 2.01541 |
| **MMP25** | 0.01229 | 0.05232 | 0.04139 | 0.11208 | 0.03160 | 0.21128 | 0.19113 |
| **MMP28** | 0.03224 | 0.00784 | 0.00988 | 0.00701 | 0.04434 | 0.00803 | 0.06585 |
| **MMP8** | 0.01874 | 0.01631 | 0.02021 | 0.01823 | 0.00921 | 0.01839 | 0.05446 |
| **MMP9** | 0.00578 | 0.00944 | 0.00000 | 0.09803 | 0.43445 | 1.92779 | 0.00940 |
| **MMRN2** | 0.16735 | 0.00000 | 0.34749 | 0.19458 | 0.04689 | 0.01242 | 0.16520 |
| **MMS19** | 8.00944 | 6.90128 | 5.34307 | 8.95394 | 1.85861 | 10.21523 | 9.52299 |
| **MMS22L** | 12.94540 | 0.94699 | 4.89925 | 5.67098 | 2.49974 | 5.05788 | 0.09782 |
| **MN1** | 0.00714 | 0.00816 | 0.01723 | 0.07604 | 0.11444 | 0.57974 | 0.03950 |
| **MNAT1** | 9.47229 | 5.49249 | 6.45942 | 3.52100 | 0.40614 | 3.34404 | 8.94299 |
| **MND1** | 6.42518 | 1.77068 | 4.26512 | 0.84471 | 0.68335 | 2.37139 | 5.59879 |
| **MNDA** | 0.01297 | 0.04677 | 0.00685 | 0.01322 | 0.00805 | 0.25416 | 0.01839 |
| **MNS1** | 0.68994 | 0.00000 | 1.74305 | 0.00000 | 1.74352 | 0.02845 | 0.85587 |
| **MNT** | 0.00000 | 0.71661 | 0.53715 | 1.02880 | 0.46706 | 0.96445 | 5.64110 |
| **MNX1** | 0.01113 | 0.05911 | 0.06593 | 0.00754 | 0.00652 | 0.04667 | 0.52408 |
| **MNX1-AS1** | 0.00932 | 0.01386 | 0.01309 | 0.01571 | 0.01018 | 0.13207 | 0.15757 |
| **MOAP1** | 3.14310 | 1.56665 | 4.93883 | 7.15842 | 4.84199 | 1.35046 | 0.00000 |
| **MOB1A** | 23.28078 | 6.59337 | 8.67773 | 11.45569 | 3.02375 | 8.60036 | 0.00000 |
| **MOB1B** | 5.23888 | 0.00000 | 3.46226 | 3.61374 | 3.44707 | 1.52840 | 9.26225 |
| **MOB2** | 0.00000 | 3.65288 | 0.90062 | 2.21670 | 1.15705 | 3.65893 | 7.92335 |
| **MOB3A** | 0.00000 | 1.40613 | 2.35081 | 1.75351 | 0.63308 | 0.65321 | 3.67181 |
| **MOB3B** | 32.88379 | 10.73136 | 0.00000 | 15.94075 | 0.70299 | 9.85463 | 10.72093 |
| **MOB3C** | 0.00585 | 0.00000 | 0.14289 | 0.17264 | 0.05603 | 0.01694 | 0.08015 |
| **MOB4.uid-3968** | 1.20390 | 1.39430 | 1.66532 | 1.87738 | 0.34489 | 0.87733 | 0.00000 |
| **MOB4.uid-3970** | 0.02574 | 0.02686 | 0.01196 | 0.03223 | 0.04340 | 0.03645 | 0.01434 |
| **MOB4.uid-3972** | 0.01496 | 0.01563 | 0.01300 | 0.02951 | 0.00774 | 0.01273 | 0.03303 |

| | | | | | | | |
|---|---|---|---|---|---|---|---|
| **MOBP** | 0.03118 | 0.00000 | 0.32345 | 0.00976 | 0.46091 | 0.43614 | 0.24774 |
| **MOCOS** | 0.11031 | 0.13164 | 0.00000 | 1.08404 | 0.37575 | 0.04477 | 0.13002 |
| **MOCS1** | 1.84030 | 1.92729 | 0.60259 | 4.98530 | 0.09561 | 1.54331 | 0.34519 |
| **MOCS2** | 11.46829 | 2.98877 | 4.38915 | 7.64336 | 0.69480 | 6.20807 | 0.00495 |
| **MOCS3** | 6.86759 | 3.64040 | 5.97317 | 3.60962 | 0.41569 | 3.21660 | 0.00000 |
| **MOG** | 0.02522 | 0.11774 | 0.15015 | 0.03237 | 0.02416 | 0.76776 | 0.16882 |
| **MOGAT1** | 0.03365 | 0.00144 | 0.02255 | 0.00756 | 0.00832 | 0.01298 | 0.12357 |
| **MOGAT2** | 0.03136 | 0.00732 | 0.17351 | 0.01907 | 0.00852 | 0.03375 | 0.01005 |
| **MOGAT3** | 0.18157 | 0.77060 | 0.00000 | 0.00000 | 0.00586 | 1.17448 | 0.08374 |
| **MOGS** | 3.39045 | 1.85598 | 3.35385 | 9.07124 | 0.05426 | 4.29037 | 11.99689 |
| **MOK** | 0.13855 | 0.97012 | 1.60846 | 1.43237 | 5.60228 | 0.37215 | 0.32675 |
| **MON1A** | 2.27639 | 5.70381 | 3.41700 | 3.95824 | 0.32231 | 2.45114 | 4.32439 |
| **MON1B** | 9.21776 | 7.03166 | 3.67798 | 14.09539 | 1.87633 | 2.96854 | 0.39745 |
| **MON2** | 6.57091 | 0.00001 | 2.21456 | 5.09022 | 2.40371 | 1.89466 | 2.82455 |
| **MORC1** | 0.04125 | 0.04753 | 0.21714 | 0.02429 | 0.09343 | 0.01850 | 0.00105 |
| **MORC2** | 3.91877 | 2.51110 | 3.31084 | 5.71966 | 0.64736 | 1.31565 | 7.02360 |
| **MORC2-AS1** | 9.28862 | 7.60601 | 2.93749 | 10.17314 | 0.90731 | 4.90140 | 0.00000 |
| **MORC3** | 13.77669 | 0.00000 | 6.13321 | 4.99707 | 3.95281 | 4.29288 | 3.86314 |
| **MORC4** | 7.44665 | 4.09336 | 1.48253 | 0.00000 | 0.29007 | 1.55642 | 13.87997 |
| **MORF4L1** | 21.08555 | 6.65987 | 11.44532 | 14.34684 | 2.40113 | 12.60300 | 11.27500 |
| **MORF4L2** | 59.61278 | 13.79436 | 17.20134 | 25.03754 | 7.31648 | 12.87189 | 8.46746 |
| **MORF4L2-AS1** | 0.62826 | 0.00000 | 0.07728 | 0.11058 | 0.07483 | 0.04482 | 0.16860 |
| **MORN1** | 0.11442 | 0.93124 | 0.46564 | 0.27283 | 0.02520 | 0.04199 | 0.00000 |
| **MORN2** | 4.47412 | 0.65503 | 10.48306 | 4.02760 | 2.33681 | 0.30905 | 1.55746 |
| **MORN3** | 0.01615 | 0.05432 | 0.77383 | 2.04402 | 0.11528 | 0.10408 | 0.00006 |
| **MORN4** | 0.24958 | 0.03405 | 0.23448 | 0.13227 | 0.02144 | 0.03511 | 0.01755 |
| **MORN5** | 0.02776 | 0.00919 | 0.02154 | 0.02851 | 0.01692 | 0.01127 | 0.02513 |
| **MOS** | 0.02184 | 0.00319 | 0.02984 | 0.01946 | 0.47947 | 0.01242 | 0.02059 |
| **MOSPD1** | 11.95711 | 1.98365 | 23.78710 | 1.43114 | 14.05808 | 8.36629 | 3.43964 |
| **MOSPD2** | 7.80194 | 0.11183 | 2.13719 | 4.50314 | 2.22449 | 0.89787 | 0.76661 |
| **MOSPD3** | 0.00000 | 0.89456 | 3.06589 | 1.11694 | 1.01305 | 1.01851 | 2.57805 |
| **MOV10** | 8.28790 | 6.41195 | 1.12723 | 9.02605 | 0.50883 | 3.85542 | 5.65813 |
| **MOV10L1** | 0.00000 | 0.09317 | 0.58554 | 0.00694 | 0.48847 | 0.00000 | 0.08610 |
| **MOXD1** | 0.05231 | 0.00671 | 0.02636 | 0.01638 | 0.00696 | 0.01029 | 0.02072 |
| **MPC1** | 12.72517 | 1.72424 | 0.69723 | 6.29841 | 1.50450 | 6.83483 | 1.47776 |

| | | | | | | | |
|---|---|---|---|---|---|---|---|
| **MPC1L** | 0.00000 | 0.63571 | 3.85396 | 0.00000 | 0.29573 | 0.00000 | 0.02148 |
| **MPC2** | 4.43635 | 7.26414 | 3.32357 | 0.36503 | 2.06239 | 7.98949 | 2.27066 |
| **MPDU1** | 21.60301 | 12.81400 | 7.64301 | 19.31723 | 0.20938 | 11.34244 | 8.35427 |
| **MPDZ** | 3.55730 | 0.00000 | 1.78840 | 1.03539 | 0.48742 | 3.21462 | 3.72462 |
| **MPG** | 5.96212 | 2.86624 | 0.00000 | 5.02729 | 5.49909 | 3.08113 | 3.44994 |
| **MPHOSPH10** | 19.17531 | 5.26826 | 15.36239 | 11.79219 | 7.98180 | 7.77338 | 12.90899 |
| **MPHOSPH6** | 20.29966 | 5.76725 | 11.66154 | 10.83276 | 1.26442 | 5.51467 | 1.97595 |
| **MPHOSPH8** | 5.82362 | 9.39988 | 7.67033 | 10.12352 | 7.85193 | 7.61296 | 17.11594 |
| **MPHOSPH9** | 5.86560 | 0.00000 | 3.41364 | 1.54400 | 1.64976 | 1.84407 | 2.32122 |
| **MPI** | 8.12978 | 12.71260 | 7.60902 | 5.31518 | 0.00000 | 7.35541 | 1.45706 |
| **MPL** | 0.03322 | 0.03161 | 0.07200 | 0.19417 | 0.09330 | 0.18695 | 0.00000 |
| **MPLKIP** | 4.78406 | 9.54760 | 13.53050 | 12.63172 | 4.84525 | 5.10737 | 9.82939 |
| **MPND** | 0.10969 | 0.88066 | 0.38235 | 0.13799 | 0.40373 | 1.10530 | 1.49989 |
| **MPO** | 0.83991 | 0.00000 | 0.00000 | 0.50852 | 0.04496 | 0.06732 | 0.52310 |
| **MPP1** | 28.28363 | 1.57586 | 0.00000 | 35.25259 | 0.36570 | 2.98719 | 5.29238 |
| **MPP2** | 0.00000 | 0.05954 | 0.00000 | 0.05220 | 2.94265 | 1.57380 | 0.08519 |
| **MPP3** | 0.06083 | 0.11729 | 0.83017 | 0.10916 | 0.62914 | 0.17206 | 0.08355 |
| **MPP4** | 0.02063 | 0.00000 | 0.07410 | 0.06546 | 0.24259 | 0.00780 | 0.03954 |
| **MPP5** | 7.71053 | 0.41154 | 5.18731 | 4.25888 | 6.46642 | 2.65661 | 8.65622 |
| **MPP6** | 2.33700 | 0.00000 | 2.13276 | 6.59872 | 0.57597 | 3.00076 | 12.90793 |
| **MPP7** | 2.69628 | 0.00000 | 3.36911 | 3.89277 | 0.64078 | 3.57778 | 1.42941 |
| **MPPE1** | 3.03980 | 0.41460 | 0.95960 | 0.64694 | 0.49548 | 0.46311 | 0.00000 |
| **MPPED2** | 0.02132 | 0.00000 | 0.03851 | 0.02819 | 0.01871 | 0.21957 | 0.02170 |
| **MPRIP** | 4.53687 | 3.25080 | 0.84699 | 4.74874 | 0.37504 | 2.90210 | 17.99201 |
| **MPST** | 2.66401 | 9.19891 | 1.64531 | 16.36603 | 0.00000 | 1.82521 | 11.09202 |
| **MPV17** | 14.48319 | 11.02573 | 8.52407 | 11.96183 | 1.52805 | 3.11876 | 0.00000 |
| **MPV17L** | 1.43713 | 0.56334 | 1.08666 | 0.00032 | 0.00000 | 0.22031 | 0.00000 |
| **MPV17L2** | 22.39817 | 28.27974 | 10.43886 | 17.44284 | 3.88532 | 11.38281 | 0.00000 |
| **MPZ** | 0.09645 | 0.00000 | 0.06972 | 0.15613 | 0.06836 | 0.06416 | 0.38481 |
| **MPZL1** | 23.88168 | 25.07561 | 13.21524 | 27.25880 | 1.92389 | 1.99449 | 41.67497 |
| **MPZL2** | 0.17201 | 0.00000 | 0.18175 | 0.04481 | 0.12975 | 0.20325 | 0.00000 |
| **MPZL3** | 0.99749 | 0.08469 | 0.63639 | 0.91546 | 1.85139 | 0.06527 | 0.00000 |
| **MR1** | 0.00703 | 0.08010 | 0.18942 | 0.08802 | 0.00934 | 0.12285 | 0.00392 |
| **MRAP** | 0.01850 | 0.04386 | 0.02881 | 0.01858 | 0.00993 | 0.03344 | 0.03795 |
| **MRAP2** | 0.37939 | 0.00000 | 0.25363 | 0.92287 | 0.08822 | 1.56993 | 0.01756 |

| | | | | | | | |
|---|---|---|---|---|---|---|---|
| **MRAS** | 0.00000 | 0.10122 | 0.68400 | 0.88961 | 0.76655 | 1.51445 | 0.73400 |
| **MRC1** | 0.02133 | 0.01928 | 0.00866 | 0.01816 | 0.11131 | 0.01520 | 0.02129 |
| **MRC2** | 0.05774 | 0.00000 | 0.01145 | 0.35429 | 0.20079 | 2.52967 | 0.00000 |
| **MRE11A** | 11.53750 | 3.09287 | 4.40014 | 5.43781 | 1.43182 | 3.15571 | 4.18321 |
| **MREG** | 1.79631 | 0.09805 | 0.35484 | 3.76903 | 0.00000 | 0.25253 | 2.40941 |
| **MRFAP1** | 52.75935 | 32.12508 | 27.50152 | 29.41955 | 9.99401 | 24.31867 | 29.13926 |
| **MRFAP1L1** | 24.89812 | 5.76719 | 15.40201 | 12.19106 | 2.35274 | 8.90228 | 6.85240 |
| **MRGBP** | 1.00677 | 0.65056 | 0.12814 | 2.18443 | 0.43081 | 0.86441 | 7.12034 |
| **MRGPRD** | 0.01874 | 0.05328 | 0.01205 | 0.02808 | 0.00752 | 0.01071 | 0.02115 |
| **MRGPRF** | 0.10670 | 0.03215 | 0.00278 | 0.00311 | 0.03477 | 0.10840 | 0.14805 |
| **MRGPRG-AS1** | 0.02784 | 0.03628 | 0.00905 | 0.07187 | 0.00803 | 0.01225 | 0.01145 |
| **MRGPRX1** | 3.39322 | 0.00000 | 0.00816 | 21.00849 | 0.00327 | 0.00425 | 0.00000 |
| **MRGPRX2** | 0.00882 | 0.01737 | 0.01558 | 0.52566 | 0.00823 | 0.01705 | 0.00179 |
| **MRGPRX3** | 0.01320 | 0.20064 | 0.20512 | 0.19195 | 0.00575 | 0.02118 | 0.00000 |
| **MRGPRX4** | 0.02459 | 0.01948 | 0.01255 | 0.03984 | 0.00811 | 0.01336 | 0.01035 |
| **MRI1** | 4.70902 | 1.81171 | 2.35257 | 5.31532 | 0.08747 | 0.99398 | 2.30754 |
| **MRM1** | 7.31734 | 2.18752 | 7.64867 | 6.72126 | 0.13355 | 2.36564 | 0.00000 |
| **MRO** | 0.03353 | 0.01390 | 0.00757 | 0.03917 | 0.11183 | 0.02867 | 0.00452 |
| **MROH1** | 1.07530 | 2.27316 | 1.49137 | 2.28333 | 9.87844 | 0.85637 | 1.59304 |
| **MROH2A** | 0.02618 | 0.02003 | 0.00251 | 0.02586 | 0.14102 | 0.04693 | 0.02522 |
| **MROH5** | 0.04126 | 0.01615 | 0.01297 | 0.02560 | 0.00799 | 0.01481 | 0.00857 |
| **MROH6** | 0.66351 | 0.00000 | 0.49775 | 1.46655 | 0.00000 | 0.00000 | 10.07524 |
| **MROH7** | 0.01684 | 0.07261 | 0.34550 | 0.01370 | 0.03085 | 0.01819 | 0.00618 |
| **MROH7-TTC4** | 0.03288 | 0.00662 | 0.03094 | 0.01876 | 0.05265 | 0.01816 | 0.01021 |
| **MROH8** | 0.05741 | 0.53364 | 0.32245 | 0.12800 | 0.07016 | 0.23167 | 0.15714 |
| **MROH9** | 0.03466 | 0.01285 | 0.00956 | 0.01374 | 0.04484 | 0.02121 | 0.02376 |
| **MRPL1** | 17.62024 | 3.98337 | 15.07659 | 0.00000 | 0.00000 | 6.27011 | 10.84949 |
| **MRPL10** | 21.79810 | 23.24976 | 10.45744 | 15.03805 | 1.35401 | 11.57302 | 8.89933 |
| **MRPL11** | 28.78029 | 25.49331 | 12.14933 | 25.88842 | 1.18361 | 17.13284 | 15.05132 |
| **MRPL12** | 23.85842 | 43.79433 | 18.48486 | 24.03707 | 5.37315 | 20.42184 | 34.50209 |
| **MRPL13** | 40.26471 | 14.42975 | 14.95123 | 17.22169 | 1.90640 | 13.72520 | 9.28438 |
| **MRPL14** | 42.06425 | 43.31379 | 21.25405 | 37.82011 | 5.92108 | 22.12092 | 17.64222 |
| **MRPL15** | 35.57232 | 24.77176 | 25.26833 | 15.63747 | 2.70991 | 19.60217 | 15.62082 |
| **MRPL16** | 38.28225 | 23.45016 | 18.92146 | 19.52604 | 10.31048 | 16.50879 | 0.98425 |
| **MRPL17** | 27.48747 | 28.26347 | 16.85070 | 20.03160 | 3.18732 | 16.33463 | 10.15793 |

| | | | | | | | |
|---|---|---|---|---|---|---|---|
| **MRPL18** | 36.06946 | 29.88642 | 18.85935 | 19.55552 | 10.27785 | 25.29067 | 5.55599 |
| **MRPL19** | 28.55395 | 12.69782 | 14.50514 | 12.13581 | 7.10562 | 12.60316 | 8.54706 |
| **MRPL2** | 28.42528 | 30.69635 | 12.06285 | 16.85890 | 6.25278 | 11.95820 | 3.69779 |
| **MRPL20** | 30.84891 | 33.40250 | 22.64056 | 26.99807 | 9.79798 | 17.78580 | 21.47738 |
| **MRPL21** | 30.04661 | 27.26520 | 18.09227 | 20.70669 | 3.16861 | 14.66169 | 15.03385 |
| **MRPL22** | 31.61379 | 14.50855 | 14.95401 | 19.90074 | 4.72698 | 12.24348 | 5.43813 |
| **MRPL23** | 0.40739 | 11.48144 | 1.17059 | 1.19021 | 0.00000 | 5.56398 | 14.73480 |
| **MRPL23-AS1** | 0.00892 | 0.09573 | 0.04611 | 0.02247 | 0.01678 | 0.00648 | 0.01717 |
| **MRPL24** | 17.49962 | 18.84727 | 9.21376 | 8.58761 | 1.84515 | 6.01689 | 6.69950 |
| **MRPL27** | 28.28016 | 37.80677 | 16.21868 | 29.42983 | 2.12703 | 12.57744 | 6.68970 |
| **MRPL28** | 33.68818 | 44.23763 | 15.65696 | 24.83463 | 1.00558 | 12.62426 | 0.54504 |
| **MRPL3** | 47.67716 | 14.43382 | 23.19015 | 21.91345 | 1.83558 | 15.27324 | 9.79200 |
| **MRPL30** | 13.79451 | 2.78655 | 11.28858 | 7.18144 | 1.64308 | 4.69102 | 8.77130 |
| **MRPL32** | 28.04539 | 17.53355 | 13.08514 | 20.66459 | 2.44733 | 12.80481 | 3.29020 |
| **MRPL33** | 27.40435 | 10.21429 | 13.67869 | 20.83227 | 0.48794 | 10.46202 | 17.04703 |
| **MRPL34** | 30.11016 | 26.17663 | 17.03459 | 21.97181 | 3.34736 | 20.28148 | 9.48352 |
| **MRPL35** | 27.59670 | 10.11679 | 16.56496 | 10.10348 | 3.29129 | 7.30466 | 5.66131 |
| **MRPL36** | 21.97326 | 27.35391 | 12.80935 | 18.87830 | 0.83349 | 8.00118 | 2.49825 |
| **MRPL37** | 41.10527 | 43.44173 | 24.52320 | 29.71012 | 4.50245 | 22.90908 | 16.50629 |
| **MRPL38** | 32.66407 | 37.90033 | 10.21781 | 27.36364 | 0.00011 | 16.43990 | 8.22223 |
| **MRPL39** | 34.33771 | 10.76803 | 16.58925 | 10.45903 | 1.51370 | 9.80088 | 0.00000 |
| **MRPL4** | 29.24907 | 39.02932 | 12.35586 | 34.00902 | 5.56769 | 17.30859 | 17.89204 |
| **MRPL40** | 23.59516 | 21.79848 | 9.62186 | 5.60263 | 2.34012 | 5.87486 | 11.90748 |
| **MRPL41** | 7.10258 | 38.64582 | 11.12052 | 15.12738 | 8.06084 | 13.40576 | 27.13151 |
| **MRPL42** | 42.51859 | 7.35745 | 12.15850 | 20.96041 | 1.65345 | 11.82179 | 3.27927 |
| **MRPL42P5** | 0.05681 | 0.01338 | 0.02700 | 0.02056 | 0.00647 | 0.00696 | 0.03098 |
| **MRPL43** | 19.51244 | 22.79734 | 14.04930 | 18.41586 | 14.34901 | 11.04570 | 5.94982 |
| **MRPL44** | 30.20366 | 7.49760 | 17.00316 | 16.27044 | 5.49718 | 10.75307 | 4.99745 |
| **MRPL45** | 31.16235 | 10.74798 | 13.28740 | 16.75969 | 1.74743 | 8.67745 | 3.14151 |
| **MRPL45P2** | 0.51858 | 0.17079 | 0.50654 | 0.40987 | 0.25998 | 0.35984 | 0.80927 |
| **MRPL46** | 25.65104 | 23.70842 | 11.99578 | 14.92393 | 2.36929 | 8.47498 | 2.84642 |
| **MRPL47** | 28.62922 | 6.66002 | 15.66345 | 21.14588 | 1.79919 | 15.15664 | 13.75766 |
| **MRPL48** | 20.74295 | 13.71937 | 18.64013 | 2.56503 | 2.46656 | 13.92712 | 7.32133 |
| **MRPL49** | 40.12619 | 24.02851 | 9.72630 | 29.43576 | 7.28414 | 13.33764 | 0.00000 |
| **MRPL50** | 35.23101 | 10.29489 | 21.57949 | 17.14568 | 6.67809 | 9.89584 | 0.00000 |

| | | | | | | | |
|---|---|---|---|---|---|---|---|
| **MRPL51** | 31.61875 | 30.73848 | 22.04351 | 26.63944 | 5.48169 | 20.19827 | 12.62721 |
| **MRPL52** | 25.13489 | 16.76356 | 14.94097 | 26.83192 | 2.44383 | 8.07403 | 25.64294 |
| **MRPL53** | 8.22453 | 19.65374 | 7.53053 | 10.95948 | 5.23803 | 8.11873 | 11.06362 |
| **MRPL54** | 12.39324 | 18.09292 | 7.14317 | 7.97202 | 0.43634 | 4.76760 | 11.36266 |
| **MRPL55** | 24.85366 | 37.79007 | 14.84765 | 20.32689 | 6.31879 | 10.96846 | 8.66794 |
| **MRPL57** | 25.54032 | 33.08275 | 15.25517 | 18.87224 | 8.41998 | 19.09000 | 19.09048 |
| **MRPL9** | 30.56566 | 23.65510 | 19.88605 | 13.41009 | 1.62447 | 16.69385 | 7.29924 |
| **MRPS10** | 15.22058 | 5.85228 | 6.97951 | 6.03689 | 0.28713 | 4.65839 | 5.04667 |
| **MRPS11** | 40.93916 | 27.97895 | 18.23925 | 26.08969 | 2.29824 | 10.71483 | 2.77840 |
| **MRPS12** | 41.31621 | 43.77069 | 13.72361 | 30.27373 | 4.44334 | 19.64953 | 9.17304 |
| **MRPS14** | 25.25913 | 13.87373 | 9.83665 | 14.78734 | 4.47037 | 8.57717 | 1.31055 |
| **MRPS15** | 36.59251 | 27.26275 | 19.69566 | 37.14785 | 2.44436 | 17.64454 | 16.89991 |
| **MRPS16** | 34.96500 | 29.53636 | 16.13294 | 20.05904 | 4.89236 | 17.18508 | 20.16747 |
| **MRPS17** | 27.90510 | 15.44076 | 9.43898 | 18.66854 | 0.69783 | 8.48747 | 7.52184 |
| **MRPS18A** | 34.30722 | 37.40846 | 13.36305 | 22.99515 | 2.23632 | 15.44128 | 2.74252 |
| **MRPS18B** | 36.97717 | 20.58868 | 18.17013 | 24.06997 | 4.78328 | 15.59454 | 2.37395 |
| **MRPS18C** | 21.00361 | 14.39663 | 11.97553 | 11.36286 | 2.63157 | 5.10631 | 3.06737 |
| **MRPS2** | 18.74387 | 30.04387 | 12.51527 | 2.83357 | 0.43502 | 11.70421 | 16.21414 |
| **MRPS21** | 31.35936 | 17.86620 | 23.01528 | 32.29387 | 9.78367 | 11.38324 | 17.93400 |
| **MRPS22** | 23.63483 | 6.92515 | 13.93182 | 15.13228 | 3.16678 | 8.67377 | 0.33407 |
| **MRPS23** | 32.95065 | 20.50145 | 26.63329 | 14.61216 | 10.23542 | 27.68705 | 5.42002 |
| **MRPS24** | 1.32983 | 3.09539 | 1.19252 | 1.44864 | 0.18755 | 0.88180 | 0.02143 |
| **MRPS25** | 32.42660 | 35.97987 | 18.99649 | 21.95597 | 7.07551 | 18.47860 | 11.80286 |
| **MRPS26** | 15.36090 | 37.01093 | 13.94076 | 19.01399 | 2.12672 | 20.21580 | 21.63901 |
| **MRPS27** | 33.42985 | 6.48816 | 7.36320 | 16.89406 | 0.38469 | 5.79349 | 0.00000 |
| **MRPS28** | 20.96941 | 8.66998 | 14.95254 | 14.25797 | 3.04112 | 7.83899 | 4.21946 |
| **MRPS30** | 32.44527 | 17.69806 | 11.80147 | 14.84628 | 4.31552 | 12.99083 | 2.66935 |
| **MRPS31** | 18.20668 | 11.89507 | 10.49732 | 4.13625 | 1.19305 | 8.71020 | 8.92677 |
| **MRPS31P5** | 1.98973 | 0.37447 | 0.69158 | 0.89342 | 0.58651 | 1.10011 | 2.16814 |
| **MRPS33** | 13.08832 | 10.94077 | 15.83105 | 5.85346 | 1.04427 | 6.26750 | 10.56590 |
| **MRPS34** | 32.04113 | 40.09657 | 14.80773 | 30.62381 | 3.08919 | 17.74241 | 17.82849 |
| **MRPS35** | 34.18309 | 6.25956 | 14.03129 | 15.24136 | 2.24765 | 16.04153 | 11.52023 |
| **MRPS36** | 13.16952 | 4.73898 | 3.20842 | 9.71290 | 0.37540 | 9.29172 | 7.30787 |
| **MRPS5** | 33.29175 | 31.59994 | 10.86804 | 16.05350 | 1.57253 | 21.08397 | 19.31015 |
| **MRPS6** | 18.44792 | 1.06813 | 6.03528 | 16.68715 | 1.30677 | 7.59058 | 9.51222 |

| | | | | | | | |
|---|---|---|---|---|---|---|---|
| **MRPS7** | 45.66471 | 33.07924 | 21.15550 | 33.05447 | 11.35058 | 19.65293 | 2.67503 |
| **MRPS9** | 29.00637 | 10.74209 | 9.37244 | 15.24970 | 1.81892 | 8.52504 | 4.53979 |
| **MRRF** | 16.34889 | 8.77200 | 10.79808 | 6.83027 | 3.04770 | 5.43065 | 6.08984 |
| **MRS2** | 28.32154 | 16.87905 | 28.76199 | 14.74002 | 0.40442 | 29.08474 | 0.00000 |
| **MRS2P2** | 1.05548 | 0.06319 | 1.80914 | 0.83375 | 0.19240 | 1.51311 | 0.58080 |
| **MRTO4** | 30.85767 | 25.11670 | 17.58461 | 22.89371 | 10.26785 | 10.57237 | 11.71324 |
| **MRVI1** | 0.15606 | 0.00613 | 0.04223 | 0.23118 | 2.50444 | 0.16676 | 0.82831 |
| **MRVI1-AS1** | 0.01356 | 0.03075 | 0.01563 | 0.03507 | 0.78844 | 0.06409 | 0.02875 |
| **MS4A1** | 0.16403 | 0.00610 | 0.03939 | 0.04361 | 0.00605 | 0.07447 | 0.00443 |
| **MS4A10** | 0.02092 | 0.01367 | 0.01642 | 0.01945 | 0.01902 | 0.06226 | 0.04182 |
| **MS4A12** | 0.04878 | 0.07014 | 0.84471 | 0.00538 | 0.00410 | 0.14564 | 0.00737 |
| **MS4A13** | 0.03667 | 0.00000 | 0.13657 | 0.05080 | 0.00445 | 0.01971 | 0.02689 |
| **MS4A14** | 0.03924 | 0.02658 | 0.03780 | 0.01472 | 0.00612 | 0.01261 | 0.01316 |
| **MS4A15** | 1.80752 | 6.53970 | 0.00766 | 1.34135 | 0.00272 | 2.62718 | 0.00000 |
| **MS4A2** | 0.31239 | 0.00000 | 0.11613 | 0.02073 | 0.00405 | 0.00362 | 0.01635 |
| **MS4A3** | 0.15326 | 0.00000 | 0.19378 | 0.17319 | 0.08490 | 0.00411 | 0.00000 |
| **MS4A4A** | 0.03426 | 0.01101 | 0.02965 | 0.03131 | 0.00786 | 0.01213 | 0.01201 |
| **MS4A5** | 0.01685 | 0.02648 | 0.03666 | 0.02224 | 0.00624 | 0.01442 | 0.02649 |
| **MS4A7** | 0.06106 | 0.00000 | 0.05609 | 0.00934 | 0.01201 | 0.03359 | 0.02197 |
| **MS4A8** | 0.07180 | 0.00329 | 0.16420 | 0.00673 | 0.08285 | 0.00000 | 0.12171 |
| **MSANTD1** | 0.00000 | 0.04409 | 0.04671 | 0.01809 | 0.00669 | 0.05881 | 0.18066 |
| **MSANTD2** | 0.75074 | 0.06365 | 0.77394 | 0.52376 | 0.63576 | 1.90890 | 2.25781 |
| **MSANTD3** | 0.71435 | 0.23034 | 0.56357 | 1.67954 | 0.05137 | 0.22054 | 2.24498 |
| **MSANTD4** | 4.42516 | 0.00000 | 0.40296 | 2.32513 | 0.36656 | 1.48325 | 0.00000 |
| **MSC** | 0.01135 | 0.01726 | 0.00850 | 0.01831 | 0.02182 | 0.21414 | 0.03130 |
| **MSH2** | 28.08628 | 11.98875 | 18.92562 | 14.03261 | 13.61425 | 15.78196 | 0.00000 |
| **MSH3** | 6.03165 | 0.44350 | 2.11098 | 2.73625 | 3.61321 | 3.58962 | 7.92164 |
| **MSH4** | 0.18093 | 0.01698 | 0.05300 | 0.01550 | 0.00591 | 0.04007 | 0.16794 |
| **MSH5-SAPCD1** | 0.06266 | 0.03557 | 0.01378 | 0.04445 | 0.00662 | 0.08638 | 0.05267 |
| **MSH5.uid-7870** | 0.03749 | 0.03911 | 0.01804 | 0.05729 | 0.00727 | 0.02012 | 0.10327 |
| **MSH5.uid-7872** | 0.03410 | 0.02423 | 0.01314 | 0.02156 | 0.01000 | 0.02441 | 0.02669 |
| **MSH6** | 18.55107 | 17.45537 | 21.75664 | 8.09849 | 10.26689 | 32.24158 | 7.47072 |
| **MSI1** | 0.74549 | 4.76718 | 0.21004 | 1.96866 | 0.23559 | 0.92707 | 1.05889 |
| **MSI2** | 0.00000 | 0.16975 | 0.38980 | 3.00049 | 1.61327 | 0.48616 | 2.23477 |
| **MSL1** | 6.08852 | 0.00000 | 3.43781 | 2.05474 | 1.77566 | 1.77540 | 10.64338 |

| | | | | | | | |
|---|---|---|---|---|---|---|---|
| **MSL2** | 2.56740 | 0.00000 | 3.91129 | 2.17256 | 6.82836 | 1.77237 | 8.07442 |
| **MSL3** | 5.82676 | 2.96417 | 5.69071 | 4.36277 | 5.40371 | 5.16427 | 7.87257 |
| **MSL3P1** | 1.29032 | 0.21457 | 8.37615 | 1.71939 | 0.47334 | 0.77609 | 0.41060 |
| **MSLN** | 0.01121 | 0.01938 | 0.00670 | 0.01351 | 0.00749 | 0.39249 | 0.18005 |
| **MSMB** | 0.96888 | 0.69034 | 0.46319 | 0.43205 | 0.98188 | 0.44776 | 0.04563 |
| **MSMO1** | 60.53020 | 5.18215 | 7.39826 | 12.21598 | 0.63925 | 14.72481 | 13.03536 |
| **MSMP** | 0.04122 | 0.01269 | 0.10424 | 0.24992 | 0.04757 | 0.05643 | 0.42370 |
| **MSN** | 1.70093 | 14.97673 | 2.90429 | 0.24618 | 0.10598 | 5.66238 | 0.00000 |
| **MSR1** | 0.04021 | 0.05906 | 0.01312 | 0.00334 | 0.17653 | 0.12053 | 0.12179 |
| **MSRA** | 0.06903 | 0.00000 | 0.46476 | 0.15027 | 3.13965 | 0.14491 | 0.51315 |
| **MSRB1** | 3.21057 | 1.27871 | 4.27063 | 11.74906 | 2.11852 | 6.15671 | 0.00000 |
| **MSRB2** | 0.00000 | 0.22837 | 0.02048 | 0.21438 | 0.09412 | 0.36441 | 0.27599 |
| **MSRB3** | 0.00000 | 0.00846 | 0.74763 | 0.05352 | 0.14030 | 0.00000 | 0.06694 |
| **MSS51** | 0.26377 | 0.16425 | 0.08181 | 0.07966 | 0.00630 | 0.00000 | 0.28612 |
| **MST1** | 0.09452 | 0.14359 | 0.07213 | 1.17622 | 0.00860 | 0.64516 | 0.67508 |
| **MST1L** | 0.02240 | 0.00877 | 0.02087 | 0.04024 | 0.01379 | 0.40653 | 0.01713 |
| **MST1P2** | 0.04185 | 0.01928 | 0.02290 | 0.09176 | 0.01088 | 0.14408 | 0.13591 |
| **MST1R** | 0.07011 | 0.49665 | 0.07779 | 0.06391 | 0.00326 | 0.04247 | 0.08107 |
| **MST4** | 5.17591 | 0.17633 | 0.63176 | 0.24432 | 0.00000 | 0.08156 | 3.09376 |
| **MSTN** | 0.02264 | 0.01502 | 0.01320 | 0.02315 | 0.00776 | 0.01310 | 0.02713 |
| **MSTO1** | 1.43835 | 0.94507 | 1.14337 | 1.52127 | 0.04075 | 2.73584 | 1.59416 |
| **MSTO2P.uid-19** | 0.03284 | 0.00787 | 0.06844 | 0.06019 | 0.00794 | 0.05766 | 0.12675 |
| **MSX1** | 0.22938 | 0.32673 | 0.05385 | 0.00000 | 0.35475 | 0.26062 | 2.55416 |
| **MSX2** | 2.59008 | 7.29287 | 1.29447 | 19.71879 | 0.60048 | 0.00000 | 25.28750 |
| **MSX2P1** | 0.00656 | 0.07498 | 0.00572 | 0.37858 | 0.11964 | 0.00262 | 0.30106 |
| **MT1A** | 0.00000 | 0.24363 | 0.18268 | 0.21546 | 0.14184 | 0.55903 | 0.02943 |
| **MT1B** | 0.01937 | 0.00129 | 0.01372 | 0.01043 | 0.06476 | 0.02686 | 0.15973 |
| **MT1DP** | 0.00390 | 0.07105 | 0.00525 | 0.02187 | 0.04944 | 0.03485 | 0.13891 |
| **MT1E** | 0.00000 | 0.75946 | 0.55224 | 0.00000 | 1.33036 | 3.60075 | 0.03139 |
| **MT1F** | 0.00000 | 0.16588 | 0.03049 | 0.00000 | 0.42032 | 3.85801 | 0.16307 |
| **MT1G** | 0.00000 | 1.51244 | 0.47245 | 0.00000 | 1.53686 | 13.61165 | 0.21665 |
| **MT1H** | 0.00000 | 0.75754 | 0.48568 | 0.00000 | 0.04919 | 6.66238 | 0.02807 |
| **MT1HL1** | 0.00000 | 0.11211 | 0.15288 | 0.00000 | 0.02576 | 1.62808 | 0.13091 |
| **MT1IP** | 0.02223 | 0.01655 | 0.03122 | 0.02179 | 0.00712 | 0.03089 | 0.01429 |
| **MT1JP** | 0.02087 | 0.01554 | 0.01364 | 0.01755 | 0.00769 | 0.01882 | 0.03441 |

| | | | | | | | |
|---|---|---|---|---|---|---|---|
| **MT1L** | 0.00000 | 0.31040 | 0.23225 | 0.01366 | 0.00619 | 0.81528 | 0.00750 |
| **MT1M** | 0.00000 | 0.48402 | 0.12564 | 0.00000 | 0.13889 | 0.86560 | 0.03300 |
| **MT1X** | 0.00000 | 13.29966 | 12.28900 | 0.00000 | 7.76992 | 24.81958 | 0.00000 |
| **MT2A** | 0.00000 | 1.75187 | 0.54976 | 0.00000 | 2.26237 | 7.10458 | 0.71786 |
| **MT3** | 0.02451 | 0.01572 | 0.01326 | 0.02050 | 0.02882 | 0.01364 | 0.01852 |
| **MT4** | 0.03713 | 0.03483 | 0.00996 | 0.01920 | 0.00811 | 0.01322 | 0.00666 |
| **MTA1** | 0.00000 | 0.07408 | 0.17976 | 0.43178 | 0.28440 | 0.30737 | 4.08527 |
| **MTA2** | 0.89078 | 15.52733 | 3.09505 | 12.81781 | 8.72848 | 4.28702 | 19.58860 |
| **MTA3** | 2.17685 | 4.77464 | 6.19735 | 3.54492 | 1.88182 | 11.40734 | 17.82206 |
| **MTAP** | 12.44215 | 0.00000 | 0.41979 | 12.73549 | 0.35983 | 0.37223 | 2.01849 |
| **MTBP** | 4.15029 | 0.00000 | 3.30207 | 1.62973 | 0.64270 | 1.02118 | 0.00000 |
| **MTCH1** | 5.45754 | 13.57058 | 7.26612 | 13.44262 | 2.77159 | 15.99475 | 28.65158 |
| **MTCH2** | 38.99917 | 17.16718 | 15.47615 | 25.74964 | 2.58139 | 18.55438 | 13.37452 |
| **MTCL1** | 0.00022 | 0.11439 | 3.24075 | 0.16444 | 0.04016 | 4.84351 | 0.42199 |
| **MTCL1-AS1** | 0.02160 | 0.06416 | 0.03814 | 0.01664 | 0.00751 | 0.02285 | 0.00898 |
| **MTCP1** | 0.21170 | 0.00896 | 0.14609 | 0.17466 | 0.25921 | 0.22089 | 0.07348 |
| **MTDH** | 14.42107 | 5.31335 | 7.67659 | 4.91198 | 4.51036 | 4.57821 | 34.79823 |
| **MTERF** | 10.75461 | 1.22912 | 7.38890 | 3.16400 | 2.98641 | 0.93623 | 0.00000 |
| **MTERFD1** | 11.76368 | 5.20180 | 12.68459 | 2.41108 | 1.65171 | 4.94762 | 0.00000 |
| **MTERFD2** | 12.25926 | 3.58495 | 9.71111 | 4.99247 | 0.39761 | 3.41680 | 3.88779 |
| **MTERFD3** | 1.62997 | 0.59639 | 1.37134 | 1.21020 | 0.98564 | 0.73653 | 0.00000 |
| **MTF1** | 7.21741 | 3.14852 | 4.21471 | 7.85649 | 4.94562 | 3.83090 | 7.71113 |
| **MTF2** | 20.09315 | 3.98824 | 12.61998 | 14.70745 | 7.76334 | 11.06129 | 12.59383 |
| **MTFMT** | 7.54096 | 6.32951 | 3.72817 | 2.60001 | 0.00000 | 1.47245 | 1.48088 |
| **MTFP1** | 18.90377 | 21.07954 | 0.98053 | 9.68020 | 0.00060 | 13.76714 | 4.37743 |
| **MTFR1** | 3.68462 | 1.22751 | 8.31617 | 2.81079 | 3.25771 | 6.79041 | 6.02136 |
| **MTFR1L** | 10.06037 | 11.46223 | 5.12376 | 7.14373 | 1.76034 | 6.42136 | 0.00000 |
| **MTFR2** | 10.62319 | 4.33383 | 9.02775 | 4.90672 | 4.35331 | 3.24542 | 3.43091 |
| **MTG1** | 19.54260 | 15.49516 | 6.70999 | 12.45589 | 0.87955 | 8.49489 | 5.18628 |
| **MTG2** | 8.62798 | 3.97507 | 6.54627 | 4.70116 | 1.64575 | 2.78043 | 2.97097 |
| **MTHFD1** | 31.10099 | 12.95366 | 6.11902 | 28.32382 | 0.60305 | 15.05243 | 4.81252 |
| **MTHFD1L** | 13.71784 | 1.14191 | 0.71265 | 8.26261 | 0.00000 | 2.84997 | 25.33241 |
| **MTHFD2** | 40.02345 | 0.01604 | 8.51231 | 22.76945 | 3.52425 | 14.69263 | 23.51045 |
| **MTHFD2L** | 0.82611 | 0.39975 | 0.36443 | 1.23106 | 0.03024 | 0.00000 | 0.44292 |
| **MTHFD2P1** | 0.14849 | 0.03129 | 0.02460 | 0.13565 | 0.01466 | 0.04627 | 0.10391 |

| | | | | | | | |
|---|---|---|---|---|---|---|---|
| **MTHFR** | 1.16737 | 0.98981 | 0.14185 | 2.28974 | 0.35867 | 1.81381 | 1.24814 |
| **MTHFS.uid-196** | 0.01998 | 0.03410 | 0.02436 | 0.03389 | 0.00791 | 0.01470 | 0.01893 |
| **MTHFS.uid-196** | 1.28772 | 2.74938 | 0.06837 | 2.05626 | 0.09048 | 1.01806 | 0.30778 |
| **MTHFSD** | 1.20043 | 1.99875 | 2.48016 | 1.07615 | 0.00000 | 0.41917 | 0.84916 |
| **MTIF2** | 24.46434 | 9.35406 | 18.26138 | 7.76548 | 1.20782 | 10.85818 | 8.44536 |
| **MTIF3** | 14.87007 | 4.48073 | 9.21267 | 4.91051 | 0.84932 | 3.71453 | 6.34161 |
| **5.00000** | 0.00000 | 0.00000 | 0.15031 | 0.31701 | 0.48890 | 0.53897 | 0.58984 |
| **MTM1** | 5.23700 | 0.57158 | 2.22091 | 0.47912 | 1.28401 | 4.99916 | 0.00000 |
| **MTMR1** | 0.71235 | 1.16495 | 0.35840 | 0.08758 | 1.35532 | 0.49040 | 4.12523 |
| **MTMR10** | 2.37523 | 1.72579 | 0.34599 | 6.54540 | 0.52751 | 1.29684 | 4.49834 |
| **MTMR11** | 0.10911 | 0.00803 | 0.07020 | 0.08623 | 0.03618 | 1.04473 | 0.42309 |
| **MTMR12** | 3.66823 | 4.07100 | 9.17571 | 3.99405 | 1.48038 | 4.59544 | 4.32170 |
| **MTMR14** | 1.85832 | 21.50393 | 36.55900 | 20.12330 | 4.65323 | 19.76647 | 1.02400 |
| **MTMR2** | 21.91529 | 5.29502 | 2.76315 | 12.83622 | 1.64194 | 4.34548 | 0.00000 |
| **MTMR3** | 0.19652 | 2.69923 | 3.42170 | 2.93501 | 10.34322 | 3.38055 | 6.75943 |
| **MTMR4** | 5.26427 | 1.95427 | 2.09016 | 5.99891 | 0.86512 | 2.00817 | 3.17554 |
| **MTMR6** | 10.35322 | 1.91884 | 2.82803 | 3.33664 | 0.03993 | 2.01292 | 5.04779 |
| **MTMR7** | 4.51046 | 0.00000 | 0.04395 | 2.09017 | 0.14880 | 0.00000 | 2.37433 |
| **MTMR8** | 0.69086 | 0.60938 | 0.40685 | 0.02689 | 0.47145 | 0.37516 | 0.00000 |
| **MTMR9** | 7.19873 | 0.16483 | 4.73770 | 1.75990 | 0.91098 | 4.85181 | 8.40370 |
| **MTMR9LP** | 0.07991 | 0.01247 | 0.16036 | 0.15709 | 0.10980 | 0.18566 | 0.02524 |
| **MTNR1B** | 0.00487 | 0.02306 | 0.01568 | 0.12572 | 0.02637 | 0.01857 | 0.02159 |
| **MTO1** | 18.19315 | 5.97767 | 7.05322 | 7.23594 | 3.25952 | 5.69818 | 0.00000 |
| **MTOR** | 9.63519 | 13.33804 | 5.25841 | 8.53658 | 2.62763 | 12.70816 | 18.45163 |
| **MTOR-AS1** | 0.01784 | 0.03499 | 0.01484 | 0.02028 | 0.01082 | 0.02371 | 0.14742 |
| **MTPAP** | 32.08703 | 16.02501 | 18.21448 | 10.68428 | 9.71245 | 7.11162 | 0.00000 |
| **MTR** | 6.62986 | 2.04601 | 3.37431 | 0.30479 | 1.83905 | 3.01869 | 7.21797 |
| **MTRF1** | 10.24216 | 1.65510 | 2.80879 | 4.55055 | 0.52619 | 2.99365 | 0.00127 |
| **MTRF1L** | 1.40133 | 1.24292 | 2.48851 | 1.84107 | 1.47544 | 1.69158 | 1.24030 |
| **MTRNR2L1** | 25.30920 | 0.00000 | 18.59200 | 24.20017 | 8.02166 | 5.89884 | 6.42816 |
| **MTRNR2L10** | 1.19644 | 0.00000 | 0.39525 | 0.27909 | 0.42175 | 0.58218 | 7.08645 |
| **MTRNR2L2** | 0.01180 | 0.36853 | 0.65757 | 0.12731 | 0.40898 | 0.86721 | 6.54615 |
| **MTRNR2L3** | 0.05356 | 0.09863 | 0.11723 | 0.11217 | 0.11591 | 0.23112 | 1.38201 |
| **MTRNR2L4** | 0.01108 | 0.04778 | 0.02520 | 0.02371 | 0.01326 | 0.09440 | 0.15868 |
| **MTRNR2L5** | 0.03017 | 0.02123 | 0.09281 | 0.02827 | 0.03442 | 0.09466 | 0.97822 |

| | | | | | | | |
|---|---|---|---|---|---|---|---|
| **MTRNR2L6** | 0.15798 | 0.00000 | 0.20121 | 0.00000 | 0.10599 | 0.22447 | 2.83173 |
| **MTRNR2L7** | 0.02409 | 0.02241 | 0.03666 | 0.01394 | 0.02817 | 0.03646 | 0.30303 |
| **MTRNR2L8** | 0.00000 | 0.25202 | 0.49444 | 0.06392 | 0.29062 | 0.35586 | 3.72484 |
| **MTRNR2L9** | 0.02101 | 0.21674 | 0.18863 | 0.04472 | 0.03973 | 0.18033 | 1.63046 |
| **MTRR** | 12.60913 | 3.10766 | 1.11919 | 5.01894 | 4.20660 | 5.33369 | 0.00000 |
| **MTSS1** | 0.36798 | 0.00000 | 0.53235 | 0.33302 | 0.76058 | 3.18260 | 0.55615 |
| **MTSS1L** | 0.00000 | 0.38657 | 0.07383 | 0.22993 | 0.60982 | 0.00000 | 0.56015 |
| **MTTP** | 0.04126 | 0.01165 | 0.03992 | 0.07944 | 0.03108 | 0.74223 | 0.00000 |
| **MTURN** | 0.00000 | 0.05804 | 0.40064 | 0.12100 | 0.30001 | 0.16203 | 0.51586 |
| **MTUS1** | 6.29549 | 0.00000 | 0.52438 | 5.57793 | 4.70722 | 2.93832 | 6.13993 |
| **MTUS2** | 3.63906 | 0.25269 | 0.02384 | 0.01476 | 1.89980 | 0.00000 | 0.73191 |
| **MTUS2-AS1** | 0.02036 | 0.01554 | 0.01149 | 0.01703 | 0.00757 | 0.01999 | 0.05578 |
| **MTX1** | 7.31436 | 8.36503 | 3.41090 | 6.02396 | 0.07496 | 2.84658 | 0.00000 |
| **MTX2** | 20.63021 | 12.21675 | 8.81780 | 9.73303 | 0.44100 | 9.67987 | 0.36469 |
| **MTX3** | 6.30715 | 0.00748 | 4.58701 | 3.24670 | 3.06207 | 0.90231 | 2.96915 |
| **MUC1** | 0.19180 | 0.02924 | 0.01135 | 0.04976 | 0.00500 | 0.14651 | 0.53265 |
| **MUC12** | 0.00840 | 0.00000 | 0.14288 | 0.08384 | 1.41459 | 0.02472 | 0.02121 |
| **MUC13** | 0.11594 | 0.01328 | 0.00250 | 0.00000 | 0.10960 | 0.16541 | 0.04956 |
| **MUC15** | 1.57312 | 0.00000 | 0.01152 | 19.63001 | 0.04432 | 0.00492 | 0.00000 |
| **MUC16** | 0.00000 | 0.00000 | 1.41211 | 0.17726 | 1.34779 | 0.11710 | 0.43789 |
| **MUC17** | 0.04250 | 0.08112 | 0.00921 | 0.04373 | 0.02841 | 0.08271 | 0.01000 |
| **MUC19** | 0.07446 | 0.02224 | 0.00000 | 0.08634 | 0.14390 | 0.00378 | 0.12630 |
| **MUC2** | 0.00575 | 0.00000 | 0.08822 | 0.03045 | 0.01153 | 0.01959 | 0.08524 |
| **MUC20** | 0.14055 | 0.02380 | 0.07612 | 0.02170 | 1.68018 | 0.06666 | 0.26639 |
| **MUC21** | 0.07022 | 0.25105 | 0.94884 | 0.00000 | 0.00021 | 0.43321 | 0.02139 |
| **MUC22** | 0.05586 | 0.01385 | 0.01655 | 0.02290 | 0.09641 | 0.02300 | 0.01053 |
| **MUC3A** | 0.11700 | 0.14070 | 0.99009 | 0.67335 | 3.35417 | 0.16479 | 1.35215 |
| **MUC4** | 0.03483 | 0.20996 | 0.03478 | 0.19608 | 0.01571 | 5.61135 | 0.00000 |
| **MUC5B** | 0.00000 | 0.50375 | 0.71543 | 0.00000 | 0.00044 | 0.54446 | 0.19189 |
| **MUC6** | 0.00000 | 0.03133 | 0.02149 | 0.07326 | 0.00622 | 0.03229 | 0.10304 |
| **MUCL1** | 0.00000 | 0.04610 | 0.02389 | 0.67806 | 0.00861 | 0.00587 | 0.00212 |
| **MUL1** | 17.04307 | 19.68925 | 10.11005 | 6.27062 | 1.42843 | 6.30661 | 0.00000 |
| **MUM1** | 0.65787 | 8.44460 | 1.98760 | 3.08221 | 0.82428 | 3.34660 | 7.43585 |
| **MUM1L1** | 0.02157 | 0.01752 | 0.00969 | 0.02542 | 0.15225 | 0.01247 | 0.02089 |
| **MURC** | 0.00000 | 0.21313 | 0.30170 | 0.30239 | 0.00670 | 0.08468 | 0.28766 |

| | | | | | | | |
|---|---|---|---|---|---|---|---|
| **MUS81** | 2.87911 | 9.51594 | 3.09072 | 4.84987 | 0.81085 | 4.16693 | 6.44613 |
| **MUSK** | 0.02761 | 0.01872 | 0.02023 | 0.01885 | 0.00726 | 0.01256 | 0.01340 |
| **MUSTN1** | 0.02091 | 0.01970 | 0.02964 | 0.13152 | 0.00874 | 0.03837 | 0.08236 |
| **MUT** | 6.43737 | 1.52333 | 2.23507 | 4.25334 | 0.85841 | 2.10357 | 0.00000 |
| **MUTYH** | 2.35083 | 4.32898 | 4.44007 | 3.43612 | 0.38192 | 2.28833 | 0.23199 |
| **MVB12A** | 2.20664 | 6.32941 | 3.04317 | 7.53753 | 1.33039 | 3.66842 | 3.93530 |
| **MVB12B** | 0.00000 | 1.47587 | 0.31561 | 0.38284 | 0.21224 | 0.32363 | 0.61536 |
| **MVD** | 24.13985 | 8.69396 | 0.00000 | 14.40184 | 0.31221 | 4.72469 | 15.70142 |
| **MVK** | 6.80398 | 4.10558 | 0.63118 | 4.60214 | 0.32860 | 1.94895 | 6.12481 |
| **MVP** | 1.54134 | 3.60139 | 1.59450 | 4.71064 | 10.05698 | 6.40368 | 1.25296 |
| **MX1** | 7.34394 | 37.78040 | 0.97390 | 0.00000 | 0.00000 | 28.75656 | 0.00000 |
| **MX2** | 0.07713 | 0.08525 | 0.06556 | 0.11759 | 0.33180 | 0.17628 | 0.00000 |
| **MXD1** | 4.37086 | 3.55808 | 0.00000 | 16.40028 | 4.24041 | 0.00000 | 0.40011 |
| **MXD3** | 0.33800 | 0.98981 | 0.13853 | 1.00241 | 1.22875 | 1.74750 | 1.52926 |
| **MXD4** | 0.00000 | 2.56834 | 0.08802 | 0.32155 | 1.60405 | 1.71835 | 2.41419 |
| **MXI1** | 0.49309 | 0.68386 | 0.95479 | 0.56508 | 0.46297 | 2.28095 | 3.21021 |
| **MXRA5** | 0.04499 | 0.02762 | 0.01738 | 0.00190 | 0.00660 | 0.16778 | 0.04731 |
| **MXRA7** | 0.87367 | 2.74885 | 0.16108 | 0.63426 | 0.06567 | 0.00000 | 3.46818 |
| **MXRA8** | 1.90056 | 0.05377 | 0.03019 | 3.52003 | 0.00000 | 2.79442 | 11.75213 |
| **MYADM** | 21.49595 | 38.26040 | 9.79953 | 2.40841 | 7.09750 | 9.91856 | 17.16500 |
| **MYADML** | 0.00351 | 0.01150 | 0.02766 | 0.00000 | 0.01008 | 0.05642 | 0.49248 |
| **MYADML2** | 0.02443 | 0.01768 | 0.01283 | 0.01923 | 0.00791 | 0.01521 | 0.01930 |
| **MYB** | 0.10875 | 0.02017 | 0.74997 | 0.00000 | 0.28263 | 0.74052 | 0.05997 |
| **MYBBP1A** | 10.03279 | 8.80559 | 8.66079 | 16.14440 | 1.14740 | 4.54888 | 9.72942 |
| **MYBL1** | 0.34525 | 0.02845 | 0.31943 | 0.01361 | 0.46014 | 0.14308 | 0.82216 |
| **MYBL2** | 19.65457 | 56.27316 | 21.34310 | 13.92933 | 5.15762 | 28.25970 | 46.18588 |
| **MYBPC1** | 0.02577 | 0.00000 | 0.10147 | 0.02516 | 0.07780 | 0.01144 | 0.01416 |
| **MYBPC2** | 0.00000 | 0.03455 | 0.07642 | 0.06024 | 0.21849 | 0.37559 | 0.04642 |
| **MYBPC3** | 0.02883 | 0.01659 | 0.00873 | 0.05338 | 0.08843 | 0.02742 | 0.00092 |
| **MYBPH** | 0.07556 | 0.00218 | 0.01482 | 0.01277 | 0.00749 | 0.03777 | 0.03661 |
| **MYBPHL** | 0.14340 | 0.01148 | 0.01365 | 0.11524 | 0.05054 | 0.27981 | 0.00000 |
| **MYC** | 43.70680 | 20.68930 | 14.58630 | 19.71140 | 22.97421 | 0.12030 | 6.86509 |
| **MYCBP** | 0.48798 | 1.03853 | 0.57949 | 0.64648 | 0.35241 | 0.14749 | 0.07457 |
| **MYCBP2** | 7.67496 | 1.00506 | 4.50857 | 4.26195 | 0.26408 | 6.74638 | 17.33403 |
| **MYCBP2-AS1** | 0.03569 | 0.01898 | 0.01348 | 0.02989 | 0.00769 | 0.01907 | 0.02297 |

| | | | | | | | |
|---|---|---|---|---|---|---|---|
| **MYCBPAP** | 0.02340 | 0.00538 | 0.01380 | 0.00713 | 0.01567 | 0.01124 | 0.14476 |
| **MYCL** | 0.00000 | 4.75522 | 4.44417 | 1.78875 | 0.08827 | 8.96216 | 2.13075 |
| **MYCN** | 0.78599 | 9.12663 | 5.53080 | 3.54679 | 0.51094 | 11.95820 | 19.84563 |
| **MYCNOS** | 0.12401 | 2.25466 | 1.44529 | 0.24741 | 0.00082 | 0.45541 | 0.02414 |
| **MYCT1** | 9.70771 | 0.00000 | 0.00000 | 7.72574 | 0.00000 | 0.00000 | 3.69194 |
| **MYD88** | 4.71184 | 0.00000 | 0.09064 | 8.88605 | 0.00000 | 6.94209 | 0.39954 |
| **MYEF2** | 0.15774 | 0.05777 | 0.00000 | 0.31298 | 0.34600 | 0.06161 | 0.33415 |
| **MYEOV** | 0.02712 | 0.01813 | 0.01285 | 0.05201 | 0.00801 | 0.01352 | 0.00859 |
| **MYEOV2** | 18.27117 | 21.18534 | 11.57404 | 14.43748 | 3.60817 | 10.47147 | 17.18165 |
| **MYF6** | 0.02359 | 0.01767 | 0.01104 | 0.04291 | 0.03248 | 0.01303 | 0.00868 |
| **MYH1** | 0.06218 | 0.31721 | 0.28460 | 0.04572 | 0.00242 | 0.01503 | 0.04743 |
| **MYH10** | 38.99176 | 14.55423 | 0.00000 | 19.49673 | 1.49482 | 11.82932 | 27.56825 |
| **MYH11** | 0.07186 | 0.21219 | 0.05373 | 0.14669 | 0.97975 | 0.00637 | 0.28888 |
| **MYH13** | 0.18296 | 0.01184 | 0.00956 | 0.08534 | 0.19444 | 0.16963 | 0.12006 |
| **MYH14** | 0.00000 | 0.02563 | 0.14085 | 0.02797 | 0.11607 | 0.28388 | 0.65068 |
| **MYH15** | 0.19582 | 0.18344 | 0.00000 | 0.12385 | 2.32539 | 0.07275 | 0.00000 |
| **MYH16** | 0.01608 | 0.02708 | 0.02109 | 0.01624 | 0.07930 | 0.03033 | 0.03155 |
| **MYH2** | 0.10238 | 0.10250 | 0.04160 | 0.03287 | 0.01047 | 0.00579 | 0.00000 |
| **MYH3** | 0.03728 | 0.45299 | 0.33388 | 0.80483 | 0.03075 | 0.41138 | 0.68769 |
| **MYH4** | 0.10305 | 0.14987 | 0.11673 | 0.01797 | 0.03253 | 0.00000 | 0.16107 |
| **MYH6** | 0.03558 | 0.00356 | 0.02910 | 0.04181 | 0.19896 | 0.02341 | 0.00000 |
| **MYH7** | 0.01562 | 0.02679 | 0.00716 | 0.00678 | 0.20812 | 0.02377 | 0.08654 |
| **MYH7B** | 0.00000 | 0.51302 | 0.10512 | 0.13480 | 0.01115 | 0.24503 | 0.17225 |
| **MYH8** | 0.04407 | 0.16847 | 0.20825 | 0.14393 | 0.01823 | 0.01880 | 0.11592 |
| **MYH9** | 17.38081 | 34.14542 | 9.22691 | 21.72542 | 1.57005 | 9.31886 | 38.91874 |
| **MYL1** | 0.02396 | 0.01699 | 0.01046 | 0.01937 | 0.03910 | 0.01253 | 0.02181 |
| **MYL10** | 0.01879 | 0.01458 | 0.01533 | 0.02221 | 0.01887 | 0.01233 | 0.02836 |
| **MYL12A** | 35.95653 | 17.74722 | 8.79009 | 28.87009 | 1.42653 | 15.35345 | 4.57779 |
| **MYL12B** | 44.80465 | 23.26589 | 10.70467 | 32.93040 | 5.68070 | 15.02268 | 14.99016 |
| **MYL2** | 0.01767 | 0.01600 | 0.01247 | 0.03900 | 0.06576 | 0.01329 | 0.01674 |
| **MYL3** | 0.00903 | 0.00917 | 0.01177 | 0.07497 | 0.00822 | 0.38385 | 0.01229 |
| **MYL4** | 0.46448 | 0.29421 | 0.11927 | 0.28857 | 0.29591 | 3.98496 | 0.00000 |
| **MYL5** | 0.92657 | 3.34207 | 0.13404 | 0.00000 | 0.03501 | 1.42304 | 1.41126 |
| **MYL6** | 47.26477 | 48.61096 | 25.59951 | 47.63093 | 7.62659 | 31.54892 | 22.42221 |
| **MYL6B** | 3.08827 | 3.59394 | 1.68017 | 12.85856 | 0.10124 | 6.80465 | 4.70959 |

| | | | | | | | |
|---|---|---|---|---|---|---|---|
| **MYL7** | 0.87707 | 0.88587 | 0.00000 | 0.00000 | 0.03499 | 0.00000 | 0.02702 |
| **MYL9** | 0.07409 | 0.03823 | 0.00000 | 1.48648 | 0.04533 | 0.00000 | 0.73491 |
| **MYLIP** | 5.92495 | 0.00000 | 0.49046 | 2.58560 | 2.08169 | 1.77656 | 16.94983 |
| **MYLK** | 0.11035 | 0.00000 | 0.84175 | 0.45005 | 1.04314 | 0.00000 | 0.58990 |
| **MYLK-AS1** | 0.32497 | 0.00000 | 0.60839 | 0.29348 | 0.02835 | 0.06617 | 0.04648 |
| **MYLK2** | 0.00000 | 0.05828 | 0.22531 | 0.22335 | 0.07062 | 0.06657 | 0.76016 |
| **MYLK3** | 0.00000 | 0.04235 | 0.00000 | 0.67131 | 1.41180 | 0.04387 | 0.24677 |
| **MYLK4** | 0.02463 | 0.01881 | 0.01357 | 0.02550 | 0.00829 | 0.01408 | 0.02539 |
| **MYLPF** | 3.82846 | 7.06367 | 0.03961 | 3.03381 | 0.09446 | 0.00000 | 1.32780 |
| **MYNN** | 10.10458 | 2.06184 | 6.07845 | 4.05154 | 3.87408 | 4.30042 | 0.00000 |
| **MYO10** | 1.95976 | 1.61966 | 5.78122 | 18.52998 | 4.21765 | 5.85854 | 4.00480 |
| **MYO15A** | 0.04997 | 0.03120 | 0.05718 | 0.17102 | 0.09617 | 0.00923 | 0.13758 |
| **MYO15B** | 0.00000 | 0.29938 | 0.04597 | 1.21348 | 0.06612 | 0.00000 | 1.10695 |
| **MYO16** | 0.08282 | 0.03594 | 0.08406 | 0.00000 | 1.42346 | 0.00979 | 0.01671 |
| **MYO18A** | 0.00000 | 0.20642 | 0.37701 | 0.97770 | 0.87932 | 2.84256 | 0.68628 |
| **MYO18B** | 0.03061 | 0.00000 | 0.11947 | 0.93642 | 0.12844 | 0.01067 | 0.05691 |
| **MYO19** | 17.52184 | 9.21778 | 5.77927 | 15.87749 | 1.73432 | 6.71114 | 10.35864 |
| **MYO1A** | 0.04828 | 0.00000 | 0.14733 | 0.01310 | 0.00343 | 0.04420 | 0.02755 |
| **MYO1B** | 11.64835 | 9.15090 | 9.42583 | 7.85783 | 1.82242 | 11.87456 | 19.00862 |
| **MYO1C** | 7.56912 | 8.00628 | 6.07120 | 13.16177 | 0.49954 | 2.13967 | 12.45266 |
| **MYO1D** | 0.51822 | 1.97578 | 0.24071 | 2.02088 | 0.16269 | 0.86279 | 1.74053 |
| **MYO1E** | 0.64747 | 2.98952 | 5.22937 | 11.21456 | 1.95680 | 5.51343 | 2.30131 |
| **MYO1F** | 0.06425 | 0.00847 | 0.19027 | 0.18808 | 0.16123 | 0.48129 | 0.20489 |
| **MYO1G** | 0.07376 | 0.74143 | 0.00000 | 0.00000 | 0.02045 | 2.20703 | 0.03876 |
| **MYO1H** | 0.03682 | 0.17249 | 0.06317 | 0.24582 | 1.44604 | 0.01088 | 0.22377 |
| **MYO3A** | 0.00000 | 0.30716 | 0.02073 | 0.00264 | 0.02837 | 0.18641 | 0.10882 |
| **MYO3B** | 0.12738 | 0.18916 | 0.06078 | 0.00000 | 0.97347 | 0.21043 | 0.00000 |
| **MYO5A** | 2.36806 | 5.20082 | 15.02009 | 3.04802 | 1.82644 | 5.02738 | 7.03211 |
| **MYO5B** | 5.08038 | 3.36978 | 0.00000 | 3.15844 | 2.03139 | 5.07475 | 10.28430 |
| **MYO5C** | 1.84163 | 0.65029 | 3.60382 | 1.22807 | 0.11808 | 0.98547 | 0.00000 |
| **MYO6** | 18.00506 | 2.58325 | 2.18462 | 20.50911 | 4.59871 | 0.00000 | 14.40723 |
| **MYO7A** | 0.40566 | 0.00000 | 0.00000 | 5.53980 | 0.56773 | 0.00000 | 4.66847 |
| **MYO7B** | 0.00167 | 0.00000 | 0.07073 | 0.02868 | 0.65338 | 0.02566 | 0.10509 |
| **MYO9A** | 1.77502 | 5.09491 | 3.87540 | 1.98483 | 4.64678 | 10.10974 | 7.11402 |
| **MYO9B** | 0.00000 | 3.05257 | 0.42113 | 5.99076 | 0.90896 | 1.45937 | 14.80750 |

| | | | | | | | |
|---|---|---|---|---|---|---|---|
| **MYOC** | 0.02556 | 0.01244 | 0.01182 | 0.01821 | 0.31296 | 0.01289 | 0.01681 |
| **MYOCD** | 0.01026 | 0.00000 | 0.01527 | 0.06293 | 0.25701 | 0.01745 | 0.17966 |
| **MYOD1** | 0.14159 | 0.15803 | 0.15543 | 0.01000 | 0.00204 | 0.00000 | 0.03945 |
| **MYOF** | 27.10862 | 17.91156 | 0.00000 | 16.33103 | 1.58688 | 0.00000 | 22.83161 |
| **MYOG** | 0.01404 | 0.02485 | 0.01133 | 0.01646 | 0.00939 | 0.01436 | 0.04878 |
| **MYOM1** | 0.27828 | 0.98140 | 0.10115 | 0.00098 | 0.05925 | 0.27757 | 0.00000 |
| **MYOM2** | 0.00870 | 0.52878 | 1.50831 | 0.17733 | 0.05069 | 0.01323 | 0.00000 |
| **MYOM3** | 0.05565 | 0.00000 | 0.27603 | 0.00784 | 0.02987 | 0.07344 | 0.03007 |
| **MYOT** | 0.03219 | 0.00000 | 0.30571 | 0.00000 | 0.02139 | 0.02940 | 0.50952 |
| **MYOZ1** | 0.02016 | 0.10791 | 0.17590 | 0.35660 | 0.00608 | 0.20846 | 0.00000 |
| **MYOZ2** | 0.02887 | 0.01262 | 0.01036 | 0.03046 | 0.05797 | 0.03271 | 0.00929 |
| **MYOZ3** | 0.08075 | 0.00000 | 0.29616 | 0.03356 | 0.10784 | 0.51473 | 0.00000 |
| **MYPN** | 0.00647 | 0.03602 | 0.09087 | 0.04886 | 0.20236 | 0.00964 | 0.05913 |
| **MYPOP** | 0.00000 | 0.27697 | 0.11487 | 0.55451 | 0.08120 | 0.22426 | 3.01707 |
| **MYRF** | 0.57528 | 0.28450 | 0.17186 | 0.00000 | 0.02940 | 1.63085 | 0.96574 |
| **MYRIP** | 0.00000 | 0.16378 | 0.01704 | 0.17639 | 1.44118 | 0.20687 | 0.22038 |
| **MYSM1** | 6.93122 | 0.00000 | 3.52604 | 1.92148 | 0.95778 | 7.22744 | 7.95230 |
| **MYT1** | 0.00090 | 0.01044 | 0.92600 | 0.03472 | 0.04042 | 0.01650 | 0.02576 |
| **MYT1L** | 0.02566 | 0.21992 | 0.69703 | 0.24503 | 0.17829 | 0.00000 | 0.00000 |
| **MYZAP** | 0.05916 | 0.04168 | 0.00000 | 0.10916 | 0.29367 | 0.21921 | 0.03705 |
| **MZB1** | 0.09858 | 0.00000 | 0.01173 | 0.12735 | 0.00704 | 0.00516 | 0.12530 |
| **MZF1** | 0.16209 | 1.48646 | 0.11211 | 2.20011 | 0.36819 | 0.21154 | 4.30114 |
| **MZT1** | 19.76092 | 0.00000 | 13.01670 | 8.61023 | 5.41569 | 5.11154 | 5.26531 |
| **MZT2A** | 0.11710 | 2.27155 | 0.31927 | 2.84626 | 0.27158 | 1.43876 | 6.83655 |
| **MZT2B** | 0.00000 | 2.76218 | 0.32835 | 2.87383 | 0.11724 | 1.67106 | 6.31106 |
| **N4BP1** | 1.95723 | 0.77044 | 2.71026 | 1.37683 | 1.20787 | 2.59163 | 6.59847 |
| **N4BP2** | 9.00023 | 6.81919 | 14.74694 | 0.00000 | 1.30579 | 14.49922 | 16.18664 |
| **N4BP2L1** | 0.13723 | 0.00000 | 0.04005 | 0.03885 | 0.08379 | 0.04323 | 0.03021 |
| **N4BP2L2** | 11.32293 | 0.00000 | 12.08130 | 7.37027 | 4.66277 | 5.22193 | 12.98804 |
| **N4BP2L2-IT2** | 0.55709 | 0.00000 | 0.14119 | 0.12587 | 0.00153 | 0.18491 | 0.41921 |
| **N4BP3** | 0.00000 | 2.21311 | 0.02634 | 1.46790 | 0.00087 | 4.25446 | 3.31227 |
| **N6AMT1** | 4.53601 | 2.45459 | 3.95828 | 1.01306 | 0.12995 | 2.00506 | 0.00001 |
| **N6AMT2** | 2.04497 | 0.69260 | 4.93734 | 0.97959 | 0.22659 | 0.00000 | 0.47720 |
| **NAA10** | 28.96187 | 25.70838 | 9.50967 | 23.62863 | 0.98828 | 12.76988 | 18.21870 |
| **NAA11** | 7.35381 | 0.00000 | 6.49765 | 5.35668 | 4.06263 | 2.92959 | 1.12691 |

| | | | | | | | |
|---|---|---|---|---|---|---|---|
| **NAA15** | 40.02517 | 8.04330 | 18.34939 | 16.61915 | 4.97512 | 14.77062 | 20.89410 |
| **NAA16** | 3.01066 | 0.00000 | 1.69305 | 3.06817 | 0.60541 | 4.81902 | 4.49807 |
| **NAA20** | 27.84443 | 12.44104 | 19.40656 | 15.84641 | 1.84641 | 8.65933 | 1.66358 |
| **NAA25** | 23.98519 | 1.93832 | 10.44241 | 13.08950 | 2.03272 | 9.53914 | 9.41018 |
| **NAA30** | 4.05630 | 0.71496 | 3.65460 | 1.39601 | 1.63790 | 3.39672 | 8.45060 |
| **NAA35** | 10.97988 | 1.70771 | 8.53087 | 8.60535 | 4.97554 | 4.30805 | 5.91429 |
| **NAA38** | 19.04896 | 27.08726 | 5.41067 | 26.04120 | 3.82318 | 9.86451 | 12.85692 |
| **NAA40** | 4.80441 | 4.75860 | 6.11575 | 5.47207 | 3.38626 | 2.88182 | 1.52023 |
| **NAA50** | 58.69204 | 20.45343 | 24.61395 | 31.57294 | 11.84903 | 18.99408 | 7.25029 |
| **NAA60** | 2.29116 | 7.34408 | 6.82756 | 15.65931 | 4.00650 | 7.58751 | 9.37376 |
| **NAAA** | 0.53534 | 0.00000 | 1.37065 | 0.08402 | 0.18007 | 0.64389 | 1.45390 |
| **NAALAD2** | 16.22561 | 0.48184 | 1.55091 | 2.54321 | 1.31640 | 0.00000 | 0.00000 |
| **NAALADL1** | 0.01442 | 0.02958 | 0.09657 | 0.02617 | 0.00516 | 0.02656 | 0.05632 |
| **NAALADL2** | 0.05025 | 0.00000 | 0.02120 | 0.00000 | 0.19368 | 0.15239 | 0.21326 |
| **NAB1** | 2.41898 | 1.14368 | 2.13566 | 0.19272 | 0.14189 | 0.00000 | 3.46560 |
| **NAB2** | 0.16083 | 0.00000 | 0.04034 | 1.01544 | 0.09746 | 0.02255 | 0.46118 |
| **NABP1** | 24.59137 | 6.13039 | 11.75694 | 36.48754 | 16.08854 | 5.68385 | 0.00000 |
| **NABP2** | 5.03034 | 0.16793 | 3.07032 | 12.67976 | 3.00618 | 0.79385 | 0.39125 |
| **NACA** | 55.22806 | 26.25308 | 31.11039 | 42.47216 | 12.82951 | 27.47887 | 32.93743 |
| **NACA2** | 20.53831 | 6.78422 | 10.87594 | 15.33658 | 3.21938 | 9.60365 | 12.25792 |
| **NACAD** | 0.00000 | 0.26256 | 0.29946 | 0.00000 | 0.04801 | 1.46209 | 0.28527 |
| **NACAP1** | 3.11534 | 1.65793 | 1.63598 | 2.16355 | 0.41814 | 1.43526 | 1.03233 |
| **NACC1** | 0.00000 | 1.05101 | 0.87901 | 0.76846 | 0.17373 | 1.94629 | 13.66778 |
| **NACC2** | 0.00000 | 4.03636 | 0.30276 | 0.00000 | 0.05169 | 0.57483 | 2.02802 |
| **NADK** | 4.53935 | 5.34018 | 0.39497 | 7.08937 | 0.04566 | 9.58000 | 8.03267 |
| **NADK2** | 1.23038 | 1.77216 | 0.64922 | 1.37303 | 0.22302 | 0.84902 | 1.80168 |
| **NADSYN1** | 5.23545 | 5.90993 | 0.63523 | 5.74901 | 0.25384 | 2.12296 | 3.94995 |
| **NAE1** | 31.95109 | 8.16150 | 14.49679 | 13.60846 | 5.46663 | 7.85307 | 7.60054 |
| **NAF1** | 2.79317 | 3.76534 | 4.22617 | 2.19506 | 4.70102 | 1.14831 | 3.54832 |
| **NAGA** | 10.49448 | 19.11633 | 0.00000 | 0.00000 | 1.31396 | 8.45626 | 0.04515 |
| **NAGK** | 11.47714 | 11.38085 | 3.54646 | 16.74075 | 1.62828 | 4.90914 | 0.31998 |
| **NAGLU** | 0.10938 | 0.61740 | 0.03674 | 0.11190 | 0.02277 | 0.87044 | 1.67810 |
| **NAGPA** | 2.47230 | 3.73519 | 2.28277 | 6.85064 | 0.55687 | 2.76706 | 9.17603 |
| **NAGPA-AS1** | 0.05666 | 0.14498 | 0.08571 | 0.06618 | 0.01507 | 0.03304 | 0.08176 |
| **NAGS** | 0.09854 | 0.00000 | 0.02200 | 0.10896 | 0.42110 | 0.25364 | 0.00000 |

| | | | | | | | |
|---|---|---|---|---|---|---|---|
| **NAIF1** | 0.11114 | 0.40552 | 1.09541 | 0.72499 | 0.85976 | 0.71917 | 3.06537 |
| **NAIP** | 0.02371 | 0.00998 | 0.01672 | 0.03133 | 0.02845 | 0.02824 | 0.02732 |
| **NALCN** | 0.02444 | 0.00560 | 0.02142 | 0.02046 | 0.16378 | 0.01346 | 0.02305 |
| **NALCN-AS1** | 0.02097 | 0.02115 | 0.02724 | 0.02101 | 0.06281 | 0.01171 | 0.02166 |
| **NAMA** | 0.02007 | 0.01435 | 0.01151 | 0.02484 | 0.02226 | 0.03377 | 0.01725 |
| **NAMPT** | 7.58371 | 0.39231 | 7.13384 | 4.51230 | 1.78038 | 4.19179 | 0.78061 |
| **NANOG** | 0.02929 | 0.16177 | 1.29062 | 0.01037 | 1.11206 | 13.78040 | 0.00000 |
| **NANOGNB** | 0.38963 | 6.30728 | 14.03079 | 0.00000 | 9.64740 | 1.07039 | 0.60420 |
| **NANOS1** | 0.96458 | 1.06001 | 0.35011 | 0.00000 | 1.29188 | 1.60866 | 6.56225 |
| **NANOS2** | 0.02486 | 0.06874 | 0.05016 | 0.02493 | 0.03359 | 0.00671 | 0.00311 |
| **NANOS3** | 0.02228 | 0.10986 | 0.00301 | 0.02373 | 0.00933 | 0.38932 | 0.00003 |
| **NANP** | 9.26865 | 0.46186 | 4.60194 | 3.74807 | 2.28701 | 1.75446 | 0.00000 |
| **NANS** | 26.70149 | 14.17683 | 8.54305 | 19.61963 | 10.65700 | 8.18606 | 0.00000 |
| **NAP1L1** | 30.69551 | 5.16447 | 13.65589 | 23.49341 | 8.23435 | 16.56359 | 41.17345 |
| **NAP1L2** | 0.06609 | 0.01457 | 0.04565 | 0.00646 | 0.01739 | 0.03498 | 0.01784 |
| **NAP1L3** | 0.25564 | 0.00000 | 0.24179 | 0.02186 | 0.82073 | 0.10403 | 0.00000 |
| **NAP1L4** | 33.89733 | 21.52652 | 10.14992 | 25.96633 | 8.97327 | 18.41820 | 11.16094 |
| **NAP1L5** | 0.10766 | 0.15415 | 0.31121 | 0.25415 | 0.88509 | 0.23183 | 0.20904 |
| **NAP1L6** | 0.03160 | 0.02984 | 0.01106 | 0.02850 | 0.00873 | 0.08187 | 0.01293 |
| **NAPA** | 20.37089 | 20.30046 | 9.14625 | 25.38201 | 8.80236 | 10.68818 | 8.10055 |
| **NAPA-AS1** | 0.05218 | 0.14597 | 0.02486 | 0.20448 | 0.13451 | 0.05159 | 0.03660 |
| **NAPB** | 5.27143 | 1.59913 | 2.07398 | 1.73833 | 1.11959 | 1.94614 | 1.25048 |
| **NAPEPLD** | 1.08247 | 0.27673 | 0.63176 | 0.48690 | 0.14260 | 1.30784 | 0.94773 |
| **NAPG** | 16.69777 | 7.58538 | 5.84757 | 8.17769 | 2.33737 | 6.00665 | 6.45606 |
| **NAPRT1** | 2.18768 | 2.12173 | 1.50789 | 0.62765 | 0.06024 | 4.34736 | 4.10433 |
| **NAPSA** | 0.00000 | 0.27064 | 0.11933 | 0.03510 | 0.07518 | 0.07874 | 0.17085 |
| **NAPSB** | 0.03793 | 0.01293 | 0.01154 | 0.01675 | 0.22535 | 0.01347 | 0.02893 |
| **NARF** | 27.09604 | 30.29931 | 12.78914 | 23.56497 | 8.43643 | 16.25799 | 0.41741 |
| **NARFL** | 7.68720 | 9.44501 | 3.28993 | 8.82001 | 0.00000 | 5.03950 | 3.44505 |
| **NARS** | 60.20966 | 22.40001 | 19.86911 | 34.34920 | 2.87548 | 22.35891 | 17.92169 |
| **NARS2** | 15.01666 | 7.59339 | 7.32096 | 11.11652 | 3.17794 | 6.33303 | 0.00000 |
| **NASP** | 38.01650 | 26.19664 | 20.23485 | 31.30402 | 10.56851 | 22.68107 | 38.69271 |
| **NAT1** | 3.18630 | 0.19542 | 0.12574 | 1.46988 | 0.24600 | 1.15579 | 0.00000 |
| **NAT10** | 18.71539 | 14.38118 | 9.83966 | 15.67297 | 3.26996 | 6.98227 | 10.70623 |
| **NAT14** | 0.00000 | 0.74859 | 0.14730 | 0.15492 | 0.28070 | 0.37468 | 2.78645 |

| | | | | | | | |
|---|---|---|---|---|---|---|---|
| **NAT16** | 0.00766 | 0.08270 | 0.05008 | 0.07486 | 0.00624 | 0.03314 | 0.06454 |
| **NAT2** | 0.01864 | 0.01147 | 0.01373 | 0.03514 | 0.15645 | 0.02470 | 0.02564 |
| **NAT6** | 1.14893 | 2.35922 | 1.20849 | 1.33356 | 1.67587 | 1.89665 | 0.50499 |
| **NAT8** | 0.02448 | 0.01558 | 0.02217 | 0.02257 | 0.00710 | 0.01738 | 0.02121 |
| **NAT8B** | 0.03104 | 0.01834 | 0.01249 | 0.01940 | 0.00787 | 0.01605 | 0.01808 |
| **NAT8L** | 0.00000 | 0.56911 | 0.08644 | 0.00000 | 0.00279 | 0.06651 | 1.71766 |
| **NAT9** | 13.64816 | 10.77906 | 4.76660 | 15.09102 | 4.38119 | 5.10571 | 0.00000 |
| **NATD1** | 0.00000 | 1.49380 | 7.97207 | 9.88075 | 0.29118 | 4.86675 | 5.86758 |
| **NAV1** | 0.00000 | 0.30392 | 0.24850 | 5.49162 | 0.12187 | 6.56374 | 5.23936 |
| **NAV2** | 2.91998 | 4.37829 | 0.54424 | 1.41471 | 2.74897 | 1.38716 | 5.07586 |
| **NAV2-AS2** | 0.02023 | 0.06032 | 0.00742 | 0.05782 | 0.01059 | 0.02188 | 0.01363 |
| **NAV2-AS4** | 0.05182 | 0.01884 | 0.00984 | 0.15254 | 0.00724 | 0.00613 | 0.08588 |
| **NAV3** | 0.00562 | 0.00050 | 0.02338 | 0.04321 | 1.93209 | 0.02917 | 0.08899 |
| **NBAS** | 3.64670 | 12.33485 | 4.19616 | 0.00000 | 1.08908 | 12.14551 | 7.27247 |
| **NBEA** | 0.13967 | 0.00000 | 0.29116 | 0.02399 | 0.57960 | 0.15812 | 0.26096 |
| **NBEAL1** | 3.54313 | 0.00000 | 3.18248 | 1.28466 | 1.61026 | 2.53056 | 4.47448 |
| **NBEAL2** | 0.00000 | 1.16549 | 0.13431 | 0.69324 | 0.00325 | 1.93645 | 3.83658 |
| **NBEAP1** | 0.02806 | 0.00000 | 0.18699 | 0.05242 | 0.78313 | 0.00585 | 0.01256 |
| **NBN** | 14.29206 | 0.88164 | 6.59986 | 2.44216 | 0.72610 | 4.76265 | 3.42427 |
| **NBPF1** | 1.56219 | 2.73070 | 2.33267 | 3.65114 | 1.92962 | 1.04318 | 1.50463 |
| **NBPF10** | 0.00000 | 0.09745 | 0.01452 | 0.49928 | 0.02218 | 0.14028 | 0.40942 |
| **NBPF11.uid-434** | 0.02060 | 0.01765 | 0.01264 | 0.01982 | 0.00854 | 0.01262 | 0.02769 |
| **NBPF12** | 0.03490 | 0.12203 | 0.05811 | 0.07837 | 0.02791 | 0.20459 | 0.00000 |
| **NBPF13P** | 0.02542 | 0.01802 | 0.01490 | 0.07489 | 0.00862 | 0.01301 | 0.01041 |
| **NBPF15** | 0.49674 | 0.54027 | 0.64175 | 0.58434 | 0.12938 | 0.50282 | 0.00000 |
| **NBPF22P** | 0.06959 | 0.00779 | 0.01767 | 0.00822 | 0.00676 | 0.01236 | 0.06632 |
| **NBPF25P.uid-43** | 0.04120 | 0.02768 | 0.01668 | 0.04121 | 0.00681 | 0.04665 | 0.05478 |
| **NBPF25P.uid-43** | 0.02534 | 0.01519 | 0.01422 | 0.02279 | 0.00789 | 0.02597 | 0.01498 |
| **NBPF3** | 0.97471 | 1.27507 | 1.20584 | 0.91338 | 0.24312 | 0.85899 | 0.00000 |
| **NBPF4** | 0.05492 | 0.03056 | 0.02579 | 0.09265 | 0.00712 | 0.00986 | 0.00000 |
| **NBPF6** | 0.03894 | 0.02037 | 0.01657 | 0.03761 | 0.00836 | 0.01303 | 0.00858 |
| **NBPF7** | 0.02584 | 0.01790 | 0.01402 | 0.02375 | 0.00826 | 0.02201 | 0.01248 |
| **NBPF8.uid-74** | 0.04139 | 0.03634 | 0.04921 | 0.05271 | 0.02088 | 0.05622 | 0.02653 |
| **NBPF8.uid-77** | 0.02891 | 0.04587 | 0.02314 | 0.01828 | 0.00827 | 0.02718 | 0.01488 |
| **NBPF9.uid-78** | 0.01866 | 0.03042 | 0.02681 | 0.20141 | 0.01866 | 0.03584 | 0.11514 |

| | | | | | | | |
|---|---|---|---|---|---|---|---|
| **NBR1** | 4.31389 | 3.95970 | 4.81716 | 5.14531 | 1.14478 | 8.87817 | 2.38773 |
| **NBR2** | 1.16911 | 0.26132 | 0.77088 | 0.94243 | 1.92435 | 0.71368 | 0.70809 |
| **NCALD** | 0.54526 | 0.34125 | 0.06965 | 0.00000 | 1.40054 | 0.10601 | 0.78402 |
| **NCAM1** | 0.02460 | 0.05098 | 0.15733 | 0.00000 | 0.33751 | 0.18167 | 0.46778 |
| **NCAM2** | 0.22953 | 0.00000 | 0.27415 | 0.11133 | 0.00986 | 0.04400 | 0.00000 |
| **NCAN** | 0.01005 | 0.06756 | 0.00000 | 0.03492 | 0.05253 | 0.00749 | 0.24739 |
| **NCAPD2** | 14.48211 | 15.35057 | 10.06619 | 14.54385 | 6.28011 | 5.39925 | 8.10368 |
| **NCAPD3** | 8.92248 | 4.87226 | 3.62211 | 8.23927 | 1.90696 | 7.54851 | 4.40088 |
| **NCAPG** | 20.27716 | 4.55175 | 11.49808 | 10.69527 | 4.50586 | 2.99442 | 6.36659 |
| **NCAPG2** | 16.36444 | 5.10207 | 3.54305 | 9.42513 | 2.90288 | 7.94750 | 1.30339 |
| **NCAPH** | 15.36768 | 9.88901 | 4.80952 | 12.64553 | 4.02936 | 5.11199 | 3.99224 |
| **NCAPH2** | 17.91044 | 22.43956 | 4.25169 | 19.28024 | 4.23197 | 7.47706 | 5.90955 |
| **NCBP1** | 27.64704 | 9.19081 | 9.31687 | 10.52171 | 5.39767 | 8.67113 | 6.75258 |
| **NCBP2** | 35.54721 | 14.85025 | 26.12669 | 23.10326 | 10.77289 | 26.71221 | 6.33359 |
| **NCBP2-AS2** | 14.41898 | 26.98535 | 12.80168 | 11.88214 | 17.29749 | 11.46414 | 0.00000 |
| **NCCRP1** | 0.00000 | 0.50770 | 0.00000 | 2.95385 | 0.58210 | 2.10950 | 0.00000 |
| **NCDN** | 0.00000 | 3.13275 | 3.81269 | 5.26934 | 0.01576 | 2.60595 | 6.99219 |
| **NCEH1** | 11.57921 | 2.67887 | 3.48839 | 0.00000 | 1.69154 | 7.99646 | 1.38014 |
| **NCF1** | 0.02265 | 0.00260 | 0.08985 | 0.01862 | 0.18523 | 0.01089 | 0.01807 |
| **NCF1B** | 0.02494 | 0.00406 | 0.07718 | 0.01492 | 0.03661 | 0.02760 | 0.02995 |
| **NCF1C** | 0.02661 | 0.01062 | 0.10016 | 0.01723 | 0.10343 | 0.00881 | 0.01549 |
| **NCF2** | 0.95202 | 0.32417 | 0.02679 | 0.00152 | 0.35428 | 0.32253 | 0.42440 |
| **NCF4** | 0.01762 | 0.02292 | 0.01063 | 0.06227 | 0.00810 | 0.06845 | 0.00941 |
| **NCK1** | 12.48249 | 0.00000 | 12.37797 | 6.32823 | 5.51127 | 5.32638 | 0.00000 |
| **NCK1-AS1** | 0.38618 | 0.00022 | 1.07457 | 0.91078 | 4.30531 | 0.13977 | 0.00000 |
| **NCK2** | 12.68669 | 6.34554 | 0.49730 | 9.16665 | 1.69114 | 5.16830 | 6.71837 |
| **NCKAP1** | 15.26533 | 0.00000 | 11.85036 | 6.49058 | 1.34977 | 8.64400 | 23.56513 |
| **NCKAP1L** | 0.03030 | 0.00385 | 0.07683 | 0.02487 | 0.00563 | 0.01433 | 0.02207 |
| **NCKAP5** | 0.01645 | 0.05071 | 0.01422 | 0.01896 | 0.05894 | 0.02995 | 0.03262 |
| **NCKAP5L** | 0.00000 | 0.00000 | 0.20682 | 1.33874 | 1.09192 | 2.62865 | 2.60151 |
| **NCKIPSD** | 3.48735 | 4.17554 | 0.99003 | 4.45327 | 0.27806 | 3.84083 | 5.13956 |
| **NCL** | 82.32643 | 49.55979 | 38.77487 | 57.51848 | 22.23526 | 37.46039 | 48.13784 |
| **NCLN** | 6.39273 | 18.78943 | 9.54049 | 15.54272 | 0.04555 | 7.72196 | 33.20806 |
| **NCMAP** | 1.42407 | 0.41247 | 0.00001 | 1.42171 | 0.03592 | 0.00000 | 0.11296 |
| **NCOA1** | 2.10109 | 7.08255 | 5.91801 | 0.71254 | 3.38501 | 12.62387 | 4.96492 |

| | | | | | | | |
|---|---|---|---|---|---|---|---|
| **NCOA2** | 0.89647 | 2.31983 | 2.70143 | 3.25701 | 2.77258 | 2.75132 | 2.46634 |
| **NCOA3** | 15.70817 | 7.82937 | 6.38699 | 19.64584 | 7.08601 | 16.21462 | 8.38693 |
| **NCOA4** | 58.22943 | 29.29263 | 21.45333 | 28.41095 | 5.75481 | 16.59658 | 8.30110 |
| **NCOA5** | 4.89185 | 6.31476 | 2.56917 | 3.31259 | 0.34447 | 2.86067 | 10.28567 |
| **NCOA6** | 6.83484 | 5.95791 | 4.04127 | 6.86053 | 6.36952 | 3.46027 | 14.12817 |
| **NCOA7** | 2.37548 | 0.00000 | 3.47788 | 2.62789 | 0.04103 | 3.20158 | 5.61292 |
| **NCOR1** | 25.53856 | 12.18066 | 11.65027 | 13.92554 | 12.88674 | 13.29689 | 19.22187 |
| **NCOR1P1** | 0.02548 | 0.01627 | 0.01701 | 0.02175 | 0.24843 | 0.01304 | 0.01573 |
| **NCOR2** | 0.00000 | 1.95415 | 0.00000 | 2.20700 | 0.55493 | 0.62150 | 19.23089 |
| **NCR1** | 0.17964 | 1.49272 | 0.26484 | 0.00693 | 0.06175 | 0.13836 | 0.28105 |
| **NCR3** | 0.05556 | 0.01132 | 0.03061 | 0.01886 | 0.01453 | 0.18383 | 0.00054 |
| **NCR3LG1** | 0.00000 | 0.00000 | 0.86757 | 0.06907 | 0.04512 | 0.00000 | 0.10518 |
| **NCRNA00185** | 0.02545 | 0.01839 | 0.01329 | 0.03042 | 0.00879 | 0.01312 | 0.01053 |
| **NCRUPAR** | 0.02728 | 0.02976 | 0.09964 | 0.01819 | 0.00397 | 0.00590 | 0.02988 |
| **NCS1** | 0.00000 | 0.00000 | 0.09029 | 0.04596 | 0.03578 | 0.09516 | 0.21875 |
| **NCSTN** | 15.68106 | 8.72747 | 10.38151 | 12.40915 | 1.79945 | 8.02448 | 0.00001 |
| **NDC1** | 32.94371 | 5.27923 | 13.96061 | 9.54375 | 1.33183 | 13.00853 | 13.85251 |
| **NDC80** | 15.25274 | 4.33604 | 10.84391 | 12.48197 | 0.39915 | 5.81029 | 0.00000 |
| **NDE1** | 0.51369 | 1.08003 | 1.03477 | 2.24948 | 3.79086 | 1.23507 | 2.87370 |
| **NDEL1** | 5.00515 | 4.41986 | 1.84715 | 9.87384 | 1.27575 | 3.88955 | 4.16903 |
| **NDFIP1** | 24.38687 | 8.55043 | 8.30944 | 9.50398 | 1.74443 | 14.45231 | 11.57813 |
| **NDFIP2** | 24.63173 | 8.93281 | 11.53602 | 14.40236 | 3.58050 | 4.99303 | 4.84381 |
| **NDFIP2-AS1** | 0.02576 | 0.02152 | 0.01297 | 0.02576 | 0.01227 | 0.01073 | 0.02000 |
| **NDN** | 0.00000 | 0.59299 | 0.82306 | 2.52178 | 0.33164 | 1.14624 | 3.46318 |
| **NDNF** | 0.05644 | 0.00000 | 0.02314 | 0.02198 | 0.37868 | 0.01763 | 0.00000 |
| **NDNL2** | 0.02566 | 1.43218 | 1.13367 | 0.85327 | 0.00293 | 1.02252 | 4.37569 |
| **NDOR1** | 0.00000 | 7.87721 | 2.90434 | 5.77089 | 0.29618 | 3.11907 | 16.54426 |
| **NDP** | 0.03149 | 0.01475 | 0.00928 | 0.01578 | 0.06100 | 0.01624 | 0.01893 |
| **NDRG1** | 0.00000 | 4.25349 | 0.00000 | 1.27526 | 0.56580 | 1.13174 | 3.38663 |
| **NDRG2** | 9.59567 | 2.80624 | 0.00000 | 0.00000 | 0.04060 | 1.55867 | 0.16375 |
| **NDRG3** | 11.84654 | 4.58348 | 10.06911 | 8.32701 | 3.12384 | 5.13883 | 8.35407 |
| **NDRG4** | 0.12750 | 0.05630 | 0.14851 | 0.07939 | 1.90713 | 0.43541 | 0.51759 |
| **NDST1** | 8.96335 | 3.92671 | 2.26399 | 4.11937 | 0.89633 | 6.81061 | 12.09099 |
| **NDST2** | 0.35255 | 0.73661 | 0.67560 | 1.20976 | 0.64374 | 2.59925 | 1.55774 |
| **NDST3** | 0.01876 | 0.03344 | 0.01132 | 0.01550 | 0.18364 | 0.13240 | 0.06244 |

| | | | | | | | |
|---|---|---|---|---|---|---|---|
| **NDST4** | 0.06342 | 0.01081 | 0.01373 | 0.03962 | 0.01518 | 0.02851 | 0.00297 |
| **NDUFA1** | 46.44188 | 29.08580 | 28.69070 | 25.00376 | 10.23774 | 26.49668 | 34.33140 |
| **NDUFA10** | 33.16549 | 25.71998 | 7.76144 | 23.10029 | 2.71045 | 15.91677 | 13.91156 |
| **NDUFA11** | 30.55917 | 38.09625 | 6.87497 | 37.00013 | 3.44420 | 20.73447 | 21.43749 |
| **NDUFA12** | 33.25301 | 21.99904 | 13.15352 | 13.62283 | 0.96547 | 23.50062 | 7.86441 |
| **NDUFA13** | 32.86783 | 37.73691 | 13.95182 | 32.22778 | 15.44138 | 27.23306 | 19.75404 |
| **NDUFA2** | 36.81128 | 20.88617 | 15.09007 | 24.00870 | 10.78067 | 21.31363 | 21.92782 |
| **NDUFA3** | 20.34797 | 22.79040 | 7.04973 | 18.84332 | 3.15894 | 15.14184 | 23.55042 |
| **NDUFA4** | 52.45795 | 21.72310 | 19.42708 | 36.07533 | 6.58822 | 30.18497 | 34.73104 |
| **NDUFA4L2** | 0.04597 | 0.00532 | 0.02478 | 0.03003 | 0.00884 | 0.01693 | 0.01918 |
| **NDUFA5** | 22.52485 | 0.07745 | 7.71564 | 7.09855 | 0.63288 | 7.83568 | 6.45231 |
| **NDUFA6** | 46.74504 | 37.71805 | 15.43642 | 32.24971 | 15.78654 | 27.73043 | 29.90547 |
| **NDUFA6-AS1** | 0.17264 | 0.13119 | 1.44865 | 0.03422 | 0.95972 | 0.00861 | 0.08031 |
| **NDUFA7** | 14.07755 | 14.75807 | 11.69142 | 10.16499 | 1.02507 | 7.81837 | 12.38821 |
| **NDUFA8** | 35.32403 | 42.19954 | 9.58933 | 24.13560 | 0.28132 | 22.76875 | 6.76324 |
| **NDUFA9** | 37.50218 | 27.69674 | 14.12755 | 20.89582 | 1.78579 | 18.19548 | 0.00006 |
| **NDUFAB1** | 50.39795 | 51.09831 | 32.69520 | 29.83230 | 8.81629 | 41.74411 | 28.71274 |
| **NDUFAF1** | 14.14289 | 4.07643 | 8.28895 | 5.46874 | 2.40342 | 3.62182 | 0.00000 |
| **NDUFAF2** | 3.97523 | 1.91822 | 2.24883 | 6.14210 | 0.49136 | 1.88378 | 4.01097 |
| **NDUFAF3** | 5.77243 | 11.37284 | 1.07288 | 2.97802 | 0.63635 | 6.03552 | 3.87529 |
| **NDUFAF4** | 27.90542 | 13.04752 | 14.45330 | 11.56373 | 2.55974 | 12.04588 | 10.13105 |
| **NDUFAF4P1** | 0.95013 | 0.00554 | 0.54191 | 0.19258 | 0.21636 | 0.41774 | 0.45835 |
| **NDUFAF5** | 7.80344 | 6.90566 | 3.94297 | 4.53965 | 3.03125 | 5.14061 | 1.60894 |
| **NDUFAF6** | 7.47532 | 5.19012 | 1.13067 | 7.86880 | 0.08046 | 3.96352 | 4.25391 |
| **NDUFAF7** | 13.98218 | 6.67401 | 7.71458 | 8.60443 | 7.29831 | 14.03212 | 0.00000 |
| **NDUFB1** | 17.37476 | 9.07124 | 5.79973 | 13.85443 | 0.60009 | 13.56611 | 22.33592 |
| **NDUFB10** | 43.29247 | 32.25316 | 6.33551 | 27.99376 | 9.11854 | 23.98967 | 20.01034 |
| **NDUFB11** | 32.00372 | 31.24931 | 20.04375 | 26.32050 | 9.25001 | 18.73967 | 30.70372 |
| **NDUFB2** | 37.20760 | 35.35445 | 10.74548 | 29.65727 | 5.72112 | 22.44700 | 24.29640 |
| **NDUFB2-AS1** | 0.33460 | 1.94460 | 0.95923 | 0.69575 | 0.16580 | 0.08816 | 1.44621 |
| **NDUFB3** | 24.56225 | 8.05516 | 15.15838 | 4.40660 | 0.60061 | 12.12347 | 13.02298 |
| **NDUFB4** | 28.70348 | 14.27400 | 8.25006 | 20.17896 | 2.82020 | 13.67305 | 16.52528 |
| **NDUFB5** | 25.71838 | 10.24360 | 10.10747 | 10.62267 | 2.43471 | 11.64176 | 5.78560 |
| **NDUFB6** | 32.13077 | 24.34379 | 16.67863 | 18.24181 | 6.57126 | 16.40348 | 13.19095 |
| **NDUFB7** | 30.16420 | 42.84264 | 12.93718 | 24.98496 | 4.51857 | 19.11386 | 23.77135 |

| | | | | | | | |
|---|---|---|---|---|---|---|---|
| **NDUFB8** | 43.39425 | 37.71172 | 8.94055 | 36.04731 | 12.72735 | 21.07662 | 7.67678 |
| **NDUFB9** | 47.54815 | 37.27412 | 20.39789 | 29.60561 | 10.40746 | 30.77656 | 33.73844 |
| **NDUFC1** | 32.68364 | 15.07164 | 8.60885 | 17.87222 | 2.17151 | 14.25829 | 23.34365 |
| **NDUFC2** | 23.00731 | 4.99118 | 11.98306 | 8.96515 | 0.67766 | 13.08368 | 20.82250 |
| **NDUFS1** | 33.94249 | 9.64877 | 8.60936 | 16.85084 | 1.88910 | 14.30794 | 0.92421 |
| **NDUFS2** | 37.12318 | 22.73589 | 12.40207 | 19.45296 | 2.64125 | 16.01866 | 3.20317 |
| **NDUFS3** | 35.73395 | 29.38692 | 8.24775 | 32.72547 | 3.24260 | 18.31035 | 9.31315 |
| **NDUFS4** | 16.04697 | 4.19574 | 5.78250 | 8.02299 | 0.42906 | 7.08412 | 10.10953 |
| **NDUFS5** | 45.91268 | 37.17012 | 17.74177 | 33.89971 | 0.89233 | 25.43521 | 32.18881 |
| **NDUFS6** | 33.64659 | 43.88199 | 9.24768 | 30.77404 | 1.10562 | 21.37103 | 26.03817 |
| **NDUFS7** | 29.85366 | 26.07624 | 2.01065 | 15.41326 | 0.68862 | 14.97314 | 9.57373 |
| **NDUFS8** | 28.30077 | 28.78760 | 7.19143 | 27.61583 | 1.58861 | 14.97823 | 8.18102 |
| **NDUFV1** | 49.47855 | 54.43496 | 12.73955 | 24.80546 | 4.97296 | 27.72983 | 8.72595 |
| **NDUFV2** | 29.13999 | 14.42725 | 13.33239 | 16.25011 | 1.02409 | 13.25405 | 9.65904 |
| **NDUFV3** | 17.20294 | 13.69753 | 6.35383 | 6.86521 | 0.72293 | 6.43054 | 11.56318 |
| **NEAT1** | 0.00000 | 1.28777 | 1.26062 | 2.55882 | 0.12649 | 3.16905 | 11.01594 |
| **NEB** | 0.89211 | 0.00000 | 0.90145 | 1.16027 | 1.20756 | 1.41758 | 1.80905 |
| **NEBL** | 0.06913 | 0.38165 | 0.00000 | 0.61095 | 0.98437 | 0.00000 | 0.53007 |
| **NEBL-AS1** | 0.02360 | 0.01525 | 0.01339 | 0.02460 | 0.00781 | 0.01214 | 0.02296 |
| **NECAB1** | 0.30293 | 0.05649 | 0.20804 | 0.19330 | 0.06041 | 0.37825 | 0.12808 |
| **NECAB2** | 0.02113 | 0.02547 | 0.09554 | 0.04066 | 0.09171 | 0.00819 | 0.02006 |
| **NECAB3** | 0.97865 | 2.86111 | 0.40172 | 1.57298 | 0.16378 | 1.42035 | 5.47014 |
| **NECAP1** | 11.44245 | 4.64591 | 6.79644 | 3.93070 | 1.88001 | 4.93520 | 0.00000 |
| **NECAP2** | 11.16213 | 10.51119 | 6.66597 | 10.14578 | 2.94015 | 4.94550 | 3.26208 |
| **NEDD1** | 12.58280 | 2.24073 | 3.85226 | 7.19030 | 4.70527 | 2.03336 | 1.83106 |
| **NEDD4** | 0.63519 | 0.00000 | 2.02274 | 0.71987 | 0.25395 | 0.83539 | 0.73374 |
| **NEDD4L** | 0.25060 | 1.72018 | 0.03999 | 6.19124 | 2.32906 | 3.45397 | 4.28408 |
| **NEDD8** | 29.38095 | 17.13004 | 17.78171 | 25.76253 | 2.61937 | 17.47155 | 16.84828 |
| **NEDD9** | 0.65708 | 5.13080 | 0.21098 | 1.83156 | 0.24118 | 0.79821 | 0.00000 |
| **NEFH** | 11.59732 | 10.78432 | 5.53923 | 0.00000 | 5.19841 | 12.80607 | 16.32162 |
| **NEFL** | 0.00000 | 7.03965 | 4.12226 | 0.00000 | 1.73114 | 5.30863 | 0.00000 |
| **NEFM** | 0.87133 | 0.02759 | 0.22582 | 0.24288 | 9.16152 | 0.44193 | 0.24651 |
| **NEGR1** | 0.02893 | 0.03302 | 0.02052 | 0.00000 | 0.14037 | 0.15787 | 0.17578 |
| **NEIL1** | 0.35190 | 0.00000 | 0.08579 | 0.72180 | 0.00209 | 1.66964 | 0.72977 |
| **NEIL2** | 5.25763 | 12.40096 | 4.35979 | 0.00000 | 2.22635 | 3.46763 | 0.02499 |

| | | | | | | | |
|---|---|---|---|---|---|---|---|
| **NEIL3** | 6.01002 | 0.51968 | 0.84061 | 2.31098 | 0.26566 | 0.94079 | 0.00000 |
| **NEK1** | 5.44801 | 3.44320 | 4.00372 | 2.33725 | 8.59497 | 3.22703 | 6.23123 |
| **NEK10** | 0.02098 | 0.02157 | 0.01255 | 0.07253 | 0.07002 | 0.01058 | 0.01864 |
| **NEK11** | 0.27673 | 0.00000 | 0.69794 | 0.47085 | 0.36682 | 0.11059 | 0.00000 |
| **NEK2** | 8.82952 | 7.17177 | 11.85529 | 5.08859 | 5.63000 | 5.88750 | 0.00000 |
| **NEK3** | 2.80102 | 2.08655 | 1.38405 | 0.26940 | 0.32809 | 3.94669 | 0.33394 |
| **NEK4** | 4.15142 | 0.93492 | 2.87914 | 0.93542 | 1.05904 | 2.53638 | 2.95701 |
| **NEK5** | 0.10498 | 0.07900 | 0.31882 | 0.03881 | 0.17818 | 0.13566 | 0.00165 |
| **NEK6** | 9.50450 | 16.19250 | 0.57920 | 5.70885 | 0.00000 | 2.07859 | 6.60870 |
| **NEK7** | 3.48300 | 0.00000 | 2.41269 | 1.90958 | 1.25366 | 0.99626 | 5.34609 |
| **NEK8** | 1.87360 | 1.47186 | 0.74587 | 1.29153 | 0.07468 | 0.57729 | 0.90745 |
| **NEK9** | 5.74079 | 2.12716 | 2.44778 | 6.46940 | 0.30335 | 3.26152 | 7.42222 |
| **NELFA** | 8.16888 | 8.39673 | 7.45326 | 9.28717 | 1.85475 | 4.63449 | 6.77369 |
| **NELFB** | 0.00000 | 11.46306 | 6.93468 | 4.55895 | 0.20367 | 4.76409 | 18.10947 |
| **NELFCD** | 21.79564 | 13.98268 | 14.56600 | 23.46628 | 1.70358 | 12.06951 | 3.89261 |
| **NELFE** | 25.41004 | 27.86141 | 14.45535 | 23.71027 | 13.92146 | 14.26193 | 2.30970 |
| **NELL1** | 0.03949 | 0.00000 | 0.04201 | 0.00785 | 2.46698 | 0.02396 | 0.03253 |
| **NELL2** | 0.00000 | 1.08938 | 1.98074 | 0.18767 | 0.10757 | 0.00930 | 0.00142 |
| **NEMF** | 17.53259 | 7.52701 | 4.62419 | 9.20907 | 4.04840 | 8.59656 | 19.28352 |
| **NENF** | 2.70924 | 1.39213 | 0.00000 | 5.97842 | 0.06317 | 0.00000 | 18.86768 |
| **NEO1** | 0.85788 | 1.83512 | 4.97467 | 0.91810 | 6.09664 | 2.44347 | 1.24997 |
| **NES** | 0.00000 | 0.00000 | 0.75843 | 0.04272 | 0.27637 | 0.69847 | 0.10331 |
| **NET1** | 7.34474 | 1.48911 | 3.70083 | 27.71643 | 3.68612 | 2.66204 | 12.57194 |
| **NETO1** | 0.03223 | 0.01555 | 0.01260 | 0.01930 | 0.00904 | 0.03821 | 0.01805 |
| **NETO2** | 0.00601 | 1.95727 | 3.54327 | 0.55392 | 0.85163 | 5.23648 | 4.81596 |
| **NEU1** | 15.30533 | 15.79614 | 16.63134 | 19.41514 | 6.27321 | 12.63263 | 5.03002 |
| **NEU3** | 0.19062 | 0.35621 | 0.32445 | 0.24855 | 0.12216 | 0.19900 | 0.00000 |
| **NEU4** | 0.02404 | 0.01918 | 0.01339 | 0.02001 | 0.00896 | 0.01627 | 0.01540 |
| **NEURL1** | 0.00000 | 0.95465 | 4.08483 | 3.22244 | 0.64344 | 0.06406 | 0.25275 |
| **NEURL1B** | 0.02337 | 0.01139 | 0.02854 | 0.04682 | 0.03811 | 0.00895 | 0.04722 |
| **NEURL2** | 0.00000 | 3.76894 | 0.00821 | 0.93612 | 0.00279 | 1.63041 | 1.49733 |
| **NEURL3** | 0.01739 | 0.01555 | 0.05452 | 0.08559 | 0.00591 | 0.00723 | 0.03680 |
| **NEURL4** | 1.58770 | 1.75414 | 1.61323 | 2.15639 | 0.00000 | 3.02454 | 9.07816 |
| **NEUROD1** | 0.02385 | 0.00000 | 0.03608 | 0.01103 | 0.01866 | 0.05436 | 0.09215 |
| **NEUROD2** | 0.04688 | 0.00000 | 0.04135 | 0.01603 | 0.02744 | 0.02961 | 0.17055 |

| | | | | | | | |
|---|---|---|---|---|---|---|---|
| **NEUROD4** | 0.02451 | 0.01387 | 0.01537 | 0.02121 | 0.05169 | 0.01342 | 0.01733 |
| **NEUROD6** | 0.02890 | 0.01462 | 0.00979 | 0.01577 | 0.04119 | 0.01150 | 0.02922 |
| **NEUROG1** | 0.00000 | 0.00000 | 0.62166 | 0.00698 | 0.00823 | 0.06431 | 0.48571 |
| **NEUROG2** | 0.02493 | 0.00000 | 0.06667 | 0.01839 | 0.10878 | 0.01639 | 0.02054 |
| **NEUROG3** | 0.06430 | 0.00457 | 0.03071 | 0.02839 | 0.00583 | 0.00125 | 0.11342 |
| **NEXN** | 0.32955 | 1.24654 | 0.43939 | 0.00000 | 0.14170 | 1.39986 | 0.68079 |
| **NEXN-AS1** | 0.00000 | 0.13759 | 0.03152 | 0.05087 | 0.14930 | 0.11423 | 0.18483 |
| **NF1** | 5.80765 | 0.00000 | 2.82476 | 6.46618 | 4.58561 | 1.48920 | 15.95505 |
| **NF2** | 36.35538 | 30.91246 | 15.92837 | 3.67572 | 5.63066 | 23.37359 | 2.92541 |
| **NFAM1** | 0.02766 | 0.01053 | 0.01176 | 0.01181 | 0.00916 | 0.00895 | 0.08384 |
| **NFASC** | 0.02941 | 0.00000 | 0.10233 | 0.01342 | 0.54361 | 0.00326 | 0.08307 |
| **NFAT5** | 3.20866 | 0.00000 | 1.22704 | 4.45003 | 2.01206 | 1.38353 | 11.76616 |
| **NFATC1** | 0.02655 | 0.52107 | 0.47532 | 0.11385 | 3.81568 | 0.20396 | 0.14641 |
| **NFATC2** | 0.73369 | 0.04930 | 0.13525 | 0.48178 | 0.09739 | 0.00000 | 1.28710 |
| **NFATC2IP** | 4.58839 | 5.54025 | 8.71482 | 6.81133 | 5.48899 | 5.61615 | 18.38723 |
| **NFATC3** | 10.68101 | 7.55545 | 4.07681 | 11.48710 | 2.22795 | 5.38982 | 11.51539 |
| **NFATC4** | 0.07992 | 0.04094 | 0.01414 | 0.11430 | 0.00816 | 0.15954 | 0.00000 |
| **NFE2** | 0.47205 | 1.11351 | 0.60296 | 3.17274 | 0.03155 | 0.21463 | 0.00000 |
| **NFE2L1** | 3.06246 | 9.24404 | 1.16931 | 2.05851 | 0.17924 | 4.61618 | 21.09853 |
| **NFE2L2** | 13.00086 | 8.65327 | 3.75070 | 10.66069 | 0.65145 | 3.63514 | 18.64049 |
| **NFE2L3** | 4.01603 | 1.84520 | 3.01403 | 15.84432 | 0.49146 | 2.84323 | 21.53638 |
| **NFIA** | 0.12131 | 0.00000 | 0.04897 | 0.00000 | 0.12616 | 0.66747 | 0.08618 |
| **NFIB** | 0.00989 | 0.05892 | 0.10645 | 0.02315 | 0.26009 | 0.01450 | 0.08872 |
| **NFIC** | 0.08562 | 1.30603 | 0.39439 | 1.01276 | 0.58528 | 0.27025 | 2.62512 |
| **NFIL3** | 0.00000 | 1.55406 | 0.85756 | 4.33015 | 0.63133 | 0.27845 | 7.32000 |
| **NFIX** | 0.01678 | 0.03019 | 0.07159 | 0.01351 | 0.15423 | 0.11562 | 0.07425 |
| **NFKB1** | 3.12092 | 0.00000 | 0.00000 | 2.25579 | 0.09682 | 0.13763 | 3.64950 |
| **NFKB2** | 0.00000 | 12.50276 | 0.97768 | 1.56589 | 0.82256 | 12.43055 | 6.99808 |
| **NFKBIA** | 2.93420 | 25.56507 | 14.86082 | 6.31353 | 15.46776 | 21.09639 | 22.87702 |
| **NFKBIB** | 1.80858 | 5.77045 | 1.52327 | 8.21073 | 0.76708 | 6.08113 | 8.46948 |
| **NFKBID** | 0.00000 | 0.01623 | 0.01137 | 0.37509 | 0.09486 | 0.55533 | 0.48187 |
| **NFKBIE** | 0.00000 | 0.76537 | 0.12454 | 2.91195 | 0.04003 | 1.13044 | 2.36365 |
| **NFKBIL1** | 0.00000 | 2.82665 | 1.82509 | 1.26307 | 0.27188 | 1.88537 | 4.96148 |
| **NFKBIZ** | 0.33819 | 0.00000 | 0.42702 | 1.48346 | 0.04863 | 0.75730 | 2.33619 |
| **NFRKB** | 12.51203 | 8.43224 | 6.20838 | 10.57333 | 5.02584 | 7.32108 | 14.73429 |

| | | | | | | | |
|---|---|---|---|---|---|---|---|
| **NFS1** | 16.61285 | 5.13783 | 4.71545 | 11.70686 | 1.06696 | 4.40001 | 3.06148 |
| **NFU1** | 8.69489 | 3.92655 | 3.08855 | 5.69391 | 1.26358 | 3.14444 | 1.26227 |
| **NFX1** | 16.83472 | 12.99421 | 9.22076 | 8.22995 | 4.26496 | 4.41786 | 1.14003 |
| **NFXL1** | 13.59335 | 2.30733 | 5.59770 | 6.65910 | 0.78186 | 6.21708 | 8.24175 |
| **NFYA** | 6.60186 | 3.31552 | 4.51507 | 4.88133 | 11.35830 | 4.35906 | 2.19156 |
| **NFYB** | 22.82748 | 6.08449 | 4.89079 | 14.74598 | 6.54451 | 8.87357 | 0.00000 |
| **NFYC** | 15.03166 | 16.73737 | 11.25739 | 10.56203 | 2.69285 | 7.68034 | 6.32216 |
| **NFYC-AS1** | 0.56057 | 0.40888 | 0.03671 | 0.33382 | 0.07582 | 0.15952 | 0.81933 |
| **NGB** | 0.01691 | 0.08317 | 0.21907 | 0.02424 | 0.06420 | 0.00282 | 0.00255 |
| **NGDN** | 25.15330 | 16.32550 | 22.20549 | 19.10271 | 12.30141 | 7.77654 | 0.00000 |
| **NGEF** | 0.00000 | 2.09296 | 2.81228 | 0.00000 | 0.09483 | 5.27059 | 0.03373 |
| **NGF** | 0.02841 | 0.01373 | 0.01507 | 0.04987 | 0.00827 | 0.02674 | 0.00274 |
| **NGFR** | 0.00000 | 0.01190 | 0.05510 | 0.01845 | 0.47759 | 0.95171 | 0.09666 |
| **NGFRAP1** | 14.08726 | 0.61385 | 6.03231 | 24.44316 | 2.30162 | 0.35868 | 3.64586 |
| **NGLY1** | 14.89705 | 6.43327 | 7.24645 | 7.27863 | 2.29012 | 4.20127 | 0.00000 |
| **NGRN** | 21.57843 | 23.54698 | 23.17911 | 21.67089 | 8.68512 | 8.71547 | 7.61111 |
| **NHEJ1** | 8.15886 | 5.35595 | 2.49554 | 5.90539 | 2.34734 | 2.34036 | 0.00000 |
| **NHLH1** | 0.08253 | 0.00000 | 0.08370 | 0.02074 | 0.00310 | 0.10542 | 0.61802 |
| **NHLH2** | 0.01993 | 0.01263 | 0.01345 | 0.04491 | 0.00777 | 0.05202 | 0.05662 |
| **NHLRC1** | 0.10582 | 0.52151 | 0.04806 | 0.00607 | 0.00454 | 0.00391 | 0.00000 |
| **NHLRC2** | 4.08683 | 1.48748 | 3.03113 | 1.35409 | 1.89555 | 2.09344 | 6.30797 |
| **NHLRC3** | 5.28145 | 0.00019 | 0.70454 | 2.26580 | 3.58004 | 0.23785 | 1.09913 |
| **NHLRC4** | 0.07242 | 0.02292 | 0.02570 | 0.00819 | 0.01472 | 0.00846 | 0.03689 |
| **NHP2** | 32.53229 | 26.97900 | 23.76348 | 32.49075 | 17.75709 | 19.19422 | 8.85586 |
| **NHP2L1** | 50.09635 | 35.00705 | 27.15610 | 45.27477 | 20.12688 | 26.27962 | 28.91732 |
| **NHS** | 0.00000 | 0.00000 | 0.17825 | 0.07757 | 0.15093 | 1.14440 | 0.32120 |
| **NHSL1** | 16.81955 | 5.36508 | 2.13292 | 5.37529 | 1.96530 | 4.33332 | 4.35298 |
| **NHSL2** | 0.00000 | 0.41338 | 0.04543 | 0.00000 | 0.09325 | 0.28564 | 0.04427 |
| **NICN1** | 1.27097 | 0.00000 | 1.56140 | 1.09930 | 1.07923 | 1.20095 | 0.00000 |
| **NID1** | 30.59142 | 7.16781 | 0.00000 | 9.31877 | 0.00000 | 16.58345 | 19.28807 |
| **NID2** | 0.23843 | 0.00000 | 0.44842 | 0.39718 | 2.66706 | 3.72430 | 0.00000 |
| **NIF3L1** | 11.89089 | 6.05899 | 9.92244 | 6.87514 | 3.47433 | 6.25724 | 0.00046 |
| **NIFK** | 34.78631 | 12.27336 | 18.25043 | 18.87673 | 8.31002 | 13.01256 | 9.49458 |
| **NIFK-AS1** | 0.70382 | 0.00000 | 1.64577 | 0.54077 | 0.48106 | 0.57847 | 1.65482 |
| **NIM1K** | 4.42747 | 0.00000 | 0.00000 | 2.27592 | 0.09205 | 0.00000 | 0.89302 |

| | | | | | | | |
|---|---|---|---|---|---|---|---|
| **NIN** | 0.95536 | 2.42827 | 0.80517 | 0.51271 | 0.52608 | 1.53857 | 4.15610 |
| **NINJ1** | 1.17703 | 2.14270 | 0.70749 | 2.18745 | 0.76992 | 0.31670 | 0.99017 |
| **NINJ2** | 0.00000 | 1.19135 | 1.05267 | 0.26188 | 0.00944 | 0.22952 | 0.01177 |
| **NINL** | 2.18016 | 7.26210 | 1.32470 | 1.46423 | 0.74457 | 3.70804 | 1.19320 |
| **NIP7** | 30.36951 | 13.59049 | 21.41750 | 17.07490 | 13.59226 | 11.53875 | 0.00159 |
| **NIPA1** | 1.01521 | 0.07507 | 0.07265 | 0.18626 | 0.95111 | 3.65471 | 1.22482 |
| **NIPA2** | 20.54978 | 5.27506 | 13.55560 | 17.61438 | 7.37493 | 15.37287 | 21.39617 |
| **NIPAL1** | 6.88588 | 5.14910 | 5.83432 | 0.97057 | 0.00000 | 0.37757 | 0.00000 |
| **NIPAL2** | 0.01723 | 0.03774 | 0.02274 | 0.02196 | 0.01115 | 0.05019 | 0.03442 |
| **NIPAL3** | 2.20698 | 2.52761 | 0.42799 | 2.41677 | 2.03688 | 0.45889 | 0.00000 |
| **NIPAL4** | 0.02828 | 0.00734 | 0.03958 | 0.03181 | 0.00924 | 0.01105 | 0.01859 |
| **NIPBL** | 11.59447 | 3.86865 | 9.64757 | 8.74227 | 5.82870 | 9.31134 | 22.14339 |
| **NIPSNAP1** | 4.78514 | 2.89439 | 2.20216 | 1.22089 | 2.15493 | 3.01254 | 2.36404 |
| **NIPSNAP3A** | 3.05949 | 0.82362 | 2.07106 | 1.74334 | 0.12396 | 0.48104 | 0.00000 |
| **NIPSNAP3B** | 0.07990 | 0.03607 | 0.02565 | 0.16458 | 0.04121 | 0.12396 | 0.00000 |
| **NISCH** | 3.96749 | 5.10056 | 3.02667 | 8.72038 | 1.67899 | 5.03414 | 10.01184 |
| **NIT1** | 4.51494 | 4.33631 | 1.76114 | 7.20068 | 0.74966 | 1.97874 | 0.03911 |
| **NIT2** | 14.20353 | 12.90867 | 3.66693 | 7.19609 | 0.05039 | 9.20497 | 7.68219 |
| **NKAIN1** | 0.00000 | 0.35515 | 0.14742 | 0.07366 | 0.02131 | 0.24240 | 0.17028 |
| **NKAIN2** | 0.02553 | 0.01553 | 0.01870 | 0.02745 | 0.06986 | 0.02993 | 0.01442 |
| **NKAIN4** | 0.77411 | 0.00000 | 0.00000 | 0.17915 | 0.00403 | 0.00000 | 0.57943 |
| **NKAP** | 9.85716 | 17.92272 | 14.31683 | 0.48236 | 2.01881 | 9.66389 | 11.20359 |
| **NKAPL** | 0.04001 | 0.02416 | 0.44577 | 0.25740 | 0.12547 | 0.07270 | 0.22288 |
| **NKAPP1** | 0.17265 | 1.00071 | 1.98775 | 0.00000 | 0.11945 | 1.18777 | 0.30639 |
| **NKD1** | 0.00000 | 0.16283 | 1.41362 | 0.28136 | 1.25394 | 0.31772 | 0.63799 |
| **NKD2** | 2.28614 | 10.54965 | 0.00000 | 1.18241 | 1.80764 | 0.00000 | 14.19135 |
| **NKG7** | 0.00000 | 0.16062 | 0.07911 | 0.00997 | 0.00929 | 0.23612 | 0.02766 |
| **NKIRAS1** | 5.46007 | 0.12902 | 3.74146 | 3.56554 | 8.07746 | 3.35429 | 0.00000 |
| **NKIRAS2** | 11.54678 | 13.37809 | 12.51511 | 10.57195 | 3.04065 | 5.17549 | 8.97489 |
| **NKPD1** | 0.02142 | 0.04202 | 0.01500 | 0.06112 | 0.04480 | 0.00000 | 0.10069 |
| **NKRF** | 9.45252 | 0.38523 | 5.08505 | 1.02268 | 2.06418 | 2.82071 | 16.60940 |
| **NKTR** | 16.44508 | 1.57383 | 10.13270 | 6.97403 | 6.79439 | 9.36186 | 21.26609 |
| **NKX1-1** | 0.02258 | 0.01965 | 0.01210 | 0.01905 | 0.01107 | 0.01246 | 0.02381 |
| **NKX1-2** | 0.02258 | 0.01863 | 0.01133 | 0.02326 | 0.02397 | 0.01550 | 0.01437 |
| **NKX2-1** | 0.02708 | 0.01401 | 0.01501 | 0.01876 | 0.04776 | 0.01323 | 0.01525 |

| | | | | | | | |
|---|---|---|---|---|---|---|---|
| **NKX2-2** | 0.01158 | 0.10961 | 0.03210 | 0.00000 | 0.05577 | 0.17097 | 0.10561 |
| **NKX2-3** | 0.04780 | 0.04261 | 0.01072 | 0.02304 | 0.00840 | 0.00000 | 0.14744 |
| **NKX2-4** | 0.01783 | 0.02678 | 0.01562 | 0.00871 | 0.02753 | 0.00963 | 0.07674 |
| **NKX2-5** | 0.01293 | 0.00000 | 0.52050 | 0.01201 | 0.32834 | 0.00249 | 0.06895 |
| **NKX2-8** | 0.01706 | 0.01680 | 0.02257 | 0.02417 | 0.00708 | 0.01469 | 0.02549 |
| **NKX3-1** | 0.00000 | 4.22916 | 0.11948 | 0.81885 | 5.21685 | 0.41790 | 0.66634 |
| **NKX3-2** | 0.01116 | 0.03369 | 0.06542 | 0.02410 | 0.05487 | 0.04749 | 0.02499 |
| **NKX6-1** | 0.02256 | 0.01123 | 0.01978 | 0.02002 | 0.03186 | 0.01353 | 0.02592 |
| **NKX6-2** | 0.02300 | 0.01699 | 0.01057 | 0.01949 | 0.05370 | 0.01780 | 0.01604 |
| **NKX6-3** | 0.02972 | 0.02579 | 0.01505 | 0.01785 | 0.00746 | 0.01590 | 0.01690 |
| **NLE1** | 11.20298 | 7.58476 | 5.43298 | 6.02686 | 5.42709 | 3.59041 | 3.44352 |
| **NLGN1** | 0.02940 | 0.00000 | 0.06841 | 0.04175 | 0.66804 | 0.23237 | 0.00000 |
| **NLGN2** | 0.02870 | 0.00000 | 0.01809 | 1.64836 | 0.13140 | 0.35984 | 2.30389 |
| **NLGN3** | 0.19963 | 0.00000 | 0.12030 | 0.03703 | 0.01366 | 0.00000 | 0.61875 |
| **NLGN4X** | 0.75604 | 0.77310 | 0.20498 | 0.49101 | 0.26643 | 0.12753 | 4.52555 |
| **NLGN4Y** | 0.02458 | 0.01778 | 0.01278 | 0.02259 | 0.01015 | 0.01296 | 0.01486 |
| **NLK** | 0.97241 | 0.79199 | 2.01522 | 2.23141 | 0.83677 | 2.33082 | 4.13141 |
| **NLN** | 17.35843 | 2.19420 | 3.22008 | 12.28530 | 0.97926 | 1.97059 | 3.81569 |
| **NLRC3** | 0.03753 | 0.28206 | 0.19250 | 0.07426 | 0.02886 | 0.19552 | 0.20662 |
| **NLRC4** | 0.05346 | 0.81471 | 1.47445 | 0.09099 | 0.40193 | 1.25745 | 0.17229 |
| **NLRC5** | 0.02304 | 0.01742 | 0.01170 | 0.04174 | 0.01845 | 0.10855 | 0.02377 |
| **NLRP1** | 0.00000 | 0.00000 | 0.15768 | 0.29356 | 0.17542 | 0.20495 | 0.03131 |
| **NLRP10** | 0.02268 | 0.01786 | 0.01251 | 0.11454 | 0.00812 | 0.02966 | 0.00585 |
| **NLRP11** | 0.00000 | 0.40458 | 0.00000 | 0.01728 | 17.22715 | 0.00000 | 0.00841 |
| **NLRP12** | 1.28930 | 10.07166 | 0.46904 | 0.67917 | 0.84006 | 6.24874 | 1.47355 |
| **NLRP13** | 0.00000 | 0.47658 | 0.00000 | 0.00000 | 20.19247 | 0.04348 | 0.02243 |
| **NLRP14** | 0.09660 | 0.00000 | 0.07323 | 0.21321 | 0.94721 | 0.00000 | 0.10166 |
| **NLRP2** | 67.22516 | 59.00335 | 31.13822 | 51.42246 | 4.18644 | 42.21738 | 44.10207 |
| **NLRP3** | 0.01078 | 0.04425 | 0.02538 | 0.02731 | 0.01248 | 0.05154 | 0.09791 |
| **NLRP4** | 0.32917 | 1.03976 | 0.87737 | 0.05956 | 27.41192 | 0.00000 | 0.00000 |
| **NLRP5** | 0.00000 | 0.42327 | 0.39095 | 0.00000 | 7.07748 | 0.00062 | 0.03769 |
| **NLRP6** | 0.02166 | 0.01544 | 0.01500 | 0.01955 | 0.01179 | 0.01243 | 0.02445 |
| **NLRP7** | 71.01583 | 76.76506 | 15.18374 | 33.22957 | 2.86742 | 46.24517 | 32.63620 |
| **NLRP8** | 0.01161 | 0.03025 | 0.00000 | 0.02117 | 4.85092 | 0.00651 | 0.01840 |
| **NLRP9** | 0.00000 | 0.70724 | 0.00000 | 0.00000 | 15.56946 | 4.57061 | 0.32503 |

| | | | | | | | |
|---|---|---|---|---|---|---|---|
| **NLRX1** | 0.00000 | 0.11977 | 0.11488 | 0.46595 | 0.01857 | 0.10814 | 0.05397 |
| **NMB** | 0.00000 | 0.56657 | 0.52523 | 1.76049 | 0.00861 | 0.27318 | 0.47695 |
| **NMBR** | 0.01991 | 0.02121 | 0.01255 | 0.04495 | 0.00781 | 0.01294 | 0.01876 |
| **NMD3** | 19.05489 | 5.76953 | 7.12087 | 4.70635 | 2.30902 | 2.25894 | 0.00000 |
| **NME1** | 37.73185 | 18.08986 | 13.10010 | 29.08995 | 0.08036 | 13.50256 | 28.24965 |
| **NME1-NME2** | 0.12742 | 0.13962 | 0.07917 | 0.27188 | 0.01289 | 0.04613 | 0.00000 |
| **NME2.uid-2204** | 8.05496 | 8.73941 | 2.46483 | 9.45385 | 0.21758 | 4.43415 | 2.76578 |
| **NME2.uid-2204** | 0.80138 | 1.72973 | 0.67311 | 0.47259 | 0.01874 | 0.80811 | 0.48372 |
| **NME2.uid-2204** | 0.05367 | 0.02596 | 0.02167 | 0.07337 | 0.01303 | 0.06228 | 0.00731 |
| **NME2.uid-2204** | 3.24319 | 5.62893 | 0.79215 | 3.61623 | 0.11729 | 2.25256 | 1.31050 |
| **NME3** | 1.35034 | 1.58014 | 0.13216 | 2.94481 | 0.37822 | 0.13995 | 2.35926 |
| **NME4** | 0.00000 | 3.09053 | 0.99176 | 3.43072 | 0.32785 | 1.00445 | 6.43682 |
| **NME5** | 0.14135 | 0.02379 | 0.34866 | 0.03118 | 0.13193 | 0.08009 | 0.01759 |
| **NME6** | 12.96162 | 8.27433 | 8.64383 | 7.61996 | 2.32403 | 5.00992 | 0.00000 |
| **NME7** | 2.07400 | 0.00000 | 1.61933 | 4.11682 | 2.14952 | 0.85885 | 0.00000 |
| **NME8** | 0.05043 | 0.00273 | 0.03318 | 0.01209 | 0.00624 | 0.00991 | 0.03397 |
| **NME9** | 0.19935 | 0.05830 | 0.12657 | 0.00000 | 0.03156 | 0.32471 | 0.67089 |
| **NMI** | 11.20344 | 0.02229 | 0.00000 | 2.67138 | 0.18115 | 2.33420 | 0.00000 |
| **NMNAT1** | 1.43638 | 1.85853 | 2.70197 | 3.08624 | 5.15395 | 1.43858 | 0.22885 |
| **NMNAT2** | 0.00000 | 0.04476 | 0.91849 | 1.85764 | 0.01236 | 2.10017 | 0.00000 |
| **NMNAT3** | 0.02534 | 0.00922 | 0.05786 | 0.01662 | 0.16225 | 0.01060 | 0.03542 |
| **NMRAL1** | 12.47794 | 9.38974 | 3.11674 | 8.77827 | 0.07320 | 4.03986 | 2.21629 |
| **NMRK1** | 0.61517 | 0.03347 | 0.85420 | 1.26536 | 0.06305 | 0.49601 | 0.00000 |
| **NMRK2** | 7.43907 | 5.88413 | 0.57673 | 1.78037 | 1.75481 | 1.08181 | 2.01769 |
| **matr** | 0.02292 | 0.01983 | 0.01066 | 0.01898 | 0.02320 | 0.01311 | 0.01809 |
| **NMT1** | 32.09540 | 28.80600 | 12.78180 | 24.67064 | 7.32358 | 7.30895 | 4.46693 |
| **NMT2** | 1.33650 | 2.88697 | 4.01057 | 0.69686 | 8.58798 | 8.55071 | 0.00000 |
| **NMU** | 1.83887 | 18.93162 | 0.86266 | 0.00000 | 0.00000 | 1.36908 | 0.00113 |
| **NMUR1** | 0.00000 | 0.38329 | 0.16921 | 0.00000 | 0.00401 | 0.42005 | 0.13242 |
| **NMUR2** | 0.05542 | 0.01957 | 0.00997 | 0.02685 | 0.10808 | 0.03128 | 0.01031 |
| **NNAT** | 0.00000 | 0.39429 | 0.06863 | 1.72152 | 0.10983 | 0.47738 | 0.00000 |
| **NNMT** | 0.07121 | 0.07246 | 0.00665 | 0.87993 | 0.01745 | 0.06252 | 0.00000 |
| **NNT** | 4.27076 | 0.71536 | 0.02941 | 0.17875 | 0.86554 | 8.91297 | 0.15612 |
| **NNT-AS1** | 2.04943 | 2.34219 | 2.73761 | 1.18499 | 0.00000 | 3.13972 | 0.49252 |
| **NOA1** | 7.26929 | 9.56362 | 6.61194 | 8.02028 | 2.41916 | 5.23450 | 11.21767 |

| | | | | | | | |
|---|---|---|---|---|---|---|---|
| **NOB1** | 21.59483 | 16.23831 | 16.45014 | 16.79106 | 5.38579 | 6.40799 | 7.30993 |
| **NOBOX** | 0.01780 | 0.02223 | 0.00304 | 0.01911 | 0.31949 | 0.03864 | 0.02846 |
| **NOC2L** | 18.80996 | 23.95521 | 8.05015 | 19.64494 | 3.17819 | 8.06036 | 0.00000 |
| **NOC3L** | 20.36257 | 0.15361 | 12.52772 | 7.37142 | 4.13768 | 2.46312 | 11.28584 |
| **NOC4L** | 3.72404 | 34.83212 | 7.76764 | 7.68992 | 1.84202 | 4.69411 | 12.57541 |
| **NOD1** | 0.18965 | 0.00000 | 0.77045 | 0.87393 | 1.61500 | 0.59772 | 0.76749 |
| **NOD2** | 0.05024 | 0.05469 | 0.04504 | 0.02450 | 0.00707 | 0.01090 | 0.00000 |
| **NODAL** | 0.00000 | 0.00000 | 0.09271 | 1.16188 | 0.21090 | 20.52200 | 0.02535 |
| **NOG** | 0.13000 | 0.00000 | 0.48451 | 0.07164 | 0.08190 | 0.72015 | 0.52091 |
| **NOL10** | 27.63662 | 9.15178 | 13.28983 | 15.23946 | 6.19775 | 6.77958 | 5.95088 |
| **NOL11** | 26.65208 | 2.90741 | 13.49781 | 15.54790 | 14.48317 | 9.03061 | 5.18694 |
| **NOL12** | 17.11722 | 23.18208 | 11.89251 | 12.37079 | 12.24005 | 4.82022 | 1.98137 |
| **NOL3** | 0.00000 | 0.66174 | 0.64288 | 0.73196 | 0.03918 | 0.72626 | 0.93087 |
| **NOL4** | 0.26472 | 0.00910 | 0.01958 | 0.00000 | 0.11786 | 0.00551 | 0.03902 |
| **NOL4L** | 1.30873 | 2.82655 | 0.60853 | 0.00000 | 2.36632 | 1.50716 | 1.84966 |
| **NOL6** | 12.23933 | 10.52989 | 12.64952 | 15.84173 | 2.24803 | 8.13154 | 18.30478 |
| **NOL7** | 5.42786 | 15.45216 | 12.29330 | 13.63162 | 3.01551 | 9.10902 | 31.23975 |
| **NOL8** | 18.86541 | 3.81788 | 12.99339 | 8.14929 | 6.61288 | 5.66467 | 14.63634 |
| **NOL9** | 11.95462 | 6.27197 | 12.49675 | 7.14615 | 5.70011 | 4.16904 | 14.34014 |
| **NOLC1** | 51.56906 | 29.66907 | 32.38034 | 37.05668 | 18.71867 | 16.69276 | 21.54662 |
| **NOM1** | 1.21902 | 0.77096 | 2.25765 | 0.89906 | 1.80055 | 3.43471 | 11.87105 |
| **NOMO1** | 1.60881 | 2.36599 | 0.31818 | 1.82715 | 0.13307 | 1.70034 | 4.46758 |
| **NOMO2** | 0.25238 | 0.27108 | 0.03761 | 0.19938 | 0.03260 | 0.20228 | 0.17166 |
| **NOMO3** | 0.11533 | 0.14801 | 0.01730 | 0.07855 | 0.02100 | 0.06907 | 0.21962 |
| **NONO** | 46.41870 | 31.02314 | 23.65252 | 25.12421 | 9.04118 | 22.68422 | 21.18797 |
| **NOP10** | 43.76766 | 30.41750 | 31.36955 | 40.57957 | 16.36512 | 16.50826 | 22.97197 |
| **NOP14** | 3.51269 | 3.66725 | 2.60157 | 2.44290 | 0.72560 | 3.86665 | 17.27150 |
| **NOP14-AS1** | 0.17900 | 1.58617 | 0.73898 | 0.28184 | 0.27731 | 0.30331 | 0.33774 |
| **NOP16** | 52.19753 | 36.48351 | 20.87698 | 28.40787 | 12.16766 | 22.06607 | 9.06711 |
| **NOP2** | 32.85829 | 24.06110 | 19.86477 | 28.55611 | 7.77435 | 13.22460 | 8.17113 |
| **NOP56** | 44.21173 | 31.48247 | 29.59145 | 34.29864 | 11.17219 | 24.01968 | 39.05522 |
| **NOP58** | 52.77430 | 21.86039 | 21.75262 | 29.54862 | 16.49005 | 19.68061 | 23.95922 |
| **NOP9** | 1.67513 | 3.65603 | 1.20800 | 2.36155 | 2.45413 | 3.39142 | 7.59418 |
| **NOS1** | 0.00000 | 0.08581 | 0.13708 | 0.01965 | 0.90904 | 0.02454 | 0.02939 |
| **NOS1AP** | 1.96719 | 1.19122 | 0.45951 | 0.00001 | 2.04654 | 0.33785 | 1.10846 |

| | | | | | | | |
|---|---|---|---|---|---|---|---|
| **NOS2** | 0.04027 | 0.02169 | 0.02509 | 8.77731 | 0.18648 | 0.02572 | 0.00243 |
| **NOS3** | 0.00000 | 1.99546 | 2.17238 | 0.81300 | 0.05280 | 0.86917 | 0.00000 |
| **NOSIP** | 29.61724 | 30.17559 | 12.48815 | 27.53753 | 19.44926 | 9.55483 | 0.45466 |
| **NOSTRIN** | 0.18497 | 0.00000 | 0.04444 | 0.23275 | 0.50716 | 0.98371 | 0.29284 |
| **NOTCH1** | 0.00000 | 1.45213 | 1.55892 | 0.18816 | 0.11230 | 1.95164 | 0.19536 |
| **NOTCH2** | 0.23063 | 1.88896 | 0.42345 | 0.79394 | 0.12305 | 0.66077 | 1.04901 |
| **NOTCH2NL** | 0.00000 | 1.28896 | 0.10279 | 0.95863 | 0.58687 | 0.97008 | 1.16452 |
| **NOTCH3** | 0.14018 | 0.02917 | 0.01710 | 0.04655 | 0.00717 | 0.00000 | 1.10978 |
| **NOTCH4** | 0.00000 | 0.02347 | 0.00756 | 0.11019 | 0.14006 | 0.09969 | 0.08066 |
| **NOTO** | 0.00000 | 0.42573 | 1.12505 | 0.00083 | 0.00948 | 0.00000 | 0.00522 |
| **NOTUM** | 0.00000 | 0.01299 | 0.01173 | 0.91655 | 0.00690 | 0.03733 | 0.19658 |
| **NOV** | 0.01066 | 0.02081 | 0.01349 | 0.04434 | 0.00872 | 0.02150 | 0.02468 |
| **NOVA1** | 0.00000 | 0.00000 | 0.61524 | 0.00000 | 0.24386 | 2.33004 | 6.55025 |
| **NOVA2** | 0.02393 | 0.01811 | 0.01269 | 0.01934 | 0.00791 | 0.01269 | 0.01891 |
| **NOX1** | 0.18006 | 0.00000 | 0.06112 | 0.01241 | 0.01058 | 0.02563 | 0.18331 |
| **NOX3** | 0.02562 | 0.01770 | 0.01615 | 0.02102 | 0.00858 | 0.01599 | 0.01801 |
| **NOX4** | 0.04379 | 0.00000 | 0.04999 | 0.02645 | 0.33447 | 0.08408 | 0.01419 |
| **NOX5** | 0.05417 | 0.00088 | 0.01165 | 0.01078 | 0.62063 | 0.01026 | 0.02312 |
| **NOXA1** | 0.02096 | 0.04816 | 0.00650 | 0.01240 | 0.02955 | 0.00920 | 0.08455 |
| **NOXO1** | 0.00000 | 0.16770 | 0.04212 | 0.49863 | 0.01017 | 0.03711 | 0.87330 |
| **NOXRED1** | 0.12238 | 0.00000 | 0.08781 | 0.05630 | 0.00358 | 0.00483 | 0.30647 |
| **NPAP1** | 0.01940 | 0.00281 | 0.05493 | 0.04409 | 0.01148 | 0.01108 | 0.03426 |
| **NPAS1** | 0.00000 | 0.72821 | 0.30234 | 0.02775 | 0.23560 | 0.18251 | 0.15233 |
| **NPAS2** | 0.00000 | 0.22358 | 0.15842 | 0.01492 | 0.17472 | 0.50979 | 0.10225 |
| **NPAS3** | 0.04614 | 0.01194 | 0.00000 | 0.03370 | 1.26437 | 0.16462 | 0.01140 |
| **NPAS4** | 0.01225 | 0.02936 | 0.14905 | 0.02807 | 0.00791 | 0.00919 | 0.04542 |
| **NPAT** | 7.68643 | 0.00000 | 4.94026 | 7.38513 | 7.32518 | 3.64779 | 5.81879 |
| **NPB** | 0.00000 | 0.29119 | 0.53638 | 0.13411 | 0.01305 | 0.06903 | 0.14699 |
| **NPBWR1** | 0.21592 | 0.00000 | 0.01198 | 0.10423 | 0.00707 | 0.00000 | 0.13173 |
| **NPC1** | 11.79109 | 6.51220 | 0.00000 | 9.88240 | 4.61326 | 3.29316 | 6.20682 |
| **NPC1L1** | 0.05649 | 0.03332 | 0.00619 | 0.16258 | 0.00925 | 0.43512 | 0.03534 |
| **NPC2** | 37.37233 | 16.52494 | 11.39435 | 18.78065 | 2.68621 | 6.12529 | 25.87632 |
| **NPDC1** | 0.00000 | 0.37828 | 0.00000 | 0.21985 | 0.12323 | 0.88428 | 0.63589 |
| **NPEPL1.uid-246** | 0.03210 | 0.01669 | 0.01247 | 0.01496 | 0.00773 | 0.01994 | 0.02751 |
| **NPEPL1.uid-246** | 0.03716 | 0.08717 | 0.00455 | 0.05116 | 0.02005 | 0.07020 | 0.00541 |

| | | | | | | | |
|---|---|---|---|---|---|---|---|
| **NPEPPS** | 22.68955 | 14.04004 | 7.92930 | 16.00400 | 3.26231 | 21.88748 | 19.86367 |
| **NPFF** | 0.28133 | 0.11653 | 0.02655 | 0.06254 | 0.06931 | 0.14257 | 0.57010 |
| **NPFFR2** | 1.33244 | 0.40552 | 0.07316 | 2.33187 | 0.00994 | 0.35904 | 0.00000 |
| **NPHP1** | 0.42162 | 0.08064 | 1.31408 | 0.30679 | 1.61936 | 0.25730 | 0.00000 |
| **NPHP3** | 0.09536 | 0.00000 | 0.03033 | 0.22729 | 0.05692 | 0.26651 | 0.46539 |
| **NPHP3-ACAD11** | 0.11780 | 0.01482 | 0.04378 | 0.01955 | 0.00582 | 0.07302 | 0.00000 |
| **NPHP3-AS1** | 0.02436 | 0.01927 | 0.04355 | 0.01813 | 0.03719 | 0.01095 | 0.01842 |
| **NPHP4** | 0.00000 | 1.44251 | 1.09410 | 0.49392 | 1.48558 | 1.22273 | 0.84210 |
| **NPHS1** | 0.00855 | 0.02592 | 0.02508 | 0.03002 | 0.08888 | 0.00922 | 0.05732 |
| **NPHS2** | 0.02296 | 0.01946 | 0.01341 | 0.01872 | 0.01004 | 0.01471 | 0.02294 |
| **NPIPA1** | 0.04671 | 0.09407 | 0.10388 | 0.18072 | 0.01962 | 0.13049 | 0.47320 |
| **NPIPA5** | 0.01056 | 0.08075 | 0.10749 | 0.09178 | 0.02108 | 0.24916 | 0.28815 |
| **NPIPB3** | 0.33999 | 0.48173 | 0.21582 | 0.30096 | 0.03787 | 0.21536 | 0.15050 |
| **NPIPB5** | 0.59111 | 0.50210 | 0.29835 | 0.43066 | 0.04739 | 0.28800 | 0.52903 |
| **NPIPB6** | 0.00569 | 0.04855 | 0.23642 | 0.03260 | 0.04068 | 0.03559 | 0.11595 |
| **NPIPB9** | 0.02680 | 0.01187 | 0.02196 | 0.03759 | 0.00917 | 0.03072 | 0.02377 |
| **NPL** | 1.01872 | 0.49907 | 4.88826 | 1.04550 | 0.49492 | 0.58628 | 0.00000 |
| **NPLOC4** | 6.04988 | 14.47355 | 6.34199 | 12.28212 | 1.10717 | 9.10820 | 23.80348 |
| **NPM1** | 56.78790 | 25.83601 | 30.93296 | 37.04032 | 12.39566 | 27.39844 | 30.83106 |
| **NPM2** | 0.00000 | 0.80908 | 0.37684 | 0.00000 | 6.72419 | 0.00000 | 0.07659 |
| **NPM3** | 20.40088 | 1.31612 | 3.16927 | 16.46351 | 8.48082 | 11.55069 | 9.47883 |
| **NPNT** | 2.23187 | 0.00000 | 0.00000 | 1.30472 | 0.04128 | 0.00000 | 0.68265 |
| **NPPA** | 0.60240 | 0.00000 | 0.01010 | 0.39312 | 0.01539 | 0.01287 | 0.07242 |
| **NPPA-AS1** | 0.46488 | 0.00039 | 0.01199 | 0.31232 | 0.03474 | 0.00799 | 0.00000 |
| **NPPB** | 0.79803 | 3.49234 | 0.00000 | 4.00472 | 0.00152 | 0.00000 | 0.31046 |
| **NPPC** | 0.00000 | 0.01389 | 0.35751 | 0.01927 | 0.14086 | 0.06408 | 0.08678 |
| **1.00000** | 0.02391 | 0.21235 | 0.05169 | 0.44228 | 0.28834 | 0.02313 | 0.00000 |
| **2.00000** | 0.01202 | 0.03725 | 0.15418 | 0.01419 | 0.05606 | 0.03685 | 0.09136 |
| **3.00000** | 0.02921 | 0.00576 | 0.03273 | 0.02537 | 0.30598 | 0.01667 | 0.01003 |
| **NPRL2** | 11.68106 | 7.46523 | 3.79931 | 9.67155 | 1.26005 | 5.07769 | 0.00000 |
| **NPRL3** | 4.39804 | 3.66901 | 6.30124 | 8.10987 | 0.87444 | 3.26495 | 14.34969 |
| **NPSR1** | 0.03281 | 0.01876 | 0.01084 | 0.01793 | 0.01514 | 0.03155 | 0.01429 |
| **NPSR1-AS1** | 0.02779 | 0.01719 | 0.03316 | 0.01670 | 0.21132 | 0.06688 | 0.01141 |
| **NPTN** | 23.82494 | 1.88329 | 8.45622 | 12.40031 | 6.76798 | 8.14774 | 7.97930 |
| **NPTN-IT1** | 0.79005 | 0.00000 | 0.42694 | 0.38543 | 0.18485 | 0.64188 | 2.97708 |

| | | | | | | | |
|---|---|---|---|---|---|---|---|
| **NPTX1** | 1.29493 | 9.22040 | 0.16783 | 0.00000 | 0.00000 | 0.11490 | 1.50879 |
| **NPTX2** | 0.01480 | 0.08220 | 1.18826 | 0.03167 | 1.78848 | 0.00000 | 0.04269 |
| **NPTXR** | 0.00000 | 0.14532 | 0.05812 | 0.01282 | 0.00377 | 0.22782 | 0.11182 |
| **NPW** | 0.00000 | 0.20893 | 0.22536 | 0.03373 | 0.06204 | 0.03715 | 0.21213 |
| **NPY** | 0.03191 | 0.00518 | 0.04753 | 0.02383 | 0.02095 | 0.05729 | 0.01177 |
| **NPY1R** | 0.04449 | 0.01210 | 0.01298 | 0.02266 | 0.00706 | 0.24668 | 0.04255 |
| **NPY4R** | 0.03555 | 0.01838 | 0.01443 | 0.02131 | 0.01217 | 0.00977 | 0.02864 |
| **NPY6R** | 0.02844 | 0.01899 | 0.01219 | 0.01694 | 0.00779 | 0.01110 | 0.02429 |
| **NQO1** | 20.30129 | 4.56189 | 0.00000 | 5.11442 | 0.56877 | 2.91401 | 3.89284 |
| **NQO2** | 7.66311 | 12.36614 | 2.51648 | 6.56019 | 1.42484 | 17.09108 | 0.00000 |
| **NR0B1** | 0.02142 | 0.00000 | 0.13999 | 0.01845 | 0.03013 | 0.01034 | 0.02194 |
| **NR0B2** | 0.02079 | 0.01295 | 0.01777 | 0.01378 | 0.09115 | 0.01168 | 0.05493 |
| **NR1D1** | 0.33509 | 0.28097 | 1.58754 | 0.25528 | 0.23587 | 0.49136 | 0.82130 |
| **NR1D2** | 1.15914 | 0.16998 | 3.61246 | 0.71059 | 0.86045 | 2.18765 | 3.53031 |
| **NR1H2** | 9.77506 | 23.56533 | 9.88522 | 12.83945 | 2.72334 | 18.13640 | 1.02174 |
| **NR1H3** | 3.68771 | 7.12708 | 2.73679 | 6.13622 | 0.13132 | 1.21691 | 0.19193 |
| **NR1H4** | 0.00789 | 0.01046 | 0.00884 | 0.92649 | 0.55490 | 0.01218 | 0.00000 |
| **NR1I2** | 0.01169 | 0.05301 | 0.02026 | 0.07624 | 0.00703 | 0.02623 | 0.03513 |
| **NR1I3** | 0.00000 | 0.03955 | 0.01699 | 0.10631 | 0.00955 | 0.16555 | 0.04931 |
| **NR2C1** | 14.18575 | 5.30323 | 14.73255 | 6.35236 | 4.22992 | 10.11007 | 3.37529 |
| **NR2C2** | 4.25422 | 0.64833 | 3.49593 | 1.87096 | 0.65614 | 1.59019 | 7.50040 |
| **NR2C2AP** | 11.24772 | 5.18423 | 3.60996 | 9.34604 | 1.84603 | 2.56411 | 1.83982 |
| **NR2E1** | 0.01206 | 0.00000 | 1.50916 | 0.00365 | 2.35863 | 0.00000 | 0.04672 |
| **NR2E3** | 0.07289 | 0.00000 | 0.02295 | 0.09554 | 0.02870 | 0.10359 | 0.15686 |
| **NR2F1** | 0.08040 | 0.05867 | 0.01381 | 0.00000 | 0.10594 | 0.02913 | 0.17320 |
| **NR2F1-AS1** | 0.04798 | 0.08428 | 0.08312 | 0.01408 | 0.12256 | 0.13099 | 0.00731 |
| **NR2F2** | 6.21413 | 0.00000 | 0.00000 | 35.83237 | 0.23113 | 0.00000 | 6.82312 |
| **NR2F2-AS1** | 0.02115 | 0.00695 | 0.03053 | 0.46895 | 0.00709 | 0.02452 | 0.00263 |
| **NR2F6** | 4.15565 | 8.65295 | 0.00000 | 10.43687 | 0.17525 | 7.06592 | 42.76752 |
| **NR3C1** | 0.17988 | 0.26814 | 0.22084 | 0.02536 | 0.74639 | 1.24289 | 0.00000 |
| **NR3C2** | 0.19278 | 0.00000 | 0.48163 | 0.08298 | 3.89919 | 0.05904 | 0.25934 |
| **NR4A1** | 0.00000 | 1.01558 | 0.38261 | 3.04091 | 0.00516 | 0.06208 | 1.74100 |
| **NR4A2** | 0.07272 | 2.19470 | 2.61551 | 0.77151 | 0.44917 | 1.56265 | 0.28161 |
| **NR4A3** | 0.00000 | 0.11100 | 0.00104 | 0.10694 | 0.27198 | 0.06050 | 0.24085 |
| **NR5A1** | 0.02584 | 0.01992 | 0.01322 | 0.05913 | 0.03255 | 0.03608 | 0.01570 |

| | | | | | | | |
|---|---|---|---|---|---|---|---|
| **NR5A2** | 2.96524 | 0.37184 | 3.96092 | 0.00000 | 0.86199 | 4.03371 | 0.69119 |
| **NR6A1** | 42.37255 | 13.53619 | 4.72209 | 49.63007 | 5.52380 | 13.50306 | 18.40454 |
| **NRADDP** | 0.12637 | 0.03987 | 0.06238 | 0.11121 | 0.00396 | 0.00544 | 0.06483 |
| **NRAP** | 0.01888 | 0.00000 | 0.41935 | 0.01317 | 2.60714 | 0.16504 | 0.01859 |
| **NRARP** | 0.00000 | 0.63744 | 0.06316 | 0.00000 | 0.04628 | 0.53968 | 0.65185 |
| **NRAS** | 22.50684 | 3.69037 | 14.30600 | 12.47416 | 6.34252 | 7.59022 | 1.14092 |
| **NRBF2** | 18.23645 | 12.76502 | 17.66343 | 6.95528 | 12.47218 | 10.79326 | 0.00051 |
| **NRBP1** | 25.39602 | 25.83779 | 14.82524 | 28.32788 | 13.15079 | 27.65999 | 0.70466 |
| **NRBP2** | 0.00000 | 1.60613 | 2.69830 | 0.00000 | 0.30500 | 9.71758 | 2.36173 |
| **NRCAM** | 0.18672 | 0.15557 | 0.45981 | 0.00000 | 0.65879 | 1.36832 | 0.00000 |
| **NRD1** | 27.30863 | 16.72031 | 12.14563 | 20.37124 | 1.31977 | 21.78562 | 10.67545 |
| **NRDE2** | 5.48944 | 2.76396 | 2.01198 | 6.33856 | 2.98638 | 0.81781 | 4.38219 |
| **NREP** | 0.31256 | 0.00000 | 0.00000 | 0.30499 | 0.71583 | 3.69387 | 0.00000 |
| **NREP-AS1** | 0.02072 | 0.01385 | 0.01096 | 0.01845 | 0.03202 | 0.01135 | 0.03609 |
| **NRF1** | 8.71378 | 7.16125 | 4.11371 | 4.46971 | 10.67361 | 4.37411 | 0.00000 |
| **NRG1** | 0.00000 | 0.28666 | 0.31782 | 0.01733 | 1.40515 | 1.53542 | 0.00229 |
| **NRG1-IT1** | 0.03286 | 0.01352 | 0.01921 | 0.03225 | 0.00935 | 0.02024 | 0.03569 |
| **NRG2** | 0.00030 | 0.00929 | 0.01139 | 0.02657 | 0.01092 | 0.16410 | 0.09719 |
| **NRG3** | 0.06190 | 0.01165 | 0.01821 | 0.01227 | 0.02414 | 0.01165 | 0.02314 |
| **NRG4** | 0.81595 | 0.00000 | 0.61361 | 0.17979 | 0.85508 | 0.40143 | 0.57757 |
| **NRGN** | 0.00000 | 0.35770 | 0.09404 | 0.00000 | 0.05678 | 4.50826 | 0.12128 |
| **NRIP1** | 0.00000 | 0.00000 | 0.17754 | 1.80856 | 0.68659 | 0.50171 | 1.39137 |
| **NRIP2** | 0.21684 | 0.14229 | 0.27680 | 0.04068 | 0.07896 | 0.11805 | 0.75419 |
| **NRIP3** | 6.03772 | 21.03686 | 6.00461 | 0.00000 | 2.33275 | 11.37198 | 0.00000 |
| **NRK** | 0.21969 | 0.00000 | 0.73526 | 0.10616 | 0.00105 | 0.17326 | 0.06820 |
| **NRL** | 0.00000 | 0.12833 | 0.11127 | 0.29656 | 0.03793 | 0.04462 | 0.04719 |
| **NRM** | 2.42980 | 0.64338 | 1.44933 | 1.87977 | 0.89170 | 0.90586 | 1.46369 |
| **NRN1** | 0.94533 | 0.94708 | 0.21829 | 0.00000 | 0.12935 | 2.40345 | 0.44118 |
| **NRN1L** | 1.09643 | 0.00000 | 0.00000 | 3.55802 | 0.00781 | 0.00000 | 1.28869 |
| **NRON** | 0.02138 | 0.01482 | 0.01829 | 0.02729 | 0.00742 | 0.02321 | 0.02028 |
| **NRP1** | 0.15756 | 0.00000 | 0.01413 | 11.36410 | 0.02897 | 0.03404 | 0.00000 |
| **NRP2** | 0.00000 | 0.09785 | 0.30593 | 0.23295 | 0.00712 | 0.10500 | 0.01896 |
| **NRROS** | 0.01522 | 0.00812 | 0.02076 | 0.01602 | 0.01469 | 0.38180 | 0.02010 |
| **NRSN1** | 0.04235 | 0.01548 | 0.01385 | 0.01891 | 0.00810 | 0.01369 | 0.00500 |
| **NRSN2** | 3.32213 | 2.75143 | 0.06396 | 2.23146 | 0.13770 | 3.99116 | 1.79394 |

| | | | | | | | |
|---|---|---|---|---|---|---|---|
| **NRTN** | 0.02374 | 0.14867 | 0.02330 | 0.05607 | 0.08139 | 0.00208 | 0.12277 |
| **NRXN1** | 0.04896 | 0.05922 | 0.00000 | 0.01276 | 0.57957 | 0.03530 | 0.09952 |
| **NRXN2** | 0.00000 | 3.04184 | 0.38611 | 0.00000 | 0.07285 | 0.85879 | 3.89189 |
| **NRXN3** | 0.19715 | 0.09940 | 0.00000 | 3.16447 | 0.47809 | 0.13839 | 1.03637 |
| **NS3BP** | 0.32438 | 1.19152 | 0.21178 | 0.34468 | 0.03898 | 0.17945 | 0.05934 |
| **NSA2** | 14.86116 | 0.00000 | 11.19754 | 8.11093 | 4.13916 | 8.03851 | 15.97174 |
| **NSD1** | 14.12470 | 9.14578 | 12.48911 | 9.69653 | 4.20381 | 14.33904 | 27.36175 |
| **NSDHL** | 30.06503 | 20.82787 | 7.94332 | 12.81547 | 7.23763 | 10.92882 | 9.34411 |
| **NSF** | 13.00562 | 4.27138 | 5.21071 | 5.98567 | 1.37134 | 5.38744 | 1.52936 |
| **NSFL1C** | 19.70093 | 26.76447 | 10.30490 | 12.17899 | 3.95738 | 12.83303 | 7.49378 |
| **NSG1** | 0.36019 | 1.52142 | 2.03080 | 0.14539 | 6.57230 | 0.00000 | 2.35968 |
| **NSL1** | 14.66602 | 4.96924 | 7.61069 | 5.08292 | 0.81250 | 9.79109 | 1.87141 |
| **NSMAF** | 9.07359 | 5.76523 | 6.53387 | 7.33968 | 0.57412 | 15.08659 | 2.80497 |
| **NSMCE1** | 5.53041 | 2.97543 | 4.72554 | 5.67275 | 0.54530 | 3.42890 | 1.63791 |
| **NSMCE2** | 13.17982 | 2.66382 | 8.87301 | 6.01787 | 3.56566 | 2.99432 | 4.39711 |
| **NSMCE4A** | 7.93210 | 7.06654 | 3.88712 | 4.82256 | 9.20479 | 5.23710 | 13.77330 |
| **NSMF** | 0.00000 | 1.70032 | 1.75312 | 0.36988 | 0.97323 | 1.50147 | 3.32468 |
| **NSRP1** | 12.31213 | 4.49054 | 10.48102 | 10.67545 | 4.15797 | 6.78152 | 18.00358 |
| **NSUN2** | 20.48663 | 16.80312 | 10.99917 | 14.85078 | 2.27925 | 9.88696 | 16.47335 |
| **NSUN3** | 1.54409 | 0.00000 | 1.34700 | 0.51808 | 0.90631 | 0.40508 | 0.00000 |
| **NSUN4** | 12.34977 | 8.97440 | 8.91203 | 6.62634 | 2.89424 | 5.16606 | 4.34119 |
| **NSUN5** | 13.71708 | 26.01173 | 10.58507 | 14.21120 | 1.56273 | 8.39917 | 4.09319 |
| **NSUN5P1** | 1.92977 | 1.68792 | 0.63122 | 3.61022 | 0.13405 | 0.74740 | 1.17458 |
| **NSUN5P2** | 0.02383 | 0.02565 | 0.01444 | 0.01878 | 0.00782 | 0.01743 | 0.02262 |
| **NSUN6** | 2.23268 | 0.36804 | 2.45748 | 1.78425 | 1.49360 | 1.26145 | 0.78511 |
| **NSUN7** | 1.78471 | 2.48407 | 1.70481 | 1.29667 | 0.39245 | 0.00000 | 0.00000 |
| **NT5C** | 7.06390 | 11.96571 | 3.39511 | 10.14356 | 1.23488 | 7.95513 | 4.04783 |
| **NT5C1A** | 0.03502 | 0.01873 | 0.01828 | 0.01486 | 0.01404 | 0.01252 | 0.01561 |
| **NT5C1B** | 0.02483 | 0.01932 | 0.01482 | 0.02127 | 0.00769 | 0.01394 | 0.01625 |
| **NT5C2** | 19.75759 | 1.92135 | 8.08284 | 11.85300 | 2.86270 | 4.70612 | 0.50224 |
| **NT5C3A** | 9.82281 | 0.04362 | 3.81329 | 4.58947 | 1.25175 | 2.74202 | 3.35536 |
| **NT5C3B** | 1.26022 | 10.96047 | 2.55763 | 9.15002 | 0.19197 | 4.99036 | 8.80109 |
| **NT5DC1** | 6.24067 | 3.45527 | 2.53352 | 2.67464 | 0.09754 | 2.99046 | 0.00006 |
| **NT5DC2** | 0.97368 | 0.00000 | 0.00000 | 7.12342 | 0.05379 | 2.01487 | 2.06419 |
| **NT5DC3** | 18.18885 | 4.67894 | 4.74770 | 6.91843 | 8.59509 | 1.63431 | 0.00000 |

| | | | | | | | |
|---|---|---|---|---|---|---|---|
| **NT5E** | 0.42968 | 0.39646 | 0.00354 | 0.01161 | 0.00613 | 1.13302 | 0.00000 |
| **NT5M** | 0.06015 | 0.00000 | 0.10835 | 0.00000 | 0.02339 | 0.12917 | 0.07433 |
| **NTAN1** | 0.38983 | 2.32697 | 0.38084 | 1.89849 | 0.72464 | 2.06691 | 1.15809 |
| **NTF3** | 0.09919 | 0.00677 | 0.00661 | 0.09429 | 0.05152 | 0.45972 | 0.00000 |
| **NTF4** | 0.00000 | 0.07961 | 0.16519 | 0.34869 | 0.12585 | 0.00000 | 0.03434 |
| **NTHL1** | 8.51886 | 10.23447 | 2.13579 | 9.38564 | 2.86175 | 6.87771 | 4.52812 |
| **NTM** | 0.05096 | 0.02206 | 0.00699 | 0.01416 | 0.07577 | 0.01260 | 0.02599 |
| **NTMT1** | 24.39864 | 34.58395 | 10.61496 | 17.57830 | 6.78133 | 8.13806 | 0.00000 |
| **NTN1** | 0.06305 | 0.02291 | 0.07618 | 0.02003 | 0.56079 | 0.11478 | 0.03974 |
| **NTN3** | 0.02325 | 0.01815 | 0.01275 | 0.02375 | 0.00795 | 0.01355 | 0.01629 |
| **NTN4** | 0.19630 | 0.63126 | 0.49299 | 0.01077 | 0.10493 | 0.89222 | 0.00000 |
| **NTN5** | 0.02357 | 0.02049 | 0.01094 | 0.02186 | 0.03275 | 0.01911 | 0.01651 |
| **NTNG1** | 0.00000 | 0.00191 | 2.10076 | 0.00000 | 0.00231 | 0.96943 | 0.28369 |
| **NTNG2** | 0.01567 | 0.01975 | 0.00842 | 0.01831 | 0.04109 | 0.02449 | 0.03332 |
| **NTPCR** | 9.19673 | 13.64155 | 4.07179 | 9.33161 | 0.72169 | 6.19612 | 0.00000 |
| **NTRK1** | 0.03372 | 0.00770 | 0.03984 | 0.03587 | 0.10947 | 0.01209 | 0.03322 |
| **NTRK2** | 0.33289 | 0.00000 | 0.00000 | 0.66850 | 0.91298 | 0.06789 | 1.24970 |
| **NTRK3** | 0.14135 | 0.28820 | 0.00000 | 0.00917 | 1.73128 | 1.82410 | 0.02717 |
| **NTRK3-AS1** | 0.02467 | 0.01735 | 0.01302 | 0.01978 | 0.01206 | 0.01274 | 0.01769 |
| **NTS** | 0.06887 | 0.00383 | 0.00737 | 0.01338 | 0.03383 | 0.04331 | 0.14221 |
| **NTSR1** | 0.02746 | 0.01890 | 0.01215 | 0.01918 | 0.00785 | 0.02115 | 0.02043 |
| **NTSR2** | 0.01535 | 0.00000 | 0.42390 | 0.01601 | 0.00247 | 0.00504 | 0.02409 |
| **NUAK1** | 0.00000 | 0.12529 | 0.04196 | 0.48479 | 0.00611 | 1.28710 | 0.69137 |
| **NUAK2** | 6.72719 | 4.58007 | 0.00000 | 16.79314 | 0.42555 | 0.00000 | 11.05171 |
| **NUB1** | 7.85704 | 6.57967 | 3.99761 | 9.72198 | 0.88237 | 2.80037 | 0.22816 |
| **NUBP1** | 9.81808 | 7.21403 | 4.90205 | 9.92266 | 0.32411 | 2.79410 | 0.00000 |
| **NUBP2** | 19.95572 | 18.41395 | 10.27630 | 17.11811 | 1.27640 | 9.59870 | 7.05286 |
| **NUBPL** | 6.97394 | 2.04013 | 3.46775 | 3.52017 | 0.08208 | 5.19930 | 1.31712 |
| **NUCB1** | 9.84250 | 39.12698 | 6.50455 | 10.75932 | 1.28197 | 19.91451 | 12.67204 |
| **NUCB1-AS1** | 0.12271 | 0.44987 | 0.11001 | 0.11646 | 0.03689 | 0.27170 | 0.00000 |
| **NUCB2** | 0.82803 | 0.10574 | 2.19328 | 10.02367 | 0.95267 | 0.66209 | 0.66271 |
| **NUCKS1** | 14.90751 | 15.69687 | 4.76568 | 2.45934 | 1.74981 | 15.18550 | 27.55246 |
| **NUDC** | 46.65098 | 39.58193 | 17.38412 | 38.06172 | 12.44157 | 20.30378 | 7.48312 |
| **NUDCD1** | 23.44792 | 0.44748 | 11.84590 | 10.08520 | 7.64812 | 8.87730 | 3.90834 |
| **NUDCD2** | 21.19522 | 9.32647 | 11.58007 | 13.28653 | 2.98419 | 10.75412 | 0.17126 |

| | | | | | | | |
|---|---|---|---|---|---|---|---|
| **NUDCD3** | 10.30208 | 6.85745 | 1.64614 | 7.33538 | 0.88342 | 4.88121 | 6.79963 |
| **NUDT1** | 6.39773 | 19.71107 | 11.00650 | 6.72252 | 0.28454 | 8.74976 | 3.86452 |
| **NUDT10** | 0.01495 | 0.01163 | 0.30521 | 0.10538 | 0.38135 | 0.26861 | 0.00000 |
| **NUDT11** | 0.06443 | 0.00000 | 0.38544 | 0.39330 | 0.43569 | 0.64982 | 0.00000 |
| **NUDT12** | 0.54536 | 0.00000 | 0.46429 | 0.20702 | 0.11409 | 0.12483 | 0.08214 |
| **NUDT13** | 0.62826 | 0.00000 | 0.71382 | 0.47017 | 0.48259 | 0.36906 | 0.77077 |
| **NUDT14** | 0.35983 | 2.79904 | 0.30425 | 1.05613 | 0.52730 | 0.51379 | 1.77733 |
| **NUDT15** | 21.29876 | 2.95713 | 7.64530 | 11.29800 | 2.78003 | 13.37388 | 2.58005 |
| **NUDT16** | 4.03404 | 3.40890 | 0.00000 | 5.63819 | 1.16610 | 2.00861 | 3.87497 |
| **NUDT16L1** | 9.13198 | 17.68304 | 4.46875 | 12.88022 | 6.72266 | 9.13967 | 0.00000 |
| **NUDT16P1** | 1.03069 | 1.16308 | 0.73957 | 1.83974 | 0.51074 | 0.40605 | 0.00000 |
| **NUDT17** | 0.22883 | 0.19320 | 0.44785 | 0.41091 | 0.26972 | 0.24650 | 0.01229 |
| **NUDT18** | 0.00000 | 0.28319 | 0.05629 | 0.05831 | 0.05805 | 0.03728 | 0.24704 |
| **NUDT19** | 1.29024 | 0.43563 | 0.19297 | 1.86561 | 0.05258 | 0.54156 | 0.00000 |
| **NUDT2** | 5.30806 | 1.30489 | 5.29441 | 6.85371 | 5.97476 | 2.15652 | 2.36337 |
| **NUDT21** | 27.67612 | 10.61973 | 18.02787 | 18.52283 | 3.97249 | 12.47121 | 11.84934 |
| **NUDT22** | 6.53123 | 2.31728 | 1.33826 | 5.28343 | 0.00000 | 5.05711 | 0.00005 |
| **NUDT3** | 0.08866 | 0.59633 | 0.15846 | 0.20013 | 0.15353 | 0.17307 | 0.05637 |
| **NUDT4** | 0.47044 | 1.24988 | 2.15896 | 0.22302 | 0.21336 | 2.56568 | 4.48135 |
| **NUDT5** | 31.83205 | 19.07888 | 15.67165 | 27.77322 | 3.97433 | 15.07746 | 3.70268 |
| **NUDT6** | 0.42035 | 0.81070 | 2.59330 | 0.44055 | 0.19950 | 0.79975 | 0.00000 |
| **NUDT7** | 0.28603 | 0.68096 | 0.84838 | 0.15639 | 0.59219 | 0.02245 | 0.01105 |
| **NUDT8** | 0.05575 | 0.80224 | 0.26482 | 0.35263 | 0.01280 | 0.74498 | 1.75713 |
| **NUDT9** | 12.15440 | 1.52168 | 5.99161 | 5.35510 | 0.63717 | 6.38920 | 1.21665 |
| **NUDT9P1** | 0.08319 | 0.01409 | 0.10817 | 0.02213 | 0.00588 | 0.06029 | 0.01445 |
| **NUF2** | 9.55728 | 1.83262 | 7.45369 | 7.30286 | 2.43414 | 3.72396 | 1.35136 |
| **NUFIP1** | 8.51427 | 6.22131 | 9.19512 | 3.62991 | 7.94318 | 4.15396 | 4.87940 |
| **NUFIP2** | 20.82378 | 3.27266 | 13.53850 | 14.34036 | 11.42323 | 4.75116 | 25.04987 |
| **NUGGC** | 0.01420 | 0.01074 | 0.01309 | 0.03893 | 0.00820 | 0.12147 | 0.03256 |
| **NUMA1** | 1.00246 | 3.00333 | 2.71486 | 12.08174 | 1.80061 | 6.65070 | 16.26413 |
| **NUMB** | 9.41151 | 6.23233 | 4.43246 | 12.52285 | 6.05760 | 8.19110 | 0.00000 |
| **NUMBL** | 0.08261 | 0.37569 | 0.75666 | 1.77962 | 0.00007 | 1.09503 | 1.61142 |
| **NUP107** | 27.48489 | 8.96495 | 14.24693 | 14.10331 | 3.30448 | 8.10795 | 0.33105 |
| **NUP133** | 17.11129 | 3.10010 | 6.71272 | 9.29753 | 7.13064 | 6.31512 | 6.87568 |
| **NUP153** | 13.67030 | 2.45871 | 15.34150 | 13.41802 | 11.34375 | 11.21518 | 25.34026 |

| | | | | | | | |
|---|---|---|---|---|---|---|---|
| **NUP155** | 17.78603 | 1.66728 | 12.03541 | 9.37032 | 4.60894 | 4.51288 | 3.74129 |
| **NUP160** | 32.94280 | 4.29585 | 12.12504 | 21.33619 | 4.35555 | 11.03316 | 3.67582 |
| **NUP188** | 18.76004 | 12.27459 | 8.23412 | 18.55665 | 1.60822 | 10.44364 | 9.53648 |
| **NUP205** | 15.21733 | 2.84308 | 11.13969 | 8.50069 | 1.46024 | 8.89073 | 12.67449 |
| **NUP210** | 0.31933 | 0.18565 | 0.04921 | 0.00000 | 0.21394 | 0.00000 | 1.19072 |
| **NUP210L** | 0.02487 | 0.00000 | 0.25739 | 0.08296 | 0.14138 | 0.04532 | 0.29397 |
| **NUP214** | 8.27136 | 9.65489 | 8.90135 | 9.73181 | 2.73741 | 7.60672 | 13.32965 |
| **NUP35** | 23.71801 | 0.98966 | 14.68137 | 8.29044 | 7.98557 | 6.90677 | 0.00000 |
| **NUP37** | 12.65630 | 5.43384 | 8.61815 | 7.13777 | 1.14083 | 3.29818 | 0.00000 |
| **NUP43** | 18.89280 | 7.02730 | 8.44024 | 8.87700 | 3.89272 | 6.41531 | 4.12123 |
| **NUP50** | 30.33832 | 19.63391 | 12.41673 | 19.98122 | 8.47664 | 9.01096 | 10.68282 |
| **NUP50-AS1** | 0.98518 | 2.74632 | 2.21904 | 1.39374 | 0.78919 | 0.38538 | 2.00219 |
| **NUP54** | 23.67726 | 1.96622 | 14.47216 | 9.96962 | 8.66208 | 11.79303 | 3.43024 |
| **NUP62** | 35.62075 | 34.26490 | 20.30010 | 33.32491 | 12.21017 | 12.22169 | 9.42848 |
| **NUP62CL** | 1.36574 | 0.56689 | 0.00000 | 0.35610 | 0.58314 | 0.57571 | 0.00000 |
| **NUP85** | 23.68087 | 13.55110 | 10.13149 | 18.17676 | 3.65687 | 11.05291 | 8.48091 |
| **NUP88** | 25.81101 | 7.29546 | 12.89959 | 12.58941 | 2.99122 | 7.05396 | 0.00000 |
| **NUP93** | 32.57295 | 23.26807 | 12.92645 | 21.15215 | 6.35149 | 9.69405 | 0.00000 |
| **NUP98** | 28.52031 | 12.67961 | 21.88186 | 15.54905 | 8.92622 | 9.73600 | 14.14646 |
| **NUPL1** | 15.08313 | 0.00000 | 8.00660 | 6.06908 | 4.82828 | 3.30670 | 7.61657 |
| **NUPL2** | 15.03817 | 2.43485 | 6.34870 | 4.40781 | 3.57558 | 3.54332 | 0.00000 |
| **NUPR1** | 0.25706 | 0.54705 | 0.00000 | 6.21622 | 0.06961 | 0.83911 | 0.00252 |
| **NUPR1L** | 0.12689 | 0.07306 | 0.03311 | 0.03111 | 0.00905 | 0.06909 | 0.08113 |
| **NUS1** | 5.46055 | 0.00000 | 2.25508 | 2.93676 | 0.31298 | 1.20019 | 6.75362 |
| **NUSAP1** | 29.26356 | 28.28414 | 23.93063 | 26.91131 | 21.53020 | 16.85113 | 6.99892 |
| **NUTF2** | 29.88713 | 21.30316 | 11.37977 | 24.98413 | 2.36942 | 17.74489 | 9.84917 |
| **NUTM1** | 0.35646 | 0.65974 | 0.57648 | 0.08570 | 0.16414 | 0.02098 | 0.19850 |
| **NUTM2A-AS1** | 2.48597 | 1.92339 | 1.35639 | 1.67064 | 1.11089 | 0.71294 | 0.62155 |
| **NUTM2B** | 0.02244 | 0.01855 | 0.01315 | 0.01826 | 0.00780 | 0.01231 | 0.02683 |
| **NUTM2D** | 0.03812 | 0.04915 | 0.01116 | 0.04143 | 0.00870 | 0.00993 | 0.00539 |
| **NUTM2F** | 0.01650 | 0.02564 | 0.01108 | 0.09296 | 0.00832 | 0.01912 | 0.01157 |
| **NUTM2G** | 0.00000 | 0.15413 | 0.05197 | 0.17015 | 0.13615 | 0.01630 | 0.24329 |
| **NVL** | 17.20912 | 6.68528 | 11.31894 | 5.21469 | 2.11436 | 5.86120 | 3.98413 |
| **NWD1** | 0.01461 | 0.03715 | 0.02648 | 0.07850 | 0.00668 | 0.14992 | 0.03922 |
| **NWD2** | 0.07762 | 0.01980 | 0.02376 | 0.07203 | 0.07844 | 0.01506 | 0.00000 |

| | | | | | | | |
|---|---|---|---|---|---|---|---|
| **NXF1** | 34.73568 | 10.91484 | 12.02539 | 26.02436 | 18.39207 | 12.47801 | 1.50878 |
| **NXF2+NXF2B.ui** | 0.17600 | 0.16554 | 0.02725 | 0.02757 | 0.01535 | 0.00065 | 0.11893 |
| **NXF2+NXF2B.ui** | 0.04679 | 0.02383 | 0.07118 | 0.02692 | 0.03687 | 0.00600 | 0.23484 |
| **NXF3** | 0.00000 | 0.23429 | 0.03398 | 0.00978 | 0.00619 | 0.23236 | 0.02210 |
| **NXF4** | 0.01735 | 0.02721 | 0.01902 | 0.02032 | 0.00701 | 0.01212 | 0.02640 |
| **NXF5** | 0.02408 | 0.01760 | 0.01293 | 0.01989 | 0.00977 | 0.01269 | 0.01856 |
| **NXN** | 0.07528 | 0.09252 | 1.77995 | 1.33723 | 0.20228 | 1.45095 | 2.60906 |
| **NXNL1** | 0.02013 | 0.00785 | 0.01755 | 0.01677 | 0.01036 | 0.04419 | 0.05652 |
| **NXNL2** | 0.31642 | 0.10496 | 0.00654 | 0.25814 | 0.04952 | 0.00000 | 0.09105 |
| **NXPE2** | 0.02266 | 0.01534 | 0.01266 | 0.02505 | 0.00795 | 0.03138 | 0.01741 |
| **NXPE3** | 0.18852 | 0.40368 | 0.61794 | 2.06586 | 0.83376 | 0.60696 | 0.00000 |
| **NXPH1** | 0.03163 | 0.01397 | 0.08603 | 0.02916 | 0.04273 | 0.00928 | 0.01042 |
| **NXPH2** | 0.08730 | 0.08537 | 0.19711 | 0.00235 | 0.00298 | 0.12178 | 0.00000 |
| **NXPH3** | 0.01665 | 0.02036 | 0.26400 | 0.15846 | 0.02015 | 0.24819 | 0.01416 |
| **NXPH4** | 0.01472 | 0.08555 | 0.02549 | 0.02814 | 0.55043 | 0.02152 | 0.03305 |
| **NXT1** | 18.24275 | 11.76174 | 11.62380 | 12.69049 | 6.00262 | 7.03765 | 2.45704 |
| **NXT2** | 7.08783 | 0.00000 | 10.92900 | 0.20686 | 1.19761 | 2.53823 | 1.28168 |
| **NYAP1** | 0.02237 | 0.01650 | 0.01396 | 0.02027 | 0.00802 | 0.03325 | 0.03489 |
| **NYAP2** | 0.01197 | 0.03214 | 0.06870 | 0.02804 | 0.35528 | 0.01258 | 0.02276 |
| **NYNRIN** | 0.44225 | 0.03672 | 0.02301 | 0.00000 | 0.01013 | 0.19499 | 0.67464 |
| **NYX** | 0.00000 | 0.19915 | 0.25796 | 0.01535 | 0.00732 | 0.02819 | 0.01482 |
| **OACYLP** | 0.02211 | 0.00768 | 0.03254 | 0.01881 | 0.07325 | 0.04035 | 0.01853 |
| **OAF** | 1.35620 | 0.00000 | 0.00000 | 0.00603 | 0.21366 | 0.00000 | 2.68704 |
| **OARD1** | 9.43866 | 2.52737 | 10.69779 | 7.38208 | 4.34625 | 5.41831 | 0.91394 |
| **OAS1** | 0.07364 | 0.00251 | 0.01080 | 0.05930 | 0.00801 | 0.66289 | 0.00000 |
| **OAS2** | 0.03284 | 0.01342 | 0.02617 | 0.05306 | 0.03776 | 0.01834 | 0.01626 |
| **OAS3** | 0.05374 | 0.03725 | 0.03042 | 0.02667 | 0.29847 | 0.01920 | 0.00000 |
| **OASL** | 0.00000 | 0.11746 | 1.67475 | 0.01288 | 0.20457 | 0.00000 | 0.01010 |
| **OAT** | 43.52099 | 29.89056 | 3.84234 | 4.29690 | 8.15342 | 11.10807 | 7.61696 |
| **OAZ1** | 70.36554 | 60.38755 | 27.34977 | 61.89844 | 22.65679 | 38.49686 | 38.41051 |
| **OAZ2** | 6.70559 | 0.49878 | 2.57574 | 13.06745 | 9.10370 | 8.32545 | 0.00000 |
| **OAZ3** | 0.45573 | 0.70241 | 1.88949 | 1.39016 | 0.58535 | 0.66749 | 0.00000 |
| **OBFC1** | 5.49365 | 0.63089 | 2.12290 | 1.06584 | 1.57756 | 3.34100 | 0.00000 |
| **OBP2A** | 0.02315 | 0.02132 | 0.01183 | 0.01854 | 0.00959 | 0.01822 | 0.01888 |
| **OBSCN** | 0.04090 | 0.36321 | 0.11142 | 0.38504 | 0.01759 | 0.51777 | 0.57882 |

| | | | | | | | |
|---|---|---|---|---|---|---|---|
| **OBSL1** | 0.10430 | 0.43770 | 0.03605 | 1.30446 | 0.72578 | 0.01010 | 0.68695 |
| **OC90** | 0.04038 | 0.00271 | 0.01159 | 0.06465 | 0.00786 | 0.73493 | 0.00000 |
| **OCA2** | 0.00000 | 0.80430 | 3.09596 | 0.00000 | 1.61172 | 0.22228 | 0.00000 |
| **OCEL1** | 0.32732 | 1.62182 | 0.35232 | 3.17088 | 0.00000 | 2.77328 | 5.73158 |
| **OCIAD1** | 22.83833 | 2.62217 | 16.97978 | 13.64734 | 1.15405 | 11.74109 | 17.86556 |
| **OCIAD2** | 0.00927 | 0.00862 | 0.00466 | 0.65684 | 0.08916 | 0.82876 | 0.00000 |
| **OCLM** | 0.51516 | 0.02852 | 0.08995 | 0.24072 | 0.00382 | 0.01959 | 0.00000 |
| **OCLN** | 10.63557 | 4.19911 | 0.50414 | 4.34532 | 0.43083 | 4.98505 | 0.00000 |
| **OCM** | 0.02304 | 0.01798 | 0.02680 | 0.01951 | 0.03020 | 0.01621 | 0.01600 |
| **OCM2** | 0.02357 | 0.01742 | 0.00891 | 0.02015 | 0.04971 | 0.01384 | 0.01677 |
| **OCRL** | 2.91213 | 2.05788 | 5.14351 | 0.00000 | 1.79320 | 1.19089 | 3.54618 |
| **ODAM** | 8.52906 | 0.00000 | 0.00399 | 12.47503 | 0.00107 | 0.00000 | 0.04023 |
| **ODC1** | 35.67965 | 33.81233 | 25.81615 | 18.76958 | 11.80589 | 4.33093 | 73.38944 |
| **ODF2** | 9.61395 | 7.22652 | 4.80366 | 9.48101 | 5.07509 | 4.72161 | 0.90477 |
| **ODF2L** | 2.61607 | 0.59211 | 0.20339 | 5.81081 | 0.23736 | 0.82879 | 1.40813 |
| **ODF3** | 0.02138 | 0.01763 | 0.01179 | 0.01198 | 0.00728 | 0.00758 | 0.08009 |
| **ODF3B** | 0.00466 | 0.05232 | 0.04155 | 0.02964 | 0.00533 | 0.00579 | 0.05512 |
| **ODF3L1** | 0.02393 | 0.00697 | 0.02549 | 0.01831 | 0.16755 | 0.01364 | 0.01933 |
| **ODF3L2** | 0.00024 | 0.02825 | 0.27856 | 0.01882 | 0.28785 | 0.00000 | 0.09032 |
| **ODF4** | 0.01236 | 0.01260 | 0.04307 | 0.03714 | 0.00708 | 0.02385 | 0.02364 |
| **OFD1** | 12.17197 | 12.16542 | 8.45872 | 10.63711 | 8.07996 | 6.02949 | 14.65868 |
| **OGDH** | 16.29484 | 12.23552 | 1.78332 | 18.22209 | 3.18863 | 10.13839 | 8.67199 |
| **OGDHL** | 0.45902 | 0.27609 | 0.24554 | 0.07160 | 0.15962 | 0.75795 | 0.89381 |
| **OGFOD1** | 24.62729 | 16.48640 | 9.28986 | 17.03719 | 6.90351 | 5.73192 | 0.00000 |
| **OGFOD2** | 6.14259 | 5.94226 | 5.98579 | 8.01553 | 2.14084 | 2.47313 | 0.00000 |
| **OGFOD3** | 0.25402 | 2.81079 | 0.41823 | 0.21505 | 0.25398 | 1.28399 | 5.18947 |
| **OGFR** | 0.00000 | 0.86273 | 0.00000 | 0.63397 | 0.00409 | 1.34230 | 6.10646 |
| **OGFR-AS1** | 0.01783 | 0.01928 | 0.02092 | 0.02332 | 0.01761 | 0.01076 | 0.04749 |
| **OGFRL1** | 0.00000 | 1.20756 | 1.07333 | 0.00000 | 0.00000 | 1.04032 | 1.27990 |
| **OGFRP1** | 0.01378 | 0.15431 | 0.02044 | 0.03687 | 0.03323 | 0.03785 | 0.00000 |
| **OGG1** | 1.58154 | 2.85291 | 3.92535 | 4.66220 | 2.90257 | 1.36171 | 0.55406 |
| **OGN** | 0.13392 | 0.00000 | 0.12485 | 0.00000 | 0.04953 | 0.21755 | 0.77995 |
| **OGT** | 29.03164 | 7.40806 | 15.33675 | 5.90814 | 3.83863 | 11.28052 | 23.56827 |
| **OIP5** | 18.68092 | 9.52752 | 14.25509 | 14.94107 | 3.87036 | 12.50910 | 1.32156 |
| **OIP5-AS1** | 9.83679 | 9.39592 | 10.24527 | 3.55221 | 0.57384 | 7.91135 | 0.68595 |

| | | | | | | | |
|---|---|---|---|---|---|---|---|
| **OIT3** | 0.10019 | 0.00000 | 0.09426 | 0.01286 | 0.00461 | 0.20218 | 0.00000 |
| **OLA1** | 16.13704 | 1.19049 | 4.37974 | 13.19568 | 0.31054 | 4.59630 | 10.06644 |
| **OLAH** | 0.00749 | 3.17590 | 1.18780 | 0.00000 | 0.00318 | 0.62174 | 0.00336 |
| **OLFM1** | 0.01730 | 0.01662 | 0.31672 | 0.00641 | 0.82279 | 0.00192 | 0.04778 |
| **OLFM2** | 0.00000 | 0.72754 | 0.01762 | 0.07813 | 0.51050 | 0.11954 | 0.00667 |
| **OLFM3** | 0.02560 | 0.01840 | 0.01076 | 0.01925 | 0.01474 | 0.01346 | 0.04318 |
| **OLFM4** | 0.03346 | 0.00131 | 0.03846 | 0.03132 | 0.86060 | 0.01223 | 0.00318 |
| **OLFML1** | 9.43765 | 4.57868 | 0.00000 | 4.80400 | 1.53652 | 0.00000 | 0.00000 |
| **OLFML2A** | 0.04606 | 0.22981 | 0.02892 | 0.02126 | 0.06814 | 0.01313 | 0.00449 |
| **OLFML2B** | 0.13536 | 0.00141 | 0.01390 | 0.09068 | 0.00885 | 0.07506 | 0.00000 |
| **OLFML3** | 1.13880 | 0.00000 | 0.01164 | 2.68737 | 0.09528 | 1.14225 | 0.00000 |
| **OLIG1** | 0.01974 | 0.01599 | 0.00797 | 0.01885 | 0.20873 | 0.02249 | 0.02216 |
| **OLIG2** | 0.01059 | 0.02552 | 0.01144 | 0.01706 | 0.00915 | 0.08908 | 0.02867 |
| **OLIG3** | 0.02134 | 0.01355 | 0.01263 | 0.01709 | 0.00784 | 0.08408 | 0.02271 |
| **OLR1** | 17.50247 | 1.37251 | 1.57178 | 10.95690 | 0.00087 | 0.00000 | 0.00000 |
| **OMA1** | 9.49374 | 2.12224 | 5.81038 | 0.58544 | 0.00000 | 2.52391 | 1.43154 |
| **OMD** | 0.08998 | 0.00000 | 0.20776 | 0.11882 | 0.02334 | 0.24204 | 0.53288 |
| **OMG** | 0.04952 | 0.00521 | 0.15715 | 0.15264 | 0.00727 | 0.01493 | 0.01091 |
| **OMP** | 0.00921 | 0.04186 | 0.02136 | 0.03625 | 0.00704 | 0.03954 | 0.03125 |
| **ONECUT1** | 0.01892 | 0.01826 | 0.01345 | 0.04708 | 0.04533 | 0.01746 | 0.03033 |
| **ONECUT2** | 0.01268 | 0.16566 | 0.08811 | 0.56974 | 0.00837 | 0.01919 | 0.25695 |
| **ONECUT3** | 0.02293 | 0.01945 | 0.01240 | 0.02127 | 0.00783 | 0.01175 | 0.02397 |
| **OOEP** | 0.34956 | 10.73230 | 14.41041 | 0.09034 | 24.21002 | 4.20725 | 0.12638 |
| **OOSP2** | 0.00658 | 0.00000 | 1.78634 | 0.00421 | 23.80519 | 0.00000 | 0.02553 |
| **OPA1** | 20.33145 | 3.10488 | 7.52636 | 6.12925 | 7.10123 | 6.18098 | 5.51950 |
| **OPA1-AS1** | 0.09751 | 0.06255 | 0.02800 | 0.03931 | 0.11281 | 0.02597 | 0.12065 |
| **OPA3** | 8.31855 | 9.98338 | 3.83841 | 10.67420 | 3.23609 | 4.79586 | 7.29732 |
| **OPALIN** | 0.02841 | 0.04858 | 0.00000 | 0.01645 | 0.27287 | 0.04226 | 0.00364 |
| **OPCML** | 0.03980 | 0.03791 | 0.01324 | 0.04453 | 0.00861 | 0.01743 | 0.00000 |
| **OPHN1** | 0.00000 | 0.00000 | 0.51298 | 0.26292 | 0.05302 | 0.24003 | 0.19270 |
| **OPLAH** | 0.09879 | 0.00000 | 0.00685 | 0.06118 | 0.00680 | 0.14206 | 0.95647 |
| **OPN1LW** | 0.02175 | 0.01797 | 0.02099 | 0.02374 | 0.07701 | 0.01483 | 0.01284 |
| **OPN1SW** | 0.05177 | 0.01169 | 0.02621 | 0.03133 | 0.19408 | 0.01478 | 0.00601 |
| **OPN3** | 0.00000 | 0.12669 | 0.11225 | 1.82095 | 0.08930 | 0.00915 | 0.56547 |
| **OPN4** | 0.03079 | 0.01592 | 0.01313 | 0.02370 | 0.00797 | 0.01315 | 0.01484 |

| | | | | | | | |
|---|---|---|---|---|---|---|---|
| **OPN5** | 0.03457 | 0.01808 | 0.01242 | 0.02073 | 0.00839 | 0.01399 | 0.01343 |
| **OPRD1** | 0.01703 | 0.01217 | 0.10988 | 0.01836 | 0.00617 | 0.01139 | 0.02615 |
| **OPRK1** | 0.04032 | 0.01655 | 0.02245 | 0.16165 | 0.00856 | 0.00877 | 0.01882 |
| **OPRL1** | 0.00000 | 0.10326 | 0.03557 | 0.03759 | 0.09328 | 0.02672 | 0.07408 |
| **OPRM1** | 0.07153 | 0.03669 | 0.00000 | 0.02034 | 0.27192 | 0.01166 | 0.10339 |
| **OPTC** | 0.01681 | 0.01473 | 0.01589 | 0.06387 | 0.00776 | 0.02304 | 0.01372 |
| **OPTN** | 0.46572 | 4.84805 | 2.31195 | 2.36160 | 4.60367 | 0.64353 | 0.57593 |
| **OR10A2** | 0.05305 | 0.03253 | 0.02278 | 0.01746 | 0.00641 | 0.00772 | 0.03790 |
| **OR10A4** | 0.02877 | 0.05408 | 0.04591 | 0.02633 | 0.00607 | 0.00744 | 0.00734 |
| **OR10A5** | 0.02261 | 0.04588 | 0.02945 | 0.01291 | 0.00583 | 0.05870 | 0.07074 |
| **OR10AD1** | 0.02468 | 0.01639 | 0.01324 | 0.02301 | 0.00789 | 0.01484 | 0.01716 |
| **OR10AG1** | 0.02633 | 0.01938 | 0.01284 | 0.01925 | 0.00806 | 0.01511 | 0.01716 |
| **OR10G3** | 0.05126 | 0.01363 | 0.01090 | 0.00391 | 0.02653 | 0.00839 | 0.04361 |
| **OR10J3** | 0.02398 | 0.01514 | 0.01349 | 0.01637 | 0.00800 | 0.02667 | 0.02661 |
| **OR11A1** | 0.02088 | 0.01192 | 0.01380 | 0.02567 | 0.00748 | 0.01229 | 0.04273 |
| **OR13A1** | 0.00532 | 0.11036 | 0.04108 | 0.02019 | 0.00594 | 0.04361 | 0.02913 |
| **OR13H1** | 0.02809 | 0.06699 | 0.01635 | 0.01787 | 0.00716 | 0.00853 | 0.00654 |
| **OR1E1** | 0.02978 | 0.01061 | 0.01966 | 0.01885 | 0.04733 | 0.01279 | 0.01902 |
| **OR1E2** | 0.02388 | 0.00901 | 0.02376 | 0.03437 | 0.04108 | 0.02780 | 0.02509 |
| **OR1F1** | 0.01566 | 0.05683 | 0.01845 | 0.04541 | 0.00688 | 0.06471 | 0.01587 |
| **OR1F2P** | 0.02185 | 0.00288 | 0.04760 | 0.01825 | 0.10676 | 0.01202 | 0.02470 |
| **OR1J4** | 0.02770 | 0.01667 | 0.01285 | 0.02552 | 0.01274 | 0.01544 | 0.01292 |
| **OR1N1** | 0.13697 | 0.06839 | 0.12288 | 0.19304 | 0.42906 | 0.00043 | 0.00000 |
| **OR1N2** | 0.03310 | 0.01139 | 0.02318 | 0.01685 | 0.01127 | 0.01380 | 0.01850 |
| **OR2A25** | 0.01891 | 0.02213 | 0.01128 | 0.01684 | 0.00769 | 0.01124 | 0.03838 |
| **OR2A7** | 0.02402 | 0.01818 | 0.01281 | 0.02291 | 0.00797 | 0.01334 | 0.01474 |
| **OR2AG1** | 0.02781 | 0.00000 | 0.07550 | 0.02440 | 0.00693 | 0.01355 | 0.01766 |
| **OR2AG2** | 0.04401 | 0.05602 | 0.02407 | 0.02952 | 0.17786 | 0.00227 | 0.02987 |
| **OR2B2** | 0.01638 | 0.01735 | 0.02105 | 0.03513 | 0.25190 | 0.01233 | 0.02207 |
| **OR2B6** | 0.07601 | 0.00000 | 0.03677 | 0.18100 | 0.40049 | 0.02133 | 0.00000 |
| **OR2C1** | 0.03180 | 0.01642 | 0.01330 | 0.01774 | 0.00994 | 0.01301 | 0.01454 |
| **OR2C3** | 0.02405 | 0.02515 | 0.01358 | 0.01709 | 0.00759 | 0.03054 | 0.01762 |
| **OR2D2** | 0.05375 | 0.00169 | 0.02823 | 0.04410 | 0.01430 | 0.02598 | 0.02004 |
| **OR2D3** | 0.05129 | 0.00000 | 0.12860 | 0.00257 | 0.08083 | 0.04162 | 0.17256 |
| **OR2F1** | 0.02451 | 0.01798 | 0.01273 | 0.03197 | 0.00797 | 0.01308 | 0.01480 |

| | | | | | | | |
|---|---|---|---|---|---|---|---|
| **OR2H1** | 0.02747 | 0.01650 | 0.01465 | 0.01784 | 0.00790 | 0.01522 | 0.01799 |
| **OR2H2** | 0.03560 | 0.01250 | 0.01456 | 0.01722 | 0.00792 | 0.03643 | 0.00807 |
| **OR2L13** | 0.02346 | 0.01617 | 0.01354 | 0.01950 | 0.02075 | 0.01458 | 0.01757 |
| **OR2M7** | 0.02459 | 0.03235 | 0.03480 | 0.01923 | 0.01235 | 0.01424 | 0.01018 |
| **OR2S2** | 0.04243 | 0.01307 | 0.01321 | 0.01112 | 0.00764 | 0.01053 | 0.03289 |
| **OR2T11** | 0.03249 | 0.00535 | 0.03428 | 0.01907 | 0.04879 | 0.01212 | 0.02063 |
| **OR2T2** | 0.02683 | 0.01894 | 0.01449 | 0.01234 | 0.01136 | 0.01617 | 0.03821 |
| **OR2T27** | 0.02147 | 0.01325 | 0.02439 | 0.02075 | 0.02403 | 0.01766 | 0.02048 |
| **OR2T29** | 0.02313 | 0.01934 | 0.01243 | 0.01854 | 0.00793 | 0.01993 | 0.01993 |
| **OR2T33** | 0.02707 | 0.02180 | 0.01693 | 0.02196 | 0.00758 | 0.01636 | 0.01086 |
| **OR2T4** | 0.03221 | 0.00141 | 0.02961 | 0.01627 | 0.26988 | 0.02130 | 0.01670 |
| **OR2T8** | 0.02732 | 0.01733 | 0.01456 | 0.01876 | 0.00798 | 0.01411 | 0.01689 |
| **OR2V1** | 0.03800 | 0.01034 | 0.01426 | 0.01614 | 0.01537 | 0.01343 | 0.03259 |
| **OR2W3** | 0.00957 | 0.01478 | 0.01155 | 0.02927 | 0.00739 | 0.07929 | 0.05848 |
| **OR2W5** | 0.01271 | 0.02409 | 0.01688 | 0.01762 | 0.00826 | 0.05408 | 0.02903 |
| **OR3A1** | 0.10521 | 0.01592 | 0.18935 | 0.10412 | 0.26222 | 0.00383 | 0.00000 |
| **OR3A2** | 0.10992 | 0.08255 | 0.29497 | 0.25694 | 0.00282 | 0.03237 | 0.00000 |
| **OR3A3** | 0.03841 | 0.04696 | 0.01605 | 0.08494 | 0.02022 | 0.01219 | 0.00000 |
| **OR3A4P** | 0.03926 | 0.02343 | 0.02438 | 0.02077 | 0.00709 | 0.00768 | 0.02267 |
| **OR4D1** | 0.04352 | 0.00000 | 0.05637 | 0.01370 | 0.02437 | 0.02822 | 0.02382 |
| **OR4F13P** | 0.01925 | 0.04388 | 0.02388 | 0.01876 | 0.03595 | 0.00898 | 0.01941 |
| **OR4F16+OR4F2** | 0.02357 | 0.01628 | 0.01380 | 0.01828 | 0.00881 | 0.01474 | 0.03262 |
| **OR4N3P** | 0.02856 | 0.01622 | 0.01453 | 0.02283 | 0.00850 | 0.01398 | 0.01417 |
| **OR51B2** | 0.05906 | 0.01022 | 0.01359 | 0.01449 | 0.00772 | 0.01257 | 0.02216 |
| **OR51B4** | 0.05443 | 0.00554 | 0.01501 | 0.07917 | 0.00707 | 0.00253 | 0.08414 |
| **OR51B5** | 1.50196 | 0.00000 | 0.64583 | 0.01285 | 0.54821 | 0.00000 | 1.55928 |
| **OR51B6** | 0.12107 | 0.01562 | 0.01725 | 0.00402 | 0.00652 | 0.01240 | 0.05601 |
| **OR51D1** | 0.02225 | 0.01693 | 0.01320 | 0.01915 | 0.00847 | 0.01995 | 0.01976 |
| **OR51E1** | 0.01360 | 0.09362 | 0.01755 | 0.01408 | 0.23168 | 0.04359 | 0.03361 |
| **OR51E2** | 0.02307 | 0.01981 | 0.01343 | 0.01986 | 0.00794 | 0.01280 | 0.01709 |
| **OR51I1** | 0.24909 | 0.00000 | 0.07996 | 0.00000 | 0.00509 | 0.00014 | 0.22844 |
| **OR51I2** | 0.01616 | 0.08879 | 0.02926 | 0.00200 | 0.00836 | 0.00457 | 0.21317 |
| **OR51L1** | 0.02310 | 0.02076 | 0.01159 | 0.01632 | 0.00773 | 0.01112 | 0.03675 |
| **OR51M1** | 0.03695 | 0.00200 | 0.05953 | 0.02077 | 0.00669 | 0.01036 | 0.02296 |
| **OR51Q1** | 0.08288 | 0.01583 | 0.01582 | 0.00019 | 0.03166 | 0.00857 | 0.04908 |

| | | | | | | | |
|---|---|---|---|---|---|---|---|
| **OR52D1** | 0.30822 | 0.00000 | 0.04285 | 0.00277 | 0.25276 | 0.00300 | 0.04709 |
| **OR52E4** | 0.04213 | 0.00000 | 0.06188 | 0.13354 | 0.00436 | 0.00336 | 0.17371 |
| **OR52E6** | 0.03513 | 0.06821 | 0.12088 | 0.05253 | 0.08473 | 0.03552 | 0.25181 |
| **OR52E8** | 0.02874 | 0.01451 | 0.02940 | 0.02774 | 0.00848 | 0.00335 | 0.11090 |
| **OR52H1** | 0.02744 | 0.01512 | 0.01309 | 0.01755 | 0.00784 | 0.01322 | 0.02294 |
| **OR52N1** | 0.01163 | 0.01303 | 0.01540 | 0.03836 | 0.00585 | 0.00199 | 0.37954 |
| **OR52N2** | 0.00755 | 0.00000 | 0.06659 | 0.01926 | 0.00488 | 0.00750 | 0.22513 |
| **OR52N4** | 0.00051 | 0.00000 | 0.07743 | 0.08667 | 0.00411 | 0.11186 | 0.49984 |
| **OR52N5** | 0.04168 | 0.00000 | 0.10303 | 0.05265 | 0.00289 | 0.00532 | 0.39320 |
| **OR52W1** | 0.02130 | 0.01821 | 0.01274 | 0.02873 | 0.00793 | 0.01256 | 0.01772 |
| **OR56A1** | 0.03808 | 0.04001 | 0.01281 | 0.03471 | 0.00958 | 0.01011 | 0.00000 |
| **OR56A3** | 0.02475 | 0.01821 | 0.01322 | 0.07224 | 0.00801 | 0.01457 | 0.00717 |
| **OR56A5** | 0.02389 | 0.01683 | 0.01493 | 0.01980 | 0.00843 | 0.01301 | 0.01692 |
| **OR56B1** | 0.01945 | 0.01293 | 0.01254 | 0.04076 | 0.00798 | 0.06311 | 0.01826 |
| **OR56B4** | 0.04314 | 0.00606 | 0.02507 | 0.02458 | 0.00699 | 0.00965 | 0.02742 |
| **OR5A1** | 0.02816 | 0.00780 | 0.03500 | 0.01787 | 0.00894 | 0.01315 | 0.01729 |
| **OR5AU1** | 0.00000 | 0.01072 | 0.01926 | 0.00000 | 0.00644 | 0.36906 | 0.61474 |
| **OR5E1P** | 0.02582 | 0.01489 | 0.00969 | 0.03377 | 0.11262 | 0.01385 | 0.01035 |
| **OR5P2** | 0.02748 | 0.01438 | 0.01672 | 0.04177 | 0.01219 | 0.01577 | 0.00770 |
| **OR5P3** | 0.01980 | 0.01898 | 0.00000 | 0.02086 | 0.27658 | 0.01514 | 0.02345 |
| **OR5W2** | 0.02419 | 0.02015 | 0.01443 | 0.01776 | 0.00875 | 0.01525 | 0.02029 |
| **OR6A2** | 0.04318 | 0.02426 | 0.01711 | 0.01954 | 0.03281 | 0.01560 | 0.03620 |
| **OR6C2** | 0.03606 | 0.05882 | 0.01257 | 0.00970 | 0.00776 | 0.12427 | 0.01166 |
| **OR6C68** | 0.02327 | 0.01873 | 0.01240 | 0.01801 | 0.00786 | 0.01689 | 0.02178 |
| **OR6C70** | 0.03697 | 0.04040 | 0.04381 | 0.01926 | 0.00721 | 0.02168 | 0.00123 |
| **OR6S1** | 0.02113 | 0.01683 | 0.00820 | 0.02342 | 0.07868 | 0.01649 | 0.01695 |
| **OR6W1P** | 0.02017 | 0.01857 | 0.01875 | 0.02621 | 0.01237 | 0.01100 | 0.03271 |
| **OR7C1** | 0.02118 | 0.02394 | 0.03097 | 0.02466 | 0.00837 | 0.01211 | 0.01534 |
| **OR7D2** | 0.03868 | 0.00615 | 0.01562 | 0.03866 | 0.00752 | 0.02500 | 0.04228 |
| **OR7E12P** | 0.09508 | 0.00000 | 0.10414 | 0.00000 | 0.16677 | 0.07959 | 0.47413 |
| **OR7E14P** | 0.00052 | 0.04315 | 0.04566 | 0.23822 | 0.10185 | 0.07533 | 0.38961 |
| **OR7E156P** | 0.02612 | 0.01616 | 0.01188 | 0.01388 | 0.00780 | 0.07987 | 0.02631 |
| **OR7E24** | 0.02664 | 0.00639 | 0.03179 | 0.01900 | 0.05332 | 0.01301 | 0.01706 |
| **OR7E2P** | 0.01303 | 0.03183 | 0.01508 | 0.01400 | 0.00659 | 0.00699 | 0.11870 |
| **OR7E47P** | 0.03919 | 0.02291 | 0.01909 | 0.04479 | 0.01150 | 0.04117 | 0.01926 |

| | | | | | | | |
|---|---|---|---|---|---|---|---|
| **OR7E5P** | 0.02440 | 0.20180 | 0.01777 | 0.02372 | 0.00904 | 0.09975 | 0.00713 |
| **OR7E91P** | 0.08327 | 0.01788 | 0.01115 | 0.54406 | 0.00778 | 0.01283 | 0.00000 |
| **OR8A1** | 0.00605 | 0.02364 | 0.01218 | 0.04011 | 0.00736 | 0.05106 | 0.08682 |
| **OR8B12** | 0.02480 | 0.02753 | 0.00984 | 0.01894 | 0.00784 | 0.04813 | 0.02528 |
| **OR8B2** | 0.04713 | 0.01369 | 0.01187 | 0.01412 | 0.00775 | 0.02798 | 0.03556 |
| **OR8B3** | 0.02442 | 0.01426 | 0.01229 | 0.01662 | 0.00744 | 0.01288 | 0.05366 |
| **OR8B4** | 0.03576 | 0.01682 | 0.01548 | 0.01755 | 0.00761 | 0.01669 | 0.02128 |
| **OR8B8** | 0.00197 | 0.02527 | 0.01088 | 0.01722 | 0.00851 | 0.01091 | 0.13587 |
| **OR8D1** | 0.06024 | 0.00000 | 0.01316 | 0.00012 | 0.00778 | 0.00343 | 0.16141 |
| **OR8D2** | 0.03284 | 0.01104 | 0.01336 | 0.01638 | 0.00760 | 0.02113 | 0.04211 |
| **OR8G1** | 0.02442 | 0.00606 | 0.02559 | 0.01722 | 0.10576 | 0.04601 | 0.04232 |
| **OR8G2** | 0.04234 | 0.03830 | 0.02130 | 0.02405 | 0.01162 | 0.01435 | 0.01367 |
| **OR8G5** | 0.03834 | 0.00000 | 0.15679 | 0.05146 | 0.37099 | 0.10467 | 0.03570 |
| **OR8J3** | 0.02229 | 0.01635 | 0.01299 | 0.01819 | 0.00784 | 0.02059 | 0.02244 |
| **OR8S1** | 0.08284 | 0.00375 | 0.01369 | 0.03252 | 0.00743 | 0.00521 | 0.04959 |
| **OR9Q1** | 0.02399 | 0.02224 | 0.01015 | 0.02009 | 0.02436 | 0.01314 | 0.01972 |
| **ORAI1** | 0.00000 | 7.37437 | 1.05120 | 2.27860 | 0.28115 | 5.62188 | 7.69776 |
| **ORAI2** | 0.24429 | 0.21062 | 0.42806 | 0.40079 | 0.01059 | 0.00000 | 0.94938 |
| **ORAI3** | 0.03885 | 0.00000 | 0.24133 | 0.20483 | 0.05056 | 0.08734 | 0.66933 |
| **ORAOV1** | 3.52234 | 3.68847 | 3.46565 | 5.05997 | 0.70794 | 2.98512 | 2.44687 |
| **ORC1** | 20.88478 | 12.36547 | 9.71702 | 14.85131 | 5.32432 | 8.89737 | 3.67598 |
| **ORC2** | 13.74353 | 0.00351 | 8.96934 | 8.29894 | 2.38686 | 6.85029 | 2.55344 |
| **ORC3** | 12.14507 | 0.00000 | 6.86848 | 7.69578 | 1.39072 | 3.84617 | 0.00006 |
| **ORC4** | 23.04825 | 2.61628 | 12.00993 | 12.79925 | 5.68591 | 7.13134 | 1.18903 |
| **ORC5** | 15.55552 | 6.63302 | 8.90662 | 5.54597 | 3.19544 | 3.45039 | 0.00000 |
| **ORC6** | 19.98258 | 8.82696 | 9.47658 | 15.01078 | 4.82061 | 13.96694 | 4.62092 |
| **ORMDL1** | 24.78018 | 1.91315 | 14.90885 | 19.42678 | 2.99402 | 7.56531 | 0.00003 |
| **ORMDL2** | 11.25736 | 19.83882 | 12.94778 | 15.76951 | 13.99787 | 7.70868 | 3.17408 |
| **ORMDL3** | 1.92628 | 2.20448 | 1.80671 | 2.86755 | 2.70106 | 2.34153 | 6.31067 |
| **OS9** | 17.20482 | 14.90381 | 7.34506 | 20.36877 | 2.62563 | 10.55666 | 2.34020 |
| **OSBP** | 7.12612 | 5.06466 | 2.90546 | 6.94604 | 4.31366 | 4.62913 | 12.47501 |
| **OSBP2** | 0.10061 | 0.99366 | 1.04994 | 0.08580 | 0.44728 | 1.41304 | 0.07402 |
| **OSBPL10** | 1.19807 | 0.00000 | 0.14149 | 1.73590 | 9.18766 | 2.32683 | 2.13044 |
| **OSBPL10-AS1** | 0.02694 | 0.01040 | 0.01325 | 0.02364 | 0.01088 | 0.02493 | 0.03507 |
| **OSBPL11** | 10.40329 | 0.65036 | 13.92700 | 8.09871 | 1.55639 | 9.33859 | 9.70200 |

| | | | | | | | |
|---|---|---|---|---|---|---|---|
| **OSBPL1A** | 1.68546 | 0.25208 | 0.68492 | 0.00000 | 1.37412 | 0.00000 | 4.07446 |
| **OSBPL2** | 9.47059 | 8.40632 | 1.12157 | 9.74121 | 1.97886 | 4.11535 | 2.39771 |
| **OSBPL3** | 7.05299 | 3.32707 | 1.84995 | 3.52914 | 0.33752 | 5.34350 | 1.46040 |
| **OSBPL5** | 0.00000 | 0.09420 | 0.10986 | 0.00000 | 0.59892 | 0.14741 | 0.32991 |
| **OSBPL6** | 9.04323 | 13.16323 | 5.82532 | 0.29372 | 5.56808 | 0.00000 | 2.21037 |
| **OSBPL7** | 0.00000 | 0.89667 | 0.00000 | 0.35187 | 0.02298 | 0.00000 | 0.11541 |
| **OSBPL8** | 8.35445 | 0.70172 | 3.95678 | 6.61482 | 4.52067 | 8.51164 | 14.67007 |
| **OSBPL9** | 19.10952 | 2.75675 | 4.39160 | 22.40861 | 3.46983 | 2.98111 | 4.10496 |
| **OSCAR** | 0.24852 | 0.11409 | 0.10145 | 0.16047 | 0.32080 | 0.08299 | 0.02488 |
| **OSCP1** | 0.09866 | 0.00000 | 0.95446 | 0.57741 | 0.78271 | 0.10432 | 0.00000 |
| **OSER1** | 23.33246 | 11.79384 | 18.33862 | 21.83913 | 18.71589 | 7.90130 | 0.00000 |
| **OSER1-AS1** | 0.41507 | 0.80771 | 2.59457 | 1.40660 | 0.53475 | 0.76670 | 0.00000 |
| **OSGEP** | 15.72917 | 12.83516 | 13.35054 | 5.76808 | 0.90655 | 12.32588 | 4.01584 |
| **OSGEPL1** | 5.17081 | 2.20028 | 5.40529 | 0.65948 | 0.09329 | 0.59858 | 1.70095 |
| **OSGEPL1-AS1** | 0.10874 | 0.08628 | 0.23624 | 0.04117 | 0.04634 | 0.00001 | 0.03711 |
| **OSGIN1** | 0.23786 | 0.01485 | 0.03252 | 1.12052 | 0.00531 | 0.00349 | 0.00000 |
| **OSGIN2** | 5.46192 | 0.00000 | 0.64884 | 1.74145 | 0.00022 | 1.34939 | 0.30317 |
| **OSM** | 0.01667 | 0.01766 | 0.01398 | 0.05059 | 0.02504 | 0.01189 | 0.02281 |
| **OSMR** | 0.00000 | 0.00000 | 0.20644 | 0.18992 | 0.55549 | 2.50236 | 0.54827 |
| **OSMR-AS1** | 0.02270 | 0.01304 | 0.01677 | 0.02303 | 0.00969 | 0.02661 | 0.02120 |
| **OSR1** | 0.00994 | 0.01129 | 0.31684 | 0.02399 | 0.15422 | 0.00305 | 0.02622 |
| **OSR2** | 0.39928 | 4.98910 | 10.03535 | 1.85392 | 3.01415 | 1.37481 | 0.00000 |
| **OST4** | 25.68023 | 27.95770 | 14.28955 | 21.08030 | 4.76263 | 15.03610 | 12.68828 |
| **OSTC** | 28.52754 | 6.64594 | 18.70199 | 15.36185 | 3.06493 | 12.11294 | 8.37880 |
| **OSTCP1** | 1.61738 | 0.27899 | 1.07821 | 0.97688 | 0.07882 | 0.46489 | 0.00599 |
| **OSTF1** | 7.89913 | 4.41554 | 2.09623 | 6.69795 | 2.79940 | 3.12990 | 0.00000 |
| **OSTM1** | 6.14671 | 0.00000 | 2.97535 | 5.34897 | 1.75698 | 0.83679 | 5.27462 |
| **OTC** | 0.04341 | 0.02801 | 0.02237 | 0.00460 | 0.05575 | 0.01796 | 0.03860 |
| **OTOA** | 0.03394 | 0.06721 | 0.01002 | 0.01782 | 0.22662 | 0.02033 | 0.02471 |
| **OTOF** | 0.20082 | 0.00000 | 0.01441 | 0.03594 | 0.00519 | 0.01808 | 0.33979 |
| **OTOG** | 0.01233 | 0.01483 | 0.01342 | 0.04841 | 0.02101 | 0.02500 | 0.05896 |
| **OTOGL** | 0.18889 | 0.00000 | 0.02737 | 0.30053 | 0.69294 | 0.01072 | 0.02090 |
| **OTOP2** | 0.00000 | 0.06480 | 0.01616 | 0.01522 | 0.00749 | 0.07448 | 0.04924 |
| **OTOP3** | 0.01598 | 0.03169 | 0.14411 | 0.02129 | 0.02132 | 0.02524 | 0.01580 |
| **OTOR** | 0.03149 | 0.01790 | 0.01224 | 0.01853 | 0.00799 | 0.01151 | 0.02149 |

| | | | | | | | |
|---|---|---|---|---|---|---|---|
| **OTOS** | 0.06557 | 0.01312 | 0.01188 | 0.01231 | 0.02568 | 0.01955 | 0.00219 |
| **OTP** | 0.08546 | 0.00841 | 0.01138 | 0.00917 | 0.01569 | 0.00156 | 0.07009 |
| **OTUB1** | 31.65689 | 25.23703 | 14.87268 | 31.71085 | 7.51663 | 18.17155 | 8.47182 |
| **OTUB2** | 0.35445 | 0.14068 | 0.12531 | 0.82508 | 0.03056 | 0.19363 | 0.06610 |
| **OTUD1** | 0.40223 | 0.92652 | 1.33196 | 0.10236 | 0.16837 | 5.61055 | 2.60430 |
| **OTUD3** | 4.87818 | 2.57960 | 0.75574 | 5.24168 | 4.21735 | 0.75294 | 1.87444 |
| **OTUD4** | 3.52164 | 0.21549 | 4.65382 | 1.49156 | 1.41910 | 4.18389 | 11.21685 |
| **OTUD5** | 0.71424 | 8.70130 | 5.16617 | 2.88894 | 4.09449 | 6.23653 | 16.97816 |
| **OTUD6A** | 0.06243 | 0.14761 | 0.62388 | 0.00000 | 0.67072 | 0.03549 | 0.00305 |
| **OTUD6B** | 12.36535 | 0.00000 | 6.29304 | 4.89454 | 1.04522 | 2.97126 | 5.26623 |
| **OTUD6B-AS1** | 9.67688 | 0.86093 | 11.49625 | 5.33377 | 1.15343 | 1.84678 | 4.09393 |
| **OTUD7A** | 0.02143 | 0.00000 | 0.08344 | 0.10921 | 0.05264 | 0.01771 | 0.01650 |
| **OTUD7B** | 0.55121 | 0.95124 | 0.72643 | 1.47844 | 0.54143 | 0.66820 | 4.42058 |
| **OTULIN** | 2.72408 | 2.58280 | 1.43861 | 2.41079 | 0.84466 | 2.24745 | 7.34683 |
| **OTX1** | 1.18624 | 0.57231 | 4.36831 | 0.00000 | 2.10472 | 1.43711 | 1.54828 |
| **OTX2** | 0.00000 | 3.60436 | 3.21638 | 0.20327 | 23.88024 | 7.34755 | 0.00000 |
| **OTX2-AS1** | 0.03673 | 0.01016 | 0.01419 | 0.01608 | 0.01022 | 0.03074 | 0.02142 |
| **OVCA2** | 0.37825 | 2.92473 | 0.87175 | 1.62081 | 0.22502 | 0.73872 | 0.43827 |
| **OVCH1** | 0.02883 | 0.02883 | 0.04297 | 0.02157 | 0.63406 | 0.02403 | 0.02798 |
| **OVCH1-AS1** | 0.12587 | 0.12700 | 0.10976 | 0.13290 | 0.15615 | 0.09384 | 0.00000 |
| **OVCH2** | 0.00000 | 0.00199 | 0.09184 | 0.05954 | 0.33588 | 0.00420 | 0.22088 |
| **OVGP1** | 0.61673 | 0.00000 | 0.00000 | 1.22282 | 0.72360 | 2.24934 | 1.64492 |
| **OVOL1** | 0.49987 | 17.28248 | 0.00000 | 36.54825 | 6.59181 | 6.36201 | 11.48598 |
| **OVOL1-AS1** | 0.01926 | 0.02275 | 0.01000 | 0.05549 | 0.02037 | 0.01579 | 0.01904 |
| **OVOL2** | 0.00000 | 4.82027 | 5.20435 | 0.00000 | 0.30836 | 3.47440 | 3.60947 |
| **OVOL3** | 0.10434 | 0.06068 | 0.08342 | 0.12169 | 0.14111 | 0.10970 | 0.02031 |
| **OXA1L** | 32.37986 | 18.66618 | 17.36755 | 16.42197 | 2.04729 | 15.71764 | 16.19739 |
| **OXCT1** | 0.20607 | 0.68936 | 3.14021 | 0.46220 | 0.54180 | 0.35503 | 0.28072 |
| **OXCT1-AS1** | 0.02281 | 0.02546 | 0.07463 | 0.01784 | 0.00910 | 0.01066 | 0.01568 |
| **OXCT2** | 0.01547 | 0.01335 | 0.01434 | 0.02533 | 0.00768 | 0.02579 | 0.03165 |
| **OXER1** | 0.01201 | 0.01535 | 0.01736 | 0.03191 | 0.00769 | 0.13729 | 0.02619 |
| **OXGR1** | 0.05627 | 0.00672 | 0.01673 | 0.09826 | 0.00797 | 0.03650 | 0.00000 |
| **OXLD1** | 0.00000 | 4.26897 | 2.37688 | 2.47933 | 0.02782 | 0.66938 | 1.75287 |
| **OXNAD1** | 10.35574 | 5.80787 | 5.86041 | 3.39848 | 1.90222 | 5.31393 | 0.00000 |
| **OXR1** | 2.51602 | 0.14449 | 0.88561 | 1.16056 | 0.18626 | 0.17486 | 4.38698 |

| | | | | | | | |
|---|---|---|---|---|---|---|---|
| **OXSM** | 8.62106 | 1.63896 | 9.30208 | 3.53093 | 0.88503 | 1.19691 | 0.56708 |
| **OXSR1** | 9.33247 | 1.65075 | 8.30056 | 6.92401 | 2.04990 | 4.28361 | 12.70908 |
| **OXT** | 2.24760 | 12.77970 | 2.77083 | 0.00000 | 6.62621 | 15.66313 | 1.39282 |
| **OXTR** | 0.31679 | 0.00000 | 0.10175 | 0.22289 | 0.00150 | 0.40541 | 1.83417 |
| **P2RX1** | 0.02099 | 0.01143 | 0.00775 | 0.03038 | 0.07483 | 0.05267 | 0.02409 |
| **P2RX2** | 0.04283 | 0.00759 | 0.01334 | 0.00561 | 0.00698 | 0.00646 | 0.12378 |
| **P2RX3** | 0.03727 | 0.00855 | 0.02949 | 0.02581 | 0.06511 | 0.01328 | 0.00405 |
| **P2RX4** | 2.58273 | 4.56025 | 0.16344 | 1.93040 | 1.01946 | 0.06092 | 2.10281 |
| **P2RX5** | 0.00000 | 2.71837 | 2.98595 | 0.23303 | 0.37097 | 0.38552 | 0.00000 |
| **P2RX5-TAX1BP3** | 0.33737 | 1.83081 | 0.67075 | 1.11099 | 0.13134 | 0.59919 | 1.18432 |
| **P2RX6** | 0.00000 | 0.01898 | 0.02357 | 0.26840 | 0.00593 | 0.06935 | 0.43388 |
| **P2RX6P** | 0.02338 | 0.01855 | 0.01253 | 0.02596 | 0.00791 | 0.01212 | 0.01830 |
| **P2RX7** | 0.11642 | 0.07495 | 0.01539 | 0.02935 | 0.03509 | 0.04671 | 0.00224 |
| **P2RY1** | 0.02456 | 0.00000 | 0.18586 | 0.07307 | 0.27166 | 0.08543 | 0.00154 |
| **P2RY11** | 0.01462 | 0.48038 | 0.11618 | 0.08227 | 0.07135 | 0.12386 | 0.72410 |
| **P2RY12** | 0.07749 | 0.00000 | 0.06497 | 0.43859 | 0.02077 | 0.02097 | 0.26536 |
| **P2RY13** | 0.00000 | 0.00000 | 0.07436 | 0.49500 | 0.00398 | 0.05216 | 0.76622 |
| **P2RY14** | 0.21523 | 0.00000 | 0.14136 | 1.34597 | 0.01307 | 0.03459 | 0.68504 |
| **P2RY2** | 0.89923 | 0.00000 | 0.00000 | 1.17598 | 0.00276 | 0.00000 | 0.93703 |
| **P2RY4** | 0.03885 | 0.00614 | 0.02993 | 0.01419 | 0.00813 | 0.01376 | 0.02359 |
| **P2RY6** | 12.53176 | 4.88861 | 0.00000 | 24.14737 | 0.00000 | 0.00000 | 0.80751 |
| **P4HA1** | 6.33604 | 0.00000 | 4.47066 | 2.72644 | 3.06630 | 5.23898 | 0.00899 |
| **P4HA2** | 4.22450 | 4.38056 | 1.18142 | 1.83724 | 0.11334 | 1.26050 | 0.00000 |
| **P4HA2-AS1** | 0.16284 | 0.00842 | 0.06014 | 0.00373 | 0.00467 | 0.03679 | 0.04226 |
| **P4HA3** | 0.00000 | 0.30889 | 0.36633 | 0.02317 | 0.09218 | 0.07614 | 0.01027 |
| **P4HB** | 40.91730 | 32.20632 | 15.84184 | 43.67320 | 5.46620 | 22.62425 | 34.25612 |
| **P4HTM** | 0.00000 | 0.01075 | 0.02142 | 0.35167 | 0.02306 | 0.10375 | 0.07861 |
| **PA2G4** | 42.66725 | 31.91297 | 21.64156 | 39.22631 | 7.10294 | 20.95284 | 37.66337 |
| **PA2G4P4** | 9.06566 | 3.82734 | 3.94634 | 6.71764 | 0.76354 | 3.45105 | 5.61024 |
| **PAAF1** | 4.96836 | 7.51942 | 2.34005 | 0.53108 | 1.86512 | 3.81741 | 0.00000 |
| **PABPC1** | 51.28949 | 42.86288 | 27.59084 | 40.50590 | 11.70020 | 31.22360 | 74.25650 |
| **PABPC1L** | 0.47820 | 0.00000 | 0.29217 | 1.72562 | 0.99715 | 1.05063 | 3.53605 |
| **PABPC1L2A** | 0.01523 | 0.01645 | 0.02335 | 0.01408 | 0.00666 | 0.04998 | 0.04267 |
| **PABPC1L2B** | 0.03337 | 0.00569 | 0.04529 | 0.01671 | 0.00641 | 0.01592 | 0.01957 |
| **PABPC1P2** | 0.13467 | 0.08647 | 0.06035 | 0.03868 | 0.08718 | 0.06342 | 0.19436 |

| | | | | | | | |
|---|---|---|---|---|---|---|---|
| **PABPC3** | 9.44092 | 6.57500 | 6.22824 | 7.45018 | 1.25725 | 7.52697 | 32.94496 |
| **PABPC4** | 30.96403 | 27.25153 | 14.24892 | 7.90370 | 2.17129 | 3.69934 | 43.60148 |
| **PABPC4L** | 0.05520 | 0.00000 | 0.29654 | 0.00813 | 0.02606 | 0.03618 | 0.02004 |
| **PABPN1** | 0.00000 | 1.24281 | 0.19733 | 0.53170 | 0.14894 | 0.46723 | 0.71431 |
| **PABPN1L** | 0.00000 | 0.15683 | 0.04357 | 0.00241 | 16.18483 | 0.00000 | 0.02200 |
| **PACRG** | 0.03016 | 0.03546 | 0.01307 | 0.02748 | 0.04405 | 0.01230 | 0.00976 |
| **PACRG-AS1** | 0.02655 | 0.01719 | 0.01302 | 0.01996 | 0.00797 | 0.01352 | 0.01453 |
| **PACRGL** | 1.90438 | 1.17824 | 2.56774 | 1.97326 | 1.39619 | 1.13088 | 1.10876 |
| **PACS1** | 0.00000 | 1.76707 | 0.00000 | 0.52177 | 0.56411 | 0.91899 | 5.67662 |
| **PACS2** | 0.00000 | 1.94530 | 0.00032 | 0.67353 | 0.80911 | 1.13651 | 1.42003 |
| **PACSIN1** | 0.50786 | 0.08073 | 0.02830 | 0.30171 | 0.00321 | 0.07700 | 0.35129 |
| **PACSIN2** | 13.70262 | 25.78270 | 3.14753 | 19.44513 | 3.33411 | 10.48117 | 0.97284 |
| **PACSIN3** | 8.94515 | 17.69233 | 1.08750 | 3.22549 | 0.11525 | 9.49468 | 7.27332 |
| **PADI1** | 0.02484 | 0.01032 | 0.04508 | 0.04737 | 0.02643 | 0.01314 | 0.01052 |
| **PADI2** | 0.23390 | 0.20519 | 0.06862 | 0.00009 | 0.08335 | 0.00102 | 0.07922 |
| **PADI3** | 0.05555 | 0.01242 | 0.02394 | 0.02257 | 0.10104 | 0.01613 | 0.00000 |
| **PADI4** | 0.02706 | 0.01284 | 0.02159 | 0.01765 | 0.02252 | 0.01291 | 0.01925 |
| **PADI6** | 0.00000 | 0.96153 | 0.09191 | 0.00000 | 13.10715 | 0.00001 | 0.01561 |
| **PAEP** | 0.00477 | 0.02315 | 0.01266 | 0.35645 | 0.00803 | 0.01199 | 0.00628 |
| **PAF1** | 15.93519 | 19.30343 | 9.98263 | 16.09712 | 5.40810 | 8.64323 | 2.24879 |
| **PAFAH1B1** | 11.62235 | 10.54224 | 10.88136 | 11.09056 | 1.81804 | 8.26949 | 18.82879 |
| **PAFAH1B2** | 41.56757 | 22.06231 | 18.99910 | 17.68834 | 17.29418 | 16.03687 | 4.16826 |
| **PAFAH1B3** | 15.89888 | 10.88627 | 9.68474 | 15.39266 | 1.25141 | 10.49776 | 7.67461 |
| **PAFAH2** | 7.62487 | 9.02431 | 5.86441 | 3.31840 | 3.13332 | 2.22525 | 0.85880 |
| **PAG1** | 0.00000 | 0.00000 | 0.15986 | 0.08644 | 2.26583 | 0.84637 | 0.10953 |
| **PAGE1** | 4.00914 | 7.21113 | 0.00000 | 0.00000 | 0.61902 | 0.00000 | 1.10546 |
| **PAGE2** | 4.11014 | 10.47298 | 4.37156 | 2.76809 | 0.00000 | 1.33685 | 0.45424 |
| **PAGE2B** | 0.82592 | 7.75443 | 6.24487 | 0.52814 | 0.03062 | 0.32058 | 0.45935 |
| **PAGE4** | 0.21938 | 0.00000 | 0.01926 | 2.39309 | 0.22187 | 0.03387 | 0.00000 |
| **PAGE5** | 2.71041 | 13.45832 | 7.23318 | 1.09132 | 0.00000 | 0.98752 | 0.75885 |
| **PAGR1** | 1.10680 | 11.84626 | 4.79695 | 13.04797 | 1.84021 | 8.85516 | 20.52065 |
| **PAICS** | 44.44898 | 13.67778 | 5.44536 | 22.85686 | 0.44152 | 14.19077 | 22.42952 |
| **PAIP1** | 7.02514 | 9.62026 | 2.90089 | 2.61317 | 8.77593 | 5.36783 | 3.29583 |
| **PAIP2** | 31.92079 | 9.74211 | 21.39379 | 28.48170 | 9.91478 | 17.34477 | 2.89922 |
| **PAIP2B** | 1.68418 | 0.11271 | 0.34778 | 7.81673 | 2.51414 | 1.32895 | 0.00000 |

| | | | | | | | |
|---|---|---|---|---|---|---|---|
| **PAK1** | 3.85976 | 3.26255 | 1.57874 | 2.37705 | 1.48434 | 9.13059 | 3.80143 |
| **PAK1IP1** | 31.84084 | 8.59434 | 20.47082 | 18.80374 | 7.51700 | 10.52658 | 8.91986 |
| **PAK2** | 10.01385 | 2.25847 | 7.33119 | 11.13774 | 1.59647 | 10.69120 | 24.51515 |
| **PAK3** | 0.01101 | 0.00684 | 0.03000 | 2.20921 | 0.02319 | 0.01291 | 0.00208 |
| **PAK4** | 0.00000 | 4.79791 | 1.60113 | 6.63275 | 0.79410 | 3.02650 | 16.01715 |
| **PAK6** | 1.10759 | 0.24129 | 0.12042 | 4.06058 | 0.02981 | 0.03236 | 1.28826 |
| **PAK7** | 0.04832 | 0.02107 | 0.00579 | 0.01159 | 0.17543 | 0.01094 | 0.01470 |
| **PALB2** | 5.47310 | 0.23921 | 3.54622 | 0.80212 | 1.96494 | 1.98627 | 1.73682 |
| **PALD1** | 0.00000 | 0.01636 | 0.01041 | 0.04068 | 0.08547 | 0.30081 | 0.11539 |
| **PALLD** | 17.69465 | 2.38148 | 3.40405 | 8.39353 | 0.39462 | 0.00000 | 25.83800 |
| **PALM** | 0.00000 | 0.05497 | 2.09722 | 0.00488 | 0.32351 | 0.00000 | 0.02089 |
| **PALM2** | 0.17104 | 0.00000 | 0.06781 | 0.00000 | 0.11928 | 0.01092 | 0.05584 |
| **PALM3** | 0.00000 | 0.00000 | 0.91068 | 1.96737 | 0.49055 | 0.00000 | 1.99364 |
| **PALMD** | 0.05808 | 0.01265 | 0.01381 | 0.02383 | 0.02292 | 0.01600 | 0.02219 |
| **PAM** | 0.41321 | 2.26628 | 0.14436 | 0.26282 | 0.12202 | 0.00000 | 0.00000 |
| **PAM16** | 0.61785 | 0.60406 | 0.96971 | 0.51306 | 0.89589 | 0.44884 | 0.34995 |
| **PAMR1** | 0.00000 | 1.00789 | 0.02048 | 0.14634 | 1.33011 | 4.54084 | 0.00000 |
| **PAN2** | 1.15871 | 0.62768 | 0.43503 | 0.43416 | 1.80829 | 3.63064 | 0.00000 |
| **PAN3** | 22.34669 | 4.51239 | 9.60982 | 4.08149 | 9.32424 | 9.99834 | 0.00000 |
| **PAN3-AS1** | 0.14245 | 0.02708 | 0.17526 | 0.06641 | 0.09208 | 0.21377 | 0.28794 |
| **PANDAR** | 2.94283 | 0.96065 | 0.79190 | 1.37913 | 0.64615 | 1.15812 | 0.00000 |
| **PANK1** | 0.76170 | 0.00000 | 0.00000 | 2.37546 | 0.09961 | 1.73114 | 1.05420 |
| **PANK2** | 3.51745 | 1.58841 | 1.14456 | 3.12246 | 1.08918 | 3.81495 | 7.02655 |
| **PANK3** | 7.34588 | 1.25521 | 6.65549 | 1.07487 | 0.44403 | 8.40791 | 9.96862 |
| **PANK4** | 4.31580 | 5.82568 | 0.58473 | 4.08037 | 0.41914 | 1.94870 | 0.00000 |
| **PANO** | 0.00000 | 0.07201 | 0.01703 | 0.04325 | 0.01682 | 0.01389 | 0.14600 |
| **PANX1** | 0.17806 | 0.32728 | 4.94349 | 2.70847 | 3.54221 | 0.13835 | 6.72082 |
| **PANX2** | 0.00000 | 0.33516 | 0.00248 | 0.14801 | 0.00603 | 0.00254 | 0.45874 |
| **PANX3** | 0.02177 | 0.01756 | 0.01274 | 0.03460 | 0.00801 | 0.02809 | 0.01406 |
| **PAOX** | 0.40437 | 0.85422 | 0.17111 | 0.00000 | 0.53273 | 0.22907 | 0.44182 |
| **PAPD4** | 15.46845 | 1.96082 | 4.36471 | 5.11005 | 3.26751 | 4.74443 | 2.98558 |
| **PAPD5** | 11.72945 | 2.45109 | 6.43509 | 4.42580 | 5.32347 | 7.20430 | 15.66188 |
| **PAPD7** | 2.95491 | 4.60974 | 2.25716 | 3.10254 | 5.16043 | 1.88087 | 12.65618 |
| **PAPL** | 0.00000 | 0.22717 | 0.14086 | 0.03075 | 0.01290 | 0.06442 | 0.01564 |
| **PAPLN** | 0.07275 | 0.40034 | 0.16148 | 0.45726 | 0.53897 | 0.37163 | 0.33076 |

| | | | | | | | |
|---|---|---|---|---|---|---|---|
| **PAPOLA** | 29.18047 | 7.33633 | 27.84696 | 52.23236 | 9.92994 | 11.58883 | 36.53900 |
| **PAPOLB** | 0.44803 | 0.03857 | 1.23717 | 3.16843 | 0.46917 | 0.20776 | 0.87284 |
| **PAPOLG** | 5.98251 | 0.00000 | 5.56087 | 3.97721 | 4.83116 | 2.86182 | 1.61385 |
| **PAPPA** | 0.30252 | 0.33966 | 0.08192 | 0.00000 | 0.14356 | 0.07828 | 0.01472 |
| **PAPPA2** | 0.08567 | 0.00000 | 0.09191 | 0.01967 | 0.18445 | 0.06535 | 0.02669 |
| **PAPSS1** | 8.48335 | 2.95873 | 3.22303 | 6.95487 | 1.45571 | 8.00997 | 0.00000 |
| **PAPSS2** | 0.16413 | 0.00000 | 0.26874 | 0.00000 | 0.14724 | 2.55676 | 0.14320 |
| **PAQR3** | 3.94100 | 0.00000 | 1.76956 | 0.76543 | 0.00000 | 4.29186 | 6.89615 |
| **PAQR4** | 0.00000 | 0.88410 | 0.17358 | 1.22943 | 0.12636 | 1.38860 | 1.75938 |
| **PAQR5** | 0.17859 | 0.08705 | 0.55287 | 0.11240 | 0.64847 | 0.15829 | 0.00000 |
| **PAQR6** | 0.31687 | 0.00000 | 0.68256 | 0.67020 | 0.00052 | 0.02353 | 0.72398 |
| **PAQR7** | 1.24117 | 2.01133 | 0.00000 | 2.57792 | 6.82730 | 0.45800 | 0.87561 |
| **PAQR8** | 0.19421 | 0.07869 | 0.15834 | 0.17417 | 0.02638 | 0.69387 | 0.43294 |
| **PAQR9** | 0.02432 | 0.01294 | 0.02282 | 0.01917 | 0.01586 | 0.01300 | 0.01858 |
| **PARD3** | 1.93314 | 7.84240 | 6.66160 | 5.11079 | 2.44717 | 13.42153 | 19.13407 |
| **PARD3-AS1** | 0.03357 | 0.01452 | 0.01411 | 0.02652 | 0.03134 | 0.01187 | 0.02201 |
| **PARD3B** | 0.00000 | 0.00957 | 0.03176 | 0.49117 | 0.96760 | 0.00000 | 0.79195 |
| **PARD6A** | 0.66337 | 0.00000 | 0.02484 | 0.93066 | 0.21333 | 0.00301 | 0.08342 |
| **PARD6B** | 41.56704 | 8.36670 | 14.86104 | 10.87185 | 10.43816 | 8.75053 | 5.15137 |
| **PARD6G** | 0.03738 | 0.03811 | 0.57892 | 0.14000 | 0.97379 | 0.41835 | 0.20420 |
| **PARD6G-AS1** | 0.10721 | 0.00735 | 0.02788 | 0.05381 | 0.05410 | 0.02685 | 0.01777 |
| **PARG** | 6.31319 | 0.82592 | 5.08791 | 1.92463 | 0.77754 | 2.71068 | 1.09411 |
| **PARK2** | 0.00122 | 1.99341 | 4.50692 | 0.00000 | 0.51275 | 2.01859 | 0.00247 |
| **PARK7** | 46.44729 | 28.52725 | 15.20023 | 36.14763 | 5.05548 | 19.64836 | 15.22242 |
| **PARL** | 15.56453 | 13.75538 | 9.59336 | 14.45722 | 22.95058 | 9.08360 | 2.20473 |
| **PARM1** | 0.00000 | 1.69123 | 1.34921 | 0.09185 | 0.00000 | 2.01604 | 0.05698 |
| **PARN** | 20.22590 | 13.40996 | 11.62920 | 10.54580 | 5.81832 | 15.69520 | 2.37152 |
| **PARP1** | 35.78971 | 29.13039 | 14.87975 | 21.54366 | 3.75832 | 31.47381 | 19.23654 |
| **PARP10** | 0.03544 | 0.01649 | 0.01299 | 0.03168 | 0.00784 | 0.01729 | 0.01621 |
| **PARP11** | 1.52967 | 0.49417 | 1.73789 | 0.46746 | 2.07928 | 0.61051 | 0.44494 |
| **PARP12** | 0.00000 | 2.97295 | 0.00000 | 0.00000 | 2.11005 | 1.41123 | 0.16138 |
| **PARP14** | 0.00000 | 0.56928 | 0.59046 | 0.00000 | 0.06730 | 1.29056 | 0.12954 |
| **PARP15** | 0.05119 | 0.01976 | 0.01732 | 0.01569 | 0.00766 | 0.02270 | 0.00000 |
| **PARP16** | 0.78015 | 0.81738 | 2.31019 | 0.14800 | 4.10260 | 3.63539 | 0.01089 |
| **PARP2** | 28.38537 | 5.20149 | 17.77559 | 10.20242 | 3.80591 | 7.70736 | 0.99806 |

| | | | | | | | |
|---|---|---|---|---|---|---|---|
| **PARP3** | 0.11588 | 0.08187 | 0.10588 | 0.22220 | 0.02160 | 0.43711 | 0.00000 |
| **PARP4** | 1.35614 | 0.26937 | 1.36612 | 0.81910 | 0.05581 | 1.01122 | 0.00000 |
| **PARP6** | 3.08813 | 0.75526 | 0.66406 | 0.84937 | 0.09254 | 2.74276 | 6.17706 |
| **PARP8** | 0.12302 | 0.00000 | 0.22725 | 0.21100 | 0.19204 | 0.35881 | 0.19569 |
| **PARP9** | 0.00000 | 0.08908 | 0.00245 | 0.03834 | 0.14140 | 1.85402 | 0.00858 |
| **PARPBP** | 13.77854 | 0.00000 | 8.00378 | 4.96581 | 3.26291 | 4.75023 | 0.48015 |
| **PARS2** | 2.92879 | 3.61534 | 4.38659 | 2.61923 | 2.21882 | 1.04342 | 0.00000 |
| **PART1** | 0.02027 | 0.02506 | 0.00694 | 0.09385 | 0.15866 | 0.04797 | 0.03751 |
| **PARVA** | 9.80098 | 5.63603 | 9.34174 | 18.18112 | 1.24327 | 4.68858 | 4.17577 |
| **PARVB** | 0.00000 | 0.06450 | 0.05100 | 0.07043 | 0.02364 | 0.32630 | 0.31648 |
| **PARVG** | 0.02067 | 0.01221 | 0.00393 | 0.01567 | 0.83163 | 0.01145 | 0.02508 |
| **PASD1** | 0.00000 | 0.09559 | 1.79147 | 0.00722 | 0.08516 | 0.00611 | 0.01620 |
| **PASK** | 5.93547 | 5.40254 | 1.85579 | 6.04827 | 0.76702 | 2.18433 | 3.76661 |
| **PATE1** | 0.02385 | 0.01621 | 0.01589 | 0.02107 | 0.00780 | 0.01289 | 0.01682 |
| **PATE2** | 0.04953 | 0.29491 | 0.24432 | 0.07525 | 0.00165 | 0.00000 | 0.03412 |
| **PATE3** | 0.02377 | 0.02022 | 0.01145 | 0.01785 | 0.00774 | 0.00896 | 0.05252 |
| **PATE4** | 0.02925 | 0.01051 | 0.02073 | 0.02877 | 0.02247 | 0.01035 | 0.02966 |
| **PATL1** | 7.21399 | 0.53482 | 5.37202 | 3.99516 | 1.99870 | 5.74022 | 12.94645 |
| **PATL2** | 0.00000 | 0.25621 | 0.00000 | 0.17357 | 8.66622 | 0.00000 | 0.09863 |
| **PATZ1** | 0.78604 | 0.10793 | 1.31041 | 2.18522 | 0.08381 | 2.25813 | 2.34181 |
| **PAWR** | 1.32060 | 0.00000 | 1.27250 | 2.66079 | 0.00134 | 0.80398 | 12.58658 |
| **PAX2** | 0.03802 | 0.01703 | 0.01728 | 0.02198 | 0.01478 | 0.01698 | 0.03273 |
| **PAX3** | 0.03600 | 0.01873 | 0.03150 | 0.07131 | 0.01967 | 0.06438 | 0.01066 |
| **PAX4** | 0.02123 | 0.01650 | 0.02262 | 0.05841 | 0.00760 | 0.01448 | 0.00814 |
| **PAX5** | 0.00000 | 0.00000 | 0.21337 | 0.11227 | 0.11142 | 0.39316 | 0.82912 |
| **PAX6** | 0.26451 | 0.00000 | 0.24320 | 0.02545 | 0.21376 | 0.23779 | 0.42037 |
| **PAX7** | 0.02352 | 0.01217 | 0.02108 | 0.01769 | 0.01464 | 0.01233 | 0.02649 |
| **PAX8** | 0.22117 | 0.25898 | 0.69976 | 0.20268 | 0.25084 | 0.18678 | 0.00000 |
| **PAX8-AS1.uid-3** | 3.11656 | 3.78944 | 1.02412 | 2.43546 | 0.00196 | 1.89435 | 1.18688 |
| **PAX8-AS1.uid-3** | 9.98326 | 7.89021 | 2.12414 | 4.42971 | 0.00000 | 1.93872 | 0.00000 |
| **PAX9** | 0.77154 | 0.00000 | 1.42156 | 0.27689 | 0.93948 | 0.01051 | 0.00000 |
| **PAXBP1** | 8.04489 | 0.38333 | 5.29822 | 1.18621 | 3.31208 | 3.49516 | 10.24069 |
| **PAXBP1-AS1** | 0.21190 | 0.00000 | 0.08846 | 0.07472 | 0.18174 | 0.02369 | 0.91779 |
| **PAXIP1** | 3.73928 | 1.26989 | 5.22042 | 1.87679 | 1.46932 | 6.18402 | 9.27905 |
| **PAXIP1-AS1** | 0.12843 | 0.56640 | 1.65514 | 1.19064 | 0.04786 | 0.35114 | 2.04485 |

| | | | | | | | |
|---|---|---|---|---|---|---|---|
| **PAXIP1OS** | 0.23646 | 0.03182 | 0.23087 | 0.13791 | 0.07099 | 0.32821 | 0.61324 |
| **PBDC1** | 23.36995 | 12.86707 | 21.49145 | 4.72156 | 2.84140 | 6.69128 | 12.57152 |
| **PBK** | 14.83820 | 3.87351 | 9.36111 | 3.34151 | 0.84689 | 2.81419 | 4.48678 |
| **PBLD** | 0.10584 | 0.28416 | 0.24877 | 0.06120 | 0.17855 | 0.01253 | 0.19185 |
| **PBOV1** | 0.02211 | 0.01713 | 0.01444 | 0.02299 | 0.01029 | 0.01416 | 0.02156 |
| **PBRM1** | 10.70290 | 1.83341 | 3.14491 | 7.34784 | 1.40160 | 6.47445 | 20.77677 |
| **PBX1** | 0.87534 | 0.00000 | 0.02615 | 0.00000 | 1.64061 | 3.42817 | 0.77971 |
| **PBX2** | 0.06174 | 0.47171 | 0.25957 | 0.75463 | 0.00000 | 1.05954 | 3.54623 |
| **PBX3** | 0.03738 | 0.84639 | 4.68884 | 0.00000 | 1.17126 | 6.44086 | 2.42876 |
| **PBX4** | 1.24670 | 1.90581 | 3.87243 | 1.17116 | 0.00000 | 2.57185 | 0.84402 |
| **PBXIP1** | 0.52580 | 2.17681 | 0.00000 | 2.41001 | 0.09271 | 0.73189 | 0.87217 |
| **PC** | 1.78535 | 1.49671 | 0.67880 | 3.36988 | 0.66621 | 1.44469 | 2.36801 |
| **PCA3** | 0.00000 | 0.01166 | 0.12692 | 0.02007 | 0.00723 | 0.30765 | 0.11916 |
| **PCAT1** | 0.06049 | 0.08256 | 0.03234 | 0.00499 | 0.01436 | 0.08714 | 0.02459 |
| **PCAT18** | 0.05434 | 0.02277 | 0.02341 | 0.20768 | 0.00901 | 0.07930 | 0.03059 |
| **PCAT19** | 0.00692 | 0.01596 | 0.04368 | 0.09733 | 0.08140 | 0.03955 | 0.02715 |
| **PCAT2** | 0.02277 | 0.01679 | 0.01301 | 0.01892 | 0.00766 | 0.01983 | 0.03103 |
| **PCAT4** | 0.36887 | 0.00000 | 0.17955 | 0.00064 | 0.45613 | 0.19230 | 0.00470 |
| **PCAT6** | 0.41325 | 1.52297 | 0.00000 | 1.34665 | 1.31179 | 0.00000 | 0.71513 |
| **PCBD1** | 8.69183 | 1.17497 | 0.13888 | 6.29018 | 0.03086 | 3.31804 | 1.96348 |
| **PCBD2** | 4.28363 | 4.25416 | 2.15762 | 0.00000 | 0.00292 | 3.59370 | 2.74658 |
| **PCBP1** | 20.18566 | 31.08087 | 16.61284 | 26.79282 | 4.66334 | 23.44198 | 54.05814 |
| **PCBP1-AS1** | 0.59991 | 0.04277 | 0.23957 | 0.27989 | 0.21297 | 0.15836 | 0.12309 |
| **PCBP2** | 17.24277 | 7.97523 | 12.16514 | 11.21805 | 5.61767 | 12.52904 | 46.36920 |
| **PCBP2-OT1** | 0.14820 | 0.17706 | 0.13970 | 0.09366 | 0.03582 | 0.25444 | 1.04753 |
| **PCBP3** | 0.00000 | 0.62486 | 0.08935 | 0.00000 | 0.02912 | 0.73972 | 0.27871 |
| **PCBP4** | 0.06294 | 0.19293 | 0.83937 | 0.98094 | 1.30795 | 0.72941 | 2.22605 |
| **PCCA** | 2.68236 | 0.49371 | 0.31602 | 2.72546 | 0.27909 | 2.71467 | 0.00000 |
| **PCCA-AS1** | 0.04960 | 0.02798 | 0.01448 | 0.04244 | 0.00752 | 0.02150 | 0.01119 |
| **PCCB** | 18.80920 | 15.00718 | 4.07278 | 16.89787 | 2.99400 | 8.35148 | 0.00000 |
| **PCDH1** | 9.78658 | 15.51241 | 0.00000 | 0.00000 | 0.00529 | 4.60593 | 10.40885 |
| **PCDH10** | 0.00712 | 0.00545 | 0.01190 | 0.15475 | 0.19226 | 0.00759 | 0.20391 |
| **PCDH11X** | 0.01606 | 0.01670 | 0.01030 | 0.00893 | 2.21344 | 0.02220 | 0.07804 |
| **PCDH11Y** | 0.02353 | 0.01661 | 0.01186 | 0.01946 | 0.02584 | 0.01376 | 0.01906 |
| **PCDH12** | 0.02777 | 0.00000 | 0.02678 | 0.08889 | 0.06203 | 0.05207 | 0.19943 |

| | | | | | | | |
|---|---|---|---|---|---|---|---|
| **PCDH15** | 0.12148 | 0.48568 | 0.74219 | 0.00000 | 2.94734 | 1.02355 | 0.00022 |
| **PCDH17** | 0.25768 | 0.00000 | 0.00332 | 0.55078 | 0.00523 | 1.04864 | 1.39796 |
| **PCDH18** | 0.06556 | 0.00078 | 0.01179 | 0.02411 | 0.01918 | 0.40071 | 0.00000 |
| **PCDH19** | 0.06918 | 0.07341 | 0.00000 | 0.00674 | 0.90271 | 0.12427 | 0.00336 |
| **PCDH20** | 0.01523 | 0.00000 | 0.15734 | 0.02354 | 0.06557 | 0.04429 | 0.02394 |
| **PCDH7** | 0.02037 | 0.02829 | 0.00679 | 0.02507 | 0.05648 | 0.22682 | 0.02717 |
| **PCDH8** | 0.03246 | 0.25818 | 0.02453 | 0.00000 | 0.35937 | 0.47919 | 0.00645 |
| **PCDH9** | 0.35907 | 0.00924 | 0.00445 | 0.02326 | 3.46601 | 0.21519 | 0.00000 |
| **PCDH9-AS3** | 0.02585 | 0.01603 | 0.01593 | 0.02353 | 0.00947 | 0.01296 | 0.01474 |
| **PCDHA1** | 0.04490 | 0.00000 | 0.30566 | 0.00708 | 0.19391 | 0.00348 | 0.03660 |
| **PCDHA10** | 0.04783 | 0.01803 | 0.13265 | 0.08465 | 0.00610 | 0.04447 | 0.00000 |
| **PCDHA11** | 0.04256 | 0.00177 | 0.11589 | 0.05845 | 0.00816 | 0.01183 | 0.00000 |
| **PCDHA12** | 0.10691 | 0.05224 | 0.03884 | 0.11469 | 0.02324 | 0.04358 | 0.00000 |
| **PCDHA13** | 0.03135 | 0.04962 | 0.04547 | 0.01540 | 0.02205 | 0.01052 | 0.01062 |
| **PCDHA2** | 0.04560 | 0.00788 | 0.03784 | 0.02789 | 0.00898 | 0.01554 | 0.00110 |
| **PCDHA3** | 0.02814 | 0.00000 | 0.09087 | 0.02047 | 0.00549 | 0.01699 | 0.02187 |
| **PCDHA4** | 0.02450 | 0.01767 | 0.01334 | 0.05034 | 0.00789 | 0.01209 | 0.01302 |
| **PCDHA5** | 0.01251 | 0.07176 | 0.02758 | 0.01902 | 0.00651 | 0.00584 | 0.03312 |
| **PCDHA6** | 0.02446 | 0.01071 | 0.02548 | 0.01696 | 0.01116 | 0.01211 | 0.02682 |
| **PCDHA8** | 0.02018 | 0.01598 | 0.01549 | 0.02018 | 0.00917 | 0.01173 | 0.02917 |
| **PCDHA9** | 0.05133 | 0.01042 | 0.01831 | 0.04131 | 0.01184 | 0.01489 | 0.03251 |
| **PCDHAC2** | 0.02200 | 0.00914 | 0.01408 | 0.01846 | 0.00767 | 0.13916 | 0.02013 |
| **PCDHB1** | 0.01723 | 0.01922 | 0.10241 | 0.02256 | 0.07383 | 0.01198 | 0.01909 |
| **PCDHB11** | 0.02562 | 0.01746 | 0.01301 | 0.02035 | 0.00896 | 0.01241 | 0.02104 |
| **PCDHB12** | 0.02456 | 0.01664 | 0.03566 | 0.02080 | 0.00777 | 0.01163 | 0.01683 |
| **PCDHB13** | 0.02099 | 0.01255 | 0.01859 | 0.01848 | 0.00727 | 0.01119 | 0.03618 |
| **PCDHB14** | 0.03865 | 0.00387 | 0.02630 | 0.02498 | 0.01462 | 0.01313 | 0.02540 |
| **PCDHB15** | 0.03635 | 0.01061 | 0.02093 | 0.01711 | 0.00740 | 0.01555 | 0.02173 |
| **PCDHB16** | 0.02148 | 0.07583 | 0.04867 | 0.01831 | 0.00625 | 0.02619 | 0.00059 |
| **PCDHB18** | 0.02279 | 0.01071 | 0.03561 | 0.01660 | 0.00882 | 0.01149 | 0.03165 |
| **PCDHB19P** | 0.01453 | 0.00000 | 0.37749 | 0.01625 | 0.32704 | 0.00596 | 0.02149 |
| **PCDHB2** | 0.00000 | 0.46485 | 1.38266 | 0.12232 | 0.13847 | 0.01190 | 0.03210 |
| **PCDHB3** | 0.17563 | 0.00000 | 1.29865 | 0.11678 | 0.06254 | 0.01260 | 0.04859 |
| **PCDHB4** | 0.03510 | 0.01449 | 0.06564 | 0.04251 | 0.03805 | 0.01089 | 0.00254 |
| **PCDHB5** | 0.05749 | 0.01085 | 0.72639 | 0.02083 | 0.01204 | 0.00324 | 0.00000 |

| | | | | | | | |
|---|---|---|---|---|---|---|---|
| **PCDHB6** | 0.01530 | 0.04343 | 0.04904 | 0.03020 | 0.00844 | 0.00928 | 0.01372 |
| **PCDHB7** | 0.01829 | 0.01675 | 0.06578 | 0.02368 | 0.02618 | 0.01869 | 0.01584 |
| **PCDHB8** | 0.03208 | 0.00240 | 0.06086 | 0.01763 | 0.00608 | 0.01724 | 0.01278 |
| **PCDHB9** | 0.02065 | 0.01073 | 0.02400 | 0.02052 | 0.00707 | 0.02140 | 0.02829 |
| **PCDHGA10** | 0.07079 | 0.00000 | 0.28638 | 0.00033 | 0.12661 | 0.00297 | 0.11054 |
| **PCDHGA11** | 0.08442 | 0.00776 | 0.07125 | 0.05136 | 0.08500 | 0.06898 | 0.06957 |
| **PCDHGA12** | 0.01854 | 0.00000 | 0.75919 | 0.01761 | 0.61743 | 0.00434 | 0.01670 |
| **PCDHGA2** | 0.01376 | 0.00000 | 0.64849 | 0.01478 | 0.00349 | 0.00332 | 0.02329 |
| **PCDHGA3** | 0.02534 | 0.03282 | 0.08791 | 0.01820 | 0.00549 | 0.00843 | 0.00830 |
| **PCDHGA4** | 0.03609 | 0.00575 | 0.29998 | 0.01860 | 0.03037 | 0.00924 | 0.00493 |
| **PCDHGA5** | 0.02720 | 0.00749 | 0.04179 | 0.01909 | 0.00745 | 0.01299 | 0.01658 |
| **PCDHGA6** | 0.01788 | 0.02780 | 0.01660 | 0.01843 | 0.06055 | 0.01134 | 0.02755 |
| **PCDHGA7** | 0.01766 | 0.00000 | 0.09082 | 0.05651 | 0.00793 | 0.00835 | 0.03067 |
| **PCDHGA8** | 0.03124 | 0.00000 | 0.11789 | 0.01458 | 0.00565 | 0.00860 | 0.02671 |
| **PCDHGA9** | 0.02571 | 0.00000 | 0.08908 | 0.02133 | 0.07487 | 0.01093 | 0.02126 |
| **PCDHGB1** | 0.03271 | 0.00168 | 0.04933 | 0.01810 | 0.00592 | 0.01601 | 0.02930 |
| **PCDHGB2** | 0.02271 | 0.00368 | 0.09826 | 0.02355 | 0.00543 | 0.01232 | 0.02062 |
| **PCDHGB4** | 0.05822 | 0.00061 | 0.06187 | 0.01894 | 0.00627 | 0.06508 | 0.00000 |
| **PCDHGB5** | 0.05595 | 0.00452 | 0.17205 | 0.07081 | 0.00397 | 0.00942 | 0.00000 |
| **PCDHGB6** | 0.03135 | 0.00564 | 0.04078 | 0.07205 | 0.02920 | 0.01252 | 0.00085 |
| **PCDHGB7** | 0.02982 | 0.01723 | 0.04065 | 0.02338 | 0.01673 | 0.01053 | 0.01392 |
| **PCDHGB8P** | 0.01734 | 0.01932 | 0.08100 | 0.03101 | 0.29585 | 0.01806 | 0.00988 |
| **PCDHGC3** | 0.03926 | 0.01447 | 0.02669 | 0.10549 | 0.00935 | 0.34191 | 0.00000 |
| **PCDHGC4** | 0.02300 | 0.00000 | 0.11921 | 0.02292 | 0.00758 | 0.01515 | 0.02078 |
| **PCDHGC5** | 0.02396 | 0.00819 | 0.02258 | 0.01635 | 0.00722 | 0.04190 | 0.02451 |
| **PCDP1** | 0.06176 | 0.03520 | 0.05242 | 0.01274 | 0.21994 | 0.12422 | 0.00000 |
| **PCED1A** | 0.00000 | 0.27720 | 0.31684 | 0.84598 | 0.01137 | 0.44351 | 0.48313 |
| **PCED1B** | 0.19371 | 0.33082 | 1.62744 | 0.48640 | 0.02657 | 4.65940 | 0.08100 |
| **PCED1B-AS1** | 0.02697 | 0.01152 | 0.03463 | 0.05778 | 0.00660 | 0.11113 | 0.01247 |
| **PCF11** | 9.82737 | 0.62754 | 8.07337 | 8.11711 | 4.59832 | 6.79541 | 16.16709 |
| **PCGF1** | 5.55060 | 4.57602 | 3.68738 | 7.22023 | 4.47794 | 1.89719 | 1.21988 |
| **PCGF2** | 0.19039 | 0.16698 | 0.00940 | 0.37825 | 0.00501 | 0.93212 | 0.87950 |
| **PCGF3** | 5.87001 | 6.83469 | 3.13589 | 8.09093 | 5.01247 | 3.25895 | 21.30496 |
| **PCGF5** | 3.87973 | 6.67619 | 7.13357 | 1.07131 | 1.77664 | 2.00985 | 3.98072 |
| **PCGF6** | 5.92401 | 0.00000 | 11.23089 | 5.86745 | 2.36137 | 4.54556 | 4.41529 |

| | | | | | | | |
|---|---|---|---|---|---|---|---|
| **PCID2** | 17.40176 | 5.59238 | 5.90694 | 3.89454 | 3.58239 | 6.07389 | 4.47601 |
| **PCIF1** | 3.31705 | 2.12526 | 1.83417 | 5.68508 | 2.29096 | 1.92439 | 0.14905 |
| **PCK1** | 0.01769 | 0.01131 | 0.03072 | 0.01280 | 0.01571 | 0.00888 | 0.06258 |
| **PCK2** | 0.21468 | 0.08648 | 0.00000 | 1.58134 | 0.00468 | 0.04258 | 1.26263 |
| **PCLO** | 0.18818 | 0.00000 | 1.37343 | 0.13201 | 1.72645 | 1.27151 | 1.11071 |
| **PCM1** | 11.44641 | 5.10728 | 9.05793 | 7.44427 | 5.34406 | 9.74330 | 17.62627 |
| **PCMT1** | 51.16042 | 28.58486 | 22.93254 | 29.45637 | 12.24061 | 24.73847 | 7.99875 |
| **PCMTD1** | 4.87927 | 0.00000 | 2.18082 | 2.06783 | 2.09228 | 1.85224 | 4.03110 |
| **PCMTD2** | 7.66688 | 0.56423 | 9.57496 | 0.46083 | 1.54085 | 8.54158 | 3.76560 |
| **PCNA** | 34.55595 | 5.57949 | 24.24057 | 21.96519 | 12.04713 | 15.13819 | 18.75478 |
| **PCNAP1** | 0.03553 | 0.03302 | 0.03314 | 0.02056 | 0.02188 | 0.01798 | 0.06680 |
| **PCNP** | 24.73787 | 3.37226 | 12.07248 | 12.76161 | 1.99310 | 12.21808 | 6.73747 |
| **PCNT** | 12.18073 | 8.23825 | 8.16362 | 7.86315 | 2.21328 | 6.78195 | 20.61548 |
| **PCNX** | 3.19336 | 1.21571 | 4.19926 | 1.67029 | 1.40236 | 2.68634 | 6.65967 |
| **PCNXL2** | 0.07993 | 0.00000 | 0.69897 | 0.02114 | 0.41913 | 0.51942 | 0.35739 |
| **PCNXL3** | 2.91564 | 2.83338 | 2.08019 | 4.31981 | 0.27764 | 5.88002 | 16.92183 |
| **PCNXL4** | 4.11811 | 0.00787 | 2.77191 | 6.08349 | 1.53805 | 1.75466 | 1.36953 |
| **PCOLCE** | 0.06946 | 0.00000 | 0.09248 | 0.31143 | 0.01123 | 0.58085 | 0.39505 |
| **PCOLCE-AS1** | 0.02562 | 0.01833 | 0.01744 | 0.03845 | 0.02483 | 0.01376 | 0.01199 |
| **PCOLCE2** | 0.34824 | 0.00000 | 1.29716 | 0.09730 | 0.61120 | 0.38274 | 0.05611 |
| **PCP2** | 0.02358 | 0.06575 | 0.02713 | 0.03117 | 0.00825 | 0.01501 | 0.03311 |
| **PCP4** | 0.36211 | 0.01105 | 0.00673 | 0.09836 | 0.08586 | 0.01945 | 0.00000 |
| **PCP4L1** | 0.27683 | 0.00000 | 1.53016 | 0.86531 | 25.01470 | 4.26465 | 0.00000 |
| **PCSK1** | 0.02619 | 0.00000 | 0.01359 | 0.07278 | 0.06979 | 0.93720 | 0.00000 |
| **PCSK1N** | 0.00000 | 0.08448 | 0.04659 | 0.12422 | 0.31630 | 1.32324 | 0.28924 |
| **PCSK2** | 0.02374 | 0.00645 | 0.04470 | 0.01140 | 0.05758 | 0.01839 | 0.05934 |
| **PCSK4** | 0.00000 | 0.21599 | 0.03051 | 0.01303 | 0.21437 | 0.34226 | 0.23816 |
| **PCSK5** | 1.24435 | 0.00000 | 0.82091 | 0.18517 | 3.63111 | 6.74455 | 0.17342 |
| **PCSK6** | 2.16095 | 0.00000 | 0.11344 | 0.50394 | 0.14548 | 0.00000 | 4.74185 |
| **PCSK7** | 2.40027 | 9.22271 | 3.12330 | 5.65088 | 2.70445 | 6.70044 | 3.29037 |
| **PCSK9** | 0.49690 | 0.26642 | 0.80845 | 0.82484 | 0.10152 | 0.98566 | 0.00000 |
| **PCTP** | 5.29830 | 1.31692 | 0.01990 | 6.20671 | 0.97579 | 4.79886 | 0.00000 |
| **PCYOX1** | 19.87385 | 11.96651 | 14.91416 | 5.97211 | 5.16102 | 15.24006 | 3.80422 |
| **PCYOX1L** | 0.10406 | 0.00000 | 0.00000 | 0.45849 | 0.74621 | 0.03677 | 0.62538 |
| **PCYT1A** | 7.38421 | 3.12490 | 4.48525 | 10.66897 | 1.62416 | 2.47676 | 1.84944 |

| | | | | | | | |
|---|---|---|---|---|---|---|---|
| **PCYT1B** | 0.05060 | 0.00000 | 0.04004 | 0.02057 | 0.52375 | 0.09901 | 0.00000 |
| **PCYT1B-AS1** | 0.02505 | 0.01733 | 0.00822 | 0.01930 | 0.06999 | 0.01459 | 0.01393 |
| **PCYT2** | 11.92390 | 12.69968 | 0.00000 | 1.34719 | 0.13156 | 1.42375 | 20.90560 |
| **PDAP1** | 12.58279 | 15.19873 | 9.41456 | 16.38358 | 2.19513 | 9.33182 | 17.25831 |
| **PDC** | 0.02558 | 0.02152 | 0.01276 | 0.02228 | 0.00774 | 0.01061 | 0.02681 |
| **PDCD1** | 0.02108 | 0.02808 | 0.01258 | 0.02923 | 0.04544 | 0.02089 | 0.03733 |
| **PDCD10** | 27.92280 | 5.19603 | 12.11074 | 12.43044 | 3.95460 | 10.09914 | 4.23722 |
| **PDCD11** | 28.86073 | 23.62583 | 15.75787 | 24.38539 | 6.91330 | 10.79398 | 26.57874 |
| **PDCD1LG2** | 0.02450 | 0.02584 | 0.01359 | 0.09652 | 0.01088 | 0.01516 | 0.00279 |
| **PDCD2** | 9.51156 | 1.26338 | 4.17494 | 8.92088 | 0.53446 | 4.81191 | 17.14300 |
| **PDCD2L** | 9.65166 | 10.02640 | 15.10989 | 8.62854 | 2.18079 | 5.12656 | 2.82001 |
| **PDCD4** | 10.90686 | 0.00000 | 2.88439 | 3.10100 | 1.75853 | 4.82288 | 15.29215 |
| **PDCD4-AS1** | 0.03068 | 0.00822 | 0.04695 | 0.02545 | 0.06073 | 0.02752 | 0.12313 |
| **PDCD5** | 21.63854 | 19.35050 | 5.96592 | 16.77043 | 1.32674 | 10.08256 | 12.75817 |
| **PDCD6** | 36.90185 | 30.07913 | 22.12887 | 21.40096 | 4.08884 | 11.68619 | 3.60711 |
| **PDCD6IP** | 25.17440 | 12.09140 | 8.00092 | 15.18703 | 1.77712 | 10.24162 | 8.08026 |
| **PDCD6IPP2** | 0.08643 | 0.29481 | 0.07723 | 0.02247 | 0.06109 | 0.03888 | 0.00574 |
| **PDCD7** | 8.30724 | 7.57556 | 4.33509 | 2.14792 | 5.95652 | 3.45648 | 4.62165 |
| **PDCL** | 14.00448 | 10.68505 | 14.37075 | 11.74430 | 14.24977 | 13.61502 | 0.00000 |
| **PDCL2** | 2.24901 | 0.36355 | 2.92222 | 0.00000 | 0.00000 | 2.45353 | 0.17773 |
| **PDCL3** | 19.19238 | 63.22454 | 30.72546 | 0.94674 | 17.42288 | 8.89554 | 3.33124 |
| **PDCL3P4** | 1.14306 | 3.42677 | 0.53219 | 0.00000 | 0.36799 | 1.87679 | 0.83400 |
| **PDDC1** | 2.04508 | 2.73619 | 2.07594 | 5.99952 | 0.01765 | 0.97382 | 6.33748 |
| **PDE10A** | 0.06411 | 0.00264 | 0.03845 | 0.01474 | 0.03849 | 0.14586 | 0.00477 |
| **PDE11A** | 0.07642 | 0.00000 | 0.90901 | 0.01592 | 1.42001 | 0.25826 | 0.03345 |
| **PDE12** | 16.30944 | 7.84993 | 5.59410 | 7.68824 | 6.28145 | 4.59113 | 0.00000 |
| **PDE1A** | 0.17790 | 0.00000 | 0.00863 | 0.02597 | 0.13945 | 0.00253 | 0.06593 |
| **PDE1B** | 0.03213 | 0.00400 | 0.03756 | 0.04792 | 0.01641 | 0.18387 | 0.00774 |
| **PDE1C** | 0.07064 | 0.01928 | 0.01776 | 0.01973 | 0.14021 | 0.23755 | 0.10898 |
| **PDE2A** | 0.54718 | 0.78264 | 0.19896 | 0.09305 | 0.00945 | 0.09704 | 0.06644 |
| **PDE3A** | 0.03878 | 0.00000 | 0.09606 | 0.00559 | 1.32684 | 0.39223 | 0.03599 |
| **PDE3B** | 0.46983 | 0.00000 | 0.12466 | 0.20900 | 0.00585 | 0.00000 | 0.65688 |
| **PDE4A** | 0.00000 | 0.63810 | 2.80277 | 0.28726 | 0.47996 | 2.46943 | 0.70936 |
| **PDE4B** | 0.02100 | 0.01712 | 0.01242 | 0.01977 | 0.01255 | 0.02872 | 0.03105 |
| **PDE4C** | 0.00000 | 1.38237 | 0.23292 | 0.03254 | 0.10371 | 0.41598 | 0.06532 |

| | | | | | | | |
|---|---|---|---|---|---|---|---|
| **PDE4D** | 0.75604 | 0.00000 | 0.74716 | 1.27287 | 0.23019 | 2.38286 | 2.91717 |
| **PDE4DIP** | 12.16793 | 1.18120 | 4.08574 | 11.34232 | 3.36772 | 7.11303 | 4.06403 |
| **PDE5A** | 1.27922 | 0.00000 | 0.66576 | 0.20379 | 0.12788 | 1.53033 | 1.65166 |
| **PDE6A** | 0.03767 | 0.19623 | 0.02382 | 0.03242 | 0.04159 | 0.02830 | 0.03785 |
| **PDE6B** | 0.06838 | 0.33065 | 0.00000 | 0.02081 | 0.97694 | 0.15509 | 0.20096 |
| **PDE6C** | 0.01596 | 0.09218 | 0.02256 | 0.01010 | 2.14878 | 0.00000 | 0.06967 |
| **PDE6D** | 6.02582 | 10.31157 | 4.61096 | 8.45956 | 1.47697 | 2.24951 | 0.00000 |
| **PDE6G** | 0.45524 | 1.83563 | 0.29239 | 0.00000 | 0.00115 | 2.26549 | 0.08066 |
| **PDE7A** | 0.84905 | 0.23861 | 0.91933 | 2.85386 | 0.24684 | 2.82011 | 7.22965 |
| **PDE7B** | 0.03228 | 0.00000 | 0.03277 | 0.02691 | 0.08194 | 0.03549 | 0.04713 |
| **PDE8A** | 3.86349 | 1.14563 | 1.00191 | 2.34104 | 0.32902 | 1.29498 | 5.07466 |
| **PDE8B** | 0.03461 | 0.29254 | 0.41701 | 0.00000 | 11.19122 | 0.20007 | 0.46014 |
| **PDE9A** | 0.01069 | 0.09285 | 0.03562 | 0.00000 | 1.02211 | 0.04561 | 0.12092 |
| **PDF** | 0.00000 | 2.51009 | 0.22659 | 0.28881 | 0.20802 | 0.72254 | 1.27769 |
| **PDGFA** | 25.98520 | 29.35694 | 0.27317 | 7.59555 | 0.90623 | 1.25590 | 20.91668 |
| **PDGFB** | 0.00665 | 0.02113 | 0.01208 | 0.21199 | 0.00931 | 0.04882 | 0.02353 |
| **PDGFC** | 0.12672 | 0.00000 | 0.00243 | 0.00000 | 0.28837 | 0.06755 | 0.31663 |
| **PDGFD** | 0.03150 | 0.09127 | 0.28200 | 0.09057 | 0.15064 | 0.12570 | 0.00000 |
| **PDGFRA** | 1.73591 | 0.74806 | 0.00000 | 0.00000 | 1.27280 | 24.32619 | 0.00000 |
| **PDGFRB** | 0.05363 | 0.00590 | 0.01544 | 0.16599 | 0.00819 | 0.06613 | 0.00000 |
| **PDGFRL** | 0.37618 | 0.00156 | 0.04147 | 0.32883 | 3.80283 | 0.01607 | 0.00000 |
| **PDHA1** | 70.27653 | 44.62474 | 24.32455 | 25.11370 | 3.76903 | 27.59823 | 23.89763 |
| **PDHA2** | 0.79140 | 0.28148 | 0.31799 | 0.13569 | 0.01969 | 0.29842 | 0.26726 |
| **PDHB** | 39.20231 | 22.70901 | 17.94049 | 23.71282 | 3.95223 | 22.23780 | 0.00701 |
| **PDHX** | 27.11806 | 8.04643 | 7.28356 | 10.79255 | 1.06116 | 10.19737 | 0.00000 |
| **PDIA2** | 0.03644 | 0.01177 | 0.01439 | 0.04343 | 0.02090 | 0.05920 | 0.01500 |
| **PDIA3** | 44.56899 | 13.57904 | 11.91734 | 34.70689 | 10.65328 | 10.77635 | 0.00000 |
| **PDIA3P1** | 20.75205 | 5.20970 | 4.68343 | 16.81648 | 5.00184 | 3.39199 | 0.00000 |
| **PDIA4** | 26.24848 | 11.24215 | 6.11009 | 24.38405 | 5.35810 | 16.78089 | 0.00000 |
| **PDIA5** | 1.71228 | 1.57106 | 0.15215 | 1.85941 | 0.06306 | 1.66154 | 6.58653 |
| **PDIA6** | 39.51844 | 18.53707 | 5.22409 | 30.08901 | 4.39954 | 18.57431 | 0.00000 |
| **PDIK1L** | 1.19727 | 0.32971 | 0.69297 | 0.51638 | 0.93532 | 0.29372 | 0.66572 |
| **PDK1** | 0.00000 | 0.00000 | 1.36039 | 1.47391 | 2.35640 | 3.31751 | 0.27565 |
| **PDK2** | 0.07873 | 0.69854 | 0.05032 | 0.07649 | 0.60895 | 0.19522 | 0.00496 |
| **PDK3** | 0.96875 | 0.20861 | 2.84014 | 0.30348 | 5.09313 | 3.44938 | 0.02021 |

| | | | | | | | |
|---|---|---|---|---|---|---|---|
| **PDK4** | 0.02424 | 0.01986 | 0.01218 | 0.01880 | 0.00796 | 0.01253 | 0.01906 |
| **PDLIM1** | 40.51915 | 28.18764 | 8.96458 | 17.15165 | 1.51283 | 3.42152 | 17.79993 |
| **PDLIM2** | 0.00000 | 0.32911 | 0.13343 | 12.30497 | 2.20590 | 0.00000 | 0.48372 |
| **PDLIM3** | 0.00002 | 0.00000 | 0.15323 | 0.17552 | 0.38694 | 0.08061 | 0.01566 |
| **PDLIM4** | 0.00001 | 0.45977 | 0.00508 | 0.11691 | 0.00619 | 0.23017 | 0.10664 |
| **PDLIM5** | 7.61586 | 3.66175 | 6.06751 | 4.38769 | 4.28573 | 5.41773 | 2.14068 |
| **PDLIM7** | 7.15723 | 10.92744 | 1.98314 | 5.65584 | 2.30132 | 10.68110 | 0.00000 |
| **PDP1** | 3.86650 | 1.68880 | 3.82780 | 0.85823 | 7.35419 | 6.17305 | 0.48205 |
| **PDP2** | 5.57222 | 3.13610 | 1.02963 | 5.94910 | 2.16693 | 0.89049 | 7.61127 |
| **PDPK1** | 3.21862 | 3.70108 | 1.84819 | 5.04650 | 2.61415 | 4.61725 | 18.93147 |
| **PDPN** | 4.46707 | 9.85371 | 1.18371 | 0.00000 | 6.27877 | 14.87710 | 0.00000 |
| **PDPR** | 2.70424 | 0.00000 | 7.58698 | 2.18848 | 3.17189 | 1.62796 | 5.12439 |
| **PDRG1** | 17.58064 | 19.32854 | 13.99722 | 14.07580 | 10.47366 | 6.34794 | 0.00000 |
| **PDS5A** | 13.27049 | 1.14538 | 13.67029 | 2.41946 | 1.15633 | 11.53581 | 27.61970 |
| **PDS5B** | 15.13233 | 0.24716 | 10.89077 | 4.97188 | 2.32955 | 7.64036 | 13.02801 |
| **PDSS1** | 2.91127 | 0.93732 | 1.28376 | 2.10271 | 0.04574 | 1.65718 | 6.68400 |
| **PDSS2** | 5.76873 | 1.50381 | 2.32345 | 3.71872 | 0.36953 | 3.54938 | 1.99432 |
| **PDX1** | 0.02614 | 0.00118 | 0.02962 | 0.02545 | 0.06266 | 0.01022 | 0.05056 |
| **PDXDC1** | 16.82759 | 7.15614 | 1.65730 | 13.73669 | 5.25082 | 8.34825 | 13.08887 |
| **PDXDC2P** | 0.81044 | 0.40605 | 0.27442 | 0.84848 | 0.25374 | 1.51860 | 1.75861 |
| **PDXK** | 19.48485 | 13.42965 | 0.00000 | 5.68770 | 0.15762 | 4.59811 | 29.79269 |
| **PDXP** | 1.17165 | 1.58737 | 2.39449 | 0.96335 | 1.41752 | 3.14652 | 8.89124 |
| **PDZD11** | 12.55738 | 15.95213 | 9.09342 | 2.70305 | 3.21628 | 8.14901 | 4.13344 |
| **PDZD2** | 1.28772 | 0.12970 | 2.74550 | 1.53802 | 0.89983 | 1.46621 | 3.80508 |
| **PDZD3** | 0.02622 | 0.02711 | 0.10237 | 0.02398 | 0.00493 | 0.00928 | 0.02017 |
| **PDZD4** | 0.02121 | 0.01879 | 0.01200 | 0.02271 | 0.13028 | 0.01285 | 0.02355 |
| **PDZD7** | 0.00000 | 0.00736 | 0.02987 | 0.12534 | 0.97050 | 0.10814 | 0.07382 |
| **PDZD8** | 1.22430 | 0.34788 | 1.68215 | 0.30030 | 1.28326 | 1.71551 | 6.14380 |
| **PDZD9** | 0.01549 | 0.02124 | 0.02112 | 0.02706 | 0.01107 | 0.01759 | 0.04063 |
| **PDZK1** | 0.32723 | 0.08252 | 0.01763 | 0.05682 | 0.13870 | 0.07790 | 0.24525 |
| **PDZK1IP1** | 0.07163 | 0.00533 | 0.01173 | 0.02213 | 0.00879 | 0.00183 | 0.07636 |
| **PDZRN3** | 0.23321 | 0.00000 | 0.11482 | 0.00000 | 0.30479 | 1.33878 | 0.02940 |
| **PDZRN3-AS1** | 0.00179 | 0.01606 | 0.00427 | 0.03502 | 0.20447 | 0.00842 | 0.08721 |
| **PDZRN4** | 0.02022 | 0.01555 | 0.01230 | 0.02650 | 0.01151 | 0.01194 | 0.03579 |
| **PEA15** | 3.01778 | 6.37369 | 1.88330 | 11.94571 | 1.66637 | 2.41815 | 12.15967 |

| | | | | | | | |
|---|---|---|---|---|---|---|---|
| **PEAK1** | 3.05686 | 0.93433 | 0.56898 | 7.59123 | 1.64478 | 0.00000 | 10.85706 |
| **PEAR1** | 0.04690 | 0.00000 | 0.01344 | 0.06154 | 0.02674 | 0.02201 | 0.13904 |
| **PEBP1** | 41.57026 | 14.13772 | 0.43715 | 40.61035 | 1.87811 | 25.53498 | 22.14177 |
| **PEBP4** | 0.02239 | 0.01805 | 0.00250 | 0.01999 | 0.59855 | 0.01259 | 0.01320 |
| **PECAM1** | 0.00000 | 0.23739 | 0.40533 | 0.06512 | 0.62908 | 1.66839 | 0.00827 |
| **PECR** | 7.60030 | 0.20114 | 0.22482 | 3.04679 | 0.78858 | 1.39988 | 1.47159 |
| **PEF1** | 20.94274 | 15.38734 | 11.30550 | 15.66764 | 1.63136 | 5.79021 | 0.00000 |
| **PEG10** | 14.60512 | 0.00000 | 0.00000 | 16.71637 | 0.09219 | 0.00000 | 3.90142 |
| **PEG3** | 1.67120 | 0.00000 | 1.79451 | 2.04690 | 0.38657 | 1.05976 | 1.19691 |
| **PELI1** | 7.35605 | 2.61058 | 2.79679 | 2.97681 | 6.27030 | 10.98378 | 0.00000 |
| **PELI2** | 0.02484 | 0.20563 | 0.47985 | 0.12820 | 0.54700 | 2.89925 | 0.09192 |
| **PELI3** | 0.31615 | 0.53483 | 0.57631 | 0.46116 | 0.10580 | 0.21202 | 0.88975 |
| **PELO** | 15.05269 | 6.07374 | 3.36842 | 28.68701 | 2.14695 | 4.00054 | 0.00000 |
| **PELP1** | 0.30104 | 6.59221 | 4.30648 | 6.09608 | 0.10439 | 3.40160 | 25.16197 |
| **PEMT** | 9.53653 | 27.27321 | 0.75486 | 3.04623 | 0.00000 | 12.51170 | 2.32330 |
| **PENK** | 0.00000 | 0.00000 | 0.57690 | 0.06075 | 0.05698 | 0.00000 | 0.24325 |
| **PEPD** | 16.13676 | 12.99538 | 4.07070 | 16.60079 | 0.00000 | 11.15582 | 0.00953 |
| **PER1** | 0.35163 | 0.75430 | 0.76582 | 3.12762 | 1.09950 | 2.16512 | 6.92963 |
| **PER2** | 9.56945 | 3.40798 | 3.52584 | 2.75605 | 1.96648 | 2.32543 | 2.77199 |
| **PER3** | 1.84464 | 1.73489 | 4.12767 | 2.19763 | 0.86963 | 0.43417 | 0.00000 |
| **PERM1** | 0.02777 | 0.00000 | 0.06320 | 0.03430 | 0.03998 | 0.01424 | 0.03777 |
| **PERP** | 35.72964 | 10.19465 | 4.36083 | 19.10507 | 0.16293 | 2.12101 | 22.91951 |
| **1.00000** | 5.32387 | 24.92843 | 8.64180 | 16.68983 | 3.00470 | 7.21259 | 17.90400 |
| **PET100** | 4.07492 | 7.34852 | 4.80775 | 7.93854 | 2.07612 | 2.63377 | 3.52153 |
| **PET112** | 16.27866 | 8.34875 | 3.41093 | 9.27492 | 0.94265 | 5.30558 | 0.00000 |
| **PET117** | 2.68061 | 6.44211 | 2.86230 | 0.19699 | 0.29924 | 1.51698 | 3.26273 |
| **PEX1** | 3.19097 | 2.39643 | 1.61689 | 0.57959 | 1.17775 | 3.28919 | 2.07486 |
| **PEX10** | 0.82764 | 2.35639 | 0.62677 | 1.75687 | 0.01425 | 0.73551 | 0.00000 |
| **PEX11A** | 9.16847 | 1.06752 | 2.27207 | 1.55657 | 1.71830 | 0.12629 | 2.47952 |
| **PEX11B** | 3.26653 | 1.29720 | 4.06067 | 1.53174 | 3.85156 | 1.68391 | 0.00000 |
| **PEX11G** | 0.24548 | 0.65576 | 0.07288 | 0.10129 | 0.09071 | 0.21820 | 0.00000 |
| **PEX12** | 2.80560 | 0.65416 | 2.43170 | 0.66889 | 4.83549 | 1.18711 | 0.47710 |
| **PEX13** | 11.48013 | 3.54938 | 6.30394 | 6.09563 | 5.98221 | 3.80025 | 0.00000 |
| **PEX14** | 0.83311 | 3.71865 | 1.37042 | 3.04998 | 1.03699 | 1.81157 | 4.04071 |
| **PEX16** | 13.45276 | 20.62729 | 3.34829 | 11.04553 | 0.66929 | 4.87299 | 0.00000 |

| | | | | | | | |
|---|---|---|---|---|---|---|---|
| **PEX19** | 19.20805 | 10.08866 | 9.23416 | 14.11472 | 1.39529 | 6.33040 | 0.00000 |
| **PEX2** | 10.54170 | 1.18889 | 7.38406 | 5.42871 | 2.52110 | 1.67852 | 1.05890 |
| **PEX26** | 0.57563 | 2.12073 | 0.59877 | 1.51354 | 0.24189 | 1.09548 | 5.42297 |
| **PEX3** | 21.93653 | 0.02609 | 14.11215 | 5.08710 | 0.41519 | 1.28997 | 0.00000 |
| **PEX5** | 4.53948 | 0.95389 | 4.67276 | 1.20815 | 0.55141 | 4.81902 | 3.77863 |
| **PEX5L** | 0.08327 | 0.00000 | 0.10689 | 0.00000 | 0.14824 | 0.03294 | 0.20639 |
| **PEX6** | 0.24946 | 1.57192 | 0.40511 | 2.86896 | 0.02208 | 1.33305 | 3.58553 |
| **PEX7** | 0.43667 | 0.24431 | 0.22326 | 0.39399 | 0.00271 | 0.83412 | 1.95810 |
| **PFAS** | 14.88399 | 9.20508 | 7.03902 | 11.14503 | 0.16936 | 7.34927 | 11.82592 |
| **PFDN1** | 22.86789 | 12.91455 | 18.97042 | 22.71636 | 6.53888 | 7.97629 | 8.23748 |
| **PFDN2** | 30.43375 | 19.29604 | 17.02516 | 30.92405 | 12.97695 | 10.85931 | 15.50345 |
| **PFDN4** | 18.39679 | 5.55300 | 7.95914 | 12.55067 | 1.10003 | 9.41774 | 11.42005 |
| **PFDN5** | 27.80031 | 21.37873 | 20.53954 | 15.57815 | 12.13792 | 18.34103 | 25.78859 |
| **PFDN6** | 2.31270 | 2.46267 | 3.95842 | 5.97732 | 1.78700 | 2.04576 | 3.18176 |
| **PFKFB1** | 0.04585 | 0.08686 | 0.07779 | 0.03548 | 0.00485 | 0.00971 | 0.00000 |
| **PFKFB2** | 6.03181 | 0.69711 | 0.93150 | 1.47349 | 2.01911 | 1.21828 | 0.66618 |
| **PFKFB3** | 0.00000 | 5.37756 | 26.51592 | 2.76971 | 9.39961 | 3.65669 | 0.00000 |
| **PFKFB4** | 12.59460 | 6.45236 | 0.02762 | 11.52985 | 0.06661 | 0.96575 | 10.73565 |
| **PFKL** | 22.02517 | 23.76168 | 2.60597 | 14.26324 | 0.09425 | 17.00416 | 17.19081 |
| **PFKM** | 5.62193 | 0.87182 | 3.12908 | 2.91765 | 0.27496 | 0.51254 | 0.58565 |
| **PFKP** | 30.48685 | 44.57959 | 13.63174 | 37.19759 | 3.83265 | 26.02988 | 15.15801 |
| **PFN1** | 70.58862 | 73.26096 | 35.51684 | 65.42479 | 26.08439 | 43.07404 | 36.81106 |
| **PFN1P2** | 0.03325 | 0.07473 | 0.07659 | 0.02305 | 0.00483 | 0.12913 | 0.08253 |
| **PFN2** | 12.57736 | 0.93650 | 5.22962 | 19.66973 | 1.86999 | 7.41592 | 5.87484 |
| **PFN3** | 0.02108 | 0.01551 | 0.02229 | 0.02509 | 0.01398 | 0.01083 | 0.03092 |
| **PFN4** | 0.04227 | 0.15062 | 0.00287 | 0.04983 | 0.00900 | 0.01350 | 0.01173 |
| **PGA3** | 0.02519 | 0.00900 | 0.02696 | 0.02311 | 0.06471 | 0.01379 | 0.01531 |
| **PGA5** | 0.00928 | 0.01548 | 0.03736 | 0.17856 | 0.01810 | 0.00629 | 0.04464 |
| **PGAM1** | 16.79672 | 3.24791 | 7.68524 | 9.63737 | 1.78264 | 11.40966 | 1.88420 |
| **PGAM1P5** | 0.13316 | 0.03585 | 0.04157 | 0.08339 | 0.01644 | 0.07006 | 0.00000 |
| **PGAM2** | 0.06338 | 0.02327 | 0.10833 | 0.13100 | 0.00784 | 0.00000 | 0.09941 |
| **PGAM4** | 0.24526 | 0.15596 | 0.08377 | 0.15070 | 0.07326 | 0.23904 | 0.00000 |
| **PGAM5** | 9.60229 | 7.46595 | 1.83172 | 7.69119 | 0.45323 | 4.92902 | 19.10933 |
| **PGAP1** | 1.55572 | 0.00000 | 0.34098 | 1.89974 | 0.75457 | 0.00000 | 3.27810 |
| **PGAP2** | 14.26904 | 4.78221 | 12.56952 | 12.08469 | 1.90671 | 3.12175 | 5.12498 |

| | | | | | | | |
|---|---|---|---|---|---|---|---|
| **PGAP3** | 0.15294 | 0.46646 | 0.49866 | 0.16830 | 0.56693 | 0.15323 | 0.34810 |
| **PGBD1** | 1.77986 | 1.24651 | 3.49965 | 0.00000 | 0.90863 | 1.54313 | 1.43344 |
| **PGBD2** | 5.00169 | 0.41975 | 4.16702 | 3.54305 | 1.64550 | 1.62136 | 0.00000 |
| **PGBD4** | 0.54240 | 0.21149 | 0.25513 | 0.32867 | 0.41949 | 0.00000 | 0.20425 |
| **PGBD5** | 0.04217 | 0.44695 | 0.21215 | 0.10436 | 0.02803 | 0.30636 | 0.00000 |
| **PGC** | 0.00000 | 0.07720 | 0.02827 | 12.09897 | 0.00650 | 0.18083 | 0.00000 |
| **PGCP1** | 0.02506 | 0.01608 | 0.01327 | 0.02029 | 0.01652 | 0.01828 | 0.01881 |
| **PGD** | 26.09108 | 19.65020 | 10.12675 | 23.90178 | 2.66440 | 25.07812 | 25.38777 |
| **PGF** | 0.48335 | 3.29996 | 0.42586 | 64.81075 | 0.02927 | 0.00228 | 7.39103 |
| **PGGT1B** | 10.22324 | 0.24961 | 4.26581 | 3.69396 | 2.65095 | 3.68539 | 0.00000 |
| **1.00000** | 71.65683 | 29.06793 | 18.17654 | 40.01960 | 0.96983 | 30.26267 | 26.76997 |
| **2.00000** | 0.06736 | 0.02779 | 0.01842 | 0.04011 | 0.00748 | 0.03176 | 0.00000 |
| **PGLS** | 4.05218 | 10.59969 | 1.24441 | 6.51737 | 0.01482 | 3.58691 | 6.58090 |
| **PGLYRP1** | 0.03850 | 0.05225 | 0.04425 | 0.05756 | 0.00587 | 0.02129 | 0.06559 |
| **PGLYRP2** | 0.02973 | 0.00000 | 0.12660 | 0.02179 | 0.07628 | 0.00961 | 0.02269 |
| **PGLYRP4** | 0.02998 | 0.02871 | 0.10304 | 0.02699 | 0.02031 | 0.00486 | 0.02266 |
| **PGM1** | 11.23423 | 7.40916 | 0.00000 | 1.99424 | 0.35484 | 4.63629 | 0.64256 |
| **PGM2** | 20.82910 | 9.82705 | 5.09065 | 2.78002 | 0.05329 | 19.01781 | 1.93162 |
| **PGM2L1** | 4.49592 | 0.00000 | 7.91790 | 2.74147 | 2.48830 | 10.88883 | 1.43815 |
| **PGM3** | 25.93783 | 6.03278 | 9.38529 | 16.32103 | 4.37822 | 4.13620 | 0.00000 |
| **PGM5** | 0.01012 | 0.06706 | 0.04026 | 0.01839 | 0.04629 | 0.00675 | 0.02519 |
| **PGM5P2** | 0.02184 | 0.00000 | 0.04952 | 0.01731 | 0.05836 | 0.04642 | 0.08667 |
| **PGP** | 0.00000 | 8.01158 | 1.34076 | 5.62155 | 0.08351 | 6.93198 | 29.26000 |
| **PGPEP1** | 0.86190 | 2.91033 | 0.14277 | 1.86000 | 2.30669 | 0.44910 | 1.94461 |
| **PGPEP1L** | 0.02303 | 0.02241 | 0.07585 | 0.01303 | 0.04062 | 0.00661 | 0.03671 |
| **PGR** | 0.02070 | 0.02049 | 0.01190 | 0.02865 | 0.00805 | 0.02198 | 0.01376 |
| **PGRMC1** | 25.36601 | 16.00071 | 3.75151 | 8.84101 | 1.97641 | 7.15527 | 9.98076 |
| **PGRMC2** | 0.22334 | 0.00000 | 4.73402 | 7.43646 | 1.43998 | 2.98513 | 14.85352 |
| **PGS1** | 7.08799 | 5.23957 | 0.96147 | 9.37407 | 1.44508 | 2.73286 | 0.92119 |
| **PHACTR1** | 0.15074 | 0.25910 | 0.42816 | 0.55383 | 0.50578 | 1.35682 | 0.00000 |
| **PHACTR2** | 11.27785 | 2.73618 | 0.00000 | 7.60742 | 0.47719 | 5.46126 | 7.23891 |
| **PHACTR3** | 0.01304 | 0.10253 | 0.16956 | 0.00000 | 5.76045 | 0.03458 | 0.02815 |
| **PHACTR4** | 4.60100 | 0.00000 | 4.50168 | 8.75111 | 5.29340 | 6.22459 | 8.61925 |
| **PHAX** | 26.20487 | 10.92497 | 19.56874 | 15.59747 | 11.93989 | 12.27532 | 3.59599 |
| **PHB** | 44.53789 | 49.68889 | 21.00488 | 33.27873 | 8.39283 | 22.39637 | 13.46973 |

| | | | | | | | |
|---|---|---|---|---|---|---|---|
| **PHB2** | 60.01130 | 47.10924 | 23.89545 | 43.29543 | 7.09278 | 29.34167 | 23.30740 |
| **PHC1** | 0.07603 | 0.02666 | 0.09890 | 0.87871 | 0.07369 | 0.73157 | 0.00790 |
| **PHC2** | 2.46705 | 0.67547 | 0.49830 | 6.07430 | 0.30170 | 0.00000 | 13.46464 |
| **PHC3** | 8.96231 | 1.89329 | 6.76312 | 7.04393 | 3.81623 | 2.50279 | 2.93442 |
| **PHEX** | 0.00000 | 0.18753 | 0.00000 | 0.00000 | 3.05316 | 0.02512 | 0.10532 |
| **PHEX-AS1** | 0.01749 | 0.02088 | 0.01141 | 0.01576 | 0.01583 | 0.02330 | 0.04315 |
| **PHF1** | 0.00000 | 0.53288 | 0.64265 | 1.02757 | 0.45370 | 2.51354 | 3.13845 |
| **PHF10** | 0.18527 | 0.43733 | 0.24994 | 0.39114 | 0.01262 | 0.16743 | 2.00475 |
| **PHF11** | 0.70304 | 0.63608 | 0.29512 | 0.58328 | 0.07818 | 1.24139 | 0.91560 |
| **PHF12** | 3.80871 | 1.79006 | 2.90921 | 2.20759 | 3.30670 | 4.32041 | 7.62502 |
| **PHF13** | 0.00000 | 3.09807 | 1.58506 | 3.33579 | 6.92588 | 4.66157 | 5.92132 |
| **PHF14** | 4.24151 | 1.10155 | 4.64610 | 5.23901 | 2.73274 | 3.43375 | 24.28870 |
| **PHF19** | 0.00000 | 0.00000 | 0.03003 | 0.13401 | 0.07918 | 0.92698 | 0.05689 |
| **PHF2** | 0.55009 | 1.06792 | 0.45316 | 1.15973 | 0.00000 | 1.46025 | 1.98848 |
| **PHF20** | 6.95152 | 3.75550 | 5.44140 | 6.67821 | 2.11416 | 3.82470 | 4.83473 |
| **PHF20L1** | 3.37951 | 2.73109 | 2.86128 | 1.91747 | 1.05923 | 1.70658 | 9.78602 |
| **PHF21A** | 1.90949 | 1.40009 | 0.57044 | 0.52935 | 0.92130 | 1.29466 | 6.19461 |
| **PHF21B** | 0.00000 | 2.42052 | 0.08353 | 0.72344 | 0.07201 | 1.02486 | 0.94256 |
| **PHF23** | 2.85439 | 5.98668 | 5.80358 | 7.77171 | 4.75299 | 6.52944 | 20.80323 |
| **PHF3** | 12.06242 | 1.16093 | 12.52882 | 7.51525 | 4.46359 | 6.76966 | 21.53064 |
| **PHF5A** | 27.79035 | 10.26439 | 12.60388 | 21.38742 | 8.50855 | 11.89340 | 2.84053 |
| **PHF6** | 9.37542 | 0.07948 | 8.09723 | 2.61280 | 6.34364 | 2.44317 | 5.64227 |
| **PHF7** | 0.39169 | 1.83695 | 1.19271 | 1.22702 | 0.83690 | 0.40599 | 1.58072 |
| **PHF8** | 6.31032 | 4.96570 | 11.54800 | 3.26535 | 6.10143 | 2.40673 | 13.56636 |
| **PHGDH** | 13.73618 | 0.00000 | 0.00000 | 18.66330 | 0.00000 | 5.91497 | 10.09316 |
| **PHGR1** | 0.03429 | 0.00714 | 0.01177 | 0.00528 | 0.00708 | 0.00487 | 0.10965 |
| **PHIP** | 10.07984 | 3.71295 | 7.90774 | 9.53891 | 1.36750 | 11.89270 | 21.74363 |
| **PHKA1** | 9.93148 | 11.27744 | 0.00000 | 0.45540 | 0.10953 | 13.17226 | 22.67664 |
| **PHKA2** | 1.56287 | 2.33540 | 1.90234 | 0.35794 | 0.86659 | 2.96857 | 2.70674 |
| **PHKA2-AS1** | 0.80813 | 1.19413 | 0.71923 | 0.00000 | 0.34147 | 3.75126 | 0.11416 |
| **PHKB** | 10.02012 | 4.42409 | 3.86853 | 4.50659 | 1.85079 | 4.81813 | 4.24086 |
| **PHKG1** | 0.01208 | 0.28792 | 0.07349 | 0.10923 | 0.33050 | 0.16590 | 0.17774 |
| **PHKG2** | 10.10007 | 11.69055 | 2.42322 | 30.62231 | 1.79867 | 7.06106 | 4.37508 |
| **PHLDA1** | 0.00000 | 5.04701 | 0.00000 | 0.24084 | 0.02623 | 17.39324 | 1.26758 |
| **PHLDA2** | 4.82683 | 19.98029 | 12.99134 | 10.29923 | 4.70888 | 7.58011 | 8.36799 |

| | | | | | | | |
|---|---|---|---|---|---|---|---|
| **PHLDA3** | 0.00000 | 10.08089 | 0.49695 | 4.53518 | 0.29594 | 8.31391 | 4.74722 |
| **PHLDB1** | 2.06550 | 2.74739 | 0.58053 | 1.87651 | 0.28959 | 2.30056 | 9.73006 |
| **PHLDB2** | 0.24219 | 0.89274 | 0.84755 | 3.24937 | 0.26319 | 5.02231 | 1.20283 |
| **PHLDB3** | 0.00000 | 5.36652 | 5.97503 | 10.74139 | 0.27124 | 1.80018 | 0.60078 |
| **PHLPP1** | 2.16262 | 4.36265 | 2.80647 | 5.43955 | 1.20980 | 3.44774 | 8.39480 |
| **PHLPP2** | 2.09072 | 2.36692 | 3.12025 | 1.95097 | 0.51431 | 1.28839 | 3.55365 |
| **PHOSPHO1** | 14.06864 | 60.26263 | 28.85221 | 0.04520 | 10.70886 | 0.00000 | 3.03540 |
| **PHOSPHO2** | 1.66922 | 0.14581 | 1.13468 | 0.73685 | 0.79588 | 0.08065 | 0.31836 |
| **PHOX2A** | 0.00000 | 0.15255 | 0.30662 | 0.01523 | 0.00344 | 0.00000 | 0.07948 |
| **PHOX2B** | 0.02329 | 0.01388 | 0.04008 | 0.01531 | 0.34618 | 0.01183 | 0.02404 |
| **PHPT1** | 18.42339 | 28.51023 | 2.19325 | 22.61669 | 2.38174 | 16.45111 | 12.55084 |
| **PHRF1** | 0.20156 | 3.48066 | 1.66125 | 3.53466 | 4.02847 | 3.31589 | 14.04643 |
| **PHTF1** | 1.02556 | 0.03270 | 1.17704 | 4.93900 | 2.07569 | 1.15623 | 3.64963 |
| **PHTF2** | 10.52983 | 0.00000 | 2.86971 | 12.47774 | 6.72234 | 4.19335 | 2.88685 |
| **PHYH** | 0.27030 | 0.04654 | 0.61715 | 0.29453 | 0.28913 | 0.47635 | 0.02682 |
| **PHYHD1** | 2.08052 | 0.10320 | 0.87489 | 0.49537 | 0.17397 | 2.96833 | 2.03878 |
| **PHYHIP** | 0.00000 | 0.05103 | 0.35005 | 0.28975 | 0.02576 | 0.00000 | 0.03716 |
| **PHYHIPL** | 0.02381 | 0.02333 | 0.19294 | 0.10882 | 0.31857 | 0.00000 | 0.22073 |
| **PHYKPL** | 1.01241 | 1.59441 | 0.92154 | 1.98774 | 0.73789 | 2.30549 | 2.86690 |
| **PI15** | 0.02767 | 0.00000 | 0.21066 | 0.00357 | 0.00275 | 0.00985 | 0.30277 |
| **PI16** | 0.00000 | 0.36739 | 0.24211 | 0.00000 | 0.00183 | 4.12223 | 0.12355 |
| **PI3** | 0.01430 | 0.01962 | 0.01287 | 0.02463 | 0.00763 | 0.02247 | 0.03656 |
| **PI4K2A** | 0.10803 | 5.39246 | 4.26031 | 4.67433 | 0.19616 | 4.51637 | 8.60853 |
| **PI4K2B** | 0.53154 | 0.40781 | 0.00000 | 0.00000 | 0.56998 | 1.80957 | 2.77333 |
| **PI4KA** | 5.50652 | 3.40436 | 0.13277 | 4.32912 | 0.80787 | 2.45502 | 9.09252 |
| **PI4KAP1** | 3.32317 | 5.10949 | 0.00000 | 1.71983 | 0.04080 | 0.56754 | 2.49286 |
| **PI4KAP2** | 3.18008 | 3.25127 | 0.09175 | 2.58047 | 0.05603 | 0.05455 | 2.23454 |
| **PI4KB** | 6.99409 | 17.52856 | 10.18909 | 11.28967 | 6.64891 | 3.93968 | 3.14676 |
| **PIANP** | 0.01834 | 0.00000 | 0.05669 | 0.02865 | 0.52152 | 0.02620 | 0.08128 |
| **PIAS1** | 16.38676 | 5.58323 | 11.23697 | 7.69521 | 11.37146 | 6.42281 | 0.00000 |
| **PIAS2** | 3.08839 | 2.52057 | 4.36299 | 1.22320 | 3.60105 | 5.41391 | 5.73575 |
| **PIAS3** | 2.14437 | 0.00000 | 2.53421 | 2.34651 | 1.81118 | 3.49221 | 1.69959 |
| **PIAS4** | 2.53437 | 13.02743 | 8.38347 | 9.66016 | 3.39078 | 5.00812 | 10.78620 |
| **PIBF1** | 9.66346 | 0.42197 | 2.03520 | 0.00000 | 1.76482 | 1.29580 | 5.98058 |
| **PICALM** | 8.84996 | 0.00000 | 9.45009 | 7.65224 | 2.70574 | 5.08638 | 18.66380 |

| | | | | | | | |
|---|---|---|---|---|---|---|---|
| **PICK1** | 0.00000 | 1.67119 | 0.21990 | 0.17715 | 0.13965 | 0.41458 | 2.71763 |
| **PID1** | 2.90026 | 0.56050 | 0.46377 | 12.40945 | 0.39930 | 0.00000 | 0.56461 |
| **PIDD1** | 0.22569 | 1.38912 | 0.13562 | 3.67668 | 0.00653 | 1.95783 | 7.80586 |
| **PIEZO1** | 2.50451 | 3.03130 | 1.78476 | 5.42568 | 1.12421 | 0.00000 | 21.41441 |
| **PIEZO2** | 0.07395 | 0.00930 | 0.04672 | 0.04054 | 0.13314 | 0.03385 | 0.05729 |
| **PIF1** | 5.68906 | 5.40827 | 1.41354 | 2.67397 | 0.31230 | 2.36433 | 6.02062 |
| **PIFO** | 0.55199 | 0.51230 | 0.94413 | 0.61833 | 0.08701 | 3.41425 | 0.00000 |
| **PIGA** | 10.33154 | 0.38629 | 5.66910 | 4.64734 | 5.51111 | 2.21543 | 0.54252 |
| **PIGB** | 4.56724 | 3.09414 | 3.80257 | 2.40663 | 1.62918 | 1.52605 | 0.00000 |
| **PIGC** | 9.01805 | 4.42294 | 8.08997 | 4.95924 | 0.40577 | 3.30377 | 0.80768 |
| **PIGF** | 4.40481 | 0.63063 | 4.10318 | 2.09820 | 1.44731 | 1.31558 | 1.13869 |
| **PIGG** | 12.51556 | 7.73708 | 3.52367 | 7.18197 | 4.07912 | 8.03714 | 6.17030 |
| **PIGH** | 9.34395 | 2.07218 | 6.30093 | 9.39194 | 4.30577 | 1.51365 | 0.00000 |
| **PIGK** | 9.29001 | 1.49601 | 6.38113 | 2.80436 | 0.31060 | 2.99132 | 0.66104 |
| **PIGL** | 5.87044 | 4.94727 | 4.66688 | 3.93988 | 0.19412 | 3.53836 | 0.57952 |
| **PIGM** | 2.25046 | 2.20745 | 2.83641 | 0.79119 | 0.80302 | 0.19908 | 1.16676 |
| **PIGN** | 6.19985 | 2.05019 | 1.35967 | 4.58042 | 0.57488 | 2.68760 | 0.00053 |
| **PIGO** | 6.61115 | 8.05205 | 3.21351 | 1.83430 | 0.06250 | 4.50777 | 2.81966 |
| **PIGP** | 4.12852 | 0.34331 | 4.46953 | 0.98645 | 0.22570 | 0.31223 | 1.43899 |
| **PIGQ** | 0.13484 | 3.65672 | 0.34810 | 2.23653 | 0.00000 | 1.04485 | 4.65500 |
| **PIGR** | 0.02992 | 0.02912 | 0.01063 | 0.04062 | 0.00852 | 0.03550 | 0.01192 |
| **PIGS** | 10.85719 | 8.10929 | 4.95109 | 8.75976 | 1.49141 | 5.45263 | 7.61452 |
| **PIGT** | 18.22234 | 8.97198 | 8.61974 | 14.13791 | 2.74113 | 6.27518 | 5.60644 |
| **PIGU** | 3.59055 | 3.54679 | 2.82409 | 3.73935 | 2.56242 | 2.26361 | 3.12092 |
| **PIGV** | 1.56895 | 1.27481 | 2.87796 | 1.07471 | 0.68081 | 0.22973 | 0.00000 |
| **PIGW** | 11.49233 | 1.15516 | 9.24106 | 5.22615 | 1.56476 | 3.56012 | 0.00000 |
| **PIGX** | 0.20299 | 1.03682 | 0.72499 | 0.10769 | 0.07687 | 2.69479 | 2.97416 |
| **PIGY+PYURF** | 23.55759 | 11.94555 | 13.00570 | 12.64993 | 5.47443 | 6.47187 | 3.54575 |
| **PIGZ** | 0.00000 | 0.05481 | 0.22809 | 0.00660 | 0.13354 | 0.29932 | 0.04662 |
| **PIH1D1** | 14.80467 | 9.18102 | 4.46698 | 21.45495 | 6.33722 | 10.44244 | 2.91636 |
| **PIH1D2** | 0.13044 | 0.03149 | 0.51079 | 1.34781 | 0.14002 | 0.12401 | 0.00000 |
| **PIH1D3** | 0.02537 | 0.01805 | 0.01270 | 0.02106 | 0.00805 | 0.01441 | 0.01630 |
| **PIK3AP1** | 0.25746 | 0.02323 | 2.71279 | 0.00000 | 0.04610 | 1.04153 | 0.02885 |
| **PIK3C2A** | 13.46959 | 0.00000 | 4.83369 | 14.39915 | 5.80000 | 6.27793 | 13.81304 |
| **PIK3C2B** | 1.47209 | 0.00000 | 0.42272 | 1.68986 | 0.59221 | 0.00000 | 4.60629 |

| | | | | | | | |
|---|---|---|---|---|---|---|---|
| **PIK3C2G** | 0.08627 | 0.01272 | 0.01129 | 0.07250 | 0.03125 | 0.02349 | 0.00000 |
| **PIK3C3** | 11.43274 | 0.15931 | 8.77646 | 7.47634 | 5.99638 | 3.84634 | 0.01592 |
| **PIK3CA** | 0.34952 | 0.15991 | 1.34647 | 0.28398 | 1.37266 | 1.14882 | 1.98645 |
| **PIK3CB** | 5.86779 | 0.90885 | 9.73647 | 8.90131 | 1.59449 | 1.95863 | 13.34619 |
| **PIK3CD** | 0.00000 | 1.25086 | 0.47793 | 0.22957 | 0.29159 | 0.37853 | 0.77369 |
| **PIK3CG** | 0.11164 | 0.00000 | 0.15762 | 0.11690 | 0.09965 | 0.04046 | 0.05461 |
| **PIK3IP1** | 1.42047 | 0.00000 | 1.34207 | 0.34580 | 0.21126 | 0.07057 | 0.18269 |
| **PIK3R1** | 0.24456 | 0.11171 | 0.68863 | 0.54542 | 1.58222 | 2.83123 | 1.12501 |
| **PIK3R2** | 0.00000 | 0.91804 | 0.38841 | 1.17161 | 0.31615 | 1.38835 | 4.18135 |
| **PIK3R3** | 13.56405 | 0.00000 | 1.57056 | 9.14507 | 5.49476 | 3.24809 | 4.48630 |
| **PIK3R4** | 10.48103 | 0.89002 | 9.92353 | 3.73558 | 6.26836 | 6.12388 | 4.18680 |
| **PIK3R5** | 0.22472 | 0.00000 | 0.05439 | 0.00748 | 0.03753 | 0.39982 | 0.00000 |
| **PIK3R6** | 0.03994 | 0.00956 | 0.00774 | 0.02174 | 0.08346 | 0.05353 | 0.02839 |
| **PIKFYVE** | 3.60606 | 0.00000 | 3.60051 | 1.81371 | 3.14975 | 5.09837 | 9.12714 |
| **PILRA** | 0.08105 | 0.44514 | 0.10499 | 0.15022 | 0.06626 | 0.04619 | 0.04965 |
| **PILRB** | 0.07261 | 0.43655 | 0.04436 | 0.07508 | 0.23208 | 0.20868 | 0.05065 |
| **PIM1** | 2.39247 | 2.68787 | 27.70939 | 30.16341 | 10.99586 | 7.60007 | 15.36011 |
| **PIM2** | 12.31197 | 6.96846 | 0.02581 | 2.97044 | 1.72702 | 35.42678 | 1.22128 |
| **PIM3** | 0.00000 | 7.81981 | 0.68864 | 6.02655 | 1.54481 | 2.72717 | 18.40727 |
| **PIN1** | 7.26132 | 25.00507 | 4.78147 | 11.91608 | 1.14495 | 7.67992 | 8.62491 |
| **PIN1P1** | 0.25210 | 1.09164 | 0.14565 | 0.47991 | 0.09092 | 0.25186 | 0.26600 |
| **PIN4** | 13.03521 | 16.13882 | 14.15125 | 7.50900 | 1.03755 | 10.79828 | 10.69575 |
| **PIN4P1** | 4.68349 | 9.55557 | 4.60238 | 2.73890 | 0.40459 | 3.68269 | 2.41652 |
| **PINK1** | 0.00000 | 6.92856 | 1.95155 | 0.00000 | 0.10928 | 4.04952 | 1.55505 |
| **PINK1-AS** | 0.00000 | 2.10144 | 0.39075 | 0.64800 | 0.00036 | 0.70141 | 0.69414 |
| **PINLYP** | 0.92707 | 0.61031 | 0.05557 | 0.38077 | 0.03899 | 0.87422 | 0.00000 |
| **PINX1** | 15.80841 | 12.28872 | 9.00066 | 8.44731 | 4.35178 | 5.66660 | 2.28864 |
| **PIP4K2A** | 28.46308 | 7.77908 | 2.71676 | 9.58141 | 4.27843 | 6.94111 | 2.02740 |
| **PIP4K2B** | 4.59600 | 1.48376 | 1.22978 | 4.58491 | 1.28079 | 4.38249 | 6.51632 |
| **PIP4K2C** | 18.53910 | 9.62486 | 7.40023 | 13.83080 | 6.48272 | 7.58577 | 0.00074 |
| **PIP5K1A** | 16.27531 | 6.12013 | 6.56864 | 17.86221 | 8.61809 | 5.93658 | 1.90240 |
| **PIP5K1B** | 7.75448 | 7.17521 | 10.24590 | 0.02294 | 0.53409 | 0.00000 | 3.54095 |
| **PIP5K1C** | 0.14511 | 3.28561 | 0.84540 | 2.70798 | 0.30160 | 1.25746 | 13.11055 |
| **PIP5K1P1** | 0.34170 | 0.04308 | 0.15529 | 0.26992 | 0.25114 | 0.09171 | 0.00000 |
| **PIP5KL1** | 0.00000 | 0.14305 | 0.01223 | 0.64711 | 0.15361 | 0.01987 | 0.15671 |

| | | | | | | | |
|---|---|---|---|---|---|---|---|
| **PIPOX** | 0.00611 | 0.21862 | 0.14584 | 0.09124 | 0.00250 | 0.64397 | 0.00000 |
| **PIPSL** | 16.55906 | 13.51824 | 5.48027 | 15.80721 | 2.83476 | 5.01577 | 0.00000 |
| **PIR** | 3.38528 | 0.50008 | 0.00000 | 2.23940 | 0.00000 | 0.17445 | 1.56909 |
| **PIR-FIGF** | 0.04738 | 0.04644 | 0.02410 | 0.02281 | 0.00691 | 0.00952 | 0.00492 |
| **PISD** | 19.25769 | 15.36263 | 3.52886 | 14.90315 | 2.00010 | 6.36633 | 0.00000 |
| **PISRT1** | 0.05633 | 0.00122 | 0.01311 | 0.01817 | 0.04592 | 0.03283 | 0.03489 |
| **PITHD1** | 9.52776 | 9.84255 | 13.22870 | 18.47733 | 9.13266 | 9.77144 | 27.94913 |
| **PITPNA** | 2.50162 | 4.64257 | 4.55749 | 6.77134 | 0.86430 | 2.17195 | 16.41622 |
| **PITPNA-AS1** | 2.07187 | 2.73541 | 7.30990 | 2.45881 | 0.60204 | 1.52843 | 1.42918 |
| **PITPNB** | 28.42111 | 9.14107 | 10.28427 | 27.18312 | 10.90540 | 14.71651 | 5.53623 |
| **PITPNC1** | 5.49945 | 5.98195 | 0.80212 | 1.13756 | 0.02574 | 8.79710 | 17.69916 |
| **PITPNM1** | 0.00000 | 0.26901 | 0.58422 | 0.81654 | 0.15805 | 1.15184 | 0.84975 |
| **PITPNM2** | 0.00000 | 0.00313 | 0.00464 | 1.06643 | 0.00575 | 3.12743 | 0.45733 |
| **PITPNM3** | 0.01825 | 0.12183 | 0.01563 | 0.03262 | 0.11208 | 0.10085 | 0.26507 |
| **PITRM1** | 16.86602 | 4.82375 | 3.98922 | 8.87722 | 7.01508 | 4.79172 | 3.11762 |
| **PITRM1-AS1** | 0.18486 | 0.22591 | 0.17600 | 0.63603 | 0.14509 | 0.73124 | 0.83754 |
| **PITX1** | 0.00000 | 4.12925 | 1.49843 | 0.39684 | 0.50436 | 3.28096 | 7.45950 |
| **PITX2** | 0.45487 | 0.26447 | 0.68440 | 0.57232 | 0.33237 | 5.58657 | 0.00000 |
| **PITX3** | 0.00000 | 0.10103 | 0.26359 | 0.04420 | 0.00595 | 0.12537 | 0.08243 |
| **PIWIL1** | 0.02043 | 0.02335 | 0.00971 | 0.02111 | 0.57697 | 0.01152 | 0.01495 |
| **PIWIL2** | 0.29017 | 2.43775 | 4.80853 | 0.10859 | 1.56547 | 0.00000 | 0.00000 |
| **PIWIL3** | 1.83953 | 0.00000 | 0.00000 | 0.62408 | 2.04143 | 0.00000 | 2.61725 |
| **PIWIL4** | 0.94642 | 0.00000 | 0.50519 | 0.68807 | 0.00090 | 0.00000 | 0.62359 |
| **PJA1** | 2.40741 | 1.89720 | 3.71230 | 0.35167 | 1.08645 | 0.94686 | 0.04715 |
| **PJA2** | 5.56401 | 0.00000 | 6.47050 | 3.70302 | 1.68687 | 5.22823 | 2.65248 |
| **PKD1** | 0.12439 | 1.79616 | 0.80654 | 0.71462 | 0.14241 | 0.93517 | 4.14156 |
| **PKD1L1** | 0.00000 | 0.46940 | 0.73589 | 0.23732 | 0.12612 | 2.97672 | 0.19669 |
| **PKD1L2** | 0.76198 | 2.54029 | 0.57733 | 0.99840 | 0.00170 | 1.14802 | 0.24981 |
| **PKD1L3** | 0.02235 | 0.01349 | 0.07369 | 0.04992 | 3.83598 | 0.00144 | 0.05928 |
| **PKD1P1** | 0.01835 | 0.12234 | 0.02473 | 0.10215 | 0.02734 | 0.04267 | 0.26987 |
| **PKD2** | 0.15273 | 0.00000 | 0.01225 | 0.00000 | 0.00754 | 0.84613 | 0.28822 |
| **PKD2L1** | 0.03338 | 0.02337 | 0.02531 | 0.03783 | 0.07374 | 0.03961 | 0.00000 |
| **PKD2L2** | 0.03313 | 0.00425 | 0.04204 | 0.03165 | 0.03138 | 0.02226 | 0.07712 |
| **PKDCC** | 0.00000 | 0.70349 | 0.95608 | 0.02397 | 0.00004 | 1.65631 | 0.02891 |
| **PKHD1** | 0.08997 | 0.39161 | 0.75033 | 0.00741 | 1.50575 | 1.04553 | 0.00000 |

| | | | | | | | |
|---|---|---|---|---|---|---|---|
| **PKHD1L1** | 0.02252 | 0.00000 | 0.60750 | 0.01195 | 0.17563 | 0.01284 | 0.08038 |
| **PKI55** | 0.27314 | 0.05923 | 0.19283 | 0.28141 | 0.00110 | 0.10980 | 0.40695 |
| **PKIA** | 0.00000 | 0.00389 | 0.00000 | 4.07298 | 0.83608 | 0.10223 | 0.14323 |
| **PKIB** | 11.15620 | 0.00000 | 0.72261 | 11.75775 | 0.33872 | 0.00000 | 1.93532 |
| **PKIG** | 1.07000 | 3.88662 | 0.51917 | 5.63014 | 0.32077 | 3.83652 | 0.85837 |
| **PKLR** | 0.03952 | 0.01102 | 0.02014 | 0.02022 | 0.18672 | 0.03269 | 0.00430 |
| **PKM** | 49.88505 | 33.86659 | 14.06011 | 58.00808 | 4.92990 | 26.60293 | 35.62360 |
| **PKMYT1** | 4.76780 | 11.85527 | 3.26363 | 12.85928 | 0.00000 | 3.87330 | 7.71277 |
| **PKN1** | 0.24811 | 0.83122 | 1.65080 | 0.82024 | 0.27544 | 0.94604 | 3.08719 |
| **PKN2** | 17.05550 | 4.47514 | 10.39419 | 9.75426 | 5.81804 | 8.58892 | 12.24138 |
| **PKN2-AS1** | 0.01812 | 0.02388 | 0.01629 | 0.01639 | 0.03298 | 0.03065 | 0.05487 |
| **PKN3** | 0.44774 | 0.66308 | 0.00000 | 1.54964 | 0.15876 | 0.15887 | 4.25577 |
| **PKNOX1** | 13.17405 | 10.83746 | 7.51892 | 7.38015 | 7.15039 | 1.20296 | 4.08587 |
| **PKNOX2** | 0.00000 | 0.14819 | 0.03901 | 0.06752 | 0.96723 | 0.00119 | 0.01706 |
| **PKP1** | 0.05486 | 0.08864 | 0.00098 | 0.05287 | 0.00708 | 0.00000 | 0.31374 |
| **PKP2** | 27.06288 | 8.21274 | 0.00000 | 20.82536 | 0.00000 | 4.37250 | 0.10037 |
| **PKP3** | 3.96844 | 0.00000 | 0.00000 | 6.88482 | 0.00000 | 0.00000 | 23.31467 |
| **PKP4** | 2.75338 | 4.59524 | 0.00000 | 2.65404 | 0.31320 | 1.77691 | 7.30323 |
| **PLA1A** | 0.05679 | 0.01691 | 0.01142 | 0.01560 | 0.02391 | 0.00957 | 0.02292 |
| **PLA2G10** | 0.25092 | 0.26220 | 0.00000 | 0.41383 | 0.00573 | 2.26987 | 1.42512 |
| **PLA2G12A** | 9.15610 | 7.34749 | 3.12829 | 4.14904 | 2.56251 | 3.81267 | 3.51372 |
| **PLA2G12B** | 0.19895 | 0.00000 | 0.00459 | 0.03027 | 0.07220 | 0.81906 | 0.00000 |
| **PLA2G15** | 3.69153 | 6.01742 | 9.62314 | 13.51030 | 3.19052 | 2.95150 | 0.00000 |
| **PLA2G16** | 4.62696 | 2.32073 | 1.26948 | 16.40551 | 11.86365 | 9.46449 | 1.75222 |
| **PLA2G1B** | 0.02634 | 0.01752 | 0.01371 | 0.01941 | 0.01280 | 0.01478 | 0.01497 |
| **PLA2G2A** | 0.10912 | 0.00000 | 0.01240 | 0.10050 | 0.17501 | 0.17259 | 0.00000 |
| **PLA2G2E** | 0.01827 | 0.01820 | 0.01308 | 0.03843 | 0.00802 | 0.01563 | 0.01587 |
| **PLA2G2F** | 0.09517 | 0.00267 | 0.01363 | 0.00923 | 0.01336 | 0.00817 | 0.03164 |
| **PLA2G3** | 0.28240 | 0.05086 | 0.04526 | 0.13758 | 0.02008 | 0.02228 | 0.00000 |
| **PLA2G4A** | 0.12352 | 0.02068 | 0.02051 | 0.01734 | 0.02135 | 0.01675 | 0.00000 |
| **PLA2G4B** | 0.02571 | 0.02099 | 0.01392 | 0.02038 | 0.00770 | 0.01295 | 0.01772 |
| **PLA2G4C** | 0.01027 | 0.10131 | 0.18121 | 0.03558 | 1.86744 | 0.00000 | 0.00596 |
| **PLA2G4D** | 0.02344 | 0.00112 | 0.09359 | 0.02250 | 0.00523 | 0.01213 | 0.02041 |
| **PLA2G4E** | 0.17765 | 0.03502 | 0.01569 | 0.00000 | 0.02313 | 0.01214 | 0.03149 |
| **PLA2G4F** | 0.83154 | 3.42909 | 3.00843 | 0.00000 | 0.00000 | 5.67859 | 0.04774 |

| | | | | | | | |
|---|---|---|---|---|---|---|---|
| **PLA2G5** | 0.02346 | 0.00258 | 0.07936 | 0.01995 | 0.00623 | 0.01310 | 0.02236 |
| **PLA2G6** | 0.29996 | 0.88402 | 0.28643 | 0.04909 | 0.19739 | 0.34471 | 0.71182 |
| **PLA2G7** | 0.02381 | 0.02568 | 0.01076 | 0.01875 | 0.00929 | 0.02687 | 0.01365 |
| **PLA2R1** | 0.03894 | 0.04465 | 0.03207 | 0.05306 | 0.00731 | 0.24997 | 0.00489 |
| **PLAA** | 19.92429 | 6.61203 | 6.70746 | 18.52241 | 2.96615 | 7.26862 | 2.58977 |
| **PLAC1** | 0.69301 | 0.00000 | 0.13434 | 0.08794 | 0.42958 | 0.00009 | 0.05642 |
| **PLAC4** | 0.00000 | 0.01038 | 0.03982 | 0.41770 | 0.02457 | 0.03242 | 0.01033 |
| **PLAC8** | 13.32928 | 4.81423 | 0.59552 | 1.22691 | 0.02091 | 0.00000 | 3.61488 |
| **PLAC8L1** | 0.38834 | 0.00000 | 0.03755 | 0.29245 | 0.00366 | 0.27406 | 0.65885 |
| **PLAC9** | 0.00000 | 0.10777 | 0.06584 | 0.92226 | 0.02672 | 0.00957 | 0.00354 |
| **PLAG1** | 1.33684 | 0.27505 | 1.09319 | 0.33274 | 4.12926 | 0.57191 | 0.18903 |
| **PLAGL1** | 0.04634 | 0.08319 | 0.04569 | 0.02735 | 0.36232 | 0.02097 | 0.00000 |
| **PLAGL2** | 3.66273 | 1.88534 | 3.39954 | 2.05891 | 3.28234 | 0.79463 | 9.90170 |
| **PLAT** | 0.05382 | 0.03040 | 0.01675 | 0.11324 | 0.09816 | 0.02351 | 0.20047 |
| **PLAU** | 0.67034 | 0.00000 | 0.17169 | 1.59879 | 0.00353 | 5.18412 | 1.41452 |
| **PLAUR** | 0.75063 | 3.32630 | 1.82477 | 0.45824 | 0.63424 | 0.67966 | 0.00000 |
| **PLB1** | 0.09725 | 0.06414 | 0.00000 | 0.75319 | 1.01554 | 0.42447 | 0.00000 |
| **PLBD1** | 10.49036 | 1.38187 | 13.77767 | 4.02450 | 0.38119 | 16.30631 | 8.29190 |
| **PLBD2** | 0.75298 | 1.38024 | 0.46401 | 2.25838 | 0.00198 | 0.50875 | 5.64385 |
| **PLCB1** | 0.24677 | 0.99795 | 1.45028 | 0.25122 | 1.67359 | 2.03312 | 0.99873 |
| **PLCB2** | 0.06250 | 0.00232 | 0.01342 | 0.02676 | 0.04010 | 0.01724 | 0.06332 |
| **PLCB3** | 0.00000 | 4.90880 | 2.42092 | 1.31966 | 0.59323 | 2.19278 | 9.29803 |
| **PLCB4** | 0.00000 | 1.13386 | 0.00771 | 0.00000 | 0.09356 | 5.19424 | 0.20368 |
| **PLCD1** | 0.00000 | 9.28358 | 8.88494 | 0.39378 | 0.46569 | 6.19887 | 0.28530 |
| **PLCD3** | 6.08834 | 8.63711 | 1.68687 | 4.37476 | 0.21690 | 0.00000 | 16.43475 |
| **PLCD4** | 0.08693 | 0.55201 | 0.26316 | 0.06489 | 0.27377 | 0.30337 | 1.91276 |
| **PLCE1** | 1.15984 | 5.57618 | 0.76473 | 0.17599 | 2.93762 | 2.88530 | 3.95191 |
| **PLCE1-AS1** | 0.05143 | 0.07421 | 0.01632 | 0.27300 | 0.01984 | 0.03610 | 0.00157 |
| **PLCG1** | 2.46599 | 1.01204 | 0.22787 | 1.99975 | 0.38530 | 1.84974 | 6.68602 |
| **PLCG1-AS1** | 0.12242 | 0.00000 | 0.16595 | 0.20305 | 0.16740 | 0.12809 | 1.02852 |
| **PLCG2** | 0.96475 | 0.77127 | 1.02721 | 0.00000 | 0.64778 | 1.44680 | 0.79957 |
| **PLCH1** | 0.07967 | 0.00149 | 0.12422 | 0.00897 | 0.59955 | 0.22719 | 0.00072 |
| **PLCH2** | 0.01550 | 0.00958 | 0.01901 | 0.02644 | 0.17986 | 0.02269 | 0.03286 |
| **PLCL1** | 0.01515 | 0.15665 | 0.00000 | 0.00476 | 2.55509 | 0.04478 | 0.01187 |
| **PLCL2** | 0.02127 | 0.09780 | 0.84442 | 0.03376 | 0.09289 | 0.40251 | 0.13832 |

| | | | | | | | |
|---|---|---|---|---|---|---|---|
| **PLCXD2** | 0.00742 | 0.06454 | 0.02320 | 0.04908 | 0.04089 | 0.00902 | 0.01336 |
| **PLCXD2-AS1** | 0.03892 | 0.02281 | 0.01302 | 0.02285 | 0.00791 | 0.01181 | 0.00997 |
| **PLCXD3** | 0.02112 | 0.01722 | 0.03118 | 0.02799 | 0.01684 | 0.04013 | 0.02468 |
| **PLCZ1** | 0.02364 | 0.01832 | 0.01174 | 0.01880 | 0.02788 | 0.01640 | 0.01844 |
| **PLD1** | 0.77795 | 0.09025 | 0.03712 | 1.04965 | 1.05217 | 0.47602 | 0.08835 |
| **PLD2** | 4.32368 | 2.51990 | 11.83106 | 8.90016 | 0.96962 | 2.04661 | 6.22730 |
| **PLD3** | 4.22266 | 9.05707 | 10.44021 | 24.02386 | 1.43899 | 2.10018 | 6.09485 |
| **PLD4** | 0.02728 | 0.01476 | 0.01586 | 0.02072 | 0.00776 | 0.03681 | 0.01079 |
| **PLD5** | 0.00000 | 0.07264 | 0.00416 | 0.10599 | 0.08229 | 0.00273 | 0.08242 |
| **PLD6** | 0.98143 | 0.00000 | 1.03744 | 1.52974 | 0.22087 | 0.40410 | 1.22624 |
| **PLEC** | 0.00000 | 22.75938 | 1.28244 | 10.19529 | 0.02416 | 11.07683 | 43.91876 |
| **PLEK** | 0.00219 | 0.01740 | 0.00757 | 0.07370 | 0.05499 | 0.01653 | 0.05814 |
| **PLEK2** | 1.13220 | 0.72099 | 0.00000 | 3.94126 | 11.04949 | 0.00000 | 2.14590 |
| **PLEKHA1** | 13.79045 | 6.62212 | 4.85076 | 3.56486 | 0.12737 | 3.35399 | 12.95421 |
| **PLEKHA2** | 0.19029 | 0.45241 | 0.15264 | 0.00000 | 0.00107 | 0.00000 | 1.28211 |
| **PLEKHA3** | 0.18834 | 0.00000 | 1.73726 | 0.40650 | 0.15956 | 1.66897 | 3.97593 |
| **PLEKHA4** | 0.00000 | 1.23671 | 0.37215 | 1.74169 | 0.93046 | 4.27975 | 0.39534 |
| **PLEKHA5** | 3.35393 | 4.14022 | 14.29691 | 1.05076 | 2.14116 | 9.37921 | 3.02539 |
| **PLEKHA6** | 0.29512 | 0.37633 | 0.00000 | 17.43386 | 0.02160 | 0.00000 | 5.76072 |
| **PLEKHA7** | 0.00000 | 3.31776 | 2.22166 | 3.06176 | 0.39056 | 2.27269 | 6.38656 |
| **PLEKHA8** | 2.10799 | 2.54417 | 2.38821 | 1.82470 | 0.00000 | 3.38061 | 6.37661 |
| **PLEKHA8P1** | 1.56748 | 1.21940 | 0.52546 | 2.47225 | 0.45437 | 0.16087 | 1.07878 |
| **PLEKHB1** | 0.00000 | 0.34471 | 1.78377 | 0.08047 | 5.01230 | 0.45081 | 0.10379 |
| **PLEKHB2** | 54.41440 | 42.05702 | 21.74340 | 33.23439 | 17.19352 | 25.66687 | 13.14313 |
| **PLEKHD1** | 0.06539 | 0.00838 | 0.01232 | 0.01830 | 0.00706 | 0.03156 | 0.06389 |
| **PLEKHF1** | 0.00000 | 0.01026 | 0.00147 | 15.34734 | 0.00379 | 1.41665 | 5.79854 |
| **PLEKHF2** | 9.63613 | 2.68866 | 0.89444 | 6.11865 | 0.30609 | 0.57138 | 5.08203 |
| **PLEKHG1** | 0.15267 | 0.26063 | 0.04405 | 0.00000 | 1.80262 | 0.04978 | 0.55666 |
| **PLEKHG2** | 0.00000 | 0.18394 | 0.54643 | 1.86163 | 0.10274 | 1.18181 | 1.91198 |
| **PLEKHG3** | 0.04404 | 4.06649 | 0.78445 | 2.22916 | 0.20580 | 0.57721 | 7.52750 |
| **PLEKHG4** | 0.00000 | 0.89173 | 0.12365 | 0.03302 | 0.09417 | 0.19921 | 0.03329 |
| **PLEKHG4B** | 1.82798 | 0.75267 | 0.24061 | 2.32357 | 0.00000 | 0.00000 | 1.89974 |
| **PLEKHG5** | 0.00000 | 0.20395 | 0.31652 | 0.11101 | 0.56393 | 1.22639 | 0.10360 |
| **PLEKHG6** | 0.00000 | 0.17801 | 0.74742 | 1.17764 | 0.00000 | 1.61502 | 0.69380 |
| **PLEKHG7** | 0.04666 | 0.00000 | 0.04809 | 0.06511 | 0.00591 | 0.00905 | 0.21652 |

| | | | | | | | |
|---|---|---|---|---|---|---|---|
| **PLEKHH1** | 1.77765 | 0.00000 | 0.05940 | 0.26301 | 0.04322 | 2.11111 | 2.01861 |
| **PLEKHH2** | 0.17596 | 0.37869 | 0.28489 | 0.00277 | 0.09663 | 0.07908 | 0.00000 |
| **PLEKHH3** | 0.00000 | 0.98121 | 0.33601 | 2.57650 | 0.47960 | 0.86125 | 5.47022 |
| **PLEKHJ1** | 19.47162 | 20.03157 | 5.42030 | 13.64574 | 0.87291 | 10.62345 | 11.32223 |
| **PLEKHM1** | 0.50584 | 1.62256 | 1.59362 | 1.66093 | 1.00593 | 0.71645 | 4.42928 |
| **PLEKHM1P** | 0.12684 | 0.11347 | 0.46866 | 0.61669 | 0.68646 | 0.30658 | 1.36096 |
| **PLEKHM2** | 2.02087 | 11.40842 | 2.24036 | 5.64473 | 1.72503 | 4.18033 | 15.67292 |
| **PLEKHM3** | 2.97330 | 0.77114 | 2.42431 | 2.90522 | 1.90925 | 0.72703 | 0.00000 |
| **PLEKHN1** | 0.00440 | 0.06454 | 0.08447 | 0.12136 | 0.11307 | 0.00977 | 0.05540 |
| **PLEKHO1** | 0.00000 | 0.13733 | 2.33579 | 0.28492 | 0.28246 | 0.33774 | 0.25364 |
| **PLEKHO2** | 0.00000 | 4.39285 | 0.00000 | 5.69212 | 1.89790 | 1.67595 | 4.97197 |
| **PLEKHS1** | 0.02338 | 0.01895 | 0.01370 | 0.02240 | 0.00773 | 0.01224 | 0.02033 |
| **PLET1** | 0.08326 | 0.03169 | 0.02240 | 0.04600 | 0.01160 | 0.00125 | 0.08383 |
| **PLG** | 0.02157 | 0.01507 | 0.01486 | 0.02302 | 0.00824 | 0.01268 | 0.02487 |
| **PLGLA** | 0.01694 | 0.02255 | 0.01654 | 0.01825 | 0.01112 | 0.03604 | 0.02266 |
| **PLGRKT** | 7.57457 | 2.59138 | 1.67316 | 3.10023 | 1.92734 | 2.66024 | 0.00000 |
| **PLIN1** | 0.21480 | 0.00000 | 0.02296 | 0.00000 | 0.00861 | 0.00563 | 0.24368 |
| **PLIN2** | 34.13642 | 22.58656 | 33.64118 | 61.01697 | 10.37703 | 9.20149 | 7.37773 |
| **PLIN3** | 19.23570 | 18.16475 | 3.92958 | 11.37916 | 10.50622 | 19.30123 | 0.00000 |
| **PLIN4** | 0.04465 | 0.30271 | 0.16996 | 0.28566 | 0.04760 | 0.03279 | 0.16532 |
| **PLIN5** | 0.04992 | 0.00000 | 0.04446 | 0.01314 | 0.63597 | 0.07472 | 0.17167 |
| **PLK1** | 36.05986 | 48.35794 | 26.24750 | 28.76594 | 11.88232 | 15.94654 | 6.83327 |
| **PLK2** | 0.00000 | 4.56693 | 0.32703 | 7.39367 | 5.66608 | 2.82474 | 0.00000 |
| **PLK3** | 0.00000 | 6.05474 | 3.09371 | 2.56938 | 2.12492 | 3.32246 | 6.04143 |
| **PLK4** | 10.62128 | 0.00002 | 5.08620 | 5.91000 | 2.70132 | 4.03579 | 0.77431 |
| **PLK5** | 0.07923 | 0.00000 | 0.04405 | 0.00000 | 0.00516 | 0.04218 | 0.23112 |
| **PLLP** | 0.00000 | 7.36353 | 9.33910 | 0.00000 | 0.35933 | 5.78705 | 0.00000 |
| **PLN** | 0.04883 | 0.00000 | 0.08010 | 0.02182 | 0.04158 | 0.01283 | 0.06267 |
| **PLOD1** | 10.27505 | 7.42455 | 0.60461 | 9.46367 | 0.17600 | 10.90297 | 3.10127 |
| **PLOD2** | 2.14704 | 0.32325 | 1.36500 | 2.14178 | 0.71320 | 10.49448 | 2.63369 |
| **PLOD3** | 6.61183 | 3.33426 | 0.58241 | 9.51121 | 1.18300 | 5.50940 | 5.39479 |
| **PLP1** | 0.50823 | 0.06897 | 0.10934 | 0.00000 | 0.03162 | 0.97900 | 0.00191 |
| **PLP2** | 0.72698 | 0.16798 | 0.08831 | 0.29572 | 0.09395 | 0.13863 | 1.05439 |
| **PLRG1** | 33.78105 | 13.71542 | 12.80275 | 18.44389 | 6.19290 | 10.32015 | 1.54559 |
| **PLS1** | 2.45109 | 0.77340 | 1.48055 | 4.38211 | 6.04694 | 0.72674 | 0.00000 |

| | | | | | | | |
|---|---|---|---|---|---|---|---|
| **PLS3** | 48.30483 | 15.11728 | 16.63645 | 23.61090 | 0.19882 | 2.39912 | 5.49356 |
| **PLS3-AS1** | 0.15123 | 0.11122 | 0.08657 | 0.25155 | 0.00386 | 0.03506 | 0.05811 |
| **PLSCR1** | 4.09976 | 4.21234 | 0.30061 | 8.98027 | 0.00000 | 15.81934 | 0.00000 |
| **PLSCR2** | 0.04986 | 0.02275 | 0.02293 | 0.03008 | 0.06141 | 0.03039 | 0.02217 |
| **PLSCR3** | 0.03076 | 0.02323 | 0.01937 | 0.04364 | 0.01085 | 0.01834 | 0.01343 |
| **PLSCR4** | 0.05031 | 0.10572 | 1.42796 | 0.03024 | 0.32199 | 1.68281 | 0.00000 |
| **PLTP** | 22.06050 | 6.07371 | 1.53860 | 16.40846 | 0.11695 | 11.06992 | 18.46276 |
| **PLVAP** | 0.00000 | 0.00248 | 2.10288 | 0.00600 | 5.79372 | 0.00000 | 0.02679 |
| **PLXDC1** | 0.05946 | 0.16175 | 0.12828 | 0.06384 | 0.38476 | 0.04334 | 0.00000 |
| **PLXDC2** | 5.03127 | 0.00889 | 0.00000 | 7.34029 | 0.08125 | 0.07311 | 0.85123 |
| **PLXNA1** | 1.35076 | 4.99035 | 0.86320 | 16.82101 | 0.53849 | 0.49762 | 21.80487 |
| **PLXNA2** | 0.47071 | 4.61568 | 0.20400 | 14.53403 | 0.02920 | 0.33665 | 8.41369 |
| **PLXNA3** | 1.76771 | 0.00000 | 0.53150 | 1.65543 | 0.31136 | 0.19802 | 9.33497 |
| **PLXNA4** | 0.05614 | 0.00000 | 0.10487 | 0.00000 | 4.06481 | 0.13410 | 0.03495 |
| **PLXNB1** | 2.86765 | 0.00000 | 0.00000 | 11.42244 | 0.00000 | 0.00000 | 4.74871 |
| **PLXNB2** | 0.00000 | 15.11104 | 1.21550 | 7.56739 | 0.00618 | 4.81577 | 5.74473 |
| **PLXNB3** | 0.02346 | 0.19549 | 0.08669 | 0.00000 | 2.74199 | 0.00000 | 0.06561 |
| **PLXNC1** | 0.44724 | 0.00000 | 0.24174 | 0.13682 | 0.71909 | 0.16286 | 0.70955 |
| **PLXND1** | 1.19597 | 7.53098 | 0.57138 | 9.48895 | 0.05453 | 11.78728 | 11.25809 |
| **PM20D1** | 0.01605 | 0.00000 | 0.09156 | 0.06236 | 0.68353 | 0.01235 | 0.02351 |
| **PM20D2** | 3.46707 | 0.00000 | 0.00000 | 2.14208 | 0.25083 | 0.66501 | 1.92736 |
| **PMAIP1** | 1.78668 | 2.98578 | 2.22214 | 0.36001 | 10.26852 | 17.33298 | 0.34094 |
| **PMCH** | 0.07127 | 0.00995 | 0.02372 | 0.04583 | 0.00780 | 0.05110 | 0.04875 |
| **PMCHL2** | 0.02172 | 0.01578 | 0.01269 | 0.02006 | 0.01075 | 0.02631 | 0.02143 |
| **PMEL** | 2.35169 | 0.04691 | 0.13584 | 9.78739 | 0.06878 | 0.00000 | 0.66997 |
| **PMEPA1** | 0.01882 | 0.00000 | 0.00826 | 0.03787 | 0.01005 | 3.43327 | 0.03442 |
| **PMF1** | 9.74629 | 8.13996 | 0.03556 | 11.34355 | 0.28576 | 4.94328 | 3.09054 |
| **PMFBP1** | 0.06562 | 0.08395 | 0.03582 | 0.09670 | 0.00481 | 0.13016 | 0.05617 |
| **PML** | 0.62216 | 1.17209 | 1.19737 | 3.50667 | 0.57885 | 1.15341 | 2.20776 |
| **PMM1** | 3.62698 | 3.26713 | 3.02164 | 7.83331 | 9.36759 | 2.46881 | 3.72683 |
| **PMM2** | 23.41654 | 26.38293 | 16.05169 | 19.27897 | 4.89623 | 11.97834 | 10.14548 |
| **PMP2** | 0.03843 | 0.02176 | 0.01143 | 0.02906 | 0.00795 | 0.01161 | 0.01179 |
| **PMP22** | 3.11828 | 6.11766 | 1.72935 | 0.74769 | 0.13380 | 7.85990 | 0.00000 |
| **PMPCA** | 23.07301 | 30.11799 | 10.88506 | 25.64021 | 3.12895 | 10.55770 | 0.00000 |
| **PMPCB** | 27.64325 | 7.45017 | 13.35059 | 15.42710 | 0.80355 | 11.29777 | 2.86273 |

| | | | | | | | |
|---|---|---|---|---|---|---|---|
| **PMS1** | 9.72700 | 0.00834 | 4.09638 | 3.03282 | 0.80432 | 2.86431 | 5.00240 |
| **PMS2** | 2.27238 | 0.60099 | 0.83443 | 0.78735 | 0.66026 | 0.95986 | 0.00000 |
| **PMS2CL** | 0.11073 | 0.07010 | 0.13250 | 0.10659 | 0.10646 | 0.09643 | 0.36220 |
| **PMS2L2** | 0.02247 | 0.03858 | 0.02275 | 0.02088 | 0.01101 | 0.01281 | 0.02228 |
| **PMS2P1** | 0.61987 | 0.70281 | 1.58876 | 0.33805 | 0.90374 | 0.69426 | 0.08173 |
| **PMS2P3** | 0.10311 | 0.13590 | 0.01756 | 0.07780 | 0.02313 | 0.03912 | 0.10418 |
| **PMS2P4** | 0.05539 | 0.47098 | 0.69865 | 0.00000 | 0.09920 | 0.21588 | 0.33534 |
| **PMS2P5.uid-92** | 0.15781 | 0.06889 | 0.27533 | 0.07310 | 0.32295 | 0.14922 | 0.04115 |
| **PMS2P9** | 0.03286 | 0.03132 | 0.04572 | 0.06293 | 0.01100 | 0.02448 | 0.05323 |
| **PMVK** | 9.88121 | 8.86704 | 1.22235 | 2.93313 | 0.16798 | 4.61896 | 2.86325 |
| **PNCK** | 0.04703 | 0.23639 | 0.10345 | 0.00000 | 0.00245 | 0.15871 | 0.17922 |
| **PNISR** | 19.74468 | 7.38547 | 16.93956 | 18.49731 | 14.45099 | 14.49098 | 22.62006 |
| **PNKD** | 9.06894 | 20.06500 | 12.70971 | 12.64832 | 6.87376 | 7.71868 | 6.02037 |
| **PNKP** | 0.39609 | 3.81740 | 0.00525 | 1.55546 | 0.35678 | 2.78050 | 5.68726 |
| **PNLDC1** | 0.91441 | 0.27802 | 0.42384 | 0.00000 | 3.95352 | 1.29961 | 0.37417 |
| **PNLIPRP1** | 0.02297 | 0.02160 | 0.01333 | 0.02236 | 0.01132 | 0.01254 | 0.01590 |
| **PNLIPRP2** | 0.02161 | 0.01838 | 0.01356 | 0.03418 | 0.00788 | 0.01552 | 0.01498 |
| **PNMA1** | 5.99123 | 9.50723 | 19.57627 | 9.73455 | 6.04893 | 8.55004 | 5.99443 |
| **PNMA2** | 0.03986 | 0.00000 | 3.47680 | 0.01666 | 0.36302 | 1.50720 | 0.01222 |
| **PNMA3** | 0.01620 | 0.02090 | 0.25239 | 0.01677 | 0.00549 | 0.00382 | 0.02476 |
| **PNMA5** | 0.00000 | 7.27026 | 13.33567 | 0.00000 | 0.46789 | 0.00000 | 0.00000 |
| **PNMAL1** | 0.76804 | 0.12172 | 0.74625 | 0.74896 | 0.24156 | 0.65486 | 1.95374 |
| **PNMAL2** | 0.00844 | 0.13707 | 0.00309 | 0.02468 | 0.01395 | 0.02347 | 0.02104 |
| **PNMT** | 0.33530 | 0.00000 | 0.00913 | 0.00000 | 0.00615 | 0.00160 | 0.09365 |
| **PNN** | 37.33650 | 6.43146 | 16.71540 | 20.84317 | 18.64728 | 13.30099 | 33.57974 |
| **PNO1** | 33.57184 | 13.79102 | 17.00721 | 21.46602 | 9.52877 | 10.94620 | 3.46922 |
| **PNOC** | 0.01544 | 0.02355 | 0.01065 | 0.01898 | 0.05344 | 0.03643 | 0.02297 |
| **PNP** | 16.08664 | 0.00000 | 2.77020 | 33.63192 | 13.63564 | 22.07622 | 6.15364 |
| **PNPLA1** | 0.13350 | 0.13050 | 0.00002 | 0.02860 | 0.07575 | 0.09940 | 0.00000 |
| **PNPLA2** | 0.00000 | 1.66698 | 0.35168 | 2.90644 | 2.85863 | 2.33472 | 9.32451 |
| **PNPLA3** | 1.08498 | 0.22218 | 0.57616 | 4.25847 | 0.17100 | 0.48912 | 3.35461 |
| **PNPLA4** | 9.35490 | 14.47541 | 0.60128 | 3.07132 | 2.89656 | 4.21675 | 0.17289 |
| **PNPLA5** | 0.01979 | 0.01856 | 0.01281 | 0.01804 | 0.00847 | 0.01237 | 0.03329 |
| **PNPLA6** | 2.64451 | 6.15363 | 0.12848 | 10.52184 | 0.08880 | 4.52606 | 13.15255 |
| **PNPLA7** | 0.00000 | 0.00917 | 0.07529 | 0.15564 | 0.03956 | 0.08535 | 0.78552 |

| | | | | | | | |
|---|---|---|---|---|---|---|---|
| **PNPLA8** | 17.53175 | 1.32542 | 3.47453 | 9.72106 | 5.04949 | 5.79205 | 3.19206 |
| **PNPO** | 5.26105 | 7.23426 | 1.34994 | 0.00000 | 0.26995 | 2.78688 | 0.80184 |
| **PNPT1** | 31.52340 | 19.61947 | 19.43048 | 7.72499 | 2.68504 | 14.01253 | 0.00000 |
| **PNRC1** | 0.00000 | 0.00000 | 8.32425 | 0.85530 | 9.11033 | 9.12842 | 5.10660 |
| **PNRC2** | 18.97312 | 3.87962 | 12.37550 | 13.59303 | 8.83050 | 5.59808 | 1.41473 |
| **POC1A** | 4.64722 | 9.34735 | 5.91545 | 3.63950 | 0.14558 | 3.28292 | 2.72467 |
| **POC1B** | 6.17418 | 1.12253 | 0.03337 | 1.29813 | 0.65405 | 2.99869 | 1.84739 |
| **POC5** | 9.90651 | 1.62101 | 4.71286 | 6.18933 | 2.70639 | 3.38152 | 0.00000 |
| **PODN** | 0.02346 | 0.01615 | 0.01519 | 0.01938 | 0.00863 | 0.01733 | 0.01984 |
| **PODNL1** | 0.07601 | 0.07429 | 0.08316 | 0.04016 | 0.01259 | 0.02104 | 0.00000 |
| **PODXL** | 0.00000 | 1.23119 | 3.57699 | 2.27200 | 1.72929 | 6.07458 | 3.05776 |
| **PODXL2** | 0.05526 | 0.08683 | 0.03218 | 0.19397 | 0.00481 | 0.12303 | 0.23390 |
| **POF1B** | 0.01645 | 0.02505 | 0.02155 | 0.02107 | 0.00688 | 0.01813 | 0.03096 |
| **POFUT1** | 9.66743 | 4.79377 | 4.36499 | 2.03809 | 0.40990 | 2.80597 | 5.39810 |
| **POFUT2** | 0.00000 | 0.99995 | 1.16496 | 1.25460 | 1.01016 | 1.31930 | 2.08160 |
| **POGK** | 1.37802 | 1.99173 | 2.86846 | 2.32033 | 0.00000 | 3.13338 | 3.98005 |
| **POGLUT1** | 3.43722 | 0.20694 | 0.00309 | 1.17812 | 0.00486 | 2.46389 | 0.09722 |
| **POGZ** | 3.90247 | 3.62894 | 3.24625 | 3.38487 | 1.65354 | 4.53182 | 8.28406 |
| **POLA1** | 9.97559 | 4.88340 | 4.73847 | 0.00000 | 4.10812 | 3.63402 | 7.88027 |
| **POLA2** | 11.76127 | 6.78540 | 2.56074 | 6.28987 | 0.78449 | 3.31403 | 0.00000 |
| **POLB** | 6.33357 | 0.21354 | 8.07195 | 6.82425 | 3.31433 | 2.68863 | 5.91724 |
| **POLD1** | 3.08519 | 20.64580 | 7.02712 | 14.54824 | 0.80964 | 14.16712 | 16.90222 |
| **POLD2** | 26.56757 | 19.97530 | 5.88706 | 15.73891 | 0.46070 | 9.18976 | 2.10456 |
| **POLD3** | 14.39600 | 7.51814 | 14.21793 | 11.03342 | 7.59842 | 5.18189 | 4.12888 |
| **POLD4** | 0.00000 | 3.10549 | 0.96210 | 1.14339 | 0.05309 | 4.71094 | 1.25075 |
| **POLDIP2** | 13.51265 | 25.75989 | 8.87559 | 11.42996 | 3.05827 | 11.99321 | 24.84530 |
| **POLDIP3** | 24.33881 | 18.75597 | 12.71064 | 21.23792 | 10.12146 | 14.14728 | 8.44255 |
| **POLE** | 5.21877 | 8.32011 | 2.83114 | 9.93920 | 1.83991 | 5.54024 | 17.95777 |
| **POLE2** | 10.78895 | 3.18612 | 3.25659 | 2.71957 | 0.00000 | 3.89888 | 1.45082 |
| **POLE3** | 28.85615 | 13.49639 | 16.56632 | 23.15395 | 16.23192 | 8.81927 | 11.25027 |
| **POLE4** | 0.00000 | 2.54165 | 3.84543 | 4.61697 | 1.63145 | 3.82419 | 8.99170 |
| **POLG** | 3.11176 | 5.20748 | 1.76760 | 3.13092 | 1.49999 | 1.80047 | 6.48745 |
| **POLG2** | 5.84491 | 1.32337 | 2.07723 | 3.83380 | 2.02245 | 2.80190 | 0.70044 |
| **POLH** | 3.12269 | 4.96469 | 7.69970 | 5.77657 | 8.14030 | 5.57348 | 2.79293 |
| **POLI** | 3.64866 | 0.00000 | 3.36817 | 2.09043 | 0.62535 | 0.26120 | 0.00000 |

| | | | | | | | |
|---|---|---|---|---|---|---|---|
| **POLK** | 3.84829 | 0.00000 | 1.14868 | 2.26912 | 1.43368 | 1.95678 | 3.17042 |
| **POLL** | 0.93263 | 4.74696 | 2.56564 | 4.13994 | 1.11157 | 2.36206 | 4.39158 |
| **POLM** | 1.45477 | 3.14135 | 0.77952 | 3.23869 | 0.24454 | 3.14485 | 3.90238 |
| **POLN** | 0.27767 | 0.00511 | 0.15246 | 0.04909 | 0.21484 | 0.12803 | 0.58067 |
| **POLQ** | 2.71755 | 3.20472 | 3.12898 | 0.00000 | 3.06961 | 3.27882 | 3.75501 |
| **POLR1A** | 7.34561 | 7.47911 | 4.16556 | 6.98489 | 4.67874 | 2.20370 | 6.68816 |
| **POLR1B** | 26.78949 | 8.79693 | 4.58156 | 12.76929 | 2.17623 | 6.00372 | 0.47100 |
| **POLR1C** | 30.33349 | 19.29613 | 17.54994 | 20.29733 | 14.04483 | 9.89790 | 0.00000 |
| **POLR1D** | 29.28370 | 6.16260 | 20.46688 | 19.11770 | 7.91943 | 13.48988 | 33.47896 |
| **POLR1E** | 9.84016 | 4.90631 | 6.45148 | 6.85195 | 4.73265 | 2.69685 | 5.44146 |
| **POLR2A** | 14.80114 | 9.06744 | 11.82764 | 30.55452 | 7.23649 | 10.89007 | 35.67669 |
| **POLR2B** | 28.48872 | 2.38822 | 15.26887 | 15.61268 | 6.42638 | 12.54910 | 15.86440 |
| **POLR2C** | 22.47141 | 13.31069 | 11.71965 | 19.30304 | 12.86348 | 7.32306 | 0.00000 |
| **POLR2D** | 31.47249 | 16.70170 | 13.12686 | 19.55545 | 3.38241 | 7.77942 | 0.00000 |
| **POLR2E** | 52.18486 | 51.09758 | 24.39621 | 43.31991 | 15.88959 | 24.03458 | 12.70039 |
| **POLR2F** | 32.87575 | 43.08627 | 22.69189 | 37.00891 | 17.57190 | 19.18495 | 17.34385 |
| **POLR2G** | 28.40813 | 15.06233 | 13.11302 | 31.53174 | 5.27326 | 13.82253 | 4.76771 |
| **POLR2H** | 28.93045 | 20.92625 | 5.48282 | 33.48167 | 2.60888 | 16.47843 | 20.24794 |
| **POLR2I** | 23.67732 | 33.32372 | 11.49309 | 16.08107 | 4.73948 | 18.22129 | 13.22101 |
| **POLR2J** | 10.98266 | 20.52490 | 8.71623 | 11.40302 | 4.78263 | 7.82252 | 11.49229 |
| **POLR2J2** | 0.10472 | 0.04871 | 0.02613 | 0.02011 | 0.02111 | 0.08474 | 0.08114 |
| **POLR2J3** | 0.21603 | 0.13291 | 0.12516 | 0.11148 | 0.01865 | 0.06226 | 0.17347 |
| **POLR2J4** | 1.05110 | 0.82875 | 1.04919 | 0.53524 | 0.35372 | 0.00000 | 0.63283 |
| **POLR2K** | 32.81973 | 1.34015 | 21.29770 | 17.59477 | 4.48148 | 10.41345 | 13.26599 |
| **POLR2L** | 43.40266 | 48.82446 | 23.72400 | 44.23815 | 14.18793 | 25.58721 | 29.07214 |
| **POLR2M** | 0.26637 | 0.39775 | 0.21809 | 0.47476 | 0.32626 | 0.05155 | 0.10302 |
| **POLR3A** | 13.91836 | 5.81211 | 6.30957 | 11.21623 | 4.92770 | 6.22250 | 4.94412 |
| **POLR3B** | 10.72907 | 3.04197 | 5.75523 | 2.66734 | 1.74297 | 2.41158 | 6.93402 |
| **POLR3C** | 18.59511 | 21.23056 | 15.37143 | 3.31659 | 9.76367 | 8.26556 | 0.00000 |
| **POLR3D** | 2.19330 | 3.83770 | 4.59003 | 5.74840 | 2.85482 | 0.80419 | 11.50892 |
| **POLR3E** | 13.06116 | 8.75653 | 6.88591 | 13.56738 | 1.72913 | 8.05605 | 17.63074 |
| **POLR3F** | 10.11539 | 1.41235 | 8.00914 | 5.35746 | 2.15468 | 4.57642 | 2.89605 |
| **POLR3G** | 12.67436 | 0.00000 | 7.11882 | 0.00021 | 1.55548 | 1.36850 | 4.67571 |
| **POLR3GL** | 0.00000 | 1.63484 | 1.76946 | 0.13647 | 3.69460 | 0.67150 | 0.00000 |
| **POLR3H** | 20.26454 | 17.36034 | 9.18998 | 14.43896 | 8.24846 | 4.81084 | 7.10930 |

| | | | | | | | |
|---|---|---|---|---|---|---|---|
| **POLR3K** | 39.81696 | 23.98824 | 11.60854 | 22.47758 | 16.30116 | 15.86759 | 3.61990 |
| **POLRMT** | 0.00000 | 7.74298 | 7.35208 | 2.60188 | 0.99038 | 3.80751 | 12.32955 |
| **POM121** | 1.56621 | 3.47957 | 2.00355 | 3.88839 | 1.91841 | 2.86788 | 10.06292 |
| **POM121C** | 1.33700 | 2.16446 | 2.52705 | 3.77129 | 1.62166 | 2.99578 | 14.15760 |
| **POM121L10P** | 0.01101 | 0.02018 | 0.01284 | 0.02692 | 0.01304 | 0.17504 | 0.02786 |
| **POM121L1P** | 0.02737 | 0.01818 | 0.01400 | 0.02322 | 0.00791 | 0.01456 | 0.01250 |
| **POM121L2** | 0.06139 | 0.01331 | 0.01398 | 0.01506 | 0.00798 | 0.01226 | 0.00198 |
| **POM121L4P** | 0.02404 | 0.02025 | 0.01297 | 0.01974 | 0.00788 | 0.01331 | 0.01563 |
| **POM121L8P** | 0.02480 | 0.01801 | 0.01316 | 0.02210 | 0.00792 | 0.01480 | 0.01550 |
| **POM121L9P** | 0.01406 | 0.03999 | 0.09503 | 0.03502 | 0.00905 | 0.01583 | 0.02847 |
| **POMC** | 9.43252 | 16.58667 | 1.61937 | 0.00000 | 5.60556 | 1.40612 | 8.60710 |
| **POMGNT1** | 4.93623 | 4.42895 | 2.94947 | 7.44252 | 0.80232 | 2.57374 | 10.33854 |
| **POMGNT2** | 0.53972 | 7.28305 | 1.98973 | 2.94932 | 0.00000 | 2.60575 | 4.79184 |
| **POMK** | 3.37149 | 0.56380 | 1.54713 | 1.71360 | 2.85449 | 1.65693 | 0.00000 |
| **POMP** | 46.08727 | 20.14626 | 31.90618 | 38.04121 | 9.54705 | 23.56787 | 27.83817 |
| **POMT1** | 3.20112 | 3.97804 | 1.02836 | 9.32435 | 0.40172 | 1.97215 | 6.65741 |
| **POMT2** | 2.02696 | 6.00546 | 1.92749 | 3.29525 | 0.20120 | 2.31699 | 12.99565 |
| **POMZP3** | 0.00000 | 0.09006 | 0.43382 | 0.20125 | 0.46868 | 0.31077 | 0.94748 |
| **PON1** | 0.03246 | 0.02746 | 0.00442 | 0.02333 | 0.17920 | 0.01415 | 0.01517 |
| **PON2** | 1.39403 | 0.29083 | 0.68557 | 1.96313 | 2.54791 | 0.26425 | 0.44345 |
| **PON3** | 0.05917 | 0.01539 | 0.00721 | 0.03259 | 0.09281 | 0.01378 | 0.00663 |
| **POP1** | 13.38433 | 8.08811 | 6.10748 | 2.62705 | 1.76831 | 3.48678 | 7.98865 |
| **POP4** | 24.91837 | 16.31199 | 17.74144 | 15.56515 | 2.36539 | 6.98493 | 1.72144 |
| **POP5** | 19.31770 | 8.53109 | 12.65692 | 17.52863 | 3.15832 | 5.01143 | 0.00000 |
| **POP7** | 32.71493 | 32.31569 | 21.47967 | 35.44107 | 11.33896 | 16.32958 | 15.53760 |
| **POPDC2** | 0.03147 | 0.00000 | 0.04043 | 0.13464 | 0.10361 | 0.01474 | 0.03940 |
| **POPDC3** | 0.06270 | 0.27166 | 0.20301 | 0.06347 | 0.14067 | 0.20347 | 0.00000 |
| **POR** | 15.97126 | 20.05684 | 5.16759 | 28.58690 | 2.56468 | 7.52958 | 7.82561 |
| **PORCN** | 2.69765 | 7.49989 | 0.00000 | 0.00000 | 0.42195 | 1.88144 | 2.44643 |
| **POT1** | 4.59601 | 0.00000 | 3.21570 | 1.43945 | 2.19982 | 1.35539 | 0.00000 |
| **POTEA** | 0.02285 | 0.02010 | 0.01283 | 0.02481 | 0.00858 | 0.01336 | 0.01688 |
| **POTEC** | 0.03637 | 0.00635 | 0.01685 | 0.01155 | 0.03578 | 0.01140 | 0.04595 |
| **POTEE** | 0.20209 | 0.00000 | 0.35644 | 0.29150 | 0.16610 | 0.06902 | 0.19883 |
| **POTEF** | 0.11148 | 0.03414 | 0.38761 | 0.22252 | 0.25427 | 0.06320 | 0.31945 |
| **POTEG** | 0.03740 | 0.00831 | 0.01928 | 0.02574 | 0.02685 | 0.01635 | 0.03293 |

| | | | | | | | |
|---|---|---|---|---|---|---|---|
| **POTEH** | 0.04044 | 0.00000 | 0.02966 | 0.02714 | 0.05085 | 0.00730 | 0.09983 |
| **POTEH-AS1.uid** | 0.02479 | 0.01763 | 0.01262 | 0.02091 | 0.00985 | 0.01299 | 0.01542 |
| **POTEI** | 0.02955 | 0.02086 | 0.02533 | 0.01797 | 0.00726 | 0.01387 | 0.05312 |
| **POTEJ** | 0.03213 | 0.03495 | 0.01247 | 0.05780 | 0.00799 | 0.01107 | 0.00066 |
| **POTEKP** | 0.05282 | 0.01007 | 0.06412 | 0.03173 | 0.00701 | 0.01710 | 0.06286 |
| **POTEM** | 0.03220 | 0.00887 | 0.01524 | 0.02570 | 0.01739 | 0.01075 | 0.05679 |
| **POU2AF1** | 0.02839 | 0.01880 | 0.01038 | 0.03496 | 0.01258 | 0.04837 | 0.03326 |
| **POU2F1** | 11.13733 | 11.49829 | 5.05451 | 10.84555 | 6.53882 | 7.09329 | 18.61838 |
| **POU2F2** | 0.00000 | 0.10038 | 0.11474 | 0.37336 | 0.07635 | 0.38855 | 0.91713 |
| **POU2F3** | 0.00000 | 0.41302 | 0.02367 | 0.92197 | 0.00664 | 0.15518 | 0.04475 |
| **POU3F1** | 0.00000 | 0.04035 | 0.06178 | 0.03708 | 0.12544 | 0.67528 | 0.03477 |
| **POU3F2** | 0.01911 | 0.00128 | 0.05228 | 0.03406 | 0.01728 | 0.08212 | 0.03139 |
| **POU3F3** | 0.02681 | 0.01836 | 0.01336 | 0.02337 | 0.00790 | 0.01261 | 0.01476 |
| **POU3F4** | 0.02446 | 0.02007 | 0.00835 | 0.02128 | 0.05038 | 0.01313 | 0.01411 |
| **POU4F1** | 0.04840 | 0.65601 | 0.66292 | 0.00000 | 0.11620 | 0.18175 | 3.07547 |
| **POU4F2** | 0.02278 | 0.01764 | 0.01319 | 0.03680 | 0.00791 | 0.01305 | 0.01532 |
| **POU4F3** | 0.02085 | 0.01956 | 0.01207 | 0.02180 | 0.00785 | 0.01297 | 0.02797 |
| **POU5F1** | 3.61862 | 21.86263 | 9.79371 | 0.00000 | 0.28753 | 26.96489 | 0.00018 |
| **POU5F1B** | 0.00000 | 0.63481 | 0.27025 | 0.00000 | 0.01245 | 1.43341 | 0.23101 |
| **POU5F1P3** | 0.37006 | 5.69442 | 2.70987 | 0.00000 | 0.04764 | 8.21723 | 0.00000 |
| **POU5F1P4** | 0.33864 | 1.86790 | 1.11976 | 0.00000 | 0.00375 | 3.47650 | 0.00000 |
| **POU5F2** | 0.02202 | 0.01675 | 0.02278 | 0.01935 | 0.00714 | 0.01904 | 0.01912 |
| **POU6F1** | 0.00000 | 0.75137 | 0.08945 | 0.47475 | 0.00275 | 0.93556 | 0.34061 |
| **POU6F2** | 0.29515 | 0.10061 | 0.11122 | 0.13931 | 0.09473 | 0.06025 | 0.00000 |
| **PP12613** | 0.00000 | 0.02158 | 0.01318 | 0.09375 | 0.00764 | 0.04150 | 0.04639 |
| **PP14571** | 0.98300 | 0.04596 | 0.00000 | 0.50880 | 0.01860 | 0.00000 | 1.92996 |
| **PP2D1** | 0.11686 | 0.00218 | 0.10471 | 0.02156 | 0.03681 | 0.02002 | 0.00918 |
| **PP7080** | 0.19049 | 14.10810 | 2.62856 | 0.79117 | 0.29390 | 3.46344 | 4.48939 |
| **PPA1** | 41.46861 | 9.95058 | 21.58737 | 19.85959 | 2.28298 | 20.54700 | 14.35602 |
| **PPA2** | 21.51399 | 2.77961 | 2.62676 | 12.50307 | 0.34144 | 7.52291 | 5.26585 |
| **PPAN** | 0.00000 | 1.46733 | 0.31401 | 0.06070 | 0.29420 | 0.45071 | 3.28208 |
| **PPAP2A** | 0.00000 | 8.75740 | 6.78186 | 0.00000 | 0.00000 | 12.95606 | 1.77541 |
| **PPAP2B** | 9.00385 | 6.76427 | 0.00000 | 21.73557 | 0.07209 | 5.40607 | 12.64672 |
| **PPAP2C** | 0.00000 | 2.06515 | 24.55942 | 0.84377 | 4.79440 | 2.93252 | 0.00000 |
| **PPAPDC1A** | 0.00000 | 0.41479 | 0.06439 | 0.00000 | 0.51020 | 0.27396 | 0.02783 |

| | | | | | | | |
|---|---|---|---|---|---|---|---|
| **PPAPDC1B** | 1.47038 | 0.00000 | 0.30404 | 4.81925 | 0.43906 | 2.41742 | 0.09505 |
| **PPAPDC2** | 0.52231 | 1.74753 | 1.17529 | 1.04875 | 0.09078 | 0.28985 | 0.43116 |
| **PPAPDC3** | 0.28215 | 0.36881 | 0.00000 | 0.31516 | 0.24778 | 0.05770 | 0.17408 |
| **PPARA** | 0.32615 | 1.22032 | 0.39475 | 0.77631 | 0.07743 | 0.05576 | 0.98959 |
| **PPARD** | 0.00000 | 4.37621 | 1.07046 | 9.65206 | 3.85587 | 2.45528 | 10.14228 |
| **PPARG** | 6.44989 | 0.00000 | 0.26602 | 18.02291 | 0.00244 | 0.00000 | 3.07702 |
| **PPARGC1A** | 0.06793 | 0.04614 | 0.00000 | 0.00062 | 0.29759 | 0.08000 | 0.04967 |
| **PPARGC1B** | 5.29198 | 16.81814 | 9.50354 | 1.02532 | 2.13455 | 8.64330 | 9.73034 |
| **PPAT** | 23.98652 | 4.45169 | 8.69589 | 7.89730 | 4.56500 | 7.69662 | 6.49828 |
| **PPCDC** | 2.60989 | 3.74231 | 2.12216 | 2.75299 | 0.21594 | 1.48772 | 1.19374 |
| **PPCS** | 20.00888 | 10.65266 | 4.17668 | 8.95018 | 0.22507 | 7.06828 | 0.88219 |
| **PPDPF** | 0.00000 | 4.61879 | 0.13597 | 0.22467 | 0.54462 | 1.37818 | 7.45514 |
| **PPEF1** | 0.01005 | 0.06978 | 0.00000 | 0.01310 | 1.98619 | 0.10060 | 0.01265 |
| **PPEF1-AS1** | 0.03547 | 0.01563 | 0.02149 | 0.01662 | 0.01139 | 0.01143 | 0.01542 |
| **PPEF2** | 0.01953 | 0.00000 | 0.27632 | 0.01703 | 0.26019 | 0.01431 | 0.02791 |
| **PPFIA1** | 6.50297 | 9.86007 | 5.96582 | 12.41620 | 3.36461 | 5.85777 | 28.31885 |
| **PPFIA2** | 0.04166 | 0.00000 | 0.03527 | 0.01965 | 0.89183 | 0.01176 | 0.03645 |
| **PPFIA3** | 0.00000 | 0.83262 | 0.33860 | 1.09437 | 1.25211 | 0.27908 | 2.20047 |
| **PPFIA4** | 0.03493 | 0.00000 | 0.14517 | 0.07908 | 0.12988 | 0.15243 | 0.05749 |
| **PPFIBP1** | 4.46428 | 1.24672 | 2.89735 | 3.51992 | 1.11827 | 2.47354 | 4.83346 |
| **PPFIBP2** | 13.92063 | 16.96617 | 5.09300 | 16.20004 | 0.74628 | 4.12620 | 4.00110 |
| **PPHLN1** | 23.68503 | 7.74148 | 17.08841 | 14.31731 | 7.16969 | 12.67473 | 7.68906 |
| **PPIA** | 29.90303 | 17.93185 | 11.83576 | 20.46278 | 4.28288 | 16.98234 | 19.59407 |
| **PPIAL4B+PPIAL** | 0.02423 | 0.01795 | 0.01355 | 0.01940 | 0.00793 | 0.01310 | 0.01622 |
| **PPIAL4G** | 0.03537 | 0.01387 | 0.01645 | 0.01894 | 0.00937 | 0.01962 | 0.05054 |
| **PPIAP30** | 0.01721 | 0.03587 | 0.00000 | 0.02056 | 1.56690 | 0.13248 | 0.03726 |
| **PPIB** | 28.41895 | 11.94658 | 8.13259 | 25.05930 | 1.21425 | 11.88094 | 3.86731 |
| **PPIC** | 14.21590 | 9.09316 | 1.56891 | 0.91305 | 1.54575 | 7.93898 | 1.49348 |
| **PPID** | 33.99274 | 9.95381 | 10.22796 | 20.39590 | 1.73928 | 8.27696 | 8.01705 |
| **PPIE** | 18.41263 | 23.01537 | 9.55402 | 13.27700 | 4.69439 | 5.51794 | 0.04948 |
| **PPIEL** | 0.18192 | 0.35956 | 0.09434 | 0.34540 | 0.06538 | 0.24024 | 0.14894 |
| **PPIF** | 45.69665 | 29.44350 | 16.36188 | 27.24679 | 5.38655 | 15.15821 | 21.42121 |
| **PPIG** | 19.14108 | 5.00746 | 17.21688 | 14.68676 | 14.21043 | 11.84309 | 28.67475 |
| **PPIH** | 11.58687 | 11.58650 | 5.68770 | 10.91674 | 0.49099 | 7.49826 | 17.72004 |
| **PPIL1** | 27.55554 | 3.94485 | 20.30999 | 17.41657 | 2.26425 | 8.45401 | 3.58379 |

| | | | | | | | |
|---|---|---|---|---|---|---|---|
| **PPIL2** | 9.81869 | 11.25626 | 3.62343 | 6.96619 | 1.23258 | 4.61436 | 1.79066 |
| **PPIL3** | 14.08013 | 0.92986 | 10.57885 | 4.68239 | 6.65013 | 4.94490 | 4.34988 |
| **PPIL4** | 17.93629 | 2.29646 | 7.28089 | 8.80035 | 1.29757 | 5.94790 | 3.44462 |
| **PPIL6** | 0.31926 | 0.15756 | 0.45797 | 0.20679 | 0.05890 | 0.00751 | 0.12246 |
| **PPIP5K1** | 0.28544 | 0.00000 | 1.18421 | 0.47976 | 0.62759 | 0.16086 | 0.43302 |
| **PPIP5K2** | 10.17031 | 0.00000 | 3.62364 | 6.45718 | 1.27674 | 3.39530 | 9.31210 |
| **PPL** | 0.00000 | 0.00000 | 0.29709 | 2.85727 | 2.30615 | 0.36134 | 2.63429 |
| **PPM1A** | 4.96560 | 0.00001 | 7.12185 | 4.86706 | 3.07203 | 5.03372 | 13.62308 |
| **PPM1B** | 4.20234 | 0.00000 | 4.26449 | 19.18349 | 2.40130 | 15.54369 | 18.73196 |
| **PPM1D** | 3.59817 | 7.39101 | 7.69275 | 8.28250 | 6.64914 | 5.53825 | 2.31162 |
| **PPM1E** | 0.00000 | 0.00000 | 2.50720 | 0.41021 | 0.80853 | 1.05613 | 0.68140 |
| **PPM1F** | 4.80284 | 6.39217 | 1.91638 | 3.67585 | 0.04448 | 0.84286 | 3.76932 |
| **PPM1G** | 27.24946 | 27.61967 | 21.61965 | 29.43945 | 5.12015 | 21.65547 | 26.94251 |
| **PPM1H** | 3.35646 | 0.47565 | 0.00000 | 0.33272 | 1.02075 | 3.53193 | 0.91687 |
| **PPM1J** | 0.01924 | 1.33040 | 0.24291 | 0.86181 | 0.00000 | 3.68619 | 0.31252 |
| **PPM1K** | 0.55285 | 0.07134 | 0.87435 | 0.28697 | 0.35430 | 1.22457 | 0.11967 |
| **PPM1L** | 0.00000 | 0.00000 | 0.05233 | 0.15075 | 0.00971 | 0.10055 | 0.39250 |
| **PPM1M** | 0.32266 | 0.39947 | 0.03978 | 0.51714 | 1.44966 | 0.53322 | 0.19052 |
| **PPM1N** | 0.20773 | 1.60919 | 7.42152 | 8.24935 | 1.37406 | 0.85414 | 1.67601 |
| **PPME1** | 21.96769 | 9.14806 | 3.21841 | 29.95297 | 6.37050 | 1.48040 | 2.30724 |
| **PPOX** | 1.62839 | 0.53975 | 0.33333 | 1.20872 | 0.74538 | 2.91772 | 0.01782 |
| **PPP1CA** | 38.64028 | 35.22573 | 11.56822 | 32.02644 | 8.95614 | 19.20820 | 0.00000 |
| **PPP1CB** | 47.51155 | 8.61481 | 15.45493 | 38.05597 | 4.05060 | 17.93419 | 8.43606 |
| **PPP1CC** | 21.34158 | 2.71010 | 16.79953 | 6.35472 | 9.52198 | 17.48469 | 49.62386 |
| **PPP1R10** | 11.78800 | 9.05335 | 11.45265 | 19.20509 | 5.72245 | 15.09332 | 21.76465 |
| **PPP1R11** | 14.32615 | 9.50079 | 11.52411 | 15.29419 | 1.84766 | 9.30331 | 1.65236 |
| **PPP1R12A** | 21.12892 | 5.90803 | 9.79349 | 12.54299 | 11.54802 | 10.66766 | 18.58071 |
| **PPP1R12B** | 0.81227 | 1.62583 | 0.94639 | 1.69465 | 1.28346 | 5.94695 | 4.81891 |
| **PPP1R12C** | 0.00000 | 1.12546 | 0.22580 | 0.19860 | 0.01596 | 1.06863 | 7.16517 |
| **PPP1R13B** | 0.00045 | 0.64584 | 4.74917 | 0.31843 | 3.12991 | 0.73323 | 2.83390 |
| **PPP1R13L** | 1.52761 | 1.48906 | 0.30631 | 4.73707 | 0.15629 | 0.51923 | 12.67378 |
| **PPP1R14A** | 17.88606 | 70.66642 | 5.29852 | 12.38710 | 0.00000 | 8.45363 | 6.63660 |
| **PPP1R14B** | 0.00000 | 2.48737 | 2.55089 | 0.82446 | 0.00000 | 2.69888 | 11.82290 |
| **PPP1R14C** | 0.01493 | 0.10254 | 0.00000 | 0.03453 | 1.83570 | 0.02151 | 0.04334 |
| **PPP1R14D** | 1.26049 | 0.41909 | 0.50976 | 1.55752 | 0.07096 | 0.00000 | 0.03021 |

| | | | | | | | |
|---|---|---|---|---|---|---|---|
| **PPP1R15A** | 0.61479 | 17.07873 | 18.93020 | 21.37223 | 16.68344 | 14.65027 | 3.03251 |
| **PPP1R15B** | 12.90572 | 1.98472 | 6.16378 | 10.61047 | 3.54746 | 2.04237 | 19.73908 |
| **PPP1R16A** | 0.74116 | 4.38619 | 0.31469 | 2.94938 | 0.42914 | 1.94083 | 9.22065 |
| **PPP1R16B** | 0.00000 | 0.49277 | 0.24557 | 0.00000 | 0.02731 | 1.66494 | 0.03416 |
| **PPP1R17** | 0.03354 | 0.01066 | 0.01356 | 0.02003 | 0.00809 | 0.03941 | 0.01821 |
| **PPP1R18** | 0.12969 | 0.06844 | 0.17990 | 1.63359 | 0.40377 | 0.92745 | 2.55846 |
| **PPP1R1A** | 0.22089 | 0.00000 | 0.00449 | 0.03490 | 0.00703 | 0.22245 | 0.24015 |
| **PPP1R1B** | 0.01379 | 0.01674 | 0.02503 | 0.01416 | 0.12416 | 0.51765 | 0.01981 |
| **PPP1R1C** | 0.07381 | 0.09234 | 0.31944 | 0.03555 | 0.00818 | 0.10611 | 0.06839 |
| **PPP1R2** | 24.50280 | 3.56225 | 10.48960 | 16.51002 | 5.58127 | 4.27364 | 2.47516 |
| **PPP1R21** | 3.58360 | 0.46911 | 1.36370 | 1.22836 | 0.51174 | 2.87737 | 6.48102 |
| **PPP1R26** | 0.00000 | 0.94494 | 3.22996 | 0.00000 | 0.01445 | 1.01394 | 0.18959 |
| **PPP1R26-AS1** | 0.00000 | 0.37327 | 0.81656 | 0.06078 | 0.00214 | 0.11825 | 0.00169 |
| **PPP1R27** | 0.00000 | 0.01798 | 0.03666 | 0.11791 | 0.01974 | 0.00286 | 0.24294 |
| **PPP1R2P3** | 1.56189 | 0.10864 | 0.78321 | 1.13645 | 0.47592 | 0.20284 | 0.00000 |
| **PPP1R2P9** | 0.07947 | 0.34808 | 0.70900 | 1.01741 | 0.29061 | 0.16294 | 0.00000 |
| **PPP1R32** | 0.28356 | 0.01433 | 0.15339 | 0.36107 | 0.00310 | 0.04864 | 0.00000 |
| **PPP1R35** | 0.51422 | 0.46833 | 0.00000 | 2.20366 | 4.25500 | 0.73412 | 2.62018 |
| **PPP1R36** | 0.00000 | 0.05664 | 0.11903 | 0.05884 | 0.06799 | 0.04534 | 0.13929 |
| **PPP1R37** | 0.00000 | 0.79059 | 0.15618 | 0.13926 | 0.71582 | 0.53465 | 3.31114 |
| **PPP1R3A** | 0.02278 | 0.02882 | 0.05562 | 0.04739 | 0.47064 | 0.00697 | 0.03377 |
| **PPP1R3B** | 11.82293 | 4.89335 | 22.43454 | 0.51584 | 9.75014 | 7.24308 | 5.63486 |
| **PPP1R3C** | 0.07717 | 0.00484 | 0.06871 | 0.00730 | 2.07587 | 0.00199 | 0.05435 |
| **PPP1R3D** | 0.06472 | 0.02781 | 0.02536 | 0.18202 | 0.82290 | 0.17396 | 0.36302 |
| **PPP1R3E** | 0.08161 | 0.10590 | 0.04367 | 0.76091 | 0.69202 | 0.00000 | 0.35090 |
| **PPP1R3F** | 0.00000 | 0.33924 | 0.00895 | 0.00000 | 0.29563 | 0.02332 | 0.11152 |
| **PPP1R3G** | 0.04998 | 0.00644 | 0.78752 | 0.00179 | 0.01388 | 0.00000 | 0.08606 |
| **PPP1R42** | 0.02804 | 0.00854 | 0.04004 | 0.02128 | 0.03836 | 0.01205 | 0.01489 |
| **PPP1R7** | 20.49527 | 17.36379 | 4.96963 | 13.78759 | 0.94634 | 8.44065 | 4.84601 |
| **PPP1R8** | 3.96860 | 1.23712 | 6.93165 | 2.79425 | 4.00438 | 4.12112 | 12.22789 |
| **PPP1R9A** | 6.51360 | 7.76300 | 1.50406 | 1.52379 | 0.05967 | 3.30685 | 6.07353 |
| **PPP1R9B** | 0.00000 | 0.07653 | 0.16628 | 0.13776 | 0.11058 | 1.34784 | 3.73211 |
| **PPP2CA** | 17.35475 | 6.07040 | 15.83653 | 20.41656 | 11.54356 | 15.51175 | 38.53141 |
| **PPP2CB** | 10.86511 | 1.10895 | 7.57330 | 6.80307 | 1.57033 | 5.88659 | 13.87917 |
| **PPP2R1A** | 31.92006 | 30.40974 | 9.17268 | 29.11788 | 6.27486 | 19.95667 | 14.76590 |

| | | | | | | | |
|---|---|---|---|---|---|---|---|
| **PPP2R1B** | 19.64526 | 5.55173 | 7.10391 | 11.64726 | 6.10205 | 4.21032 | 0.00000 |
| **PPP2R2A** | 8.86086 | 1.41970 | 9.40754 | 6.23184 | 1.48547 | 4.64846 | 22.01996 |
| **PPP2R2B** | 0.07881 | 0.02221 | 0.00000 | 0.03454 | 1.30617 | 1.17242 | 0.01383 |
| **PPP2R2C** | 0.02705 | 0.11430 | 0.05787 | 0.00507 | 0.76189 | 0.00295 | 0.03134 |
| **PPP2R2D** | 1.10783 | 4.38825 | 2.53203 | 2.70682 | 0.83037 | 4.96784 | 11.56709 |
| **PPP2R3A** | 4.13506 | 0.00000 | 1.16041 | 8.77672 | 1.02055 | 0.64923 | 3.54564 |
| **PPP2R3C** | 10.65243 | 6.65003 | 15.86307 | 11.63017 | 3.82901 | 5.30956 | 7.20445 |
| **PPP2R4** | 3.16339 | 7.40899 | 3.16528 | 15.80372 | 0.49689 | 5.64693 | 19.22413 |
| **PPP2R5A** | 9.31786 | 5.56772 | 10.77288 | 4.83197 | 0.06949 | 4.68876 | 25.40683 |
| **PPP2R5B** | 0.06510 | 1.79251 | 3.75269 | 0.17687 | 0.00019 | 0.38436 | 1.94528 |
| **PPP2R5C** | 21.34605 | 5.11448 | 7.87925 | 11.27755 | 3.05244 | 5.56755 | 2.82724 |
| **PPP2R5D** | 9.98734 | 6.87355 | 2.81892 | 10.76196 | 2.64880 | 6.91112 | 5.91010 |
| **PPP2R5E** | 3.04361 | 1.02396 | 4.52545 | 9.21296 | 1.67459 | 5.53334 | 22.03539 |
| **PPP3CA** | 1.11419 | 0.25863 | 2.67946 | 1.57835 | 3.32383 | 1.30464 | 0.29757 |
| **PPP3CB** | 1.12362 | 0.20162 | 1.00250 | 1.36057 | 0.54785 | 1.81751 | 7.53380 |
| **PPP3CC** | 0.59976 | 0.68210 | 0.41616 | 0.44165 | 0.31712 | 0.26991 | 2.84556 |
| **PPP3R1** | 2.01190 | 0.00000 | 2.29639 | 0.83851 | 1.35168 | 1.39681 | 5.99295 |
| **PPP4C** | 3.98141 | 22.00761 | 5.91521 | 13.98576 | 5.04856 | 12.43272 | 33.68922 |
| **PPP4R1** | 8.90845 | 4.49029 | 2.24959 | 8.62745 | 1.72544 | 4.19834 | 11.30654 |
| **PPP4R1L** | 0.24159 | 0.00000 | 0.79851 | 0.41118 | 1.08138 | 0.43990 | 0.00000 |
| **PPP4R2** | 2.55465 | 1.27582 | 2.44089 | 1.22889 | 1.47280 | 2.38119 | 7.29522 |
| **PPP4R4** | 0.03426 | 0.00354 | 0.02071 | 0.02470 | 1.61788 | 0.05564 | 0.01330 |
| **PPP5C** | 5.17218 | 13.23887 | 4.11763 | 12.66134 | 0.00000 | 9.26132 | 10.50511 |
| **PPP5D1** | 0.05041 | 0.23081 | 0.09065 | 0.21952 | 0.05455 | 0.04251 | 0.00000 |
| **PPP6C** | 17.31237 | 1.97117 | 15.96956 | 13.81089 | 5.36082 | 9.26043 | 9.67626 |
| **PPP6R1** | 1.54746 | 8.70971 | 2.74177 | 8.70960 | 1.02700 | 5.49042 | 22.12962 |
| **PPP6R2** | 1.78345 | 5.54556 | 1.73752 | 4.66225 | 0.78459 | 2.52857 | 7.57078 |
| **PPP6R3** | 26.80852 | 12.02508 | 11.60661 | 15.22065 | 6.70389 | 10.35701 | 6.69311 |
| **PPRC1** | 7.80622 | 10.80681 | 13.84165 | 10.23841 | 6.06133 | 10.65030 | 31.73049 |
| **PPT1** | 79.37055 | 23.47270 | 10.27862 | 30.59958 | 2.83787 | 1.10888 | 7.35410 |
| **PPT2** | 5.75570 | 19.80515 | 9.28449 | 2.73980 | 1.74267 | 5.87071 | 5.36680 |
| **PPT2-EGFL8** | 0.17489 | 0.93505 | 0.53531 | 0.01806 | 0.00039 | 0.09280 | 0.01245 |
| **PPTC7** | 1.48832 | 0.92670 | 2.44331 | 0.96652 | 2.82340 | 3.26421 | 5.26148 |
| **PPWD1** | 17.39563 | 1.64401 | 9.53029 | 11.93381 | 5.85707 | 5.80178 | 0.00000 |
| **PPY** | 0.08883 | 0.02096 | 0.00889 | 0.18790 | 0.52356 | 0.03171 | 0.01596 |

| | | | | | | | |
|---|---|---|---|---|---|---|---|
| **PPY2** | 0.01320 | 0.02171 | 0.16413 | 0.16203 | 0.00665 | 0.13341 | 0.00000 |
| **PQBP1** | 24.59030 | 21.69898 | 26.08005 | 11.86252 | 17.29573 | 13.76201 | 12.09670 |
| **PQLC1** | 0.00000 | 6.90133 | 5.60757 | 0.22963 | 0.28110 | 9.25134 | 6.28432 |
| **PQLC2** | 4.02819 | 2.70120 | 2.18206 | 11.92560 | 0.35841 | 0.65612 | 5.12663 |
| **PQLC3** | 9.72700 | 17.32956 | 6.85549 | 0.47053 | 1.67940 | 18.34551 | 2.10804 |
| **PRAC1** | 0.01267 | 0.02400 | 0.01474 | 0.08455 | 0.00755 | 0.09720 | 0.00732 |
| **PRAC2** | 0.02880 | 0.00170 | 0.05567 | 0.03698 | 0.00656 | 0.11600 | 0.01327 |
| **PRADC1** | 5.05551 | 7.38080 | 0.00000 | 1.11841 | 1.71022 | 5.25103 | 4.66097 |
| **PRAF2** | 1.86909 | 1.21268 | 3.34119 | 1.98313 | 1.63095 | 1.09528 | 0.22767 |
| **PRAM1** | 0.01163 | 0.00000 | 0.10963 | 0.05972 | 0.15371 | 0.09322 | 0.03688 |
| **PRAME** | 2.27189 | 1.88933 | 6.77462 | 0.00000 | 6.88159 | 0.00000 | 0.00000 |
| **PRAMEF1** | 0.01096 | 0.03958 | 0.00000 | 0.00529 | 5.49104 | 0.01527 | 0.04248 |
| **PRAMEF11** | 0.01110 | 0.00683 | 0.02691 | 0.00998 | 10.25139 | 0.01950 | 0.02652 |
| **PRAMEF12** | 0.06790 | 0.35137 | 0.21939 | 0.00660 | 19.20244 | 0.00000 | 0.01656 |
| **PRAMEF15.uid-** | 0.02356 | 0.01762 | 0.01194 | 0.01925 | 0.01685 | 0.01305 | 0.01718 |
| **PRAMEF17.uid-** | 0.01439 | 0.02030 | 0.72100 | 0.00959 | 0.90817 | 0.00000 | 0.01631 |
| **PRAMEF19.uid-** | 0.76339 | 2.60437 | 2.76901 | 0.24475 | 3.86971 | 0.00000 | 0.00000 |
| **PRAMEF19.uid-** | 0.04926 | 0.43366 | 0.55224 | 0.00000 | 0.55221 | 0.00000 | 0.00000 |
| **PRAMEF2** | 0.00000 | 0.11923 | 0.00000 | 0.00967 | 8.48819 | 0.04589 | 0.01561 |
| **PRAMEF20.uid-** | 0.01659 | 0.04172 | 0.03641 | 0.01945 | 0.20410 | 0.00939 | 0.01303 |
| **PRAMEF20.uid-** | 0.00886 | 0.08748 | 0.19061 | 0.01019 | 2.40140 | 0.00051 | 0.00947 |
| **PRAMEF4** | 0.00730 | 0.01462 | 0.00000 | 0.01012 | 11.98078 | 0.01635 | 0.02643 |
| **PRAMEF5.uid-8** | 0.02042 | 0.00746 | 0.00614 | 0.01685 | 1.44183 | 0.01237 | 0.01594 |
| **PRAMEF6.uid-8** | 0.01692 | 0.02040 | 0.06625 | 0.01194 | 2.86738 | 0.01202 | 0.01761 |
| **PRAMEF7+PRA** | 0.02257 | 0.04887 | 0.00000 | 0.01307 | 0.63041 | 0.01299 | 0.01817 |
| **PRAP1** | 15.91891 | 53.05685 | 0.00000 | 0.00000 | 4.00245 | 3.71761 | 3.57494 |
| **PRB3** | 0.01814 | 0.22383 | 0.36091 | 0.01045 | 0.03978 | 0.03629 | 0.01417 |
| **PRC1** | 21.83407 | 6.99754 | 10.96137 | 17.85529 | 6.62340 | 3.20351 | 12.74955 |
| **PRC1-AS1** | 0.07958 | 0.00000 | 0.04172 | 0.05929 | 0.07098 | 0.02082 | 0.08773 |
| **PRCC** | 9.05487 | 13.19543 | 9.74529 | 20.53857 | 4.63450 | 8.01885 | 14.97313 |
| **PRCD** | 0.02681 | 0.01555 | 0.03711 | 0.02344 | 0.02123 | 0.02777 | 0.01536 |
| **PRCP** | 7.49914 | 2.21762 | 0.44602 | 5.56031 | 0.12775 | 10.71817 | 0.00000 |
| **PRDM1** | 0.32081 | 0.29238 | 0.07945 | 0.03392 | 0.06646 | 4.85294 | 0.00000 |
| **PRDM10** | 2.18818 | 1.78656 | 0.09470 | 1.43310 | 4.71437 | 0.96764 | 3.77120 |
| **PRDM11** | 0.08720 | 0.00000 | 0.03774 | 0.42154 | 0.02727 | 1.31341 | 0.00000 |

| | | | | | | | |
|---|---|---|---|---|---|---|---|
| **PRDM12** | 0.02276 | 0.01989 | 0.01004 | 0.01925 | 0.02321 | 0.01518 | 0.02488 |
| **PRDM13** | 0.05358 | 0.08785 | 0.07987 | 0.02270 | 0.00592 | 0.01276 | 0.17262 |
| **PRDM14** | 0.00000 | 1.65284 | 4.58639 | 0.00000 | 0.20434 | 15.18473 | 0.00000 |
| **PRDM15** | 1.98310 | 1.07156 | 1.63302 | 3.30134 | 0.53598 | 0.71667 | 2.30226 |
| **PRDM16** | 0.00957 | 0.10831 | 0.00974 | 0.00881 | 0.92395 | 0.01193 | 0.10713 |
| **PRDM2** | 5.40738 | 0.70221 | 4.83621 | 7.05165 | 4.45463 | 1.80122 | 15.23978 |
| **PRDM4** | 1.92589 | 1.10147 | 0.44279 | 2.78128 | 2.51670 | 1.51842 | 6.35793 |
| **PRDM5** | 0.01634 | 0.01678 | 0.07276 | 0.24907 | 0.04535 | 0.01114 | 0.00863 |
| **PRDM6** | 0.02175 | 0.33173 | 0.03000 | 0.12718 | 0.06178 | 0.05695 | 0.10454 |
| **PRDM7** | 0.02177 | 0.01885 | 0.00577 | 0.02351 | 0.11788 | 0.01566 | 0.01691 |
| **PRDM8** | 0.02257 | 0.01522 | 0.01359 | 0.01690 | 0.22028 | 0.01188 | 0.02768 |
| **PRDM9** | 0.02485 | 0.02060 | 0.01092 | 0.24016 | 0.03179 | 0.01693 | 0.00000 |
| **PRDX1** | 85.28627 | 74.79603 | 33.18134 | 61.16254 | 13.98694 | 52.94613 | 56.05439 |
| **PRDX2** | 50.61568 | 34.50921 | 1.05764 | 45.17763 | 6.11348 | 24.32144 | 9.74309 |
| **PRDX3** | 57.21257 | 19.11704 | 21.46082 | 29.23260 | 5.00041 | 29.27208 | 8.13534 |
| **PRDX4** | 17.75806 | 29.12435 | 12.35300 | 10.50430 | 0.44104 | 24.92968 | 3.17730 |
| **PRDX5** | 33.04411 | 19.77907 | 9.17890 | 35.94665 | 7.38256 | 20.78152 | 14.18489 |
| **PRDX6** | 26.12297 | 11.65167 | 4.87987 | 36.07558 | 1.73316 | 22.38717 | 19.25514 |
| **PREB** | 5.57410 | 16.15216 | 7.45231 | 13.37230 | 1.25377 | 12.30921 | 18.66882 |
| **PRELID1** | 13.52141 | 21.61184 | 4.47234 | 14.23293 | 2.13927 | 7.04383 | 18.31753 |
| **PRELID2** | 0.82728 | 0.00509 | 0.00000 | 0.05013 | 3.76464 | 0.07497 | 0.19567 |
| **PRELP** | 0.08570 | 0.00000 | 0.02206 | 0.23420 | 0.01262 | 0.62211 | 0.26176 |
| **PREP** | 5.32538 | 2.66365 | 0.77624 | 3.26886 | 2.07787 | 2.06833 | 5.18258 |
| **PREPL** | 10.25384 | 3.44697 | 6.65057 | 2.65682 | 7.26403 | 2.33341 | 4.50119 |
| **PREX1** | 1.50729 | 0.90891 | 2.90707 | 2.31504 | 1.08180 | 2.95146 | 2.63532 |
| **PREX2** | 0.02208 | 0.00786 | 0.00779 | 0.01936 | 0.06510 | 0.17886 | 0.01833 |
| **PRF1** | 0.00000 | 0.04183 | 0.00277 | 0.00000 | 0.01765 | 1.21272 | 0.29722 |
| **PRG2** | 0.00000 | 0.38106 | 0.32730 | 0.03318 | 0.00316 | 0.00000 | 0.00621 |
| **PRG3** | 0.02522 | 0.00000 | 0.05282 | 0.01724 | 0.34698 | 0.01200 | 0.01933 |
| **PRG4** | 0.07138 | 0.00000 | 0.05022 | 0.07904 | 0.01950 | 0.00000 | 0.21799 |
| **PRH1** | 0.03578 | 0.02276 | 0.01956 | 0.06966 | 0.01045 | 0.01748 | 0.06315 |
| **PRH1-PRR4** | 0.05089 | 0.01648 | 0.02001 | 0.04810 | 0.00757 | 0.03685 | 0.00000 |
| **PRH2** | 0.15542 | 0.00000 | 0.03314 | 0.00011 | 0.00457 | 0.07423 | 0.25402 |
| **PRICKLE1** | 0.20596 | 0.70855 | 1.61642 | 0.00000 | 0.30405 | 7.31447 | 0.00000 |
| **PRICKLE2** | 0.05845 | 0.02570 | 0.02580 | 0.00849 | 0.23723 | 0.06097 | 0.35163 |

| | | | | | | | |
|---|---|---|---|---|---|---|---|
| **PRICKLE2-AS1** | 0.02338 | 0.01650 | 0.01515 | 0.01963 | 0.00768 | 0.01525 | 0.02004 |
| **PRICKLE2-AS3** | 0.02730 | 0.02200 | 0.01230 | 0.01711 | 0.00904 | 0.01349 | 0.03273 |
| **PRICKLE3** | 1.51116 | 1.90446 | 0.46655 | 0.44079 | 0.06265 | 0.00000 | 0.42785 |
| **PRICKLE4** | 1.08164 | 0.46708 | 0.17959 | 1.88557 | 0.64837 | 0.08681 | 0.48347 |
| **PRIM1** | 17.84666 | 7.23863 | 11.67480 | 7.55486 | 0.00000 | 9.57092 | 5.14019 |
| **PRIM2** | 5.90154 | 1.22946 | 4.80319 | 4.28186 | 2.51896 | 4.02360 | 0.65841 |
| **PRIMA1** | 0.07705 | 0.07824 | 0.07585 | 0.00000 | 0.00306 | 0.08486 | 0.11443 |
| **PRIMPOL** | 0.96903 | 0.16134 | 0.25741 | 0.44488 | 0.25955 | 0.23344 | 0.23329 |
| **PRKAA1** | 7.60095 | 0.00000 | 8.02663 | 3.95916 | 4.76176 | 3.85276 | 1.69189 |
| **PRKAA2** | 1.21676 | 0.00000 | 0.40149 | 3.91538 | 0.03755 | 0.00000 | 1.27818 |
| **PRKAB1** | 8.66315 | 10.17379 | 5.23147 | 15.17939 | 3.11189 | 6.67009 | 0.00000 |
| **PRKAB2** | 2.34298 | 2.34345 | 7.26061 | 0.76847 | 1.11808 | 8.87069 | 2.89622 |
| **PRKACA** | 0.00000 | 1.32793 | 1.09212 | 0.85426 | 0.38008 | 0.74077 | 4.08604 |
| **PRKACB** | 0.42698 | 0.00000 | 0.44276 | 0.80495 | 0.18497 | 1.05357 | 0.00000 |
| **PRKACG** | 0.03587 | 0.03896 | 0.10559 | 0.01602 | 0.00875 | 0.00708 | 0.02068 |
| **PRKAG1** | 21.14779 | 6.90280 | 12.84852 | 11.79066 | 5.41737 | 5.34085 | 0.00000 |
| **PRKAG2** | 0.26908 | 0.07997 | 0.44624 | 1.12155 | 0.11867 | 0.18695 | 0.21913 |
| **PRKAG2-AS1** | 0.02046 | 0.02395 | 0.01250 | 0.01960 | 0.01447 | 0.02296 | 0.01687 |
| **PRKAG3** | 0.01268 | 0.01475 | 0.01211 | 0.02599 | 0.02654 | 0.05200 | 0.02797 |
| **PRKAR1A** | 22.78681 | 8.21978 | 6.73110 | 12.36325 | 5.10382 | 9.22328 | 1.05232 |
| **PRKAR1B** | 4.87840 | 6.41258 | 0.00000 | 7.25671 | 0.54408 | 0.31046 | 1.31165 |
| **PRKAR2A** | 2.48312 | 1.58862 | 1.33906 | 3.24995 | 5.48505 | 1.55225 | 2.61431 |
| **PRKAR2A-AS1** | 0.08832 | 0.33879 | 0.02173 | 0.03177 | 0.12475 | 0.02957 | 0.00000 |
| **PRKAR2B** | 1.64363 | 0.00000 | 0.51706 | 0.00000 | 0.33008 | 0.86193 | 1.99730 |
| **PRKCA** | 2.34866 | 2.03837 | 1.06132 | 0.00000 | 0.26802 | 0.82931 | 2.16599 |
| **PRKCB** | 0.00000 | 0.36109 | 0.33030 | 0.12221 | 0.11554 | 0.05211 | 0.01348 |
| **PRKCD** | 0.87483 | 5.88091 | 0.39300 | 2.80709 | 0.44342 | 5.09719 | 5.67023 |
| **PRKCDBP** | 0.00153 | 0.69336 | 1.76231 | 1.38598 | 1.95971 | 0.84221 | 0.43926 |
| **PRKCE** | 0.00000 | 0.02928 | 0.00000 | 0.11013 | 0.94427 | 0.07047 | 0.12921 |
| **PRKCG** | 0.00000 | 0.12468 | 0.32531 | 0.07436 | 4.80901 | 0.26151 | 0.08684 |
| **PRKCH** | 0.00000 | 0.04436 | 0.04217 | 7.58587 | 0.06810 | 0.00293 | 0.00749 |
| **PRKCI** | 4.31702 | 2.10904 | 4.30660 | 4.58193 | 1.06185 | 4.51141 | 11.86295 |
| **PRKCQ** | 0.00000 | 1.10455 | 1.28528 | 0.00000 | 0.87566 | 0.54866 | 0.00351 |
| **PRKCQ-AS1** | 0.09610 | 0.06921 | 1.52775 | 0.01096 | 0.07523 | 0.69386 | 0.82923 |
| **PRKCSH** | 4.08670 | 8.30381 | 5.34104 | 13.75494 | 0.80151 | 6.84259 | 14.10214 |

| | | | | | | | |
|---|---|---|---|---|---|---|---|
| **PRKCZ** | 0.00000 | 0.89566 | 0.00000 | 3.06417 | 0.61759 | 0.69119 | 0.79843 |
| **PRKD1** | 0.00000 | 0.25040 | 0.18126 | 0.01386 | 0.33517 | 0.02246 | 0.03649 |
| **PRKD2** | 2.71280 | 1.12261 | 11.44890 | 5.49240 | 11.40379 | 2.91303 | 3.13271 |
| **PRKD3** | 2.66160 | 1.12334 | 4.77090 | 4.06849 | 1.78473 | 5.34650 | 5.81147 |
| **PRKDC** | 39.62265 | 13.66831 | 20.87485 | 21.91908 | 1.05439 | 20.48657 | 45.64074 |
| **PRKG1** | 0.00000 | 0.00000 | 0.13872 | 0.10278 | 1.00355 | 0.00315 | 0.18012 |
| **PRKG2** | 0.03904 | 0.00000 | 0.06522 | 0.08294 | 0.18510 | 0.11109 | 0.04944 |
| **PRKRA** | 12.54675 | 2.84961 | 5.14774 | 9.27272 | 2.79569 | 6.86159 | 0.00000 |
| **PRKRIP1** | 1.09870 | 1.80885 | 1.93721 | 0.92478 | 1.35221 | 0.34905 | 0.20101 |
| **PRKRIR** | 2.21452 | 0.41391 | 2.57596 | 0.87131 | 1.73762 | 2.22036 | 6.69316 |
| **PRKX** | 4.84230 | 4.98733 | 1.76196 | 4.60662 | 0.29999 | 3.07330 | 10.53738 |
| **PRKX-AS1** | 0.02874 | 0.02605 | 0.02488 | 0.01708 | 0.00968 | 0.01127 | 0.05208 |
| **PRKXP1** | 0.49864 | 0.00000 | 0.55426 | 0.77576 | 0.38624 | 0.00000 | 1.43631 |
| **PRKY** | 0.02318 | 0.03846 | 0.02483 | 0.05535 | 0.01155 | 0.02029 | 0.00785 |
| **PRL** | 0.02328 | 0.01921 | 0.00656 | 0.02014 | 0.08751 | 0.01331 | 0.01531 |
| **PRLHR** | 0.01478 | 0.00000 | 0.19851 | 0.03368 | 0.00290 | 0.46827 | 0.00859 |
| **PRLR** | 1.16386 | 0.00000 | 0.00000 | 1.70543 | 0.02257 | 0.00000 | 1.07733 |
| **PRM1** | 0.01301 | 0.02283 | 0.01158 | 0.03416 | 0.00802 | 0.02818 | 0.02130 |
| **PRM3** | 0.03559 | 0.01573 | 0.01395 | 0.01164 | 0.00719 | 0.00729 | 0.05028 |
| **PRMT1** | 31.71600 | 26.64862 | 14.54047 | 33.66617 | 8.18201 | 17.29580 | 19.44954 |
| **PRMT2** | 4.83339 | 7.18375 | 1.05573 | 5.92094 | 1.94811 | 3.73026 | 0.00000 |
| **PRMT3** | 18.09378 | 3.29913 | 7.60359 | 4.01224 | 0.08996 | 4.71655 | 0.00000 |
| **PRMT5** | 34.44631 | 15.92236 | 12.71585 | 20.95650 | 2.54946 | 8.91355 | 0.00000 |
| **PRMT5-AS1** | 0.18228 | 0.32028 | 0.11185 | 0.24263 | 0.06456 | 0.16904 | 0.06385 |
| **PRMT6** | 9.05250 | 5.55705 | 4.50944 | 8.18844 | 0.73866 | 3.99860 | 0.00000 |
| **PRMT7** | 14.16035 | 16.41538 | 5.89828 | 12.40845 | 1.87772 | 4.46255 | 0.76589 |
| **PRMT8** | 0.02839 | 0.01504 | 0.01317 | 0.01999 | 0.00790 | 0.02823 | 0.01760 |
| **PRMT9** | 4.21873 | 3.31213 | 1.17872 | 3.45876 | 0.97676 | 1.74812 | 0.00000 |
| **PRNCR1** | 2.27880 | 1.15080 | 0.59285 | 0.96853 | 0.46878 | 0.63897 | 0.80948 |
| **PRND** | 0.03038 | 0.01385 | 0.01598 | 0.01863 | 0.03434 | 0.01332 | 0.02457 |
| **PRNP** | 3.44055 | 3.30693 | 12.46997 | 0.53100 | 2.72229 | 4.18810 | 2.81346 |
| **PRNT** | 0.05383 | 0.01872 | 0.01169 | 0.00927 | 0.00782 | 0.01850 | 0.01807 |
| **PROB1** | 0.00296 | 0.02222 | 0.01558 | 0.08084 | 0.02569 | 0.05788 | 0.05995 |
| **PROC** | 0.00000 | 0.04870 | 0.01760 | 0.05107 | 0.00853 | 0.49788 | 0.05799 |
| **PROCA1** | 0.37648 | 0.47262 | 0.23948 | 0.20782 | 1.23837 | 0.91404 | 0.55607 |

| | | | | | | | |
|---|---|---|---|---|---|---|---|
| **PROCR** | 6.00880 | 10.36256 | 4.49589 | 9.21496 | 4.25859 | 15.73468 | 0.15552 |
| **PRODH** | 12.10398 | 6.44847 | 0.28064 | 0.27530 | 0.05118 | 5.43738 | 12.84189 |
| **PRODH2** | 0.01061 | 0.00000 | 0.07432 | 0.02125 | 0.02871 | 0.13996 | 0.04336 |
| **PROK1** | 0.06338 | 0.02152 | 0.01075 | 0.00381 | 0.00889 | 0.00706 | 0.03156 |
| **PROK2** | 0.02655 | 0.01117 | 0.01293 | 0.01479 | 0.01028 | 0.11026 | 0.01782 |
| **PROM1** | 57.55474 | 12.30329 | 6.54442 | 8.59108 | 0.21557 | 16.55487 | 13.47747 |
| **PROM2** | 0.06235 | 0.00376 | 0.02389 | 0.11839 | 0.01143 | 0.11714 | 0.13675 |
| **PROP1** | 0.01889 | 0.02003 | 0.01205 | 0.01750 | 0.01420 | 0.02882 | 0.03616 |
| **PRORSD1P** | 0.25909 | 0.36799 | 2.36073 | 0.25430 | 0.37889 | 1.51400 | 0.02945 |
| **PROS1** | 0.85521 | 0.15214 | 0.02729 | 0.92620 | 0.00397 | 4.96370 | 0.00000 |
| **PROSC** | 20.32726 | 5.56928 | 6.33486 | 12.03289 | 1.62173 | 4.17558 | 0.00000 |
| **PROSER1** | 10.85035 | 4.17396 | 4.67774 | 16.81069 | 15.90624 | 4.93404 | 37.46006 |
| **PROSER2** | 0.00000 | 0.89549 | 0.85266 | 0.92012 | 0.22196 | 0.12642 | 4.87738 |
| **PROSER2-AS1** | 2.27937 | 4.76404 | 3.85163 | 8.14068 | 2.71838 | 0.00000 | 3.26189 |
| **PROSER3** | 0.01174 | 0.09511 | 0.60815 | 0.48121 | 0.59555 | 0.25177 | 0.57270 |
| **PROX1** | 0.03578 | 0.00000 | 0.00100 | 0.01485 | 0.20875 | 0.86820 | 0.07374 |
| **PROX2** | 0.14443 | 0.09542 | 0.14079 | 0.02730 | 0.15055 | 0.05982 | 0.00661 |
| **PROZ** | 0.16315 | 0.07312 | 0.07941 | 0.15795 | 0.00256 | 0.12789 | 0.39651 |
| **PRPF18** | 9.65100 | 2.67255 | 5.68115 | 4.84849 | 8.51398 | 3.91748 | 0.76029 |
| **PRPF19** | 0.07426 | 5.44751 | 7.95145 | 8.16923 | 2.53359 | 1.90577 | 31.24991 |
| **PRPF3** | 20.05071 | 10.79535 | 9.34031 | 13.63587 | 6.36931 | 6.12505 | 3.88869 |
| **PRPF31** | 25.43818 | 26.38146 | 15.29873 | 22.47547 | 3.62609 | 11.23780 | 2.75418 |
| **PRPF38A** | 23.31276 | 9.44186 | 20.96553 | 16.76488 | 14.53751 | 8.85500 | 2.82275 |
| **PRPF38B** | 18.92575 | 10.72069 | 9.35954 | 14.85383 | 12.43675 | 12.34171 | 26.23519 |
| **PRPF39** | 11.79484 | 0.00000 | 5.26632 | 3.75680 | 4.78243 | 4.15226 | 3.00882 |
| **PRPF4** | 25.28220 | 18.07204 | 10.73376 | 15.78955 | 4.86245 | 7.99214 | 4.44335 |
| **PRPF40A** | 41.18742 | 19.28428 | 26.69377 | 32.16680 | 16.59811 | 19.02206 | 23.84648 |
| **PRPF40B** | 0.68420 | 0.31848 | 0.29898 | 0.58165 | 0.00219 | 0.34241 | 0.05882 |
| **PRPF4B** | 20.94494 | 1.06131 | 9.29545 | 7.63121 | 6.64677 | 7.03744 | 29.01973 |
| **PRPF6** | 18.01803 | 17.34764 | 10.55916 | 17.09760 | 4.24030 | 8.89208 | 6.66922 |
| **PRPF8** | 26.02336 | 29.21410 | 21.54682 | 25.43165 | 9.88166 | 19.81212 | 30.25556 |
| **PRPH** | 0.04288 | 0.29567 | 4.49714 | 0.01045 | 2.74951 | 0.17345 | 0.00569 |
| **PRPH2** | 0.00000 | 0.21817 | 0.44150 | 0.10969 | 0.99462 | 0.27718 | 0.06544 |
| **PRPS1** | 24.69439 | 43.54670 | 14.67109 | 7.07366 | 2.76996 | 5.67198 | 0.45380 |
| **PRPS1L1** | 1.00598 | 2.67349 | 0.85165 | 0.11798 | 0.16632 | 0.05056 | 0.00000 |

| | | | | | | | |
|---|---|---|---|---|---|---|---|
| **PRPS2** | 37.49130 | 7.66834 | 5.71981 | 11.45271 | 2.73885 | 4.69141 | 0.00000 |
| **PRPSAP1** | 9.34992 | 19.29695 | 9.19844 | 0.00000 | 5.26837 | 12.07562 | 3.17926 |
| **PRPSAP2** | 15.25278 | 6.85613 | 9.13385 | 8.89132 | 1.43541 | 4.64658 | 1.91019 |
| **PRR11** | 6.21714 | 5.24234 | 5.94692 | 8.29217 | 1.28994 | 5.06170 | 12.99203 |
| **PRR12** | 0.00000 | 0.39395 | 0.03185 | 0.41515 | 0.16429 | 0.64382 | 4.47059 |
| **PRR13** | 19.72994 | 42.42582 | 27.77281 | 9.70744 | 8.25696 | 26.51660 | 3.63505 |
| **PRR14** | 0.87694 | 0.66483 | 1.60672 | 5.50526 | 3.04479 | 1.55540 | 6.54884 |
| **PRR14L** | 11.37512 | 6.20878 | 5.12406 | 12.39988 | 6.57673 | 4.72839 | 16.40115 |
| **PRR15** | 0.00000 | 0.39629 | 0.00000 | 0.00000 | 4.79557 | 0.87789 | 0.10490 |
| **PRR15L** | 0.92274 | 1.13585 | 0.13547 | 0.00000 | 0.00051 | 0.74936 | 0.01546 |
| **PRR16** | 0.02661 | 0.00000 | 0.18173 | 0.00534 | 0.28876 | 0.05786 | 0.05645 |
| **PRR18** | 0.03669 | 0.02124 | 0.01191 | 0.01999 | 0.00809 | 0.01372 | 0.00941 |
| **PRR19** | 2.28925 | 0.23940 | 7.62228 | 4.98339 | 0.42755 | 1.94355 | 0.88278 |
| **PRR22** | 0.00000 | 0.78333 | 0.68540 | 0.51581 | 0.00000 | 0.00000 | 4.14864 |
| **PRR23A** | 0.02589 | 0.50402 | 1.27027 | 0.88240 | 2.11466 | 0.00000 | 0.44876 |
| **PRR23B** | 0.00000 | 0.00000 | 1.74939 | 0.05325 | 0.73713 | 0.14702 | 0.43157 |
| **PRR23C** | 0.06451 | 0.04724 | 0.87238 | 0.12386 | 0.98599 | 0.04656 | 0.14487 |
| **PRR24** | 0.00000 | 0.21923 | 0.08472 | 0.27542 | 0.15149 | 0.32052 | 1.67353 |
| **PRR25** | 0.02383 | 0.02492 | 0.01363 | 0.02434 | 0.00767 | 0.01223 | 0.01749 |
| **PRR26** | 0.00000 | 0.03701 | 0.21747 | 0.11071 | 0.00115 | 0.14899 | 0.94558 |
| **PRR27** | 0.02612 | 0.01105 | 0.01395 | 0.02353 | 0.00756 | 0.00972 | 0.04175 |
| **PRR29** | 0.01749 | 0.00921 | 0.03898 | 0.03964 | 0.02228 | 0.00989 | 0.03381 |
| **PRR3** | 2.91440 | 0.62015 | 3.79033 | 3.84452 | 1.24310 | 1.20745 | 2.37960 |
| **PRR30** | 0.01974 | 0.01522 | 0.01689 | 0.01703 | 0.00835 | 0.05365 | 0.02549 |
| **PRR32** | 0.02321 | 0.02410 | 0.00000 | 0.01865 | 0.41301 | 0.01373 | 0.01329 |
| **PRR34** | 0.03024 | 0.01718 | 0.01509 | 0.01896 | 0.00788 | 0.01277 | 0.01496 |
| **PRR35** | 0.01444 | 0.03599 | 0.00881 | 0.02600 | 0.00809 | 0.03047 | 0.01954 |
| **PRR4** | 0.04102 | 0.01611 | 0.01399 | 0.01946 | 0.00775 | 0.01118 | 0.01706 |
| **PRR5** | 3.28986 | 13.69609 | 0.02578 | 12.49626 | 0.51613 | 0.00000 | 28.28501 |
| **PRR5L** | 0.01191 | 0.08243 | 0.00000 | 0.03967 | 3.51431 | 0.01167 | 0.03209 |
| **PRR7** | 0.00000 | 0.21688 | 0.01749 | 0.36982 | 0.43088 | 0.06499 | 0.43042 |
| **PRR7-AS1** | 0.04526 | 0.01922 | 0.01306 | 0.03269 | 0.00727 | 0.01055 | 0.04020 |
| **PRR9** | 0.00000 | 0.01674 | 0.01102 | 14.76177 | 0.00655 | 0.00699 | 0.00000 |
| **PRRC1** | 15.51145 | 1.75156 | 7.60758 | 10.21763 | 1.75912 | 6.88120 | 1.63506 |
| **PRRC2A** | 0.00000 | 5.42268 | 6.46140 | 11.20273 | 0.49457 | 7.80575 | 33.48533 |

| | | | | | | | |
|---|---|---|---|---|---|---|---|
| **PRRC2B** | 3.82306 | 15.34418 | 9.55360 | 14.30801 | 2.10923 | 17.81239 | 45.19791 |
| **PRRC2C** | 31.88516 | 17.11461 | 17.54709 | 30.62290 | 9.84033 | 17.68822 | 50.30714 |
| **PRRG1** | 2.67841 | 0.00000 | 6.76444 | 1.31495 | 3.22556 | 0.34162 | 1.84549 |
| **PRRG2** | 0.08244 | 0.38403 | 0.16932 | 1.08998 | 0.59962 | 0.44617 | 0.41363 |
| **PRRG3** | 0.02023 | 0.05497 | 0.01650 | 0.02106 | 0.20527 | 0.01041 | 0.01673 |
| **PRRG4** | 0.00000 | 0.02322 | 0.36593 | 0.66057 | 0.68796 | 0.00000 | 0.10864 |
| **PRRT1** | 0.00000 | 0.51190 | 1.19000 | 0.01160 | 0.00166 | 0.12365 | 0.10610 |
| **PRRT2** | 0.07249 | 0.00000 | 0.15502 | 0.20296 | 0.18742 | 0.19108 | 1.08682 |
| **PRRT3** | 0.25776 | 1.42468 | 0.41938 | 0.11121 | 0.18881 | 0.45577 | 0.82502 |
| **PRRT3-AS1** | 0.06696 | 0.06382 | 0.02673 | 0.24465 | 0.01773 | 0.13545 | 0.00000 |
| **PRRT4** | 0.00000 | 0.23775 | 0.00000 | 0.01911 | 0.12633 | 0.13524 | 0.04347 |
| **PRRX1** | 0.03369 | 0.01034 | 0.02234 | 0.01880 | 0.01329 | 0.01825 | 0.01795 |
| **PRRX2** | 0.00000 | 0.38366 | 0.01185 | 0.03361 | 0.15450 | 0.28450 | 0.02098 |
| **PRSS1** | 0.14101 | 0.00000 | 0.14105 | 0.01698 | 0.48866 | 0.03501 | 0.01538 |
| **PRSS12** | 1.73813 | 0.00000 | 0.05133 | 0.65913 | 0.00000 | 3.41735 | 4.15530 |
| **PRSS16** | 1.50375 | 0.00000 | 0.84767 | 3.02449 | 1.01225 | 0.22291 | 0.59550 |
| **PRSS21** | 0.40628 | 0.10309 | 0.22769 | 0.39820 | 0.09406 | 0.31765 | 0.00000 |
| **PRSS22** | 0.00000 | 0.19944 | 0.11055 | 0.12255 | 0.00789 | 0.07628 | 0.10468 |
| **PRSS23** | 21.02834 | 38.37792 | 21.32477 | 0.00000 | 0.00000 | 15.77220 | 0.73324 |
| **PRSS27** | 0.00739 | 0.10713 | 0.09069 | 0.05114 | 0.06016 | 0.01238 | 0.01697 |
| **PRSS3** | 0.00000 | 0.69190 | 0.12990 | 0.00000 | 0.42064 | 5.46315 | 0.01040 |
| **PRSS30P** | 0.05710 | 0.02067 | 0.08246 | 0.79433 | 0.00541 | 0.03264 | 0.08888 |
| **PRSS33** | 0.00000 | 0.04857 | 0.05508 | 0.03061 | 0.05026 | 0.08749 | 0.04539 |
| **PRSS35** | 0.09255 | 0.00000 | 0.06152 | 0.01027 | 0.00435 | 1.00921 | 0.00000 |
| **PRSS36** | 0.00000 | 0.29134 | 0.75615 | 0.21873 | 0.08183 | 0.27835 | 0.08936 |
| **PRSS37** | 0.11408 | 0.01214 | 0.03045 | 0.02940 | 0.02504 | 0.01716 | 0.05234 |
| **PRSS38** | 0.01954 | 0.08056 | 0.01128 | 0.03004 | 0.00761 | 0.01201 | 0.00800 |
| **PRSS3P2** | 0.02528 | 0.01793 | 0.05501 | 0.01548 | 0.22556 | 0.00977 | 0.02226 |
| **PRSS41** | 0.00000 | 1.89701 | 1.39578 | 0.00000 | 0.25872 | 0.79324 | 0.00256 |
| **PRSS45** | 0.02059 | 0.00395 | 0.06479 | 0.02756 | 0.00830 | 0.01249 | 0.02091 |
| **PRSS48** | 0.05976 | 0.07622 | 0.05433 | 0.00000 | 0.00346 | 0.05770 | 0.29703 |
| **PRSS50** | 0.02217 | 0.47110 | 0.19843 | 0.00182 | 0.00329 | 0.02787 | 0.00610 |
| **PRSS53** | 0.00008 | 0.42427 | 0.56423 | 0.87219 | 0.23456 | 0.08952 | 1.60418 |
| **PRSS57** | 0.02380 | 0.01932 | 0.00988 | 0.01994 | 0.03081 | 0.01292 | 0.01706 |
| **PRSS58** | 0.02197 | 0.01048 | 0.03528 | 0.01860 | 0.40342 | 0.01157 | 0.01553 |

| | | | | | | | |
|---|---|---|---|---|---|---|---|
| **PRSS8** | 49.31097 | 10.02205 | 0.00000 | 32.40230 | 0.00000 | 0.00000 | 26.24809 |
| **PRTFDC1** | 1.77299 | 0.00000 | 0.08942 | 0.30494 | 0.01382 | 0.28692 | 1.22132 |
| **PRTG** | 3.67391 | 0.00000 | 0.00000 | 6.48836 | 4.98754 | 0.00000 | 10.43090 |
| **PRTN3** | 0.01451 | 0.01451 | 0.25020 | 0.02072 | 0.04306 | 0.00495 | 0.01869 |
| **PRUNE** | 5.68650 | 0.24866 | 3.85790 | 2.29368 | 0.98539 | 2.66761 | 0.71438 |
| **PRUNE2** | 0.11064 | 1.62785 | 0.76482 | 0.00000 | 1.26654 | 7.96421 | 0.00000 |
| **PRX** | 0.00000 | 0.15754 | 0.24730 | 0.09491 | 0.78352 | 0.00000 | 0.41401 |
| **PSAP** | 81.75926 | 82.45848 | 26.84335 | 57.67918 | 12.78650 | 40.85655 | 34.16647 |
| **PSAPL1** | 0.00000 | 0.19755 | 0.05083 | 0.03688 | 0.00556 | 0.02387 | 0.00661 |
| **PSAT1** | 37.93074 | 0.00000 | 1.01594 | 27.97651 | 0.00000 | 3.98518 | 7.76804 |
| **PSCA** | 0.08253 | 0.01153 | 0.00808 | 0.02672 | 0.07660 | 0.00898 | 0.01653 |
| **PSD** | 0.00000 | 0.38903 | 0.02451 | 0.01232 | 0.01092 | 0.22084 | 0.21855 |
| **PSD2** | 0.01338 | 0.05169 | 0.01320 | 0.02538 | 0.00719 | 0.01564 | 0.02846 |
| **PSD3** | 1.05170 | 0.00000 | 1.53221 | 0.00000 | 4.40107 | 0.59539 | 2.45223 |
| **PSD4** | 0.00000 | 1.39353 | 0.98821 | 1.65448 | 0.08720 | 0.04417 | 0.25271 |
| **PSEN1** | 15.48651 | 6.45253 | 9.86533 | 17.23403 | 3.82817 | 12.14248 | 0.00000 |
| **PSEN2** | 0.54151 | 0.14023 | 0.00000 | 1.46672 | 2.90414 | 2.63360 | 0.38532 |
| **PSENEN** | 12.44799 | 15.18967 | 7.98866 | 15.80184 | 1.07207 | 12.67951 | 7.97081 |
| **PSG1** | 0.02245 | 0.01740 | 0.01327 | 0.03342 | 0.00796 | 0.01732 | 0.01615 |
| **PSG10P** | 0.01674 | 0.01247 | 0.02269 | 0.01721 | 0.02712 | 0.02498 | 0.03418 |
| **PSG11** | 0.01692 | 0.02029 | 0.01291 | 0.02707 | 0.00777 | 0.02060 | 0.02281 |
| **PSG2** | 0.01361 | 0.01907 | 0.01328 | 0.12011 | 0.00860 | 0.01298 | 0.00551 |
| **PSG3** | 0.03292 | 0.01867 | 0.01202 | 0.01806 | 0.01055 | 0.01265 | 0.02753 |
| **PSG4** | 0.00000 | 0.02042 | 0.01248 | 0.24598 | 0.01003 | 0.01517 | 0.02331 |
| **PSG5** | 0.03146 | 0.01512 | 0.01689 | 0.03038 | 0.00777 | 0.01273 | 0.00904 |
| **PSG6** | 0.01711 | 0.02214 | 0.01114 | 0.06271 | 0.00801 | 0.02995 | 0.02565 |
| **PSG7** | 0.02249 | 0.01800 | 0.01279 | 0.02596 | 0.00799 | 0.01272 | 0.01706 |
| **PSG9** | 0.02769 | 0.02781 | 0.01243 | 0.02374 | 0.00777 | 0.03213 | 0.01690 |
| **PSIP1** | 5.18616 | 0.00000 | 1.63552 | 5.51791 | 3.24442 | 6.65855 | 17.15460 |
| **PSKH1** | 0.00000 | 1.63447 | 1.19442 | 0.97065 | 0.22967 | 0.72118 | 1.52456 |
| **PSMA1** | 52.26912 | 14.92143 | 25.50993 | 33.70692 | 0.21993 | 22.59110 | 21.45708 |
| **PSMA2** | 51.11020 | 16.18137 | 22.81101 | 40.40789 | 9.24784 | 24.39365 | 13.23602 |
| **PSMA3** | 49.32530 | 11.41722 | 24.07896 | 35.39099 | 6.67423 | 19.06625 | 18.50281 |
| **PSMA4** | 51.58647 | 19.89362 | 23.17226 | 37.85142 | 2.12068 | 19.63605 | 17.67740 |
| **PSMA5** | 50.13572 | 20.39396 | 18.47211 | 41.14269 | 9.49959 | 20.35759 | 19.70044 |

| | | | | | | | |
|---|---|---|---|---|---|---|---|
| **PSMA6** | 42.36136 | 20.38078 | 21.25877 | 33.36711 | 11.63866 | 19.20627 | 11.34507 |
| **PSMA7** | 46.91604 | 46.99099 | 16.51992 | 51.62799 | 11.31585 | 27.51070 | 36.18998 |
| **PSMA8** | 0.22468 | 0.24394 | 0.27115 | 0.14293 | 0.03806 | 0.02214 | 0.00000 |
| **PSMB1** | 56.27268 | 37.80394 | 27.64750 | 45.29654 | 8.60367 | 25.34597 | 19.26870 |
| **PSMB10** | 14.73233 | 6.51446 | 0.67905 | 10.75919 | 0.00000 | 6.38477 | 2.63879 |
| **PSMB11** | 0.02070 | 0.02014 | 0.01072 | 0.03440 | 0.03087 | 0.01414 | 0.01554 |
| **PSMB2** | 45.36871 | 27.47930 | 27.48016 | 46.93112 | 9.03237 | 20.88569 | 18.91914 |
| **PSMB3** | 48.29250 | 39.94500 | 19.80889 | 49.38055 | 11.50554 | 26.64761 | 19.20391 |
| **PSMB4** | 42.31982 | 40.76635 | 21.16453 | 48.76503 | 10.42911 | 23.44240 | 20.33185 |
| **PSMB5** | 62.13362 | 37.88923 | 26.90974 | 57.60396 | 4.42844 | 25.89947 | 8.71664 |
| **PSMB6** | 47.88214 | 48.04432 | 30.38250 | 49.19596 | 13.33286 | 24.67479 | 16.72032 |
| **PSMB7** | 47.15860 | 31.18275 | 21.81320 | 41.08075 | 5.32911 | 16.63962 | 0.00000 |
| **PSMB8** | 0.02665 | 0.04345 | 0.78598 | 0.02996 | 0.39623 | 0.00000 | 0.07878 |
| **PSMB9** | 0.00000 | 0.19871 | 1.02936 | 0.01712 | 0.01021 | 0.00698 | 0.00553 |
| **PSMC1** | 0.57692 | 0.85986 | 0.22381 | 1.02842 | 0.12808 | 0.14922 | 0.89774 |
| **PSMC2** | 45.65600 | 21.54899 | 13.81082 | 37.99490 | 3.38448 | 13.16584 | 0.00331 |
| **PSMC3** | 40.01087 | 36.91720 | 13.56972 | 38.75148 | 5.32717 | 17.71607 | 3.06158 |
| **PSMC3IP** | 4.80454 | 4.37595 | 1.41998 | 5.47575 | 4.33577 | 1.56455 | 0.00000 |
| **PSMC4** | 44.55668 | 38.95055 | 23.90146 | 49.56111 | 20.32830 | 25.24416 | 0.00000 |
| **PSMC5** | 41.57195 | 32.75401 | 18.17949 | 39.12840 | 7.10547 | 19.94132 | 9.86902 |
| **PSMC6** | 23.24475 | 4.13526 | 7.49269 | 17.08418 | 0.92855 | 7.06614 | 2.65893 |
| **PSMD1** | 50.98481 | 24.37405 | 15.90232 | 39.19238 | 9.82543 | 20.76062 | 13.82408 |
| **PSMD10** | 30.34374 | 7.49461 | 15.39213 | 10.92106 | 4.16934 | 9.10170 | 0.00000 |
| **PSMD11** | 55.53751 | 34.21134 | 16.68998 | 49.13941 | 7.90122 | 21.66257 | 11.85373 |
| **PSMD12** | 50.69418 | 20.02569 | 22.08523 | 28.72529 | 6.39227 | 18.60815 | 6.46898 |
| **PSMD13** | 34.13513 | 27.95106 | 13.53027 | 37.75434 | 3.57357 | 13.11909 | 8.23529 |
| **PSMD14** | 45.04025 | 19.10664 | 13.00899 | 42.64459 | 5.79903 | 16.61649 | 10.06539 |
| **PSMD2** | 21.74678 | 29.25290 | 12.86053 | 32.17729 | 15.49117 | 19.12497 | 32.46118 |
| **PSMD3** | 3.05291 | 16.86807 | 3.77983 | 14.17219 | 1.62599 | 9.93210 | 34.45939 |
| **PSMD4** | 42.67442 | 34.05360 | 16.91643 | 38.19295 | 7.00244 | 15.87218 | 4.36785 |
| **PSMD5** | 12.95377 | 0.13108 | 1.73225 | 7.53939 | 0.88858 | 3.40042 | 0.00000 |
| **PSMD5-AS1** | 0.51450 | 0.52715 | 0.28669 | 0.87898 | 0.27549 | 0.40401 | 1.50196 |
| **PSMD6** | 40.97132 | 19.24492 | 20.82419 | 31.46942 | 5.89647 | 16.24093 | 4.43513 |
| **PSMD6-AS2** | 1.44361 | 0.05717 | 0.53640 | 0.67805 | 0.23924 | 0.03638 | 0.24038 |
| **PSMD7** | 40.09025 | 32.88408 | 15.90166 | 29.54305 | 3.28961 | 18.15786 | 26.05999 |

| | | | | | | | |
|---|---|---|---|---|---|---|---|
| **PSMD8** | 40.16678 | 34.32831 | 21.00254 | 37.45192 | 10.13795 | 21.52259 | 11.91416 |
| **PSMD9** | 27.51040 | 28.20041 | 14.78195 | 27.22348 | 16.85728 | 12.33547 | 0.00000 |
| **PSME1** | 2.58660 | 0.00000 | 1.32278 | 4.73078 | 1.34915 | 5.79265 | 0.42999 |
| **PSME2** | 7.24123 | 3.11607 | 1.55890 | 7.44419 | 0.40207 | 8.55855 | 1.98218 |
| **PSME3** | 50.60235 | 30.72030 | 18.62670 | 33.08171 | 11.00026 | 19.54464 | 5.64294 |
| **PSME4** | 11.06944 | 10.24469 | 13.46600 | 12.23213 | 1.83799 | 17.66988 | 24.22845 |
| **PSMF1** | 23.69097 | 20.35316 | 13.92501 | 16.27758 | 3.53438 | 7.76469 | 4.51593 |
| **PSMG1** | 31.66317 | 10.24740 | 12.96574 | 15.90041 | 6.72396 | 12.70609 | 1.60713 |
| **PSMG2** | 22.76308 | 7.38480 | 15.32074 | 15.15209 | 0.11460 | 9.82457 | 10.36142 |
| **PSMG3** | 24.27814 | 32.35451 | 7.67816 | 13.57973 | 2.42916 | 15.94141 | 7.05922 |
| **PSMG3-AS1** | 0.21905 | 0.64102 | 0.00000 | 0.09431 | 0.10807 | 0.66124 | 0.00000 |
| **PSMG4** | 2.56747 | 11.16910 | 5.52951 | 3.05484 | 0.92450 | 3.10458 | 6.73779 |
| **PSORS1C1** | 0.00000 | 0.08992 | 0.79993 | 0.00445 | 0.00405 | 0.35418 | 0.03364 |
| **PSORS1C2** | 0.00000 | 1.79991 | 18.34298 | 0.00000 | 0.00000 | 7.42576 | 0.00000 |
| **PSORS1C3** | 0.02321 | 0.00486 | 0.05135 | 0.01734 | 0.00971 | 0.01155 | 0.02666 |
| **PSPC1** | 14.20730 | 16.72004 | 18.17740 | 16.96502 | 12.48078 | 6.32811 | 15.38974 |
| **PSPH** | 13.76918 | 0.00000 | 1.16096 | 6.01283 | 0.38692 | 4.46475 | 2.62312 |
| **PSPN** | 0.00287 | 0.05471 | 0.01388 | 0.09687 | 0.02025 | 0.02047 | 0.06685 |
| **PSRC1** | 2.62150 | 7.63418 | 2.97191 | 6.58281 | 0.45042 | 18.74675 | 2.19652 |
| **PSTK** | 4.03270 | 1.74207 | 5.15367 | 2.07051 | 0.13488 | 0.00000 | 0.00000 |
| **PSTPIP1** | 0.00000 | 0.02332 | 0.03842 | 0.05324 | 1.48267 | 0.03673 | 0.14752 |
| **PSTPIP2** | 1.85497 | 0.06675 | 0.25073 | 0.00000 | 0.26371 | 0.00000 | 2.89215 |
| **PTAFR** | 0.76693 | 0.20635 | 0.62193 | 0.41184 | 0.52555 | 0.01146 | 0.03651 |
| **PTAR1** | 7.04685 | 0.00000 | 8.18134 | 1.07494 | 4.21774 | 4.50076 | 10.46579 |
| **PTBP1** | 13.65136 | 25.60891 | 23.74187 | 29.48935 | 6.35372 | 19.70753 | 71.88579 |
| **PTBP2** | 4.39293 | 0.00000 | 1.22232 | 2.07376 | 2.97988 | 5.18517 | 2.16868 |
| **PTBP3** | 22.29960 | 5.09757 | 3.44090 | 15.87697 | 1.05531 | 8.83194 | 14.72343 |
| **PTCD1** | 0.02305 | 0.25502 | 0.39688 | 0.27440 | 0.03327 | 0.24549 | 0.29867 |
| **PTCD2** | 5.37095 | 0.31825 | 2.61563 | 1.91163 | 0.36782 | 1.77740 | 0.00000 |
| **PTCD3** | 30.52182 | 13.42997 | 12.48237 | 18.15938 | 5.68679 | 13.42805 | 9.22867 |
| **PTCH1** | 0.14269 | 0.00000 | 1.24040 | 0.96340 | 0.05944 | 0.46528 | 1.41319 |
| **PTCH2** | 0.05280 | 0.19041 | 0.72465 | 0.11701 | 0.07273 | 0.16269 | 0.51371 |
| **PTCHD1** | 0.01662 | 0.24421 | 0.42397 | 0.00000 | 0.00286 | 1.29554 | 0.00140 |
| **PTCHD1-AS** | 0.02869 | 0.02888 | 0.01204 | 0.01880 | 0.01804 | 0.09406 | 0.02730 |
| **PTCHD2** | 0.17890 | 0.06488 | 0.03506 | 0.50980 | 0.36010 | 0.01750 | 0.13725 |

| | | | | | | | |
|---|---|---|---|---|---|---|---|
| **PTCHD3** | 0.01181 | 0.03204 | 0.03564 | 0.01722 | 0.09997 | 0.01380 | 0.04860 |
| **PTCHD4** | 0.01074 | 0.01629 | 0.01184 | 0.11636 | 0.00749 | 0.01693 | 0.08041 |
| **PTCRA** | 0.01506 | 0.03305 | 0.00764 | 0.01781 | 0.03280 | 0.15214 | 0.01603 |
| **PTDSS1** | 13.00561 | 9.41152 | 13.74391 | 17.23553 | 2.85547 | 10.28418 | 17.55395 |
| **PTDSS2** | 3.14603 | 11.29552 | 0.64199 | 7.89803 | 0.53561 | 5.98563 | 22.27845 |
| **PTEN** | 14.42931 | 3.75813 | 5.32432 | 3.92641 | 5.75194 | 3.75225 | 1.12305 |
| **PTENP1** | 0.44546 | 0.00000 | 0.67281 | 0.49081 | 0.18158 | 0.34643 | 1.08655 |
| **PTENP1-AS** | 0.01490 | 0.00454 | 0.27734 | 0.02048 | 0.11256 | 0.01312 | 0.02791 |
| **PTER** | 6.73960 | 5.34561 | 2.41612 | 3.38315 | 0.22534 | 0.16223 | 0.00000 |
| **PTGDR** | 0.02251 | 0.01782 | 0.01557 | 0.01905 | 0.07359 | 0.01284 | 0.01969 |
| **PTGDR2** | 0.04849 | 0.00003 | 0.10132 | 0.00000 | 0.23335 | 0.02657 | 0.20175 |
| **PTGDS** | 0.48430 | 0.15511 | 0.00000 | 0.25945 | 0.01221 | 0.00000 | 0.19505 |
| **PTGER2** | 0.02934 | 0.01659 | 0.02013 | 0.04665 | 0.00811 | 0.01231 | 0.01026 |
| **PTGER3** | 0.02408 | 0.01778 | 0.00000 | 0.02769 | 0.90141 | 0.03236 | 0.00931 |
| **PTGER4** | 0.00695 | 0.13334 | 0.26750 | 0.03278 | 0.25852 | 0.00662 | 0.02507 |
| **PTGER4P2-CDK** | 0.02253 | 0.02638 | 0.01268 | 0.02084 | 0.02767 | 0.02951 | 0.01209 |
| **PTGES** | 41.84543 | 70.31924 | 0.00000 | 20.39141 | 0.00000 | 0.00000 | 13.80336 |
| **PTGES2** | 1.45245 | 14.03969 | 4.24432 | 0.28393 | 0.06766 | 6.92278 | 9.78989 |
| **PTGES2-AS1** | 0.03543 | 0.01221 | 0.04175 | 0.03370 | 0.03260 | 0.01824 | 0.01487 |
| **PTGES3** | 51.22813 | 11.98865 | 22.24539 | 40.86507 | 7.92753 | 21.67715 | 35.84505 |
| **PTGES3L** | 0.11569 | 0.07698 | 0.17494 | 0.15117 | 0.02753 | 0.42288 | 0.00000 |
| **PTGES3L-AARS** | 0.02066 | 0.01793 | 0.01341 | 0.02612 | 0.00783 | 0.02126 | 0.01832 |
| **PTGFR** | 0.02534 | 0.01857 | 0.01189 | 0.01888 | 0.01837 | 0.01363 | 0.01540 |
| **PTGFRN** | 2.80651 | 7.09477 | 3.35565 | 0.05802 | 1.73648 | 5.65591 | 0.33604 |
| **PTGIS** | 0.08201 | 0.00000 | 0.02211 | 0.12098 | 0.01797 | 0.00000 | 0.22606 |
| **PTGR1** | 20.27967 | 11.98708 | 0.00000 | 2.36675 | 0.08438 | 19.68539 | 5.87591 |
| **PTGR2** | 0.96438 | 0.08967 | 1.60215 | 0.62897 | 0.23290 | 0.65556 | 1.05923 |
| **PTGS1** | 0.06757 | 0.02314 | 0.02979 | 0.01822 | 0.07162 | 0.02872 | 0.05806 |
| **PTGS2** | 0.02005 | 0.04735 | 0.03099 | 0.07155 | 0.02582 | 0.04259 | 0.05496 |
| **PTH1R** | 0.00000 | 0.23807 | 0.57999 | 0.00000 | 0.28523 | 0.52342 | 0.17597 |
| **PTH2** | 0.02216 | 0.02295 | 0.00651 | 0.02004 | 0.10502 | 0.02149 | 0.01246 |
| **PTH2R** | 0.11362 | 0.00002 | 0.02510 | 0.00034 | 0.07351 | 0.06103 | 0.01874 |
| **PTHLH** | 0.18388 | 1.07042 | 0.08570 | 0.00000 | 0.02590 | 0.46345 | 0.76245 |
| **PTK2** | 11.71709 | 4.68546 | 4.84343 | 12.92212 | 0.44185 | 4.75143 | 9.85280 |
| **PTK2B** | 0.11884 | 0.34729 | 0.25685 | 1.54215 | 2.47675 | 0.71182 | 0.47906 |

| | | | | | | | |
|---|---|---|---|---|---|---|---|
| **PTK6** | 0.03756 | 0.14541 | 0.00878 | 0.05755 | 0.02851 | 0.04411 | 0.00001 |
| **PTK7** | 1.42118 | 0.94752 | 1.63369 | 5.12734 | 0.46773 | 5.28770 | 2.84879 |
| **PTMA** | 45.65142 | 46.61700 | 21.70388 | 40.32567 | 14.12437 | 31.94274 | 48.90213 |
| **PTMS** | 0.00000 | 2.33655 | 0.10890 | 1.41577 | 0.08355 | 5.64208 | 14.23820 |
| **PTN** | 15.72046 | 0.00000 | 0.00000 | 55.53989 | 3.35134 | 0.00000 | 16.69963 |
| **PTOV1** | 0.31947 | 3.99363 | 0.00000 | 5.48855 | 1.75457 | 4.30236 | 21.37846 |
| **PTOV1-AS1** | 0.59694 | 0.28355 | 0.22317 | 1.30780 | 0.02751 | 0.98044 | 3.00229 |
| **PTP4A1** | 70.22807 | 18.84421 | 31.75131 | 31.51736 | 23.17706 | 16.83961 | 16.94679 |
| **PTP4A2** | 13.36691 | 5.82067 | 4.33942 | 11.43144 | 3.46005 | 8.99368 | 24.91242 |
| **PTP4A3** | 0.02060 | 0.27297 | 0.01673 | 0.01874 | 0.00732 | 0.00986 | 0.00816 |
| **PTPDC1** | 0.14490 | 0.18874 | 0.24539 | 0.12797 | 0.00127 | 0.50399 | 0.60687 |
| **PTPLA** | 0.74460 | 0.30191 | 0.10253 | 0.26372 | 0.03889 | 1.07552 | 0.91943 |
| **PTPLAD1** | 34.91575 | 15.57644 | 30.50283 | 33.86562 | 2.89944 | 9.80804 | 12.71862 |
| **PTPLAD2** | 0.03872 | 0.12755 | 0.44111 | 0.04053 | 0.02628 | 0.01305 | 0.24254 |
| **PTPLB** | 0.88879 | 0.00000 | 0.95025 | 1.47223 | 0.10354 | 0.71857 | 2.33978 |
| **PTPMT1** | 6.70522 | 11.19930 | 2.51778 | 9.29845 | 2.87349 | 0.68453 | 9.48213 |
| **PTPN1** | 7.03395 | 1.67045 | 5.31556 | 9.96339 | 3.01490 | 4.38406 | 7.12360 |
| **PTPN11** | 15.82766 | 5.67462 | 2.77589 | 8.10877 | 1.88568 | 3.79669 | 10.70178 |
| **PTPN12** | 1.80267 | 0.00000 | 7.93723 | 6.45467 | 1.19405 | 13.28802 | 9.15215 |
| **PTPN13** | 0.22441 | 0.00000 | 0.35677 | 0.00000 | 2.09738 | 0.09519 | 0.48181 |
| **PTPN14** | 8.32705 | 6.53267 | 2.57353 | 12.59804 | 1.98239 | 0.00000 | 16.28469 |
| **PTPN18** | 0.35404 | 0.31885 | 0.21102 | 2.58960 | 0.00000 | 0.00000 | 2.22312 |
| **PTPN2** | 13.02833 | 8.62056 | 12.32007 | 6.88344 | 6.72923 | 13.09040 | 5.36671 |
| **PTPN20B.uid-13** | 0.03993 | 0.00621 | 0.02184 | 0.01597 | 0.00712 | 0.01231 | 0.05368 |
| **PTPN20B.uid-14** | 0.02733 | 0.02010 | 0.01217 | 0.01883 | 0.00800 | 0.01482 | 0.01530 |
| **PTPN21** | 2.88067 | 0.68797 | 3.32547 | 6.08810 | 3.91400 | 0.00000 | 4.07590 |
| **PTPN22** | 0.06057 | 0.00007 | 0.01589 | 0.04119 | 1.57685 | 0.01420 | 0.01894 |
| **PTPN23** | 0.00440 | 1.79230 | 0.90964 | 2.65691 | 0.57988 | 2.36622 | 10.13214 |
| **PTPN3** | 2.43068 | 3.40046 | 0.71389 | 3.67536 | 4.56481 | 1.35015 | 6.57115 |
| **PTPN4** | 2.83522 | 0.00000 | 0.88358 | 1.44079 | 0.05478 | 1.98432 | 4.96763 |
| **PTPN5** | 0.00000 | 0.25258 | 0.77048 | 0.51591 | 0.85423 | 2.14373 | 0.00000 |
| **PTPN6** | 2.09566 | 0.49043 | 0.02376 | 3.60874 | 0.63822 | 0.37086 | 1.13363 |
| **PTPN7** | 0.03195 | 0.00905 | 0.01271 | 0.01918 | 0.01834 | 0.19194 | 0.01638 |
| **PTPN9** | 2.20367 | 2.91748 | 6.10768 | 1.12092 | 3.05255 | 3.84402 | 4.45014 |
| **PTPRA** | 1.05483 | 2.28212 | 2.23776 | 2.15946 | 1.06300 | 3.65297 | 7.05475 |

| | | | | | | | |
|---|---|---|---|---|---|---|---|
| **PTPRB** | 1.27832 | 1.13922 | 1.66613 | 0.16741 | 0.29677 | 0.52413 | 0.00000 |
| **PTPRC** | 0.01259 | 0.02317 | 0.00968 | 0.00931 | 0.00854 | 0.01346 | 0.12634 |
| **PTPRCAP** | 0.09764 | 0.00000 | 0.29276 | 0.14652 | 0.19200 | 0.43281 | 0.11190 |
| **PTPRD** | 0.04185 | 0.00000 | 0.01912 | 0.04070 | 2.72245 | 2.11326 | 0.02145 |
| **PTPRD-AS1** | 0.01519 | 0.00281 | 0.01678 | 0.01706 | 0.10501 | 0.08099 | 0.11288 |
| **PTPRE** | 0.09280 | 0.00000 | 0.00000 | 5.35753 | 0.47783 | 0.08460 | 0.22854 |
| **PTPRF** | 19.08132 | 11.49294 | 1.16014 | 15.90805 | 0.46661 | 9.02173 | 31.30772 |
| **PTPRG** | 0.72226 | 0.46523 | 0.91414 | 0.46618 | 1.79006 | 4.29616 | 1.31933 |
| **PTPRG-AS1** | 1.41556 | 3.49178 | 2.81224 | 0.73574 | 0.42098 | 0.41244 | 0.23297 |
| **PTPRH** | 0.01309 | 0.03713 | 0.06864 | 0.09974 | 2.05149 | 0.01955 | 0.00000 |
| **PTPRJ** | 0.00000 | 0.76535 | 4.59471 | 0.35800 | 1.48418 | 2.54595 | 0.94437 |
| **PTPRK** | 2.45040 | 0.00000 | 0.00000 | 1.10408 | 2.09238 | 0.69680 | 2.55334 |
| **PTPRM** | 0.06353 | 0.37011 | 0.00000 | 1.12913 | 0.44667 | 0.14709 | 0.27970 |
| **PTPRN** | 0.02362 | 0.01805 | 0.02111 | 0.04351 | 0.00833 | 0.01119 | 0.01667 |
| **PTPRN2** | 0.00566 | 0.15501 | 0.07041 | 0.02326 | 1.33366 | 0.00610 | 0.00579 |
| **PTPRO** | 0.02533 | 0.09665 | 0.11467 | 0.09796 | 0.22175 | 0.00735 | 0.09224 |
| **PTPRQ** | 0.02452 | 0.01541 | 0.01065 | 0.02232 | 0.06246 | 0.01682 | 0.01909 |
| **PTPRR** | 0.09745 | 0.01202 | 0.01997 | 0.02922 | 0.38716 | 0.28217 | 0.00000 |
| **PTPRS** | 1.45982 | 5.52943 | 4.85593 | 0.00000 | 2.17562 | 2.81702 | 0.84228 |
| **PTPRT** | 0.00952 | 0.07267 | 0.00228 | 0.03651 | 0.27824 | 0.01218 | 0.09948 |
| **PTPRU** | 6.57545 | 12.71759 | 0.12963 | 6.89693 | 0.00000 | 12.00677 | 17.26485 |
| **PTPRVP** | 0.01574 | 0.02994 | 0.01734 | 0.02407 | 0.02448 | 0.04576 | 0.01961 |
| **PTPRZ1** | 0.12803 | 0.06770 | 0.07447 | 0.02686 | 0.18290 | 1.56224 | 0.05667 |
| **PTRF** | 0.00000 | 16.58752 | 0.77920 | 1.52563 | 1.86948 | 1.43533 | 0.00000 |
| **PTRH1** | 0.30963 | 4.32206 | 1.81984 | 1.20231 | 0.37275 | 1.33196 | 1.24074 |
| **PTRH2** | 26.09106 | 15.04331 | 13.60105 | 19.42458 | 7.29808 | 7.29161 | 10.04262 |
| **PTRHD1** | 13.73870 | 8.53079 | 3.03360 | 12.24160 | 5.19255 | 3.52890 | 2.50290 |
| **PTS** | 18.39793 | 11.11226 | 11.00043 | 24.16156 | 0.43614 | 4.24762 | 3.74238 |
| **PTTG1** | 20.64152 | 12.55474 | 20.74908 | 24.16419 | 37.98721 | 10.76127 | 5.32829 |
| **PTTG1IP** | 43.53698 | 36.90832 | 11.22251 | 25.95649 | 5.39932 | 16.77160 | 18.41013 |
| **PTTG2** | 0.41666 | 0.00105 | 0.58945 | 0.47150 | 6.33336 | 0.32425 | 0.40437 |
| **PTTG3P** | 4.24246 | 1.43913 | 4.99365 | 5.29410 | 20.50935 | 2.08427 | 0.00000 |
| **PUF60** | 44.20221 | 43.22239 | 28.79380 | 31.43520 | 12.66139 | 23.18910 | 11.20651 |
| **PUM1** | 23.93329 | 12.19086 | 8.81941 | 22.04806 | 2.00043 | 9.57807 | 20.64364 |
| **PUM2** | 13.92972 | 0.00000 | 7.16771 | 7.92982 | 5.38173 | 6.44545 | 14.71812 |

| | | | | | | | |
|---|---|---|---|---|---|---|---|
| **PURA** | 2.05857 | 0.95687 | 0.84649 | 2.39912 | 0.02766 | 1.61423 | 9.63967 |
| **PURB** | 0.00001 | 0.00000 | 3.81996 | 1.96377 | 0.48911 | 0.41044 | 5.90119 |
| **PURG** | 0.03616 | 0.01565 | 0.01704 | 0.02375 | 0.03183 | 0.03055 | 0.01422 |
| **PUS1** | 3.79275 | 11.63568 | 3.39765 | 6.30244 | 0.21789 | 6.37157 | 26.10395 |
| **PUS10** | 1.16122 | 0.28749 | 0.65970 | 0.70966 | 1.20484 | 0.23141 | 0.23986 |
| **PUS3** | 12.50598 | 0.92121 | 8.98380 | 8.03415 | 1.45938 | 5.03297 | 5.93769 |
| **PUS7** | 22.13490 | 1.05965 | 4.01143 | 11.25323 | 0.95799 | 2.05436 | 1.98238 |
| **PUS7L** | 3.89462 | 0.00000 | 3.19210 | 4.51971 | 0.40140 | 2.13100 | 3.39119 |
| **PUSL1** | 8.72392 | 10.69477 | 0.00000 | 8.43508 | 0.01078 | 5.14772 | 5.49751 |
| **PVR** | 3.19584 | 10.04670 | 5.12578 | 6.39678 | 0.68197 | 10.02290 | 5.30812 |
| **PVRIG** | 0.00000 | 0.14010 | 0.04268 | 0.21997 | 0.01930 | 0.40398 | 0.40852 |
| **PVRL1** | 0.00000 | 3.36440 | 0.39899 | 15.09626 | 0.54716 | 3.68334 | 20.13795 |
| **PVRL2** | 1.76277 | 27.85369 | 5.70844 | 5.81430 | 0.29016 | 18.73352 | 24.15583 |
| **PVRL3** | 0.00000 | 0.00000 | 1.39560 | 2.87672 | 1.05210 | 0.41696 | 1.50869 |
| **PVRL3-AS1** | 0.03129 | 0.00887 | 0.02123 | 0.03012 | 0.05548 | 0.01784 | 0.00784 |
| **PVRL4** | 5.14487 | 1.87564 | 6.01208 | 18.21413 | 0.00000 | 8.48154 | 4.16030 |
| **PVT1** | 1.38870 | 3.27241 | 1.59911 | 3.78345 | 0.32926 | 0.56940 | 7.57664 |
| **PWAR1** | 0.00801 | 0.05515 | 0.01842 | 0.00751 | 0.01260 | 0.08056 | 0.08181 |
| **PWAR5** | 0.48288 | 0.00000 | 0.39671 | 0.03633 | 0.33040 | 0.99844 | 0.51386 |
| **PWARSN** | 3.50025 | 2.21941 | 3.46966 | 0.76223 | 0.24659 | 5.83648 | 2.28832 |
| **PWP1** | 36.09419 | 12.25841 | 19.23882 | 24.00857 | 11.00607 | 7.87073 | 13.26134 |
| **PWP2** | 21.18030 | 15.23650 | 8.74732 | 15.35784 | 2.40475 | 8.94517 | 3.02589 |
| **PWRN1** | 0.02017 | 0.00983 | 0.02052 | 0.01863 | 0.03839 | 0.02892 | 0.03202 |
| **PWWP2A** | 2.75325 | 0.48112 | 2.90672 | 1.12727 | 1.62577 | 5.79521 | 7.42933 |
| **PWWP2B** | 1.92583 | 1.46696 | 0.00000 | 9.78438 | 0.00000 | 0.00000 | 12.02232 |
| **PXDC1** | 0.00000 | 0.00000 | 0.03310 | 0.16127 | 0.03205 | 0.21816 | 0.25449 |
| **PXDN** | 0.00000 | 0.04729 | 0.00000 | 0.02422 | 1.62561 | 12.66087 | 0.40867 |
| **PXDNL** | 0.13666 | 0.07511 | 0.00655 | 0.00734 | 0.15911 | 0.00119 | 0.01478 |
| **PXK** | 2.69530 | 1.39832 | 0.32907 | 0.00000 | 5.13843 | 1.65186 | 3.12643 |
| **PXMP2** | 0.35095 | 2.23750 | 0.43539 | 1.21470 | 0.04761 | 0.85699 | 3.92139 |
| **PXMP4** | 4.56642 | 8.18634 | 7.17586 | 1.52719 | 1.48756 | 3.60522 | 0.00028 |
| **PXN** | 6.97215 | 7.01023 | 6.27228 | 15.41519 | 0.40527 | 7.07326 | 24.82107 |
| **PXN-AS1** | 6.70804 | 4.33515 | 0.76853 | 4.89563 | 0.63228 | 0.31799 | 0.00000 |
| **PXT1** | 0.21957 | 0.07006 | 0.01138 | 0.04101 | 0.33402 | 0.01363 | 0.00000 |
| **PXYLP1** | 1.61114 | 1.60585 | 0.19266 | 0.00000 | 2.51763 | 0.00000 | 0.52372 |

| | | | | | | | |
|---|---|---|---|---|---|---|---|
| **PYCARD** | 0.08402 | 0.49378 | 0.00000 | 0.00000 | 0.04663 | 1.40590 | 0.13674 |
| **PYCARDOS** | 0.00000 | 0.19249 | 0.06715 | 0.06157 | 0.32710 | 0.19759 | 0.16837 |
| **PYCR1** | 11.33906 | 1.63636 | 7.21595 | 8.39473 | 0.39224 | 5.25253 | 10.94611 |
| **PYCR2** | 5.55822 | 5.52429 | 2.89636 | 8.62313 | 0.53195 | 3.96644 | 5.78129 |
| **PYCRL** | 1.02568 | 4.28829 | 2.45586 | 2.32917 | 0.00000 | 0.82320 | 0.00000 |
| **PYDC1** | 0.01576 | 0.33102 | 0.01552 | 0.00000 | 0.07192 | 0.25718 | 0.08628 |
| **PYDC2** | 0.02548 | 0.02425 | 0.01446 | 0.02000 | 0.00790 | 0.01846 | 0.01461 |
| **PYGB** | 7.11340 | 45.96710 | 14.78765 | 1.89055 | 0.25572 | 39.26282 | 10.44017 |
| **PYGL** | 8.20287 | 11.12741 | 0.66599 | 16.29030 | 0.00000 | 12.86779 | 0.00000 |
| **PYGM** | 0.23957 | 0.05798 | 0.11502 | 0.36671 | 0.19217 | 0.00000 | 0.18432 |
| **PYGO1** | 4.16604 | 1.03622 | 1.33752 | 1.43798 | 0.65428 | 0.00000 | 0.67611 |
| **PYGO2** | 1.70697 | 5.49327 | 2.62376 | 5.44923 | 0.36853 | 2.48519 | 13.17708 |
| **PYHIN1** | 0.02011 | 0.02060 | 0.01088 | 0.01848 | 0.00767 | 0.06829 | 0.05005 |
| **PYROXD1** | 2.13600 | 0.09619 | 1.86813 | 2.74088 | 0.17362 | 0.10277 | 1.11480 |
| **PYROXD2** | 0.14221 | 0.13170 | 0.03386 | 0.02596 | 0.00594 | 0.00740 | 0.02147 |
| **PYY** | 0.00000 | 0.16614 | 1.08815 | 0.00000 | 0.34115 | 0.38685 | 0.06391 |
| **PYY2** | 0.00000 | 2.19905 | 1.93538 | 0.85183 | 0.48274 | 3.28711 | 1.19737 |
| **PZP** | 0.06093 | 0.00630 | 0.03232 | 0.05372 | 0.00776 | 0.01388 | 0.00000 |
| **QARS** | 47.09162 | 33.08333 | 2.28072 | 28.32653 | 2.80740 | 29.29696 | 19.37731 |
| **QDPR** | 5.49592 | 12.76073 | 3.39774 | 0.00000 | 2.47199 | 4.06565 | 5.28581 |
| **QKI** | 15.82664 | 1.86382 | 4.61214 | 8.19240 | 3.02520 | 7.64287 | 25.05627 |
| **QPCT** | 0.00000 | 0.54649 | 0.88233 | 0.10794 | 0.00000 | 1.80842 | 0.00044 |
| **QPCTL** | 2.85158 | 2.62784 | 1.10575 | 2.52761 | 0.48470 | 0.60754 | 1.27660 |
| **QPRT** | 19.66450 | 0.35494 | 0.00000 | 26.88038 | 0.00000 | 4.64079 | 7.28863 |
| **QRFP** | 0.00612 | 0.13654 | 0.05170 | 0.08278 | 0.03241 | 0.00256 | 0.06647 |
| **QRFPR** | 0.03543 | 0.01408 | 0.01369 | 0.03228 | 0.00917 | 0.02835 | 0.01537 |
| **QRICH1** | 10.01177 | 6.72283 | 9.64869 | 13.31291 | 7.77402 | 8.99440 | 17.33513 |
| **QRICH2** | 0.33913 | 3.35418 | 1.31049 | 0.00000 | 0.01748 | 1.30960 | 1.65691 |
| **QRSL1** | 17.93246 | 6.32849 | 6.82371 | 7.85467 | 1.28130 | 5.17475 | 0.00000 |
| **QSER1** | 1.15304 | 0.00000 | 0.77398 | 3.03203 | 0.06129 | 1.46781 | 4.33713 |
| **QSOX1** | 0.00000 | 2.65199 | 3.19450 | 13.83725 | 0.79138 | 4.70967 | 2.01626 |
| **QSOX2** | 0.76177 | 0.49134 | 0.98551 | 1.06576 | 0.43502 | 0.36313 | 5.78432 |
| **QTRT1** | 12.47977 | 10.39066 | 5.14605 | 9.74078 | 3.55867 | 7.67575 | 0.00000 |
| **QTRTD1** | 19.99854 | 7.41162 | 12.10633 | 11.23639 | 4.76799 | 7.47265 | 0.68530 |
| **R3HCC1** | 4.33543 | 12.93410 | 1.78421 | 0.00000 | 0.37048 | 1.30933 | 1.14169 |

| | | | | | | | |
|---|---|---|---|---|---|---|---|
| **R3HCC1L** | 9.34577 | 4.21150 | 6.98297 | 5.24154 | 6.31196 | 6.45979 | 1.03537 |
| **R3HDM1** | 6.79631 | 0.59181 | 4.63835 | 4.69820 | 1.79795 | 5.30559 | 6.74195 |
| **R3HDM2** | 7.21767 | 5.49894 | 1.45029 | 3.24478 | 1.55419 | 3.77886 | 5.55142 |
| **R3HDM4** | 0.00000 | 3.04414 | 1.89353 | 4.29311 | 3.76427 | 2.99923 | 16.17941 |
| **R3HDML** | 0.00548 | 0.08600 | 0.85097 | 0.00657 | 0.50216 | 0.01840 | 0.02127 |
| **RAB10** | 21.06878 | 16.39943 | 15.17235 | 20.91673 | 10.11359 | 13.76152 | 18.09733 |
| **RAB11A** | 43.13520 | 4.97094 | 23.97921 | 31.35391 | 12.89041 | 19.24746 | 33.70663 |
| **RAB11B** | 0.00000 | 2.96362 | 2.18112 | 2.68950 | 0.37176 | 5.20196 | 9.13974 |
| **RAB11B-AS1** | 0.05638 | 0.52196 | 0.07853 | 0.30583 | 0.39995 | 0.11103 | 0.08804 |
| **RAB11FIP1** | 25.17942 | 8.52821 | 1.69760 | 25.80002 | 2.15469 | 1.30541 | 33.98223 |
| **RAB11FIP2** | 6.28708 | 0.00000 | 4.01952 | 1.11620 | 0.71401 | 3.14135 | 9.63085 |
| **RAB11FIP3** | 0.36295 | 2.38408 | 0.54759 | 0.54264 | 0.33621 | 0.73270 | 3.86359 |
| **RAB11FIP4** | 17.63324 | 9.08047 | 0.97176 | 14.54378 | 0.00000 | 0.00000 | 28.64799 |
| **RAB11FIP5** | 0.51275 | 0.00000 | 0.34346 | 4.28684 | 0.00947 | 0.00051 | 9.19469 |
| **RAB12** | 0.41115 | 0.11187 | 1.17299 | 0.00000 | 0.83817 | 1.98640 | 4.80368 |
| **RAB13** | 0.10470 | 0.63775 | 0.60526 | 1.42773 | 0.20102 | 1.23965 | 1.38840 |
| **RAB14** | 20.33710 | 8.41195 | 9.67294 | 13.85462 | 1.50918 | 6.89168 | 15.16209 |
| **RAB15** | 4.29070 | 20.27351 | 7.67842 | 0.57362 | 0.03249 | 31.37739 | 13.23503 |
| **RAB17** | 0.54609 | 0.34826 | 0.00243 | 4.32187 | 0.03301 | 0.00000 | 0.05334 |
| **RAB18** | 40.72902 | 17.84329 | 15.08158 | 12.72706 | 0.00000 | 10.78524 | 4.58396 |
| **RAB19** | 0.02345 | 0.00000 | 0.10131 | 0.02799 | 0.01426 | 0.12631 | 0.04440 |
| **RAB1A** | 46.01995 | 15.29632 | 22.47159 | 29.92339 | 15.51272 | 19.63174 | 6.79112 |
| **RAB1B** | 19.19833 | 22.86264 | 7.43993 | 21.39308 | 2.92107 | 12.18403 | 6.75691 |
| **RAB20** | 4.01194 | 3.32686 | 2.09196 | 6.82598 | 0.92411 | 0.00000 | 12.38821 |
| **RAB21** | 3.72213 | 1.19682 | 4.22368 | 1.81433 | 1.51904 | 2.73567 | 8.21622 |
| **RAB22A** | 1.10351 | 0.88320 | 2.47896 | 1.94384 | 2.42839 | 1.83202 | 3.29375 |
| **RAB23** | 3.10149 | 0.00000 | 3.81197 | 2.20475 | 0.31832 | 2.89408 | 4.32308 |
| **RAB24** | 5.95846 | 6.22634 | 2.10530 | 7.67415 | 2.80605 | 3.60116 | 1.82504 |
| **RAB25** | 27.41938 | 7.88634 | 0.00000 | 6.16693 | 0.59461 | 0.00000 | 19.18198 |
| **RAB26** | 0.08426 | 0.22551 | 0.10972 | 0.74270 | 0.04092 | 0.52866 | 1.39067 |
| **RAB27A** | 0.35254 | 1.46193 | 0.97231 | 1.05963 | 2.02678 | 1.86128 | 0.21393 |
| **RAB27B** | 0.03766 | 0.00000 | 0.08912 | 0.01391 | 0.47936 | 0.00552 | 0.09698 |
| **RAB28** | 3.80059 | 2.09540 | 2.64478 | 2.44265 | 0.72634 | 2.30986 | 0.43945 |
| **RAB29** | 0.66435 | 0.00000 | 12.18973 | 2.91552 | 1.50408 | 4.19277 | 0.92687 |
| **RAB2A** | 8.22214 | 7.81669 | 12.15382 | 9.94762 | 3.61017 | 12.29504 | 29.90978 |

| | | | | | | | |
|---|---|---|---|---|---|---|---|
| **RAB2B** | 4.89211 | 0.00000 | 4.40438 | 3.33859 | 3.75786 | 1.21254 | 2.96321 |
| **RAB30** | 4.71892 | 0.89824 | 6.06335 | 1.08926 | 1.11702 | 0.94208 | 1.24580 |
| **RAB30-AS1** | 3.04638 | 1.58653 | 9.75312 | 3.90957 | 0.01531 | 0.43227 | 1.53959 |
| **RAB31** | 11.19140 | 0.00000 | 0.00000 | 17.89146 | 0.00000 | 0.00000 | 14.56752 |
| **RAB32** | 0.02047 | 0.00000 | 0.02483 | 0.12606 | 0.00539 | 0.09474 | 0.18352 |
| **RAB33A** | 0.00113 | 0.05538 | 0.00627 | 0.04280 | 0.00774 | 0.00466 | 0.10383 |
| **RAB33B** | 5.13501 | 0.28121 | 1.70284 | 2.24725 | 3.54383 | 1.06013 | 0.74900 |
| **RAB34** | 0.62603 | 0.00000 | 0.03061 | 11.62691 | 0.42965 | 0.84106 | 0.00032 |
| **RAB35** | 11.00063 | 18.36539 | 3.17842 | 14.31748 | 4.44625 | 8.33616 | 10.80155 |
| **RAB36** | 1.74862 | 0.17922 | 0.00000 | 5.82410 | 4.33203 | 0.00000 | 1.25383 |
| **RAB37** | 0.50802 | 0.80935 | 0.15223 | 0.19716 | 0.02044 | 0.15541 | 0.00000 |
| **RAB38** | 0.50935 | 0.00000 | 0.10848 | 0.68744 | 0.33690 | 0.96783 | 0.17388 |
| **RAB39A** | 0.45344 | 0.29525 | 0.74552 | 2.33385 | 0.56631 | 0.37828 | 0.20830 |
| **RAB39B** | 0.15738 | 0.05371 | 0.11327 | 0.86860 | 0.26717 | 0.32454 | 0.00000 |
| **RAB3A** | 0.20647 | 0.58090 | 0.01318 | 0.00000 | 2.44630 | 0.40551 | 0.28016 |
| **RAB3B** | 13.32455 | 0.00000 | 0.00000 | 7.06537 | 0.04173 | 17.23575 | 2.88568 |
| **RAB3C** | 0.02908 | 0.01620 | 0.04965 | 0.16452 | 0.04827 | 0.14291 | 0.00000 |
| **RAB3D** | 2.75660 | 6.63607 | 7.65043 | 1.80668 | 5.13072 | 5.91502 | 0.00000 |
| **RAB3GAP1** | 5.08123 | 1.68632 | 3.64363 | 6.27388 | 0.76569 | 6.16958 | 5.82774 |
| **RAB3GAP2** | 6.60477 | 0.65598 | 3.22558 | 4.32699 | 3.40880 | 2.91754 | 4.00881 |
| **RAB3IL1** | 0.00000 | 0.29550 | 0.25603 | 0.88984 | 0.10402 | 0.41051 | 1.50911 |
| **RAB3IP** | 8.47267 | 7.36069 | 7.18278 | 5.50620 | 1.52492 | 8.62890 | 3.58268 |
| **RAB40A** | 0.04097 | 0.11581 | 0.05930 | 0.12443 | 0.02017 | 0.05407 | 0.02908 |
| **RAB40AL** | 0.09500 | 0.06375 | 0.03774 | 0.04448 | 0.00492 | 0.02506 | 0.07012 |
| **RAB40B** | 1.48705 | 0.00000 | 1.63936 | 0.00000 | 0.19334 | 0.14266 | 4.37021 |
| **RAB40C** | 0.07714 | 2.65036 | 0.74456 | 2.59280 | 0.43840 | 2.09710 | 5.58667 |
| **RAB41** | 0.06214 | 0.01895 | 0.45513 | 0.08547 | 0.35662 | 0.04167 | 0.04909 |
| **RAB42** | 0.00000 | 2.00218 | 1.77351 | 0.18883 | 0.37322 | 0.47604 | 0.00000 |
| **RAB43** | 0.03058 | 1.56446 | 1.18772 | 0.00901 | 0.02678 | 0.15717 | 0.06620 |
| **RAB44** | 0.02242 | 0.01712 | 0.01118 | 0.02050 | 0.02848 | 0.01320 | 0.01889 |
| **RAB4A** | 3.70280 | 0.10223 | 2.11026 | 3.84383 | 0.41124 | 2.62077 | 7.26807 |
| **RAB5A** | 13.67899 | 1.45501 | 20.42623 | 7.84291 | 5.32769 | 10.41477 | 6.17842 |
| **RAB5B** | 2.76089 | 1.63076 | 2.88763 | 7.60520 | 2.95794 | 4.00024 | 8.39232 |
| **RAB5C** | 33.72382 | 32.83954 | 14.90618 | 25.87835 | 9.44776 | 11.74549 | 11.36216 |
| **RAB6A** | 26.18795 | 11.92636 | 21.16128 | 21.87832 | 11.63055 | 14.43113 | 0.00000 |

| | | | | | | | |
|---|---|---|---|---|---|---|---|
| **RAB6B** | 0.18597 | 1.01779 | 0.38011 | 0.64729 | 0.84757 | 0.57952 | 0.00000 |
| **RAB6C** | 0.02372 | 0.02349 | 0.02322 | 0.03161 | 0.02269 | 0.03744 | 0.04405 |
| **RAB6C-AS1** | 0.03712 | 0.01537 | 0.02083 | 0.02066 | 0.02564 | 0.03107 | 0.00738 |
| **RAB7A** | 41.54004 | 23.30466 | 24.55605 | 30.35279 | 9.69701 | 24.09504 | 12.86575 |
| **RAB8A** | 41.70819 | 20.23394 | 4.83930 | 27.02608 | 4.70273 | 12.34835 | 4.72207 |
| **RAB8B** | 2.48887 | 0.00000 | 3.20707 | 9.67976 | 6.37077 | 1.06613 | 0.00000 |
| **RAB9A** | 16.30821 | 4.37607 | 19.18171 | 12.03258 | 2.14699 | 6.10075 | 2.31733 |
| **RAB9B** | 0.03861 | 0.01092 | 0.04987 | 0.02352 | 0.00571 | 0.02190 | 0.01746 |
| **RAB9BP1** | 0.07463 | 0.03310 | 0.02032 | 0.02341 | 0.01820 | 0.07260 | 0.03504 |
| **RABAC1** | 7.12794 | 18.35090 | 8.68403 | 11.82956 | 4.69001 | 16.88330 | 10.44074 |
| **RABEP1** | 6.27279 | 4.62968 | 13.60020 | 6.83621 | 5.49501 | 3.24636 | 16.47337 |
| **RABEP2** | 0.05102 | 1.05320 | 1.09028 | 2.44820 | 0.73650 | 1.17971 | 2.76109 |
| **RABEPK** | 21.93105 | 19.47404 | 5.54997 | 21.36727 | 1.30491 | 6.92805 | 0.00000 |
| **RABGAP1** | 1.58515 | 0.13602 | 1.63402 | 3.27313 | 2.50328 | 2.36536 | 6.50651 |
| **RABGAP1L** | 1.90915 | 0.00000 | 1.70191 | 1.21702 | 1.17345 | 1.06799 | 1.67055 |
| **RABGEF1** | 10.99316 | 13.50958 | 14.75338 | 11.76827 | 17.22221 | 5.10902 | 0.00000 |
| **RABGGTA** | 6.88369 | 2.60953 | 1.18708 | 11.21718 | 0.20524 | 3.04119 | 1.27955 |
| **RABGGTB** | 33.68138 | 4.70559 | 15.43691 | 13.03391 | 1.71581 | 16.40706 | 27.93571 |
| **RABIF** | 15.66205 | 19.16298 | 15.90075 | 13.53639 | 6.25841 | 7.04864 | 3.34203 |
| **RABL2A** | 0.16329 | 0.69154 | 0.20921 | 0.27510 | 0.43404 | 0.03044 | 0.00000 |
| **RABL2B** | 1.80369 | 1.82922 | 0.51210 | 0.94818 | 0.42306 | 0.29629 | 0.00004 |
| **RABL3** | 8.80817 | 0.86022 | 4.65905 | 6.45925 | 1.25563 | 1.75732 | 0.00000 |
| **RABL6** | 0.00000 | 3.87869 | 0.04236 | 1.98303 | 0.06284 | 2.62568 | 20.49109 |
| **RAC1** | 5.34335 | 4.72340 | 3.00443 | 8.20366 | 2.16861 | 9.22154 | 38.94938 |
| **RAC2** | 0.29480 | 2.49107 | 0.00000 | 0.00000 | 2.04448 | 1.05358 | 0.20210 |
| **RAC3** | 0.00000 | 0.02875 | 0.01471 | 0.97046 | 0.03400 | 0.80064 | 1.35839 |
| **RACGAP1** | 19.31940 | 7.14262 | 8.47606 | 15.63099 | 14.50629 | 4.85149 | 2.99259 |
| **RACGAP1P** | 1.20428 | 0.59392 | 0.46517 | 0.95364 | 1.79495 | 0.12948 | 0.00000 |
| **RAD1** | 17.78546 | 7.18246 | 11.59042 | 9.48108 | 5.34115 | 5.69250 | 5.98262 |
| **RAD17** | 11.47478 | 0.75053 | 6.24463 | 2.38385 | 2.44861 | 4.39862 | 3.80635 |
| **RAD18** | 10.11319 | 2.12955 | 7.56958 | 5.61868 | 1.72528 | 2.47152 | 4.84725 |
| **RAD21** | 15.41415 | 3.45390 | 16.48413 | 10.65199 | 5.67766 | 10.20350 | 32.97723 |
| **RAD21-AS1** | 0.05892 | 0.07764 | 0.06844 | 0.05730 | 0.01780 | 0.05676 | 0.05342 |
| **RAD21L1** | 0.05645 | 0.00671 | 0.02988 | 0.03065 | 0.08194 | 0.04217 | 0.00233 |
| **RAD23A** | 19.41222 | 30.47073 | 9.76831 | 26.58187 | 1.49178 | 14.35766 | 11.45448 |

| | | | | | | | |
|---|---|---|---|---|---|---|---|
| **RAD23B** | 4.16369 | 2.99759 | 4.42333 | 5.32182 | 0.12227 | 5.72108 | 29.77000 |
| **RAD50** | 8.53213 | 1.30975 | 5.37128 | 3.47219 | 6.54078 | 5.01419 | 8.15043 |
| **RAD51** | 26.69158 | 28.31660 | 16.43914 | 13.62775 | 6.74841 | 21.66213 | 0.00000 |
| **RAD51-AS1** | 0.17620 | 1.28868 | 1.23317 | 0.88335 | 0.54757 | 0.36150 | 1.17830 |
| **RAD51AP1** | 11.73153 | 2.33939 | 9.80435 | 7.37825 | 4.08274 | 4.65802 | 7.55662 |
| **RAD51B** | 1.64682 | 0.44608 | 0.92259 | 0.50151 | 0.39046 | 0.96635 | 0.00000 |
| **RAD51C** | 19.30375 | 10.25323 | 15.58797 | 12.26302 | 3.57168 | 4.76954 | 0.00000 |
| **RAD51D** | 1.86793 | 2.66689 | 1.86225 | 1.94689 | 0.19555 | 1.29842 | 1.47338 |
| **RAD51L3-RFFL** | 0.05146 | 0.01356 | 0.01421 | 0.02930 | 0.00788 | 0.01176 | 0.00819 |
| **RAD52** | 4.65657 | 2.79433 | 4.80287 | 0.54400 | 2.03244 | 1.26269 | 1.13208 |
| **RAD54B** | 6.64238 | 0.00000 | 3.51280 | 1.41946 | 1.50261 | 1.94149 | 2.01236 |
| **RAD54L** | 9.01151 | 10.24207 | 3.25807 | 7.86035 | 7.45006 | 9.84152 | 5.61376 |
| **RAD54L2** | 2.09365 | 7.19352 | 6.07079 | 3.17669 | 3.05926 | 6.89999 | 14.78651 |
| **RAD9A** | 0.58920 | 2.61809 | 0.49239 | 2.89754 | 0.58615 | 1.24023 | 3.12730 |
| **RAD9B** | 0.09269 | 0.00000 | 0.29890 | 0.07454 | 0.22090 | 0.11517 | 0.11552 |
| **RADIL** | 0.00000 | 0.83972 | 0.23158 | 0.03041 | 1.50694 | 1.10782 | 0.30804 |
| **RAE1** | 27.06574 | 19.37948 | 19.42820 | 24.85354 | 9.01639 | 8.18904 | 0.00000 |
| **RAET1E** | 0.03049 | 0.01717 | 0.01345 | 0.02282 | 0.01210 | 0.03105 | 0.01173 |
| **RAET1K** | 0.05349 | 0.01238 | 0.01293 | 0.01808 | 0.09792 | 0.04269 | 0.00648 |
| **RAF1** | 26.62805 | 22.73383 | 11.26532 | 26.95917 | 4.30766 | 15.77055 | 0.64725 |
| **RAG1** | 0.06429 | 0.00000 | 0.30708 | 0.03683 | 0.12693 | 0.04770 | 0.02509 |
| **RAG2** | 0.04400 | 0.01296 | 0.02140 | 0.02644 | 0.00744 | 0.01499 | 0.00727 |
| **RAI1** | 0.00000 | 0.03059 | 0.24559 | 0.01450 | 0.07071 | 0.15536 | 0.28443 |
| **RAI14** | 16.98810 | 6.22492 | 0.00000 | 9.33969 | 2.02011 | 1.68017 | 15.57579 |
| **RAI2** | 0.00000 | 0.10681 | 0.14044 | 0.00996 | 0.00252 | 0.10397 | 0.07731 |
| **RALA** | 16.57380 | 12.61391 | 5.72182 | 16.74206 | 3.03587 | 11.12506 | 8.36964 |
| **RALB** | 4.54428 | 0.46036 | 0.88722 | 2.85714 | 2.51795 | 1.39784 | 0.25166 |
| **RALBP1** | 22.07682 | 20.96649 | 4.98714 | 27.89255 | 5.40763 | 6.57898 | 26.24482 |
| **RALGAPA1** | 0.16747 | 0.00000 | 0.18686 | 0.00000 | 0.00912 | 0.15049 | 0.59304 |
| **RALGAPA1P** | 0.02274 | 0.01796 | 0.01372 | 0.01936 | 0.00779 | 0.01486 | 0.02270 |
| **RALGAPA2** | 9.84142 | 1.16103 | 0.05548 | 7.77062 | 0.27767 | 1.66025 | 14.84862 |
| **RALGAPB** | 4.82129 | 1.95357 | 4.64202 | 3.21764 | 2.99482 | 6.06058 | 9.29490 |
| **RALGDS** | 1.15639 | 0.67648 | 0.01534 | 0.70516 | 0.22022 | 0.00000 | 4.26555 |
| **RALGPS1** | 0.73073 | 0.15762 | 0.00000 | 1.40186 | 0.57573 | 0.64858 | 4.16144 |
| **RALGPS2** | 2.76543 | 2.75614 | 3.22396 | 0.73676 | 0.95216 | 5.48943 | 4.38754 |

| | | | | | | | |
|---|---|---|---|---|---|---|---|
| **RALY** | 0.00000 | 11.21937 | 5.49357 | 10.82917 | 6.34813 | 8.19674 | 23.81355 |
| **RALY-AS1** | 0.85301 | 0.36865 | 1.12820 | 0.61643 | 1.06858 | 0.00000 | 0.47822 |
| **RALYL** | 0.03020 | 0.00490 | 0.07355 | 0.04906 | 0.26582 | 0.01914 | 0.01133 |
| **RAMP1** | 0.00000 | 0.97272 | 1.78963 | 0.00000 | 0.43587 | 1.03088 | 0.00089 |
| **RAMP2** | 0.08303 | 0.03849 | 0.04850 | 0.03068 | 0.02299 | 0.05921 | 0.00000 |
| **RAMP2-AS1** | 0.05604 | 0.00821 | 0.00931 | 0.03821 | 0.05878 | 0.03235 | 0.07841 |
| **RAN** | 63.52528 | 37.74029 | 30.71899 | 48.38882 | 17.81793 | 29.98950 | 20.44792 |
| **RANBP1** | 22.97927 | 28.93306 | 11.26517 | 21.44836 | 9.97860 | 13.34834 | 15.77604 |
| **RANBP10** | 2.63853 | 3.68952 | 1.43368 | 4.66228 | 0.93348 | 2.11199 | 6.97290 |
| **RANBP17** | 3.06187 | 1.29571 | 1.07121 | 0.56943 | 0.18656 | 1.32100 | 2.37852 |
| **RANBP2** | 15.01215 | 9.50725 | 16.47067 | 6.56966 | 5.57892 | 14.26171 | 28.72544 |
| **RANBP3** | 12.47817 | 12.64240 | 7.50870 | 13.96409 | 4.29130 | 5.85055 | 3.35803 |
| **RANBP3L** | 0.02086 | 0.01777 | 0.00979 | 0.04193 | 0.04334 | 0.01209 | 0.01713 |
| **RANBP6** | 2.90033 | 0.00000 | 3.39134 | 1.19258 | 0.84871 | 0.39596 | 0.00000 |
| **RANBP9** | 3.16807 | 1.14289 | 2.92941 | 5.20111 | 2.22033 | 4.82634 | 11.75844 |
| **RANGAP1** | 15.46077 | 40.08743 | 17.90678 | 28.21100 | 0.70691 | 13.48218 | 34.59444 |
| **RANGRF** | 7.57017 | 1.49376 | 0.38166 | 6.72469 | 0.21207 | 2.58799 | 2.42084 |
| **RAP1A** | 1.91614 | 4.96810 | 5.83493 | 0.42245 | 0.37069 | 10.00268 | 12.57750 |
| **RAP1B** | 8.77262 | 1.33669 | 11.97860 | 6.97512 | 8.12397 | 4.32863 | 1.98415 |
| **RAP1GAP** | 0.00927 | 0.16519 | 0.09733 | 0.10633 | 0.19797 | 0.18249 | 0.23341 |
| **RAP1GAP2** | 0.00000 | 2.25469 | 0.46925 | 0.74849 | 0.20491 | 2.74268 | 2.15992 |
| **RAP1GDS1** | 20.60916 | 0.34593 | 9.93074 | 10.10368 | 4.24791 | 2.86468 | 3.45049 |
| **RAP2A** | 2.57550 | 0.00000 | 2.72268 | 2.22629 | 1.88859 | 2.28152 | 5.82195 |
| **RAP2B** | 29.94985 | 24.33668 | 23.20938 | 18.16217 | 2.68273 | 17.41434 | 4.68878 |
| **RAP2C** | 5.93226 | 0.85608 | 2.15092 | 1.15848 | 0.57649 | 4.47163 | 5.29599 |
| **RAP2C-AS1** | 0.07138 | 0.00544 | 0.02784 | 0.03917 | 0.01328 | 0.05186 | 0.00000 |
| **RAPGEF1** | 1.07981 | 3.58928 | 0.31820 | 11.18044 | 0.08195 | 6.63182 | 20.80678 |
| **RAPGEF2** | 11.82461 | 11.47524 | 2.49740 | 7.58368 | 1.17194 | 3.50421 | 22.07781 |
| **RAPGEF3** | 0.15961 | 0.05501 | 0.00244 | 0.03858 | 0.06928 | 0.00000 | 0.19693 |
| **RAPGEF4** | 0.01459 | 0.29317 | 0.03889 | 0.08331 | 0.01640 | 0.21090 | 0.02438 |
| **RAPGEF4-AS1** | 0.02994 | 0.01544 | 0.01397 | 0.01956 | 0.00787 | 0.02149 | 0.01360 |
| **RAPGEF5** | 0.00000 | 0.15748 | 0.02388 | 0.03331 | 0.04539 | 0.06200 | 0.02428 |
| **RAPGEF6** | 8.56185 | 1.18349 | 1.75787 | 3.63870 | 1.38800 | 1.72635 | 4.68773 |
| **RAPGEFL1** | 0.17381 | 0.13940 | 0.42616 | 0.72920 | 0.12249 | 0.45756 | 2.62821 |
| **RAPH1** | 3.30348 | 0.81054 | 4.16943 | 2.63253 | 3.84484 | 2.70408 | 8.00620 |

| | | | | | | | |
|---|---|---|---|---|---|---|---|
| **RAPSN** | 0.08709 | 0.00959 | 0.02221 | 0.07809 | 0.00662 | 0.00843 | 0.05259 |
| **RARA** | 0.02654 | 0.00000 | 0.23925 | 0.92680 | 0.87309 | 0.07376 | 2.91409 |
| **RARA-AS1** | 0.03658 | 0.02069 | 0.00930 | 0.05510 | 0.01417 | 0.00749 | 0.05058 |
| **RARB** | 0.09355 | 0.48002 | 0.14234 | 0.03510 | 0.90838 | 0.06917 | 0.01502 |
| **RARG** | 0.23061 | 0.75933 | 0.67708 | 0.10084 | 0.10620 | 1.11417 | 0.86054 |
| **RARRES1** | 0.29374 | 0.03397 | 0.03731 | 0.39689 | 0.00754 | 0.00000 | 1.32331 |
| **RARRES2** | 0.00000 | 0.37925 | 2.78890 | 0.00000 | 10.78014 | 0.00000 | 0.01882 |
| **RARRES3** | 0.02895 | 0.26017 | 0.08594 | 0.00103 | 0.00404 | 0.12612 | 0.01176 |
| **RARS** | 43.36357 | 11.32945 | 15.88693 | 22.13382 | 0.82874 | 11.39802 | 10.03105 |
| **RARS2** | 14.37942 | 4.10032 | 12.31133 | 13.49429 | 4.62232 | 7.08852 | 0.00000 |
| **RASA1** | 1.59862 | 0.00000 | 2.44581 | 3.83433 | 1.08722 | 2.55501 | 5.56606 |
| **RASA2** | 1.77027 | 0.00000 | 1.90918 | 3.19211 | 0.74879 | 0.93453 | 1.65073 |
| **RASA3** | 0.01151 | 0.25488 | 0.02136 | 0.00679 | 2.48323 | 0.32937 | 0.00017 |
| **RASA4** | 0.02600 | 0.01335 | 0.01443 | 0.01900 | 0.00766 | 0.01976 | 0.02693 |
| **RASA4B.uid-999** | 0.02680 | 0.01882 | 0.01606 | 0.02167 | 0.00761 | 0.01988 | 0.01667 |
| **RASA4CP** | 0.03030 | 0.03327 | 0.01349 | 0.01131 | 0.12254 | 0.02991 | 0.05980 |
| **RASAL1** | 0.13227 | 0.00492 | 0.01150 | 0.01690 | 0.07505 | 0.01625 | 0.00000 |
| **RASAL2** | 0.61827 | 0.85326 | 0.90779 | 7.13694 | 0.13919 | 2.41975 | 2.81343 |
| **RASAL2-AS1** | 0.00634 | 0.08389 | 0.02144 | 0.23105 | 0.00813 | 0.05641 | 0.00889 |
| **RASAL3** | 0.01641 | 0.04805 | 0.00000 | 0.01780 | 0.23029 | 0.01004 | 0.03720 |
| **RASD1** | 0.00000 | 8.96469 | 0.02514 | 0.22363 | 1.16566 | 9.58657 | 0.00000 |
| **RASEF** | 0.03932 | 0.13381 | 0.01280 | 0.00000 | 0.02569 | 0.04155 | 0.60641 |
| **RASGEF1A** | 1.78794 | 13.75511 | 4.74049 | 0.00000 | 0.79346 | 6.38978 | 0.00000 |
| **RASGEF1B** | 0.00000 | 0.18056 | 0.00000 | 0.01885 | 0.09665 | 0.41801 | 0.17329 |
| **RASGEF1C** | 0.05137 | 0.08403 | 0.11915 | 0.00839 | 0.01493 | 0.00272 | 0.00959 |
| **RASGRF1** | 1.01591 | 0.00000 | 0.00000 | 0.46570 | 0.83189 | 0.00000 | 2.32481 |
| **RASGRF2** | 0.06500 | 0.47311 | 0.09620 | 0.67514 | 0.70402 | 1.30809 | 1.11041 |
| **RASGRF2-AS1** | 0.02153 | 0.02053 | 0.02843 | 0.03084 | 0.00682 | 0.08062 | 0.01471 |
| **RASGRP1** | 0.07367 | 0.00000 | 0.07213 | 0.14127 | 0.17130 | 0.00452 | 0.87169 |
| **RASGRP2** | 9.25308 | 9.38307 | 1.34676 | 9.85022 | 0.00000 | 2.46865 | 20.09396 |
| **RASGRP3** | 0.10901 | 0.00166 | 0.06362 | 0.06390 | 0.85103 | 0.06702 | 0.24175 |
| **RASGRP4** | 0.01438 | 0.22263 | 0.17370 | 0.00000 | 0.11945 | 0.00603 | 0.13962 |
| **RASIP1** | 0.12719 | 0.00000 | 0.37126 | 0.14522 | 0.40991 | 0.25097 | 1.69110 |
| **RASL10A** | 0.16931 | 0.35985 | 0.01945 | 0.23648 | 0.30000 | 0.53377 | 0.25098 |
| **RASL10B** | 0.02805 | 0.02965 | 0.00923 | 0.12414 | 0.02213 | 0.00865 | 0.00170 |

| | | | | | | | |
|---|---|---|---|---|---|---|---|
| **RASL11A** | 0.60544 | 0.81566 | 1.37031 | 0.13992 | 1.27732 | 1.14812 | 0.05447 |
| **RASL11B** | 2.26344 | 0.00000 | 1.30081 | 3.65314 | 0.72311 | 0.97567 | 0.00000 |
| **RASL12** | 0.01721 | 0.01579 | 0.01298 | 0.04167 | 0.01732 | 0.01811 | 0.03539 |
| **RASSF1** | 8.18032 | 11.01177 | 5.37501 | 15.45059 | 0.17578 | 3.97754 | 0.00000 |
| **RASSF1-AS1** | 0.05628 | 0.22049 | 0.04751 | 0.06065 | 0.03299 | 0.04141 | 0.02014 |
| **RASSF10** | 0.00597 | 0.01377 | 0.01948 | 0.04205 | 0.00766 | 0.00959 | 0.06094 |
| **RASSF2** | 6.25103 | 0.00000 | 0.14832 | 5.36888 | 0.14662 | 0.46196 | 2.51005 |
| **RASSF3** | 6.28200 | 1.02933 | 2.31365 | 2.96751 | 2.83418 | 2.98986 | 7.25563 |
| **RASSF4** | 2.57609 | 0.06634 | 0.00000 | 0.84887 | 0.09800 | 1.29698 | 0.00000 |
| **RASSF5** | 0.01638 | 0.17340 | 0.01193 | 0.01114 | 0.42951 | 0.04427 | 0.01034 |
| **RASSF6** | 0.05891 | 0.00000 | 0.06942 | 0.03401 | 0.02371 | 0.28692 | 0.01967 |
| **RASSF7** | 3.66541 | 7.21298 | 1.27462 | 17.54487 | 0.00000 | 4.01784 | 5.71481 |
| **RASSF8** | 2.59123 | 0.00000 | 6.40035 | 0.18046 | 0.93406 | 6.99949 | 8.36703 |
| **RASSF8-AS1** | 0.49045 | 0.00000 | 2.17982 | 0.04904 | 0.72215 | 0.82413 | 0.91771 |
| **RASSF9** | 0.07649 | 0.00000 | 0.03491 | 0.01617 | 0.02238 | 0.01317 | 0.01890 |
| **RAVER1** | 2.64392 | 3.48550 | 3.41932 | 6.43721 | 0.78436 | 3.27365 | 20.08259 |
| **RAVER2** | 0.12358 | 0.03255 | 0.06691 | 0.00000 | 0.28738 | 0.66717 | 0.65157 |
| **RAX** | 0.02841 | 0.01243 | 0.01031 | 0.01393 | 0.03183 | 0.01811 | 0.03648 |
| **RAX2** | 0.01070 | 0.08684 | 0.00000 | 0.01955 | 0.49232 | 0.00673 | 0.02150 |
| **RB1** | 1.84611 | 0.00000 | 1.78766 | 2.45942 | 0.53960 | 0.69445 | 2.18128 |
| **RB1CC1** | 13.51856 | 1.28548 | 30.25335 | 11.49498 | 14.57776 | 9.78641 | 22.16037 |
| **RBAK** | 4.12867 | 0.00000 | 4.56781 | 0.99778 | 0.20821 | 0.77701 | 7.79486 |
| **RBAKDN** | 0.01941 | 0.05315 | 0.02867 | 0.02269 | 0.08852 | 0.01508 | 0.00847 |
| **RBBP4** | 20.64092 | 1.62756 | 10.65284 | 11.00356 | 3.10311 | 12.81421 | 6.04789 |
| **RBBP5** | 16.63929 | 10.17140 | 5.40054 | 8.75138 | 10.26207 | 6.09118 | 0.25488 |
| **RBBP6** | 12.39731 | 3.66745 | 11.07857 | 5.26533 | 11.11580 | 9.02451 | 35.89664 |
| **RBBP7** | 30.38870 | 21.69698 | 25.12238 | 20.06473 | 14.06444 | 8.69559 | 6.07952 |
| **RBBP8** | 9.64832 | 2.13958 | 5.74857 | 3.56461 | 1.95498 | 2.28323 | 5.97507 |
| **RBBP8NL** | 0.54821 | 0.00000 | 0.00000 | 1.89651 | 0.00187 | 0.00000 | 4.04236 |
| **RBBP9** | 2.64265 | 1.75637 | 8.60719 | 1.14429 | 0.25859 | 3.45546 | 0.19537 |
| **RBCK1** | 4.36715 | 7.51768 | 2.37477 | 5.28699 | 0.72793 | 1.66930 | 3.87557 |
| **RBFA** | 10.13277 | 5.02297 | 3.70179 | 9.87192 | 1.11840 | 3.45496 | 6.17913 |
| **RBFADN** | 0.07553 | 0.00000 | 0.06064 | 0.02799 | 0.00705 | 0.03427 | 0.15741 |
| **RBFOX1** | 0.11087 | 0.07678 | 0.07520 | 0.00000 | 0.45105 | 0.06243 | 0.03294 |
| **RBFOX2** | 7.23302 | 1.59521 | 1.96519 | 10.65398 | 0.41612 | 7.49940 | 14.85717 |

| | | | | | | | |
|---|---|---|---|---|---|---|---|
| **RBFOX3** | 0.00000 | 0.04698 | 0.02575 | 0.03461 | 0.32564 | 0.01390 | 0.05954 |
| **RBKS** | 0.46456 | 1.73236 | 0.46663 | 0.31804 | 0.03420 | 0.21334 | 0.00000 |
| **RBL1** | 3.41951 | 0.38461 | 2.03926 | 1.72548 | 1.05203 | 1.83912 | 3.51492 |
| **RBL2** | 1.10878 | 1.44977 | 1.53214 | 0.50158 | 0.80713 | 0.75534 | 1.76850 |
| **RBM10** | 13.81808 | 19.67372 | 7.13804 | 17.12354 | 2.52929 | 7.92737 | 17.89713 |
| **RBM11** | 0.20123 | 0.00000 | 0.34830 | 0.27577 | 0.06454 | 0.09843 | 0.21934 |
| **RBM12** | 20.61222 | 9.24015 | 9.68814 | 12.70135 | 2.43504 | 8.66869 | 9.21055 |
| **RBM12B** | 6.54062 | 0.00000 | 5.94627 | 2.36631 | 2.17121 | 1.72515 | 2.70628 |
| **RBM12B-AS1** | 0.12113 | 0.00000 | 0.21595 | 0.35755 | 0.03188 | 0.40700 | 0.37943 |
| **RBM14** | 22.63156 | 18.91900 | 15.79167 | 21.78168 | 6.95182 | 12.33216 | 13.52686 |
| **RBM15** | 8.00186 | 1.90837 | 7.19497 | 6.89068 | 3.42873 | 3.09629 | 6.15577 |
| **RBM15B** | 3.48301 | 3.00834 | 5.12744 | 8.59297 | 0.71396 | 4.50492 | 14.26417 |
| **RBM17** | 18.15810 | 27.08116 | 16.13244 | 19.34407 | 7.17211 | 14.61477 | 25.12944 |
| **RBM18** | 18.19708 | 7.38103 | 9.32259 | 12.08111 | 5.05901 | 7.26457 | 0.82158 |
| **RBM19** | 10.49095 | 10.72823 | 7.94781 | 12.83486 | 4.77692 | 7.05567 | 6.42106 |
| **RBM20** | 0.00229 | 0.07102 | 0.00895 | 0.01337 | 0.00889 | 0.02522 | 0.35477 |
| **RBM22** | 19.19417 | 7.78056 | 10.74355 | 15.91341 | 6.21382 | 8.26759 | 7.24936 |
| **RBM23** | 16.90672 | 6.35171 | 6.52315 | 11.55763 | 5.57297 | 11.38113 | 0.80165 |
| **RBM24** | 0.00834 | 0.05243 | 1.69454 | 0.01434 | 0.32878 | 0.00000 | 0.01854 |
| **RBM25** | 29.09556 | 15.58485 | 20.29132 | 19.10555 | 15.43486 | 17.13615 | 39.17270 |
| **RBM26** | 7.85176 | 0.00000 | 4.82570 | 1.84283 | 0.98992 | 4.22959 | 15.66958 |
| **RBM26-AS1** | 0.28440 | 0.66758 | 0.01591 | 0.01642 | 0.04614 | 0.04663 | 0.04244 |
| **RBM27** | 11.43824 | 1.84788 | 7.05624 | 10.37565 | 4.58643 | 8.07256 | 21.90598 |
| **RBM28** | 19.58469 | 11.79866 | 12.38970 | 12.27683 | 4.71269 | 6.88337 | 7.90502 |
| **RBM3** | 55.79918 | 17.49769 | 29.62539 | 27.96567 | 8.77428 | 25.14000 | 34.84996 |
| **RBM33** | 6.28912 | 3.16355 | 7.34722 | 6.46073 | 3.07295 | 2.83298 | 21.05826 |
| **RBM34** | 24.44635 | 12.89769 | 14.45539 | 17.13133 | 9.13419 | 7.72690 | 7.37929 |
| **RBM38** | 6.19942 | 22.26571 | 2.93238 | 7.26789 | 3.06600 | 8.41376 | 29.04497 |
| **RBM39** | 40.48823 | 24.96019 | 20.66632 | 26.98815 | 20.66131 | 17.27293 | 22.17650 |
| **RBM4** | 12.01039 | 5.14931 | 6.49525 | 7.47555 | 4.96319 | 5.33387 | 0.00000 |
| **RBM41** | 11.37068 | 3.15075 | 3.59823 | 0.00000 | 0.72672 | 3.08292 | 6.78005 |
| **RBM42** | 2.31860 | 25.71947 | 11.00472 | 17.45440 | 6.33973 | 9.69432 | 25.00532 |
| **RBM43** | 0.04916 | 0.03227 | 0.01399 | 0.01642 | 0.01131 | 0.03203 | 0.00119 |
| **RBM44** | 0.10821 | 0.00000 | 3.88273 | 0.00000 | 2.29140 | 0.00000 | 0.63424 |
| **RBM45** | 4.47135 | 2.61386 | 1.41023 | 0.89594 | 1.17639 | 1.29114 | 0.00000 |

| | | | | | | | |
|---|---|---|---|---|---|---|---|
| **RBM46** | 0.32486 | 0.00000 | 0.69808 | 1.04726 | 5.38404 | 0.04100 | 0.59851 |
| **RBM47** | 38.63393 | 26.10396 | 8.04656 | 24.25246 | 0.00000 | 7.56117 | 29.60926 |
| **RBM48** | 6.09269 | 1.18578 | 10.21193 | 4.64554 | 10.67064 | 2.06456 | 0.00000 |
| **RBM4B** | 7.56945 | 2.64903 | 5.72051 | 7.54403 | 3.32862 | 1.23524 | 0.00000 |
| **RBM5** | 11.75870 | 9.03382 | 9.65127 | 11.97885 | 5.37191 | 11.80761 | 6.45699 |
| **RBM5-AS1** | 0.51317 | 0.15685 | 0.51585 | 0.16650 | 0.07970 | 0.35511 | 0.59404 |
| **RBM6** | 20.38050 | 12.55789 | 7.95321 | 11.33273 | 7.62988 | 10.30572 | 8.05222 |
| **RBM7** | 18.06580 | 0.36139 | 7.36126 | 14.37948 | 6.94850 | 6.61649 | 0.00000 |
| **RBM8A** | 26.79284 | 25.31302 | 23.04404 | 25.38582 | 16.19799 | 21.79674 | 4.95728 |
| **RBMS1** | 2.24368 | 2.84936 | 3.46347 | 4.04460 | 0.35357 | 7.57223 | 16.13150 |
| **RBMS2** | 0.30843 | 0.13789 | 0.25940 | 0.75439 | 0.98354 | 0.59380 | 0.51885 |
| **RBMS3** | 0.04076 | 0.04189 | 0.02553 | 0.00895 | 0.31947 | 0.02173 | 0.02114 |
| **RBMS3-AS3** | 0.02351 | 0.02070 | 0.01443 | 0.02063 | 0.00975 | 0.01254 | 0.01981 |
| **RBMX** | 42.70507 | 20.80859 | 20.17115 | 18.28741 | 18.45342 | 18.07197 | 28.26552 |
| **RBMX2** | 7.23197 | 8.21318 | 6.49808 | 4.02536 | 8.17351 | 1.49841 | 7.19444 |
| **RBMXL1** | 14.62448 | 4.44642 | 8.24296 | 7.73106 | 2.19316 | 4.53991 | 6.20130 |
| **RBMXL2** | 0.02449 | 0.00000 | 0.11713 | 0.17150 | 0.00705 | 0.00927 | 0.18971 |
| **RBMXL3** | 0.07239 | 0.00000 | 2.28011 | 0.25812 | 0.70983 | 0.07045 | 0.00000 |
| **RBMY2FP** | 0.02600 | 0.01060 | 0.02988 | 0.01914 | 0.00887 | 0.01275 | 0.01654 |
| **RBP1** | 2.52153 | 0.00000 | 0.06075 | 0.00000 | 1.07871 | 0.79814 | 1.06027 |
| **RBP2** | 0.03158 | 0.01666 | 0.01843 | 0.04285 | 0.00768 | 0.05146 | 0.00000 |
| **RBP3** | 0.01620 | 0.01422 | 0.01307 | 0.10961 | 0.00900 | 0.00768 | 0.03587 |
| **RBP4** | 0.03382 | 0.00234 | 0.04435 | 0.07502 | 0.19904 | 0.01181 | 0.01413 |
| **RBP5** | 0.01897 | 0.01198 | 0.02082 | 0.02836 | 0.00710 | 0.21544 | 0.01735 |
| **RBP7** | 0.00000 | 5.68512 | 20.68914 | 0.00000 | 2.71373 | 0.14372 | 0.00000 |
| **RBPJ** | 23.48456 | 9.42659 | 10.19560 | 11.49469 | 6.88364 | 11.29394 | 23.03425 |
| **RBPJL** | 0.00000 | 0.11670 | 0.99823 | 0.02747 | 0.00315 | 0.03129 | 0.01423 |
| **RBPMS** | 1.20663 | 3.64215 | 1.60801 | 6.63676 | 0.60959 | 6.21955 | 23.89505 |
| **RBPMS-AS1** | 0.22891 | 0.53073 | 0.49930 | 0.69851 | 0.03470 | 0.02960 | 0.10808 |
| **RBPMS2** | 3.05350 | 24.94045 | 13.57225 | 0.97083 | 2.66650 | 12.70747 | 18.03255 |
| **RBX1** | 28.25010 | 16.22197 | 16.26636 | 24.70705 | 6.44118 | 15.40576 | 18.44100 |
| **RC3H1** | 14.50587 | 0.78839 | 3.10481 | 6.77888 | 8.08684 | 4.25496 | 7.38762 |
| **RC3H2** | 3.09575 | 5.13694 | 3.12473 | 4.45732 | 2.05226 | 5.41005 | 4.66260 |
| **RCAN1** | 1.02000 | 0.00000 | 0.68763 | 0.08329 | 0.55710 | 1.13700 | 3.72192 |
| **RCAN2** | 0.00000 | 0.26335 | 0.00000 | 0.01554 | 0.49494 | 0.01377 | 0.02313 |

| | | | | | | | |
|---|---|---|---|---|---|---|---|
| **RCAN3** | 1.01542 | 1.78557 | 1.83063 | 0.51120 | 0.27995 | 0.18174 | 3.08378 |
| **RCAN3AS** | 0.02293 | 0.01602 | 0.02263 | 0.02162 | 0.02449 | 0.01350 | 0.01783 |
| **RCBTB1** | 4.85499 | 5.05858 | 10.14830 | 0.42988 | 1.61283 | 18.33810 | 0.70340 |
| **RCBTB2** | 0.12125 | 0.12059 | 0.13377 | 0.15843 | 0.58974 | 0.39915 | 0.00000 |
| **RCC1** | 52.32851 | 25.83246 | 15.85171 | 35.52632 | 4.13611 | 15.53854 | 0.00010 |
| **RCC2** | 15.37702 | 16.96979 | 10.95947 | 12.13760 | 1.41367 | 20.44324 | 31.61859 |
| **RCCD1** | 1.32580 | 1.70172 | 1.00691 | 1.52626 | 0.50869 | 1.18254 | 5.91571 |
| **RCE1** | 4.24726 | 9.53015 | 4.61806 | 7.14576 | 4.01619 | 3.99880 | 11.46129 |
| **RCHY1** | 15.28306 | 3.65454 | 13.63661 | 4.91595 | 0.41039 | 7.13495 | 2.49217 |
| **RCL1** | 25.45048 | 10.80964 | 13.66859 | 12.64659 | 4.45610 | 5.12745 | 2.60047 |
| **RCN1** | 10.84672 | 0.01318 | 0.00000 | 12.78804 | 0.00000 | 13.61926 | 14.76561 |
| **RCN2** | 5.33121 | 0.00000 | 1.55528 | 10.74794 | 0.00000 | 8.70356 | 23.55544 |
| **RCN3** | 0.00000 | 1.30942 | 1.59001 | 0.25347 | 0.70074 | 0.26312 | 0.00000 |
| **RCOR1** | 5.08167 | 2.61377 | 7.40032 | 5.50961 | 3.73838 | 3.61145 | 20.63752 |
| **RCOR2** | 0.00209 | 0.03218 | 0.17320 | 0.15958 | 0.00126 | 0.42453 | 3.51702 |
| **RCOR3** | 4.26115 | 2.70327 | 5.18932 | 1.49652 | 1.89417 | 6.21814 | 4.96855 |
| **RCSD1** | 0.00000 | 0.10990 | 0.02520 | 0.03410 | 0.00569 | 0.58348 | 0.02559 |
| **RCVRN** | 0.00341 | 0.10446 | 0.01545 | 0.02839 | 0.02201 | 0.01452 | 0.10597 |
| **RD3** | 0.00439 | 0.03087 | 0.07192 | 0.03477 | 0.01756 | 0.02555 | 0.03924 |
| **RD3L** | 0.02890 | 0.00694 | 0.03021 | 0.02062 | 0.01510 | 0.01212 | 0.02245 |
| **RDH10** | 9.23674 | 1.01573 | 2.10590 | 2.91313 | 4.07506 | 4.69189 | 5.90424 |
| **RDH11** | 42.50398 | 9.34393 | 7.22038 | 29.14790 | 0.00000 | 14.03871 | 2.73102 |
| **RDH12** | 0.09502 | 0.00000 | 0.08191 | 0.03271 | 0.00464 | 0.02691 | 0.05561 |
| **RDH13** | 17.46096 | 14.40359 | 2.91733 | 8.35341 | 2.04529 | 4.39110 | 2.99743 |
| **RDH14** | 0.00000 | 0.43848 | 0.08065 | 0.25893 | 0.01062 | 0.48993 | 0.57322 |
| **RDH16** | 0.01245 | 0.00910 | 0.03244 | 0.12571 | 0.02102 | 0.11914 | 0.08327 |
| **RDH5** | 0.04052 | 0.05811 | 0.04297 | 0.07434 | 0.01019 | 0.00754 | 0.00000 |
| **RDM1** | 1.28891 | 0.68464 | 0.51628 | 0.00000 | 0.28080 | 0.03077 | 0.00000 |
| **RDX** | 26.48802 | 12.64266 | 10.87393 | 17.97029 | 5.74740 | 10.70529 | 21.77750 |
| **REC114** | 0.02664 | 0.31916 | 0.00000 | 0.00429 | 1.88229 | 0.09793 | 0.10172 |
| **REC8** | 0.29587 | 0.14253 | 0.17696 | 0.98081 | 0.00860 | 4.82302 | 1.21696 |
| **RECK** | 1.10507 | 0.53004 | 0.19388 | 0.26822 | 0.28420 | 0.21561 | 0.16375 |
| **RECQL** | 7.65914 | 3.76407 | 2.90608 | 5.98340 | 1.19167 | 2.58035 | 0.00000 |
| **RECQL4** | 3.77352 | 6.20966 | 1.20170 | 4.40830 | 0.73173 | 4.89415 | 22.78733 |
| **RECQL5** | 0.90072 | 4.12214 | 3.45147 | 3.25407 | 1.26460 | 0.73734 | 3.34815 |

| | | | | | | | |
|---|---|---|---|---|---|---|---|
| **REEP1** | 18.26018 | 5.43324 | 0.65293 | 23.36841 | 0.00000 | 1.67362 | 19.15162 |
| **REEP2** | 0.03073 | 0.16164 | 0.06278 | 0.03181 | 1.93891 | 0.00032 | 0.01073 |
| **REEP3** | 6.23630 | 3.38387 | 0.90434 | 5.47730 | 0.07730 | 2.88159 | 13.06117 |
| **REEP4** | 8.11172 | 10.79380 | 8.55803 | 11.15662 | 0.30048 | 4.48249 | 13.07632 |
| **REEP5** | 32.68779 | 19.52734 | 7.88564 | 24.06407 | 1.89588 | 20.34552 | 14.00258 |
| **REEP6** | 1.05911 | 4.53267 | 0.12105 | 2.21123 | 0.82966 | 5.13150 | 0.00000 |
| **REG1A** | 0.00000 | 0.10465 | 0.03380 | 0.02977 | 0.02646 | 1.19383 | 0.01195 |
| **REG1B** | 0.02842 | 0.02069 | 0.00826 | 0.02110 | 0.04494 | 0.01347 | 0.01351 |
| **REG1P** | 0.02283 | 0.01760 | 0.00857 | 0.01953 | 0.06159 | 0.01334 | 0.01779 |
| **REG4** | 0.02444 | 0.02248 | 0.01074 | 0.01823 | 0.00900 | 0.01772 | 0.02926 |
| **REL** | 1.32814 | 0.92119 | 1.75737 | 2.17715 | 0.91354 | 2.81267 | 6.30058 |
| **RELA** | 2.02938 | 5.02894 | 5.32551 | 7.90439 | 1.17463 | 3.68135 | 8.14325 |
| **RELB** | 0.02684 | 0.26715 | 0.09989 | 0.28491 | 2.38932 | 0.56627 | 0.12214 |
| **RELL1** | 34.11936 | 18.39042 | 14.40916 | 7.88092 | 1.07836 | 9.82119 | 5.41508 |
| **RELL2** | 0.00216 | 0.01525 | 0.04689 | 0.43990 | 0.28930 | 0.10728 | 0.08405 |
| **RELN** | 0.04286 | 0.03479 | 0.06875 | 0.01502 | 0.10604 | 0.00552 | 0.02150 |
| **RELT** | 0.00000 | 0.76051 | 0.00000 | 4.94816 | 0.12462 | 0.17269 | 4.05120 |
| **REM1** | 0.00000 | 0.46424 | 2.37096 | 0.00602 | 0.16986 | 0.92835 | 0.00000 |
| **REM2** | 0.07840 | 0.00000 | 0.09504 | 0.04010 | 0.03311 | 0.14337 | 0.00925 |
| **RENBP** | 0.00519 | 0.06628 | 0.67926 | 0.01991 | 0.00399 | 0.10421 | 0.07439 |
| **REP15** | 0.09828 | 0.00000 | 0.74229 | 0.07273 | 0.08316 | 0.06280 | 0.27570 |
| **REPIN1** | 1.00805 | 3.95468 | 0.00000 | 3.13205 | 0.00000 | 0.63374 | 9.77351 |
| **REPS1** | 4.32155 | 7.92917 | 7.19283 | 3.08181 | 0.29177 | 3.93497 | 15.20312 |
| **REPS2** | 0.00000 | 1.00291 | 0.87292 | 0.00000 | 0.47957 | 1.83193 | 0.10357 |
| **RER1** | 24.62809 | 28.99215 | 15.52092 | 20.29479 | 1.66479 | 11.54327 | 20.07178 |
| **RERE** | 0.03158 | 5.15269 | 3.01715 | 5.88564 | 2.92703 | 6.98786 | 20.65636 |
| **REREP3** | 0.00000 | 0.04436 | 0.05683 | 0.00470 | 0.24460 | 0.20499 | 0.12647 |
| **RERG** | 2.69500 | 0.00835 | 0.48343 | 1.97773 | 0.06126 | 0.57593 | 0.00000 |
| **RERG-AS1** | 0.01675 | 0.02565 | 0.00570 | 0.02565 | 0.16390 | 0.01221 | 0.02117 |
| **RERGL** | 0.11120 | 0.03202 | 0.02526 | 0.00000 | 0.37905 | 0.14211 | 0.03216 |
| **RESP18** | 0.02017 | 0.01778 | 0.00481 | 0.02832 | 0.21797 | 0.01295 | 0.01548 |
| **REST** | 13.29237 | 8.29888 | 20.07755 | 4.77524 | 6.35636 | 28.19580 | 32.74579 |
| **RET** | 0.00000 | 6.89653 | 1.24953 | 0.00000 | 0.04981 | 2.50229 | 0.31523 |
| **RETSAT** | 4.30525 | 1.14463 | 2.21424 | 3.29362 | 1.74217 | 0.49698 | 0.60696 |
| **REV1** | 2.55080 | 1.45600 | 1.55464 | 2.97099 | 2.72747 | 2.68143 | 10.81015 |

| | | | | | | | |
|---|---|---|---|---|---|---|---|
| **REV3L** | 11.32540 | 0.00000 | 5.19136 | 3.86829 | 4.10776 | 3.79494 | 15.53578 |
| **REXO1** | 0.00000 | 1.16847 | 0.72810 | 1.86657 | 0.48519 | 1.77621 | 15.86156 |
| **REXO2** | 18.67417 | 10.26477 | 4.46976 | 25.52321 | 5.71819 | 10.97328 | 6.84903 |
| **REXO4** | 5.54739 | 18.68649 | 4.95286 | 5.22293 | 1.31248 | 5.89524 | 12.75277 |
| **RFC1** | 10.17520 | 1.09003 | 10.05537 | 6.43377 | 6.44647 | 7.98016 | 12.77812 |
| **RFC2** | 26.16780 | 16.94336 | 9.88821 | 14.84024 | 5.63923 | 7.03592 | 0.00000 |
| **RFC3** | 29.64002 | 1.43804 | 19.21481 | 10.88890 | 7.89826 | 6.62510 | 6.18470 |
| **RFC4** | 19.68750 | 7.11212 | 14.10151 | 15.27670 | 3.77468 | 6.99554 | 0.00000 |
| **RFC5** | 26.51452 | 10.04975 | 5.88519 | 14.45064 | 2.68958 | 6.08016 | 0.00000 |
| **RFESD** | 0.84015 | 0.00000 | 0.76926 | 0.09382 | 0.22641 | 0.02273 | 0.02900 |
| **RFFL** | 0.48095 | 0.26555 | 0.07740 | 0.24977 | 0.52136 | 0.09455 | 0.00000 |
| **RFK** | 42.64231 | 6.08160 | 19.83887 | 10.28859 | 15.98341 | 2.90227 | 12.35036 |
| **RFNG** | 0.22359 | 4.08762 | 0.12836 | 2.39895 | 0.72886 | 2.29522 | 8.70364 |
| **RFPL1** | 0.01786 | 0.01079 | 0.00000 | 0.01662 | 2.95834 | 0.01041 | 0.01780 |
| **RFPL1S** | 0.13727 | 0.04473 | 0.00430 | 0.00000 | 0.81337 | 0.12879 | 0.08758 |
| **RFPL2** | 0.00660 | 0.00851 | 0.00000 | 0.00905 | 12.94245 | 0.02395 | 0.02982 |
| **RFPL3** | 0.02275 | 0.01627 | 0.00000 | 0.02029 | 0.72520 | 0.01294 | 0.01257 |
| **RFPL3S** | 0.33540 | 0.00000 | 0.08826 | 0.36647 | 0.03901 | 0.07380 | 0.43138 |
| **RFPL4A** | 0.01742 | 0.01373 | 0.00000 | 0.01367 | 6.46008 | 0.03156 | 0.01431 |
| **RFPL4AL1** | 0.01668 | 0.01317 | 0.00000 | 0.01607 | 3.91504 | 0.02533 | 0.01473 |
| **RFPL4B** | 5.01011 | 0.14319 | 8.08808 | 11.54893 | 11.19086 | 0.00000 | 0.00000 |
| **RFT1** | 7.60806 | 10.45483 | 4.39866 | 4.90378 | 1.75658 | 4.47052 | 0.00000 |
| **RFTN1** | 0.04440 | 0.22982 | 0.11384 | 0.14765 | 0.42946 | 0.16969 | 0.55970 |
| **RFTN2** | 0.12659 | 0.00000 | 1.51313 | 0.26921 | 0.29496 | 0.38482 | 0.10816 |
| **RFWD2** | 1.87378 | 1.73008 | 2.28139 | 2.64997 | 0.28027 | 5.28624 | 11.81946 |
| **RFWD3** | 19.67277 | 8.54539 | 13.56372 | 13.90780 | 8.70566 | 10.03827 | 1.99322 |
| **RFX1** | 0.00000 | 0.74569 | 0.92115 | 0.97969 | 0.61338 | 0.57971 | 6.11557 |
| **RFX2** | 0.48160 | 0.00000 | 0.65487 | 0.70642 | 0.25632 | 0.49687 | 0.70010 |
| **RFX3** | 0.35857 | 0.09409 | 0.17312 | 1.28549 | 0.44460 | 0.44599 | 1.64721 |
| **RFX4** | 0.03324 | 0.00000 | 0.17572 | 0.03899 | 0.14260 | 0.01398 | 0.00813 |
| **RFX5** | 1.31358 | 0.00000 | 0.26833 | 1.34985 | 0.52686 | 2.31542 | 0.30501 |
| **RFX6** | 0.05123 | 0.00000 | 0.06898 | 0.02198 | 0.04315 | 0.00663 | 0.04412 |
| **RFX7** | 1.66554 | 0.00000 | 1.04588 | 0.84767 | 0.57787 | 2.08888 | 2.97212 |
| **RFXANK** | 6.33199 | 3.37614 | 1.07049 | 9.03621 | 0.70963 | 3.04741 | 3.16906 |
| **RFXAP** | 0.49046 | 0.06706 | 0.02799 | 0.22772 | 0.00178 | 0.27245 | 2.37822 |

| | | | | | | | |
|---|---|---|---|---|---|---|---|
| **RGAG1** | 0.01072 | 0.02885 | 0.00547 | 0.09947 | 0.04421 | 0.13894 | 0.01903 |
| **RGAG4** | 0.00000 | 0.30773 | 0.19845 | 0.01412 | 0.04193 | 0.00356 | 0.02309 |
| **RGCC** | 1.04364 | 7.50727 | 0.12459 | 0.00000 | 0.02450 | 8.81669 | 9.44684 |
| **RGL1** | 0.95631 | 0.00000 | 0.00000 | 4.11594 | 0.42523 | 0.00000 | 1.19439 |
| **RGL2** | 3.32888 | 0.06269 | 0.82460 | 16.93628 | 1.16860 | 0.96891 | 8.61125 |
| **RGL3** | 0.00000 | 0.00000 | 1.16258 | 1.01465 | 0.18432 | 0.37072 | 0.48853 |
| **RGL4** | 0.06672 | 0.10146 | 0.11057 | 0.50956 | 0.05736 | 0.11309 | 0.90890 |
| **RGMA** | 0.00000 | 8.85998 | 0.17545 | 0.53190 | 1.03415 | 0.00000 | 4.79723 |
| **RGMB** | 1.09854 | 0.73621 | 1.36254 | 1.78375 | 1.52663 | 2.22008 | 5.27376 |
| **RGMB-AS1** | 0.03735 | 0.05103 | 0.04588 | 0.04067 | 0.22270 | 0.14179 | 0.05758 |
| **RGN** | 0.12330 | 0.08140 | 0.01450 | 0.12238 | 0.26286 | 0.00541 | 0.13704 |
| **RGP1** | 0.53433 | 1.03794 | 0.54254 | 1.51738 | 0.35914 | 0.54719 | 0.20664 |
| **RGPD1.uid-2613** | 0.04476 | 0.01165 | 0.01391 | 0.02451 | 0.01065 | 0.01989 | 0.03282 |
| **RGPD1.uid-2866** | 0.04855 | 0.01105 | 0.01889 | 0.02133 | 0.00720 | 0.01669 | 0.03522 |
| **RGPD2.uid-2611** | 0.02083 | 0.07622 | 0.01476 | 0.04475 | 0.03508 | 0.02035 | 0.06276 |
| **RGPD2.uid-2863** | 0.02315 | 0.02675 | 0.01264 | 0.02346 | 0.00808 | 0.01706 | 0.02402 |
| **RGPD3** | 0.03185 | 0.01830 | 0.03759 | 0.02403 | 0.13901 | 0.02934 | 0.02507 |
| **RGPD4** | 0.05569 | 0.00862 | 0.03872 | 0.05970 | 0.12970 | 0.02966 | 0.07012 |
| **RGPD8** | 0.02100 | 0.02811 | 0.01089 | 0.02143 | 0.00794 | 0.01497 | 0.02212 |
| **RGS10** | 5.62464 | 10.62519 | 7.38258 | 0.00000 | 0.65298 | 8.39824 | 2.47361 |
| **RGS11** | 0.00000 | 0.08155 | 0.10679 | 0.01341 | 0.07927 | 0.09710 | 0.04989 |
| **RGS12** | 0.00000 | 0.32801 | 0.19776 | 0.83429 | 0.01117 | 0.91039 | 0.83781 |
| **RGS13** | 21.07472 | 0.00108 | 0.00759 | 0.00000 | 0.00185 | 0.08558 | 0.04945 |
| **RGS14** | 0.84502 | 0.17385 | 0.97341 | 0.61921 | 2.12562 | 1.76074 | 0.00000 |
| **RGS16** | 8.02791 | 12.44634 | 0.00000 | 34.69200 | 2.49130 | 9.64401 | 2.20664 |
| **RGS17** | 0.08407 | 0.04382 | 0.95576 | 0.10817 | 0.33163 | 0.18183 | 0.00010 |
| **RGS18** | 0.02371 | 0.01659 | 0.01026 | 0.01952 | 0.06127 | 0.01323 | 0.01633 |
| **RGS19** | 1.71684 | 1.38944 | 0.01865 | 2.23934 | 0.07056 | 1.87782 | 1.73141 |
| **RGS2** | 5.42542 | 5.00184 | 27.74995 | 1.61704 | 19.22284 | 5.94650 | 0.00000 |
| **RGS20** | 0.04370 | 0.06285 | 0.34315 | 0.03763 | 0.24064 | 0.29323 | 0.02331 |
| **RGS22** | 0.03498 | 0.01006 | 0.03936 | 0.02068 | 0.14540 | 0.02405 | 0.01993 |
| **RGS3** | 1.92003 | 0.00000 | 0.16561 | 7.25200 | 0.03015 | 0.00000 | 0.41344 |
| **RGS4** | 0.02178 | 0.02647 | 0.01008 | 0.02283 | 0.00832 | 0.04780 | 0.01086 |
| **RGS5** | 1.18685 | 0.37652 | 1.20549 | 0.14498 | 0.05539 | 1.81023 | 0.15333 |
| **RGS6** | 0.05416 | 0.01727 | 0.01413 | 0.02179 | 0.14774 | 0.00672 | 0.37347 |

| | | | | | | | |
|---|---|---|---|---|---|---|---|
| **RGS7** | 0.12686 | 0.02210 | 0.11134 | 0.09598 | 0.46290 | 0.01025 | 0.00000 |
| **RGS7BP** | 0.02186 | 0.02651 | 0.01059 | 0.04856 | 0.00840 | 0.06113 | 0.00135 |
| **RGS8** | 0.04106 | 0.00000 | 0.04065 | 0.02836 | 0.10012 | 0.02392 | 0.00263 |
| **RGS9** | 0.00000 | 0.46003 | 0.00868 | 0.05320 | 0.01299 | 0.11075 | 0.03593 |
| **RGS9BP** | 0.02715 | 0.01212 | 0.01185 | 0.00975 | 0.09180 | 0.00824 | 0.06965 |
| **RGSL1** | 0.02719 | 0.01010 | 0.09796 | 0.01739 | 0.13569 | 0.02698 | 0.01386 |
| **RHBDD1** | 2.80496 | 0.00000 | 3.02296 | 1.21502 | 3.03015 | 0.61156 | 1.68836 |
| **RHBDD2** | 7.57591 | 9.36010 | 2.78582 | 8.44682 | 1.72383 | 6.50553 | 8.00319 |
| **RHBDD3** | 7.00247 | 8.40283 | 1.17225 | 19.20492 | 0.23021 | 4.37707 | 6.76925 |
| **RHBDF1** | 0.02902 | 1.23456 | 0.13598 | 1.35573 | 0.15606 | 0.79355 | 6.01394 |
| **RHBDF2** | 0.03587 | 1.23450 | 1.54457 | 2.04620 | 0.93396 | 0.37368 | 1.77363 |
| **RHBDL1** | 0.00214 | 0.02842 | 0.03339 | 0.23445 | 0.03017 | 0.13315 | 1.07516 |
| **RHBDL2** | 0.00000 | 0.95067 | 0.29407 | 0.15729 | 0.11170 | 0.00000 | 0.03420 |
| **RHBDL3** | 0.02897 | 0.02219 | 0.02750 | 0.01618 | 0.08554 | 0.01260 | 0.04878 |
| **RHBG** | 0.12595 | 0.00000 | 0.18030 | 0.14057 | 0.45652 | 0.06912 | 0.19419 |
| **RHCE** | 0.01085 | 0.02089 | 0.00945 | 0.04590 | 0.06119 | 0.02532 | 0.02085 |
| **RHCG** | 0.01863 | 0.01884 | 0.01213 | 0.01852 | 0.01167 | 0.01568 | 0.02850 |
| **RHD** | 0.06141 | 0.02080 | 0.04231 | 0.05522 | 0.02487 | 0.02378 | 0.08107 |
| **RHEB** | 13.16123 | 8.22979 | 5.07144 | 13.35299 | 1.67133 | 6.25288 | 4.66013 |
| **RHEBL1** | 0.37134 | 0.00000 | 6.06620 | 1.85710 | 10.82732 | 2.10021 | 1.24700 |
| **RHNO1** | 8.10013 | 3.22521 | 9.00586 | 4.56540 | 0.10640 | 2.03369 | 1.26289 |
| **RHO** | 0.00000 | 0.02446 | 0.04000 | 0.03445 | 0.00625 | 0.03687 | 0.06557 |
| **RHOA** | 55.28907 | 24.45140 | 32.38662 | 40.01487 | 18.81785 | 25.84348 | 30.70048 |
| **RHOB** | 0.74202 | 5.01172 | 9.12944 | 2.51573 | 0.37146 | 1.52568 | 11.03940 |
| **RHOBTB1** | 3.49876 | 0.00000 | 0.00000 | 19.90392 | 0.51067 | 0.00000 | 0.43797 |
| **RHOBTB2** | 3.90863 | 0.10618 | 0.32268 | 5.37782 | 0.36564 | 0.00000 | 5.57965 |
| **RHOBTB3** | 1.47479 | 0.00000 | 0.06336 | 0.57200 | 1.59330 | 3.01319 | 3.38198 |
| **RHOC** | 43.15413 | 59.77128 | 0.61146 | 37.38037 | 0.74809 | 21.23501 | 18.66053 |
| **RHOD** | 0.00000 | 0.00000 | 0.27513 | 1.87884 | 0.24665 | 0.00000 | 0.96649 |
| **RHOF** | 0.32752 | 0.51838 | 0.02969 | 0.15583 | 0.03023 | 0.26879 | 0.06095 |
| **RHOG** | 7.95637 | 10.26930 | 4.65820 | 15.14430 | 0.32024 | 10.82580 | 16.82231 |
| **RHOH** | 0.16908 | 0.19219 | 0.01243 | 0.03272 | 0.01142 | 0.41514 | 0.00000 |
| **RHOJ** | 0.02152 | 0.02576 | 0.01984 | 0.02099 | 0.00709 | 0.01202 | 0.01569 |
| **RHOQ** | 0.52553 | 0.00000 | 0.42064 | 0.18860 | 0.07391 | 0.69402 | 0.33275 |
| **RHOT1** | 1.59237 | 0.00000 | 0.75912 | 0.99242 | 0.82827 | 2.32647 | 5.96296 |

| | | | | | | | |
|---|---|---|---|---|---|---|---|
| **RHOT2** | 15.32761 | 23.40859 | 2.99062 | 15.54123 | 0.82831 | 12.44360 | 18.03667 |
| **RHOU** | 10.77750 | 0.25497 | 0.00000 | 11.03718 | 0.00000 | 0.00000 | 25.23709 |
| **RHOV** | 1.62932 | 11.77971 | 0.00000 | 2.40998 | 0.00000 | 0.30967 | 4.01676 |
| **RHOXF1** | 0.00000 | 0.69236 | 0.81793 | 0.00000 | 0.12743 | 0.00000 | 0.11994 |
| **RHPN1** | 0.01142 | 0.04746 | 0.10590 | 0.08177 | 0.00999 | 0.02121 | 0.29000 |
| **RHPN1-AS1** | 0.02887 | 0.02605 | 0.00635 | 0.15330 | 0.12602 | 0.00697 | 0.03201 |
| **RHPN2** | 9.56661 | 9.86865 | 7.63113 | 3.20946 | 0.85715 | 1.40877 | 11.43749 |
| **RIBC1** | 0.00000 | 0.45113 | 0.35541 | 0.41858 | 0.42911 | 0.42200 | 0.43382 |
| **RIBC2** | 1.04620 | 0.92110 | 1.43474 | 0.61503 | 1.50446 | 0.00000 | 0.00000 |
| **RIC3** | 0.48807 | 0.70631 | 0.88000 | 0.41995 | 0.07642 | 0.10531 | 0.65121 |
| **RIC8A** | 12.60114 | 9.34886 | 5.35748 | 15.38795 | 8.50606 | 7.97159 | 3.85679 |
| **RIC8B** | 2.55619 | 0.00000 | 2.91089 | 2.25824 | 0.55866 | 0.86825 | 0.45658 |
| **RICTOR** | 5.19524 | 0.00000 | 0.72391 | 2.79945 | 2.05011 | 1.43794 | 4.14731 |
| **RIF1** | 22.86796 | 2.37770 | 19.19164 | 7.35606 | 11.41834 | 16.67027 | 33.74881 |
| **RIIAD1** | 0.01908 | 0.03090 | 0.04717 | 0.01460 | 2.10621 | 0.00602 | 0.01111 |
| **RILP** | 0.06581 | 2.48649 | 0.26632 | 0.00000 | 0.08332 | 0.49851 | 2.90352 |
| **RILPL1** | 0.00000 | 0.30802 | 0.01983 | 3.08681 | 0.04026 | 0.52637 | 1.15968 |
| **RILPL2** | 0.00000 | 0.03370 | 0.12577 | 6.55807 | 0.04908 | 0.30834 | 0.68292 |
| **RIMBP2** | 0.02876 | 0.00000 | 0.03850 | 0.01930 | 0.53373 | 0.01422 | 0.03263 |
| **RIMBP3** | 0.01068 | 0.06286 | 0.02469 | 0.03316 | 0.01599 | 0.01134 | 0.01647 |
| **RIMKLA** | 0.04006 | 0.08753 | 0.15010 | 0.11732 | 2.72816 | 0.00000 | 0.36176 |
| **RIMKLB** | 7.31075 | 10.21696 | 14.64709 | 0.00000 | 1.37476 | 8.18822 | 4.70367 |
| **RIMS1** | 0.00000 | 0.00000 | 0.26624 | 0.00166 | 0.41273 | 0.04800 | 0.44319 |
| **RIMS2** | 0.22013 | 0.46364 | 0.00000 | 0.30670 | 0.50923 | 0.24393 | 0.31306 |
| **RIMS3** | 0.11126 | 0.31600 | 0.04651 | 0.10672 | 0.31042 | 0.30726 | 0.09111 |
| **RIMS4** | 0.00000 | 0.34409 | 1.77854 | 0.00000 | 0.03703 | 0.00000 | 0.06808 |
| **RIN1** | 0.06790 | 0.00000 | 0.06418 | 0.06300 | 0.01283 | 0.04198 | 0.31633 |
| **RIN2** | 3.62959 | 6.13261 | 1.79111 | 0.58507 | 0.07621 | 3.92844 | 0.77739 |
| **RIN3** | 0.36175 | 0.00000 | 0.00000 | 4.45161 | 0.11250 | 0.00000 | 6.26872 |
| **RING1** | 0.00000 | 0.72544 | 0.63328 | 0.77889 | 0.44901 | 0.53327 | 4.70260 |
| **RINL** | 0.26728 | 0.06123 | 0.27045 | 1.20629 | 0.00112 | 1.50062 | 0.79397 |
| **RINT1** | 15.37052 | 1.41672 | 6.73555 | 3.25060 | 3.84442 | 3.98882 | 0.00000 |
| **RIOK1** | 22.23674 | 8.97388 | 14.51194 | 16.03307 | 5.91491 | 4.17426 | 6.38725 |
| **RIOK2** | 16.03822 | 3.28720 | 15.03891 | 3.05651 | 3.00579 | 6.85934 | 2.70924 |
| **RIOK3** | 17.36511 | 10.04475 | 9.96605 | 10.76299 | 8.98519 | 6.46610 | 11.79685 |

| | | | | | | | |
|---|---|---|---|---|---|---|---|
| **RIPK1** | 1.06817 | 0.84442 | 1.62229 | 1.20328 | 0.87458 | 0.68389 | 3.43786 |
| **RIPK2** | 9.37249 | 5.76574 | 2.92012 | 7.57030 | 0.90615 | 1.95739 | 0.51754 |
| **RIPK4** | 1.85337 | 4.22777 | 1.28729 | 9.14511 | 0.67966 | 5.81673 | 2.82303 |
| **RIPPLY2** | 0.02455 | 0.01886 | 0.00593 | 0.01991 | 0.14975 | 0.01788 | 0.01182 |
| **RIPPLY3** | 0.01729 | 0.01948 | 0.01491 | 0.05670 | 0.00835 | 0.01486 | 0.02588 |
| **RIT1** | 4.07901 | 1.67636 | 1.82153 | 5.65608 | 6.22466 | 2.89740 | 0.00000 |
| **RIT2** | 0.93052 | 0.00000 | 0.04265 | 0.23142 | 0.23739 | 0.00000 | 0.55500 |
| **RITA1** | 6.60230 | 8.89154 | 3.72984 | 11.41185 | 1.64952 | 4.72867 | 5.85571 |
| **RLBP1** | 0.00871 | 0.02512 | 0.01169 | 0.05684 | 0.00771 | 0.00987 | 0.04090 |
| **RLF** | 12.97890 | 0.60071 | 8.47621 | 8.70424 | 10.85737 | 3.89473 | 7.92407 |
| **RLIM** | 7.83813 | 3.82468 | 12.71796 | 1.89114 | 14.26888 | 9.21926 | 23.28134 |
| **RLN1** | 0.04137 | 0.05073 | 0.00547 | 0.03990 | 0.06623 | 0.00838 | 0.02050 |
| **RLN2** | 0.04233 | 0.01171 | 0.01647 | 0.08447 | 0.00762 | 0.04817 | 0.00507 |
| **RLN3** | 0.03348 | 0.17240 | 0.08789 | 0.02066 | 0.00633 | 0.01851 | 0.02010 |
| **RLTPR** | 0.01802 | 0.09986 | 0.00742 | 0.28104 | 0.05407 | 0.13275 | 0.10630 |
| **RMDN1** | 13.98672 | 9.89443 | 4.47784 | 3.10498 | 0.16889 | 4.99063 | 1.55536 |
| **RMDN2** | 0.38956 | 0.00000 | 1.43987 | 0.04092 | 0.07118 | 0.78598 | 0.04344 |
| **RMDN2-AS1** | 0.02165 | 0.04365 | 0.06795 | 0.02389 | 0.00502 | 0.01197 | 0.00913 |
| **RMDN3** | 18.69017 | 9.60146 | 6.79196 | 11.74701 | 1.83074 | 4.76942 | 1.26821 |
| **RMI1** | 9.80366 | 0.00000 | 5.28224 | 1.92403 | 5.75320 | 3.12449 | 1.14068 |
| **RMI2** | 0.02309 | 1.80941 | 1.18265 | 0.46768 | 0.08063 | 2.11447 | 3.27768 |
| **RMND1** | 21.17250 | 6.64759 | 22.48354 | 11.55643 | 5.23014 | 11.27621 | 0.00000 |
| **RMND5A** | 27.02367 | 8.53809 | 6.80980 | 17.08490 | 11.86102 | 10.96502 | 3.45981 |
| **RMND5B** | 3.26197 | 5.61543 | 0.92854 | 4.24502 | 0.74597 | 3.91724 | 2.59309 |
| **RMRP** | 0.00000 | 0.99931 | 0.04972 | 0.00000 | 0.42894 | 2.29449 | 2.51485 |
| **RMST** | 0.03096 | 0.04100 | 0.01091 | 0.01838 | 0.02567 | 0.01094 | 0.00658 |
| **RNASE1** | 12.58989 | 0.00000 | 0.00000 | 0.00000 | 0.11854 | 5.35305 | 2.41409 |
| **RNASE10** | 0.00000 | 0.18992 | 1.31326 | 0.00303 | 0.08573 | 0.88590 | 0.05475 |
| **RNASE11** | 0.02238 | 0.01026 | 0.03183 | 0.02268 | 0.03508 | 0.01261 | 0.01895 |
| **RNASE4** | 0.10082 | 0.00000 | 0.30500 | 0.07640 | 0.21053 | 0.20287 | 0.00000 |
| **RNASE7** | 0.02478 | 0.01671 | 0.01446 | 0.05046 | 0.00819 | 0.01270 | 0.00781 |
| **RNASEH1** | 16.59559 | 15.41836 | 8.18714 | 8.90118 | 2.06844 | 6.38732 | 7.52468 |
| **RNASEH1-AS1** | 2.86926 | 3.08748 | 3.04445 | 2.39263 | 1.83356 | 1.83771 | 1.02946 |
| **RNASEH2A** | 19.40799 | 16.74699 | 8.50716 | 17.43482 | 1.24554 | 11.70791 | 4.34841 |
| **RNASEH2B** | 3.74055 | 1.08824 | 2.91314 | 1.67822 | 0.00000 | 1.29355 | 6.62073 |

| | | | | | | | |
|---|---|---|---|---|---|---|---|
| **RNASEH2B-AS1** | 0.02459 | 0.05546 | 0.00826 | 0.04207 | 0.01659 | 0.00602 | 0.05129 |
| **RNASEH2C** | 12.51029 | 21.61646 | 9.10832 | 12.95394 | 7.00515 | 10.24746 | 8.66062 |
| **RNASEK-C17or** | 0.06917 | 0.10017 | 0.05817 | 0.06439 | 0.01067 | 0.03791 | 0.00000 |
| **RNASEK.uid-21** | 1.10546 | 0.00000 | 0.11492 | 1.48377 | 0.01833 | 0.06119 | 0.53149 |
| **RNASEL** | 0.77053 | 0.00000 | 0.30609 | 1.62610 | 0.02538 | 0.73296 | 0.00000 |
| **RNASET2** | 0.20733 | 4.39073 | 0.65726 | 0.73326 | 3.04545 | 9.24079 | 3.47619 |
| **RND1** | 0.06527 | 0.00676 | 0.92902 | 0.13091 | 7.78815 | 0.38263 | 0.00260 |
| **RND2** | 2.46887 | 7.23186 | 1.43165 | 0.00000 | 0.16605 | 8.68257 | 1.36601 |
| **RND3** | 0.06688 | 0.00000 | 0.00861 | 0.44883 | 1.67559 | 0.10131 | 0.00386 |
| **RNF10** | 9.97973 | 14.68887 | 7.17105 | 16.45920 | 6.70922 | 9.17126 | 23.59353 |
| **RNF103** | 0.97733 | 0.00000 | 1.11465 | 3.95469 | 0.55090 | 1.02695 | 2.78603 |
| **RNF103-CHMP3** | 0.02029 | 0.02565 | 0.01484 | 0.25447 | 0.01730 | 0.03209 | 0.00931 |
| **RNF11** | 2.95865 | 0.00000 | 22.35670 | 4.68475 | 7.42492 | 4.20272 | 9.81627 |
| **RNF111** | 8.26399 | 2.01122 | 6.35946 | 10.54805 | 4.72773 | 3.20451 | 8.91907 |
| **RNF112** | 0.00000 | 0.05226 | 0.01153 | 0.01533 | 0.00655 | 0.11371 | 0.22454 |
| **RNF113A** | 14.09498 | 12.80356 | 14.76727 | 6.18441 | 12.91216 | 7.64785 | 2.91321 |
| **RNF113B** | 0.00000 | 0.00000 | 0.56981 | 0.03954 | 0.26738 | 0.03559 | 0.49913 |
| **RNF114** | 29.24927 | 26.85101 | 20.59561 | 20.73111 | 10.39869 | 16.72002 | 4.22709 |
| **RNF115** | 4.23543 | 5.84873 | 10.84474 | 0.00000 | 2.46398 | 6.33931 | 10.44050 |
| **RNF121** | 7.66811 | 5.28808 | 4.99676 | 5.86905 | 3.25981 | 4.16240 | 1.14071 |
| **RNF122** | 0.00000 | 0.06643 | 0.58713 | 0.09092 | 6.93040 | 0.55568 | 0.28091 |
| **RNF123** | 1.81161 | 3.60675 | 2.14806 | 6.13625 | 0.26638 | 1.67958 | 7.75497 |
| **RNF125** | 7.36181 | 6.39468 | 19.39615 | 1.35277 | 9.35769 | 14.58617 | 3.32397 |
| **RNF126** | 0.00000 | 14.23362 | 6.48113 | 5.43234 | 1.24884 | 5.27136 | 27.24106 |
| **RNF126P1** | 0.00000 | 0.99103 | 0.35162 | 0.24115 | 0.04395 | 0.37542 | 1.05279 |
| **RNF128** | 0.02209 | 0.01415 | 0.14065 | 0.02278 | 0.05319 | 0.00837 | 0.01698 |
| **RNF13** | 11.25988 | 0.00000 | 8.42953 | 5.07291 | 1.31675 | 1.57713 | 0.80986 |
| **RNF130** | 0.27986 | 0.49069 | 0.24776 | 0.48495 | 1.21000 | 1.10413 | 0.45157 |
| **RNF135** | 0.73977 | 6.99215 | 3.15495 | 4.45197 | 0.56554 | 5.17929 | 5.65974 |
| **RNF138** | 21.83955 | 0.00000 | 15.44933 | 3.50700 | 10.25241 | 8.99090 | 3.69657 |
| **RNF138P1** | 1.96919 | 0.00000 | 2.90215 | 0.35612 | 1.80626 | 2.48427 | 0.00000 |
| **RNF139** | 5.43663 | 0.38611 | 4.02371 | 1.29249 | 2.19691 | 4.77617 | 11.09219 |
| **RNF139-AS1** | 0.30305 | 0.17142 | 0.24882 | 0.17521 | 0.18780 | 0.19507 | 0.00005 |
| **RNF14** | 15.64995 | 5.62298 | 4.23635 | 3.03170 | 3.06265 | 7.81999 | 0.40986 |
| **RNF141** | 4.68655 | 5.80200 | 3.09359 | 0.45609 | 0.06517 | 3.04643 | 0.87028 |

| | | | | | | | |
|---|---|---|---|---|---|---|---|
| **RNF144A** | 0.00000 | 0.07389 | 0.14942 | 0.20894 | 0.06924 | 0.00796 | 0.32473 |
| **RNF144A-AS1** | 0.01736 | 0.10624 | 0.01202 | 0.02004 | 0.00811 | 0.00823 | 0.00938 |
| **RNF144B** | 0.43552 | 0.04740 | 0.45394 | 0.13115 | 1.33512 | 0.05146 | 0.18233 |
| **RNF145** | 7.03235 | 0.95323 | 7.96721 | 2.96563 | 2.34341 | 3.42325 | 8.92421 |
| **RNF146** | 2.58515 | 1.91314 | 1.19036 | 1.86837 | 0.07631 | 0.90683 | 0.00000 |
| **RNF148** | 0.03072 | 0.01094 | 0.00753 | 0.00852 | 0.05598 | 0.08245 | 0.04224 |
| **RNF149** | 2.66390 | 0.57961 | 2.11377 | 4.21864 | 0.53406 | 1.08609 | 9.92327 |
| **RNF150** | 0.03330 | 0.00087 | 0.01157 | 0.07946 | 0.32983 | 0.42914 | 0.00000 |
| **RNF151** | 0.07099 | 0.13799 | 0.02568 | 0.04203 | 0.09069 | 0.00823 | 0.00000 |
| **RNF152** | 0.01016 | 0.05928 | 0.01163 | 0.02593 | 0.13545 | 0.19588 | 0.01438 |
| **RNF157** | 0.00000 | 0.01619 | 0.00320 | 0.30007 | 0.25699 | 0.34414 | 0.63289 |
| **RNF157-AS1** | 0.02252 | 0.01526 | 0.02580 | 0.03036 | 0.01981 | 0.02139 | 0.01743 |
| **RNF165** | 0.14621 | 0.26294 | 0.21624 | 0.11488 | 0.19126 | 0.03156 | 0.03908 |
| **RNF166** | 0.00000 | 3.76931 | 0.10476 | 0.57572 | 0.01568 | 0.69936 | 3.74346 |
| **RNF167** | 14.14393 | 13.43019 | 15.50350 | 20.04957 | 3.48637 | 13.23117 | 16.57178 |
| **RNF168** | 14.06270 | 2.12585 | 11.01515 | 12.95511 | 22.51123 | 9.16173 | 4.45990 |
| **RNF169** | 1.19043 | 0.28976 | 0.78084 | 0.55865 | 0.39145 | 0.50569 | 1.61156 |
| **RNF17** | 3.68997 | 0.00000 | 3.99766 | 0.03330 | 2.32436 | 1.69049 | 2.58183 |
| **RNF170** | 4.76603 | 2.39362 | 4.33336 | 1.01409 | 0.60925 | 3.62197 | 0.00263 |
| **RNF180** | 0.14292 | 0.01438 | 0.28908 | 0.00000 | 0.47931 | 0.04008 | 0.08656 |
| **RNF181** | 18.97491 | 26.89810 | 7.60412 | 27.80194 | 8.73374 | 13.07902 | 5.35079 |
| **RNF182** | 0.67308 | 0.00000 | 0.07806 | 0.05003 | 0.12548 | 0.00000 | 0.10682 |
| **RNF183** | 0.22278 | 0.74678 | 0.15613 | 0.00000 | 0.30008 | 0.17281 | 0.26073 |
| **RNF185** | 8.48334 | 3.70544 | 3.49957 | 7.81501 | 1.55672 | 3.05161 | 0.00000 |
| **RNF187** | 1.68855 | 8.08289 | 1.24132 | 1.79360 | 1.33258 | 5.88950 | 25.06709 |
| **RNF19A** | 4.46264 | 0.00000 | 2.73366 | 2.82164 | 2.45183 | 4.60807 | 3.42380 |
| **RNF19B** | 0.00000 | 8.26604 | 11.44615 | 7.18942 | 5.54412 | 8.47560 | 9.39368 |
| **RNF2** | 20.81549 | 2.16338 | 11.22731 | 9.15953 | 7.25364 | 9.13949 | 2.34582 |
| **RNF20** | 7.51785 | 5.00997 | 9.43441 | 6.24798 | 6.04961 | 7.21516 | 6.49077 |
| **RNF207** | 1.21231 | 0.52943 | 0.02189 | 0.93549 | 0.10461 | 0.00000 | 1.44314 |
| **RNF208** | 0.02359 | 0.13758 | 0.00824 | 0.03970 | 0.00922 | 0.07284 | 0.30608 |
| **RNF212** | 4.50968 | 3.12248 | 3.21794 | 0.15413 | 1.61947 | 0.38670 | 0.12597 |
| **RNF212B** | 0.38390 | 0.27079 | 0.27397 | 0.47532 | 0.21614 | 0.05139 | 0.23124 |
| **RNF213** | 0.00000 | 2.79384 | 2.13680 | 0.95657 | 1.56879 | 2.22451 | 1.27072 |
| **RNF214** | 1.31897 | 2.42388 | 4.41718 | 0.84623 | 0.49327 | 8.26876 | 8.44199 |

| | | | | | | | |
|---|---|---|---|---|---|---|---|
| **RNF215** | 0.04027 | 0.08603 | 0.04839 | 0.03567 | 0.00677 | 0.21654 | 0.43779 |
| **RNF216** | 9.94106 | 12.13153 | 7.55820 | 6.29041 | 3.56731 | 7.89280 | 5.69258 |
| **RNF216-IT1** | 0.03158 | 0.05368 | 0.01660 | 0.01992 | 0.00709 | 0.01134 | 0.02098 |
| **RNF216P1** | 8.79780 | 9.01483 | 10.30674 | 9.84598 | 6.42142 | 4.87765 | 2.14789 |
| **RNF217** | 0.03526 | 0.00000 | 0.01469 | 0.71496 | 0.01178 | 0.10396 | 0.07070 |
| **RNF219** | 6.82369 | 0.00000 | 10.90147 | 2.31482 | 3.69942 | 4.39441 | 5.39374 |
| **RNF219-AS1** | 0.02605 | 0.01122 | 0.02822 | 0.01940 | 0.01485 | 0.05127 | 0.04261 |
| **RNF220** | 1.51175 | 3.83661 | 4.92882 | 4.43276 | 0.53515 | 4.91265 | 9.34034 |
| **RNF222** | 0.00000 | 0.01278 | 0.03838 | 0.11416 | 0.00768 | 0.00947 | 0.12643 |
| **RNF223** | 0.02453 | 0.02114 | 0.01234 | 0.04594 | 0.00807 | 0.01250 | 0.00780 |
| **RNF24** | 1.84855 | 3.38116 | 1.03750 | 1.21165 | 2.10245 | 11.34623 | 3.10880 |
| **RNF25** | 4.16605 | 8.53190 | 6.17593 | 8.43941 | 2.72284 | 4.02534 | 0.00000 |
| **RNF26** | 9.72381 | 20.14387 | 10.76465 | 16.04158 | 0.00000 | 6.69082 | 8.18089 |
| **RNF31** | 1.30102 | 7.05298 | 3.22311 | 1.31553 | 0.86745 | 5.41607 | 7.82303 |
| **RNF32** | 0.02810 | 0.06998 | 0.50932 | 0.17819 | 0.07589 | 0.16134 | 0.15096 |
| **RNF34** | 28.08177 | 15.11938 | 13.29118 | 15.80448 | 16.28441 | 8.56918 | 0.00000 |
| **RNF38** | 14.29953 | 4.09108 | 7.67177 | 11.57120 | 6.61472 | 7.94081 | 9.87612 |
| **RNF39** | 0.34200 | 2.70891 | 0.30556 | 0.66672 | 0.21963 | 1.02383 | 2.08802 |
| **RNF4** | 25.23734 | 16.55409 | 12.30058 | 15.26498 | 12.74154 | 11.65279 | 8.72248 |
| **RNF40** | 7.00193 | 10.87302 | 3.87673 | 13.95012 | 1.11293 | 7.05176 | 13.00873 |
| **RNF41** | 9.48988 | 1.55537 | 3.01587 | 9.44447 | 3.05187 | 2.79017 | 6.58625 |
| **RNF43** | 2.22005 | 0.20639 | 0.00000 | 6.28917 | 0.13206 | 0.00000 | 1.50225 |
| **RNF44** | 1.02662 | 7.94989 | 7.43000 | 7.38522 | 0.35486 | 14.78469 | 19.90679 |
| **RNF5** | 0.06183 | 0.05000 | 0.06006 | 0.07059 | 0.00658 | 0.06246 | 0.01705 |
| **RNF5P1.uid-107** | 0.78236 | 0.55860 | 1.22586 | 1.35724 | 0.00000 | 0.93907 | 0.10012 |
| **RNF6** | 14.65817 | 2.77361 | 9.21838 | 5.32749 | 2.18053 | 3.11877 | 0.47920 |
| **RNF7** | 18.20433 | 7.07978 | 13.07552 | 17.60721 | 4.02414 | 9.67717 | 10.09536 |
| **RNF8** | 6.91828 | 2.90617 | 5.64424 | 6.98951 | 3.78326 | 7.85945 | 1.58468 |
| **RNFT1** | 3.23447 | 0.00000 | 4.20577 | 1.82004 | 1.38599 | 0.19014 | 0.00000 |
| **RNFT2** | 1.98844 | 3.20377 | 0.54209 | 0.00000 | 0.06960 | 0.44939 | 1.33502 |
| **RNGTT** | 13.34809 | 1.31451 | 9.52947 | 5.97934 | 3.72707 | 2.66825 | 0.92829 |
| **RNH1** | 22.46942 | 27.67827 | 11.65433 | 19.92078 | 1.05488 | 9.01303 | 7.02693 |
| **RNLS** | 1.82855 | 0.26224 | 0.73120 | 0.64326 | 0.04828 | 0.72841 | 0.00000 |
| **RNMT** | 13.19398 | 0.69973 | 9.61213 | 7.08361 | 4.56405 | 4.70521 | 13.52322 |
| **RNMTL1** | 24.60174 | 19.43044 | 13.48915 | 18.83980 | 4.62722 | 7.05465 | 0.00000 |

| | | | | | | | |
|---|---|---|---|---|---|---|---|
| **RNPC3** | 2.61968 | 0.43278 | 1.38795 | 1.58941 | 1.94039 | 0.82142 | 3.64695 |
| **RNPEP** | 5.02819 | 5.08107 | 0.97898 | 1.10429 | 0.00000 | 4.19154 | 11.73568 |
| **RNPEPL1** | 1.27090 | 1.49683 | 0.05592 | 0.84261 | 0.00000 | 1.02881 | 7.82367 |
| **RNPS1** | 35.66885 | 28.66999 | 21.13214 | 27.66160 | 9.30156 | 18.73770 | 9.73364 |
| **RNU11** | 0.00000 | 0.00652 | 0.02394 | 0.04690 | 0.02688 | 0.07824 | 0.25045 |
| **RNU12** | 0.00000 | 0.04851 | 0.12491 | 0.09350 | 0.20278 | 0.13073 | 0.79659 |
| **RNU4ATAC** | 0.00000 | 0.00000 | 0.12618 | 0.02730 | 0.03477 | 0.27540 | 3.95145 |
| **RNU5F-1** | 0.02329 | 0.01502 | 0.01877 | 0.02059 | 0.00747 | 0.01262 | 0.01896 |
| **RNU6-19P** | 0.02191 | 0.01476 | 0.01309 | 0.01749 | 0.00772 | 0.02131 | 0.02886 |
| **RNU6-28P.uid-1** | 0.02368 | 0.01816 | 0.01321 | 0.01956 | 0.00784 | 0.01524 | 0.01872 |
| **RNU6-71P** | 0.02371 | 0.01839 | 0.01319 | 0.02055 | 0.00790 | 0.01312 | 0.01588 |
| **RNU6-78P** | 0.02537 | 0.01518 | 0.02092 | 0.01895 | 0.00740 | 0.02377 | 0.02722 |
| **RNU6-83P.uid-1** | 0.02464 | 0.01789 | 0.01263 | 0.01954 | 0.00797 | 0.01583 | 0.01606 |
| **RNU6ATAC** | 0.02119 | 0.05377 | 0.01719 | 0.04138 | 0.01722 | 0.02876 | 0.08141 |
| **RNU86** | 0.37421 | 0.41363 | 0.19528 | 0.80698 | 0.21854 | 0.79791 | 1.20676 |
| **RNVU1-19.uid-1** | 0.01072 | 0.00406 | 0.01434 | 0.02200 | 0.00862 | 0.05220 | 0.31230 |
| **RNVU1-19.uid-1** | 0.00244 | 0.00729 | 0.01733 | 0.03035 | 0.00684 | 0.04182 | 0.15275 |
| **RNVU1-8** | 0.00000 | 0.03762 | 0.01548 | 0.03081 | 0.00638 | 0.19731 | 0.22700 |
| **ROBO1** | 0.13007 | 0.55874 | 0.24681 | 0.73835 | 0.44587 | 4.16066 | 0.59176 |
| **ROBO2** | 0.15623 | 0.00000 | 0.01047 | 0.03203 | 0.54208 | 0.42347 | 0.00000 |
| **ROBO3** | 0.05556 | 0.00000 | 0.07771 | 0.29588 | 1.35115 | 0.30637 | 0.00000 |
| **ROBO4** | 0.00000 | 0.26234 | 0.01209 | 0.00000 | 0.78380 | 0.04904 | 0.12684 |
| **ROCK1** | 4.46585 | 0.00000 | 8.85707 | 5.00396 | 3.31513 | 7.92073 | 23.52025 |
| **ROCK1P1** | 0.06399 | 0.00000 | 0.03559 | 0.03032 | 0.01474 | 0.07146 | 0.21921 |
| **ROCK2** | 1.73465 | 1.43913 | 3.20725 | 2.84883 | 0.62733 | 2.45206 | 15.31775 |
| **ROGDI** | 2.82053 | 4.83037 | 0.01033 | 7.52481 | 0.55069 | 6.24421 | 8.69028 |
| **ROM1** | 0.66184 | 0.53958 | 0.02948 | 0.88628 | 0.08699 | 0.12834 | 0.36206 |
| **ROMO1** | 28.28586 | 43.30733 | 14.11493 | 36.84416 | 4.64332 | 20.33087 | 24.84200 |
| **ROPN1B** | 0.03390 | 0.08818 | 0.15440 | 0.01516 | 0.00494 | 0.00522 | 0.00000 |
| **ROPN1L** | 0.02268 | 0.11339 | 0.28117 | 0.03700 | 0.25904 | 0.00170 | 0.00000 |
| **ROPN1L-AS1** | 0.01458 | 0.01928 | 0.02422 | 0.04102 | 0.03119 | 0.01143 | 0.01978 |
| **ROR1** | 0.12003 | 0.39118 | 0.00000 | 0.11985 | 0.08084 | 1.34952 | 0.26489 |
| **ROR1-AS1** | 0.02130 | 0.02735 | 0.01101 | 0.02415 | 0.00785 | 0.01952 | 0.02641 |
| **ROR2** | 0.00000 | 0.00000 | 0.01804 | 0.11032 | 0.84405 | 0.66490 | 0.23978 |
| **RORA** | 0.00000 | 0.00000 | 1.86850 | 0.10658 | 1.28225 | 2.81805 | 9.09071 |

| | | | | | | | |
|---|---|---|---|---|---|---|---|
| **RORB** | 0.01698 | 0.00629 | 0.10502 | 0.00902 | 0.16160 | 0.00000 | 0.22640 |
| **RORC** | 0.01854 | 0.01199 | 0.01444 | 0.02122 | 0.00728 | 0.01461 | 0.05448 |
| **ROS1** | 0.01318 | 0.06027 | 0.00000 | 0.01082 | 3.57159 | 0.00482 | 0.02625 |
| **RP1** | 0.01293 | 0.00000 | 0.06670 | 0.01266 | 0.05686 | 0.06188 | 0.05893 |
| **RP11-87M18.2** | 0.02265 | 0.01103 | 0.04187 | 0.01881 | 0.16636 | 0.01192 | 0.01687 |
| **RP1L1** | 0.01332 | 0.01148 | 0.01190 | 0.01959 | 0.00736 | 0.10563 | 0.14301 |
| **RP2** | 14.72481 | 1.86149 | 3.18821 | 23.83762 | 5.04804 | 0.00292 | 2.15816 |
| **RP9** | 0.27201 | 4.06717 | 3.23090 | 0.80448 | 0.53374 | 2.87776 | 7.02526 |
| **RP9P** | 0.06423 | 0.00000 | 0.04011 | 0.09905 | 0.04801 | 0.12111 | 0.00000 |
| **RPA1** | 25.80324 | 8.69307 | 6.74069 | 24.90306 | 7.78445 | 7.95308 | 2.77026 |
| **RPA2** | 34.88626 | 10.93602 | 18.15264 | 27.93785 | 6.50549 | 12.86145 | 0.00000 |
| **RPA3** | 14.77078 | 7.82712 | 8.70339 | 9.57248 | 3.65612 | 9.99202 | 9.87756 |
| **RPA3OS** | 8.37061 | 0.28921 | 4.87128 | 4.01253 | 2.51482 | 1.55557 | 0.00000 |
| **RPA4** | 0.00000 | 0.00533 | 0.03636 | 0.13952 | 0.40034 | 0.12179 | 0.32852 |
| **RPAIN** | 19.73586 | 6.73051 | 7.57776 | 16.04012 | 0.39206 | 7.37304 | 12.38702 |
| **RPAP1** | 6.48840 | 2.15233 | 9.62841 | 12.62302 | 1.39249 | 3.36239 | 16.34303 |
| **RPAP2** | 7.58158 | 0.07289 | 3.74027 | 3.04104 | 0.65510 | 2.31813 | 2.37971 |
| **RPAP3** | 18.73719 | 2.86594 | 5.33093 | 9.37856 | 2.72756 | 4.31795 | 7.62553 |
| **RPARP-AS1** | 8.52487 | 13.58932 | 5.69342 | 11.52741 | 0.80075 | 2.14618 | 2.13495 |
| **RPE** | 16.18697 | 4.27598 | 9.03427 | 7.28329 | 2.38881 | 4.58842 | 0.00000 |
| **RPE65** | 0.02133 | 0.03374 | 0.00832 | 0.03015 | 0.03415 | 0.05228 | 0.01985 |
| **RPEL1** | 2.07869 | 1.32653 | 1.14121 | 1.34635 | 0.27162 | 0.46843 | 0.00000 |
| **RPF1** | 24.18275 | 9.51845 | 18.30744 | 13.38710 | 10.88296 | 5.17511 | 0.35959 |
| **RPF2** | 30.90317 | 2.22511 | 19.94610 | 14.99852 | 12.73188 | 7.32805 | 10.08657 |
| **RPGR** | 4.03023 | 0.00000 | 4.25421 | 0.90409 | 5.01094 | 1.20783 | 1.56826 |
| **RPGRIP1** | 0.24068 | 0.00000 | 1.93027 | 0.09965 | 1.64756 | 0.02207 | 0.00000 |
| **RPGRIP1L** | 1.61314 | 0.80881 | 1.52673 | 1.26531 | 1.48743 | 1.74745 | 1.34114 |
| **RPH3A** | 0.01481 | 0.00234 | 0.29684 | 0.02257 | 3.44310 | 0.00185 | 0.02749 |
| **RPH3AL** | 0.01170 | 0.27068 | 0.00901 | 0.04588 | 0.71577 | 0.11504 | 0.00124 |
| **RPIA** | 3.29744 | 2.38677 | 1.82872 | 6.00291 | 1.04217 | 2.41748 | 5.17809 |
| **RPL10** | 31.18440 | 21.24187 | 17.48825 | 19.22747 | 6.41551 | 16.65546 | 23.26039 |
| **RPL10A** | 36.23848 | 27.40371 | 20.94966 | 28.68389 | 8.62252 | 19.15773 | 25.72931 |
| **RPL10L** | 0.43781 | 1.42008 | 4.70459 | 0.77342 | 8.14640 | 0.11108 | 0.00000 |
| **RPL11** | 49.27616 | 36.50331 | 36.09547 | 46.24882 | 9.32272 | 35.49438 | 64.37670 |
| **RPL12** | 15.55957 | 18.38099 | 14.59560 | 21.52779 | 1.56516 | 13.36180 | 45.95147 |

| | | | | | | | |
|---|---|---|---|---|---|---|---|
| **RPL13** | 38.85668 | 46.04179 | 24.81728 | 48.76760 | 6.78584 | 27.34703 | 44.60315 |
| **RPL13A** | 30.59786 | 10.54545 | 15.77836 | 41.33412 | 11.22064 | 20.94458 | 42.43830 |
| **RPL13AP17** | 0.02109 | 0.01171 | 0.01547 | 0.01960 | 0.00717 | 0.00924 | 0.07264 |
| **RPL13AP20** | 1.18283 | 0.63826 | 0.61996 | 0.35849 | 0.06326 | 0.45930 | 2.46848 |
| **RPL13AP3** | 0.19007 | 0.07099 | 0.10157 | 0.17631 | 0.11123 | 0.12245 | 0.27220 |
| **RPL13AP6** | 4.98657 | 5.38527 | 2.15492 | 7.46658 | 0.96203 | 2.96103 | 2.12999 |
| **RPL13P5** | 0.00000 | 3.46648 | 1.51164 | 0.18657 | 0.03387 | 4.56575 | 0.40220 |
| **RPL14** | 22.96920 | 18.45078 | 18.56003 | 26.96112 | 3.75544 | 12.73672 | 41.68353 |
| **RPL15** | 41.68680 | 21.80205 | 26.64464 | 35.61327 | 10.91519 | 26.55717 | 37.43816 |
| **RPL17** | 15.63320 | 10.00380 | 6.08711 | 13.14162 | 0.79354 | 7.20827 | 5.38899 |
| **RPL18** | 50.28711 | 44.21343 | 33.21446 | 55.07000 | 14.48970 | 34.70264 | 57.12760 |
| **RPL18A** | 13.47537 | 20.04692 | 7.75058 | 17.21703 | 4.11047 | 9.02306 | 5.64351 |
| **RPL19** | 52.20476 | 37.11810 | 31.28201 | 51.29205 | 7.50327 | 31.85914 | 54.54824 |
| **RPL19P12** | 9.66415 | 5.99115 | 5.24143 | 11.47768 | 0.35810 | 5.56176 | 7.76855 |
| **RPL21P44** | 0.04036 | 0.10919 | 0.00777 | 0.08975 | 0.11229 | 0.12226 | 0.29670 |
| **RPL22** | 10.74731 | 5.85546 | 7.01204 | 10.69730 | 0.65111 | 5.92905 | 18.21098 |
| **RPL22L1** | 14.95024 | 0.00000 | 13.88147 | 1.40566 | 0.97675 | 10.48590 | 34.84772 |
| **RPL23** | 42.12920 | 41.95169 | 29.01374 | 45.45531 | 11.05493 | 29.26942 | 52.53646 |
| **RPL23A** | 31.96454 | 8.82657 | 19.92447 | 31.99476 | 8.00466 | 17.33850 | 33.53217 |
| **RPL23AP32** | 0.16624 | 0.06029 | 0.50203 | 0.02324 | 0.07889 | 0.24679 | 0.40979 |
| **RPL23AP53** | 0.92538 | 0.00000 | 0.69909 | 0.57281 | 2.32027 | 0.05622 | 0.50809 |
| **RPL23AP64** | 0.03695 | 0.00000 | 0.08243 | 0.09504 | 0.00758 | 0.16223 | 0.30145 |
| **RPL23AP7** | 0.77411 | 1.11663 | 0.85901 | 1.03640 | 1.01469 | 0.01132 | 0.00000 |
| **RPL23AP82** | 1.69238 | 1.93681 | 2.02007 | 1.24088 | 0.72302 | 0.34443 | 0.00000 |
| **RPL23P8** | 2.14395 | 1.17267 | 1.97547 | 2.83813 | 0.34186 | 1.58452 | 3.11323 |
| **RPL24** | 20.57831 | 15.33662 | 18.35087 | 20.93972 | 3.94896 | 14.51020 | 41.10199 |
| **RPL26** | 32.31554 | 17.35877 | 24.82338 | 29.35927 | 8.12612 | 22.44300 | 46.32818 |
| **RPL26L1** | 8.98232 | 9.63252 | 6.99601 | 3.16447 | 6.13907 | 2.92742 | 1.86017 |
| **RPL27** | 50.59207 | 39.99231 | 29.11348 | 48.56947 | 11.54745 | 29.82773 | 45.42999 |
| **RPL27A** | 27.24671 | 24.98155 | 17.78547 | 35.74548 | 6.60006 | 19.77874 | 50.41484 |
| **RPL28** | 20.89353 | 35.10899 | 21.31643 | 37.19712 | 14.26988 | 22.47323 | 64.05622 |
| **RPL29** | 27.17875 | 26.51638 | 21.52966 | 30.16401 | 5.69672 | 18.17071 | 37.15817 |
| **RPL29P2** | 0.02994 | 0.02900 | 0.02226 | 0.01745 | 0.01021 | 0.02569 | 0.09250 |
| **RPL3** | 60.33020 | 49.33010 | 31.81922 | 48.93106 | 11.61307 | 32.41701 | 33.30497 |
| **RPL30** | 51.34524 | 25.94594 | 37.22958 | 46.34595 | 18.42408 | 32.92513 | 57.94175 |

| | | | | | | | |
|---|---|---|---|---|---|---|---|
| **RPL31** | 43.89360 | 27.57732 | 28.29455 | 42.70069 | 7.47691 | 29.04947 | 48.79319 |
| **RPL31P11** | 1.00750 | 0.28154 | 0.79197 | 0.71607 | 0.08787 | 0.75033 | 1.21620 |
| **RPL32** | 36.06398 | 29.18974 | 25.85002 | 36.60693 | 7.38696 | 24.44780 | 47.92924 |
| **RPL32P3** | 2.04072 | 0.24659 | 0.94125 | 2.25269 | 1.68299 | 0.91348 | 2.38790 |
| **RPL34** | 34.28467 | 19.89288 | 20.71314 | 27.47078 | 3.69679 | 19.96069 | 37.40937 |
| **RPL34-AS1** | 0.15901 | 0.03001 | 0.27765 | 0.25478 | 0.62782 | 0.02979 | 0.12885 |
| **RPL35** | 34.72692 | 48.70883 | 23.26737 | 40.18689 | 9.81630 | 24.37681 | 45.80671 |
| **RPL35A** | 46.84512 | 30.56834 | 33.42231 | 42.31364 | 9.42079 | 29.71029 | 51.46601 |
| **RPL36** | 50.18139 | 50.71100 | 34.22221 | 53.14880 | 19.59022 | 34.52518 | 55.10775 |
| **RPL36A.uid-130** | 0.10938 | 0.00000 | 0.17581 | 0.02548 | 0.03977 | 0.05416 | 0.36335 |
| **RPL36A.uid-130** | 2.11406 | 0.00000 | 2.07484 | 0.00000 | 0.37249 | 1.64206 | 6.97058 |
| **RPL36AL** | 37.25146 | 25.74868 | 22.54717 | 28.50876 | 3.71063 | 16.84270 | 18.17644 |
| **RPL37** | 39.65051 | 30.00108 | 29.13838 | 41.48371 | 11.12030 | 28.78272 | 56.19567 |
| **RPL37A** | 40.00683 | 34.19789 | 30.27047 | 45.90896 | 10.77185 | 31.32522 | 68.59443 |
| **RPL38** | 45.27462 | 39.06028 | 30.52546 | 44.64569 | 10.10491 | 30.41587 | 53.31179 |
| **RPL39** | 8.66361 | 0.00000 | 10.10580 | 5.98996 | 1.33618 | 7.87449 | 29.23466 |
| **RPL39L** | 7.37745 | 9.60003 | 5.71484 | 7.94178 | 1.22534 | 6.41637 | 3.52324 |
| **RPL3L** | 0.04725 | 0.02090 | 0.03354 | 0.09768 | 0.11795 | 0.19351 | 0.06746 |
| **RPL4** | 51.41635 | 31.26178 | 29.30098 | 40.18205 | 7.10537 | 25.27769 | 42.62314 |
| **RPL41** | 8.59068 | 7.25237 | 9.54230 | 7.98927 | 3.34591 | 6.22807 | 9.24293 |
| **RPL5** | 60.84895 | 25.97742 | 34.37395 | 48.27856 | 12.77940 | 31.83048 | 45.64715 |
| **RPL6** | 28.75582 | 15.73199 | 18.26092 | 19.84337 | 4.03299 | 13.21708 | 16.91342 |
| **RPL7** | 0.94636 | 0.35442 | 0.56047 | 0.21438 | 0.29577 | 0.48257 | 0.49346 |
| **RPL7A** | 19.52549 | 18.86711 | 16.45075 | 23.65718 | 6.51407 | 15.80133 | 33.81029 |
| **RPL7L1** | 14.67071 | 6.22121 | 8.29643 | 7.16583 | 1.14656 | 5.24047 | 3.99787 |
| **RPL8** | 67.45952 | 71.00082 | 35.85991 | 64.45115 | 23.70631 | 48.67051 | 59.50401 |
| **RPL9** | 16.45605 | 6.54815 | 9.87685 | 11.46443 | 2.52592 | 6.26100 | 4.61266 |
| **RPLP0** | 44.77513 | 41.75700 | 23.19152 | 44.63967 | 12.61401 | 27.34246 | 41.34739 |
| **RPLP0P2** | 0.40637 | 6.55816 | 1.99911 | 1.41868 | 0.62982 | 3.39659 | 4.65215 |
| **RPLP1** | 57.80374 | 44.99645 | 32.95482 | 65.99160 | 15.07810 | 35.73000 | 67.77417 |
| **RPLP2** | 43.78603 | 43.31597 | 34.23183 | 58.00644 | 13.20725 | 31.75552 | 77.90721 |
| **RPN1** | 46.63161 | 29.33971 | 26.74825 | 35.66524 | 6.86205 | 23.06898 | 8.93903 |
| **RPN2** | 46.44352 | 41.62033 | 24.27337 | 38.58355 | 5.18389 | 30.82375 | 17.77362 |
| **RPP14** | 18.68269 | 11.95475 | 15.81188 | 9.50049 | 5.21099 | 10.05352 | 3.74057 |
| **RPP21** | 1.70330 | 4.32501 | 1.00406 | 1.73447 | 1.13209 | 1.17943 | 0.22543 |

| | | | | | | | |
|---|---|---|---|---|---|---|---|
| **RPP25** | 2.63785 | 12.84707 | 9.99883 | 7.08857 | 1.87195 | 5.70353 | 20.68745 |
| **RPP25L** | 3.66553 | 11.59210 | 7.95265 | 7.27171 | 0.44620 | 3.33851 | 0.64271 |
| **RPP30** | 20.34975 | 11.46700 | 12.37752 | 15.40206 | 6.65658 | 7.79061 | 2.51644 |
| **RPP38** | 9.93037 | 13.11922 | 10.66970 | 7.69442 | 1.39426 | 6.29887 | 11.36481 |
| **RPP40** | 15.26601 | 3.06532 | 10.68225 | 10.14296 | 2.39936 | 2.66951 | 0.00000 |
| **RPPH1** | 0.00004 | 2.32433 | 1.82548 | 0.69840 | 3.59322 | 0.37740 | 1.87972 |
| **RPRD1A** | 14.57805 | 2.05594 | 11.24690 | 5.67776 | 4.59969 | 7.60615 | 9.00210 |
| **RPRD1B** | 12.99510 | 4.27664 | 5.53592 | 6.47494 | 4.03698 | 3.44460 | 2.93110 |
| **RPRD2** | 5.07090 | 1.64742 | 4.05145 | 4.98221 | 1.59587 | 3.35507 | 12.31875 |
| **RPRM** | 0.00000 | 0.26837 | 0.45279 | 0.18056 | 0.01882 | 1.54848 | 0.01525 |
| **RPRML** | 0.00000 | 0.03702 | 0.05157 | 0.10348 | 0.00609 | 0.06120 | 0.06868 |
| **RPS10** | 7.49660 | 8.14161 | 3.21571 | 8.34250 | 2.31221 | 4.08662 | 0.00000 |
| **RPS10P7** | 0.02111 | 0.04020 | 0.02784 | 0.08056 | 0.02933 | 0.02289 | 0.07203 |
| **RPS11** | 43.71697 | 46.98615 | 32.48552 | 52.97083 | 18.46653 | 30.54636 | 55.88853 |
| **RPS12** | 22.48359 | 10.92150 | 22.89575 | 33.24263 | 5.67374 | 19.60796 | 62.87873 |
| **RPS13** | 46.11806 | 22.27606 | 32.53442 | 44.36290 | 14.53693 | 29.58263 | 54.37835 |
| **RPS14** | 69.45031 | 55.97236 | 32.72340 | 66.76807 | 20.03652 | 40.73716 | 43.80474 |
| **RPS14P3** | 13.71966 | 11.87575 | 6.59654 | 16.45250 | 3.61438 | 9.10725 | 11.25122 |
| **RPS15** | 23.29294 | 34.10928 | 16.16312 | 34.36093 | 6.48647 | 17.52897 | 30.15023 |
| **RPS15A** | 28.20982 | 18.88490 | 19.84489 | 29.13744 | 5.43619 | 19.24243 | 30.00058 |
| **RPS15AP10** | 0.00000 | 0.41428 | 0.38866 | 0.22608 | 0.34885 | 0.24541 | 1.40433 |
| **RPS16** | 45.43591 | 47.04401 | 28.20443 | 50.52394 | 12.89978 | 30.99173 | 60.88305 |
| **RPS16P5** | 0.00000 | 0.00000 | 0.09813 | 0.18135 | 0.13053 | 0.04570 | 0.30671 |
| **RPS17.uid-1969** | 0.02089 | 0.03421 | 0.01228 | 0.03447 | 0.01287 | 0.02861 | 0.07141 |
| **RPS17.uid-1970** | 0.02506 | 0.05282 | 0.02250 | 0.01887 | 0.00805 | 0.01218 | 0.00779 |
| **RPS18** | 37.82555 | 30.61468 | 26.69601 | 37.65251 | 10.39970 | 24.61670 | 31.52671 |
| **RPS18P9** | 2.77060 | 3.65639 | 2.67616 | 4.00552 | 0.91483 | 2.57392 | 3.61642 |
| **RPS19** | 57.74133 | 59.88156 | 35.72698 | 68.18713 | 22.93427 | 38.34635 | 53.66855 |
| **RPS19BP1** | 20.28039 | 31.61985 | 12.44005 | 30.16724 | 2.75176 | 15.14569 | 25.34546 |
| **RPS2** | 1.01680 | 17.23002 | 7.04178 | 14.67264 | 3.86926 | 13.31019 | 58.44724 |
| **RPS20** | 41.47959 | 34.16659 | 27.65204 | 39.80333 | 12.66864 | 22.10278 | 47.32605 |
| **RPS21** | 29.93730 | 30.13851 | 23.67168 | 39.39702 | 9.48556 | 23.65119 | 60.58897 |
| **RPS23** | 50.29896 | 24.36792 | 31.21498 | 47.21415 | 7.83273 | 30.10874 | 47.03074 |
| **RPS24** | 43.91212 | 25.49131 | 30.32848 | 38.12748 | 11.87820 | 28.85978 | 52.25057 |
| **RPS25** | 41.89518 | 32.06787 | 28.89250 | 37.04163 | 11.94780 | 22.24250 | 32.48859 |

| | | | | | | | |
|---|---|---|---|---|---|---|---|
| **RPS26** | 9.32162 | 3.20658 | 7.40285 | 12.98886 | 1.43351 | 3.71181 | 0.00000 |
| **RPS26P11** | 0.08441 | 0.31691 | 0.08724 | 0.05932 | 0.04691 | 0.08104 | 0.11745 |
| **RPS27** | 2.84960 | 3.12704 | 2.95058 | 1.86898 | 1.70042 | 2.42748 | 4.40661 |
| **RPS27A** | 28.36837 | 16.36123 | 21.50398 | 26.47345 | 4.11675 | 20.32170 | 40.59321 |
| **RPS27L** | 18.93104 | 20.09372 | 16.41809 | 16.89851 | 10.88006 | 16.17853 | 14.09520 |
| **RPS28** | 4.65977 | 2.73258 | 6.01589 | 0.00000 | 1.60686 | 4.30789 | 20.93599 |
| **RPS29** | 32.59109 | 14.83646 | 28.43059 | 31.58237 | 8.69371 | 21.31167 | 56.13449 |
| **RPS2P32** | 0.00000 | 0.10108 | 0.18608 | 0.28521 | 0.06877 | 0.28942 | 0.15256 |
| **RPS3** | 61.11195 | 48.13073 | 35.68964 | 62.56491 | 14.51430 | 39.94944 | 62.18007 |
| **RPS3A** | 5.03335 | 1.26796 | 3.24690 | 2.24136 | 0.87559 | 4.36144 | 9.47616 |
| **RPS4X** | 55.16348 | 36.36895 | 36.01021 | 52.96195 | 11.84800 | 36.03084 | 66.98295 |
| **RPS4Y1** | 0.00000 | 0.05504 | 0.11883 | 0.35222 | 0.00649 | 0.06726 | 0.10749 |
| **RPS4Y2** | 0.01859 | 0.01298 | 0.01232 | 0.01819 | 0.00742 | 0.01003 | 0.06410 |
| **RPS5** | 59.65741 | 50.86495 | 35.21718 | 57.72327 | 20.08057 | 37.10986 | 57.75392 |
| **RPS6** | 66.71061 | 50.28031 | 43.65245 | 59.20627 | 19.64627 | 38.51133 | 51.71142 |
| **RPS6KA1** | 3.36797 | 11.73660 | 2.49067 | 4.13278 | 0.59888 | 8.48211 | 8.14222 |
| **RPS6KA2** | 0.38339 | 0.10534 | 0.00000 | 0.05995 | 0.24803 | 2.38946 | 0.44487 |
| **RPS6KA2-AS1** | 0.02047 | 0.01960 | 0.01107 | 0.02022 | 0.02674 | 0.01657 | 0.02067 |
| **RPS6KA3** | 2.29993 | 0.61565 | 3.27305 | 1.94969 | 1.46690 | 2.58419 | 5.92023 |
| **RPS6KA4** | 0.38972 | 0.89301 | 0.00000 | 1.43689 | 0.00114 | 0.00000 | 15.46964 |
| **RPS6KA5** | 2.91330 | 4.49430 | 1.45528 | 3.95269 | 2.55099 | 2.33763 | 5.33709 |
| **RPS6KA6** | 0.98175 | 0.00000 | 0.00000 | 0.70918 | 0.27418 | 0.23900 | 1.82239 |
| **RPS6KB1** | 15.02115 | 2.10249 | 4.25658 | 10.61086 | 0.70724 | 5.14105 | 1.39923 |
| **RPS6KB2** | 9.65004 | 15.68979 | 10.66031 | 12.21512 | 1.84574 | 8.25225 | 8.58851 |
| **RPS6KC1** | 5.27937 | 0.74789 | 7.15523 | 2.59828 | 3.57851 | 4.42993 | 0.69704 |
| **RPS6KL1** | 0.00000 | 1.55135 | 1.79172 | 0.35199 | 1.92492 | 0.54263 | 0.62867 |
| **RPS7** | 7.30233 | 10.96305 | 7.37447 | 9.47464 | 1.53155 | 5.33957 | 12.93879 |
| **RPS7P5** | 0.02549 | 0.01251 | 0.01943 | 0.02077 | 0.00810 | 0.00754 | 0.09899 |
| **RPS8** | 43.48869 | 39.02886 | 29.39622 | 46.48998 | 7.49165 | 28.91045 | 54.45289 |
| **RPS9** | 35.52998 | 38.98533 | 20.32171 | 36.34093 | 8.61235 | 24.00718 | 49.63117 |
| **RPSA** | 16.84961 | 16.63057 | 7.39651 | 21.44327 | 1.71269 | 18.85596 | 31.83762 |
| **RPSAP52** | 0.08854 | 0.07209 | 0.02769 | 0.01215 | 0.01607 | 0.06090 | 0.05034 |
| **RPSAP58** | 5.68132 | 1.86846 | 2.64069 | 5.21224 | 2.01032 | 4.71257 | 2.53264 |
| **RPSAP9** | 0.12643 | 0.05720 | 0.02752 | 0.12207 | 0.33111 | 0.10636 | 0.06939 |
| **RPTN** | 0.02794 | 0.01755 | 0.01283 | 0.02022 | 0.00823 | 0.01829 | 0.01567 |

| | | | | | | | |
|---|---|---|---|---|---|---|---|
| **RPTOR** | 1.20011 | 7.24584 | 5.57987 | 3.19629 | 4.16684 | 1.69819 | 8.99363 |
| **RPUSD1** | 0.00000 | 14.09718 | 2.18154 | 9.98348 | 0.08927 | 8.13640 | 28.35739 |
| **RPUSD2** | 4.90082 | 7.35128 | 9.75540 | 6.84904 | 1.38710 | 5.22536 | 8.27182 |
| **RPUSD3** | 27.03926 | 29.58732 | 18.66445 | 18.82860 | 1.37665 | 16.75504 | 1.33311 |
| **RPUSD4** | 17.35270 | 2.86524 | 7.14643 | 10.94007 | 5.72045 | 4.53517 | 6.92663 |
| **RQCD1** | 43.65121 | 21.45171 | 12.61854 | 23.11990 | 9.55602 | 12.63576 | 0.69122 |
| **RRAD** | 0.12078 | 12.78378 | 0.00000 | 0.00000 | 0.14029 | 11.75346 | 0.12851 |
| **RRAGA** | 6.77924 | 8.07104 | 5.98042 | 14.07409 | 1.64319 | 6.53761 | 5.05787 |
| **RRAGB** | 1.65138 | 0.58916 | 3.72254 | 0.34046 | 2.46391 | 2.11446 | 0.26213 |
| **RRAGC** | 1.41837 | 2.23374 | 2.51865 | 8.77317 | 1.64089 | 4.65336 | 12.33583 |
| **RRAGD** | 0.03060 | 0.09908 | 0.03627 | 0.08754 | 0.11899 | 0.06435 | 0.20682 |
| **RRAS** | 0.19916 | 0.05742 | 1.24916 | 0.19178 | 1.16652 | 0.12804 | 0.42487 |
| **RRAS2** | 2.20563 | 1.31483 | 3.35422 | 1.65591 | 0.13326 | 4.37035 | 12.63407 |
| **RRBP1** | 11.62013 | 34.14080 | 3.56735 | 17.67699 | 3.54294 | 13.06051 | 30.58688 |
| **RREB1** | 12.59103 | 13.77378 | 14.68839 | 8.59416 | 6.99215 | 6.26859 | 28.34308 |
| **RRH** | 0.13019 | 0.00000 | 0.09013 | 0.08873 | 0.03378 | 0.04685 | 0.04320 |
| **RRM1** | 34.87929 | 16.26939 | 26.52520 | 27.84103 | 5.07940 | 13.00401 | 8.04904 |
| **RRM2** | 46.68706 | 26.21195 | 37.67201 | 52.75842 | 20.07608 | 18.03941 | 9.51736 |
| **RRM2B** | 6.84479 | 2.01459 | 4.25920 | 3.24705 | 5.67540 | 1.96861 | 0.00000 |
| **RRN3** | 12.68811 | 3.35079 | 4.89217 | 5.06293 | 7.94324 | 4.91896 | 1.30992 |
| **RRN3P1** | 0.03163 | 0.01904 | 0.09960 | 0.01465 | 0.11250 | 0.16090 | 0.02087 |
| **RRN3P2** | 0.03135 | 0.04740 | 0.17988 | 0.04547 | 0.08673 | 0.01391 | 0.00000 |
| **RRN3P3** | 0.00000 | 0.71941 | 0.46771 | 0.82959 | 0.12287 | 1.13516 | 1.48554 |
| **RRNAD1** | 1.51951 | 2.03141 | 1.70203 | 2.23003 | 0.12881 | 1.18052 | 1.12529 |
| **RRP1** | 4.35702 | 16.63762 | 7.36531 | 11.72644 | 0.85504 | 4.83337 | 20.10408 |
| **RRP12** | 13.56723 | 13.69059 | 16.97653 | 21.90782 | 4.95421 | 10.88304 | 10.01010 |
| **RRP15** | 20.74465 | 3.19078 | 12.76674 | 10.21839 | 3.80593 | 8.15923 | 11.30681 |
| **RRP1B** | 25.25891 | 8.85702 | 12.24479 | 12.50134 | 3.63682 | 8.09976 | 19.58430 |
| **RRP36** | 1.37674 | 4.41048 | 6.12095 | 5.87109 | 0.18251 | 4.84404 | 22.74567 |
| **RRP7A** | 25.22103 | 34.56501 | 12.51586 | 23.60328 | 3.76454 | 10.18623 | 12.22627 |
| **RRP7B** | 4.28603 | 11.72656 | 1.29870 | 4.66397 | 1.40084 | 1.22002 | 2.28184 |
| **RRP8** | 9.49707 | 12.22387 | 7.16413 | 3.69794 | 4.49902 | 2.43732 | 3.34251 |
| **RRP9** | 12.88087 | 24.22800 | 12.31415 | 16.28721 | 5.11781 | 11.21407 | 13.41501 |
| **RRS1** | 14.12573 | 9.93634 | 18.27300 | 16.25331 | 7.77992 | 4.26771 | 7.77857 |
| **RRS1-AS1** | 0.08895 | 0.04747 | 0.13887 | 0.05952 | 0.36697 | 0.01403 | 0.00000 |

| | | | | | | | |
|---|---|---|---|---|---|---|---|
| **RS1** | 0.02753 | 0.00000 | 0.04311 | 0.00881 | 2.45704 | 0.01550 | 0.06542 |
| **RSAD1** | 1.56206 | 6.13944 | 1.99051 | 0.00000 | 0.00000 | 4.05825 | 8.27008 |
| **RSAD2** | 0.04463 | 0.04424 | 0.01153 | 0.01245 | 0.01314 | 0.01119 | 0.01793 |
| **RSBN1** | 3.81482 | 3.45753 | 3.23519 | 6.03432 | 5.20468 | 4.24523 | 5.59519 |
| **RSBN1L** | 1.32645 | 0.00000 | 2.19148 | 1.67643 | 1.49634 | 1.48692 | 2.90396 |
| **RSC1A1** | 1.94056 | 0.32717 | 1.53049 | 0.68501 | 1.58763 | 0.66602 | 2.60379 |
| **RSF1** | 2.86794 | 4.48160 | 4.76046 | 3.35777 | 6.39917 | 7.42826 | 8.10947 |
| **RSG1** | 0.00000 | 0.82944 | 0.04015 | 0.23299 | 0.00536 | 0.06111 | 0.35690 |
| **RSL1D1** | 61.44814 | 25.83101 | 29.06954 | 35.39296 | 14.53347 | 19.87806 | 20.68539 |
| **RSL24D1** | 30.39270 | 0.00000 | 17.54580 | 16.54784 | 6.50997 | 9.39532 | 17.00981 |
| **RSPH1** | 0.07229 | 0.15134 | 0.04341 | 0.00000 | 0.22738 | 0.23223 | 0.01466 |
| **RSPH10B+RSPI** | 0.00653 | 0.13046 | 0.24195 | 0.01329 | 0.13746 | 0.00708 | 0.01396 |
| **RSPH10B+RSPI** | 0.00000 | 0.18364 | 0.52818 | 0.01068 | 0.25032 | 0.01818 | 0.02758 |
| **RSPH3** | 0.97795 | 1.69275 | 2.11055 | 0.70336 | 0.56826 | 0.76629 | 0.00000 |
| **RSPH4A** | 0.13260 | 0.02525 | 0.06924 | 0.09966 | 0.51604 | 0.08363 | 0.19903 |
| **RSPH6A** | 0.01997 | 0.03380 | 0.01422 | 0.03423 | 0.00733 | 0.00829 | 0.02985 |
| **RSPH9** | 0.36126 | 0.54699 | 2.14033 | 0.13549 | 3.88453 | 0.07597 | 0.00000 |
| **RSPO1** | 0.02202 | 0.00000 | 0.26800 | 0.01835 | 0.13794 | 0.00572 | 0.02331 |
| **RSPO2** | 0.00000 | 0.21704 | 0.13584 | 0.00446 | 2.60179 | 0.02974 | 0.02292 |
| **RSPO3** | 0.07338 | 0.00000 | 0.02360 | 0.82413 | 0.02093 | 7.34684 | 0.02069 |
| **RSPO4** | 0.03296 | 0.01500 | 0.01928 | 0.02539 | 0.00756 | 0.01609 | 0.01069 |
| **RSPRY1** | 10.08372 | 12.69317 | 8.81185 | 7.82315 | 2.15906 | 8.85422 | 1.05255 |
| **RSRC1** | 8.03984 | 5.40831 | 6.34294 | 1.33202 | 1.35916 | 4.09840 | 9.83838 |
| **RSRC2** | 22.74017 | 12.19134 | 15.85965 | 14.95434 | 13.55755 | 11.91682 | 16.68919 |
| **RSRP1** | 16.37089 | 8.45387 | 6.11041 | 9.24932 | 21.52234 | 3.48349 | 0.84885 |
| **RSU1** | 16.47763 | 7.71489 | 3.07064 | 11.44520 | 1.73568 | 9.34807 | 0.00000 |
| **RSU1P2** | 0.02700 | 0.01737 | 0.01416 | 0.02110 | 0.00815 | 0.01808 | 0.01289 |
| **RTBDN** | 0.00047 | 0.71739 | 1.76248 | 0.32833 | 0.27278 | 0.44275 | 0.03510 |
| **RTCA** | 14.25977 | 5.74265 | 1.54062 | 12.61364 | 0.22271 | 8.22167 | 2.38240 |
| **RTCA-AS1** | 0.04083 | 0.04061 | 0.01193 | 0.07179 | 0.00751 | 0.01310 | 0.06605 |
| **RTCB** | 41.16706 | 28.62795 | 15.22325 | 25.22229 | 3.73781 | 12.88739 | 0.00000 |
| **RTDR1** | 0.01306 | 0.04870 | 0.01596 | 0.01854 | 0.14855 | 0.00839 | 0.02789 |
| **RTEL1** | 0.00000 | 1.04208 | 0.41004 | 0.23309 | 0.00545 | 0.61795 | 2.01796 |
| **RTF1** | 5.12549 | 9.90905 | 4.16880 | 10.85142 | 7.30268 | 8.30071 | 19.51682 |
| **RTFDC1** | 27.05532 | 19.04789 | 12.87026 | 21.12946 | 6.60384 | 9.54856 | 0.00000 |

| | | | | | | | |
|---|---|---|---|---|---|---|---|
| **RTKN** | 13.50662 | 18.23872 | 0.00000 | 13.12796 | 0.00000 | 9.81734 | 12.02301 |
| **RTKN2** | 5.97735 | 0.00000 | 1.15828 | 5.82766 | 0.86632 | 0.59994 | 4.75794 |
| **RTL1** | 0.00568 | 0.01358 | 0.00889 | 0.01478 | 0.00759 | 0.82624 | 0.02655 |
| **RTN1** | 0.02759 | 0.00000 | 0.06396 | 0.01830 | 0.06200 | 0.06860 | 0.04099 |
| **RTN2** | 0.00000 | 0.55939 | 0.80849 | 1.45963 | 1.00209 | 0.63583 | 2.01677 |
| **RTN3** | 24.72750 | 4.74004 | 12.79742 | 18.71783 | 6.21962 | 6.26135 | 6.65526 |
| **RTN4** | 4.28738 | 0.00000 | 2.87835 | 9.71500 | 2.02376 | 15.19929 | 24.13508 |
| **RTN4IP1** | 8.62926 | 3.34597 | 3.57480 | 5.38999 | 2.79217 | 1.00790 | 0.00000 |
| **RTN4R** | 0.00000 | 0.88485 | 0.10249 | 0.32509 | 0.40764 | 0.00000 | 0.80709 |
| **RTN4RL1** | 0.02209 | 0.01718 | 0.02945 | 0.01103 | 0.01026 | 0.12895 | 0.03466 |
| **RTN4RL2** | 0.01967 | 0.00000 | 0.04899 | 0.11943 | 0.08740 | 0.02125 | 0.24109 |
| **RTP1** | 0.18543 | 0.80644 | 0.68262 | 0.00000 | 0.54581 | 1.35259 | 0.00000 |
| **RTP2** | 0.02594 | 0.04821 | 0.01459 | 0.02165 | 0.01827 | 0.02644 | 0.00656 |
| **RTP3** | 0.06251 | 4.23956 | 4.19379 | 0.00000 | 0.00000 | 4.89225 | 0.00000 |
| **RTTN** | 2.51053 | 0.38570 | 1.46882 | 2.48704 | 2.11773 | 1.16669 | 4.67124 |
| **RUFY1** | 6.41334 | 6.74626 | 7.17790 | 2.82151 | 1.50289 | 5.46180 | 4.95664 |
| **RUFY2** | 2.42990 | 0.00000 | 2.67972 | 0.63882 | 1.30926 | 1.24924 | 3.88893 |
| **RUFY3** | 7.81974 | 3.06280 | 6.22370 | 3.55292 | 0.14412 | 6.85397 | 8.12154 |
| **RUNDC1** | 1.68446 | 5.27083 | 2.04815 | 3.07306 | 0.43738 | 2.09675 | 2.18363 |
| **RUNDC3A** | 0.03263 | 0.36888 | 0.04973 | 0.15813 | 0.27538 | 0.34512 | 0.09027 |
| **RUNDC3B** | 0.05695 | 0.00832 | 0.04712 | 0.11343 | 0.69194 | 0.02820 | 0.01000 |
| **RUNX1** | 0.12838 | 0.61114 | 0.01538 | 4.37259 | 0.07655 | 2.56628 | 7.54635 |
| **RUNX1T1** | 0.00000 | 0.94484 | 4.15099 | 0.00000 | 3.12437 | 5.90229 | 0.37233 |
| **RUNX2** | 0.00786 | 0.02054 | 0.02643 | 0.02452 | 0.57803 | 0.02346 | 0.17216 |
| **RUNX3** | 0.03453 | 0.01133 | 0.01006 | 0.01001 | 0.03158 | 0.20074 | 0.03364 |
| **RUSC1** | 9.24700 | 6.50576 | 4.85350 | 14.18339 | 0.10709 | 8.96094 | 8.34906 |
| **RUSC1-AS1** | 0.03666 | 0.01865 | 0.02351 | 0.03632 | 0.00720 | 0.02577 | 0.00162 |
| **RUSC2** | 0.00000 | 0.00000 | 1.98263 | 0.26556 | 0.44766 | 1.63017 | 0.19924 |
| **RUVBL1** | 30.35048 | 19.28896 | 14.50929 | 23.35101 | 0.33281 | 13.27371 | 3.31219 |
| **RUVBL2** | 28.53047 | 34.32191 | 9.80455 | 25.19035 | 2.70939 | 13.61343 | 7.68650 |
| **RWDD1** | 14.10027 | 5.48051 | 6.86128 | 7.95981 | 2.68642 | 7.26630 | 18.77115 |
| **RWDD2A** | 2.04516 | 0.00000 | 1.64601 | 1.35108 | 0.09802 | 0.32636 | 0.00000 |
| **RWDD2B** | 1.06858 | 0.23344 | 0.68285 | 0.70285 | 0.10453 | 0.47524 | 0.03477 |
| **RWDD3** | 3.79728 | 0.66288 | 3.22649 | 2.40006 | 0.06869 | 1.04518 | 0.00000 |
| **RWDD4** | 2.49890 | 0.00000 | 2.46782 | 1.11828 | 0.03184 | 1.08325 | 2.14245 |

| | | | | | | | |
|---|---|---|---|---|---|---|---|
| **RXFP1** | 0.03533 | 0.00000 | 0.09021 | 0.00605 | 0.01554 | 0.04057 | 0.06960 |
| **RXRA** | 0.75584 | 0.96564 | 0.00634 | 10.85341 | 0.00000 | 0.00000 | 8.70612 |
| **RXRB** | 0.77287 | 1.38737 | 0.77786 | 2.95813 | 0.31472 | 0.76272 | 4.80353 |
| **RXRG** | 0.03175 | 0.01081 | 0.01346 | 0.01378 | 0.00759 | 0.01317 | 0.03833 |
| **RYBP** | 10.26127 | 2.60609 | 5.15000 | 11.72744 | 3.03902 | 6.91846 | 30.07862 |
| **RYK** | 1.74793 | 1.08001 | 3.04779 | 2.20504 | 0.00000 | 1.28176 | 7.26016 |
| **RYR1** | 0.91087 | 0.86685 | 2.22626 | 0.00000 | 0.34618 | 1.06360 | 6.93004 |
| **RYR2** | 0.02145 | 0.57027 | 0.41156 | 0.04880 | 1.12026 | 1.08039 | 0.01640 |
| **RYR3** | 0.42037 | 0.06881 | 0.08019 | 0.44700 | 1.27053 | 0.34328 | 0.26296 |
| **S100A1** | 0.04273 | 0.07048 | 0.04673 | 0.02707 | 0.00434 | 0.07167 | 0.19047 |
| **S100A10** | 60.87378 | 38.24355 | 9.05422 | 31.30443 | 0.00000 | 14.79232 | 48.99441 |
| **S100A11** | 34.55481 | 25.91786 | 21.56637 | 57.66568 | 0.76728 | 23.52352 | 22.32859 |
| **S100A12** | 0.21944 | 0.00505 | 0.02161 | 0.05228 | 0.00689 | 0.00789 | 0.00000 |
| **S100A13** | 53.20777 | 15.03083 | 0.00000 | 24.99844 | 0.07193 | 25.33265 | 62.95224 |
| **S100A14** | 33.60430 | 47.17105 | 0.00000 | 26.71351 | 0.00000 | 28.96890 | 11.63817 |
| **S100A16** | 71.71291 | 64.37875 | 2.64491 | 20.08038 | 0.00000 | 13.69721 | 31.91813 |
| **S100A2** | 0.68522 | 0.83446 | 0.02952 | 0.01259 | 0.01818 | 0.04687 | 0.37398 |
| **S100A3** | 0.07981 | 0.01593 | 0.09858 | 0.03814 | 0.00513 | 0.02940 | 0.00000 |
| **S100A4** | 1.06603 | 0.00000 | 0.09154 | 0.89405 | 0.33896 | 0.00000 | 0.23612 |
| **S100A5** | 0.04226 | 0.00000 | 0.12484 | 0.03023 | 0.01707 | 0.01154 | 0.00399 |
| **S100A6** | 47.46352 | 0.20483 | 0.00000 | 30.86745 | 0.00000 | 0.00000 | 28.08472 |
| **S100A7** | 0.02074 | 0.01963 | 0.01362 | 0.03056 | 0.00788 | 0.01579 | 0.01612 |
| **S100A8** | 0.01212 | 0.04607 | 0.02411 | 0.02484 | 0.00924 | 0.03191 | 0.02134 |
| **S100A9** | 0.19496 | 0.00810 | 0.01415 | 0.18239 | 0.00622 | 0.00870 | 0.20870 |
| **S100B** | 0.04969 | 0.00808 | 0.03740 | 0.01619 | 0.04212 | 0.01334 | 0.00414 |
| **S100P** | 15.93729 | 0.00000 | 0.00000 | 12.36786 | 0.00000 | 5.12687 | 16.02001 |
| **S100PBP** | 9.31035 | 4.32195 | 6.98088 | 3.13618 | 2.71463 | 6.27106 | 2.02518 |
| **S100Z** | 0.01027 | 0.16265 | 0.41719 | 0.00000 | 0.45242 | 1.34076 | 0.00920 |
| **S1PR1** | 0.00581 | 0.01378 | 0.03027 | 0.38022 | 0.04625 | 0.01458 | 0.00156 |
| **S1PR2** | 2.12849 | 0.34599 | 0.34910 | 27.56955 | 0.11194 | 0.01722 | 0.00000 |
| **S1PR3** | 0.70924 | 0.32413 | 0.98515 | 0.00000 | 0.07744 | 1.30768 | 0.49512 |
| **S1PR5** | 0.00000 | 0.37096 | 0.11503 | 0.00898 | 0.03029 | 0.22345 | 0.02675 |
| **SAA1** | 0.00000 | 0.00228 | 0.35542 | 0.57277 | 0.72059 | 0.00000 | 0.03234 |
| **SAA2.uid-15279** | 0.02885 | 0.01688 | 0.01365 | 0.01951 | 0.00793 | 0.01362 | 0.01338 |
| **SAA2.uid-15282** | 0.00134 | 0.01278 | 0.01622 | 0.11131 | 0.01245 | 0.00953 | 0.05284 |

| | | | | | | | |
|---|---|---|---|---|---|---|---|
| **SAA3P** | 0.01751 | 0.01406 | 0.02008 | 0.03359 | 0.01489 | 0.01301 | 0.02489 |
| **SAAL1** | 14.56254 | 6.37184 | 8.03947 | 5.30319 | 2.30264 | 5.13512 | 0.00000 |
| **SAC3D1** | 0.28065 | 1.73818 | 2.80966 | 2.48610 | 1.29931 | 1.00512 | 3.79180 |
| **SACM1L** | 11.70234 | 2.80463 | 2.70406 | 3.50681 | 1.83571 | 4.55956 | 4.88264 |
| **SACS** | 3.05057 | 0.87794 | 5.65988 | 2.23420 | 2.06110 | 0.70193 | 9.44412 |
| **SACS-AS1** | 0.00000 | 0.44268 | 1.53881 | 0.54407 | 0.43965 | 0.00000 | 0.00000 |
| **SAE1** | 32.17657 | 16.18152 | 11.66007 | 29.07141 | 8.61828 | 14.56143 | 11.55217 |
| **SAFB** | 21.33490 | 27.73124 | 11.45799 | 23.57876 | 7.62139 | 12.26029 | 10.33650 |
| **SAFB2** | 13.13199 | 12.99657 | 7.17158 | 12.49589 | 3.60506 | 7.56461 | 18.61483 |
| **SAG** | 0.17030 | 0.18348 | 0.12458 | 0.01741 | 0.06433 | 0.01120 | 0.00000 |
| **SAGE1** | 0.03588 | 0.01405 | 0.02059 | 0.01133 | 0.00737 | 0.02626 | 0.03378 |
| **SALL1** | 0.00878 | 0.00000 | 0.33797 | 0.06673 | 0.09783 | 1.38611 | 0.01715 |
| **SALL2** | 0.00000 | 0.00000 | 0.51859 | 0.01104 | 0.20351 | 0.34192 | 0.07622 |
| **SALL3** | 0.29722 | 0.95902 | 0.34790 | 0.00000 | 0.80434 | 1.46667 | 0.00000 |
| **SALL4** | 24.89507 | 32.50915 | 18.02418 | 25.39979 | 8.46166 | 15.06396 | 8.26864 |
| **SAMD1** | 0.58813 | 4.99446 | 1.38221 | 1.40666 | 0.74244 | 3.50272 | 9.54383 |
| **SAMD10** | 0.08841 | 0.00000 | 0.08003 | 0.19039 | 0.01660 | 0.00000 | 0.12906 |
| **SAMD11** | 0.00000 | 0.07285 | 0.07608 | 0.06386 | 0.03889 | 0.86975 | 0.21966 |
| **SAMD12** | 0.57938 | 0.01112 | 0.00000 | 0.46054 | 3.19974 | 0.33205 | 0.25010 |
| **SAMD12-AS1** | 0.02042 | 0.02068 | 0.00848 | 0.03608 | 0.04155 | 0.02881 | 0.01421 |
| **SAMD13** | 0.05197 | 0.20654 | 0.04449 | 0.01802 | 0.02545 | 0.03711 | 0.00000 |
| **SAMD14** | 0.01382 | 0.01637 | 0.01229 | 0.02058 | 0.20134 | 0.06763 | 0.05445 |
| **SAMD15** | 0.31439 | 0.12598 | 0.58590 | 0.09340 | 0.17201 | 0.01417 | 0.09945 |
| **SAMD3** | 0.03984 | 0.01500 | 0.00000 | 0.01164 | 0.92784 | 0.00948 | 0.02590 |
| **SAMD4A** | 0.07251 | 0.03988 | 0.00000 | 0.10249 | 1.13970 | 0.00000 | 0.18649 |
| **SAMD4B** | 1.49588 | 3.35277 | 3.60313 | 10.40715 | 0.35793 | 5.20271 | 14.91168 |
| **SAMD5** | 0.02140 | 0.00000 | 0.29723 | 0.06788 | 0.00423 | 0.72034 | 0.08731 |
| **SAMD7** | 0.03744 | 0.08140 | 0.00766 | 0.01339 | 0.08789 | 0.00412 | 0.02986 |
| **SAMD8** | 6.67732 | 0.72573 | 23.42443 | 9.27443 | 7.83431 | 4.86489 | 7.89178 |
| **SAMD9** | 0.01449 | 0.01695 | 0.00620 | 0.03426 | 0.10045 | 0.01486 | 0.02445 |
| **SAMD9L** | 0.01536 | 0.02339 | 0.01122 | 0.01882 | 0.00794 | 0.10078 | 0.01871 |
| **SAMHD1** | 7.66201 | 4.24526 | 0.41087 | 2.46532 | 4.77343 | 14.28523 | 0.00000 |
| **SAMM50** | 17.26918 | 19.42917 | 7.53433 | 15.07987 | 0.13053 | 10.47523 | 7.89083 |
| **SAMSN1** | 0.02779 | 0.01570 | 0.01320 | 0.02257 | 0.00794 | 0.01434 | 0.01611 |
| **SAP130** | 9.40050 | 6.83314 | 4.98645 | 16.00568 | 3.81709 | 7.12100 | 11.33447 |

| | | | | | | | |
|---|---|---|---|---|---|---|---|
| **SAP18** | 43.50034 | 23.62367 | 28.58788 | 29.72382 | 20.36914 | 26.41001 | 25.98600 |
| **SAP25** | 0.00000 | 1.47758 | 1.39077 | 0.11475 | 0.07022 | 0.32312 | 1.03548 |
| **SAP30** | 0.00000 | 0.06867 | 1.38148 | 0.02457 | 0.33629 | 1.12525 | 2.18338 |
| **SAP30BP** | 23.77598 | 22.24774 | 12.20340 | 25.68633 | 5.44588 | 7.80931 | 5.38693 |
| **SAP30L** | 0.74885 | 0.00000 | 0.63001 | 1.27407 | 0.02252 | 1.72823 | 8.03188 |
| **SAP30L-AS1** | 2.19878 | 0.93669 | 0.00000 | 0.00000 | 0.02577 | 0.00000 | 2.71089 |
| **SAPCD2** | 4.26362 | 9.21664 | 0.00000 | 0.49205 | 0.00000 | 5.86186 | 29.45766 |
| **SAR1A** | 46.65469 | 19.28713 | 21.66119 | 29.96238 | 12.05141 | 15.68155 | 12.54706 |
| **SAR1B** | 17.36869 | 6.10894 | 8.92702 | 8.48617 | 2.13488 | 7.33114 | 2.69470 |
| **SARDH** | 0.00000 | 0.13821 | 0.00324 | 0.05712 | 0.77051 | 0.16478 | 0.00000 |
| **SARM1** | 0.10388 | 0.07927 | 0.10344 | 0.18314 | 0.21074 | 0.06333 | 0.31980 |
| **SARNP** | 18.79256 | 8.96916 | 13.33883 | 15.29390 | 6.11694 | 10.08983 | 9.95858 |
| **SARS** | 39.90188 | 25.31483 | 15.33985 | 41.45929 | 13.90910 | 16.40317 | 12.60010 |
| **SARS2** | 9.04932 | 7.04453 | 2.35905 | 6.95047 | 0.04437 | 4.85599 | 5.78037 |
| **SART1** | 2.04462 | 9.56759 | 2.89762 | 7.57933 | 1.62306 | 5.32164 | 12.66317 |
| **SART3** | 20.01429 | 9.92589 | 8.67254 | 12.92437 | 3.11458 | 7.96461 | 6.34192 |
| **SASH1** | 0.68885 | 6.56899 | 4.44010 | 0.62280 | 0.17714 | 5.27050 | 3.52537 |
| **SASH3** | 0.00486 | 0.02392 | 0.02624 | 0.01141 | 0.00780 | 0.04104 | 0.10180 |
| **SASS6** | 9.07060 | 0.00095 | 3.15947 | 5.68890 | 8.40039 | 2.48258 | 9.26019 |
| **SAT1** | 3.85601 | 3.85657 | 14.28706 | 1.47801 | 0.74442 | 31.53368 | 0.00000 |
| **SAT2** | 0.40632 | 0.00000 | 1.96522 | 2.27281 | 0.45819 | 2.95916 | 0.12103 |
| **SATB1** | 1.29100 | 0.00000 | 0.17100 | 4.69780 | 0.43143 | 0.00000 | 1.79685 |
| **SATB2** | 6.82604 | 1.72429 | 2.79000 | 0.00000 | 5.06919 | 0.00000 | 3.90857 |
| **SATB2-AS1** | 0.02989 | 0.00061 | 0.07420 | 0.01655 | 0.18221 | 0.01029 | 0.02106 |
| **SATL1** | 0.03759 | 0.01214 | 0.01146 | 0.02511 | 0.02539 | 0.02527 | 0.01451 |
| **SAV1** | 3.63406 | 5.65559 | 2.96500 | 1.85461 | 0.68536 | 10.19015 | 15.89892 |
| **SAYSD1** | 3.14425 | 3.69791 | 3.96454 | 3.39820 | 1.50358 | 2.87979 | 0.00000 |
| **SBDS** | 27.43514 | 15.73929 | 9.79601 | 15.98489 | 11.43317 | 9.75266 | 3.26646 |
| **SBDSP1** | 13.66862 | 7.98173 | 8.52315 | 7.89940 | 8.71289 | 5.05398 | 1.30100 |
| **SBF1** | 0.00000 | 7.24603 | 0.49157 | 4.81798 | 1.15724 | 3.16052 | 14.42710 |
| **SBF1P1** | 0.00885 | 0.34386 | 0.03151 | 0.08652 | 0.04021 | 0.08014 | 0.63593 |
| **SBF2** | 2.21059 | 1.08800 | 1.66184 | 2.56535 | 0.96680 | 1.28502 | 2.32307 |
| **SBF2-AS1** | 3.41043 | 3.37927 | 1.30006 | 0.71477 | 0.05695 | 0.00000 | 1.32797 |
| **SBK1** | 0.00000 | 0.26228 | 0.00864 | 0.10093 | 0.31367 | 0.09603 | 0.10961 |
| **SBK3** | 0.02245 | 0.01381 | 0.00790 | 0.01832 | 0.21309 | 0.01308 | 0.01940 |

| | | | | | | | |
|---|---|---|---|---|---|---|---|
| **SBNO1** | 35.22034 | 11.15627 | 18.28656 | 25.47274 | 14.68663 | 11.30200 | 20.34925 |
| **SBNO2** | 0.00000 | 0.50356 | 0.07096 | 0.36712 | 0.29429 | 0.64839 | 4.12787 |
| **SBSN** | 0.01998 | 0.00910 | 0.03209 | 0.27975 | 0.23204 | 0.01954 | 0.00000 |
| **SBSPON** | 1.16907 | 0.00000 | 0.05524 | 2.12593 | 0.07438 | 0.92859 | 0.00000 |
| **SC5D** | 23.48859 | 0.00000 | 0.76425 | 15.22806 | 1.00505 | 2.49091 | 0.66054 |
| **SCAF1** | 0.00000 | 0.19788 | 0.86135 | 0.93904 | 0.28175 | 0.84565 | 10.30440 |
| **SCAF11** | 21.07345 | 14.10212 | 14.43025 | 12.85596 | 6.69202 | 9.15735 | 17.72094 |
| **SCAF4** | 10.54865 | 8.90744 | 5.11205 | 9.55062 | 8.16427 | 8.24662 | 12.74349 |
| **SCAF8** | 7.26325 | 0.95273 | 6.82808 | 1.10296 | 4.21428 | 5.94808 | 8.20569 |
| **SCAI** | 3.10219 | 0.00000 | 1.66825 | 0.68665 | 2.30242 | 0.86191 | 3.74483 |
| **SCAMP1** | 11.84672 | 0.87865 | 11.04582 | 0.00000 | 2.04179 | 15.31331 | 5.32629 |
| **SCAMP1-AS1** | 0.47552 | 1.06597 | 0.74093 | 0.37008 | 0.46147 | 0.19389 | 0.00000 |
| **SCAMP2** | 8.53448 | 11.40708 | 4.12640 | 6.47435 | 1.24898 | 3.74332 | 1.71240 |
| **SCAMP3** | 23.93675 | 13.72321 | 5.67117 | 24.41493 | 2.42921 | 9.44745 | 0.93929 |
| **SCAMP4** | 0.00000 | 4.86301 | 4.72476 | 6.21551 | 0.14970 | 1.64528 | 15.58325 |
| **SCAMP5** | 0.00000 | 0.66676 | 0.54542 | 3.26978 | 0.26404 | 2.47593 | 0.30936 |
| **SCAND1** | 2.48586 | 13.75232 | 6.31330 | 13.95115 | 6.03331 | 5.89028 | 17.06232 |
| **SCAND2P** | 0.79196 | 0.00000 | 0.90705 | 1.66038 | 2.08544 | 0.95636 | 1.47831 |
| **SCAP** | 0.37935 | 13.08972 | 3.66861 | 9.63413 | 3.88455 | 8.74920 | 20.91683 |
| **SCAPER** | 1.55560 | 1.24861 | 1.84559 | 2.30366 | 1.89265 | 3.39268 | 4.83745 |
| **SCARA3** | 0.99866 | 0.45274 | 0.00000 | 0.17085 | 0.00900 | 0.00000 | 1.46992 |
| **SCARA5** | 0.78458 | 0.00000 | 0.00190 | 0.00000 | 1.00633 | 0.79208 | 0.04098 |
| **SCARB1** | 3.39801 | 0.00000 | 0.10714 | 2.64723 | 0.03181 | 2.15765 | 4.26652 |
| **SCARB2** | 8.47707 | 0.74007 | 2.05663 | 1.04199 | 1.11263 | 2.97296 | 12.55005 |
| **SCARF1** | 0.08937 | 0.00000 | 0.03568 | 0.00000 | 0.00462 | 0.02216 | 0.29398 |
| **SCARF2** | 0.00000 | 0.12288 | 0.07057 | 0.04738 | 0.32254 | 0.06417 | 0.04012 |
| **SCARNA1** | 0.05290 | 0.00000 | 0.13040 | 0.12732 | 0.04507 | 0.06450 | 0.35541 |
| **SCARNA10** | 0.00000 | 0.01209 | 0.02851 | 0.01081 | 0.15513 | 0.08047 | 0.18584 |
| **SCARNA11** | 0.01916 | 0.01628 | 0.01366 | 0.03759 | 0.00783 | 0.01212 | 0.02461 |
| **SCARNA12** | 0.61242 | 1.44213 | 0.23027 | 1.56799 | 0.68945 | 1.49817 | 1.74693 |
| **SCARNA14** | 0.01802 | 0.00338 | 0.02546 | 0.07632 | 0.02593 | 0.01751 | 0.13018 |
| **SCARNA16** | 0.07236 | 0.15794 | 0.04341 | 0.03247 | 0.03969 | 0.02207 | 0.06869 |
| **SCARNA17** | 0.06362 | 0.17515 | 0.00000 | 0.00000 | 0.10472 | 0.00461 | 0.38497 |
| **SCARNA18** | 0.00289 | 0.01347 | 0.06921 | 0.10054 | 0.08403 | 0.02977 | 0.15029 |
| **SCARNA2** | 0.01284 | 0.14203 | 0.04260 | 0.00818 | 0.27587 | 0.08155 | 1.24419 |

| | | | | | | | |
|---|---|---|---|---|---|---|---|
| **SCARNA20** | 0.02436 | 0.01859 | 0.02867 | 0.04582 | 0.01211 | 0.03334 | 0.03928 |
| **SCARNA21** | 0.03472 | 0.01175 | 0.00947 | 0.00771 | 0.05734 | 0.02752 | 0.05778 |
| **SCARNA22** | 0.00000 | 1.49235 | 1.05127 | 0.22850 | 2.50947 | 0.00000 | 1.94340 |
| **SCARNA23** | 0.01997 | 0.01612 | 0.03153 | 0.03472 | 0.03888 | 0.02698 | 0.09388 |
| **SCARNA27** | 0.03501 | 0.00808 | 0.03117 | 0.03792 | 0.01097 | 0.05444 | 0.07949 |
| **SCARNA3** | 0.00000 | 0.04062 | 0.03014 | 0.05001 | 0.01497 | 0.07574 | 0.11233 |
| **SCARNA4** | 0.00587 | 0.02725 | 0.02174 | 0.09485 | 0.02423 | 0.06910 | 0.05752 |
| **SCARNA5** | 0.03644 | 0.08029 | 0.15708 | 0.00000 | 0.29881 | 0.16229 | 0.42243 |
| **SCARNA6** | 0.03268 | 0.02104 | 0.02125 | 0.02221 | 0.21755 | 0.02210 | 0.01134 |
| **SCARNA7** | 0.02099 | 0.01013 | 0.01587 | 0.02559 | 0.01779 | 0.01509 | 0.05653 |
| **SCARNA8** | 0.03018 | 0.01173 | 0.04052 | 0.06561 | 0.02625 | 0.00814 | 0.08405 |
| **SCARNA9** | 0.07306 | 0.00000 | 0.02299 | 0.00000 | 0.01967 | 0.12541 | 0.71687 |
| **SCARNA9L** | 0.02140 | 0.04306 | 0.05402 | 0.03132 | 0.08729 | 0.15195 | 0.46191 |
| **SCART1** | 0.10370 | 0.10080 | 0.02937 | 0.14091 | 0.06148 | 0.03918 | 0.05997 |
| **SCCPDH** | 1.04286 | 0.17021 | 0.26121 | 3.04275 | 0.33775 | 0.99325 | 0.93303 |
| **SCD** | 67.41373 | 12.00810 | 25.84178 | 56.50248 | 2.63669 | 18.44284 | 41.52859 |
| **SCD5** | 0.01042 | 0.02874 | 0.05444 | 0.37992 | 0.01191 | 0.03477 | 0.42788 |
| **SCEL** | 0.03548 | 0.00000 | 0.04214 | 0.01768 | 0.04588 | 0.02227 | 0.01600 |
| **SCFD1** | 18.99526 | 2.96004 | 7.11559 | 7.46821 | 1.05245 | 4.65480 | 0.00000 |
| **SCFD2** | 6.76442 | 2.49116 | 2.34098 | 3.48709 | 1.86733 | 2.83068 | 1.33941 |
| **SCG2** | 0.02443 | 0.00368 | 0.40019 | 0.04239 | 1.31506 | 0.01434 | 0.00093 |
| **SCG3** | 0.08499 | 0.02062 | 0.37951 | 0.04763 | 0.00538 | 0.21723 | 0.01769 |
| **SCG5** | 0.08284 | 0.08326 | 0.01851 | 0.14260 | 0.06433 | 0.35065 | 0.20630 |
| **SCGB1A1** | 0.02331 | 0.08160 | 0.00978 | 0.01929 | 0.00862 | 0.06307 | 0.01113 |
| **SCGB1B2P** | 0.02760 | 0.05581 | 0.05785 | 0.06152 | 0.02473 | 0.04472 | 0.00000 |
| **SCGB1D1** | 0.26589 | 0.17601 | 0.53199 | 0.08317 | 0.00144 | 0.00182 | 0.02290 |
| **SCGB1D2** | 0.01024 | 0.05322 | 0.00857 | 0.03774 | 0.00831 | 0.17989 | 0.01228 |
| **SCGB1D4** | 0.00974 | 0.06471 | 0.04905 | 0.01620 | 0.00516 | 0.08162 | 0.01719 |
| **SCGB2A1** | 0.02751 | 0.06590 | 0.00965 | 0.00644 | 0.01931 | 0.04953 | 0.04920 |
| **SCGB2A2** | 0.02090 | 0.01856 | 0.01309 | 0.09969 | 0.00853 | 0.01311 | 0.00038 |
| **SCGB3A2** | 3.05685 | 19.42428 | 6.23170 | 0.00000 | 2.04322 | 15.56046 | 0.00000 |
| **SCGN** | 0.01344 | 0.01430 | 0.03344 | 0.02398 | 0.03912 | 0.01517 | 0.03410 |
| **SCHIP1.uid-546** | 0.03062 | 0.01978 | 0.02438 | 0.02820 | 0.00701 | 0.08448 | 0.01357 |
| **SCHLAP1** | 0.17258 | 0.02947 | 0.01399 | 0.08829 | 0.01231 | 0.06988 | 0.03705 |
| **SCIMP** | 0.09529 | 0.10556 | 0.08434 | 0.00086 | 0.01140 | 0.00125 | 0.01170 |

| | | | | | | | |
|---|---|---|---|---|---|---|---|
| **SCIN** | 0.90711 | 0.40886 | 0.03374 | 0.03709 | 0.76239 | 0.06173 | 0.00000 |
| **SCLT1** | 3.69026 | 2.24484 | 1.02081 | 1.70246 | 1.66332 | 0.96904 | 1.67865 |
| **SCLY** | 0.06644 | 0.05130 | 0.04177 | 0.17476 | 0.02781 | 0.11552 | 0.00000 |
| **SCMH1** | 0.26374 | 0.18271 | 0.12352 | 0.70012 | 0.06260 | 0.32895 | 0.47205 |
| **SCML1** | 0.99977 | 1.81062 | 0.28147 | 0.00000 | 3.41012 | 1.22183 | 0.28328 |
| **SCML2** | 3.97518 | 0.67036 | 9.84083 | 0.35088 | 2.23045 | 1.40244 | 0.72340 |
| **SCML4** | 0.02887 | 0.01233 | 0.13686 | 0.00661 | 0.03248 | 0.62697 | 0.04149 |
| **SCN10A** | 0.03778 | 0.03119 | 0.04780 | 0.01679 | 0.00546 | 0.03984 | 0.01739 |
| **SCN11A** | 0.05353 | 0.14113 | 0.03810 | 0.00986 | 0.10600 | 0.23194 | 0.08796 |
| **SCN1A** | 0.03919 | 0.00000 | 0.13675 | 0.01424 | 0.59744 | 0.09412 | 0.02341 |
| **SCN1B** | 0.00000 | 0.02805 | 0.05121 | 0.06222 | 0.03412 | 0.10745 | 0.05939 |
| **SCN2A** | 0.57204 | 0.16409 | 0.35441 | 0.21174 | 0.05516 | 0.54056 | 0.68232 |
| **SCN2B** | 0.04733 | 0.01335 | 0.01729 | 0.01727 | 0.00742 | 0.02722 | 0.01582 |
| **SCN3A** | 0.35405 | 0.31317 | 1.07165 | 0.19172 | 0.13189 | 0.34409 | 0.24416 |
| **SCN3B** | 0.40278 | 0.72329 | 0.09496 | 0.14473 | 0.00565 | 0.00000 | 1.24118 |
| **SCN4A** | 0.04846 | 0.00281 | 0.01586 | 0.07711 | 0.07949 | 0.03606 | 0.01626 |
| **SCN4B** | 0.03049 | 0.05848 | 0.09919 | 0.00000 | 0.00553 | 0.00000 | 0.14156 |
| **SCN5A** | 0.01897 | 0.26135 | 0.00861 | 0.25670 | 0.80242 | 0.64185 | 0.02392 |
| **SCN7A** | 0.04289 | 0.00000 | 0.03974 | 0.01347 | 0.28726 | 0.01547 | 0.04405 |
| **SCN8A** | 0.49116 | 0.43234 | 0.85428 | 1.49077 | 0.58107 | 1.65629 | 2.20342 |
| **SCN9A** | 0.02050 | 0.02572 | 0.00512 | 0.02550 | 0.16793 | 0.04596 | 0.02264 |
| **SCNM1** | 4.18604 | 2.89397 | 2.95788 | 4.35739 | 5.51611 | 1.54537 | 0.00000 |
| **SCNN1A** | 0.81088 | 0.18637 | 0.05663 | 0.00000 | 0.00154 | 5.74893 | 0.70609 |
| **SCNN1B** | 0.02412 | 0.02949 | 0.00928 | 0.02385 | 0.04552 | 0.01140 | 0.02847 |
| **SCNN1D** | 0.00985 | 0.19507 | 0.00823 | 0.14494 | 0.01292 | 0.01287 | 0.02777 |
| **SCNN1G** | 0.03465 | 0.00621 | 0.03016 | 0.02945 | 0.01337 | 0.05083 | 0.01021 |
| **SCO1** | 29.12683 | 10.88933 | 14.44846 | 15.04821 | 7.37308 | 9.07002 | 0.00501 |
| **SCO2.uid-26107** | 1.26606 | 2.36838 | 0.32813 | 0.64680 | 0.04335 | 0.40512 | 0.00000 |
| **SCOC** | 9.45185 | 0.00000 | 8.29565 | 2.64162 | 1.78679 | 5.43418 | 2.56834 |
| **SCOC-AS1** | 0.12721 | 0.39660 | 0.54856 | 0.00000 | 0.06432 | 0.14758 | 0.00782 |
| **SCP2** | 20.81315 | 2.67777 | 6.59070 | 4.98200 | 0.93756 | 7.09263 | 4.66931 |
| **SCPEP1** | 26.91572 | 15.34234 | 8.46366 | 1.11932 | 0.00000 | 6.93133 | 0.00000 |
| **SCRG1** | 0.02159 | 0.01417 | 0.04227 | 0.02010 | 0.02415 | 0.01187 | 0.01850 |
| **SCRIB** | 0.00000 | 3.31941 | 0.48053 | 0.74387 | 0.09418 | 3.14978 | 13.08937 |
| **SCRN1** | 0.24459 | 2.07827 | 0.00000 | 0.48106 | 0.02104 | 6.11164 | 0.08003 |

| | | | | | | | |
|---|---|---|---|---|---|---|---|
| **SCRN2** | 2.01704 | 1.11182 | 0.00000 | 3.60885 | 0.08912 | 0.57105 | 0.63294 |
| **SCRN3** | 2.47206 | 0.37290 | 1.78758 | 0.57460 | 0.18048 | 1.02199 | 0.13087 |
| **SCRT1** | 0.01593 | 0.04548 | 0.02317 | 0.01887 | 0.02098 | 0.06777 | 0.03288 |
| **SCRT2** | 0.02274 | 0.02447 | 0.02093 | 0.02093 | 0.03039 | 0.01225 | 0.01285 |
| **SCTR** | 0.01428 | 0.22534 | 0.02384 | 0.06389 | 0.00742 | 0.04203 | 0.00632 |
| **SCUBE1** | 0.00000 | 0.13517 | 0.02666 | 0.01592 | 0.42542 | 0.00358 | 0.05599 |
| **SCUBE2** | 0.00000 | 1.91204 | 0.66613 | 0.00000 | 0.15183 | 1.66757 | 0.40513 |
| **SCUBE3** | 0.33413 | 0.00000 | 0.05523 | 1.16111 | 0.00218 | 0.40951 | 1.34363 |
| **SCYL1** | 6.20704 | 10.23056 | 5.52180 | 11.88138 | 2.02735 | 5.93902 | 3.02379 |
| **SCYL2** | 7.83542 | 0.00000 | 5.80287 | 6.35522 | 1.66050 | 4.41554 | 6.76170 |
| **SCYL3** | 2.70877 | 0.64102 | 1.32170 | 1.87471 | 2.87721 | 0.72871 | 0.46429 |
| **SDAD1** | 19.35153 | 4.32776 | 13.13338 | 7.11533 | 4.78924 | 7.00920 | 11.46230 |
| **SDC1** | 0.88612 | 0.88754 | 0.00000 | 24.53686 | 0.00105 | 0.00000 | 1.88112 |
| **SDC2** | 0.00000 | 0.31480 | 1.37888 | 0.00000 | 0.55665 | 0.58265 | 0.02663 |
| **SDC3** | 0.00000 | 0.04242 | 0.04510 | 0.47620 | 0.07569 | 0.21993 | 0.50131 |
| **SDC4** | 42.64384 | 35.79330 | 14.09234 | 24.79241 | 2.71543 | 16.27729 | 22.63152 |
| **SDC4P** | 0.02626 | 0.01747 | 0.01383 | 0.02156 | 0.00783 | 0.01326 | 0.01612 |
| **SDCBP** | 18.07483 | 8.90783 | 10.51821 | 19.64095 | 2.19567 | 24.34566 | 0.00000 |
| **SDCBP2-AS1** | 0.52876 | 0.00000 | 0.33899 | 0.37861 | 0.02505 | 0.42573 | 0.81937 |
| **SDCBP2.uid-24** | 0.02181 | 0.03628 | 0.01285 | 0.02105 | 0.00804 | 0.01097 | 0.01436 |
| **SDCCAG3** | 1.09998 | 4.71798 | 2.35654 | 5.77272 | 0.06951 | 4.15985 | 14.34249 |
| **SDCCAG8** | 2.90401 | 2.26857 | 0.63038 | 3.71802 | 3.94373 | 2.29860 | 1.85519 |
| **SDE2** | 11.95115 | 3.61684 | 8.88883 | 9.53399 | 12.74756 | 5.56445 | 0.98671 |
| **SDF2** | 10.21458 | 6.79153 | 6.85731 | 14.94746 | 1.44985 | 6.07656 | 0.44020 |
| **SDF2L1** | 16.45051 | 14.79779 | 1.84302 | 29.05435 | 1.52928 | 9.66660 | 13.69046 |
| **SDF4** | 0.00000 | 5.91226 | 1.60675 | 3.65535 | 0.66458 | 5.48394 | 15.09358 |
| **SDHA** | 24.65494 | 22.53152 | 6.05962 | 11.44767 | 2.93569 | 9.19921 | 0.00000 |
| **SDHAF1** | 0.73846 | 7.41829 | 2.49235 | 6.56881 | 3.15552 | 2.62319 | 6.52722 |
| **SDHAF2** | 20.49757 | 12.53456 | 8.77427 | 17.05985 | 1.68576 | 7.54254 | 4.30271 |
| **SDHAP1** | 1.88731 | 2.14989 | 0.54958 | 1.28445 | 1.15384 | 1.24531 | 0.42874 |
| **SDHAP2** | 1.09752 | 0.98004 | 0.31303 | 0.66476 | 0.39717 | 0.42152 | 0.00000 |
| **SDHAP3** | 0.74839 | 1.51420 | 1.27892 | 0.10395 | 0.12484 | 1.72577 | 0.15522 |
| **SDHB** | 48.82315 | 23.18537 | 16.84197 | 28.41470 | 6.12770 | 18.18606 | 11.68693 |
| **SDHC** | 13.85375 | 3.13764 | 5.11448 | 9.85361 | 0.28935 | 6.63273 | 0.00000 |
| **SDHD** | 33.30673 | 9.00354 | 10.78704 | 12.07000 | 1.05737 | 13.57779 | 0.00000 |

| | | | | | | | |
|---|---|---|---|---|---|---|---|
| **SDK1** | 0.00741 | 0.00000 | 0.01139 | 0.24918 | 0.89638 | 0.23965 | 0.61583 |
| **SDK2** | 0.00000 | 0.45136 | 0.04935 | 0.13393 | 2.41770 | 0.49238 | 0.33564 |
| **SDPR** | 0.00000 | 0.03075 | 0.00477 | 0.02775 | 0.25034 | 0.16262 | 0.04268 |
| **SDR16C5** | 0.03224 | 0.13709 | 0.41180 | 0.00982 | 0.00407 | 0.09704 | 0.00000 |
| **SDR39U1** | 3.08232 | 1.06541 | 0.40987 | 3.80021 | 0.11319 | 0.89401 | 0.00000 |
| **SDR42E1** | 1.00991 | 1.66068 | 1.12569 | 0.00000 | 0.56682 | 1.12554 | 0.00000 |
| **SDS** | 0.01805 | 0.02781 | 0.00680 | 0.05610 | 0.65252 | 0.04106 | 0.01369 |
| **SDSL** | 0.03697 | 0.57813 | 0.00000 | 0.00072 | 0.18378 | 0.01522 | 0.00709 |
| **SEBOX** | 0.00979 | 0.02691 | 0.01875 | 0.01438 | 0.01711 | 0.00836 | 0.12570 |
| **SEC11A** | 29.01533 | 7.89874 | 11.95250 | 30.20503 | 2.16608 | 7.07424 | 22.74836 |
| **SEC11C** | 14.34742 | 6.98338 | 2.36107 | 9.33291 | 0.03623 | 5.96676 | 5.26512 |
| **SEC13** | 24.52184 | 25.21852 | 16.35702 | 27.88686 | 9.57277 | 12.13531 | 5.28876 |
| **SEC14L1** | 1.23655 | 3.41094 | 4.26681 | 3.11489 | 5.52913 | 5.39053 | 2.84724 |
| **SEC14L1P1** | 0.12796 | 0.24894 | 0.28768 | 0.14620 | 0.54848 | 0.15864 | 0.18230 |
| **SEC14L2** | 0.09961 | 0.13887 | 0.33067 | 0.39793 | 0.16536 | 0.38294 | 0.48699 |
| **SEC14L4** | 0.11643 | 0.00000 | 0.63122 | 0.65510 | 0.00189 | 0.00000 | 0.55861 |
| **SEC14L5** | 0.03215 | 0.04986 | 0.03893 | 0.01018 | 0.46116 | 0.01626 | 0.23488 |
| **SEC14L6** | 0.02171 | 0.00837 | 0.10338 | 0.02369 | 0.03523 | 0.01277 | 0.01811 |
| **SEC16A** | 6.25436 | 9.00695 | 2.42084 | 7.39921 | 1.07292 | 2.52307 | 19.69868 |
| **SEC16B** | 0.06337 | 0.02503 | 0.01596 | 0.01327 | 0.45978 | 0.00338 | 0.06206 |
| **SEC1P** | 0.03714 | 0.00000 | 0.21063 | 0.02183 | 0.01745 | 0.01345 | 0.01609 |
| **SEC22A** | 6.43243 | 3.04821 | 1.77599 | 3.40179 | 2.80365 | 1.09048 | 0.00000 |
| **SEC22B** | 25.60499 | 8.16540 | 9.93258 | 13.69339 | 3.53086 | 11.76407 | 1.56513 |
| **SEC22C** | 4.53651 | 0.00000 | 0.52777 | 4.38876 | 0.41252 | 2.34648 | 1.43273 |
| **SEC23A** | 8.02094 | 2.31379 | 2.35471 | 3.79461 | 2.11239 | 10.96739 | 0.00000 |
| **SEC23B** | 38.66634 | 25.73742 | 14.09204 | 18.32988 | 4.67650 | 13.49090 | 0.00000 |
| **SEC23IP** | 13.54299 | 2.84317 | 6.11244 | 11.04093 | 1.77336 | 3.96103 | 0.00000 |
| **SEC24A** | 13.06211 | 0.15143 | 5.80294 | 7.19736 | 5.50630 | 5.11055 | 5.88409 |
| **SEC24B** | 7.65472 | 0.28300 | 7.42141 | 3.37234 | 2.78190 | 2.76178 | 11.65037 |
| **SEC24B-AS1** | 0.13960 | 0.00000 | 0.26206 | 0.16712 | 0.05120 | 0.09250 | 0.10397 |
| **SEC24C** | 10.95226 | 7.15085 | 11.33754 | 16.67281 | 4.15152 | 6.98517 | 8.83796 |
| **SEC24D** | 1.85638 | 0.00000 | 1.04093 | 2.49750 | 0.62431 | 2.97362 | 0.12996 |
| **SEC31A** | 25.72892 | 11.53278 | 8.75117 | 15.56335 | 3.69810 | 8.65823 | 8.31806 |
| **SEC31B** | 0.11318 | 0.17685 | 0.41313 | 1.31984 | 0.06345 | 0.41976 | 0.87322 |
| **SEC61A1** | 34.21741 | 27.81038 | 20.06426 | 34.71260 | 12.82277 | 15.98567 | 12.09244 |

| | | | | | | | |
|---|---|---|---|---|---|---|---|
| **SEC61A2** | 3.01917 | 1.11823 | 8.11312 | 1.68599 | 4.82601 | 1.95381 | 0.98602 |
| **SEC61B** | 23.04835 | 17.16916 | 18.26315 | 26.52438 | 6.58042 | 14.84641 | 18.38742 |
| **SEC61G** | 25.01713 | 10.87388 | 18.57338 | 21.45332 | 6.53275 | 24.41022 | 18.89171 |
| **SEC62** | 19.97376 | 6.57410 | 16.55252 | 13.11470 | 5.11070 | 15.06477 | 19.77903 |
| **SEC63** | 16.44671 | 10.92709 | 9.99016 | 7.55477 | 2.81222 | 18.32588 | 19.89285 |
| **SECISBP2** | 6.62431 | 6.24411 | 5.04569 | 5.42572 | 3.72802 | 6.33878 | 9.18918 |
| **SECISBP2L** | 4.90958 | 1.02577 | 2.33957 | 3.18951 | 2.20238 | 1.60912 | 7.36948 |
| **SECTM1** | 0.03365 | 0.01951 | 0.01207 | 0.01555 | 0.00954 | 0.01334 | 0.01545 |
| **SEH1L** | 44.68926 | 13.86317 | 15.15832 | 22.39639 | 10.04735 | 13.12829 | 5.11751 |
| **SEL1L** | 12.37520 | 4.85712 | 4.29236 | 7.00792 | 8.58399 | 4.17693 | 0.23434 |
| **SEL1L2** | 0.02701 | 0.06472 | 0.04351 | 0.01299 | 0.04191 | 0.00569 | 0.02560 |
| **SEL1L3** | 0.15373 | 0.06774 | 0.07999 | 0.08901 | 0.07401 | 0.08042 | 0.16059 |
| **SELENBP1** | 0.32427 | 0.00000 | 0.00261 | 0.28857 | 0.00552 | 0.13684 | 0.31701 |
| **SELK** | 15.06834 | 14.36114 | 9.45137 | 23.69889 | 7.83013 | 11.19361 | 3.22710 |
| **SELL** | 0.11060 | 0.00554 | 0.01762 | 0.01305 | 0.00814 | 0.01723 | 0.00000 |
| **SELM** | 3.64618 | 0.27306 | 1.68215 | 4.57161 | 9.24822 | 1.51761 | 0.00010 |
| **SELO** | 0.10599 | 1.52034 | 0.54093 | 2.19381 | 0.00000 | 1.13294 | 6.14101 |
| **SELP** | 0.02941 | 0.01906 | 0.01224 | 0.02057 | 0.00818 | 0.01277 | 0.01882 |
| **SELPLG** | 0.56673 | 0.00296 | 0.17914 | 0.12643 | 0.13761 | 0.20048 | 0.07699 |
| **SELT** | 46.94073 | 17.54774 | 25.86822 | 19.46763 | 1.16080 | 23.82167 | 2.06960 |
| **SELV** | 0.00503 | 0.00000 | 0.07968 | 0.16187 | 0.00404 | 0.00608 | 0.15754 |
| **SEMA3A** | 0.11048 | 0.00000 | 0.16357 | 0.08015 | 0.12775 | 0.08455 | 0.00000 |
| **SEMA3B** | 0.00000 | 0.46465 | 0.41613 | 0.05520 | 0.09405 | 0.14033 | 0.06046 |
| **SEMA3C** | 0.02584 | 0.01386 | 0.01204 | 0.02975 | 0.07421 | 0.01091 | 0.03058 |
| **SEMA3D** | 0.08168 | 0.00617 | 0.01042 | 0.00583 | 0.38529 | 0.01066 | 0.02198 |
| **SEMA3E** | 0.03706 | 0.00000 | 0.15571 | 0.01682 | 0.20992 | 0.24980 | 0.04696 |
| **SEMA3F** | 0.49497 | 0.07535 | 0.09510 | 0.27848 | 0.06866 | 0.00000 | 3.79279 |
| **SEMA3G** | 0.00475 | 0.01497 | 0.01231 | 0.06118 | 0.01885 | 0.02090 | 0.04509 |
| **SEMA4A** | 0.41269 | 1.07697 | 0.02277 | 0.00000 | 0.09389 | 0.86833 | 0.29423 |
| **SEMA4B** | 0.11257 | 3.66530 | 5.87336 | 3.26620 | 1.69091 | 6.12291 | 6.98195 |
| **SEMA4C** | 1.52669 | 4.67207 | 4.20086 | 13.67294 | 1.10133 | 2.95686 | 27.94169 |
| **SEMA4D** | 1.24510 | 1.15456 | 0.15340 | 1.25054 | 0.63601 | 2.47990 | 4.01669 |
| **SEMA4F** | 0.39027 | 0.00000 | 0.54905 | 0.26399 | 0.26660 | 0.31995 | 0.70081 |
| **SEMA4G** | 0.13629 | 0.11081 | 0.07260 | 0.23037 | 0.19821 | 2.64347 | 0.43668 |
| **SEMA5A** | 0.02005 | 0.02656 | 0.00000 | 0.10470 | 1.36083 | 0.01164 | 0.03794 |

| | | | | | | | |
|---|---|---|---|---|---|---|---|
| **SEMA5B** | 0.00015 | 0.00609 | 0.02555 | 0.03181 | 0.05918 | 0.03050 | 0.10970 |
| **SEMA6A** | 29.74654 | 19.77891 | 0.52597 | 41.95749 | 0.80663 | 23.85555 | 39.86492 |
| **SEMA6B** | 0.06876 | 0.07552 | 0.01009 | 0.03094 | 0.02332 | 0.01117 | 0.00000 |
| **SEMA6C** | 0.00000 | 0.03725 | 0.03810 | 0.21514 | 0.86865 | 0.15237 | 0.29580 |
| **SEMA6D** | 12.29620 | 2.41128 | 0.00000 | 1.02987 | 0.00000 | 0.00000 | 4.56259 |
| **SEMA7A** | 0.00000 | 0.00000 | 0.17226 | 5.53032 | 0.04010 | 0.00000 | 0.90421 |
| **SENCR** | 0.02337 | 0.02471 | 0.01339 | 0.02852 | 0.01640 | 0.01186 | 0.01570 |
| **SENP1** | 5.55409 | 1.55528 | 3.51415 | 3.74995 | 1.39420 | 2.32504 | 2.62352 |
| **SENP2** | 14.52465 | 6.36832 | 10.53929 | 4.71766 | 1.22029 | 10.71120 | 7.77930 |
| **SENP3** | 0.08716 | 0.03522 | 0.06573 | 0.01538 | 0.06828 | 0.30313 | 0.29492 |
| **SENP3-EIF4A1** | 0.03049 | 0.03937 | 0.04256 | 0.16011 | 0.05939 | 0.02078 | 0.00800 |
| **SENP5** | 21.24180 | 14.95408 | 11.56870 | 12.94391 | 16.63167 | 6.77960 | 3.01543 |
| **SENP6** | 5.61438 | 0.00000 | 7.13488 | 5.37531 | 1.77156 | 5.36293 | 16.02101 |
| **SENP7** | 1.22411 | 0.00000 | 0.90737 | 0.90686 | 1.41612 | 2.03400 | 5.81688 |
| **SENP8** | 0.99155 | 1.37336 | 1.50195 | 0.53361 | 0.83558 | 0.89296 | 0.19679 |
| **SEP15** | 33.88513 | 8.23064 | 21.37773 | 18.42368 | 6.68896 | 18.17833 | 14.54047 |
| **SEPHS1** | 13.82068 | 11.58012 | 8.09148 | 8.18547 | 0.87370 | 9.66014 | 7.29751 |
| **SEPHS2** | 4.32347 | 8.92530 | 8.46793 | 6.66464 | 0.00000 | 23.50164 | 28.13509 |
| **SEPN1** | 0.00000 | 1.43875 | 0.00340 | 1.23613 | 0.69483 | 2.14047 | 6.25741 |
| **SEPP1** | 12.71736 | 0.00000 | 0.00000 | 2.26108 | 0.08198 | 2.47971 | 0.36985 |
| **SEPSECS** | 1.93711 | 1.52889 | 2.99700 | 0.59803 | 0.09396 | 0.78818 | 2.02622 |
| **SEPSECS-AS1** | 0.33780 | 0.49920 | 0.16196 | 0.26279 | 0.03243 | 0.14127 | 0.27843 |
| **SEPT1** | 0.00000 | 1.02113 | 0.62646 | 0.05856 | 0.42612 | 0.52460 | 0.63390 |
| **SEPT10** | 2.53721 | 0.00000 | 2.63894 | 1.86204 | 1.92401 | 2.11741 | 0.70357 |
| **SEPT11** | 3.59737 | 17.36279 | 17.82675 | 9.69521 | 5.24939 | 28.06653 | 21.80065 |
| **SEPT12** | 0.00000 | 2.20050 | 1.63158 | 0.22813 | 0.91758 | 0.25021 | 0.00000 |
| **SEPT14** | 0.04268 | 0.03451 | 0.12478 | 0.00137 | 0.22519 | 0.01607 | 0.10197 |
| **SEPT2** | 47.84646 | 20.80427 | 14.32224 | 26.29702 | 9.58070 | 19.86922 | 9.30485 |
| **SEPT3** | 0.00000 | 0.00000 | 0.18460 | 0.39351 | 0.20470 | 0.03943 | 0.08815 |
| **SEPT4** | 0.01852 | 0.03468 | 0.05955 | 0.03250 | 0.00498 | 0.02293 | 0.02623 |
| **SEPT5-GP1BB** | 0.02251 | 0.02534 | 0.01185 | 0.01630 | 0.01259 | 0.01036 | 0.03007 |
| **SEPT6** | 0.08600 | 0.00000 | 0.00000 | 0.07124 | 0.29760 | 3.94860 | 0.64853 |
| **SEPT7** | 23.11977 | 2.69922 | 14.13321 | 15.08494 | 3.16443 | 9.50031 | 6.14894 |
| **SEPT7P2** | 2.46039 | 0.00000 | 0.71069 | 1.30231 | 0.50611 | 1.57619 | 1.30206 |
| **SEPT7P9** | 0.03350 | 0.02059 | 0.01521 | 0.02371 | 0.00956 | 0.01291 | 0.03675 |

| | | | | | | | |
|---|---|---|---|---|---|---|---|
| **SEPT8** | 7.90934 | 6.24543 | 0.42094 | 0.08043 | 0.55174 | 0.81588 | 7.62677 |
| **SEPT9** | 0.00000 | 0.85115 | 0.88389 | 9.88923 | 1.91999 | 8.81715 | 10.89730 |
| **SEPW1** | 19.58605 | 43.28482 | 17.35298 | 10.36494 | 3.72318 | 18.62629 | 14.14765 |
| **SERAC1** | 6.70521 | 7.88975 | 3.18784 | 3.87206 | 0.64827 | 2.01748 | 0.87746 |
| **SERBP1** | 40.99974 | 27.46772 | 23.74678 | 27.01255 | 3.74162 | 20.37453 | 34.05683 |
| **SERF2** | 40.52095 | 53.24454 | 31.48992 | 39.19388 | 16.43527 | 17.77411 | 29.74885 |
| **SERGEF** | 5.83776 | 2.41681 | 0.28630 | 5.19522 | 0.35659 | 2.21234 | 0.88532 |
| **SERHL** | 0.01959 | 0.01929 | 0.00992 | 0.03033 | 0.04473 | 0.01302 | 0.02758 |
| **SERHL2** | 0.48197 | 1.17154 | 0.05399 | 0.52830 | 1.21567 | 0.12346 | 1.09715 |
| **SERINC1** | 19.15481 | 0.00000 | 14.32393 | 14.77367 | 11.36831 | 5.48606 | 1.73800 |
| **SERINC2** | 0.00053 | 0.01292 | 0.00000 | 3.63023 | 0.73481 | 0.14620 | 0.93827 |
| **SERINC3** | 15.96568 | 8.77857 | 17.16379 | 18.96480 | 4.22927 | 12.10439 | 0.19666 |
| **SERINC4** | 0.10903 | 0.33438 | 0.21434 | 0.19025 | 0.23221 | 0.03104 | 0.32688 |
| **SERINC5** | 19.24244 | 8.34050 | 3.27866 | 8.51319 | 8.24130 | 24.76906 | 0.00000 |
| **SERP1** | 25.40997 | 28.76258 | 15.94856 | 35.13912 | 5.71931 | 16.81063 | 25.31864 |
| **SERP2** | 0.00000 | 0.00000 | 0.11474 | 0.19330 | 0.01747 | 0.20376 | 0.21341 |
| **SERPINA1** | 0.02301 | 0.02966 | 0.00483 | 0.01711 | 0.07154 | 0.01129 | 0.02262 |
| **SERPINA5** | 0.25398 | 0.00000 | 0.01314 | 0.00000 | 0.17013 | 0.25075 | 0.20232 |
| **SERPINB1** | 2.16191 | 0.22724 | 0.10520 | 3.30096 | 0.00259 | 2.79168 | 0.00000 |
| **SERPINB12** | 0.01544 | 0.01844 | 0.01265 | 0.06865 | 0.00916 | 0.01120 | 0.01991 |
| **SERPINB13** | 0.02504 | 0.01816 | 0.01276 | 0.02236 | 0.00806 | 0.01343 | 0.01604 |
| **SERPINB2** | 0.00000 | 0.01759 | 0.01367 | 0.18585 | 0.00841 | 0.02510 | 0.01814 |
| **SERPINB3** | 0.01898 | 0.03674 | 0.02082 | 0.01887 | 0.01163 | 0.01103 | 0.01836 |
| **SERPINB5** | 0.05775 | 0.00000 | 6.99327 | 0.15452 | 1.39075 | 1.77905 | 0.02871 |
| **SERPINB6** | 1.55738 | 12.27173 | 12.72088 | 4.03854 | 0.50485 | 18.77604 | 0.00000 |
| **SERPINB7** | 0.02271 | 0.01853 | 0.01608 | 0.02711 | 0.00774 | 0.01330 | 0.01478 |
| **SERPINB8** | 0.87945 | 0.07845 | 0.07113 | 2.95196 | 1.51621 | 0.00000 | 0.02992 |
| **SERPINB9** | 11.09336 | 12.69971 | 0.00656 | 52.08026 | 0.06834 | 28.33662 | 0.00668 |
| **SERPINB9P1** | 0.03787 | 0.01186 | 0.01710 | 0.04855 | 0.00761 | 0.01187 | 0.01594 |
| **SERPINC1** | 1.95465 | 0.00000 | 0.00000 | 2.69989 | 0.00153 | 0.00000 | 0.29016 |
| **SERPIND1** | 0.00000 | 0.14831 | 0.00032 | 0.13479 | 0.03285 | 0.04569 | 0.08976 |
| **SERPINE1** | 0.00000 | 9.31300 | 2.15702 | 2.76380 | 0.00000 | 5.38629 | 0.00000 |
| **SERPINE2** | 0.56271 | 0.84276 | 6.59086 | 2.77118 | 4.04418 | 10.16329 | 0.00000 |
| **SERPINE3** | 0.03254 | 0.01928 | 0.00610 | 0.02939 | 0.07930 | 0.01239 | 0.06377 |
| **SERPINF1** | 0.07816 | 1.25904 | 2.66053 | 0.16980 | 11.32421 | 0.74433 | 0.59773 |

| | | | | | | | |
|---|---|---|---|---|---|---|---|
| **SERPINF2** | 0.00000 | 0.02987 | 0.06463 | 0.02479 | 0.40118 | 0.03074 | 0.14493 |
| **SERPING1** | 0.00383 | 0.00511 | 0.08093 | 0.21166 | 0.06218 | 0.75964 | 0.00316 |
| **SERPINH1** | 0.97951 | 1.38948 | 0.06406 | 0.11792 | 0.00000 | 21.69992 | 0.20797 |
| **SERPINI1** | 1.52792 | 0.55387 | 0.82944 | 0.11646 | 0.16806 | 0.60901 | 0.05451 |
| **SERPINI2** | 0.04385 | 0.00084 | 0.02232 | 0.00894 | 0.05025 | 0.00944 | 0.04331 |
| **SERTAD1** | 4.56186 | 2.14451 | 17.07815 | 12.90596 | 18.89514 | 3.53395 | 3.28712 |
| **SERTAD2** | 8.65666 | 4.78221 | 1.19243 | 6.05351 | 1.63972 | 0.49531 | 17.81859 |
| **SERTAD3** | 14.06233 | 9.73602 | 21.03394 | 12.22722 | 22.21403 | 9.25268 | 5.04745 |
| **SERTAD4** | 0.08802 | 0.00000 | 0.31633 | 3.94573 | 0.00270 | 0.00000 | 1.34925 |
| **SERTAD4-AS1** | 0.00000 | 0.06477 | 0.14336 | 0.38516 | 0.00351 | 0.11684 | 0.01144 |
| **SERTM1** | 0.03352 | 0.01197 | 0.01150 | 0.01256 | 0.13858 | 0.01062 | 0.02842 |
| **SESN1** | 6.24560 | 5.12750 | 10.04831 | 3.81274 | 1.80457 | 8.50368 | 0.00000 |
| **SESN2** | 8.86228 | 18.45395 | 5.37803 | 11.48569 | 7.70656 | 6.69622 | 20.70846 |
| **SESN3** | 0.39152 | 0.24608 | 4.73809 | 0.00000 | 0.66802 | 5.04402 | 2.59303 |
| **SESTD1** | 2.41642 | 0.64105 | 5.05719 | 3.81380 | 0.81263 | 7.09664 | 8.42842 |
| **SET** | 60.52166 | 35.25734 | 27.12282 | 37.17081 | 21.50316 | 26.55281 | 30.66283 |
| **SETBP1** | 2.43613 | 0.14790 | 0.00000 | 0.95953 | 0.08180 | 0.36346 | 1.60703 |
| **SETD1A** | 0.05341 | 1.80494 | 3.00558 | 1.70998 | 0.84882 | 4.06991 | 17.25802 |
| **SETD1B** | 6.60541 | 2.28787 | 3.45692 | 3.94106 | 2.20603 | 5.08981 | 11.91645 |
| **SETD2** | 8.44601 | 6.83732 | 8.10774 | 4.71558 | 1.99432 | 11.00653 | 20.42791 |
| **SETD3** | 29.60187 | 10.87621 | 5.84486 | 14.95171 | 1.43834 | 4.58290 | 15.86407 |
| **SETD4** | 6.36346 | 0.85292 | 4.54218 | 6.71296 | 2.87174 | 1.47887 | 0.00000 |
| **SETD5** | 7.30996 | 3.42707 | 4.16729 | 10.58330 | 4.04174 | 8.27017 | 29.51073 |
| **SETD6** | 5.41138 | 1.01366 | 0.86314 | 6.65831 | 2.03554 | 0.99769 | 3.79828 |
| **SETD7** | 6.06407 | 0.00000 | 0.00000 | 4.41157 | 0.03564 | 0.00000 | 4.87513 |
| **SETD8** | 3.91548 | 11.92456 | 2.12112 | 8.97204 | 6.80581 | 2.39186 | 6.91685 |
| **SETD9** | 2.21643 | 1.22800 | 0.58024 | 1.45171 | 0.11298 | 1.10805 | 0.22230 |
| **SETDB1** | 5.26151 | 2.50251 | 4.36224 | 4.68582 | 1.10208 | 3.78659 | 5.97533 |
| **SETDB2** | 0.96825 | 1.10230 | 1.96884 | 0.47112 | 1.19535 | 1.79359 | 0.84990 |
| **SETMAR** | 3.32178 | 0.74412 | 1.07920 | 2.22686 | 1.12727 | 0.59429 | 0.63004 |
| **SETSIP** | 3.55419 | 0.49277 | 1.77135 | 1.47276 | 0.70409 | 1.29502 | 1.29546 |
| **SETX** | 8.62067 | 4.04382 | 9.82882 | 5.13432 | 8.06516 | 8.86353 | 11.03336 |
| **SEZ6** | 0.06273 | 0.00000 | 0.07634 | 0.05649 | 0.02007 | 0.00210 | 0.25966 |
| **SEZ6L** | 0.02797 | 0.01132 | 0.01362 | 0.01807 | 0.03735 | 0.03719 | 0.02225 |
| **SEZ6L2** | 3.62103 | 4.72389 | 2.73180 | 2.30376 | 2.02576 | 4.26844 | 0.48027 |

| | | | | | | | |
|---|---|---|---|---|---|---|---|
| **SF1** | 7.64698 | 12.46521 | 11.35479 | 13.46333 | 5.93293 | 14.10855 | 51.62287 |
| **SF3A1** | 9.35608 | 12.54025 | 7.08863 | 16.36773 | 1.54799 | 7.91964 | 26.91794 |
| **SF3A2** | 0.00000 | 5.22238 | 3.36987 | 3.20960 | 0.06050 | 5.23533 | 24.64725 |
| **SF3A3** | 39.22342 | 20.48785 | 20.24590 | 26.49735 | 11.31079 | 12.67665 | 4.62871 |
| **SF3B1** | 45.42571 | 16.38684 | 20.12313 | 27.92698 | 10.00158 | 18.60114 | 19.43804 |
| **SF3B2** | 30.72162 | 35.77054 | 14.80626 | 29.51083 | 7.01326 | 18.23978 | 20.53400 |
| **SF3B3** | 40.70088 | 18.24456 | 19.09222 | 24.68096 | 3.13624 | 18.41201 | 12.34051 |
| **SF3B4** | 19.27338 | 18.05257 | 13.24522 | 24.05584 | 5.86578 | 11.77372 | 8.52826 |
| **SF3B5** | 27.45891 | 30.06124 | 16.63438 | 26.22512 | 10.48169 | 14.22578 | 20.76418 |
| **SF3B6** | 34.76483 | 8.28962 | 21.97635 | 27.18400 | 13.62837 | 15.86418 | 19.47082 |
| **SFI1** | 1.06932 | 3.10251 | 0.96398 | 0.78091 | 1.16984 | 0.76993 | 0.23210 |
| **SFMBT1** | 2.52782 | 2.95005 | 2.50326 | 3.11289 | 0.85948 | 0.81965 | 2.63234 |
| **SFMBT2** | 0.00333 | 0.33509 | 0.69245 | 0.06827 | 0.00002 | 1.26837 | 0.61880 |
| **SFN** | 2.51372 | 0.78911 | 0.00000 | 10.26676 | 0.03783 | 0.60794 | 5.42423 |
| **SFPQ** | 19.52013 | 10.96803 | 13.39786 | 15.97606 | 5.38400 | 14.49828 | 50.28378 |
| **SFR1** | 3.45511 | 2.37885 | 5.40095 | 0.36052 | 0.52983 | 0.81785 | 0.00000 |
| **SFRP1** | 0.00000 | 0.02845 | 0.00027 | 0.08301 | 0.03671 | 6.06943 | 0.55448 |
| **SFRP2** | 0.02190 | 0.01177 | 0.01060 | 0.05929 | 0.00775 | 0.74635 | 0.00000 |
| **SFRP5** | 0.02572 | 0.01895 | 0.01237 | 0.02696 | 0.00815 | 0.03161 | 0.00823 |
| **SFSWAP** | 2.17736 | 5.93286 | 6.06949 | 6.71035 | 1.97926 | 3.87400 | 13.93748 |
| **SFT2D1** | 19.13847 | 7.62427 | 12.64886 | 12.00478 | 0.41959 | 8.49197 | 2.11909 |
| **SFT2D2** | 9.16281 | 12.31088 | 2.48835 | 1.71882 | 2.13698 | 3.96617 | 0.00000 |
| **SFT2D3** | 0.00000 | 0.07341 | 0.32867 | 0.57528 | 0.08158 | 0.39854 | 1.87325 |
| **SFTA2** | 0.07941 | 0.01244 | 0.01594 | 0.01003 | 0.01973 | 0.01730 | 0.00469 |
| **SFTA3** | 0.02370 | 0.01717 | 0.00929 | 0.01932 | 0.06313 | 0.01339 | 0.01585 |
| **SFTPA2** | 0.02147 | 0.01641 | 0.01288 | 0.02087 | 0.00777 | 0.01197 | 0.02663 |
| **SFTPB** | 0.03429 | 0.00722 | 0.01637 | 0.01162 | 0.01064 | 0.00553 | 0.13559 |
| **SFTPD** | 0.00000 | 0.38944 | 3.98156 | 0.00000 | 1.59752 | 0.36127 | 0.01391 |
| **SFXN1** | 33.26947 | 10.73084 | 9.01397 | 22.43281 | 0.55471 | 2.48494 | 0.55564 |
| **SFXN2** | 4.56675 | 2.94079 | 0.59515 | 2.71121 | 0.11135 | 0.21347 | 0.00000 |
| **SFXN3** | 0.01540 | 0.01294 | 0.02279 | 0.18523 | 0.01166 | 0.01299 | 0.00888 |
| **SFXN4** | 15.08097 | 10.51391 | 0.00000 | 6.24281 | 0.46992 | 0.97748 | 8.68017 |
| **SFXN5** | 3.24673 | 4.74181 | 0.19494 | 0.90862 | 0.25209 | 3.41419 | 2.01632 |
| **SGCB** | 3.22254 | 4.93195 | 12.83386 | 9.39791 | 1.09086 | 7.13160 | 0.00000 |
| **SGCD** | 0.03277 | 0.01288 | 0.00000 | 0.03778 | 0.84026 | 0.04057 | 0.00532 |

| | | | | | | | |
|---|---|---|---|---|---|---|---|
| **SGCE** | 1.66596 | 0.00000 | 0.00752 | 2.75245 | 0.17115 | 0.17766 | 0.00000 |
| **SGCG** | 0.07658 | 0.06851 | 0.09335 | 0.01063 | 0.44690 | 0.00000 | 0.07355 |
| **SGCZ** | 0.03649 | 0.01445 | 0.00285 | 0.02990 | 0.48025 | 0.01790 | 0.01009 |
| **SGIP1** | 0.02891 | 0.01598 | 0.01262 | 0.03969 | 0.01370 | 0.01930 | 0.00843 |
| **SGK1** | 17.00869 | 23.56367 | 11.95441 | 27.43967 | 16.95512 | 17.23000 | 0.00000 |
| **SGK2** | 0.07010 | 0.04303 | 0.02379 | 0.02899 | 0.02019 | 0.07610 | 0.00000 |
| **SGK223** | 0.48187 | 0.78649 | 0.23846 | 0.11452 | 0.12611 | 0.77053 | 7.15800 |
| **SGK3.uid-10446** | 0.09527 | 0.05220 | 0.07917 | 0.11180 | 0.12336 | 0.04733 | 0.00000 |
| **SGK3.uid-10450** | 0.03863 | 0.02521 | 0.01155 | 0.04699 | 0.00842 | 0.02516 | 0.01145 |
| **SGK494** | 3.27789 | 3.59413 | 2.17990 | 2.31777 | 0.34915 | 0.79754 | 1.77882 |
| **SGMS1** | 8.38984 | 0.58922 | 1.78295 | 6.76893 | 0.63986 | 0.54577 | 9.63964 |
| **SGMS2** | 0.02900 | 0.03505 | 0.07173 | 0.00329 | 1.61020 | 0.03624 | 0.05403 |
| **SGOL1** | 9.78394 | 1.18761 | 8.46503 | 7.97279 | 4.07487 | 4.08983 | 5.22485 |
| **SGOL1-AS1** | 0.03355 | 0.03165 | 0.03657 | 0.04174 | 0.01534 | 0.02389 | 0.02461 |
| **SGOL2** | 12.59498 | 0.00000 | 10.74459 | 10.53380 | 9.18941 | 6.13782 | 10.18959 |
| **SGPL1** | 52.46522 | 22.51885 | 3.30163 | 20.62886 | 3.71914 | 10.61656 | 0.00000 |
| **SGPP1** | 8.35788 | 0.00000 | 3.02905 | 7.51815 | 2.27493 | 0.00000 | 14.91402 |
| **SGPP2** | 0.32628 | 0.23603 | 0.98940 | 0.00000 | 0.00000 | 0.00000 | 0.75862 |
| **SGSH** | 1.53042 | 0.69710 | 0.29666 | 1.37403 | 0.04273 | 1.04038 | 3.61570 |
| **SGSM1** | 0.14961 | 0.19068 | 0.00000 | 0.17677 | 0.16970 | 0.15761 | 0.19304 |
| **SGSM2** | 0.47453 | 0.00000 | 0.28129 | 2.16556 | 0.15792 | 1.65185 | 3.22665 |
| **SGSM3** | 4.34553 | 3.15927 | 1.56459 | 9.19102 | 0.54368 | 4.08792 | 1.92506 |
| **SGTA** | 33.64144 | 29.40129 | 10.91773 | 28.94124 | 4.86681 | 10.97540 | 6.11216 |
| **SGTB** | 5.69862 | 0.00000 | 4.54707 | 3.30242 | 0.55838 | 2.50178 | 0.00000 |
| **SH2B1** | 0.00000 | 1.85993 | 0.77059 | 1.13720 | 0.66924 | 3.04467 | 6.46233 |
| **SH2B2** | 0.00000 | 0.00998 | 0.02169 | 0.31830 | 0.00546 | 0.01064 | 0.63164 |
| **SH2B3** | 3.27881 | 0.85122 | 2.48599 | 0.13459 | 1.57196 | 7.70229 | 5.81123 |
| **SH2D1A** | 0.09460 | 0.00350 | 0.02590 | 0.00111 | 0.08766 | 0.02186 | 0.01952 |
| **SH2D1B** | 0.37106 | 0.32340 | 0.00000 | 0.00000 | 0.59203 | 2.66074 | 0.00000 |
| **SH2D2A** | 0.06635 | 0.13132 | 0.13874 | 0.01894 | 0.00752 | 0.07455 | 0.00000 |
| **SH2D3A** | 0.36065 | 4.94212 | 0.18751 | 1.65454 | 0.10547 | 1.42058 | 4.44779 |
| **SH2D3C** | 0.01642 | 0.03973 | 0.00000 | 0.01693 | 0.37112 | 0.06904 | 0.06116 |
| **SH2D4A** | 16.70834 | 0.08570 | 0.00000 | 25.43557 | 8.99745 | 0.00000 | 19.16793 |
| **SH2D4B** | 0.14825 | 0.45564 | 0.39326 | 0.00000 | 0.72439 | 0.47689 | 0.00000 |
| **SH2D5** | 0.49649 | 1.62553 | 0.39249 | 2.05198 | 0.00000 | 2.48444 | 4.66107 |

| | | | | | | | |
|---|---|---|---|---|---|---|---|
| **SH2D6** | 0.01714 | 0.02309 | 0.02532 | 0.02304 | 0.00662 | 0.02357 | 0.02080 |
| **SH2D7** | 0.10846 | 0.03490 | 0.00947 | 0.02937 | 0.00695 | 0.02752 | 0.08400 |
| **SH3BGR** | 0.33142 | 0.17444 | 1.00637 | 0.12738 | 0.27233 | 0.07410 | 0.00000 |
| **SH3BGRL** | 0.17555 | 0.00000 | 0.00679 | 0.02292 | 2.91145 | 0.46346 | 0.00000 |
| **SH3BGRL2** | 15.00731 | 0.00000 | 0.68894 | 16.01435 | 1.69075 | 0.00000 | 4.23035 |
| **SH3BGRL3** | 16.40079 | 17.96831 | 18.21504 | 17.86866 | 12.28185 | 9.19207 | 3.20568 |
| **SH3BP1** | 0.00000 | 0.00690 | 0.00955 | 0.64274 | 0.03627 | 0.34782 | 3.31586 |
| **SH3BP2** | 0.09372 | 0.00000 | 0.06762 | 1.32057 | 0.01377 | 0.00000 | 4.32366 |
| **SH3BP4** | 6.70620 | 9.27416 | 4.42675 | 2.89509 | 0.00000 | 3.85008 | 14.02888 |
| **SH3BP5** | 0.24856 | 0.52236 | 0.00000 | 3.59500 | 0.19699 | 0.03287 | 0.60357 |
| **SH3BP5-AS1** | 0.20420 | 0.00000 | 0.07656 | 0.98921 | 0.04431 | 0.00000 | 1.44721 |
| **SH3BP5L** | 0.33266 | 1.43878 | 1.27608 | 1.62625 | 1.96163 | 0.87118 | 3.05274 |
| **SH3D19** | 2.24576 | 1.26350 | 0.01030 | 1.17349 | 0.49056 | 3.35339 | 2.57215 |
| **SH3D21** | 0.51810 | 2.93612 | 0.73950 | 0.58481 | 0.45650 | 0.00000 | 3.84573 |
| **SH3GL1** | 2.53618 | 7.96575 | 1.26235 | 8.27828 | 0.67788 | 3.06264 | 27.20477 |
| **SH3GL1P1** | 0.06925 | 0.14781 | 0.00000 | 0.07590 | 0.03765 | 0.14349 | 0.04297 |
| **SH3GL1P2** | 0.00000 | 0.16792 | 0.02241 | 0.01283 | 0.06778 | 0.09294 | 0.14778 |
| **SH3GL2** | 0.00000 | 0.02633 | 0.08098 | 0.05554 | 0.70623 | 0.06920 | 0.04261 |
| **SH3GL3** | 5.50253 | 0.00000 | 0.00000 | 6.29527 | 0.05494 | 0.00000 | 9.31840 |
| **SH3GLB1** | 9.52627 | 5.34870 | 10.30694 | 11.98046 | 5.27717 | 5.87580 | 1.22942 |
| **SH3GLB2** | 2.12014 | 6.55612 | 0.00000 | 0.00000 | 0.16481 | 1.50504 | 19.57363 |
| **SH3KBP1** | 13.46445 | 4.32879 | 0.40556 | 12.25242 | 2.83850 | 0.00000 | 26.68582 |
| **SH3PXD2A** | 0.00000 | 0.00000 | 5.52229 | 0.00000 | 0.12452 | 7.42249 | 2.34689 |
| **SH3PXD2A-AS1** | 0.00000 | 0.17289 | 0.02201 | 0.03195 | 0.00683 | 0.10070 | 0.02271 |
| **SH3PXD2B** | 0.96998 | 0.31537 | 0.92523 | 1.71638 | 0.34099 | 1.57511 | 0.20495 |
| **SH3RF1** | 3.32202 | 9.04306 | 3.89127 | 2.81227 | 0.00000 | 4.84767 | 3.52322 |
| **SH3RF2** | 2.79457 | 1.20236 | 0.00000 | 0.00000 | 0.00265 | 0.01331 | 0.01863 |
| **SH3RF3** | 0.01436 | 0.02464 | 0.02802 | 0.11273 | 0.00690 | 0.09992 | 0.00130 |
| **SH3TC1** | 2.55042 | 3.65955 | 0.00000 | 1.27748 | 0.00000 | 0.35341 | 13.20760 |
| **SH3TC2** | 0.01633 | 0.25990 | 0.11739 | 1.22670 | 0.00246 | 0.10415 | 0.12469 |
| **SH3YL1** | 1.06033 | 0.09348 | 0.69546 | 1.46893 | 0.15403 | 0.83666 | 0.04465 |
| **SHANK1** | 0.02129 | 0.01925 | 0.01627 | 0.02457 | 0.00796 | 0.04357 | 0.01909 |
| **SHANK2** | 0.45352 | 1.98238 | 0.04033 | 0.02392 | 0.82941 | 3.50330 | 1.38948 |
| **SHANK2-AS1** | 0.01695 | 0.02029 | 0.01166 | 0.01740 | 0.00802 | 0.05738 | 0.02870 |
| **SHANK2-AS3** | 0.04973 | 0.04562 | 0.00502 | 0.00809 | 0.00742 | 0.07645 | 0.10003 |

| | | | | | | | |
|---|---|---|---|---|---|---|---|
| **SHANK3** | 0.00000 | 0.56729 | 0.00392 | 0.00000 | 0.37102 | 0.33288 | 0.99931 |
| **SHARPIN** | 1.01459 | 8.33812 | 1.19284 | 1.07204 | 0.01656 | 3.14856 | 5.19644 |
| **SHB** | 0.00000 | 6.13119 | 2.06698 | 0.40012 | 0.21063 | 6.34704 | 6.76565 |
| **SHBG** | 0.02421 | 0.01845 | 0.15411 | 0.15547 | 0.02291 | 0.00456 | 0.00204 |
| **SHC1** | 8.19428 | 12.26850 | 6.97561 | 23.89964 | 3.87898 | 8.05063 | 3.53680 |
| **SHC2** | 0.00000 | 0.06679 | 0.23979 | 0.01211 | 0.31292 | 0.23066 | 0.04859 |
| **SHC3** | 0.00208 | 0.00677 | 0.06538 | 0.03935 | 0.10648 | 0.19153 | 0.05687 |
| **SHC4** | 0.02570 | 0.01659 | 0.01480 | 0.02487 | 0.12187 | 0.01166 | 0.02285 |
| **SHCBP1** | 18.67257 | 12.31452 | 8.41806 | 7.19220 | 5.23347 | 8.22317 | 0.00000 |
| **SHCBP1L** | 0.14793 | 0.05555 | 0.01503 | 0.17096 | 0.00679 | 0.04938 | 0.00000 |
| **SHD** | 0.28342 | 1.46662 | 0.54161 | 0.21878 | 1.18242 | 2.58903 | 0.00000 |
| **SHE** | 0.02949 | 0.00890 | 0.00000 | 0.00000 | 0.93087 | 0.66242 | 0.07025 |
| **SHF** | 0.00000 | 0.00000 | 0.61470 | 0.50195 | 0.31011 | 1.07501 | 0.21770 |
| **SHFM1** | 30.77860 | 20.85481 | 9.00657 | 28.39566 | 3.45421 | 15.74511 | 22.50010 |
| **SHH** | 0.02392 | 0.00174 | 0.04004 | 0.01923 | 0.27966 | 0.01330 | 0.01724 |
| **SHISA2** | 0.22372 | 0.06154 | 0.13960 | 0.00603 | 0.18566 | 0.51493 | 0.27457 |
| **SHISA3** | 0.00000 | 0.07402 | 0.17190 | 0.00000 | 0.00683 | 0.29898 | 0.10083 |
| **SHISA4** | 0.32306 | 0.41582 | 0.00081 | 1.69741 | 0.00325 | 0.00000 | 1.19954 |
| **SHISA5** | 51.06553 | 61.68461 | 0.00000 | 37.66513 | 1.26232 | 31.32828 | 31.33512 |
| **SHISA6** | 0.02714 | 0.00527 | 0.03002 | 0.11251 | 0.01152 | 0.01545 | 0.04138 |
| **SHISA7** | 0.05133 | 0.05825 | 0.00981 | 0.02475 | 0.13498 | 0.02182 | 0.00000 |
| **SHISA8** | 0.00444 | 0.00782 | 0.01125 | 0.04566 | 0.09512 | 0.05443 | 0.15507 |
| **SHISA9** | 0.00777 | 0.03281 | 0.01142 | 0.01794 | 0.02414 | 0.17279 | 0.03020 |
| **SHKBP1** | 7.33932 | 6.58225 | 0.49146 | 10.39673 | 0.00000 | 5.86088 | 2.89163 |
| **SHMT1** | 21.34472 | 16.18645 | 6.78938 | 14.68276 | 0.84801 | 9.81282 | 0.00001 |
| **SHMT2** | 31.36791 | 8.59766 | 4.83026 | 31.11212 | 3.55263 | 12.78445 | 7.84304 |
| **SHOC2** | 12.27732 | 1.14395 | 4.92017 | 6.69643 | 8.22735 | 4.33604 | 1.53124 |
| **SHOX2** | 0.03836 | 0.00361 | 0.08473 | 0.02888 | 0.21295 | 0.00290 | 0.07248 |
| **SHPK** | 3.80376 | 17.17000 | 12.13618 | 1.81739 | 2.17422 | 17.43522 | 1.68906 |
| **SHPRH** | 6.06140 | 1.39574 | 3.26111 | 2.64134 | 1.58723 | 2.42164 | 4.21351 |
| **SHQ1** | 14.69080 | 1.48322 | 8.71064 | 6.22419 | 12.58061 | 4.03145 | 0.00000 |
| **SHROOM1** | 0.11097 | 0.22989 | 0.01471 | 0.29395 | 0.00513 | 0.00000 | 0.61531 |
| **SHROOM2** | 5.94816 | 4.34293 | 0.47770 | 0.00451 | 0.51061 | 0.31038 | 9.09397 |
| **SHROOM3** | 6.54540 | 6.75076 | 2.39982 | 0.00000 | 0.81070 | 4.79894 | 8.74658 |
| **SHROOM4** | 0.05071 | 0.05406 | 0.02178 | 0.03960 | 0.02702 | 0.15594 | 0.00000 |

| | | | | | | | |
|---|---|---|---|---|---|---|---|
| **SI** | 0.03111 | 0.01963 | 0.01276 | 0.02576 | 0.00828 | 0.01475 | 0.01388 |
| **SIAE** | 5.69992 | 5.42270 | 0.31927 | 1.18151 | 0.01309 | 6.24373 | 0.00000 |
| **SIAH1** | 1.22185 | 0.00000 | 7.02177 | 2.40005 | 13.72322 | 6.54068 | 6.51849 |
| **SIAH2** | 0.54117 | 0.00000 | 0.02964 | 4.55268 | 0.40940 | 0.00000 | 3.16104 |
| **SIAH3** | 0.00508 | 0.03518 | 0.04553 | 0.03425 | 0.23378 | 0.07753 | 0.03929 |
| **SIDT1** | 0.00000 | 0.13331 | 0.17789 | 0.03289 | 0.00395 | 0.01454 | 0.02451 |
| **SIDT2** | 1.03911 | 0.28075 | 2.51871 | 2.98501 | 1.52129 | 1.00393 | 3.51060 |
| **SIGIRR** | 8.21138 | 4.30238 | 0.45705 | 5.82793 | 0.67459 | 6.51921 | 10.94090 |
| **SIGLEC1** | 0.02838 | 0.02104 | 0.00578 | 0.01595 | 0.06181 | 0.01646 | 0.03415 |
| **SIGLEC10** | 0.01587 | 0.00000 | 0.09149 | 0.08373 | 2.77060 | 0.02339 | 0.00001 |
| **SIGLEC11** | 0.01186 | 0.06676 | 0.02558 | 0.06051 | 0.02747 | 0.25994 | 0.00961 |
| **SIGLEC12** | 0.01904 | 0.02221 | 0.00880 | 0.03067 | 0.04614 | 0.01421 | 0.01481 |
| **SIGLEC14** | 0.02339 | 0.04363 | 0.02463 | 0.37686 | 0.00667 | 0.01013 | 0.00000 |
| **SIGLEC15** | 0.03639 | 0.01795 | 0.02274 | 0.01796 | 0.00657 | 0.00803 | 0.03895 |
| **SIGLEC16** | 0.01423 | 0.02825 | 0.01027 | 0.04048 | 0.00792 | 0.05574 | 0.01756 |
| **SIGLEC17P** | 0.03483 | 0.01252 | 0.02030 | 0.01774 | 0.02236 | 0.01416 | 0.01163 |
| **SIGLEC5** | 0.18285 | 0.06639 | 0.00683 | 0.25388 | 0.01426 | 0.00790 | 0.00000 |
| **SIGLEC6** | 0.02084 | 0.01511 | 0.01175 | 0.13912 | 0.02732 | 0.02273 | 0.01106 |
| **SIGLEC7** | 0.02363 | 0.01356 | 0.01532 | 0.01909 | 0.09065 | 0.01310 | 0.01771 |
| **SIGLEC8** | 0.02593 | 0.01887 | 0.01296 | 0.01923 | 0.00919 | 0.01268 | 0.01585 |
| **SIGLEC9** | 0.01538 | 0.01246 | 0.01354 | 0.03237 | 0.00794 | 0.01016 | 0.04719 |
| **SIGLECL1** | 0.05875 | 0.01863 | 0.00795 | 0.00846 | 0.04261 | 0.04875 | 0.02269 |
| **SIGMAR1** | 3.34420 | 5.50421 | 5.90240 | 12.42532 | 1.78664 | 3.85962 | 2.16213 |
| **SIK1** | 1.69323 | 0.91780 | 1.00105 | 0.73643 | 0.70915 | 1.85318 | 7.08156 |
| **SIK2** | 0.76628 | 1.86270 | 0.88659 | 1.23961 | 1.01739 | 2.04543 | 0.59033 |
| **SIK3** | 0.00000 | 0.99789 | 1.34172 | 2.90669 | 0.21526 | 2.49117 | 5.67332 |
| **SIKE1** | 20.35490 | 13.09842 | 18.44778 | 16.88082 | 9.75455 | 12.04716 | 4.13969 |
| **SIL1** | 9.19599 | 13.09940 | 7.15530 | 13.41490 | 0.64871 | 5.17793 | 3.60294 |
| **SIM1** | 0.03295 | 0.02581 | 0.01761 | 0.03660 | 0.01600 | 0.04120 | 0.00000 |
| **SIM2** | 0.07903 | 0.00000 | 0.03913 | 0.04973 | 0.00884 | 0.00000 | 0.31218 |
| **SIMC1** | 1.08344 | 4.75873 | 2.97744 | 0.49068 | 0.58863 | 4.06642 | 1.94573 |
| **SIN3A** | 10.93232 | 9.92248 | 12.62574 | 8.91027 | 11.56134 | 9.15205 | 12.12996 |
| **SIN3B** | 0.00000 | 5.87784 | 1.92094 | 7.95421 | 1.19247 | 2.29879 | 12.56741 |
| **SIPA1** | 0.18490 | 0.26554 | 0.02943 | 1.22998 | 0.21201 | 1.85059 | 2.02045 |
| **SIPA1L1** | 4.85591 | 2.84974 | 2.94988 | 7.86478 | 2.90778 | 2.17409 | 11.10534 |

| | | | | | | | |
|---|---|---|---|---|---|---|---|
| **SIPA1L2** | 0.00000 | 1.17183 | 2.26864 | 0.27462 | 3.81285 | 7.23966 | 0.17777 |
| **SIPA1L3** | 0.00686 | 2.74210 | 0.23164 | 1.21367 | 0.31248 | 0.84492 | 10.34610 |
| **SIRPA** | 0.00000 | 0.01672 | 0.00000 | 0.29471 | 0.40593 | 0.12096 | 0.05735 |
| **SIRPB2** | 0.02435 | 0.01720 | 0.01310 | 0.02701 | 0.00812 | 0.01345 | 0.01848 |
| **SIRT1** | 4.99446 | 0.00000 | 4.89098 | 2.67956 | 5.27475 | 5.65464 | 6.50928 |
| **SIRT2** | 4.16656 | 2.89089 | 0.73655 | 7.18357 | 0.53369 | 3.38875 | 0.00000 |
| **SIRT3** | 0.40971 | 1.52569 | 2.74094 | 1.01109 | 0.80837 | 0.69577 | 0.73355 |
| **SIRT4** | 1.15713 | 0.57886 | 0.61847 | 1.37983 | 0.45445 | 0.05873 | 0.39428 |
| **SIRT5** | 4.16632 | 4.38409 | 1.42705 | 0.95645 | 0.73526 | 2.79318 | 0.00000 |
| **SIRT6** | 1.41136 | 6.18031 | 3.67131 | 9.09930 | 0.60709 | 1.88920 | 4.73435 |
| **SIRT7** | 1.03771 | 5.34967 | 1.45614 | 3.40233 | 0.72421 | 2.51150 | 10.75018 |
| **1.00000** | 0.02916 | 0.01682 | 0.01238 | 0.01739 | 0.01050 | 0.01715 | 0.01790 |
| **SIVA1** | 21.04077 | 33.80445 | 10.40306 | 30.59258 | 7.93642 | 15.25502 | 18.36892 |
| **SIX1** | 0.28180 | 0.09422 | 0.05330 | 0.00089 | 0.17305 | 0.64581 | 0.00000 |
| **SIX2** | 0.00000 | 1.91114 | 0.56672 | 0.00000 | 0.64885 | 1.38328 | 0.06632 |
| **SIX3** | 0.03923 | 0.00229 | 0.01977 | 0.01310 | 0.03768 | 0.09110 | 0.03581 |
| **SIX3-AS1** | 0.02314 | 0.01911 | 0.00679 | 0.01921 | 0.07640 | 0.01300 | 0.01754 |
| **SIX4** | 1.85304 | 0.00000 | 0.88113 | 1.67854 | 0.62986 | 0.09201 | 2.33247 |
| **SIX5** | 0.00000 | 2.34464 | 0.02471 | 1.64270 | 0.01734 | 0.50571 | 4.01973 |
| **SKA1** | 11.66777 | 3.01478 | 7.36400 | 12.44112 | 3.02404 | 4.17793 | 2.46492 |
| **SKA2** | 14.45324 | 4.77059 | 16.98592 | 10.12293 | 11.21737 | 6.65606 | 7.78765 |
| **SKA3** | 29.41417 | 7.80601 | 15.98011 | 12.29312 | 9.82476 | 10.88286 | 2.78979 |
| **SKAP1** | 0.02325 | 0.01782 | 0.00869 | 0.02181 | 0.04767 | 0.01182 | 0.02614 |
| **SKAP2** | 8.23077 | 3.47071 | 2.84635 | 3.21700 | 3.26557 | 1.87616 | 4.19349 |
| **SKI** | 0.14021 | 1.27553 | 1.06916 | 0.80379 | 0.96917 | 0.64077 | 8.52107 |
| **SKIDA1** | 2.14851 | 2.80295 | 0.25341 | 5.11154 | 0.05083 | 1.26040 | 2.16311 |
| **SKIL** | 12.90648 | 0.00000 | 11.59861 | 0.64787 | 2.53529 | 11.17040 | 5.26532 |
| **SKINTL** | 0.02156 | 0.00000 | 0.06186 | 0.01719 | 0.00674 | 0.01626 | 0.04094 |
| **SKIV2L** | 4.05180 | 4.23370 | 2.72005 | 7.43171 | 0.51598 | 4.01515 | 4.12894 |
| **SKIV2L2** | 18.53542 | 0.52671 | 9.86107 | 11.68170 | 1.95552 | 4.59547 | 9.98897 |
| **SKOR1** | 0.00790 | 0.07458 | 0.00000 | 0.01496 | 1.34603 | 0.00633 | 0.02232 |
| **SKOR2** | 0.01561 | 0.01376 | 0.01218 | 0.01759 | 0.00903 | 0.03109 | 0.04579 |
| **SKP1** | 45.43011 | 14.67311 | 17.27109 | 33.19369 | 8.74474 | 21.44300 | 15.15329 |
| **SKP1P2** | 9.62459 | 2.02747 | 2.88930 | 6.92697 | 1.46933 | 3.55172 | 0.00000 |
| **SKP2** | 32.59877 | 13.79092 | 3.93992 | 18.49284 | 6.42324 | 3.19324 | 0.00000 |

| | | | | | | | |
|---|---|---|---|---|---|---|---|
| **SLA** | 0.03990 | 0.01509 | 0.01417 | 0.01800 | 0.00800 | 0.02070 | 0.00663 |
| **SLA2** | 0.00000 | 0.08806 | 0.00000 | 0.08047 | 1.01372 | 0.08377 | 0.24877 |
| **SLAIN1** | 0.04549 | 0.00000 | 0.06739 | 0.07515 | 0.07318 | 0.34089 | 0.21518 |
| **SLAIN2** | 1.87608 | 6.61075 | 4.27018 | 2.87987 | 2.47289 | 10.00840 | 13.15743 |
| **SLAMF1** | 0.02499 | 0.01481 | 0.03636 | 0.04244 | 0.00681 | 0.01417 | 0.01462 |
| **SLAMF6** | 0.04894 | 0.01588 | 0.01560 | 0.01659 | 0.00946 | 0.02031 | 0.00528 |
| **SLAMF7** | 0.02408 | 0.01523 | 0.01766 | 0.08163 | 0.00790 | 0.01818 | 0.00365 |
| **SLAMF8** | 0.02311 | 0.01936 | 0.01311 | 0.01932 | 0.00883 | 0.01288 | 0.01819 |
| **SLAMF9** | 0.03539 | 0.01619 | 0.01488 | 0.02246 | 0.00788 | 0.01638 | 0.00789 |
| **SLBP** | 19.05591 | 16.80705 | 13.19540 | 6.59537 | 7.40302 | 6.99955 | 31.92850 |
| **SLC10A1** | 0.06678 | 0.00249 | 0.08268 | 0.06348 | 0.09964 | 0.01641 | 0.00995 |
| **SLC10A3** | 4.19351 | 3.62252 | 0.58843 | 2.08680 | 0.52311 | 0.67106 | 5.73841 |
| **SLC10A4** | 0.00000 | 0.04779 | 0.07086 | 0.13464 | 0.32317 | 0.25371 | 0.36520 |
| **SLC10A5** | 0.06271 | 0.04821 | 0.00491 | 0.01320 | 0.02982 | 0.01286 | 0.03272 |
| **SLC10A6** | 0.01034 | 0.02257 | 0.01042 | 0.01915 | 0.00758 | 0.00825 | 0.07378 |
| **SLC10A7** | 4.27905 | 0.00000 | 0.93451 | 3.01025 | 0.51665 | 0.98612 | 0.14857 |
| **SLC11A1** | 0.00466 | 0.41696 | 0.24943 | 0.06237 | 0.00297 | 0.00000 | 0.39598 |
| **SLC11A2** | 29.28479 | 0.03253 | 0.31750 | 19.11379 | 0.92487 | 1.48975 | 4.77631 |
| **SLC12A1** | 0.02161 | 0.01381 | 0.02031 | 0.02069 | 0.05885 | 0.01203 | 0.02666 |
| **SLC12A2** | 6.14148 | 0.44986 | 2.01869 | 3.98958 | 5.18801 | 3.67553 | 1.68989 |
| **SLC12A3** | 28.98654 | 22.87352 | 0.00000 | 0.00000 | 0.00000 | 0.00000 | 21.61660 |
| **SLC12A4** | 0.99095 | 2.70944 | 0.47164 | 3.77520 | 0.08502 | 1.42511 | 2.60745 |
| **SLC12A5** | 0.01074 | 0.03140 | 0.12985 | 0.02103 | 1.09355 | 0.07170 | 0.15383 |
| **SLC12A6** | 2.04062 | 0.33032 | 0.38997 | 0.98538 | 1.40557 | 2.24268 | 0.68780 |
| **SLC12A7** | 0.00000 | 13.72822 | 2.41768 | 0.82141 | 0.11748 | 1.40700 | 3.18816 |
| **SLC12A8** | 0.25687 | 0.19841 | 0.00000 | 0.36752 | 1.37423 | 0.02497 | 0.00000 |
| **SLC12A9** | 0.25027 | 1.12130 | 0.24550 | 1.37346 | 0.03106 | 0.46636 | 9.94231 |
| **SLC13A1** | 0.03167 | 0.01571 | 0.01017 | 0.02004 | 0.03608 | 0.01255 | 0.01719 |
| **SLC13A2** | 0.01919 | 0.01422 | 0.01285 | 0.03070 | 0.00818 | 0.04383 | 0.01755 |
| **SLC13A3** | 0.08531 | 0.05655 | 0.00000 | 0.59003 | 1.18120 | 0.06052 | 0.22883 |
| **SLC13A4** | 0.00000 | 0.10151 | 0.02987 | 7.77573 | 1.39138 | 0.13603 | 0.00000 |
| **SLC13A5** | 0.07560 | 0.01581 | 0.01185 | 0.00000 | 0.00917 | 0.00000 | 0.11044 |
| **SLC14A1** | 0.04001 | 0.05924 | 0.00294 | 0.01359 | 0.07508 | 0.00409 | 0.04161 |
| **SLC14A2** | 0.26816 | 0.07519 | 0.00000 | 0.00000 | 0.17616 | 0.00000 | 0.03126 |
| **SLC15A1** | 0.18139 | 0.00652 | 0.01717 | 0.00000 | 0.04253 | 0.01006 | 0.07855 |

| | | | | | | | |
|---|---|---|---|---|---|---|---|
| **SLC15A2** | 8.63010 | 0.00000 | 0.02354 | 0.72903 | 0.01340 | 0.00000 | 1.06178 |
| **SLC15A3** | 0.00000 | 0.21545 | 1.79762 | 0.00000 | 0.08667 | 0.00000 | 0.22416 |
| **SLC15A4** | 3.95680 | 2.74819 | 0.54461 | 0.75317 | 0.26401 | 4.44270 | 7.14605 |
| **SLC15A5** | 0.08801 | 0.00560 | 0.02430 | 0.08013 | 0.00728 | 0.01624 | 0.00000 |
| **SLC16A1** | 70.69655 | 29.30915 | 39.78421 | 38.55820 | 22.62442 | 19.45453 | 21.43254 |
| **SLC16A1-AS1** | 1.41355 | 0.81510 | 1.14345 | 1.09682 | 1.01828 | 0.28310 | 0.99866 |
| **SLC16A10** | 1.72671 | 0.00000 | 0.11403 | 0.39899 | 0.53516 | 4.61702 | 1.97741 |
| **SLC16A11** | 0.00000 | 0.01326 | 0.01098 | 0.36041 | 0.05309 | 0.02938 | 0.03528 |
| **SLC16A12** | 0.00966 | 0.00670 | 0.03215 | 0.02597 | 0.12942 | 0.03141 | 0.12475 |
| **SLC16A13** | 0.38772 | 0.00000 | 0.09011 | 1.07001 | 0.23373 | 0.01304 | 0.00000 |
| **SLC16A14** | 0.18839 | 0.21026 | 0.24381 | 0.00000 | 0.10382 | 0.18394 | 0.06698 |
| **SLC16A2** | 0.00000 | 1.91240 | 0.19537 | 0.00000 | 0.04067 | 8.15708 | 0.13238 |
| **SLC16A3** | 1.70866 | 0.15297 | 0.00000 | 2.57867 | 0.00000 | 0.00000 | 3.18416 |
| **SLC16A4** | 0.17431 | 0.00000 | 0.19751 | 0.53287 | 0.34567 | 0.27651 | 0.77234 |
| **SLC16A5** | 1.63564 | 0.57219 | 0.00000 | 1.02070 | 1.10599 | 0.54155 | 1.07306 |
| **SLC16A6** | 0.13166 | 0.06458 | 0.69610 | 0.58992 | 9.74787 | 0.00000 | 0.53755 |
| **SLC16A7** | 0.02974 | 0.05112 | 0.04030 | 0.03061 | 0.00623 | 0.03302 | 0.00038 |
| **SLC16A8** | 0.01415 | 0.06573 | 0.01430 | 0.03695 | 0.08115 | 0.01030 | 0.05467 |
| **SLC16A9** | 0.08448 | 0.00000 | 5.34485 | 0.11940 | 1.26559 | 8.89273 | 0.21579 |
| **SLC17A2** | 0.07744 | 0.02056 | 0.07882 | 0.00852 | 0.05643 | 0.00454 | 0.02649 |
| **SLC17A4** | 0.03068 | 0.01653 | 0.01381 | 0.06321 | 0.00817 | 0.03360 | 0.00000 |
| **SLC17A5** | 9.94301 | 26.41838 | 3.41749 | 0.62326 | 0.24755 | 5.46967 | 0.45780 |
| **SLC17A6** | 0.01955 | 0.01749 | 0.01318 | 0.02845 | 0.00783 | 0.01189 | 0.02380 |
| **SLC17A7** | 0.02741 | 0.00000 | 0.45160 | 0.01878 | 1.33184 | 0.22948 | 0.24182 |
| **SLC17A8** | 0.00526 | 0.02246 | 0.01050 | 0.03070 | 0.08009 | 0.08089 | 0.05912 |
| **SLC17A9** | 0.01375 | 0.00693 | 0.01012 | 0.11593 | 0.03657 | 0.24510 | 0.03574 |
| **SLC18A2** | 0.95331 | 0.04891 | 0.29288 | 0.54377 | 1.99732 | 1.34318 | 0.69737 |
| **SLC18A3** | 0.02466 | 0.01947 | 0.00928 | 0.02651 | 0.03546 | 0.01248 | 0.01548 |
| **SLC18B1** | 3.57455 | 0.00000 | 0.02925 | 1.92374 | 0.33923 | 0.29731 | 0.00000 |
| **SLC19A1** | 0.57091 | 0.29530 | 0.15248 | 3.37460 | 0.16731 | 0.25035 | 1.64960 |
| **SLC19A2** | 12.01446 | 0.00000 | 0.00000 | 8.31778 | 1.67484 | 0.00000 | 6.38599 |
| **SLC19A3** | 7.16319 | 3.14053 | 0.15296 | 3.43641 | 0.08942 | 0.00000 | 3.47162 |
| **SLC1A1** | 0.04651 | 0.02656 | 0.31201 | 0.04497 | 0.19911 | 0.07163 | 0.00000 |
| **SLC1A2** | 0.10664 | 0.00000 | 0.02580 | 0.05132 | 0.10757 | 1.08527 | 0.00548 |
| **SLC1A3** | 36.95716 | 0.00000 | 0.00000 | 23.41862 | 0.00000 | 5.67539 | 10.82564 |

| | | | | | | | |
|---|---|---|---|---|---|---|---|
| **SLC1A4** | 0.28471 | 0.00000 | 0.04584 | 0.40193 | 0.12963 | 0.00000 | 0.91427 |
| **SLC1A5** | 17.80743 | 0.00000 | 8.10461 | 41.13606 | 4.61628 | 4.10425 | 28.76895 |
| **SLC1A6** | 0.00000 | 0.31111 | 0.07825 | 0.46060 | 1.88503 | 0.00432 | 0.00000 |
| **SLC1A7** | 0.00000 | 0.13295 | 0.08177 | 0.02985 | 0.03543 | 0.00000 | 0.13399 |
| **SLC20A1** | 14.77932 | 1.90585 | 7.58554 | 18.04980 | 4.14640 | 5.85000 | 0.50549 |
| **SLC20A2** | 2.90711 | 1.99670 | 0.84027 | 15.27542 | 0.01576 | 2.00055 | 3.60993 |
| **SLC22A1** | 0.06624 | 0.07270 | 0.02082 | 0.08854 | 0.00507 | 0.11421 | 0.14425 |
| **SLC22A10** | 0.03222 | 0.00000 | 0.07868 | 0.01214 | 0.00488 | 0.05906 | 0.02845 |
| **SLC22A11** | 0.46244 | 0.11975 | 0.04361 | 5.94243 | 0.01978 | 0.00000 | 0.24616 |
| **SLC22A12** | 0.03316 | 0.01647 | 0.01219 | 0.03980 | 0.01038 | 0.01045 | 0.01608 |
| **SLC22A13** | 0.05252 | 0.02491 | 0.07975 | 0.01908 | 0.01672 | 0.01518 | 0.05754 |
| **SLC22A14** | 0.01647 | 0.00961 | 0.08737 | 0.05261 | 0.00495 | 0.01829 | 0.03951 |
| **SLC22A15** | 0.07180 | 0.09411 | 0.04587 | 0.00000 | 0.04079 | 0.34609 | 0.07433 |
| **SLC22A16** | 0.02080 | 0.00896 | 0.05964 | 0.16204 | 0.05438 | 0.01310 | 0.00678 |
| **SLC22A17** | 0.02412 | 0.00634 | 0.01296 | 0.16263 | 0.02159 | 0.00000 | 0.32273 |
| **SLC22A18** | 0.58050 | 0.94780 | 0.01237 | 0.17214 | 0.00681 | 1.44682 | 0.62324 |
| **SLC22A18AS** | 0.06922 | 0.00000 | 0.02275 | 0.01268 | 0.00789 | 0.12324 | 0.11635 |
| **SLC22A2** | 0.06223 | 0.09861 | 0.02831 | 0.02634 | 0.26490 | 0.00779 | 0.00000 |
| **SLC22A20** | 0.00000 | 0.00000 | 3.69978 | 0.20329 | 0.18653 | 1.28983 | 0.23168 |
| **SLC22A23** | 1.55394 | 1.37438 | 0.39318 | 5.26974 | 0.80216 | 0.60790 | 4.07364 |
| **SLC22A24** | 0.09762 | 0.03040 | 0.03684 | 0.00577 | 0.00585 | 0.00103 | 0.04628 |
| **SLC22A3** | 0.00933 | 0.02930 | 0.01145 | 0.08539 | 0.00733 | 0.01133 | 0.04868 |
| **SLC22A31** | 0.00000 | 0.50231 | 0.03422 | 0.15679 | 0.00348 | 0.35710 | 0.18599 |
| **SLC22A4** | 0.08384 | 0.05871 | 0.01212 | 0.02278 | 0.12387 | 0.10183 | 0.00000 |
| **SLC22A5** | 2.27475 | 2.02254 | 0.01700 | 2.11480 | 0.12094 | 0.46527 | 0.26849 |
| **SLC22A7** | 0.00000 | 0.09208 | 0.01604 | 0.00925 | 0.01366 | 0.48304 | 0.11869 |
| **SLC22A8** | 0.02671 | 0.03086 | 0.02293 | 0.02098 | 0.01659 | 0.01085 | 0.01486 |
| **SLC22A9** | 0.02809 | 0.02689 | 0.01632 | 0.01848 | 0.01063 | 0.01931 | 0.01782 |
| **SLC23A1** | 0.30112 | 0.00000 | 0.06953 | 0.32088 | 0.07038 | 0.04539 | 0.00000 |
| **SLC23A2** | 8.78419 | 12.55397 | 4.81806 | 3.41923 | 0.00000 | 10.52989 | 5.75842 |
| **SLC23A3** | 0.07434 | 0.04324 | 0.02668 | 0.35099 | 0.00477 | 0.00000 | 0.48307 |
| **SLC24A1** | 0.02097 | 0.49368 | 0.04559 | 0.20143 | 0.03347 | 0.69824 | 0.65869 |
| **SLC24A2** | 0.08740 | 0.05705 | 0.27417 | 0.09038 | 0.42790 | 0.06910 | 0.00000 |
| **SLC24A3** | 0.04670 | 0.04892 | 0.00797 | 0.00455 | 0.02057 | 0.10063 | 0.03330 |
| **SLC24A4** | 0.00000 | 0.11172 | 0.03550 | 0.03615 | 0.07469 | 0.01738 | 0.09750 |

| | | | | | | | |
|---|---|---|---|---|---|---|---|
| **SLC24A5** | 0.03674 | 0.01414 | 0.00615 | 0.01799 | 0.11571 | 0.01130 | 0.02626 |
| **SLC25A1** | 16.57324 | 9.66745 | 0.01592 | 18.44801 | 0.55356 | 12.63597 | 32.95446 |
| **SLC25A10** | 4.13510 | 0.98044 | 0.29043 | 5.52258 | 0.00372 | 3.05873 | 6.87114 |
| **SLC25A11** | 23.98893 | 26.07882 | 6.07527 | 15.08225 | 2.08563 | 14.44687 | 3.23295 |
| **SLC25A12** | 0.15558 | 0.90565 | 5.25919 | 0.02744 | 2.10316 | 6.56691 | 0.00000 |
| **SLC25A13** | 15.13846 | 3.33159 | 0.08567 | 17.91453 | 1.04014 | 5.96719 | 7.05654 |
| **SLC25A14** | 4.89350 | 1.25287 | 4.94895 | 3.56075 | 4.29733 | 1.70517 | 0.00000 |
| **SLC25A15** | 0.32129 | 0.97353 | 0.79469 | 0.55681 | 0.00000 | 1.50413 | 1.40687 |
| **SLC25A16** | 3.41882 | 8.81824 | 10.24671 | 2.18237 | 0.40018 | 11.16513 | 12.94472 |
| **SLC25A17** | 14.79962 | 2.69340 | 8.02480 | 13.89731 | 3.62792 | 4.59639 | 1.12157 |
| **SLC25A18** | 0.00000 | 1.81269 | 0.61512 | 0.20644 | 0.17465 | 2.02110 | 0.00000 |
| **SLC25A19** | 13.36600 | 7.92778 | 6.41956 | 9.31185 | 0.72776 | 1.21618 | 0.00000 |
| **SLC25A2** | 0.02463 | 0.02436 | 0.01488 | 0.02568 | 0.00768 | 0.01476 | 0.01051 |
| **SLC25A20** | 1.21693 | 2.05917 | 3.12945 | 0.88799 | 0.73771 | 0.86753 | 2.12896 |
| **SLC25A21** | 0.46692 | 0.00000 | 0.23909 | 0.01370 | 0.12407 | 0.01935 | 0.00000 |
| **SLC25A21-AS1** | 0.09541 | 0.05190 | 0.06840 | 0.00000 | 0.00592 | 0.03777 | 0.01090 |
| **SLC25A22** | 0.96452 | 2.76696 | 0.00000 | 0.54965 | 0.00524 | 4.43789 | 12.45076 |
| **SLC25A23** | 2.67248 | 2.56562 | 0.92861 | 0.00000 | 0.61434 | 4.19926 | 8.65532 |
| **SLC25A24** | 2.64670 | 0.48558 | 0.63031 | 2.11680 | 0.00000 | 0.85033 | 0.06486 |
| **SLC25A25** | 0.00000 | 2.46571 | 1.77593 | 3.08503 | 2.43625 | 3.12120 | 3.28638 |
| **SLC25A26** | 10.12756 | 4.75026 | 3.27251 | 16.41567 | 3.75181 | 5.45618 | 6.07524 |
| **SLC25A27** | 0.00000 | 0.02365 | 0.01980 | 0.08239 | 0.01753 | 0.02006 | 0.05849 |
| **SLC25A28** | 0.02746 | 2.42760 | 1.24936 | 1.23100 | 0.13453 | 3.35613 | 6.02559 |
| **SLC25A29** | 0.00000 | 9.07371 | 0.14259 | 1.70644 | 0.08477 | 9.45738 | 13.66541 |
| **SLC25A3** | 68.48173 | 35.44642 | 32.28789 | 46.24715 | 9.99808 | 32.62228 | 48.68768 |
| **SLC25A30** | 0.79959 | 0.25194 | 0.13841 | 0.20979 | 0.65279 | 0.16250 | 1.16152 |
| **SLC25A30-AS1** | 0.02049 | 0.06304 | 0.00594 | 0.01514 | 0.01629 | 0.02921 | 0.03758 |
| **SLC25A31** | 0.06070 | 0.00000 | 0.50343 | 0.03112 | 0.17094 | 0.03245 | 0.00000 |
| **SLC25A32** | 17.70780 | 0.00000 | 9.07949 | 7.22465 | 5.63053 | 4.18934 | 0.00000 |
| **SLC25A33** | 0.54336 | 0.28727 | 1.38740 | 1.11689 | 1.00692 | 1.93194 | 8.75504 |
| **SLC25A34** | 0.13910 | 0.06626 | 0.14319 | 0.19532 | 0.00333 | 0.00000 | 0.44113 |
| **SLC25A35** | 2.44050 | 1.05261 | 0.04841 | 1.65317 | 0.02460 | 1.84145 | 0.67720 |
| **SLC25A36** | 20.92931 | 4.35353 | 9.02408 | 8.25059 | 8.28833 | 7.34305 | 10.48626 |
| **SLC25A37** | 3.17792 | 5.62741 | 1.26122 | 7.40675 | 0.18876 | 0.28828 | 5.83071 |
| **SLC25A38** | 12.78107 | 18.19205 | 9.45455 | 14.54202 | 3.92918 | 16.54330 | 2.51261 |

| | | | | | | | |
|---|---|---|---|---|---|---|---|
| **SLC25A39** | 48.87840 | 53.63987 | 24.25166 | 23.97891 | 7.01939 | 29.01877 | 15.62456 |
| **SLC25A3P1** | 0.02030 | 0.00255 | 0.01908 | 0.07079 | 0.38335 | 0.12236 | 0.01213 |
| **SLC25A4** | 3.23037 | 4.21085 | 2.01636 | 3.01329 | 0.42952 | 5.25580 | 0.00000 |
| **SLC25A40** | 2.67331 | 0.00000 | 1.08285 | 0.90321 | 0.05191 | 0.77647 | 0.95770 |
| **SLC25A41** | 0.06289 | 0.06603 | 0.01688 | 0.02684 | 0.01158 | 0.00307 | 0.16224 |
| **SLC25A42** | 0.00000 | 0.86189 | 0.20315 | 0.98411 | 0.37321 | 1.26408 | 0.44885 |
| **SLC25A43** | 3.53660 | 2.12519 | 2.80337 | 0.00000 | 0.31615 | 1.72910 | 6.66043 |
| **SLC25A44** | 36.22771 | 19.49505 | 14.05958 | 10.61352 | 16.20396 | 22.26691 | 5.37311 |
| **SLC25A45** | 0.04757 | 0.18714 | 0.14848 | 0.30059 | 0.26087 | 0.01368 | 0.00515 |
| **SLC25A46** | 11.11228 | 1.37111 | 2.88091 | 6.76116 | 6.43171 | 3.68291 | 0.00000 |
| **SLC25A47** | 0.00000 | 0.25541 | 0.08223 | 0.03610 | 0.00562 | 0.02486 | 0.01858 |
| **SLC25A48** | 0.05022 | 0.04843 | 0.01126 | 0.03164 | 0.41985 | 0.01180 | 0.00575 |
| **SLC25A5** | 89.31810 | 58.77088 | 21.10637 | 47.31127 | 23.27461 | 44.86405 | 31.83973 |
| **SLC25A5-AS1** | 0.22478 | 0.21307 | 0.18509 | 0.13265 | 0.32217 | 0.41190 | 0.31486 |
| **SLC25A51** | 1.75783 | 2.35112 | 2.03578 | 1.28885 | 0.08394 | 1.00167 | 1.31868 |
| **SLC25A51P1** | 0.02223 | 0.02445 | 0.01826 | 0.02200 | 0.00720 | 0.01482 | 0.02214 |
| **SLC25A52** | 0.07782 | 0.05197 | 0.13137 | 0.02677 | 0.00457 | 0.00829 | 0.00000 |
| **SLC25A53** | 0.24422 | 0.31930 | 0.23523 | 0.42805 | 0.69406 | 0.09310 | 0.01669 |
| **SLC26A1** | 0.02893 | 0.01660 | 0.00898 | 0.01847 | 0.06253 | 0.01285 | 0.01733 |
| **SLC26A10** | 0.01880 | 0.04312 | 0.13290 | 0.06327 | 0.02296 | 0.00934 | 0.02694 |
| **SLC26A11** | 2.35235 | 1.54321 | 0.71104 | 1.70996 | 0.00000 | 3.14439 | 3.26688 |
| **SLC26A2** | 0.74521 | 0.00000 | 0.07572 | 1.30537 | 0.39924 | 0.00890 | 0.01720 |
| **SLC26A3** | 0.02784 | 0.00370 | 0.03572 | 0.02172 | 0.08709 | 0.01282 | 0.02027 |
| **SLC26A4** | 0.08826 | 0.04527 | 0.08757 | 0.08552 | 0.00367 | 0.03384 | 0.17376 |
| **SLC26A4-AS1** | 0.05765 | 0.06778 | 0.10371 | 0.00989 | 0.00733 | 0.00444 | 0.03525 |
| **SLC26A5** | 0.07784 | 0.03933 | 0.01890 | 0.11527 | 0.05478 | 0.01412 | 0.00000 |
| **SLC26A6** | 9.74379 | 2.65860 | 1.12547 | 7.52841 | 3.23901 | 5.49543 | 5.45923 |
| **SLC26A7** | 0.00000 | 0.00000 | 0.04469 | 0.05817 | 0.15747 | 0.02138 | 0.18759 |
| **SLC26A8** | 0.22040 | 1.88635 | 0.08823 | 0.00011 | 1.00428 | 0.00000 | 0.40256 |
| **SLC26A9** | 0.03518 | 0.00000 | 0.09194 | 0.02930 | 0.00527 | 0.01345 | 0.01468 |
| **SLC27A1** | 0.00000 | 0.37114 | 0.98311 | 0.37003 | 0.94428 | 0.24952 | 0.48206 |
| **SLC27A2** | 2.63803 | 0.46906 | 0.00000 | 5.30073 | 0.29130 | 0.65313 | 2.00213 |
| **SLC27A3** | 0.14208 | 0.79165 | 0.62009 | 0.40645 | 0.00000 | 1.99973 | 1.60650 |
| **SLC27A4** | 3.86441 | 26.37382 | 6.99075 | 6.89352 | 0.00000 | 7.18045 | 17.39233 |
| **SLC27A5** | 1.67438 | 0.32682 | 0.40575 | 1.50249 | 0.26092 | 0.44239 | 0.65141 |

| | | | | | | | |
|---|---|---|---|---|---|---|---|
| **SLC27A6** | 0.04469 | 0.00266 | 0.07132 | 0.88772 | 2.35235 | 0.00292 | 0.02825 |
| **SLC28A1** | 0.02195 | 0.04646 | 0.03425 | 0.04643 | 0.01304 | 0.00734 | 0.01836 |
| **SLC28A2** | 0.15421 | 0.03839 | 0.02264 | 0.04987 | 0.00959 | 0.24099 | 0.00000 |
| **SLC28A3** | 28.88103 | 3.43119 | 4.97810 | 7.89478 | 0.70470 | 0.00000 | 8.28224 |
| **SLC29A1** | 18.33956 | 25.96015 | 4.34391 | 27.67823 | 6.01432 | 10.98633 | 8.48556 |
| **SLC29A2** | 7.02822 | 3.07597 | 1.18238 | 8.47730 | 0.00000 | 3.19655 | 12.82776 |
| **SLC29A3** | 3.73969 | 2.38634 | 2.59111 | 7.49776 | 0.93907 | 1.12638 | 0.50149 |
| **SLC29A4** | 0.02474 | 0.25214 | 0.04890 | 0.09215 | 2.10058 | 0.01696 | 0.18637 |
| **SLC2A1** | 50.84079 | 65.36069 | 2.05911 | 59.17591 | 1.68991 | 17.43083 | 17.14581 |
| **SLC2A1-AS1** | 0.00000 | 0.18629 | 0.00629 | 0.25517 | 0.00910 | 0.01001 | 0.16523 |
| **SLC2A10** | 0.01240 | 0.15292 | 0.15229 | 0.02220 | 0.02291 | 0.09263 | 0.02119 |
| **SLC2A11** | 2.80276 | 3.63507 | 0.15624 | 3.47220 | 0.00000 | 6.21087 | 0.84457 |
| **SLC2A12** | 0.75972 | 0.00000 | 0.51083 | 0.10082 | 0.44907 | 0.26415 | 0.68529 |
| **SLC2A13** | 0.15333 | 0.04645 | 0.00463 | 0.36662 | 0.47209 | 0.85951 | 0.85015 |
| **SLC2A14** | 1.24310 | 0.00000 | 4.39305 | 1.69057 | 1.21407 | 0.59265 | 0.85422 |
| **SLC2A2** | 0.05405 | 0.00000 | 0.01908 | 0.02341 | 0.15081 | 0.05156 | 0.00000 |
| **SLC2A3** | 51.95607 | 20.48409 | 50.19301 | 53.05297 | 18.54501 | 10.01161 | 32.62173 |
| **SLC2A4** | 0.76034 | 0.30852 | 1.06900 | 0.61195 | 0.78668 | 1.32388 | 1.41714 |
| **SLC2A4RG** | 0.00000 | 0.57063 | 0.12904 | 0.31991 | 0.14488 | 0.55212 | 2.18730 |
| **SLC2A5** | 0.05519 | 0.18686 | 1.17257 | 0.04084 | 0.39948 | 0.27323 | 0.00000 |
| **SLC2A6** | 0.00611 | 0.00216 | 0.04698 | 0.16338 | 0.01742 | 0.23345 | 0.02395 |
| **SLC2A7** | 0.02742 | 0.01572 | 0.01150 | 0.01806 | 0.02581 | 0.01753 | 0.01632 |
| **SLC2A8** | 2.36019 | 0.12880 | 0.51184 | 4.04795 | 0.84503 | 0.68947 | 6.94008 |
| **SLC2A9** | 0.16405 | 0.16635 | 0.07787 | 0.12155 | 0.02075 | 0.26990 | 0.00000 |
| **SLC30A1** | 0.00000 | 2.96790 | 1.98651 | 2.97422 | 1.15122 | 2.30479 | 3.37430 |
| **SLC30A10** | 0.02449 | 0.01822 | 0.00756 | 0.01928 | 0.06941 | 0.01340 | 0.01598 |
| **SLC30A2** | 1.46321 | 5.58949 | 0.00000 | 0.00000 | 0.00000 | 0.00000 | 2.70401 |
| **SLC30A3** | 0.09313 | 0.03401 | 0.07108 | 0.06015 | 0.36607 | 0.01457 | 0.02064 |
| **SLC30A4** | 0.04729 | 0.15013 | 0.16022 | 0.02063 | 0.37215 | 0.04095 | 0.00219 |
| **SLC30A5** | 7.30412 | 0.00000 | 3.78174 | 3.74775 | 0.51552 | 4.89683 | 13.24685 |
| **SLC30A6** | 10.68771 | 1.58774 | 7.77113 | 4.12045 | 1.69935 | 3.73895 | 2.78810 |
| **SLC30A7** | 15.83870 | 4.08400 | 12.48103 | 10.76006 | 6.64528 | 4.74407 | 0.00000 |
| **SLC30A8** | 0.29335 | 0.00000 | 0.60207 | 0.04806 | 0.53744 | 0.11960 | 0.00000 |
| **SLC30A9** | 13.94703 | 5.03898 | 2.39155 | 3.05221 | 1.24341 | 5.37857 | 1.75353 |
| **SLC31A1** | 26.53902 | 14.59405 | 2.45124 | 16.31562 | 3.08781 | 4.23871 | 0.00000 |

| | | | | | | | |
|---|---|---|---|---|---|---|---|
| **SLC31A2** | 2.79216 | 1.75874 | 3.30655 | 5.05764 | 1.48296 | 0.50083 | 0.00000 |
| **SLC32A1** | 0.03033 | 0.00839 | 0.02521 | 0.02390 | 0.00735 | 0.04391 | 0.01331 |
| **SLC33A1** | 10.30017 | 2.62772 | 6.48918 | 5.26577 | 1.52887 | 3.60217 | 0.00000 |
| **SLC34A1** | 0.02677 | 0.01834 | 0.01118 | 0.01679 | 0.01380 | 0.01750 | 0.02470 |
| **SLC34A2** | 12.99070 | 22.70884 | 0.53082 | 4.65047 | 30.29409 | 0.00000 | 0.19790 |
| **SLC34A3** | 0.02106 | 0.01709 | 0.01335 | 0.01846 | 0.00775 | 0.02942 | 0.02225 |
| **SLC35A1** | 12.14935 | 2.90025 | 2.31528 | 0.36021 | 1.10856 | 5.09126 | 4.06497 |
| **SLC35A2** | 18.17110 | 8.33199 | 4.70075 | 12.00289 | 0.30971 | 6.62843 | 2.88526 |
| **SLC35A3** | 3.50643 | 0.53372 | 0.75626 | 3.02584 | 0.00000 | 2.36282 | 5.92773 |
| **SLC35A4** | 20.49438 | 12.04611 | 15.37796 | 31.22546 | 1.69663 | 8.48583 | 0.00000 |
| **SLC35A5** | 8.03305 | 0.00000 | 14.97153 | 3.05187 | 4.92480 | 8.08489 | 0.00000 |
| **SLC35B1** | 20.53240 | 15.10569 | 19.31188 | 14.83059 | 6.13740 | 11.80935 | 7.67645 |
| **SLC35B2** | 5.94152 | 8.20531 | 3.32622 | 6.67385 | 0.34232 | 5.00823 | 5.58515 |
| **SLC35B3** | 5.40231 | 1.39455 | 2.39315 | 3.03553 | 0.36240 | 1.61383 | 0.62438 |
| **SLC35B4** | 25.46793 | 8.88109 | 4.45011 | 5.79386 | 0.72489 | 5.73177 | 3.78695 |
| **SLC35C1** | 7.77732 | 3.71514 | 1.92572 | 7.44499 | 2.25166 | 0.25357 | 12.09611 |
| **SLC35C2** | 3.90204 | 6.14866 | 5.94611 | 12.30200 | 1.00824 | 2.01021 | 1.16008 |
| **SLC35D1** | 5.80688 | 0.01714 | 1.00294 | 5.46973 | 3.08811 | 2.17069 | 0.83756 |
| **SLC35D2** | 0.47198 | 0.83615 | 0.26782 | 0.52732 | 0.11243 | 1.03743 | 4.76911 |
| **SLC35E1** | 5.99935 | 4.74332 | 3.10862 | 5.92433 | 1.76191 | 1.88740 | 14.17680 |
| **SLC35E2** | 0.16340 | 0.38516 | 0.13260 | 0.52252 | 0.08636 | 0.05622 | 0.69647 |
| **SLC35E2B** | 0.36471 | 1.05962 | 0.29070 | 1.37605 | 0.14148 | 0.77579 | 1.48705 |
| **SLC35E3** | 7.40511 | 1.48567 | 2.57125 | 5.56998 | 0.23060 | 2.87988 | 0.00000 |
| **SLC35E4** | 0.00000 | 13.78403 | 16.45480 | 0.00000 | 3.46650 | 6.40220 | 1.41661 |
| **SLC35F1** | 0.32761 | 0.09742 | 0.07516 | 0.14848 | 0.03494 | 0.31398 | 0.00000 |
| **SLC35F2** | 27.16256 | 17.60602 | 29.93628 | 12.32796 | 11.23504 | 9.75887 | 0.00000 |
| **SLC35F3** | 0.00000 | 0.14164 | 0.36059 | 0.00320 | 0.50184 | 0.01676 | 0.04382 |
| **SLC35F4** | 0.03026 | 0.01961 | 0.04615 | 0.02337 | 0.24825 | 0.00981 | 0.01729 |
| **SLC35F5** | 16.18166 | 2.20506 | 3.34121 | 6.85335 | 2.87483 | 1.64656 | 0.00000 |
| **SLC35F6** | 18.88911 | 36.82617 | 10.92249 | 12.65499 | 0.52371 | 26.86709 | 0.00000 |
| **SLC35G1** | 3.69098 | 0.66906 | 3.11796 | 2.47389 | 1.51601 | 0.46838 | 0.92404 |
| **SLC35G2** | 0.00000 | 0.03034 | 0.40145 | 0.11737 | 0.17179 | 0.11272 | 0.01317 |
| **SLC35G3** | 0.00000 | 0.67647 | 0.85927 | 0.00000 | 0.11708 | 0.33302 | 0.00777 |
| **SLC35G4P** | 0.02410 | 0.02954 | 0.04773 | 0.01803 | 0.09432 | 0.01258 | 0.01103 |
| **SLC35G5** | 0.00000 | 0.17756 | 0.09568 | 0.00000 | 0.11781 | 0.24800 | 0.12198 |

| | | | | | | | |
|---|---|---|---|---|---|---|---|
| **SLC35G6** | 0.03019 | 0.01785 | 0.00954 | 0.02886 | 0.02507 | 0.00777 | 0.05148 |
| **SLC36A1** | 5.84048 | 0.00000 | 0.92931 | 4.19196 | 0.34966 | 1.61198 | 5.72140 |
| **SLC36A2** | 1.13421 | 0.50054 | 0.12688 | 0.02646 | 0.52667 | 0.00000 | 2.35588 |
| **SLC36A3** | 0.09668 | 0.06283 | 0.00088 | 0.02536 | 0.06744 | 0.01337 | 0.03770 |
| **SLC36A4** | 0.00000 | 0.06948 | 0.64594 | 3.79757 | 0.00125 | 0.37834 | 0.98875 |
| **SLC37A1** | 0.93436 | 0.89279 | 0.00000 | 0.00000 | 0.68558 | 0.81169 | 1.45084 |
| **SLC37A2** | 0.00888 | 0.05995 | 0.09063 | 0.02611 | 0.41932 | 0.01790 | 0.04970 |
| **SLC37A3** | 19.52003 | 11.99157 | 9.98069 | 11.54842 | 0.93664 | 12.96253 | 0.90758 |
| **SLC37A4** | 6.37551 | 12.11106 | 7.37893 | 6.43304 | 0.00000 | 14.11499 | 11.04468 |
| **SLC38A1** | 26.22265 | 0.00000 | 0.00000 | 53.01224 | 0.00000 | 0.00000 | 6.45220 |
| **SLC38A10** | 0.00000 | 2.89766 | 1.10183 | 2.06939 | 0.02186 | 1.55537 | 10.05818 |
| **SLC38A11** | 0.04817 | 0.01252 | 0.01325 | 0.01822 | 0.00717 | 0.00485 | 0.08147 |
| **SLC38A2** | 27.74900 | 0.00000 | 1.33636 | 32.05966 | 1.59333 | 12.29889 | 10.36077 |
| **SLC38A3** | 0.00000 | 0.01924 | 1.14409 | 4.55836 | 0.01026 | 0.00000 | 0.00000 |
| **SLC38A4** | 0.06736 | 0.01777 | 0.01072 | 0.42651 | 0.01839 | 0.00773 | 0.00000 |
| **SLC38A5** | 0.09139 | 0.14700 | 0.01350 | 0.00000 | 0.04322 | 0.04732 | 0.34763 |
| **SLC38A6** | 1.90530 | 0.99295 | 1.08055 | 5.04684 | 1.52778 | 0.31105 | 0.00000 |
| **SLC38A7** | 7.49422 | 7.82951 | 2.95375 | 8.87389 | 1.77035 | 4.26210 | 0.00000 |
| **SLC38A8** | 0.00000 | 0.10750 | 0.33927 | 0.22118 | 4.26050 | 0.00000 | 0.19245 |
| **SLC38A9** | 14.30544 | 3.73328 | 0.59563 | 7.80604 | 0.10938 | 7.21730 | 0.00000 |
| **SLC39A1** | 38.78325 | 30.70373 | 14.64821 | 39.47552 | 2.14253 | 16.36010 | 0.00054 |
| **SLC39A10** | 9.75687 | 0.00000 | 9.05369 | 2.37365 | 6.88968 | 23.53983 | 7.49796 |
| **SLC39A11** | 5.24811 | 3.94173 | 1.08902 | 0.00000 | 0.14173 | 1.45641 | 1.74151 |
| **SLC39A12** | 0.02055 | 0.06715 | 0.03405 | 0.02677 | 0.07348 | 0.00801 | 0.01113 |
| **SLC39A13** | 0.82722 | 0.48225 | 0.26526 | 1.54946 | 0.03620 | 1.68084 | 0.41572 |
| **SLC39A14** | 26.82583 | 3.48636 | 11.35538 | 12.73110 | 1.33658 | 3.13714 | 14.34582 |
| **SLC39A2** | 0.42138 | 0.00000 | 0.03409 | 0.01909 | 0.04077 | 0.01382 | 0.00000 |
| **SLC39A3** | 1.97540 | 13.82909 | 9.74766 | 10.58117 | 1.79608 | 3.25312 | 14.42305 |
| **SLC39A4** | 7.52115 | 9.19650 | 1.40822 | 3.30775 | 1.02319 | 6.59273 | 3.26596 |
| **SLC39A5** | 0.00093 | 0.01074 | 0.01271 | 0.04046 | 0.00870 | 0.15748 | 0.08619 |
| **SLC39A6** | 9.23286 | 0.00000 | 9.27112 | 9.16918 | 5.96856 | 2.77502 | 0.00000 |
| **SLC39A7** | 12.18371 | 7.08325 | 6.33509 | 14.55152 | 0.29759 | 7.17573 | 2.61010 |
| **SLC39A8** | 0.08145 | 0.04833 | 0.00506 | 0.24571 | 0.01238 | 0.77376 | 0.00000 |
| **SLC39A9** | 19.63554 | 4.28014 | 6.76807 | 13.22006 | 5.65368 | 2.80422 | 4.71700 |
| **SLC3A1** | 0.00952 | 0.00000 | 0.04309 | 0.18841 | 4.80604 | 0.40100 | 0.00148 |

| | | | | | | | |
|---|---|---|---|---|---|---|---|
| **SLC3A2** | 78.35873 | 55.67575 | 36.02088 | 67.37143 | 19.05631 | 25.34539 | 18.14076 |
| **SLC40A1** | 4.54069 | 1.21277 | 0.00000 | 26.06979 | 0.04828 | 0.76335 | 0.00000 |
| **SLC41A1** | 0.40484 | 0.00000 | 6.40081 | 1.51741 | 2.34350 | 1.66163 | 1.28859 |
| **SLC41A2** | 1.79538 | 0.08787 | 0.59004 | 0.32534 | 1.40736 | 0.10402 | 0.00000 |
| **SLC41A3** | 3.76734 | 0.00000 | 0.78077 | 8.79022 | 0.38250 | 1.46167 | 5.18559 |
| **SLC43A1** | 0.14702 | 0.01718 | 0.10265 | 0.38048 | 0.16771 | 1.24192 | 0.01010 |
| **SLC43A2** | 0.00000 | 9.11622 | 8.23503 | 6.25473 | 0.66970 | 0.00000 | 7.37499 |
| **SLC43A3** | 0.07749 | 0.83324 | 3.38349 | 0.11987 | 2.59852 | 0.00000 | 0.00000 |
| **SLC44A1** | 2.90805 | 1.72894 | 1.43622 | 1.45623 | 0.21851 | 9.09469 | 5.33193 |
| **SLC44A2** | 13.07170 | 34.35063 | 4.65983 | 20.46067 | 0.00000 | 16.02004 | 30.68907 |
| **SLC44A3** | 0.39319 | 0.05930 | 0.01666 | 0.08171 | 0.03566 | 0.72033 | 0.41150 |
| **SLC44A4** | 0.01123 | 0.05039 | 3.50462 | 0.13064 | 0.00116 | 0.00000 | 0.02124 |
| **SLC44A5** | 0.21320 | 0.00000 | 0.78232 | 0.34987 | 0.07710 | 0.29841 | 0.31214 |
| **SLC45A1** | 0.03909 | 0.01859 | 0.02449 | 0.00191 | 0.11792 | 0.03795 | 0.09012 |
| **SLC45A3** | 0.00000 | 0.02762 | 0.04091 | 0.12327 | 0.54887 | 0.18936 | 0.08897 |
| **SLC45A4** | 0.00000 | 0.61252 | 0.96302 | 3.91493 | 0.50357 | 5.32165 | 3.14214 |
| **SLC46A1** | 0.60002 | 4.25629 | 0.54812 | 0.68897 | 0.00000 | 0.00000 | 1.83425 |
| **SLC46A2** | 0.11596 | 0.08251 | 0.04101 | 0.00000 | 0.00433 | 0.02368 | 0.23792 |
| **SLC46A3** | 0.10180 | 0.00996 | 0.01548 | 0.00259 | 0.05556 | 0.03885 | 0.06511 |
| **SLC47A1** | 0.00000 | 3.56793 | 6.40585 | 0.00000 | 0.06162 | 7.52622 | 0.00000 |
| **SLC47A2** | 0.00000 | 0.36639 | 0.06946 | 0.03460 | 0.07799 | 0.05025 | 0.00610 |
| **SLC48A1** | 3.02114 | 7.73191 | 0.01006 | 0.00000 | 0.37172 | 0.00000 | 1.22382 |
| **SLC4A1** | 0.00000 | 0.29973 | 0.11763 | 0.26717 | 0.04451 | 0.08856 | 0.00000 |
| **SLC4A10** | 0.02939 | 0.01369 | 0.01344 | 0.02205 | 0.00809 | 0.01599 | 0.02849 |
| **SLC4A11** | 8.13589 | 41.72874 | 9.41065 | 0.00010 | 0.15607 | 6.71270 | 18.19590 |
| **SLC4A1AP** | 14.96985 | 10.99615 | 9.73424 | 8.10365 | 4.61904 | 6.37936 | 1.49689 |
| **SLC4A2** | 3.73729 | 11.85860 | 3.09376 | 19.03890 | 0.00000 | 5.15624 | 30.79892 |
| **SLC4A3** | 0.00000 | 0.01831 | 0.04387 | 0.21578 | 0.16961 | 0.07274 | 0.17900 |
| **SLC4A4** | 0.07680 | 0.00000 | 0.05777 | 0.00212 | 0.06510 | 0.00618 | 0.05200 |
| **SLC4A5** | 5.33904 | 0.00000 | 0.48322 | 2.13774 | 0.31305 | 0.88689 | 6.66673 |
| **SLC4A7** | 1.44448 | 0.00000 | 0.71112 | 1.01781 | 6.79345 | 0.67329 | 3.95087 |
| **SLC4A8** | 1.11787 | 2.44706 | 1.66554 | 0.01470 | 0.15868 | 11.36339 | 0.00000 |
| **SLC4A9** | 0.03211 | 0.02996 | 0.03303 | 0.05769 | 0.00816 | 0.01240 | 0.00600 |
| **SLC50A1** | 6.02115 | 2.18659 | 3.77580 | 6.90264 | 2.20090 | 1.28133 | 0.00000 |
| **SLC51A** | 0.02762 | 0.01443 | 0.02907 | 0.04569 | 0.23605 | 0.02682 | 0.10951 |

| | | | | | | | |
|---|---|---|---|---|---|---|---|
| **SLC52A1** | 0.56685 | 4.55695 | 3.18900 | 3.57186 | 0.56559 | 4.03668 | 2.04221 |
| **SLC52A2** | 1.83075 | 20.91485 | 7.90578 | 9.75823 | 0.35012 | 8.28637 | 19.02191 |
| **SLC52A3** | 2.64167 | 4.68634 | 2.67254 | 1.95437 | 0.00000 | 0.73431 | 0.00000 |
| **SLC5A1** | 0.15231 | 0.00000 | 0.00988 | 0.12755 | 0.12255 | 0.11145 | 0.00000 |
| **SLC5A10** | 0.44243 | 0.93469 | 0.00000 | 1.57886 | 0.00483 | 0.26698 | 1.52055 |
| **SLC5A11** | 0.04806 | 0.26523 | 0.00000 | 0.26274 | 3.27899 | 0.07135 | 0.15627 |
| **SLC5A12** | 0.05279 | 0.01787 | 0.01346 | 0.02058 | 0.06126 | 0.01640 | 0.07221 |
| **SLC5A2** | 0.03972 | 0.02561 | 0.02146 | 0.01257 | 0.02615 | 0.00943 | 0.01974 |
| **SLC5A3** | 5.06882 | 0.00000 | 1.99481 | 2.89053 | 0.74656 | 1.62885 | 8.66024 |
| **SLC5A4** | 0.02003 | 0.01707 | 0.02031 | 0.02753 | 0.02617 | 0.01229 | 0.01782 |
| **SLC5A5** | 2.07548 | 5.23490 | 10.46062 | 0.41980 | 4.33681 | 6.28577 | 0.00000 |
| **SLC5A6** | 50.35569 | 11.38352 | 11.06808 | 39.56879 | 2.43642 | 12.98259 | 20.19036 |
| **SLC5A7** | 0.02070 | 0.03018 | 0.01649 | 0.01903 | 0.00735 | 0.01168 | 0.01964 |
| **SLC5A9** | 0.02792 | 0.04166 | 0.02363 | 0.02754 | 0.07524 | 0.33245 | 0.00000 |
| **SLC6A1** | 0.01012 | 0.01715 | 0.01213 | 0.05394 | 0.00754 | 0.00952 | 0.04866 |
| **SLC6A10P** | 0.00000 | 0.08251 | 0.04918 | 0.04093 | 0.01247 | 0.01706 | 0.11298 |
| **SLC6A11** | 0.05697 | 0.01844 | 0.10563 | 0.03359 | 0.17514 | 0.00000 | 0.12533 |
| **SLC6A12** | 0.07934 | 0.00000 | 0.11319 | 0.02681 | 0.07496 | 0.01785 | 0.00617 |
| **SLC6A13** | 0.02923 | 0.06390 | 0.02758 | 0.02651 | 0.24807 | 0.07449 | 0.16735 |
| **SLC6A14** | 0.02599 | 0.00793 | 0.02886 | 0.02700 | 0.05764 | 0.01614 | 0.01477 |
| **SLC6A15** | 0.03377 | 0.00000 | 0.01260 | 0.35415 | 0.14411 | 0.02752 | 0.07254 |
| **SLC6A16** | 0.01821 | 0.00000 | 0.32792 | 0.25784 | 0.75305 | 0.09860 | 0.89026 |
| **SLC6A17** | 0.04011 | 0.00000 | 0.04868 | 0.03376 | 0.22512 | 0.11938 | 0.02938 |
| **SLC6A18** | 0.00636 | 0.01421 | 0.01374 | 0.05160 | 0.00954 | 0.02128 | 0.06313 |
| **SLC6A19** | 22.30793 | 26.62058 | 4.74636 | 3.80953 | 0.00000 | 11.15764 | 17.98726 |
| **SLC6A2** | 0.00003 | 0.01389 | 0.08440 | 0.08776 | 0.00403 | 0.00000 | 0.18655 |
| **SLC6A20** | 0.00000 | 0.03254 | 0.03350 | 0.08778 | 0.50311 | 0.14159 | 0.06992 |
| **SLC6A3** | 0.19771 | 0.70409 | 0.09524 | 0.81489 | 1.60303 | 0.82460 | 0.00000 |
| **SLC6A4** | 0.04937 | 0.04529 | 0.13110 | 0.00000 | 0.01249 | 0.00000 | 0.70472 |
| **SLC6A5** | 0.00342 | 0.00044 | 1.73479 | 0.00866 | 6.33035 | 0.00000 | 0.02540 |
| **SLC6A6** | 4.87456 | 1.85349 | 0.00000 | 10.26431 | 0.33555 | 16.99934 | 8.72879 |
| **SLC6A7** | 0.01405 | 0.03322 | 0.02806 | 0.02669 | 0.00614 | 0.01233 | 0.03578 |
| **SLC6A8** | 0.00000 | 37.93060 | 27.03833 | 0.00000 | 1.21087 | 14.60716 | 13.39618 |
| **SLC6A9** | 1.56395 | 0.00000 | 0.00762 | 4.70076 | 1.30233 | 0.20529 | 3.39292 |
| **SLC7A1** | 0.72654 | 0.00000 | 0.02137 | 0.77228 | 0.00552 | 0.00000 | 3.11998 |

| | | | | | | | |
|---|---|---|---|---|---|---|---|
| **SLC7A10** | 0.00000 | 0.04673 | 0.01070 | 0.14243 | 0.01911 | 0.06339 | 0.05184 |
| **SLC7A11** | 9.28946 | 0.00000 | 0.00000 | 9.12702 | 0.81217 | 0.00000 | 12.09957 |
| **SLC7A11-AS1** | 0.29061 | 0.00000 | 0.04168 | 0.00000 | 0.56338 | 1.25546 | 0.46129 |
| **SLC7A14** | 0.01966 | 0.01924 | 0.00779 | 0.02883 | 0.12599 | 0.08013 | 0.01803 |
| **SLC7A2** | 46.24378 | 3.94548 | 0.75586 | 27.93589 | 0.29666 | 0.00000 | 46.27281 |
| **SLC7A3** | 7.03123 | 25.34856 | 2.35080 | 0.00000 | 0.00000 | 13.11536 | 0.00000 |
| **SLC7A4** | 37.69462 | 3.69406 | 0.00000 | 23.32380 | 0.00000 | 0.00000 | 5.92388 |
| **SLC7A5** | 27.32474 | 24.24437 | 0.17741 | 33.28057 | 0.23020 | 0.00000 | 63.16560 |
| **SLC7A5P1** | 0.00000 | 0.45475 | 0.00000 | 0.16096 | 0.05970 | 0.09463 | 0.29900 |
| **SLC7A5P2** | 0.00000 | 0.75098 | 0.06057 | 0.04023 | 0.08967 | 0.12755 | 2.25357 |
| **SLC7A6** | 7.49947 | 2.04895 | 4.02018 | 19.09374 | 5.36213 | 5.69275 | 11.28947 |
| **SLC7A6OS** | 6.44167 | 6.81928 | 2.71245 | 4.72402 | 11.60056 | 1.82799 | 0.00000 |
| **SLC7A7** | 44.94776 | 21.18586 | 3.59707 | 19.33802 | 5.28853 | 6.57319 | 15.30792 |
| **SLC7A8** | 71.04963 | 21.64154 | 0.00000 | 60.36231 | 6.96907 | 14.21642 | 47.46254 |
| **SLC7A9** | 0.05677 | 0.08221 | 0.00000 | 0.04556 | 3.22630 | 0.00439 | 0.00000 |
| **SLC8A1** | 0.73539 | 0.00000 | 0.61725 | 0.01845 | 2.47831 | 0.04009 | 0.61837 |
| **SLC8A1-AS1** | 0.02243 | 0.00750 | 0.00368 | 0.00922 | 0.20139 | 0.00968 | 0.15444 |
| **SLC8A2** | 0.00000 | 0.27301 | 0.07169 | 0.07669 | 2.69420 | 0.09372 | 0.13203 |
| **SLC8A3** | 0.01623 | 0.00842 | 0.01329 | 0.36354 | 0.00709 | 0.01169 | 0.08336 |
| **SLC8B1** | 6.07768 | 29.24310 | 1.42980 | 1.77623 | 0.00000 | 8.07428 | 3.81404 |
| **SLC9A1** | 0.83980 | 12.67421 | 2.52257 | 6.04306 | 2.74055 | 9.51554 | 26.11683 |
| **SLC9A2** | 0.10169 | 0.12903 | 0.00462 | 0.00000 | 0.01109 | 0.00000 | 0.23198 |
| **SLC9A3** | 0.00000 | 0.46243 | 0.31191 | 0.04843 | 0.08475 | 0.33942 | 0.66289 |
| **SLC9A3R1** | 17.43469 | 38.41796 | 2.25482 | 26.18392 | 1.06432 | 6.87687 | 43.94509 |
| **SLC9A3R2** | 0.00000 | 3.08419 | 2.99084 | 0.00000 | 0.06369 | 2.70340 | 0.65310 |
| **SLC9A4** | 0.02918 | 0.02074 | 0.00792 | 0.03300 | 0.04923 | 0.02338 | 0.02795 |
| **SLC9A5** | 0.00000 | 0.01712 | 0.23175 | 0.24093 | 0.01778 | 0.07931 | 0.29722 |
| **SLC9A6** | 5.05203 | 4.03855 | 2.01602 | 3.85143 | 2.67862 | 0.13685 | 4.08504 |
| **SLC9A7** | 0.13065 | 1.18693 | 0.45654 | 0.59042 | 0.54672 | 0.51375 | 1.55034 |
| **SLC9A7P1** | 0.07446 | 0.01928 | 0.02434 | 0.03680 | 0.01664 | 0.00963 | 0.02881 |
| **SLC9A8** | 0.82356 | 2.85637 | 2.66871 | 1.33742 | 0.31345 | 0.71705 | 3.34096 |
| **SLC9A9** | 0.04378 | 0.02272 | 0.00000 | 0.01633 | 0.82203 | 0.02878 | 0.00752 |
| **SLC9B1** | 0.63407 | 0.08840 | 0.15607 | 0.44509 | 0.04427 | 0.20826 | 0.00000 |
| **SLC9B2** | 0.26897 | 0.10449 | 0.12439 | 0.42722 | 0.29260 | 0.14516 | 0.20961 |
| **SLC9C1** | 0.05011 | 0.02283 | 0.02118 | 0.04406 | 0.04371 | 0.03641 | 0.12390 |

| | | | | | | | |
|---|---|---|---|---|---|---|---|
| **SLC9C2** | 0.02789 | 0.01474 | 0.01294 | 0.02151 | 0.00912 | 0.10025 | 0.01634 |
| **SLCO1A2** | 1.14555 | 0.00000 | 0.68152 | 0.03848 | 0.00065 | 1.30911 | 0.00000 |
| **SLCO1B1** | 0.09796 | 0.00000 | 0.05991 | 0.00000 | 0.07072 | 0.09110 | 0.17571 |
| **SLCO1B3** | 0.08309 | 0.00000 | 0.02542 | 0.07321 | 0.42522 | 0.20049 | 0.00373 |
| **SLCO1B7** | 0.01159 | 0.01607 | 0.01665 | 0.01663 | 0.03943 | 0.04170 | 0.04213 |
| **SLCO1C1** | 0.03842 | 0.00000 | 0.03412 | 0.02398 | 0.01178 | 0.21335 | 0.02395 |
| **SLCO2A1** | 0.99780 | 0.03240 | 0.00000 | 4.30250 | 0.09371 | 3.22774 | 1.80521 |
| **SLCO2B1** | 1.67603 | 2.84427 | 0.59380 | 0.00000 | 0.00000 | 0.00000 | 2.76354 |
| **SLCO3A1** | 0.30081 | 0.36451 | 0.41021 | 0.00000 | 1.90450 | 0.05900 | 0.00000 |
| **SLCO4A1** | 0.00000 | 0.01839 | 0.01745 | 12.70917 | 1.26105 | 0.03157 | 0.39477 |
| **SLCO4A1-AS1** | 0.02071 | 0.01941 | 0.01260 | 0.05814 | 0.01777 | 0.01318 | 0.00914 |
| **SLCO4C1** | 7.56570 | 1.33661 | 7.04561 | 0.00000 | 0.05065 | 4.01507 | 0.87336 |
| **SLCO5A1** | 0.02362 | 0.13534 | 0.01261 | 0.00460 | 0.00928 | 1.12958 | 0.00706 |
| **SLCO6A1** | 0.05945 | 0.00017 | 0.02459 | 0.11384 | 0.00880 | 0.01072 | 0.03255 |
| **SLFN11** | 0.02440 | 0.01684 | 0.01260 | 0.01939 | 0.00953 | 0.01918 | 0.01931 |
| **SLFN12** | 0.00004 | 0.28822 | 0.03975 | 0.51196 | 0.18445 | 0.18156 | 0.05342 |
| **SLFN12L** | 0.02462 | 0.00000 | 0.06940 | 0.01857 | 0.24332 | 0.01247 | 0.01597 |
| **SLFN13** | 0.08555 | 0.00000 | 0.30727 | 0.06525 | 0.45434 | 0.08762 | 0.00000 |
| **SLFN14** | 0.02475 | 0.01667 | 0.00462 | 0.01812 | 0.24312 | 0.01310 | 0.01575 |
| **SLFN5** | 0.04800 | 0.00000 | 0.24643 | 0.09180 | 0.00225 | 0.03932 | 0.18584 |
| **SLFNL1** | 0.04977 | 0.05454 | 0.09831 | 0.14543 | 0.00420 | 0.00246 | 0.01205 |
| **SLFNL1-AS1** | 0.08414 | 0.00000 | 0.38718 | 0.32022 | 0.00836 | 0.00000 | 0.52620 |
| **SLIRP** | 66.02811 | 57.85106 | 30.89903 | 47.89535 | 11.04427 | 35.81031 | 30.96435 |
| **SLIT1** | 0.04655 | 0.02389 | 0.06283 | 0.03407 | 0.00710 | 0.06091 | 0.03966 |
| **SLIT2** | 0.07176 | 0.54507 | 0.05254 | 0.03693 | 0.06953 | 0.21835 | 0.09329 |
| **SLIT3** | 0.03777 | 0.03884 | 0.10125 | 0.01953 | 4.31142 | 0.15611 | 0.00283 |
| **SLITRK1** | 0.37239 | 0.13355 | 0.08150 | 0.22959 | 0.27158 | 0.20187 | 0.19789 |
| **SLITRK2** | 0.00000 | 0.77796 | 0.43549 | 0.00000 | 0.00802 | 1.10324 | 0.33327 |
| **SLITRK4** | 0.04177 | 0.00000 | 0.03543 | 0.00000 | 0.03860 | 0.04470 | 0.40185 |
| **SLITRK5** | 0.01150 | 0.03578 | 0.00958 | 0.01976 | 0.00853 | 0.07752 | 0.03650 |
| **SLK** | 14.21868 | 3.73745 | 2.38243 | 1.39165 | 0.00000 | 12.12901 | 27.24369 |
| **SLMAP** | 6.14170 | 0.01665 | 5.21504 | 3.28943 | 0.92023 | 6.16389 | 12.62385 |
| **SLMO1** | 0.04023 | 0.02674 | 0.26166 | 0.18180 | 0.00424 | 0.00000 | 0.55264 |
| **SLMO2** | 39.04765 | 7.76710 | 4.28801 | 10.09301 | 0.52371 | 11.84388 | 5.06290 |
| **SLN** | 0.26846 | 0.20985 | 0.14400 | 0.67268 | 0.07598 | 0.22631 | 0.00760 |

| | | | | | | | |
|---|---|---|---|---|---|---|---|
| **SLPI** | 3.46923 | 0.50179 | 0.00000 | 0.67467 | 0.01113 | 0.00000 | 1.69402 |
| **SLTM** | 16.90042 | 6.59233 | 14.60585 | 24.14386 | 3.04961 | 13.99568 | 30.98720 |
| **SLU7** | 16.76074 | 3.25038 | 17.32649 | 13.15729 | 11.05424 | 7.61494 | 3.05625 |
| **SLX4** | 0.49042 | 1.04170 | 2.17456 | 1.32510 | 0.56217 | 0.77719 | 4.40877 |
| **SLX4IP** | 0.93491 | 0.74236 | 1.86879 | 0.63987 | 0.88419 | 1.19869 | 0.73092 |
| **SMA4** | 0.02260 | 0.01702 | 0.01340 | 0.01860 | 0.00778 | 0.01240 | 0.02524 |
| **SMA5** | 0.02068 | 0.01944 | 0.01447 | 0.02172 | 0.00760 | 0.01387 | 0.02214 |
| **SMAD1** | 2.54917 | 0.34594 | 0.48800 | 1.09247 | 1.19022 | 0.59157 | 3.79894 |
| **SMAD1-AS2** | 0.05468 | 0.00735 | 0.01414 | 0.01922 | 0.00759 | 0.00960 | 0.03311 |
| **SMAD2** | 13.18401 | 0.97241 | 9.19962 | 6.29863 | 2.87489 | 8.16814 | 11.92754 |
| **SMAD3** | 0.00000 | 3.55799 | 0.75404 | 0.52396 | 0.51707 | 4.11238 | 0.40619 |
| **SMAD4** | 6.97627 | 0.86555 | 2.38698 | 4.22492 | 0.83428 | 4.32334 | 10.70314 |
| **SMAD5** | 24.03908 | 13.55793 | 4.88046 | 13.96052 | 3.11227 | 4.66327 | 17.81281 |
| **SMAD5-AS1** | 0.19487 | 1.11253 | 0.03685 | 0.62791 | 0.02912 | 0.03806 | 0.87181 |
| **SMAD6** | 0.00000 | 0.05964 | 0.00779 | 0.00000 | 0.04288 | 1.62507 | 0.27026 |
| **SMAD7** | 1.92019 | 0.00000 | 0.00000 | 1.96438 | 0.00000 | 5.21650 | 7.22320 |
| **SMAD9** | 1.08976 | 1.18797 | 0.95286 | 0.37772 | 0.15270 | 1.49879 | 2.68352 |
| **SMAGP** | 17.25386 | 6.23587 | 0.00000 | 27.88601 | 0.12999 | 0.16122 | 13.19856 |
| **SMAP1** | 12.37951 | 12.73988 | 6.48940 | 10.76258 | 1.08486 | 6.10627 | 15.82406 |
| **SMAP2** | 0.77141 | 0.78996 | 0.43422 | 1.45950 | 0.34804 | 1.04739 | 4.31752 |
| **SMARCA1** | 0.48901 | 0.00649 | 0.05239 | 0.29956 | 0.36089 | 0.02574 | 0.06929 |
| **SMARCA2** | 3.26974 | 0.00000 | 0.89310 | 12.88409 | 4.42293 | 2.44144 | 9.85941 |
| **SMARCA4** | 13.48265 | 17.10256 | 6.60069 | 17.39237 | 2.08607 | 12.54924 | 33.98098 |
| **SMARCA5** | 30.33332 | 6.94159 | 13.46396 | 13.79196 | 7.87823 | 12.80259 | 16.56761 |
| **SMARCA5-AS1** | 0.00000 | 0.02064 | 0.05003 | 0.05147 | 0.12791 | 0.03399 | 0.07098 |
| **SMARCAD1** | 18.35942 | 0.00000 | 10.36888 | 9.18198 | 0.89127 | 11.43648 | 12.81442 |
| **SMARCAL1** | 7.61167 | 3.54294 | 1.60615 | 5.52847 | 0.20932 | 2.37514 | 0.00000 |
| **SMARCB1** | 17.84800 | 23.89512 | 6.40974 | 19.45996 | 7.81069 | 9.70831 | 6.03822 |
| **SMARCC1** | 17.05632 | 8.41592 | 7.74412 | 18.70253 | 1.34235 | 11.38131 | 45.27365 |
| **SMARCC2** | 1.35354 | 5.61774 | 4.98526 | 7.73362 | 3.93237 | 5.42717 | 12.43555 |
| **SMARCD1** | 0.86950 | 1.38378 | 1.22372 | 1.39902 | 1.31731 | 2.49643 | 11.43184 |
| **SMARCD2** | 9.73904 | 8.60615 | 1.21077 | 7.62381 | 2.31803 | 7.89476 | 19.30602 |
| **SMARCD3** | 0.52123 | 0.29386 | 0.00000 | 0.28231 | 0.03042 | 0.00000 | 0.55624 |
| **SMARCE1** | 21.86333 | 14.12755 | 10.25001 | 9.71582 | 0.56606 | 8.07160 | 10.87144 |
| **SMC1A** | 35.42492 | 15.05509 | 13.90858 | 16.80777 | 3.79711 | 19.19210 | 30.20114 |

| | | | | | | | |
|---|---|---|---|---|---|---|---|
| **SMC1B** | 0.52594 | 0.18689 | 0.73889 | 0.06611 | 0.78680 | 0.07012 | 0.60871 |
| **SMC2** | 21.64526 | 11.84212 | 14.66095 | 14.04486 | 7.72947 | 10.66043 | 18.25418 |
| **SMC3** | 29.35491 | 7.76236 | 20.30122 | 17.94112 | 11.43070 | 12.53340 | 21.92909 |
| **SMC4** | 26.44184 | 9.53059 | 20.63338 | 16.37429 | 9.57593 | 13.10695 | 30.43562 |
| **SMC5** | 9.92003 | 3.37840 | 7.22770 | 7.55996 | 4.36610 | 6.05348 | 9.50787 |
| **SMC5-AS1** | 0.03357 | 0.05763 | 0.07336 | 0.04633 | 0.48154 | 0.01966 | 0.02347 |
| **SMC6** | 11.91351 | 2.85487 | 5.66218 | 4.13108 | 0.72631 | 9.08634 | 16.84521 |
| **SMCHD1** | 2.11441 | 2.74600 | 2.58266 | 2.62931 | 0.91975 | 3.70026 | 15.23049 |
| **SMCO2** | 0.01821 | 0.01718 | 0.01224 | 0.01774 | 0.00878 | 0.06781 | 0.02875 |
| **SMCO3** | 0.10127 | 0.07040 | 0.00000 | 0.03821 | 0.32453 | 0.03828 | 0.00000 |
| **SMCO4** | 0.34811 | 3.29162 | 0.78929 | 0.80254 | 0.68565 | 5.08457 | 0.00000 |
| **SMCR5** | 0.01375 | 0.00700 | 0.01303 | 0.01836 | 0.00745 | 0.06998 | 0.06821 |
| **SMCR8** | 0.23858 | 1.16586 | 2.08970 | 2.34463 | 0.49233 | 0.18464 | 2.16543 |
| **SMCR9** | 0.02480 | 0.01943 | 0.01250 | 0.02082 | 0.00797 | 0.01277 | 0.01744 |
| **SMDT1** | 0.57230 | 0.78316 | 0.96201 | 4.48062 | 0.37028 | 1.35321 | 2.68014 |
| **SMEK1** | 11.82540 | 1.21759 | 14.11640 | 9.79421 | 8.15811 | 7.70607 | 24.83701 |
| **SMEK2** | 16.82210 | 4.46394 | 11.51468 | 11.20125 | 8.01188 | 7.29308 | 0.58131 |
| **SMEK3P** | 0.72895 | 0.40625 | 1.03883 | 1.11927 | 0.45026 | 0.63307 | 0.00000 |
| **SMG1** | 20.53032 | 6.19785 | 10.00098 | 9.41045 | 10.26298 | 9.68488 | 30.87771 |
| **SMG1P1** | 0.01074 | 0.02539 | 0.02918 | 0.04419 | 0.02421 | 0.04411 | 0.19179 |
| **SMG1P2.uid-20** | 0.05009 | 0.09938 | 0.11846 | 0.24828 | 0.08921 | 0.20250 | 0.46158 |
| **SMG1P3** | 0.04223 | 0.47990 | 0.01671 | 0.12356 | 2.62229 | 0.32588 | 0.31490 |
| **SMG1P5** | 0.01555 | 0.19382 | 0.00252 | 0.12805 | 0.26170 | 0.08468 | 0.02800 |
| **SMG1P7** | 0.02782 | 0.15165 | 5.99628 | 0.07566 | 0.05660 | 2.12414 | 0.28693 |
| **SMG5** | 4.10210 | 12.05673 | 5.36496 | 14.80373 | 1.91022 | 9.65004 | 18.90923 |
| **SMG6** | 1.56228 | 2.24013 | 3.56363 | 2.97096 | 4.33306 | 3.05548 | 11.16723 |
| **SMG7** | 8.68862 | 3.84384 | 6.12928 | 16.56608 | 3.85724 | 5.28210 | 17.90736 |
| **SMG7-AS1** | 0.06520 | 0.00954 | 0.09939 | 0.39215 | 0.10728 | 0.07394 | 0.25044 |
| **SMG8** | 13.11521 | 3.61358 | 9.28882 | 6.19196 | 3.28249 | 5.24184 | 0.00000 |
| **SMG9** | 2.88572 | 4.01797 | 2.17353 | 7.77586 | 0.34791 | 3.03288 | 4.55650 |
| **SMIM1** | 2.69977 | 0.20847 | 0.00000 | 0.08982 | 0.00699 | 0.02161 | 0.69717 |
| **SMIM10** | 0.26533 | 0.08189 | 0.04847 | 0.00000 | 0.10686 | 0.00000 | 0.12248 |
| **SMIM11** | 3.71106 | 1.62132 | 2.11489 | 2.93319 | 0.66700 | 0.62435 | 1.40930 |
| **SMIM12** | 20.74370 | 25.37100 | 13.57394 | 12.78964 | 2.82722 | 13.67397 | 3.73706 |
| **SMIM13** | 2.59266 | 2.11365 | 6.57613 | 6.82688 | 2.05242 | 1.12181 | 9.90566 |

| | | | | | | | |
|---|---|---|---|---|---|---|---|
| **SMIM14** | 0.77749 | 0.53502 | 0.31998 | 2.11561 | 1.16120 | 0.71348 | 0.00000 |
| **SMIM15** | 21.46990 | 1.11939 | 15.01490 | 11.49821 | 4.70830 | 4.33860 | 0.00000 |
| **SMIM17** | 0.06092 | 0.02510 | 0.02760 | 0.01783 | 0.04065 | 0.00000 | 0.26238 |
| **SMIM18** | 0.02816 | 0.02818 | 0.09571 | 0.05109 | 0.10080 | 0.07604 | 0.14766 |
| **SMIM19** | 4.36461 | 5.14312 | 7.94499 | 0.87876 | 0.00000 | 3.59024 | 3.46328 |
| **SMIM2** | 0.04280 | 0.01026 | 0.02479 | 0.02766 | 0.00751 | 0.02609 | 0.01239 |
| **SMIM2-AS1** | 0.14828 | 0.01965 | 0.23254 | 0.06770 | 0.00709 | 0.09142 | 0.00000 |
| **SMIM20** | 9.33879 | 8.01134 | 7.51648 | 2.19567 | 0.47843 | 5.04033 | 7.64073 |
| **SMIM21** | 0.02102 | 0.02071 | 0.01670 | 0.01908 | 0.00826 | 0.01540 | 0.01945 |
| **SMIM22** | 1.17335 | 0.00610 | 0.00000 | 0.60479 | 0.00249 | 0.33976 | 1.10200 |
| **SMIM23** | 0.03507 | 0.00982 | 0.01463 | 0.01295 | 0.00768 | 0.01792 | 0.03308 |
| **SMIM24** | 0.03203 | 0.06021 | 0.11251 | 0.00000 | 0.05706 | 0.00000 | 0.10020 |
| **SMIM3** | 0.22678 | 0.00062 | 0.01319 | 0.43624 | 0.00716 | 0.08693 | 0.00000 |
| **SMIM4** | 8.35667 | 11.34734 | 2.26396 | 10.78049 | 1.00053 | 3.54302 | 5.31947 |
| **SMIM5** | 0.00000 | 0.06463 | 0.02457 | 0.41858 | 0.00547 | 0.00000 | 0.58943 |
| **SMIM6** | 0.01606 | 0.01773 | 0.01025 | 0.04969 | 0.02812 | 0.01114 | 0.02532 |
| **SMIM7** | 12.34234 | 7.98793 | 12.51911 | 10.96956 | 1.30505 | 6.17050 | 6.97916 |
| **SMIM8** | 3.71368 | 1.24599 | 3.24641 | 0.65432 | 0.41528 | 1.44802 | 0.76433 |
| **SMIM9** | 0.00000 | 0.01760 | 0.01912 | 0.46117 | 0.01559 | 0.01030 | 0.01505 |
| **SMKR1** | 0.01192 | 0.04220 | 0.05407 | 0.08030 | 1.33242 | 0.10084 | 0.03413 |
| **SMLR1** | 0.10045 | 0.04637 | 0.00354 | 0.00215 | 0.03224 | 1.40478 | 0.00000 |
| **SMNDC1** | 10.94944 | 0.00000 | 9.79616 | 10.65138 | 1.88974 | 7.68805 | 16.86127 |
| **SMO** | 0.08784 | 0.00000 | 0.00627 | 0.11354 | 0.00598 | 0.20934 | 0.69740 |
| **SMOC1** | 0.02501 | 0.02276 | 0.02126 | 0.02320 | 0.05057 | 0.02402 | 0.01718 |
| **SMOC2** | 0.46864 | 0.00000 | 0.08272 | 0.13321 | 0.00721 | 0.15279 | 0.14685 |
| **SMOX** | 6.53967 | 29.64355 | 7.79807 | 12.61516 | 5.32344 | 6.25503 | 27.41857 |
| **SMPD1** | 3.66926 | 3.47794 | 0.91692 | 0.00000 | 0.40929 | 0.36776 | 9.11059 |
| **SMPD2** | 1.33016 | 4.39502 | 1.36601 | 4.60525 | 0.21932 | 1.48095 | 7.21079 |
| **SMPD3** | 0.00000 | 1.85385 | 0.41184 | 0.88619 | 3.30419 | 0.98964 | 0.97396 |
| **SMPD4** | 10.67242 | 5.68156 | 8.45777 | 13.66587 | 0.80822 | 6.46849 | 4.37195 |
| **SMPDL3A** | 35.38682 | 7.56322 | 4.33830 | 7.73250 | 0.84293 | 3.29467 | 12.76210 |
| **SMPDL3B** | 0.00000 | 0.02234 | 0.13457 | 9.65101 | 0.07153 | 0.00066 | 0.00000 |
| **SMPX** | 0.04169 | 0.01340 | 0.01790 | 0.02342 | 0.00910 | 0.01311 | 0.00597 |
| **SMS** | 11.03076 | 1.58594 | 0.36753 | 3.80910 | 3.28976 | 3.74575 | 6.94710 |
| **SMTN** | 2.12567 | 4.53820 | 0.43752 | 8.15258 | 0.08099 | 1.59386 | 4.00133 |

| | | | | | | | |
|---|---|---|---|---|---|---|---|
| **SMTNL1** | 0.06427 | 0.00954 | 0.01354 | 0.01576 | 0.00768 | 0.00914 | 0.02286 |
| **SMTNL2** | 2.17256 | 0.00000 | 0.00000 | 1.53978 | 0.01653 | 0.00185 | 1.22863 |
| **SMU1** | 20.94270 | 11.11992 | 6.60227 | 14.82258 | 7.42995 | 6.94340 | 0.00000 |
| **SMUG1** | 10.23491 | 14.59639 | 7.88786 | 9.63228 | 0.00257 | 3.49417 | 0.00000 |
| **SMURF1** | 1.15351 | 1.86098 | 0.71247 | 3.68711 | 3.50673 | 2.05452 | 5.74325 |
| **SMURF2** | 1.11050 | 5.10229 | 2.48553 | 1.87177 | 1.76656 | 2.90661 | 2.52635 |
| **SMYD1** | 0.00000 | 0.00000 | 0.08979 | 0.09182 | 0.13457 | 0.00625 | 0.06180 |
| **SMYD2** | 9.93371 | 5.32431 | 6.11801 | 9.13360 | 0.07425 | 0.83073 | 6.28169 |
| **SMYD3** | 4.93980 | 1.56520 | 4.79571 | 3.81996 | 1.54483 | 1.70257 | 0.00000 |
| **SMYD4** | 2.23922 | 2.97058 | 1.71746 | 2.15151 | 1.09932 | 1.80821 | 2.41179 |
| **SMYD5** | 8.39633 | 5.79033 | 5.80922 | 10.27532 | 0.15965 | 6.50701 | 6.29709 |
| **SNAI1** | 1.15416 | 1.67117 | 9.96318 | 7.29509 | 8.62093 | 3.38845 | 3.26328 |
| **SNAI2** | 0.07529 | 0.00316 | 0.01371 | 0.03789 | 0.00894 | 0.01708 | 0.05578 |
| **SNAI3** | 0.06582 | 0.01976 | 0.19619 | 0.11952 | 0.52744 | 0.00809 | 0.21439 |
| **SNAI3-AS1** | 0.01653 | 0.09002 | 0.22298 | 0.31605 | 0.03388 | 0.11129 | 0.82220 |
| **SNAP23** | 36.80719 | 11.90752 | 14.81433 | 17.31482 | 0.66016 | 12.00279 | 2.45262 |
| **SNAP25** | 0.00000 | 0.00000 | 0.23239 | 0.26782 | 0.01100 | 0.29852 | 0.03053 |
| **SNAP25-AS1** | 0.02493 | 0.02033 | 0.01834 | 0.08874 | 0.00826 | 0.14912 | 0.00000 |
| **SNAP29** | 4.40750 | 6.91539 | 1.68352 | 5.59032 | 0.64578 | 1.33354 | 9.67227 |
| **SNAP47** | 17.23448 | 16.99703 | 12.37483 | 14.44540 | 1.81692 | 6.03676 | 0.00000 |
| **SNAP91** | 0.00004 | 0.01541 | 0.02547 | 0.11010 | 0.28274 | 0.01868 | 0.19029 |
| **SNAPC1** | 4.74343 | 0.00000 | 8.86158 | 3.12507 | 8.35898 | 2.26466 | 2.83394 |
| **SNAPC2** | 0.10815 | 0.80980 | 12.99097 | 1.50787 | 11.49261 | 1.94814 | 5.95959 |
| **SNAPC3** | 6.57769 | 6.09036 | 5.93707 | 2.63066 | 3.67752 | 10.09287 | 1.35818 |
| **SNAPC4** | 0.07439 | 3.06415 | 1.04862 | 2.49885 | 1.91731 | 1.30421 | 14.28234 |
| **SNAPC5** | 3.53885 | 5.08224 | 1.96330 | 3.28493 | 1.71447 | 1.73730 | 0.00000 |
| **SNAPIN** | 7.72660 | 8.55004 | 4.61426 | 5.56203 | 2.77083 | 4.25893 | 0.16651 |
| **SNAR-B1+SNAF** | 0.03208 | 0.02998 | 0.00989 | 0.01869 | 0.01703 | 0.01826 | 0.02043 |
| **SNAR-C3** | 0.02978 | 0.10525 | 0.03442 | 0.03069 | 0.01424 | 0.01767 | 0.09525 |
| **SNAR-D** | 0.00000 | 1.52901 | 0.13390 | 0.14525 | 0.00684 | 0.26083 | 0.46593 |
| **SNAR-E** | 26.58528 | 15.98326 | 11.68694 | 6.64809 | 2.69027 | 13.40498 | 27.14647 |
| **SNAR-F** | 3.21025 | 2.74635 | 0.91568 | 0.59846 | 0.03847 | 0.10888 | 0.00000 |
| **SNAR-G1** | 0.65185 | 17.39719 | 0.38987 | 0.12582 | 0.00000 | 3.93158 | 2.05431 |
| **SNAR-G2** | 0.14959 | 0.19071 | 0.05595 | 0.09244 | 0.01625 | 0.13665 | 0.60880 |
| **SNAR-H** | 3.92621 | 11.94779 | 0.00000 | 3.90107 | 0.00000 | 2.15486 | 8.04478 |

| | | | | | | | |
|---|---|---|---|---|---|---|---|
| **SNAR-I** | 0.77102 | 18.13223 | 0.51396 | 1.71545 | 0.21163 | 2.88492 | 1.84159 |
| **SNCA** | 0.02141 | 0.10795 | 0.00000 | 0.03278 | 0.14641 | 0.08522 | 0.00695 |
| **SNCAIP** | 0.22564 | 0.12104 | 0.01777 | 0.00000 | 0.01971 | 0.40113 | 0.08537 |
| **SNCB** | 0.01949 | 0.02549 | 0.00000 | 0.01412 | 0.46927 | 0.02627 | 0.02749 |
| **SNCG** | 0.00818 | 0.00531 | 0.00652 | 0.43472 | 0.07588 | 0.36856 | 0.02113 |
| **SND1** | 24.47642 | 13.75054 | 4.69131 | 26.25990 | 1.53246 | 11.47430 | 16.11348 |
| **SND1-IT1** | 1.14748 | 0.00000 | 0.85908 | 0.60257 | 0.02167 | 0.98943 | 6.38440 |
| **SNED1** | 0.14514 | 0.05808 | 0.25816 | 0.00000 | 0.00595 | 0.02433 | 0.23332 |
| **SNF8** | 1.95808 | 15.36840 | 11.00388 | 10.52340 | 5.89084 | 5.00135 | 22.19020 |
| **SNHG1** | 19.09137 | 6.74323 | 17.14507 | 21.68997 | 10.68650 | 20.39574 | 42.43275 |
| **SNHG10** | 5.88111 | 5.51954 | 6.37991 | 5.08473 | 2.56597 | 0.46273 | 0.70630 |
| **SNHG11** | 2.99758 | 6.35756 | 2.29720 | 1.87438 | 4.15619 | 3.89553 | 1.30798 |
| **SNHG12** | 10.83357 | 5.24800 | 5.67075 | 11.47898 | 3.17803 | 7.82188 | 25.95166 |
| **SNHG15** | 10.40850 | 30.14012 | 30.18511 | 12.28298 | 14.43576 | 24.43444 | 9.21746 |
| **SNHG16** | 35.72702 | 19.70829 | 20.85322 | 21.01072 | 8.42266 | 13.05773 | 23.33614 |
| **SNHG17** | 10.55999 | 18.38983 | 7.36832 | 12.99150 | 6.92615 | 17.40154 | 13.86624 |
| **SNHG18** | 0.10295 | 0.01273 | 0.00000 | 0.41539 | 0.85739 | 0.00627 | 0.00000 |
| **SNHG20** | 2.38808 | 1.83181 | 1.33444 | 0.01690 | 0.66170 | 0.05620 | 0.38731 |
| **SNHG21** | 5.37907 | 1.27807 | 1.88655 | 2.08744 | 3.16238 | 0.41895 | 0.00000 |
| **SNHG22** | 0.02127 | 0.01041 | 0.01209 | 0.03813 | 0.00807 | 0.00698 | 0.09462 |
| **SNHG24** | 0.01396 | 0.00999 | 0.01039 | 0.01281 | 0.00947 | 0.47253 | 0.02884 |
| **SNHG3** | 14.38598 | 10.87751 | 10.45168 | 12.40085 | 7.46235 | 10.66216 | 25.81130 |
| **SNHG4.uid-743[illegible]** | 0.63781 | 0.00000 | 0.16171 | 0.19946 | 0.08544 | 0.17015 | 0.00000 |
| **SNHG4.uid-743[illegible]** | 1.25088 | 0.00000 | 0.26989 | 0.46878 | 0.06238 | 0.26447 | 1.35853 |
| **SNHG5** | 34.60585 | 0.00000 | 9.44785 | 9.51258 | 3.70837 | 9.06783 | 30.30952 |
| **SNHG6** | 12.23705 | 0.50201 | 9.05695 | 8.67972 | 2.89546 | 5.36586 | 28.68600 |
| **SNHG7** | 0.53867 | 7.08943 | 1.56384 | 4.71782 | 1.82142 | 3.64014 | 19.78448 |
| **SNHG8** | 21.49326 | 11.27666 | 11.39050 | 20.25668 | 6.73231 | 11.40844 | 21.14251 |
| **SNHG9** | 6.37659 | 6.44995 | 2.81352 | 4.34530 | 2.50468 | 0.58953 | 10.29053 |
| **SNIP1** | 4.07315 | 1.06373 | 3.33024 | 4.22926 | 7.45211 | 1.65479 | 3.20351 |
| **SNN** | 0.00000 | 1.18926 | 0.59139 | 0.67128 | 0.19326 | 0.50381 | 1.24552 |
| **SNORA1** | 0.13095 | 0.00000 | 0.08248 | 0.06799 | 0.03196 | 0.13288 | 0.19728 |
| **SNORA10** | 0.00019 | 0.42877 | 0.12193 | 0.21628 | 0.13050 | 0.32001 | 1.89232 |
| **SNORA11B** | 0.01964 | 0.10189 | 0.02377 | 0.01791 | 0.00822 | 0.00887 | 0.01936 |
| **SNORA11C** | 0.01871 | 0.01602 | 0.01699 | 0.01552 | 0.04370 | 0.03058 | 0.03295 |

| | | | | | | | |
|---|---|---|---|---|---|---|---|
| **SNORA12** | 0.04412 | 0.00000 | 0.26872 | 0.01975 | 0.55690 | 0.00000 | 0.33303 |
| **SNORA13** | 0.06682 | 0.00777 | 0.04776 | 0.12276 | 0.07927 | 0.11470 | 0.24562 |
| **SNORA14A** | 0.04204 | 0.00510 | 0.01667 | 0.04555 | 0.03413 | 0.01891 | 0.03932 |
| **SNORA14B** | 0.00000 | 0.01330 | 0.10867 | 0.09149 | 0.01798 | 0.09091 | 0.19716 |
| **SNORA15** | 0.02426 | 0.01757 | 0.01328 | 0.02634 | 0.00800 | 0.01278 | 0.02357 |
| **SNORA16A** | 0.02686 | 0.02245 | 0.07627 | 0.08159 | 0.05399 | 0.12232 | 1.15185 |
| **SNORA16B** | 0.03977 | 0.00891 | 0.02345 | 0.01502 | 0.00692 | 0.02919 | 0.06552 |
| **SNORA18** | 0.60119 | 0.09616 | 0.46823 | 0.29176 | 0.12899 | 0.39100 | 1.47145 |
| **SNORA19** | 0.00000 | 0.00000 | 0.12458 | 0.18929 | 0.31322 | 0.12976 | 0.52094 |
| **SNORA20** | 0.06042 | 0.00000 | 0.05649 | 0.09721 | 0.05311 | 0.15251 | 0.44811 |
| **SNORA21** | 0.22128 | 0.00000 | 0.25522 | 0.21656 | 0.21653 | 0.26719 | 0.53568 |
| **SNORA22** | 0.03803 | 0.01092 | 0.01753 | 0.02204 | 0.00752 | 0.04622 | 0.02102 |
| **SNORA23** | 0.00000 | 0.00029 | 0.01415 | 0.22157 | 0.49787 | 0.06810 | 0.56213 |
| **SNORA24** | 0.62625 | 0.14255 | 0.19928 | 0.15141 | 0.07784 | 0.29811 | 0.85616 |
| **SNORA25** | 0.44836 | 0.00000 | 0.18350 | 0.19543 | 0.01635 | 0.65460 | 1.86010 |
| **SNORA26** | 0.22385 | 0.00000 | 0.19075 | 0.35767 | 0.05832 | 0.00875 | 0.69387 |
| **SNORA27** | 0.60511 | 0.00000 | 0.15487 | 0.17615 | 0.25014 | 0.35888 | 0.62523 |
| **SNORA28** | 0.01479 | 0.03023 | 0.11158 | 0.09142 | 0.13479 | 0.04052 | 0.34322 |
| **SNORA29** | 0.13322 | 0.03806 | 0.06199 | 0.06540 | 0.04739 | 0.14762 | 0.00465 |
| **SNORA2A** | 0.00130 | 0.01749 | 0.01692 | 0.01939 | 0.17494 | 0.06201 | 0.13039 |
| **SNORA2B** | 0.03410 | 0.01251 | 0.08626 | 0.03552 | 0.40552 | 0.04783 | 0.06986 |
| **SNORA30** | 0.01483 | 0.01583 | 0.01401 | 0.03287 | 0.01772 | 0.01028 | 0.04217 |
| **SNORA31** | 0.04887 | 0.00000 | 0.06121 | 0.00926 | 0.00733 | 0.13566 | 0.48441 |
| **SNORA32** | 0.24123 | 0.00000 | 0.21013 | 0.04047 | 0.02533 | 0.17364 | 0.82829 |
| **SNORA33** | 0.10906 | 0.00000 | 0.11652 | 0.28449 | 0.08465 | 0.44136 | 1.63079 |
| **SNORA34** | 0.02683 | 0.07365 | 0.01207 | 0.04343 | 0.01354 | 0.04766 | 0.03045 |
| **SNORA36C** | 0.03625 | 0.01453 | 0.01291 | 0.01684 | 0.00753 | 0.01167 | 0.03443 |
| **SNORA37** | 0.01728 | 0.00311 | 0.02613 | 0.01679 | 0.01346 | 0.03381 | 0.19684 |
| **SNORA38** | 0.02525 | 0.01911 | 0.01488 | 0.01728 | 0.00832 | 0.01238 | 0.03113 |
| **SNORA38B** | 0.01926 | 0.01916 | 0.01541 | 0.01561 | 0.01592 | 0.01695 | 0.03782 |
| **SNORA4** | 0.36680 | 0.10352 | 0.30928 | 0.13009 | 0.29946 | 0.59904 | 0.84509 |
| **SNORA40** | 0.33258 | 0.00000 | 0.33060 | 0.17492 | 0.06139 | 0.27241 | 1.01486 |
| **SNORA41** | 0.08313 | 0.06375 | 0.03474 | 0.09199 | 0.05963 | 0.11415 | 0.25277 |
| **SNORA44** | 0.02910 | 0.30411 | 0.08525 | 0.09335 | 0.11856 | 0.27700 | 0.94141 |
| **SNORA45A** | 0.12836 | 0.18321 | 0.16174 | 0.23991 | 0.06491 | 0.36326 | 0.40948 |

| | | | | | | | |
|---|---|---|---|---|---|---|---|
| **SNORA45B** | 0.16197 | 0.00000 | 0.15956 | 0.17628 | 0.02699 | 0.24655 | 1.54749 |
| **SNORA46** | 0.01312 | 0.02358 | 0.00661 | 0.02849 | 0.06586 | 0.03917 | 0.05599 |
| **SNORA47** | 0.01331 | 0.01384 | 0.01189 | 0.01662 | 0.01009 | 0.08780 | 0.03561 |
| **SNORA48** | 0.22206 | 0.31137 | 0.01179 | 0.41736 | 1.40685 | 0.57818 | 0.72089 |
| **SNORA49** | 0.01031 | 0.01371 | 0.03120 | 0.02100 | 0.02867 | 0.03604 | 0.06728 |
| **SNORA51** | 0.00000 | 0.15126 | 0.18241 | 0.08950 | 1.17425 | 0.25653 | 0.37960 |
| **SNORA52** | 0.26826 | 0.31506 | 0.10481 | 0.23254 | 0.06770 | 0.23834 | 0.43195 |
| **SNORA53** | 0.09593 | 0.00000 | 0.03494 | 0.12590 | 0.04491 | 0.09115 | 0.63728 |
| **SNORA54** | 0.00000 | 0.04364 | 0.02471 | 0.03556 | 0.01173 | 0.04887 | 0.11108 |
| **SNORA55** | 0.00950 | 0.02358 | 0.03007 | 0.05826 | 0.01477 | 0.04595 | 0.42370 |
| **SNORA57** | 0.96837 | 0.90157 | 0.05630 | 0.96892 | 0.87428 | 0.91070 | 3.82343 |
| **SNORA58** | 0.09077 | 0.23156 | 0.51976 | 0.11435 | 0.09126 | 0.37215 | 1.59248 |
| **SNORA5A** | 0.05385 | 0.03980 | 0.05378 | 0.06366 | 0.01122 | 0.05561 | 0.12285 |
| **SNORA5B** | 0.03406 | 0.02854 | 0.02648 | 0.02350 | 0.00784 | 0.01698 | 0.00432 |
| **SNORA5C** | 0.01584 | 0.12630 | 0.04100 | 0.03995 | 0.00968 | 0.07108 | 0.05124 |
| **SNORA6** | 0.09944 | 0.11106 | 0.13576 | 0.08447 | 0.22599 | 0.31376 | 0.84339 |
| **SNORA60** | 0.03030 | 0.02608 | 0.01400 | 0.03762 | 0.04336 | 0.09208 | 0.02288 |
| **SNORA61** | 1.67133 | 0.00002 | 0.88684 | 1.54743 | 0.37439 | 1.59518 | 5.48806 |
| **SNORA62** | 0.19217 | 0.39558 | 0.11648 | 0.17496 | 0.30116 | 0.66651 | 0.59039 |
| **SNORA63** | 0.27490 | 0.15392 | 0.58503 | 0.54215 | 1.06671 | 0.42101 | 0.86022 |
| **SNORA64** | 0.02302 | 0.44220 | 0.06875 | 0.29496 | 0.03227 | 0.65489 | 1.17639 |
| **SNORA65** | 0.12346 | 0.03491 | 0.14771 | 0.09141 | 0.06416 | 0.05610 | 0.32859 |
| **SNORA66** | 0.35108 | 0.00002 | 0.08710 | 0.17820 | 0.05124 | 0.20608 | 0.41235 |
| **SNORA67** | 0.83324 | 1.92285 | 0.60351 | 1.48321 | 1.11301 | 1.76647 | 3.46579 |
| **SNORA68** | 0.10870 | 0.06880 | 0.02901 | 0.14494 | 0.56818 | 0.11273 | 0.06991 |
| **SNORA69** | 0.03726 | 0.00000 | 0.03895 | 0.02544 | 0.02514 | 0.01099 | 0.14115 |
| **SNORA70** | 1.08173 | 1.35344 | 0.67123 | 0.00000 | 0.15663 | 1.60598 | 3.20553 |
| **SNORA70B** | 0.01180 | 0.01552 | 0.02052 | 0.02597 | 0.00695 | 0.03139 | 0.08094 |
| **SNORA70D** | 0.02036 | 0.01814 | 0.01790 | 0.02342 | 0.00723 | 0.02843 | 0.02434 |
| **SNORA70F** | 0.03685 | 0.02521 | 0.01159 | 0.05903 | 0.00661 | 0.05615 | 0.10907 |
| **SNORA70G** | 0.15240 | 0.00000 | 0.16705 | 0.18457 | 0.02580 | 0.21803 | 0.20223 |
| **SNORA71A** | 0.00222 | 0.05568 | 0.04789 | 0.06529 | 0.01691 | 0.04855 | 0.20737 |
| **SNORA71B** | 0.15120 | 0.00000 | 0.06681 | 0.13812 | 0.00720 | 0.13329 | 0.06974 |
| **SNORA71C** | 0.02265 | 0.08636 | 0.07804 | 0.07733 | 0.00493 | 0.04571 | 0.17184 |
| **SNORA71D** | 0.00000 | 0.08454 | 0.02086 | 0.08119 | 0.00952 | 0.04478 | 0.05366 |

| | | | | | | | |
|---|---|---|---|---|---|---|---|
| **SNORA71E** | 0.05298 | 0.02992 | 0.04008 | 0.02391 | 0.00782 | 0.06293 | 0.00000 |
| **SNORA72** | 0.13368 | 0.00000 | 0.06457 | 0.14010 | 0.03215 | 0.16554 | 0.35830 |
| **SNORA74A** | 0.00000 | 0.03325 | 0.01381 | 0.05197 | 0.03762 | 0.11527 | 0.17038 |
| **SNORA74B** | 0.05013 | 0.00000 | 0.03278 | 0.02571 | 0.08820 | 0.02920 | 0.25609 |
| **SNORA75** | 0.01367 | 0.01535 | 0.05872 | 0.05515 | 0.29596 | 0.14773 | 0.47453 |
| **SNORA76A** | 0.04652 | 0.00683 | 0.02200 | 0.02865 | 0.01075 | 0.03243 | 0.07329 |
| **SNORA76C** | 0.00000 | 0.00000 | 0.12327 | 0.16499 | 0.07021 | 0.08073 | 1.96347 |
| **SNORA77** | 0.12622 | 0.17096 | 0.01269 | 0.08716 | 0.01875 | 0.05291 | 0.17104 |
| **SNORA78** | 0.29044 | 0.34686 | 0.04539 | 0.31091 | 0.13785 | 0.02775 | 0.29420 |
| **SNORA79** | 0.01439 | 0.00000 | 0.06960 | 0.05956 | 0.07685 | 0.01015 | 0.04150 |
| **SNORA7B** | 0.07146 | 0.11133 | 0.04743 | 0.06229 | 0.09372 | 0.09997 | 0.12171 |
| **SNORA8** | 0.44798 | 0.00000 | 0.27783 | 0.13826 | 0.10437 | 0.46966 | 1.67486 |
| **SNORA80A** | 0.03299 | 0.08234 | 0.15611 | 0.02783 | 0.17475 | 0.00000 | 0.46677 |
| **SNORA80B** | 0.00245 | 0.19167 | 0.15579 | 0.02206 | 0.23292 | 0.00127 | 1.03069 |
| **SNORA80E** | 0.00000 | 0.29434 | 0.12050 | 0.23856 | 0.83780 | 0.10781 | 0.87340 |
| **SNORA81** | 0.13153 | 0.04526 | 0.07970 | 0.07139 | 0.02063 | 0.13702 | 0.10407 |
| **SNORA84** | 0.08876 | 0.15651 | 0.07232 | 0.11645 | 0.00849 | 0.09111 | 0.15301 |
| **SNORA9** | 0.47192 | 0.55794 | 0.68136 | 0.11880 | 0.15729 | 0.51219 | 0.18294 |
| **SNORD10** | 0.43477 | 0.12484 | 0.17579 | 0.68417 | 0.14515 | 0.84350 | 0.91467 |
| **SNORD100** | 0.14481 | 0.00000 | 0.04331 | 0.08032 | 0.08139 | 0.12326 | 0.41925 |
| **SNORD101** | 0.06822 | 0.00002 | 0.06509 | 0.05051 | 0.02029 | 0.08328 | 0.14908 |
| **SNORD102** | 0.78458 | 0.00000 | 0.09886 | 0.05731 | 0.12170 | 0.38996 | 0.11434 |
| **SNORD104** | 0.00000 | 0.51380 | 0.37683 | 0.31542 | 0.09937 | 1.06901 | 3.67589 |
| **SNORD105** | 0.03523 | 0.01910 | 0.01445 | 0.02123 | 0.00729 | 0.01497 | 0.03762 |
| **SNORD105B** | 0.02234 | 0.01441 | 0.01317 | 0.02181 | 0.00885 | 0.02359 | 0.02925 |
| **SNORD109A+SI** | 0.01490 | 0.02181 | 0.01914 | 0.01591 | 0.00886 | 0.06047 | 0.09594 |
| **SNORD11** | 0.04198 | 0.00000 | 0.15200 | 0.08301 | 0.18642 | 0.21674 | 0.92428 |
| **SNORD110** | 0.21800 | 0.07279 | 0.14605 | 0.14137 | 0.03228 | 0.32944 | 0.43732 |
| **SNORD111** | 0.02349 | 0.01487 | 0.01602 | 0.01836 | 0.00740 | 0.03466 | 0.03464 |
| **SNORD111B** | 0.02913 | 0.00990 | 0.01959 | 0.01483 | 0.00744 | 0.01475 | 0.03873 |
| **SNORD113-2** | 0.02472 | 0.01545 | 0.01312 | 0.01877 | 0.00797 | 0.02308 | 0.01612 |
| **SNORD113-4** | 0.02478 | 0.01711 | 0.01256 | 0.01862 | 0.00858 | 0.01748 | 0.01789 |
| **SNORD114-1** | 0.02331 | 0.01473 | 0.01202 | 0.01889 | 0.01227 | 0.07327 | 0.01617 |
| **SNORD114-10** | 0.02606 | 0.01809 | 0.01253 | 0.01912 | 0.00801 | 0.02146 | 0.01360 |
| **SNORD114-13** | 0.01944 | 0.01389 | 0.01243 | 0.01586 | 0.00932 | 0.12455 | 0.03137 |

| | | | | | | | |
|---|---|---|---|---|---|---|---|
| **SNORD114-2** | 0.02376 | 0.01444 | 0.01315 | 0.01931 | 0.00791 | 0.03497 | 0.01749 |
| **SNORD114-20** | 0.03067 | 0.01454 | 0.01248 | 0.01235 | 0.00743 | 0.08987 | 0.04794 |
| **SNORD114-21** | 0.02237 | 0.01719 | 0.01335 | 0.01958 | 0.00813 | 0.02095 | 0.01901 |
| **SNORD114-29** | 0.01748 | 0.01339 | 0.01312 | 0.02517 | 0.00806 | 0.05222 | 0.01890 |
| **SNORD114-3** | 0.00450 | 0.03520 | 0.00961 | 0.01532 | 0.00789 | 0.20701 | 0.05143 |
| **SNORD114-6** | 0.02301 | 0.01546 | 0.01306 | 0.01886 | 0.00795 | 0.03308 | 0.01607 |
| **SNORD114-7** | 0.02472 | 0.01744 | 0.01271 | 0.01956 | 0.01080 | 0.01871 | 0.01676 |
| **SNORD114-9** | 0.02286 | 0.01709 | 0.01275 | 0.02101 | 0.00805 | 0.02058 | 0.01634 |
| **SNORD116-1** | 0.02262 | 0.01941 | 0.01297 | 0.01861 | 0.02284 | 0.02036 | 0.02413 |
| **SNORD116-11** | 0.02471 | 0.01736 | 0.01289 | 0.01878 | 0.00903 | 0.01658 | 0.01880 |
| **SNORD116-12** | 0.02527 | 0.01763 | 0.01338 | 0.01866 | 0.01006 | 0.01330 | 0.01789 |
| **SNORD116-13** | 0.03303 | 0.01815 | 0.01585 | 0.01705 | 0.01257 | 0.02123 | 0.02345 |
| **SNORD116-14** | 0.02454 | 0.01858 | 0.01208 | 0.01802 | 0.01390 | 0.01623 | 0.02095 |
| **SNORD116-15** | 0.02389 | 0.02025 | 0.01216 | 0.01972 | 0.01507 | 0.01288 | 0.01543 |
| **SNORD116-16** | 0.02270 | 0.01722 | 0.01172 | 0.01855 | 0.01586 | 0.01681 | 0.02024 |
| **SNORD116-17+** | 0.02360 | 0.01703 | 0.01408 | 0.02477 | 0.00798 | 0.01432 | 0.01428 |
| **SNORD116-18** | 0.02272 | 0.02810 | 0.01071 | 0.02532 | 0.01091 | 0.01242 | 0.01954 |
| **SNORD116-2** | 0.01888 | 0.01897 | 0.01566 | 0.01671 | 0.01486 | 0.02473 | 0.05047 |
| **SNORD116-2+S** | 0.01775 | 0.02012 | 0.01141 | 0.00917 | 0.01096 | 0.01725 | 0.13774 |
| **SNORD116-20** | 0.02570 | 0.01816 | 0.01290 | 0.02068 | 0.01190 | 0.01474 | 0.02739 |
| **SNORD116-21** | 0.02835 | 0.01699 | 0.01348 | 0.01731 | 0.01158 | 0.01283 | 0.01976 |
| **SNORD116-22** | 0.02776 | 0.01934 | 0.01328 | 0.02027 | 0.01049 | 0.01732 | 0.01859 |
| **SNORD116-23** | 0.02581 | 0.02175 | 0.01126 | 0.02247 | 0.03324 | 0.01505 | 0.01223 |
| **SNORD116-24** | 0.03034 | 0.01117 | 0.02209 | 0.02075 | 0.05792 | 0.02674 | 0.02123 |
| **SNORD116-25** | 0.03834 | 0.01491 | 0.02065 | 0.02734 | 0.01822 | 0.04709 | 0.01152 |
| **SNORD116-26** | 0.13947 | 0.04044 | 0.16324 | 0.05295 | 0.02354 | 0.35649 | 0.12789 |
| **SNORD116-27** | 0.02268 | 0.01875 | 0.01671 | 0.02014 | 0.01473 | 0.03324 | 0.01520 |
| **SNORD116-28** | 0.02394 | 0.01809 | 0.01162 | 0.02182 | 0.02506 | 0.01432 | 0.01431 |
| **SNORD116-29** | 0.01920 | 0.02078 | 0.01947 | 0.01639 | 0.01609 | 0.01919 | 0.08961 |
| **SNORD116-4** | 0.02950 | 0.04920 | 0.01744 | 0.03638 | 0.01052 | 0.01896 | 0.00607 |
| **SNORD116-8** | 0.02113 | 0.01567 | 0.01221 | 0.01852 | 0.02191 | 0.02199 | 0.02950 |
| **SNORD117** | 0.02125 | 0.00818 | 0.01803 | 0.02519 | 0.02018 | 0.04287 | 0.04629 |
| **SNORD119** | 0.02931 | 0.03174 | 0.01941 | 0.05367 | 0.04116 | 0.05672 | 0.05101 |
| **SNORD11B** | 0.01639 | 0.00000 | 0.04348 | 0.01651 | 0.05534 | 0.02981 | 0.31890 |
| **SNORD12** | 0.10268 | 0.06930 | 0.08450 | 0.09329 | 0.04437 | 0.06941 | 0.13924 |

| | | | | | | | |
|---|---|---|---|---|---|---|---|
| **SNORD121A** | 0.03960 | 0.04118 | 0.07508 | 0.02739 | 0.01305 | 0.03031 | 0.02323 |
| **SNORD124** | 0.01367 | 0.01314 | 0.01582 | 0.01925 | 0.01157 | 0.01141 | 0.06898 |
| **SNORD126** | 0.02868 | 0.01505 | 0.02193 | 0.02491 | 0.00878 | 0.01836 | 0.02198 |
| **SNORD127** | 0.01867 | 0.01409 | 0.01943 | 0.02034 | 0.00949 | 0.03978 | 0.02987 |
| **SNORD12B** | 0.16466 | 0.06350 | 0.07006 | 0.10282 | 0.02001 | 0.06465 | 0.25785 |
| **SNORD12C** | 0.22784 | 0.01931 | 0.15458 | 0.29593 | 0.10113 | 0.35055 | 1.27170 |
| **SNORD15A** | 0.20606 | 0.33796 | 0.19741 | 0.11591 | 0.01161 | 0.18695 | 0.04479 |
| **SNORD15B** | 0.14552 | 0.04007 | 0.05533 | 0.06295 | 0.02457 | 0.30120 | 0.53810 |
| **SNORD16** | 0.17206 | 0.05108 | 0.08460 | 0.09607 | 0.40356 | 0.15932 | 0.25668 |
| **SNORD17** | 4.06956 | 8.25970 | 1.22700 | 1.40020 | 0.36823 | 2.09801 | 1.61276 |
| **SNORD18A** | 0.05177 | 0.01900 | 0.02002 | 0.05120 | 0.02865 | 0.04701 | 0.02859 |
| **SNORD18B** | 0.04496 | 0.00508 | 0.02145 | 0.01798 | 0.03099 | 0.02255 | 0.09419 |
| **SNORD18C** | 0.07726 | 0.01375 | 0.05803 | 0.06208 | 0.01670 | 0.09418 | 0.06862 |
| **SNORD19** | 0.05384 | 0.04820 | 0.05311 | 0.07828 | 0.01082 | 0.15637 | 0.24682 |
| **SNORD19B** | 0.39317 | 0.00000 | 0.22887 | 0.05599 | 0.01746 | 0.11590 | 0.36336 |
| **SNORD1A** | 0.05046 | 0.05348 | 0.05444 | 0.06640 | 0.01297 | 0.04254 | 0.30786 |
| **SNORD1B** | 0.03183 | 0.00853 | 0.02104 | 0.01799 | 0.03508 | 0.01628 | 0.10840 |
| **SNORD1C** | 0.09520 | 0.00000 | 0.05477 | 0.07978 | 0.01273 | 0.03529 | 0.07851 |
| **SNORD2** | 0.38180 | 0.18276 | 0.12586 | 0.11157 | 0.07909 | 0.06388 | 0.03832 |
| **SNORD20** | 0.05688 | 0.00000 | 0.03927 | 0.01289 | 0.01641 | 0.04863 | 0.18141 |
| **SNORD21** | 0.01151 | 0.01006 | 0.05672 | 0.02706 | 0.02888 | 0.11172 | 0.31354 |
| **SNORD22** | 0.56395 | 0.00000 | 0.45861 | 0.33959 | 0.02782 | 1.10823 | 1.47857 |
| **SNORD23** | 0.09002 | 0.29598 | 0.15110 | 0.35222 | 0.06579 | 0.12151 | 0.21539 |
| **SNORD24** | 0.02366 | 0.02149 | 0.02785 | 0.06096 | 0.12281 | 0.06683 | 0.07189 |
| **SNORD25** | 0.30999 | 0.16326 | 0.11186 | 0.18929 | 0.09676 | 0.28126 | 0.00000 |
| **SNORD26** | 0.09284 | 0.05068 | 0.06746 | 0.07939 | 0.03250 | 0.26795 | 0.29913 |
| **SNORD27** | 0.20477 | 0.02743 | 0.14695 | 0.13295 | 0.18090 | 0.49949 | 0.85027 |
| **SNORD28** | 0.33258 | 0.00000 | 0.18041 | 0.09509 | 0.00516 | 0.48482 | 0.43603 |
| **SNORD29** | 0.26171 | 0.03311 | 0.18077 | 0.15784 | 0.06373 | 0.44982 | 0.39802 |
| **SNORD30** | 0.20379 | 0.07721 | 0.15373 | 0.12547 | 0.10768 | 0.26874 | 0.19111 |
| **SNORD31** | 0.19249 | 0.10934 | 0.15085 | 0.14300 | 0.07513 | 0.45943 | 0.29558 |
| **SNORD32A** | 0.59704 | 0.86236 | 0.14725 | 0.33339 | 0.18668 | 0.50105 | 0.65690 |
| **SNORD33** | 0.12983 | 0.18368 | 0.05934 | 0.11519 | 0.04163 | 0.09226 | 0.16249 |
| **SNORD34** | 0.21191 | 0.03438 | 0.05663 | 0.19297 | 0.13663 | 0.13403 | 0.37651 |
| **SNORD35A** | 0.14987 | 0.17781 | 0.03999 | 0.17400 | 0.40388 | 0.25053 | 0.54897 |

| | | | | | | | |
|---|---|---|---|---|---|---|---|
| **SNORD35B** | 0.58325 | 0.49015 | 0.22388 | 0.61016 | 0.23573 | 0.46614 | 0.54972 |
| **SNORD36A** | 0.29832 | 0.03377 | 0.24027 | 0.26003 | 0.24006 | 0.62520 | 1.18723 |
| **SNORD36B** | 0.08136 | 0.00199 | 0.03809 | 0.06818 | 0.02323 | 0.19926 | 0.22915 |
| **SNORD36C** | 0.03929 | 0.00835 | 0.02951 | 0.07076 | 0.02536 | 0.18440 | 0.55319 |
| **SNORD37** | 0.01960 | 0.02820 | 0.01208 | 0.02753 | 0.00800 | 0.01578 | 0.04030 |
| **SNORD38A** | 0.02469 | 0.04983 | 0.05081 | 0.06161 | 0.01969 | 0.26050 | 0.84057 |
| **SNORD38B** | 0.02964 | 0.08027 | 0.02850 | 0.07383 | 0.18104 | 0.14945 | 0.51102 |
| **SNORD41** | 0.02173 | 0.02979 | 0.01144 | 0.01906 | 0.02447 | 0.01677 | 0.04893 |
| **SNORD42A** | 0.16616 | 0.00000 | 0.10560 | 0.08975 | 0.08920 | 0.16288 | 1.64145 |
| **SNORD42B** | 0.12765 | 0.00000 | 0.06080 | 0.05690 | 0.05971 | 0.11545 | 0.34377 |
| **SNORD43** | 0.06088 | 0.00859 | 0.08004 | 0.06287 | 0.05458 | 0.09490 | 0.36433 |
| **SNORD44** | 0.25591 | 0.00000 | 0.07600 | 0.09494 | 0.03300 | 0.24614 | 0.27358 |
| **SNORD45A** | 0.18942 | 0.00000 | 0.15376 | 0.01031 | 0.00585 | 0.04680 | 0.11257 |
| **SNORD45B** | 0.40890 | 0.00000 | 0.12481 | 0.00488 | 0.00889 | 0.12619 | 0.09415 |
| **SNORD45C** | 0.46645 | 0.03451 | 0.16486 | 0.01744 | 0.00469 | 0.11705 | 0.12274 |
| **SNORD46** | 0.00000 | 0.25767 | 0.01065 | 0.18455 | 0.25773 | 0.19384 | 0.54222 |
| **SNORD47** | 0.98472 | 0.00000 | 0.50949 | 0.17384 | 0.02537 | 1.34098 | 5.08464 |
| **SNORD48** | 0.24958 | 0.40743 | 0.13498 | 0.12605 | 0.04162 | 0.24551 | 0.18388 |
| **SNORD49A** | 0.54446 | 0.00000 | 0.23654 | 0.19228 | 0.00527 | 0.30478 | 1.80353 |
| **SNORD49B** | 0.48212 | 0.22883 | 0.32490 | 0.52816 | 0.02165 | 0.36061 | 0.71946 |
| **SNORD4A** | 0.16997 | 0.07054 | 0.06099 | 0.10725 | 0.06643 | 0.14089 | 0.39623 |
| **SNORD4B** | 0.07349 | 0.00000 | 0.03964 | 0.02884 | 0.01818 | 0.02216 | 0.19005 |
| **SNORD5** | 0.15288 | 0.00000 | 0.11827 | 0.02618 | 0.03156 | 0.11880 | 0.19826 |
| **SNORD50A** | 0.06527 | 0.00000 | 0.18700 | 0.44802 | 0.06017 | 0.42218 | 5.67293 |
| **SNORD50B** | 0.29127 | 0.00000 | 0.11743 | 0.08100 | 0.00793 | 0.18307 | 1.17650 |
| **SNORD51** | 0.06185 | 0.02189 | 0.03283 | 0.01455 | 0.03984 | 0.02964 | 0.09748 |
| **SNORD52** | 0.95358 | 0.21377 | 0.24107 | 0.10567 | 0.04776 | 0.45905 | 0.17971 |
| **SNORD53** | 0.03252 | 0.00609 | 0.03063 | 0.01851 | 0.01676 | 0.00446 | 0.10007 |
| **SNORD54** | 0.18622 | 0.04580 | 0.05643 | 0.03307 | 0.01626 | 0.08820 | 0.12042 |
| **SNORD55** | 0.00000 | 0.26337 | 0.17583 | 0.23117 | 0.45906 | 0.46369 | 0.86206 |
| **SNORD56** | 0.09474 | 0.20963 | 0.03299 | 0.21174 | 0.06034 | 0.59129 | 0.79576 |
| **SNORD56B** | 0.03162 | 0.02341 | 0.01365 | 0.02025 | 0.01477 | 0.01747 | 0.01818 |
| **SNORD57** | 0.11254 | 0.02275 | 0.26997 | 0.10282 | 0.05600 | 0.31296 | 0.94571 |
| **SNORD58A** | 0.05783 | 0.02551 | 0.03007 | 0.02900 | 0.01490 | 0.08028 | 0.10870 |
| **SNORD58C** | 0.14963 | 0.00000 | 0.10993 | 0.07381 | 0.03464 | 0.19818 | 0.81636 |

| | | | | | | | |
|---|---|---|---|---|---|---|---|
| **SNORD59A** | 0.03839 | 0.02036 | 0.01424 | 0.02631 | 0.00886 | 0.12509 | 0.12527 |
| **SNORD59B** | 0.04240 | 0.04454 | 0.00590 | 0.04981 | 0.05016 | 0.05318 | 0.05197 |
| **SNORD6** | 0.12877 | 0.00000 | 0.06576 | 0.01688 | 0.01069 | 0.06231 | 0.13866 |
| **SNORD60** | 0.14622 | 0.04374 | 0.05861 | 0.00002 | 0.04680 | 0.34040 | 1.00888 |
| **SNORD61** | 0.03236 | 0.01262 | 0.01767 | 0.01836 | 0.01436 | 0.02311 | 0.01581 |
| **SNORD62A+SN** | 0.02057 | 0.02274 | 0.01955 | 0.03884 | 0.01377 | 0.01683 | 0.02913 |
| **SNORD62A+SN** | 0.03766 | 0.04945 | 0.03304 | 0.02443 | 0.01493 | 0.02719 | 0.10919 |
| **SNORD63** | 0.11540 | 0.00000 | 0.18455 | 0.10007 | 0.09421 | 0.51617 | 1.38705 |
| **SNORD65** | 0.28067 | 0.00000 | 0.14336 | 0.09920 | 0.01139 | 0.25381 | 0.88136 |
| **SNORD66** | 0.02344 | 0.01231 | 0.01328 | 0.02317 | 0.01680 | 0.01655 | 0.04620 |
| **SNORD67** | 0.06102 | 0.08668 | 0.04152 | 0.03662 | 0.07404 | 0.06689 | 0.00659 |
| **SNORD68** | 0.14841 | 0.31028 | 0.10442 | 0.32997 | 0.18107 | 0.42601 | 0.51754 |
| **SNORD69** | 0.91391 | 0.19004 | 0.16463 | 0.10288 | 0.16888 | 0.04141 | 0.00000 |
| **SNORD7** | 0.10094 | 0.00089 | 0.03007 | 0.04527 | 0.03147 | 0.04513 | 0.13041 |
| **SNORD70** | 0.02111 | 0.06574 | 0.05473 | 0.00643 | 0.00958 | 0.00996 | 0.34077 |
| **SNORD71** | 0.02609 | 0.02739 | 0.01896 | 0.02432 | 0.16073 | 0.01235 | 0.01917 |
| **SNORD72** | 0.03748 | 0.00000 | 0.03267 | 0.02534 | 0.02098 | 0.04543 | 0.14928 |
| **SNORD74** | 0.18134 | 0.08204 | 0.07476 | 0.20155 | 0.05622 | 0.25633 | 0.12774 |
| **SNORD75** | 0.05975 | 0.02087 | 0.02923 | 0.02845 | 0.01104 | 0.05849 | 0.05478 |
| **SNORD76** | 0.41884 | 0.00000 | 0.10192 | 0.12246 | 0.23936 | 0.59162 | 0.60346 |
| **SNORD77** | 0.19228 | 0.01295 | 0.11025 | 0.04679 | 0.07456 | 0.19433 | 0.18641 |
| **SNORD78** | 0.23916 | 0.03065 | 0.07151 | 0.03867 | 0.07325 | 0.15884 | 0.26223 |
| **SNORD79** | 0.67894 | 0.00000 | 0.16478 | 0.15007 | 0.17799 | 0.55897 | 1.13309 |
| **SNORD8** | 0.01811 | 0.02648 | 0.04783 | 0.06247 | 0.09249 | 0.07075 | 0.19699 |
| **SNORD80** | 1.58485 | 0.00000 | 0.48855 | 0.31937 | 0.18184 | 1.26464 | 1.59674 |
| **SNORD81** | 0.47636 | 0.00000 | 0.21891 | 0.08997 | 0.51427 | 0.55964 | 1.79540 |
| **SNORD82** | 0.03195 | 0.01873 | 0.01832 | 0.02322 | 0.03029 | 0.01483 | 0.05466 |
| **SNORD83A** | 0.03264 | 0.22996 | 0.01559 | 0.04771 | 0.05000 | 0.20398 | 0.18245 |
| **SNORD83B** | 0.05979 | 0.12181 | 0.08969 | 0.12558 | 0.09869 | 0.17917 | 0.58238 |
| **SNORD84** | 0.03788 | 0.00617 | 0.01964 | 0.02230 | 0.02279 | 0.03423 | 0.10223 |
| **SNORD85** | 0.01479 | 0.02279 | 0.01027 | 0.02645 | 0.00935 | 0.03141 | 0.03320 |
| **SNORD87** | 0.39161 | 0.00000 | 0.11278 | 0.00000 | 0.14854 | 0.08036 | 0.86495 |
| **SNORD88A** | 0.02006 | 0.02889 | 0.02711 | 0.02173 | 0.00774 | 0.05778 | 0.05676 |
| **SNORD88B** | 1.06193 | 1.12018 | 0.63494 | 0.57181 | 0.13445 | 0.30621 | 0.00000 |
| **SNORD88C** | 0.02566 | 0.13677 | 0.06889 | 0.03412 | 0.03420 | 0.07398 | 0.21487 |

| | | | | | | | |
|---|---|---|---|---|---|---|---|
| **SNORD89** | 0.13310 | 0.00000 | 0.09686 | 0.07817 | 0.05639 | 0.07948 | 0.20536 |
| **SNORD9** | 0.02604 | 0.01737 | 0.01935 | 0.01763 | 0.03234 | 0.01474 | 0.02458 |
| **SNORD90** | 0.01932 | 0.02118 | 0.00888 | 0.01941 | 0.03411 | 0.02274 | 0.03978 |
| **SNORD91A** | 0.04908 | 0.09421 | 0.07826 | 0.06017 | 0.00981 | 0.10894 | 0.44718 |
| **SNORD91B** | 0.02831 | 0.02097 | 0.01505 | 0.00731 | 0.01429 | 0.02537 | 0.10248 |
| **SNORD92** | 0.02513 | 0.01273 | 0.02853 | 0.02601 | 0.01912 | 0.01444 | 0.03137 |
| **SNORD93** | 0.01472 | 0.01421 | 0.01718 | 0.01186 | 0.01336 | 0.03852 | 0.11464 |
| **SNORD94** | 0.10361 | 0.13972 | 0.00155 | 0.05714 | 0.23022 | 0.09008 | 0.00000 |
| **SNORD95** | 0.04390 | 0.13017 | 0.03582 | 0.05004 | 0.09027 | 0.05067 | 0.00673 |
| **SNORD96A** | 0.28901 | 0.12085 | 0.18789 | 0.12017 | 0.05765 | 0.52424 | 0.30696 |
| **SNORD96B** | 0.01679 | 0.01384 | 0.01426 | 0.01964 | 0.00731 | 0.02363 | 0.04788 |
| **SNORD97** | 0.02135 | 0.01490 | 0.01692 | 0.03342 | 0.01919 | 0.00776 | 0.14195 |
| **SNORD98** | 0.27806 | 0.06536 | 0.13285 | 0.07873 | 0.05365 | 0.07947 | 0.23377 |
| **SNORD99** | 3.20090 | 1.67597 | 0.53180 | 0.93120 | 1.00952 | 1.53317 | 1.28891 |
| **SNPH** | 0.00000 | 0.06039 | 0.01719 | 0.00571 | 0.61729 | 1.20095 | 0.04479 |
| **SNRK** | 1.76415 | 0.00000 | 2.30548 | 1.68901 | 0.70744 | 1.95751 | 3.51404 |
| **SNRK-AS1** | 0.05017 | 0.01223 | 0.06146 | 0.08157 | 0.00956 | 0.02158 | 0.06708 |
| **SNRNP200** | 25.91799 | 19.38983 | 11.97909 | 23.95218 | 5.33026 | 14.87149 | 23.65709 |
| **SNRNP25** | 20.46768 | 20.85005 | 5.49785 | 20.11023 | 8.68507 | 11.80055 | 6.26320 |
| **SNRNP27** | 13.59251 | 1.01615 | 13.40783 | 9.25854 | 4.36839 | 4.29270 | 6.28114 |
| **SNRNP35** | 6.93730 | 7.68549 | 11.10201 | 5.08645 | 7.04601 | 3.75219 | 3.14964 |
| **SNRNP40** | 32.82639 | 14.93526 | 11.52611 | 22.80587 | 6.95710 | 10.53780 | 0.00000 |
| **SNRNP48** | 5.53732 | 0.00000 | 1.90195 | 4.52659 | 3.74456 | 2.30774 | 6.15012 |
| **SNRNP70** | 0.00000 | 6.01620 | 0.97333 | 1.96586 | 1.66148 | 4.20152 | 20.82657 |
| **SNRPA** | 16.82915 | 25.87553 | 13.30566 | 21.92974 | 2.02668 | 10.92369 | 20.68318 |
| **SNRPA1** | 12.70132 | 7.91480 | 5.15953 | 9.20330 | 4.17627 | 3.90607 | 0.76768 |
| **SNRPB** | 53.84915 | 45.08641 | 30.44106 | 55.76237 | 18.11709 | 34.06970 | 29.24350 |
| **SNRPB2** | 23.98875 | 12.05082 | 18.47150 | 19.28343 | 10.79697 | 10.57416 | 21.68525 |
| **SNRPC** | 43.05215 | 31.37386 | 32.23221 | 31.98066 | 16.80305 | 21.32894 | 17.45192 |
| **SNRPD1** | 41.28366 | 19.33647 | 22.79037 | 26.73015 | 9.16904 | 16.41496 | 22.11689 |
| **SNRPD2** | 46.01047 | 46.38662 | 33.22859 | 46.06335 | 9.88040 | 26.15178 | 28.50282 |
| **SNRPD2P2** | 0.02599 | 0.02995 | 0.01317 | 0.02295 | 0.03339 | 0.02072 | 0.02030 |
| **SNRPD3** | 37.27089 | 24.11088 | 20.01515 | 31.59422 | 13.68014 | 22.50393 | 28.59700 |
| **SNRPE** | 24.54876 | 7.29943 | 15.65428 | 17.56553 | 4.32131 | 12.42307 | 23.13732 |
| **SNRPF** | 28.44423 | 18.76439 | 17.41529 | 25.87750 | 8.53810 | 17.06251 | 26.96232 |

| | | | | | | | |
|---|---|---|---|---|---|---|---|
| **SNRPG** | 25.34306 | 15.35353 | 13.28780 | 20.16966 | 5.64645 | 11.66890 | 13.61881 |
| **SNRPN.uid-189** | 0.07861 | 0.01650 | 0.01039 | 0.03661 | 0.02814 | 0.04196 | 0.00000 |
| **SNTA1** | 0.06634 | 0.76085 | 0.47708 | 0.06566 | 0.39310 | 0.25448 | 0.30033 |
| **SNTB1** | 2.09101 | 0.00000 | 0.02096 | 0.00000 | 0.09097 | 0.00000 | 0.88527 |
| **SNTB2** | 2.20255 | 1.58514 | 3.02465 | 5.47952 | 1.30010 | 2.45783 | 8.54339 |
| **SNTG1** | 0.02740 | 0.01619 | 0.01239 | 0.01694 | 0.00948 | 0.02320 | 0.01747 |
| **SNTG2** | 0.02729 | 0.00864 | 0.02387 | 0.01936 | 0.07130 | 0.06154 | 0.01283 |
| **SNTN** | 0.02916 | 0.02792 | 0.01320 | 0.01910 | 0.00991 | 0.01391 | 0.00855 |
| **SNUPN** | 7.85784 | 3.89847 | 17.44001 | 3.35795 | 10.21874 | 5.77654 | 1.52233 |
| **SNURF.uid-1921** | 0.11392 | 0.15064 | 0.13031 | 0.09468 | 0.00712 | 0.16471 | 0.00000 |
| **SNW1** | 32.42942 | 20.17472 | 19.73613 | 22.32201 | 9.18152 | 16.53333 | 7.73205 |
| **SNX1** | 8.30345 | 3.35210 | 5.50504 | 5.76748 | 3.09228 | 0.43408 | 1.96254 |
| **SNX10** | 1.41439 | 0.51180 | 1.22410 | 1.64513 | 0.31181 | 1.04530 | 1.24975 |
| **SNX11** | 8.56594 | 5.04676 | 3.34389 | 7.44536 | 0.68675 | 3.17662 | 0.00000 |
| **SNX12** | 5.19966 | 4.40033 | 3.46123 | 0.08937 | 1.17956 | 0.44151 | 13.50967 |
| **SNX13** | 6.30960 | 0.00000 | 3.79215 | 7.23655 | 2.61156 | 2.15725 | 2.76427 |
| **SNX14** | 7.41468 | 0.00000 | 1.53237 | 6.87268 | 7.28753 | 2.53613 | 2.20786 |
| **SNX16** | 2.05504 | 0.00000 | 5.07250 | 0.64346 | 0.58147 | 1.21945 | 0.63374 |
| **SNX17** | 19.74666 | 16.78147 | 4.08823 | 11.76987 | 0.12908 | 11.30146 | 3.08697 |
| **SNX18** | 4.00543 | 0.73880 | 2.31016 | 1.32605 | 0.24027 | 2.18803 | 12.54639 |
| **SNX19** | 0.10143 | 1.11275 | 4.84635 | 1.37589 | 0.78528 | 0.10886 | 2.06756 |
| **SNX2** | 73.87629 | 26.22243 | 17.75116 | 33.32929 | 2.63202 | 16.76601 | 1.37017 |
| **SNX20** | 0.02177 | 0.01732 | 0.01180 | 0.01978 | 0.01663 | 0.01309 | 0.02133 |
| **SNX21** | 0.00000 | 6.31064 | 0.91982 | 0.00000 | 1.07518 | 2.08953 | 1.89253 |
| **SNX22** | 2.84652 | 2.07634 | 0.93863 | 4.34439 | 0.15509 | 1.67626 | 1.46682 |
| **SNX24** | 9.75493 | 4.45014 | 11.60776 | 3.21249 | 0.56220 | 6.38478 | 9.61320 |
| **SNX25** | 0.29084 | 0.32517 | 1.31122 | 0.00000 | 1.72911 | 1.23462 | 0.57633 |
| **SNX27** | 15.38965 | 14.61264 | 3.19620 | 18.05208 | 3.71661 | 13.80438 | 2.03462 |
| **SNX29** | 3.14242 | 4.47106 | 0.80585 | 2.40232 | 1.35429 | 0.00000 | 8.19494 |
| **SNX29P1** | 0.01894 | 0.02901 | 0.01623 | 0.02043 | 0.00739 | 0.01346 | 0.02874 |
| **SNX29P2** | 0.02614 | 0.02506 | 0.01906 | 0.01981 | 0.00818 | 0.01356 | 0.02558 |
| **SNX3** | 18.94122 | 5.77660 | 7.34474 | 10.56932 | 2.60062 | 10.56355 | 33.21206 |
| **SNX30** | 6.52452 | 2.55478 | 1.20651 | 1.34086 | 1.36157 | 3.88178 | 4.61867 |
| **SNX31** | 0.02818 | 0.00638 | 0.03468 | 0.02510 | 0.06694 | 0.03990 | 0.01734 |
| **SNX32** | 0.11970 | 0.00000 | 0.01189 | 0.00341 | 0.01218 | 0.01908 | 0.13224 |

| | | | | | | | |
|---|---|---|---|---|---|---|---|
| **SNX33** | 0.22977 | 0.14417 | 0.00000 | 1.74163 | 0.00294 | 0.22298 | 3.64846 |
| **SNX4** | 18.46624 | 4.20768 | 8.95839 | 9.18014 | 1.30119 | 7.61059 | 1.03950 |
| **SNX5** | 29.35079 | 15.83521 | 10.80759 | 14.19957 | 3.21826 | 13.23986 | 30.99617 |
| **SNX6** | 36.92503 | 7.20143 | 16.84037 | 20.82777 | 3.95535 | 16.69078 | 14.54755 |
| **SNX7** | 1.21099 | 0.80511 | 0.32499 | 0.00000 | 0.04573 | 3.51111 | 3.34003 |
| **SNX8** | 31.36157 | 34.91847 | 9.52301 | 3.18622 | 3.86518 | 7.95702 | 8.71425 |
| **SNX9** | 4.65855 | 3.02465 | 0.65564 | 4.51204 | 0.15846 | 1.22830 | 10.99544 |
| **SOAT1** | 52.35656 | 23.63836 | 35.74368 | 18.09542 | 5.84454 | 7.02039 | 16.23661 |
| **SOAT2** | 0.57934 | 3.40706 | 15.11879 | 0.00000 | 1.64420 | 2.96557 | 0.00000 |
| **SOBP** | 0.00186 | 0.00000 | 0.31382 | 0.00409 | 0.32645 | 0.34851 | 0.12889 |
| **SOCS1** | 0.00000 | 2.22579 | 0.00000 | 3.41142 | 0.18009 | 2.34622 | 1.44624 |
| **SOCS2** | 1.30400 | 0.00000 | 1.96276 | 0.72180 | 1.18624 | 3.82260 | 2.79916 |
| **SOCS2-AS1** | 0.01869 | 0.01551 | 0.36382 | 0.11132 | 0.04205 | 0.26306 | 0.03939 |
| **SOCS3** | 0.00000 | 3.60166 | 0.00000 | 2.99631 | 0.01353 | 9.67489 | 3.26537 |
| **SOCS4** | 9.13471 | 3.42680 | 10.84960 | 6.42132 | 5.20593 | 10.43733 | 5.82314 |
| **SOCS5** | 1.53006 | 0.00000 | 0.94182 | 0.45102 | 1.55683 | 3.03547 | 2.51191 |
| **SOCS6** | 8.23182 | 3.35269 | 0.74706 | 11.24651 | 0.66196 | 1.19445 | 3.10774 |
| **SOCS7** | 3.98572 | 1.39967 | 4.81777 | 4.66288 | 2.97954 | 3.31677 | 8.28551 |
| **SOD1** | 47.65360 | 27.78379 | 18.09371 | 39.53441 | 2.89408 | 16.67695 | 29.33538 |
| **SOD2** | 18.85609 | 20.95987 | 4.28078 | 18.90804 | 1.35882 | 12.94962 | 0.00000 |
| **SOD3** | 0.00000 | 0.01222 | 0.01184 | 0.05573 | 0.00699 | 0.00663 | 0.13001 |
| **SOGA1** | 0.00000 | 1.10642 | 0.22646 | 0.81096 | 0.00000 | 1.04875 | 1.89287 |
| **SOGA3** | 0.02112 | 0.01481 | 0.00922 | 0.02205 | 0.27750 | 0.03319 | 0.01983 |
| **SOHLH1** | 0.05611 | 0.01217 | 0.02722 | 0.02001 | 0.09480 | 0.00621 | 0.03291 |
| **SOHLH2** | 0.50405 | 0.43195 | 0.22488 | 0.50432 | 0.87460 | 0.13231 | 0.71891 |
| **SON** | 38.68867 | 18.88331 | 22.67281 | 25.13876 | 15.09379 | 18.74486 | 38.06385 |
| **SORBS1** | 11.50894 | 27.51448 | 8.88393 | 0.57789 | 3.28393 | 3.13563 | 1.33653 |
| **SORBS2** | 0.05554 | 0.22760 | 0.00671 | 0.41971 | 0.17459 | 0.01119 | 0.00000 |
| **SORBS3** | 0.00000 | 1.60921 | 0.04706 | 0.84580 | 0.04150 | 0.54996 | 2.09870 |
| **SORCS1** | 0.12670 | 0.00000 | 0.04529 | 0.00000 | 0.17855 | 0.03592 | 0.04255 |
| **SORCS2** | 0.03544 | 0.04364 | 0.00000 | 0.00482 | 0.16515 | 0.00300 | 0.07022 |
| **SORCS3** | 0.00000 | 0.06628 | 0.00000 | 0.05891 | 0.57310 | 0.00883 | 0.10509 |
| **SORD** | 0.29109 | 0.17065 | 0.87125 | 1.08295 | 0.02619 | 0.00000 | 0.00000 |
| **SORL1** | 0.86509 | 1.04621 | 0.17768 | 0.00000 | 0.57088 | 1.70937 | 0.26975 |
| **SORT1** | 4.41836 | 5.49250 | 0.00000 | 9.90955 | 2.24395 | 6.89900 | 11.16956 |

| | | | | | | | |
|---|---|---|---|---|---|---|---|
| **1.00000** | 5.39318 | 1.45640 | 2.88155 | 2.67822 | 0.05870 | 2.55700 | 10.04564 |
| **2.00000** | 0.78713 | 0.20418 | 1.25040 | 0.68106 | 0.31522 | 0.69608 | 3.40010 |
| **SOST** | 0.02304 | 0.01735 | 0.00456 | 0.01841 | 0.16376 | 0.01326 | 0.01890 |
| **SOSTDC1** | 0.05439 | 0.01180 | 0.01634 | 0.05778 | 0.07847 | 0.02927 | 0.00000 |
| **SOWAHA** | 0.46365 | 0.03116 | 0.00319 | 0.00000 | 0.00513 | 0.00000 | 1.20151 |
| **SOWAHB** | 0.00020 | 0.27997 | 0.09284 | 0.17039 | 0.00281 | 0.19633 | 0.05043 |
| **SOWAHC** | 1.82977 | 0.29764 | 0.32675 | 2.13797 | 0.16546 | 1.26848 | 4.02939 |
| **SOWAHD** | 0.00000 | 0.06132 | 0.02922 | 0.07586 | 0.06905 | 0.71431 | 0.27879 |
| **SOX1** | 0.01757 | 0.04012 | 0.01300 | 0.00345 | 0.09284 | 0.01884 | 0.11164 |
| **SOX10** | 0.01855 | 0.08210 | 0.06479 | 0.00983 | 0.01524 | 0.00000 | 0.10445 |
| **SOX11** | 0.02653 | 0.11636 | 0.00000 | 0.03243 | 3.28803 | 0.08304 | 0.01894 |
| **SOX12** | 0.00000 | 0.21134 | 2.19993 | 0.22303 | 0.15788 | 0.68182 | 2.01571 |
| **SOX13** | 0.00000 | 0.32627 | 3.70868 | 11.54114 | 3.90664 | 2.93091 | 12.26270 |
| **SOX14** | 0.01753 | 0.01588 | 0.01337 | 0.06782 | 0.00746 | 0.00883 | 0.04092 |
| **SOX15** | 6.19478 | 18.13434 | 31.08204 | 20.71414 | 19.29830 | 10.13698 | 0.00000 |
| **SOX17** | 0.00000 | 0.04043 | 0.01528 | 0.05409 | 0.00668 | 5.16411 | 0.16624 |
| **SOX18** | 0.02219 | 0.01472 | 0.01314 | 0.02449 | 0.00777 | 0.04065 | 0.02207 |
| **SOX2** | 0.00000 | 0.25916 | 7.61962 | 0.00000 | 1.20699 | 9.64411 | 0.16628 |
| **SOX2-OT** | 0.04333 | 0.00000 | 0.31598 | 0.01809 | 0.18147 | 0.82758 | 0.06714 |
| **SOX21** | 0.08506 | 0.00000 | 0.01437 | 0.00000 | 0.05602 | 0.05984 | 0.07327 |
| **SOX21-AS1** | 0.10120 | 0.00000 | 0.13965 | 0.00767 | 0.06086 | 0.12922 | 0.02101 |
| **SOX3** | 0.01838 | 0.01964 | 0.01145 | 0.02616 | 0.04350 | 0.03656 | 0.01850 |
| **SOX30** | 0.00000 | 0.05473 | 0.82664 | 0.07162 | 0.60421 | 0.02114 | 0.03247 |
| **SOX4** | 46.96163 | 28.47983 | 5.12426 | 43.17666 | 4.69605 | 21.71323 | 38.65113 |
| **SOX5** | 0.00000 | 0.08838 | 0.01412 | 0.00748 | 0.24869 | 0.20930 | 0.10307 |
| **SOX6** | 0.08145 | 0.00000 | 0.15648 | 0.09008 | 0.09899 | 0.02302 | 0.01211 |
| **SOX7** | 0.03568 | 0.01826 | 0.14733 | 0.06524 | 0.04879 | 0.17859 | 0.25397 |
| **SOX8** | 0.01735 | 0.01273 | 0.01053 | 0.02587 | 0.00948 | 0.02381 | 0.10364 |
| **SOX9** | 0.03060 | 0.00000 | 0.01074 | 0.04172 | 0.01004 | 3.73960 | 0.05660 |
| **SOX9-AS1** | 0.03852 | 0.02383 | 0.00741 | 0.02166 | 0.04961 | 0.01100 | 0.01923 |
| **SP1** | 11.34720 | 6.03316 | 13.75613 | 10.42756 | 7.47193 | 7.82748 | 17.61110 |
| **SP100** | 0.51721 | 0.02568 | 0.10701 | 0.05755 | 0.00443 | 0.41398 | 0.05169 |
| **SP110** | 0.09738 | 0.03775 | 0.10303 | 0.01399 | 0.61677 | 0.07919 | 0.00000 |
| **SP140** | 0.05188 | 0.12424 | 0.00776 | 0.01618 | 0.01094 | 0.02348 | 0.00000 |
| **SP140L** | 0.85503 | 0.00000 | 0.83415 | 1.80099 | 0.10582 | 0.32564 | 0.02620 |

| | | | | | | | |
|---|---|---|---|---|---|---|---|
| **SP2** | 2.83991 | 4.02721 | 3.63212 | 6.80080 | 2.03009 | 2.02013 | 4.48452 |
| **SP3** | 9.34344 | 0.00000 | 7.24354 | 5.20693 | 3.26029 | 4.82320 | 14.05559 |
| **SP4** | 5.00441 | 0.23448 | 0.19112 | 2.54604 | 2.49020 | 2.03373 | 0.00000 |
| **SP5** | 0.00000 | 0.10427 | 0.08716 | 0.01216 | 0.11108 | 1.00009 | 0.76288 |
| **SP6** | 5.74328 | 0.00000 | 0.00000 | 32.71043 | 0.04250 | 0.00000 | 8.19119 |
| **SP7** | 0.00151 | 0.08920 | 0.00690 | 1.82426 | 0.16282 | 0.13672 | 0.00000 |
| **SP8** | 0.00000 | 0.00000 | 1.21864 | 0.27504 | 0.00003 | 3.49752 | 1.24965 |
| **SP9** | 0.02253 | 0.02200 | 0.01110 | 0.01009 | 0.01089 | 0.04551 | 0.05041 |
| **SPA17** | 2.07660 | 1.27812 | 0.96725 | 1.72214 | 0.01068 | 1.97246 | 0.54523 |
| **SPACA1** | 0.02210 | 0.00000 | 0.11238 | 0.02009 | 0.22882 | 0.01018 | 0.01774 |
| **SPACA3** | 0.02315 | 0.01707 | 0.01702 | 0.04118 | 0.01032 | 0.02556 | 0.00844 |
| **SPACA4** | 0.03575 | 0.00373 | 0.03885 | 0.06226 | 0.00864 | 0.01502 | 0.01503 |
| **SPACA6P** | 0.08170 | 0.15811 | 0.02792 | 0.07160 | 0.09800 | 0.26154 | 0.13028 |
| **SPACA6P-AS** | 0.00001 | 0.00846 | 0.03408 | 0.03599 | 0.06970 | 0.05633 | 0.10059 |
| **SPAG1** | 0.23119 | 0.05695 | 0.35094 | 0.24486 | 0.62653 | 0.40803 | 1.50173 |
| **SPAG16** | 1.20542 | 0.24477 | 0.43389 | 0.79036 | 0.94834 | 0.63646 | 0.04597 |
| **SPAG17** | 0.05251 | 0.00000 | 0.17255 | 0.04124 | 0.08626 | 0.01670 | 0.30925 |
| **SPAG4** | 0.00659 | 0.03964 | 0.15490 | 0.16574 | 1.22492 | 0.19305 | 0.00000 |
| **SPAG5** | 26.26971 | 23.23062 | 17.23903 | 16.87037 | 2.50103 | 13.51666 | 0.00000 |
| **SPAG5-AS1** | 0.36251 | 0.14271 | 0.14296 | 0.16848 | 0.05895 | 0.32844 | 0.14962 |
| **SPAG6** | 0.34033 | 0.01950 | 0.79285 | 0.11877 | 1.15752 | 0.00000 | 0.00000 |
| **SPAG7** | 14.30961 | 8.76098 | 3.89400 | 11.61698 | 2.09954 | 8.07275 | 2.15615 |
| **SPAG8** | 0.00918 | 0.07347 | 0.10265 | 0.01534 | 0.04248 | 0.00596 | 0.03045 |
| **SPAG9** | 6.67432 | 2.84489 | 3.25072 | 7.61364 | 0.80623 | 6.84898 | 16.89533 |
| **SPAM1** | 0.13102 | 0.01461 | 0.02120 | 0.01841 | 0.08529 | 0.00423 | 0.03676 |
| **SPANXA2-OT1** | 0.09092 | 0.03365 | 0.02459 | 0.06924 | 0.08039 | 0.04599 | 0.00000 |
| **SPANXD** | 0.02403 | 0.02193 | 0.01216 | 0.02515 | 0.00795 | 0.01220 | 0.01534 |
| **SPANXN3** | 0.02562 | 0.01695 | 0.00691 | 0.02436 | 0.09877 | 0.03346 | 0.00728 |
| **SPARC** | 26.12018 | 23.67274 | 3.35002 | 5.19307 | 0.04237 | 40.04563 | 3.75334 |
| **SPARCL1** | 0.08573 | 0.00000 | 0.06164 | 0.00015 | 0.01102 | 0.34050 | 0.05266 |
| **SPAST** | 5.22239 | 0.00000 | 2.86246 | 4.64598 | 1.15372 | 2.61527 | 4.11278 |
| **SPATA12** | 0.08751 | 0.07305 | 0.02842 | 0.00000 | 0.00564 | 0.23381 | 0.04608 |
| **SPATA13** | 0.98793 | 0.25327 | 0.77528 | 7.48034 | 0.41088 | 1.43129 | 1.62983 |
| **SPATA13-AS1** | 0.02334 | 0.01749 | 0.01563 | 0.02302 | 0.00778 | 0.01449 | 0.01524 |
| **SPATA16** | 0.02357 | 0.00965 | 0.02281 | 0.02178 | 0.04088 | 0.01471 | 0.02149 |

| | | | | | | | |
|---|---|---|---|---|---|---|---|
| **SPATA17** | 1.01175 | 0.00000 | 1.77585 | 0.06622 | 0.07624 | 0.02195 | 0.12474 |
| **SPATA18** | 0.45729 | 0.70007 | 1.60102 | 0.48284 | 0.01239 | 0.61202 | 0.62156 |
| **SPATA2** | 0.00033 | 7.00864 | 8.46571 | 5.12136 | 6.67227 | 3.44107 | 6.26912 |
| **SPATA20** | 0.06436 | 0.12856 | 0.02203 | 0.00000 | 0.01847 | 0.15216 | 0.01951 |
| **SPATA21** | 0.01718 | 0.18687 | 0.01560 | 0.03110 | 0.00845 | 0.01811 | 0.00987 |
| **SPATA22** | 1.06078 | 0.00000 | 0.66780 | 0.69115 | 0.60143 | 0.49683 | 0.65325 |
| **SPATA24** | 0.41674 | 0.25513 | 0.21486 | 0.63374 | 0.10431 | 0.47190 | 0.00000 |
| **SPATA25** | 0.01260 | 0.12763 | 0.00220 | 0.03608 | 0.01582 | 0.00000 | 0.19907 |
| **SPATA2L** | 0.07000 | 1.62642 | 0.30655 | 1.99282 | 0.06098 | 1.43082 | 2.10333 |
| **SPATA3** | 0.02021 | 0.03301 | 0.01054 | 0.01646 | 0.00961 | 0.01463 | 0.02790 |
| **SPATA3-AS1** | 0.05651 | 0.03156 | 0.04505 | 0.01401 | 0.06108 | 0.01171 | 0.00537 |
| **SPATA31A6** | 0.01980 | 0.02894 | 0.01285 | 0.01851 | 0.00822 | 0.01130 | 0.02197 |
| **SPATA31C1** | 0.04186 | 0.09981 | 0.01921 | 0.01142 | 0.01546 | 0.10630 | 0.03470 |
| **SPATA31C2** | 0.08322 | 0.00660 | 0.11496 | 0.03285 | 0.00804 | 0.04499 | 0.05244 |
| **SPATA31D1** | 0.12749 | 0.01659 | 0.18581 | 0.17670 | 0.01314 | 0.00275 | 0.00000 |
| **SPATA31D5P** | 0.02639 | 0.01615 | 0.01256 | 0.01719 | 0.01960 | 0.01200 | 0.02414 |
| **SPATA31E1** | 0.01756 | 0.01020 | 0.06623 | 0.03283 | 0.01510 | 0.01574 | 0.11508 |
| **SPATA32** | 0.02368 | 0.01216 | 0.02112 | 0.02176 | 0.01443 | 0.01498 | 0.01852 |
| **SPATA33** | 2.25189 | 1.26880 | 3.05171 | 3.23175 | 5.88782 | 1.94365 | 1.14118 |
| **SPATA4** | 0.03577 | 0.01191 | 0.02151 | 0.03910 | 0.02426 | 0.01256 | 0.00975 |
| **SPATA41** | 0.02938 | 0.01550 | 0.02658 | 0.03930 | 0.03511 | 0.04614 | 0.03062 |
| **SPATA42** | 0.02259 | 0.01541 | 0.01269 | 0.03611 | 0.05034 | 0.01469 | 0.01077 |
| **SPATA45** | 0.05406 | 0.03124 | 0.02485 | 0.02132 | 0.00798 | 0.02262 | 0.00848 |
| **SPATA5** | 4.80296 | 0.00127 | 4.50706 | 3.89079 | 4.63050 | 4.14255 | 4.98633 |
| **SPATA5L1** | 1.16054 | 1.29633 | 2.73170 | 2.36810 | 0.76435 | 2.24630 | 7.27232 |
| **SPATA6** | 0.00000 | 0.15827 | 1.28642 | 0.12407 | 0.17786 | 2.75191 | 0.88241 |
| **SPATA6L** | 0.00749 | 0.15998 | 0.14685 | 0.15439 | 1.92167 | 0.07701 | 0.03980 |
| **SPATA7** | 0.91326 | 0.64936 | 1.31590 | 0.38384 | 0.72758 | 1.09460 | 0.14454 |
| **SPATA8** | 0.03000 | 0.01258 | 0.01312 | 0.05933 | 0.00768 | 0.02209 | 0.02508 |
| **SPATA8-AS1** | 0.02738 | 0.01579 | 0.01321 | 0.03405 | 0.00797 | 0.01862 | 0.01281 |
| **SPATA9** | 0.04636 | 0.01258 | 0.01973 | 0.02000 | 0.01882 | 0.01502 | 0.01090 |
| **SPATC1** | 0.01440 | 0.01613 | 0.01098 | 0.03491 | 0.02523 | 0.02833 | 0.02592 |
| **SPATC1L** | 0.00000 | 0.22601 | 0.04605 | 0.70658 | 0.29914 | 0.20822 | 0.73714 |
| **SPATS1** | 0.01246 | 0.02026 | 0.00950 | 0.09286 | 0.02665 | 0.11196 | 0.00779 |
| **SPATS2** | 5.46766 | 5.04704 | 2.73524 | 4.90992 | 0.44223 | 6.62830 | 3.69822 |

| | | | | | | | |
|---|---|---|---|---|---|---|---|
| **SPATS2L** | 0.00000 | 4.38193 | 0.23287 | 1.21533 | 0.01221 | 7.42450 | 0.52647 |
| **SPC24** | 1.41519 | 1.66500 | 0.38788 | 1.11644 | 0.08983 | 0.52262 | 0.00000 |
| **SPC25** | 10.23682 | 3.16139 | 4.06606 | 8.27180 | 0.27323 | 6.57177 | 1.96558 |
| **SPCS1** | 35.13611 | 12.39123 | 21.37191 | 24.85368 | 2.52338 | 16.48647 | 17.51342 |
| **SPCS2** | 6.34572 | 5.21598 | 3.12937 | 5.24918 | 1.45833 | 2.98821 | 1.17136 |
| **SPCS3** | 21.64939 | 6.66736 | 18.32894 | 19.93363 | 3.06872 | 4.42595 | 10.17664 |
| **SPDEF** | 0.11122 | 0.02678 | 0.02205 | 0.05586 | 0.02426 | 0.06620 | 0.02293 |
| **SPDL1** | 19.39364 | 6.13311 | 8.93628 | 12.03394 | 13.94215 | 2.57970 | 1.15685 |
| **SPDYA** | 0.29794 | 0.00000 | 0.10667 | 0.50889 | 0.14254 | 0.14202 | 0.00000 |
| **SPDYC** | 0.88595 | 0.00953 | 4.50772 | 1.16425 | 5.86798 | 0.00000 | 0.00000 |
| **SPDYE1** | 0.04231 | 0.01848 | 0.10616 | 0.05854 | 0.04096 | 0.06150 | 0.33120 |
| **SPDYE2+SPDYI** | 0.02331 | 0.02242 | 0.01342 | 0.01904 | 0.00779 | 0.01407 | 0.01692 |
| **SPDYE2+SPDYI** | 0.02569 | 0.01606 | 0.01589 | 0.02106 | 0.00771 | 0.01353 | 0.01525 |
| **SPDYE3** | 0.04251 | 0.02628 | 0.04742 | 0.06057 | 0.02124 | 0.03575 | 0.03110 |
| **SPDYE5** | 0.09180 | 0.11482 | 0.04750 | 0.12110 | 0.00405 | 0.05222 | 0.43081 |
| **SPDYE6** | 0.03935 | 0.03916 | 0.03718 | 0.03553 | 0.01128 | 0.00929 | 0.01402 |
| **SPDYE7P** | 0.02528 | 0.01780 | 0.01380 | 0.01870 | 0.00784 | 0.01252 | 0.01894 |
| **SPDYE8P.uid-92** | 0.02516 | 0.01793 | 0.01303 | 0.02366 | 0.00788 | 0.01220 | 0.01756 |
| **SPECC1** | 8.63949 | 5.91250 | 5.21310 | 11.71378 | 5.31171 | 6.37207 | 11.73417 |
| **SPECC1L** | 1.95336 | 7.00155 | 2.45100 | 2.25786 | 0.30507 | 2.99779 | 8.12038 |
| **SPECC1L-ADOI** | 0.01888 | 0.01594 | 0.01336 | 0.01747 | 0.00740 | 0.01088 | 0.04847 |
| **SPEF1** | 0.00000 | 0.03705 | 0.02266 | 0.07147 | 0.04973 | 0.07952 | 0.05120 |
| **SPEF2** | 0.05218 | 0.62318 | 0.03029 | 0.00000 | 0.00977 | 0.00753 | 0.41523 |
| **SPEG** | 0.09883 | 0.00000 | 0.02533 | 0.03685 | 0.01291 | 0.05532 | 0.10180 |
| **SPEM1** | 0.01478 | 0.01865 | 0.01192 | 0.03799 | 0.00757 | 0.01022 | 0.04229 |
| **SPEN** | 9.00977 | 5.57260 | 1.80278 | 7.57038 | 0.37040 | 9.29008 | 35.52699 |
| **SPERT** | 0.05335 | 0.02204 | 0.04201 | 0.01616 | 0.06164 | 0.07377 | 0.02805 |
| **SPESP1** | 1.41218 | 1.22996 | 5.63676 | 0.21323 | 1.19012 | 0.34746 | 0.00000 |
| **SPG11** | 3.73198 | 1.62081 | 2.60468 | 1.79026 | 1.18855 | 2.42513 | 4.75882 |
| **SPG20** | 16.48350 | 3.35765 | 3.57043 | 8.01662 | 0.92257 | 0.75259 | 0.66681 |
| **SPG20OS** | 0.01739 | 0.00179 | 0.02753 | 0.05334 | 0.00700 | 0.01181 | 0.16232 |
| **SPG21** | 8.31925 | 3.78624 | 1.67035 | 10.96122 | 0.56448 | 3.44559 | 15.23764 |
| **SPG7** | 5.79285 | 5.45548 | 3.69025 | 5.42747 | 0.92685 | 5.10084 | 9.28362 |
| **SPHK1** | 0.00000 | 12.46264 | 1.34979 | 0.00000 | 0.40190 | 5.75160 | 2.21441 |
| **SPHK2** | 1.76751 | 0.64600 | 2.10026 | 13.27512 | 1.74289 | 0.50032 | 2.63118 |

| | | | | | | | |
|---|---|---|---|---|---|---|---|
| **SPHKAP** | 0.03443 | 0.00000 | 0.08720 | 0.00694 | 0.13621 | 0.01346 | 0.06570 |
| **SPI1** | 0.02325 | 0.01862 | 0.01268 | 0.01919 | 0.00793 | 0.01557 | 0.01711 |
| **SPIB** | 0.00000 | 0.90502 | 1.40863 | 0.76757 | 0.00000 | 0.76461 | 0.69676 |
| **SPIC** | 0.00000 | 0.23947 | 1.07773 | 0.00000 | 0.00000 | 4.51531 | 0.00444 |
| **SPICE1** | 4.88593 | 0.67356 | 5.34541 | 2.83299 | 1.96321 | 2.66828 | 0.00000 |
| **SPIDR** | 4.89158 | 0.98928 | 1.23386 | 4.29658 | 1.38246 | 2.19533 | 3.43150 |
| **SPIN1** | 22.00987 | 3.64321 | 14.32037 | 8.55110 | 3.39402 | 14.22986 | 18.26581 |
| **SPIN2B** | 0.10447 | 0.52593 | 0.30770 | 0.06507 | 0.29453 | 0.03786 | 0.00000 |
| **SPIN3** | 1.64209 | 0.14408 | 1.05220 | 0.73658 | 2.20923 | 0.00000 | 0.14726 |
| **SPIN4** | 16.07654 | 7.45059 | 2.97051 | 11.07447 | 0.06461 | 0.78973 | 5.29737 |
| **SPINK1** | 0.28470 | 0.27110 | 0.78694 | 0.00000 | 0.02116 | 1.37529 | 0.00173 |
| **SPINK13** | 0.02284 | 0.01818 | 0.01402 | 0.01874 | 0.00776 | 0.02508 | 0.01803 |
| **SPINK14** | 0.02641 | 0.01986 | 0.02374 | 0.02046 | 0.00826 | 0.01624 | 0.01624 |
| **SPINK2** | 0.11389 | 0.13267 | 0.69049 | 0.00465 | 0.25040 | 0.30004 | 0.00000 |
| **SPINK4** | 0.00610 | 0.05156 | 0.04756 | 0.03670 | 0.00472 | 0.00911 | 0.25666 |
| **SPINK5** | 0.21630 | 0.54270 | 1.69383 | 0.02812 | 0.28673 | 1.57920 | 1.11607 |
| **SPINK6** | 0.02786 | 0.01312 | 0.02256 | 0.01579 | 0.00708 | 0.03171 | 0.02025 |
| **SPINK7** | 0.02385 | 0.16009 | 0.00809 | 0.00000 | 1.71674 | 0.02727 | 0.02087 |
| **SPINK8** | 0.02106 | 0.01650 | 0.00955 | 0.03488 | 0.04474 | 0.01554 | 0.02493 |
| **SPINT1** | 8.24175 | 25.36758 | 6.61579 | 29.15777 | 1.38409 | 9.76871 | 35.26108 |
| **SPINT2** | 45.82762 | 56.61675 | 16.45161 | 49.67669 | 14.77780 | 25.98923 | 22.58140 |
| **SPIRE1** | 1.48239 | 3.14018 | 0.13054 | 0.58723 | 0.49369 | 0.44002 | 1.79390 |
| **SPIRE2** | 0.19531 | 0.84987 | 0.00000 | 1.98724 | 0.04286 | 0.52120 | 1.19768 |
| **SPN** | 0.00000 | 0.05015 | 0.01878 | 0.23838 | 0.00891 | 0.09790 | 0.02934 |
| **SPNS1** | 1.97193 | 10.02527 | 1.60914 | 15.40103 | 0.43000 | 7.86903 | 23.86135 |
| **SPNS2** | 0.00000 | 0.06997 | 0.11936 | 0.16568 | 0.07744 | 0.00000 | 0.86049 |
| **SPNS3** | 0.34099 | 0.99375 | 0.51648 | 0.00000 | 0.04763 | 0.16998 | 0.46908 |
| **SPO11** | 0.02525 | 0.00691 | 0.03210 | 0.01900 | 0.07604 | 0.01298 | 0.01663 |
| **SPOCD1** | 0.14004 | 0.00000 | 0.06711 | 0.05547 | 0.00446 | 0.00000 | 0.13723 |
| **SPOCK1** | 0.01519 | 0.00000 | 1.02977 | 0.00000 | 0.83432 | 0.31081 | 0.10652 |
| **SPOCK2** | 0.13517 | 1.68486 | 1.73333 | 0.00000 | 0.00070 | 0.32828 | 0.14110 |
| **SPOCK3** | 0.08689 | 0.03352 | 0.00684 | 0.04408 | 0.41375 | 0.07706 | 0.00776 |
| **SPON1** | 0.31176 | 0.17563 | 0.08337 | 0.15249 | 0.75533 | 0.11316 | 0.20368 |
| **SPON2** | 0.63422 | 0.00000 | 0.08740 | 0.64782 | 0.01030 | 1.88569 | 0.39990 |
| **SPOP** | 8.78304 | 4.27525 | 4.08011 | 8.92040 | 4.37390 | 3.11172 | 1.66761 |

| | | | | | | | |
|---|---|---|---|---|---|---|---|
| **SPOPL** | 7.61176 | 0.38858 | 5.90602 | 7.38403 | 0.12076 | 1.06510 | 6.91797 |
| **SPP1** | 1.20009 | 0.00000 | 0.00409 | 0.05279 | 0.01241 | 12.13413 | 0.00000 |
| **SPPL2A** | 16.27640 | 0.54082 | 10.69575 | 16.58096 | 1.13750 | 3.89324 | 19.46313 |
| **SPPL2B** | 0.00000 | 1.27452 | 0.93161 | 3.12616 | 0.22574 | 1.45134 | 6.52925 |
| **SPPL3** | 0.51888 | 3.51470 | 12.73803 | 2.68755 | 3.43895 | 5.16453 | 14.50838 |
| **SPR** | 1.46560 | 7.19971 | 8.46750 | 2.93926 | 0.26704 | 0.43026 | 5.63861 |
| **SPRED1** | 0.98792 | 0.00000 | 0.17063 | 0.00000 | 0.15849 | 3.25944 | 1.85546 |
| **SPRED2** | 1.46288 | 0.80931 | 5.58278 | 2.05559 | 1.44904 | 10.48446 | 11.50758 |
| **SPRED3** | 0.00691 | 0.09634 | 0.03136 | 0.25146 | 0.17186 | 0.23314 | 0.22020 |
| **SPRN** | 0.79759 | 0.14951 | 0.84870 | 0.94972 | 0.00187 | 0.68379 | 1.29270 |
| **SPRNP1** | 0.02070 | 0.01594 | 0.01424 | 0.01993 | 0.00760 | 0.02863 | 0.02622 |
| **SPRR2A** | 0.08392 | 0.04077 | 0.00755 | 0.00340 | 0.00805 | 0.01194 | 0.01477 |
| **SPRR2D** | 0.01499 | 0.01820 | 0.01462 | 0.06071 | 0.00769 | 0.01615 | 0.01735 |
| **SPRR2G** | 0.02440 | 0.01634 | 0.01326 | 0.04395 | 0.00793 | 0.01862 | 0.01296 |
| **SPRR3** | 0.02289 | 0.02041 | 0.01125 | 0.02214 | 0.02250 | 0.01289 | 0.01480 |
| **SPRR4** | 0.03520 | 0.01819 | 0.01280 | 0.01837 | 0.00783 | 0.01995 | 0.01492 |
| **SPRTN** | 8.97535 | 3.15983 | 4.47615 | 6.82624 | 5.00609 | 0.88771 | 1.48497 |
| **SPRY1** | 0.67210 | 0.00000 | 0.00000 | 0.18768 | 0.07227 | 1.01069 | 1.26481 |
| **SPRY2** | 0.00000 | 0.00000 | 1.43906 | 0.00000 | 0.03886 | 10.61178 | 1.27801 |
| **SPRY4** | 0.00000 | 0.88645 | 0.78914 | 1.93380 | 4.61130 | 7.70682 | 4.16726 |
| **SPRYD3** | 1.90141 | 1.06552 | 1.97021 | 3.98252 | 0.35528 | 1.84625 | 2.19995 |
| **SPRYD4** | 3.31618 | 2.63627 | 2.87660 | 5.98478 | 0.55114 | 1.87754 | 0.00000 |
| **SPRYD7** | 10.27557 | 7.15502 | 1.92899 | 4.76728 | 0.91042 | 1.82209 | 3.54885 |
| **SPSB1** | 0.00000 | 0.98605 | 0.21049 | 3.04907 | 0.12774 | 0.39554 | 8.69619 |
| **SPSB2** | 0.51395 | 8.02279 | 6.15276 | 2.47223 | 0.07443 | 5.18355 | 0.79532 |
| **SPSB3** | 9.43363 | 24.03604 | 10.73418 | 16.14489 | 1.35695 | 5.70309 | 3.42581 |
| **SPSB4** | 5.87651 | 0.55210 | 0.01761 | 0.63958 | 0.27535 | 0.12462 | 12.27922 |
| **SPTA1** | 0.03661 | 0.01803 | 0.01162 | 0.01146 | 0.00832 | 0.01997 | 0.03124 |
| **SPTAN1** | 16.31290 | 28.56449 | 4.74489 | 10.08600 | 4.29224 | 3.77437 | 22.72833 |
| **SPTB** | 0.00000 | 5.29622 | 0.65640 | 1.81003 | 1.40196 | 0.30955 | 1.77517 |
| **SPTBN1** | 26.03862 | 33.48607 | 8.52111 | 5.50198 | 4.17949 | 15.99172 | 23.67559 |
| **SPTBN2** | 0.00000 | 0.18998 | 2.12795 | 0.46659 | 0.32012 | 0.00000 | 1.32645 |
| **SPTBN4** | 0.00000 | 0.00000 | 0.60042 | 0.03785 | 0.33465 | 0.24428 | 0.09001 |
| **SPTBN5** | 0.09020 | 0.33668 | 0.32633 | 0.53880 | 0.00000 | 0.00000 | 1.71674 |
| **SPTLC1** | 28.88900 | 5.01293 | 10.17275 | 17.10626 | 2.28344 | 10.81659 | 0.75082 |

| | | | | | | | |
|---|---|---|---|---|---|---|---|
| **SPTLC2** | 20.15575 | 28.70655 | 5.14335 | 0.00000 | 0.00000 | 16.84654 | 6.53901 |
| **SPTLC3** | 0.84858 | 0.16303 | 0.05776 | 0.75783 | 0.00309 | 0.11858 | 0.04791 |
| **SPTSSA** | 18.60635 | 13.93596 | 10.59270 | 9.10063 | 0.77986 | 16.74060 | 6.50634 |
| **SPTSSB** | 0.07228 | 0.00000 | 0.49082 | 0.01942 | 0.06197 | 0.00571 | 0.00000 |
| **SPTY2D1** | 7.82886 | 4.02468 | 3.90078 | 8.88358 | 9.31920 | 5.04146 | 6.44103 |
| **SPTY2D1-AS1** | 0.21996 | 0.02622 | 0.11793 | 0.15917 | 0.08206 | 0.11944 | 0.30708 |
| **SPX** | 0.04684 | 0.01681 | 0.01149 | 0.02454 | 0.03293 | 0.01507 | 0.00428 |
| **SPZ1** | 0.07264 | 0.00000 | 0.15533 | 0.01108 | 0.01506 | 0.42895 | 0.00021 |
| **SQLE** | 53.09211 | 8.99387 | 0.00000 | 19.32997 | 1.63360 | 15.34994 | 16.16777 |
| **SQRDL** | 0.03639 | 0.09973 | 0.02901 | 0.00581 | 5.23469 | 0.00001 | 0.02853 |
| **SQSTM1** | 18.10562 | 21.44785 | 6.53820 | 22.43327 | 3.64088 | 12.30952 | 8.35460 |
| **SRA1** | 24.75111 | 19.52673 | 8.22551 | 22.75262 | 6.69233 | 11.38413 | 3.16559 |
| **SRBD1** | 6.25050 | 0.37247 | 4.81814 | 0.89028 | 1.65234 | 2.59663 | 2.26921 |
| **SRC** | 0.00000 | 0.00000 | 0.33064 | 7.68628 | 0.00050 | 0.55178 | 2.99659 |
| **SRCAP** | 6.64123 | 5.60445 | 11.03685 | 17.50892 | 3.29060 | 11.95321 | 42.34003 |
| **SRCIN1** | 0.00000 | 0.09807 | 0.03745 | 0.05190 | 0.02391 | 0.06018 | 0.03479 |
| **SRCRB4D** | 0.13708 | 0.17491 | 0.11130 | 0.05724 | 0.13932 | 0.00506 | 0.00636 |
| **SRD5A1** | 2.37499 | 0.98544 | 0.26857 | 2.23156 | 0.40145 | 1.41957 | 1.26543 |
| **SRD5A1P1** | 0.12550 | 0.03852 | 0.01814 | 0.09561 | 0.02906 | 0.05611 | 0.00000 |
| **SRD5A2** | 0.03270 | 0.00000 | 0.13734 | 0.01534 | 0.19674 | 0.01086 | 0.01780 |
| **SRD5A3** | 5.33038 | 6.99844 | 0.88235 | 0.00000 | 0.00000 | 3.37418 | 6.44088 |
| **SRD5A3-AS1** | 0.06255 | 0.02715 | 0.02016 | 0.01222 | 0.05363 | 0.01752 | 0.01355 |
| **SREBF1** | 1.06936 | 0.10779 | 0.00000 | 5.73078 | 0.16627 | 2.20826 | 21.53657 |
| **SREBF2** | 3.84477 | 3.56466 | 0.00000 | 6.64952 | 1.53690 | 6.97816 | 36.29595 |
| **SREK1** | 19.19516 | 6.71992 | 5.04098 | 13.73378 | 6.43899 | 8.14343 | 20.01414 |
| **SREK1IP1** | 7.80748 | 0.00000 | 3.21947 | 7.72802 | 2.26753 | 4.64492 | 8.05999 |
| **SRF** | 1.47178 | 3.64680 | 1.62806 | 2.36118 | 0.43297 | 5.86198 | 16.10761 |
| **SRFBP1** | 11.66059 | 0.00000 | 7.32544 | 3.14655 | 1.73313 | 2.81914 | 8.30225 |
| **SRGAP1** | 0.51310 | 4.29160 | 3.13305 | 0.01334 | 0.60243 | 9.66454 | 0.67804 |
| **SRGAP2** | 0.15985 | 1.16329 | 1.39522 | 4.05330 | 0.05969 | 4.92736 | 3.40598 |
| **SRGAP2-AS1** | 0.02402 | 0.02041 | 0.01276 | 0.02255 | 0.00936 | 0.01527 | 0.01571 |
| **SRGAP2D.uid-4** | 0.69056 | 0.00000 | 0.55901 | 0.48338 | 0.18193 | 1.56905 | 0.35030 |
| **SRGAP3** | 0.00000 | 0.06878 | 0.00000 | 0.05962 | 0.21936 | 0.04899 | 0.33354 |
| **SRGN** | 0.01282 | 0.04846 | 0.08939 | 0.01489 | 0.02059 | 0.01379 | 0.09003 |
| **SRI** | 12.95268 | 8.83061 | 2.20363 | 17.26358 | 2.06687 | 8.33978 | 1.38957 |

| | | | | | | | |
|---|---|---|---|---|---|---|---|
| **SRL** | 0.01455 | 0.01755 | 0.01288 | 0.03111 | 0.00884 | 0.01334 | 0.04468 |
| **SRM** | 21.11912 | 46.80532 | 18.75123 | 10.52651 | 2.74491 | 9.98964 | 76.39485 |
| **SRMS** | 0.00000 | 0.02188 | 0.01014 | 0.01377 | 0.00729 | 0.05921 | 0.10842 |
| **SRP14** | 47.93583 | 26.87872 | 27.75652 | 40.23878 | 18.72161 | 20.59686 | 19.40386 |
| **SRP14-AS1** | 1.81731 | 1.13194 | 0.74439 | 2.38871 | 0.13516 | 0.67056 | 0.03871 |
| **SRP19** | 22.98146 | 4.57221 | 14.58762 | 24.25846 | 8.00920 | 9.80660 | 11.79346 |
| **SRP54** | 30.37402 | 5.36882 | 14.38496 | 17.71489 | 6.96511 | 9.22345 | 1.61548 |
| **SRP68** | 9.60266 | 14.04508 | 1.58380 | 8.31574 | 2.66590 | 9.63919 | 21.20601 |
| **SRP72** | 24.74355 | 8.00477 | 15.18773 | 16.07113 | 3.61929 | 10.84236 | 22.51619 |
| **SRP9** | 28.44730 | 0.00000 | 19.14382 | 16.77674 | 5.80530 | 11.16108 | 5.02136 |
| **SRPK1** | 41.41345 | 16.40402 | 16.76963 | 23.92348 | 10.54454 | 9.30166 | 21.12033 |
| **SRPK2** | 10.91630 | 16.96364 | 15.89446 | 6.87655 | 3.28912 | 14.45554 | 8.33618 |
| **SRPK3** | 0.01355 | 0.02527 | 0.04801 | 0.01651 | 0.04004 | 0.00485 | 0.08737 |
| **SRPR** | 21.77119 | 12.08071 | 10.34627 | 20.95564 | 4.33171 | 7.13532 | 7.41231 |
| **SRPRB** | 26.43273 | 17.29119 | 11.90730 | 21.32333 | 0.18227 | 14.82625 | 4.29013 |
| **SRPX** | 0.01822 | 0.00000 | 0.28677 | 0.02983 | 0.04812 | 0.05051 | 0.01949 |
| **SRPX2** | 0.02447 | 0.26184 | 0.02181 | 0.01350 | 0.13859 | 0.05161 | 0.00000 |
| **SRR** | 1.54988 | 0.15281 | 1.16880 | 0.98781 | 0.19380 | 0.75928 | 0.58909 |
| **SRRD** | 2.40086 | 2.54933 | 2.55445 | 2.70963 | 1.15584 | 5.35452 | 14.14873 |
| **SRRM1** | 34.05171 | 16.75849 | 15.26746 | 20.21163 | 8.19170 | 13.33581 | 27.64081 |
| **SRRM2** | 12.91617 | 33.62737 | 18.48126 | 21.11235 | 12.96752 | 17.65611 | 58.74507 |
| **SRRM2-AS1** | 0.02836 | 0.43494 | 0.57839 | 0.12158 | 0.08103 | 0.00000 | 0.02921 |
| **SRRM3** | 0.03546 | 0.23727 | 0.00515 | 0.54793 | 0.23405 | 0.00011 | 0.44469 |
| **SRRM4** | 0.09454 | 0.04422 | 0.00000 | 0.38966 | 0.06276 | 0.31525 | 0.97378 |
| **SRRM5** | 0.02000 | 0.10307 | 0.16397 | 0.10636 | 1.88134 | 0.36621 | 0.25397 |
| **SRRT** | 18.35008 | 19.79387 | 8.37240 | 24.55284 | 10.83556 | 13.99622 | 27.75555 |
| **SRSF1** | 58.60393 | 20.32731 | 31.22158 | 38.29570 | 15.16081 | 20.42440 | 27.35142 |
| **SRSF10.uid-100** | 43.72730 | 11.31575 | 21.27127 | 18.90902 | 13.18146 | 20.16153 | 13.21510 |
| **SRSF11** | 29.18078 | 14.29020 | 19.52428 | 26.55391 | 11.93959 | 16.05956 | 41.22867 |
| **SRSF12** | 0.27606 | 1.09157 | 1.88639 | 0.67736 | 1.37234 | 3.12766 | 0.00000 |
| **SRSF2** | 31.88692 | 14.46108 | 25.21194 | 27.58328 | 13.74679 | 23.79129 | 53.60706 |
| **SRSF3** | 56.68101 | 22.63156 | 31.06592 | 37.11413 | 16.23668 | 24.52154 | 29.15018 |
| **SRSF4** | 23.02419 | 14.80527 | 8.20561 | 18.10527 | 7.56610 | 11.91705 | 12.80539 |
| **SRSF5** | 30.66380 | 9.66012 | 19.09289 | 25.60193 | 15.39947 | 9.33404 | 6.44999 |
| **SRSF6** | 46.60324 | 12.68790 | 17.71547 | 21.80791 | 8.80577 | 13.85495 | 5.96108 |

| | | | | | | | |
|---|---|---|---|---|---|---|---|
| **SRSF7** | 60.07108 | 25.54397 | 31.22779 | 35.33370 | 14.90990 | 29.11521 | 20.61913 |
| **SRSF8** | 3.13837 | 0.00000 | 0.16090 | 4.19605 | 2.18905 | 0.00000 | 4.94054 |
| **SRSF9** | 2.54977 | 10.29826 | 3.81269 | 6.49824 | 4.20018 | 5.37463 | 12.25419 |
| **SRXN1** | 7.05862 | 8.03100 | 0.42885 | 11.39142 | 0.01389 | 0.34403 | 16.23668 |
| **SS18** | 25.34905 | 7.30327 | 13.38831 | 16.98760 | 6.30362 | 11.63106 | 0.00000 |
| **SS18L1** | 2.49305 | 0.19395 | 0.93100 | 3.98170 | 1.38634 | 1.09945 | 5.96092 |
| **SS18L2** | 22.03046 | 3.36808 | 9.60155 | 18.37040 | 3.44662 | 8.84988 | 5.31816 |
| **SSB** | 31.38793 | 7.41659 | 13.73942 | 20.86354 | 6.82528 | 13.32705 | 21.14545 |
| **SSBP1** | 48.22352 | 20.67622 | 26.75383 | 35.32615 | 1.92486 | 22.56371 | 23.92857 |
| **SSBP2** | 0.85973 | 0.07105 | 0.93941 | 0.00037 | 0.81421 | 1.57844 | 0.00000 |
| **SSBP3** | 2.47001 | 21.81361 | 1.35982 | 6.68582 | 1.05504 | 3.37608 | 18.37534 |
| **SSBP3-AS1** | 0.00001 | 0.47227 | 0.74410 | 0.32907 | 0.09079 | 0.36608 | 4.27522 |
| **SSBP4** | 0.00000 | 0.54493 | 0.00059 | 0.17428 | 0.03543 | 0.69184 | 1.99054 |
| **SSC5D** | 0.02834 | 0.00000 | 0.09944 | 0.01345 | 0.03333 | 0.01076 | 0.04295 |
| **SSFA2** | 0.46513 | 0.00000 | 1.56006 | 1.40669 | 3.23494 | 1.42679 | 1.93271 |
| **SSH1** | 3.74452 | 2.15914 | 1.19087 | 2.53969 | 1.82261 | 1.55928 | 8.86669 |
| **SSH2** | 3.39464 | 1.11349 | 1.46461 | 2.55954 | 2.30629 | 1.27959 | 5.29245 |
| **SSH3** | 0.70633 | 1.97396 | 0.37993 | 1.66418 | 0.03505 | 0.00000 | 2.30203 |
| **SSMEM1** | 0.02551 | 0.02924 | 0.00660 | 0.02732 | 0.22614 | 0.01128 | 0.03684 |
| **SSNA1** | 11.30250 | 24.74574 | 11.11795 | 12.81135 | 2.14684 | 9.70817 | 9.76918 |
| **SSPN** | 0.06568 | 0.03432 | 0.02359 | 0.01588 | 0.08897 | 0.01598 | 0.04320 |
| **SSPO** | 0.01220 | 0.02547 | 0.07168 | 0.09754 | 0.09684 | 0.03171 | 0.09148 |
| **SSR1** | 28.04750 | 1.76684 | 15.28468 | 16.41460 | 10.02456 | 12.09546 | 14.19509 |
| **SSR2** | 32.89375 | 18.57866 | 14.65775 | 37.49350 | 9.14215 | 15.80858 | 18.38649 |
| **SSR3** | 29.81422 | 7.46347 | 18.40022 | 20.89155 | 3.37291 | 10.93947 | 24.01628 |
| **SSR4** | 23.13460 | 28.97949 | 15.43802 | 18.60854 | 8.41890 | 15.75644 | 26.72762 |
| **SSR4P1** | 0.04910 | 0.03802 | 0.00975 | 0.01916 | 0.00668 | 0.00000 | 0.11325 |
| **SSRP1** | 15.31236 | 21.39050 | 9.08552 | 2.33082 | 0.83032 | 10.92154 | 42.14096 |
| **SSSCA1** | 17.73906 | 22.67362 | 17.02541 | 14.89883 | 7.79752 | 5.97102 | 3.49418 |
| **SSSCA1-AS1** | 0.17073 | 0.41326 | 0.63568 | 0.31986 | 0.42252 | 0.24102 | 0.00000 |
| **SSTR1** | 0.03863 | 0.02461 | 0.06457 | 0.01220 | 0.00492 | 0.16947 | 0.01613 |
| **SSTR2** | 0.07513 | 0.00000 | 0.12208 | 0.24215 | 0.00367 | 0.42057 | 0.00000 |
| **SSTR3** | 0.05171 | 0.00538 | 0.02367 | 0.00311 | 0.17640 | 0.01465 | 0.04881 |
| **SSTR5-AS1** | 0.00000 | 0.25314 | 0.42555 | 0.01568 | 7.58490 | 0.03202 | 0.11935 |
| **SSU72** | 14.51343 | 35.97463 | 21.29417 | 24.81960 | 5.65017 | 21.09270 | 26.75350 |

| | | | | | | | |
|---|---|---|---|---|---|---|---|
| **SSUH2** | 0.07501 | 0.00000 | 0.06301 | 0.11862 | 0.03980 | 0.21129 | 0.19572 |
| **SSX1** | 0.00000 | 0.15896 | 1.09562 | 0.17587 | 1.54027 | 0.15031 | 0.04719 |
| **SSX2IP** | 9.68765 | 0.00000 | 10.07983 | 0.70752 | 1.94911 | 0.86158 | 3.49343 |
| **SSX3** | 0.02521 | 0.45439 | 1.99197 | 0.00000 | 0.21782 | 0.06121 | 0.41057 |
| **SSX5** | 0.00000 | 0.08947 | 1.62233 | 0.00086 | 0.00102 | 0.03421 | 0.04187 |
| **SSX6** | 0.00000 | 0.21711 | 1.73296 | 0.00000 | 1.52367 | 0.15579 | 0.00639 |
| **SSX7** | 0.03125 | 0.01516 | 0.01505 | 0.01424 | 0.00836 | 0.01165 | 0.02717 |
| **SSX8** | 0.02487 | 0.01531 | 0.03761 | 0.02030 | 0.03534 | 0.01593 | 0.01773 |
| **SSX9** | 0.02889 | 0.01659 | 0.01209 | 0.01862 | 0.03835 | 0.01401 | 0.01355 |
| **ST13** | 20.57860 | 5.29908 | 5.43931 | 13.35112 | 2.07601 | 8.16346 | 13.91417 |
| **ST13P4** | 0.05492 | 0.02481 | 0.01567 | 0.03088 | 0.00794 | 0.02822 | 0.08763 |
| **ST14** | 7.43307 | 8.25287 | 0.00000 | 0.00473 | 3.56255 | 0.42877 | 18.19766 |
| **ST18** | 0.02128 | 0.01906 | 0.01674 | 0.01232 | 0.05813 | 0.02810 | 0.04192 |
| **ST20** | 1.28172 | 0.11525 | 1.22038 | 1.86506 | 0.00359 | 1.17387 | 1.23179 |
| **ST3GAL1** | 3.89178 | 17.54912 | 5.22889 | 6.34763 | 1.26273 | 15.27762 | 14.78092 |
| **ST3GAL2** | 2.61612 | 6.30672 | 6.03318 | 1.76358 | 0.69418 | 0.80355 | 12.29688 |
| **ST3GAL3** | 0.15452 | 0.11596 | 0.00562 | 0.01101 | 0.02775 | 0.07462 | 0.00000 |
| **ST3GAL4** | 3.02867 | 1.99030 | 0.31891 | 10.43021 | 2.94317 | 2.75712 | 1.01630 |
| **ST3GAL4-AS1** | 0.58303 | 0.13767 | 0.53636 | 1.07887 | 0.01088 | 0.06069 | 0.72736 |
| **ST3GAL5** | 0.00389 | 0.01989 | 0.00753 | 0.12564 | 0.06383 | 0.01647 | 0.07498 |
| **ST3GAL6** | 2.01761 | 0.00000 | 4.29888 | 2.53486 | 0.00000 | 1.08069 | 3.44200 |
| **ST3GAL6-AS1** | 0.02054 | 0.02164 | 0.27307 | 0.64778 | 0.00227 | 0.00000 | 0.41336 |
| **ST5** | 0.00000 | 7.69762 | 3.43032 | 0.33566 | 0.61790 | 2.42030 | 4.29984 |
| **ST6GAL1** | 26.33461 | 16.99113 | 12.38028 | 7.82434 | 0.00000 | 15.27003 | 16.86743 |
| **ST6GAL2** | 0.03185 | 0.00884 | 0.02391 | 0.02697 | 0.14267 | 0.01229 | 0.02315 |
| **ST6GALNAC1** | 8.82644 | 8.02979 | 0.00034 | 0.00000 | 0.00103 | 2.84738 | 0.09251 |
| **ST6GALNAC2** | 0.58479 | 1.62526 | 0.40716 | 1.34527 | 0.00000 | 0.00000 | 2.21985 |
| **ST6GALNAC3** | 0.00000 | 0.45104 | 0.07524 | 0.00310 | 0.46259 | 4.10288 | 0.01487 |
| **ST6GALNAC4** | 0.00944 | 0.01060 | 0.06196 | 0.24067 | 0.05787 | 0.03429 | 0.02983 |
| **ST6GALNAC5** | 0.06386 | 0.09607 | 0.02096 | 0.02690 | 0.19032 | 0.34842 | 0.00000 |
| **ST6GALNAC6** | 0.00000 | 0.30684 | 0.21224 | 0.41203 | 0.30654 | 0.73107 | 0.14506 |
| **ST7** | 3.23090 | 0.67709 | 1.96307 | 3.28203 | 0.34880 | 3.43656 | 1.30570 |
| **ST7-AS1** | 0.04062 | 0.03662 | 0.01190 | 0.02111 | 0.01361 | 0.03325 | 0.02160 |
| **ST7-AS2.uid-10** | 0.02073 | 0.02480 | 0.01337 | 0.03524 | 0.04170 | 0.02618 | 0.03380 |
| **ST7-AS2.uid-93** | 0.02092 | 0.03054 | 0.02783 | 0.03647 | 0.00994 | 0.03228 | 0.03604 |

| | | | | | | | |
|---|---|---|---|---|---|---|---|
| **ST7-OT3** | 0.02870 | 0.00975 | 0.03894 | 0.03060 | 0.03193 | 0.03491 | 0.02656 |
| **ST7-OT4** | 0.05951 | 0.01556 | 0.15853 | 0.08488 | 0.06057 | 0.12104 | 0.07669 |
| **ST7L** | 4.01190 | 1.94812 | 1.28523 | 0.94660 | 0.37716 | 1.70584 | 0.41895 |
| **ST8SIA1** | 0.01868 | 0.00000 | 0.18858 | 0.28080 | 0.03787 | 0.00502 | 0.05637 |
| **ST8SIA2** | 0.00000 | 0.10113 | 0.08428 | 0.05915 | 0.04532 | 0.51450 | 0.10662 |
| **ST8SIA3** | 0.02732 | 0.01737 | 0.01311 | 0.08479 | 0.00877 | 0.01499 | 0.00000 |
| **ST8SIA4** | 0.37900 | 0.08254 | 0.02389 | 6.72501 | 0.00435 | 0.33654 | 0.04236 |
| **ST8SIA5** | 0.03889 | 0.03068 | 0.51550 | 0.08864 | 0.57353 | 0.76160 | 0.00000 |
| **ST8SIA6** | 0.03660 | 0.00000 | 0.01244 | 0.00937 | 0.01023 | 0.07101 | 0.14842 |
| **ST8SIA6-AS1** | 2.31788 | 0.26596 | 0.81144 | 2.42181 | 0.08359 | 0.57441 | 0.77566 |
| **STAB1** | 0.00107 | 0.02379 | 0.02690 | 0.04864 | 0.09125 | 0.10781 | 0.14221 |
| **STAB2** | 0.16977 | 0.08961 | 0.17738 | 0.08134 | 0.69438 | 0.00000 | 0.33985 |
| **STAC** | 0.66691 | 1.54361 | 0.41529 | 0.00000 | 0.27903 | 0.00000 | 0.35467 |
| **STAC2** | 0.00000 | 0.01670 | 0.01646 | 0.04210 | 0.09965 | 0.14125 | 0.06863 |
| **STAC3** | 0.10234 | 0.02557 | 0.02935 | 0.04783 | 0.04124 | 0.06564 | 0.02102 |
| **STAG1** | 10.90990 | 1.45041 | 3.65087 | 7.37194 | 3.05611 | 2.59315 | 7.25560 |
| **STAG2** | 21.30310 | 1.28825 | 23.60342 | 11.66899 | 5.02173 | 13.77430 | 17.35827 |
| **STAG3** | 4.35214 | 1.83199 | 0.86036 | 2.71402 | 5.75421 | 0.00000 | 6.29462 |
| **STAG3L1.uid-97** | 0.02444 | 0.01206 | 0.06255 | 0.03495 | 0.10388 | 0.02089 | 0.07435 |
| **STAG3L2** | 0.25365 | 0.93677 | 0.56882 | 0.52513 | 2.59174 | 0.24806 | 0.00000 |
| **STAG3L3** | 0.02852 | 0.04661 | 0.01264 | 0.02807 | 0.08757 | 0.03318 | 0.02941 |
| **STAG3L4** | 5.53920 | 6.83074 | 3.58661 | 2.77263 | 4.85319 | 3.14908 | 0.00000 |
| **STAG3L5P** | 0.12322 | 0.00000 | 0.38404 | 0.12242 | 0.27096 | 0.00000 | 0.20728 |
| **STAG3L5P-PVR** | 0.78106 | 0.19789 | 0.20417 | 0.81067 | 0.60803 | 0.55769 | 0.36413 |
| **STAM** | 14.99386 | 1.92499 | 7.36553 | 9.63276 | 8.23177 | 7.90077 | 0.92334 |
| **STAM-AS1** | 0.34303 | 0.35139 | 0.52145 | 1.26943 | 0.54363 | 0.36855 | 0.76407 |
| **STAM2** | 10.74191 | 5.75769 | 8.75201 | 5.02222 | 1.44760 | 4.31441 | 0.00000 |
| **STAMBP** | 10.47553 | 5.80049 | 7.75810 | 8.54266 | 3.57582 | 3.08042 | 0.09346 |
| **STAMBPL1** | 0.77913 | 0.00000 | 0.16230 | 0.15305 | 0.00405 | 0.28338 | 0.00000 |
| **STAP1** | 0.02376 | 0.01908 | 0.01853 | 0.01907 | 0.01187 | 0.01578 | 0.01746 |
| **STAP2** | 0.25084 | 0.49681 | 1.17253 | 0.05200 | 1.91032 | 0.12395 | 0.00000 |
| **STAR** | 0.13579 | 0.31315 | 0.08767 | 0.00000 | 0.07337 | 0.55904 | 0.00893 |
| **STARD10** | 6.07023 | 4.47937 | 0.00000 | 11.59687 | 0.09989 | 0.00000 | 30.01211 |
| **STARD13** | 0.14311 | 0.01135 | 0.00505 | 0.19307 | 0.01498 | 0.00000 | 0.16395 |
| **STARD13-AS** | 0.02034 | 0.01681 | 0.01481 | 0.03259 | 0.00791 | 0.01260 | 0.01878 |

| | | | | | | | |
|---|---|---|---|---|---|---|---|
| **STARD3** | 2.82498 | 7.95494 | 4.39998 | 11.36646 | 1.79678 | 5.09993 | 4.88975 |
| **STARD3NL** | 7.05875 | 0.62226 | 2.49453 | 5.62527 | 1.07344 | 2.38915 | 0.12023 |
| **STARD4** | 5.20254 | 0.00000 | 0.00008 | 3.08838 | 0.03541 | 1.33752 | 0.06502 |
| **STARD4-AS1** | 0.21409 | 0.00000 | 0.01503 | 0.26493 | 0.18524 | 0.28061 | 0.10288 |
| **STARD5** | 0.97691 | 0.40511 | 0.00000 | 0.65075 | 0.61202 | 0.21900 | 0.00000 |
| **STARD6** | 0.05259 | 0.00207 | 0.08176 | 0.03355 | 0.00678 | 0.01422 | 0.00000 |
| **STARD7** | 26.79715 | 15.61004 | 14.92362 | 20.21116 | 4.73124 | 18.32509 | 32.01423 |
| **STARD7-AS1** | 0.25011 | 0.58952 | 0.05911 | 0.43065 | 0.05967 | 0.18469 | 0.04342 |
| **STARD8** | 0.02101 | 0.02695 | 0.01080 | 0.05473 | 0.00797 | 0.07549 | 0.00821 |
| **STARD9** | 0.00000 | 0.34367 | 0.51498 | 0.19894 | 2.32331 | 1.56661 | 0.66606 |
| **STAT1** | 9.43970 | 3.89802 | 2.38353 | 6.14736 | 0.61386 | 1.31649 | 0.71884 |
| **STAT2** | 2.01289 | 1.32716 | 1.35994 | 5.54432 | 0.58916 | 4.91848 | 0.00000 |
| **STAT3** | 5.27868 | 14.31112 | 21.32201 | 9.35551 | 1.47027 | 19.43249 | 0.00000 |
| **STAT4** | 0.00674 | 0.00000 | 0.03635 | 0.02601 | 0.00690 | 0.11067 | 0.21761 |
| **STAT5A** | 0.00000 | 0.00000 | 0.17114 | 0.01550 | 0.36433 | 0.18606 | 0.32913 |
| **STAT5B** | 0.13599 | 0.82306 | 0.27188 | 1.78791 | 0.08459 | 0.50916 | 1.51398 |
| **STAT6** | 0.23754 | 0.00000 | 0.00111 | 0.55896 | 0.00609 | 1.38073 | 0.37925 |
| **STAU1** | 29.51695 | 10.17906 | 19.89105 | 27.83261 | 10.70832 | 13.49835 | 28.97078 |
| **STAU2** | 9.65635 | 4.13924 | 3.76293 | 1.28466 | 1.13521 | 2.47791 | 4.43870 |
| **STAU2-AS1** | 0.05339 | 0.01457 | 0.00200 | 0.02377 | 0.26724 | 0.01502 | 0.01110 |
| **STBD1** | 0.01900 | 0.09503 | 0.03043 | 0.04935 | 0.00615 | 0.03711 | 0.00000 |
| **STC1** | 0.06808 | 0.00096 | 0.01075 | 0.00219 | 0.01738 | 0.00415 | 0.13632 |
| **STC2** | 20.21287 | 0.00000 | 0.00000 | 9.10807 | 0.03967 | 3.71894 | 18.69339 |
| **STEAP1** | 0.73192 | 0.21027 | 6.51894 | 0.21750 | 0.00000 | 0.97174 | 0.00000 |
| **STEAP1B** | 0.00817 | 0.00000 | 0.71184 | 0.63141 | 0.00252 | 0.22629 | 0.43661 |
| **STEAP2** | 0.03827 | 0.00000 | 0.03339 | 0.00957 | 0.02333 | 0.06161 | 0.03801 |
| **STEAP3** | 5.72225 | 0.00000 | 0.00000 | 6.58903 | 0.00000 | 1.10165 | 5.25539 |
| **STEAP4** | 0.15048 | 0.16666 | 0.01700 | 0.01857 | 0.02821 | 0.09373 | 0.00000 |
| **STIL** | 13.87811 | 0.39367 | 8.87313 | 7.87462 | 8.51933 | 5.29083 | 2.72349 |
| **STIM1** | 1.69796 | 0.30600 | 5.01940 | 3.91425 | 1.51750 | 0.10490 | 4.41519 |
| **STIM2** | 2.20301 | 0.00000 | 2.69311 | 0.71456 | 0.89536 | 0.93944 | 5.46879 |
| **STIP1** | 54.09565 | 43.10359 | 17.35388 | 36.82557 | 10.14275 | 22.89552 | 11.57621 |
| **STK10** | 6.98531 | 8.31695 | 3.33076 | 6.40183 | 0.80504 | 5.96303 | 14.34790 |
| **STK11** | 0.00000 | 1.44283 | 0.16221 | 0.28162 | 0.04315 | 1.13140 | 4.70410 |
| **STK11IP** | 0.83708 | 0.62546 | 1.39288 | 2.50095 | 0.18811 | 0.54762 | 2.15754 |

| | | | | | | | |
|---|---|---|---|---|---|---|---|
| **STK16** | 1.98558 | 0.80818 | 0.99462 | 0.95696 | 0.55083 | 2.18435 | 1.21631 |
| **STK17A** | 1.11963 | 2.40671 | 5.15536 | 3.34912 | 0.13243 | 3.01266 | 10.72578 |
| **STK17B** | 12.53720 | 0.00020 | 1.46048 | 2.82241 | 4.86261 | 1.47322 | 1.74326 |
| **STK19.uid-8457** | 1.32410 | 1.95060 | 1.56501 | 2.52018 | 2.57857 | 0.45852 | 0.00000 |
| **STK24** | 4.44353 | 7.88038 | 1.56043 | 3.14594 | 4.09454 | 4.62312 | 23.20547 |
| **STK25** | 6.02457 | 9.71317 | 2.22504 | 9.80731 | 0.53630 | 5.85676 | 21.34224 |
| **STK3** | 8.17929 | 1.13604 | 1.36763 | 5.20607 | 0.68335 | 1.65366 | 3.04001 |
| **STK31** | 0.00000 | 0.17271 | 5.24848 | 0.00384 | 2.95906 | 0.25090 | 0.01970 |
| **STK32A** | 0.03457 | 0.02910 | 0.00593 | 0.03514 | 0.07064 | 0.11048 | 0.00000 |
| **STK32B** | 0.00000 | 0.04523 | 0.60805 | 0.03127 | 4.60824 | 0.00000 | 0.08052 |
| **STK32C** | 0.00000 | 0.20589 | 0.11718 | 0.04488 | 0.14182 | 0.02596 | 0.12714 |
| **STK33** | 0.04944 | 0.44277 | 0.49609 | 0.00000 | 0.35951 | 0.60826 | 0.00760 |
| **STK35** | 6.08889 | 9.44305 | 3.56261 | 7.79307 | 5.00716 | 8.24192 | 14.04882 |
| **STK36** | 0.76589 | 1.75576 | 1.98693 | 0.80296 | 0.92323 | 4.38150 | 1.70052 |
| **STK38** | 8.34929 | 3.70684 | 2.66528 | 6.91306 | 0.96763 | 1.66147 | 4.52402 |
| **STK38L** | 8.39704 | 0.00000 | 2.10752 | 5.37853 | 1.40109 | 2.30468 | 0.72330 |
| **STK39** | 0.00000 | 0.07465 | 0.00000 | 0.00000 | 1.19409 | 0.73394 | 0.38809 |
| **STK4** | 10.53264 | 2.17493 | 3.28906 | 9.49367 | 3.74930 | 3.92152 | 7.92934 |
| **STK4-AS1** | 0.00796 | 0.02143 | 0.27754 | 0.09899 | 0.25273 | 0.00471 | 0.05133 |
| **STK40** | 0.00000 | 1.42501 | 1.03706 | 5.45774 | 1.36500 | 0.81361 | 6.37965 |
| **STKLD1** | 0.00000 | 0.55190 | 0.16633 | 0.05278 | 0.00740 | 0.35677 | 0.05963 |
| **STL** | 0.03861 | 0.01543 | 0.01289 | 0.04248 | 0.00800 | 0.01239 | 0.01913 |
| **STMN1** | 63.99928 | 37.96170 | 44.62260 | 30.04214 | 6.98145 | 44.87061 | 27.19510 |
| **STMN2** | 0.03789 | 0.00700 | 0.03068 | 0.02551 | 0.05175 | 0.02801 | 0.01220 |
| **STMN3** | 0.00000 | 0.74645 | 0.69039 | 0.05782 | 0.52672 | 0.58866 | 0.07825 |
| **STMN4** | 0.02051 | 0.02118 | 0.01201 | 0.01817 | 0.00909 | 0.04399 | 0.04797 |
| **STMND1** | 0.02898 | 0.00516 | 0.04077 | 0.02261 | 0.02612 | 0.01520 | 0.01507 |
| **STOM** | 9.90381 | 0.00836 | 12.75013 | 6.84896 | 8.55880 | 17.77542 | 0.00000 |
| **STOML1** | 0.82173 | 3.53637 | 0.22188 | 0.08358 | 0.07458 | 0.74310 | 0.67429 |
| **STOML2** | 45.06058 | 43.36878 | 24.78190 | 41.08551 | 14.40113 | 24.52661 | 11.77043 |
| **STOML3** | 0.00510 | 0.01047 | 0.03574 | 0.20600 | 0.01724 | 0.01350 | 0.01084 |
| **STON1** | 2.61456 | 0.07041 | 0.04015 | 0.75676 | 0.08745 | 0.15545 | 1.63035 |
| **STON1-GTF2A1** | 0.03030 | 0.02335 | 0.01120 | 0.01737 | 0.00827 | 0.02352 | 0.02471 |
| **STON2** | 23.53105 | 6.58117 | 1.09849 | 8.31447 | 0.00000 | 6.70338 | 5.12167 |
| **STOX1** | 0.00000 | 0.45378 | 0.00000 | 0.08073 | 0.38415 | 0.72714 | 0.19441 |

| | | | | | | | |
|---|---|---|---|---|---|---|---|
| **STOX2** | 0.00000 | 0.21047 | 0.17893 | 0.00000 | 0.05834 | 0.71889 | 0.03111 |
| **STPG1** | 2.20315 | 1.89836 | 0.19678 | 1.87816 | 1.58351 | 0.14696 | 0.00000 |
| **STPG2** | 0.04233 | 0.00891 | 0.02925 | 0.01664 | 0.09146 | 0.02553 | 0.02754 |
| **STPG2-AS1** | 0.02506 | 0.01990 | 0.01514 | 0.01935 | 0.00773 | 0.01924 | 0.01304 |
| **STRA13** | 22.25885 | 19.82170 | 2.51933 | 27.45048 | 1.43187 | 8.75777 | 2.00319 |
| **STRA6** | 0.39162 | 0.00000 | 0.00000 | 6.90476 | 0.01002 | 0.00000 | 2.64897 |
| **STRA8** | 0.02376 | 0.02813 | 0.00701 | 0.22908 | 0.12373 | 0.01383 | 0.00000 |
| **STRADA** | 1.00587 | 2.48987 | 1.59339 | 1.35589 | 0.24948 | 2.16720 | 2.27052 |
| **STRADB** | 1.44637 | 1.67281 | 2.97005 | 0.55838 | 0.24448 | 2.12617 | 1.64759 |
| **STRAP** | 49.87149 | 16.38466 | 24.87428 | 35.01305 | 10.76483 | 19.95928 | 14.55109 |
| **STRBP** | 7.45333 | 8.35410 | 4.27358 | 3.88502 | 1.78516 | 4.36193 | 20.98189 |
| **STRC** | 0.01935 | 0.01469 | 0.02049 | 0.04959 | 0.03402 | 0.01753 | 0.01466 |
| **STRIP1** | 3.57262 | 4.32389 | 1.98228 | 5.87837 | 1.17010 | 5.46453 | 4.33501 |
| **STRIP2** | 2.58823 | 1.41685 | 0.76490 | 3.95365 | 0.74949 | 1.23889 | 0.34364 |
| **STRN** | 2.20366 | 0.00000 | 3.37829 | 1.65943 | 0.45280 | 1.83298 | 8.90849 |
| **STRN3** | 24.80081 | 0.00000 | 10.64691 | 6.22133 | 5.76052 | 8.26466 | 5.42371 |
| **STRN4** | 1.89908 | 4.41378 | 1.61189 | 6.13108 | 0.00000 | 7.36909 | 18.69016 |
| **STS** | 32.60028 | 0.00000 | 0.00000 | 34.12261 | 0.00000 | 1.36116 | 17.70040 |
| **STT3A** | 36.49612 | 8.88840 | 10.51761 | 17.71054 | 8.98668 | 14.08215 | 4.80591 |
| **STT3B** | 11.98741 | 0.00000 | 7.43971 | 7.66028 | 1.62832 | 7.75096 | 23.30047 |
| **STUB1** | 0.47512 | 4.16284 | 0.27164 | 1.18930 | 0.07816 | 1.33218 | 3.45147 |
| **STX10** | 2.11644 | 4.22360 | 0.69751 | 10.34244 | 0.63563 | 2.12230 | 5.76269 |
| **STX11** | 0.00000 | 0.75696 | 3.81344 | 1.26291 | 0.73923 | 0.00000 | 0.14483 |
| **STX12** | 10.09749 | 0.00000 | 6.68917 | 9.34709 | 5.13822 | 2.50052 | 3.08523 |
| **STX16** | 6.13246 | 5.66667 | 4.03582 | 3.40127 | 0.33825 | 4.61495 | 11.52771 |
| **STX16-NPEPL1** | 0.04736 | 0.06395 | 0.00782 | 0.01934 | 0.01352 | 0.04936 | 0.01950 |
| **STX17** | 1.07657 | 0.75773 | 2.34485 | 1.58545 | 1.92827 | 1.28508 | 2.18511 |
| **STX18** | 12.83958 | 7.80794 | 4.81260 | 4.29436 | 1.53180 | 3.50260 | 1.72153 |
| **STX18-AS1** | 1.46344 | 1.29248 | 0.19065 | 0.04255 | 0.17865 | 0.64142 | 0.27938 |
| **STX19** | 0.04804 | 0.00000 | 0.05628 | 0.01344 | 0.00578 | 0.02828 | 0.04882 |
| **STX1A** | 0.88060 | 1.14108 | 0.59547 | 2.67302 | 0.17633 | 0.23407 | 2.76182 |
| **STX1B** | 0.01405 | 0.00000 | 0.19249 | 0.03062 | 0.31933 | 0.09632 | 0.02720 |
| **STX2** | 1.92834 | 0.00000 | 0.94563 | 0.00000 | 1.68700 | 3.96906 | 0.25285 |
| **STX3** | 12.77941 | 7.60284 | 9.32004 | 10.31451 | 3.37128 | 6.97973 | 21.64576 |
| **STX4** | 0.00000 | 4.16974 | 1.16297 | 2.80269 | 0.35940 | 3.44826 | 2.46530 |

| | | | | | | | |
|---|---|---|---|---|---|---|---|
| **STX5** | 0.00000 | 3.55584 | 1.39671 | 3.42621 | 2.48085 | 1.88273 | 4.99206 |
| **STX6** | 1.36462 | 0.33161 | 1.29411 | 2.37373 | 5.01782 | 2.23485 | 11.32638 |
| **STX7** | 21.89133 | 2.38226 | 7.65075 | 6.89122 | 1.07208 | 3.01259 | 2.51148 |
| **STX8** | 7.03650 | 3.99727 | 3.00939 | 4.69098 | 6.90614 | 3.51760 | 4.27280 |
| **STXBP1** | 0.91003 | 0.25463 | 0.57617 | 2.34081 | 1.64213 | 0.20314 | 3.25655 |
| **STXBP2** | 3.62966 | 6.98536 | 0.18650 | 7.38369 | 0.94781 | 1.68147 | 2.35098 |
| **STXBP3** | 9.02854 | 0.00000 | 9.15603 | 4.86772 | 3.31360 | 4.39930 | 0.00000 |
| **STXBP4** | 6.24836 | 0.00000 | 0.99641 | 3.01002 | 0.57180 | 0.80559 | 3.12701 |
| **STXBP5** | 2.98488 | 1.16124 | 0.45491 | 1.59473 | 0.02714 | 3.16625 | 5.10501 |
| **STXBP5-AS1** | 0.03705 | 0.00980 | 0.01255 | 0.01450 | 0.07509 | 0.01930 | 0.03471 |
| **STXBP5L** | 0.11110 | 0.00000 | 3.45136 | 0.00000 | 0.36241 | 0.98553 | 0.22931 |
| **STXBP6** | 10.82718 | 5.32227 | 1.36537 | 0.00000 | 0.48511 | 0.28506 | 1.27183 |
| **STYK1** | 1.28568 | 0.44601 | 0.77513 | 0.25542 | 0.10127 | 0.00926 | 0.00000 |
| **STYX** | 12.65062 | 0.97352 | 7.21730 | 5.71400 | 7.14652 | 4.56197 | 6.61714 |
| **STYXL1** | 3.48136 | 3.91759 | 0.50224 | 3.87801 | 0.17082 | 3.82202 | 1.34272 |
| **SUB1** | 32.69765 | 7.39092 | 26.50291 | 21.56187 | 11.34589 | 14.48504 | 19.88998 |
| **SUCLA2** | 20.62315 | 2.18795 | 5.80814 | 7.21565 | 0.97275 | 13.69189 | 0.00000 |
| **SUCLG1** | 30.41262 | 18.86819 | 4.91653 | 17.50980 | 0.87656 | 17.74793 | 17.01513 |
| **SUCLG2** | 4.52745 | 1.53702 | 0.68705 | 0.11257 | 0.07548 | 2.05895 | 2.62824 |
| **SUCLG2-AS1** | 0.06289 | 0.16158 | 0.05479 | 0.00000 | 0.06761 | 0.12016 | 0.07877 |
| **SUCNR1** | 0.00747 | 0.02744 | 0.02458 | 0.07634 | 0.00605 | 0.02178 | 0.24722 |
| **SUCO** | 3.53026 | 1.36475 | 3.84468 | 9.47195 | 1.80160 | 6.83035 | 10.58377 |
| **SUDS3** | 8.68182 | 8.61842 | 6.47873 | 7.41040 | 6.64671 | 8.30349 | 13.08454 |
| **SUFU** | 2.64695 | 2.45219 | 1.47565 | 2.79229 | 0.78233 | 1.46700 | 5.07437 |
| **SUGCT** | 0.52601 | 1.09937 | 0.39845 | 0.44788 | 2.66622 | 0.46012 | 0.77257 |
| **SUGP1** | 13.81125 | 12.10306 | 7.21611 | 10.13687 | 3.84384 | 4.37238 | 0.26999 |
| **SUGP2** | 1.90155 | 5.03552 | 3.65202 | 4.06525 | 1.29473 | 5.41963 | 8.28899 |
| **SUGT1** | 15.16924 | 2.19345 | 8.87867 | 9.96797 | 3.39616 | 7.17840 | 12.94229 |
| **SUGT1P1** | 0.02600 | 0.01794 | 0.01283 | 0.02295 | 0.00777 | 0.01915 | 0.02146 |
| **SUGT1P3** | 0.04728 | 0.04258 | 0.04032 | 0.02917 | 0.02519 | 0.05567 | 0.06321 |
| **SULF1** | 0.00797 | 0.00670 | 0.02017 | 0.01910 | 0.16914 | 0.03877 | 0.05665 |
| **SULF2** | 0.80412 | 2.80513 | 0.00000 | 1.46824 | 1.79803 | 1.45276 | 3.12997 |
| **SULT1A1** | 0.24276 | 0.89311 | 0.00092 | 0.23323 | 0.01081 | 1.80511 | 0.04541 |
| **SULT1A2** | 0.07260 | 0.12324 | 0.01495 | 0.02622 | 0.00891 | 0.25633 | 0.03810 |
| **SULT1B1** | 0.03592 | 0.03388 | 0.00709 | 0.06010 | 0.10000 | 0.05467 | 0.00000 |

| | | | | | | | |
|---|---|---|---|---|---|---|---|
| **SULT1C2** | 0.02612 | 0.01742 | 0.01418 | 0.02668 | 0.00810 | 0.01705 | 0.01488 |
| **SULT1C2P1** | 0.02122 | 0.01215 | 0.01419 | 0.01862 | 0.00754 | 0.03492 | 0.03320 |
| **SULT1C4** | 0.02616 | 0.02040 | 0.01625 | 0.01610 | 0.00777 | 0.01083 | 0.02639 |
| **SULT1E1** | 0.02854 | 0.01743 | 0.01634 | 0.01968 | 0.01145 | 0.01470 | 0.01317 |
| **SULT2A1** | 0.61672 | 0.00000 | 0.02176 | 1.15222 | 0.34158 | 0.00525 | 0.01711 |
| **SULT2B1** | 0.48179 | 2.16932 | 0.52951 | 0.12485 | 2.26509 | 0.00000 | 0.00000 |
| **SULT4A1** | 0.00567 | 0.03751 | 0.78290 | 0.00000 | 0.34551 | 0.25059 | 0.12461 |
| **SUMF1** | 6.85814 | 4.12895 | 0.24635 | 1.57217 | 0.00000 | 1.89214 | 2.89985 |
| **SUMF2** | 10.26706 | 8.75228 | 2.92878 | 3.96431 | 0.31159 | 4.89538 | 0.00001 |
| **SUMO1** | 26.72463 | 8.51830 | 11.69038 | 18.76536 | 6.42981 | 13.02358 | 0.79396 |
| **SUMO1P1** | 0.01705 | 0.44833 | 0.28153 | 0.00688 | 0.00269 | 0.10733 | 0.07819 |
| **SUMO1P3** | 4.48416 | 0.57951 | 1.85581 | 2.47583 | 0.78639 | 2.06045 | 0.00000 |
| **SUMO2** | 16.81701 | 3.47386 | 14.96997 | 8.12477 | 5.82780 | 9.35393 | 11.05458 |
| **SUMO3** | 37.56709 | 16.67842 | 8.73737 | 13.85373 | 0.54082 | 8.94203 | 8.99632 |
| **SUMO4** | 0.03630 | 0.00885 | 0.08545 | 0.03213 | 0.06351 | 0.12105 | 0.26612 |
| **SUN1** | 13.54916 | 7.54542 | 3.25621 | 5.03059 | 7.33117 | 7.64251 | 1.53653 |
| **SUN2** | 0.60211 | 3.02784 | 0.00000 | 2.69623 | 0.16068 | 0.08972 | 8.45917 |
| **SUN3** | 20.10863 | 3.80067 | 0.22132 | 13.38758 | 0.00000 | 2.11229 | 4.73971 |
| **SUOX** | 2.98291 | 0.45181 | 0.41778 | 4.89835 | 0.07923 | 1.13499 | 0.00000 |
| **SUPT16H** | 31.47429 | 11.44970 | 16.98335 | 24.72535 | 5.51952 | 11.86153 | 20.37391 |
| **SUPT20H** | 9.78846 | 3.41010 | 7.20345 | 6.26344 | 2.21102 | 3.61671 | 1.25277 |
| **SUPT20HL1** | 0.02235 | 0.03838 | 0.01191 | 0.02539 | 0.00962 | 0.01352 | 0.02008 |
| **SUPT20HL2** | 0.01990 | 0.21946 | 0.07746 | 0.02803 | 0.50696 | 0.00000 | 0.02929 |
| **SUPT3H** | 1.29039 | 1.18097 | 1.98488 | 1.90776 | 1.90321 | 0.98495 | 2.69422 |
| **SUPT4H1** | 27.57690 | 8.81099 | 15.69341 | 26.17389 | 10.37488 | 10.53060 | 13.47267 |
| **SUPT5H** | 12.73883 | 12.81488 | 12.74415 | 26.07836 | 3.88536 | 13.26846 | 28.92988 |
| **SUPT6H** | 6.85072 | 10.15085 | 6.67287 | 12.72727 | 3.46474 | 8.41564 | 15.68544 |
| **SUPT7L** | 14.77680 | 16.08849 | 15.11070 | 10.68583 | 7.87254 | 8.14361 | 1.72672 |
| **SUPV3L1** | 26.82258 | 7.05081 | 12.09518 | 15.77634 | 5.55574 | 9.08604 | 2.65235 |
| **SURF1** | 0.19211 | 5.12178 | 1.21234 | 2.74216 | 0.22428 | 5.62678 | 13.02658 |
| **SURF2** | 19.24551 | 26.78014 | 11.62487 | 21.13603 | 3.34647 | 9.98577 | 2.13036 |
| **SURF4** | 27.95104 | 26.27787 | 12.59759 | 30.75754 | 5.88870 | 25.90337 | 15.61286 |
| **SURF6** | 7.92433 | 23.14569 | 9.49485 | 13.13420 | 5.40871 | 11.32677 | 20.33549 |
| **SUSD1** | 2.70542 | 0.61512 | 0.11341 | 1.41135 | 0.12250 | 0.00000 | 3.44040 |
| **SUSD2** | 8.77542 | 28.97442 | 5.79629 | 0.45805 | 0.00000 | 27.68071 | 5.91433 |

| | | | | | | | |
|---|---|---|---|---|---|---|---|
| **SUSD3** | 0.00000 | 0.64976 | 0.00000 | 0.00000 | 2.35408 | 2.27522 | 0.08793 |
| **SUSD4** | 0.00000 | 0.09387 | 0.06333 | 0.01935 | 0.09150 | 0.03664 | 0.06374 |
| **SUSD5** | 0.01267 | 0.02557 | 0.61437 | 0.00000 | 0.05132 | 0.99667 | 0.14416 |
| **SUV39H1** | 3.74244 | 7.64022 | 3.30195 | 3.08063 | 2.13435 | 0.71027 | 4.95748 |
| **SUV39H2** | 11.66675 | 1.19831 | 5.81745 | 6.20544 | 1.93422 | 2.04357 | 1.40655 |
| **SUV420H1** | 1.72227 | 4.61673 | 4.65404 | 3.19444 | 2.99037 | 1.78466 | 10.43561 |
| **SUV420H2** | 0.00000 | 0.12960 | 0.15489 | 0.43475 | 0.17614 | 0.67090 | 2.41076 |
| **SUZ12** | 11.63751 | 0.00000 | 4.25143 | 5.78299 | 0.54486 | 4.01742 | 25.57155 |
| **SUZ12P1** | 0.00000 | 0.85252 | 0.96330 | 0.00000 | 0.63606 | 1.55275 | 1.16910 |
| **SV2A** | 0.21277 | 0.02758 | 0.04137 | 0.81770 | 0.04140 | 0.00000 | 0.43639 |
| **SV2B** | 0.00211 | 0.03012 | 0.00000 | 0.04739 | 1.61179 | 0.03185 | 0.03482 |
| **SV2C** | 1.21471 | 0.36573 | 0.16036 | 0.12260 | 0.06724 | 0.44492 | 0.19426 |
| **SVEP1** | 0.08054 | 0.04834 | 0.30533 | 0.08057 | 1.03105 | 0.26514 | 0.00000 |
| **SVIL** | 1.12638 | 3.82706 | 1.31718 | 0.81152 | 0.03869 | 2.97668 | 1.78537 |
| **SVILP1** | 0.02744 | 0.04539 | 0.01062 | 0.01992 | 0.01012 | 0.02052 | 0.00957 |
| **SVIP** | 2.07725 | 0.76015 | 0.66745 | 3.83088 | 1.75905 | 0.00000 | 3.39750 |
| **SVOP** | 0.70316 | 0.04642 | 0.08056 | 0.06244 | 0.00667 | 0.00000 | 0.07978 |
| **SVOPL** | 0.88938 | 0.52686 | 1.40690 | 0.00000 | 2.83829 | 0.44355 | 0.52893 |
| **SWAP70** | 9.66779 | 9.10462 | 7.47817 | 8.65850 | 2.44340 | 2.69128 | 10.51768 |
| **SWI5** | 7.56355 | 11.44102 | 2.44134 | 15.38781 | 0.88648 | 3.07929 | 2.55146 |
| **SWSAP1** | 2.23611 | 5.65080 | 2.69358 | 3.79675 | 0.83119 | 3.06171 | 0.00000 |
| **SWT1** | 1.41387 | 0.40546 | 1.91970 | 3.41105 | 3.54108 | 1.63010 | 1.47769 |
| **SYAP1** | 17.28449 | 18.91244 | 10.77898 | 9.74234 | 7.56339 | 10.27822 | 11.51049 |
| **SYBU** | 7.68400 | 0.00000 | 0.06573 | 1.80293 | 0.05651 | 0.00000 | 3.00359 |
| **SYCE1** | 0.13847 | 3.25524 | 3.38650 | 0.44870 | 0.30926 | 0.88867 | 0.00000 |
| **SYCE1L** | 0.00000 | 0.18319 | 0.04965 | 0.09982 | 0.15737 | 0.08550 | 0.01985 |
| **SYCE2** | 0.04548 | 0.33201 | 0.40636 | 0.25335 | 0.16203 | 0.21697 | 0.06314 |
| **SYCE3** | 0.02197 | 0.17720 | 0.01764 | 0.01229 | 0.43086 | 0.00499 | 0.00003 |
| **SYCN** | 0.00603 | 0.14936 | 0.03824 | 0.01877 | 0.00550 | 0.00603 | 0.01327 |
| **SYCP1** | 0.00286 | 0.00000 | 0.94757 | 0.01405 | 0.07290 | 0.00954 | 0.31027 |
| **SYCP2** | 0.05626 | 0.05578 | 1.12538 | 0.00000 | 4.83104 | 0.24039 | 0.28564 |
| **SYCP2L** | 1.62758 | 0.00000 | 2.17630 | 0.00000 | 1.74356 | 0.74934 | 2.56779 |
| **SYCP3** | 20.58459 | 12.03164 | 14.02474 | 17.39900 | 7.06417 | 9.76368 | 0.00000 |
| **SYDE1** | 0.08651 | 0.10836 | 0.02690 | 0.19305 | 0.03645 | 0.00000 | 0.13882 |
| **SYDE2** | 2.12293 | 0.20417 | 0.00000 | 1.93035 | 0.10550 | 0.00000 | 6.93439 |

| | | | | | | | |
|---|---|---|---|---|---|---|---|
| **SYF2** | 15.24375 | 11.57328 | 15.39135 | 14.23511 | 6.96277 | 8.00948 | 10.74003 |
| **SYK** | 0.00000 | 6.07414 | 0.91816 | 0.65844 | 0.16779 | 2.80481 | 0.92126 |
| **SYMPK** | 0.87623 | 2.57168 | 2.14715 | 4.41800 | 0.58586 | 3.31092 | 5.51344 |
| **SYN1** | 0.00951 | 0.06346 | 0.00310 | 0.01582 | 0.02546 | 0.01538 | 0.09709 |
| **SYN2** | 0.12369 | 0.21586 | 0.09017 | 0.43986 | 1.67116 | 0.58528 | 0.01367 |
| **SYN3** | 0.09306 | 0.13195 | 0.28792 | 0.01476 | 0.54118 | 0.64335 | 0.08327 |
| **SYNC** | 0.06111 | 0.39849 | 0.54153 | 0.00000 | 0.03417 | 0.00000 | 0.07083 |
| **SYNCRIP** | 34.65551 | 14.60051 | 17.06594 | 19.45914 | 3.73839 | 13.46413 | 31.76708 |
| **SYNDIG1** | 0.02338 | 0.01102 | 0.02731 | 0.02204 | 0.00828 | 0.01523 | 0.01796 |
| **SYNDIG1L** | 0.07481 | 0.01892 | 0.01940 | 0.04550 | 0.01929 | 0.03947 | 0.02048 |
| **SYNE1** | 1.76126 | 5.37808 | 1.39664 | 0.42688 | 6.86396 | 0.69434 | 0.31724 |
| **SYNE2** | 9.51400 | 9.37573 | 15.23812 | 20.64955 | 8.90420 | 8.69796 | 31.85447 |
| **SYNE3** | 0.10013 | 0.05603 | 0.00225 | 0.96757 | 0.02814 | 0.00000 | 0.09553 |
| **SYNE4** | 0.00000 | 0.00000 | 0.02446 | 1.68843 | 0.05557 | 0.18526 | 0.47848 |
| **SYNGAP1** | 0.00000 | 0.01000 | 0.05678 | 0.06081 | 0.01752 | 0.13822 | 0.17869 |
| **SYNGR1** | 0.16731 | 0.19032 | 0.49335 | 1.28595 | 0.01083 | 0.00000 | 0.10823 |
| **SYNGR2** | 27.44970 | 38.66505 | 12.43563 | 16.14471 | 1.25036 | 7.45303 | 0.94675 |
| **SYNGR3** | 7.53863 | 9.53721 | 0.99332 | 0.00000 | 0.60699 | 2.66952 | 3.72640 |
| **SYNGR4** | 0.22712 | 0.22488 | 1.99133 | 0.00134 | 0.14062 | 0.10174 | 0.00000 |
| **SYNJ1** | 4.32153 | 0.00000 | 4.22732 | 3.03471 | 4.40020 | 0.09761 | 4.19318 |
| **SYNJ2** | 2.61769 | 3.78512 | 0.57196 | 3.10246 | 0.36109 | 2.22270 | 8.35113 |
| **SYNJ2-IT1** | 0.00520 | 0.01488 | 0.01765 | 0.09390 | 0.00732 | 0.03937 | 0.03953 |
| **SYNJ2BP** | 12.24215 | 7.02018 | 3.54350 | 7.07609 | 1.53808 | 7.49131 | 3.68646 |
| **SYNM** | 0.03748 | 1.01571 | 0.20009 | 0.00000 | 1.54839 | 0.15842 | 0.01293 |
| **SYNPO** | 0.00847 | 0.00330 | 0.00171 | 7.61892 | 0.00439 | 0.00000 | 2.46323 |
| **SYNPO2** | 0.03226 | 0.00000 | 0.06306 | 0.03024 | 0.00662 | 0.02590 | 0.06872 |
| **SYNPO2L** | 0.00000 | 0.18050 | 0.10461 | 0.19267 | 0.01781 | 0.00000 | 0.37864 |
| **SYNPR** | 0.02563 | 0.01532 | 0.00899 | 0.02494 | 0.06927 | 0.01337 | 0.01942 |
| **SYNPR-AS1** | 0.04476 | 0.01247 | 0.01229 | 0.02307 | 0.01711 | 0.04958 | 0.00274 |
| **SYNRG** | 1.84677 | 5.18572 | 2.47233 | 3.91792 | 3.28814 | 4.81231 | 8.95039 |
| **SYP** | 0.04361 | 0.02033 | 0.25480 | 0.02462 | 0.41920 | 0.03698 | 0.18181 |
| **SYP-AS1** | 0.02119 | 0.02477 | 0.01776 | 0.02401 | 0.02776 | 0.01190 | 0.02109 |
| **SYPL1** | 21.38245 | 0.00000 | 10.72887 | 11.52884 | 3.40068 | 10.29603 | 6.96705 |
| **SYPL2** | 0.05487 | 0.21202 | 0.00000 | 0.00000 | 0.17354 | 0.28811 | 0.03207 |
| **SYS1** | 12.32109 | 1.19365 | 13.32617 | 12.52724 | 4.48122 | 2.47688 | 5.56757 |

| | | | | | | | |
|---|---|---|---|---|---|---|---|
| **SYT1** | 0.35122 | 0.00000 | 0.47590 | 0.32489 | 0.88449 | 0.45244 | 0.48080 |
| **SYT10** | 0.02578 | 0.00888 | 0.02791 | 0.01885 | 0.11181 | 0.01423 | 0.01504 |
| **SYT11** | 0.02769 | 0.09500 | 0.00000 | 0.09694 | 0.81265 | 0.02198 | 0.02149 |
| **SYT12** | 0.00000 | 0.01017 | 0.07814 | 0.07054 | 0.00385 | 0.05783 | 0.24739 |
| **SYT13** | 0.00000 | 0.00894 | 0.01816 | 0.27050 | 0.04230 | 1.51777 | 0.05871 |
| **SYT14** | 0.00000 | 0.06182 | 0.73717 | 0.06575 | 0.22412 | 0.29822 | 0.08169 |
| **SYT14L** | 0.02203 | 0.01891 | 0.01974 | 0.01928 | 0.01407 | 0.01536 | 0.01848 |
| **SYT15** | 0.17714 | 0.00017 | 0.06139 | 0.03831 | 0.04619 | 0.00000 | 0.24762 |
| **SYT16** | 0.00000 | 0.00488 | 0.00000 | 0.05359 | 1.34612 | 0.22954 | 0.07596 |
| **SYT17** | 0.00000 | 0.59525 | 0.53036 | 0.43999 | 1.49849 | 0.02929 | 0.43373 |
| **SYT2** | 1.93148 | 3.28269 | 1.07204 | 0.00000 | 5.69645 | 6.42108 | 7.09848 |
| **SYT3** | 0.01444 | 0.00413 | 0.05306 | 0.07239 | 0.01942 | 0.06522 | 0.03092 |
| **SYT5** | 0.02323 | 0.00926 | 0.02860 | 0.01721 | 0.15524 | 0.04292 | 0.04066 |
| **SYT6** | 0.00000 | 0.12713 | 0.05401 | 0.34943 | 0.29951 | 0.37106 | 0.00000 |
| **SYT7** | 0.30200 | 0.56720 | 0.00000 | 0.69171 | 0.00676 | 0.40913 | 3.24546 |
| **SYT8** | 0.01466 | 0.02228 | 0.01154 | 0.12272 | 0.00782 | 0.00940 | 0.01956 |
| **SYT9** | 0.02322 | 0.00000 | 0.11408 | 0.01625 | 0.80473 | 0.02383 | 0.01688 |
| **SYTL1** | 0.07389 | 0.00000 | 0.02142 | 0.02224 | 0.01131 | 0.22222 | 0.05442 |
| **SYTL2** | 0.00000 | 0.10849 | 0.67375 | 0.00000 | 2.53175 | 1.07351 | 0.18901 |
| **SYTL3** | 0.83938 | 0.35965 | 0.27116 | 0.18861 | 6.00486 | 0.00000 | 0.22920 |
| **SYTL4** | 0.00000 | 1.84860 | 2.34512 | 0.00000 | 0.49416 | 0.57420 | 0.00000 |
| **SYTL5** | 0.02056 | 0.00651 | 0.04098 | 0.02945 | 1.76958 | 0.00387 | 0.08221 |
| **SYVN1** | 3.92827 | 0.00000 | 2.05395 | 10.33915 | 0.98900 | 5.15920 | 9.29207 |
| **SZRD1** | 28.66289 | 32.28740 | 14.95546 | 26.02604 | 7.36135 | 12.61636 | 5.39764 |
| **SZT2** | 1.44276 | 0.47006 | 1.07223 | 1.80870 | 0.24254 | 0.91315 | 4.95439 |
| **T** | 0.03817 | 0.01234 | 0.01456 | 0.02913 | 0.00778 | 0.01803 | 0.01683 |
| **TAAR5** | 0.00000 | 0.03737 | 0.24275 | 0.00000 | 0.08623 | 0.22354 | 1.36170 |
| **TAAR6** | 0.01939 | 0.01889 | 0.00294 | 0.02017 | 0.16608 | 0.01258 | 0.02313 |
| **TAB1** | 0.55030 | 3.68441 | 1.05225 | 1.50649 | 0.59685 | 1.26086 | 3.15004 |
| **TAB2** | 4.69437 | 0.00000 | 4.79461 | 3.37710 | 1.50885 | 4.63800 | 10.55446 |
| **TAB3** | 2.75945 | 0.00000 | 1.01620 | 1.96010 | 0.26148 | 1.15706 | 7.59123 |
| **TAC1** | 0.02295 | 0.00000 | 0.02112 | 0.02621 | 1.05497 | 0.01267 | 0.01611 |
| **TAC3** | 0.05404 | 0.00382 | 0.01516 | 0.09882 | 0.08318 | 0.07939 | 0.02663 |
| **TAC4** | 0.34645 | 0.00000 | 0.01205 | 0.11680 | 0.00681 | 0.00000 | 0.37322 |
| **TACC1** | 24.27076 | 11.95019 | 4.79548 | 18.67732 | 3.12090 | 0.00000 | 34.75158 |

| | | | | | | | |
|---|---|---|---|---|---|---|---|
| **TACC2** | 3.17678 | 2.22886 | 0.63992 | 4.18360 | 0.35335 | 1.85195 | 9.69986 |
| **TACC3** | 16.91044 | 28.34264 | 12.27092 | 10.73288 | 5.09160 | 9.32063 | 20.11396 |
| **TACO1** | 0.00000 | 10.60340 | 3.36175 | 0.48338 | 0.00000 | 5.05291 | 11.16074 |
| **TACR1** | 0.00000 | 0.52042 | 0.08202 | 0.08943 | 0.05497 | 0.00609 | 0.10467 |
| **TACR2** | 0.04765 | 0.00000 | 0.04885 | 0.10408 | 0.03342 | 0.00640 | 0.13717 |
| **TACR3** | 0.03338 | 0.00000 | 0.03094 | 0.01930 | 0.15953 | 0.03528 | 0.01431 |
| **TACSTD2** | 27.08736 | 3.89158 | 0.00000 | 25.00109 | 0.00000 | 0.00000 | 12.46577 |
| **TADA1** | 7.55914 | 0.00000 | 11.58964 | 4.60344 | 0.84721 | 4.27642 | 0.00000 |
| **TADA2A** | 4.52714 | 3.06605 | 4.06743 | 2.37054 | 2.05708 | 1.39937 | 0.00000 |
| **TADA2B** | 1.68097 | 5.73577 | 1.01368 | 0.75979 | 0.75288 | 3.11181 | 5.55838 |
| **TADA3** | 2.91621 | 0.82864 | 2.95382 | 9.18391 | 1.27605 | 5.74748 | 4.39363 |
| **TAF1** | 2.69198 | 1.13327 | 4.36423 | 0.00000 | 2.68634 | 4.31989 | 6.92540 |
| **TAF10** | 0.00000 | 5.45007 | 3.12636 | 2.19266 | 0.30036 | 4.99830 | 16.53101 |
| **TAF11** | 17.91470 | 5.24460 | 9.51554 | 23.54140 | 7.89053 | 5.76649 | 10.67685 |
| **TAF12** | 17.25316 | 10.55873 | 18.32554 | 17.39974 | 5.23346 | 4.44458 | 2.24598 |
| **TAF13** | 10.58902 | 1.84357 | 10.21530 | 6.17310 | 9.07272 | 5.38643 | 0.00000 |
| **TAF15** | 2.38844 | 0.00000 | 2.33451 | 2.59673 | 0.09310 | 1.24583 | 8.83584 |
| **TAF1A** | 4.62393 | 0.00000 | 4.87463 | 2.78365 | 1.19047 | 1.31017 | 0.00000 |
| **TAF1A-AS1** | 0.12427 | 0.39027 | 1.86141 | 0.45547 | 0.30208 | 0.00000 | 0.29693 |
| **TAF1B** | 12.83781 | 2.37756 | 4.21198 | 9.64383 | 2.32562 | 4.40360 | 6.44253 |
| **TAF1C** | 0.74965 | 3.13416 | 3.98676 | 5.95761 | 0.63326 | 2.64665 | 12.82372 |
| **TAF1D** | 12.72568 | 2.21430 | 10.58103 | 11.02764 | 3.76843 | 12.69266 | 37.09015 |
| **TAF1L** | 0.19102 | 0.18241 | 0.34607 | 0.08422 | 0.76183 | 0.20945 | 0.38909 |
| **TAF2** | 13.60981 | 2.16614 | 4.17572 | 8.04796 | 1.81127 | 5.59785 | 5.17501 |
| **TAF3** | 5.42549 | 6.61369 | 5.71419 | 9.87666 | 9.28868 | 4.41729 | 8.94559 |
| **TAF4** | 0.20373 | 0.18636 | 1.38935 | 0.43901 | 1.03896 | 0.69693 | 6.54680 |
| **TAF4B** | 1.11041 | 0.00000 | 6.12214 | 4.34879 | 1.45367 | 4.83136 | 10.51823 |
| **TAF5** | 1.25853 | 0.00000 | 3.52851 | 2.74013 | 4.00344 | 1.05712 | 8.21229 |
| **TAF5L** | 2.63999 | 0.00000 | 7.65764 | 0.84031 | 6.15573 | 0.90816 | 10.75371 |
| **TAF6** | 1.78011 | 7.03368 | 3.50340 | 9.38020 | 1.26953 | 5.65748 | 8.26092 |
| **TAF6L** | 0.00000 | 1.06244 | 0.24595 | 0.50623 | 0.38232 | 0.55276 | 2.53061 |
| **TAF7** | 14.91645 | 10.72770 | 24.11455 | 14.36752 | 13.82433 | 4.51850 | 4.18960 |
| **TAF7L** | 0.10620 | 0.00000 | 0.58805 | 0.14161 | 0.69070 | 0.41242 | 0.36162 |
| **TAF8** | 5.45604 | 1.62438 | 3.85949 | 5.81161 | 2.75430 | 1.51367 | 3.24243 |
| **TAF9** | 38.24964 | 3.41377 | 17.03539 | 21.41774 | 5.64102 | 11.82610 | 6.72455 |

| | | | | | | | |
|---|---|---|---|---|---|---|---|
| **TAF9B** | 3.63140 | 0.00000 | 3.73836 | 3.16445 | 1.08919 | 1.57309 | 0.00469 |
| **TAGAP** | 0.00003 | 0.00000 | 0.05452 | 0.00576 | 0.00805 | 0.10965 | 0.39048 |
| **TAGLN** | 0.00000 | 0.75749 | 1.27430 | 1.05046 | 0.07944 | 0.13349 | 4.77226 |
| **TAGLN2** | 64.22853 | 43.23518 | 12.56318 | 29.88344 | 2.60716 | 8.57251 | 33.68245 |
| **TAGLN3** | 0.00444 | 0.01852 | 0.01115 | 0.04176 | 0.00725 | 0.00950 | 0.08346 |
| **TAL1** | 0.03675 | 0.02595 | 0.01062 | 0.01405 | 0.00746 | 0.00673 | 0.04360 |
| **TAL2** | 0.02976 | 0.00955 | 0.02692 | 0.01558 | 0.00927 | 0.04946 | 0.01732 |
| **TALDO1** | 61.61528 | 44.27070 | 6.69306 | 44.40669 | 10.95817 | 25.54229 | 24.65183 |
| **TAMM41** | 6.60934 | 3.53419 | 5.72316 | 0.04262 | 1.77199 | 1.47516 | 5.30670 |
| **TANC1** | 0.45904 | 0.41715 | 4.53992 | 1.39137 | 2.17092 | 0.61758 | 3.42893 |
| **TANC2** | 0.00000 | 0.24221 | 1.21833 | 6.58947 | 1.48383 | 1.27103 | 3.86656 |
| **TANGO2** | 7.18718 | 4.60086 | 1.34311 | 5.36198 | 0.06892 | 2.01534 | 0.03716 |
| **TANGO6** | 3.10006 | 2.58329 | 3.70171 | 1.93645 | 0.25259 | 1.21506 | 0.00000 |
| **TANK** | 8.64365 | 0.00000 | 2.36125 | 7.97135 | 2.04942 | 2.32858 | 0.00000 |
| **TAOK1** | 8.27790 | 2.73587 | 8.73308 | 7.17046 | 2.87078 | 5.68264 | 24.60076 |
| **TAOK2** | 0.00000 | 5.70207 | 2.05518 | 5.67551 | 0.61813 | 2.56170 | 16.19430 |
| **TAOK3** | 9.55227 | 9.63429 | 7.72439 | 4.61848 | 8.89760 | 6.55084 | 18.06212 |
| **TAP1** | 0.00612 | 0.11297 | 0.55802 | 1.14812 | 0.00220 | 0.00000 | 0.00000 |
| **TAP2** | 0.09055 | 0.00000 | 0.36141 | 0.07030 | 0.20532 | 0.10969 | 0.02835 |
| **TAPBP** | 0.24874 | 0.55475 | 0.76770 | 1.08402 | 0.64122 | 0.72401 | 0.47321 |
| **TAPBPL** | 0.04975 | 0.23685 | 0.06533 | 0.04017 | 0.27417 | 0.01562 | 0.09179 |
| **TAPT1** | 0.22245 | 0.22782 | 0.84766 | 0.36643 | 0.59816 | 0.81090 | 2.47815 |
| **TAPT1-AS1** | 4.57762 | 9.42712 | 5.54091 | 11.12724 | 1.91086 | 6.23079 | 21.05122 |
| **TARBP1** | 1.91636 | 1.13856 | 0.31630 | 0.00000 | 0.63539 | 1.88887 | 2.38035 |
| **TARBP2** | 12.27467 | 14.84292 | 7.51204 | 10.47219 | 1.32994 | 8.50303 | 8.53257 |
| **TARDBP** | 40.53555 | 22.38614 | 18.10289 | 31.05577 | 12.36748 | 16.40546 | 2.34855 |
| **TARS** | 53.83001 | 12.37260 | 2.98495 | 27.17414 | 3.82816 | 20.42885 | 19.00341 |
| **TARS2** | 9.50140 | 7.29512 | 3.73957 | 3.58976 | 1.39646 | 5.78729 | 0.00000 |
| **TARSL2** | 1.02222 | 0.63169 | 1.29376 | 1.05932 | 0.75718 | 0.92202 | 1.18242 |
| **TAS1R1** | 0.00000 | 0.00000 | 5.48918 | 0.28373 | 2.01799 | 0.00458 | 0.00636 |
| **TAS1R2** | 0.02646 | 0.01766 | 0.00990 | 0.01613 | 0.01713 | 0.00593 | 0.08439 |
| **TAS1R3** | 0.00140 | 0.02819 | 0.00764 | 0.01274 | 0.00715 | 0.00400 | 0.24369 |
| **TAS2R1** | 0.03015 | 0.01742 | 0.01280 | 0.01947 | 0.00799 | 0.01243 | 0.01446 |
| **TAS2R10** | 0.02398 | 0.01654 | 0.01292 | 0.01875 | 0.00764 | 0.01334 | 0.03156 |
| **TAS2R14** | 0.18724 | 0.00000 | 0.03109 | 0.01750 | 0.00615 | 0.04657 | 0.06074 |

| | | | | | | | |
|---|---|---|---|---|---|---|---|
| **TAS2R19** | 0.05788 | 0.02149 | 0.04112 | 0.02562 | 0.00563 | 0.01467 | 0.02765 |
| **TAS2R20** | 0.28693 | 0.08588 | 0.14157 | 0.38378 | 0.00156 | 0.16435 | 0.13108 |
| **TAS2R3** | 0.03653 | 0.01493 | 0.01538 | 0.02306 | 0.00769 | 0.01270 | 0.01506 |
| **TAS2R30** | 0.01076 | 0.00989 | 0.01533 | 0.05150 | 0.00742 | 0.02465 | 0.10893 |
| **TAS2R31** | 0.02391 | 0.05272 | 0.01316 | 0.03869 | 0.00737 | 0.01652 | 0.08868 |
| **TAS2R4** | 0.01098 | 0.02470 | 0.02442 | 0.04198 | 0.00547 | 0.15732 | 0.19184 |
| **TAS2R42** | 0.03088 | 0.02760 | 0.01709 | 0.02041 | 0.00734 | 0.01512 | 0.01069 |
| **TAS2R43** | 0.03520 | 0.01473 | 0.02940 | 0.02189 | 0.00644 | 0.02394 | 0.05378 |
| **TAS2R46** | 0.02903 | 0.01979 | 0.02699 | 0.01946 | 0.00671 | 0.01534 | 0.04597 |
| **TAS2R5** | 0.37440 | 0.00000 | 0.14880 | 0.10242 | 0.01675 | 0.00000 | 0.48319 |
| **TAS2R50** | 0.02898 | 0.01918 | 0.01442 | 0.01797 | 0.00749 | 0.01018 | 0.03539 |
| **TAS2R60** | 0.04509 | 0.03581 | 0.00977 | 0.02219 | 0.00804 | 0.01638 | 0.00473 |
| **TASP1** | 6.89630 | 3.09141 | 5.67627 | 3.35349 | 2.32353 | 1.97075 | 0.00000 |
| **TAT** | 0.01907 | 0.05918 | 0.04943 | 0.01668 | 0.20454 | 0.01525 | 0.02055 |
| **TATDN1** | 1.05783 | 0.00000 | 3.03950 | 2.84445 | 0.56694 | 0.50477 | 1.43071 |
| **TATDN2** | 2.12782 | 3.02239 | 0.45137 | 3.12744 | 1.52448 | 2.07602 | 8.41788 |
| **TATDN3** | 6.51288 | 4.02170 | 4.55212 | 1.69320 | 0.77268 | 2.05682 | 1.65926 |
| **TAX1BP1** | 31.10938 | 18.54504 | 27.18012 | 24.36096 | 13.42527 | 17.78611 | 15.78029 |
| **TAX1BP3** | 2.02183 | 2.82118 | 0.00000 | 0.93505 | 0.02419 | 1.23304 | 0.67078 |
| **TAZ** | 0.00000 | 1.96451 | 1.58783 | 0.00000 | 0.00000 | 0.85984 | 3.03873 |
| **TBC1D1** | 13.60791 | 7.28206 | 0.00000 | 0.00000 | 5.35641 | 4.26304 | 22.54658 |
| **TBC1D10A** | 0.00000 | 1.74293 | 0.61932 | 8.24316 | 1.26235 | 1.47767 | 1.61798 |
| **TBC1D10B** | 0.75381 | 3.52448 | 1.71692 | 2.23846 | 1.74975 | 5.11582 | 19.66057 |
| **TBC1D10C** | 0.02416 | 0.02186 | 0.01177 | 0.02698 | 0.00812 | 0.01909 | 0.01112 |
| **TBC1D12** | 1.56352 | 0.00000 | 0.66857 | 1.93062 | 1.05965 | 1.25121 | 3.17494 |
| **TBC1D13** | 3.02505 | 7.28617 | 8.96876 | 2.37041 | 2.04527 | 4.04553 | 10.18415 |
| **TBC1D14** | 2.81230 | 2.84256 | 0.54104 | 0.78136 | 2.65147 | 2.42967 | 0.72051 |
| **TBC1D15** | 9.13354 | 0.00000 | 6.89161 | 12.83954 | 5.34785 | 7.55050 | 2.86713 |
| **TBC1D16** | 0.04643 | 0.18150 | 0.93242 | 0.16323 | 0.00092 | 4.75831 | 1.32574 |
| **TBC1D17** | 0.30293 | 1.42442 | 0.25403 | 4.33163 | 4.98223 | 1.62615 | 5.18910 |
| **TBC1D19** | 0.48075 | 0.00000 | 0.78624 | 0.10105 | 0.12354 | 0.11387 | 0.10706 |
| **TBC1D2** | 0.80715 | 3.17706 | 0.29677 | 3.40429 | 0.24065 | 0.05255 | 0.97729 |
| **TBC1D20** | 1.92472 | 3.08089 | 3.34971 | 4.05784 | 0.75390 | 2.49882 | 10.59443 |
| **TBC1D21** | 0.02016 | 0.01606 | 0.01295 | 0.02072 | 0.00770 | 0.01420 | 0.03038 |
| **TBC1D22A** | 1.52084 | 3.13874 | 0.62944 | 3.25223 | 0.33305 | 1.93267 | 5.08505 |

| | | | | | | | |
|---|---|---|---|---|---|---|---|
| **TBC1D22B** | 2.22051 | 2.03921 | 0.96816 | 3.09362 | 0.45943 | 2.07772 | 1.71441 |
| **TBC1D23** | 16.13621 | 5.70602 | 27.29508 | 7.19387 | 8.67645 | 21.98577 | 0.00000 |
| **TBC1D24** | 0.25595 | 0.65143 | 1.92132 | 3.19110 | 0.00000 | 1.53920 | 1.96182 |
| **TBC1D25** | 2.40815 | 3.06394 | 3.04337 | 1.93323 | 1.16650 | 4.89845 | 4.45272 |
| **TBC1D26** | 0.02870 | 0.02027 | 0.02010 | 0.05505 | 0.00788 | 0.01196 | 0.00420 |
| **TBC1D29** | 0.02611 | 0.02911 | 0.02545 | 0.03311 | 0.00692 | 0.01825 | 0.01692 |
| **TBC1D2B** | 0.72168 | 0.04036 | 0.28789 | 1.99627 | 0.39637 | 0.31188 | 2.12693 |
| **TBC1D30** | 0.38422 | 0.00000 | 0.89455 | 0.03184 | 0.21083 | 0.07577 | 0.21406 |
| **TBC1D31** | 2.91827 | 1.29866 | 1.16833 | 1.39616 | 1.16225 | 1.91507 | 0.68766 |
| **TBC1D32** | 1.08002 | 0.00000 | 0.86254 | 1.43043 | 0.31201 | 0.32646 | 1.98519 |
| **TBC1D3P1-DHX** | 0.02444 | 0.01791 | 0.01304 | 0.02190 | 0.00985 | 0.01354 | 0.01584 |
| **TBC1D3P2** | 0.02150 | 0.01849 | 0.01344 | 0.12936 | 0.00818 | 0.01924 | 0.00000 |
| **TBC1D3P5** | 0.03267 | 0.01595 | 0.01565 | 0.03110 | 0.00784 | 0.01246 | 0.00714 |
| **TBC1D4** | 0.66446 | 0.14651 | 0.21636 | 0.94542 | 0.00826 | 0.00000 | 1.47147 |
| **TBC1D5** | 2.00609 | 2.30826 | 1.21867 | 3.19932 | 0.09896 | 4.94521 | 6.72161 |
| **TBC1D7** | 10.59478 | 7.89902 | 8.38614 | 5.28578 | 5.71194 | 6.78236 | 0.00076 |
| **TBC1D8** | 1.12377 | 1.74716 | 1.83163 | 2.89122 | 0.41261 | 2.76126 | 5.19687 |
| **TBC1D8B** | 0.16976 | 0.00000 | 0.09807 | 0.04279 | 0.04971 | 0.06016 | 0.02392 |
| **TBC1D9** | 0.35391 | 0.00000 | 2.22724 | 0.33629 | 0.65383 | 0.00000 | 0.53857 |
| **TBC1D9B** | 6.99321 | 6.26558 | 2.63694 | 6.90633 | 0.21036 | 4.88075 | 14.79125 |
| **TBCA** | 19.00467 | 11.31701 | 7.83870 | 11.35053 | 1.85525 | 20.87139 | 26.73956 |
| **TBCB** | 10.75304 | 11.38088 | 5.16237 | 20.23228 | 10.88262 | 16.36962 | 6.17525 |
| **TBCC** | 15.53500 | 14.70549 | 15.68694 | 7.80634 | 15.68402 | 9.05407 | 0.00000 |
| **TBCCD1** | 6.59742 | 1.66218 | 4.95485 | 1.53940 | 1.76276 | 3.16837 | 0.00000 |
| **TBCD** | 6.15543 | 23.17556 | 5.22430 | 0.00000 | 0.38407 | 17.25483 | 13.24054 |
| **TBCE** | 10.19663 | 1.48891 | 6.23689 | 7.21642 | 3.23634 | 2.26683 | 0.00000 |
| **TBCEL** | 0.76844 | 0.00000 | 0.67518 | 0.26380 | 0.90556 | 0.11411 | 0.80893 |
| **TBCK** | 2.26228 | 1.31567 | 1.07148 | 0.89970 | 0.00000 | 2.25618 | 1.79281 |
| **TBK1** | 19.24297 | 1.26217 | 3.29394 | 12.85693 | 1.59815 | 5.44943 | 0.00000 |
| **TBKBP1** | 0.00000 | 0.94805 | 1.06632 | 0.00000 | 0.08543 | 0.61850 | 0.58374 |
| **TBL1X** | 0.50368 | 1.91594 | 2.07779 | 8.83206 | 0.40372 | 0.00000 | 19.28524 |
| **TBL1XR1** | 39.96843 | 9.70049 | 22.44489 | 12.82190 | 5.34731 | 15.06424 | 16.41598 |
| **TBL1Y** | 0.02267 | 0.01797 | 0.01284 | 0.01978 | 0.00782 | 0.01287 | 0.02259 |
| **TBL2** | 7.90364 | 3.40007 | 1.01972 | 8.45766 | 0.02692 | 0.17075 | 7.87620 |
| **TBL3** | 17.43280 | 24.03086 | 7.68753 | 13.03973 | 2.94484 | 5.01203 | 2.58242 |

| | | | | | | | |
|---|---|---|---|---|---|---|---|
| **TBP** | 15.03508 | 10.98077 | 9.38499 | 7.45936 | 9.53399 | 2.25690 | 0.00000 |
| **TBPL1** | 21.51047 | 3.08006 | 19.29335 | 6.41123 | 16.43674 | 3.92735 | 4.29501 |
| **TBPL2** | 0.19827 | 0.51085 | 0.79669 | 0.00000 | 0.29384 | 0.12046 | 0.06783 |
| **TBR1** | 0.01919 | 0.02616 | 0.00733 | 0.02416 | 0.04059 | 0.02867 | 0.02618 |
| **TBRG1** | 11.95642 | 10.76568 | 8.58400 | 10.89271 | 5.79600 | 8.39923 | 4.88971 |
| **TBRG4** | 34.74095 | 34.54186 | 15.22251 | 27.06672 | 3.02494 | 19.15320 | 0.48221 |
| **TBX1** | 0.02987 | 0.02151 | 0.01867 | 0.11673 | 0.05058 | 0.00392 | 0.15296 |
| **TBX10** | 0.02276 | 0.03098 | 0.01060 | 0.04350 | 0.00814 | 0.02959 | 0.00693 |
| **TBX15** | 0.03568 | 0.03624 | 0.00000 | 0.03631 | 0.21796 | 0.02323 | 0.00000 |
| **TBX18** | 0.18034 | 0.06763 | 0.09440 | 0.00793 | 0.02070 | 0.11383 | 0.15852 |
| **TBX19** | 0.86549 | 1.92941 | 0.16078 | 0.16959 | 0.18470 | 0.06009 | 0.67995 |
| **TBX2** | 0.00054 | 0.25386 | 0.00000 | 2.96227 | 0.00597 | 0.00000 | 4.57186 |
| **TBX20** | 0.12588 | 0.21721 | 0.01236 | 0.23264 | 0.33151 | 0.01245 | 0.20677 |
| **TBX21** | 0.03178 | 0.01375 | 0.01302 | 0.01775 | 0.00930 | 0.03008 | 0.01297 |
| **TBX22** | 0.03745 | 0.02080 | 0.04833 | 0.01971 | 0.01029 | 0.05471 | 0.00119 |
| **TBX3** | 16.10890 | 24.49678 | 0.23077 | 7.38951 | 0.51109 | 6.10383 | 28.62380 |
| **TBX4** | 0.01895 | 0.01966 | 0.03196 | 0.02222 | 0.00648 | 0.01079 | 0.02404 |
| **TBX5** | 0.03062 | 0.10020 | 0.00276 | 0.03621 | 1.28047 | 0.00652 | 0.06305 |
| **TBX6** | 0.01413 | 0.02266 | 0.01512 | 0.06451 | 0.00738 | 0.06334 | 0.02940 |
| **TBXA2R** | 0.14184 | 0.17379 | 0.06623 | 0.15068 | 0.00665 | 0.43183 | 0.32681 |
| **TBXAS1** | 0.02353 | 0.02151 | 0.01304 | 0.01575 | 0.08445 | 0.01485 | 0.03295 |
| **TC2N** | 1.84713 | 0.00000 | 1.29167 | 0.66167 | 4.44394 | 0.00000 | 1.96035 |
| **TCAIM** | 4.86440 | 0.00000 | 2.70995 | 5.58493 | 0.61289 | 1.06675 | 0.76524 |
| **TCAM1P** | 0.07438 | 0.03197 | 0.01139 | 0.00609 | 0.04067 | 0.01407 | 0.04637 |
| **TCAP** | 0.01357 | 0.11755 | 0.00877 | 0.13555 | 0.11299 | 0.03550 | 0.01105 |
| **TCEA1** | 7.24474 | 2.21978 | 11.11194 | 1.55296 | 2.14984 | 15.97692 | 9.05692 |
| **TCEA2** | 0.00000 | 0.17747 | 0.01227 | 0.23987 | 0.21472 | 0.36359 | 1.23594 |
| **TCEA3** | 0.42019 | 0.03910 | 0.02221 | 0.21376 | 0.00757 | 0.16460 | 0.02270 |
| **TCEAL1** | 0.57951 | 0.60293 | 0.70935 | 0.00000 | 0.04367 | 0.28572 | 0.00292 |
| **TCEAL2** | 0.02869 | 0.01130 | 0.06659 | 0.01267 | 0.02013 | 0.00935 | 0.03343 |
| **TCEAL3** | 0.00059 | 0.02255 | 0.20663 | 0.02456 | 0.03069 | 0.07770 | 0.05349 |
| **TCEAL4** | 1.16983 | 1.86892 | 3.50634 | 0.00000 | 0.00000 | 0.29406 | 0.42200 |
| **TCEAL7** | 0.01292 | 0.00820 | 0.02882 | 0.33888 | 0.03949 | 0.01459 | 0.00761 |
| **TCEAL8** | 9.59953 | 0.00000 | 6.26647 | 0.59925 | 0.00008 | 4.43463 | 5.77339 |
| **TCEANC** | 0.63823 | 0.13202 | 1.00265 | 0.10042 | 1.32338 | 0.18439 | 0.00000 |

| | | | | | | | |
|---|---|---|---|---|---|---|---|
| **TCEANC2** | 4.75578 | 2.70464 | 2.35856 | 3.20280 | 4.11827 | 1.33507 | 0.00000 |
| **TCEB1** | 22.42893 | 11.91387 | 21.45463 | 17.65301 | 15.01200 | 11.06916 | 15.06632 |
| **TCEB2** | 24.41005 | 53.83873 | 31.21498 | 25.35157 | 10.05290 | 22.67222 | 16.66376 |
| **TCEB3** | 32.22210 | 25.89145 | 17.95145 | 20.21179 | 14.08738 | 10.92708 | 7.79266 |
| **TCEB3-AS1** | 0.49904 | 0.75032 | 1.15035 | 0.63457 | 0.44262 | 0.21422 | 0.00000 |
| **TCEB3B** | 0.02286 | 0.01795 | 0.00843 | 0.01929 | 0.07614 | 0.01323 | 0.02329 |
| **TCERG1** | 16.43111 | 6.03055 | 12.50176 | 15.38631 | 5.15609 | 14.87757 | 34.10368 |
| **TCERG1L** | 0.98536 | 1.32106 | 0.01832 | 0.55254 | 0.01965 | 0.81691 | 3.77626 |
| **TCERG1L-AS1** | 0.01520 | 0.02543 | 0.01790 | 0.05876 | 0.00756 | 0.01263 | 0.01708 |
| **TCF12** | 7.82117 | 1.56989 | 1.01681 | 3.95129 | 0.75109 | 3.92007 | 2.48894 |
| **TCF15** | 0.00000 | 0.77481 | 0.57667 | 0.08806 | 0.25464 | 0.00000 | 1.46446 |
| **TCF19** | 1.76645 | 3.30003 | 1.91375 | 2.18432 | 0.24879 | 5.58445 | 0.00000 |
| **TCF20** | 0.24347 | 0.00000 | 2.30769 | 1.31907 | 2.36211 | 1.81619 | 12.05904 |
| **TCF21** | 0.02224 | 0.01772 | 0.01249 | 0.01914 | 0.01070 | 0.01485 | 0.02254 |
| **TCF23** | 0.02150 | 0.02377 | 0.01134 | 0.02115 | 0.00792 | 0.01847 | 0.02135 |
| **TCF24** | 0.40442 | 0.00000 | 1.33317 | 0.07130 | 0.96202 | 0.12990 | 1.26858 |
| **TCF25** | 13.99175 | 10.95767 | 4.00470 | 7.56176 | 2.97412 | 13.85820 | 19.76070 |
| **TCF3** | 2.47210 | 6.43919 | 2.83073 | 6.15468 | 1.29819 | 2.95933 | 16.87208 |
| **TCF4** | 1.12487 | 0.00000 | 0.47598 | 0.01708 | 0.03409 | 1.46919 | 1.73169 |
| **TCF7** | 0.19238 | 2.96541 | 0.74771 | 0.51246 | 1.46139 | 4.48751 | 4.58460 |
| **TCF7L1** | 4.51583 | 8.15071 | 0.00000 | 2.54071 | 0.36508 | 0.00000 | 19.63562 |
| **TCF7L2** | 10.98280 | 3.69867 | 0.09207 | 25.80562 | 0.55851 | 0.00000 | 20.47331 |
| **TCFL5** | 0.66348 | 0.46284 | 0.28252 | 0.15761 | 0.14495 | 0.38511 | 1.30348 |
| **TCHH** | 0.10392 | 0.00548 | 1.38305 | 0.17012 | 1.64968 | 0.00000 | 0.40892 |
| **TCHHL1** | 0.03881 | 0.01427 | 0.01474 | 0.11133 | 0.00848 | 0.01179 | 0.00000 |
| **TCHP** | 3.65184 | 5.70581 | 4.42180 | 1.84722 | 2.30087 | 0.58791 | 4.04754 |
| **TCIRG1** | 0.71450 | 1.71334 | 0.21754 | 3.84792 | 0.00000 | 1.62031 | 5.53486 |
| **TCL1A** | 0.00000 | 1.27799 | 12.50419 | 0.00000 | 35.42673 | 0.00000 | 0.00000 |
| **TCL1B** | 0.64817 | 2.56560 | 10.96022 | 0.00000 | 4.54116 | 3.73126 | 0.00000 |
| **TCL6** | 0.16902 | 0.00000 | 0.01475 | 0.49283 | 0.00799 | 0.10358 | 0.34125 |
| **TCN1** | 0.02343 | 0.02211 | 0.02572 | 0.05137 | 0.02142 | 0.01162 | 0.01882 |
| **TCN2** | 0.33137 | 0.00000 | 0.76514 | 0.16455 | 1.71757 | 1.98359 | 0.21676 |
| **TCOF1** | 20.60904 | 25.59955 | 12.85179 | 25.31092 | 10.50224 | 17.45374 | 27.23991 |
| **TCP1** | 61.81342 | 29.73899 | 24.71332 | 37.18495 | 6.48196 | 28.44642 | 11.39436 |
| **TCP10L** | 0.31769 | 0.28663 | 0.16416 | 0.49412 | 0.03842 | 0.02678 | 0.09908 |

| | | | | | | | |
|---|---|---|---|---|---|---|---|
| **TCP11** | 0.00810 | 0.01177 | 0.01920 | 0.04368 | 0.50593 | 0.06477 | 0.10234 |
| **TCP11L1** | 0.60182 | 0.00000 | 0.75702 | 1.11709 | 0.38040 | 1.30773 | 0.00000 |
| **TCP11L2** | 0.16432 | 0.36038 | 0.11295 | 0.60713 | 0.57607 | 0.04700 | 0.03798 |
| **TCTA** | 2.92203 | 4.56840 | 4.07779 | 3.20544 | 1.42211 | 1.76273 | 0.00000 |
| **TCTE1** | 0.01586 | 0.16203 | 0.36851 | 0.02936 | 0.01501 | 0.03485 | 0.01857 |
| **TCTE3** | 0.07811 | 0.15932 | 0.06096 | 0.10306 | 0.06495 | 0.03510 | 0.11806 |
| **TCTEX1D2** | 0.97475 | 0.32325 | 0.03182 | 2.08777 | 0.06748 | 0.81123 | 0.03950 |
| **TCTEX1D4** | 0.01318 | 0.01750 | 0.01568 | 0.02650 | 0.00741 | 0.01428 | 0.05225 |
| **TCTN1** | 0.22347 | 0.15363 | 0.48859 | 0.14009 | 0.77490 | 0.10145 | 0.03611 |
| **TCTN2** | 0.73251 | 0.29829 | 1.69098 | 1.90107 | 2.95289 | 0.66150 | 1.13662 |
| **TCTN3** | 13.43038 | 2.60730 | 0.77354 | 9.64117 | 0.47394 | 4.29351 | 0.00000 |
| **TDG** | 16.37793 | 7.61463 | 0.00000 | 14.55377 | 0.00000 | 3.22096 | 8.24614 |
| **TDGF1** | 3.29485 | 0.09201 | 0.00000 | 0.00010 | 0.06617 | 23.31041 | 0.00000 |
| **TDGF1P3** | 0.81402 | 0.11481 | 0.00000 | 0.00000 | 0.08384 | 8.48886 | 0.00001 |
| **TDH** | 0.35663 | 0.34886 | 0.46834 | 0.06572 | 0.20826 | 1.19926 | 0.00000 |
| **TDO2** | 0.27113 | 0.00000 | 0.04999 | 0.24497 | 0.03708 | 0.10510 | 0.00000 |
| **TDP1** | 9.92328 | 4.90466 | 4.41338 | 6.79086 | 4.45893 | 4.79492 | 0.00000 |
| **TDP2** | 10.87749 | 1.03561 | 9.53839 | 9.06290 | 2.93686 | 6.61596 | 2.95570 |
| **TDRD1** | 0.31098 | 0.00000 | 0.15179 | 0.04455 | 2.29393 | 0.08442 | 0.00000 |
| **TDRD10** | 0.02176 | 0.00260 | 0.06303 | 0.02866 | 0.33385 | 0.28506 | 0.00321 |
| **TDRD12** | 0.00000 | 0.02000 | 0.13243 | 0.03798 | 0.04966 | 0.01519 | 0.11651 |
| **TDRD3** | 2.10989 | 0.76323 | 1.42706 | 1.06073 | 0.05474 | 2.58400 | 3.75003 |
| **TDRD5** | 0.08671 | 0.00000 | 0.02819 | 0.01145 | 0.17523 | 0.11567 | 0.08918 |
| **TDRD6** | 0.15249 | 0.00000 | 1.49231 | 0.39161 | 2.65037 | 0.33325 | 0.64978 |
| **TDRD7** | 0.46320 | 0.63292 | 1.60623 | 0.10791 | 0.62850 | 0.91320 | 0.00000 |
| **TDRD9** | 0.06144 | 0.00000 | 0.34111 | 0.00387 | 0.95829 | 0.26772 | 0.38002 |
| **TDRG1** | 0.01971 | 0.04863 | 0.01066 | 0.02318 | 0.71482 | 0.03996 | 0.00072 |
| **TDRKH** | 2.70936 | 0.29440 | 1.99672 | 2.81957 | 1.02906 | 3.83411 | 0.00000 |
| **TDRP** | 1.56212 | 2.74776 | 0.00000 | 9.82640 | 0.31409 | 0.00000 | 13.45959 |
| **TEAD1** | 19.09486 | 3.85947 | 5.48767 | 21.43964 | 0.66794 | 0.00000 | 38.84919 |
| **TEAD2** | 0.71148 | 1.40229 | 1.70169 | 0.94827 | 0.66212 | 4.74819 | 8.50703 |
| **TEAD3** | 2.83148 | 0.90756 | 0.00000 | 24.29240 | 0.77844 | 0.00000 | 9.62785 |
| **TEAD4** | 23.02827 | 34.30450 | 19.77070 | 18.97632 | 1.22965 | 17.20439 | 15.11243 |
| **TEC** | 1.37823 | 1.44495 | 1.96177 | 0.00000 | 0.68797 | 5.98143 | 0.00000 |
| **TECPR1** | 0.63873 | 3.54439 | 0.59513 | 2.50803 | 0.05693 | 1.32047 | 2.47739 |

| | | | | | | | |
|---|---|---|---|---|---|---|---|
| **TECPR2** | 0.00000 | 3.53381 | 1.18918 | 1.13009 | 2.72925 | 3.15481 | 4.25096 |
| **TECR** | 26.82483 | 20.93393 | 9.13139 | 16.24191 | 3.11372 | 11.87803 | 0.00000 |
| **TECRL** | 0.02468 | 0.01710 | 0.01296 | 0.02486 | 0.00800 | 0.01936 | 0.01278 |
| **TECTA** | 0.00376 | 0.08649 | 0.00000 | 0.02352 | 5.92407 | 0.00243 | 0.05430 |
| **TECTB** | 0.03028 | 0.01955 | 0.01325 | 0.01663 | 0.01086 | 0.00932 | 0.03966 |
| **TEDDM1** | 0.01907 | 0.03879 | 0.03929 | 0.02911 | 0.00770 | 0.01344 | 0.00729 |
| **TEF** | 0.71884 | 0.73556 | 0.77408 | 1.04716 | 0.25924 | 0.55826 | 2.31886 |
| **TEFM** | 12.97044 | 4.29711 | 9.89736 | 7.23166 | 7.70278 | 4.74082 | 1.09489 |
| **TEK** | 0.29516 | 0.59110 | 0.00000 | 0.59023 | 1.14875 | 0.00000 | 0.25208 |
| **TEKT2** | 0.36469 | 0.00000 | 0.34468 | 0.06459 | 0.04561 | 0.03587 | 0.06958 |
| **TEKT3** | 0.04093 | 0.09328 | 0.07053 | 0.20198 | 0.01498 | 0.32494 | 0.00000 |
| **TEKT4** | 0.13738 | 0.11735 | 0.03858 | 0.10241 | 0.01748 | 0.01971 | 0.06861 |
| **TEKT4P2** | 5.17041 | 2.84856 | 3.36019 | 1.92300 | 0.77413 | 0.77615 | 1.94886 |
| **TEKT5** | 0.61065 | 0.00000 | 0.04449 | 0.80148 | 0.20858 | 0.00000 | 0.04498 |
| **TELO2** | 0.80269 | 8.37195 | 1.46812 | 5.10743 | 0.21393 | 4.42804 | 11.19789 |
| **TEN1-CDK3** | 0.00000 | 0.29282 | 0.01986 | 0.04010 | 0.00728 | 0.05799 | 0.01698 |
| **TENC1** | 0.07500 | 0.00000 | 0.16736 | 0.11759 | 0.07220 | 0.07967 | 0.25335 |
| **TENM1** | 0.00698 | 0.00000 | 0.09847 | 0.13335 | 0.18052 | 0.07123 | 0.85382 |
| **TENM2** | 0.00000 | 0.10555 | 0.12712 | 0.02533 | 0.39503 | 0.35403 | 0.08324 |
| **TENM3** | 1.69691 | 0.00000 | 0.03752 | 19.16076 | 0.06261 | 0.00000 | 9.45981 |
| **TENM4** | 0.00000 | 0.01252 | 0.07275 | 0.01131 | 0.30442 | 0.66088 | 0.15175 |
| **TEP1** | 0.00000 | 0.18208 | 1.62338 | 0.40864 | 0.11418 | 1.60324 | 0.71960 |
| **TEPP** | 0.01467 | 0.04937 | 0.00294 | 0.14128 | 0.19203 | 0.10981 | 0.00002 |
| **TERC** | 0.00000 | 0.12075 | 0.18720 | 0.24444 | 0.10032 | 0.06385 | 1.48735 |
| **TERF1** | 0.62929 | 0.15691 | 1.52595 | 1.52716 | 0.64196 | 3.83955 | 6.77746 |
| **TERF2** | 6.43771 | 2.68301 | 2.05996 | 3.31893 | 1.04598 | 2.28518 | 3.66327 |
| **TERF2IP** | 0.00000 | 0.36336 | 6.79208 | 0.96423 | 7.16964 | 2.02970 | 5.78073 |
| **TERT** | 2.07827 | 1.72426 | 2.43500 | 0.00000 | 0.00000 | 0.00000 | 2.43509 |
| **TES** | 24.92044 | 7.35657 | 1.07733 | 17.00943 | 1.56175 | 4.05553 | 11.28204 |
| **TESC** | 0.00000 | 0.10975 | 0.45745 | 0.00937 | 1.04610 | 0.03423 | 0.01747 |
| **TESK1** | 0.00000 | 3.34917 | 4.72563 | 1.14636 | 1.40254 | 2.01065 | 4.23020 |
| **TESK2** | 1.84083 | 0.18244 | 2.34446 | 2.16416 | 4.55513 | 2.62862 | 0.01757 |
| **TESPA1** | 0.02380 | 0.01875 | 0.01142 | 0.01966 | 0.01748 | 0.01355 | 0.01663 |
| **TET1** | 22.16422 | 20.84833 | 2.57675 | 11.80114 | 0.94638 | 22.58168 | 45.42325 |
| **TET2** | 33.48393 | 37.79541 | 1.31756 | 15.37371 | 1.81091 | 0.00000 | 27.02313 |

| | | | | | | | |
|---|---|---|---|---|---|---|---|
| **TET3** | 3.95366 | 2.20480 | 3.97698 | 6.41970 | 2.10374 | 1.70779 | 17.08162 |
| **TEX10** | 24.05132 | 14.35784 | 12.16937 | 17.32381 | 3.03636 | 5.11772 | 0.00002 |
| **TEX101** | 0.04160 | 0.00000 | 0.17773 | 0.05896 | 0.07549 | 0.01620 | 0.01075 |
| **TEX11** | 2.70501 | 4.87121 | 6.50467 | 0.00000 | 0.14683 | 3.62837 | 0.00000 |
| **TEX12** | 0.12969 | 0.00000 | 0.41775 | 0.01664 | 0.23792 | 0.04905 | 0.13804 |
| **TEX13A** | 0.02439 | 0.00000 | 0.11031 | 0.02402 | 0.00776 | 0.01743 | 0.01554 |
| **TEX13B** | 0.01743 | 0.00903 | 0.01243 | 0.01477 | 0.01534 | 0.07580 | 0.05073 |
| **TEX14** | 0.04951 | 0.30402 | 0.48270 | 0.48698 | 1.54570 | 0.00000 | 0.00000 |
| **TEX15** | 0.04677 | 0.00000 | 0.34599 | 0.00000 | 1.66031 | 0.13701 | 0.02920 |
| **TEX19** | 0.03902 | 0.09375 | 5.39478 | 0.01761 | 2.98512 | 0.05839 | 0.00000 |
| **TEX2** | 3.21685 | 5.13483 | 2.12612 | 3.86591 | 3.56523 | 1.06446 | 5.93328 |
| **TEX21P** | 0.04381 | 0.00837 | 0.11227 | 0.04012 | 0.02916 | 0.01768 | 0.04889 |
| **TEX22** | 0.03011 | 0.01383 | 0.07473 | 0.01630 | 0.02473 | 0.00942 | 0.02094 |
| **TEX26** | 0.06647 | 0.01302 | 0.01160 | 0.02308 | 0.04557 | 0.03030 | 0.00002 |
| **TEX26-AS1** | 0.05507 | 0.18527 | 0.09492 | 0.00000 | 0.02244 | 0.00000 | 0.08627 |
| **TEX261** | 6.77916 | 5.82203 | 4.50021 | 4.36800 | 0.39752 | 1.42735 | 12.39000 |
| **TEX264** | 10.91297 | 18.80245 | 8.23350 | 6.63892 | 0.00000 | 6.60382 | 1.18707 |
| **TEX29** | 0.08129 | 0.00000 | 0.05293 | 0.02429 | 0.00742 | 0.00000 | 0.27884 |
| **TEX30** | 9.97160 | 0.00000 | 12.42240 | 5.93915 | 1.22698 | 0.83246 | 1.78953 |
| **TEX35** | 0.47553 | 0.34244 | 0.00130 | 0.08582 | 0.00807 | 0.00076 | 0.00000 |
| **TEX37** | 0.55095 | 1.67241 | 0.08255 | 1.29212 | 5.58558 | 0.04191 | 0.00000 |
| **TEX38** | 0.07863 | 0.00000 | 0.04330 | 0.02283 | 0.06835 | 0.07076 | 0.11554 |
| **TEX40** | 1.97723 | 1.03182 | 1.09289 | 0.23397 | 0.05266 | 0.22705 | 0.97743 |
| **TEX41** | 0.23376 | 0.03391 | 0.94701 | 0.03056 | 0.47373 | 0.05039 | 0.00000 |
| **TEX43** | 0.02558 | 0.00639 | 0.02954 | 0.02016 | 0.03027 | 0.01442 | 0.04128 |
| **TEX9** | 0.37669 | 0.00000 | 0.35648 | 0.27950 | 0.66871 | 0.08027 | 0.00000 |
| **TF** | 0.00000 | 0.33845 | 0.01629 | 0.00350 | 0.00602 | 1.00218 | 0.02410 |
| **TFAM** | 31.11494 | 10.81327 | 19.22344 | 19.27332 | 12.43105 | 11.30094 | 19.08463 |
| **TFAMP1** | 0.49445 | 0.00000 | 0.50948 | 0.34273 | 0.31934 | 0.25296 | 0.78089 |
| **TFAP2A** | 5.39582 | 0.67095 | 0.14438 | 11.31119 | 0.51047 | 0.00000 | 6.20508 |
| **TFAP2A-AS1** | 0.16538 | 0.02182 | 0.02061 | 0.13034 | 0.01755 | 0.00000 | 0.04100 |
| **TFAP2B** | 0.00000 | 0.20233 | 0.05697 | 0.06051 | 0.87483 | 0.22699 | 0.02586 |
| **TFAP2C** | 39.40591 | 18.35630 | 17.55939 | 31.63000 | 19.78113 | 7.49277 | 21.02117 |
| **TFAP2D** | 0.02353 | 0.01155 | 0.01868 | 0.01755 | 0.47327 | 0.01173 | 0.02232 |
| **TFAP2E** | 0.00831 | 0.05989 | 0.02146 | 0.08401 | 0.01920 | 0.06632 | 0.07826 |

| | | | | | | | |
|---|---|---|---|---|---|---|---|
| **TFAP4** | 0.00000 | 0.88350 | 0.09435 | 0.03838 | 1.05685 | 0.54313 | 0.99020 |
| **TFB1M** | 9.75040 | 6.78911 | 9.05350 | 4.73532 | 3.02443 | 4.37915 | 0.00000 |
| **TFB2M** | 23.97504 | 2.58423 | 19.70873 | 12.80542 | 13.27316 | 4.79394 | 0.00000 |
| **TFCP2** | 9.48457 | 2.12134 | 4.44937 | 9.13709 | 1.88839 | 3.58641 | 7.02374 |
| **TFCP2L1** | 33.66737 | 35.21407 | 6.62324 | 12.77378 | 0.00000 | 20.31136 | 32.40413 |
| **TFDP1** | 10.80382 | 6.78308 | 4.83609 | 3.77354 | 0.35963 | 4.53723 | 18.83765 |
| **TFDP2** | 8.77996 | 15.08465 | 20.37623 | 14.17030 | 4.26349 | 2.08027 | 7.13287 |
| **TFDP3** | 0.67318 | 0.28844 | 1.04141 | 0.62796 | 0.45520 | 0.21738 | 0.00000 |
| **TFE3** | 3.90216 | 7.38642 | 0.00000 | 0.71710 | 0.90388 | 2.34652 | 18.92392 |
| **TFEB** | 0.00000 | 7.07311 | 1.83514 | 18.88345 | 0.14740 | 0.19655 | 2.68680 |
| **TFEC** | 0.03113 | 0.00357 | 0.06044 | 0.12305 | 0.01381 | 0.01278 | 0.00651 |
| **TFF1** | 0.26358 | 0.00000 | 0.00000 | 1.79272 | 0.00457 | 0.00000 | 0.74568 |
| **TFF2** | 0.00798 | 0.01754 | 0.01245 | 0.25293 | 0.01604 | 0.02583 | 0.00377 |
| **TFG** | 38.64476 | 12.84659 | 12.76766 | 26.91328 | 5.38683 | 12.74282 | 4.21648 |
| **TFIP11** | 13.86871 | 14.58308 | 9.33498 | 14.24352 | 13.71115 | 6.40757 | 0.00000 |
| **TFPI** | 1.32445 | 2.47124 | 0.05579 | 2.56988 | 0.07036 | 2.98203 | 0.00000 |
| **TFPI2** | 0.00737 | 0.00004 | 0.04452 | 0.06378 | 0.17452 | 1.27834 | 0.02764 |
| **TFPT** | 5.70316 | 10.29240 | 1.85535 | 6.53119 | 10.25738 | 4.14541 | 0.14461 |
| **TFR2** | 0.02860 | 0.00000 | 0.05495 | 0.10160 | 0.43293 | 0.18758 | 0.07449 |
| **TFRC** | 60.73258 | 8.50941 | 9.16588 | 31.53507 | 2.26425 | 4.21565 | 30.58649 |
| **TG** | 0.04587 | 0.01301 | 0.08893 | 0.04630 | 0.08696 | 0.03471 | 0.00023 |
| **TGDS** | 14.59254 | 0.69065 | 5.26107 | 1.83468 | 0.35613 | 2.88145 | 0.00000 |
| **TGFA** | 0.01829 | 0.01386 | 0.04094 | 0.01759 | 0.91156 | 0.01021 | 0.01810 |
| **TGFB1** | 0.00000 | 0.15228 | 0.25252 | 0.17518 | 0.00094 | 0.84900 | 0.60088 |
| **TGFB1I1** | 0.02493 | 0.01766 | 0.00000 | 0.03439 | 0.40918 | 0.02980 | 0.00599 |
| **TGFB2** | 0.03470 | 0.01907 | 0.02744 | 0.04304 | 0.06703 | 0.00222 | 0.06946 |
| **TGFB2-AS1** | 0.03553 | 0.01765 | 0.01262 | 0.01842 | 0.00783 | 0.01217 | 0.01963 |
| **TGFB3** | 0.00000 | 1.76677 | 3.12134 | 0.58159 | 0.00000 | 5.02678 | 0.16736 |
| **TGFBI** | 0.52575 | 2.05875 | 0.13541 | 0.00000 | 0.22866 | 0.00000 | 0.13095 |
| **TGFBR1** | 3.88390 | 4.23390 | 3.53499 | 3.29563 | 2.41245 | 2.60619 | 0.05210 |
| **TGFBR2** | 0.03490 | 0.15865 | 0.01586 | 5.86328 | 0.00481 | 0.06679 | 0.07188 |
| **TGFBR3** | 24.61481 | 15.82196 | 2.78553 | 8.96580 | 0.28946 | 0.00000 | 29.03694 |
| **TGFBR3L** | 0.00016 | 0.06509 | 0.42414 | 0.04674 | 0.40930 | 0.03875 | 0.15419 |
| **TGFBRAP1** | 5.77807 | 3.30905 | 0.00000 | 5.62091 | 0.18769 | 0.37074 | 9.87782 |
| **TGIF1** | 17.23719 | 12.24058 | 24.07481 | 25.25901 | 6.03098 | 18.65924 | 2.17809 |

| | | | | | | | |
|---|---|---|---|---|---|---|---|
| **TGIF2** | 4.95531 | 4.29931 | 5.28850 | 3.72113 | 2.58967 | 6.02985 | 22.19742 |
| **TGIF2LX** | 0.04797 | 0.01554 | 0.06190 | 0.04351 | 0.17193 | 0.00812 | 0.00964 |
| **TGM1** | 0.34652 | 0.24317 | 0.07265 | 1.78803 | 0.00118 | 0.00000 | 0.70850 |
| **TGM2** | 14.79043 | 0.00000 | 0.00000 | 9.53564 | 0.00000 | 0.00000 | 3.95348 |
| **TGM3** | 0.01447 | 0.01162 | 0.05822 | 0.01998 | 0.05658 | 0.03852 | 0.02500 |
| **TGM4** | 0.02790 | 0.07497 | 0.05746 | 0.01322 | 0.70548 | 0.05111 | 0.07303 |
| **TGM5** | 0.01822 | 0.01724 | 0.00768 | 0.01769 | 0.05073 | 0.13966 | 0.02009 |
| **TGM6** | 0.03990 | 0.02431 | 0.01764 | 0.02295 | 0.00849 | 0.00868 | 0.02883 |
| **TGM7** | 0.01929 | 0.00000 | 0.03991 | 0.01810 | 1.36712 | 0.09014 | 0.01873 |
| **TGOLN2** | 15.95727 | 7.37001 | 7.75427 | 12.11574 | 12.11940 | 7.92008 | 15.32349 |
| **TGS1** | 12.53117 | 0.00000 | 19.75071 | 5.61504 | 7.64186 | 9.49641 | 18.96433 |
| **TH** | 0.06886 | 0.04230 | 0.00503 | 0.01138 | 0.03541 | 0.00602 | 0.01150 |
| **THADA** | 8.09220 | 2.44590 | 6.01891 | 4.63279 | 1.27878 | 2.81682 | 6.33665 |
| **THAP1** | 9.53472 | 0.00000 | 8.69090 | 6.02179 | 9.41162 | 3.58060 | 0.00000 |
| **THAP10** | 0.88884 | 0.37884 | 0.39513 | 0.00004 | 0.12543 | 0.53016 | 0.12579 |
| **THAP11** | 1.75125 | 3.02883 | 3.90794 | 7.00546 | 6.32805 | 8.76038 | 13.80561 |
| **THAP2** | 1.19128 | 0.57763 | 0.50171 | 2.02678 | 0.84891 | 0.14238 | 0.00000 |
| **THAP3** | 2.90829 | 4.10990 | 1.80186 | 4.36930 | 0.18777 | 1.16797 | 0.75471 |
| **THAP4** | 1.31472 | 14.67447 | 4.05932 | 6.67568 | 0.74585 | 5.45387 | 17.49970 |
| **THAP5** | 4.98597 | 1.54065 | 2.96983 | 2.38806 | 3.50360 | 1.24014 | 0.00000 |
| **THAP6** | 3.93606 | 0.00000 | 3.91589 | 0.89124 | 2.56053 | 0.53425 | 0.00000 |
| **THAP7** | 7.22339 | 12.84735 | 6.75562 | 7.67065 | 1.12103 | 2.58419 | 2.73230 |
| **THAP7-AS1** | 0.21144 | 1.36393 | 0.23509 | 0.40683 | 0.10753 | 0.00000 | 0.19780 |
| **THAP8** | 0.00000 | 2.20251 | 0.77683 | 1.08672 | 0.58058 | 1.11055 | 0.33724 |
| **THAP9** | 5.28991 | 0.50093 | 5.25838 | 1.18903 | 1.82833 | 1.44551 | 0.86423 |
| **THAP9-AS1** | 10.71416 | 0.00003 | 2.83663 | 5.26319 | 0.51408 | 4.95674 | 4.74845 |
| **THBD** | 0.00375 | 0.01166 | 0.01142 | 0.02375 | 0.00745 | 0.05790 | 0.09400 |
| **THBS1** | 0.05771 | 0.05605 | 0.05695 | 0.19714 | 0.00440 | 0.00436 | 0.07829 |
| **THBS2** | 0.01418 | 0.04770 | 0.03641 | 0.05933 | 0.63747 | 0.26850 | 0.00460 |
| **THBS3** | 0.04798 | 0.26843 | 0.04865 | 0.18802 | 0.01572 | 0.13720 | 0.15305 |
| **THBS4** | 0.05930 | 0.04562 | 0.00000 | 0.04792 | 4.08171 | 0.33344 | 0.00000 |
| **THEG** | 0.00000 | 0.00504 | 0.60550 | 0.01104 | 1.40476 | 0.19914 | 0.02997 |
| **THEGL** | 0.02623 | 0.01308 | 0.01765 | 0.03058 | 0.62553 | 0.01475 | 0.02441 |
| **THEM4** | 5.44489 | 0.46413 | 1.26270 | 2.83620 | 0.00837 | 0.20021 | 4.10128 |
| **THEM5** | 0.03859 | 0.02110 | 0.00725 | 0.30053 | 0.04823 | 0.04254 | 0.01818 |

| | | | | | | | |
|---|---|---|---|---|---|---|---|
| **THEM6** | 0.35415 | 0.00000 | 1.55122 | 0.69486 | 0.41350 | 0.02925 | 0.37532 |
| **THEMIS** | 0.08561 | 0.01267 | 0.15926 | 0.00000 | 0.68502 | 0.00494 | 0.02345 |
| **THEMIS2** | 0.80191 | 0.00000 | 2.83666 | 0.82844 | 0.66102 | 1.81560 | 0.00000 |
| **THG1L** | 5.08697 | 2.27673 | 6.45118 | 0.12325 | 2.48174 | 3.01843 | 1.01827 |
| **THNSL1** | 0.61523 | 0.00000 | 0.06167 | 0.50778 | 0.93559 | 0.35596 | 0.00000 |
| **THNSL2** | 0.12312 | 0.11431 | 0.44676 | 0.14866 | 0.07422 | 0.11520 | 0.36636 |
| **THOC1** | 18.78525 | 3.27116 | 7.71245 | 8.60609 | 1.90160 | 4.15076 | 1.10234 |
| **THOC2** | 33.01119 | 7.73914 | 21.82503 | 14.35272 | 10.95346 | 19.90128 | 33.15119 |
| **THOC3** | 3.61446 | 5.37320 | 1.66338 | 3.86937 | 2.16047 | 2.51974 | 0.00000 |
| **THOC5** | 19.02856 | 14.55755 | 15.05575 | 12.18231 | 6.13165 | 12.83590 | 2.91920 |
| **THOC6** | 16.45829 | 17.50889 | 2.93860 | 16.98235 | 1.87045 | 6.92922 | 2.60901 |
| **THOC7** | 12.15875 | 0.00000 | 8.38539 | 6.27929 | 1.23058 | 5.18111 | 13.86308 |
| **THOC7-AS1** | 0.02856 | 0.01361 | 0.02051 | 0.01326 | 0.00885 | 0.05601 | 0.09246 |
| **THOP1** | 3.19683 | 9.79351 | 0.82356 | 7.96697 | 0.00000 | 3.01572 | 20.37679 |
| **THPO** | 0.09371 | 0.06585 | 0.02482 | 0.02411 | 0.00888 | 0.50354 | 0.00000 |
| **THRA** | 0.40390 | 0.06428 | 0.06526 | 0.04533 | 0.28013 | 0.83303 | 1.05145 |
| **THRAP3** | 43.90729 | 29.82936 | 18.34733 | 24.26534 | 9.49462 | 17.29940 | 16.25885 |
| **THRB** | 0.00000 | 0.00000 | 0.06817 | 0.17753 | 2.15320 | 0.35288 | 0.53447 |
| **THRIL** | 1.75674 | 0.10973 | 0.08916 | 0.20705 | 0.23802 | 0.00000 | 1.91433 |
| **THRSP** | 0.40742 | 0.00000 | 0.07781 | 0.43087 | 0.18285 | 0.01765 | 0.00000 |
| **THSD1** | 1.27906 | 1.05797 | 0.78625 | 1.50458 | 0.54455 | 0.00119 | 0.00000 |
| **THSD4** | 0.00000 | 0.00000 | 0.32237 | 0.15969 | 1.43851 | 0.23588 | 0.10566 |
| **THSD7A** | 0.00000 | 0.15737 | 0.02299 | 0.01007 | 2.58118 | 0.07443 | 0.04582 |
| **THSD7B** | 0.01622 | 0.05409 | 0.01675 | 0.02720 | 0.35830 | 0.01134 | 0.04086 |
| **THTPA** | 1.43978 | 3.65524 | 1.90012 | 2.04237 | 0.57964 | 0.72514 | 0.19420 |
| **THUMPD1** | 27.71528 | 7.77260 | 16.91568 | 14.71485 | 11.87214 | 9.46738 | 6.99122 |
| **THUMPD2** | 4.83650 | 0.46793 | 3.40715 | 4.17669 | 0.25917 | 0.89066 | 0.00000 |
| **THUMPD3** | 15.38848 | 9.01225 | 12.19339 | 7.53158 | 1.74708 | 5.81777 | 5.83732 |
| **THUMPD3-AS1** | 6.41869 | 0.86722 | 4.96946 | 4.07348 | 2.19571 | 1.58063 | 4.21645 |
| **THY1** | 0.03765 | 0.00000 | 0.78273 | 0.06888 | 0.40833 | 3.27408 | 0.00000 |
| **THYN1** | 9.79264 | 7.32531 | 4.31087 | 11.55445 | 7.70939 | 3.79744 | 0.00000 |
| **TIA1** | 17.51304 | 1.33980 | 10.97861 | 11.30994 | 3.20629 | 6.35369 | 3.95380 |
| **TIAF1** | 0.01234 | 0.11110 | 0.00560 | 0.04879 | 0.05187 | 0.19299 | 0.02814 |
| **TIAL1** | 12.06668 | 3.91460 | 9.45934 | 6.70614 | 6.39233 | 8.02669 | 25.84802 |
| **TIAM1** | 12.77583 | 6.84324 | 2.89673 | 10.40717 | 4.88414 | 4.93324 | 13.85102 |

| | | | | | | | |
|---|---|---|---|---|---|---|---|
| **TIAM2** | 0.13896 | 0.27946 | 1.85011 | 0.00000 | 0.95092 | 0.62177 | 0.00000 |
| **TICAM1** | 0.05616 | 0.00000 | 0.45771 | 0.01213 | 0.25441 | 0.00105 | 0.04576 |
| **TICAM2** | 0.02406 | 0.01869 | 0.01516 | 0.01924 | 0.00772 | 0.01491 | 0.01700 |
| **TICRR** | 3.37473 | 3.18449 | 1.71106 | 3.12581 | 0.97994 | 4.30504 | 11.17060 |
| **TIE1** | 0.02394 | 0.01957 | 0.01196 | 0.01972 | 0.00796 | 0.01587 | 0.02183 |
| **TIFA** | 0.01769 | 0.02047 | 0.13342 | 0.02606 | 1.82582 | 0.00375 | 0.01496 |
| **TIGD1** | 1.25232 | 0.71322 | 2.42953 | 1.08960 | 0.05814 | 0.43399 | 0.43722 |
| **TIGD2** | 0.00000 | 0.00000 | 1.39707 | 0.59055 | 0.33589 | 0.07522 | 1.24098 |
| **TIGD3** | 0.00000 | 0.10933 | 0.07078 | 0.32059 | 0.01911 | 0.00921 | 0.37762 |
| **TIGD4** | 0.09729 | 0.09881 | 0.12375 | 0.07224 | 0.25860 | 0.60310 | 0.00000 |
| **TIGD5** | 0.16014 | 1.57028 | 0.05830 | 0.00000 | 0.00898 | 0.65087 | 2.77992 |
| **TIGD6** | 0.48683 | 0.67331 | 0.88615 | 0.63950 | 1.15688 | 0.19529 | 0.00000 |
| **TIGD7** | 1.68879 | 0.48396 | 2.99968 | 1.56883 | 0.84699 | 0.73503 | 0.45509 |
| **TIGIT** | 0.01964 | 0.02392 | 0.00695 | 0.01491 | 1.95533 | 0.01123 | 0.01552 |
| **TIMD4** | 1.88400 | 0.33032 | 0.16801 | 0.80886 | 1.55404 | 0.10917 | 0.72955 |
| **TIMELESS** | 10.30368 | 4.74083 | 8.11892 | 8.55268 | 0.61080 | 9.79781 | 8.81819 |
| **TIMM10** | 32.13737 | 32.99005 | 22.56971 | 24.25038 | 9.77655 | 17.01805 | 21.03150 |
| **TIMM10B** | 23.71582 | 13.13156 | 13.16030 | 17.76491 | 17.49872 | 6.73558 | 0.00000 |
| **TIMM13** | 43.87496 | 52.79794 | 22.69246 | 39.58321 | 9.63012 | 26.17763 | 26.30313 |
| **TIMM17A** | 54.53429 | 20.40880 | 21.54686 | 27.26711 | 7.89368 | 19.09283 | 4.57306 |
| **TIMM17B** | 12.51911 | 18.33965 | 8.11107 | 6.60472 | 1.68620 | 11.99822 | 8.37864 |
| **TIMM21** | 23.52722 | 17.24943 | 11.44213 | 10.60662 | 0.25207 | 6.21891 | 0.00045 |
| **TIMM22** | 20.72970 | 26.13943 | 13.07587 | 9.49754 | 7.04133 | 12.09910 | 0.00000 |
| **TIMM23** | 29.66433 | 18.21320 | 19.38145 | 14.92068 | 6.12509 | 10.78295 | 7.75929 |
| **TIMM23B** | 0.62904 | 0.20348 | 0.35001 | 0.30444 | 0.61342 | 0.31594 | 0.14384 |
| **TIMM44** | 25.24163 | 32.00977 | 8.58754 | 17.81909 | 5.09323 | 10.34596 | 20.81116 |
| **TIMM50** | 34.85526 | 36.72985 | 29.60347 | 22.37440 | 10.69294 | 20.96426 | 6.26156 |
| **TIMM8A** | 23.12008 | 19.52248 | 17.39019 | 6.33347 | 12.93799 | 8.62274 | 8.84579 |
| **TIMM8B** | 26.94774 | 25.18691 | 14.55368 | 22.08617 | 4.61412 | 13.41343 | 7.57386 |
| **TIMM9** | 14.61221 | 7.81863 | 7.07260 | 3.88854 | 0.02699 | 6.35843 | 14.02855 |
| **TIMMDC1** | 33.43611 | 21.70908 | 16.32675 | 19.41050 | 6.22276 | 11.22289 | 0.00000 |
| **TIMP1** | 0.00065 | 1.04112 | 2.93507 | 0.00000 | 1.95606 | 0.89628 | 0.00000 |
| **TIMP2** | 0.00000 | 0.57560 | 0.32708 | 0.89650 | 1.19572 | 0.00000 | 2.08251 |
| **TIMP3** | 0.25689 | 0.00000 | 0.20835 | 0.29279 | 0.01935 | 0.13461 | 1.41703 |
| **TIMP4** | 1.44441 | 0.00000 | 0.00833 | 8.73569 | 0.00373 | 0.53010 | 0.00000 |

| | | | | | | | |
|---|---|---|---|---|---|---|---|
| **TINAGL1** | 0.00000 | 0.02224 | 0.00138 | 11.21184 | 0.00740 | 0.00000 | 0.69324 |
| **TINCR** | 7.07344 | 5.54817 | 2.06409 | 12.28542 | 0.00097 | 0.00000 | 10.60946 |
| **TINF2** | 5.08356 | 3.86220 | 1.35513 | 7.87629 | 3.41134 | 4.76672 | 0.00000 |
| **TIPARP** | 0.75460 | 0.81656 | 0.12678 | 3.50947 | 9.62466 | 1.02158 | 2.35415 |
| **TIPARP-AS1** | 0.08788 | 0.13945 | 0.00000 | 0.16186 | 0.37814 | 0.00527 | 0.11393 |
| **TIPIN** | 28.87204 | 3.35595 | 6.22221 | 19.47043 | 7.53150 | 2.54331 | 7.08486 |
| **TIPRL** | 18.59636 | 13.49403 | 15.11657 | 9.18770 | 1.72402 | 10.69969 | 4.01891 |
| **TIRAP** | 0.47614 | 0.00000 | 0.44636 | 1.61270 | 0.10755 | 0.42974 | 0.72378 |
| **TJAP1** | 2.77165 | 3.79017 | 4.45841 | 4.35317 | 2.80301 | 2.07848 | 5.64384 |
| **TJP1** | 20.33023 | 11.00127 | 5.89956 | 12.12324 | 5.00847 | 7.49650 | 34.86913 |
| **TJP2** | 16.15170 | 15.81291 | 0.00000 | 10.32335 | 0.79150 | 12.75149 | 23.19819 |
| **TJP3** | 6.51323 | 2.01387 | 0.00000 | 5.74713 | 0.16353 | 0.94520 | 18.44144 |
| **TK1** | 20.33315 | 34.88566 | 7.13305 | 14.99433 | 4.88742 | 14.05890 | 0.00000 |
| **TK2** | 0.00000 | 1.92504 | 0.45357 | 0.08477 | 0.54768 | 1.24650 | 0.03631 |
| **TKT** | 40.13831 | 21.65391 | 5.25990 | 43.35208 | 2.88202 | 23.55863 | 39.69813 |
| **TKTL1** | 12.37591 | 45.60512 | 11.64746 | 0.00000 | 4.50982 | 12.87786 | 0.00000 |
| **TKTL2** | 0.05533 | 0.03730 | 0.03380 | 0.06950 | 0.01022 | 0.02431 | 0.00000 |
| **TLCD1** | 2.43413 | 8.14392 | 0.11720 | 2.91300 | 0.49625 | 4.14981 | 10.07725 |
| **TLCD2** | 0.00000 | 0.86594 | 0.91770 | 0.03496 | 0.00378 | 0.28518 | 0.54653 |
| **TLDC1** | 2.76490 | 0.81588 | 0.00000 | 3.96656 | 0.15873 | 0.04203 | 0.00537 |
| **TLDC2** | 0.01938 | 0.05243 | 0.00000 | 0.00990 | 0.15285 | 0.21776 | 0.09374 |
| **TLE1** | 17.31476 | 15.32185 | 1.52874 | 0.79435 | 0.02589 | 0.00000 | 30.54228 |
| **TLE2** | 0.00000 | 0.97576 | 0.71424 | 0.09960 | 1.11217 | 0.51269 | 0.09146 |
| **TLE3** | 19.51789 | 33.77581 | 0.33980 | 28.96019 | 0.00000 | 10.71199 | 15.59109 |
| **TLE4** | 0.81812 | 0.38381 | 1.94161 | 5.43600 | 3.01684 | 1.37756 | 5.97675 |
| **TLE6** | 0.00000 | 0.67888 | 0.80993 | 0.00475 | 10.27691 | 0.07421 | 0.00355 |
| **TLK1** | 17.28388 | 0.00000 | 10.17333 | 10.33643 | 7.11693 | 6.59333 | 1.23287 |
| **TLK2** | 1.82042 | 0.91305 | 1.17671 | 0.87008 | 1.78893 | 3.15549 | 6.63229 |
| **TLL1** | 0.03306 | 0.02235 | 0.00000 | 0.02122 | 0.40383 | 0.01383 | 0.02906 |
| **TLL2** | 0.00000 | 0.00000 | 0.53914 | 0.16141 | 0.19456 | 1.93346 | 0.25941 |
| **TLN1** | 0.00000 | 3.01418 | 0.88463 | 9.55876 | 0.08295 | 5.30442 | 11.70864 |
| **TLN2** | 5.04146 | 1.97650 | 0.00000 | 1.61173 | 1.44067 | 6.47649 | 13.90318 |
| **TLR10** | 0.03635 | 0.01589 | 0.01626 | 0.01686 | 0.00955 | 0.01688 | 0.00882 |
| **TLR3** | 0.02320 | 0.01836 | 0.01207 | 0.01794 | 0.01525 | 0.01311 | 0.02217 |
| **TLR4** | 0.01683 | 0.01757 | 0.01040 | 0.01476 | 0.00783 | 0.39395 | 0.01317 |

| | | | | | | | |
|---|---|---|---|---|---|---|---|
| **TLR5** | 0.12771 | 0.03247 | 0.06564 | 0.12815 | 0.11041 | 0.01433 | 0.00000 |
| **TLR6** | 0.00704 | 0.02721 | 0.02271 | 0.02421 | 0.00631 | 0.02826 | 0.06033 |
| **TLR7** | 1.65935 | 0.00000 | 0.02775 | 0.42096 | 0.00324 | 0.00000 | 0.81342 |
| **TLR8** | 0.05285 | 0.01181 | 0.00901 | 0.00419 | 0.03880 | 0.02216 | 0.06691 |
| **TLR8-AS1** | 0.13506 | 0.06322 | 0.02994 | 0.02164 | 0.29236 | 0.19761 | 0.00000 |
| **TLR9** | 0.01212 | 0.02121 | 0.00941 | 0.01866 | 0.02372 | 0.06559 | 0.06800 |
| **TLX1** | 0.01873 | 0.11143 | 0.01594 | 0.01965 | 0.00737 | 0.00890 | 0.00249 |
| **TLX2** | 0.04914 | 0.20004 | 0.11149 | 0.00036 | 0.04311 | 0.01682 | 0.02499 |
| **TLX3** | 0.01115 | 0.01919 | 0.02807 | 0.01864 | 0.02477 | 0.07159 | 0.03582 |
| **TM2D1** | 9.36359 | 0.57521 | 6.56890 | 4.39492 | 1.33743 | 2.83498 | 3.07890 |
| **TM2D2** | 24.91428 | 5.81801 | 5.64730 | 17.50144 | 2.49999 | 8.31137 | 3.90844 |
| **TM2D3** | 13.04453 | 2.47003 | 10.08678 | 8.09984 | 2.54434 | 3.52037 | 0.00000 |
| **TM4SF1** | 0.01168 | 0.00000 | 0.44020 | 0.02778 | 0.18547 | 0.33702 | 0.07209 |
| **TM4SF1-AS1** | 0.01785 | 0.40862 | 0.37848 | 0.00104 | 0.15687 | 0.03995 | 0.01123 |
| **TM4SF18** | 0.08486 | 0.00000 | 0.01464 | 0.01634 | 0.00746 | 0.22526 | 0.02027 |
| **TM4SF19** | 0.02560 | 0.02909 | 0.04102 | 0.02096 | 0.00963 | 0.01205 | 0.03207 |
| **TM4SF19-AS1** | 0.02092 | 0.01413 | 0.02153 | 0.02910 | 0.02934 | 0.02745 | 0.03059 |
| **TM4SF19-TCTE** | 0.02372 | 0.01716 | 0.01298 | 0.02256 | 0.00791 | 0.01325 | 0.01745 |
| **TM4SF20** | 0.02337 | 0.02591 | 0.01349 | 0.03498 | 0.00789 | 0.01156 | 0.01646 |
| **TM4SF4** | 0.02040 | 0.02687 | 0.05862 | 0.01720 | 0.00541 | 0.01233 | 0.02451 |
| **TM4SF5** | 0.00000 | 3.83096 | 4.03498 | 0.03457 | 0.00038 | 0.23166 | 0.06144 |
| **TM6SF1** | 0.04869 | 0.09546 | 0.05136 | 0.01808 | 0.08428 | 0.22888 | 0.01827 |
| **TM6SF2** | 0.00000 | 0.04287 | 0.01725 | 0.04042 | 0.00641 | 0.49058 | 0.02507 |
| **TM7SF2** | 3.09825 | 0.14531 | 0.25070 | 5.21648 | 0.00000 | 0.20011 | 4.83348 |
| **TM7SF3** | 4.98442 | 2.30771 | 0.16935 | 2.82881 | 0.07332 | 1.97020 | 7.47571 |
| **TM9SF1** | 14.04738 | 5.20540 | 5.25793 | 12.27385 | 0.20010 | 8.86264 | 4.39542 |
| **TM9SF2** | 33.78720 | 11.71145 | 11.00624 | 15.57535 | 4.44531 | 13.87497 | 4.03415 |
| **TM9SF3** | 17.50694 | 0.00000 | 13.11145 | 9.58893 | 2.57380 | 10.77738 | 34.02786 |
| **TM9SF4** | 10.40882 | 7.36697 | 6.53050 | 9.69645 | 2.05973 | 5.65908 | 2.90085 |
| **TMA16** | 13.99814 | 11.93792 | 10.14380 | 2.93131 | 5.66258 | 4.99752 | 14.74739 |
| **TMA7** | 9.91059 | 8.07275 | 8.23431 | 8.91136 | 1.97189 | 4.16331 | 9.30115 |
| **TMBIM1** | 7.52808 | 34.70914 | 6.13816 | 12.60837 | 2.51210 | 28.54495 | 0.00000 |
| **TMBIM4** | 17.55027 | 7.36754 | 8.77216 | 4.14465 | 0.08971 | 10.60670 | 7.00896 |
| **TMBIM6** | 61.25555 | 35.33280 | 30.05344 | 47.18663 | 22.49376 | 34.06187 | 24.17544 |
| **TMC1** | 0.09103 | 0.47987 | 0.36887 | 0.36577 | 0.06719 | 0.36756 | 0.84351 |

| | | | | | | | |
|---|---|---|---|---|---|---|---|
| **TMC2** | 0.02614 | 0.01737 | 0.01479 | 0.01741 | 0.05770 | 0.01335 | 0.01822 |
| **TMC3** | 0.09092 | 0.01386 | 0.16969 | 0.12664 | 0.39044 | 0.00179 | 0.00000 |
| **TMC4** | 0.00000 | 0.24929 | 0.04738 | 0.59186 | 0.07032 | 0.67350 | 0.26961 |
| **TMC5** | 4.32923 | 0.25884 | 0.74453 | 3.79999 | 1.82786 | 0.00000 | 3.87865 |
| **TMC6** | 0.00000 | 0.11260 | 0.04268 | 0.78203 | 0.00348 | 1.17014 | 1.76087 |
| **TMC7** | 3.49914 | 11.04247 | 5.02214 | 0.00000 | 0.05454 | 6.48178 | 3.47429 |
| **TMC8** | 0.05747 | 0.13543 | 0.27794 | 0.00338 | 0.04879 | 0.00000 | 0.01919 |
| **TMCC1** | 0.34349 | 1.32940 | 0.76260 | 0.92450 | 2.22979 | 1.26913 | 1.41466 |
| **TMCC1-AS1** | 0.17129 | 0.12806 | 0.29452 | 0.05387 | 0.00622 | 0.00000 | 0.00009 |
| **TMCC2** | 0.03124 | 0.17726 | 0.00000 | 0.00000 | 1.16095 | 0.10717 | 0.37102 |
| **TMCC3** | 6.37005 | 0.00000 | 0.34809 | 2.94738 | 0.29267 | 0.00000 | 0.73843 |
| **TMCO1** | 33.91420 | 9.53878 | 23.40294 | 20.84551 | 11.45353 | 9.27206 | 7.45720 |
| **TMCO2** | 0.02123 | 0.01685 | 0.01344 | 0.01955 | 0.04594 | 0.01708 | 0.02601 |
| **TMCO3** | 5.41844 | 0.00000 | 0.18436 | 4.19862 | 0.38303 | 1.77443 | 3.30260 |
| **TMCO4** | 0.22533 | 2.91846 | 0.89191 | 0.00439 | 0.02389 | 1.98671 | 0.00000 |
| **TMCO5B** | 0.02211 | 0.03158 | 0.01570 | 0.03107 | 0.00762 | 0.01484 | 0.01003 |
| **TMCO6** | 0.84973 | 0.23537 | 0.99486 | 2.37360 | 0.19069 | 0.90045 | 2.62152 |
| **TMED1** | 8.78977 | 14.45687 | 3.82588 | 5.06103 | 1.98855 | 9.21647 | 4.36946 |
| **TMED10** | 32.82764 | 14.84592 | 15.02033 | 25.47609 | 1.83317 | 13.44716 | 7.65050 |
| **TMED10P1** | 1.69028 | 0.75990 | 1.02821 | 1.38243 | 0.33620 | 0.64015 | 0.07529 |
| **TMED11P** | 0.14872 | 0.00000 | 0.08108 | 0.00000 | 0.02891 | 0.02400 | 0.17376 |
| **TMED2** | 57.59243 | 19.92379 | 20.73997 | 41.69219 | 9.15541 | 20.46323 | 3.92488 |
| **TMED3** | 24.78227 | 19.41796 | 7.68247 | 17.64189 | 0.51433 | 4.97045 | 5.88422 |
| **TMED4** | 9.07113 | 7.18031 | 0.88991 | 7.77442 | 0.88209 | 3.67618 | 0.00000 |
| **TMED5** | 19.28560 | 0.45019 | 15.58692 | 11.63846 | 3.44013 | 4.93067 | 8.04395 |
| **TMED6** | 0.25070 | 0.00000 | 0.31411 | 0.25692 | 0.03741 | 0.61441 | 0.50230 |
| **TMED7** | 20.02065 | 0.00000 | 8.20665 | 9.79691 | 0.80931 | 4.58894 | 5.81066 |
| **TMED8** | 0.00000 | 1.07359 | 0.35995 | 0.55859 | 0.06227 | 0.29263 | 1.07348 |
| **TMED9** | 43.66672 | 31.95556 | 11.35975 | 34.16114 | 3.95687 | 28.70178 | 0.00000 |
| **TMEFF1** | 0.01383 | 0.04943 | 0.02084 | 0.04561 | 0.00771 | 0.01554 | 0.01428 |
| **TMEFF2** | 0.08609 | 0.00000 | 0.00218 | 0.10061 | 1.85832 | 0.32802 | 0.00000 |
| **TMEM101** | 4.06502 | 5.02876 | 6.22298 | 6.23788 | 3.76247 | 1.58473 | 0.00000 |
| **TMEM102** | 0.00000 | 0.63619 | 0.16196 | 0.25367 | 0.19427 | 0.19027 | 0.91463 |
| **TMEM104** | 3.53237 | 1.83553 | 2.38879 | 4.58428 | 1.12284 | 0.73800 | 4.13796 |
| **TMEM105** | 0.04023 | 0.02650 | 0.01833 | 0.05877 | 0.00892 | 0.02262 | 0.17723 |

| | | | | | | | |
|---|---|---|---|---|---|---|---|
| **TMEM106A** | 0.13372 | 0.00000 | 0.06536 | 0.19547 | 0.25352 | 1.09904 | 0.00079 |
| **TMEM106B** | 4.58629 | 0.00000 | 2.31842 | 2.23499 | 0.07245 | 1.91542 | 2.80885 |
| **TMEM106C** | 15.51281 | 8.10811 | 3.32772 | 24.82596 | 1.86917 | 0.00000 | 0.68203 |
| **TMEM107** | 1.23161 | 0.23025 | 0.32238 | 2.38735 | 1.35577 | 1.16652 | 0.22675 |
| **TMEM108** | 0.09202 | 0.28818 | 1.03177 | 0.24323 | 2.02482 | 1.24258 | 0.00000 |
| **TMEM108-AS1** | 0.02756 | 0.05327 | 0.04268 | 0.02312 | 0.02832 | 0.05236 | 0.00082 |
| **TMEM109** | 16.94347 | 23.33386 | 3.72273 | 16.73775 | 2.69747 | 15.06483 | 12.10772 |
| **TMEM11** | 26.14637 | 22.14569 | 18.86991 | 22.01146 | 10.78601 | 13.83217 | 0.00000 |
| **TMEM110** | 0.52769 | 1.40009 | 0.73452 | 1.38147 | 0.07207 | 1.67057 | 4.29842 |
| **TMEM115** | 0.00000 | 12.20168 | 4.00739 | 12.03735 | 0.88911 | 4.45183 | 9.02884 |
| **TMEM116** | 2.00353 | 0.34505 | 1.71810 | 0.94310 | 0.29886 | 0.80292 | 0.68704 |
| **TMEM117** | 0.63682 | 0.01221 | 0.10743 | 0.06455 | 0.00127 | 1.02363 | 0.38571 |
| **TMEM119** | 0.01785 | 0.01947 | 0.01524 | 0.05136 | 0.00794 | 0.01658 | 0.01937 |
| **TMEM120A** | 6.99201 | 6.58747 | 0.18525 | 5.77481 | 0.00000 | 3.48078 | 2.80357 |
| **TMEM120B** | 3.24616 | 3.99603 | 0.93134 | 1.73116 | 1.17413 | 2.44905 | 6.88987 |
| **TMEM123** | 2.59307 | 0.00000 | 1.50589 | 7.79612 | 1.95730 | 10.22621 | 7.50511 |
| **TMEM125** | 0.42986 | 0.00000 | 0.00000 | 2.52387 | 0.22494 | 0.01746 | 0.08819 |
| **TMEM126A** | 15.26192 | 0.74293 | 13.46393 | 9.82854 | 5.06318 | 5.02758 | 4.15992 |
| **TMEM126B** | 16.78290 | 0.00000 | 8.50397 | 13.69652 | 3.63123 | 6.19440 | 3.29050 |
| **TMEM127** | 2.88408 | 6.00124 | 2.27478 | 2.53062 | 0.94771 | 3.23550 | 11.29323 |
| **TMEM128** | 8.44063 | 5.10093 | 3.17839 | 0.00000 | 2.69560 | 2.67229 | 0.26500 |
| **TMEM129** | 0.52163 | 2.34657 | 0.69858 | 0.76575 | 0.13180 | 2.17468 | 3.14979 |
| **TMEM130** | 0.02548 | 0.00921 | 0.01699 | 0.01922 | 0.14215 | 0.01396 | 0.02054 |
| **TMEM131** | 3.73733 | 2.63087 | 1.91946 | 4.88614 | 1.39442 | 4.50939 | 17.04512 |
| **TMEM132A** | 0.00000 | 0.16957 | 0.01302 | 0.05904 | 0.00491 | 0.91771 | 2.37246 |
| **TMEM132B** | 0.02358 | 0.17192 | 0.00000 | 0.01544 | 2.15013 | 0.00000 | 0.04402 |
| **TMEM132C** | 0.03614 | 0.00570 | 0.00971 | 0.01703 | 0.01168 | 0.98442 | 0.02094 |
| **TMEM132D** | 0.00915 | 0.06833 | 0.00000 | 0.02407 | 1.87671 | 0.00637 | 0.04956 |
| **TMEM132E** | 0.02436 | 0.00997 | 0.01047 | 0.01023 | 0.00658 | 0.00562 | 0.24628 |
| **TMEM133** | 0.87997 | 0.05966 | 0.00731 | 0.05779 | 0.01163 | 1.86239 | 0.43032 |
| **TMEM134** | 1.34568 | 5.66307 | 9.21218 | 1.42137 | 2.37319 | 2.97299 | 0.00000 |
| **TMEM135** | 7.50644 | 0.00000 | 2.18638 | 7.40019 | 0.22870 | 1.90986 | 3.30303 |
| **TMEM136** | 0.47517 | 1.61652 | 0.48245 | 0.00000 | 0.21516 | 1.04740 | 0.38451 |
| **TMEM138** | 12.23454 | 6.29158 | 14.11755 | 10.82852 | 6.12478 | 1.00980 | 0.00000 |
| **TMEM139** | 1.54380 | 0.22299 | 0.34398 | 2.71357 | 0.27011 | 1.40213 | 0.00000 |

| | | | | | | | |
|---|---|---|---|---|---|---|---|
| **TMEM140** | 0.71781 | 0.82979 | 0.38758 | 1.28114 | 0.00226 | 0.31758 | 0.29404 |
| **TMEM141** | 3.33119 | 13.72237 | 3.78224 | 1.81643 | 0.11986 | 6.29407 | 0.16223 |
| **TMEM143** | 1.30484 | 2.14316 | 1.08620 | 1.39221 | 1.58787 | 0.70772 | 0.00000 |
| **TMEM144** | 1.17859 | 0.28294 | 0.06690 | 0.26980 | 0.00146 | 2.41155 | 0.00000 |
| **TMEM145** | 0.03797 | 0.01458 | 0.01847 | 0.02182 | 0.02747 | 0.03759 | 0.02940 |
| **TMEM147** | 22.35118 | 19.48789 | 3.31203 | 21.51821 | 1.41319 | 16.26369 | 24.23399 |
| **TMEM14A** | 10.23841 | 6.91081 | 5.64912 | 3.02040 | 0.81610 | 3.53230 | 0.53291 |
| **TMEM14B** | 16.25210 | 5.53630 | 4.45435 | 8.10531 | 1.28928 | 7.56680 | 4.26384 |
| **TMEM14C** | 14.51594 | 5.08566 | 2.51330 | 8.89043 | 0.42258 | 11.41484 | 6.74291 |
| **TMEM14E** | 0.02319 | 0.00828 | 0.02098 | 0.01454 | 0.01465 | 0.00413 | 0.11213 |
| **TMEM150A** | 2.77101 | 5.58617 | 1.32268 | 7.77755 | 0.37830 | 2.78084 | 6.19009 |
| **TMEM150B** | 0.02543 | 0.01700 | 0.00000 | 0.01177 | 0.69696 | 0.00853 | 0.04726 |
| **TMEM150C** | 0.00000 | 1.32734 | 0.81162 | 0.26685 | 0.00000 | 0.95419 | 0.70231 |
| **TMEM151A** | 0.02723 | 0.03035 | 0.02450 | 0.02017 | 0.00672 | 0.01121 | 0.01480 |
| **TMEM151B** | 0.01899 | 0.05108 | 0.07869 | 0.08731 | 0.03425 | 0.00000 | 0.07255 |
| **TMEM154** | 0.58122 | 0.00000 | 0.00057 | 8.34964 | 0.58113 | 1.60140 | 0.00000 |
| **TMEM155** | 0.00862 | 0.00908 | 0.02270 | 0.57731 | 0.00760 | 0.12591 | 0.00000 |
| **TMEM156** | 0.02478 | 0.01722 | 0.01198 | 0.01924 | 0.04002 | 0.01721 | 0.01860 |
| **TMEM158** | 0.00000 | 0.00000 | 0.60612 | 0.07498 | 0.07592 | 0.30265 | 0.70800 |
| **TMEM159** | 0.92703 | 0.46511 | 4.68757 | 0.36133 | 3.92038 | 6.07710 | 0.00000 |
| **TMEM160** | 0.00000 | 9.46497 | 0.00574 | 1.17059 | 1.78679 | 5.36088 | 12.95295 |
| **TMEM161A** | 7.21585 | 6.25379 | 5.03321 | 11.06567 | 0.26790 | 3.76800 | 3.88773 |
| **TMEM161B** | 7.71692 | 0.40826 | 2.96823 | 2.01456 | 0.68173 | 2.08769 | 5.36108 |
| **TMEM161B-AS1** | 4.19949 | 2.14650 | 2.28601 | 2.09368 | 0.08197 | 1.40522 | 1.36433 |
| **TMEM163** | 0.00652 | 0.08575 | 0.03185 | 0.01831 | 2.17132 | 0.01784 | 0.06816 |
| **TMEM164** | 8.44604 | 1.73561 | 3.01262 | 1.27618 | 0.65549 | 5.59527 | 9.58271 |
| **TMEM165** | 8.73993 | 0.00000 | 3.73509 | 3.58987 | 0.08799 | 5.44693 | 21.12814 |
| **TMEM167A** | 37.98605 | 0.15592 | 19.69368 | 22.34482 | 9.60394 | 10.25428 | 13.51519 |
| **TMEM167B** | 5.14077 | 2.52841 | 8.17884 | 5.48206 | 6.94822 | 4.13586 | 6.22239 |
| **TMEM168** | 12.44346 | 2.61426 | 8.99314 | 3.64419 | 2.76892 | 1.17828 | 1.84938 |
| **TMEM169** | 0.33642 | 0.00000 | 0.00000 | 0.45996 | 0.23837 | 0.00000 | 0.67733 |
| **TMEM17** | 0.33462 | 0.18654 | 0.02793 | 0.12017 | 0.13048 | 0.15819 | 0.00000 |
| **TMEM170A** | 13.18764 | 7.59338 | 7.17663 | 11.95195 | 0.48340 | 7.54004 | 5.26008 |
| **TMEM170B** | 0.08007 | 0.00000 | 0.00661 | 0.07363 | 0.01219 | 0.00000 | 0.36332 |
| **TMEM171** | 2.28046 | 1.11879 | 0.04997 | 1.65882 | 0.30815 | 0.00000 | 1.82062 |

| | | | | | | | |
|---|---|---|---|---|---|---|---|
| **TMEM173** | 0.02897 | 0.02231 | 0.00767 | 0.01280 | 0.04170 | 0.02233 | 0.03121 |
| **TMEM174** | 0.02808 | 0.02285 | 0.01202 | 0.04619 | 0.00826 | 0.01370 | 0.00001 |
| **TMEM175** | 0.56359 | 2.04800 | 0.45970 | 0.24108 | 0.53878 | 0.38639 | 0.44506 |
| **TMEM176A** | 0.02354 | 0.00950 | 0.01353 | 0.02091 | 0.10025 | 0.11702 | 0.00743 |
| **TMEM176B** | 0.01623 | 0.02245 | 0.01030 | 0.01811 | 0.02161 | 0.04177 | 0.03875 |
| **TMEM177** | 10.94731 | 7.05217 | 5.06888 | 2.33806 | 0.44193 | 2.46869 | 0.00000 |
| **TMEM178A** | 0.01741 | 0.01745 | 0.00873 | 0.02821 | 0.04856 | 0.03838 | 0.04599 |
| **TMEM178B** | 0.08643 | 0.03602 | 0.00000 | 0.02751 | 0.16852 | 0.00000 | 0.09831 |
| **TMEM179** | 0.00000 | 0.36054 | 0.75421 | 0.00000 | 0.17404 | 0.00000 | 0.31129 |
| **TMEM179B** | 17.08602 | 19.12683 | 11.44520 | 8.54451 | 2.81359 | 9.22221 | 2.67306 |
| **TMEM18** | 12.93547 | 11.18278 | 3.42722 | 5.25046 | 1.89124 | 8.36579 | 3.50652 |
| **TMEM180** | 4.92562 | 5.87022 | 8.81547 | 7.64032 | 0.30544 | 3.36731 | 3.58852 |
| **TMEM181** | 9.26343 | 0.00000 | 3.49906 | 2.80947 | 0.59232 | 2.38162 | 16.72627 |
| **TMEM182** | 0.62528 | 0.36210 | 0.39399 | 0.53512 | 0.00113 | 0.17535 | 0.07232 |
| **TMEM183A** | 0.30516 | 0.00000 | 0.09826 | 0.02619 | 0.02329 | 0.03704 | 0.48115 |
| **TMEM184A** | 0.00000 | 0.02408 | 0.00974 | 0.02528 | 0.01479 | 0.94538 | 0.02868 |
| **TMEM184B** | 0.00000 | 3.31318 | 6.24088 | 7.00100 | 0.97030 | 4.25850 | 12.90664 |
| **TMEM184C** | 19.32789 | 5.58791 | 10.71822 | 6.27109 | 3.89634 | 2.18870 | 0.00000 |
| **TMEM185A** | 0.49859 | 1.71287 | 4.32231 | 0.34541 | 3.62002 | 0.09447 | 0.00000 |
| **TMEM185B** | 0.16358 | 0.00000 | 1.73269 | 0.32998 | 0.75416 | 0.00000 | 2.42909 |
| **TMEM186** | 8.81585 | 4.67168 | 6.88559 | 7.89215 | 2.08847 | 2.28484 | 0.00000 |
| **TMEM187** | 0.38548 | 0.49280 | 0.16357 | 0.07829 | 0.02415 | 0.00000 | 0.09258 |
| **TMEM189** | 5.89363 | 4.19736 | 1.81673 | 9.52019 | 2.39146 | 1.86996 | 15.65880 |
| **TMEM19** | 6.08360 | 2.77311 | 5.62839 | 5.18241 | 0.23219 | 1.89566 | 0.00000 |
| **TMEM191A** | 0.26190 | 0.31543 | 1.05767 | 0.39869 | 0.06370 | 0.39816 | 1.28798 |
| **TMEM191B** | 0.06076 | 0.09089 | 0.01306 | 0.11096 | 0.00937 | 0.06849 | 0.21773 |
| **TMEM191C** | 0.03668 | 0.09394 | 0.04359 | 0.33184 | 0.01936 | 0.06961 | 0.72945 |
| **TMEM192** | 7.16932 | 2.45819 | 2.53905 | 3.68459 | 0.31238 | 2.22066 | 0.15943 |
| **TMEM194A** | 5.44053 | 0.00000 | 5.31117 | 2.49057 | 1.37504 | 2.07569 | 0.00000 |
| **TMEM194B** | 0.40597 | 0.23917 | 0.39665 | 0.12725 | 0.45752 | 0.09902 | 0.16797 |
| **TMEM198** | 0.02365 | 0.00651 | 0.02228 | 0.63722 | 0.14924 | 0.06813 | 0.06123 |
| **TMEM198B** | 0.00000 | 0.14688 | 0.81155 | 1.53733 | 0.27684 | 0.21080 | 0.86091 |
| **TMEM199** | 14.89467 | 11.28669 | 10.03389 | 11.32779 | 4.99997 | 7.95313 | 0.00000 |
| **TMEM2** | 4.58027 | 1.63113 | 1.51288 | 7.88256 | 11.81310 | 4.22112 | 5.78363 |
| **TMEM200A** | 0.00000 | 0.03163 | 0.00000 | 0.75242 | 0.72763 | 0.00000 | 0.23042 |

| | | | | | | | |
|---|---|---|---|---|---|---|---|
| **TMEM200B** | 0.00000 | 1.39167 | 0.50051 | 0.00000 | 0.03983 | 0.13940 | 0.10137 |
| **TMEM200C** | 0.01279 | 0.00000 | 0.11010 | 0.02025 | 0.00524 | 0.04041 | 0.05297 |
| **TMEM201** | 1.16981 | 9.65594 | 3.98681 | 3.09913 | 1.45097 | 3.37594 | 11.62957 |
| **TMEM202** | 0.01048 | 0.10209 | 0.22647 | 0.01929 | 0.00253 | 0.06667 | 0.01894 |
| **TMEM203** | 0.56019 | 5.07978 | 2.06694 | 2.64533 | 0.12830 | 1.52075 | 8.33914 |
| **TMEM204** | 0.06142 | 0.12473 | 0.53213 | 0.03213 | 0.00482 | 0.00000 | 0.03101 |
| **TMEM205** | 12.69698 | 9.80062 | 1.39222 | 10.90178 | 0.04999 | 7.36728 | 5.18612 |
| **TMEM206** | 9.15646 | 3.12895 | 0.59931 | 7.52271 | 1.61593 | 3.20342 | 0.00000 |
| **TMEM207** | 0.02029 | 0.01752 | 0.02451 | 0.02205 | 0.00683 | 0.03682 | 0.02580 |
| **TMEM208** | 21.37844 | 25.96884 | 18.98692 | 24.75893 | 8.89717 | 13.73006 | 7.36948 |
| **TMEM209** | 16.63774 | 0.00000 | 16.65274 | 3.63828 | 6.54261 | 2.65335 | 5.25674 |
| **TMEM210** | 0.01983 | 0.01109 | 0.03675 | 0.02155 | 0.02837 | 0.01232 | 0.02475 |
| **TMEM211** | 0.05762 | 0.42115 | 0.00316 | 0.00201 | 0.00827 | 0.05048 | 0.00000 |
| **TMEM213** | 0.08713 | 0.01254 | 0.01297 | 0.00734 | 0.00786 | 0.06188 | 0.01777 |
| **TMEM214** | 1.56935 | 4.14029 | 1.35372 | 5.91676 | 0.15407 | 3.82984 | 14.95923 |
| **TMEM215** | 0.01543 | 0.01741 | 0.01124 | 0.02735 | 0.00698 | 0.00833 | 0.12912 |
| **TMEM216** | 3.12757 | 2.26615 | 0.92617 | 2.80808 | 0.99801 | 1.13006 | 0.27264 |
| **TMEM217** | 0.03270 | 0.01662 | 0.01336 | 0.26263 | 0.00855 | 0.00000 | 0.39139 |
| **TMEM218** | 1.22355 | 0.67953 | 3.26907 | 2.91191 | 1.20240 | 0.85771 | 0.00000 |
| **TMEM219** | 4.67020 | 2.77948 | 3.48597 | 9.05303 | 0.18974 | 4.20776 | 6.62845 |
| **TMEM220** | 0.57788 | 0.00000 | 1.61513 | 0.06632 | 0.07981 | 1.31455 | 0.14919 |
| **TMEM220-AS1** | 0.02322 | 0.03313 | 0.03441 | 0.02084 | 0.00738 | 0.03547 | 0.01341 |
| **TMEM221** | 0.04890 | 0.01848 | 0.03816 | 0.05443 | 0.01550 | 0.06534 | 0.03823 |
| **TMEM222** | 13.32099 | 9.55978 | 4.52481 | 13.09784 | 2.07219 | 5.70468 | 3.55097 |
| **TMEM223** | 6.61341 | 9.26602 | 2.18311 | 9.22576 | 0.33576 | 2.53716 | 2.09922 |
| **TMEM229A** | 0.02339 | 0.01884 | 0.01169 | 0.03123 | 0.00837 | 0.03496 | 0.01832 |
| **TMEM229B** | 0.05758 | 0.00000 | 4.26404 | 0.00000 | 1.75278 | 2.48447 | 0.33181 |
| **TMEM230** | 14.09121 | 4.59373 | 9.11349 | 8.39833 | 0.75599 | 7.55003 | 3.81212 |
| **TMEM231** | 0.35261 | 0.29389 | 0.14489 | 0.00000 | 0.57636 | 0.32501 | 0.07745 |
| **TMEM232** | 0.03744 | 0.01757 | 0.05974 | 0.04741 | 0.42603 | 0.01953 | 0.00602 |
| **TMEM234** | 1.51663 | 1.95608 | 0.83577 | 2.49651 | 1.53163 | 1.10980 | 0.00000 |
| **TMEM236.uid-1** | 0.02576 | 0.01713 | 0.01261 | 0.01928 | 0.00882 | 0.01274 | 0.01916 |
| **TMEM236.uid-1** | 0.04137 | 0.00072 | 0.02537 | 0.02453 | 0.03328 | 0.00806 | 0.06978 |
| **TMEM237** | 7.16584 | 1.05633 | 3.44239 | 5.31817 | 0.42634 | 1.94164 | 0.00000 |
| **TMEM238** | 0.00274 | 0.10720 | 0.01079 | 0.01874 | 0.02205 | 0.00545 | 0.08574 |

| | | | | | | | |
|---|---|---|---|---|---|---|---|
| **TMEM241** | 3.59364 | 0.00000 | 0.99041 | 1.64803 | 0.06327 | 0.34947 | 0.57125 |
| **TMEM242** | 8.34966 | 7.16665 | 5.25542 | 3.63846 | 2.20074 | 2.37836 | 1.12367 |
| **TMEM243** | 0.32513 | 0.00000 | 0.70708 | 2.47926 | 0.50835 | 0.40651 | 0.49800 |
| **TMEM245** | 13.94769 | 2.20131 | 5.36438 | 9.45498 | 2.00523 | 4.12767 | 19.71594 |
| **TMEM246** | 0.01894 | 0.17935 | 0.02289 | 0.01912 | 0.00556 | 0.48554 | 0.00345 |
| **TMEM247** | 0.13408 | 0.22873 | 0.04053 | 1.44693 | 0.14788 | 0.16347 | 0.00000 |
| **TMEM248** | 3.15787 | 6.25404 | 4.81132 | 6.43914 | 6.07725 | 8.70043 | 21.45898 |
| **TMEM249** | 0.07331 | 0.00000 | 0.08889 | 0.13669 | 0.03179 | 0.08756 | 0.17116 |
| **TMEM25** | 1.46879 | 0.46045 | 0.19100 | 0.00000 | 0.93999 | 1.40334 | 0.02612 |
| **TMEM251** | 10.22827 | 5.50529 | 10.60204 | 6.31740 | 1.91214 | 2.58676 | 0.00000 |
| **TMEM252** | 0.01914 | 0.01303 | 0.00000 | 0.01683 | 1.52124 | 0.01195 | 0.01550 |
| **TMEM253** | 0.05189 | 0.00000 | 0.08527 | 0.18586 | 0.21284 | 0.02919 | 0.00073 |
| **TMEM254** | 4.82635 | 2.29991 | 4.56613 | 4.45401 | 0.23619 | 0.87872 | 0.24262 |
| **TMEM254-AS1** | 1.01313 | 0.00000 | 0.82119 | 2.75329 | 5.83417 | 0.38780 | 0.63149 |
| **TMEM255A** | 0.02343 | 0.04343 | 0.05912 | 0.02013 | 0.02288 | 0.01298 | 0.01961 |
| **TMEM255B** | 0.03513 | 0.20780 | 0.00363 | 0.00000 | 0.30271 | 0.00406 | 0.18359 |
| **TMEM256** | 5.44372 | 0.00000 | 1.30165 | 6.62419 | 3.73798 | 4.73074 | 7.59471 |
| **TMEM256-PLSC** | 0.03113 | 0.02041 | 0.01258 | 0.01970 | 0.00763 | 0.02304 | 0.02977 |
| **TMEM257** | 0.01874 | 0.01294 | 0.11364 | 0.01484 | 0.00378 | 0.04104 | 0.02915 |
| **TMEM258** | 26.60756 | 21.42088 | 12.21136 | 22.95530 | 3.41695 | 19.53083 | 23.53490 |
| **TMEM259** | 0.00000 | 5.16700 | 0.85994 | 1.27257 | 0.04666 | 4.09362 | 17.61056 |
| **TMEM260** | 2.75436 | 0.00000 | 1.13947 | 0.32450 | 0.33357 | 2.16433 | 2.94883 |
| **TMEM261** | 19.27709 | 33.24762 | 11.51479 | 12.41477 | 0.04643 | 13.79507 | 11.00714 |
| **TMEM262** | 0.04285 | 0.01397 | 0.01444 | 0.04650 | 0.00760 | 0.00818 | 0.02763 |
| **TMEM263** | 1.57442 | 0.54138 | 0.79521 | 1.08100 | 1.27866 | 1.57348 | 0.00000 |
| **TMEM27** | 0.66827 | 0.55380 | 0.33468 | 0.44828 | 0.07881 | 0.31962 | 0.37423 |
| **TMEM30A** | 26.39315 | 0.00041 | 14.35338 | 12.74593 | 8.28979 | 6.83363 | 3.34108 |
| **TMEM30B** | 0.05862 | 0.00000 | 0.00000 | 1.80840 | 0.04139 | 0.41861 | 1.28181 |
| **TMEM30C** | 0.01401 | 0.01356 | 0.01305 | 0.01639 | 0.00763 | 0.07709 | 0.03289 |
| **TMEM31** | 1.38142 | 0.93386 | 0.00000 | 0.00000 | 0.86919 | 0.25124 | 0.25797 |
| **TMEM33** | 45.92566 | 8.20738 | 23.57193 | 14.41976 | 4.96737 | 9.16514 | 13.02451 |
| **TMEM35** | 0.02082 | 0.02179 | 0.05405 | 0.02473 | 0.03923 | 0.01966 | 0.01600 |
| **TMEM37** | 3.17378 | 33.11374 | 6.18253 | 0.00000 | 0.00000 | 4.89017 | 0.00000 |
| **TMEM38A** | 0.19891 | 0.00000 | 0.01217 | 0.17076 | 0.01920 | 0.26750 | 0.00721 |
| **TMEM38B** | 17.27080 | 4.98761 | 5.23530 | 7.76577 | 0.80063 | 4.99053 | 2.84773 |

| | | | | | | | |
|---|---|---|---|---|---|---|---|
| **TMEM39A** | 9.84133 | 2.18438 | 6.76307 | 9.75143 | 2.49754 | 4.89584 | 0.00000 |
| **TMEM39B** | 6.13307 | 4.49127 | 1.14178 | 4.52938 | 0.94121 | 2.97539 | 0.35215 |
| **TMEM40** | 0.00000 | 0.22238 | 0.15583 | 0.93923 | 0.03931 | 0.00118 | 0.00000 |
| **TMEM41A** | 20.37613 | 16.47199 | 6.67823 | 8.52441 | 3.80427 | 9.61796 | 12.57035 |
| **TMEM41B** | 32.64175 | 5.38989 | 14.58086 | 18.13550 | 12.54464 | 15.06320 | 2.28167 |
| **TMEM42** | 0.29150 | 0.00007 | 0.86741 | 1.15485 | 0.34784 | 0.20291 | 0.00000 |
| **TMEM43** | 15.94833 | 8.05695 | 4.00793 | 15.39729 | 1.39536 | 6.13224 | 0.00000 |
| **TMEM44** | 0.10607 | 0.47359 | 0.10245 | 0.93626 | 0.07754 | 0.25463 | 1.23176 |
| **TMEM44-AS1** | 0.51976 | 0.79785 | 0.84662 | 0.37434 | 0.30783 | 0.00000 | 0.03305 |
| **TMEM45A** | 0.12648 | 0.00000 | 0.15246 | 0.17211 | 0.12702 | 0.28110 | 0.00000 |
| **TMEM45B** | 3.58646 | 3.84647 | 6.08134 | 1.98163 | 0.01180 | 0.22805 | 0.23840 |
| **TMEM47** | 0.02286 | 0.02518 | 0.23306 | 0.05063 | 0.06386 | 0.01607 | 0.00302 |
| **TMEM5** | 3.45261 | 1.82121 | 3.42311 | 1.04200 | 0.09500 | 2.20885 | 6.59335 |
| **TMEM50A** | 24.82363 | 13.25956 | 19.85421 | 21.55760 | 6.01801 | 14.01664 | 6.32883 |
| **TMEM50B** | 8.89170 | 2.19710 | 8.05444 | 2.29560 | 5.61418 | 2.94179 | 0.00000 |
| **TMEM51** | 0.00000 | 5.84020 | 2.03112 | 0.41415 | 0.11614 | 5.96597 | 1.93960 |
| **TMEM51-AS1** | 0.00000 | 0.36888 | 0.03293 | 0.01407 | 0.00530 | 0.30788 | 0.10272 |
| **TMEM52** | 1.67589 | 0.11988 | 0.18235 | 4.49398 | 1.71981 | 0.48217 | 0.99352 |
| **TMEM52B** | 4.23932 | 0.00000 | 0.62922 | 5.50147 | 0.04464 | 0.00000 | 0.40252 |
| **TMEM53** | 0.61201 | 1.03091 | 0.16657 | 2.02574 | 0.59499 | 1.88588 | 0.00000 |
| **TMEM54** | 1.56597 | 9.62167 | 2.04186 | 4.76048 | 0.36953 | 0.00000 | 15.90552 |
| **TMEM55A** | 1.42240 | 0.17614 | 0.14362 | 2.68877 | 0.32436 | 0.19975 | 0.69877 |
| **TMEM55B** | 0.23775 | 2.86083 | 2.48996 | 2.68526 | 2.52339 | 4.13801 | 9.25690 |
| **TMEM56** | 3.24573 | 13.85613 | 6.00442 | 0.00000 | 0.00000 | 0.00000 | 5.09231 |
| **TMEM56-RWDD** | 0.13700 | 0.18139 | 0.14942 | 0.00711 | 0.03093 | 0.00399 | 0.05693 |
| **TMEM57** | 0.39598 | 2.03145 | 0.00000 | 2.42764 | 1.22138 | 1.84310 | 8.07062 |
| **TMEM59** | 29.26360 | 11.95402 | 11.02956 | 20.23323 | 10.50085 | 15.35696 | 1.07500 |
| **TMEM59L** | 0.03960 | 0.01031 | 0.01690 | 0.02065 | 0.00813 | 0.01362 | 0.02906 |
| **TMEM60** | 2.27616 | 1.10988 | 1.04808 | 4.34253 | 2.51740 | 1.29582 | 0.00000 |
| **TMEM61** | 0.06536 | 0.00000 | 0.03438 | 0.00826 | 0.44408 | 0.03407 | 0.00732 |
| **TMEM62** | 5.46103 | 0.85420 | 0.83204 | 9.44596 | 0.09565 | 1.29437 | 10.59619 |
| **TMEM63A** | 0.25127 | 3.01451 | 0.73895 | 4.32017 | 0.25032 | 7.06148 | 2.03780 |
| **TMEM63B** | 0.00000 | 1.24927 | 0.56738 | 1.63615 | 0.59080 | 0.61118 | 4.24087 |
| **TMEM63C** | 0.02922 | 0.07503 | 0.05585 | 0.02052 | 0.32411 | 0.03249 | 0.01541 |
| **TMEM64** | 9.02887 | 1.80199 | 2.50143 | 0.03260 | 0.04731 | 0.13371 | 8.59305 |

| | | | | | | | |
|---|---|---|---|---|---|---|---|
| **TMEM65** | 1.11475 | 0.00000 | 1.40428 | 0.00000 | 0.32057 | 0.13022 | 3.50275 |
| **TMEM66** | 21.96697 | 5.66943 | 20.33496 | 10.52730 | 4.55357 | 12.72500 | 10.05796 |
| **TMEM67** | 1.36847 | 0.00000 | 0.39842 | 0.67006 | 0.37161 | 0.57715 | 0.07541 |
| **TMEM68** | 9.87819 | 0.00000 | 14.77894 | 10.39035 | 3.82881 | 2.05771 | 4.37763 |
| **TMEM69** | 22.99993 | 10.56752 | 12.03568 | 13.28926 | 2.79496 | 5.54223 | 0.00000 |
| **TMEM70** | 32.10590 | 9.65747 | 17.97357 | 22.88077 | 4.75302 | 15.98669 | 6.60283 |
| **TMEM71** | 0.03803 | 0.01543 | 0.01917 | 0.03015 | 0.03040 | 0.01201 | 0.01394 |
| **TMEM74** | 0.31794 | 0.00000 | 1.02236 | 0.26407 | 1.36389 | 0.08978 | 0.06262 |
| **TMEM74B** | 0.00002 | 0.43352 | 2.42023 | 0.10174 | 0.12107 | 0.28569 | 0.76961 |
| **TMEM75** | 0.06467 | 0.06276 | 0.12918 | 0.12373 | 0.00422 | 0.00000 | 0.48618 |
| **TMEM79** | 5.65066 | 0.46159 | 1.21429 | 6.56044 | 0.68684 | 0.01108 | 1.28719 |
| **TMEM80** | 1.85402 | 0.99216 | 0.49725 | 3.11612 | 0.20596 | 0.15448 | 2.97867 |
| **TMEM81** | 0.17402 | 0.15232 | 0.06387 | 0.65131 | 0.19644 | 0.34939 | 0.62639 |
| **TMEM82** | 0.02750 | 0.05208 | 0.01959 | 0.01691 | 0.04654 | 0.02064 | 0.01974 |
| **TMEM86A** | 0.66409 | 2.00769 | 0.00000 | 0.24262 | 0.16598 | 3.47364 | 0.32704 |
| **TMEM86B** | 0.01059 | 0.00000 | 0.11381 | 0.63056 | 0.00462 | 0.34222 | 1.54629 |
| **TMEM87A** | 19.45526 | 3.90593 | 13.10792 | 16.60099 | 4.98977 | 7.74336 | 3.20705 |
| **TMEM87B** | 4.69621 | 1.90751 | 3.00213 | 1.77375 | 0.00000 | 2.65458 | 1.95598 |
| **TMEM88** | 1.44536 | 0.41632 | 0.00916 | 2.51255 | 0.01317 | 0.67993 | 0.00000 |
| **TMEM88B** | 0.02335 | 0.01729 | 0.01870 | 0.02118 | 0.01341 | 0.01498 | 0.01483 |
| **TMEM89** | 0.01917 | 0.00351 | 0.02355 | 0.05595 | 0.01089 | 0.01724 | 0.31905 |
| **TMEM8A** | 1.38682 | 4.88805 | 0.34103 | 1.34206 | 0.79543 | 8.42665 | 7.66125 |
| **TMEM8B** | 0.00000 | 0.08550 | 0.10380 | 0.21897 | 0.15505 | 0.11016 | 0.09610 |
| **TMEM9** | 18.46221 | 11.49987 | 8.59621 | 12.77300 | 7.67996 | 7.73312 | 1.72712 |
| **TMEM91** | 0.00000 | 0.42313 | 0.36990 | 0.35403 | 0.02704 | 0.03989 | 0.13732 |
| **TMEM92** | 0.04514 | 0.64767 | 13.47498 | 0.32137 | 8.36005 | 0.35768 | 0.34907 |
| **TMEM95** | 0.02004 | 0.01330 | 0.01139 | 0.02475 | 0.02805 | 0.01094 | 0.03663 |
| **TMEM97** | 30.87798 | 1.46921 | 0.52642 | 17.33836 | 2.59762 | 3.28354 | 14.30155 |
| **TMEM98** | 0.15231 | 0.08541 | 0.00427 | 0.25573 | 0.00719 | 0.71110 | 0.00000 |
| **TMEM99** | 4.12971 | 1.70243 | 14.81870 | 3.15936 | 0.55083 | 0.00000 | 1.42077 |
| **TMEM9B** | 17.56526 | 15.93004 | 12.31367 | 8.61583 | 3.61640 | 9.87621 | 4.35899 |
| **TMEM9B-AS1** | 0.00000 | 0.49150 | 0.08464 | 0.19864 | 0.12258 | 0.05975 | 0.10707 |
| **TMF1** | 15.56616 | 2.70290 | 6.40895 | 9.89700 | 7.78671 | 4.91332 | 7.27183 |
| **TMIE** | 0.02359 | 0.01887 | 0.01137 | 0.05873 | 0.01995 | 0.01296 | 0.00764 |
| **TMIGD2** | 0.01831 | 0.02963 | 0.01508 | 0.05475 | 0.01150 | 0.01265 | 0.01199 |

| | | | | | | | |
|---|---|---|---|---|---|---|---|
| **TMLHE** | 2.03592 | 0.44861 | 0.27292 | 1.19237 | 0.03141 | 0.13328 | 0.42643 |
| **TMOD1** | 2.86347 | 6.41185 | 1.11774 | 0.00000 | 0.37957 | 0.00000 | 0.67525 |
| **TMOD2** | 0.04377 | 0.00610 | 0.05366 | 0.06046 | 0.71311 | 0.08522 | 0.00000 |
| **TMOD3** | 19.32105 | 6.34575 | 7.12096 | 8.57075 | 4.87431 | 6.92694 | 7.42365 |
| **TMOD4** | 0.05787 | 0.10702 | 0.03982 | 0.17098 | 0.04715 | 0.00000 | 0.11529 |
| **TMPO** | 31.41073 | 2.12007 | 18.84910 | 8.48215 | 6.71110 | 14.42200 | 25.36920 |
| **TMPO-AS1** | 0.11327 | 0.18309 | 0.62505 | 0.22340 | 0.63347 | 0.30162 | 0.45568 |
| **TMPPE** | 2.40972 | 0.58326 | 1.90377 | 0.73352 | 0.33175 | 1.31689 | 2.12313 |
| **TMPRSS11A** | 0.01552 | 0.02307 | 0.01298 | 0.02573 | 0.15967 | 0.01058 | 0.03003 |
| **TMPRSS11B** | 0.02754 | 0.01927 | 0.01527 | 0.01846 | 0.00831 | 0.01834 | 0.02102 |
| **TMPRSS11E** | 0.87263 | 0.00000 | 0.05106 | 0.12290 | 0.00592 | 0.69468 | 0.00000 |
| **TMPRSS11F** | 0.00746 | 0.06020 | 0.01651 | 0.08432 | 0.00676 | 0.01102 | 0.02668 |
| **TMPRSS12** | 0.03751 | 0.01138 | 0.01200 | 0.04667 | 0.02585 | 0.00887 | 0.02803 |
| **TMPRSS13** | 10.83219 | 0.00000 | 0.00000 | 6.99892 | 0.00000 | 0.00000 | 4.11195 |
| **TMPRSS15** | 0.08202 | 0.00000 | 0.16317 | 0.00234 | 0.17181 | 0.00154 | 0.09783 |
| **TMPRSS2** | 12.52559 | 0.00000 | 0.00000 | 2.31215 | 0.00000 | 0.00000 | 8.59344 |
| **TMPRSS3** | 0.05661 | 0.01124 | 0.01509 | 0.01998 | 0.01010 | 0.03025 | 0.00922 |
| **TMPRSS4** | 1.40281 | 0.00000 | 0.00000 | 0.17427 | 0.00323 | 1.04322 | 1.58212 |
| **TMPRSS4-AS1** | 0.01708 | 0.01353 | 0.02519 | 0.02819 | 0.01002 | 0.01023 | 0.03579 |
| **TMPRSS5** | 0.02769 | 0.03988 | 0.03077 | 0.11436 | 0.04140 | 0.10761 | 0.13620 |
| **TMPRSS6** | 0.01598 | 0.26757 | 0.01273 | 0.04033 | 0.05072 | 0.02469 | 0.00267 |
| **TMPRSS7** | 0.00000 | 0.08243 | 0.02227 | 0.06144 | 0.02026 | 0.00762 | 0.03078 |
| **TMPRSS9** | 0.01516 | 0.20068 | 0.01211 | 0.02247 | 0.05371 | 0.04880 | 0.26835 |
| **TMSB10** | 26.40707 | 38.25732 | 3.90575 | 76.17359 | 7.17476 | 41.19010 | 48.63237 |
| **TMSB15A** | 0.04051 | 0.08298 | 0.41642 | 0.00000 | 0.54161 | 0.14721 | 0.10587 |
| **TMSB15B** | 0.15013 | 0.08365 | 0.80637 | 0.02209 | 0.01017 | 0.21854 | 0.00000 |
| **TMSB4X** | 36.87405 | 13.38743 | 10.47166 | 41.88683 | 23.83048 | 27.70281 | 13.51709 |
| **TMSB4Y** | 0.02738 | 0.00306 | 0.01614 | 0.03956 | 0.05165 | 0.07187 | 0.10589 |
| **TMTC1** | 0.00000 | 0.01354 | 0.00698 | 0.90718 | 0.49272 | 0.36043 | 0.05020 |
| **TMTC2** | 0.13710 | 0.00000 | 0.09679 | 0.75603 | 0.84594 | 0.64779 | 1.08443 |
| **TMTC3** | 40.75716 | 1.54601 | 9.42653 | 15.88487 | 0.96197 | 3.96238 | 15.12760 |
| **TMTC4** | 3.70615 | 0.28607 | 0.19932 | 0.21693 | 0.00000 | 1.41743 | 3.97327 |
| **TMUB1** | 0.32718 | 0.99833 | 0.13714 | 9.04622 | 0.10348 | 4.10824 | 12.09082 |
| **TMUB2** | 3.59118 | 7.30500 | 6.71513 | 12.61324 | 7.34984 | 3.38075 | 2.00163 |
| **TMX1** | 41.06547 | 5.18886 | 19.75924 | 17.39897 | 4.29533 | 16.13195 | 0.00000 |

| | | | | | | | |
|---|---|---|---|---|---|---|---|
| **TMX2** | 32.31887 | 18.08264 | 22.42598 | 22.59550 | 2.80303 | 14.71075 | 9.17258 |
| **TMX3** | 8.16907 | 0.00000 | 3.94316 | 4.79985 | 2.65307 | 2.35229 | 2.75277 |
| **TMX4** | 0.00000 | 1.99049 | 0.08026 | 0.24171 | 1.28253 | 6.12053 | 1.68443 |
| **TNC** | 0.02016 | 0.01719 | 0.00944 | 0.07818 | 0.03734 | 0.02688 | 0.03089 |
| **TNF** | 0.15097 | 0.30511 | 0.00293 | 0.04169 | 0.00599 | 0.02779 | 0.10435 |
| **TNFAIP1** | 5.20699 | 9.02297 | 9.06294 | 11.74216 | 5.16894 | 3.95152 | 0.00000 |
| **TNFAIP2** | 0.09879 | 0.07567 | 0.02174 | 4.36765 | 0.17465 | 0.01822 | 0.00000 |
| **TNFAIP3** | 0.00000 | 0.95150 | 1.21111 | 6.58526 | 0.00000 | 2.96028 | 0.91697 |
| **TNFAIP6** | 0.01821 | 0.01040 | 0.01533 | 0.05339 | 0.27243 | 0.01605 | 0.01142 |
| **TNFAIP8** | 7.78323 | 0.00000 | 0.23175 | 10.31960 | 0.36448 | 1.80506 | 0.00000 |
| **TNFAIP8L1** | 0.17719 | 0.78182 | 0.07179 | 0.25574 | 1.27779 | 0.02623 | 1.93875 |
| **TNFAIP8L2** | 0.14277 | 0.00000 | 0.07497 | 0.12260 | 0.14866 | 0.10620 | 0.00420 |
| **TNFAIP8L3** | 0.04493 | 0.00000 | 0.03696 | 0.05521 | 0.00574 | 0.05988 | 0.15768 |
| **TNFRSF10A** | 0.01317 | 0.00919 | 0.01638 | 0.04540 | 0.02014 | 0.00697 | 0.14634 |
| **TNFRSF10B** | 0.00000 | 2.58913 | 0.15209 | 6.10772 | 0.49160 | 1.05615 | 2.60380 |
| **TNFRSF10C** | 0.08823 | 0.14609 | 0.01515 | 0.39397 | 0.00498 | 0.00000 | 0.16368 |
| **TNFRSF10D** | 0.00000 | 0.49205 | 0.19888 | 0.06477 | 0.89770 | 0.59245 | 0.25722 |
| **TNFRSF11A** | 0.03651 | 0.16797 | 0.14437 | 0.00000 | 0.04258 | 0.02155 | 0.02685 |
| **TNFRSF11B** | 0.03000 | 0.01240 | 0.01894 | 0.01755 | 0.03029 | 0.01376 | 0.01636 |
| **TNFRSF12A** | 10.84469 | 21.00976 | 3.64391 | 14.74324 | 11.39035 | 4.38365 | 5.11297 |
| **TNFRSF13C** | 0.03338 | 0.02847 | 0.01823 | 0.01533 | 0.00829 | 0.01471 | 0.01751 |
| **TNFRSF14** | 0.00707 | 0.02637 | 0.01744 | 0.06505 | 0.00836 | 0.01076 | 0.02634 |
| **TNFRSF18** | 0.00000 | 0.18319 | 0.55575 | 0.00940 | 0.02631 | 0.18450 | 0.00836 |
| **TNFRSF19** | 0.10366 | 0.00000 | 0.04415 | 0.12754 | 0.32944 | 0.03011 | 0.00000 |
| **TNFRSF1A** | 0.83087 | 1.15724 | 0.00000 | 1.38631 | 0.01222 | 9.09699 | 2.61617 |
| **TNFRSF1B** | 0.00000 | 0.06001 | 0.06018 | 3.30927 | 0.33450 | 0.00350 | 0.00000 |
| **TNFRSF21** | 2.70712 | 0.00000 | 4.06163 | 16.30011 | 0.17468 | 4.26204 | 3.31608 |
| **TNFRSF25** | 0.00000 | 1.66624 | 0.26706 | 0.00000 | 0.00181 | 1.91685 | 0.20280 |
| **TNFRSF4** | 0.02279 | 0.02605 | 0.01626 | 0.02323 | 0.00750 | 0.01483 | 0.01342 |
| **TNFRSF6B** | 0.02476 | 0.01846 | 0.01541 | 0.02624 | 0.00776 | 0.01859 | 0.01698 |
| **TNFRSF8** | 0.00000 | 0.05876 | 1.50627 | 0.14203 | 0.35348 | 3.40666 | 0.00015 |
| **TNFRSF9** | 0.03478 | 0.00000 | 0.08155 | 0.03368 | 0.02750 | 0.01189 | 0.02638 |
| **TNFSF10** | 0.03541 | 0.00793 | 0.01317 | 0.00832 | 0.00704 | 0.01090 | 0.13744 |
| **TNFSF11** | 0.02686 | 0.02634 | 0.00986 | 0.01962 | 0.02429 | 0.01641 | 0.01280 |
| **TNFSF12** | 0.17910 | 0.05022 | 0.02437 | 1.06566 | 0.03093 | 0.00000 | 2.73412 |

| | | | | | | | |
|---|---|---|---|---|---|---|---|
| **TNFSF13** | 0.00106 | 0.01323 | 0.05682 | 0.15193 | 0.34611 | 0.00294 | 0.17674 |
| **TNFSF13B** | 0.00000 | 0.00000 | 0.18903 | 0.04818 | 0.02444 | 0.59482 | 2.82895 |
| **TNFSF14** | 0.02395 | 0.01389 | 0.05878 | 0.07420 | 0.00584 | 0.01291 | 0.00643 |
| **TNFSF15** | 0.11742 | 0.16154 | 0.00000 | 0.00000 | 0.00989 | 0.00000 | 0.53876 |
| **TNFSF4** | 0.01653 | 0.01812 | 0.00870 | 0.11679 | 0.03566 | 0.09636 | 0.00214 |
| **TNFSF8** | 0.06074 | 0.04454 | 0.00731 | 0.14816 | 0.01887 | 0.00196 | 0.03528 |
| **TNFSF9** | 0.15218 | 6.59719 | 1.20050 | 0.00000 | 4.41005 | 6.86495 | 0.00000 |
| **TNIK** | 2.46105 | 0.00000 | 0.00000 | 1.93047 | 4.58771 | 0.00000 | 5.04167 |
| **TNIP1** | 0.00000 | 2.55544 | 0.33620 | 1.19269 | 0.02691 | 1.35086 | 5.69073 |
| **TNIP2** | 2.84312 | 9.80747 | 1.18916 | 10.39826 | 0.47076 | 5.01384 | 19.95076 |
| **TNIP3** | 0.07611 | 0.00000 | 0.04152 | 0.09557 | 0.01886 | 0.97053 | 0.00000 |
| **TNK1** | 0.38367 | 0.00000 | 0.14952 | 0.45197 | 0.18886 | 0.10658 | 0.25234 |
| **TNK2** | 0.00000 | 1.72035 | 0.78043 | 3.75597 | 0.49268 | 4.70832 | 11.41326 |
| **TNKS** | 2.66796 | 1.41208 | 2.21404 | 3.52832 | 0.87001 | 1.43514 | 6.55343 |
| **TNKS1BP1** | 0.00000 | 3.74845 | 0.78569 | 3.73482 | 0.41854 | 3.19817 | 13.96850 |
| **TNKS2** | 4.15549 | 0.00000 | 3.66420 | 4.96300 | 1.17232 | 3.25980 | 12.73297 |
| **TNKS2-AS1** | 0.05646 | 0.03589 | 0.26306 | 0.07386 | 0.11392 | 0.01211 | 0.00167 |
| **TNMD** | 0.01465 | 0.05239 | 0.02737 | 0.02308 | 0.00623 | 0.01628 | 0.02272 |
| **TNN** | 0.21612 | 0.00815 | 0.03294 | 0.01099 | 0.00799 | 0.00735 | 0.01428 |
| **TNNC1** | 2.97720 | 0.55887 | 0.00000 | 7.39947 | 0.17946 | 0.00000 | 0.47218 |
| **TNNC2** | 0.72842 | 4.36628 | 0.00000 | 0.00000 | 1.06413 | 0.46415 | 0.49441 |
| **TNNI1** | 0.00000 | 0.06504 | 0.00000 | 0.05045 | 0.15117 | 0.01222 | 0.13801 |
| **TNNI2** | 0.02476 | 0.01186 | 0.01372 | 0.03509 | 0.00753 | 0.01352 | 0.03660 |
| **TNNI3** | 0.00893 | 0.18017 | 0.83742 | 0.00870 | 5.13264 | 0.16552 | 0.00133 |
| **TNNT1** | 0.00000 | 1.27504 | 2.59136 | 0.00000 | 3.40704 | 0.93587 | 0.02816 |
| **TNNT2** | 0.31604 | 0.28380 | 0.00000 | 0.64569 | 0.01693 | 0.00037 | 0.00000 |
| **TNNT3** | 0.02593 | 0.00793 | 0.01194 | 0.01281 | 0.00739 | 0.06757 | 0.06437 |
| **TNPO1** | 36.17436 | 6.52779 | 15.28313 | 21.02659 | 5.46986 | 11.61847 | 20.83465 |
| **TNPO2** | 8.08150 | 15.07951 | 9.11880 | 7.11491 | 5.40826 | 3.11839 | 23.02751 |
| **TNPO3** | 25.96163 | 9.81411 | 9.54837 | 19.56188 | 13.40278 | 11.04187 | 3.57298 |
| **TNR** | 0.08856 | 0.06346 | 0.01389 | 0.13926 | 0.00727 | 0.01598 | 0.00003 |
| **TNRC18** | 0.00000 | 1.22672 | 0.78310 | 1.79838 | 0.38299 | 1.18422 | 4.78415 |
| **TNRC18P1** | 0.00066 | 0.03024 | 0.03054 | 0.05387 | 0.08750 | 0.03768 | 0.09074 |
| **TNRC6A** | 7.13429 | 4.11465 | 5.31054 | 5.33751 | 3.63498 | 6.77952 | 19.14205 |
| **TNRC6B** | 11.74222 | 3.90679 | 4.31772 | 5.56513 | 4.32382 | 8.90350 | 12.29843 |

| | | | | | | | |
|---|---|---|---|---|---|---|---|
| **TNRC6C** | 0.00000 | 1.86472 | 1.31572 | 0.56738 | 1.16649 | 6.35702 | 2.71967 |
| **TNRC6C-AS1** | 0.02601 | 0.02277 | 0.02077 | 0.02841 | 0.00605 | 0.00282 | 0.21035 |
| **TNS1** | 0.18964 | 0.26477 | 0.18252 | 1.29772 | 0.00273 | 0.00000 | 1.74297 |
| **TNS3** | 6.58727 | 16.53848 | 0.30537 | 13.49524 | 0.96124 | 1.77708 | 12.71490 |
| **TNS4** | 0.01520 | 0.02643 | 0.01362 | 0.01930 | 0.00778 | 0.04664 | 0.03177 |
| **TNXB.uid-8455** | 0.02388 | 0.01861 | 0.01333 | 0.02179 | 0.00788 | 0.01289 | 0.01582 |
| **TNXB.uid-8469** | 0.06029 | 0.21084 | 0.04011 | 0.05029 | 0.00373 | 0.06348 | 0.42642 |
| **TOB1** | 5.11860 | 2.97968 | 0.31123 | 7.46479 | 3.41316 | 3.32302 | 0.00000 |
| **TOB1-AS1** | 0.02954 | 0.01593 | 0.01567 | 0.04563 | 0.00900 | 0.02474 | 0.01106 |
| **TOB2** | 10.41312 | 9.84915 | 1.80097 | 11.88281 | 6.27336 | 4.17550 | 18.57145 |
| **TOB2P1** | 0.04061 | 0.06573 | 0.01785 | 0.30767 | 0.00747 | 0.00222 | 0.12100 |
| **TOE1** | 18.86468 | 16.55890 | 9.90911 | 14.29452 | 4.91750 | 5.33273 | 0.59308 |
| **TOLLIP** | 9.13228 | 17.16035 | 4.98889 | 9.55797 | 1.80353 | 5.31119 | 2.85987 |
| **TOLLIP-AS1** | 0.04600 | 0.32389 | 0.26512 | 0.33429 | 0.18612 | 0.22726 | 0.11632 |
| **TOM1** | 4.01861 | 6.18739 | 0.63036 | 13.87986 | 0.12957 | 5.26000 | 1.84526 |
| **TOM1L1** | 2.32723 | 0.72603 | 1.38001 | 0.53260 | 1.52934 | 0.50356 | 0.00000 |
| **TOM1L2** | 0.64059 | 2.59390 | 0.48946 | 1.20176 | 0.46900 | 0.79558 | 0.00000 |
| **TOMM20** | 26.86747 | 8.10685 | 8.63867 | 13.30242 | 5.38462 | 10.11062 | 22.34701 |
| **TOMM20L** | 0.39255 | 0.09796 | 0.20731 | 0.05877 | 0.02564 | 0.13099 | 0.42858 |
| **TOMM22** | 42.00741 | 32.64426 | 20.75306 | 29.69631 | 7.76115 | 22.92810 | 23.19457 |
| **TOMM34** | 21.53071 | 10.52487 | 2.27026 | 23.59407 | 3.96613 | 5.98827 | 5.70908 |
| **TOMM40** | 2.48205 | 26.27210 | 6.21845 | 6.50947 | 2.12346 | 12.04683 | 40.05190 |
| **TOMM40L** | 0.50410 | 0.89039 | 0.25659 | 3.61570 | 0.26187 | 3.03465 | 1.10394 |
| **TOMM5** | 45.08337 | 27.87068 | 30.68845 | 33.88600 | 17.75052 | 12.86942 | 19.25033 |
| **TOMM6** | 42.74820 | 27.13652 | 32.10544 | 30.42143 | 9.02960 | 21.59243 | 34.02936 |
| **TOMM7** | 34.55603 | 29.98961 | 25.11610 | 30.07269 | 5.36116 | 27.34460 | 32.51610 |
| **TOMM70A** | 16.68543 | 2.19383 | 10.37945 | 3.31193 | 2.57675 | 8.41138 | 31.91577 |
| **TONSL** | 2.56113 | 6.26495 | 1.85684 | 2.00687 | 0.00000 | 2.85036 | 14.09054 |
| **1.00000** | 21.08560 | 11.38441 | 10.88874 | 27.59270 | 9.21696 | 8.14366 | 36.78153 |
| **TOP1MT** | 11.51591 | 16.56910 | 13.73022 | 6.98324 | 0.56707 | 5.98740 | 4.73080 |
| **TOP1P1** | 1.37423 | 0.35406 | 0.52695 | 0.99855 | 0.53196 | 0.32743 | 0.59804 |
| **TOP1P2** | 0.15540 | 0.00002 | 0.01846 | 0.06176 | 0.00662 | 0.00000 | 0.32282 |
| **TOP2A** | 35.32722 | 18.10398 | 18.36453 | 24.13108 | 6.97961 | 11.35276 | 24.88412 |
| **TOP2B** | 10.23311 | 4.03937 | 0.00000 | 5.66356 | 5.02197 | 6.73342 | 13.38668 |
| **TOP3A** | 3.15833 | 5.57268 | 6.24767 | 6.00768 | 4.35687 | 6.43570 | 2.44248 |

| | | | | | | | |
|---|---|---|---|---|---|---|---|
| **TOP3B** | 2.51502 | 5.60432 | 2.31437 | 2.61907 | 0.00000 | 0.99897 | 1.05111 |
| **TOPAZ1** | 0.01666 | 0.00000 | 0.81485 | 0.08023 | 0.40847 | 0.00000 | 0.29949 |
| **TOPBP1** | 9.46372 | 2.64151 | 7.57877 | 5.32515 | 1.69555 | 7.47051 | 13.32711 |
| **TOPORS** | 10.64126 | 1.96442 | 15.91852 | 9.45333 | 21.10374 | 5.90637 | 4.48753 |
| **TOPORS-AS1** | 0.23605 | 1.61242 | 2.00878 | 1.68022 | 3.58796 | 0.81460 | 0.51249 |
| **TOR1A** | 19.37095 | 13.34962 | 17.41172 | 20.96531 | 9.47349 | 7.56823 | 0.00000 |
| **TOR1AIP1** | 17.16589 | 0.42910 | 6.62712 | 10.98793 | 5.15389 | 4.10069 | 0.00000 |
| **TOR1AIP2** | 34.49098 | 20.15529 | 12.25033 | 27.35585 | 17.06156 | 6.40726 | 12.14390 |
| **TOR1B** | 15.10411 | 5.57987 | 18.99200 | 12.90075 | 12.58946 | 3.82282 | 2.49660 |
| **TOR2A** | 3.04138 | 7.79496 | 1.33668 | 1.58591 | 2.15917 | 1.27045 | 0.40014 |
| **TOR3A** | 6.12247 | 10.78536 | 0.00000 | 9.10540 | 3.26265 | 12.87321 | 10.12550 |
| **TOR4A** | 0.00000 | 0.32012 | 0.10844 | 0.02521 | 0.02027 | 0.07342 | 0.06498 |
| **TOX** | 0.01434 | 0.04876 | 0.00000 | 0.04588 | 0.71135 | 0.12411 | 0.01495 |
| **TOX2** | 0.00000 | 0.16266 | 0.60232 | 0.03289 | 5.23563 | 0.18241 | 0.05619 |
| **TOX3** | 0.02244 | 0.09492 | 0.00300 | 0.01861 | 0.02778 | 0.36441 | 0.00000 |
| **TOX4** | 16.75977 | 14.98814 | 13.93161 | 14.79523 | 20.37166 | 7.34989 | 2.93560 |
| **TP53** | 2.10649 | 6.88910 | 1.94387 | 8.16126 | 0.22228 | 5.07483 | 3.22519 |
| **TP53AIP1** | 0.02887 | 0.01940 | 0.06669 | 0.18517 | 0.00551 | 0.00813 | 0.00000 |
| **TP53BP1** | 3.19078 | 0.93167 | 3.96997 | 5.04177 | 1.14630 | 2.44303 | 3.36299 |
| **TP53BP2** | 6.33435 | 0.75733 | 3.36147 | 4.02034 | 1.75053 | 2.39259 | 11.51173 |
| **TP53I11** | 0.00000 | 0.02882 | 0.41074 | 0.06030 | 0.00858 | 0.01038 | 0.05866 |
| **TP53I13** | 0.00000 | 4.87254 | 0.79539 | 0.45836 | 0.00000 | 1.20807 | 3.89995 |
| **TP53I3** | 0.59324 | 0.02977 | 0.00000 | 0.89458 | 0.15091 | 0.39029 | 0.37146 |
| **TP53INP1** | 2.25302 | 0.06594 | 0.37927 | 5.71164 | 1.55473 | 1.53086 | 7.51727 |
| **TP53INP2** | 0.00000 | 2.62657 | 2.55941 | 0.55333 | 1.19574 | 3.08781 | 2.03895 |
| **TP53RK** | 8.92923 | 11.60827 | 5.05567 | 11.31117 | 5.55287 | 5.77692 | 9.17645 |
| **TP53TG1** | 0.24554 | 0.18343 | 0.05112 | 0.37801 | 0.19953 | 0.29155 | 0.00000 |
| **TP53TG3+TP53T** | 0.02444 | 0.01503 | 0.01571 | 0.01917 | 0.02979 | 0.01338 | 0.01636 |
| **TP53TG3D** | 0.05085 | 0.00000 | 0.40754 | 0.09636 | 1.43164 | 0.00404 | 0.04144 |
| **TP53TG5** | 0.02402 | 0.02348 | 0.01823 | 0.02090 | 0.01154 | 0.00824 | 0.04023 |
| **TP63** | 4.05675 | 0.00000 | 0.00000 | 6.81459 | 1.71364 | 0.00000 | 0.64011 |
| **TP73** | 0.00000 | 0.21316 | 0.76725 | 2.17664 | 0.15022 | 0.33731 | 0.78392 |
| **TP73-AS1** | 0.51994 | 1.64087 | 0.07990 | 0.85398 | 1.61551 | 1.19873 | 2.36077 |
| **TPBG** | 3.10215 | 0.00000 | 1.57654 | 3.17498 | 0.55711 | 0.65733 | 7.49520 |
| **TPBGL** | 0.01020 | 0.01189 | 0.01586 | 0.01335 | 0.03682 | 0.04992 | 0.07035 |

| | | | | | | | |
|---|---|---|---|---|---|---|---|
| **TPCN1** | 2.38920 | 2.77710 | 0.07317 | 2.43735 | 0.13988 | 0.47720 | 4.03433 |
| **TPCN2** | 4.36275 | 1.45303 | 0.23934 | 1.04551 | 0.00000 | 2.18517 | 9.02317 |
| **TPD52** | 12.04734 | 0.00000 | 2.23363 | 6.82027 | 4.62262 | 5.48556 | 5.64958 |
| **TPD52L1** | 5.96218 | 0.46944 | 0.00000 | 7.54307 | 0.00000 | 0.39029 | 0.31917 |
| **TPD52L2** | 33.51458 | 22.65478 | 13.98203 | 22.37188 | 12.04680 | 10.96449 | 5.26854 |
| **TPGS1** | 0.00000 | 0.30004 | 0.05680 | 0.08986 | 0.00783 | 0.22006 | 0.23341 |
| **TPGS2** | 7.34522 | 2.38497 | 0.86828 | 7.17747 | 1.04771 | 1.02797 | 0.00000 |
| **TPH1** | 0.21379 | 0.05118 | 0.05283 | 0.01350 | 0.00503 | 0.00984 | 0.00000 |
| **TPH2** | 0.01740 | 0.06266 | 0.00131 | 0.01467 | 0.31256 | 0.00612 | 0.03326 |
| **TPI1** | 63.79966 | 59.78666 | 9.75283 | 51.77510 | 7.44819 | 42.37226 | 19.89926 |
| **TPI1P2** | 3.12119 | 4.15032 | 0.27064 | 2.46001 | 0.33036 | 2.82568 | 0.00000 |
| **TPI1P3** | 0.27268 | 0.25402 | 0.01587 | 0.13967 | 0.01794 | 0.28665 | 0.00000 |
| **TPK1** | 2.82137 | 0.00000 | 2.33421 | 1.76962 | 1.06527 | 0.00226 | 0.00000 |
| **TPM1** | 9.46028 | 19.20718 | 0.90411 | 34.06528 | 2.53743 | 15.73333 | 18.53760 |
| **TPM2** | 3.12560 | 13.83113 | 15.96629 | 3.27674 | 2.80595 | 8.99776 | 5.04894 |
| **TPM3** | 57.83344 | 38.15149 | 18.69603 | 30.76890 | 16.64829 | 25.68550 | 10.05105 |
| **TPM3P9** | 3.21663 | 3.66741 | 1.03928 | 4.43863 | 2.91017 | 1.14741 | 0.63672 |
| **TPM4** | 47.53101 | 2.39633 | 0.00000 | 39.19324 | 0.00000 | 8.53051 | 21.74391 |
| **TPMT** | 12.79399 | 2.59567 | 6.07434 | 5.64597 | 0.38713 | 4.62011 | 0.00000 |
| **TPO** | 0.00974 | 0.03192 | 0.24187 | 0.01830 | 0.14506 | 0.01454 | 0.03086 |
| **TPP1** | 10.48722 | 0.00002 | 0.12526 | 3.38924 | 0.00000 | 4.94048 | 1.78312 |
| **TPP2** | 14.25573 | 1.58153 | 2.83348 | 5.25399 | 2.67616 | 5.65512 | 11.67992 |
| **TPPP** | 0.00000 | 1.35783 | 2.76853 | 0.00000 | 0.80881 | 0.97758 | 0.74969 |
| **TPPP2** | 0.05869 | 0.01345 | 0.01343 | 0.01481 | 0.00804 | 0.02512 | 0.00254 |
| **TPPP3** | 0.38185 | 0.02670 | 0.07462 | 0.17127 | 0.27391 | 0.04789 | 0.00000 |
| **TPR** | 23.99407 | 15.84837 | 10.74081 | 25.27126 | 4.73283 | 14.35217 | 27.72487 |
| **TPRA1** | 0.38208 | 9.09433 | 0.99504 | 5.31654 | 0.25725 | 5.86376 | 14.53949 |
| **TPRG1** | 0.22211 | 0.23549 | 0.67910 | 0.00000 | 0.93111 | 0.15583 | 0.00000 |
| **TPRG1-AS1** | 0.04651 | 0.06343 | 0.13508 | 0.03766 | 0.01029 | 0.05013 | 0.00125 |
| **TPRG1-AS2** | 0.02147 | 0.03294 | 0.04343 | 0.01784 | 0.01909 | 0.00906 | 0.01793 |
| **TPRG1L** | 1.46397 | 7.84775 | 2.90202 | 2.60547 | 1.91336 | 3.91308 | 4.15379 |
| **TPRKB** | 16.63203 | 1.59224 | 20.12867 | 13.13886 | 4.95965 | 5.78071 | 3.24146 |
| **TPRN** | 1.34611 | 3.01509 | 0.00000 | 3.19987 | 0.00401 | 2.87455 | 10.38319 |
| **TPRX1** | 0.00000 | 0.00000 | 1.93862 | 0.00371 | 28.46189 | 0.00000 | 0.02979 |
| **TPRXL** | 0.07958 | 1.53454 | 1.01373 | 0.28752 | 16.50084 | 0.07248 | 0.00000 |

| | | | | | | | |
|---|---|---|---|---|---|---|---|
| **TPSG1** | 0.01708 | 0.02891 | 0.01563 | 0.01963 | 0.04613 | 0.01578 | 0.02576 |
| **TPST1** | 0.86766 | 0.51564 | 1.64671 | 2.72222 | 6.42045 | 2.41408 | 2.74347 |
| **TPST2** | 8.20866 | 9.74589 | 3.78684 | 9.32281 | 3.57184 | 25.03005 | 3.92555 |
| **TPT1** | 37.01651 | 29.82513 | 34.34045 | 31.66178 | 13.46140 | 29.88170 | 79.35027 |
| **TPT1-AS1** | 0.75596 | 0.15088 | 0.02766 | 1.44166 | 0.05212 | 1.02488 | 1.31537 |
| **TPTE** | 2.34920 | 0.00000 | 2.34518 | 1.15640 | 4.37764 | 1.95771 | 0.30556 |
| **TPTE2** | 0.30036 | 0.03110 | 0.32770 | 0.04250 | 0.09436 | 0.12181 | 0.16486 |
| **TPTE2P1** | 0.01523 | 0.52991 | 0.20179 | 0.00000 | 0.20421 | 0.80455 | 0.01043 |
| **TPTE2P3** | 0.10762 | 0.00000 | 0.20315 | 0.07610 | 0.02080 | 0.07779 | 0.19788 |
| **TPTE2P5** | 0.03607 | 0.16039 | 0.09962 | 0.01222 | 0.00231 | 0.05304 | 0.27859 |
| **TPTE2P6** | 0.08797 | 0.03977 | 0.95980 | 0.00000 | 0.00053 | 1.11312 | 0.07800 |
| **TPTEP1** | 4.25784 | 7.01870 | 6.10705 | 2.18785 | 0.21984 | 3.51065 | 4.44126 |
| **TPX2** | 35.03739 | 19.02943 | 21.74138 | 26.32874 | 14.94294 | 10.11728 | 10.47659 |
| **TRA2A** | 38.82499 | 4.07697 | 20.97890 | 17.85934 | 17.75408 | 16.10075 | 9.84195 |
| **TRA2B** | 37.81726 | 15.82594 | 20.88013 | 23.85359 | 14.64127 | 18.04362 | 20.14332 |
| **TRABD** | 3.45249 | 10.94868 | 0.71272 | 12.80148 | 0.00000 | 3.67222 | 19.44809 |
| **TRABD2A** | 0.00000 | 0.00990 | 0.01728 | 2.22128 | 0.00728 | 0.40487 | 0.61688 |
| **TRABD2B** | 0.00000 | 0.05663 | 0.18088 | 0.01620 | 0.00179 | 1.42998 | 0.06150 |
| **TRADD** | 1.08312 | 1.76355 | 1.51703 | 2.75291 | 0.00000 | 0.46505 | 0.59360 |
| **TRAF1** | 0.02258 | 0.01442 | 0.00887 | 0.06514 | 0.11692 | 0.05917 | 0.01843 |
| **TRAF2** | 10.88132 | 13.78762 | 4.81939 | 12.38235 | 2.49052 | 5.95020 | 3.67160 |
| **TRAF3** | 3.50517 | 0.00000 | 0.36641 | 5.41153 | 0.33261 | 0.11660 | 10.34185 |
| **TRAF3IP1** | 0.69827 | 1.58770 | 0.53443 | 1.22672 | 1.55765 | 1.05868 | 3.59293 |
| **TRAF3IP2** | 2.57320 | 1.18057 | 0.00000 | 1.89890 | 1.87822 | 1.51033 | 2.12523 |
| **TRAF3IP2-AS1** | 0.96267 | 1.04063 | 1.37946 | 0.95697 | 0.47183 | 1.08668 | 1.73425 |
| **TRAF3IP3** | 0.04443 | 0.12812 | 0.14996 | 0.19016 | 0.03172 | 0.00550 | 0.00891 |
| **TRAF4** | 2.43031 | 10.33681 | 1.97961 | 13.26113 | 1.21749 | 8.52358 | 31.60819 |
| **TRAF5** | 0.47171 | 0.00000 | 5.78045 | 0.04734 | 0.32643 | 0.52623 | 0.17804 |
| **TRAF6** | 5.97274 | 3.09501 | 3.45078 | 1.57227 | 6.20551 | 2.25699 | 0.00000 |
| **TRAF7** | 9.10670 | 6.67014 | 1.71366 | 10.46414 | 0.88446 | 6.77735 | 9.57024 |
| **TRAFD1** | 9.98279 | 7.73912 | 3.62926 | 7.18248 | 6.72391 | 3.63542 | 0.00000 |
| **TRAIP** | 7.88993 | 4.74997 | 5.72296 | 5.20795 | 3.87039 | 2.67508 | 0.00000 |
| **TRAK1** | 2.41159 | 8.39434 | 6.81145 | 7.39915 | 2.34096 | 8.21920 | 15.00111 |
| **TRAK2** | 4.32432 | 4.91480 | 4.36836 | 3.69621 | 5.32356 | 4.93346 | 2.81903 |
| **TRAM1** | 20.17113 | 8.57740 | 10.10129 | 11.30576 | 2.85275 | 9.27901 | 12.69179 |

| | | | | | | | |
|---|---|---|---|---|---|---|---|
| **TRAM1L1** | 0.43071 | 0.08167 | 0.67011 | 0.20374 | 0.03880 | 0.26464 | 0.00000 |
| **TRAM2** | 3.23762 | 4.38147 | 10.71103 | 12.32278 | 5.40647 | 6.91580 | 14.27236 |
| **TRAM2-AS1** | 0.11980 | 0.00000 | 0.07122 | 0.07430 | 0.04731 | 0.31931 | 0.16338 |
| **TRANK1** | 0.06564 | 0.01970 | 0.09070 | 0.03693 | 0.14795 | 0.57038 | 0.01009 |
| **TRAP1** | 41.75331 | 24.55908 | 6.86018 | 24.84797 | 2.95051 | 12.46515 | 12.07014 |
| **TRAPPC1** | 9.95098 | 9.32235 | 5.78654 | 13.09205 | 3.16424 | 7.59501 | 4.78031 |
| **TRAPPC10** | 4.90540 | 3.02948 | 2.07317 | 2.60454 | 0.72535 | 3.50930 | 13.25918 |
| **TRAPPC11** | 7.31788 | 2.49939 | 3.77039 | 2.02452 | 0.74807 | 3.99535 | 1.39336 |
| **TRAPPC12** | 0.69388 | 1.93385 | 0.24324 | 0.38406 | 0.14947 | 1.45158 | 4.64784 |
| **TRAPPC13** | 14.25998 | 1.80589 | 4.32463 | 8.39613 | 0.86341 | 2.75462 | 0.00000 |
| **TRAPPC2** | 7.84188 | 0.99119 | 1.85246 | 5.40396 | 0.82835 | 1.88891 | 3.14285 |
| **TRAPPC2L** | 31.88907 | 28.02697 | 10.22291 | 23.51653 | 0.16012 | 17.64024 | 12.22663 |
| **TRAPPC2P1** | 3.98650 | 0.87784 | 4.11157 | 3.34127 | 3.07715 | 0.78153 | 0.41477 |
| **TRAPPC3** | 24.39512 | 13.18501 | 15.19864 | 25.30187 | 3.76748 | 8.29699 | 2.76096 |
| **TRAPPC3L** | 0.01730 | 0.01629 | 0.01423 | 0.02625 | 0.03366 | 0.03156 | 0.04991 |
| **TRAPPC4** | 26.71966 | 20.59206 | 20.90657 | 19.50204 | 9.98913 | 8.61499 | 1.49824 |
| **TRAPPC5** | 2.72699 | 22.18480 | 7.51697 | 17.32383 | 1.79498 | 5.10201 | 9.38253 |
| **TRAPPC6A** | 1.67089 | 1.34372 | 0.23587 | 1.81935 | 0.53104 | 1.03469 | 0.43070 |
| **TRAPPC6B** | 8.17775 | 0.36662 | 7.64818 | 4.47430 | 5.66582 | 3.94620 | 5.13304 |
| **TRAPPC8** | 12.42356 | 0.00000 | 11.18433 | 4.81170 | 2.52955 | 9.07862 | 15.96525 |
| **TRAPPC9** | 0.61603 | 0.87285 | 0.05655 | 0.70436 | 0.60066 | 1.05558 | 1.44536 |
| **TRDMT1** | 2.77542 | 0.27684 | 2.47235 | 0.00837 | 0.29548 | 0.92378 | 0.28362 |
| **TRDN** | 0.02186 | 0.00000 | 0.08804 | 0.00000 | 0.02808 | 0.08550 | 0.49432 |
| **TREH** | 0.05465 | 0.01891 | 0.01049 | 0.06299 | 0.02262 | 0.00347 | 0.05884 |
| **TREM1** | 0.00000 | 0.00877 | 0.11503 | 8.18005 | 0.00691 | 0.00472 | 0.00000 |
| **TREM2** | 0.01795 | 0.02032 | 0.01085 | 0.28203 | 0.02991 | 0.00988 | 0.00000 |
| **TREML2** | 0.00000 | 0.01424 | 0.01112 | 4.31706 | 0.06712 | 0.03679 | 0.00000 |
| **TREML3P** | 0.00000 | 0.02528 | 0.01289 | 0.74053 | 0.00781 | 0.01036 | 0.00712 |
| **TREML4** | 0.01573 | 0.02319 | 0.01147 | 0.04847 | 0.00821 | 0.02967 | 0.01111 |
| **TRERF1** | 6.12364 | 0.00533 | 0.81774 | 6.23278 | 0.00000 | 1.69157 | 13.60310 |
| **TREX1** | 1.27027 | 0.94010 | 0.09935 | 2.23859 | 0.05835 | 0.00000 | 0.54494 |
| **TREX2** | 0.00000 | 0.01971 | 0.04193 | 0.06823 | 0.00689 | 0.11027 | 0.08369 |
| **TRG-AS1** | 0.62681 | 0.00000 | 0.00372 | 0.00000 | 0.00949 | 0.00000 | 0.22006 |
| **TRH** | 0.00000 | 0.01870 | 0.04175 | 0.04713 | 0.00771 | 1.04538 | 0.01912 |
| **TRHDE** | 0.04711 | 0.00000 | 0.02779 | 0.05427 | 0.32790 | 0.05009 | 0.04308 |

| | | | | | | | |
|---|---|---|---|---|---|---|---|
| **TRHDE-AS1** | 0.05160 | 0.01094 | 0.00788 | 0.02445 | 0.10208 | 0.02569 | 0.00646 |
| **TRHR** | 0.01644 | 0.01268 | 0.03604 | 0.03875 | 0.00560 | 0.00660 | 0.08304 |
| **TRIAP1** | 35.80926 | 24.27664 | 21.28201 | 29.70414 | 8.43177 | 14.32685 | 0.00000 |
| **TRIB1** | 0.00000 | 1.59806 | 2.91038 | 2.29826 | 0.15758 | 2.18201 | 4.75190 |
| **TRIB2** | 0.11939 | 0.00000 | 0.01771 | 0.09580 | 0.06916 | 1.57742 | 0.00000 |
| **TRIB3** | 1.40080 | 2.33744 | 0.34531 | 11.53509 | 0.00000 | 3.65954 | 9.81865 |
| **TRIL** | 0.05266 | 0.00255 | 0.01849 | 0.02372 | 0.13768 | 0.01028 | 0.05796 |
| **TRIM10** | 0.01504 | 0.01854 | 0.01243 | 0.05669 | 0.00779 | 0.01017 | 0.03052 |
| **TRIM11** | 2.14837 | 8.58272 | 2.21533 | 8.01940 | 0.53788 | 2.89102 | 16.25582 |
| **TRIM13** | 20.15953 | 4.68496 | 10.02339 | 11.44597 | 4.31130 | 6.87178 | 6.83322 |
| **TRIM14** | 0.32129 | 1.83314 | 0.32975 | 0.17539 | 0.11207 | 4.18671 | 0.29341 |
| **TRIM16** | 3.30585 | 1.95680 | 1.44519 | 0.08097 | 0.02428 | 2.89295 | 0.89639 |
| **TRIM16L** | 0.22066 | 0.23290 | 0.38962 | 0.38226 | 0.02356 | 0.00685 | 0.09606 |
| **TRIM17** | 0.00000 | 0.13171 | 0.01049 | 0.36146 | 0.00573 | 0.32963 | 0.49507 |
| **TRIM2** | 7.64492 | 4.91880 | 1.02649 | 0.00000 | 0.00000 | 11.63290 | 2.49542 |
| **TRIM21** | 1.21491 | 0.63517 | 1.53573 | 0.34283 | 0.33161 | 0.16486 | 0.00000 |
| **TRIM22** | 0.21387 | 0.14114 | 0.35026 | 0.41308 | 0.00000 | 1.14575 | 0.58860 |
| **TRIM23** | 5.02701 | 0.00215 | 3.05259 | 2.59939 | 0.71093 | 2.03637 | 1.16334 |
| **TRIM24** | 3.96232 | 1.51434 | 1.87948 | 5.41889 | 0.16419 | 5.06850 | 11.15178 |
| **TRIM25** | 0.14759 | 0.00000 | 0.01532 | 0.17428 | 0.00705 | 0.47741 | 0.14320 |
| **TRIM26** | 3.60088 | 3.35144 | 1.53872 | 1.46201 | 1.76572 | 10.43064 | 5.06005 |
| **TRIM27** | 13.42175 | 13.84112 | 12.50531 | 17.57901 | 4.11362 | 7.79536 | 13.77324 |
| **TRIM28** | 23.50710 | 43.23190 | 22.33062 | 18.92207 | 7.51738 | 28.57824 | 63.77343 |
| **TRIM29** | 0.46255 | 0.00000 | 0.40336 | 3.62242 | 0.12927 | 0.00000 | 0.29367 |
| **TRIM3** | 0.65034 | 1.13476 | 0.58719 | 2.28389 | 0.54861 | 0.85771 | 0.55948 |
| **TRIM31** | 0.01068 | 0.00313 | 0.04575 | 0.02673 | 0.03671 | 0.04146 | 0.07576 |
| **TRIM32** | 11.45745 | 4.34021 | 3.00499 | 3.71049 | 2.18836 | 1.15033 | 0.00000 |
| **TRIM33** | 11.85850 | 2.85895 | 9.44177 | 13.95157 | 3.16073 | 8.34223 | 33.82912 |
| **TRIM34.uid-151** | 0.02951 | 0.01134 | 0.02044 | 0.01928 | 0.00732 | 0.01169 | 0.03727 |
| **TRIM34.uid-151** | 0.02527 | 0.02089 | 0.01619 | 0.03680 | 0.00772 | 0.02302 | 0.00899 |
| **TRIM34.uid-151** | 0.03490 | 0.02311 | 0.01396 | 0.06208 | 0.00887 | 0.01904 | 0.00814 |
| **TRIM35** | 5.79651 | 5.56267 | 3.54330 | 5.21416 | 1.20978 | 3.83860 | 0.00000 |
| **TRIM36** | 0.31895 | 0.00000 | 0.27449 | 0.00957 | 6.26929 | 0.13749 | 0.00865 |
| **TRIM37** | 11.75716 | 2.28025 | 3.41961 | 5.25062 | 1.43732 | 4.96039 | 6.83936 |
| **TRIM38** | 0.88737 | 1.03057 | 3.09170 | 0.00000 | 2.51114 | 2.98515 | 0.10043 |

| | | | | | | | |
|---|---|---|---|---|---|---|---|
| **TRIM39** | 1.51242 | 0.84270 | 2.44738 | 3.00110 | 1.95955 | 1.66308 | 2.91399 |
| **TRIM4** | 1.22287 | 0.00000 | 0.41926 | 3.24813 | 0.19499 | 0.16529 | 0.53927 |
| **TRIM40** | 0.11118 | 0.00564 | 0.01253 | 0.00000 | 0.01005 | 0.06857 | 0.03510 |
| **TRIM41** | 0.00006 | 1.65666 | 1.49481 | 2.48997 | 1.28052 | 1.98887 | 8.53363 |
| **TRIM42** | 0.02255 | 0.00460 | 0.02104 | 0.02790 | 0.52664 | 0.01382 | 0.02898 |
| **TRIM43** | 0.01958 | 0.14072 | 0.00000 | 0.00420 | 1.32683 | 0.05090 | 0.01757 |
| **TRIM43B** | 0.03508 | 0.17185 | 0.00000 | 0.00225 | 0.93533 | 0.03468 | 0.01438 |
| **TRIM44** | 0.14588 | 2.89527 | 2.40602 | 0.00000 | 1.03804 | 2.76135 | 3.81391 |
| **TRIM45** | 1.42083 | 0.14518 | 0.08899 | 0.64360 | 0.62647 | 0.19460 | 0.69053 |
| **TRIM46** | 0.24677 | 1.81316 | 0.00000 | 0.00000 | 0.14470 | 0.70020 | 0.40514 |
| **TRIM47** | 0.00000 | 0.03064 | 0.12319 | 0.26127 | 0.00425 | 0.30221 | 0.05618 |
| **TRIM48** | 0.02275 | 0.02005 | 0.00000 | 0.01600 | 0.97873 | 0.01406 | 0.01541 |
| **TRIM49** | 0.03136 | 0.10114 | 0.00000 | 0.00395 | 0.39990 | 0.01180 | 0.02919 |
| **TRIM49B** | 0.02936 | 0.04746 | 0.00000 | 0.01490 | 0.70795 | 0.00883 | 0.01277 |
| **TRIM49C** | 0.02503 | 0.10836 | 0.00000 | 0.00121 | 0.48581 | 0.01811 | 0.04981 |
| **TRIM5** | 4.51697 | 0.90210 | 2.03124 | 4.70906 | 0.08325 | 1.08736 | 2.03923 |
| **TRIM50** | 0.00000 | 0.00000 | 0.81488 | 0.01099 | 0.26511 | 0.01648 | 0.08871 |
| **TRIM51** | 0.02376 | 0.03125 | 0.00000 | 0.01327 | 0.69605 | 0.00890 | 0.02784 |
| **TRIM51HP** | 0.02255 | 0.02138 | 0.00413 | 0.01915 | 0.18271 | 0.01318 | 0.01466 |
| **TRIM52** | 0.85865 | 0.53562 | 0.40614 | 0.67713 | 1.55717 | 1.24145 | 0.32209 |
| **TRIM52-AS1** | 0.40913 | 3.00094 | 2.54304 | 0.18086 | 0.36815 | 0.91322 | 1.18364 |
| **TRIM53AP.uid-1** | 0.00000 | 0.17052 | 0.17142 | 0.00669 | 0.70471 | 0.04114 | 0.02457 |
| **TRIM53AP.uid-1** | 0.00000 | 0.09600 | 0.31284 | 0.00204 | 1.24043 | 0.12475 | 0.03488 |
| **TRIM54** | 0.10542 | 0.27357 | 1.10595 | 0.00000 | 2.22206 | 0.07905 | 0.14226 |
| **TRIM55** | 0.05593 | 0.01540 | 0.01640 | 0.17550 | 0.00633 | 0.00000 | 0.08737 |
| **TRIM56** | 0.00000 | 2.99581 | 0.24116 | 0.25378 | 0.00000 | 4.35233 | 0.44290 |
| **TRIM58** | 0.40791 | 0.35410 | 0.09504 | 0.06743 | 0.04148 | 0.56508 | 1.06157 |
| **TRIM59** | 30.04198 | 9.54887 | 17.79854 | 15.39260 | 4.00064 | 3.00563 | 0.00000 |
| **TRIM6** | 13.17758 | 0.44198 | 1.79098 | 3.63268 | 1.09469 | 5.02420 | 1.95992 |
| **TRIM60** | 13.47642 | 10.09411 | 23.45398 | 4.57128 | 9.93021 | 8.81512 | 0.00000 |
| **TRIM61** | 2.67841 | 7.22333 | 11.11129 | 0.42254 | 2.28430 | 4.14009 | 0.00000 |
| **TRIM62** | 0.75552 | 0.10900 | 0.00000 | 0.93516 | 1.18038 | 0.10521 | 1.16788 |
| **TRIM63** | 0.36887 | 2.05797 | 0.00000 | 0.00000 | 0.08323 | 0.08858 | 0.98188 |
| **TRIM64** | 0.02351 | 0.01794 | 0.00400 | 0.01852 | 0.14747 | 0.01403 | 0.01694 |
| **TRIM64B** | 0.03809 | 0.06240 | 0.00156 | 0.01324 | 0.13038 | 0.05192 | 0.00257 |

| | | | | | | | |
|---|---|---|---|---|---|---|---|
| **TRIM64C** | 0.02244 | 0.01843 | 0.00242 | 0.01880 | 0.20959 | 0.01343 | 0.01635 |
| **TRIM65** | 0.31281 | 1.06126 | 0.06186 | 0.81834 | 0.00000 | 0.70874 | 2.56665 |
| **TRIM66** | 0.27632 | 0.03650 | 0.68421 | 0.49399 | 0.00308 | 0.00000 | 1.30735 |
| **TRIM67** | 0.00665 | 0.09246 | 0.00619 | 0.02447 | 0.00794 | 0.02156 | 0.03105 |
| **TRIM68** | 1.59894 | 0.37656 | 1.59392 | 1.69455 | 0.15518 | 1.35626 | 1.52245 |
| **TRIM7** | 0.04921 | 0.19427 | 0.16987 | 0.02790 | 0.24750 | 0.12302 | 0.09276 |
| **TRIM71** | 2.22548 | 9.73173 | 6.30096 | 7.12629 | 1.54381 | 14.85474 | 18.16193 |
| **TRIM72** | 0.01817 | 0.02792 | 0.01992 | 0.02067 | 0.04187 | 0.04428 | 0.02494 |
| **TRIM77** | 0.01640 | 0.21681 | 0.00000 | 0.00662 | 3.37052 | 0.01398 | 0.01047 |
| **TRIM8** | 0.00000 | 11.92070 | 0.17788 | 0.00000 | 0.13881 | 7.49831 | 4.99369 |
| **TRIM9** | 0.03197 | 0.01860 | 0.06545 | 0.03405 | 0.14803 | 0.05462 | 0.01435 |
| **TRIML1** | 8.89931 | 0.32898 | 0.04944 | 8.69305 | 0.00033 | 0.00000 | 0.15693 |
| **TRIML2** | 13.39935 | 13.47204 | 8.21009 | 11.77553 | 3.08492 | 10.40167 | 6.24904 |
| **TRIO** | 0.00000 | 1.26216 | 0.38010 | 4.41025 | 3.75153 | 0.05437 | 6.98254 |
| **TRIOBP** | 0.00000 | 2.16929 | 0.24764 | 2.58776 | 0.49867 | 3.54225 | 3.63250 |
| **TRIP10** | 13.38250 | 30.78612 | 0.35896 | 14.21635 | 3.27391 | 13.41633 | 11.75039 |
| **TRIP11** | 7.21307 | 4.65952 | 7.72096 | 5.74740 | 3.66851 | 4.61452 | 11.87444 |
| **TRIP12** | 21.39173 | 12.33815 | 21.15563 | 13.63590 | 12.68867 | 9.86316 | 21.30944 |
| **TRIP13** | 19.05142 | 12.19024 | 8.21767 | 12.64990 | 1.12806 | 5.35792 | 1.93279 |
| **TRIP4** | 20.25086 | 7.04377 | 7.16620 | 11.15604 | 0.97959 | 0.98892 | 0.85264 |
| **TRIP6** | 0.73780 | 0.00000 | 0.00000 | 10.37135 | 0.11210 | 0.00000 | 7.59677 |
| **TRIQK** | 0.87762 | 0.05694 | 0.14375 | 0.38190 | 0.13211 | 1.21282 | 0.00000 |
| **TRIT1** | 7.91877 | 4.26880 | 6.53816 | 7.69088 | 2.40340 | 6.78528 | 1.20367 |
| **TRMT1** | 12.47060 | 13.56663 | 10.86499 | 15.80771 | 1.32935 | 9.85960 | 5.29870 |
| **TRMT10A** | 7.94066 | 1.58603 | 6.92842 | 3.39197 | 2.87134 | 0.21011 | 3.50366 |
| **TRMT10B** | 1.17175 | 0.98925 | 0.57751 | 0.91810 | 0.12776 | 0.39747 | 1.91623 |
| **TRMT10C** | 36.58675 | 2.17457 | 22.39618 | 19.13821 | 7.15989 | 10.35689 | 17.08847 |
| **TRMT11** | 11.83253 | 0.00000 | 2.41624 | 7.85754 | 1.07517 | 3.69822 | 1.47997 |
| **TRMT112** | 23.98093 | 17.29305 | 23.32592 | 26.75328 | 19.58147 | 12.48720 | 16.47760 |
| **TRMT12** | 3.93070 | 1.86254 | 5.08835 | 4.17915 | 1.56187 | 1.15054 | 0.49887 |
| **TRMT13** | 5.67270 | 0.87214 | 4.33976 | 5.00253 | 1.66947 | 2.97849 | 2.28844 |
| **TRMT1L** | 4.30302 | 3.30425 | 6.51666 | 2.20423 | 0.98890 | 4.35065 | 3.79098 |
| **TRMT2A** | 7.70908 | 7.56413 | 1.46338 | 12.22352 | 0.12616 | 2.95789 | 10.65439 |
| **TRMT2B** | 6.71703 | 4.97833 | 0.18465 | 2.66323 | 0.51595 | 0.71832 | 1.45027 |
| **TRMT44** | 0.55965 | 1.59836 | 0.95715 | 1.27749 | 4.44283 | 0.58031 | 4.37439 |

| | | | | | | | |
|---|---|---|---|---|---|---|---|
| **TRMT5** | 7.56135 | 3.63891 | 9.93556 | 4.51239 | 2.11331 | 1.83985 | 0.07435 |
| **TRMT6** | 19.53615 | 11.82328 | 11.07792 | 16.59000 | 5.07757 | 5.28426 | 8.95357 |
| **TRMT61A** | 3.52220 | 16.48443 | 7.91556 | 11.19205 | 2.01101 | 3.52511 | 9.42566 |
| **TRMT61B** | 7.52001 | 3.14995 | 6.54134 | 6.42838 | 2.48333 | 3.26311 | 0.00000 |
| **TRMU** | 24.19199 | 14.00208 | 3.25319 | 18.39723 | 1.75742 | 6.51930 | 0.00000 |
| **TRNAU1AP** | 9.67970 | 4.27930 | 6.88130 | 8.45127 | 4.38372 | 5.30534 | 5.21261 |
| **TRNP1** | 0.00002 | 0.20960 | 0.03398 | 0.09533 | 0.00854 | 1.02413 | 1.16277 |
| **TRNT1** | 15.45948 | 1.29961 | 9.52931 | 4.47113 | 2.63839 | 2.26722 | 0.00000 |
| **TRO** | 0.07264 | 1.54907 | 2.07820 | 0.00000 | 1.03544 | 0.88890 | 0.09535 |
| **TROAP** | 17.64791 | 13.08265 | 9.62471 | 14.36549 | 7.50155 | 7.80952 | 1.65785 |
| **TROVE2** | 11.21069 | 0.70699 | 6.00694 | 8.29066 | 5.13331 | 5.27251 | 3.99264 |
| **TRPA1** | 0.01568 | 0.02456 | 0.00731 | 0.01740 | 1.68508 | 0.01528 | 0.02909 |
| **TRPC1** | 0.10694 | 0.00000 | 0.44632 | 0.10179 | 0.14558 | 0.53255 | 0.37519 |
| **TRPC3** | 0.02939 | 0.01538 | 0.00000 | 0.01478 | 0.59995 | 0.01401 | 0.02115 |
| **TRPC4AP** | 2.18301 | 9.36921 | 3.24228 | 9.28559 | 3.34215 | 8.79060 | 17.71933 |
| **TRPC5** | 0.02680 | 0.00000 | 0.06288 | 0.04093 | 0.04930 | 0.17829 | 0.01616 |
| **TRPC5OS** | 0.03344 | 0.02122 | 0.00744 | 0.04490 | 0.05888 | 0.02843 | 0.00000 |
| **TRPC6** | 0.02371 | 0.02735 | 0.01529 | 0.01379 | 0.07783 | 0.01302 | 0.04613 |
| **TRPC7** | 0.29592 | 0.00000 | 2.06341 | 0.01270 | 0.66147 | 0.35792 | 0.01439 |
| **TRPM1** | 0.18841 | 0.39101 | 0.07888 | 0.24243 | 0.18097 | 0.00629 | 0.04275 |
| **TRPM2** | 0.18724 | 0.50688 | 0.03318 | 0.00000 | 0.22675 | 0.19511 | 0.04044 |
| **TRPM2-AS** | 0.03755 | 0.01285 | 0.02882 | 0.00678 | 0.00685 | 0.08791 | 0.05025 |
| **TRPM3** | 0.02300 | 0.02515 | 0.05680 | 0.00413 | 0.15935 | 0.00445 | 0.08777 |
| **TRPM4** | 2.21948 | 2.22095 | 2.04991 | 1.46640 | 0.79733 | 1.10411 | 5.91944 |
| **TRPM5** | 0.00000 | 0.06538 | 0.05856 | 0.08584 | 0.90864 | 0.00945 | 0.09745 |
| **TRPM6** | 8.08780 | 4.67685 | 7.99150 | 2.28173 | 1.69777 | 0.00000 | 3.19401 |
| **TRPM7** | 9.40766 | 0.00000 | 5.61826 | 9.83781 | 3.91231 | 2.87711 | 14.58468 |
| **TRPM8** | 0.02720 | 0.01894 | 0.01016 | 0.02578 | 0.02925 | 0.01348 | 0.01174 |
| **TRPS1** | 1.43123 | 0.03556 | 0.50010 | 0.00000 | 0.07206 | 0.50823 | 0.69261 |
| **TRPT1** | 3.46545 | 2.09091 | 1.46736 | 4.28974 | 0.07730 | 4.01215 | 2.53267 |
| **TRPV1** | 0.00000 | 4.05363 | 2.50131 | 1.76932 | 0.75653 | 2.55797 | 1.32491 |
| **TRPV2** | 1.02917 | 0.00000 | 0.10691 | 13.71656 | 0.02862 | 0.30213 | 0.00000 |
| **TRPV3** | 3.31696 | 6.42722 | 0.00000 | 5.88055 | 0.43763 | 2.73077 | 5.01236 |
| **TRPV4** | 0.00000 | 0.44523 | 0.10960 | 0.32321 | 0.98907 | 0.00000 | 1.78105 |
| **TRPV5** | 0.02725 | 0.01795 | 0.01557 | 0.01236 | 0.00724 | 0.05772 | 0.02721 |

| | | | | | | | |
|---|---|---|---|---|---|---|---|
| **TRPV6** | 0.00000 | 0.00000 | 0.02828 | 0.19734 | 0.53719 | 0.01233 | 0.10161 |
| **TRRAP** | 4.01023 | 7.39075 | 7.54630 | 4.43832 | 5.38138 | 7.39722 | 12.49377 |
| **TRUB1** | 9.71260 | 1.74434 | 4.11274 | 1.25260 | 0.39343 | 2.43758 | 1.35263 |
| **TRUB2** | 24.68515 | 22.92418 | 13.81602 | 19.32448 | 2.64951 | 9.27172 | 0.00000 |
| **TRY2P** | 0.01883 | 0.00000 | 0.25253 | 0.01785 | 0.07605 | 0.00710 | 0.01967 |
| **TSACC** | 1.52108 | 1.76093 | 0.71746 | 1.01737 | 0.21057 | 0.62671 | 0.75066 |
| **TSC1** | 9.59166 | 5.95750 | 4.67921 | 7.73453 | 6.97657 | 4.36279 | 0.41638 |
| **TSC2** | 1.74854 | 3.59572 | 2.26199 | 4.45442 | 0.37721 | 3.92818 | 7.33268 |
| **TSC22D1** | 19.55377 | 0.00000 | 2.31601 | 20.17106 | 4.07266 | 10.03880 | 5.89058 |
| **TSC22D1-AS1** | 0.06892 | 0.00000 | 0.02504 | 0.02974 | 0.05804 | 0.00286 | 0.15138 |
| **TSC22D2** | 5.11586 | 0.11667 | 2.26841 | 7.25512 | 2.61359 | 1.61317 | 12.32084 |
| **TSC22D3** | 0.24970 | 0.00000 | 0.54167 | 0.02519 | 0.79395 | 0.30306 | 0.00000 |
| **TSC22D4** | 0.00000 | 0.78931 | 0.00000 | 0.68809 | 0.05774 | 0.87857 | 2.98912 |
| **TSEN15** | 13.76250 | 3.22571 | 5.50794 | 0.33796 | 0.00000 | 0.23501 | 4.60725 |
| **TSEN2** | 8.98682 | 2.24360 | 5.49530 | 3.08774 | 0.17492 | 1.70401 | 0.00000 |
| **TSEN34** | 0.14855 | 3.61758 | 1.08878 | 4.51713 | 2.71181 | 4.06224 | 9.72687 |
| **TSEN54** | 12.77038 | 16.89668 | 4.91155 | 6.78018 | 0.17821 | 4.06591 | 16.64737 |
| **TSFM** | 31.01944 | 15.53535 | 16.00061 | 15.98034 | 12.67201 | 11.12921 | 2.34638 |
| **TSG1** | 0.03212 | 0.01614 | 0.00858 | 0.01853 | 0.06432 | 0.01303 | 0.01389 |
| **TSG101** | 16.22779 | 9.42376 | 11.29282 | 13.30266 | 2.80421 | 11.47814 | 0.00000 |
| **TSGA10** | 0.62321 | 0.89045 | 2.21251 | 0.44033 | 0.67277 | 1.03198 | 1.59691 |
| **TSGA10IP** | 0.01541 | 0.00506 | 0.03973 | 0.02581 | 0.00653 | 0.14599 | 0.02112 |
| **TSHR** | 0.02162 | 0.07835 | 0.02250 | 0.02067 | 0.13819 | 0.00000 | 0.11711 |
| **TSHZ1** | 0.00000 | 0.23678 | 0.37070 | 0.03718 | 0.55872 | 0.35297 | 0.05943 |
| **TSHZ2** | 0.01831 | 0.03851 | 0.02037 | 0.02239 | 0.20236 | 0.04775 | 0.04584 |
| **TSHZ3** | 0.00000 | 0.12781 | 0.00000 | 0.03004 | 0.83615 | 0.96842 | 0.21392 |
| **TSIX** | 0.40519 | 0.04674 | 1.00134 | 0.30963 | 0.22095 | 0.29079 | 0.52673 |
| **TSKS** | 0.19277 | 0.17981 | 0.36756 | 0.00000 | 0.84541 | 0.20151 | 0.79467 |
| **TSKU** | 1.14207 | 3.13021 | 0.88452 | 6.11464 | 0.58869 | 4.36583 | 4.91420 |
| **TSLP** | 0.02896 | 0.01444 | 0.06263 | 0.02713 | 0.00605 | 0.01333 | 0.01691 |
| **TSN** | 32.62975 | 9.81858 | 16.21995 | 13.18818 | 5.80199 | 9.45731 | 2.57228 |
| **TSNARE1** | 0.09232 | 0.79018 | 0.37882 | 0.67064 | 0.07977 | 0.63896 | 0.52892 |
| **TSNAX** | 10.68776 | 0.00000 | 5.55650 | 3.96076 | 1.76208 | 3.02212 | 3.72799 |
| **TSNAXIP1** | 0.00000 | 0.01048 | 0.08019 | 0.13843 | 0.24905 | 0.01960 | 0.16482 |
| **TSPAN1** | 3.20299 | 0.34769 | 0.02607 | 1.53900 | 0.04426 | 0.00000 | 0.01660 |

| | | | | | | | |
|---|---|---|---|---|---|---|---|
| **TSPAN10** | 0.00000 | 0.24688 | 0.50608 | 0.03029 | 0.03505 | 0.00203 | 0.02031 |
| **TSPAN11** | 0.02711 | 0.02109 | 0.01256 | 0.01877 | 0.00862 | 0.01257 | 0.01747 |
| **TSPAN12** | 0.35863 | 0.25331 | 0.17782 | 0.46121 | 1.41387 | 0.55487 | 0.00000 |
| **TSPAN13** | 18.10140 | 5.31580 | 4.41183 | 5.10133 | 2.63506 | 22.40386 | 1.87545 |
| **TSPAN14** | 19.87309 | 22.71987 | 11.56404 | 14.52656 | 0.98535 | 7.28955 | 26.99497 |
| **TSPAN15** | 19.51873 | 0.00000 | 0.00000 | 13.70506 | 0.40004 | 0.00000 | 10.77824 |
| **TSPAN16** | 0.00000 | 0.14641 | 0.14203 | 0.02120 | 0.00631 | 0.06931 | 0.03576 |
| **TSPAN17** | 0.00000 | 1.48868 | 2.36064 | 0.55268 | 0.04447 | 0.56275 | 0.60357 |
| **TSPAN18** | 0.00000 | 5.75251 | 0.00000 | 0.21584 | 1.11325 | 12.51259 | 0.72886 |
| **TSPAN2** | 0.00000 | 0.08785 | 0.02933 | 0.05222 | 0.00672 | 0.00585 | 0.03101 |
| **TSPAN3** | 19.27281 | 5.92278 | 1.83140 | 13.11887 | 0.30324 | 17.25776 | 3.08902 |
| **TSPAN31** | 1.03986 | 0.06042 | 0.51952 | 0.35548 | 0.03062 | 0.48925 | 0.00000 |
| **TSPAN32** | 0.02622 | 0.01401 | 0.01342 | 0.01678 | 0.00789 | 0.02942 | 0.01902 |
| **TSPAN33** | 0.06233 | 0.00000 | 0.01609 | 0.27625 | 0.12831 | 0.46953 | 0.45470 |
| **TSPAN4** | 3.25075 | 3.69943 | 0.00000 | 4.95911 | 0.33454 | 2.38525 | 2.03675 |
| **TSPAN5** | 0.17719 | 0.00000 | 0.69439 | 0.71211 | 1.97344 | 0.09566 | 1.53088 |
| **TSPAN6** | 29.91326 | 11.63848 | 12.64008 | 9.98303 | 0.25643 | 8.49159 | 6.55708 |
| **TSPAN7** | 0.14619 | 0.00000 | 0.45449 | 0.04125 | 1.08820 | 0.07127 | 0.00000 |
| **TSPAN8** | 0.07744 | 0.00431 | 0.01567 | 0.12031 | 0.00744 | 0.00602 | 0.02588 |
| **TSPAN9** | 0.00000 | 3.44732 | 3.74322 | 6.97662 | 1.02217 | 6.06109 | 0.75420 |
| **TSPEAR** | 0.05740 | 0.01355 | 0.01113 | 0.06580 | 0.35687 | 0.01211 | 0.00000 |
| **TSPEAR-AS1** | 0.04627 | 0.04005 | 0.01057 | 0.00854 | 0.05451 | 0.01917 | 0.03467 |
| **TSPO** | 0.06720 | 0.62454 | 0.29151 | 5.13027 | 0.17390 | 0.00000 | 0.00000 |
| **TSPO2** | 0.03971 | 0.01706 | 0.01296 | 0.03794 | 0.00794 | 0.01170 | 0.01137 |
| **TSPYL1** | 25.67243 | 6.17567 | 7.68297 | 12.17670 | 7.31098 | 6.00391 | 0.00000 |
| **TSPYL2** | 1.19002 | 1.46939 | 2.33984 | 0.79091 | 3.01787 | 0.70089 | 2.01365 |
| **TSPYL4** | 1.82270 | 0.00000 | 1.71532 | 5.74121 | 1.57817 | 1.14441 | 1.46693 |
| **TSPYL5** | 1.05681 | 0.00000 | 2.30307 | 1.11710 | 0.35304 | 0.00000 | 1.24720 |
| **TSPYL6** | 1.03781 | 0.96804 | 3.19531 | 2.39101 | 1.35770 | 0.38789 | 0.00000 |
| **TSR1** | 42.27955 | 16.90312 | 18.26169 | 22.17395 | 7.33302 | 10.88302 | 0.68286 |
| **TSR2** | 23.39387 | 22.35657 | 12.38454 | 8.50293 | 3.32156 | 10.60335 | 1.19772 |
| **TSR3** | 0.26809 | 8.06828 | 0.55231 | 1.29374 | 0.00000 | 1.90480 | 9.48737 |
| **TSSC1** | 10.20144 | 9.51198 | 1.91253 | 5.76139 | 1.56294 | 6.88948 | 0.00000 |
| **TSSC4** | 1.01715 | 16.01206 | 5.30540 | 11.23028 | 1.25978 | 5.03119 | 14.97379 |
| **TSSK1B** | 0.01622 | 0.01719 | 0.01341 | 0.02984 | 0.00791 | 0.01204 | 0.03437 |

| | | | | | | | |
|---|---|---|---|---|---|---|---|
| **TSSK3** | 0.04541 | 0.01316 | 0.06910 | 0.18356 | 0.00699 | 0.00000 | 0.16578 |
| **TSSK4** | 0.08987 | 0.35822 | 0.05447 | 0.13073 | 0.00942 | 0.10128 | 0.02653 |
| **TSSK6** | 0.03198 | 0.00000 | 0.16634 | 0.23218 | 0.65043 | 0.04391 | 0.08681 |
| **TST** | 1.02117 | 1.26841 | 0.13188 | 2.77978 | 0.00000 | 0.55518 | 2.54723 |
| **TSTA3** | 21.26271 | 11.73951 | 3.13470 | 12.87643 | 0.52303 | 6.99471 | 5.43312 |
| **TSTD1** | 3.70868 | 5.64605 | 3.14901 | 2.77664 | 4.06740 | 3.40826 | 1.01067 |
| **TSTD2** | 3.70211 | 3.31219 | 2.76420 | 3.41874 | 3.38090 | 1.36617 | 0.00000 |
| **TSTD3** | 0.81778 | 0.31327 | 0.29106 | 0.95323 | 0.14175 | 0.22079 | 0.76503 |
| **TTBK1** | 0.01898 | 0.00000 | 0.06782 | 0.03046 | 0.01460 | 0.01942 | 0.06397 |
| **TTBK2** | 0.28125 | 0.25890 | 0.54027 | 0.00000 | 0.87215 | 0.68411 | 1.87187 |
| **TTC1** | 27.59888 | 10.42596 | 14.91662 | 25.99022 | 7.58877 | 9.97508 | 8.04741 |
| **TTC12** | 0.71237 | 1.55409 | 0.87773 | 0.47219 | 1.51220 | 0.63894 | 0.01107 |
| **TTC13** | 0.20828 | 0.12812 | 0.01520 | 0.38750 | 0.01862 | 0.30217 | 1.04835 |
| **TTC14** | 7.54805 | 0.24492 | 8.08196 | 5.74382 | 3.35086 | 2.19782 | 10.95418 |
| **TTC16** | 0.00240 | 0.00000 | 0.16425 | 0.02722 | 0.10856 | 0.02099 | 0.07845 |
| **TTC17** | 8.01989 | 2.58388 | 6.53731 | 5.36165 | 2.26880 | 2.48473 | 4.81651 |
| **TTC18** | 0.44514 | 0.28857 | 0.96196 | 0.51067 | 0.88554 | 0.15753 | 0.04013 |
| **TTC19** | 5.51921 | 3.10463 | 2.83442 | 0.90182 | 0.77651 | 8.92259 | 10.81000 |
| **TTC21A** | 0.07238 | 0.00000 | 0.26728 | 0.21656 | 0.18101 | 0.41777 | 0.74888 |
| **TTC21B** | 1.38857 | 0.59970 | 1.89958 | 0.36037 | 0.58473 | 1.73716 | 0.90879 |
| **TTC21B-AS1** | 0.03038 | 0.08768 | 0.02056 | 0.02015 | 0.00933 | 0.01781 | 0.00000 |
| **TTC22** | 0.02079 | 0.04257 | 0.01259 | 0.10890 | 0.00772 | 0.02340 | 0.00000 |
| **TTC23** | 0.56234 | 1.01964 | 0.36718 | 0.07434 | 0.20854 | 0.32754 | 0.02802 |
| **TTC23L** | 0.05956 | 0.00000 | 0.07522 | 0.30692 | 0.40542 | 0.00000 | 0.15234 |
| **TTC24** | 0.01732 | 0.00894 | 0.01205 | 0.01954 | 0.05808 | 0.03619 | 0.04178 |
| **TTC25** | 0.04021 | 0.45392 | 0.08599 | 0.21853 | 0.32104 | 0.00000 | 0.43633 |
| **TTC26** | 1.48794 | 0.46747 | 3.42229 | 0.03341 | 0.58882 | 1.78308 | 0.00001 |
| **TTC27** | 12.97056 | 4.35391 | 11.79360 | 8.67222 | 2.09812 | 3.87846 | 0.46836 |
| **TTC28** | 4.60132 | 1.90235 | 0.00000 | 0.65076 | 1.75336 | 0.00000 | 2.62538 |
| **TTC28-AS1** | 1.61011 | 1.64042 | 2.54702 | 3.86926 | 3.56851 | 3.13355 | 2.87298 |
| **TTC29** | 0.01447 | 0.00410 | 0.05093 | 0.01644 | 0.29117 | 0.00908 | 0.04790 |
| **TTC3** | 7.40322 | 0.25865 | 1.55138 | 3.43452 | 2.56720 | 1.39472 | 15.36351 |
| **TTC30A** | 0.25502 | 0.06429 | 0.53267 | 0.23399 | 0.21435 | 0.06667 | 0.00000 |
| **TTC30B** | 0.16845 | 0.29398 | 0.22233 | 0.14545 | 0.17292 | 0.01371 | 0.00000 |
| **TTC31** | 5.20528 | 3.26861 | 3.90239 | 4.46798 | 0.60210 | 3.75137 | 0.00000 |

| | | | | | | | |
|---|---|---|---|---|---|---|---|
| **TTC32** | 3.37208 | 0.00000 | 2.50741 | 2.00401 | 0.33063 | 1.19209 | 0.00000 |
| **TTC33** | 3.51981 | 0.06108 | 2.60320 | 1.39404 | 1.58664 | 1.05821 | 0.00000 |
| **TTC34** | 0.02721 | 0.01711 | 0.01807 | 0.02791 | 0.01246 | 0.03012 | 0.00583 |
| **TTC36** | 0.02390 | 0.03208 | 0.01180 | 0.01981 | 0.00773 | 0.03076 | 0.01719 |
| **TTC37** | 14.51186 | 0.28965 | 1.84856 | 3.41336 | 0.72947 | 4.19699 | 8.70330 |
| **TTC38** | 2.55687 | 0.70553 | 0.02164 | 4.56947 | 0.31297 | 0.30242 | 0.00000 |
| **TTC39A** | 0.90964 | 0.00000 | 0.23616 | 2.25209 | 0.18199 | 0.08295 | 0.00000 |
| **TTC39B** | 0.66628 | 0.20864 | 1.34231 | 0.09293 | 0.30987 | 6.16022 | 0.00000 |
| **TTC39C** | 0.35694 | 0.00000 | 1.85143 | 0.18983 | 2.45787 | 0.06663 | 0.08414 |
| **TTC3P1** | 0.18336 | 0.00000 | 0.07795 | 0.05866 | 0.12277 | 0.14330 | 1.34043 |
| **TTC4** | 1.30867 | 1.49766 | 1.76687 | 0.34692 | 0.92729 | 0.39421 | 0.00000 |
| **TTC40** | 0.23143 | 0.56643 | 0.03894 | 0.00000 | 0.09426 | 0.67147 | 0.01106 |
| **TTC5** | 6.54917 | 2.42122 | 5.32838 | 0.11613 | 2.58834 | 2.95563 | 0.33913 |
| **TTC7A** | 1.79920 | 5.23980 | 1.28707 | 2.50899 | 0.07949 | 0.43069 | 5.71244 |
| **TTC7B** | 4.19245 | 2.12121 | 2.28254 | 5.14601 | 0.08032 | 0.00000 | 11.86123 |
| **TTC8** | 1.06951 | 0.20858 | 1.16110 | 0.50563 | 0.00000 | 1.03044 | 0.73518 |
| **TTC9** | 0.38796 | 0.04858 | 0.01675 | 0.85913 | 0.04689 | 0.00000 | 3.81665 |
| **TTC9B** | 0.02400 | 0.11248 | 0.15808 | 0.02929 | 0.28373 | 0.01917 | 0.00000 |
| **TTC9C** | 10.78649 | 3.31744 | 11.54430 | 7.39455 | 11.54374 | 4.22180 | 2.58643 |
| **TTF1** | 9.45205 | 7.60347 | 11.26025 | 5.84152 | 3.81306 | 3.96598 | 8.56170 |
| **TTF2** | 11.76110 | 2.74535 | 2.45477 | 4.56713 | 3.29953 | 5.11879 | 5.92961 |
| **TTI1** | 14.94876 | 7.37568 | 7.25545 | 9.72633 | 3.83559 | 3.14223 | 0.00000 |
| **TTI2** | 8.45058 | 3.67850 | 6.17808 | 4.00147 | 3.61548 | 1.27192 | 0.00000 |
| **TTK** | 19.34837 | 0.00000 | 14.77544 | 9.06147 | 6.45431 | 3.18562 | 1.44933 |
| **TTL** | 5.67287 | 2.92959 | 2.67711 | 0.60508 | 1.15642 | 2.56929 | 9.72906 |
| **TTLL1** | 0.11084 | 0.00000 | 0.37809 | 0.09314 | 1.30724 | 0.08553 | 0.14337 |
| **TTLL10** | 0.01839 | 0.01629 | 0.02126 | 0.02120 | 0.06916 | 0.03078 | 0.02123 |
| **TTLL11** | 0.00000 | 0.28020 | 0.00069 | 0.21656 | 0.16428 | 0.01230 | 0.55561 |
| **TTLL12** | 2.21416 | 12.62715 | 2.19494 | 7.41212 | 0.22758 | 4.49124 | 22.53067 |
| **TTLL13** | 0.05955 | 0.06115 | 0.12260 | 0.11154 | 0.00751 | 0.02058 | 0.00000 |
| **TTLL3** | 0.15588 | 0.00000 | 0.30985 | 0.58123 | 0.00604 | 0.11246 | 0.53973 |
| **TTLL4** | 13.73030 | 7.93739 | 5.52216 | 12.99745 | 6.06172 | 11.60856 | 12.49935 |
| **TTLL5** | 7.03696 | 0.71822 | 5.68265 | 7.45821 | 5.61489 | 4.28799 | 6.09056 |
| **TTLL6** | 0.03778 | 0.01712 | 0.01247 | 0.03259 | 0.01244 | 0.01259 | 0.00539 |
| **TTLL7** | 0.05895 | 0.00803 | 0.00903 | 0.34133 | 0.24590 | 0.03181 | 0.75996 |

| | | | | | | | |
|---|---|---|---|---|---|---|---|
| **TTLL9** | 0.21988 | 0.58269 | 0.19655 | 0.03670 | 0.10914 | 0.04357 | 0.00000 |
| **TTN** | 5.46780 | 0.00000 | 3.17225 | 6.65775 | 2.76425 | 1.68170 | 16.07459 |
| **TTN-AS1** | 0.83773 | 0.67836 | 1.06836 | 1.35624 | 2.07703 | 0.94355 | 0.00000 |
| **TTPA** | 0.02153 | 0.06313 | 0.08970 | 0.02356 | 0.03521 | 0.00585 | 0.00512 |
| **TTPAL** | 3.05564 | 2.72392 | 3.33624 | 3.00011 | 3.07888 | 1.90781 | 1.93137 |
| **TTR** | 0.06102 | 0.00366 | 0.02007 | 0.06295 | 0.03512 | 0.12746 | 0.00747 |
| **TTTY15** | 0.00000 | 0.01020 | 0.14599 | 0.04473 | 0.00566 | 0.05116 | 0.10332 |
| **TTYH1** | 0.49446 | 0.00000 | 0.02876 | 0.00000 | 0.00246 | 0.88393 | 0.87767 |
| **TTYH2** | 0.00000 | 0.32243 | 0.15181 | 0.00730 | 1.45935 | 0.07044 | 0.10340 |
| **TTYH3** | 0.06887 | 0.00000 | 0.04973 | 1.30190 | 0.00504 | 1.83669 | 1.72088 |
| **TUB** | 0.75734 | 0.24477 | 0.01211 | 0.09664 | 0.30130 | 0.28616 | 0.04428 |
| **TUBA1A** | 2.73139 | 0.18006 | 1.26614 | 0.91715 | 0.09853 | 4.97848 | 1.57743 |
| **TUBA1B** | 65.80162 | 36.29701 | 15.51651 | 48.15516 | 8.12056 | 28.61190 | 36.10030 |
| **TUBA1C** | 62.22591 | 54.40803 | 25.68464 | 37.94510 | 15.19833 | 34.46261 | 61.41502 |
| **TUBA3C** | 0.00000 | 3.63830 | 2.87590 | 0.00000 | 1.84116 | 0.22452 | 0.00125 |
| **TUBA3D** | 1.54055 | 5.68702 | 2.53075 | 1.14312 | 4.44298 | 2.79911 | 0.00000 |
| **TUBA3E** | 0.29025 | 1.04699 | 0.29569 | 0.32049 | 3.84234 | 0.84981 | 0.00000 |
| **TUBA3FP** | 0.11430 | 0.00000 | 0.04117 | 0.07910 | 0.02761 | 0.10577 | 0.07389 |
| **TUBA4A** | 0.34483 | 0.46086 | 0.08595 | 0.51267 | 2.52318 | 0.42390 | 0.16205 |
| **TUBA4B** | 0.05240 | 0.01329 | 0.14582 | 0.03642 | 4.54649 | 0.01764 | 0.04750 |
| **TUBA8** | 0.24599 | 0.42654 | 0.01676 | 0.45094 | 0.01008 | 0.30668 | 0.68671 |
| **TUBAL3** | 0.00000 | 0.05315 | 0.02024 | 0.15486 | 1.31216 | 0.03748 | 0.00363 |
| **TUBB** | 69.90009 | 29.71990 | 20.22534 | 44.81948 | 2.89916 | 34.20156 | 36.94573 |
| **TUBB1** | 0.23028 | 0.00000 | 0.49188 | 0.50473 | 0.57296 | 0.17247 | 0.43530 |
| **TUBB2A** | 5.53763 | 3.33773 | 0.94897 | 0.94094 | 0.46006 | 14.25389 | 1.05140 |
| **TUBB2B** | 5.43283 | 0.00000 | 0.00056 | 0.00000 | 0.00127 | 11.67838 | 3.00681 |
| **TUBB3** | 8.81677 | 13.13810 | 0.18927 | 7.02760 | 0.63771 | 17.23830 | 0.31987 |
| **TUBB4A** | 61.38158 | 86.12334 | 17.43260 | 0.00000 | 0.00000 | 52.10471 | 17.86736 |
| **TUBB4B** | 52.75108 | 58.09786 | 28.07993 | 45.26484 | 14.48274 | 26.63781 | 20.04653 |
| **TUBB6** | 24.19438 | 19.20455 | 3.84311 | 25.02293 | 2.12088 | 7.39747 | 4.63107 |
| **TUBB8** | 0.00000 | 0.47397 | 0.65592 | 0.00000 | 19.36754 | 0.00000 | 0.00670 |
| **TUBBP5** | 0.00000 | 0.72494 | 1.98684 | 0.09719 | 10.85098 | 0.70156 | 0.08922 |
| **TUBD1** | 10.47204 | 0.82516 | 6.63260 | 6.58985 | 0.22432 | 2.72578 | 0.31195 |
| **TUBE1** | 6.32074 | 0.79376 | 4.22430 | 1.31072 | 0.10903 | 1.94163 | 1.31677 |
| **TUBG1** | 16.93215 | 15.36296 | 3.47894 | 20.04747 | 1.32380 | 10.96923 | 0.00000 |

| | | | | | | | |
|---|---|---|---|---|---|---|---|
| **TUBG2** | 1.04213 | 0.91979 | 0.56008 | 1.61540 | 1.79162 | 1.62513 | 0.00000 |
| **TUBGCP2** | 5.69716 | 11.43221 | 3.08626 | 4.67461 | 0.36054 | 6.83193 | 12.37652 |
| **TUBGCP3** | 5.62360 | 3.12868 | 4.82574 | 2.21968 | 0.72543 | 1.87619 | 8.68509 |
| **TUBGCP4** | 10.65814 | 4.56827 | 2.37493 | 5.72805 | 1.23317 | 1.94542 | 1.39893 |
| **TUBGCP5** | 3.93243 | 1.15824 | 3.21984 | 1.30297 | 0.33243 | 1.11493 | 3.30267 |
| **TUBGCP6** | 0.34219 | 2.65459 | 1.48269 | 2.45735 | 0.69596 | 1.19369 | 7.77697 |
| **TUFM** | 50.66925 | 42.82987 | 23.47299 | 45.35864 | 3.54784 | 27.90156 | 21.71582 |
| **TUFT1** | 6.80892 | 13.75585 | 2.02929 | 16.94721 | 1.82398 | 0.81575 | 3.11929 |
| **TUG1** | 19.31198 | 7.52789 | 14.19952 | 11.11607 | 5.92146 | 4.53433 | 16.59024 |
| **TULP1** | 0.00000 | 0.01049 | 0.06459 | 1.54863 | 0.19898 | 0.00849 | 0.00853 |
| **TULP2** | 0.49919 | 1.93007 | 1.09191 | 0.21290 | 0.29046 | 1.30264 | 0.00000 |
| **TULP3** | 5.74689 | 6.25773 | 9.52691 | 6.51065 | 10.23051 | 4.30686 | 3.01522 |
| **TULP4** | 23.21950 | 19.17447 | 8.20493 | 12.94521 | 2.97302 | 6.30458 | 6.77657 |
| **TUNAR** | 0.02247 | 0.01651 | 0.00719 | 0.02824 | 0.17238 | 0.02285 | 0.03021 |
| **TUSC1** | 0.00000 | 0.86748 | 0.57154 | 0.97682 | 0.15927 | 1.02000 | 2.98036 |
| **TUSC2** | 8.97364 | 18.93420 | 10.45842 | 19.03098 | 6.37537 | 5.03316 | 7.38314 |
| **TUSC3** | 0.12408 | 0.00000 | 0.80245 | 6.60302 | 0.48597 | 4.31948 | 6.67343 |
| **TUSC5** | 0.03576 | 0.01058 | 0.02325 | 0.01464 | 0.70153 | 0.01078 | 0.01128 |
| **TUSC8** | 0.03258 | 0.01190 | 0.01030 | 0.01889 | 0.13689 | 0.03305 | 0.01329 |
| **TUT1** | 5.50154 | 4.68908 | 4.22351 | 4.89115 | 2.81784 | 2.55285 | 2.41795 |
| **TVP23A** | 0.16643 | 0.12883 | 0.09832 | 0.26260 | 0.13960 | 0.01994 | 0.24921 |
| **TVP23B** | 4.51438 | 0.19620 | 3.43141 | 2.47219 | 2.58167 | 1.58628 | 0.61263 |
| **TVP23C.uid-209** | 0.30062 | 0.00000 | 0.57319 | 0.09525 | 0.41889 | 0.26600 | 0.17316 |
| **TVP23C.uid-209** | 0.45856 | 0.79497 | 0.48652 | 0.31843 | 0.44331 | 0.19065 | 1.48607 |
| **TWF1** | 12.58120 | 0.00000 | 4.53444 | 3.42969 | 0.80141 | 2.84385 | 2.05681 |
| **TWF2** | 14.57015 | 22.39241 | 3.19214 | 9.71421 | 0.89026 | 17.03807 | 4.56305 |
| **TWIST1** | 0.00000 | 0.03375 | 0.00000 | 0.27471 | 0.13498 | 1.45599 | 0.81238 |
| **TWIST2** | 0.00638 | 0.03046 | 0.00936 | 0.01837 | 0.00737 | 0.14356 | 0.05700 |
| **TWISTNB** | 21.25216 | 9.02069 | 10.27984 | 12.33839 | 5.97139 | 10.01119 | 14.32200 |
| **TWSG1** | 1.09353 | 0.00000 | 1.43290 | 5.01831 | 2.28311 | 0.42580 | 0.63274 |
| **TXK** | 0.18346 | 1.11670 | 0.33635 | 0.00986 | 0.05727 | 1.34338 | 0.00000 |
| **TXLNA** | 23.56980 | 15.58550 | 7.49300 | 18.40764 | 3.94845 | 8.95758 | 2.97839 |
| **TXLNB** | 0.03043 | 0.00192 | 0.04831 | 0.02769 | 0.04812 | 0.02312 | 0.01692 |
| **TXLNG** | 16.94525 | 5.24351 | 7.53553 | 8.47168 | 5.23958 | 9.13822 | 24.53212 |
| **TXLNGY** | 0.02259 | 0.01871 | 0.01282 | 0.06199 | 0.00788 | 0.01359 | 0.01408 |

| | | | | | | | |
|---|---|---|---|---|---|---|---|
| **TXN** | 49.63833 | 41.70237 | 33.28267 | 38.90618 | 5.19250 | 27.28750 | 25.88033 |
| **TXN2** | 30.27559 | 29.11610 | 17.11327 | 19.78525 | 10.55456 | 10.75184 | 11.49031 |
| **TXNDC11** | 2.23703 | 3.61528 | 1.52341 | 1.21633 | 1.62577 | 1.91938 | 4.43343 |
| **TXNDC12** | 28.81294 | 23.05896 | 17.38538 | 19.91669 | 0.52803 | 21.98384 | 5.63147 |
| **TXNDC15** | 5.88342 | 4.43989 | 6.09399 | 6.45071 | 0.00000 | 7.00495 | 0.03848 |
| **TXNDC16** | 0.90322 | 0.23294 | 0.48198 | 0.52616 | 1.15575 | 0.74745 | 0.81692 |
| **TXNDC17** | 29.29600 | 18.53068 | 5.54590 | 21.75751 | 13.73311 | 16.02345 | 19.13820 |
| **TXNDC2** | 0.00352 | 0.00922 | 0.01428 | 0.09558 | 0.00715 | 0.00912 | 0.09147 |
| **TXNDC5.uid-81** | 0.02377 | 0.01732 | 0.01286 | 0.02973 | 0.00872 | 0.01431 | 0.01435 |
| **TXNDC5.uid-81** | 0.01395 | 0.12039 | 0.01682 | 0.03944 | 0.01273 | 0.07790 | 0.14347 |
| **TXNDC8** | 0.06232 | 0.01878 | 0.02572 | 0.00467 | 0.00664 | 0.01088 | 0.02504 |
| **TXNDC9** | 25.60755 | 3.12574 | 10.19260 | 11.67496 | 5.14722 | 6.53859 | 1.12386 |
| **TXNIP** | 14.68075 | 5.66062 | 5.35308 | 3.78007 | 0.00000 | 1.73782 | 4.49028 |
| **TXNL1** | 35.48017 | 20.10908 | 9.96865 | 29.24775 | 6.30011 | 11.19062 | 1.02329 |
| **TXNL4A** | 4.26070 | 6.28522 | 3.79052 | 8.95364 | 0.14801 | 8.54429 | 30.74164 |
| **TXNL4B** | 7.92464 | 6.07351 | 7.01254 | 10.30604 | 9.40583 | 2.12483 | 0.71762 |
| **TXNRD1** | 51.84282 | 29.33756 | 29.62473 | 28.30807 | 0.93239 | 19.84291 | 23.46216 |
| **TXNRD2** | 0.99164 | 6.67552 | 0.07003 | 0.00000 | 0.05621 | 1.15535 | 2.90138 |
| **TXNRD3** | 0.00838 | 0.03807 | 0.02620 | 0.01949 | 0.00546 | 0.00264 | 0.11901 |
| **TYK2** | 3.19539 | 2.20663 | 1.88588 | 3.10495 | 0.00069 | 2.42993 | 7.11108 |
| **TYMP** | 0.00000 | 0.12897 | 0.00841 | 0.06902 | 0.00743 | 0.05931 | 0.70418 |
| **TYMS** | 3.71092 | 8.34994 | 6.44665 | 8.26577 | 0.31545 | 2.94047 | 0.00274 |
| **TYMSOS** | 0.24689 | 1.47314 | 0.85635 | 0.67336 | 0.27846 | 0.82863 | 0.85643 |
| **TYR** | 0.02432 | 0.01352 | 0.01633 | 0.02051 | 0.00756 | 0.01849 | 0.02237 |
| **TYRO3** | 3.78704 | 9.94180 | 5.27862 | 20.14734 | 0.00000 | 5.48033 | 25.27381 |
| **TYRO3P** | 0.11248 | 0.02196 | 0.08262 | 0.17313 | 0.00498 | 0.04475 | 0.10557 |
| **TYROBP** | 0.04000 | 0.00000 | 0.14928 | 0.02603 | 0.11740 | 0.02368 | 0.00612 |
| **TYRP1** | 0.01706 | 0.04040 | 0.00057 | 0.03756 | 0.13576 | 0.02123 | 0.01446 |
| **TYSND1** | 0.82542 | 0.00000 | 1.80566 | 4.54181 | 1.19568 | 1.45345 | 3.58963 |
| **TYW1** | 5.30888 | 1.53966 | 2.89862 | 3.40517 | 2.98691 | 3.52360 | 3.35273 |
| **TYW1B** | 0.53356 | 0.02813 | 0.55566 | 0.60450 | 3.58614 | 0.71361 | 1.75898 |
| **TYW3** | 9.13280 | 3.01349 | 6.59397 | 3.55729 | 0.82666 | 5.61852 | 6.79359 |
| **TYW5** | 9.06287 | 2.00521 | 4.22012 | 3.34552 | 1.17861 | 3.13242 | 0.00000 |
| **U2AF1** | 46.21763 | 39.91857 | 18.93801 | 39.49605 | 10.52171 | 18.28519 | 10.88305 |
| **U2AF1L4** | 4.12185 | 5.50901 | 1.67008 | 2.15532 | 3.15029 | 6.13366 | 0.00000 |

| | | | | | | | |
|---|---|---|---|---|---|---|---|
| **U2AF2** | 2.85571 | 3.97072 | 7.93906 | 10.87847 | 1.45750 | 5.37547 | 27.47152 |
| **U2SURP** | 29.85234 | 6.15215 | 21.84474 | 11.26277 | 6.95400 | 15.15955 | 24.69639 |
| **UACA** | 6.06705 | 34.14209 | 5.96833 | 0.68534 | 4.05226 | 48.22486 | 2.95379 |
| **UAP1** | 12.25964 | 9.54697 | 9.05071 | 5.26788 | 4.51561 | 5.87841 | 0.00000 |
| **UAP1L1** | 3.53459 | 12.92968 | 0.00000 | 0.00000 | 0.03665 | 6.41927 | 5.97342 |
| **UBA1** | 22.68392 | 30.56239 | 21.82807 | 20.76802 | 5.72127 | 14.46175 | 35.22361 |
| **UBA2** | 9.16222 | 1.90421 | 7.41848 | 6.35071 | 0.45445 | 7.79837 | 21.79750 |
| **UBA3** | 23.63997 | 1.64754 | 8.47480 | 6.62508 | 0.87672 | 5.72176 | 2.02883 |
| **UBA5** | 4.34493 | 3.93226 | 4.16028 | 2.11099 | 0.95592 | 4.73901 | 10.59278 |
| **UBA52** | 44.17798 | 44.46920 | 31.08483 | 48.95046 | 14.06822 | 29.18435 | 53.62441 |
| **UBA6** | 12.79667 | 0.48195 | 8.01161 | 7.13895 | 3.87482 | 8.36310 | 3.02973 |
| **UBA6-AS1** | 1.39892 | 0.43269 | 0.56899 | 2.04774 | 0.00997 | 0.67556 | 0.15172 |
| **UBA7** | 0.00000 | 0.11894 | 0.01189 | 0.07476 | 0.00601 | 0.07085 | 0.22550 |
| **UBAC1** | 10.44581 | 17.96891 | 1.78962 | 11.50809 | 3.96114 | 5.46183 | 13.55814 |
| **UBAC2** | 13.23127 | 18.31164 | 6.66907 | 15.00003 | 0.02001 | 10.61424 | 6.35258 |
| **UBAC2-AS1** | 0.24910 | 1.19208 | 0.89166 | 0.01919 | 0.02029 | 0.47446 | 0.27836 |
| **UBALD1** | 0.00000 | 2.21520 | 1.10285 | 1.77947 | 0.08642 | 0.92549 | 5.32258 |
| **UBALD2** | 0.00000 | 3.19607 | 0.42327 | 4.40604 | 0.17461 | 2.41644 | 13.14481 |
| **UBAP1** | 12.46162 | 11.42279 | 7.01506 | 16.07421 | 6.84397 | 9.53832 | 2.73793 |
| **UBAP1L** | 0.05889 | 0.03856 | 0.14531 | 0.12708 | 0.25197 | 0.02275 | 0.83178 |
| **UBAP2** | 18.34085 | 18.19587 | 21.27141 | 17.76074 | 3.52177 | 19.13771 | 20.79162 |
| **UBAP2L** | 29.12370 | 20.12475 | 13.63269 | 23.73416 | 5.65194 | 12.29661 | 12.37668 |
| **UBASH3B** | 1.98854 | 17.94709 | 5.90558 | 1.71627 | 0.23274 | 19.44808 | 0.48982 |
| **UBB** | 68.29475 | 36.50828 | 25.46333 | 57.30098 | 23.22899 | 30.12440 | 20.03797 |
| **UBC** | 32.82367 | 10.66573 | 15.03904 | 35.08059 | 8.01177 | 16.00848 | 18.54097 |
| **UBE2A** | 30.80595 | 19.12543 | 2.86857 | 28.43489 | 3.09653 | 15.96466 | 8.22536 |
| **UBE2B** | 20.28816 | 5.21513 | 10.18572 | 10.98398 | 11.94861 | 8.53569 | 4.84214 |
| **UBE2C** | 36.77265 | 43.91429 | 40.79151 | 30.90331 | 24.40456 | 18.60899 | 16.10555 |
| **UBE2D1** | 3.94136 | 0.00000 | 6.39369 | 3.26768 | 0.26311 | 3.64522 | 11.71280 |
| **UBE2D2** | 4.95139 | 4.04375 | 3.00673 | 11.30472 | 1.25315 | 9.80787 | 29.23303 |
| **UBE2D3** | 41.96116 | 16.51910 | 22.55420 | 27.08061 | 12.97564 | 17.70212 | 20.14645 |
| **UBE2D4** | 2.02361 | 2.38359 | 1.79849 | 0.63688 | 0.00000 | 0.00000 | 1.59548 |
| **UBE2DNL** | 0.03251 | 0.04199 | 0.00000 | 0.13211 | 0.36615 | 0.10593 | 0.00000 |
| **UBE2E1** | 2.85270 | 0.00000 | 4.45250 | 2.78747 | 0.44673 | 3.11082 | 13.67609 |
| **UBE2E1-AS1** | 0.04901 | 0.16859 | 0.03070 | 0.00662 | 0.04806 | 0.01036 | 0.01186 |

| | | | | | | | |
|---|---|---|---|---|---|---|---|
| **UBE2E2** | 4.11558 | 0.30924 | 5.31467 | 2.88447 | 1.11954 | 1.81267 | 5.81925 |
| **UBE2E3** | 8.12011 | 0.62157 | 3.45567 | 6.08979 | 0.11654 | 2.36870 | 10.33142 |
| **UBE2E4P** | 1.28878 | 0.26491 | 0.33380 | 0.91753 | 0.02588 | 0.30142 | 0.22555 |
| **UBE2F** | 15.73710 | 5.25581 | 6.45690 | 8.85366 | 1.13901 | 7.17074 | 10.18499 |
| **UBE2F-SCLY** | 0.02292 | 0.01621 | 0.01602 | 0.01958 | 0.01159 | 0.01238 | 0.02214 |
| **UBE2G1** | 20.21422 | 3.72317 | 30.42116 | 16.24018 | 11.76552 | 13.46523 | 18.51870 |
| **UBE2G2** | 26.38847 | 5.48104 | 4.77014 | 18.27910 | 0.19934 | 1.80643 | 6.13597 |
| **UBE2H** | 0.62749 | 4.19298 | 3.02534 | 12.57675 | 2.33198 | 5.21895 | 12.83011 |
| **UBE2I** | 44.29091 | 37.94880 | 10.28802 | 29.86483 | 7.19853 | 18.88960 | 17.93736 |
| **UBE2J1** | 5.39874 | 0.00000 | 0.00000 | 14.71726 | 0.04709 | 0.00000 | 11.39178 |
| **UBE2J2** | 11.85439 | 18.73718 | 8.76853 | 13.68577 | 1.83810 | 8.31406 | 15.81792 |
| **UBE2K** | 28.21788 | 15.17242 | 14.16782 | 21.37298 | 6.76694 | 13.03761 | 16.85131 |
| **UBE2L3** | 27.36638 | 18.89538 | 9.60411 | 22.34388 | 4.08551 | 11.69899 | 7.17648 |
| **UBE2L6** | 0.05505 | 0.00058 | 0.15831 | 0.74194 | 0.51906 | 0.00243 | 0.00000 |
| **UBE2M** | 0.80211 | 7.86780 | 1.04150 | 3.84573 | 1.34934 | 3.56269 | 20.12989 |
| **UBE2MP1** | 0.34220 | 0.78268 | 0.12400 | 0.50037 | 0.25339 | 0.44905 | 1.00732 |
| **UBE2N** | 40.00006 | 13.91571 | 16.47064 | 24.67456 | 6.26464 | 15.93370 | 3.34394 |
| **UBE2NL** | 1.00693 | 0.08850 | 0.35800 | 0.49098 | 0.11122 | 0.29843 | 0.00000 |
| **UBE2O** | 2.29934 | 10.64413 | 3.29860 | 7.63393 | 2.06682 | 7.50050 | 16.55253 |
| **UBE2Q1** | 2.15515 | 1.45833 | 4.85687 | 4.49831 | 0.29621 | 3.99815 | 10.57842 |
| **UBE2Q2** | 4.71336 | 6.92054 | 3.49268 | 3.82914 | 7.19711 | 8.83319 | 17.92479 |
| **UBE2Q2P1** | 0.04785 | 0.17723 | 0.04752 | 0.00000 | 0.07554 | 0.17405 | 0.48456 |
| **UBE2QL1** | 0.02015 | 0.01580 | 0.01019 | 0.02152 | 0.02222 | 0.01076 | 0.04914 |
| **UBE2R2** | 3.56371 | 6.09535 | 5.81864 | 5.42336 | 1.15762 | 5.90330 | 26.15492 |
| **UBE2S** | 0.00000 | 3.03278 | 0.70610 | 0.43755 | 0.56378 | 1.12104 | 4.60303 |
| **UBE2T** | 30.37058 | 13.37734 | 14.89953 | 14.97349 | 5.88736 | 16.12410 | 0.00008 |
| **UBE2U** | 0.01974 | 0.06132 | 0.52097 | 0.00005 | 1.21455 | 0.33030 | 0.02856 |
| **UBE2V1** | 3.53758 | 1.95505 | 2.96196 | 2.55490 | 1.66150 | 1.52316 | 0.00010 |
| **UBE2V2** | 29.99932 | 2.79125 | 15.04834 | 14.33879 | 1.85644 | 8.31376 | 5.58966 |
| **UBE2W** | 12.41363 | 0.00000 | 7.92179 | 4.67179 | 5.06172 | 3.28172 | 0.53993 |
| **UBE2Z** | 5.56689 | 2.78437 | 6.35139 | 6.99867 | 2.04135 | 10.87814 | 24.92205 |
| **UBE3A** | 8.30610 | 0.84179 | 6.30621 | 4.57193 | 4.09705 | 5.76666 | 24.53166 |
| **UBE3B** | 4.06107 | 1.36540 | 5.15132 | 2.13128 | 0.50416 | 5.21882 | 11.26224 |
| **UBE3C** | 12.83537 | 5.27387 | 7.15945 | 7.22969 | 2.51335 | 5.61971 | 13.91149 |
| **UBE3D** | 6.80554 | 0.78753 | 1.25622 | 3.17573 | 0.21939 | 1.20554 | 0.00002 |

| | | | | | | | |
|---|---|---|---|---|---|---|---|
| **UBE4A** | 18.91139 | 3.49981 | 12.01166 | 9.16975 | 5.13625 | 7.26055 | 7.87802 |
| **UBE4B** | 9.72141 | 9.48965 | 6.41994 | 15.65850 | 5.05530 | 4.76517 | 8.49367 |
| **UBFD1** | 8.05692 | 6.44617 | 3.98478 | 13.04713 | 2.75792 | 9.44016 | 19.74831 |
| **UBIAD1** | 14.46243 | 14.24908 | 10.08941 | 13.86350 | 4.43206 | 7.26961 | 0.00000 |
| **UBL3** | 5.53677 | 1.18914 | 0.43544 | 1.22035 | 1.00668 | 0.62757 | 10.99891 |
| **UBL4A** | 23.76090 | 13.65842 | 2.38821 | 11.55128 | 0.00000 | 7.73218 | 11.32153 |
| **UBL4B** | 0.02903 | 0.01103 | 0.02030 | 0.01773 | 0.00748 | 0.01355 | 0.02602 |
| **UBL5** | 33.13085 | 30.89404 | 26.10506 | 27.98802 | 17.22493 | 16.35316 | 27.28706 |
| **UBL7** | 16.56038 | 11.82739 | 10.42348 | 16.09197 | 8.57790 | 8.07996 | 3.79940 |
| **UBL7-AS1** | 0.53005 | 1.05220 | 1.37607 | 2.46004 | 1.03030 | 0.76835 | 1.05750 |
| **UBLCP1** | 7.07083 | 0.00000 | 3.93204 | 4.07109 | 0.77312 | 3.00545 | 2.99045 |
| **UBN1** | 5.18669 | 4.71674 | 3.21241 | 5.53749 | 1.98008 | 8.34361 | 17.50551 |
| **UBN2** | 3.31278 | 0.00000 | 7.23422 | 3.21377 | 5.12985 | 5.70909 | 12.88482 |
| **UBOX5** | 2.12828 | 2.01726 | 1.40667 | 1.83723 | 0.39590 | 1.63097 | 0.00000 |
| **UBOX5-AS1** | 0.05097 | 0.02530 | 0.03408 | 0.01011 | 0.02739 | 0.04918 | 0.15161 |
| **UBP1** | 6.09244 | 0.00000 | 5.37038 | 3.14579 | 2.64165 | 4.26522 | 12.70520 |
| **UBQLN1** | 30.44158 | 30.85216 | 20.17938 | 26.83343 | 12.63267 | 16.96682 | 13.85380 |
| **UBQLN2** | 9.95812 | 5.02357 | 8.08530 | 4.30738 | 1.35791 | 0.29003 | 2.18710 |
| **UBQLN4** | 1.33109 | 2.74221 | 1.90407 | 1.97198 | 0.00000 | 0.84059 | 6.28845 |
| **UBQLNL** | 0.06126 | 0.00753 | 0.01322 | 0.10398 | 0.10944 | 0.01141 | 0.13143 |
| **UBR1** | 9.50089 | 0.37907 | 7.14707 | 8.28291 | 1.99964 | 5.59661 | 12.92153 |
| **UBR2** | 10.93226 | 6.59587 | 9.47808 | 7.79582 | 1.60953 | 8.79930 | 7.94289 |
| **UBR3** | 3.99451 | 0.27183 | 0.97126 | 2.68719 | 0.51139 | 3.73895 | 10.55286 |
| **UBR4** | 8.78870 | 17.00463 | 7.57289 | 21.54705 | 1.74166 | 11.75831 | 27.05579 |
| **UBR5** | 6.30613 | 0.85189 | 11.24334 | 7.96298 | 3.66043 | 6.91260 | 21.44716 |
| **UBR7** | 14.88399 | 5.89100 | 6.66303 | 7.69131 | 12.57247 | 3.41563 | 2.41453 |
| **UBTD1** | 0.00000 | 2.37842 | 4.86087 | 0.76253 | 1.62985 | 7.65337 | 0.57489 |
| **UBTD2** | 6.64696 | 0.42786 | 3.33758 | 2.01651 | 2.00201 | 3.17300 | 10.94566 |
| **UBTF** | 5.38631 | 13.18137 | 8.51177 | 7.37341 | 2.73041 | 9.92918 | 16.76796 |
| **UBTFL1** | 0.02450 | 0.01209 | 0.00000 | 0.01501 | 1.05610 | 0.01153 | 0.02030 |
| **UBXN1** | 20.75685 | 28.76012 | 19.26224 | 27.66891 | 17.78324 | 13.56809 | 10.53080 |
| **UBXN10** | 0.02679 | 0.00944 | 0.01476 | 0.01478 | 0.30453 | 0.07402 | 0.02219 |
| **UBXN11** | 3.43592 | 5.96229 | 2.69887 | 3.40609 | 14.84673 | 0.84587 | 0.01942 |
| **UBXN2A** | 6.11616 | 2.62322 | 5.86999 | 4.40733 | 6.31093 | 3.13316 | 2.74957 |
| **UBXN2B** | 10.42465 | 2.72495 | 2.53428 | 6.08530 | 0.53105 | 3.54466 | 0.62296 |

| | | | | | | | |
|---|---|---|---|---|---|---|---|
| **UBXN4** | 19.68044 | 14.53852 | 9.73703 | 20.76170 | 7.39069 | 14.02607 | 17.19006 |
| **UBXN6** | 0.00000 | 6.22338 | 2.78969 | 1.16293 | 1.37556 | 3.25529 | 8.18595 |
| **UBXN7** | 9.32182 | 0.79677 | 13.35769 | 7.56918 | 9.68771 | 3.78709 | 6.09037 |
| **UBXN8** | 9.24907 | 0.00000 | 12.18792 | 4.04098 | 1.81355 | 2.44961 | 5.04225 |
| **UCA1** | 2.65449 | 0.00000 | 0.00000 | 7.69734 | 0.00000 | 0.00000 | 1.40616 |
| **UCHL1** | 1.35491 | 0.00000 | 0.87985 | 14.23455 | 9.70788 | 0.69612 | 0.35549 |
| **UCHL3** | 23.33935 | 9.01243 | 5.87027 | 12.43577 | 1.77064 | 8.43116 | 6.66935 |
| **UCHL5** | 24.07882 | 6.19543 | 4.22383 | 7.46155 | 0.36393 | 5.53026 | 4.89835 |
| **UCK1** | 2.70040 | 12.66968 | 3.20740 | 14.26858 | 2.64108 | 5.40241 | 1.74349 |
| **UCK2** | 8.07295 | 9.40665 | 7.68936 | 10.38921 | 0.20534 | 1.98882 | 14.04041 |
| **UCKL1** | 9.34742 | 9.72715 | 0.75778 | 7.89726 | 1.56619 | 6.39188 | 4.39088 |
| **UCKL1-AS1** | 0.00000 | 0.20277 | 0.06918 | 0.18064 | 0.04609 | 0.38900 | 0.95236 |
| **UCMA** | 0.01845 | 0.00000 | 0.66632 | 0.01915 | 0.60592 | 0.00550 | 0.02188 |
| **UCN** | 0.01395 | 0.00484 | 0.03183 | 0.06675 | 0.02007 | 0.00589 | 0.16324 |
| **UCN2** | 0.05954 | 0.00000 | 0.05735 | 0.12974 | 0.00448 | 0.00000 | 0.23091 |
| **UCP1** | 0.12975 | 0.02404 | 0.04985 | 0.01895 | 0.01568 | 0.00938 | 0.00000 |
| **UCP2** | 4.94141 | 7.38200 | 3.48359 | 7.10255 | 7.69664 | 8.95540 | 0.00000 |
| **UCP3** | 0.12181 | 0.01635 | 0.06400 | 0.08069 | 0.00509 | 0.07039 | 0.10552 |
| **UEVLD** | 3.43343 | 0.52401 | 2.42639 | 1.28947 | 0.63997 | 0.26752 | 0.41579 |
| **UFC1** | 36.02053 | 6.95527 | 17.09964 | 42.33130 | 2.16248 | 12.03733 | 11.53065 |
| **UFD1L** | 23.22437 | 17.67256 | 6.42567 | 28.57454 | 2.06986 | 9.65783 | 0.97113 |
| **UFL1** | 7.91191 | 1.15936 | 2.83615 | 5.28288 | 1.26989 | 3.05870 | 5.96923 |
| **UFM1** | 17.42201 | 10.88331 | 13.80947 | 9.38540 | 2.40473 | 11.27106 | 4.40781 |
| **UFSP1** | 0.00000 | 0.67291 | 0.34354 | 1.08401 | 0.16624 | 0.31188 | 0.33310 |
| **UFSP2** | 8.49961 | 1.95556 | 4.84995 | 4.38891 | 1.02039 | 2.47149 | 0.33523 |
| **UG0898H09** | 0.02432 | 0.01920 | 0.01292 | 0.02037 | 0.00793 | 0.01418 | 0.01661 |
| **UGCG** | 11.17044 | 13.74354 | 0.13567 | 0.00000 | 0.42492 | 8.17651 | 13.42716 |
| **UGDH** | 12.73897 | 0.00000 | 2.39432 | 7.78629 | 4.76460 | 4.85568 | 0.00000 |
| **UGDH-AS1** | 0.04476 | 0.00000 | 0.03367 | 0.09779 | 0.01125 | 0.01735 | 0.13801 |
| **UGGT1** | 16.91702 | 2.29839 | 6.33594 | 9.35575 | 3.41915 | 6.88234 | 11.88567 |
| **UGGT2** | 4.24261 | 1.40136 | 1.20157 | 1.08872 | 1.05259 | 1.21302 | 4.31709 |
| **UGP2** | 32.08322 | 10.23481 | 10.42945 | 16.33125 | 2.53138 | 25.91663 | 9.80529 |
| **UGT2A1** | 0.02210 | 0.00575 | 0.02877 | 0.02799 | 0.06303 | 0.01847 | 0.01944 |
| **UGT2B11** | 0.02939 | 0.01750 | 0.01364 | 0.06090 | 0.00810 | 0.01210 | 0.00452 |
| **UGT2B15** | 0.03227 | 0.01570 | 0.01383 | 0.02312 | 0.00803 | 0.01751 | 0.00836 |

| | | | | | | | |
|---|---|---|---|---|---|---|---|
| **UGT2B28** | 0.03120 | 0.01531 | 0.01418 | 0.18372 | 0.00819 | 0.01170 | 0.00000 |
| **UGT2B4** | 0.02617 | 0.01879 | 0.01336 | 0.06521 | 0.00833 | 0.01393 | 0.00144 |
| **UGT2B7** | 0.02623 | 0.01493 | 0.01278 | 0.04086 | 0.00907 | 0.01177 | 0.02690 |
| **UGT3A1** | 0.16182 | 0.00214 | 0.00968 | 0.41113 | 0.12247 | 0.00857 | 0.00000 |
| **UGT3A2** | 0.18454 | 0.00743 | 0.01336 | 0.09554 | 0.00735 | 0.04641 | 0.00000 |
| **UGT8** | 1.01000 | 0.25331 | 1.78561 | 0.00000 | 0.75168 | 0.82309 | 0.00819 |
| **UHMK1** | 14.73906 | 5.97577 | 10.76347 | 4.79785 | 8.82555 | 13.25512 | 10.60442 |
| **UHRF1** | 6.26031 | 3.66077 | 0.00000 | 5.04361 | 5.39897 | 2.00997 | 0.00000 |
| **UHRF1BP1** | 7.22285 | 1.49338 | 4.86927 | 2.98624 | 1.22909 | 2.84062 | 9.84363 |
| **UHRF1BP1L** | 5.02573 | 0.78351 | 3.08383 | 2.32949 | 4.43902 | 5.14910 | 0.58118 |
| **UHRF2** | 3.11501 | 0.90530 | 2.17549 | 2.62929 | 2.78985 | 1.41772 | 9.82712 |
| **UIMC1** | 7.86080 | 1.26923 | 5.63636 | 6.09055 | 3.98266 | 5.04668 | 2.12521 |
| **ULBP1** | 0.30628 | 0.00000 | 0.00233 | 0.04442 | 0.00594 | 0.06016 | 0.50551 |
| **ULBP2** | 0.04553 | 0.02091 | 0.30715 | 0.03145 | 0.08612 | 0.04350 | 0.00000 |
| **ULBP3** | 0.12585 | 0.01598 | 0.01601 | 0.02208 | 0.04515 | 0.01308 | 0.00000 |
| **ULK1** | 4.14770 | 14.80046 | 5.50303 | 9.69461 | 1.09033 | 8.18421 | 29.32182 |
| **ULK2** | 0.35414 | 0.67839 | 15.80841 | 0.00001 | 7.83093 | 1.54869 | 0.57797 |
| **ULK3** | 1.59145 | 1.33684 | 0.75779 | 4.05023 | 0.00000 | 2.75302 | 4.94787 |
| **ULK4** | 1.94793 | 0.73025 | 3.20715 | 2.11144 | 2.94226 | 3.70795 | 2.01032 |
| **ULK4P3.uid-189** | 0.01685 | 0.03578 | 0.02058 | 0.01785 | 0.01777 | 0.00949 | 0.02792 |
| **UMOD** | 0.01807 | 0.08571 | 0.00994 | 0.03112 | 0.00762 | 0.01876 | 0.01010 |
| **UMODL1** | 0.02353 | 0.00267 | 0.00079 | 0.01684 | 0.31431 | 0.24918 | 0.04829 |
| **UMPS** | 40.93181 | 27.07651 | 16.52889 | 17.29681 | 3.02724 | 14.36609 | 0.00000 |
| **UNC119** | 0.09543 | 2.43840 | 0.43068 | 1.39730 | 2.04297 | 5.76846 | 3.53132 |
| **UNC119B** | 2.07027 | 2.65909 | 1.86857 | 1.33284 | 0.00000 | 1.79218 | 2.09193 |
| **UNC13A** | 0.00000 | 0.00000 | 0.16830 | 0.02856 | 0.19401 | 0.10882 | 0.21756 |
| **UNC13B** | 7.20001 | 3.36062 | 2.22695 | 1.69426 | 0.33950 | 5.11075 | 6.63008 |
| **UNC13C** | 0.02109 | 0.06391 | 0.18109 | 0.00000 | 11.16816 | 0.09609 | 0.13172 |
| **UNC13D** | 0.85103 | 0.83028 | 0.00000 | 1.19539 | 0.94105 | 1.52386 | 0.42914 |
| **UNC45A** | 8.10177 | 4.33595 | 0.58410 | 14.58866 | 0.06145 | 5.57918 | 2.26217 |
| **UNC45B** | 0.04620 | 0.03797 | 0.11881 | 0.01535 | 0.01328 | 0.01838 | 0.01252 |
| **UNC50** | 17.29146 | 7.29914 | 6.20291 | 10.48249 | 0.50781 | 5.40292 | 0.00000 |
| **UNC5A** | 0.04352 | 0.07811 | 0.03373 | 0.01502 | 0.00722 | 0.00737 | 0.04360 |
| **UNC5B** | 0.00000 | 0.66186 | 0.00000 | 0.25201 | 0.02026 | 13.19943 | 1.43952 |
| **UNC5B-AS1** | 0.00000 | 0.07130 | 0.00153 | 0.03172 | 0.12795 | 0.64593 | 0.01650 |

| | | | | | | | |
|---|---|---|---|---|---|---|---|
| **UNC5C** | 0.13403 | 0.12206 | 0.00000 | 0.42967 | 3.00194 | 0.00000 | 0.00000 |
| **UNC5CL** | 0.00729 | 0.24308 | 0.11804 | 0.22881 | 0.00336 | 0.04812 | 0.39547 |
| **UNC5D** | 0.01545 | 0.02303 | 0.03586 | 0.01748 | 0.32616 | 0.21587 | 0.03627 |
| **UNC79** | 0.00000 | 0.05530 | 0.03769 | 0.13570 | 0.17077 | 0.12685 | 0.05559 |
| **UNC80** | 0.02464 | 0.01730 | 0.00000 | 0.02281 | 0.68123 | 0.02394 | 0.01723 |
| **UNC93A** | 0.10553 | 0.32441 | 0.00000 | 0.00276 | 2.07153 | 1.80838 | 0.00000 |
| **UNC93B1** | 0.00000 | 0.03305 | 0.00516 | 0.24102 | 0.00637 | 0.00171 | 0.82717 |
| **UNCX** | 0.02381 | 0.02063 | 0.01079 | 0.02009 | 0.03398 | 0.01286 | 0.01635 |
| **UNG** | 15.83446 | 11.22459 | 16.07428 | 16.35515 | 8.77548 | 14.67935 | 12.42696 |
| **UNK** | 5.42290 | 3.54058 | 6.08549 | 5.51155 | 1.80633 | 2.73520 | 0.17193 |
| **UNKL** | 2.96038 | 7.33362 | 2.23735 | 7.31005 | 3.78007 | 0.59190 | 8.08444 |
| **UNQ6494** | 0.00000 | 0.01983 | 0.00994 | 0.10666 | 0.00783 | 0.33072 | 0.02411 |
| **UOX** | 0.25790 | 0.01341 | 0.02423 | 0.00000 | 0.06574 | 0.05149 | 0.01010 |
| **UPB1** | 0.00459 | 0.15554 | 0.00000 | 0.01961 | 0.94603 | 0.00412 | 0.02625 |
| **UPF1** | 4.75868 | 10.28285 | 5.71957 | 5.92934 | 1.71235 | 5.91875 | 22.13152 |
| **UPF2** | 11.17950 | 6.46854 | 7.93525 | 11.98242 | 6.39063 | 5.63140 | 17.36340 |
| **UPF3A** | 1.64992 | 1.81466 | 0.61930 | 1.76271 | 0.72956 | 2.01045 | 5.13115 |
| **UPF3B** | 11.01788 | 4.56574 | 6.80580 | 10.44664 | 6.81113 | 9.00296 | 15.06208 |
| **UPK1A** | 0.01175 | 0.30201 | 0.00833 | 0.05670 | 0.01796 | 0.03298 | 0.36626 |
| **UPK1A-AS1** | 0.02042 | 0.03132 | 0.01727 | 0.02616 | 0.02908 | 0.01262 | 0.01158 |
| **UPK1B** | 0.02544 | 0.02052 | 0.01236 | 0.02916 | 0.00838 | 0.01755 | 0.01211 |
| **UPK2** | 0.37872 | 0.00000 | 0.00000 | 1.23219 | 0.02122 | 0.00000 | 1.40131 |
| **UPK3B** | 0.02124 | 0.01638 | 0.01953 | 0.01992 | 0.00740 | 0.01769 | 0.01974 |
| **UPP1** | 17.62200 | 51.10458 | 13.70891 | 25.50739 | 0.19483 | 41.57291 | 24.67045 |
| **UPP2** | 0.01641 | 0.02589 | 0.00725 | 0.04001 | 0.03931 | 0.04043 | 0.02222 |
| **UPRT** | 4.44477 | 0.94565 | 0.42115 | 0.00480 | 0.36685 | 1.66866 | 0.00000 |
| **UQCC1** | 17.41087 | 13.17703 | 5.68238 | 16.86492 | 0.40430 | 7.11983 | 0.00000 |
| **UQCC2** | 7.14945 | 12.33139 | 4.59361 | 2.12638 | 0.60400 | 3.99153 | 0.75720 |
| **UQCR10** | 43.84678 | 46.50695 | 16.50759 | 34.55377 | 7.13424 | 27.51812 | 22.43929 |
| **UQCR11** | 47.34632 | 36.05683 | 10.10834 | 37.91576 | 0.43483 | 27.90014 | 25.92035 |
| **UQCRB** | 30.14974 | 4.45095 | 17.77653 | 16.27366 | 0.98662 | 17.08058 | 25.73870 |
| **UQCRBP1** | 2.74731 | 0.79647 | 1.25094 | 1.30964 | 0.04064 | 1.35808 | 1.63425 |
| **UQCRC1** | 63.07155 | 50.39770 | 13.09420 | 42.48086 | 3.05129 | 30.43087 | 8.18251 |
| **UQCRC2** | 47.86928 | 13.21378 | 11.22080 | 22.98186 | 7.64766 | 17.77357 | 8.25246 |
| **UQCRFS1** | 31.70073 | 20.82036 | 13.72406 | 22.36849 | 3.18935 | 19.43163 | 12.97386 |

| | | | | | | | |
|---|---|---|---|---|---|---|---|
| **UQCRH** | 50.80532 | 30.46417 | 26.00657 | 37.08212 | 6.69972 | 41.28929 | 36.20850 |
| **UQCRHL** | 25.87863 | 18.08717 | 10.90330 | 19.32382 | 1.99473 | 22.46744 | 8.40818 |
| **UQCRQ** | 49.81964 | 31.21963 | 15.09458 | 35.44490 | 8.26932 | 22.00759 | 33.78312 |
| **URAD** | 0.41042 | 0.15497 | 0.00000 | 0.08771 | 0.00411 | 0.00000 | 0.70949 |
| **URAHP** | 0.04671 | 0.06596 | 0.04326 | 0.03238 | 0.04654 | 0.03692 | 0.01676 |
| **URB1** | 8.97017 | 3.97328 | 6.21538 | 3.41639 | 1.99015 | 1.97145 | 6.59506 |
| **URB2** | 11.15217 | 4.06575 | 6.07950 | 8.36444 | 1.25508 | 3.27046 | 7.43990 |
| **URGCP** | 0.17657 | 2.15618 | 6.95121 | 0.75950 | 0.67103 | 2.76294 | 1.75910 |
| **URI1** | 4.22586 | 0.00105 | 6.42106 | 3.45726 | 0.52169 | 3.03717 | 16.34607 |
| **URM1** | 6.77730 | 14.11546 | 9.08102 | 15.65287 | 0.64120 | 6.32077 | 13.59526 |
| **UROC1** | 0.02360 | 0.07631 | 0.00900 | 0.01996 | 0.00808 | 0.00970 | 0.00240 |
| **UROD** | 22.36300 | 15.42270 | 7.95024 | 18.99866 | 8.12050 | 10.45561 | 0.00000 |
| **UROS** | 1.50072 | 2.04280 | 0.52917 | 0.28791 | 0.02091 | 1.55060 | 5.12654 |
| **USB1** | 7.76346 | 2.62769 | 5.09265 | 11.37344 | 3.11424 | 2.25380 | 1.26850 |
| **USE1** | 4.34019 | 3.36637 | 1.84726 | 5.01150 | 0.81450 | 1.49551 | 0.67114 |
| **USF1** | 6.06133 | 8.72603 | 3.72142 | 8.62680 | 3.63777 | 5.87086 | 1.69794 |
| **USF2** | 0.00000 | 12.84165 | 1.67035 | 4.72558 | 0.52796 | 4.94205 | 17.28162 |
| **USH1C** | 0.19585 | 0.00000 | 0.36634 | 0.95453 | 0.00618 | 0.00000 | 0.31313 |
| **USH1G** | 0.02653 | 0.02568 | 0.02111 | 0.02093 | 0.00915 | 0.01547 | 0.02757 |
| **USH2A** | 0.09849 | 0.00000 | 0.01824 | 0.00000 | 0.79733 | 0.07804 | 0.18151 |
| **USHBP1** | 0.23928 | 0.06574 | 0.03599 | 0.93474 | 0.11678 | 0.12678 | 0.14778 |
| **USMG5** | 43.05707 | 17.13484 | 18.01953 | 26.98093 | 6.11829 | 19.61163 | 30.45914 |
| **USO1** | 12.92734 | 2.48038 | 5.71774 | 6.43997 | 0.80806 | 7.03150 | 9.84313 |
| **USP1** | 22.65482 | 4.61684 | 15.72620 | 9.33301 | 13.31980 | 14.22498 | 18.72934 |
| **USP10** | 34.45258 | 19.96920 | 15.48754 | 25.23300 | 6.68284 | 13.63809 | 4.93486 |
| **USP11** | 3.90017 | 5.91020 | 6.73636 | 2.29121 | 5.33477 | 3.69934 | 0.00000 |
| **USP12** | 0.69089 | 0.00000 | 2.16064 | 0.00000 | 0.23255 | 1.34145 | 2.02271 |
| **USP12-AS1** | 0.01549 | 0.01648 | 0.02191 | 0.01022 | 0.00906 | 0.07532 | 0.10838 |
| **USP12-AS2** | 0.02285 | 0.01559 | 0.01513 | 0.01931 | 0.03940 | 0.01561 | 0.01831 |
| **USP13** | 0.45600 | 0.67641 | 1.20360 | 0.19434 | 4.48100 | 3.28089 | 0.40385 |
| **USP14** | 34.46921 | 5.97276 | 13.29448 | 22.80573 | 2.00659 | 11.71153 | 14.28726 |
| **USP15** | 22.47983 | 4.04800 | 9.68174 | 11.05246 | 3.31188 | 4.19617 | 2.50333 |
| **USP16** | 20.26471 | 4.65057 | 18.96647 | 9.80601 | 7.87820 | 8.24460 | 15.04374 |
| **USP18** | 0.38369 | 0.09264 | 0.16152 | 0.12060 | 0.01856 | 0.07278 | 0.00000 |
| **USP19** | 1.97055 | 5.87513 | 3.40733 | 4.57406 | 0.71697 | 4.00193 | 9.19017 |

| | | | | | | | |
|---|---|---|---|---|---|---|---|
| **USP2** | 1.37478 | 4.67858 | 1.01314 | 0.92638 | 11.99131 | 1.62956 | 0.00000 |
| **USP2-AS1** | 0.35547 | 0.00000 | 2.52106 | 0.58464 | 1.42349 | 0.05759 | 0.16122 |
| **USP20** | 0.00000 | 2.59105 | 0.21790 | 1.36656 | 0.97789 | 0.44579 | 1.36600 |
| **USP21** | 0.55160 | 1.49786 | 0.74217 | 1.66253 | 0.63781 | 0.92594 | 2.16431 |
| **USP22** | 37.29097 | 33.70995 | 9.95658 | 23.56463 | 2.21370 | 19.70338 | 18.58401 |
| **USP24** | 9.21988 | 3.87435 | 3.25273 | 5.05842 | 3.64642 | 3.08052 | 13.65822 |
| **USP25** | 3.91898 | 0.00000 | 1.52873 | 2.36060 | 0.57657 | 1.47741 | 10.83416 |
| **USP26** | 0.03298 | 0.00000 | 0.06788 | 0.02048 | 0.08948 | 0.01172 | 0.01529 |
| **USP27X** | 0.00000 | 0.11834 | 0.07708 | 0.24248 | 0.04720 | 0.02351 | 1.20036 |
| **USP27X-AS1** | 0.04289 | 0.21395 | 0.09594 | 0.09302 | 0.09426 | 0.01597 | 0.00000 |
| **USP28** | 18.15224 | 5.78853 | 7.81603 | 3.54906 | 0.21804 | 29.51766 | 5.38010 |
| **USP29** | 0.02718 | 0.00000 | 0.03721 | 0.01804 | 1.74755 | 0.01490 | 0.01647 |
| **USP3** | 13.18924 | 0.02780 | 1.04158 | 10.42690 | 2.55307 | 0.00000 | 6.56261 |
| **USP3-AS1** | 0.66703 | 0.00000 | 0.11669 | 0.18683 | 0.92431 | 0.00000 | 0.62753 |
| **USP30** | 3.25272 | 2.54219 | 3.24683 | 2.77631 | 1.00976 | 0.95348 | 0.23030 |
| **USP30-AS1** | 0.02745 | 0.03045 | 0.02362 | 0.03944 | 0.00965 | 0.02348 | 0.01529 |
| **USP31** | 3.24420 | 2.73969 | 1.72242 | 0.87064 | 0.44442 | 2.13089 | 5.88812 |
| **USP32** | 4.41544 | 0.75710 | 1.87637 | 2.82499 | 1.18226 | 3.70964 | 4.58476 |
| **USP32P1** | 0.08371 | 0.00000 | 0.67617 | 0.02694 | 0.09966 | 0.08246 | 0.21990 |
| **USP32P2** | 0.04788 | 0.01658 | 0.09543 | 0.03589 | 0.06245 | 0.06324 | 0.02861 |
| **USP33** | 10.36714 | 4.11735 | 4.79887 | 10.08879 | 1.58819 | 5.23964 | 3.82326 |
| **USP34** | 9.97123 | 1.15733 | 8.79913 | 7.40875 | 12.37776 | 4.71471 | 25.00006 |
| **USP35** | 0.00000 | 0.01353 | 0.00000 | 0.50473 | 0.74023 | 0.56476 | 1.03373 |
| **USP36** | 13.42054 | 11.87577 | 6.68024 | 10.82520 | 4.87756 | 6.54855 | 16.73878 |
| **USP37** | 10.24468 | 1.42821 | 6.06338 | 4.79520 | 4.61872 | 3.50212 | 4.83301 |
| **USP38** | 6.43010 | 3.09522 | 5.18148 | 4.99705 | 4.65226 | 3.28289 | 3.46594 |
| **USP39** | 26.78128 | 15.44395 | 10.18738 | 18.10341 | 4.18257 | 13.02817 | 5.99852 |
| **USP4** | 8.63225 | 5.68819 | 4.03690 | 1.45799 | 1.37358 | 0.00000 | 3.31337 |
| **USP40** | 3.20088 | 3.07703 | 2.83940 | 1.64454 | 1.31158 | 0.46462 | 2.23268 |
| **USP42** | 1.91474 | 0.70379 | 1.13851 | 4.72471 | 1.83296 | 2.10676 | 6.56232 |
| **USP43** | 1.40316 | 5.64041 | 3.57166 | 1.03014 | 0.28963 | 0.83303 | 5.89269 |
| **USP44** | 0.10932 | 0.00000 | 0.15860 | 0.34570 | 1.22109 | 0.38295 | 0.00000 |
| **USP45** | 3.66094 | 0.00000 | 2.33132 | 1.13156 | 0.69230 | 1.11060 | 2.20500 |
| **USP46** | 3.68665 | 0.41482 | 3.32457 | 2.46763 | 0.42477 | 2.89871 | 5.03131 |
| **USP47** | 13.96615 | 4.17071 | 5.06618 | 12.98632 | 4.50216 | 3.75947 | 12.26452 |

| | | | | | | | |
|---|---|---|---|---|---|---|---|
| **USP48** | 4.36555 | 4.38712 | 6.42042 | 3.30739 | 1.44286 | 8.84434 | 21.23835 |
| **USP49** | 0.32875 | 0.00000 | 1.73870 | 3.03132 | 0.37116 | 0.29462 | 6.00982 |
| **USP5** | 5.91169 | 13.48292 | 1.95773 | 15.78930 | 2.16927 | 6.63056 | 7.59021 |
| **USP50** | 0.02547 | 0.02962 | 0.02277 | 0.03025 | 0.15527 | 0.02580 | 0.02048 |
| **USP51** | 0.03612 | 0.01002 | 0.29645 | 0.01143 | 0.03779 | 0.23006 | 0.01899 |
| **USP53** | 6.41446 | 0.21951 | 0.60063 | 5.54011 | 1.05925 | 2.49903 | 10.33843 |
| **USP54** | 2.61120 | 0.00000 | 4.29058 | 2.69644 | 3.80141 | 1.89808 | 3.03673 |
| **USP6** | 0.06261 | 0.33999 | 0.35012 | 0.04319 | 0.16132 | 0.35202 | 0.28771 |
| **USP6NL** | 1.74508 | 0.06104 | 1.64025 | 1.23055 | 0.52987 | 0.42527 | 3.86353 |
| **USP7** | 17.86175 | 16.81760 | 37.70778 | 7.63467 | 16.51930 | 13.11656 | 35.38532 |
| **USP8** | 16.68989 | 0.73395 | 7.62987 | 9.95420 | 3.68208 | 6.86613 | 14.20513 |
| **USP9X** | 13.93489 | 8.62011 | 12.36188 | 2.87234 | 7.02601 | 13.59537 | 26.97929 |
| **USP9Y** | 0.00139 | 0.01890 | 0.01740 | 0.02217 | 0.00657 | 0.08074 | 0.09557 |
| **USPL1** | 6.86302 | 0.00000 | 4.93606 | 3.23761 | 5.83397 | 1.87345 | 3.41810 |
| **UST** | 1.52494 | 0.22564 | 0.00000 | 0.00000 | 0.25622 | 1.41528 | 2.91326 |
| **UTF1** | 0.00000 | 0.01080 | 0.00442 | 0.01987 | 0.01510 | 5.87639 | 0.07460 |
| **UTP11L** | 29.17486 | 12.37057 | 13.69226 | 18.52067 | 11.90641 | 9.16751 | 1.49498 |
| **UTP14A** | 24.18422 | 13.70862 | 14.69677 | 6.17605 | 7.62676 | 7.22573 | 7.32129 |
| **UTP14C** | 3.76483 | 1.50012 | 2.21175 | 2.63515 | 0.19754 | 2.34447 | 5.18453 |
| **UTP15** | 19.97867 | 0.25043 | 14.64178 | 8.13118 | 2.31017 | 4.99480 | 2.34298 |
| **UTP18** | 23.29633 | 8.56978 | 15.97743 | 17.69075 | 4.18778 | 9.21881 | 4.69559 |
| **UTP20** | 12.94478 | 3.03618 | 9.56120 | 9.82661 | 2.20918 | 6.63537 | 15.65527 |
| **UTP23** | 11.19584 | 3.65128 | 9.11594 | 7.46281 | 4.88945 | 4.84456 | 4.10132 |
| **UTP3** | 4.34718 | 0.18680 | 9.85756 | 2.71311 | 3.20686 | 3.34382 | 9.88770 |
| **UTP6** | 38.66393 | 17.04364 | 13.03951 | 23.72413 | 11.30500 | 8.45635 | 2.29170 |
| **UTRN** | 12.73709 | 6.20237 | 3.81612 | 5.63206 | 0.18799 | 0.81149 | 27.97312 |
| **UTS2** | 0.02562 | 0.00780 | 0.13604 | 0.02034 | 0.34672 | 0.02207 | 0.01164 |
| **UTS2B** | 0.02000 | 0.00002 | 0.04492 | 0.01728 | 1.06963 | 0.09442 | 0.01430 |
| **UTS2R** | 0.02892 | 0.02512 | 0.01716 | 0.01020 | 0.00692 | 0.00750 | 0.04462 |
| **UTY** | 0.02809 | 0.02424 | 0.02548 | 0.03975 | 0.00680 | 0.01415 | 0.01599 |
| **UVRAG** | 1.39676 | 1.33335 | 2.99647 | 0.50620 | 0.05914 | 2.78716 | 4.28248 |
| **UVSSA** | 0.49558 | 0.65364 | 0.38002 | 0.00000 | 0.14076 | 0.10286 | 1.75585 |
| **UXS1** | 13.82295 | 6.69237 | 3.72803 | 7.47281 | 4.22470 | 4.32599 | 3.49611 |
| **UXT** | 22.05299 | 14.38983 | 18.29940 | 16.22102 | 9.24114 | 14.18704 | 18.92095 |
| **UXT-AS1** | 0.03863 | 0.01920 | 0.03216 | 0.03606 | 0.00924 | 0.01084 | 0.00736 |

| | | | | | | | |
|---|---|---|---|---|---|---|---|
| **VAC14** | 2.93240 | 3.76366 | 1.21313 | 4.09628 | 0.52600 | 4.75852 | 10.72601 |
| **VAC14-AS1** | 0.11582 | 0.00000 | 0.12235 | 0.07952 | 0.20619 | 0.12523 | 1.10996 |
| **VAMP1** | 1.88223 | 2.62960 | 2.48230 | 0.52479 | 6.86215 | 1.53998 | 0.00000 |
| **VAMP2** | 0.76545 | 1.38937 | 2.16636 | 5.98624 | 2.46289 | 1.25565 | 2.01835 |
| **VAMP3** | 15.10132 | 6.05225 | 13.22147 | 11.35638 | 2.30852 | 7.62270 | 9.87856 |
| **VAMP4** | 1.45211 | 0.22433 | 2.06819 | 0.09488 | 0.31065 | 0.54935 | 0.43128 |
| **VAMP5** | 0.51046 | 1.24704 | 0.35015 | 1.66901 | 0.05165 | 0.10934 | 0.46386 |
| **VAMP8** | 27.70109 | 36.10275 | 10.23182 | 26.94189 | 0.12937 | 3.79401 | 14.27126 |
| **VANGL1** | 2.01357 | 3.65262 | 1.80832 | 1.33709 | 0.05240 | 3.35886 | 4.01551 |
| **VANGL2** | 0.24800 | 0.00000 | 0.03661 | 0.00000 | 0.00720 | 1.61681 | 0.86332 |
| **VAPA** | 51.66898 | 25.30434 | 21.69561 | 26.49585 | 8.25890 | 14.93263 | 9.76773 |
| **VAPB** | 7.39964 | 10.36593 | 7.58905 | 6.86469 | 0.79647 | 6.93885 | 11.31712 |
| **VARS** | 10.04103 | 13.60987 | 5.12007 | 10.22087 | 0.01776 | 8.59522 | 26.42270 |
| **VARS2** | 0.12922 | 0.90226 | 1.30989 | 0.74816 | 0.02140 | 1.85369 | 5.53421 |
| **VASH1** | 0.00000 | 1.09893 | 0.97743 | 0.00000 | 0.41514 | 1.49751 | 0.23444 |
| **VASH2** | 0.15334 | 0.00000 | 0.01326 | 0.14924 | 0.39884 | 5.31422 | 0.00000 |
| **VASN** | 0.00000 | 0.00004 | 0.11401 | 0.69968 | 0.06738 | 0.36314 | 0.66477 |
| **VASP** | 0.49506 | 5.12176 | 4.77218 | 5.52393 | 0.80620 | 1.66294 | 22.36969 |
| **VAT1** | 17.15021 | 9.21136 | 8.22491 | 34.32105 | 6.51893 | 18.74698 | 26.15979 |
| **VAT1L** | 0.29827 | 0.17325 | 0.00000 | 0.00000 | 0.03261 | 0.00000 | 0.39096 |
| **VAV1** | 0.38441 | 0.28132 | 0.00051 | 1.57779 | 0.02199 | 0.21849 | 0.10890 |
| **VAV2** | 1.01510 | 14.35382 | 3.02832 | 2.31839 | 0.00361 | 9.95541 | 8.65112 |
| **VAV3** | 0.13188 | 0.00000 | 0.10395 | 0.52045 | 2.89634 | 0.00000 | 1.05462 |
| **VAV3-AS1** | 0.02197 | 0.02226 | 0.00797 | 0.02265 | 0.17967 | 0.01257 | 0.01722 |
| **VAX1** | 0.02944 | 0.00000 | 0.04548 | 0.03949 | 0.20082 | 0.01130 | 0.02724 |
| **VAX2** | 0.01008 | 0.07101 | 0.02910 | 0.02070 | 0.00532 | 0.00781 | 0.08057 |
| **VBP1** | 40.04388 | 4.37757 | 21.22950 | 18.66350 | 3.62434 | 16.71283 | 4.67864 |
| **VCAN** | 0.00000 | 0.20788 | 2.59338 | 0.27177 | 0.00000 | 22.25510 | 0.00812 |
| **VCL** | 12.04813 | 8.30383 | 2.19608 | 5.16664 | 0.82695 | 14.28258 | 11.54595 |
| **VCP** | 59.38762 | 50.45747 | 17.69930 | 59.25047 | 11.85998 | 28.49085 | 11.75558 |
| **VCPIP1** | 6.72534 | 0.00000 | 4.57073 | 3.07678 | 4.44886 | 2.67364 | 5.55736 |
| **VCPKMT** | 4.74348 | 0.94764 | 2.46648 | 6.36961 | 2.69203 | 2.36914 | 0.74271 |
| **VCX** | 0.21394 | 9.20828 | 0.00000 | 0.13804 | 0.70047 | 1.52659 | 0.00000 |
| **VCX2** | 0.37338 | 2.00507 | 0.00000 | 0.22796 | 0.72131 | 1.47218 | 0.00000 |
| **VCX3A** | 0.25807 | 1.47245 | 0.00000 | 0.41782 | 0.15425 | 1.80621 | 0.62247 |

| | | | | | | | |
|---|---|---|---|---|---|---|---|
| **VCX3B** | 0.15367 | 2.07670 | 0.05057 | 0.77171 | 0.53548 | 0.38229 | 0.00000 |
| **VDAC1** | 38.67095 | 14.87633 | 7.31726 | 20.47848 | 2.13296 | 16.59720 | 20.25414 |
| **VDAC2** | 28.36836 | 16.94749 | 11.58657 | 14.33931 | 3.05095 | 12.74412 | 10.48973 |
| **VDAC3** | 45.81840 | 22.34127 | 11.88237 | 23.44328 | 1.82883 | 19.51453 | 10.55756 |
| **VDR** | 0.03490 | 0.08536 | 0.18817 | 0.00340 | 0.15203 | 0.00000 | 0.06671 |
| **VEGFA** | 1.06349 | 0.00000 | 0.00000 | 4.78671 | 0.01932 | 0.00000 | 6.33688 |
| **VEGFB** | 0.01317 | 0.15283 | 0.07274 | 0.92675 | 0.01214 | 0.44672 | 1.29879 |
| **VEGFC** | 0.02163 | 0.01043 | 0.11465 | 0.03937 | 0.28173 | 0.01253 | 0.04548 |
| **VENTX** | 0.05555 | 0.10721 | 0.00220 | 0.02702 | 0.00823 | 14.29665 | 0.00000 |
| **VENTXP1** | 0.02103 | 0.01667 | 0.01263 | 0.02199 | 0.01112 | 0.01785 | 0.03155 |
| **VENTXP7** | 0.02964 | 0.00764 | 0.02252 | 0.01464 | 0.01068 | 0.12184 | 0.05680 |
| **VEPH1** | 1.42722 | 0.14171 | 0.00000 | 0.00000 | 0.79596 | 0.00000 | 0.09564 |
| **VEZF1** | 3.34750 | 0.00000 | 3.03708 | 5.72972 | 1.25468 | 3.50386 | 7.54958 |
| **VEZT** | 7.15866 | 0.57745 | 3.89341 | 10.25631 | 1.11270 | 6.76067 | 0.23284 |
| **VGF** | 0.00000 | 0.24603 | 0.31591 | 0.15699 | 0.01516 | 0.45218 | 0.88278 |
| **VGLL1** | 7.20270 | 0.00000 | 0.00000 | 27.60062 | 0.00000 | 0.00000 | 5.06744 |
| **VGLL3** | 0.01791 | 0.01405 | 0.01777 | 0.13883 | 0.00896 | 0.00992 | 0.02648 |
| **VGLL4** | 9.81767 | 15.82850 | 4.31651 | 9.10477 | 0.81911 | 0.00000 | 2.71277 |
| **VHL** | 2.09180 | 1.50806 | 1.72499 | 4.76641 | 1.47948 | 5.02752 | 18.76329 |
| **VHLL** | 0.02684 | 0.01713 | 0.01358 | 0.02784 | 0.00797 | 0.01307 | 0.01121 |
| **VIL1** | 0.27997 | 0.03746 | 0.00000 | 0.03728 | 0.53181 | 1.48694 | 0.00000 |
| **VILL** | 0.11145 | 0.27213 | 0.27312 | 0.22571 | 0.00908 | 0.05992 | 0.16018 |
| **VIM** | 0.11677 | 0.29685 | 0.08784 | 0.00817 | 0.14213 | 0.16257 | 0.00022 |
| **VIM-AS1** | 0.03584 | 0.02420 | 0.01399 | 0.02912 | 0.00992 | 0.02092 | 0.02891 |
| **VIMP** | 17.61979 | 17.80999 | 8.26578 | 18.97996 | 7.13002 | 9.44177 | 6.82705 |
| **VIP** | 0.03657 | 0.01546 | 0.01498 | 0.03927 | 0.01065 | 0.01254 | 0.00800 |
| **VIPAS39** | 3.24285 | 4.99371 | 2.87735 | 5.31870 | 1.05725 | 2.05758 | 1.87619 |
| **VIPR1** | 0.01024 | 0.02042 | 0.01221 | 0.08749 | 0.00795 | 0.10167 | 0.02062 |
| **VIPR2** | 0.02404 | 0.01945 | 0.01673 | 0.01907 | 0.09408 | 0.01727 | 0.04091 |
| **VIT** | 0.09652 | 0.13640 | 0.00007 | 0.02598 | 0.04341 | 0.07536 | 0.00000 |
| **VKORC1** | 22.94672 | 10.98110 | 23.40398 | 23.02850 | 7.80782 | 8.23634 | 10.71326 |
| **VKORC1L1** | 4.43452 | 3.68055 | 5.43724 | 4.85640 | 0.92199 | 2.25421 | 12.22062 |
| **VLDLR** | 0.40054 | 0.00000 | 0.10929 | 0.23036 | 1.33857 | 0.79915 | 1.68094 |
| **VLDLR-AS1** | 0.01503 | 0.00988 | 0.00000 | 0.06658 | 0.64034 | 0.04763 | 0.06077 |
| **VMA21** | 18.40584 | 9.16737 | 17.41111 | 13.28493 | 2.11774 | 3.76701 | 17.65424 |

| | | | | | | | |
|---|---|---|---|---|---|---|---|
| **VMAC** | 0.05125 | 0.59036 | 0.00000 | 0.27494 | 0.38073 | 0.15561 | 0.09410 |
| **VMO1** | 0.00000 | 2.32396 | 5.86382 | 0.11508 | 2.28173 | 0.00000 | 0.00000 |
| **VMP1** | 31.06418 | 5.53946 | 15.59109 | 24.23543 | 9.68612 | 12.50922 | 0.00000 |
| **VN1R1** | 0.05050 | 0.08456 | 0.04149 | 0.07708 | 0.05210 | 0.15010 | 0.24914 |
| **VN1R2** | 0.04140 | 0.03624 | 0.01807 | 0.01866 | 0.02199 | 0.01639 | 0.00000 |
| **VN1R4** | 0.03602 | 0.01361 | 0.02976 | 0.01892 | 0.01711 | 0.01507 | 0.00706 |
| **VN1R5** | 0.02878 | 0.01754 | 0.01340 | 0.04185 | 0.00771 | 0.02199 | 0.01931 |
| **1.00000** | 0.03056 | 0.01775 | 0.00564 | 0.03246 | 0.13291 | 0.01469 | 0.00576 |
| **2.00000** | 0.07306 | 0.00523 | 0.01551 | 0.03190 | 0.00740 | 0.01015 | 0.02878 |
| **3.00000** | 0.02404 | 0.00000 | 0.04590 | 0.01848 | 0.34992 | 0.01260 | 0.01711 |
| **VOPP1** | 0.69318 | 0.74758 | 0.27241 | 0.36652 | 0.03225 | 0.47218 | 3.19853 |
| **VPRBP** | 16.63775 | 0.56742 | 5.38121 | 13.77287 | 3.27397 | 6.64191 | 5.36664 |
| **VPREB1** | 0.02326 | 0.00309 | 0.05427 | 0.02196 | 0.01257 | 0.01225 | 0.02067 |
| **VPREB3** | 0.01523 | 0.01731 | 0.05932 | 0.07468 | 0.00641 | 0.00904 | 0.15806 |
| **VPS11** | 9.21696 | 7.67873 | 3.84922 | 9.27136 | 2.06998 | 5.39891 | 2.37531 |
| **VPS13A** | 3.87315 | 0.00000 | 1.39305 | 1.83804 | 1.68906 | 2.84827 | 11.42621 |
| **VPS13A-AS1** | 0.02050 | 0.01664 | 0.01341 | 0.03811 | 0.00779 | 0.02193 | 0.02396 |
| **VPS13B** | 1.97144 | 1.74858 | 2.15956 | 1.68351 | 2.70807 | 3.32841 | 6.81339 |
| **VPS13C** | 0.92393 | 0.00000 | 0.51097 | 0.00000 | 0.18607 | 1.88162 | 3.22281 |
| **VPS13D** | 3.03198 | 3.86940 | 3.23820 | 1.01584 | 1.46684 | 3.18095 | 9.67205 |
| **VPS16** | 4.51128 | 3.61029 | 2.13343 | 4.46745 | 0.19638 | 3.49103 | 2.67450 |
| **VPS18** | 2.25923 | 6.14234 | 3.31470 | 8.71651 | 0.14651 | 2.83837 | 15.34900 |
| **VPS25** | 27.96572 | 22.92645 | 14.34242 | 14.82335 | 2.20293 | 12.85035 | 5.27819 |
| **VPS26A** | 33.53355 | 3.06650 | 14.34097 | 17.96598 | 6.64037 | 9.07808 | 0.00000 |
| **VPS26B** | 2.26499 | 2.34265 | 1.54037 | 2.15368 | 0.00000 | 3.87914 | 8.02842 |
| **VPS28** | 9.64011 | 7.72915 | 4.30775 | 8.50574 | 0.84814 | 11.61396 | 11.36761 |
| **VPS29** | 34.79510 | 13.64051 | 9.68995 | 22.50378 | 1.69026 | 14.51678 | 6.10275 |
| **VPS33A** | 26.17228 | 16.07498 | 9.70430 | 13.69592 | 0.64451 | 7.60486 | 0.00000 |
| **VPS33B** | 4.18844 | 3.79147 | 1.43069 | 3.07145 | 0.45787 | 0.80032 | 0.00000 |
| **VPS35** | 29.38764 | 15.43223 | 8.67863 | 18.06895 | 3.26751 | 15.37757 | 7.73950 |
| **VPS36** | 15.57182 | 0.50799 | 3.94006 | 6.09097 | 0.71457 | 4.44303 | 5.97394 |
| **VPS37A** | 14.87372 | 4.45074 | 6.80265 | 12.63920 | 2.26976 | 4.49843 | 2.95192 |
| **VPS37B** | 0.91689 | 3.91940 | 1.59113 | 5.79382 | 2.05317 | 3.30264 | 8.91043 |
| **VPS37C** | 0.47715 | 4.07989 | 3.11434 | 4.08233 | 0.76500 | 3.27981 | 15.46640 |
| **VPS37D** | 0.03724 | 0.00768 | 0.03515 | 0.02710 | 0.00631 | 0.00941 | 0.06789 |

| | | | | | | | |
|---|---|---|---|---|---|---|---|
| **VPS39** | 6.21228 | 3.82646 | 4.98519 | 5.18695 | 0.33344 | 3.69563 | 2.50562 |
| **VPS41** | 9.42999 | 1.54124 | 1.91294 | 4.87537 | 0.44397 | 1.81514 | 3.67151 |
| **VPS45** | 7.95595 | 2.76879 | 4.09341 | 6.01558 | 1.44994 | 1.59460 | 0.00000 |
| **VPS4A** | 1.98896 | 14.68917 | 4.49807 | 2.87837 | 1.89356 | 7.19730 | 17.18202 |
| **VPS4B** | 21.82625 | 9.63752 | 10.28498 | 13.66898 | 10.50656 | 7.44962 | 7.57427 |
| **VPS51** | 0.33834 | 4.08439 | 0.86914 | 1.93242 | 1.86163 | 5.60835 | 12.79036 |
| **VPS52** | 9.15528 | 7.98130 | 4.37954 | 7.05367 | 1.80529 | 5.98896 | 0.96868 |
| **VPS53** | 7.69324 | 9.02149 | 2.09219 | 3.92626 | 5.20587 | 4.76329 | 3.72949 |
| **VPS54** | 3.10328 | 0.00000 | 2.53952 | 0.32739 | 2.81222 | 2.69838 | 4.21236 |
| **VPS72** | 17.23696 | 10.17499 | 8.34047 | 15.82976 | 3.18135 | 6.88495 | 7.00655 |
| **VPS8** | 2.95106 | 0.55532 | 1.28305 | 2.04378 | 0.92058 | 1.87647 | 2.38651 |
| **VPS9D1** | 0.15755 | 0.61429 | 0.16539 | 0.08824 | 0.08177 | 0.33133 | 0.43478 |
| **VPS9D1-AS1** | 0.00000 | 4.90662 | 0.77562 | 1.48616 | 0.05832 | 1.18402 | 4.40568 |
| **VRK1** | 26.39438 | 9.37236 | 15.78446 | 13.38593 | 2.56441 | 9.14234 | 4.82621 |
| **VRK2** | 12.87456 | 10.77715 | 4.36575 | 2.58526 | 0.92180 | 4.77481 | 0.00000 |
| **VRK3** | 10.30505 | 10.28343 | 6.75036 | 9.35422 | 5.04694 | 4.10128 | 0.00000 |
| **VRTN** | 0.00000 | 0.00000 | 7.78573 | 0.00596 | 0.59565 | 1.47060 | 0.15112 |
| **VSIG1** | 0.02306 | 0.00518 | 0.05858 | 0.01799 | 0.10843 | 0.01884 | 0.03042 |
| **VSIG10** | 16.30431 | 30.02266 | 10.18258 | 0.00000 | 3.55889 | 25.49685 | 2.16421 |
| **VSIG10L** | 0.05946 | 0.00000 | 0.10120 | 0.06650 | 0.19505 | 0.11481 | 0.01062 |
| **VSIG2** | 0.00000 | 0.03170 | 0.01378 | 0.04140 | 0.01291 | 0.61056 | 0.06029 |
| **VSIG8** | 0.02098 | 0.01953 | 0.01203 | 0.02080 | 0.00796 | 0.01500 | 0.02454 |
| **VSNL1** | 0.05318 | 0.00000 | 0.00842 | 0.27995 | 0.00782 | 4.24799 | 0.00000 |
| **VSTM1** | 0.02407 | 0.00935 | 0.04248 | 0.02423 | 0.01044 | 0.01267 | 0.01578 |
| **VSTM2A** | 0.01090 | 0.01939 | 0.02320 | 0.01697 | 0.00727 | 0.14937 | 0.03468 |
| **VSTM2A-OT1** | 0.03218 | 0.02579 | 0.01635 | 0.01596 | 0.00748 | 0.02440 | 0.01138 |
| **VSTM2B** | 0.01180 | 0.01271 | 0.01887 | 0.01994 | 0.01334 | 0.11566 | 0.02808 |
| **VSTM2L** | 0.05542 | 0.03552 | 0.00935 | 0.14497 | 0.00599 | 0.00000 | 0.44712 |
| **VSTM4** | 0.02206 | 0.02964 | 0.05326 | 0.01816 | 0.00524 | 0.03764 | 0.04089 |
| **VSTM5** | 0.08266 | 0.00935 | 0.01378 | 0.01215 | 0.00771 | 0.01296 | 0.01260 |
| **VSX1** | 0.04085 | 0.01797 | 0.01290 | 0.01632 | 0.00868 | 0.03327 | 0.00898 |
| **VSX2** | 0.01525 | 0.00200 | 0.08617 | 0.02699 | 0.05023 | 0.01129 | 0.03643 |
| **VTA1** | 37.72704 | 9.41668 | 15.93259 | 18.79768 | 4.21962 | 11.33988 | 0.00000 |
| **VTCN1** | 3.79175 | 0.00000 | 0.02360 | 2.96429 | 1.15923 | 0.00034 | 0.00831 |
| **VTI1A** | 6.98480 | 5.38254 | 3.50141 | 6.53672 | 1.90283 | 3.87428 | 2.07184 |

| | | | | | | | |
|---|---|---|---|---|---|---|---|
| **VTI1B** | 13.08362 | 18.61885 | 7.00041 | 5.89407 | 0.62198 | 6.96372 | 1.05701 |
| **VTN** | 0.00000 | 0.03690 | 0.01980 | 0.20317 | 0.00657 | 0.02919 | 0.06856 |
| **VTRNA1-1** | 0.01636 | 0.10190 | 0.00740 | 0.02909 | 0.02833 | 0.07783 | 0.08597 |
| **VTRNA1-2** | 0.08737 | 0.74966 | 0.09561 | 0.15300 | 0.03941 | 0.07223 | 0.00000 |
| **VTRNA1-3** | 0.03438 | 0.13395 | 0.01553 | 0.67711 | 0.00488 | 0.27723 | 0.63435 |
| **VTRNA2-1** | 0.04019 | 0.03770 | 0.00878 | 0.02373 | 0.00791 | 0.03384 | 0.02008 |
| **VWA1** | 0.00000 | 2.70184 | 0.11825 | 4.42060 | 0.00000 | 7.01385 | 13.09499 |
| **VWA2** | 0.02162 | 0.00391 | 0.06337 | 0.02639 | 0.03059 | 0.00261 | 0.12262 |
| **VWA3A** | 0.00000 | 0.01665 | 0.00561 | 0.08109 | 0.29145 | 0.11974 | 0.03726 |
| **VWA3B** | 0.03338 | 0.20937 | 0.06549 | 0.02827 | 0.01423 | 0.00320 | 0.00000 |
| **VWA5A** | 0.13672 | 1.20181 | 0.81743 | 0.00000 | 3.32235 | 4.68142 | 0.00000 |
| **VWA5B2** | 0.01649 | 0.02966 | 0.01093 | 0.03550 | 0.00921 | 0.06748 | 0.03375 |
| **VWA7** | 0.00000 | 0.07028 | 0.26546 | 0.22053 | 0.19315 | 0.00873 | 0.00688 |
| **VWA8** | 2.96107 | 2.79796 | 1.31990 | 0.00000 | 0.48091 | 2.10845 | 4.79703 |
| **VWA8-AS1** | 0.01317 | 0.11174 | 0.03469 | 0.02930 | 0.00595 | 0.00840 | 0.01246 |
| **VWA9** | 17.46041 | 3.80343 | 7.15983 | 15.46507 | 1.52386 | 4.76806 | 0.00000 |
| **VWC2** | 0.00494 | 0.00000 | 0.48468 | 0.01304 | 0.00253 | 0.16143 | 0.03143 |
| **VWC2L-IT1** | 0.02943 | 0.02199 | 0.01079 | 0.01575 | 0.01995 | 0.01239 | 0.02002 |
| **VWCE** | 0.00000 | 0.24559 | 0.41817 | 0.96191 | 0.29687 | 0.57008 | 0.23493 |
| **VWDE** | 0.22987 | 0.00000 | 0.05453 | 0.08730 | 0.00425 | 0.33587 | 0.16964 |
| **VWF** | 0.10068 | 0.00000 | 0.03929 | 0.09665 | 0.33170 | 0.03199 | 0.34356 |
| **WAC** | 16.00948 | 4.72350 | 14.99904 | 15.27134 | 5.54920 | 11.02389 | 30.84198 |
| **WAC-AS1** | 0.49488 | 0.49931 | 0.65604 | 0.23824 | 0.13804 | 0.20079 | 1.87312 |
| **WAPAL** | 17.42334 | 6.34132 | 13.60483 | 11.20994 | 6.05048 | 12.15343 | 25.28333 |
| **WARS** | 36.39421 | 43.58129 | 19.81177 | 15.24825 | 3.49485 | 35.88076 | 7.04445 |
| **WARS2** | 6.08604 | 1.18766 | 2.52453 | 4.87740 | 0.88119 | 1.95046 | 0.79145 |
| **WAS** | 0.01149 | 0.06521 | 0.01988 | 0.03343 | 0.04277 | 0.04093 | 0.11991 |
| **WASF1** | 3.03618 | 0.00000 | 3.03978 | 5.34228 | 6.98940 | 1.07161 | 5.61388 |
| **WASF2** | 12.44768 | 11.08700 | 0.67760 | 15.61525 | 1.96222 | 6.64679 | 19.40394 |
| **WASF3** | 0.00000 | 0.50592 | 4.98053 | 0.16543 | 1.55410 | 0.46194 | 0.80962 |
| **WASH1** | 0.02522 | 0.26976 | 0.23497 | 0.09223 | 0.06000 | 0.13947 | 0.56633 |
| **WASH2P** | 0.05935 | 0.47247 | 0.20069 | 0.09367 | 0.02411 | 0.07703 | 0.30908 |
| **WASH3P** | 0.02937 | 0.17229 | 0.19715 | 0.00623 | 0.01558 | 0.04377 | 0.23867 |
| **WASH5P** | 0.02441 | 0.01816 | 0.01675 | 0.01939 | 0.00823 | 0.01382 | 0.01617 |
| **WASH7P** | 0.02280 | 0.01960 | 0.01724 | 0.01916 | 0.00732 | 0.01247 | 0.02326 |

| | | | | | | | |
|---|---|---|---|---|---|---|---|
| **WASL** | 16.24717 | 9.33023 | 8.53017 | 9.42111 | 2.51598 | 15.61233 | 17.83584 |
| **WBP1** | 0.05154 | 0.08122 | 0.03825 | 0.14495 | 0.00784 | 0.09777 | 0.05949 |
| **WBP11** | 15.99163 | 9.73392 | 10.98376 | 16.85334 | 9.78353 | 8.67230 | 15.88543 |
| **WBP11P1** | 0.94217 | 0.23161 | 0.63703 | 0.84874 | 0.79067 | 0.32518 | 0.98592 |
| **WBP1L** | 8.67960 | 11.84245 | 2.52794 | 16.43648 | 1.96434 | 2.99117 | 6.48896 |
| **WBP2** | 6.80280 | 5.16689 | 3.79369 | 21.80603 | 1.74764 | 6.99326 | 3.37193 |
| **WBP2NL** | 0.03491 | 0.00853 | 0.06750 | 0.01457 | 0.07452 | 0.01514 | 0.06605 |
| **WBP4** | 8.75195 | 5.28280 | 8.85174 | 6.03422 | 3.61706 | 6.53423 | 12.14435 |
| **WBP5** | 16.28830 | 0.00000 | 17.91731 | 14.24639 | 10.31650 | 0.00000 | 12.97411 |
| **WBSCR16** | 0.20288 | 1.31146 | 0.20257 | 0.77421 | 0.13378 | 0.54822 | 1.73292 |
| **WBSCR17** | 0.13731 | 0.06204 | 0.04179 | 0.00000 | 0.06251 | 0.14166 | 0.48916 |
| **WBSCR22** | 34.45168 | 23.70055 | 14.97422 | 30.19346 | 13.68227 | 10.49818 | 3.93775 |
| **WBSCR27** | 0.00000 | 0.69800 | 0.09370 | 0.20835 | 0.02570 | 0.56541 | 0.62070 |
| **WDFY1** | 6.80108 | 1.56440 | 0.21338 | 5.12469 | 10.45593 | 3.44769 | 0.90519 |
| **WDFY2** | 4.93650 | 3.40147 | 2.19872 | 6.08787 | 8.30659 | 2.42801 | 0.20571 |
| **WDFY3** | 3.73593 | 0.94299 | 3.40246 | 2.54219 | 2.07625 | 3.44592 | 8.26207 |
| **WDFY3-AS2** | 0.07360 | 0.02681 | 0.18140 | 0.01866 | 0.03805 | 0.05122 | 0.19068 |
| **WDFY4** | 0.02460 | 0.03959 | 0.00781 | 0.04251 | 0.02529 | 0.00918 | 0.05297 |
| **WDHD1** | 14.28232 | 5.14768 | 13.41104 | 6.89155 | 4.17533 | 11.72698 | 1.53837 |
| **WDPCP** | 1.27221 | 0.97824 | 0.39284 | 0.72609 | 0.22190 | 0.67596 | 0.40560 |
| **WDR1** | 24.63754 | 43.10829 | 9.78340 | 26.07386 | 6.15089 | 25.81281 | 16.24126 |
| **WDR11** | 9.83704 | 6.18155 | 7.02183 | 0.00000 | 2.28483 | 9.76510 | 2.63887 |
| **WDR11-AS1** | 0.02458 | 1.42013 | 2.59795 | 0.00000 | 0.08687 | 2.08777 | 0.00000 |
| **WDR12** | 31.27151 | 8.85708 | 13.96494 | 10.82930 | 4.13097 | 8.83329 | 0.89228 |
| **WDR13** | 4.18755 | 9.24522 | 1.46570 | 1.68005 | 0.63661 | 6.05788 | 2.39212 |
| **WDR16** | 0.06919 | 0.17954 | 0.21064 | 0.00000 | 0.11095 | 0.03653 | 0.05791 |
| **WDR17** | 0.03455 | 0.01500 | 0.02332 | 0.24138 | 0.01752 | 0.01470 | 0.01584 |
| **WDR18** | 17.24932 | 22.99955 | 9.88588 | 27.81592 | 0.00000 | 9.20423 | 11.53444 |
| **WDR19** | 1.95445 | 0.00000 | 0.90349 | 0.68455 | 2.02102 | 0.58426 | 0.68786 |
| **WDR20** | 13.32490 | 8.60345 | 7.85188 | 9.15027 | 12.19287 | 5.15501 | 0.00000 |
| **WDR24** | 3.20305 | 4.35187 | 3.47751 | 3.60759 | 0.03659 | 1.29901 | 4.19610 |
| **WDR25** | 2.05135 | 11.64736 | 7.14119 | 0.29996 | 2.30875 | 2.40377 | 0.50674 |
| **WDR26** | 5.75872 | 1.90004 | 8.29989 | 8.27087 | 3.28001 | 5.54670 | 16.28372 |
| **WDR27** | 0.08990 | 0.31164 | 0.52833 | 0.57014 | 0.22172 | 0.27070 | 1.59569 |
| **WDR3** | 29.35284 | 4.99009 | 14.10360 | 14.41263 | 7.07119 | 5.45702 | 0.38070 |

| | | | | | | | |
|---|---|---|---|---|---|---|---|
| **WDR31** | 0.02822 | 0.01942 | 0.09111 | 0.05952 | 0.20291 | 0.11446 | 0.15929 |
| **WDR33** | 19.77060 | 9.43631 | 9.48930 | 14.68352 | 2.88739 | 11.59394 | 12.41165 |
| **WDR34** | 6.73937 | 27.32493 | 8.98189 | 15.21907 | 0.76933 | 6.99281 | 20.85097 |
| **WDR35** | 0.42396 | 0.00000 | 0.43607 | 0.50880 | 1.16522 | 0.03643 | 0.20290 |
| **WDR36** | 23.63313 | 0.37038 | 10.33165 | 9.08588 | 3.50897 | 5.17363 | 4.98769 |
| **WDR37** | 1.73967 | 0.19751 | 1.57021 | 2.48071 | 4.03677 | 1.10351 | 1.85911 |
| **WDR38** | 0.00844 | 0.05979 | 0.02792 | 0.03630 | 0.65677 | 0.01652 | 0.11539 |
| **WDR4** | 14.94632 | 18.00389 | 7.79128 | 5.96623 | 2.78251 | 8.26187 | 3.65132 |
| **WDR41** | 7.52461 | 4.28994 | 4.80899 | 4.18399 | 1.84990 | 4.16785 | 0.00000 |
| **WDR43** | 43.62807 | 23.24178 | 23.96481 | 25.89331 | 12.15187 | 14.42593 | 20.97015 |
| **WDR44** | 4.44177 | 0.00000 | 4.59261 | 1.44741 | 2.41445 | 4.90834 | 2.09240 |
| **WDR45** | 9.72621 | 22.56970 | 31.01945 | 12.29254 | 8.68818 | 5.68400 | 2.49976 |
| **WDR45B** | 26.34842 | 20.45352 | 12.68190 | 18.65445 | 15.50681 | 12.48102 | 9.75820 |
| **WDR46** | 31.13632 | 25.54751 | 17.27926 | 28.30522 | 7.46918 | 13.93196 | 0.01567 |
| **WDR47** | 6.78091 | 0.85461 | 15.35368 | 4.96687 | 20.95318 | 7.49047 | 0.54321 |
| **WDR48** | 14.84966 | 4.27707 | 9.33688 | 10.59420 | 4.12679 | 5.51281 | 0.00000 |
| **WDR49** | 0.05975 | 0.00116 | 0.03783 | 0.00734 | 0.03681 | 0.02001 | 0.04285 |
| **WDR5** | 11.21701 | 22.85918 | 14.80222 | 7.98112 | 3.13572 | 14.90899 | 35.57483 |
| **WDR52** | 0.58079 | 0.00000 | 0.49957 | 0.35607 | 0.13740 | 0.02293 | 2.28482 |
| **WDR52-AS1** | 0.02625 | 0.01266 | 0.02236 | 0.02095 | 0.00777 | 0.01272 | 0.01664 |
| **WDR53** | 4.27731 | 4.02611 | 5.56159 | 4.48576 | 1.15497 | 2.69962 | 0.00000 |
| **WDR54** | 1.52351 | 1.22870 | 0.37868 | 1.70297 | 0.22849 | 1.96270 | 1.13712 |
| **WDR55** | 25.76476 | 9.40407 | 7.68848 | 14.36419 | 5.97586 | 9.39554 | 0.00000 |
| **WDR59** | 2.48787 | 3.36656 | 2.33310 | 4.08421 | 0.75619 | 3.20261 | 10.49352 |
| **WDR5B** | 1.18586 | 0.56862 | 1.27146 | 0.81064 | 0.03571 | 0.00000 | 0.63077 |
| **WDR6** | 7.31429 | 1.73436 | 2.08653 | 18.54869 | 0.54151 | 11.34666 | 4.34664 |
| **WDR60** | 0.64877 | 1.09538 | 1.37417 | 0.16556 | 2.10725 | 1.33126 | 4.74791 |
| **WDR61** | 20.67243 | 6.03884 | 9.83958 | 9.37905 | 3.02083 | 5.32165 | 2.23407 |
| **WDR62** | 0.27540 | 2.49795 | 2.14224 | 5.37105 | 0.55518 | 5.56586 | 5.64932 |
| **WDR63** | 0.03966 | 0.00000 | 0.01402 | 0.03438 | 0.34434 | 0.01527 | 0.10338 |
| **WDR64** | 0.04047 | 0.13012 | 0.02301 | 0.12829 | 0.01679 | 0.00000 | 0.03887 |
| **WDR65** | 0.00000 | 0.04011 | 0.06696 | 0.37106 | 0.81916 | 0.00678 | 0.13497 |
| **WDR66** | 4.16553 | 10.30682 | 10.84524 | 0.20608 | 3.19412 | 10.21576 | 0.20338 |
| **WDR7** | 2.09507 | 1.72493 | 1.67397 | 0.17855 | 2.33389 | 1.27534 | 3.15090 |
| **WDR70** | 9.52033 | 5.19103 | 6.15325 | 3.18911 | 3.41945 | 2.96420 | 8.60494 |

| | | | | | | | |
|---|---|---|---|---|---|---|---|
| **WDR72** | 1.45400 | 0.51949 | 1.09228 | 0.17940 | 0.76663 | 1.60025 | 0.88027 |
| **WDR73** | 12.65698 | 5.38637 | 8.54912 | 10.66702 | 3.86490 | 3.56834 | 2.65531 |
| **WDR74** | 35.19642 | 23.95025 | 22.18007 | 28.78338 | 5.72571 | 11.28487 | 6.81735 |
| **WDR75** | 27.99519 | 5.13475 | 14.74475 | 10.24208 | 7.05767 | 8.23933 | 3.68370 |
| **WDR76** | 4.95382 | 2.10721 | 4.08823 | 3.03249 | 5.82312 | 0.48781 | 2.66345 |
| **WDR77** | 40.42817 | 24.22299 | 17.52976 | 30.67075 | 2.77901 | 17.12938 | 4.10316 |
| **WDR78** | 0.06266 | 0.14930 | 0.12988 | 0.53644 | 0.21113 | 0.06684 | 0.09131 |
| **WDR81** | 0.15061 | 0.00000 | 1.06482 | 0.91929 | 0.00644 | 0.79976 | 2.16587 |
| **WDR82** | 8.64688 | 5.79518 | 6.95000 | 5.59599 | 2.64618 | 7.63053 | 20.59184 |
| **WDR83** | 9.19884 | 11.56559 | 4.21394 | 7.26637 | 8.52572 | 3.26339 | 0.00000 |
| **WDR83OS** | 21.52085 | 27.27661 | 12.87933 | 23.29944 | 3.35805 | 14.67727 | 18.39645 |
| **WDR86-AS1** | 0.02509 | 0.01213 | 0.01680 | 0.01918 | 0.09131 | 0.01336 | 0.01496 |
| **WDR87** | 0.00000 | 0.26234 | 0.16987 | 0.15404 | 0.06806 | 0.00000 | 1.27398 |
| **WDR88** | 0.02457 | 0.02806 | 0.03162 | 0.04356 | 0.05501 | 0.02115 | 0.00000 |
| **WDR89** | 14.09872 | 0.00000 | 6.26833 | 11.40043 | 5.08463 | 2.97867 | 2.65898 |
| **WDR90** | 0.38736 | 0.36993 | 0.93694 | 0.91273 | 0.24075 | 0.86211 | 1.50064 |
| **WDR91** | 1.11051 | 0.00000 | 1.26341 | 1.77007 | 0.44558 | 0.60885 | 0.00000 |
| **WDR92** | 4.18610 | 1.13806 | 5.78548 | 2.74571 | 0.63763 | 2.42218 | 0.91431 |
| **WDR93** | 0.19904 | 0.09080 | 0.16363 | 0.01690 | 0.47955 | 0.01016 | 0.12357 |
| **WDR96** | 0.04700 | 0.17664 | 0.41457 | 0.07491 | 0.03182 | 0.19611 | 0.00000 |
| **WDSUB1** | 8.32256 | 1.74319 | 1.32176 | 3.85115 | 0.08699 | 3.65043 | 1.95501 |
| **WDTC1** | 0.00000 | 7.46645 | 8.58859 | 0.73652 | 2.52903 | 3.69423 | 5.26563 |
| **WDYHV1** | 4.82972 | 1.63882 | 3.18816 | 1.51340 | 0.48334 | 2.45905 | 1.24177 |
| **WEE1** | 3.18628 | 0.32914 | 4.22742 | 11.65138 | 0.06891 | 3.84997 | 14.20200 |
| **WEE2** | 0.00000 | 1.34827 | 1.40173 | 0.00000 | 32.90582 | 0.00000 | 0.00000 |
| **WEE2-AS1** | 0.01525 | 0.14783 | 0.00000 | 0.29188 | 1.41041 | 0.19749 | 0.01571 |
| **WFDC10A** | 0.02455 | 0.00135 | 0.04780 | 0.01934 | 0.07560 | 0.01242 | 0.02176 |
| **WFDC10B** | 0.01839 | 0.01689 | 0.02035 | 0.01754 | 0.05032 | 0.02392 | 0.03183 |
| **WFDC13** | 0.02495 | 0.01271 | 0.02011 | 0.01870 | 0.02954 | 0.01286 | 0.01827 |
| **WFDC2** | 0.37401 | 0.00000 | 3.41806 | 3.20402 | 7.06096 | 0.00000 | 4.45740 |
| **WFDC21P** | 0.03839 | 0.01363 | 0.01677 | 0.01712 | 0.00793 | 0.01726 | 0.01146 |
| **WFDC3** | 0.02054 | 0.00000 | 0.29041 | 0.03698 | 0.01909 | 0.04488 | 0.02372 |
| **WFDC6** | 0.04859 | 0.02662 | 0.08929 | 0.02332 | 0.02216 | 0.00202 | 0.04364 |
| **WFDC8** | 0.03735 | 0.00127 | 0.05718 | 0.01618 | 0.01656 | 0.01672 | 0.05463 |
| **WFDC9** | 0.02544 | 0.01320 | 0.02110 | 0.01880 | 0.01035 | 0.01320 | 0.01703 |

| | | | | | | | |
|---|---|---|---|---|---|---|---|
| **WFIKKN1** | 0.01326 | 0.00000 | 0.03730 | 0.02364 | 0.00510 | 0.06737 | 0.18823 |
| **WFIKKN2** | 0.01642 | 0.00708 | 0.02584 | 0.02423 | 0.01538 | 0.00818 | 0.07405 |
| **WFS1** | 1.84650 | 5.83376 | 0.60701 | 3.12007 | 5.16286 | 3.13223 | 8.86825 |
| **WHAMM** | 0.55860 | 1.34260 | 0.12321 | 1.14099 | 1.61542 | 1.20867 | 3.74115 |
| **WHAMMP1** | 0.04949 | 0.01515 | 0.02054 | 0.01351 | 0.03688 | 0.00589 | 0.05328 |
| **WHAMMP2** | 0.03101 | 0.01845 | 0.01606 | 0.01769 | 0.01177 | 0.01490 | 0.01421 |
| **WHAMMP3** | 0.03230 | 0.02440 | 0.01407 | 0.01988 | 0.02415 | 0.01605 | 0.01955 |
| **WHSC1** | 2.12522 | 7.39553 | 3.03272 | 2.33154 | 4.11184 | 5.02438 | 21.08302 |
| **WHSC1L1** | 3.09312 | 2.04332 | 0.38199 | 7.35300 | 0.54264 | 2.63566 | 16.50273 |
| **WIBG** | 21.99797 | 19.92556 | 6.59749 | 15.31715 | 2.14389 | 8.06897 | 0.51064 |
| **WIF1** | 0.02402 | 0.01386 | 0.01764 | 0.01985 | 0.04531 | 0.01313 | 0.01669 |
| **WIPF1** | 0.15111 | 0.00000 | 0.08827 | 0.41740 | 0.05987 | 0.12793 | 0.85981 |
| **WIPF2** | 3.21987 | 6.04345 | 3.44616 | 6.22725 | 5.82436 | 4.05948 | 15.23835 |
| **WIPF3** | 1.22272 | 0.16035 | 0.34818 | 0.00000 | 0.00349 | 2.08488 | 1.37218 |
| **WIPI1** | 0.42105 | 0.74360 | 0.31082 | 5.56517 | 0.32773 | 0.11713 | 0.00000 |
| **WIPI2** | 2.15487 | 11.96477 | 10.92595 | 9.73643 | 6.41318 | 9.27311 | 19.18723 |
| **WISP1** | 0.02640 | 0.01798 | 0.01269 | 0.01845 | 0.00801 | 0.01307 | 0.01674 |
| **WISP3** | 0.03088 | 0.01179 | 0.02185 | 0.01386 | 0.00934 | 0.02746 | 0.03615 |
| **WIZ** | 1.11374 | 1.62344 | 2.03218 | 1.75253 | 0.46693 | 1.41671 | 7.79269 |
| **WLS** | 5.82963 | 0.00000 | 0.92233 | 12.45570 | 3.89711 | 0.00000 | 3.29414 |
| **WNK1** | 7.57858 | 3.34792 | 11.59565 | 7.96359 | 2.95208 | 7.14896 | 25.46466 |
| **WNK2** | 0.17553 | 0.22603 | 0.24793 | 1.72873 | 0.00431 | 0.22602 | 2.29662 |
| **WNK3** | 1.34677 | 0.00000 | 9.41118 | 0.30367 | 6.62085 | 2.09252 | 1.62440 |
| **WNK4** | 0.23699 | 0.03060 | 0.57997 | 0.30437 | 0.03522 | 0.31065 | 0.00000 |
| **WNT1** | 0.00000 | 0.03185 | 0.02684 | 0.49333 | 0.01270 | 0.00881 | 0.01314 |
| **WNT10A** | 0.00536 | 0.03404 | 0.02274 | 0.03943 | 0.01284 | 0.00634 | 0.06934 |
| **WNT10B** | 0.11812 | 0.16249 | 0.38022 | 2.75648 | 0.00170 | 0.33048 | 0.00000 |
| **WNT11** | 0.87806 | 13.10305 | 0.72900 | 5.27657 | 0.05537 | 2.09421 | 0.88819 |
| **WNT16** | 1.78376 | 0.00531 | 0.11519 | 0.00000 | 0.05380 | 0.01884 | 0.42905 |
| **WNT2** | 0.89079 | 0.01347 | 0.01898 | 0.00002 | 0.00477 | 0.01788 | 0.03284 |
| **WNT2B** | 0.16600 | 0.13079 | 0.02056 | 0.07226 | 0.04183 | 0.06316 | 0.00373 |
| **WNT3** | 0.00000 | 0.98235 | 1.61008 | 1.03638 | 0.14853 | 7.50045 | 0.00000 |
| **WNT3A** | 0.13311 | 0.08823 | 0.00000 | 0.00000 | 0.10549 | 0.00000 | 1.22691 |
| **WNT4** | 0.02819 | 0.05647 | 0.01347 | 0.06675 | 0.00985 | 0.00603 | 0.01186 |
| **WNT5A** | 0.26419 | 0.00000 | 0.03089 | 0.21248 | 0.08476 | 0.19328 | 0.57217 |

| | | | | | | | |
|---|---|---|---|---|---|---|---|
| **WNT5B** | 1.48215 | 8.12003 | 1.41936 | 1.98243 | 0.00029 | 1.61461 | 1.06479 |
| **WNT6** | 0.09266 | 0.00000 | 0.00000 | 2.78216 | 0.00448 | 0.00000 | 6.12745 |
| **WNT7A** | 2.30220 | 0.00000 | 0.00000 | 1.01495 | 0.00000 | 0.00000 | 6.38670 |
| **WNT7B** | 1.16290 | 0.00000 | 0.00000 | 1.86664 | 0.00029 | 0.00000 | 7.34273 |
| **WNT8A** | 0.01377 | 0.03144 | 0.04361 | 0.02635 | 0.08174 | 0.10839 | 0.00969 |
| **WNT8B** | 0.03217 | 0.01196 | 0.02081 | 0.01950 | 0.04627 | 0.01447 | 0.01864 |
| **WNT9A** | 0.24094 | 1.49532 | 0.16455 | 0.09888 | 0.16114 | 0.07453 | 0.00148 |
| **WNT9B** | 0.02556 | 0.02108 | 0.01182 | 0.01667 | 0.00787 | 0.01691 | 0.02087 |
| **WRAP53** | 14.93170 | 4.75827 | 8.04863 | 16.43925 | 7.06547 | 3.93797 | 1.86461 |
| **WRAP73** | 9.50346 | 6.85495 | 3.22338 | 12.52308 | 0.28146 | 4.69180 | 3.09774 |
| **WRB** | 13.15518 | 0.24741 | 4.66512 | 7.15258 | 2.04211 | 1.17547 | 0.00000 |
| **WRN** | 3.45470 | 0.00000 | 2.81105 | 3.62128 | 1.85565 | 2.47237 | 4.70587 |
| **WRNIP1** | 3.23458 | 2.50259 | 1.37432 | 3.77589 | 0.42185 | 2.91401 | 11.24626 |
| **WSB1** | 33.93530 | 9.90462 | 16.19561 | 31.41289 | 10.39405 | 22.10927 | 18.01970 |
| **WSB2** | 3.58756 | 0.04499 | 1.13348 | 5.33970 | 0.47327 | 2.63898 | 3.77694 |
| **WSCD1** | 0.17887 | 0.23035 | 0.00000 | 0.00000 | 0.53339 | 0.10695 | 0.55190 |
| **WSCD2** | 0.02008 | 0.01120 | 0.02087 | 0.02429 | 0.04033 | 0.02787 | 0.02238 |
| **WT1** | 0.02050 | 0.03828 | 0.01961 | 0.02305 | 0.00724 | 0.01476 | 0.01439 |
| **WTAP** | 28.09860 | 8.13649 | 17.21539 | 13.43354 | 7.89480 | 13.47536 | 11.72285 |
| **WTAPP1** | 0.02706 | 0.01661 | 0.01321 | 0.01816 | 0.00787 | 0.01250 | 0.01844 |
| **WTH3DI** | 0.86945 | 0.08893 | 1.03568 | 0.71064 | 0.76749 | 0.45305 | 0.00000 |
| **WTIP** | 0.00000 | 2.25792 | 0.00000 | 0.59097 | 0.06379 | 0.21239 | 3.15732 |
| **WWC1** | 4.16704 | 8.89359 | 3.36672 | 6.98533 | 0.00000 | 0.00000 | 14.55130 |
| **WWC2** | 13.92732 | 5.13444 | 2.51116 | 4.27640 | 0.00000 | 3.26333 | 17.92169 |
| **WWC2-AS2** | 0.13737 | 0.18225 | 0.00000 | 0.33248 | 0.08481 | 0.38440 | 0.41314 |
| **WWC3** | 3.07749 | 2.46602 | 0.09687 | 4.88899 | 1.09382 | 0.00000 | 8.01955 |
| **WWOX** | 3.23806 | 4.07289 | 0.06199 | 0.93138 | 1.45322 | 1.01757 | 1.36883 |
| **WWP1** | 6.63655 | 0.00000 | 1.36626 | 1.57282 | 0.15514 | 2.54536 | 10.79149 |
| **WWP2** | 5.93288 | 3.96088 | 0.92399 | 8.08705 | 4.75050 | 3.14040 | 7.32352 |
| **WWTR1** | 20.18986 | 13.29922 | 18.40121 | 24.48616 | 0.40177 | 14.66617 | 7.08837 |
| **WWTR1-AS1** | 0.09224 | 0.03792 | 0.18443 | 0.08495 | 0.02856 | 0.00000 | 0.21465 |
| **XAB2** | 3.60343 | 8.33544 | 8.18287 | 13.93594 | 1.68048 | 4.97567 | 11.51544 |
| **XAF1** | 0.04606 | 0.01323 | 0.01141 | 0.00871 | 0.00750 | 0.01191 | 0.05386 |
| **XAGE2.uid-1286** | 0.03588 | 0.03051 | 0.01115 | 0.02347 | 0.00797 | 0.00867 | 0.02254 |
| **XAGE3** | 1.45011 | 11.00634 | 1.92974 | 0.00000 | 0.00135 | 0.00000 | 0.00910 |

| | | | | | | | |
|---|---|---|---|---|---|---|---|
| **XAGE5** | 0.00000 | 0.19812 | 1.07433 | 0.26311 | 0.00513 | 0.04924 | 0.09880 |
| **XBP1** | 11.08384 | 3.97381 | 2.32984 | 13.71569 | 0.00000 | 11.14709 | 28.52576 |
| **XCL1** | 0.20034 | 0.00399 | 0.01560 | 0.00000 | 0.07180 | 0.04538 | 0.21079 |
| **XCL2** | 0.03252 | 0.01078 | 0.02015 | 0.00104 | 0.00765 | 0.01627 | 0.13568 |
| **XCR1** | 0.01994 | 0.03095 | 0.00519 | 0.17359 | 0.30661 | 0.01523 | 0.00039 |
| **XDH** | 0.03809 | 0.06090 | 0.19270 | 0.05841 | 0.00263 | 0.09085 | 0.06428 |
| **XG** | 0.13530 | 0.17068 | 0.00156 | 0.16771 | 0.09942 | 0.00835 | 0.02156 |
| **XIAP** | 4.69371 | 0.00000 | 9.25753 | 5.54456 | 0.05076 | 7.19856 | 17.37163 |
| **XIRP2** | 0.10923 | 0.00000 | 0.09269 | 0.11943 | 0.50710 | 0.01768 | 0.00000 |
| **XIST** | 16.28833 | 0.00000 | 3.80505 | 16.86019 | 0.00000 | 10.96748 | 31.27040 |
| **XK** | 0.08370 | 0.18961 | 0.03010 | 0.00360 | 0.04901 | 0.03063 | 0.00000 |
| **XKR4** | 0.03969 | 0.09251 | 0.02078 | 0.01203 | 0.02580 | 0.00881 | 0.00821 |
| **XKR5** | 0.02003 | 0.01782 | 0.01335 | 0.02021 | 0.00819 | 0.01929 | 0.02987 |
| **XKR6** | 0.42385 | 0.00000 | 0.05705 | 0.05772 | 0.00413 | 0.01332 | 0.10027 |
| **XKR7** | 0.02452 | 0.02577 | 0.01521 | 0.02170 | 0.00759 | 0.01377 | 0.01251 |
| **XKR8** | 1.12340 | 1.33348 | 0.04180 | 7.30520 | 1.18363 | 3.52574 | 4.55286 |
| **XKR9** | 0.05902 | 0.10883 | 0.00519 | 0.00908 | 0.03028 | 0.05448 | 0.00000 |
| **XKRX** | 0.04875 | 0.00000 | 1.61834 | 0.00000 | 0.01465 | 0.12955 | 0.14487 |
| **XPA** | 3.08231 | 2.22823 | 1.03865 | 3.32288 | 0.00000 | 0.54164 | 2.81711 |
| **XPC** | 6.48224 | 10.87269 | 4.09905 | 8.97942 | 1.62359 | 4.60471 | 0.97642 |
| **XPNPEP1** | 12.67211 | 10.60337 | 1.12465 | 11.75092 | 0.19907 | 0.00000 | 3.18885 |
| **XPNPEP2** | 0.03304 | 0.02465 | 0.01169 | 0.01773 | 0.00776 | 0.01059 | 0.02218 |
| **XPNPEP3** | 7.03812 | 2.17207 | 7.69607 | 3.75955 | 0.89757 | 2.76779 | 8.09570 |
| **XPO1** | 32.62434 | 0.97207 | 22.15571 | 15.13752 | 6.81318 | 16.81109 | 33.66360 |
| **XPO4** | 14.43197 | 2.75343 | 7.35180 | 5.16216 | 2.14456 | 6.82837 | 15.19052 |
| **XPO5** | 22.84779 | 14.00035 | 11.76405 | 15.07934 | 3.71661 | 8.55911 | 13.33285 |
| **XPO6** | 11.89599 | 13.98480 | 4.76807 | 7.86725 | 1.20944 | 8.22444 | 24.48166 |
| **XPO7** | 10.13706 | 4.22391 | 11.74137 | 7.83178 | 2.23274 | 7.68415 | 15.84595 |
| **XPOT** | 25.73785 | 3.24079 | 8.56201 | 14.52661 | 3.84143 | 7.86195 | 7.88825 |
| **XPR1** | 4.43730 | 0.00000 | 3.44773 | 3.12034 | 3.45186 | 4.04079 | 2.78131 |
| **XRCC1** | 2.51887 | 3.38406 | 2.62658 | 2.94818 | 2.52553 | 4.03423 | 4.11349 |
| **XRCC2** | 4.56235 | 3.21456 | 5.29633 | 3.16221 | 0.58930 | 0.86834 | 2.22423 |
| **XRCC3** | 0.25210 | 2.36150 | 1.92960 | 3.70152 | 0.61365 | 1.38418 | 5.70066 |
| **XRCC4** | 12.52667 | 0.08165 | 6.18098 | 4.26072 | 2.26509 | 2.64526 | 2.33523 |
| **XRCC5** | 61.30628 | 24.42786 | 27.45577 | 40.69486 | 6.19098 | 23.94486 | 20.74968 |

| | | | | | | | |
|---|---|---|---|---|---|---|---|
| **XRCC6** | 55.76164 | 37.08496 | 21.69670 | 38.35758 | 9.17139 | 21.37667 | 8.59839 |
| **XRCC6BP1** | 2.30372 | 2.27659 | 4.33782 | 0.15210 | 0.59685 | 3.36171 | 0.00000 |
| **XRCC6P5** | 2.42392 | 1.24461 | 0.66465 | 1.22661 | 0.23145 | 0.64799 | 0.00000 |
| **XRN1** | 2.55052 | 0.97262 | 2.99887 | 1.50059 | 2.81557 | 1.93253 | 7.07811 |
| **XRN2** | 25.41138 | 9.86816 | 12.41983 | 18.82225 | 3.81093 | 15.00628 | 18.23170 |
| **XRRA1** | 1.67299 | 2.11161 | 0.64721 | 1.99574 | 1.73770 | 0.86448 | 2.53733 |
| **XXYLT1** | 1.48549 | 1.89284 | 0.38387 | 0.53457 | 0.59288 | 2.20864 | 2.86450 |
| **XXYLT1-AS2** | 0.01646 | 0.01870 | 0.01707 | 0.01494 | 0.00695 | 0.01130 | 0.05284 |
| **XYLB** | 1.88806 | 1.37742 | 0.81733 | 3.18112 | 0.03838 | 1.22006 | 1.67482 |
| **XYLT1** | 0.00000 | 0.13968 | 0.12015 | 0.08345 | 1.41699 | 0.46846 | 0.28928 |
| **XYLT2** | 0.61467 | 5.89625 | 6.16901 | 1.34404 | 0.67572 | 2.10451 | 1.99223 |
| **YAE1D1** | 8.31069 | 0.00000 | 4.99915 | 6.69836 | 0.37415 | 1.46731 | 0.00000 |
| **YAF2** | 3.14935 | 4.02338 | 3.27190 | 3.34829 | 4.48135 | 3.03461 | 4.72536 |
| **YAP1** | 4.88482 | 0.00000 | 2.43032 | 10.06533 | 1.36649 | 1.85760 | 25.21494 |
| **YARS** | 43.34018 | 24.09833 | 9.19194 | 20.95095 | 0.86660 | 19.06396 | 19.39711 |
| **YARS2** | 15.42267 | 6.67831 | 11.92512 | 10.51870 | 5.48889 | 6.36679 | 2.46442 |
| **YBEY** | 2.05333 | 5.49491 | 6.54403 | 3.15638 | 0.15612 | 1.01198 | 5.53179 |
| **YBX1** | 8.53053 | 13.32714 | 7.29638 | 9.18879 | 4.34095 | 7.12565 | 32.25687 |
| **YBX2** | 0.00000 | 1.30212 | 0.31600 | 0.00000 | 1.89945 | 0.58498 | 0.76635 |
| **YBX3** | 0.00000 | 6.62313 | 4.28413 | 1.59499 | 0.35289 | 8.86643 | 9.75547 |
| **YBX3P1** | 0.00000 | 0.74963 | 0.23780 | 0.13808 | 0.12385 | 0.60826 | 0.42636 |
| **YDJC** | 13.14399 | 14.24505 | 0.76982 | 16.26789 | 0.00000 | 3.84964 | 12.63579 |
| **YEATS2** | 6.85060 | 4.07212 | 8.17979 | 4.11982 | 5.56856 | 1.42429 | 8.21892 |
| **YEATS4** | 19.12101 | 3.68728 | 11.06575 | 3.47930 | 1.35837 | 8.01391 | 5.68035 |
| **YES1** | 15.50972 | 1.36926 | 3.93007 | 10.18957 | 1.46632 | 2.59142 | 16.73520 |
| **YIF1A** | 20.34179 | 22.90358 | 7.12088 | 20.06022 | 0.29900 | 10.37485 | 2.89901 |
| **YIF1B** | 19.81714 | 27.94666 | 13.95665 | 10.37568 | 4.01909 | 14.05226 | 0.00000 |
| **YIPF1** | 14.22861 | 4.31103 | 3.28156 | 7.85498 | 1.91538 | 3.22252 | 1.46607 |
| **YIPF2** | 4.17590 | 8.89355 | 2.27841 | 8.92063 | 2.47744 | 4.62094 | 9.31697 |
| **YIPF3** | 11.49388 | 8.15899 | 8.61014 | 21.43249 | 1.62841 | 10.77323 | 5.42620 |
| **YIPF4** | 10.81532 | 10.66303 | 27.83125 | 8.18941 | 1.86042 | 18.01655 | 0.51357 |
| **YIPF5** | 16.08768 | 3.61840 | 11.23423 | 11.68883 | 4.07763 | 5.80917 | 0.00000 |
| **YIPF6** | 20.17015 | 8.60199 | 7.33377 | 12.20663 | 2.42735 | 6.62091 | 13.37601 |
| **YIPF7** | 0.02718 | 0.00721 | 0.04618 | 0.01906 | 0.01189 | 0.01567 | 0.02260 |
| **YJEFN3** | 0.45082 | 1.99933 | 0.42197 | 3.44857 | 0.22521 | 0.37001 | 0.90367 |

| | | | | | | | |
|---|---|---|---|---|---|---|---|
| **YKT6** | 3.15718 | 12.91266 | 4.91063 | 10.74275 | 6.58024 | 6.35288 | 23.82571 |
| **YLPM1** | 8.19260 | 4.07096 | 6.67962 | 6.90324 | 5.63396 | 5.14461 | 20.00073 |
| **YME1L1** | 26.15327 | 15.00900 | 16.71814 | 17.07518 | 9.53604 | 11.59607 | 16.54700 |
| **YOD1** | 5.98116 | 0.00000 | 2.20244 | 2.61799 | 2.53586 | 1.16882 | 3.26586 |
| **YPEL1** | 0.21434 | 0.40561 | 0.17715 | 0.16770 | 1.42650 | 0.16815 | 0.71411 |
| **YPEL2** | 10.25200 | 33.57466 | 26.67768 | 0.00000 | 11.73284 | 19.85802 | 10.85482 |
| **YPEL3** | 0.00000 | 2.69860 | 0.63330 | 1.80704 | 0.50515 | 1.87637 | 5.81494 |
| **YPEL4** | 0.01768 | 0.04713 | 0.03760 | 0.04587 | 0.00555 | 0.05654 | 0.08258 |
| **YPEL5** | 8.03633 | 9.65083 | 22.22733 | 20.13627 | 27.24909 | 7.68375 | 0.00000 |
| **YRDC** | 8.64623 | 2.91512 | 9.91149 | 9.12405 | 7.30703 | 5.09872 | 15.13459 |
| **YTHDC1** | 11.06317 | 9.99477 | 6.44612 | 14.87989 | 4.20023 | 11.51677 | 25.56996 |
| **YTHDC2** | 3.75045 | 0.99475 | 1.64471 | 1.79545 | 0.20672 | 2.53759 | 9.59880 |
| **YTHDF1** | 12.11067 | 18.60984 | 10.31264 | 6.33808 | 3.00337 | 10.85880 | 32.18296 |
| **YTHDF2** | 36.76476 | 17.50300 | 14.98340 | 33.62057 | 14.77305 | 14.51543 | 8.09044 |
| **YTHDF3** | 27.48000 | 4.78688 | 16.13447 | 10.29459 | 12.18605 | 10.04811 | 3.49226 |
| **YTHDF3-AS1** | 0.16429 | 0.88127 | 0.73276 | 0.18686 | 0.87712 | 0.11717 | 0.00000 |
| **YWHAB** | 40.36961 | 25.87541 | 20.80926 | 32.13406 | 14.09158 | 23.09311 | 37.98536 |
| **YWHAE** | 52.65117 | 36.25336 | 33.44120 | 41.42188 | 16.23327 | 30.48012 | 26.03099 |
| **YWHAEP1** | 0.10688 | 0.00000 | 0.20931 | 0.13245 | 0.71308 | 0.07028 | 0.02164 |
| **YWHAEP7** | 0.01948 | 0.01034 | 0.02346 | 0.02074 | 0.02584 | 0.03033 | 0.02704 |
| **YWHAG** | 4.51741 | 0.00000 | 6.12422 | 7.23472 | 5.35008 | 7.27076 | 25.42574 |
| **YWHAH** | 11.94968 | 12.28927 | 5.04612 | 13.27224 | 2.37399 | 13.59965 | 38.88985 |
| **YWHAQ** | 55.31313 | 25.21902 | 19.04458 | 33.83755 | 14.07801 | 19.27995 | 0.00000 |
| **YWHAZ** | 53.50024 | 11.13269 | 20.21563 | 32.07420 | 4.84439 | 21.45437 | 22.67297 |
| **YY1** | 12.87975 | 8.04522 | 12.95606 | 7.66859 | 6.95174 | 12.61487 | 28.32160 |
| **YY1AP1** | 7.52472 | 14.99141 | 13.39235 | 13.53369 | 5.46978 | 9.60104 | 9.37745 |
| **YY1P2** | 0.02639 | 0.02009 | 0.01649 | 0.02464 | 0.01060 | 0.01858 | 0.02635 |
| **YY2** | 0.21738 | 0.04218 | 0.09257 | 0.00000 | 1.33330 | 0.08243 | 0.49147 |
| **ZACN** | 0.02991 | 0.02864 | 0.00917 | 0.03721 | 0.01254 | 0.00869 | 0.02438 |
| **ZADH2** | 0.01623 | 0.40106 | 0.29649 | 0.09777 | 0.00096 | 0.21068 | 0.36957 |
| **ZAK** | 0.45950 | 0.00000 | 1.80891 | 1.96924 | 1.49605 | 0.82366 | 1.30145 |
| **ZAN** | 0.01976 | 0.17885 | 0.08286 | 0.06361 | 0.18971 | 0.06691 | 0.04967 |
| **ZAP70** | 0.00000 | 0.28161 | 0.00042 | 0.05779 | 0.01000 | 0.16910 | 0.04632 |
| **1.00000** | 0.00493 | 0.06435 | 0.00000 | 0.02166 | 0.67234 | 0.01938 | 0.12802 |
| **ZAR1L** | 0.00000 | 0.08294 | 0.34923 | 0.00001 | 4.32519 | 0.02884 | 0.06295 |

| | | | | | | | |
|---|---|---|---|---|---|---|---|
| **ZBBX** | 0.01693 | 0.00000 | 0.11105 | 0.01357 | 0.09187 | 0.02955 | 0.04661 |
| **ZBED3** | 0.00000 | 0.06471 | 0.02262 | 0.00000 | 0.03089 | 0.67483 | 0.50384 |
| **ZBED3-AS1** | 0.01550 | 0.00000 | 0.03146 | 0.01083 | 0.07224 | 0.26456 | 0.16263 |
| **ZBED4** | 3.79333 | 2.27018 | 0.66904 | 2.83959 | 1.37132 | 0.52150 | 6.64181 |
| **ZBED5** | 2.22089 | 0.00000 | 3.81184 | 6.69660 | 1.84895 | 4.37222 | 2.32472 |
| **ZBED5-AS1** | 0.09135 | 0.29862 | 2.34001 | 0.65060 | 1.25780 | 0.34682 | 0.09404 |
| **ZBED6** | 0.67061 | 0.00000 | 0.49793 | 3.40771 | 0.52777 | 1.47744 | 2.96596 |
| **ZBED6CL** | 0.81321 | 1.41899 | 0.25938 | 2.45429 | 0.00000 | 1.18151 | 1.57670 |
| **ZBED8** | 0.05836 | 0.00000 | 0.33293 | 0.19585 | 0.12640 | 0.27963 | 0.00000 |
| **ZBED9** | 1.87695 | 0.36064 | 1.32990 | 0.23163 | 0.38182 | 0.29859 | 0.00000 |
| **ZBP1** | 0.02515 | 0.01690 | 0.01167 | 0.01838 | 0.01440 | 0.01329 | 0.03047 |
| **ZBTB1** | 8.77211 | 0.00000 | 5.74625 | 3.92113 | 5.05460 | 3.12046 | 0.32929 |
| **ZBTB10** | 4.94502 | 0.00000 | 7.98531 | 5.69536 | 4.16562 | 4.97158 | 17.78500 |
| **ZBTB11** | 6.19922 | 0.00000 | 7.79494 | 2.92419 | 7.26746 | 3.23364 | 5.50495 |
| **ZBTB11-AS1** | 0.37398 | 1.67002 | 2.59612 | 0.78235 | 1.18837 | 1.18650 | 2.59178 |
| **ZBTB12** | 0.01256 | 0.01246 | 0.02208 | 0.02115 | 0.00772 | 0.04804 | 0.06406 |
| **ZBTB14** | 4.86033 | 2.22188 | 3.22454 | 1.55411 | 0.98995 | 1.95578 | 0.00000 |
| **ZBTB16** | 0.00000 | 0.03184 | 0.33426 | 0.02860 | 5.87404 | 0.00000 | 0.02423 |
| **ZBTB17** | 2.22078 | 4.32740 | 2.53290 | 5.71584 | 3.43348 | 1.91328 | 3.90065 |
| **ZBTB18** | 0.10126 | 0.15134 | 0.22694 | 0.01755 | 0.54607 | 0.77668 | 1.95019 |
| **ZBTB2** | 6.31044 | 1.27504 | 6.37797 | 3.29198 | 2.74445 | 5.99573 | 5.67809 |
| **ZBTB20** | 0.19108 | 0.00000 | 0.28045 | 0.17053 | 0.13383 | 0.94307 | 1.63718 |
| **ZBTB20-AS4** | 0.02718 | 0.02001 | 0.04312 | 0.06342 | 0.00686 | 0.01390 | 0.00000 |
| **ZBTB21** | 4.42440 | 1.27500 | 5.31197 | 2.01215 | 5.18347 | 1.21259 | 1.12781 |
| **ZBTB22** | 0.00000 | 0.99621 | 0.29817 | 0.93384 | 0.19285 | 0.38353 | 0.46724 |
| **ZBTB24** | 2.46259 | 0.87729 | 2.78526 | 0.10680 | 4.95660 | 2.60952 | 7.07807 |
| **ZBTB25** | 0.70973 | 0.00000 | 0.74534 | 0.74780 | 0.27730 | 0.94794 | 0.35203 |
| **ZBTB26** | 0.60837 | 0.57874 | 0.36272 | 0.68347 | 0.17867 | 0.07504 | 0.28875 |
| **ZBTB3** | 1.06112 | 1.57526 | 1.62044 | 2.15846 | 1.14724 | 1.06948 | 0.48432 |
| **ZBTB32** | 0.01577 | 0.05885 | 0.01055 | 0.10368 | 0.05212 | 0.08854 | 0.01303 |
| **ZBTB33** | 6.74104 | 0.00000 | 5.71795 | 0.90525 | 4.44787 | 1.23082 | 4.28593 |
| **ZBTB34** | 2.44274 | 1.26551 | 1.15826 | 2.96043 | 0.42256 | 0.35840 | 4.76574 |
| **ZBTB37** | 0.40309 | 0.00000 | 1.62377 | 0.65789 | 1.74432 | 0.00000 | 0.31258 |
| **ZBTB38** | 0.97098 | 0.22288 | 0.32364 | 2.34366 | 1.35942 | 0.14109 | 0.59333 |
| **ZBTB39** | 0.99557 | 1.69488 | 1.51826 | 0.50641 | 1.38204 | 1.16798 | 3.45919 |

| | | | | | | | |
|---|---|---|---|---|---|---|---|
| **ZBTB4** | 0.16972 | 0.00000 | 0.52758 | 1.68553 | 0.35343 | 0.48126 | 2.82625 |
| **ZBTB40** | 6.45536 | 6.04503 | 6.00798 | 10.89182 | 1.96511 | 3.83255 | 10.86583 |
| **ZBTB41** | 2.05382 | 0.00000 | 1.66479 | 0.58825 | 0.41913 | 0.45293 | 1.18927 |
| **ZBTB42** | 0.00000 | 0.11656 | 0.00330 | 0.24159 | 0.01447 | 0.04940 | 0.46868 |
| **ZBTB43** | 8.42641 | 5.35203 | 4.76994 | 6.59864 | 3.72269 | 2.97384 | 8.68503 |
| **ZBTB44** | 6.91229 | 0.22753 | 3.52660 | 2.73637 | 0.87824 | 6.87005 | 15.04506 |
| **ZBTB45** | 0.73903 | 3.01755 | 0.80853 | 1.66406 | 0.08645 | 1.41249 | 5.76426 |
| **ZBTB46** | 0.00000 | 0.45140 | 0.40989 | 0.25369 | 0.02012 | 0.53279 | 1.01634 |
| **ZBTB46-AS1** | 0.06825 | 1.05198 | 0.21515 | 0.00818 | 0.00281 | 0.00000 | 0.00311 |
| **ZBTB47** | 0.02159 | 0.02708 | 0.02715 | 0.03127 | 0.00701 | 0.01328 | 0.03322 |
| **ZBTB48** | 1.40633 | 4.12068 | 2.74600 | 3.04913 | 0.50636 | 0.65226 | 1.08475 |
| **ZBTB49** | 1.27507 | 1.85697 | 0.84795 | 0.56163 | 2.55161 | 0.93606 | 0.00000 |
| **ZBTB5** | 0.97338 | 0.49028 | 1.40666 | 1.70135 | 1.72386 | 0.97248 | 4.57876 |
| **ZBTB6** | 1.62431 | 0.00000 | 2.70246 | 1.47764 | 0.82059 | 1.21686 | 0.63771 |
| **ZBTB7A** | 0.00000 | 0.34925 | 0.07299 | 0.53995 | 0.03205 | 0.56348 | 8.10181 |
| **ZBTB7B** | 0.00000 | 0.48594 | 0.01947 | 2.07466 | 0.08765 | 0.55460 | 6.43069 |
| **ZBTB7C** | 0.60845 | 0.00000 | 0.00000 | 0.20092 | 0.09893 | 0.00000 | 3.50723 |
| **ZBTB8A** | 0.79383 | 0.00000 | 7.29008 | 0.54671 | 0.68634 | 1.78563 | 2.33563 |
| **ZBTB8B** | 0.11146 | 0.04873 | 0.14537 | 0.21415 | 0.50910 | 0.37562 | 0.00000 |
| **ZBTB8OS** | 7.95975 | 3.45326 | 8.52279 | 5.70117 | 0.90575 | 8.69842 | 2.05525 |
| **ZBTB9** | 4.81874 | 0.81183 | 5.09135 | 5.36092 | 0.48125 | 2.41041 | 0.00000 |
| **ZC2HC1A** | 2.37256 | 0.21574 | 2.03046 | 4.19516 | 1.42783 | 0.27281 | 1.20772 |
| **ZC2HC1B** | 0.11712 | 0.05186 | 0.00000 | 0.02309 | 2.45695 | 0.12104 | 0.04212 |
| **ZC2HC1C** | 0.00000 | 1.90047 | 0.00136 | 7.85051 | 0.21366 | 0.15563 | 0.00000 |
| **ZC3H10** | 0.37240 | 2.52555 | 1.29006 | 1.84439 | 2.90830 | 1.40057 | 0.00000 |
| **ZC3H11A** | 6.72028 | 2.27762 | 4.19490 | 8.57326 | 3.51992 | 3.79770 | 6.88859 |
| **ZC3H12A** | 1.67544 | 1.52220 | 0.32899 | 4.38938 | 0.32675 | 1.67171 | 4.36469 |
| **ZC3H12B** | 1.67174 | 1.25640 | 0.61024 | 0.59199 | 0.44098 | 1.19589 | 4.24308 |
| **ZC3H12C** | 0.47383 | 0.00000 | 1.02438 | 0.00000 | 0.00030 | 0.82243 | 0.68591 |
| **ZC3H12D** | 0.03098 | 0.00858 | 0.02160 | 0.01207 | 0.00916 | 0.01955 | 0.05094 |
| **ZC3H13** | 16.46718 | 7.31207 | 12.45138 | 10.94952 | 9.76517 | 9.60551 | 26.14826 |
| **ZC3H14** | 23.18134 | 10.39775 | 7.55314 | 13.64371 | 3.63870 | 4.67731 | 4.04833 |
| **ZC3H15** | 35.40395 | 10.94904 | 23.22940 | 23.27627 | 7.82786 | 20.30435 | 23.46866 |
| **ZC3H18** | 2.06507 | 5.22936 | 3.73169 | 5.51758 | 6.89696 | 4.56491 | 14.32801 |
| **ZC3H3** | 0.51617 | 5.76728 | 4.24549 | 3.92730 | 1.04570 | 2.45643 | 7.11701 |

| | | | | | | | |
|---|---|---|---|---|---|---|---|
| **ZC3H4** | 12.40462 | 4.36196 | 8.60941 | 7.54174 | 7.59915 | 4.86149 | 8.30188 |
| **ZC3H6** | 0.03630 | 0.00881 | 0.17243 | 0.05766 | 0.63074 | 0.02030 | 0.17149 |
| **ZC3H7A** | 9.29824 | 3.18456 | 1.30147 | 3.04256 | 1.53672 | 2.06949 | 5.52796 |
| **ZC3H7B** | 0.27169 | 1.69770 | 0.08435 | 1.11865 | 0.02933 | 2.17216 | 4.70925 |
| **ZC3H8** | 4.68925 | 0.26630 | 3.97119 | 5.75800 | 0.84997 | 1.04203 | 5.64854 |
| **ZC3HAV1** | 1.92428 | 8.76563 | 0.40505 | 0.00000 | 0.14693 | 31.36552 | 0.25785 |
| **ZC3HAV1L** | 0.01758 | 0.03165 | 0.01852 | 0.08718 | 0.01844 | 0.07211 | 0.04672 |
| **ZC3HC1** | 17.86304 | 8.17589 | 9.05453 | 13.64906 | 3.32941 | 4.88508 | 0.00000 |
| **ZC4H2** | 5.44091 | 0.09590 | 2.45308 | 0.08611 | 0.09375 | 3.44457 | 0.00000 |
| **ZCCHC10** | 11.96149 | 0.00003 | 6.84685 | 8.01628 | 7.84178 | 6.05360 | 4.26013 |
| **ZCCHC11** | 3.28013 | 0.11509 | 2.05714 | 4.84495 | 0.47023 | 3.62367 | 5.22997 |
| **ZCCHC12** | 0.03998 | 0.90831 | 0.00000 | 0.02264 | 0.43791 | 0.00162 | 0.00000 |
| **ZCCHC13** | 0.02627 | 0.00358 | 0.04066 | 0.02663 | 0.02690 | 0.03299 | 0.01590 |
| **ZCCHC14** | 0.98756 | 2.03685 | 1.60614 | 0.43256 | 3.75763 | 1.53882 | 5.58033 |
| **ZCCHC16** | 0.02461 | 0.01751 | 0.01259 | 0.01888 | 0.01114 | 0.01329 | 0.01824 |
| **ZCCHC17** | 22.41559 | 11.89089 | 10.45334 | 21.08526 | 9.95989 | 10.94043 | 6.06701 |
| **ZCCHC18** | 0.05111 | 0.01355 | 0.01479 | 0.01614 | 0.00938 | 0.01403 | 0.00506 |
| **ZCCHC2** | 2.11770 | 0.87192 | 0.27633 | 0.66092 | 3.60896 | 1.37004 | 3.06055 |
| **ZCCHC24** | 0.25585 | 0.47687 | 0.00000 | 0.10274 | 0.08103 | 1.05217 | 0.00000 |
| **ZCCHC3** | 1.53728 | 0.22462 | 3.61084 | 2.75990 | 0.40329 | 0.00000 | 9.23715 |
| **ZCCHC4** | 3.13124 | 2.01839 | 3.54268 | 1.30781 | 0.71323 | 1.54008 | 0.50854 |
| **ZCCHC6** | 6.31781 | 2.42402 | 3.92063 | 7.49467 | 7.91581 | 2.77056 | 3.68792 |
| **ZCCHC7** | 9.62188 | 3.78152 | 14.66717 | 5.11203 | 4.90478 | 4.04952 | 11.57529 |
| **ZCCHC8** | 15.46987 | 5.72549 | 9.44786 | 18.74574 | 5.36480 | 8.84605 | 1.20001 |
| **ZCCHC9** | 9.31080 | 0.59919 | 7.58872 | 7.20245 | 2.42575 | 5.17818 | 4.68352 |
| **ZCRB1** | 16.70932 | 11.26861 | 13.43770 | 10.63731 | 6.94282 | 9.27811 | 12.49335 |
| **ZCWPW1** | 0.43687 | 0.00000 | 0.39936 | 0.04902 | 2.41051 | 0.60150 | 0.08179 |
| **ZCWPW2** | 0.05818 | 0.01720 | 0.28885 | 0.04976 | 0.03078 | 0.00363 | 0.00540 |
| **ZDBF2** | 3.35497 | 0.00000 | 1.63922 | 1.93335 | 1.01906 | 1.98255 | 6.37768 |
| **ZDHHC1** | 0.01485 | 0.01642 | 0.00912 | 0.04316 | 0.01648 | 0.04691 | 0.09679 |
| **ZDHHC11** | 1.56368 | 0.26071 | 0.68004 | 1.49311 | 0.00000 | 0.90420 | 5.70148 |
| **ZDHHC12** | 8.98345 | 14.93939 | 1.93071 | 10.16855 | 0.22821 | 6.88891 | 2.41510 |
| **ZDHHC13** | 3.35589 | 0.72815 | 1.35707 | 5.05671 | 0.40778 | 1.74331 | 0.00000 |
| **ZDHHC14** | 0.55684 | 0.00000 | 0.01279 | 2.42655 | 0.17065 | 0.00000 | 3.75357 |
| **ZDHHC15** | 0.01414 | 0.05400 | 0.03699 | 0.01985 | 0.02311 | 0.03269 | 0.04672 |

| | | | | | | | |
|---|---|---|---|---|---|---|---|
| **ZDHHC16** | 0.00000 | 5.23083 | 3.91411 | 6.33464 | 0.12815 | 6.60772 | 7.76592 |
| **ZDHHC17** | 8.21884 | 0.00000 | 5.40031 | 1.64011 | 2.37462 | 1.67637 | 6.23803 |
| **ZDHHC18** | 1.57063 | 7.18981 | 3.47773 | 4.80309 | 0.55579 | 2.29182 | 11.78856 |
| **ZDHHC19** | 0.00578 | 0.32550 | 0.06624 | 0.12221 | 0.07419 | 0.03613 | 0.00000 |
| **ZDHHC2** | 0.42420 | 0.00000 | 0.45525 | 0.40409 | 0.27444 | 0.00000 | 1.23914 |
| **ZDHHC20** | 7.73902 | 0.00000 | 3.41387 | 5.37219 | 0.64119 | 7.00136 | 14.10811 |
| **ZDHHC21** | 1.90842 | 0.00238 | 0.93228 | 1.57318 | 0.08982 | 0.12529 | 2.87215 |
| **ZDHHC22** | 0.01980 | 0.06832 | 0.03368 | 0.01831 | 0.02256 | 0.01175 | 0.01441 |
| **ZDHHC23** | 0.21737 | 1.19816 | 0.71434 | 0.00000 | 0.00000 | 1.53913 | 1.04638 |
| **ZDHHC24** | 0.21658 | 1.79349 | 0.71533 | 0.62621 | 0.00000 | 1.52302 | 3.38437 |
| **ZDHHC3** | 4.12074 | 5.23306 | 5.04020 | 9.85234 | 1.36011 | 7.96458 | 21.21818 |
| **ZDHHC4** | 8.08786 | 3.38606 | 3.07748 | 5.59190 | 1.08151 | 4.39783 | 4.07654 |
| **ZDHHC5** | 8.71426 | 12.67699 | 6.72153 | 13.79886 | 5.09470 | 6.45866 | 10.84418 |
| **ZDHHC6** | 23.48395 | 3.06966 | 6.72596 | 14.79855 | 1.59532 | 7.10714 | 0.00000 |
| **ZDHHC7** | 6.40672 | 6.27253 | 3.12104 | 4.11968 | 1.01133 | 5.96770 | 14.69783 |
| **ZDHHC8** | 0.00000 | 0.00218 | 0.00000 | 0.21448 | 0.38107 | 0.96935 | 2.94856 |
| **ZDHHC8P1** | 0.11058 | 0.00000 | 0.09417 | 0.29060 | 0.00690 | 0.25270 | 0.06555 |
| **ZDHHC9** | 8.30326 | 16.90412 | 9.30800 | 1.03440 | 0.67673 | 11.23327 | 6.05024 |
| **ZEB1** | 0.01895 | 0.07252 | 0.26334 | 0.11938 | 1.39492 | 0.00000 | 0.05290 |
| **ZEB1-AS1** | 0.02431 | 0.27275 | 1.19544 | 0.16983 | 0.13239 | 0.03725 | 0.00000 |
| **ZEB2** | 0.07366 | 0.00000 | 0.13512 | 0.01643 | 0.23755 | 0.28673 | 0.03752 |
| **ZER1** | 0.41810 | 7.16376 | 2.94540 | 1.83015 | 0.67883 | 1.19879 | 3.91581 |
| **ZFAND1** | 10.50536 | 0.67642 | 3.46133 | 4.12760 | 0.38502 | 4.03491 | 1.48146 |
| **ZFAND2A** | 2.71678 | 14.23305 | 6.76705 | 16.56580 | 15.02273 | 6.22541 | 0.00000 |
| **ZFAND2B** | 1.95207 | 6.57900 | 1.45470 | 2.90914 | 0.64422 | 4.43077 | 1.37475 |
| **ZFAND3** | 0.00000 | 2.48052 | 1.37156 | 2.58313 | 1.28976 | 3.19591 | 8.28518 |
| **ZFAND4** | 1.14348 | 0.18265 | 1.73316 | 0.30916 | 1.19731 | 0.35364 | 0.15566 |
| **ZFAND5** | 3.07222 | 6.84241 | 30.29326 | 10.44530 | 15.48226 | 9.93681 | 27.63956 |
| **ZFAND6** | 23.66968 | 3.29532 | 10.28875 | 15.47215 | 7.34827 | 9.37671 | 2.26188 |
| **ZFAS1** | 19.42942 | 2.10075 | 15.35597 | 28.29427 | 6.57645 | 20.01889 | 45.66311 |
| **ZFAT** | 0.83354 | 0.33528 | 1.25323 | 8.04387 | 0.43408 | 1.04707 | 1.60730 |
| **ZFAT-AS1** | 0.02665 | 0.01634 | 0.01973 | 0.07294 | 0.00826 | 0.01472 | 0.00920 |
| **ZFC3H1** | 7.66325 | 0.33940 | 3.68145 | 8.86487 | 3.30338 | 2.75018 | 13.57893 |
| **ZFHX2** | 0.00000 | 0.01257 | 0.01619 | 0.39974 | 0.16945 | 0.90402 | 0.44255 |
| **ZFHX3** | 9.36511 | 1.51418 | 0.51691 | 15.40252 | 1.00174 | 0.00000 | 35.08766 |

| | | | | | | | |
|---|---|---|---|---|---|---|---|
| **ZFHX4** | 0.08342 | 0.00000 | 0.02719 | 0.05210 | 0.86681 | 0.09853 | 0.00000 |
| **ZFP1** | 2.87577 | 0.00000 | 2.00359 | 1.57072 | 0.31410 | 1.09030 | 0.08733 |
| **ZFP14** | 0.19288 | 0.00000 | 0.31878 | 0.41417 | 0.31094 | 0.30749 | 0.46227 |
| **ZFP2** | 0.18422 | 0.14006 | 0.10003 | 0.00000 | 0.01244 | 0.11079 | 0.10616 |
| **ZFP28** | 0.64876 | 0.34564 | 0.75498 | 0.18601 | 4.93465 | 0.32098 | 0.00000 |
| **ZFP3** | 0.80740 | 0.00000 | 1.93204 | 0.14452 | 2.37645 | 0.30677 | 0.00000 |
| **ZFP30** | 0.52094 | 0.00000 | 1.18182 | 0.15107 | 0.02289 | 0.87881 | 0.97841 |
| **ZFP36** | 0.04265 | 0.16141 | 0.24114 | 3.23700 | 0.19540 | 1.31915 | 2.92183 |
| **ZFP36L1** | 7.01125 | 4.72805 | 3.38398 | 23.71071 | 1.63561 | 10.65824 | 11.86008 |
| **ZFP36L2** | 13.89771 | 11.46702 | 2.75455 | 2.64375 | 0.58702 | 16.08098 | 25.37298 |
| **ZFP37** | 0.64376 | 0.00000 | 0.71651 | 1.02436 | 2.61404 | 0.34012 | 0.08199 |
| **ZFP41** | 0.00000 | 0.70302 | 1.23196 | 0.01320 | 0.26346 | 0.15496 | 1.03055 |
| **ZFP42** | 48.71197 | 25.94087 | 19.52149 | 21.97301 | 4.49258 | 20.06610 | 16.88882 |
| **ZFP57** | 8.21453 | 14.51269 | 4.29137 | 4.41998 | 0.00000 | 11.02128 | 0.00000 |
| **ZFP62** | 2.91673 | 0.00000 | 3.70657 | 2.75601 | 2.13417 | 0.17099 | 1.72116 |
| **ZFP64** | 10.95525 | 6.40041 | 1.62480 | 8.10521 | 0.06863 | 2.14046 | 4.31375 |
| **ZFP69** | 0.46510 | 0.37104 | 0.29631 | 0.23062 | 0.21214 | 0.16318 | 0.90958 |
| **ZFP69B** | 0.92062 | 0.00000 | 1.14296 | 0.38390 | 0.59070 | 0.12823 | 0.69019 |
| **ZFP82** | 8.27761 | 0.60476 | 4.82603 | 2.59961 | 2.23617 | 1.43468 | 0.00000 |
| **ZFP90** | 0.07547 | 0.10696 | 0.92832 | 0.12072 | 0.87123 | 0.13146 | 0.71538 |
| **ZFP91** | 3.52019 | 0.00000 | 2.73351 | 2.97211 | 1.28487 | 1.78172 | 12.84675 |
| **ZFPL1** | 7.43057 | 11.63794 | 3.39686 | 9.13773 | 0.56152 | 3.29715 | 1.86367 |
| **ZFPM1** | 0.00000 | 0.15096 | 0.06481 | 0.00000 | 0.00715 | 0.11361 | 0.42124 |
| **ZFPM2** | 0.01775 | 0.00220 | 0.03042 | 0.01725 | 2.44400 | 0.01157 | 0.03745 |
| **ZFR** | 23.41483 | 15.96339 | 14.09637 | 16.90486 | 5.43030 | 8.94126 | 10.56017 |
| **ZFR2** | 0.00000 | 0.15594 | 5.08993 | 0.00000 | 1.30913 | 0.36582 | 0.15303 |
| **ZFX** | 9.99252 | 1.66806 | 4.30072 | 2.88811 | 1.86092 | 3.08398 | 3.67511 |
| **ZFX-AS1** | 0.02202 | 0.01685 | 0.01760 | 0.02405 | 0.01473 | 0.01684 | 0.02000 |
| **ZFY** | 0.03266 | 0.00620 | 0.03508 | 0.02445 | 0.00668 | 0.01373 | 0.02305 |
| **ZFYVE1** | 1.07149 | 11.13266 | 19.13225 | 5.53165 | 9.85499 | 4.55039 | 0.00000 |
| **ZFYVE16** | 4.92237 | 0.00000 | 2.26525 | 2.42530 | 1.01085 | 2.42500 | 5.44187 |
| **ZFYVE19** | 4.12801 | 7.26622 | 14.10174 | 9.78326 | 5.74516 | 1.47537 | 3.55854 |
| **ZFYVE20** | 7.39699 | 5.89558 | 2.43838 | 5.57950 | 0.40808 | 4.49434 | 5.09818 |
| **ZFYVE21** | 27.30126 | 20.79665 | 0.99174 | 12.70840 | 2.71670 | 7.58698 | 3.63936 |
| **ZFYVE26** | 0.94767 | 1.20241 | 0.27666 | 3.00103 | 0.00123 | 2.86522 | 3.87471 |

| | | | | | | | |
|---|---|---|---|---|---|---|---|
| **ZFYVE27** | 3.26861 | 3.93391 | 3.69577 | 7.52753 | 2.09156 | 2.23684 | 2.16091 |
| **ZFYVE28** | 0.06113 | 0.04980 | 0.05689 | 0.02793 | 0.03471 | 0.02685 | 0.21735 |
| **ZFYVE9** | 0.00108 | 1.42875 | 2.07800 | 0.23502 | 0.56171 | 3.02385 | 5.55423 |
| **ZG16** | 0.32588 | 0.57710 | 0.03162 | 0.20016 | 0.00615 | 0.00000 | 0.00201 |
| **ZG16B** | 0.03176 | 0.02485 | 0.00928 | 0.02212 | 0.02103 | 0.03719 | 0.01635 |
| **ZGLP1** | 0.56396 | 0.71210 | 0.43022 | 0.46816 | 0.77450 | 0.00000 | 0.02858 |
| **ZGPAT** | 5.54041 | 16.17692 | 5.74275 | 10.29355 | 4.31242 | 4.44294 | 0.64810 |
| **ZGRF1** | 4.19530 | 1.19186 | 5.98637 | 0.50257 | 1.35497 | 3.99069 | 4.86580 |
| **ZHX1** | 1.54038 | 2.06184 | 2.93735 | 0.56354 | 3.84104 | 4.20707 | 2.18681 |
| **ZHX1-C8orf76** | 0.39455 | 0.22702 | 0.76679 | 0.49406 | 0.56571 | 0.44201 | 0.59107 |
| **ZHX2** | 0.00000 | 0.10371 | 0.28903 | 0.51147 | 0.74614 | 0.12325 | 0.00000 |
| **ZHX3** | 0.19762 | 1.04812 | 0.07515 | 0.00000 | 8.53194 | 0.64646 | 2.71538 |
| **ZIC1** | 0.02605 | 0.01013 | 0.03698 | 0.02286 | 0.03392 | 0.01775 | 0.02580 |
| **ZIC2** | 0.00000 | 0.24620 | 0.18009 | 0.01872 | 0.25100 | 1.47230 | 0.02926 |
| **ZIC3** | 0.03550 | 0.00000 | 0.00538 | 0.05077 | 0.06457 | 2.19661 | 0.00016 |
| **ZIC4** | 0.02083 | 0.01188 | 0.01238 | 0.01739 | 0.02404 | 0.01273 | 0.04652 |
| **ZIC5** | 0.51976 | 0.66328 | 0.14895 | 0.00000 | 0.27135 | 0.42946 | 0.04231 |
| **ZIK1** | 2.93247 | 0.14363 | 9.62006 | 1.79221 | 8.41976 | 2.65579 | 0.00000 |
| **ZIM2** | 0.13475 | 0.00000 | 1.27112 | 0.09117 | 1.47594 | 0.00211 | 0.00000 |
| **ZIM3** | 0.00714 | 0.00000 | 4.72132 | 0.00298 | 17.19701 | 0.00000 | 0.03348 |
| **ZKSCAN1** | 5.80205 | 1.36879 | 1.95304 | 4.68554 | 1.82529 | 0.00000 | 3.39560 |
| **ZKSCAN2** | 0.59522 | 1.16994 | 1.45235 | 2.26952 | 1.63042 | 0.80913 | 3.88858 |
| **ZKSCAN3** | 1.18325 | 0.59744 | 1.09765 | 0.20361 | 0.00845 | 0.09563 | 0.05839 |
| **ZKSCAN4** | 0.30789 | 0.22456 | 4.15015 | 0.00000 | 1.00845 | 0.63609 | 0.28706 |
| **ZKSCAN5** | 4.50521 | 1.50999 | 3.03006 | 1.89785 | 6.00291 | 1.97149 | 1.92371 |
| **ZKSCAN7** | 0.09626 | 0.01425 | 0.04520 | 0.10102 | 0.14192 | 0.01419 | 0.00493 |
| **ZKSCAN8** | 3.92050 | 0.00000 | 2.19824 | 0.37900 | 2.59661 | 1.14206 | 3.54595 |
| **ZMAT1** | 0.05327 | 0.00152 | 0.03178 | 0.01610 | 0.02448 | 0.04388 | 0.02095 |
| **ZMAT2** | 10.26330 | 3.11697 | 9.00431 | 14.29669 | 6.00712 | 8.62778 | 16.77024 |
| **ZMAT3** | 2.29154 | 2.95386 | 3.54575 | 1.58645 | 0.74611 | 6.95772 | 9.21318 |
| **ZMAT4** | 0.01888 | 0.01475 | 0.01195 | 0.05510 | 0.09643 | 0.06694 | 0.04426 |
| **ZMAT5** | 3.05039 | 9.46881 | 5.24713 | 6.97513 | 10.10328 | 0.45259 | 0.00000 |
| **ZMIZ1** | 11.61557 | 8.06556 | 2.71498 | 3.04977 | 0.01256 | 8.52525 | 20.27727 |
| **ZMIZ1-AS1** | 0.02999 | 0.01184 | 0.01505 | 0.01778 | 0.02321 | 0.02513 | 0.01818 |
| **ZMIZ2** | 0.47289 | 2.72739 | 0.00000 | 0.81741 | 0.07423 | 4.08770 | 6.31279 |

| | | | | | | | |
|---|---|---|---|---|---|---|---|
| **ZMPSTE24** | 35.08332 | 2.92210 | 12.11368 | 11.67282 | 4.70439 | 11.24415 | 8.24797 |
| **ZMYM1** | 8.54327 | 0.78151 | 13.61055 | 2.12538 | 1.05922 | 2.56598 | 6.47741 |
| **ZMYM2** | 9.97244 | 0.00000 | 9.00133 | 3.01622 | 4.51393 | 8.39216 | 11.90939 |
| **ZMYM3** | 2.83292 | 1.13112 | 3.45419 | 0.00000 | 0.66700 | 5.07692 | 4.79684 |
| **ZMYM4** | 4.25623 | 1.66048 | 4.05787 | 3.29848 | 1.32265 | 3.62205 | 16.28718 |
| **ZMYM5** | 5.95768 | 0.69225 | 6.34695 | 1.95719 | 2.23515 | 3.54653 | 1.56835 |
| **ZMYM6** | 3.27449 | 0.02212 | 2.95002 | 1.52951 | 2.46548 | 1.06077 | 1.27105 |
| **ZMYM6NB** | 0.59328 | 3.01831 | 7.79683 | 0.43133 | 0.17676 | 2.70689 | 0.00000 |
| **ZMYND10** | 0.04994 | 0.08043 | 0.05909 | 0.43293 | 0.00975 | 0.08117 | 0.24605 |
| **ZMYND11** | 8.13056 | 0.08321 | 1.21413 | 1.62237 | 0.48317 | 0.00000 | 8.52944 |
| **ZMYND12** | 0.24512 | 0.05702 | 0.00000 | 0.04685 | 0.27744 | 0.08332 | 0.00000 |
| **ZMYND15** | 0.01884 | 0.00000 | 0.05355 | 0.01757 | 0.03064 | 0.01537 | 0.03966 |
| **ZMYND19** | 0.33346 | 5.30417 | 2.45388 | 1.98317 | 1.71381 | 2.52836 | 12.84462 |
| **ZMYND8** | 13.81085 | 7.34828 | 6.22721 | 13.63172 | 10.21306 | 16.54915 | 12.48494 |
| **ZNF10** | 1.60744 | 0.00000 | 2.08467 | 1.47812 | 1.55708 | 0.82301 | 0.00000 |
| **ZNF100** | 6.56038 | 0.00000 | 8.44615 | 1.82306 | 5.34076 | 4.74681 | 5.17878 |
| **ZNF101** | 4.26344 | 1.18529 | 7.51271 | 0.81415 | 1.71498 | 3.03093 | 0.79814 |
| **ZNF106** | 29.62572 | 16.42193 | 23.88740 | 15.21948 | 5.27017 | 12.00712 | 31.95556 |
| **ZNF107** | 4.08937 | 0.00000 | 3.94089 | 1.67585 | 5.24965 | 2.07972 | 6.05938 |
| **ZNF112** | 1.13141 | 0.21099 | 0.25830 | 0.47691 | 0.33646 | 0.43847 | 0.10029 |
| **ZNF114** | 11.91714 | 7.40383 | 4.04514 | 12.37254 | 0.27050 | 1.40687 | 0.00000 |
| **ZNF117** | 0.50278 | 0.00000 | 0.27180 | 1.25775 | 0.06867 | 0.07508 | 1.39354 |
| **ZNF12** | 2.06658 | 0.00000 | 2.79242 | 0.88435 | 0.86913 | 2.11377 | 5.45738 |
| **ZNF121** | 3.96244 | 0.00000 | 1.58251 | 0.87673 | 1.11405 | 0.65139 | 1.30650 |
| **ZNF124** | 6.67527 | 0.00000 | 3.87703 | 2.87477 | 0.80568 | 1.57445 | 2.00163 |
| **ZNF131** | 7.03300 | 0.39848 | 5.31268 | 3.04983 | 7.60736 | 5.65551 | 9.47125 |
| **ZNF132** | 0.13478 | 0.10293 | 2.07432 | 0.37875 | 2.01523 | 0.14925 | 0.00000 |
| **ZNF133** | 3.38349 | 2.36020 | 1.18072 | 3.01477 | 0.48796 | 1.74086 | 1.29846 |
| **ZNF134** | 6.75584 | 3.80338 | 15.29506 | 2.99891 | 6.74006 | 1.98565 | 0.00000 |
| **ZNF135** | 0.10975 | 0.62683 | 0.19336 | 0.02495 | 0.07484 | 0.06904 | 0.03710 |
| **ZNF136** | 5.95754 | 0.10541 | 4.42393 | 2.04147 | 9.50553 | 1.95469 | 0.76149 |
| **ZNF137P** | 0.47637 | 0.00000 | 0.54891 | 0.26883 | 0.28348 | 0.29262 | 1.17267 |
| **ZNF138** | 6.60333 | 0.21937 | 9.37177 | 2.77399 | 2.11399 | 3.38082 | 1.02294 |
| **ZNF14** | 2.82842 | 0.39784 | 1.00004 | 0.98199 | 0.38074 | 1.21453 | 0.59508 |
| **ZNF140** | 4.65557 | 0.54980 | 4.05007 | 2.34211 | 6.13572 | 2.42273 | 0.00000 |

| | | | | | | | |
|---|---|---|---|---|---|---|---|
| **ZNF141** | 2.27372 | 0.21054 | 2.08382 | 0.21653 | 0.69346 | 0.44209 | 0.53242 |
| **ZNF142** | 3.75831 | 4.95632 | 5.23383 | 0.00000 | 5.73390 | 3.77682 | 10.49068 |
| **ZNF143** | 14.78109 | 1.68686 | 5.16706 | 18.20711 | 5.96260 | 4.81569 | 0.00000 |
| **ZNF146** | 36.66440 | 7.66482 | 12.83190 | 19.77807 | 2.68701 | 14.41047 | 10.37349 |
| **ZNF148** | 8.24665 | 0.00000 | 5.56937 | 3.10006 | 4.08326 | 4.13790 | 10.16041 |
| **ZNF154** | 0.27538 | 0.10629 | 0.21384 | 0.14271 | 0.03835 | 1.02722 | 3.04731 |
| **ZNF155** | 1.04236 | 0.41418 | 1.00720 | 0.27064 | 2.95726 | 0.00000 | 0.00000 |
| **ZNF157** | 0.02788 | 0.00000 | 0.55698 | 0.02509 | 3.52554 | 0.17893 | 0.00000 |
| **ZNF16** | 1.08884 | 2.02536 | 4.65915 | 0.24319 | 1.44689 | 0.62978 | 0.00000 |
| **ZNF160** | 4.79339 | 2.21524 | 2.24351 | 2.03892 | 2.55325 | 1.78604 | 0.00000 |
| **ZNF165** | 20.92592 | 5.28568 | 1.81387 | 19.92439 | 0.62958 | 0.00556 | 0.00000 |
| **ZNF169** | 0.60104 | 0.39457 | 0.93745 | 0.41662 | 0.63737 | 0.31342 | 0.61870 |
| **ZNF17** | 2.22397 | 0.27607 | 2.75008 | 0.70833 | 2.04828 | 1.70301 | 0.64381 |
| **ZNF174** | 3.99526 | 2.60581 | 4.99187 | 2.18398 | 4.01903 | 2.25915 | 1.34858 |
| **ZNF175** | 1.10345 | 0.18923 | 0.80286 | 0.45525 | 0.45126 | 0.44595 | 0.45581 |
| **ZNF177.uid-232** | 0.02875 | 0.02069 | 0.01617 | 0.01844 | 0.00943 | 0.01333 | 0.01279 |
| **ZNF18** | 0.96160 | 2.11592 | 2.76482 | 0.35354 | 2.23881 | 2.48184 | 0.00000 |
| **ZNF180** | 4.95388 | 0.00000 | 3.23195 | 1.70086 | 1.69343 | 1.48083 | 0.56708 |
| **ZNF181** | 0.26916 | 0.06135 | 0.36611 | 0.21663 | 0.19894 | 0.70841 | 0.13491 |
| **ZNF182** | 1.80689 | 0.57638 | 1.92316 | 0.04477 | 2.06313 | 1.71294 | 0.31413 |
| **ZNF184** | 2.19302 | 0.00000 | 2.18688 | 0.75930 | 3.69218 | 0.52325 | 0.00000 |
| **ZNF185** | 0.46115 | 0.14937 | 0.10970 | 0.12233 | 0.14704 | 0.14328 | 0.00000 |
| **ZNF189** | 0.34292 | 0.00000 | 1.69378 | 0.85425 | 1.26555 | 0.92697 | 1.67203 |
| **ZNF19** | 0.15665 | 0.32987 | 2.76448 | 0.00000 | 0.36530 | 0.09031 | 0.00000 |
| **ZNF192P1** | 0.02665 | 0.06392 | 0.10667 | 0.12324 | 0.20704 | 0.00000 | 0.06284 |
| **ZNF195** | 18.82479 | 1.11606 | 10.60000 | 11.72006 | 2.81277 | 8.48808 | 8.98216 |
| **ZNF197** | 0.19351 | 0.20022 | 0.57683 | 0.00000 | 0.07332 | 0.01868 | 0.41088 |
| **ZNF197-AS1** | 0.03167 | 0.05797 | 0.01194 | 0.01746 | 0.00781 | 0.00910 | 0.00425 |
| **ZNF2** | 1.44941 | 0.48758 | 1.13556 | 0.78599 | 1.67992 | 0.21499 | 0.00000 |
| **ZNF20** | 0.03126 | 0.03244 | 0.01314 | 0.02116 | 0.01142 | 0.01541 | 0.00766 |
| **ZNF200** | 10.65833 | 2.91744 | 16.07212 | 8.23277 | 8.21708 | 6.26333 | 0.00000 |
| **ZNF202** | 3.43150 | 1.36807 | 2.36326 | 1.92819 | 3.20804 | 1.69915 | 1.42267 |
| **ZNF204P** | 0.55292 | 0.00000 | 0.00150 | 0.92719 | 0.15440 | 0.29092 | 0.02464 |
| **ZNF205** | 0.00000 | 0.66132 | 0.05837 | 0.03910 | 0.00154 | 0.60525 | 2.15936 |
| **ZNF205-AS1** | 0.05107 | 0.08167 | 0.12138 | 0.03249 | 0.01206 | 0.00295 | 0.02640 |

| | | | | | | | |
|---|---|---|---|---|---|---|---|
| **ZNF207** | 43.07216 | 19.49545 | 23.46227 | 24.06040 | 11.60221 | 15.81856 | 12.13948 |
| **ZNF208** | 7.96907 | 0.00000 | 4.01697 | 0.40928 | 1.67628 | 6.74212 | 16.33773 |
| **ZNF211** | 3.63330 | 2.08198 | 2.05514 | 4.67890 | 6.28306 | 1.46909 | 0.00000 |
| **ZNF212** | 4.14093 | 3.93696 | 7.34551 | 3.50982 | 2.71205 | 2.13512 | 1.81120 |
| **ZNF213** | 0.30054 | 0.23578 | 0.08999 | 0.23017 | 0.05896 | 0.19705 | 0.19798 |
| **ZNF213-AS1** | 0.97644 | 0.44922 | 0.96554 | 0.62065 | 0.14557 | 0.00000 | 0.00000 |
| **ZNF214** | 0.51786 | 0.00000 | 0.53098 | 0.39197 | 1.42780 | 0.16250 | 0.00000 |
| **ZNF215** | 4.16619 | 0.00000 | 5.08814 | 3.33158 | 4.59983 | 1.95192 | 1.53014 |
| **ZNF217** | 2.60887 | 0.00000 | 7.73091 | 3.83539 | 1.68524 | 7.46189 | 13.37653 |
| **ZNF219** | 0.00000 | 0.31799 | 0.02253 | 0.46174 | 0.01396 | 0.39226 | 0.88320 |
| **ZNF22** | 5.08929 | 4.28708 | 12.23181 | 0.00000 | 3.27156 | 3.00055 | 1.36092 |
| **ZNF221** | 0.25369 | 0.00000 | 0.28272 | 0.20540 | 1.27032 | 0.11989 | 0.00000 |
| **ZNF222** | 0.99722 | 0.46558 | 1.52691 | 2.27605 | 0.76679 | 0.21640 | 0.00000 |
| **ZNF223** | 1.37520 | 0.20126 | 0.44599 | 0.26724 | 0.28458 | 0.38401 | 0.00000 |
| **ZNF224** | 0.35018 | 0.13849 | 0.15429 | 0.23370 | 0.93977 | 0.00530 | 0.00000 |
| **ZNF225** | 1.41295 | 0.74829 | 2.19863 | 0.19097 | 0.50022 | 0.24624 | 0.00000 |
| **ZNF226** | 3.17492 | 0.80010 | 6.73297 | 1.53662 | 1.92755 | 0.37829 | 0.00000 |
| **ZNF227** | 5.73978 | 0.00000 | 5.83293 | 4.30966 | 1.82574 | 1.15674 | 0.00000 |
| **ZNF229** | 2.12544 | 0.56834 | 3.43200 | 1.85965 | 5.61988 | 0.57795 | 0.00000 |
| **ZNF23** | 2.71167 | 0.58096 | 5.48422 | 0.69061 | 2.50121 | 1.21688 | 0.00000 |
| **ZNF230** | 1.35622 | 0.00000 | 2.99448 | 1.01594 | 2.12718 | 0.32817 | 0.00000 |
| **ZNF232** | 10.33966 | 2.04919 | 14.87821 | 8.99529 | 1.07887 | 6.87466 | 0.00000 |
| **ZNF233** | 0.16562 | 0.20201 | 0.54297 | 0.34313 | 0.04874 | 0.00000 | 0.66830 |
| **ZNF234** | 3.44971 | 0.29754 | 3.15284 | 2.22276 | 0.35319 | 1.65110 | 0.19190 |
| **ZNF235** | 1.58501 | 0.20670 | 2.12660 | 1.77035 | 0.31571 | 0.57770 | 0.00000 |
| **ZNF236** | 3.82275 | 1.26189 | 2.57742 | 2.04009 | 0.49330 | 2.59588 | 7.42826 |
| **ZNF239** | 0.47517 | 0.00000 | 1.42243 | 0.30696 | 0.59074 | 0.43225 | 0.87463 |
| **ZNF24** | 11.72216 | 3.83309 | 10.85191 | 8.21202 | 3.12376 | 6.85673 | 10.35491 |
| **ZNF248** | 1.83934 | 0.67701 | 0.28690 | 1.36674 | 0.01558 | 0.44479 | 0.12459 |
| **ZNF25** | 0.03563 | 0.06436 | 0.00000 | 0.11273 | 0.16184 | 0.01403 | 0.02599 |
| **ZNF250** | 1.36715 | 1.22245 | 1.09725 | 0.22934 | 1.83227 | 0.98652 | 0.00000 |
| **ZNF251** | 2.76653 | 2.20125 | 1.30496 | 1.17818 | 1.34712 | 1.28242 | 0.48511 |
| **ZNF252P** | 1.14862 | 0.00000 | 3.67532 | 0.27843 | 2.11276 | 0.87261 | 0.37194 |
| **ZNF252P-AS1** | 0.03845 | 0.00000 | 0.11859 | 0.03202 | 0.36477 | 0.01748 | 0.01781 |
| **ZNF253** | 4.60943 | 0.00000 | 1.57421 | 1.43756 | 1.27949 | 0.54938 | 0.00000 |

| | | | | | | | |
|---|---|---|---|---|---|---|---|
| **ZNF254** | 9.79963 | 0.00000 | 5.89461 | 4.75124 | 3.42085 | 1.93594 | 1.83935 |
| **ZNF256** | 4.18615 | 2.86658 | 9.96168 | 1.70903 | 9.28667 | 0.95225 | 0.00000 |
| **ZNF257** | 7.68270 | 0.00000 | 5.27088 | 1.77857 | 0.34529 | 4.56226 | 7.34521 |
| **ZNF26.uid-1752** | 3.10494 | 1.47906 | 2.79401 | 1.52669 | 2.33352 | 1.24225 | 0.00000 |
| **ZNF260** | 6.78231 | 0.00000 | 11.30549 | 4.46957 | 2.59229 | 2.51902 | 3.61931 |
| **ZNF263** | 4.70141 | 4.65221 | 3.32858 | 1.89311 | 2.67155 | 6.32530 | 8.12519 |
| **ZNF264** | 2.09000 | 0.00000 | 1.81504 | 1.03584 | 1.61778 | 1.40219 | 4.31211 |
| **ZNF266** | 3.67390 | 1.02286 | 6.70215 | 2.83059 | 4.39565 | 1.05847 | 3.32995 |
| **ZNF267** | 13.17310 | 0.00000 | 11.53004 | 5.11950 | 5.72220 | 3.98148 | 1.76155 |
| **ZNF268** | 5.20501 | 0.00000 | 14.03674 | 2.98860 | 4.95955 | 3.44643 | 1.53525 |
| **ZNF271** | 9.76053 | 3.57416 | 6.50802 | 6.29914 | 2.24907 | 3.15779 | 2.63203 |
| **ZNF273** | 4.56145 | 0.00000 | 7.29613 | 1.69077 | 2.30985 | 3.18934 | 0.72300 |
| **ZNF274** | 5.86486 | 3.94128 | 5.20487 | 4.48381 | 1.16873 | 1.63493 | 0.00000 |
| **ZNF275** | 0.00000 | 1.37357 | 0.37639 | 0.33190 | 0.64287 | 0.40302 | 0.44396 |
| **ZNF276** | 0.78875 | 2.16075 | 0.38498 | 0.98268 | 0.35547 | 1.52405 | 4.14962 |
| **ZNF277** | 5.78529 | 1.20842 | 2.45740 | 12.52677 | 0.64887 | 2.46043 | 1.38973 |
| **ZNF28** | 15.18484 | 5.60050 | 13.19371 | 6.25586 | 2.99849 | 5.63934 | 0.00000 |
| **ZNF280A** | 1.39579 | 1.43190 | 20.37432 | 0.00000 | 24.10883 | 0.99856 | 0.00000 |
| **ZNF280B** | 1.91629 | 0.24214 | 4.87821 | 0.37212 | 18.46096 | 2.19793 | 0.10379 |
| **ZNF280C** | 1.25866 | 0.00000 | 0.85760 | 0.30935 | 3.12915 | 0.12367 | 0.42690 |
| **ZNF280D** | 0.23934 | 0.00000 | 0.52495 | 0.33932 | 0.42322 | 0.22321 | 0.57161 |
| **ZNF281** | 15.98213 | 1.55774 | 14.66028 | 8.38166 | 9.88149 | 11.64687 | 17.11349 |
| **ZNF282** | 0.17733 | 1.37649 | 0.21937 | 0.98450 | 0.04274 | 0.87758 | 5.59482 |
| **ZNF283** | 2.32684 | 0.03906 | 1.29346 | 0.96977 | 0.29770 | 0.26005 | 0.00000 |
| **ZNF284** | 0.28055 | 0.27911 | 0.50133 | 0.25161 | 1.24969 | 0.05356 | 0.00000 |
| **ZNF285** | 0.30808 | 0.62334 | 1.50317 | 0.54602 | 0.80248 | 0.09686 | 0.00000 |
| **ZNF286A** | 6.14903 | 1.40556 | 1.63892 | 5.72922 | 3.26198 | 2.58930 | 0.00000 |
| **ZNF286B** | 1.16056 | 0.22640 | 0.16529 | 1.29461 | 1.18362 | 0.50551 | 0.00000 |
| **ZNF287** | 0.08739 | 0.00559 | 0.66061 | 0.13214 | 0.55418 | 0.08883 | 0.00000 |
| **ZNF292** | 5.25521 | 0.00000 | 5.98452 | 20.73812 | 3.46277 | 8.25711 | 11.44307 |
| **ZNF295-AS1** | 0.08698 | 0.05769 | 0.07635 | 0.13527 | 0.04242 | 0.00204 | 0.02030 |
| **ZNF296** | 2.95492 | 23.45700 | 18.74124 | 2.53054 | 15.47061 | 13.32853 | 13.78105 |
| **ZNF3** | 10.09287 | 7.27494 | 2.86455 | 5.98485 | 0.98621 | 6.59913 | 6.25946 |
| **ZNF30** | 0.46330 | 0.78474 | 3.61087 | 0.00175 | 0.88331 | 1.70633 | 0.00000 |
| **ZNF300** | 2.86554 | 0.96531 | 1.01168 | 4.21606 | 0.96196 | 1.86191 | 0.10514 |

| | | | | | | | |
|---|---|---|---|---|---|---|---|
| **ZNF300P1** | 1.57881 | 0.27076 | 0.26431 | 1.39761 | 0.25840 | 0.43431 | 0.27604 |
| **ZNF302** | 1.74354 | 0.96547 | 3.22733 | 1.93930 | 0.61942 | 1.31982 | 1.11803 |
| **ZNF304** | 4.64804 | 2.35476 | 5.00310 | 1.34797 | 5.02027 | 2.27739 | 0.00000 |
| **ZNF311** | 0.00000 | 0.31214 | 1.23160 | 0.00000 | 1.60604 | 0.22209 | 0.12594 |
| **ZNF316** | 0.13006 | 0.23340 | 0.04061 | 0.57952 | 0.04493 | 0.00000 | 1.83033 |
| **ZNF317** | 8.67332 | 5.54712 | 1.64685 | 7.80504 | 0.97790 | 4.00101 | 2.88378 |
| **ZNF318** | 1.53921 | 1.68200 | 6.21154 | 0.88433 | 1.61354 | 7.08379 | 9.47829 |
| **ZNF319** | 0.02653 | 0.14366 | 0.00719 | 0.10414 | 0.00978 | 0.00000 | 0.26198 |
| **ZNF32** | 0.47523 | 0.60870 | 0.58066 | 0.89992 | 0.00000 | 0.85499 | 1.26836 |
| **ZNF32-AS1** | 0.02283 | 0.01824 | 0.01310 | 0.02343 | 0.00765 | 0.01424 | 0.02729 |
| **ZNF32-AS2** | 0.05819 | 0.05384 | 0.11329 | 0.04250 | 0.00790 | 0.06781 | 0.00524 |
| **ZNF320** | 6.30504 | 2.45970 | 3.18594 | 3.29147 | 1.04878 | 3.50254 | 6.52176 |
| **ZNF321P** | 0.04063 | 0.11633 | 0.05363 | 0.13052 | 0.10225 | 0.02660 | 0.08833 |
| **ZNF322** | 0.53126 | 0.29075 | 0.40458 | 0.40371 | 0.42848 | 0.24551 | 0.08955 |
| **ZNF324** | 3.09041 | 5.22312 | 3.68453 | 2.38018 | 2.39258 | 1.66360 | 3.19693 |
| **ZNF324B** | 0.11549 | 0.31897 | 0.63565 | 0.37111 | 0.04502 | 0.08825 | 0.19408 |
| **ZNF326** | 13.84562 | 3.76839 | 11.15125 | 14.51281 | 1.17458 | 6.24409 | 11.36247 |
| **ZNF329** | 0.30372 | 0.00000 | 1.83083 | 0.33552 | 0.67795 | 0.47156 | 0.17845 |
| **ZNF330** | 14.44014 | 8.17685 | 10.49073 | 5.81707 | 7.20340 | 6.10734 | 1.51358 |
| **ZNF331** | 9.07206 | 6.24950 | 4.20872 | 20.98089 | 0.52318 | 5.27192 | 8.94502 |
| **ZNF333** | 0.87406 | 0.00000 | 0.30263 | 0.87975 | 0.04143 | 0.93097 | 0.48112 |
| **ZNF334** | 0.39379 | 0.12697 | 1.59002 | 0.55934 | 0.04791 | 0.21549 | 0.00000 |
| **ZNF335** | 2.75118 | 10.04960 | 4.59618 | 10.47116 | 2.73030 | 13.19272 | 12.50499 |
| **ZNF337** | 2.76360 | 1.32138 | 3.84409 | 1.67984 | 0.98938 | 0.91655 | 0.73140 |
| **ZNF33A** | 5.14294 | 0.52832 | 3.78830 | 1.60575 | 0.14573 | 1.98589 | 3.96816 |
| **ZNF33B** | 1.63046 | 0.00000 | 0.83243 | 1.27992 | 0.43629 | 0.42448 | 0.81443 |
| **ZNF33BP1** | 0.02105 | 0.01932 | 0.01346 | 0.02049 | 0.00810 | 0.01711 | 0.02171 |
| **ZNF34** | 0.58989 | 1.58280 | 1.49928 | 0.42345 | 1.23435 | 2.04121 | 1.43166 |
| **ZNF341** | 0.49622 | 0.55275 | 0.54810 | 4.19744 | 0.21333 | 0.88074 | 3.07951 |
| **ZNF341-AS1** | 0.03366 | 0.02651 | 0.01174 | 0.06839 | 0.01364 | 0.01554 | 0.00032 |
| **ZNF343** | 2.85327 | 4.59306 | 5.51292 | 1.13662 | 3.51318 | 2.97009 | 0.00000 |
| **ZNF345** | 1.44942 | 0.00000 | 1.90238 | 1.68246 | 1.36947 | 0.65000 | 0.43364 |
| **ZNF346** | 5.05220 | 4.44575 | 0.93680 | 5.58785 | 2.51133 | 1.91150 | 0.87367 |
| **ZNF347** | 1.35829 | 0.71865 | 2.06874 | 1.47679 | 1.27671 | 0.70560 | 0.38767 |
| **ZNF35** | 0.35843 | 0.28901 | 0.00000 | 0.54318 | 0.61911 | 0.01475 | 0.37515 |

| | | | | | | | |
|---|---|---|---|---|---|---|---|
| **ZNF350** | 3.62725 | 4.80127 | 11.09644 | 1.15564 | 9.05706 | 3.39291 | 0.00000 |
| **ZNF350-AS1** | 0.21807 | 0.63755 | 0.35979 | 0.18669 | 0.41800 | 0.02003 | 0.00000 |
| **ZNF354A** | 0.90044 | 0.70435 | 3.58352 | 2.31334 | 1.65813 | 1.42114 | 0.85299 |
| **ZNF354B** | 0.03104 | 0.00000 | 0.99392 | 0.41195 | 0.11461 | 0.12106 | 0.72291 |
| **ZNF354C** | 0.15070 | 0.00000 | 0.13851 | 0.06118 | 0.05292 | 0.02178 | 0.00000 |
| **ZNF358** | 0.00000 | 0.45116 | 0.06888 | 0.24367 | 0.00384 | 0.12663 | 1.54740 |
| **ZNF362** | 0.03940 | 0.60901 | 0.00000 | 3.60483 | 0.45259 | 0.00000 | 2.61438 |
| **ZNF365** | 0.02237 | 0.00000 | 0.21203 | 0.07433 | 0.21359 | 0.01733 | 0.01735 |
| **ZNF366** | 0.04454 | 0.01291 | 0.01362 | 0.03396 | 0.00787 | 0.00981 | 0.01903 |
| **ZNF367** | 1.11303 | 0.00004 | 0.05622 | 0.26732 | 0.00040 | 1.05547 | 1.83244 |
| **ZNF37A** | 6.60514 | 1.48978 | 1.94886 | 6.49540 | 0.34490 | 2.56264 | 5.65291 |
| **ZNF37BP** | 3.40068 | 0.60675 | 1.70896 | 2.87749 | 0.45328 | 1.08417 | 1.32235 |
| **ZNF382** | 0.86247 | 0.13626 | 1.23722 | 0.27151 | 0.26661 | 0.00000 | 0.00000 |
| **ZNF383** | 2.79711 | 0.00000 | 1.90617 | 0.56238 | 0.02982 | 0.62813 | 0.57120 |
| **ZNF384** | 8.22690 | 7.42138 | 6.94234 | 7.72639 | 3.19126 | 5.28700 | 9.30372 |
| **ZNF385A** | 2.50427 | 0.62346 | 0.00000 | 6.93529 | 0.02477 | 0.00000 | 10.33758 |
| **ZNF385B** | 0.10926 | 0.19557 | 0.16117 | 0.03381 | 0.27337 | 0.00987 | 0.00000 |
| **ZNF385C** | 0.07060 | 0.09332 | 0.03840 | 0.19379 | 0.03700 | 0.00000 | 0.04335 |
| **ZNF385D** | 0.04459 | 0.00997 | 0.02271 | 0.01158 | 0.03635 | 0.03850 | 0.04035 |
| **ZNF385D-AS1** | 1.24364 | 0.15016 | 0.55135 | 0.35596 | 0.64671 | 1.05034 | 0.00000 |
| **ZNF391** | 0.66294 | 0.50034 | 1.19315 | 0.36051 | 0.24993 | 0.37352 | 0.00000 |
| **ZNF394** | 6.53293 | 9.73969 | 9.29677 | 6.14907 | 18.06950 | 3.01962 | 0.00000 |
| **ZNF395** | 0.74298 | 2.02722 | 0.40350 | 0.29007 | 0.86174 | 2.47414 | 3.94694 |
| **ZNF396** | 0.04545 | 0.17794 | 0.08352 | 0.18277 | 0.01092 | 0.00097 | 0.00000 |
| **ZNF397** | 1.00845 | 0.00000 | 4.02563 | 1.21005 | 1.81870 | 1.38893 | 3.17020 |
| **ZNF398** | 1.81125 | 5.32404 | 5.85169 | 0.79852 | 0.58124 | 5.43556 | 4.64448 |
| **ZNF404** | 0.30505 | 0.17921 | 0.32004 | 0.31396 | 0.10960 | 0.13482 | 0.00000 |
| **ZNF407** | 1.43351 | 0.00000 | 3.03797 | 2.30566 | 1.16773 | 0.42266 | 4.36350 |
| **ZNF408** | 3.09801 | 10.81889 | 7.07360 | 5.29206 | 8.11270 | 6.88980 | 2.13049 |
| **ZNF41** | 2.40212 | 1.12516 | 1.07721 | 0.42931 | 0.06742 | 0.00000 | 0.07402 |
| **ZNF410** | 24.47582 | 5.23146 | 9.21444 | 18.41203 | 13.10299 | 3.55975 | 2.53248 |
| **ZNF414** | 0.00000 | 0.92617 | 0.42605 | 0.16525 | 0.21602 | 0.19714 | 0.86334 |
| **ZNF415** | 2.46335 | 0.77344 | 4.23144 | 0.92114 | 1.16429 | 0.01525 | 0.00000 |
| **ZNF416** | 2.88308 | 2.56263 | 9.68375 | 0.92399 | 6.93226 | 0.83324 | 0.00000 |
| **ZNF417** | 1.31745 | 0.24109 | 0.92593 | 0.70373 | 0.86261 | 0.05856 | 1.02692 |

| | | | | | | | |
|---|---|---|---|---|---|---|---|
| **ZNF418** | 1.12535 | 0.10957 | 0.58785 | 0.28834 | 0.35387 | 0.44534 | 0.39946 |
| **ZNF419** | 3.43039 | 2.00402 | 3.92699 | 2.36951 | 4.02672 | 0.57773 | 0.21293 |
| **ZNF420** | 5.49937 | 0.00038 | 1.99840 | 2.65565 | 0.32828 | 1.23833 | 0.19458 |
| **ZNF423** | 0.01338 | 0.00667 | 0.00634 | 0.06935 | 0.04488 | 0.35741 | 0.08040 |
| **ZNF425** | 0.17392 | 0.29367 | 0.39622 | 0.21464 | 0.50608 | 0.04177 | 0.00000 |
| **ZNF426** | 5.58966 | 1.89530 | 5.42107 | 4.60303 | 2.66271 | 2.67074 | 0.43076 |
| **ZNF428** | 0.00000 | 2.40868 | 1.68028 | 0.43718 | 0.22998 | 5.12761 | 2.91729 |
| **ZNF429** | 1.47727 | 0.00000 | 2.58224 | 0.44626 | 0.26231 | 1.49725 | 0.12749 |
| **ZNF43** | 8.76536 | 0.00000 | 6.80711 | 3.33032 | 3.45761 | 6.02171 | 9.00776 |
| **ZNF430** | 9.82525 | 1.12120 | 2.65768 | 3.16766 | 1.56657 | 4.00340 | 2.83909 |
| **ZNF431** | 3.61116 | 0.00000 | 2.12341 | 2.12882 | 0.59472 | 2.03486 | 1.93580 |
| **ZNF432** | 1.06226 | 0.91825 | 0.81437 | 1.76345 | 1.57766 | 1.47853 | 0.00000 |
| **ZNF433** | 1.91559 | 0.19533 | 1.23486 | 1.28242 | 1.34383 | 0.04663 | 0.16334 |
| **ZNF436** | 2.42090 | 0.27707 | 0.88975 | 5.33417 | 4.20343 | 0.00000 | 0.00000 |
| **ZNF438** | 0.42039 | 0.11218 | 0.73847 | 0.41108 | 0.50397 | 0.08388 | 0.00000 |
| **ZNF439** | 0.14036 | 0.00000 | 0.26329 | 1.62366 | 0.23995 | 0.00000 | 0.18896 |
| **ZNF44** | 5.43752 | 2.74708 | 3.31400 | 2.31771 | 1.92533 | 3.51854 | 4.16814 |
| **ZNF440** | 3.21082 | 0.00000 | 0.00867 | 3.01984 | 0.01173 | 0.42066 | 2.50179 |
| **ZNF441** | 0.60281 | 0.00000 | 1.23628 | 0.30455 | 0.55257 | 0.41328 | 0.48920 |
| **ZNF442** | 0.03588 | 0.00000 | 0.17283 | 0.04427 | 0.23667 | 0.04140 | 0.00435 |
| **ZNF443** | 1.64169 | 0.06835 | 2.31924 | 0.61509 | 1.38601 | 0.76979 | 0.52177 |
| **ZNF444** | 0.00000 | 1.93948 | 0.56309 | 1.22101 | 0.02640 | 1.08117 | 9.32253 |
| **ZNF445** | 4.93384 | 0.53391 | 2.57528 | 1.05212 | 0.68877 | 1.17945 | 3.63743 |
| **ZNF446** | 0.11739 | 0.73295 | 0.41799 | 1.95575 | 0.37869 | 1.52442 | 1.08681 |
| **ZNF449** | 1.75351 | 0.46198 | 3.50404 | 0.00000 | 1.20547 | 1.72754 | 0.84901 |
| **ZNF45** | 3.96727 | 0.01132 | 3.15116 | 1.49476 | 1.54913 | 0.94465 | 0.00000 |
| **ZNF451** | 6.96157 | 0.00000 | 11.63998 | 6.93918 | 4.13421 | 9.26012 | 2.04931 |
| **ZNF454** | 0.11315 | 0.00000 | 0.27679 | 0.11685 | 0.07073 | 0.13561 | 0.07874 |
| **ZNF460** | 3.88475 | 0.00000 | 1.74979 | 2.77434 | 1.54364 | 0.87643 | 2.28254 |
| **ZNF461** | 1.85079 | 0.20245 | 1.88784 | 1.82541 | 0.89073 | 0.50587 | 0.00000 |
| **ZNF462** | 0.00760 | 0.14547 | 0.72135 | 0.05668 | 1.36362 | 4.04204 | 0.00000 |
| **ZNF467** | 0.00097 | 2.00894 | 0.00034 | 0.00000 | 0.00840 | 0.46903 | 0.66522 |
| **ZNF468** | 27.29090 | 9.02144 | 10.62656 | 16.44032 | 5.13937 | 4.93012 | 0.00000 |
| **ZNF469** | 0.01119 | 0.04330 | 0.02868 | 0.01840 | 0.00603 | 0.02427 | 0.05292 |
| **ZNF470** | 0.75677 | 0.50336 | 0.85878 | 1.50034 | 0.66537 | 0.27849 | 0.46362 |

| | | | | | | | |
|---|---|---|---|---|---|---|---|
| **ZNF471** | 0.62292 | 0.25394 | 0.45598 | 0.66214 | 0.11690 | 0.07365 | 0.00000 |
| **ZNF473** | 8.86032 | 5.26891 | 6.86504 | 6.16583 | 14.93028 | 4.94079 | 0.00000 |
| **ZNF474** | 0.02240 | 0.00000 | 0.04305 | 0.04165 | 0.05888 | 0.00799 | 0.07165 |
| **ZNF479** | 0.02357 | 0.01418 | 0.01304 | 0.01680 | 0.33685 | 0.01532 | 0.02090 |
| **ZNF48** | 0.00772 | 2.34847 | 0.78157 | 0.16818 | 0.83997 | 3.74270 | 2.96088 |
| **ZNF480** | 20.36106 | 3.37375 | 8.34735 | 3.67676 | 0.14241 | 7.04484 | 8.25715 |
| **ZNF483** | 0.43161 | 0.00000 | 0.03531 | 0.56574 | 0.52347 | 0.79465 | 0.24361 |
| **ZNF484** | 1.68551 | 0.00000 | 1.51000 | 0.59419 | 0.38052 | 0.78410 | 0.00000 |
| **ZNF485** | 1.63441 | 1.07073 | 3.60538 | 0.60148 | 0.37576 | 0.07626 | 0.00000 |
| **ZNF486** | 6.01415 | 0.15513 | 5.60455 | 0.76787 | 2.31791 | 7.53592 | 7.55670 |
| **ZNF487** | 0.36102 | 0.16908 | 3.02208 | 0.49078 | 1.77347 | 0.14557 | 0.00000 |
| **ZNF488** | 0.02748 | 0.03328 | 0.01115 | 0.64198 | 0.00794 | 0.33638 | 0.06777 |
| **ZNF490** | 1.27616 | 4.14980 | 3.68884 | 1.41494 | 6.01684 | 5.31616 | 0.00000 |
| **ZNF491** | 0.14001 | 0.00264 | 0.05741 | 0.02304 | 0.02837 | 0.01334 | 0.01075 |
| **ZNF492** | 1.21419 | 0.00000 | 0.70333 | 0.48722 | 0.19427 | 0.57924 | 1.13809 |
| **ZNF493** | 3.15153 | 0.00000 | 2.06813 | 1.44024 | 0.73644 | 0.79240 | 2.81321 |
| **ZNF496** | 0.00000 | 6.51028 | 0.00940 | 2.41588 | 1.30219 | 0.76411 | 6.55257 |
| **ZNF497** | 0.00000 | 0.15205 | 0.11697 | 0.10827 | 0.08162 | 0.04198 | 0.20712 |
| **ZNF500** | 0.67197 | 2.37544 | 0.97854 | 2.19984 | 0.08035 | 1.25913 | 6.46435 |
| **ZNF501** | 0.41921 | 0.01901 | 2.42082 | 0.13236 | 3.25445 | 0.33753 | 0.08036 |
| **ZNF502** | 1.09777 | 0.00000 | 3.36607 | 0.41902 | 3.54949 | 0.68164 | 0.24457 |
| **ZNF503** | 0.00000 | 0.06818 | 0.12436 | 0.00950 | 0.29297 | 1.02116 | 0.11215 |
| **ZNF503-AS1** | 0.06528 | 0.00000 | 0.04301 | 0.01181 | 0.06801 | 0.01433 | 0.01041 |
| **ZNF503-AS2** | 0.03847 | 0.03033 | 0.02930 | 0.02640 | 0.15672 | 0.27977 | 0.13115 |
| **ZNF506** | 4.07712 | 0.00000 | 4.39156 | 1.78270 | 7.29274 | 0.97057 | 3.07233 |
| **ZNF507** | 3.93165 | 2.99491 | 4.67811 | 2.05608 | 0.43962 | 1.67215 | 2.87059 |
| **ZNF510** | 1.53397 | 1.02286 | 1.69989 | 1.02261 | 0.28563 | 0.48242 | 1.67682 |
| **ZNF511** | 6.64962 | 12.30237 | 2.68401 | 7.43144 | 0.34333 | 6.02303 | 10.33309 |
| **ZNF512** | 4.50962 | 0.74651 | 1.91450 | 2.56929 | 2.36487 | 1.13220 | 0.67235 |
| **ZNF512B** | 0.00000 | 1.00215 | 0.22080 | 0.49217 | 0.15936 | 0.82335 | 2.00001 |
| **ZNF513** | 0.40203 | 1.33956 | 0.59255 | 1.69698 | 0.34504 | 0.95175 | 1.98478 |
| **ZNF514** | 0.33796 | 0.00000 | 0.21991 | 0.63723 | 0.04473 | 0.32318 | 0.15672 |
| **ZNF516** | 1.11012 | 0.33054 | 2.07394 | 0.31540 | 0.38003 | 1.27313 | 0.72449 |
| **ZNF517** | 0.03349 | 0.05706 | 0.10534 | 0.07688 | 0.06950 | 0.12780 | 0.19838 |
| **ZNF518A** | 11.47809 | 0.00000 | 14.47033 | 7.33857 | 3.23675 | 7.64988 | 10.56468 |

| | | | | | | | |
|---|---|---|---|---|---|---|---|
| **ZNF518B** | 7.00807 | 0.07676 | 5.43401 | 5.77823 | 1.17943 | 2.23198 | 7.05877 |
| **ZNF519** | 3.71645 | 0.00000 | 0.94440 | 1.26239 | 0.66247 | 0.53999 | 1.01045 |
| **ZNF521** | 0.06141 | 0.01623 | 0.02960 | 0.00927 | 0.15120 | 0.02900 | 0.02435 |
| **ZNF524** | 0.04853 | 0.45462 | 0.47973 | 0.80501 | 0.00000 | 0.06532 | 1.55587 |
| **ZNF525** | 5.40099 | 1.16346 | 5.46016 | 2.98610 | 1.41381 | 1.05740 | 3.19281 |
| **ZNF526** | 2.45607 | 5.78629 | 7.50248 | 3.20732 | 1.09757 | 1.78977 | 3.62951 |
| **ZNF527** | 1.76195 | 0.87336 | 0.25441 | 0.72308 | 0.62701 | 0.95580 | 0.00000 |
| **ZNF528** | 1.01996 | 0.68595 | 7.31868 | 0.34828 | 1.90030 | 0.43386 | 0.09149 |
| **ZNF529** | 4.37750 | 0.22123 | 2.72705 | 1.73224 | 1.55800 | 1.66553 | 4.05581 |
| **ZNF530** | 2.60762 | 2.02683 | 1.58613 | 2.02859 | 6.27760 | 0.74007 | 0.00003 |
| **ZNF532** | 6.12455 | 3.41887 | 1.24244 | 4.40487 | 1.19347 | 2.73614 | 8.07418 |
| **ZNF534** | 3.48072 | 0.89305 | 13.39480 | 1.55965 | 3.30867 | 2.93312 | 0.23525 |
| **ZNF536** | 0.01949 | 0.03195 | 0.01392 | 0.02290 | 0.18448 | 0.01122 | 0.01824 |
| **ZNF540** | 0.27278 | 0.00000 | 0.49870 | 0.14879 | 0.09411 | 0.24575 | 0.05187 |
| **ZNF541** | 0.00645 | 0.00000 | 0.36094 | 0.02313 | 0.41514 | 0.01213 | 0.04852 |
| **ZNF542P** | 4.17966 | 0.76065 | 5.18287 | 1.44606 | 3.34095 | 0.88221 | 0.00000 |
| **ZNF543** | 2.41058 | 1.21910 | 1.52660 | 0.22582 | 1.02787 | 1.27168 | 0.00000 |
| **ZNF544** | 7.40238 | 0.26836 | 3.54705 | 7.85985 | 5.38619 | 1.81735 | 0.00000 |
| **ZNF546** | 0.08771 | 0.00000 | 0.27107 | 0.12451 | 0.22253 | 0.37021 | 0.11063 |
| **ZNF547** | 1.87913 | 0.18119 | 2.08497 | 0.14143 | 1.04361 | 0.00000 | 0.09513 |
| **ZNF548** | 3.44931 | 0.42784 | 2.21823 | 0.97112 | 3.28009 | 0.93116 | 0.00000 |
| **ZNF549** | 4.63588 | 1.47458 | 4.20504 | 1.33457 | 3.60550 | 0.67480 | 0.00000 |
| **ZNF550** | 3.89156 | 1.19730 | 2.97178 | 1.92243 | 1.59337 | 2.51995 | 0.00000 |
| **ZNF551** | 8.05725 | 3.95343 | 2.27121 | 2.68936 | 1.14994 | 1.18381 | 0.00000 |
| **ZNF552** | 5.16224 | 6.79542 | 8.67495 | 3.20678 | 0.83685 | 3.36930 | 0.35153 |
| **ZNF554** | 0.00000 | 1.84589 | 0.11455 | 0.15181 | 1.04396 | 0.20517 | 0.56745 |
| **ZNF555** | 1.06116 | 0.00000 | 1.41613 | 0.13877 | 2.97353 | 0.62500 | 0.35327 |
| **ZNF556** | 0.40608 | 0.00000 | 0.00853 | 0.00000 | 0.06286 | 0.00440 | 0.06136 |
| **ZNF557** | 6.59690 | 4.56950 | 6.93799 | 2.46816 | 5.30075 | 1.74382 | 2.41543 |
| **ZNF558** | 0.87824 | 0.23057 | 0.56622 | 0.39470 | 0.22027 | 0.46995 | 0.94788 |
| **ZNF559** | 2.74023 | 0.00000 | 4.63908 | 0.73167 | 1.27685 | 1.37743 | 0.00000 |
| **ZNF559-ZNF177** | 0.03489 | 0.01929 | 0.01706 | 0.02218 | 0.02370 | 0.03526 | 0.01011 |
| **ZNF559-ZNF177** | 0.03619 | 0.00241 | 0.04806 | 0.02408 | 0.02700 | 0.02172 | 0.01967 |
| **ZNF560** | 3.18910 | 0.00000 | 3.94528 | 2.87055 | 1.34104 | 0.81385 | 0.00000 |
| **ZNF561** | 3.69215 | 0.68866 | 6.76896 | 0.94144 | 3.60014 | 0.54461 | 1.02584 |

| | | | | | | | |
|---|---|---|---|---|---|---|---|
| **ZNF562** | 8.03432 | 3.06903 | 5.32391 | 6.04999 | 1.75060 | 1.86753 | 0.00000 |
| **ZNF563** | 0.01746 | 0.00160 | 0.07529 | 0.03645 | 0.05931 | 0.01057 | 0.04374 |
| **ZNF564** | 4.02679 | 1.14780 | 7.18639 | 1.16348 | 0.43195 | 1.87805 | 0.00425 |
| **ZNF565** | 1.49034 | 2.12043 | 3.77786 | 2.22851 | 0.80780 | 0.09826 | 0.02038 |
| **ZNF566** | 8.92307 | 0.00000 | 12.04553 | 2.41750 | 1.80912 | 2.33230 | 3.75168 |
| **ZNF567** | 6.53672 | 0.15483 | 3.61150 | 5.49530 | 1.14350 | 2.79845 | 1.36958 |
| **ZNF568** | 2.81620 | 0.57315 | 1.44521 | 1.23330 | 4.46861 | 1.76536 | 0.26332 |
| **ZNF569** | 1.95822 | 0.87966 | 4.84744 | 0.53187 | 1.53485 | 1.60754 | 0.75626 |
| **ZNF57** | 12.33951 | 19.32992 | 9.02913 | 2.62131 | 2.33615 | 4.16043 | 0.00000 |
| **ZNF570** | 0.55530 | 0.31150 | 1.82158 | 1.30501 | 0.50045 | 0.96241 | 0.00000 |
| **ZNF571** | 1.40547 | 0.00000 | 2.08419 | 0.24157 | 1.10172 | 1.26773 | 0.00000 |
| **ZNF571-AS1** | 0.52243 | 0.10414 | 0.22753 | 0.72713 | 0.19062 | 0.19359 | 0.00000 |
| **ZNF572** | 0.10359 | 0.21962 | 0.22824 | 0.00000 | 0.17111 | 0.18334 | 0.00000 |
| **ZNF573** | 1.88984 | 0.00000 | 2.48040 | 0.72646 | 1.23493 | 0.70188 | 1.27757 |
| **ZNF574** | 4.47018 | 15.85460 | 13.46480 | 6.98300 | 6.98318 | 6.16468 | 9.39712 |
| **ZNF575** | 0.00000 | 0.01428 | 0.11218 | 0.02686 | 0.18596 | 0.07474 | 0.16054 |
| **ZNF576** | 13.96749 | 24.28595 | 9.24045 | 7.15369 | 7.63811 | 6.73885 | 0.00000 |
| **ZNF577** | 1.22789 | 2.15969 | 2.86821 | 1.11187 | 1.04669 | 0.87521 | 0.44488 |
| **ZNF578** | 2.13524 | 2.83469 | 7.33886 | 0.67707 | 7.86328 | 1.46387 | 0.09975 |
| **ZNF579** | 0.00000 | 0.23396 | 0.08148 | 0.01039 | 0.00671 | 0.00154 | 0.06578 |
| **ZNF580** | 0.05482 | 0.08995 | 0.16714 | 0.21732 | 0.08238 | 0.31355 | 3.17960 |
| **ZNF581** | 8.16569 | 19.19821 | 18.73632 | 4.67842 | 3.69593 | 9.40459 | 13.02079 |
| **ZNF582** | 0.89178 | 0.47772 | 1.18891 | 0.29785 | 0.73054 | 0.18289 | 0.00000 |
| **ZNF582-AS1** | 0.85997 | 1.01541 | 1.22688 | 1.26563 | 0.07050 | 0.34641 | 0.00000 |
| **ZNF583** | 0.73317 | 0.94686 | 0.70170 | 0.41668 | 0.12015 | 0.05232 | 0.18836 |
| **ZNF584** | 0.89728 | 2.23823 | 7.61430 | 2.01131 | 5.09128 | 0.40517 | 0.00000 |
| **ZNF585A** | 6.05308 | 4.76885 | 2.66670 | 3.54161 | 1.29174 | 2.14187 | 0.28749 |
| **ZNF585B** | 2.01167 | 0.00000 | 2.15865 | 0.80957 | 0.49294 | 0.93045 | 0.23053 |
| **ZNF586** | 8.40776 | 3.85763 | 11.88833 | 3.95831 | 8.65757 | 5.69658 | 0.00000 |
| **ZNF587** | 2.62336 | 1.19422 | 3.77109 | 3.63743 | 1.68915 | 4.75558 | 5.70973 |
| **ZNF587B** | 5.06459 | 4.54686 | 5.06130 | 2.35257 | 1.34543 | 2.63088 | 2.38700 |
| **ZNF589** | 16.52998 | 9.45910 | 10.98760 | 9.29536 | 6.34035 | 7.50472 | 0.01339 |
| **ZNF592** | 5.40640 | 5.49187 | 6.93309 | 5.11708 | 0.00000 | 1.06156 | 10.84338 |
| **ZNF593** | 11.11738 | 27.36341 | 18.16612 | 25.50449 | 3.60372 | 13.85856 | 15.46404 |
| **ZNF594** | 0.33452 | 0.00000 | 0.19510 | 0.48228 | 0.00201 | 0.15787 | 0.68731 |

| | | | | | | | |
|---|---|---|---|---|---|---|---|
| **ZNF595** | 4.43416 | 0.00000 | 6.03519 | 0.55207 | 3.38377 | 1.15610 | 0.52572 |
| **ZNF596** | 0.89834 | 0.00000 | 1.47334 | 0.14192 | 2.68968 | 0.00000 | 0.37896 |
| **ZNF597** | 1.93967 | 0.00000 | 5.89690 | 2.71592 | 9.63914 | 0.98654 | 0.00000 |
| **ZNF598** | 0.46572 | 5.32121 | 0.69028 | 1.96555 | 0.38957 | 2.37287 | 17.66500 |
| **ZNF599** | 0.28539 | 0.27249 | 0.71920 | 0.38169 | 0.42377 | 0.00000 | 0.11404 |
| **ZNF600** | 7.60379 | 7.17907 | 7.44720 | 10.68902 | 2.14831 | 8.10247 | 0.14729 |
| **ZNF605** | 1.27040 | 0.00000 | 2.48000 | 1.37158 | 1.25193 | 1.04280 | 1.15290 |
| **ZNF606** | 0.64583 | 0.62176 | 1.80921 | 0.52801 | 4.09691 | 0.68967 | 0.58439 |
| **ZNF607** | 0.98029 | 3.45137 | 1.81809 | 1.37100 | 0.46793 | 1.45050 | 2.54223 |
| **ZNF608** | 0.77792 | 0.00000 | 0.84196 | 0.27374 | 0.36771 | 2.17534 | 0.00000 |
| **ZNF609** | 1.45810 | 1.25529 | 1.81778 | 3.12145 | 0.44706 | 2.50636 | 12.83675 |
| **ZNF610** | 0.18268 | 0.00000 | 3.20641 | 1.08613 | 0.58760 | 0.54081 | 0.29096 |
| **ZNF611** | 2.99722 | 3.57426 | 4.52032 | 3.04718 | 3.18524 | 2.94791 | 3.59468 |
| **ZNF613** | 1.47179 | 2.22587 | 4.92265 | 0.61976 | 1.04780 | 1.48066 | 0.19152 |
| **ZNF614** | 4.41920 | 1.48049 | 9.84616 | 2.44229 | 1.36956 | 4.61503 | 0.52307 |
| **ZNF615** | 1.87235 | 0.58558 | 2.86093 | 1.36475 | 0.70546 | 1.43686 | 0.00000 |
| **ZNF616** | 1.20831 | 0.25044 | 2.47943 | 1.69018 | 3.85217 | 1.00206 | 0.00000 |
| **ZNF618** | 0.00000 | 0.85610 | 0.54596 | 0.00000 | 5.26954 | 0.00000 | 0.33147 |
| **ZNF619** | 0.46016 | 0.33258 | 0.97832 | 0.00000 | 0.15073 | 0.56119 | 0.49615 |
| **ZNF620** | 1.37148 | 1.48023 | 1.40199 | 0.20057 | 0.54111 | 0.27672 | 1.34332 |
| **ZNF621** | 2.34951 | 0.81665 | 0.90852 | 1.51758 | 2.05392 | 0.42495 | 2.55660 |
| **ZNF622** | 21.91150 | 25.17145 | 10.92643 | 16.55707 | 13.98073 | 6.96499 | 0.00000 |
| **ZNF623** | 1.13427 | 1.66110 | 3.77967 | 0.40803 | 1.79481 | 1.56787 | 2.71710 |
| **ZNF624** | 0.94978 | 0.00000 | 0.64358 | 0.13862 | 0.33834 | 0.67698 | 0.30958 |
| **ZNF625** | 1.58554 | 0.22500 | 0.91067 | 0.47667 | 0.66445 | 0.25818 | 0.00000 |
| **ZNF625-ZNF20** | 0.97681 | 0.53788 | 0.58794 | 0.18799 | 0.26965 | 0.63411 | 0.35004 |
| **ZNF626** | 10.12320 | 0.00000 | 6.84967 | 3.25088 | 1.37031 | 0.94481 | 3.46940 |
| **ZNF627** | 2.23899 | 0.80831 | 0.36239 | 1.36357 | 0.44770 | 0.00000 | 0.74080 |
| **ZNF628** | 0.00000 | 1.38411 | 0.25309 | 0.43397 | 0.39308 | 0.85508 | 3.94094 |
| **ZNF629** | 0.00000 | 0.08956 | 0.05086 | 0.12836 | 0.15004 | 0.45311 | 0.07014 |
| **ZNF630** | 0.21807 | 0.00000 | 0.63960 | 0.04644 | 0.38167 | 0.03622 | 0.00000 |
| **ZNF630-AS1** | 0.01981 | 0.03686 | 0.01258 | 0.01952 | 0.02805 | 0.01130 | 0.01682 |
| **ZNF638** | 18.63550 | 1.09196 | 12.94895 | 10.52189 | 6.73504 | 8.42159 | 15.89693 |
| **ZNF639** | 4.00216 | 0.00000 | 8.06499 | 0.44407 | 6.15347 | 4.67978 | 12.76433 |
| **ZNF641** | 1.92271 | 0.95410 | 1.24457 | 0.31230 | 0.53637 | 0.47887 | 2.30749 |

| | | | | | | | |
|---|---|---|---|---|---|---|---|
| **ZNF644** | 5.01045 | 0.00000 | 3.24743 | 6.36627 | 0.84112 | 3.70025 | 5.22459 |
| **ZNF645** | 0.02355 | 0.01796 | 0.01386 | 0.01779 | 0.00877 | 0.01214 | 0.02484 |
| **ZNF646** | 0.00000 | 1.19109 | 1.56028 | 0.94319 | 0.34920 | 0.59489 | 2.29155 |
| **ZNF648** | 0.03386 | 0.18397 | 0.04138 | 0.00569 | 0.00870 | 0.00418 | 0.03808 |
| **ZNF649** | 3.55887 | 2.60978 | 7.17628 | 1.35676 | 0.43085 | 4.43498 | 0.00000 |
| **ZNF652** | 9.91620 | 2.47862 | 4.21806 | 2.45867 | 6.14272 | 3.46966 | 7.52357 |
| **ZNF653** | 0.00000 | 0.02796 | 0.04846 | 0.06172 | 0.01416 | 0.14607 | 0.27460 |
| **ZNF654** | 2.20205 | 0.00000 | 3.01221 | 1.56425 | 4.62701 | 1.68121 | 2.12597 |
| **ZNF655** | 2.08810 | 0.00000 | 3.89001 | 9.83434 | 5.67955 | 2.62343 | 2.94288 |
| **ZNF658** | 0.04949 | 0.00791 | 0.02249 | 0.03187 | 0.00695 | 0.00763 | 0.04521 |
| **ZNF660** | 0.02907 | 0.01264 | 0.01396 | 0.02592 | 0.04854 | 0.01432 | 0.01688 |
| **ZNF662** | 0.17601 | 1.09802 | 0.92894 | 0.15858 | 0.96017 | 0.15001 | 0.45431 |
| **ZNF663P** | 0.05544 | 0.01793 | 0.01449 | 0.02402 | 0.01168 | 0.01287 | 0.03285 |
| **ZNF664** | 4.36672 | 0.51735 | 5.40262 | 3.32628 | 0.58937 | 1.03236 | 11.89189 |
| **ZNF664-FAM10** | 0.07919 | 0.01456 | 0.07673 | 0.02942 | 0.01324 | 0.08637 | 0.00000 |
| **ZNF665** | 0.84208 | 0.00014 | 2.07383 | 0.49879 | 0.71412 | 0.25355 | 0.79110 |
| **ZNF667** | 1.21325 | 1.22664 | 0.92945 | 2.07662 | 0.86449 | 0.86984 | 0.00000 |
| **ZNF667-AS1** | 2.28452 | 1.49391 | 9.66701 | 4.61363 | 3.55860 | 0.32700 | 0.00000 |
| **ZNF668** | 2.06072 | 2.98495 | 5.44577 | 6.27242 | 5.04343 | 2.02322 | 3.59953 |
| **ZNF669** | 1.50231 | 0.00000 | 3.58831 | 1.62400 | 4.90107 | 0.74270 | 0.31694 |
| **ZNF670** | 7.73651 | 0.00000 | 7.29534 | 4.40973 | 4.11278 | 1.68175 | 0.29945 |
| **ZNF670-ZNF69** | 0.09451 | 0.01726 | 0.01797 | 0.05145 | 0.00711 | 0.04078 | 0.03654 |
| **ZNF671** | 2.17574 | 0.22670 | 1.36388 | 0.76923 | 0.06626 | 0.35907 | 0.00000 |
| **ZNF672** | 1.43469 | 2.75170 | 0.94041 | 4.17519 | 0.18813 | 1.44557 | 7.61934 |
| **ZNF674** | 0.35828 | 0.00000 | 1.29571 | 0.00000 | 0.28609 | 0.54650 | 0.49568 |
| **ZNF674-AS1** | 1.43771 | 1.16609 | 4.15329 | 0.50771 | 2.01869 | 0.37396 | 0.00000 |
| **ZNF675** | 4.20379 | 0.00000 | 14.67698 | 1.64317 | 3.70395 | 4.16297 | 1.69082 |
| **ZNF676** | 4.03339 | 0.00000 | 4.07392 | 1.03936 | 0.94461 | 1.72020 | 2.70445 |
| **ZNF677** | 2.67856 | 0.65348 | 5.98433 | 1.14303 | 3.12132 | 0.50961 | 0.79360 |
| **ZNF678** | 1.68061 | 0.00000 | 0.96756 | 0.48282 | 0.42128 | 0.76575 | 0.46346 |
| **ZNF679** | 0.00540 | 0.02171 | 0.00000 | 0.00247 | 14.64553 | 0.01066 | 0.07841 |
| **ZNF680** | 8.70725 | 0.00000 | 6.86280 | 1.53848 | 1.65148 | 3.66498 | 2.32929 |
| **ZNF681** | 3.65433 | 0.00000 | 4.26237 | 0.89283 | 0.28586 | 1.86070 | 1.74367 |
| **ZNF682** | 6.07055 | 0.00000 | 2.42702 | 2.66601 | 4.79090 | 3.32062 | 2.27958 |
| **ZNF683** | 0.02468 | 0.00901 | 0.03544 | 0.02727 | 0.05337 | 0.01196 | 0.01753 |

| | | | | | | | |
|---|---|---|---|---|---|---|---|
| **ZNF684** | 2.13586 | 0.14260 | 7.67261 | 0.45522 | 3.45371 | 2.28806 | 0.11827 |
| **ZNF687** | 0.62933 | 0.00000 | 1.15313 | 1.85241 | 0.46034 | 1.60354 | 3.97575 |
| **ZNF688** | 0.09766 | 0.02890 | 0.47773 | 0.00000 | 0.00048 | 0.26450 | 0.56295 |
| **ZNF689** | 4.86460 | 5.60609 | 6.59277 | 3.87450 | 7.32667 | 4.04272 | 0.73205 |
| **ZNF69** | 0.14578 | 0.00000 | 0.07132 | 0.95732 | 0.03690 | 0.48767 | 0.51855 |
| **ZNF691** | 5.23197 | 6.24725 | 5.27624 | 2.81128 | 2.80112 | 0.66563 | 0.00000 |
| **ZNF692** | 5.08843 | 3.89533 | 2.23911 | 5.89377 | 0.35399 | 0.67816 | 7.91696 |
| **ZNF695** | 3.17853 | 1.01358 | 0.96845 | 1.94049 | 0.07013 | 1.50963 | 3.61169 |
| **ZNF696** | 0.44521 | 1.45330 | 1.95142 | 0.00000 | 0.04092 | 1.54166 | 3.63338 |
| **ZNF697** | 1.48436 | 0.00000 | 0.42407 | 2.67073 | 0.01774 | 1.19729 | 4.93429 |
| **ZNF699** | 0.14543 | 0.00000 | 0.20222 | 0.07169 | 0.07085 | 0.08052 | 0.00000 |
| **ZNF7** | 7.11099 | 3.15527 | 9.17344 | 1.36332 | 1.43759 | 3.40604 | 0.36694 |
| **ZNF70** | 0.19431 | 0.13912 | 0.47747 | 0.10896 | 0.25901 | 0.00000 | 0.31702 |
| **ZNF700** | 3.46515 | 0.00000 | 2.66591 | 1.65423 | 1.28985 | 0.93935 | 0.28681 |
| **ZNF701** | 2.35354 | 3.05425 | 5.54531 | 1.54585 | 6.86169 | 1.77507 | 1.36945 |
| **ZNF702P** | 2.73621 | 0.61725 | 0.62056 | 4.20155 | 0.51999 | 0.00824 | 0.01359 |
| **ZNF703** | 0.00000 | 8.94837 | 0.00000 | 0.73890 | 0.27363 | 0.16847 | 6.67931 |
| **ZNF704** | 4.36357 | 0.07458 | 0.00000 | 1.73549 | 0.09927 | 0.00000 | 6.22805 |
| **ZNF705A** | 0.01783 | 0.01232 | 0.00000 | 0.01600 | 2.20346 | 0.01695 | 0.01662 |
| **ZNF705B** | 0.02404 | 0.01763 | 0.01196 | 0.01942 | 0.01952 | 0.01316 | 0.01586 |
| **ZNF705D** | 0.02343 | 0.01376 | 0.03008 | 0.01931 | 0.02730 | 0.01595 | 0.01828 |
| **ZNF705E** | 0.02465 | 0.00200 | 0.01221 | 0.02689 | 2.20129 | 0.02460 | 0.01424 |
| **ZNF705G** | 0.02262 | 0.01764 | 0.00000 | 0.01818 | 0.42940 | 0.01334 | 0.01509 |
| **ZNF706** | 16.16466 | 16.88554 | 7.56711 | 13.64202 | 1.40623 | 8.63444 | 4.51333 |
| **ZNF707** | 0.42787 | 0.90992 | 1.51672 | 0.58143 | 5.09612 | 1.26804 | 1.13511 |
| **ZNF708** | 2.07097 | 0.00000 | 0.36649 | 2.16325 | 0.12857 | 0.60438 | 1.00562 |
| **ZNF709** | 1.97309 | 0.25996 | 2.41913 | 1.47565 | 1.26992 | 1.37477 | 0.00000 |
| **ZNF71** | 1.11903 | 2.82266 | 0.15393 | 2.15602 | 0.29387 | 0.59603 | 1.31193 |
| **ZNF710** | 0.02180 | 0.00000 | 0.95689 | 0.13203 | 0.06121 | 1.25447 | 0.93175 |
| **ZNF711** | 0.20329 | 0.00000 | 0.58508 | 0.00000 | 0.55716 | 0.17705 | 0.14988 |
| **ZNF713** | 0.18046 | 0.00000 | 0.13231 | 0.15079 | 0.00223 | 0.08552 | 0.27744 |
| **ZNF714** | 4.90578 | 0.00000 | 0.21254 | 3.73403 | 0.37428 | 0.89394 | 5.58364 |
| **ZNF716** | 0.41563 | 0.00000 | 2.81047 | 0.00000 | 5.61750 | 0.05671 | 0.01065 |
| **ZNF717** | 1.13076 | 0.05059 | 1.39761 | 0.91203 | 0.53813 | 0.72181 | 0.26541 |
| **ZNF718** | 4.94854 | 0.39883 | 3.72969 | 1.22294 | 2.47648 | 0.50522 | 1.85944 |

| | | | | | | | |
|---|---|---|---|---|---|---|---|
| **ZNF720** | 0.93390 | 0.03860 | 0.42959 | 0.84110 | 0.16617 | 0.21523 | 0.74081 |
| **ZNF721** | 8.49153 | 0.05148 | 5.78045 | 1.51626 | 1.46697 | 4.20623 | 6.85665 |
| **ZNF724P** | 1.49241 | 0.00000 | 1.70016 | 0.31042 | 0.77342 | 0.60023 | 2.48368 |
| **ZNF726** | 0.78761 | 0.36165 | 0.15226 | 0.57814 | 0.04164 | 0.27980 | 0.45581 |
| **ZNF727P** | 0.14461 | 0.10407 | 0.30753 | 0.22403 | 2.29577 | 0.10631 | 0.00000 |
| **ZNF728** | 5.24679 | 0.18460 | 5.59364 | 1.57852 | 6.70768 | 0.83466 | 0.27300 |
| **ZNF729** | 2.27352 | 0.49074 | 5.87628 | 0.00000 | 1.57470 | 3.58681 | 1.57672 |
| **ZNF730** | 2.07925 | 0.00000 | 0.90306 | 0.91597 | 0.54481 | 1.07181 | 2.11669 |
| **ZNF732** | 1.70165 | 0.00000 | 1.44509 | 0.44917 | 0.69561 | 0.38095 | 0.57951 |
| **ZNF733P** | 0.02868 | 0.00205 | 0.02712 | 0.02294 | 0.33960 | 0.01302 | 0.01311 |
| **ZNF735P** | 0.00000 | 0.00000 | 0.92542 | 0.00753 | 12.78632 | 0.00000 | 0.02851 |
| **ZNF736** | 0.30678 | 0.00923 | 0.61584 | 0.36235 | 4.38109 | 0.31395 | 0.00000 |
| **ZNF737** | 2.95278 | 0.00000 | 2.74101 | 1.03564 | 0.79657 | 1.52885 | 1.90301 |
| **ZNF738** | 12.32160 | 2.53665 | 6.92491 | 8.03878 | 4.08545 | 4.81952 | 6.62729 |
| **ZNF74** | 0.89006 | 2.07411 | 0.32062 | 3.04062 | 0.00187 | 0.86432 | 5.28378 |
| **ZNF740** | 1.36308 | 0.84089 | 0.48324 | 0.30494 | 0.00000 | 0.50720 | 6.88838 |
| **ZNF746** | 0.20790 | 1.73397 | 1.33096 | 1.40465 | 0.40003 | 0.82352 | 5.48424 |
| **ZNF747** | 0.00000 | 0.54086 | 0.10252 | 0.11925 | 0.08323 | 0.47946 | 0.00338 |
| **ZNF749** | 3.14990 | 4.62494 | 1.95850 | 1.05433 | 1.50842 | 0.77387 | 0.00000 |
| **ZNF750** | 5.52274 | 0.00000 | 0.00000 | 12.24864 | 0.00000 | 0.00000 | 3.23969 |
| **ZNF75A** | 3.38535 | 0.89459 | 4.55841 | 2.02328 | 0.24346 | 0.82546 | 1.41119 |
| **ZNF75D** | 3.03065 | 1.31782 | 2.11941 | 1.19369 | 1.90772 | 5.42301 | 2.13481 |
| **ZNF76** | 3.69604 | 5.58219 | 1.71567 | 8.20800 | 1.62508 | 3.18108 | 2.96220 |
| **ZNF761** | 12.60144 | 2.26000 | 5.02041 | 7.55413 | 2.50116 | 4.81765 | 0.00000 |
| **ZNF763** | 0.09542 | 0.23610 | 0.07062 | 0.20053 | 0.12132 | 0.00000 | 0.01713 |
| **ZNF764** | 0.88106 | 1.04163 | 3.51670 | 1.04906 | 0.30621 | 1.63232 | 3.97708 |
| **ZNF765** | 8.49078 | 2.89911 | 6.07578 | 2.97695 | 2.78891 | 3.01043 | 0.00000 |
| **ZNF766** | 6.32009 | 4.78748 | 11.84784 | 2.10410 | 5.55802 | 2.63771 | 0.00000 |
| **ZNF767P** | 0.60741 | 0.29764 | 0.43120 | 0.35522 | 0.12085 | 0.24391 | 1.80417 |
| **ZNF768** | 0.00000 | 1.36931 | 0.36790 | 0.08785 | 0.19033 | 1.39851 | 0.93596 |
| **ZNF77** | 3.50123 | 0.00000 | 1.35666 | 0.47977 | 1.50051 | 0.68340 | 0.12934 |
| **ZNF770** | 10.95000 | 0.00000 | 10.60903 | 4.97956 | 2.44817 | 4.42916 | 18.93929 |
| **ZNF771** | 0.00000 | 0.82307 | 1.68784 | 1.37144 | 0.04119 | 0.55126 | 0.39694 |
| **ZNF772** | 2.01106 | 0.50796 | 2.03640 | 0.49867 | 0.94020 | 0.24344 | 0.13250 |
| **ZNF773** | 3.18252 | 2.96069 | 2.36903 | 2.78857 | 3.61652 | 0.26859 | 0.00000 |

| | | | | | | | |
|---|---|---|---|---|---|---|---|
| **ZNF774** | 0.18816 | 0.03384 | 0.15451 | 0.43667 | 0.03152 | 0.05739 | 0.00000 |
| **ZNF775** | 0.00000 | 0.34541 | 0.00806 | 0.03416 | 0.00814 | 0.26677 | 0.40128 |
| **ZNF776** | 5.28702 | 0.29615 | 3.29406 | 1.66406 | 3.90572 | 2.30370 | 0.00000 |
| **ZNF777** | 0.00000 | 0.00000 | 1.04238 | 0.14242 | 0.67075 | 0.22649 | 3.27772 |
| **ZNF778** | 2.93992 | 1.47074 | 2.22456 | 0.94894 | 2.98981 | 0.53204 | 0.00000 |
| **ZNF780A** | 2.52043 | 1.04249 | 0.24727 | 1.43391 | 0.00000 | 0.77494 | 1.46963 |
| **ZNF780B** | 0.53910 | 0.00000 | 0.18039 | 0.37917 | 0.02239 | 0.61457 | 0.87770 |
| **ZNF781** | 0.36004 | 0.00000 | 0.54489 | 0.55404 | 0.03094 | 0.23975 | 0.00000 |
| **ZNF782** | 0.46203 | 0.00000 | 0.39324 | 0.02657 | 0.00199 | 0.13135 | 1.33289 |
| **ZNF783** | 2.20558 | 4.78905 | 1.70869 | 0.00000 | 0.00000 | 1.01929 | 0.83219 |
| **ZNF784** | 0.07382 | 0.11166 | 0.05740 | 0.12762 | 0.00330 | 0.17814 | 0.37134 |
| **ZNF785** | 0.11529 | 0.15198 | 0.16403 | 0.02868 | 0.03447 | 0.28818 | 0.17426 |
| **ZNF786** | 1.40540 | 0.23714 | 0.91656 | 1.68187 | 0.29584 | 0.00000 | 1.44504 |
| **ZNF787** | 0.00000 | 1.65453 | 0.00989 | 0.15001 | 0.49565 | 0.26973 | 4.64555 |
| **ZNF788** | 4.33970 | 0.35289 | 1.23927 | 1.06991 | 2.50340 | 0.81468 | 0.59966 |
| **ZNF789** | 5.48432 | 0.23843 | 2.14750 | 2.99710 | 2.73721 | 1.50814 | 2.13953 |
| **ZNF79** | 2.07529 | 3.17457 | 3.37358 | 2.56699 | 3.23949 | 0.79104 | 0.00000 |
| **ZNF790** | 1.33176 | 0.00000 | 0.51147 | 0.72186 | 0.29423 | 0.73325 | 0.84740 |
| **ZNF790-AS1** | 1.06634 | 0.00000 | 1.06261 | 1.44807 | 2.86441 | 0.86202 | 0.28789 |
| **ZNF791** | 8.46394 | 3.79751 | 6.14888 | 4.20796 | 7.13545 | 4.58694 | 0.00000 |
| **ZNF792** | 0.01820 | 0.27318 | 0.41401 | 0.07005 | 0.24563 | 0.23229 | 0.06442 |
| **ZNF793** | 0.72743 | 0.00000 | 1.74182 | 0.87364 | 0.40273 | 1.78400 | 0.23739 |
| **ZNF799** | 0.86992 | 0.00000 | 1.51798 | 0.69104 | 0.88972 | 0.49315 | 0.21585 |
| **ZNF8** | 1.52291 | 4.19874 | 3.62379 | 2.48803 | 1.11676 | 1.75407 | 0.88511 |
| **ZNF80** | 0.01354 | 0.01489 | 0.01561 | 0.02930 | 0.00921 | 0.04158 | 0.03563 |
| **ZNF800** | 1.52556 | 1.26004 | 2.03844 | 2.70930 | 0.44426 | 1.06449 | 12.28758 |
| **ZNF804A** | 0.02018 | 0.01112 | 0.01352 | 0.04123 | 0.01157 | 0.01032 | 0.06244 |
| **ZNF805** | 6.86218 | 1.45436 | 2.84206 | 2.18920 | 2.08161 | 2.42977 | 1.44317 |
| **ZNF808** | 1.61777 | 0.55374 | 1.75504 | 0.70119 | 1.69025 | 0.50069 | 0.82391 |
| **ZNF81** | 1.26855 | 0.00000 | 0.46035 | 0.07277 | 0.02456 | 0.89482 | 1.04876 |
| **ZNF812** | 0.03708 | 0.00000 | 0.14955 | 0.10377 | 0.17628 | 0.01000 | 0.00000 |
| **ZNF813** | 3.09655 | 0.68775 | 1.72028 | 0.77289 | 0.97180 | 1.00530 | 0.00000 |
| **ZNF814** | 14.59900 | 12.05078 | 14.43928 | 7.40513 | 6.56968 | 4.36495 | 1.06379 |
| **ZNF815P** | 0.19188 | 0.00000 | 0.77126 | 0.13116 | 0.59868 | 0.12148 | 0.27447 |
| **ZNF816** | 7.83700 | 0.35374 | 8.30041 | 3.00035 | 2.65515 | 1.29454 | 0.00000 |

| | | | | | | | |
|---|---|---|---|---|---|---|---|
| **ZNF818P** | 0.21943 | 0.26201 | 1.22628 | 0.20326 | 0.20169 | 0.08395 | 0.95161 |
| **ZNF821** | 0.26629 | 0.09219 | 0.71815 | 0.75380 | 1.91541 | 0.86716 | 1.76606 |
| **ZNF823** | 10.09080 | 0.81222 | 3.38492 | 4.13314 | 6.00431 | 2.63475 | 2.97518 |
| **ZNF826P** | 0.55200 | 0.62029 | 2.38154 | 0.55318 | 0.61694 | 1.64862 | 0.77208 |
| **ZNF827** | 0.07647 | 0.10524 | 0.02064 | 0.01573 | 0.45548 | 0.97407 | 0.37285 |
| **ZNF829** | 4.12584 | 0.00000 | 4.75238 | 1.08497 | 4.16164 | 1.60480 | 2.17216 |
| **ZNF83** | 6.21854 | 0.00000 | 7.32423 | 3.62662 | 1.44025 | 3.30101 | 0.70859 |
| **ZNF830** | 7.93918 | 1.08281 | 10.53142 | 7.02905 | 7.43463 | 3.09864 | 3.31749 |
| **ZNF831** | 0.04233 | 0.01480 | 0.01349 | 0.03863 | 0.00796 | 0.03129 | 0.00427 |
| **ZNF833P** | 0.10683 | 0.00000 | 0.12533 | 0.43442 | 0.90503 | 0.23532 | 0.41496 |
| **ZNF835** | 0.76399 | 0.94523 | 0.00000 | 0.40127 | 0.11624 | 0.00000 | 0.02526 |
| **ZNF836** | 1.51031 | 0.00000 | 0.93948 | 0.75678 | 0.06986 | 0.03495 | 0.16294 |
| **ZNF837** | 0.01155 | 0.01415 | 0.20768 | 0.03200 | 0.06065 | 0.07953 | 0.06660 |
| **ZNF839** | 0.67213 | 1.16638 | 2.33707 | 0.06627 | 0.16350 | 0.48644 | 1.35151 |
| **ZNF84.uid-1752** | 5.04922 | 0.59217 | 4.48784 | 2.91380 | 3.33293 | 2.20921 | 1.21875 |
| **ZNF841** | 2.61034 | 0.89548 | 1.92446 | 1.37643 | 2.75466 | 1.07663 | 0.25843 |
| **ZNF843** | 0.00000 | 0.06131 | 0.44964 | 0.01670 | 0.26950 | 0.00000 | 0.03239 |
| **ZNF844** | 0.61805 | 0.03788 | 1.16152 | 0.64356 | 3.98196 | 1.16460 | 0.55787 |
| **ZNF845** | 5.01409 | 0.00000 | 7.09910 | 2.45172 | 4.29253 | 1.41720 | 0.00000 |
| **ZNF846** | 0.02324 | 0.00000 | 0.27076 | 0.02602 | 0.27063 | 0.04244 | 0.06766 |
| **ZNF85** | 3.15648 | 0.00000 | 2.66075 | 1.36147 | 0.99484 | 1.40885 | 1.20798 |
| **ZNF850** | 8.97019 | 2.38906 | 7.34912 | 4.16197 | 2.36827 | 2.46678 | 7.87488 |
| **ZNF852** | 1.29748 | 0.00000 | 1.38078 | 0.47463 | 0.82457 | 0.09412 | 0.00000 |
| **ZNF853** | 0.02559 | 0.02974 | 0.02305 | 0.02244 | 0.01186 | 0.01608 | 0.03088 |
| **ZNF860** | 0.61497 | 0.00000 | 0.04981 | 0.78635 | 0.16983 | 0.08543 | 0.07221 |
| **ZNF862** | 0.03191 | 0.04831 | 0.13170 | 0.06193 | 0.04141 | 0.18208 | 0.22951 |
| **ZNF865** | 0.00000 | 0.25249 | 0.17317 | 0.00000 | 0.27260 | 0.06692 | 0.53303 |
| **ZNF876P** | 4.96055 | 0.01762 | 5.14257 | 1.35816 | 0.78861 | 0.89206 | 0.79951 |
| **ZNF878** | 1.22481 | 0.00000 | 0.58164 | 0.85480 | 4.08730 | 0.32930 | 0.00000 |
| **ZNF879** | 0.49928 | 0.24932 | 0.69152 | 0.48401 | 0.17706 | 0.06290 | 0.00000 |
| **ZNF880** | 2.01576 | 1.82052 | 6.39861 | 1.78513 | 2.03461 | 1.57744 | 0.00000 |
| **ZNF883** | 0.14112 | 0.00000 | 0.13352 | 0.00000 | 0.22228 | 0.08950 | 0.37033 |
| **ZNF890P** | 0.02472 | 0.03172 | 0.02487 | 0.02099 | 0.02783 | 0.01642 | 0.00881 |
| **ZNF891** | 1.66292 | 0.00000 | 2.87963 | 1.22316 | 1.98679 | 0.10182 | 0.00000 |
| **ZNF90** | 10.21816 | 0.00000 | 15.97813 | 1.80594 | 2.83268 | 13.08428 | 10.33224 |

| | | | | | | | |
|---|---|---|---|---|---|---|---|
| **ZNF91** | 3.99174 | 0.00000 | 2.81942 | 3.80874 | 1.26158 | 1.82948 | 4.83081 |
| **ZNF92** | 11.44293 | 0.00000 | 11.38286 | 1.90708 | 5.82921 | 3.72528 | 3.49010 |
| **ZNF93** | 9.72270 | 0.00000 | 5.83317 | 5.71979 | 7.54664 | 3.31798 | 0.91728 |
| **ZNF98** | 1.65578 | 0.00000 | 0.81279 | 0.11205 | 0.32682 | 0.63095 | 0.37811 |
| **ZNF99** | 1.91932 | 0.43895 | 1.14339 | 0.00000 | 0.55102 | 0.90887 | 0.33912 |
| **ZNFX1** | 6.15911 | 4.98675 | 8.90333 | 8.44585 | 5.54836 | 3.79951 | 7.87964 |
| **ZNHIT1** | 19.97982 | 18.29674 | 15.87522 | 24.30014 | 10.31672 | 17.49906 | 10.27992 |
| **ZNHIT2** | 1.09816 | 5.17035 | 5.50527 | 6.09859 | 2.72432 | 2.46844 | 3.01249 |
| **ZNHIT3** | 19.50305 | 7.96931 | 14.88767 | 16.26017 | 15.43949 | 11.07695 | 9.54273 |
| **ZNHIT6** | 9.17714 | 0.00000 | 9.56120 | 6.15620 | 4.61038 | 3.95896 | 11.67230 |
| **ZNRD1** | 7.57424 | 11.76931 | 9.08783 | 6.38510 | 4.38763 | 5.41837 | 7.18457 |
| **ZNRD1-AS1** | 0.25595 | 0.49707 | 0.40666 | 0.00000 | 0.34486 | 0.02474 | 0.00795 |
| **ZNRF1** | 0.11699 | 1.56667 | 2.42269 | 6.24736 | 0.06942 | 0.97576 | 22.13094 |
| **ZNRF2** | 0.88850 | 0.00000 | 0.05311 | 0.71439 | 0.06035 | 0.39035 | 2.27195 |
| **ZNRF2P1** | 0.00376 | 0.02329 | 0.01185 | 0.06488 | 0.01746 | 0.07215 | 0.12317 |
| **ZNRF3** | 0.71156 | 2.16760 | 2.25778 | 3.72717 | 0.06245 | 4.11758 | 14.34599 |
| **ZNRF3-AS1** | 0.01859 | 0.00000 | 0.14987 | 0.04553 | 0.03052 | 0.10449 | 0.57747 |
| **ZP1** | 0.03518 | 0.01171 | 1.69094 | 0.00000 | 8.90058 | 0.04041 | 0.12908 |
| **ZP2** | 0.00000 | 0.43187 | 4.77380 | 0.00000 | 20.05280 | 0.00000 | 0.09793 |
| **ZP3** | 0.00000 | 0.17608 | 7.81922 | 0.00000 | 23.89462 | 0.00000 | 0.27973 |
| **ZP4** | 0.00000 | 0.02314 | 1.86188 | 0.00000 | 15.42761 | 0.00000 | 0.05289 |
| **ZPBP** | 0.01643 | 0.02406 | 0.18695 | 0.02896 | 0.06246 | 0.03087 | 0.02101 |
| **ZPBP2** | 0.01539 | 0.00000 | 0.45296 | 0.01599 | 0.23970 | 0.00436 | 0.01964 |
| **ZPLD1** | 0.02212 | 0.00000 | 0.02500 | 0.02499 | 0.40049 | 0.00830 | 0.05676 |
| **ZPR1** | 9.75011 | 11.34718 | 4.43955 | 10.14305 | 0.34957 | 3.48490 | 22.32488 |
| **ZRANB1** | 4.11983 | 1.89217 | 3.02380 | 2.32275 | 4.27366 | 4.10888 | 11.19035 |
| **ZRANB2** | 27.35598 | 4.61566 | 20.35196 | 14.92732 | 5.98930 | 6.80196 | 13.70803 |
| **ZRANB2-AS1** | 0.11290 | 0.02392 | 0.06309 | 0.05333 | 0.01642 | 0.09304 | 0.08202 |
| **ZRANB2-AS2** | 0.11951 | 0.03661 | 0.41600 | 0.07301 | 0.15424 | 0.01934 | 0.02833 |
| **ZRANB3** | 3.52347 | 0.42619 | 2.35435 | 1.37554 | 0.62894 | 1.67045 | 1.00859 |
| **ZRSR2** | 0.14225 | 0.08853 | 1.49108 | 0.62166 | 0.85049 | 0.57148 | 3.00394 |
| **ZSCAN1** | 0.00000 | 0.15808 | 0.28320 | 0.04546 | 0.00352 | 0.01659 | 0.03822 |
| **ZSCAN10** | 0.59739 | 0.00000 | 0.00000 | 0.97156 | 0.14055 | 6.33162 | 0.08645 |
| **ZSCAN12** | 1.68308 | 0.22144 | 0.47417 | 0.00000 | 0.26225 | 0.04938 | 0.21741 |
| **ZSCAN12P1** | 1.02551 | 0.00000 | 0.22708 | 2.20139 | 0.00328 | 0.00135 | 0.00000 |

| | | | | | | | |
|---|---|---|---|---|---|---|---|
| **ZSCAN16** | 1.68493 | 1.08709 | 0.62291 | 0.00000 | 0.68344 | 0.26073 | 0.53712 |
| **ZSCAN16-AS1** | 0.90970 | 0.63247 | 1.07743 | 0.10097 | 0.19154 | 0.21827 | 1.28563 |
| **ZSCAN18** | 0.00000 | 0.29493 | 1.53734 | 0.50472 | 1.33269 | 0.10408 | 0.56832 |
| **ZSCAN2** | 1.48194 | 0.82262 | 1.97905 | 2.12080 | 1.58925 | 3.57472 | 0.37141 |
| **ZSCAN20** | 0.27217 | 0.41362 | 0.35454 | 0.49862 | 0.24593 | 0.31365 | 0.06664 |
| **ZSCAN21** | 2.64915 | 2.49232 | 4.13221 | 1.13614 | 1.09967 | 1.07328 | 0.00000 |
| **ZSCAN22** | 1.22435 | 2.61001 | 0.93476 | 2.14612 | 1.24816 | 0.05133 | 0.48052 |
| **ZSCAN23** | 0.17383 | 0.01189 | 0.15111 | 0.00000 | 0.01612 | 0.02057 | 0.03855 |
| **ZSCAN25** | 4.86236 | 1.31343 | 3.91176 | 1.67119 | 1.38412 | 1.97790 | 3.78569 |
| **ZSCAN26** | 0.77839 | 0.19450 | 1.88267 | 0.00000 | 0.77703 | 0.44586 | 0.00000 |
| **ZSCAN29** | 1.51229 | 0.92084 | 3.33617 | 3.40911 | 1.44493 | 1.92246 | 8.74620 |
| **ZSCAN30** | 0.29281 | 0.49686 | 0.50063 | 0.20692 | 0.35452 | 0.30278 | 0.26477 |
| **ZSCAN31** | 1.00360 | 0.29248 | 0.09105 | 0.47603 | 0.14091 | 0.15401 | 0.00000 |
| **ZSCAN32** | 4.46583 | 0.05664 | 4.36497 | 3.92150 | 4.14157 | 3.48614 | 0.18377 |
| **ZSCAN4** | 0.00000 | 0.00000 | 1.65053 | 0.08810 | 29.59116 | 0.00000 | 0.03610 |
| **ZSCAN5A** | 2.79431 | 2.30470 | 1.98143 | 1.71258 | 6.85192 | 0.49853 | 0.00000 |
| **ZSCAN5B** | 0.02483 | 0.01586 | 3.85656 | 0.04567 | 20.83562 | 0.00000 | 0.00000 |
| **ZSCAN9** | 2.02187 | 0.30673 | 3.17188 | 0.32965 | 0.78455 | 0.67929 | 0.28520 |
| **ZSWIM1** | 2.71232 | 1.87737 | 1.53771 | 2.80628 | 0.13838 | 1.97750 | 0.00000 |
| **ZSWIM3** | 0.57217 | 3.25825 | 2.94232 | 0.71231 | 1.72396 | 3.98076 | 0.42452 |
| **ZSWIM4** | 0.00000 | 0.00000 | 0.15044 | 1.02814 | 0.40861 | 0.06317 | 0.78544 |
| **ZSWIM5** | 0.04467 | 0.33832 | 0.24785 | 0.01189 | 0.23145 | 0.33890 | 0.16707 |
| **ZSWIM6** | 0.14371 | 0.00000 | 0.03819 | 0.17039 | 0.26923 | 0.60747 | 0.28825 |
| **ZSWIM7** | 12.29194 | 11.36982 | 5.74962 | 7.44525 | 2.04641 | 9.26010 | 5.43602 |
| **ZSWIM8** | 1.16729 | 1.91618 | 0.60672 | 6.11687 | 0.54931 | 1.00496 | 6.62959 |
| **ZSWIM8-AS1** | 0.12692 | 0.07497 | 0.21621 | 0.11189 | 0.09363 | 0.15923 | 0.40133 |
| **ZUFSP** | 9.08282 | 0.65305 | 7.48722 | 3.47183 | 1.64661 | 2.35426 | 0.00000 |
| **ZW10** | 12.46588 | 3.98198 | 4.55285 | 6.80654 | 3.27319 | 4.90239 | 0.00000 |
| **ZWILCH** | 26.38108 | 0.38195 | 12.36134 | 12.19934 | 1.98336 | 8.89355 | 3.11446 |
| **ZWINT** | 46.69718 | 20.95559 | 21.12738 | 30.44427 | 10.67140 | 13.56871 | 0.00000 |
| **ZXDA** | 0.06147 | 0.05465 | 0.01594 | 0.00000 | 0.00686 | 0.00000 | 0.12490 |
| **ZXDB** | 0.37798 | 0.11531 | 0.00303 | 0.03128 | 0.00355 | 0.45703 | 1.28123 |
| **ZXDC** | 2.64611 | 3.17306 | 1.28513 | 4.28814 | 0.19392 | 3.09098 | 8.82355 |
| **ZYG11A** | 4.29219 | 4.10293 | 3.69672 | 1.84070 | 2.48050 | 13.89562 | 2.85946 |
| **ZYG11B** | 2.25965 | 0.00000 | 0.64890 | 1.87116 | 0.39322 | 1.56133 | 7.64767 |

| | | | | | | | |
|---|---|---|---|---|---|---|---|
| **ZYX** | 2.60588 | 17.04812 | 2.90649 | 8.37723 | 2.21060 | 11.62496 | 20.51014 |
| **ZZEF1** | 2.79515 | 4.64940 | 1.58622 | 4.55599 | 1.16368 | 4.05005 | 9.15947 |
| **ZZZ3** | 12.99931 | 1.30554 | 4.75710 | 5.43833 | 7.97458 | 4.12900 | 9.52713 |
